# Rapid effects of valproic acid on the fetal brain transcriptome: Implications for brain development and autism

**DOI:** 10.1101/2023.05.01.538959

**Authors:** Susan G. Dorsey, Evelina Mocci, Malcolm V. Lane, Bruce K. Krueger

## Abstract

There is an increased incidence of autism among the children of women who take the anti-epileptic, mood stabilizing drug, valproic acid (VPA) during pregnancy; moreover, exposure to VPA *in utero* causes autistic-like symptoms in rodents and non-human primates. Analysis of RNA-seq data obtained from E12.5 fetal mouse brains 3 hours after VPA administration revealed that VPA significantly increased or decreased the expression of approximately 7,300 genes. No significant sex differences in VPA-induced gene expression were observed. Expression of genes associated with neurodevelopmental disorders (NDDs) such as autism as well as neurogenesis, axon growth and synaptogenesis, GABAergic, glutaminergic and dopaminergic synaptic transmission, perineuronal nets, and circadian rhythms was dysregulated by VPA. Moreover, expression of 399 autism risk genes was significantly altered by VPA as was expression of 252 genes that have been reported to play fundamental roles in the development of the nervous system but are not otherwise linked to autism. The goal of this study was to identify mouse genes that are: **(a) significantly up- or down-regulated by VPA in the fetal brain** and **(b) known to be associated with autism and/or to play a role in embryonic neurodevelopmental processes, perturbation of which has the potential to alter brain connectivity in the postnatal and adult brain.** The set of genes meeting these criteria provides potential targets for future hypothesis-driven approaches to elucidating the proximal underlying causes of defective brain connectivity in NDDs such as autism.

## INTRODUCTION

Autism is a neurodevelopmental disorder (NDD) characterized by social interaction deficits including language, and repetitive, stereotyped behavior with restricted interests (American Psychiatric Association, 2022). Autistic individuals may also display an increased incidence of anxiety, intellectual disability and seizures. The reported incidence of autism has increased dramatically over the last two decades (Zeidan et al., 2022) with some estimates now as high as 1:44 (Maenner et al., 2021). The severity of autism varies widely, ranging from high-functioning with minimal disability, to severely afflicted, where aggressive, self-destructive repetitive behaviors pose a threat to the safety of the patient, caregivers, and family. The term “autism spectrum disorders” (ASDs) reflects this symptomatic variability. The neurological mechanisms underlying ASDs are not understood nor is it known whether autism is a single disorder or multiple disorders sharing common core features.

Twin studies have revealed that 60 to 88% of autism cases are inherited (Hallmayer et al., 2011; Rosenberg et al., 2009), however, many cases have been linked to *in utero* exposure to environmental factors such as pharmaceuticals, air pollution, chemical insecticides, and maternal infection (Landrigan, 2010; Rosenberg et al., 2009; Karimi et al., 2017).

The diagnosis of autism typically is made at around two years of age when the child fails to meet normal milestones for social development and language. However, it has become increasingly evident that autism arises before birth, with infants failing to show normal attention to faces (Macari et al., 2021). Epidemiological studies indicate that the onset of autism occurs during fetal development; one such study found that folate supplementation for pregnant women reduced the incidence of autism, but only when administered between two weeks before and four weeks after conception (Surén et al., 2013). Exposure to a variety of environmental factors, notably the anti-epileptic, mood stabilizing drug, valproic acid (VPA; Depakote®) (Christensen et al., 2013) and the organophosphate insecticide, chlorpyrifos (Burke et al., 2017), increases the incidence of autism and other NDDs in the offspring of women who are exposed during pregnancy. Another environmental cause of autism is maternal immune activation (MIA) in which pregnant women who have systemic bacterial or viral infections with a hyper-immune response have an increased incidence of autism in their children (Bilbo et al., 2018; Careaga et al., 2017; Zawadzka et al., 2021).

Studying the etiology of autism in humans is limited to epidemiological approaches implicating genetic variants (single nucleotide polymorphisms, copy number variants, and single-gene syndromic mutations). The Simons Foundation Autism Research Initiative (SFARI) has compiled a list of 1,115 human genes (https://gene.sfari.org/database/human-gene/) for which there is evidence for association with autism based on genome-wide association studies (GWAS), referred to as the “SFARI List” below.

Hypothesis-driven experiments to determine cause and effect need to be done in animal models, which display a range of behaviors that are remarkably similar to the core symptoms of autism (Silverman et al., 2010; Lan et al., 2017; Nicolini and Fahnestock, 2018). These autistic-like behaviors have been reported in transgenic animals lacking ASD-linked genes such as *Cntnap2* and *Shank2* (Peñagarikano and Geschwind, 2012; Eltokhi et al., 2018), supporting the conclusion that pathogenic mutations in these genes are causative for syndromic autism and leading to the widespread use of these transgenic mice as animal models (Bey and Jiang, 2014; Kazdoba et al., 2016). Systemic administration of VPA (Nicolini and Fahnestock, 2018), chlorpyrifos (Lan et al., 2017) or induction of MIA (Careaga et al., 2017) in genetically normal pregnant rodents also leads to increased autistic-like behaviors in their offspring. Unlike transgenic mouse models, these environmental toxicity models of autism provide the opportunity to control the timing of exposure of the fetus to the toxic stressor, as will be discussed below. Gene expression analyses of the brains of mice in the VPA and MIA models have been conducted using RNA microarray analysis or next-generation RNA sequencing (RNA-seq). In the majority of those animal studies, the fetus was exposed to the drug during mid-gestation, but RNA expression was analyzed in postnatal animals, usually following behavioral testing (e.g., see Qian et al., 2018; Zhao et al., 2019; Matsuo et al., 2022; Zhang et al., 2018; Guerra et al., 2023). This approach has the potential to explain behavior in terms of contemporaneous gene expression but does not inform as to the initial molecular events occurring at the time the environmental stressor is present.

There are two competing, but not mutually exclusive, theories about the role of differential gene expression in autism. In one, abnormal gene expression in the child or adult at the time of behavioral testing, is responsible for autistic-like behavior. The second posits that abnormal gene expression, due to genomic variants or *in utero* environmental stressors, interferes with one or more critical steps in the “program” controlling fetal brain development. Such errors could carry forward throughout life leading to a brain with subtle anatomical or connectivity defects that underlie the abnormal development of an autistic brain. This has been referred to as a developmental “signature” (Ben-Ari, 2008). The present study focused on the second hypothesis by examining altered gene expression in fetal brains when the environmental stressor (VPA) is still present; these genes need not be continuously dysregulated throughout life. Consequently, genes associated with critical steps in early brain development such as neurogenesis, neuron fate specification, axon and dendrite growth, and synaptogenesis are of particular interest.

The incidence of autism is about 4-times higher in males than in females (Ferri et al., 2018). Sex differences in neural function are generally considered to be mediated by sex hormones acting from late prenatal brain development through to the adult. Since the time of VPA administration in the present study (E12.5) is just prior to the maturation of gonads and production of sex hormones (Ross and Capel, 2005), any sex differences in gene expression observed likely would be due to sexually-dimorphic gene expression rather than to the effects of sex hormones.

The complexity and variability of the behavioral symptoms of autism together with identification of over 1,100 ASD-associated gene variants make a mechanistic understanding of the causes of the disorder a daunting task. One approach to this problem is to compare the results of multiple studies using GWAS results from human patients and gene expression results from animal models of autism, looking for common differentially expressed genes. The subset of genes in common from both types of studies can provide a short(er) list of candidate genes that could be subjected to future hypothesis-driven studies to establish causal relationships between gene dysregulation and ASD symptoms.

The goal of the present study was to address this question using the VPA mouse model in which pregnant mice receive a single i.p. injection of VPA at gestational day 12.5 (E12.5). Pharmacokinetic studies have demonstrated that injected VPA dissipates within 3−5 hours due to metabolism of the drug by the maternal liver (Nau and Löscher, 1982; Nau, 1986a); VPA levels in the fetal brain follow a similar trajectory (Nau, 1986b; Dickinson et al., 1980). Fetal brains were processed for RNA-seq three hours after VPA administration to the pregnant dam; male and female fetuses were analyzed separately. We propose that VPA-induced molecular events occurring on E12.5 are both necessary and sufficient for the autistic-like behaviors in VPA-treated animals assessed 5−6 weeks later. Analysis of these data revealed that VPA induced a significant increase or decrease in the expression of approximately 7,300 genes, of which 399 are among the 1,115 genes on the SFARI List and at least 252 additional genes are known to play fundamental roles in the development and function of the nervous system.

## METHODS

Timed-pregnant C57BL6 mice were generated by the University of Maryland School of Medicine Division of Veterinary Resources. All animal procedures were approved by the Institutional Animal Care and Use Committee. VPA was obtained from Sigma.

One male was paired overnight with two females; the day of separation was designated E0.5. On E12.5 pregnant females received i.p injections of VPA (400 mg/kg) in sterile saline or saline alone. 7 pregnant females received VPA and 7 received saline. Three hours after VPA administration, pregnant females were euthanized by cervical dislocation and decapitation. Fetuses were transferred to ice-cold saline, decapitated and the entire brain was removed to Trizol and disrupted for 60 sec with a Bead Beater using 0.2 mm beads. The 105 individual fetal brain samples were frozen on dry ice and stored at −80°C prior to RNA extraction and quality control (RIN = 10 for all samples).

Sex was determined by analyzing *Sry* RNA derived from the torso of each fetus by PCR. Males were identified by the presence of both *Sry* and *Gapdh* RNA determined; *Gapdh* but not *Sry* RNA was expressed in females (Konopko et al., 2017). One male and one female brain from each of the 7 pregnancies was processed for RNA-seq by the University of Maryland Institute for Genome Sciences.

Libraries were prepared from 25ng of RNA using the NEB Ultra II Directional RNA kit. Samples were sequenced on an Illumina NovaSeq 6000 with a 150bp paired-end read configuration. The quality of sequences was evaluated by using FastQC (Wingett and Andrews, 2018). The alignment was performed using HiSat (version HISAT2-2.0.4) (Kim et al., 2019) and *Mus musculus* GRCm38 as reference genome and annotation (version 102). Aligned bam files were used to determine number of reads by gene using HTSeq (Anders et al., 2015). On average, we sequenced 187,000,000 reads. 96.6% of them properly mapped the reference: 87% mapped exons, 2.4% mapped introns and the remaining 10.6% mapped intergenic regions.

Differential gene expression between mice treated with VPA and saline controls was conducted using DESeq2 (Love et al., 2014), which models gene counts using the negative binomial distribution. P-values were corrected for false discovery rate (FDR) using the Benjamini-Hochberg procedure; p_FDR_ ≤ 0.025 and log_2_-fold change ≥0.32 was used as the criterion for significance. Differentially expressed genes due to VPA treatment were tested for enrichment in SFARI autism risk genes (https://gene.sfari.org/database/human-gene/) using Fisher’s Exact Test.

Significance of sex-differences in FC was analyzed by 2-way ANOVA using the Limma-Voom tool (Law et al., 2014).

## RESULTS

### RNA-seq

19,721 individual genes were analyzed in fetal mouse brain by RNA-seq. Male and female brains were analyzed separately. The raw data showing base counts (mean of male and female counts) for each gene in each of seven independent fetal brain samples of each sex, each from a different pregnancy, are shown in Supplemental Table 1 (S1), i.e., n = 7 fetal brains from 7 different pregnancies for each condition. Only genes with base counts ≥100 were analyzed further. Genes that increased or decreased in response to VPA by < 1.25-fold and > −1.25-fold and those with adjusted p-values (p_FDR_) > 0.025 were also filtered out; a gene was included if VPA increased or decreased its expression by ≤ 1.25-fold and ≥ −1.25-fold in only one sex. Shown in Supplemental Table 4 (S2) are the 6516 genes significantly affected by VPA (p_FDR_ ≤ 0.025) in both sexes, 498 in females only and 280 in males only (listed in three separate sections in Table S2). These apparent sex differences were due to variability in the replicates of one of the sexes (i.e., p_FDR_ > 0.025).

Throughout this report, a positive fold-change (FC) indicates that VPA increased gene expression; a negative FC indicates that VPA decreased expression such that the FC is the ratio of control to VPA gene expression. For example, FC = −3.0 indicates that V/C = 0.33, a 67% reduction by VPA.

### Curating the mouse genes dysregulated by VPA in the fetal brain

The 7294 genes that were significantly altered by VPA (Table S2) were further analyzed in two ways. First, they were merged with the genes on the SFARI List identifying 399 common genes (Table S3). It should be noted that inclusion on the SFARI List is based on GWAS; in many cases, a mechanistic role in neither brain development nor the etiology of autism is known.

Second, we identified 252 genes (Table S4) significantly altered by VPA not on the list of SFARI risk genes but having a documented role in brain development including neurogenesis, synaptogenesis, and axon or dendrite growth. As will be discussed below, although these latter genes have not been associated with autism in GWAS, it is plausible that changes (either positive or negative) in the expression of their gene products in the fetal brain might adversely affect the trajectory of brain development leaving a permanent “signature” (Ben-Ari, 2008) on brain structure and connectivity leading to autistic-like behavior in the adult.

### No significant sexually-dimorphic effects of VPA

Figure 1 is a plot of the log_2_ fold-change (FC) of males versus females for the 651 genes in Tables S3 and S4. The gap in the data reflects the ±1.25- fold FC cut-off. Deviations from the regression line (r^2^=0.973) indicate possible sex differences; however, in no case was the effect of VPA found to be significantly different between males and females. 20 genes (magenta) were upregulated by > 5-fold or downregulated > 80% by VPA (average of males and females

**Figure 1.**
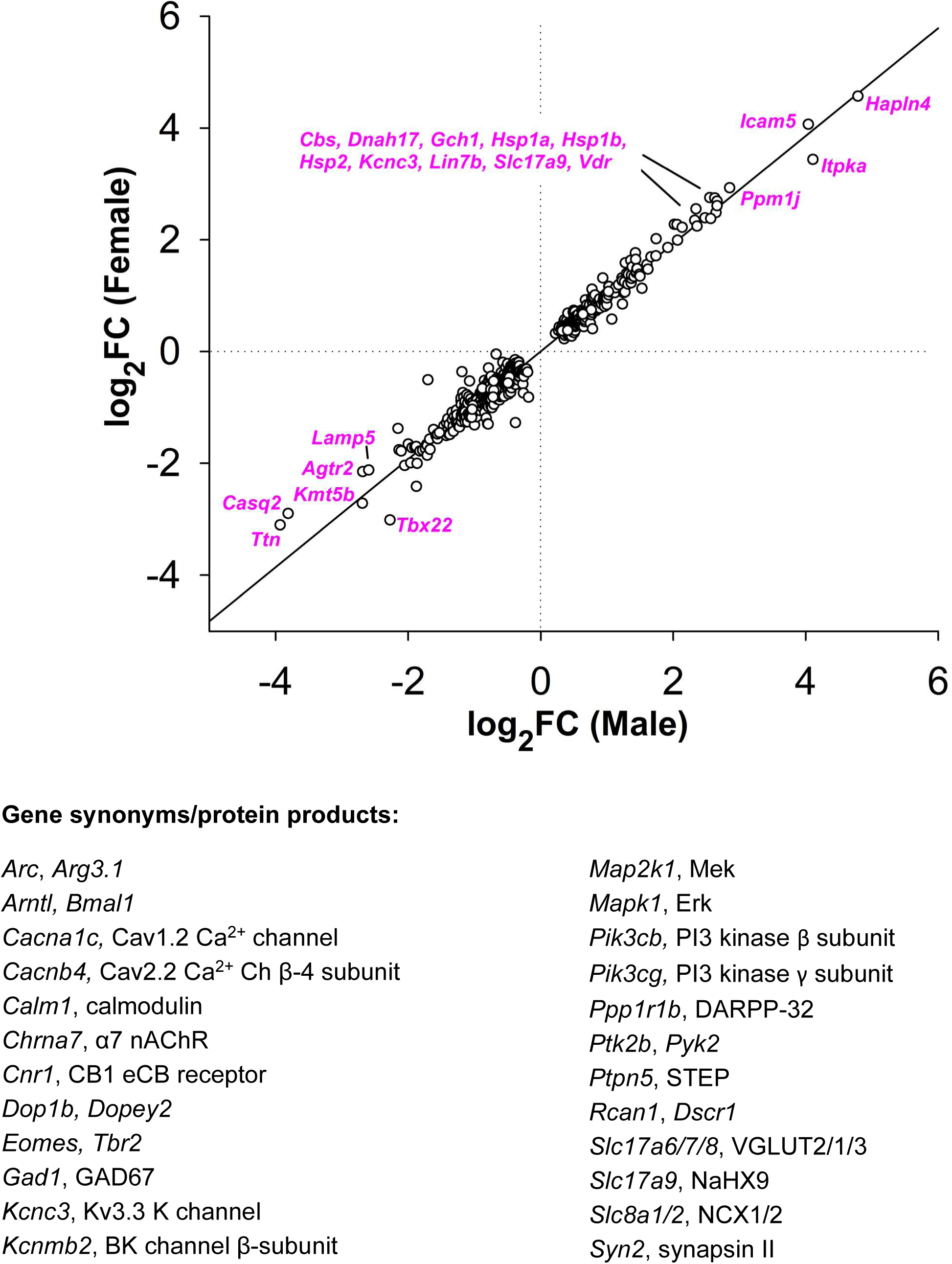

## DISCUSSION

The overarching hypothesis for this research is that NDDs involving intellectual disability (ID) such as ASDs are the result of abnormal connections among multiple brain regions. Normal connectivity is established beginning during fetal brain development as the various brain regions become populated with specific classes of neurons that subsequently connect with other neurons to establish the complex neuronal networks that underlie cognition and behavior. Although these networks are refined during late pre- and postnatal brain development, it is likely that certain early prenatal developmental epochs are particularly vulnerable to alterations in the neurodevelopmental “program” (Ben-Ari, 2008). Although it is by no means clear that the fetal rodent brain accurately recapitulates the autistic human brain during prenatal development, the VPA model enables experimental control of the timing of exposure to an environmental factor that causes abnormal behavior resembling autism. In the present experimental paradigm, VPA is administered to the pregnant dam at E12.5, a time when multiple early neurodevelopmental events critical for brain organization and connectivity are occurring. These include proliferation of neural progenitors (NPs), determination of the neuronal fate of these NPs, migration of newborn neurons away from the proliferative ventricular zones, differentiation of NPs to establish a mature neuronal phenotype, extension and branching of axons and dendrites and establishment of synaptic connections.

Approximately 7,300 genes were significantly up- or downregulated in the E12.5 fetal brain 3 hr after exposure to VPA (Table S2). 399 of those genes have been linked to autism in GWAS (Table S3), although in many cases the mechanism of action is unknown. In addition, at least 252 of the genes dysregulated by VPA but not on the SFARI Autism Risk list (Table S4), are known to play a mechanistic role in mediating one or more of the neurodevelopmental processes discussed above. Therefore, the goal of this study was to identify genes that are:

a. **significantly up- or down-regulated by VPA in the fetal mouse brain (Table S2) AND**
b. **known to be linked to autism in GWAS (Table S3) or to play a role in embryonic neurodevelopmental processes, perturbation of which has the potential to alter brain connectivity in the postnatal and adult brain (Table S4).**

The set of genes meeting these criteria would provide potential targets for future hypothesis-driven approaches to understanding the underlying proximal causes of defective brain connectivity in NDDs such as autism.

It was not practical to vet each of the nearly 7,300 VPA-dysregulated genes for a potential role in abnormal brain development. However, as described below, the 252 genes in Table S4 and some of the genes in Table S3 have been reported to play substantial roles in specific aspects of fetal brain development. Consequently, it is plausible that the up- or downregulation of one or more of those genes could contribute to the abnormal connectivity in subjects with NDDs. The validity of this potential contribution could be tested in animals by determining the effect of manipulating the expression of individual (or combinations of) genes in the fetal brain on subsequent behavior.

### A common subset of genes in GWAS and VPA-fetal mouse brain studies

Several GWAS have identified genes linked to autism (see SFARI Autism list). Here we tabulated the findings of four GWAS studies (Satterstrom et al., 2020; Yuen et al., 2017; Stessman et al., 2017; Ruzzo et al., 2019) and one report categorizing autism risk genes potentially involved in neurogenesis (Garcia-Forn et al., 2020). As shown in Table 1, many of the genes identified in these reports are also dysregulated by VPA in the fetal mouse brain (Tables S3 and S4). Genes discussed in the respective papers are listed in each column in Table 1; genes that are also dysregulated by VPA in the present study are shown in **bold**. Five VPA-regulated genes in Tables S3 and S4 that also appear in all five of the published reports are in **red**. (*Arid1b*, *Dyrk1a*, *Pogz*, *Pten*, *Tbr1*). Six genes in Tables S3 and S4 that also appear in four of the five other studies are in **blue**. (*Adnp*, *Ash1l*, *Chd2*, *Kmt5b*, *Tcf7l2*, *Wac*). There were four genes (yellow) identified in all five previously published studies that were not affected by VPA in the fetal mouse brain. *Chd2*, which is listed in three of the four published studies, and *Chd3*, (50% reduced by VPA), are both SFARI genes and have been reported to have similar functions to *Chd8*. Of the 11 **red** and **blue** genes in Table 1, three (*Adnp*, *Dyrk1a*, *Pten*) appear in Table 2 as VPA-regulated, autism-associated genes involved with the structural stability of neurons (Lin et al., 2016).

**Table 1.** (previous page). Autism-risk genes identified in five published studies (Satterstrom, et al., 2020; Garcia-Forn, et al., 2020; Yuen et. al., 2017; Stessman et al., 2017; Ruzzo et al., 2019). Genes dysregulated by VPA in fetal brain in the present study (Table S3 and 2) are indicated in ***bold***. The percentage of VPA-regulated genes is shown for each study. ***Red***: genes identified in all five published papers and **were** affected by VPA the present study; ***blue***: genes identified in four of the five published papers and **were** affected by VPA the present study; ***yellow***: autism-associated genes identified in all five published papers but **were not** affected by VPA in the present study.

| Satterstrom | Garcia-Forn | Yuen | Stessman | Ruzzo |
| --- | --- | --- | --- | --- |
| **41%** | **50%** | **37%** | **41%** | **44%** |
|  |  | ADCY3 | ADCY5 |  |
| **ADNP** |  | **ADNP** | **ADNP** | **ADNP** |
|  |  | AFF2 |  |  |
|  |  | AGAP2 |  | **AKAP9** |
| **ANK2** | **ANK2** | **ANK2** | **ANK2** | **ANK2** |
| ANKRD11 |  | ANKRD11 | ANKRD11 |  |
| AP2S1 |  |  |  |  |
| **ARID1B** | **ARID1B** | **ARID1B** | **ARID1B** | **ARID1B** |
| **ASH1L** |  | **ASH1L** | **ASH1L** | **ASH1L** |
| **ASXL3** |  | **ASXL3** | **ASXL3** |  |
| BABRB3 |  |  |  |  |
| **BCL11A** | **BCL11A** |  |  | **BCL11A** |
| BRIN2B |  |  |  | BTRC |
| CACNA1E |  |  |  | C16orf13 |
| CELF4 |  | CACNA2D3 | CACNA2D3 | CACNA2D3 |
|  |  |  | **CASK** | CAPN12 |
|  |  |  | **CDC42BPB** | CCSER1 |
| **CHD2** |  | **CHD2** | **CHD2** | **CHD2** |
| **CHD8** | **CHD8** | **CHD8** | **CHD8** | **CHD8** |
|  |  | CIC |  | **CMPK2** |
|  |  | CLASP1 | COL4A3BP |  |
|  |  | CNOT3 |  |  |
| **CREBBP** |  |  |  |  |
| CTNNB1 | CTNNB1 |  | CTNNB1 | CTTNBP2 |
|  |  | CUL3 | CUL3 | CUL3 |
| **DEAF1** | **DDX3X** | **DIP2C** | **DDX3X** | **DDX3X** |
| **DNMT3A** |  | **DNMT3A** | DIP2A | **DNMT3A** |
| DPYSL2 |  |  | DLG4 |  |
|  |  |  | DLGAP1 |  |
| DSCAM |  | DSCAM | **DOCK8** | DSCAM |
|  |  | DYNC1H1 | DSCAM |  |
| **DYRK1A** | **DYRK1A** | **DYRK1A** | **DYRK1A** | **DYRK1A** |
| EIF3G | FMR1 | FAM47A |  | **ERBIN** |
|  |  |  |  | ETFB |
|  |  |  |  | **FAM98C** |
| **FOXP1** | **FOXP1** | **FOXP1** | **FOXP1** | **FOXP1** |
| **FOXP2** |  |  |  | GABRB3 |
| GFAP |  |  |  |  |
| GIGYF1 |  | GIGYF1 | GIGYF2 | GIGYF1 |
| GNAI1 |  | GRIN2B | GRIN2B | GRIA1 |
| IRF2BPL |  | KDM6A | HIVEP3 | GRIN2B |
| **KCNQ3** |  |  | ILF2 | ILF2 |
| KDM5B |  |  | ITPR1 | **INTS6** |
| **KDM6B** |  | **KDM6B** |  | **KDM6B** |
| KIAA0232 |  | KIAA2022 | KATNAL2 | KATNAL2 |
|  |  | KMT2A | KMT2A | KDM5B |
| KMT2C |  | KMT2C | KMT2C | KMT2C |
|  |  |  | **KMT2E** | **KMT2E** |
| **KMT5B** |  | **KMT5B** | **KMT5B** | **KMT5B** |
| **LDB1** |  |  | LAMC3 | MFRP |
| **MAP1A** |  | **MECP2** | **MECP2** | MLANA |
| MBD5 |  |  |  |  |
| MED13L |  | MED13 | MED13L |  |

\*
\*
\*

| Satterstrom | Garcia-Forn | Yuen | Stessman | Ruzzo |
| --- | --- | --- | --- | --- |
| MKX |  | **MYO5A** |  | **MYO5A** |
| MYT1L |  | MYT1L | MYT1L | MYT1L |
| NRXN1 |  | NAA15 | NAA15 | NCKAP1 |
| **NSD1** |  | NLGN3 | NCKAP1 | NRXN1 |
| PAX5 |  | NLGN4X | NRXN1 | P2RX5 |
| PHF12 |  | PAX5 | PARD3B | PCM1 |
| PHF21A |  | PCDH11X | PAX5 |  |
|  |  | **PHF2** | **PHF2** | **PHF2** |
|  |  | PHF3 | **PHIP** |  |
|  |  |  | **PLXNB1** |  |
| **POGZ** | **POGZ** | **POGZ** | **POGZ** | **POGZ** |
|  |  |  | PPM1D | PRKAR1B |
| **PPP2R5D** |  |  | **PPP2R5D** |  |
| **PRR12** |  |  |  |  |
| **PTEN** | **PTEN** | **PTEN** | **PTEN** | **PTEN** |
| **RAI1** | SCN2A | SCN2A | **PTK7** | **RANBP17** |
| RFX3 |  |  | **PTPN11** | RAPGEF4 |
| RORB |  |  | RELN | SCN2A |
| SATB1 |  |  | RIMS1 |  |
|  |  |  | SCN2A |  |
| SCN2A |  |  | **SETBP1** |  |
|  |  |  | SETD2 |  |
| SETD5 |  |  | SETD5 | SETD5 |
| **SHANK2** |  | **SHANK2** |  | **SHANK2** |
| SHANK3 | SHANK3 | SHANK3 |  | SHANK3 |
| **SIN3A** |  |  | SLC6A1 | SLC6A1 |
| SKI |  | SLC6A1 | SMC3 | SMURF1 |
| SLC6A1 |  |  |  |  |
| **SMARCC2** |  | **SMARCC2** |  |  |
| SPAST |  | SPAST | SPAST | SPAST |
| SRPRA |  |  |  |  |
|  |  | SRSF11 | **SRCAP** |  |
|  |  |  | SRGAP3 |  |
| STXBP1 |  |  | STXBP1 |  |
| **SYNGAP1** | **SYNGAP1** | **SYNGAP1** | **SYNGAP1** | **SYNGAP1** |
|  |  |  | TANC2 |  |
| **TBL1XR1** |  | **TAF6** | **TBL1XR1** |  |
| **TBR1** | **TBR1** | **TBR1** | **TBR1** | **TBR1** |
| TCF20 |  |  |  |  |
| **TCF4** | **TCF4** |  | **TCF4** |  |
| **TCF7L2** |  | **TCF7L2** | **TCF7L2** | **TCF7L2** |
| **TLK2** |  |  |  | **TMEM39B** |
| TM9SF4 |  | **TNRC6B** | **TNRC6B** | **TNRC6B** |
|  |  | UBN2 | **TRIO** |  |
| TRIP12 |  |  | TRIP12 | TRIP12 |
| **VEZF1** |  | UPF3B | UNC80 | **TSPAN4** |
|  |  |  |  | UIMC1 |
|  |  |  |  | **USP45** |
| **WAC** |  | **WAC** | **WAC** | **WAC** |
| ZYMND8 |  | **WDFY3** | **WDFY3** | **WDFY3** |
|  |  |  | WDR45 | ZNF559 |
|  |  |  | **ZC3H4** |  |
|  |  |  | ZNF292 |  |

\*Not affected by VPA
\*

**Table 2.** Genes linked to autism in GWAS that have also been reported to regulate “neurite outgrowth”, “synapse/spine formation” or “synaptic plasticity” (Lin et al., 2016). Genes in red ***bold*** are also up- or down-regulated by VPA in the fetal brain (Table S3). Lin et al. (2016) did not distinguish among the multiple known isoforms of cadherins (*Cdh’*s); *Cdh11* is downregulated 60% by VPA in the fetal brain (c.f., Table S3).

| Neurites | Synapse | Plasticity |
| --- | --- | --- |
| *CTTNBP2* | **ADNP** | **CDH(S)** |
| **CDH(S)** | **CDH(S)** | **CDKL5** |
| **CDKL5** | **CDKL5** | *DLGAP2* |
| *CNTN* | *CNTN* | *FMR1* |
| **CNTNAP2** | **CNTNAP2** | *GRIK2* |
| **DYRK1A** | *CTTNBP2* | **GRIK4** |
| *ELMO1* | *DLGAP2* | *GRIN2B* |
| *FMR1* | **DYRK1A** | **MECP2** |
| *GRIN2B* | *ELMO1* | *NLGN* |
| **MECP2** | *FMR1* | *NRXN* |
| *MYO16* | *GRIK2* | **PTEN** |
| *NCAM2* | **GRIK4** | *SHANK3* |
| *NLGN* | *GRIN2B* | **STXBP5** |
| *NRXN* | **MECP2** | *SYNGAP1* |
| *NTRK2* | *NLGN* | *UBE3A* |
| *PCDH* | *NRXN* |  |
| **PTEN** | *NTRK2* |  |
| *SPRICKLE1* | *PCDH* |  |
| **STXBP5** | **PTEN** |  |
| *TSC1* | *SHANK3* |  |
| **TSC2** | *SYNGAP1* |  |
| *UBE3A* | *TSC1* |  |
|  | **TSC2** |  |
|  | *UBE3A* |  |

### VPA dysregulation of high-confidence (hc)ASD genes in mid-fetal layer 5/6 cortical projection neurons

(Willsey et al., 2013) identified 9 hcASD genes that are expressed in layer 5/6 projection neurons in the fetal human brain. Three of these genes (*Dyrk1a*, *Pogz*, *Tbr1*) are downregulated in fetal mouse brain following VPA administration to the pregnant dam (Table S3). Willsey et al. (2013) used the nine hcASD genes as seed genes for co-expression network analysis which revealed 10 probable ASD genes, of which two (*Bcl11a*, *Nfia*) were down-regulated and one (*Aph1a*) was up-regulated by VPA (Table S4). All of these genes are on the SFARI autism risk gene list (Table S3) and the three seed genes were identified in all 5 reports analyzed in this study (Table 1). These findings raise the possibility that dysregulation of one or more of these genes in developing layer 5/6 projection neurons in cortical plate of the fetal brain contributes to permanent connectivity defects underlying autistic-like behaviors.

### Autism-associated genes and structural stability of neurons

Lin et al. (2016) tabulated genes linked to autism in GWAS that have also been reported to be involved in the structural stability of neurons. These genes were further sorted into three categories, viz., “neurite outgrowth”, “spine/synapse formation” and “synaptic plasticity”. 61 genes were assigned to these three categories although some genes appeared in two or all three categories, resulting in 29 different genes among the three categories (Table 2). Shown in red ***bold*** are 10 of these genes (35%) that were significantly up- or downregulated by VPA (i.e., in Table S3). As discussed in Lin et al. (2016), dysregulation of one or more of these genes (which could be due to gene variants or fetal exposure to VPA) could alter the structure and function of synapses throughout brain development.

Examination of Tables 1 and 2 revealed that *Pten* and *Dyrk1a* are common to the Lin et al., (2016) study and to all five of the GWAS. Consequently, these two genes may be good candidates for investigating the molecular basis for the proximal causes of autism.

### Biological roles of genes dysregulated by VPA

Table 3 (next page) shows the biological function of genes that were significantly dysregulated by VPA in the E12.5 fetal brain. One or more developmental errors caused by these changes could induce a “signature” of abnormal brain circuitry resulting in abnormal behavior in juveniles and adults. In this discussion, the change in gene expression induced by VPA at E12.5 (expressed as fold-change) is shown in parentheses (mean of males and females). Following Table 3, the rationale for inclusion of the genes in the 18 categories is provided together with references to the literature.

**Table 3.**
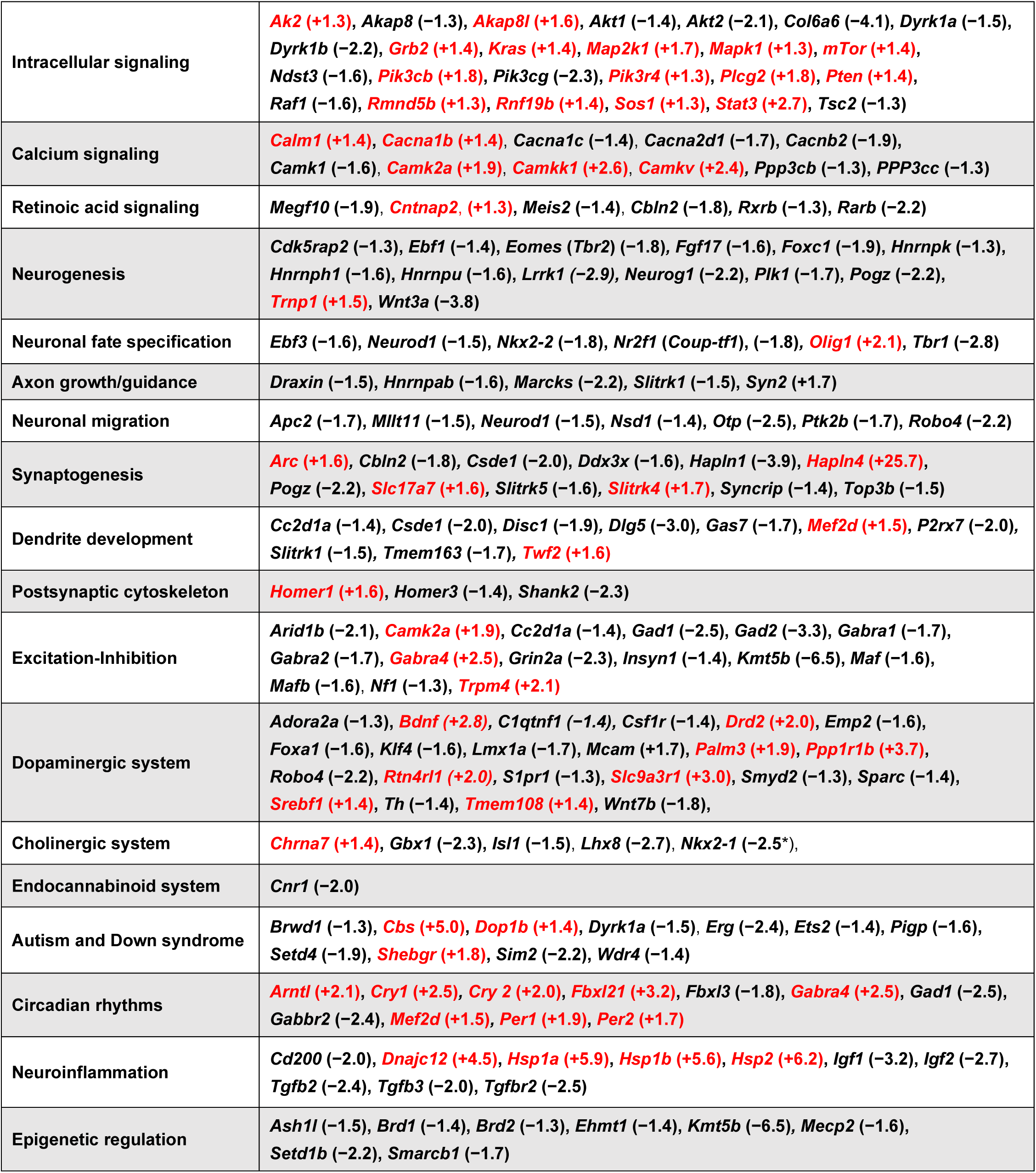
The biological roles of genes that are dysregulated by VPA in the fetal brain and are associated with autism or prenatal neurodevelopmental processes (from Tables S3 and S4). The number in parentheses shows the FC induced by VPA (mean of males and females); **red** text indicates that VPA increased gene expression. Note that some genes are included in more than one category. The basis for this categorization, with references, is given below.

### Intracellular signaling

As summarized in Figure 2, the genes encoding 10 members of the canonical signaling pathways downstream from receptor tyrosine kinase receptors are significantly up-regulated by VPA in the fetal brain (green); four are down-regulated (red). ***Pten* (+1.4)** (Phosphatase and tensin homolog), a negative regulator of the PI3K-AKT-mTor signaling pathway, has a particularly strong association with autism, appearing in all GWAS (Tables 1, 3) and associated with the structural stability of neurons (Table 2) (Lin et al., 2016).

**Figure 2.**
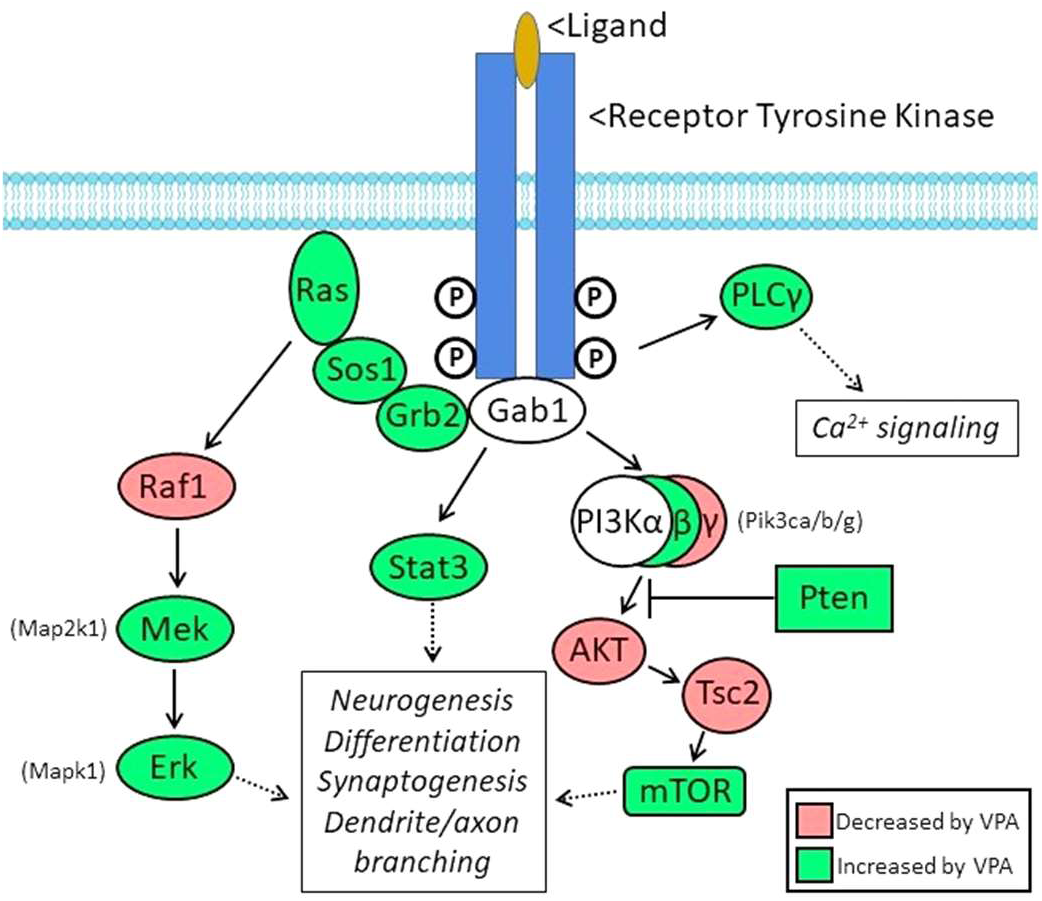

Twelve additional genes associated with intracellular signaling are included in Table 3. Thus, abnormal signaling via these pathways is a potential mechanism by which VPA could interfere with the establishment of normal brain connectivity. Meta-analysis of GWAS and copy number variant studies of autism-related genes revealed that three signaling networks, regulating steroidogenesis, neurite outgrowth and excitatory synaptic function, were enriched (Poelmans et al., 2013). A-kinase anchoring proteins (Akaps) functionally integrate signaling cascades within and among these networks. ***Akap8* (−1.3)** and ***Akap8l* (+1.6)** was decreased and increased, respectively, in fetal mouse brains exposed to VPA. ***Dyrk1b* (−2.2)** (which encodes dual specificity tyrosine phosphorylation regulated kinase 1B) is an early mediator of double-stranded DNA break repair (Dong et al., 2020) and regulates hedgehog signaling through activation of the mTor/AKT pathway (Singh et al., 2017; Upadhyay et al., 2021; Singh and Lauth, 2017).

*Dyrk1b* has been linked to metabolic syndrome and autism (Orenstein et al., 2022). The 55% reduction in *Dyrk1b* expression induced by VPA could contribute to autistic-like behavior by dysregulating hedgehog or mTor/AKT signaling. ***Ppp1r1b* (+3.7)**, encodes the dopamine- and cAMP-regulated neuronal phosphoprotein (DARPP-32). DARPP-32 amplifies and/or mediates many actions of cyclic AMP-dependent protein kinase at the plasma membrane and in the cytoplasm, with a broad spectrum of potential targets and functions (Yger and Girault, 2011).

### Calcium signaling

Ca^2+^ is an intracellular messenger in a wide variety of cellular responses; calmodulin, encoded by ***Calm1* (+1.4)**, acts as the Ca^2+^-sensor in many responses including Ca^2+^-calmodulin dependent kinases [***Camk1* (−1.6)**, ***Camk2a* (+1.9)**, ***Camkk1* (+2.6)**, ***Camkv* (+2.4)**], calcineurin, [***Ppp3cb* (−1.3)** and ***PPP3cc* (−1.3)**]. Four genes encoding subunits of voltage-gated Ca^2+^ channels on the SFARI list (Liao and Li, 2020; Breitenkamp et al., 2014) (Table S3) were dysregulated by VPA in the fetal brain: ***Cacna1b* (+1.4)**, ***Cacna1c* (−1.4)**, ***Cacna2d1* (−1.7)** and ***Cacnb2* (−1.9)**. Like *Cacnb2*, ***Cacnb4* (−1.5)** is an auxiliary subunit, however, it is not linked to autism by GWAS. *Cacna1c* is associated with Timothy syndrome, an ASD characterized by ID as well as long-QT syndrome, a frequently lethal heart rhythm defect (Herold et al., 2023; Endres et al., 2020).

### Retinoic acid signaling

Retinoic acid (RA) is a diffusible derivative of vitamin A that has been reported to regulate the fetal development of the prefrontal cortex and the dopaminergic system in the striatum; dysregulation of RA signaling in mice can lead to autistic-like behavior. ***Rarb* (−2.2)**, which encodes the RA receptor, RARβ, is downregulated by more than 50% by VPA in the E12.5 mouse brain. In addition, four additional genes dysregulated by VPA **[*Megf10* (−1.9)**, ***Cntnap2***, **(+1.3) *Meis2* (−1.4)**, ***Cbln2* (−1.8)]** are involved in RA signaling; three of these (*Megf10*, *Cntnap2*, *Meis2*) are on the SFARI list. RA-dependent developmental processes in the fetal brain generation of GABAergic striatal projection neurons and interneurons (Chatzi et al., 2013), expression of D2 dopamine receptors in the ventral striatum (Krezel et al., 1998) and regulation of cortical neurogenesis and cortical plate thickness (Haushalter et al., 2017). Shibata et al. (2021) reported that genetic deletion of *Rarb* disrupts proper molecular patterning of prefrontal and motor areas, development of prefrontal cortex (PFC)-mediodorsal thalamus connectivity and development of intra-PFC dendritic spinogenesis. RA signaling occurs through the RA receptors, RXRB and RARB, and genetic deletions of ***Rxrb* (−1.3)** and *Rarb* demonstrated that these molecules are involved in the proper molecular patterning of the prefrontal and motor cortices, the development of cortico-thalamic connectivity and cortical spinogenesis (Shibata et al., 2021). RA signaling may also regulate the development of the dopamine (DA) system (see below), (Krezel et al., 1998); deletion of *Rxrb* and *Rarb* resulted in locomotor defects related to dysfunction of the mesolimbic DA signaling pathway, as well as expression of D1 and D2 DA receptors in the ventral striatum. VPA exposure reduced expression of *Rarb* (55%) and *Rxrb* (23%) in the fetal mouse brain. Interestingly, VPA reduced the expression the D2 DA receptor, ***Drd2* (−1.9)** by nearly half. Thus, these findings suggest that VPA could reduce the strength of RA signaling in the fetal brain, particularly in the dopaminergic system, resulting in altered numbers and/or connectivity of cortical or striatal neurons.

### Neurogenesis

Excitatory neurons in the mammalian cortex are generated from proliferating radial glia cells (RGs) which undergo mitosis at the ventricular surface to expand the pool of neuroblasts and consequently determine the final number of neurons (Silbereis et al., 2016). Neurogenesis proceeds according to a “program” that is tightly-regulated by intercellular signals from a variety of brain cells as well as the meninges. Disturbances in this regulation, for example by environmental factors such as VPA or gene mutations that disrupt the expression of the genes encoding the relevant intercellular signals, have the potential to alter the configuration and neuronal population of the developing brain leading to permanent defects in connectivity that could lead to altered behavior.

As discussed above, RA plays a number of roles in the regulation of cortical neurogenesis (Haushalter et al., 2017; Shibata et al., 2021). The RA has been reported to be produced in the dorsal forebrain meninges and to act at receptors on the end-feet of RGs (Siegenthaler et al., 2009). This group used a transgenic mouse with reduced expression of ***Foxc1* (−1.9)** leading to defects in forebrain meningeal formation and loss of neuron progenitor production due to failure of proliferating RGs to exit the cell cycle. This would be predicted to lead to a delay in the generation of postmitotic neurons and consequently, an expansion of the proliferating RG pool neuroprogenitor pool. ***Eomes* (*Tbr2*) (−1.8)** is also implicated in the regulation of cortical neurogenesis (Englund et al., 2005; Baala et al., 2007; Arnold et al., 2008; Elsen et al., 2013; Sessa et al., 2017), while ***Neurog1* (−2.2)** has been reported to play a role as a negative regulator of cortical neurogenesis (Han et al., 2018).

The conversion from symmetrical to asymmetrical mitoses associated with the generation of postmitotic neurons is regulated by the ***Plk1* (−1.7)** – ***Lrrk1* (−2.9)** – ***Cdk5rap2* (−1.3)** cascade (Hanafusa et al., 2015). Suliman-Lavie et al. (2020) reported that ***Pogz* (−2.2)** is a negative regulator of transcription and *Pogz* deficiency upregulates expression of genes associated with ASD resulting in disruption of embryonic neurogenesis; *Pogz* was found to be associated with autism in all five GWAS (Table 1). Several members of the heterogeneous nuclear ribonucleoprotein family have been linked to NDDs including ***Hnrnph1* (−1.6), *Hnrnpk* (−1.3), *Hnrnpu* (−1.6);** expression of these genes in radial glia has been reported to be critical during neurodevelopment (Gillentine et al., 2021).

Comparison of the mechanisms controlling neurogenesis in rodents and humans has revealed that gyrification of the human brain is driven by changes in the expression of regulatory genes and growth factors (Borrell and Götz, 2014). Stahl et al. (2013) reported that, over the period E12 to E16, decreased expression of the DNA-associated protein, ***Trnp1* (+1.5)**, leads to radial expansion of the cortex and the appearance of gyri-like folding of the mouse brain (Stahl et al., 2013). In the present study, VPA induced a 50% increase in the expression of *Trnp1* in the E12.5 fetal mouse brain; this would be predicted to inhibit or delay the shift to radial growth of the cortex. Fibroblast growth factor 2 (FGF2) has also been reported to induce gyrification in the mouse brain, having an effect opposite to that of *Trnp1* (Rash et al., 2013). In the present study, VPA caused a 65% reduction on the expression of ***Fgf2* (−2.9)** (Table S4). This would also be predicted to bias cortical neurogenesis toward tangential and away from radial growth that would lead to gyrification. These findings together with the present results lead to a hypothesis that increased *Trnp1* and decreased *Fgf2* expression, induced by VPA in the fetal brain at E12.5, cause a delayed shift in radial growth resulting in autism-like behavioral defects. This hypothesis could be tested by experimentally decreasing *Trnp1* and increasing *Fgf2* expression in the mouse fetal brain exposed to VPA; these maneuvers would be predicted to normalize behavior.

WNT/β-catenin signaling plays a number of roles in regulating neurogenesis in the developing neocortex (Da Silva et al., 2021). ***Wnt3a* (−3.8)** regulates the timing of differentiation of neocortical intermediate progenitors into neurons (Munji et al., 2011). The 74% reduction in *Wnt3a* expression induced by VPA has the potential to disrupt the normal timing of cortical neurogenesis in the fetal brain.

### Neuronal fate specification

***Tbr1* (−2.8)** encodes T-brain-1, a brain-specific T-box transcription factor that has been linked to autism in multiple GWAS (see Table 1). VPA reduced *Tbr1* expression by 63% in the fetal mouse brain (Table S3). *Tbr1* has been described as a “master regulator of ASDs” because it regulates the expression of multiple genes with links to autism (Chuang et al., 2015), thereby controlling brain connectivity in ASDs (Huang and Hsueh, 2015; Notwell et al., 2016). *Tbr1* has been reported to regulate differentiation of the preplate and layer 6 of the fetal cortex (Hevner et al., 2001) as well as the connectivity of cortical layer 6 (Fazel Darbandi et al., 2018). Recently, different *Tbr1* mutations found in individuals with ASD, intellectual disability and developmental delay had differing effects on cortical development (Co et al., 2022); one such mutation (K228E) caused significant upregulation of *Tbr1* but similar behavioral phenotypes to those of *Tbr1* KO mice (Yook et al., 2019). Thus, it appears that both positive and negative deviations from an optimal level of *Tbr1* expression are associated with an autism-like behavioral phenotype. Considering its links to autism in GWAS and its central role in regulating fetal brain development, VPA-induced changes in *Tbr1* expression in the fetal brain could contribute to the behavioral abnormalities observed in animals exposed to VPA *in utero*.

***Nr2f1* (*Coup-tf1*) (−1.8)** encodes an orphan nuclear receptor that has been implicated in a range of neurodevelopmental functions including coordinating cortical patterning, neurogenesis, and laminar fate (Armentano et al., 2007; Faedo et al., 2008; Tocco et al., 2021). *Nr2f1* has also been reported to control subtype and laminar identity of cortical interneurons derived from the medial ganglionic eminence (Hu et al., 2017) as well as regional dynamics of neuroprogenitors in the cortex, possibly being responsible for abnormal gyrification (Bertacchi et al., 2020). *Nr2f1*, together with *Nr1f2*, regulates cell migration in the basal forebrain (Tripodi et al., 2004) and is required for forebrain commissural projections (Armentano et al., 2006). As discussed below, Engeln et al. (2021) reported that *Nr2f1* is a common regulator for multiple genes involved in spine morphology of D1-MSNs in the striatum.

Neuronal differentiation and regional specification in the CNS are regulated by the *Ebf* family of genes including ***Ebf3* (−1.6)** (Garel et al., 1997). Variants of *EBF3* are associated with hypotonia, developmental delay, intellectual disability, and autism (Padhi et al., 2021; Tanaka et al., 2017). The *Olig* genes (*Olig1/2/3*), which encode members of the basic helix-loop-helix (bHLH) family of transcription factors, were originally identified as regulators of oligodendrocyte production in the CNS. More recently, the *Olig* genes have been found to regulate the specification and differentiation of neuronal phenotype during fetal brain development (Chakrabarti et al., 2010; Szu et al., 2021). ***Olig1* (+2.1**, significant in females only) expression was more than doubled in fetal brains exposed to VPA. The basic Helix-loop-Helix gene, ***Neurod1* (−1.5)**, regulates multiple steps in cortical development including neuronal fate specification, differentiation, and migration (Tutukova et al., 2021).

### Axon growth/guidance

Myristoylated Alanine-Rich C-kinase Substrate, encoded by ***Marcks* (−2.2),** plays a wide variety of roles in brain development and function (El Amri et al., 2018; Stumpo et al., 1995). These may be due at least in part to the regulation of axon growth in which MARCKS mediates membrane targeting of plasmalemmal precursor vesicles during axon development (Xu et al., 2014). Although total deletion of *Marcks* in mice is embryonic lethal, the 55% reduction in *Marcks* expression induced by VPA may be less severe and cause subtle changes in brain connectivity that could lead to autistic-like behavior. ***Syn2* (+1.7)** (synapsin-2) has been reported to be required for normal axon formation (Ferreira et al., 1998). ***Draxin* (−1.5)** (dorsal inhibitory axon guidance protein) is essential for the axon guidance underlying development of the corpus callosum and thalamocortical projections (Liu et al., 2018; Ahmed and Shinmyo, 2021; Morcom et al., 2021). Heterogeneous nuclear ribonucleoprotein A/B encoded by ***Hnrnpab* (−1.6)** facilitates olfactory sensory neuron maturation and axon projections by regulating the local expression of its target genes at axon terminals (Fukuda et al., 2023).

### Neuronal migration

***Robo4* (−2.2)** has been reported to regulate the radial migration of newborn neurons in the developing neocortex (Zheng et al., 2012; Gonda et al., 2020); *Robo4* expression was reduced by more than half in the fetal brain by VPA. Dysregulation of *Robo4* expression at E12.5 could affect the developing connectivity of the fetal brain leading to behavioral abnormalities in the adult. ***Mllt11* (−1.5)** regulates migration and neurite outgrowth of cortical projection neurons during development (Stanton-Turcotte et al., 2022). ***Ptk2b* (−1.7)** (*Pyk2*) is a non-receptor cell-adhesion kinase and scaffold protein which, if either overexpressed or knocked down, impairs cortical neuron migration by altering growth cone dynamics (Fan et al., 2018). *Ptk2b* is regulated by ***Ptpn5* (−1.5)** (STEP), a tyrosine phosphatase (Chatterjee et al., 2020). None of these three genes has been linked to autism in GWAS but downregulation of *Ptk2b*, *Ptpn5* and/or *Mllt11a* by VPA has the potential to alter neuronal migration at a critical time when circuits are forming in the fetal brain. ***Otp* (−2.5)** encodes a homeodomain transcription factor that is associated with the development of the hypothalamus in vertebrates. *Otp* is necessary for the migration of diencephalic neurons to the amygdala (García-Moreno et al., 2010) and affects neuropeptide switching in oxytocin neurons (Wircer et al., 2017; Acampora et al., 1999; Ryu et al., 2007). A loss of function mutation in ***Apc2* (−1.7)** causes Sotos syndrome which is characterized by ID and characteristic facial features; this is because *Apc2* is a downstream effector of ***Nsd1* (−1.4)** in regulating the migration and laminar positioning of cortical neurons (Almuriekhi et al., 2015). Thus, VPA inhibits this critical developmental pathway at two steps.

### Synaptogenesis

***Slc17a7* (+1.6)**, which encodes the vesicular glutamate transporter VGLUT1, promotes the development of cortical presynaptic terminals (Berry et al., 2012). The VPA-induced increase in *Slc17a7* would be consistent with the abnormal acceleration or premature initiation of this process. *SLTRKs* are single-pass transmembrane proteins of the leucine-rich repeat (LRR) superfamily that have been proposed as candidate genes for neuropsychiatric disorders (Proenca et al., 2011). There are 6 members of this family in humans and mice, encoded by *Slitrk1−6*. ***Slitrk1* (−1.5)** has been shown to modulate neurite outgrowth (Aruga and Mikoshiba, 2003) and to promote the development of excitatory synapses (Beaubien et al., 2016), *Slitrk2* (not significantly affected by VPA) is on the SFARI list and has been linked to schizophrenia and X-linked NDDs (El Chehadeh et al., 2022) and ***Slitrk5* (−1.6)** deficiency impairs corticostriatal circuitry (Shmelkov et al., 2010) and synaptogenesis (Song et al., 2017). *Slitrk1* and *Slitrk5* expression in the fetal brain was reduced 33% and 37% by VPA; ***Slitrk4* (+1.7)** expression was increased 67% by VPA (Table S4).

Activity-dependent immediate early genes (IEGs) have been shown to regulate synaptogenesis (Kim et al., 2018). The IEG, ***Arc* (+1.6)** (*Arg 3.1*), has been described as a flexible hub for synaptic plasticity and cognition (Nikolaienko et al., 2018) and has been reported to mediate activity dependent synapse elimination in the developing cerebellum (Mikuni et al., 2013).The cerebellins (CBLNs) are secreted glycoproteins that link presynaptic neurexins with postsynaptic δ1 glutamate receptors to form transsynaptic adhesion complexes and promote the formation and stability of excitatory synapses (Seigneur et al., 2021). The formation of these complexes is deficient in the ***Cbln2* (−1.8)** knockout mouse (Tao et al., 2018) resulting in the destabilization of excitatory synapses. Consequently, the 45% reduction in *Cbln* expression induced by VPA in the fetal mouse brain would be predicted to compromise excitatory synaptogenesis.

***Top3b* (−1.5)** encodes an RNA topoisomerase that works with FMRP to promote synapse formation (Xu et al., 2013) and is a high-confidence gene on the SFARI list (Table S3). *Top* family members have been reported to facilitate transcription of long genes linked to autism (King et al., 2013). ***Syncrip* (−1.4)** (SFARI list) (also known as *Hnrnpg*) encodes the synaptotagmin-binding cytoplasmic RNA-interacting protein, which is a candidate gene for ASD and ID (Rauch et al., 2012; Lelieveld et al., 2016; Semino et al., 2021). Bannai et al. (2004) reported that *Syncrip* is a component of mRNA-containing granules in dendrites, possibly regulating local protein synthesis in developing dendritic spines.

Disruption of *POGZ* is associated with ID, ASDs (Stessman et al., 2016, 2017) and impaired cortical development (Matsumura et al., 2020). Markenscoff-Papadimitriou et al. (2021) reported that ***Pogz* (−2.2)** promotes chromatin accessibility and expression of clustered synaptic gene; moreover, brain-specific conditional knockout of *Pogz* results in gene expression changes associated with synaptic function as well as an autistic-like behavioral phenotype (Suliman-Lavie et al., 2020). The X-linked gene, ***Ddx3x* (−1.6),** encodes an RNA helicase that functions in corticogenesis and synaptogenesis (Lennox et al., 2020; (Hoye et al., 2022). Mutations in *DDX3X* account for approximately 2% of intellectual disability in females (DDX3X syndrome) (Boitnott et al., 2021; Weil et al., 2020). Knockdown of the RNA binding protein, ***Csde1* (−2.0)**, has been reported to cause abnormal dendritic spine morphology and synapse formation (Guo et al., 2019).

### Dendrite development

***P2rx7* (−2.0)** (ionotrophic P2X7 receptor, which is activated by extracellular ATP) has been described as the “central hub of brain diseases” (Andrejew et al., 2020). Inhibition of P2X7 receptors ameliorated dendritic spine pathology under pathological conditions (in *Mecp2*-deficient mice) (Garré et al., 2020), while P2rx7 deficiency caused dendritic branching deficits on otherwise normal mice (Mut-Arbona et al., 2023). Interestingly, ***Tmem163* (−1.7)** is required for full function of P2X7 receptors (Salm et al., 2020). ***Sema3a* (−2.0)** regulates dendritic complexity via ***Farp1* (−1.7)** in an activity dependent manner (Cheadle and Biederer, 2014). *Farp1* links postsynaptic cytoskeletal dynamics and transsynaptic organization to coordinate synaptic development (Cheadle and Biederer, 2012). In contrast, ***Itpka* (+14.1)**, which has been reported to regulate dendrite morphology (Windhorst et al., 2012), was overexpressed by 14-fold in fetal brain exposed to VPA. Thus, the reduction in expression of *P2rx7*, *Tmem163*, *Sema3a* and *Farp1* and the increase in *Itpka* induced by VPA may work together to alter dendritic branching of developing neurons in the E12.5 brain. ***Icam5* (+16.6)** (intercellular adhesion molecule 5) is overexpressed in the brain of the *Fmr1* knockout mouse (which models fragile X syndrome, an ASD), leading to dendritic spine abnormalities. The massive increase in *Icam5* expression induced by VPA has the potential to severely disrupt the connectivity of the developing brain leading to autism-like behavior. ***Gas7* (−1.7)** encodes a spine initiation factor that responds to neuronal activity; *Gas7* knockdown decreased spine density in hippocampal neurons (Khanal et al., 2023). ***Dlg5* (−3.0)** encodes a membrane-associated guanylate kinase (MAGUK) protein, which is required for dendritic spine formation and synaptogenesis (Wang et al., 2014), raising the possibility that synaptic development and organization may be impacted by the 67% reduction *Dlg5* expression in fetal brain exposed to VPA at E12.5. Normal expression levels of ***Disc1* (−1.9)** (Disrupted in schizophrenia-1) are required for microtubule function and depletion of *Disc1* or dominant negative *Disc1* constructs impairs neurite outgrowth *in vitro* and proper development of the cortex *in vivo* (Kamiya et al., 2005). Increased expression of Twinfillin-2 which is encoded by ***Twf2* (+1.6)**, increases of thin dendritic spine length in hippocampal neurons (Walker et al., 2023).

### Postsynaptic cytoskeleton

The organization and assembly of glutamate receptors in the postsynaptic membrane are dependent on scaffold proteins including members of the HOMER and SHANK families (Guang et al., 2018). ***Homer1* (+1.6)** and ***Shank2* (−2.3)** are associated with autism by GWAS (Table S3) and are responsible for rare syndromic cases of autism. ***Homer3* (−1.4)** is also dysregulated in the fetal brain. (Eltokhi et al., 2018; Ronesi et al., 2012; Zaslavsky et al., 2019; Yoon et al., 2021). Disruption of *Homer* and *Shank* gene expression by VPA would be expected to alter synaptic function throughout development.

### Excitation-inhibition

A widely discussed hypothesis is that autism is caused by an imbalance between neuronal excitation and inhibition due to increased excitation, reduced inhibition, or both (Rubenstein and Merzenich, 2003; Nelson and Valakh, 2015). Consequently, alterations in the developmental program in the fetal brain that result in too many excitatory (glutamatergic) or too few inhibitory (GABAergic) neurons (or synapses) could contribute to autistic behavior. Two related genes, ***Maf* (−1.6)** and ***Mafb* (−1.6)**, which are both downregulated by about 40% by VPA, have been reported to play redundant roles in the generation of interneurons from the fetal median ganglionic eminence; their deletion results in decreased numbers of cortical somatostatin-releasing, inhibitory interneurons (Pai et al., 2019). ***Gad1* (−2.5)** and ***Gad2* (−3.3)** encode isoforms of the enzyme that synthesizes the inhibitory transmitter, GABA*; Gad1* and *Gad2* were reduced by 60% and 70%, respectively; however, reduced *Gad* expression may be a consequence of a deficit in the number inhibitory GABAergic neurons. ***Insyn1* (−1.4)** encodes a component of the dystroglycan complex at inhibitory synapses; loss of *Insyn1* alters the composition of the GABAergic synapses, excitatory/inhibitory balance, and cognitive behavior (Uezu et al., 2016, 2019) Taken together, these findings are consistent with VPA increasing net brain excitation by decreasing the number of inhibitory interneurons generated during fetal brain development. These reductions in gene expression may contribute to a decrease in inhibition in mice exposed to VPA *in utero* (Markram et al., 2008). ***Nf1* (−1.3)** encodes NF1, which is mutated in neurofibromatosis type 1, an inherited neurocutaneous disorder associated with NDDs including autism. *Nf1* deletion results in the specific loss of parvalbumin-expressing inhibitory cortical interneurons (Angara et al., 2020). In the present study, *Nf1* was downregulated by about 25% in fetal mouse brains exposed to VPA.

***Hapln1* (−3.7)** and ***Hapln4* (+25.7)**, encode extracellular matrix proteins that are components of perineuronal nets (PNNs) (Bosiacki et al., 2019; Nojima et al., 2021). Ramsaran et al. (2023) recently reported that *Hapln1* mediates the functional maturation of hippocampal parvalbumin neurons through assembly of PNNs; this mechanism mediates the development of memory precision during early childhood. *Hapln4* is a selective regulator of the formation of GABAergic synapses between Purkinje and deep cerebellar nuclei neurons (Edamatsu et al., 2018); the cerebellum is one of many brain regions implicated in the etiology of autism (Mapelli et al., 2022). The 70% decrease in *Hapln1* expression and the massive, 25-fold increase in *Hapln4* induced by VPA would be expected to alter PNN density which could disrupt the connectivity of neuronal circuitry leading to cognitive behavioral deficits.

### Dopaminergic system

Several lines of evidence link autistic symptoms with dysfunction in the mesencephalic DA system (Langen et al., 2014; Subramanian et al., 2017; DiCarlo and Wallace, 2022). This can be explained by abnormal circuitry (Golden et al., 2018), notably in the indirect pathway downstream from D2-receptor expressing medium spiny neurons (MSNs). The subthalamic nucleus (STN), an excitatory nucleus in the indirect pathway downstream from the D2-MSNs, is a key factor in the normal functioning of the basal ganglia. ***Foxa1* (−1.6)**, which is reduced by 38% by VPA, has been reported to be essential for the development and functional integrity of the STN (Ferri et al., 2007; Lee et al., 2010; Gasser et al., 2016). *Foxa1* and ***Lmx1a* (−1.7)**, which are both reduced by about 40% in the E12.5 brain exposed to VPA (Table S4), have been shown to promote mesencephalic DA neuron development and diversity (Yan et al., 2011; Doucet-Beaupré et al., 2015; Lin et al., 2009; Deng et al., 2011; Bodea and Blaess, 2015). The basic helix-loop-helix transcription factor ***Srebf1* (+1.4** significant in males only) has been shown to be both necessary and sufficient for midbrain dopaminergic neurogenesis (Toledo et al., 2020). Engeln et al. (2021) reported that transgenic mice with the BDNF receptor, trkB, knocked out specifically in D1 MSNs, displayed autism-like stereotypic behavior that was reversed by chemogenetic inhibition of D1-MSNs. Using RNA-seq of ribosome-associated mRNA, the authors found that 26 genes were dysregulated in the transgenic mouse D1-MSNs. Of these 17 were significantly dysregulated by VPA in the present study [***Slc9a3r1* (+3.0)**, ***Csf1r* (−1.4)**, ***Sparc* (−1.4)**, ***Mcam* (+1.7)**, ***Robo4* (−2.2)**, ***Emp2* (−1.6)**, ***Klf4* (−1.6)**, ***Rtn4rl1* (+2.0)**, ***Bdnf* (+2.8)**, ***C1qtnf1* (−1.4)**, ***Tmem108* (+1.4)**, ***Smyd2* (−1.3)**, ***Wnt7b* (−1.8)**, ***Palm3* (+1.9)**, ***Adora2a* (−1.3)**, ***Drd2* (+2.0)**, ***S1pr1* (−1.3)**]. As discussed above, another gene linked to the development of the DA system is ***Nr2f1* (−1.8)**, which was downregulated by 45% by VPA. It is of interest that all three DA receptor subtypes (*Drd1*, *Drd2*, *Drd3*) are linked to autism by GWAS (SFARI List) and that partial antagonists of the D2 DA receptor (encoded by *Drd2*) have demonstrated utility in reducing stereotypic behaviors in human subjects (Mandic-Maravic et al., 2022); ***Drd2* (+2.2)** expression was doubled by VPA in the present study. Tyrosine hydroxylase encoded by the ***Th* (−1.4)** gene is the rate limiting step in the biosynthesis of catecholamines including DA; *Th* is downregulated by about 35% in fetal brain exposed to VPA.

***Ppp1r1b* (+3.7)**, encodes the dopamine- and cAMP-regulated neuronal phosphoprotein (DARPP-32). DARPP-32 amplifies and/or mediates many actions of cyclic AMP-dependent protein kinase at the plasma membrane and in the cytoplasm, with a broad spectrum of potential targets and functions within the dopaminergic system and throughout the brain (Yger and Girault, 2011).

### Cholinergic system

Cholinergic projection neurons in the nucleus basalis and cholinergic interneurons in the striatum are specified in the embryonic forebrain beginning at about E10 in mouse (Allaway and Machold, 2017). This process is regulated by several transcription factors including ***Lhx8* (−2.7)**, ***Isl1* (−1.5)**, and ***Gbx1* (−2.3)** all of which are downregulated in response to VPA in the fetal mouse brain. In the developing forebrain, *Isl1*-*Lhx8* hexameric complexes promote cholinergic identity (Cho et al., 2014). Conditional deletion of *Isl1* depletes cholinergic projection- and interneurons resulting in abolition of cholinergic innervation of the cortex (Elshatory and Gan, 2008). ***Nkx2-1* (−2.5**, significant in females only), a homeodomain transcription factor, is required for the development of cholinergic septohippocampal neurons and large subsets of basal forebrain cholinergic neurons (Magno et al., 2017). The 33−67% decrease in the expression of these transcription factors induced in the fetal brain by VPA has the potential to alter the proper development of circuits critical for normal brain function. Indeed, the loss of these neurons due to lack of *Nkx2-1* causes alterations in hippocampal theta rhythms and learning and memory defects (Magno et al., 2017). In addition, *Nkx2-1* acting together with *Lhx6* (not affected by VPA), is necessary to specify pallidal projection neurons and forebrain interneurons in the medial ganglionic eminence (Sandberg et al., 2016).

### Endocannabinoid system

***Cnr1* (−2.0)**, encodes the primary endocannabinoid (eCB) receptor in the brain; *Cnr1* has been linked to autism in GWAS. eCBs have been reported to regulate a wide range of embryonic neurodevelopmental processes (Harkany et al., 2007; Gomes et al., 2020) including neural progenitor proliferation (Aguado et al., 2005), long-range axon patterning (Mulder et al., 2008), interneuron migration and morphogenesis (Berghuis et al., 2005), and neuronal fate specification (Harkany et al., 2008). A 50% decrease in expression of *Cnr1* would be predicted to alter the cellular responses to eCB and could disrupt normal fetal brain development (Chakrabarti et al., 2015).

### Autism and Down syndrome (DS)

While the cause of autism is unknown, DS is caused by triplication of human chromosome 21 (Hsa21), leading to the hypothesis that a 50% increase in gene dosage of one or more Hsa21 genes is the cause of DS symptoms. ASDs and DS share ID as a common feature; moreover, DS individuals are diagnosed with ASD at a higher frequency than the general population (Hepburn et al., 2008; Rachubinski et al., 2017). In fact, several genes triplicated in DS that have been liked to autism or to prenatal brain development are up- or downregulated by VPA in the fetal brain; these include ***Dyrk1a* (−1.5)**, ***Brwd1* (−1.3)**, ***Cbs* (+5.0)** and ***Wdr4* (−1.4)** (Park and Chung, 2013; Duchon and Herault, 2016; Fulton et al., 2022; Marechal et al., 2019). ID in both ASDs and DS has been attributed to overexpression of *Cbs* (cystathionine beta synthase) (Conan et al., 2023; Tisato et al., 2021); dysregulation of *Cbs* and dihydrofolate reductase **(*Dhfr*, +1.5)** contributes to inborn errors of amino acid metabolism and is linked to ASDs (Zigman et al., 2021; Hoxha et al., 2021). *Dyrk1a* (which encodes dual specificity tyrosine phosphorylation regulated kinase 1A) is associated with autism in all five GWAS (Table 1) and is believed to underlie intellectual disability in DS due to its increased gene dosage in DS (Duchon and Herault, 2016). In contrast, haploinsufficiency of *DYRK1A* results in a (non-DS) syndrome characterized by intellectual disability including impaired speech development, autism spectrum disorder including anxious and/or stereotypic behavior problems, and microcephaly (Levy et al., 2021; Dang et al., 2018).

### Circadian rhythms

Eleven genes involved in generating circadian rhythms were significantly affected by VPA in the fetal mouse brain (Table 3). Two of these (*Per1* and *Per1*) are SFARI risk genes. While ***Per1* (+1.9), *Per2* +1.7), *Cry1* (+2.5)*, Cry 2* (+2.0), *Arntl* (+2.0) and *Mef2d* (+1.5)** (Geoffray et al., 2016; Mohawk et al., 2019; Lorsung et al., 2021) were upregulated by VPA, *Clock* was not affected. ***Fbxl3* (−1.8)** a negative regulator of *Cry1* and *Cry2*, is downregulated by VPA whereas ***Fbxl21* (+3.2)**, which is up-regulated by VPA, down-regulates *Fbxl3* (Hirano et al., 2013). Thus, these VPA-induced changes lead to increased *Per* and *Cry* gene expression. ***Arntl* (*Bmal1*) (+2.1)**, together with *Clock*, activates rhythmic transcription of *Per* and *Cry* genes; knockout of *Arntl* (*Bmal1*) induces autistic-like behavior and cerebellar dysfunction which was ameliorated by mTORC1 inhibition (Liu et al., 2022).

Daily rhythms in the fetal brain are thought to be entrained by the mother, however, whether there is a functional *Per/Cry/Clock* cycle in the fetal brain has not been explored. It is possible that disrupting the expression of these genes could alter early embryonic brain development through a mechanism unrelated to sleep. For example, (Noda et al., 2019) reported that *in utero* knockdown of *Per3* caused abnormal positioning of cortical excitatory neurons as well as impaired axon extension and dendritic arbor formation.

If VPA-induced changes in circadian rhythm gene expression were to persist postnatally, they could cause disrupted sleep patterns, a common occurrence in autism. Indeed, exposure of fetal rats to VPA on E12.5 caused sleep disturbances that were similar to those reported in autistic children (Cusmano and Mong, 2014); moreover, a VPA-induced reduction in GAD-67 (*Gad1*) was also found in the juvenile rat brains. In the present study, ***Gad1* (−2.5)** was reduced by 60% while RNAs encoding GABA-A and -B receptors subunits, encoded by ***Gabra4* (+2.5)** and ***Gabbr2* (−2.4),** were found to be upregulated 2.5-fold and downregulated 60% by VPA, respectively.

### Neuroinflammation

Immune dysregulation is believed to play a key role in the pathogenesis of autism. Neuroinflammatory molecules such as insulin-like growth factors [***Igf1* (−3.2), *Igf2* (−2.7)**], transforming growth factor-β isoforms [***Tgfb2* (−2.4), *Tgfb3* (−2.0)**], and the TGF-β receptor, ***Tgfbr2* (−2.5)**, are decreased in fetal brain in response to VPA exposure. Expression of several heat-shock proteins [***Hsp1a* (+5.9), *Hsp1b* (+5.6), *Hsp2* (+6.2) and *Dnajc12* (+4.5)**] was dramatically increased by VPA exposure; this may be a response to immune dysregulation following the administration of a high dose of VPA. *DNAJC12* encodes a chaperone protein responsible for the proper folding of phenylalanine hydrolase and *DNAJC12* deficiency causes hyperphenylalaninemia leading symptoms ranging from mild autistic features or hyperactivity to severe intellectual disability (Blau et al., 2018; Anikster et al., 2017). ***Cd200* (−2.0)** is a surface glycoprotein expressed by neurons and its receptor, *Cd200r*, is expressed on microglia. Disturbances in this signaling pathway result in microglial activation and can lead to neuroinflammation (Zawadzka et al., 2021) as is observed in MIA, a cause of autism.

### Epigenetic regulation of gene expression in the action of VPA

The mechanism(s) by which VPA alters gene expression in the fetal brain is not known; however, VPA is a Class I histone deacetyase (HDAC) inhibitor, suggesting that many of the changes reported here may involve increased histone acetylation (Bourin, 2020; Gurvich et al., 2004). Indeed, Konopko et al. (2017) reported that VPA increased the acetylation of several histones at the promoter regions of *Bdnf* 5’ untranslated exons 1, 4, and 6 and that this was positively correlated with increased expression of exon 9, the protein coding region.

It is also likely that many of the genes targeted by VPA are themselves epigenetic writers, erasers, or readers; moreover, there is well-established crosstalk among epigenetic marks (Berger, 2007), including between DNA CpG methylation and covalent histone modifications, which could explain why HDAC inhibition could both increase and decrease gene expression. ***Setd1b* (−2.2)** encodes a lysine-specific methyltransferase which has been linked to autism in GWAS studies (Table S3). De novo variants of *SET1B* are associated with intellectual disability, ASD and epilepsy (Hiraide et al., 2019; Roston et al., 2021). ***Kmt5b* (−6.5)** encodes the epigenetic writer, lysine methyl transferase 5B, which methylates lysine residues on histones, resulting in gene activation or silencing, depending on the histone and lysine residue. Loss of function mutations in *KMT5B* lead to developmental delay and ASD (Eliyahu et al., 2022) (SFARI List). *Kmt5b* expression is reduced by 85% in fetal mouse brains exposed to VPA (Table S3). Pathogenic mutations in ***Ehmt1* (−1.4)** (SFARI List), a lysine methyltransferase, and ***Smarcb2* (−1.7)**, an actin-dependent regulator of chromatin, are causative for Kleefstra Syndrome Spectrum, an ASD; reduced expression of these genes leads to increased neuronal excitability (Kleefstra et al., 2012; Frega et al., 2020).

***Mecp2* (−1.6)** is an X-linked gene that encodes the epigenetic reader, methyl cytosine binding protein 2. Mutations in *MECP2* cause Rett syndrome, an ASD that affects mostly girls (Chahrour and Zoghbi, 2007); *MECP2* is on the SFARI autism list. MeCP2 binds to methylated CpGs in DNA mediating gene repression (Ip et al., 2018, Gonzales and LaSalle, 2010; Guy et al., 2011). Either increased or decreased expression of MeCP2 can cause developmental brain disorders. MeCP2 binds to CpGs throughout the genome; reduced MeCP2 would be expected to alter the expression of many genes. One such gene is ***Bdnf* (+2.8)**. MeCP2 binds to CpGs in the 5’UTR of exon 4 of *Bdnf* leading to suppression of BDNF expression (Martinowich et al., 2003). Reduction of MeCP2 as observed with VPA (Table S3), would be predicted to decrease normal epigenetic suppression of CpG methylated genes. Although reduced *Mecp2* expression induced by VPA would be expected to increase BDNF expression, VPA did not alter CpG methylation in the regulatory regions of *Bdnf*; instead, increased *Bdnf* expression was due to increased histone acetylation and methylation (Konopko et al., 2017).

Paulsen et al. (2022) reported that haploinsufficiency of the chromatin remodelers, *Kmt5b, Arid1b* and *Chd8*, increased the numbers of inhibitory GABAergic neurons in organoid models of human cortex leading to reduced spontaneous circuit activity. In the present study, ***Kmt5b* (−6.5)** and ***Arid1b* (−2.1)** were downregulated in the fetal brain by 85% and 50% respectively, in response to VPA. Although *Chd8* expression was unaffected by VPA, ***Chd2***, ***Chd3*** and ***Chd7***, which have similar chromatin remodeling functions (Marfella and Imbalzano, 2007), were reduced by 23%, 50% and 23% respectively.

### Limitations

1. The results reported here are restricted to VPA-induced changes in RNA levels; the extent to which these changes reflect commensurate changes in the expression levels of the proteins encoded by these genes is not known. Although in many cases, RNA and protein levels are correlated, the extent to which RNA levels of a given gene track its corresponding protein levels may vary.
2. These results represent VPA-induced changes in gene expression at a single point in time (3 hr after VPA administration on E12.5). Whether the changes are transient and reverse as the VPA levels dissipate, as has been reported for *Bdnf* (Almeida et al., 2014), or are persistent, is not known. It is also not known whether the same VPA induced changes would be observed after VPA administration on a different gestational day.
3. There are many genes for which expression is increased or decreased by ≥ 1.25-fold or ≤ −1.25-fold in response to VPA but the FDR-corrected p-value did not reach the p_FDR_ ≤ 0.025 criterion applied in this study (Supplemental Table S1). Higher statistical power would be needed to determine whether these changes are significant.
4. The ± 1.25 FC cutoff used here is arbitrary; it is possible that smaller changes in the expression of one or more critical genes have profound effects on fetal brain development and these would not be identified in this analysis.
5. Only a fraction of the approximately 7,300 genes whose expression is significantly affected by VPA were curated in this study. It is likely that other genes, in addition to those shown in Tables S3 and S4, could play a role in mediating the effects of VPA on fetal brain development.

### Reproducibility

The stimulation by VPA of *Bdnf* expression (Table S4) has been independently confirmed by quantitative RT-PCR (Almeida et al., 2014; Konopko et al., 2017).

By separately analyzing gene expression in male and female brains, the results of this study were effectively replicated with independent biological samples (7 of each sex ± VPA, each from a different pregnancy). With very few exceptions, male and female gene expression levels were similar (c.f., Figure 1) demonstrating reproducibility across independent samples.

## SUMMARY

The approach for this study can be described as “hypothesis generating” in that we began without any preconceived idea (hypothesis) as to the mechanism by which a brief, transient dose of VPA, administered to the pregnant mouse at E12.5, can cause abnormal, autistic-like behavior in her offspring many weeks later. The analysis identified approximately 7,300 genes, expression of which was significantly affected by VPA. This set of genes was narrowed down in two ways, viz., identifying those genes significantly affected by VPA that were a) also linked to autism by GWAS (SFARI List; Table S3) or b) not on the SFARI list (Table S4), but known to play a role in critical steps of early brain development such as neuroprogenitor proliferation and fate determination, axon and dendrite growth, and synaptogenesis. Interference with one or more of these steps in the fetal brain has the potential to interfere with the “program” directing brain development to create a persistent pathological “signature” that leads to abnormal neuronal circuitry in the adult, long after the increase in VPA in the fetal brain and the initial changes in gene expression have dissipated. Of these 651 genes, about half have known mechanisms of action associated with brain development or function; none showed significant sexually-dimorphic effects of VPA. Several genes affected by VPA in the fetal mouse brain appear in multiple GWAS studies (c.f., Tables 1 and 2) including *Adnp*, *Arid1b*, *Ash1l*, *Cdh2*, *Cdk5l*, *Dyrk1a*, *Kmt5b*, *Mecp2*, *Nr2f1*, *Pogz*, *Pten*, *Tbr1*, *Tcf7l2*, and *Wac*. This “short list” is a potential starting point for future hypothesis-driven studies to determine whether dysregulation of one or more of these genes by VPA is a proximal cause of the behavioral abnormalities identified in the adult animals. Nevertheless, it remains possible that dysregulation of one or more of the other genes altered by VPA contributes to the proximal cause of the autistic-like behavior. This hypothesis could be tested by determining the effect of manipulating gene expression (singly or in combination) on VPA-induced behavioral abnormalities.

## Acknowledgements

We thank Tom Abrams, Edna Pereira Albuquerque, Brad Alger, Seth Ament, Tarik Haydar and Alex Poulopoulos for helpful discussions and comments on the manuscript.

## REFERENCES

Acampora D, Postiglione MP, Avantaggiato V, Bonito M Di, Vaccarino FM, Michaud J, Simeone A (1999) Progressive impairment of developing neuroendocrine cell lineages in the hypothalamus of mice lacking the Orthopedia gene. Genes Dev 13:2787–2800 Available at: www.genesdev.org.

Aguado T, Monory K, Palazuelos J, Stella N, Cravatt B, Lutz B, Marsicano G, Kokaia Z, Guzmán M, Galve-Roperh I (2005) The endocannabinoid system drives neural progenitor proliferation. FASEB J 19:1704–1706.

Ahmed G, Shinmyo Y (2021) Multiple Functions of Draxin/Netrin-1 Signaling in the Development of Neural Circuits in the Spinal Cord and the Brain. Front Neuroanat 15:766911.

Allaway KC, Machold R (2017) Developmental specification of forebrain cholinergic neurons. Dev Biol 421:1–7.

Almuriekhi M, Shintani T, Fahiminiya S, Fujikawa A, Kuboyama K, Takeuchi Y, Nawaz Z, Nadaf J, Kamel H, Kitam AK, Samiha Z, Mahmoud L, Ben-Omran T, Majewski J, Noda M (2015) Loss-of-function mutation in APC2 causes Sotos syndrome features. Cell Rep 10:1585–1598.

American Psychiatric Association (2022) Diagnostic and Statistical Manual of Mental Disorders (5th ed., text rev.).

Anders S, Pyl PT, Huber W (2015) HTSeq-A Python framework to work with high-throughput sequencing data. Bioinformatics 31:166–169.

Andrejew R, Oliveira-Giacomelli Á, Ribeiro DE, Glaser T, Arnaud-Sampaio VF, Lameu C, Ulrich H (2020) The P2X7 Receptor: Central Hub of Brain Diseases. Front Mol Neurosci 13:Article 124.

Angara K, Ling-Lin Pai E, Bilinovich SM, Stafford AM, Nguyen JT, Li KX, Paul A, Rubenstein JL, Vogt D (2020) Nf1 deletion results in depletion of the Lhx6 transcription factor and a specific loss of parvalbumin + cortical interneurons. Proceedings of the National Academy of Sciences, USA 117:6189–6195 Available at: https://www.pnas.org.

Anikster Y et al. (2017) Biallelic Mutations in DNAJC12 Cause Hyperphenylalaninemia, Dystonia, and Intellectual Disability. Am J Hum Genet 100:257–266.

Armentano M, Chou SJ, Srubek Tomassy G, Leingärtner A, O’Leary DDM, Studer M (2007) COUP-TFI regulates the balance of cortical patterning between frontal/motor and sensory areas. Nat Neurosci 10:1277–1286.

Armentano M, Filosa A, Andolfi G, Studer M (2006) COUP-TFI is required for the formation of commissural projections in the forebrain by regulating axonal growth. Development 133:4151–4162.

Arnold SJ, Huang GJ, Cheung AFP, Era T, Nishikawa SI, Bikoff EK, Molnár Z, Robertson EJ, Groszer M (2008) The T-box transcription factor Eomes/Tbr2 regulates neurogenesis in the cortical subventricular zone. Genes Dev 22:2479–2484.

Aruga J, Mikoshiba K (2003) Identification and characterization of Slitrk, a novel neuronal transmembrane protein family controlling neurite outgrowth. Mol Cell Neurosci 24:117–129.

Baala L, Briault S, Etchevers HC, Laumonnier F, Natiq A, Amiel J, Boddaert N, Picard C, Sbiti A, Asermouh A, Attié-Bitach T, Encha-Razavi F, Munnich A, Sefiani A, Lyonnet S (2007) Homozygous silencing of T-box transcription factor EOMES leads to microcephaly with polymicrogyria and corpus callosum agenesis. Nat Genet 39:454–456.

Bannai H, Fukatsu K, Mizutani A, Natsume T, Iemura SI, Ikegami T, Inoue T, Mikoshiba K (2004) An RNA-interacting protein, SYNCRIP (heterogeneous nuclear ribonuclear protein Q1/NSAP1) is a component of mRNA granule transported with inositol 1,4,5-trisphosphate receptor type 1 mRNA in neuronal dendrites. Journal of Biological Chemistry 279:53427–53434.

Beaubien F, Raja R, Kennedy TE, Fournier AE, Cloutier JF (2016) Slitrk1 is localized to excitatory synapses and promotes their development. Sci Rep 6:27343.

Ben-Ari Y (2008) Neuro-archaeology: pre-symptomatic architecture and signature of neurological disorders. Trends Neurosci 31:626–636.

Berger SL (2007) The complex language of chromatin regulation during transcription. Nature 447:407–412.

Berghuis P, Dobszay MB, Wang X, Spano S, Ledda F, Sousa KM, Schulte G, Ernfors P, Mackie K, Paratcha G, Hurd YL, Harkany T (2005) Endocannabinoids regulate interneuron migration and morphogenesis by transactivating the TrkB receptor. Proc Natl Acad Sci U S A 102:19115– 19120.

Berry CT, Sceniak MP, Zhou L, Sabo SL (2012) Developmental Up-Regulation of Vesicular Glutamate Transporter-1 Promotes Neocortical Presynaptic Terminal Development. PLoS One 7:e50911.

Bertacchi M et al. (2020) NR2F1 regulates regional progenitor dynamics in the mouse neocortex and cortical gyrification in BBSOAS patients. EMBO J 39:e104163.

Bey AL, Jiang Y hui (2014) Overview of mouse models of autism spectrum disorders. Curr Protoc Pharmacol 2014:5.66.1–5.66.26.

Bilbo SD, Block CL, Bolton JL, Hanamsagar R, Tran PK (2018) Beyond infection - Maternal immune activation by environmental factors, microglial development, and relevance for autism spectrum disorders. Exp Neurol 299:241–251.

Blau N, Martinez A, Hoffmann GF, Thöny B (2018) DNAJC12 deficiency: A new strategy in the diagnosis of hyperphenylalaninemias. Mol Genet Metab 123:1–5.

Bodea GO, Blaess S (2015) Establishing diversity in the dopaminergic system. FEBS Lett 589:3773– 3785.

Boitnott A, Garcia-Forn M, Ung DC, Niblo K, Mendonca D, Park Y, Flores M, Maxwell S, Ellegood J, Qiu LR, Grice DE, Lerch JP, Rasin MR, Buxbaum JD, Drapeau E, De Rubeis S (2021) Developmental and Behavioral Phenotypes in a Mouse Model of DDX3X Syndrome. Biol Psychiatry 90:742–755.

Borrell V, Götz M (2014) Role of radial glial cells in cerebral cortex folding. Curr Opin Neurobiol 27:39–46.

Bosiacki M, Gąssowska-Dobrowolska M, Kojder K, Fabiańska M, Jeżewski D, Gutowska I, Lubkowska A (2019) Perineuronal nets and their role in synaptic homeostasis. Int J Mol Sci 20:4108.

Bourin M (2020) Mechanism of Action of Valproic Acid and Its Derivatives. SOJ Pharm Pharm Sci 7:1–4.

Breitenkamp AFS, Matthes J, Nass RD, Sinzig J, Lehmkuhl G, Nürnberg P, Herzig S (2014) Rare mutations of CACNB2 found in autism spectrum disease-affected families alter calcium channel function. PLoS One 9.

Buddell T, Friedman V, Drozd CJ, Quinn CC (2019) An autism-causing calcium channel variant functions with selective autophagy to alter axon targeting and behavior. PLoS Genet 15.

Burke RD, Todd SW, Lumsden E, Mullins RJ, Mamczarz J, Fawcett WP, Gullapalli RP, Randall WR, Pereira EFR, Albuquerque EX (2017) Developmental neurotoxicity of the organophosphorus insecticide chlorpyrifos: from clinical findings to preclinical models and potential mechanisms. J Neurochem 142 Suppl 2:162–177.

Careaga M, Murai T, Bauman MD (2017) Maternal Immune Activation and Autism Spectrum Disorder: From Rodents to Nonhuman and Human Primates. Biol Psychiatry 81:391–401.

Chahrour M, Zoghbi HY (2007) The story of Rett syndrome: from clinic to neurobiology. Neuron 56:422–437.

Chakrabarti B, Persico A, Battista N, Maccarrone M (2015) Endocannabinoid Signaling in Autism. Neurotherapeutics 12:837–847.

Chakrabarti L, Best TK, Cramer NP, Carney RSE, Isaac JTR, Galdzicki Z, Haydar TF (2010) Olig1 and Olig2 triplication causes developmental brain defects in Down syndrome. Nat Neurosci 13:927–934.

Chatterjee M, Singh P, Xu J, Lombroso PJ, Kurup PK (2020) Inhibition of striatal-enriched protein tyrosine phosphatase (STEP) activity reverses behavioral deficits in a rodent model of autism. Behavioural Brain Research 391:112713.

Chatzi C, Cunningham TJ, Duester G (2013) Investigation of retinoic acid function during embryonic brain development using retinaldehyde-rescued Rdh10 knockout mice. Developmental Dynamics 242:1056–1065.

Cheadle L, Biederer T (2012) The novel synaptogenic protein farp1 links postsynaptic cytoskeletal dynamics and transsynaptic organization. Journal of Cell Biology 199:985–1001.

Cheadle L, Biederer T (2014) Activity-dependent regulation of dendritic complexity by Semaphorin 3A through Farp1. Journal of Neuroscience 34:7999–8009.

Cho H-H, Cargnin F, Kim Y, Lee B, Kwon R-J, Nam H, Shen R, Barnes AP, Lee JW, Lee S, Lee S-K (2014) Isl1 directly controls a cholinergic neuronal identity in the developing forebrain and spinal cord by forming cell type-specific complexes. PLoS Genet 10:e1004280.

Christensen J, Grønborg TK, Sørensen MJ, Schendel D, Parner ET, Pedersen LH, Vestergaard M (2013) Prenatal Valproate Exposure and Risk of Autism Spectrum Disorders and Childhood Autism. JAMA 309:1696–1703.

Chuang HC, Huang TN, Hsueh YP (2015) T-Brain-1 - A Potential Master Regulator in Autism Spectrum Disorders. Autism Research 8:412–426.

Co M, Barnard RA, Jahncke JN, Grindstaff S, Fedorov LM, Adey AC, Wright KM, O’Roak BJ (2022) Shared and distinct functional effects of patient-specific Tbr1 mutations on cortical development. The Journal of Neuroscience:JN-RM-0409–22.

Conan P, Léon A, Caroff N, Rollet C, Chaïr L, Martin J, Bihel F, Mignen O, Voisset C, Friocourt G (2023) New insights into the regulation of Cystathionine beta synthase (CBS), an enzyme involved in intellectual deficiency in Down syndrome. Front Neurosci 16:1110163.

Cusmano DM, Mong JA (2014) In Utero exposure to valproic acid changes sleep in juvenile rats: A model for sleep disturbances in autism. Sleep 37:1489–1499.

Da Silva F, Zhang K, Pinson A, Fatti E, Wilsch-Bräuninger M, Herbst J, Vidal V, Schedl A, Huttner WB, Niehrs C (2021) Mitotic WNT signalling orchestrates neurogenesis in the developing neocortex. EMBO J 40.

Dang T, Duan WY, Yu B, Tong DL, Cheng C, Zhang YF, Wu W, Ye K, Zhang WX, Wu M, Wu BB, An Y, Qiu ZL, Wu BL (2018) Autism-associated Dyrk1a truncation mutants impair neuronal dendritic and spine growth and interfere with postnatal cortical development. Mol Psychiatry 23:747–758.

Deng Q, Andersson E, Hedlund E, Alekseenko Z, Coppola E, Panman L, Millonig JH, Brunet JF, Ericson J, Perlmann T (2011) Specific and integrated roles of Lmx1a, Lmx1b and Phox2a in ventral midbrain development. Development 138:3399–3408.

DiCarlo GE, Wallace MT (2022) Modeling dopamine dysfunction in autism spectrum disorder: From invertebrates to vertebrates. Neurosci Biobehav Rev 133:104494.

Dickinson R G, Lawyer C H, Kaufman S N, Lynn RK, Gerber N, Novy M J, Cook M J (1980) Materno-fetal pharmacokinetics and fetal distribution of valproic acid in a pregnant rhesus monkey. Pediatric Pharmacology (New York) 1:71–83.

Dong C, West KL, Tan XY, Li J, Ishibashi T, Yu C, Sy SMH, Leung JWC, Huen MSY (2020) Screen identifies DYRK1B network as mediator of transcription repression on damaged chromatin. Proceedings of the National Academy of Sciences 117:17019–17030.

Doucet-Beaupré H, Ang SL, Lévesque M (2015) Cell fate determination, neuronal maintenance and disease state: The emerging role of transcription factors Lmx1a and Lmx1b. FEBS Lett 589:3727–3738.

Duchon A, Herault Y (2016) DYRK1A, a dosage-sensitive gene involved in neurodevelopmental disorders, Is a target for drug development in down syndrome. Front Behav Neurosci 10:104.

Edamatsu M, Miyano R, Fujikawa A, Fujii F, Hori T, Sakaba T, Oohashi T (2018) Hapln4/Bral2 is a selective regulator for formation and transmission of GABAergic synapses between Purkinje and deep cerebellar nuclei neurons. J Neurochem 147:748–763.

el Amri M, Fitzgerald U, Schlosser G (2018) MARCKS and MARCKS-like proteins in development and regeneration. J Biomed Sci 25:43.

el Chehadeh S, et al. (2022) SLITRK2 variants associated with neurodevelopmental disorders impair excitatory synaptic function and cognition in mice. Nat Commun 13:4112.

Eliyahu A, Barel O, Greenbaum L, Zaks Hoffer G, Goldberg Y, Raas-Rothschild A, Singer A, Bar-Joseph I, Kunik V, Javasky E, Staretz-Chacham O, Pode-Shakked N, Bazak L, Ruhrman-Shahar N, Pras E, Frydman M, Shohat M, Pode-Shakked B (2022) Refining the Phenotypic Spectrum of KMT5B-Associated Developmental Delay. Front Pediatr 10:Article 844845.

Elsen GE, Hodge RD, Bedogni F, Daza RAM, Nelson BR, Shiba N, Reiner SL, Hevner RF (2013) The protomap is propagated to cortical plate neurons through an Eomes-dependent intermediate map. Proc Natl Acad Sci U S A 110:4081–4086.

Elshatory Y, Gan L (2008) The LIM-homeobox gene Islet-1 is required for the development of restricted forebrain cholinergic neurons. J Neurosci 28:3291–3297.

Eltokhi A, Rappold G, Sprengel R (2018) Distinct Phenotypes of Shank2 Mouse Models Reflect Neuropsychiatric Spectrum Disorders of Human Patients With SHANK2 Variants. Front Mol Neurosci 11:240.

Endres D, Decher N, Röhr I, Vowinkel K, Domschke K, Komlosi K, Tzschach A, Gläser B, Schiele MA, Runge K, Süß P, Schuchardt F, Nickel K, Stallmeyer B, Rinné S, Schulze-Bahr E, van Elst LT (2020) New CaV1.2 channelopathy with high-functioning autism, affective disorder, severe dental enamel defects, a short QT interval, and a novel cacna1c loss-of-function mutation. Int J Mol Sci 21:1–8.

Engeln M, Song Y, Chandra R, La A, Fox ME, Evans B, Turner MD, Thomas S, Francis TC, Hertzano R, Lobo MK (2021) Individual differences in stereotypy and neuron subtype translatome with TrkB deletion. Mol Psychiatry 26:1846–1859.

Englund C, Fink A, Lau C, Pham D, Daza RAM, Bulfone A, Kowalczyk T, Hevner RF (2005) Pax6, Tbr2, and Tbr1 are expressed sequentially by radial glia, intermediate progenitor cells, and postmitotic neurons in developing neocortex. Journal of Neuroscience 25:247–251.

Faedo A, Tomassy GS, Ruan Y, Teichmann H, Krauss S, Pleasure SJ, Tsai SY, Tsai MJ, Studer M, Rubenstein JLR (2008) COUP-TFI coordinates cortical patterning, neurogenesis, and laminar fate and modulates MAPK/ERK, AKT, and β-catenin signaling. Cerebral Cortex 18:2117–2131.

Fan L, Lu Y, Shen X, Shao H, Suo L, Wu Q (2018) Alpha protocadherins and Pyk2 kinase regulate cortical neuron migration and cytoskeletal dynamics via Rac1 GTPase and WAVE complex in mice. Elife 7:e35242.

Fazel Darbandi S, Robinson Schwartz SE, Qi Q, Catta-Preta R, Pai ELL, Mandell JD, Everitt A, Rubin A, Krasnoff RA, Katzman S, Tastad D, Nord AS, Willsey AJ, Chen B, State MW, Sohal VS, Rubenstein JLR (2018) Neonatal Tbr1 Dosage Controls Cortical Layer 6 Connectivity. Neuron 100:831–845.e7.

Ferreira A, Chin L-S, Lian L, Lanier LM, Kosik KS, Greengard P (1998) Distinct roles of synapsin I an synapsin II during neuronal development. Molecular Medicine 4:2228.

Ferri ALM, Lin W, Mavromatakis YE, Wang JC, Sasaki H, Whitsett JA, Ang SL (2007) Foxa1 and Foxa2 regulate multiple phases of midbrain dopaminergic neuron development in a dosage-dependent manner. Development 134:2761–2769.

Ferri SL, Abel T, Brodkin ES (2018) Sex Differences in Autism Spectrum Disorder: a Review. Curr Psychiatry Rep 20:9.

Frega M, Selten M, Mossink B, Keller JM, Linda K, Moerschen R, Qu J, Koerner P, Jansen S, Oudakker A, Kleefstra T, van Bokhoven H, Zhou H, Schubert D, Nadif Kasri N (2020) Distinct Pathogenic Genes Causing Intellectual Disability and Autism Exhibit a Common Neuronal Network Hyperactivity Phenotype. Cell Rep 30:173–186.e6.

Fukuda N, Fukuda T, Percipalle P, Oda K, Takei N, Czaplinski K, Touhara K, Yoshihara Y, Sasaoka T (2023) Axonal mRNA binding of hnRNP A/B is crucial for axon targeting and maturation of olfactory sensory neurons. Cell Rep 42.

Fulton SL et al. (2022) Rescue of deficits by Brwd1 copy number restoration in the Ts65Dn mouse model of Down syndrome. Nat Commun 13:6384.

Garcia-Forn M, Boitnott A, Akpinar Z, de Rubeis S (2020) Linking autism risk genes to disruption of cortical development. Cells 9:1–24.

García-Moreno F, Pedraza M, Di Giovannantonio LG, Di Salvio M, López-Mascaraque L, Simeone A, De Carlos JA (2010) A neuronal migratory pathway crossing from diencephalon to telencephalon populates amygdala nuclei. Nat Neurosci 13:680–689.

Garel S, Marin F, Marin M, Genevie M-G, Matte G, Vesque C, Vincent A, Charnay P (1997) Family of Ebf/Olf-1-Related Genes Potentially Involved in Neuronal Differentiation and Regional Specification in the Central Nervous System. Dev Dyn 210:191–205.

Garré JM, Silva HM, Lafaille JJ, Yang G (2020) P2X7 receptor inhibition ameliorates dendritic spine pathology and social behavioral deficits in Rett syndrome mice. Nat Commun 11:1784.

Gasser E, Johannssen HC, Rülicke T, Zeilhofer HU, Stoffel M (2016) Foxa1 is essential for development and functional integrity of the subthalamic nucleus. Sci Rep 6:38611.

Geoffray MM, Nicolas A, Speranza M, Georgieff N (2016) Are circadian rhythms new pathways to understand Autism Spectrum Disorder? J Physiol Paris 110:434–438.

Gillentine MA et al. (2021) Rare deleterious mutations of HNRNP genes result in shared neurodevelopmental disorders. Genome Med 13.

Golden CE, Buxbaum JD, de Rubeis S (2018) Disrupted circuits in mouse models of autism spectrum disorder and intellectual disability. Curr Opin Neurobiol 48:106–112.

Gomes TM, Dias da Silva D, Carmo H, Carvalho F, Silva JP (2020) Epigenetics and the endocannabinoid system signaling: An intricate interplay modulating neurodevelopment. Pharmacol Res 162:105237.

Gonda Y, Namba T, Hanashima C (2020) Beyond Axon Guidance: Roles of Slit-Robo Signaling in Neocortical Formation. Front Cell Dev Biol 8:Article 607415.

Gonzales ML, LaSalle JM (2010) The role of MeCP2 in brain development and neurodevelopmental disorders. Curr Psychiatry Rep 12:127–134.

Guang S, Pang N, Deng X, Yang L, He F, Wu L, Chen C, Yin F, Peng J (2018) Synaptopathology involved in autism spectrum disorder. Front Cell Neurosci 12.

Guerra M, Medici V, Weatheritt R, Corvino V, Palacios D, Geloso MC, Farini D, Sette C (2023) Fetal exposure to valproic acid dysregulates the expression of autism-linked genes in the developing cerebellum. Transl Psychiatry 13:114.

Guo H et al. (2019) Disruptive variants of CSDE1 associate with autism and interfere with neuronal development and synaptic transmission. Sci Adv 5:eaax2166 Available at: https://gene.sfari.org/.

Gurvich N, Tsygankova OM, Meinkoth JL, Klein PS (2004) Histone Deacetylase Is a Target of Valproic Acid-Mediated Cellular Differentiation. Cancer Res 64:1079–1086 Available at: http://aacrjournals.org/cancerres/article-pdf/64/3/1079/2522598/zch00304001079.pdf.

Guy J, Cheval H, Selfridge J, Bird A (2011) The role of MeCP2 in the brain. Annu Rev Cell Dev Biol 27:631–652.

Hallmayer J, Cleveland S, Torres A, Phillips J, Cohen B, Torigoe T, Miller J, Fedele A, Collins J, Smith K, Lotspeich L, Croen LA, Ozonoff S, Lajonchere C, Grether JK, Risch N (2011) Genetic heritability and shared environmental factors among twin pairs with autism. Arch Gen Psychiatry 68:1095–1102.

Han S, Dennis DJ, Balakrishnan A, Dixit R, Britz O, Zinyk D, Touahri Y, Olender T, Brand M, Guillemot F, Kurrasch D, Schuurmans C (2018) A non-canonical role for the proneural gene neurog1 as a negative regulator of neocortical neurogenesis. Development (Cambridge) 145:dev157719.

Hanafusa H, Kedashiro S, Tezuka M, Funatsu M, Usami S, Toyoshima F, Matsumoto K (2015) PLK1-dependent activation of LRRK1 regulates spindle orientation by phosphorylating CDK5RAP2. Nat Cell Biol 17:1024–1035.

Harkany T, Guzmán M, Galve-Roperh I, Berghuis P, Devi LA, Mackie K (2007) The emerging functions of endocannabinoid signaling during CNS development. Trends Pharmacol Sci 28:83–92.

Harkany T, Keimpema E, Barabás K, Mulder J (2008) Endocannabinoid functions controlling neuronal specification during brain development. Mol Cell Endocrinol 286S:S84–S90.

Haushalter C, Asselin L, Fraulob V, Dollé P, Rhinn M (2017) Retinoic acid controls early neurogenesis in the developing mouse cerebral cortex. Dev Biol 430:129–141.

Hepburn S, Philofsky A, Fidler DJ, Rogers S (2008) Autism symptoms in toddlers with Down syndrome: A descriptive study. Journal of Applied Research in Intellectual Disabilities 21:48–57.

Herold KG, Hussey JW, Dick IE (2023) CACNA1C-Related Channelopathies. Springer Nature Switzerland.

Hevner RF, Shi L, Justice N, Bulfone A, Goffinet AM, Campagnoni AT, Rubenstein JLR (2001) Tbr1 regulates differentiation of the preplate and layer 6. Neuron 29:353–366.

Hiraide T, Hattori A, Ieda D, Hori I, Saitoh S, Nakashima M, Saitsu H (2019) De novo variants in SETD1B cause intellectual disability, autism spectrum disorder, and epilepsy with myoclonic absences. Epilepsia Open 4:476–481.

Hirano A, Yumimoto K, Tsunematsu R, Matsumoto M, Oyama M, Kozuka-Hata H, Nakagawa T, Lanjakornsiripan D, Nakayama KI, Fukada Y (2013) FBXL21 regulates oscillation of the circadian clock through ubiquitination and stabilization of cryptochromes. Cell 152:1106–1118.

Hoxha B, Hoxha M, Zappacosta B, Domi E, Gervasoni J, Persichilli S, Malaj V (2021) Folic acid and autism: A systematic review of the current state of knowledge. Cells 10:1976.

Hoye ML, Calviello L, Poff AJ, Ejimogu N-E, Newman CR, Montgomery MD, Ou J, Floor SN, Silver DL (2022) Aberrant cortical development is driven by impaired cell cycle and translational control in a DDX3X syndrome model. Elife 11:e78203.

Hu JS, Vogt D, Lindtner S, Sandberg M, Silberberg SN, Rubenstein JLR (2017) Coup-TF1 and coup-TF2 control subtype and laminar identity of mge-derived neocortical interneurons. Development (Cambridge) 144:2837–2851.

Huang TN, Hsueh YP (2015) Brain-specific transcriptional regulator T-brain-1 controls brain wiring and neuronal activity in autism spectrum disorders. Front Neurosci 9:Article 406.

Ip JPK, Mellios N, Sur M (2018) Rett syndrome: Insights into genetic, molecular and circuit mechanisms. Nat Rev Neurosci 19:368–382.

Kamiya A, Kubo K, Tomoda T, Takaki M, Youn R, Ozeki Y, Sawamura N, Park U, Kudo C, Okawa M, Ross CA, Hatten ME, Nakajima K, Sawa A (2005) A schizophrenia-associated mutation of DISC1 perturbs cerebral cortex development. Nat Cell Biol 7:1167–1178.

Karimi P, Kamali E, Mousavi SM, Karahmadi M (2017) Environmental factors influencing the risk of autism. Journal of Research in Medical Sciences 22:27.

Kazdoba TM, Leach PT, Yang M, Silverman JL, Solomon M, Crawley JN (2016) Translational mouse models of autism: Advancing toward pharmacological therapeutics. Curr Top Behav Neurosci 28:1–52.

Khanal P, Boskovic Z, Lahti L, Ghimire A, Minkeviciene R, Opazo P, Hotulainen P (2023) Gas7 Is a Novel Dendritic Spine Initiation Factor. eNeuro 10:0344–22.

Kim D, Paggi JM, Park C, Bennett C, Salzberg SL (2019) Graph-based genome alignment and genotyping with HISAT2 and HISAT-genotype. Nat Biotechnol 37:907–915.

Kim S, Kim H, Um JW (2018) Synapse development organized by neuronal activity-regulated immediate-early genes. Exp Mol Med 50:11.

King IF, Yandava CN, Mabb AM, Hsiao JS, Huang HS, Pearson BL, Calabrese JM, Starmer J, Parker JS, Magnuson T, Chamberlain SJ, Philpot BD, Zylka MJ (2013) Topoisomerases facilitate transcription of long genes linked to autism. Nature 501:58–62.

Kleefstra T et al. (2012) Disruption of an EHMT1-associated chromatin-modification module causes intellectual disability. Am J Hum Genet 91:73–82.

Krezel W, Ghyselinck N, Samad TA, Dupé V, Kastner P, Borrelli E, Chambon P (1998) Impaired locomotion and dopamine signaling in retinoid receptor mutant mice. Science 279:863–867.

Lan A, Kalimian M, Amram B, Kofman O (2017) Prenatal chlorpyrifos leads to autism-like deficits in C57Bl6/J mice. Environ Health 16:43.

Landrigan PJ (2010) What causes autism? Exploring the environmental contribution. Curr Opin Pediatr 22:219–225.

Langen M, Bos D, Noordermeer SDS, Nederveen H, van Engeland H, Durston S (2014) Changes in the development of striatum are involved in repetitive behavior in autism. Biol Psychiatry 76:405– 411.

Law CW, Chen Y, Shi W, Smyth GK (2014) voom: precision weights unlock linear model analysis tools for RNA-seq read counts. Genome Biol 15:R29 Available at: http://genomebiology.com/2014/15/2/R29.

Lee HS, Bae EJ, Yi SH, Shim JW, Jo AY, Kang JS, Yoon EH, Rhee YH, Park CH, Koh HC, Kim HJ, Choi HS, Han JW, Lee YS, Kim J, Li JY, Brundin P, Lee SH (2010) Foxa2 and Nurr1 synergistically yield A9 nigral dopamine neurons exhibiting improved differentiation, function, and cell survival. Stem Cells 28:501–512.

Lelieveld SH et al. (2016) Meta-analysis of 2,104 trios provides support for 10 new genes for intellectual disability. Nat Neurosci 19:1194–1196.

Lennox AL et al. (2020) Pathogenic DDX3X Mutations Impair RNA Metabolism and Neurogenesis during Fetal Cortical Development. Neuron 106:404–420.e8.

Levy JA, LaFlamme CW, Tsaprailis G, Crynen G, Page DT (2021) Dyrk1a Mutations Cause Undergrowth of Cortical Pyramidal Neurons via Dysregulated Growth Factor Signaling. Biol Psychiatry 90:295–306.

Liao X, Li Y (2020) Genetic associations between voltage-gated calcium channels and autism spectrum disorder: A systematic review. Mol Brain 13:96.

Lin W, Metzakopian E, Mavromatakis YE, Gao N, Balaskas N, Sasaki H, Briscoe J, Whitsett JA, Goulding M, Kaestner KH, Ang SL (2009) Foxa1 and Foxa2 function both upstream of and cooperatively with Lmx1a and Lmx1b in a feedforward loop promoting mesodiencephalic dopaminergic neuron development. Dev Biol 333:386–396.

Lin YC, Frei JA, Kilander MBC, Shen W, Blatt GJ (2016) A subset of autism-associated genes regulate the structural stability of neurons. Front Cell Neurosci 10:Article 263.

Liu D, Nanclares C, Simbriger K, Fang K, Lorsung E, Le N, Amorim IS, Chalkiadaki K, Pathak SS, Li J, Gewirtz JC, Jin VX, Kofuji P, Araque A, Orr HT, Gkogkas CG, Cao R (2022) Autistic-like behavior and cerebellar dysfunction in Bmal1 mutant mice ameliorated by mTORC1 inhibition. Mol Psychiatry:s41380.

Liu Y, Bhowmick T, Liu Y, Gao X, Mertens HDT, Svergun DI, Xiao J, Zhang Y, Wang J huai, Meijers R (2018) Structural Basis for Draxin-Modulated Axon Guidance and Fasciculation by Netrin-1 through DCC. Neuron 97:1261–1267.e4.

Lorsung E, Karthikeyan R, Cao R (2021) Biological Timing and Neurodevelopmental Disorders: A Role for Circadian Dysfunction in Autism Spectrum Disorders. Front Neurosci 15:642745.

Love MI, Huber W, Anders S (2014) Moderated estimation of fold change and dispersion for RNA-seq data with DESeq2. Genome Biol 15:550.

Macari S, Milgramm A, Reed J, Shic F, Powell KK, Macris D, Chawarska K (2021) Context-Specific Dyadic Attention Vulnerabilities During the First Year in Infants Later Developing Autism Spectrum Disorder. J Am Acad Child Adolesc Psychiatry 60:166–175.

Maenner MJ et al. (2021) Prevalence and Characteristics of Autism Spectrum Disorder Among Children Aged 8 Years — Autism and Developmental Disabilities Monitoring Network, 11 Sites, United States, 2018. MMWR Surveillance Summary 70:1–16 Available at: https://stacks.cdc.

Magno L, Barry C, Schmidt-Hieber C, Theodotou P, Häusser M, Kessaris N (2017) NKX2-1 Is Required in the Embryonic Septum for Cholinergic System Development, Learning, and Memory. Cell Rep 20:1572–1584.

Mandic-Maravic V, Grujicic R, Milutinovic L, Munjiza-Jovanovic A, Pejovic-Milovancevic M (2022) Dopamine in Autism Spectrum Disorders—Focus on D2/D3 Partial Agonists and Their Possible Use in Treatment. Front Psychiatry 12.

Mapelli L, Soda T, D’Angelo E, Prestori F (2022) The Cerebellar Involvement in Autism Spectrum Disorders: From the Social Brain to Mouse Models. Int J Mol Sci 23:3894.

Marechal D, Brault V, Leon A, Martin D, Pereira PL, Loaëc N, Birling MC, Friocourt G, Blondel M, Herault Y (2019) Cbs overdosage is necessary and sufficient to induce cognitive phenotypes in mouse models of Down syndrome and interacts genetically with Dyrk1a. Hum Mol Genet 28:1561–1577.

Marfella CGA, Imbalzano AN (2007) The Chd family of chromatin remodelers. Mutation Research - Fundamental and Molecular Mechanisms of Mutagenesis 618:30–40.

Markenscoff-Papadimitriou E, Binyameen F, Whalen S, Price J, Lim K, Ypsilanti AR, Catta-Preta R, Pai ELL, Mu X, Xu D, Pollard KS, Nord AS, State MW, Rubenstein JL (2021) Autism risk gene POGZ promotes chromatin accessibility and expression of clustered synaptic genes. Cell Rep 37:110089.

Markram K, Rinaldi T, la Mendola D, Sandi C, Markram H (2008) Abnormal fear conditioning and amygdala processing in an animal model of autism. Neuropsychopharmacology 33:901–912.

Martinowich K, Hattori D, Wu H, Fouse S, He F, Hu Y, Fan G, Sun YE (2003) DNA methylation-related chromatin remodeling in activity-dependent BDNF gene regulation. Science 302:890–893.

Matsumura K et al. (2020) Pathogenic POGZ mutation causes impaired cortical development and reversible autism-like phenotypes. Nat Commun 11:859.

Matsuo K, Shinoda Y, Abolhassani N, Nakabeppu Y, Fukunaga K (2022) Transcriptome Analysis in Hippocampus of Rats Prenatally Exposed to Valproic Acid and Effects of Intranasal Treatment of Oxytocin. Front Psychiatry 13:Article 859198.

Mikuni T, Uesaka N, Okuno H, Hirai H, Deisseroth K, Bito H, Kano M (2013) Arc/Arg3.1 Is a Postsynaptic Mediator of Activity-Dependent Synapse Elimination in the Developing Cerebellum. Neuron 78:1024–1035.

Mohawk JA, Cox KH, Sato M, Yoo S-H, Yanagisawa M, Olson EN, Takahashi JS (2019) Neuronal Myocyte-Specific Enhancer Factor 2D (MEF2D) Is Required for Normal Circadian and Sleep Behavior in Mice. The Journal of Neuroscience 39:7958–7967.

Morcom L, Edwards TJ, Rider E, Jones-Davis D, Lim JWC, Chen KS, Dean RJ, Bunt J, Ye Y, Gobius I, Suárez R, Mandelstam S, Sherr EH, Richards LJ (2021) Draxin regulates interhemispheric fissure remodelling to influence the extent of corpus callosum formation. Elife 10:e61618.

Munji RN, Choe Y, Li G, Siegenthaler JA, Pleasure SJ (2011) Wnt signaling regulates neuronal differentiation of cortical intermediate progenitors. Journal of Neuroscience 31:1676–1687.

Mut-Arbona P, Huang L, Baranyi M, Tod P, Iring A, Calzaferri F, de Los Ríos C, Sperlágh B (2023) Dual role of the P2X7 receptor in dendritic outgrowth during physiological and pathological brain development. The Journal of neuroscience : 43:1125–1142 Available at: http://www.ncbi.nlm.nih.gov/pubmed/36732073.

Nau H (1986a) Species Differences in Pharmacokinetics and Drug Teratogenesis. Environ Health Perspect 70:113–129 Available at: https://about.jstor.org/terms.

Nau H (1986b) Transfer of valproic acid and its main active unsaturated metabolite to the gestational tissue: Correlation with neural tube defect formation in the mouse. Teratology 33:21–27.

Nau H, Löscher W (1982) Valproic acid: brain and plasma levels of the drug and its metabolites, anticonvulsant effects and gamma-aminobutyric acid (GABA) metabolism in the mouse. J Pharmacol Exp Ther 220:654–659.

Nelson SB, Valakh V (2015) Excitatory/Inhibitory Balance and Circuit Homeostasis in Autism Spectrum Disorders. Neuron 87:684–698.

Nicolini C, Fahnestock M (2018) The valproic acid-induced rodent model of autism. Exp Neurol 299:217–227.

Nikolaienko O, Patil S, Eriksen MS, Bramham CR (2018) Arc protein: a flexible hub for synaptic plasticity and cognition. Semin Cell Dev Biol 77:33–42.

Noda M, Iwamoto I, Tabata H, Yamagata T, Ito H, Nagata K ichi (2019) Role of Per3, a circadian clock gene, in embryonic development of mouse cerebral cortex. Sci Rep 9:5874.

Nojima K, Miyazaki H, Hori T, Vargova L, Oohashi T (2021) Assessment of Possible Contributions of Hyaluronan and Proteoglycan Binding Link Protein 4 to Differential Perineuronal Net Formation at the Calyx of Held. Front Cell Dev Biol 9:730550.

Notwell JH, Heavner WE, Darbandi SF, Katzman S, McKenna WL, Ortiz-Londono CF, Tastad D, Eckler MJ, Rubenstei JLR, McConnell SK, Chen B, Bejerano G (2016) TBR1 regulates autism risk genes in the developing neocortex. Genome Res 26:1013–1022.

Orenstein N, Gofin Y, Shomron N, Ruhrman-Shahar N, Magal N, Hagari O, Azulay N, Bazak L, Goldberg Y, Basel-Salmon L (2022) DYRK1B haploinsufficiency in a family with metabolic syndrome and abnormal cognition. Clin Genet 101:265–266.

Pai ELL, Vogt D, Clemente-Perez A, McKinsey GL, Cho FS, Hu JS, Wimer M, Paul A, Fazel Darbandi S, Pla R, Nowakowski TJ, Goodrich L v., Paz JT, Rubenstein JLR (2019) Mafb and c-Maf Have Prenatal Compensatory and Postnatal Antagonistic Roles in Cortical Interneuron Fate and Function. Cell Rep 26:1157–1173.e5.

Park J, Chung KC (2013) New Perspectives of Dyrk1A Role in Neurogenesis and Neuropathologic Features of Down Syndrome. Exp Neurobiol 22:244–248.

Paulsen B et al. (2022) Autism genes converge on asynchronous development of shared neuron classes. Nature 602:268–273.

Peñagarikano O, Geschwind DH (2012) What does CNTNAP2 reveal about autism spectrum disorder? Trends Mol Med 18:156–163.

Poelmans G, Franke B, Pauls DL, Glennon JC, Buitelaar JK (2013) AKAPs integrate genetic findings for autism spectrum disorders. Transl Psychiatry 3:e270.

Proenca CC, Gao KP, Shmelkov S v., Rafii S, Lee FS (2011) Slitrks as emerging candidate genes involved in neuropsychiatric disorders. Trends Neurosci 34:143–153.

Qian Z, Zhang R, Zhou J, Sun S, Di Y, Ren W, Tian Y (2018) RNA-Seq data on prefrontal cortex in valproic acid model of autism and control rats. Data Brief 18:787–789.

Rachubinski AL, Hepburn S, Elias ER, Gardiner K, Shaikh TH (2017) The co-occurrence of Down syndrome and autism spectrum disorder: is it because of additional genetic variations? Prenat Diagn 37:31–36.

Ramsaran AI et al. (2023) A shift in the mechanisms controlling hippocampal engram formation during brain maturation. Science 380:543–551.

Rash BG, Tomasi S, Lim HD, Suh CY, Vaccarino FM (2013) Cortical gyrification induced by fibroblast growth factor 2 in the mouse brain. Journal of Neuroscience 33:10802–10814.

Rauch A et al. (2012) Range of genetic mutations associated with severe non-syndromic sporadic intellectual disability: An exome sequencing study. The Lancet 380:1674–1682.

Ronesi JA, Collins KA, Hays SA, Tsai NP, Guo W, Birnbaum SG, Hu JH, Worley PF, Gibson JR, Huber KM (2012) Disrupted Homer scaffolds mediate abnormal mGluR5 function in a mouse model of fragile X syndrome. Nat Neurosci 15:431–440.

Rosenberg RE, Law JK, Yenokyan G, Mcgready J, Kaufmann WE, Law PA (2009) Characteristics and Concordance of Autism Spectrum Disorders Among 277 Twin Pairs. Arch Pediatr Adolesc Med 163:907–914.

Ross AJ, Capel B (2005) Signaling at the crossroads of gonad development. Trends in Endocrinology and Metabolism 16:19–25.

Roston A, Evans D, Gill H, McKinnon M, Isidor B, Cogné B, Mwenifumbo J, van Karnebeek C, An J, Jones SJM, Farrer M, Demos M, Connolly M, Gibson WT (2021) SETD1B -associated neurodevelopmental disorder. J Med Genet 58:196–204.

Rubenstein JLR, Merzenich MM (2003) Model of autism: increased ratio of excitation/ inhibition in key neural systems. Genes Brain Behav 2:255–267.

Ruzzo EK, Pérez-Cano L, Jung JY, Wang L kai, Kashef-Haghighi D, Hartl C, Singh C, Xu J, Hoekstra JN, Leventhal O, Leppä VM, Gandal MJ, Paskov K, Stockham N, Polioudakis D, Lowe JK, Prober DA, Geschwind DH, Wall DP (2019) Inherited and De Novo Genetic Risk for Autism Impacts Shared Networks. Cell 178:850–866.e26.

Ryu S, Mahler J, Acampora D, Holzschuh J, Erhardt S, Omodei D, Simeone A, Driever W (2007) Orthopedia Homeodomain Protein Is Essential for Diencephalic Dopaminergic Neuron Development. Current Biology 17:873–880.

Salm EJ, Dunn PJ, Shan L, Yamasaki M, Malewicz NM, Miyazaki T, Park J, Sumioka A, Hamer RRL, He WW, Morimoto-Tomita M, LaMotte RH, Tomita S (2020) TMEM163 Regulates ATP-Gated P2X Receptor and Behavior. Cell Rep 31.

Sandberg M, Flandin P, Silberberg S, Su-Feher L, Price JD, Hu JS, Kim C, Visel A, Nord AS, Rubenstein JLR (2016) Transcriptional Networks Controlled by NKX2-1 in the Development of Forebrain GABAergic Neurons. Neuron 91:1260–1275.

Satterstrom FK et al. (2020) Large-Scale Exome Sequencing Study Implicates Both Developmental and Functional Changes in the Neurobiology of Autism. Cell 180:568–584.e23.

Seigneur E, Wang J, Dai J, Polepalli J, Südhof TC (2021) Cerebellin-2 regulates a serotonergic dorsal raphe circuit that controls compulsive behaviors. Mol Psychiatry 26:7509–7521.

Semino F, Schröter J, Willemsen MH, Bast T, Biskup S, Beck-Woedl S, Brennenstuhl H, Schaaf CP, Kölker S, Hoffmann GF, Haack TB, Syrbe S (2021) Further evidence for de novo variants in SYNCRIP as the cause of a neurodevelopmental disorder. Hum Mutat 42:1094–1100.

Sessa A, Ciabatti E, Drechsel D, Massimino L, Colasante G, Giannelli S, Satoh T, Akira S, Guillemot F, Broccoli V (2017) The Tbr2 Molecular Network Controls Cortical Neuronal Differentiation Through Complementary Genetic and Epigenetic Pathways. Cerebral Cortex 27:3378–3396.

Shibata M, Pattabiraman K, Lorente-Galdos B, Andrijevic D, Kim SK, Kaur N, Muchnik SK, Xing X, Santpere G, Sousa AMM, Sestan N (2021) Regulation of prefrontal patterning and connectivity by retinoic acid. Nature 598:483–488.

Shiraishi-Yamaguchi Y, Furuichi T (2007) The Homer family proteins. Genome Biol 8:206.

Shmelkov S v, Hormigo A, Jing D, Proenca CC, Bath KG, Milde T, Shmelkov E, Kushner JS, Baljevic M, Dincheva I, Murphy AJ, Valenzuela DM, Gale NW, Yancopoulos GD, Ninan I, Lee FS, Rafii S (2010) Slitrk5 deficiency impairs corticostriatal circuitry and leads to obsessive-compulsive-like behaviors in mice. Nat Med 16:598–602, 1p following 602.

Siegenthaler JA, Ashique AM, Zarbalis K, Patterson KP, Hecht JH, Kane MA, Folias AE, Choe Y, May SR, Kume T, Napoli JL, Peterson AS, Pleasure SJ (2009) Retinoic Acid from the Meninges Regulates Cortical Neuron Generation. Cell 139:597–609.

Silbereis JC, Pochareddy S, Zhu Y, Li M, Sestan N (2016) The Cellular and Molecular Landscapes of the Developing Human Central Nervous System. Neuron 89:248–268.

Silverman JL, Yang M, Lord C, Crawley JN (2010) Behavioural phenotyping assays for mouse models of autism. Nat Rev Neurosci 11:490–502.

Singh R, Kumar Dhanyamraju P, Lauth M (2017) Oncotarget 833 www.impactjournals.com/oncotarget DYRK1B blocks canonical and promotes non-canonical Hedgehog signaling through activation of the mTOR/AKT pathway. Oncotarget 8:833–845 Available at: www.impactjournals.com/oncotarget/.

Singh R, Lauth M (2017) Emerging roles of DYRK kinases in embryogenesis and Hedgehog pathway control. J Dev Biol 5:13.

Song M, Mathews CA, Stewart SE, Shmelkov S v., Mezey JG, Rodriguez-Flores JL, Rasmussen SA, Britton JC, Oh YS, Walkup JT, Lee FS, Glatt CE (2017) Rare synaptogenesis-impairing mutations in SLITRK5 are associated with obsessive compulsive disorder. PLoS One 12.

Stahl R, Walcher T, de Juan Romero C, Pilz GA, Cappello S, Irmler M, Sanz-Aquela JM, Beckers J, Blum R, Borrell V, Götz M (2013) Trnp1 regulates expansion and folding of the mammalian cerebral cortex by control of radial glial fate. Cell 153:535–549.

Stanton-Turcotte D, Hsu K, Moore SA, Yamada M, Fawcett JP, Iulianella A (2022) Mllt11 regulates migration and neurite outgrowth of cortical projection neurons during development. The Journal of Neuroscience 42:3931–3948 Available at: https://www.jneurosci.org/lookup/doi/10.1523/JNEUROSCI.0124-22.2022.

Stessman HAF et al. (2016) Disruption of POGZ Is Associated with Intellectual Disability and Autism Spectrum Disorders. Am J Hum Genet 98:541–552.

Stessman HAF et al. (2017) Targeted sequencing identifies 91 neurodevelopmental-disorder risk genes with autism and developmental-disability biases. Nat Genet 49:515–526.

Stumpo DJ, Bockt CB, Tuttle JS, Blackshear PJ (1995) MARCKS deficiency in mice leads to abnormal brain development and perinatal death (myristoylated, alanine-rich C kinase substrate/protein kinase C/Macs gene). Dev Biol 92:944–948.

Subramanian K, Brandenburg C, Orsati F, Soghomonian JJ, Hussman JP, Blatt GJ (2017) Basal ganglia and autism – a translational perspective. Autism Research 10:1751–1775.

Suliman-Lavie R, Title B, Cohen Y, Hamada N, Tal M, Tal N, Monderer-Rothkoff G, Gudmundsdottir B, Gudmundsson KO, Keller JR, Huang GJ, Nagata K ichi, Yarom Y, Shifman S (2020) Pogz deficiency leads to transcription dysregulation and impaired cerebellar activity underlying autism-like behavior in mice. Nat Commun 11:5836.

Surén P, Roth C, Bresnahan M, Haugen M, Hornig M, Hirtz D, Lie KK, Lipkin WI, Reichborn-Kjennerud T, Schjølberg S, George M, Smith D, Øyen A-S, Susser E, Stoltenberg C (2013) Association Between Maternal Use of Folic Acid Supplements and Risk of Autism Spectrum Disorders in Children. J Am Med Assoc 309:570–577 Available at: www.jama.com.

Szu J, Wojcinski A, Jiang P, Kesari S (2021) Impact of the Olig Family on Neurodevelopmental Disorders. Front Neurosci 15:659601.

Tanaka AJ, Cho MT, Willaert R, Retterer K, Zarate YA, Bosanko K, Stefans V, Oishi K, Williamson A, Wilson GN, Basinger A, Barbaro-Dieber T, Ortega L, Sorrentino S, Gabriel MK, Anderson IJ, Sacoto MJG, Schnur RE, Chung WK (2017) De novo variants in EBF3 are associated with hypotonia, developmental delay, intellectual disability, and autism. Cold Spring Harb Mol Case Stud 3:a002097.

Tao W, Díaz-Alonso J, Sheng N, Nicoll RA (2018) Postsynaptic δ1 glutamate receptor assembles and maintains hippocampal synapses via Cbln2 and neurexin. Proceedings of the National Academy of Sciences USA 115:E5373–E5381.

Tisato V, Silva JA, Longo G, Gallo I, Singh A V., Milani D, Gemmati D (2021) Genetics and epigenetics of one-carbon metabolism pathway in autism spectrum disorder: A sex-specific brain epigenome? Genes (Basel) 12:782.

Tocco C, Bertacchi M, Studer M (2021) Structural and Functional Aspects of the Neurodevelopmental Gene NR2F1: From Animal Models to Human Pathology. Front Mol Neurosci 14:767965.

Toledo EM, Yang S, Gyllborg D, van Wijk KE, Sinha I, Varas-Godoy M, Grigsby CL, Lönnerberg P, Islam S, Steffensen KR, Linnarsson S, Arenas E (2020) Srebf1 Controls Midbrain Dopaminergic Neurogenesis. Cell Rep 31:107607.

Tripodi M, Filosa A, Armentano M, Studer M (2004) The COUP-TF nuclear receptors regulate cell migration in the mammalian basal forebrain. Development 131:6119–6129.

Tutukova S, Tarabykin V, Hernandez-Miranda LR (2021) The Role of Neurod Genes in Brain Development, Function, and Disease. Front Mol Neurosci 14:662774.

Uezu A, Hisey E, Kobayashi Y, Gao Y, Bradshaw TWA, Devlin P, Rodriguiz R, Tata PR, Soderling SH (2019) Essential role for insyn1 in dystroglycan complex integrity and cognitive behaviors in mice. Elife 8.

Uezu A, Kanak DJ, Bradshaw TWA, Soderblom EJ, Catavero CM, Burette AC, Weinberg RJ, Soderling SH (2016) Identification of an elaborate complex mediating postsynaptic inhibition. Science (1979) 353:1123–1129.

Walker CK, Greathouse KM, Tuscher JJ, Dammer EB, Weber AJ, Liu E, Curtis KA, Boros BD, Freeman CD, Seo JV, Ramdas R, Hurst C, Duong DM, Gearing M, Murchison CF, Day JJ, Seyfried NT, Herskowitz JH (2023) Cross-Platform Synaptic Network Analysis of Human Entorhinal Cortex Identifies TWF2 as a Modulator of Dendritic Spine Length. The Journal of Neuroscience:JN-RM-2102–22 Available at:autism spectrum disorderss https://www.jneurosci.org/lookup/doi/10.1523/JNEUROSCI.2102-22.2023.

Wang SHJ, Celic I, Choi SY, Riccomagno M, Wang Q, Sun LO, Mitchell SP, Vasioukhin V, Huganir RL, Kolodkin AL (2014) Dlg5 regulates dendritic spine formation and synaptogenesis by controlling subcellular N-cadherin localization. Journal of Neuroscience 34:12745–12761.

Weil D, Piton A, Lessel D, Standart N (2020) Mutations in genes encoding regulators of mRNA decapping and translation initiation: Links to intellectual disability. Biochem Soc Trans 48:1199– 1211.

Willsey AJ et al. (2013) Coexpression networks implicate human midfetal deep cortical projection neurons in the pathogenesis of autism. Cell 155:997–1007.

Windhorst S, Minge D, Bähring R, Hüser S, Schob C, Blechner C, Lin HY, Mayr GW, Kindler S (2012) Inositol-1,4,5-trisphosphate 3-kinase A regulates dendritic morphology and shapes synaptic Ca 2+ transients. Cell Signal 24:750–757.

Wingett SW, Andrews S (2018) Fastq screen: A tool for multi-genome mapping and quality control. F1000Res 7:1–14.

Wircer E, Blechman J, Borodovsky N, Tsoory M, Nunes AR, Oliveira RF, Levkowitz G (2017) Homeodomain protein Otp affects developmental neuropeptide switching in oxytocin neurons associated with a long-term effect on social behavior. Elife 6:e22170.

Xu D et al. (2013) Top3β is an RNA topoisomerase that works with fragile X syndrome protein to promote synapse formation. Nat Neurosci 16:1238–1247.

Xu XH, Deng CY, Liu Y, He M, Peng J, Wang T, Yuan L, Zheng ZS, Blackshear PJ, Luo ZG (2014) MARCKS regulates membrane targeting of Rab10 vesicles to promote axon development. Cell Res 24:576–594.

Yan CH, Levesque M, Claxton S, Johnson RL, Ang SL (2011) Lmx1a and Lmx1b function cooperatively to regulate proliferation, specification, and differentiation of midbrain dopaminergic progenitors. Journal of Neuroscience 31:12413–12425.

Yger M, Girault JA (2011) DARPP-32, jack of all trades…master of which? Front Behav Neurosci 5:Article 56.

Yook C, Kim K, Kim D, Kang H, Kim SG, Kim E, Kim SY (2019) A TBR1-K228E Mutation Induces Tbr1 Upregulation, Altered Cortical Distribution of Interneurons, Increased Inhibitory Synaptic Transmission, and Autistic-Like Behavioral Deficits in Mice. Front Mol Neurosci 12:241.

Yoon S, Piguel NH, Khalatyan N, Dionisio LE, Savas JN, Penzes P (2021) Homer1 promotes dendritic spine growth through ankyrin-G and its loss reshapes the synaptic proteome. Mol Psychiatry 26:1775–1789.

Yuen RKC et al. (2017) Whole genome sequencing resource identifies 18 new candidate genes for autism spectrum disorder. Nat Neurosci 20:602–611.

Zaslavsky K et al. (2019) SHANK2 mutations associated with autism spectrum disorder cause hyper-connectivity of human neurons. Nat Neurosci 22:556–564.

Zawadzka A, Cieślik M, Adamczyk A (2021) The role of maternal immune activation in the pathogenesis of autism: A review of the evidence, proposed mechanisms and implications for treatment. Int J Mol Sci 22:11516.

Zeidan J, Fombonne E, Scorah J, Ibrahim A, Durkin MS, Saxena S, Yusuf A, Shih A, Elsabbagh M (2022) Global prevalence of autism: A systematic review update. Autism Research 15:778–790.

Zhang R, Zhou J, Ren J, Sun S, Di Y, Wang H, An X, Zhang K, Zhang J, Qian Z, Shi M, Qiao Y, Ren W, Tian Y (2018) Transcriptional and splicing dysregulation in the prefrontal cortex in valproic acid rat model of autism. Reproductive Toxicology 77:53–61.

Zhao H, Wang Q, Yan T, Zhang Y, Xu H juan, Yu H peng, Tu Z, Guo X, Jiang Y hui, Li X jiang, Zhou H, Zhang YQ (2019) Maternal valproic acid exposure leads to neurogenesis defects and autism-like behaviors in non-human primates. Transl Psychiatry 9:267.

Zheng W, Geng AQ, Li PF, Wang Y, Yuan XB (2012) Robo4 regulates the radial migration of newborn neurons in developing neocortex. Cerebral Cortex 22:2587–2601.

Zigman T, Petković Ramadža D, Šimić G, Barić I (2021) Inborn Errors of Metabolism Associated With Autism Spectrum Disorders: Approaches to Intervention. Front Neurosci 15:673600.

