## Supplemental Table S1 for "Rapid effects of valproic acid on the fetal brain transcriptome: Implications for brain development and autism"

| | VPA | VPA | VPA | VPA | VPA | VPA | VPA | Control | Control | Control | Control | Control | Control | Control | VPA | VPA | VPA | VPA | VPA | VPA | VPA | Control | Control | Control | Control | Control | Control | Control | |
|---|---|---|---|---|---|---|---|---|---|---|---|---|---|---|---|---|---|---|---|---|---|---|---|---|---|---|---|---|---|
| | male | male | male | male | male | male | male | male | male | male | male | male | male | male | female | female | female | female | female | female | female | female | female | female | female | female | female | female | |
| Gene | V1A | V2B | V3B | V4C | V5A | V6D | V7A | C1D | C2A | C3A | C4C | C5C | C6B | C7A | V1B | V2A | V3A | V4A | V5F | V6A | V7C | C1A | C2C | C3B | C4A | C5A | C6A | C7C | Gene |
| 0610009B22Rik | 889 | 916 | 1136 | 1004 | 1019 | 818 | 1186 | 1299 | 1121 | 1125 | 1088 | 1047 | 1159 | 1361 | 1085 | 935 | 1272 | 1159 | 973 | 1312 | 1147 | 954 | 1213 | 1193 | 1083 | 1315 | 1189 | 1386 | 0610009B22Rik |
| 0610010K14Rik | 161 | 201 | 247 | 240 | 225 | 229 | 235 | 255 | 220 | 289 | 272 | 235 | 257 | 303 | 170 | 141 | 240 | 222 | 214 | 178 | 268 | 249 | 195 | 284 | 213 | 294 | 235 | 285 | 0610010K14Rik |
| 0610012G03Rik | 1219 | 1420 | 1049 | 1073 | 1100 | 1032 | 1244 | 1369 | 1296 | 1490 | 1337 | 1334 | 1323 | 1464 | 1131 | 975 | 1258 | 1158 | 1292 | 1358 | 1244 | 1390 | 1335 | 1381 | 1416 | 1561 | 1281 | 1362 | 0610012G03Rik |
| 0610030E20Rik | 114 | 41 | 39 | 82 | 66 | 34 | 48 | 85 | 49 | 52 | 57 | 88 | 67 | 64 | 70 | 56 | 42 | 42 | 55 | 31 | 32 | 77 | 65 | 60 | 50 | 56 | 66 | 64 | 0610030E20Rik |
| 0610040J01Rik | 114 | 124 | 90 | 90 | 141 | 114 | 112 | 103 | 129 | 115 | 149 | 59 | 90 | 81 | 104 | 144 | 82 | 108 | 115 | 93 | 117 | 100 | 158 | 102 | 93 | 86 | 60 | 71 | 0610040J01Rik |
| 1110002E22Rik | 37 | 39 | 12 | 64 | 55 | 64 | 41 | 59 | 105 | 58 | 555 | 36 | 87 | 21 | 47 | 61 | 44 | 72 | 45 | 35 | 37 | 119 | 374 | 57 | 55 | 17 | 26 | 15 | 1110002E22Rik |
| 1110004F10Rik | 8880 | 8767 | 10225 | 8794 | 8571 | 8678 | 9571 | 10492 | 10156 | 10830 | 10106 | 11692 | 10727 | 13064 | 8923 | 8181 | 9616 | 8556 | 8933 | 10116 | 9400 | 9535 | 9716 | 11709 | 11517 | 13661 | 12535 | 13396 | 1110004F10Rik |
| 1110017D15Rik | 378 | 536 | 364 | 737 | 653 | 571 | 607 | 139 | 152 | 161 | 113 | 167 | 221 | 130 | 375 | 508 | 561 | 810 | 659 | 573 | 676 | 199 | 207 | 148 | 152 | 148 | 143 | 159 | 1110017D15Rik |
| 1110025L11Rik | 0 | 0 | 0 | 0 | 0 | 4 | 0 | 0 | 0 | 0 | 0 | 0 | 0 | 0 | 0 | 0 | 0 | 0 | 0 | 0 | 0 | 0 | 0 | 0 | 0 | 0 | 0 | 0 | 1110025L11Rik |
| 1110032A03Rik | 1526 | 1495 | 1335 | 1422 | 1508 | 1352 | 1501 | 1387 | 1308 | 1290 | 1173 | 1218 | 1439 | 1397 | 1683 | 1391 | 1428 | 1305 | 1340 | 1629 | 1547 | 1365 | 1394 | 1242 | 1281 | 1401 | 1399 | 1310 | 1110032A03Rik |
| 1110032F04Rik | 223 | 177 | 220 | 271 | 316 | 279 | 286 | 201 | 265 | 247 | 253 | 250 | 204 | 208 | 257 | 178 | 301 | 282 | 334 | 294 | 311 | 284 | 273 | 246 | 301 | 187 | 207 | 179 | 1110032F04Rik |
| 1110038F14Rik | 1068 | 1168 | 1434 | 1223 | 1211 | 1069 | 1091 | 1480 | 1795 | 1828 | 1883 | 1977 | 1907 | 1876 | 984 | 1148 | 1333 | 1125 | 1297 | 1254 | 963 | 1529 | 1772 | 2106 | 1763 | 2534 | 2164 | 2143 | 1110038F14Rik |
| 1110051M20Rik | 3398 | 3971 | 2817 | 3503 | 3466 | 3307 | 3832 | 3677 | 3426 | 3345 | 3121 | 3515 | 3272 | 3859 | 3611 | 3168 | 3231 | 3374 | 3607 | 3762 | 3751 | 3740 | 3447 | 3495 | 3605 | 3709 | 3475 | 3893 | 1110051M20Rik |
| 1110059E24Rik | 4479 | 3648 | 3903 | 3464 | 3004 | 3331 | 3923 | 3303 | 2708 | 2307 | 2282 | 2830 | 2733 | 3069 | 4718 | 4158 | 3668 | 3320 | 3385 | 4227 | 3649 | 2364 | 2706 | 2606 | 2601 | 3058 | 3051 | 3207 | 1110059E24Rik |
| 1110059G10Rik | 867 | 633 | 827 | 865 | 818 | 751 | 896 | 1489 | 1233 | 1346 | 1231 | 1328 | 1296 | 1586 | 853 | 643 | 819 | 710 | 871 | 1030 | 820 | 1193 | 1311 | 1469 | 1297 | 1459 | 1474 | 1455 | 1110059G10Rik |
| 1110065P20Rik | 751 | 875 | 640 | 875 | 832 | 829 | 790 | 901 | 887 | 883 | 824 | 974 | 900 | 867 | 736 | 833 | 885 | 763 | 780 | 873 | 837 | 714 | 830 | 893 | 887 | 915 | 809 | 946 | 1110065P20Rik |
| 1190005I06Rik | 1 | 10 | 12 | 9 | 6 | 10 | 8 | 4 | 3 | 8 | 3 | 0 | 1 | 3 | 3 | 11 | 12 | 12 | 0 | 7 | 10 | 2 | 5 | 2 | 2 | 2 | 6 | 4 | 1190005I06Rik |
| 1190007I07Rik | 344 | 300 | 409 | 286 | 391 | 379 | 399 | 491 | 438 | 616 | 474 | 579 | 556 | 613 | 330 | 276 | 341 | 314 | 316 | 377 | 338 | 528 | 440 | 614 | 616 | 566 | 539 | 622 | 1190007I07Rik |
| 1300017J02Rik | 82 | 141 | 166 | 58 | 29 | 52 | 21 | 38 | 43 | 43 | 36 | 71 | 322 | 192 | 83 | 129 | 81 | 39 | 54 | 37 | 55 | 23 | 30 | 38 | 12 | 89 | 141 | 354 | 1300017J02Rik |
| 1500009C09Rik | 1323 | 1836 | 1364 | 2377 | 2398 | 2307 | 2767 | 1305 | 1200 | 1518 | 1149 | 1338 | 792 | 1484 | 1669 | 934 | 2168 | 2206 | 2906 | 2534 | 3084 | 1369 | 1177 | 1398 | 1705 | 1138 | 1088 | 1298 | 1500009C09Rik |
| 1500009L16Rik | 1430 | 1353 | 1500 | 1454 | 1334 | 1292 | 1655 | 880 | 773 | 827 | 753 | 770 | 814 | 935 | 1547 | 1439 | 1534 | 1384 | 1280 | 1422 | 1482 | 819 | 858 | 805 | 871 | 893 | 823 | 1013 | 1500009L16Rik |
| 1500011B03Rik | 3831 | 3873 | 3538 | 4049 | 4017 | 3838 | 4421 | 4070 | 3508 | 4004 | 3484 | 3696 | 3498 | 4078 | 4237 | 3294 | 4172 | 3954 | 3919 | 4287 | 4412 | 4111 | 3358 | 3809 | 4131 | 3838 | 3691 | 3983 | 1500011B03Rik |
| 1520401A03Rik | 33 | 10 | 13 | 41 | 43 | 17 | 35 | 22 | 44 | 27 | 184 | 14 | 16 | 27 | 20 | 19 | 35 | 22 | 25 | 20 | 25 | 43 | 111 | 19 | 23 | 18 | 9 | 12 | 1520401A03Rik |
| 1600012H06Rik | 2230 | 2017 | 1955 | 2029 | 1621 | 1754 | 2006 | 2176 | 2165 | 2010 | 2152 | 2038 | 2203 | 2087 | 2341 | 2245 | 1920 | 2001 | 1718 | 2071 | 1779 | 2095 | 2273 | 1995 | 1886 | 2494 | 2172 | 2238 | 1600012H06Rik |
| 1600014C10Rik | 685 | 747 | 749 | 653 | 853 | 726 | 670 | 1465 | 1303 | 1409 | 1358 | 1159 | 1275 | 1180 | 819 | 776 | 781 | 637 | 655 | 781 | 679 | 1480 | 1405 | 1332 | 1243 | 1188 | 1226 | 1280 | 1600014C10Rik |
| 1600014C23Rik | 1 | 0 | 1 | 0 | 0 | 0 | 0 | 0 | 1 | 1 | 3 | 0 | 2 | 3 | 7 | 2 | 0 | 7 | 1 | 3 | 2 | 0 | 3 | 3 | 1 | 4 | 1 | 7 | 1600014C23Rik |
| 1700001C19Rik | 3 | 0 | 2 | 0 | 1 | 3 | 4 | 0 | 4 | 2 | 0 | 1 | 5 | 4 | 0 | 0 | 3 | 1 | 2 | 4 | 3 | 3 | 3 | 1 | 1 | 3 | 0 | 2 | 1700001C19Rik |
| 1700001F09Rik | 0 | 0 | 0 | 0 | 0 | 0 | 0 | 1 | 1 | 0 | 8 | 0 | 0 | 3 | 3 | 1 | 0 | 0 | 0 | 2 | 0 | 5 | 1 | 2 | 0 | 0 | 0 | 0 | 1700001F09Rik |
| 1700001J03Rik | 24 | 34 | 52 | 72 | 74 | 54 | 50 | 10 | 12 | 14 | 28 | 24 | 5 | 21 | 59 | 25 | 57 | 80 | 57 | 30 | 48 | 22 | 13 | 16 | 10 | 28 | 28 | 14 | 1700001J03Rik |
| 1700001K19Rik | 0 | 1 | 0 | 1 | 0 | 0 | 1 | 0 | 0 | 0 | 0 | 0 | 0 | 0 | 0 | 0 | 0 | 0 | 0 | 0 | 0 | 0 | 0 | 0 | 0 | 0 | 0 | 0 | 1700001K19Rik |
| 1700001L19Rik | 104 | 96 | 127 | 167 | 160 | 116 | 113 | 56 | 53 | 63 | 57 | 53 | 70 | 48 | 137 | 109 | 123 | 109 | 137 | 108 | 132 | 77 | 54 | 40 | 62 | 58 | 40 | 49 | 1700001L19Rik |
| 1700001O22Rik | 251 | 309 | 294 | 300 | 243 | 242 | 269 | 64 | 121 | 80 | 76 | 139 | 90 | 99 | 227 | 248 | 329 | 281 | 288 | 247 | 244 | 72 | 87 | 122 | 79 | 171 | 94 | 107 | 1700001O22Rik |
| 1700001P01Rik | 54 | 64 | 88 | 88 | 89 | 78 | 58 | 7 | 30 | 23 | 27 | 62 | 33 | 37 | 36 | 49 | 81 | 107 | 83 | 64 | 51 | 40 | 32 | 40 | 41 | 49 | 27 | 25 | 1700001P01Rik |
| 1700003E16Rik | 294 | 516 | 370 | 450 | 369 | 357 | 326 | 76 | 86 | 118 | 77 | 78 | 103 | 48 | 286 | 368 | 454 | 494 | 402 | 349 | 374 | 80 | 139 | 98 | 90 | 122 | 66 | 76 | 1700003E16Rik |
| 1700003F12Rik | 10 | 22 | 23 | 18 | 16 | 26 | 17 | 7 | 4 | 12 | 10 | 21 | 17 | 16 | 17 | 30 | 12 | 20 | 21 | 27 | 10 | 12 | 10 | 17 | 6 | 12 | 16 | 11 | 1700003F12Rik |
| 1700006A11Rik | 0 | 0 | 0 | 0 | 0 | 0 | 0 | 0 | 0 | 0 | 0 | 0 | 0 | 0 | 0 | 0 | 0 | 1 | 0 | 0 | 0 | 0 | 0 | 0 | 0 | 0 | 0 | 0 | 1700006A11Rik |
| 1700007K13Rik | 81 | 78 | 71 | 80 | 66 | 87 | 64 | 38 | 40 | 57 | 32 | 54 | 60 | 64 | 102 | 100 | 51 | 64 | 58 | 77 | 79 | 35 | 78 | 47 | 40 | 49 | 46 | 51 | 1700007K13Rik |
| 1700008O03Rik | 33 | 38 | 48 | 42 | 44 | 68 | 30 | 31 | 59 | 25 | 48 | 56 | 53 | 39 | 24 | 28 | 48 | 23 | 37 | 40 | 29 | 23 | 49 | 33 | 47 | 70 | 36 | 69 | 1700008O03Rik |
| 1700008P02Rik | 0 | 0 | 0 | 0 | 0 | 0 | 0 | 1 | 0 | 0 | 0 | 0 | 0 | 0 | 0 | 0 | 0 | 0 | 0 | 0 | 0 | 2 | 0 | 0 | 0 | 0 | 0 | 0 | 1700008P02Rik |
| 1700009N14Rik | 0 | 1 | 0 | 6 | 0 | 0 | 0 | 0 | 0 | 0 | 4 | 3 | 3 | 1 | 5 | 4 | 3 | 0 | 0 | 0 | 0 | 2 | 5 | 0 | 2 | 5 | 13 | 1 | 1700009N14Rik |
| 1700010B08Rik | 1 | 1 | 1 | 0 | 2 | 4 | 3 | 1 | 0 | 0 | 0 | 2 | 0 | 0 | 3 | 2 | 0 | 0 | 0 | 3 | 1 | 0 | 1 | 0 | 1 | 0 | 0 | 0 | 1700010B08Rik |
| 1700010I14Rik | 197 | 192 | 236 | 188 | 160 | 195 | 149 | 175 | 170 | 187 | 161 | 237 | 228 | 233 | 227 | 183 | 245 | 183 | 146 | 196 | 151 | 180 | 161 | 194 | 191 | 218 | 275 | 258 | 1700010I14Rik |
| 1700011L22Rik | 1 | 3 | 1 | 2 | 7 | 26 | 4 | 4 | 4 | 0 | 1 | 0 | 6 | 11 | 5 | 6 | 3 | 3 | 6 | 5 | 13 | 8 | 3 | 2 | 6 | 1 | 5 | 0 | 1700011L22Rik |
| 1700011M02Rik | 0 | 0 | 0 | 0 | 0 | 0 | 0 | 0 | 0 | 0 | 1 | 0 | 0 | 0 | 0 | 0 | 0 | 0 | 0 | 0 | 0 | 0 | 0 | 0 | 0 | 0 | 0 | 0 | 1700011M02Rik |
| 1700012B07Rik | 0 | 0 | 0 | 0 | 0 | 0 | 0 | 1 | 0 | 0 | 0 | 1 | 0 | 0 | 0 | 0 | 0 | 1 | 0 | 0 | 0 | 0 | 0 | 0 | 0 | 1 | 0 | 0 | 1700012B07Rik |
| 1700012B09Rik | 15 | 21 | 3 | 8 | 22 | 35 | 25 | 28 | 16 | 21 | 18 | 26 | 43 | 23 | 36 | 31 | 2 | 18 | 10 | 24 | 19 | 35 | 41 | 28 | 17 | 36 | 13 | 14 | 1700012B09Rik |
| 1700012P22Rik | 1 | 2 | 1 | 0 | 0 | 7 | 0 | 1 | 3 | 5 | 3 | 7 | 15 | 0 | 1 | 2 | 0 | 0 | 0 | 2 | 0 | 3 | 8 | 0 | 0 | 0 | 0 | 0 | 1700012P22Rik |
| 1700013F07Rik | 38 | 37 | 35 | 59 | 52 | 63 | 45 | 37 | 53 | 38 | 40 | 30 | 45 | 43 | 26 | 41 | 46 | 36 | 44 | 45 | 47 | 25 | 56 | 24 | 31 | 20 | 35 | 26 | 1700013F07Rik |
| 1700013G24Rik | 0 | 0 | 0 | 0 | 0 | 2 | 0 | 0 | 1 | 0 | 1 | 0 | 0 | 0 | 0 | 0 | 0 | 0 | 0 | 0 | 0 | 0 | 0 | 0 | 0 | 0 | 0 | 0 | 1700013G24Rik |
| 1700013H16Rik | 5 | 1 | 6 | 11 | 0 | 10 | 6 | 0 | 2 | 3 | 5 | 10 | 2 | 1 | 4 | 5 | 7 | 0 | 1 | 0 | 2 | 2 | 4 | 1 | 6 | 1 | 6 | 2 | 1700013H16Rik |
| 1700014B07Rik | 6 | 5 | 3 | 0 | 0 | 0 | 3 | 1 | 7 | 5 | 4 | 2 | 11 | 9 | 4 | 5 | 8 | 1 | 3 | 2 | 1 | 3 | 6 | 2 | 15 | 8 | 5 | 11 | 1700014B07Rik |
| 1700014D04Rik | 38 | 35 | 17 | 24 | 25 | 33 | 37 | 18 | 16 | 17 | 20 | 16 | 7 | 9 | 42 | 31 | 12 | 36 | 26 | 29 | 28 | 8 | 15 | 16 | 5 | 8 | 0 | 13 | 1700014D04Rik |
| 1700015G11Rik | 0 | 0 | 0 | 0 | 1 | 0 | 0 | 0 | 0 | 0 | 0 | 0 | 0 | 0 | 0 | 0 | 1 | 0 | 0 | 0 | 0 | 0 | 0 | 0 | 0 | 0 | 0 | 0 | 1700015G11Rik |
| 1700016C15Rik | 0 | 0 | 0 | 0 | 0 | 0 | 0 | 1 | 1 | 0 | 0 | 2 | 0 | 1 | 0 | 0 | 0 | 0 | 0 | 0 | 0 | 2 | 2 | 0 | 0 | 0 | 1 | 0 | 1700016C15Rik |
| 1700016D06Rik | 1 | 0 | 0 | 1 | 0 | 0 | 0 | 0 | 2 | 0 | 6 | 0 | 0 | 1 | 0 | 0 | 1 | 0 | 0 | 0 | 1 | 0 | 1 | 0 | 2 | 0 | 0 | 0 | 1700016D06Rik |
| 1700016H13Rik | 1 | 9 | 8 | 10 | 15 | 1 | 6 | 11 | 14 | 0 | 6 | 12 | 20 | 7 | 8 | 10 | 20 | 11 | 9 | 16 | 8 | 13 | 14 | 15 | 8 | 19 | 7 | 12 | 1700016H13Rik |
| 1700016K19Rik | 24 | 39 | 36 | 42 | 47 | 35 | 42 | 35 | 42 | 52 | 37 | 56 | 60 | 22 | 45 | 53 | 57 | 24 | 31 | 41 | 30 | 37 | 63 | 25 | 16 | 15 | 23 | 31 | 1700016K19Rik |
| 1700017B05Rik | 1441 | 1646 | 1757 | 1629 | 1227 | 1325 | 1073 | 664 | 904 | 966 | 1109 | 999 | 966 | 874 | 1230 | 1829 | 1654 | 1547 | 1488 | 1023 | 1093 | 705 | 908 | 1101 | 748 | 1450 | 923 | 1010 | 1700017B05Rik |
| 1700017N19Rik | 0 | 0 | 0 | 3 | 0 | 2 | 0 | 1 | 1 | 1 | 0 | 0 | 1 | 0 | 0 | 0 | 0 | 1 | 2 | 0 | 3 | 0 | 0 | 2 | 0 | 0 | 0 | 1 | 1700017N19Rik |
| 1700018B08Rik | 7 | 25 | 7 | 35 | 25 | 13 | 21 | 0 | 1 | 3 | 0 | 2 | 0 | 0 | 17 | 25 | 14 | 26 | 9 | 10 | 17 | 2 | 5 | 1 | 5 | 2 | 3 | 1 | 1700018B08Rik |
| 1700018F24Rik | 0 | 0 | 0 | 0 | 0 | 0 | 0 | 0 | 1 | 1 | 0 | 0 | 0 | 0 | 0 | 0 | 0 | 0 | 0 | 0 | 0 | 0 | 0 | 0 | 0 | 0 | 0 | 0 | 1700018F24Rik |
| 1700019A02Rik | 78 | 42 | 90 | 73 | 107 | 106 | 110 | 44 | 44 | 61 | 63 | 34 | 41 | 46 | 58 | 66 | 67 | 58 | 79 | 108 | 98 | 42 | 36 | 38 | 49 | 58 | 37 | 43 | 1700019A02Rik |
| 1700019D03Rik | 820 | 625 | 799 | 509 | 448 | 484 | 374 | 139 | 123 | 101 | 172 | 184 | 262 | 178 | 851 | 877 | 532 | 445 | 339 | 396 | 318 | 182 | 154 | 104 | 71 | 156 | 202 | 234 | 1700019D03Rik |
| 1700019N19Rik | 2 | 3 | 1 | 4 | 4 | 1 | 0 | 0 | 2 | 1 | 2 | 7 | 0 | 0 | 0 | 2 | 0 | 0 | 4 | 0 | 1 | 3 | 2 | 2 | 1 | 0 | 0 | 0 | 1700019N19Rik |
| 1700020A23Rik | 0 | 0 | 0 | 0 | 0 | 0 | 0 | 0 | 0 | 0 | 0 | 0 | 0 | 0 | 0 | 0 | 0 | 0 | 0 | 0 | 1 | 0 | 0 | 0 | 0 | 0 | 1 | 0 | 1700020A23Rik |
| 1700020L13Rik | 7 | 4 | 2 | 12 | 7 | 12 | 10 | 5 | 11 | 17 | 10 | 9 | 7 | 16 | 8 | 8 | 0 | 3 | 6 | 7 | 11 | 5 | 9 | 15 | 20 | 5 | 5 | 9 | 1700020L13Rik |
| 1700020L24Rik | 2 | 3 | 10 | 5 | 5 | 2 | 1 | 4 | 7 | 5 | 14 | 14 | 14 | 9 | 1 | 1 | 3 | 1 | 3 | 0 | 1 | 10 | 8 | 7 | 5 | 11 | 2 | 0 | 1700020L24Rik |
| 1700020N01Rik | 3 | 0 | 0 | 2 | 0 | 0 | 0 | 1 | 1 | 0 | 0 | 0 | 0 | 0 | 0 | 0 | 0 | 0 | 4 | 3 | 0 | 0 | 0 | 0 | 0 | 0 | 0 | 0 | 1700020N01Rik |
| 1700020N15Rik | 1 | 0 | 0 | 4 | 1 | 5 | 1 | 0 | 3 | 1 | 0 | 1 | 0 | 0 | 3 | 1 | 0 | 2 | 2 | 4 | 0 | 0 | 1 | 2 | 0 | 3 | 1 | 0 | 1700020N15Rik |
| 1700021F07Rik | 0 | 0 | 0 | 0 | 0 | 0 | 0 | 0 | 2 | 0 | 0 | 0 | 0 | 0 | 0 | 0 | 0 | 0 | 0 | 0 | 0 | 0 | 0 | 0 | 0 | 0 | 0 | 0 | 1700021F07Rik |
| 1700022I11Rik | 12 | 22 | 10 | 12 | 10 | 14 | 4 | 15 | 3 | 1 | 2 | 17 | 12 | 10 | 9 | 7 | 12 | 4 | 16 | 7 | 13 | 5 | 6 | 21 | 5 | 22 | 19 | 1 | 1700022I11Rik |
| 1700024G13Rik | 1 | 4 | 9 | 4 | 9 | 0 | 4 | 12 | 8 | 7 | 7 | 11 | 7 | 3 | 8 | 12 | 3 | 12 | 0 | 5 | 12 | 7 | 10 | 5 | 2 | 2 | 5 | 2 | 1700024G13Rik |
| 1700025G04Rik | 7124 | 6994 | 6160 | 7478 | 8260 | 8459 | 8383 | 8619 | 7669 | 8651 | 7690 | 8073 | 8172 | 8371 | 7644 | 7020 | 7408 | 7390 | 8191 | 7495 | 8192 | 8905 | 7976 | 7823 | 9112 | 6082 | 8370 | 8005 | 1700025G04Rik |
| 1700028J19Rik | 21 | 25 | 24 | 15 | 20 | 25 | 18 | 39 | 44 | 73 | 49 | 65 | 45 | 65 | 8 | 33 | 19 | 12 | 15 | 24 | 14 | 65 | 46 | 35 | 53 | 67 | 52 | 62 | 1700028J19Rik |
| 1700028K03Rik | 13 | 6 | 4 | 15 | 15 | 26 | 13 | 12 | 4 | 8 | 10 | 12 | 6 | 7 | 7 | 3 | 22 | 14 | 23 | 12 | 17 | 5 | 8 | 12 | 11 | 2 | 6 | 0 | 1700028K03Rik |

| | | | | | | | | | | | | | | | | | | | | | | | | | | | | | |
|---|---|---|---|---|---|---|---|---|---|---|---|---|---|---|---|---|---|---|---|---|---|---|---|---|---|---|---|---|---|
| 1700028P14Rik | 35 | 27 | 8 | 23 | 21 | 42 | 31 | 35 | 35 | 19 | 27 | 24 | 37 | 28 | 41 | 30 | 7 | 33 | 26 | 33 | 31 | 42 | 50 | 24 | 32 | 20 | 35 | 25 | 1700028P14Rik |
| 1700029H14Rik | 9 | 16 | 6 | 3 | 2 | 12 | 4 | 1 | 6 | 1 | 0 | 4 | 1 | 0 | 13 | 10 | 6 | 4 | 2 | 4 | 2 | 2 | 1 | 0 | 3 | 3 | 6 | 2 | 1700029H14Rik |
| 1700029I15Rik | 102 | 117 | 137 | 148 | 124 | 108 | 89 | 82 | 83 | 90 | 86 | 115 | 90 | 81 | 104 | 103 | 138 | 122 | 138 | 94 | 99 | 107 | 87 | 92 | 101 | 90 | 80 | 106 | 1700029I15Rik |
| 1700029J07Rik | 208 | 226 | 224 | 248 | 262 | 244 | 274 | 506 | 446 | 493 | 517 | 383 | 387 | 419 | 265 | 217 | 305 | 263 | 260 | 255 | 259 | 536 | 495 | 434 | 479 | 375 | 483 | 392 | 1700029J07Rik |
| 1700030J22Rik | 599 | 848 | 707 | 735 | 664 | 703 | 651 | 372 | 356 | 425 | 334 | 346 | 400 | 349 | 608 | 720 | 703 | 792 | 711 | 603 | 703 | 438 | 368 | 376 | 449 | 260 | 381 | 323 | 1700030J22Rik |
| 1700030K09Rik | 176 | 169 | 226 | 231 | 328 | 253 | 251 | 1303 | 1388 | 1474 | 1504 | 1340 | 1337 | 1344 | 200 | 160 | 255 | 206 | 330 | 293 | 264 | 1419 | 1298 | 1455 | 1632 | 1405 | 1446 | 1356 | 1700030K09Rik |
| 1700031F05Rik | 0 | 0 | 4 | 2 | 0 | 1 | 2 | 1 | 0 | 0 | 0 | 0 | 2 | 0 | 0 | 1 | 0 | 0 | 0 | 0 | 2 | 0 | 1 | 0 | 0 | 0 | 0 | 0 | 1700031F05Rik |
| 1700031L13Rik | 0 | 0 | 0 | 0 | 0 | 0 | 0 | 1 | 0 | 0 | 0 | 0 | 2 | 3 | 0 | 1 | 0 | 0 | 0 | 0 | 1 | 2 | 0 | 0 | 0 | 4 | 0 | 0 | 1700031L13Rik |
| 1700034E13Rik | 0 | 0 | 0 | 0 | 0 | 0 | 0 | 0 | 7 | 0 | 0 | 0 | 0 | 0 | 0 | 0 | 0 | 0 | 0 | 0 | 0 | 0 | 0 | 0 | 0 | 0 | 0 | 0 | 1700034E13Rik |
| 1700034I23Rik | 0 | 0 | 0 | 1 | 1 | 0 | 0 | 0 | 0 | 0 | 0 | 0 | 1 | 0 | 0 | 0 | 0 | 2 | 0 | 0 | 0 | 0 | 0 | 1 | 0 | 0 | 0 | 0 | 1700034I23Rik |
| 1700037C18Rik | 63 | 59 | 46 | 49 | 70 | 89 | 80 | 104 | 198 | 164 | 194 | 159 | 154 | 172 | 55 | 59 | 66 | 53 | 66 | 81 | 59 | 135 | 170 | 178 | 214 | 162 | 206 | 147 | 1700037C18Rik |
| 1700037H04Rik | 4339 | 5272 | 4530 | 4150 | 2963 | 2968 | 3862 | 1977 | 1973 | 1855 | 1846 | 2221 | 2487 | 2461 | 4158 | 4505 | 4460 | 3935 | 3507 | 3726 | 3744 | 1933 | 1776 | 2225 | 2157 | 2783 | 2715 | 2839 | 1700037H04Rik |
| 1700056E22Rik | 33 | 49 | 60 | 29 | 21 | 25 | 29 | 7 | 24 | 30 | 27 | 48 | 35 | 46 | 51 | 26 | 32 | 21 | 23 | 20 | 29 | 20 | 19 | 30 | 20 | 45 | 30 | 44 | 1700056E22Rik |
| 1700057G04Rik | 0 | 0 | 0 | 0 | 0 | 0 | 0 | 0 | 0 | 3 | 0 | 0 | 0 | 0 | 0 | 0 | 0 | 0 | 2 | 0 | 0 | 0 | 0 | 0 | 0 | 0 | 0 | 0 | 1700057G04Rik |
| 1700064H15Rik | 0 | 1 | 1 | 9 | 0 | 3 | 1 | 0 | 5 | 4 | 3 | 3 | 6 | 3 | 3 | 0 | 1 | 0 | 1 | 0 | 3 | 2 | 3 | 1 | 0 | 3 | 0 | 0 | 1700064H15Rik |
| 1700066B19Rik | 2 | 8 | 6 | 11 | 12 | 17 | 9 | 2 | 16 | 5 | 9 | 0 | 14 | 11 | 17 | 9 | 7 | 4 | 4 | 14 | 4 | 12 | 11 | 7 | 8 | 11 | 5 | 19 | 1700066B19Rik |
| 1700066M21Rik | 1924 | 1415 | 1365 | 1898 | 2032 | 2057 | 2567 | 1721 | 1479 | 1387 | 1362 | 1345 | 1194 | 1315 | 2018 | 1704 | 1409 | 2058 | 1843 | 2353 | 2327 | 1489 | 1595 | 1477 | 1501 | 1403 | 1167 | 1174 | 1700066M21Rik |
| 1700067K01Rik | 60 | 73 | 84 | 85 | 75 | 77 | 63 | 51 | 41 | 64 | 63 | 67 | 66 | 73 | 47 | 67 | 115 | 83 | 103 | 63 | 84 | 63 | 35 | 59 | 60 | 51 | 76 | 63 | 1700067K01Rik |
| 1700069L16Rik | 3 | 1 | 2 | 0 | 1 | 4 | 3 | 2 | 0 | 0 | 1 | 0 | 3 | 1 | 7 | 0 | 0 | 1 | 0 | 1 | 2 | 0 | 5 | 0 | 1 | 2 | 2 | 4 | 1700069L16Rik |
| 1700084M14Rik | 0 | 0 | 0 | 0 | 0 | 0 | 0 | 1 | 0 | 0 | 0 | 0 | 0 | 0 | 0 | 0 | 0 | 0 | 0 | 0 | 0 | 0 | 0 | 0 | 0 | 0 | 0 | 0 | 1700084M14Rik |
| 1700086D15Rik | 0 | 2 | 0 | 0 | 1 | 0 | 2 | 2 | 2 | 0 | 8 | 1 | 2 | 2 | 0 | 4 | 2 | 0 | 0 | 1 | 1 | 2 | 5 | 3 | 1 | 0 | 0 | 0 | 1700086D15Rik |
| 1700088E04Rik | 617 | 980 | 741 | 809 | 657 | 690 | 673 | 232 | 308 | 313 | 290 | 260 | 359 | 253 | 729 | 827 | 811 | 780 | 771 | 749 | 757 | 312 | 330 | 279 | 260 | 247 | 265 | 292 | 1700088E04Rik |
| 1700092M07Rik | 57 | 71 | 42 | 105 | 61 | 55 | 63 | 22 | 40 | 44 | 37 | 33 | 22 | 38 | 33 | 54 | 139 | 81 | 65 | 33 | 85 | 47 | 36 | 39 | 42 | 24 | 42 | 53 | 1700092M07Rik |
| 1700093K21Rik | 118 | 128 | 87 | 283 | 238 | 242 | 263 | 34 | 21 | 45 | 45 | 55 | 67 | 38 | 175 | 143 | 117 | 259 | 188 | 191 | 352 | 35 | 21 | 27 | 47 | 42 | 30 | 26 | 1700093K21Rik |
| 1700094D03Rik | 340 | 399 | 456 | 441 | 543 | 447 | 503 | 243 | 235 | 258 | 255 | 278 | 186 | 234 | 436 | 356 | 524 | 501 | 503 | 484 | 513 | 316 | 186 | 277 | 282 | 270 | 258 | 254 | 1700094D03Rik |
| 1700102P08Rik | 258 | 349 | 306 | 376 | 430 | 365 | 381 | 181 | 174 | 203 | 176 | 204 | 181 | 197 | 244 | 286 | 480 | 403 | 413 | 308 | 417 | 254 | 160 | 214 | 213 | 173 | 178 | 157 | 1700102P08Rik |
| 1700109H08Rik | 912 | 764 | 400 | 633 | 628 | 872 | 813 | 735 | 569 | 590 | 659 | 591 | 634 | 610 | 862 | 787 | 521 | 530 | 631 | 939 | 791 | 546 | 592 | 441 | 602 | 350 | 717 | 573 | 1700109H08Rik |
| 1700113H08Rik | 6 | 3 | 3 | 2 | 2 | 1 | 4 | 6 | 7 | 5 | 12 | 3 | 8 | 11 | 11 | 1 | 3 | 0 | 2 | 1 | 4 | 13 | 4 | 11 | 5 | 2 | 9 | 5 | 1700113H08Rik |
| 1700123K08Rik | 0 | 6 | 1 | 8 | 2 | 3 | 7 | 0 | 1 | 0 | 1 | 0 | 2 | 3 | 9 | 0 | 5 | 6 | 2 | 7 | 2 | 2 | 1 | 0 | 0 | 7 | 1 | 4 | 1700123K08Rik |
| 1700123O20Rik | 3012 | 2774 | 2880 | 2762 | 2294 | 2375 | 2369 | 2317 | 2137 | 2184 | 2033 | 2393 | 2266 | 2250 | 3026 | 2797 | 2773 | 2497 | 2565 | 2946 | 2420 | 1998 | 2263 | 2201 | 2316 | 2572 | 2429 | 2388 | 1700123O20Rik |
| 1700125H20Rik | 21 | 21 | 7 | 39 | 20 | 27 | 23 | 12 | 7 | 23 | 16 | 7 | 18 | 14 | 22 | 12 | 20 | 25 | 16 | 15 | 14 | 7 | 10 | 9 | 11 | 19 | 8 | 11 | 1700125H20Rik |
| 1810009A15Rik | 23 | 20 | 34 | 50 | 20 | 25 | 34 | 27 | 26 | 31 | 22 | 13 | 25 | 35 | 13 | 19 | 41 | 30 | 13 | 32 | 38 | 23 | 23 | 38 | 23 | 29 | 27 | 43 | 1810009A15Rik |
| 1810010H24Rik | 224 | 192 | 304 | 207 | 291 | 254 | 274 | 132 | 136 | 157 | 140 | 108 | 125 | 177 | 214 | 182 | 309 | 243 | 299 | 229 | 279 | 192 | 122 | 121 | 149 | 124 | 110 | 96 | 1810010H24Rik |
| 1810013L24Rik | 4834 | 4375 | 4572 | 5947 | 5659 | 5465 | 5336 | 6138 | 5258 | 5281 | 5434 | 5613 | 5719 | 5752 | 5331 | 4942 | 5062 | 5619 | 5322 | 4878 | 5398 | 6462 | 5160 | 5112 | 6047 | 4685 | 5697 | 5673 | 1810013L24Rik |
| 1810020O05Rik | 22 | 40 | 19 | 46 | 61 | 32 | 39 | 35 | 43 | 14 | 40 | 56 | 53 | 22 | 55 | 46 | 23 | 37 | 26 | 22 | 43 | 40 | 56 | 16 | 36 | 8 | 37 | 26 | 1810020O05Rik |
| 1810024B03Rik | 89 | 87 | 58 | 58 | 46 | 75 | 74 | 191 | 168 | 155 | 136 | 163 | 196 | 239 | 82 | 73 | 60 | 50 | 62 | 89 | 56 | 134 | 135 | 170 | 146 | 205 | 198 | 248 | 1810024B03Rik |
| 1810030O07Rik | 2733 | 2607 | 2602 | 2841 | 2752 | 2698 | 3051 | 3120 | 3057 | 2930 | 3068 | 3277 | 2960 | 3388 | 2681 | 2502 | 2662 | 2849 | 2939 | 2731 | 2895 | 3257 | 2958 | 3280 | 3164 | 3304 | 3083 | 3452 | 1810030O07Rik |
| 1810037I17Rik | 2195 | 2493 | 3115 | 2888 | 2278 | 1895 | 2609 | 1954 | 1824 | 2476 | 2005 | 2167 | 2084 | 2842 | 2339 | 1903 | 2789 | 2299 | 2188 | 2793 | 2561 | 1587 | 1886 | 2589 | 2244 | 2659 | 2255 | 2785 | 1810037I17Rik |
| 1810046K07Rik | 0 | 0 | 0 | 0 | 0 | 0 | 2 | 0 | 1 | 0 | 0 | 0 | 0 | 0 | 0 | 0 | 0 | 0 | 0 | 0 | 0 | 0 | 3 | 0 | 0 | 0 | 0 | 0 | 1810046K07Rik |
| 1810049J17Rik | 3 | 1 | 5 | 3 | 4 | 6 | 6 | 4 | 3 | 1 | 1 | 3 | 2 | 3 | 3 | 3 | 2 | 4 | 5 | 3 | 4 | 2 | 5 | 3 | 2 | 4 | 1 | 1 | 1810049J17Rik |
| 1810055G02Rik | 2676 | 3327 | 2739 | 2989 | 2257 | 2497 | 1833 | 1352 | 1590 | 1748 | 1807 | 1640 | 1635 | 1307 | 2622 | 3200 | 2971 | 2878 | 2354 | 2088 | 2011 | 1536 | 1679 | 1623 | 1618 | 1791 | 1624 | 1426 | 1810055G02Rik |
| 1810058I24Rik | 3969 | 4564 | 3953 | 3758 | 3161 | 3504 | 3461 | 2646 | 3104 | 2737 | 3415 | 3282 | 3785 | 3395 | 3955 | 4288 | 4128 | 3934 | 3500 | 3857 | 3477 | 2727 | 3342 | 3006 | 3025 | 3872 | 3655 | 3779 | 1810058I24Rik |
| 1810062G17Rik | 40 | 22 | 23 | 10 | 18 | 25 | 20 | 11 | 12 | 21 | 10 | 9 | 23 | 21 | 38 | 29 | 13 | 14 | 19 | 50 | 17 | 3 | 25 | 12 | 5 | 20 | 14 | 17 | 1810062G17Rik |
| 1810065E05Rik | 0 | 0 | 0 | 0 | 0 | 0 | 0 | 0 | 0 | 0 | 0 | 1 | 0 | 0 | 0 | 0 | 0 | 0 | 0 | 0 | 0 | 0 | 0 | 0 | 0 | 2 | 0 | 0 | 1810065E05Rik |
| 2010003K11Rik | 1 | 2 | 0 | 3 | 0 | 0 | 0 | 0 | 2 | 0 | 5 | 4 | 6 | 0 | 4 | 0 | 0 | 1 | 2 | 0 | 0 | 2 | 1 | 0 | 0 | 0 | 1 | 1 | 2010003K11Rik |
| 2010106E10Rik | 5 | 0 | 0 | 1 | 5 | 0 | 2 | 1 | 3 | 0 | 0 | 0 | 1 | 1 | 1 | 2 | 8 | 0 | 0 | 1 | 0 | 0 | 0 | 2 | 0 | 0 | 0 | 0 | 2010106E10Rik |
| 2010109A12Rik | 16 | 16 | 11 | 5 | 21 | 6 | 10 | 32 | 33 | 40 | 25 | 42 | 36 | 38 | 9 | 6 | 14 | 4 | 6 | 16 | 17 | 17 | 30 | 34 | 29 | 39 | 28 | 27 | 2010109A12Rik |
| 2010315B03Rik | 0 | 1 | 1 | 0 | 0 | 0 | 0 | 0 | 0 | 0 | 0 | 0 | 1 | 0 | 0 | 0 | 0 | 0 | 0 | 0 | 0 | 0 | 0 | 0 | 8 | 0 | 0 | 0 | 2010315B03Rik |
| 2200002D01Rik | 23 | 27 | 34 | 13 | 11 | 18 | 4 | 17 | 19 | 27 | 21 | 11 | 33 | 18 | 17 | 30 | 17 | 24 | 26 | 10 | 13 | 27 | 13 | 13 | 23 | 25 | 31 | 23 | 2200002D01Rik |
| 2200002J24Rik | 1 | 0 | 0 | 0 | 0 | 0 | 0 | 0 | 0 | 0 | 1 | 0 | 0 | 0 | 0 | 2 | 0 | 3 | 0 | 0 | 0 | 0 | 1 | 0 | 0 | 2 | 0 | 0 | 2200002J24Rik |
| 2210016F16Rik | 1336 | 1404 | 1530 | 1361 | 1396 | 1255 | 1370 | 1301 | 1241 | 1233 | 1076 | 1453 | 1353 | 1320 | 1432 | 1237 | 1570 | 1282 | 1470 | 1308 | 1449 | 1151 | 1357 | 1401 | 1221 | 1509 | 1541 | 1539 | 2210016F16Rik |
| 2210016L21Rik | 1584 | 1812 | 1816 | 1758 | 2008 | 1766 | 1894 | 2077 | 2360 | 2528 | 2454 | 2249 | 2193 | 2447 | 1548 | 1625 | 1795 | 1862 | 1958 | 1806 | 1972 | 2401 | 2333 | 2551 | 2491 | 2407 | 2071 | 2258 | 2210016L21Rik |
| 2210408I21Rik | 1283 | 1133 | 916 | 946 | 991 | 1175 | 1284 | 1319 | 1049 | 745 | 850 | 908 | 917 | 1158 | 1527 | 1177 | 891 | 1017 | 983 | 1395 | 1165 | 874 | 1071 | 888 | 990 | 939 | 1026 | 1099 | 2210408I21Rik |
| 2210418O10Rik | 98 | 56 | 107 | 43 | 49 | 72 | 60 | 172 | 124 | 427 | 79 | 73 | 175 | 64 | 127 | 63 | 35 | 25 | 42 | 70 | 67 | 92 | 145 | 84 | 169 | 62 | 88 | 55 | 2210418O10Rik |
| 2300002M23Rik | 0 | 0 | 4 | 0 | 0 | 0 | 0 | 1 | 1 | 0 | 1 | 0 | 1 | 0 | 3 | 1 | 0 | 1 | 1 | 0 | 0 | 0 | 1 | 4 | 0 | 2 | 0 | 0 | 2300002M23Rik |
| 2300009A05Rik | 672 | 576 | 595 | 747 | 597 | 642 | 820 | 766 | 658 | 583 | 689 | 677 | 668 | 776 | 696 | 628 | 716 | 581 | 601 | 705 | 710 | 613 | 587 | 697 | 638 | 565 | 646 | 718 | 2300009A05Rik |
| 2310002L09Rik | 6 | 0 | 0 | 0 | 1 | 0 | 0 | 5 | 22 | 3 | 151 | 0 | 15 | 0 | 3 | 2 | 0 | 0 | 0 | 0 | 0 | 23 | 78 | 0 | 7 | 0 | 0 | 2 | 2310002L09Rik |
| 2310003L06Rik | 0 | 0 | 0 | 0 | 0 | 0 | 0 | 0 | 1 | 0 | 0 | 0 | 1 | 1 | 0 | 0 | 0 | 0 | 0 | 0 | 0 | 0 | 0 | 0 | 0 | 0 | 0 | 0 | 2310003L06Rik |
| 2310009B15Rik | 443 | 511 | 526 | 511 | 452 | 430 | 560 | 396 | 424 | 452 | 353 | 526 | 483 | 622 | 484 | 372 | 557 | 452 | 501 | 596 | 568 | 359 | 399 | 564 | 537 | 583 | 589 | 559 | 2310009B15Rik |
| 2310011J03Rik | 2151 | 2684 | 2301 | 2290 | 1620 | 1971 | 1789 | 1025 | 1187 | 1125 | 1099 | 1393 | 1367 | 1192 | 1999 | 2432 | 2226 | 2108 | 1883 | 1973 | 1751 | 894 | 1299 | 1316 | 1107 | 1715 | 1363 | 1329 | 2310011J03Rik |
| 2310022A10Rik | 1576 | 1804 | 2338 | 2213 | 1767 | 1649 | 1626 | 1414 | 1681 | 1892 | 1788 | 2084 | 1666 | 1894 | 1404 | 1682 | 2152 | 2006 | 2127 | 1426 | 1672 | 1741 | 1604 | 2088 | 1801 | 2386 | 1685 | 1838 | 2310022A10Rik |
| 2310022B05Rik | 6150 | 7188 | 7244 | 7629 | 8080 | 7391 | 6651 | 4695 | 6082 | 6435 | 6081 | 5718 | 5319 | 5007 | 5943 | 7432 | 7540 | 8625 | 7971 | 5874 | 7362 | 5850 | 6798 | 6133 | 6368 | 6052 | 4871 | 5079 | 2310022B05Rik |
| 2310030G06Rik | 68 | 16 | 23 | 13 | 25 | 28 | 19 | 81 | 87 | 74 | 87 | 48 | 77 | 32 | 51 | 32 | 24 | 31 | 12 | 32 | 13 | 42 | 87 | 70 | 13 | 73 | 46 | 38 | 2310030G06Rik |
| 2310033P09Rik | 1423 | 1681 | 1881 | 1414 | 1356 | 1471 | 1246 | 1126 | 1210 | 1238 | 1192 | 1462 | 1429 | 1336 | 1431 | 1399 | 1889 | 1405 | 1563 | 1524 | 1247 | 1168 | 1120 | 1358 | 1208 | 1761 | 1534 | 1335 | 2310033P09Rik |
| 2310034C09Rik | 0 | 0 | 0 | 0 | 0 | 0 | 0 | 0 | 0 | 0 | 0 | 0 | 0 | 0 | 0 | 0 | 0 | 0 | 0 | 0 | 0 | 0 | 3 | 0 | 0 | 0 | 0 | 0 | 2310034C09Rik |
| 2310034G01Rik | 38 | 35 | 19 | 39 | 43 | 14 | 45 | 43 | 38 | 29 | 45 | 24 | 26 | 32 | 34 | 24 | 25 | 22 | 24 | 30 | 52 | 35 | 27 | 25 | 27 | 28 | 21 | 28 | 2310034G01Rik |
| 2310039H08Rik | 505 | 601 | 545 | 572 | 395 | 480 | 444 | 432 | 416 | 366 | 431 | 376 | 384 | 462 | 460 | 445 | 492 | 404 | 436 | 533 | 415 | 262 | 416 | 485 | 384 | 530 | 450 | 379 | 2310039H08Rik |
| 2310057J18Rik | 0 | 2 | 0 | 0 | 2 | 2 | 0 | 0 | 0 | 0 | 1 | 0 | 3 | 1 | 0 | 0 | 0 | 0 | 0 | 0 | 3 | 5 | 1 | 2 | 0 | 1 | 0 | 0 | 2310057J18Rik |
| 2310057M21Rik | 1343 | 1195 | 1134 | 1230 | 1387 | 1412 | 1376 | 1487 | 1444 | 1301 | 1504 | 1366 | 1373 | 1513 | 1345 | 1343 | 1145 | 1205 | 1268 | 1454 | 1460 | 1447 | 1390 | 1242 | 1520 | 1156 | 1449 | 1367 | 2310057M21Rik |
| 2310061I04Rik | 1009 | 1209 | 1223 | 1325 | 1569 | 1575 | 1315 | 1249 | 1382 | 1627 | 1424 | 1471 | 1444 | 1388 | 1000 | 970 | 1254 | 1351 | 1450 | 1289 | 1386 | 1810 | 1296 | 1537 | 1708 | 1490 | 1343 | 1316 | 2310061I04Rik |
| 2410002F23Rik | 4986 | 5121 | 4682 | 5061 | 5080 | 4894 | 5123 | 7016 | 7354 | 7292 | 6898 | 7258 | 6569 | 7497 | 4820 | 4298 | 4804 | 4633 | 4964 | 5100 | 5138 | 6839 | 6728 | 7153 | 7090 | 7479 | 6922 | 7262 | 2410002F23Rik |
| 2410004B18Rik | 2298 | 2470 | 2437 | 2542 | 2183 | 2178 | 2554 | 1768 | 1554 | 1645 | 1409 | 1523 | 1396 | 1755 | 2498 | 2262 | 2351 | 2521 | 2322 | 2768 | 2624 | 1708 | 1641 | 1622 | 1650 | 1609 | 1575 | 1749 | 2410004B18Rik |
| 2410004P03Rik | 59 | 62 | 57 | 36 | 77 | 78 | 48 | 77 | 96 | 124 | 58 | 123 | 126 | 57 | 47 | 72 | 37 | 24 | 58 | 61 | 78 | 102 | 159 | 60 | 70 | 55 | 37 | 53 | 2410004P03Rik |
| 2410137M14Rik | 9 | 12 | 20 | 15 | 15 | 12 | 18 | 20 | 20 | 31 | 19 | 30 | 34 | 40 | 8 | 5 | 19 | 17 | 18 | 12 | 23 | 35 | 16 | 20 | 26 | 18 | 23 | 28 | 2410137M14Rik |
| 2510002D24Rik | 391 | 355 | 433 | 441 | 436 | 384 | 470 | 579 | 628 | 673 | 589 | 579 | 566 | 553 | 442 | 375 | 421 | 350 | 408 | 411 | 474 | 620 | 560 | 717 | 584 | 638 | 507 | 615 | 2510002D24Rik |
| 2510009E07Rik | 7717 | 7390 | 10741 | 9087 | 10316 | 10660 | 9011 | 9636 | 7640 | 9776 | 8874 | 12432 | 12010 | 12075 | 8590 | 6731 | 10222 | 8857 | 10839 | 9763 | 9903 | 10955 | 8638 | 10024 | 10778 | 7950 | 15036 | 14952 | 2510009E07Rik |
| 2510039O18Rik | 3120 | 3911 | 3412 | 3598 | 2671 | 2683 | 2353 | 1782 | 2144 | 2067 | 2130 | 2029 | 1982 | 1821 | 2755 | 3731 | 3348 | 3374 | 2833 | 2288 | 2351 | 2079 | 2119 | 2053 | 2133 | 2387 | 1871 | 1849 | 2510039O18Rik |
| 2610001J05Rik | 3703 | 3114 | 3940 | 3551 | 3458 | 3502 | 3879 | 2947 | 2608 | 2583 | 2733 | 3052 | 3149 | 3285 | 4117 | 2872 | 3474 | 3504 | 3643 | 4262 | 3616 | 2239 | 2785 | 3139 | 2650 | 3523 | 3477 | 3368 | 2610001J05Rik |
| 2610002M06Rik | 2523 | 1970 | 1933 | 2330 | 3044 | 2668 | 2808 | 3389 | 3119 | 3222 | 2964 | 2614 | 2624 | 2804 | 2449 | 2094 | 2194 | 2470 | 2613 | 2666 | 2784 | 3479 | 3260 | 2760 | 2978 | 2366 | 2579 | 2483 | 2610002M06Rik |

| | | | | | | | | | | | | | | | | | | | | | | | | | | | | | |
|---|---|---|---|---|---|---|---|---|---|---|---|---|---|---|---|---|---|---|---|---|---|---|---|---|---|---|---|---|---|
| 2610008E11Rik | 6243 | 5056 | 6511 | 6341 | 6827 | 6583 | 7123 | 11225 | 9010 | 9041 | 8831 | 8883 | 8552 | 9447 | 7216 | 6292 | 6435 | 6236 | 6465 | 7421 | 6585 | 10597 | 8729 | 9183 | 9872 | 8024 | 8613 | 8967 | 2610008E11Rik |
| 2610021A01Rik | 3294 | 2930 | 2893 | 3009 | 3136 | 3267 | 3357 | 3175 | 2678 | 2271 | 2166 | 2378 | 2384 | 2594 | 3473 | 3127 | 3008 | 2982 | 2996 | 3802 | 3342 | 2565 | 2530 | 2266 | 2419 | 2390 | 2587 | 2483 | 2610021A01Rik |
| 2610028H24Rik | 0 | 0 | 0 | 0 | 0 | 0 | 0 | 0 | 0 | 0 | 1 | 0 | 2 | 0 | 0 | 0 | 0 | 0 | 1 | 0 | 0 | 0 | 1 | 2 | 0 | 0 | 5 | 1 | 2610028H24Rik |
| 2610042L04Rik | 31 | 10 | 31 | 36 | 56 | 70 | 73 | 154 | 144 | 168 | 121 | 124 | 78 | 122 | 22 | 26 | 38 | 29 | 50 | 56 | 59 | 172 | 146 | 147 | 164 | 156 | 112 | 148 | 2610042L04Rik |
| 2610044O15Rik8 | 1795 | 1364 | 2136 | 1844 | 1860 | 2003 | 1853 | 2660 | 2274 | 2074 | 2266 | 2207 | 2274 | 2433 | 1878 | 1714 | 1874 | 1788 | 1898 | 2203 | 1807 | 2359 | 2322 | 2147 | 2426 | 2489 | 2453 | 2343 | 2610044O15Rik8 |
| 2610301B20Rik | 4291 | 3877 | 3492 | 3815 | 3913 | 3997 | 4524 | 3499 | 2731 | 2600 | 2442 | 2841 | 2835 | 2972 | 4465 | 3996 | 3837 | 3934 | 3966 | 4852 | 4357 | 3048 | 3016 | 2631 | 2623 | 2835 | 3087 | 3267 | 2610301B20Rik |
| 2610318N02Rik | 2741 | 3284 | 3304 | 2482 | 1770 | 2002 | 2211 | 1346 | 1391 | 1329 | 1204 | 2456 | 1992 | 2473 | 2421 | 2675 | 3266 | 2154 | 2302 | 2166 | 2059 | 1063 | 1096 | 2028 | 1450 | 2885 | 2697 | 2460 | 2610318N02Rik |
| 2610507B11Rik | 25301 | 25880 | 17856 | 21699 | 21293 | 22676 | 22329 | 17227 | 17988 | 16293 | 16838 | 17220 | 17580 | 16609 | 23409 | 22675 | 19019 | 21742 | 21522 | 21495 | 22919 | 15714 | 17224 | 15330 | 16997 | 12889 | 18174 | 16884 | 2610507B11Rik |
| 2610528A11Rik | 9 | 6 | 9 | 20 | 7 | 3 | 7 | 7 | 9 | 9 | 6 | 1 | 5 | 4 | 12 | 8 | 6 | 12 | 12 | 5 | 14 | 2 | 7 | 2 | 8 | 4 | 2 | 5 | 2610528A11Rik |
| 2610528J11Rik | 28 | 13 | 16 | 6 | 0 | 6 | 3 | 10 | 16 | 8 | 20 | 4 | 9 | 12 | 13 | 23 | 2 | 7 | 0 | 1 | 1 | 8 | 19 | 27 | 0 | 13 | 7 | 5 | 2610528J11Rik |
| 2700049A03Rik | 4302 | 4056 | 3462 | 3485 | 3519 | 3650 | 4082 | 3402 | 3145 | 2740 | 2631 | 3414 | 2825 | 3347 | 4371 | 4075 | 3621 | 3572 | 3472 | 4167 | 3885 | 2755 | 2961 | 2861 | 3122 | 2488 | 3106 | 2973 | 2700049A03Rik |
| 2700062C07Rik | 513 | 489 | 660 | 431 | 512 | 581 | 533 | 971 | 894 | 829 | 832 | 1032 | 865 | 1004 | 534 | 411 | 562 | 357 | 517 | 623 | 547 | 954 | 966 | 985 | 928 | 989 | 997 | 900 | 2700062C07Rik |
| 2700097O09Rik | 2903 | 2272 | 2196 | 2679 | 2274 | 2400 | 2747 | 1948 | 1694 | 1797 | 1962 | 1963 | 1776 | 2071 | 2556 | 2695 | 2437 | 2709 | 2337 | 2619 | 2723 | 1888 | 1666 | 1804 | 1938 | 1789 | 1901 | 2065 | 2700097O09Rik |
| 2810004N23Rik | 3715 | 3645 | 3687 | 3158 | 2747 | 3378 | 3372 | 3684 | 3527 | 3459 | 3435 | 3793 | 3932 | 4174 | 3695 | 3382 | 3575 | 2954 | 3025 | 3328 | 3228 | 2765 | 3603 | 3750 | 3629 | 4425 | 4345 | 4004 | 2810004N23Rik |
| 2810021J22Rik | 1722 | 1593 | 1492 | 1611 | 1582 | 1595 | 1568 | 1966 | 1664 | 1441 | 1578 | 1729 | 1692 | 1755 | 1751 | 1439 | 1607 | 1461 | 1531 | 1638 | 1490 | 1882 | 1608 | 1664 | 1847 | 1668 | 1658 | 1780 | 2810021J22Rik |
| 2810408A11Rik | 1232 | 1164 | 1237 | 1101 | 1055 | 1149 | 1181 | 1031 | 1130 | 1239 | 1111 | 1638 | 1423 | 1755 | 1093 | 943 | 1263 | 1016 | 1210 | 1126 | 1118 | 1026 | 989 | 1575 | 1354 | 1875 | 1628 | 1676 | 2810408A11Rik |
| 2810459M11Rik | 45 | 51 | 48 | 79 | 266 | 212 | 181 | 172 | 274 | 263 | 198 | 49 | 50 | 74 | 46 | 37 | 29 | 120 | 214 | 128 | 210 | 196 | 339 | 217 | 370 | 61 | 21 | 26 | 2810459M11Rik |
| 2900026A02Rik | 3537 | 4374 | 4330 | 5546 | 4241 | 4103 | 4181 | 1662 | 2280 | 2377 | 2410 | 2321 | 2006 | 2260 | 3435 | 4000 | 4923 | 5410 | 4630 | 3432 | 4398 | 1878 | 2324 | 2708 | 2198 | 2182 | 1807 | 1991 | 2900026A02Rik |
| 2900092C05Rik | 0 | 1 | 0 | 0 | 0 | 0 | 0 | 0 | 0 | 0 | 0 | 2 | 0 | 0 | 9 | 2 | 1 | 0 | 4 | 0 | 0 | 0 | 2 | 0 | 1 | 0 | 0 | 1 | 2900092C05Rik |
| 3110001I22Rik | 356 | 416 | 363 | 369 | 314 | 408 | 371 | 541 | 412 | 366 | 328 | 400 | 429 | 420 | 409 | 396 | 377 | 281 | 308 | 443 | 340 | 434 | 370 | 337 | 426 | 355 | 460 | 409 | 3110001I22Rik |
| 3110009E18Rik | 104 | 94 | 81 | 81 | 90 | 48 | 133 | 156 | 129 | 133 | 125 | 153 | 128 | 204 | 96 | 65 | 90 | 46 | 96 | 113 | 93 | 147 | 136 | 181 | 160 | 221 | 172 | 199 | 3110009E18Rik |
| 3110021N24Rik | 16 | 10 | 11 | 7 | 13 | 9 | 20 | 33 | 16 | 23 | 24 | 9 | 18 | 29 | 24 | 16 | 13 | 11 | 19 | 20 | 25 | 13 | 18 | 15 | 9 | 20 | 24 | 11 | 3110021N24Rik |
| 3110040M04Rik | 0 | 0 | 0 | 1 | 0 | 0 | 0 | 0 | 1 | 0 | 0 | 0 | 5 | 0 | 0 | 0 | 0 | 0 | 0 | 0 | 0 | 0 | 0 | 0 | 0 | 0 | 0 | 0 | 3110040M04Rik |
| 3110040N11Rik | 415 | 498 | 601 | 417 | 425 | 497 | 453 | 894 | 886 | 870 | 937 | 860 | 933 | 877 | 452 | 457 | 477 | 395 | 508 | 518 | 470 | 722 | 878 | 962 | 879 | 957 | 1001 | 834 | 3110040N11Rik |
| 3110082I17Rik | 1310 | 1749 | 2078 | 1444 | 1059 | 1153 | 1079 | 675 | 789 | 845 | 862 | 1059 | 990 | 1032 | 1288 | 1490 | 1661 | 1224 | 1272 | 1346 | 1087 | 633 | 845 | 1072 | 931 | 1486 | 1098 | 1092 | 3110082I17Rik |
| 3110082J24Rik | 53 | 54 | 82 | 71 | 87 | 88 | 101 | 209 | 123 | 160 | 180 | 223 | 218 | 275 | 87 | 44 | 102 | 89 | 96 | 76 | 75 | 206 | 133 | 193 | 185 | 223 | 189 | 288 | 3110082J24Rik |
| 3300002I08Rik | 258 | 263 | 147 | 132 | 225 | 275 | 228 | 355 | 339 | 216 | 274 | 226 | 266 | 263 | 239 | 217 | 162 | 164 | 203 | 356 | 239 | 172 | 293 | 171 | 291 | 99 | 237 | 217 | 3300002I08Rik |
| 3425401B19Rik | 3 | 2 | 0 | 2 | 7 | 9 | 0 | 9 | 34 | 5 | 186 | 1 | 7 | 0 | 3 | 4 | 0 | 0 | 2 | 0 | 2 | 13 | 92 | 7 | 0 | 2 | 0 | 1 | 3425401B19Rik |
| 3830403N18Rik | 53 | 22 | 24 | 59 | 16 | 36 | 18 | 18 | 22 | 41 | 31 | 12 | 45 | 21 | 12 | 40 | 15 | 20 | 11 | 15 | 14 | 52 | 57 | 7 | 23 | 18 | 38 | 37 | 3830403N18Rik |
| 3830406C13Rik | 739 | 504 | 548 | 684 | 704 | 771 | 963 | 1131 | 1028 | 893 | 780 | 834 | 814 | 861 | 764 | 604 | 611 | 661 | 648 | 872 | 816 | 991 | 1045 | 927 | 843 | 991 | 826 | 879 | 3830406C13Rik |
| 4833420G17Rik | 9691 | 8910 | 8182 | 9624 | 10082 | 10537 | 11508 | 10169 | 8981 | 9651 | 8641 | 9544 | 9183 | 10312 | 10059 | 8539 | 9042 | 9355 | 9841 | 5219 | 11076 | 10134 | 8334 | 9081 | 10221 | 8612 | 9541 | 9566 | 4833420G17Rik |
| 4833439L19Rik | 9939 | 10312 | 10126 | 10596 | 10153 | 9994 | 11048 | 8970 | 8880 | 9502 | 8864 | 11061 | 9531 | 11375 | 9849 | 9761 | 10549 | 10445 | 10833 | 9414 | 10871 | 10592 | 8121 | 10493 | 10723 | 10919 | 10163 | 11363 | 4833439L19Rik |
| 4921504E06Rik | 0 | 6 | 0 | 1 | 7 | 4 | 0 | 16 | 9 | 3 | 2 | 0 | 34 | 1 | 9 | 22 | 0 | 0 | 0 | 5 | 0 | 7 | 6 | 1 | 8 | 0 | 0 | 0 | 4921504E06Rik |
| 4921507P07Rik | 219 | 191 | 187 | 251 | 348 | 307 | 232 | 183 | 185 | 199 | 132 | 165 | 206 | 177 | 262 | 227 | 245 | 250 | 280 | 276 | 248 | 227 | 200 | 141 | 210 | 138 | 171 | 170 | 4921507P07Rik |
| 4921509C19Rik | 1 | 0 | 0 | 0 | 0 | 0 | 0 | 0 | 0 | 0 | 0 | 0 | 0 | 0 | 0 | 0 | 0 | 0 | 0 | 0 | 0 | 0 | 0 | 0 | 0 | 0 | 0 | 0 | 4921509C19Rik |
| 4921517D22Rik | 0 | 1 | 0 | 1 | 1 | 0 | 1 | 0 | 0 | 0 | 0 | 0 | 0 | 1 | 0 | 1 | 0 | 4 | 0 | 0 | 1 | 0 | 0 | 0 | 0 | 0 | 0 | 0 | 4921517D22Rik |
| 4921524J17Rik | 2014 | 1871 | 1835 | 2150 | 2041 | 1877 | 2368 | 1481 | 1152 | 1003 | 1119 | 1157 | 1141 | 1309 | 2385 | 2101 | 1684 | 2088 | 1916 | 2452 | 2150 | 1258 | 1265 | 1264 | 1235 | 1343 | 1126 | 1280 | 4921524J17Rik |
| 4921524L21Rik | 0 | 0 | 0 | 0 | 0 | 0 | 0 | 0 | 1 | 0 | 0 | 0 | 0 | 0 | 0 | 0 | 0 | 0 | 0 | 0 | 0 | 0 | 0 | 0 | 0 | 0 | 0 | 0 | 4921524L21Rik |
| 4921536K21Rik | 213 | 288 | 261 | 271 | 247 | 223 | 292 | 154 | 147 | 172 | 105 | 226 | 196 | 202 | 193 | 228 | 257 | 248 | 235 | 211 | 248 | 144 | 117 | 192 | 169 | 215 | 196 | 229 | 4921536K21Rik |
| 4921539E11Rik | 0 | 0 | 0 | 0 | 0 | 0 | 0 | 0 | 0 | 0 | 0 | 0 | 0 | 0 | 0 | 0 | 0 | 0 | 1 | 0 | 0 | 0 | 0 | 0 | 0 | 0 | 0 | 0 | 4921539E11Rik |
| 4930402F06Rik | 0 | 3 | 3 | 0 | 2 | 0 | 0 | 0 | 0 | 0 | 0 | 0 | 0 | 0 | 0 | 0 | 0 | 0 | 0 | 0 | 0 | 0 | 0 | 0 | 0 | 0 | 0 | 0 | 4930402F06Rik |
| 4930404H24Rik | 0 | 0 | 0 | 0 | 0 | 0 | 3 | 0 | 0 | 0 | 0 | 1 | 0 | 0 | 1 | 0 | 1 | 0 | 1 | 1 | 2 | 2 | 2 | 0 | 0 | 0 | 0 | 2 | 4930404H24Rik |
| 4930404N11Rik | 17 | 15 | 21 | 15 | 9 | 21 | 8 | 2 | 6 | 6 | 6 | 2 | 6 | 5 | 7 | 12 | 21 | 15 | 12 | 15 | 11 | 12 | 4 | 7 | 12 | 11 | 7 | 14 | 4930404N11Rik |
| 4930407I10Rik | 2 | 2 | 2 | 9 | 1 | 0 | 1 | 1 | 4 | 1 | 1 | 9 | 7 | 1 | 5 | 5 | 0 | 0 | 2 | 0 | 0 | 3 | 2 | 3 | 1 | 4 | 0 | 0 | 4930407I10Rik |
| 4930412O13Rik | 130 | 145 | 197 | 182 | 148 | 177 | 102 | 147 | 109 | 133 | 122 | 138 | 231 | 165 | 119 | 168 | 185 | 154 | 182 | 106 | 172 | 164 | 122 | 162 | 142 | 50 | 201 | 126 | 4930412O13Rik |
| 4930430F08Rik | 1649 | 1494 | 1634 | 1627 | 1413 | 1549 | 1666 | 1261 | 980 | 906 | 925 | 951 | 1015 | 1022 | 1999 | 1665 | 1608 | 1487 | 1522 | 1781 | 1709 | 999 | 1038 | 934 | 1016 | 1047 | 951 | 1187 | 4930430F08Rik |
| 4930432M17Rik | 0 | 0 | 0 | 0 | 0 | 0 | 0 | 0 | 0 | 0 | 0 | 0 | 0 | 0 | 0 | 0 | 0 | 1 | 0 | 0 | 0 | 0 | 0 | 0 | 0 | 0 | 0 | 0 | 4930432M17Rik |
| 4930438A08Rik | 2 | 7 | 6 | 2 | 3 | 0 | 6 | 0 | 7 | 4 | 5 | 4 | 1 | 1 | 13 | 7 | 6 | 2 | 5 | 8 | 4 | 0 | 1 | 5 | 2 | 4 | 3 | 2 | 4930438A08Rik |
| 4930444P10Rik | 47 | 51 | 44 | 62 | 65 | 61 | 75 | 132 | 133 | 178 | 130 | 119 | 127 | 153 | 40 | 37 | 52 | 67 | 60 | 74 | 78 | 169 | 148 | 116 | 164 | 93 | 141 | 125 | 4930444P10Rik |
| 4930447C04Rik | 1060 | 557 | 670 | 535 | 677 | 766 | 472 | 863 | 947 | 791 | 1317 | 618 | 1099 | 727 | 737 | 1106 | 604 | 819 | 366 | 440 | 496 | 1180 | 950 | 728 | 873 | 624 | 780 | 605 | 4930447C04Rik |
| 4930451I11Rik | 1 | 5 | 0 | 0 | 0 | 0 | 0 | 0 | 1 | 1 | 0 | 0 | 0 | 0 | 0 | 1 | 0 | 0 | 0 | 0 | 2 | 0 | 1 | 2 | 2 | 1 | 9 | 1 | 4930451I11Rik |
| 4930453N24Rik | 4195 | 3576 | 3859 | 3688 | 3134 | 3506 | 3866 | 3831 | 2979 | 2817 | 2892 | 3255 | 3106 | 3575 | 4385 | 3758 | 3585 | 3190 | 3208 | 3802 | 3780 | 2958 | 3032 | 3229 | 3035 | 3705 | 3421 | 3847 | 4930453N24Rik |
| 4930455H04Rik | 0 | 0 | 0 | 0 | 0 | 0 | 0 | 0 | 0 | 0 | 0 | 0 | 0 | 0 | 0 | 0 | 0 | 0 | 0 | 0 | 0 | 0 | 0 | 0 | 0 | 0 | 1 | 0 | 4930455H04Rik |
| 4930467E23Rik | 6 | 3 | 0 | 0 | 3 | 3 | 1 | 0 | 5 | 0 | 0 | 1 | 0 | 0 | 5 | 3 | 0 | 0 | 0 | 5 | 4 | 0 | 2 | 0 | 0 | 0 | 0 | 2 | 4930467E23Rik |
| 4930469K13Rik | 2 | 0 | 0 | 0 | 65 | 0 | 0 | 20 | 16 | 38 | 67 | 0 | 11 | 0 | 3 | 0 | 3 | 0 | 2 | 0 | 0 | 5 | 62 | 4 | 4 | 1 | 0 | 0 | 4930469K13Rik |
| 4930470P17Rik | 0 | 0 | 0 | 0 | 0 | 0 | 0 | 0 | 1 | 1 | 0 | 0 | 2 | 0 | 0 | 2 | 0 | 0 | 0 | 0 | 0 | 0 | 1 | 0 | 0 | 1 | 0 | 2 | 4930470P17Rik |
| 4930474N05Rik | 0 | 0 | 0 | 0 | 0 | 0 | 0 | 0 | 0 | 0 | 0 | 0 | 0 | 1 | 0 | 0 | 0 | 0 | 0 | 0 | 0 | 2 | 0 | 0 | 0 | 0 | 0 | 0 | 4930474N05Rik |
| 4930503B20Rik | 8 | 3 | 10 | 13 | 2 | 7 | 3 | 4 | 5 | 1 | 5 | 3 | 5 | 2 | 7 | 7 | 3 | 3 | 5 | 1 | 4 | 0 | 5 | 4 | 2 | 2 | 2 | 5 | 4930503B20Rik |
| 4930503E14Rik | 0 | 0 | 0 | 0 | 0 | 0 | 0 | 0 | 0 | 0 | 2 | 0 | 0 | 0 | 0 | 0 | 1 | 2 | 0 | 0 | 0 | 0 | 1 | 2 | 0 | 2 | 0 | 1 | 4930503E14Rik |
| 4930503L19Rik | 4943 | 3569 | 3903 | 3589 | 3715 | 4461 | 4584 | 5013 | 4187 | 3152 | 3672 | 4692 | 4511 | 4763 | 4896 | 4533 | 3684 | 3455 | 3777 | 5196 | 4261 | 3350 | 3750 | 3797 | 3684 | 4662 | 4962 | 4752 | 4930503L19Rik |
| 4930505A04Rik | 13 | 3 | 8 | 2 | 6 | 8 | 4 | 2 | 7 | 5 | 4 | 11 | 14 | 5 | 4 | 5 | 1 | 0 | 2 | 3 | 0 | 8 | 8 | 3 | 10 | 6 | 7 | 2 | 4930505A04Rik |
| 4930513O06Rik | 0 | 0 | 0 | 0 | 0 | 0 | 0 | 0 | 0 | 0 | 0 | 0 | 0 | 0 | 0 | 0 | 0 | 0 | 0 | 0 | 0 | 0 | 0 | 2 | 0 | 1 | 0 | 0 | 4930513O06Rik |
| 4930516K23Rik | 0 | 0 | 0 | 0 | 0 | 0 | 0 | 0 | 0 | 0 | 0 | 0 | 0 | 0 | 0 | 0 | 0 | 0 | 0 | 0 | 0 | 0 | 3 | 0 | 0 | 0 | 0 | 0 | 4930516K23Rik |
| 4930519G04Rik | 0 | 4 | 2 | 0 | 0 | 0 | 1 | 0 | 0 | 0 | 0 | 0 | 0 | 0 | 4 | 2 | 6 | 1 | 0 | 0 | 1 | 0 | 0 | 1 | 0 | 0 | 1 | 0 | 4930519G04Rik |
| 4930522H14Rik | 1 | 2 | 0 | 0 | 0 | 0 | 3 | 0 | 0 | 0 | 0 | 0 | 0 | 0 | 0 | 1 | 0 | 1 | 0 | 5 | 0 | 0 | 0 | 0 | 0 | 0 | 0 | 0 | 4930522H14Rik |
| 4930522L14Rik | 482 | 398 | 464 | 450 | 398 | 497 | 438 | 292 | 286 | 204 | 246 | 288 | 316 | 245 | 522 | 593 | 534 | 469 | 516 | 642 | 421 | 239 | 313 | 273 | 297 | 346 | 298 | 282 | 4930522L14Rik |
| 4930523C07Rik | 195 | 154 | 204 | 112 | 141 | 208 | 127 | 132 | 176 | 198 | 240 | 115 | 175 | 101 | 185 | 198 | 144 | 176 | 147 | 146 | 117 | 137 | 299 | 125 | 115 | 107 | 120 | 147 | 4930523C07Rik |
| 4930524B15Rik | 15 | 16 | 9 | 13 | 21 | 16 | 17 | 23 | 24 | 29 | 28 | 21 | 51 | 24 | 26 | 27 | 10 | 3 | 12 | 21 | 9 | 27 | 28 | 32 | 30 | 20 | 15 | 31 | 4930524B15Rik |
| 4930544D05Rik | 1 | 4 | 2 | 0 | 4 | 8 | 6 | 0 | 1 | 3 | 0 | 0 | 2 | 1 | 4 | 2 | 1 | 2 | 0 | 0 | 3 | 0 | 1 | 0 | 0 | 0 | 2 | 0 | 4930544D05Rik |
| 4930548H24Rik | 28 | 18 | 31 | 27 | 12 | 16 | 32 | 4 | 10 | 9 | 6 | 16 | 14 | 15 | 25 | 26 | 24 | 29 | 30 | 19 | 15 | 25 | 11 | 17 | 15 | 22 | 8 | 21 | 4930548H24Rik |
| 4930550C14Rik | 24 | 19 | 25 | 13 | 11 | 14 | 17 | 33 | 33 | 9 | 36 | 29 | 25 | 24 | 25 | 19 | 23 | 17 | 9 | 7 | 16 | 15 | 57 | 11 | 15 | 17 | 33 | 17 | 4930550C14Rik |
| 4930556J24Rik | 6 | 0 | 1 | 1 | 2 | 2 | 0 | 0 | 1 | 1 | 4 | 0 | 1 | 2 | 8 | 1 | 1 | 4 | 0 | 1 | 3 | 7 | 0 | 6 | 4 | 4 | 3 | 0 | 4930556J24Rik |
| 4930558C23Rik | 2 | 0 | 0 | 0 | 0 | 0 | 0 | 2 | 0 | 0 | 0 | 0 | 0 | 0 | 0 | 0 | 0 | 0 | 0 | 0 | 0 | 0 | 2 | 2 | 0 | 0 | 0 | 2 | 4930558C23Rik |
| 4930558K02Rik | 3 | 3 | 6 | 2 | 7 | 1 | 6 | 1 | 4 | 1 | 8 | 1 | 2 | 2 | 0 | 2 | 12 | 11 | 2 | 4 | 6 | 10 | 3 | 2 | 1 | 3 | 0 | 1 | 4930558K02Rik |
| 4930562C15Rik | 1 | 5 | 5 | 4 | 5 | 2 | 8 | 0 | 1 | 0 | 1 | 0 | 0 | 2 | 8 | 2 | 1 | 6 | 8 | 2 | 1 | 0 | 0 | 2 | 0 | 0 | 0 | 0 | 4930562C15Rik |
| 4930563E22Rik | 299 | 247 | 347 | 375 | 354 | 316 | 384 | 504 | 480 | 583 | 521 | 526 | 458 | 496 | 349 | 264 | 358 | 410 | 376 | 371 | 387 | 622 | 471 | 499 | 547 | 498 | 532 | 422 | 4930563E22Rik |
| 4930563M21Rik | 0 | 1 | 0 | 0 | 1 | 1 | 0 | 0 | 0 | 0 | 0 | 0 | 0 | 1 | 0 | 2 | 1 | 0 | 3 | 0 | 2 | 0 | 0 | 2 | 0 | 0 | 0 | 0 | 4930563M21Rik |
| 4930564D02Rik | 0 | 0 | 0 | 0 | 0 | 0 | 0 | 0 | 0 | 0 | 0 | 1 | 0 | 0 | 0 | 0 | 0 | 0 | 0 | 0 | 0 | 0 | 0 | 0 | 0 | 0 | 0 | 0 | 4930564D02Rik |
| 4930578G10Rik | 18 | 14 | 23 | 16 | 17 | 48 | 15 | 10 | 5 | 2 | 8 | 2 | 14 | 2 | 24 | 21 | 43 | 26 | 12 | 17 | 28 | 10 | 6 | 19 | 5 | 4 | 7 | 4 | 4930578G10Rik |

| | | | | | | | | | | | | | | | | | | | | | | | | | | | | | |
|---|---|---|---|---|---|---|---|---|---|---|---|---|---|---|---|---|---|---|---|---|---|---|---|---|---|---|---|---|---|
| 4930579F01Rik | 5 | 3 | 0 | 2 | 0 | 0 | 0 | 4 | 4 | 8 | 5 | 3 | 3 | 0 | 1 | 2 | 0 | 1 | 1 | 3 | 0 | 0 | 5 | 10 | 0 | 2 | 0 | 1 | 4930579F01Rik |
| 4930579G24Rik | 1257 | 956 | 1068 | 959 | 648 | 714 | 701 | 510 | 492 | 431 | 520 | 507 | 568 | 528 | 1068 | 1164 | 972 | 873 | 678 | 779 | 710 | 386 | 507 | 506 | 368 | 632 | 581 | 597 | 4930579G24Rik |
| 4930583I09Rik | 0 | 1 | 5 | 0 | 0 | 0 | 0 | 0 | 0 | 0 | 0 | 0 | 0 | 0 | 0 | 2 | 0 | 0 | 0 | 0 | 0 | 0 | 0 | 0 | 0 | 0 | 0 | 0 | 4930583I09Rik |
| 4930590J08Rik | 499 | 526 | 282 | 442 | 548 | 539 | 561 | 368 | 289 | 343 | 275 | 321 | 339 | 286 | 562 | 479 | 393 | 555 | 494 | 562 | 527 | 416 | 244 | 261 | 340 | 222 | 343 | 274 | 4930590J08Rik |
| 4930595D18Rik | 2 | 3 | 0 | 0 | 9 | 6 | 6 | 5 | 9 | 9 | 8 | 5 | 3 | 13 | 4 | 2 | 0 | 7 | 9 | 8 | 3 | 8 | 8 | 7 | 9 | 12 | 12 | 4 | 4930595D18Rik |
| 4930595M18Rik | 0 | 0 | 0 | 0 | 0 | 0 | 1 | 0 | 0 | 0 | 0 | 0 | 0 | 0 | 0 | 0 | 0 | 0 | 0 | 0 | 0 | 0 | 0 | 0 | 0 | 0 | 0 | 0 | 4930595M18Rik |
| 4931406B18Rik | 0 | 0 | 0 | 0 | 0 | 0 | 0 | 0 | 0 | 3 | 1 | 1 | 0 | 0 | 0 | 0 | 0 | 2 | 0 | 0 | 0 | 0 | 3 | 2 | 0 | 1 | 0 | 0 | 4931406B18Rik |
| 4931406C07Rik | 2226 | 2101 | 2245 | 2181 | 2016 | 2312 | 2204 | 3070 | 2740 | 2746 | 2943 | 2603 | 2662 | 2855 | 2552 | 2083 | 2080 | 2167 | 1978 | 2303 | 2218 | 2892 | 2928 | 2545 | 2624 | 2566 | 2571 | 2667 | 4931406C07Rik |
| 4931408C20Rik | 0 | 0 | 0 | 0 | 0 | 0 | 0 | 0 | 0 | 0 | 0 | 0 | 0 | 0 | 0 | 0 | 0 | 2 | 0 | 0 | 0 | 0 | 0 | 1 | 0 | 0 | 0 | 0 | 4931408C20Rik |
| 4931414P19Rik | 836 | 1001 | 1094 | 1009 | 860 | 772 | 932 | 702 | 817 | 925 | 810 | 923 | 792 | 852 | 857 | 814 | 1132 | 873 | 1039 | 889 | 904 | 742 | 761 | 1102 | 901 | 986 | 870 | 875 | 4931414P19Rik |
| 4931422A03Rik | 6 | 2 | 11 | 5 | 2 | 2 | 3 | 0 | 2 | 3 | 0 | 2 | 1 | 2 | 1 | 0 | 3 | 2 | 5 | 3 | 2 | 3 | 3 | 2 | 2 | 5 | 7 | 2 | 4931422A03Rik |
| 4931428F04Rik | 6664 | 8587 | 8274 | 10986 | 13070 | 10823 | 11255 | 7880 | 8971 | 10951 | 9455 | 9601 | 8800 | 9691 | 7543 | 7567 | 10164 | 11537 | 12123 | 9640 | 11879 | 12509 | 8066 | 10416 | 12589 | 7992 | 8807 | 8822 | 4931428F04Rik |
| 4931428L18Rik | 0 | 0 | 0 | 0 | 0 | 0 | 0 | 0 | 0 | 0 | 0 | 0 | 0 | 0 | 0 | 0 | 0 | 0 | 0 | 0 | 0 | 0 | 0 | 2 | 0 | 0 | 0 | 0 | 4931428L18Rik |
| 4931429L15Rik | 6 | 6 | 6 | 6 | 10 | 2 | 7 | 7 | 6 | 3 | 2 | 8 | 12 | 7 | 8 | 6 | 6 | 2 | 4 | 8 | 3 | 5 | 6 | 10 | 5 | 8 | 2 | 6 | 4931429L15Rik |
| 4932414N04Rik | 0 | 0 | 0 | 0 | 0 | 1 | 0 | 0 | 2 | 0 | 0 | 0 | 0 | 0 | 0 | 0 | 5 | 0 | 0 | 0 | 0 | 0 | 0 | 0 | 0 | 0 | 0 | 0 | 4932414N04Rik |
| 4932415D10Rik | 0 | 0 | 0 | 3 | 0 | 4 | 2 | 5 | 3 | 2 | 2 | 1 | 7 | 3 | 4 | 3 | 2 | 0 | 0 | 0 | 2 | 0 | 3 | 0 | 1 | 0 | 0 | 2 | 4932415D10Rik |
| 4932429P05Rik | 0 | 0 | 0 | 0 | 0 | 0 | 0 | 0 | 0 | 0 | 0 | 1 | 0 | 0 | 0 | 0 | 0 | 0 | 0 | 0 | 0 | 0 | 0 | 0 | 2 | 0 | 0 | 0 | 4932429P05Rik |
| 4932438A13Rik | 19144 | 16218 | 6762 | 9590 | 10934 | 14644 | 13370 | 15344 | 14505 | 7911 | 9022 | 6863 | 7655 | 7313 | 19317 | 17879 | 7398 | 10162 | 9143 | 18521 | 14974 | 8241 | 13271 | 5887 | 8427 | 4150 | 8400 | 6246 | 4932438A13Rik |
| 4932438H23Rik | 29 | 26 | 30 | 24 | 45 | 16 | 15 | 38 | 50 | 49 | 27 | 40 | 44 | 19 | 30 | 36 | 23 | 22 | 33 | 42 | 30 | 27 | 58 | 27 | 23 | 31 | 26 | 25 | 4932438H23Rik |
| 4933400A11Rik | 0 | 0 | 0 | 0 | 0 | 0 | 0 | 1 | 0 | 0 | 0 | 0 | 0 | 0 | 0 | 0 | 0 | 0 | 0 | 0 | 0 | 0 | 0 | 0 | 0 | 0 | 0 | 0 | 4933400A11Rik |
| 4933402D24Rik | 1 | 4 | 6 | 4 | 8 | 4 | 4 | 7 | 9 | 5 | 0 | 1 | 1 | 0 | 7 | 2 | 10 | 4 | 9 | 1 | 3 | 10 | 3 | 1 | 8 | 8 | 8 | 2 | 4933402D24Rik |
| 4933402J07Rik | 17 | 18 | 4 | 23 | 20 | 14 | 22 | 0 | 0 | 1 | 4 | 2 | 1 | 2 | 17 | 12 | 6 | 36 | 12 | 18 | 22 | 0 | 3 | 0 | 2 | 4 | 2 | 2 | 4933402J07Rik |
| 4933402N22Rik | 0 | 0 | 0 | 0 | 0 | 0 | 0 | 0 | 0 | 0 | 2 | 0 | 0 | 0 | 0 | 0 | 0 | 0 | 0 | 0 | 0 | 0 | 0 | 0 | 0 | 0 | 0 | 0 | 4933402N22Rik |
| 4933403O08Rik | 97 | 71 | 9 | 41 | 69 | 66 | 67 | 62 | 44 | 21 | 28 | 39 | 15 | 35 | 76 | 58 | 22 | 32 | 37 | 79 | 60 | 23 | 35 | 20 | 45 | 2 | 55 | 33 | 4933403O08Rik |
| 4933405L10Rik | 0 | 4 | 2 | 5 | 4 | 13 | 0 | 4 | 2 | 3 | 1 | 8 | 7 | 1 | 1 | 1 | 1 | 4 | 9 | 5 | 4 | 2 | 5 | 8 | 5 | 11 | 2 | 2 | 4933405L10Rik |
| 4933405O20Rik | 0 | 0 | 0 | 0 | 2 | 0 | 0 | 4 | 0 | 0 | 0 | 0 | 0 | 0 | 3 | 0 | 2 | 0 | 0 | 0 | 0 | 5 | 0 | 0 | 0 | 0 | 0 | 0 | 4933405O20Rik |
| 4933407L21Rik | 72 | 86 | 61 | 109 | 132 | 123 | 177 | 184 | 198 | 203 | 177 | 212 | 161 | 185 | 88 | 55 | 82 | 86 | 170 | 170 | 183 | 212 | 194 | 251 | 225 | 203 | 164 | 190 | 4933407L21Rik |
| 4933409G03Rik | 0 | 0 | 0 | 0 | 0 | 0 | 0 | 1 | 0 | 2 | 5 | 0 | 0 | 0 | 1 | 0 | 0 | 0 | 0 | 0 | 0 | 3 | 1 | 0 | 0 | 0 | 0 | 0 | 4933409G03Rik |
| 4933411K16Rik | 14 | 17 | 6 | 8 | 6 | 18 | 13 | 6 | 16 | 12 | 21 | 8 | 19 | 5 | 15 | 23 | 6 | 10 | 2 | 6 | 8 | 15 | 14 | 3 | 3 | 0 | 3 | 5 | 4933411K16Rik |
| 4933412E24Rik | 0 | 0 | 0 | 0 | 0 | 0 | 0 | 0 | 0 | 0 | 0 | 0 | 0 | 0 | 0 | 0 | 0 | 0 | 0 | 0 | 0 | 0 | 0 | 0 | 0 | 0 | 3 | 0 | 4933412E24Rik |
| 4933414I15Rik | 0 | 0 | 0 | 0 | 0 | 0 | 0 | 0 | 1 | 0 | 0 | 0 | 0 | 0 | 0 | 0 | 0 | 0 | 0 | 0 | 0 | 0 | 0 | 0 | 0 | 2 | 0 | 0 | 4933414I15Rik |
| 4933417A18Rik | 0 | 0 | 0 | 0 | 0 | 0 | 1 | 0 | 0 | 0 | 1 | 0 | 0 | 0 | 0 | 0 | 0 | 0 | 0 | 0 | 0 | 0 | 1 | 0 | 0 | 0 | 0 | 0 | 4933417A18Rik |
| 4933421I07Rik | 0 | 0 | 0 | 5 | 0 | 2 | 0 | 0 | 0 | 0 | 1 | 0 | 0 | 1 | 0 | 0 | 0 | 0 | 0 | 0 | 0 | 0 | 0 | 0 | 0 | 1 | 0 | 0 | 4933421I07Rik |
| 4933424G06Rik | 6 | 8 | 0 | 4 | 2 | 2 | 8 | 17 | 16 | 10 | 9 | 27 | 5 | 9 | 5 | 3 | 5 | 1 | 2 | 7 | 7 | 18 | 10 | 13 | 9 | 16 | 5 | 9 | 4933424G06Rik |
| 4933427D14Rik | 8 | 10 | 2 | 6 | 13 | 12 | 13 | 21 | 7 | 8 | 13 | 16 | 12 | 15 | 12 | 8 | 6 | 3 | 5 | 17 | 13 | 7 | 15 | 10 | 13 | 4 | 14 | 19 | 4933427D14Rik |
| 4933428G20Rik | 25 | 16 | 23 | 39 | 21 | 36 | 49 | 2 | 11 | 9 | 22 | 9 | 6 | 10 | 18 | 16 | 24 | 51 | 37 | 35 | 42 | 8 | 8 | 5 | 12 | 11 | 7 | 11 | 4933428G20Rik |
| 4933430I17Rik | 325 | 241 | 228 | 223 | 259 | 232 | 259 | 147 | 149 | 168 | 142 | 237 | 219 | 223 | 316 | 224 | 242 | 224 | 236 | 304 | 232 | 182 | 144 | 188 | 232 | 189 | 285 | 236 | 4933430I17Rik |
| 4933434E20Rik | 945 | 788 | 743 | 836 | 867 | 845 | 1079 | 1258 | 871 | 909 | 843 | 1054 | 886 | 1229 | 903 | 722 | 866 | 844 | 910 | 932 | 1184 | 1101 | 830 | 1070 | 1026 | 968 | 949 | 1195 | 4933434E20Rik |
| 5031439G07Rik | 19305 | 21054 | 21182 | 24009 | 22863 | 21756 | 21566 | 10893 | 11576 | 13424 | 12898 | 11961 | 11677 | 11414 | 18923 | 20586 | 23063 | 25281 | 24277 | 17625 | 22858 | 15241 | 11296 | 12593 | 14446 | 9776 | 11460 | 11317 | 5031439G07Rik |
| 5430403G16Rik | 298 | 224 | 297 | 332 | 328 | 289 | 355 | 192 | 208 | 254 | 215 | 211 | 234 | 202 | 355 | 283 | 316 | 252 | 292 | 364 | 312 | 194 | 226 | 179 | 146 | 169 | 185 | 203 | 5430403G16Rik |
| 5530401A14Rik | 3 | 4 | 0 | 0 | 1 | 3 | 0 | 5 | 3 | 2 | 5 | 4 | 11 | 2 | 4 | 3 | 1 | 0 | 0 | 0 | 1 | 0 | 6 | 7 | 2 | 5 | 2 | 4 | 5530401A14Rik |
| 5730409E04Rik | 210 | 253 | 226 | 289 | 727 | 586 | 351 | 1044 | 1302 | 1265 | 1447 | 787 | 921 | 860 | 347 | 298 | 350 | 457 | 408 | 419 | 422 | 1399 | 1337 | 1055 | 1096 | 931 | 895 | 809 | 5730409E04Rik |
| 5730455P16Rik | 2986 | 2394 | 2926 | 2592 | 2814 | 2918 | 2841 | 3861 | 3571 | 3582 | 3583 | 3973 | 3814 | 4316 | 3054 | 2442 | 2605 | 2535 | 2656 | 2824 | 2623 | 3686 | 3359 | 3460 | 3935 | 3855 | 4073 | 4190 | 5730455P16Rik |
| 5730480H06Rik | 1307 | 1099 | 1122 | 1004 | 929 | 904 | 1134 | 1340 | 1182 | 1148 | 1137 | 1232 | 1221 | 1395 | 1325 | 1210 | 1092 | 1002 | 913 | 1190 | 1077 | 1081 | 1224 | 1168 | 1215 | 1351 | 1391 | 1533 | 5730480H06Rik |
| 5730507C01Rik | 68 | 48 | 151 | 53 | 83 | 53 | 36 | 54 | 95 | 85 | 58 | 53 | 120 | 77 | 91 | 118 | 87 | 51 | 72 | 76 | 56 | 114 | 121 | 69 | 73 | 105 | 103 | 141 | 5730507C01Rik |
| 5730522E02Rik | 248 | 157 | 96 | 170 | 239 | 235 | 325 | 302 | 307 | 245 | 259 | 234 | 170 | 318 | 206 | 140 | 162 | 217 | 216 | 320 | 311 | 353 | 282 | 274 | 307 | 269 | 230 | 214 | 5730522E02Rik |
| 6030458C11Rik | 3157 | 2981 | 2908 | 3043 | 2897 | 2963 | 3018 | 3776 | 3745 | 3664 | 4077 | 4051 | 4024 | 3984 | 3299 | 3062 | 3038 | 3035 | 2950 | 3081 | 2918 | 3613 | 3737 | 3755 | 3724 | 3397 | 4026 | 3796 | 6030458C11Rik |
| 6030468B19Rik | 2 | 4 | 14 | 2 | 2 | 1 | 1 | 4 | 3 | 9 | 8 | 18 | 0 | 4 | 4 | 7 | 3 | 1 | 2 | 2 | 1 | 7 | 3 | 2 | 2 | 6 | 0 | 31 | 6030468B19Rik |
| 6030498E09Rik | 1 | 0 | 7 | 3 | 4 | 2 | 0 | 4 | 5 | 3 | 10 | 4 | 3 | 3 | 1 | 2 | 2 | 1 | 0 | 3 | 1 | 3 | 3 | 2 | 1 | 1 | 0 | 5 | 6030498E09Rik |
| 6330403K07Rik | 9054 | 10612 | 9745 | 10964 | 13042 | 12339 | 12081 | 17027 | 15021 | 15826 | 14860 | 15459 | 15618 | 16153 | 10362 | 8565 | 10855 | 10871 | 13339 | 13169 | 12545 | 16404 | 14914 | 16444 | 17285 | 14974 | 16467 | 16014 | 6330403K07Rik |
| 6330409D20Rik | 179 | 231 | 167 | 251 | 209 | 209 | 173 | 110 | 98 | 152 | 112 | 165 | 118 | 123 | 202 | 160 | 251 | 253 | 254 | 193 | 201 | 144 | 119 | 152 | 147 | 166 | 180 | 150 | 6330409D20Rik |
| 6330411D24Rik | 1 | 0 | 0 | 6 | 0 | 3 | 0 | 1 | 1 | 0 | 0 | 0 | 0 | 0 | 1 | 1 | 0 | 0 | 0 | 1 | 0 | 2 | 5 | 0 | 2 | 0 | 1 | 0 | 6330411D24Rik |
| 6430548M08Rik | 4955 | 6823 | 5476 | 7846 | 7234 | 6833 | 6737 | 2522 | 2645 | 3098 | 2709 | 2845 | 2595 | 2600 | 5548 | 5896 | 6609 | 7947 | 7662 | 6248 | 7502 | 3755 | 2576 | 2707 | 3305 | 1978 | 2782 | 2693 | 6430548M08Rik |
| 6430550D23Rik | 324 | 355 | 194 | 331 | 256 | 341 | 323 | 137 | 124 | 139 | 159 | 135 | 141 | 127 | 324 | 347 | 291 | 322 | 313 | 318 | 320 | 172 | 124 | 118 | 124 | 100 | 132 | 160 | 6430550D23Rik |
| 6430571L13Rik | 287 | 358 | 398 | 454 | 541 | 366 | 405 | 282 | 323 | 319 | 268 | 312 | 324 | 287 | 342 | 268 | 446 | 495 | 517 | 418 | 399 | 364 | 271 | 297 | 357 | 191 | 268 | 311 | 6430571L13Rik |
| 6430628N08Rik | 2 | 7 | 6 | 6 | 6 | 1 | 3 | 2 | 2 | 0 | 6 | 0 | 5 | 4 | 1 | 0 | 3 | 3 | 8 | 13 | 5 | 2 | 1 | 1 | 0 | 0 | 2 | 1 | 6430628N08Rik |
| 6720489N17Rik | 1261 | 832 | 1053 | 936 | 975 | 1074 | 1130 | 1348 | 1063 | 891 | 991 | 1049 | 1099 | 1018 | 1184 | 985 | 1093 | 886 | 1024 | 1204 | 893 | 964 | 1131 | 1058 | 932 | 1320 | 1238 | 1260 | 6720489N17Rik |
| 6820408C15Rik | 42 | 86 | 84 | 90 | 109 | 73 | 83 | 98 | 84 | 106 | 105 | 113 | 147 | 97 | 73 | 86 | 73 | 74 | 98 | 87 | 69 | 119 | 122 | 98 | 72 | 96 | 119 | 87 | 6820408C15Rik |
| 7420426K07Rik | 0 | 0 | 0 | 0 | 0 | 0 | 0 | 0 | 0 | 0 | 0 | 0 | 0 | 0 | 0 | 1 | 0 | 0 | 0 | 0 | 1 | 0 | 0 | 0 | 0 | 0 | 0 | 0 | 7420426K07Rik |
| 7530416G11Rik | 1 | 0 | 5 | 0 | 2 | 0 | 1 | 1 | 6 | 3 | 0 | 3 | 3 | 3 | 1 | 0 | 0 | 0 | 6 | 1 | 5 | 5 | 3 | 2 | 0 | 1 | 6 | 1 | 7530416G11Rik |
| 8030462N17Rik | 155 | 89 | 106 | 156 | 138 | 90 | 87 | 89 | 84 | 54 | 56 | 103 | 92 | 54 | 186 | 129 | 125 | 122 | 97 | 160 | 81 | 107 | 52 | 61 | 109 | 55 | 146 | 209 | 8030462N17Rik |
| 8030474K03Rik | 2 | 0 | 6 | 1 | 0 | 8 | 2 | 6 | 4 | 11 | 9 | 5 | 11 | 10 | 0 | 2 | 2 | 0 | 0 | 1 | 0 | 12 | 13 | 4 | 5 | 4 | 2 | 4 | 8030474K03Rik |
| 9030025P20Rik | 87 | 88 | 78 | 92 | 76 | 88 | 115 | 166 | 161 | 155 | 140 | 148 | 151 | 165 | 116 | 92 | 70 | 70 | 86 | 121 | 120 | 202 | 148 | 176 | 166 | 146 | 175 | 167 | 9030025P20Rik |
| 9030624G23Rik | 276 | 200 | 168 | 147 | 213 | 165 | 101 | 171 | 182 | 200 | 123 | 135 | 144 | 143 | 195 | 287 | 162 | 142 | 170 | 141 | 157 | 185 | 177 | 130 | 150 | 129 | 209 | 127 | 9030624G23Rik |
| 9130008F23Rik | 107 | 54 | 81 | 35 | 91 | 76 | 30 | 54 | 82 | 34 | 69 | 60 | 53 | 43 | 105 | 83 | 88 | 63 | 29 | 60 | 35 | 32 | 83 | 29 | 36 | 64 | 58 | 43 | 9130008F23Rik |
| 9130019O22Rik | 787 | 1042 | 685 | 810 | 641 | 817 | 771 | 482 | 594 | 511 | 568 | 537 | 551 | 432 | 778 | 762 | 681 | 724 | 644 | 861 | 771 | 369 | 462 | 400 | 539 | 368 | 502 | 483 | 9130019O22Rik |
| 9130023H24Rik | 1061 | 1055 | 1165 | 1081 | 999 | 911 | 961 | 852 | 887 | 932 | 971 | 1069 | 1112 | 1029 | 1018 | 980 | 1073 | 967 | 969 | 1098 | 920 | 1088 | 925 | 964 | 981 | 1236 | 1103 | 1146 | 9130023H24Rik |
| 9130230L23Rik | 1 | 1 | 0 | 0 | 4 | 2 | 1 | 0 | 4 | 4 | 2 | 11 | 8 | 3 | 4 | 0 | 3 | 0 | 2 | 3 | 0 | 0 | 3 | 2 | 2 | 0 | 1 | 2 | 9130230L23Rik |
| 9130401M01Rik | 0 | 0 | 0 | 0 | 0 | 0 | 0 | 0 | 0 | 0 | 0 | 0 | 0 | 0 | 0 | 0 | 0 | 0 | 0 | 0 | 0 | 0 | 0 | 0 | 0 | 0 | 2 | 0 | 9130401M01Rik |
| 9130409I23Rik | 0 | 0 | 0 | 0 | 0 | 4 | 0 | 0 | 0 | 0 | 0 | 0 | 0 | 0 | 0 | 0 | 0 | 0 | 0 | 0 | 0 | 0 | 0 | 0 | 0 | 0 | 0 | 0 | 9130409I23Rik |
| 9230102O04Rik | 23 | 33 | 39 | 39 | 51 | 38 | 37 | 48 | 33 | 72 | 48 | 41 | 70 | 54 | 37 | 33 | 51 | 50 | 50 | 44 | 53 | 74 | 25 | 29 | 47 | 5 | 96 | 52 | 9230102O04Rik |
| 9330159F19Rik | 3038 | 2820 | 3920 | 5824 | 8233 | 7171 | 8767 | 7330 | 6108 | 8835 | 7356 | 7642 | 6067 | 8738 | 3700 | 2864 | 5353 | 6866 | 7896 | 6104 | 8525 | 12626 | 5725 | 8763 | 10615 | 6107 | 6312 | 7155 | 9330159F19Rik |
| 9330182O14Rik | 3 | 0 | 0 | 0 | 2 | 0 | 0 | 0 | 0 | 0 | 0 | 0 | 0 | 0 | 0 | 0 | 0 | 0 | 0 | 1 | 0 | 0 | 2 | 0 | 0 | 1 | 0 | 0 | 9330182O14Rik |
| 9430015G10Rik | 4932 | 5279 | 4348 | 4383 | 4223 | 4160 | 4703 | 5153 | 4820 | 4749 | 4352 | 5159 | 4699 | 5233 | 4523 | 4501 | 4592 | 4451 | 4376 | 4389 | 4525 | 4490 | 4448 | 5212 | 4983 | 4976 | 5226 | 5158 | 9430015G10Rik |
| 9430038I01Rik | 548 | 499 | 418 | 380 | 413 | 430 | 427 | 435 | 491 | 380 | 504 | 393 | 502 | 479 | 589 | 554 | 409 | 417 | 295 | 451 | 406 | 279 | 529 | 300 | 342 | 504 | 521 | 394 | 9430038I01Rik |
| 9430069I07Rik | 0 | 2 | 0 | 1 | 2 | 3 | 0 | 0 | 2 | 3 | 0 | 0 | 0 | 0 | 5 | 0 | 3 | 0 | 4 | 4 | 1 | 0 | 0 | 0 | 0 | 0 | 0 | 0 | 9430069I07Rik |
| 9530003J23Rik | 2 | 0 | 0 | 0 | 0 | 0 | 0 | 0 | 0 | 0 | 0 | 2 | 0 | 0 | 0 | 0 | 3 | 0 | 0 | 0 | 0 | 0 | 1 | 0 | 0 | 0 | 0 | 0 | 9530003J23Rik |
| 9530053A07Rik | 7 | 3 | 5 | 3 | 11 | 15 | 3 | 20 | 24 | 32 | 14 | 4 | 15 | 10 | 4 | 6 | 5 | 7 | 12 | 7 | 10 | 10 | 30 | 19 | 9 | 13 | 10 | 6 | 9530053A07Rik |
| 9530068E07Rik | 7214 | 6935 | 8678 | 6604 | 6257 | 6865 | 6099 | 8033 | 8242 | 7942 | 9031 | 8901 | 9062 | 8651 | 7346 | 7821 | 7326 | 6644 | 6152 | 5463 | 6147 | 8535 | 9528 | 8436 | 7523 | 9839 | 8428 | 9282 | 9530068E07Rik |

| | | | | | | | | | | | | | | | | | | | | | | | | | | | | | |
|---|---|---|---|---|---|---|---|---|---|---|---|---|---|---|---|---|---|---|---|---|---|---|---|---|---|---|---|---|---|
| 9530077C05Rik | 536 | 573 | 605 | 680 | 592 | 605 | 655 | 354 | 367 | 440 | 333 | 496 | 383 | 486 | 579 | 492 | 630 | 559 | 660 | 613 | 559 | 374 | 392 | 423 | 460 | 450 | 562 | 503 | 9530077C05Rik |
| 9630041A04Rik | 0 | 0 | 0 | 0 | 0 | 0 | 0 | 0 | 0 | 0 | 0 | 0 | 0 | 2 | 0 | 0 | 0 | 0 | 1 | 0 | 2 | 0 | 0 | 0 | 0 | 1 | 0 | 0 | 9630041A04Rik |
| 9930012K11Rik | 564 | 564 | 567 | 417 | 367 | 374 | 312 | 353 | 388 | 389 | 335 | 375 | 438 | 313 | 549 | 643 | 526 | 333 | 374 | 381 | 293 | 349 | 410 | 347 | 300 | 448 | 350 | 333 | 9930012K11Rik |
| 9930021J03Rik | 5341 | 4844 | 3607 | 4836 | 4822 | 5317 | 5338 | 7684 | 6550 | 6104 | 6031 | 6448 | 5692 | 6297 | 5181 | 4910 | 4088 | 4754 | 4836 | 5224 | 5214 | 6924 | 6269 | 5447 | 6501 | 4730 | 6242 | 5985 | 9930021J03Rik |
| 9930022D16Rik | 2 | 0 | 0 | 0 | 1 | 0 | 0 | 1 | 2 | 0 | 3 | 0 | 9 | 0 | 0 | 0 | 0 | 2 | 0 | 0 | 0 | 3 | 1 | 0 | 0 | 0 | 0 | 1 | 9930022D16Rik |
| 9930104L06Rik | 2501 | 2738 | 2778 | 2930 | 2628 | 2667 | 2580 | 1890 | 1800 | 1989 | 1991 | 2077 | 2045 | 2228 | 2352 | 2618 | 2880 | 2859 | 2622 | 2412 | 2531 | 1838 | 1754 | 2062 | 2187 | 2051 | 2251 | 2187 | 9930104L06Rik |
| 9930111J21Rik1 | 0 | 0 | 0 | 0 | 2 | 0 | 0 | 2 | 2 | 1 | 8 | 3 | 2 | 5 | 4 | 1 | 0 | 1 | 2 | 0 | 0 | 2 | 8 | 4 | 1 | 0 | 10 | 2 | 9930111J21Rik1 |
| 9930111J21Rik2 | 16 | 9 | 7 | 12 | 16 | 23 | 11 | 51 | 55 | 42 | 80 | 39 | 32 | 29 | 11 | 14 | 3 | 1 | 10 | 12 | 13 | 67 | 74 | 25 | 32 | 25 | 41 | 21 | 9930111J21Rik2 |
| a | 59 | 10 | 28 | 20 | 19 | 22 | 16 | 76 | 69 | 60 | 44 | 10 | 39 | 22 | 21 | 24 | 6 | 25 | 12 | 32 | 29 | 15 | 78 | 48 | 38 | 40 | 31 | 28 | a |
| A030003K21Rik | 0 | 0 | 0 | 0 | 0 | 0 | 0 | 0 | 0 | 0 | 0 | 0 | 0 | 0 | 0 | 0 | 0 | 0 | 0 | 0 | 0 | 0 | 3 | 0 | 0 | 0 | 0 | 0 | A030003K21Rik |
| A030005L19Rik | 47 | 73 | 57 | 93 | 59 | 69 | 75 | 22 | 23 | 28 | 31 | 39 | 43 | 42 | 40 | 60 | 66 | 73 | 84 | 51 | 71 | 28 | 26 | 33 | 46 | 48 | 26 | 24 | A030005L19Rik |
| A030014E15Rik | 6 | 3 | 5 | 6 | 11 | 22 | 10 | 7 | 1 | 5 | 1 | 0 | 0 | 3 | 1 | 2 | 9 | 15 | 11 | 12 | 8 | 3 | 1 | 4 | 0 | 1 | 3 | 2 | A030014E15Rik |
| A130010J15Rik | 825 | 680 | 905 | 785 | 856 | 918 | 898 | 1022 | 972 | 954 | 938 | 1084 | 1042 | 1043 | 832 | 783 | 819 | 826 | 812 | 914 | 831 | 894 | 1065 | 1043 | 981 | 1118 | 1070 | 1140 | A130010J15Rik |
| A1bg | 3 | 2 | 0 | 1 | 0 | 0 | 0 | 1 | 2 | 0 | 0 | 1 | 0 | 1 | 1 | 2 | 0 | 0 | 0 | 0 | 0 | 0 | 0 | 0 | 0 | 0 | 1 | 0 | A1bg |
| A1cf | 0 | 1 | 0 | 0 | 2 | 0 | 0 | 2 | 0 | 0 | 0 | 1 | 0 | 0 | 0 | 1 | 0 | 0 | 0 | 0 | 0 | 0 | 1 | 0 | 0 | 0 | 0 | 0 | A1cf |
| A230072I06Rik | 0 | 0 | 0 | 0 | 2 | 0 | 3 | 0 | 0 | 0 | 2 | 0 | 0 | 3 | 0 | 0 | 0 | 1 | 0 | 0 | 0 | 0 | 1 | 0 | 2 | 0 | 3 | 1 | A230072I06Rik |
| A2m | 578 | 741 | 764 | 391 | 790 | 701 | 391 | 854 | 821 | 767 | 784 | 821 | 1060 | 887 | 624 | 705 | 497 | 529 | 477 | 546 | 514 | 1004 | 984 | 600 | 707 | 508 | 1047 | 985 | A2m |
| A2ml1 | 14 | 3 | 2 | 16 | 11 | 19 | 19 | 17 | 25 | 41 | 20 | 10 | 7 | 14 | 4 | 3 | 2 | 4 | 16 | 15 | 14 | 7 | 32 | 29 | 19 | 14 | 2 | 5 | A2ml1 |
| A330070K13Rik | 0 | 0 | 0 | 0 | 0 | 0 | 1 | 1 | 1 | 0 | 0 | 0 | 0 | 0 | 0 | 0 | 0 | 0 | 0 | 0 | 0 | 0 | 1 | 0 | 0 | 0 | 1 | 0 | A330070K13Rik |
| A3galt2 | 2284 | 3340 | 1796 | 3757 | 2959 | 2819 | 3161 | 386 | 362 | 480 | 414 | 422 | 380 | 442 | 2300 | 3026 | 2715 | 4085 | 3189 | 2679 | 3327 | 548 | 354 | 433 | 485 | 250 | 356 | 373 | A3galt2 |
| A430005L14Rik | 1224 | 1498 | 1430 | 1220 | 986 | 1018 | 1101 | 1017 | 1165 | 1122 | 1058 | 1333 | 1287 | 1276 | 1188 | 1261 | 1381 | 1164 | 1206 | 1315 | 1046 | 984 | 1177 | 1224 | 1024 | 1680 | 1527 | 1494 | A430005L14Rik |
| A430033K04Rik | 2264 | 2274 | 1991 | 2336 | 2166 | 2107 | 2320 | 1885 | 1766 | 1788 | 1657 | 1486 | 1477 | 1605 | 2378 | 2233 | 2141 | 2226 | 2166 | 2570 | 2360 | 1873 | 1821 | 1588 | 1899 | 1640 | 1374 | 1488 | A430033K04Rik |
| A4galt | 172 | 164 | 191 | 162 | 164 | 176 | 112 | 112 | 137 | 126 | 173 | 97 | 168 | 105 | 116 | 205 | 204 | 196 | 121 | 149 | 125 | 157 | 147 | 176 | 111 | 149 | 170 | 103 | A4galt |
| A4gnt | 0 | 0 | 0 | 0 | 1 | 0 | 0 | 0 | 0 | 0 | 3 | 0 | 0 | 0 | 0 | 1 | 0 | 0 | 0 | 0 | 0 | 0 | 0 | 0 | 0 | 1 | 0 | 0 | A4gnt |
| A530016L24Rik | 5 | 0 | 1 | 0 | 0 | 0 | 0 | 4 | 7 | 4 | 5 | 3 | 1 | 1 | 0 | 1 | 0 | 0 | 0 | 0 | 0 | 10 | 10 | 5 | 3 | 0 | 1 | 1 | A530016L24Rik |
| A530032D15Rik | 1 | 0 | 0 | 1 | 0 | 0 | 0 | 0 | 1 | 0 | 0 | 0 | 0 | 0 | 0 | 0 | 0 | 0 | 0 | 1 | 0 | 2 | 0 | 3 | 0 | 0 | 0 | 0 | A530032D15Rik |
| A530084C06Rik | 23 | 22 | 37 | 48 | 56 | 22 | 46 | 31 | 55 | 33 | 73 | 27 | 35 | 30 | 22 | 39 | 43 | 14 | 38 | 41 | 38 | 50 | 47 | 61 | 49 | 49 | 48 | 50 | A530084C06Rik |
| A630001G21Rik | 59 | 42 | 66 | 55 | 60 | 66 | 34 | 139 | 128 | 160 | 220 | 164 | 231 | 132 | 46 | 48 | 61 | 48 | 44 | 52 | 25 | 139 | 198 | 117 | 128 | 168 | 177 | 145 | A630001G21Rik |
| A630010A05Rik | 0 | 0 | 0 | 0 | 0 | 4 | 0 | 0 | 0 | 0 | 0 | 0 | 0 | 0 | 0 | 0 | 0 | 0 | 0 | 0 | 0 | 0 | 0 | 2 | 0 | 0 | 0 | 0 | A630010A05Rik |
| A630023A22Rik | 0 | 0 | 0 | 3 | 0 | 3 | 0 | 0 | 0 | 0 | 1 | 1 | 0 | 3 | 0 | 0 | 0 | 0 | 0 | 0 | 1 | 0 | 0 | 1 | 2 | 0 | 0 | 0 | A630023A22Rik |
| A630076J17Rik | 0 | 0 | 0 | 0 | 0 | 1 | 0 | 0 | 1 | 1 | 0 | 0 | 0 | 0 | 0 | 0 | 0 | 3 | 0 | 0 | 1 | 0 | 0 | 0 | 0 | 1 | 0 | 0 | A630076J17Rik |
| A630089N07Rik | 929 | 818 | 724 | 694 | 757 | 771 | 702 | 1357 | 1178 | 927 | 909 | 1102 | 980 | 1015 | 955 | 1003 | 721 | 593 | 788 | 999 | 666 | 839 | 1176 | 807 | 950 | 867 | 1140 | 1123 | A630089N07Rik |
| A630095N17Rik | 1 | 3 | 0 | 0 | 2 | 2 | 0 | 0 | 0 | 3 | 0 | 1 | 3 | 2 | 1 | 1 | 1 | 0 | 0 | 0 | 1 | 0 | 2 | 0 | 0 | 0 | 5 | 0 | A630095N17Rik |
| A730009L09Rik | 1 | 0 | 0 | 0 | 0 | 0 | 0 | 0 | 0 | 4 | 3 | 0 | 1 | 0 | 0 | 1 | 0 | 0 | 0 | 0 | 2 | 0 | 1 | 0 | 5 | 0 | 3 | 0 | A730009L09Rik |
| A730046J19Rik | 52 | 37 | 31 | 32 | 89 | 65 | 73 | 43 | 46 | 52 | 116 | 13 | 14 | 15 | 50 | 47 | 20 | 65 | 43 | 63 | 63 | 75 | 52 | 24 | 30 | 14 | 31 | 18 | A730046J19Rik |
| A730049H05Rik | 28 | 19 | 23 | 26 | 25 | 33 | 25 | 45 | 27 | 12 | 25 | 35 | 29 | 28 | 30 | 40 | 25 | 32 | 30 | 29 | 27 | 50 | 46 | 25 | 23 | 27 | 30 | 27 | A730049H05Rik |
| A730071L15Rik | 6 | 3 | 7 | 7 | 6 | 3 | 7 | 29 | 9 | 14 | 8 | 25 | 2 | 22 | 13 | 8 | 3 | 7 | 3 | 7 | 6 | 8 | 7 | 15 | 15 | 15 | 9 | 24 | A730071L15Rik |
| A830005F24Rik | 62 | 74 | 69 | 47 | 48 | 72 | 61 | 48 | 56 | 64 | 41 | 40 | 76 | 52 | 78 | 53 | 67 | 61 | 70 | 71 | 60 | 69 | 63 | 52 | 58 | 77 | 50 | 55 | A830005F24Rik |
| A830018L16Rik | 143 | 89 | 75 | 88 | 187 | 86 | 87 | 282 | 347 | 278 | 341 | 149 | 220 | 185 | 92 | 95 | 63 | 109 | 62 | 104 | 122 | 294 | 328 | 202 | 206 | 175 | 138 | 176 | A830018L16Rik |
| A830031A19Rik | 22 | 17 | 12 | 10 | 16 | 23 | 32 | 13 | 14 | 15 | 8 | 8 | 23 | 13 | 11 | 5 | 32 | 29 | 30 | 22 | 14 | 12 | 14 | 8 | 21 | 8 | 12 | 17 | A830031A19Rik |
| A930002H24Rik | 37 | 42 | 46 | 39 | 34 | 30 | 19 | 23 | 20 | 33 | 16 | 25 | 25 | 19 | 28 | 33 | 42 | 52 | 28 | 36 | 22 | 17 | 21 | 18 | 11 | 28 | 31 | 18 | A930002H24Rik |
| A930003A15Rik | 0 | 0 | 2 | 0 | 2 | 0 | 1 | 4 | 2 | 2 | 8 | 7 | 0 | 4 | 1 | 2 | 0 | 0 | 2 | 5 | 1 | 5 | 8 | 0 | 0 | 1 | 0 | 2 | A930003A15Rik |
| A930004D18Rik | 694 | 755 | 998 | 785 | 704 | 799 | 811 | 1097 | 1189 | 1129 | 1153 | 1410 | 1424 | 1430 | 708 | 681 | 1065 | 722 | 891 | 793 | 738 | 964 | 1071 | 1504 | 1336 | 1547 | 1262 | 1504 | A930004D18Rik |
| A930007A09Rik | 389 | 363 | 355 | 323 | 445 | 431 | 445 | 443 | 354 | 366 | 395 | 405 | 388 | 449 | 397 | 313 | 322 | 387 | 372 | 445 | 463 | 329 | 401 | 381 | 379 | 315 | 360 | 319 | A930007A09Rik |
| A930009A15Rik | 3 | 15 | 10 | 11 | 9 | 5 | 3 | 33 | 21 | 25 | 34 | 29 | 34 | 21 | 7 | 5 | 9 | 13 | 13 | 8 | 10 | 45 | 21 | 21 | 20 | 4 | 29 | 43 | A930009A15Rik |
| A930017K11Rik | 28 | 37 | 12 | 22 | 27 | 33 | 15 | 13 | 22 | 26 | 17 | 10 | 7 | 6 | 13 | 21 | 19 | 33 | 41 | 25 | 25 | 28 | 27 | 7 | 25 | 6 | 3 | 7 | A930017K11Rik |
| A930018P22Rik | 3 | 0 | 4 | 0 | 1 | 0 | 1 | 0 | 0 | 0 | 0 | 0 | 0 | 1 | 0 | 3 | 0 | 1 | 2 | 1 | 0 | 0 | 1 | 0 | 0 | 0 | 3 | 1 | A930018P22Rik |
| A930033H14Rik | 95 | 71 | 82 | 61 | 83 | 89 | 78 | 166 | 224 | 187 | 251 | 219 | 208 | 210 | 94 | 51 | 75 | 85 | 79 | 90 | 70 | 202 | 241 | 227 | 194 | 261 | 138 | 201 | A930033H14Rik |
| AA792892 | 0 | 0 | 0 | 0 | 0 | 0 | 0 | 0 | 0 | 1 | 0 | 0 | 0 | 0 | 0 | 0 | 0 | 0 | 0 | 0 | 0 | 0 | 0 | 0 | 0 | 0 | 0 | 0 | AA792892 |
| AA986860 | 163 | 233 | 170 | 243 | 140 | 133 | 196 | 70 | 123 | 99 | 115 | 54 | 99 | 78 | 165 | 205 | 232 | 281 | 200 | 180 | 205 | 63 | 73 | 87 | 68 | 75 | 73 | 76 | AA986860 |
| Aaas | 4305 | 4956 | 4615 | 3883 | 2670 | 3011 | 3472 | 2033 | 2344 | 2089 | 2078 | 2725 | 2620 | 2797 | 3572 | 4364 | 4228 | 3449 | 3441 | 3231 | 3117 | 1775 | 2136 | 2645 | 2174 | 3400 | 2815 | 3175 | Aaas |
| Aacs | 4346 | 6385 | 5343 | 5043 | 4678 | 4440 | 4170 | 2118 | 2399 | 2645 | 2283 | 2550 | 2573 | 2464 | 4478 | 5195 | 5349 | 5025 | 4756 | 4552 | 4104 | 2389 | 2439 | 2346 | 2593 | 2759 | 2717 | 2744 | Aacs |
| Aadac | 0 | 0 | 0 | 0 | 2 | 0 | 0 | 0 | 6 | 2 | 3 | 0 | 0 | 0 | 0 | 0 | 0 | 0 | 0 | 0 | 0 | 0 | 5 | 0 | 0 | 0 | 0 | 0 | Aadac |
| Aadacl2 | 0 | 0 | 0 | 0 | 0 | 0 | 0 | 0 | 3 | 0 | 0 | 0 | 0 | 0 | 0 | 0 | 0 | 0 | 0 | 0 | 0 | 0 | 2 | 0 | 0 | 0 | 0 | 0 | Aadacl2 |
| Aadacl2fm1 | 0 | 0 | 0 | 0 | 0 | 0 | 0 | 0 | 1 | 1 | 0 | 0 | 0 | 0 | 0 | 0 | 0 | 0 | 0 | 0 | 0 | 0 | 0 | 0 | 0 | 0 | 0 | 0 | Aadacl2fm1 |
| Aadacl2fm2 | 1 | 0 | 0 | 0 | 0 | 6 | 0 | 2 | 0 | 0 | 0 | 0 | 2 | 0 | 0 | 0 | 0 | 2 | 0 | 0 | 0 | 0 | 0 | 0 | 0 | 0 | 0 | 0 | Aadacl2fm2 |
| Aadacl2fm3 | 0 | 0 | 0 | 0 | 0 | 0 | 0 | 0 | 0 | 0 | 0 | 0 | 0 | 0 | 0 | 0 | 0 | 0 | 0 | 0 | 0 | 0 | 1 | 0 | 0 | 0 | 0 | 0 | Aadacl2fm3 |
| AAdacl4fm3 | 0 | 0 | 0 | 1 | 0 | 0 | 0 | 0 | 0 | 0 | 0 | 0 | 0 | 0 | 0 | 0 | 0 | 0 | 0 | 0 | 0 | 0 | 0 | 0 | 0 | 0 | 0 | 0 | AAdacl4fm3 |
| Aadacl4fm5 | 0 | 0 | 0 | 0 | 0 | 0 | 0 | 0 | 0 | 6 | 0 | 0 | 0 | 0 | 0 | 1 | 0 | 0 | 0 | 0 | 0 | 0 | 0 | 0 | 0 | 0 | 0 | 0 | Aadacl4fm5 |
| Aadat | 23 | 22 | 35 | 23 | 38 | 23 | 18 | 51 | 41 | 31 | 21 | 45 | 34 | 53 | 33 | 45 | 17 | 18 | 36 | 41 | 29 | 33 | 34 | 9 | 44 | 53 | 37 | 47 | Aadat |
| Aagab | 7436 | 9065 | 7944 | 7947 | 7358 | 7706 | 7029 | 4682 | 4421 | 4569 | 3906 | 4586 | 4390 | 4365 | 8192 | 7762 | 8383 | 7827 | 7436 | 8569 | 7184 | 4441 | 4601 | 4255 | 4671 | 4465 | 4726 | 4457 | Aagab |
| Aak1 | 19224 | 18726 | 12354 | 17866 | 21216 | 22449 | 22096 | 14724 | 13642 | 12704 | 14109 | 12092 | 12470 | 12172 | 19711 | 18541 | 14581 | 19099 | 18997 | 21201 | 23565 | 14171 | 13253 | 10565 | 14077 | 7284 | 13115 | 11097 | Aak1 |
| Aamdc | 1129 | 1109 | 1068 | 1092 | 1114 | 1146 | 1130 | 1044 | 962 | 907 | 1037 | 823 | 908 | 893 | 1179 | 1056 | 1082 | 1120 | 1079 | 1189 | 1143 | 871 | 1107 | 909 | 918 | 840 | 867 | 897 | Aamdc |
| Aamp | 7842 | 9950 | 9453 | 9118 | 7800 | 7567 | 7021 | 6575 | 7687 | 7369 | 7141 | 7352 | 7435 | 6949 | 7713 | 8756 | 9452 | 8435 | 8941 | 7690 | 7446 | 6585 | 7857 | 7861 | 7618 | 8692 | 7435 | 7288 | Aamp |
| Aanat | 3 | 4 | 24 | 15 | 15 | 3 | 10 | 22 | 23 | 19 | 33 | 23 | 37 | 32 | 5 | 3 | 7 | 2 | 9 | 3 | 21 | 52 | 26 | 44 | 47 | 22 | 51 | 28 | Aanat |
| Aar2 | 2705 | 2972 | 2802 | 2663 | 2608 | 2640 | 2664 | 2853 | 2841 | 2901 | 2641 | 3038 | 2845 | 2800 | 2653 | 2456 | 2802 | 2579 | 3024 | 2745 | 2694 | 2936 | 2877 | 2822 | 3021 | 3246 | 3011 | 3197 | Aar2 |
| Aard | 170 | 162 | 206 | 150 | 195 | 176 | 107 | 250 | 251 | 249 | 294 | 224 | 320 | 230 | 166 | 202 | 214 | 179 | 100 | 154 | 113 | 252 | 320 | 249 | 274 | 269 | 274 | 279 | Aard |
| Aars | 11388 | 13132 | 12172 | 12412 | 11797 | 11432 | 10811 | 11032 | 11739 | 11487 | 10826 | 12975 | 11827 | 12631 | 10863 | 11096 | 12063 | 11554 | 12595 | 11131 | 11315 | 10560 | 10683 | 11695 | 12025 | 13368 | 12794 | 13655 | Aars |
| Aars2 | 1066 | 1347 | 1441 | 1273 | 1118 | 1183 | 1025 | 1241 | 1519 | 1778 | 1482 | 1983 | 1819 | 1623 | 905 | 1238 | 1418 | 1161 | 1296 | 967 | 962 | 1504 | 1425 | 1751 | 1605 | 1994 | 1943 | 1610 | Aars2 |
| Aarsd1 | 102 | 94 | 86 | 135 | 104 | 67 | 121 | 117 | 90 | 131 | 77 | 114 | 105 | 121 | 115 | 96 | 132 | 106 | 107 | 65 | 80 | 105 | 71 | 101 | 94 | 91 | 77 | 142 | Aarsd1 |
| Aasdh | 1170 | 1132 | 901 | 1188 | 1341 | 1305 | 1359 | 2071 | 2029 | 1946 | 2022 | 2004 | 1796 | 1952 | 1114 | 1123 | 1172 | 1153 | 1248 | 1300 | 1329 | 1820 | 1977 | 1949 | 2309 | 1550 | 2076 | 1682 | Aasdh |
| Aasdhppt | 3666 | 2860 | 3241 | 3339 | 3058 | 3240 | 3585 | 2952 | 2625 | 2270 | 2351 | 2473 | 2404 | 2464 | 4038 | 3724 | 2993 | 3252 | 3053 | 3791 | 3253 | 2491 | 2624 | 2482 | 2495 | 2622 | 2471 | 2630 | Aasdhppt |
| Aass | 1020 | 779 | 559 | 359 | 285 | 489 | 387 | 195 | 332 | 189 | 317 | 228 | 285 | 195 | 872 | 864 | 552 | 521 | 348 | 339 | 313 | 174 | 463 | 209 | 166 | 267 | 224 | 161 | Aass |
| Aatf | 2843 | 3201 | 3226 | 2957 | 2609 | 2638 | 2826 | 2491 | 2207 | 2317 | 2262 | 2374 | 2446 | 2637 | 2995 | 2593 | 3038 | 2674 | 2644 | 2817 | 2932 | 2184 | 2182 | 2383 | 2340 | 2712 | 2801 | 2690 | Aatf |
| Aatk | 547 | 580 | 715 | 995 | 2136 | 1099 | 1178 | 2004 | 2582 | 2963 | 2894 | 1729 | 1740 | 1327 | 505 | 598 | 944 | 1172 | 1584 | 1131 | 1247 | 3840 | 2219 | 2260 | 2917 | 1323 | 1389 | 1175 | Aatk |
| AB124611 | 48 | 56 | 37 | 80 | 61 | 71 | 65 | 27 | 48 | 31 | 69 | 40 | 26 | 29 | 58 | 58 | 68 | 98 | 66 | 40 | 47 | 40 | 77 | 27 | 37 | 57 | 43 | 58 | AB124611 |
| Abat | 9913 | 11745 | 7447 | 10529 | 11968 | 10367 | 11317 | 7121 | 7503 | 7347 | 6912 | 6047 | 5583 | 5876 | 10969 | 10300 | 9961 | 11251 | 10498 | 9918 | 11154 | 8585 | 7171 | 5802 | 6898 | 5121 | 5445 | 4987 | Abat |
| Abca1 | 4427 | 4234 | 2789 | 4017 | 5789 | 5770 | 5450 | 6138 | 7644 | 6516 | 8511 | 5591 | 6192 | 5492 | 4756 | 4760 | 3342 | 5052 | 4682 | 5281 | 5812 | 7621 | 9190 | 4921 | 7062 | 3344 | 5624 | 4199 | Abca1 |
| Abca12 | 8 | 3 | 15 | 1 | 5 | 8 | 4 | 21 | 26 | 19 | 21 | 14 | 29 | 7 | 12 | 6 | 2 | 1 | 9 | 7 | 10 | 8 | 45 | 9 | 12 | 4 | 16 | 19 | Abca12 |

| | | | | | | | | | | | | | | | | | | | | | | | | | | | | | |
|---|---|---|---|---|---|---|---|---|---|---|---|---|---|---|---|---|---|---|---|---|---|---|---|---|---|---|---|---|---|
| Abca13 | 50 | 39 | 5 | 22 | 8 | 22 | 23 | 49 | 61 | 41 | 35 | 23 | 25 | 18 | 40 | 58 | 8 | 9 | 12 | 17 | 13 | 8 | 52 | 41 | 19 | 17 | 13 | 15 | Abca13 |
| Abca14 | 0 | 0 | 0 | 4 | 0 | 0 | 0 | 0 | 2 | 3 | 3 | 0 | 0 | 0 | 3 | 1 | 0 | 4 | 0 | 0 | 0 | 3 | 0 | 0 | 5 | 3 | 1 | 0 | Abca14 |
| Abca15 | 0 | 0 | 0 | 2 | 0 | 0 | 2 | 2 | 2 | 0 | 0 | 0 | 0 | 0 | 0 | 1 | 0 | 0 | 2 | 4 | 0 | 0 | 0 | 0 | 0 | 0 | 0 | 0 | Abca15 |
| Abca16 | 0 | 3 | 0 | 0 | 0 | 0 | 0 | 0 | 2 | 3 | 1 | 0 | 0 | 0 | 0 | 5 | 0 | 0 | 0 | 0 | 0 | 3 | 2 | 1 | 1 | 1 | 0 | 0 | Abca16 |
| Abca17 | 21 | 20 | 27 | 38 | 71 | 31 | 30 | 31 | 40 | 38 | 61 | 8 | 28 | 17 | 18 | 36 | 24 | 24 | 42 | 33 | 55 | 33 | 32 | 23 | 29 | 25 | 12 | 25 | Abca17 |
| Abca2 | 2228 | 2956 | 2773 | 4069 | 4996 | 4295 | 3996 | 3852 | 5116 | 5442 | 4631 | 5409 | 4510 | 4494 | 2313 | 2482 | 3344 | 4260 | 5051 | 3783 | 4378 | 6191 | 4442 | 5213 | 6148 | 3217 | 4530 | 4273 | Abca2 |
| Abca3 | 5737 | 6639 | 6250 | 8079 | 9516 | 8101 | 8189 | 5072 | 5532 | 6441 | 5856 | 5806 | 5635 | 5461 | 6125 | 6368 | 7414 | 8408 | 9037 | 7365 | 8660 | 7222 | 5323 | 5821 | 7027 | 3853 | 5327 | 5169 | Abca3 |
| Abca4 | 352 | 349 | 202 | 182 | 352 | 274 | 192 | 412 | 500 | 368 | 415 | 471 | 564 | 325 | 287 | 433 | 189 | 223 | 235 | 288 | 213 | 510 | 693 | 223 | 276 | 274 | 446 | 332 | Abca4 |
| Abca5 | 5580 | 4043 | 3135 | 4100 | 4030 | 4643 | 4419 | 2551 | 2476 | 2088 | 2527 | 1685 | 1809 | 1657 | 5458 | 5230 | 3279 | 4459 | 3675 | 5191 | 4293 | 2067 | 2769 | 1530 | 1909 | 1102 | 1641 | 1510 | Abca5 |
| Abca6 | 196 | 83 | 38 | 60 | 74 | 84 | 56 | 110 | 146 | 78 | 130 | 31 | 57 | 37 | 76 | 113 | 50 | 120 | 49 | 98 | 81 | 70 | 239 | 72 | 77 | 12 | 71 | 40 | Abca6 |
| Abca7 | 4140 | 6116 | 6580 | 7409 | 6827 | 6051 | 6092 | 2026 | 2150 | 2796 | 2351 | 2622 | 2477 | 2259 | 4387 | 5341 | 7539 | 7500 | 7790 | 5765 | 6142 | 2923 | 2181 | 2699 | 2586 | 1749 | 2597 | 2531 | Abca7 |
| Abca8a | 31 | 27 | 8 | 39 | 120 | 82 | 110 | 106 | 192 | 107 | 678 | 30 | 61 | 42 | 37 | 32 | 10 | 91 | 63 | 72 | 129 | 129 | 616 | 44 | 90 | 24 | 36 | 31 | Abca8a |
| Abca8b | 1001 | 855 | 533 | 794 | 1159 | 1090 | 1153 | 884 | 1023 | 862 | 1259 | 588 | 727 | 612 | 973 | 927 | 637 | 923 | 856 | 1204 | 1037 | 1008 | 1220 | 637 | 817 | 380 | 530 | 469 | Abca8b |
| Abca9 | 1318 | 1267 | 1105 | 1053 | 1189 | 1552 | 977 | 1859 | 2226 | 1986 | 3296 | 1952 | 2487 | 1938 | 1447 | 1456 | 955 | 1401 | 848 | 1376 | 1087 | 2074 | 3736 | 1927 | 2053 | 1089 | 2086 | 1709 | Abca9 |
| Abcb10 | 2060 | 2169 | 2224 | 1946 | 2060 | 1951 | 1707 | 2296 | 2360 | 2249 | 2371 | 2600 | 3083 | 2489 | 2151 | 2233 | 1954 | 1767 | 1926 | 1782 | 1631 | 2410 | 2348 | 2268 | 2364 | 2561 | 2751 | 3190 | Abcb10 |
| Abcb11 | 0 | 0 | 13 | 2 | 9 | 9 | 6 | 6 | 5 | 3 | 4 | 5 | 9 | 3 | 1 | 3 | 0 | 3 | 5 | 3 | 2 | 5 | 1 | 1 | 11 | 4 | 1 | 2 | Abcb11 |
| Abcb1a | 211 | 196 | 272 | 284 | 416 | 328 | 335 | 369 | 302 | 410 | 357 | 382 | 364 | 380 | 269 | 221 | 250 | 338 | 391 | 392 | 374 | 431 | 403 | 336 | 483 | 295 | 320 | 345 | Abcb1a |
| Abcb1b | 146 | 106 | 153 | 80 | 79 | 113 | 98 | 85 | 110 | 86 | 121 | 77 | 74 | 69 | 167 | 164 | 93 | 94 | 110 | 115 | 90 | 60 | 117 | 59 | 60 | 63 | 92 | 80 | Abcb1b |
| Abcb4 | 102 | 105 | 76 | 93 | 94 | 140 | 132 | 82 | 74 | 66 | 55 | 99 | 71 | 85 | 84 | 75 | 123 | 76 | 147 | 195 | 144 | 60 | 63 | 57 | 86 | 106 | 91 | 81 | Abcb4 |
| Abcb5 | 0 | 0 | 1 | 2 | 0 | 0 | 0 | 2 | 0 | 0 | 0 | 3 | 0 | 0 | 1 | 0 | 0 | 0 | 0 | 0 | 0 | 0 | 3 | 0 | 0 | 1 | 0 | 1 | Abcb5 |
| Abcb6 | 191 | 147 | 139 | 155 | 177 | 87 | 227 | 96 | 70 | 148 | 100 | 110 | 99 | 116 | 156 | 93 | 172 | 142 | 154 | 129 | 172 | 135 | 73 | 145 | 89 | 117 | 115 | 76 | Abcb6 |
| Abcb7 | 3840 | 2953 | 3252 | 3202 | 3513 | 3728 | 4119 | 6298 | 4578 | 4175 | 4548 | 5470 | 4880 | 6076 | 4174 | 3297 | 3416 | 3420 | 3391 | 4027 | 3813 | 4719 | 4687 | 4867 | 5072 | 5028 | 5896 | 5841 | Abcb7 |
| Abcb8 | 815 | 1000 | 891 | 900 | 974 | 998 | 786 | 2256 | 2723 | 2602 | 2383 | 2987 | 2956 | 2756 | 874 | 914 | 972 | 853 | 958 | 946 | 803 | 2804 | 2650 | 2818 | 2825 | 3016 | 2990 | 2963 | Abcb8 |
| Abcb9 | 324 | 435 | 498 | 510 | 512 | 410 | 337 | 182 | 291 | 280 | 220 | 251 | 257 | 214 | 376 | 537 | 396 | 552 | 444 | 358 | 345 | 294 | 285 | 240 | 293 | 213 | 222 | 189 | Abcb9 |
| Abcc1 | 2473 | 2437 | 1890 | 1932 | 2627 | 2508 | 2286 | 2074 | 2108 | 2016 | 1891 | 1807 | 2009 | 1737 | 2254 | 2308 | 1862 | 1996 | 2390 | 2394 | 2224 | 1756 | 1946 | 1624 | 1893 | 1453 | 1682 | 1661 | Abcc1 |
| Abcc10 | 1113 | 1424 | 1614 | 1784 | 1740 | 1562 | 1246 | 1102 | 1313 | 1615 | 1600 | 1575 | 1436 | 1377 | 1015 | 1460 | 1629 | 1597 | 1698 | 1369 | 1304 | 1664 | 1289 | 1560 | 1575 | 1275 | 1497 | 1334 | Abcc10 |
| Abcc12 | 2 | 10 | 10 | 9 | 9 | 0 | 17 | 24 | 21 | 21 | 24 | 7 | 26 | 17 | 5 | 15 | 3 | 9 | 6 | 16 | 11 | 10 | 16 | 16 | 10 | 7 | 23 | 20 | Abcc12 |
| Abcc2 | 3 | 5 | 2 | 2 | 6 | 3 | 0 | 15 | 21 | 2 | 28 | 7 | 6 | 0 | 5 | 7 | 1 | 10 | 7 | 3 | 4 | 5 | 19 | 3 | 2 | 5 | 12 | 12 | Abcc2 |
| Abcc3 | 90 | 82 | 82 | 69 | 102 | 77 | 58 | 88 | 125 | 98 | 147 | 86 | 122 | 101 | 80 | 106 | 67 | 65 | 58 | 59 | 64 | 100 | 177 | 126 | 105 | 100 | 124 | 68 | Abcc3 |
| Abcc4 | 854 | 733 | 772 | 557 | 488 | 542 | 368 | 503 | 506 | 552 | 469 | 630 | 640 | 482 | 756 | 993 | 673 | 515 | 504 | 432 | 448 | 558 | 498 | 498 | 447 | 596 | 635 | 555 | Abcc4 |
| Abcc5 | 10366 | 10726 | 6959 | 10307 | 14122 | 13236 | 13120 | 19702 | 19924 | 17736 | 19298 | 14905 | 15331 | 15100 | 11565 | 10288 | 8895 | 11410 | 11911 | 13369 | 13767 | 18376 | 18863 | 13477 | 19047 | 8446 | 14955 | 12991 | Abcc5 |
| Abcc6 | 20 | 13 | 24 | 13 | 28 | 14 | 18 | 48 | 71 | 60 | 74 | 50 | 81 | 71 | 22 | 28 | 27 | 35 | 26 | 30 | 21 | 62 | 81 | 65 | 82 | 58 | 58 | 46 | Abcc6 |
| Abcc8 | 219 | 246 | 216 | 403 | 826 | 529 | 626 | 310 | 503 | 488 | 476 | 219 | 166 | 186 | 218 | 252 | 320 | 623 | 686 | 440 | 775 | 378 | 331 | 365 | 560 | 112 | 139 | 133 | Abcc8 |
| Abcc9 | 1019 | 944 | 830 | 1132 | 1481 | 1461 | 1334 | 1633 | 1755 | 1844 | 2431 | 1783 | 1672 | 1744 | 1079 | 990 | 970 | 1252 | 1218 | 1376 | 1418 | 2257 | 2337 | 1727 | 1908 | 1138 | 1585 | 1612 | Abcc9 |
| Abcd1 | 1729 | 1991 | 1889 | 1889 | 1769 | 1846 | 1574 | 1258 | 1645 | 1656 | 1585 | 1543 | 1522 | 1410 | 1667 | 1932 | 1966 | 1972 | 1894 | 1582 | 1625 | 1526 | 1604 | 1515 | 1601 | 1615 | 1543 | 1396 | Abcd1 |
| Abcd2 | 6025 | 5835 | 6243 | 6083 | 5352 | 5797 | 7109 | 3756 | 2800 | 2787 | 2444 | 4129 | 3157 | 4283 | 6837 | 4846 | 6103 | 5801 | 5954 | 6816 | 6717 | 2679 | 2726 | 3700 | 3678 | 3631 | 4442 | 4131 | Abcd2 |
| Abcd3 | 8560 | 6995 | 6788 | 7573 | 7738 | 7656 | 8262 | 8095 | 7421 | 6834 | 6835 | 7038 | 6644 | 7292 | 8586 | 6983 | 7178 | 7741 | 7525 | 7552 | 7626 | 7588 | 7394 | 7038 | 7063 | 7419 | 6871 | 7270 | Abcd3 |
| Abcd4 | 1669 | 1934 | 2184 | 2496 | 2783 | 2587 | 2689 | 1353 | 1563 | 1888 | 1747 | 1840 | 1774 | 1799 | 1658 | 1999 | 2300 | 3002 | 2638 | 2270 | 2548 | 1803 | 1480 | 1934 | 2036 | 1579 | 1722 | 1773 | Abcd4 |
| Abce1 | 15819 | 13141 | 13794 | 11543 | 11102 | 11803 | 12424 | 19273 | 15969 | 13966 | 14176 | 16249 | 15816 | 16298 | 16441 | 14717 | 12507 | 11451 | 11322 | 14686 | 11651 | 14574 | 16261 | 14515 | 14239 | 17364 | 17460 | 16981 | Abce1 |
| Abcf1 | 13951 | 15931 | 17241 | 15867 | 12738 | 12679 | 13711 | 7587 | 8345 | 9666 | 8880 | 10735 | 9683 | 10592 | 13378 | 14562 | 16045 | 14835 | 14022 | 12078 | 13683 | 8318 | 8129 | 10172 | 9377 | 11228 | 10388 | 10841 | Abcf1 |
| Abcf2 | 9574 | 10838 | 9537 | 9363 | 8030 | 8937 | 8771 | 7563 | 7676 | 7012 | 6723 | 8492 | 7764 | 7857 | 9200 | 9003 | 8998 | 8900 | 9555 | 9632 | 9177 | 6234 | 7440 | 7893 | 7592 | 9553 | 9267 | 8534 | Abcf2 |
| Abcf3 | 6601 | 7825 | 5952 | 6337 | 5911 | 6140 | 5991 | 4476 | 4630 | 3973 | 4014 | 4463 | 4415 | 4021 | 6879 | 6520 | 6071 | 6324 | 6263 | 6647 | 6134 | 4049 | 4689 | 4225 | 4349 | 4643 | 4396 | 4448 | Abcf3 |
| Abcg1 | 1496 | 1626 | 1652 | 2043 | 1999 | 1693 | 1861 | 1320 | 1463 | 1704 | 1324 | 1516 | 1305 | 1438 | 1435 | 1843 | 1760 | 1810 | 1908 | 1593 | 1722 | 1594 | 1345 | 1714 | 1518 | 1719 | 1214 | 1283 | Abcg1 |
| Abcg2 | 3477 | 2695 | 2948 | 2164 | 2587 | 2579 | 2249 | 3051 | 2776 | 2345 | 2737 | 2602 | 2871 | 2541 | 3390 | 3432 | 2691 | 2452 | 2208 | 2815 | 2188 | 2613 | 3026 | 2174 | 2218 | 2832 | 2667 | 2623 | Abcg2 |
| Abcg3 | 39 | 38 | 37 | 26 | 27 | 43 | 39 | 94 | 77 | 66 | 111 | 51 | 84 | 71 | 44 | 48 | 30 | 24 | 35 | 40 | 22 | 94 | 104 | 80 | 57 | 103 | 77 | 58 | Abcg3 |
| Abcg4 | 1275 | 1576 | 1656 | 1748 | 2505 | 2007 | 1853 | 1277 | 1563 | 1622 | 1532 | 1585 | 1403 | 1533 | 1209 | 1287 | 1905 | 1826 | 2229 | 1899 | 1956 | 1978 | 1296 | 1517 | 1823 | 1098 | 1352 | 1425 | Abcg4 |
| Abcg5 | 121 | 99 | 111 | 62 | 103 | 80 | 87 | 216 | 176 | 188 | 148 | 252 | 264 | 161 | 112 | 134 | 83 | 56 | 103 | 95 | 91 | 246 | 322 | 150 | 146 | 199 | 164 | 182 | Abcg5 |
| Abcg8 | 42 | 54 | 43 | 23 | 54 | 40 | 40 | 64 | 49 | 43 | 56 | 106 | 104 | 55 | 41 | 54 | 57 | 39 | 31 | 58 | 30 | 50 | 109 | 38 | 46 | 69 | 38 | 76 | Abcg8 |
| Abhd10 | 1919 | 1591 | 1862 | 1843 | 1663 | 1730 | 1978 | 2573 | 2105 | 2203 | 2225 | 2714 | 2379 | 2579 | 2023 | 1632 | 1846 | 1712 | 1679 | 1870 | 1920 | 2555 | 2164 | 2414 | 2215 | 2383 | 2603 | 2750 | Abhd10 |
| Abhd11 | 1601 | 1956 | 1734 | 1769 | 1720 | 1569 | 1697 | 1229 | 1428 | 1475 | 1393 | 1221 | 1208 | 1213 | 1606 | 1670 | 1692 | 1727 | 1785 | 1827 | 1589 | 1228 | 1372 | 1322 | 1401 | 1369 | 1295 | 1213 | Abhd11 |
| Abhd12 | 3673 | 3699 | 3014 | 3620 | 3654 | 3520 | 3557 | 2602 | 2838 | 2816 | 2952 | 2544 | 2480 | 2587 | 3320 | 3389 | 3376 | 3706 | 3741 | 3295 | 3895 | 3029 | 2928 | 2746 | 2670 | 2650 | 2408 | 2510 | Abhd12 |
| Abhd12b | 5 | 12 | 11 | 18 | 27 | 42 | 9 | 24 | 15 | 30 | 19 | 18 | 45 | 35 | 8 | 11 | 9 | 13 | 27 | 22 | 21 | 33 | 32 | 12 | 20 | 12 | 12 | 14 | Abhd12b |
| Abhd13 | 5163 | 4401 | 3969 | 4335 | 3850 | 4290 | 4478 | 4198 | 3449 | 3240 | 3275 | 3484 | 3207 | 3451 | 5230 | 4535 | 3869 | 4097 | 3741 | 4493 | 4337 | 3546 | 3617 | 3148 | 3380 | 3509 | 3519 | 3516 | Abhd13 |
| Abhd14a | 1916 | 2249 | 1934 | 2283 | 1991 | 1976 | 1863 | 1207 | 1534 | 1568 | 1530 | 1316 | 1382 | 1204 | 1721 | 2117 | 2138 | 2278 | 2165 | 1855 | 1999 | 1410 | 1713 | 1442 | 1522 | 1459 | 1334 | 1270 | Abhd14a |
| Abhd14b | 1321 | 1400 | 1109 | 1102 | 914 | 1048 | 898 | 488 | 699 | 609 | 742 | 482 | 566 | 404 | 1197 | 1600 | 1114 | 1241 | 909 | 869 | 827 | 605 | 757 | 632 | 450 | 492 | 325 | 362 | Abhd14b |
| Abhd15 | 582 | 764 | 628 | 608 | 475 | 493 | 429 | 93 | 81 | 85 | 94 | 98 | 70 | 88 | 562 | 654 | 585 | 582 | 425 | 507 | 478 | 89 | 89 | 96 | 68 | 60 | 101 | 107 | Abhd15 |
| Abhd16a | 914 | 1155 | 1185 | 1236 | 1384 | 1286 | 1057 | 2100 | 2631 | 2375 | 2529 | 2223 | 2448 | 2147 | 836 | 1048 | 1203 | 1135 | 1295 | 1181 | 1013 | 2461 | 2723 | 2510 | 2583 | 2383 | 2246 | 2106 | Abhd16a |
| Abhd16b | 2 | 0 | 0 | 0 | 0 | 0 | 0 | 1 | 0 | 0 | 0 | 2 | 0 | 0 | 1 | 2 | 0 | 0 | 0 | 0 | 0 | 0 | 0 | 0 | 0 | 0 | 0 | 0 | Abhd16b |
| Abhd17a | 3435 | 4876 | 3881 | 4485 | 3601 | 3622 | 3180 | 2179 | 2544 | 2746 | 2573 | 2563 | 2622 | 2212 | 3303 | 3992 | 4209 | 4102 | 4204 | 3288 | 3403 | 2214 | 2731 | 2679 | 2655 | 2887 | 2535 | 2254 | Abhd17a |
| Abhd17b | 4013 | 2952 | 3998 | 4008 | 3984 | 4125 | 4555 | 3414 | 2840 | 2905 | 2864 | 3446 | 3226 | 3449 | 4245 | 3658 | 3770 | 4118 | 4013 | 4143 | 4302 | 3332 | 2972 | 3125 | 3322 | 3714 | 3461 | 3710 | Abhd17b |
| Abhd17c | 3620 | 3354 | 4155 | 4627 | 4663 | 4243 | 4568 | 4068 | 4002 | 4531 | 4265 | 4979 | 4183 | 4755 | 3690 | 3460 | 4486 | 4579 | 4669 | 3672 | 4612 | 4771 | 3726 | 4872 | 4800 | 5574 | 4130 | 4671 | Abhd17c |
| Abhd18 | 255 | 137 | 69 | 247 | 126 | 150 | 227 | 51 | 85 | 116 | 22 | 67 | 79 | 26 | 215 | 98 | 104 | 292 | 301 | 189 | 85 | 75 | 33 | 68 | 163 | 20 | 62 | 106 | Abhd18 |
| Abhd2 | 7784 | 8509 | 7195 | 9364 | 10037 | 9714 | 10096 | 6598 | 7517 | 7644 | 6925 | 7354 | 6855 | 6720 | 7942 | 8787 | 8039 | 10371 | 9707 | 8636 | 9882 | 7860 | 8011 | 6692 | 8342 | 5356 | 6517 | 6285 | Abhd2 |
| Abhd3 | 424 | 440 | 579 | 477 | 376 | 308 | 344 | 128 | 118 | 123 | 97 | 101 | 108 | 93 | 460 | 490 | 439 | 439 | 342 | 385 | 343 | 120 | 168 | 135 | 111 | 72 | 80 | 75 | Abhd3 |
| Abhd4 | 3796 | 4583 | 3431 | 3366 | 3366 | 3296 | 3323 | 2536 | 3355 | 2392 | 2631 | 2694 | 2740 | 2470 | 3789 | 4070 | 3524 | 3606 | 3440 | 3206 | 3137 | 2764 | 3107 | 2480 | 2495 | 2899 | 2528 | 2643 | Abhd4 |
| Abhd5 | 1241 | 1130 | 1120 | 958 | 1026 | 1044 | 916 | 1619 | 1548 | 1324 | 1427 | 1225 | 1300 | 1304 | 1324 | 1284 | 931 | 1042 | 884 | 832 | 867 | 1577 | 1709 | 1256 | 1229 | 1442 | 1267 | 1238 | Abhd5 |
| Abhd6 | 1465 | 1610 | 1329 | 1571 | 1696 | 1691 | 1729 | 951 | 957 | 946 | 800 | 773 | 755 | 772 | 1468 | 1526 | 1400 | 1680 | 1594 | 1906 | 1774 | 779 | 1039 | 785 | 810 | 893 | 816 | 752 | Abhd6 |
| Abhd8 | 5727 | 7518 | 6323 | 7134 | 6883 | 6417 | 6215 | 3468 | 3628 | 3604 | 3476 | 3795 | 3104 | 3264 | 5618 | 5759 | 7608 | 7186 | 7938 | 6338 | 6853 | 3654 | 3489 | 3893 | 3980 | 3821 | 3643 | 3330 | Abhd8 |
| Abi1 | 7279 | 5836 | 6904 | 7203 | 7046 | 7538 | 7747 | 7135 | 6344 | 6384 | 6353 | 6359 | 6094 | 6737 | 7433 | 6572 | 6913 | 7096 | 7097 | 7081 | 7687 | 7035 | 6396 | 6599 | 6596 | 6921 | 6192 | 6768 | Abi1 |
| Abi2 | 8474 | 7272 | 8984 | 11056 | 13169 | 12803 | 13284 | 10960 | 9652 | 11142 | 9972 | 11596 | 10570 | 12184 | 9265 | 8200 | 10218 | 11396 | 12119 | 11085 | 12790 | 13767 | 9078 | 10755 | 12944 | 10123 | 11433 | 11237 | Abi2 |
| Abi3 | 102 | 124 | 84 | 86 | 82 | 75 | 51 | 93 | 145 | 128 | 148 | 117 | 129 | 98 | 66 | 116 | 96 | 118 | 53 | 64 | 71 | 87 | 148 | 123 | 135 | 122 | 102 | 113 | Abi3 |
| Abi3bp | 223 | 174 | 200 | 197 | 79 | 169 | 47 | 231 | 506 | 245 | 1054 | 146 | 188 | 119 | 187 | 345 | 126 | 173 | 47 | 64 | 37 | 277 | 900 | 218 | 135 | 147 | 156 | 110 | Abi3bp |
| Abitram | 3622 | 3199 | 2760 | 2727 | 2392 | 2654 | 3030 | 3002 | 3016 | 2891 | 2782 | 3121 | 3201 | 3074 | 3202 | 2923 | 2613 | 2739 | 2728 | 2939 | 2802 | 2958 | 2831 | 2872 | 2952 | 3298 | 3133 | 3370 | Abitram |
| Abl1 | 12499 | 13599 | 14346 | 12883 | 12188 | 12429 | 11336 | 10643 | 12691 | 12260 | 12634 | 14320 | 13554 | 12572 | 10835 | 12995 | 14330 | 12489 | 13902 | 10912 | 11542 | 11085 | 11756 | 13366 | 12022 | 13089 | 13113 | 12914 | Abl1 |
| Abl2 | 5216 | 4795 | 4320 | 5653 | 6856 | 6529 | 6165 | 7188 | 7142 | 7243 | 8040 | 7525 | 7493 | 8122 | 5112 | 4725 | 4707 | 5674 | 6495 | 5472 | 6718 | 7219 | 6677 | 6452 | 8675 | 4464 | 8190 | 7106 | Abl2 |
| Ablim1 | 4319 | 3607 | 3416 | 3451 | 4956 | 3999 | 3867 | 5709 | 6064 | 5859 | 5864 | 4139 | 5256 | 4189 | 3990 | 4421 | 3219 | 3913 | 3775 | 3842 | 4061 | 6749 | 5657 | 4717 | 5659 | 3411 | 3975 | 3972 | Ablim1 |
| Ablim2 | 212 | 340 | 162 | 380 | 601 | 404 | 355 | 204 | 213 | 265 | 252 | 59 | 110 | 60 | 253 | 263 | 222 | 407 | 350 | 407 | 443 | 271 | 274 | 137 | 190 | 62 | 44 | 63 | Ablim2 |
| Ablim3 | 501 | 657 | 473 | 1481 | 3610 | 2170 | 2071 | 2420 | 3192 | 3385 | 4666 | 1225 | 1564 | 1208 | 848 | 716 | 1004 | 2216 | 2509 | 1845 | 2620 | 4909 | 3360 | 1986 | 3350 | 725 | 923 | 879 | Ablim3 |
| Abo | 0 | 0 | 0 | 0 | 0 | 0 | 0 | 0 | 0 | 0 | 0 | 0 | 0 | 0 | 1 | 0 | 0 | 0 | 0 | 0 | 0 | 0 | 0 | 0 | 0 | 0 | 0 | 0 | Abo |

| | | | | | | | | | | | | | | | | | | | | | | | | | | | | | |
|---|---|---|---|---|---|---|---|---|---|---|---|---|---|---|---|---|---|---|---|---|---|---|---|---|---|---|---|---|---|
| Abr | 7264 | 7425 | 7436 | 9361 | 12839 | 11553 | 10760 | 14854 | 15540 | 17738 | 15224 | 15637 | 12686 | 15197 | 7481 | 6497 | 9469 | 9729 | 11995 | 9963 | 11465 | 19453 | 14861 | 16208 | 18938 | 13520 | 13444 | 13824 | Abr |
| Abra | 0 | 0 | 0 | 0 | 1 | 0 | 0 | 4 | 1 | 0 | 42 | 0 | 2 | 0 | 3 | 0 | 0 | 0 | 0 | 0 | 0 | 5 | 32 | 0 | 0 | 0 | 0 | 0 | Abra |
| Abracl | 2969 | 2093 | 2780 | 2269 | 2248 | 2473 | 3508 | 4384 | 3681 | 3747 | 3543 | 4712 | 3782 | 5991 | 2896 | 2263 | 2743 | 2180 | 2693 | 3126 | 3040 | 3126 | 3216 | 5144 | 4733 | 5854 | 4819 | 5712 | Abracl |
| Abraxas1 | 796 | 590 | 724 | 625 | 558 | 668 | 650 | 871 | 747 | 749 | 807 | 786 | 832 | 915 | 746 | 709 | 669 | 621 | 617 | 668 | 650 | 632 | 746 | 868 | 707 | 993 | 952 | 971 | Abraxas1 |
| Abraxas2 | 3588 | 3427 | 3290 | 3280 | 2970 | 3063 | 3289 | 4021 | 3400 | 3176 | 3167 | 3459 | 3215 | 3461 | 3645 | 3396 | 3370 | 3142 | 3103 | 3524 | 3111 | 3232 | 3500 | 3318 | 3452 | 3702 | 3433 | 3531 | Abraxas2 |
| Abt1 | 1609 | 1969 | 1674 | 1826 | 1193 | 1230 | 1402 | 1078 | 1094 | 1096 | 1046 | 1336 | 1134 | 1225 | 1474 | 1578 | 1610 | 1407 | 1412 | 1568 | 1351 | 840 | 1115 | 1176 | 1123 | 1346 | 1287 | 1240 | Abt1 |
| Abtb1 | 2009 | 2712 | 2275 | 2576 | 2338 | 2244 | 2181 | 1741 | 2043 | 2031 | 1818 | 2017 | 2038 | 1694 | 2089 | 2353 | 2429 | 2487 | 2408 | 2300 | 2052 | 2279 | 2015 | 1942 | 2153 | 1797 | 2007 | 1800 | Abtb1 |
| Abtb2 | 2858 | 3855 | 3667 | 3880 | 3581 | 3246 | 2874 | 1294 | 1651 | 1634 | 1596 | 1617 | 1693 | 1397 | 3049 | 3574 | 3673 | 4008 | 3556 | 2821 | 3125 | 1659 | 1538 | 1551 | 1396 | 1574 | 1626 | 1543 | Abtb2 |
| Acaa1a | 3488 | 4310 | 3999 | 4692 | 4583 | 4166 | 3743 | 3588 | 4407 | 4048 | 4534 | 4125 | 3963 | 3588 | 3446 | 3967 | 4266 | 4477 | 4514 | 3916 | 3830 | 4248 | 4278 | 4469 | 4615 | 4076 | 4098 | 3660 | Acaa1a |
| Acaa1b | 189 | 207 | 231 | 232 | 205 | 188 | 145 | 83 | 103 | 72 | 69 | 66 | 73 | 72 | 166 | 220 | 279 | 207 | 210 | 169 | 128 | 79 | 84 | 87 | 41 | 73 | 64 | 45 | Acaa1b |
| Acaa2 | 3060 | 3064 | 2699 | 2256 | 1993 | 2336 | 2019 | 1718 | 2289 | 1690 | 1803 | 1590 | 1775 | 1402 | 2760 | 2768 | 2506 | 2194 | 2292 | 2337 | 2038 | 1382 | 2456 | 1717 | 1498 | 1860 | 1513 | 1525 | Acaa2 |
| Acaca | 15165 | 16056 | 9412 | 13504 | 14441 | 16301 | 14225 | 12767 | 13215 | 12262 | 11716 | 13396 | 13730 | 12573 | 15381 | 15075 | 10785 | 13795 | 14611 | 14687 | 16001 | 10599 | 11378 | 9798 | 13124 | 7997 | 14177 | 12352 | Acaca |
| Acacb | 758 | 1088 | 825 | 1161 | 901 | 958 | 962 | 205 | 264 | 263 | 260 | 277 | 311 | 221 | 703 | 1030 | 1005 | 1191 | 846 | 865 | 895 | 249 | 334 | 244 | 236 | 203 | 277 | 249 | Acacb |
| Acad10 | 188 | 225 | 276 | 284 | 236 | 247 | 215 | 201 | 236 | 249 | 233 | 269 | 287 | 203 | 190 | 213 | 286 | 265 | 251 | 197 | 208 | 234 | 224 | 299 | 218 | 265 | 246 | 275 | Acad10 |
| Acad11 | 1410 | 1386 | 1087 | 893 | 920 | 896 | 878 | 1071 | 1022 | 871 | 825 | 794 | 906 | 784 | 1466 | 1462 | 1148 | 991 | 916 | 1026 | 886 | 775 | 1166 | 856 | 722 | 801 | 854 | 888 | Acad11 |
| Acad12 | 128 | 156 | 135 | 142 | 112 | 145 | 120 | 173 | 185 | 196 | 190 | 153 | 186 | 180 | 152 | 150 | 123 | 128 | 104 | 147 | 121 | 177 | 233 | 208 | 188 | 257 | 184 | 159 | Acad12 |
| Acad8 | 2024 | 1861 | 2272 | 1926 | 2007 | 1868 | 2092 | 2359 | 2375 | 2255 | 2502 | 2656 | 2369 | 2488 | 1909 | 1802 | 2071 | 2082 | 2156 | 2096 | 1901 | 2304 | 2485 | 2411 | 2281 | 2806 | 2398 | 2427 | Acad8 |
| Acad9 | 3624 | 3725 | 2898 | 3281 | 2947 | 3182 | 3235 | 3120 | 2942 | 2637 | 2863 | 2780 | 2860 | 2927 | 3700 | 3587 | 2968 | 3198 | 3028 | 3484 | 3334 | 2511 | 2996 | 2661 | 2960 | 2372 | 3169 | 2873 | Acad9 |
| Acadl | 2711 | 2444 | 3124 | 1976 | 2083 | 2078 | 1863 | 2181 | 2445 | 2040 | 2703 | 2274 | 2557 | 2374 | 2848 | 2923 | 2369 | 2004 | 1997 | 1901 | 1893 | 2326 | 2889 | 1943 | 1942 | 2445 | 2266 | 2546 | Acadl |
| Acadm | 5180 | 5453 | 5199 | 5342 | 4669 | 5070 | 5222 | 2955 | 3009 | 2656 | 2974 | 2696 | 3063 | 2710 | 5662 | 5541 | 5244 | 5194 | 4950 | 5680 | 5067 | 2685 | 3224 | 2589 | 2493 | 2792 | 2888 | 2791 | Acadm |
| Acads | 1638 | 1978 | 2299 | 1745 | 1623 | 1610 | 1222 | 1172 | 1373 | 1341 | 1316 | 1516 | 1729 | 1403 | 1526 | 2030 | 1876 | 1665 | 1714 | 1436 | 1361 | 1068 | 1467 | 1454 | 1342 | 1870 | 1614 | 1561 | Acads |
| Acadsb | 6615 | 5532 | 5368 | 5716 | 5905 | 6157 | 6011 | 7017 | 6287 | 6143 | 6088 | 5561 | 5846 | 5577 | 6998 | 6406 | 5198 | 6009 | 5556 | 6895 | 6293 | 6211 | 6585 | 5286 | 5994 | 4814 | 5684 | 5617 | Acadsb |
| Acadvl | 2523 | 2949 | 2480 | 2568 | 2310 | 2407 | 2151 | 2651 | 2881 | 2536 | 2795 | 2474 | 2758 | 2263 | 2457 | 2783 | 2554 | 2531 | 2311 | 2383 | 2129 | 2521 | 3106 | 2632 | 2587 | 2617 | 2538 | 2489 | Acadvl |
| Acan | 144 | 200 | 51 | 424 | 155 | 1141 | 31 | 441 | 2586 | 3469 | 7662 | 137 | 342 | 180 | 150 | 262 | 102 | 932 | 422 | 36 | 30 | 3945 | 1416 | 1880 | 2452 | 397 | 193 | 87 | Acan |
| Acap1 | 92 | 111 | 125 | 131 | 93 | 70 | 107 | 70 | 64 | 89 | 109 | 119 | 93 | 133 | 80 | 79 | 165 | 123 | 111 | 68 | 107 | 72 | 81 | 106 | 76 | 121 | 131 | 112 | Acap1 |
| Acap2 | 11798 | 10121 | 8894 | 9790 | 9583 | 11028 | 11225 | 6398 | 6068 | 5475 | 6130 | 6221 | 6135 | 6641 | 11224 | 11147 | 9199 | 10202 | 8951 | 10803 | 11046 | 5544 | 6092 | 5801 | 5790 | 4789 | 6478 | 5986 | Acap2 |
| Acap3 | 6031 | 8022 | 6833 | 9766 | 9133 | 8509 | 7725 | 5112 | 5464 | 6472 | 5764 | 6118 | 5285 | 5337 | 6029 | 7016 | 8343 | 9928 | 9163 | 6924 | 8619 | 7603 | 4910 | 6110 | 7660 | 4828 | 5498 | 5001 | Acap3 |
| Acat1 | 7111 | 6316 | 5043 | 7415 | 8231 | 8129 | 8494 | 8981 | 9459 | 7815 | 9269 | 6178 | 6347 | 6685 | 7632 | 6185 | 5828 | 8015 | 7910 | 9187 | 8706 | 8224 | 9930 | 7577 | 8350 | 6683 | 5850 | 6139 | Acat1 |
| Acat2 | 12041 | 12278 | 13143 | 11982 | 10740 | 11301 | 11135 | 13184 | 10149 | 10123 | 8912 | 12914 | 12853 | 13534 | 12657 | 11055 | 13128 | 11748 | 12150 | 11753 | 11827 | 10843 | 9215 | 11739 | 12137 | 13984 | 14655 | 16301 | Acat2 |
| Acat3 | 130 | 97 | 115 | 116 | 90 | 144 | 87 | 72 | 41 | 70 | 74 | 55 | 50 | 66 | 127 | 98 | 109 | 109 | 86 | 86 | 80 | 84 | 32 | 59 | 76 | 67 | 60 | 75 | Acat3 |
| Acbd3 | 4520 | 4085 | 4702 | 4810 | 4696 | 4483 | 4559 | 4500 | 4157 | 4230 | 4870 | 4452 | 4280 | 4548 | 4585 | 4358 | 4731 | 4866 | 4507 | 4126 | 4442 | 5021 | 4162 | 4610 | 4514 | 4335 | 4256 | 4322 | Acbd3 |
| Acbd4 | 738 | 739 | 879 | 694 | 623 | 591 | 596 | 151 | 255 | 256 | 246 | 197 | 264 | 184 | 633 | 819 | 716 | 759 | 660 | 585 | 533 | 202 | 315 | 226 | 180 | 213 | 216 | 165 | Acbd4 |
| Acbd5 | 5812 | 4564 | 5729 | 6305 | 6884 | 6445 | 7634 | 6977 | 5820 | 5855 | 5709 | 5813 | 5736 | 6608 | 6651 | 5599 | 5981 | 6234 | 6338 | 7193 | 7073 | 6986 | 5875 | 5934 | 6163 | 5458 | 5957 | 6051 | Acbd5 |
| Acbd6 | 2549 | 2555 | 2801 | 2897 | 2914 | 2876 | 3064 | 2105 | 2104 | 2313 | 2283 | 2439 | 2087 | 2580 | 2643 | 2239 | 2932 | 2845 | 3123 | 2423 | 3228 | 2418 | 2186 | 2531 | 2605 | 2734 | 2269 | 2492 | Acbd6 |
| Acbd7 | 39 | 29 | 32 | 76 | 33 | 27 | 54 | 17 | 13 | 3 | 14 | 7 | 3 | 11 | 32 | 31 | 27 | 35 | 35 | 25 | 31 | 15 | 16 | 18 | 16 | 5 | 2 | 14 | Acbd7 |
| Accs | 1381 | 1300 | 1197 | 1342 | 1623 | 1706 | 1643 | 1646 | 1798 | 1960 | 1801 | 2180 | 2082 | 2162 | 1366 | 1224 | 1403 | 1393 | 1572 | 1598 | 1524 | 1731 | 1581 | 1963 | 1975 | 1790 | 2141 | 1995 | Accs |
| Accsl | 16 | 18 | 13 | 27 | 41 | 29 | 54 | 13 | 18 | 22 | 24 | 22 | 14 | 22 | 38 | 16 | 3 | 39 | 36 | 33 | 37 | 30 | 17 | 11 | 16 | 19 | 35 | 28 | Accsl |
| Acd | 4361 | 5911 | 4451 | 5299 | 4773 | 4367 | 4784 | 2198 | 2454 | 2561 | 2426 | 2822 | 2584 | 2736 | 4350 | 4641 | 5201 | 4998 | 5121 | 4424 | 4693 | 2602 | 2238 | 2734 | 2695 | 2995 | 2535 | 2578 | Acd |
| Ace | 193 | 270 | 300 | 338 | 449 | 353 | 250 | 307 | 383 | 392 | 348 | 262 | 330 | 176 | 249 | 325 | 253 | 371 | 352 | 292 | 356 | 541 | 441 | 336 | 389 | 235 | 279 | 246 | Ace |
| Ace2 | 22 | 10 | 12 | 0 | 16 | 0 | 0 | 40 | 59 | 43 | 16 | 1 | 11 | 6 | 15 | 18 | 2 | 10 | 2 | 9 | 3 | 17 | 104 | 14 | 23 | 9 | 3 | 11 | Ace2 |
| Ace3 | 0 | 0 | 0 | 0 | 0 | 0 | 0 | 0 | 0 | 0 | 0 | 0 | 0 | 0 | 1 | 0 | 0 | 0 | 0 | 0 | 0 | 0 | 0 | 0 | 0 | 0 | 0 | 0 | Ace3 |
| Acer1 | 0 | 0 | 0 | 0 | 0 | 0 | 0 | 4 | 3 | 5 | 0 | 0 | 0 | 0 | 0 | 0 | 0 | 0 | 1 | 1 | 1 | 0 | 0 | 0 | 0 | 0 | 0 | 0 | Acer1 |
| Acer2 | 1051 | 929 | 1116 | 1025 | 1034 | 1164 | 870 | 1025 | 1071 | 1404 | 1191 | 930 | 1076 | 994 | 1043 | 1099 | 1049 | 941 | 974 | 1021 | 895 | 1327 | 1165 | 1192 | 1123 | 1055 | 954 | 983 | Acer2 |
| Acer3 | 1959 | 1377 | 1596 | 1372 | 1651 | 1803 | 1700 | 1253 | 1217 | 952 | 1077 | 948 | 1117 | 1126 | 1941 | 1835 | 1399 | 1608 | 1380 | 1806 | 1616 | 1121 | 1206 | 852 | 935 | 1013 | 1190 | 1093 | Acer3 |
| Ache | 767 | 903 | 1124 | 1343 | 1602 | 1151 | 1196 | 579 | 681 | 740 | 1202 | 401 | 625 | 454 | 790 | 938 | 1356 | 1267 | 1364 | 995 | 1167 | 1034 | 841 | 499 | 616 | 431 | 399 | 500 | Ache |
| Acin1 | 16594 | 18709 | 24755 | 23426 | 21893 | 20676 | 21007 | 23913 | 25047 | 31593 | 29256 | 33202 | 29290 | 34805 | 16220 | 16707 | 24501 | 21332 | 24206 | 18135 | 21715 | 27686 | 23532 | 35565 | 32445 | 32849 | 31337 | 33146 | Acin1 |
| Ackr1 | 1048 | 1580 | 1514 | 2308 | 2434 | 1967 | 2485 | 824 | 819 | 1051 | 861 | 869 | 797 | 835 | 1288 | 1307 | 1882 | 2641 | 2371 | 1830 | 2330 | 1369 | 741 | 862 | 1100 | 676 | 764 | 724 | Ackr1 |
| Ackr2 | 29 | 45 | 18 | 53 | 77 | 66 | 78 | 103 | 105 | 80 | 66 | 104 | 60 | 82 | 54 | 21 | 59 | 61 | 48 | 86 | 60 | 122 | 87 | 101 | 118 | 63 | 99 | 76 | Ackr2 |
| Ackr3 | 2651 | 2579 | 2217 | 1833 | 1532 | 1784 | 1683 | 3116 | 3016 | 2315 | 2081 | 2728 | 2734 | 2688 | 2531 | 2334 | 1856 | 1684 | 1675 | 1888 | 1652 | 2045 | 2922 | 2687 | 2193 | 3144 | 2920 | 3185 | Ackr3 |
| Ackr4 | 1 | 0 | 0 | 1 | 1 | 0 | 1 | 11 | 29 | 10 | 28 | 13 | 17 | 10 | 3 | 1 | 3 | 1 | 1 | 1 | 0 | 5 | 25 | 11 | 4 | 17 | 5 | 4 | Ackr4 |
| Acly | 49147 | 60885 | 50005 | 50667 | 49300 | 48603 | 49087 | 32756 | 32114 | 31561 | 28605 | 37367 | 36105 | 36966 | 51922 | 51970 | 53101 | 50485 | 52039 | 49065 | 50443 | 31222 | 29573 | 33052 | 34481 | 34087 | 39139 | 39010 | Acly |
| Acmsd | 12 | 11 | 5 | 3 | 2 | 4 | 3 | 12 | 9 | 4 | 2 | 7 | 6 | 9 | 15 | 6 | 3 | 1 | 9 | 13 | 9 | 2 | 8 | 4 | 4 | 8 | 2 | 12 | Acmsd |
| Acnat1 | 10 | 6 | 8 | 2 | 6 | 1 | 1 | 9 | 16 | 2 | 0 | 8 | 1 | 2 | 5 | 9 | 5 | 2 | 3 | 3 | 4 | 2 | 14 | 5 | 2 | 8 | 2 | 1 | Acnat1 |
| Acnat2 | 0 | 0 | 0 | 0 | 0 | 0 | 0 | 0 | 0 | 0 | 2 | 0 | 0 | 0 | 0 | 0 | 0 | 0 | 0 | 0 | 0 | 0 | 0 | 0 | 0 | 0 | 0 | 0 | Acnat2 |
| Aco1 | 13441 | 14928 | 11344 | 11466 | 9925 | 10609 | 11561 | 6876 | 6947 | 5811 | 5830 | 6958 | 6643 | 6970 | 13018 | 12172 | 11522 | 11225 | 11357 | 10728 | 11749 | 6301 | 6514 | 5946 | 6766 | 7136 | 7078 | 7347 | Aco1 |
| Aco2 | 13705 | 17109 | 12743 | 13797 | 14101 | 14153 | 13348 | 14663 | 16434 | 15545 | 13904 | 15412 | 14341 | 15051 | 14249 | 13093 | 13612 | 13461 | 15197 | 15219 | 14635 | 14492 | 15888 | 15071 | 15574 | 15656 | 15966 | 14808 | Aco2 |
| Acod1 | 2 | 0 | 0 | 0 | 0 | 0 | 1 | 0 | 1 | 0 | 0 | 1 | 0 | 0 | 0 | 0 | 0 | 2 | 0 | 0 | 0 | 0 | 1 | 2 | 0 | 2 | 0 | 1 | Acod1 |
| Acot1 | 1227 | 1183 | 1243 | 1119 | 1519 | 1378 | 1387 | 1682 | 1811 | 1589 | 1661 | 1481 | 1573 | 1646 | 1236 | 1088 | 1343 | 1192 | 1523 | 1519 | 1451 | 1823 | 1785 | 1618 | 1582 | 1668 | 1514 | 1640 | Acot1 |
| Acot10 | 12 | 8 | 3 | 4 | 4 | 6 | 4 | 21 | 7 | 5 | 5 | 11 | 8 | 9 | 12 | 6 | 3 | 6 | 7 | 7 | 5 | 18 | 18 | 3 | 8 | 9 | 15 | 8 | Acot10 |
| Acot11 | 429 | 496 | 314 | 299 | 275 | 305 | 197 | 250 | 356 | 363 | 470 | 310 | 353 | 275 | 405 | 557 | 316 | 401 | 230 | 225 | 182 | 515 | 493 | 315 | 351 | 330 | 342 | 351 | Acot11 |
| Acot12 | 54 | 65 | 14 | 83 | 51 | 91 | 79 | 12 | 11 | 3 | 5 | 14 | 5 | 12 | 59 | 40 | 35 | 84 | 53 | 44 | 78 | 8 | 16 | 7 | 5 | 1 | 2 | 1 | Acot12 |
| Acot13 | 1105 | 1149 | 962 | 967 | 776 | 829 | 1059 | 706 | 675 | 604 | 674 | 726 | 716 | 805 | 1131 | 967 | 969 | 1051 | 867 | 1093 | 1038 | 550 | 715 | 791 | 537 | 812 | 741 | 830 | Acot13 |
| Acot2 | 89 | 87 | 133 | 80 | 142 | 129 | 83 | 276 | 325 | 261 | 329 | 335 | 373 | 302 | 128 | 122 | 127 | 76 | 117 | 116 | 111 | 262 | 338 | 296 | 236 | 371 | 334 | 332 | Acot2 |
| Acot3 | 3 | 3 | 8 | 7 | 2 | 11 | 6 | 10 | 10 | 19 | 12 | 3 | 6 | 10 | 3 | 3 | 0 | 1 | 2 | 2 | 3 | 15 | 3 | 9 | 14 | 4 | 8 | 8 | Acot3 |
| Acot4 | 18 | 4 | 16 | 7 | 11 | 16 | 6 | 27 | 30 | 19 | 35 | 22 | 49 | 23 | 16 | 9 | 2 | 7 | 2 | 4 | 10 | 28 | 32 | 21 | 23 | 18 | 30 | 27 | Acot4 |
| Acot5 | 0 | 0 | 0 | 4 | 0 | 0 | 0 | 1 | 0 | 0 | 3 | 0 | 2 | 3 | 1 | 1 | 0 | 0 | 0 | 0 | 0 | 2 | 0 | 5 | 0 | 0 | 0 | 0 | Acot5 |
| Acot6 | 111 | 75 | 124 | 76 | 132 | 111 | 148 | 363 | 442 | 355 | 446 | 365 | 348 | 403 | 120 | 98 | 78 | 84 | 148 | 135 | 109 | 398 | 415 | 343 | 368 | 453 | 372 | 459 | Acot6 |
| Acot7 | 3842 | 5577 | 4530 | 5106 | 6012 | 5002 | 4613 | 4340 | 4810 | 4701 | 4142 | 3534 | 3809 | 3144 | 4764 | 4583 | 5081 | 5131 | 5963 | 5346 | 5146 | 5123 | 4819 | 3943 | 4514 | 3235 | 3724 | 3100 | Acot7 |
| Acot8 | 1017 | 1363 | 1137 | 1414 | 1158 | 1190 | 1133 | 609 | 777 | 753 | 867 | 878 | 820 | 662 | 997 | 1242 | 1213 | 1449 | 1230 | 1163 | 1089 | 752 | 735 | 883 | 825 | 779 | 837 | 736 | Acot8 |
| Acot9 | 1171 | 860 | 876 | 749 | 1074 | 893 | 921 | 1712 | 1587 | 1243 | 1449 | 1111 | 1310 | 1170 | 1209 | 1173 | 773 | 844 | 857 | 1016 | 871 | 1591 | 1750 | 1153 | 1216 | 1154 | 1121 | 1263 | Acot9 |
| Acox1 | 7919 | 9210 | 8167 | 9433 | 9677 | 8861 | 9588 | 6459 | 6620 | 6964 | 6908 | 6856 | 6532 | 6918 | 8326 | 8569 | 8998 | 9811 | 9202 | 8750 | 9476 | 7556 | 6633 | 6977 | 7855 | 5814 | 6548 | 6539 | Acox1 |
| Acox2 | 0 | 0 | 0 | 0 | 2 | 0 | 0 | 0 | 3 | 3 | 0 | 0 | 3 | 0 | 0 | 2 | 0 | 0 | 0 | 0 | 0 | 0 | 1 | 0 | 0 | 0 | 0 | 1 | Acox2 |
| Acox3 | 2099 | 2320 | 2255 | 2068 | 2396 | 2344 | 2029 | 2118 | 2442 | 2236 | 2224 | 2402 | 2533 | 2388 | 2064 | 2188 | 2140 | 2076 | 2180 | 2213 | 2068 | 2421 | 2177 | 2283 | 2531 | 2036 | 2444 | 2410 | Acox3 |
| Acoxl | 0 | 0 | 0 | 3 | 4 | 4 | 4 | 0 | 2 | 3 | 5 | 0 | 5 | 0 | 0 | 2 | 0 | 0 | 2 | 0 | 0 | 0 | 9 | 2 | 5 | 3 | 0 | 4 | Acoxl |
| Acp1 | 6547 | 6052 | 7153 | 5301 | 4629 | 5103 | 5561 | 5435 | 5325 | 4972 | 4712 | 6096 | 5702 | 6348 | 6803 | 7002 | 5867 | 4859 | 4753 | 6505 | 4787 | 4106 | 5203 | 5311 | 5010 | 7395 | 6603 | 7030 | Acp1 |
| Acp2 | 3654 | 3470 | 3192 | 3105 | 3778 | 3862 | 3555 | 6624 | 6228 | 5922 | 5862 | 5971 | 6073 | 6155 | 3964 | 3430 | 3014 | 3012 | 3448 | 3821 | 3716 | 6497 | 6266 | 5384 | 5994 | 5491 | 6195 | 6309 | Acp2 |
| Acp4 | 0 | 1 | 0 | 0 | 0 | 0 | 0 | 0 | 0 | 0 | 2 | 0 | 0 | 0 | 0 | 0 | 0 | 0 | 0 | 0 | 0 | 0 | 1 | 0 | 0 | 0 | 0 | 0 | Acp4 |
| Acp5 | 133 | 192 | 308 | 79 | 32 | 38 | 28 | 64 | 110 | 103 | 84 | 200 | 592 | 334 | 125 | 227 | 129 | 43 | 31 | 41 | 25 | 53 | 133 | 65 | 69 | 260 | 306 | 797 | Acp5 |
| Acp6 | 2355 | 2675 | 2017 | 2220 | 2152 | 2172 | 2226 | 1224 | 1242 | 1032 | 1138 | 974 | 1018 | 1072 | 2135 | 2355 | 2277 | 2223 | 2146 | 2093 | 2165 | 847 | 1111 | 1016 | 991 | 1142 | 1110 | 1097 | Acp6 |

| | | | | | | | | | | | | | | | | | | | | | | | | | | | | | |
|---|---|---|---|---|---|---|---|---|---|---|---|---|---|---|---|---|---|---|---|---|---|---|---|---|---|---|---|---|---|
| Acp7 | 2 | 1 | 0 | 0 | 6 | 0 | 0 | 1 | 2 | 3 | 4 | 0 | 1 | 1 | 3 | 0 | 3 | 6 | 1 | 2 | 1 | 2 | 5 | 0 | 1 | 0 | 0 | 0 | Acp7 |
| Acpp | 153 | 51 | 17 | 32 | 70 | 24 | 21 | 110 | 138 | 123 | 67 | 17 | 73 | 28 | 66 | 81 | 23 | 30 | 28 | 37 | 31 | 50 | 146 | 67 | 50 | 48 | 16 | 21 | Acpp |
| Acr | 120 | 144 | 117 | 188 | 310 | 260 | 238 | 147 | 229 | 267 | 233 | 230 | 238 | 269 | 109 | 129 | 136 | 248 | 275 | 234 | 238 | 287 | 174 | 254 | 302 | 203 | 210 | 185 | Acr |
| Acrbp | 408 | 509 | 519 | 579 | 670 | 564 | 571 | 723 | 666 | 949 | 845 | 773 | 820 | 838 | 410 | 412 | 687 | 683 | 703 | 479 | 620 | 944 | 653 | 995 | 940 | 784 | 757 | 803 | Acrbp |
| Acrv1 | 0 | 0 | 0 | 0 | 0 | 0 | 0 | 0 | 0 | 0 | 0 | 0 | 0 | 0 | 0 | 0 | 0 | 0 | 0 | 0 | 0 | 0 | 0 | 0 | 0 | 1 | 0 | 0 | Acrv1 |
| Acsbg1 | 496 | 563 | 496 | 447 | 544 | 560 | 539 | 524 | 725 | 715 | 568 | 1006 | 598 | 930 | 406 | 305 | 539 | 494 | 592 | 433 | 553 | 585 | 506 | 684 | 772 | 976 | 983 | 892 | Acsbg1 |
| Acsbg2 | 3 | 1 | 0 | 0 | 1 | 0 | 2 | 0 | 1 | 0 | 4 | 3 | 2 | 2 | 1 | 0 | 0 | 0 | 2 | 1 | 0 | 0 | 1 | 0 | 0 | 0 | 0 | 0 | Acsbg2 |
| Acsbg3 | 0 | 2 | 1 | 0 | 4 | 1 | 4 | 1 | 2 | 3 | 1 | 5 | 8 | 1 | 0 | 2 | 5 | 3 | 1 | 0 | 4 | 2 | 6 | 2 | 2 | 1 | 10 | 4 | Acsbg3 |
| Acsf2 | 1404 | 1516 | 1823 | 1291 | 1126 | 1073 | 780 | 1123 | 1584 | 1517 | 1720 | 1623 | 1505 | 1527 | 1193 | 1819 | 1440 | 1372 | 1089 | 841 | 862 | 1267 | 1606 | 1539 | 1176 | 2151 | 1447 | 1412 | Acsf2 |
| Acsf3 | 339 | 394 | 447 | 448 | 494 | 444 | 371 | 913 | 1162 | 1155 | 1313 | 1388 | 1246 | 1194 | 324 | 410 | 429 | 381 | 489 | 407 | 406 | 1158 | 1166 | 1283 | 1204 | 1364 | 1339 | 1258 | Acsf3 |
| Acsl1 | 3638 | 3496 | 3261 | 3325 | 3227 | 3042 | 3196 | 1106 | 1174 | 1064 | 1309 | 816 | 1195 | 791 | 3617 | 3658 | 3182 | 3258 | 2923 | 2865 | 3161 | 1347 | 1273 | 833 | 927 | 752 | 884 | 892 | Acsl1 |
| Acsl3 | 12334 | 11245 | 11381 | 10783 | 9401 | 9360 | 10715 | 8067 | 6401 | 6525 | 6514 | 7450 | 6931 | 8020 | 12634 | 11843 | 10888 | 10564 | 9432 | 11137 | 9605 | 7102 | 6319 | 7106 | 6549 | 8579 | 7845 | 8428 | Acsl3 |
| Acsl4 | 5368 | 4427 | 5374 | 4848 | 4813 | 4959 | 5322 | 3229 | 2687 | 2575 | 3048 | 3170 | 3561 | 3460 | 5840 | 5455 | 4943 | 4412 | 4470 | 4697 | 4851 | 3101 | 2500 | 2430 | 2846 | 2814 | 3519 | 3666 | Acsl4 |
| Acsl5 | 1066 | 809 | 924 | 603 | 766 | 733 | 593 | 1657 | 1631 | 1371 | 1518 | 1426 | 1626 | 1344 | 1048 | 1012 | 713 | 580 | 631 | 773 | 599 | 1343 | 1792 | 1409 | 1117 | 1665 | 1419 | 1540 | Acsl5 |
| Acsl6 | 2543 | 2756 | 3000 | 3436 | 5501 | 4665 | 4597 | 5046 | 4440 | 5305 | 4352 | 5158 | 4540 | 5154 | 3003 | 2196 | 3554 | 3487 | 4955 | 4375 | 4899 | 6154 | 4267 | 4760 | 6057 | 3395 | 4918 | 4771 | Acsl6 |
| Acsm1 | 0 | 0 | 0 | 0 | 0 | 0 | 0 | 4 | 6 | 7 | 2 | 0 | 0 | 1 | 0 | 0 | 0 | 0 | 0 | 0 | 0 | 0 | 1 | 0 | 0 | 6 | 0 | 1 | Acsm1 |
| Acsm2 | 1 | 0 | 0 | 0 | 0 | 0 | 0 | 0 | 0 | 1 | 0 | 0 | 0 | 0 | 0 | 0 | 0 | 0 | 0 | 0 | 0 | 0 | 1 | 0 | 0 | 0 | 0 | 0 | Acsm2 |
| Acsm3 | 3 | 2 | 0 | 0 | 1 | 1 | 1 | 1 | 7 | 0 | 2 | 5 | 3 | 2 | 0 | 0 | 0 | 0 | 2 | 0 | 0 | 7 | 3 | 7 | 6 | 1 | 2 | 1 | Acsm3 |
| Acsm4 | 338 | 215 | 0 | 302 | 0 | 2 | 1 | 375 | 235 | 486 | 320 | 1 | 228 | 103 | 235 | 275 | 64 | 0 | 1 | 0 | 0 | 0 | 401 | 713 | 1 | 911 | 0 | 0 | Acsm4 |
| Acsm5 | 0 | 0 | 0 | 0 | 0 | 0 | 0 | 0 | 0 | 0 | 0 | 0 | 0 | 0 | 0 | 0 | 0 | 0 | 0 | 0 | 2 | 0 | 0 | 0 | 0 | 1 | 0 | 0 | Acsm5 |
| Acss1 | 2331 | 2769 | 2438 | 2676 | 2468 | 2789 | 2870 | 3116 | 3030 | 3119 | 2427 | 4586 | 3224 | 4373 | 2669 | 2257 | 3115 | 2490 | 2748 | 2370 | 2481 | 3018 | 2023 | 3511 | 3531 | 4507 | 3707 | 3734 | Acss1 |
| Acss2 | 3869 | 4668 | 3511 | 4143 | 3629 | 3711 | 3791 | 2105 | 1919 | 1933 | 1760 | 1912 | 1948 | 1808 | 4221 | 4045 | 3819 | 3649 | 3896 | 4009 | 3870 | 2012 | 1889 | 1844 | 1914 | 1929 | 1983 | 1875 | Acss2 |
| Acss3 | 2211 | 1394 | 1648 | 1054 | 807 | 992 | 705 | 977 | 1037 | 828 | 1083 | 683 | 1093 | 650 | 1692 | 2500 | 1105 | 1075 | 723 | 808 | 634 | 800 | 1175 | 779 | 453 | 637 | 586 | 646 | Acss3 |
| Acta1 | 123 | 41 | 23 | 50 | 61 | 47 | 47 | 211 | 586 | 120 | 4987 | 41 | 108 | 36 | 59 | 109 | 75 | 41 | 55 | 37 | 45 | 386 | 3415 | 100 | 86 | 21 | 17 | 20 | Acta1 |
| Acta2 | 3151 | 3653 | 2124 | 2072 | 411 | 2947 | 231 | 2805 | 3707 | 1808 | 6515 | 3818 | 4560 | 3873 | 3993 | 3579 | 2525 | 2845 | 445 | 400 | 277 | 2115 | 6405 | 2903 | 1743 | 2212 | 4283 | 3607 | Acta2 |
| Actb | 88182 | 99792 | 96469 | 87355 | 82410 | 87040 | 84310 | 99147 | 120896 | 118443 | 111570 | 132637 | 135651 | 131534 | 89394 | 94787 | 91605 | 83304 | 86617 | 84993 | 83424 | 108985 | 118458 | 129768 | 121549 | 157428 | 134215 | 139806 | Actb |
| Actbl2 | 0 | 0 | 0 | 0 | 0 | 0 | 0 | 1 | 2 | 0 | 1 | 0 | 3 | 3 | 0 | 2 | 0 | 0 | 0 | 0 | 1 | 5 | 5 | 2 | 3 | 6 | 0 | 0 | Actbl2 |
| Actc1 | 503 | 38 | 18 | 9 | 65 | 28 | 22 | 662 | 1804 | 190 | 11268 | 70 | 500 | 16 | 174 | 459 | 38 | 35 | 8 | 13 | 9 | 1138 | 7861 | 181 | 189 | 39 | 16 | 26 | Actc1 |
| Acte1 | 0 | 3 | 2 | 1 | 4 | 3 | 0 | 0 | 1 | 0 | 0 | 0 | 0 | 0 | 0 | 2 | 0 | 1 | 3 | 0 | 0 | 0 | 0 | 0 | 0 | 0 | 0 | 0 | Acte1 |
| Actg1 | 79836 | 98697 | 83982 | 85665 | 83983 | 83203 | 77385 | 69487 | 84269 | 74917 | 77123 | 82592 | 86579 | 80138 | 83632 | 95706 | 85787 | 85956 | 88860 | 78917 | 79013 | 84895 | 81019 | 78078 | 84443 | 91024 | 84703 | 82380 | Actg1 |
| Actg2 | 174 | 188 | 111 | 141 | 31 | 187 | 27 | 183 | 204 | 137 | 255 | 247 | 225 | 295 | 227 | 231 | 109 | 163 | 29 | 38 | 32 | 142 | 262 | 201 | 152 | 187 | 301 | 249 | Actg2 |
| Actl10 | 1 | 1 | 5 | 3 | 2 | 0 | 4 | 0 | 0 | 0 | 0 | 1 | 0 | 3 | 3 | 2 | 0 | 1 | 2 | 2 | 5 | 3 | 3 | 4 | 3 | 0 | 6 | 1 | Actl10 |
| Actl6a | 6106 | 6011 | 6076 | 4973 | 4084 | 4652 | 5085 | 7214 | 6266 | 5831 | 5093 | 8018 | 6975 | 8125 | 5850 | 5226 | 5270 | 4340 | 5129 | 5550 | 4749 | 5237 | 5752 | 6957 | 6375 | 9177 | 8625 | 8780 | Actl6a |
| Actl6b | 2293 | 3655 | 2814 | 5733 | 7100 | 5995 | 6120 | 4217 | 4364 | 5592 | 4393 | 4121 | 3670 | 4339 | 3257 | 2636 | 4824 | 6101 | 7028 | 5994 | 6579 | 6041 | 4121 | 4769 | 6329 | 3201 | 3683 | 3521 | Actl6b |
| Actl7a | 0 | 0 | 0 | 0 | 0 | 0 | 0 | 1 | 0 | 2 | 0 | 0 | 1 | 0 | 0 | 2 | 0 | 0 | 0 | 0 | 0 | 0 | 1 | 3 | 1 | 0 | 0 | 0 | Actl7a |
| Actl7b | 9 | 8 | 28 | 15 | 6 | 3 | 7 | 1 | 3 | 6 | 2 | 0 | 1 | 3 | 3 | 5 | 6 | 9 | 5 | 4 | 9 | 3 | 3 | 2 | 9 | 3 | 1 | 4 | Actl7b |
| Actl9 | 0 | 0 | 0 | 0 | 2 | 5 | 0 | 4 | 5 | 0 | 4 | 5 | 0 | 0 | 0 | 0 | 0 | 0 | 1 | 2 | 2 | 0 | 1 | 4 | 4 | 1 | 2 | 4 | Actl9 |
| Actn1 | 7173 | 7113 | 8457 | 6478 | 5312 | 5446 | 4894 | 4684 | 5712 | 5517 | 6364 | 6042 | 6526 | 5829 | 6328 | 8073 | 7103 | 6430 | 5822 | 4060 | 5278 | 4640 | 5744 | 6450 | 4876 | 7871 | 6163 | 6101 | Actn1 |
| Actn2 | 345 | 250 | 279 | 253 | 304 | 203 | 264 | 250 | 652 | 190 | 4473 | 98 | 222 | 67 | 291 | 386 | 268 | 232 | 246 | 232 | 202 | 456 | 2617 | 101 | 150 | 65 | 80 | 81 | Actn2 |
| Actn3 | 161 | 229 | 126 | 160 | 99 | 151 | 73 | 140 | 230 | 165 | 347 | 122 | 171 | 136 | 187 | 244 | 102 | 133 | 107 | 90 | 60 | 152 | 329 | 147 | 129 | 114 | 126 | 108 | Actn3 |
| Actn4 | 20617 | 23736 | 21591 | 18818 | 15203 | 16826 | 14434 | 10805 | 14190 | 12556 | 14129 | 13240 | 14078 | 12638 | 19229 | 24464 | 18426 | 19167 | 16113 | 13634 | 15026 | 11294 | 14378 | 13421 | 11896 | 14412 | 13179 | 13111 | Actn4 |
| Actr10 | 10961 | 9848 | 9394 | 10914 | 11103 | 11210 | 11939 | 10378 | 8296 | 8382 | 7885 | 8090 | 8219 | 8699 | 11772 | 9997 | 9843 | 11008 | 10996 | 11912 | 12141 | 9454 | 8589 | 8360 | 8838 | 8169 | 8306 | 8748 | Actr10 |
| Actr1a | 16815 | 20136 | 15878 | 16374 | 18213 | 18447 | 18739 | 19925 | 19462 | 17411 | 16132 | 18236 | 17047 | 18729 | 17845 | 15524 | 17207 | 16753 | 19113 | 19550 | 19272 | 19233 | 18716 | 18208 | 20023 | 18153 | 18599 | 18465 | Actr1a |
| Actr1b | 11066 | 12628 | 12735 | 12295 | 11596 | 11287 | 11238 | 7344 | 8424 | 8946 | 9378 | 9236 | 9118 | 9357 | 10568 | 11858 | 12466 | 12097 | 11697 | 10204 | 11268 | 8808 | 8181 | 9246 | 9149 | 10302 | 9104 | 9193 | Actr1b |
| Actr2 | 24592 | 20397 | 25541 | 23148 | 23131 | 22421 | 24825 | 26036 | 21523 | 21461 | 21509 | 22685 | 22531 | 24778 | 26074 | 22292 | 23330 | 23140 | 22152 | 26104 | 23670 | 23078 | 21591 | 22352 | 24372 | 23051 | 24033 | 24519 | Actr2 |
| Actr3 | 18497 | 14489 | 18612 | 17581 | 18803 | 18251 | 20131 | 21704 | 19411 | 19438 | 18742 | 20996 | 19490 | 21506 | 18834 | 16026 | 18052 | 17460 | 18741 | 17820 | 19147 | 21158 | 19935 | 20090 | 20544 | 21844 | 20529 | 21367 | Actr3 |
| Actr3b | 1142 | 1192 | 988 | 1386 | 1416 | 1287 | 1627 | 617 | 752 | 671 | 518 | 792 | 540 | 667 | 1096 | 856 | 1286 | 1480 | 1604 | 1554 | 1557 | 658 | 592 | 704 | 871 | 934 | 588 | 640 | Actr3b |
| Actr5 | 1739 | 2126 | 1631 | 2317 | 1887 | 1963 | 1867 | 1189 | 1233 | 1169 | 1305 | 1310 | 1312 | 1330 | 1721 | 1611 | 2045 | 2166 | 2138 | 1829 | 2075 | 1103 | 1155 | 1295 | 1315 | 1434 | 1354 | 1296 | Actr5 |
| Actr6 | 2608 | 2029 | 2278 | 2382 | 2059 | 2318 | 2730 | 1986 | 1811 | 1668 | 1454 | 1650 | 1716 | 1806 | 2660 | 2420 | 2267 | 2367 | 2349 | 2916 | 2440 | 1611 | 1869 | 1777 | 1631 | 1926 | 1758 | 1782 | Actr6 |
| Actr8 | 2540 | 2415 | 2344 | 2176 | 2564 | 2339 | 2430 | 3918 | 3536 | 3655 | 3381 | 3922 | 3880 | 3978 | 2349 | 2090 | 2458 | 2139 | 2717 | 2489 | 2456 | 3706 | 3456 | 3771 | 3790 | 4071 | 4063 | 3961 | Actr8 |
| Actrt3 | 46 | 41 | 33 | 37 | 83 | 69 | 80 | 11 | 7 | 7 | 9 | 20 | 15 | 14 | 70 | 46 | 67 | 57 | 56 | 72 | 49 | 2 | 10 | 9 | 19 | 8 | 7 | 11 | Actrt3 |
| Acvr1 | 1991 | 1912 | 2349 | 2387 | 2491 | 2277 | 2435 | 2097 | 2186 | 2221 | 2652 | 2035 | 2101 | 2051 | 1960 | 1958 | 2239 | 2461 | 2547 | 1843 | 2564 | 2393 | 2242 | 2345 | 2287 | 2197 | 1981 | 2061 | Acvr1 |
| Acvr1b | 6511 | 7055 | 8204 | 9204 | 10775 | 9215 | 9819 | 6638 | 6991 | 8142 | 7404 | 8115 | 7317 | 8168 | 6422 | 6527 | 9494 | 9663 | 10629 | 8295 | 10017 | 8895 | 6895 | 8688 | 9367 | 7365 | 7268 | 7592 | Acvr1b |
| Acvr1c | 194 | 169 | 121 | 212 | 250 | 290 | 341 | 82 | 95 | 96 | 70 | 115 | 71 | 76 | 178 | 119 | 137 | 249 | 231 | 328 | 300 | 67 | 96 | 98 | 115 | 74 | 79 | 84 | Acvr1c |
| Acvr2a | 7671 | 6214 | 7053 | 8691 | 10185 | 9463 | 10766 | 10710 | 8941 | 9084 | 9058 | 8502 | 8088 | 9158 | 8715 | 6911 | 7665 | 8842 | 8947 | 9415 | 10222 | 11855 | 9137 | 8389 | 10118 | 7767 | 8171 | 8810 | Acvr2a |
| Acvr2b | 1549 | 2005 | 812 | 2122 | 2485 | 2473 | 2126 | 3258 | 3347 | 4079 | 3667 | 4868 | 4314 | 4990 | 1483 | 1551 | 1599 | 1765 | 2416 | 2128 | 2481 | 2966 | 2834 | 3064 | 4681 | 1071 | 5206 | 4385 | Acvr2b |
| Acvrl1 | 834 | 1032 | 1026 | 759 | 722 | 749 | 587 | 719 | 894 | 961 | 1092 | 991 | 1125 | 802 | 844 | 1125 | 834 | 817 | 631 | 623 | 626 | 936 | 1077 | 942 | 871 | 1117 | 949 | 884 | Acvrl1 |
| Acy1 | 2182 | 2682 | 2369 | 2337 | 1852 | 2119 | 1990 | 1108 | 1349 | 1275 | 1302 | 1396 | 1401 | 1297 | 2091 | 2352 | 2460 | 2229 | 2277 | 2213 | 2003 | 964 | 1313 | 1319 | 1259 | 1517 | 1498 | 1409 | Acy1 |
| Acy3 | 10 | 20 | 31 | 8 | 22 | 14 | 8 | 39 | 66 | 41 | 63 | 41 | 54 | 27 | 15 | 19 | 24 | 22 | 6 | 14 | 8 | 67 | 82 | 47 | 38 | 38 | 22 | 27 | Acy3 |
| Acyp1 | 789 | 949 | 1089 | 1042 | 947 | 859 | 1106 | 728 | 684 | 713 | 729 | 806 | 762 | 955 | 870 | 867 | 1150 | 856 | 915 | 1038 | 1092 | 630 | 742 | 885 | 765 | 890 | 830 | 971 | Acyp1 |
| Acyp2 | 37 | 52 | 42 | 42 | 90 | 46 | 48 | 104 | 125 | 113 | 139 | 110 | 142 | 166 | 59 | 51 | 53 | 53 | 65 | 50 | 71 | 120 | 170 | 122 | 128 | 128 | 143 | 144 | Acyp2 |
| Ada | 398 | 389 | 322 | 330 | 255 | 310 | 255 | 188 | 251 | 197 | 256 | 261 | 271 | 308 | 411 | 357 | 312 | 316 | 331 | 300 | 263 | 170 | 244 | 266 | 167 | 339 | 268 | 230 | Ada |
| Adad1 | 28 | 16 | 22 | 16 | 7 | 4 | 8 | 16 | 22 | 19 | 9 | 14 | 26 | 14 | 25 | 28 | 10 | 19 | 7 | 10 | 4 | 33 | 38 | 12 | 19 | 16 | 9 | 13 | Adad1 |
| Adad2 | 5 | 9 | 10 | 3 | 2 | 3 | 8 | 10 | 12 | 10 | 13 | 7 | 3 | 9 | 3 | 7 | 3 | 2 | 1 | 4 | 6 | 13 | 7 | 5 | 11 | 9 | 10 | 8 | Adad2 |
| Adal | 1395 | 1438 | 1367 | 1290 | 1394 | 1509 | 1652 | 2577 | 2300 | 2176 | 2114 | 2108 | 2038 | 2168 | 1701 | 1312 | 1380 | 1382 | 1496 | 1755 | 1608 | 2179 | 2291 | 2102 | 2157 | 1849 | 2172 | 2172 | Adal |
| Adam10 | 16366 | 13306 | 17365 | 16208 | 17039 | 16556 | 17826 | 15410 | 13234 | 14868 | 15521 | 15101 | 16887 | 16136 | 17096 | 15943 | 16023 | 17038 | 15656 | 15158 | 16826 | 16872 | 13948 | 14644 | 15936 | 14966 | 15737 | 16463 | Adam10 |
| Adam11 | 1551 | 2071 | 2118 | 2042 | 3567 | 2121 | 1868 | 2812 | 3541 | 3620 | 3741 | 2902 | 4182 | 2562 | 1870 | 2505 | 2531 | 2656 | 2289 | 1819 | 1994 | 4879 | 2342 | 2621 | 3858 | 2566 | 2626 | 2642 | Adam11 |
| Adam12 | 615 | 596 | 523 | 428 | 497 | 675 | 310 | 1047 | 1459 | 1321 | 2450 | 898 | 1571 | 830 | 591 | 748 | 388 | 574 | 346 | 412 | 349 | 1597 | 2458 | 1069 | 1164 | 568 | 1206 | 807 | Adam12 |
| Adam15 | 4562 | 6034 | 5474 | 6159 | 5182 | 4980 | 4742 | 1441 | 2008 | 1898 | 2049 | 1728 | 1854 | 1412 | 4361 | 5620 | 5724 | 6451 | 5739 | 4854 | 4900 | 1913 | 2227 | 1887 | 1710 | 1884 | 1694 | 1542 | Adam15 |
| Adam17 | 6286 | 5667 | 5467 | 4918 | 4347 | 4620 | 4317 | 4978 | 4902 | 4289 | 4822 | 4777 | 5086 | 4845 | 5948 | 6344 | 4990 | 4880 | 4097 | 4894 | 4251 | 4299 | 5227 | 4455 | 4151 | 4665 | 5351 | 5069 | Adam17 |
| Adam18 | 2 | 2 | 0 | 2 | 5 | 5 | 7 | 1 | 1 | 3 | 5 | 2 | 2 | 2 | 7 | 2 | 0 | 2 | 9 | 5 | 4 | 2 | 4 | 0 | 2 | 0 | 5 | 0 | Adam18 |
| Adam19 | 3061 | 3102 | 3146 | 2811 | 3712 | 3063 | 2554 | 3866 | 4586 | 4722 | 6316 | 4266 | 5796 | 4168 | 3171 | 4073 | 3044 | 3381 | 2939 | 2325 | 2543 | 5265 | 4890 | 4176 | 4499 | 3958 | 4281 | 4283 | Adam19 |
| Adam1a | 420 | 430 | 439 | 547 | 772 | 657 | 613 | 608 | 541 | 737 | 678 | 595 | 491 | 515 | 456 | 371 | 568 | 697 | 707 | 654 | 655 | 745 | 522 | 606 | 724 | 366 | 588 | 460 | Adam1a |
| Adam1b | 0 | 5 | 0 | 1 | 1 | 4 | 4 | 0 | 0 | 0 | 0 | 2 | 1 | 0 | 3 | 2 | 6 | 3 | 4 | 2 | 2 | 0 | 1 | 0 | 0 | 0 | 0 | 0 | Adam1b |
| Adam2 | 5 | 0 | 1 | 0 | 1 | 0 | 0 | 1 | 5 | 3 | 12 | 1 | 3 | 3 | 4 | 2 | 7 | 0 | 5 | 0 | 1 | 3 | 7 | 2 | 7 | 1 | 1 | 1 | Adam2 |
| Adam21 | 23 | 21 | 37 | 32 | 57 | 37 | 45 | 32 | 42 | 54 | 31 | 41 | 24 | 39 | 29 | 29 | 30 | 45 | 50 | 27 | 46 | 50 | 34 | 34 | 35 | 34 | 34 | 52 | Adam21 |
| Adam22 | 2741 | 2496 | 1945 | 2963 | 4595 | 4023 | 4063 | 5983 | 5641 | 5586 | 5415 | 4582 | 4348 | 4889 | 3145 | 2733 | 2452 | 3531 | 3497 | 3840 | 4031 | 6602 | 5069 | 4678 | 5980 | 3251 | 4286 | 3934 | Adam22 |
| Adam23 | 3744 | 3609 | 2637 | 3562 | 4311 | 4169 | 4168 | 3985 | 3921 | 3106 | 3784 | 3194 | 2807 | 3610 | 4209 | 3318 | 3303 | 3450 | 3461 | 4266 | 4301 | 4328 | 3427 | 2808 | 3781 | 2668 | 3297 | 3269 | Adam23 |
| Adam26a | 0 | 0 | 0 | 3 | 0 | 2 | 1 | 2 | 0 | 0 | 0 | 0 | 0 | 1 | 0 | 0 | 0 | 0 | 0 | 0 | 0 | 2 | 0 | 0 | 0 | 0 | 0 | 0 | Adam26a |

| | | | | | | | | | | | | | | | | | | | | | | | | | | | | | |
|---|---|---|---|---|---|---|---|---|---|---|---|---|---|---|---|---|---|---|---|---|---|---|---|---|---|---|---|---|---|
| Adam26b | 0 | 0 | 0 | 0 | 0 | 1 | 0 | 0 | 0 | 0 | 0 | 0 | 0 | 0 | 0 | 0 | 0 | 0 | 0 | 0 | 0 | 0 | 0 | 0 | 0 | 0 | 0 | 0 | Adam26b |
| Adam28 | 6 | 4 | 0 | 8 | 2 | 0 | 0 | 74 | 96 | 59 | 76 | 1 | 9 | 2 | 4 | 5 | 0 | 0 | 0 | 7 | 4 | 12 | 126 | 52 | 3 | 14 | 0 | 2 | Adam28 |
| Adam3 | 7 | 4 | 5 | 8 | 9 | 11 | 6 | 7 | 11 | 8 | 10 | 10 | 3 | 9 | 5 | 5 | 0 | 3 | 9 | 7 | 4 | 12 | 8 | 4 | 7 | 2 | 22 | 11 | Adam3 |
| Adam30 | 6 | 3 | 2 | 4 | 4 | 7 | 9 | 1 | 0 | 0 | 2 | 2 | 0 | 0 | 4 | 1 | 1 | 9 | 3 | 7 | 5 | 5 | 0 | 1 | 0 | 2 | 0 | 0 | Adam30 |
| Adam32 | 88 | 87 | 88 | 132 | 110 | 141 | 126 | 21 | 16 | 32 | 14 | 31 | 29 | 27 | 88 | 72 | 96 | 174 | 109 | 87 | 121 | 38 | 22 | 16 | 20 | 28 | 31 | 27 | Adam32 |
| Adam33 | 179 | 150 | 189 | 132 | 50 | 223 | 51 | 215 | 400 | 366 | 579 | 254 | 286 | 189 | 150 | 234 | 154 | 202 | 79 | 54 | 44 | 266 | 600 | 419 | 338 | 176 | 237 | 244 | Adam33 |
| Adam34 | 1 | 1 | 0 | 0 | 2 | 6 | 2 | 2 | 0 | 1 | 0 | 1 | 3 | 0 | 1 | 1 | 0 | 2 | 0 | 3 | 0 | 2 | 1 | 1 | 4 | 1 | 0 | 0 | Adam34 |
| Adam34l | 0 | 1 | 0 | 0 | 0 | 0 | 0 | 1 | 0 | 0 | 0 | 0 | 2 | 0 | 0 | 0 | 0 | 1 | 2 | 0 | 0 | 0 | 3 | 0 | 1 | 2 | 0 | 1 | Adam34l |
| Adam4 | 35 | 31 | 44 | 49 | 63 | 42 | 56 | 59 | 48 | 47 | 62 | 64 | 59 | 72 | 36 | 33 | 49 | 41 | 55 | 32 | 33 | 74 | 35 | 48 | 72 | 46 | 64 | 78 | Adam4 |
| Adam5 | 29 | 41 | 8 | 35 | 48 | 34 | 31 | 35 | 45 | 28 | 26 | 21 | 64 | 21 | 53 | 44 | 15 | 55 | 28 | 26 | 43 | 47 | 63 | 41 | 20 | 36 | 31 | 23 | Adam5 |
| Adam7 | 1 | 1 | 0 | 2 | 1 | 0 | 1 | 4 | 1 | 0 | 4 | 0 | 2 | 0 | 0 | 2 | 0 | 1 | 0 | 0 | 0 | 0 | 1 | 2 | 0 | 1 | 0 | 0 | Adam7 |
| Adam8 | 1284 | 2014 | 1881 | 2779 | 2311 | 2151 | 2239 | 811 | 808 | 932 | 887 | 973 | 870 | 880 | 1443 | 1779 | 2311 | 2882 | 2498 | 1857 | 2220 | 1185 | 755 | 1075 | 1147 | 872 | 850 | 910 | Adam8 |
| Adam9 | 6542 | 6088 | 6857 | 5752 | 5660 | 5768 | 5775 | 3729 | 3608 | 3454 | 3778 | 3630 | 3712 | 3693 | 6526 | 6548 | 6185 | 5790 | 5448 | 5721 | 5590 | 3803 | 3707 | 3389 | 3455 | 3606 | 3719 | 3659 | Adam9 |
| Adamdec1 | 2 | 0 | 0 | 0 | 1 | 2 | 0 | 4 | 2 | 9 | 3 | 1 | 1 | 0 | 5 | 1 | 3 | 2 | 5 | 0 | 0 | 0 | 12 | 6 | 0 | 2 | 0 | 1 | Adamdec1 |
| Adamts1 | 2040 | 1298 | 1614 | 1212 | 1638 | 1374 | 1008 | 2292 | 3136 | 2576 | 4240 | 1714 | 2456 | 1611 | 1501 | 1808 | 1336 | 1497 | 1195 | 1200 | 1120 | 2864 | 4276 | 2181 | 1998 | 2507 | 1482 | 1496 | Adamts1 |
| Adamts10 | 7067 | 7774 | 7680 | 7859 | 7495 | 7697 | 6609 | 5675 | 7553 | 8293 | 8934 | 7064 | 7547 | 6575 | 6098 | 8138 | 7701 | 7885 | 7613 | 6322 | 6540 | 7752 | 7446 | 8100 | 7034 | 6738 | 7137 | 6382 | Adamts10 |
| Adamts12 | 1982 | 1792 | 1431 | 1809 | 1653 | 2608 | 1141 | 2535 | 3315 | 2918 | 4156 | 2786 | 3562 | 2728 | 2417 | 2429 | 1292 | 2433 | 1043 | 1664 | 1340 | 3482 | 4781 | 2748 | 2860 | 1711 | 2918 | 2333 | Adamts12 |
| Adamts13 | 10 | 2 | 2 | 0 | 0 | 0 | 23 | 5 | 2 | 1 | 3 | 1 | 0 | 2 | 21 | 3 | 5 | 0 | 1 | 2 | 1 | 5 | 1 | 3 | 12 | 4 | 0 | 1 | Adamts13 |
| Adamts14 | 213 | 251 | 320 | 223 | 162 | 249 | 86 | 201 | 381 | 343 | 662 | 281 | 441 | 261 | 182 | 321 | 229 | 304 | 115 | 75 | 62 | 376 | 485 | 262 | 377 | 243 | 278 | 228 | Adamts14 |
| Adamts15 | 1604 | 1702 | 1675 | 1245 | 1020 | 1371 | 601 | 1469 | 2159 | 1859 | 3793 | 2169 | 2457 | 1751 | 1478 | 2461 | 1159 | 1796 | 677 | 721 | 609 | 2114 | 3449 | 1990 | 1694 | 2188 | 2141 | 2124 | Adamts15 |
| Adamts16 | 116 | 149 | 94 | 167 | 282 | 255 | 79 | 350 | 323 | 339 | 607 | 179 | 378 | 172 | 280 | 246 | 83 | 226 | 138 | 158 | 124 | 353 | 555 | 173 | 196 | 101 | 237 | 148 | Adamts16 |
| Adamts17 | 1096 | 838 | 578 | 630 | 620 | 818 | 429 | 1188 | 1895 | 1439 | 2286 | 610 | 982 | 587 | 815 | 1187 | 489 | 835 | 375 | 356 | 518 | 957 | 2298 | 1146 | 878 | 698 | 638 | 540 | Adamts17 |
| Adamts18 | 260 | 169 | 116 | 189 | 337 | 246 | 321 | 730 | 1085 | 869 | 807 | 607 | 415 | 663 | 214 | 184 | 210 | 208 | 373 | 310 | 345 | 585 | 1065 | 767 | 809 | 624 | 545 | 495 | Adamts18 |
| Adamts19 | 1052 | 646 | 787 | 463 | 366 | 606 | 393 | 849 | 971 | 734 | 920 | 816 | 972 | 791 | 770 | 1138 | 448 | 539 | 442 | 499 | 372 | 403 | 753 | 753 | 658 | 850 | 853 | 915 | Adamts19 |
| Adamts2 | 536 | 739 | 453 | 710 | 571 | 1019 | 202 | 850 | 1523 | 1400 | 2255 | 797 | 1289 | 726 | 616 | 785 | 484 | 1093 | 362 | 272 | 231 | 1584 | 2080 | 1176 | 1377 | 571 | 723 | 583 | Adamts2 |
| Adamts20 | 3790 | 3477 | 2888 | 3223 | 2872 | 3414 | 3461 | 1745 | 1784 | 1582 | 1873 | 2227 | 2243 | 2207 | 3401 | 3909 | 2970 | 3529 | 2855 | 3353 | 3262 | 947 | 1608 | 1678 | 1394 | 2470 | 1792 | 2263 | Adamts20 |
| Adamts3 | 752 | 732 | 451 | 563 | 839 | 847 | 749 | 977 | 1387 | 1068 | 1132 | 774 | 770 | 763 | 634 | 677 | 443 | 688 | 673 | 686 | 804 | 922 | 1182 | 802 | 1120 | 478 | 815 | 763 | Adamts3 |
| Adamts4 | 694 | 733 | 911 | 831 | 890 | 887 | 827 | 896 | 1044 | 1196 | 1031 | 912 | 1040 | 913 | 622 | 780 | 911 | 813 | 1086 | 657 | 913 | 978 | 1098 | 1177 | 1127 | 779 | 1066 | 959 | Adamts4 |
| Adamts5 | 438 | 351 | 385 | 475 | 801 | 645 | 589 | 1017 | 1542 | 1175 | 1898 | 604 | 882 | 599 | 468 | 454 | 476 | 626 | 611 | 500 | 619 | 1577 | 2111 | 960 | 1286 | 677 | 526 | 526 | Adamts5 |
| Adamts6 | 2254 | 1802 | 1589 | 1750 | 1784 | 1960 | 1915 | 1900 | 2174 | 1832 | 2360 | 1858 | 2453 | 1871 | 2362 | 2136 | 1651 | 1536 | 1636 | 1996 | 1792 | 1669 | 2344 | 1503 | 1454 | 1307 | 2156 | 1932 | Adamts6 |
| Adamts7 | 2874 | 3457 | 3976 | 2888 | 3050 | 3343 | 2350 | 3004 | 3718 | 3305 | 3940 | 4089 | 3980 | 3499 | 2788 | 3359 | 3437 | 2990 | 3402 | 2320 | 2452 | 3701 | 3647 | 4006 | 3730 | 3829 | 3920 | 3621 | Adamts7 |
| Adamts8 | 73 | 89 | 72 | 109 | 127 | 142 | 70 | 137 | 180 | 165 | 325 | 154 | 209 | 128 | 92 | 104 | 73 | 135 | 95 | 58 | 82 | 180 | 295 | 153 | 174 | 181 | 137 | 128 | Adamts8 |
| Adamts9 | 2062 | 1569 | 1068 | 1150 | 1169 | 1789 | 946 | 2014 | 3302 | 2374 | 4426 | 1435 | 2234 | 1426 | 1826 | 2100 | 1042 | 1498 | 1113 | 1040 | 947 | 2010 | 4082 | 2005 | 1630 | 1378 | 1535 | 1301 | Adamts9 |
| Adamtsl1 | 911 | 1180 | 822 | 780 | 723 | 1223 | 314 | 985 | 1936 | 1771 | 2572 | 1174 | 1844 | 1198 | 948 | 1301 | 717 | 1171 | 537 | 350 | 391 | 1464 | 2284 | 1347 | 1229 | 656 | 1319 | 952 | Adamtsl1 |
| Adamtsl2 | 295 | 446 | 483 | 464 | 499 | 524 | 473 | 479 | 613 | 645 | 826 | 706 | 854 | 680 | 340 | 399 | 519 | 504 | 560 | 427 | 434 | 790 | 717 | 717 | 777 | 693 | 781 | 727 | Adamtsl2 |
| Adamtsl3 | 110 | 81 | 44 | 109 | 115 | 176 | 88 | 327 | 489 | 434 | 597 | 98 | 200 | 78 | 119 | 121 | 51 | 110 | 98 | 67 | 113 | 341 | 438 | 250 | 272 | 123 | 100 | 58 | Adamtsl3 |
| Adamtsl4 | 48 | 73 | 69 | 25 | 66 | 48 | 45 | 27 | 34 | 30 | 55 | 27 | 37 | 17 | 28 | 79 | 56 | 81 | 48 | 51 | 48 | 60 | 65 | 35 | 20 | 51 | 24 | 21 | Adamtsl4 |
| Adamtsl5 | 111 | 96 | 109 | 85 | 58 | 71 | 57 | 37 | 93 | 78 | 119 | 40 | 57 | 34 | 80 | 86 | 111 | 125 | 59 | 73 | 49 | 74 | 148 | 98 | 74 | 28 | 43 | 33 | Adamtsl5 |
| Adap1 | 243 | 308 | 167 | 542 | 1294 | 748 | 628 | 736 | 1083 | 1119 | 1116 | 391 | 499 | 366 | 332 | 315 | 342 | 817 | 886 | 696 | 853 | 1348 | 1159 | 592 | 1137 | 279 | 321 | 294 | Adap1 |
| Adap2 | 96 | 101 | 113 | 132 | 100 | 116 | 101 | 128 | 159 | 149 | 233 | 152 | 141 | 160 | 116 | 117 | 102 | 109 | 123 | 115 | 102 | 196 | 174 | 157 | 187 | 168 | 182 | 132 | Adap2 |
| Adar | 7414 | 8840 | 8227 | 10367 | 11009 | 10227 | 10594 | 5928 | 7018 | 8171 | 7047 | 7721 | 7042 | 7730 | 7654 | 8517 | 9642 | 10334 | 10968 | 9056 | 10876 | 8091 | 6231 | 7343 | 8500 | 5791 | 7165 | 7225 | Adar |
| Adarb1 | 2646 | 3040 | 2851 | 3401 | 3665 | 3301 | 3234 | 1395 | 1642 | 1874 | 1886 | 1673 | 1648 | 1566 | 2637 | 3165 | 3086 | 3476 | 3297 | 3096 | 3409 | 1897 | 1673 | 1560 | 2240 | 1234 | 1607 | 1487 | Adarb1 |
| Adarb2 | 1300 | 1811 | 862 | 1771 | 2133 | 2598 | 2052 | 1914 | 1795 | 1480 | 1459 | 1150 | 1385 | 1200 | 1882 | 1675 | 1305 | 1716 | 1626 | 2598 | 2419 | 1943 | 1347 | 945 | 1736 | 581 | 1171 | 917 | Adarb2 |
| Adat1 | 1384 | 1620 | 1569 | 1510 | 1249 | 1267 | 1332 | 1334 | 1301 | 1385 | 1411 | 1527 | 1480 | 1465 | 1353 | 1472 | 1516 | 1431 | 1477 | 1373 | 1406 | 1089 | 1315 | 1410 | 1382 | 1486 | 1518 | 1444 | Adat1 |
| Adat2 | 339 | 361 | 405 | 436 | 398 | 389 | 307 | 330 | 387 | 319 | 443 | 425 | 403 | 445 | 285 | 334 | 386 | 325 | 356 | 352 | 312 | 234 | 388 | 345 | 356 | 453 | 465 | 469 | Adat2 |
| Adck1 | 760 | 955 | 785 | 837 | 932 | 907 | 900 | 1602 | 1605 | 1345 | 1271 | 1243 | 1353 | 1187 | 861 | 816 | 746 | 854 | 941 | 868 | 827 | 1384 | 1529 | 1285 | 1196 | 1326 | 1253 | 1158 | Adck1 |
| Adck2 | 824 | 872 | 971 | 1035 | 1015 | 984 | 930 | 973 | 1118 | 1257 | 1162 | 1189 | 962 | 1033 | 786 | 850 | 951 | 1061 | 996 | 853 | 948 | 1111 | 1051 | 1059 | 1091 | 959 | 977 | 1085 | Adck2 |
| Adck5 | 1060 | 1215 | 1217 | 1294 | 1285 | 1289 | 1214 | 1148 | 1293 | 1426 | 1360 | 1404 | 1323 | 1326 | 1079 | 1104 | 1276 | 1367 | 1378 | 1340 | 1239 | 1240 | 1324 | 1587 | 1426 | 1764 | 1341 | 1296 | Adck5 |
| Adcy1 | 2329 | 2700 | 1785 | 2817 | 5896 | 4026 | 3529 | 3444 | 3921 | 4078 | 5138 | 2712 | 3442 | 2586 | 2649 | 3063 | 1970 | 3511 | 3490 | 3480 | 3983 | 4213 | 4339 | 2511 | 4522 | 1135 | 2840 | 2550 | Adcy1 |
| Adcy10 | 27 | 28 | 14 | 15 | 12 | 25 | 23 | 12 | 7 | 12 | 14 | 13 | 10 | 1 | 45 | 19 | 19 | 21 | 18 | 10 | 13 | 5 | 4 | 2 | 7 | 2 | 0 | 1 | Adcy10 |
| Adcy2 | 588 | 598 | 410 | 506 | 797 | 615 | 546 | 909 | 1089 | 987 | 1106 | 666 | 705 | 606 | 644 | 653 | 442 | 646 | 535 | 556 | 572 | 1118 | 1133 | 633 | 781 | 434 | 572 | 436 | Adcy2 |
| Adcy3 | 705 | 701 | 860 | 751 | 902 | 908 | 741 | 1142 | 1555 | 1627 | 1688 | 1222 | 1166 | 1010 | 649 | 833 | 847 | 861 | 745 | 606 | 661 | 1434 | 1620 | 1620 | 1365 | 1252 | 1059 | 1073 | Adcy3 |
| Adcy4 | 693 | 818 | 719 | 553 | 475 | 569 | 330 | 604 | 968 | 920 | 1238 | 748 | 1076 | 610 | 647 | 950 | 627 | 605 | 391 | 347 | 332 | 886 | 997 | 806 | 616 | 998 | 750 | 678 | Adcy4 |
| Adcy5 | 2519 | 3689 | 3168 | 3889 | 5106 | 4115 | 3669 | 2882 | 3297 | 3987 | 3326 | 2686 | 3142 | 2460 | 2900 | 3328 | 3744 | 4504 | 4818 | 3867 | 4132 | 3942 | 3159 | 3139 | 3649 | 2212 | 2510 | 2449 | Adcy5 |
| Adcy6 | 5861 | 6874 | 9202 | 8118 | 8078 | 7739 | 6355 | 5787 | 6695 | 8204 | 8713 | 8246 | 7845 | 7543 | 5418 | 7035 | 8263 | 8000 | 8537 | 5953 | 7015 | 8390 | 6440 | 8594 | 8462 | 7159 | 7932 | 7551 | Adcy6 |
| Adcy7 | 374 | 368 | 385 | 239 | 367 | 403 | 253 | 438 | 613 | 456 | 696 | 530 | 533 | 484 | 377 | 484 | 273 | 376 | 211 | 315 | 297 | 495 | 758 | 430 | 326 | 474 | 435 | 487 | Adcy7 |
| Adcy8 | 431 | 542 | 539 | 486 | 1013 | 584 | 513 | 480 | 477 | 430 | 509 | 403 | 504 | 330 | 579 | 545 | 504 | 694 | 674 | 569 | 638 | 647 | 668 | 333 | 453 | 158 | 343 | 377 | Adcy8 |
| Adcy9 | 806 | 977 | 839 | 1201 | 1558 | 1333 | 1183 | 541 | 621 | 599 | 665 | 544 | 604 | 478 | 799 | 861 | 905 | 1251 | 1278 | 1217 | 1168 | 744 | 612 | 556 | 645 | 167 | 481 | 458 | Adcy9 |
| Adcyap1 | 291 | 276 | 416 | 305 | 615 | 434 | 361 | 1330 | 976 | 888 | 883 | 1021 | 1075 | 1060 | 386 | 298 | 375 | 306 | 411 | 565 | 402 | 1367 | 1153 | 851 | 1096 | 691 | 1166 | 1314 | Adcyap1 |
| Adcyap1r1 | 3422 | 4977 | 4637 | 6768 | 10666 | 8146 | 7780 | 7111 | 6508 | 7680 | 6774 | 6179 | 8557 | 5701 | 6274 | 4464 | 6171 | 7810 | 8386 | 6230 | 8658 | 8920 | 6352 | 5816 | 7789 | 3134 | 5949 | 5644 | Adcyap1r1 |
| Add1 | 22636 | 24288 | 22719 | 24368 | 24813 | 24598 | 23978 | 21461 | 23960 | 24016 | 22498 | 24654 | 23524 | 24028 | 22887 | 22771 | 23600 | 25638 | 24751 | 21072 | 25023 | 25064 | 23680 | 24191 | 25239 | 23524 | 22936 | 24303 | Add1 |
| Add2 | 7271 | 8661 | 7018 | 11014 | 15462 | 14195 | 13339 | 20645 | 19724 | 23217 | 18825 | 22261 | 19376 | 21550 | 8951 | 6805 | 9761 | 11427 | 14847 | 12774 | 14502 | 26040 | 17081 | 20679 | 28450 | 13236 | 21514 | 20598 | Add2 |
| Add3 | 13723 | 12239 | 11548 | 10302 | 8808 | 9426 | 9729 | 5355 | 6395 | 5029 | 6164 | 5016 | 5643 | 5260 | 12369 | 13890 | 10472 | 10801 | 8694 | 9521 | 9358 | 4293 | 7504 | 5214 | 4454 | 6213 | 5168 | 5523 | Add3 |
| Adgb | 66 | 66 | 11 | 31 | 77 | 45 | 41 | 60 | 77 | 17 | 44 | 50 | 51 | 35 | 88 | 97 | 22 | 47 | 32 | 55 | 36 | 30 | 82 | 35 | 44 | 33 | 36 | 14 | Adgb |
| Adgra1 | 230 | 354 | 391 | 316 | 550 | 463 | 306 | 592 | 532 | 750 | 529 | 806 | 925 | 700 | 347 | 320 | 510 | 392 | 482 | 360 | 402 | 861 | 469 | 649 | 752 | 445 | 847 | 803 | Adgra1 |
| Adgra2 | 4548 | 5167 | 5160 | 3492 | 2363 | 3617 | 1495 | 3788 | 5732 | 5903 | 7631 | 4114 | 6032 | 3661 | 4003 | 6414 | 3839 | 4074 | 2118 | 1509 | 1595 | 4468 | 6310 | 5796 | 3905 | 5188 | 4663 | 4175 | Adgra2 |
| Adgra3 | 7750 | 7467 | 7023 | 6396 | 6260 | 6504 | 6186 | 8210 | 8938 | 9222 | 9018 | 9532 | 9224 | 8869 | 6680 | 7249 | 6558 | 6004 | 6736 | 5959 | 5965 | 7606 | 8334 | 9131 | 8097 | 10296 | 9135 | 9562 | Adgra3 |
| Adgrb1 | 1926 | 2764 | 3313 | 4301 | 5987 | 4688 | 4076 | 4686 | 5054 | 6669 | 4830 | 6742 | 5315 | 5435 | 2251 | 1734 | 4194 | 4508 | 5842 | 4107 | 4529 | 7416 | 4257 | 6617 | 7446 | 4731 | 5619 | 5095 | Adgrb1 |
| Adgrb2 | 5932 | 8180 | 8632 | 10713 | 11468 | 10427 | 8605 | 4774 | 6048 | 7438 | 6192 | 6045 | 5845 | 4598 | 6154 | 7018 | 10690 | 10903 | 12603 | 7990 | 9743 | 7581 | 5539 | 6916 | 7108 | 4659 | 4980 | 4397 | Adgrb2 |
| Adgrb3 | 1806 | 1854 | 1808 | 2508 | 3157 | 2929 | 3109 | 3859 | 3350 | 3204 | 3030 | 2887 | 2588 | 3202 | 2238 | 1611 | 2046 | 2786 | 2950 | 3024 | 3282 | 4578 | 3471 | 2854 | 4139 | 2175 | 2834 | 2758 | Adgrb3 |
| Adgrd1 | 8 | 11 | 11 | 7 | 6 | 21 | 7 | 20 | 41 | 29 | 66 | 26 | 26 | 16 | 11 | 12 | 6 | 9 | 14 | 6 | 6 | 27 | 89 | 17 | 39 | 10 | 16 | 28 | Adgrd1 |
| Adgre1 | 471 | 475 | 472 | 404 | 439 | 455 | 323 | 575 | 620 | 440 | 738 | 413 | 525 | 413 | 576 | 545 | 393 | 474 | 329 | 430 | 394 | 595 | 922 | 437 | 474 | 476 | 423 | 427 | Adgre1 |
| Adgre4 | 6 | 2 | 6 | 0 | 2 | 5 | 3 | 6 | 7 | 1 | 8 | 5 | 5 | 9 | 4 | 8 | 0 | 2 | 5 | 0 | 2 | 3 | 8 | 2 | 2 | 2 | 2 | 1 | Adgre4 |
| Adgre5 | 1132 | 1139 | 1157 | 800 | 772 | 929 | 606 | 1203 | 1304 | 1327 | 1877 | 1344 | 1694 | 1228 | 1056 | 1323 | 944 | 877 | 697 | 659 | 659 | 1245 | 1863 | 1416 | 1234 | 1455 | 1506 | 1332 | Adgre5 |
| Adgrf1 | 0 | 1 | 0 | 9 | 2 | 1 | 3 | 5 | 3 | 5 | 11 | 0 | 6 | 0 | 4 | 1 | 2 | 1 | 1 | 1 | 4 | 2 | 1 | 0 | 0 | 0 | 0 | 2 | Adgrf1 |
| Adgrf2 | 35 | 46 | 10 | 44 | 43 | 57 | 39 | 42 | 39 | 14 | 18 | 25 | 28 | 24 | 36 | 50 | 17 | 46 | 27 | 41 | 63 | 13 | 34 | 16 | 40 | 8 | 16 | 23 | Adgrf2 |
| Adgrf3 | 0 | 0 | 0 | 1 | 0 | 1 | 0 | 0 | 1 | 0 | 5 | 0 | 1 | 0 | 0 | 2 | 0 | 0 | 0 | 0 | 3 | 0 | 0 | 0 | 0 | 0 | 0 | 0 | Adgrf3 |
| Adgrf4 | 89 | 73 | 50 | 78 | 100 | 83 | 105 | 54 | 46 | 30 | 48 | 54 | 44 | 33 | 73 | 60 | 67 | 84 | 81 | 89 | 91 | 52 | 59 | 27 | 68 | 28 | 37 | 21 | Adgrf4 |
| Adgrf5 | 2192 | 2037 | 1694 | 1691 | 2251 | 2237 | 1904 | 2622 | 2555 | 2776 | 3346 | 2492 | 2785 | 2569 | 2194 | 2016 | 1683 | 2025 | 1929 | 2297 | 2222 | 2988 | 3278 | 2466 | 2632 | 1969 | 2550 | 2355 | Adgrf5 |

| | | | | | | | | | | | | | | | | | | | | | | | | | | | | | |
|---|---|---|---|---|---|---|---|---|---|---|---|---|---|---|---|---|---|---|---|---|---|---|---|---|---|---|---|---|---|
| Adgrg1 | 5118 | 6560 | 7787 | 7524 | 9803 | 8340 | 8419 | 14842 | 15384 | 18766 | 14160 | 23079 | 19251 | 20783 | 6247 | 5415 | 9115 | 7204 | 10491 | 7386 | 8292 | 21564 | 13414 | 19932 | 21367 | 20443 | 18986 | 20610 | Adgrg1 |
| Adgrg2 | 218 | 160 | 119 | 201 | 336 | 189 | 182 | 583 | 694 | 526 | 683 | 343 | 359 | 377 | 260 | 236 | 149 | 216 | 210 | 198 | 175 | 769 | 530 | 327 | 450 | 226 | 370 | 320 | Adgrg2 |
| Adgrg3 | 193 | 253 | 213 | 185 | 204 | 151 | 159 | 131 | 159 | 175 | 219 | 196 | 296 | 154 | 198 | 287 | 257 | 199 | 186 | 175 | 187 | 192 | 186 | 134 | 173 | 216 | 191 | 165 | Adgrg3 |
| Adgrg4 | 0 | 1 | 0 | 0 | 0 | 0 | 0 | 1 | 0 | 0 | 1 | 1 | 0 | 2 | 0 | 0 | 0 | 0 | 0 | 0 | 0 | 0 | 2 | 0 | 0 | 0 | 0 | 0 | Adgrg4 |
| Adgrg5 | 14 | 11 | 18 | 6 | 11 | 10 | 6 | 15 | 18 | 6 | 19 | 9 | 14 | 12 | 13 | 15 | 2 | 17 | 13 | 13 | 7 | 12 | 13 | 15 | 10 | 21 | 19 | 18 | Adgrg5 |
| Adgrg6 | 989 | 872 | 656 | 569 | 584 | 876 | 253 | 1409 | 1817 | 1329 | 2698 | 1132 | 1683 | 1109 | 1100 | 1062 | 663 | 914 | 337 | 303 | 273 | 1641 | 2177 | 1295 | 1198 | 723 | 1298 | 990 | Adgrg6 |
| Adgrg7 | 0 | 0 | 0 | 0 | 0 | 0 | 0 | 0 | 2 | 0 | 0 | 0 | 0 | 0 | 0 | 0 | 0 | 0 | 0 | 2 | 0 | 0 | 1 | 0 | 2 | 0 | 0 | 0 | Adgrg7 |
| Adgrl1 | 22026 | 27298 | 33710 | 37764 | 36918 | 32634 | 33245 | 19501 | 22673 | 29683 | 27019 | 26426 | 23534 | 25059 | 20276 | 24974 | 37521 | 37243 | 39123 | 26165 | 33738 | 32078 | 20186 | 30291 | 32104 | 22372 | 22048 | 22947 | Adgrl1 |
| Adgrl2 | 13086 | 12558 | 12906 | 13449 | 17290 | 16140 | 15488 | 17215 | 17932 | 20191 | 19705 | 20299 | 20080 | 19987 | 12869 | 11628 | 13254 | 13804 | 15716 | 13925 | 16050 | 19380 | 17204 | 18033 | 20236 | 12994 | 20161 | 20302 | Adgrl2 |
| Adgrl3 | 4591 | 4450 | 3096 | 5177 | 6379 | 6165 | 5744 | 6567 | 6377 | 6030 | 6034 | 4712 | 5101 | 4582 | 4956 | 4445 | 4162 | 5491 | 5421 | 6143 | 6046 | 7294 | 6215 | 4606 | 6169 | 3023 | 4614 | 4599 | Adgrl3 |
| Adgrl4 | 865 | 751 | 861 | 871 | 939 | 1015 | 914 | 1065 | 916 | 844 | 865 | 901 | 875 | 913 | 1030 | 834 | 732 | 963 | 801 | 1064 | 912 | 1081 | 1088 | 830 | 980 | 940 | 930 | 814 | Adgrl4 |
| Adgrv1 | 14366 | 17598 | 8258 | 8507 | 7751 | 11084 | 10211 | 10159 | 7659 | 4731 | 4291 | 6928 | 6108 | 7123 | 17275 | 14299 | 9347 | 8106 | 8679 | 15189 | 11757 | 5396 | 7587 | 4481 | 5399 | 4529 | 9621 | 6903 | Adgrv1 |
| Adh1 | 14 | 17 | 5 | 15 | 16 | 78 | 4 | 43 | 80 | 80 | 100 | 29 | 18 | 37 | 24 | 12 | 8 | 30 | 19 | 9 | 9 | 15 | 133 | 85 | 46 | 19 | 27 | 19 | Adh1 |
| Adh4 | 5 | 1 | 1 | 0 | 0 | 0 | 2 | 0 | 0 | 0 | 1 | 1 | 0 | 0 | 0 | 2 | 0 | 0 | 2 | 0 | 1 | 2 | 0 | 0 | 0 | 0 | 1 | 0 | Adh4 |
| Adh5 | 12505 | 13235 | 11535 | 11381 | 10158 | 10836 | 11824 | 11364 | 10539 | 9566 | 8971 | 10540 | 9679 | 10867 | 13043 | 11799 | 11778 | 10636 | 11974 | 13382 | 11750 | 9052 | 10485 | 10358 | 10043 | 11906 | 11390 | 11472 | Adh5 |
| Adh6a | 2 | 0 | 0 | 0 | 1 | 0 | 0 | 2 | 0 | 3 | 0 | 0 | 0 | 0 | 0 | 0 | 0 | 0 | 0 | 0 | 0 | 0 | 3 | 1 | 0 | 0 | 0 | 0 | Adh6a |
| Adh6b | 15 | 10 | 36 | 2 | 5 | 10 | 11 | 23 | 26 | 11 | 22 | 20 | 27 | 28 | 11 | 18 | 28 | 11 | 18 | 7 | 8 | 20 | 34 | 25 | 20 | 9 | 41 | 24 | Adh6b |
| Adh7 | 13 | 21 | 24 | 53 | 33 | 64 | 16 | 78 | 163 | 150 | 112 | 42 | 25 | 47 | 21 | 21 | 19 | 58 | 26 | 17 | 12 | 55 | 170 | 108 | 69 | 7 | 33 | 18 | Adh7 |
| Adhfe1 | 116 | 124 | 69 | 114 | 129 | 136 | 125 | 115 | 125 | 102 | 120 | 28 | 43 | 36 | 111 | 132 | 67 | 151 | 86 | 101 | 119 | 125 | 116 | 79 | 52 | 46 | 45 | 36 | Adhfe1 |
| Adi1 | 871 | 613 | 708 | 560 | 573 | 675 | 631 | 1396 | 1384 | 1325 | 1249 | 1287 | 1405 | 1396 | 662 | 711 | 590 | 573 | 627 | 575 | 654 | 1151 | 1427 | 1479 | 1292 | 1958 | 1319 | 1525 | Adi1 |
| Adig | 5 | 7 | 8 | 7 | 8 | 5 | 2 | 0 | 4 | 0 | 6 | 0 | 5 | 3 | 4 | 3 | 9 | 11 | 10 | 6 | 4 | 2 | 5 | 2 | 0 | 2 | 10 | 4 | Adig |
| Adipoq | 3 | 3 | 0 | 1 | 4 | 3 | 2 | 5 | 3 | 0 | 1 | 0 | 0 | 4 | 5 | 2 | 5 | 0 | 0 | 0 | 0 | 3 | 9 | 2 | 0 | 2 | 2 | 4 | Adipoq |
| Adipor1 | 6091 | 6643 | 6350 | 6536 | 5645 | 5712 | 5611 | 4726 | 4823 | 4658 | 5199 | 4701 | 5530 | 5058 | 5690 | 6420 | 6351 | 6417 | 5685 | 5416 | 5728 | 4547 | 5192 | 4811 | 4796 | 5043 | 4923 | 5365 | Adipor1 |
| Adipor2 | 10492 | 11980 | 11361 | 12182 | 10321 | 10523 | 10644 | 7424 | 6863 | 7238 | 7182 | 7782 | 7353 | 7744 | 10589 | 10684 | 11333 | 11999 | 11187 | 10186 | 11124 | 7215 | 6996 | 7723 | 7756 | 7645 | 7936 | 8464 | Adipor2 |
| Adk | 4866 | 3723 | 4364 | 3125 | 2819 | 3279 | 3985 | 3010 | 2880 | 2298 | 2446 | 2999 | 2969 | 3583 | 4432 | 4210 | 3505 | 3335 | 3350 | 4212 | 3478 | 1838 | 2687 | 2938 | 2515 | 3824 | 3127 | 3489 | Adk |
| Adm | 98 | 109 | 99 | 85 | 128 | 134 | 115 | 73 | 87 | 70 | 87 | 56 | 52 | 79 | 75 | 121 | 141 | 111 | 118 | 132 | 101 | 72 | 99 | 70 | 114 | 76 | 51 | 81 | Adm |
| Adm2 | 2 | 3 | 0 | 0 | 0 | 0 | 4 | 1 | 2 | 0 | 0 | 4 | 0 | 4 | 0 | 5 | 0 | 4 | 1 | 0 | 0 | 0 | 0 | 1 | 0 | 3 | 3 | 1 | Adm2 |
| Adnp | 225 | 226 | 156 | 157 | 205 | 185 | 127 | 338 | 351 | 269 | 288 | 291 | 299 | 256 | 173 | 181 | 159 | 142 | 194 | 222 | 148 | 247 | 276 | 300 | 337 | 217 | 313 | 233 | Adnp |
| Adnp2 | 4116 | 4156 | 4414 | 4160 | 3800 | 3451 | 3848 | 3251 | 3310 | 3549 | 3125 | 3926 | 3724 | 3704 | 3572 | 3798 | 4117 | 4010 | 4004 | 3693 | 3724 | 3380 | 3088 | 3443 | 3326 | 3912 | 3833 | 3916 | Adnp2 |
| Ado | 0 | 0 | 0 | 0 | 0 | 0 | 0 | 0 | 0 | 0 | 1 | 0 | 0 | 0 | 0 | 0 | 0 | 0 | 0 | 0 | 0 | 0 | 0 | 0 | 0 | 0 | 0 | 0 | Ado |
| Adora1 | 250 | 307 | 213 | 441 | 1012 | 667 | 586 | 391 | 566 | 618 | 601 | 260 | 252 | 237 | 302 | 258 | 227 | 638 | 643 | 524 | 667 | 675 | 518 | 381 | 638 | 222 | 149 | 131 | Adora1 |
| Adora2a | 163 | 221 | 268 | 288 | 313 | 239 | 274 | 287 | 280 | 313 | 413 | 304 | 290 | 319 | 186 | 204 | 294 | 297 | 351 | 217 | 271 | 341 | 340 | 403 | 409 | 392 | 325 | 272 | Adora2a |
| Adora2b | 316 | 265 | 257 | 169 | 171 | 202 | 122 | 160 | 188 | 148 | 205 | 109 | 164 | 139 | 215 | 301 | 209 | 190 | 181 | 176 | 122 | 125 | 260 | 192 | 135 | 206 | 128 | 152 | Adora2b |
| Adora3 | 1 | 6 | 0 | 0 | 7 | 4 | 2 | 7 | 11 | 12 | 6 | 4 | 6 | 5 | 0 | 3 | 3 | 7 | 0 | 1 | 2 | 2 | 9 | 7 | 0 | 6 | 1 | 2 | Adora3 |
| Adpgk | 4151 | 4027 | 4555 | 4036 | 3548 | 3657 | 3463 | 3003 | 3143 | 2998 | 3602 | 3568 | 3839 | 3474 | 4187 | 4469 | 3999 | 4030 | 3634 | 3324 | 3532 | 3108 | 3133 | 3459 | 3224 | 3673 | 3571 | 3525 | Adpgk |
| Adprh | 4929 | 5616 | 6650 | 5599 | 5433 | 4953 | 5068 | 4818 | 5596 | 6039 | 5690 | 6453 | 6256 | 6497 | 4846 | 5203 | 6398 | 5168 | 5674 | 4578 | 5010 | 5618 | 5622 | 6275 | 6122 | 7272 | 5739 | 6867 | Adprh |
| Adprhl1 | 2 | 10 | 1 | 6 | 2 | 9 | 3 | 9 | 18 | 4 | 36 | 3 | 5 | 2 | 8 | 2 | 2 | 0 | 5 | 4 | 1 | 2 | 32 | 2 | 5 | 8 | 3 | 2 | Adprhl1 |
| Adprm | 1215 | 1266 | 1330 | 1249 | 1268 | 1168 | 1265 | 1602 | 1355 | 1391 | 1380 | 1334 | 1458 | 1553 | 1334 | 1125 | 1389 | 1122 | 1281 | 1363 | 1268 | 1420 | 1425 | 1520 | 1351 | 1541 | 1460 | 1511 | Adprm |
| Adprs | 1564 | 1871 | 1912 | 1825 | 1572 | 1695 | 1463 | 2066 | 2472 | 2228 | 2230 | 2386 | 2138 | 2059 | 1491 | 1693 | 1808 | 1751 | 1778 | 1652 | 1525 | 1995 | 2464 | 2424 | 2164 | 2764 | 2085 | 2012 | Adprs |
| Adra1a | 416 | 509 | 210 | 326 | 551 | 655 | 396 | 399 | 317 | 221 | 269 | 273 | 486 | 269 | 500 | 483 | 299 | 292 | 387 | 635 | 508 | 199 | 285 | 133 | 313 | 62 | 352 | 223 | Adra1a |
| Adra1b | 48 | 46 | 56 | 29 | 81 | 52 | 34 | 95 | 112 | 172 | 133 | 100 | 98 | 60 | 57 | 51 | 37 | 55 | 39 | 30 | 46 | 132 | 124 | 109 | 98 | 85 | 70 | 58 | Adra1b |
| Adra1d | 38 | 18 | 60 | 20 | 26 | 18 | 10 | 13 | 21 | 16 | 34 | 14 | 23 | 3 | 16 | 40 | 36 | 25 | 28 | 9 | 12 | 12 | 38 | 8 | 19 | 13 | 14 | 1 | Adra1d |
| Adra2a | 433 | 466 | 541 | 573 | 811 | 618 | 489 | 818 | 839 | 1207 | 1004 | 926 | 854 | 891 | 462 | 491 | 510 | 523 | 586 | 446 | 455 | 1019 | 1052 | 1065 | 930 | 1161 | 843 | 917 | Adra2a |
| Adra2b | 62 | 112 | 114 | 57 | 39 | 41 | 29 | 10 | 18 | 12 | 30 | 70 | 112 | 92 | 113 | 152 | 79 | 30 | 23 | 28 | 27 | 18 | 27 | 4 | 8 | 81 | 58 | 250 | Adra2b |
| Adra2c | 45 | 51 | 31 | 63 | 153 | 89 | 63 | 85 | 76 | 65 | 51 | 35 | 26 | 33 | 50 | 51 | 75 | 88 | 93 | 73 | 76 | 94 | 89 | 80 | 108 | 27 | 17 | 18 | Adra2c |
| Adrb1 | 185 | 188 | 186 | 235 | 206 | 214 | 204 | 66 | 76 | 74 | 81 | 72 | 49 | 60 | 169 | 163 | 192 | 206 | 226 | 188 | 234 | 80 | 92 | 101 | 60 | 111 | 37 | 51 | Adrb1 |
| Adrb2 | 116 | 140 | 152 | 77 | 65 | 106 | 58 | 56 | 106 | 42 | 63 | 54 | 118 | 63 | 102 | 170 | 108 | 110 | 52 | 106 | 71 | 60 | 106 | 51 | 47 | 85 | 45 | 93 | Adrb2 |
| Adrb3 | 70 | 117 | 80 | 46 | 34 | 44 | 35 | 15 | 28 | 14 | 21 | 22 | 33 | 13 | 59 | 105 | 73 | 67 | 41 | 40 | 40 | 30 | 30 | 14 | 15 | 4 | 7 | 13 | Adrb3 |
| Adrm1 | 4184 | 5056 | 5503 | 5118 | 4662 | 4415 | 4132 | 3705 | 4148 | 4415 | 4288 | 5111 | 4538 | 4694 | 3898 | 4251 | 5443 | 4918 | 5398 | 4064 | 4426 | 4253 | 3965 | 4807 | 4670 | 6058 | 4907 | 4991 | Adrm1 |
| Adsl | 3217 | 3720 | 3679 | 3043 | 2453 | 2886 | 2789 | 2882 | 3170 | 2812 | 3024 | 3480 | 3316 | 3629 | 2825 | 3185 | 3110 | 2839 | 2877 | 2837 | 2599 | 2608 | 3325 | 3261 | 2787 | 4335 | 3853 | 3731 | Adsl |
| Adss | 12370 | 9403 | 11066 | 9598 | 10245 | 10723 | 11605 | 9191 | 8339 | 7317 | 7469 | 8149 | 8056 | 8354 | 12775 | 11180 | 9922 | 9702 | 10016 | 11723 | 11026 | 7865 | 8406 | 7397 | 7380 | 8734 | 8870 | 8759 | Adss |
| Adssl1 | 96 | 120 | 87 | 79 | 99 | 68 | 86 | 76 | 107 | 73 | 190 | 51 | 58 | 67 | 103 | 101 | 125 | 108 | 111 | 66 | 79 | 99 | 196 | 81 | 74 | 84 | 53 | 87 | Adssl1 |
| Adtrp | 9 | 8 | 0 | 4 | 5 | 8 | 3 | 9 | 16 | 13 | 21 | 1 | 10 | 5 | 7 | 5 | 3 | 0 | 1 | 0 | 4 | 13 | 24 | 11 | 3 | 8 | 12 | 2 | Adtrp |
| Aebp1 | 2840 | 5413 | 5544 | 4266 | 2351 | 4200 | 963 | 2358 | 3789 | 4340 | 5531 | 4771 | 6482 | 4493 | 3599 | 5674 | 4384 | 4883 | 1419 | 1257 | 1239 | 4814 | 5168 | 4573 | 4326 | 4333 | 5571 | 4475 | Aebp1 |
| Aebp2 | 6981 | 5250 | 9221 | 7991 | 7438 | 7541 | 7561 | 6462 | 5857 | 7001 | 7207 | 8300 | 8101 | 8778 | 6703 | 6408 | 8193 | 7280 | 7647 | 6455 | 7370 | 7192 | 6134 | 7764 | 7539 | 7748 | 9010 | 9334 | Aebp2 |
| Aen | 5790 | 6753 | 6186 | 6249 | 5852 | 5851 | 5604 | 2782 | 2944 | 2881 | 2877 | 3565 | 3145 | 3297 | 5625 | 5808 | 6108 | 6212 | 6385 | 5883 | 6041 | 2720 | 2727 | 3078 | 3195 | 3377 | 3491 | 3561 | Aen |
| AF067063 | 7 | 9 | 24 | 7 | 12 | 13 | 13 | 0 | 0 | 0 | 0 | 1 | 1 | 4 | 7 | 9 | 10 | 8 | 11 | 7 | 8 | 3 | 3 | 0 | 4 | 0 | 0 | 4 | AF067063 |
| AF366264 | 0 | 0 | 0 | 0 | 0 | 0 | 0 | 0 | 0 | 0 | 0 | 0 | 0 | 0 | 0 | 0 | 0 | 0 | 0 | 0 | 1 | 0 | 0 | 0 | 0 | 0 | 0 | 0 | AF366264 |
| Afap1 | 4638 | 5316 | 5365 | 7228 | 8556 | 7176 | 6486 | 6732 | 7077 | 8655 | 8618 | 7911 | 9438 | 7801 | 5856 | 5832 | 6846 | 7556 | 7289 | 5833 | 6870 | 10093 | 7025 | 7701 | 9397 | 5238 | 7499 | 7722 | Afap1 |
| Afap1l1 | 1038 | 1021 | 977 | 769 | 859 | 952 | 765 | 1250 | 1511 | 1233 | 1621 | 1106 | 1065 | 954 | 1122 | 1231 | 815 | 824 | 825 | 868 | 760 | 1071 | 1785 | 1149 | 964 | 1098 | 1160 | 1162 | Afap1l1 |
| Afap1l2 | 565 | 553 | 633 | 460 | 658 | 534 | 363 | 628 | 826 | 717 | 844 | 446 | 769 | 444 | 607 | 746 | 508 | 494 | 395 | 385 | 402 | 830 | 917 | 495 | 576 | 493 | 542 | 477 | Afap1l2 |
| Afdn | 17446 | 16383 | 14739 | 19582 | 21809 | 22758 | 23692 | 20668 | 19074 | 21947 | 19730 | 22931 | 19057 | 23495 | 16672 | 15003 | 16382 | 19391 | 20543 | 20006 | 23429 | 20402 | 17906 | 19590 | 23449 | 14305 | 20632 | 19574 | Afdn |
| Aff1 | 3414 | 3486 | 3714 | 3653 | 3182 | 3461 | 2708 | 1281 | 1556 | 1604 | 1729 | 1661 | 2220 | 1740 | 3203 | 3768 | 3444 | 3661 | 3150 | 2288 | 2924 | 1519 | 1549 | 1534 | 1497 | 1152 | 1798 | 1695 | Aff1 |
| Aff2 | 1849 | 1823 | 531 | 1607 | 2295 | 2475 | 2106 | 2192 | 2624 | 2037 | 2087 | 1872 | 1692 | 1613 | 2023 | 1733 | 1118 | 1766 | 1882 | 2344 | 2417 | 1830 | 2302 | 1244 | 2271 | 418 | 1799 | 1244 | Aff2 |
| Aff3 | 3553 | 3594 | 3826 | 4863 | 5541 | 4487 | 4235 | 5370 | 6133 | 7170 | 6810 | 5960 | 5632 | 5322 | 3676 | 4089 | 4539 | 5019 | 4903 | 3731 | 4635 | 6722 | 5860 | 6627 | 6883 | 5340 | 4998 | 4912 | Aff3 |
| Aff4 | 31041 | 27723 | 18708 | 26651 | 30507 | 35167 | 33814 | 33974 | 30639 | 27313 | 27743 | 29145 | 28183 | 30248 | 31537 | 26799 | 22404 | 27362 | 28454 | 32554 | 34932 | 28069 | 29060 | 23703 | 30727 | 17593 | 31220 | 27510 | Aff4 |
| Afg1l | 209 | 166 | 226 | 207 | 171 | 172 | 225 | 376 | 395 | 331 | 363 | 312 | 276 | 299 | 195 | 188 | 253 | 201 | 218 | 222 | 205 | 368 | 416 | 333 | 294 | 322 | 325 | 288 | Afg1l |
| Afg3l1 | 2899 | 3196 | 3424 | 3391 | 3365 | 3382 | 2932 | 3833 | 4508 | 4933 | 5138 | 5935 | 5544 | 5323 | 2682 | 2848 | 3110 | 3133 | 3535 | 3035 | 3041 | 4376 | 4508 | 5322 | 5573 | 5834 | 6112 | 5638 | Afg3l1 |
| Afg3l2 | 8973 | 9392 | 7659 | 7548 | 6553 | 7176 | 7211 | 4135 | 4662 | 4282 | 4884 | 4127 | 4576 | 4244 | 8466 | 9322 | 7368 | 7637 | 6798 | 7326 | 7229 | 4072 | 4697 | 4075 | 4180 | 4229 | 4172 | 4378 | Afg3l2 |
| Afm | 0 | 3 | 0 | 4 | 0 | 3 | 4 | 11 | 4 | 16 | 7 | 4 | 2 | 3 | 7 | 3 | 5 | 4 | 5 | 0 | 4 | 20 | 4 | 4 | 6 | 1 | 8 | 4 | Afm |
| Afmid | 317 | 381 | 299 | 430 | 263 | 306 | 318 | 83 | 152 | 139 | 149 | 184 | 120 | 158 | 266 | 422 | 344 | 355 | 293 | 247 | 299 | 97 | 137 | 163 | 137 | 154 | 169 | 158 | Afmid |
| Afp | 2 | 129 | 2 | 4 | 118 | 1 | 12 | 4 | 758 | 9 | 5 | 4 | 1 | 13 | 4 | 283 | 0 | 14 | 3 | 22 | 4 | 3 | 166 | 4 | 0 | 14 | 3 | 24 | Afp |
| Aftph | 8709 | 7975 | 6786 | 8076 | 8102 | 8337 | 8929 | 7074 | 5575 | 5048 | 4687 | 5158 | 4907 | 5347 | 9432 | 7949 | 7529 | 8004 | 7833 | 9649 | 8769 | 5990 | 5424 | 4617 | 5374 | 4212 | 5283 | 5199 | Aftph |
| Aga | 349 | 328 | 347 | 248 | 311 | 254 | 292 | 556 | 636 | 502 | 618 | 593 | 662 | 605 | 286 | 323 | 374 | 278 | 298 | 311 | 279 | 540 | 720 | 554 | 513 | 625 | 608 | 564 | Aga |
| Agap1 | 8105 | 9360 | 12189 | 15422 | 17703 | 14244 | 14351 | 9557 | 10111 | 14715 | 12698 | 13023 | 12440 | 12887 | 8678 | 8883 | 14922 | 15654 | 16286 | 11696 | 15273 | 17241 | 10132 | 13643 | 15930 | 9372 | 11721 | 12631 | Agap1 |
| Agap2 | 5580 | 7769 | 8247 | 13582 | 16282 | 12725 | 12366 | 6735 | 7731 | 10688 | 8346 | 7437 | 5896 | 7037 | 5825 | 5587 | 11802 | 13745 | 16440 | 11841 | 13918 | 11762 | 6715 | 9542 | 12544 | 5317 | 6141 | 5980 | Agap2 |
| Agap3 | 6274 | 8283 | 8160 | 9573 | 10539 | 8969 | 8075 | 6367 | 7341 | 8559 | 7649 | 8618 | 6913 | 7487 | 6289 | 6857 | 9391 | 9377 | 11044 | 7934 | 8841 | 8532 | 7016 | 8807 | 9049 | 8401 | 7513 | 7416 | Agap3 |
| Agbl1 | 14 | 6 | 0 | 7 | 11 | 17 | 6 | 12 | 12 | 5 | 14 | 14 | 7 | 12 | 8 | 12 | 6 | 17 | 14 | 18 | 19 | 10 | 19 | 6 | 13 | 7 | 5 | 2 | Agbl1 |
| Agbl2 | 209 | 161 | 126 | 64 | 139 | 137 | 72 | 197 | 194 | 178 | 150 | 159 | 248 | 112 | 189 | 232 | 131 | 142 | 69 | 91 | 90 | 192 | 300 | 132 | 85 | 173 | 135 | 150 | Agbl2 |

| | | | | | | | | | | | | | | | | | | | | | | | | | | | | | |
|---|---|---|---|---|---|---|---|---|---|---|---|---|---|---|---|---|---|---|---|---|---|---|---|---|---|---|---|---|---|
| Agbl3 | 713 | 441 | 484 | 556 | 702 | 697 | 792 | 832 | 852 | 886 | 776 | 876 | 741 | 827 | 724 | 657 | 484 | 634 | 629 | 842 | 777 | 884 | 872 | 797 | 718 | 665 | 782 | 717 | Agbl3 |
| Agbl4 | 108 | 115 | 80 | 111 | 102 | 152 | 142 | 128 | 102 | 56 | 80 | 71 | 129 | 70 | 140 | 118 | 96 | 99 | 99 | 152 | 155 | 105 | 116 | 77 | 68 | 85 | 93 | 101 | Agbl4 |
| Agbl5 | 2834 | 3113 | 3256 | 3324 | 3832 | 3253 | 3390 | 4273 | 4870 | 5393 | 4949 | 5545 | 5202 | 5213 | 2654 | 2709 | 3631 | 3187 | 3837 | 3060 | 3443 | 5384 | 4480 | 5431 | 5792 | 5086 | 4960 | 4995 | Agbl5 |
| Ager | 369 | 455 | 496 | 450 | 491 | 400 | 439 | 316 | 342 | 357 | 332 | 481 | 465 | 405 | 331 | 418 | 512 | 483 | 512 | 394 | 386 | 396 | 273 | 455 | 368 | 586 | 424 | 440 | Ager |
| Agfg1 | 14020 | 11430 | 10286 | 11566 | 12616 | 13287 | 14122 | 11574 | 10159 | 8878 | 8761 | 9190 | 8955 | 9558 | 14652 | 12784 | 10741 | 11755 | 11610 | 14364 | 13626 | 9929 | 9537 | 8450 | 10159 | 7785 | 9178 | 9196 | Agfg1 |
| Agfg2 | 3149 | 4089 | 3742 | 3903 | 2539 | 2991 | 2995 | 1165 | 1290 | 1360 | 1112 | 1685 | 1506 | 1504 | 3003 | 3518 | 3605 | 3548 | 2801 | 2906 | 2881 | 1235 | 1182 | 1481 | 1375 | 1912 | 1680 | 1746 | Agfg2 |
| Aggf1 | 6854 | 7503 | 7979 | 7885 | 6650 | 7268 | 7840 | 5625 | 5324 | 5620 | 5403 | 6650 | 6199 | 6509 | 6940 | 6869 | 7840 | 7574 | 7271 | 7372 | 7258 | 5329 | 5344 | 6274 | 6231 | 6881 | 6720 | 6806 | Aggf1 |
| Agk | 956 | 959 | 956 | 856 | 1100 | 1029 | 1076 | 1769 | 1970 | 1591 | 1817 | 1778 | 1645 | 1735 | 953 | 907 | 951 | 978 | 1139 | 1101 | 932 | 1678 | 1885 | 1709 | 1666 | 1828 | 1807 | 1659 | Agk |
| Agl | 8298 | 7649 | 5692 | 7641 | 7684 | 8280 | 8209 | 6064 | 6034 | 5487 | 6562 | 5867 | 6032 | 5588 | 8327 | 7538 | 6780 | 7401 | 7246 | 8055 | 8242 | 5726 | 6237 | 4847 | 5676 | 4366 | 6020 | 5565 | Agl |
| Agmat | 0 | 1 | 1 | 0 | 1 | 0 | 0 | 0 | 4 | 0 | 0 | 0 | 2 | 0 | 3 | 1 | 0 | 0 | 0 | 0 | 0 | 2 | 5 | 1 | 0 | 0 | 0 | 0 | Agmat |
| Agmo | 17 | 15 | 9 | 15 | 34 | 27 | 10 | 24 | 34 | 35 | 28 | 14 | 23 | 13 | 12 | 21 | 12 | 14 | 13 | 36 | 13 | 27 | 47 | 19 | 23 | 22 | 16 | 21 | Agmo |
| Ago1 | 21176 | 21870 | 24315 | 26991 | 27803 | 27819 | 28927 | 25368 | 22649 | 28174 | 24000 | 40453 | 35578 | 38395 | 21511 | 20034 | 27063 | 26027 | 29418 | 23184 | 28150 | 29731 | 20853 | 31545 | 34313 | 26485 | 39094 | 40282 | Ago1 |
| Ago2 | 11468 | 11710 | 12405 | 14007 | 15252 | 14590 | 14717 | 9879 | 11182 | 12324 | 13977 | 14122 | 12990 | 14160 | 9928 | 12141 | 12617 | 13848 | 13866 | 10728 | 14179 | 11488 | 10411 | 11302 | 12873 | 11114 | 13503 | 13306 | Ago2 |
| Ago3 | 7969 | 7330 | 4836 | 6624 | 6273 | 8120 | 7727 | 5119 | 4694 | 4056 | 4362 | 3976 | 4048 | 4317 | 7770 | 8276 | 5388 | 6834 | 5808 | 7882 | 8744 | 3975 | 4383 | 3363 | 4280 | 2629 | 4408 | 3685 | Ago3 |
| Ago4 | 2744 | 2712 | 2513 | 3292 | 3881 | 3557 | 3658 | 3659 | 3377 | 3796 | 3090 | 3140 | 2829 | 2978 | 2831 | 2594 | 3114 | 3365 | 3620 | 3580 | 3714 | 4219 | 3056 | 3313 | 4070 | 2065 | 2957 | 2605 | Ago4 |
| Agpat1 | 5855 | 8558 | 7930 | 8128 | 7283 | 6993 | 7011 | 4476 | 5156 | 5589 | 5060 | 5996 | 5311 | 5512 | 5785 | 7055 | 8522 | 7936 | 7741 | 6390 | 7185 | 5479 | 4882 | 5875 | 5564 | 6057 | 5231 | 5622 | Agpat1 |
| Agpat2 | 112 | 147 | 188 | 79 | 97 | 146 | 73 | 250 | 303 | 200 | 284 | 193 | 257 | 164 | 145 | 181 | 108 | 122 | 102 | 88 | 71 | 219 | 377 | 230 | 172 | 273 | 229 | 197 | Agpat2 |
| Agpat3 | 2734 | 3220 | 2984 | 3820 | 4835 | 3931 | 4286 | 4840 | 5501 | 5919 | 5692 | 5248 | 4747 | 5674 | 2970 | 2527 | 3453 | 3765 | 4568 | 3844 | 4564 | 5477 | 5466 | 5873 | 6202 | 5038 | 4974 | 5357 | Agpat3 |
| Agpat4 | 1774 | 2476 | 2258 | 2005 | 2042 | 1864 | 1707 | 1877 | 1917 | 2050 | 1877 | 2011 | 2161 | 1852 | 1800 | 2189 | 2110 | 1946 | 2187 | 1996 | 1798 | 1868 | 2093 | 1906 | 2130 | 2244 | 2091 | 2026 | Agpat4 |
| Agpat5 | 5301 | 4959 | 5115 | 6341 | 6037 | 5842 | 6440 | 4803 | 4115 | 4206 | 4167 | 3965 | 3787 | 4156 | 6103 | 5264 | 5604 | 6189 | 6086 | 6439 | 6184 | 4712 | 4177 | 4217 | 4516 | 4131 | 4043 | 3900 | Agpat5 |
| Agps | 12974 | 9244 | 10605 | 8750 | 9205 | 9818 | 9243 | 10360 | 9266 | 8220 | 9195 | 9481 | 10016 | 9404 | 12580 | 12244 | 9398 | 8961 | 8412 | 9070 | 8765 | 8918 | 9652 | 8139 | 8333 | 8114 | 9976 | 9731 | Agps |
| Agr2 | 3 | 4 | 0 | 6 | 1 | 4 | 3 | 7 | 8 | 3 | 9 | 2 | 2 | 1 | 0 | 4 | 8 | 0 | 2 | 2 | 1 | 5 | 12 | 10 | 0 | 4 | 0 | 1 | Agr2 |
| Agr3 | 22 | 6 | 0 | 11 | 0 | 0 | 0 | 46 | 33 | 30 | 46 | 2 | 14 | 4 | 17 | 7 | 2 | 0 | 0 | 0 | 1 | 22 | 41 | 44 | 8 | 64 | 2 | 1 | Agr3 |
| Agrn | 41303 | 53980 | 54794 | 61923 | 62404 | 56823 | 49910 | 39346 | 45374 | 52753 | 47621 | 57315 | 55651 | 48439 | 42288 | 51007 | 62644 | 63301 | 62782 | 45675 | 52707 | 56972 | 41328 | 53333 | 54363 | 38134 | 51276 | 48562 | Agrn |
| Agrp | 6 | 8 | 8 | 5 | 9 | 6 | 2 | 12 | 16 | 16 | 25 | 9 | 8 | 15 | 5 | 12 | 2 | 1 | 4 | 13 | 5 | 18 | 16 | 26 | 17 | 14 | 10 | 12 | Agrp |
| Agt | 1 | 2 | 0 | 4 | 4 | 0 | 1 | 0 | 14 | 8 | 16 | 2 | 14 | 0 | 4 | 4 | 0 | 0 | 4 | 0 | 0 | 8 | 8 | 0 | 5 | 0 | 0 | 1 | Agt |
| Agtpbp1 | 6739 | 7066 | 4327 | 7010 | 9067 | 8295 | 9269 | 7584 | 7420 | 6681 | 7071 | 5775 | 5693 | 5959 | 7710 | 6720 | 5696 | 8041 | 7559 | 8915 | 9275 | 7478 | 6967 | 5363 | 7753 | 3563 | 5753 | 5034 | Agtpbp1 |
| Agtr1a | 384 | 308 | 400 | 232 | 175 | 242 | 180 | 137 | 136 | 185 | 229 | 155 | 269 | 186 | 314 | 399 | 247 | 257 | 169 | 118 | 153 | 154 | 211 | 137 | 108 | 174 | 248 | 174 | Agtr1a |
| Agtr1b | 0 | 4 | 0 | 0 | 11 | 4 | 0 | 0 | 2 | 3 | 24 | 0 | 0 | 0 | 4 | 6 | 2 | 0 | 2 | 1 | 0 | 8 | 6 | 0 | 9 | 2 | 0 | 1 | Agtr1b |
| Agtr2 | 386 | 173 | 101 | 135 | 218 | 374 | 115 | 752 | 2456 | 899 | 4654 | 153 | 607 | 154 | 355 | 333 | 123 | 295 | 298 | 153 | 92 | 1444 | 4223 | 473 | 591 | 269 | 188 | 150 | Agtr2 |
| Agtrap | 2320 | 2496 | 3321 | 2558 | 2008 | 1951 | 1933 | 839 | 968 | 871 | 1118 | 1072 | 1209 | 1140 | 2265 | 2576 | 2765 | 2275 | 2175 | 1844 | 1896 | 805 | 1087 | 974 | 894 | 1436 | 1206 | 1169 | Agtrap |
| Agxt | 0 | 0 | 0 | 0 | 0 | 0 | 0 | 0 | 0 | 0 | 0 | 0 | 0 | 0 | 0 | 0 | 0 | 1 | 0 | 0 | 0 | 0 | 0 | 0 | 0 | 0 | 0 | 0 | Agxt |
| Agxt2 | 0 | 0 | 0 | 0 | 0 | 0 | 1 | 0 | 0 | 0 | 0 | 0 | 0 | 0 | 0 | 0 | 0 | 0 | 0 | 0 | 0 | 0 | 0 | 0 | 0 | 0 | 0 | 0 | Agxt2 |
| Ahctf1 | 14451 | 12816 | 8194 | 9809 | 9578 | 12100 | 11521 | 14097 | 12307 | 10088 | 10936 | 12117 | 11780 | 12314 | 13767 | 12591 | 9009 | 9762 | 9626 | 12542 | 11303 | 9519 | 11690 | 9226 | 10698 | 8228 | 13233 | 11345 | Ahctf1 |
| Ahcy | 7545 | 7565 | 7694 | 4892 | 3918 | 4816 | 4132 | 5525 | 6155 | 4792 | 4881 | 6402 | 6055 | 5829 | 6090 | 7034 | 5990 | 4309 | 5090 | 4925 | 4299 | 4080 | 5891 | 5454 | 5040 | 8273 | 7197 | 6941 | Ahcy |
| Ahcyl1 | 20458 | 17113 | 17979 | 19019 | 19751 | 19503 | 21210 | 19468 | 16796 | 17250 | 16720 | 18616 | 17759 | 19700 | 20376 | 17406 | 18294 | 18901 | 18834 | 19111 | 20788 | 18372 | 17046 | 17637 | 18491 | 17861 | 19500 | 19306 | Ahcyl1 |
| Ahcyl2 | 5881 | 5356 | 5379 | 5293 | 5623 | 5699 | 5839 | 3915 | 3564 | 3800 | 3709 | 3579 | 4009 | 3680 | 5838 | 5583 | 5621 | 5515 | 5496 | 4991 | 5638 | 4365 | 3730 | 3458 | 3460 | 3564 | 3333 | 3761 | Ahcyl2 |
| Ahdc1 | 2709 | 3891 | 4147 | 4560 | 4975 | 3840 | 3302 | 2771 | 3621 | 4370 | 4322 | 3551 | 3251 | 2994 | 2544 | 3568 | 4661 | 4916 | 4848 | 3150 | 3677 | 4684 | 3770 | 4198 | 4448 | 3900 | 3001 | 2924 | Ahdc1 |
| Ahi1 | 10944 | 11471 | 8701 | 13544 | 14557 | 13930 | 15100 | 10343 | 8938 | 9105 | 7268 | 7234 | 5745 | 7632 | 11939 | 9302 | 10364 | 14045 | 14544 | 15000 | 15411 | 11344 | 8336 | 8008 | 9960 | 5657 | 6839 | 6585 | Ahi1 |
| Ahnak | 7931 | 6319 | 4689 | 3348 | 3025 | 6435 | 2272 | 16055 | 20286 | 12795 | 20881 | 9751 | 14247 | 10314 | 7228 | 8049 | 3529 | 4933 | 2424 | 3353 | 2744 | 9969 | 23484 | 11105 | 10446 | 7181 | 12459 | 10439 | Ahnak |
| Ahr | 984 | 916 | 773 | 811 | 700 | 698 | 584 | 850 | 1083 | 776 | 1137 | 463 | 604 | 471 | 938 | 1298 | 578 | 974 | 465 | 616 | 568 | 792 | 1263 | 561 | 493 | 413 | 381 | 436 | Ahr |
| Ahrr | 116 | 83 | 78 | 64 | 130 | 125 | 106 | 303 | 350 | 285 | 263 | 373 | 336 | 333 | 127 | 93 | 95 | 76 | 111 | 119 | 115 | 204 | 265 | 240 | 238 | 300 | 364 | 335 | Ahrr |
| Ahsa1 | 6211 | 6643 | 6738 | 5901 | 5308 | 5038 | 5776 | 6009 | 5718 | 5578 | 5384 | 5734 | 5482 | 6009 | 6128 | 6179 | 6362 | 5647 | 5916 | 6108 | 5546 | 5375 | 5921 | 5880 | 5217 | 6588 | 5767 | 6164 | Ahsa1 |
| Ahsa2 | 7128 | 6286 | 6550 | 7260 | 8362 | 8071 | 9332 | 10761 | 8668 | 9348 | 8732 | 10452 | 8370 | 10499 | 7822 | 6225 | 7599 | 7221 | 8345 | 8923 | 8951 | 10537 | 8512 | 9523 | 10114 | 9580 | 9265 | 9920 | Ahsa2 |
| Ahsg | 1 | 3 | 0 | 0 | 0 | 0 | 2 | 4 | 4 | 0 | 10 | 3 | 0 | 5 | 3 | 2 | 0 | 0 | 0 | 3 | 0 | 0 | 5 | 2 | 2 | 2 | 1 | 4 | Ahsg |
| AI182371 | 35 | 29 | 30 | 44 | 62 | 49 | 46 | 54 | 78 | 81 | 77 | 66 | 77 | 68 | 55 | 39 | 32 | 35 | 58 | 39 | 53 | 77 | 69 | 69 | 57 | 82 | 71 | 68 | AI182371 |
| AI413582 | 705 | 941 | 849 | 1020 | 848 | 780 | 771 | 305 | 335 | 362 | 324 | 278 | 379 | 284 | 781 | 880 | 976 | 928 | 824 | 871 | 806 | 314 | 361 | 327 | 309 | 324 | 281 | 258 | AI413582 |
| AI429214 | 438 | 450 | 399 | 440 | 407 | 337 | 448 | 330 | 297 | 349 | 251 | 275 | 306 | 289 | 468 | 445 | 476 | 371 | 365 | 482 | 446 | 322 | 314 | 279 | 307 | 341 | 272 | 286 | AI429214 |
| AI467606 | 42 | 35 | 37 | 26 | 25 | 45 | 30 | 78 | 73 | 93 | 88 | 53 | 51 | 35 | 36 | 40 | 44 | 41 | 28 | 42 | 38 | 87 | 101 | 46 | 48 | 48 | 49 | 38 | AI467606 |
| AI593442 | 490 | 476 | 531 | 855 | 1414 | 1024 | 1049 | 813 | 724 | 845 | 814 | 405 | 634 | 458 | 704 | 614 | 728 | 1027 | 991 | 1060 | 1212 | 1208 | 732 | 534 | 918 | 263 | 408 | 299 | AI593442 |
| AI597479 | 5241 | 4717 | 4656 | 4728 | 4242 | 4553 | 4960 | 5367 | 4440 | 4048 | 3831 | 4539 | 4175 | 4412 | 5660 | 4699 | 4807 | 4978 | 4459 | 5180 | 4858 | 4906 | 4491 | 4426 | 4328 | 4623 | 4549 | 4552 | AI597479 |
| AI661453 | 190 | 76 | 91 | 71 | 65 | 81 | 25 | 193 | 294 | 212 | 261 | 108 | 108 | 110 | 98 | 149 | 76 | 91 | 76 | 50 | 42 | 112 | 332 | 207 | 119 | 157 | 109 | 144 | AI661453 |
| AI837181 | 864 | 1133 | 1214 | 977 | 1173 | 1020 | 891 | 1502 | 1879 | 1832 | 1920 | 1760 | 1844 | 1639 | 857 | 924 | 1145 | 921 | 1247 | 865 | 1044 | 1584 | 2046 | 1910 | 1816 | 1994 | 1799 | 1832 | AI837181 |
| AI854703 | 255 | 374 | 327 | 454 | 695 | 472 | 488 | 363 | 470 | 520 | 333 | 222 | 238 | 196 | 323 | 338 | 533 | 536 | 567 | 441 | 451 | 592 | 425 | 286 | 408 | 188 | 175 | 171 | AI854703 |
| AI987944 | 1323 | 1126 | 1386 | 1211 | 995 | 1150 | 1138 | 717 | 624 | 655 | 682 | 802 | 762 | 672 | 1489 | 1502 | 1202 | 1209 | 969 | 1361 | 1079 | 690 | 770 | 623 | 694 | 754 | 646 | 752 | AI987944 |
| Aicda | 14 | 8 | 16 | 13 | 22 | 2 | 9 | 21 | 17 | 19 | 17 | 16 | 18 | 13 | 13 | 7 | 27 | 6 | 12 | 27 | 10 | 17 | 20 | 12 | 15 | 9 | 5 | 11 | Aicda |
| Aida | 10062 | 8639 | 9148 | 8452 | 7885 | 8126 | 8812 | 6222 | 5843 | 5769 | 6173 | 6217 | 6561 | 6294 | 10261 | 9686 | 8843 | 8412 | 8082 | 8633 | 8595 | 5793 | 6176 | 5664 | 5452 | 6397 | 6544 | 6913 | Aida |
| Aif1 | 98 | 88 | 109 | 96 | 109 | 93 | 75 | 126 | 123 | 168 | 186 | 141 | 117 | 132 | 104 | 134 | 90 | 110 | 63 | 109 | 93 | 110 | 144 | 121 | 95 | 111 | 117 | 107 | Aif1 |
| Aif1l | 4406 | 4121 | 4002 | 3332 | 3229 | 3797 | 3738 | 4078 | 4135 | 3881 | 3626 | 4728 | 4110 | 4849 | 4039 | 3696 | 3611 | 3219 | 3691 | 3613 | 3780 | 3090 | 3164 | 4448 | 4020 | 5792 | 4365 | 4781 | Aif1l |
| Aifm1 | 6552 | 6271 | 7522 | 6221 | 5074 | 5343 | 5993 | 3979 | 3559 | 3892 | 3787 | 4215 | 4082 | 4514 | 6253 | 6259 | 6449 | 5760 | 5333 | 6108 | 5404 | 3210 | 3478 | 4138 | 3814 | 5015 | 4341 | 4432 | Aifm1 |
| Aifm2 | 1188 | 1272 | 1252 | 1011 | 904 | 954 | 689 | 380 | 495 | 342 | 518 | 421 | 440 | 351 | 973 | 1398 | 1035 | 1080 | 786 | 813 | 738 | 463 | 577 | 450 | 373 | 499 | 467 | 420 | Aifm2 |
| Aifm3 | 368 | 452 | 506 | 705 | 678 | 663 | 692 | 733 | 766 | 893 | 705 | 896 | 765 | 866 | 393 | 438 | 535 | 609 | 754 | 627 | 645 | 1026 | 830 | 1043 | 1139 | 883 | 801 | 695 | Aifm3 |
| Aig1 | 997 | 996 | 851 | 989 | 1206 | 1073 | 1086 | 1070 | 1136 | 1016 | 1055 | 807 | 1008 | 909 | 1226 | 998 | 1104 | 1029 | 1093 | 1067 | 1130 | 1196 | 1159 | 988 | 1055 | 867 | 801 | 854 | Aig1 |
| Aim2 | 884 | 863 | 1017 | 887 | 758 | 1008 | 1031 | 1286 | 1128 | 1202 | 876 | 1893 | 1334 | 1969 | 985 | 745 | 953 | 755 | 873 | 876 | 920 | 1141 | 915 | 1369 | 1433 | 2031 | 2062 | 1989 | Aim2 |
| Aimp1 | 4711 | 4726 | 4897 | 4228 | 3535 | 3484 | 4471 | 3968 | 3834 | 3405 | 3447 | 3785 | 3534 | 4222 | 4800 | 4694 | 4327 | 3987 | 4102 | 4747 | 4115 | 3071 | 3822 | 3562 | 3539 | 4532 | 4054 | 4436 | Aimp1 |
| Aimp2 | 1673 | 2022 | 1633 | 1485 | 1291 | 1331 | 1405 | 1788 | 2033 | 1966 | 1709 | 1855 | 1945 | 2071 | 1515 | 1574 | 1612 | 1436 | 1465 | 1509 | 1343 | 1679 | 1914 | 1943 | 1816 | 2212 | 2133 | 2097 | Aimp2 |
| Aip | 0 | 0 | 0 | 0 | 0 | 0 | 0 | 0 | 0 | 0 | 0 | 0 | 0 | 0 | 0 | 0 | 0 | 0 | 0 | 0 | 0 | 0 | 0 | 0 | 0 | 0 | 0 | 1 | Aip |
| Aipl1 | 42 | 43 | 36 | 31 | 47 | 58 | 44 | 32 | 34 | 29 | 30 | 47 | 32 | 27 | 22 | 33 | 51 | 59 | 37 | 28 | 28 | 42 | 24 | 39 | 37 | 24 | 30 | 52 | Aipl1 |
| Aire | 14 | 18 | 21 | 12 | 13 | 24 | 12 | 29 | 39 | 65 | 62 | 47 | 36 | 19 | 13 | 12 | 16 | 14 | 12 | 2 | 15 | 32 | 36 | 44 | 33 | 53 | 33 | 39 | Aire |
| Ajap1 | 329 | 503 | 517 | 605 | 885 | 653 | 631 | 567 | 534 | 822 | 766 | 546 | 474 | 502 | 496 | 432 | 672 | 544 | 708 | 699 | 639 | 715 | 687 | 532 | 612 | 461 | 440 | 470 | Ajap1 |
| Ajm1 | 216 | 308 | 249 | 535 | 883 | 590 | 542 | 376 | 477 | 610 | 492 | 252 | 303 | 258 | 248 | 267 | 399 | 615 | 700 | 527 | 568 | 710 | 486 | 439 | 608 | 176 | 249 | 205 | Ajm1 |
| Ajuba | 3959 | 4784 | 5387 | 4467 | 3107 | 3624 | 3207 | 2854 | 3354 | 2810 | 3038 | 3703 | 3699 | 3293 | 3621 | 4089 | 4649 | 3890 | 3822 | 3266 | 3092 | 2144 | 3256 | 3300 | 2606 | 4697 | 3885 | 3996 | Ajuba |
| Ak1 | 799 | 910 | 1000 | 1086 | 1352 | 1104 | 1152 | 1393 | 1768 | 1681 | 1903 | 1360 | 1348 | 1341 | 891 | 949 | 1034 | 1192 | 1248 | 1317 | 1156 | 1729 | 2061 | 1637 | 1905 | 1302 | 1188 | 1186 | Ak1 |
| AK157302 | 89 | 61 | 68 | 72 | 49 | 72 | 44 | 57 | 39 | 45 | 38 | 34 | 40 | 42 | 117 | 61 | 79 | 79 | 67 | 73 | 62 | 48 | 40 | 51 | 45 | 39 | 41 | 52 | AK157302 |
| Ak2 | 8428 | 8447 | 9899 | 8626 | 6730 | 7062 | 8641 | 4952 | 5311 | 5054 | 5522 | 6720 | 5963 | 7632 | 7400 | 7790 | 9143 | 7926 | 7991 | 7349 | 8460 | 4231 | 5185 | 6642 | 5503 | 8017 | 6800 | 7837 | Ak2 |
| Ak3 | 4570 | 3932 | 4014 | 2813 | 3499 | 3651 | 3620 | 5412 | 4982 | 4254 | 4592 | 4672 | 4808 | 4632 | 4597 | 3926 | 3418 | 3175 | 3702 | 3730 | 3576 | 5073 | 5306 | 4524 | 4545 | 5161 | 4940 | 5522 | Ak3 |
| Ak4 | 2173 | 1782 | 2247 | 1523 | 1501 | 1759 | 1568 | 1845 | 1600 | 1703 | 1773 | 1706 | 1765 | 2058 | 1875 | 1794 | 2044 | 1658 | 1600 | 1750 | 1448 | 1343 | 1886 | 1806 | 1454 | 2011 | 2138 | 2238 | Ak4 |
| Ak5 | 110 | 111 | 82 | 97 | 189 | 157 | 117 | 198 | 211 | 222 | 249 | 100 | 124 | 134 | 154 | 165 | 73 | 160 | 134 | 89 | 118 | 259 | 302 | 160 | 225 | 125 | 102 | 141 | Ak5 |
| Ak6 | 1637 | 1744 | 1855 | 1690 | 1414 | 1383 | 2008 | 1189 | 1046 | 1073 | 950 | 1115 | 1026 | 1439 | 1817 | 1606 | 1933 | 1650 | 1576 | 1868 | 1720 | 964 | 967 | 1255 | 1016 | 1355 | 1230 | 1429 | Ak6 |

| | | | | | | | | | | | | | | | | | | | | | | | | | | | | | |
|---|---|---|---|---|---|---|---|---|---|---|---|---|---|---|---|---|---|---|---|---|---|---|---|---|---|---|---|---|---|
| Ak7 | 231 | 343 | 210 | 381 | 457 | 406 | 348 | 264 | 261 | 297 | 204 | 345 | 381 | 165 | 305 | 446 | 284 | 360 | 323 | 356 | 375 | 399 | 441 | 194 | 250 | 152 | 200 | 159 | Ak7 |
| Ak8 | 39 | 46 | 24 | 41 | 37 | 36 | 36 | 51 | 59 | 46 | 43 | 47 | 56 | 30 | 54 | 64 | 39 | 25 | 39 | 39 | 57 | 53 | 75 | 49 | 35 | 45 | 35 | 36 | Ak8 |
| Ak9 | 54 | 68 | 41 | 36 | 71 | 65 | 40 | 114 | 82 | 70 | 63 | 98 | 134 | 42 | 58 | 83 | 28 | 55 | 43 | 65 | 54 | 89 | 144 | 56 | 51 | 43 | 55 | 33 | Ak9 |
| Akain1 | 15 | 16 | 34 | 41 | 36 | 21 | 31 | 15 | 9 | 6 | 15 | 8 | 5 | 3 | 24 | 31 | 23 | 34 | 35 | 29 | 36 | 15 | 16 | 5 | 7 | 2 | 1 | 1 | Akain1 |
| Akap1 | 5707 | 6632 | 6315 | 7522 | 6958 | 6893 | 6673 | 3138 | 3591 | 4132 | 3815 | 3981 | 3743 | 3901 | 5851 | 5849 | 6657 | 7471 | 7267 | 6126 | 6868 | 4044 | 3429 | 4193 | 4335 | 3625 | 3946 | 3858 | Akap1 |
| Akap10 | 1615 | 1340 | 1187 | 1353 | 1430 | 1531 | 1576 | 2447 | 2268 | 2203 | 2320 | 2149 | 1934 | 2121 | 1538 | 1421 | 1292 | 1297 | 1283 | 1477 | 1590 | 2003 | 2333 | 1977 | 2246 | 1666 | 2220 | 2071 | Akap10 |
| Akap11 | 20767 | 18289 | 10818 | 16221 | 17722 | 20731 | 20496 | 21366 | 18465 | 14467 | 15368 | 14831 | 14632 | 15604 | 21441 | 18806 | 12660 | 15483 | 15610 | 21802 | 20598 | 15321 | 17553 | 11485 | 16707 | 8566 | 15657 | 12977 | Akap11 |
| Akap12 | 19210 | 23437 | 21992 | 16078 | 12291 | 18954 | 7257 | 15536 | 18458 | 16036 | 21144 | 18252 | 24913 | 18227 | 24571 | 30626 | 15052 | 20164 | 8156 | 9888 | 7973 | 17981 | 24601 | 16493 | 16247 | 10931 | 20811 | 18112 | Akap12 |
| Akap13 | 6953 | 7315 | 4431 | 6865 | 7337 | 7682 | 7411 | 4956 | 5605 | 5137 | 6190 | 5300 | 5997 | 5282 | 6844 | 6977 | 5485 | 6804 | 6823 | 6467 | 8192 | 4166 | 5700 | 4119 | 4847 | 3212 | 5772 | 4789 | Akap13 |
| Akap14 | 5 | 10 | 2 | 13 | 13 | 3 | 9 | 11 | 13 | 11 | 8 | 14 | 23 | 6 | 4 | 6 | 1 | 2 | 9 | 6 | 6 | 20 | 26 | 2 | 14 | 8 | 3 | 8 | Akap14 |
| Akap17b | 2646 | 2515 | 2664 | 2998 | 3084 | 3281 | 3217 | 2352 | 1863 | 2231 | 1986 | 1947 | 2302 | 2108 | 2897 | 2669 | 2861 | 2797 | 2825 | 3034 | 3118 | 2550 | 1823 | 2052 | 2102 | 1526 | 2074 | 2032 | Akap17b |
| Akap3 | 22 | 14 | 8 | 19 | 16 | 22 | 21 | 12 | 12 | 17 | 15 | 20 | 19 | 13 | 22 | 22 | 13 | 8 | 21 | 20 | 14 | 12 | 14 | 2 | 12 | 9 | 22 | 7 | Akap3 |
| Akap4 | 0 | 2 | 0 | 0 | 0 | 0 | 0 | 0 | 1 | 0 | 0 | 0 | 0 | 0 | 0 | 0 | 0 | 0 | 0 | 0 | 0 | 2 | 0 | 0 | 0 | 0 | 0 | 0 | Akap4 |
| Akap5 | 1468 | 1550 | 1462 | 1907 | 2207 | 2065 | 2016 | 1264 | 1181 | 1229 | 1162 | 1120 | 1098 | 1220 | 1669 | 1493 | 1641 | 1755 | 1882 | 2025 | 2133 | 1527 | 1131 | 1143 | 1406 | 882 | 1132 | 1031 | Akap5 |
| Akap6 | 11284 | 13068 | 7510 | 12344 | 15514 | 14913 | 14698 | 15091 | 12505 | 11671 | 11635 | 10224 | 13013 | 11274 | 15813 | 12777 | 10871 | 11786 | 12599 | 15816 | 15313 | 13936 | 12062 | 8505 | 12758 | 5758 | 11767 | 10295 | Akap6 |
| Akap7 | 709 | 475 | 680 | 630 | 849 | 658 | 591 | 1436 | 1216 | 1207 | 1442 | 1184 | 1340 | 1305 | 723 | 695 | 541 | 584 | 553 | 607 | 600 | 1591 | 1337 | 1154 | 1269 | 1474 | 1201 | 1276 | Akap7 |
| Akap8 | 13625 | 13687 | 15320 | 15979 | 16376 | 15832 | 17118 | 18520 | 17206 | 19461 | 18596 | 20770 | 19605 | 22207 | 13448 | 12600 | 15712 | 16140 | 16480 | 15411 | 16613 | 19061 | 15827 | 20162 | 20656 | 19808 | 20798 | 20810 | Akap8 |
| Akap8l | 8298 | 10523 | 9434 | 12179 | 11112 | 10327 | 10922 | 5886 | 6122 | 7467 | 6748 | 6811 | 6278 | 6315 | 8694 | 9532 | 11208 | 12120 | 11706 | 9549 | 11349 | 7782 | 5697 | 7204 | 7460 | 5565 | 6405 | 6022 | Akap8l |
| Akap9 | 11708 | 10535 | 6198 | 9547 | 14017 | 16327 | 15094 | 35960 | 30764 | 26260 | 24642 | 24050 | 23161 | 26949 | 13092 | 10386 | 7934 | 10135 | 12043 | 18534 | 15867 | 23200 | 27606 | 19763 | 30301 | 13456 | 27674 | 20515 | Akap9 |
| Akip1 | 361 | 308 | 356 | 270 | 246 | 345 | 303 | 352 | 399 | 352 | 449 | 366 | 463 | 379 | 332 | 251 | 302 | 249 | 244 | 259 | 217 | 312 | 466 | 346 | 279 | 467 | 442 | 455 | Akip1 |
| Akirin1 | 10101 | 9272 | 9149 | 9491 | 8057 | 8235 | 9404 | 6923 | 5361 | 5134 | 4763 | 5868 | 5799 | 6270 | 10789 | 9517 | 9170 | 9037 | 8097 | 9892 | 8560 | 6246 | 5575 | 5581 | 5606 | 6419 | 6012 | 6788 | Akirin1 |
| Akirin2 | 6439 | 5174 | 6550 | 6642 | 6699 | 6098 | 6773 | 8570 | 6914 | 7671 | 6835 | 7838 | 6870 | 8022 | 6697 | 5136 | 6614 | 6299 | 7106 | 6600 | 7024 | 8467 | 7013 | 8092 | 8139 | 8732 | 7280 | 7956 | Akirin2 |
| Akna | 2578 | 3549 | 4071 | 3963 | 3679 | 3255 | 3286 | 1778 | 2178 | 2565 | 2315 | 4173 | 2282 | 3402 | 2370 | 2763 | 4352 | 3893 | 4089 | 2574 | 3206 | 2259 | 1730 | 3646 | 2988 | 4125 | 2994 | 3064 | Akna |
| Aknad1 | 12 | 8 | 8 | 7 | 1 | 4 | 0 | 29 | 37 | 12 | 186 | 4 | 19 | 6 | 5 | 17 | 0 | 1 | 0 | 3 | 3 | 25 | 129 | 16 | 2 | 11 | 16 | 2 | Aknad1 |
| Akp3 | 2 | 2 | 0 | 2 | 3 | 2 | 2 | 0 | 0 | 0 | 2 | 0 | 0 | 0 | 0 | 7 | 8 | 13 | 5 | 7 | 4 | 0 | 1 | 0 | 0 | 0 | 0 | 0 | Akp3 |
| Akr1a1 | 21534 | 22709 | 18138 | 18686 | 18623 | 19068 | 19750 | 20402 | 22177 | 19438 | 19261 | 18418 | 18764 | 18867 | 21228 | 20067 | 18056 | 19052 | 20437 | 20905 | 20235 | 18188 | 23014 | 19342 | 19318 | 22657 | 19029 | 20144 | Akr1a1 |
| Akr1b10 | 917 | 884 | 740 | 600 | 569 | 626 | 512 | 409 | 367 | 300 | 358 | 462 | 451 | 368 | 847 | 897 | 725 | 578 | 548 | 642 | 550 | 346 | 462 | 295 | 283 | 409 | 485 | 535 | Akr1b10 |
| Akr1b3 | 5319 | 6247 | 5404 | 4162 | 3395 | 3760 | 3358 | 3132 | 3624 | 2573 | 2926 | 3310 | 3464 | 3118 | 5093 | 6092 | 4758 | 3657 | 4054 | 4685 | 3341 | 2645 | 3625 | 2835 | 2711 | 4173 | 4064 | 3527 | Akr1b3 |
| Akr1b7 | 2 | 0 | 1 | 0 | 0 | 4 | 4 | 2 | 7 | 1 | 0 | 0 | 0 | 0 | 1 | 3 | 0 | 2 | 2 | 0 | 1 | 0 | 5 | 0 | 0 | 4 | 0 | 1 | Akr1b7 |
| Akr1b8 | 706 | 872 | 808 | 396 | 351 | 432 | 339 | 579 | 476 | 379 | 491 | 658 | 782 | 653 | 893 | 949 | 689 | 410 | 359 | 499 | 263 | 488 | 568 | 446 | 468 | 943 | 916 | 932 | Akr1b8 |
| Akr1c12 | 9 | 2 | 6 | 0 | 4 | 3 | 4 | 12 | 8 | 5 | 10 | 1 | 12 | 10 | 3 | 6 | 9 | 4 | 4 | 1 | 4 | 3 | 6 | 6 | 11 | 6 | 10 | 0 | Akr1c12 |
| Akr1c13 | 22 | 16 | 11 | 22 | 74 | 50 | 31 | 76 | 77 | 78 | 79 | 47 | 41 | 32 | 28 | 29 | 20 | 33 | 41 | 41 | 41 | 75 | 89 | 48 | 46 | 57 | 29 | 33 | Akr1c13 |
| Akr1c14 | 43 | 44 | 79 | 69 | 101 | 80 | 76 | 48 | 72 | 54 | 72 | 38 | 34 | 26 | 79 | 60 | 59 | 66 | 65 | 105 | 68 | 85 | 115 | 60 | 54 | 57 | 29 | 45 | Akr1c14 |
| Akr1c18 | 0 | 0 | 0 | 0 | 2 | 0 | 2 | 0 | 1 | 3 | 0 | 0 | 0 | 0 | 0 | 0 | 0 | 2 | 0 | 0 | 0 | 0 | 3 | 0 | 0 | 0 | 0 | 0 | Akr1c18 |
| Akr1c19 | 20 | 18 | 8 | 26 | 51 | 66 | 58 | 154 | 171 | 122 | 122 | 61 | 71 | 55 | 26 | 23 | 32 | 41 | 65 | 63 | 62 | 117 | 160 | 74 | 101 | 13 | 53 | 44 | Akr1c19 |
| Akr1c21 | 0 | 2 | 0 | 1 | 0 | 0 | 0 | 0 | 1 | 0 | 0 | 0 | 0 | 0 | 0 | 0 | 0 | 0 | 0 | 0 | 0 | 0 | 1 | 0 | 0 | 0 | 0 | 0 | Akr1c21 |
| Akr1cl | 8 | 8 | 17 | 6 | 2 | 6 | 9 | 6 | 3 | 0 | 2 | 20 | 11 | 7 | 16 | 12 | 7 | 6 | 2 | 7 | 7 | 5 | 5 | 3 | 10 | 11 | 9 | 20 | Akr1cl |
| Akr1d1 | 0 | 0 | 0 | 0 | 0 | 0 | 0 | 0 | 2 | 0 | 0 | 0 | 1 | 0 | 0 | 0 | 0 | 0 | 0 | 0 | 0 | 0 | 0 | 0 | 0 | 0 | 0 | 0 | Akr1d1 |
| Akr1e1 | 2521 | 2465 | 2654 | 2292 | 2324 | 2387 | 2335 | 2886 | 2456 | 2158 | 2132 | 2367 | 2793 | 2369 | 3270 | 2842 | 2596 | 2353 | 2236 | 2828 | 2324 | 2568 | 2808 | 2344 | 2309 | 2596 | 2838 | 2579 | Akr1e1 |
| Akr7a5 | 1939 | 2364 | 1889 | 1767 | 1423 | 1580 | 1484 | 1055 | 1239 | 1123 | 1181 | 1203 | 1328 | 1103 | 1766 | 1910 | 1946 | 1763 | 1680 | 1666 | 1560 | 917 | 1308 | 1294 | 1024 | 1565 | 1284 | 1284 | Akr7a5 |
| Akt1 | 11130 | 12766 | 13259 | 13612 | 14912 | 14017 | 12635 | 15591 | 18217 | 19763 | 18835 | 19943 | 18160 | 18337 | 10730 | 11567 | 14236 | 14187 | 15503 | 11798 | 13851 | 17483 | 18261 | 20040 | 20136 | 20901 | 18245 | 18509 | Akt1 |
| Akt1s1 | 1400 | 1647 | 1655 | 1558 | 1378 | 1280 | 1233 | 2864 | 3159 | 2930 | 2985 | 3080 | 2860 | 2903 | 1403 | 1372 | 1815 | 1428 | 1558 | 1334 | 1324 | 2780 | 3271 | 3516 | 3037 | 3668 | 3134 | 3072 | Akt1s1 |
| Akt2 | 3066 | 3587 | 3142 | 3357 | 3169 | 3342 | 2911 | 5505 | 7049 | 6744 | 7024 | 6900 | 6565 | 5982 | 2774 | 3250 | 3333 | 3219 | 3702 | 3033 | 3089 | 5944 | 7103 | 6857 | 6866 | 7205 | 6398 | 6259 | Akt2 |
| Akt3 | 10883 | 10316 | 8917 | 11236 | 13700 | 14120 | 14852 | 17824 | 15464 | 15151 | 13469 | 16026 | 14413 | 16644 | 12384 | 9650 | 10755 | 11683 | 12846 | 14655 | 15111 | 16188 | 14641 | 14001 | 17845 | 9931 | 16192 | 15040 | Akt3 |
| Aktip | 6443 | 6847 | 6457 | 6718 | 5938 | 5781 | 6519 | 5266 | 3986 | 4219 | 3742 | 4340 | 4065 | 4787 | 7007 | 6362 | 6647 | 6421 | 6136 | 7321 | 6349 | 4451 | 4180 | 4408 | 4732 | 4504 | 4581 | 4664 | Aktip |
| Alad | 1866 | 2183 | 2003 | 1769 | 1580 | 1765 | 1609 | 1098 | 1441 | 1398 | 1426 | 1624 | 1627 | 1500 | 1652 | 2111 | 1931 | 1778 | 1848 | 1426 | 1524 | 1158 | 1363 | 1554 | 1387 | 1834 | 1611 | 1747 | Alad |
| Alas1 | 2175 | 2484 | 2250 | 2199 | 2246 | 2131 | 1912 | 2236 | 2511 | 2399 | 2416 | 2382 | 2422 | 2257 | 2086 | 2367 | 2418 | 2188 | 2326 | 2151 | 2126 | 2257 | 2482 | 2425 | 2343 | 2583 | 2341 | 2392 | Alas1 |
| Alas2 | 1403 | 2496 | 1687 | 1291 | 794 | 1271 | 1646 | 1481 | 2897 | 1453 | 1222 | 1233 | 3882 | 1894 | 1733 | 1945 | 1130 | 1272 | 1615 | 2136 | 1073 | 926 | 3008 | 985 | 1581 | 1441 | 1599 | 3859 | Alas2 |
| Alb | 0 | 1 | 4 | 0 | 6 | 1 | 2 | 60 | 63 | 0 | 9 | 0 | 0 | 4 | 11 | 9 | 0 | 0 | 3 | 3 | 0 | 2 | 39 | 8 | 7 | 0 | 0 | 2 | Alb |
| Albfm1 | 6 | 3 | 6 | 3 | 2 | 10 | 1 | 1 | 2 | 1 | 1 | 4 | 0 | 2 | 4 | 3 | 9 | 2 | 5 | 0 | 3 | 0 | 1 | 3 | 3 | 0 | 6 | 0 | Albfm1 |
| Alcam | 17355 | 15494 | 18738 | 13467 | 10817 | 14449 | 10260 | 11495 | 12952 | 13391 | 13816 | 14684 | 21455 | 15701 | 19998 | 23809 | 12848 | 14990 | 8698 | 9173 | 9553 | 13251 | 13562 | 13477 | 12685 | 12452 | 16178 | 16676 | Alcam |
| Aldh16a1 | 1068 | 1313 | 1284 | 1103 | 974 | 1064 | 848 | 1453 | 1819 | 1703 | 1637 | 2139 | 2074 | 1762 | 998 | 1226 | 1355 | 930 | 1126 | 911 | 916 | 1318 | 1720 | 1935 | 1749 | 2430 | 2117 | 1902 | Aldh16a1 |
| Aldh18a1 | 11627 | 13385 | 12491 | 12592 | 11016 | 11689 | 12022 | 5979 | 7052 | 7423 | 7112 | 8937 | 8166 | 8956 | 10640 | 11294 | 12865 | 12003 | 12927 | 10502 | 11903 | 6560 | 6578 | 8067 | 7903 | 9093 | 8795 | 9509 | Aldh18a1 |
| Aldh1a1 | 75 | 62 | 37 | 24 | 52 | 40 | 30 | 51 | 82 | 60 | 54 | 31 | 90 | 31 | 62 | 74 | 32 | 37 | 62 | 44 | 41 | 75 | 113 | 56 | 47 | 21 | 17 | 33 | Aldh1a1 |
| Aldh1a2 | 1613 | 549 | 145 | 654 | 719 | 302 | 280 | 1257 | 1574 | 1088 | 2300 | 319 | 640 | 403 | 836 | 1102 | 285 | 678 | 191 | 372 | 374 | 1242 | 1862 | 1157 | 532 | 1242 | 268 | 271 | Aldh1a2 |
| Aldh1a3 | 1277 | 375 | 178 | 420 | 256 | 254 | 293 | 1006 | 988 | 1134 | 1160 | 252 | 346 | 277 | 472 | 625 | 221 | 249 | 245 | 89 | 256 | 475 | 1045 | 1211 | 289 | 1803 | 101 | 138 | Aldh1a3 |
| Aldh1a7 | 172 | 136 | 100 | 39 | 92 | 51 | 10 | 127 | 122 | 116 | 164 | 27 | 107 | 31 | 132 | 229 | 50 | 42 | 11 | 22 | 12 | 72 | 240 | 65 | 52 | 53 | 42 | 51 | Aldh1a7 |
| Aldh1b1 | 671 | 917 | 713 | 879 | 1100 | 810 | 889 | 390 | 485 | 443 | 454 | 399 | 260 | 392 | 723 | 752 | 841 | 1034 | 1026 | 958 | 929 | 478 | 458 | 445 | 502 | 381 | 332 | 345 | Aldh1b1 |
| Aldh1l1 | 328 | 647 | 425 | 822 | 1737 | 1070 | 790 | 1277 | 1200 | 1224 | 1242 | 613 | 1043 | 549 | 702 | 566 | 689 | 1252 | 1319 | 1011 | 1292 | 1566 | 1340 | 854 | 1347 | 214 | 725 | 529 | Aldh1l1 |
| Aldh1l2 | 2315 | 2236 | 1435 | 2144 | 2817 | 2486 | 2870 | 3105 | 2800 | 2672 | 2511 | 2561 | 2266 | 2770 | 2446 | 2001 | 2009 | 2078 | 2449 | 2659 | 2806 | 3273 | 2494 | 2259 | 3266 | 1489 | 2510 | 2332 | Aldh1l2 |
| Aldh2 | 7135 | 7978 | 8770 | 7908 | 6570 | 6539 | 5611 | 4264 | 5766 | 5842 | 6447 | 5017 | 5370 | 4832 | 6195 | 8422 | 8499 | 7582 | 7419 | 5413 | 6100 | 4639 | 6099 | 5865 | 4829 | 7104 | 4732 | 4970 | Aldh2 |
| Aldh3a1 | 1 | 0 | 1 | 2 | 0 | 2 | 0 | 0 | 5 | 3 | 3 | 0 | 1 | 0 | 4 | 0 | 0 | 8 | 1 | 2 | 0 | 0 | 5 | 2 | 5 | 2 | 0 | 0 | Aldh3a1 |
| Aldh3a2 | 1692 | 1596 | 1604 | 1531 | 1705 | 1726 | 1655 | 2357 | 2409 | 2195 | 2233 | 2151 | 2236 | 2228 | 1834 | 1594 | 1530 | 1493 | 1650 | 1790 | 1610 | 2274 | 2593 | 2176 | 2158 | 2038 | 2210 | 2305 | Aldh3a2 |
| Aldh3b1 | 17 | 26 | 26 | 23 | 51 | 48 | 42 | 27 | 84 | 69 | 83 | 33 | 54 | 26 | 15 | 39 | 39 | 40 | 39 | 44 | 33 | 57 | 103 | 31 | 62 | 50 | 36 | 52 | Aldh3b1 |
| Aldh3b2 | 55 | 51 | 44 | 3 | 7 | 38 | 1 | 64 | 103 | 52 | 60 | 43 | 75 | 30 | 47 | 53 | 13 | 36 | 6 | 6 | 0 | 12 | 122 | 65 | 15 | 14 | 53 | 55 | Aldh3b2 |
| Aldh3b3 | 1 | 0 | 0 | 0 | 0 | 0 | 0 | 1 | 1 | 3 | 3 | 0 | 0 | 1 | 0 | 0 | 0 | 2 | 0 | 0 | 0 | 0 | 3 | 0 | 0 | 0 | 0 | 0 | Aldh3b3 |
| Aldh4a1 | 2163 | 2515 | 2282 | 2725 | 2263 | 2180 | 2541 | 939 | 1098 | 1160 | 1160 | 1027 | 1006 | 1075 | 2052 | 2392 | 2554 | 2819 | 2303 | 1853 | 2474 | 1034 | 1132 | 1226 | 1119 | 1161 | 959 | 978 | Aldh4a1 |
| Aldh5a1 | 2957 | 2594 | 3400 | 3005 | 3379 | 2989 | 3146 | 3059 | 3019 | 3235 | 2980 | 3789 | 3458 | 3657 | 2801 | 2585 | 3299 | 2836 | 3257 | 2815 | 3003 | 3255 | 2818 | 2950 | 3226 | 3493 | 3839 | 3793 | Aldh5a1 |
| Aldh6a1 | 5080 | 4390 | 3551 | 4069 | 3485 | 3790 | 4208 | 2919 | 3059 | 2532 | 2800 | 2287 | 2303 | 2412 | 4741 | 4753 | 3700 | 4097 | 3322 | 4047 | 3897 | 2821 | 3243 | 2353 | 2432 | 2280 | 2158 | 2223 | Aldh6a1 |
| Aldh7a1 | 7194 | 7520 | 7960 | 6661 | 5670 | 5911 | 5195 | 4670 | 5555 | 4939 | 5691 | 5126 | 5475 | 4549 | 6676 | 7580 | 6992 | 6346 | 6091 | 5724 | 5392 | 4480 | 5908 | 5087 | 4638 | 5842 | 5259 | 4989 | Aldh7a1 |
| Aldh8a1 | 33 | 24 | 23 | 48 | 31 | 29 | 36 | 9 | 7 | 19 | 5 | 4 | 11 | 6 | 24 | 37 | 37 | 43 | 31 | 41 | 40 | 7 | 3 | 11 | 5 | 9 | 22 | 5 | Aldh8a1 |
| Aldh9a1 | 3353 | 3373 | 3609 | 3052 | 2836 | 2877 | 2743 | 2260 | 2623 | 2486 | 2489 | 2545 | 3010 | 2450 | 3367 | 3625 | 3089 | 3103 | 2909 | 2657 | 2718 | 2460 | 2941 | 2610 | 2226 | 2985 | 2682 | 2843 | Aldh9a1 |
| Aldoa | 25504 | 29280 | 23609 | 22027 | 18489 | 19886 | 16797 | 14967 | 19993 | 16870 | 18675 | 15169 | 16687 | 15306 | 23667 | 26321 | 22633 | 21387 | 20472 | 20240 | 17957 | 14735 | 21924 | 16793 | 14915 | 19835 | 17146 | 17025 | Aldoa |
| Aldoa | 261 | 324 | 280 | 435 | 282 | 290 | 321 | 122 | 168 | 249 | 158 | 217 | 194 | 227 | 237 | 292 | 353 | 360 | 321 | 298 | 248 | 157 | 171 | 208 | 196 | 217 | 242 | 211 | Aldoa |
| Aldoart1 | 8 | 3 | 2 | 0 | 2 | 3 | 0 | 1 | 2 | 8 | 2 | 0 | 0 | 1 | 1 | 2 | 1 | 10 | 8 | 0 | 4 | 3 | 3 | 3 | 1 | 1 | 1 | 0 | Aldoart1 |
| Aldoart2 | 1 | 1 | 6 | 2 | 2 | 3 | 2 | 0 | 2 | 2 | 2 | 4 | 0 | 2 | 1 | 2 | 3 | 1 | 3 | 1 | 2 | 5 | 1 | 1 | 1 | 1 | 1 | 0 | Aldoart2 |
| Aldob | 15 | 11 | 13 | 13 | 12 | 27 | 26 | 6 | 30 | 21 | 19 | 12 | 10 | 6 | 11 | 7 | 10 | 26 | 23 | 18 | 24 | 3 | 22 | 7 | 14 | 4 | 3 | 7 | Aldob |
| Aldoc | 940 | 999 | 551 | 1963 | 3669 | 2631 | 3648 | 2414 | 3697 | 3672 | 3203 | 1962 | 1758 | 2348 | 931 | 639 | 910 | 2804 | 3225 | 3048 | 3590 | 1880 | 3818 | 3157 | 3972 | 1777 | 1323 | 1572 | Aldoc |
| Alg1 | 964 | 1131 | 1036 | 975 | 1035 | 1004 | 906 | 964 | 1077 | 935 | 1122 | 983 | 968 | 916 | 856 | 1052 | 1006 | 946 | 987 | 963 | 834 | 962 | 1056 | 949 | 1047 | 956 | 1001 | 847 | Alg1 |

| | | | | | | | | | | | | | | | | | | | | | | | | | | | | | |
|---|---|---|---|---|---|---|---|---|---|---|---|---|---|---|---|---|---|---|---|---|---|---|---|---|---|---|---|---|---|
| Alg10b | 15478 | 12006 | 12553 | 13036 | 14642 | 15613 | 16023 | 19718 | 16645 | 15817 | 16492 | 17973 | 17786 | 18793 | 16043 | 13549 | 13149 | 13625 | 13508 | 15624 | 15076 | 17597 | 16041 | 16053 | 17858 | 13734 | 18708 | 18573 | Alg10b |
| Alg11 | 4666 | 3750 | 3326 | 3311 | 2933 | 3179 | 3263 | 4081 | 3761 | 3325 | 3560 | 3252 | 3449 | 3336 | 4340 | 4191 | 3006 | 3335 | 2648 | 3368 | 3238 | 3823 | 3850 | 3166 | 3227 | 3244 | 3611 | 3522 | Alg11 |
| Alg12 | 795 | 855 | 871 | 898 | 729 | 829 | 710 | 782 | 985 | 838 | 974 | 981 | 985 | 810 | 678 | 811 | 891 | 815 | 818 | 648 | 689 | 754 | 891 | 973 | 927 | 958 | 938 | 832 | Alg12 |
| Alg13 | 5811 | 4596 | 4988 | 4628 | 3816 | 4519 | 5227 | 2331 | 2117 | 1919 | 2274 | 2607 | 2415 | 2607 | 5196 | 5288 | 4534 | 4709 | 4303 | 4246 | 4774 | 1922 | 1958 | 2486 | 2273 | 2688 | 2738 | 2741 | Alg13 |
| Alg14 | 1278 | 1216 | 1296 | 1089 | 1005 | 1034 | 1174 | 1237 | 1060 | 974 | 1134 | 1229 | 1308 | 1277 | 1290 | 1106 | 1204 | 1111 | 994 | 1273 | 1034 | 1156 | 1221 | 1130 | 977 | 1402 | 1282 | 1359 | Alg14 |
| Alg2 | 9202 | 9875 | 9129 | 13081 | 15700 | 14088 | 13017 | 12187 | 11661 | 14080 | 13075 | 11456 | 12922 | 10745 | 10624 | 9492 | 11098 | 13247 | 13814 | 12719 | 13688 | 16434 | 11172 | 11776 | 14656 | 10127 | 11167 | 10443 | Alg2 |
| Alg3 | 946 | 1076 | 843 | 852 | 778 | 778 | 813 | 996 | 1093 | 958 | 1044 | 1064 | 1186 | 1142 | 895 | 844 | 936 | 820 | 814 | 889 | 752 | 1008 | 1076 | 1114 | 1017 | 1314 | 1216 | 1167 | Alg3 |
| Alg5 | 1378 | 1148 | 1208 | 1196 | 1218 | 1116 | 1309 | 1627 | 1478 | 1283 | 1464 | 1434 | 1475 | 1413 | 1476 | 1301 | 1100 | 1203 | 1164 | 1405 | 1320 | 1377 | 1484 | 1350 | 1271 | 1496 | 1369 | 1413 | Alg5 |
| Alg6 | 122 | 86 | 77 | 52 | 74 | 41 | 37 | 155 | 87 | 130 | 200 | 58 | 98 | 153 | 124 | 249 | 107 | 79 | 107 | 94 | 53 | 105 | 139 | 153 | 83 | 165 | 222 | 97 | Alg6 |
| Alg8 | 1398 | 1174 | 1251 | 916 | 908 | 934 | 1016 | 1867 | 1804 | 1640 | 1812 | 2135 | 1871 | 2105 | 1184 | 1194 | 1119 | 920 | 970 | 1024 | 935 | 1521 | 1741 | 1757 | 1658 | 2323 | 1987 | 2092 | Alg8 |
| Alg9 | 1723 | 1732 | 1569 | 1680 | 1776 | 1699 | 1562 | 2236 | 2193 | 2043 | 2047 | 1899 | 1824 | 1818 | 1854 | 1692 | 1716 | 1520 | 1698 | 1624 | 1684 | 2481 | 2306 | 1933 | 2000 | 2001 | 2030 | 1867 | Alg9 |
| Alk | 131 | 93 | 106 | 248 | 366 | 309 | 226 | 194 | 260 | 318 | 402 | 153 | 172 | 149 | 104 | 118 | 144 | 223 | 377 | 217 | 306 | 384 | 303 | 208 | 289 | 87 | 116 | 96 | Alk |
| Alkal1 | 6 | 0 | 4 | 0 | 0 | 0 | 6 | 0 | 1 | 0 | 0 | 0 | 0 | 2 | 3 | 1 | 0 | 2 | 2 | 2 | 0 | 0 | 1 | 2 | 2 | 3 | 5 | 4 | Alkal1 |
| Alkal2 | 99 | 51 | 63 | 26 | 10 | 37 | 11 | 84 | 98 | 71 | 78 | 62 | 74 | 43 | 50 | 123 | 35 | 17 | 5 | 17 | 9 | 35 | 110 | 78 | 18 | 204 | 36 | 74 | Alkal2 |
| Alkbh1 | 2693 | 2286 | 2566 | 2560 | 2347 | 2376 | 2356 | 3375 | 3200 | 2992 | 3401 | 3731 | 3497 | 3428 | 2491 | 2287 | 2433 | 2250 | 2355 | 2316 | 2335 | 3182 | 3172 | 3311 | 3175 | 3403 | 3631 | 3246 | Alkbh1 |
| Alkbh2 | 315 | 398 | 448 | 392 | 355 | 350 | 350 | 210 | 280 | 229 | 257 | 336 | 325 | 274 | 284 | 322 | 357 | 365 | 406 | 367 | 315 | 254 | 268 | 331 | 279 | 376 | 291 | 294 | Alkbh2 |
| Alkbh3 | 772 | 655 | 478 | 540 | 531 | 546 | 591 | 1084 | 1123 | 892 | 950 | 986 | 949 | 966 | 708 | 655 | 550 | 545 | 563 | 676 | 605 | 810 | 1117 | 831 | 970 | 844 | 1016 | 932 | Alkbh3 |
| Alkbh4 | 926 | 1182 | 987 | 998 | 830 | 842 | 924 | 644 | 746 | 574 | 692 | 749 | 680 | 662 | 876 | 1058 | 983 | 876 | 920 | 939 | 777 | 702 | 640 | 724 | 746 | 846 | 857 | 738 | Alkbh4 |
| Alkbh5 | 8079 | 8255 | 9807 | 10251 | 7679 | 7539 | 7746 | 5621 | 5407 | 6711 | 6589 | 7790 | 6998 | 7308 | 7295 | 8195 | 10143 | 9782 | 8253 | 6481 | 7754 | 6520 | 5676 | 7408 | 6737 | 7865 | 6907 | 8002 | Alkbh5 |
| Alkbh6 | 1226 | 1637 | 1146 | 1284 | 1266 | 1184 | 1073 | 833 | 934 | 831 | 872 | 854 | 907 | 804 | 1211 | 1465 | 1133 | 1491 | 1319 | 1329 | 1116 | 916 | 896 | 976 | 959 | 1084 | 1012 | 796 | Alkbh6 |
| Alkbh7 | 419 | 653 | 585 | 658 | 579 | 532 | 622 | 371 | 462 | 505 | 418 | 533 | 492 | 469 | 469 | 469 | 538 | 582 | 598 | 571 | 590 | 456 | 437 | 487 | 472 | 554 | 494 | 474 | Alkbh7 |
| Alkbh8 | 7683 | 6380 | 5510 | 6756 | 7864 | 8327 | 8674 | 9810 | 8202 | 7706 | 8010 | 8238 | 8130 | 8548 | 8130 | 6644 | 6161 | 7072 | 7367 | 8504 | 8327 | 8380 | 7855 | 7290 | 8781 | 5498 | 8516 | 8272 | Alkbh8 |
| Allc | 0 | 1 | 0 | 1 | 0 | 2 | 0 | 0 | 3 | 0 | 4 | 4 | 0 | 0 | 1 | 0 | 0 | 0 | 0 | 1 | 4 | 0 | 1 | 0 | 0 | 0 | 0 | 1 | Allc |
| Alms1 | 9130 | 7591 | 3336 | 4977 | 5259 | 6989 | 7271 | 6682 | 5370 | 3727 | 3981 | 4662 | 4332 | 4760 | 8847 | 7227 | 3613 | 4427 | 5157 | 8660 | 7063 | 2902 | 4916 | 3241 | 4007 | 2418 | 5306 | 4125 | Alms1 |
| Alox12 | 168 | 125 | 108 | 58 | 93 | 67 | 40 | 178 | 195 | 173 | 148 | 125 | 215 | 95 | 102 | 176 | 104 | 91 | 43 | 103 | 66 | 114 | 208 | 99 | 73 | 210 | 173 | 102 | Alox12 |
| Alox12b | 5 | 3 | 3 | 2 | 7 | 0 | 1 | 4 | 4 | 19 | 11 | 9 | 6 | 2 | 7 | 9 | 10 | 4 | 2 | 5 | 3 | 3 | 13 | 7 | 6 | 5 | 0 | 1 | Alox12b |
| Alox12e | 2 | 1 | 0 | 0 | 0 | 0 | 0 | 5 | 8 | 0 | 3 | 0 | 0 | 0 | 1 | 1 | 1 | 0 | 0 | 0 | 0 | 0 | 6 | 4 | 0 | 5 | 0 | 0 | Alox12e |
| Alox15 | 46 | 16 | 28 | 24 | 15 | 19 | 16 | 12 | 12 | 21 | 14 | 10 | 15 | 17 | 24 | 28 | 20 | 18 | 14 | 10 | 18 | 12 | 21 | 20 | 5 | 18 | 9 | 18 | Alox15 |
| Alox5 | 69 | 72 | 84 | 48 | 55 | 43 | 37 | 18 | 28 | 24 | 54 | 23 | 35 | 28 | 63 | 78 | 80 | 59 | 44 | 45 | 34 | 42 | 54 | 38 | 18 | 47 | 36 | 27 | Alox5 |
| Alox5ap | 83 | 116 | 92 | 76 | 106 | 79 | 108 | 98 | 109 | 80 | 131 | 73 | 91 | 101 | 140 | 130 | 79 | 129 | 66 | 92 | 85 | 114 | 154 | 121 | 93 | 138 | 121 | 107 | Alox5ap |
| Alox8 | 7 | 11 | 13 | 3 | 2 | 8 | 2 | 2 | 15 | 14 | 22 | 10 | 5 | 9 | 7 | 12 | 6 | 13 | 2 | 2 | 4 | 3 | 10 | 11 | 5 | 8 | 12 | 1 | Alox8 |
| Aloxe3 | 68 | 67 | 78 | 75 | 110 | 106 | 75 | 12 | 27 | 22 | 32 | 13 | 19 | 15 | 57 | 77 | 79 | 79 | 96 | 74 | 90 | 30 | 30 | 15 | 23 | 10 | 7 | 24 | Aloxe3 |
| Alpk1 | 738 | 725 | 455 | 571 | 650 | 776 | 822 | 689 | 798 | 721 | 704 | 818 | 668 | 830 | 757 | 632 | 534 | 752 | 628 | 852 | 773 | 503 | 858 | 644 | 692 | 481 | 809 | 602 | Alpk1 |
| Alpk2 | 3 | 10 | 13 | 8 | 1 | 5 | 1 | 22 | 46 | 9 | 210 | 14 | 34 | 9 | 7 | 8 | 3 | 6 | 0 | 14 | 3 | 40 | 139 | 14 | 14 | 8 | 9 | 4 | Alpk2 |
| Alpk3 | 23 | 22 | 43 | 30 | 47 | 27 | 35 | 53 | 115 | 38 | 627 | 43 | 96 | 29 | 21 | 40 | 16 | 42 | 48 | 28 | 35 | 95 | 356 | 41 | 21 | 41 | 15 | 39 | Alpk3 |
| Alpl | 640 | 711 | 737 | 675 | 484 | 658 | 398 | 1015 | 1195 | 1183 | 865 | 1072 | 961 | 850 | 568 | 729 | 765 | 800 | 598 | 393 | 413 | 881 | 1000 | 1288 | 913 | 1143 | 906 | 862 | Alpl |
| Als2 | 7996 | 8958 | 7271 | 8036 | 7744 | 7502 | 8149 | 4957 | 4606 | 4805 | 4740 | 5137 | 4948 | 5061 | 8372 | 8408 | 7619 | 8205 | 7821 | 8023 | 8466 | 4888 | 4664 | 4557 | 5190 | 3716 | 5165 | 5003 | Als2 |
| Als2cl | 422 | 456 | 601 | 383 | 454 | 458 | 304 | 228 | 373 | 414 | 320 | 328 | 277 | 234 | 392 | 597 | 474 | 525 | 400 | 247 | 332 | 312 | 324 | 306 | 218 | 314 | 260 | 206 | Als2cl |
| Alx1 | 2128 | 626 | 880 | 510 | 74 | 231 | 57 | 2886 | 1473 | 1882 | 727 | 926 | 1088 | 847 | 936 | 1604 | 521 | 373 | 48 | 75 | 55 | 219 | 1352 | 2271 | 472 | 3442 | 797 | 1118 | Alx1 |
| Alx3 | 392 | 244 | 266 | 200 | 78 | 134 | 69 | 462 | 292 | 344 | 200 | 299 | 370 | 312 | 278 | 437 | 145 | 143 | 72 | 81 | 80 | 74 | 326 | 443 | 178 | 861 | 277 | 344 | Alx3 |
| Alx4 | 2392 | 1589 | 2197 | 1044 | 386 | 1002 | 280 | 3886 | 3454 | 3874 | 2976 | 3263 | 4276 | 2961 | 1593 | 2397 | 1254 | 989 | 321 | 369 | 256 | 2401 | 3224 | 4296 | 2105 | 5245 | 3459 | 3806 | Alx4 |
| Alyref | 5847 | 5697 | 5619 | 5078 | 4463 | 4853 | 4875 | 6829 | 6211 | 5711 | 5951 | 7702 | 6698 | 7767 | 5416 | 4836 | 5355 | 4581 | 4941 | 5441 | 4965 | 5355 | 5896 | 6929 | 6399 | 8252 | 8270 | 8148 | Alyref |
| Alyref2 | 205 | 267 | 289 | 374 | 475 | 348 | 416 | 381 | 451 | 506 | 482 | 474 | 361 | 466 | 278 | 216 | 460 | 369 | 485 | 430 | 403 | 448 | 471 | 512 | 557 | 446 | 475 | 436 | Alyref2 |
| Amacr | 248 | 260 | 246 | 248 | 239 | 215 | 217 | 495 | 536 | 418 | 569 | 433 | 502 | 456 | 266 | 287 | 224 | 206 | 207 | 176 | 186 | 428 | 587 | 413 | 444 | 461 | 374 | 452 | Amacr |
| Ambn | 10 | 4 | 0 | 0 | 0 | 0 | 0 | 15 | 27 | 14 | 4 | 4 | 0 | 0 | 3 | 1 | 0 | 4 | 3 | 3 | 1 | 3 | 12 | 1 | 0 | 0 | 1 | 0 | Ambn |
| Ambp | 0 | 0 | 0 | 0 | 0 | 0 | 1 | 0 | 2 | 0 | 0 | 0 | 0 | 1 | 0 | 1 | 0 | 3 | 0 | 0 | 0 | 0 | 1 | 0 | 0 | 3 | 0 | 0 | Ambp |
| Ambra1 | 4991 | 5664 | 6454 | 6638 | 6516 | 6469 | 5625 | 5101 | 5759 | 6671 | 6300 | 6746 | 6021 | 6156 | 4739 | 5314 | 6789 | 6632 | 7027 | 4904 | 6439 | 6697 | 5489 | 6375 | 6841 | 5849 | 5753 | 6108 | Ambra1 |
| Amd1 | 9883 | 8783 | 8401 | 7749 | 6889 | 8032 | 7996 | 8107 | 6958 | 5631 | 5864 | 7142 | 7164 | 7329 | 9965 | 10243 | 8105 | 8140 | 7033 | 9279 | 7346 | 5798 | 6932 | 6146 | 6072 | 8618 | 7706 | 7570 | Amd1 |
| Amd2 | 294 | 203 | 226 | 141 | 137 | 159 | 180 | 236 | 176 | 123 | 124 | 147 | 160 | 130 | 314 | 254 | 177 | 176 | 127 | 223 | 161 | 140 | 158 | 127 | 138 | 194 | 187 | 178 | Amd2 |
| Amdhd1 | 17 | 22 | 33 | 22 | 8 | 13 | 3 | 12 | 16 | 9 | 8 | 23 | 12 | 10 | 22 | 31 | 8 | 19 | 6 | 15 | 10 | 10 | 16 | 13 | 7 | 10 | 21 | 8 | Amdhd1 |
| Amdhd2 | 864 | 1122 | 1040 | 1097 | 1150 | 1084 | 991 | 801 | 1041 | 1050 | 1094 | 1101 | 1125 | 1004 | 891 | 1026 | 1176 | 1124 | 1091 | 1008 | 906 | 881 | 1050 | 1092 | 1045 | 1068 | 1059 | 978 | Amdhd2 |
| Amelx | 0 | 0 | 0 | 0 | 0 | 0 | 0 | 0 | 0 | 2 | 0 | 0 | 0 | 0 | 0 | 0 | 0 | 0 | 0 | 0 | 0 | 0 | 1 | 0 | 0 | 0 | 0 | 0 | Amelx |
| Amer1 | 6965 | 5679 | 3737 | 4330 | 4922 | 5374 | 5111 | 5205 | 4696 | 4114 | 4206 | 4076 | 4288 | 4261 | 6163 | 5449 | 4067 | 4749 | 4634 | 4973 | 5330 | 4578 | 4565 | 3336 | 3977 | 2761 | 4680 | 4069 | Amer1 |
| Amer2 | 5624 | 5091 | 4573 | 7463 | 8372 | 8230 | 9817 | 8569 | 7236 | 8251 | 6682 | 8726 | 6103 | 9569 | 6268 | 4525 | 5887 | 7005 | 8061 | 8159 | 9642 | 8898 | 6986 | 8063 | 10192 | 6255 | 8414 | 8179 | Amer2 |
| Amer3 | 325 | 404 | 435 | 736 | 1339 | 1013 | 939 | 1717 | 1865 | 2470 | 1817 | 1724 | 1261 | 1541 | 398 | 290 | 648 | 771 | 1243 | 919 | 1074 | 2842 | 1788 | 1905 | 2529 | 1205 | 1248 | 1314 | Amer3 |
| Amfr | 5356 | 5389 | 5916 | 5979 | 7079 | 6568 | 6502 | 8400 | 8630 | 8646 | 8448 | 9328 | 8481 | 9240 | 5577 | 5261 | 5727 | 5804 | 7310 | 5917 | 6919 | 9003 | 8624 | 9025 | 9681 | 9269 | 8745 | 9419 | Amfr |
| Amh | 42 | 96 | 99 | 85 | 63 | 87 | 60 | 38 | 59 | 75 | 47 | 60 | 66 | 32 | 36 | 81 | 101 | 106 | 97 | 61 | 73 | 85 | 42 | 39 | 60 | 68 | 59 | 49 | Amh |
| Amhr2 | 25 | 41 | 58 | 78 | 82 | 63 | 49 | 20 | 31 | 34 | 30 | 31 | 37 | 30 | 36 | 37 | 61 | 72 | 66 | 42 | 72 | 33 | 30 | 37 | 23 | 25 | 27 | 24 | Amhr2 |
| Amigo1 | 3002 | 3405 | 3192 | 4256 | 4522 | 4006 | 4009 | 2298 | 2338 | 2548 | 2527 | 1977 | 2107 | 1916 | 3353 | 3137 | 3822 | 4580 | 4363 | 3365 | 4224 | 2963 | 2196 | 2146 | 2621 | 1719 | 1677 | 1907 | Amigo1 |
| Amigo2 | 433 | 454 | 561 | 540 | 810 | 577 | 627 | 739 | 673 | 740 | 621 | 556 | 641 | 471 | 489 | 533 | 603 | 627 | 598 | 594 | 651 | 852 | 659 | 556 | 752 | 529 | 582 | 584 | Amigo2 |
| Amigo3 | 483 | 560 | 399 | 501 | 437 | 463 | 439 | 177 | 220 | 245 | 248 | 231 | 217 | 201 | 369 | 495 | 428 | 616 | 443 | 359 | 423 | 202 | 195 | 239 | 211 | 244 | 210 | 220 | Amigo3 |
| Ammecr1 | 2327 | 1475 | 2165 | 1672 | 1635 | 1934 | 2123 | 2319 | 2254 | 2111 | 2889 | 2401 | 2366 | 2417 | 2047 | 2136 | 1845 | 1885 | 1785 | 1608 | 1935 | 2181 | 2374 | 2218 | 2138 | 2712 | 2260 | 2504 | Ammecr1 |
| Ammecr1l | 4505 | 3987 | 4609 | 4731 | 5548 | 5329 | 5282 | 5882 | 5420 | 5673 | 5979 | 6312 | 6111 | 6078 | 4272 | 3864 | 4629 | 4789 | 5484 | 4437 | 5543 | 6351 | 5641 | 5739 | 5960 | 5423 | 6240 | 6389 | Ammecr1l |
| Amn | 63 | 65 | 40 | 92 | 88 | 69 | 53 | 21 | 28 | 30 | 23 | 11 | 16 | 17 | 46 | 62 | 41 | 64 | 69 | 60 | 46 | 22 | 21 | 23 | 26 | 8 | 6 | 11 | Amn |
| Amn1 | 2117 | 1532 | 1467 | 1929 | 1967 | 1964 | 2266 | 3475 | 2880 | 2780 | 2676 | 2868 | 2625 | 3256 | 2190 | 1690 | 1640 | 1677 | 1926 | 2305 | 2207 | 2755 | 2867 | 2590 | 3023 | 2317 | 2914 | 2883 | Amn1 |
| Amot | 15845 | 12023 | 10172 | 10259 | 9001 | 10343 | 9424 | 7399 | 9385 | 7749 | 13849 | 7375 | 9324 | 7513 | 13940 | 15566 | 9757 | 10397 | 8840 | 8375 | 9136 | 7917 | 10591 | 6977 | 6839 | 7261 | 7796 | 7266 | Amot |
| Amotl1 | 8786 | 7113 | 5967 | 6123 | 6731 | 7258 | 5595 | 9523 | 11246 | 10846 | 12664 | 10474 | 11535 | 9813 | 7010 | 8738 | 6178 | 6437 | 6243 | 5395 | 5580 | 8905 | 11491 | 9759 | 9821 | 6807 | 10951 | 9552 | Amotl1 |
| Amotl2 | 3925 | 4667 | 6173 | 4955 | 5677 | 5057 | 4037 | 6892 | 7119 | 7997 | 7987 | 8973 | 8280 | 7925 | 3899 | 4556 | 6006 | 4678 | 5753 | 4246 | 4577 | 8293 | 7384 | 8904 | 8248 | 8347 | 8207 | 8681 | Amotl2 |
| Ampd1 | 14 | 18 | 17 | 24 | 11 | 10 | 10 | 10 | 15 | 2 | 155 | 8 | 3 | 4 | 8 | 13 | 15 | 11 | 3 | 7 | 8 | 18 | 157 | 2 | 10 | 5 | 0 | 9 | Ampd1 |
| Ampd2 | 4497 | 5690 | 5654 | 5322 | 4855 | 4605 | 4235 | 3242 | 4033 | 4219 | 4291 | 4355 | 4161 | 4107 | 4137 | 5313 | 5413 | 4981 | 5103 | 4376 | 4261 | 4380 | 3839 | 4347 | 4316 | 4464 | 4162 | 4053 | Ampd2 |
| Ampd3 | 106 | 82 | 52 | 50 | 146 | 71 | 79 | 221 | 365 | 356 | 325 | 167 | 265 | 171 | 83 | 107 | 58 | 99 | 91 | 86 | 74 | 388 | 372 | 218 | 253 | 250 | 146 | 196 | Ampd3 |
| Amph | 1867 | 2442 | 1740 | 2662 | 4024 | 3043 | 3401 | 3261 | 3127 | 3076 | 2667 | 2046 | 2303 | 2158 | 2632 | 1959 | 2406 | 2981 | 3259 | 3482 | 3613 | 3688 | 3015 | 2138 | 2995 | 1454 | 1968 | 1979 | Amph |
| Amt | 2410 | 3002 | 2993 | 3097 | 3256 | 2652 | 3027 | 2394 | 2245 | 2707 | 2281 | 2470 | 2388 | 2502 | 2678 | 2493 | 3219 | 3026 | 3422 | 2907 | 3014 | 2912 | 2106 | 2715 | 2138 | 2811 | 2308 | 2607 | 2580 | Amt |
| Amtn | 1 | 0 | 0 | 0 | 0 | 5 | 0 | 1 | 7 | 2 | 6 | 0 | 1 | 5 | 1 | 2 | 0 | 1 | 0 | 0 | 0 | 0 | 6 | 0 | 0 | 1 | 0 | 2 | Amtn |
| Amy1 | 376 | 417 | 338 | 752 | 1014 | 899 | 1016 | 763 | 632 | 640 | 653 | 438 | 442 | 500 | 541 | 417 | 539 | 905 | 843 | 998 | 943 | 1026 | 590 | 473 | 747 | 289 | 335 | 363 | Amy1 |
| Amy2a1 | 1 | 0 | 0 | 0 | 2 | 0 | 0 | 0 | 0 | 0 | 0 | 0 | 0 | 0 | 0 | 1 | 0 | 0 | 0 | 0 | 2 | 0 | 0 | 0 | 0 | 0 | 0 | 0 | Amy2a1 |
| Amy2a2 | 0 | 0 | 0 | 0 | 0 | 0 | 0 | 0 | 0 | 0 | 0 | 0 | 0 | 0 | 0 | 0 | 0 | 0 | 2 | 0 | 1 | 0 | 0 | 0 | 0 | 0 | 0 | 0 | Amy2a2 |
| Amy2a3 | 0 | 0 | 0 | 0 | 1 | 0 | 1 | 0 | 0 | 0 | 0 | 0 | 0 | 0 | 0 | 0 | 7 | 0 | 0 | 0 | 1 | 0 | 0 | 0 | 0 | 0 | 0 | 0 | Amy2a3 |
| Amy2a4 | 0 | 0 | 0 | 0 | 0 | 0 | 0 | 0 | 0 | 0 | 0 | 0 | 0 | 0 | 0 | 0 | 0 | 0 | 0 | 0 | 0 | 0 | 0 | 0 | 1 | 0 | 0 | 0 | Amy2a4 |

| | | | | | | | | | | | | | | | | | | | | | | | | | | | | | |
|---|---|---|---|---|---|---|---|---|---|---|---|---|---|---|---|---|---|---|---|---|---|---|---|---|---|---|---|---|---|
| Amy2a5 | 0 | 0 | 0 | 0 | 0 | 0 | 0 | 0 | 0 | 0 | 0 | 0 | 0 | 0 | 0 | 0 | 0 | 0 | 5 | 0 | 0 | 0 | 1 | 0 | 0 | 0 | 0 | 0 | Amy2a5 |
| Amz1 | 104 | 98 | 82 | 190 | 137 | 122 | 151 | 109 | 143 | 109 | 116 | 104 | 130 | 104 | 107 | 114 | 143 | 142 | 163 | 123 | 143 | 157 | 131 | 133 | 192 | 100 | 114 | 101 | Amz1 |
| Amz2 | 7989 | 8080 | 7719 | 9703 | 10748 | 9777 | 10756 | 11138 | 10299 | 11131 | 9648 | 10085 | 9097 | 10105 | 8380 | 6755 | 8887 | 9928 | 10516 | 9465 | 10944 | 12288 | 10015 | 10226 | 11631 | 9571 | 9603 | 9892 | Amz2 |
| Anapc1 | 21644 | 19671 | 13283 | 14506 | 14872 | 17814 | 16694 | 25565 | 23511 | 19863 | 20910 | 24411 | 23537 | 23851 | 20031 | 18714 | 13753 | 14339 | 14822 | 17297 | 16974 | 19248 | 22411 | 18442 | 21680 | 16778 | 25782 | 23265 | Anapc1 |
| Anapc10 | 2170 | 1841 | 2066 | 1981 | 1856 | 1815 | 2066 | 2019 | 1738 | 1657 | 1529 | 1707 | 1659 | 1919 | 2399 | 1981 | 1892 | 1915 | 1835 | 2282 | 1980 | 1656 | 1834 | 1718 | 1729 | 1913 | 1802 | 1958 | Anapc10 |
| Anapc11 | 2643 | 3090 | 3783 | 3301 | 2925 | 2985 | 2874 | 3104 | 3365 | 4024 | 3848 | 3989 | 3656 | 3880 | 2529 | 2767 | 3505 | 3213 | 3375 | 2923 | 2952 | 3322 | 3539 | 4269 | 4039 | 4557 | 3976 | 4162 | Anapc11 |
| Anapc13 | 1540 | 1815 | 2425 | 2159 | 1584 | 1379 | 2018 | 1774 | 1568 | 2542 | 2220 | 2418 | 2138 | 3575 | 1571 | 1037 | 2228 | 1604 | 1654 | 1732 | 2154 | 1484 | 1647 | 3029 | 2398 | 2870 | 2500 | 3504 | Anapc13 |
| Anapc15 | 1690 | 1804 | 1769 | 1690 | 1672 | 1612 | 1738 | 3219 | 2891 | 3080 | 2685 | 3396 | 3215 | 3632 | 1734 | 1511 | 1819 | 1608 | 1854 | 2113 | 1770 | 2854 | 2812 | 3242 | 3312 | 3654 | 3484 | 3640 | Anapc15 |
| Anapc15-ps | 24 | 28 | 19 | 24 | 16 | 23 | 18 | 33 | 19 | 24 | 14 | 33 | 33 | 45 | 33 | 14 | 27 | 8 | 15 | 20 | 11 | 22 | 28 | 24 | 19 | 40 | 31 | 31 | Anapc15-ps |
| Anapc16 | 2340 | 2455 | 2360 | 2243 | 2049 | 2135 | 2161 | 2998 | 2674 | 2668 | 2574 | 2853 | 2667 | 2878 | 2320 | 2022 | 2335 | 1869 | 2232 | 2274 | 2157 | 2724 | 2721 | 2716 | 2602 | 3021 | 2933 | 2963 | Anapc16 |
| Anapc2 | 11374 | 12103 | 12582 | 11551 | 10341 | 10425 | 10124 | 9431 | 10463 | 11100 | 10209 | 11942 | 11480 | 11059 | 10489 | 10731 | 11925 | 11526 | 11916 | 9436 | 10860 | 10347 | 10480 | 11416 | 11586 | 12308 | 11443 | 11720 | Anapc2 |
| Anapc4 | 10531 | 9021 | 9069 | 8841 | 9171 | 9274 | 9452 | 12038 | 10899 | 9531 | 9747 | 10833 | 10664 | 11043 | 11230 | 9156 | 8857 | 8526 | 9100 | 10435 | 8951 | 11134 | 10528 | 10152 | 10316 | 10873 | 10838 | 11272 | Anapc4 |
| Anapc5 | 25617 | 28587 | 22453 | 24453 | 25641 | 25521 | 26980 | 31614 | 31232 | 30610 | 29206 | 30401 | 28388 | 30544 | 26212 | 23521 | 24469 | 24448 | 26752 | 27729 | 27910 | 30239 | 30958 | 29179 | 31352 | 29673 | 30477 | 30688 | Anapc5 |
| Anapc7 | 6815 | 7975 | 6371 | 7215 | 6065 | 6596 | 5948 | 5186 | 5053 | 5127 | 5020 | 5805 | 5407 | 5435 | 6679 | 6611 | 6575 | 6540 | 6955 | 6651 | 6683 | 4610 | 5087 | 5247 | 5421 | 6049 | 5555 | 5700 | Anapc7 |
| Ang | 85 | 40 | 51 | 34 | 44 | 40 | 30 | 66 | 54 | 71 | 81 | 76 | 85 | 40 | 62 | 79 | 61 | 56 | 47 | 56 | 38 | 58 | 73 | 60 | 44 | 44 | 45 | 51 | Ang |
| Ang2 | 0 | 0 | 0 | 0 | 0 | 0 | 0 | 0 | 2 | 0 | 0 | 0 | 0 | 0 | 0 | 0 | 0 | 0 | 0 | 0 | 0 | 0 | 0 | 0 | 0 | 0 | 0 | 0 | Ang2 |
| Ang4 | 0 | 0 | 0 | 0 | 0 | 0 | 0 | 0 | 0 | 0 | 0 | 0 | 0 | 0 | 1 | 0 | 0 | 0 | 0 | 0 | 0 | 8 | 0 | 0 | 0 | 1 | 0 | 0 | Ang4 |
| Angel1 | 2006 | 2626 | 2656 | 2714 | 2305 | 2199 | 2185 | 1010 | 1148 | 1128 | 1184 | 1206 | 1243 | 1040 | 2073 | 2437 | 2615 | 2770 | 2485 | 2111 | 2242 | 1078 | 1062 | 1091 | 1237 | 1193 | 1074 | 1182 | Angel1 |
| Angel2 | 3233 | 2750 | 3404 | 2726 | 3232 | 3076 | 3374 | 7586 | 6372 | 6073 | 6076 | 7035 | 6815 | 7422 | 3332 | 2770 | 2950 | 2677 | 3239 | 3618 | 3202 | 6371 | 6333 | 6336 | 6765 | 7053 | 7678 | 7177 | Angel2 |
| Angpt1 | 1606 | 1137 | 1229 | 1273 | 1028 | 1501 | 972 | 2136 | 2286 | 1749 | 2214 | 1615 | 1971 | 1799 | 1822 | 1451 | 1305 | 1327 | 955 | 1230 | 993 | 2222 | 2799 | 1674 | 1774 | 1345 | 1864 | 1720 | Angpt1 |
| Angpt2 | 108 | 112 | 135 | 110 | 146 | 165 | 112 | 228 | 325 | 278 | 411 | 265 | 265 | 249 | 185 | 126 | 114 | 145 | 122 | 173 | 136 | 373 | 579 | 282 | 311 | 308 | 265 | 206 | Angpt2 |
| Angpt4 | 3 | 2 | 0 | 0 | 0 | 3 | 2 | 6 | 12 | 1 | 10 | 4 | 5 | 7 | 11 | 1 | 1 | 0 | 1 | 13 | 4 | 2 | 8 | 2 | 1 | 4 | 7 | 2 | Angpt4 |
| Angptl1 | 200 | 92 | 45 | 64 | 55 | 98 | 19 | 264 | 709 | 419 | 2129 | 110 | 305 | 101 | 156 | 191 | 50 | 100 | 34 | 47 | 22 | 516 | 1709 | 185 | 231 | 218 | 134 | 120 | Angptl1 |
| Angptl2 | 3453 | 3506 | 3030 | 2356 | 1943 | 2526 | 1446 | 2071 | 2918 | 2377 | 3513 | 2544 | 3312 | 2260 | 3208 | 4009 | 2730 | 2647 | 1673 | 1903 | 1460 | 2739 | 3547 | 2397 | 2173 | 2778 | 3029 | 2454 | Angptl2 |
| Angptl3 | 0 | 0 | 0 | 0 | 0 | 0 | 1 | 4 | 2 | 1 | 0 | 0 | 0 | 1 | 0 | 1 | 0 | 0 | 0 | 2 | 0 | 2 | 2 | 4 | 0 | 0 | 6 | 4 | Angptl3 |
| Angptl4 | 535 | 617 | 473 | 416 | 390 | 459 | 253 | 1102 | 1589 | 1070 | 1455 | 1006 | 1306 | 991 | 554 | 681 | 367 | 395 | 305 | 338 | 275 | 939 | 1996 | 993 | 957 | 1113 | 1076 | 1009 | Angptl4 |
| Angptl6 | 69 | 121 | 117 | 190 | 150 | 142 | 156 | 178 | 176 | 243 | 202 | 149 | 136 | 179 | 112 | 96 | 129 | 160 | 191 | 187 | 170 | 172 | 166 | 217 | 202 | 162 | 185 | 153 | Angptl6 |
| Angptl7 | 0 | 0 | 0 | 0 | 2 | 0 | 0 | 1 | 8 | 5 | 60 | 0 | 0 | 0 | 3 | 0 | 0 | 1 | 0 | 0 | 0 | 5 | 29 | 3 | 0 | 5 | 0 | 1 | Angptl7 |
| Angptl8 | 0 | 0 | 0 | 0 | 0 | 0 | 0 | 0 | 4 | 0 | 0 | 0 | 0 | 2 | 0 | 0 | 2 | 0 | 0 | 0 | 0 | 0 | 0 | 0 | 0 | 0 | 0 | 1 | Angptl8 |
| Ank | 1239 | 1228 | 1163 | 1409 | 1753 | 1400 | 1450 | 1655 | 1858 | 1938 | 1840 | 1538 | 1608 | 1469 | 1304 | 1357 | 1377 | 1462 | 1563 | 1370 | 1516 | 1932 | 1676 | 1894 | 1822 | 1599 | 1434 | 1507 | Ank |
| Ank1 | 413 | 656 | 698 | 544 | 1169 | 656 | 632 | 819 | 1182 | 1146 | 1716 | 945 | 1531 | 990 | 463 | 642 | 545 | 664 | 630 | 569 | 677 | 1390 | 1166 | 763 | 1288 | 568 | 908 | 1523 | Ank1 |
| Ank2 | 14229 | 16363 | 10938 | 17197 | 23275 | 25030 | 21908 | 23517 | 21994 | 22819 | 21392 | 21191 | 21020 | 22545 | 17921 | 15707 | 13568 | 17249 | 18839 | 21832 | 25782 | 24488 | 19935 | 17791 | 26681 | 11898 | 23336 | 18574 | Ank2 |
| Ank3 | 17162 | 19675 | 14300 | 24127 | 31675 | 35256 | 29195 | 28107 | 28517 | 31611 | 30881 | 24085 | 29227 | 26338 | 21726 | 20386 | 19668 | 24202 | 25563 | 28952 | 35887 | 31528 | 24886 | 21885 | 36426 | 12392 | 28259 | 22533 | Ank3 |
| Ankar | 0 | 0 | 4 | 0 | 0 | 0 | 0 | 0 | 5 | 0 | 1 | 4 | 0 | 3 | 3 | 0 | 3 | 7 | 0 | 0 | 1 | 0 | 0 | 2 | 1 | 3 | 1 | 2 | Ankar |
| Ankdd1a | 8 | 29 | 9 | 13 | 56 | 40 | 26 | 32 | 69 | 30 | 64 | 54 | 52 | 39 | 15 | 44 | 27 | 30 | 38 | 14 | 20 | 62 | 46 | 30 | 63 | 4 | 31 | 14 | Ankdd1a |
| Ankdd1b | 7 | 6 | 0 | 5 | 5 | 2 | 3 | 7 | 2 | 10 | 7 | 2 | 9 | 0 | 4 | 21 | 8 | 19 | 7 | 4 | 5 | 3 | 4 | 5 | 5 | 0 | 2 | 0 | Ankdd1b |
| Ankef1 | 104 | 105 | 62 | 90 | 71 | 91 | 101 | 72 | 67 | 81 | 74 | 73 | 53 | 92 | 83 | 92 | 82 | 78 | 69 | 95 | 97 | 60 | 93 | 58 | 41 | 74 | 70 | 56 | Ankef1 |
| Ankfn1 | 215 | 253 | 62 | 174 | 223 | 271 | 236 | 289 | 326 | 243 | 228 | 261 | 254 | 259 | 218 | 188 | 130 | 171 | 217 | 312 | 258 | 261 | 231 | 200 | 372 | 74 | 311 | 211 | Ankfn1 |
| Ankfy1 | 10548 | 10331 | 6767 | 8343 | 8719 | 9114 | 9526 | 11242 | 11554 | 9400 | 9672 | 10083 | 9722 | 9748 | 10358 | 9786 | 7211 | 8680 | 8436 | 9860 | 9473 | 9347 | 10989 | 8615 | 9954 | 6585 | 10035 | 9130 | Ankfy1 |
| Ankhd1 | 14947 | 13377 | 6698 | 10678 | 10819 | 13262 | 12647 | 10926 | 9327 | 7684 | 7713 | 8484 | 8452 | 9003 | 14778 | 12376 | 8472 | 10272 | 9938 | 13674 | 13856 | 7010 | 8714 | 6409 | 8059 | 3677 | 9334 | 8062 | Ankhd1 |
| Ankib1 | 7874 | 7496 | 5739 | 8165 | 8771 | 8941 | 9184 | 6489 | 6044 | 5932 | 5873 | 5918 | 5697 | 6000 | 8375 | 7329 | 6981 | 8554 | 8419 | 8659 | 9388 | 6462 | 5929 | 5552 | 6591 | 4562 | 5975 | 5914 | Ankib1 |
| Ankk1 | 14 | 14 | 10 | 21 | 18 | 14 | 13 | 0 | 3 | 0 | 1 | 2 | 0 | 1 | 11 | 20 | 21 | 18 | 16 | 1 | 13 | 0 | 1 | 2 | 0 | 0 | 0 | 1 | Ankk1 |
| Ankle1 | 1336 | 1511 | 1606 | 1206 | 964 | 1154 | 1020 | 1111 | 1300 | 1375 | 1334 | 1930 | 1655 | 1583 | 1065 | 1284 | 1552 | 1080 | 1200 | 1166 | 916 | 1028 | 1243 | 1673 | 1286 | 2204 | 2133 | 1656 | Ankle1 |
| Ankle2 | 3973 | 3636 | 4218 | 3714 | 3724 | 3938 | 3998 | 5959 | 5113 | 5290 | 5209 | 5815 | 5349 | 5794 | 3939 | 3542 | 3815 | 3319 | 3973 | 3800 | 3707 | 5451 | 5095 | 5349 | 5285 | 5526 | 5715 | 5719 | Ankle2 |
| Ankmy1 | 54 | 56 | 53 | 82 | 56 | 87 | 45 | 34 | 36 | 30 | 46 | 28 | 27 | 31 | 51 | 62 | 44 | 47 | 49 | 55 | 61 | 25 | 37 | 14 | 18 | 24 | 23 | 37 | Ankmy1 |
| Ankmy2 | 3102 | 2906 | 2713 | 3102 | 3718 | 3420 | 3851 | 3604 | 3207 | 3024 | 2873 | 2448 | 2474 | 2763 | 3299 | 2666 | 2941 | 3387 | 3547 | 4011 | 3980 | 3430 | 3323 | 2960 | 2962 | 2618 | 2529 | 2534 | Ankmy2 |
| Ankra2 | 2726 | 2522 | 2370 | 2911 | 3160 | 2797 | 3403 | 3734 | 3232 | 3372 | 2882 | 2996 | 2735 | 3351 | 3119 | 2407 | 2716 | 2914 | 2982 | 3329 | 3091 | 3882 | 3180 | 3371 | 3543 | 2869 | 2826 | 3084 | Ankra2 |
| Ankrd1 | 10 | 3 | 6 | 0 | 0 | 3 | 0 | 17 | 21 | 8 | 151 | 0 | 6 | 3 | 4 | 2 | 0 | 7 | 2 | 0 | 0 | 7 | 93 | 6 | 12 | 5 | 0 | 4 | Ankrd1 |
| Ankrd10 | 11642 | 10547 | 9184 | 11545 | 11027 | 11616 | 12259 | 13575 | 12545 | 11309 | 12158 | 12716 | 12284 | 12983 | 10787 | 11386 | 9750 | 11825 | 11567 | 12140 | 12405 | 11135 | 11721 | 11537 | 12739 | 11740 | 13142 | 12043 | Ankrd10 |
| Ankrd11 | 20274 | 21298 | 18418 | 24519 | 23948 | 24098 | 21369 | 16505 | 18839 | 21186 | 21375 | 22525 | 20995 | 21141 | 19077 | 21284 | 22155 | 22734 | 23974 | 19675 | 23172 | 18945 | 17384 | 19603 | 21995 | 12437 | 21372 | 19737 | Ankrd11 |
| Ankrd12 | 9983 | 8487 | 6489 | 10980 | 12284 | 13789 | 12990 | 9767 | 8406 | 7731 | 7780 | 6078 | 6199 | 6367 | 11493 | 9128 | 8049 | 10848 | 10428 | 14378 | 13793 | 8555 | 7824 | 6159 | 8566 | 3813 | 5918 | 5425 | Ankrd12 |
| Ankrd13a | 13320 | 16199 | 14568 | 14440 | 10933 | 11795 | 11624 | 5603 | 5698 | 5866 | 5936 | 6342 | 6205 | 6289 | 13270 | 14759 | 14173 | 13479 | 11441 | 10468 | 11804 | 5830 | 5615 | 5719 | 6113 | 6494 | 6307 | 6815 | Ankrd13a |
| Ankrd13b | 5856 | 7883 | 8156 | 9823 | 11060 | 9174 | 8819 | 7638 | 8472 | 10735 | 8938 | 10306 | 8522 | 8892 | 5705 | 6247 | 9560 | 10029 | 11766 | 8026 | 9512 | 10432 | 7405 | 10971 | 11545 | 10544 | 8427 | 8057 | Ankrd13b |
| Ankrd13c | 3735 | 3187 | 4543 | 4988 | 4756 | 4395 | 5163 | 3936 | 3419 | 4019 | 3678 | 4182 | 3857 | 4725 | 3853 | 3419 | 4525 | 4876 | 4440 | 3989 | 4929 | 4640 | 3450 | 4130 | 4486 | 4361 | 4233 | 4645 | Ankrd13c |
| Ankrd13d | 1465 | 1830 | 1637 | 2051 | 2316 | 2065 | 1656 | 1152 | 1402 | 1496 | 1416 | 1312 | 1354 | 1076 | 1440 | 1734 | 2002 | 1944 | 1986 | 1838 | 1761 | 1688 | 1333 | 1307 | 1535 | 932 | 1239 | 1019 | Ankrd13d |
| Ankrd16 | 6138 | 5319 | 4849 | 5431 | 5737 | 6489 | 5843 | 6637 | 6197 | 6435 | 7178 | 7493 | 7274 | 7451 | 5724 | 5480 | 4835 | 5150 | 5241 | 6141 | 5711 | 6691 | 5731 | 6355 | 7030 | 5466 | 7193 | 6945 | Ankrd16 |
| Ankrd17 | 23584 | 20913 | 13390 | 18450 | 19571 | 21587 | 20385 | 21648 | 19378 | 16435 | 17676 | 17514 | 17346 | 17375 | 23186 | 19893 | 15796 | 17854 | 18537 | 20663 | 21185 | 17426 | 18698 | 14479 | 17962 | 11348 | 18525 | 16610 | Ankrd17 |
| Ankrd2 | 12 | 14 | 15 | 27 | 25 | 35 | 25 | 7 | 8 | 18 | 24 | 9 | 9 | 5 | 11 | 28 | 13 | 47 | 23 | 30 | 14 | 8 | 23 | 19 | 18 | 8 | 5 | 12 | Ankrd2 |
| Ankrd22 | 6 | 3 | 0 | 1 | 0 | 0 | 0 | 21 | 18 | 6 | 9 | 3 | 5 | 0 | 3 | 3 | 0 | 1 | 0 | 0 | 0 | 2 | 34 | 11 | 0 | 24 | 3 | 0 | Ankrd22 |
| Ankrd23 | 106 | 131 | 127 | 141 | 114 | 127 | 116 | 55 | 47 | 72 | 76 | 55 | 81 | 76 | 55 | 136 | 129 | 125 | 132 | 151 | 150 | 75 | 45 | 60 | 47 | 35 | 73 | 45 | Ankrd23 |
| Ankrd24 | 1380 | 1456 | 1810 | 1900 | 2056 | 1693 | 1662 | 1756 | 2136 | 2601 | 2894 | 1875 | 2096 | 1815 | 1303 | 1710 | 1790 | 2024 | 1889 | 1540 | 1667 | 2592 | 2083 | 2584 | 2205 | 1894 | 1796 | 1703 | Ankrd24 |
| Ankrd26 | 4171 | 3963 | 3107 | 3656 | 3920 | 3973 | 4183 | 4927 | 4180 | 4412 | 3909 | 5047 | 4352 | 5121 | 4116 | 3861 | 3669 | 3502 | 3967 | 4813 | 4312 | 3360 | 3796 | 3940 | 4614 | 3204 | 4804 | 4644 | Ankrd26 |
| Ankrd27 | 4665 | 5167 | 4622 | 4409 | 4140 | 4345 | 4484 | 2610 | 3243 | 3289 | 3384 | 3017 | 2959 | 3061 | 4291 | 4860 | 4512 | 4855 | 4397 | 4158 | 4385 | 2617 | 3031 | 3222 | 3270 | 2938 | 3041 | 2959 | Ankrd27 |
| Ankrd28 | 11300 | 8657 | 7227 | 9334 | 8411 | 9884 | 10505 | 7700 | 7113 | 6450 | 6636 | 7482 | 7269 | 7661 | 10610 | 8834 | 7232 | 8899 | 8179 | 9468 | 10013 | 6681 | 6922 | 6131 | 7103 | 5801 | 7448 | 7334 | Ankrd28 |
| Ankrd29 | 589 | 495 | 539 | 491 | 655 | 597 | 516 | 618 | 619 | 570 | 549 | 445 | 577 | 394 | 636 | 597 | 524 | 460 | 573 | 521 | 503 | 662 | 617 | 427 | 454 | 426 | 450 | 456 | Ankrd29 |
| Ankrd31 | 3 | 0 | 5 | 0 | 3 | 4 | 6 | 1 | 1 | 1 | 6 | 1 | 1 | 3 | 1 | 2 | 6 | 0 | 2 | 5 | 3 | 5 | 3 | 0 | 3 | 0 | 3 | 8 | Ankrd31 |
| Ankrd33 | 0 | 2 | 0 | 1 | 0 | 3 | 1 | 0 | 1 | 0 | 2 | 0 | 0 | 0 | 0 | 1 | 0 | 0 | 0 | 2 | 0 | 3 | 1 | 0 | 0 | 0 | 0 | 0 | Ankrd33 |
| Ankrd33b | 194 | 135 | 237 | 183 | 177 | 151 | 135 | 84 | 66 | 89 | 132 | 97 | 107 | 95 | 173 | 175 | 169 | 180 | 125 | 116 | 173 | 99 | 93 | 82 | 77 | 55 | 117 | 101 | Ankrd33b |
| Ankrd34a | 538 | 799 | 643 | 813 | 899 | 709 | 735 | 238 | 352 | 399 | 340 | 248 | 168 | 244 | 491 | 598 | 811 | 938 | 981 | 705 | 879 | 388 | 277 | 354 | 371 | 225 | 195 | 174 | Ankrd34a |
| Ankrd34b | 532 | 658 | 629 | 735 | 931 | 800 | 942 | 176 | 132 | 184 | 201 | 74 | 98 | 102 | 666 | 598 | 661 | 870 | 888 | 942 | 965 | 237 | 152 | 118 | 117 | 40 | 91 | 66 | Ankrd34b |
| Ankrd34c | 6 | 5 | 13 | 11 | 43 | 40 | 20 | 7 | 8 | 0 | 5 | 0 | 2 | 0 | 24 | 9 | 8 | 14 | 6 | 16 | 13 | 8 | 3 | 0 | 9 | 5 | 1 | 0 | Ankrd34c |
| Ankrd35 | 489 | 543 | 552 | 587 | 369 | 391 | 367 | 101 | 131 | 137 | 157 | 108 | 69 | 109 | 491 | 702 | 446 | 621 | 336 | 328 | 323 | 119 | 167 | 181 | 135 | 121 | 79 | 89 | Ankrd35 |
| Ankrd36 | 5 | 5 | 10 | 7 | 7 | 8 | 1 | 1 | 4 | 0 | 1 | 1 | 7 | 6 | 3 | 5 | 3 | 2 | 2 | 6 | 2 | 0 | 1 | 2 | 4 | 3 | 27 | 13 | Ankrd36 |
| Ankrd37 | 264 | 269 | 225 | 415 | 336 | 410 | 379 | 409 | 403 | 412 | 462 | 298 | 339 | 401 | 314 | 238 | 290 | 342 | 373 | 468 | 366 | 389 | 524 | 519 | 537 | 315 | 358 | 349 | Ankrd37 |
| Ankrd39 | 1001 | 1080 | 870 | 1147 | 1045 | 956 | 980 | 525 | 530 | 644 | 611 | 553 | 579 | 549 | 997 | 1012 | 1052 | 1004 | 997 | 937 | 993 | 657 | 622 | 525 | 615 | 385 | 538 | 586 | Ankrd39 |
| Ankrd40 | 3894 | 4038 | 5409 | 4128 | 3868 | 3649 | 3709 | 3742 | 3928 | 4483 | 4700 | 5060 | 4452 | 5196 | 3492 | 4012 | 4967 | 3637 | 4122 | 2969 | 3806 | 4187 | 3968 | 5046 | 4287 | 6106 | 4788 | 5328 | Ankrd40 |
| Ankrd42 | 2177 | 2201 | 1741 | 2122 | 2131 | 2128 | 2320 | 1180 | 1067 | 1102 | 972 | 988 | 930 | 1037 | 2140 | 2107 | 2072 | 2154 | 1963 | 2131 | 2206 | 1034 | 1039 | 877 | 1035 | 651 | 1011 | 914 | Ankrd42 |
| Ankrd44 | 2420 | 1757 | 1328 | 1665 | 2027 | 2245 | 1894 | 3778 | 3976 | 3699 | 4081 | 3125 | 3229 | 3196 | 2014 | 2373 | 1353 | 1830 | 1785 | 1966 | 1961 | 3955 | 3752 | 3174 | 3872 | 2469 | 3067 | 2659 | Ankrd44 |
| Ankrd45 | 912 | 750 | 704 | 934 | 1316 | 1182 | 1440 | 1318 | 1160 | 1160 | 926 | 1098 | 915 | 1037 | 1019 | 748 | 950 | 1119 | 1360 | 1368 | 1288 | 1464 | 1099 | 1146 | 1441 | 1000 | 1008 | 1053 | Ankrd45 |

| | | | | | | | | | | | | | | | | | | | | | | | | | | | | | |
|---|---|---|---|---|---|---|---|---|---|---|---|---|---|---|---|---|---|---|---|---|---|---|---|---|---|---|---|---|---|
| Ankrd46 | 7448 | 7144 | 6629 | 6942 | 6732 | 7041 | 6612 | 7588 | 7373 | 7021 | 6970 | 6934 | 6783 | 6556 | 7622 | 6988 | 6782 | 6748 | 7279 | 7409 | 6864 | 7329 | 7502 | 6892 | 7313 | 6932 | 7494 | 6824 | Ankrd46 |
| Ankrd49 | 3332 | 2458 | 2541 | 2257 | 2217 | 2652 | 2972 | 3626 | 3284 | 2798 | 2645 | 3404 | 3208 | 3473 | 3225 | 2762 | 2730 | 2303 | 2484 | 3199 | 2683 | 2983 | 3146 | 3282 | 2969 | 3843 | 3749 | 3755 | Ankrd49 |
| Ankrd50 | 9646 | 11033 | 8929 | 12029 | 10596 | 10820 | 10047 | 9782 | 9009 | 9599 | 9424 | 10567 | 11253 | 10120 | 9965 | 10543 | 10339 | 11180 | 10823 | 9649 | 10881 | 10368 | 8857 | 9157 | 10141 | 6784 | 10838 | 10272 | Ankrd50 |
| Ankrd52 | 5890 | 6499 | 7634 | 7709 | 8759 | 7618 | 7777 | 6881 | 8098 | 9336 | 9136 | 10622 | 9191 | 10245 | 5087 | 5756 | 8379 | 8033 | 9006 | 6045 | 8258 | 9914 | 7175 | 10203 | 9983 | 6505 | 9209 | 9360 | Ankrd52 |
| Ankrd53 | 0 | 0 | 1 | 0 | 1 | 0 | 1 | 0 | 0 | 1 | 1 | 0 | 1 | 1 | 0 | 1 | 0 | 0 | 0 | 0 | 3 | 2 | 0 | 2 | 0 | 5 | 0 | 0 | Ankrd53 |
| Ankrd54 | 2754 | 3687 | 2646 | 3303 | 2477 | 2559 | 2407 | 1655 | 1921 | 1799 | 1650 | 1972 | 1871 | 1969 | 2557 | 2984 | 3005 | 2916 | 3060 | 2586 | 2364 | 1751 | 1826 | 1932 | 1771 | 2261 | 2079 | 2044 | Ankrd54 |
| Ankrd55 | 118 | 149 | 124 | 175 | 226 | 221 | 179 | 203 | 232 | 213 | 301 | 200 | 239 | 165 | 146 | 222 | 156 | 195 | 192 | 226 | 219 | 259 | 296 | 166 | 244 | 161 | 210 | 196 | Ankrd55 |
| Ankrd6 | 4888 | 5381 | 5538 | 5749 | 5866 | 6102 | 5637 | 4741 | 4107 | 4494 | 3702 | 4854 | 4727 | 4871 | 5401 | 5182 | 5655 | 5657 | 5799 | 5677 | 6205 | 4836 | 3770 | 4529 | 5216 | 3486 | 5421 | 5141 | Ankrd6 |
| Ankrd60 | 5 | 3 | 0 | 1 | 2 | 0 | 2 | 0 | 0 | 1 | 0 | 1 | 3 | 2 | 1 | 1 | 8 | 11 | 3 | 1 | 1 | 0 | 1 | 0 | 0 | 0 | 0 | 0 | Ankrd60 |
| Ankrd61 | 59 | 71 | 35 | 51 | 47 | 55 | 60 | 18 | 37 | 33 | 27 | 40 | 45 | 39 | 32 | 41 | 32 | 70 | 48 | 53 | 57 | 48 | 39 | 20 | 43 | 39 | 43 | 38 | Ankrd61 |
| Ankrd63 | 48 | 54 | 81 | 46 | 65 | 42 | 72 | 23 | 22 | 25 | 27 | 9 | 2 | 9 | 36 | 43 | 101 | 46 | 58 | 65 | 57 | 10 | 6 | 39 | 22 | 16 | 7 | 12 | Ankrd63 |
| Ankrd65 | 0 | 0 | 1 | 0 | 3 | 2 | 0 | 7 | 1 | 0 | 3 | 5 | 2 | 2 | 0 | 0 | 2 | 0 | 0 | 0 | 0 | 2 | 1 | 2 | 2 | 2 | 0 | 0 | Ankrd65 |
| Ankrd66 | 20 | 34 | 33 | 9 | 57 | 18 | 21 | 26 | 42 | 32 | 28 | 72 | 75 | 5 | 37 | 41 | 1 | 9 | 26 | 15 | 17 | 62 | 81 | 20 | 15 | 11 | 12 | 11 | Ankrd66 |
| Ankrd7 | 17 | 13 | 11 | 20 | 20 | 17 | 23 | 13 | 12 | 4 | 15 | 25 | 27 | 27 | 20 | 21 | 37 | 29 | 13 | 21 | 35 | 22 | 17 | 19 | 20 | 21 | 10 | 17 | Ankrd7 |
| Ankrd9 | 353 | 460 | 461 | 412 | 293 | 310 | 315 | 172 | 181 | 164 | 213 | 216 | 243 | 245 | 322 | 396 | 462 | 382 | 354 | 319 | 311 | 204 | 206 | 182 | 169 | 267 | 259 | 261 | Ankrd9 |
| Anks1 | 5619 | 7667 | 8448 | 8520 | 8732 | 7656 | 7156 | 6151 | 6650 | 8112 | 7181 | 8653 | 7186 | 7142 | 5902 | 7051 | 8757 | 8665 | 9411 | 6742 | 7774 | 8502 | 5737 | 8344 | 8853 | 7133 | 7102 | 7188 | Anks1 |
| Anks1b | 741 | 902 | 788 | 1657 | 2686 | 2087 | 2338 | 2424 | 2453 | 2888 | 2403 | 1995 | 1676 | 2081 | 1196 | 696 | 1311 | 2100 | 2297 | 2155 | 2373 | 3863 | 2340 | 2483 | 3555 | 1488 | 1646 | 1690 | Anks1b |
| Anks3 | 4398 | 5385 | 3941 | 4993 | 4381 | 4672 | 4403 | 3848 | 4294 | 4194 | 4090 | 4060 | 4073 | 3660 | 4383 | 5063 | 4457 | 4933 | 4634 | 4870 | 4423 | 4207 | 4161 | 4007 | 4432 | 3643 | 4350 | 3648 | Anks3 |
| Anks4b | 9 | 9 | 2 | 13 | 11 | 10 | 3 | 7 | 8 | 1 | 3 | 3 | 17 | 6 | 12 | 2 | 7 | 8 | 6 | 3 | 7 | 7 | 9 | 7 | 4 | 7 | 5 | 8 | Anks4b |
| Anks6 | 1130 | 1437 | 1274 | 1359 | 1508 | 1388 | 1052 | 723 | 971 | 891 | 1078 | 817 | 754 | 701 | 1062 | 1380 | 1329 | 1354 | 1333 | 1087 | 1105 | 912 | 981 | 910 | 961 | 862 | 752 | 716 | Anks6 |
| Ankub1 | 0 | 0 | 0 | 6 | 0 | 0 | 1 | 6 | 3 | 3 | 1 | 0 | 11 | 3 | 0 | 3 | 0 | 2 | 0 | 3 | 0 | 17 | 6 | 0 | 1 | 0 | 0 | 0 | Ankub1 |
| Ankzf1 | 2663 | 3141 | 2921 | 3397 | 2858 | 2881 | 2966 | 2094 | 2247 | 2356 | 2323 | 2139 | 2236 | 2206 | 2673 | 2769 | 3046 | 3264 | 3097 | 2903 | 2788 | 1920 | 2120 | 2300 | 2336 | 2058 | 2237 | 2052 | Ankzf1 |
| Anln | 4802 | 3937 | 3982 | 2746 | 2466 | 2984 | 3209 | 5235 | 5037 | 4059 | 4136 | 5459 | 5059 | 5509 | 4145 | 4084 | 3509 | 2848 | 3024 | 3288 | 2847 | 3462 | 4842 | 4609 | 3788 | 5605 | 5610 | 5715 | Anln |
| Ano1 | 922 | 832 | 858 | 769 | 821 | 903 | 701 | 1495 | 1443 | 1539 | 1261 | 2322 | 1630 | 2256 | 655 | 814 | 969 | 670 | 978 | 635 | 754 | 1437 | 1034 | 1758 | 1620 | 2035 | 2449 | 2235 | Ano1 |
| Ano10 | 4201 | 4521 | 3796 | 3361 | 3671 | 3650 | 3447 | 2836 | 2780 | 2443 | 2709 | 2618 | 3020 | 2500 | 4441 | 4587 | 3896 | 3778 | 3582 | 3475 | 3514 | 3013 | 2896 | 2402 | 2547 | 2370 | 2707 | 2626 | Ano10 |
| Ano2 | 8 | 10 | 10 | 4 | 9 | 10 | 3 | 27 | 28 | 8 | 28 | 15 | 37 | 6 | 16 | 26 | 2 | 6 | 0 | 1 | 6 | 10 | 27 | 29 | 3 | 42 | 13 | 14 | Ano2 |
| Ano3 | 243 | 246 | 304 | 248 | 268 | 170 | 229 | 313 | 269 | 248 | 234 | 220 | 270 | 211 | 301 | 312 | 309 | 187 | 182 | 275 | 220 | 297 | 263 | 154 | 194 | 161 | 224 | 260 | Ano3 |
| Ano4 | 236 | 269 | 189 | 316 | 482 | 397 | 388 | 712 | 620 | 517 | 437 | 403 | 371 | 495 | 318 | 210 | 246 | 362 | 336 | 494 | 412 | 749 | 469 | 362 | 631 | 290 | 475 | 389 | Ano4 |
| Ano5 | 130 | 84 | 112 | 83 | 244 | 215 | 170 | 110 | 92 | 80 | 283 | 64 | 88 | 72 | 178 | 114 | 114 | 112 | 144 | 214 | 189 | 125 | 197 | 72 | 83 | 37 | 87 | 61 | Ano5 |
| Ano6 | 6502 | 6271 | 4609 | 4786 | 5304 | 5567 | 4875 | 5409 | 6023 | 5520 | 5796 | 4835 | 5640 | 4799 | 6076 | 6600 | 4713 | 5122 | 4752 | 5322 | 4894 | 5098 | 6117 | 4877 | 4926 | 4125 | 5063 | 4753 | Ano6 |
| Ano7 | 5 | 9 | 9 | 4 | 9 | 1 | 2 | 15 | 22 | 13 | 32 | 4 | 9 | 26 | 3 | 11 | 9 | 8 | 3 | 7 | 2 | 10 | 21 | 8 | 5 | 5 | 8 | 8 | Ano7 |
| Ano8 | 3308 | 3940 | 5339 | 5534 | 6245 | 5411 | 4954 | 6557 | 6844 | 9267 | 7546 | 9455 | 7666 | 8601 | 3319 | 3454 | 6070 | 5497 | 6820 | 4882 | 5290 | 9346 | 5812 | 9982 | 10296 | 9089 | 8450 | 8001 | Ano8 |
| Ano9 | 168 | 75 | 31 | 31 | 8 | 19 | 4 | 104 | 88 | 71 | 60 | 5 | 24 | 14 | 57 | 78 | 12 | 59 | 9 | 4 | 4 | 28 | 124 | 81 | 16 | 64 | 8 | 7 | Ano9 |
| Anp32a | 29109 | 25734 | 36041 | 29703 | 24818 | 26510 | 30388 | 29771 | 26028 | 30219 | 25946 | 33827 | 31207 | 38626 | 29204 | 24789 | 31217 | 26106 | 28442 | 31604 | 28774 | 22020 | 25161 | 36038 | 30912 | 43961 | 36101 | 39303 | Anp32a |
| Anp32b | 12664 | 10028 | 15697 | 10851 | 8966 | 9646 | 10876 | 15520 | 15765 | 18696 | 17815 | 23248 | 20155 | 26787 | 11409 | 11011 | 12913 | 9127 | 10299 | 9043 | 10085 | 14051 | 15312 | 22514 | 17304 | 30072 | 23423 | 28188 | Anp32b |
| Anp32e | 25535 | 21153 | 30084 | 23791 | 22054 | 23199 | 26418 | 25457 | 21626 | 23966 | 22410 | 32727 | 27857 | 36316 | 24857 | 20119 | 27310 | 21454 | 23927 | 25784 | 23582 | 20568 | 20152 | 29882 | 17314 | 39452 | 35805 | 35719 | Anp32e |
| Anpep | 629 | 785 | 916 | 734 | 707 | 810 | 402 | 545 | 864 | 884 | 1206 | 706 | 864 | 607 | 636 | 967 | 676 | 977 | 545 | 393 | 449 | 782 | 1157 | 854 | 722 | 667 | 652 | 547 | Anpep |
| Antkmt | 1816 | 2452 | 3031 | 2584 | 1755 | 1567 | 1796 | 996 | 1090 | 1339 | 1193 | 1634 | 1352 | 1503 | 1723 | 1870 | 2972 | 2187 | 2219 | 1662 | 1692 | 993 | 1013 | 1548 | 1306 | 1920 | 1656 | 1567 | Antkmt |
| Antxr1 | 2653 | 1789 | 1917 | 1514 | 1411 | 1889 | 1019 | 4068 | 5065 | 4586 | 6666 | 3683 | 4262 | 3688 | 2002 | 2475 | 1658 | 1589 | 1340 | 1037 | 1146 | 4030 | 6512 | 4322 | 3220 | 4288 | 3453 | 3668 | Antxr1 |
| Antxr2 | 910 | 772 | 894 | 530 | 610 | 706 | 577 | 1215 | 1487 | 1061 | 1965 | 1149 | 1406 | 1296 | 881 | 910 | 727 | 623 | 565 | 546 | 597 | 1332 | 1681 | 1137 | 1023 | 1447 | 1564 | 1330 | Antxr2 |
| Anxa1 | 366 | 210 | 205 | 129 | 142 | 366 | 56 | 1106 | 1397 | 1014 | 1023 | 332 | 555 | 460 | 348 | 326 | 110 | 382 | 91 | 120 | 86 | 540 | 1770 | 779 | 428 | 306 | 368 | 432 | Anxa1 |
| Anxa10 | 0 | 0 | 0 | 0 | 1 | 0 | 0 | 0 | 0 | 0 | 0 | 0 | 0 | 0 | 0 | 0 | 0 | 0 | 0 | 1 | 0 | 0 | 1 | 0 | 3 | 1 | 1 | 2 | Anxa10 |
| Anxa11 | 476 | 473 | 562 | 374 | 277 | 321 | 170 | 172 | 227 | 287 | 333 | 153 | 198 | 113 | 426 | 600 | 370 | 308 | 238 | 261 | 161 | 157 | 353 | 250 | 144 | 330 | 177 | 135 | Anxa11 |
| Anxa13 | 0 | 0 | 0 | 0 | 0 | 0 | 0 | 0 | 0 | 0 | 0 | 0 | 0 | 0 | 0 | 0 | 0 | 0 | 0 | 0 | 0 | 0 | 1 | 0 | 0 | 0 | 0 | 0 | Anxa13 |
| Anxa2 | 14492 | 11804 | 8836 | 5853 | 5109 | 7996 | 3123 | 13219 | 18123 | 10420 | 14597 | 8647 | 12895 | 8822 | 13383 | 14843 | 6464 | 8981 | 3905 | 4625 | 3471 | 10260 | 19634 | 10263 | 7892 | 9074 | 9590 | 9721 | Anxa2 |
| Anxa3 | 1873 | 1760 | 2116 | 1200 | 1015 | 1339 | 965 | 1236 | 1346 | 1230 | 1476 | 1491 | 1969 | 1743 | 2015 | 2251 | 1418 | 1273 | 939 | 1290 | 925 | 1502 | 1774 | 1314 | 1247 | 1483 | 1995 | 1798 | Anxa3 |
| Anxa4 | 679 | 557 | 537 | 520 | 354 | 626 | 269 | 641 | 741 | 574 | 814 | 437 | 606 | 456 | 715 | 836 | 533 | 604 | 317 | 372 | 269 | 548 | 989 | 602 | 422 | 606 | 439 | 475 | Anxa4 |
| Anxa5 | 7125 | 5820 | 5511 | 3915 | 3576 | 3841 | 2992 | 5177 | 6215 | 4605 | 6420 | 3532 | 4626 | 3857 | 6893 | 7153 | 4511 | 4767 | 2969 | 3506 | 3117 | 4958 | 7182 | 4209 | 3796 | 4831 | 4192 | 4048 | Anxa5 |
| Anxa6 | 10823 | 10127 | 8340 | 5949 | 4896 | 6981 | 3284 | 7097 | 10711 | 7337 | 12520 | 6485 | 9027 | 5773 | 10334 | 13025 | 6189 | 7383 | 3893 | 4065 | 3604 | 6863 | 13389 | 6588 | 5722 | 6847 | 7203 | 6503 | Anxa6 |
| Anxa7 | 2651 | 2129 | 2131 | 1540 | 1417 | 1714 | 1306 | 1834 | 2197 | 1769 | 2293 | 1499 | 1979 | 1463 | 2399 | 2784 | 1665 | 1776 | 1301 | 1211 | 1225 | 1616 | 2559 | 1609 | 1378 | 1723 | 1419 | 1562 | Anxa7 |
| Anxa8 | 91 | 35 | 36 | 6 | 7 | 38 | 0 | 132 | 217 | 71 | 159 | 27 | 95 | 70 | 63 | 89 | 1 | 28 | 0 | 1 | 0 | 22 | 271 | 52 | 19 | 16 | 28 | 68 | Anxa8 |
| Anxa9 | 50 | 48 | 34 | 39 | 38 | 42 | 42 | 92 | 110 | 108 | 112 | 134 | 101 | 104 | 49 | 55 | 60 | 26 | 44 | 24 | 40 | 114 | 107 | 132 | 107 | 116 | 117 | 132 | Anxa9 |
| Aoah | 44 | 46 | 51 | 32 | 30 | 45 | 28 | 40 | 54 | 44 | 52 | 65 | 71 | 37 | 59 | 37 | 35 | 47 | 22 | 41 | 37 | 69 | 98 | 71 | 49 | 50 | 72 | 65 | Aoah |
| Aoc1 | 0 | 0 | 0 | 0 | 0 | 4 | 0 | 0 | 1 | 1 | 0 | 7 | 3 | 1 | 0 | 2 | 0 | 0 | 0 | 0 | 0 | 0 | 0 | 2 | 0 | 6 | 0 | 1 | Aoc1 |
| Aoc2 | 1294 | 1773 | 1584 | 2041 | 1955 | 1653 | 1943 | 815 | 765 | 972 | 894 | 943 | 1008 | 1051 | 1238 | 1653 | 1815 | 2255 | 1935 | 1570 | 1859 | 988 | 752 | 902 | 1022 | 916 | 923 | 939 | Aoc2 |
| Aoc3 | 100 | 153 | 109 | 178 | 106 | 149 | 121 | 23 | 39 | 61 | 104 | 25 | 41 | 49 | 78 | 161 | 198 | 185 | 132 | 89 | 122 | 53 | 112 | 29 | 20 | 28 | 27 | 34 | Aoc3 |
| Aopep | 2945 | 1943 | 1397 | 1342 | 1739 | 1340 | 1364 | 2166 | 2450 | 1773 | 2236 | 1151 | 1973 | 1067 | 1957 | 2261 | 1578 | 1526 | 1284 | 1506 | 1427 | 1404 | 2394 | 1413 | 1323 | 1609 | 1247 | 1129 | Aopep |
| Aox1 | 131 | 79 | 67 | 81 | 65 | 113 | 46 | 70 | 76 | 75 | 105 | 85 | 103 | 88 | 125 | 124 | 71 | 76 | 50 | 61 | 55 | 94 | 139 | 82 | 65 | 82 | 85 | 59 | Aox1 |
| Aox2 | 15 | 5 | 4 | 25 | 3 | 5 | 1 | 90 | 105 | 173 | 126 | 1 | 12 | 10 | 17 | 19 | 5 | 0 | 0 | 1 | 2 | 10 | 166 | 223 | 4 | 73 | 6 | 1 | Aox2 |
| Aox3 | 46 | 15 | 4 | 5 | 6 | 6 | 8 | 34 | 42 | 50 | 78 | 4 | 12 | 5 | 30 | 40 | 6 | 9 | 4 | 4 | 13 | 30 | 83 | 59 | 17 | 54 | 3 | 5 | Aox3 |
| Aox4 | 2 | 7 | 1 | 5 | 11 | 21 | 9 | 31 | 41 | 27 | 21 | 0 | 3 | 3 | 0 | 3 | 0 | 17 | 13 | 7 | 18 | 7 | 71 | 34 | 22 | 1 | 5 | 9 | Aox4 |
| Ap1ar | 8172 | 7775 | 8274 | 9260 | 8081 | 7962 | 9231 | 6491 | 5789 | 6374 | 6157 | 6626 | 6437 | 6884 | 8313 | 7431 | 8073 | 8781 | 8349 | 8000 | 8636 | 7008 | 5579 | 6561 | 6853 | 6979 | 6415 | 6705 | Ap1ar |
| Ap1b1 | 6125 | 7464 | 6504 | 7123 | 7588 | 7255 | 6757 | 6095 | 7669 | 7493 | 6980 | 7817 | 7612 | 7066 | 6233 | 6624 | 7429 | 7192 | 7838 | 6506 | 7108 | 7895 | 7414 | 7380 | 8060 | 6987 | 7422 | 7068 | Ap1b1 |
| Ap1g1 | 14827 | 13291 | 12460 | 12876 | 13854 | 14102 | 14565 | 17801 | 15199 | 14622 | 14714 | 15465 | 14514 | 15935 | 14852 | 12721 | 13001 | 13558 | 13362 | 14183 | 14534 | 16668 | 14539 | 14111 | 15809 | 13774 | 15208 | 14968 | Ap1g1 |
| Ap1g2 | 464 | 509 | 536 | 660 | 686 | 605 | 622 | 512 | 659 | 644 | 513 | 634 | 484 | 549 | 471 | 469 | 699 | 687 | 734 | 573 | 654 | 581 | 615 | 676 | 705 | 702 | 538 | 522 | Ap1g2 |
| Ap1m1 | 4828 | 6140 | 5167 | 5523 | 6148 | 5647 | 5378 | 8122 | 8662 | 8734 | 8168 | 9722 | 8564 | 9041 | 5259 | 4980 | 5791 | 5357 | 6347 | 5639 | 5936 | 9404 | 8271 | 9004 | 10011 | 9640 | 9528 | 9477 | Ap1m1 |
| Ap1m2 | 535 | 216 | 107 | 108 | 50 | 82 | 8 | 304 | 363 | 338 | 213 | 21 | 116 | 72 | 245 | 282 | 42 | 124 | 18 | 14 | 8 | 89 | 386 | 299 | 39 | 336 | 43 | 75 | Ap1m2 |
| Ap1s1 | 5087 | 5807 | 5009 | 5423 | 5086 | 5118 | 6111 | 5752 | 5800 | 5567 | 5175 | 6217 | 5500 | 6807 | 5119 | 4910 | 5760 | 5045 | 5746 | 5750 | 5905 | 5394 | 5814 | 6308 | 6001 | 6858 | 6083 | 6596 | Ap1s1 |
| Ap1s2 | 5526 | 4172 | 4799 | 5842 | 7337 | 6948 | 8381 | 5130 | 4608 | 4489 | 4720 | 4340 | 3800 | 4803 | 6062 | 5060 | 5416 | 6634 | 6629 | 6583 | 7971 | 5825 | 4838 | 4456 | 5123 | 4228 | 4312 | 4600 | Ap1s2 |
| Ap1s3 | 865 | 501 | 736 | 546 | 428 | 435 | 472 | 187 | 202 | 203 | 196 | 127 | 178 | 162 | 632 | 811 | 494 | 538 | 392 | 338 | 396 | 85 | 205 | 205 | 97 | 310 | 102 | 201 | Ap1s3 |
| Ap2a1 | 8007 | 10425 | 11190 | 12073 | 11472 | 10028 | 8924 | 5709 | 6924 | 8728 | 7821 | 8905 | 7351 | 7508 | 7473 | 9547 | 12269 | 12307 | 12130 | 7945 | 10146 | 8293 | 6873 | 8633 | 8492 | 9155 | 7226 | 7060 | Ap2a1 |
| Ap2a2 | 16182 | 18865 | 17987 | 18406 | 18131 | 17709 | 16199 | 11483 | 12970 | 13320 | 12908 | 13700 | 13012 | 12190 | 15555 | 17010 | 18337 | 18330 | 19163 | 15552 | 17141 | 12938 | 12688 | 12977 | 13857 | 13216 | 12955 | 12876 | Ap2a2 |
| Ap2b1 | 24666 | 24247 | 21449 | 24142 | 26421 | 25891 | 26273 | 24814 | 24542 | 23689 | 21768 | 23343 | 22970 | 22663 | 24958 | 22821 | 23526 | 25546 | 26814 | 24050 | 27255 | 28340 | 22912 | 22229 | 25253 | 20451 | 22634 | 23250 | Ap2b1 |
| Ap2m1 | 17602 | 22473 | 20203 | 23861 | 21486 | 22146 | 21839 | 17265 | 19218 | 18634 | 18264 | 19262 | 19224 | 18713 | 18092 | 19706 | 20892 | 21289 | 25983 | 21447 | 21963 | 17248 | 19476 | 20138 | 21207 | 21470 | 20299 | 18910 | Ap2m1 |
| Ap2s1 | 4516 | 5738 | 4080 | 4791 | 4649 | 4668 | 4646 | 4513 | 4930 | 4397 | 4172 | 4281 | 4426 | 4415 | 4721 | 4484 | 4691 | 4640 | 4966 | 5145 | 4908 | 4151 | 4858 | 4640 | 4716 | 4680 | 4389 | 4609 | Ap2s1 |
| Ap3b1 | 6649 | 6075 | 5814 | 5189 | 5183 | 5571 | 5615 | 5217 | 5127 | 4654 | 4980 | 4474 | 4609 | 4299 | 6269 | 6122 | 5386 | 5555 | 5382 | 5270 | 5621 | 4563 | 5465 | 4776 | 4615 | 4408 | 4825 | 4798 | Ap3b1 |
| Ap3b2 | 4331 | 6372 | 4739 | 7464 | 9650 | 8351 | 7904 | 7888 | 7666 | 8917 | 7066 | 6795 | 6289 | 6738 | 5689 | 4582 | 6592 | 7717 | 9021 | 8009 | 8930 | 9895 | 6754 | 7442 | 9720 | 4927 | 6651 | 5903 | Ap3b2 |
| Ap3d1 | 15560 | 18438 | 15855 | 18559 | 16235 | 15870 | 15545 | 11237 | 13104 | 14026 | 12992 | 13725 | 12737 | 12641 | 15004 | 16588 | 16932 | 17758 | 17572 | 14213 | 16429 | 14432 | 12796 | 13607 | 13666 | 12519 | 12526 | 12848 | Ap3d1 |
| Ap3m1 | 14275 | 12737 | 11720 | 10900 | 9970 | 10487 | 12608 | 10807 | 8980 | 7671 | 7520 | 9588 | 8659 | 9383 | 15840 | 12985 | 11346 | 10793 | 10514 | 13429 | 11516 | 8582 | 8890 | 8173 | 8275 | 9292 | 9737 | 9995 | Ap3m1 |

| | | | | | | | | | | | | | | | | | | | | | | | | | | | | | |
|---|---|---|---|---|---|---|---|---|---|---|---|---|---|---|---|---|---|---|---|---|---|---|---|---|---|---|---|---|---|
| Ap3m2 | 3787 | 5232 | 3661 | 5345 | 5506 | 5051 | 5006 | 3780 | 3681 | 3786 | 3300 | 3355 | 3399 | 3244 | 4713 | 4237 | 4595 | 5507 | 5354 | 5538 | 5319 | 4351 | 3500 | 3289 | 3774 | 2874 | 3266 | 3218 | Ap3m2 |
| Ap3s1 | 8975 | 6913 | 8029 | 9063 | 8631 | 8939 | 11394 | 7192 | 6018 | 6287 | 5775 | 6604 | 6243 | 7331 | 9370 | 8018 | 8496 | 8900 | 8837 | 10031 | 10367 | 6662 | 6256 | 6497 | 6695 | 6881 | 6386 | 7113 | Ap3s1 |
| Ap3s2 | 9388 | 10268 | 10508 | 11392 | 10736 | 10294 | 10983 | 8280 | 8039 | 8883 | 7933 | 9747 | 8566 | 9851 | 9566 | 8783 | 11356 | 11184 | 10841 | 9574 | 11422 | 8834 | 7549 | 9077 | 10273 | 9167 | 9376 | 9653 | Ap3s2 |
| Ap4b1 | 68 | 55 | 32 | 50 | 44 | 43 | 50 | 93 | 80 | 64 | 51 | 105 | 50 | 96 | 79 | 43 | 29 | 37 | 37 | 47 | 60 | 63 | 63 | 83 | 62 | 46 | 50 | 62 | Ap4b1 |
| Ap4e1 | 2624 | 2128 | 1706 | 2256 | 2274 | 2671 | 2616 | 3329 | 2970 | 2886 | 3213 | 2888 | 2999 | 3141 | 2520 | 2445 | 1815 | 2248 | 2222 | 2461 | 2602 | 2844 | 2862 | 2589 | 2916 | 1681 | 3045 | 2763 | Ap4e1 |
| Ap4m1 | 2475 | 2858 | 2786 | 2751 | 2959 | 2587 | 2913 | 5385 | 5401 | 5642 | 5849 | 5908 | 5389 | 6030 | 2447 | 2430 | 2741 | 2641 | 3006 | 2664 | 2811 | 5513 | 5256 | 6235 | 5951 | 5853 | 5736 | 5681 | Ap4m1 |
| Ap4s1 | 615 | 567 | 677 | 584 | 668 | 659 | 739 | 1440 | 1069 | 1137 | 1089 | 1397 | 1266 | 1561 | 674 | 579 | 663 | 530 | 634 | 790 | 630 | 1228 | 1062 | 1245 | 1314 | 1297 | 1460 | 1600 | Ap4s1 |
| Ap5b1 | 175 | 216 | 219 | 145 | 215 | 192 | 161 | 156 | 238 | 265 | 221 | 214 | 223 | 170 | 131 | 206 | 212 | 163 | 202 | 178 | 163 | 209 | 242 | 186 | 205 | 170 | 184 | 161 | Ap5b1 |
| Ap5m1 | 5666 | 4939 | 4735 | 4587 | 3738 | 4634 | 4407 | 4798 | 3869 | 3410 | 3452 | 3751 | 3920 | 3939 | 5792 | 5251 | 4216 | 4040 | 3876 | 5013 | 4452 | 3604 | 3871 | 3140 | 3328 | 3085 | 4210 | 4160 | Ap5m1 |
| Ap5s1 | 952 | 1137 | 1123 | 1047 | 1017 | 1002 | 1077 | 568 | 766 | 745 | 768 | 799 | 745 | 685 | 969 | 961 | 1304 | 1052 | 1159 | 1045 | 1032 | 690 | 728 | 727 | 806 | 838 | 788 | 777 | Ap5s1 |
| Ap5z1 | 1413 | 2008 | 1953 | 2069 | 1726 | 1777 | 1820 | 1185 | 1430 | 1714 | 1518 | 1658 | 1583 | 1619 | 1412 | 1840 | 2221 | 2172 | 1908 | 1511 | 1757 | 1529 | 1489 | 1647 | 1624 | 1559 | 1456 | 1499 | Ap5z1 |
| Apaf1 | 11988 | 10066 | 8804 | 10459 | 10366 | 11179 | 11515 | 6303 | 5883 | 5956 | 6531 | 6570 | 6427 | 6292 | 11404 | 10728 | 10050 | 11077 | 10202 | 10194 | 10879 | 6219 | 5972 | 5643 | 6293 | 5341 | 6393 | 6267 | Apaf1 |
| Apba1 | 1826 | 2390 | 2419 | 3936 | 5600 | 4040 | 3636 | 3280 | 3515 | 4895 | 4081 | 2723 | 2899 | 2635 | 2259 | 2098 | 3461 | 4709 | 4472 | 3637 | 4291 | 5738 | 3407 | 3481 | 4946 | 1931 | 2310 | 2300 | Apba1 |
| Apba2 | 3372 | 4331 | 3887 | 5507 | 7221 | 6487 | 6446 | 7199 | 7928 | 9585 | 7119 | 9621 | 7921 | 8961 | 3938 | 3572 | 5030 | 5852 | 6982 | 6218 | 6593 | 9900 | 7445 | 9014 | 11029 | 6991 | 8214 | 8268 | Apba2 |
| Apba3 | 915 | 1249 | 1205 | 1131 | 979 | 943 | 794 | 718 | 973 | 956 | 1026 | 891 | 959 | 765 | 881 | 1136 | 1172 | 995 | 966 | 886 | 799 | 891 | 1018 | 1084 | 874 | 1047 | 898 | 808 | Apba3 |
| Apbb1 | 6485 | 8702 | 8517 | 10303 | 11274 | 10481 | 9198 | 9358 | 9923 | 11356 | 8922 | 10276 | 9300 | 9618 | 7234 | 7131 | 9430 | 9584 | 12127 | 10200 | 10585 | 10079 | 9688 | 10525 | 11532 | 9938 | 9358 | 9088 | Apbb1 |
| Apbb1ip | 133 | 122 | 132 | 71 | 105 | 112 | 60 | 160 | 194 | 144 | 212 | 185 | 217 | 159 | 153 | 158 | 79 | 107 | 113 | 73 | 84 | 175 | 223 | 159 | 137 | 206 | 164 | 139 | Apbb1ip |
| Apbb2 | 3829 | 3785 | 3264 | 4538 | 5492 | 5221 | 4733 | 4488 | 4827 | 5423 | 5093 | 4016 | 4388 | 3858 | 4092 | 3931 | 4130 | 4914 | 4841 | 4824 | 4837 | 5683 | 4577 | 4427 | 5133 | 3141 | 3975 | 3836 | Apbb2 |
| Apbb3 | 2774 | 2902 | 2502 | 3362 | 3283 | 3278 | 3084 | 2640 | 2795 | 2836 | 3146 | 2950 | 3004 | 2872 | 2759 | 2697 | 2878 | 3175 | 3171 | 3110 | 3073 | 2889 | 2544 | 2938 | 2942 | 2275 | 3022 | 2655 | Apbb3 |
| Apc | 26404 | 24306 | 14195 | 23055 | 32127 | 36727 | 34134 | 44512 | 38458 | 33076 | 31744 | 30316 | 30481 | 32290 | 30005 | 23140 | 18463 | 24434 | 26772 | 37361 | 36282 | 33175 | 34783 | 24988 | 38261 | 17763 | 32847 | 26743 | Apc |
| Apc2 | 10576 | 16116 | 18029 | 25548 | 28939 | 24028 | 21499 | 22769 | 26060 | 35717 | 26925 | 35575 | 26055 | 30702 | 12481 | 11576 | 24087 | 24384 | 30677 | 20128 | 24929 | 40494 | 22340 | 36222 | 42782 | 21437 | 28123 | 28106 | Apc2 |
| Apcdd1 | 10785 | 8746 | 11312 | 8627 | 10014 | 9721 | 8670 | 10479 | 12286 | 10506 | 9699 | 11499 | 11509 | 11346 | 8097 | 8069 | 9123 | 9441 | 10031 | 9066 | 9509 | 9115 | 13966 | 10260 | 10235 | 9981 | 12420 | 12142 | Apcdd1 |
| Apeh | 5317 | 6565 | 5711 | 5178 | 4379 | 4527 | 3961 | 3373 | 3313 | 3314 | 3133 | 3627 | 3955 | 3312 | 5164 | 5524 | 5738 | 4833 | 4998 | 4895 | 4290 | 3046 | 3366 | 3340 | 3353 | 3603 | 3760 | 3748 | Apeh |
| Apela | 172 | 105 | 84 | 102 | 71 | 77 | 99 | 65 | 63 | 74 | 45 | 38 | 10 | 80 | 144 | 99 | 145 | 88 | 100 | 54 | 92 | 40 | 39 | 60 | 46 | 61 | 71 | 55 | Apela |
| Apex1 | 7782 | 7565 | 6918 | 6205 | 6139 | 6535 | 7099 | 8387 | 6982 | 6462 | 5950 | 7979 | 7274 | 8886 | 7805 | 6382 | 6968 | 5832 | 7006 | 8102 | 7022 | 6082 | 6614 | 7778 | 7040 | 9615 | 8548 | 9165 | Apex1 |
| Apex2 | 1076 | 1119 | 909 | 896 | 991 | 1075 | 903 | 1694 | 1594 | 1528 | 1422 | 1552 | 1489 | 1522 | 1033 | 1132 | 980 | 754 | 1090 | 1097 | 1009 | 1424 | 1551 | 1461 | 1517 | 1821 | 1621 | 1594 | Apex2 |
| Aph1a | 5360 | 5876 | 7108 | 6091 | 5273 | 5474 | 4911 | 3555 | 3882 | 3763 | 4081 | 4483 | 4308 | 3965 | 5035 | 5672 | 6592 | 6125 | 6208 | 5114 | 4900 | 3798 | 3959 | 4306 | 4331 | 5357 | 4245 | 4154 | Aph1a |
| Aph1b | 5051 | 4380 | 4083 | 5032 | 5829 | 5756 | 6369 | 4426 | 3870 | 3640 | 3307 | 3781 | 3053 | 3653 | 5426 | 4153 | 4552 | 5316 | 5573 | 5806 | 6157 | 4029 | 3519 | 3620 | 4140 | 3217 | 3524 | 3648 | Aph1b |
| Aph1c | 1539 | 1462 | 1189 | 1428 | 1233 | 1344 | 1393 | 252 | 278 | 213 | 240 | 211 | 230 | 227 | 1593 | 1542 | 1270 | 1508 | 1266 | 1290 | 1399 | 264 | 320 | 210 | 247 | 183 | 182 | 208 | Aph1c |
| Api5 | 24314 | 20707 | 21704 | 19385 | 16176 | 17762 | 19791 | 16133 | 13479 | 12715 | 12439 | 15610 | 15221 | 15618 | 23512 | 21925 | 20067 | 18610 | 16892 | 19783 | 18291 | 12953 | 13286 | 13838 | 12818 | 17484 | 16095 | 16798 | Api5 |
| Apip | 1584 | 1386 | 1466 | 1294 | 1278 | 1289 | 1569 | 1274 | 1388 | 1365 | 1452 | 1237 | 1326 | 1457 | 1468 | 1418 | 1523 | 1383 | 1474 | 1437 | 1447 | 1150 | 1310 | 1403 | 1233 | 1472 | 1353 | 1477 | Apip |
| Aplf | 2540 | 2571 | 2674 | 2864 | 2306 | 2377 | 2439 | 950 | 991 | 998 | 1078 | 974 | 1009 | 1053 | 2669 | 2356 | 2824 | 2674 | 2451 | 2408 | 2436 | 866 | 999 | 1068 | 965 | 1201 | 965 | 1000 | Aplf |
| Apln | 843 | 677 | 889 | 849 | 883 | 1111 | 800 | 943 | 1027 | 1063 | 1084 | 829 | 793 | 805 | 905 | 710 | 841 | 972 | 1009 | 667 | 819 | 993 | 1081 | 1026 | 834 | 869 | 716 | 820 | Apln |
| Aplnr | 1043 | 985 | 1611 | 824 | 1240 | 994 | 1014 | 1977 | 2529 | 2297 | 2890 | 2426 | 3072 | 2414 | 981 | 1104 | 1125 | 1037 | 1018 | 1016 | 970 | 2558 | 2897 | 2147 | 2508 | 2693 | 2712 | 2300 | Aplnr |
| Aplp1 | 9724 | 13331 | 10301 | 17768 | 21559 | 18477 | 17021 | 16918 | 17409 | 19396 | 15529 | 14949 | 13981 | 14089 | 11890 | 10615 | 15286 | 18421 | 21860 | 18415 | 19793 | 21763 | 17421 | 16865 | 20495 | 12226 | 13093 | 13309 | Aplp1 |
| Aplp2 | 28305 | 29933 | 26283 | 29138 | 27513 | 26115 | 27436 | 17594 | 17386 | 17497 | 16637 | 15329 | 16484 | 15439 | 29024 | 28778 | 25910 | 31094 | 26135 | 25661 | 28188 | 17615 | 17656 | 15953 | 16330 | 14693 | 15137 | 15338 | Aplp2 |
| Apmap | 3516 | 3742 | 3377 | 3501 | 2994 | 3172 | 3254 | 2497 | 2262 | 2133 | 1934 | 2303 | 2250 | 2384 | 3840 | 3348 | 3117 | 3184 | 3125 | 3531 | 3118 | 2261 | 2305 | 2262 | 2160 | 2546 | 2256 | 2366 | Apmap |
| Apoa1 | 8 | 21 | 10 | 8 | 26 | 6 | 9 | 10 | 86 | 13 | 5 | 0 | 6 | 14 | 8 | 23 | 3 | 13 | 5 | 10 | 6 | 10 | 28 | 11 | 11 | 6 | 9 | 20 | Apoa1 |
| Apoa2 | 32 | 48 | 62 | 49 | 43 | 42 | 26 | 29 | 65 | 30 | 40 | 48 | 56 | 50 | 32 | 51 | 44 | 36 | 37 | 60 | 29 | 43 | 45 | 41 | 33 | 52 | 53 | 56 | Apoa2 |
| Apoa4 | 1 | 7 | 0 | 0 | 9 | 0 | 0 | 0 | 42 | 1 | 2 | 0 | 0 | 2 | 0 | 19 | 1 | 4 | 0 | 1 | 0 | 8 | 14 | 0 | 0 | 0 | 0 | 1 | Apoa4 |
| Apoa5 | 0 | 0 | 0 | 0 | 0 | 0 | 0 | 0 | 2 | 0 | 0 | 0 | 5 | 0 | 0 | 0 | 0 | 0 | 0 | 0 | 0 | 0 | 0 | 0 | 0 | 0 | 0 | 0 | Apoa5 |
| Apob | 8 | 42 | 25 | 2 | 24 | 16 | 8 | 44 | 119 | 36 | 64 | 41 | 48 | 18 | 20 | 48 | 20 | 10 | 5 | 4 | 4 | 75 | 72 | 7 | 21 | 10 | 31 | 13 | Apob |
| Apobec1 | 130 | 113 | 124 | 134 | 185 | 190 | 200 | 145 | 187 | 160 | 149 | 160 | 137 | 152 | 146 | 116 | 123 | 124 | 182 | 205 | 205 | 139 | 170 | 127 | 177 | 180 | 153 | 148 | Apobec1 |
| Apobec2 | 1 | 0 | 0 | 0 | 1 | 0 | 0 | 0 | 2 | 0 | 11 | 0 | 0 | 0 | 1 | 0 | 0 | 3 | 2 | 0 | 0 | 0 | 6 | 0 | 0 | 0 | 0 | 1 | Apobec2 |
| Apobec3 | 200 | 203 | 192 | 174 | 147 | 163 | 164 | 242 | 377 | 355 | 415 | 298 | 266 | 321 | 174 | 202 | 172 | 178 | 186 | 168 | 165 | 261 | 355 | 313 | 272 | 358 | 280 | 306 | Apobec3 |
| Apobec4 | 0 | 0 | 0 | 0 | 0 | 0 | 0 | 6 | 2 | 4 | 0 | 1 | 0 | 0 | 0 | 0 | 1 | 0 | 1 | 0 | 0 | 0 | 0 | 1 | 0 | 0 | 0 | 1 | Apobec4 |
| Apobr | 22 | 21 | 36 | 36 | 16 | 29 | 8 | 48 | 47 | 38 | 80 | 65 | 45 | 49 | 15 | 27 | 30 | 20 | 16 | 14 | 16 | 43 | 89 | 72 | 43 | 62 | 48 | 68 | Apobr |
| Apoc1 | 14 | 13 | 10 | 9 | 11 | 7 | 7 | 11 | 16 | 19 | 20 | 5 | 25 | 14 | 12 | 14 | 12 | 9 | 13 | 15 | 7 | 7 | 27 | 35 | 8 | 11 | 9 | 19 | Apoc1 |
| Apoc3 | 0 | 0 | 0 | 0 | 0 | 0 | 0 | 0 | 0 | 0 | 0 | 0 | 0 | 0 | 0 | 0 | 0 | 0 | 0 | 0 | 1 | 2 | 0 | 1 | 0 | 0 | 0 | 0 | Apoc3 |
| Apoc4 | 0 | 0 | 0 | 0 | 0 | 0 | 0 | 0 | 0 | 0 | 0 | 1 | 0 | 0 | 0 | 0 | 0 | 0 | 0 | 0 | 0 | 0 | 0 | 0 | 0 | 0 | 0 | 0 | Apoc4 |
| Apod | 1279 | 1791 | 1983 | 1198 | 1172 | 1707 | 1045 | 1410 | 1868 | 1579 | 1754 | 1751 | 2631 | 1925 | 1623 | 1962 | 1155 | 1431 | 1146 | 1507 | 1176 | 1719 | 2566 | 1588 | 1812 | 1547 | 1963 | 2082 | Apod |
| Apoe | 3772 | 5430 | 5934 | 4790 | 4767 | 4588 | 3265 | 4078 | 5402 | 5378 | 7263 | 4625 | 5245 | 4534 | 4468 | 5666 | 4744 | 5226 | 3621 | 4426 | 3677 | 5070 | 7898 | 5352 | 5222 | 5709 | 5102 | 4586 | Apoe |
| Apof | 0 | 0 | 0 | 0 | 0 | 0 | 0 | 0 | 1 | 0 | 4 | 0 | 0 | 0 | 0 | 0 | 0 | 0 | 0 | 0 | 1 | 2 | 0 | 0 | 0 | 0 | 0 | 0 | Apof |
| Apoh | 0 | 1 | 0 | 1 | 8 | 0 | 2 | 2 | 1 | 8 | 2 | 0 | 5 | 2 | 0 | 6 | 0 | 1 | 0 | 0 | 0 | 2 | 7 | 7 | 0 | 6 | 0 | 4 | Apoh |
| Apol10a | 0 | 0 | 0 | 0 | 0 | 0 | 0 | 1 | 0 | 0 | 0 | 0 | 0 | 0 | 0 | 0 | 0 | 0 | 0 | 0 | 0 | 2 | 0 | 0 | 0 | 0 | 3 | 0 | Apol10a |
| Apol10b | 1 | 1 | 7 | 0 | 2 | 0 | 0 | 2 | 6 | 10 | 12 | 7 | 6 | 3 | 1 | 4 | 9 | 0 | 2 | 2 | 2 | 7 | 6 | 11 | 3 | 4 | 13 | 11 | Apol10b |
| Apol11a | 0 | 0 | 0 | 0 | 0 | 0 | 0 | 0 | 0 | 0 | 0 | 4 | 0 | 0 | 0 | 0 | 0 | 0 | 0 | 0 | 0 | 2 | 0 | 0 | 0 | 0 | 0 | 0 | Apol11a |
| Apol11b | 0 | 0 | 0 | 0 | 1 | 0 | 0 | 2 | 5 | 0 | 2 | 0 | 0 | 0 | 1 | 0 | 0 | 0 | 1 | 3 | 0 | 2 | 5 | 5 | 3 | 0 | 0 | 4 | Apol11b |
| Apol6 | 2 | 0 | 0 | 0 | 2 | 0 | 1 | 9 | 7 | 10 | 9 | 2 | 3 | 1 | 0 | 0 | 0 | 0 | 0 | 0 | 1 | 5 | 12 | 2 | 5 | 0 | 0 | 4 | Apol6 |
| Apol7a | 0 | 2 | 0 | 0 | 0 | 0 | 0 | 0 | 6 | 0 | 8 | 0 | 0 | 0 | 0 | 1 | 0 | 0 | 0 | 0 | 0 | 3 | 3 | 0 | 0 | 0 | 0 | 0 | Apol7a |
| Apol7b | 0 | 0 | 0 | 0 | 0 | 0 | 0 | 0 | 0 | 0 | 0 | 0 | 0 | 1 | 0 | 0 | 0 | 0 | 0 | 0 | 0 | 0 | 1 | 0 | 0 | 0 | 0 | 0 | Apol7b |
| Apol7c | 1 | 0 | 0 | 0 | 0 | 0 | 0 | 0 | 0 | 0 | 0 | 0 | 0 | 0 | 0 | 0 | 0 | 0 | 0 | 0 | 0 | 0 | 0 | 0 | 0 | 0 | 0 | 0 | Apol7c |
| Apol7e | 0 | 0 | 0 | 1 | 1 | 0 | 0 | 0 | 0 | 0 | 2 | 0 | 0 | 0 | 0 | 0 | 0 | 0 | 0 | 0 | 0 | 0 | 2 | 0 | 1 | 0 | 0 | 0 | Apol7e |
| Apol8 | 48 | 54 | 38 | 64 | 50 | 41 | 59 | 23 | 12 | 27 | 14 | 26 | 42 | 23 | 59 | 57 | 50 | 39 | 35 | 36 | 31 | 22 | 28 | 11 | 13 | 37 | 52 | 74 | Apol8 |
| Apol9a | 1 | 0 | 0 | 0 | 0 | 0 | 0 | 1 | 1 | 0 | 0 | 0 | 0 | 0 | 4 | 0 | 0 | 0 | 0 | 0 | 0 | 7 | 2 | 0 | 0 | 0 | 0 | 0 | Apol9a |
| Apol9b | 0 | 0 | 0 | 0 | 0 | 2 | 1 | 0 | 1 | 1 | 1 | 0 | 0 | 1 | 0 | 1 | 0 | 0 | 0 | 0 | 0 | 0 | 1 | 1 | 0 | 1 | 0 | 0 | Apol9b |
| Apold1 | 705 | 501 | 589 | 763 | 628 | 647 | 646 | 352 | 367 | 414 | 488 | 405 | 414 | 514 | 790 | 672 | 688 | 662 | 572 | 531 | 620 | 326 | 425 | 410 | 507 | 381 | 410 | 510 | Apold1 |
| Apom | 29 | 35 | 41 | 20 | 39 | 21 | 30 | 40 | 54 | 96 | 46 | 59 | 62 | 82 | 21 | 19 | 61 | 64 | 50 | 30 | 26 | 55 | 45 | 83 | 61 | 71 | 71 | 95 | Apom |
| Apon | 32 | 34 | 51 | 45 | 9 | 54 | 35 | 38 | 29 | 45 | 26 | 28 | 36 | 46 | 15 | 33 | 38 | 25 | 23 | 12 | 37 | 23 | 30 | 29 | 39 | 77 | 52 | 39 | Apon |
| Apoo | 2512 | 2356 | 2326 | 2535 | 2248 | 2333 | 2991 | 2315 | 1728 | 1701 | 1620 | 1896 | 1948 | 2425 | 2742 | 2420 | 2300 | 2266 | 2596 | 2947 | 2885 | 1716 | 1821 | 1845 | 1939 | 2077 | 2108 | 2166 | Apoo |
| Apool | 1736 | 1395 | 1780 | 1277 | 1004 | 1160 | 1221 | 1861 | 1525 | 1329 | 1386 | 1721 | 1835 | 1839 | 1811 | 1689 | 1559 | 1034 | 1123 | 1558 | 1166 | 1340 | 1623 | 1527 | 1242 | 2130 | 1942 | 2016 | Apool |
| App | 20977 | 23177 | 24626 | 25608 | 30232 | 26992 | 23732 | 24101 | 28906 | 34155 | 33750 | 27234 | 30563 | 25798 | 23189 | 25906 | 24676 | 29477 | 25337 | 21644 | 24982 | 33352 | 30462 | 30269 | 30356 | 26613 | 25451 | 25763 | App |
| Appbp2 | 11595 | 10497 | 10295 | 10673 | 10992 | 11383 | 11722 | 11333 | 10141 | 10058 | 10404 | 11753 | 11174 | 11935 | 11590 | 10653 | 10593 | 10848 | 10898 | 11480 | 11284 | 11349 | 10167 | 10408 | 11375 | 10186 | 12146 | 11947 | Appbp2 |
| Appl1 | 11227 | 8652 | 7885 | 9620 | 10381 | 10787 | 11210 | 11383 | 9846 | 9249 | 9721 | 9114 | 8780 | 9269 | 11168 | 10175 | 8509 | 9930 | 9484 | 11357 | 10697 | 9625 | 9668 | 8112 | 9737 | 6993 | 9052 | 8492 | Appl1 |
| Appl2 | 5595 | 5383 | 4707 | 5311 | 5488 | 5322 | 5494 | 4513 | 4485 | 4568 | 4711 | 4098 | 4308 | 4068 | 5495 | 5840 | 4917 | 5547 | 5143 | 5360 | 5367 | 4712 | 4527 | 4158 | 4246 | 3862 | 4075 | 3980 | Appl2 |
| Aprt | 3295 | 4386 | 3468 | 2995 | 2365 | 2558 | 2434 | 2379 | 3190 | 2600 | 2844 | 2557 | 2989 | 2517 | 3011 | 3685 | 3394 | 2837 | 2737 | 3032 | 2318 | 1815 | 3236 | 2707 | 2272 | 3552 | 3025 | 2897 | Aprt |
| Aptx | 2241 | 2263 | 2426 | 2615 | 2822 | 2743 | 2853 | 2972 | 3129 | 3359 | 3004 | 3510 | 3309 | 3766 | 2275 | 2065 | 2694 | 2550 | 2643 | 2759 | 2817 | 3355 | 2864 | 3407 | 3711 | 3191 | 3525 | 3692 | Aptx |
| Aqp1 | 1167 | 1487 | 841 | 1007 | 1179 | 1878 | 329 | 1789 | 2721 | 2614 | 2946 | 1630 | 2256 | 1024 | 1176 | 1930 | 727 | 1567 | 517 | 463 | 474 | 2605 | 4495 | 2110 | 1690 | 1144 | 982 | 1036 | Aqp1 |
| Aqp11 | 731 | 840 | 758 | 800 | 816 | 648 | 838 | 425 | 362 | 388 | 380 | 268 | 308 | 320 | 827 | 774 | 700 | 796 | 707 | 894 | 795 | 498 | 448 | 341 | 392 | 291 | 332 | 279 | Aqp11 |

| | | | | | | | | | | | | | | | | | | | | | | | | | | | | | |
|---|---|---|---|---|---|---|---|---|---|---|---|---|---|---|---|---|---|---|---|---|---|---|---|---|---|---|---|---|---|
| Aqp12 | 0 | 0 | 0 | 1 | 0 | 0 | 0 | 0 | 0 | 0 | 0 | 0 | 2 | 0 | 0 | 0 | 0 | 0 | 0 | 0 | 0 | 0 | 0 | 0 | 0 | 0 | 0 | 0 | Aqp12 |
| Aqp2 | 0 | 0 | 0 | 0 | 0 | 0 | 1 | 0 | 0 | 0 | 1 | 0 | 0 | 0 | 0 | 0 | 0 | 0 | 0 | 0 | 0 | 0 | 0 | 0 | 0 | 0 | 0 | 0 | Aqp2 |
| Aqp3 | 72 | 41 | 25 | 53 | 16 | 22 | 11 | 134 | 198 | 206 | 100 | 4 | 42 | 35 | 41 | 72 | 20 | 19 | 16 | 16 | 13 | 28 | 212 | 190 | 21 | 121 | 21 | 82 | Aqp3 |
| Aqp4 | 186 | 164 | 135 | 122 | 232 | 121 | 134 | 265 | 242 | 193 | 197 | 72 | 211 | 77 | 269 | 205 | 88 | 176 | 124 | 146 | 102 | 227 | 216 | 135 | 112 | 62 | 83 | 74 | Aqp4 |
| Aqp5 | 17 | 34 | 42 | 40 | 57 | 49 | 21 | 21 | 32 | 38 | 35 | 14 | 22 | 7 | 21 | 23 | 46 | 52 | 42 | 30 | 32 | 27 | 37 | 10 | 12 | 6 | 13 | 20 | Aqp5 |
| Aqp6 | 16 | 30 | 38 | 42 | 155 | 82 | 2 | 182 | 157 | 243 | 174 | 73 | 254 | 39 | 44 | 35 | 68 | 74 | 70 | 45 | 42 | 306 | 111 | 78 | 118 | 7 | 86 | 46 | Aqp6 |
| Aqp7 | 0 | 0 | 0 | 0 | 0 | 0 | 2 | 0 | 0 | 3 | 3 | 0 | 0 | 0 | 0 | 0 | 0 | 0 | 0 | 0 | 1 | 0 | 1 | 0 | 0 | 3 | 0 | 0 | Aqp7 |
| Aqp8 | 1 | 5 | 12 | 5 | 12 | 2 | 4 | 2 | 12 | 0 | 0 | 7 | 28 | 9 | 3 | 8 | 2 | 2 | 0 | 3 | 2 | 2 | 4 | 0 | 0 | 7 | 19 | 40 | Aqp8 |
| Aqp9 | 3 | 3 | 0 | 12 | 20 | 22 | 12 | 13 | 23 | 25 | 31 | 41 | 41 | 28 | 3 | 2 | 8 | 8 | 9 | 20 | 18 | 17 | 49 | 51 | 31 | 21 | 14 | 20 | Aqp9 |
| Aqr | 11258 | 11866 | 8944 | 8479 | 7976 | 8672 | 9323 | 9639 | 9198 | 7793 | 8262 | 9192 | 8864 | 8969 | 11011 | 9899 | 8841 | 8350 | 8522 | 10202 | 8841 | 7778 | 9082 | 7840 | 8225 | 8631 | 9459 | 9146 | Aqr |
| Ar | 174 | 119 | 129 | 141 | 108 | 183 | 108 | 34 | 40 | 40 | 43 | 14 | 27 | 27 | 166 | 157 | 129 | 138 | 76 | 114 | 117 | 30 | 60 | 20 | 23 | 23 | 27 | 20 | Ar |
| Araf | 6744 | 6578 | 6122 | 7345 | 8264 | 7742 | 7707 | 9548 | 9212 | 10078 | 10208 | 9394 | 9292 | 9279 | 6810 | 7183 | 6780 | 7860 | 7759 | 7545 | 7814 | 11028 | 9366 | 9653 | 10643 | 7927 | 9554 | 9137 | Araf |
| Arap1 | 1275 | 1569 | 1512 | 1349 | 1141 | 1522 | 816 | 1288 | 2066 | 1441 | 2352 | 1295 | 1556 | 1191 | 1245 | 1595 | 1471 | 1497 | 976 | 772 | 868 | 1310 | 2539 | 1660 | 1363 | 1341 | 1429 | 1358 | Arap1 |
| Arap2 | 290 | 235 | 158 | 223 | 355 | 359 | 298 | 611 | 587 | 446 | 457 | 377 | 418 | 364 | 371 | 314 | 189 | 244 | 267 | 334 | 298 | 470 | 662 | 413 | 397 | 223 | 394 | 338 | Arap2 |
| Arap3 | 1059 | 1376 | 1622 | 1164 | 1272 | 1266 | 912 | 1387 | 1820 | 2102 | 2431 | 2139 | 2307 | 1772 | 963 | 1573 | 1292 | 1271 | 1139 | 835 | 945 | 2154 | 2187 | 1961 | 1824 | 2099 | 2173 | 1715 | Arap3 |
| Arc | 144 | 209 | 164 | 309 | 250 | 175 | 191 | 95 | 131 | 156 | 103 | 161 | 88 | 143 | 166 | 156 | 258 | 252 | 251 | 185 | 248 | 139 | 124 | 149 | 156 | 138 | 95 | 115 | Arc |
| Arcn1 | 20738 | 18519 | 17255 | 15218 | 14605 | 15124 | 16346 | 19380 | 17766 | 16282 | 17380 | 16531 | 16146 | 17043 | 20591 | 18856 | 15988 | 15506 | 14601 | 16852 | 15418 | 16987 | 18402 | 16201 | 15569 | 16785 | 16665 | 17488 | Arcn1 |
| Areg | 8 | 12 | 8 | 9 | 17 | 11 | 15 | 16 | 5 | 3 | 4 | 1 | 7 | 0 | 17 | 21 | 9 | 12 | 2 | 6 | 5 | 2 | 3 | 1 | 3 | 6 | 2 | 0 | Areg |
| Arel1 | 6457 | 6996 | 5106 | 6434 | 6907 | 6697 | 6486 | 7023 | 6779 | 6506 | 6034 | 6337 | 6121 | 6241 | 6721 | 5972 | 5838 | 6464 | 6371 | 6676 | 6797 | 6796 | 6368 | 6231 | 7055 | 5313 | 6357 | 6018 | Arel1 |
| Arf1 | 20528 | 24618 | 21136 | 21222 | 20440 | 20088 | 20397 | 19593 | 21774 | 19815 | 20438 | 19481 | 20204 | 19485 | 21153 | 21320 | 21998 | 20632 | 21407 | 22168 | 21099 | 17498 | 22880 | 20695 | 20041 | 22269 | 20812 | 20950 | Arf1 |
| Arf2 | 12713 | 14148 | 12909 | 12756 | 11924 | 12082 | 13319 | 9758 | 8188 | 7486 | 7024 | 8309 | 7891 | 9330 | 14412 | 12513 | 12776 | 13119 | 12180 | 15208 | 13069 | 8809 | 7942 | 8389 | 9230 | 8073 | 9108 | 9353 | Arf2 |
| Arf3 | 12074 | 12720 | 13847 | 14314 | 14575 | 13511 | 12361 | 10422 | 10890 | 10982 | 10187 | 10911 | 10354 | 9679 | 12129 | 12270 | 14457 | 14845 | 15055 | 12159 | 12876 | 13491 | 10802 | 11266 | 11862 | 11577 | 9936 | 9511 | Arf3 |
| Arf4 | 0 | 1 | 0 | 0 | 4 | 0 | 0 | 0 | 0 | 0 | 0 | 1 | 0 | 1 | 7 | 0 | 1 | 0 | 0 | 0 | 0 | 2 | 1 | 0 | 0 | 1 | 0 | 1 | Arf4 |
| Arf5 | 5252 | 5918 | 5491 | 5717 | 5136 | 4788 | 4414 | 5300 | 5912 | 5343 | 5530 | 5330 | 5266 | 4749 | 4833 | 5038 | 5639 | 5390 | 5668 | 4969 | 4866 | 5015 | 6114 | 5677 | 5264 | 5992 | 5396 | 5009 | Arf5 |
| Arf6 | 3381 | 3748 | 4232 | 4039 | 3109 | 3128 | 3020 | 3178 | 3083 | 3323 | 3216 | 3208 | 3168 | 3408 | 3189 | 3289 | 3948 | 3763 | 3234 | 3112 | 3218 | 2968 | 3159 | 3380 | 3178 | 3913 | 3284 | 3328 | Arf6 |
| Arfgap1 | 5343 | 6190 | 5525 | 5857 | 5870 | 5604 | 5268 | 4945 | 5975 | 5642 | 6709 | 5646 | 5429 | 5313 | 5301 | 5506 | 5691 | 5784 | 5940 | 5079 | 5435 | 5817 | 5987 | 5869 | 5951 | 5110 | 5621 | 5167 | Arfgap1 |
| Arfgap2 | 5168 | 5008 | 4670 | 4284 | 4211 | 4810 | 4165 | 7437 | 7481 | 6629 | 6437 | 7904 | 7598 | 6784 | 4853 | 4372 | 4546 | 4122 | 4957 | 4489 | 4210 | 6644 | 7214 | 7090 | 7244 | 8540 | 8435 | 7417 | Arfgap2 |
| Arfgap3 | 3094 | 3519 | 3326 | 3143 | 2987 | 3080 | 2918 | 1927 | 2296 | 2162 | 2389 | 2240 | 2210 | 2165 | 3025 | 3322 | 3063 | 2983 | 2990 | 2975 | 2831 | 2211 | 2402 | 2259 | 2322 | 2173 | 2253 | 2222 | Arfgap3 |
| Arfgef1 | 11156 | 9471 | 7596 | 9299 | 10601 | 11663 | 11194 | 13085 | 11398 | 10328 | 10716 | 10060 | 10394 | 10215 | 11313 | 10054 | 8619 | 9153 | 9744 | 10342 | 11314 | 10830 | 10912 | 8579 | 10767 | 6603 | 10397 | 9842 | Arfgef1 |
| Arfgef2 | 4848 | 4803 | 4011 | 4563 | 4439 | 4865 | 4328 | 2318 | 2639 | 2660 | 2786 | 2684 | 2984 | 2615 | 4557 | 5153 | 4006 | 4308 | 4210 | 3934 | 4443 | 2548 | 2668 | 2204 | 2569 | 1804 | 2905 | 2554 | Arfgef2 |
| Arfgef3 | 1592 | 1571 | 1147 | 1839 | 3228 | 2961 | 2778 | 2751 | 2760 | 2494 | 2782 | 1869 | 2110 | 1984 | 2016 | 1850 | 1637 | 2062 | 2053 | 2730 | 2995 | 3171 | 2549 | 1708 | 2784 | 1007 | 2158 | 1767 | Arfgef3 |
| Arfip1 | 295 | 302 | 265 | 225 | 191 | 170 | 179 | 221 | 165 | 204 | 223 | 172 | 147 | 212 | 284 | 170 | 283 | 190 | 125 | 205 | 213 | 147 | 146 | 212 | 139 | 186 | 144 | 229 | Arfip1 |
| Arfip2 | 7186 | 8404 | 6538 | 8008 | 7310 | 7424 | 7795 | 5290 | 5515 | 5381 | 5523 | 5055 | 5151 | 5283 | 7225 | 7190 | 7004 | 7852 | 7681 | 7704 | 7701 | 4984 | 5811 | 5275 | 5216 | 5376 | 5192 | 5343 | Arfip2 |
| Arfrp1 | 2884 | 3213 | 3359 | 3465 | 3168 | 3195 | 3546 | 3903 | 3729 | 4175 | 3998 | 4568 | 4078 | 4494 | 2981 | 2712 | 3603 | 3491 | 3684 | 3184 | 3446 | 4343 | 3488 | 4445 | 4359 | 4897 | 4389 | 4523 | Arfrp1 |
| Arg1 | 100 | 27 | 7 | 21 | 20 | 7 | 8 | 46 | 60 | 74 | 66 | 31 | 59 | 23 | 22 | 63 | 9 | 11 | 7 | 15 | 9 | 12 | 89 | 57 | 10 | 218 | 23 | 32 | Arg1 |
| Arg2 | 328 | 296 | 209 | 295 | 458 | 365 | 441 | 520 | 456 | 476 | 409 | 308 | 306 | 327 | 342 | 274 | 260 | 355 | 479 | 471 | 362 | 597 | 449 | 364 | 459 | 327 | 286 | 345 | Arg2 |
| Arglu1 | 21879 | 15314 | 23086 | 24570 | 25540 | 24814 | 28426 | 27426 | 22647 | 27775 | 27815 | 30364 | 28907 | 33233 | 21844 | 17700 | 22651 | 23165 | 24701 | 23475 | 27245 | 29895 | 22474 | 29174 | 30350 | 29554 | 28760 | 31105 | Arglu1 |
| Arhgap1 | 5351 | 6473 | 5210 | 4968 | 4921 | 4854 | 4766 | 4460 | 5587 | 4831 | 5317 | 4632 | 4814 | 4395 | 5283 | 6070 | 5077 | 5094 | 4846 | 4687 | 4475 | 5157 | 5552 | 4697 | 4819 | 4813 | 4326 | 4273 | Arhgap1 |
| Arhgap10 | 1706 | 1430 | 1471 | 862 | 740 | 775 | 644 | 1091 | 1189 | 948 | 1169 | 1020 | 1115 | 870 | 1408 | 1823 | 1205 | 907 | 727 | 702 | 563 | 817 | 1342 | 1015 | 678 | 1256 | 1065 | 1000 | Arhgap10 |
| Arhgap11a | 10520 | 9461 | 12696 | 12102 | 8749 | 10186 | 11674 | 7377 | 7230 | 7901 | 7409 | 11870 | 9885 | 12552 | 9701 | 10768 | 11350 | 10423 | 10136 | 10477 | 10573 | 6191 | 6200 | 10316 | 9376 | 11333 | 11390 | 11223 | Arhgap11a |
| Arhgap12 | 8705 | 7411 | 7147 | 7883 | 8341 | 8111 | 8634 | 8730 | 7121 | 6653 | 6243 | 7303 | 7035 | 7496 | 9320 | 7671 | 7502 | 7449 | 7674 | 9576 | 8059 | 7297 | 6860 | 6302 | 7039 | 6301 | 7227 | 7246 | Arhgap12 |
| Arhgap15 | 32 | 35 | 45 | 39 | 72 | 37 | 30 | 57 | 85 | 56 | 58 | 26 | 59 | 26 | 69 | 84 | 54 | 37 | 37 | 69 | 37 | 45 | 107 | 40 | 40 | 48 | 24 | 28 | Arhgap15 |
| Arhgap17 | 3393 | 3582 | 3460 | 3400 | 3030 | 3096 | 2803 | 2602 | 2642 | 2737 | 3116 | 3393 | 3172 | 3172 | 3096 | 3609 | 3531 | 3285 | 2759 | 2469 | 2876 | 2931 | 2639 | 2821 | 2884 | 3317 | 3204 | 3120 | Arhgap17 |
| Arhgap18 | 3616 | 2928 | 3200 | 2873 | 2918 | 3286 | 3601 | 2832 | 2396 | 2198 | 2437 | 2578 | 2225 | 2522 | 3777 | 3083 | 3089 | 2987 | 3048 | 3690 | 3283 | 2526 | 2802 | 2404 | 2134 | 2502 | 2546 | 2300 | Arhgap18 |
| Arhgap19 | 5954 | 5299 | 5840 | 4984 | 3928 | 4534 | 4380 | 4285 | 3874 | 3510 | 3561 | 5377 | 4637 | 5064 | 5545 | 4900 | 5283 | 4584 | 4687 | 4414 | 4412 | 3128 | 3562 | 4488 | 4006 | 5452 | 5517 | 5085 | Arhgap19 |
| Arhgap20 | 3104 | 3132 | 2541 | 3412 | 3892 | 3722 | 3557 | 1969 | 2121 | 1903 | 2402 | 1847 | 2083 | 1785 | 3476 | 3211 | 2870 | 3310 | 3168 | 3770 | 3753 | 2338 | 2882 | 1530 | 2011 | 1383 | 1965 | 1707 | Arhgap20 |
| Arhgap21 | 11167 | 12225 | 9303 | 12370 | 15678 | 16033 | 14292 | 19484 | 17681 | 18267 | 16043 | 20014 | 19006 | 19177 | 13003 | 10027 | 11868 | 12427 | 14850 | 15035 | 14948 | 19026 | 15503 | 16141 | 20951 | 11850 | 20024 | 17655 | Arhgap21 |
| Arhgap22 | 280 | 306 | 351 | 246 | 241 | 168 | 132 | 117 | 176 | 124 | 151 | 60 | 121 | 63 | 244 | 332 | 269 | 228 | 224 | 198 | 134 | 129 | 151 | 64 | 90 | 60 | 55 | 57 | Arhgap22 |
| Arhgap23 | 3320 | 4318 | 4341 | 4989 | 5243 | 4411 | 3986 | 2386 | 3066 | 3481 | 3538 | 2914 | 3178 | 2661 | 3254 | 3724 | 4679 | 5151 | 5085 | 3644 | 4530 | 3527 | 3089 | 3232 | 3389 | 2830 | 2501 | 2712 | Arhgap23 |
| Arhgap24 | 335 | 321 | 316 | 208 | 365 | 301 | 220 | 567 | 802 | 550 | 696 | 304 | 524 | 279 | 364 | 371 | 218 | 302 | 258 | 265 | 230 | 540 | 838 | 512 | 388 | 430 | 342 | 306 | Arhgap24 |
| Arhgap25 | 208 | 163 | 198 | 140 | 128 | 128 | 116 | 154 | 212 | 159 | 218 | 146 | 220 | 146 | 179 | 191 | 166 | 178 | 141 | 124 | 116 | 150 | 261 | 172 | 163 | 198 | 175 | 145 | Arhgap25 |
| Arhgap26 | 900 | 817 | 1020 | 1076 | 1424 | 1044 | 1219 | 542 | 538 | 642 | 771 | 475 | 668 | 513 | 940 | 932 | 1093 | 1319 | 1138 | 771 | 1096 | 760 | 605 | 492 | 661 | 381 | 510 | 541 | Arhgap26 |
| Arhgap27 | 290 | 368 | 387 | 406 | 392 | 316 | 280 | 381 | 611 | 482 | 625 | 575 | 599 | 447 | 236 | 322 | 392 | 420 | 318 | 278 | 292 | 553 | 606 | 514 | 446 | 626 | 507 | 436 | Arhgap27 |
| Arhgap28 | 6697 | 5410 | 5090 | 3456 | 2950 | 3478 | 2243 | 4286 | 5466 | 4432 | 6622 | 3529 | 5650 | 3101 | 5888 | 7423 | 3828 | 3847 | 2318 | 2482 | 2187 | 3708 | 6252 | 3674 | 2936 | 4036 | 3897 | 3519 | Arhgap28 |
| Arhgap29 | 10691 | 8315 | 7783 | 5728 | 5642 | 6327 | 4838 | 6361 | 6536 | 4952 | 7429 | 5641 | 6860 | 5169 | 10026 | 11871 | 6045 | 7157 | 4243 | 6325 | 4796 | 5482 | 9092 | 5165 | 4584 | 4909 | 5892 | 5217 | Arhgap29 |
| Arhgap30 | 67 | 68 | 97 | 79 | 78 | 49 | 44 | 171 | 191 | 153 | 262 | 196 | 213 | 209 | 70 | 98 | 81 | 74 | 72 | 62 | 49 | 211 | 299 | 212 | 160 | 191 | 224 | 182 | Arhgap30 |
| Arhgap31 | 1914 | 2126 | 2249 | 1661 | 1891 | 2104 | 1397 | 2588 | 3175 | 3198 | 4282 | 3022 | 4231 | 2863 | 2008 | 2567 | 1734 | 1885 | 1508 | 1248 | 1377 | 3168 | 3828 | 3042 | 2927 | 2504 | 3145 | 2987 | Arhgap31 |
| Arhgap32 | 7120 | 6505 | 6387 | 9114 | 11729 | 10339 | 10397 | 7344 | 7598 | 8723 | 8520 | 8114 | 7617 | 8137 | 7301 | 7099 | 7570 | 9837 | 10286 | 8512 | 11074 | 8970 | 6985 | 7297 | 9850 | 5363 | 7612 | 7022 | Arhgap32 |
| Arhgap33 | 8170 | 11391 | 12782 | 15364 | 17108 | 14354 | 14006 | 10861 | 13238 | 16963 | 13788 | 19513 | 14166 | 17011 | 8442 | 8914 | 16010 | 14926 | 17409 | 11991 | 14680 | 19258 | 10797 | 17970 | 20127 | 17766 | 15515 | 15088 | Arhgap33 |
| Arhgap33os | 5 | 3 | 1 | 6 | 11 | 10 | 6 | 2 | 2 | 10 | 9 | 15 | 8 | 9 | 13 | 12 | 8 | 6 | 4 | 0 | 4 | 12 | 3 | 12 | 16 | 10 | 6 | 6 | Arhgap33os |
| Arhgap35 | 16542 | 16468 | 13953 | 17606 | 20998 | 19856 | 19837 | 23706 | 23913 | 24706 | 21996 | 24582 | 22068 | 23390 | 16483 | 14954 | 16383 | 18378 | 21322 | 17320 | 21198 | 26911 | 21219 | 23125 | 26861 | 16440 | 22635 | 22735 | Arhgap35 |
| Arhgap36 | 166 | 181 | 124 | 139 | 227 | 146 | 116 | 187 | 159 | 165 | 367 | 116 | 101 | 119 | 232 | 157 | 151 | 150 | 133 | 98 | 165 | 199 | 256 | 139 | 118 | 122 | 126 | 113 | Arhgap36 |
| Arhgap39 | 5006 | 6385 | 5602 | 6716 | 6428 | 6312 | 5721 | 3363 | 3914 | 4402 | 4300 | 4247 | 3898 | 3849 | 4685 | 5611 | 6059 | 6778 | 6606 | 5210 | 6068 | 4293 | 3671 | 3878 | 4582 | 3390 | 3721 | 3597 | Arhgap39 |
| Arhgap4 | 581 | 775 | 560 | 789 | 834 | 699 | 816 | 353 | 448 | 483 | 508 | 426 | 394 | 442 | 662 | 662 | 713 | 821 | 730 | 723 | 773 | 428 | 431 | 473 | 375 | 355 | 370 | 377 | Arhgap4 |
| Arhgap40 | 21 | 26 | 19 | 32 | 25 | 16 | 19 | 28 | 37 | 43 | 18 | 24 | 42 | 27 | 15 | 14 | 21 | 14 | 34 | 26 | 17 | 23 | 50 | 45 | 14 | 30 | 28 | 7 | Arhgap40 |
| Arhgap42 | 993 | 709 | 729 | 583 | 843 | 947 | 736 | 3389 | 3171 | 2902 | 3172 | 2839 | 3155 | 2751 | 997 | 1008 | 625 | 802 | 694 | 664 | 770 | 2441 | 3240 | 2667 | 2664 | 2547 | 2692 | 2719 | Arhgap42 |
| Arhgap44 | 1081 | 1277 | 1123 | 1366 | 1667 | 1210 | 1298 | 1064 | 1129 | 1347 | 1171 | 946 | 980 | 888 | 1058 | 1128 | 1150 | 1359 | 1409 | 1216 | 1322 | 1220 | 1135 | 1109 | 1177 | 1003 | 879 | 974 | Arhgap44 |
| Arhgap45 | 510 | 666 | 846 | 513 | 405 | 468 | 257 | 382 | 619 | 593 | 830 | 613 | 839 | 526 | 475 | 809 | 693 | 583 | 390 | 287 | 263 | 518 | 743 | 515 | 466 | 737 | 795 | 699 | Arhgap45 |
| Arhgap5 | 14422 | 9297 | 9268 | 10036 | 10441 | 12879 | 12752 | 17637 | 14989 | 11127 | 12273 | 14901 | 14257 | 15226 | 15207 | 12359 | 9418 | 9310 | 10630 | 14153 | 12219 | 11974 | 14683 | 11837 | 12068 | 11298 | 15748 | 15264 | Arhgap5 |
| Arhgap6 | 166 | 140 | 191 | 114 | 107 | 104 | 84 | 173 | 254 | 211 | 364 | 197 | 230 | 178 | 157 | 204 | 141 | 147 | 108 | 90 | 94 | 194 | 342 | 210 | 148 | 201 | 222 | 196 | Arhgap6 |
| Arhgap8 | 170 | 126 | 112 | 56 | 81 | 73 | 25 | 56 | 105 | 54 | 87 | 49 | 65 | 47 | 120 | 183 | 85 | 95 | 47 | 29 | 31 | 35 | 115 | 81 | 39 | 57 | 51 | 57 | Arhgap8 |
| Arhgap9 | 115 | 84 | 97 | 62 | 65 | 54 | 23 | 53 | 68 | 71 | 70 | 54 | 60 | 48 | 57 | 130 | 70 | 83 | 28 | 45 | 33 | 57 | 113 | 69 | 75 | 99 | 62 | 57 | Arhgap9 |
| Arhgdia | 16842 | 21126 | 21503 | 18093 | 16216 | 16426 | 14898 | 15014 | 19860 | 19616 | 19833 | 21518 | 21629 | 20518 | 16038 | 19644 | 20321 | 16901 | 17961 | 15078 | 15521 | 17679 | 19355 | 20300 | 19311 | 24591 | 20661 | 21385 | Arhgdia |
| Arhgdib | 1044 | 807 | 740 | 536 | 353 | 723 | 334 | 912 | 1285 | 994 | 1106 | 676 | 1000 | 722 | 906 | 1124 | 539 | 748 | 374 | 430 | 337 | 720 | 1478 | 1130 | 662 | 1116 | 691 | 763 | Arhgdib |
| Arhgdig | 149 | 118 | 36 | 173 | 281 | 117 | 124 | 308 | 325 | 441 | 401 | 90 | 185 | 123 | 136 | 121 | 66 | 122 | 127 | 120 | 141 | 358 | 344 | 346 | 172 | 326 | 70 | 90 | Arhgdig |
| Arhgef1 | 5359 | 7078 | 7570 | 7524 | 7491 | 6942 | 6619 | 4805 | 6508 | 7155 | 7376 | 6952 | 6537 | 6481 | 4942 | 6630 | 8287 | 7911 | 7962 | 5539 | 6727 | 7003 | 5977 | 7603 | 7393 | 6921 | 6244 | 6035 | Arhgef1 |
| Arhgef10 | 2122 | 2656 | 2811 | 2360 | 2226 | 2273 | 1741 | 1779 | 2352 | 2118 | 2527 | 2191 | 2624 | 1926 | 2285 | 2681 | 2289 | 2477 | 1994 | 1803 | 1698 | 1937 | 2454 | 2248 | 1944 | 2103 | 2316 | 2078 | Arhgef10 |
| Arhgef10l | 1599 | 2236 | 2409 | 2244 | 2532 | 2118 | 1886 | 2689 | 3057 | 3574 | 3255 | 4003 | 3461 | 3577 | 1615 | 1920 | 2447 | 2277 | 2632 | 1909 | 2109 | 3427 | 2807 | 3915 | 4010 | 3543 | 3737 | 3757 | Arhgef10l |
| Arhgef11 | 8003 | 8445 | 7073 | 9894 | 10157 | 10039 | 9205 | 5388 | 5892 | 6511 | 6421 | 6620 | 6008 | 5794 | 7668 | 8404 | 8199 | 10566 | 9496 | 8240 | 9359 | 7010 | 5341 | 6501 | 6999 | 4366 | 5977 | 5738 | Arhgef11 |

| | | | | | | | | | | | | | | | | | | | | | | | | | | | | | |
|---|---|---|---|---|---|---|---|---|---|---|---|---|---|---|---|---|---|---|---|---|---|---|---|---|---|---|---|---|---|
| Arhgef12 | 16387 | 15836 | 11286 | 16874 | 18261 | 18708 | 18676 | 16201 | 16665 | 17145 | 17062 | 15699 | 15686 | 15915 | 15427 | 15411 | 13418 | 17288 | 17093 | 15344 | 19827 | 16753 | 15392 | 14442 | 18367 | 10241 | 15869 | 14445 | Arhgef12 |
| Arhgef15 | 352 | 430 | 545 | 458 | 620 | 593 | 465 | 965 | 1233 | 1272 | 1453 | 1017 | 1298 | 1141 | 398 | 385 | 467 | 407 | 570 | 469 | 517 | 1497 | 1286 | 1119 | 1456 | 1177 | 1168 | 1069 | Arhgef15 |
| Arhgef16 | 269 | 207 | 122 | 75 | 77 | 75 | 23 | 98 | 193 | 97 | 148 | 47 | 118 | 33 | 169 | 255 | 111 | 97 | 44 | 15 | 23 | 87 | 179 | 86 | 52 | 103 | 45 | 46 | Arhgef16 |
| Arhgef17 | 4954 | 5390 | 5657 | 4692 | 4808 | 4564 | 3741 | 4176 | 5437 | 5811 | 5945 | 5427 | 5533 | 4423 | 4044 | 6049 | 5247 | 4642 | 4957 | 3303 | 3771 | 4749 | 5554 | 5675 | 4365 | 5065 | 4983 | 4707 | Arhgef17 |
| Arhgef18 | 4863 | 5942 | 5223 | 6074 | 5332 | 5458 | 5158 | 3444 | 3634 | 3911 | 3678 | 3924 | 3983 | 3523 | 4684 | 5197 | 5651 | 5986 | 5841 | 4730 | 5495 | 4080 | 3638 | 3801 | 4002 | 3318 | 3665 | 3767 | Arhgef18 |
| Arhgef19 | 682 | 637 | 533 | 654 | 489 | 564 | 374 | 668 | 1070 | 832 | 805 | 262 | 498 | 275 | 537 | 717 | 608 | 712 | 507 | 371 | 406 | 541 | 1041 | 707 | 287 | 708 | 257 | 240 | Arhgef19 |
| Arhgef2 | 5173 | 5282 | 6523 | 6244 | 8635 | 7479 | 7169 | 10811 | 11603 | 13952 | 13765 | 13760 | 12911 | 13659 | 5041 | 5277 | 6667 | 6366 | 8219 | 6061 | 7144 | 13892 | 10928 | 14711 | 15807 | 12132 | 12723 | 13010 | Arhgef2 |
| Arhgef25 | 6004 | 6167 | 8029 | 6548 | 6080 | 6290 | 5291 | 6377 | 7345 | 8048 | 9005 | 7300 | 7652 | 6839 | 5731 | 6329 | 7793 | 6433 | 6314 | 4839 | 5426 | 6794 | 7580 | 8544 | 7087 | 8762 | 7143 | 7031 | Arhgef25 |
| Arhgef26 | 1563 | 1553 | 1525 | 1584 | 2356 | 2030 | 2128 | 1652 | 1881 | 1845 | 1828 | 1508 | 1659 | 1417 | 1841 | 1598 | 1757 | 1973 | 2007 | 1631 | 2067 | 1890 | 1803 | 1454 | 1635 | 1369 | 1306 | 1318 | Arhgef26 |
| Arhgef28 | 641 | 563 | 484 | 481 | 763 | 496 | 460 | 1016 | 1167 | 1021 | 1157 | 626 | 1061 | 664 | 661 | 700 | 523 | 567 | 447 | 557 | 481 | 1084 | 1154 | 782 | 940 | 598 | 669 | 672 | Arhgef28 |
| Arhgef3 | 706 | 695 | 456 | 481 | 526 | 529 | 417 | 441 | 726 | 638 | 770 | 285 | 535 | 276 | 796 | 806 | 395 | 550 | 360 | 456 | 380 | 421 | 854 | 429 | 286 | 258 | 227 | 243 | Arhgef3 |
| Arhgef33 | 103 | 97 | 52 | 88 | 160 | 105 | 153 | 46 | 60 | 74 | 86 | 49 | 57 | 61 | 123 | 104 | 86 | 101 | 80 | 111 | 122 | 70 | 96 | 40 | 63 | 45 | 80 | 49 | Arhgef33 |
| Arhgef37 | 193 | 193 | 137 | 180 | 241 | 277 | 202 | 217 | 278 | 273 | 272 | 177 | 229 | 186 | 208 | 210 | 134 | 254 | 199 | 208 | 178 | 247 | 266 | 242 | 295 | 118 | 188 | 186 | Arhgef37 |
| Arhgef38 | 55 | 21 | 7 | 39 | 9 | 5 | 7 | 84 | 69 | 112 | 56 | 10 | 45 | 14 | 36 | 26 | 7 | 14 | 7 | 6 | 3 | 22 | 98 | 91 | 11 | 49 | 13 | 1 | Arhgef38 |
| Arhgef39 | 698 | 807 | 907 | 615 | 509 | 497 | 618 | 1012 | 1091 | 1114 | 1028 | 1571 | 1315 | 1325 | 575 | 623 | 856 | 525 | 678 | 612 | 549 | 745 | 952 | 1359 | 1071 | 1860 | 1510 | 1489 | Arhgef39 |
| Arhgef4 | 806 | 1256 | 868 | 1410 | 2306 | 1773 | 1401 | 1771 | 2007 | 2026 | 1824 | 1349 | 2075 | 1359 | 1331 | 1347 | 1337 | 1556 | 1591 | 1496 | 1457 | 2064 | 2048 | 1285 | 1875 | 837 | 1422 | 1247 | Arhgef4 |
| Arhgef40 | 6597 | 7280 | 7278 | 8063 | 8162 | 7850 | 7507 | 7040 | 8581 | 9117 | 10284 | 9295 | 9311 | 8312 | 5906 | 6696 | 7930 | 8128 | 8584 | 6349 | 7426 | 8457 | 8420 | 9268 | 8546 | 8443 | 8681 | 8285 | Arhgef40 |
| Arhgef5 | 627 | 350 | 361 | 218 | 362 | 376 | 207 | 1442 | 1624 | 1396 | 1480 | 910 | 1204 | 832 | 477 | 470 | 271 | 299 | 259 | 229 | 193 | 993 | 1764 | 1177 | 877 | 935 | 943 | 934 | Arhgef5 |
| Arhgef6 | 784 | 733 | 459 | 611 | 743 | 844 | 685 | 1172 | 1241 | 1053 | 1228 | 804 | 895 | 867 | 965 | 866 | 708 | 676 | 640 | 806 | 672 | 1206 | 1263 | 743 | 961 | 682 | 906 | 739 | Arhgef6 |
| Arhgef7 | 9513 | 10782 | 9719 | 10898 | 11295 | 11062 | 10335 | 12121 | 13422 | 14563 | 13282 | 15278 | 12647 | 14440 | 9461 | 9804 | 10331 | 11203 | 11389 | 9935 | 10873 | 14437 | 12234 | 14679 | 16790 | 13360 | 13702 | 13362 | Arhgef7 |
| Arhgef9 | 4507 | 4204 | 3729 | 4815 | 6057 | 5694 | 6234 | 7607 | 6990 | 6606 | 6530 | 5376 | 5611 | 6024 | 5151 | 3772 | 4226 | 5197 | 5206 | 6071 | 6355 | 7579 | 6597 | 5492 | 6873 | 3801 | 5441 | 4977 | Arhgef9 |
| Arid1a | 11439 | 13015 | 18195 | 18090 | 19874 | 16530 | 15969 | 17795 | 20033 | 25602 | 22177 | 30939 | 24214 | 27842 | 10962 | 12352 | 19873 | 17130 | 21369 | 12812 | 17399 | 27317 | 17807 | 28500 | 27447 | 24255 | 24181 | 27033 | Arid1a |
| Arid1b | 2629 | 2852 | 3635 | 3920 | 4934 | 3998 | 3461 | 6079 | 7121 | 8755 | 8117 | 8439 | 8062 | 7659 | 2612 | 2875 | 3694 | 3933 | 4574 | 3335 | 3850 | 8341 | 6702 | 8540 | 8845 | 7043 | 6925 | 7510 | Arid1b |
| Arid2 | 8293 | 7351 | 7498 | 8116 | 9654 | 9748 | 10023 | 14153 | 12941 | 13873 | 12552 | 15108 | 14630 | 15310 | 8044 | 7443 | 7932 | 7866 | 9590 | 9193 | 9692 | 13167 | 11934 | 13394 | 14391 | 11777 | 15418 | 14972 | Arid2 |
| Arid3a | 1617 | 1518 | 2550 | 1781 | 1829 | 1455 | 1521 | 1360 | 1411 | 1878 | 1771 | 1831 | 2259 | 2071 | 1377 | 1648 | 2056 | 1756 | 1678 | 1256 | 1490 | 1607 | 1597 | 1957 | 1703 | 2166 | 1951 | 2212 | Arid3a |
| Arid3b | 2578 | 3043 | 3386 | 2441 | 1999 | 2069 | 1950 | 1252 | 1215 | 1259 | 1191 | 1782 | 1731 | 1770 | 2532 | 2786 | 2977 | 2292 | 2141 | 1911 | 2068 | 1507 | 1154 | 1398 | 1463 | 1892 | 1977 | 1981 | Arid3b |
| Arid3c | 77 | 74 | 68 | 73 | 70 | 40 | 85 | 23 | 28 | 67 | 26 | 30 | 40 | 40 | 58 | 79 | 81 | 81 | 73 | 75 | 63 | 23 | 19 | 34 | 28 | 40 | 38 | 55 | Arid3c |
| Arid4a | 4384 | 3876 | 4075 | 4501 | 4022 | 4406 | 4317 | 3791 | 3506 | 3437 | 3475 | 3327 | 2930 | 3561 | 4381 | 4184 | 3964 | 4582 | 3779 | 4386 | 4061 | 3414 | 3168 | 3310 | 3816 | 3006 | 3208 | 3081 | Arid4a |
| Arid4b | 11661 | 8547 | 8624 | 10338 | 10636 | 11679 | 12040 | 13361 | 11224 | 11042 | 10782 | 12348 | 11649 | 13256 | 11885 | 9463 | 9332 | 10248 | 10244 | 11653 | 11605 | 11496 | 10705 | 10740 | 11783 | 9011 | 12700 | 12100 | Arid4b |
| Arid5a | 468 | 684 | 578 | 635 | 459 | 457 | 407 | 161 | 232 | 217 | 293 | 147 | 196 | 142 | 447 | 660 | 659 | 672 | 503 | 425 | 473 | 192 | 273 | 204 | 186 | 179 | 145 | 120 | Arid5a |
| Arid5b | 5501 | 4986 | 5274 | 6157 | 6347 | 5827 | 5575 | 3565 | 3821 | 3849 | 4099 | 3036 | 3763 | 2841 | 5190 | 5343 | 5836 | 6238 | 6347 | 5034 | 5613 | 3474 | 3767 | 3394 | 2898 | 2682 | 2825 | 2851 | Arid5b |
| Arih1 | 18790 | 15979 | 16748 | 18061 | 16503 | 17002 | 18979 | 16546 | 14026 | 14343 | 13743 | 14847 | 14159 | 15564 | 19035 | 17108 | 16811 | 17595 | 16536 | 16900 | 18043 | 16379 | 14038 | 14588 | 15206 | 14790 | 14573 | 16009 | Arih1 |
| Arih2 | 6359 | 6632 | 5601 | 6299 | 5453 | 5865 | 5981 | 4513 | 4000 | 4225 | 4350 | 4811 | 4517 | 4739 | 6095 | 5889 | 5806 | 5935 | 5866 | 5890 | 6057 | 3992 | 4099 | 4294 | 4747 | 4820 | 4810 | 4844 | Arih2 |
| Arl1 | 10776 | 8349 | 9337 | 8536 | 7976 | 8506 | 8956 | 8526 | 7922 | 6639 | 8278 | 7342 | 7404 | 7776 | 11108 | 10006 | 8659 | 8940 | 7606 | 9254 | 8245 | 7469 | 8527 | 7444 | 7327 | 8293 | 7689 | 7938 | Arl1 |
| Arl10 | 843 | 1142 | 1066 | 1200 | 1505 | 1347 | 1163 | 1451 | 1725 | 1615 | 1398 | 1503 | 1450 | 1285 | 944 | 963 | 1333 | 1270 | 1559 | 1450 | 1266 | 1790 | 1689 | 1631 | 1733 | 1389 | 1477 | 1343 | Arl10 |
| Arl11 | 7 | 9 | 13 | 9 | 6 | 4 | 16 | 15 | 21 | 21 | 22 | 16 | 14 | 14 | 13 | 5 | 7 | 6 | 9 | 12 | 13 | 18 | 33 | 24 | 17 | 10 | 15 | 23 | Arl11 |
| Arl13b | 5659 | 4509 | 4801 | 5007 | 4322 | 4666 | 4707 | 2532 | 2494 | 2174 | 2431 | 2173 | 2391 | 2392 | 5343 | 5625 | 4317 | 5188 | 4080 | 4412 | 4487 | 2333 | 2397 | 2201 | 2158 | 2318 | 2074 | 2058 | Arl13b |
| Arl14 | 0 | 0 | 0 | 0 | 0 | 0 | 0 | 0 | 0 | 0 | 0 | 0 | 0 | 0 | 0 | 0 | 0 | 0 | 0 | 0 | 0 | 0 | 1 | 0 | 0 | 0 | 0 | 0 | Arl14 |
| Arl14ep | 5383 | 4294 | 4911 | 4621 | 4795 | 5115 | 5315 | 5639 | 5281 | 5059 | 5362 | 5709 | 5633 | 5986 | 4862 | 4823 | 4946 | 4556 | 4708 | 4991 | 5059 | 5065 | 5288 | 5398 | 4996 | 5437 | 5751 | 6162 | Arl14ep |
| Arl14epl | 58 | 22 | 10 | 73 | 9 | 5 | 15 | 150 | 80 | 212 | 143 | 0 | 22 | 27 | 26 | 31 | 20 | 25 | 4 | 8 | 8 | 3 | 113 | 305 | 2 | 257 | 0 | 1 | Arl14epl |
| Arl15 | 4054 | 3805 | 3636 | 4588 | 4781 | 4233 | 5623 | 4045 | 3145 | 3405 | 3361 | 3438 | 3144 | 3779 | 4366 | 3338 | 4522 | 4661 | 4810 | 5203 | 5093 | 3514 | 3206 | 3636 | 3915 | 3466 | 3385 | 3731 | Arl15 |
| Arl16 | 1546 | 1595 | 1589 | 1424 | 1318 | 1298 | 1491 | 999 | 948 | 968 | 902 | 1017 | 1040 | 1014 | 1578 | 1604 | 1510 | 1394 | 1419 | 1427 | 1368 | 1014 | 1041 | 844 | 879 | 1089 | 1119 | 1155 | Arl16 |
| Arl2 | 1286 | 1558 | 1641 | 1422 | 1417 | 1410 | 1391 | 2066 | 2348 | 2462 | 2290 | 2300 | 2412 | 2397 | 1291 | 1374 | 1606 | 1370 | 1641 | 1447 | 1468 | 2259 | 2330 | 2455 | 2341 | 2460 | 2515 | 2477 | Arl2 |
| Arl2bp | 8352 | 8796 | 9227 | 8713 | 8309 | 7925 | 9581 | 8206 | 7811 | 8574 | 7579 | 10342 | 9216 | 11541 | 8244 | 7138 | 9327 | 7870 | 8777 | 8220 | 9623 | 8475 | 7497 | 9653 | 9369 | 10996 | 10263 | 11412 | Arl2bp |
| Arl3 | 6082 | 6871 | 7360 | 7469 | 6648 | 6680 | 7649 | 3994 | 4019 | 4723 | 4156 | 4873 | 4258 | 5385 | 6127 | 5816 | 7590 | 6995 | 7425 | 7399 | 7509 | 3945 | 4223 | 5003 | 4923 | 5587 | 4931 | 5438 | Arl3 |
| Arl4a | 6569 | 5780 | 6636 | 6440 | 6620 | 6506 | 7192 | 5733 | 4926 | 4678 | 5372 | 5335 | 4910 | 5393 | 8053 | 6109 | 6091 | 6581 | 5436 | 8375 | 6814 | 6087 | 6470 | 4527 | 5174 | 4474 | 5760 | 5599 | Arl4a |
| Arl4c | 4688 | 4895 | 5641 | 5854 | 6740 | 6514 | 6094 | 6179 | 5944 | 6920 | 5964 | 6631 | 5905 | 6106 | 5045 | 4424 | 5971 | 5871 | 6937 | 6260 | 6208 | 8069 | 6029 | 6539 | 7454 | 5744 | 6285 | 6325 | Arl4c |
| Arl4d | 535 | 815 | 436 | 922 | 865 | 715 | 934 | 646 | 824 | 852 | 681 | 823 | 431 | 907 | 624 | 536 | 925 | 968 | 965 | 586 | 894 | 810 | 413 | 1049 | 1042 | 1019 | 602 | 711 | Arl4d |
| Arl5a | 13334 | 9545 | 10928 | 11954 | 12558 | 13032 | 14546 | 11764 | 10000 | 9673 | 9948 | 10618 | 10181 | 11035 | 13129 | 11397 | 11436 | 12304 | 12499 | 12843 | 13740 | 10873 | 10158 | 9816 | 10310 | 9404 | 11029 | 11331 | Arl5a |
| Arl5b | 12930 | 10439 | 10244 | 10637 | 10830 | 11896 | 11516 | 12698 | 9957 | 9348 | 10146 | 11795 | 12187 | 12633 | 12798 | 12631 | 10719 | 10564 | 9853 | 11502 | 11291 | 10773 | 9589 | 9524 | 10494 | 8561 | 13143 | 12577 | Arl5b |
| Arl5c | 382 | 384 | 488 | 307 | 344 | 353 | 325 | 62 | 86 | 101 | 96 | 99 | 86 | 70 | 286 | 400 | 363 | 333 | 318 | 365 | 278 | 69 | 93 | 89 | 47 | 87 | 88 | 96 | Arl5c |
| Arl6 | 3652 | 3301 | 3418 | 3323 | 3000 | 3106 | 3724 | 2077 | 1835 | 1578 | 1464 | 1978 | 1765 | 2059 | 4279 | 3745 | 3188 | 3065 | 3060 | 4045 | 3514 | 1780 | 1827 | 1673 | 1654 | 2117 | 1991 | 2058 | Arl6 |
| Arl6ip1 | 34835 | 33657 | 33169 | 31197 | 25706 | 27147 | 30437 | 24260 | 21147 | 19766 | 18230 | 23561 | 21950 | 23513 | 36476 | 32992 | 31831 | 30042 | 28687 | 34173 | 29020 | 20347 | 21665 | 21906 | 21053 | 26716 | 24199 | 25144 | Arl6ip1 |
| Arl6ip4 | 1730 | 2139 | 2376 | 2263 | 1871 | 1896 | 1693 | 1909 | 1952 | 2219 | 2109 | 2387 | 2224 | 2321 | 1656 | 1934 | 2394 | 1795 | 1985 | 2020 | 1658 | 1681 | 1970 | 2369 | 2198 | 2888 | 2553 | 2538 | Arl6ip4 |
| Arl6ip5 | 4956 | 5326 | 5451 | 5851 | 6135 | 5708 | 5686 | 4504 | 3992 | 4355 | 4538 | 3852 | 3793 | 3622 | 5718 | 5016 | 5669 | 5968 | 5992 | 6544 | 6138 | 4298 | 4599 | 4068 | 4211 | 4131 | 3774 | 3773 | Arl6ip5 |
| Arl6ip6 | 7060 | 5392 | 6690 | 6380 | 4788 | 5539 | 6962 | 3976 | 3269 | 3005 | 2917 | 4494 | 3739 | 4554 | 6877 | 6053 | 6106 | 5827 | 5632 | 6030 | 6257 | 3360 | 3159 | 3790 | 3602 | 5448 | 4549 | 4931 | Arl6ip6 |
| Arl8a | 3775 | 4291 | 4629 | 6355 | 7888 | 6329 | 6772 | 6364 | 6849 | 8768 | 7368 | 6573 | 5956 | 7004 | 4114 | 3912 | 5687 | 6619 | 7336 | 5659 | 7588 | 8973 | 6680 | 7789 | 8947 | 6096 | 5412 | 5925 | Arl8a |
| Arl8b | 10484 | 8517 | 9293 | 9204 | 8908 | 9042 | 10409 | 8154 | 7064 | 7216 | 6941 | 7061 | 6957 | 7415 | 10407 | 9051 | 9026 | 9466 | 8733 | 9260 | 9746 | 8261 | 7284 | 7088 | 7469 | 7439 | 6734 | 7531 | Arl8b |
| Arl9 | 137 | 154 | 111 | 128 | 124 | 87 | 91 | 39 | 28 | 30 | 39 | 43 | 22 | 44 | 119 | 168 | 126 | 173 | 117 | 93 | 121 | 33 | 34 | 34 | 30 | 50 | 50 | 36 | Arl9 |
| Armc1 | 8507 | 7380 | 7233 | 7623 | 6517 | 6726 | 7794 | 6803 | 5518 | 5095 | 5104 | 5913 | 5523 | 5934 | 9022 | 7963 | 6942 | 7396 | 6770 | 7723 | 7494 | 6024 | 5644 | 5560 | 5650 | 6307 | 6101 | 6224 | Armc1 |
| Armc10 | 8612 | 7889 | 8485 | 7110 | 6470 | 6799 | 7403 | 8063 | 6679 | 6895 | 6360 | 8533 | 7503 | 8842 | 8530 | 7475 | 7814 | 6762 | 7161 | 7321 | 7372 | 7501 | 6934 | 7671 | 7779 | 8992 | 8323 | 9176 | Armc10 |
| Armc12 | 2 | 5 | 5 | 1 | 5 | 2 | 6 | 0 | 0 | 0 | 0 | 0 | 0 | 0 | 0 | 3 | 6 | 3 | 2 | 0 | 3 | 0 | 1 | 2 | 0 | 1 | 5 | 0 | Armc12 |
| Armc2 | 611 | 504 | 418 | 619 | 596 | 598 | 668 | 372 | 493 | 483 | 440 | 320 | 299 | 271 | 556 | 583 | 513 | 714 | 597 | 531 | 665 | 506 | 551 | 378 | 403 | 256 | 270 | 247 | Armc2 |
| Armc3 | 122 | 133 | 82 | 65 | 130 | 105 | 135 | 116 | 117 | 131 | 97 | 125 | 126 | 74 | 137 | 111 | 53 | 89 | 129 | 139 | 103 | 155 | 195 | 78 | 123 | 71 | 80 | 75 | Armc3 |
| Armc5 | 521 | 552 | 705 | 612 | 806 | 652 | 559 | 901 | 1210 | 1190 | 1260 | 1310 | 1303 | 1122 | 521 | 541 | 846 | 681 | 806 | 540 | 631 | 1125 | 1245 | 1204 | 1163 | 1511 | 1310 | 1146 | Armc5 |
| Armc6 | 914 | 1486 | 1061 | 1687 | 1612 | 1337 | 1286 | 1925 | 2228 | 2236 | 2022 | 1742 | 1811 | 1838 | 1162 | 1181 | 1357 | 1404 | 1553 | 1541 | 1470 | 2226 | 2104 | 2005 | 2347 | 1702 | 1668 | 1733 | Armc6 |
| Armc7 | 639 | 954 | 664 | 983 | 759 | 757 | 725 | 503 | 564 | 640 | 609 | 615 | 624 | 544 | 683 | 906 | 754 | 964 | 672 | 632 | 759 | 673 | 559 | 610 | 638 | 554 | 502 | 610 | Armc7 |
| Armc8 | 4879 | 4693 | 4146 | 5066 | 5285 | 5345 | 5605 | 5670 | 5126 | 4646 | 4694 | 5158 | 4689 | 5287 | 5343 | 4463 | 4813 | 5133 | 4977 | 5914 | 5726 | 5668 | 5024 | 4941 | 5256 | 4786 | 4938 | 4907 | Armc8 |
| Armc9 | 4565 | 5206 | 3930 | 4528 | 3880 | 3925 | 4215 | 1931 | 1932 | 1841 | 1745 | 1927 | 1951 | 1888 | 4230 | 4757 | 4092 | 4339 | 4089 | 4033 | 4281 | 1898 | 1985 | 1714 | 1938 | 1776 | 1885 | 1990 | Armc9 |
| Armcx1 | 2445 | 2157 | 2870 | 2556 | 2993 | 2634 | 3006 | 5754 | 5244 | 5658 | 6211 | 5363 | 5329 | 6408 | 2575 | 2391 | 2825 | 2609 | 3015 | 2910 | 2806 | 6161 | 5386 | 5724 | 5999 | 5793 | 5403 | 6019 | Armcx1 |
| Armcx2 | 20777 | 21453 | 19773 | 17941 | 16389 | 18191 | 15871 | 16855 | 16858 | 14807 | 16119 | 16927 | 16796 | 15910 | 20598 | 20843 | 18227 | 17082 | 17549 | 17166 | 16093 | 17338 | 17531 | 15591 | 15907 | 16876 | 17482 | 16559 | Armcx2 |
| Armcx3 | 5261 | 5140 | 5343 | 5005 | 5606 | 5585 | 5757 | 3700 | 3867 | 3444 | 4310 | 3517 | 3390 | 3625 | 5738 | 5443 | 4985 | 5532 | 5056 | 6426 | 5099 | 3778 | 4185 | 3541 | 3677 | 3801 | 3540 | 3569 | Armcx3 |
| Armcx4 | 17411 | 16554 | 14926 | 17568 | 18236 | 19515 | 18476 | 19148 | 18742 | 21151 | 20996 | 23840 | 22567 | 24544 | 16376 | 16748 | 16742 | 17035 | 17775 | 18151 | 17662 | 19920 | 17896 | 21064 | 24353 | 17000 | 24846 | 21499 | Armcx4 |
| Armcx5 | 1069 | 644 | 615 | 640 | 676 | 713 | 877 | 826 | 768 | 863 | 821 | 791 | 753 | 951 | 986 | 661 | 564 | 576 | 599 | 758 | 747 | 692 | 771 | 686 | 721 | 818 | 712 | 808 | Armcx5 |
| Armcx6 | 1291 | 1348 | 1290 | 1351 | 1486 | 1436 | 1433 | 1533 | 1610 | 1609 | 1742 | 1454 | 1422 | 1480 | 1276 | 1255 | 1341 | 1354 | 1542 | 1445 | 1470 | 1539 | 1737 | 1514 | 1439 | 1486 | 1381 | 1459 | Armcx6 |
| Armh1 | 141 | 187 | 151 | 166 | 97 | 142 | 113 | 40 | 46 | 71 | 47 | 35 | 39 | 16 | 119 | 172 | 156 | 190 | 111 | 92 | 113 | 38 | 63 | 34 | 32 | 40 | 13 | 28 | Armh1 |
| Armh2 | 6 | 3 | 2 | 18 | 6 | 10 | 9 | 4 | 5 | 8 | 1 | 4 | 0 | 2 | 4 | 2 | 5 | 18 | 8 | 5 | 11 | 7 | 3 | 2 | 0 | 0 | 2 | 1 | Armh2 |
| Armh3 | 1295 | 1196 | 1314 | 1417 | 1473 | 1321 | 1407 | 2004 | 2033 | 2126 | 2030 | 2227 | 2090 | 2200 | 1250 | 1151 | 1314 | 1218 | 1483 | 1140 | 1398 | 2296 | 1935 | 2157 | 2250 | 2277 | 2165 | 2111 | Armh3 |
| Armh4 | 5685 | 5374 | 5357 | 3613 | 4296 | 3679 | 3448 | 3969 | 3623 | 2758 | 3080 | 2635 | 4051 | 2670 | 6281 | 6128 | 4344 | 3608 | 3426 | 4824 | 3471 | 3669 | 3570 | 1965 | 2576 | 1960 | 3531 | 3555 | Armh4 |

| | | | | | | | | | | | | | | | | | | | | | | | | | | | | | |
|---|---|---|---|---|---|---|---|---|---|---|---|---|---|---|---|---|---|---|---|---|---|---|---|---|---|---|---|---|---|
| Armt1 | 2242 | 2070 | 2439 | 2255 | 1800 | 1944 | 2065 | 1890 | 1776 | 1729 | 1880 | 2317 | 2093 | 2172 | 2171 | 2027 | 2298 | 1998 | 2147 | 2035 | 1844 | 1577 | 1716 | 1861 | 1633 | 2321 | 2259 | 2327 | Armt1 |
| Arnt | 6730 | 5452 | 6318 | 5439 | 5204 | 5730 | 5446 | 5741 | 5777 | 5992 | 6597 | 5729 | 6233 | 6134 | 5938 | 6571 | 5702 | 5850 | 5046 | 4629 | 5275 | 5566 | 5899 | 5994 | 5647 | 5725 | 5891 | 5930 | Arnt |
| Arnt2 | 12029 | 13167 | 12569 | 16917 | 19964 | 19067 | 18405 | 19048 | 19466 | 22997 | 19081 | 24969 | 20422 | 24440 | 13264 | 10543 | 15140 | 17006 | 19412 | 15922 | 19126 | 26309 | 16519 | 22282 | 26930 | 18632 | 22086 | 22921 | Arnt2 |
| Arntl | 1468 | 1716 | 1328 | 1723 | 1587 | 1664 | 1753 | 775 | 843 | 735 | 741 | 682 | 618 | 769 | 1630 | 1593 | 1627 | 1829 | 1509 | 1684 | 1785 | 800 | 827 | 830 | 897 | 886 | 712 | 667 | Arntl |
| Arntl2 | 664 | 729 | 444 | 755 | 695 | 700 | 706 | 353 | 445 | 442 | 544 | 294 | 405 | 255 | 603 | 707 | 613 | 800 | 604 | 642 | 732 | 361 | 415 | 339 | 342 | 212 | 270 | 230 | Arntl2 |
| Arpc1a | 7627 | 8841 | 7823 | 7874 | 8034 | 7668 | 7782 | 8139 | 8398 | 8415 | 7671 | 8160 | 8218 | 8491 | 7635 | 7675 | 8226 | 7970 | 8391 | 7753 | 8123 | 8158 | 8633 | 8414 | 9027 | 8438 | 8505 | 8537 | Arpc1a |
| Arpc1b | 1735 | 2171 | 2014 | 1468 | 1370 | 1659 | 1018 | 1304 | 1881 | 1472 | 2134 | 1264 | 1771 | 1284 | 1793 | 2390 | 1657 | 1658 | 1342 | 1364 | 1164 | 1394 | 2441 | 1560 | 1431 | 1672 | 1517 | 1400 | Arpc1b |
| Arpc2 | 15763 | 14369 | 14577 | 13849 | 13052 | 13092 | 14661 | 12086 | 11147 | 11214 | 10651 | 11391 | 11075 | 11785 | 15730 | 14319 | 13965 | 13760 | 13499 | 14216 | 14582 | 11256 | 11595 | 10883 | 11226 | 12409 | 11037 | 12045 | Arpc2 |
| Arpc3 | 6712 | 8986 | 8296 | 8394 | 7511 | 7940 | 7365 | 6847 | 7262 | 6626 | 6391 | 6130 | 6843 | 6483 | 7882 | 8063 | 8013 | 8013 | 8208 | 10064 | 7497 | 5708 | 7025 | 6823 | 7169 | 7268 | 7181 | 6608 | Arpc3 |
| Arpc4 | 5758 | 5852 | 6857 | 6426 | 5618 | 5246 | 6124 | 4250 | 4359 | 5024 | 4891 | 5070 | 4974 | 5644 | 5430 | 5312 | 6343 | 5984 | 5910 | 4760 | 6379 | 4625 | 4765 | 5245 | 5032 | 5911 | 5033 | 5899 | Arpc4 |
| Arpc5 | 11344 | 9862 | 11630 | 11507 | 12245 | 11227 | 13255 | 10235 | 9059 | 9535 | 9050 | 9538 | 9656 | 10261 | 12554 | 11416 | 12135 | 11959 | 12293 | 12026 | 12940 | 11084 | 9565 | 9771 | 10642 | 10181 | 9932 | 10325 | Arpc5 |
| Arpc5l | 6035 | 5740 | 5949 | 5894 | 5215 | 5268 | 5989 | 4471 | 3733 | 4106 | 3896 | 4434 | 4082 | 4494 | 5809 | 5079 | 6042 | 5701 | 5319 | 5366 | 5676 | 4259 | 3691 | 4389 | 4319 | 4840 | 4240 | 4586 | Arpc5l |
| Arpin | 826 | 844 | 978 | 627 | 740 | 692 | 593 | 1243 | 1225 | 991 | 1151 | 1312 | 1272 | 1408 | 928 | 745 | 853 | 616 | 711 | 657 | 618 | 1173 | 1185 | 1178 | 1121 | 1365 | 1481 | 1525 | Arpin |
| Arpp19 | 20455 | 16397 | 17905 | 17166 | 16368 | 17230 | 19081 | 15430 | 13055 | 12818 | 12624 | 13484 | 12959 | 14725 | 21089 | 17284 | 17796 | 16766 | 16523 | 19373 | 17963 | 13050 | 12960 | 12923 | 13507 | 14189 | 14207 | 14597 | Arpp19 |
| Arpp21 | 158 | 139 | 108 | 245 | 542 | 365 | 421 | 935 | 1149 | 1119 | 1658 | 540 | 561 | 589 | 212 | 159 | 243 | 320 | 388 | 394 | 405 | 1153 | 1478 | 780 | 1149 | 433 | 431 | 409 | Arpp21 |
| Arr3 | 1 | 7 | 0 | 2 | 0 | 8 | 0 | 5 | 2 | 5 | 6 | 9 | 10 | 0 | 0 | 5 | 2 | 0 | 2 | 5 | 3 | 0 | 8 | 0 | 4 | 0 | 0 | 0 | Arr3 |
| Arrb1 | 4865 | 5352 | 4862 | 5852 | 8759 | 7151 | 6044 | 7780 | 7953 | 10173 | 9079 | 7871 | 8930 | 7564 | 5507 | 5963 | 6057 | 6336 | 7148 | 6051 | 6982 | 11242 | 7822 | 8017 | 11051 | 4936 | 7838 | 7860 | Arrb1 |
| Arrb2 | 3372 | 4591 | 4677 | 5032 | 4689 | 4169 | 4480 | 2961 | 2887 | 3422 | 3234 | 3362 | 3266 | 3506 | 3483 | 3555 | 5043 | 4812 | 5104 | 4037 | 5025 | 3399 | 2869 | 3702 | 3493 | 3729 | 3230 | 3595 | Arrb2 |
| Arrdc1 | 1475 | 1821 | 1578 | 1683 | 1752 | 1615 | 1591 | 652 | 778 | 766 | 635 | 577 | 575 | 533 | 1393 | 1611 | 1739 | 1700 | 1833 | 1575 | 1434 | 702 | 801 | 691 | 608 | 813 | 552 | 526 | Arrdc1 |
| Arrdc2 | 61 | 30 | 76 | 17 | 59 | 15 | 18 | 43 | 9 | 41 | 5 | 12 | 28 | 4 | 83 | 63 | 17 | 48 | 13 | 90 | 34 | 13 | 11 | 52 | 38 | 39 | 31 | 8 | Arrdc2 |
| Arrdc3 | 6481 | 3972 | 4939 | 4956 | 4489 | 4792 | 5555 | 6900 | 8759 | 6776 | 7628 | 7667 | 7083 | 7070 | 7243 | 4946 | 4723 | 4981 | 3848 | 5160 | 4555 | 6087 | 8889 | 7073 | 6318 | 7499 | 6130 | 6490 | Arrdc3 |
| Arrdc4 | 3184 | 2878 | 3225 | 3230 | 3077 | 2834 | 3073 | 3718 | 3544 | 3097 | 3097 | 2598 | 2799 | 2429 | 3299 | 2969 | 3113 | 3411 | 2951 | 3040 | 3095 | 3840 | 3722 | 2929 | 3152 | 2718 | 2781 | 2263 | Arrdc4 |
| Arrdc5 | 0 | 1 | 0 | 1 | 0 | 0 | 2 | 0 | 0 | 0 | 0 | 0 | 0 | 0 | 8 | 1 | 0 | 0 | 0 | 1 | 0 | 0 | 0 | 2 | 0 | 0 | 0 | 0 | Arrdc5 |
| Arsa | 1004 | 1449 | 1470 | 1375 | 1520 | 1301 | 1248 | 1149 | 1424 | 1532 | 1548 | 1222 | 1202 | 1195 | 1205 | 1333 | 1483 | 1477 | 1441 | 1188 | 1277 | 1474 | 1645 | 1459 | 1206 | 1288 | 1016 | 1043 | Arsa |
| Arsb | 1363 | 1201 | 1319 | 1136 | 1072 | 1370 | 976 | 1762 | 2363 | 1929 | 2442 | 1933 | 1812 | 1796 | 1245 | 1367 | 1162 | 1269 | 1048 | 1002 | 1071 | 2151 | 2487 | 1865 | 1833 | 1865 | 1819 | 1729 | Arsb |
| Arsg | 482 | 540 | 453 | 346 | 414 | 435 | 332 | 430 | 405 | 414 | 458 | 348 | 463 | 331 | 475 | 523 | 381 | 386 | 292 | 370 | 379 | 463 | 651 | 340 | 302 | 397 | 368 | 348 | Arsg |
| Arsi | 187 | 205 | 231 | 303 | 253 | 467 | 131 | 402 | 715 | 667 | 1138 | 452 | 583 | 416 | 214 | 183 | 260 | 470 | 227 | 192 | 170 | 959 | 999 | 659 | 680 | 372 | 451 | 350 | Arsi |
| Arsj | 23 | 19 | 16 | 32 | 5 | 32 | 6 | 62 | 149 | 56 | 146 | 45 | 68 | 39 | 46 | 44 | 28 | 37 | 16 | 14 | 4 | 95 | 131 | 63 | 47 | 45 | 29 | 43 | Arsj |
| Arsk | 862 | 619 | 744 | 689 | 585 | 652 | 721 | 905 | 976 | 769 | 959 | 833 | 921 | 750 | 769 | 748 | 782 | 683 | 625 | 782 | 634 | 829 | 1074 | 757 | 800 | 818 | 824 | 807 | Arsk |
| Art1 | 8 | 3 | 6 | 1 | 8 | 1 | 7 | 24 | 46 | 27 | 194 | 8 | 28 | 18 | 5 | 9 | 0 | 2 | 4 | 5 | 4 | 40 | 140 | 12 | 4 | 24 | 19 | 23 | Art1 |
| Art3 | 16 | 25 | 27 | 28 | 70 | 45 | 37 | 59 | 82 | 58 | 89 | 41 | 65 | 60 | 33 | 42 | 30 | 46 | 44 | 42 | 42 | 104 | 93 | 50 | 90 | 47 | 62 | 38 | Art3 |
| Art4 | 33 | 24 | 28 | 9 | 6 | 16 | 3 | 81 | 101 | 57 | 117 | 80 | 115 | 72 | 32 | 41 | 7 | 13 | 2 | 2 | 3 | 18 | 125 | 92 | 15 | 60 | 56 | 133 | Art4 |
| Art5 | 78 | 71 | 67 | 39 | 52 | 62 | 63 | 49 | 60 | 41 | 84 | 71 | 50 | 68 | 69 | 49 | 74 | 34 | 48 | 55 | 52 | 38 | 60 | 56 | 80 | 79 | 78 | 68 | Art5 |
| Artn | 24 | 58 | 42 | 42 | 25 | 34 | 27 | 17 | 49 | 26 | 47 | 12 | 23 | 20 | 22 | 45 | 57 | 36 | 41 | 41 | 29 | 35 | 32 | 29 | 9 | 20 | 17 | 21 | Artn |
| Arv1 | 1296 | 1384 | 1012 | 1235 | 1222 | 1446 | 1274 | 1116 | 1131 | 1142 | 1085 | 1315 | 1143 | 1278 | 1213 | 1251 | 1137 | 1350 | 1276 | 1255 | 1304 | 1285 | 1069 | 1209 | 1356 | 1029 | 1421 | 1206 | Arv1 |
| Arx | 2141 | 2113 | 2436 | 3383 | 3050 | 2844 | 4383 | 2514 | 2927 | 3497 | 2320 | 5153 | 1039 | 5803 | 1760 | 875 | 3153 | 2981 | 3843 | 2612 | 3866 | 3335 | 1582 | 5184 | 4580 | 6255 | 3270 | 4535 | Arx |
| Arxes1 | 1423 | 1102 | 1299 | 1563 | 1963 | 1503 | 2094 | 1553 | 1299 | 1310 | 1295 | 1095 | 1201 | 1525 | 1569 | 1153 | 1451 | 1670 | 1753 | 1721 | 2003 | 1753 | 1392 | 1392 | 1490 | 1288 | 1197 | 1327 | Arxes1 |
| Arxes2 | 3460 | 2602 | 3148 | 3490 | 4368 | 4117 | 4567 | 4087 | 3095 | 2997 | 2676 | 3120 | 2862 | 3242 | 4188 | 3169 | 3494 | 3779 | 4068 | 5111 | 4382 | 4244 | 3312 | 3215 | 3252 | 3052 | 3116 | 3305 | Arxes2 |
| As3mt | 623 | 676 | 706 | 529 | 632 | 585 | 497 | 1194 | 1137 | 1007 | 1171 | 1097 | 1252 | 1065 | 696 | 660 | 651 | 551 | 553 | 732 | 583 | 1071 | 1074 | 945 | 1102 | 1038 | 1167 | 1031 | As3mt |
| Asah1 | 11003 | 9193 | 9765 | 8545 | 7839 | 8637 | 8726 | 5951 | 5128 | 4533 | 4931 | 5175 | 5316 | 5467 | 12012 | 10770 | 8700 | 8484 | 7819 | 9716 | 8557 | 5210 | 5545 | 4739 | 4840 | 5631 | 5657 | 5844 | Asah1 |
| Asah2 | 266 | 129 | 250 | 125 | 243 | 282 | 188 | 446 | 616 | 427 | 738 | 363 | 460 | 314 | 280 | 312 | 155 | 197 | 147 | 185 | 157 | 528 | 690 | 399 | 382 | 348 | 344 | 299 | Asah2 |
| Asap1 | 6177 | 6308 | 6483 | 6480 | 7514 | 6480 | 6462 | 5937 | 6283 | 6979 | 7160 | 6879 | 7523 | 6642 | 6783 | 7380 | 7050 | 6958 | 6772 | 5534 | 6343 | 7780 | 6097 | 6414 | 6592 | 6539 | 6430 | 6741 | Asap1 |
| Asap2 | 3853 | 3733 | 4920 | 5106 | 5811 | 5041 | 5136 | 4324 | 4504 | 5785 | 5169 | 6571 | 5420 | 6388 | 3669 | 3820 | 4582 | 4911 | 5666 | 3939 | 5199 | 5504 | 4106 | 5979 | 5817 | 5737 | 5531 | 5893 | Asap2 |
| Asap3 | 2505 | 2831 | 2921 | 2633 | 2461 | 2284 | 1943 | 1047 | 1336 | 1440 | 1831 | 1099 | 1184 | 948 | 2326 | 2836 | 2870 | 2732 | 2436 | 2011 | 2175 | 1358 | 1730 | 1151 | 1010 | 1155 | 1147 | 1008 | Asap3 |
| Asb1 | 5150 | 5770 | 3988 | 4611 | 4377 | 4597 | 4539 | 3306 | 3149 | 2830 | 2915 | 3061 | 3083 | 2921 | 5283 | 5263 | 4465 | 4873 | 4526 | 4729 | 4574 | 3402 | 2915 | 2626 | 3322 | 2545 | 3221 | 2924 | Asb1 |
| Asb10 | 1 | 4 | 6 | 5 | 4 | 0 | 6 | 18 | 27 | 10 | 31 | 17 | 16 | 10 | 5 | 8 | 0 | 0 | 5 | 5 | 3 | 10 | 39 | 12 | 4 | 7 | 3 | 13 | Asb10 |
| Asb11 | 3 | 3 | 3 | 1 | 2 | 2 | 2 | 1 | 2 | 0 | 1 | 3 | 2 | 3 | 4 | 2 | 3 | 12 | 4 | 9 | 7 | 3 | 1 | 2 | 3 | 4 | 8 | 1 | Asb11 |
| Asb12 | 3 | 11 | 3 | 1 | 11 | 2 | 12 | 5 | 4 | 4 | 2 | 5 | 5 | 7 | 8 | 5 | 12 | 9 | 13 | 7 | 6 | 0 | 2 | 9 | 3 | 8 | 2 | 9 | Asb12 |
| Asb13 | 1136 | 1037 | 1109 | 1194 | 1412 | 1215 | 1388 | 1600 | 1383 | 1728 | 1563 | 1769 | 1673 | 1893 | 990 | 942 | 1187 | 1196 | 1365 | 1229 | 1191 | 1718 | 1284 | 1896 | 1846 | 2030 | 1898 | 1761 | Asb13 |
| Asb14 | 2 | 0 | 1 | 2 | 0 | 1 | 1 | 0 | 1 | 0 | 1 | 1 | 3 | 0 | 0 | 0 | 10 | 0 | 2 | 0 | 0 | 2 | 1 | 1 | 3 | 2 | 5 | 0 | Asb14 |
| Asb15 | 6 | 8 | 0 | 0 | 8 | 1 | 3 | 6 | 2 | 0 | 6 | 0 | 0 | 1 | 0 | 2 | 8 | 4 | 6 | 5 | 5 | 12 | 5 | 1 | 1 | 5 | 3 | 2 | Asb15 |
| Asb16 | 16 | 17 | 10 | 7 | 3 | 10 | 8 | 11 | 34 | 38 | 35 | 23 | 28 | 17 | 11 | 16 | 12 | 13 | 6 | 5 | 3 | 27 | 35 | 21 | 17 | 24 | 19 | 32 | Asb16 |
| Asb17 | 0 | 3 | 4 | 0 | 5 | 5 | 0 | 2 | 6 | 4 | 0 | 3 | 10 | 2 | 1 | 9 | 3 | 10 | 4 | 0 | 2 | 2 | 1 | 0 | 0 | 7 | 0 | 7 | Asb17 |
| Asb18 | 50 | 41 | 39 | 42 | 74 | 70 | 87 | 27 | 34 | 38 | 50 | 24 | 25 | 28 | 66 | 59 | 66 | 40 | 57 | 54 | 79 | 32 | 41 | 17 | 40 | 28 | 22 | 14 | Asb18 |
| Asb2 | 18 | 33 | 71 | 38 | 32 | 26 | 13 | 24 | 72 | 38 | 257 | 37 | 57 | 26 | 26 | 35 | 39 | 18 | 20 | 20 | 21 | 69 | 179 | 31 | 22 | 27 | 35 | 36 | Asb2 |
| Asb3 | 1283 | 1234 | 1076 | 1065 | 1173 | 1120 | 1380 | 2194 | 1968 | 1474 | 1653 | 1708 | 1616 | 1730 | 1452 | 1239 | 1167 | 1097 | 1155 | 1527 | 1160 | 1761 | 1995 | 1711 | 1800 | 1769 | 1648 | 1767 | Asb3 |
| Asb4 | 2828 | 2268 | 1931 | 2239 | 2030 | 2043 | 2326 | 2889 | 3091 | 2366 | 3366 | 2872 | 1593 | 2902 | 2438 | 2521 | 2377 | 2163 | 2169 | 2484 | 2213 | 2782 | 2583 | 2490 | 2472 | 3887 | 2735 | 2479 | Asb4 |
| Asb5 | 29 | 32 | 12 | 25 | 36 | 29 | 38 | 37 | 21 | 23 | 20 | 22 | 36 | 32 | 30 | 38 | 37 | 18 | 19 | 38 | 49 | 15 | 33 | 39 | 32 | 19 | 31 | 20 | Asb5 |
| Asb6 | 1937 | 2491 | 2065 | 2268 | 1944 | 2127 | 1713 | 1502 | 1684 | 1669 | 1544 | 1694 | 1788 | 1569 | 1745 | 2044 | 2383 | 2024 | 2312 | 2164 | 1826 | 1551 | 1643 | 1937 | 1675 | 2117 | 1832 | 1722 | Asb6 |
| Asb7 | 3122 | 2775 | 3164 | 3773 | 3656 | 3748 | 3818 | 2130 | 2135 | 2410 | 2421 | 2602 | 2545 | 2537 | 3158 | 2963 | 3468 | 3668 | 3733 | 3152 | 3769 | 2516 | 2089 | 2412 | 2582 | 2531 | 2492 | 2690 | Asb7 |
| Asb8 | 2270 | 2494 | 2476 | 2174 | 2152 | 2022 | 2309 | 2229 | 2245 | 2148 | 2075 | 2061 | 2144 | 2399 | 2352 | 2312 | 2278 | 2082 | 2030 | 2143 | 2132 | 2137 | 2186 | 2223 | 2143 | 2351 | 2105 | 2267 | Asb8 |
| Asb9 | 22 | 22 | 16 | 1 | 1 | 18 | 0 | 7 | 18 | 32 | 36 | 10 | 37 | 12 | 13 | 21 | 10 | 7 | 2 | 0 | 1 | 17 | 29 | 14 | 16 | 36 | 14 | 27 | Asb9 |
| Ascc1 | 1204 | 1353 | 1163 | 1133 | 1281 | 1341 | 1260 | 1579 | 1707 | 1736 | 1536 | 1661 | 1555 | 1609 | 1269 | 1131 | 1329 | 1215 | 1324 | 1308 | 1358 | 1582 | 1672 | 1797 | 1576 | 1841 | 1694 | 1691 | Ascc1 |
| Ascc2 | 4977 | 5526 | 4983 | 5355 | 4867 | 4842 | 4928 | 4262 | 4683 | 4934 | 4696 | 5317 | 5219 | 5213 | 4762 | 5337 | 4964 | 5546 | 5315 | 4401 | 4691 | 4746 | 4495 | 4969 | 4888 | 4725 | 5031 | 5103 | Ascc2 |
| Ascc3 | 5598 | 5090 | 3329 | 4292 | 3863 | 4236 | 4756 | 5438 | 4761 | 3646 | 3913 | 3789 | 3799 | 3894 | 5722 | 4839 | 3787 | 4187 | 3942 | 5271 | 4686 | 3907 | 4667 | 3479 | 3701 | 2683 | 4052 | 3891 | Ascc3 |
| Ascl1 | 2640 | 3118 | 6099 | 3981 | 4173 | 3962 | 4207 | 4301 | 3200 | 4555 | 3186 | 7083 | 4540 | 7568 | 3165 | 1842 | 6382 | 3456 | 5378 | 3421 | 4627 | 5060 | 2676 | 6477 | 5303 | 6243 | 6752 | 7729 | Ascl1 |
| Ascl2 | 87 | 145 | 98 | 83 | 72 | 53 | 60 | 5 | 12 | 12 | 18 | 14 | 6 | 10 | 61 | 111 | 86 | 105 | 65 | 38 | 55 | 13 | 17 | 16 | 6 | 6 | 12 | 8 | Ascl2 |
| Ascl3 | 2 | 11 | 7 | 3 | 1 | 7 | 3 | 0 | 3 | 1 | 2 | 2 | 1 | 2 | 3 | 5 | 7 | 8 | 9 | 1 | 5 | 2 | 2 | 2 | 0 | 0 | 0 | 0 | Ascl3 |
| Ascl4 | 77 | 70 | 94 | 88 | 103 | 85 | 93 | 35 | 22 | 16 | 16 | 26 | 25 | 28 | 78 | 53 | 80 | 57 | 97 | 86 | 99 | 15 | 32 | 29 | 18 | 29 | 45 | 32 | Ascl4 |
| Ascl5 | 15 | 21 | 0 | 4 | 11 | 14 | 7 | 1 | 2 | 0 | 1 | 2 | 2 | 1 | 8 | 17 | 7 | 17 | 7 | 12 | 7 | 3 | 3 | 1 | 0 | 2 | 0 | 2 | Ascl5 |
| Asf1a | 4861 | 3879 | 3931 | 3943 | 3635 | 4246 | 4548 | 4527 | 3629 | 3128 | 3011 | 4111 | 4007 | 4724 | 5144 | 3961 | 3800 | 3481 | 3821 | 4880 | 4203 | 3228 | 3541 | 3475 | 3673 | 4353 | 4471 | 4530 | Asf1a |
| Asf1b | 4840 | 4933 | 4792 | 3675 | 2065 | 2781 | 2806 | 1327 | 1207 | 1045 | 1090 | 1599 | 1465 | 1609 | 4109 | 4431 | 4051 | 3256 | 2852 | 3033 | 2660 | 897 | 1216 | 1539 | 1189 | 2056 | 1753 | 1921 | Asf1b |
| Asgr1 | 77 | 98 | 51 | 126 | 114 | 69 | 137 | 33 | 40 | 41 | 46 | 26 | 34 | 38 | 58 | 69 | 127 | 133 | 116 | 87 | 87 | 53 | 50 | 42 | 43 | 50 | 35 | 34 | Asgr1 |
| Asgr2 | 1 | 3 | 0 | 2 | 0 | 0 | 0 | 4 | 0 | 0 | 2 | 0 | 3 | 0 | 1 | 4 | 3 | 1 | 1 | 1 | 2 | 0 | 3 | 2 | 2 | 2 | 3 | 1 | Asgr2 |
| Ash1l | 7894 | 7226 | 4089 | 6188 | 7863 | 9120 | 8694 | 13913 | 13951 | 10898 | 12765 | 11147 | 11200 | 12228 | 7888 | 7293 | 4687 | 6259 | 6717 | 8924 | 9173 | 9154 | 12596 | 8603 | 12658 | 6453 | 11949 | 9871 | Ash1l |
| Ash2l | 7260 | 6305 | 6582 | 6502 | 6775 | 6750 | 6850 | 8971 | 8504 | 9071 | 8291 | 10111 | 9482 | 10116 | 7030 | 6194 | 6448 | 5954 | 6464 | 6757 | 6725 | 8691 | 8400 | 9363 | 9129 | 10840 | 9909 | 10207 | Ash2l |
| Asic1 | 1658 | 2540 | 2412 | 3670 | 6358 | 4149 | 3800 | 4415 | 4693 | 7032 | 5602 | 4876 | 5715 | 4501 | 2399 | 2122 | 3614 | 4178 | 5481 | 3697 | 4685 | 7673 | 4510 | 5454 | 6544 | 2687 | 4095 | 4593 | Asic1 |
| Asic2 | 221 | 352 | 388 | 488 | 917 | 668 | 545 | 570 | 714 | 669 | 581 | 422 | 557 | 440 | 361 | 341 | 447 | 529 | 771 | 571 | 619 | 907 | 613 | 621 | 778 | 478 | 406 | 388 | Asic2 |
| Asic3 | 44 | 41 | 15 | 43 | 55 | 30 | 34 | 115 | 126 | 100 | 116 | 33 | 73 | 46 | 38 | 55 | 46 | 56 | 30 | 33 | 32 | 134 | 104 | 49 | 74 | 55 | 38 | 26 | Asic3 |

| | | | | | | | | | | | | | | | | | | | | | | | | | | | | | |
|---|---|---|---|---|---|---|---|---|---|---|---|---|---|---|---|---|---|---|---|---|---|---|---|---|---|---|---|---|---|
| Asic4 | 959 | 1284 | 1844 | 1970 | 2242 | 1805 | 1530 | 1882 | 1364 | 2350 | 1761 | 2751 | 2824 | 2674 | 1143 | 1009 | 1876 | 1629 | 2332 | 1882 | 2120 | 2687 | 1520 | 2815 | 2700 | 721 | 2987 | 2886 | Asic4 |
| Asic5 | 0 | 3 | 0 | 0 | 2 | 6 | 3 | 1 | 4 | 0 | 3 | 7 | 3 | 3 | 0 | 2 | 0 | 2 | 1 | 7 | 3 | 3 | 0 | 4 | 0 | 3 | 1 | 1 | Asic5 |
| Asl | 1801 | 2515 | 2225 | 2203 | 2018 | 2120 | 1686 | 978 | 959 | 864 | 1030 | 1056 | 1133 | 958 | 1875 | 2225 | 1991 | 2086 | 2092 | 2091 | 1928 | 984 | 1072 | 978 | 1156 | 1142 | 1069 | 985 | Asl |
| Asmt | 1 | 1 | 1 | 1 | 2 | 3 | 3 | 4 | 8 | 5 | 5 | 9 | 10 | 6 | 0 | 1 | 5 | 2 | 8 | 1 | 1 | 10 | 8 | 13 | 9 | 17 | 3 | 9 | Asmt |
| Asns | 3922 | 3746 | 2922 | 2785 | 2910 | 2910 | 3042 | 2356 | 2349 | 1960 | 1922 | 1631 | 1958 | 2033 | 3893 | 3733 | 2751 | 2960 | 2907 | 3232 | 2917 | 2067 | 2263 | 1712 | 1803 | 2070 | 1873 | 2445 | Asns |
| Asnsd1 | 290 | 251 | 242 | 195 | 145 | 135 | 287 | 316 | 216 | 204 | 188 | 239 | 248 | 469 | 351 | 204 | 218 | 190 | 139 | 205 | 269 | 221 | 226 | 258 | 224 | 238 | 229 | 399 | Asnsd1 |
| Aspa | 15 | 17 | 0 | 0 | 34 | 11 | 12 | 27 | 30 | 36 | 50 | 17 | 28 | 13 | 17 | 16 | 6 | 25 | 4 | 12 | 17 | 52 | 62 | 11 | 19 | 6 | 3 | 6 | Aspa |
| Aspdh | 163 | 282 | 327 | 373 | 417 | 455 | 360 | 402 | 430 | 530 | 364 | 603 | 493 | 568 | 202 | 205 | 431 | 483 | 528 | 384 | 407 | 732 | 366 | 575 | 707 | 409 | 502 | 539 | Aspdh |
| Aspg | 96 | 111 | 129 | 72 | 52 | 75 | 34 | 27 | 47 | 47 | 44 | 37 | 77 | 40 | 58 | 124 | 136 | 80 | 55 | 47 | 37 | 45 | 69 | 43 | 38 | 58 | 66 | 63 | Aspg |
| Asph | 9357 | 7381 | 8114 | 6933 | 6243 | 7096 | 5727 | 6283 | 6793 | 5965 | 7673 | 5689 | 7775 | 5873 | 9330 | 9965 | 7174 | 6974 | 5611 | 6170 | 5560 | 5633 | 7453 | 6071 | 5175 | 6258 | 5999 | 5936 | Asph |
| Asphd1 | 143 | 168 | 195 | 294 | 457 | 349 | 312 | 316 | 358 | 500 | 444 | 399 | 298 | 328 | 161 | 110 | 336 | 292 | 403 | 284 | 338 | 536 | 339 | 494 | 569 | 306 | 356 | 399 | Asphd1 |
| Asphd2 | 2941 | 4165 | 3178 | 3937 | 4130 | 3964 | 3921 | 2317 | 2448 | 2454 | 1989 | 2648 | 2216 | 2255 | 3334 | 3067 | 3664 | 3772 | 4508 | 4354 | 4219 | 2490 | 2094 | 2466 | 2654 | 2154 | 2227 | 2070 | Asphd2 |
| Aspm | 11770 | 10811 | 6872 | 7073 | 5932 | 7877 | 8335 | 12822 | 11016 | 7804 | 7720 | 12511 | 10819 | 12666 | 11020 | 9495 | 6749 | 6231 | 6816 | 10298 | 8340 | 5715 | 9183 | 8322 | 8926 | 8736 | 14346 | 11459 | Aspm |
| Aspn | 167 | 105 | 41 | 279 | 67 | 571 | 15 | 1480 | 2016 | 1585 | 2814 | 563 | 569 | 622 | 237 | 178 | 123 | 556 | 75 | 46 | 24 | 1482 | 3565 | 1366 | 1322 | 545 | 471 | 482 | Aspn |
| Asprv1 | 43 | 45 | 32 | 48 | 49 | 46 | 45 | 83 | 131 | 97 | 83 | 50 | 61 | 81 | 26 | 43 | 29 | 50 | 37 | 41 | 54 | 33 | 143 | 96 | 69 | 62 | 52 | 55 | Asprv1 |
| Aspscr1 | 2366 | 2909 | 2868 | 2980 | 2792 | 2852 | 2522 | 3366 | 3888 | 3709 | 3530 | 4029 | 3867 | 3888 | 2554 | 2512 | 3034 | 2717 | 2920 | 2848 | 2791 | 3710 | 3735 | 3948 | 3832 | 4282 | 4285 | 3890 | Aspscr1 |
| Asrgl1 | 3914 | 4205 | 3600 | 3950 | 4358 | 4010 | 4626 | 2463 | 2389 | 2313 | 2112 | 2474 | 1889 | 2380 | 3823 | 3160 | 4222 | 4127 | 4251 | 4206 | 4408 | 2174 | 1910 | 2344 | 2400 | 2701 | 2160 | 2265 | Asrgl1 |
| Ass1 | 272 | 298 | 189 | 166 | 174 | 120 | 107 | 177 | 213 | 151 | 179 | 174 | 204 | 151 | 270 | 322 | 192 | 168 | 121 | 100 | 120 | 154 | 247 | 183 | 144 | 164 | 185 | 161 | Ass1 |
| Aste1 | 1318 | 1325 | 984 | 1211 | 973 | 1100 | 1117 | 1033 | 907 | 828 | 707 | 818 | 790 | 852 | 1189 | 1255 | 1056 | 1188 | 996 | 1293 | 1194 | 703 | 845 | 727 | 761 | 699 | 945 | 833 | Aste1 |
| Astl | 1 | 1 | 1 | 0 | 0 | 0 | 1 | 5 | 5 | 3 | 2 | 0 | 1 | 2 | 0 | 2 | 1 | 0 | 0 | 0 | 0 | 0 | 5 | 0 | 0 | 0 | 1 | 1 | Astl |
| Astn1 | 6577 | 6385 | 5452 | 8195 | 10762 | 10748 | 10523 | 12193 | 11769 | 12611 | 10500 | 12055 | 10735 | 11819 | 7671 | 5752 | 7011 | 8817 | 9919 | 9823 | 11366 | 15024 | 9976 | 11272 | 14350 | 8224 | 11702 | 10595 | Astn1 |
| Astn2 | 1152 | 1248 | 1100 | 1816 | 2389 | 1927 | 1672 | 1685 | 1851 | 2254 | 2076 | 1624 | 1909 | 1578 | 1304 | 1238 | 1387 | 2168 | 1896 | 1402 | 1936 | 2665 | 1916 | 1707 | 2055 | 1339 | 1502 | 1475 | Astn2 |
| Asxl1 | 7019 | 6966 | 7952 | 8339 | 9025 | 8504 | 8555 | 9308 | 9863 | 10991 | 10362 | 12761 | 12507 | 12832 | 6691 | 6514 | 8314 | 8263 | 9262 | 6630 | 9092 | 11294 | 8999 | 10729 | 11283 | 10371 | 12776 | 12589 | Asxl1 |
| Asxl2 | 4926 | 3736 | 3185 | 3938 | 4234 | 4699 | 4662 | 4539 | 4352 | 4215 | 4478 | 4622 | 4695 | 4846 | 4741 | 3989 | 3537 | 3796 | 3970 | 4345 | 4482 | 3634 | 4078 | 3715 | 4437 | 3328 | 4855 | 4186 | Asxl2 |
| Asxl3 | 4106 | 3654 | 2710 | 3583 | 4624 | 4785 | 4324 | 7342 | 7217 | 7223 | 7228 | 5975 | 7347 | 6220 | 4146 | 4094 | 3027 | 3434 | 3867 | 3888 | 4539 | 7158 | 6499 | 5279 | 7092 | 4211 | 6627 | 5626 | Asxl3 |
| Asz1 | 0 | 0 | 0 | 2 | 0 | 0 | 0 | 0 | 0 | 0 | 1 | 0 | 0 | 0 | 0 | 0 | 0 | 0 | 0 | 2 | 0 | 0 | 0 | 0 | 0 | 9 | 0 | 0 | Asz1 |
| Atad1 | 11344 | 9192 | 10403 | 10340 | 10531 | 10665 | 11957 | 13065 | 10411 | 9579 | 9159 | 10457 | 9992 | 10863 | 12922 | 10622 | 10384 | 10210 | 10395 | 13414 | 11042 | 11486 | 10545 | 9988 | 10462 | 10699 | 10928 | 11136 | Atad1 |
| Atad2 | 0 | 0 | 0 | 0 | 0 | 0 | 0 | 0 | 0 | 0 | 0 | 1 | 0 | 0 | 0 | 0 | 0 | 1 | 0 | 0 | 0 | 0 | 0 | 0 | 0 | 0 | 1 | 0 | Atad2 |
| Atad2b | 5708 | 4333 | 4632 | 5158 | 5985 | 6136 | 5995 | 10555 | 8932 | 9856 | 9745 | 10736 | 10350 | 10895 | 5570 | 5068 | 4894 | 5056 | 5699 | 5420 | 5819 | 9875 | 8484 | 8921 | 10416 | 7648 | 11152 | 10153 | Atad2b |
| Atad3a | 4883 | 5859 | 5712 | 6040 | 5114 | 4928 | 5207 | 3105 | 3798 | 3938 | 3831 | 4607 | 4230 | 4202 | 4512 | 5531 | 5651 | 5760 | 5400 | 4625 | 5182 | 3534 | 3412 | 4276 | 4050 | 4789 | 4105 | 4453 | Atad3a |
| Atad5 | 9281 | 7264 | 6084 | 6214 | 5175 | 6832 | 7585 | 5619 | 5087 | 4272 | 4251 | 6211 | 5324 | 6308 | 8784 | 7547 | 5917 | 6265 | 5541 | 7766 | 6783 | 3805 | 4343 | 4529 | 4674 | 5097 | 6528 | 5873 | Atad5 |
| Atat1 | 9806 | 11901 | 11974 | 17282 | 18858 | 16302 | 17763 | 13166 | 13428 | 18658 | 14533 | 17706 | 14149 | 17440 | 10172 | 10248 | 16171 | 17511 | 19601 | 13372 | 18743 | 22257 | 12393 | 17714 | 19842 | 14584 | 13933 | 15599 | Atat1 |
| Atcay | 3809 | 5495 | 5246 | 7212 | 9348 | 8133 | 7625 | 7507 | 8131 | 9542 | 7418 | 8292 | 6761 | 8240 | 4909 | 4201 | 7013 | 7542 | 9524 | 7916 | 8519 | 10591 | 7431 | 8865 | 11342 | 6919 | 7637 | 7430 | Atcay |
| Ate1 | 5301 | 5135 | 5092 | 5704 | 6181 | 6029 | 6279 | 6017 | 5236 | 5243 | 5252 | 5303 | 5034 | 5443 | 5768 | 5312 | 5475 | 6195 | 5690 | 6091 | 6255 | 6204 | 5239 | 5138 | 5705 | 4453 | 5120 | 5234 | Ate1 |
| Atf1 | 870 | 714 | 1332 | 913 | 889 | 946 | 853 | 1911 | 1879 | 2168 | 2209 | 2621 | 2456 | 2543 | 815 | 761 | 1109 | 852 | 1111 | 798 | 870 | 1980 | 1895 | 2510 | 2276 | 3458 | 2551 | 2715 | Atf1 |
| Atf2 | 7978 | 6488 | 8180 | 7907 | 8609 | 8376 | 8982 | 9274 | 8239 | 8341 | 7904 | 9231 | 8727 | 9678 | 7975 | 7727 | 8327 | 8187 | 8348 | 8312 | 8368 | 9984 | 7996 | 8697 | 9484 | 9093 | 9136 | 9280 | Atf2 |
| Atf3 | 300 | 343 | 432 | 266 | 124 | 195 | 159 | 94 | 101 | 82 | 107 | 73 | 87 | 76 | 266 | 322 | 275 | 173 | 132 | 166 | 177 | 72 | 115 | 95 | 88 | 124 | 89 | 88 | Atf3 |
| Atf4 | 6149 | 7525 | 6927 | 6421 | 5731 | 6070 | 6216 | 8249 | 7949 | 6932 | 7611 | 8711 | 8387 | 9296 | 6158 | 6171 | 6743 | 5900 | 6373 | 6508 | 5947 | 7563 | 7596 | 8373 | 8272 | 9723 | 8947 | 9556 | Atf4 |
| Atf5 | 1888 | 2085 | 1981 | 2790 | 1667 | 1586 | 1466 | 2512 | 2600 | 3545 | 3137 | 2435 | 2233 | 2521 | 1610 | 1640 | 2066 | 1681 | 2091 | 1613 | 1643 | 1706 | 2758 | 4554 | 2212 | 4853 | 2249 | 2478 | Atf5 |
| Atf6 | 7709 | 7482 | 5565 | 6413 | 5933 | 6419 | 6358 | 3689 | 3138 | 3242 | 3302 | 3214 | 3385 | 3318 | 7473 | 7212 | 5733 | 6147 | 5837 | 6170 | 6478 | 3183 | 3329 | 2787 | 3215 | 2468 | 3375 | 3232 | Atf6 |
| Atf6b | 4534 | 5948 | 6145 | 6075 | 5509 | 5121 | 4785 | 4408 | 5188 | 5674 | 5235 | 5941 | 5177 | 5455 | 4415 | 4984 | 6198 | 5766 | 5991 | 4514 | 5079 | 5217 | 5138 | 6069 | 5528 | 6517 | 5227 | 5478 | Atf6b |
| Atf7 | 3179 | 2983 | 2187 | 3040 | 3287 | 3164 | 3067 | 3770 | 4022 | 3655 | 4779 | 3128 | 3757 | 3294 | 3117 | 3209 | 2301 | 3226 | 2834 | 2766 | 3202 | 3611 | 4132 | 3009 | 3614 | 2536 | 3470 | 3023 | Atf7 |
| Atf7ip | 9806 | 8881 | 9030 | 10906 | 12029 | 12023 | 12187 | 14883 | 15935 | 17520 | 16736 | 18558 | 18394 | 17945 | 9394 | 9829 | 10247 | 11280 | 12368 | 10196 | 11976 | 17776 | 14595 | 17084 | 18945 | 13826 | 17845 | 17514 | Atf7ip |
| Atf7ip2 | 360 | 189 | 231 | 99 | 88 | 107 | 108 | 93 | 95 | 60 | 76 | 83 | 122 | 99 | 235 | 271 | 173 | 106 | 60 | 107 | 96 | 43 | 92 | 83 | 49 | 112 | 130 | 101 | Atf7ip2 |
| Atg10 | 298 | 257 | 245 | 265 | 293 | 305 | 311 | 503 | 421 | 423 | 428 | 414 | 424 | 516 | 348 | 266 | 297 | 222 | 219 | 302 | 259 | 436 | 447 | 378 | 400 | 531 | 447 | 472 | Atg10 |
| Atg101 | 1983 | 2857 | 2159 | 2434 | 1793 | 1903 | 1743 | 1501 | 1562 | 1477 | 1383 | 1517 | 1441 | 1327 | 1905 | 2361 | 2385 | 2253 | 2037 | 2140 | 1772 | 1308 | 1467 | 1729 | 1576 | 1708 | 1539 | 1392 | Atg101 |
| Atg12 | 10242 | 10819 | 9197 | 10354 | 9076 | 8670 | 10457 | 7891 | 6329 | 5916 | 5561 | 6584 | 6203 | 6508 | 12063 | 10834 | 9775 | 10463 | 8747 | 11090 | 9507 | 6876 | 6297 | 5932 | 6680 | 6088 | 6607 | 6722 | Atg12 |
| Atg13 | 4767 | 5425 | 4898 | 5549 | 5196 | 5235 | 5342 | 3497 | 3683 | 3984 | 3791 | 3937 | 3943 | 3856 | 4847 | 5079 | 5208 | 5564 | 5329 | 4827 | 5491 | 4025 | 3586 | 3765 | 4358 | 3476 | 3746 | 3889 | Atg13 |
| Atg14 | 1761 | 2033 | 2003 | 2049 | 1910 | 1893 | 2035 | 2030 | 1980 | 1840 | 1792 | 1983 | 1870 | 1965 | 1826 | 1791 | 1940 | 1791 | 1997 | 1962 | 1951 | 1835 | 1831 | 1923 | 1979 | 1909 | 2000 | 1932 | Atg14 |
| Atg16l1 | 4278 | 4391 | 3440 | 4211 | 4211 | 4357 | 4004 | 4835 | 4674 | 4243 | 4354 | 4086 | 4149 | 3929 | 4184 | 3847 | 3735 | 4011 | 3997 | 4715 | 4012 | 4326 | 4271 | 3957 | 4581 | 3731 | 4230 | 3801 | Atg16l1 |
| Atg16l2 | 1208 | 1442 | 1436 | 1236 | 1103 | 1163 | 1088 | 652 | 718 | 680 | 701 | 629 | 620 | 722 | 1100 | 1246 | 1415 | 1219 | 1334 | 1162 | 1024 | 774 | 659 | 751 | 722 | 632 | 733 | 744 | Atg16l2 |
| Atg2a | 0 | 0 | 0 | 0 | 0 | 0 | 0 | 0 | 0 | 0 | 0 | 0 | 0 | 0 | 0 | 0 | 0 | 0 | 0 | 0 | 0 | 2 | 0 | 0 | 0 | 0 | 0 | 0 | Atg2a |
| Atg2b | 6014 | 6251 | 4328 | 6397 | 6950 | 6988 | 6852 | 4824 | 4721 | 4781 | 5052 | 4845 | 4600 | 4768 | 5822 | 5955 | 5467 | 6522 | 6808 | 6106 | 7322 | 4627 | 4496 | 4268 | 5392 | 3062 | 4577 | 4174 | Atg2b |
| Atg3 | 3216 | 2811 | 2959 | 2823 | 2756 | 2602 | 3168 | 3040 | 2614 | 2708 | 2397 | 2497 | 2476 | 2869 | 3456 | 2788 | 2714 | 2825 | 2622 | 2703 | 2749 | 2836 | 2599 | 2540 | 2696 | 2801 | 2494 | 2808 | Atg3 |
| Atg4a | 23 | 166 | 117 | 10 | 11 | 142 | 47 | 13 | 166 | 12 | 12 | 42 | 15 | 24 | 30 | 35 | 21 | 152 | 26 | 13 | 43 | 3 | 15 | 6 | 22 | 110 | 9 | 8 | Atg4a |
| Atg4a-ps | 269 | 193 | 204 | 141 | 135 | 185 | 137 | 219 | 202 | 146 | 180 | 127 | 125 | 130 | 252 | 276 | 175 | 188 | 164 | 163 | 128 | 172 | 214 | 149 | 137 | 159 | 143 | 142 | Atg4a-ps |
| Atg4b | 9450 | 10635 | 8414 | 9077 | 8655 | 8870 | 8656 | 7314 | 6741 | 6291 | 6070 | 6507 | 6597 | 6431 | 9737 | 9269 | 9035 | 8856 | 9130 | 9336 | 9161 | 6644 | 6658 | 6117 | 6777 | 6068 | 6503 | 6548 | Atg4b |
| Atg4c | 1204 | 910 | 960 | 1108 | 1481 | 1253 | 1454 | 1238 | 1206 | 1183 | 1250 | 1074 | 1111 | 1209 | 1146 | 1139 | 1101 | 1271 | 1320 | 1291 | 1326 | 1275 | 1197 | 1160 | 1110 | 1188 | 1088 | 1137 | Atg4c |
| Atg4d | 2136 | 2838 | 2551 | 2924 | 2675 | 2564 | 2486 | 1529 | 1734 | 1867 | 1722 | 2066 | 1707 | 1874 | 2123 | 2312 | 2643 | 2921 | 3033 | 2407 | 2507 | 1798 | 1701 | 2079 | 2026 | 2164 | 1991 | 1937 | Atg4d |
| Atg5 | 3369 | 2998 | 3219 | 2651 | 2417 | 2549 | 2864 | 2684 | 2389 | 2067 | 1953 | 2183 | 2369 | 2409 | 3568 | 3443 | 3009 | 2810 | 2632 | 3364 | 2766 | 2114 | 2573 | 2120 | 2134 | 2471 | 2312 | 2575 | Atg5 |
| Atg7 | 915 | 1234 | 1006 | 1043 | 1019 | 1083 | 946 | 929 | 1182 | 1117 | 985 | 1143 | 1030 | 1115 | 1007 | 977 | 975 | 1039 | 966 | 916 | 986 | 967 | 1069 | 997 | 1092 | 1224 | 1044 | 1165 | Atg7 |
| Atg9a | 1395 | 1557 | 1522 | 1656 | 1681 | 1528 | 1504 | 1481 | 1739 | 1890 | 1684 | 1536 | 1664 | 1618 | 1249 | 1571 | 1684 | 1717 | 1904 | 1284 | 1659 | 2075 | 1750 | 1561 | 1858 | 1374 | 1435 | 1576 | Atg9a |
| Atg9b | 33 | 40 | 66 | 27 | 49 | 45 | 31 | 38 | 44 | 49 | 42 | 34 | 37 | 29 | 41 | 40 | 52 | 34 | 57 | 46 | 25 | 22 | 53 | 49 | 34 | 30 | 22 | 23 | Atg9b |
| Atic | 6527 | 6659 | 6298 | 5291 | 5142 | 5417 | 4755 | 5583 | 6162 | 5242 | 5856 | 5566 | 5865 | 5139 | 5970 | 6456 | 5778 | 5440 | 5563 | 5258 | 4956 | 4328 | 5847 | 5622 | 4935 | 6963 | 5875 | 5849 | Atic |
| Atl1 | 1137 | 1106 | 1060 | 1196 | 2247 | 1635 | 1868 | 5075 | 3760 | 3764 | 3484 | 3394 | 3481 | 3962 | 1404 | 984 | 1158 | 1264 | 1802 | 1911 | 1998 | 5111 | 3882 | 3662 | 4460 | 2757 | 3903 | 3742 | Atl1 |
| Atl2 | 5823 | 4981 | 5176 | 5930 | 5377 | 5578 | 6279 | 4921 | 4359 | 4131 | 4316 | 4687 | 4568 | 4696 | 6156 | 5583 | 5171 | 5293 | 5375 | 5937 | 6180 | 4717 | 4467 | 4250 | 4471 | 4733 | 4783 | 4988 | Atl2 |
| Atl3 | 4349 | 2925 | 2812 | 3365 | 3748 | 4275 | 3786 | 6372 | 6991 | 5646 | 7489 | 5641 | 6458 | 5612 | 4092 | 3774 | 2880 | 3729 | 2964 | 3693 | 3632 | 5349 | 7298 | 5544 | 5681 | 4594 | 5798 | 5272 | Atl3 |
| Atm | 11311 | 10115 | 3626 | 5059 | 5468 | 8603 | 7551 | 8402 | 7317 | 4612 | 5348 | 5762 | 5864 | 6127 | 11086 | 10335 | 3939 | 4933 | 4866 | 9872 | 8418 | 3078 | 6725 | 3224 | 5159 | 2410 | 7698 | 5035 | Atm |
| Atmin | 5493 | 6222 | 4610 | 5432 | 5411 | 5607 | 5579 | 3858 | 3942 | 3939 | 3566 | 3894 | 3548 | 3927 | 5474 | 5440 | 5057 | 5613 | 5501 | 5425 | 5591 | 4084 | 3559 | 3750 | 4047 | 3615 | 3655 | 3826 | Atmin |
| Atn1 | 266 | 309 | 444 | 584 | 560 | 377 | 794 | 507 | 458 | 1013 | 754 | 659 | 634 | 1212 | 239 | 167 | 581 | 568 | 496 | 350 | 836 | 683 | 456 | 1149 | 889 | 900 | 637 | 1053 | Atn1 |
| Atoh1 | 32 | 162 | 29 | 92 | 155 | 69 | 9 | 167 | 190 | 124 | 139 | 25 | 305 | 2 | 145 | 255 | 207 | 72 | 30 | 99 | 18 | 179 | 222 | 0 | 75 | 1 | 5 | 8 | Atoh1 |
| Atoh7 | 36 | 13 | 23 | 13 | 34 | 37 | 32 | 12 | 17 | 14 | 16 | 18 | 20 | 6 | 28 | 32 | 44 | 17 | 23 | 38 | 33 | 25 | 21 | 124 | 22 | 38 | 33 | 12 | Atoh7 |
| Atoh8 | 246 | 325 | 247 | 219 | 158 | 229 | 158 | 221 | 359 | 345 | 422 | 317 | 359 | 280 | 229 | 312 | 264 | 220 | 228 | 214 | 125 | 211 | 426 | 299 | 237 | 392 | 300 | 272 | Atoh8 |
| Atox1 | 2330 | 2832 | 2392 | 2536 | 2059 | 1940 | 2573 | 1507 | 1571 | 1619 | 1904 | 1367 | 1819 | 1763 | 2153 | 2153 | 2452 | 2331 | 2264 | 2375 | 2405 | 1188 | 1674 | 1711 | 1640 | 1771 | 1621 | 1782 | Atox1 |
| Atp10a | 374 | 426 | 251 | 286 | 372 | 320 | 326 | 546 | 644 | 569 | 755 | 728 | 552 | 652 | 353 | 340 | 278 | 317 | 351 | 283 | 305 | 491 | 605 | 555 | 585 | 608 | 724 | 638 | Atp10a |
| Atp10b | 82 | 51 | 6 | 26 | 74 | 25 | 13 | 254 | 321 | 162 | 292 | 48 | 217 | 43 | 71 | 113 | 24 | 24 | 17 | 18 | 25 | 277 | 289 | 60 | 135 | 51 | 35 | 37 | Atp10b |
| Atp11a | 10697 | 10534 | 7018 | 9025 | 9406 | 9745 | 9288 | 8190 | 9314 | 8556 | 9076 | 8085 | 7660 | 7294 | 8909 | 10253 | 7356 | 9255 | 8503 | 9051 | 9442 | 6919 | 8361 | 7197 | 8243 | 5986 | 8279 | 6518 | Atp11a |
| Atp11b | 8494 | 6130 | 6609 | 7078 | 8102 | 8085 | 8544 | 7139 | 6586 | 6418 | 7282 | 6297 | 6514 | 6528 | 8024 | 7238 | 6806 | 7545 | 7187 | 7506 | 8416 | 6697 | 6514 | 6025 | 6650 | 5286 | 6403 | 5929 | Atp11b |
| Atp11c | 5922 | 4037 | 4827 | 5596 | 5920 | 6751 | 7158 | 7640 | 6254 | 5789 | 5929 | 7031 | 6600 | 7261 | 6311 | 5133 | 5167 | 5407 | 5781 | 6929 | 6807 | 6958 | 5941 | 5831 | 6797 | 6216 | 7089 | 7049 | Atp11c |

| | | | | | | | | | | | | | | | | | | | | | | | | | | | | | |
|---|---|---|---|---|---|---|---|---|---|---|---|---|---|---|---|---|---|---|---|---|---|---|---|---|---|---|---|---|---|
| Atp12a | 16 | 6 | 10 | 2 | 5 | 1 | 7 | 4 | 8 | 8 | 13 | 3 | 9 | 9 | 11 | 3 | 7 | 4 | 4 | 5 | 6 | 7 | 14 | 14 | 11 | 3 | 2 | 17 | Atp12a |
| Atp13a1 | 5520 | 6672 | 6242 | 6638 | 6431 | 6050 | 6112 | 5297 | 6164 | 6013 | 5792 | 6637 | 6038 | 5801 | 5329 | 6154 | 6439 | 6489 | 6666 | 5669 | 5917 | 6315 | 5834 | 6283 | 6418 | 6206 | 6186 | 6107 | Atp13a1 |
| Atp13a2 | 7041 | 9898 | 9504 | 11670 | 12851 | 10421 | 9137 | 5907 | 7158 | 8181 | 7444 | 7342 | 6484 | 5958 | 7508 | 8759 | 10856 | 11975 | 12478 | 8707 | 9849 | 8676 | 6594 | 7620 | 8289 | 6693 | 6288 | 5671 | Atp13a2 |
| Atp13a3 | 13755 | 10025 | 10101 | 12354 | 12084 | 13184 | 14067 | 7558 | 7009 | 6601 | 8098 | 7379 | 7005 | 7731 | 12957 | 12606 | 11038 | 12682 | 11461 | 11309 | 13495 | 7331 | 7237 | 6998 | 7158 | 6244 | 7252 | 7444 | Atp13a3 |
| Atp13a4 | 3 | 7 | 5 | 4 | 13 | 25 | 15 | 50 | 41 | 35 | 31 | 29 | 18 | 42 | 11 | 6 | 6 | 17 | 16 | 21 | 20 | 42 | 47 | 23 | 48 | 47 | 27 | 34 | Atp13a4 |
| Atp13a5 | 17 | 22 | 1 | 15 | 43 | 25 | 20 | 59 | 41 | 63 | 101 | 35 | 27 | 20 | 24 | 14 | 12 | 9 | 19 | 20 | 34 | 69 | 76 | 38 | 42 | 44 | 19 | 24 | Atp13a5 |
| Atp1a1 | 12978 | 15129 | 15311 | 13874 | 14087 | 13037 | 11778 | 11375 | 12687 | 12541 | 13407 | 12211 | 13021 | 11082 | 13239 | 15118 | 14394 | 14425 | 13881 | 12221 | 12528 | 12226 | 14350 | 11807 | 10904 | 13615 | 12455 | 12208 | Atp1a1 |
| Atp1a2 | 1843 | 2401 | 2094 | 3279 | 6250 | 4441 | 4012 | 5616 | 6708 | 7133 | 7276 | 4670 | 4829 | 4611 | 2529 | 2076 | 2613 | 4192 | 4959 | 4476 | 4773 | 7758 | 8643 | 6037 | 7576 | 3966 | 4241 | 3628 | Atp1a2 |
| Atp1a3 | 6921 | 10726 | 10939 | 20554 | 25269 | 19270 | 17242 | 9714 | 11536 | 17266 | 13050 | 11752 | 9538 | 10095 | 8426 | 8168 | 17685 | 22503 | 24695 | 14939 | 21688 | 20100 | 10519 | 13817 | 18114 | 9554 | 8459 | 9207 | Atp1a3 |
| Atp1a4 | 7 | 12 | 4 | 8 | 18 | 14 | 9 | 18 | 11 | 8 | 9 | 8 | 3 | 11 | 16 | 9 | 7 | 8 | 8 | 14 | 13 | 8 | 6 | 10 | 14 | 6 | 22 | 6 | Atp1a4 |
| Atp1b1 | 3073 | 3519 | 4356 | 5523 | 7480 | 5653 | 5396 | 4316 | 3541 | 4606 | 4377 | 3269 | 3763 | 3154 | 4138 | 3404 | 5087 | 5591 | 5885 | 5369 | 5870 | 6041 | 4148 | 3653 | 4609 | 2874 | 3129 | 3185 | Atp1b1 |
| Atp1b2 | 2829 | 3276 | 3973 | 4105 | 4739 | 3917 | 4138 | 1754 | 2142 | 2703 | 2387 | 2088 | 1793 | 2004 | 3090 | 3216 | 4325 | 4469 | 4567 | 3293 | 4366 | 3293 | 2551 | 2344 | 2756 | 1872 | 1541 | 1861 | Atp1b2 |
| Atp1b3 | 10555 | 9764 | 10927 | 10026 | 9685 | 9417 | 10778 | 8007 | 6921 | 7407 | 7271 | 7983 | 7922 | 8328 | 10797 | 9944 | 10298 | 10121 | 9949 | 10429 | 10154 | 8154 | 7607 | 7976 | 7587 | 8793 | 8318 | 8578 | Atp1b3 |
| Atp1b4 | 5 | 2 | 1 | 4 | 2 | 1 | 4 | 13 | 21 | 3 | 420 | 3 | 0 | 0 | 4 | 2 | 17 | 9 | 0 | 0 | 5 | 13 | 276 | 3 | 2 | 0 | 0 | 0 | Atp1b4 |
| Atp23 | 1129 | 1160 | 1164 | 1095 | 917 | 964 | 980 | 466 | 444 | 486 | 388 | 527 | 462 | 449 | 1077 | 1218 | 1169 | 1268 | 890 | 1046 | 932 | 434 | 432 | 458 | 387 | 609 | 510 | 484 | Atp23 |
| Atp2a1 | 138 | 57 | 50 | 60 | 37 | 46 | 36 | 131 | 340 | 45 | 2125 | 47 | 92 | 23 | 84 | 181 | 63 | 73 | 34 | 52 | 37 | 262 | 1296 | 61 | 36 | 49 | 29 | 26 | Atp2a1 |
| Atp2a2 | 21533 | 21343 | 24257 | 26415 | 30089 | 26180 | 29059 | 25563 | 23850 | 28010 | 28334 | 27908 | 26584 | 27636 | 21605 | 20049 | 26438 | 28196 | 30035 | 23009 | 30068 | 30350 | 24013 | 28606 | 28779 | 23412 | 26877 | 28769 | Atp2a2 |
| Atp2a3 | 156 | 197 | 308 | 178 | 226 | 143 | 164 | 95 | 129 | 175 | 177 | 101 | 247 | 97 | 182 | 283 | 192 | 208 | 161 | 96 | 169 | 157 | 191 | 110 | 119 | 155 | 167 | 103 | Atp2a3 |
| Atp2b1 | 14056 | 12065 | 11314 | 13232 | 16540 | 16115 | 16394 | 14824 | 13830 | 13323 | 15327 | 10690 | 13126 | 11636 | 15116 | 13434 | 12512 | 14296 | 14159 | 15918 | 16370 | 14290 | 13889 | 10418 | 13600 | 7603 | 11381 | 10833 | Atp2b1 |
| Atp2b2 | 894 | 1167 | 956 | 1703 | 3116 | 2181 | 2276 | 1969 | 2496 | 2835 | 2413 | 1951 | 1929 | 1869 | 1246 | 1111 | 1517 | 2167 | 2239 | 1852 | 2254 | 3116 | 1982 | 2116 | 3260 | 1165 | 1413 | 1508 | Atp2b2 |
| Atp2b3 | 108 | 194 | 108 | 278 | 718 | 451 | 329 | 652 | 920 | 742 | 817 | 360 | 533 | 366 | 183 | 189 | 147 | 316 | 481 | 438 | 532 | 1076 | 904 | 446 | 910 | 144 | 273 | 218 | Atp2b3 |
| Atp2b4 | 11479 | 11041 | 8885 | 9748 | 10698 | 10488 | 9246 | 6637 | 7697 | 7707 | 8442 | 6040 | 6754 | 5983 | 11176 | 13105 | 9587 | 10879 | 8462 | 8285 | 9643 | 7969 | 7749 | 6155 | 6866 | 4260 | 5736 | 5782 | Atp2b4 |
| Atp2c1 | 11118 | 9891 | 9643 | 11460 | 12905 | 11779 | 13047 | 11894 | 9866 | 9452 | 9754 | 9042 | 9036 | 9433 | 12150 | 10063 | 10326 | 11637 | 11672 | 12781 | 12804 | 11498 | 9849 | 9207 | 10030 | 8133 | 9164 | 8984 | Atp2c1 |
| Atp2c2 | 100 | 74 | 83 | 6 | 7 | 58 | 3 | 55 | 96 | 49 | 40 | 62 | 92 | 73 | 80 | 118 | 21 | 73 | 12 | 3 | 4 | 12 | 136 | 61 | 14 | 19 | 56 | 69 | Atp2c2 |
| Atp4a | 234 | 229 | 273 | 224 | 241 | 188 | 213 | 71 | 60 | 49 | 48 | 48 | 65 | 65 | 167 | 187 | 228 | 204 | 200 | 195 | 162 | 48 | 47 | 44 | 60 | 47 | 80 | 53 | Atp4a |
| Atp5a1 | 2476 | 1692 | 1335 | 1438 | 1121 | 1532 | 962 | 2161 | 1418 | 1205 | 1344 | 1479 | 1489 | 1070 | 2412 | 1832 | 1966 | 1716 | 1416 | 1493 | 1626 | 1828 | 1486 | 1333 | 1981 | 1387 | 1326 | 1226 | Atp5a1 |
| Atp5b | 72391 | 72958 | 65096 | 63884 | 56231 | 58574 | 60728 | 67402 | 63809 | 59043 | 56772 | 65794 | 60617 | 65868 | 71596 | 62678 | 65077 | 60841 | 65607 | 62191 | 64589 | 59486 | 63966 | 63502 | 62393 | 74417 | 67877 | 70324 | Atp5b |
| Atp5c1 | 12340 | 12030 | 10494 | 11238 | 11169 | 11138 | 12692 | 12986 | 12049 | 11267 | 10663 | 11879 | 11533 | 11947 | 12675 | 11660 | 10653 | 10780 | 11908 | 13758 | 12419 | 9638 | 12787 | 11443 | 11350 | 12839 | 12261 | 12931 | Atp5c1 |
| Atp5d | 5128 | 6962 | 5757 | 5993 | 5128 | 5108 | 5079 | 4009 | 4641 | 4810 | 4681 | 4742 | 4603 | 4702 | 4776 | 5118 | 6236 | 5695 | 5727 | 5156 | 5465 | 3907 | 4687 | 5075 | 4800 | 5366 | 4767 | 4975 | Atp5d |
| Atp5e | 4083 | 5274 | 4516 | 5088 | 4693 | 3304 | 6762 | 4821 | 4427 | 6057 | 5972 | 5452 | 5147 | 8436 | 3789 | 2973 | 5254 | 4215 | 4327 | 4390 | 6520 | 3832 | 4634 | 6952 | 5626 | 5929 | 5467 | 8572 | Atp5e |
| Atp5g1 | 3835 | 4506 | 3940 | 4311 | 3720 | 3601 | 3986 | 3075 | 3369 | 3078 | 3434 | 3180 | 3433 | 3351 | 3662 | 3961 | 3819 | 4035 | 3930 | 4142 | 3933 | 2981 | 3536 | 3277 | 3207 | 3704 | 3366 | 3571 | Atp5g1 |
| Atp5g2 | 4674 | 6417 | 5163 | 4541 | 4130 | 4472 | 4957 | 5034 | 5927 | 4673 | 5336 | 5837 | 6317 | 7354 | 4603 | 4926 | 4908 | 4078 | 4979 | 5783 | 4944 | 4204 | 5705 | 6141 | 5568 | 7117 | 6846 | 7380 | Atp5g2 |
| Atp5g3 | 12003 | 11593 | 13605 | 12285 | 11440 | 9635 | 11360 | 10534 | 9203 | 11131 | 10473 | 11288 | 10566 | 11143 | 11248 | 10129 | 13166 | 11316 | 12157 | 11241 | 11237 | 9424 | 9756 | 11674 | 10482 | 12771 | 11545 | 12290 | Atp5g3 |
| Atp5h | 10299 | 11657 | 11920 | 11139 | 9720 | 9265 | 11589 | 9638 | 8883 | 9258 | 8964 | 9512 | 9512 | 12270 | 10900 | 9285 | 11425 | 10160 | 10615 | 12082 | 11751 | 7618 | 9041 | 9883 | 9265 | 11055 | 10494 | 12463 | Atp5h |
| Atp5j | 8030 | 6854 | 9685 | 8483 | 7515 | 6830 | 10097 | 8287 | 6600 | 7940 | 7895 | 8748 | 7869 | 10987 | 8024 | 5992 | 9048 | 7557 | 7719 | 9091 | 9260 | 6929 | 6644 | 9289 | 8696 | 10048 | 8835 | 10684 | Atp5j |
| Atp5j2 | 7209 | 8468 | 8893 | 9051 | 7082 | 6003 | 10153 | 8372 | 8362 | 9287 | 9589 | 8582 | 8899 | 12025 | 7348 | 5861 | 8846 | 7491 | 7922 | 8845 | 9224 | 5955 | 8390 | 9927 | 9437 | 9933 | 8971 | 12506 | Atp5j2 |
| Atp5k | 2170 | 2663 | 2041 | 2665 | 2140 | 1485 | 4052 | 2302 | 1989 | 3089 | 2620 | 2399 | 2367 | 5998 | 1826 | 1159 | 2650 | 1949 | 1767 | 2244 | 4093 | 1726 | 1931 | 3795 | 2707 | 2795 | 2696 | 5436 | Atp5k |
| Atp5l | 6536 | 6880 | 7372 | 8498 | 7238 | 7247 | 9249 | 5561 | 6053 | 6780 | 7282 | 6672 | 6775 | 8737 | 6835 | 6551 | 7723 | 6899 | 8078 | 9613 | 8779 | 4926 | 6291 | 6861 | 7197 | 7047 | 7224 | 8039 | Atp5l |
| Atp5md | 7239 | 6389 | 6831 | 8268 | 6887 | 5249 | 11843 | 7679 | 5846 | 8026 | 7832 | 5987 | 6262 | 11884 | 7465 | 4371 | 8055 | 6676 | 6657 | 7582 | 11545 | 5793 | 6034 | 8554 | 7883 | 6958 | 6314 | 11066 | Atp5md |
| Atp5mpl | 4370 | 3808 | 4147 | 4444 | 3822 | 2573 | 6297 | 4737 | 3571 | 5166 | 4553 | 4213 | 3864 | 8324 | 4398 | 2593 | 4329 | 3506 | 3471 | 4154 | 5999 | 3407 | 3628 | 5346 | 4690 | 4574 | 3963 | 7433 | Atp5mpl |
| Atp5o | 1067 | 1293 | 842 | 886 | 877 | 729 | 1159 | 1106 | 803 | 1237 | 857 | 967 | 872 | 1712 | 985 | 578 | 1204 | 795 | 761 | 1011 | 1356 | 937 | 874 | 1680 | 945 | 1048 | 1001 | 1616 | Atp5o |
| Atp5pb | 17946 | 16626 | 16472 | 15832 | 14977 | 14838 | 19525 | 19131 | 16174 | 14950 | 14083 | 16167 | 15442 | 18310 | 19937 | 16055 | 16026 | 15056 | 15359 | 18917 | 17648 | 15871 | 16243 | 16487 | 15386 | 17663 | 16490 | 19522 | Atp5pb |
| Atp6ap1 | 8729 | 9624 | 7718 | 9042 | 9836 | 8881 | 9341 | 9226 | 8894 | 8757 | 8282 | 7348 | 7851 | 7059 | 9525 | 8833 | 8577 | 9894 | 9321 | 9603 | 9341 | 10644 | 9207 | 7500 | 8636 | 7262 | 7434 | 7092 | Atp6ap1 |
| Atp6ap1l | 72 | 68 | 14 | 66 | 131 | 71 | 97 | 71 | 87 | 65 | 48 | 37 | 56 | 22 | 124 | 111 | 50 | 128 | 69 | 79 | 92 | 65 | 61 | 27 | 58 | 30 | 9 | 17 | Atp6ap1l |
| Atp6ap2 | 8053 | 6855 | 7072 | 7525 | 8253 | 7589 | 8961 | 6575 | 5361 | 5132 | 5089 | 5310 | 5087 | 5403 | 9053 | 7675 | 7667 | 8156 | 7867 | 9179 | 8376 | 6323 | 5603 | 5252 | 5555 | 5205 | 5244 | 5459 | Atp6ap2 |
| Atp6v0a1 | 6163 | 7165 | 6400 | 9608 | 12329 | 9606 | 9904 | 8206 | 8871 | 10356 | 10179 | 8750 | 8731 | 8371 | 6883 | 7128 | 8241 | 10786 | 10505 | 8460 | 10652 | 12131 | 8532 | 9058 | 11165 | 6344 | 7835 | 7790 | Atp6v0a1 |
| Atp6v0a2 | 7632 | 8491 | 7097 | 7131 | 6330 | 6963 | 6959 | 4756 | 4821 | 5119 | 5152 | 5621 | 5145 | 5009 | 7087 | 8169 | 7204 | 7547 | 6480 | 6274 | 7170 | 5207 | 4981 | 4752 | 5387 | 4608 | 5265 | 5448 | Atp6v0a2 |
| Atp6v0a4 | 68 | 69 | 105 | 129 | 145 | 100 | 135 | 44 | 84 | 124 | 67 | 33 | 39 | 15 | 88 | 75 | 105 | 155 | 107 | 103 | 145 | 57 | 71 | 51 | 59 | 23 | 22 | 17 | Atp6v0a4 |
| Atp6v0b | 6665 | 8633 | 6930 | 8795 | 8101 | 7825 | 7644 | 5026 | 4895 | 5116 | 4719 | 4412 | 4307 | 4116 | 7371 | 7454 | 8236 | 9037 | 8766 | 8590 | 8182 | 5474 | 5122 | 5147 | 5246 | 4432 | 4230 | 4177 | Atp6v0b |
| Atp6v0c | 4484 | 7203 | 6255 | 6607 | 7173 | 6350 | 5662 | 4996 | 6377 | 6491 | 6474 | 5955 | 5937 | 5337 | 5003 | 5734 | 6839 | 6743 | 7423 | 6766 | 6254 | 5870 | 6546 | 6106 | 6939 | 6094 | 5996 | 5404 | Atp6v0c |
| Atp6v0d1 | 7098 | 8891 | 7716 | 8696 | 9009 | 8566 | 9447 | 6431 | 6676 | 6682 | 6536 | 6583 | 6507 | 6676 | 7494 | 7706 | 8632 | 9014 | 9354 | 7684 | 9646 | 7833 | 7048 | 6704 | 7271 | 6475 | 6212 | 6749 | Atp6v0d1 |
| Atp6v0d2 | 2 | 4 | 0 | 0 | 3 | 0 | 0 | 0 | 7 | 0 | 5 | 3 | 2 | 3 | 3 | 2 | 0 | 6 | 1 | 0 | 3 | 0 | 7 | 1 | 0 | 4 | 0 | 5 | Atp6v0d2 |
| Atp6v0e | 2487 | 2602 | 2259 | 2644 | 2212 | 2311 | 2487 | 2107 | 2273 | 2080 | 2088 | 1690 | 1954 | 1776 | 2516 | 2474 | 2305 | 2386 | 2595 | 2960 | 2422 | 1781 | 2388 | 1986 | 1961 | 1961 | 1820 | 1848 | Atp6v0e |
| Atp6v0e2 | 3874 | 5123 | 4003 | 4969 | 6352 | 5587 | 5535 | 4947 | 4949 | 4889 | 4226 | 4784 | 4266 | 4670 | 4640 | 3712 | 4826 | 5513 | 6148 | 6141 | 5930 | 5945 | 4647 | 4593 | 5643 | 4124 | 4747 | 4435 | Atp6v0e2 |
| Atp6v1a | 19314 | 20096 | 17440 | 18293 | 18038 | 17546 | 19206 | 12411 | 11364 | 10709 | 10152 | 10236 | 9827 | 10161 | 21074 | 19087 | 17369 | 18812 | 17803 | 20812 | 18443 | 11336 | 11467 | 9985 | 11192 | 9275 | 10322 | 10268 | Atp6v1a |
| Atp6v1b1 | 12 | 5 | 1 | 14 | 14 | 3 | 7 | 53 | 82 | 82 | 21 | 4 | 5 | 1 | 11 | 12 | 3 | 12 | 13 | 15 | 8 | 5 | 82 | 74 | 0 | 13 | 1 | 1 | Atp6v1b1 |
| Atp6v1b2 | 14081 | 15689 | 12590 | 14219 | 15061 | 13876 | 14852 | 14803 | 12421 | 12231 | 10759 | 12138 | 11237 | 12058 | 15956 | 13101 | 13614 | 14544 | 14908 | 15597 | 14895 | 14710 | 12486 | 11476 | 13615 | 11510 | 12437 | 12479 | Atp6v1b2 |
| Atp6v1c1 | 294 | 202 | 527 | 1437 | 158 | 214 | 194 | 1794 | 127 | 722 | 114 | 1352 | 1736 | 164 | 1245 | 821 | 2319 | 1539 | 638 | 960 | 166 | 1367 | 131 | 235 | 373 | 478 | 862 | 905 | Atp6v1c1 |
| Atp6v1c2 | 48 | 22 | 20 | 41 | 16 | 10 | 10 | 64 | 44 | 71 | 62 | 5 | 10 | 15 | 30 | 35 | 20 | 0 | 12 | 14 | 8 | 13 | 78 | 77 | 14 | 41 | 9 | 11 | Atp6v1c2 |
| Atp6v1d | 7775 | 7723 | 7839 | 8402 | 7601 | 7045 | 8718 | 6421 | 5391 | 5561 | 5034 | 5437 | 4933 | 6038 | 8759 | 7362 | 7873 | 8061 | 7948 | 8949 | 8651 | 5914 | 5418 | 5529 | 5761 | 5481 | 5339 | 5889 | Atp6v1d |
| Atp6v1e1 | 7343 | 9061 | 7408 | 8541 | 8534 | 7904 | 9113 | 7513 | 7172 | 6627 | 6456 | 6541 | 6420 | 7061 | 8553 | 7594 | 8281 | 8682 | 8573 | 9748 | 9256 | 7056 | 7315 | 6521 | 7479 | 6490 | 6412 | 6857 | Atp6v1e1 |
| Atp6v1e2 | 5 | 5 | 0 | 22 | 11 | 8 | 4 | 6 | 3 | 3 | 0 | 18 | 1 | 6 | 7 | 12 | 17 | 11 | 6 | 16 | 13 | 5 | 3 | 5 | 5 | 0 | 17 | 6 | Atp6v1e2 |
| Atp6v1f | 2137 | 2677 | 2571 | 2569 | 2603 | 2380 | 2731 | 2776 | 2982 | 2652 | 2831 | 2582 | 2863 | 3009 | 2293 | 2201 | 2630 | 2321 | 2746 | 3001 | 2696 | 2603 | 3018 | 3166 | 2914 | 2835 | 2814 | 2867 | Atp6v1f |
| Atp6v1fnb | 2 | 3 | 0 | 0 | 0 | 7 | 0 | 1 | 1 | 0 | 4 | 2 | 0 | 3 | 5 | 0 | 0 | 0 | 4 | 0 | 4 | 2 | 3 | 0 | 0 | 0 | 2 | 0 | Atp6v1fnb |
| Atp6v1g1 | 9193 | 8960 | 9330 | 9541 | 9409 | 8843 | 10992 | 9652 | 8369 | 7932 | 7970 | 8550 | 8108 | 10104 | 10345 | 8957 | 9612 | 9228 | 9376 | 11585 | 10781 | 8194 | 8626 | 9005 | 8830 | 9096 | 8612 | 10174 | Atp6v1g1 |
| Atp6v1g2 | 576 | 672 | 830 | 1081 | 1476 | 1162 | 1258 | 1082 | 1171 | 1545 | 1344 | 1324 | 1294 | 1323 | 696 | 669 | 1074 | 1229 | 1294 | 1099 | 1348 | 1688 | 1156 | 1366 | 1609 | 1160 | 1283 | 1345 | Atp6v1g2 |
| Atp6v1g3 | 2 | 1 | 0 | 15 | 0 | 0 | 0 | 20 | 6 | 11 | 6 | 1 | 7 | 2 | 7 | 0 | 0 | 0 | 2 | 0 | 0 | 3 | 10 | 11 | 0 | 14 | 0 | 0 | Atp6v1g3 |
| Atp6v1h | 8588 | 8651 | 7613 | 9127 | 9352 | 9398 | 10715 | 9432 | 7454 | 7040 | 6330 | 7028 | 6927 | 7430 | 10221 | 7683 | 8693 | 9112 | 9884 | 11162 | 10402 | 8575 | 7127 | 6890 | 7819 | 6876 | 6962 | 7312 | Atp6v1h |
| Atp7a | 6960 | 5787 | 2322 | 3712 | 3954 | 4869 | 4013 | 4958 | 4508 | 3185 | 3954 | 3488 | 3938 | 3553 | 6570 | 6152 | 3018 | 4034 | 3310 | 5155 | 4737 | 2974 | 4599 | 2580 | 3140 | 1185 | 3593 | 3147 | Atp7a |
| Atp7b | 818 | 953 | 1086 | 981 | 947 | 846 | 872 | 405 | 475 | 489 | 493 | 587 | 486 | 484 | 765 | 819 | 1068 | 837 | 858 | 744 | 733 | 429 | 474 | 465 | 460 | 381 | 497 | 503 | Atp7b |
| Atp8a1 | 10933 | 10785 | 8732 | 12552 | 14723 | 13376 | 14501 | 7476 | 6660 | 7052 | 6886 | 5998 | 6554 | 5993 | 12635 | 10909 | 10880 | 12531 | 12849 | 13970 | 14086 | 8748 | 6277 | 5645 | 6966 | 3746 | 6449 | 6165 | Atp8a1 |
| Atp8a2 | 2262 | 2430 | 1363 | 3045 | 5100 | 4324 | 3873 | 4250 | 4270 | 4498 | 4042 | 2959 | 3370 | 2855 | 2966 | 2593 | 2321 | 3502 | 3522 | 3878 | 4541 | 4821 | 3484 | 2665 | 4980 | 1573 | 2940 | 2314 | Atp8a2 |
| Atp8b1 | 90 | 50 | 38 | 85 | 65 | 34 | 46 | 133 | 167 | 107 | 230 | 56 | 43 | 77 | 66 | 73 | 38 | 55 | 26 | 38 | 28 | 89 | 176 | 99 | 73 | 118 | 50 | 58 | Atp8b1 |
| Atp8b2 | 7739 | 8660 | 7466 | 8186 | 7665 | 7671 | 7066 | 4936 | 5693 | 5395 | 6074 | 5076 | 5505 | 5178 | 7226 | 8885 | 7402 | 8225 | 7339 | 6572 | 7049 | 6017 | 5588 | 5178 | 5767 | 4267 | 5244 | 5033 | Atp8b2 |
| Atp8b3 | 135 | 149 | 120 | 165 | 136 | 117 | 103 | 89 | 113 | 99 | 109 | 76 | 124 | 78 | 144 | 151 | 153 | 111 | 170 | 158 | 125 | 137 | 95 | 109 | 124 | 66 | 113 | 150 | Atp8b3 |
| Atp8b4 | 206 | 80 | 11 | 122 | 11 | 17 | 7 | 414 | 230 | 398 | 290 | 22 | 103 | 68 | 103 | 137 | 56 | 18 | 7 | 12 | 5 | 22 | 344 | 526 | 18 | 569 | 21 | 25 | Atp8b4 |
| Atp8b5 | 37 | 65 | 40 | 46 | 71 | 75 | 89 | 142 | 88 | 100 | 63 | 76 | 92 | 98 | 62 | 37 | 44 | 45 | 55 | 59 | 81 | 70 | 82 | 87 | 77 | 59 | 99 | 102 | Atp8b5 |
| Atp9a | 4282 | 5553 | 5333 | 7258 | 10357 | 8330 | 7097 | 8350 | 10930 | 11746 | 10864 | 8878 | 8074 | 7565 | 4717 | 4841 | 6400 | 8217 | 9690 | 6984 | 8740 | 12850 | 10777 | 10128 | 12344 | 7166 | 7433 | 7437 | Atp9a |
| Atp9b | 5135 | 5422 | 4470 | 4707 | 4764 | 4958 | 4817 | 5487 | 5682 | 5418 | 5292 | 5343 | 5232 | 5231 | 5236 | 5044 | 4605 | 4589 | 4664 | 4962 | 4846 | 5096 | 5553 | 4882 | 5333 | 4714 | 5460 | 5168 | Atp9b |

| | | | | | | | | | | | | | | | | | | | | | | | | | | | | | |
|---|---|---|---|---|---|---|---|---|---|---|---|---|---|---|---|---|---|---|---|---|---|---|---|---|---|---|---|---|---|
| Atpaf1 | 767 | 608 | 811 | 605 | 722 | 660 | 705 | 1080 | 1022 | 1011 | 1069 | 974 | 1140 | 981 | 803 | 750 | 694 | 622 | 683 | 652 | 684 | 1011 | 1128 | 1101 | 904 | 1280 | 1118 | 1089 | Atpaf1 |
| Atpaf2 | 1541 | 1589 | 1440 | 1556 | 1275 | 1363 | 1514 | 1296 | 1477 | 1332 | 1425 | 1501 | 1454 | 1494 | 1440 | 1654 | 1471 | 1506 | 1418 | 1415 | 1438 | 1308 | 1368 | 1431 | 1459 | 1743 | 1502 | 1457 | Atpaf2 |
| Atpif1 | 5727 | 6692 | 6441 | 6646 | 6653 | 5993 | 7967 | 6949 | 7191 | 8008 | 7097 | 6938 | 6924 | 9486 | 5919 | 5063 | 6633 | 6345 | 6668 | 7993 | 8057 | 6433 | 7163 | 8643 | 7657 | 7738 | 7510 | 9121 | Atpif1 |
| Atpsckmt | 1027 | 1012 | 545 | 746 | 879 | 1029 | 830 | 1702 | 1590 | 1488 | 1600 | 1497 | 1503 | 1549 | 938 | 819 | 661 | 638 | 783 | 1065 | 972 | 1180 | 1507 | 1251 | 1761 | 868 | 1897 | 1388 | Atpsckmt |
| Atr | 10028 | 7276 | 4963 | 5801 | 5150 | 6976 | 7137 | 6879 | 6339 | 4408 | 5173 | 5435 | 5502 | 5778 | 9806 | 8482 | 4940 | 5399 | 5054 | 7887 | 6764 | 4137 | 6122 | 4054 | 4849 | 4007 | 6199 | 5446 | Atr |
| Atraid | 2712 | 2772 | 2387 | 2542 | 2372 | 2472 | 2744 | 2773 | 2562 | 2310 | 2480 | 2336 | 2304 | 2467 | 2933 | 2322 | 2464 | 2479 | 2455 | 2820 | 2683 | 2762 | 2668 | 2478 | 2435 | 2520 | 2375 | 2592 | Atraid |
| Atrip | 2339 | 2641 | 2232 | 2691 | 2653 | 2485 | 2486 | 2301 | 2160 | 2294 | 2037 | 2271 | 2258 | 2156 | 2229 | 2454 | 2370 | 2599 | 2442 | 2547 | 2515 | 2252 | 1948 | 2237 | 2411 | 1991 | 2308 | 2161 | Atrip |
| Atrn | 9844 | 9392 | 6282 | 8403 | 10033 | 10007 | 9505 | 8664 | 8497 | 7899 | 7880 | 7556 | 7460 | 7469 | 9936 | 9404 | 7044 | 8630 | 8503 | 9477 | 9798 | 8281 | 7882 | 6405 | 8306 | 4764 | 7216 | 6584 | Atrn |
| Atrnl1 | 9870 | 8712 | 7968 | 9870 | 10964 | 10628 | 11173 | 8131 | 7968 | 8528 | 8906 | 7698 | 8377 | 7954 | 9955 | 9667 | 9251 | 10630 | 10243 | 9582 | 11481 | 9389 | 7963 | 7379 | 8501 | 5242 | 8062 | 7563 | Atrnl1 |
| Atrx | 27726 | 18643 | 13408 | 18713 | 22418 | 27626 | 27765 | 51321 | 44406 | 33897 | 36596 | 37263 | 36807 | 41311 | 27960 | 23977 | 14865 | 18368 | 20715 | 30479 | 27708 | 31900 | 41520 | 28239 | 37603 | 22086 | 41154 | 33484 | Atrx |
| Atxn1 | 1330 | 1165 | 1754 | 1738 | 2140 | 2015 | 2292 | 1535 | 1562 | 2054 | 1925 | 1692 | 1349 | 1682 | 1342 | 1138 | 1776 | 1951 | 2172 | 1437 | 2096 | 2354 | 1515 | 1897 | 2082 | 1414 | 1435 | 1467 | Atxn1 |
| Atxn10 | 21224 | 21376 | 17491 | 17923 | 19014 | 18145 | 20770 | 22167 | 20034 | 20181 | 17283 | 20191 | 18872 | 20411 | 22618 | 17397 | 18310 | 18308 | 19739 | 19415 | 21294 | 23064 | 19792 | 19242 | 20318 | 20596 | 19717 | 21687 | Atxn10 |
| Atxn1l | 5515 | 5459 | 5394 | 5101 | 4490 | 4917 | 4679 | 4295 | 4517 | 4384 | 5114 | 5002 | 4911 | 4908 | 4863 | 5345 | 5116 | 4932 | 4602 | 4175 | 4971 | 4134 | 4399 | 4668 | 4262 | 4083 | 4747 | 4938 | Atxn1l |
| Atxn2 | 5982 | 6137 | 6862 | 7634 | 8069 | 7566 | 7171 | 6763 | 7064 | 8018 | 8024 | 8114 | 7635 | 7853 | 5544 | 6068 | 7158 | 7718 | 7697 | 6046 | 7638 | 8245 | 6470 | 7798 | 8458 | 6949 | 7191 | 7316 | Atxn2 |
| Atxn2l | 9183 | 11749 | 14364 | 15070 | 15194 | 12697 | 12956 | 12383 | 14225 | 17767 | 15882 | 18678 | 15305 | 17617 | 9028 | 10210 | 15807 | 14345 | 15665 | 10724 | 13989 | 18735 | 12863 | 19503 | 18097 | 20524 | 15894 | 16887 | Atxn2l |
| Atxn3 | 4119 | 3568 | 4238 | 4321 | 3831 | 4386 | 4376 | 3631 | 2857 | 3166 | 3007 | 3593 | 3543 | 3838 | 4233 | 3875 | 3957 | 4275 | 3783 | 4188 | 4364 | 3350 | 2790 | 3037 | 3427 | 2910 | 3607 | 3677 | Atxn3 |
| Atxn7 | 1688 | 1552 | 1693 | 1950 | 1935 | 2215 | 2305 | 2997 | 2956 | 3095 | 3302 | 4034 | 3699 | 4003 | 1543 | 1577 | 1655 | 1837 | 2084 | 2035 | 2098 | 2966 | 2933 | 3392 | 3735 | 2995 | 3946 | 3757 | Atxn7 |
| Atxn7l1 | 1129 | 1253 | 930 | 1145 | 1567 | 1383 | 1284 | 1536 | 1566 | 1527 | 1432 | 1389 | 1463 | 1288 | 1197 | 1194 | 1090 | 1224 | 1369 | 1367 | 1325 | 1551 | 1422 | 1277 | 1527 | 1091 | 1330 | 1250 | Atxn7l1 |
| Atxn7l2 | 698 | 917 | 1007 | 1069 | 1450 | 1207 | 1241 | 2249 | 2802 | 3171 | 2995 | 3338 | 2913 | 3257 | 719 | 767 | 1169 | 1084 | 1354 | 1079 | 1223 | 3213 | 2429 | 3544 | 3410 | 3064 | 2919 | 3068 | Atxn7l2 |
| Atxn7l3 | 8780 | 10613 | 10770 | 12426 | 13331 | 12057 | 11567 | 11048 | 11977 | 13151 | 12112 | 12912 | 11936 | 11974 | 8761 | 8816 | 12302 | 12171 | 13695 | 11167 | 12343 | 13257 | 11408 | 12921 | 13305 | 13247 | 12183 | 11941 | Atxn7l3 |
| Atxn7l3b | 33410 | 28170 | 35636 | 36935 | 38454 | 36630 | 39191 | 39886 | 35398 | 39873 | 38714 | 41586 | 40335 | 42465 | 34598 | 29116 | 37066 | 36542 | 38899 | 33510 | 39523 | 45609 | 35529 | 41607 | 42188 | 41695 | 41032 | 44699 | Atxn7l3b |
| AU015836 | 14 | 14 | 16 | 4 | 0 | 8 | 0 | 9 | 18 | 3 | 5 | 10 | 19 | 15 | 33 | 17 | 15 | 1 | 3 | 0 | 1 | 10 | 12 | 9 | 7 | 12 | 16 | 14 | AU015836 |
| AU018091 | 0 | 0 | 0 | 0 | 0 | 0 | 0 | 5 | 1 | 0 | 6 | 7 | 1 | 0 | 0 | 0 | 0 | 0 | 0 | 0 | 0 | 0 | 0 | 1 | 0 | 0 | 1 | 1 | AU018091 |
| AU021092 | 153 | 177 | 106 | 161 | 217 | 218 | 184 | 253 | 354 | 268 | 235 | 290 | 244 | 270 | 158 | 100 | 153 | 195 | 241 | 211 | 193 | 237 | 357 | 266 | 299 | 280 | 285 | 202 | AU021092 |
| AU022252 | 11086 | 11970 | 9426 | 11796 | 9452 | 10407 | 9731 | 5454 | 4823 | 5251 | 5329 | 6083 | 6042 | 5815 | 11044 | 10982 | 10515 | 11055 | 10211 | 9373 | 10158 | 5238 | 4692 | 5260 | 5958 | 4655 | 6422 | 6076 | AU022252 |
| AU040320 | 2896 | 3448 | 3772 | 4052 | 3979 | 3727 | 3518 | 3121 | 3438 | 4215 | 3994 | 4191 | 3903 | 3687 | 2926 | 3203 | 3793 | 3744 | 4074 | 3260 | 3651 | 4311 | 3617 | 4028 | 4166 | 3928 | 3550 | 3542 | AU040320 |
| AU041133 | 467 | 325 | 311 | 236 | 279 | 414 | 313 | 788 | 831 | 592 | 499 | 558 | 627 | 582 | 452 | 395 | 287 | 242 | 346 | 530 | 375 | 620 | 819 | 471 | 523 | 570 | 595 | 547 | AU041133 |
| Auh | 1090 | 1220 | 1083 | 1228 | 1422 | 1319 | 1329 | 1490 | 1518 | 1512 | 1472 | 1552 | 1653 | 1697 | 1143 | 1224 | 1214 | 1285 | 1242 | 1307 | 1349 | 1494 | 1594 | 1599 | 1693 | 1514 | 1684 | 1665 | Auh |
| Aunip | 603 | 628 | 747 | 478 | 318 | 442 | 419 | 442 | 392 | 373 | 384 | 604 | 515 | 622 | 496 | 557 | 567 | 410 | 435 | 513 | 417 | 231 | 368 | 440 | 399 | 824 | 682 | 608 | Aunip |
| Aup1 | 2748 | 3346 | 3314 | 3177 | 2752 | 2697 | 2892 | 2865 | 3043 | 3018 | 3103 | 3186 | 2963 | 3227 | 2683 | 3131 | 3227 | 3101 | 3113 | 2619 | 2786 | 3365 | 3053 | 3398 | 3316 | 3399 | 3049 | 3065 | Aup1 |
| Aurka | 3194 | 3647 | 3914 | 2690 | 1972 | 2269 | 2425 | 2958 | 3117 | 2865 | 2566 | 4056 | 3622 | 3780 | 2557 | 3072 | 2938 | 2212 | 2780 | 2719 | 2314 | 1861 | 2782 | 3274 | 2780 | 5277 | 4541 | 4208 | Aurka |
| Aurkaip1 | 2476 | 2859 | 2443 | 2417 | 2088 | 2012 | 2232 | 2185 | 2310 | 2030 | 2074 | 2192 | 2131 | 2594 | 2502 | 2348 | 2537 | 2082 | 2355 | 2561 | 2334 | 1761 | 2335 | 2303 | 2245 | 2727 | 2298 | 2552 | Aurkaip1 |
| Aurkb | 4327 | 4505 | 5390 | 4112 | 2955 | 3198 | 3696 | 2848 | 3451 | 3374 | 2955 | 5370 | 4425 | 5201 | 3477 | 4111 | 4990 | 3562 | 4072 | 3202 | 3344 | 2430 | 2913 | 4371 | 3774 | 6809 | 4811 | 5381 | Aurkb |
| Aurkc | 0 | 2 | 4 | 4 | 6 | 1 | 2 | 4 | 2 | 0 | 4 | 0 | 3 | 3 | 5 | 2 | 1 | 0 | 5 | 2 | 4 | 2 | 3 | 2 | 1 | 1 | 0 | 1 | Aurkc |
| Auts2 | 373 | 1391 | 863 | 1700 | 1498 | 748 | 896 | 835 | 1041 | 1795 | 1436 | 1727 | 1233 | 1258 | 974 | 560 | 1144 | 824 | 776 | 1694 | 1225 | 1480 | 1072 | 1075 | 2743 | 1262 | 1710 | 914 | Auts2 |
| AV320801 | 0 | 0 | 0 | 0 | 0 | 0 | 0 | 0 | 0 | 0 | 0 | 1 | 0 | 0 | 0 | 0 | 0 | 0 | 0 | 0 | 0 | 0 | 3 | 0 | 0 | 0 | 0 | 0 | AV320801 |
| Aven | 264 | 203 | 255 | 180 | 171 | 155 | 171 | 688 | 613 | 600 | 708 | 760 | 689 | 744 | 255 | 238 | 236 | 145 | 168 | 189 | 155 | 673 | 579 | 691 | 539 | 850 | 618 | 768 | Aven |
| Avil | 273 | 452 | 325 | 404 | 893 | 307 | 226 | 458 | 701 | 391 | 723 | 159 | 485 | 161 | 394 | 498 | 387 | 748 | 428 | 302 | 305 | 896 | 415 | 149 | 533 | 95 | 193 | 239 | Avil |
| Avl9 | 9397 | 8302 | 6530 | 8755 | 10452 | 10501 | 11622 | 12331 | 9768 | 9880 | 8804 | 9820 | 9285 | 10194 | 10098 | 8292 | 7697 | 9016 | 9693 | 11027 | 11434 | 10921 | 9405 | 8887 | 10921 | 6687 | 9975 | 9437 | Avl9 |
| Avp | 1 | 0 | 5 | 1 | 0 | 2 | 1 | 0 | 2 | 1 | 0 | 0 | 3 | 0 | 0 | 0 | 0 | 1 | 0 | 2 | 1 | 2 | 0 | 0 | 0 | 0 | 0 | 0 | Avp |
| Avpi1 | 541 | 550 | 478 | 450 | 274 | 405 | 273 | 161 | 275 | 201 | 238 | 194 | 209 | 175 | 460 | 452 | 426 | 404 | 369 | 400 | 284 | 147 | 287 | 236 | 165 | 316 | 199 | 211 | Avpi1 |
| Avpr1a | 141 | 24 | 35 | 27 | 23 | 16 | 28 | 216 | 170 | 116 | 74 | 33 | 24 | 29 | 22 | 24 | 16 | 32 | 19 | 33 | 27 | 85 | 260 | 94 | 50 | 52 | 24 | 24 | Avpr1a |
| Avpr1b | 0 | 0 | 0 | 0 | 0 | 3 | 1 | 0 | 1 | 0 | 0 | 0 | 2 | 0 | 1 | 3 | 0 | 0 | 2 | 0 | 1 | 0 | 1 | 5 | 0 | 0 | 1 | 0 | Avpr1b |
| Avpr2 | 1 | 0 | 0 | 0 | 2 | 0 | 1 | 0 | 0 | 0 | 0 | 2 | 0 | 1 | 0 | 0 | 0 | 0 | 0 | 0 | 3 | 0 | 0 | 1 | 0 | 0 | 0 | 0 | Avpr2 |
| AW146154 | 622 | 474 | 754 | 576 | 451 | 563 | 517 | 526 | 442 | 490 | 510 | 628 | 589 | 506 | 605 | 700 | 648 | 560 | 555 | 485 | 466 | 560 | 557 | 488 | 426 | 600 | 522 | 605 | AW146154 |
| AW209491 | 1051 | 1003 | 1111 | 996 | 1014 | 1035 | 1009 | 1557 | 1361 | 1134 | 1244 | 1229 | 1280 | 1270 | 1146 | 976 | 1137 | 905 | 1026 | 1268 | 1075 | 1364 | 1408 | 1228 | 1275 | 1365 | 1334 | 1285 | AW209491 |
| AW549877 | 12017 | 9508 | 9412 | 9373 | 10369 | 10904 | 11616 | 15107 | 12612 | 10726 | 11572 | 12843 | 12938 | 13193 | 12552 | 10160 | 9822 | 9683 | 9825 | 11925 | 11004 | 12724 | 12376 | 11087 | 11722 | 10222 | 13536 | 13027 | AW549877 |
| AW551984 | 6342 | 5294 | 4590 | 4591 | 6114 | 6202 | 5822 | 7486 | 7778 | 6130 | 8234 | 4770 | 5237 | 4782 | 6603 | 5626 | 4664 | 5508 | 5293 | 6304 | 5625 | 7549 | 8827 | 5087 | 5788 | 4646 | 4412 | 4656 | AW551984 |
| AW554918 | 2136 | 1764 | 1468 | 1861 | 1899 | 1853 | 2223 | 2638 | 2360 | 2208 | 2119 | 2293 | 2108 | 2405 | 1876 | 1798 | 1707 | 1852 | 1961 | 1945 | 2081 | 2438 | 2157 | 2134 | 2257 | 2092 | 2235 | 2302 | AW554918 |
| Awat2 | 28 | 30 | 38 | 25 | 19 | 19 | 28 | 35 | 40 | 18 | 51 | 52 | 39 | 31 | 40 | 36 | 42 | 45 | 22 | 29 | 17 | 20 | 28 | 25 | 36 | 29 | 35 | 34 | Awat2 |
| Axdnd1 | 189 | 154 | 129 | 173 | 139 | 114 | 164 | 39 | 37 | 39 | 37 | 26 | 45 | 42 | 156 | 152 | 129 | 174 | 127 | 136 | 189 | 28 | 38 | 25 | 50 | 24 | 52 | 40 | Axdnd1 |
| Axin1 | 3563 | 4148 | 5106 | 4524 | 4007 | 4194 | 3834 | 3740 | 4374 | 4509 | 4498 | 4977 | 4804 | 4747 | 3330 | 3946 | 4578 | 4359 | 4495 | 3455 | 3892 | 4361 | 4352 | 5124 | 4714 | 5793 | 4866 | 5006 | Axin1 |
| Axin2 | 4963 | 5480 | 8092 | 5507 | 5105 | 5047 | 4685 | 3666 | 3926 | 4520 | 3976 | 6064 | 6357 | 6024 | 4544 | 5223 | 6569 | 5244 | 5648 | 4455 | 4864 | 4089 | 4590 | 5430 | 4421 | 5750 | 5914 | 6957 | Axin2 |
| Axl | 6592 | 5394 | 5260 | 4210 | 3934 | 4853 | 3434 | 4517 | 6060 | 5214 | 7486 | 4791 | 5358 | 4625 | 5016 | 6442 | 4406 | 4874 | 3566 | 3493 | 3057 | 4582 | 6953 | 5056 | 4192 | 5648 | 4736 | 4488 | Axl |
| AY074887 | 5 | 3 | 1 | 0 | 2 | 6 | 3 | 2 | 7 | 12 | 6 | 10 | 15 | 12 | 1 | 4 | 0 | 1 | 5 | 7 | 5 | 8 | 8 | 7 | 4 | 26 | 6 | 13 | AY074887 |
| AY358078 | 0 | 0 | 0 | 0 | 0 | 0 | 0 | 0 | 0 | 0 | 0 | 0 | 0 | 0 | 0 | 0 | 0 | 0 | 0 | 0 | 0 | 0 | 1 | 0 | 0 | 0 | 0 | 0 | AY358078 |
| Aym1 | 2 | 0 | 0 | 0 | 1 | 1 | 0 | 0 | 0 | 0 | 0 | 0 | 0 | 0 | 1 | 1 | 0 | 1 | 0 | 1 | 0 | 0 | 0 | 0 | 0 | 0 | 0 | 0 | Aym1 |
| Azgp1 | 0 | 0 | 0 | 0 | 0 | 0 | 0 | 0 | 0 | 4 | 1 | 0 | 0 | 0 | 0 | 0 | 0 | 0 | 0 | 0 | 0 | 2 | 1 | 2 | 0 | 0 | 0 | 1 | Azgp1 |
| Azi2 | 6467 | 5237 | 5034 | 5176 | 5761 | 5733 | 6515 | 8241 | 6951 | 6114 | 6162 | 6610 | 6159 | 6737 | 6746 | 5819 | 5047 | 5093 | 5559 | 6239 | 6205 | 6906 | 6880 | 6212 | 6500 | 6083 | 6215 | 6274 | Azi2 |
| Azin1 | 27289 | 22476 | 23710 | 26072 | 26751 | 28228 | 30792 | 22740 | 19415 | 18021 | 18678 | 20256 | 19390 | 20898 | 28825 | 22869 | 24663 | 26844 | 26603 | 29764 | 29258 | 20936 | 18990 | 18353 | 21225 | 19720 | 20811 | 20458 | Azin1 |
| Azin2 | 1980 | 2230 | 1616 | 2097 | 1594 | 1759 | 1351 | 561 | 752 | 598 | 667 | 646 | 676 | 517 | 1772 | 2365 | 1951 | 1969 | 1612 | 1603 | 1383 | 535 | 819 | 599 | 511 | 597 | 533 | 535 | Azin2 |
| B020031M17Rik | 0 | 1 | 0 | 0 | 0 | 0 | 0 | 0 | 0 | 0 | 0 | 0 | 0 | 0 | 0 | 0 | 0 | 0 | 0 | 0 | 0 | 0 | 0 | 0 | 0 | 0 | 0 | 0 | B020031M17Rik |
| B230217C12Rik | 820 | 797 | 777 | 916 | 1074 | 926 | 940 | 797 | 751 | 689 | 718 | 737 | 699 | 748 | 894 | 713 | 918 | 820 | 848 | 986 | 911 | 767 | 730 | 751 | 697 | 843 | 812 | 659 | B230217C12Rik |
| B230219D22Rik | 10650 | 8180 | 10199 | 10537 | 12153 | 11485 | 12290 | 14521 | 13359 | 13292 | 14017 | 13599 | 13463 | 13894 | 11040 | 9775 | 10371 | 10638 | 11288 | 10690 | 12007 | 14955 | 14474 | 13838 | 13853 | 12714 | 13238 | 13623 | B230219D22Rik |
| B230307C23Rik | 668 | 639 | 427 | 512 | 612 | 716 | 653 | 562 | 571 | 450 | 530 | 454 | 469 | 535 | 655 | 695 | 430 | 584 | 561 | 779 | 723 | 431 | 571 | 417 | 454 | 253 | 524 | 449 | B230307C23Rik |
| B2m | 2955 | 2339 | 2845 | 2138 | 1950 | 2099 | 1885 | 2235 | 2333 | 1927 | 2559 | 1729 | 2078 | 1912 | 3091 | 2890 | 2161 | 2249 | 1772 | 2457 | 1919 | 2276 | 3019 | 1975 | 1759 | 2296 | 1983 | 2171 | B2m |
| B3galnt1 | 2054 | 1860 | 1973 | 2218 | 2732 | 2475 | 2426 | 2596 | 2543 | 2274 | 2275 | 2066 | 2121 | 2063 | 2397 | 1962 | 2145 | 2235 | 2463 | 2569 | 2346 | 2617 | 2621 | 2127 | 2393 | 2038 | 2031 | 2123 | B3galnt1 |
| B3galnt2 | 6981 | 5524 | 5627 | 5097 | 4672 | 5218 | 5327 | 3965 | 3580 | 3359 | 3659 | 4027 | 3936 | 4247 | 6154 | 5972 | 5273 | 4910 | 4901 | 5553 | 5020 | 3459 | 3634 | 3666 | 3879 | 4230 | 4101 | 4462 | B3galnt2 |
| B3galt1 | 1870 | 1906 | 1564 | 2513 | 3418 | 3083 | 3319 | 3204 | 2689 | 2727 | 2588 | 2008 | 2011 | 2075 | 2409 | 1878 | 1997 | 2573 | 2786 | 3392 | 3186 | 3288 | 2497 | 2120 | 3095 | 1434 | 2092 | 1899 | B3galt1 |
| B3galt2 | 465 | 390 | 328 | 581 | 917 | 847 | 834 | 1601 | 1382 | 969 | 1063 | 915 | 680 | 1027 | 733 | 362 | 444 | 614 | 829 | 1056 | 977 | 1455 | 1330 | 949 | 1352 | 681 | 915 | 845 | B3galt2 |
| B3galt4 | 80 | 112 | 117 | 100 | 130 | 132 | 86 | 227 | 274 | 297 | 231 | 235 | 242 | 246 | 102 | 71 | 153 | 94 | 130 | 118 | 111 | 284 | 341 | 212 | 278 | 278 | 234 | 177 | B3galt4 |
| B3galt5 | 197 | 202 | 222 | 179 | 400 | 245 | 241 | 203 | 220 | 172 | 185 | 128 | 180 | 146 | 227 | 269 | 192 | 196 | 223 | 213 | 225 | 247 | 183 | 138 | 226 | 104 | 102 | 142 | B3galt5 |
| B3galt6 | 611 | 618 | 619 | 678 | 835 | 683 | 759 | 940 | 934 | 974 | 1023 | 971 | 853 | 895 | 616 | 575 | 674 | 688 | 666 | 747 | 732 | 962 | 1014 | 988 | 952 | 888 | 888 | 996 | B3galt6 |
| B3gat1 | 3091 | 4099 | 3561 | 7036 | 10538 | 7991 | 8535 | 5930 | 6547 | 8526 | 7133 | 6309 | 5023 | 6015 | 3974 | 3091 | 5792 | 8210 | 9139 | 7045 | 9097 | 10195 | 5689 | 7202 | 9279 | 4501 | 4489 | 4793 | B3gat1 |
| B3gat2 | 396 | 414 | 368 | 807 | 1819 | 1275 | 1305 | 987 | 1029 | 1286 | 1126 | 605 | 582 | 626 | 555 | 374 | 639 | 1008 | 1259 | 1100 | 1331 | 1425 | 998 | 902 | 1360 | 436 | 495 | 428 | B3gat2 |
| B3gat3 | 3825 | 5181 | 4932 | 5480 | 5064 | 5179 | 4773 | 3461 | 3789 | 3909 | 3976 | 4121 | 4210 | 4177 | 4044 | 4335 | 5380 | 5146 | 5517 | 4770 | 5058 | 3750 | 3698 | 4323 | 4216 | 3834 | 3901 | 3849 | B3gat3 |
| B3glct | 2263 | 2024 | 2226 | 2443 | 3151 | 3023 | 3004 | 3179 | 3166 | 3178 | 3004 | 3592 | 3574 | 3704 | 2531 | 2220 | 2315 | 2717 | 3014 | 2876 | 2929 | 3810 | 2963 | 3203 | 3687 | 3573 | 3226 | 3630 | B3glct |
| B3gnt2 | 14 | 9 | 8 | 6 | 9 | 10 | 7 | 0 | 3 | 5 | 4 | 2 | 2 | 2 | 12 | 6 | 9 | 6 | 2 | 2 | 9 | 7 | 4 | 6 | 4 | 8 | 1 | 2 | B3gnt2 |
| B3gnt3 | 73 | 76 | 103 | 61 | 89 | 75 | 83 | 42 | 46 | 60 | 65 | 48 | 69 | 63 | 102 | 79 | 103 | 68 | 67 | 54 | 71 | 53 | 79 | 58 | 39 | 63 | 23 | 70 | B3gnt3 |

| | | | | | | | | | | | | | | | | | | | | | | | | | | | | | |
|---|---|---|---|---|---|---|---|---|---|---|---|---|---|---|---|---|---|---|---|---|---|---|---|---|---|---|---|---|---|
| B3gnt4 | 118 | 124 | 116 | 121 | 124 | 149 | 121 | 82 | 79 | 114 | 72 | 100 | 82 | 80 | 90 | 146 | 124 | 131 | 142 | 140 | 106 | 82 | 85 | 111 | 94 | 123 | 121 | 81 | B3gnt4 |
| B3gnt5 | 6649 | 4749 | 5457 | 4823 | 5424 | 5412 | 5955 | 8961 | 7510 | 6766 | 6292 | 8043 | 7489 | 8171 | 7116 | 5687 | 5241 | 4767 | 5116 | 6360 | 5384 | 7384 | 6772 | 6917 | 7388 | 7590 | 8358 | 8562 | B3gnt5 |
| B3gnt6 | 12 | 13 | 16 | 16 | 25 | 8 | 15 | 1 | 6 | 8 | 5 | 5 | 3 | 4 | 12 | 11 | 25 | 30 | 5 | 12 | 12 | 8 | 5 | 5 | 2 | 2 | 5 | 0 | B3gnt6 |
| B3gnt7 | 376 | 332 | 396 | 292 | 264 | 319 | 188 | 201 | 241 | 275 | 242 | 164 | 201 | 145 | 301 | 447 | 309 | 264 | 316 | 202 | 201 | 221 | 256 | 212 | 136 | 279 | 157 | 122 | B3gnt7 |
| B3gnt8 | 35 | 21 | 28 | 9 | 17 | 43 | 12 | 46 | 93 | 55 | 75 | 27 | 41 | 28 | 24 | 33 | 22 | 24 | 15 | 9 | 18 | 35 | 118 | 72 | 33 | 54 | 41 | 59 | B3gnt8 |
| B3gnt9 | 194 | 202 | 252 | 177 | 135 | 247 | 105 | 291 | 559 | 485 | 656 | 405 | 523 | 333 | 173 | 231 | 234 | 151 | 164 | 162 | 104 | 455 | 744 | 424 | 377 | 312 | 456 | 394 | B3gnt9 |
| B3gntl1 | 746 | 759 | 569 | 516 | 600 | 583 | 574 | 663 | 840 | 713 | 848 | 663 | 807 | 643 | 655 | 741 | 557 | 521 | 516 | 634 | 591 | 647 | 794 | 608 | 645 | 566 | 737 | 743 | B3gntl1 |
| B430305J03Rik | 12 | 18 | 17 | 11 | 24 | 9 | 21 | 24 | 34 | 36 | 23 | 43 | 27 | 51 | 20 | 18 | 13 | 18 | 5 | 20 | 9 | 35 | 34 | 23 | 28 | 8 | 30 | 17 | B430305J03Rik |
| B430306N03Rik | 0 | 0 | 0 | 0 | 1 | 0 | 0 | 1 | 0 | 0 | 1 | 0 | 3 | 0 | 3 | 0 | 0 | 0 | 0 | 0 | 0 | 0 | 1 | 0 | 3 | 0 | 0 | 0 | B430306N03Rik |
| B4galnt1 | 804 | 930 | 961 | 1289 | 1919 | 1489 | 1372 | 1436 | 1645 | 2058 | 1683 | 1346 | 1326 | 1344 | 822 | 854 | 1399 | 1402 | 1905 | 1473 | 1516 | 2281 | 1568 | 1753 | 2230 | 1060 | 1027 | 1105 | B4galnt1 |
| B4galnt2 | 3 | 4 | 1 | 0 | 0 | 0 | 1 | 1 | 5 | 0 | 0 | 1 | 1 | 1 | 1 | 2 | 2 | 1 | 1 | 0 | 3 | 3 | 2 | 4 | 0 | 0 | 0 | 1 | B4galnt2 |
| B4galnt3 | 219 | 149 | 115 | 116 | 116 | 117 | 72 | 193 | 205 | 297 | 308 | 68 | 142 | 74 | 127 | 193 | 95 | 132 | 63 | 73 | 60 | 79 | 200 | 279 | 88 | 173 | 44 | 55 | B4galnt3 |
| B4galnt4 | 2846 | 3620 | 3499 | 3713 | 4232 | 3677 | 3384 | 5010 | 5656 | 6085 | 5143 | 6113 | 5687 | 5660 | 2740 | 3203 | 3853 | 3688 | 4277 | 3420 | 3503 | 6025 | 5016 | 6001 | 6354 | 5982 | 5635 | 5466 | B4galnt4 |
| B4galt1 | 3864 | 3170 | 3474 | 3023 | 2300 | 2911 | 2359 | 2277 | 2709 | 2434 | 3578 | 2488 | 3041 | 2349 | 3307 | 4077 | 2881 | 3273 | 2290 | 2187 | 2218 | 2445 | 3093 | 2549 | 2139 | 2954 | 2508 | 2301 | B4galt1 |
| B4galt2 | 2645 | 3208 | 3140 | 3350 | 3856 | 3705 | 3175 | 4306 | 5221 | 5429 | 5350 | 5225 | 5038 | 4791 | 2698 | 2842 | 3409 | 3367 | 3878 | 3089 | 3347 | 5127 | 4718 | 5291 | 5744 | 5075 | 4968 | 4678 | B4galt2 |
| B4galt3 | 3212 | 4156 | 3704 | 4102 | 3697 | 3420 | 3795 | 2699 | 3306 | 3494 | 3503 | 3505 | 3127 | 3193 | 3218 | 3785 | 3987 | 3949 | 4058 | 3439 | 3772 | 3547 | 3170 | 3553 | 3630 | 3496 | 3213 | 3143 | B4galt3 |
| B4galt4 | 1531 | 1607 | 1679 | 1281 | 1346 | 1443 | 1259 | 1504 | 1533 | 1492 | 1555 | 1418 | 1699 | 1475 | 1642 | 1850 | 1386 | 1290 | 1340 | 1311 | 1283 | 1569 | 1682 | 1444 | 1351 | 1659 | 1464 | 1689 | B4galt4 |
| B4galt5 | 3688 | 4341 | 5207 | 5257 | 4609 | 4505 | 4137 | 2182 | 2684 | 2929 | 3305 | 3191 | 3061 | 2902 | 3575 | 4365 | 4934 | 5208 | 4947 | 3615 | 4218 | 3136 | 2539 | 3175 | 3243 | 3347 | 2957 | 3021 | B4galt5 |
| B4galt6 | 4475 | 4117 | 5376 | 5859 | 6752 | 5948 | 5882 | 3789 | 3639 | 4332 | 4477 | 4182 | 4393 | 4077 | 4771 | 4995 | 5696 | 5904 | 5901 | 4874 | 6010 | 5013 | 3852 | 3791 | 4414 | 3486 | 3807 | 4030 | B4galt6 |
| B4galt7 | 1926 | 2277 | 2471 | 2464 | 2371 | 2276 | 2171 | 1602 | 1888 | 1785 | 2028 | 1868 | 1867 | 1940 | 1991 | 2273 | 2516 | 2313 | 2328 | 2114 | 2157 | 1977 | 1794 | 1974 | 2043 | 1981 | 1789 | 1876 | B4galt7 |
| B4gat1 | 9 | 11 | 6 | 8 | 0 | 8 | 7 | 4 | 2 | 8 | 1 | 8 | 6 | 6 | 11 | 2 | 2 | 4 | 2 | 0 | 18 | 3 | 3 | 9 | 6 | 5 | 8 | 5 | B4gat1 |
| B630019K06Rik | 882 | 814 | 1177 | 1596 | 2248 | 1780 | 1856 | 1914 | 1619 | 2208 | 1958 | 1686 | 1572 | 1632 | 1017 | 801 | 1234 | 1613 | 1849 | 1508 | 1860 | 2784 | 1641 | 1957 | 2333 | 1502 | 1410 | 1530 | B630019K06Rik |
| B830017H08Rik | 8 | 3 | 7 | 6 | 20 | 9 | 7 | 28 | 51 | 63 | 46 | 17 | 35 | 19 | 4 | 7 | 3 | 2 | 13 | 4 | 5 | 47 | 44 | 25 | 56 | 14 | 12 | 12 | B830017H08Rik |
| B930094E09Rik | 1 | 2 | 1 | 0 | 2 | 3 | 8 | 6 | 7 | 7 | 0 | 8 | 9 | 11 | 0 | 5 | 3 | 0 | 2 | 3 | 3 | 8 | 12 | 10 | 5 | 12 | 19 | 15 | B930094E09Rik |
| B9d1 | 678 | 752 | 558 | 569 | 603 | 590 | 575 | 1036 | 997 | 787 | 834 | 958 | 889 | 932 | 654 | 719 | 545 | 521 | 596 | 798 | 577 | 735 | 1075 | 869 | 935 | 1018 | 1082 | 929 | B9d1 |
| B9d2 | 328 | 514 | 381 | 495 | 457 | 409 | 420 | 670 | 684 | 750 | 676 | 751 | 746 | 706 | 367 | 405 | 526 | 474 | 561 | 557 | 443 | 653 | 726 | 786 | 758 | 840 | 782 | 765 | B9d2 |
| Baalc | 3 | 4 | 2 | 4 | 3 | 10 | 1 | 12 | 16 | 7 | 19 | 3 | 7 | 5 | 9 | 5 | 3 | 4 | 2 | 6 | 4 | 17 | 14 | 7 | 8 | 8 | 5 | 11 | Baalc |
| Baat | 59 | 60 | 46 | 69 | 32 | 63 | 34 | 31 | 12 | 14 | 0 | 31 | 16 | 30 | 55 | 52 | 54 | 56 | 28 | 31 | 41 | 13 | 17 | 20 | 28 | 34 | 31 | 26 | Baat |
| Babam1 | 3309 | 3952 | 3520 | 3399 | 3101 | 3286 | 3123 | 3098 | 3518 | 3516 | 3320 | 3721 | 3506 | 3655 | 2989 | 3252 | 3500 | 3232 | 3608 | 3222 | 3467 | 2882 | 3389 | 3704 | 3522 | 4302 | 3850 | 3751 | Babam1 |
| Babam2 | 1179 | 1313 | 1108 | 966 | 1038 | 1146 | 1106 | 1362 | 1285 | 1108 | 1028 | 1300 | 1171 | 1189 | 1218 | 1093 | 1116 | 925 | 1053 | 1171 | 1104 | 1305 | 1229 | 1188 | 1251 | 1396 | 1287 | 1283 | Babam2 |
| Bace1 | 8382 | 9051 | 7518 | 8347 | 7800 | 8022 | 7704 | 4531 | 4631 | 4712 | 5161 | 3872 | 4088 | 3975 | 8631 | 9263 | 8017 | 9248 | 7341 | 7000 | 7983 | 5370 | 4706 | 4381 | 4679 | 3643 | 3805 | 3730 | Bace1 |
| Bace2 | 281 | 230 | 211 | 241 | 263 | 232 | 175 | 167 | 212 | 150 | 279 | 110 | 192 | 122 | 251 | 241 | 211 | 233 | 192 | 209 | 164 | 157 | 296 | 101 | 129 | 93 | 96 | 129 | Bace2 |
| Bach1 | 8767 | 8084 | 8428 | 8128 | 7996 | 8633 | 8537 | 8994 | 8374 | 7887 | 8400 | 9049 | 9443 | 9983 | 8980 | 8506 | 8383 | 7964 | 7956 | 8555 | 8518 | 8621 | 7706 | 8317 | 9152 | 7699 | 9401 | 10159 | Bach1 |
| Bach2 | 2713 | 2494 | 3259 | 5135 | 5774 | 4856 | 5626 | 4125 | 4582 | 6233 | 5415 | 5840 | 5601 | 5739 | 2888 | 2941 | 4516 | 5086 | 5656 | 4018 | 5605 | 5987 | 4311 | 6178 | 7430 | 5196 | 4814 | 5287 | Bach2 |
| Bad | 1603 | 2328 | 1837 | 1981 | 1771 | 1741 | 1654 | 1430 | 1617 | 1551 | 1598 | 1440 | 1345 | 1537 | 1567 | 1794 | 1905 | 1960 | 2069 | 1877 | 1845 | 1399 | 1681 | 1526 | 1573 | 1596 | 1388 | 1489 | Bad |
| Bag1 | 8815 | 8124 | 8494 | 8055 | 7473 | 6723 | 8896 | 6945 | 6185 | 6620 | 6280 | 6414 | 6262 | 7735 | 8594 | 7430 | 8423 | 8166 | 7684 | 7479 | 8887 | 6475 | 6530 | 7063 | 6504 | 7124 | 6503 | 7516 | Bag1 |
| Bag2 | 850 | 833 | 871 | 803 | 838 | 769 | 820 | 686 | 739 | 809 | 865 | 810 | 772 | 758 | 798 | 957 | 889 | 799 | 921 | 703 | 701 | 592 | 819 | 855 | 706 | 958 | 748 | 833 | Bag2 |
| Bag3 | 1006 | 1168 | 1315 | 913 | 808 | 831 | 496 | 459 | 685 | 714 | 973 | 663 | 629 | 475 | 885 | 1290 | 1168 | 1002 | 780 | 556 | 544 | 585 | 918 | 650 | 572 | 754 | 655 | 521 | Bag3 |
| Bag4 | 5553 | 4763 | 4348 | 5324 | 5247 | 5570 | 5882 | 4098 | 3601 | 3412 | 3261 | 3934 | 3569 | 3761 | 5470 | 4600 | 4766 | 5151 | 5165 | 5759 | 5837 | 3659 | 3344 | 3546 | 3859 | 3710 | 3640 | 3806 | Bag4 |
| Bag5 | 2754 | 3149 | 3007 | 3254 | 2879 | 2919 | 3034 | 3171 | 2970 | 2874 | 2926 | 3178 | 2972 | 3216 | 2883 | 2845 | 3041 | 2996 | 2891 | 3186 | 2954 | 2896 | 2986 | 3079 | 3149 | 3606 | 3196 | 3345 | Bag5 |
| Bag6 | 14464 | 18134 | 22552 | 23439 | 22056 | 19499 | 18093 | 17059 | 20937 | 24624 | 22523 | 27432 | 21814 | 22923 | 14424 | 16173 | 23417 | 22237 | 24654 | 15622 | 19879 | 24465 | 19285 | 27651 | 26114 | 29111 | 21633 | 22840 | Bag6 |
| Bahcc1 | 6575 | 8867 | 10953 | 11109 | 9009 | 8800 | 7832 | 6533 | 7531 | 10639 | 9065 | 12820 | 9781 | 10570 | 6070 | 7728 | 10810 | 10258 | 9694 | 6498 | 8040 | 10833 | 6370 | 11535 | 11126 | 9104 | 10620 | 10170 | Bahcc1 |
| Bahd1 | 4832 | 6908 | 8665 | 8813 | 7688 | 6596 | 6777 | 2281 | 2540 | 3198 | 2794 | 3389 | 3005 | 3316 | 4767 | 5748 | 9170 | 8650 | 8505 | 5555 | 7511 | 3110 | 2552 | 3656 | 3548 | 3515 | 3159 | 3406 | Bahd1 |
| Baiap2 | 2535 | 3078 | 2281 | 3048 | 2694 | 2764 | 2741 | 895 | 1149 | 1065 | 991 | 957 | 891 | 912 | 2561 | 2924 | 2620 | 3177 | 2911 | 2976 | 2871 | 922 | 1136 | 1035 | 1017 | 920 | 807 | 882 | Baiap2 |
| Baiap2l1 | 938 | 752 | 840 | 659 | 384 | 435 | 377 | 160 | 197 | 159 | 274 | 124 | 153 | 93 | 738 | 1077 | 618 | 547 | 422 | 433 | 296 | 189 | 329 | 174 | 69 | 178 | 86 | 110 | Baiap2l1 |
| Baiap2l2 | 158 | 156 | 181 | 144 | 153 | 134 | 91 | 45 | 66 | 77 | 63 | 53 | 29 | 32 | 123 | 190 | 166 | 134 | 159 | 99 | 88 | 72 | 78 | 54 | 51 | 94 | 41 | 65 | Baiap2l2 |
| Baiap3 | 511 | 809 | 768 | 911 | 961 | 836 | 658 | 342 | 444 | 491 | 360 | 305 | 345 | 346 | 558 | 706 | 769 | 894 | 809 | 843 | 762 | 610 | 398 | 409 | 465 | 244 | 257 | 241 | Baiap3 |
| Bak1 | 1369 | 1519 | 1679 | 1046 | 1031 | 1140 | 864 | 1780 | 2298 | 2094 | 2118 | 2274 | 2434 | 2036 | 1321 | 1576 | 1306 | 965 | 1146 | 1106 | 932 | 1616 | 2508 | 2066 | 1891 | 2810 | 2423 | 2325 | Bak1 |
| Bambi | 1010 | 822 | 714 | 555 | 449 | 513 | 443 | 927 | 775 | 590 | 623 | 484 | 625 | 447 | 779 | 913 | 487 | 512 | 410 | 542 | 406 | 607 | 863 | 600 | 409 | 705 | 547 | 551 | Bambi |
| Banf1 | 3951 | 4694 | 4371 | 3887 | 3090 | 3147 | 4122 | 4776 | 4619 | 4675 | 4413 | 5607 | 5033 | 6829 | 3653 | 3588 | 4311 | 3317 | 3635 | 4038 | 3857 | 4055 | 4566 | 5629 | 4942 | 7047 | 6237 | 7028 | Banf1 |
| Bank1 | 23 | 24 | 25 | 24 | 31 | 24 | 20 | 31 | 54 | 19 | 39 | 37 | 44 | 30 | 37 | 36 | 19 | 36 | 10 | 39 | 22 | 45 | 65 | 43 | 41 | 41 | 35 | 15 | Bank1 |
| Banp | 1867 | 2780 | 3235 | 3379 | 2838 | 2275 | 2562 | 3065 | 2913 | 3314 | 3205 | 2858 | 2715 | 2680 | 1726 | 2406 | 3072 | 3298 | 3297 | 2563 | 2648 | 3233 | 2801 | 3494 | 3486 | 2860 | 2574 | 2683 | Banp |
| Bap1 | 9724 | 11384 | 11140 | 10957 | 9971 | 9985 | 9700 | 9393 | 9823 | 10758 | 9673 | 12396 | 11222 | 11514 | 9535 | 9460 | 10940 | 10150 | 11178 | 10091 | 10011 | 9707 | 9388 | 11581 | 11400 | 12412 | 12033 | 12252 | Bap1 |
| Bard1 | 1846 | 1373 | 1994 | 1765 | 1292 | 1384 | 1590 | 1387 | 1314 | 1356 | 1307 | 2026 | 1678 | 1963 | 1311 | 1386 | 1728 | 1472 | 1519 | 1152 | 1427 | 1153 | 1087 | 1772 | 1382 | 2316 | 1911 | 2057 | Bard1 |
| Barhl1 | 321 | 436 | 898 | 699 | 922 | 894 | 517 | 983 | 723 | 1296 | 931 | 1399 | 1056 | 1376 | 315 | 270 | 782 | 769 | 1198 | 718 | 876 | 1724 | 1191 | 1544 | 1571 | 511 | 1524 | 1559 | Barhl1 |
| Barhl2 | 927 | 1146 | 2306 | 1470 | 1010 | 986 | 1011 | 883 | 476 | 879 | 667 | 1849 | 945 | 1926 | 1009 | 525 | 1820 | 1010 | 1435 | 1129 | 1221 | 1235 | 698 | 1573 | 1273 | 1701 | 2145 | 2479 | Barhl2 |
| Barx1 | 607 | 92 | 57 | 135 | 33 | 35 | 18 | 217 | 531 | 197 | 519 | 10 | 120 | 28 | 100 | 291 | 73 | 35 | 27 | 22 | 16 | 142 | 222 | 194 | 10 | 234 | 1 | 17 | Barx1 |
| Barx2 | 52 | 18 | 9 | 24 | 20 | 15 | 17 | 74 | 149 | 78 | 239 | 52 | 43 | 44 | 21 | 29 | 30 | 19 | 27 | 16 | 24 | 43 | 190 | 59 | 29 | 43 | 30 | 38 | Barx2 |
| Basp1 | 21454 | 21552 | 25581 | 29841 | 36057 | 31220 | 37253 | 38747 | 32902 | 43977 | 38332 | 40166 | 39977 | 49631 | 24667 | 18897 | 28636 | 31177 | 34144 | 32224 | 38254 | 45893 | 34058 | 46401 | 49426 | 38996 | 41416 | 47508 | Basp1 |
| Batf | 39 | 24 | 55 | 17 | 20 | 46 | 7 | 24 | 54 | 52 | 36 | 17 | 49 | 29 | 46 | 39 | 28 | 20 | 20 | 26 | 10 | 37 | 63 | 43 | 14 | 57 | 50 | 31 | Batf |
| Batf2 | 2 | 2 | 3 | 4 | 7 | 1 | 2 | 9 | 8 | 8 | 8 | 8 | 8 | 5 | 4 | 5 | 0 | 2 | 4 | 1 | 5 | 7 | 6 | 13 | 3 | 16 | 0 | 2 | Batf2 |
| Batf3 | 100 | 92 | 103 | 64 | 43 | 74 | 41 | 59 | 69 | 69 | 52 | 61 | 61 | 70 | 94 | 116 | 110 | 63 | 48 | 41 | 42 | 40 | 68 | 64 | 50 | 64 | 51 | 45 | Batf3 |
| Bax | 1962 | 2425 | 2180 | 2547 | 1965 | 1632 | 2133 | 1494 | 1364 | 1662 | 1429 | 1605 | 1380 | 1936 | 1808 | 1489 | 2316 | 2169 | 2134 | 1968 | 2259 | 1278 | 1338 | 1788 | 1820 | 1698 | 1706 | 1728 | Bax |
| Baz1a | 3917 | 3261 | 3639 | 3069 | 3679 | 3775 | 3639 | 4575 | 3583 | 3646 | 3497 | 4889 | 4752 | 5104 | 3396 | 3558 | 3504 | 2831 | 3926 | 3583 | 3601 | 2842 | 3536 | 4617 | 3882 | 5041 | 5371 | 5404 | Baz1a |
| Baz1b | 27437 | 25937 | 27610 | 28252 | 27222 | 28704 | 29795 | 29278 | 28044 | 28682 | 29868 | 35742 | 32515 | 37293 | 25847 | 24725 | 27712 | 27535 | 29454 | 27153 | 28969 | 27121 | 25572 | 32501 | 32481 | 28441 | 35490 | 35532 | Baz1b |
| Baz2a | 15175 | 17807 | 16769 | 20903 | 18002 | 18025 | 18628 | 8490 | 8375 | 10253 | 9449 | 11838 | 11248 | 11447 | 14600 | 16439 | 19244 | 20720 | 18827 | 14551 | 18573 | 10492 | 8132 | 10426 | 11291 | 7084 | 11028 | 11027 | Baz2a |
| Baz2b | 8450 | 8168 | 5510 | 7713 | 9647 | 10129 | 9600 | 14865 | 13144 | 12844 | 12416 | 13685 | 13306 | 14380 | 9279 | 7551 | 6905 | 7690 | 8423 | 9928 | 9994 | 11734 | 12176 | 10978 | 12789 | 8539 | 15376 | 12705 | Baz2b |
| BB014433 | 0 | 0 | 0 | 0 | 0 | 0 | 0 | 0 | 0 | 0 | 3 | 0 | 0 | 0 | 0 | 0 | 0 | 0 | 0 | 0 | 0 | 2 | 0 | 0 | 0 | 0 | 0 | 0 | BB014433 |
| Bbc3 | 708 | 1178 | 988 | 1252 | 1198 | 1061 | 945 | 423 | 476 | 517 | 455 | 475 | 518 | 446 | 723 | 814 | 1163 | 1250 | 1275 | 1042 | 1031 | 434 | 445 | 535 | 541 | 543 | 490 | 418 | Bbc3 |
| Bbip1 | 3130 | 2826 | 2906 | 2945 | 3121 | 3076 | 3587 | 4351 | 3709 | 3792 | 3450 | 4165 | 3919 | 4560 | 3498 | 2766 | 2992 | 2867 | 3198 | 3881 | 3561 | 3897 | 3623 | 4005 | 3990 | 4209 | 4122 | 4558 | Bbip1 |
| Bbln | 982 | 1372 | 1083 | 1096 | 1106 | 952 | 893 | 664 | 627 | 637 | 683 | 565 | 632 | 577 | 932 | 1085 | 1092 | 1059 | 992 | 1021 | 1020 | 688 | 709 | 599 | 656 | 638 | 672 | 648 | Bbln |
| Bbof1 | 549 | 484 | 329 | 579 | 609 | 755 | 707 | 475 | 485 | 483 | 496 | 312 | 278 | 378 | 597 | 445 | 444 | 567 | 496 | 742 | 701 | 331 | 476 | 339 | 477 | 255 | 310 | 292 | Bbof1 |
| Bbox1 | 14 | 2 | 5 | 8 | 2 | 7 | 0 | 10 | 23 | 9 | 34 | 4 | 24 | 4 | 12 | 14 | 5 | 10 | 6 | 0 | 4 | 15 | 30 | 7 | 13 | 8 | 2 | 11 | Bbox1 |
| Bbs1 | 2007 | 2320 | 2181 | 2418 | 2432 | 2427 | 2430 | 2590 | 2678 | 3095 | 2752 | 3382 | 3057 | 3077 | 2169 | 2092 | 2453 | 2326 | 2612 | 2430 | 2408 | 3429 | 2586 | 3143 | 3268 | 2615 | 3312 | 3312 | Bbs1 |
| Bbs10 | 521 | 548 | 633 | 375 | 464 | 359 | 470 | 692 | 626 | 582 | 468 | 651 | 555 | 650 | 538 | 439 | 563 | 393 | 462 | 546 | 374 | 602 | 673 | 599 | 590 | 754 | 662 | 726 | Bbs10 |
| Bbs12 | 1353 | 965 | 1086 | 898 | 788 | 960 | 1014 | 1095 | 1011 | 829 | 760 | 973 | 880 | 762 | 1257 | 1192 | 978 | 841 | 864 | 1136 | 842 | 724 | 967 | 799 | 731 | 1007 | 839 | 970 | Bbs12 |
| Bbs2 | 1931 | 2040 | 1793 | 1902 | 1892 | 1849 | 1906 | 2659 | 2408 | 2456 | 2242 | 2616 | 2583 | 2677 | 2194 | 1932 | 2107 | 1853 | 2034 | 1954 | 1976 | 2550 | 2331 | 2281 | 2680 | 2428 | 2766 | 2559 | Bbs2 |
| Bbs4 | 2346 | 2315 | 1872 | 2173 | 2288 | 2294 | 2605 | 2948 | 2411 | 2251 | 2346 | 2273 | 2236 | 2579 | 2496 | 2221 | 2146 | 2161 | 2356 | 2597 | 2420 | 2725 | 2523 | 2333 | 2465 | 2317 | 2420 | 2382 | Bbs4 |
| Bbs5 | 1190 | 1004 | 1017 | 1134 | 1003 | 1082 | 1150 | 983 | 881 | 903 | 936 | 849 | 866 | 843 | 1096 | 1070 | 1012 | 1136 | 1029 | 1155 | 1188 | 984 | 896 | 824 | 943 | 756 | 723 | 892 | Bbs5 |

| | | | | | | | | | | | | | | | | | | | | | | | | | | | | | |
|---|---|---|---|---|---|---|---|---|---|---|---|---|---|---|---|---|---|---|---|---|---|---|---|---|---|---|---|---|---|
| Bbs7 | 2184 | 2031 | 2121 | 1914 | 1906 | 1981 | 2047 | 1871 | 1775 | 1734 | 1827 | 1962 | 1917 | 1993 | 2302 | 1946 | 2103 | 1772 | 1802 | 2340 | 2084 | 1644 | 1704 | 1758 | 1781 | 1891 | 1956 | 1923 | Bbs7 |
| Bbs9 | 2089 | 2179 | 1971 | 1828 | 1782 | 1618 | 1698 | 1754 | 1752 | 1546 | 1315 | 1535 | 1398 | 1480 | 2148 | 1927 | 2102 | 1699 | 1760 | 1930 | 1707 | 1534 | 1726 | 1547 | 1536 | 1653 | 1518 | 1423 | Bbs9 |
| Bbx | 6757 | 5675 | 4706 | 5713 | 6332 | 6788 | 6265 | 4505 | 4253 | 3503 | 4435 | 3823 | 4107 | 3949 | 6915 | 5868 | 5341 | 5718 | 6081 | 7094 | 6509 | 3432 | 4203 | 3372 | 3709 | 2874 | 4023 | 3370 | Bbx |
| BC003965 | 2679 | 2493 | 2347 | 2553 | 2176 | 2146 | 2443 | 2119 | 1953 | 1886 | 1847 | 2218 | 1905 | 2117 | 2454 | 2353 | 2327 | 2428 | 2302 | 2640 | 2336 | 1866 | 1884 | 2255 | 2035 | 2299 | 2200 | 2203 | BC003965 |
| BC004004 | 1885 | 2069 | 1666 | 1750 | 1873 | 2121 | 2015 | 2462 | 2485 | 2250 | 2251 | 2296 | 2311 | 2232 | 1967 | 1826 | 1751 | 1727 | 1843 | 2125 | 2182 | 2421 | 2650 | 2130 | 2318 | 2184 | 2332 | 2220 | BC004004 |
| BC005537 | 18239 | 16677 | 22677 | 25017 | 24399 | 22250 | 26691 | 19883 | 18712 | 22952 | 22569 | 23889 | 21463 | 24148 | 19183 | 17442 | 24591 | 26060 | 25011 | 19345 | 26059 | 25267 | 18430 | 24644 | 24947 | 24486 | 21065 | 23482 | BC005537 |
| BC005624 | 3275 | 3970 | 4092 | 3935 | 3267 | 3294 | 3593 | 3390 | 3343 | 3675 | 3417 | 3682 | 3730 | 3942 | 3291 | 3527 | 3845 | 3639 | 3444 | 3688 | 3550 | 3322 | 3353 | 3858 | 3786 | 3922 | 3704 | 3878 | BC005624 |
| BC016579 | 27 | 31 | 43 | 68 | 50 | 47 | 46 | 31 | 11 | 14 | 16 | 22 | 23 | 22 | 26 | 31 | 54 | 70 | 69 | 46 | 46 | 42 | 12 | 22 | 23 | 18 | 14 | 18 | BC016579 |
| BC024063 | 465 | 392 | 368 | 366 | 457 | 589 | 420 | 838 | 774 | 585 | 626 | 562 | 619 | 592 | 531 | 505 | 512 | 375 | 460 | 551 | 448 | 655 | 717 | 489 | 594 | 525 | 562 | 541 | BC024063 |
| BC024139 | 632 | 1267 | 1469 | 1033 | 1532 | 1155 | 802 | 1850 | 1769 | 1933 | 1792 | 2013 | 3433 | 1814 | 1250 | 1475 | 1578 | 991 | 1134 | 1329 | 977 | 2288 | 1761 | 1490 | 1983 | 922 | 2310 | 2015 | BC024139 |
| BC024978 | 299 | 309 | 447 | 438 | 465 | 433 | 456 | 772 | 826 | 1170 | 976 | 1038 | 1095 | 1213 | 274 | 247 | 399 | 375 | 544 | 396 | 438 | 1043 | 773 | 1106 | 1017 | 1053 | 1017 | 1161 | BC024978 |
| BC025920 | 520 | 463 | 508 | 420 | 419 | 565 | 534 | 872 | 823 | 743 | 676 | 939 | 743 | 898 | 568 | 411 | 557 | 372 | 550 | 573 | 504 | 667 | 782 | 865 | 917 | 907 | 931 | 885 | BC025920 |
| BC028528 | 146 | 109 | 191 | 140 | 158 | 147 | 159 | 244 | 233 | 301 | 419 | 215 | 290 | 228 | 171 | 158 | 110 | 157 | 139 | 160 | 166 | 307 | 342 | 279 | 251 | 292 | 201 | 256 | BC028528 |
| BC029722 | 504 | 562 | 501 | 545 | 571 | 597 | 565 | 635 | 689 | 793 | 806 | 688 | 724 | 763 | 566 | 557 | 586 | 576 | 564 | 533 | 640 | 757 | 747 | 744 | 759 | 667 | 652 | 757 | BC029722 |
| BC030500 | 134 | 148 | 141 | 180 | 219 | 190 | 200 | 176 | 233 | 240 | 267 | 163 | 138 | 155 | 161 | 129 | 218 | 183 | 192 | 193 | 191 | 261 | 219 | 174 | 215 | 118 | 105 | 157 | BC030500 |
| BC031181 | 5009 | 5107 | 5266 | 5710 | 5123 | 5162 | 5525 | 4892 | 4495 | 4728 | 4341 | 4782 | 4433 | 5014 | 5639 | 5139 | 5496 | 5108 | 5491 | 6165 | 5623 | 4674 | 4678 | 5048 | 5124 | 5049 | 5090 | 4951 | BC031181 |
| BC034090 | 270 | 274 | 386 | 318 | 380 | 373 | 261 | 856 | 1038 | 1090 | 1196 | 1111 | 1218 | 875 | 284 | 333 | 326 | 288 | 394 | 350 | 264 | 904 | 1120 | 985 | 947 | 766 | 1026 | 929 | BC034090 |
| BC035044 | 2 | 6 | 20 | 6 | 5 | 24 | 9 | 22 | 23 | 14 | 18 | 4 | 7 | 24 | 11 | 15 | 7 | 10 | 9 | 14 | 8 | 13 | 38 | 12 | 21 | 23 | 13 | 13 | BC035044 |
| BC035947 | 62 | 33 | 48 | 68 | 101 | 88 | 70 | 178 | 156 | 258 | 204 | 139 | 138 | 135 | 61 | 51 | 41 | 69 | 58 | 35 | 55 | 206 | 162 | 190 | 164 | 240 | 71 | 103 | BC035947 |
| BC048403 | 374 | 308 | 371 | 328 | 353 | 322 | 343 | 248 | 331 | 423 | 402 | 380 | 350 | 409 | 280 | 310 | 371 | 364 | 338 | 283 | 343 | 346 | 342 | 396 | 389 | 436 | 407 | 398 | BC048403 |
| BC048507 | 1 | 5 | 4 | 7 | 2 | 3 | 0 | 0 | 1 | 3 | 0 | 0 | 3 | 2 | 1 | 3 | 7 | 8 | 6 | 5 | 4 | 2 | 2 | 2 | 0 | 0 | 1 | 1 | BC048507 |
| BC048562 | 0 | 3 | 0 | 0 | 0 | 0 | 0 | 0 | 0 | 0 | 2 | 0 | 0 | 0 | 0 | 0 | 0 | 0 | 0 | 0 | 0 | 0 | 0 | 0 | 0 | 0 | 0 | 0 | BC048562 |
| BC048671 | 0 | 0 | 0 | 0 | 0 | 0 | 1 | 0 | 0 | 0 | 0 | 0 | 0 | 0 | 0 | 0 | 0 | 0 | 0 | 0 | 0 | 0 | 0 | 0 | 0 | 0 | 0 | 0 | BC048671 |
| BC048679 | 0 | 0 | 0 | 0 | 0 | 0 | 0 | 0 | 0 | 6 | 0 | 0 | 0 | 1 | 0 | 1 | 5 | 0 | 0 | 0 | 0 | 0 | 0 | 0 | 0 | 0 | 0 | 0 | BC048679 |
| BC049352 | 3 | 4 | 2 | 12 | 5 | 2 | 2 | 32 | 25 | 45 | 38 | 1 | 17 | 13 | 9 | 2 | 3 | 7 | 3 | 2 | 6 | 0 | 30 | 56 | 4 | 33 | 1 | 2 | BC049352 |
| BC049715 | 17 | 25 | 9 | 24 | 17 | 44 | 19 | 34 | 52 | 38 | 31 | 55 | 37 | 54 | 26 | 26 | 22 | 8 | 14 | 15 | 19 | 37 | 32 | 33 | 23 | 21 | 42 | 23 | BC049715 |
| BC049762 | 8 | 14 | 8 | 4 | 10 | 5 | 3 | 6 | 11 | 12 | 6 | 8 | 15 | 11 | 7 | 2 | 8 | 3 | 6 | 6 | 7 | 7 | 11 | 9 | 7 | 5 | 7 | 7 | BC049762 |
| BC051019 | 6 | 2 | 0 | 8 | 0 | 8 | 4 | 6 | 10 | 8 | 5 | 18 | 12 | 6 | 4 | 9 | 2 | 3 | 11 | 3 | 5 | 8 | 16 | 5 | 6 | 8 | 6 | 4 | BC051019 |
| BC051142 | 227 | 271 | 250 | 305 | 369 | 351 | 285 | 285 | 221 | 232 | 238 | 289 | 333 | 397 | 290 | 272 | 260 | 351 | 387 | 360 | 411 | 406 | 224 | 271 | 342 | 243 | 410 | 398 | BC051142 |
| BC055324 | 4478 | 3691 | 3056 | 2783 | 2019 | 2592 | 2894 | 2756 | 2446 | 1788 | 1936 | 2850 | 2456 | 2719 | 3914 | 3728 | 2817 | 2565 | 2551 | 3090 | 2748 | 1696 | 2145 | 2265 | 2080 | 3009 | 2764 | 2703 | BC055324 |
| BC061237 | 0 | 0 | 0 | 1 | 4 | 0 | 2 | 0 | 1 | 0 | 3 | 0 | 0 | 2 | 1 | 0 | 5 | 3 | 0 | 0 | 0 | 0 | 3 | 0 | 1 | 0 | 0 | 0 | BC061237 |
| BC107364 | 36 | 31 | 22 | 21 | 13 | 6 | 9 | 4 | 7 | 0 | 3 | 1 | 1 | 0 | 42 | 42 | 9 | 9 | 23 | 16 | 5 | 5 | 2 | 1 | 7 | 3 | 3 | 0 | BC107364 |
| BC147527 | 2 | 0 | 1 | 0 | 2 | 1 | 4 | 5 | 7 | 12 | 15 | 11 | 2 | 1 | 0 | 0 | 0 | 0 | 2 | 4 | 1 | 5 | 8 | 5 | 7 | 1 | 7 | 0 | BC147527 |
| Bcam | 1658 | 1220 | 1180 | 573 | 626 | 725 | 403 | 1210 | 2092 | 1172 | 1583 | 818 | 1162 | 686 | 1042 | 1496 | 738 | 924 | 581 | 464 | 358 | 590 | 2450 | 1007 | 574 | 1012 | 798 | 871 | Bcam |
| Bcan | 300 | 387 | 480 | 759 | 1581 | 942 | 1120 | 481 | 828 | 953 | 850 | 270 | 134 | 362 | 307 | 382 | 561 | 950 | 1244 | 820 | 1238 | 1014 | 898 | 655 | 1108 | 256 | 179 | 141 | Bcan |
| Bcap29 | 1316 | 897 | 1243 | 1159 | 1334 | 1394 | 1330 | 1625 | 1454 | 1349 | 1570 | 1370 | 1633 | 1520 | 1342 | 1252 | 1093 | 1159 | 1192 | 1564 | 1226 | 1337 | 1480 | 1369 | 1377 | 1726 | 1445 | 1637 | Bcap29 |
| Bcap31 | 5193 | 4773 | 5061 | 4448 | 3961 | 3881 | 4166 | 3665 | 3394 | 3165 | 3388 | 3150 | 3383 | 3434 | 5239 | 4815 | 4687 | 4266 | 4013 | 4531 | 3984 | 3412 | 3930 | 3369 | 2939 | 3827 | 3385 | 3704 | Bcap31 |
| Bcar1 | 4399 | 6088 | 5884 | 6818 | 6089 | 5268 | 4608 | 3488 | 4215 | 4623 | 4159 | 5142 | 4490 | 4296 | 4412 | 5555 | 6693 | 6568 | 6497 | 4603 | 5303 | 4645 | 3838 | 5027 | 4923 | 5051 | 4210 | 4206 | Bcar1 |
| Bcar3 | 532 | 487 | 831 | 484 | 579 | 458 | 445 | 565 | 577 | 459 | 531 | 615 | 632 | 640 | 506 | 586 | 605 | 359 | 482 | 436 | 458 | 588 | 606 | 527 | 524 | 703 | 791 | 705 | Bcar3 |
| Bcas1 | 135 | 136 | 215 | 172 | 297 | 252 | 162 | 310 | 310 | 487 | 473 | 222 | 411 | 219 | 181 | 177 | 225 | 295 | 235 | 223 | 256 | 548 | 433 | 276 | 330 | 135 | 404 | 297 | Bcas1 |
| Bcas2 | 5913 | 5048 | 6280 | 6058 | 5875 | 5454 | 6661 | 6736 | 6135 | 6520 | 6793 | 7780 | 6794 | 8310 | 5608 | 5404 | 6096 | 5892 | 6062 | 5653 | 6279 | 6649 | 6098 | 7844 | 7313 | 8329 | 7583 | 8413 | Bcas2 |
| Bcas3 | 1681 | 1773 | 1408 | 1500 | 1695 | 1682 | 1466 | 1694 | 1895 | 1842 | 1587 | 1651 | 1846 | 1549 | 1706 | 1835 | 1494 | 1557 | 1585 | 1591 | 1560 | 1900 | 1774 | 1640 | 1816 | 1564 | 1611 | 1508 | Bcas3 |
| Bcat1 | 4161 | 4472 | 4568 | 4156 | 5031 | 5127 | 4984 | 6364 | 5469 | 5845 | 5189 | 6789 | 6327 | 6857 | 4524 | 3699 | 5202 | 4131 | 5577 | 5350 | 5142 | 6493 | 5047 | 6271 | 6965 | 6387 | 7703 | 7276 | Bcat1 |
| Bcat2 | 1194 | 1202 | 1368 | 1179 | 1136 | 1054 | 917 | 1570 | 2192 | 1811 | 2105 | 1716 | 1931 | 1618 | 1073 | 1405 | 1212 | 1018 | 1113 | 878 | 862 | 1820 | 2073 | 1946 | 1850 | 2068 | 1923 | 1724 | Bcat2 |
| Bccip | 4820 | 4324 | 5536 | 4708 | 3862 | 3926 | 4510 | 4099 | 3585 | 4437 | 4242 | 4672 | 4622 | 4956 | 4593 | 4310 | 4828 | 3979 | 4417 | 4591 | 4169 | 3417 | 3838 | 4732 | 4238 | 6015 | 4821 | 5301 | Bccip |
| Bcdin3d | 860 | 782 | 852 | 881 | 837 | 984 | 1011 | 983 | 1040 | 1027 | 965 | 1231 | 1068 | 1322 | 843 | 655 | 895 | 675 | 903 | 883 | 883 | 1004 | 950 | 1232 | 1086 | 1307 | 1285 | 1187 | Bcdin3d |
| Bche | 965 | 611 | 628 | 474 | 589 | 540 | 446 | 907 | 977 | 629 | 899 | 671 | 869 | 668 | 1005 | 896 | 546 | 486 | 448 | 721 | 490 | 774 | 955 | 487 | 536 | 631 | 805 | 739 | Bche |
| Bckdha | 2402 | 2864 | 2119 | 2290 | 1810 | 2025 | 1760 | 1638 | 2060 | 1747 | 1696 | 1730 | 1770 | 1432 | 2024 | 2513 | 2132 | 2111 | 2320 | 2144 | 1903 | 1516 | 2065 | 1796 | 1750 | 2018 | 1704 | 1604 | Bckdha |
| Bckdhb | 1751 | 1602 | 1795 | 1550 | 1268 | 1243 | 1532 | 1094 | 1089 | 1117 | 1103 | 1212 | 1222 | 1306 | 1714 | 1733 | 1615 | 1505 | 1382 | 1400 | 1392 | 1026 | 1028 | 1218 | 1122 | 1487 | 1302 | 1347 | Bckdhb |
| Bckdk | 2480 | 2763 | 2292 | 2065 | 1915 | 2187 | 1979 | 2549 | 3093 | 2448 | 2424 | 2993 | 2887 | 2845 | 2367 | 2424 | 2113 | 1921 | 2095 | 2294 | 2013 | 2283 | 2948 | 2709 | 2632 | 3608 | 3224 | 3038 | Bckdk |
| Bcl10 | 3079 | 3336 | 3176 | 3303 | 2335 | 2454 | 2842 | 1768 | 1759 | 1612 | 1674 | 1749 | 1804 | 1826 | 3112 | 3191 | 2967 | 3064 | 2682 | 2492 | 2600 | 1626 | 1752 | 1709 | 1671 | 2019 | 1836 | 2012 | Bcl10 |
| Bcl11a | 5783 | 5238 | 6471 | 8418 | 9568 | 8559 | 10042 | 11369 | 11957 | 13989 | 11659 | 13050 | 9740 | 14336 | 5964 | 5002 | 7778 | 7593 | 9354 | 7502 | 9492 | 12950 | 9897 | 15033 | 16872 | 12970 | 9939 | 11661 | Bcl11a |
| Bcl11b | 3849 | 2745 | 3264 | 6062 | 7743 | 6249 | 7881 | 8051 | 9628 | 10817 | 8400 | 8527 | 5116 | 9337 | 3432 | 2623 | 5251 | 6424 | 7921 | 4721 | 7687 | 9302 | 6695 | 12596 | 12871 | 9674 | 5877 | 6381 | Bcl11b |
| Bcl2 | 1668 | 1380 | 1762 | 1977 | 2728 | 2402 | 2340 | 2457 | 2888 | 3027 | 2930 | 2496 | 2588 | 2427 | 1665 | 1735 | 1829 | 2429 | 2438 | 1938 | 2490 | 3053 | 2944 | 2531 | 2990 | 2379 | 2487 | 2441 | Bcl2 |
| Bcl2a1a | 2 | 0 | 0 | 0 | 0 | 6 | 0 | 0 | 0 | 0 | 2 | 0 | 0 | 2 | 0 | 1 | 0 | 0 | 0 | 0 | 0 | 0 | 0 | 0 | 3 | 0 | 0 | 0 | Bcl2a1a |
| Bcl2a1b | 0 | 0 | 1 | 0 | 0 | 0 | 0 | 0 | 1 | 3 | 0 | 4 | 0 | 1 | 0 | 0 | 0 | 0 | 0 | 0 | 2 | 2 | 0 | 1 | 0 | 0 | 0 | 0 | Bcl2a1b |
| Bcl2a1d | 0 | 0 | 0 | 1 | 0 | 0 | 1 | 0 | 2 | 0 | 1 | 0 | 3 | 0 | 0 | 2 | 0 | 0 | 2 | 3 | 0 | 0 | 3 | 5 | 0 | 1 | 0 | 0 | Bcl2a1d |
| Bcl2l1 | 3102 | 3539 | 3265 | 3772 | 4637 | 3972 | 4098 | 6134 | 6625 | 6944 | 5746 | 7282 | 6509 | 7170 | 3169 | 3024 | 3712 | 3572 | 4622 | 3987 | 4024 | 7021 | 6188 | 7544 | 7362 | 7770 | 6517 | 7401 | Bcl2l1 |
| Bcl2l10 | 5 | 3 | 5 | 4 | 5 | 1 | 3 | 0 | 0 | 0 | 0 | 2 | 0 | 0 | 3 | 5 | 3 | 2 | 4 | 15 | 9 | 2 | 0 | 0 | 1 | 0 | 0 | 0 | Bcl2l10 |
| Bcl2l11 | 2731 | 2603 | 2898 | 3312 | 2813 | 2645 | 2496 | 1860 | 1932 | 1904 | 2252 | 2134 | 2339 | 2080 | 2604 | 3103 | 2848 | 3557 | 2544 | 2267 | 2727 | 2062 | 2153 | 1990 | 1814 | 2450 | 1941 | 2190 | Bcl2l11 |
| Bcl2l12 | 1188 | 1150 | 1516 | 1090 | 817 | 971 | 926 | 1233 | 1519 | 1348 | 1424 | 1731 | 1921 | 1798 | 984 | 1209 | 1167 | 872 | 1042 | 1018 | 810 | 1011 | 1354 | 1639 | 1292 | 2203 | 1972 | 1886 | Bcl2l12 |
| Bcl2l13 | 7356 | 7272 | 6259 | 6164 | 5758 | 6220 | 6150 | 6730 | 6406 | 5680 | 5916 | 7547 | 6777 | 7598 | 6949 | 6433 | 6288 | 5642 | 6254 | 5999 | 6026 | 5770 | 5956 | 6427 | 6541 | 6795 | 7661 | 7792 | Bcl2l13 |
| Bcl2l14 | 29 | 15 | 4 | 7 | 1 | 1 | 2 | 23 | 21 | 18 | 34 | 9 | 14 | 7 | 17 | 14 | 5 | 6 | 0 | 0 | 1 | 2 | 24 | 14 | 11 | 25 | 7 | 13 | Bcl2l14 |
| Bcl2l15 | 77 | 83 | 99 | 128 | 108 | 108 | 113 | 70 | 43 | 73 | 67 | 71 | 54 | 67 | 79 | 86 | 110 | 101 | 91 | 83 | 88 | 75 | 50 | 53 | 64 | 49 | 53 | 55 | Bcl2l15 |
| Bcl2l2 | 3167 | 3048 | 2824 | 3511 | 4061 | 3706 | 3511 | 3363 | 3058 | 3553 | 3604 | 3406 | 3487 | 3319 | 3191 | 2872 | 3373 | 3470 | 3883 | 3711 | 3790 | 3350 | 3158 | 3420 | 3744 | 2940 | 3581 | 3414 | Bcl2l2 |
| Bcl3 | 18 | 18 | 11 | 26 | 29 | 9 | 12 | 33 | 40 | 79 | 88 | 11 | 24 | 11 | 24 | 38 | 21 | 22 | 21 | 7 | 16 | 30 | 46 | 69 | 32 | 50 | 13 | 25 | Bcl3 |
| Bcl6 | 354 | 326 | 244 | 369 | 293 | 290 | 372 | 242 | 376 | 375 | 423 | 147 | 161 | 165 | 203 | 329 | 326 | 443 | 309 | 222 | 383 | 209 | 348 | 399 | 240 | 279 | 156 | 136 | Bcl6 |
| Bcl6b | 963 | 899 | 968 | 882 | 835 | 806 | 721 | 583 | 673 | 787 | 875 | 697 | 771 | 675 | 707 | 910 | 873 | 778 | 754 | 596 | 752 | 742 | 794 | 722 | 694 | 879 | 669 | 662 | Bcl6b |
| Bcl7a | 4014 | 4596 | 6379 | 6408 | 7801 | 6459 | 7030 | 9128 | 10043 | 12442 | 10716 | 13276 | 11535 | 13359 | 4184 | 4063 | 7204 | 6214 | 8044 | 5623 | 7340 | 10317 | 8470 | 13956 | 13701 | 12889 | 11443 | 12650 | Bcl7a |
| Bcl7b | 423 | 542 | 690 | 577 | 590 | 559 | 523 | 888 | 1133 | 1245 | 1064 | 1339 | 1186 | 1379 | 404 | 414 | 644 | 495 | 622 | 620 | 553 | 1121 | 1003 | 1316 | 1260 | 1450 | 1264 | 1307 | Bcl7b |
| Bcl7c | 5214 | 5641 | 6002 | 6208 | 7053 | 6147 | 6030 | 6174 | 6461 | 7536 | 7211 | 6631 | 7009 | 6615 | 5111 | 5280 | 6079 | 6081 | 6823 | 5957 | 6549 | 7386 | 6183 | 6657 | 7449 | 5933 | 6472 | 6286 | Bcl7c |
| Bcl9 | 6584 | 7478 | 9367 | 9899 | 11095 | 9544 | 8841 | 9966 | 11731 | 13940 | 12929 | 14787 | 13159 | 13686 | 6393 | 7035 | 10430 | 10000 | 11099 | 8073 | 9181 | 13709 | 10352 | 15371 | 14833 | 14462 | 12561 | 13016 | Bcl9 |
| Bcl9l | 5939 | 8329 | 8373 | 9941 | 8639 | 7944 | 6726 | 4532 | 6181 | 7126 | 6710 | 6521 | 5632 | 5359 | 5239 | 7588 | 8854 | 9918 | 8963 | 6308 | 7306 | 6786 | 5799 | 7194 | 6915 | 7064 | 5241 | 4891 | Bcl9l |
| Bclaf1 | 24568 | 18711 | 20791 | 20783 | 21781 | 23836 | 25636 | 38373 | 32470 | 30416 | 30964 | 35228 | 34551 | 36725 | 25648 | 20194 | 20614 | 20030 | 22343 | 27791 | 23792 | 30178 | 31153 | 32496 | 32955 | 32355 | 37058 | 35243 | Bclaf1 |
| Bclaf3 | 3390 | 2497 | 3135 | 3231 | 3370 | 3659 | 4053 | 3421 | 2860 | 2869 | 2851 | 3194 | 3314 | 3284 | 3629 | 2945 | 3131 | 3180 | 3293 | 3541 | 3477 | 3203 | 2856 | 2826 | 3163 | 3085 | 3277 | 3322 | Bclaf3 |
| Bco1 | 16 | 16 | 13 | 5 | 9 | 15 | 12 | 13 | 21 | 15 | 26 | 34 | 32 | 7 | 18 | 30 | 3 | 7 | 5 | 4 | 6 | 13 | 14 | 19 | 14 | 13 | 10 | 13 | Bco1 |
| Bco2 | 29 | 13 | 19 | 8 | 14 | 16 | 26 | 44 | 44 | 58 | 52 | 53 | 44 | 50 | 33 | 30 | 35 | 15 | 23 | 28 | 25 | 25 | 57 | 52 | 56 | 36 | 57 | 53 | Bco2 |
| Bcor | 6230 | 6574 | 6772 | 7751 | 8465 | 7900 | 7743 | 5693 | 6125 | 7223 | 6534 | 8389 | 7410 | 7711 | 6302 | 6385 | 7575 | 7246 | 8698 | 7060 | 7787 | 7386 | 5625 | 7443 | 7587 | 5939 | 7659 | 7685 | Bcor |
| Bcorl1 | 4565 | 5684 | 6813 | 7820 | 7435 | 6734 | 6089 | 2652 | 2957 | 3582 | 3648 | 3933 | 3517 | 3561 | 4089 | 5233 | 7428 | 7985 | 7790 | 4775 | 6853 | 3825 | 2893 | 3853 | 3842 | 3339 | 3284 | 3531 | Bcorl1 |

| | | | | | | | | | | | | | | | | | | | | | | | | | | | | | |
|---|---|---|---|---|---|---|---|---|---|---|---|---|---|---|---|---|---|---|---|---|---|---|---|---|---|---|---|---|---|
| Bcr | 7809 | 9322 | 10016 | 12754 | 13542 | 11168 | 10758 | 6450 | 7463 | 9862 | 8135 | 8509 | 7443 | 7717 | 7920 | 8793 | 11800 | 13132 | 13712 | 8800 | 11901 | 10009 | 6575 | 8907 | 9913 | 7082 | 6579 | 7294 | Bcr |
| Bcs1l | 1210 | 1424 | 1355 | 1445 | 1229 | 1194 | 1326 | 1169 | 1213 | 1321 | 1268 | 1257 | 1075 | 1212 | 1312 | 1301 | 1304 | 1395 | 1453 | 1306 | 1349 | 1353 | 1234 | 1395 | 1321 | 1315 | 1343 | 1247 | Bcs1l |
| Bdh1 | 1703 | 2046 | 2014 | 2117 | 2990 | 2522 | 2472 | 3362 | 3873 | 4330 | 3538 | 3346 | 2774 | 3385 | 1829 | 1755 | 2031 | 2216 | 2826 | 2571 | 2552 | 3935 | 3446 | 3982 | 4582 | 3205 | 2960 | 3153 | Bdh1 |
| Bdh2 | 209 | 135 | 127 | 102 | 101 | 113 | 95 | 236 | 307 | 252 | 269 | 190 | 186 | 162 | 175 | 166 | 109 | 109 | 82 | 123 | 88 | 100 | 292 | 228 | 143 | 316 | 182 | 145 | Bdh2 |
| Bdkrb1 | 1 | 2 | 0 | 0 | 0 | 0 | 0 | 1 | 1 | 3 | 0 | 0 | 0 | 0 | 0 | 0 | 2 | 0 | 1 | 0 | 0 | 2 | 2 | 0 | 0 | 0 | 0 | 0 | Bdkrb1 |
| Bdkrb2 | 263 | 282 | 289 | 203 | 248 | 223 | 132 | 145 | 213 | 157 | 209 | 64 | 185 | 67 | 280 | 339 | 228 | 245 | 169 | 152 | 136 | 201 | 185 | 83 | 128 | 76 | 92 | 89 | Bdkrb2 |
| Bdnf | 744 | 570 | 775 | 634 | 597 | 595 | 521 | 285 | 252 | 245 | 291 | 150 | 346 | 158 | 669 | 685 | 650 | 616 | 548 | 551 | 571 | 232 | 346 | 175 | 210 | 106 | 143 | 170 | Bdnf |
| Bdp1 | 8726 | 7605 | 4675 | 6894 | 5931 | 7473 | 7176 | 6633 | 5821 | 4859 | 5407 | 5340 | 5372 | 5873 | 8321 | 8139 | 4879 | 6254 | 5578 | 7986 | 7429 | 4672 | 5487 | 4088 | 5366 | 3385 | 6047 | 5028 | Bdp1 |
| Bean1 | 40 | 44 | 44 | 54 | 160 | 73 | 36 | 197 | 335 | 324 | 255 | 136 | 263 | 142 | 51 | 73 | 56 | 81 | 74 | 48 | 63 | 393 | 273 | 198 | 256 | 112 | 114 | 109 | Bean1 |
| Becn1 | 8349 | 8254 | 9051 | 8870 | 7860 | 7880 | 8503 | 7701 | 7053 | 7534 | 7234 | 8464 | 7700 | 8422 | 8331 | 7855 | 8528 | 8733 | 8117 | 7929 | 8675 | 8118 | 6852 | 7941 | 8451 | 8394 | 8481 | 8781 | Becn1 |
| Becn2 | 33 | 19 | 47 | 48 | 16 | 27 | 36 | 5 | 2 | 8 | 11 | 12 | 1 | 9 | 28 | 30 | 34 | 24 | 48 | 17 | 19 | 3 | 6 | 11 | 9 | 13 | 7 | 9 | Becn2 |
| Begain | 641 | 902 | 912 | 1242 | 1764 | 1325 | 1184 | 1362 | 1692 | 2002 | 1918 | 1396 | 1518 | 1329 | 803 | 800 | 1293 | 1445 | 1608 | 1143 | 1352 | 2262 | 1545 | 1722 | 1983 | 1321 | 1247 | 1218 | Begain |
| Bend3 | 634 | 593 | 898 | 810 | 1012 | 846 | 873 | 2100 | 2147 | 2992 | 2415 | 4015 | 3266 | 3472 | 669 | 600 | 1026 | 771 | 1029 | 700 | 834 | 2714 | 2073 | 3438 | 2986 | 3962 | 3579 | 3879 | Bend3 |
| Bend4 | 3519 | 4094 | 3842 | 3918 | 4125 | 3936 | 3587 | 2425 | 2043 | 2333 | 2510 | 2312 | 2262 | 2206 | 4000 | 4231 | 3999 | 4255 | 4034 | 3827 | 3947 | 2603 | 1976 | 2074 | 2539 | 1589 | 2539 | 2218 | Bend4 |
| Bend5 | 870 | 910 | 760 | 996 | 1222 | 988 | 988 | 1661 | 1990 | 1946 | 1829 | 1563 | 1682 | 1692 | 885 | 793 | 966 | 928 | 1101 | 982 | 1031 | 1920 | 1920 | 1948 | 1897 | 1620 | 1404 | 1502 | Bend5 |
| Bend6 | 431 | 416 | 417 | 590 | 1186 | 772 | 988 | 725 | 779 | 750 | 573 | 311 | 307 | 353 | 547 | 382 | 425 | 774 | 849 | 973 | 1113 | 760 | 779 | 455 | 700 | 288 | 278 | 241 | Bend6 |
| Bend7 | 422 | 460 | 644 | 523 | 650 | 486 | 582 | 579 | 412 | 439 | 405 | 443 | 509 | 443 | 485 | 344 | 532 | 469 | 658 | 510 | 570 | 565 | 390 | 442 | 444 | 420 | 480 | 452 | Bend7 |
| Best1 | 60 | 59 | 79 | 44 | 35 | 57 | 38 | 90 | 101 | 119 | 108 | 93 | 94 | 79 | 53 | 64 | 75 | 70 | 51 | 64 | 37 | 87 | 111 | 74 | 79 | 103 | 138 | 91 | Best1 |
| Best2 | 5 | 9 | 5 | 13 | 8 | 4 | 6 | 2 | 12 | 5 | 3 | 4 | 0 | 2 | 8 | 9 | 12 | 17 | 5 | 10 | 13 | 3 | 9 | 10 | 1 | 0 | 5 | 2 | Best2 |
| Best3 | 13 | 3 | 17 | 4 | 16 | 13 | 6 | 16 | 24 | 14 | 130 | 10 | 6 | 24 | 7 | 10 | 10 | 8 | 9 | 4 | 3 | 33 | 60 | 8 | 16 | 10 | 10 | 11 | Best3 |
| Bet1 | 3573 | 2701 | 3565 | 2999 | 2453 | 2525 | 2978 | 3041 | 2749 | 2493 | 2891 | 2802 | 2919 | 3007 | 3603 | 3572 | 3269 | 2956 | 2519 | 3358 | 2734 | 2533 | 2972 | 2849 | 2328 | 3055 | 2853 | 3000 | Bet1 |
| Bet1l | 736 | 856 | 934 | 833 | 748 | 780 | 738 | 865 | 996 | 907 | 1037 | 836 | 847 | 760 | 737 | 853 | 787 | 768 | 775 | 867 | 752 | 827 | 1035 | 838 | 861 | 871 | 862 | 826 | Bet1l |
| Bex1 | 3218 | 4098 | 4547 | 4712 | 4279 | 3827 | 4672 | 2395 | 2473 | 3363 | 3087 | 2942 | 2719 | 3549 | 3360 | 3006 | 4591 | 4196 | 4417 | 4825 | 4444 | 2247 | 2634 | 3375 | 3333 | 3384 | 2765 | 3538 | Bex1 |
| Bex2 | 3985 | 4790 | 6044 | 6611 | 6662 | 5808 | 7013 | 5678 | 5496 | 7664 | 6055 | 7086 | 6008 | 8689 | 4208 | 3624 | 6567 | 5874 | 6798 | 6608 | 6874 | 6171 | 5093 | 8216 | 8325 | 7997 | 6992 | 7716 | Bex2 |
| Bex3 | 9718 | 11355 | 9383 | 10033 | 8614 | 8588 | 9379 | 9255 | 8669 | 9625 | 7950 | 9105 | 9170 | 9915 | 9923 | 8479 | 10022 | 8647 | 9510 | 10316 | 9499 | 8251 | 8818 | 9597 | 9112 | 10791 | 9538 | 10460 | Bex3 |
| Bex4 | 3663 | 3484 | 3935 | 3806 | 3308 | 2804 | 4278 | 2328 | 2126 | 2456 | 1961 | 2338 | 2291 | 3179 | 3546 | 3009 | 3452 | 3478 | 3428 | 4012 | 3909 | 1973 | 2030 | 2665 | 2377 | 3252 | 2547 | 2932 | Bex4 |
| Bex6 | 2 | 1 | 0 | 0 | 0 | 0 | 0 | 0 | 0 | 0 | 10 | 4 | 1 | 1 | 0 | 1 | 0 | 0 | 0 | 0 | 0 | 0 | 2 | 0 | 0 | 1 | 2 | 0 | Bex6 |
| Bfar | 3442 | 3531 | 3637 | 3295 | 2944 | 3303 | 3303 | 3949 | 3395 | 3498 | 3681 | 3480 | 3502 | 3498 | 3542 | 3301 | 3646 | 3232 | 3138 | 3500 | 3022 | 3557 | 3694 | 3739 | 4090 | 3795 | 3536 | 3489 | Bfar |
| Bfsp1 | 42 | 32 | 21 | 27 | 37 | 25 | 16 | 94 | 134 | 133 | 153 | 192 | 105 | 163 | 28 | 44 | 53 | 21 | 17 | 31 | 30 | 155 | 152 | 111 | 135 | 130 | 164 | 153 | Bfsp1 |
| Bfsp2 | 32 | 12 | 29 | 34 | 24 | 22 | 15 | 96 | 76 | 78 | 67 | 91 | 75 | 80 | 30 | 28 | 38 | 26 | 23 | 48 | 18 | 32 | 94 | 132 | 88 | 103 | 72 | 76 | Bfsp2 |
| Bglap | 32 | 31 | 24 | 38 | 26 | 21 | 34 | 18 | 26 | 13 | 6 | 28 | 25 | 32 | 13 | 18 | 16 | 33 | 36 | 22 | 34 | 13 | 16 | 20 | 29 | 38 | 35 | 18 | Bglap |
| Bglap2 | 2 | 3 | 1 | 8 | 0 | 6 | 3 | 1 | 2 | 0 | 2 | 1 | 1 | 1 | 3 | 3 | 0 | 0 | 1 | 2 | 4 | 0 | 2 | 2 | 1 | 4 | 0 | 6 | Bglap2 |
| Bglap3 | 2 | 2 | 8 | 1 | 2 | 8 | 1 | 2 | 5 | 8 | 3 | 8 | 0 | 2 | 1 | 4 | 0 | 3 | 1 | 0 | 0 | 5 | 1 | 6 | 3 | 2 | 3 | 2 | Bglap3 |
| Bgn | 5617 | 6863 | 6658 | 5663 | 3750 | 6852 | 2942 | 6243 | 8463 | 8048 | 11257 | 6858 | 9308 | 6036 | 6261 | 8051 | 5475 | 6934 | 3665 | 3469 | 3181 | 8644 | 10974 | 7891 | 7243 | 7754 | 7403 | 6978 | Bgn |
| Bhlha15 | 21 | 30 | 12 | 18 | 16 | 30 | 16 | 11 | 11 | 29 | 21 | 9 | 3 | 3 | 15 | 19 | 14 | 21 | 16 | 16 | 14 | 13 | 6 | 11 | 10 | 20 | 13 | 17 | Bhlha15 |
| Bhlha9 | 1 | 7 | 1 | 5 | 6 | 4 | 3 | 6 | 5 | 7 | 3 | 2 | 1 | 0 | 4 | 2 | 12 | 10 | 3 | 8 | 4 | 2 | 3 | 3 | 1 | 1 | 13 | 0 | Bhlha9 |
| Bhlhb9 | 7951 | 6896 | 7747 | 7373 | 9011 | 8720 | 9832 | 10794 | 9489 | 9374 | 8699 | 9558 | 8245 | 9887 | 8681 | 6442 | 7880 | 7791 | 8642 | 9443 | 9094 | 10305 | 9532 | 9560 | 9754 | 9872 | 9254 | 9311 | Bhlhb9 |
| Bhlhe22 | 348 | 275 | 326 | 772 | 1838 | 1225 | 1740 | 2578 | 2106 | 2616 | 2017 | 2064 | 1472 | 2337 | 543 | 227 | 644 | 917 | 1956 | 1453 | 1982 | 2079 | 2240 | 2822 | 3609 | 1825 | 1395 | 1794 | Bhlhe22 |
| Bhlhe23 | 173 | 343 | 512 | 304 | 252 | 247 | 304 | 215 | 68 | 227 | 165 | 457 | 313 | 416 | 260 | 275 | 418 | 231 | 321 | 211 | 322 | 317 | 167 | 280 | 294 | 154 | 459 | 540 | Bhlhe23 |
| Bhlhe40 | 400 | 358 | 312 | 278 | 456 | 425 | 432 | 423 | 608 | 493 | 654 | 548 | 501 | 612 | 376 | 312 | 396 | 445 | 548 | 440 | 397 | 528 | 659 | 596 | 643 | 664 | 507 | 614 | Bhlhe40 |
| Bhlhe41 | 362 | 462 | 508 | 533 | 453 | 407 | 375 | 206 | 241 | 259 | 349 | 199 | 184 | 154 | 439 | 438 | 612 | 533 | 418 | 346 | 413 | 301 | 343 | 232 | 230 | 204 | 158 | 183 | Bhlhe41 |
| Bhmt | 5 | 15 | 0 | 4 | 11 | 11 | 1 | 26 | 21 | 6 | 11 | 12 | 27 | 1 | 1 | 15 | 6 | 3 | 4 | 7 | 4 | 22 | 35 | 6 | 13 | 5 | 8 | 5 | Bhmt |
| Bhmt2 | 2 | 6 | 0 | 3 | 6 | 1 | 2 | 0 | 8 | 3 | 2 | 10 | 5 | 3 | 4 | 3 | 2 | 1 | 0 | 0 | 4 | 3 | 6 | 0 | 0 | 1 | 2 | 0 | Bhmt2 |
| Bicc1 | 3376 | 3156 | 3267 | 2474 | 1834 | 2643 | 1253 | 2749 | 3774 | 3573 | 5526 | 3106 | 3694 | 2901 | 3320 | 4187 | 2561 | 3028 | 1444 | 1552 | 1375 | 3262 | 4804 | 3434 | 2913 | 2832 | 3477 | 2905 | Bicc1 |
| Bicd1 | 2541 | 2045 | 2420 | 3714 | 4777 | 4434 | 4879 | 4296 | 3409 | 3540 | 3538 | 3667 | 3433 | 4035 | 3264 | 2220 | 2885 | 3572 | 4154 | 4548 | 5010 | 4284 | 3079 | 3265 | 4553 | 2500 | 3543 | 3624 | Bicd1 |
| Bicd2 | 7792 | 9078 | 7898 | 8467 | 8885 | 8220 | 8168 | 7427 | 7180 | 7117 | 7329 | 7519 | 7911 | 7804 | 8257 | 8634 | 8503 | 8623 | 8722 | 8263 | 8634 | 7681 | 6530 | 6920 | 7820 | 6293 | 7594 | 7517 | Bicd2 |
| Bicdl1 | 1356 | 2024 | 1923 | 2317 | 2746 | 1978 | 2202 | 1628 | 1636 | 2039 | 1852 | 1581 | 1461 | 1485 | 1551 | 1597 | 2371 | 2389 | 2471 | 2036 | 2357 | 2378 | 1704 | 1718 | 2234 | 1311 | 1429 | 1444 | Bicdl1 |
| Bicdl2 | 43 | 33 | 20 | 27 | 14 | 8 | 3 | 34 | 40 | 29 | 29 | 9 | 11 | 7 | 41 | 52 | 10 | 39 | 7 | 2 | 1 | 5 | 35 | 30 | 9 | 11 | 7 | 7 | Bicdl2 |
| Bicra | 1985 | 2685 | 3126 | 3264 | 3067 | 2779 | 2247 | 2484 | 2768 | 3574 | 2992 | 4012 | 2935 | 3019 | 1826 | 2182 | 3150 | 3091 | 3250 | 2039 | 2666 | 3708 | 2317 | 3746 | 3574 | 3522 | 3099 | 2893 | Bicra |
| Bicral | 1637 | 1692 | 2198 | 2419 | 2855 | 2319 | 2600 | 2372 | 2656 | 3317 | 3091 | 2840 | 2739 | 2869 | 1642 | 1712 | 2496 | 2585 | 3010 | 1971 | 2586 | 3507 | 2382 | 3245 | 3463 | 2145 | 2295 | 2820 | Bicral |
| Bid | 2043 | 2236 | 1945 | 1995 | 1891 | 1922 | 1766 | 768 | 844 | 772 | 674 | 627 | 685 | 635 | 2035 | 2110 | 1840 | 1808 | 1997 | 2173 | 1759 | 789 | 755 | 626 | 730 | 651 | 693 | 560 | Bid |
| Bik | 171 | 215 | 217 | 167 | 140 | 92 | 130 | 28 | 26 | 19 | 29 | 21 | 27 | 33 | 144 | 168 | 183 | 122 | 143 | 147 | 115 | 18 | 41 | 40 | 23 | 61 | 44 | 50 | Bik |
| Bin1 | 3746 | 4368 | 4523 | 4443 | 5918 | 5007 | 4210 | 5820 | 6626 | 7001 | 8243 | 5924 | 6157 | 5546 | 3930 | 4223 | 4694 | 4870 | 5368 | 4654 | 4775 | 7441 | 7223 | 6339 | 6953 | 4796 | 5487 | 5365 | Bin1 |
| Bin2 | 99 | 139 | 199 | 89 | 79 | 72 | 96 | 100 | 120 | 135 | 126 | 139 | 170 | 101 | 92 | 186 | 105 | 84 | 77 | 88 | 63 | 132 | 145 | 125 | 73 | 224 | 146 | 112 | Bin2 |
| Bin3 | 2354 | 2631 | 2027 | 2054 | 1779 | 1978 | 1911 | 1337 | 1488 | 1252 | 1141 | 1413 | 1464 | 1462 | 2312 | 2444 | 2013 | 2192 | 1772 | 2165 | 1873 | 1208 | 1510 | 1377 | 1359 | 1489 | 1668 | 1538 | Bin3 |
| Birc2 | 4864 | 3930 | 3932 | 4504 | 4889 | 4732 | 5003 | 6920 | 5891 | 5063 | 5271 | 5418 | 5037 | 5427 | 5091 | 4379 | 4354 | 4358 | 4581 | 5505 | 4861 | 5282 | 5746 | 5240 | 5347 | 5086 | 5291 | 5022 | Birc2 |
| Birc3 | 211 | 143 | 163 | 71 | 84 | 153 | 54 | 391 | 518 | 366 | 555 | 300 | 452 | 358 | 187 | 171 | 144 | 94 | 122 | 103 | 57 | 338 | 572 | 359 | 261 | 330 | 374 | 360 | Birc3 |
| Birc5 | 4128 | 3728 | 4248 | 2918 | 2162 | 2706 | 3024 | 3607 | 3564 | 3191 | 3012 | 4810 | 4400 | 5163 | 3724 | 3613 | 3698 | 2744 | 2833 | 3053 | 2707 | 2381 | 3165 | 4256 | 3290 | 6386 | 5298 | 5585 | Birc5 |
| Birc6 | 25230 | 23301 | 10421 | 13537 | 14388 | 21014 | 18676 | 24892 | 23322 | 13594 | 16292 | 14524 | 15863 | 16182 | 24031 | 22513 | 11946 | 13458 | 13585 | 23582 | 21498 | 13017 | 21055 | 12193 | 15529 | 9669 | 18661 | 14572 | Birc6 |
| Birc7 | 1 | 6 | 0 | 3 | 2 | 10 | 2 | 5 | 7 | 0 | 6 | 13 | 10 | 3 | 4 | 4 | 0 | 2 | 0 | 0 | 4 | 2 | 6 | 7 | 6 | 6 | 14 | 6 | Birc7 |
| Bivm | 2458 | 1999 | 2182 | 1966 | 2500 | 2114 | 2319 | 4185 | 3665 | 3562 | 3331 | 3602 | 3407 | 3408 | 2545 | 2069 | 2312 | 2043 | 2325 | 2374 | 2382 | 3885 | 3468 | 3384 | 3568 | 3480 | 3452 | 3666 | Bivm |
| Blcap | 7307 | 8556 | 7182 | 8213 | 8419 | 8140 | 8206 | 7503 | 7419 | 6979 | 6553 | 7217 | 5849 | 6846 | 7568 | 7059 | 8194 | 8516 | 8690 | 8774 | 8698 | 8779 | 6870 | 7234 | 8261 | 7124 | 6893 | 6813 | Blcap |
| Blk | 0 | 0 | 0 | 0 | 0 | 0 | 0 | 0 | 0 | 0 | 0 | 0 | 0 | 0 | 0 | 1 | 0 | 0 | 0 | 0 | 1 | 0 | 1 | 0 | 0 | 0 | 1 | 0 | Blk |
| Blm | 7353 | 6023 | 5762 | 5071 | 4298 | 5311 | 6024 | 4534 | 4191 | 3518 | 3591 | 5534 | 4887 | 5806 | 6931 | 6311 | 5691 | 4863 | 4733 | 6005 | 5445 | 2887 | 3624 | 4101 | 3933 | 5101 | 5851 | 5705 | Blm |
| Blmh | 12156 | 11790 | 13260 | 10325 | 9337 | 9887 | 9708 | 11748 | 10542 | 10252 | 10043 | 11704 | 11614 | 11471 | 12812 | 11915 | 11931 | 9573 | 9688 | 10926 | 9499 | 9648 | 10840 | 10403 | 10222 | 13167 | 12279 | 12250 | Blmh |
| Blnk | 48 | 23 | 3 | 53 | 10 | 12 | 7 | 68 | 44 | 99 | 59 | 14 | 24 | 18 | 33 | 32 | 6 | 9 | 5 | 6 | 10 | 30 | 63 | 85 | 10 | 103 | 12 | 13 | Blnk |
| Bloc1s1 | 28 | 38 | 27 | 21 | 22 | 14 | 23 | 20 | 21 | 33 | 29 | 25 | 19 | 36 | 15 | 16 | 49 | 22 | 22 | 31 | 25 | 37 | 33 | 47 | 27 | 21 | 20 | 40 | Bloc1s1 |
| Bloc1s2 | 1835 | 1958 | 2030 | 1967 | 1982 | 1821 | 2233 | 1934 | 1932 | 1930 | 1790 | 2014 | 1792 | 2039 | 1954 | 1833 | 1893 | 1842 | 2162 | 2173 | 1960 | 2039 | 1976 | 1963 | 2078 | 1941 | 2023 | 2077 | Bloc1s2 |
| Bloc1s3 | 659 | 621 | 601 | 603 | 587 | 606 | 590 | 683 | 673 | 653 | 628 | 688 | 691 | 722 | 549 | 589 | 695 | 614 | 748 | 598 | 617 | 688 | 695 | 752 | 754 | 811 | 669 | 686 | Bloc1s3 |
| Bloc1s4 | 540 | 541 | 537 | 610 | 524 | 504 | 527 | 477 | 453 | 535 | 524 | 480 | 442 | 442 | 555 | 534 | 588 | 546 | 493 | 489 | 478 | 461 | 458 | 574 | 377 | 546 | 496 | 443 | Bloc1s4 |
| Bloc1s5 | 1651 | 1735 | 1522 | 1566 | 1464 | 1416 | 1591 | 1796 | 1692 | 1559 | 1521 | 1647 | 1703 | 1850 | 1739 | 1629 | 1524 | 1486 | 1492 | 1533 | 1550 | 1589 | 1689 | 1681 | 1695 | 1802 | 1712 | 1887 | Bloc1s5 |
| Bloc1s6 | 2873 | 2972 | 2919 | 2566 | 2737 | 2714 | 3115 | 4200 | 3474 | 3424 | 3287 | 3920 | 3797 | 4393 | 2997 | 2681 | 2917 | 2569 | 2604 | 2964 | 2755 | 4040 | 3546 | 3745 | 3820 | 4146 | 4076 | 4126 | Bloc1s6 |
| Blvra | 489 | 452 | 451 | 416 | 540 | 500 | 480 | 692 | 838 | 767 | 758 | 627 | 692 | 670 | 546 | 466 | 442 | 523 | 486 | 510 | 533 | 712 | 878 | 712 | 591 | 721 | 674 | 643 | Blvra |
| Blvrb | 1107 | 1354 | 1169 | 1119 | 722 | 823 | 657 | 388 | 623 | 436 | 454 | 369 | 727 | 470 | 922 | 1167 | 955 | 978 | 797 | 935 | 630 | 336 | 657 | 434 | 334 | 479 | 540 | 667 | Blvrb |
| Blzf1 | 2228 | 2096 | 2045 | 2072 | 1967 | 1914 | 2056 | 2474 | 2146 | 2254 | 2181 | 2182 | 2248 | 2351 | 2374 | 2187 | 2028 | 1918 | 1750 | 2157 | 1885 | 2276 | 2206 | 2132 | 2211 | 2156 | 2335 | 2391 | Blzf1 |
| Bmerb1 | 1589 | 2085 | 1499 | 2687 | 4118 | 3136 | 3399 | 4399 | 4200 | 4458 | 3749 | 3418 | 3045 | 3590 | 2142 | 1530 | 2090 | 2992 | 3472 | 3461 | 3476 | 5247 | 3827 | 3837 | 4692 | 2849 | 3095 | 3145 | Bmerb1 |
| Bmf | 7957 | 9284 | 9933 | 9937 | 6524 | 7517 | 6909 | 2574 | 3258 | 3168 | 3925 | 2374 | 3616 | 2489 | 7766 | 9623 | 9247 | 8162 | 8112 | 6121 | 7766 | 3053 | 3346 | 3101 | 2408 | 3460 | 2216 | 2613 | Bmf |
| Bmi1 | 4628 | 3556 | 4748 | 4451 | 5003 | 4675 | 5871 | 11604 | 9275 | 9141 | 8686 | 10262 | 8902 | 10476 | 4723 | 3755 | 4618 | 4392 | 5312 | 5229 | 5484 | 10400 | 9196 | 10096 | 10735 | 10422 | 9770 | 10270 | Bmi1 |
| Bmp1 | 5043 | 4991 | 4838 | 4309 | 4903 | 4993 | 3104 | 6829 | 8856 | 9080 | 10905 | 7428 | 8597 | 6220 | 4650 | 6261 | 4539 | 4854 | 4454 | 3466 | 3310 | 8109 | 9057 | 8004 | 7366 | 7596 | 7102 | 6488 | Bmp1 |

| | | | | | | | | | | | | | | | | | | | | | | | | | | | | | |
|---|---|---|---|---|---|---|---|---|---|---|---|---|---|---|---|---|---|---|---|---|---|---|---|---|---|---|---|---|---|
| Bmp10 | 0 | 0 | 0 | 0 | 0 | 0 | 0 | 0 | 1 | 0 | 0 | 0 | 0 | 0 | 0 | 1 | 0 | 0 | 0 | 0 | 0 | 0 | 3 | 1 | 0 | 0 | 0 | 0 | Bmp10 |
| Bmp15 | 7 | 3 | 18 | 1 | 11 | 10 | 11 | 17 | 19 | 8 | 19 | 17 | 31 | 26 | 7 | 16 | 2 | 12 | 2 | 7 | 5 | 28 | 22 | 13 | 19 | 13 | 21 | 24 | Bmp15 |
| Bmp2 | 441 | 229 | 258 | 242 | 172 | 231 | 144 | 388 | 443 | 381 | 562 | 267 | 311 | 209 | 257 | 293 | 226 | 269 | 136 | 119 | 148 | 426 | 498 | 400 | 251 | 332 | 265 | 250 | Bmp2 |
| Bmp2k | 3065 | 2732 | 2881 | 3204 | 2976 | 2971 | 3085 | 1757 | 1963 | 1830 | 2004 | 1800 | 2099 | 2060 | 3128 | 3020 | 3096 | 3081 | 2829 | 2826 | 3075 | 1945 | 1948 | 1781 | 1967 | 1654 | 1977 | 2293 | Bmp2k |
| Bmp3 | 351 | 207 | 212 | 269 | 320 | 291 | 292 | 829 | 984 | 601 | 851 | 318 | 381 | 343 | 256 | 244 | 173 | 284 | 244 | 250 | 276 | 690 | 954 | 591 | 562 | 378 | 311 | 252 | Bmp3 |
| Bmp4 | 1021 | 1044 | 1050 | 599 | 331 | 541 | 265 | 818 | 839 | 727 | 790 | 555 | 814 | 488 | 725 | 1419 | 707 | 661 | 344 | 352 | 268 | 495 | 1070 | 873 | 440 | 1130 | 723 | 655 | Bmp4 |
| Bmp5 | 1427 | 977 | 1415 | 752 | 722 | 913 | 672 | 800 | 1114 | 860 | 1691 | 630 | 879 | 663 | 1369 | 1706 | 909 | 749 | 634 | 912 | 680 | 971 | 1492 | 778 | 653 | 792 | 625 | 667 | Bmp5 |
| Bmp6 | 742 | 774 | 1019 | 648 | 794 | 619 | 486 | 519 | 630 | 686 | 778 | 481 | 642 | 397 | 800 | 861 | 731 | 759 | 641 | 667 | 566 | 675 | 962 | 521 | 561 | 524 | 475 | 502 | Bmp6 |
| Bmp7 | 2450 | 2089 | 2666 | 1831 | 1483 | 1708 | 1249 | 1998 | 2551 | 2238 | 2220 | 2207 | 2578 | 1794 | 1800 | 2764 | 1730 | 1845 | 1515 | 1664 | 1307 | 1796 | 2972 | 2355 | 1617 | 2303 | 2265 | 2129 | Bmp7 |
| Bmp8a | 81 | 130 | 118 | 95 | 56 | 55 | 48 | 17 | 10 | 7 | 6 | 7 | 10 | 10 | 98 | 118 | 96 | 127 | 79 | 67 | 72 | 2 | 16 | 9 | 5 | 4 | 5 | 2 | Bmp8a |
| Bmp8b | 106 | 77 | 71 | 84 | 58 | 62 | 68 | 50 | 74 | 45 | 33 | 20 | 16 | 32 | 92 | 121 | 45 | 62 | 46 | 56 | 53 | 32 | 43 | 22 | 10 | 13 | 19 | 17 | Bmp8b |
| Bmper | 659 | 530 | 292 | 550 | 720 | 646 | 656 | 1299 | 1819 | 1431 | 1770 | 1222 | 781 | 1261 | 568 | 426 | 523 | 665 | 726 | 745 | 722 | 1823 | 1427 | 1548 | 1391 | 1634 | 1006 | 876 | Bmper |
| Bmpr1a | 15050 | 10284 | 9454 | 9133 | 8980 | 10034 | 10503 | 14124 | 13035 | 9862 | 11618 | 11034 | 11609 | 11425 | 13653 | 11875 | 8960 | 9428 | 9054 | 11077 | 9553 | 9655 | 13140 | 10465 | 9310 | 11500 | 11354 | 12113 | Bmpr1a |
| Bmpr1b | 1052 | 845 | 666 | 754 | 728 | 808 | 839 | 1540 | 1474 | 1412 | 1784 | 1510 | 1660 | 1665 | 963 | 992 | 787 | 724 | 754 | 816 | 795 | 1031 | 1515 | 1658 | 1003 | 1851 | 1564 | 1337 | Bmpr1b |
| Bmpr2 | 12664 | 9413 | 9337 | 12129 | 14080 | 15719 | 15539 | 16004 | 14660 | 13448 | 15442 | 14062 | 15485 | 15398 | 12920 | 11662 | 10050 | 12545 | 12828 | 12689 | 15931 | 14952 | 14019 | 12684 | 15918 | 11218 | 15402 | 14878 | Bmpr2 |
| Bms1 | 8409 | 7789 | 9034 | 7718 | 6596 | 6759 | 7059 | 7601 | 7680 | 7946 | 7906 | 9097 | 8245 | 9105 | 7544 | 8090 | 8028 | 6957 | 6756 | 6945 | 6612 | 7005 | 7187 | 8208 | 7786 | 8856 | 8906 | 8548 | Bms1 |
| Bmt2 | 6485 | 5072 | 6083 | 6381 | 5617 | 6020 | 6911 | 4266 | 3746 | 3184 | 3362 | 3681 | 3582 | 3863 | 6587 | 5572 | 5862 | 6206 | 6192 | 5886 | 7038 | 4042 | 3698 | 3623 | 3747 | 3718 | 3719 | 3935 | Bmt2 |
| Bmx | 15 | 16 | 24 | 19 | 31 | 32 | 18 | 48 | 58 | 37 | 54 | 84 | 77 | 62 | 32 | 20 | 22 | 14 | 19 | 23 | 24 | 87 | 87 | 66 | 71 | 80 | 76 | 65 | Bmx |
| Bmyc | 622 | 709 | 747 | 731 | 728 | 618 | 618 | 677 | 680 | 750 | 769 | 645 | 665 | 666 | 560 | 678 | 757 | 715 | 615 | 598 | 604 | 719 | 719 | 769 | 656 | 782 | 583 | 642 | Bmyc |
| Bnc1 | 165 | 144 | 168 | 54 | 98 | 67 | 30 | 193 | 293 | 142 | 296 | 93 | 286 | 82 | 120 | 229 | 81 | 135 | 30 | 32 | 20 | 180 | 220 | 123 | 108 | 92 | 53 | 102 | Bnc1 |
| Bnc2 | 629 | 616 | 215 | 269 | 434 | 588 | 180 | 1362 | 1771 | 1286 | 1929 | 803 | 1757 | 813 | 541 | 815 | 194 | 305 | 224 | 484 | 349 | 901 | 1763 | 619 | 1121 | 352 | 1335 | 714 | Bnc2 |
| Bnip1 | 1337 | 1261 | 1095 | 1444 | 1493 | 1416 | 1406 | 1846 | 1931 | 2001 | 1991 | 2027 | 2088 | 2183 | 1386 | 1164 | 1236 | 1312 | 1477 | 1418 | 1398 | 2075 | 1900 | 1968 | 2040 | 1963 | 2260 | 2092 | Bnip1 |
| Bnip2 | 8222 | 7228 | 7627 | 6777 | 7325 | 7715 | 7785 | 8899 | 7559 | 7066 | 7736 | 8374 | 7743 | 8779 | 8140 | 7506 | 7582 | 6919 | 7136 | 8298 | 7750 | 7322 | 7722 | 7530 | 7748 | 7912 | 8839 | 8603 | Bnip2 |
| Bnip3 | 6590 | 5031 | 4957 | 5031 | 4635 | 4957 | 5204 | 3289 | 3108 | 3123 | 3118 | 2210 | 2851 | 2661 | 6878 | 5998 | 5129 | 5037 | 4641 | 5741 | 4711 | 2760 | 3543 | 2705 | 2556 | 2656 | 2432 | 2624 | Bnip3 |
| Bnip3l | 15132 | 12696 | 12804 | 13090 | 13061 | 13510 | 15672 | 14972 | 13182 | 12130 | 11493 | 12182 | 13113 | 13148 | 16200 | 13064 | 13428 | 13304 | 13532 | 15946 | 14765 | 12861 | 13804 | 12012 | 13075 | 12685 | 12829 | 13680 | Bnip3l |
| Bnip5 | 31 | 33 | 75 | 31 | 35 | 48 | 38 | 35 | 44 | 42 | 25 | 34 | 33 | 18 | 50 | 69 | 46 | 51 | 49 | 62 | 38 | 48 | 43 | 25 | 41 | 18 | 33 | 18 | Bnip5 |
| Bnipl | 58 | 51 | 39 | 48 | 20 | 26 | 31 | 111 | 131 | 117 | 95 | 54 | 66 | 32 | 21 | 32 | 42 | 29 | 20 | 35 | 24 | 43 | 170 | 109 | 24 | 92 | 26 | 49 | Bnipl |
| Boc | 2831 | 3220 | 3818 | 3342 | 3076 | 3317 | 2421 | 4347 | 5081 | 5200 | 5303 | 5905 | 6366 | 4987 | 2656 | 3061 | 3692 | 2797 | 3813 | 2724 | 2627 | 4024 | 5288 | 5511 | 4823 | 4819 | 5617 | 5547 | Boc |
| Bod1 | 6428 | 7622 | 6900 | 6696 | 6144 | 5982 | 6317 | 5769 | 5338 | 5194 | 4987 | 6095 | 5645 | 5885 | 6783 | 6108 | 6815 | 6342 | 6831 | 6923 | 6395 | 5041 | 5439 | 5875 | 5757 | 6730 | 6115 | 6000 | Bod1 |
| Bod1l | 13339 | 12127 | 7899 | 10767 | 11093 | 13331 | 11852 | 12016 | 10513 | 8769 | 9271 | 8807 | 9064 | 9104 | 13837 | 13009 | 8962 | 10051 | 10137 | 13496 | 12583 | 8126 | 9939 | 7339 | 8993 | 5542 | 9508 | 7913 | Bod1l |
| Bok | 508 | 523 | 603 | 517 | 781 | 851 | 629 | 1155 | 1277 | 1294 | 1101 | 1518 | 1213 | 1185 | 558 | 477 | 584 | 610 | 808 | 881 | 583 | 1125 | 1294 | 1409 | 1451 | 1396 | 1374 | 1235 | Bok |
| Bola1 | 483 | 721 | 640 | 553 | 574 | 626 | 548 | 727 | 828 | 713 | 737 | 867 | 821 | 899 | 559 | 476 | 637 | 574 | 743 | 651 | 547 | 481 | 681 | 826 | 842 | 1099 | 1052 | 916 | Bola1 |
| Bola2 | 2519 | 3509 | 3549 | 3602 | 2712 | 2114 | 3529 | 1680 | 1560 | 1983 | 1845 | 1991 | 1898 | 2545 | 2600 | 2155 | 3545 | 3044 | 2809 | 2957 | 3356 | 1328 | 1470 | 2312 | 1909 | 2246 | 1928 | 2652 | Bola2 |
| Bola3 | 608 | 619 | 562 | 536 | 371 | 411 | 485 | 564 | 558 | 591 | 514 | 654 | 603 | 817 | 553 | 508 | 564 | 406 | 414 | 494 | 450 | 520 | 568 | 711 | 626 | 887 | 737 | 860 | Bola3 |
| Boll | 17 | 9 | 20 | 18 | 13 | 14 | 9 | 12 | 7 | 8 | 10 | 4 | 7 | 10 | 25 | 13 | 19 | 7 | 5 | 15 | 9 | 12 | 8 | 3 | 5 | 10 | 10 | 5 | Boll |
| Bop1 | 6245 | 8466 | 6508 | 6875 | 4997 | 5402 | 5455 | 2997 | 3447 | 3037 | 2996 | 3541 | 3328 | 3036 | 5763 | 6948 | 6582 | 6249 | 6399 | 5584 | 5704 | 2896 | 3300 | 3360 | 3342 | 3933 | 3297 | 3356 | Bop1 |
| Bora | 5575 | 5021 | 5186 | 5225 | 3428 | 4496 | 5082 | 2267 | 2343 | 2203 | 2262 | 3419 | 3070 | 3457 | 5338 | 4995 | 4815 | 4757 | 4007 | 4319 | 4563 | 1855 | 2078 | 2887 | 2392 | 3613 | 3519 | 3447 | Bora |
| Borcs5 | 957 | 1111 | 1116 | 1192 | 1071 | 995 | 1224 | 753 | 836 | 873 | 790 | 859 | 819 | 889 | 897 | 913 | 1185 | 1021 | 1253 | 966 | 1029 | 876 | 755 | 914 | 893 | 968 | 876 | 821 | Borcs5 |
| Borcs6 | 1126 | 1593 | 1362 | 1651 | 1521 | 1366 | 1144 | 603 | 664 | 633 | 651 | 591 | 595 | 546 | 1141 | 1336 | 1548 | 1613 | 1518 | 1239 | 1320 | 675 | 719 | 588 | 674 | 575 | 605 | 591 | Borcs6 |
| Borcs7 | 234 | 232 | 359 | 291 | 326 | 299 | 315 | 988 | 1068 | 1281 | 1159 | 1344 | 1306 | 1361 | 236 | 276 | 302 | 283 | 314 | 289 | 296 | 1213 | 1075 | 1168 | 1215 | 1336 | 1320 | 1422 | Borcs7 |
| Borcs8 | 1279 | 1564 | 1358 | 1361 | 1303 | 1202 | 1255 | 1110 | 1265 | 1218 | 1176 | 1212 | 1119 | 1148 | 1217 | 1363 | 1188 | 1343 | 1443 | 1450 | 1241 | 1024 | 1170 | 1244 | 1255 | 1211 | 1176 | 1158 | Borcs8 |
| Bpgm | 4684 | 5675 | 5188 | 5141 | 4926 | 4726 | 5399 | 4567 | 4413 | 4309 | 3667 | 4732 | 5882 | 5336 | 5226 | 4858 | 5550 | 4693 | 5125 | 5975 | 5220 | 4197 | 4578 | 4367 | 4690 | 5032 | 5539 | 6703 | Bpgm |
| Bphl | 932 | 857 | 725 | 670 | 699 | 676 | 780 | 973 | 1047 | 886 | 1056 | 876 | 919 | 1014 | 862 | 747 | 775 | 761 | 635 | 775 | 691 | 832 | 1115 | 932 | 903 | 1033 | 923 | 862 | Bphl |
| Bpi | 0 | 3 | 0 | 0 | 2 | 3 | 1 | 0 | 0 | 0 | 0 | 0 | 1 | 0 | 0 | 4 | 0 | 0 | 5 | 5 | 2 | 2 | 1 | 0 | 0 | 1 | 0 | 0 | Bpi |
| Bpifa1 | 0 | 0 | 0 | 0 | 0 | 0 | 0 | 2 | 13 | 2 | 2 | 0 | 0 | 0 | 0 | 0 | 0 | 0 | 0 | 0 | 0 | 0 | 17 | 2 | 0 | 0 | 0 | 0 | Bpifa1 |
| Bpifa6 | 0 | 0 | 0 | 0 | 0 | 0 | 0 | 0 | 0 | 0 | 0 | 0 | 0 | 0 | 0 | 0 | 0 | 0 | 0 | 0 | 0 | 0 | 1 | 0 | 0 | 0 | 0 | 0 | Bpifa6 |
| Bpifb1 | 0 | 2 | 0 | 0 | 5 | 0 | 7 | 1 | 2 | 0 | 7 | 3 | 0 | 2 | 0 | 0 | 0 | 0 | 5 | 13 | 5 | 8 | 2 | 3 | 1 | 3 | 1 | 0 | Bpifb1 |
| Bpifb2 | 1 | 1 | 0 | 1 | 0 | 0 | 2 | 0 | 0 | 0 | 1 | 0 | 0 | 0 | 0 | 1 | 0 | 0 | 2 | 0 | 0 | 0 | 5 | 0 | 0 | 0 | 0 | 0 | Bpifb2 |
| Bpifb3 | 0 | 1 | 0 | 0 | 1 | 0 | 0 | 0 | 0 | 4 | 2 | 1 | 1 | 2 | 3 | 0 | 1 | 0 | 2 | 1 | 1 | 3 | 2 | 0 | 0 | 3 | 0 | 6 | Bpifb3 |
| Bpifb4 | 0 | 1 | 0 | 0 | 0 | 0 | 0 | 1 | 0 | 0 | 0 | 0 | 0 | 0 | 0 | 1 | 2 | 0 | 0 | 0 | 0 | 0 | 0 | 0 | 0 | 4 | 3 | 1 | Bpifb4 |
| Bpifb5 | 6 | 2 | 6 | 1 | 7 | 6 | 0 | 4 | 16 | 8 | 13 | 13 | 8 | 6 | 12 | 12 | 0 | 0 | 8 | 6 | 9 | 12 | 8 | 0 | 0 | 3 | 1 | 4 | Bpifb5 |
| Bpifb6 | 0 | 0 | 2 | 0 | 1 | 0 | 0 | 0 | 0 | 0 | 0 | 0 | 0 | 0 | 0 | 1 | 0 | 1 | 0 | 0 | 0 | 0 | 0 | 0 | 1 | 0 | 0 | 1 | Bpifb6 |
| Bpifc | 1 | 3 | 1 | 6 | 2 | 0 | 0 | 1 | 4 | 0 | 5 | 4 | 0 | 0 | 8 | 0 | 7 | 0 | 3 | 2 | 0 | 0 | 5 | 2 | 0 | 5 | 0 | 0 | Bpifc |
| Bpnt1 | 1967 | 2118 | 2171 | 2519 | 2196 | 1925 | 2451 | 1690 | 1537 | 1718 | 1657 | 1614 | 1577 | 1646 | 2012 | 1919 | 2213 | 2369 | 2247 | 2210 | 2401 | 1793 | 1575 | 1749 | 1638 | 1718 | 1544 | 1717 | Bpnt1 |
| Bpnt2 | 23156 | 15607 | 19577 | 20407 | 19235 | 22165 | 22256 | 16938 | 15200 | 14166 | 14872 | 16301 | 15348 | 15976 | 22131 | 19531 | 19207 | 20159 | 19410 | 20837 | 21326 | 15771 | 14749 | 14449 | 15621 | 15517 | 16274 | 16393 | Bpnt2 |
| Bptf | 9561 | 9214 | 7175 | 9361 | 11178 | 11274 | 10594 | 18120 | 17991 | 18630 | 18249 | 20298 | 18741 | 19966 | 9113 | 9402 | 8241 | 9118 | 10454 | 9895 | 11207 | 17134 | 16394 | 15307 | 19424 | 10999 | 19969 | 17626 | Bptf |
| Braf | 8470 | 6882 | 5923 | 7608 | 8434 | 9452 | 9603 | 8988 | 8460 | 7794 | 8040 | 8082 | 8340 | 8804 | 8817 | 7270 | 6526 | 7633 | 7661 | 8876 | 9405 | 8335 | 7839 | 6914 | 8237 | 5861 | 8803 | 7881 | Braf |
| Brap | 5429 | 5860 | 5494 | 6250 | 5258 | 5025 | 5510 | 3799 | 3893 | 4076 | 3822 | 4136 | 3847 | 4274 | 5044 | 5294 | 5614 | 6039 | 5259 | 4860 | 5544 | 3957 | 3522 | 4150 | 4496 | 4123 | 3936 | 4195 | Brap |
| Brat1 | 1989 | 2814 | 2442 | 2994 | 2493 | 2541 | 2387 | 1603 | 1908 | 2080 | 1956 | 1840 | 1928 | 1861 | 2165 | 2393 | 2743 | 3023 | 2563 | 2147 | 2506 | 1945 | 1844 | 2037 | 2050 | 2001 | 1926 | 1824 | Brat1 |
| Brca1 | 8824 | 7895 | 5364 | 5208 | 4011 | 5161 | 5821 | 4562 | 4221 | 3416 | 3378 | 4848 | 4166 | 4934 | 7713 | 6907 | 5499 | 4835 | 4744 | 6382 | 5321 | 2543 | 3661 | 3787 | 3578 | 4083 | 5414 | 4931 | Brca1 |
| Brca2 | 4672 | 3299 | 2458 | 2449 | 2369 | 3292 | 3310 | 3711 | 3963 | 2960 | 2949 | 4186 | 4215 | 4310 | 4176 | 4093 | 2461 | 2185 | 2400 | 3453 | 3144 | 2020 | 3478 | 2685 | 2941 | 2917 | 5464 | 3923 | Brca2 |
| Brcc3 | 4019 | 2984 | 3193 | 3664 | 3697 | 3805 | 4472 | 3925 | 3434 | 3241 | 3311 | 4097 | 3943 | 4416 | 4373 | 3400 | 3480 | 3148 | 3695 | 4329 | 4141 | 3594 | 3318 | 3657 | 3557 | 4046 | 4047 | 4386 | Brcc3 |
| Brd1 | 4538 | 4951 | 6524 | 6477 | 6678 | 6353 | 5870 | 6499 | 7022 | 8663 | 8253 | 9585 | 8245 | 9460 | 4246 | 4616 | 6354 | 6358 | 6911 | 5243 | 6383 | 7436 | 6443 | 8964 | 9370 | 8843 | 8537 | 9223 | Brd1 |
| Brd2 | 13979 | 16101 | 16916 | 19280 | 17436 | 16442 | 16689 | 19199 | 19773 | 21261 | 20687 | 22719 | 20550 | 23000 | 13876 | 13511 | 17527 | 18483 | 18446 | 14892 | 17773 | 22541 | 18254 | 22558 | 22440 | 24544 | 21330 | 22383 | Brd2 |
| Brd3 | 11633 | 11944 | 16462 | 16186 | 15610 | 14444 | 14975 | 13446 | 14698 | 19545 | 17747 | 20549 | 18223 | 19764 | 10644 | 12916 | 16799 | 15825 | 16689 | 12138 | 15036 | 17448 | 13566 | 20976 | 20598 | 20970 | 18128 | 20013 | Brd3 |
| Brd3os | 844 | 1037 | 813 | 939 | 1054 | 1051 | 1055 | 813 | 909 | 965 | 831 | 853 | 769 | 860 | 847 | 770 | 965 | 876 | 1146 | 1012 | 1083 | 993 | 879 | 804 | 899 | 809 | 796 | 764 | Brd3os |
| Brd4 | 6952 | 7800 | 11919 | 10953 | 10019 | 8914 | 7610 | 7478 | 8485 | 10888 | 10499 | 12688 | 9927 | 10224 | 6295 | 7500 | 11366 | 10811 | 11070 | 6441 | 8371 | 11650 | 7860 | 12564 | 11156 | 13331 | 10118 | 10391 | Brd4 |
| Brd7 | 9436 | 8516 | 9124 | 8421 | 7260 | 7780 | 8506 | 8474 | 7618 | 8051 | 7536 | 8462 | 8163 | 9346 | 9245 | 8077 | 8552 | 7892 | 7876 | 8367 | 7954 | 7548 | 7556 | 8528 | 7729 | 9336 | 8885 | 9053 | Brd7 |
| Brd8 | 9523 | 8356 | 8002 | 8350 | 8881 | 9276 | 9626 | 16394 | 15279 | 15877 | 15892 | 18178 | 16709 | 19168 | 9302 | 7701 | 8450 | 7578 | 9121 | 9573 | 9467 | 15679 | 14728 | 16306 | 16893 | 15224 | 18675 | 17232 | Brd8 |
| Brd8dc | 399 | 399 | 459 | 618 | 513 | 518 | 591 | 515 | 330 | 472 | 477 | 479 | 480 | 460 | 450 | 425 | 612 | 618 | 597 | 542 | 596 | 677 | 374 | 507 | 535 | 455 | 438 | 605 | Brd8dc |
| Brd9 | 5184 | 5223 | 4648 | 4783 | 5722 | 5463 | 5845 | 10498 | 9587 | 9021 | 8691 | 9313 | 9003 | 10003 | 5471 | 4896 | 5120 | 4951 | 5460 | 5969 | 5733 | 10056 | 9180 | 9180 | 9718 | 8035 | 9469 | 9327 | Brd9 |
| Brdt | 1113 | 845 | 939 | 1016 | 967 | 932 | 910 | 387 | 392 | 439 | 471 | 321 | 399 | 317 | 981 | 1036 | 976 | 994 | 860 | 847 | 856 | 394 | 471 | 378 | 307 | 315 | 338 | 347 | Brdt |
| Brf1 | 5727 | 6879 | 7144 | 7586 | 6813 | 6678 | 6411 | 3826 | 4239 | 5030 | 4812 | 5605 | 4899 | 5215 | 5261 | 6414 | 7221 | 7240 | 7088 | 5592 | 6665 | 4360 | 4206 | 5100 | 5157 | 4837 | 5464 | 4977 | Brf1 |
| Brf2 | 676 | 804 | 874 | 840 | 808 | 822 | 789 | 763 | 929 | 926 | 869 | 900 | 965 | 901 | 728 | 755 | 767 | 796 | 853 | 754 | 822 | 817 | 840 | 985 | 932 | 1117 | 895 | 1016 | Brf2 |
| Bri3 | 1625 | 1878 | 2012 | 1958 | 1785 | 1653 | 1538 | 1443 | 1301 | 1749 | 1489 | 1727 | 1560 | 1689 | 1597 | 1510 | 2195 | 1646 | 1855 | 1610 | 1768 | 1392 | 1411 | 1681 | 1564 | 2029 | 1732 | 1678 | Bri3 |
| Bri3bp | 6757 | 6920 | 7190 | 8283 | 7968 | 7801 | 7897 | 4224 | 4299 | 4775 | 4529 | 4995 | 4696 | 4947 | 6666 | 6627 | 8090 | 8621 | 8216 | 6788 | 8204 | 4949 | 3927 | 5006 | 5208 | 4669 | 4911 | 5209 | Bri3bp |
| Bricd5 | 69 | 103 | 66 | 70 | 65 | 46 | 51 | 18 | 22 | 15 | 26 | 20 | 11 | 28 | 47 | 67 | 76 | 89 | 51 | 38 | 71 | 17 | 7 | 15 | 7 | 28 | 31 | 30 | Bricd5 |
| Brinp1 | 1305 | 1482 | 1208 | 1551 | 2320 | 1749 | 1649 | 1614 | 2018 | 2200 | 2077 | 1341 | 1500 | 1331 | 1447 | 1449 | 1384 | 1806 | 1760 | 1762 | 1809 | 2321 | 2256 | 1463 | 1905 | 887 | 1221 | 1280 | Brinp1 |
| Brinp2 | 2775 | 3698 | 2787 | 4489 | 5308 | 4948 | 5470 | 1424 | 1764 | 1829 | 1469 | 1636 | 776 | 1562 | 3160 | 2511 | 3967 | 4451 | 5499 | 5108 | 5627 | 2015 | 1486 | 1852 | 2411 | 1462 | 1216 | 1200 | Brinp2 |

| | | | | | | | | | | | | | | | | | | | | | | | | | | | | | |
|---|---|---|---|---|---|---|---|---|---|---|---|---|---|---|---|---|---|---|---|---|---|---|---|---|---|---|---|---|---|
| Brinp3 | 133 | 133 | 89 | 178 | 360 | 240 | 244 | 449 | 468 | 413 | 398 | 185 | 166 | 214 | 181 | 91 | 161 | 204 | 235 | 304 | 252 | 551 | 523 | 270 | 495 | 190 | 222 | 230 | Brinp3 |
| Brip1 | 2378 | 1859 | 1645 | 1296 | 1000 | 1271 | 1535 | 1525 | 1345 | 1056 | 1245 | 1750 | 1516 | 1665 | 1909 | 1934 | 1619 | 1224 | 1319 | 1512 | 1393 | 988 | 1274 | 1267 | 1119 | 1484 | 1687 | 1635 | Brip1 |
| Brix1 | 4103 | 3439 | 4366 | 3624 | 3382 | 3355 | 3770 | 4617 | 3954 | 4226 | 3889 | 4833 | 4453 | 4833 | 4146 | 3895 | 3894 | 3296 | 3550 | 3924 | 3447 | 4172 | 4105 | 4432 | 4066 | 5533 | 4891 | 5161 | Brix1 |
| Brk1 | 5531 | 6236 | 5601 | 5973 | 5284 | 5279 | 6200 | 5250 | 5440 | 5157 | 5384 | 5078 | 5124 | 5503 | 5943 | 5848 | 5948 | 5607 | 5576 | 6372 | 5792 | 4787 | 5311 | 5409 | 4985 | 6125 | 5542 | 6014 | Brk1 |
| Brme1 | 165 | 183 | 147 | 173 | 147 | 185 | 129 | 153 | 153 | 159 | 128 | 135 | 149 | 129 | 162 | 152 | 140 | 133 | 147 | 174 | 140 | 109 | 142 | 173 | 138 | 121 | 152 | 121 | Brme1 |
| Brms1 | 24 | 22 | 15 | 28 | 22 | 22 | 22 | 21 | 17 | 3 | 21 | 18 | 16 | 20 | 21 | 19 | 16 | 17 | 17 | 13 | 22 | 22 | 15 | 10 | 23 | 11 | 10 | 23 | Brms1 |
| Brms1l | 4058 | 3731 | 3653 | 3687 | 3730 | 3632 | 4168 | 4478 | 3821 | 3703 | 3596 | 3964 | 3644 | 3945 | 4120 | 3351 | 4164 | 3805 | 3879 | 4130 | 3897 | 4393 | 3917 | 4049 | 3967 | 3859 | 3994 | 4050 | Brms1l |
| Brox | 6394 | 6438 | 5677 | 6857 | 8382 | 7818 | 8134 | 9987 | 8563 | 8187 | 7795 | 7783 | 7979 | 7783 | 7402 | 6591 | 6383 | 7170 | 7624 | 8963 | 8258 | 9244 | 8484 | 7616 | 8982 | 6805 | 7850 | 7741 | Brox |
| Brpf1 | 5621 | 6862 | 6466 | 7296 | 5702 | 6078 | 5909 | 3764 | 4062 | 4432 | 4147 | 5294 | 4572 | 4859 | 5003 | 6073 | 6825 | 6729 | 6278 | 5176 | 5882 | 3918 | 3686 | 4831 | 4608 | 5131 | 5001 | 4907 | Brpf1 |
| Brpf3 | 791 | 955 | 1225 | 939 | 1033 | 926 | 836 | 3207 | 3973 | 4139 | 4109 | 4776 | 4196 | 4580 | 740 | 862 | 1045 | 847 | 993 | 830 | 838 | 3830 | 3480 | 4349 | 4561 | 4098 | 4579 | 4767 | Brpf3 |
| Brs3 | 0 | 0 | 0 | 0 | 2 | 3 | 1 | 4 | 1 | 6 | 2 | 0 | 2 | 0 | 0 | 0 | 0 | 7 | 4 | 0 | 1 | 13 | 10 | 0 | 1 | 0 | 0 | 1 | Brs3 |
| Brsk1 | 2993 | 3814 | 3750 | 5829 | 7543 | 6027 | 5757 | 6329 | 7207 | 9252 | 7540 | 8009 | 6527 | 7437 | 3285 | 2959 | 5131 | 6216 | 7492 | 5304 | 6738 | 9324 | 6218 | 8665 | 9719 | 6558 | 6632 | 6793 | Brsk1 |
| Brsk2 | 2751 | 3748 | 4310 | 5401 | 7468 | 6234 | 5852 | 7140 | 7329 | 9708 | 7559 | 11149 | 9213 | 10182 | 3225 | 2570 | 5533 | 5328 | 7181 | 5424 | 6404 | 10462 | 6471 | 9859 | 11819 | 8155 | 9789 | 9667 | Brsk2 |
| Brwd1 | 12637 | 11229 | 9218 | 10707 | 12422 | 13210 | 12883 | 19882 | 16813 | 14625 | 14779 | 15234 | 15018 | 16077 | 12774 | 10865 | 10128 | 10355 | 11497 | 13920 | 13408 | 16188 | 16402 | 13594 | 16317 | 12982 | 16586 | 15061 | Brwd1 |
| Brwd3 | 9777 | 7179 | 5364 | 6688 | 6945 | 8836 | 8666 | 7182 | 5694 | 4296 | 4841 | 5023 | 5180 | 5544 | 9873 | 7999 | 5421 | 6253 | 6716 | 9166 | 8847 | 4251 | 5501 | 4017 | 4959 | 3215 | 5808 | 4974 | Brwd3 |
| Bscl2 | 1744 | 2199 | 1977 | 2248 | 2197 | 2201 | 2149 | 1820 | 1999 | 2210 | 1885 | 2149 | 1977 | 2139 | 1859 | 1765 | 1971 | 1926 | 2253 | 2106 | 2229 | 2072 | 1832 | 2164 | 2316 | 1986 | 2199 | 2280 | Bscl2 |
| Bsdc1 | 4241 | 5707 | 4324 | 4946 | 4054 | 4219 | 4138 | 3115 | 3597 | 3152 | 3130 | 3167 | 2894 | 3279 | 4482 | 5189 | 4570 | 4944 | 4214 | 4150 | 4118 | 3332 | 3306 | 3321 | 3242 | 3479 | 3066 | 3262 | Bsdc1 |
| Bsg | 13115 | 14610 | 12697 | 12380 | 11437 | 11024 | 10911 | 13140 | 14817 | 13930 | 14038 | 11771 | 12445 | 14000 | 12744 | 12905 | 12127 | 11660 | 11200 | 11328 | 11937 | 12788 | 16309 | 13676 | 13061 | 14130 | 12213 | 14036 | Bsg |
| Bsn | 2809 | 4179 | 4175 | 7814 | 10560 | 7941 | 6687 | 4608 | 5217 | 7954 | 6222 | 6574 | 5320 | 5277 | 3247 | 3374 | 6859 | 8649 | 9641 | 6026 | 8263 | 10148 | 4330 | 6673 | 9052 | 2809 | 5014 | 4766 | Bsn |
| Bsnd | 2 | 0 | 0 | 0 | 0 | 0 | 0 | 0 | 2 | 0 | 1 | 3 | 2 | 0 | 1 | 1 | 0 | 1 | 0 | 0 | 0 | 0 | 2 | 0 | 0 | 1 | 2 | 2 | Bsnd |
| Bsph1 | 0 | 0 | 0 | 6 | 0 | 2 | 3 | 0 | 0 | 0 | 0 | 0 | 0 | 0 | 0 | 2 | 0 | 0 | 0 | 1 | 0 | 0 | 0 | 2 | 0 | 0 | 0 | 0 | Bsph1 |
| Bspry | 288 | 166 | 112 | 105 | 83 | 56 | 28 | 74 | 85 | 113 | 93 | 20 | 53 | 37 | 175 | 165 | 67 | 96 | 33 | 51 | 39 | 32 | 140 | 80 | 24 | 97 | 23 | 37 | Bspry |
| Bst1 | 3 | 4 | 24 | 8 | 4 | 9 | 4 | 5 | 6 | 7 | 11 | 20 | 8 | 5 | 4 | 11 | 2 | 11 | 0 | 2 | 4 | 8 | 5 | 5 | 10 | 5 | 9 | 5 | Bst1 |
| Bst2 | 346 | 486 | 575 | 426 | 374 | 310 | 226 | 245 | 251 | 376 | 425 | 330 | 314 | 255 | 427 | 622 | 396 | 399 | 296 | 460 | 321 | 316 | 406 | 341 | 340 | 421 | 425 | 316 | Bst2 |
| Bsx | 103 | 105 | 108 | 129 | 103 | 78 | 145 | 101 | 96 | 100 | 94 | 171 | 12 | 199 | 69 | 50 | 115 | 94 | 111 | 106 | 137 | 137 | 89 | 179 | 150 | 230 | 148 | 176 | Bsx |
| Btaf1 | 19615 | 12995 | 7997 | 9879 | 8971 | 13074 | 13766 | 12123 | 10239 | 6551 | 9027 | 7816 | 7294 | 8283 | 19524 | 12705 | 8524 | 9840 | 8879 | 11892 | 13371 | 6029 | 9354 | 5989 | 9007 | 6425 | 7963 | 7444 | Btaf1 |
| Btbd1 | 11224 | 9192 | 9932 | 11307 | 10722 | 10813 | 12659 | 11696 | 8882 | 9111 | 9020 | 9645 | 8614 | 10361 | 12293 | 9608 | 10331 | 10692 | 11011 | 12056 | 12283 | 10527 | 9578 | 9242 | 9620 | 9698 | 9534 | 10371 | Btbd1 |
| Btbd10 | 3224 | 2689 | 3235 | 3218 | 3457 | 3192 | 3581 | 3504 | 3149 | 3451 | 3016 | 3276 | 2951 | 3423 | 3574 | 3027 | 3063 | 3378 | 3235 | 3204 | 3276 | 3883 | 2987 | 3151 | 3566 | 3231 | 3003 | 3294 | Btbd10 |
| Btbd11 | 1933 | 2105 | 2321 | 2372 | 2969 | 2388 | 2271 | 1362 | 1386 | 1480 | 1483 | 1276 | 1357 | 1146 | 2064 | 1981 | 2292 | 2626 | 2648 | 2064 | 2360 | 1733 | 1473 | 1119 | 1407 | 1000 | 1154 | 1241 | Btbd11 |
| Btbd16 | 1 | 2 | 2 | 4 | 2 | 5 | 0 | 0 | 2 | 1 | 1 | 9 | 1 | 1 | 4 | 5 | 0 | 0 | 2 | 3 | 4 | 2 | 3 | 2 | 1 | 1 | 0 | 1 | Btbd16 |
| Btbd17 | 201 | 258 | 378 | 193 | 237 | 177 | 238 | 893 | 765 | 1016 | 740 | 1662 | 1387 | 1534 | 235 | 173 | 292 | 182 | 270 | 231 | 200 | 916 | 739 | 1229 | 1139 | 1671 | 1859 | 1736 | Btbd17 |
| Btbd18 | 5 | 1 | 0 | 2 | 0 | 0 | 0 | 2 | 2 | 8 | 0 | 2 | 0 | 4 | 0 | 2 | 2 | 0 | 2 | 0 | 4 | 2 | 2 | 3 | 0 | 0 | 3 | 5 | Btbd18 |
| Btbd19 | 91 | 85 | 162 | 75 | 110 | 57 | 116 | 192 | 208 | 204 | 316 | 214 | 213 | 231 | 90 | 101 | 122 | 84 | 98 | 84 | 97 | 190 | 254 | 239 | 214 | 245 | 200 | 214 | Btbd19 |
| Btbd2 | 4133 | 5290 | 5307 | 4924 | 4266 | 3932 | 3565 | 3471 | 4372 | 4685 | 4552 | 4954 | 4542 | 4262 | 3666 | 4866 | 5110 | 4626 | 4747 | 3429 | 3782 | 4428 | 4623 | 4939 | 4649 | 5561 | 4321 | 4302 | Btbd2 |
| Btbd3 | 5531 | 5166 | 5308 | 6180 | 7956 | 7007 | 7554 | 6533 | 6223 | 6551 | 6553 | 5217 | 5532 | 5466 | 5975 | 5210 | 5589 | 6929 | 7267 | 7168 | 7204 | 7720 | 6655 | 5573 | 6745 | 4350 | 5263 | 5260 | Btbd3 |
| Btbd6 | 336 | 408 | 354 | 407 | 572 | 503 | 459 | 1039 | 1196 | 1059 | 1060 | 963 | 972 | 924 | 376 | 374 | 467 | 497 | 611 | 534 | 494 | 1158 | 1105 | 1073 | 1303 | 1064 | 947 | 989 | Btbd6 |
| Btbd7 | 11204 | 9779 | 8780 | 10327 | 10352 | 10913 | 11129 | 11546 | 10686 | 10001 | 10311 | 11081 | 10715 | 11213 | 10661 | 10678 | 9304 | 9828 | 10154 | 10582 | 10930 | 10387 | 10494 | 9713 | 10578 | 8557 | 10658 | 10615 | Btbd7 |
| Btbd8 | 1015 | 874 | 781 | 1029 | 1322 | 1286 | 1468 | 1548 | 1377 | 1401 | 1269 | 1322 | 1234 | 1525 | 1208 | 763 | 897 | 982 | 1195 | 1369 | 1375 | 1447 | 1361 | 1237 | 1459 | 932 | 1502 | 1267 | Btbd8 |
| Btbd9 | 2560 | 2998 | 2953 | 3873 | 4489 | 4000 | 3862 | 3726 | 4077 | 4736 | 4371 | 4636 | 4127 | 3951 | 2644 | 2634 | 3617 | 4118 | 4660 | 3487 | 4245 | 5310 | 3941 | 4548 | 4949 | 4136 | 3978 | 3977 | Btbd9 |
| Btc | 9 | 4 | 8 | 19 | 5 | 6 | 2 | 43 | 28 | 33 | 25 | 12 | 23 | 5 | 15 | 4 | 2 | 11 | 3 | 10 | 5 | 0 | 45 | 27 | 9 | 40 | 10 | 14 | Btc |
| Btd | 535 | 596 | 485 | 448 | 508 | 402 | 400 | 877 | 1159 | 859 | 1058 | 845 | 1015 | 911 | 483 | 613 | 423 | 470 | 528 | 494 | 439 | 988 | 1240 | 994 | 847 | 1014 | 893 | 1006 | Btd |
| Btf3 | 25704 | 22521 | 25567 | 22946 | 20478 | 20517 | 24558 | 23455 | 21091 | 21836 | 21332 | 22450 | 22628 | 25683 | 26008 | 21555 | 24180 | 20585 | 22735 | 24414 | 23926 | 19199 | 22433 | 24362 | 21959 | 27466 | 24767 | 27911 | Btf3 |
| Btf3l4 | 7177 | 5542 | 5986 | 5491 | 5858 | 5831 | 6955 | 11479 | 9208 | 8084 | 7726 | 10038 | 9180 | 11014 | 7668 | 5802 | 5850 | 5186 | 5834 | 7458 | 6496 | 9910 | 9064 | 9172 | 9076 | 10951 | 10634 | 11145 | Btf3l4 |
| Btg1 | 9818 | 8053 | 8597 | 9479 | 8028 | 8483 | 9075 | 9572 | 8694 | 9194 | 8476 | 8871 | 9196 | 8846 | 9373 | 8642 | 8684 | 9382 | 8475 | 9435 | 8934 | 9332 | 9017 | 8995 | 9380 | 9598 | 8564 | 9347 | Btg1 |
| Btg2 | 4191 | 5412 | 6247 | 5015 | 4404 | 4493 | 4866 | 5195 | 4397 | 4689 | 4154 | 7128 | 6569 | 7921 | 4656 | 4630 | 6145 | 4754 | 5099 | 5002 | 4673 | 4299 | 4260 | 6454 | 5733 | 7106 | 7804 | 8336 | Btg2 |
| Btg3 | 1825 | 1792 | 2255 | 2061 | 1734 | 1851 | 2140 | 942 | 942 | 905 | 1019 | 1093 | 1056 | 1173 | 1875 | 1679 | 1992 | 1942 | 1946 | 1845 | 1885 | 784 | 1043 | 969 | 941 | 1252 | 1201 | 1320 | Btg3 |
| Btg4 | 0 | 0 | 0 | 0 | 0 | 0 | 0 | 1 | 1 | 1 | 0 | 0 | 1 | 0 | 0 | 0 | 0 | 4 | 1 | 1 | 2 | 0 | 0 | 0 | 0 | 0 | 1 | 0 | Btg4 |
| Btk | 58 | 55 | 95 | 46 | 62 | 51 | 39 | 105 | 139 | 93 | 130 | 130 | 141 | 80 | 69 | 89 | 35 | 57 | 34 | 63 | 37 | 100 | 140 | 140 | 59 | 144 | 112 | 112 | Btk |
| Btla | 47 | 54 | 81 | 122 | 115 | 92 | 97 | 51 | 24 | 24 | 31 | 51 | 39 | 38 | 88 | 45 | 108 | 105 | 55 | 86 | 94 | 37 | 43 | 38 | 29 | 62 | 106 | 77 | Btla |
| Btn1a1 | 240 | 187 | 155 | 198 | 164 | 171 | 170 | 7 | 17 | 14 | 9 | 18 | 12 | 12 | 222 | 222 | 232 | 160 | 154 | 144 | 186 | 13 | 4 | 6 | 6 | 3 | 9 | 19 | Btn1a1 |
| Btn2a2 | 518 | 338 | 242 | 327 | 271 | 251 | 283 | 51 | 62 | 34 | 50 | 65 | 39 | 20 | 368 | 450 | 318 | 333 | 333 | 303 | 286 | 25 | 52 | 45 | 28 | 92 | 28 | 25 | Btn2a2 |
| Btnl10 | 2 | 2 | 5 | 1 | 1 | 6 | 3 | 9 | 3 | 9 | 4 | 16 | 9 | 9 | 1 | 3 | 5 | 1 | 2 | 3 | 5 | 5 | 5 | 11 | 5 | 13 | 10 | 9 | Btnl10 |
| Btnl2 | 2 | 3 | 0 | 0 | 1 | 3 | 1 | 2 | 0 | 6 | 9 | 3 | 6 | 1 | 0 | 5 | 3 | 4 | 0 | 1 | 0 | 0 | 4 | 1 | 0 | 8 | 1 | 0 | Btnl2 |
| Btnl4 | 1 | 0 | 0 | 0 | 0 | 0 | 0 | 0 | 0 | 0 | 0 | 0 | 0 | 0 | 0 | 0 | 0 | 0 | 0 | 0 | 0 | 0 | 0 | 0 | 0 | 0 | 0 | 0 | Btnl4 |
| Btnl6 | 0 | 0 | 0 | 0 | 0 | 0 | 0 | 0 | 0 | 0 | 0 | 0 | 0 | 0 | 0 | 0 | 0 | 0 | 0 | 0 | 3 | 0 | 0 | 0 | 0 | 0 | 0 | 0 | Btnl6 |
| Btnl9 | 22 | 10 | 17 | 11 | 16 | 10 | 11 | 45 | 55 | 58 | 63 | 28 | 24 | 16 | 7 | 15 | 13 | 18 | 6 | 5 | 5 | 35 | 50 | 29 | 19 | 33 | 22 | 17 | Btnl9 |
| Btrc | 3863 | 3851 | 2413 | 3605 | 4313 | 4224 | 3846 | 5169 | 5184 | 4680 | 5120 | 4626 | 4794 | 4868 | 3765 | 3765 | 2920 | 3466 | 3732 | 4060 | 4014 | 4010 | 5117 | 3615 | 5118 | 3124 | 5026 | 4436 | Btrc |
| Bub1 | 11858 | 9415 | 10646 | 7866 | 5207 | 6783 | 8425 | 7627 | 6568 | 5782 | 5505 | 8748 | 7720 | 9048 | 10847 | 9981 | 8583 | 6741 | 7024 | 8669 | 7232 | 4821 | 5860 | 6994 | 6122 | 9936 | 9775 | 9405 | Bub1 |
| Bub1b | 12545 | 11574 | 11249 | 8978 | 6105 | 8039 | 8384 | 9101 | 8428 | 7891 | 7250 | 11619 | 9813 | 12023 | 10613 | 10465 | 9879 | 7496 | 8819 | 8358 | 8132 | 5882 | 7413 | 9774 | 8235 | 13564 | 12257 | 12244 | Bub1b |
| Bub3 | 7865 | 8949 | 8289 | 7350 | 6733 | 6434 | 6464 | 9431 | 8983 | 7981 | 7555 | 9658 | 8671 | 9444 | 8027 | 7235 | 7684 | 6516 | 7238 | 7871 | 6990 | 7536 | 8471 | 8854 | 8275 | 11024 | 9950 | 9937 | Bub3 |
| Bud13 | 1551 | 1540 | 1726 | 1427 | 1409 | 1382 | 1328 | 1286 | 1530 | 1600 | 1516 | 1839 | 1543 | 1665 | 1474 | 1576 | 1678 | 1360 | 1467 | 1272 | 1254 | 1688 | 1451 | 1705 | 1514 | 2056 | 1526 | 1805 | Bud13 |
| Bud23 | 2645 | 3459 | 3263 | 2880 | 2291 | 2370 | 2372 | 2082 | 2273 | 2199 | 2239 | 2560 | 2490 | 2554 | 2512 | 3016 | 3059 | 2685 | 2621 | 2446 | 2326 | 1942 | 2218 | 2616 | 2280 | 2949 | 2639 | 2528 | Bud23 |
| Bud31 | 6813 | 6488 | 6681 | 6284 | 5370 | 5693 | 6426 | 6631 | 5819 | 5972 | 5273 | 6093 | 6053 | 7102 | 7242 | 6402 | 6445 | 6097 | 5992 | 7374 | 6336 | 5233 | 6034 | 5833 | 5782 | 7005 | 6560 | 7425 | Bud31 |
| Bves | 383 | 370 | 540 | 415 | 545 | 327 | 352 | 255 | 255 | 333 | 392 | 244 | 356 | 213 | 448 | 531 | 519 | 448 | 432 | 312 | 405 | 339 | 289 | 219 | 201 | 204 | 208 | 212 | Bves |
| Bysl | 2730 | 3225 | 3225 | 3253 | 2758 | 2755 | 2979 | 1949 | 2225 | 2524 | 2299 | 2747 | 2689 | 2823 | 2743 | 2903 | 3373 | 3174 | 3084 | 2420 | 2995 | 2526 | 2149 | 2669 | 2709 | 3182 | 2581 | 3007 | Bysl |
| Bzw1 | 59673 | 49187 | 54116 | 47231 | 44408 | 47840 | 55752 | 48852 | 40411 | 36097 | 36597 | 43034 | 41364 | 46659 | 62556 | 53968 | 50234 | 47878 | 46043 | 57484 | 49732 | 39401 | 40788 | 39361 | 40076 | 46247 | 46015 | 48406 | Bzw1 |
| Bzw2 | 20031 | 21836 | 18741 | 20873 | 20570 | 19590 | 24033 | 15216 | 14314 | 14018 | 12219 | 13131 | 12034 | 13915 | 20484 | 20115 | 19901 | 19873 | 22298 | 24468 | 23894 | 13017 | 14325 | 13996 | 14925 | 14047 | 12809 | 13754 | Bzw2 |
| C030006K11Rik | 498 | 621 | 747 | 610 | 632 | 628 | 666 | 366 | 504 | 569 | 513 | 597 | 434 | 568 | 538 | 406 | 641 | 588 | 651 | 548 | 558 | 408 | 465 | 525 | 582 | 632 | 558 | 653 | C030006K11Rik |
| C130026I21Rik | 0 | 0 | 1 | 0 | 0 | 0 | 0 | 7 | 4 | 0 | 0 | 2 | 0 | 0 | 3 | 5 | 0 | 0 | 0 | 0 | 0 | 3 | 2 | 5 | 3 | 2 | 0 | 0 | C130026I21Rik |
| C130050O18Rik | 13 | 6 | 3 | 11 | 7 | 7 | 8 | 17 | 49 | 37 | 56 | 14 | 23 | 9 | 11 | 19 | 0 | 13 | 19 | 8 | 9 | 47 | 47 | 32 | 19 | 11 | 17 | 6 | C130050O18Rik |
| C130073F10Rik | 0 | 0 | 0 | 0 | 0 | 0 | 0 | 0 | 2 | 0 | 0 | 0 | 0 | 0 | 0 | 0 | 0 | 0 | 0 | 0 | 0 | 0 | 0 | 0 | 0 | 0 | 0 | 0 | C130073F10Rik |
| C130074G19Rik | 493 | 543 | 725 | 579 | 783 | 734 | 615 | 912 | 1069 | 1393 | 1419 | 1275 | 1445 | 1293 | 526 | 569 | 606 | 639 | 643 | 616 | 723 | 1180 | 1309 | 1382 | 1333 | 1404 | 1117 | 1207 | C130074G19Rik |
| C1d | 3992 | 2661 | 3471 | 3365 | 3941 | 3861 | 4094 | 6563 | 5960 | 4973 | 5246 | 5355 | 5668 | 5810 | 4261 | 3667 | 3267 | 3299 | 3677 | 4904 | 3835 | 6034 | 6160 | 5357 | 5638 | 5920 | 5812 | 5981 | C1d |
| C1galt1 | 5654 | 3720 | 3605 | 3795 | 3382 | 3991 | 3884 | 3331 | 3056 | 2660 | 3339 | 2429 | 2862 | 2267 | 5471 | 4989 | 3542 | 3781 | 3275 | 4123 | 3593 | 2946 | 3298 | 2462 | 2381 | 2232 | 2488 | 2508 | C1galt1 |
| C1galt1c1 | 734 | 696 | 713 | 621 | 755 | 685 | 758 | 1216 | 1101 | 901 | 928 | 841 | 973 | 1034 | 807 | 769 | 627 | 638 | 786 | 879 | 726 | 994 | 1139 | 865 | 932 | 985 | 922 | 1022 | C1galt1c1 |
| C1qa | 206 | 292 | 227 | 208 | 290 | 307 | 193 | 327 | 412 | 379 | 508 | 328 | 344 | 288 | 281 | 330 | 212 | 286 | 202 | 312 | 219 | 279 | 656 | 412 | 322 | 400 | 367 | 350 | C1qa |
| C1qb | 482 | 559 | 562 | 476 | 467 | 504 | 354 | 574 | 778 | 651 | 791 | 573 | 567 | 459 | 543 | 650 | 433 | 593 | 355 | 627 | 388 | 583 | 955 | 601 | 574 | 684 | 674 | 609 | C1qb |
| C1qbp | 6209 | 5596 | 6866 | 5922 | 4889 | 5126 | 5499 | 4727 | 4378 | 4887 | 4784 | 5502 | 4711 | 6162 | 5443 | 5102 | 6179 | 5283 | 5533 | 4370 | 5747 | 4841 | 4393 | 5435 | 4829 | 6782 | 4981 | 6246 | C1qbp |
| C1qc | 256 | 379 | 375 | 321 | 369 | 320 | 238 | 414 | 463 | 434 | 547 | 383 | 375 | 399 | 316 | 445 | 337 | 324 | 275 | 373 | 278 | 398 | 728 | 492 | 399 | 486 | 404 | 376 | C1qc |

| | | | | | | | | | | | | | | | | | | | | | | | | | | | | | |
|---|---|---|---|---|---|---|---|---|---|---|---|---|---|---|---|---|---|---|---|---|---|---|---|---|---|---|---|---|---|
| C1ql1 | 465 | 697 | 846 | 637 | 458 | 437 | 445 | 320 | 281 | 254 | 227 | 357 | 471 | 305 | 502 | 558 | 740 | 472 | 519 | 563 | 394 | 301 | 186 | 306 | 271 | 310 | 452 | 522 | C1ql1 |
| C1ql2 | 5 | 22 | 16 | 38 | 40 | 41 | 53 | 68 | 62 | 114 | 60 | 77 | 60 | 65 | 24 | 12 | 34 | 30 | 58 | 29 | 46 | 109 | 61 | 108 | 109 | 14 | 40 | 40 | C1ql2 |
| C1ql3 | 98 | 95 | 68 | 144 | 284 | 179 | 156 | 376 | 360 | 396 | 441 | 163 | 381 | 140 | 185 | 133 | 159 | 155 | 220 | 202 | 194 | 439 | 382 | 206 | 241 | 133 | 96 | 104 | C1ql3 |
| C1ql4 | 210 | 528 | 453 | 648 | 567 | 614 | 358 | 408 | 330 | 438 | 311 | 543 | 821 | 408 | 438 | 466 | 912 | 563 | 709 | 678 | 673 | 553 | 284 | 454 | 494 | 63 | 500 | 475 | C1ql4 |
| C1qtnf1 | 366 | 349 | 282 | 413 | 385 | 443 | 317 | 488 | 597 | 570 | 664 | 417 | 551 | 405 | 291 | 387 | 357 | 504 | 334 | 340 | 336 | 575 | 703 | 519 | 500 | 451 | 478 | 419 | C1qtnf1 |
| C1qtnf12 | 711 | 761 | 713 | 520 | 455 | 536 | 380 | 509 | 634 | 502 | 626 | 520 | 534 | 428 | 585 | 745 | 698 | 605 | 587 | 378 | 390 | 498 | 588 | 514 | 493 | 579 | 508 | 458 | C1qtnf12 |
| C1qtnf2 | 83 | 88 | 75 | 87 | 128 | 143 | 74 | 159 | 208 | 179 | 278 | 149 | 238 | 136 | 67 | 114 | 70 | 116 | 82 | 87 | 58 | 152 | 358 | 170 | 192 | 98 | 171 | 122 | C1qtnf2 |
| C1qtnf3 | 148 | 166 | 106 | 28 | 47 | 460 | 9 | 256 | 846 | 699 | 3081 | 191 | 513 | 223 | 256 | 181 | 72 | 324 | 178 | 20 | 4 | 947 | 2941 | 322 | 596 | 41 | 182 | 261 | C1qtnf3 |
| C1qtnf6 | 706 | 784 | 825 | 602 | 508 | 768 | 310 | 1287 | 1696 | 1406 | 1752 | 1086 | 1477 | 1008 | 712 | 991 | 592 | 772 | 438 | 368 | 353 | 1308 | 2071 | 1480 | 1102 | 1415 | 1111 | 1056 | C1qtnf6 |
| C1qtnf7 | 82 | 33 | 13 | 23 | 22 | 83 | 15 | 149 | 224 | 96 | 404 | 112 | 143 | 62 | 75 | 53 | 27 | 36 | 41 | 30 | 24 | 115 | 353 | 67 | 107 | 49 | 69 | 62 | C1qtnf7 |
| C1qtnf9 | 65 | 102 | 62 | 146 | 91 | 72 | 127 | 24 | 32 | 38 | 50 | 42 | 41 | 43 | 70 | 77 | 83 | 147 | 101 | 84 | 145 | 30 | 58 | 46 | 35 | 57 | 27 | 38 | C1qtnf9 |
| C1ra | 12 | 23 | 14 | 22 | 17 | 21 | 9 | 33 | 26 | 38 | 42 | 9 | 26 | 23 | 8 | 16 | 16 | 14 | 12 | 13 | 12 | 27 | 52 | 35 | 29 | 17 | 21 | 20 | C1ra |
| C1rb | 0 | 0 | 0 | 0 | 0 | 0 | 0 | 0 | 0 | 3 | 0 | 0 | 0 | 0 | 0 | 0 | 0 | 0 | 0 | 0 | 0 | 2 | 0 | 0 | 0 | 0 | 0 | 1 | C1rb |
| C1rl | 7 | 15 | 9 | 13 | 11 | 6 | 8 | 17 | 30 | 24 | 24 | 4 | 15 | 15 | 7 | 15 | 22 | 8 | 10 | 4 | 2 | 17 | 19 | 18 | 12 | 12 | 22 | 8 | C1rl |
| C1s1 | 45 | 44 | 50 | 51 | 30 | 31 | 18 | 76 | 81 | 75 | 145 | 34 | 48 | 34 | 58 | 55 | 32 | 41 | 52 | 18 | 23 | 60 | 128 | 75 | 73 | 49 | 45 | 40 | C1s1 |
| C1s2 | 16 | 14 | 13 | 12 | 33 | 29 | 25 | 23 | 19 | 33 | 24 | 46 | 35 | 36 | 16 | 11 | 8 | 23 | 20 | 24 | 31 | 40 | 30 | 15 | 31 | 30 | 33 | 24 | C1s2 |
| C2 | 60 | 88 | 101 | 91 | 97 | 84 | 82 | 43 | 45 | 43 | 58 | 68 | 51 | 80 | 67 | 61 | 105 | 96 | 81 | 71 | 67 | 53 | 40 | 83 | 54 | 65 | 73 | 75 | C2 |
| C2cd2 | 2921 | 2702 | 2622 | 2424 | 2673 | 2600 | 2386 | 1749 | 1780 | 1725 | 1977 | 1807 | 2039 | 1717 | 2710 | 2866 | 2473 | 2512 | 2287 | 2280 | 2298 | 1731 | 1896 | 1545 | 1626 | 1747 | 1848 | 1832 | C2cd2 |
| C2cd2l | 2718 | 4120 | 3493 | 5235 | 4359 | 4088 | 3711 | 1498 | 1760 | 1997 | 1820 | 1922 | 1751 | 1689 | 2789 | 3326 | 4320 | 5148 | 4577 | 3310 | 4158 | 2450 | 1734 | 2133 | 2349 | 1865 | 1714 | 1652 | C2cd2l |
| C2cd3 | 4381 | 4494 | 3025 | 3554 | 3887 | 3952 | 4091 | 7708 | 7977 | 7624 | 6690 | 7982 | 7420 | 8099 | 4051 | 3922 | 3529 | 3627 | 3968 | 4114 | 4043 | 6754 | 6824 | 7242 | 7529 | 5159 | 7771 | 7335 | C2cd3 |
| C2cd4a | 12 | 3 | 10 | 10 | 4 | 5 | 3 | 5 | 4 | 4 | 9 | 1 | 5 | 1 | 15 | 10 | 8 | 8 | 7 | 7 | 1 | 3 | 5 | 2 | 5 | 0 | 0 | 1 | C2cd4a |
| C2cd4b | 15 | 12 | 3 | 20 | 8 | 4 | 7 | 1 | 9 | 8 | 6 | 5 | 0 | 7 | 9 | 4 | 5 | 13 | 3 | 6 | 6 | 8 | 14 | 16 | 10 | 5 | 5 | 6 | C2cd4b |
| C2cd4c | 557 | 825 | 727 | 1035 | 906 | 805 | 829 | 209 | 201 | 289 | 235 | 189 | 162 | 173 | 545 | 752 | 805 | 1034 | 867 | 702 | 864 | 329 | 247 | 240 | 249 | 146 | 138 | 134 | C2cd4c |
| C2cd4d | 7 | 4 | 0 | 7 | 5 | 3 | 2 | 9 | 2 | 15 | 23 | 7 | 18 | 4 | 9 | 5 | 1 | 7 | 1 | 4 | 1 | 5 | 16 | 15 | 8 | 18 | 0 | 9 | C2cd4d |
| C2cd5 | 4496 | 4442 | 4494 | 4669 | 5351 | 5420 | 5069 | 7110 | 7168 | 7631 | 7998 | 7628 | 7097 | 7648 | 4655 | 4663 | 4526 | 4708 | 5146 | 4771 | 5330 | 8002 | 6590 | 7174 | 8423 | 6275 | 7512 | 6807 | C2cd5 |
| C2cd6 | 6 | 5 | 2 | 12 | 10 | 8 | 8 | 7 | 6 | 7 | 8 | 7 | 6 | 13 | 11 | 7 | 7 | 14 | 7 | 18 | 21 | 3 | 7 | 7 | 2 | 3 | 6 | 11 | C2cd6 |
| C3 | 30 | 27 | 30 | 23 | 65 | 47 | 42 | 27 | 81 | 57 | 90 | 24 | 44 | 43 | 32 | 41 | 34 | 58 | 33 | 18 | 33 | 80 | 85 | 78 | 41 | 24 | 48 | 31 | C3 |
| C330018D20Rik | 947 | 804 | 1053 | 880 | 875 | 899 | 1015 | 1558 | 1447 | 1356 | 1322 | 1447 | 1391 | 1669 | 1013 | 883 | 956 | 818 | 959 | 1070 | 979 | 1404 | 1370 | 1527 | 1377 | 1701 | 1596 | 1521 | C330018D20Rik |
| C3ar1 | 80 | 53 | 82 | 39 | 74 | 87 | 53 | 199 | 254 | 175 | 335 | 219 | 252 | 171 | 69 | 75 | 50 | 42 | 53 | 76 | 47 | 214 | 397 | 209 | 164 | 267 | 265 | 199 | C3ar1 |
| C4b | 0 | 0 | 0 | 4 | 6 | 10 | 1 | 0 | 7 | 1 | 32 | 1 | 2 | 12 | 1 | 2 | 6 | 2 | 0 | 8 | 1 | 3 | 8 | 2 | 5 | 0 | 5 | 15 | C4b |
| C4bp | 5 | 2 | 9 | 13 | 5 | 3 | 2 | 0 | 2 | 0 | 7 | 0 | 1 | 1 | 5 | 6 | 22 | 9 | 2 | 4 | 4 | 8 | 2 | 0 | 3 | 0 | 0 | 1 | C4bp |
| C5ar1 | 30 | 18 | 37 | 31 | 24 | 22 | 27 | 40 | 39 | 54 | 65 | 31 | 75 | 51 | 34 | 29 | 15 | 39 | 19 | 17 | 28 | 75 | 83 | 78 | 49 | 38 | 42 | 55 | C5ar1 |
| C5ar2 | 0 | 5 | 16 | 3 | 2 | 4 | 2 | 6 | 6 | 6 | 18 | 12 | 5 | 13 | 4 | 6 | 16 | 11 | 2 | 1 | 3 | 10 | 21 | 13 | 15 | 13 | 3 | 20 | C5ar2 |
| C6 | 0 | 0 | 0 | 0 | 2 | 0 | 0 | 0 | 0 | 0 | 3 | 0 | 0 | 0 | 1 | 0 | 0 | 0 | 2 | 0 | 4 | 0 | 1 | 0 | 0 | 0 | 0 | 0 | C6 |
| C7 | 2 | 9 | 7 | 4 | 16 | 10 | 11 | 23 | 18 | 10 | 19 | 5 | 10 | 11 | 16 | 13 | 15 | 9 | 12 | 17 | 24 | 28 | 32 | 6 | 29 | 11 | 8 | 15 | C7 |
| C77080 | 2455 | 3105 | 3226 | 2874 | 2396 | 2304 | 1971 | 1081 | 1416 | 1567 | 1485 | 1449 | 1518 | 1274 | 2080 | 2929 | 2943 | 2760 | 2486 | 1936 | 2078 | 1644 | 1658 | 1449 | 1379 | 1521 | 1227 | 1290 | C77080 |
| C87436 | 804 | 887 | 787 | 794 | 821 | 723 | 819 | 1034 | 1057 | 981 | 834 | 1024 | 899 | 1075 | 833 | 718 | 822 | 793 | 830 | 918 | 819 | 986 | 1010 | 1037 | 984 | 1110 | 941 | 984 | C87436 |
| C8a | 2 | 3 | 0 | 4 | 0 | 0 | 0 | 1 | 1 | 0 | 2 | 7 | 0 | 0 | 3 | 5 | 2 | 0 | 2 | 1 | 0 | 0 | 1 | 0 | 1 | 0 | 0 | 0 | C8a |
| C8b | 17 | 22 | 11 | 15 | 20 | 27 | 41 | 5 | 3 | 0 | 7 | 7 | 8 | 5 | 17 | 9 | 3 | 18 | 7 | 29 | 32 | 8 | 3 | 4 | 16 | 6 | 7 | 9 | C8b |
| C8g | 30 | 32 | 72 | 20 | 20 | 28 | 36 | 51 | 75 | 50 | 63 | 60 | 85 | 55 | 32 | 26 | 19 | 40 | 35 | 28 | 29 | 47 | 83 | 51 | 66 | 68 | 89 | 46 | C8g |
| C9 | 1 | 1 | 0 | 3 | 0 | 0 | 0 | 0 | 1 | 0 | 4 | 0 | 1 | 1 | 3 | 3 | 6 | 1 | 0 | 0 | 3 | 0 | 2 | 0 | 0 | 1 | 7 | 1 | C9 |
| C9orf72 | 5779 | 5363 | 4780 | 6295 | 6317 | 5881 | 6931 | 4400 | 3443 | 3271 | 2908 | 3288 | 3047 | 3398 | 6862 | 5358 | 5406 | 6218 | 5828 | 7264 | 6815 | 3967 | 3422 | 3380 | 3734 | 3130 | 3349 | 3533 | C9orf72 |
| Caap1 | 2077 | 1295 | 1941 | 1737 | 1577 | 1653 | 2011 | 1702 | 1388 | 1393 | 1373 | 1844 | 1660 | 1975 | 1954 | 1580 | 1729 | 1672 | 1457 | 1756 | 1657 | 1516 | 1499 | 1667 | 1664 | 2003 | 1801 | 1996 | Caap1 |
| Cab39 | 18108 | 15724 | 16020 | 16360 | 17184 | 17158 | 19300 | 17507 | 14362 | 13824 | 12109 | 14997 | 13667 | 14690 | 19524 | 15701 | 16238 | 15983 | 17206 | 19810 | 18233 | 15842 | 13992 | 13626 | 14679 | 13687 | 14614 | 15257 | Cab39 |
| Cab39l | 3475 | 2983 | 3131 | 2439 | 2365 | 2414 | 2428 | 2635 | 2490 | 2294 | 2220 | 2543 | 2709 | 2383 | 3280 | 3237 | 2678 | 2396 | 2413 | 2665 | 2509 | 2309 | 2636 | 2400 | 2275 | 2756 | 2613 | 2699 | Cab39l |
| Cabcoco1 | 33 | 33 | 18 | 11 | 25 | 14 | 23 | 34 | 23 | 17 | 14 | 13 | 34 | 16 | 21 | 20 | 15 | 36 | 16 | 30 | 19 | 17 | 34 | 14 | 20 | 24 | 15 | 13 | Cabcoco1 |
| Cabin1 | 8548 | 10591 | 9745 | 11007 | 10521 | 10375 | 9219 | 5964 | 7358 | 8036 | 7557 | 8721 | 8030 | 7466 | 8147 | 10097 | 10707 | 10814 | 10961 | 8609 | 9744 | 7703 | 6950 | 7835 | 8184 | 5965 | 7620 | 7525 | Cabin1 |
| Cables1 | 329 | 314 | 313 | 277 | 420 | 350 | 310 | 454 | 480 | 584 | 463 | 458 | 439 | 357 | 318 | 361 | 321 | 326 | 418 | 369 | 289 | 563 | 508 | 458 | 498 | 440 | 408 | 367 | Cables1 |
| Cables2 | 2729 | 3017 | 2841 | 3413 | 3859 | 3377 | 3429 | 2601 | 2747 | 3481 | 3080 | 3433 | 2875 | 3114 | 2640 | 2741 | 3354 | 3735 | 3720 | 2952 | 3749 | 3674 | 2634 | 3083 | 3370 | 3188 | 2731 | 2998 | Cables2 |
| Cabp1 | 1714 | 2314 | 1949 | 3231 | 3833 | 3044 | 3479 | 843 | 1099 | 1498 | 1178 | 933 | 852 | 887 | 1833 | 1781 | 2559 | 3433 | 3775 | 2741 | 3648 | 946 | 843 | 1175 | 1373 | 545 | 590 | 565 | Cabp1 |
| Cabp4 | 0 | 0 | 0 | 1 | 0 | 0 | 0 | 0 | 0 | 0 | 0 | 0 | 0 | 0 | 0 | 0 | 0 | 0 | 0 | 0 | 0 | 0 | 0 | 0 | 0 | 0 | 0 | 0 | Cabp4 |
| Cabp5 | 48 | 70 | 97 | 123 | 86 | 73 | 96 | 20 | 24 | 41 | 27 | 25 | 17 | 26 | 62 | 78 | 130 | 139 | 80 | 67 | 121 | 43 | 27 | 33 | 37 | 26 | 27 | 19 | Cabp5 |
| Cabp7 | 126 | 191 | 163 | 273 | 419 | 263 | 225 | 164 | 139 | 146 | 181 | 136 | 177 | 102 | 153 | 119 | 234 | 255 | 333 | 302 | 313 | 194 | 126 | 159 | 197 | 130 | 132 | 138 | Cabp7 |
| Cabs1 | 0 | 0 | 0 | 0 | 0 | 0 | 0 | 0 | 1 | 0 | 0 | 0 | 0 | 0 | 0 | 0 | 0 | 0 | 0 | 0 | 0 | 0 | 0 | 0 | 0 | 0 | 0 | 0 | Cabs1 |
| Cabyr | 91 | 84 | 56 | 93 | 119 | 89 | 130 | 56 | 77 | 105 | 78 | 67 | 84 | 51 | 102 | 79 | 92 | 111 | 96 | 98 | 112 | 89 | 53 | 83 | 80 | 47 | 77 | 58 | Cabyr |
| Cacfd1 | 5881 | 6125 | 9642 | 7307 | 7314 | 6627 | 6291 | 6609 | 6961 | 7803 | 7656 | 9994 | 8839 | 8514 | 5979 | 6681 | 8608 | 7162 | 7220 | 6087 | 5647 | 8684 | 6706 | 9561 | 8890 | 11184 | 9420 | 7946 | Cacfd1 |
| Cachd1 | 7408 | 7139 | 9226 | 7370 | 8118 | 8079 | 7152 | 7546 | 7021 | 9260 | 7135 | 10340 | 10751 | 9536 | 6556 | 7813 | 8253 | 7814 | 9182 | 7419 | 8062 | 7874 | 8231 | 9220 | 8962 | 7787 | 10013 | 10763 | Cachd1 |
| Cacna1a | 1299 | 1671 | 1603 | 2308 | 3427 | 2509 | 2252 | 2387 | 2690 | 3560 | 3103 | 2855 | 2810 | 2503 | 1416 | 1515 | 2148 | 2357 | 2530 | 2183 | 2517 | 3989 | 2192 | 2696 | 3577 | 1589 | 2573 | 2204 | Cacna1a |
| Cacna1b | 5413 | 6206 | 5445 | 7599 | 10145 | 8228 | 7616 | 5283 | 5667 | 6999 | 5992 | 4726 | 4627 | 4453 | 5921 | 6403 | 6575 | 7967 | 8080 | 7205 | 8044 | 8181 | 5029 | 4658 | 7306 | 2419 | 4097 | 3701 | Cacna1b |
| Cacna1c | 2668 | 2449 | 1967 | 2779 | 3726 | 3626 | 3181 | 3407 | 4299 | 5141 | 5765 | 3537 | 4306 | 3489 | 2371 | 3279 | 2130 | 3417 | 2716 | 2170 | 3287 | 5168 | 3963 | 4086 | 4894 | 2383 | 3388 | 2887 | Cacna1c |
| Cacna1d | 979 | 866 | 555 | 1409 | 2469 | 1829 | 1781 | 2277 | 2604 | 2390 | 2726 | 1219 | 1441 | 1251 | 1155 | 1035 | 854 | 1642 | 1585 | 1805 | 2222 | 2533 | 2238 | 1617 | 2488 | 666 | 1239 | 959 | Cacna1d |
| Cacna1e | 2398 | 2572 | 1395 | 3518 | 7522 | 6231 | 7055 | 6676 | 7444 | 6472 | 6999 | 3138 | 4241 | 3665 | 3429 | 3304 | 2283 | 4872 | 4923 | 6266 | 7947 | 8335 | 5813 | 4081 | 7676 | 1864 | 3204 | 2772 | Cacna1e |
| Cacna1f | 2 | 6 | 3 | 1 | 7 | 22 | 6 | 4 | 15 | 3 | 10 | 8 | 7 | 1 | 11 | 2 | 2 | 9 | 7 | 14 | 9 | 8 | 12 | 15 | 6 | 2 | 9 | 6 | Cacna1f |
| Cacna1g | 10735 | 12867 | 15217 | 13930 | 14725 | 13405 | 10799 | 9252 | 11359 | 13389 | 14314 | 10600 | 13250 | 9483 | 9767 | 13652 | 14100 | 14861 | 13616 | 9559 | 11145 | 12071 | 12514 | 12679 | 10533 | 7939 | 10162 | 9919 | Cacna1g |
| Cacna1h | 5881 | 7167 | 7936 | 9080 | 10492 | 8475 | 7799 | 5912 | 7158 | 8566 | 7508 | 7481 | 6798 | 6559 | 5695 | 7753 | 8993 | 9305 | 9650 | 7145 | 8236 | 9909 | 5888 | 8380 | 8751 | 5270 | 6335 | 6038 | Cacna1h |
| Cacna1i | 175 | 213 | 264 | 467 | 815 | 474 | 558 | 424 | 464 | 767 | 537 | 409 | 313 | 468 | 178 | 185 | 488 | 554 | 721 | 500 | 625 | 966 | 452 | 668 | 813 | 392 | 363 | 398 | Cacna1i |
| Cacna1s | 31 | 49 | 23 | 8 | 5 | 27 | 7 | 51 | 142 | 27 | 594 | 23 | 56 | 16 | 28 | 51 | 10 | 25 | 10 | 4 | 6 | 85 | 381 | 25 | 28 | 13 | 42 | 26 | Cacna1s |
| Cacna2d1 | 6103 | 4983 | 3962 | 6072 | 8272 | 8588 | 8633 | 13998 | 11492 | 11492 | 12861 | 11693 | 11669 | 12454 | 7358 | 6280 | 5714 | 6733 | 6897 | 8902 | 8571 | 14125 | 11697 | 10342 | 12733 | 7493 | 12378 | 11506 | Cacna2d1 |
| Cacna2d2 | 3269 | 4432 | 5121 | 7196 | 9710 | 7225 | 6383 | 4893 | 5260 | 7258 | 5961 | 6827 | 6704 | 5706 | 4043 | 4087 | 6932 | 7583 | 8299 | 5926 | 6994 | 8930 | 4362 | 6474 | 7856 | 4513 | 5521 | 5257 | Cacna2d2 |
| Cacna2d3 | 1230 | 1252 | 1230 | 1476 | 2346 | 1860 | 1856 | 2220 | 2107 | 1918 | 2138 | 1426 | 1589 | 1428 | 1507 | 1102 | 1285 | 1847 | 1682 | 1550 | 1822 | 2503 | 2273 | 1382 | 1846 | 883 | 1353 | 1400 | Cacna2d3 |
| Cacna2d4 | 13 | 13 | 14 | 27 | 30 | 12 | 23 | 20 | 26 | 9 | 9 | 8 | 15 | 17 | 11 | 7 | 14 | 23 | 30 | 23 | 31 | 37 | 36 | 21 | 18 | 7 | 5 | 12 | Cacna2d4 |
| Cacnb1 | 2882 | 4249 | 4662 | 5763 | 6307 | 5503 | 5116 | 3378 | 3542 | 4974 | 4544 | 4213 | 3937 | 3829 | 3160 | 3396 | 5592 | 5686 | 6448 | 4692 | 5394 | 5386 | 3620 | 4447 | 5298 | 3589 | 3649 | 3595 | Cacnb1 |
| Cacnb2 | 276 | 288 | 193 | 297 | 474 | 395 | 352 | 727 | 700 | 695 | 688 | 608 | 671 | 645 | 355 | 322 | 286 | 254 | 369 | 484 | 406 | 665 | 709 | 577 | 690 | 584 | 608 | 637 | Cacnb2 |
| Cacnb3 | 6431 | 7976 | 8564 | 10080 | 11119 | 9323 | 9215 | 7496 | 8267 | 10750 | 8907 | 9921 | 8320 | 9078 | 6368 | 7259 | 10189 | 9894 | 11838 | 8449 | 10606 | 11939 | 8202 | 10744 | 11317 | 9150 | 8510 | 8761 | Cacnb3 |
| Cacnb4 | 1111 | 1018 | 790 | 1403 | 2390 | 2127 | 2295 | 3364 | 2692 | 2910 | 2610 | 2247 | 2489 | 2373 | 1568 | 1010 | 1321 | 1625 | 2084 | 2336 | 2314 | 3349 | 2576 | 2181 | 3216 | 1099 | 2210 | 2106 | Cacnb4 |
| Cacng1 | 22 | 21 | 5 | 37 | 36 | 42 | 44 | 10 | 34 | 21 | 113 | 21 | 11 | 13 | 11 | 19 | 8 | 28 | 32 | 30 | 38 | 38 | 72 | 26 | 18 | 18 | 23 | 27 | Cacng1 |
| Cacng2 | 320 | 460 | 711 | 1242 | 2318 | 1552 | 1393 | 1432 | 1533 | 2431 | 1960 | 1569 | 1579 | 1679 | 518 | 450 | 1043 | 1416 | 1754 | 1325 | 1650 | 2787 | 1421 | 2017 | 2746 | 1182 | 1416 | 1468 | Cacng2 |
| Cacng3 | 46 | 76 | 89 | 108 | 303 | 198 | 191 | 227 | 180 | 300 | 190 | 152 | 139 | 138 | 103 | 83 | 152 | 206 | 207 | 207 | 230 | 333 | 212 | 198 | 282 | 130 | 156 | 126 | Cacng3 |
| Cacng4 | 4685 | 6246 | 7648 | 7979 | 9955 | 8084 | 7454 | 8956 | 9465 | 12284 | 9718 | 11185 | 11133 | 10750 | 5701 | 6876 | 8230 | 8041 | 9619 | 8624 | 8254 | 13535 | 9740 | 11176 | 13012 | 9838 | 9522 | 10383 | Cacng4 |

| | | | | | | | | | | | | | | | | | | | | | | | | | | | | | |
|---|---|---|---|---|---|---|---|---|---|---|---|---|---|---|---|---|---|---|---|---|---|---|---|---|---|---|---|---|---|
| Cacng5 | 326 | 455 | 417 | 556 | 829 | 541 | 505 | 177 | 197 | 271 | 249 | 128 | 181 | 136 | 364 | 378 | 492 | 650 | 578 | 487 | 651 | 307 | 174 | 168 | 252 | 92 | 113 | 120 | Cacng5 |
| Cacng7 | 0 | 0 | 0 | 0 | 0 | 0 | 0 | 0 | 0 | 0 | 0 | 0 | 0 | 0 | 0 | 0 | 0 | 0 | 0 | 0 | 1 | 0 | 0 | 0 | 0 | 0 | 0 | 0 | Cacng7 |
| Cactin | 2754 | 3747 | 4003 | 3696 | 3001 | 2934 | 2665 | 2383 | 2784 | 2971 | 2903 | 3744 | 3109 | 3211 | 2515 | 3315 | 4117 | 3360 | 3197 | 2680 | 2725 | 2585 | 2639 | 3491 | 3032 | 4220 | 3483 | 3216 | Cactin |
| Cacul1 | 11064 | 8630 | 9912 | 9346 | 9548 | 9854 | 10874 | 8796 | 7168 | 6887 | 7366 | 8189 | 8151 | 8654 | 10704 | 9361 | 8941 | 9444 | 9082 | 9521 | 9975 | 8779 | 7210 | 7234 | 7877 | 7821 | 8312 | 8808 | Cacul1 |
| Cacybp | 14397 | 11839 | 14114 | 13560 | 12103 | 12040 | 15908 | 11080 | 8469 | 8906 | 7918 | 9588 | 8882 | 10320 | 15564 | 13276 | 13726 | 12490 | 13535 | 16054 | 14674 | 8662 | 9420 | 9743 | 9010 | 10914 | 9775 | 10788 | Cacybp |
| Cad | 7216 | 9052 | 10653 | 9247 | 7682 | 7412 | 6800 | 5427 | 7636 | 7783 | 7875 | 11901 | 9378 | 9797 | 6319 | 8636 | 10288 | 8854 | 9040 | 6065 | 7261 | 6657 | 7217 | 9448 | 8082 | 9267 | 9417 | 10326 | Cad |
| Cadm1 | 12572 | 11460 | 10769 | 14468 | 16299 | 14472 | 17126 | 12742 | 11998 | 12264 | 10775 | 10176 | 9804 | 10253 | 13919 | 11721 | 13446 | 14889 | 15306 | 15348 | 16479 | 14250 | 10558 | 11921 | 12974 | 11885 | 8707 | 8733 | Cadm1 |
| Cadm2 | 1187 | 936 | 850 | 2356 | 4178 | 3482 | 3572 | 2804 | 2920 | 2790 | 2684 | 1561 | 1530 | 1640 | 1626 | 1089 | 1343 | 3003 | 2983 | 3050 | 4085 | 3858 | 2523 | 1946 | 3259 | 1039 | 1205 | 1234 | Cadm2 |
| Cadm3 | 8945 | 11470 | 10300 | 17474 | 21940 | 18117 | 17252 | 10353 | 10167 | 13866 | 11585 | 9464 | 9891 | 9294 | 10961 | 9512 | 13833 | 19070 | 19112 | 15530 | 20251 | 16123 | 9950 | 10761 | 14449 | 5848 | 8178 | 8667 | Cadm3 |
| Cadm4 | 7097 | 9736 | 10222 | 14971 | 14996 | 12508 | 12496 | 8498 | 8692 | 11999 | 10014 | 11844 | 9473 | 11413 | 7694 | 6748 | 13663 | 13921 | 15257 | 11146 | 14317 | 11232 | 8141 | 12719 | 14066 | 10965 | 10282 | 11132 | Cadm4 |
| Cadps | 5508 | 6374 | 4757 | 7979 | 12210 | 10002 | 10793 | 11871 | 10223 | 11164 | 9699 | 8430 | 8940 | 8495 | 7477 | 5991 | 6686 | 8745 | 9700 | 10323 | 11256 | 12790 | 10087 | 8391 | 11065 | 5293 | 7877 | 7888 | Cadps |
| Cadps2 | 734 | 584 | 507 | 679 | 918 | 710 | 940 | 1056 | 1018 | 858 | 759 | 658 | 575 | 594 | 754 | 574 | 517 | 800 | 709 | 866 | 879 | 1029 | 891 | 812 | 731 | 787 | 629 | 551 | Cadps2 |
| Cage1 | 247 | 255 | 171 | 206 | 264 | 242 | 275 | 234 | 259 | 196 | 239 | 180 | 163 | 180 | 274 | 269 | 198 | 220 | 211 | 297 | 237 | 197 | 229 | 163 | 227 | 137 | 209 | 126 | Cage1 |
| Calb1 | 901 | 744 | 723 | 1162 | 1256 | 1159 | 1755 | 1272 | 926 | 1042 | 923 | 1106 | 633 | 1394 | 843 | 642 | 942 | 1195 | 1273 | 1309 | 1545 | 1482 | 1009 | 1283 | 1272 | 1305 | 986 | 1131 | Calb1 |
| Calb2 | 386 | 429 | 327 | 789 | 855 | 756 | 1044 | 790 | 928 | 875 | 715 | 647 | 499 | 792 | 555 | 414 | 446 | 724 | 897 | 819 | 919 | 797 | 847 | 973 | 994 | 909 | 519 | 634 | Calb2 |
| Calca | 869 | 959 | 1022 | 596 | 461 | 576 | 258 | 469 | 607 | 444 | 502 | 711 | 956 | 752 | 986 | 1197 | 936 | 854 | 295 | 268 | 274 | 690 | 535 | 533 | 519 | 852 | 774 | 703 | Calca |
| Calcb | 45 | 43 | 36 | 38 | 40 | 19 | 11 | 18 | 26 | 25 | 36 | 24 | 52 | 22 | 47 | 73 | 36 | 40 | 4 | 12 | 7 | 25 | 21 | 20 | 19 | 32 | 24 | 31 | Calcb |
| Calcoco1 | 6522 | 8065 | 7677 | 8654 | 7203 | 7127 | 7375 | 4212 | 4838 | 4627 | 4632 | 4546 | 4223 | 4142 | 6444 | 7498 | 8115 | 8392 | 7711 | 6814 | 7398 | 4583 | 4926 | 4903 | 4886 | 4563 | 3871 | 4059 | Calcoco1 |
| Calcoco2 | 0 | 0 | 0 | 0 | 0 | 0 | 0 | 2 | 2 | 1 | 2 | 0 | 2 | 0 | 0 | 4 | 1 | 0 | 0 | 0 | 1 | 2 | 5 | 1 | 1 | 5 | 0 | 7 | Calcoco2 |
| Calcr | 241 | 297 | 156 | 369 | 506 | 440 | 444 | 554 | 372 | 411 | 319 | 232 | 256 | 213 | 406 | 240 | 367 | 434 | 350 | 431 | 430 | 623 | 354 | 323 | 475 | 248 | 230 | 166 | Calcr |
| Calcrl | 1321 | 678 | 1033 | 746 | 767 | 751 | 781 | 1227 | 1244 | 1086 | 1354 | 981 | 1338 | 1106 | 1100 | 1200 | 782 | 768 | 637 | 999 | 731 | 1113 | 1737 | 1090 | 919 | 1427 | 1207 | 1218 | Calcrl |
| Cald1 | 19890 | 13998 | 15608 | 9899 | 7967 | 11347 | 7573 | 16248 | 17153 | 15150 | 17891 | 14588 | 18191 | 15123 | 17795 | 20845 | 12067 | 10830 | 7635 | 8269 | 7617 | 13231 | 19181 | 15641 | 11406 | 17808 | 16837 | 16555 | Cald1 |
| Calhm1 | 0 | 0 | 0 | 0 | 0 | 2 | 0 | 0 | 0 | 0 | 0 | 0 | 0 | 0 | 0 | 0 | 0 | 0 | 0 | 0 | 0 | 0 | 0 | 0 | 0 | 0 | 0 | 0 | Calhm1 |
| Calhm2 | 50 | 24 | 45 | 35 | 37 | 40 | 25 | 103 | 150 | 134 | 199 | 114 | 139 | 103 | 33 | 40 | 32 | 24 | 40 | 30 | 29 | 94 | 201 | 168 | 124 | 134 | 126 | 116 | Calhm2 |
| Calhm3 | 0 | 1 | 0 | 9 | 2 | 0 | 4 | 0 | 0 | 0 | 0 | 0 | 0 | 0 | 0 | 0 | 0 | 2 | 0 | 0 | 2 | 0 | 0 | 0 | 0 | 1 | 0 | 0 | Calhm3 |
| Calhm4 | 0 | 0 | 0 | 0 | 1 | 0 | 3 | 0 | 0 | 0 | 0 | 1 | 0 | 0 | 0 | 1 | 3 | 1 | 0 | 3 | 3 | 0 | 1 | 1 | 0 | 2 | 0 | 0 | Calhm4 |
| Calhm5 | 42 | 40 | 78 | 26 | 43 | 62 | 31 | 87 | 87 | 87 | 97 | 54 | 68 | 82 | 57 | 59 | 27 | 45 | 20 | 39 | 31 | 75 | 116 | 80 | 83 | 57 | 84 | 87 | Calhm5 |
| Calhm6 | 21 | 9 | 9 | 9 | 9 | 1 | 9 | 10 | 17 | 7 | 12 | 10 | 17 | 11 | 12 | 12 | 7 | 8 | 4 | 4 | 14 | 13 | 8 | 7 | 4 | 10 | 10 | 4 | Calhm6 |
| Calm1 | 60584 | 61190 | 56252 | 61473 | 60296 | 58434 | 65067 | 48778 | 43218 | 42320 | 39332 | 40999 | 38519 | 41751 | 68880 | 57545 | 58961 | 62704 | 59539 | 67913 | 64068 | 46023 | 44067 | 42225 | 44378 | 42459 | 42424 | 43151 | Calm1 |
| Calm2 | 33846 | 30751 | 29806 | 35630 | 39606 | 35452 | 42102 | 49515 | 41525 | 38327 | 36988 | 37074 | 35722 | 42258 | 42475 | 34158 | 32609 | 35588 | 36563 | 46097 | 40712 | 43387 | 43328 | 38888 | 41183 | 42711 | 37829 | 40937 | Calm2 |
| Calm3 | 27084 | 34922 | 31593 | 35397 | 36231 | 31764 | 31969 | 28556 | 31883 | 32871 | 30411 | 27710 | 27535 | 28838 | 28537 | 29845 | 33642 | 34297 | 35011 | 34006 | 34731 | 29711 | 32402 | 31530 | 32427 | 29904 | 27444 | 28802 | Calm3 |
| Calm4 | 1 | 0 | 3 | 3 | 0 | 0 | 0 | 1 | 13 | 0 | 0 | 0 | 0 | 0 | 0 | 0 | 0 | 0 | 0 | 0 | 0 | 0 | 8 | 5 | 3 | 0 | 0 | 1 | Calm4 |
| Calm5 | 0 | 0 | 0 | 0 | 1 | 0 | 0 | 0 | 0 | 0 | 0 | 0 | 0 | 0 | 0 | 0 | 0 | 0 | 0 | 0 | 0 | 0 | 0 | 0 | 0 | 0 | 0 | 0 | Calm5 |
| Calml3 | 7 | 5 | 4 | 2 | 0 | 0 | 0 | 26 | 43 | 30 | 22 | 12 | 14 | 6 | 3 | 5 | 8 | 11 | 1 | 0 | 0 | 0 | 49 | 18 | 0 | 3 | 6 | 6 | Calml3 |
| Calml4 | 68 | 110 | 73 | 89 | 218 | 117 | 118 | 200 | 251 | 227 | 261 | 191 | 280 | 99 | 83 | 132 | 54 | 108 | 89 | 107 | 139 | 247 | 355 | 156 | 176 | 64 | 114 | 104 | Calml4 |
| Caln1 | 204 | 162 | 58 | 349 | 780 | 425 | 431 | 617 | 758 | 753 | 771 | 230 | 325 | 243 | 203 | 186 | 144 | 347 | 381 | 354 | 570 | 1046 | 742 | 478 | 611 | 225 | 202 | 220 | Caln1 |
| Calr | 52246 | 51227 | 57541 | 48965 | 37642 | 38456 | 45306 | 42140 | 41861 | 43328 | 47184 | 37726 | 41374 | 43296 | 51696 | 51648 | 51481 | 49052 | 44063 | 39032 | 44721 | 42902 | 43713 | 45555 | 39847 | 43180 | 39574 | 45525 | Calr |
| Calr3 | 31 | 35 | 33 | 19 | 25 | 45 | 36 | 55 | 50 | 46 | 35 | 53 | 48 | 76 | 42 | 32 | 20 | 19 | 24 | 44 | 27 | 55 | 58 | 65 | 40 | 51 | 73 | 61 | Calr3 |
| Calr4 | 8 | 22 | 6 | 24 | 30 | 21 | 20 | 85 | 50 | 58 | 66 | 28 | 77 | 29 | 36 | 39 | 7 | 12 | 18 | 14 | 21 | 82 | 82 | 38 | 50 | 40 | 37 | 31 | Calr4 |
| Calu | 39431 | 33251 | 32998 | 27539 | 22602 | 26668 | 26024 | 27612 | 27964 | 23261 | 29051 | 22718 | 24267 | 23126 | 39655 | 38167 | 29080 | 29366 | 23474 | 28821 | 24643 | 24533 | 31400 | 23308 | 21981 | 24273 | 23474 | 24318 | Calu |
| Caly | 368 | 580 | 354 | 797 | 975 | 807 | 886 | 506 | 643 | 622 | 563 | 363 | 310 | 393 | 426 | 328 | 584 | 731 | 899 | 882 | 904 | 799 | 596 | 579 | 708 | 379 | 367 | 290 | Caly |
| Camk1 | 1521 | 1409 | 1364 | 1371 | 1750 | 1613 | 1444 | 2431 | 2611 | 2403 | 2691 | 2228 | 2180 | 2331 | 1527 | 1604 | 1462 | 1559 | 1531 | 1555 | 1490 | 2734 | 2678 | 2589 | 2445 | 2197 | 2157 | 2324 | Camk1 |
| Camk1d | 833 | 751 | 743 | 1071 | 1598 | 1496 | 1678 | 1373 | 1710 | 1716 | 1498 | 1259 | 1046 | 1211 | 819 | 834 | 864 | 1399 | 1524 | 1238 | 1661 | 1731 | 1709 | 1428 | 1682 | 726 | 981 | 909 | Camk1d |
| Camk1g | 1633 | 1694 | 1352 | 1764 | 2171 | 1937 | 1979 | 1728 | 1768 | 1672 | 1418 | 1592 | 1318 | 1530 | 1671 | 1424 | 1572 | 1797 | 1973 | 2109 | 2071 | 2125 | 1600 | 1696 | 1937 | 1344 | 1434 | 1387 | Camk1g |
| Camk2a | 3832 | 5135 | 3623 | 6042 | 5788 | 6003 | 6943 | 2292 | 2703 | 2835 | 2689 | 3535 | 1918 | 3331 | 3872 | 3009 | 5493 | 6501 | 6680 | 5976 | 6922 | 3228 | 2626 | 3219 | 3391 | 3363 | 2500 | 2524 | Camk2a |
| Camk2b | 1020 | 1354 | 921 | 3099 | 5747 | 3765 | 3974 | 2766 | 3760 | 4438 | 3704 | 1934 | 1804 | 1865 | 1303 | 989 | 1977 | 3853 | 4498 | 3889 | 4492 | 5103 | 3305 | 3356 | 4835 | 1377 | 1457 | 1413 | Camk2b |
| Camk2d | 2819 | 2257 | 2290 | 3145 | 4407 | 3719 | 3662 | 3368 | 3545 | 3585 | 3938 | 2351 | 2418 | 2287 | 3237 | 2696 | 2586 | 3239 | 3388 | 3502 | 3684 | 4114 | 3863 | 2827 | 3299 | 2256 | 1977 | 2029 | Camk2d |
| Camk2g | 6605 | 6989 | 5505 | 6899 | 8303 | 8304 | 7632 | 5873 | 6562 | 6134 | 6259 | 5970 | 5929 | 5751 | 6637 | 6809 | 6743 | 7559 | 7635 | 7576 | 7973 | 6378 | 6195 | 5641 | 6581 | 5767 | 6030 | 5389 | Camk2g |
| Camk2n1 | 3233 | 3388 | 5158 | 5227 | 6679 | 5752 | 6214 | 5722 | 4102 | 5790 | 5131 | 4729 | 4995 | 5698 | 4007 | 3309 | 5599 | 5447 | 6620 | 5807 | 6821 | 6667 | 4840 | 5999 | 6598 | 4213 | 5112 | 5797 | Camk2n1 |
| Camk2n2 | 962 | 1210 | 1206 | 1762 | 2096 | 1643 | 1829 | 822 | 845 | 1303 | 1150 | 921 | 743 | 973 | 965 | 891 | 1554 | 1858 | 2091 | 1384 | 2003 | 1342 | 783 | 1125 | 1318 | 853 | 687 | 750 | Camk2n2 |
| Camk4 | 221 | 257 | 102 | 151 | 351 | 371 | 296 | 465 | 504 | 352 | 492 | 206 | 243 | 241 | 266 | 306 | 152 | 213 | 176 | 373 | 427 | 269 | 537 | 193 | 412 | 135 | 264 | 211 | Camk4 |
| Camkk1 | 2624 | 2929 | 2904 | 3093 | 3302 | 3167 | 2959 | 1081 | 1172 | 1395 | 1276 | 1013 | 1017 | 926 | 2413 | 2854 | 3037 | 3327 | 3404 | 2965 | 2994 | 1489 | 1249 | 1175 | 1384 | 907 | 1069 | 1031 | Camkk1 |
| Camkk2 | 5462 | 6253 | 6512 | 7713 | 9042 | 7889 | 7768 | 3761 | 4539 | 5111 | 4914 | 4491 | 4419 | 4306 | 5521 | 5724 | 7258 | 7989 | 8785 | 7287 | 8566 | 5148 | 4214 | 4908 | 5549 | 4139 | 4171 | 4015 | Camkk2 |
| Camkmt | 544 | 560 | 514 | 512 | 499 | 418 | 551 | 432 | 445 | 498 | 440 | 445 | 447 | 490 | 479 | 426 | 486 | 513 | 498 | 504 | 474 | 521 | 449 | 476 | 501 | 506 | 449 | 509 | Camkmt |
| Camkv | 2476 | 2670 | 2714 | 2668 | 2869 | 2785 | 3109 | 1077 | 1137 | 1235 | 1081 | 1199 | 804 | 1195 | 2066 | 2097 | 2942 | 2850 | 3474 | 3025 | 3369 | 1310 | 1123 | 1377 | 1543 | 1079 | 1094 | 1079 | Camkv |
| Caml | 2817 | 2974 | 2854 | 3272 | 3129 | 3138 | 3384 | 2941 | 2630 | 2964 | 2815 | 2813 | 2604 | 3064 | 3020 | 2705 | 2965 | 3076 | 3156 | 3266 | 3271 | 2709 | 2596 | 2776 | 3078 | 2872 | 2524 | 2944 | Caml |
| Camp | 3 | 1 | 7 | 2 | 7 | 4 | 4 | 5 | 7 | 5 | 23 | 7 | 14 | 5 | 3 | 9 | 7 | 7 | 2 | 0 | 3 | 5 | 17 | 23 | 6 | 3 | 14 | 6 | Camp |
| Camsap1 | 9027 | 10235 | 9574 | 12371 | 14336 | 13215 | 13110 | 13202 | 13411 | 15230 | 13467 | 16507 | 14233 | 16701 | 9539 | 9087 | 11138 | 12557 | 13835 | 11658 | 13718 | 16158 | 11739 | 14775 | 18352 | 11301 | 15439 | 16175 | Camsap1 |
| Camsap2 | 17767 | 17460 | 12906 | 18377 | 21158 | 21988 | 21009 | 21659 | 17412 | 16857 | 15622 | 15993 | 15528 | 16402 | 19788 | 16248 | 15764 | 19515 | 19591 | 22250 | 22219 | 19696 | 15810 | 14196 | 19384 | 10375 | 16563 | 15320 | Camsap2 |
| Camsap3 | 1942 | 2296 | 2453 | 3101 | 4246 | 3325 | 3115 | 3069 | 3432 | 4484 | 3413 | 3826 | 3181 | 3391 | 1940 | 2046 | 2960 | 2992 | 3842 | 2829 | 3114 | 4953 | 3133 | 3924 | 4661 | 3158 | 2815 | 3183 | Camsap3 |
| Camta1 | 6038 | 5596 | 8017 | 9182 | 10798 | 9055 | 10339 | 8731 | 8416 | 10954 | 9247 | 10447 | 8461 | 10851 | 5872 | 5220 | 8500 | 8825 | 9919 | 7985 | 10182 | 12459 | 7781 | 10763 | 12028 | 9157 | 9705 | 10187 | Camta1 |
| Camta2 | 1480 | 2054 | 2190 | 2670 | 2878 | 2126 | 2256 | 1605 | 2099 | 2689 | 2529 | 2326 | 1952 | 2024 | 1365 | 1907 | 2556 | 2749 | 2766 | 1976 | 2381 | 2730 | 1933 | 2314 | 2677 | 1953 | 1964 | 1869 | Camta2 |
| Cand1 | 16654 | 14569 | 16700 | 15627 | 16586 | 16245 | 17040 | 19141 | 18133 | 18266 | 18049 | 20726 | 20226 | 21268 | 15427 | 14843 | 15907 | 14911 | 16850 | 14162 | 16905 | 19011 | 18028 | 18334 | 18757 | 20686 | 20779 | 21517 | Cand1 |
| Cand2 | 5982 | 6516 | 6908 | 5295 | 5066 | 5181 | 4771 | 4578 | 5978 | 5653 | 5737 | 7018 | 6581 | 6425 | 5275 | 6524 | 6656 | 5450 | 5674 | 4411 | 4839 | 4687 | 5537 | 6323 | 5210 | 7930 | 6575 | 6713 | Cand2 |
| Cant1 | 2721 | 3474 | 3673 | 3763 | 3310 | 3287 | 3085 | 3065 | 3679 | 4209 | 3980 | 4063 | 3848 | 3924 | 2735 | 3269 | 3746 | 3653 | 3747 | 2759 | 3280 | 3974 | 3725 | 4175 | 4041 | 4274 | 3720 | 3922 | Cant1 |
| Canx | 43198 | 35706 | 43164 | 35348 | 31606 | 33485 | 35995 | 38532 | 34948 | 32577 | 36173 | 32923 | 34417 | 35516 | 44224 | 41170 | 38020 | 35424 | 32573 | 36575 | 34042 | 34597 | 36748 | 34585 | 32718 | 36352 | 35395 | 37400 | Canx |
| Cap1 | 16150 | 17248 | 17462 | 16755 | 14885 | 15092 | 14817 | 12414 | 13523 | 13020 | 12625 | 14585 | 13793 | 13748 | 16064 | 16034 | 17121 | 16058 | 16108 | 14850 | 15448 | 12711 | 13455 | 12836 | 13285 | 15071 | 13498 | 14281 | Cap1 |
| Cap2 | 937 | 1030 | 1110 | 912 | 911 | 970 | 1282 | 1076 | 1135 | 1022 | 1564 | 1165 | 944 | 1231 | 1004 | 758 | 1055 | 883 | 1045 | 1120 | 1037 | 998 | 1482 | 1249 | 1067 | 1205 | 1240 | 1197 | Cap2 |
| Capg | 281 | 182 | 204 | 146 | 164 | 152 | 121 | 231 | 450 | 325 | 293 | 179 | 248 | 158 | 189 | 293 | 159 | 149 | 141 | 170 | 134 | 162 | 418 | 235 | 183 | 251 | 171 | 153 | Capg |
| Capn1 | 376 | 477 | 507 | 272 | 346 | 394 | 153 | 572 | 835 | 644 | 867 | 524 | 691 | 410 | 400 | 588 | 384 | 343 | 260 | 193 | 164 | 650 | 1045 | 471 | 497 | 473 | 579 | 586 | Capn1 |
| Capn10 | 2720 | 3696 | 3130 | 3646 | 3454 | 3482 | 3056 | 2497 | 2792 | 2803 | 2521 | 2791 | 2766 | 2607 | 2884 | 3125 | 3687 | 3476 | 3734 | 3271 | 3340 | 2757 | 2695 | 2707 | 3124 | 2795 | 2721 | 2909 | Capn10 |
| Capn11 | 1058 | 96 | 772 | 69 | 83 | 72 | 733 | 17 | 14 | 32 | 284 | 250 | 12 | 81 | 62 | 44 | 49 | 265 | 114 | 75 | 84 | 177 | 65 | 757 | 24 | 285 | 13 | 665 | Capn11 |
| Capn12 | 10 | 11 | 15 | 20 | 14 | 19 | 4 | 10 | 8 | 0 | 20 | 25 | 14 | 9 | 16 | 9 | 29 | 17 | 9 | 17 | 20 | 8 | 8 | 15 | 9 | 14 | 6 | 9 | Capn12 |
| Capn13 | 8 | 3 | 3 | 6 | 7 | 1 | 1 | 0 | 6 | 2 | 12 | 3 | 7 | 15 | 7 | 3 | 1 | 2 | 3 | 3 | 2 | 2 | 3 | 4 | 4 | 1 | 3 | 12 | Capn13 |
| Capn15 | 1944 | 2611 | 3620 | 3596 | 3600 | 3027 | 2871 | 2545 | 3325 | 3973 | 3864 | 4238 | 3718 | 3450 | 1820 | 2497 | 3978 | 3439 | 3712 | 2349 | 2706 | 4186 | 3062 | 4376 | 4207 | 4449 | 3787 | 3609 | Capn15 |
| Capn2 | 6006 | 5532 | 6011 | 4558 | 5508 | 5679 | 4270 | 8422 | 9358 | 8913 | 9742 | 9040 | 9776 | 9096 | 6258 | 6350 | 5587 | 5070 | 4802 | 4320 | 4491 | 8768 | 10087 | 8798 | 8191 | 9784 | 8721 | 9337 | Capn2 |
| Capn3 | 2 | 11 | 6 | 2 | 5 | 12 | 13 | 9 | 17 | 6 | 43 | 17 | 18 | 20 | 3 | 2 | 6 | 4 | 7 | 13 | 4 | 17 | 62 | 20 | 5 | 21 | 28 | 25 | Capn3 |
| Capn5 | 1945 | 2733 | 2121 | 2659 | 3216 | 2490 | 2582 | 1458 | 1943 | 1839 | 1804 | 1522 | 1346 | 1454 | 2405 | 2482 | 2518 | 2830 | 3008 | 2358 | 2646 | 1975 | 1867 | 1545 | 1761 | 1342 | 1392 | 1227 | Capn5 |
| Capn6 | 8477 | 3279 | 2059 | 2084 | 1200 | 2165 | 1047 | 4829 | 7517 | 4451 | 12222 | 1562 | 2931 | 1559 | 4232 | 6045 | 1801 | 1961 | 1320 | 980 | 1013 | 4385 | 9774 | 3193 | 1796 | 3692 | 1361 | 1621 | Capn6 |

| | | | | | | | | | | | | | | | | | | | | | | | | | | | | | |
|---|---|---|---|---|---|---|---|---|---|---|---|---|---|---|---|---|---|---|---|---|---|---|---|---|---|---|---|---|---|
| Capn7 | 9666 | 8543 | 7461 | 9228 | 9585 | 9775 | 10321 | 9493 | 7495 | 6979 | 7362 | 6960 | 6546 | 7009 | 10578 | 8614 | 7922 | 9044 | 8827 | 10815 | 10226 | 8641 | 8036 | 6713 | 7612 | 5906 | 6833 | 6577 | Capn7 |
| Capn8 | 10 | 14 | 0 | 13 | 30 | 12 | 15 | 29 | 30 | 19 | 27 | 30 | 12 | 17 | 17 | 9 | 17 | 10 | 5 | 25 | 9 | 15 | 10 | 12 | 18 | 10 | 14 | 12 | Capn8 |
| Capn9 | 5 | 3 | 1 | 5 | 3 | 0 | 7 | 0 | 2 | 8 | 4 | 1 | 0 | 5 | 3 | 2 | 2 | 6 | 2 | 0 | 4 | 2 | 1 | 0 | 4 | 2 | 1 | 6 | Capn9 |
| Capns1 | 10691 | 12461 | 10100 | 9885 | 9045 | 9352 | 8574 | 7351 | 8397 | 7304 | 8005 | 6769 | 7764 | 6771 | 10437 | 11157 | 9959 | 9765 | 9840 | 8758 | 8854 | 7636 | 8989 | 7205 | 6970 | 8047 | 7348 | 7382 | Capns1 |
| Capns2 | 69 | 108 | 7 | 37 | 62 | 83 | 54 | 85 | 86 | 47 | 93 | 43 | 25 | 36 | 99 | 114 | 14 | 39 | 41 | 91 | 70 | 15 | 93 | 29 | 26 | 9 | 33 | 17 | Capns2 |
| Caprin1 | 45382 | 39523 | 43057 | 38687 | 43228 | 41527 | 46241 | 78053 | 64320 | 61547 | 57519 | 71664 | 65022 | 71738 | 47329 | 38871 | 41773 | 38660 | 43302 | 47056 | 44590 | 65702 | 62239 | 65716 | 68530 | 71335 | 72471 | 74656 | Caprin1 |
| Caprin2 | 4610 | 3655 | 3553 | 3310 | 2971 | 3673 | 3463 | 2943 | 2700 | 2668 | 2713 | 3023 | 3136 | 2981 | 4301 | 3981 | 3364 | 3361 | 2823 | 3651 | 3342 | 2650 | 2700 | 2393 | 2574 | 2389 | 3274 | 2962 | Caprin2 |
| Caps2 | 44 | 31 | 34 | 53 | 84 | 61 | 72 | 57 | 82 | 98 | 91 | 75 | 76 | 90 | 49 | 50 | 58 | 39 | 48 | 62 | 48 | 104 | 98 | 69 | 38 | 95 | 59 | 71 | Caps2 |
| Capsl | 13 | 21 | 9 | 26 | 28 | 67 | 36 | 43 | 42 | 43 | 38 | 38 | 65 | 22 | 33 | 33 | 15 | 14 | 32 | 31 | 22 | 43 | 64 | 35 | 35 | 53 | 16 | 28 | Capsl |
| Capza1 | 12998 | 10804 | 11208 | 9490 | 9673 | 10077 | 11263 | 14396 | 12529 | 10343 | 10774 | 12029 | 12169 | 12192 | 14122 | 12136 | 10339 | 9515 | 9735 | 12426 | 10144 | 11882 | 12908 | 11037 | 11277 | 13013 | 12740 | 12892 | Capza1 |
| Capza2 | 13434 | 10285 | 13062 | 11852 | 12390 | 12046 | 13957 | 15358 | 12824 | 12677 | 13291 | 14685 | 13674 | 15403 | 14029 | 11998 | 12399 | 12394 | 12249 | 12968 | 12635 | 14297 | 13496 | 13410 | 13538 | 15687 | 14587 | 16021 | Capza2 |
| Capzb | 13666 | 14251 | 13981 | 12856 | 11952 | 11836 | 11824 | 10953 | 11602 | 11763 | 11486 | 12041 | 12301 | 12588 | 12879 | 13403 | 13786 | 12660 | 12436 | 11173 | 12871 | 10572 | 12435 | 12412 | 11111 | 13622 | 11993 | 13190 | Capzb |
| Car1 | 1 | 4 | 5 | 3 | 0 | 4 | 3 | 4 | 1 | 0 | 1 | 0 | 2 | 3 | 1 | 3 | 0 | 0 | 0 | 0 | 1 | 2 | 1 | 0 | 0 | 0 | 1 | 5 | Car1 |
| Car10 | 262 | 377 | 341 | 524 | 1455 | 700 | 711 | 812 | 844 | 1058 | 953 | 390 | 611 | 380 | 508 | 437 | 440 | 669 | 736 | 684 | 808 | 1497 | 987 | 538 | 921 | 222 | 397 | 354 | Car10 |
| Car11 | 572 | 853 | 692 | 944 | 1397 | 981 | 926 | 1208 | 1245 | 1601 | 1421 | 1021 | 1234 | 1047 | 731 | 810 | 1014 | 1120 | 1206 | 1060 | 1082 | 1699 | 1253 | 1313 | 1561 | 774 | 1027 | 864 | Car11 |
| Car12 | 582 | 401 | 271 | 389 | 522 | 378 | 383 | 648 | 796 | 686 | 693 | 239 | 342 | 194 | 429 | 511 | 251 | 335 | 390 | 383 | 415 | 545 | 1244 | 473 | 286 | 372 | 139 | 179 | Car12 |
| Car13 | 495 | 369 | 404 | 195 | 209 | 234 | 162 | 293 | 325 | 263 | 337 | 217 | 308 | 211 | 378 | 476 | 214 | 284 | 133 | 163 | 127 | 302 | 302 | 169 | 132 | 222 | 242 | 223 | Car13 |
| Car14 | 841 | 673 | 797 | 524 | 586 | 629 | 578 | 898 | 803 | 842 | 614 | 1069 | 1178 | 1196 | 674 | 720 | 581 | 346 | 645 | 739 | 642 | 453 | 869 | 1034 | 870 | 1461 | 1147 | 1342 | Car14 |
| Car15 | 129 | 138 | 197 | 179 | 186 | 97 | 222 | 106 | 77 | 98 | 94 | 111 | 59 | 126 | 127 | 97 | 188 | 177 | 231 | 117 | 198 | 152 | 65 | 122 | 113 | 73 | 83 | 91 | Car15 |
| Car2 | 1263 | 1701 | 1579 | 696 | 637 | 684 | 741 | 873 | 1194 | 890 | 797 | 1126 | 2658 | 1877 | 1557 | 1668 | 947 | 673 | 677 | 1037 | 687 | 675 | 1229 | 815 | 695 | 1323 | 1502 | 3593 | Car2 |
| Car3 | 75 | 64 | 115 | 39 | 63 | 46 | 36 | 76 | 150 | 109 | 246 | 68 | 136 | 119 | 69 | 95 | 67 | 52 | 53 | 62 | 41 | 115 | 179 | 76 | 140 | 38 | 99 | 109 | Car3 |
| Car4 | 99 | 106 | 171 | 89 | 108 | 93 | 114 | 84 | 73 | 102 | 68 | 79 | 93 | 90 | 100 | 130 | 103 | 119 | 94 | 129 | 100 | 149 | 60 | 60 | 103 | 93 | 131 | 108 | Car4 |
| Car5a | 9 | 3 | 2 | 3 | 7 | 7 | 4 | 4 | 2 | 11 | 9 | 7 | 3 | 2 | 7 | 3 | 9 | 8 | 10 | 9 | 5 | 5 | 6 | 7 | 8 | 4 | 6 | 12 | Car5a |
| Car5b | 572 | 418 | 457 | 328 | 289 | 308 | 327 | 77 | 92 | 65 | 80 | 49 | 58 | 63 | 493 | 562 | 440 | 386 | 244 | 308 | 297 | 92 | 91 | 47 | 51 | 58 | 67 | 52 | Car5b |
| Car6 | 0 | 1 | 1 | 0 | 1 | 0 | 1 | 7 | 4 | 3 | 9 | 2 | 11 | 4 | 5 | 2 | 2 | 2 | 2 | 2 | 1 | 3 | 3 | 7 | 1 | 11 | 8 | 4 | Car6 |
| Car7 | 120 | 161 | 97 | 156 | 142 | 131 | 120 | 44 | 61 | 63 | 41 | 55 | 51 | 54 | 127 | 118 | 160 | 136 | 132 | 186 | 150 | 45 | 48 | 71 | 63 | 47 | 43 | 58 | Car7 |
| Car8 | 648 | 473 | 458 | 448 | 486 | 403 | 468 | 191 | 222 | 175 | 249 | 185 | 163 | 152 | 534 | 563 | 448 | 479 | 396 | 382 | 431 | 217 | 291 | 179 | 141 | 186 | 172 | 157 | Car8 |
| Car9 | 40 | 20 | 40 | 27 | 25 | 18 | 22 | 23 | 35 | 13 | 44 | 20 | 20 | 24 | 29 | 39 | 31 | 39 | 19 | 26 | 17 | 25 | 37 | 31 | 39 | 47 | 22 | 36 | Car9 |
| Card10 | 1093 | 1447 | 1523 | 1232 | 1078 | 1050 | 839 | 333 | 575 | 475 | 659 | 436 | 532 | 399 | 1031 | 1448 | 1360 | 1217 | 1083 | 855 | 809 | 404 | 622 | 500 | 380 | 556 | 485 | 379 | Card10 |
| Card11 | 7 | 6 | 1 | 1 | 10 | 9 | 2 | 10 | 16 | 14 | 35 | 18 | 5 | 9 | 1 | 9 | 6 | 12 | 3 | 4 | 8 | 33 | 17 | 11 | 3 | 2 | 5 | 9 | Card11 |
| Card14 | 12 | 6 | 9 | 3 | 2 | 4 | 0 | 24 | 48 | 30 | 21 | 16 | 16 | 24 | 9 | 10 | 5 | 11 | 5 | 0 | 2 | 3 | 65 | 21 | 5 | 17 | 10 | 27 | Card14 |
| Card19 | 625 | 717 | 606 | 670 | 567 | 553 | 551 | 233 | 349 | 365 | 321 | 191 | 304 | 184 | 580 | 720 | 586 | 679 | 568 | 565 | 506 | 299 | 353 | 236 | 226 | 242 | 201 | 171 | Card19 |
| Card6 | 164 | 158 | 196 | 164 | 151 | 159 | 162 | 198 | 211 | 195 | 322 | 209 | 249 | 179 | 182 | 167 | 191 | 208 | 176 | 112 | 147 | 202 | 261 | 227 | 208 | 267 | 171 | 157 | Card6 |
| Card9 | 60 | 99 | 77 | 96 | 97 | 86 | 66 | 62 | 63 | 57 | 72 | 62 | 82 | 57 | 63 | 92 | 90 | 106 | 77 | 83 | 53 | 70 | 86 | 69 | 68 | 86 | 86 | 66 | Card9 |
| Carf | 1300 | 920 | 905 | 1250 | 1297 | 1295 | 1440 | 1557 | 1563 | 1509 | 1808 | 1477 | 1558 | 1623 | 1273 | 1136 | 1125 | 1170 | 1114 | 1233 | 1412 | 1644 | 1598 | 1389 | 1573 | 1193 | 1404 | 1456 | Carf |
| Carhsp1 | 23286 | 26984 | 29445 | 27421 | 25830 | 25518 | 26492 | 13423 | 12114 | 13709 | 13375 | 16725 | 15234 | 16316 | 24331 | 23637 | 28971 | 26053 | 28409 | 23326 | 26311 | 15944 | 12101 | 15143 | 15919 | 15967 | 16784 | 17764 | Carhsp1 |
| Carlr | 29 | 46 | 40 | 78 | 48 | 42 | 51 | 55 | 39 | 40 | 66 | 79 | 44 | 83 | 62 | 23 | 83 | 36 | 51 | 73 | 47 | 55 | 27 | 48 | 52 | 69 | 80 | 72 | Carlr |
| Carm1 | 5188 | 5935 | 6800 | 5606 | 5808 | 5574 | 4708 | 6322 | 7531 | 8246 | 8204 | 9131 | 8196 | 7665 | 4584 | 5692 | 6406 | 5212 | 6201 | 4368 | 5223 | 8056 | 7526 | 8436 | 7789 | 10080 | 7458 | 7990 | Carm1 |
| Carmil1 | 1583 | 1535 | 1384 | 1307 | 1541 | 1554 | 1571 | 1917 | 2357 | 2061 | 2161 | 1858 | 1845 | 1932 | 1341 | 1562 | 1289 | 1454 | 1504 | 1139 | 1481 | 2082 | 2062 | 2052 | 1972 | 1843 | 1714 | 1683 | Carmil1 |
| Carmil2 | 329 | 475 | 702 | 665 | 1015 | 750 | 624 | 789 | 817 | 1040 | 947 | 831 | 838 | 738 | 413 | 463 | 662 | 725 | 863 | 624 | 742 | 1247 | 750 | 928 | 1012 | 692 | 722 | 672 | Carmil2 |
| Carmil3 | 4673 | 6541 | 6404 | 9002 | 10541 | 9193 | 9426 | 6379 | 8137 | 10074 | 7378 | 9394 | 7043 | 8212 | 4894 | 5127 | 8248 | 9590 | 11176 | 8306 | 9487 | 9449 | 6601 | 10245 | 10968 | 8024 | 7124 | 6636 | Carmil3 |
| Carnmt1 | 2655 | 1815 | 2444 | 1902 | 1837 | 2137 | 2090 | 1785 | 1523 | 1458 | 1413 | 1824 | 2016 | 1994 | 2371 | 2364 | 1897 | 1966 | 1940 | 1981 | 1920 | 1589 | 2043 | 1722 | 1492 | 1928 | 1643 | 1816 | Carnmt1 |
| Carns1 | 8 | 4 | 5 | 4 | 2 | 2 | 6 | 10 | 5 | 3 | 5 | 5 | 9 | 11 | 7 | 5 | 0 | 6 | 4 | 4 | 2 | 8 | 3 | 5 | 1 | 5 | 5 | 6 | Carns1 |
| Cars | 4585 | 4958 | 4691 | 4387 | 3989 | 4110 | 4053 | 2937 | 3264 | 3354 | 3210 | 3653 | 3327 | 4110 | 4159 | 4330 | 4354 | 4252 | 4120 | 4119 | 4081 | 3080 | 3039 | 3396 | 3538 | 4011 | 3656 | 4086 | Cars |
| Cars2 | 1551 | 1573 | 1342 | 1631 | 1342 | 1509 | 1455 | 1473 | 1828 | 1745 | 1970 | 1714 | 1566 | 1487 | 1274 | 1492 | 1516 | 1536 | 1472 | 1488 | 1339 | 1721 | 1745 | 1726 | 1872 | 1647 | 1531 | 1402 | Cars2 |
| Cartpt | 187 | 227 | 260 | 174 | 185 | 155 | 207 | 216 | 171 | 173 | 198 | 301 | 310 | 366 | 202 | 181 | 272 | 160 | 164 | 264 | 158 | 212 | 210 | 228 | 161 | 234 | 535 | 503 | Cartpt |
| Casc1 | 849 | 730 | 512 | 856 | 626 | 753 | 873 | 256 | 266 | 203 | 209 | 202 | 228 | 211 | 872 | 813 | 629 | 914 | 670 | 764 | 891 | 212 | 227 | 186 | 173 | 168 | 237 | 222 | Casc1 |
| Casc3 | 6660 | 7157 | 8544 | 9467 | 8968 | 7969 | 8495 | 7052 | 7381 | 9070 | 8132 | 10230 | 8544 | 10146 | 6389 | 6857 | 8985 | 9591 | 8917 | 6739 | 8631 | 9514 | 6827 | 10059 | 9366 | 10424 | 8875 | 9597 | Casc3 |
| Casd1 | 2531 | 2115 | 2075 | 2045 | 2576 | 2401 | 2481 | 7897 | 6990 | 6020 | 6302 | 6217 | 6366 | 6092 | 2815 | 2368 | 2121 | 1954 | 2299 | 2833 | 2299 | 6976 | 6757 | 5811 | 6215 | 6205 | 6564 | 6037 | Casd1 |
| Cask | 7874 | 7083 | 6854 | 8785 | 11105 | 10483 | 10625 | 12422 | 11330 | 12451 | 11909 | 10875 | 10957 | 11446 | 8420 | 7100 | 7737 | 9018 | 10063 | 9580 | 10979 | 14059 | 11021 | 10726 | 12897 | 8001 | 10766 | 10318 | Cask |
| Caskin1 | 2625 | 3938 | 4637 | 5479 | 7027 | 5576 | 4783 | 4021 | 4542 | 6358 | 5087 | 5122 | 4356 | 4226 | 2971 | 3289 | 5346 | 5795 | 6869 | 4684 | 5229 | 7140 | 3922 | 5257 | 6290 | 3143 | 4173 | 3773 | Caskin1 |
| Caskin2 | 2701 | 3886 | 3983 | 3665 | 3615 | 3262 | 2720 | 2612 | 3123 | 3230 | 3138 | 3623 | 3319 | 2881 | 2622 | 3572 | 4074 | 4001 | 3839 | 2726 | 3004 | 3039 | 3112 | 3457 | 3054 | 4268 | 3242 | 2942 | Caskin2 |
| Casp1 | 32 | 22 | 27 | 39 | 16 | 35 | 35 | 61 | 60 | 43 | 59 | 26 | 58 | 40 | 26 | 44 | 24 | 41 | 11 | 27 | 32 | 43 | 78 | 35 | 37 | 72 | 71 | 58 | Casp1 |
| Casp12 | 0 | 3 | 10 | 0 | 8 | 3 | 10 | 18 | 20 | 6 | 69 | 15 | 9 | 7 | 8 | 3 | 2 | 6 | 6 | 12 | 12 | 18 | 61 | 15 | 16 | 7 | 15 | 9 | Casp12 |
| Casp14 | 2 | 1 | 0 | 0 | 0 | 0 | 1 | 1 | 12 | 4 | 8 | 2 | 5 | 2 | 1 | 2 | 3 | 1 | 0 | 0 | 3 | 0 | 17 | 0 | 1 | 0 | 0 | 5 | Casp14 |
| Casp2 | 6516 | 5837 | 5278 | 5029 | 4711 | 4988 | 5441 | 7367 | 6128 | 5339 | 5098 | 6391 | 5769 | 6535 | 6158 | 5521 | 5233 | 4616 | 5201 | 5754 | 5352 | 6039 | 5738 | 5750 | 5769 | 6476 | 6495 | 6776 | Casp2 |
| Casp3 | 18837 | 16973 | 18576 | 18521 | 18929 | 17656 | 21154 | 15990 | 13767 | 13370 | 12467 | 13131 | 12576 | 14365 | 21418 | 19052 | 18199 | 18772 | 18687 | 21951 | 20498 | 14440 | 13396 | 13800 | 14568 | 14836 | 13527 | 14383 | Casp3 |
| Casp4 | 3 | 5 | 0 | 12 | 7 | 15 | 4 | 13 | 15 | 12 | 24 | 9 | 27 | 13 | 11 | 9 | 1 | 6 | 5 | 12 | 21 | 7 | 20 | 16 | 23 | 15 | 9 | 9 | Casp4 |
| Casp6 | 2526 | 2427 | 2644 | 2212 | 1801 | 1832 | 1934 | 1603 | 1612 | 1422 | 1605 | 1763 | 1906 | 1767 | 2279 | 2451 | 2365 | 2117 | 1934 | 2087 | 1826 | 1302 | 1749 | 1858 | 1413 | 2272 | 1861 | 1968 | Casp6 |
| Casp7 | 320 | 262 | 422 | 380 | 359 | 343 | 323 | 1011 | 1077 | 963 | 1139 | 1118 | 1120 | 1127 | 319 | 338 | 350 | 316 | 344 | 381 | 319 | 907 | 1148 | 1008 | 966 | 1304 | 1085 | 1050 | Casp7 |
| Casp8 | 511 | 320 | 489 | 296 | 295 | 306 | 184 | 812 | 795 | 702 | 941 | 724 | 915 | 649 | 390 | 490 | 366 | 329 | 254 | 284 | 221 | 690 | 1070 | 873 | 567 | 979 | 758 | 741 | Casp8 |
| Casp8ap2 | 12439 | 10096 | 8224 | 9638 | 8132 | 9441 | 10547 | 8651 | 7273 | 6083 | 6326 | 7923 | 7282 | 8477 | 12812 | 11203 | 8360 | 9361 | 8648 | 11857 | 9778 | 5753 | 6496 | 6377 | 6447 | 6126 | 8562 | 8025 | Casp8ap2 |
| Casp9 | 3480 | 3962 | 3832 | 3919 | 3507 | 3497 | 3486 | 2292 | 2237 | 2287 | 2395 | 2530 | 2606 | 2428 | 3632 | 3727 | 3950 | 3977 | 3707 | 3367 | 3461 | 2563 | 2278 | 2252 | 2271 | 2390 | 2420 | 2406 | Casp9 |
| Casq1 | 58 | 78 | 56 | 128 | 135 | 128 | 135 | 143 | 210 | 161 | 364 | 136 | 137 | 154 | 74 | 79 | 81 | 129 | 104 | 146 | 160 | 147 | 262 | 160 | 204 | 122 | 137 | 108 | Casq1 |
| Casq2 | 67 | 16 | 0 | 3 | 18 | 27 | 12 | 112 | 257 | 47 | 1478 | 31 | 66 | 16 | 61 | 73 | 10 | 20 | 6 | 4 | 8 | 132 | 1104 | 42 | 28 | 10 | 26 | 21 | Casq2 |
| Casr | 39 | 17 | 12 | 12 | 11 | 5 | 4 | 57 | 42 | 56 | 47 | 11 | 29 | 12 | 18 | 17 | 7 | 3 | 12 | 10 | 4 | 7 | 71 | 74 | 6 | 32 | 13 | 8 | Casr |
| Cass4 | 17 | 29 | 16 | 19 | 9 | 12 | 17 | 9 | 13 | 12 | 9 | 10 | 26 | 13 | 12 | 23 | 20 | 17 | 22 | 20 | 7 | 17 | 14 | 16 | 7 | 12 | 14 | 14 | Cass4 |
| Cast | 2587 | 1970 | 2414 | 1420 | 1549 | 1787 | 1177 | 1823 | 2299 | 1750 | 2640 | 1892 | 2589 | 1853 | 2560 | 2896 | 1687 | 1648 | 1226 | 1309 | 1114 | 1781 | 3020 | 1727 | 1484 | 1922 | 2101 | 2074 | Cast |
| Castor1 | 174 | 236 | 220 | 200 | 177 | 144 | 115 | 73 | 128 | 71 | 125 | 81 | 91 | 67 | 145 | 258 | 182 | 200 | 141 | 111 | 113 | 62 | 128 | 65 | 74 | 74 | 71 | 80 | Castor1 |
| Castor2 | 9732 | 11168 | 13424 | 15100 | 17207 | 14532 | 14266 | 5764 | 6063 | 8168 | 6846 | 7618 | 6098 | 7002 | 9679 | 10312 | 15288 | 15639 | 17480 | 12170 | 15227 | 8729 | 5794 | 8216 | 8930 | 5926 | 6199 | 6677 | Castor2 |
| Casz1 | 1348 | 2147 | 2264 | 2573 | 1938 | 1661 | 1431 | 576 | 752 | 867 | 1240 | 524 | 1042 | 503 | 1386 | 2116 | 2610 | 2291 | 1928 | 1392 | 1565 | 1008 | 764 | 571 | 731 | 412 | 442 | 498 | Casz1 |
| Cat | 8433 | 9113 | 8703 | 7461 | 6960 | 7499 | 7671 | 6295 | 6345 | 5838 | 5252 | 7211 | 7208 | 6870 | 8732 | 8446 | 8460 | 7458 | 7582 | 8068 | 7385 | 5882 | 6642 | 6341 | 6023 | 7849 | 7286 | 7831 | Cat |
| Catip | 45 | 52 | 55 | 62 | 79 | 74 | 50 | 62 | 88 | 95 | 96 | 88 | 132 | 92 | 74 | 61 | 38 | 97 | 57 | 56 | 67 | 125 | 149 | 32 | 75 | 67 | 78 | 68 | Catip |
| Catsper2 | 127 | 145 | 145 | 169 | 224 | 241 | 185 | 416 | 386 | 436 | 534 | 446 | 404 | 456 | 181 | 131 | 214 | 193 | 203 | 193 | 212 | 543 | 400 | 477 | 487 | 346 | 388 | 432 | Catsper2 |
| Catsper3 | 1 | 9 | 4 | 12 | 8 | 12 | 11 | 5 | 3 | 1 | 12 | 5 | 7 | 11 | 3 | 11 | 3 | 12 | 4 | 4 | 7 | 2 | 4 | 11 | 16 | 7 | 1 | 5 | Catsper3 |
| Catsper4 | 18 | 14 | 18 | 27 | 11 | 13 | 7 | 9 | 7 | 9 | 7 | 0 | 7 | 4 | 11 | 17 | 7 | 26 | 4 | 16 | 15 | 2 | 8 | 6 | 2 | 0 | 2 | 2 | Catsper4 |
| Catsperb | 0 | 0 | 0 | 0 | 0 | 0 | 0 | 0 | 0 | 0 | 0 | 1 | 0 | 0 | 0 | 0 | 0 | 0 | 0 | 0 | 0 | 0 | 0 | 0 | 0 | 0 | 3 | 0 | Catsperb |
| Catsperd | 116 | 130 | 86 | 194 | 256 | 233 | 228 | 192 | 219 | 204 | 196 | 179 | 202 | 146 | 160 | 138 | 257 | 260 | 257 | 279 | 223 | 331 | 189 | 168 | 252 | 91 | 158 | 176 | Catsperd |
| Catspere1 | 0 | 0 | 0 | 0 | 0 | 0 | 2 | 0 | 0 | 0 | 0 | 0 | 0 | 1 | 3 | 2 | 0 | 0 | 0 | 2 | 0 | 0 | 3 | 0 | 0 | 0 | 0 | 1 | Catspere1 |

| | | | | | | | | | | | | | | | | | | | | | | | | | | | | | |
|---|---|---|---|---|---|---|---|---|---|---|---|---|---|---|---|---|---|---|---|---|---|---|---|---|---|---|---|---|---|
| Catspere2 | 45 | 58 | 31 | 39 | 43 | 40 | 34 | 24 | 32 | 34 | 31 | 25 | 44 | 20 | 30 | 43 | 22 | 53 | 33 | 33 | 29 | 40 | 38 | 33 | 20 | 17 | 27 | 17 | Catspere2 |
| Catsperg1 | 531 | 534 | 484 | 691 | 830 | 793 | 653 | 374 | 353 | 484 | 443 | 300 | 293 | 308 | 555 | 657 | 626 | 747 | 675 | 724 | 742 | 555 | 304 | 305 | 509 | 196 | 293 | 241 | Catsperg1 |
| Catsperg2 | 14 | 13 | 6 | 21 | 28 | 47 | 19 | 22 | 27 | 29 | 29 | 41 | 37 | 35 | 29 | 15 | 16 | 25 | 25 | 45 | 50 | 90 | 46 | 42 | 71 | 16 | 22 | 32 | Catsperg2 |
| Catsperz | 20 | 25 | 15 | 14 | 17 | 14 | 31 | 7 | 11 | 3 | 13 | 7 | 3 | 7 | 17 | 41 | 17 | 15 | 14 | 26 | 13 | 10 | 8 | 7 | 2 | 0 | 7 | 4 | Catsperz |
| Cav1 | 829 | 572 | 583 | 641 | 929 | 975 | 894 | 1660 | 2380 | 1709 | 2727 | 1138 | 1276 | 1215 | 802 | 700 | 568 | 927 | 936 | 1186 | 886 | 1826 | 3839 | 1825 | 1711 | 1142 | 953 | 1161 | Cav1 |
| Cav2 | 337 | 225 | 314 | 285 | 394 | 405 | 420 | 606 | 669 | 511 | 724 | 482 | 600 | 533 | 372 | 336 | 330 | 339 | 360 | 431 | 375 | 460 | 909 | 611 | 547 | 552 | 437 | 538 | Cav2 |
| Cav3 | 6 | 3 | 0 | 0 | 1 | 0 | 0 | 10 | 19 | 15 | 173 | 0 | 11 | 0 | 4 | 2 | 1 | 2 | 0 | 0 | 0 | 15 | 133 | 7 | 3 | 0 | 0 | 2 | Cav3 |
| Cavin1 | 3481 | 3704 | 3570 | 2997 | 2356 | 3564 | 1581 | 3416 | 4747 | 4205 | 5644 | 3081 | 3928 | 3281 | 3658 | 4003 | 3071 | 4210 | 1963 | 1730 | 1810 | 3989 | 6468 | 4679 | 3667 | 3324 | 3819 | 3298 | Cavin1 |
| Cavin2 | 260 | 159 | 214 | 222 | 350 | 352 | 202 | 556 | 958 | 653 | 1623 | 439 | 598 | 372 | 293 | 267 | 240 | 305 | 238 | 300 | 214 | 725 | 1572 | 519 | 552 | 419 | 382 | 362 | Cavin2 |
| Cavin3 | 356 | 455 | 435 | 337 | 295 | 490 | 232 | 793 | 1020 | 803 | 949 | 775 | 881 | 797 | 355 | 403 | 292 | 386 | 276 | 289 | 232 | 690 | 1197 | 842 | 774 | 782 | 764 | 793 | Cavin3 |
| Cavin4 | 242 | 260 | 264 | 383 | 368 | 398 | 318 | 397 | 321 | 404 | 465 | 532 | 437 | 486 | 237 | 251 | 337 | 278 | 365 | 341 | 348 | 471 | 375 | 397 | 530 | 308 | 522 | 447 | Cavin4 |
| Cbarp | 3073 | 4238 | 3963 | 5596 | 6135 | 4913 | 4199 | 3032 | 3872 | 5012 | 4042 | 3754 | 3241 | 3193 | 3066 | 3544 | 5277 | 5735 | 5634 | 4216 | 4951 | 5798 | 3425 | 4185 | 4987 | 3484 | 2990 | 2872 | Cbarp |
| Cbfa2t2 | 8431 | 8102 | 7159 | 8424 | 8376 | 8482 | 8713 | 8054 | 7192 | 7183 | 6785 | 8692 | 8326 | 8617 | 8329 | 7633 | 8108 | 8349 | 8449 | 7995 | 8577 | 7646 | 6988 | 7825 | 8518 | 7341 | 8932 | 9070 | Cbfa2t2 |
| Cbfa2t3 | 735 | 863 | 1208 | 1697 | 3201 | 2059 | 1861 | 2560 | 3223 | 4998 | 4192 | 3884 | 3244 | 3672 | 873 | 810 | 1820 | 2023 | 2559 | 1563 | 2096 | 5249 | 2937 | 4090 | 4766 | 2770 | 2884 | 3205 | Cbfa2t3 |
| Cbfb | 7350 | 4727 | 6220 | 5602 | 5906 | 5793 | 6459 | 8902 | 8319 | 7186 | 9156 | 7635 | 8868 | 8546 | 7023 | 6460 | 5976 | 6309 | 5721 | 5354 | 5855 | 8119 | 8337 | 8060 | 7116 | 10304 | 7548 | 9169 | Cbfb |
| Cbl | 13937 | 12606 | 9214 | 12623 | 13630 | 15202 | 14377 | 14510 | 13401 | 12803 | 12512 | 14104 | 13622 | 14329 | 13497 | 12832 | 10929 | 12455 | 12839 | 13489 | 15093 | 12352 | 12145 | 10874 | 14568 | 8911 | 14674 | 12709 | Cbl |
| Cblb | 4946 | 4491 | 6737 | 6727 | 6465 | 5819 | 6268 | 5434 | 5010 | 6435 | 6155 | 7619 | 8781 | 7349 | 5070 | 5474 | 6730 | 6185 | 6154 | 4837 | 5919 | 7439 | 5218 | 6655 | 6591 | 7037 | 7284 | 8146 | Cblb |
| Cblc | 53 | 14 | 11 | 5 | 6 | 6 | 0 | 29 | 51 | 34 | 29 | 2 | 8 | 10 | 22 | 25 | 1 | 9 | 7 | 1 | 0 | 2 | 50 | 21 | 1 | 24 | 6 | 4 | Cblc |
| Cbll1 | 3509 | 2975 | 4154 | 4393 | 4260 | 4143 | 4159 | 3352 | 3222 | 3579 | 3394 | 3985 | 3751 | 4009 | 3526 | 3519 | 4342 | 4264 | 4101 | 3141 | 4362 | 4062 | 2976 | 3717 | 3947 | 3783 | 3860 | 4120 | Cbll1 |
| Cbln1 | 1719 | 2499 | 2574 | 2845 | 3165 | 2400 | 2254 | 2171 | 2277 | 2685 | 2113 | 1990 | 2743 | 1985 | 2474 | 2647 | 2525 | 2684 | 2286 | 2414 | 2317 | 2874 | 2349 | 1584 | 2159 | 2094 | 1772 | 2105 | Cbln1 |
| Cbln2 | 976 | 1073 | 1072 | 1273 | 1897 | 1697 | 1923 | 3412 | 2657 | 2826 | 2594 | 2055 | 2235 | 2281 | 1487 | 840 | 1347 | 1351 | 1827 | 1831 | 1724 | 3628 | 2955 | 2250 | 3169 | 1884 | 2034 | 1924 | Cbln2 |
| Cbln3 | 63 | 51 | 73 | 52 | 57 | 37 | 76 | 71 | 74 | 57 | 85 | 72 | 87 | 82 | 54 | 55 | 66 | 55 | 43 | 47 | 57 | 72 | 67 | 78 | 60 | 76 | 67 | 42 | Cbln3 |
| Cbln4 | 113 | 114 | 135 | 199 | 557 | 269 | 311 | 475 | 435 | 501 | 485 | 237 | 260 | 230 | 190 | 127 | 140 | 265 | 319 | 286 | 328 | 647 | 407 | 313 | 483 | 243 | 198 | 193 | Cbln4 |
| Cbr1 | 3462 | 3675 | 3176 | 3541 | 3661 | 3317 | 3499 | 3447 | 3330 | 3579 | 3250 | 3247 | 3326 | 3412 | 3600 | 3217 | 3759 | 3417 | 3709 | 3873 | 3417 | 3280 | 3524 | 3656 | 3546 | 3600 | 3403 | 3368 | Cbr1 |
| Cbr2 | 25 | 21 | 21 | 19 | 6 | 14 | 1 | 114 | 131 | 128 | 116 | 30 | 17 | 18 | 28 | 18 | 14 | 8 | 9 | 7 | 5 | 27 | 207 | 98 | 26 | 24 | 27 | 18 | Cbr2 |
| Cbr3 | 73 | 77 | 85 | 34 | 57 | 66 | 65 | 23 | 56 | 36 | 43 | 10 | 29 | 12 | 62 | 83 | 95 | 77 | 50 | 66 | 53 | 17 | 54 | 29 | 14 | 14 | 15 | 23 | Cbr3 |
| Cbr4 | 882 | 713 | 752 | 730 | 694 | 749 | 821 | 1241 | 1163 | 1047 | 1033 | 1382 | 1349 | 1473 | 887 | 650 | 737 | 585 | 635 | 865 | 702 | 1103 | 1224 | 1166 | 1098 | 1306 | 1397 | 1216 | Cbr4 |
| Cbs | 959 | 1179 | 920 | 1602 | 1310 | 1196 | 1452 | 227 | 297 | 313 | 265 | 193 | 196 | 221 | 873 | 1008 | 972 | 1552 | 1428 | 1360 | 1401 | 182 | 285 | 314 | 343 | 206 | 166 | 188 | Cbs |
| Cbwd1 | 2337 | 2261 | 2718 | 2342 | 2007 | 1930 | 2242 | 1561 | 1277 | 1198 | 1215 | 1550 | 1469 | 1495 | 2546 | 2336 | 2381 | 2161 | 2051 | 2502 | 2112 | 1218 | 1309 | 1476 | 1327 | 1644 | 1599 | 1600 | Cbwd1 |
| Cbx1 | 24267 | 19925 | 31416 | 24772 | 19136 | 18713 | 23253 | 22252 | 18166 | 21820 | 20437 | 25296 | 23661 | 31207 | 22759 | 19972 | 27148 | 21308 | 21707 | 22207 | 22305 | 18989 | 17387 | 26934 | 22675 | 31953 | 27923 | 32155 | Cbx1 |
| Cbx2 | 5969 | 7032 | 8209 | 6607 | 4933 | 5426 | 5014 | 4564 | 4863 | 5130 | 4683 | 7491 | 6399 | 6491 | 5163 | 6798 | 7581 | 6092 | 5944 | 4595 | 5126 | 4553 | 4322 | 6397 | 5246 | 9454 | 6876 | 7244 | Cbx2 |
| Cbx3 | 11262 | 8921 | 13899 | 11312 | 11365 | 11343 | 13238 | 18221 | 15522 | 16113 | 15416 | 20398 | 17736 | 22395 | 11628 | 9837 | 12747 | 10991 | 12204 | 14296 | 12591 | 14551 | 15262 | 19998 | 18717 | 24004 | 21611 | 23099 | Cbx3 |
| Cbx4 | 1902 | 2248 | 2180 | 2553 | 2173 | 2086 | 1977 | 1441 | 1546 | 1854 | 1576 | 1653 | 1354 | 1381 | 1734 | 1997 | 2420 | 2479 | 2187 | 1763 | 2153 | 1668 | 1451 | 1831 | 1772 | 1619 | 1424 | 1355 | Cbx4 |
| Cbx5 | 44010 | 36234 | 35705 | 34661 | 35249 | 37874 | 39026 | 59348 | 49479 | 45998 | 45675 | 57731 | 50574 | 58458 | 42072 | 33619 | 35829 | 33355 | 36359 | 40104 | 38459 | 46500 | 47750 | 46610 | 51860 | 49862 | 62277 | 58955 | Cbx5 |
| Cbx6 | 7116 | 8909 | 11517 | 11891 | 11981 | 10962 | 10299 | 8695 | 9101 | 12256 | 12412 | 11906 | 10494 | 12034 | 6674 | 7476 | 12422 | 12389 | 12725 | 8592 | 11241 | 12039 | 9982 | 12621 | 12857 | 9515 | 10775 | 11763 | Cbx6 |
| Cbx7 | 2480 | 3090 | 3565 | 3564 | 2860 | 3106 | 2870 | 501 | 620 | 640 | 572 | 693 | 679 | 659 | 2273 | 2591 | 3775 | 3565 | 3307 | 2500 | 2891 | 612 | 592 | 662 | 675 | 630 | 595 | 666 | Cbx7 |
| Cbx8 | 1639 | 2380 | 2631 | 2896 | 1816 | 1959 | 1788 | 1232 | 1308 | 1574 | 1349 | 1667 | 1483 | 1521 | 1617 | 2068 | 2911 | 2642 | 2208 | 1837 | 1900 | 1491 | 1218 | 1749 | 1731 | 1671 | 1601 | 1450 | Cbx8 |
| Cby1 | 2600 | 3380 | 3442 | 3416 | 2872 | 2864 | 3057 | 2318 | 2341 | 2666 | 2120 | 2717 | 2277 | 2817 | 2798 | 2509 | 3390 | 3199 | 3291 | 3307 | 3335 | 2044 | 2375 | 2876 | 2696 | 3308 | 2725 | 2870 | Cby1 |
| Cby2 | 0 | 0 | 0 | 0 | 0 | 2 | 0 | 0 | 0 | 0 | 0 | 0 | 0 | 0 | 0 | 0 | 0 | 0 | 0 | 0 | 0 | 0 | 0 | 0 | 0 | 0 | 0 | 0 | Cby2 |
| Cby3 | 0 | 3 | 3 | 0 | 1 | 3 | 7 | 1 | 2 | 5 | 4 | 5 | 0 | 4 | 1 | 1 | 1 | 0 | 9 | 0 | 1 | 2 | 2 | 7 | 1 | 3 | 0 | 0 | Cby3 |
| Cc2d1a | 1130 | 1457 | 1560 | 1873 | 2194 | 1850 | 1581 | 1800 | 2488 | 2795 | 2605 | 2311 | 2182 | 1942 | 1015 | 1453 | 1763 | 1931 | 2260 | 1482 | 1835 | 2647 | 2313 | 2690 | 2698 | 2536 | 1947 | 2040 | Cc2d1a |
| Cc2d1b | 2398 | 2588 | 2567 | 2928 | 2823 | 2583 | 2594 | 2977 | 3849 | 3990 | 3892 | 3974 | 3809 | 3811 | 2422 | 2423 | 2863 | 2799 | 2828 | 2566 | 2607 | 3559 | 3464 | 3921 | 3860 | 3944 | 3847 | 3526 | Cc2d1b |
| Cc2d2a | 1007 | 985 | 898 | 836 | 1078 | 1033 | 948 | 2319 | 2374 | 2190 | 2395 | 2467 | 2496 | 2493 | 1022 | 935 | 912 | 840 | 925 | 1107 | 949 | 2025 | 2434 | 2112 | 2044 | 1900 | 2778 | 2409 | Cc2d2a |
| Cc2d2b | 18 | 17 | 18 | 11 | 14 | 24 | 22 | 31 | 22 | 31 | 23 | 21 | 11 | 17 | 18 | 14 | 14 | 25 | 26 | 25 | 21 | 15 | 36 | 23 | 44 | 18 | 16 | 26 | Cc2d2b |
| Ccar1 | 14534 | 11104 | 14240 | 12561 | 13191 | 14329 | 14460 | 24277 | 18322 | 19616 | 18275 | 22499 | 20889 | 24640 | 14249 | 12116 | 13216 | 11853 | 14412 | 15657 | 13809 | 20397 | 17831 | 21809 | 21604 | 23411 | 23424 | 24301 | Ccar1 |
| Ccar2 | 9878 | 10984 | 12672 | 12444 | 11413 | 11197 | 11153 | 11339 | 12307 | 13753 | 12291 | 16011 | 14020 | 15088 | 9082 | 9628 | 12843 | 11482 | 12684 | 10564 | 11659 | 13293 | 11235 | 14954 | 15102 | 16184 | 15271 | 15578 | Ccar2 |
| Ccbe1 | 728 | 673 | 849 | 417 | 209 | 437 | 137 | 430 | 363 | 477 | 502 | 829 | 1109 | 810 | 645 | 1040 | 557 | 427 | 115 | 131 | 130 | 456 | 550 | 497 | 486 | 543 | 1027 | 987 | Ccbe1 |
| Ccdc102a | 413 | 459 | 550 | 418 | 361 | 452 | 288 | 879 | 1261 | 1169 | 1526 | 1033 | 1287 | 983 | 327 | 565 | 516 | 424 | 388 | 287 | 265 | 934 | 1420 | 1289 | 939 | 1582 | 1127 | 1076 | Ccdc102a |
| Ccdc103 | 212 | 278 | 225 | 173 | 169 | 220 | 163 | 90 | 91 | 104 | 81 | 112 | 93 | 67 | 248 | 204 | 219 | 184 | 177 | 211 | 181 | 97 | 102 | 87 | 100 | 113 | 110 | 109 | Ccdc103 |
| Ccdc105 | 15 | 22 | 13 | 13 | 16 | 11 | 19 | 5 | 7 | 10 | 4 | 7 | 5 | 6 | 17 | 20 | 30 | 14 | 15 | 21 | 21 | 13 | 5 | 4 | 5 | 3 | 1 | 7 | Ccdc105 |
| Ccdc106 | 482 | 404 | 446 | 527 | 799 | 556 | 523 | 659 | 922 | 884 | 929 | 497 | 496 | 501 | 351 | 450 | 489 | 610 | 613 | 492 | 554 | 762 | 1012 | 753 | 815 | 430 | 427 | 472 | Ccdc106 |
| Ccdc107 | 1192 | 1187 | 1083 | 1183 | 1033 | 1056 | 1189 | 815 | 761 | 844 | 788 | 759 | 759 | 833 | 1182 | 1118 | 1133 | 1206 | 1179 | 1141 | 1223 | 755 | 843 | 817 | 750 | 845 | 765 | 806 | Ccdc107 |
| Ccdc110 | 7 | 12 | 6 | 20 | 16 | 2 | 15 | 34 | 19 | 21 | 8 | 23 | 28 | 29 | 16 | 14 | 16 | 11 | 25 | 20 | 13 | 27 | 19 | 33 | 22 | 20 | 30 | 26 | Ccdc110 |
| Ccdc112 | 3945 | 3195 | 4334 | 3946 | 3589 | 3736 | 4063 | 2561 | 2178 | 2189 | 2198 | 2164 | 2042 | 2292 | 3828 | 3642 | 3715 | 3772 | 3546 | 3854 | 3957 | 2358 | 2229 | 2317 | 2303 | 2490 | 2121 | 2184 | Ccdc112 |
| Ccdc113 | 54 | 51 | 35 | 35 | 68 | 46 | 70 | 50 | 50 | 74 | 39 | 62 | 85 | 30 | 53 | 58 | 44 | 52 | 51 | 87 | 77 | 53 | 129 | 42 | 50 | 31 | 24 | 36 | Ccdc113 |
| Ccdc115 | 2659 | 3155 | 3320 | 2928 | 2464 | 2562 | 3028 | 2280 | 2235 | 2423 | 2203 | 2599 | 2498 | 2931 | 2789 | 2590 | 3086 | 2656 | 2786 | 3069 | 2930 | 2015 | 2283 | 2610 | 2676 | 2785 | 2615 | 2946 | Ccdc115 |
| Ccdc116 | 155 | 171 | 118 | 192 | 200 | 202 | 172 | 150 | 168 | 256 | 226 | 151 | 170 | 133 | 206 | 166 | 174 | 168 | 182 | 184 | 176 | 239 | 153 | 153 | 182 | 92 | 134 | 138 | Ccdc116 |
| Ccdc117 | 4334 | 4120 | 4885 | 4156 | 3794 | 3520 | 4202 | 3624 | 2990 | 3062 | 3234 | 3456 | 3078 | 3308 | 4040 | 4234 | 4673 | 4382 | 4379 | 3919 | 4018 | 3370 | 3016 | 3388 | 3458 | 3854 | 3540 | 3509 | Ccdc117 |
| Ccdc12 | 666 | 924 | 967 | 837 | 798 | 738 | 815 | 1515 | 1648 | 1960 | 1677 | 2204 | 2127 | 2381 | 673 | 807 | 1043 | 745 | 902 | 1019 | 752 | 1338 | 1680 | 2318 | 2016 | 2963 | 2181 | 2346 | Ccdc12 |
| Ccdc120 | 1691 | 1994 | 1959 | 2348 | 2577 | 2286 | 2215 | 1624 | 1714 | 1979 | 1716 | 1757 | 1503 | 1577 | 1427 | 1590 | 2176 | 2476 | 2581 | 1992 | 2409 | 2159 | 1563 | 1889 | 2063 | 1775 | 1362 | 1558 | Ccdc120 |
| Ccdc121 | 0 | 0 | 0 | 8 | 10 | 0 | 2 | 5 | 4 | 3 | 4 | 1 | 6 | 2 | 0 | 4 | 0 | 1 | 1 | 2 | 0 | 0 | 4 | 4 | 0 | 3 | 3 | 2 | Ccdc121 |
| Ccdc122 | 128 | 55 | 90 | 62 | 47 | 63 | 50 | 79 | 73 | 69 | 70 | 78 | 78 | 46 | 113 | 102 | 51 | 64 | 56 | 76 | 55 | 48 | 99 | 47 | 56 | 72 | 88 | 74 | Ccdc122 |
| Ccdc124 | 3254 | 4363 | 4902 | 4589 | 3928 | 3890 | 3779 | 3337 | 3612 | 4081 | 3568 | 4328 | 4025 | 4634 | 3233 | 3131 | 4706 | 3842 | 4722 | 4147 | 4278 | 3013 | 3346 | 4842 | 4171 | 5049 | 4812 | 4946 | Ccdc124 |
| Ccdc125 | 82 | 79 | 128 | 113 | 92 | 96 | 112 | 142 | 204 | 174 | 252 | 204 | 276 | 194 | 78 | 75 | 112 | 73 | 108 | 77 | 101 | 164 | 194 | 253 | 145 | 251 | 202 | 192 | Ccdc125 |
| Ccdc126 | 325 | 298 | 346 | 334 | 369 | 350 | 386 | 423 | 352 | 346 | 368 | 315 | 308 | 339 | 313 | 407 | 336 | 370 | 344 | 332 | 364 | 343 | 380 | 293 | 305 | 322 | 291 | 311 | Ccdc126 |
| Ccdc127 | 8079 | 7534 | 6602 | 7421 | 7023 | 7176 | 7947 | 6205 | 5722 | 5555 | 5775 | 5784 | 5590 | 6314 | 7751 | 7807 | 7163 | 7805 | 6844 | 7444 | 7861 | 5497 | 5744 | 5534 | 5783 | 5437 | 5943 | 6064 | Ccdc127 |
| Ccdc13 | 308 | 471 | 411 | 445 | 355 | 455 | 409 | 206 | 222 | 259 | 140 | 311 | 296 | 254 | 356 | 409 | 447 | 395 | 465 | 423 | 362 | 256 | 205 | 233 | 252 | 287 | 273 | 268 | Ccdc13 |
| Ccdc130 | 549 | 694 | 622 | 714 | 558 | 604 | 551 | 674 | 787 | 672 | 613 | 752 | 691 | 752 | 530 | 576 | 650 | 632 | 595 | 653 | 554 | 760 | 732 | 693 | 809 | 757 | 777 | 689 | Ccdc130 |
| Ccdc134 | 3224 | 3799 | 3143 | 3551 | 2768 | 2918 | 2914 | 1534 | 1658 | 1580 | 2064 | 1813 | 1653 | 1699 | 3086 | 3417 | 3097 | 3565 | 2952 | 2814 | 3015 | 1546 | 1752 | 1711 | 1923 | 1579 | 1773 | 1820 | Ccdc134 |
| Ccdc136 | 4547 | 4752 | 2724 | 3464 | 4983 | 5925 | 4411 | 3952 | 4306 | 3872 | 4654 | 3390 | 4231 | 3680 | 4229 | 5027 | 2964 | 3758 | 4140 | 5562 | 5345 | 2984 | 4433 | 3158 | 4586 | 2519 | 4357 | 3288 | Ccdc136 |
| Ccdc137 | 1930 | 2024 | 2108 | 1862 | 1876 | 1878 | 1900 | 1881 | 1835 | 1794 | 2026 | 2041 | 1931 | 1949 | 1871 | 2005 | 2176 | 1729 | 2018 | 1822 | 1912 | 1878 | 1877 | 2062 | 1925 | 2171 | 1966 | 2104 | Ccdc137 |
| Ccdc138 | 1167 | 992 | 1179 | 1086 | 935 | 1049 | 1158 | 1016 | 892 | 883 | 811 | 1183 | 917 | 1216 | 1147 | 1011 | 1194 | 1035 | 1000 | 1180 | 1040 | 857 | 862 | 1077 | 962 | 1240 | 1170 | 1130 | Ccdc138 |
| Ccdc14 | 2664 | 2305 | 2199 | 1989 | 1874 | 1989 | 2200 | 1800 | 1641 | 1391 | 1453 | 1933 | 1691 | 1939 | 2676 | 2466 | 1991 | 1873 | 1942 | 2302 | 2090 | 1268 | 1523 | 1532 | 1273 | 1890 | 1968 | 1895 | Ccdc14 |
| Ccdc141 | 526 | 498 | 407 | 519 | 808 | 542 | 582 | 793 | 929 | 747 | 1822 | 587 | 889 | 655 | 630 | 760 | 483 | 550 | 529 | 637 | 542 | 947 | 1524 | 519 | 779 | 377 | 531 | 705 | Ccdc141 |
| Ccdc142 | 5 | 4 | 6 | 3 | 13 | 8 | 2 | 4 | 2 | 0 | 2 | 7 | 0 | 5 | 4 | 3 | 3 | 2 | 0 | 3 | 2 | 2 | 2 | 0 | 3 | 2 | 5 | 2 | Ccdc142 |
| Ccdc144b | 0 | 0 | 0 | 0 | 0 | 0 | 0 | 0 | 0 | 0 | 0 | 0 | 0 | 0 | 0 | 0 | 0 | 0 | 1 | 1 | 0 | 0 | 0 | 0 | 3 | 0 | 0 | 0 | Ccdc144b |
| Ccdc146 | 66 | 103 | 47 | 74 | 119 | 115 | 56 | 79 | 85 | 71 | 45 | 100 | 122 | 50 | 96 | 137 | 78 | 52 | 86 | 89 | 68 | 100 | 155 | 64 | 53 | 56 | 51 | 40 | Ccdc146 |
| Ccdc148 | 408 | 439 | 358 | 630 | 628 | 557 | 648 | 354 | 288 | 262 | 254 | 294 | 205 | 292 | 504 | 398 | 498 | 632 | 531 | 637 | 619 | 401 | 333 | 276 | 339 | 233 | 210 | 267 | Ccdc148 |
| Ccdc149 | 137 | 146 | 176 | 214 | 380 | 255 | 247 | 756 | 707 | 846 | 732 | 745 | 733 | 722 | 160 | 136 | 191 | 238 | 269 | 249 | 242 | 952 | 669 | 754 | 906 | 676 | 708 | 663 | Ccdc149 |

| | | | | | | | | | | | | | | | | | | | | | | | | | | | | | |
|---|---|---|---|---|---|---|---|---|---|---|---|---|---|---|---|---|---|---|---|---|---|---|---|---|---|---|---|---|---|
| Ccdc15 | 609 | 474 | 282 | 440 | 415 | 460 | 470 | 652 | 588 | 522 | 522 | 529 | 549 | 550 | 556 | 504 | 395 | 359 | 370 | 530 | 462 | 471 | 583 | 395 | 498 | 335 | 673 | 553 | Ccdc15 |
| Ccdc150 | 77 | 86 | 129 | 75 | 56 | 107 | 74 | 101 | 79 | 71 | 110 | 159 | 136 | 151 | 109 | 108 | 98 | 62 | 47 | 56 | 52 | 74 | 91 | 102 | 94 | 122 | 160 | 154 | Ccdc150 |
| Ccdc152 | 1 | 1 | 0 | 1 | 4 | 2 | 0 | 0 | 0 | 1 | 1 | 1 | 0 | 2 | 1 | 1 | 5 | 2 | 0 | 1 | 0 | 12 | 1 | 1 | 0 | 0 | 5 | 0 | Ccdc152 |
| Ccdc153 | 1 | 5 | 0 | 8 | 7 | 0 | 0 | 4 | 1 | 0 | 8 | 1 | 1 | 2 | 4 | 2 | 0 | 3 | 1 | 10 | 0 | 0 | 1 | 0 | 2 | 3 | 2 | 1 | Ccdc153 |
| Ccdc154 | 22 | 31 | 24 | 22 | 3 | 12 | 21 | 1 | 7 | 6 | 6 | 0 | 5 | 9 | 21 | 14 | 7 | 34 | 22 | 10 | 14 | 10 | 5 | 9 | 3 | 4 | 6 | 4 | Ccdc154 |
| Ccdc157 | 2396 | 2618 | 2436 | 2469 | 2582 | 2440 | 2111 | 2121 | 2185 | 2416 | 2870 | 2292 | 2420 | 2072 | 2256 | 2477 | 2501 | 2357 | 2445 | 2117 | 2171 | 2483 | 2368 | 2293 | 2146 | 1951 | 2101 | 2106 | Ccdc157 |
| Ccdc158 | 865 | 798 | 777 | 722 | 558 | 667 | 687 | 217 | 198 | 201 | 193 | 185 | 147 | 160 | 854 | 861 | 588 | 741 | 537 | 701 | 625 | 159 | 210 | 198 | 117 | 156 | 185 | 208 | Ccdc158 |
| Ccdc159 | 338 | 437 | 363 | 612 | 645 | 678 | 521 | 536 | 531 | 606 | 696 | 559 | 560 | 536 | 396 | 407 | 531 | 584 | 584 | 533 | 546 | 782 | 514 | 599 | 656 | 362 | 573 | 504 | Ccdc159 |
| Ccdc160 | 311 | 355 | 386 | 330 | 367 | 274 | 363 | 183 | 160 | 199 | 167 | 215 | 218 | 213 | 436 | 389 | 346 | 300 | 287 | 443 | 343 | 264 | 179 | 172 | 156 | 203 | 229 | 211 | Ccdc160 |
| Ccdc162 | 119 | 156 | 128 | 137 | 169 | 174 | 141 | 118 | 222 | 153 | 195 | 131 | 221 | 115 | 158 | 211 | 100 | 134 | 179 | 217 | 162 | 196 | 320 | 114 | 114 | 118 | 174 | 107 | Ccdc162 |
| Ccdc163 | 1459 | 1580 | 1662 | 1516 | 1067 | 1174 | 1435 | 1269 | 1114 | 1265 | 1179 | 1561 | 1406 | 1750 | 1298 | 1261 | 1618 | 1307 | 1277 | 1288 | 1326 | 1135 | 1043 | 1545 | 1346 | 1676 | 1619 | 1746 | Ccdc163 |
| Ccdc166 | 367 | 392 | 379 | 443 | 452 | 451 | 441 | 502 | 559 | 664 | 542 | 559 | 500 | 474 | 394 | 313 | 423 | 383 | 515 | 409 | 466 | 592 | 537 | 538 | 557 | 596 | 471 | 506 | Ccdc166 |
| Ccdc167 | 1145 | 1116 | 1004 | 1190 | 1312 | 1359 | 1506 | 2150 | 2227 | 2214 | 2108 | 2356 | 2013 | 2480 | 1155 | 861 | 1107 | 1264 | 1305 | 1345 | 1496 | 1955 | 2017 | 2357 | 2280 | 1936 | 2239 | 2416 | Ccdc167 |
| Ccdc169 | 36 | 30 | 28 | 54 | 44 | 48 | 49 | 38 | 17 | 24 | 32 | 15 | 10 | 15 | 44 | 46 | 54 | 54 | 62 | 41 | 46 | 55 | 17 | 29 | 26 | 23 | 12 | 19 | Ccdc169 |
| Ccdc17 | 145 | 171 | 159 | 167 | 135 | 179 | 160 | 165 | 161 | 132 | 182 | 167 | 187 | 167 | 179 | 161 | 154 | 171 | 141 | 142 | 165 | 187 | 161 | 141 | 161 | 177 | 148 | 150 | Ccdc17 |
| Ccdc170 | 52 | 62 | 50 | 80 | 121 | 89 | 75 | 76 | 103 | 120 | 75 | 124 | 90 | 44 | 73 | 121 | 81 | 96 | 93 | 87 | 97 | 100 | 155 | 43 | 56 | 35 | 37 | 38 | Ccdc170 |
| Ccdc171 | 1370 | 1301 | 834 | 1327 | 1178 | 1216 | 1422 | 876 | 810 | 812 | 614 | 694 | 636 | 698 | 1349 | 1255 | 1194 | 1096 | 1149 | 1411 | 1304 | 645 | 826 | 713 | 667 | 524 | 740 | 503 | Ccdc171 |
| Ccdc172 | 1 | 0 | 0 | 0 | 0 | 0 | 0 | 0 | 0 | 0 | 0 | 0 | 0 | 0 | 0 | 0 | 0 | 0 | 0 | 0 | 1 | 2 | 1 | 0 | 1 | 0 | 0 | 2 | Ccdc172 |
| Ccdc173 | 739 | 717 | 1004 | 924 | 663 | 764 | 837 | 587 | 490 | 561 | 539 | 634 | 626 | 784 | 853 | 759 | 1000 | 801 | 701 | 845 | 768 | 622 | 532 | 674 | 558 | 543 | 636 | 717 | Ccdc173 |
| Ccdc174 | 2461 | 2342 | 2570 | 2582 | 2391 | 2442 | 2760 | 3108 | 2636 | 2796 | 2522 | 3085 | 2854 | 3385 | 2512 | 2326 | 2459 | 2448 | 2448 | 2867 | 2495 | 2729 | 2709 | 3167 | 3135 | 3370 | 3387 | 3355 | Ccdc174 |
| Ccdc175 | 10 | 10 | 12 | 13 | 6 | 12 | 22 | 4 | 4 | 6 | 9 | 10 | 11 | 12 | 20 | 6 | 5 | 17 | 9 | 9 | 22 | 3 | 3 | 11 | 5 | 9 | 15 | 6 | Ccdc175 |
| Ccdc177 | 1262 | 1203 | 1650 | 1715 | 2401 | 2004 | 2273 | 2691 | 2516 | 3610 | 2596 | 3894 | 2826 | 3971 | 1394 | 981 | 1866 | 1735 | 2590 | 1940 | 2247 | 3424 | 2181 | 3731 | 3825 | 3363 | 3527 | 3652 | Ccdc177 |
| Ccdc178 | 2 | 6 | 4 | 0 | 4 | 0 | 1 | 1 | 10 | 0 | 5 | 2 | 7 | 1 | 11 | 6 | 1 | 2 | 5 | 4 | 2 | 0 | 6 | 4 | 3 | 4 | 1 | 1 | Ccdc178 |
| Ccdc18 | 2343 | 1493 | 1518 | 1655 | 1129 | 1537 | 1734 | 1237 | 1176 | 1009 | 934 | 1382 | 1195 | 1439 | 1995 | 1965 | 1582 | 1402 | 1407 | 1927 | 1543 | 620 | 1001 | 1052 | 844 | 1057 | 1490 | 1214 | Ccdc18 |
| Ccdc180 | 98 | 153 | 43 | 70 | 132 | 121 | 116 | 82 | 107 | 93 | 59 | 76 | 108 | 21 | 86 | 170 | 79 | 90 | 116 | 108 | 113 | 85 | 148 | 47 | 65 | 59 | 42 | 37 | Ccdc180 |
| Ccdc181 | 1002 | 991 | 1060 | 1108 | 1073 | 1110 | 1193 | 1547 | 1344 | 1350 | 1215 | 1308 | 1339 | 1505 | 1046 | 1021 | 1001 | 1017 | 1057 | 1154 | 1053 | 1405 | 1418 | 1340 | 1486 | 1321 | 1453 | 1434 | Ccdc181 |
| Ccdc182 | 0 | 2 | 5 | 0 | 0 | 0 | 0 | 2 | 1 | 0 | 1 | 1 | 0 | 1 | 0 | 0 | 0 | 0 | 0 | 0 | 0 | 0 | 1 | 1 | 0 | 0 | 2 | 0 | Ccdc182 |
| Ccdc183 | 0 | 4 | 10 | 5 | 2 | 2 | 3 | 6 | 3 | 0 | 3 | 2 | 0 | 0 | 7 | 3 | 1 | 7 | 14 | 0 | 0 | 0 | 7 | 0 | 4 | 0 | 0 | 2 | Ccdc183 |
| Ccdc184 | 1441 | 2247 | 1699 | 2906 | 3278 | 2728 | 2914 | 1399 | 1422 | 1806 | 1443 | 1166 | 1076 | 1206 | 1756 | 1502 | 2671 | 2896 | 3236 | 2865 | 3201 | 2152 | 1385 | 1543 | 1849 | 1095 | 1019 | 1200 | Ccdc184 |
| Ccdc185 | 0 | 0 | 0 | 0 | 0 | 0 | 0 | 0 | 0 | 0 | 0 | 0 | 0 | 1 | 0 | 0 | 0 | 0 | 0 | 0 | 0 | 0 | 0 | 0 | 0 | 4 | 0 | 0 | Ccdc185 |
| Ccdc186 | 7947 | 7414 | 5386 | 7634 | 7298 | 7651 | 7894 | 6646 | 5687 | 5240 | 5487 | 4888 | 5117 | 5067 | 8219 | 7464 | 6215 | 7655 | 6674 | 8623 | 7967 | 5452 | 5567 | 4788 | 5745 | 3259 | 5125 | 4472 | Ccdc186 |
| Ccdc187 | 226 | 249 | 88 | 329 | 416 | 432 | 318 | 800 | 690 | 877 | 691 | 777 | 902 | 721 | 225 | 334 | 196 | 275 | 330 | 396 | 331 | 680 | 896 | 955 | 683 | 650 | 644 | 602 | Ccdc187 |
| Ccdc189 | 664 | 777 | 643 | 669 | 626 | 569 | 702 | 348 | 400 | 374 | 355 | 403 | 466 | 435 | 785 | 742 | 629 | 683 | 620 | 735 | 676 | 343 | 414 | 365 | 419 | 450 | 442 | 433 | Ccdc189 |
| Ccdc190 | 47 | 58 | 65 | 86 | 117 | 78 | 56 | 217 | 207 | 281 | 199 | 127 | 177 | 168 | 73 | 44 | 72 | 47 | 68 | 68 | 116 | 297 | 280 | 187 | 244 | 203 | 188 | 167 | Ccdc190 |
| Ccdc191 | 558 | 477 | 613 | 488 | 482 | 448 | 525 | 1222 | 1173 | 1200 | 1089 | 1247 | 1252 | 1285 | 520 | 486 | 579 | 410 | 457 | 517 | 482 | 896 | 1098 | 1133 | 1177 | 1309 | 1361 | 1375 | Ccdc191 |
| Ccdc192 | 0 | 3 | 0 | 0 | 1 | 6 | 0 | 0 | 1 | 0 | 4 | 0 | 0 | 1 | 4 | 0 | 0 | 0 | 0 | 12 | 1 | 2 | 4 | 2 | 5 | 3 | 0 | 0 | Ccdc192 |
| Ccdc194 | 5 | 10 | 8 | 7 | 10 | 9 | 3 | 13 | 25 | 31 | 19 | 12 | 23 | 16 | 7 | 6 | 7 | 1 | 2 | 4 | 3 | 12 | 21 | 10 | 6 | 17 | 3 | 13 | Ccdc194 |
| ccdc198 | 2 | 1 | 1 | 0 | 3 | 3 | 2 | 1 | 5 | 10 | 6 | 7 | 0 | 0 | 1 | 1 | 2 | 0 | 1 | 5 | 0 | 5 | 10 | 4 | 9 | 2 | 9 | 4 | ccdc198 |
| Ccdc201 | 2 | 0 | 0 | 0 | 0 | 0 | 0 | 0 | 0 | 0 | 0 | 1 | 0 | 0 | 0 | 0 | 0 | 0 | 0 | 0 | 0 | 0 | 2 | 1 | 0 | 0 | 0 | 4 | Ccdc201 |
| Ccdc22 | 15 | 19 | 19 | 28 | 22 | 18 | 19 | 34 | 13 | 26 | 19 | 15 | 22 | 38 | 25 | 7 | 22 | 11 | 16 | 21 | 38 | 27 | 10 | 33 | 20 | 14 | 26 | 19 | Ccdc22 |
| Ccdc24 | 54 | 85 | 115 | 137 | 169 | 121 | 145 | 116 | 86 | 112 | 105 | 150 | 141 | 148 | 67 | 75 | 136 | 180 | 139 | 97 | 142 | 159 | 78 | 150 | 174 | 122 | 89 | 127 | Ccdc24 |
| Ccdc25 | 4760 | 4362 | 5575 | 4697 | 3931 | 4276 | 4542 | 3118 | 2899 | 3029 | 2856 | 3836 | 3339 | 3888 | 4781 | 4341 | 4748 | 4416 | 4313 | 4418 | 4265 | 3312 | 2876 | 3568 | 3478 | 4272 | 3799 | 3987 | Ccdc25 |
| Ccdc27 | 0 | 2 | 5 | 0 | 2 | 0 | 3 | 0 | 1 | 3 | 4 | 3 | 0 | 0 | 0 | 0 | 0 | 0 | 3 | 2 | 0 | 0 | 4 | 2 | 0 | 4 | 0 | 0 | Ccdc27 |
| Ccdc28a | 873 | 932 | 760 | 912 | 673 | 644 | 753 | 397 | 397 | 374 | 386 | 283 | 418 | 309 | 847 | 1025 | 710 | 801 | 643 | 798 | 729 | 351 | 370 | 373 | 378 | 338 | 331 | 312 | Ccdc28a |
| Ccdc28b | 2152 | 2449 | 2441 | 2622 | 2639 | 2473 | 2504 | 1810 | 1893 | 1949 | 1796 | 1987 | 1835 | 1903 | 2230 | 2228 | 2608 | 2555 | 2662 | 2859 | 2583 | 1945 | 1866 | 2062 | 2131 | 1859 | 2059 | 1844 | Ccdc28b |
| Ccdc3 | 421 | 556 | 619 | 519 | 620 | 596 | 360 | 1044 | 1118 | 1123 | 937 | 622 | 1093 | 620 | 873 | 700 | 523 | 647 | 455 | 519 | 383 | 964 | 1484 | 842 | 782 | 495 | 668 | 587 | Ccdc3 |
| Ccdc30 | 1395 | 1548 | 1252 | 1687 | 1694 | 1708 | 1776 | 1064 | 936 | 1066 | 843 | 989 | 976 | 1000 | 1534 | 1292 | 1596 | 1684 | 1626 | 1759 | 1867 | 1226 | 908 | 909 | 997 | 830 | 1011 | 980 | Ccdc30 |
| Ccdc32 | 2551 | 3167 | 2857 | 2902 | 2498 | 2500 | 2937 | 1406 | 1392 | 1463 | 1318 | 1420 | 1396 | 1487 | 2859 | 2768 | 2787 | 2945 | 2699 | 2973 | 2937 | 1379 | 1383 | 1473 | 1508 | 1503 | 1411 | 1388 | Ccdc32 |
| Ccdc33 | 59 | 64 | 68 | 65 | 65 | 89 | 68 | 72 | 112 | 68 | 52 | 131 | 87 | 93 | 58 | 54 | 74 | 62 | 38 | 65 | 67 | 79 | 72 | 78 | 72 | 81 | 72 | 75 | Ccdc33 |
| Ccdc34 | 5165 | 4572 | 6291 | 4658 | 3696 | 3767 | 4417 | 5569 | 4458 | 5253 | 4205 | 5976 | 5306 | 6465 | 5418 | 4667 | 5762 | 3710 | 4317 | 4983 | 4278 | 4288 | 4499 | 5936 | 5007 | 6627 | 6294 | 7104 | Ccdc34 |
| Ccdc38 | 5 | 7 | 6 | 16 | 3 | 14 | 3 | 39 | 26 | 46 | 15 | 14 | 34 | 21 | 11 | 12 | 23 | 3 | 7 | 7 | 8 | 33 | 39 | 25 | 31 | 16 | 19 | 30 | Ccdc38 |
| Ccdc39 | 2204 | 2029 | 1932 | 2015 | 2157 | 2455 | 2402 | 1987 | 1749 | 1727 | 1731 | 1683 | 1929 | 1728 | 2540 | 2326 | 1920 | 2012 | 2131 | 2591 | 2552 | 2137 | 2034 | 1414 | 1739 | 1061 | 1617 | 1421 | Ccdc39 |
| Ccdc40 | 1232 | 1538 | 1208 | 1368 | 1581 | 1480 | 1367 | 1744 | 1645 | 1759 | 1376 | 2287 | 2090 | 1973 | 1367 | 1305 | 1393 | 1544 | 1548 | 1500 | 1422 | 1813 | 1657 | 1674 | 1791 | 1898 | 2236 | 2136 | Ccdc40 |
| Ccdc42 | 17 | 24 | 27 | 4 | 8 | 17 | 10 | 12 | 8 | 8 | 19 | 17 | 34 | 20 | 8 | 18 | 7 | 4 | 4 | 4 | 2 | 13 | 17 | 11 | 9 | 18 | 8 | 23 | Ccdc42 |
| Ccdc43 | 2879 | 2374 | 2894 | 2429 | 2184 | 2430 | 2557 | 2949 | 2555 | 2402 | 2569 | 2925 | 2717 | 3194 | 2884 | 2313 | 2844 | 2222 | 2522 | 2684 | 2426 | 2602 | 2705 | 2877 | 2491 | 3355 | 3194 | 3310 | Ccdc43 |
| Ccdc47 | 9597 | 8264 | 9797 | 9390 | 8926 | 8761 | 9843 | 8633 | 7358 | 7496 | 7708 | 7497 | 7329 | 8326 | 9810 | 8700 | 9387 | 9666 | 8360 | 9852 | 9815 | 8074 | 7347 | 7956 | 8142 | 7585 | 7482 | 7753 | Ccdc47 |
| Ccdc50 | 14966 | 12185 | 11982 | 12405 | 14174 | 14442 | 14504 | 20932 | 17328 | 16760 | 17328 | 18372 | 17568 | 18953 | 15604 | 14146 | 12588 | 12764 | 12623 | 15070 | 14673 | 18036 | 17139 | 15646 | 18328 | 12636 | 18543 | 18157 | Ccdc50 |
| Ccdc51 | 701 | 841 | 817 | 893 | 758 | 653 | 697 | 716 | 676 | 716 | 695 | 767 | 724 | 757 | 761 | 750 | 785 | 810 | 841 | 777 | 684 | 632 | 625 | 704 | 805 | 850 | 737 | 614 | Ccdc51 |
| Ccdc54 | 1 | 0 | 8 | 3 | 0 | 0 | 1 | 0 | 0 | 0 | 0 | 0 | 0 | 0 | 0 | 0 | 1 | 2 | 0 | 0 | 1 | 0 | 1 | 0 | 0 | 0 | 0 | 0 | Ccdc54 |
| Ccdc57 | 563 | 580 | 502 | 685 | 605 | 586 | 617 | 576 | 783 | 733 | 738 | 800 | 761 | 774 | 513 | 576 | 616 | 567 | 590 | 612 | 556 | 655 | 713 | 829 | 768 | 728 | 743 | 712 | Ccdc57 |
| Ccdc58 | 1546 | 1177 | 1520 | 1147 | 1006 | 1066 | 1196 | 1922 | 1897 | 1846 | 1829 | 1925 | 2100 | 2555 | 1360 | 1223 | 1352 | 954 | 1059 | 1382 | 1205 | 1335 | 1757 | 1915 | 1873 | 2332 | 2137 | 2605 | Ccdc58 |
| Ccdc59 | 2495 | 2195 | 2572 | 2477 | 2257 | 2242 | 2635 | 3125 | 2862 | 2939 | 2922 | 3006 | 2759 | 3309 | 2505 | 2221 | 2604 | 2226 | 2502 | 2479 | 2541 | 2909 | 2880 | 3291 | 2850 | 3488 | 3105 | 3160 | Ccdc59 |
| Ccdc6 | 6165 | 5887 | 8482 | 8202 | 9178 | 8003 | 8203 | 6016 | 6007 | 7509 | 7601 | 7628 | 7058 | 7491 | 6039 | 6222 | 8703 | 8341 | 8514 | 6575 | 8283 | 7640 | 5838 | 7250 | 7881 | 7067 | 6721 | 7547 | Ccdc6 |
| Ccdc60 | 204 | 326 | 348 | 213 | 324 | 269 | 235 | 247 | 269 | 239 | 254 | 285 | 325 | 197 | 306 | 385 | 265 | 267 | 252 | 345 | 235 | 237 | 273 | 199 | 250 | 204 | 272 | 217 | Ccdc60 |
| Ccdc61 | 2467 | 3291 | 3205 | 3460 | 2216 | 2361 | 2342 | 1028 | 1159 | 1253 | 1133 | 1538 | 1401 | 1429 | 2425 | 2719 | 3342 | 3133 | 2749 | 2408 | 2254 | 906 | 1109 | 1440 | 1288 | 1744 | 1476 | 1375 | Ccdc61 |
| Ccdc62 | 389 | 343 | 364 | 477 | 431 | 457 | 477 | 460 | 447 | 612 | 590 | 572 | 534 | 549 | 381 | 328 | 411 | 379 | 422 | 389 | 410 | 625 | 472 | 561 | 561 | 442 | 483 | 489 | Ccdc62 |
| Ccdc63 | 2 | 8 | 9 | 6 | 12 | 5 | 2 | 9 | 6 | 5 | 7 | 7 | 7 | 4 | 3 | 4 | 19 | 12 | 8 | 6 | 8 | 12 | 10 | 2 | 2 | 0 | 2 | 7 | Ccdc63 |
| Ccdc65 | 129 | 179 | 168 | 180 | 205 | 136 | 140 | 101 | 113 | 121 | 124 | 118 | 112 | 82 | 144 | 173 | 162 | 255 | 188 | 166 | 184 | 134 | 145 | 92 | 110 | 109 | 72 | 77 | Ccdc65 |
| Ccdc66 | 3543 | 2763 | 2889 | 3014 | 2773 | 2860 | 3140 | 3675 | 3292 | 3001 | 2946 | 3415 | 3200 | 3530 | 3430 | 3178 | 2887 | 2863 | 2947 | 3363 | 2940 | 3088 | 3135 | 3347 | 3110 | 3530 | 3438 | 3310 | Ccdc66 |
| Ccdc68 | 48 | 32 | 24 | 13 | 27 | 15 | 11 | 72 | 97 | 66 | 48 | 47 | 64 | 50 | 34 | 40 | 16 | 33 | 21 | 19 | 15 | 77 | 96 | 47 | 35 | 43 | 55 | 40 | Ccdc68 |
| Ccdc69 | 21 | 10 | 25 | 8 | 16 | 15 | 9 | 15 | 7 | 19 | 14 | 22 | 18 | 6 | 18 | 16 | 19 | 7 | 30 | 12 | 6 | 15 | 27 | 16 | 13 | 18 | 3 | 11 | Ccdc69 |
| Ccdc71 | 2413 | 2611 | 2739 | 2627 | 2297 | 2470 | 2031 | 2525 | 2725 | 3024 | 2618 | 2789 | 2676 | 2680 | 2238 | 2220 | 2651 | 2487 | 2727 | 2724 | 2307 | 2670 | 2809 | 2955 | 2763 | 3239 | 2789 | 2690 | Ccdc71 |
| Ccdc71l | 1535 | 1014 | 1448 | 1575 | 1813 | 1696 | 2073 | 2390 | 2010 | 1995 | 2016 | 2252 | 2067 | 2273 | 1456 | 1302 | 1623 | 1768 | 1878 | 1797 | 1853 | 2435 | 2112 | 2264 | 2208 | 2229 | 2056 | 2299 | Ccdc71l |
| Ccdc73 | 22 | 37 | 51 | 53 | 29 | 32 | 47 | 57 | 30 | 47 | 36 | 37 | 39 | 99 | 11 | 19 | 28 | 23 | 30 | 24 | 41 | 28 | 40 | 56 | 39 | 39 | 43 | 61 | Ccdc73 |
| Ccdc74a | 172 | 185 | 167 | 157 | 203 | 201 | 134 | 194 | 221 | 191 | 223 | 205 | 198 | 181 | 194 | 171 | 127 | 149 | 162 | 145 | 140 | 244 | 246 | 191 | 194 | 260 | 177 | 214 | Ccdc74a |
| Ccdc77 | 2012 | 1887 | 1900 | 1698 | 1274 | 1401 | 1583 | 2091 | 1877 | 1895 | 1982 | 2252 | 2104 | 2549 | 1956 | 1876 | 1836 | 1664 | 1454 | 1637 | 1430 | 1644 | 1753 | 2080 | 1907 | 2455 | 2449 | 2470 | Ccdc77 |
| Ccdc78 | 9 | 10 | 22 | 3 | 7 | 0 | 4 | 7 | 7 | 6 | 3 | 4 | 18 | 5 | 7 | 7 | 1 | 8 | 7 | 5 | 6 | 7 | 6 | 11 | 8 | 3 | 6 | 9 | Ccdc78 |
| Ccdc7a | 0 | 1 | 0 | 1 | 0 | 0 | 0 | 0 | 0 | 0 | 0 | 1 | 0 | 1 | 0 | 0 | 0 | 1 | 0 | 0 | 2 | 2 | 1 | 0 | 0 | 0 | 0 | 0 | Ccdc7a |
| Ccdc8 | 2606 | 2750 | 3113 | 2349 | 2090 | 2396 | 2019 | 2916 | 3950 | 3252 | 4743 | 3779 | 4039 | 3725 | 2251 | 2954 | 2574 | 2395 | 2113 | 2023 | 1772 | 2806 | 4475 | 3899 | 2983 | 4844 | 3897 | 3836 | Ccdc8 |

| | | | | | | | | | | | | | | | | | | | | | | | | | | | | | |
|---|---|---|---|---|---|---|---|---|---|---|---|---|---|---|---|---|---|---|---|---|---|---|---|---|---|---|---|---|---|
| Ccdc80 | 1284 | 1317 | 1649 | 661 | 417 | 875 | 255 | 1248 | 1832 | 1570 | 2609 | 1290 | 1914 | 1337 | 1497 | 1810 | 917 | 784 | 320 | 367 | 293 | 1375 | 2935 | 1472 | 1115 | 1359 | 1825 | 1496 | Ccdc80 |
| Ccdc81 | 100 | 111 | 97 | 170 | 157 | 137 | 166 | 93 | 74 | 94 | 78 | 84 | 82 | 43 | 84 | 119 | 131 | 211 | 164 | 140 | 156 | 112 | 89 | 57 | 61 | 25 | 56 | 38 | Ccdc81 |
| Ccdc82 | 5574 | 4887 | 4842 | 6111 | 5757 | 6228 | 6206 | 3692 | 3170 | 2831 | 3048 | 3294 | 3098 | 3303 | 5939 | 5576 | 4968 | 5789 | 5294 | 5954 | 6169 | 3380 | 2993 | 2669 | 3260 | 2464 | 3269 | 3082 | Ccdc82 |
| Ccdc83 | 5 | 3 | 0 | 1 | 1 | 1 | 0 | 2 | 1 | 3 | 8 | 1 | 0 | 2 | 0 | 2 | 0 | 0 | 4 | 1 | 0 | 3 | 3 | 2 | 0 | 0 | 1 | 1 | Ccdc83 |
| Ccdc84 | 2151 | 2369 | 2121 | 2482 | 2034 | 1935 | 2221 | 1324 | 1191 | 1216 | 1208 | 1191 | 1178 | 1387 | 2219 | 2110 | 2080 | 2395 | 2027 | 2015 | 2199 | 1375 | 1044 | 1224 | 1188 | 1264 | 1269 | 1224 | Ccdc84 |
| Ccdc85a | 264 | 268 | 266 | 430 | 831 | 527 | 579 | 1025 | 1069 | 1050 | 1161 | 714 | 992 | 815 | 309 | 315 | 364 | 485 | 542 | 514 | 574 | 1507 | 1018 | 762 | 1130 | 658 | 732 | 634 | Ccdc85a |
| Ccdc85b | 1893 | 2255 | 2244 | 2516 | 2384 | 2313 | 2088 | 1239 | 1197 | 1452 | 1421 | 1412 | 1391 | 1318 | 1859 | 1771 | 2484 | 2388 | 2563 | 2285 | 2247 | 1440 | 1293 | 1493 | 1532 | 1506 | 1453 | 1275 | Ccdc85b |
| Ccdc85c | 8105 | 11606 | 12905 | 13993 | 11079 | 10397 | 10832 | 4295 | 4576 | 6118 | 5029 | 7729 | 6419 | 7393 | 8087 | 8859 | 13520 | 12532 | 12595 | 9641 | 11877 | 5733 | 4446 | 7141 | 6725 | 7895 | 6707 | 7517 | Ccdc85c |
| Ccdc86 | 1446 | 1587 | 1627 | 1541 | 1091 | 1158 | 1105 | 918 | 1008 | 986 | 1129 | 1185 | 1168 | 1252 | 1354 | 1782 | 1467 | 1427 | 1228 | 995 | 1095 | 971 | 1029 | 1258 | 946 | 1465 | 1051 | 1200 | Ccdc86 |
| Ccdc87 | 116 | 124 | 54 | 119 | 84 | 68 | 88 | 28 | 40 | 21 | 35 | 18 | 42 | 27 | 132 | 121 | 104 | 98 | 58 | 77 | 85 | 37 | 53 | 30 | 22 | 33 | 23 | 28 | Ccdc87 |
| Ccdc88a | 19947 | 14076 | 10392 | 16274 | 19241 | 23213 | 22822 | 24342 | 20436 | 17007 | 16671 | 17931 | 16056 | 18183 | 21129 | 16179 | 13151 | 15978 | 18122 | 23351 | 22868 | 17584 | 18274 | 14686 | 18664 | 11520 | 19000 | 16306 | Ccdc88a |
| Ccdc88b | 80 | 114 | 136 | 94 | 107 | 84 | 36 | 61 | 82 | 96 | 121 | 113 | 90 | 72 | 98 | 152 | 125 | 89 | 58 | 39 | 53 | 70 | 123 | 129 | 83 | 79 | 110 | 107 | Ccdc88b |
| Ccdc88c | 6463 | 7916 | 6987 | 10017 | 11470 | 10641 | 11149 | 6892 | 7177 | 8622 | 6862 | 10256 | 8089 | 9982 | 6936 | 6312 | 9525 | 10128 | 12094 | 10094 | 11415 | 7364 | 6144 | 9820 | 10055 | 6900 | 8613 | 9265 | Ccdc88c |
| Ccdc89 | 454 | 478 | 504 | 377 | 370 | 398 | 389 | 171 | 166 | 155 | 131 | 226 | 259 | 203 | 618 | 543 | 452 | 439 | 358 | 500 | 357 | 164 | 206 | 184 | 181 | 219 | 218 | 216 | Ccdc89 |
| Ccdc9 | 2616 | 3194 | 4176 | 4050 | 3058 | 3127 | 2882 | 2916 | 3423 | 4599 | 4290 | 4664 | 4344 | 4564 | 2421 | 2669 | 4108 | 3597 | 3509 | 2636 | 2985 | 3845 | 3339 | 4899 | 4620 | 5043 | 4365 | 4524 | Ccdc9 |
| Ccdc90b | 1975 | 1786 | 2350 | 1867 | 1718 | 1668 | 2045 | 2674 | 2603 | 2318 | 2510 | 2757 | 2855 | 3025 | 2159 | 1898 | 2035 | 1681 | 1831 | 2316 | 1822 | 2520 | 2742 | 2678 | 2541 | 2987 | 2972 | 3144 | Ccdc90b |
| Ccdc91 | 1380 | 1456 | 1209 | 1179 | 1285 | 1278 | 1238 | 2811 | 2769 | 2579 | 2381 | 2386 | 2335 | 2320 | 1462 | 1206 | 1301 | 1224 | 1288 | 1451 | 1368 | 2384 | 2727 | 2430 | 2248 | 2243 | 2474 | 2442 | Ccdc91 |
| Ccdc92 | 1077 | 1520 | 1454 | 1847 | 1843 | 1354 | 1516 | 692 | 906 | 1037 | 967 | 943 | 869 | 796 | 1047 | 1195 | 1892 | 1802 | 1801 | 1367 | 1658 | 1140 | 824 | 970 | 1150 | 915 | 657 | 805 | Ccdc92 |
| Ccdc92b | 422 | 549 | 423 | 712 | 819 | 628 | 730 | 416 | 424 | 524 | 445 | 466 | 706 | 528 | 455 | 418 | 743 | 791 | 795 | 529 | 687 | 637 | 390 | 489 | 535 | 501 | 407 | 777 | Ccdc92b |
| Ccdc93 | 6489 | 6369 | 4849 | 6010 | 6403 | 6657 | 5845 | 4653 | 4587 | 4344 | 4745 | 4013 | 4124 | 3842 | 6414 | 6539 | 5394 | 6217 | 5925 | 6301 | 6244 | 4114 | 4575 | 3974 | 4328 | 2588 | 3956 | 3863 | Ccdc93 |
| Ccdc96 | 1166 | 1454 | 1340 | 1439 | 1180 | 1193 | 1344 | 492 | 459 | 528 | 388 | 659 | 661 | 673 | 1224 | 995 | 1393 | 1269 | 1287 | 1345 | 1268 | 451 | 464 | 579 | 516 | 670 | 625 | 587 | Ccdc96 |
| Ccdc97 | 3073 | 3974 | 4517 | 4303 | 4076 | 3706 | 3505 | 3038 | 3606 | 4252 | 3963 | 4795 | 3962 | 4401 | 2893 | 3455 | 4612 | 4146 | 4377 | 3116 | 3700 | 4328 | 3309 | 4772 | 4401 | 4799 | 3901 | 4277 | Ccdc97 |
| Ccdc9b | 218 | 186 | 214 | 159 | 247 | 268 | 289 | 565 | 583 | 677 | 559 | 810 | 721 | 900 | 215 | 138 | 199 | 161 | 302 | 254 | 241 | 744 | 455 | 778 | 702 | 853 | 891 | 870 | Ccdc9b |
| Ccer1 | 2 | 2 | 5 | 1 | 0 | 0 | 0 | 0 | 3 | 0 | 0 | 0 | 0 | 0 | 1 | 0 | 0 | 0 | 0 | 0 | 3 | 0 | 1 | 0 | 0 | 3 | 0 | 2 | Ccer1 |
| Ccer2 | 20 | 54 | 7 | 7 | 54 | 5 | 4 | 89 | 100 | 74 | 68 | 5 | 108 | 7 | 63 | 80 | 45 | 25 | 2 | 2 | 3 | 85 | 35 | 2 | 48 | 4 | 12 | 2 | Ccer2 |
| Cchcr1 | 2580 | 3086 | 3865 | 3421 | 2683 | 2854 | 2658 | 1418 | 1846 | 1844 | 1995 | 2392 | 2259 | 2265 | 2329 | 3026 | 3494 | 3305 | 3310 | 2476 | 2641 | 1462 | 1732 | 2310 | 1869 | 2537 | 2437 | 2271 | Cchcr1 |
| Ccin | 42 | 40 | 29 | 49 | 43 | 37 | 23 | 16 | 24 | 13 | 26 | 12 | 26 | 13 | 44 | 52 | 45 | 47 | 42 | 44 | 31 | 15 | 21 | 16 | 23 | 8 | 33 | 9 | Ccin |
| Cck | 72 | 94 | 67 | 84 | 123 | 123 | 182 | 133 | 149 | 97 | 102 | 139 | 34 | 175 | 69 | 72 | 109 | 110 | 160 | 221 | 185 | 177 | 151 | 181 | 130 | 126 | 102 | 127 | Cck |
| Cckar | 106 | 130 | 120 | 120 | 173 | 170 | 194 | 198 | 182 | 144 | 149 | 201 | 239 | 199 | 167 | 125 | 100 | 157 | 146 | 211 | 188 | 140 | 179 | 199 | 165 | 229 | 164 | 172 | Cckar |
| Cckbr | 32 | 19 | 27 | 28 | 58 | 27 | 27 | 22 | 49 | 38 | 33 | 18 | 22 | 10 | 16 | 16 | 19 | 35 | 32 | 28 | 22 | 35 | 30 | 39 | 30 | 51 | 15 | 19 | Cckbr |
| Ccl1 | 0 | 0 | 0 | 0 | 1 | 0 | 0 | 0 | 2 | 2 | 0 | 0 | 1 | 2 | 4 | 0 | 0 | 1 | 0 | 1 | 2 | 2 | 0 | 0 | 0 | 0 | 0 | 5 | Ccl1 |
| Ccl11 | 0 | 0 | 0 | 0 | 7 | 3 | 0 | 4 | 16 | 16 | 99 | 0 | 0 | 0 | 1 | 0 | 0 | 3 | 0 | 0 | 0 | 30 | 97 | 11 | 2 | 0 | 0 | 0 | Ccl11 |
| Ccl12 | 2 | 3 | 0 | 4 | 3 | 0 | 4 | 16 | 26 | 22 | 24 | 17 | 16 | 26 | 1 | 2 | 1 | 2 | 2 | 3 | 1 | 12 | 25 | 16 | 20 | 14 | 8 | 30 | Ccl12 |
| Ccl17 | 3 | 3 | 1 | 5 | 7 | 10 | 9 | 0 | 2 | 0 | 0 | 1 | 0 | 1 | 7 | 2 | 5 | 2 | 0 | 0 | 3 | 2 | 1 | 0 | 5 | 5 | 1 | 0 | Ccl17 |
| Ccl2 | 2 | 4 | 2 | 6 | 7 | 14 | 0 | 12 | 40 | 5 | 69 | 25 | 19 | 19 | 22 | 5 | 0 | 3 | 3 | 1 | 4 | 22 | 51 | 24 | 32 | 16 | 9 | 11 | Ccl2 |
| Ccl20 | 0 | 0 | 0 | 0 | 0 | 2 | 0 | 0 | 1 | 0 | 0 | 0 | 0 | 1 | 0 | 0 | 0 | 0 | 1 | 0 | 0 | 0 | 2 | 0 | 0 | 4 | 1 | 1 | Ccl20 |
| Ccl21a | 12 | 0 | 0 | 0 | 0 | 0 | 0 | 0 | 0 | 0 | 0 | 0 | 0 | 0 | 0 | 0 | 0 | 0 | 0 | 0 | 0 | 0 | 0 | 0 | 27 | 0 | 0 | 0 | Ccl21a |
| Ccl21b | 10 | 10 | 4 | 12 | 7 | 3 | 2 | 2 | 0 | 2 | 0 | 1 | 0 | 0 | 3 | 6 | 1 | 4 | 3 | 8 | 10 | 0 | 0 | 1 | 3 | 2 | 0 | 1 | Ccl21b |
| Ccl21d | 0 | 3 | 4 | 2 | 0 | 3 | 0 | 0 | 0 | 1 | 0 | 1 | 3 | 1 | 0 | 4 | 1 | 0 | 5 | 1 | 0 | 0 | 0 | 0 | 0 | 0 | 0 | 0 | Ccl21d |
| Ccl22 | 0 | 3 | 0 | 0 | 0 | 0 | 0 | 0 | 0 | 0 | 0 | 0 | 0 | 0 | 0 | 0 | 0 | 0 | 1 | 0 | 0 | 0 | 0 | 0 | 0 | 0 | 0 | 0 | Ccl22 |
| Ccl24 | 22 | 24 | 49 | 35 | 36 | 40 | 13 | 61 | 46 | 43 | 63 | 27 | 56 | 38 | 21 | 33 | 36 | 36 | 24 | 32 | 20 | 53 | 62 | 44 | 51 | 43 | 79 | 36 | Ccl24 |
| Ccl25 | 1344 | 1166 | 935 | 1115 | 1100 | 1198 | 1151 | 1468 | 1243 | 1044 | 1232 | 985 | 1235 | 1062 | 1189 | 1137 | 1010 | 921 | 901 | 1354 | 1144 | 929 | 1111 | 985 | 1092 | 812 | 1219 | 954 | Ccl25 |
| Ccl26 | 0 | 0 | 0 | 8 | 1 | 0 | 0 | 0 | 0 | 0 | 0 | 0 | 0 | 0 | 0 | 0 | 0 | 0 | 3 | 0 | 0 | 0 | 3 | 0 | 0 | 0 | 0 | 0 | Ccl26 |
| Ccl27a | 2160 | 2599 | 2610 | 3580 | 3932 | 3529 | 3832 | 2811 | 2696 | 2937 | 2831 | 2656 | 2427 | 2659 | 2471 | 2301 | 3304 | 3793 | 4203 | 3823 | 3904 | 3240 | 2597 | 2839 | 3307 | 2712 | 2539 | 2455 | Ccl27a |
| Ccl27b | 7 | 4 | 7 | 11 | 2 | 9 | 2 | 4 | 7 | 3 | 6 | 3 | 6 | 0 | 3 | 9 | 2 | 3 | 13 | 5 | 4 | 10 | 6 | 3 | 10 | 1 | 5 | 8 | Ccl27b |
| Ccl28 | 125 | 87 | 99 | 159 | 167 | 246 | 226 | 89 | 133 | 126 | 121 | 168 | 118 | 105 | 185 | 81 | 180 | 68 | 104 | 203 | 273 | 244 | 198 | 230 | 86 | 138 | 228 | 161 | Ccl28 |
| Ccl3 | 9 | 10 | 7 | 16 | 11 | 11 | 15 | 15 | 12 | 21 | 16 | 10 | 10 | 9 | 5 | 10 | 9 | 23 | 5 | 22 | 13 | 22 | 21 | 11 | 17 | 15 | 10 | 13 | Ccl3 |
| Ccl4 | 0 | 0 | 2 | 0 | 1 | 1 | 0 | 5 | 5 | 3 | 13 | 3 | 2 | 2 | 3 | 7 | 2 | 6 | 1 | 0 | 2 | 2 | 5 | 1 | 0 | 9 | 0 | 2 | Ccl4 |
| Ccl5 | 0 | 0 | 3 | 0 | 0 | 0 | 0 | 1 | 2 | 0 | 8 | 0 | 1 | 0 | 3 | 0 | 0 | 0 | 0 | 0 | 0 | 0 | 3 | 2 | 0 | 0 | 2 | 0 | Ccl5 |
| Ccl6 | 25 | 20 | 25 | 13 | 28 | 22 | 18 | 49 | 36 | 23 | 50 | 22 | 33 | 47 | 13 | 37 | 22 | 33 | 19 | 36 | 26 | 32 | 41 | 33 | 31 | 38 | 44 | 51 | Ccl6 |
| Ccl7 | 3 | 0 | 5 | 6 | 4 | 3 | 1 | 11 | 36 | 15 | 48 | 15 | 11 | 7 | 4 | 2 | 2 | 9 | 5 | 4 | 3 | 20 | 44 | 28 | 44 | 17 | 7 | 9 | Ccl7 |
| Ccl8 | 0 | 0 | 0 | 0 | 0 | 0 | 0 | 0 | 0 | 0 | 1 | 0 | 0 | 0 | 0 | 0 | 0 | 0 | 0 | 0 | 0 | 0 | 0 | 0 | 0 | 0 | 0 | 0 | Ccl8 |
| Ccl9 | 57 | 35 | 52 | 42 | 56 | 50 | 51 | 64 | 84 | 69 | 90 | 51 | 61 | 65 | 55 | 70 | 37 | 70 | 37 | 64 | 65 | 80 | 113 | 75 | 60 | 108 | 83 | 107 | Ccl9 |
| Ccm2 | 3054 | 3858 | 3754 | 3966 | 3458 | 3405 | 3489 | 2251 | 2348 | 2541 | 2616 | 2620 | 2624 | 2572 | 2949 | 3343 | 3854 | 3712 | 4053 | 3525 | 3694 | 2400 | 2695 | 2604 | 2681 | 2781 | 2621 | 2718 | Ccm2 |
| Ccm2l | 81 | 94 | 141 | 125 | 140 | 125 | 122 | 198 | 221 | 281 | 334 | 298 | 361 | 319 | 107 | 125 | 141 | 134 | 149 | 94 | 113 | 296 | 307 | 309 | 298 | 355 | 387 | 292 | Ccm2l |
| Ccn1 | 964 | 1488 | 1307 | 1042 | 637 | 796 | 683 | 834 | 1118 | 1070 | 1582 | 1168 | 860 | 1055 | 1199 | 1488 | 1094 | 1083 | 785 | 601 | 668 | 921 | 1196 | 1161 | 1055 | 1402 | 1025 | 1061 | Ccn1 |
| Ccn2 | 636 | 770 | 1076 | 829 | 580 | 680 | 377 | 774 | 1100 | 1353 | 1660 | 618 | 768 | 582 | 822 | 918 | 884 | 925 | 527 | 418 | 400 | 1054 | 1169 | 1235 | 647 | 903 | 666 | 564 | Ccn2 |
| Ccn3 | 136 | 98 | 84 | 101 | 111 | 148 | 88 | 211 | 300 | 276 | 411 | 100 | 108 | 106 | 108 | 114 | 82 | 145 | 69 | 100 | 98 | 234 | 490 | 260 | 170 | 128 | 85 | 74 | Ccn3 |
| Ccn4 | 850 | 835 | 829 | 554 | 522 | 793 | 240 | 1456 | 2298 | 1592 | 2922 | 1298 | 1837 | 1284 | 981 | 1139 | 464 | 878 | 231 | 372 | 261 | 2054 | 3183 | 1325 | 1615 | 1027 | 1331 | 1233 | Ccn4 |
| Ccn5 | 39 | 54 | 30 | 35 | 30 | 37 | 35 | 10 | 14 | 10 | 19 | 9 | 10 | 17 | 28 | 44 | 49 | 28 | 39 | 22 | 24 | 8 | 14 | 11 | 9 | 17 | 2 | 4 | Ccn5 |
| Ccn6 | 12 | 16 | 11 | 36 | 20 | 28 | 23 | 16 | 19 | 24 | 43 | 31 | 18 | 14 | 17 | 19 | 22 | 40 | 30 | 20 | 37 | 27 | 16 | 20 | 11 | 10 | 12 | 19 | Ccn6 |
| Ccna1 | 77 | 163 | 109 | 120 | 80 | 101 | 121 | 110 | 96 | 93 | 94 | 137 | 124 | 112 | 141 | 142 | 174 | 118 | 101 | 109 | 105 | 94 | 102 | 118 | 93 | 128 | 138 | 90 | Ccna1 |
| Ccna2 | 14927 | 11768 | 13406 | 9288 | 7292 | 9259 | 11210 | 15387 | 12987 | 11008 | 10345 | 17831 | 14369 | 18087 | 13949 | 10931 | 11594 | 8130 | 9903 | 11264 | 9690 | 10029 | 11519 | 14407 | 12305 | 21150 | 18173 | 19832 | Ccna2 |
| Ccnb1 | 11146 | 9902 | 11040 | 7332 | 5297 | 6675 | 8149 | 12284 | 10489 | 8369 | 8121 | 14212 | 11907 | 15535 | 10291 | 9090 | 9011 | 6349 | 7566 | 9044 | 7120 | 7277 | 9506 | 11432 | 9850 | 17402 | 15619 | 16290 | Ccnb1 |
| Ccnb1ip1 | 3 | 5 | 5 | 0 | 7 | 4 | 0 | 4 | 2 | 3 | 0 | 0 | 0 | 7 | 3 | 3 | 0 | 0 | 2 | 4 | 3 | 2 | 1 | 5 | 0 | 5 | 0 | 1 | Ccnb1ip1 |
| Ccnb2 | 2746 | 2808 | 2916 | 1968 | 1429 | 1806 | 1905 | 3014 | 3044 | 2721 | 2585 | 4256 | 3598 | 4335 | 2712 | 2535 | 2408 | 1662 | 1954 | 2008 | 1672 | 2206 | 2834 | 3492 | 2815 | 5003 | 4475 | 4524 | Ccnb2 |
| Ccnb3 | 0 | 0 | 0 | 0 | 0 | 0 | 0 | 0 | 2 | 0 | 19 | 0 | 0 | 0 | 3 | 5 | 0 | 0 | 0 | 2 | 0 | 0 | 5 | 0 | 0 | 0 | 0 | 0 | Ccnb3 |
| Ccnc | 3230 | 2328 | 3108 | 2864 | 2990 | 3063 | 3342 | 4038 | 3240 | 2896 | 2861 | 3509 | 3569 | 3905 | 3750 | 2998 | 3116 | 2763 | 2976 | 3920 | 3138 | 3377 | 3174 | 3189 | 3449 | 3604 | 3647 | 3913 | Ccnc |
| Ccnd1 | 14813 | 15079 | 20809 | 12035 | 8897 | 11064 | 9286 | 16695 | 15512 | 16206 | 16194 | 27860 | 21899 | 27545 | 14227 | 12036 | 16964 | 9216 | 11461 | 11846 | 10058 | 14920 | 15950 | 21922 | 16483 | 31982 | 31231 | 32918 | Ccnd1 |
| Ccnd2 | 66220 | 61543 | 69998 | 67071 | 51837 | 64705 | 75203 | 51550 | 49998 | 51328 | 47082 | 74500 | 62754 | 78329 | 60825 | 63180 | 64067 | 61760 | 68059 | 68394 | 71708 | 34487 | 44473 | 56044 | 66390 | 69422 | 75828 | 76419 | Ccnd2 |
| Ccnd3 | 8962 | 10517 | 8882 | 7170 | 5570 | 6487 | 5412 | 4930 | 6494 | 5092 | 7358 | 5122 | 5917 | 4672 | 7734 | 9691 | 8466 | 7263 | 6050 | 5817 | 5403 | 4553 | 7834 | 5373 | 4276 | 7048 | 5710 | 5491 | Ccnd3 |
| Ccndbp1 | 2750 | 2743 | 2353 | 2920 | 2623 | 2485 | 2991 | 2858 | 2596 | 2355 | 2256 | 2528 | 2566 | 2754 | 2836 | 2651 | 2700 | 2492 | 2546 | 3037 | 2984 | 2617 | 2393 | 2527 | 2705 | 2369 | 2586 | 2604 | Ccndbp1 |
| Ccne1 | 3298 | 3078 | 2913 | 2400 | 2050 | 2004 | 1999 | 1291 | 1282 | 1013 | 1511 | 1576 | 1515 | 1398 | 2909 | 3011 | 2612 | 2313 | 2159 | 2140 | 2036 | 1106 | 1407 | 1253 | 1071 | 2040 | 1692 | 1710 | Ccne1 |
| Ccne2 | 2467 | 1668 | 2335 | 1641 | 1358 | 1807 | 1831 | 2643 | 2389 | 1831 | 2271 | 2777 | 2437 | 2963 | 2395 | 2203 | 1803 | 1378 | 1527 | 1991 | 1613 | 1952 | 2381 | 2442 | 2182 | 3407 | 3019 | 3455 | Ccne2 |
| Ccnf | 3902 | 4739 | 4508 | 3530 | 2819 | 3177 | 3032 | 3308 | 3490 | 3490 | 2946 | 5146 | 4691 | 4930 | 3574 | 3882 | 4113 | 3111 | 3613 | 3253 | 3160 | 2501 | 3279 | 4302 | 3375 | 6413 | 5433 | 5629 | Ccnf |
| Ccng1 | 7789 | 6650 | 7056 | 5914 | 5705 | 5585 | 6462 | 5195 | 4663 | 3950 | 4098 | 4050 | 4456 | 3985 | 8317 | 7506 | 6684 | 6243 | 5678 | 6822 | 6120 | 4655 | 5339 | 3743 | 3770 | 3894 | 4047 | 4581 | Ccng1 |
| Ccng2 | 6993 | 6108 | 6900 | 7525 | 8021 | 7719 | 8463 | 12305 | 9836 | 10170 | 9532 | 11158 | 9584 | 11483 | 6907 | 5888 | 7405 | 7326 | 8337 | 8268 | 8020 | 11062 | 9335 | 11116 | 11784 | 10347 | 10548 | 10865 | Ccng2 |
| Ccnh | 5610 | 4812 | 5037 | 5269 | 4173 | 4539 | 5413 | 3529 | 2945 | 2576 | 2730 | 3254 | 3027 | 3527 | 5980 | 5363 | 4815 | 4799 | 4494 | 5442 | 4966 | 2824 | 3014 | 2963 | 2792 | 3496 | 3332 | 3637 | Ccnh |
| Ccni | 27347 | 22138 | 28215 | 30187 | 30062 | 30080 | 35303 | 24391 | 21476 | 25570 | 22655 | 26162 | 22470 | 27590 | 27022 | 22807 | 28957 | 31457 | 32740 | 26092 | 35349 | 27389 | 21647 | 27061 | 26937 | 28647 | 23882 | 27687 | Ccni |

| | | | | | | | | | | | | | | | | | | | | | | | | | | | | | |
|---|---|---|---|---|---|---|---|---|---|---|---|---|---|---|---|---|---|---|---|---|---|---|---|---|---|---|---|---|---|
| Ccnj | 1888 | 1700 | 2759 | 2041 | 2284 | 2095 | 2292 | 3466 | 2998 | 3504 | 3054 | 4830 | 4204 | 5110 | 1843 | 1606 | 2544 | 1789 | 2352 | 1929 | 2154 | 3487 | 2795 | 4258 | 4052 | 5020 | 4865 | 5242 | Ccnj |
| Ccnjl | 1329 | 1640 | 1994 | 1500 | 1540 | 1386 | 1345 | 1072 | 938 | 998 | 953 | 1434 | 1446 | 1276 | 1506 | 1561 | 1846 | 1513 | 1626 | 1371 | 1369 | 1126 | 870 | 1057 | 1030 | 1531 | 1470 | 1511 | Ccnjl |
| Ccnk | 5257 | 6026 | 7490 | 6933 | 5530 | 5166 | 5428 | 4114 | 4298 | 5096 | 4743 | 6225 | 4970 | 5918 | 4986 | 5589 | 7123 | 6516 | 6400 | 4431 | 5940 | 4782 | 3935 | 5600 | 5528 | 6968 | 5087 | 5994 | Ccnk |
| Ccnl1 | 6124 | 4950 | 6118 | 6088 | 5543 | 5760 | 5841 | 9867 | 8057 | 7466 | 8381 | 9104 | 8579 | 8542 | 6224 | 5667 | 5786 | 5529 | 5531 | 6369 | 5525 | 8808 | 8471 | 8004 | 7812 | 8338 | 8836 | 8579 | Ccnl1 |
| Ccnl2 | 10117 | 9709 | 11112 | 11726 | 12584 | 11460 | 11587 | 17484 | 16200 | 17309 | 19171 | 16993 | 16082 | 16814 | 9973 | 10110 | 11304 | 11855 | 12338 | 11298 | 11491 | 19298 | 16371 | 18271 | 18075 | 16575 | 16518 | 17165 | Ccnl2 |
| Ccno | 153 | 170 | 171 | 162 | 133 | 137 | 154 | 105 | 130 | 96 | 95 | 276 | 229 | 229 | 150 | 183 | 177 | 160 | 148 | 138 | 140 | 87 | 185 | 173 | 160 | 335 | 218 | 259 | Ccno |
| Ccnq | 683 | 775 | 691 | 628 | 561 | 490 | 723 | 685 | 737 | 655 | 648 | 893 | 716 | 795 | 644 | 616 | 712 | 601 | 624 | 645 | 616 | 530 | 782 | 772 | 634 | 916 | 816 | 890 | Ccnq |
| Ccnt1 | 10146 | 10051 | 9055 | 10382 | 10137 | 10140 | 10464 | 10297 | 8576 | 8909 | 8393 | 9046 | 8781 | 9010 | 10199 | 9749 | 10031 | 10657 | 10137 | 10619 | 10613 | 8928 | 8418 | 8773 | 9717 | 7083 | 9547 | 9227 | Ccnt1 |
| Ccnt2 | 15733 | 13408 | 12520 | 15191 | 15273 | 17087 | 16955 | 18516 | 15599 | 15490 | 16776 | 17512 | 16894 | 17969 | 15380 | 13374 | 13257 | 14712 | 14758 | 17409 | 16753 | 15797 | 14756 | 15185 | 17505 | 12678 | 18500 | 16185 | Ccnt2 |
| Ccny | 12708 | 10739 | 11763 | 12142 | 10998 | 12038 | 12262 | 11337 | 9921 | 9919 | 10415 | 10315 | 10287 | 10722 | 13102 | 10789 | 11581 | 11483 | 11681 | 11950 | 11717 | 10475 | 10270 | 10216 | 10452 | 10519 | 10322 | 10863 | Ccny |
| Ccnyl1 | 5783 | 5023 | 5542 | 5154 | 4330 | 4573 | 4975 | 2911 | 2741 | 2978 | 2932 | 3420 | 3333 | 3274 | 5625 | 5453 | 5016 | 5113 | 4472 | 4403 | 4513 | 3220 | 2875 | 3121 | 3078 | 3415 | 3362 | 3635 | Ccnyl1 |
| Ccp110 | 16827 | 15260 | 13025 | 14181 | 13689 | 14474 | 16584 | 14550 | 12125 | 12453 | 11991 | 13421 | 12465 | 14458 | 16143 | 14702 | 14205 | 13596 | 14613 | 16703 | 16009 | 10415 | 11549 | 12951 | 12046 | 11728 | 14181 | 13459 | Ccp110 |
| Ccpg1 | 335 | 293 | 383 | 489 | 441 | 456 | 480 | 383 | 315 | 346 | 301 | 286 | 287 | 331 | 365 | 442 | 380 | 484 | 392 | 387 | 406 | 414 | 328 | 322 | 385 | 294 | 228 | 228 | Ccpg1 |
| Ccpg1os | 364 | 437 | 414 | 430 | 402 | 297 | 398 | 353 | 434 | 366 | 336 | 370 | 390 | 411 | 496 | 379 | 469 | 369 | 459 | 487 | 407 | 418 | 417 | 463 | 345 | 416 | 378 | 452 | Ccpg1os |
| Ccr1 | 44 | 16 | 56 | 19 | 28 | 42 | 29 | 100 | 101 | 75 | 154 | 90 | 113 | 54 | 29 | 35 | 45 | 42 | 38 | 40 | 18 | 124 | 201 | 123 | 101 | 138 | 98 | 103 | Ccr1 |
| Ccr10 | 1 | 5 | 7 | 3 | 3 | 0 | 8 | 1 | 2 | 1 | 6 | 2 | 0 | 3 | 5 | 2 | 12 | 11 | 8 | 6 | 9 | 5 | 6 | 11 | 4 | 1 | 1 | 2 | Ccr10 |
| Ccr1l1 | 0 | 0 | 0 | 0 | 0 | 0 | 0 | 0 | 2 | 2 | 1 | 0 | 1 | 0 | 0 | 1 | 0 | 0 | 0 | 0 | 1 | 0 | 1 | 0 | 0 | 0 | 0 | 0 | Ccr1l1 |
| Ccr2 | 7 | 3 | 16 | 1 | 3 | 36 | 10 | 68 | 118 | 94 | 185 | 35 | 31 | 28 | 11 | 3 | 3 | 6 | 8 | 7 | 5 | 57 | 159 | 56 | 52 | 73 | 36 | 30 | Ccr2 |
| Ccr3 | 1 | 0 | 5 | 0 | 1 | 2 | 1 | 0 | 4 | 3 | 3 | 3 | 3 | 0 | 0 | 0 | 0 | 0 | 0 | 0 | 0 | 5 | 3 | 0 | 4 | 6 | 0 | 1 | Ccr3 |
| Ccr4 | 6 | 5 | 14 | 3 | 3 | 7 | 1 | 0 | 1 | 0 | 0 | 0 | 3 | 0 | 4 | 4 | 12 | 22 | 3 | 13 | 4 | 0 | 3 | 0 | 1 | 6 | 0 | 0 | Ccr4 |
| Ccr5 | 21 | 11 | 5 | 20 | 26 | 9 | 6 | 73 | 60 | 47 | 93 | 64 | 42 | 40 | 5 | 12 | 0 | 12 | 13 | 17 | 17 | 95 | 120 | 42 | 43 | 53 | 42 | 58 | Ccr5 |
| Ccr6 | 0 | 0 | 0 | 0 | 0 | 0 | 0 | 0 | 0 | 0 | 2 | 0 | 0 | 0 | 0 | 0 | 0 | 0 | 0 | 1 | 0 | 0 | 0 | 0 | 2 | 0 | 0 | 0 | Ccr6 |
| Ccr7 | 12 | 14 | 11 | 4 | 2 | 4 | 3 | 9 | 5 | 14 | 7 | 1 | 12 | 6 | 5 | 11 | 10 | 1 | 0 | 2 | 6 | 7 | 4 | 7 | 0 | 11 | 2 | 9 | Ccr7 |
| Ccr8 | 0 | 0 | 0 | 0 | 0 | 0 | 0 | 0 | 6 | 1 | 4 | 0 | 0 | 0 | 0 | 1 | 0 | 1 | 1 | 0 | 0 | 0 | 5 | 0 | 0 | 0 | 0 | 0 | Ccr8 |
| Ccr9 | 9 | 6 | 3 | 3 | 11 | 4 | 7 | 32 | 7 | 17 | 18 | 8 | 16 | 13 | 0 | 5 | 0 | 8 | 9 | 12 | 15 | 20 | 23 | 10 | 19 | 11 | 15 | 9 | Ccr9 |
| Ccrl2 | 39 | 32 | 26 | 31 | 26 | 36 | 20 | 31 | 45 | 52 | 20 | 29 | 71 | 68 | 41 | 44 | 36 | 18 | 38 | 33 | 38 | 33 | 38 | 45 | 46 | 65 | 38 | 80 | Ccrl2 |
| Ccs | 1547 | 1837 | 1669 | 1933 | 1767 | 1905 | 1541 | 1216 | 1408 | 1378 | 1656 | 1478 | 1625 | 1436 | 1548 | 1801 | 1865 | 1875 | 1786 | 1645 | 1529 | 1253 | 1383 | 1626 | 1526 | 1531 | 1828 | 1533 | Ccs |
| Ccsap | 1051 | 1153 | 1142 | 1332 | 1156 | 1026 | 1218 | 497 | 590 | 630 | 387 | 709 | 504 | 681 | 1129 | 1110 | 1279 | 1105 | 1337 | 1097 | 1202 | 461 | 492 | 652 | 645 | 786 | 585 | 616 | Ccsap |
| Ccser1 | 1380 | 1278 | 832 | 2053 | 2570 | 2422 | 2856 | 2356 | 2252 | 1959 | 1751 | 1813 | 1398 | 1793 | 1731 | 1115 | 1460 | 2128 | 2371 | 2430 | 2673 | 2461 | 1927 | 1946 | 2522 | 1578 | 1407 | 1568 | Ccser1 |
| Ccser2 | 9197 | 7540 | 5331 | 8273 | 11207 | 10006 | 10178 | 9514 | 9094 | 8129 | 8283 | 6510 | 7629 | 6634 | 10247 | 8781 | 6697 | 8757 | 8358 | 9709 | 9623 | 9733 | 8303 | 6182 | 8422 | 4386 | 6334 | 5954 | Ccser2 |
| Cct2 | 33894 | 34854 | 34401 | 30603 | 30018 | 29696 | 34274 | 29916 | 27376 | 25849 | 25514 | 29915 | 28200 | 31556 | 34867 | 30368 | 33695 | 30660 | 31316 | 34247 | 32914 | 26174 | 26669 | 29138 | 28763 | 34081 | 31817 | 33993 | Cct2 |
| Cct3 | 27117 | 28249 | 26068 | 21694 | 20994 | 21132 | 24462 | 26668 | 24080 | 21493 | 20905 | 26998 | 24524 | 26631 | 27447 | 23934 | 24877 | 21183 | 22673 | 26058 | 23055 | 21343 | 24205 | 24405 | 22603 | 29546 | 28130 | 28968 | Cct3 |
| Cct4 | 28894 | 25568 | 25540 | 23545 | 22154 | 21703 | 26060 | 27352 | 22302 | 20660 | 19170 | 23734 | 22484 | 23571 | 30995 | 25932 | 24455 | 22774 | 22614 | 26341 | 24272 | 21672 | 22867 | 21470 | 20378 | 24943 | 24659 | 25891 | Cct4 |
| Cct5 | 30222 | 34389 | 28298 | 26053 | 24648 | 25584 | 27968 | 28924 | 27897 | 24217 | 23355 | 27591 | 25420 | 27860 | 32220 | 29491 | 27786 | 25608 | 27841 | 31129 | 27539 | 23118 | 27754 | 26644 | 25131 | 31004 | 28483 | 30197 | Cct5 |
| Cct6a | 32156 | 29365 | 30582 | 26896 | 26684 | 27453 | 30979 | 35067 | 29967 | 27740 | 27938 | 32630 | 31926 | 33738 | 33712 | 28200 | 28821 | 26387 | 28906 | 33190 | 29104 | 27693 | 29613 | 30918 | 31106 | 37314 | 35404 | 36032 | Cct6a |
| Cct6b | 52 | 68 | 50 | 89 | 57 | 49 | 80 | 24 | 25 | 19 | 21 | 37 | 37 | 36 | 66 | 49 | 44 | 62 | 54 | 50 | 41 | 18 | 27 | 28 | 22 | 21 | 34 | 37 | Cct6b |
| Cct7 | 28884 | 32204 | 28726 | 28604 | 26333 | 27507 | 28008 | 25690 | 24320 | 22314 | 21792 | 26729 | 24581 | 26648 | 29050 | 27493 | 28983 | 27467 | 30684 | 30385 | 28982 | 22262 | 24320 | 25411 | 25258 | 29113 | 27663 | 27303 | Cct7 |
| Cct8 | 39167 | 35510 | 37913 | 31819 | 28643 | 29026 | 34025 | 34221 | 27882 | 24544 | 25130 | 30987 | 28776 | 32231 | 40856 | 35392 | 35178 | 30941 | 30339 | 36214 | 31614 | 27299 | 27661 | 28282 | 27246 | 34409 | 31752 | 33583 | Cct8 |
| Cct8l1 | 1 | 0 | 8 | 6 | 6 | 0 | 0 | 0 | 0 | 0 | 0 | 0 | 0 | 0 | 3 | 4 | 2 | 9 | 4 | 2 | 3 | 0 | 0 | 3 | 0 | 0 | 0 | 0 | Cct8l1 |
| Ccz1 | 3842 | 3540 | 3648 | 3263 | 3360 | 3232 | 3835 | 5069 | 4234 | 4363 | 4271 | 4997 | 4589 | 5255 | 4092 | 3372 | 3585 | 3244 | 3438 | 3756 | 3706 | 4288 | 4360 | 4639 | 4584 | 5280 | 5210 | 5479 | Ccz1 |
| Cd101 | 37 | 61 | 14 | 49 | 51 | 48 | 53 | 15 | 16 | 16 | 16 | 24 | 16 | 19 | 30 | 44 | 34 | 51 | 63 | 43 | 56 | 10 | 8 | 25 | 23 | 20 | 10 | 13 | Cd101 |
| Cd109 | 1518 | 1104 | 1164 | 756 | 743 | 1161 | 686 | 1384 | 1448 | 1248 | 1359 | 1278 | 1346 | 1254 | 1282 | 1408 | 915 | 1081 | 756 | 879 | 721 | 1036 | 1515 | 1218 | 1121 | 1191 | 1399 | 1299 | Cd109 |
| Cd14 | 20 | 20 | 20 | 8 | 16 | 16 | 13 | 16 | 26 | 22 | 39 | 23 | 27 | 22 | 21 | 9 | 8 | 12 | 25 | 17 | 9 | 17 | 39 | 27 | 23 | 34 | 37 | 28 | Cd14 |
| Cd151 | 2668 | 2720 | 2736 | 2309 | 2083 | 2284 | 2119 | 2546 | 2639 | 2478 | 2362 | 2467 | 2608 | 2485 | 2466 | 2646 | 2605 | 2176 | 2356 | 2414 | 2232 | 2289 | 2875 | 2491 | 2100 | 2706 | 2692 | 2662 | Cd151 |
| Cd160 | 0 | 1 | 3 | 0 | 2 | 0 | 1 | 0 | 0 | 0 | 2 | 0 | 2 | 0 | 0 | 0 | 0 | 0 | 0 | 0 | 1 | 3 | 1 | 0 | 0 | 0 | 0 | 0 | Cd160 |
| Cd163 | 2 | 3 | 2 | 9 | 4 | 5 | 10 | 5 | 16 | 7 | 36 | 4 | 9 | 11 | 1 | 4 | 0 | 14 | 8 | 5 | 9 | 35 | 44 | 20 | 24 | 5 | 7 | 1 | Cd163 |
| Cd164 | 16492 | 11153 | 12691 | 11211 | 9692 | 11084 | 10694 | 12003 | 11293 | 10113 | 12617 | 10461 | 11128 | 10099 | 14600 | 15492 | 10973 | 12123 | 9809 | 10787 | 10383 | 10559 | 13230 | 10578 | 9168 | 12943 | 10479 | 10498 | Cd164 |
| Cd164l2 | 80 | 67 | 51 | 41 | 56 | 33 | 25 | 9 | 36 | 12 | 21 | 14 | 18 | 15 | 61 | 80 | 48 | 53 | 39 | 27 | 14 | 10 | 35 | 20 | 9 | 32 | 22 | 14 | Cd164l2 |
| Cd177 | 1 | 2 | 10 | 9 | 2 | 0 | 7 | 15 | 11 | 13 | 14 | 8 | 11 | 7 | 7 | 3 | 3 | 3 | 1 | 8 | 4 | 7 | 19 | 20 | 7 | 18 | 5 | 9 | Cd177 |
| Cd180 | 36 | 43 | 38 | 21 | 47 | 49 | 38 | 127 | 123 | 140 | 162 | 143 | 129 | 131 | 38 | 44 | 52 | 26 | 30 | 51 | 40 | 139 | 166 | 136 | 97 | 130 | 145 | 107 | Cd180 |
| Cd19 | 8 | 3 | 9 | 3 | 7 | 2 | 0 | 1 | 8 | 7 | 3 | 12 | 0 | 5 | 4 | 5 | 5 | 0 | 5 | 4 | 3 | 2 | 3 | 15 | 4 | 8 | 9 | 5 | Cd19 |
| Cd1d1 | 679 | 612 | 584 | 847 | 1046 | 958 | 876 | 2307 | 2100 | 2223 | 2101 | 2278 | 2227 | 2485 | 798 | 513 | 720 | 840 | 966 | 902 | 861 | 2166 | 2028 | 2304 | 2407 | 2269 | 2179 | 2038 | Cd1d1 |
| Cd2 | 1 | 1 | 0 | 0 | 2 | 3 | 1 | 2 | 2 | 14 | 0 | 0 | 7 | 0 | 0 | 6 | 0 | 1 | 0 | 3 | 1 | 0 | 3 | 5 | 2 | 0 | 0 | 2 | Cd2 |
| Cd200 | 1561 | 1571 | 1584 | 2607 | 4398 | 3497 | 3835 | 6936 | 5866 | 6652 | 5828 | 4366 | 4428 | 4935 | 2068 | 1441 | 2118 | 2906 | 3563 | 3738 | 3838 | 9123 | 6165 | 5654 | 7393 | 3535 | 4122 | 4419 | Cd200 |
| Cd200r1 | 23 | 24 | 27 | 24 | 25 | 31 | 13 | 60 | 58 | 52 | 65 | 55 | 54 | 38 | 46 | 24 | 44 | 23 | 21 | 19 | 30 | 63 | 76 | 52 | 44 | 61 | 55 | 33 | Cd200r1 |
| Cd200r2 | 0 | 1 | 0 | 0 | 3 | 0 | 0 | 0 | 2 | 3 | 0 | 0 | 0 | 0 | 0 | 1 | 1 | 0 | 0 | 0 | 1 | 0 | 2 | 0 | 5 | 0 | 0 | 0 | Cd200r2 |
| Cd200r3 | 0 | 0 | 1 | 0 | 0 | 0 | 0 | 0 | 6 | 0 | 4 | 0 | 3 | 0 | 0 | 0 | 0 | 1 | 0 | 0 | 2 | 5 | 6 | 2 | 1 | 3 | 0 | 2 | Cd200r3 |
| Cd200r4 | 9 | 9 | 10 | 0 | 0 | 7 | 0 | 7 | 3 | 2 | 10 | 2 | 1 | 5 | 7 | 3 | 0 | 3 | 4 | 2 | 1 | 5 | 5 | 0 | 15 | 1 | 6 | 9 | Cd200r4 |
| Cd207 | 0 | 4 | 3 | 6 | 2 | 2 | 6 | 1 | 0 | 0 | 2 | 0 | 0 | 0 | 0 | 0 | 0 | 2 | 1 | 0 | 3 | 3 | 0 | 0 | 0 | 1 | 0 | 0 | Cd207 |
| Cd209a | 0 | 0 | 0 | 0 | 0 | 0 | 0 | 0 | 6 | 3 | 3 | 0 | 0 | 0 | 0 | 0 | 0 | 0 | 0 | 1 | 0 | 3 | 5 | 0 | 0 | 0 | 2 | 1 | Cd209a |
| Cd209b | 0 | 0 | 0 | 0 | 0 | 0 | 0 | 0 | 2 | 0 | 2 | 0 | 0 | 0 | 0 | 0 | 0 | 0 | 0 | 0 | 0 | 0 | 0 | 0 | 0 | 0 | 0 | 0 | Cd209b |
| Cd209c | 0 | 2 | 0 | 0 | 3 | 0 | 0 | 5 | 1 | 1 | 0 | 0 | 0 | 0 | 0 | 2 | 0 | 0 | 2 | 3 | 0 | 0 | 0 | 0 | 1 | 4 | 0 | 1 | Cd209c |
| Cd209f | 0 | 0 | 0 | 0 | 0 | 1 | 0 | 0 | 0 | 0 | 0 | 0 | 0 | 0 | 0 | 0 | 0 | 0 | 1 | 0 | 0 | 0 | 0 | 1 | 0 | 0 | 0 | 0 | Cd209f |
| Cd209g | 0 | 0 | 0 | 0 | 0 | 0 | 0 | 0 | 0 | 0 | 0 | 0 | 0 | 0 | 0 | 0 | 0 | 0 | 0 | 0 | 0 | 0 | 0 | 2 | 0 | 0 | 0 | 0 | Cd209g |
| Cd22 | 6 | 5 | 0 | 1 | 2 | 2 | 0 | 4 | 2 | 0 | 6 | 0 | 3 | 0 | 0 | 3 | 3 | 2 | 1 | 2 | 1 | 5 | 2 | 2 | 8 | 2 | 9 | 0 | Cd22 |
| Cd226 | 15 | 10 | 8 | 13 | 12 | 5 | 15 | 34 | 29 | 38 | 21 | 50 | 56 | 17 | 8 | 26 | 2 | 7 | 13 | 7 | 3 | 27 | 35 | 27 | 15 | 52 | 57 | 36 | Cd226 |
| Cd244a | 2 | 5 | 5 | 1 | 0 | 0 | 3 | 5 | 4 | 3 | 2 | 2 | 3 | 1 | 1 | 1 | 0 | 1 | 0 | 1 | 1 | 0 | 2 | 0 | 0 | 0 | 7 | 4 | Cd244a |
| Cd247 | 66 | 81 | 32 | 68 | 76 | 84 | 69 | 56 | 54 | 36 | 39 | 77 | 48 | 99 | 82 | 59 | 27 | 75 | 80 | 62 | 88 | 37 | 32 | 58 | 47 | 79 | 81 | 62 | Cd247 |
| Cd248 | 2824 | 3632 | 3286 | 2508 | 1814 | 2312 | 1049 | 2058 | 3092 | 2789 | 3472 | 2436 | 2964 | 1694 | 2527 | 4243 | 2621 | 2720 | 1559 | 1390 | 1088 | 2256 | 3916 | 2721 | 2117 | 3330 | 2576 | 1911 | Cd248 |
| Cd24a | 29085 | 24789 | 22854 | 26609 | 35730 | 34252 | 36755 | 66762 | 51913 | 51852 | 46252 | 45566 | 45455 | 50888 | 34666 | 25108 | 25451 | 28182 | 32803 | 38868 | 37045 | 63127 | 52637 | 47922 | 57171 | 43178 | 46424 | 49529 | Cd24a |
| Cd27 | 1 | 3 | 0 | 2 | 1 | 2 | 6 | 9 | 6 | 2 | 4 | 1 | 7 | 3 | 1 | 1 | 2 | 0 | 0 | 1 | 0 | 5 | 8 | 7 | 5 | 0 | 7 | 5 | Cd27 |
| Cd274 | 96 | 108 | 111 | 127 | 100 | 98 | 98 | 110 | 105 | 112 | 98 | 125 | 90 | 90 | 94 | 107 | 76 | 117 | 114 | 95 | 86 | 112 | 95 | 126 | 92 | 108 | 127 | 109 | Cd274 |
| Cd276 | 2028 | 1947 | 2070 | 1796 | 2439 | 2094 | 1609 | 3595 | 4386 | 4579 | 4614 | 4402 | 4307 | 3865 | 1822 | 2216 | 2177 | 1919 | 2191 | 1818 | 1741 | 4500 | 4302 | 4608 | 4267 | 4897 | 3734 | 3985 | Cd276 |
| Cd28 | 1 | 0 | 0 | 2 | 1 | 1 | 2 | 2 | 6 | 0 | 14 | 9 | 6 | 3 | 4 | 5 | 0 | 0 | 2 | 0 | 3 | 5 | 2 | 2 | 6 | 8 | 5 | 2 | Cd28 |
| Cd2ap | 5075 | 3147 | 4432 | 4104 | 4358 | 4854 | 4631 | 4579 | 5218 | 4858 | 5754 | 4905 | 5252 | 5169 | 4644 | 4494 | 3957 | 4150 | 3635 | 3796 | 4320 | 5023 | 5174 | 4661 | 4602 | 4433 | 4665 | 4597 | Cd2ap |
| Cd2bp2 | 6258 | 6517 | 7598 | 6643 | 6758 | 6418 | 6902 | 8183 | 8860 | 9142 | 8670 | 10694 | 9939 | 11078 | 6138 | 5908 | 7488 | 6395 | 7241 | 6166 | 6757 | 8833 | 8267 | 10100 | 9897 | 11572 | 10858 | 11726 | Cd2bp2 |
| Cd300a | 209 | 218 | 207 | 221 | 170 | 210 | 125 | 154 | 158 | 129 | 216 | 181 | 168 | 114 | 223 | 294 | 173 | 188 | 141 | 134 | 115 | 164 | 221 | 127 | 139 | 155 | 136 | 144 | Cd300a |
| Cd300c | 0 | 0 | 0 | 2 | 0 | 0 | 0 | 0 | 0 | 0 | 0 | 0 | 0 | 0 | 1 | 0 | 0 | 0 | 0 | 0 | 0 | 0 | 0 | 1 | 1 | 4 | 1 | 0 | Cd300c |
| Cd300c2 | 42 | 35 | 34 | 29 | 25 | 25 | 20 | 46 | 54 | 38 | 85 | 41 | 53 | 31 | 26 | 44 | 24 | 22 | 26 | 33 | 19 | 37 | 71 | 29 | 28 | 50 | 41 | 52 | Cd300c2 |

| | | | | | | | | | | | | | | | | | | | | | | | | | | | | | |
|---|---|---|---|---|---|---|---|---|---|---|---|---|---|---|---|---|---|---|---|---|---|---|---|---|---|---|---|---|---|
| Cd300e | 0 | 0 | 0 | 1 | 0 | 0 | 0 | 1 | 3 | 0 | 3 | 1 | 1 | 2 | 1 | 0 | 0 | 0 | 0 | 0 | 1 | 2 | 1 | 8 | 0 | 0 | 0 | 0 | Cd300e |
| Cd300lb | 2 | 6 | 2 | 6 | 2 | 1 | 1 | 5 | 7 | 1 | 11 | 3 | 1 | 3 | 4 | 3 | 0 | 1 | 0 | 0 | 0 | 13 | 6 | 5 | 3 | 9 | 2 | 4 | Cd300lb |
| Cd300ld | 3 | 10 | 10 | 3 | 8 | 8 | 3 | 16 | 5 | 5 | 5 | 10 | 1 | 10 | 11 | 14 | 13 | 7 | 9 | 9 | 8 | 13 | 21 | 13 | 6 | 10 | 6 | 11 | Cd300ld |
| Cd300ld2 | 0 | 0 | 0 | 1 | 1 | 0 | 0 | 0 | 0 | 0 | 1 | 0 | 0 | 0 | 0 | 2 | 0 | 0 | 0 | 0 | 0 | 2 | 0 | 4 | 0 | 0 | 0 | 0 | Cd300ld2 |
| Cd300ld3 | 0 | 0 | 0 | 0 | 0 | 0 | 0 | 0 | 0 | 0 | 0 | 4 | 1 | 3 | 0 | 0 | 0 | 2 | 0 | 0 | 1 | 0 | 1 | 0 | 0 | 0 | 2 | 5 | Cd300ld3 |
| Cd300ld4 | 0 | 0 | 5 | 1 | 0 | 0 | 1 | 0 | 2 | 0 | 3 | 0 | 0 | 0 | 1 | 1 | 0 | 3 | 0 | 2 | 1 | 0 | 3 | 1 | 1 | 0 | 0 | 1 | Cd300ld4 |
| Cd300ld5 | 0 | 0 | 1 | 0 | 0 | 0 | 0 | 0 | 0 | 0 | 0 | 1 | 0 | 0 | 0 | 1 | 0 | 0 | 0 | 0 | 0 | 0 | 1 | 1 | 3 | 0 | 0 | 0 | Cd300ld5 |
| Cd300lf | 30 | 29 | 41 | 42 | 12 | 13 | 22 | 2 | 12 | 20 | 29 | 5 | 9 | 18 | 17 | 32 | 39 | 26 | 12 | 31 | 14 | 10 | 8 | 18 | 12 | 15 | 9 | 17 | Cd300lf |
| Cd300lg | 10 | 7 | 3 | 1 | 5 | 2 | 6 | 11 | 16 | 6 | 15 | 5 | 1 | 12 | 20 | 11 | 7 | 3 | 2 | 1 | 4 | 12 | 9 | 8 | 7 | 18 | 10 | 4 | Cd300lg |
| Cd302 | 549 | 365 | 480 | 280 | 331 | 342 | 273 | 529 | 556 | 438 | 629 | 318 | 417 | 360 | 545 | 548 | 403 | 397 | 240 | 341 | 238 | 399 | 733 | 429 | 329 | 505 | 395 | 320 | Cd302 |
| Cd320 | 2179 | 2493 | 1832 | 1814 | 1532 | 1670 | 1542 | 1468 | 1451 | 1076 | 1237 | 1334 | 1352 | 1305 | 2065 | 2308 | 1776 | 1777 | 1812 | 2188 | 1693 | 1280 | 1377 | 1309 | 1268 | 1369 | 1331 | 1418 | Cd320 |
| Cd33 | 115 | 138 | 104 | 70 | 140 | 91 | 111 | 201 | 228 | 207 | 304 | 230 | 225 | 282 | 123 | 118 | 122 | 131 | 122 | 149 | 123 | 282 | 299 | 284 | 254 | 199 | 259 | 249 | Cd33 |
| Cd34 | 3766 | 3357 | 3624 | 2622 | 2774 | 3410 | 2789 | 3731 | 3975 | 3473 | 4291 | 3476 | 4003 | 3388 | 3753 | 4030 | 2845 | 3140 | 2930 | 3462 | 2904 | 3519 | 5009 | 3359 | 3289 | 4311 | 3669 | 3637 | Cd34 |
| Cd36 | 153 | 160 | 178 | 170 | 277 | 231 | 150 | 416 | 439 | 352 | 718 | 234 | 330 | 230 | 261 | 231 | 185 | 253 | 121 | 257 | 167 | 429 | 871 | 289 | 275 | 176 | 252 | 160 | Cd36 |
| Cd37 | 25 | 31 | 37 | 33 | 23 | 30 | 25 | 38 | 51 | 38 | 64 | 35 | 33 | 22 | 18 | 38 | 32 | 36 | 30 | 25 | 25 | 48 | 39 | 41 | 44 | 45 | 36 | 21 | Cd37 |
| Cd38 | 208 | 135 | 274 | 122 | 143 | 181 | 124 | 296 | 255 | 270 | 419 | 331 | 358 | 248 | 203 | 240 | 124 | 133 | 131 | 124 | 124 | 241 | 389 | 281 | 183 | 415 | 235 | 305 | Cd38 |
| Cd3d | 1 | 0 | 0 | 0 | 0 | 0 | 0 | 0 | 0 | 0 | 0 | 0 | 0 | 1 | 0 | 0 | 0 | 0 | 0 | 0 | 0 | 2 | 1 | 0 | 0 | 0 | 0 | 0 | Cd3d |
| Cd3e | 0 | 0 | 0 | 0 | 0 | 0 | 2 | 0 | 0 | 0 | 2 | 1 | 0 | 0 | 0 | 0 | 0 | 0 | 0 | 0 | 0 | 2 | 0 | 0 | 0 | 1 | 0 | 1 | Cd3e |
| Cd3eap | 2500 | 2951 | 2885 | 2941 | 2136 | 2378 | 2492 | 1237 | 1486 | 1354 | 1594 | 1634 | 1524 | 1611 | 2446 | 2756 | 2581 | 2680 | 2251 | 2377 | 2342 | 1168 | 1425 | 1593 | 1534 | 1880 | 1526 | 1689 | Cd3eap |
| Cd3g | 0 | 0 | 3 | 0 | 0 | 0 | 0 | 0 | 2 | 2 | 7 | 1 | 2 | 2 | 4 | 0 | 0 | 1 | 0 | 0 | 0 | 0 | 8 | 0 | 0 | 4 | 0 | 0 | Cd3g |
| Cd4 | 8 | 10 | 4 | 7 | 15 | 13 | 13 | 12 | 11 | 3 | 14 | 3 | 15 | 4 | 12 | 8 | 0 | 23 | 10 | 9 | 7 | 8 | 21 | 15 | 8 | 10 | 7 | 4 | Cd4 |
| Cd40 | 273 | 343 | 339 | 257 | 221 | 236 | 183 | 195 | 204 | 166 | 197 | 176 | 253 | 194 | 287 | 347 | 256 | 299 | 237 | 227 | 247 | 206 | 208 | 225 | 225 | 237 | 241 | 216 | Cd40 |
| Cd40lg | 0 | 0 | 0 | 0 | 0 | 0 | 0 | 0 | 0 | 0 | 0 | 1 | 0 | 0 | 0 | 0 | 0 | 0 | 0 | 0 | 0 | 0 | 0 | 0 | 0 | 0 | 0 | 0 | Cd40lg |
| Cd44 | 655 | 451 | 582 | 320 | 292 | 455 | 284 | 826 | 1100 | 681 | 1266 | 481 | 595 | 488 | 657 | 715 | 351 | 541 | 323 | 303 | 225 | 692 | 1618 | 731 | 543 | 539 | 516 | 484 | Cd44 |
| Cd46 | 326 | 189 | 274 | 365 | 533 | 405 | 486 | 601 | 627 | 562 | 632 | 598 | 671 | 700 | 357 | 265 | 278 | 331 | 344 | 359 | 459 | 672 | 574 | 550 | 586 | 433 | 724 | 564 | Cd46 |
| Cd47 | 3167 | 2718 | 4244 | 4882 | 5902 | 4773 | 5264 | 3495 | 3299 | 4129 | 4202 | 3247 | 3317 | 3293 | 3687 | 2965 | 4855 | 5292 | 5416 | 4145 | 5332 | 5016 | 3428 | 4010 | 4599 | 3381 | 3225 | 3264 | Cd47 |
| Cd48 | 8 | 14 | 17 | 10 | 16 | 16 | 4 | 20 | 30 | 27 | 53 | 10 | 18 | 15 | 9 | 15 | 14 | 18 | 10 | 20 | 14 | 35 | 56 | 25 | 17 | 28 | 6 | 13 | Cd48 |
| Cd5 | 2 | 2 | 1 | 7 | 11 | 1 | 0 | 5 | 8 | 2 | 8 | 20 | 6 | 5 | 0 | 6 | 8 | 7 | 9 | 8 | 1 | 5 | 21 | 5 | 13 | 12 | 7 | 12 | Cd5 |
| Cd52 | 38 | 56 | 29 | 37 | 39 | 48 | 44 | 29 | 70 | 46 | 86 | 39 | 53 | 60 | 33 | 44 | 48 | 33 | 43 | 30 | 31 | 40 | 91 | 38 | 28 | 71 | 42 | 36 | Cd52 |
| Cd53 | 170 | 135 | 137 | 193 | 134 | 213 | 142 | 217 | 264 | 160 | 259 | 180 | 179 | 156 | 195 | 190 | 151 | 145 | 158 | 174 | 154 | 190 | 333 | 179 | 208 | 207 | 195 | 177 | Cd53 |
| Cd55 | 226 | 143 | 147 | 73 | 193 | 136 | 95 | 585 | 499 | 451 | 423 | 175 | 289 | 179 | 186 | 258 | 114 | 121 | 113 | 160 | 101 | 378 | 673 | 390 | 265 | 276 | 214 | 148 | Cd55 |
| Cd55b | 1 | 4 | 0 | 0 | 0 | 1 | 0 | 2 | 1 | 0 | 0 | 1 | 0 | 0 | 1 | 1 | 0 | 0 | 0 | 0 | 2 | 0 | 1 | 2 | 0 | 0 | 0 | 1 | Cd55b |
| Cd59a | 603 | 449 | 584 | 274 | 245 | 282 | 401 | 413 | 339 | 256 | 373 | 316 | 725 | 510 | 757 | 558 | 470 | 302 | 279 | 339 | 278 | 428 | 514 | 351 | 260 | 482 | 625 | 736 | Cd59a |
| Cd59b | 96 | 53 | 56 | 61 | 100 | 106 | 64 | 61 | 89 | 52 | 74 | 46 | 100 | 57 | 87 | 80 | 71 | 111 | 39 | 89 | 84 | 47 | 72 | 43 | 60 | 55 | 69 | 58 | Cd59b |
| Cd5l | 7 | 17 | 10 | 29 | 8 | 23 | 1 | 1 | 11 | 14 | 11 | 3 | 11 | 3 | 3 | 24 | 15 | 12 | 4 | 22 | 4 | 8 | 21 | 25 | 8 | 16 | 13 | 33 | Cd5l |
| Cd6 | 3 | 1 | 0 | 2 | 2 | 1 | 0 | 0 | 2 | 3 | 3 | 0 | 0 | 1 | 3 | 3 | 2 | 4 | 0 | 0 | 0 | 0 | 4 | 0 | 8 | 1 | 0 | 1 | Cd6 |
| Cd63 | 5246 | 5110 | 7105 | 5197 | 4866 | 5550 | 3355 | 4484 | 6136 | 5405 | 8095 | 4513 | 6102 | 3617 | 5037 | 5907 | 4880 | 5460 | 4875 | 4839 | 3623 | 4234 | 7441 | 5475 | 4660 | 6157 | 5590 | 4332 | Cd63 |
| Cd68 | 362 | 441 | 442 | 445 | 430 | 403 | 355 | 431 | 523 | 510 | 494 | 565 | 559 | 529 | 348 | 428 | 381 | 481 | 394 | 335 | 332 | 546 | 612 | 552 | 525 | 680 | 568 | 520 | Cd68 |
| Cd69 | 3 | 3 | 3 | 6 | 2 | 0 | 3 | 1 | 4 | 3 | 1 | 0 | 0 | 1 | 0 | 5 | 6 | 0 | 0 | 0 | 0 | 2 | 4 | 2 | 0 | 1 | 0 | 0 | Cd69 |
| Cd7 | 2 | 3 | 0 | 2 | 2 | 0 | 0 | 1 | 4 | 3 | 5 | 0 | 0 | 1 | 3 | 1 | 0 | 1 | 0 | 0 | 0 | 7 | 10 | 0 | 0 | 3 | 12 | 0 | Cd7 |
| Cd70 | 3 | 2 | 2 | 0 | 1 | 1 | 1 | 5 | 3 | 3 | 2 | 3 | 7 | 4 | 0 | 2 | 2 | 0 | 2 | 2 | 2 | 5 | 3 | 7 | 0 | 2 | 3 | 4 | Cd70 |
| Cd72 | 18 | 11 | 27 | 4 | 18 | 8 | 11 | 5 | 7 | 3 | 20 | 11 | 9 | 7 | 17 | 12 | 23 | 14 | 14 | 23 | 12 | 10 | 8 | 7 | 3 | 21 | 14 | 8 | Cd72 |
| Cd74 | 7 | 12 | 14 | 4 | 9 | 12 | 15 | 27 | 32 | 38 | 57 | 25 | 12 | 13 | 9 | 14 | 14 | 18 | 18 | 7 | 12 | 27 | 62 | 20 | 20 | 16 | 17 | 9 | Cd74 |
| Cd79a | 24 | 55 | 40 | 40 | 35 | 35 | 46 | 10 | 23 | 17 | 29 | 5 | 15 | 9 | 21 | 44 | 78 | 62 | 55 | 47 | 39 | 15 | 24 | 25 | 26 | 18 | 16 | 17 | Cd79a |
| Cd79b | 16 | 12 | 29 | 17 | 11 | 9 | 8 | 26 | 44 | 26 | 28 | 24 | 48 | 16 | 17 | 19 | 13 | 17 | 12 | 14 | 16 | 15 | 44 | 32 | 16 | 43 | 20 | 30 | Cd79b |
| Cd80 | 24 | 16 | 16 | 17 | 56 | 21 | 16 | 32 | 23 | 38 | 44 | 26 | 31 | 19 | 22 | 26 | 14 | 6 | 39 | 27 | 21 | 27 | 32 | 32 | 20 | 15 | 14 | 19 | Cd80 |
| Cd81 | 7375 | 7906 | 7721 | 7428 | 7326 | 7003 | 6541 | 6397 | 7474 | 6057 | 7707 | 5532 | 5998 | 5498 | 7526 | 8164 | 7714 | 7472 | 7266 | 7718 | 6882 | 6221 | 8951 | 6427 | 6037 | 6666 | 5602 | 5658 | Cd81 |
| Cd82 | 977 | 1011 | 823 | 848 | 870 | 801 | 639 | 642 | 862 | 809 | 1045 | 547 | 745 | 479 | 928 | 1145 | 773 | 862 | 745 | 815 | 670 | 742 | 1216 | 762 | 606 | 754 | 579 | 538 | Cd82 |
| Cd83 | 196 | 151 | 165 | 122 | 276 | 166 | 173 | 421 | 402 | 319 | 511 | 215 | 325 | 237 | 212 | 189 | 156 | 179 | 176 | 225 | 177 | 416 | 602 | 257 | 304 | 295 | 255 | 260 | Cd83 |
| Cd84 | 77 | 79 | 83 | 66 | 60 | 54 | 61 | 167 | 125 | 106 | 124 | 90 | 95 | 84 | 100 | 98 | 52 | 90 | 59 | 71 | 73 | 84 | 205 | 89 | 103 | 89 | 109 | 115 | Cd84 |
| Cd86 | 144 | 148 | 96 | 116 | 117 | 111 | 87 | 142 | 124 | 96 | 140 | 141 | 117 | 126 | 136 | 165 | 71 | 134 | 79 | 133 | 90 | 114 | 190 | 106 | 129 | 134 | 100 | 103 | Cd86 |
| Cd8a | 0 | 0 | 0 | 0 | 0 | 0 | 0 | 0 | 0 | 0 | 0 | 0 | 3 | 1 | 0 | 0 | 0 | 0 | 0 | 0 | 0 | 0 | 3 | 0 | 6 | 0 | 0 | 0 | Cd8a |
| Cd9 | 1855 | 1805 | 1669 | 1538 | 1669 | 1560 | 1437 | 1469 | 1807 | 1433 | 1713 | 1017 | 1111 | 1181 | 1764 | 1976 | 1484 | 1791 | 1442 | 1538 | 1457 | 1148 | 2084 | 1475 | 1204 | 1398 | 992 | 1130 | Cd9 |
| Cd93 | 2599 | 2362 | 2871 | 2396 | 3187 | 3204 | 3030 | 3683 | 3939 | 4675 | 5195 | 4362 | 4754 | 4123 | 2624 | 2642 | 2817 | 2793 | 2897 | 2831 | 3170 | 5068 | 4644 | 4377 | 4608 | 4076 | 4402 | 4321 | Cd93 |
| Cd96 | 2 | 3 | 0 | 1 | 0 | 0 | 3 | 2 | 4 | 3 | 9 | 5 | 1 | 2 | 1 | 2 | 0 | 0 | 2 | 0 | 3 | 12 | 3 | 2 | 1 | 3 | 0 | 2 | Cd96 |
| Cd99l2 | 5261 | 5790 | 4679 | 5751 | 6518 | 6291 | 6086 | 7192 | 6830 | 6356 | 5900 | 5579 | 5710 | 5604 | 6142 | 5103 | 5342 | 5877 | 6226 | 6519 | 6668 | 7045 | 6798 | 5715 | 6559 | 5473 | 5552 | 5323 | Cd99l2 |
| Cda | 55 | 55 | 52 | 30 | 60 | 48 | 21 | 28 | 62 | 55 | 58 | 35 | 46 | 45 | 53 | 76 | 45 | 39 | 35 | 39 | 30 | 62 | 63 | 39 | 35 | 53 | 31 | 36 | Cda |
| Cdadc1 | 2789 | 2782 | 2232 | 2576 | 2459 | 2415 | 2874 | 2847 | 2695 | 2278 | 2290 | 2153 | 2117 | 2363 | 2954 | 2815 | 2338 | 2633 | 2381 | 3058 | 2903 | 2415 | 2465 | 2263 | 2409 | 2171 | 2226 | 2376 | Cdadc1 |
| Cdan1 | 3691 | 4621 | 3821 | 4778 | 4393 | 4602 | 4543 | 2927 | 3133 | 3159 | 3334 | 3492 | 3170 | 3471 | 3517 | 4101 | 4285 | 4700 | 4459 | 3984 | 4579 | 3205 | 2929 | 3277 | 3508 | 3144 | 3196 | 3425 | Cdan1 |
| Cdc123 | 8919 | 7992 | 8952 | 7572 | 7013 | 7178 | 8110 | 9186 | 8331 | 7775 | 7521 | 9094 | 8774 | 9540 | 9220 | 7666 | 8331 | 7352 | 7604 | 8872 | 7468 | 7864 | 7734 | 8398 | 8388 | 10693 | 9545 | 9975 | Cdc123 |
| Cdc14a | 2367 | 1940 | 2433 | 2336 | 2563 | 2490 | 2548 | 1942 | 1800 | 1804 | 1715 | 2647 | 2095 | 2525 | 2207 | 2016 | 2509 | 2218 | 2371 | 2366 | 2434 | 1790 | 1576 | 2248 | 1902 | 2799 | 2420 | 2559 | Cdc14a |
| Cdc14b | 4412 | 3014 | 2978 | 3230 | 2834 | 3173 | 2976 | 1706 | 1736 | 1698 | 1944 | 1637 | 2117 | 1730 | 3421 | 3932 | 2892 | 3009 | 2707 | 2499 | 2733 | 1275 | 1772 | 1823 | 1430 | 1839 | 1850 | 1714 | Cdc14b |
| Cdc16 | 13103 | 12088 | 11308 | 10065 | 10386 | 10801 | 11733 | 13242 | 12117 | 10676 | 11215 | 11858 | 11870 | 11765 | 13442 | 11721 | 11053 | 10248 | 10505 | 11760 | 10730 | 11446 | 12821 | 10883 | 10857 | 12814 | 12966 | 12303 | Cdc16 |
| Cdc20 | 4582 | 4812 | 4914 | 3730 | 2942 | 3361 | 3075 | 4418 | 4746 | 4564 | 4052 | 7393 | 5932 | 6664 | 3861 | 3759 | 4494 | 3082 | 4100 | 3508 | 3180 | 3556 | 4215 | 6089 | 5186 | 9527 | 7036 | 6871 | Cdc20 |
| Cdc20b | 202 | 225 | 192 | 135 | 157 | 221 | 142 | 164 | 134 | 130 | 104 | 295 | 321 | 204 | 210 | 310 | 192 | 145 | 191 | 234 | 161 | 119 | 324 | 123 | 130 | 239 | 322 | 230 | Cdc20b |
| Cdc23 | 10536 | 9253 | 9449 | 8310 | 8274 | 8855 | 8790 | 11964 | 11379 | 11042 | 10780 | 12438 | 11382 | 12481 | 9577 | 8758 | 9218 | 8053 | 9137 | 8364 | 8745 | 11266 | 10958 | 11463 | 11408 | 13129 | 12031 | 12411 | Cdc23 |
| Cdc25a | 5768 | 5161 | 6326 | 5556 | 4995 | 5156 | 5264 | 5032 | 4742 | 4557 | 4809 | 5762 | 5527 | 6284 | 5149 | 5036 | 5884 | 5119 | 5392 | 4609 | 5206 | 4624 | 4317 | 5296 | 4703 | 6884 | 6032 | 6502 | Cdc25a |
| Cdc25b | 5191 | 6793 | 7756 | 5762 | 4664 | 4983 | 4805 | 4349 | 4225 | 4371 | 4232 | 6906 | 6166 | 6257 | 5455 | 5576 | 7218 | 5085 | 5651 | 4617 | 4904 | 4363 | 4110 | 5543 | 4882 | 6814 | 7295 | 7365 | Cdc25b |
| Cdc25c | 4896 | 4232 | 4273 | 3692 | 2343 | 3161 | 3693 | 2573 | 2388 | 2034 | 1831 | 3340 | 2814 | 3263 | 4210 | 3745 | 3708 | 3125 | 3317 | 3602 | 3158 | 1683 | 1913 | 2548 | 2186 | 4117 | 3356 | 3521 | Cdc25c |
| Cdc26 | 4209 | 3846 | 4842 | 3948 | 4380 | 4360 | 4887 | 5438 | 5515 | 5275 | 5175 | 6429 | 6064 | 6085 | 4307 | 4029 | 4408 | 3813 | 4715 | 5062 | 4637 | 4592 | 5318 | 5919 | 5716 | 7442 | 6592 | 6272 | Cdc26 |
| Cdc27 | 12102 | 9506 | 9925 | 9217 | 9857 | 10669 | 11491 | 13048 | 10989 | 9397 | 10139 | 11653 | 11284 | 12074 | 12484 | 10114 | 9501 | 9313 | 9632 | 11391 | 10238 | 10701 | 10839 | 10302 | 10182 | 10433 | 12253 | 12082 | Cdc27 |
| Cdc34 | 3960 | 5099 | 5188 | 3809 | 2936 | 3049 | 3049 | 2655 | 2966 | 2785 | 2591 | 3343 | 3209 | 3258 | 3753 | 3970 | 4488 | 3146 | 3473 | 3395 | 2966 | 2064 | 2735 | 3020 | 2455 | 4084 | 3638 | 3879 | Cdc34 |
| Cdc34b | 43 | 46 | 41 | 17 | 29 | 19 | 17 | 26 | 31 | 24 | 28 | 21 | 16 | 23 | 29 | 25 | 23 | 13 | 23 | 16 | 28 | 43 | 22 | 19 | 31 | 24 | 13 | 31 | Cdc34b |
| Cdc37 | 14350 | 18007 | 16042 | 13838 | 10869 | 10998 | 10859 | 9812 | 10794 | 9995 | 10098 | 11041 | 10452 | 10248 | 13796 | 14934 | 14753 | 12641 | 12731 | 13067 | 11634 | 8355 | 10701 | 11034 | 9638 | 13058 | 11988 | 11727 | Cdc37 |
| Cdc37l1 | 13929 | 12133 | 6992 | 8887 | 9191 | 11835 | 10968 | 12070 | 10084 | 7793 | 7905 | 7948 | 8252 | 8642 | 14144 | 12286 | 7817 | 9260 | 9007 | 15062 | 12496 | 7954 | 9805 | 7133 | 8284 | 5995 | 8956 | 7806 | Cdc37l1 |
| Cdc40 | 6068 | 5985 | 5494 | 5237 | 4719 | 5033 | 5553 | 4557 | 3789 | 3530 | 3553 | 4083 | 3561 | 3987 | 6245 | 5842 | 5412 | 4966 | 5023 | 6512 | 5259 | 3434 | 3735 | 3675 | 3778 | 4055 | 4198 | 4170 | Cdc40 |
| Cdc42 | 35029 | 27293 | 30815 | 27057 | 27479 | 28155 | 30100 | 39813 | 33927 | 29495 | 33328 | 32065 | 33481 | 33718 | 36077 | 30016 | 28314 | 28388 | 27550 | 31934 | 28249 | 35805 | 36804 | 32232 | 31970 | 36453 | 34810 | 35014 | Cdc42 |
| Cdc42bpa | 10051 | 9547 | 7909 | 10659 | 13573 | 13435 | 12824 | 10616 | 10916 | 11500 | 11503 | 10836 | 10992 | 11151 | 10879 | 10319 | 9644 | 12127 | 11552 | 11279 | 12885 | 11866 | 10306 | 10195 | 13002 | 7136 | 10808 | 10123 | Cdc42bpa |
| Cdc42bpb | 12957 | 15988 | 18261 | 19189 | 18585 | 17534 | 15365 | 9594 | 11752 | 13439 | 12978 | 14907 | 12522 | 12639 | 12845 | 15210 | 18770 | 18768 | 19304 | 13451 | 16906 | 14014 | 11066 | 13585 | 14243 | 10710 | 12405 | 12003 | Cdc42bpb |

| | | | | | | | | | | | | | | | | | | | | | | | | | | | | | |
|---|---|---|---|---|---|---|---|---|---|---|---|---|---|---|---|---|---|---|---|---|---|---|---|---|---|---|---|---|---|
| Cdc42ep1 | 279 | 439 | 421 | 293 | 320 | 323 | 244 | 512 | 613 | 636 | 766 | 535 | 660 | 512 | 364 | 468 | 343 | 255 | 236 | 224 | 234 | 560 | 771 | 526 | 451 | 716 | 536 | 542 | Cdc42ep1 |
| Cdc42ep2 | 588 | 769 | 577 | 627 | 749 | 659 | 660 | 600 | 649 | 561 | 601 | 281 | 432 | 351 | 636 | 717 | 521 | 721 | 734 | 722 | 708 | 623 | 754 | 491 | 477 | 334 | 351 | 320 | Cdc42ep2 |
| Cdc42ep3 | 1228 | 1061 | 913 | 1197 | 1222 | 1295 | 1132 | 1642 | 1846 | 1737 | 1798 | 1424 | 1479 | 1513 | 1246 | 1082 | 1043 | 1252 | 1111 | 984 | 1084 | 1890 | 1940 | 1642 | 1885 | 1292 | 1290 | 1301 | Cdc42ep3 |
| Cdc42ep4 | 6144 | 7519 | 7306 | 7063 | 6748 | 6490 | 5946 | 6232 | 6869 | 6341 | 5977 | 7857 | 7479 | 7499 | 5573 | 6486 | 7234 | 6645 | 7581 | 6712 | 6264 | 5648 | 6613 | 7407 | 6710 | 9464 | 7696 | 7903 | Cdc42ep4 |
| Cdc42ep5 | 423 | 459 | 452 | 293 | 205 | 297 | 177 | 328 | 481 | 283 | 486 | 331 | 424 | 297 | 352 | 454 | 352 | 294 | 210 | 177 | 130 | 244 | 504 | 411 | 219 | 422 | 345 | 374 | Cdc42ep5 |
| Cdc42se1 | 4637 | 5115 | 5334 | 5002 | 4743 | 4809 | 4516 | 4510 | 5157 | 4914 | 4826 | 5903 | 5211 | 5486 | 4195 | 4738 | 5443 | 4716 | 5299 | 4681 | 4464 | 4443 | 4947 | 5587 | 5411 | 6492 | 5713 | 5638 | Cdc42se1 |
| Cdc42se2 | 12629 | 13985 | 14191 | 12746 | 11900 | 12353 | 12976 | 12678 | 10508 | 10825 | 9642 | 14641 | 13189 | 14976 | 13362 | 11840 | 13655 | 12301 | 12705 | 13952 | 12720 | 11478 | 10584 | 12599 | 13023 | 14342 | 15287 | 15688 | Cdc42se2 |
| Cdc45 | 2550 | 2634 | 2545 | 1778 | 1315 | 1704 | 1918 | 2141 | 1988 | 1637 | 1564 | 2709 | 2113 | 2574 | 2346 | 2220 | 2051 | 1692 | 1905 | 2036 | 1652 | 1332 | 1827 | 2134 | 1842 | 3098 | 2638 | 2633 | Cdc45 |
| Cdc5l | 9682 | 9450 | 9222 | 8681 | 8207 | 8286 | 9269 | 8551 | 7659 | 7434 | 7323 | 7638 | 7244 | 8560 | 9508 | 9290 | 8747 | 8445 | 8608 | 9382 | 8930 | 6990 | 7262 | 7264 | 7175 | 8348 | 7741 | 8221 | Cdc5l |
| Cdc6 | 4265 | 4329 | 4117 | 3122 | 1971 | 2504 | 2759 | 2209 | 1945 | 1687 | 1593 | 2775 | 2251 | 2761 | 3394 | 3768 | 3407 | 2738 | 2593 | 2825 | 2556 | 1534 | 1746 | 2180 | 1758 | 3398 | 2874 | 3012 | Cdc6 |
| Cdc7 | 6428 | 5417 | 5494 | 5220 | 5121 | 5063 | 5515 | 5973 | 5338 | 5115 | 5002 | 5295 | 4909 | 5487 | 6120 | 6227 | 5782 | 5728 | 5368 | 6181 | 5235 | 4936 | 5094 | 5284 | 5222 | 5381 | 5433 | 5282 | Cdc7 |
| Cdc73 | 7600 | 5587 | 7695 | 8176 | 7324 | 8020 | 9095 | 6277 | 5704 | 6152 | 5942 | 7075 | 6338 | 7649 | 7366 | 6481 | 7660 | 7913 | 7772 | 7279 | 8691 | 6276 | 5608 | 6482 | 6548 | 6976 | 7108 | 7772 | Cdc73 |
| Cdca2 | 3956 | 3664 | 3666 | 2963 | 2339 | 2869 | 3060 | 4415 | 4360 | 3918 | 3583 | 5944 | 5008 | 6058 | 3418 | 3480 | 3272 | 2686 | 3069 | 3112 | 2829 | 2747 | 3619 | 4720 | 4233 | 6137 | 5972 | 5877 | Cdca2 |
| Cdca3 | 7401 | 7311 | 6865 | 4928 | 3679 | 4281 | 5296 | 4827 | 4287 | 3859 | 3249 | 6051 | 5069 | 6311 | 6572 | 5614 | 6079 | 4112 | 4754 | 5630 | 4653 | 3098 | 3680 | 5141 | 4160 | 7475 | 6107 | 6378 | Cdca3 |
| Cdca4 | 4135 | 3947 | 4204 | 3373 | 2962 | 3585 | 3378 | 4047 | 4341 | 4159 | 3714 | 5749 | 4735 | 5181 | 3535 | 3647 | 3854 | 2965 | 3777 | 3765 | 3092 | 3402 | 3893 | 4803 | 4120 | 6615 | 5972 | 5847 | Cdca4 |
| Cdca5 | 2241 | 2044 | 2247 | 1560 | 1244 | 1354 | 1658 | 1717 | 1622 | 1373 | 1212 | 2282 | 1982 | 2331 | 1834 | 1739 | 1985 | 1346 | 1771 | 1626 | 1489 | 1163 | 1339 | 1767 | 1582 | 2714 | 2529 | 2522 | Cdca5 |
| Cdca7 | 7118 | 7012 | 7906 | 6854 | 5381 | 6088 | 7156 | 6133 | 6661 | 6539 | 5773 | 9321 | 6482 | 9602 | 6484 | 6023 | 7379 | 6400 | 7291 | 6263 | 6814 | 4700 | 5400 | 8606 | 7365 | 12012 | 8911 | 9311 | Cdca7 |
| Cdca7l | 2382 | 1970 | 2403 | 1781 | 1138 | 1577 | 1604 | 1419 | 1521 | 1521 | 1270 | 1836 | 1685 | 1915 | 2000 | 1867 | 1817 | 1478 | 1414 | 1531 | 1381 | 884 | 1392 | 1579 | 1254 | 2407 | 1935 | 2059 | Cdca7l |
| Cdca8 | 5501 | 5967 | 7255 | 4947 | 3506 | 3985 | 4340 | 4258 | 4558 | 4517 | 4190 | 7404 | 6254 | 7276 | 4759 | 4939 | 5934 | 4163 | 4769 | 4208 | 3975 | 3484 | 3879 | 6068 | 4938 | 9320 | 7475 | 7847 | Cdca8 |
| Cdcp1 | 253 | 85 | 44 | 27 | 20 | 26 | 2 | 346 | 447 | 302 | 268 | 39 | 137 | 51 | 99 | 118 | 20 | 35 | 16 | 8 | 2 | 65 | 474 | 242 | 45 | 217 | 27 | 45 | Cdcp1 |
| Cdcp2 | 15 | 18 | 0 | 13 | 14 | 2 | 6 | 5 | 4 | 1 | 11 | 16 | 11 | 10 | 15 | 19 | 6 | 14 | 1 | 5 | 17 | 0 | 3 | 0 | 3 | 0 | 6 | 5 | Cdcp2 |
| Cdcp3 | 0 | 1 | 0 | 0 | 2 | 0 | 2 | 2 | 4 | 0 | 0 | 0 | 1 | 0 | 0 | 1 | 2 | 0 | 0 | 1 | 0 | 2 | 3 | 2 | 5 | 0 | 0 | 0 | Cdcp3 |
| Cdh1 | 2157 | 1012 | 725 | 535 | 336 | 685 | 110 | 1469 | 1976 | 1213 | 1263 | 563 | 878 | 523 | 1154 | 1592 | 384 | 749 | 224 | 138 | 183 | 555 | 2298 | 1271 | 365 | 906 | 549 | 571 | Cdh1 |
| Cdh10 | 1059 | 1012 | 845 | 915 | 1262 | 1127 | 1282 | 1484 | 1285 | 1356 | 1166 | 988 | 1057 | 968 | 1153 | 925 | 934 | 1011 | 1152 | 1312 | 1145 | 1248 | 1260 | 973 | 1275 | 914 | 977 | 885 | Cdh10 |
| Cdh11 | 12425 | 9485 | 9768 | 5788 | 4775 | 8203 | 3943 | 18434 | 22696 | 16449 | 21192 | 15555 | 20889 | 15409 | 11732 | 13161 | 7154 | 7284 | 4056 | 4713 | 3562 | 16456 | 24192 | 16944 | 13629 | 17939 | 17103 | 16025 | Cdh11 |
| Cdh12 | 156 | 111 | 71 | 143 | 250 | 246 | 225 | 448 | 381 | 292 | 330 | 156 | 227 | 231 | 232 | 195 | 147 | 215 | 231 | 268 | 248 | 344 | 278 | 258 | 274 | 106 | 188 | 177 | Cdh12 |
| Cdh13 | 992 | 1411 | 829 | 2174 | 3459 | 2683 | 2666 | 2368 | 2936 | 2568 | 2419 | 1330 | 1501 | 1247 | 1375 | 1121 | 1209 | 2775 | 2865 | 2701 | 3543 | 2909 | 2511 | 1914 | 2906 | 802 | 1188 | 1015 | Cdh13 |
| Cdh15 | 80 | 105 | 82 | 42 | 47 | 31 | 9 | 92 | 276 | 64 | 1144 | 49 | 90 | 52 | 87 | 173 | 56 | 53 | 26 | 32 | 29 | 242 | 677 | 62 | 63 | 51 | 44 | 46 | Cdh15 |
| Cdh16 | 0 | 1 | 0 | 0 | 0 | 5 | 0 | 0 | 0 | 0 | 1 | 0 | 0 | 0 | 0 | 0 | 0 | 4 | 2 | 2 | 0 | 0 | 1 | 0 | 0 | 0 | 0 | 0 | Cdh16 |
| Cdh17 | 27 | 15 | 51 | 35 | 38 | 28 | 18 | 56 | 63 | 37 | 93 | 39 | 61 | 45 | 26 | 27 | 46 | 34 | 27 | 25 | 47 | 45 | 82 | 50 | 41 | 46 | 72 | 33 | Cdh17 |
| Cdh18 | 203 | 204 | 142 | 242 | 464 | 274 | 275 | 320 | 349 | 277 | 316 | 109 | 203 | 110 | 307 | 254 | 191 | 314 | 307 | 394 | 297 | 309 | 281 | 150 | 214 | 101 | 116 | 85 | Cdh18 |
| Cdh19 | 87 | 32 | 19 | 25 | 61 | 15 | 12 | 143 | 188 | 135 | 251 | 39 | 177 | 46 | 70 | 119 | 54 | 39 | 2 | 19 | 9 | 231 | 168 | 70 | 79 | 57 | 53 | 47 | Cdh19 |
| Cdh2 | 13664 | 13388 | 15876 | 15107 | 15855 | 16530 | 17393 | 19784 | 19385 | 21323 | 18254 | 25640 | 22502 | 26413 | 14350 | 12076 | 15432 | 14201 | 16879 | 16215 | 17057 | 21624 | 17395 | 23141 | 24745 | 25630 | 24786 | 25501 | Cdh2 |
| Cdh20 | 306 | 345 | 579 | 315 | 372 | 323 | 340 | 332 | 354 | 339 | 317 | 396 | 418 | 393 | 382 | 311 | 404 | 263 | 380 | 343 | 340 | 381 | 250 | 298 | 353 | 307 | 346 | 412 | Cdh20 |
| Cdh22 | 625 | 916 | 986 | 1129 | 1220 | 993 | 885 | 861 | 825 | 1045 | 851 | 1199 | 840 | 1025 | 613 | 586 | 1306 | 1073 | 1395 | 964 | 1142 | 1277 | 819 | 1179 | 1131 | 1118 | 1004 | 972 | Cdh22 |
| Cdh23 | 212 | 203 | 196 | 389 | 501 | 439 | 459 | 342 | 316 | 390 | 324 | 390 | 197 | 362 | 177 | 222 | 249 | 413 | 514 | 419 | 442 | 560 | 216 | 333 | 494 | 140 | 393 | 261 | Cdh23 |
| Cdh24 | 3902 | 4569 | 4776 | 5329 | 5380 | 5139 | 4526 | 3602 | 4168 | 4932 | 4242 | 5580 | 4732 | 4736 | 3251 | 4430 | 5341 | 5584 | 6009 | 4291 | 4454 | 5237 | 3875 | 5290 | 5576 | 5520 | 4996 | 4446 | Cdh24 |
| Cdh26 | 21 | 3 | 0 | 14 | 5 | 15 | 2 | 66 | 59 | 90 | 80 | 12 | 17 | 18 | 20 | 10 | 0 | 13 | 9 | 6 | 4 | 12 | 93 | 75 | 9 | 53 | 8 | 7 | Cdh26 |
| Cdh3 | 1398 | 1026 | 892 | 407 | 216 | 630 | 105 | 888 | 1371 | 1020 | 1460 | 730 | 1144 | 677 | 861 | 1505 | 534 | 573 | 235 | 105 | 103 | 762 | 1566 | 988 | 593 | 965 | 781 | 868 | Cdh3 |
| Cdh4 | 3217 | 4343 | 4383 | 5997 | 6925 | 6453 | 6084 | 4224 | 5656 | 5945 | 5619 | 5605 | 4009 | 5300 | 3575 | 3104 | 5326 | 6437 | 6922 | 5272 | 6629 | 6759 | 5403 | 5554 | 7080 | 4853 | 4095 | 4976 | Cdh4 |
| Cdh5 | 3057 | 3410 | 4136 | 2976 | 3164 | 3445 | 2457 | 3264 | 4837 | 4790 | 5871 | 4479 | 5451 | 3963 | 2979 | 3816 | 3170 | 3398 | 2608 | 2516 | 2729 | 4921 | 5923 | 4494 | 4364 | 5393 | 4138 | 4075 | Cdh5 |
| Cdh6 | 4379 | 3114 | 3010 | 2853 | 3288 | 3426 | 3226 | 5097 | 4811 | 4385 | 4798 | 3806 | 3706 | 4074 | 4229 | 3719 | 2993 | 3042 | 3046 | 3422 | 2962 | 3870 | 4472 | 3668 | 3712 | 3767 | 3884 | 3496 | Cdh6 |
| Cdh7 | 886 | 712 | 704 | 1002 | 1219 | 1274 | 1310 | 1652 | 1397 | 1257 | 1111 | 1490 | 1001 | 1721 | 1018 | 724 | 842 | 985 | 1245 | 1181 | 1229 | 1233 | 1120 | 1341 | 1473 | 1305 | 1197 | 1310 | Cdh7 |
| Cdh8 | 2025 | 2317 | 2215 | 2344 | 3118 | 3226 | 3390 | 4216 | 2788 | 2930 | 2448 | 2730 | 2454 | 2947 | 2819 | 1694 | 2376 | 2729 | 3179 | 3954 | 3484 | 4012 | 2689 | 2558 | 3485 | 1926 | 2986 | 2976 | Cdh8 |
| Cdh9 | 106 | 96 | 100 | 112 | 217 | 172 | 166 | 169 | 216 | 188 | 232 | 94 | 163 | 125 | 119 | 133 | 126 | 173 | 175 | 209 | 204 | 229 | 172 | 136 | 208 | 114 | 112 | 115 | Cdh9 |
| Cdhr1 | 323 | 195 | 31 | 155 | 317 | 135 | 115 | 596 | 539 | 517 | 558 | 101 | 607 | 118 | 398 | 466 | 115 | 173 | 110 | 129 | 74 | 525 | 394 | 360 | 303 | 525 | 58 | 63 | Cdhr1 |
| Cdhr2 | 5 | 3 | 0 | 2 | 1 | 1 | 2 | 5 | 6 | 9 | 0 | 0 | 2 | 3 | 3 | 2 | 5 | 8 | 5 | 0 | 5 | 7 | 7 | 4 | 3 | 2 | 0 | 0 | Cdhr2 |
| Cdhr3 | 51 | 42 | 31 | 80 | 99 | 61 | 75 | 13 | 20 | 16 | 55 | 20 | 34 | 10 | 47 | 50 | 44 | 87 | 59 | 70 | 109 | 18 | 38 | 10 | 20 | 10 | 9 | 9 | Cdhr3 |
| Cdhr4 | 54 | 52 | 51 | 49 | 38 | 26 | 11 | 89 | 113 | 102 | 207 | 58 | 88 | 55 | 50 | 72 | 31 | 66 | 18 | 22 | 20 | 94 | 132 | 130 | 52 | 92 | 84 | 83 | Cdhr4 |
| Cdhr5 | 1 | 7 | 0 | 1 | 4 | 1 | 0 | 4 | 1 | 3 | 13 | 4 | 0 | 0 | 5 | 8 | 7 | 0 | 0 | 3 | 0 | 3 | 1 | 0 | 4 | 3 | 1 | 0 | Cdhr5 |
| Cdin1 | 1378 | 1450 | 1403 | 1225 | 1321 | 1166 | 1531 | 1574 | 1602 | 1634 | 1458 | 1857 | 1738 | 1920 | 1457 | 1214 | 1531 | 1350 | 1478 | 1256 | 1284 | 1445 | 1510 | 1760 | 1582 | 2026 | 1736 | 1889 | Cdin1 |
| Cdip1 | 9830 | 10272 | 8527 | 8726 | 8882 | 8623 | 9157 | 6696 | 6632 | 6245 | 6391 | 7128 | 6747 | 6893 | 9428 | 8799 | 8991 | 8714 | 8590 | 8646 | 9059 | 7108 | 6480 | 6290 | 6454 | 6737 | 6549 | 6995 | Cdip1 |
| Cdipt | 3388 | 3944 | 3403 | 3332 | 3655 | 3463 | 3482 | 3615 | 3854 | 3626 | 3605 | 3473 | 3621 | 3440 | 3270 | 3619 | 3566 | 3432 | 3600 | 3803 | 3156 | 3621 | 3901 | 3697 | 3525 | 3816 | 3581 | 3509 | Cdipt |
| Cdiptos | 22 | 16 | 12 | 13 | 18 | 23 | 16 | 21 | 30 | 24 | 25 | 15 | 28 | 45 | 20 | 12 | 10 | 13 | 19 | 25 | 18 | 25 | 45 | 25 | 21 | 18 | 27 | 28 | Cdiptos |
| Cdk1 | 6248 | 5742 | 5379 | 4288 | 3282 | 3878 | 4513 | 6307 | 6076 | 4924 | 4995 | 6525 | 6051 | 7421 | 6021 | 5425 | 4729 | 3779 | 4123 | 5402 | 3951 | 4039 | 5935 | 5976 | 4885 | 8552 | 7079 | 7392 | Cdk1 |
| Cdk10 | 2553 | 2819 | 3051 | 3048 | 2576 | 2752 | 2856 | 2119 | 2458 | 2790 | 2757 | 2862 | 2545 | 2901 | 2421 | 2504 | 3247 | 2961 | 2822 | 2408 | 2866 | 2575 | 2413 | 2789 | 3032 | 2834 | 2780 | 2938 | Cdk10 |
| Cdk11b | 10265 | 9937 | 13300 | 11060 | 9037 | 9322 | 10163 | 7592 | 7450 | 9308 | 8585 | 10494 | 9474 | 10997 | 9879 | 9753 | 12188 | 10192 | 10113 | 8854 | 9653 | 8507 | 7374 | 10291 | 9692 | 12090 | 10384 | 10892 | Cdk11b |
| Cdk12 | 4200 | 4228 | 3917 | 4732 | 4272 | 4293 | 4103 | 5994 | 5472 | 5436 | 5254 | 6536 | 5781 | 6209 | 4204 | 3932 | 4015 | 4366 | 4168 | 3966 | 4343 | 5513 | 5049 | 5766 | 5885 | 4715 | 5964 | 6035 | Cdk12 |
| Cdk13 | 9034 | 7830 | 8584 | 9966 | 9137 | 9657 | 9950 | 8322 | 7514 | 8475 | 8174 | 9385 | 8814 | 9177 | 8598 | 7843 | 8886 | 9149 | 9217 | 8148 | 9811 | 9498 | 7348 | 8204 | 8732 | 8695 | 8574 | 9278 | Cdk13 |
| Cdk14 | 4353 | 4104 | 3646 | 4617 | 5229 | 5186 | 4967 | 5968 | 5863 | 5864 | 5485 | 5060 | 5641 | 5343 | 4772 | 4710 | 4267 | 4791 | 4543 | 4871 | 5051 | 6422 | 5537 | 5078 | 5747 | 4446 | 4977 | 5091 | Cdk14 |
| Cdk15 | 37 | 44 | 42 | 11 | 43 | 30 | 12 | 33 | 44 | 51 | 50 | 40 | 52 | 16 | 50 | 58 | 37 | 25 | 21 | 28 | 25 | 48 | 52 | 34 | 38 | 29 | 27 | 37 | Cdk15 |
| Cdk16 | 8756 | 10242 | 10166 | 10225 | 10088 | 9670 | 9472 | 8767 | 10322 | 10989 | 10663 | 11248 | 10233 | 10495 | 8739 | 9164 | 10787 | 10582 | 10737 | 8304 | 9998 | 10724 | 10437 | 10786 | 10901 | 12177 | 9946 | 10682 | Cdk16 |
| Cdk17 | 4699 | 4287 | 5349 | 6140 | 7367 | 6306 | 6898 | 7315 | 6239 | 6642 | 6460 | 6528 | 6119 | 6639 | 5496 | 4354 | 5879 | 5969 | 7040 | 6277 | 7046 | 8113 | 6028 | 6903 | 7812 | 5886 | 6410 | 6576 | Cdk17 |
| Cdk18 | 322 | 317 | 322 | 184 | 133 | 180 | 118 | 84 | 112 | 104 | 135 | 61 | 71 | 42 | 239 | 467 | 231 | 211 | 127 | 163 | 128 | 63 | 118 | 128 | 42 | 179 | 42 | 46 | Cdk18 |
| Cdk19 | 7375 | 6271 | 5178 | 7556 | 7673 | 7494 | 7986 | 6892 | 6255 | 5905 | 5954 | 5822 | 5709 | 5930 | 7613 | 6422 | 6297 | 7472 | 7135 | 7455 | 8110 | 5905 | 5864 | 5922 | 6607 | 5213 | 5585 | 5352 | Cdk19 |
| Cdk2 | 9579 | 8933 | 8741 | 7973 | 5807 | 6643 | 6954 | 5332 | 5408 | 4551 | 5779 | 5929 | 6009 | 5830 | 8453 | 8365 | 7930 | 7472 | 6665 | 7302 | 6485 | 3813 | 5367 | 5420 | 4461 | 7666 | 6208 | 6355 | Cdk2 |
| Cdk20 | 744 | 872 | 858 | 1031 | 1305 | 1037 | 1071 | 576 | 690 | 699 | 625 | 667 | 598 | 729 | 729 | 729 | 936 | 1152 | 1185 | 1003 | 1085 | 770 | 715 | 765 | 802 | 677 | 676 | 597 | Cdk20 |
| Cdk2ap1 | 4364 | 4350 | 4412 | 3575 | 3939 | 4294 | 4532 | 7276 | 7785 | 8183 | 7278 | 10387 | 9689 | 9633 | 3684 | 3748 | 4401 | 3371 | 4550 | 4582 | 3833 | 6662 | 6908 | 9184 | 8865 | 11722 | 10501 | 10529 | Cdk2ap1 |
| Cdk4 | 26576 | 29103 | 26293 | 20580 | 18867 | 21096 | 22173 | 31069 | 31360 | 27783 | 26332 | 36992 | 32485 | 38489 | 24927 | 24356 | 24118 | 19301 | 22170 | 23459 | 21337 | 24364 | 29337 | 34151 | 30073 | 45211 | 39115 | 39990 | Cdk4 |
| Cdk5 | 4144 | 5107 | 4426 | 4753 | 4595 | 4123 | 4637 | 3483 | 3664 | 3450 | 3101 | 3414 | 3062 | 3978 | 4327 | 4215 | 4852 | 4376 | 4665 | 4418 | 4754 | 3654 | 3440 | 3555 | 3745 | 3488 | 3281 | 3519 | Cdk5 |
| Cdk5r1 | 4586 | 5696 | 5696 | 10488 | 13768 | 10251 | 11695 | 12998 | 12893 | 16113 | 13380 | 13751 | 10350 | 13629 | 5984 | 4321 | 8920 | 11217 | 12821 | 10149 | 12777 | 18623 | 11948 | 15993 | 19124 | 11796 | 10959 | 11997 | Cdk5r1 |
| Cdk5r2 | 1764 | 2781 | 3348 | 3750 | 4293 | 3416 | 2752 | 1556 | 1491 | 2289 | 1742 | 1786 | 1894 | 1635 | 2268 | 2060 | 3830 | 3888 | 3956 | 3058 | 3464 | 2695 | 1476 | 1874 | 2360 | 1055 | 1754 | 1892 | Cdk5r2 |
| Cdk5rap1 | 845 | 424 | 474 | 379 | 422 | 406 | 516 | 742 | 706 | 546 | 606 | 648 | 655 | 630 | 574 | 556 | 445 | 353 | 383 | 419 | 457 | 617 | 718 | 727 | 627 | 722 | 920 | 635 | Cdk5rap1 |
| Cdk5rap2 | 8364 | 8347 | 4967 | 5922 | 6886 | 7651 | 7212 | 9994 | 9728 | 8793 | 9006 | 10032 | 9299 | 10555 | 8049 | 7791 | 5149 | 5548 | 6050 | 7594 | 7368 | 7060 | 9090 | 7577 | 9452 | 6951 | 10734 | 9293 | Cdk5rap2 |
| Cdk5rap3 | 3547 | 4769 | 4197 | 4155 | 3969 | 4046 | 3940 | 3629 | 4257 | 4052 | 3936 | 4575 | 4314 | 4143 | 3989 | 3954 | 4651 | 3993 | 4182 | 4086 | 3875 | 3643 | 3893 | 4215 | 4250 | 4785 | 4682 | 4290 | Cdk5rap3 |
| Cdk6 | 2149 | 1664 | 2271 | 2534 | 2781 | 2903 | 3127 | 3707 | 4327 | 4372 | 4223 | 4757 | 3447 | 4490 | 1841 | 1667 | 2555 | 2542 | 3174 | 2350 | 2976 | 3679 | 3264 | 4850 | 5084 | 4513 | 4028 | 3936 | Cdk6 |
| Cdk7 | 3137 | 2916 | 2559 | 2420 | 2169 | 2682 | 2505 | 2987 | 2890 | 2331 | 2909 | 2649 | 2846 | 2595 | 3025 | 2779 | 2483 | 2310 | 2199 | 2743 | 2498 | 2181 | 3079 | 2460 | 2466 | 2611 | 2975 | 2928 | Cdk7 |
| Cdk8 | 8601 | 7724 | 8972 | 8181 | 7416 | 8197 | 8356 | 7008 | 5850 | 6233 | 6236 | 7652 | 6955 | 7492 | 8449 | 7974 | 8492 | 7682 | 7800 | 7851 | 7961 | 6193 | 5504 | 6755 | 6834 | 7389 | 7992 | 7711 | Cdk8 |
| Cdk9 | 5043 | 5447 | 6814 | 6554 | 5470 | 5314 | 5444 | 4112 | 4449 | 4970 | 4721 | 5447 | 4799 | 5173 | 4750 | 5127 | 6489 | 5994 | 6212 | 4960 | 5488 | 4829 | 4570 | 5460 | 5141 | 6073 | 4996 | 5167 | Cdk9 |

| | | | | | | | | | | | | | | | | | | | | | | | | | | | | | |
|---|---|---|---|---|---|---|---|---|---|---|---|---|---|---|---|---|---|---|---|---|---|---|---|---|---|---|---|---|---|
| Cdkal1 | 1028 | 984 | 926 | 819 | 762 | 810 | 895 | 1252 | 1229 | 1067 | 1045 | 1098 | 1149 | 1166 | 1040 | 907 | 860 | 828 | 810 | 878 | 874 | 1069 | 1165 | 1088 | 1183 | 1339 | 1219 | 1243 | Cdkal1 |
| Cdkl1 | 98 | 53 | 66 | 68 | 124 | 124 | 82 | 145 | 154 | 116 | 190 | 118 | 128 | 80 | 104 | 76 | 86 | 59 | 81 | 118 | 103 | 155 | 206 | 109 | 131 | 106 | 107 | 98 | Cdkl1 |
| Cdkl2 | 210 | 209 | 158 | 296 | 542 | 316 | 412 | 325 | 394 | 322 | 440 | 137 | 204 | 180 | 274 | 260 | 178 | 373 | 316 | 312 | 397 | 393 | 445 | 261 | 343 | 150 | 119 | 132 | Cdkl2 |
| Cdkl3 | 1539 | 1379 | 1087 | 1609 | 1573 | 1690 | 1659 | 1260 | 1124 | 1136 | 1014 | 1206 | 1102 | 1243 | 1630 | 1310 | 1318 | 1510 | 1451 | 1640 | 1549 | 1113 | 1108 | 1019 | 1206 | 908 | 1192 | 1097 | Cdkl3 |
| Cdkl4 | 255 | 246 | 234 | 241 | 276 | 232 | 218 | 208 | 248 | 228 | 177 | 191 | 311 | 168 | 342 | 336 | 228 | 227 | 218 | 246 | 216 | 344 | 274 | 176 | 176 | 167 | 160 | 164 | Cdkl4 |
| Cdkl5 | 4241 | 4058 | 2358 | 3594 | 3842 | 4338 | 4323 | 2905 | 2479 | 2058 | 1943 | 2140 | 1929 | 2196 | 4509 | 3591 | 2987 | 3975 | 3726 | 4994 | 4757 | 2052 | 2407 | 1950 | 2326 | 1492 | 2429 | 1978 | Cdkl5 |
| Cdkn1a | 2794 | 3080 | 2937 | 3193 | 3144 | 2931 | 2601 | 1183 | 1430 | 1257 | 2123 | 1232 | 1252 | 983 | 2689 | 2970 | 3204 | 3212 | 3013 | 2952 | 2700 | 1340 | 1930 | 1207 | 1219 | 1350 | 986 | 1098 | Cdkn1a |
| Cdkn1b | 9503 | 8466 | 11592 | 13211 | 12159 | 12965 | 14976 | 10437 | 8978 | 10340 | 9991 | 12019 | 10645 | 14387 | 9941 | 8666 | 11754 | 12715 | 13609 | 13127 | 14715 | 9892 | 8576 | 13108 | 13201 | 13116 | 11904 | 13953 | Cdkn1b |
| Cdkn1c | 21223 | 22849 | 24579 | 19918 | 16123 | 17986 | 11096 | 24337 | 26595 | 24050 | 44729 | 22615 | 28535 | 22095 | 21803 | 23179 | 22403 | 19387 | 15284 | 14327 | 13729 | 26199 | 37600 | 25715 | 22115 | 21893 | 27054 | 25280 | Cdkn1c |
| Cdkn2a | 5 | 11 | 12 | 5 | 1 | 12 | 6 | 0 | 0 | 3 | 0 | 1 | 0 | 2 | 7 | 5 | 19 | 1 | 9 | 0 | 6 | 0 | 1 | 2 | 0 | 0 | 1 | 0 | Cdkn2a |
| Cdkn2aip | 3200 | 3038 | 3523 | 3776 | 3028 | 3316 | 3552 | 3747 | 2800 | 2904 | 2965 | 3598 | 3250 | 3792 | 3487 | 3091 | 3435 | 3461 | 3204 | 3618 | 3433 | 3287 | 2763 | 3452 | 3491 | 3683 | 3750 | 3762 | Cdkn2aip |
| Cdkn2aipnl | 4951 | 4705 | 4856 | 4616 | 4339 | 4511 | 4702 | 4164 | 3978 | 3736 | 3463 | 3945 | 3747 | 4199 | 4971 | 4739 | 4695 | 4408 | 4597 | 5019 | 4640 | 3574 | 4112 | 3865 | 3691 | 4563 | 4162 | 4455 | Cdkn2aipnl |
| Cdkn2b | 44 | 48 | 29 | 37 | 34 | 36 | 29 | 21 | 18 | 12 | 32 | 21 | 11 | 6 | 54 | 42 | 17 | 33 | 17 | 20 | 21 | 5 | 28 | 18 | 0 | 5 | 10 | 5 | Cdkn2b |
| Cdkn2c | 1022 | 962 | 1056 | 1024 | 814 | 972 | 1141 | 796 | 904 | 784 | 737 | 1025 | 824 | 1096 | 885 | 798 | 1027 | 983 | 1101 | 1056 | 1153 | 642 | 785 | 1069 | 1033 | 1113 | 889 | 996 | Cdkn2c |
| Cdkn2d | 2345 | 3048 | 2430 | 2685 | 2646 | 2554 | 2455 | 1552 | 1631 | 1448 | 1324 | 1598 | 1495 | 1388 | 2465 | 2641 | 2805 | 2773 | 2956 | 3295 | 2568 | 1318 | 1546 | 1557 | 1531 | 1810 | 1592 | 1423 | Cdkn2d |
| Cdkn3 | 1277 | 1036 | 1325 | 1167 | 729 | 897 | 929 | 487 | 494 | 567 | 569 | 783 | 776 | 845 | 1179 | 1107 | 1139 | 867 | 781 | 1062 | 838 | 319 | 480 | 609 | 488 | 920 | 843 | 828 | Cdkn3 |
| Cdnf | 65 | 56 | 38 | 37 | 39 | 61 | 54 | 50 | 39 | 34 | 43 | 42 | 67 | 42 | 62 | 51 | 60 | 33 | 31 | 61 | 41 | 38 | 45 | 47 | 30 | 37 | 45 | 53 | Cdnf |
| Cdo1 | 3673 | 2699 | 2897 | 2792 | 2408 | 3333 | 2445 | 4924 | 4776 | 4510 | 4705 | 3190 | 4425 | 3962 | 3580 | 3404 | 2445 | 3191 | 2424 | 2528 | 2555 | 4293 | 5489 | 4056 | 3920 | 3608 | 3467 | 4165 | Cdo1 |
| Cdon | 13768 | 12142 | 11272 | 11978 | 10258 | 13593 | 11916 | 13713 | 15627 | 14750 | 16821 | 17955 | 19891 | 17539 | 12286 | 13387 | 10136 | 12172 | 11838 | 11423 | 11950 | 8154 | 14851 | 16535 | 16172 | 11923 | 16580 | 15991 | Cdon |
| Cdpf1 | 558 | 792 | 590 | 604 | 511 | 525 | 523 | 333 | 442 | 421 | 405 | 431 | 432 | 425 | 504 | 659 | 628 | 596 | 528 | 608 | 508 | 376 | 441 | 452 | 322 | 492 | 450 | 394 | Cdpf1 |
| Cdr2 | 4254 | 4831 | 4288 | 4397 | 4087 | 4013 | 4069 | 2936 | 2700 | 2481 | 2395 | 2405 | 2963 | 2717 | 4659 | 4432 | 4154 | 3940 | 3935 | 4238 | 3982 | 2702 | 2621 | 2107 | 2440 | 2673 | 2669 | 3059 | Cdr2 |
| Cdr2l | 3227 | 4234 | 4578 | 4360 | 4046 | 3631 | 3677 | 2786 | 2730 | 3341 | 2905 | 3579 | 3299 | 3798 | 3306 | 3636 | 4662 | 4363 | 4352 | 3437 | 3802 | 3761 | 2618 | 3654 | 3734 | 4108 | 3560 | 3762 | Cdr2l |
| Cdrt4 | 18 | 12 | 2 | 10 | 7 | 17 | 11 | 18 | 11 | 12 | 26 | 9 | 26 | 4 | 13 | 16 | 6 | 22 | 23 | 8 | 17 | 8 | 18 | 7 | 20 | 9 | 19 | 5 | Cdrt4 |
| Cds1 | 754 | 588 | 513 | 553 | 709 | 587 | 456 | 432 | 564 | 601 | 440 | 321 | 396 | 284 | 543 | 743 | 563 | 763 | 586 | 491 | 567 | 568 | 663 | 440 | 428 | 258 | 177 | 223 | Cds1 |
| Cds2 | 13455 | 14176 | 12476 | 17458 | 19477 | 17033 | 16554 | 10003 | 10387 | 11105 | 10666 | 10453 | 10142 | 9903 | 13580 | 13769 | 15308 | 18359 | 18343 | 15439 | 17465 | 12221 | 9716 | 10363 | 12641 | 7145 | 9770 | 9131 | Cds2 |
| Cdsn | 5 | 12 | 13 | 8 | 20 | 6 | 9 | 4 | 12 | 3 | 1 | 7 | 5 | 2 | 8 | 9 | 10 | 17 | 7 | 10 | 6 | 2 | 5 | 1 | 2 | 7 | 7 | 0 | Cdsn |
| Cdt1 | 2050 | 2389 | 2947 | 2478 | 1684 | 1768 | 1773 | 1320 | 1598 | 1528 | 1678 | 2499 | 1884 | 2254 | 1609 | 2170 | 2736 | 2140 | 2244 | 1540 | 1775 | 1332 | 1453 | 2040 | 1559 | 3499 | 2076 | 2293 | Cdt1 |
| Cdv3 | 20362 | 16848 | 20907 | 18165 | 16278 | 16929 | 19023 | 20290 | 17994 | 17949 | 18866 | 23076 | 21430 | 23428 | 19460 | 17987 | 19200 | 17649 | 17386 | 17750 | 17388 | 17739 | 18215 | 20314 | 19105 | 26659 | 23372 | 26248 | Cdv3 |
| Cdx1 | 0 | 2 | 3 | 2 | 2 | 2 | 2 | 0 | 0 | 0 | 4 | 0 | 1 | 0 | 1 | 2 | 0 | 1 | 4 | 0 | 1 | 3 | 0 | 1 | 0 | 5 | 1 | 1 | Cdx1 |
| Cdx2 | 0 | 1 | 0 | 0 | 0 | 0 | 0 | 0 | 2 | 0 | 0 | 0 | 0 | 0 | 0 | 0 | 0 | 3 | 0 | 0 | 0 | 0 | 1 | 3 | 1 | 0 | 0 | 0 | Cdx2 |
| Cdx4 | 0 | 0 | 0 | 0 | 7 | 0 | 0 | 0 | 0 | 0 | 0 | 0 | 0 | 1 | 0 | 0 | 0 | 0 | 0 | 0 | 0 | 2 | 0 | 0 | 0 | 0 | 0 | 0 | Cdx4 |
| Cdyl | 1498 | 1294 | 1854 | 1620 | 1447 | 1474 | 1399 | 1144 | 1270 | 1233 | 1416 | 1551 | 1572 | 1424 | 1321 | 1504 | 1679 | 1469 | 1637 | 1186 | 1378 | 1108 | 1434 | 1433 | 1141 | 1791 | 1364 | 1622 | Cdyl |
| Cdyl2 | 194 | 195 | 147 | 174 | 283 | 236 | 213 | 321 | 381 | 359 | 374 | 277 | 291 | 247 | 160 | 212 | 129 | 200 | 234 | 205 | 224 | 224 | 357 | 321 | 291 | 145 | 273 | 235 | Cdyl2 |
| Ceacam1 | 36 | 36 | 45 | 40 | 44 | 55 | 60 | 57 | 68 | 65 | 110 | 61 | 70 | 45 | 50 | 58 | 59 | 37 | 48 | 44 | 58 | 70 | 93 | 63 | 73 | 86 | 72 | 65 | Ceacam1 |
| Ceacam10 | 5 | 4 | 4 | 6 | 1 | 0 | 0 | 1 | 2 | 0 | 4 | 4 | 1 | 3 | 3 | 6 | 0 | 2 | 1 | 1 | 0 | 5 | 5 | 7 | 1 | 13 | 0 | 4 | Ceacam10 |
| Ceacam11 | 0 | 0 | 0 | 0 | 0 | 0 | 0 | 0 | 16 | 0 | 0 | 0 | 0 | 1 | 0 | 0 | 0 | 0 | 0 | 0 | 0 | 0 | 5 | 0 | 0 | 0 | 0 | 0 | Ceacam11 |
| Ceacam12 | 0 | 0 | 0 | 0 | 0 | 0 | 0 | 0 | 11 | 0 | 0 | 0 | 0 | 0 | 0 | 0 | 0 | 0 | 0 | 0 | 0 | 0 | 0 | 0 | 0 | 0 | 0 | 0 | Ceacam12 |
| Ceacam13 | 0 | 0 | 0 | 0 | 0 | 0 | 0 | 0 | 0 | 0 | 0 | 0 | 0 | 0 | 0 | 0 | 0 | 0 | 0 | 0 | 0 | 0 | 1 | 0 | 0 | 0 | 0 | 0 | Ceacam13 |
| Ceacam14 | 0 | 0 | 0 | 0 | 0 | 0 | 0 | 0 | 19 | 0 | 0 | 0 | 0 | 0 | 0 | 0 | 0 | 0 | 0 | 0 | 0 | 0 | 4 | 0 | 0 | 0 | 0 | 0 | Ceacam14 |
| Ceacam15 | 0 | 0 | 0 | 0 | 1 | 0 | 0 | 0 | 1 | 0 | 0 | 0 | 0 | 0 | 0 | 2 | 0 | 0 | 0 | 0 | 0 | 0 | 0 | 0 | 0 | 1 | 0 | 0 | Ceacam15 |
| Ceacam16 | 0 | 0 | 0 | 0 | 0 | 0 | 0 | 0 | 0 | 0 | 0 | 0 | 0 | 1 | 0 | 0 | 0 | 0 | 0 | 0 | 0 | 0 | 0 | 2 | 4 | 0 | 0 | 0 | Ceacam16 |
| Ceacam18 | 7 | 5 | 4 | 4 | 3 | 0 | 2 | 0 | 1 | 0 | 4 | 3 | 0 | 1 | 1 | 9 | 3 | 4 | 5 | 9 | 4 | 2 | 3 | 0 | 2 | 3 | 1 | 1 | Ceacam18 |
| Ceacam19 | 0 | 1 | 0 | 0 | 0 | 0 | 0 | 0 | 6 | 5 | 1 | 0 | 0 | 0 | 0 | 0 | 0 | 0 | 0 | 0 | 0 | 0 | 9 | 1 | 0 | 0 | 0 | 0 | Ceacam19 |
| Ceacam2 | 0 | 0 | 0 | 0 | 0 | 1 | 0 | 1 | 0 | 0 | 0 | 0 | 0 | 0 | 1 | 0 | 1 | 0 | 0 | 1 | 0 | 0 | 1 | 0 | 0 | 1 | 0 | 0 | Ceacam2 |
| Ceacam20 | 0 | 0 | 0 | 0 | 0 | 0 | 0 | 0 | 0 | 0 | 0 | 0 | 0 | 0 | 0 | 0 | 0 | 0 | 0 | 0 | 1 | 0 | 0 | 0 | 0 | 0 | 0 | 0 | Ceacam20 |
| Ceacam3 | 0 | 0 | 0 | 0 | 2 | 0 | 0 | 1 | 9 | 4 | 1 | 0 | 1 | 2 | 4 | 2 | 0 | 0 | 5 | 0 | 0 | 2 | 1 | 6 | 1 | 1 | 0 | 0 | Ceacam3 |
| Ceacam5 | 0 | 0 | 0 | 0 | 0 | 0 | 0 | 0 | 11 | 0 | 0 | 0 | 0 | 0 | 0 | 0 | 0 | 0 | 0 | 0 | 0 | 0 | 2 | 0 | 0 | 0 | 0 | 0 | Ceacam5 |
| Ceacam9 | 0 | 0 | 0 | 0 | 2 | 0 | 0 | 0 | 0 | 0 | 0 | 0 | 0 | 0 | 0 | 0 | 0 | 0 | 0 | 0 | 0 | 2 | 0 | 0 | 2 | 0 | 0 | 0 | Ceacam9 |
| Cebpa | 495 | 460 | 538 | 496 | 360 | 388 | 284 | 210 | 328 | 242 | 227 | 139 | 129 | 84 | 298 | 503 | 532 | 519 | 415 | 391 | 326 | 119 | 414 | 197 | 125 | 202 | 102 | 125 | Cebpa |
| Cebpb | 76 | 81 | 70 | 57 | 47 | 46 | 40 | 72 | 87 | 102 | 105 | 51 | 56 | 38 | 46 | 74 | 101 | 74 | 42 | 46 | 46 | 40 | 87 | 72 | 49 | 61 | 24 | 21 | Cebpb |
| Cebpd | 63 | 49 | 46 | 51 | 57 | 62 | 22 | 96 | 114 | 78 | 108 | 49 | 59 | 40 | 51 | 47 | 44 | 32 | 64 | 66 | 34 | 74 | 102 | 86 | 74 | 73 | 81 | 53 | Cebpd |
| Cebpe | 1 | 1 | 3 | 2 | 0 | 5 | 0 | 2 | 1 | 13 | 5 | 4 | 0 | 1 | 1 | 0 | 0 | 3 | 3 | 0 | 4 | 0 | 4 | 5 | 12 | 3 | 1 | 2 | Cebpe |
| Cebpg | 7138 | 6544 | 9103 | 8586 | 7852 | 7740 | 8336 | 7331 | 6051 | 6606 | 6424 | 7399 | 6471 | 7620 | 7089 | 7061 | 8867 | 8249 | 8445 | 8049 | 8164 | 5905 | 5831 | 7484 | 7365 | 8681 | 7340 | 7672 | Cebpg |
| Cebpz | 6846 | 6329 | 6346 | 5414 | 5009 | 5459 | 6330 | 7135 | 6119 | 5971 | 5573 | 6649 | 6143 | 6779 | 7146 | 6208 | 5789 | 5702 | 5594 | 6459 | 5790 | 5822 | 6111 | 6393 | 6137 | 7166 | 6643 | 7096 | Cebpz |
| Cebpzos | 1152 | 1054 | 1317 | 1138 | 1058 | 1113 | 1286 | 1232 | 1093 | 1141 | 1113 | 1202 | 1246 | 1434 | 1164 | 1070 | 1309 | 1207 | 1148 | 1383 | 1231 | 947 | 1204 | 1284 | 1110 | 1418 | 1283 | 1474 | Cebpzos |
| Cecr2 | 3404 | 3103 | 4292 | 3912 | 4028 | 3755 | 4108 | 3617 | 3273 | 3633 | 3288 | 4620 | 5240 | 4817 | 3451 | 3298 | 4058 | 3573 | 3982 | 3464 | 4018 | 3552 | 2934 | 4184 | 4279 | 3837 | 5084 | 5000 | Cecr2 |
| Cel | 0 | 9 | 12 | 10 | 3 | 8 | 6 | 2 | 2 | 1 | 4 | 3 | 2 | 4 | 4 | 4 | 7 | 7 | 5 | 6 | 2 | 0 | 1 | 9 | 11 | 3 | 0 | 2 | Cel |
| Cela1 | 27 | 32 | 28 | 20 | 16 | 24 | 3 | 20 | 28 | 10 | 12 | 16 | 40 | 4 | 16 | 28 | 17 | 13 | 5 | 18 | 4 | 15 | 29 | 12 | 18 | 21 | 16 | 20 | Cela1 |
| Cela3a | 0 | 0 | 0 | 0 | 0 | 0 | 0 | 0 | 0 | 0 | 0 | 0 | 0 | 0 | 0 | 0 | 0 | 0 | 0 | 0 | 0 | 0 | 1 | 0 | 0 | 0 | 0 | 0 | Cela3a |
| Cela3b | 0 | 0 | 0 | 0 | 0 | 0 | 0 | 0 | 1 | 0 | 5 | 0 | 0 | 0 | 0 | 0 | 0 | 0 | 0 | 0 | 0 | 0 | 2 | 0 | 0 | 0 | 0 | 0 | Cela3b |
| Celf1 | 0 | 1 | 0 | 0 | 1 | 1 | 0 | 0 | 0 | 0 | 0 | 0 | 0 | 3 | 1 | 2 | 0 | 0 | 0 | 0 | 1 | 5 | 1 | 0 | 0 | 0 | 1 | 0 | Celf1 |
| Celf2 | 16200 | 13996 | 14017 | 21477 | 24004 | 24648 | 30736 | 26253 | 25480 | 25210 | 23403 | 27722 | 18761 | 29982 | 17578 | 12295 | 18729 | 22331 | 24949 | 24354 | 29054 | 29061 | 23173 | 28312 | 35053 | 21892 | 23984 | 23930 | Celf2 |
| Celf3 | 7120 | 11149 | 11700 | 15036 | 16431 | 14018 | 14680 | 12263 | 12765 | 17439 | 12949 | 19917 | 14835 | 19179 | 8549 | 7790 | 15630 | 14239 | 17275 | 13057 | 15053 | 19714 | 11301 | 19063 | 20425 | 17294 | 17552 | 17478 | Celf3 |
| Celf4 | 6710 | 7684 | 9911 | 14823 | 17504 | 15449 | 15816 | 13021 | 12311 | 18699 | 14084 | 15374 | 12374 | 15664 | 8250 | 6725 | 12929 | 14705 | 16737 | 15019 | 16613 | 20384 | 11105 | 17039 | 21090 | 11569 | 14100 | 14527 | Celf4 |
| Celf5 | 4045 | 5022 | 6276 | 10422 | 13085 | 9943 | 11120 | 8140 | 8892 | 14314 | 10255 | 12083 | 8150 | 11820 | 4337 | 3842 | 9426 | 10747 | 12666 | 8776 | 12019 | 15329 | 7825 | 14045 | 16178 | 10098 | 9230 | 9911 | Celf5 |
| Celf6 | 1081 | 1563 | 1362 | 1980 | 2404 | 1983 | 2048 | 1171 | 1114 | 1290 | 1018 | 1353 | 1055 | 1198 | 1291 | 1145 | 1980 | 1922 | 2212 | 1913 | 2221 | 1796 | 976 | 1235 | 1550 | 1013 | 1090 | 1159 | Celf6 |
| Celsr1 | 4654 | 6010 | 6078 | 7134 | 7072 | 7243 | 6900 | 4992 | 7099 | 5345 | 4142 | 9521 | 6524 | 7662 | 4272 | 5003 | 7021 | 7678 | 8540 | 6414 | 7310 | 5020 | 6263 | 9674 | 6104 | 5983 | 6788 | 6831 | Celsr1 |
| Celsr2 | 9350 | 15104 | 18963 | 20801 | 19211 | 16700 | 15542 | 10862 | 12091 | 15249 | 11876 | 20927 | 18590 | 17566 | 11107 | 12989 | 22288 | 19387 | 20193 | 14012 | 17291 | 17034 | 10911 | 17359 | 17942 | 11069 | 17332 | 17932 | Celsr2 |
| Celsr3 | 0 | 0 | 0 | 0 | 1 | 0 | 0 | 0 | 0 | 0 | 0 | 0 | 0 | 0 | 0 | 0 | 0 | 0 | 0 | 0 | 0 | 0 | 0 | 0 | 0 | 0 | 0 | 0 | Celsr3 |
| Cemip | 642 | 937 | 753 | 711 | 856 | 1002 | 299 | 860 | 1594 | 1341 | 2033 | 1309 | 1988 | 1142 | 786 | 994 | 1029 | 1298 | 568 | 298 | 319 | 1866 | 2096 | 1464 | 1770 | 1000 | 1926 | 943 | Cemip |
| Cemip2 | 14739 | 15493 | 14451 | 15224 | 14071 | 15736 | 14550 | 14141 | 11799 | 13520 | 12274 | 17637 | 16306 | 17285 | 15976 | 14715 | 15487 | 14519 | 14294 | 13978 | 14556 | 13692 | 11013 | 14354 | 15520 | 12511 | 19009 | 17443 | Cemip2 |
| Cend1 | 911 | 1362 | 1029 | 1325 | 1994 | 1390 | 1364 | 1312 | 1589 | 1734 | 1530 | 1205 | 1274 | 1272 | 1067 | 942 | 1349 | 1481 | 1616 | 1540 | 1414 | 1744 | 1461 | 1396 | 1689 | 1168 | 1126 | 1074 | Cend1 |
| Cenpa | 6223 | 5635 | 5663 | 4111 | 3507 | 4189 | 4613 | 6036 | 5882 | 5844 | 5532 | 8442 | 7538 | 8617 | 5159 | 5332 | 5138 | 3981 | 4504 | 4519 | 4375 | 4341 | 5350 | 7134 | 6142 | 8937 | 8992 | 8813 | Cenpa |
| Cenpb | 15694 | 17553 | 18332 | 20465 | 18177 | 18197 | 18323 | 11941 | 14351 | 16202 | 14602 | 17961 | 16195 | 17608 | 15424 | 16351 | 19695 | 18830 | 20211 | 16914 | 19410 | 14639 | 14358 | 17767 | 17582 | 16779 | 17576 | 17253 | Cenpb |
| Cenpc1 | 7394 | 6461 | 7136 | 6908 | 6991 | 7332 | 8097 | 6690 | 5809 | 5449 | 5435 | 6778 | 6466 | 6841 | 8209 | 6639 | 7263 | 6859 | 7377 | 8977 | 7586 | 5656 | 5608 | 5976 | 5912 | 6560 | 6759 | 6680 | Cenpc1 |
| Cenpe | 15632 | 12246 | 10128 | 9143 | 6858 | 9666 | 10235 | 12206 | 10727 | 8986 | 8530 | 14092 | 12292 | 14581 | 13652 | 13188 | 9783 | 7864 | 8509 | 11800 | 9165 | 6592 | 9288 | 10128 | 9471 | 10531 | 15468 | 13511 | Cenpe |
| Cenpf | 25478 | 19763 | 16495 | 15062 | 11584 | 17105 | 17757 | 20695 | 17641 | 14585 | 13209 | 22981 | 20674 | 24094 | 23619 | 21557 | 15521 | 12503 | 13705 | 21030 | 15853 | 9825 | 15479 | 15427 | 15145 | 16439 | 26818 | 21660 | Cenpf |
| Cenph | 2970 | 2371 | 2806 | 2040 | 1420 | 2057 | 2333 | 1441 | 1386 | 1298 | 1229 | 2225 | 1706 | 2117 | 2639 | 2572 | 2388 | 1856 | 1835 | 2322 | 1895 | 877 | 1309 | 1691 | 1422 | 2520 | 2116 | 2106 | Cenph |
| Cenpi | 2280 | 1762 | 1830 | 1667 | 1413 | 1490 | 1854 | 2146 | 1851 | 1455 | 1553 | 2236 | 1809 | 2204 | 2032 | 1805 | 1714 | 1539 | 1609 | 1943 | 1684 | 1377 | 1733 | 1901 | 1776 | 2668 | 2430 | 2137 | Cenpi |

| | | | | | | | | | | | | | | | | | | | | | | | | | | | | | |
|---|---|---|---|---|---|---|---|---|---|---|---|---|---|---|---|---|---|---|---|---|---|---|---|---|---|---|---|---|---|
| Cenpj | 5187 | 4820 | 4615 | 4162 | 3318 | 3860 | 4375 | 3131 | 2640 | 2580 | 2142 | 3220 | 2565 | 3458 | 5196 | 4533 | 4212 | 3919 | 3654 | 4729 | 4169 | 2154 | 2308 | 3035 | 2598 | 3236 | 3385 | 3353 | Cenpj |
| Cenpk | 1902 | 1168 | 1801 | 1292 | 972 | 1436 | 1581 | 1768 | 1579 | 1433 | 1426 | 1991 | 1735 | 2382 | 1753 | 1592 | 1465 | 1120 | 1286 | 1752 | 1342 | 1066 | 1452 | 1811 | 1474 | 2552 | 2095 | 2256 | Cenpk |
| Cenpl | 2914 | 2347 | 2327 | 2139 | 1810 | 2076 | 2283 | 1890 | 1648 | 1530 | 1335 | 2043 | 1817 | 2256 | 2516 | 2427 | 2138 | 2076 | 2033 | 2463 | 2115 | 1265 | 1589 | 1843 | 1646 | 2597 | 2436 | 2403 | Cenpl |
| Cenpm | 775 | 899 | 734 | 648 | 510 | 643 | 629 | 563 | 638 | 486 | 523 | 881 | 797 | 822 | 589 | 712 | 800 | 558 | 634 | 675 | 612 | 393 | 559 | 772 | 657 | 1128 | 881 | 961 | Cenpm |
| Cenpn | 2211 | 1863 | 1950 | 1505 | 1271 | 1456 | 1614 | 1790 | 1830 | 1740 | 1532 | 2349 | 2065 | 2450 | 1998 | 1786 | 1847 | 1450 | 1547 | 1602 | 1447 | 1242 | 1542 | 1986 | 1681 | 2772 | 2468 | 2540 | Cenpn |
| Cenpo | 1544 | 1390 | 1308 | 1212 | 922 | 1174 | 1129 | 1420 | 1250 | 1170 | 1262 | 1433 | 1369 | 1479 | 1412 | 1289 | 1207 | 1044 | 1174 | 1326 | 1183 | 1014 | 1205 | 1259 | 1276 | 1438 | 1633 | 1525 | Cenpo |
| Cenpp | 1480 | 1225 | 1194 | 1067 | 725 | 845 | 1161 | 730 | 630 | 494 | 484 | 818 | 793 | 900 | 1407 | 1379 | 1213 | 855 | 976 | 1227 | 1003 | 419 | 499 | 710 | 582 | 1024 | 851 | 978 | Cenpp |
| Cenpq | 2271 | 1456 | 2010 | 1548 | 1174 | 1575 | 1757 | 1961 | 1756 | 1446 | 1337 | 2058 | 1772 | 2288 | 2054 | 1808 | 1607 | 1247 | 1600 | 2005 | 1457 | 1215 | 1464 | 1763 | 1468 | 2510 | 2166 | 2404 | Cenpq |
| Cenps | 414 | 381 | 541 | 409 | 394 | 452 | 386 | 679 | 819 | 798 | 681 | 989 | 1012 | 995 | 385 | 390 | 489 | 348 | 471 | 449 | 333 | 521 | 772 | 949 | 743 | 1311 | 1169 | 1030 | Cenps |
| Cenpt | 3222 | 3703 | 3694 | 3658 | 3240 | 3121 | 3193 | 3251 | 3392 | 3643 | 3524 | 4060 | 4073 | 4054 | 3055 | 3314 | 3489 | 3442 | 3288 | 2938 | 3085 | 3589 | 3082 | 3868 | 3856 | 4442 | 4215 | 4086 | Cenpt |
| Cenpu | 2584 | 2447 | 2337 | 2021 | 1187 | 1665 | 1932 | 1130 | 1032 | 899 | 802 | 1382 | 1311 | 1503 | 2531 | 2229 | 2103 | 1717 | 1700 | 2023 | 1770 | 695 | 830 | 1265 | 1064 | 1681 | 1508 | 1505 | Cenpu |
| Cenpv | 1502 | 1681 | 1569 | 1691 | 1793 | 1845 | 2168 | 1728 | 1874 | 1911 | 1709 | 2018 | 1681 | 2333 | 1490 | 1071 | 1779 | 1742 | 2021 | 1988 | 2011 | 1693 | 1702 | 2416 | 2285 | 2323 | 2186 | 2296 | Cenpv |
| Cenpw | 1425 | 938 | 1655 | 1104 | 714 | 870 | 1272 | 1092 | 840 | 754 | 761 | 1458 | 1051 | 1632 | 1311 | 1182 | 1270 | 797 | 854 | 1263 | 1004 | 735 | 933 | 1200 | 935 | 1788 | 1549 | 1869 | Cenpw |
| Cenpx | 1700 | 2048 | 1841 | 1757 | 1454 | 1442 | 1645 | 1159 | 1224 | 1145 | 1236 | 1325 | 1212 | 1467 | 1545 | 1710 | 1877 | 1532 | 1657 | 1664 | 1772 | 1036 | 1264 | 1397 | 1280 | 1535 | 1388 | 1480 | Cenpx |
| Cep104 | 2668 | 3175 | 2735 | 3009 | 3271 | 3140 | 2866 | 2709 | 3078 | 3285 | 2851 | 3539 | 3251 | 3083 | 2595 | 2867 | 2853 | 3214 | 3326 | 2738 | 2931 | 3227 | 2915 | 3312 | 3506 | 2992 | 3175 | 3348 | Cep104 |
| Cep112 | 1509 | 1214 | 1732 | 1303 | 1104 | 1335 | 1256 | 1418 | 1206 | 1101 | 1222 | 1575 | 1430 | 1723 | 1524 | 1326 | 1335 | 1080 | 1150 | 1593 | 1129 | 1243 | 1369 | 1415 | 1412 | 1486 | 1758 | 1680 | Cep112 |
| Cep120 | 7170 | 7545 | 7050 | 7907 | 8489 | 7951 | 8548 | 7498 | 6698 | 7395 | 6545 | 7215 | 6578 | 7488 | 7780 | 6893 | 7365 | 7650 | 7911 | 7731 | 8384 | 7536 | 6515 | 6925 | 7371 | 6719 | 6887 | 7157 | Cep120 |
| Cep126 | 2673 | 2432 | 2091 | 2604 | 2422 | 2564 | 2452 | 1601 | 1332 | 1375 | 1302 | 1512 | 1600 | 1334 | 2763 | 2706 | 2493 | 2498 | 2605 | 2607 | 2611 | 1532 | 1340 | 1220 | 1292 | 1115 | 1457 | 1449 | Cep126 |
| Cep128 | 2280 | 2191 | 2245 | 1917 | 1711 | 1781 | 2065 | 2031 | 1682 | 1643 | 1463 | 2209 | 1852 | 2247 | 2201 | 1942 | 2088 | 1876 | 1945 | 2154 | 1921 | 1414 | 1665 | 1803 | 1677 | 2142 | 2328 | 2083 | Cep128 |
| Cep131 | 3791 | 4884 | 6150 | 5899 | 4733 | 4775 | 4474 | 2657 | 3222 | 3986 | 3607 | 4574 | 3877 | 4358 | 3666 | 4388 | 6197 | 5222 | 5340 | 4045 | 4654 | 3218 | 2897 | 4821 | 4108 | 5087 | 4357 | 4365 | Cep131 |
| Cep135 | 2369 | 2005 | 1664 | 1674 | 1763 | 1959 | 2045 | 2920 | 2959 | 2505 | 2570 | 3048 | 3106 | 3253 | 2094 | 2076 | 1775 | 1599 | 1794 | 2235 | 1964 | 2070 | 2526 | 2624 | 2498 | 2614 | 3273 | 3045 | Cep135 |
| Cep152 | 4109 | 4155 | 2502 | 3173 | 2167 | 2629 | 2894 | 2437 | 2136 | 1886 | 1912 | 2199 | 2115 | 2183 | 3600 | 4097 | 2702 | 2677 | 2223 | 3281 | 2788 | 1226 | 2090 | 2145 | 1679 | 2318 | 2494 | 2068 | Cep152 |
| Cep162 | 5765 | 6102 | 4310 | 4631 | 4485 | 5026 | 5152 | 4096 | 3599 | 2991 | 3014 | 3696 | 3444 | 3624 | 5930 | 5939 | 4485 | 4540 | 4393 | 5811 | 5107 | 2964 | 3410 | 2994 | 3236 | 2617 | 3866 | 3510 | Cep162 |
| Cep164 | 5740 | 7007 | 7534 | 7571 | 6792 | 6802 | 6587 | 2682 | 3540 | 3700 | 3839 | 4324 | 4123 | 3928 | 5333 | 6499 | 7497 | 7381 | 7115 | 5443 | 6672 | 3288 | 3372 | 3805 | 3847 | 3506 | 3864 | 3793 | Cep164 |
| Cep170 | 16137 | 14906 | 12123 | 17350 | 21164 | 22139 | 22969 | 33556 | 28541 | 26324 | 25390 | 25470 | 23170 | 27042 | 18407 | 14651 | 15592 | 17621 | 19716 | 23056 | 23054 | 28899 | 26903 | 22823 | 30216 | 15570 | 25104 | 23346 | Cep170 |
| Cep170b | 474 | 1199 | 536 | 1485 | 463 | 466 | 885 | 319 | 981 | 345 | 167 | 381 | 633 | 241 | 479 | 940 | 1263 | 708 | 596 | 451 | 553 | 404 | 230 | 668 | 571 | 154 | 550 | 319 | Cep170b |
| Cep19 | 2025 | 1881 | 1944 | 2054 | 1843 | 1866 | 2044 | 1988 | 1716 | 1674 | 1554 | 1743 | 1677 | 1912 | 2206 | 1828 | 2163 | 2085 | 1889 | 2468 | 2069 | 1861 | 1676 | 1754 | 1708 | 1815 | 1863 | 1844 | Cep19 |
| Cep192 | 15503 | 13659 | 7785 | 9532 | 8789 | 10918 | 11512 | 8692 | 7662 | 6022 | 6504 | 7878 | 7011 | 7994 | 14486 | 13582 | 8890 | 9414 | 8480 | 12533 | 11157 | 5133 | 6921 | 5712 | 6386 | 5395 | 8938 | 7605 | Cep192 |
| Cep20 | 2590 | 1995 | 2485 | 2283 | 2525 | 2397 | 2671 | 3926 | 3573 | 3485 | 3240 | 3764 | 3485 | 4010 | 2786 | 2290 | 2218 | 2213 | 2261 | 2815 | 2475 | 3474 | 3721 | 3675 | 3629 | 4102 | 3672 | 3999 | Cep20 |
| Cep250 | 5978 | 6222 | 7403 | 7457 | 7841 | 7536 | 7061 | 7310 | 8539 | 10610 | 10130 | 10862 | 10501 | 11127 | 5387 | 6724 | 7379 | 7236 | 7516 | 6048 | 6976 | 8407 | 7591 | 10893 | 10340 | 7840 | 11011 | 10021 | Cep250 |
| Cep290 | 4045 | 3032 | 2210 | 2976 | 2884 | 3831 | 3617 | 3729 | 3238 | 2495 | 2292 | 2868 | 2790 | 3075 | 4547 | 3662 | 2484 | 2829 | 2822 | 4427 | 3572 | 2075 | 3078 | 2352 | 2540 | 1824 | 3177 | 2593 | Cep290 |
| Cep295 | 9220 | 7613 | 5067 | 6403 | 6256 | 7810 | 8035 | 9873 | 8579 | 6858 | 6781 | 8270 | 7843 | 8420 | 9115 | 8028 | 5471 | 6298 | 5951 | 8365 | 7742 | 6278 | 7802 | 6289 | 7317 | 5624 | 8671 | 7408 | Cep295 |
| Cep295nl | 16 | 21 | 13 | 14 | 12 | 8 | 7 | 35 | 34 | 6 | 37 | 11 | 16 | 9 | 18 | 26 | 9 | 10 | 12 | 25 | 11 | 13 | 22 | 22 | 32 | 15 | 19 | 25 | Cep295nl |
| Cep350 | 10638 | 9673 | 5477 | 7444 | 7621 | 10271 | 9231 | 10022 | 9387 | 7568 | 8941 | 7467 | 7787 | 7986 | 10544 | 10333 | 5937 | 7086 | 6650 | 10379 | 9908 | 6682 | 8875 | 5829 | 8031 | 4673 | 8656 | 6773 | Cep350 |
| Cep41 | 1066 | 1016 | 1013 | 1011 | 1159 | 1067 | 1269 | 2044 | 1980 | 1931 | 2302 | 2095 | 2108 | 2066 | 1068 | 917 | 1145 | 971 | 1124 | 1118 | 1170 | 1903 | 2203 | 2006 | 1904 | 2098 | 2173 | 2302 | Cep41 |
| Cep43 | 3496 | 2817 | 3275 | 2662 | 2716 | 2851 | 2932 | 3645 | 3088 | 2720 | 2735 | 3296 | 3515 | 3460 | 3351 | 3235 | 2745 | 2759 | 2710 | 3303 | 2785 | 2891 | 2915 | 2951 | 2824 | 3642 | 3500 | 3783 | Cep43 |
| Cep44 | 3129 | 2717 | 2568 | 2707 | 2537 | 2716 | 2860 | 2488 | 2175 | 1923 | 2101 | 2056 | 2081 | 2225 | 2913 | 2914 | 2629 | 2598 | 2556 | 3305 | 2891 | 1873 | 2051 | 2089 | 2092 | 2124 | 2298 | 2082 | Cep44 |
| Cep55 | 4193 | 3820 | 4584 | 3270 | 2425 | 2873 | 3339 | 3154 | 2661 | 2615 | 2123 | 3908 | 3151 | 4169 | 3894 | 3509 | 3672 | 2873 | 2887 | 3658 | 2917 | 1995 | 2325 | 3230 | 2786 | 4675 | 4040 | 4243 | Cep55 |
| Cep57 | 5469 | 4336 | 4818 | 4439 | 4014 | 4458 | 5197 | 5357 | 4765 | 4125 | 4319 | 5798 | 5077 | 6247 | 5428 | 4543 | 4558 | 4040 | 4209 | 5532 | 4400 | 4264 | 4401 | 4764 | 4560 | 6313 | 5933 | 5993 | Cep57 |
| Cep57l1 | 624 | 600 | 584 | 625 | 511 | 513 | 552 | 1247 | 1080 | 988 | 1085 | 1213 | 1188 | 1199 | 686 | 521 | 540 | 497 | 583 | 657 | 563 | 922 | 1186 | 1135 | 935 | 1198 | 1218 | 1245 | Cep57l1 |
| Cep63 | 1884 | 1635 | 1760 | 1929 | 1879 | 1795 | 1990 | 3577 | 3253 | 3566 | 3410 | 3877 | 3632 | 4070 | 1842 | 1699 | 1976 | 1702 | 1880 | 1850 | 1952 | 3347 | 3178 | 3583 | 3477 | 3834 | 3972 | 3830 | Cep63 |
| Cep68 | 5021 | 4122 | 4330 | 3999 | 4275 | 4516 | 4645 | 5699 | 5694 | 5187 | 5343 | 5910 | 5968 | 6149 | 4631 | 4261 | 4239 | 4080 | 4284 | 4230 | 4414 | 4995 | 5320 | 5586 | 5409 | 5535 | 5879 | 5878 | Cep68 |
| Cep70 | 3723 | 2853 | 2844 | 3192 | 3530 | 3763 | 4046 | 4557 | 3892 | 3690 | 3286 | 3645 | 3307 | 3964 | 3729 | 3029 | 2934 | 3143 | 3300 | 4330 | 3909 | 3883 | 3544 | 3408 | 3812 | 3491 | 3585 | 3434 | Cep70 |
| Cep72 | 1867 | 1692 | 1666 | 1476 | 1459 | 1492 | 1505 | 2216 | 2253 | 1997 | 1959 | 2787 | 2411 | 2639 | 1975 | 1632 | 1628 | 1274 | 1400 | 1685 | 1350 | 1733 | 2034 | 2055 | 1900 | 2391 | 2726 | 2613 | Cep72 |
| Cep76 | 2877 | 2642 | 3018 | 2775 | 2347 | 2603 | 2808 | 2407 | 2029 | 1943 | 1906 | 2881 | 2652 | 3002 | 2711 | 2537 | 2756 | 2511 | 2571 | 2678 | 2586 | 1892 | 1817 | 2446 | 2361 | 3020 | 2932 | 3129 | Cep76 |
| Cep78 | 5376 | 5269 | 4473 | 5106 | 4765 | 5257 | 5707 | 3910 | 3305 | 3415 | 3073 | 4100 | 3800 | 4410 | 5156 | 4644 | 4955 | 4673 | 4938 | 5717 | 5269 | 3322 | 3064 | 3685 | 3671 | 4401 | 4250 | 4232 | Cep78 |
| Cep83 | 7089 | 6094 | 6105 | 6681 | 6091 | 6746 | 7025 | 4623 | 4330 | 3997 | 3770 | 4337 | 4199 | 4516 | 7217 | 6492 | 5855 | 6321 | 6283 | 7863 | 6819 | 3897 | 4305 | 4121 | 4118 | 4755 | 4443 | 4291 | Cep83 |
| Cep85 | 4314 | 4656 | 3919 | 4167 | 3766 | 4044 | 4188 | 2260 | 2117 | 2332 | 2090 | 2452 | 2371 | 2582 | 4167 | 4217 | 3991 | 4222 | 3997 | 4137 | 4211 | 2019 | 2002 | 2152 | 2338 | 2537 | 2650 | 2521 | Cep85 |
| Cep85l | 2791 | 1954 | 1884 | 2457 | 3114 | 2906 | 3067 | 2963 | 2786 | 2461 | 2955 | 2183 | 2611 | 2468 | 2867 | 2623 | 2077 | 2796 | 2674 | 2977 | 2967 | 2834 | 2700 | 2264 | 2885 | 1564 | 2482 | 2144 | Cep85l |
| Cep89 | 3069 | 3143 | 3222 | 2840 | 2326 | 2599 | 2701 | 2807 | 2927 | 2930 | 2820 | 3691 | 3239 | 3746 | 2648 | 2736 | 2888 | 2597 | 2623 | 2665 | 2753 | 2557 | 2746 | 3335 | 3045 | 3861 | 3717 | 3808 | Cep89 |
| Cep95 | 2896 | 2824 | 2742 | 3361 | 2806 | 3084 | 3399 | 2893 | 2668 | 2741 | 2683 | 2683 | 2567 | 2647 | 2843 | 2609 | 2866 | 3204 | 3180 | 3230 | 3294 | 2812 | 2545 | 2748 | 2652 | 2633 | 2551 | 2553 | Cep95 |
| Cep97 | 2911 | 2392 | 2770 | 3150 | 3956 | 3866 | 3722 | 4857 | 3859 | 4286 | 4137 | 4549 | 4306 | 4705 | 3066 | 2729 | 3048 | 3250 | 3927 | 3885 | 3824 | 4171 | 3700 | 4250 | 4825 | 3859 | 4757 | 4427 | Cep97 |
| Cept1 | 2268 | 1726 | 2055 | 2045 | 1897 | 2042 | 1998 | 2981 | 2742 | 2574 | 2659 | 2463 | 2699 | 2466 | 2258 | 2206 | 2031 | 2093 | 1971 | 2131 | 1981 | 2994 | 2954 | 2658 | 2319 | 2951 | 2542 | 2754 | Cept1 |
| Cer1 | 7 | 10 | 8 | 13 | 12 | 9 | 13 | 15 | 15 | 22 | 27 | 16 | 15 | 6 | 8 | 12 | 12 | 6 | 7 | 14 | 10 | 30 | 22 | 11 | 35 | 20 | 10 | 9 | Cer1 |
| Cercam | 1517 | 1968 | 2389 | 1867 | 1834 | 1797 | 1336 | 1298 | 1433 | 1932 | 1904 | 1555 | 2262 | 1439 | 1723 | 2098 | 2298 | 1820 | 1720 | 1475 | 1406 | 1841 | 1763 | 1443 | 1603 | 1098 | 1596 | 1664 | Cercam |
| Cerk | 1847 | 2070 | 2366 | 2411 | 3256 | 2599 | 2662 | 2739 | 3087 | 3499 | 3083 | 3123 | 3173 | 3147 | 2016 | 1862 | 2506 | 2542 | 3215 | 2571 | 2851 | 3223 | 2671 | 3209 | 3399 | 3349 | 3008 | 3185 | Cerk |
| Cerkl | 153 | 144 | 89 | 110 | 106 | 107 | 108 | 111 | 80 | 109 | 88 | 86 | 124 | 71 | 193 | 188 | 82 | 75 | 84 | 133 | 147 | 77 | 82 | 62 | 46 | 49 | 92 | 75 | Cerkl |
| Cers1 | 513 | 739 | 657 | 836 | 908 | 768 | 733 | 498 | 560 | 747 | 589 | 604 | 473 | 531 | 434 | 575 | 811 | 878 | 927 | 726 | 827 | 842 | 577 | 636 | 777 | 480 | 443 | 434 | Cers1 |
| Cers2 | 2366 | 2103 | 2221 | 1598 | 2066 | 2026 | 1900 | 4976 | 5054 | 4587 | 5068 | 4872 | 4908 | 5036 | 2189 | 2256 | 1888 | 1775 | 2004 | 2019 | 1872 | 5003 | 5124 | 4619 | 4634 | 5650 | 4760 | 5090 | Cers2 |
| Cers3 | 118 | 96 | 44 | 69 | 64 | 110 | 96 | 131 | 218 | 112 | 93 | 76 | 95 | 92 | 87 | 97 | 71 | 105 | 67 | 62 | 88 | 47 | 195 | 96 | 69 | 89 | 73 | 90 | Cers3 |
| Cers4 | 4387 | 3787 | 3972 | 4597 | 6140 | 5932 | 5776 | 4903 | 4806 | 5374 | 5016 | 4615 | 4543 | 4636 | 4191 | 3814 | 4140 | 4439 | 5163 | 5276 | 5703 | 5675 | 4686 | 4306 | 5439 | 3395 | 4416 | 3895 | Cers4 |
| Cers5 | 3885 | 3047 | 3024 | 3083 | 3881 | 4005 | 3690 | 5332 | 5001 | 4856 | 5221 | 4876 | 4722 | 4997 | 3974 | 3347 | 3269 | 3274 | 3776 | 3956 | 3700 | 5280 | 4921 | 4651 | 4874 | 5313 | 4793 | 4894 | Cers5 |
| Cers6 | 6627 | 6753 | 4962 | 7163 | 9593 | 8668 | 8401 | 7797 | 7076 | 7274 | 6981 | 5420 | 6035 | 5472 | 7901 | 6900 | 6643 | 7889 | 8281 | 8531 | 8842 | 8362 | 6923 | 5554 | 7488 | 3643 | 5048 | 4469 | Cers6 |
| Cert1 | 7263 | 5056 | 6094 | 6175 | 6787 | 7219 | 7279 | 6042 | 5551 | 5748 | 6176 | 5635 | 5992 | 6064 | 6638 | 5994 | 6289 | 6695 | 6585 | 6380 | 7143 | 6243 | 5711 | 5302 | 5788 | 4944 | 5754 | 5775 | Cert1 |
| Ces1a | 0 | 0 | 0 | 0 | 0 | 0 | 1 | 0 | 2 | 4 | 0 | 0 | 0 | 0 | 0 | 0 | 0 | 3 | 0 | 1 | 0 | 0 | 0 | 0 | 3 | 1 | 0 | 0 | Ces1a |
| Ces1b | 0 | 0 | 0 | 0 | 0 | 0 | 0 | 0 | 2 | 0 | 0 | 0 | 0 | 0 | 0 | 0 | 0 | 0 | 0 | 0 | 0 | 0 | 0 | 0 | 0 | 0 | 0 | 0 | Ces1b |
| Ces1c | 0 | 1 | 0 | 0 | 0 | 0 | 0 | 0 | 0 | 0 | 0 | 0 | 0 | 1 | 0 | 1 | 0 | 0 | 0 | 0 | 0 | 0 | 1 | 0 | 0 | 0 | 0 | 0 | Ces1c |
| Ces1d | 36 | 20 | 16 | 23 | 8 | 4 | 11 | 98 | 73 | 93 | 52 | 13 | 28 | 11 | 13 | 33 | 12 | 1 | 2 | 1 | 12 | 20 | 110 | 78 | 20 | 70 | 26 | 17 | Ces1d |
| Ces1e | 0 | 0 | 0 | 0 | 0 | 0 | 0 | 0 | 2 | 0 | 1 | 0 | 0 | 0 | 0 | 0 | 0 | 0 | 0 | 0 | 0 | 0 | 2 | 0 | 0 | 1 | 0 | 0 | Ces1e |
| Ces1f | 0 | 1 | 0 | 0 | 0 | 0 | 2 | 0 | 2 | 1 | 1 | 0 | 0 | 2 | 3 | 1 | 0 | 0 | 2 | 0 | 1 | 0 | 0 | 0 | 0 | 2 | 0 | 1 | Ces1f |
| Ces1g | 0 | 0 | 0 | 0 | 0 | 0 | 0 | 2 | 1 | 0 | 0 | 0 | 0 | 0 | 1 | 0 | 0 | 0 | 0 | 0 | 0 | 0 | 1 | 0 | 0 | 0 | 0 | 1 | Ces1g |
| Ces1h | 0 | 0 | 0 | 0 | 0 | 0 | 0 | 0 | 0 | 0 | 0 | 0 | 0 | 0 | 1 | 0 | 0 | 0 | 2 | 0 | 0 | 0 | 1 | 0 | 0 | 0 | 0 | 0 | Ces1h |
| Ces2b | 1 | 8 | 16 | 2 | 10 | 7 | 13 | 13 | 21 | 16 | 9 | 17 | 35 | 15 | 18 | 9 | 2 | 10 | 12 | 9 | 11 | 15 | 16 | 16 | 20 | 21 | 16 | 19 | Ces2b |
| Ces2c | 0 | 0 | 0 | 0 | 0 | 0 | 0 | 0 | 1 | 0 | 0 | 0 | 0 | 0 | 0 | 0 | 0 | 0 | 0 | 0 | 0 | 0 | 0 | 0 | 0 | 0 | 0 | 0 | Ces2c |
| Ces2e | 0 | 0 | 0 | 0 | 0 | 0 | 0 | 0 | 2 | 6 | 5 | 0 | 3 | 1 | 0 | 0 | 0 | 1 | 0 | 1 | 0 | 5 | 1 | 0 | 0 | 1 | 0 | 0 | Ces2e |
| Ces2f | 2 | 1 | 0 | 0 | 0 | 0 | 0 | 6 | 12 | 6 | 11 | 11 | 11 | 7 | 4 | 4 | 0 | 3 | 0 | 0 | 0 | 0 | 11 | 2 | 0 | 0 | 10 | 6 | Ces2f |
| Ces2g | 0 | 2 | 0 | 0 | 0 | 0 | 0 | 4 | 6 | 3 | 10 | 8 | 8 | 3 | 1 | 1 | 0 | 0 | 0 | 0 | 0 | 3 | 6 | 7 | 3 | 2 | 0 | 1 | Ces2g |
| Ces4a | 1 | 1 | 2 | 2 | 2 | 0 | 2 | 0 | 1 | 0 | 1 | 0 | 0 | 1 | 0 | 0 | 2 | 3 | 2 | 0 | 0 | 0 | 2 | 1 | 3 | 4 | 0 | 0 | Ces4a |
| Ces5a | 8 | 15 | 6 | 14 | 25 | 14 | 23 | 17 | 21 | 17 | 18 | 17 | 33 | 14 | 25 | 9 | 6 | 12 | 23 | 14 | 24 | 22 | 20 | 27 | 14 | 11 | 9 | 7 | Ces5a |

| | | | | | | | | | | | | | | | | | | | | | | | | | | | | | |
|---|---|---|---|---|---|---|---|---|---|---|---|---|---|---|---|---|---|---|---|---|---|---|---|---|---|---|---|---|---|
| Cetn2 | 4295 | 3712 | 4226 | 3605 | 3371 | 3510 | 4261 | 3538 | 3011 | 2781 | 2597 | 3356 | 3128 | 3670 | 4515 | 4231 | 4078 | 3552 | 3532 | 4773 | 3799 | 2801 | 3079 | 3139 | 2794 | 3848 | 3636 | 3888 | Cetn2 |
| Cetn3 | 5892 | 4622 | 6205 | 6237 | 6070 | 6384 | 8042 | 9541 | 8372 | 7835 | 7534 | 8978 | 8143 | 10093 | 7052 | 5371 | 5997 | 5812 | 6901 | 8242 | 7660 | 7935 | 8228 | 9260 | 8784 | 9990 | 9166 | 10842 | Cetn3 |
| Cetn4 | 168 | 131 | 154 | 157 | 157 | 142 | 169 | 161 | 150 | 115 | 190 | 113 | 171 | 151 | 165 | 147 | 118 | 141 | 129 | 145 | 125 | 157 | 194 | 182 | 126 | 177 | 124 | 159 | Cetn4 |
| Cfap100 | 189 | 234 | 179 | 363 | 510 | 440 | 345 | 420 | 414 | 505 | 435 | 398 | 378 | 309 | 203 | 214 | 280 | 351 | 444 | 408 | 449 | 505 | 401 | 381 | 449 | 245 | 404 | 328 | Cfap100 |
| Cfap126 | 126 | 181 | 108 | 88 | 151 | 162 | 93 | 170 | 190 | 214 | 208 | 263 | 345 | 160 | 148 | 170 | 105 | 118 | 93 | 175 | 103 | 182 | 379 | 173 | 193 | 186 | 194 | 132 | Cfap126 |
| Cfap157 | 43 | 71 | 33 | 82 | 77 | 78 | 86 | 34 | 29 | 43 | 19 | 26 | 41 | 26 | 45 | 63 | 71 | 97 | 74 | 58 | 90 | 32 | 36 | 29 | 15 | 13 | 21 | 17 | Cfap157 |
| Cfap161 | 36 | 48 | 46 | 23 | 47 | 38 | 23 | 20 | 35 | 30 | 20 | 31 | 41 | 17 | 41 | 65 | 17 | 41 | 21 | 56 | 29 | 30 | 41 | 37 | 38 | 24 | 22 | 11 | Cfap161 |
| Cfap20 | 5868 | 6439 | 5319 | 6209 | 4988 | 5309 | 6014 | 3711 | 3612 | 3729 | 3662 | 4178 | 3625 | 3951 | 5942 | 5412 | 5244 | 5702 | 5293 | 5675 | 5574 | 3798 | 3719 | 4124 | 3840 | 4240 | 4045 | 4029 | Cfap20 |
| Cfap206 | 448 | 601 | 388 | 448 | 385 | 356 | 333 | 205 | 182 | 172 | 125 | 162 | 215 | 116 | 520 | 626 | 450 | 383 | 330 | 439 | 320 | 159 | 282 | 105 | 125 | 152 | 151 | 101 | Cfap206 |
| Cfap20dc | 756 | 663 | 631 | 622 | 443 | 544 | 594 | 683 | 714 | 622 | 658 | 817 | 674 | 769 | 683 | 660 | 621 | 518 | 534 | 583 | 580 | 486 | 634 | 757 | 623 | 1002 | 683 | 793 | Cfap20dc |
| Cfap221 | 39 | 34 | 38 | 33 | 30 | 17 | 21 | 33 | 21 | 26 | 20 | 33 | 42 | 23 | 33 | 44 | 36 | 36 | 27 | 19 | 32 | 20 | 43 | 7 | 13 | 16 | 6 | 17 | Cfap221 |
| Cfap298 | 619 | 733 | 751 | 658 | 755 | 713 | 704 | 918 | 1042 | 1192 | 1036 | 1203 | 1109 | 1208 | 667 | 619 | 674 | 657 | 716 | 799 | 762 | 882 | 1107 | 1146 | 1092 | 1402 | 1332 | 1158 | Cfap298 |
| Cfap299 | 20 | 16 | 9 | 14 | 31 | 24 | 26 | 24 | 29 | 19 | 24 | 25 | 26 | 12 | 22 | 23 | 24 | 13 | 24 | 28 | 28 | 20 | 29 | 26 | 23 | 20 | 17 | 13 | Cfap299 |
| Cfap300 | 2114 | 1741 | 1673 | 1861 | 1895 | 1773 | 2085 | 1286 | 1200 | 1051 | 1064 | 1132 | 1185 | 1006 | 2153 | 1967 | 1679 | 2052 | 1877 | 2108 | 2044 | 1297 | 1271 | 1079 | 1044 | 1130 | 1123 | 1216 | Cfap300 |
| Cfap36 | 5592 | 5142 | 5321 | 5610 | 5348 | 5031 | 6272 | 4485 | 3570 | 3547 | 3259 | 3755 | 3624 | 3995 | 6140 | 5224 | 5650 | 5461 | 4948 | 6524 | 5890 | 4149 | 3743 | 3754 | 3764 | 4010 | 3720 | 3944 | Cfap36 |
| Cfap410 | 1959 | 2354 | 1901 | 1906 | 1721 | 1752 | 1721 | 1097 | 1308 | 1275 | 1153 | 1269 | 1377 | 1200 | 1850 | 2047 | 1949 | 1778 | 1792 | 1745 | 1700 | 1170 | 1236 | 1279 | 1200 | 1341 | 1299 | 1277 | Cfap410 |
| Cfap43 | 129 | 159 | 35 | 108 | 306 | 239 | 150 | 180 | 354 | 269 | 220 | 266 | 335 | 83 | 160 | 291 | 48 | 133 | 162 | 222 | 227 | 336 | 570 | 92 | 199 | 51 | 74 | 52 | Cfap43 |
| Cfap44 | 847 | 987 | 327 | 617 | 1064 | 1160 | 836 | 1122 | 1091 | 853 | 706 | 690 | 1055 | 485 | 1096 | 1264 | 402 | 665 | 703 | 1255 | 1041 | 764 | 1583 | 385 | 671 | 170 | 504 | 360 | Cfap44 |
| Cfap45 | 475 | 447 | 420 | 513 | 412 | 315 | 391 | 115 | 141 | 183 | 132 | 163 | 171 | 121 | 439 | 514 | 416 | 393 | 357 | 385 | 357 | 160 | 219 | 144 | 111 | 140 | 143 | 109 | Cfap45 |
| Cfap46 | 185 | 258 | 135 | 195 | 349 | 244 | 198 | 231 | 297 | 302 | 256 | 287 | 256 | 218 | 223 | 283 | 188 | 216 | 252 | 286 | 284 | 309 | 357 | 181 | 296 | 98 | 155 | 154 | Cfap46 |
| Cfap47 | 126 | 140 | 88 | 105 | 231 | 223 | 189 | 272 | 252 | 184 | 176 | 186 | 285 | 151 | 220 | 199 | 105 | 135 | 78 | 206 | 197 | 224 | 307 | 154 | 193 | 86 | 202 | 132 | Cfap47 |
| Cfap52 | 168 | 256 | 135 | 167 | 216 | 204 | 142 | 161 | 130 | 123 | 138 | 173 | 226 | 84 | 212 | 266 | 155 | 166 | 150 | 200 | 179 | 175 | 302 | 94 | 78 | 142 | 102 | 125 | Cfap52 |
| Cfap53 | 92 | 115 | 96 | 175 | 144 | 136 | 154 | 73 | 69 | 63 | 74 | 68 | 92 | 71 | 138 | 103 | 88 | 167 | 140 | 114 | 147 | 99 | 106 | 60 | 79 | 69 | 45 | 44 | Cfap53 |
| Cfap54 | 549 | 666 | 332 | 568 | 607 | 685 | 613 | 473 | 422 | 349 | 304 | 355 | 439 | 255 | 686 | 656 | 399 | 470 | 491 | 806 | 677 | 393 | 583 | 218 | 352 | 173 | 341 | 255 | Cfap54 |
| Cfap57 | 103 | 161 | 141 | 97 | 169 | 127 | 117 | 201 | 184 | 187 | 188 | 169 | 249 | 146 | 144 | 171 | 102 | 147 | 159 | 151 | 139 | 229 | 213 | 112 | 167 | 143 | 141 | 204 | Cfap57 |
| Cfap58 | 30 | 48 | 22 | 66 | 44 | 47 | 47 | 46 | 30 | 32 | 32 | 49 | 41 | 18 | 47 | 40 | 74 | 52 | 37 | 65 | 43 | 42 | 40 | 27 | 27 | 23 | 16 | 31 | Cfap58 |
| Cfap61 | 131 | 198 | 150 | 152 | 156 | 162 | 131 | 144 | 119 | 131 | 117 | 116 | 129 | 71 | 202 | 207 | 161 | 157 | 213 | 149 | 145 | 184 | 196 | 129 | 92 | 76 | 93 | 100 | Cfap61 |
| Cfap65 | 213 | 336 | 136 | 270 | 488 | 305 | 359 | 374 | 410 | 429 | 352 | 507 | 643 | 287 | 306 | 530 | 271 | 297 | 402 | 402 | 328 | 536 | 741 | 275 | 369 | 210 | 339 | 275 | Cfap65 |
| Cfap69 | 3712 | 3143 | 2991 | 3728 | 3948 | 4022 | 4369 | 1371 | 1192 | 1292 | 1224 | 1270 | 1402 | 1128 | 3948 | 3704 | 3174 | 3800 | 3663 | 4187 | 4305 | 1379 | 1445 | 987 | 1079 | 803 | 1039 | 1113 | Cfap69 |
| Cfap70 | 324 | 295 | 248 | 253 | 277 | 303 | 352 | 154 | 136 | 91 | 114 | 88 | 144 | 96 | 287 | 349 | 183 | 302 | 316 | 288 | 327 | 147 | 184 | 108 | 120 | 75 | 94 | 107 | Cfap70 |
| Cfap73 | 3 | 5 | 0 | 8 | 11 | 4 | 7 | 11 | 18 | 21 | 12 | 25 | 8 | 12 | 9 | 13 | 2 | 7 | 2 | 9 | 5 | 28 | 31 | 11 | 14 | 20 | 14 | 18 | Cfap73 |
| Cfap74 | 442 | 473 | 396 | 493 | 567 | 465 | 468 | 562 | 600 | 650 | 591 | 797 | 741 | 681 | 463 | 522 | 399 | 431 | 515 | 545 | 462 | 622 | 695 | 583 | 602 | 625 | 788 | 733 | Cfap74 |
| Cfap77 | 174 | 215 | 248 | 219 | 193 | 182 | 125 | 51 | 53 | 61 | 55 | 61 | 82 | 60 | 145 | 159 | 217 | 200 | 199 | 127 | 180 | 69 | 69 | 51 | 62 | 55 | 52 | 50 | Cfap77 |
| Cfap91 | 85 | 107 | 125 | 82 | 101 | 80 | 86 | 123 | 104 | 157 | 122 | 106 | 144 | 98 | 100 | 97 | 102 | 74 | 88 | 69 | 93 | 157 | 248 | 92 | 112 | 110 | 85 | 103 | Cfap91 |
| Cfap97 | 2171 | 1831 | 2045 | 1792 | 1924 | 1991 | 1949 | 3305 | 3009 | 2831 | 2783 | 2790 | 2658 | 2569 | 2345 | 2022 | 1999 | 1895 | 1986 | 2241 | 1959 | 3098 | 3036 | 2708 | 2565 | 2676 | 3030 | 2649 | Cfap97 |
| Cfap97d1 | 1 | 1 | 0 | 7 | 0 | 3 | 1 | 6 | 1 | 0 | 4 | 7 | 0 | 1 | 3 | 2 | 0 | 0 | 0 | 1 | 1 | 0 | 0 | 0 | 0 | 0 | 3 | 0 | Cfap97d1 |
| Cfap97d2 | 61 | 72 | 64 | 77 | 124 | 75 | 103 | 100 | 59 | 96 | 87 | 127 | 110 | 139 | 87 | 69 | 78 | 70 | 91 | 129 | 89 | 77 | 116 | 87 | 107 | 90 | 126 | 123 | Cfap97d2 |
| Cfap99 | 10 | 22 | 17 | 4 | 20 | 7 | 7 | 1 | 8 | 8 | 9 | 7 | 17 | 9 | 7 | 17 | 5 | 7 | 12 | 10 | 8 | 5 | 25 | 17 | 17 | 6 | 2 | 4 | Cfap99 |
| Cfb | 0 | 0 | 0 | 0 | 0 | 0 | 0 | 0 | 0 | 0 | 0 | 7 | 1 | 1 | 0 | 0 | 6 | 0 | 0 | 2 | 0 | 0 | 0 | 0 | 0 | 2 | 0 | 0 | Cfb |
| Cfc1 | 5 | 3 | 4 | 3 | 2 | 0 | 0 | 7 | 4 | 4 | 9 | 1 | 0 | 0 | 1 | 1 | 0 | 0 | 0 | 0 | 0 | 0 | 2 | 0 | 1 | 0 | 0 | 1 | Cfc1 |
| Cfd | 1 | 0 | 0 | 0 | 0 | 0 | 0 | 0 | 0 | 3 | 0 | 0 | 0 | 0 | 0 | 0 | 0 | 0 | 0 | 0 | 0 | 2 | 0 | 0 | 0 | 0 | 0 | 0 | Cfd |
| Cfdp1 | 10558 | 9539 | 11294 | 9493 | 7761 | 7540 | 9857 | 9318 | 8511 | 7866 | 8624 | 8095 | 8255 | 10294 | 10760 | 10137 | 9812 | 8730 | 8114 | 9883 | 8851 | 7735 | 8517 | 8880 | 7845 | 9936 | 8886 | 10350 | Cfdp1 |
| Cfh | 600 | 482 | 645 | 673 | 1430 | 1154 | 1017 | 1099 | 1284 | 1117 | 1416 | 949 | 870 | 956 | 684 | 587 | 607 | 1133 | 1051 | 1505 | 1261 | 1708 | 2181 | 1138 | 1262 | 771 | 867 | 786 | Cfh |
| Cfhr2 | 0 | 1 | 0 | 0 | 3 | 2 | 0 | 0 | 0 | 0 | 2 | 0 | 5 | 2 | 0 | 2 | 0 | 0 | 6 | 1 | 4 | 2 | 3 | 2 | 1 | 4 | 7 | 1 | Cfhr2 |
| Cfhr4 | 0 | 0 | 0 | 0 | 0 | 0 | 1 | 0 | 1 | 0 | 0 | 0 | 0 | 1 | 0 | 1 | 1 | 0 | 1 | 1 | 0 | 0 | 1 | 0 | 0 | 0 | 0 | 0 | Cfhr4 |
| Cfi | 7 | 17 | 12 | 5 | 9 | 7 | 7 | 11 | 22 | 19 | 21 | 8 | 16 | 9 | 16 | 9 | 13 | 9 | 8 | 5 | 5 | 18 | 19 | 17 | 9 | 17 | 7 | 6 | Cfi |
| Cfl1 | 60557 | 74060 | 67946 | 64624 | 61080 | 60887 | 68659 | 61766 | 60556 | 63517 | 58450 | 66749 | 68241 | 78647 | 62621 | 57323 | 67525 | 62055 | 65365 | 69699 | 68116 | 55469 | 59383 | 68768 | 68644 | 74737 | 72430 | 80184 | Cfl1 |
| Cfl2 | 9260 | 6900 | 8451 | 9157 | 10394 | 9736 | 12291 | 13597 | 11044 | 10655 | 11375 | 12250 | 11154 | 13892 | 10315 | 7990 | 8948 | 9118 | 10476 | 10125 | 11566 | 13237 | 11958 | 11988 | 13014 | 13157 | 12529 | 13887 | Cfl2 |
| Cflar | 1688 | 1424 | 1707 | 1323 | 1422 | 1414 | 1219 | 1756 | 1912 | 1942 | 2359 | 1801 | 2080 | 1926 | 1640 | 1733 | 1570 | 1304 | 1367 | 1379 | 1324 | 1781 | 2141 | 1843 | 1630 | 1842 | 1929 | 1864 | Cflar |
| Cfp | 1628 | 2382 | 2237 | 2620 | 2441 | 2223 | 2400 | 1323 | 1244 | 1652 | 1482 | 1492 | 1373 | 1365 | 1664 | 2154 | 2695 | 2685 | 2667 | 2146 | 2285 | 1823 | 1220 | 1568 | 1620 | 1373 | 1541 | 1355 | Cfp |
| Cftr | 88 | 67 | 27 | 47 | 46 | 72 | 58 | 123 | 147 | 119 | 123 | 41 | 77 | 45 | 86 | 93 | 45 | 45 | 43 | 85 | 68 | 58 | 169 | 69 | 68 | 73 | 63 | 42 | Cftr |
| Cga | 57 | 37 | 1 | 2 | 34 | 0 | 0 | 42 | 42 | 8 | 29 | 1 | 1 | 0 | 47 | 17 | 2 | 48 | 1 | 1 | 1 | 60 | 45 | 10 | 0 | 30 | 0 | 0 | Cga |
| Cgas | 146 | 98 | 178 | 140 | 173 | 141 | 141 | 139 | 204 | 195 | 261 | 129 | 121 | 120 | 140 | 153 | 112 | 122 | 116 | 119 | 119 | 177 | 183 | 166 | 128 | 172 | 109 | 126 | Cgas |
| Cggbp1 | 15892 | 13306 | 14937 | 16247 | 13902 | 14419 | 16455 | 13694 | 11674 | 11566 | 12506 | 12888 | 12420 | 13696 | 15762 | 15560 | 16004 | 17087 | 14675 | 14031 | 15781 | 13578 | 11829 | 12754 | 13463 | 13347 | 13108 | 14291 | Cggbp1 |
| Cgn | 3234 | 3704 | 3297 | 4321 | 3228 | 3248 | 3272 | 900 | 1049 | 1180 | 1087 | 803 | 958 | 804 | 2934 | 3849 | 3627 | 4171 | 3252 | 2988 | 3316 | 864 | 967 | 1121 | 888 | 958 | 787 | 782 | Cgn |
| Cgnl1 | 3917 | 4081 | 4320 | 2937 | 2950 | 2962 | 2048 | 2363 | 2880 | 3206 | 3937 | 2778 | 3628 | 2788 | 3683 | 5086 | 3078 | 3258 | 2123 | 2441 | 2208 | 2642 | 3511 | 2926 | 2505 | 2710 | 2897 | 2942 | Cgnl1 |
| Cgref1 | 136 | 149 | 105 | 155 | 331 | 255 | 207 | 454 | 511 | 480 | 443 | 313 | 352 | 329 | 156 | 147 | 136 | 249 | 256 | 260 | 246 | 612 | 556 | 360 | 408 | 297 | 315 | 337 | Cgref1 |
| Cgrrf1 | 1610 | 1513 | 1335 | 1480 | 1060 | 1131 | 1347 | 758 | 605 | 526 | 573 | 641 | 619 | 608 | 1577 | 1393 | 1262 | 1208 | 1079 | 1384 | 1213 | 581 | 616 | 596 | 629 | 643 | 801 | 674 | Cgrrf1 |
| Ch25h | 55 | 64 | 39 | 41 | 13 | 34 | 10 | 67 | 140 | 120 | 94 | 64 | 104 | 97 | 55 | 58 | 29 | 58 | 30 | 7 | 5 | 63 | 178 | 131 | 68 | 60 | 52 | 66 | Ch25h |
| Chac1 | 134 | 168 | 98 | 222 | 187 | 174 | 149 | 56 | 98 | 91 | 98 | 39 | 49 | 65 | 117 | 158 | 149 | 208 | 215 | 203 | 181 | 84 | 100 | 92 | 102 | 83 | 53 | 52 | Chac1 |
| Chac2 | 569 | 496 | 679 | 646 | 665 | 682 | 783 | 523 | 459 | 484 | 421 | 386 | 429 | 531 | 658 | 535 | 665 | 682 | 586 | 748 | 771 | 481 | 509 | 492 | 478 | 412 | 399 | 564 | Chac2 |
| Chad | 3 | 10 | 6 | 9 | 5 | 9 | 1 | 21 | 71 | 93 | 167 | 0 | 5 | 9 | 3 | 18 | 2 | 8 | 9 | 2 | 3 | 70 | 37 | 69 | 30 | 50 | 2 | 4 | Chad |
| Chadl | 35 | 54 | 47 | 80 | 128 | 119 | 69 | 209 | 395 | 376 | 626 | 156 | 179 | 114 | 42 | 39 | 49 | 105 | 118 | 76 | 65 | 506 | 293 | 267 | 297 | 170 | 122 | 95 | Chadl |
| Chaf1a | 9141 | 9688 | 10848 | 8622 | 5870 | 7099 | 7049 | 4419 | 5115 | 5236 | 5189 | 7988 | 6312 | 7628 | 7643 | 8730 | 9123 | 7770 | 7561 | 6265 | 6966 | 3920 | 4807 | 6551 | 5360 | 8970 | 7645 | 8243 | Chaf1a |
| Chaf1b | 5563 | 5793 | 5229 | 4675 | 3472 | 4217 | 4242 | 2142 | 2311 | 1968 | 1860 | 2903 | 2584 | 2762 | 4923 | 5298 | 4713 | 4173 | 4285 | 4464 | 4133 | 1532 | 2045 | 2314 | 2004 | 3781 | 3277 | 3083 | Chaf1b |
| Champ1 | 5789 | 5530 | 5762 | 5254 | 5519 | 5325 | 5782 | 6848 | 6357 | 6403 | 6040 | 6977 | 6304 | 7112 | 5736 | 5084 | 5614 | 5130 | 5472 | 5648 | 5489 | 7165 | 5954 | 6887 | 6733 | 7102 | 6979 | 7027 | Champ1 |
| Chat | 5 | 31 | 4 | 22 | 92 | 29 | 20 | 35 | 68 | 65 | 92 | 13 | 62 | 12 | 44 | 26 | 41 | 34 | 23 | 28 | 27 | 74 | 40 | 11 | 21 | 6 | 23 | 1 | Chat |
| Chchd1 | 2144 | 1962 | 1959 | 2029 | 1849 | 1920 | 2312 | 3063 | 2633 | 2703 | 2683 | 2737 | 2926 | 3502 | 2116 | 1700 | 2015 | 1753 | 2011 | 2388 | 2169 | 2284 | 2582 | 3108 | 2818 | 3409 | 3211 | 3465 | Chchd1 |
| Chchd10 | 493 | 649 | 563 | 660 | 575 | 512 | 492 | 149 | 198 | 154 | 231 | 165 | 125 | 136 | 496 | 526 | 651 | 587 | 588 | 505 | 503 | 169 | 223 | 163 | 188 | 160 | 108 | 139 | Chchd10 |
| Chchd2 | 11664 | 12770 | 11949 | 12052 | 10627 | 9827 | 10699 | 11573 | 10926 | 10598 | 10483 | 11354 | 10796 | 11610 | 11238 | 10009 | 12553 | 10568 | 11719 | 12213 | 10759 | 8539 | 11079 | 12751 | 11433 | 13060 | 12022 | 11859 | Chchd2 |
| Chchd3 | 6436 | 6088 | 5999 | 5275 | 4720 | 4924 | 5595 | 5254 | 4571 | 4502 | 4049 | 4708 | 4460 | 5362 | 6636 | 5345 | 5895 | 5006 | 5092 | 5967 | 5525 | 3999 | 4432 | 4696 | 4236 | 5407 | 4659 | 5486 | Chchd3 |
| Chchd4 | 1442 | 1557 | 1562 | 1514 | 1370 | 1348 | 1526 | 1612 | 1385 | 1310 | 1241 | 1428 | 1411 | 1612 | 1468 | 1363 | 1422 | 1324 | 1348 | 1650 | 1521 | 1282 | 1383 | 1447 | 1443 | 1660 | 1569 | 1593 | Chchd4 |
| Chchd5 | 533 | 746 | 682 | 551 | 603 | 670 | 658 | 718 | 826 | 800 | 801 | 840 | 821 | 933 | 628 | 628 | 681 | 593 | 651 | 618 | 612 | 620 | 825 | 791 | 725 | 978 | 976 | 846 | Chchd5 |
| Chchd6 | 1607 | 2109 | 1710 | 1836 | 1865 | 1778 | 1922 | 1561 | 1790 | 1599 | 1424 | 1602 | 1501 | 1823 | 1625 | 1516 | 1809 | 1738 | 1919 | 2024 | 1920 | 1477 | 1706 | 1607 | 1563 | 1628 | 1677 | 1768 | Chchd6 |
| Chchd7 | 6036 | 5644 | 5491 | 6878 | 4936 | 5536 | 6349 | 1840 | 1603 | 1745 | 1778 | 1968 | 2140 | 2083 | 6125 | 5799 | 5749 | 6343 | 4932 | 5376 | 5938 | 1661 | 1570 | 1816 | 1699 | 1949 | 2045 | 2048 | Chchd7 |
| Chd1 | 11277 | 9627 | 9530 | 9352 | 8881 | 10136 | 10208 | 9149 | 7796 | 7472 | 7679 | 9545 | 9319 | 9819 | 11206 | 10629 | 8998 | 9110 | 9019 | 10246 | 9980 | 6737 | 7335 | 7560 | 7725 | 7189 | 9897 | 9635 | Chd1 |
| Chd1l | 437 | 384 | 396 | 294 | 243 | 352 | 247 | 1027 | 1114 | 1046 | 1212 | 1410 | 1428 | 1362 | 402 | 357 | 315 | 230 | 282 | 290 | 264 | 862 | 1188 | 1176 | 1043 | 1650 | 1513 | 1425 | Chd1l |
| Chd2 | 13720 | 12132 | 9045 | 14390 | 14606 | 15980 | 16692 | 11199 | 10561 | 10490 | 11596 | 11638 | 11420 | 12668 | 14054 | 11854 | 11201 | 13871 | 13492 | 15748 | 16156 | 8983 | 10330 | 10360 | 11825 | 7803 | 11525 | 10558 | Chd2 |
| Chd3 | 22277 | 23213 | 24520 | 28099 | 30480 | 30003 | 26755 | 43125 | 52822 | 61025 | 56215 | 54513 | 47194 | 52810 | 20661 | 23523 | 27195 | 29792 | 30543 | 23738 | 29064 | 54507 | 48029 | 63480 | 61642 | 46624 | 47283 | 47148 | Chd3 |

| | | | | | | | | | | | | | | | | | | | | | | | | | | | | | |
|---|---|---|---|---|---|---|---|---|---|---|---|---|---|---|---|---|---|---|---|---|---|---|---|---|---|---|---|---|---|
| Chd4 | 59991 | 62514 | 77083 | 73453 | 64138 | 65894 | 67182 | 40729 | 43039 | 53945 | 48610 | 65694 | 60461 | 67897 | 57430 | 64001 | 75668 | 69290 | 71565 | 51208 | 68563 | 49526 | 40417 | 58667 | 58480 | 54820 | 60837 | 66987 | Chd4 |
| Chd5 | 1266 | 1487 | 1300 | 3764 | 6545 | 4497 | 4618 | 3573 | 5049 | 6530 | 5401 | 3457 | 1999 | 3424 | 1377 | 1177 | 2481 | 4413 | 5209 | 3813 | 5068 | 7519 | 4023 | 5233 | 7673 | 1772 | 1964 | 2243 | Chd5 |
| Chd6 | 14681 | 14653 | 7758 | 11198 | 13361 | 14749 | 14872 | 17574 | 17145 | 14267 | 15068 | 14468 | 14351 | 15055 | 14308 | 13404 | 9741 | 11342 | 12645 | 15799 | 15656 | 12088 | 15367 | 11655 | 15616 | 8872 | 15689 | 13013 | Chd6 |
| Chd7 | 8405 | 9439 | 10139 | 11745 | 13565 | 12361 | 12524 | 11407 | 10492 | 14666 | 12099 | 19492 | 17939 | 19623 | 8931 | 9495 | 11471 | 11214 | 12655 | 10695 | 12835 | 12539 | 10200 | 14348 | 16453 | 11160 | 19866 | 18579 | Chd7 |
| Chd8 | 23469 | 23688 | 16737 | 22043 | 21251 | 23413 | 23309 | 22351 | 22373 | 21249 | 20405 | 23177 | 21675 | 23202 | 23475 | 21235 | 18967 | 20710 | 21394 | 22383 | 24479 | 19281 | 20235 | 18854 | 22662 | 14809 | 23373 | 22341 | Chd8 |
| Chd9 | 10250 | 9236 | 5561 | 7843 | 9749 | 10385 | 9736 | 13050 | 12006 | 10361 | 11032 | 9760 | 10453 | 10224 | 11003 | 10971 | 6598 | 7828 | 7893 | 11157 | 10601 | 9807 | 11103 | 7620 | 11145 | 5313 | 10796 | 8551 | Chd9 |
| Chdh | 9 | 9 | 20 | 8 | 23 | 16 | 9 | 22 | 54 | 45 | 57 | 5 | 18 | 9 | 8 | 16 | 8 | 11 | 18 | 10 | 12 | 42 | 63 | 25 | 26 | 7 | 7 | 4 | Chdh |
| Chek1 | 3229 | 2216 | 3311 | 2253 | 1694 | 2009 | 2465 | 3313 | 2760 | 2435 | 2564 | 3716 | 3302 | 3794 | 2647 | 2539 | 2630 | 2060 | 2220 | 2452 | 1952 | 2122 | 2462 | 3179 | 2738 | 4762 | 4094 | 4225 | Chek1 |
| Chek2 | 1662 | 1563 | 1998 | 1709 | 1225 | 1373 | 1640 | 1175 | 1018 | 1057 | 1002 | 1587 | 1413 | 1635 | 1551 | 1703 | 1656 | 1631 | 1406 | 1440 | 1364 | 872 | 972 | 1387 | 1122 | 1758 | 1727 | 1730 | Chek2 |
| Cherp | 5244 | 6861 | 8218 | 7244 | 6275 | 5936 | 5668 | 4980 | 6513 | 7148 | 6793 | 8720 | 7077 | 7714 | 4944 | 6309 | 7642 | 6584 | 7119 | 4818 | 5580 | 6660 | 5935 | 7970 | 7187 | 9477 | 7204 | 7418 | Cherp |
| Chfr | 8707 | 9288 | 8491 | 9308 | 9327 | 9376 | 9947 | 7778 | 6587 | 7150 | 6299 | 7346 | 6704 | 7232 | 9521 | 8216 | 8841 | 9263 | 9524 | 9818 | 9967 | 7663 | 6631 | 7027 | 7676 | 6347 | 7137 | 7056 | Chfr |
| Chga | 2390 | 3580 | 3324 | 5280 | 5654 | 4697 | 4997 | 2144 | 2460 | 3038 | 2738 | 2523 | 2155 | 2404 | 3050 | 2792 | 4545 | 5588 | 5594 | 4460 | 5694 | 3178 | 2557 | 2853 | 3291 | 2044 | 1951 | 2144 | Chga |
| Chgb | 1768 | 2329 | 2913 | 3421 | 4644 | 3530 | 3384 | 3473 | 3036 | 4050 | 3051 | 3236 | 3701 | 3509 | 2942 | 2217 | 3427 | 3311 | 3657 | 4057 | 3620 | 4806 | 3048 | 3273 | 4256 | 1873 | 3301 | 3202 | Chgb |
| Chia1 | 16 | 43 | 39 | 25 | 75 | 46 | 56 | 31 | 35 | 69 | 44 | 48 | 69 | 38 | 37 | 40 | 46 | 55 | 37 | 51 | 46 | 50 | 69 | 47 | 28 | 48 | 42 | 44 | Chia1 |
| Chic1 | 2526 | 1948 | 2258 | 2731 | 3288 | 3166 | 3521 | 3179 | 2444 | 2774 | 2961 | 2785 | 2858 | 3160 | 2730 | 2159 | 2538 | 2621 | 2991 | 2993 | 3413 | 3004 | 2477 | 2534 | 3006 | 1974 | 3052 | 3006 | Chic1 |
| Chic2 | 3035 | 2224 | 2324 | 2911 | 2975 | 2936 | 3322 | 2821 | 2383 | 2337 | 2405 | 2429 | 2116 | 2749 | 2921 | 2425 | 2589 | 2779 | 2783 | 2899 | 3186 | 2622 | 2275 | 2390 | 2541 | 2611 | 2451 | 2493 | Chic2 |
| Chid1 | 3313 | 3602 | 2374 | 2897 | 2926 | 3303 | 2968 | 2931 | 3185 | 2865 | 2948 | 2469 | 2609 | 2439 | 3570 | 3267 | 2526 | 3192 | 2846 | 3328 | 3157 | 2796 | 3105 | 2527 | 2628 | 2146 | 2719 | 2372 | Chid1 |
| Chil1 | 17 | 26 | 13 | 21 | 26 | 29 | 26 | 40 | 67 | 64 | 136 | 53 | 40 | 47 | 46 | 31 | 31 | 13 | 16 | 36 | 21 | 53 | 74 | 46 | 53 | 65 | 57 | 52 | Chil1 |
| Chil3 | 0 | 0 | 0 | 0 | 0 | 0 | 0 | 0 | 1 | 0 | 1 | 0 | 3 | 0 | 1 | 0 | 0 | 0 | 0 | 0 | 0 | 0 | 0 | 0 | 0 | 2 | 0 | 0 | Chil3 |
| Chil4 | 0 | 0 | 0 | 0 | 0 | 0 | 0 | 0 | 0 | 0 | 0 | 0 | 0 | 0 | 0 | 0 | 0 | 0 | 0 | 0 | 0 | 0 | 0 | 2 | 0 | 0 | 0 | 0 | Chil4 |
| Chil5 | 1 | 4 | 4 | 0 | 1 | 2 | 0 | 0 | 3 | 4 | 1 | 0 | 0 | 1 | 3 | 1 | 0 | 0 | 1 | 0 | 1 | 0 | 2 | 0 | 0 | 1 | 0 | 2 | Chil5 |
| Chil6 | 0 | 0 | 0 | 0 | 0 | 0 | 0 | 0 | 0 | 0 | 0 | 0 | 1 | 0 | 0 | 0 | 0 | 2 | 0 | 0 | 0 | 0 | 0 | 0 | 0 | 0 | 0 | 0 | Chil6 |
| Chit1 | 0 | 0 | 2 | 5 | 2 | 0 | 3 | 6 | 9 | 0 | 1 | 7 | 1 | 1 | 4 | 1 | 0 | 0 | 1 | 1 | 2 | 0 | 6 | 2 | 8 | 0 | 0 | 0 | Chit1 |
| Chka | 9421 | 8789 | 7402 | 7983 | 8413 | 9019 | 8942 | 5897 | 5538 | 4803 | 5092 | 5166 | 5082 | 5137 | 9013 | 9195 | 7422 | 8029 | 8064 | 8574 | 9149 | 4680 | 5052 | 4430 | 4952 | 4721 | 5555 | 4989 | Chka |
| Chkb | 104 | 108 | 152 | 193 | 139 | 144 | 172 | 106 | 83 | 123 | 113 | 92 | 115 | 184 | 102 | 62 | 153 | 121 | 136 | 111 | 192 | 140 | 78 | 204 | 144 | 117 | 115 | 165 | Chkb |
| Chl1 | 3865 | 4270 | 3099 | 6875 | 13874 | 10771 | 10561 | 20655 | 17015 | 16526 | 13727 | 8804 | 10231 | 9572 | 7530 | 4691 | 5293 | 8612 | 9575 | 12620 | 11773 | 21731 | 16453 | 10293 | 17959 | 4679 | 8519 | 8133 | Chl1 |
| Chm | 3691 | 2640 | 2724 | 2663 | 3009 | 3279 | 3295 | 4786 | 4308 | 3537 | 3641 | 3966 | 3821 | 3987 | 4212 | 3370 | 2960 | 2709 | 2856 | 3917 | 3270 | 3920 | 4262 | 3382 | 3646 | 3523 | 4033 | 4101 | Chm |
| Chml | 3489 | 2747 | 3083 | 2768 | 3034 | 3226 | 3257 | 3160 | 2248 | 2292 | 2072 | 2723 | 2954 | 2995 | 3679 | 3287 | 2858 | 2625 | 3067 | 3306 | 3001 | 2526 | 2214 | 2062 | 2459 | 1994 | 3211 | 3170 | Chml |
| Chmp1a | 6204 | 7832 | 5749 | 6623 | 5867 | 6118 | 5632 | 5726 | 6521 | 5646 | 5596 | 6228 | 5810 | 5234 | 6247 | 6400 | 6280 | 6279 | 6760 | 6439 | 5961 | 5576 | 6559 | 5970 | 6381 | 6408 | 6276 | 5745 | Chmp1a |
| Chmp1b | 4095 | 3982 | 4067 | 3945 | 3238 | 3544 | 3865 | 3975 | 3023 | 2924 | 2769 | 3448 | 3463 | 3782 | 4192 | 3963 | 3691 | 3438 | 3468 | 3910 | 3840 | 3262 | 3280 | 3362 | 3356 | 3569 | 3956 | 4170 | Chmp1b |
| Chmp2a | 2841 | 3220 | 3236 | 3239 | 3299 | 3232 | 2998 | 4113 | 3980 | 4249 | 4032 | 4126 | 4009 | 4398 | 3152 | 2748 | 3540 | 3215 | 3574 | 3540 | 3133 | 4182 | 4166 | 4398 | 4702 | 4425 | 4148 | 4264 | Chmp2a |
| Chmp2b | 6436 | 4216 | 5803 | 5541 | 6235 | 5536 | 5479 | 3585 | 3598 | 3519 | 4287 | 2257 | 2925 | 2431 | 5921 | 5590 | 5225 | 5746 | 5252 | 4949 | 5460 | 3347 | 4538 | 3024 | 2802 | 3381 | 2292 | 2671 | Chmp2b |
| Chmp3 | 1780 | 1564 | 1892 | 1712 | 1804 | 1902 | 1721 | 5135 | 5085 | 4853 | 4856 | 4857 | 4612 | 5033 | 2031 | 1672 | 1822 | 1742 | 1864 | 1931 | 1726 | 4752 | 5237 | 4778 | 4673 | 5093 | 4660 | 5026 | Chmp3 |
| Chmp4b | 7867 | 8784 | 9617 | 9108 | 7945 | 7603 | 8006 | 7297 | 7105 | 7619 | 7488 | 7715 | 7325 | 7999 | 7820 | 7705 | 9288 | 8383 | 8605 | 7789 | 8325 | 6909 | 7609 | 8208 | 7820 | 8862 | 8118 | 8293 | Chmp4b |
| Chmp4c | 65 | 55 | 56 | 11 | 29 | 32 | 3 | 35 | 77 | 47 | 55 | 31 | 34 | 14 | 75 | 58 | 21 | 17 | 13 | 9 | 7 | 40 | 71 | 29 | 17 | 23 | 40 | 24 | Chmp4c |
| Chmp5 | 6515 | 6143 | 6850 | 6717 | 6268 | 6130 | 7006 | 7767 | 6675 | 6957 | 6407 | 6903 | 6639 | 7517 | 7580 | 6699 | 7165 | 6346 | 6653 | 8460 | 6758 | 6679 | 6864 | 7118 | 7191 | 7233 | 7161 | 7244 | Chmp5 |
| Chmp6 | 1569 | 1746 | 2022 | 1684 | 1568 | 1588 | 1669 | 1729 | 2185 | 2400 | 2067 | 2452 | 2188 | 2429 | 1429 | 1526 | 1936 | 1554 | 1813 | 1600 | 1780 | 1836 | 1986 | 2396 | 2122 | 3031 | 2407 | 2662 | Chmp6 |
| Chmp7 | 8139 | 8499 | 8265 | 9072 | 8819 | 8798 | 9233 | 7565 | 6967 | 7306 | 6884 | 7643 | 6717 | 7195 | 8335 | 7432 | 8759 | 9178 | 9902 | 8562 | 9343 | 8133 | 6914 | 7701 | 7937 | 7543 | 7458 | 7541 | Chmp7 |
| Chn1 | 5144 | 5527 | 5210 | 6057 | 7487 | 6266 | 7476 | 7610 | 6567 | 7355 | 5887 | 6975 | 6542 | 7278 | 6191 | 4886 | 6101 | 6185 | 6952 | 7155 | 7325 | 8221 | 6193 | 7030 | 7731 | 6323 | 6949 | 7206 | Chn1 |
| Chn2 | 1100 | 1164 | 1166 | 1268 | 1652 | 1479 | 1669 | 1854 | 1834 | 1933 | 1575 | 1791 | 1278 | 1782 | 1238 | 1109 | 1286 | 1409 | 1630 | 1618 | 1522 | 2421 | 1652 | 1849 | 2220 | 1746 | 1713 | 1625 | Chn2 |
| Chodl | 391 | 155 | 144 | 344 | 441 | 277 | 306 | 552 | 785 | 688 | 1255 | 276 | 358 | 275 | 339 | 273 | 251 | 308 | 298 | 260 | 317 | 620 | 1020 | 463 | 485 | 392 | 212 | 280 | Chodl |
| Chordc1 | 9788 | 7461 | 9410 | 9255 | 9160 | 9174 | 10173 | 12875 | 10369 | 9978 | 10685 | 10861 | 10746 | 11262 | 10291 | 9322 | 9295 | 8488 | 9043 | 10020 | 9311 | 11593 | 10255 | 10832 | 10614 | 10251 | 11293 | 11322 | Chordc1 |
| Chp1 | 15141 | 12989 | 11696 | 12141 | 11331 | 11353 | 12752 | 6736 | 6180 | 5760 | 5656 | 6169 | 6004 | 6365 | 14181 | 12753 | 12133 | 12676 | 12212 | 11348 | 12776 | 6490 | 6437 | 5940 | 5974 | 6718 | 6136 | 6276 | Chp1 |
| Chp2 | 5 | 10 | 21 | 5 | 38 | 45 | 22 | 116 | 104 | 124 | 107 | 99 | 85 | 103 | 11 | 7 | 20 | 23 | 26 | 25 | 20 | 112 | 91 | 85 | 95 | 65 | 85 | 90 | Chp2 |
| Chpf | 2876 | 4039 | 3367 | 3828 | 3195 | 3345 | 2670 | 1577 | 1973 | 2023 | 2279 | 1766 | 1692 | 1488 | 2835 | 3434 | 3548 | 3825 | 3427 | 2597 | 2876 | 2361 | 2151 | 1803 | 1943 | 1724 | 1707 | 1518 | Chpf |
| Chpf2 | 2832 | 3490 | 3463 | 3285 | 3050 | 3068 | 2668 | 1856 | 2506 | 2563 | 2927 | 2725 | 2853 | 2473 | 2672 | 3277 | 3427 | 3341 | 3207 | 2455 | 2686 | 2605 | 2401 | 2690 | 2803 | 2666 | 2797 | 2515 | Chpf2 |
| Chpt1 | 3599 | 2064 | 2911 | 2451 | 2068 | 2814 | 2362 | 2348 | 2431 | 2012 | 2426 | 2105 | 2367 | 2169 | 3233 | 3340 | 2342 | 2469 | 2005 | 2353 | 2334 | 1910 | 2504 | 1968 | 1873 | 2073 | 1985 | 2219 | Chpt1 |
| Chrac1 | 438 | 439 | 544 | 353 | 387 | 382 | 432 | 1095 | 1083 | 1053 | 1032 | 1122 | 1168 | 1262 | 483 | 400 | 400 | 374 | 369 | 459 | 383 | 837 | 1111 | 1098 | 985 | 1404 | 1228 | 1207 | Chrac1 |
| Chrd | 905 | 1134 | 1293 | 1340 | 1371 | 1432 | 1094 | 793 | 871 | 964 | 1426 | 942 | 1058 | 902 | 832 | 1122 | 1328 | 1332 | 1318 | 1086 | 1093 | 1200 | 946 | 1016 | 936 | 828 | 924 | 813 | Chrd |
| Chrdl1 | 1054 | 915 | 839 | 718 | 772 | 1071 | 716 | 891 | 1016 | 743 | 1515 | 784 | 1063 | 767 | 1240 | 1249 | 929 | 807 | 853 | 940 | 838 | 1044 | 1316 | 728 | 769 | 595 | 854 | 761 | Chrdl1 |
| Chrdl2 | 35 | 14 | 11 | 16 | 7 | 10 | 7 | 10 | 8 | 16 | 40 | 4 | 10 | 6 | 20 | 32 | 3 | 8 | 13 | 7 | 2 | 3 | 26 | 6 | 9 | 28 | 2 | 6 | Chrdl2 |
| Chrm1 | 21 | 44 | 27 | 38 | 97 | 100 | 84 | 112 | 151 | 188 | 159 | 110 | 95 | 88 | 24 | 36 | 23 | 77 | 82 | 69 | 94 | 170 | 129 | 141 | 132 | 94 | 99 | 76 | Chrm1 |
| Chrm2 | 188 | 161 | 116 | 242 | 546 | 333 | 335 | 328 | 364 | 389 | 376 | 124 | 151 | 158 | 260 | 135 | 180 | 284 | 355 | 284 | 432 | 418 | 324 | 192 | 284 | 108 | 130 | 109 | Chrm2 |
| Chrm3 | 82 | 118 | 54 | 126 | 300 | 174 | 169 | 336 | 294 | 322 | 333 | 173 | 167 | 145 | 138 | 80 | 87 | 176 | 196 | 243 | 247 | 439 | 326 | 206 | 345 | 119 | 117 | 139 | Chrm3 |
| Chrm4 | 305 | 395 | 472 | 454 | 365 | 438 | 263 | 32 | 44 | 38 | 36 | 24 | 34 | 28 | 240 | 410 | 472 | 439 | 397 | 335 | 289 | 94 | 37 | 39 | 69 | 26 | 27 | 26 | Chrm4 |
| Chrm5 | 6 | 9 | 8 | 2 | 20 | 16 | 7 | 9 | 15 | 16 | 15 | 5 | 8 | 9 | 9 | 9 | 1 | 0 | 13 | 5 | 11 | 15 | 7 | 10 | 21 | 1 | 10 | 17 | Chrm5 |
| Chrna1 | 15 | 3 | 6 | 1 | 5 | 2 | 4 | 35 | 105 | 26 | 580 | 9 | 25 | 6 | 4 | 9 | 0 | 0 | 0 | 5 | 0 | 80 | 363 | 18 | 20 | 6 | 6 | 8 | Chrna1 |
| Chrna10 | 5 | 7 | 8 | 6 | 3 | 2 | 3 | 4 | 2 | 2 | 1 | 4 | 7 | 5 | 4 | 5 | 2 | 4 | 1 | 4 | 4 | 2 | 4 | 4 | 1 | 6 | 1 | 0 | Chrna10 |
| Chrna2 | 3 | 3 | 0 | 1 | 6 | 5 | 16 | 9 | 15 | 10 | 16 | 11 | 0 | 7 | 0 | 2 | 6 | 7 | 15 | 12 | 13 | 23 | 12 | 10 | 6 | 16 | 8 | 6 | Chrna2 |
| Chrna3 | 273 | 244 | 342 | 239 | 538 | 373 | 332 | 1125 | 971 | 1117 | 976 | 935 | 1203 | 792 | 446 | 296 | 422 | 244 | 373 | 411 | 290 | 1295 | 1047 | 664 | 881 | 720 | 852 | 886 | Chrna3 |
| Chrna4 | 1559 | 1952 | 3025 | 2843 | 3954 | 3051 | 2972 | 2649 | 2232 | 3572 | 2899 | 3882 | 3689 | 3431 | 2085 | 1960 | 3539 | 2772 | 3700 | 2420 | 3050 | 5100 | 2421 | 3153 | 3877 | 2566 | 3462 | 3935 | Chrna4 |
| Chrna5 | 933 | 973 | 912 | 1015 | 1094 | 1121 | 730 | 349 | 290 | 321 | 236 | 198 | 490 | 191 | 1106 | 1160 | 1041 | 928 | 824 | 1009 | 878 | 364 | 254 | 111 | 358 | 99 | 208 | 258 | Chrna5 |
| Chrna6 | 1 | 4 | 7 | 6 | 6 | 10 | 13 | 27 | 37 | 28 | 24 | 12 | 6 | 9 | 4 | 5 | 8 | 6 | 5 | 10 | 11 | 65 | 53 | 35 | 41 | 0 | 12 | 4 | Chrna6 |
| Chrna7 | 496 | 530 | 420 | 455 | 485 | 570 | 418 | 316 | 405 | 372 | 339 | 374 | 420 | 402 | 464 | 455 | 422 | 495 | 436 | 473 | 437 | 297 | 338 | 306 | 368 | 162 | 393 | 374 | Chrna7 |
| Chrna9 | 0 | 0 | 0 | 0 | 1 | 0 | 0 | 5 | 2 | 2 | 7 | 0 | 1 | 2 | 0 | 1 | 3 | 1 | 0 | 1 | 2 | 7 | 7 | 7 | 0 | 15 | 3 | 0 | Chrna9 |
| Chrnb1 | 246 | 307 | 261 | 354 | 272 | 222 | 244 | 148 | 242 | 155 | 800 | 125 | 162 | 97 | 212 | 329 | 312 | 358 | 318 | 243 | 259 | 267 | 540 | 144 | 109 | 150 | 105 | 90 | Chrnb1 |
| Chrnb2 | 2554 | 3209 | 2775 | 4105 | 5703 | 4858 | 4920 | 5837 | 5214 | 5797 | 4816 | 5736 | 5289 | 5539 | 3448 | 2549 | 3916 | 4338 | 5427 | 4674 | 5177 | 7359 | 4583 | 5336 | 6902 | 3879 | 5244 | 5399 | Chrnb2 |
| Chrnb3 | 9 | 14 | 12 | 4 | 38 | 21 | 17 | 12 | 20 | 6 | 7 | 7 | 5 | 10 | 24 | 15 | 13 | 36 | 5 | 16 | 18 | 35 | 14 | 7 | 14 | 1 | 0 | 5 | Chrnb3 |
| Chrnb4 | 224 | 255 | 277 | 253 | 270 | 210 | 190 | 107 | 87 | 99 | 93 | 169 | 166 | 128 | 302 | 320 | 285 | 261 | 207 | 224 | 179 | 89 | 96 | 99 | 94 | 139 | 158 | 170 | Chrnb4 |
| Chrnd | 6 | 4 | 0 | 4 | 2 | 0 | 0 | 29 | 61 | 17 | 260 | 2 | 11 | 2 | 4 | 3 | 0 | 4 | 0 | 0 | 0 | 30 | 188 | 12 | 4 | 12 | 0 | 0 | Chrnd |
| Chrne | 2 | 0 | 4 | 1 | 2 | 0 | 1 | 2 | 2 | 1 | 2 | 0 | 3 | 1 | 0 | 2 | 0 | 9 | 1 | 0 | 2 | 2 | 4 | 0 | 1 | 0 | 5 | 0 | Chrne |
| Chrng | 22 | 4 | 0 | 5 | 5 | 10 | 1 | 39 | 132 | 14 | 677 | 2 | 45 | 0 | 4 | 17 | 3 | 0 | 0 | 0 | 0 | 90 | 368 | 7 | 4 | 3 | 5 | 1 | Chrng |
| Chst1 | 819 | 891 | 1291 | 1199 | 1655 | 1459 | 1354 | 1049 | 1195 | 1759 | 1549 | 1418 | 878 | 1305 | 778 | 858 | 1289 | 1245 | 1751 | 1094 | 1596 | 2149 | 1288 | 1942 | 2221 | 1356 | 1148 | 1186 | Chst1 |
| Chst10 | 2031 | 2460 | 2070 | 2802 | 3639 | 3322 | 3556 | 3287 | 2998 | 3587 | 2621 | 3034 | 2572 | 2995 | 2596 | 1996 | 2691 | 2915 | 3442 | 3254 | 3432 | 3980 | 2992 | 3209 | 3827 | 2586 | 2481 | 2817 | Chst10 |
| Chst11 | 2864 | 2899 | 3519 | 3953 | 3692 | 3597 | 2243 | 2611 | 3042 | 3176 | 2358 | 2195 | 2259 | 2875 | 2849 | 3407 | 3664 | 3635 | 3185 | 3644 | 3133 | 2581 | 2456 | 2576 | 1908 | 1987 | 2023 | Chst11 |
| Chst12 | 568 | 832 | 916 | 846 | 644 | 642 | 474 | 294 | 480 | 519 | 568 | 396 | 483 | 392 | 554 | 890 | 851 | 925 | 690 | 435 | 549 | 505 | 528 | 547 | 459 | 445 | 424 | 389 | Chst12 |
| Chst13 | 0 | 0 | 0 | 0 | 0 | 0 | 0 | 2 | 2 | 0 | 5 | 1 | 1 | 2 | 1 | 1 | 0 | 2 | 0 | 0 | 0 | 5 | 6 | 1 | 0 | 0 | 0 | 0 | Chst13 |
| Chst14 | 766 | 777 | 734 | 601 | 403 | 544 | 279 | 755 | 946 | 772 | 1023 | 812 | 1076 | 742 | 691 | 947 | 605 | 590 | 337 | 293 | 295 | 772 | 1164 | 828 | 577 | 1045 | 804 | 849 | Chst14 |
| Chst15 | 1668 | 1533 | 1691 | 1431 | 1933 | 1945 | 1435 | 3013 | 3363 | 3326 | 3152 | 3063 | 4052 | 2817 | 1590 | 1537 | 1592 | 1305 | 1810 | 1590 | 1608 | 3041 | 3738 | 2997 | 3308 | 2465 | 3574 | 3297 | Chst15 |

| | | | | | | | | | | | | | | | | | | | | | | | | | | | | | |
|---|---|---|---|---|---|---|---|---|---|---|---|---|---|---|---|---|---|---|---|---|---|---|---|---|---|---|---|---|---|
| Chst2 | 2911 | 3005 | 4442 | 3913 | 3800 | 3898 | 3590 | 2757 | 2980 | 3279 | 3497 | 3645 | 3527 | 3634 | 2863 | 3128 | 3965 | 3649 | 3872 | 3272 | 3569 | 3888 | 3116 | 3608 | 3706 | 4439 | 3893 | 3896 | Chst2 |
| Chst3 | 1006 | 1024 | 1215 | 1059 | 1135 | 1119 | 1053 | 1270 | 1297 | 1478 | 1524 | 1617 | 1535 | 1603 | 982 | 876 | 1389 | 1042 | 1279 | 888 | 1077 | 1512 | 1181 | 1614 | 1378 | 1755 | 1529 | 1815 | Chst3 |
| Chst4 | 0 | 0 | 0 | 0 | 0 | 0 | 1 | 0 | 0 | 0 | 1 | 0 | 0 | 0 | 0 | 0 | 0 | 0 | 3 | 1 | 0 | 2 | 0 | 0 | 0 | 0 | 0 | 0 | Chst4 |
| Chst5 | 28 | 29 | 44 | 7 | 38 | 26 | 15 | 100 | 134 | 74 | 102 | 78 | 102 | 98 | 24 | 46 | 9 | 26 | 18 | 22 | 22 | 72 | 141 | 85 | 72 | 58 | 71 | 123 | Chst5 |
| Chst7 | 249 | 265 | 286 | 229 | 272 | 270 | 221 | 101 | 124 | 93 | 131 | 105 | 87 | 100 | 215 | 269 | 177 | 304 | 275 | 266 | 264 | 84 | 140 | 111 | 104 | 141 | 93 | 78 | Chst7 |
| Chst8 | 571 | 863 | 728 | 737 | 744 | 696 | 594 | 423 | 487 | 591 | 523 | 631 | 574 | 514 | 683 | 783 | 873 | 655 | 749 | 760 | 686 | 698 | 498 | 499 | 667 | 561 | 551 | 559 | Chst8 |
| Chst9 | 45 | 46 | 33 | 40 | 53 | 24 | 51 | 40 | 26 | 32 | 20 | 20 | 25 | 32 | 78 | 71 | 25 | 53 | 26 | 37 | 47 | 28 | 26 | 32 | 27 | 7 | 22 | 34 | Chst9 |
| Chsy1 | 4535 | 4006 | 5390 | 4860 | 4214 | 4268 | 3845 | 3864 | 4353 | 4570 | 5633 | 4787 | 5009 | 4619 | 4160 | 4834 | 4702 | 4566 | 4072 | 3402 | 3808 | 4542 | 4625 | 4931 | 4446 | 5658 | 4571 | 4806 | Chsy1 |
| Chsy3 | 305 | 251 | 204 | 307 | 309 | 343 | 227 | 343 | 417 | 446 | 548 | 241 | 324 | 296 | 278 | 368 | 256 | 318 | 222 | 217 | 239 | 506 | 511 | 332 | 344 | 283 | 255 | 275 | Chsy3 |
| Chtf18 | 2756 | 3400 | 3819 | 3054 | 1864 | 2195 | 2240 | 1238 | 1520 | 1550 | 1444 | 2391 | 1825 | 2027 | 2229 | 2760 | 3227 | 2676 | 2643 | 2015 | 2163 | 1043 | 1335 | 1964 | 1545 | 2975 | 2137 | 2035 | Chtf18 |
| Chtf8 | 82 | 103 | 93 | 93 | 84 | 123 | 93 | 137 | 127 | 120 | 101 | 94 | 112 | 134 | 82 | 79 | 80 | 102 | 104 | 90 | 104 | 99 | 83 | 126 | 151 | 106 | 150 | 115 | Chtf8 |
| Chtop | 14820 | 13582 | 14905 | 14555 | 14941 | 14653 | 15836 | 22514 | 20910 | 20379 | 19576 | 22761 | 20478 | 22211 | 15154 | 13105 | 15515 | 14046 | 16105 | 15714 | 15715 | 19916 | 20335 | 21570 | 21202 | 23777 | 22636 | 22869 | Chtop |
| Chuk | 9292 | 7429 | 7521 | 8317 | 8144 | 8334 | 9213 | 8636 | 7055 | 6695 | 7481 | 8051 | 7343 | 8262 | 9216 | 8450 | 7601 | 8099 | 7701 | 8431 | 8798 | 8335 | 7011 | 7173 | 7874 | 6634 | 8056 | 7900 | Chuk |
| Churc1 | 176 | 134 | 111 | 116 | 110 | 146 | 148 | 194 | 162 | 199 | 151 | 121 | 170 | 232 | 177 | 122 | 112 | 139 | 129 | 107 | 164 | 165 | 150 | 126 | 170 | 183 | 189 | 218 | Churc1 |
| Ciao1 | 3804 | 4190 | 3717 | 4000 | 3306 | 3404 | 3604 | 3042 | 2917 | 2883 | 2708 | 2631 | 2773 | 2750 | 3956 | 3696 | 3640 | 3995 | 3625 | 3998 | 3669 | 2709 | 3065 | 2913 | 2898 | 2843 | 2840 | 2773 | Ciao1 |
| Ciao2a | 4134 | 3874 | 4810 | 4011 | 3269 | 3461 | 4205 | 3497 | 3125 | 3080 | 3372 | 3596 | 3664 | 3898 | 4338 | 4154 | 4376 | 3612 | 3650 | 4062 | 3615 | 2543 | 3399 | 3663 | 3144 | 4587 | 3989 | 4196 | Ciao2a |
| Ciao2b | 1121 | 1544 | 1404 | 1539 | 1280 | 1252 | 1521 | 968 | 1003 | 1007 | 1041 | 1177 | 904 | 1213 | 1054 | 1164 | 1531 | 1442 | 1452 | 1361 | 1437 | 973 | 1006 | 1165 | 1166 | 1318 | 1149 | 1222 | Ciao2b |
| Ciao3 | 1177 | 1443 | 1377 | 1230 | 1126 | 1073 | 1142 | 1194 | 1418 | 1426 | 1314 | 1332 | 1331 | 1281 | 1087 | 1213 | 1168 | 1163 | 1302 | 1136 | 1257 | 1233 | 1411 | 1352 | 1422 | 1514 | 1345 | 1326 | Ciao3 |
| Ciapin1 | 5180 | 6230 | 5619 | 6418 | 5791 | 5733 | 6010 | 5131 | 5094 | 5027 | 4577 | 5370 | 5200 | 5630 | 5441 | 5612 | 5898 | 5746 | 6160 | 6362 | 6112 | 4868 | 4916 | 5270 | 5339 | 5703 | 5801 | 5700 | Ciapin1 |
| Ciart | 185 | 239 | 161 | 237 | 124 | 119 | 110 | 134 | 132 | 114 | 162 | 131 | 121 | 147 | 214 | 271 | 191 | 216 | 118 | 108 | 149 | 92 | 169 | 150 | 118 | 154 | 114 | 164 | Ciart |
| Cib1 | 1599 | 1913 | 1989 | 1844 | 1280 | 1234 | 1304 | 677 | 784 | 756 | 689 | 775 | 812 | 764 | 1478 | 1874 | 1728 | 1557 | 1469 | 1371 | 1322 | 683 | 832 | 778 | 720 | 1017 | 859 | 853 | Cib1 |
| Cib2 | 772 | 772 | 625 | 603 | 862 | 702 | 713 | 700 | 753 | 855 | 730 | 626 | 586 | 573 | 807 | 805 | 689 | 797 | 724 | 761 | 767 | 742 | 806 | 754 | 667 | 621 | 623 | 590 | Cib2 |
| Cib3 | 1 | 0 | 0 | 1 | 0 | 0 | 0 | 1 | 0 | 4 | 0 | 0 | 0 | 0 | 0 | 3 | 0 | 0 | 3 | 0 | 0 | 0 | 0 | 0 | 0 | 0 | 0 | 2 | Cib3 |
| Cib4 | 0 | 0 | 0 | 1 | 0 | 0 | 0 | 0 | 3 | 0 | 3 | 0 | 7 | 0 | 0 | 0 | 0 | 2 | 0 | 0 | 0 | 2 | 0 | 1 | 0 | 3 | 0 | 0 | Cib4 |
| Cibar1 | 2633 | 1873 | 2142 | 2143 | 2445 | 2534 | 2489 | 4435 | 3891 | 3650 | 3857 | 3917 | 3951 | 3889 | 2699 | 2241 | 2036 | 1935 | 2187 | 2958 | 2469 | 3656 | 4129 | 3712 | 3741 | 4137 | 4006 | 3963 | Cibar1 |
| Cibar2 | 38 | 57 | 23 | 42 | 39 | 51 | 34 | 82 | 59 | 65 | 58 | 104 | 69 | 64 | 47 | 66 | 45 | 29 | 41 | 56 | 41 | 62 | 85 | 88 | 53 | 107 | 72 | 72 | Cibar2 |
| Cic | 10893 | 15393 | 17072 | 19149 | 18026 | 15567 | 14313 | 11917 | 15050 | 18506 | 17123 | 19457 | 16037 | 16514 | 10261 | 12499 | 19047 | 18324 | 19221 | 12574 | 15326 | 17749 | 13802 | 20349 | 18403 | 19358 | 15955 | 15664 | Cic |
| Cidea | 20 | 29 | 8 | 19 | 11 | 10 | 6 | 15 | 19 | 7 | 16 | 0 | 5 | 0 | 18 | 40 | 20 | 9 | 8 | 15 | 2 | 23 | 18 | 3 | 6 | 7 | 0 | 1 | Cidea |
| Cideb | 9 | 10 | 4 | 7 | 10 | 11 | 10 | 6 | 29 | 19 | 26 | 9 | 10 | 5 | 11 | 9 | 12 | 2 | 4 | 6 | 4 | 12 | 15 | 19 | 6 | 16 | 9 | 2 | Cideb |
| Cidec | 0 | 0 | 0 | 1 | 0 | 1 | 0 | 0 | 0 | 0 | 0 | 0 | 0 | 0 | 0 | 0 | 0 | 0 | 1 | 3 | 0 | 0 | 0 | 0 | 0 | 0 | 0 | 2 | Cidec |
| Ciita | 6 | 3 | 1 | 3 | 3 | 15 | 7 | 4 | 7 | 8 | 15 | 3 | 0 | 1 | 0 | 6 | 0 | 0 | 4 | 1 | 3 | 17 | 12 | 1 | 6 | 1 | 5 | 0 | Ciita |
| Cilk1 | 4124 | 3947 | 4942 | 5085 | 6428 | 5670 | 5794 | 5755 | 5487 | 6817 | 6177 | 6703 | 6702 | 7021 | 4178 | 4109 | 5283 | 5399 | 5919 | 4575 | 6118 | 7427 | 5474 | 6823 | 7384 | 5978 | 6422 | 6694 | Cilk1 |
| Cilp | 43 | 48 | 43 | 26 | 35 | 52 | 44 | 61 | 91 | 53 | 239 | 106 | 83 | 79 | 49 | 44 | 34 | 39 | 42 | 37 | 31 | 95 | 139 | 65 | 61 | 140 | 91 | 83 | Cilp |
| Cilp2 | 138 | 181 | 202 | 171 | 263 | 179 | 143 | 93 | 101 | 161 | 144 | 71 | 44 | 42 | 166 | 131 | 205 | 206 | 185 | 170 | 145 | 89 | 135 | 78 | 76 | 59 | 59 | 55 | Cilp2 |
| Cinp | 3712 | 4000 | 3458 | 3566 | 2737 | 2909 | 3315 | 2284 | 2077 | 1972 | 2137 | 2257 | 2142 | 2422 | 3701 | 3619 | 3291 | 3431 | 2907 | 3377 | 3450 | 2000 | 2124 | 2134 | 2147 | 2524 | 2215 | 2606 | Cinp |
| Cip2a | 10092 | 7053 | 8315 | 6542 | 4968 | 6475 | 7478 | 8527 | 6896 | 5877 | 5819 | 9260 | 7482 | 9231 | 9480 | 8208 | 7339 | 6107 | 6434 | 8350 | 6595 | 5230 | 6323 | 7591 | 6580 | 9711 | 9423 | 8994 | Cip2a |
| Cipc | 3393 | 3701 | 4108 | 4729 | 5454 | 4933 | 5311 | 5689 | 5291 | 6402 | 5325 | 6576 | 5539 | 5989 | 3712 | 3248 | 4715 | 4664 | 5393 | 5043 | 4774 | 6211 | 4793 | 6971 | 6948 | 6693 | 5861 | 6132 | Cipc |
| Cir1 | 3234 | 2808 | 3115 | 3131 | 2543 | 2680 | 2972 | 2811 | 2544 | 2766 | 2495 | 2628 | 2745 | 2502 | 2920 | 3231 | 3400 | 2921 | 3133 | 2902 | 3045 | 2251 | 2341 | 2899 | 2322 | 2534 | 2379 | 2376 | Cir1 |
| Cirbp | 6534 | 6969 | 8069 | 10445 | 9647 | 9372 | 8734 | 6746 | 7950 | 9625 | 8765 | 10593 | 9122 | 10530 | 7031 | 5487 | 8902 | 9270 | 9238 | 9111 | 9119 | 8072 | 7309 | 10748 | 11353 | 11043 | 9825 | 9413 | Cirbp |
| Cisd1 | 2000 | 2717 | 2073 | 2357 | 2477 | 2305 | 2952 | 2000 | 2066 | 2071 | 1813 | 2028 | 1859 | 2177 | 2288 | 1946 | 2459 | 2324 | 2686 | 2933 | 2825 | 1860 | 1888 | 2121 | 2225 | 2132 | 2060 | 2156 | Cisd1 |
| Cisd2 | 4124 | 3409 | 4728 | 4210 | 3983 | 4473 | 5183 | 5897 | 4974 | 5339 | 5706 | 6843 | 6430 | 7257 | 4250 | 3863 | 4589 | 4113 | 4373 | 4165 | 4506 | 6211 | 5075 | 6235 | 6139 | 7229 | 7275 | 7484 | Cisd2 |
| Cisd3 | 581 | 741 | 704 | 686 | 625 | 509 | 543 | 300 | 308 | 295 | 357 | 287 | 355 | 289 | 658 | 710 | 574 | 686 | 620 | 721 | 575 | 294 | 340 | 369 | 380 | 361 | 354 | 369 | Cisd3 |
| Cish | 612 | 720 | 732 | 786 | 947 | 742 | 713 | 421 | 553 | 613 | 656 | 520 | 593 | 516 | 541 | 671 | 897 | 824 | 884 | 605 | 737 | 570 | 584 | 567 | 620 | 656 | 446 | 573 | Cish |
| Cit | 3325 | 3814 | 4378 | 4537 | 3797 | 3686 | 3432 | 2459 | 2838 | 3192 | 3047 | 5010 | 3978 | 4549 | 2860 | 3288 | 4631 | 4116 | 4260 | 2937 | 3656 | 2928 | 2319 | 3974 | 3577 | 3039 | 4267 | 4284 | Cit |
| Cited1 | 309 | 420 | 393 | 399 | 346 | 282 | 413 | 133 | 182 | 149 | 219 | 159 | 117 | 154 | 322 | 405 | 397 | 395 | 370 | 364 | 323 | 174 | 179 | 191 | 142 | 201 | 156 | 158 | Cited1 |
| Cited2 | 3246 | 3909 | 5509 | 4599 | 4770 | 3845 | 4010 | 4245 | 3804 | 5090 | 4297 | 5640 | 5400 | 5343 | 3847 | 3706 | 5186 | 4293 | 4597 | 3312 | 4097 | 6254 | 4068 | 5243 | 5434 | 5157 | 5204 | 5784 | Cited2 |
| Cited4 | 106 | 196 | 172 | 122 | 52 | 69 | 79 | 17 | 43 | 26 | 31 | 42 | 162 | 102 | 142 | 154 | 134 | 102 | 79 | 84 | 68 | 12 | 34 | 25 | 27 | 77 | 89 | 230 | Cited4 |
| Ciz1 | 5705 | 6721 | 7261 | 7429 | 6390 | 6538 | 6117 | 4833 | 5595 | 6738 | 6228 | 6932 | 6321 | 6163 | 5549 | 6204 | 7421 | 7092 | 7463 | 6196 | 6354 | 6433 | 5354 | 6918 | 6554 | 6726 | 6151 | 5864 | Ciz1 |
| CK137956 | 23 | 17 | 10 | 17 | 32 | 25 | 8 | 32 | 30 | 27 | 25 | 14 | 34 | 32 | 9 | 12 | 24 | 17 | 26 | 22 | 19 | 45 | 23 | 27 | 30 | 28 | 40 | 18 | CK137956 |
| Ckap2 | 4429 | 3975 | 4206 | 3755 | 3175 | 3605 | 3730 | 4858 | 4376 | 3874 | 3521 | 5600 | 5051 | 6001 | 4316 | 3837 | 3872 | 3163 | 3681 | 4257 | 3531 | 3312 | 3878 | 4900 | 4170 | 6425 | 5928 | 5704 | Ckap2 |
| Ckap2l | 4338 | 3823 | 4174 | 2965 | 2527 | 3059 | 2960 | 5819 | 5250 | 4570 | 4156 | 6543 | 5544 | 6465 | 3944 | 3373 | 3385 | 2708 | 3162 | 3680 | 2641 | 3893 | 4711 | 5794 | 5036 | 7939 | 6825 | 6284 | Ckap2l |
| Ckap4 | 5667 | 6214 | 8007 | 5582 | 5688 | 6135 | 4479 | 6445 | 7841 | 8218 | 10646 | 8789 | 9248 | 7880 | 5798 | 7273 | 6800 | 6450 | 5638 | 4146 | 4500 | 8809 | 9237 | 8380 | 7823 | 9809 | 8727 | 8825 | Ckap4 |
| Ckap5 | 35104 | 32005 | 23113 | 25473 | 26965 | 30169 | 32184 | 39042 | 32398 | 28122 | 25908 | 35052 | 30798 | 35676 | 36385 | 30298 | 26226 | 25118 | 27565 | 36253 | 30854 | 27111 | 29884 | 27012 | 30196 | 23324 | 36653 | 33246 | Ckap5 |
| Ckb | 31487 | 48214 | 42179 | 40065 | 32030 | 32104 | 32284 | 23549 | 24183 | 27050 | 20051 | 40747 | 29055 | 36201 | 33693 | 28056 | 44739 | 32515 | 41427 | 34460 | 32887 | 24869 | 20909 | 32869 | 30593 | 42606 | 40245 | 39311 | Ckb |
| Cklf | 3396 | 3091 | 2305 | 2549 | 2506 | 2830 | 2562 | 2356 | 2307 | 2013 | 2407 | 2259 | 2429 | 2246 | 3158 | 3244 | 2379 | 2702 | 2391 | 2997 | 2703 | 2212 | 2368 | 2054 | 2173 | 1503 | 2374 | 2305 | Cklf |
| Ckm | 8 | 2 | 0 | 4 | 2 | 0 | 1 | 0 | 3 | 1 | 85 | 0 | 5 | 3 | 1 | 4 | 0 | 3 | 2 | 0 | 0 | 3 | 107 | 0 | 0 | 1 | 2 | 0 | Ckm |
| Ckmt1 | 627 | 493 | 302 | 589 | 851 | 499 | 568 | 876 | 890 | 958 | 926 | 327 | 542 | 330 | 605 | 538 | 448 | 567 | 564 | 656 | 621 | 730 | 988 | 856 | 537 | 826 | 245 | 278 | Ckmt1 |
| Ckmt2 | 1 | 0 | 0 | 0 | 1 | 0 | 1 | 0 | 1 | 0 | 2 | 0 | 0 | 1 | 0 | 0 | 0 | 0 | 0 | 0 | 0 | 2 | 6 | 0 | 0 | 3 | 0 | 0 | Ckmt2 |
| Cks1b | 3519 | 3953 | 4056 | 2932 | 2283 | 2719 | 2887 | 2461 | 2815 | 2679 | 2669 | 3555 | 3324 | 3465 | 3094 | 3471 | 3422 | 2664 | 2932 | 3360 | 2708 | 1582 | 2640 | 3261 | 2443 | 4733 | 3552 | 3796 | Cks1b |
| Cks2 | 4244 | 4042 | 4325 | 3351 | 2139 | 2450 | 3711 | 3009 | 2568 | 2340 | 2523 | 3466 | 3132 | 4697 | 3799 | 3377 | 3670 | 2804 | 2751 | 3430 | 3207 | 1597 | 2438 | 3505 | 2604 | 4610 | 3869 | 4726 | Cks2 |
| Clasp1 | 18476 | 18496 | 14472 | 18567 | 20145 | 20738 | 20830 | 21184 | 20992 | 22922 | 20296 | 24632 | 21048 | 24212 | 18480 | 16622 | 16819 | 18407 | 20406 | 19624 | 21713 | 23484 | 18880 | 21172 | 25644 | 15806 | 23933 | 22731 | Clasp1 |
| Clasp2 | 15927 | 15654 | 11885 | 17654 | 23373 | 20964 | 24637 | 22990 | 21296 | 20426 | 18257 | 17614 | 15883 | 19083 | 18456 | 14304 | 15230 | 19203 | 20066 | 22846 | 23901 | 23494 | 19477 | 16837 | 22086 | 12541 | 16858 | 16308 | Clasp2 |
| Clasrp | 4503 | 5754 | 6852 | 6782 | 6090 | 5853 | 5417 | 4799 | 5245 | 6755 | 6264 | 6364 | 6040 | 6125 | 4019 | 5121 | 7472 | 6936 | 6733 | 4783 | 5666 | 6159 | 4868 | 6969 | 6592 | 6604 | 6029 | 5947 | Clasrp |
| Clba1 | 510 | 504 | 437 | 524 | 442 | 432 | 539 | 162 | 140 | 215 | 181 | 192 | 168 | 177 | 487 | 516 | 527 | 556 | 461 | 525 | 495 | 190 | 185 | 198 | 228 | 196 | 187 | 228 | Clba1 |
| Clca1 | 0 | 3 | 0 | 0 | 0 | 0 | 0 | 2 | 2 | 4 | 4 | 1 | 0 | 0 | 1 | 2 | 1 | 0 | 0 | 0 | 0 | 0 | 2 | 1 | 0 | 2 | 0 | 2 | Clca1 |
| Clca2 | 29 | 9 | 21 | 5 | 12 | 23 | 10 | 26 | 30 | 27 | 43 | 24 | 27 | 21 | 32 | 16 | 10 | 35 | 16 | 8 | 12 | 30 | 36 | 20 | 38 | 25 | 24 | 24 | Clca2 |
| Clca3a1 | 15 | 15 | 12 | 19 | 21 | 25 | 11 | 32 | 47 | 69 | 102 | 15 | 37 | 27 | 25 | 19 | 5 | 24 | 12 | 15 | 21 | 70 | 82 | 69 | 41 | 42 | 28 | 19 | Clca3a1 |
| Clca3a2 | 2 | 1 | 1 | 1 | 1 | 0 | 1 | 4 | 10 | 3 | 5 | 4 | 0 | 0 | 5 | 4 | 0 | 2 | 1 | 0 | 1 | 5 | 2 | 6 | 0 | 5 | 1 | 4 | Clca3a2 |
| Clca3b | 0 | 0 | 0 | 0 | 0 | 0 | 0 | 0 | 2 | 0 | 6 | 0 | 0 | 0 | 0 | 0 | 0 | 0 | 0 | 0 | 0 | 0 | 1 | 0 | 1 | 0 | 0 | 4 | Clca3b |
| Clca4a | 0 | 0 | 0 | 0 | 0 | 0 | 0 | 2 | 0 | 0 | 0 | 0 | 0 | 0 | 1 | 0 | 0 | 0 | 0 | 0 | 0 | 0 | 1 | 2 | 0 | 1 | 0 | 0 | Clca4a |
| Clca4b | 1 | 0 | 0 | 0 | 0 | 0 | 0 | 0 | 0 | 0 | 0 | 0 | 5 | 0 | 0 | 0 | 0 | 0 | 0 | 0 | 0 | 0 | 1 | 1 | 2 | 0 | 0 | 0 | Clca4b |
| Clcc1 | 3829 | 3642 | 3835 | 3431 | 2904 | 3375 | 3433 | 3036 | 2966 | 2798 | 2685 | 3168 | 2905 | 3094 | 3696 | 3815 | 3741 | 3496 | 3416 | 3820 | 3240 | 2582 | 2798 | 2920 | 2740 | 3428 | 3366 | 3215 | Clcc1 |
| Clcf1 | 5 | 3 | 2 | 2 | 2 | 5 | 0 | 6 | 7 | 5 | 2 | 4 | 2 | 6 | 4 | 5 | 2 | 1 | 6 | 3 | 1 | 7 | 4 | 2 | 3 | 6 | 0 | 4 | Clcf1 |
| Clcn1 | 1489 | 1575 | 1132 | 1793 | 1779 | 1760 | 2094 | 852 | 769 | 877 | 780 | 1227 | 1005 | 1204 | 1811 | 1239 | 1562 | 1760 | 1747 | 1840 | 2005 | 951 | 712 | 932 | 942 | 944 | 1298 | 1087 | Clcn1 |
| Clcn2 | 3911 | 4941 | 5023 | 5145 | 5827 | 5307 | 4567 | 2950 | 3453 | 3839 | 3617 | 3672 | 3908 | 3212 | 3926 | 5031 | 5358 | 5447 | 5463 | 4514 | 4530 | 3969 | 3310 | 3621 | 3494 | 3602 | 3619 | 3229 | Clcn2 |
| Clcn3 | 13313 | 12292 | 10217 | 13106 | 12576 | 13671 | 14577 | 10741 | 9252 | 8174 | 7976 | 9330 | 8840 | 9797 | 14446 | 11924 | 11079 | 13048 | 11798 | 14883 | 13952 | 9521 | 8714 | 7961 | 9838 | 7292 | 9706 | 9467 | Clcn3 |
| Clcn4 | 6684 | 6492 | 6144 | 8390 | 10512 | 9722 | 9793 | 14302 | 12880 | 14378 | 12900 | 12347 | 11587 | 12249 | 7913 | 6252 | 7546 | 8872 | 9313 | 10324 | 9795 | 14952 | 12888 | 12570 | 15213 | 9899 | 11721 | 11407 | Clcn4 |
| Clcn5 | 7989 | 6932 | 4850 | 5007 | 5709 | 6213 | 5644 | 6196 | 6676 | 5333 | 9901 | 5602 | 6089 | 5366 | 7652 | 7645 | 4799 | 5102 | 4874 | 5558 | 5560 | 5431 | 7684 | 4406 | 4973 | 3703 | 5911 | 5178 | Clcn5 |
| Clcn6 | 7891 | 9619 | 8392 | 9864 | 9848 | 9352 | 9105 | 6119 | 6251 | 7014 | 6282 | 7704 | 7285 | 7208 | 8447 | 8638 | 9531 | 9720 | 9655 | 8487 | 9287 | 7737 | 5551 | 6995 | 8193 | 5696 | 7517 | 6855 | Clcn6 |

| | | | | | | | | | | | | | | | | | | | | | | | | | | | | | |
|---|---|---|---|---|---|---|---|---|---|---|---|---|---|---|---|---|---|---|---|---|---|---|---|---|---|---|---|---|---|
| Clcn7 | 4583 | 5343 | 4323 | 4948 | 4625 | 4878 | 4604 | 3445 | 3827 | 3974 | 3776 | 3926 | 3938 | 3821 | 4100 | 5005 | 4509 | 5302 | 4697 | 4251 | 4636 | 3950 | 3551 | 3619 | 4158 | 3415 | 3849 | 3452 | Clcn7 |
| Clcnka | 13 | 3 | 5 | 2 | 0 | 7 | 2 | 5 | 26 | 7 | 27 | 1 | 15 | 1 | 5 | 7 | 6 | 2 | 0 | 0 | 4 | 15 | 13 | 9 | 0 | 12 | 1 | 2 | Clcnka |
| Clcnkb | 0 | 0 | 0 | 2 | 2 | 0 | 0 | 5 | 2 | 1 | 0 | 0 | 2 | 0 | 0 | 0 | 0 | 2 | 0 | 0 | 1 | 3 | 3 | 0 | 0 | 0 | 0 | 2 | Clcnkb |
| Cldn1 | 727 | 517 | 580 | 337 | 583 | 446 | 303 | 683 | 888 | 490 | 642 | 480 | 818 | 305 | 800 | 980 | 406 | 403 | 312 | 458 | 328 | 759 | 1211 | 372 | 312 | 412 | 363 | 417 | Cldn1 |
| Cldn10 | 127 | 92 | 115 | 105 | 70 | 87 | 47 | 16 | 38 | 6 | 53 | 27 | 37 | 10 | 116 | 136 | 93 | 88 | 48 | 59 | 65 | 10 | 34 | 17 | 13 | 17 | 22 | 12 | Cldn10 |
| Cldn11 | 1617 | 1263 | 1310 | 1942 | 1818 | 1517 | 1541 | 1859 | 2343 | 2215 | 2523 | 1642 | 1698 | 1869 | 1419 | 1670 | 1203 | 2161 | 1234 | 1468 | 1739 | 2137 | 3423 | 2653 | 2152 | 2032 | 1267 | 1381 | Cldn11 |
| Cldn12 | 5225 | 4991 | 4696 | 4340 | 3494 | 4338 | 4227 | 4279 | 3680 | 3217 | 3151 | 3865 | 3675 | 3942 | 5436 | 4647 | 4679 | 4291 | 4157 | 4919 | 4068 | 3437 | 3589 | 3643 | 3450 | 4358 | 4071 | 4315 | Cldn12 |
| Cldn13 | 13 | 22 | 18 | 16 | 0 | 10 | 2 | 15 | 16 | 21 | 9 | 15 | 70 | 51 | 21 | 30 | 16 | 3 | 3 | 17 | 5 | 3 | 13 | 14 | 2 | 39 | 36 | 147 | Cldn13 |
| Cldn14 | 1 | 1 | 0 | 3 | 0 | 0 | 2 | 4 | 4 | 3 | 0 | 0 | 6 | 0 | 3 | 2 | 0 | 0 | 0 | 0 | 3 | 0 | 1 | 2 | 0 | 0 | 0 | 4 | Cldn14 |
| Cldn15 | 179 | 206 | 173 | 204 | 187 | 179 | 189 | 60 | 49 | 36 | 45 | 61 | 77 | 81 | 124 | 199 | 251 | 230 | 226 | 132 | 204 | 53 | 42 | 45 | 60 | 59 | 94 | 55 | Cldn15 |
| Cldn16 | 0 | 0 | 0 | 0 | 0 | 0 | 0 | 0 | 2 | 0 | 0 | 0 | 0 | 0 | 0 | 0 | 0 | 0 | 0 | 0 | 0 | 0 | 0 | 0 | 0 | 0 | 0 | 0 | Cldn16 |
| Cldn18 | 0 | 0 | 0 | 0 | 0 | 0 | 0 | 0 | 1 | 0 | 0 | 0 | 0 | 0 | 0 | 0 | 0 | 0 | 0 | 0 | 0 | 0 | 0 | 0 | 0 | 0 | 0 | 0 | Cldn18 |
| Cldn19 | 6 | 10 | 6 | 4 | 18 | 12 | 6 | 33 | 71 | 91 | 59 | 34 | 59 | 64 | 12 | 9 | 2 | 6 | 14 | 16 | 17 | 57 | 125 | 47 | 82 | 49 | 53 | 58 | Cldn19 |
| Cldn2 | 15 | 15 | 16 | 6 | 29 | 22 | 11 | 44 | 54 | 46 | 72 | 28 | 57 | 22 | 13 | 41 | 7 | 4 | 5 | 14 | 14 | 72 | 126 | 24 | 23 | 15 | 22 | 25 | Cldn2 |
| Cldn22 | 0 | 0 | 0 | 0 | 0 | 0 | 0 | 9 | 12 | 8 | 17 | 0 | 11 | 0 | 0 | 0 | 0 | 0 | 0 | 0 | 0 | 15 | 8 | 2 | 0 | 0 | 0 | 0 | Cldn22 |
| Cldn23 | 84 | 58 | 27 | 62 | 20 | 41 | 36 | 110 | 114 | 117 | 89 | 67 | 73 | 66 | 54 | 60 | 45 | 44 | 70 | 30 | 45 | 42 | 145 | 159 | 46 | 139 | 70 | 70 | Cldn23 |
| Cldn24 | 0 | 0 | 0 | 0 | 0 | 0 | 0 | 13 | 6 | 3 | 13 | 0 | 3 | 0 | 0 | 0 | 0 | 0 | 0 | 0 | 0 | 3 | 2 | 0 | 0 | 1 | 1 | 0 | Cldn24 |
| Cldn3 | 125 | 85 | 63 | 61 | 43 | 63 | 19 | 161 | 180 | 234 | 148 | 40 | 102 | 42 | 84 | 103 | 39 | 35 | 48 | 39 | 29 | 90 | 192 | 235 | 19 | 256 | 41 | 46 | Cldn3 |
| Cldn34b3 | 0 | 0 | 0 | 0 | 0 | 0 | 0 | 0 | 0 | 0 | 0 | 3 | 0 | 0 | 0 | 0 | 0 | 0 | 0 | 0 | 0 | 0 | 0 | 2 | 0 | 0 | 0 | 0 | Cldn34b3 |
| Cldn34c1 | 519 | 357 | 390 | 515 | 589 | 649 | 620 | 950 | 846 | 759 | 707 | 704 | 785 | 861 | 553 | 466 | 446 | 451 | 569 | 654 | 590 | 966 | 822 | 756 | 785 | 708 | 995 | 770 | Cldn34c1 |
| Cldn34c2 | 0 | 1 | 0 | 1 | 1 | 2 | 0 | 0 | 1 | 0 | 0 | 1 | 1 | 2 | 0 | 0 | 5 | 1 | 1 | 0 | 0 | 0 | 2 | 0 | 2 | 1 | 2 | 0 | Cldn34c2 |
| Cldn34d | 0 | 0 | 0 | 0 | 0 | 0 | 0 | 0 | 0 | 0 | 0 | 0 | 0 | 0 | 0 | 0 | 0 | 0 | 2 | 0 | 0 | 0 | 0 | 0 | 0 | 0 | 0 | 0 | Cldn34d |
| Cldn4 | 451 | 275 | 194 | 59 | 20 | 180 | 2 | 360 | 563 | 333 | 384 | 161 | 262 | 143 | 306 | 357 | 58 | 119 | 15 | 17 | 3 | 75 | 603 | 366 | 39 | 364 | 151 | 218 | Cldn4 |
| Cldn5 | 702 | 816 | 1080 | 938 | 1280 | 934 | 888 | 1253 | 1390 | 1814 | 1881 | 1538 | 1540 | 1277 | 728 | 794 | 909 | 844 | 1149 | 920 | 1012 | 1479 | 1732 | 1659 | 2002 | 1645 | 1429 | 1245 | Cldn5 |
| Cldn6 | 1232 | 602 | 268 | 303 | 119 | 187 | 53 | 767 | 770 | 635 | 655 | 98 | 299 | 125 | 687 | 734 | 156 | 213 | 51 | 85 | 51 | 279 | 759 | 759 | 57 | 994 | 110 | 120 | Cldn6 |
| Cldn7 | 189 | 119 | 37 | 97 | 29 | 24 | 2 | 173 | 185 | 236 | 147 | 41 | 112 | 43 | 103 | 145 | 17 | 35 | 12 | 0 | 3 | 57 | 216 | 225 | 17 | 319 | 35 | 52 | Cldn7 |
| Cldn8 | 68 | 24 | 16 | 3 | 5 | 29 | 1 | 109 | 180 | 80 | 116 | 25 | 65 | 45 | 71 | 52 | 6 | 43 | 11 | 0 | 0 | 35 | 170 | 78 | 14 | 28 | 23 | 38 | Cldn8 |
| Cldn9 | 321 | 226 | 126 | 223 | 117 | 140 | 174 | 167 | 206 | 261 | 235 | 73 | 111 | 77 | 206 | 238 | 155 | 178 | 132 | 169 | 139 | 104 | 192 | 279 | 60 | 366 | 56 | 69 | Cldn9 |
| Cldnd1 | 6770 | 6325 | 7357 | 6809 | 6172 | 6197 | 6797 | 5254 | 5006 | 4699 | 5006 | 4819 | 5036 | 5136 | 6898 | 6615 | 7065 | 6796 | 6316 | 6742 | 6320 | 4881 | 5066 | 5065 | 4719 | 5544 | 4965 | 5256 | Cldnd1 |
| Cldnd2 | 5 | 3 | 7 | 3 | 1 | 11 | 1 | 10 | 5 | 6 | 12 | 11 | 11 | 3 | 5 | 10 | 2 | 8 | 1 | 2 | 4 | 15 | 23 | 5 | 14 | 8 | 5 | 13 | Cldnd2 |
| Clec10a | 12 | 21 | 29 | 24 | 14 | 26 | 9 | 31 | 38 | 32 | 42 | 43 | 36 | 44 | 20 | 17 | 14 | 17 | 16 | 9 | 13 | 57 | 45 | 32 | 22 | 33 | 43 | 32 | Clec10a |
| Clec11a | 102 | 155 | 210 | 144 | 123 | 168 | 116 | 105 | 140 | 194 | 174 | 126 | 69 | 113 | 138 | 154 | 152 | 139 | 134 | 126 | 125 | 135 | 227 | 158 | 177 | 126 | 103 | 83 | Clec11a |
| Clec12a | 17 | 5 | 6 | 6 | 11 | 2 | 0 | 34 | 39 | 66 | 33 | 4 | 17 | 11 | 1 | 9 | 2 | 1 | 1 | 0 | 1 | 18 | 51 | 30 | 21 | 45 | 9 | 7 | Clec12a |
| Clec12b | 2 | 0 | 0 | 1 | 3 | 8 | 3 | 2 | 2 | 10 | 0 | 3 | 5 | 2 | 5 | 2 | 0 | 0 | 2 | 0 | 1 | 3 | 5 | 0 | 3 | 1 | 0 | 5 | Clec12b |
| Clec14a | 361 | 256 | 419 | 195 | 452 | 457 | 323 | 1003 | 1147 | 1088 | 1319 | 1044 | 1410 | 1105 | 384 | 319 | 336 | 246 | 349 | 439 | 335 | 1141 | 1587 | 1039 | 1094 | 1109 | 1295 | 991 | Clec14a |
| Clec16a | 6150 | 6574 | 5736 | 7107 | 6883 | 6404 | 6740 | 3993 | 4024 | 4283 | 3779 | 4023 | 3374 | 3688 | 6091 | 5886 | 6720 | 7080 | 7058 | 5764 | 6921 | 4859 | 3831 | 3883 | 4358 | 3069 | 3449 | 3436 | Clec16a |
| Clec18a | 20 | 76 | 74 | 34 | 87 | 55 | 27 | 95 | 125 | 92 | 124 | 58 | 105 | 43 | 66 | 82 | 37 | 64 | 43 | 24 | 32 | 154 | 101 | 40 | 100 | 45 | 59 | 58 | Clec18a |
| Clec1a | 110 | 66 | 46 | 57 | 92 | 78 | 98 | 181 | 152 | 157 | 148 | 140 | 147 | 131 | 95 | 62 | 51 | 107 | 70 | 137 | 100 | 162 | 208 | 97 | 117 | 193 | 152 | 135 | Clec1a |
| Clec1b | 250 | 190 | 252 | 140 | 126 | 129 | 162 | 198 | 141 | 142 | 109 | 225 | 340 | 218 | 193 | 199 | 209 | 125 | 150 | 250 | 189 | 94 | 155 | 181 | 129 | 240 | 308 | 252 | Clec1b |
| Clec2d | 84 | 66 | 92 | 90 | 67 | 46 | 75 | 180 | 175 | 250 | 346 | 89 | 153 | 226 | 82 | 49 | 54 | 81 | 71 | 73 | 86 | 155 | 238 | 296 | 201 | 205 | 138 | 186 | Clec2d |
| Clec2e | 1 | 0 | 3 | 0 | 0 | 0 | 1 | 0 | 0 | 0 | 0 | 4 | 1 | 0 | 0 | 0 | 0 | 0 | 0 | 0 | 1 | 0 | 1 | 0 | 0 | 4 | 0 | 0 | Clec2e |
| Clec2f | 1 | 1 | 0 | 0 | 0 | 1 | 0 | 0 | 0 | 0 | 3 | 0 | 0 | 0 | 0 | 2 | 0 | 0 | 0 | 0 | 0 | 0 | 1 | 0 | 0 | 0 | 0 | 0 | Clec2f |
| Clec2g | 0 | 0 | 1 | 0 | 0 | 0 | 0 | 0 | 3 | 1 | 0 | 0 | 3 | 0 | 0 | 1 | 0 | 0 | 0 | 0 | 0 | 2 | 2 | 1 | 1 | 0 | 0 | 0 | Clec2g |
| Clec2h | 0 | 0 | 0 | 0 | 0 | 0 | 0 | 0 | 2 | 0 | 0 | 0 | 0 | 0 | 0 | 1 | 0 | 0 | 0 | 0 | 0 | 0 | 0 | 0 | 0 | 0 | 0 | 0 | Clec2h |
| Clec2i | 27 | 10 | 30 | 13 | 7 | 8 | 11 | 4 | 2 | 6 | 4 | 0 | 2 | 6 | 7 | 12 | 21 | 3 | 5 | 5 | 7 | 10 | 6 | 4 | 5 | 7 | 1 | 2 | Clec2i |
| Clec2l | 193 | 185 | 174 | 228 | 382 | 286 | 266 | 416 | 493 | 485 | 463 | 317 | 327 | 268 | 193 | 192 | 187 | 217 | 309 | 234 | 289 | 465 | 476 | 380 | 493 | 300 | 298 | 216 | Clec2l |
| Clec2m | 1 | 0 | 1 | 2 | 0 | 1 | 0 | 7 | 2 | 0 | 1 | 0 | 0 | 0 | 0 | 4 | 0 | 0 | 2 | 0 | 2 | 0 | 1 | 0 | 0 | 0 | 3 | 0 | Clec2m |
| Clec3a | 1 | 0 | 0 | 0 | 0 | 0 | 0 | 5 | 3 | 3 | 60 | 0 | 5 | 0 | 3 | 2 | 0 | 0 | 0 | 0 | 0 | 3 | 5 | 0 | 0 | 0 | 6 | 4 | Clec3a |
| Clec3b | 29 | 29 | 54 | 47 | 55 | 32 | 39 | 16 | 21 | 47 | 29 | 23 | 33 | 44 | 24 | 22 | 63 | 53 | 44 | 42 | 36 | 38 | 36 | 44 | 33 | 37 | 22 | 19 | Clec3b |
| Clec4a1 | 46 | 47 | 52 | 65 | 44 | 52 | 66 | 85 | 101 | 98 | 106 | 99 | 128 | 83 | 49 | 58 | 64 | 55 | 36 | 54 | 48 | 115 | 144 | 78 | 88 | 99 | 116 | 108 | Clec4a1 |
| Clec4a2 | 17 | 19 | 22 | 10 | 7 | 21 | 17 | 60 | 65 | 45 | 57 | 43 | 35 | 53 | 32 | 16 | 9 | 19 | 18 | 25 | 14 | 80 | 91 | 59 | 56 | 68 | 56 | 52 | Clec4a2 |
| Clec4a3 | 10 | 16 | 6 | 8 | 10 | 22 | 16 | 32 | 36 | 14 | 46 | 18 | 22 | 22 | 13 | 19 | 8 | 13 | 16 | 9 | 7 | 42 | 48 | 27 | 34 | 10 | 38 | 25 | Clec4a3 |
| Clec4a4 | 0 | 0 | 0 | 0 | 0 | 3 | 0 | 0 | 0 | 0 | 5 | 0 | 0 | 0 | 0 | 1 | 0 | 0 | 0 | 0 | 0 | 0 | 2 | 0 | 0 | 6 | 0 | 0 | Clec4a4 |
| Clec4b1 | 0 | 0 | 0 | 0 | 0 | 0 | 0 | 0 | 1 | 0 | 0 | 0 | 0 | 1 | 0 | 0 | 0 | 0 | 0 | 0 | 0 | 2 | 1 | 0 | 0 | 0 | 0 | 0 | Clec4b1 |
| Clec4d | 1 | 2 | 1 | 1 | 1 | 1 | 0 | 0 | 2 | 0 | 1 | 1 | 2 | 1 | 0 | 1 | 0 | 2 | 0 | 0 | 0 | 5 | 10 | 0 | 0 | 3 | 2 | 1 | Clec4d |
| Clec4e | 0 | 0 | 0 | 0 | 0 | 0 | 0 | 0 | 1 | 0 | 4 | 1 | 0 | 0 | 0 | 2 | 0 | 1 | 0 | 0 | 0 | 0 | 3 | 0 | 0 | 0 | 2 | 0 | Clec4e |
| Clec4f | 1 | 1 | 0 | 0 | 2 | 1 | 2 | 0 | 1 | 0 | 0 | 0 | 3 | 0 | 0 | 1 | 0 | 4 | 0 | 1 | 1 | 0 | 0 | 0 | 0 | 0 | 0 | 0 | Clec4f |
| Clec4g | 1 | 1 | 6 | 0 | 3 | 0 | 0 | 1 | 5 | 0 | 0 | 1 | 9 | 4 | 1 | 2 | 0 | 1 | 2 | 1 | 0 | 2 | 7 | 1 | 4 | 6 | 5 | 1 | Clec4g |
| Clec4n | 15 | 12 | 4 | 15 | 8 | 30 | 19 | 40 | 34 | 25 | 53 | 16 | 11 | 26 | 12 | 17 | 7 | 23 | 12 | 14 | 20 | 38 | 56 | 25 | 46 | 21 | 28 | 40 | Clec4n |
| Clec5a | 14 | 18 | 33 | 10 | 19 | 13 | 12 | 23 | 40 | 13 | 39 | 20 | 32 | 31 | 24 | 27 | 14 | 21 | 20 | 18 | 18 | 42 | 40 | 25 | 13 | 36 | 28 | 31 | Clec5a |
| Clec9a | 39 | 37 | 109 | 55 | 45 | 73 | 54 | 50 | 35 | 30 | 37 | 77 | 88 | 74 | 66 | 51 | 78 | 45 | 39 | 63 | 60 | 48 | 47 | 51 | 47 | 83 | 74 | 50 | Clec9a |
| Clgn | 791 | 641 | 732 | 836 | 1167 | 795 | 850 | 987 | 942 | 919 | 924 | 813 | 852 | 913 | 841 | 681 | 714 | 775 | 831 | 800 | 852 | 939 | 818 | 954 | 1018 | 806 | 780 | 809 | Clgn |
| Clhc1 | 38 | 41 | 47 | 31 | 34 | 35 | 31 | 34 | 21 | 19 | 31 | 30 | 24 | 29 | 30 | 44 | 43 | 18 | 40 | 52 | 26 | 35 | 28 | 19 | 19 | 38 | 15 | 17 | Clhc1 |
| Clic1 | 6061 | 6153 | 6299 | 4785 | 4253 | 4841 | 4137 | 5564 | 6302 | 5183 | 5852 | 6110 | 6404 | 5425 | 5796 | 6260 | 5663 | 4506 | 5179 | 5431 | 4154 | 4259 | 6851 | 5799 | 5150 | 7938 | 6751 | 6356 | Clic1 |
| Clic3 | 2 | 1 | 5 | 6 | 4 | 5 | 4 | 5 | 6 | 6 | 7 | 12 | 11 | 4 | 3 | 3 | 0 | 9 | 8 | 3 | 12 | 7 | 8 | 4 | 0 | 4 | 9 | 4 | Clic3 |
| Clic4 | 18477 | 17271 | 20547 | 18667 | 16927 | 17182 | 18815 | 14450 | 15474 | 16539 | 15316 | 21963 | 18901 | 21539 | 17818 | 17986 | 20148 | 18293 | 18922 | 15997 | 17940 | 15015 | 14426 | 19369 | 17226 | 23795 | 20642 | 23159 | Clic4 |
| Clic5 | 331 | 219 | 227 | 162 | 111 | 172 | 92 | 176 | 118 | 184 | 163 | 219 | 235 | 203 | 257 | 269 | 165 | 167 | 90 | 107 | 111 | 167 | 135 | 170 | 128 | 99 | 352 | 273 | Clic5 |
| Clic6 | 922 | 1158 | 764 | 812 | 1699 | 1197 | 1015 | 1721 | 2004 | 2142 | 1494 | 2217 | 2707 | 1324 | 1209 | 1631 | 878 | 782 | 1154 | 1366 | 1149 | 2099 | 3579 | 1635 | 1387 | 1688 | 1226 | 1289 | Clic6 |
| Clint1 | 8188 | 7728 | 8926 | 8066 | 6568 | 7113 | 6880 | 6914 | 7345 | 7179 | 8222 | 7769 | 7756 | 7633 | 7944 | 8580 | 7747 | 7664 | 6886 | 5919 | 6580 | 7274 | 7359 | 7803 | 7239 | 8739 | 7382 | 8049 | Clint1 |
| Clip1 | 5933 | 5667 | 5252 | 5921 | 7860 | 7300 | 7513 | 11190 | 10598 | 11884 | 11127 | 11724 | 11157 | 12260 | 6224 | 5516 | 5916 | 6371 | 6832 | 6884 | 7355 | 12146 | 9803 | 11021 | 12622 | 8573 | 11230 | 11068 | Clip1 |
| Clip2 | 5330 | 6975 | 9016 | 9674 | 10981 | 9076 | 7653 | 8120 | 9666 | 12576 | 10591 | 10728 | 9090 | 9517 | 5472 | 6361 | 9779 | 9799 | 11300 | 7052 | 8943 | 13145 | 9301 | 11696 | 13786 | 9905 | 8667 | 9108 | Clip2 |
| Clip3 | 10079 | 12294 | 14991 | 15381 | 17223 | 14119 | 12559 | 11269 | 13532 | 17083 | 14623 | 18027 | 13488 | 14947 | 9988 | 11210 | 16179 | 15962 | 18915 | 11070 | 15064 | 18603 | 12221 | 18875 | 18031 | 18651 | 12788 | 14148 | Clip3 |
| Clip4 | 1028 | 924 | 1016 | 1243 | 1491 | 1344 | 1547 | 1119 | 1251 | 1155 | 1260 | 1262 | 1135 | 1255 | 1097 | 893 | 1042 | 1310 | 1415 | 1234 | 1475 | 1285 | 1207 | 1097 | 1207 | 1077 | 911 | 1235 | Clip4 |
| Clk1 | 10154 | 8739 | 11789 | 11838 | 11261 | 11586 | 11112 | 19649 | 15966 | 15037 | 18270 | 15780 | 16669 | 17131 | 10916 | 11334 | 11490 | 11977 | 10757 | 12895 | 11088 | 19537 | 16454 | 16480 | 17237 | 15975 | 16314 | 16486 | Clk1 |
| Clk2 | 5568 | 6415 | 6680 | 6714 | 5846 | 6192 | 5760 | 5239 | 5487 | 6035 | 5748 | 6553 | 6150 | 6478 | 5341 | 6107 | 6906 | 6413 | 6460 | 5509 | 5882 | 5778 | 5222 | 6293 | 6036 | 7094 | 6408 | 6389 | Clk2 |
| Clk3 | 5753 | 6973 | 6514 | 6703 | 6297 | 6270 | 6013 | 6868 | 6846 | 6818 | 6335 | 7263 | 6975 | 7045 | 5611 | 5949 | 6697 | 6721 | 6906 | 6581 | 6142 | 7160 | 6637 | 6928 | 7366 | 7151 | 7126 | 7333 | Clk3 |
| Clk4 | 5760 | 4627 | 6323 | 6674 | 7090 | 6775 | 6802 | 10277 | 8211 | 8074 | 9148 | 8361 | 8517 | 9364 | 6030 | 5893 | 6124 | 6336 | 6752 | 7589 | 6678 | 9713 | 8045 | 8770 | 9335 | 8145 | 8610 | 8285 | Clk4 |
| Clmn | 1732 | 1612 | 1389 | 1285 | 1267 | 1346 | 911 | 1815 | 1985 | 2308 | 2564 | 1871 | 2462 | 1687 | 1445 | 2305 | 1321 | 1330 | 1099 | 895 | 896 | 1708 | 2190 | 1973 | 1679 | 1693 | 1936 | 1724 | Clmn |
| Clmp | 3898 | 3100 | 3400 | 3199 | 4370 | 4382 | 3733 | 9828 | 9442 | 10263 | 10692 | 8119 | 10030 | 8255 | 4083 | 4125 | 2998 | 3935 | 3831 | 3303 | 3891 | 11463 | 9852 | 9127 | 9773 | 7999 | 8215 | 8181 | Clmp |

| | | | | | | | | | | | | | | | | | | | | | | | | | | | | | |
|---|---|---|---|---|---|---|---|---|---|---|---|---|---|---|---|---|---|---|---|---|---|---|---|---|---|---|---|---|---|
| Cln3 | 758 | 893 | 851 | 999 | 1059 | 922 | 925 | 1036 | 1122 | 1198 | 1099 | 1073 | 1014 | 1155 | 719 | 762 | 1058 | 933 | 1072 | 830 | 944 | 1118 | 1126 | 1087 | 1200 | 1111 | 1149 | 1046 | Cln3 |
| Cln5 | 1965 | 1763 | 2021 | 1624 | 1280 | 1437 | 1285 | 1401 | 1470 | 1324 | 1613 | 1168 | 1644 | 1266 | 2149 | 2222 | 1592 | 1512 | 1167 | 1387 | 1227 | 1252 | 1666 | 1278 | 1087 | 1565 | 1519 | 1524 | Cln5 |
| Cln6 | 544 | 640 | 623 | 599 | 533 | 549 | 449 | 593 | 867 | 737 | 999 | 910 | 920 | 796 | 512 | 639 | 679 | 535 | 489 | 462 | 487 | 730 | 944 | 909 | 677 | 1215 | 831 | 845 | Cln6 |
| Cln8 | 2937 | 3734 | 3125 | 3788 | 2912 | 3201 | 2825 | 1719 | 1879 | 1821 | 1883 | 1783 | 1895 | 1760 | 2803 | 3363 | 2992 | 3690 | 2929 | 2926 | 3168 | 1855 | 1952 | 1875 | 1914 | 1844 | 1728 | 1858 | Cln8 |
| Clnk | 2 | 5 | 0 | 4 | 3 | 6 | 1 | 15 | 13 | 11 | 4 | 11 | 20 | 18 | 5 | 4 | 12 | 1 | 0 | 3 | 3 | 22 | 17 | 8 | 12 | 23 | 26 | 23 | Clnk |
| Clns1a | 8990 | 8032 | 8006 | 6830 | 6365 | 6328 | 7285 | 7061 | 6070 | 6133 | 5747 | 6862 | 6497 | 7261 | 8707 | 7843 | 7501 | 6713 | 6876 | 7618 | 7037 | 5955 | 5954 | 6513 | 6089 | 7906 | 6565 | 7729 | Clns1a |
| Clock | 5430 | 4171 | 2977 | 4183 | 5096 | 5282 | 5547 | 5766 | 5073 | 4367 | 4620 | 4154 | 4156 | 4457 | 5234 | 4644 | 3598 | 4299 | 4712 | 5562 | 5454 | 4642 | 5055 | 3721 | 4788 | 2868 | 4350 | 3966 | Clock |
| Clp1 | 2473 | 2466 | 2438 | 2039 | 1276 | 1406 | 1646 | 1683 | 1341 | 1299 | 1342 | 1585 | 1597 | 1609 | 2325 | 2461 | 1910 | 1854 | 1509 | 1777 | 1417 | 1275 | 1287 | 1450 | 1279 | 1966 | 1770 | 1829 | Clp1 |
| Clpb | 2352 | 2611 | 2093 | 2171 | 2279 | 2167 | 2178 | 2789 | 2985 | 2979 | 2739 | 2883 | 2538 | 2679 | 2284 | 2453 | 2308 | 2032 | 2288 | 2149 | 2247 | 2811 | 2779 | 2658 | 2810 | 2665 | 2869 | 2772 | Clpb |
| Clpp | 1851 | 2358 | 2083 | 2138 | 2050 | 1986 | 1928 | 1893 | 2258 | 2036 | 1856 | 2155 | 2109 | 2099 | 1875 | 1997 | 2313 | 1880 | 2214 | 2374 | 1877 | 1713 | 2106 | 2272 | 2211 | 2493 | 2296 | 2181 | Clpp |
| Clps | 0 | 1 | 0 | 0 | 0 | 1 | 0 | 0 | 0 | 0 | 0 | 0 | 0 | 0 | 0 | 0 | 0 | 2 | 0 | 0 | 0 | 0 | 3 | 0 | 0 | 0 | 0 | 0 | Clps |
| Clpsl2 | 0 | 2 | 0 | 2 | 0 | 0 | 0 | 0 | 0 | 0 | 0 | 0 | 0 | 0 | 0 | 0 | 0 | 0 | 0 | 0 | 0 | 0 | 0 | 0 | 0 | 0 | 0 | 0 | Clpsl2 |
| Clptm1 | 7100 | 8629 | 6870 | 7910 | 7273 | 7390 | 6374 | 5591 | 6336 | 6072 | 5674 | 6046 | 5528 | 5174 | 6989 | 7650 | 7268 | 7579 | 7837 | 6316 | 7081 | 6782 | 6287 | 5661 | 6017 | 6094 | 5719 | 5563 | Clptm1 |
| Clptm1l | 5222 | 5537 | 5448 | 5074 | 5015 | 4981 | 5044 | 6089 | 6278 | 5900 | 6349 | 5983 | 6007 | 5649 | 5209 | 5288 | 5065 | 5214 | 5165 | 4336 | 5197 | 6400 | 6571 | 5983 | 6123 | 6161 | 6007 | 5866 | Clptm1l |
| Clpx | 4136 | 3649 | 4035 | 3695 | 3285 | 3458 | 3836 | 3521 | 3167 | 3111 | 3195 | 3373 | 3125 | 3473 | 4246 | 3764 | 3804 | 3719 | 3500 | 3719 | 3578 | 3118 | 3293 | 3300 | 3042 | 3809 | 3253 | 3501 | Clpx |
| Clrn1 | 57 | 46 | 16 | 17 | 77 | 23 | 18 | 122 | 98 | 60 | 88 | 17 | 56 | 20 | 55 | 57 | 20 | 34 | 26 | 28 | 35 | 90 | 130 | 27 | 58 | 32 | 13 | 11 | Clrn1 |
| Clrn2 | 3 | 3 | 9 | 4 | 6 | 5 | 2 | 9 | 7 | 6 | 3 | 5 | 16 | 6 | 9 | 2 | 7 | 9 | 5 | 3 | 2 | 12 | 3 | 16 | 3 | 12 | 7 | 1 | Clrn2 |
| Clrn3 | 0 | 0 | 0 | 0 | 2 | 0 | 0 | 0 | 0 | 0 | 0 | 0 | 0 | 0 | 3 | 1 | 0 | 0 | 0 | 0 | 0 | 2 | 0 | 0 | 0 | 0 | 0 | 0 | Clrn3 |
| Clspn | 3749 | 3372 | 3801 | 3046 | 2528 | 3068 | 3310 | 4079 | 3934 | 3813 | 3514 | 5996 | 4576 | 6114 | 3186 | 2913 | 3582 | 2472 | 3024 | 3128 | 2833 | 2682 | 3300 | 4734 | 3931 | 6382 | 6340 | 6160 | Clspn |
| Clstn1 | 10247 | 12899 | 15649 | 18159 | 21729 | 17609 | 15343 | 13454 | 16228 | 20324 | 18420 | 17017 | 16276 | 15539 | 11107 | 11983 | 17468 | 18190 | 20186 | 14193 | 17326 | 19019 | 16079 | 18694 | 19654 | 16557 | 14386 | 15075 | Clstn1 |
| Clstn2 | 912 | 727 | 414 | 972 | 2083 | 1459 | 1551 | 1536 | 1987 | 1821 | 2559 | 892 | 1054 | 899 | 936 | 938 | 680 | 1215 | 1120 | 1129 | 1568 | 1751 | 2016 | 879 | 1694 | 621 | 815 | 674 | Clstn2 |
| Clstn3 | 3405 | 4552 | 4353 | 6398 | 7497 | 6448 | 5803 | 4218 | 4799 | 5807 | 4858 | 4190 | 4288 | 3899 | 4013 | 3634 | 5677 | 6260 | 7128 | 5655 | 6398 | 6452 | 4306 | 4710 | 5752 | 3386 | 3828 | 3771 | Clstn3 |
| Clta | 9322 | 10466 | 9715 | 10425 | 9116 | 8563 | 10026 | 7862 | 7721 | 8357 | 8339 | 7735 | 7608 | 8163 | 9357 | 8938 | 10109 | 9554 | 9504 | 9260 | 9906 | 7356 | 8220 | 8679 | 8178 | 8441 | 7459 | 8198 | Clta |
| Cltb | 2376 | 2826 | 2411 | 4032 | 4695 | 3386 | 3509 | 2158 | 2457 | 2796 | 2615 | 1742 | 1757 | 1840 | 2862 | 2622 | 3299 | 4260 | 3826 | 3498 | 4075 | 2981 | 2404 | 2200 | 2927 | 1514 | 1553 | 1652 | Cltb |
| Cltc | 54765 | 48405 | 40083 | 46393 | 48058 | 52562 | 53902 | 46179 | 42110 | 38983 | 40357 | 40135 | 41155 | 40781 | 55260 | 48678 | 43775 | 47852 | 47308 | 49103 | 54614 | 44623 | 42988 | 35688 | 41449 | 30696 | 41287 | 40282 | Cltc |
| Cltrn | 12 | 3 | 8 | 6 | 12 | 2 | 0 | 13 | 9 | 5 | 8 | 3 | 23 | 10 | 12 | 16 | 8 | 1 | 9 | 5 | 4 | 10 | 25 | 6 | 0 | 4 | 2 | 15 | Cltrn |
| Clu | 1123 | 1096 | 889 | 847 | 1102 | 1200 | 980 | 1196 | 1464 | 1538 | 1525 | 835 | 858 | 724 | 890 | 1226 | 788 | 1069 | 1130 | 1004 | 1146 | 1277 | 2049 | 1202 | 1108 | 1309 | 791 | 786 | Clu |
| Cluap1 | 1070 | 1045 | 903 | 1082 | 1141 | 978 | 1245 | 1459 | 1299 | 1527 | 1245 | 1361 | 1253 | 1523 | 1109 | 829 | 956 | 989 | 1012 | 993 | 1179 | 1350 | 1242 | 1425 | 1349 | 1373 | 1296 | 1320 | Cluap1 |
| Cluh | 6574 | 8167 | 8421 | 8251 | 7236 | 6713 | 6655 | 3057 | 4032 | 3881 | 4424 | 4183 | 4246 | 3960 | 6200 | 8753 | 8137 | 8009 | 7169 | 5410 | 6606 | 3810 | 4059 | 3934 | 3845 | 4100 | 3692 | 3907 | Cluh |
| Clvs1 | 1529 | 1562 | 1660 | 1967 | 3063 | 2255 | 2646 | 3537 | 2812 | 3156 | 2625 | 3002 | 2893 | 3108 | 2164 | 1442 | 2049 | 2168 | 2593 | 2606 | 2859 | 4075 | 2719 | 2961 | 3561 | 2138 | 2959 | 3204 | Clvs1 |
| Clvs2 | 156 | 168 | 242 | 254 | 391 | 320 | 313 | 677 | 512 | 605 | 482 | 502 | 598 | 585 | 269 | 162 | 264 | 274 | 376 | 334 | 289 | 825 | 548 | 521 | 581 | 310 | 608 | 585 | Clvs2 |
| Clybl | 1099 | 1088 | 1338 | 925 | 709 | 859 | 910 | 1109 | 849 | 851 | 901 | 1166 | 1183 | 1278 | 1200 | 1131 | 1015 | 777 | 874 | 1175 | 906 | 1048 | 1117 | 984 | 948 | 1427 | 1525 | 1572 | Clybl |
| Cma1 | 0 | 0 | 0 | 2 | 7 | 4 | 0 | 17 | 37 | 47 | 25 | 8 | 1 | 5 | 0 | 2 | 0 | 14 | 7 | 1 | 0 | 7 | 61 | 26 | 19 | 4 | 2 | 1 | Cma1 |
| Cma2 | 0 | 0 | 0 | 4 | 0 | 0 | 0 | 0 | 1 | 0 | 0 | 0 | 0 | 0 | 0 | 0 | 0 | 1 | 0 | 0 | 0 | 0 | 2 | 1 | 0 | 0 | 0 | 0 | Cma2 |
| Cmah | 23 | 47 | 35 | 20 | 35 | 41 | 36 | 73 | 93 | 74 | 109 | 66 | 86 | 73 | 45 | 53 | 29 | 65 | 28 | 59 | 29 | 53 | 119 | 53 | 62 | 56 | 71 | 75 | Cmah |
| Cmas | 6868 | 6322 | 6957 | 7169 | 7526 | 6842 | 8126 | 7458 | 6300 | 6422 | 6054 | 6331 | 5968 | 7473 | 7535 | 6329 | 7473 | 7080 | 7575 | 7981 | 7973 | 6963 | 6324 | 6738 | 7212 | 6429 | 6568 | 7226 | Cmas |
| Cmbl | 55 | 79 | 75 | 62 | 76 | 78 | 76 | 68 | 96 | 109 | 90 | 71 | 94 | 65 | 75 | 73 | 74 | 70 | 58 | 89 | 69 | 115 | 155 | 99 | 102 | 86 | 83 | 64 | Cmbl |
| Cmc1 | 783 | 682 | 750 | 660 | 669 | 592 | 963 | 797 | 688 | 692 | 667 | 760 | 711 | 1013 | 779 | 544 | 721 | 752 | 701 | 829 | 884 | 702 | 758 | 820 | 677 | 792 | 705 | 922 | Cmc1 |
| Cmc2 | 1310 | 1477 | 1447 | 1327 | 1028 | 1018 | 1283 | 782 | 709 | 616 | 704 | 789 | 764 | 928 | 1240 | 1382 | 1293 | 1339 | 1234 | 1279 | 1160 | 521 | 674 | 722 | 684 | 970 | 758 | 953 | Cmc2 |
| Cmc4 | 376 | 276 | 410 | 367 | 403 | 432 | 487 | 482 | 447 | 484 | 562 | 508 | 464 | 530 | 410 | 347 | 368 | 410 | 451 | 430 | 461 | 521 | 419 | 610 | 519 | 607 | 536 | 566 | Cmc4 |
| Cmip | 5124 | 6594 | 7887 | 9478 | 10890 | 8236 | 7657 | 5555 | 6354 | 9297 | 7530 | 7681 | 6382 | 6671 | 5012 | 5706 | 9258 | 10034 | 10731 | 6995 | 9044 | 9800 | 6121 | 8592 | 9474 | 7002 | 5799 | 6055 | Cmip |
| Cmklr1 | 116 | 118 | 123 | 144 | 202 | 164 | 134 | 155 | 206 | 240 | 335 | 213 | 183 | 180 | 103 | 125 | 151 | 136 | 157 | 119 | 188 | 219 | 292 | 230 | 252 | 163 | 205 | 203 | Cmklr1 |
| Cmpk1 | 5248 | 4061 | 5095 | 5046 | 5872 | 5371 | 6080 | 9889 | 8106 | 7775 | 7949 | 8174 | 7905 | 8805 | 5918 | 4793 | 4869 | 4924 | 5217 | 5876 | 5672 | 9820 | 8515 | 8077 | 8623 | 8791 | 8125 | 9003 | Cmpk1 |
| Cmpk2 | 707 | 669 | 649 | 695 | 1225 | 941 | 1056 | 791 | 663 | 703 | 633 | 644 | 642 | 650 | 903 | 614 | 797 | 956 | 976 | 1080 | 972 | 820 | 724 | 623 | 694 | 734 | 660 | 744 | Cmpk2 |
| Cmss1 | 820 | 806 | 742 | 657 | 513 | 610 | 612 | 849 | 764 | 791 | 753 | 869 | 872 | 846 | 750 | 678 | 641 | 508 | 583 | 659 | 626 | 595 | 716 | 806 | 744 | 952 | 887 | 939 | Cmss1 |
| Cmtm1 | 0 | 0 | 0 | 0 | 0 | 0 | 0 | 0 | 0 | 1 | 0 | 0 | 0 | 0 | 1 | 0 | 0 | 0 | 0 | 0 | 0 | 0 | 0 | 0 | 0 | 0 | 0 | 0 | Cmtm1 |
| Cmtm2a | 0 | 1 | 0 | 0 | 0 | 0 | 3 | 0 | 0 | 0 | 0 | 0 | 0 | 0 | 1 | 0 | 0 | 0 | 1 | 0 | 1 | 0 | 0 | 0 | 0 | 0 | 0 | 0 | Cmtm2a |
| Cmtm2b | 0 | 0 | 0 | 0 | 0 | 0 | 0 | 0 | 0 | 0 | 0 | 1 | 0 | 0 | 0 | 0 | 2 | 0 | 0 | 0 | 0 | 0 | 0 | 0 | 0 | 0 | 0 | 0 | Cmtm2b |
| Cmtm3 | 1242 | 1245 | 1254 | 742 | 828 | 916 | 620 | 2252 | 3146 | 2381 | 3320 | 2222 | 2928 | 2122 | 1033 | 1402 | 1029 | 1030 | 697 | 667 | 659 | 1977 | 3543 | 2337 | 1902 | 3000 | 2574 | 2417 | Cmtm3 |
| Cmtm4 | 13487 | 12239 | 10277 | 12903 | 12191 | 13102 | 12867 | 7210 | 7209 | 7735 | 7157 | 7634 | 7316 | 7238 | 12597 | 12411 | 11356 | 12743 | 12772 | 11592 | 12628 | 5905 | 6916 | 7691 | 7036 | 5696 | 7499 | 7027 | Cmtm4 |
| Cmtm5 | 30 | 43 | 5 | 13 | 60 | 27 | 29 | 121 | 181 | 140 | 241 | 26 | 61 | 15 | 24 | 32 | 17 | 30 | 20 | 17 | 18 | 160 | 131 | 71 | 80 | 62 | 19 | 37 | Cmtm5 |
| Cmtm6 | 3697 | 3390 | 3701 | 3452 | 2987 | 3212 | 2873 | 1983 | 1874 | 1915 | 2026 | 2042 | 2009 | 1893 | 3421 | 3882 | 3402 | 3686 | 2839 | 2919 | 2834 | 1729 | 2067 | 2218 | 1993 | 2252 | 1897 | 1875 | Cmtm6 |
| Cmtm7 | 767 | 770 | 979 | 605 | 513 | 578 | 426 | 646 | 644 | 652 | 761 | 680 | 827 | 611 | 767 | 861 | 749 | 551 | 512 | 586 | 469 | 515 | 786 | 644 | 468 | 796 | 769 | 735 | Cmtm7 |
| Cmtm8 | 495 | 553 | 606 | 405 | 416 | 307 | 396 | 256 | 283 | 201 | 235 | 225 | 332 | 261 | 496 | 485 | 437 | 336 | 421 | 521 | 363 | 140 | 186 | 255 | 196 | 277 | 196 | 228 | Cmtm8 |
| Cmtr1 | 1271 | 1615 | 1416 | 1782 | 1628 | 1689 | 1724 | 1109 | 1084 | 1156 | 1079 | 1121 | 1115 | 1131 | 1349 | 1314 | 1702 | 1688 | 1598 | 1500 | 1851 | 1257 | 1000 | 1155 | 1309 | 1078 | 1119 | 1086 | Cmtr1 |
| Cmtr2 | 662 | 721 | 620 | 810 | 838 | 685 | 710 | 1445 | 1177 | 1101 | 919 | 1196 | 1152 | 1133 | 688 | 721 | 658 | 769 | 831 | 858 | 800 | 1253 | 1190 | 1034 | 1100 | 1210 | 1171 | 1181 | Cmtr2 |
| Cmya5 | 63 | 100 | 31 | 50 | 79 | 93 | 56 | 116 | 138 | 103 | 249 | 62 | 86 | 64 | 74 | 96 | 41 | 86 | 51 | 105 | 81 | 75 | 210 | 73 | 104 | 31 | 45 | 45 | Cmya5 |
| CN725425 | 0 | 0 | 0 | 2 | 1 | 8 | 0 | 4 | 0 | 0 | 6 | 0 | 2 | 7 | 4 | 3 | 0 | 1 | 1 | 9 | 1 | 3 | 1 | 0 | 2 | 0 | 0 | 1 | CN725425 |
| Cnbd1 | 0 | 0 | 0 | 0 | 0 | 0 | 0 | 0 | 0 | 0 | 0 | 0 | 0 | 1 | 0 | 0 | 0 | 0 | 0 | 0 | 0 | 0 | 2 | 0 | 2 | 1 | 0 | 0 | Cnbd1 |
| Cnbd2 | 233 | 301 | 237 | 396 | 322 | 274 | 324 | 228 | 220 | 240 | 239 | 253 | 231 | 236 | 222 | 248 | 312 | 322 | 321 | 282 | 348 | 272 | 240 | 288 | 298 | 186 | 201 | 204 | Cnbd2 |
| Cnbp | 32688 | 27707 | 32073 | 26301 | 23005 | 25351 | 27717 | 30759 | 27903 | 24532 | 25813 | 30890 | 29181 | 31346 | 32741 | 29862 | 27505 | 24249 | 26577 | 32123 | 25624 | 21841 | 28346 | 27881 | 26138 | 37948 | 32641 | 34868 | Cnbp |
| Cndp1 | 12 | 1 | 0 | 0 | 6 | 3 | 1 | 18 | 20 | 10 | 4 | 9 | 0 | 1 | 3 | 4 | 5 | 11 | 4 | 0 | 6 | 3 | 24 | 2 | 4 | 8 | 0 | 1 | Cndp1 |
| Cndp2 | 6868 | 8133 | 7691 | 7245 | 6103 | 6225 | 6750 | 3711 | 3676 | 3795 | 3559 | 4313 | 4369 | 4340 | 6810 | 6783 | 7350 | 7109 | 6749 | 6577 | 6792 | 3278 | 3752 | 3841 | 3789 | 4837 | 4622 | 4705 | Cndp2 |
| Cnep1r1 | 5304 | 4177 | 4493 | 4518 | 4043 | 4250 | 4438 | 4274 | 3809 | 3547 | 3870 | 4341 | 4301 | 4224 | 5231 | 4916 | 4321 | 4355 | 4006 | 4713 | 4268 | 4092 | 3983 | 3996 | 3700 | 4495 | 4320 | 4597 | Cnep1r1 |
| Cnfn | 1 | 0 | 0 | 0 | 0 | 0 | 1 | 1 | 5 | 8 | 4 | 0 | 1 | 0 | 0 | 0 | 0 | 0 | 0 | 0 | 0 | 0 | 7 | 2 | 0 | 0 | 0 | 0 | Cnfn |
| Cnga1 | 0 | 1 | 0 | 0 | 1 | 0 | 0 | 2 | 1 | 0 | 3 | 1 | 0 | 0 | 1 | 0 | 3 | 0 | 1 | 0 | 0 | 0 | 0 | 1 | 2 | 3 | 0 | 1 | Cnga1 |
| Cnga2 | 42 | 29 | 17 | 37 | 23 | 42 | 31 | 53 | 44 | 56 | 37 | 13 | 22 | 20 | 30 | 33 | 28 | 31 | 30 | 32 | 33 | 37 | 51 | 49 | 17 | 34 | 9 | 13 | Cnga2 |
| Cnga3 | 12 | 14 | 6 | 5 | 21 | 10 | 8 | 13 | 12 | 12 | 8 | 10 | 3 | 4 | 12 | 10 | 10 | 14 | 15 | 16 | 16 | 8 | 10 | 6 | 7 | 1 | 12 | 11 | Cnga3 |
| Cnga4 | 25 | 42 | 40 | 38 | 25 | 50 | 29 | 15 | 16 | 12 | 25 | 20 | 16 | 14 | 40 | 35 | 52 | 36 | 41 | 25 | 44 | 15 | 9 | 8 | 5 | 8 | 15 | 13 | Cnga4 |
| Cngb1 | 18 | 35 | 17 | 23 | 14 | 40 | 38 | 73 | 53 | 60 | 81 | 99 | 110 | 81 | 28 | 23 | 24 | 24 | 19 | 45 | 21 | 63 | 65 | 79 | 60 | 84 | 66 | 80 | Cngb1 |
| Cngb3 | 7 | 2 | 0 | 2 | 3 | 5 | 4 | 7 | 12 | 4 | 3 | 8 | 1 | 6 | 5 | 2 | 1 | 3 | 14 | 8 | 4 | 10 | 9 | 9 | 4 | 8 | 6 | 4 | Cngb3 |
| Cnih1 | 6183 | 5148 | 6043 | 5438 | 5168 | 5160 | 5826 | 6206 | 5635 | 5308 | 5536 | 5931 | 5762 | 6131 | 6551 | 5904 | 5904 | 5425 | 5301 | 5427 | 5439 | 5892 | 6026 | 5458 | 5377 | 6844 | 5596 | 6686 | Cnih1 |
| Cnih2 | 2744 | 3625 | 3039 | 4592 | 5880 | 4821 | 5228 | 4229 | 4294 | 5475 | 4413 | 4935 | 3607 | 4530 | 2712 | 2597 | 4306 | 4948 | 5887 | 4582 | 5693 | 5910 | 4045 | 5543 | 5913 | 4329 | 3980 | 4004 | Cnih2 |
| Cnih3 | 140 | 145 | 93 | 169 | 470 | 291 | 391 | 220 | 293 | 286 | 280 | 211 | 181 | 205 | 212 | 98 | 173 | 270 | 317 | 382 | 464 | 222 | 306 | 275 | 367 | 215 | 201 | 198 | Cnih3 |
| Cnih4 | 4380 | 4009 | 4600 | 4328 | 3662 | 3754 | 4226 | 3007 | 2615 | 2900 | 2905 | 3051 | 2967 | 3530 | 4396 | 3982 | 4339 | 4037 | 3865 | 4150 | 4264 | 2682 | 2691 | 3036 | 3063 | 3722 | 3310 | 3616 | Cnih4 |
| Cnksr1 | 134 | 111 | 37 | 113 | 64 | 75 | 57 | 71 | 97 | 76 | 61 | 18 | 26 | 21 | 84 | 127 | 61 | 98 | 59 | 60 | 68 | 18 | 89 | 61 | 21 | 41 | 17 | 33 | Cnksr1 |
| Cnksr2 | 1482 | 1089 | 737 | 1338 | 1714 | 1781 | 1677 | 2064 | 1917 | 1978 | 1816 | 1606 | 1801 | 1790 | 1403 | 1319 | 922 | 1202 | 1468 | 1429 | 1861 | 1863 | 1717 | 1383 | 2169 | 924 | 1806 | 1421 | Cnksr2 |
| Cnksr3 | 1088 | 1197 | 1263 | 1264 | 1108 | 940 | 1001 | 623 | 717 | 666 | 827 | 751 | 714 | 685 | 973 | 1211 | 1181 | 1165 | 1011 | 887 | 1002 | 533 | 803 | 707 | 596 | 978 | 759 | 768 | Cnksr3 |
| Cnmd | 414 | 606 | 333 | 423 | 131 | 778 | 15 | 844 | 2191 | 2157 | 4181 | 446 | 972 | 436 | 585 | 654 | 349 | 602 | 295 | 37 | 33 | 2236 | 2182 | 1385 | 1351 | 629 | 693 | 503 | Cnmd |

| | | | | | | | | | | | | | | | | | | | | | | | | | | | | | |
|---|---|---|---|---|---|---|---|---|---|---|---|---|---|---|---|---|---|---|---|---|---|---|---|---|---|---|---|---|---|
| Cnn1 | 120 | 140 | 92 | 89 | 23 | 101 | 11 | 73 | 83 | 50 | 77 | 104 | 170 | 111 | 135 | 185 | 108 | 84 | 16 | 7 | 13 | 57 | 131 | 96 | 35 | 66 | 196 | 135 | Cnn1 |
| Cnn2 | 10395 | 11297 | 9563 | 4770 | 2808 | 5138 | 1743 | 4956 | 6983 | 5317 | 6932 | 6838 | 9835 | 5932 | 9264 | 14072 | 6809 | 6141 | 2474 | 2264 | 1873 | 3865 | 8031 | 6167 | 3674 | 8613 | 8458 | 7204 | Cnn2 |
| Cnn3 | 30336 | 31797 | 29305 | 25899 | 23498 | 25156 | 24620 | 30616 | 29683 | 26665 | 27988 | 30573 | 30598 | 30126 | 31389 | 30463 | 28238 | 24678 | 25287 | 29520 | 24206 | 25716 | 31088 | 28602 | 25630 | 33862 | 32189 | 32551 | Cnn3 |
| Cnnm1 | 197 | 141 | 179 | 222 | 399 | 313 | 248 | 247 | 261 | 403 | 323 | 150 | 172 | 150 | 175 | 169 | 210 | 263 | 308 | 275 | 335 | 236 | 260 | 250 | 247 | 206 | 136 | 104 | Cnnm1 |
| Cnnm2 | 1723 | 2158 | 1881 | 2237 | 2247 | 1951 | 1943 | 884 | 892 | 1145 | 1102 | 980 | 962 | 899 | 1863 | 2081 | 2063 | 2409 | 2158 | 1638 | 2131 | 1125 | 909 | 919 | 1079 | 543 | 946 | 870 | Cnnm2 |
| Cnnm3 | 2317 | 2813 | 3127 | 3540 | 3413 | 3158 | 3272 | 2231 | 2304 | 2746 | 2651 | 2811 | 2565 | 2951 | 2354 | 2571 | 3356 | 3537 | 3397 | 2603 | 3461 | 2864 | 2288 | 2931 | 2971 | 2888 | 2583 | 2705 | Cnnm3 |
| Cnnm4 | 2103 | 2439 | 2649 | 3295 | 2684 | 2443 | 2567 | 655 | 833 | 972 | 1004 | 960 | 889 | 927 | 2022 | 2412 | 2936 | 3379 | 2689 | 1813 | 2765 | 876 | 772 | 1004 | 899 | 1121 | 850 | 926 | Cnnm4 |
| Cnot1 | 34800 | 32915 | 20175 | 23345 | 23762 | 27625 | 27598 | 33352 | 29722 | 23800 | 24496 | 28063 | 26771 | 28166 | 33230 | 29918 | 21667 | 23343 | 23605 | 30135 | 27605 | 22264 | 27965 | 22081 | 24916 | 19537 | 30386 | 27098 | Cnot1 |
| Cnot10 | 8980 | 9456 | 6900 | 7561 | 6895 | 7114 | 7445 | 7058 | 7215 | 6700 | 6910 | 6962 | 6645 | 6867 | 8625 | 8553 | 6831 | 7311 | 7297 | 8060 | 7256 | 6557 | 6963 | 6365 | 6312 | 6352 | 7358 | 6986 | Cnot10 |
| Cnot11 | 3809 | 3313 | 3532 | 3362 | 3256 | 3030 | 3252 | 2989 | 2974 | 2909 | 2775 | 3410 | 2865 | 3065 | 3429 | 3576 | 3466 | 3311 | 3282 | 2991 | 3220 | 3034 | 2960 | 3383 | 3066 | 3905 | 2936 | 3348 | Cnot11 |
| Cnot2 | 6905 | 5686 | 6444 | 5884 | 5800 | 5959 | 6620 | 11057 | 9323 | 8902 | 8564 | 9245 | 9208 | 9718 | 7100 | 6309 | 6434 | 5384 | 6010 | 6282 | 6156 | 9621 | 9088 | 9277 | 8689 | 10408 | 9618 | 10030 | Cnot2 |
| Cnot3 | 4850 | 5630 | 5863 | 6353 | 5565 | 5172 | 5108 | 5191 | 5494 | 6488 | 6016 | 7311 | 6016 | 6846 | 4334 | 4787 | 6146 | 6015 | 6318 | 4229 | 5366 | 6316 | 5141 | 6865 | 6817 | 7557 | 6250 | 6457 | Cnot3 |
| Cnot4 | 3160 | 3105 | 3191 | 3313 | 3272 | 2869 | 3351 | 3665 | 3349 | 3262 | 3014 | 3169 | 3159 | 3317 | 3265 | 2984 | 3189 | 3285 | 3074 | 3195 | 3138 | 3711 | 3147 | 3255 | 3329 | 3558 | 3353 | 3440 | Cnot4 |
| Cnot6 | 18589 | 12591 | 18242 | 17563 | 19046 | 20496 | 21071 | 24119 | 21204 | 21939 | 22894 | 28279 | 25975 | 28884 | 18213 | 15389 | 17712 | 17850 | 18937 | 17477 | 19842 | 24401 | 20542 | 23883 | 25141 | 25972 | 28023 | 28544 | Cnot6 |
| Cnot6l | 6273 | 4384 | 5427 | 6261 | 6905 | 7340 | 7420 | 6625 | 5906 | 5598 | 6733 | 5870 | 6078 | 6308 | 5649 | 5355 | 5457 | 6560 | 6551 | 5730 | 7621 | 6291 | 5998 | 5583 | 6085 | 5227 | 5751 | 5868 | Cnot6l |
| Cnot7 | 12209 | 9094 | 12021 | 12095 | 12470 | 12165 | 14797 | 16203 | 13030 | 14309 | 13840 | 16706 | 14561 | 18191 | 12242 | 10129 | 12388 | 12481 | 12608 | 12756 | 13643 | 16476 | 13024 | 16096 | 16280 | 17055 | 16338 | 17463 | Cnot7 |
| Cnot8 | 3676 | 3715 | 3846 | 3764 | 3769 | 3526 | 4051 | 4995 | 4598 | 4470 | 4135 | 4985 | 4329 | 5191 | 3790 | 3434 | 3913 | 3715 | 4102 | 3864 | 3896 | 4978 | 4275 | 5125 | 4707 | 5285 | 4724 | 5067 | Cnot8 |
| Cnot9 | 6771 | 6265 | 6404 | 6068 | 5658 | 5920 | 6715 | 7068 | 6446 | 6410 | 6170 | 7941 | 7250 | 8522 | 6166 | 5698 | 6528 | 5838 | 6483 | 6152 | 6482 | 6490 | 6097 | 7414 | 6820 | 9150 | 8085 | 9044 | Cnot9 |
| Cnp | 1189 | 1217 | 1213 | 933 | 1195 | 1119 | 949 | 1031 | 1180 | 1114 | 1069 | 921 | 987 | 840 | 1299 | 1218 | 1061 | 1040 | 1034 | 1003 | 998 | 1056 | 1197 | 885 | 1016 | 1005 | 933 | 795 | Cnp |
| Cnppd1 | 3403 | 4125 | 4361 | 4651 | 4808 | 4094 | 4305 | 3318 | 3435 | 3883 | 3636 | 3863 | 3936 | 3612 | 3603 | 3713 | 4731 | 4774 | 4955 | 4028 | 4370 | 4127 | 3476 | 3992 | 4131 | 3984 | 3763 | 3853 | Cnppd1 |
| Cnpy1 | 3520 | 3487 | 5250 | 3905 | 3023 | 4056 | 855 | 5019 | 3428 | 3837 | 2756 | 5757 | 8869 | 5141 | 2972 | 5658 | 4890 | 2588 | 3869 | 4661 | 2761 | 5401 | 3744 | 2589 | 5233 | 404 | 7792 | 6451 | Cnpy1 |
| Cnpy2 | 2580 | 2492 | 2925 | 2281 | 2141 | 2167 | 2086 | 3673 | 3633 | 3446 | 3734 | 3467 | 3801 | 3963 | 2920 | 2558 | 2624 | 2025 | 2243 | 2460 | 2298 | 3016 | 3880 | 3826 | 3251 | 4230 | 3918 | 4143 | Cnpy2 |
| Cnpy3 | 2182 | 2555 | 2186 | 2329 | 2053 | 2222 | 2058 | 1804 | 2118 | 1900 | 1983 | 2057 | 2064 | 1852 | 2103 | 2226 | 2168 | 2113 | 2221 | 2149 | 2090 | 1905 | 2289 | 1995 | 1893 | 2115 | 1952 | 1946 | Cnpy3 |
| Cnpy4 | 2343 | 2009 | 2877 | 1990 | 1881 | 1974 | 1975 | 3270 | 2801 | 2825 | 3111 | 3245 | 3185 | 3444 | 2197 | 2044 | 2458 | 1808 | 2057 | 2172 | 1657 | 2938 | 3010 | 3322 | 2934 | 3948 | 3362 | 3458 | Cnpy4 |
| Cnr1 | 2506 | 2371 | 1934 | 3927 | 6633 | 6138 | 6850 | 10453 | 9711 | 9293 | 8161 | 7960 | 9217 | 8730 | 3642 | 2294 | 3242 | 4250 | 5760 | 6469 | 6293 | 9725 | 8587 | 8798 | 11677 | 6132 | 7030 | 6912 | Cnr1 |
| Cnr2 | 29 | 33 | 23 | 18 | 26 | 34 | 21 | 18 | 27 | 10 | 23 | 27 | 39 | 26 | 33 | 36 | 25 | 22 | 21 | 26 | 23 | 12 | 34 | 17 | 19 | 35 | 44 | 28 | Cnr2 |
| Cnrip1 | 741 | 627 | 582 | 850 | 1547 | 1306 | 1354 | 2020 | 2053 | 1917 | 1809 | 1413 | 1433 | 1558 | 909 | 681 | 753 | 1062 | 1258 | 1307 | 1359 | 2388 | 2062 | 1714 | 2119 | 1385 | 1282 | 1388 | Cnrip1 |
| Cnst | 4036 | 4179 | 4129 | 4966 | 5088 | 4813 | 4630 | 3965 | 3881 | 4212 | 4234 | 4000 | 3949 | 4032 | 4187 | 4199 | 4486 | 4938 | 5160 | 4280 | 4871 | 4660 | 3633 | 4299 | 4347 | 3816 | 3909 | 4020 | Cnst |
| Cntd1 | 115 | 95 | 140 | 90 | 63 | 125 | 77 | 137 | 120 | 115 | 138 | 141 | 149 | 159 | 99 | 111 | 74 | 84 | 83 | 73 | 85 | 115 | 109 | 127 | 108 | 133 | 163 | 114 | Cntd1 |
| Cntf | 39 | 41 | 56 | 42 | 41 | 45 | 48 | 39 | 30 | 41 | 39 | 42 | 57 | 53 | 54 | 21 | 50 | 29 | 43 | 55 | 43 | 23 | 34 | 38 | 42 | 40 | 50 | 42 | Cntf |
| Cntfr | 3676 | 5288 | 6469 | 5920 | 5691 | 5356 | 3956 | 3209 | 3687 | 4360 | 3896 | 5353 | 4780 | 4367 | 3940 | 4879 | 6470 | 5919 | 6353 | 4069 | 4885 | 4234 | 3451 | 4854 | 4354 | 4789 | 4635 | 4379 | Cntfr |
| Cntln | 2493 | 1861 | 1941 | 1741 | 1662 | 2223 | 1914 | 3791 | 3515 | 2991 | 3234 | 3621 | 3523 | 3672 | 2393 | 2340 | 1735 | 1768 | 1538 | 2068 | 1699 | 2714 | 3425 | 3003 | 2991 | 3386 | 3844 | 3445 | Cntln |
| Cntn1 | 1223 | 1318 | 1209 | 1405 | 2955 | 1942 | 2075 | 2146 | 2273 | 2058 | 1874 | 1115 | 1471 | 1133 | 1776 | 1407 | 1274 | 2012 | 1879 | 2199 | 2132 | 2784 | 2455 | 1285 | 2141 | 627 | 1103 | 1037 | Cntn1 |
| Cntn2 | 7623 | 10766 | 14810 | 16384 | 23316 | 18631 | 14626 | 17154 | 19033 | 26298 | 21619 | 25417 | 30991 | 23814 | 11036 | 12196 | 17985 | 18343 | 21111 | 12991 | 19262 | 29335 | 17416 | 24329 | 32969 | 10181 | 24575 | 26285 | Cntn2 |
| Cntn3 | 308 | 310 | 284 | 499 | 740 | 550 | 579 | 451 | 671 | 570 | 751 | 470 | 412 | 434 | 398 | 340 | 337 | 614 | 538 | 448 | 614 | 730 | 724 | 525 | 680 | 288 | 359 | 411 | Cntn3 |
| Cntn4 | 498 | 416 | 240 | 486 | 1023 | 532 | 593 | 1081 | 928 | 889 | 989 | 346 | 469 | 327 | 491 | 469 | 304 | 600 | 535 | 715 | 641 | 1063 | 998 | 586 | 614 | 348 | 352 | 341 | Cntn4 |
| Cntn5 | 311 | 360 | 247 | 368 | 516 | 471 | 479 | 576 | 570 | 490 | 485 | 324 | 313 | 331 | 398 | 276 | 315 | 418 | 372 | 520 | 459 | 488 | 557 | 384 | 591 | 159 | 334 | 306 | Cntn5 |
| Cntn6 | 381 | 308 | 294 | 333 | 434 | 401 | 409 | 488 | 492 | 392 | 406 | 342 | 269 | 397 | 339 | 295 | 309 | 380 | 365 | 492 | 448 | 560 | 456 | 415 | 493 | 321 | 373 | 451 | Cntn6 |
| Cntnap1 | 346 | 433 | 519 | 664 | 1027 | 705 | 549 | 339 | 444 | 549 | 657 | 326 | 361 | 272 | 434 | 404 | 601 | 908 | 626 | 552 | 599 | 712 | 519 | 350 | 452 | 246 | 317 | 228 | Cntnap1 |
| Cntnap2 | 11870 | 11604 | 11365 | 13527 | 12827 | 13701 | 16290 | 9449 | 8559 | 8190 | 6828 | 11486 | 7418 | 12118 | 12890 | 8542 | 12614 | 12775 | 14572 | 15949 | 14449 | 8968 | 7587 | 9797 | 10191 | 13329 | 11321 | 11796 | Cntnap2 |
| Cntnap3 | 51 | 48 | 9 | 80 | 106 | 65 | 131 | 126 | 227 | 142 | 194 | 148 | 2 | 181 | 54 | 15 | 49 | 64 | 108 | 98 | 167 | 155 | 153 | 200 | 249 | 136 | 117 | 100 | Cntnap3 |
| Cntnap4 | 486 | 523 | 357 | 734 | 1488 | 1131 | 1229 | 1071 | 1343 | 1169 | 1107 | 640 | 561 | 674 | 618 | 363 | 574 | 1061 | 1144 | 1448 | 1404 | 1419 | 1418 | 768 | 1311 | 440 | 531 | 502 | Cntnap4 |
| Cntnap5a | 468 | 549 | 271 | 634 | 1211 | 1027 | 898 | 1114 | 968 | 921 | 936 | 471 | 644 | 465 | 704 | 558 | 480 | 832 | 777 | 1137 | 1073 | 1340 | 1040 | 426 | 963 | 152 | 525 | 478 | Cntnap5a |
| Cntnap5b | 194 | 172 | 57 | 166 | 311 | 330 | 276 | 350 | 301 | 289 | 273 | 156 | 184 | 162 | 237 | 188 | 126 | 173 | 215 | 338 | 400 | 256 | 318 | 158 | 272 | 31 | 187 | 136 | Cntnap5b |
| Cntnap5c | 54 | 68 | 13 | 74 | 142 | 154 | 145 | 143 | 101 | 129 | 115 | 70 | 95 | 95 | 76 | 72 | 35 | 70 | 86 | 146 | 186 | 95 | 111 | 50 | 135 | 7 | 89 | 57 | Cntnap5c |
| Cntrl | 7108 | 6740 | 5205 | 5165 | 4607 | 5713 | 5547 | 7866 | 7434 | 6393 | 6597 | 8681 | 7622 | 8944 | 6622 | 6432 | 5501 | 4638 | 4737 | 6107 | 5449 | 5294 | 6359 | 6698 | 6676 | 6313 | 9885 | 7637 | Cntrl |
| Cntrob | 3700 | 4311 | 4901 | 5124 | 4576 | 4307 | 4638 | 3811 | 4120 | 4948 | 4392 | 6059 | 4910 | 5832 | 3572 | 3794 | 5475 | 4916 | 4942 | 4112 | 4459 | 4223 | 3708 | 5522 | 5043 | 5896 | 5765 | 5480 | Cntrob |
| Coa3 | 1929 | 2240 | 2329 | 2088 | 1954 | 1958 | 2002 | 2339 | 2532 | 2629 | 2382 | 2685 | 2683 | 2790 | 2028 | 2127 | 2291 | 2065 | 2134 | 2128 | 2132 | 2239 | 2660 | 2775 | 2336 | 2954 | 2703 | 2858 | Coa3 |
| Coa4 | 391 | 411 | 485 | 412 | 317 | 302 | 359 | 429 | 358 | 455 | 366 | 416 | 471 | 461 | 425 | 329 | 400 | 335 | 372 | 384 | 324 | 366 | 439 | 416 | 371 | 459 | 453 | 453 | Coa4 |
| Coa5 | 6351 | 5933 | 5692 | 5050 | 5087 | 5229 | 4808 | 6581 | 6133 | 5697 | 5642 | 6241 | 5736 | 5862 | 6535 | 5837 | 5320 | 4932 | 4894 | 5259 | 4826 | 6296 | 6180 | 5605 | 5909 | 6091 | 6150 | 5967 | Coa5 |
| Coa6 | 578 | 587 | 571 | 640 | 483 | 515 | 586 | 465 | 433 | 472 | 496 | 454 | 416 | 508 | 576 | 571 | 612 | 578 | 532 | 661 | 616 | 343 | 417 | 478 | 419 | 499 | 467 | 462 | Coa6 |
| Coa7 | 1100 | 1207 | 1327 | 1159 | 1177 | 1179 | 1162 | 1685 | 1643 | 1695 | 1553 | 2019 | 1700 | 1961 | 1138 | 1110 | 1302 | 1052 | 1266 | 1195 | 1203 | 1551 | 1698 | 1692 | 1828 | 1936 | 2077 | 2055 | Coa7 |
| Coa8 | 1020 | 1100 | 1075 | 990 | 906 | 1020 | 1049 | 1015 | 1082 | 1132 | 1019 | 1168 | 1169 | 1197 | 1058 | 949 | 1001 | 1010 | 1057 | 999 | 978 | 966 | 1001 | 1219 | 1200 | 1358 | 1268 | 1246 | Coa8 |
| Coasy | 2016 | 2499 | 2350 | 2357 | 1901 | 1938 | 2030 | 2169 | 2267 | 2311 | 2088 | 2510 | 2278 | 2237 | 2048 | 2287 | 2271 | 2010 | 2179 | 2073 | 1902 | 2125 | 2230 | 2409 | 2291 | 2567 | 2595 | 2465 | Coasy |
| Cobl | 751 | 666 | 571 | 755 | 1170 | 855 | 702 | 1067 | 1213 | 1453 | 1531 | 882 | 1015 | 837 | 738 | 955 | 627 | 774 | 849 | 713 | 852 | 1744 | 1654 | 994 | 1182 | 849 | 712 | 786 | Cobl |
| Cobll1 | 594 | 532 | 658 | 608 | 756 | 619 | 675 | 546 | 604 | 665 | 738 | 714 | 721 | 728 | 563 | 524 | 597 | 691 | 659 | 569 | 732 | 769 | 638 | 669 | 704 | 658 | 532 | 595 | Cobll1 |
| Coch | 803 | 855 | 787 | 739 | 955 | 797 | 844 | 547 | 660 | 587 | 530 | 484 | 553 | 412 | 1007 | 868 | 676 | 879 | 747 | 959 | 782 | 722 | 806 | 443 | 523 | 318 | 408 | 450 | Coch |
| Cog1 | 1705 | 2109 | 1951 | 1680 | 1987 | 1857 | 1717 | 3053 | 3549 | 3593 | 3060 | 3493 | 3618 | 3297 | 1825 | 1776 | 1882 | 1779 | 2098 | 1911 | 1863 | 3484 | 3319 | 3517 | 3618 | 3741 | 3277 | 3357 | Cog1 |
| Cog2 | 3346 | 3616 | 2596 | 2976 | 2580 | 3012 | 2914 | 3086 | 3330 | 3018 | 2880 | 3120 | 2834 | 2866 | 3390 | 3348 | 2705 | 2821 | 2776 | 2987 | 2978 | 2862 | 3342 | 2766 | 2907 | 2974 | 3226 | 3069 | Cog2 |
| Cog3 | 5429 | 5140 | 4470 | 4762 | 4545 | 4907 | 5064 | 4537 | 4298 | 3995 | 4049 | 4189 | 3944 | 4191 | 5261 | 4998 | 4704 | 4846 | 4488 | 5072 | 4824 | 4040 | 4240 | 3952 | 4130 | 3673 | 4188 | 3912 | Cog3 |
| Cog4 | 4094 | 4171 | 3520 | 3423 | 3126 | 3126 | 3019 | 4866 | 4796 | 4108 | 4165 | 4018 | 4322 | 3709 | 3883 | 3897 | 3135 | 3036 | 3080 | 3364 | 3130 | 4483 | 5099 | 4192 | 4094 | 4277 | 4477 | 4269 | Cog4 |
| Cog5 | 6403 | 5858 | 4132 | 4229 | 4313 | 5022 | 4676 | 6872 | 5986 | 4629 | 4988 | 4651 | 5104 | 4934 | 6490 | 5810 | 4212 | 4118 | 4308 | 5688 | 4905 | 4670 | 6188 | 3879 | 4268 | 3937 | 4966 | 4747 | Cog5 |
| Cog6 | 4087 | 3729 | 3346 | 3373 | 3557 | 3620 | 3675 | 5444 | 4635 | 4465 | 4304 | 4369 | 3814 | 4623 | 4152 | 3720 | 3487 | 3416 | 3451 | 3822 | 3728 | 4771 | 4697 | 4465 | 4653 | 4021 | 4159 | 4289 | Cog6 |
| Cog7 | 1192 | 1306 | 1238 | 1250 | 1231 | 1182 | 1225 | 1308 | 1556 | 1474 | 1498 | 1599 | 1590 | 1632 | 1204 | 1249 | 1260 | 1195 | 1276 | 1199 | 1127 | 1512 | 1492 | 1511 | 1424 | 1698 | 1735 | 1515 | Cog7 |
| Cog8 | 1633 | 2093 | 1688 | 1659 | 1692 | 1549 | 1711 | 1595 | 1819 | 1913 | 1807 | 1807 | 1732 | 1837 | 1598 | 1591 | 1819 | 1652 | 1595 | 1526 | 1663 | 1846 | 1848 | 1989 | 1797 | 1920 | 1768 | 2021 | Cog8 |
| Coil | 1135 | 1254 | 1446 | 1140 | 1415 | 1181 | 1273 | 2677 | 2704 | 2746 | 2575 | 3018 | 2837 | 3094 | 1230 | 1121 | 1557 | 1144 | 1319 | 1218 | 1214 | 2617 | 2628 | 2952 | 2821 | 3269 | 3240 | 3227 | Coil |
| Col10a1 | 3 | 11 | 1 | 10 | 4 | 7 | 9 | 6 | 8 | 12 | 177 | 0 | 2 | 5 | 4 | 4 | 5 | 14 | 7 | 15 | 10 | 8 | 12 | 28 | 7 | 2 | 6 | 1 | Col10a1 |
| Col11a1 | 13095 | 13735 | 13672 | 11927 | 9390 | 14188 | 8642 | 7660 | 18273 | 18332 | 33598 | 10264 | 14108 | 10631 | 13123 | 16168 | 12928 | 14011 | 9405 | 8424 | 8191 | 20879 | 14516 | 14671 | 14568 | 9312 | 12434 | 11191 | Col11a1 |
| Col11a2 | 1458 | 2147 | 2314 | 2947 | 2608 | 3068 | 1888 | 1503 | 4771 | 5667 | 11509 | 1628 | 1751 | 1488 | 1462 | 1953 | 2737 | 3734 | 3246 | 1854 | 1922 | 6829 | 2871 | 3789 | 4692 | 1525 | 1582 | 1396 | Col11a2 |
| Col12a1 | 9687 | 11399 | 5194 | 8716 | 3431 | 16154 | 1191 | 14919 | 26139 | 23824 | 37634 | 14828 | 22304 | 16156 | 11706 | 13225 | 5766 | 14105 | 2840 | 1654 | 1433 | 21398 | 30485 | 17236 | 22885 | 6615 | 17882 | 11915 | Col12a1 |
| Col13a1 | 305 | 253 | 360 | 212 | 241 | 194 | 126 | 214 | 285 | 272 | 423 | 305 | 361 | 291 | 225 | 382 | 217 | 272 | 127 | 209 | 159 | 247 | 360 | 311 | 210 | 348 | 343 | 250 | Col13a1 |
| Col14a1 | 1284 | 1494 | 636 | 1489 | 763 | 2177 | 120 | 1569 | 3618 | 3641 | 7234 | 1348 | 2629 | 1588 | 1515 | 1604 | 864 | 2333 | 316 | 169 | 154 | 3905 | 5067 | 2562 | 2804 | 949 | 1727 | 1221 | Col14a1 |
| Col15a1 | 1218 | 1386 | 1465 | 1129 | 993 | 1150 | 646 | 1374 | 2328 | 2799 | 3850 | 1854 | 2625 | 1592 | 947 | 1748 | 1306 | 1130 | 825 | 522 | 699 | 2125 | 2414 | 2442 | 1755 | 2747 | 1701 | 1448 | Col15a1 |
| Col16a1 | 2094 | 2644 | 2466 | 3184 | 1415 | 3979 | 582 | 2683 | 5271 | 6252 | 9649 | 3616 | 4643 | 3335 | 2041 | 3191 | 2313 | 4832 | 856 | 661 | 587 | 6318 | 6957 | 5891 | 5744 | 3205 | 4049 | 3135 | Col16a1 |
| Col17a1 | 482 | 134 | 89 | 29 | 4 | 132 | 6 | 785 | 1491 | 663 | 425 | 169 | 294 | 138 | 141 | 178 | 60 | 117 | 22 | 21 | 5 | 53 | 1583 | 511 | 108 | 169 | 123 | 163 | Col17a1 |
| Col18a1 | 10385 | 12437 | 15157 | 9407 | 9316 | 8698 | 6746 | 6467 | 8874 | 8953 | 10491 | 11157 | 12582 | 9872 | 9014 | 13621 | 11143 | 9945 | 8376 | 6693 | 7010 | 7800 | 10110 | 10015 | 7662 | 13831 | 10528 | 10666 | Col18a1 |
| Col19a1 | 46 | 50 | 11 | 36 | 61 | 70 | 55 | 110 | 121 | 68 | 206 | 22 | 41 | 51 | 57 | 54 | 25 | 30 | 48 | 64 | 63 | 37 | 155 | 38 | 78 | 15 | 27 | 37 | Col19a1 |
| Col1a1 | 35047 | 60361 | 55735 | 50426 | 16218 | 61595 | 5728 | 30874 | 48667 | 64571 | 90332 | 62167 | 71531 | 53408 | 38272 | 61070 | 46494 | 79382 | 11744 | 6960 | 6217 | 82898 | 80208 | 73387 | 76557 | 48638 | 67642 | 55085 | Col1a1 |
| Col1a2 | 46540 | 60563 | 63980 | 50123 | 21138 | 65340 | 9466 | 41947 | 67891 | 73802 | 113459 | 62289 | 83284 | 56528 | 47957 | 73398 | 45590 | 74783 | 17211 | 11728 | 10198 | 78717 | 102764 | 77592 | 75383 | 59905 | 67727 | 57463 | Col1a2 |

| | | | | | | | | | | | | | | | | | | | | | | | | | | | | | |
|---|---|---|---|---|---|---|---|---|---|---|---|---|---|---|---|---|---|---|---|---|---|---|---|---|---|---|---|---|---|
| Col20a1 | 465 | 590 | 432 | 701 | 555 | 429 | 466 | 227 | 301 | 287 | 517 | 192 | 235 | 184 | 498 | 612 | 535 | 757 | 475 | 390 | 443 | 324 | 390 | 199 | 177 | 122 | 149 | 182 | Col20a1 |
| Col22a1 | 885 | 1288 | 1419 | 1226 | 1640 | 1205 | 1417 | 1439 | 1711 | 1380 | 1643 | 1502 | 1225 | 1467 | 1233 | 985 | 1315 | 1510 | 1411 | 1708 | 1408 | 2015 | 1867 | 1516 | 1757 | 1567 | 1507 | 1544 | Col22a1 |
| Col23a1 | 3147 | 2911 | 3491 | 2053 | 1799 | 1996 | 1154 | 2120 | 3470 | 2677 | 3689 | 2862 | 3780 | 2517 | 2614 | 4038 | 2300 | 2126 | 1249 | 1735 | 1362 | 2794 | 3585 | 2385 | 2077 | 2543 | 3142 | 2516 | Col23a1 |
| Col24a1 | 435 | 375 | 319 | 202 | 207 | 275 | 113 | 342 | 628 | 521 | 1120 | 292 | 401 | 243 | 378 | 570 | 243 | 332 | 125 | 119 | 121 | 386 | 851 | 397 | 368 | 253 | 243 | 244 | Col24a1 |
| Col25a1 | 2535 | 1285 | 1203 | 1375 | 1773 | 1825 | 1836 | 2265 | 2782 | 1972 | 3397 | 1320 | 1649 | 1348 | 1891 | 2476 | 1271 | 1730 | 1657 | 1432 | 1972 | 2525 | 2487 | 1648 | 1847 | 1098 | 1549 | 1297 | Col25a1 |
| Col26a1 | 8202 | 8528 | 7983 | 5144 | 3165 | 5206 | 1892 | 6593 | 10317 | 7136 | 11136 | 6096 | 10020 | 5723 | 6649 | 10895 | 5268 | 5975 | 2664 | 2406 | 2007 | 5782 | 11567 | 7863 | 5173 | 9106 | 7180 | 6713 | Col26a1 |
| Col27a1 | 2014 | 1849 | 2232 | 1666 | 1159 | 1674 | 875 | 2359 | 4567 | 5179 | 7943 | 2039 | 3161 | 1814 | 1557 | 2524 | 1837 | 1805 | 1217 | 958 | 839 | 4002 | 3875 | 4135 | 3025 | 3003 | 2274 | 1951 | Col27a1 |
| Col28a1 | 36 | 24 | 17 | 13 | 160 | 44 | 30 | 129 | 201 | 104 | 228 | 15 | 62 | 17 | 37 | 37 | 20 | 86 | 31 | 44 | 55 | 257 | 137 | 29 | 139 | 31 | 21 | 23 | Col28a1 |
| Col2a1 | 13352 | 19669 | 21029 | 16046 | 9945 | 17987 | 6002 | 14480 | 34820 | 38353 | 70390 | 19930 | 28354 | 15384 | 14068 | 24464 | 16807 | 18225 | 11980 | 6905 | 6211 | 45388 | 24324 | 29218 | 29322 | 28321 | 22486 | 17899 | Col2a1 |
| Col3a1 | 50497 | 54706 | 63777 | 42514 | 24020 | 49470 | 15678 | 50006 | 69478 | 65298 | 119396 | 65751 | 84662 | 59196 | 49538 | 71555 | 41354 | 56667 | 20287 | 21227 | 15598 | 57824 | 109383 | 70063 | 57427 | 70628 | 69910 | 61638 | Col3a1 |
| Col4a1 | 20677 | 27288 | 37681 | 25302 | 26934 | 25885 | 19708 | 18309 | 24531 | 31014 | 36331 | 34944 | 39551 | 30744 | 19697 | 31263 | 28154 | 28380 | 23944 | 17187 | 21260 | 27616 | 29005 | 34689 | 29513 | 36887 | 33484 | 30769 | Col4a1 |
| Col4a2 | 14626 | 20236 | 26074 | 17774 | 17677 | 17554 | 12103 | 11453 | 15748 | 20479 | 22787 | 22237 | 25175 | 18817 | 13440 | 22534 | 19675 | 20400 | 16172 | 11517 | 13274 | 18539 | 18258 | 22470 | 19362 | 24931 | 22430 | 19630 | Col4a2 |
| Col4a3 | 14 | 13 | 7 | 5 | 34 | 29 | 20 | 29 | 39 | 38 | 47 | 30 | 50 | 37 | 25 | 36 | 9 | 33 | 24 | 20 | 17 | 35 | 63 | 20 | 47 | 11 | 30 | 21 | Col4a3 |
| Col4a4 | 104 | 67 | 87 | 63 | 88 | 101 | 104 | 84 | 101 | 135 | 115 | 129 | 109 | 114 | 94 | 82 | 85 | 99 | 88 | 75 | 121 | 58 | 109 | 86 | 132 | 70 | 66 | 80 | Col4a4 |
| Col4a5 | 3425 | 3571 | 3592 | 2783 | 3500 | 3512 | 2704 | 3243 | 3728 | 3296 | 3962 | 3568 | 4295 | 3397 | 3371 | 3923 | 3195 | 3422 | 2980 | 3627 | 2798 | 3008 | 4501 | 3262 | 3243 | 2561 | 3741 | 3300 | Col4a5 |
| Col4a6 | 29 | 58 | 47 | 59 | 84 | 85 | 63 | 46 | 62 | 35 | 84 | 43 | 75 | 54 | 53 | 65 | 56 | 51 | 51 | 76 | 64 | 30 | 52 | 47 | 68 | 41 | 67 | 62 | Col4a6 |
| Col5a1 | 11037 | 13826 | 14287 | 9842 | 6380 | 11818 | 3596 | 8783 | 15708 | 14625 | 27671 | 12926 | 17352 | 11672 | 9877 | 16756 | 11283 | 13804 | 5358 | 3942 | 3841 | 13994 | 22489 | 14931 | 12418 | 12243 | 13462 | 11576 | Col5a1 |
| Col5a2 | 16246 | 16326 | 17238 | 12721 | 7162 | 16616 | 5307 | 15807 | 22209 | 19266 | 29006 | 19956 | 25776 | 19349 | 16456 | 21431 | 13041 | 17241 | 7029 | 7446 | 5322 | 18478 | 31732 | 21919 | 19243 | 19813 | 22718 | 20425 | Col5a2 |
| Col5a3 | 205 | 290 | 247 | 176 | 179 | 201 | 68 | 116 | 259 | 179 | 406 | 100 | 248 | 131 | 212 | 347 | 234 | 193 | 99 | 92 | 92 | 284 | 389 | 127 | 168 | 117 | 151 | 129 | Col5a3 |
| Col6a1 | 4532 | 5133 | 4507 | 3807 | 2701 | 5952 | 1371 | 4711 | 8970 | 6726 | 15189 | 5162 | 7406 | 4585 | 4163 | 6122 | 3456 | 6090 | 2307 | 1953 | 1667 | 6129 | 13674 | 6174 | 5470 | 3735 | 5326 | 4466 | Col6a1 |
| Col6a2 | 5210 | 6200 | 5366 | 4570 | 3201 | 6314 | 1373 | 5337 | 9516 | 7611 | 14729 | 5925 | 8405 | 5281 | 4981 | 6918 | 4304 | 6707 | 2614 | 1808 | 1864 | 6868 | 13986 | 7391 | 6015 | 4767 | 6596 | 5478 | Col6a2 |
| Col6a3 | 7273 | 7275 | 4627 | 6196 | 4073 | 8355 | 2094 | 7334 | 11926 | 12547 | 22805 | 8287 | 11083 | 7785 | 5458 | 8564 | 4784 | 10071 | 2865 | 2461 | 2372 | 10288 | 17026 | 10292 | 9760 | 4862 | 8164 | 7355 | Col6a3 |
| Col6a4 | 58 | 61 | 34 | 53 | 47 | 71 | 83 | 92 | 119 | 81 | 95 | 96 | 68 | 74 | 40 | 30 | 45 | 56 | 60 | 75 | 77 | 117 | 72 | 86 | 79 | 76 | 91 | 74 | Col6a4 |
| Col6a5 | 18 | 18 | 10 | 48 | 42 | 66 | 51 | 48 | 59 | 77 | 91 | 47 | 23 | 53 | 26 | 18 | 35 | 26 | 29 | 81 | 46 | 77 | 65 | 65 | 81 | 33 | 29 | 27 | Col6a5 |
| Col6a6 | 112 | 110 | 93 | 109 | 61 | 66 | 38 | 112 | 294 | 260 | 1255 | 136 | 252 | 133 | 90 | 139 | 81 | 84 | 47 | 73 | 52 | 197 | 1300 | 192 | 85 | 174 | 196 | 184 | Col6a6 |
| Col7a1 | 1374 | 1231 | 1594 | 1113 | 936 | 1270 | 773 | 1165 | 1908 | 1453 | 1370 | 1114 | 1514 | 1057 | 960 | 1423 | 1220 | 1468 | 1067 | 702 | 774 | 652 | 1718 | 1651 | 757 | 993 | 1124 | 1104 | Col7a1 |
| Col8a1 | 1209 | 991 | 683 | 741 | 456 | 1887 | 127 | 2804 | 3703 | 2651 | 4232 | 2204 | 2484 | 2157 | 1312 | 1496 | 671 | 1900 | 254 | 152 | 113 | 2070 | 4554 | 3046 | 2014 | 1833 | 1918 | 1943 | Col8a1 |
| Col8a2 | 5090 | 7224 | 6187 | 3943 | 1699 | 4519 | 497 | 3480 | 5264 | 5712 | 7671 | 5148 | 7400 | 4740 | 5625 | 8844 | 4704 | 5439 | 898 | 607 | 496 | 6184 | 7449 | 5348 | 5324 | 5852 | 5919 | 5294 | Col8a2 |
| Col9a1 | 2148 | 2204 | 2010 | 1460 | 541 | 2395 | 177 | 2788 | 9372 | 7335 | 15830 | 1429 | 3309 | 1327 | 2342 | 3883 | 1197 | 2013 | 760 | 156 | 145 | 8300 | 6091 | 4707 | 4529 | 3171 | 1755 | 1345 | Col9a1 |
| Col9a2 | 1419 | 1779 | 1318 | 1562 | 1201 | 1693 | 534 | 2312 | 5530 | 5610 | 10252 | 1420 | 2665 | 1332 | 1576 | 2244 | 1224 | 1958 | 935 | 585 | 558 | 5700 | 4293 | 3840 | 3534 | 2711 | 1936 | 1335 | Col9a2 |
| Col9a3 | 1550 | 2135 | 1147 | 2354 | 2027 | 3771 | 973 | 3109 | 9709 | 8793 | 15504 | 2246 | 4018 | 1910 | 1880 | 2223 | 1418 | 3057 | 2040 | 771 | 958 | 9690 | 7288 | 5307 | 6362 | 2918 | 2121 | 1508 | Col9a3 |
| Colca2 | 31 | 40 | 40 | 17 | 26 | 22 | 20 | 92 | 76 | 87 | 82 | 56 | 64 | 55 | 24 | 26 | 32 | 14 | 16 | 22 | 28 | 28 | 113 | 87 | 44 | 80 | 50 | 44 | Colca2 |
| Colec10 | 13 | 0 | 0 | 2 | 1 | 6 | 1 | 9 | 18 | 8 | 44 | 0 | 3 | 0 | 1 | 5 | 1 | 0 | 0 | 0 | 2 | 8 | 12 | 10 | 0 | 12 | 8 | 2 | Colec10 |
| Colec11 | 14 | 22 | 20 | 18 | 29 | 28 | 37 | 16 | 14 | 15 | 28 | 11 | 19 | 14 | 20 | 22 | 14 | 37 | 28 | 35 | 36 | 15 | 43 | 19 | 18 | 18 | 34 | 11 | Colec11 |
| Colec12 | 5771 | 5924 | 8200 | 3375 | 2972 | 4235 | 2306 | 4680 | 5555 | 4649 | 6035 | 6072 | 7699 | 5675 | 6504 | 8417 | 4543 | 4145 | 2518 | 3914 | 2573 | 5571 | 7684 | 4735 | 4490 | 5263 | 7349 | 6273 | Colec12 |
| Colgalt1 | 7450 | 7746 | 6975 | 6701 | 6718 | 6458 | 5680 | 6057 | 7204 | 6250 | 7858 | 5990 | 6624 | 5605 | 7280 | 7797 | 6630 | 7038 | 6050 | 5930 | 5874 | 6600 | 7567 | 6096 | 6421 | 6324 | 6333 | 5822 | Colgalt1 |
| Colgalt2 | 392 | 416 | 334 | 326 | 442 | 421 | 346 | 414 | 468 | 468 | 641 | 285 | 380 | 229 | 434 | 477 | 350 | 377 | 310 | 446 | 323 | 578 | 513 | 355 | 400 | 321 | 302 | 278 | Colgalt2 |
| Colq | 27 | 22 | 9 | 19 | 29 | 12 | 13 | 21 | 23 | 23 | 116 | 10 | 17 | 6 | 12 | 20 | 8 | 21 | 10 | 5 | 5 | 15 | 106 | 22 | 14 | 8 | 9 | 7 | Colq |
| Commd1 | 471 | 472 | 339 | 466 | 324 | 250 | 496 | 479 | 300 | 355 | 329 | 395 | 374 | 709 | 451 | 329 | 392 | 294 | 327 | 403 | 505 | 373 | 351 | 583 | 397 | 428 | 347 | 636 | Commd1 |
| Commd10 | 2459 | 2153 | 2504 | 1885 | 1627 | 1873 | 1884 | 2276 | 1956 | 1960 | 1856 | 2414 | 2417 | 2582 | 2385 | 2150 | 2015 | 1666 | 1707 | 1987 | 1894 | 2015 | 2001 | 2081 | 1981 | 2782 | 2556 | 3021 | Commd10 |
| Commd2 | 2347 | 2348 | 2053 | 2171 | 2297 | 2284 | 2407 | 2555 | 2235 | 1894 | 2073 | 2342 | 2333 | 2299 | 2523 | 2252 | 1805 | 2065 | 2227 | 2874 | 2429 | 1937 | 2234 | 2049 | 2204 | 2442 | 2407 | 2373 | Commd2 |
| Commd3 | 2718 | 2425 | 2322 | 2543 | 2708 | 2618 | 3302 | 3856 | 3629 | 3475 | 3373 | 3116 | 2901 | 3513 | 2908 | 2523 | 2456 | 2563 | 2632 | 3001 | 3107 | 3243 | 3719 | 3452 | 3471 | 3414 | 2937 | 3258 | Commd3 |
| Commd4 | 1679 | 2153 | 2018 | 2208 | 2470 | 2198 | 2267 | 2519 | 2809 | 2869 | 2835 | 2899 | 2649 | 2792 | 1854 | 1767 | 2293 | 2141 | 2427 | 2704 | 2257 | 2652 | 2818 | 3132 | 3207 | 2999 | 2975 | 2804 | Commd4 |
| Commd5 | 25 | 19 | 40 | 23 | 11 | 23 | 23 | 11 | 16 | 15 | 11 | 21 | 8 | 42 | 25 | 13 | 30 | 25 | 18 | 27 | 29 | 23 | 23 | 29 | 12 | 16 | 3 | 31 | Commd5 |
| Commd6 | 2275 | 2284 | 2888 | 2584 | 2235 | 2125 | 2645 | 2010 | 1864 | 1925 | 1920 | 2096 | 2121 | 2507 | 2407 | 2317 | 2602 | 2403 | 2370 | 2649 | 2489 | 1811 | 1933 | 2368 | 2037 | 2390 | 2188 | 2447 | Commd6 |
| Commd7 | 1709 | 1888 | 1938 | 1938 | 2160 | 1989 | 2053 | 2751 | 2661 | 2696 | 2613 | 2713 | 2505 | 2703 | 1738 | 1678 | 1960 | 1841 | 2170 | 2202 | 2107 | 2695 | 2697 | 2842 | 2885 | 2921 | 2632 | 2824 | Commd7 |
| Commd8 | 3821 | 3471 | 4022 | 3629 | 3499 | 3301 | 4446 | 4010 | 3360 | 2996 | 3086 | 3531 | 3458 | 4106 | 4230 | 3565 | 4055 | 3758 | 3550 | 4378 | 3936 | 3166 | 3390 | 3760 | 3237 | 4072 | 3801 | 4230 | Commd8 |
| Commd9 | 1436 | 1846 | 1298 | 1543 | 1474 | 1503 | 1621 | 1752 | 1897 | 1578 | 1671 | 1850 | 1783 | 1854 | 1536 | 1584 | 1408 | 1495 | 1509 | 1637 | 1493 | 1531 | 1879 | 1673 | 1719 | 1803 | 1902 | 1799 | Commd9 |
| Comp | 6 | 22 | 32 | 25 | 7 | 14 | 11 | 10 | 26 | 32 | 431 | 12 | 15 | 4 | 8 | 26 | 22 | 13 | 16 | 8 | 5 | 55 | 36 | 8 | 18 | 7 | 2 | 6 | Comp |
| Comt | 38 | 30 | 45 | 72 | 37 | 64 | 22 | 67 | 71 | 49 | 56 | 53 | 64 | 38 | 54 | 42 | 37 | 48 | 31 | 134 | 24 | 58 | 119 | 188 | 73 | 56 | 58 | 51 | Comt |
| Comtd1 | 242 | 307 | 296 | 279 | 344 | 318 | 331 | 264 | 310 | 291 | 239 | 204 | 210 | 241 | 244 | 285 | 322 | 326 | 331 | 313 | 315 | 232 | 268 | 235 | 300 | 223 | 210 | 244 | Comtd1 |
| Cop1 | 8607 | 7433 | 7509 | 7729 | 7718 | 8022 | 9207 | 10636 | 9047 | 8653 | 8426 | 9461 | 8690 | 9340 | 8938 | 7559 | 7354 | 7704 | 7852 | 8278 | 9109 | 9359 | 9038 | 8915 | 9282 | 9847 | 9026 | 10318 | Cop1 |
| Copa | 49161 | 49413 | 35005 | 39004 | 36151 | 39556 | 38777 | 38443 | 35667 | 30367 | 32476 | 29238 | 29660 | 28897 | 49375 | 42225 | 36967 | 39025 | 39735 | 40938 | 40865 | 33849 | 36112 | 29162 | 31639 | 27028 | 31125 | 29169 | Copa |
| Copb1 | 15870 | 13596 | 12143 | 11530 | 11044 | 11336 | 12972 | 15432 | 13740 | 11636 | 12116 | 11807 | 11904 | 12238 | 16168 | 14171 | 12254 | 11820 | 11528 | 13287 | 12058 | 12855 | 14246 | 11716 | 11437 | 12567 | 12156 | 12509 | Copb1 |
| Copb2 | 24323 | 21503 | 19264 | 17949 | 16226 | 17523 | 18501 | 19403 | 17917 | 15001 | 16139 | 15420 | 15912 | 16117 | 24750 | 22100 | 18208 | 18306 | 16984 | 20055 | 17788 | 15993 | 18433 | 15082 | 15097 | 16819 | 16404 | 16734 | Copb2 |
| Cope | 4367 | 5887 | 4735 | 4907 | 4241 | 4524 | 4375 | 4584 | 4988 | 4487 | 4964 | 4575 | 4664 | 4553 | 4444 | 4842 | 5064 | 4747 | 4982 | 4951 | 4363 | 4074 | 5054 | 4911 | 5137 | 5266 | 4934 | 4675 | Cope |
| Copg1 | 12475 | 13392 | 10843 | 11160 | 11257 | 11583 | 10802 | 13801 | 14962 | 13986 | 14712 | 13484 | 13394 | 13166 | 11947 | 11899 | 10933 | 11772 | 11450 | 11205 | 11363 | 14076 | 14859 | 12977 | 13884 | 13256 | 13496 | 13212 | Copg1 |
| Copg2 | 8491 | 7874 | 7455 | 6479 | 6296 | 6354 | 7002 | 10084 | 7935 | 7414 | 6365 | 8934 | 7935 | 8992 | 9206 | 7027 | 6852 | 5973 | 6555 | 7583 | 6869 | 7585 | 7658 | 7464 | 7641 | 9128 | 9753 | 9417 | Copg2 |
| Coprs | 1720 | 1769 | 1756 | 1930 | 1781 | 1558 | 1791 | 1410 | 1153 | 1257 | 1126 | 1180 | 1243 | 1204 | 1716 | 1706 | 1814 | 1797 | 1802 | 1905 | 1762 | 1285 | 1142 | 1250 | 1322 | 1243 | 1326 | 1320 | Coprs |
| Cops2 | 15855 | 11441 | 13825 | 11610 | 10468 | 11389 | 12905 | 14100 | 12587 | 10395 | 10839 | 11869 | 12649 | 12929 | 17085 | 14436 | 12595 | 11052 | 11218 | 14326 | 11866 | 12231 | 13075 | 11237 | 10889 | 13790 | 13359 | 13657 | Cops2 |
| Cops3 | 6670 | 6125 | 5808 | 5076 | 5370 | 5539 | 6013 | 9951 | 8815 | 7578 | 7432 | 8987 | 8534 | 9533 | 7034 | 5780 | 5627 | 4666 | 5529 | 6721 | 5639 | 8057 | 8813 | 7971 | 8076 | 9880 | 9462 | 9810 | Cops3 |
| Cops4 | 6871 | 6835 | 6335 | 5865 | 5641 | 5671 | 6108 | 7790 | 7323 | 6994 | 6475 | 7239 | 7097 | 7713 | 6908 | 6037 | 6011 | 5290 | 6113 | 6969 | 6560 | 6395 | 7267 | 6969 | 6780 | 8010 | 7986 | 7971 | Cops4 |
| Cops5 | 7300 | 6734 | 7395 | 6436 | 6171 | 6470 | 6798 | 8413 | 7626 | 6837 | 6804 | 7590 | 7807 | 8243 | 8039 | 6727 | 7058 | 6214 | 6467 | 8034 | 6673 | 7043 | 7695 | 7591 | 7507 | 8760 | 8474 | 8762 | Cops5 |
| Cops6 | 7516 | 8059 | 8620 | 8019 | 6972 | 7458 | 7175 | 6430 | 6375 | 6944 | 6826 | 7290 | 7495 | 7595 | 7404 | 7359 | 8414 | 7001 | 7749 | 7411 | 7126 | 5972 | 6910 | 7572 | 6991 | 8560 | 8136 | 8530 | Cops6 |
| Cops7a | 2380 | 2561 | 2474 | 2003 | 1852 | 1958 | 1907 | 2993 | 3224 | 2951 | 3021 | 3052 | 3006 | 3242 | 2192 | 2364 | 2271 | 2075 | 2050 | 2218 | 1839 | 2938 | 3365 | 3093 | 2945 | 3690 | 3299 | 3236 | Cops7a |
| Cops7b | 3560 | 3760 | 3462 | 3129 | 2828 | 2873 | 2930 | 3656 | 3269 | 3062 | 3033 | 3466 | 3212 | 3597 | 3357 | 3153 | 3202 | 2984 | 3105 | 3143 | 2908 | 3496 | 3148 | 3373 | 3329 | 4011 | 3561 | 3813 | Cops7b |
| Cops8 | 5849 | 6049 | 5381 | 5084 | 4525 | 4927 | 5168 | 5843 | 5531 | 5209 | 5272 | 5991 | 5568 | 6040 | 5738 | 5406 | 5157 | 4642 | 4833 | 5354 | 4897 | 5137 | 5594 | 5625 | 5349 | 6518 | 6151 | 6241 | Cops8 |
| Cops9 | 2062 | 2612 | 2685 | 3334 | 2707 | 2219 | 3352 | 2508 | 2294 | 3285 | 3001 | 2603 | 2429 | 3804 | 2380 | 1755 | 3038 | 2707 | 2540 | 2789 | 3566 | 2007 | 2451 | 3562 | 3232 | 3083 | 2623 | 3604 | Cops9 |
| Copz1 | 12397 | 11863 | 10289 | 9379 | 8375 | 8790 | 9810 | 10248 | 9220 | 7787 | 7919 | 8718 | 8670 | 8674 | 11962 | 10675 | 9915 | 8943 | 9475 | 10013 | 9100 | 8497 | 9165 | 8544 | 7886 | 9833 | 9155 | 9304 | Copz1 |
| Copz2 | 1115 | 1041 | 1152 | 719 | 558 | 703 | 436 | 681 | 885 | 706 | 1300 | 558 | 802 | 564 | 1080 | 1160 | 817 | 781 | 424 | 526 | 439 | 901 | 1139 | 744 | 608 | 742 | 680 | 663 | Copz2 |
| Coq10a | 1411 | 1472 | 1290 | 1651 | 1771 | 1551 | 1835 | 1313 | 1178 | 1314 | 1299 | 1340 | 1165 | 1315 | 1501 | 1224 | 1611 | 1597 | 1720 | 1712 | 1824 | 1547 | 1268 | 1264 | 1440 | 1283 | 1232 | 1372 | Coq10a |
| Coq10b | 2079 | 2011 | 2146 | 2038 | 1762 | 1835 | 2110 | 1729 | 1285 | 1268 | 1247 | 1428 | 1330 | 1457 | 2396 | 2091 | 2109 | 2037 | 1850 | 2132 | 1959 | 1556 | 1373 | 1329 | 1409 | 1499 | 1496 | 1462 | Coq10b |
| Coq2 | 1891 | 1967 | 1769 | 1754 | 1797 | 1830 | 1891 | 2656 | 2857 | 2637 | 2746 | 2805 | 2963 | 2782 | 1847 | 1811 | 2024 | 1702 | 1788 | 1882 | 1754 | 2363 | 2730 | 2831 | 2728 | 3067 | 2918 | 2849 | Coq2 |
| Coq3 | 745 | 573 | 701 | 651 | 747 | 592 | 794 | 933 | 793 | 801 | 714 | 867 | 818 | 839 | 709 | 586 | 581 | 741 | 696 | 764 | 777 | 805 | 904 | 825 | 938 | 915 | 876 | 862 | Coq3 |
| Coq4 | 1100 | 1384 | 1191 | 1078 | 884 | 800 | 952 | 750 | 903 | 833 | 849 | 870 | 945 | 897 | 997 | 1094 | 1205 | 1021 | 1033 | 1005 | 953 | 800 | 803 | 880 | 849 | 1022 | 906 | 985 | Coq4 |
| Coq5 | 2167 | 2552 | 2427 | 2151 | 1786 | 1789 | 1968 | 1677 | 1690 | 1673 | 1540 | 1821 | 1889 | 1857 | 2081 | 2067 | 2102 | 1852 | 1804 | 2014 | 1928 | 1664 | 1698 | 1820 | 1867 | 2074 | 1857 | 1970 | Coq5 |
| Coq6 | 1894 | 2075 | 1780 | 1997 | 1638 | 1626 | 1669 | 1678 | 1763 | 1563 | 1549 | 1620 | 1769 | 1759 | 1772 | 1790 | 1648 | 1725 | 1864 | 1825 | 1717 | 1579 | 1720 | 1816 | 1709 | 2046 | 1864 | 1876 | Coq6 |
| Coq7 | 536 | 629 | 723 | 532 | 595 | 589 | 616 | 1305 | 1339 | 1422 | 1321 | 1660 | 1532 | 1763 | 614 | 492 | 618 | 414 | 512 | 711 | 548 | 1086 | 1244 | 1602 | 1561 | 2047 | 1754 | 1993 | Coq7 |
| Coq8a | 338 | 403 | 198 | 310 | 214 | 236 | 234 | 104 | 164 | 126 | 204 | 73 | 115 | 83 | 347 | 445 | 271 | 360 | 178 | 228 | 289 | 99 | 203 | 107 | 128 | 129 | 102 | 88 | Coq8a |

| | | | | | | | | | | | | | | | | | | | | | | | | | | | | | |
|---|---|---|---|---|---|---|---|---|---|---|---|---|---|---|---|---|---|---|---|---|---|---|---|---|---|---|---|---|---|
| Coq8b | 1076 | 1437 | 1363 | 1428 | 1436 | 1247 | 1336 | 1071 | 1158 | 1237 | 1112 | 1224 | 1134 | 1129 | 1026 | 1199 | 1505 | 1499 | 1624 | 1165 | 1350 | 1208 | 1139 | 1237 | 1257 | 1267 | 1146 | 1238 | Coq8b |
| Coq9 | 1834 | 1927 | 1845 | 1810 | 1565 | 1720 | 1667 | 1801 | 1959 | 1809 | 1987 | 2065 | 1920 | 2003 | 1698 | 1777 | 1957 | 1813 | 1657 | 1557 | 1671 | 1870 | 2141 | 2070 | 1781 | 2071 | 2007 | 1920 | Coq9 |
| Corin | 414 | 569 | 550 | 371 | 393 | 346 | 184 | 749 | 649 | 525 | 1124 | 525 | 1317 | 388 | 909 | 971 | 408 | 358 | 197 | 343 | 132 | 764 | 935 | 286 | 361 | 169 | 643 | 606 | Corin |
| Coro1a | 1732 | 1984 | 1808 | 1923 | 2392 | 2337 | 1844 | 1949 | 2214 | 1927 | 1844 | 1866 | 1776 | 1717 | 1611 | 1872 | 2133 | 1919 | 2707 | 2327 | 1992 | 2252 | 2273 | 1950 | 2325 | 1994 | 1909 | 1730 | Coro1a |
| Coro1b | 291 | 307 | 187 | 534 | 101 | 139 | 108 | 296 | 145 | 137 | 98 | 165 | 151 | 151 | 215 | 208 | 407 | 154 | 149 | 146 | 166 | 185 | 210 | 161 | 287 | 124 | 166 | 123 | Coro1b |
| Coro1c | 15756 | 20132 | 21668 | 22884 | 18125 | 18744 | 19042 | 9976 | 11567 | 12673 | 10946 | 15918 | 12462 | 15006 | 16313 | 17618 | 22540 | 21328 | 21039 | 16405 | 19526 | 12096 | 10389 | 14481 | 14746 | 16346 | 14440 | 15267 | Coro1c |
| Coro2a | 610 | 642 | 651 | 597 | 1037 | 657 | 603 | 706 | 704 | 838 | 900 | 344 | 771 | 360 | 692 | 889 | 674 | 720 | 633 | 496 | 603 | 1079 | 620 | 598 | 630 | 555 | 288 | 331 | Coro2a |
| Coro2b | 1692 | 2242 | 2350 | 2936 | 3723 | 3170 | 3113 | 3751 | 4911 | 5780 | 5099 | 6317 | 4852 | 6207 | 2011 | 1970 | 3179 | 3083 | 3627 | 2473 | 3228 | 5833 | 4139 | 6035 | 6571 | 5888 | 4710 | 5269 | Coro2b |
| Coro6 | 111 | 115 | 120 | 109 | 120 | 105 | 102 | 51 | 59 | 57 | 185 | 59 | 65 | 45 | 94 | 132 | 149 | 180 | 108 | 106 | 92 | 112 | 107 | 57 | 74 | 47 | 84 | 46 | Coro6 |
| Coro7 | 4783 | 5598 | 5467 | 6351 | 6316 | 6053 | 5589 | 2172 | 2773 | 2819 | 2770 | 2803 | 2759 | 2638 | 4371 | 5076 | 5985 | 6390 | 6416 | 4980 | 6042 | 2968 | 2490 | 2688 | 3165 | 2621 | 2603 | 2403 | Coro7 |
| Cort | 20 | 10 | 24 | 8 | 26 | 22 | 18 | 21 | 14 | 37 | 17 | 27 | 32 | 28 | 12 | 12 | 5 | 14 | 31 | 10 | 10 | 27 | 33 | 25 | 25 | 23 | 37 | 31 | Cort |
| Cotl1 | 9029 | 13357 | 13557 | 13064 | 11293 | 10397 | 11222 | 5775 | 7270 | 8389 | 7196 | 7304 | 7583 | 7793 | 9559 | 11725 | 13490 | 13016 | 12601 | 10135 | 11605 | 6797 | 7851 | 8809 | 8090 | 9412 | 6886 | 7602 | Cotl1 |
| Cox10 | 828 | 969 | 870 | 674 | 650 | 679 | 587 | 1037 | 1202 | 1114 | 1041 | 1351 | 1330 | 1232 | 778 | 804 | 957 | 649 | 787 | 600 | 626 | 1068 | 1232 | 1136 | 1169 | 1423 | 1253 | 1337 | Cox10 |
| Cox11 | 1299 | 1021 | 1102 | 1299 | 1108 | 1063 | 1258 | 1152 | 1072 | 1114 | 995 | 1139 | 1050 | 1048 | 1124 | 1008 | 1137 | 1208 | 1120 | 1170 | 1141 | 1086 | 1045 | 1106 | 1084 | 1212 | 1196 | 1149 | Cox11 |
| Cox14 | 1821 | 2311 | 2155 | 2391 | 1960 | 2053 | 2292 | 1892 | 1946 | 2006 | 1959 | 1918 | 1928 | 2048 | 1998 | 1749 | 2502 | 2204 | 2207 | 2405 | 2084 | 1574 | 1970 | 2064 | 1910 | 2464 | 2038 | 2130 | Cox14 |
| Cox15 | 1505 | 1582 | 1762 | 1480 | 1519 | 1440 | 1530 | 2175 | 2264 | 2328 | 2115 | 2627 | 2427 | 2495 | 1486 | 1423 | 1486 | 1412 | 1696 | 1570 | 1426 | 2186 | 2192 | 2404 | 2260 | 2708 | 2703 | 2711 | Cox15 |
| Cox16 | 78 | 68 | 73 | 92 | 87 | 146 | 94 | 104 | 110 | 151 | 121 | 135 | 128 | 106 | 66 | 90 | 122 | 99 | 107 | 130 | 92 | 74 | 108 | 129 | 106 | 159 | 106 | 110 | Cox16 |
| Cox17 | 657 | 607 | 583 | 650 | 488 | 597 | 658 | 666 | 687 | 693 | 664 | 679 | 596 | 696 | 634 | 619 | 655 | 657 | 564 | 554 | 630 | 652 | 638 | 695 | 654 | 642 | 696 | 724 | Cox17 |
| Cox18 | 543 | 612 | 614 | 609 | 605 | 584 | 596 | 568 | 681 | 628 | 656 | 693 | 666 | 628 | 570 | 641 | 503 | 629 | 539 | 559 | 576 | 615 | 673 | 612 | 612 | 713 | 682 | 624 | Cox18 |
| Cox19 | 2656 | 3422 | 3212 | 3740 | 3790 | 3598 | 3709 | 3672 | 3867 | 4318 | 3416 | 4529 | 4155 | 4872 | 3150 | 3020 | 3564 | 3425 | 3926 | 4072 | 3925 | 4067 | 3653 | 4428 | 4693 | 4266 | 4649 | 4535 | Cox19 |
| Cox20 | 75 | 29 | 65 | 36 | 21 | 39 | 32 | 95 | 111 | 249 | 59 | 464 | 27 | 59 | 276 | 143 | 64 | 29 | 40 | 361 | 33 | 90 | 104 | 83 | 64 | 209 | 209 | 856 | Cox20 |
| Cox4i1 | 15009 | 17729 | 19085 | 19082 | 16055 | 15173 | 16924 | 13802 | 13988 | 16211 | 15397 | 15998 | 15993 | 17965 | 14973 | 14195 | 19413 | 17475 | 17919 | 15912 | 18408 | 12689 | 15310 | 17293 | 16420 | 18474 | 17445 | 18856 | Cox4i1 |
| Cox4i2 | 114 | 124 | 63 | 107 | 88 | 70 | 58 | 90 | 103 | 106 | 107 | 78 | 65 | 67 | 113 | 111 | 98 | 90 | 84 | 79 | 76 | 87 | 126 | 81 | 55 | 93 | 64 | 74 | Cox4i2 |
| Cox5a | 6869 | 6918 | 6848 | 7606 | 6542 | 5495 | 8337 | 5170 | 4420 | 5458 | 4813 | 5173 | 4580 | 6628 | 6712 | 4968 | 7634 | 6664 | 7007 | 6219 | 8268 | 5075 | 4245 | 5833 | 5443 | 5808 | 4979 | 6732 | Cox5a |
| Cox5b | 3449 | 3950 | 4149 | 3545 | 3184 | 2859 | 3952 | 3430 | 3219 | 3650 | 3466 | 3383 | 3476 | 4907 | 3608 | 2812 | 4069 | 3219 | 3154 | 3595 | 3940 | 3001 | 3317 | 3979 | 3602 | 3920 | 3669 | 4624 | Cox5b |
| Cox5b-ps | 310 | 289 | 281 | 259 | 230 | 152 | 273 | 318 | 253 | 262 | 244 | 216 | 236 | 301 | 324 | 188 | 258 | 228 | 199 | 206 | 269 | 190 | 226 | 276 | 231 | 239 | 255 | 377 | Cox5b-ps |
| Cox6a1 | 5867 | 7769 | 6310 | 6468 | 5533 | 4891 | 6240 | 4375 | 4062 | 4393 | 4390 | 4712 | 4390 | 5518 | 5909 | 4867 | 6852 | 5897 | 5936 | 5930 | 6275 | 4209 | 4166 | 5276 | 4682 | 5080 | 4991 | 5421 | Cox6a1 |
| Cox6a2 | 2 | 1 | 0 | 0 | 1 | 1 | 2 | 5 | 18 | 0 | 183 | 3 | 3 | 0 | 0 | 1 | 0 | 0 | 0 | 0 | 1 | 10 | 105 | 3 | 4 | 0 | 1 | 0 | Cox6a2 |
| Cox6b1 | 8731 | 10018 | 9205 | 9849 | 9430 | 9374 | 10690 | 8672 | 8363 | 8732 | 8859 | 8918 | 9517 | 9524 | 8941 | 8000 | 9711 | 9130 | 10248 | 12308 | 10158 | 7212 | 8644 | 9195 | 9529 | 9925 | 9785 | 9862 | Cox6b1 |
| Cox6b2 | 1374 | 1820 | 2759 | 2191 | 1328 | 1224 | 1324 | 419 | 447 | 616 | 620 | 581 | 645 | 777 | 1120 | 1210 | 2490 | 1670 | 1527 | 1182 | 1508 | 359 | 541 | 768 | 455 | 845 | 632 | 882 | Cox6b2 |
| Cox6c | 7689 | 7594 | 8450 | 8806 | 8372 | 7371 | 9975 | 7489 | 7427 | 8245 | 8269 | 7707 | 8191 | 9260 | 7750 | 6612 | 8510 | 7839 | 8447 | 10104 | 9316 | 5909 | 7398 | 8688 | 8506 | 8926 | 8549 | 9952 | Cox6c |
| Cox7a1 | 23 | 40 | 41 | 43 | 38 | 29 | 42 | 46 | 30 | 40 | 33 | 42 | 52 | 43 | 40 | 33 | 54 | 36 | 58 | 42 | 45 | 52 | 42 | 60 | 53 | 38 | 33 | 57 | Cox7a1 |
| Cox7a2 | 5906 | 6005 | 5897 | 5976 | 5821 | 5467 | 8075 | 6521 | 6136 | 5433 | 5867 | 5723 | 6100 | 6685 | 6231 | 5325 | 5769 | 5880 | 6676 | 7632 | 7184 | 5519 | 6209 | 6287 | 6527 | 6659 | 6826 | 7645 | Cox7a2 |
| Cox7a2l | 9329 | 10605 | 9430 | 9994 | 8937 | 8880 | 9839 | 8267 | 8062 | 7785 | 7939 | 7767 | 7887 | 8182 | 9802 | 10270 | 9741 | 9257 | 9751 | 11032 | 9493 | 7138 | 8559 | 7912 | 8199 | 8687 | 8262 | 8452 | Cox7a2l |
| Cox7b | 5667 | 5137 | 5887 | 6122 | 5701 | 5411 | 6994 | 5958 | 4957 | 5422 | 6387 | 5168 | 6056 | 6837 | 6134 | 4780 | 5808 | 5592 | 5502 | 7335 | 6209 | 4916 | 5284 | 6289 | 6135 | 6387 | 6003 | 6817 | Cox7b |
| Cox7b2 | 0 | 0 | 0 | 0 | 4 | 5 | 8 | 0 | 0 | 0 | 5 | 2 | 0 | 3 | 0 | 0 | 2 | 1 | 0 | 1 | 5 | 3 | 4 | 0 | 0 | 0 | 1 | 0 | Cox7b2 |
| Cox7c | 4783 | 5073 | 6675 | 7364 | 6716 | 5178 | 8324 | 6809 | 6518 | 8561 | 8506 | 7698 | 7720 | 11574 | 5139 | 4018 | 6960 | 6060 | 6243 | 7394 | 8218 | 5243 | 6581 | 9893 | 8445 | 8772 | 8463 | 11284 | Cox7c |
| Cox8a | 5464 | 6862 | 5762 | 5621 | 5066 | 4230 | 5851 | 4813 | 5101 | 4956 | 5500 | 4941 | 5294 | 6452 | 5343 | 4551 | 6127 | 5110 | 5191 | 5798 | 5786 | 4450 | 5036 | 5538 | 5483 | 5844 | 5533 | 6467 | Cox8a |
| Cox8b | 0 | 0 | 0 | 0 | 0 | 0 | 0 | 2 | 1 | 0 | 16 | 0 | 0 | 0 | 0 | 0 | 0 | 0 | 0 | 0 | 0 | 5 | 9 | 1 | 0 | 0 | 0 | 0 | Cox8b |
| Cox8c | 0 | 0 | 0 | 0 | 0 | 0 | 0 | 0 | 0 | 0 | 0 | 0 | 0 | 1 | 0 | 0 | 0 | 1 | 0 | 0 | 0 | 0 | 0 | 0 | 0 | 0 | 0 | 0 | Cox8c |
| Cp | 584 | 537 | 682 | 450 | 746 | 652 | 462 | 797 | 850 | 663 | 755 | 697 | 802 | 552 | 667 | 817 | 461 | 483 | 478 | 865 | 565 | 956 | 1116 | 526 | 716 | 580 | 719 | 739 | Cp |
| Cpa1 | 10 | 9 | 15 | 9 | 27 | 23 | 21 | 12 | 18 | 32 | 17 | 15 | 6 | 4 | 7 | 13 | 10 | 13 | 22 | 14 | 18 | 10 | 17 | 12 | 12 | 18 | 5 | 9 | Cpa1 |
| Cpa2 | 72 | 67 | 104 | 69 | 54 | 63 | 38 | 118 | 87 | 76 | 86 | 136 | 126 | 98 | 111 | 88 | 51 | 63 | 72 | 70 | 43 | 85 | 74 | 102 | 80 | 149 | 130 | 102 | Cpa2 |
| Cpa3 | 52 | 17 | 37 | 24 | 14 | 48 | 6 | 87 | 199 | 116 | 204 | 40 | 68 | 22 | 53 | 40 | 17 | 78 | 37 | 18 | 1 | 69 | 293 | 116 | 68 | 41 | 14 | 27 | Cpa3 |
| Cpa4 | 110 | 104 | 82 | 70 | 65 | 106 | 99 | 276 | 260 | 181 | 236 | 401 | 146 | 304 | 124 | 53 | 102 | 87 | 92 | 107 | 64 | 229 | 198 | 295 | 251 | 456 | 311 | 306 | Cpa4 |
| Cpa5 | 12 | 16 | 2 | 10 | 2 | 5 | 3 | 7 | 9 | 14 | 21 | 0 | 2 | 9 | 12 | 14 | 19 | 15 | 4 | 7 | 7 | 10 | 24 | 2 | 2 | 9 | 9 | 12 | Cpa5 |
| Cpa6 | 15 | 12 | 6 | 11 | 7 | 21 | 2 | 20 | 41 | 25 | 28 | 9 | 18 | 17 | 9 | 7 | 6 | 17 | 2 | 5 | 0 | 40 | 49 | 38 | 18 | 11 | 14 | 17 | Cpa6 |
| Cpb1 | 7 | 3 | 0 | 2 | 7 | 0 | 0 | 5 | 4 | 0 | 3 | 2 | 1 | 2 | 3 | 3 | 0 | 2 | 0 | 0 | 0 | 2 | 12 | 1 | 4 | 0 | 1 | 0 | Cpb1 |
| Cpb2 | 1 | 0 | 0 | 0 | 2 | 4 | 1 | 1 | 1 | 0 | 3 | 2 | 0 | 1 | 0 | 0 | 0 | 0 | 0 | 2 | 0 | 2 | 1 | 0 | 0 | 0 | 1 | 0 | Cpb2 |
| Cpd | 17426 | 14933 | 10895 | 11848 | 10444 | 12161 | 11577 | 8529 | 8429 | 7063 | 8448 | 7973 | 9044 | 8143 | 16200 | 17002 | 10849 | 12218 | 10048 | 11874 | 11865 | 7478 | 8609 | 6743 | 6889 | 6469 | 8961 | 8138 | Cpd |
| Cpe | 16206 | 15281 | 16106 | 21246 | 25840 | 22787 | 25538 | 25374 | 20328 | 25014 | 22728 | 24348 | 19739 | 24841 | 18714 | 12272 | 19060 | 22393 | 25872 | 23171 | 26237 | 31930 | 20903 | 25335 | 27529 | 23295 | 23717 | 25159 | Cpe |
| Cpeb1 | 528 | 499 | 436 | 560 | 790 | 610 | 617 | 369 | 293 | 355 | 322 | 156 | 213 | 177 | 556 | 519 | 499 | 601 | 613 | 615 | 640 | 269 | 323 | 298 | 191 | 268 | 151 | 153 | Cpeb1 |
| Cpeb2 | 1632 | 1434 | 2054 | 3141 | 4339 | 3273 | 3239 | 4109 | 3270 | 4033 | 4212 | 3146 | 3785 | 3414 | 2078 | 1741 | 2551 | 3180 | 3281 | 2597 | 3541 | 6355 | 3472 | 3177 | 4661 | 2050 | 3087 | 3260 | Cpeb2 |
| Cpeb3 | 1038 | 1019 | 1376 | 2083 | 2628 | 1868 | 2094 | 1042 | 905 | 1146 | 1240 | 982 | 1016 | 1090 | 1182 | 1069 | 1729 | 2216 | 2119 | 1551 | 2305 | 1751 | 887 | 1025 | 1512 | 593 | 796 | 929 | Cpeb3 |
| Cpeb4 | 7202 | 6607 | 7497 | 11573 | 12315 | 11855 | 12588 | 6152 | 5352 | 6610 | 6211 | 6015 | 6662 | 6490 | 8143 | 7409 | 9282 | 12340 | 10514 | 9119 | 12033 | 8682 | 5323 | 5846 | 7727 | 3949 | 5939 | 6608 | Cpeb4 |
| Cped1 | 3291 | 2093 | 2189 | 1232 | 951 | 1583 | 801 | 2676 | 3059 | 2347 | 2645 | 2269 | 3351 | 2268 | 2540 | 3741 | 1218 | 1478 | 741 | 1089 | 788 | 1601 | 2915 | 2280 | 1577 | 2795 | 1947 | 2165 | Cped1 |
| Cplane1 | 6395 | 5750 | 3762 | 5466 | 6916 | 7725 | 7152 | 8784 | 8946 | 8089 | 8279 | 8044 | 7749 | 8207 | 6231 | 6213 | 4638 | 6023 | 6124 | 6775 | 7686 | 7359 | 7896 | 6532 | 9100 | 4787 | 8170 | 7154 | Cplane1 |
| Cplane2 | 269 | 313 | 316 | 310 | 280 | 347 | 288 | 177 | 161 | 181 | 214 | 232 | 229 | 192 | 210 | 319 | 318 | 390 | 323 | 334 | 312 | 209 | 195 | 203 | 246 | 171 | 229 | 199 | Cplane2 |
| Cplx1 | 281 | 453 | 284 | 605 | 1564 | 750 | 850 | 1004 | 1405 | 1647 | 1394 | 680 | 829 | 765 | 481 | 417 | 570 | 944 | 949 | 702 | 984 | 1485 | 1176 | 950 | 1424 | 566 | 573 | 572 | Cplx1 |
| Cplx2 | 4851 | 5656 | 5906 | 7004 | 8318 | 8017 | 7431 | 9268 | 9578 | 11037 | 8658 | 12128 | 9943 | 12130 | 5972 | 5565 | 7050 | 7092 | 8969 | 8715 | 8750 | 11854 | 9570 | 12754 | 13799 | 10127 | 11910 | 11776 | Cplx2 |
| Cplx3 | 21 | 45 | 44 | 41 | 30 | 39 | 11 | 18 | 10 | 26 | 19 | 54 | 56 | 61 | 42 | 26 | 32 | 28 | 30 | 33 | 27 | 25 | 27 | 32 | 18 | 4 | 87 | 95 | Cplx3 |
| Cplx4 | 0 | 1 | 4 | 6 | 0 | 3 | 0 | 1 | 2 | 9 | 3 | 1 | 3 | 1 | 3 | 5 | 3 | 2 | 0 | 1 | 4 | 5 | 6 | 2 | 0 | 1 | 1 | 0 | Cplx4 |
| Cpm | 1458 | 854 | 836 | 733 | 655 | 774 | 649 | 1051 | 1009 | 1031 | 1081 | 377 | 501 | 423 | 1001 | 1159 | 728 | 697 | 606 | 616 | 605 | 730 | 1254 | 945 | 483 | 798 | 356 | 421 | Cpm |
| Cpn1 | 2 | 6 | 5 | 2 | 2 | 5 | 4 | 5 | 4 | 3 | 4 | 8 | 6 | 2 | 3 | 11 | 1 | 3 | 8 | 7 | 4 | 3 | 6 | 0 | 6 | 3 | 1 | 4 | Cpn1 |
| Cpn2 | 5 | 1 | 2 | 2 | 1 | 1 | 1 | 4 | 1 | 0 | 0 | 2 | 1 | 0 | 1 | 4 | 2 | 0 | 0 | 0 | 0 | 0 | 1 | 2 | 0 | 0 | 7 | 1 | Cpn2 |
| Cpne1 | 28 | 37 | 34 | 61 | 53 | 70 | 56 | 53 | 47 | 58 | 55 | 55 | 54 | 72 | 25 | 37 | 64 | 62 | 53 | 39 | 54 | 57 | 38 | 87 | 38 | 40 | 72 | 68 | Cpne1 |
| Cpne2 | 362 | 438 | 574 | 465 | 525 | 579 | 470 | 708 | 881 | 967 | 809 | 1010 | 699 | 911 | 407 | 375 | 508 | 459 | 567 | 478 | 513 | 984 | 1030 | 836 | 853 | 1110 | 807 | 952 | Cpne2 |
| Cpne3 | 9988 | 7383 | 7061 | 6643 | 6086 | 7265 | 7232 | 4708 | 4503 | 3280 | 4069 | 4032 | 4183 | 3920 | 9403 | 9065 | 6450 | 7209 | 6100 | 7304 | 7083 | 3352 | 4612 | 3778 | 3428 | 3297 | 4181 | 3940 | Cpne3 |
| Cpne4 | 302 | 301 | 287 | 485 | 987 | 704 | 782 | 613 | 610 | 669 | 538 | 331 | 388 | 317 | 360 | 286 | 341 | 725 | 770 | 668 | 795 | 971 | 507 | 512 | 852 | 160 | 287 | 244 | Cpne4 |
| Cpne5 | 803 | 660 | 760 | 837 | 1031 | 918 | 896 | 731 | 1050 | 1065 | 831 | 827 | 480 | 750 | 513 | 593 | 889 | 911 | 1114 | 907 | 991 | 1056 | 887 | 970 | 981 | 702 | 612 | 698 | Cpne5 |
| Cpne6 | 104 | 187 | 137 | 341 | 405 | 226 | 257 | 159 | 245 | 212 | 201 | 83 | 167 | 106 | 144 | 136 | 254 | 379 | 399 | 230 | 335 | 289 | 175 | 176 | 244 | 90 | 88 | 76 | Cpne6 |
| Cpne7 | 137 | 129 | 152 | 111 | 131 | 98 | 65 | 59 | 76 | 63 | 113 | 45 | 39 | 40 | 109 | 178 | 87 | 114 | 86 | 95 | 90 | 74 | 71 | 74 | 68 | 28 | 19 | 28 | Cpne7 |
| Cpne8 | 431 | 318 | 368 | 378 | 375 | 558 | 327 | 570 | 568 | 505 | 595 | 376 | 450 | 393 | 417 | 448 | 311 | 373 | 338 | 420 | 364 | 598 | 766 | 415 | 458 | 398 | 467 | 400 | Cpne8 |
| Cpne9 | 82 | 125 | 82 | 145 | 140 | 79 | 114 | 22 | 29 | 52 | 26 | 23 | 17 | 27 | 104 | 94 | 89 | 149 | 118 | 85 | 125 | 58 | 61 | 29 | 26 | 19 | 36 | 13 | Cpne9 |
| Cpox | 2175 | 2059 | 2916 | 2465 | 2220 | 2138 | 2128 | 1446 | 1519 | 1452 | 1781 | 1550 | 1726 | 1819 | 1974 | 2454 | 2387 | 2429 | 2120 | 1854 | 2155 | 1572 | 1447 | 1558 | 1503 | 1810 | 1456 | 2388 | Cpox |
| Cpped1 | 761 | 704 | 566 | 553 | 731 | 755 | 748 | 1602 | 1854 | 1571 | 1528 | 1485 | 1550 | 1527 | 816 | 681 | 649 | 574 | 705 | 664 | 684 | 1417 | 1728 | 1466 | 1461 | 1651 | 1551 | 1674 | Cpped1 |
| Cpq | 920 | 735 | 739 | 403 | 485 | 625 | 382 | 805 | 881 | 651 | 905 | 658 | 896 | 695 | 889 | 1033 | 555 | 584 | 363 | 448 | 383 | 852 | 1109 | 640 | 645 | 693 | 823 | 774 | Cpq |
| Cps1 | 10 | 10 | 10 | 6 | 13 | 15 | 22 | 35 | 42 | 11 | 34 | 24 | 28 | 27 | 11 | 12 | 1 | 13 | 12 | 10 | 19 | 50 | 34 | 25 | 27 | 34 | 26 | 19 | Cps1 |
| Cpsf1 | 6181 | 7560 | 8104 | 7489 | 7031 | 6816 | 6535 | 5360 | 7130 | 7760 | 7124 | 9285 | 8315 | 8145 | 5818 | 7389 | 7893 | 7348 | 8146 | 5824 | 6779 | 7180 | 6717 | 8622 | 8162 | 9245 | 8032 | 8469 | Cpsf1 |

| | | | | | | | | | | | | | | | | | | | | | | | | | | | | | |
|---|---|---|---|---|---|---|---|---|---|---|---|---|---|---|---|---|---|---|---|---|---|---|---|---|---|---|---|---|---|
| Cpsf2 | 7262 | 6613 | 6563 | 5905 | 6532 | 6833 | 7244 | 9228 | 8074 | 7509 | 7049 | 8690 | 7971 | 9027 | 7224 | 6350 | 6372 | 5888 | 6315 | 7525 | 6781 | 7678 | 7868 | 7546 | 7625 | 8526 | 8750 | 9095 | Cpsf2 |
| Cpsf3 | 6815 | 7056 | 6330 | 5661 | 5152 | 5551 | 5732 | 7338 | 7093 | 6902 | 6465 | 8119 | 7445 | 7903 | 6521 | 6189 | 5660 | 5297 | 5747 | 6230 | 5741 | 6483 | 6810 | 6774 | 6981 | 8134 | 7819 | 7999 | Cpsf3 |
| Cpsf4 | 4151 | 4969 | 4149 | 4623 | 3855 | 3998 | 4117 | 2920 | 3263 | 3145 | 3060 | 3451 | 3386 | 3461 | 4060 | 4326 | 4283 | 4376 | 4190 | 4325 | 3962 | 2928 | 2792 | 3403 | 3366 | 3359 | 3583 | 3370 | Cpsf4 |
| Cpsf4l | 2 | 5 | 5 | 1 | 7 | 5 | 0 | 0 | 19 | 6 | 9 | 2 | 5 | 4 | 0 | 6 | 2 | 0 | 8 | 2 | 3 | 2 | 12 | 8 | 7 | 0 | 6 | 2 | Cpsf4l |
| Cpsf6 | 28905 | 20571 | 23357 | 26793 | 27312 | 31660 | 33479 | 38098 | 31150 | 28634 | 30140 | 35131 | 32114 | 37587 | 29048 | 25486 | 24696 | 25708 | 28061 | 31243 | 32549 | 31834 | 28950 | 30275 | 33101 | 29116 | 36157 | 35614 | Cpsf6 |
| Cpsf7 | 7750 | 7478 | 9377 | 8974 | 9814 | 9212 | 10299 | 12737 | 12038 | 13406 | 12669 | 14901 | 13011 | 15525 | 7555 | 6634 | 9143 | 8707 | 10474 | 8486 | 9972 | 13917 | 11138 | 14153 | 14260 | 15384 | 14000 | 15225 | Cpsf7 |
| Cpt1a | 9612 | 9342 | 8436 | 8195 | 7812 | 8632 | 7851 | 2789 | 3507 | 2869 | 3405 | 2918 | 3072 | 2977 | 8913 | 8385 | 7705 | 8055 | 7838 | 7242 | 7818 | 2734 | 3716 | 2755 | 2787 | 2551 | 2744 | 2753 | Cpt1a |
| Cpt1b | 59 | 131 | 97 | 151 | 151 | 119 | 117 | 54 | 66 | 85 | 43 | 48 | 77 | 57 | 95 | 85 | 119 | 113 | 146 | 120 | 105 | 80 | 46 | 57 | 68 | 45 | 85 | 59 | Cpt1b |
| Cpt1c | 1817 | 2197 | 2134 | 2735 | 3153 | 2892 | 2501 | 2969 | 3475 | 3962 | 3610 | 3715 | 3495 | 3386 | 1928 | 1903 | 2677 | 2645 | 3125 | 2477 | 2693 | 3995 | 3163 | 4030 | 4077 | 3322 | 3442 | 3528 | Cpt1c |
| Cpt2 | 1641 | 1980 | 1647 | 1445 | 1199 | 1247 | 1187 | 1089 | 1116 | 945 | 953 | 1083 | 1237 | 1095 | 1692 | 1674 | 1520 | 1391 | 1163 | 1387 | 1170 | 914 | 1265 | 976 | 884 | 1230 | 1217 | 1228 | Cpt2 |
| Cptp | 1806 | 1982 | 1782 | 1742 | 1591 | 1522 | 1669 | 1093 | 1083 | 1088 | 987 | 1199 | 962 | 1200 | 1717 | 1747 | 1752 | 1582 | 1689 | 1533 | 1576 | 1226 | 1043 | 1081 | 1262 | 1149 | 1227 | 1208 | Cptp |
| Cpvl | 6 | 4 | 13 | 2 | 5 | 0 | 4 | 6 | 4 | 6 | 0 | 11 | 9 | 7 | 1 | 10 | 2 | 4 | 3 | 0 | 0 | 3 | 2 | 5 | 2 | 1 | 6 | 4 | Cpvl |
| Cpxcr1 | 0 | 0 | 0 | 0 | 0 | 0 | 0 | 0 | 1 | 0 | 0 | 0 | 2 | 0 | 0 | 0 | 0 | 0 | 0 | 0 | 0 | 0 | 0 | 0 | 0 | 0 | 0 | 0 | Cpxcr1 |
| Cpxm1 | 2002 | 2098 | 1866 | 1440 | 1218 | 1616 | 681 | 2253 | 3126 | 2484 | 3257 | 2097 | 2871 | 1870 | 1899 | 2571 | 1560 | 1948 | 833 | 735 | 725 | 2737 | 3194 | 2524 | 2229 | 2472 | 2679 | 2039 | Cpxm1 |
| Cpxm2 | 472 | 381 | 247 | 271 | 80 | 311 | 44 | 593 | 738 | 745 | 460 | 382 | 587 | 353 | 405 | 625 | 152 | 342 | 56 | 58 | 58 | 369 | 677 | 668 | 345 | 513 | 381 | 406 | Cpxm2 |
| Cpz | 226 | 364 | 355 | 139 | 56 | 212 | 28 | 128 | 210 | 188 | 237 | 264 | 305 | 222 | 229 | 341 | 224 | 217 | 58 | 45 | 29 | 180 | 267 | 236 | 202 | 219 | 308 | 310 | Cpz |
| Cr1l | 1185 | 955 | 1210 | 885 | 1136 | 1303 | 1055 | 2880 | 2747 | 2502 | 2838 | 2525 | 2883 | 2692 | 1196 | 1064 | 943 | 977 | 1154 | 1147 | 1079 | 2475 | 2904 | 2567 | 2524 | 2763 | 2681 | 2843 | Cr1l |
| Cr2 | 9 | 3 | 16 | 5 | 17 | 10 | 9 | 31 | 33 | 22 | 23 | 40 | 32 | 29 | 22 | 12 | 0 | 8 | 10 | 31 | 10 | 17 | 25 | 20 | 29 | 21 | 37 | 36 | Cr2 |
| Crabp1 | 5727 | 6133 | 6429 | 2475 | 1823 | 2247 | 780 | 3234 | 4183 | 3481 | 4040 | 2918 | 5243 | 3397 | 5020 | 6796 | 4033 | 2468 | 1207 | 1637 | 1039 | 2117 | 4783 | 3054 | 1878 | 3034 | 3936 | 4815 | Crabp1 |
| Crabp2 | 9816 | 10090 | 11417 | 6219 | 3867 | 4939 | 4055 | 4559 | 3807 | 3447 | 3863 | 4425 | 5684 | 4813 | 10112 | 10222 | 9663 | 4935 | 4878 | 6127 | 4110 | 2971 | 4718 | 4220 | 2527 | 6644 | 5606 | 5844 | Crabp2 |
| Cracd | 3421 | 3870 | 4622 | 5981 | 7567 | 6365 | 6360 | 11431 | 12102 | 16759 | 13422 | 16140 | 14197 | 16431 | 4011 | 3731 | 6118 | 5862 | 7212 | 5435 | 6762 | 16967 | 10412 | 16307 | 17980 | 10809 | 14189 | 14739 | Cracd |
| Cracdl | 213 | 247 | 285 | 248 | 311 | 274 | 149 | 134 | 282 | 239 | 236 | 124 | 142 | 92 | 193 | 287 | 226 | 307 | 282 | 206 | 208 | 221 | 267 | 183 | 188 | 182 | 95 | 113 | Cracdl |
| Cracr2a | 114 | 111 | 113 | 90 | 117 | 110 | 65 | 132 | 154 | 162 | 152 | 146 | 171 | 127 | 117 | 129 | 103 | 116 | 77 | 128 | 82 | 125 | 135 | 159 | 105 | 122 | 135 | 155 | Cracr2a |
| Cracr2b | 107 | 140 | 179 | 126 | 92 | 122 | 89 | 103 | 163 | 136 | 183 | 129 | 146 | 101 | 99 | 172 | 165 | 125 | 117 | 90 | 104 | 100 | 159 | 141 | 139 | 193 | 134 | 93 | Cracr2b |
| Cradd | 211 | 209 | 221 | 169 | 177 | 184 | 181 | 454 | 419 | 443 | 435 | 438 | 504 | 567 | 171 | 156 | 184 | 156 | 235 | 226 | 186 | 431 | 419 | 522 | 471 | 646 | 518 | 510 | Cradd |
| Cramp1 | 7083 | 7311 | 6740 | 8623 | 8057 | 7877 | 8094 | 6348 | 6292 | 7093 | 6981 | 7951 | 7528 | 7885 | 6355 | 7013 | 7736 | 8597 | 8034 | 6429 | 8052 | 7802 | 5675 | 7349 | 7695 | 5722 | 7487 | 7771 | Cramp1 |
| Crat | 2253 | 2661 | 2385 | 2282 | 2484 | 2107 | 2076 | 1883 | 2177 | 2026 | 2046 | 2006 | 2051 | 1825 | 2218 | 2685 | 2433 | 2335 | 2383 | 1905 | 2045 | 2085 | 2093 | 2092 | 2104 | 2034 | 1912 | 1925 | Crat |
| Crb1 | 1613 | 1721 | 1371 | 1898 | 1892 | 2028 | 2188 | 1865 | 1324 | 1536 | 1011 | 1804 | 1232 | 1693 | 1975 | 1219 | 1849 | 1909 | 2146 | 2226 | 2244 | 1908 | 1269 | 1325 | 1562 | 1242 | 1844 | 1635 | Crb1 |
| Crb2 | 3245 | 4384 | 5580 | 4019 | 3397 | 3545 | 3472 | 2414 | 2273 | 2625 | 1975 | 5058 | 3978 | 4189 | 3400 | 3123 | 5115 | 3498 | 4464 | 3588 | 3413 | 2821 | 2447 | 3602 | 3260 | 4878 | 4794 | 4881 | Crb2 |
| Crb3 | 143 | 81 | 42 | 111 | 42 | 41 | 32 | 71 | 96 | 109 | 57 | 26 | 51 | 33 | 82 | 92 | 36 | 70 | 37 | 31 | 28 | 47 | 109 | 83 | 24 | 84 | 22 | 30 | Crb3 |
| Crbn | 3983 | 3601 | 2777 | 3171 | 3244 | 3405 | 3474 | 4855 | 4156 | 3347 | 3341 | 3332 | 3341 | 3528 | 4354 | 3796 | 3063 | 3126 | 3137 | 4149 | 3680 | 3720 | 4374 | 3148 | 3543 | 3082 | 3691 | 3553 | Crbn |
| Crcp | 1765 | 1910 | 2283 | 1742 | 1844 | 1760 | 2029 | 2974 | 2512 | 2566 | 2301 | 2685 | 2656 | 3112 | 1970 | 1649 | 1912 | 1698 | 1952 | 2207 | 1979 | 2764 | 2471 | 2686 | 2785 | 2872 | 2862 | 2963 | Crcp |
| Crct1 | 5 | 2 | 6 | 1 | 6 | 4 | 3 | 7 | 11 | 3 | 5 | 1 | 1 | 1 | 3 | 1 | 1 | 0 | 0 | 2 | 4 | 3 | 7 | 7 | 5 | 1 | 0 | 2 | Crct1 |
| Creb1 | 12312 | 9428 | 10103 | 12121 | 13377 | 13911 | 14679 | 15778 | 13180 | 12435 | 12231 | 14101 | 12893 | 14661 | 13097 | 10999 | 10710 | 12068 | 12807 | 15201 | 13926 | 13334 | 12510 | 13081 | 14016 | 11924 | 14534 | 14320 | Creb1 |
| Creb3 | 2941 | 3178 | 2606 | 3405 | 2761 | 2898 | 2694 | 2405 | 2094 | 2138 | 2028 | 1896 | 2011 | 1853 | 3029 | 3045 | 2844 | 3043 | 2894 | 3270 | 2865 | 2184 | 2183 | 2047 | 2225 | 2034 | 2092 | 1901 | Creb3 |
| Creb3l1 | 783 | 977 | 886 | 766 | 745 | 807 | 422 | 466 | 784 | 763 | 1124 | 555 | 625 | 459 | 775 | 1122 | 914 | 965 | 613 | 430 | 483 | 747 | 1131 | 772 | 669 | 518 | 582 | 469 | Creb3l1 |
| Creb3l2 | 6819 | 6199 | 6192 | 5324 | 4075 | 4717 | 3831 | 3707 | 5662 | 5241 | 8571 | 3049 | 4610 | 2929 | 5813 | 7959 | 5795 | 6294 | 4032 | 2888 | 3733 | 5616 | 5977 | 4499 | 3374 | 3797 | 2995 | 2982 | Creb3l2 |
| Creb3l3 | 10 | 13 | 8 | 6 | 1 | 0 | 1 | 20 | 18 | 5 | 21 | 18 | 28 | 10 | 8 | 12 | 9 | 1 | 2 | 10 | 8 | 17 | 16 | 14 | 8 | 8 | 9 | 9 | Creb3l3 |
| Creb3l4 | 245 | 251 | 272 | 219 | 173 | 206 | 196 | 182 | 239 | 233 | 258 | 213 | 227 | 187 | 256 | 204 | 267 | 185 | 208 | 219 | 208 | 174 | 260 | 289 | 160 | 298 | 244 | 273 | Creb3l4 |
| Creb5 | 1562 | 1167 | 1354 | 1880 | 1723 | 1521 | 2367 | 1345 | 1732 | 1678 | 2414 | 2232 | 1631 | 2388 | 1402 | 1304 | 1494 | 1852 | 1962 | 1556 | 1934 | 1268 | 1617 | 1832 | 1981 | 2019 | 1741 | 1965 | Creb5 |
| Crebbp | 2643 | 2750 | 3828 | 4313 | 4556 | 3917 | 3652 | 5470 | 6267 | 8040 | 7299 | 8016 | 6998 | 7438 | 2658 | 2829 | 4424 | 4298 | 4669 | 3024 | 3829 | 8348 | 5513 | 8328 | 8684 | 7781 | 6800 | 7505 | Crebbp |
| Crebl2 | 767 | 772 | 797 | 976 | 1079 | 936 | 876 | 598 | 527 | 751 | 699 | 509 | 525 | 533 | 711 | 798 | 882 | 1114 | 887 | 881 | 964 | 692 | 571 | 596 | 707 | 444 | 509 | 501 | Crebl2 |
| Crebrf | 2726 | 2094 | 1458 | 2556 | 2468 | 2785 | 2904 | 3337 | 2624 | 2603 | 2427 | 2274 | 2471 | 2317 | 2846 | 2437 | 2028 | 2776 | 2392 | 2946 | 3091 | 3075 | 2673 | 2382 | 2826 | 1373 | 2136 | 2267 | Crebrf |
| Crebzf | 7653 | 6157 | 7034 | 8253 | 7298 | 7448 | 8214 | 8385 | 7217 | 6683 | 7326 | 7777 | 7244 | 7601 | 7404 | 6685 | 6955 | 7617 | 7512 | 8082 | 8027 | 7620 | 7207 | 7609 | 7424 | 7788 | 7503 | 7887 | Crebzf |
| Creg1 | 2435 | 2549 | 1887 | 1956 | 1559 | 1805 | 1798 | 813 | 940 | 681 | 851 | 779 | 1227 | 816 | 2327 | 2708 | 1871 | 2134 | 1629 | 1979 | 1751 | 703 | 1117 | 776 | 695 | 988 | 809 | 1099 | Creg1 |
| Creg2 | 602 | 629 | 809 | 895 | 1269 | 1111 | 860 | 462 | 381 | 509 | 479 | 332 | 443 | 267 | 847 | 830 | 801 | 873 | 1001 | 975 | 897 | 662 | 419 | 300 | 426 | 101 | 266 | 325 | Creg2 |
| Creld1 | 1119 | 1465 | 1920 | 1768 | 2014 | 1667 | 1724 | 1774 | 2115 | 2438 | 2420 | 2332 | 2299 | 2202 | 1221 | 1368 | 1972 | 1816 | 2055 | 1669 | 1709 | 2283 | 2060 | 2621 | 2599 | 2417 | 2181 | 2277 | Creld1 |
| Creld2 | 2874 | 3584 | 3364 | 3161 | 2222 | 2298 | 2358 | 1799 | 1860 | 1878 | 2135 | 1916 | 1754 | 1800 | 2620 | 3086 | 3315 | 3127 | 2724 | 2486 | 2420 | 1775 | 1867 | 2216 | 1834 | 2262 | 1998 | 1953 | Creld2 |
| Crem | 538 | 480 | 616 | 551 | 578 | 549 | 613 | 937 | 815 | 729 | 865 | 732 | 840 | 823 | 626 | 572 | 560 | 530 | 563 | 714 | 566 | 854 | 904 | 659 | 798 | 774 | 779 | 727 | Crem |
| Crh | 5 | 6 | 9 | 13 | 15 | 13 | 12 | 28 | 51 | 23 | 31 | 2 | 32 | 1 | 7 | 5 | 8 | 17 | 2 | 8 | 21 | 62 | 54 | 8 | 11 | 0 | 0 | 1 | Crh |
| Crhbp | 20 | 35 | 36 | 23 | 41 | 23 | 28 | 72 | 72 | 97 | 101 | 21 | 128 | 32 | 57 | 44 | 21 | 23 | 41 | 29 | 24 | 92 | 109 | 31 | 36 | 42 | 41 | 46 | Crhbp |
| Crhr1 | 54 | 97 | 85 | 126 | 295 | 163 | 161 | 118 | 184 | 203 | 174 | 88 | 132 | 60 | 95 | 67 | 141 | 183 | 187 | 162 | 178 | 204 | 151 | 97 | 158 | 60 | 62 | 49 | Crhr1 |
| Crhr2 | 68 | 98 | 90 | 89 | 102 | 88 | 75 | 16 | 32 | 19 | 31 | 11 | 18 | 23 | 71 | 87 | 95 | 90 | 89 | 77 | 87 | 28 | 19 | 14 | 15 | 24 | 14 | 24 | Crhr2 |
| Crim1 | 2892 | 2851 | 3448 | 3599 | 4272 | 3948 | 3919 | 2827 | 2925 | 3023 | 3777 | 2620 | 3384 | 2993 | 3492 | 3422 | 3279 | 3884 | 3628 | 3130 | 3934 | 3725 | 2945 | 2871 | 2965 | 2082 | 2803 | 3021 | Crim1 |
| Crip1 | 594 | 732 | 760 | 554 | 384 | 471 | 427 | 397 | 491 | 477 | 588 | 387 | 619 | 579 | 574 | 564 | 548 | 617 | 367 | 416 | 382 | 354 | 645 | 605 | 480 | 484 | 569 | 642 | Crip1 |
| Crip2 | 5060 | 8040 | 8214 | 8638 | 8603 | 7204 | 8089 | 3496 | 4274 | 4823 | 4190 | 5045 | 4451 | 4864 | 5751 | 6185 | 9199 | 7905 | 9232 | 7674 | 8309 | 4639 | 4232 | 5107 | 5431 | 5048 | 4656 | 4758 | Crip2 |
| Crip3 | 110 | 140 | 212 | 309 | 340 | 194 | 215 | 105 | 145 | 234 | 163 | 112 | 121 | 98 | 116 | 143 | 294 | 335 | 301 | 215 | 293 | 212 | 114 | 147 | 163 | 116 | 106 | 103 | Crip3 |
| Cript | 4868 | 4442 | 4543 | 4672 | 4234 | 3743 | 5545 | 4451 | 3405 | 3522 | 3167 | 3925 | 3677 | 4575 | 5487 | 4602 | 4678 | 4718 | 4514 | 5112 | 5443 | 3903 | 3457 | 3858 | 3830 | 3955 | 3890 | 4482 | Cript |
| Crisp1 | 1 | 0 | 0 | 0 | 2 | 0 | 0 | 7 | 6 | 0 | 0 | 0 | 0 | 0 | 0 | 0 | 0 | 0 | 0 | 0 | 0 | 0 | 6 | 2 | 0 | 2 | 0 | 0 | Crisp1 |
| Crisp2 | 1 | 0 | 0 | 0 | 0 | 1 | 0 | 0 | 0 | 0 | 0 | 0 | 0 | 0 | 0 | 0 | 1 | 8 | 2 | 0 | 0 | 0 | 0 | 0 | 3 | 1 | 0 | 1 | Crisp2 |
| Crisp3 | 0 | 0 | 0 | 0 | 0 | 0 | 0 | 0 | 0 | 0 | 0 | 0 | 0 | 0 | 0 | 0 | 0 | 0 | 0 | 0 | 0 | 0 | 1 | 0 | 0 | 0 | 0 | 0 | Crisp3 |
| Crisp4 | 0 | 0 | 0 | 0 | 0 | 0 | 1 | 0 | 0 | 1 | 0 | 1 | 0 | 0 | 0 | 0 | 0 | 0 | 0 | 0 | 1 | 0 | 1 | 2 | 0 | 0 | 0 | 0 | Crisp4 |
| Crispld1 | 1494 | 1076 | 1143 | 900 | 594 | 1178 | 413 | 2002 | 2368 | 1821 | 2562 | 1543 | 2193 | 1760 | 1476 | 1813 | 830 | 1127 | 433 | 631 | 419 | 2074 | 2666 | 1901 | 1643 | 2015 | 1858 | 1665 | Crispld1 |
| Crispld2 | 798 | 952 | 920 | 587 | 320 | 537 | 203 | 324 | 437 | 355 | 605 | 472 | 651 | 402 | 844 | 1326 | 610 | 680 | 280 | 315 | 249 | 391 | 638 | 494 | 371 | 527 | 549 | 487 | Crispld2 |
| Crk | 23348 | 20496 | 21444 | 20361 | 18462 | 19380 | 20740 | 20309 | 16967 | 15080 | 15690 | 17767 | 16276 | 17404 | 24564 | 21436 | 20715 | 20447 | 19240 | 21960 | 19825 | 17998 | 17114 | 16829 | 16491 | 18940 | 17680 | 19015 | Crk |
| Crkl | 10873 | 10401 | 10131 | 11030 | 10555 | 10929 | 10956 | 11116 | 10055 | 10359 | 9552 | 11796 | 10579 | 11406 | 10363 | 9851 | 10729 | 11114 | 11291 | 9804 | 11348 | 11251 | 9228 | 10288 | 11643 | 11396 | 10716 | 11489 | Crkl |
| Crlf1 | 436 | 100 | 80 | 125 | 34 | 48 | 36 | 391 | 380 | 286 | 210 | 12 | 59 | 19 | 78 | 217 | 63 | 99 | 57 | 49 | 58 | 72 | 366 | 195 | 20 | 316 | 12 | 39 | Crlf1 |
| Crlf2 | 63 | 123 | 127 | 102 | 165 | 133 | 77 | 227 | 327 | 320 | 348 | 243 | 325 | 180 | 140 | 107 | 130 | 138 | 156 | 119 | 130 | 424 | 364 | 311 | 300 | 231 | 235 | 183 | Crlf2 |
| Crlf3 | 3054 | 2736 | 2422 | 2612 | 2167 | 2397 | 2611 | 1895 | 1849 | 1642 | 2005 | 1868 | 2025 | 2002 | 2598 | 2766 | 2447 | 2422 | 2313 | 2392 | 2393 | 1419 | 1928 | 1947 | 1716 | 2335 | 2108 | 2054 | Crlf3 |
| Crls1 | 2540 | 2097 | 2501 | 2124 | 2163 | 2074 | 2597 | 2522 | 2259 | 2059 | 2012 | 2320 | 2110 | 2604 | 2567 | 2213 | 2088 | 1948 | 2261 | 2489 | 2329 | 2176 | 2167 | 2343 | 2179 | 2697 | 2273 | 2678 | Crls1 |
| Crmp1 | 19152 | 25829 | 23134 | 35115 | 46921 | 38119 | 39016 | 29692 | 33858 | 42986 | 32991 | 38198 | 31495 | 34895 | 24539 | 22179 | 32123 | 37873 | 44222 | 33270 | 41213 | 47775 | 31625 | 38705 | 46439 | 31476 | 30355 | 33375 | Crmp1 |
| Crnkl1 | 6482 | 6816 | 6198 | 6018 | 4565 | 4969 | 5726 | 4684 | 4185 | 4044 | 3497 | 4528 | 4373 | 4750 | 6861 | 6286 | 5782 | 5442 | 5060 | 5921 | 5500 | 3531 | 4071 | 4315 | 3997 | 4714 | 4478 | 4685 | Crnkl1 |
| Crnn | 0 | 0 | 0 | 0 | 0 | 0 | 0 | 1 | 4 | 5 | 0 | 0 | 5 | 0 | 0 | 0 | 0 | 0 | 0 | 0 | 0 | 0 | 4 | 0 | 0 | 0 | 0 | 0 | Crnn |
| Crocc | 3617 | 4632 | 6080 | 5981 | 4962 | 4784 | 4497 | 3230 | 3884 | 4799 | 4385 | 5956 | 5173 | 5450 | 3435 | 4352 | 6437 | 5315 | 5563 | 4117 | 4676 | 4373 | 3613 | 5639 | 4812 | 5042 | 5444 | 4997 | Crocc |
| Crocc2 | 87 | 82 | 36 | 59 | 110 | 145 | 60 | 106 | 119 | 90 | 117 | 122 | 172 | 83 | 74 | 119 | 58 | 86 | 77 | 113 | 97 | 124 | 234 | 118 | 122 | 114 | 120 | 82 | Crocc2 |
| Crot | 3838 | 3218 | 3345 | 2619 | 2648 | 2778 | 3044 | 3011 | 3030 | 2208 | 2433 | 2483 | 2415 | 2589 | 4032 | 3544 | 2891 | 2641 | 2565 | 3604 | 2801 | 2214 | 3072 | 2381 | 2182 | 2957 | 2674 | 2676 | Crot |
| Crp | 0 | 1 | 0 | 1 | 3 | 1 | 0 | 0 | 0 | 0 | 13 | 0 | 0 | 0 | 0 | 1 | 0 | 0 | 0 | 0 | 0 | 2 | 1 | 1 | 0 | 2 | 0 | 0 | Crp |
| Crppa | 155 | 134 | 130 | 94 | 170 | 111 | 121 | 672 | 528 | 484 | 517 | 518 | 583 | 509 | 186 | 168 | 94 | 72 | 98 | 152 | 113 | 580 | 621 | 530 | 521 | 499 | 600 | 540 | Crppa |
| Crtac1 | 577 | 716 | 483 | 936 | 1256 | 904 | 935 | 573 | 672 | 840 | 717 | 494 | 522 | 470 | 646 | 664 | 762 | 1029 | 1071 | 930 | 938 | 765 | 702 | 687 | 696 | 610 | 472 | 385 | Crtac1 |

| | | | | | | | | | | | | | | | | | | | | | | | | | | | | | |
|---|---|---|---|---|---|---|---|---|---|---|---|---|---|---|---|---|---|---|---|---|---|---|---|---|---|---|---|---|---|
| Crtam | 18 | 16 | 42 | 41 | 19 | 24 | 23 | 12 | 14 | 22 | 16 | 12 | 31 | 6 | 42 | 32 | 24 | 44 | 15 | 27 | 31 | 8 | 11 | 15 | 17 | 12 | 8 | 12 | Crtam |
| Crtap | 1256 | 1430 | 1376 | 941 | 638 | 929 | 502 | 1462 | 2035 | 1761 | 2527 | 1374 | 1955 | 1409 | 1255 | 1631 | 907 | 1102 | 542 | 588 | 518 | 1848 | 2620 | 1555 | 1316 | 1663 | 1548 | 1501 | Crtap |
| Crtc1 | 1752 | 2185 | 2332 | 3255 | 3823 | 3174 | 2568 | 3793 | 4941 | 6194 | 5791 | 5370 | 4820 | 4535 | 1755 | 2029 | 2786 | 3229 | 3406 | 2387 | 2874 | 6697 | 4603 | 5799 | 6375 | 4711 | 4458 | 4194 | Crtc1 |
| Crtc2 | 3133 | 3778 | 3511 | 3833 | 3328 | 3355 | 3343 | 2546 | 2888 | 2840 | 2985 | 3070 | 3024 | 2899 | 2755 | 3453 | 3486 | 3683 | 3647 | 3090 | 3075 | 3096 | 2657 | 3107 | 3036 | 3147 | 2968 | 2885 | Crtc2 |
| Crtc3 | 514 | 628 | 771 | 580 | 873 | 680 | 532 | 1160 | 1681 | 1635 | 1968 | 1402 | 1504 | 1271 | 454 | 720 | 700 | 765 | 628 | 428 | 557 | 1355 | 1670 | 1833 | 1465 | 1631 | 1128 | 1299 | Crtc3 |
| Crx | 1 | 3 | 1 | 0 | 3 | 12 | 0 | 7 | 9 | 26 | 5 | 23 | 23 | 21 | 4 | 2 | 10 | 3 | 3 | 4 | 5 | 18 | 12 | 32 | 15 | 3 | 34 | 23 | Crx |
| Crxos | 6 | 3 | 10 | 3 | 1 | 5 | 0 | 4 | 0 | 0 | 0 | 3 | 7 | 5 | 5 | 2 | 1 | 3 | 0 | 4 | 4 | 0 | 1 | 2 | 4 | 0 | 2 | 6 | Crxos |
| Cry1 | 5116 | 4922 | 5605 | 5751 | 4494 | 5131 | 5268 | 2037 | 2015 | 1848 | 2002 | 2162 | 1983 | 2103 | 4754 | 4682 | 5401 | 5642 | 5075 | 4418 | 5413 | 2030 | 1919 | 2094 | 2091 | 2326 | 1954 | 2274 | Cry1 |
| Cry2 | 4872 | 6217 | 4823 | 6147 | 4709 | 4781 | 4602 | 2268 | 2454 | 2403 | 2750 | 2773 | 2961 | 2413 | 4899 | 5844 | 4960 | 6276 | 4742 | 4377 | 4845 | 3066 | 2629 | 2549 | 2880 | 2517 | 2459 | 2455 | Cry2 |
| Cryaa | 0 | 2 | 0 | 0 | 1 | 0 | 0 | 0 | 1 | 0 | 5 | 5 | 1 | 0 | 0 | 0 | 0 | 0 | 2 | 5 | 0 | 10 | 2 | 0 | 2 | 0 | 0 | 0 | Cryaa |
| Cryab | 92 | 72 | 51 | 48 | 52 | 53 | 46 | 115 | 222 | 104 | 679 | 106 | 120 | 87 | 73 | 100 | 32 | 61 | 44 | 22 | 42 | 135 | 535 | 136 | 102 | 98 | 95 | 106 | Cryab |
| Cryba1 | 20 | 26 | 10 | 4 | 17 | 4 | 4 | 31 | 15 | 18 | 30 | 18 | 43 | 11 | 45 | 32 | 24 | 6 | 4 | 23 | 5 | 28 | 20 | 22 | 27 | 15 | 16 | 40 | Cryba1 |
| Cryba2 | 2 | 3 | 17 | 3 | 18 | 15 | 4 | 21 | 17 | 9 | 12 | 13 | 42 | 7 | 5 | 11 | 8 | 3 | 9 | 6 | 3 | 13 | 14 | 10 | 7 | 0 | 8 | 6 | Cryba2 |
| Cryba4 | 18 | 22 | 16 | 6 | 19 | 31 | 17 | 28 | 31 | 33 | 18 | 27 | 20 | 27 | 16 | 30 | 15 | 8 | 27 | 22 | 14 | 20 | 37 | 20 | 27 | 24 | 19 | 28 | Cryba4 |
| Crybb1 | 18 | 12 | 9 | 22 | 30 | 15 | 25 | 29 | 32 | 28 | 58 | 16 | 19 | 14 | 22 | 25 | 13 | 23 | 25 | 25 | 28 | 77 | 44 | 21 | 32 | 36 | 53 | 9 | Crybb1 |
| Crybb2 | 0 | 0 | 0 | 1 | 0 | 0 | 0 | 0 | 0 | 0 | 0 | 0 | 0 | 0 | 0 | 0 | 0 | 0 | 0 | 0 | 0 | 0 | 0 | 0 | 0 | 0 | 0 | 0 | Crybb2 |
| Crybb3 | 210 | 339 | 363 | 476 | 283 | 335 | 283 | 34 | 43 | 74 | 60 | 64 | 46 | 54 | 166 | 314 | 382 | 491 | 376 | 223 | 273 | 77 | 61 | 72 | 90 | 104 | 26 | 51 | Crybb3 |
| Crybg1 | 211 | 121 | 137 | 83 | 61 | 77 | 57 | 167 | 325 | 160 | 192 | 96 | 111 | 80 | 128 | 172 | 82 | 86 | 66 | 82 | 73 | 117 | 304 | 177 | 85 | 93 | 135 | 107 | Crybg1 |
| Crybg2 | 206 | 155 | 187 | 90 | 89 | 153 | 79 | 101 | 147 | 93 | 104 | 63 | 109 | 56 | 162 | 178 | 97 | 141 | 96 | 108 | 80 | 15 | 191 | 114 | 29 | 35 | 73 | 95 | Crybg2 |
| Crybg3 | 2223 | 1791 | 1171 | 1109 | 1226 | 1587 | 1142 | 2710 | 2756 | 1979 | 2751 | 2036 | 2679 | 2037 | 2163 | 2303 | 1118 | 1279 | 948 | 1257 | 1205 | 1566 | 2784 | 1450 | 1624 | 1583 | 2623 | 2042 | Crybg3 |
| Cryga | 0 | 0 | 0 | 0 | 0 | 0 | 0 | 2 | 0 | 0 | 0 | 0 | 0 | 1 | 0 | 0 | 0 | 0 | 0 | 0 | 0 | 0 | 0 | 0 | 0 | 0 | 0 | 0 | Cryga |
| Crygb | 0 | 1 | 0 | 0 | 2 | 0 | 0 | 0 | 1 | 1 | 1 | 1 | 2 | 1 | 1 | 1 | 1 | 1 | 1 | 0 | 0 | 2 | 1 | 1 | 5 | 0 | 2 | 1 | Crygb |
| Crygc | 0 | 0 | 0 | 3 | 0 | 4 | 0 | 0 | 0 | 0 | 2 | 0 | 1 | 0 | 1 | 0 | 2 | 0 | 0 | 0 | 0 | 2 | 3 | 1 | 0 | 3 | 2 | 0 | Crygc |
| Crygd | 0 | 0 | 1 | 0 | 2 | 2 | 0 | 0 | 0 | 3 | 1 | 1 | 0 | 1 | 0 | 2 | 0 | 0 | 1 | 1 | 0 | 0 | 0 | 3 | 0 | 0 | 9 | 0 | Crygd |
| Cryge | 2 | 0 | 2 | 0 | 1 | 0 | 1 | 0 | 0 | 0 | 0 | 3 | 0 | 0 | 0 | 0 | 2 | 0 | 0 | 2 | 2 | 0 | 0 | 0 | 1 | 3 | 0 | 4 | Cryge |
| Crygf | 0 | 1 | 0 | 0 | 0 | 0 | 0 | 0 | 0 | 0 | 0 | 0 | 0 | 0 | 0 | 0 | 0 | 0 | 0 | 0 | 0 | 0 | 1 | 0 | 0 | 0 | 0 | 0 | Crygf |
| Crygn | 2 | 1 | 1 | 4 | 2 | 3 | 0 | 1 | 2 | 3 | 2 | 0 | 5 | 1 | 3 | 2 | 2 | 4 | 2 | 2 | 4 | 8 | 6 | 7 | 5 | 7 | 1 | 5 | Crygn |
| Crygs | 1 | 1 | 0 | 5 | 0 | 0 | 0 | 0 | 0 | 0 | 0 | 0 | 1 | 0 | 3 | 1 | 0 | 3 | 1 | 1 | 2 | 3 | 2 | 4 | 0 | 4 | 3 | 1 | Crygs |
| Cryl1 | 456 | 381 | 412 | 313 | 261 | 318 | 217 | 464 | 536 | 453 | 489 | 292 | 491 | 328 | 393 | 537 | 335 | 370 | 235 | 278 | 234 | 341 | 676 | 458 | 314 | 498 | 298 | 335 | Cryl1 |
| Crym | 1338 | 1028 | 896 | 656 | 461 | 512 | 337 | 1208 | 1043 | 1217 | 1321 | 903 | 1458 | 938 | 1030 | 1450 | 670 | 479 | 493 | 361 | 486 | 927 | 1006 | 1128 | 768 | 1478 | 1120 | 897 | Crym |
| Cryz | 1294 | 1156 | 1213 | 921 | 904 | 1073 | 941 | 1224 | 1381 | 1145 | 1228 | 1329 | 1401 | 1378 | 1294 | 1281 | 1077 | 867 | 949 | 1243 | 902 | 988 | 1405 | 1326 | 1061 | 1541 | 1357 | 1597 | Cryz |
| Cryzl1 | 4118 | 3507 | 3323 | 3689 | 3906 | 4091 | 4725 | 4137 | 3569 | 3526 | 3366 | 3438 | 3392 | 3870 | 4352 | 3816 | 3716 | 3918 | 3995 | 4357 | 4484 | 3999 | 3620 | 3661 | 3727 | 3469 | 3800 | 3614 | Cryzl1 |
| Cryzl2 | 378 | 441 | 411 | 374 | 319 | 362 | 315 | 430 | 475 | 433 | 489 | 450 | 524 | 423 | 405 | 397 | 364 | 318 | 331 | 320 | 322 | 394 | 456 | 502 | 421 | 544 | 423 | 394 | Cryzl2 |
| Cs | 29251 | 29864 | 27486 | 25264 | 23734 | 24307 | 26521 | 18429 | 18211 | 17243 | 16677 | 20329 | 18718 | 20182 | 27903 | 26279 | 27186 | 25062 | 26197 | 24020 | 25555 | 18180 | 16823 | 17871 | 17926 | 21695 | 19955 | 20843 | Cs |
| Csad | 895 | 885 | 918 | 923 | 906 | 919 | 829 | 1920 | 2290 | 2535 | 2481 | 2310 | 2350 | 2031 | 719 | 924 | 900 | 1000 | 909 | 773 | 866 | 2341 | 2425 | 2562 | 2292 | 2748 | 1984 | 2067 | Csad |
| Csdc2 | 2483 | 3327 | 2916 | 4761 | 5886 | 4574 | 4522 | 2146 | 2569 | 3134 | 2582 | 2295 | 2055 | 2024 | 2545 | 2989 | 3996 | 5187 | 5896 | 4113 | 4952 | 3816 | 2563 | 2656 | 3225 | 1900 | 1617 | 1735 | Csdc2 |
| Csde1 | 22255 | 17278 | 23402 | 18888 | 21066 | 21640 | 22669 | 47006 | 38495 | 37993 | 37433 | 43637 | 42056 | 45350 | 23546 | 19756 | 21436 | 19275 | 21575 | 23696 | 21741 | 40818 | 39535 | 40773 | 41570 | 46275 | 47197 | 46873 | Csde1 |
| Cse1l | 17338 | 14410 | 14257 | 11214 | 10486 | 12086 | 13103 | 19441 | 16614 | 13534 | 12679 | 16727 | 15752 | 16987 | 17368 | 14971 | 12524 | 10588 | 11758 | 14956 | 11950 | 13552 | 15755 | 14515 | 14316 | 18099 | 18053 | 18188 | Cse1l |
| Csf1 | 429 | 456 | 755 | 468 | 691 | 580 | 534 | 657 | 772 | 852 | 1079 | 1002 | 1054 | 872 | 446 | 441 | 578 | 512 | 579 | 550 | 567 | 869 | 1032 | 955 | 914 | 882 | 1022 | 926 | Csf1 |
| Csf1r | 1271 | 1526 | 1267 | 1098 | 1152 | 1361 | 929 | 1575 | 1851 | 1573 | 2064 | 1698 | 1726 | 1454 | 1328 | 1619 | 1031 | 1332 | 1004 | 1365 | 1066 | 1835 | 2737 | 1602 | 1594 | 1664 | 1722 | 1571 | Csf1r |
| Csf2 | 0 | 0 | 0 | 0 | 0 | 0 | 0 | 0 | 0 | 0 | 2 | 1 | 0 | 1 | 0 | 0 | 0 | 0 | 0 | 0 | 0 | 0 | 0 | 0 | 0 | 0 | 0 | 0 | Csf2 |
| Csf2ra | 556 | 764 | 801 | 909 | 1267 | 832 | 843 | 1335 | 1581 | 1882 | 1932 | 1557 | 1569 | 1326 | 592 | 648 | 985 | 1056 | 1146 | 933 | 831 | 2035 | 1521 | 1816 | 1673 | 1415 | 1325 | 1294 | Csf2ra |
| Csf2rb | 63 | 107 | 133 | 60 | 69 | 77 | 37 | 76 | 129 | 67 | 182 | 129 | 167 | 147 | 63 | 138 | 82 | 70 | 48 | 47 | 60 | 92 | 162 | 83 | 114 | 144 | 134 | 451 | Csf2rb |
| Csf2rb2 | 14 | 21 | 58 | 16 | 20 | 15 | 15 | 16 | 41 | 12 | 63 | 37 | 44 | 63 | 16 | 47 | 19 | 33 | 12 | 15 | 22 | 30 | 69 | 39 | 22 | 53 | 38 | 146 | Csf2rb2 |
| Csf3 | 6 | 2 | 1 | 1 | 3 | 2 | 2 | 1 | 4 | 1 | 1 | 2 | 3 | 3 | 1 | 2 | 0 | 4 | 6 | 7 | 1 | 7 | 3 | 5 | 4 | 3 | 1 | 1 | Csf3 |
| Csf3r | 22 | 43 | 32 | 20 | 43 | 47 | 34 | 53 | 82 | 60 | 83 | 59 | 59 | 45 | 30 | 51 | 34 | 24 | 31 | 36 | 32 | 90 | 124 | 69 | 92 | 31 | 51 | 78 | Csf3r |
| Csgalnact1 | 75 | 51 | 57 | 57 | 104 | 101 | 57 | 216 | 438 | 379 | 729 | 150 | 204 | 89 | 79 | 79 | 73 | 89 | 70 | 51 | 82 | 408 | 474 | 236 | 309 | 157 | 92 | 104 | Csgalnact1 |
| Csgalnact2 | 1850 | 1274 | 1357 | 1326 | 1392 | 1627 | 1533 | 1448 | 1451 | 1275 | 1597 | 1102 | 1344 | 1295 | 1785 | 1682 | 1401 | 1571 | 1370 | 1585 | 1531 | 1190 | 1539 | 1310 | 1331 | 1214 | 1219 | 1183 | Csgalnact2 |
| Csk | 4107 | 5156 | 5870 | 5873 | 6052 | 5430 | 4809 | 5046 | 6002 | 6746 | 6403 | 7080 | 6149 | 6242 | 4162 | 4653 | 5858 | 5664 | 6731 | 4368 | 5748 | 6716 | 5900 | 7122 | 7205 | 7841 | 6144 | 6509 | Csk |
| Csl | 7 | 7 | 2 | 1 | 5 | 4 | 6 | 4 | 7 | 2 | 1 | 1 | 1 | 3 | 7 | 2 | 2 | 2 | 2 | 1 | 4 | 5 | 2 | 2 | 5 | 3 | 1 | 8 | Csl |
| Csmd1 | 187 | 247 | 107 | 432 | 1079 | 822 | 810 | 666 | 823 | 739 | 771 | 313 | 295 | 321 | 249 | 232 | 211 | 617 | 549 | 790 | 981 | 785 | 827 | 378 | 825 | 162 | 220 | 228 | Csmd1 |
| Csmd2 | 1232 | 1944 | 1371 | 3095 | 4831 | 3893 | 3641 | 2624 | 2864 | 3664 | 3194 | 2695 | 2908 | 2584 | 1760 | 1781 | 2356 | 3760 | 4023 | 3187 | 4116 | 4659 | 2476 | 2593 | 4063 | 981 | 2414 | 2148 | Csmd2 |
| Csmd3 | 521 | 682 | 212 | 691 | 1148 | 1264 | 1294 | 1771 | 1430 | 918 | 1026 | 509 | 650 | 642 | 825 | 652 | 337 | 935 | 895 | 1663 | 1413 | 1255 | 1232 | 579 | 1076 | 179 | 587 | 456 | Csmd3 |
| Csn1s1 | 0 | 0 | 0 | 0 | 0 | 0 | 0 | 2 | 0 | 0 | 1 | 0 | 0 | 0 | 0 | 0 | 0 | 0 | 0 | 0 | 0 | 0 | 3 | 0 | 0 | 0 | 0 | 0 | Csn1s1 |
| Csn1s2a | 0 | 0 | 0 | 0 | 0 | 3 | 0 | 1 | 0 | 8 | 4 | 0 | 0 | 0 | 3 | 0 | 0 | 0 | 0 | 0 | 0 | 0 | 0 | 4 | 0 | 0 | 0 | 0 | Csn1s2a |
| Csn1s2b | 0 | 0 | 0 | 0 | 4 | 0 | 0 | 0 | 0 | 0 | 4 | 0 | 1 | 0 | 0 | 0 | 0 | 0 | 0 | 0 | 0 | 0 | 2 | 0 | 0 | 0 | 0 | 0 | Csn1s2b |
| Csn2 | 0 | 0 | 0 | 0 | 0 | 0 | 0 | 0 | 0 | 0 | 0 | 0 | 0 | 0 | 0 | 1 | 0 | 0 | 0 | 0 | 0 | 0 | 0 | 0 | 0 | 0 | 0 | 0 | Csn2 |
| Csn3 | 0 | 1 | 0 | 7 | 0 | 0 | 0 | 1 | 0 | 3 | 5 | 3 | 0 | 0 | 0 | 2 | 0 | 1 | 0 | 0 | 0 | 0 | 3 | 7 | 4 | 5 | 2 | 0 | Csn3 |
| Csnk1a1 | 26094 | 20135 | 23611 | 23728 | 26149 | 26117 | 28362 | 36359 | 32548 | 32441 | 33921 | 35882 | 35458 | 36598 | 25862 | 22857 | 23607 | 23300 | 25295 | 24519 | 26911 | 35130 | 32393 | 32135 | 34157 | 34098 | 35143 | 36177 | Csnk1a1 |
| Csnk1d | 11912 | 14213 | 15916 | 16202 | 16537 | 15007 | 14565 | 14554 | 16487 | 17433 | 17120 | 19434 | 17937 | 18310 | 11433 | 13000 | 16734 | 16120 | 17564 | 13915 | 15204 | 17450 | 15571 | 19222 | 18983 | 19790 | 18352 | 18448 | Csnk1d |
| Csnk1e | 8989 | 10946 | 12098 | 11878 | 13094 | 11686 | 11872 | 14108 | 15361 | 17852 | 15229 | 19478 | 15399 | 18354 | 9051 | 9169 | 12975 | 11717 | 13895 | 11104 | 12418 | 17619 | 14057 | 18815 | 19272 | 19180 | 16693 | 17932 | Csnk1e |
| Csnk1g1 | 4930 | 4566 | 4493 | 5553 | 6073 | 5739 | 6086 | 4677 | 4244 | 4753 | 4918 | 4987 | 4775 | 5247 | 4920 | 4403 | 5255 | 5531 | 5795 | 5205 | 6032 | 5394 | 4311 | 5019 | 5417 | 3923 | 5094 | 4946 | Csnk1g1 |
| Csnk1g2 | 7882 | 9723 | 10287 | 9788 | 7598 | 7785 | 6746 | 5362 | 6871 | 7293 | 7487 | 7714 | 7244 | 6838 | 6790 | 8770 | 10062 | 8749 | 8906 | 6946 | 7168 | 6251 | 6965 | 7951 | 6974 | 9148 | 7504 | 7168 | Csnk1g2 |
| Csnk1g3 | 7049 | 5890 | 7427 | 8209 | 7811 | 8635 | 8344 | 6708 | 5630 | 5845 | 5968 | 6081 | 5723 | 6141 | 7766 | 6824 | 7350 | 7524 | 8314 | 7702 | 8220 | 6231 | 5735 | 5992 | 6024 | 5954 | 5810 | 6146 | Csnk1g3 |
| Csnk2a1 | 15166 | 13196 | 15330 | 14374 | 13674 | 13620 | 15081 | 19825 | 16871 | 16741 | 16431 | 18773 | 17823 | 20695 | 15343 | 13420 | 15119 | 13685 | 14587 | 15426 | 14547 | 16852 | 16509 | 18631 | 18649 | 18497 | 19494 | 20340 | Csnk2a1 |
| Csnk2a2 | 7552 | 7379 | 8609 | 8699 | 7198 | 7611 | 8084 | 5195 | 5365 | 6281 | 6315 | 6852 | 5970 | 6818 | 7192 | 7284 | 8358 | 8199 | 8016 | 6573 | 8016 | 6163 | 5461 | 6630 | 6479 | 6953 | 6393 | 6997 | Csnk2a2 |
| Csnk2b | 6369 | 7872 | 6678 | 6967 | 5862 | 6032 | 6402 | 4352 | 4657 | 4651 | 4448 | 5085 | 4702 | 5229 | 6335 | 6514 | 6941 | 6459 | 6856 | 5997 | 6894 | 4309 | 4528 | 5111 | 4724 | 5670 | 5043 | 5401 | Csnk2b |
| Csnka2ip | 7 | 3 | 0 | 3 | 0 | 3 | 2 | 2 | 1 | 0 | 0 | 2 | 0 | 0 | 3 | 1 | 1 | 0 | 12 | 0 | 0 | 0 | 3 | 2 | 4 | 0 | 0 | 0 | Csnka2ip |
| Cspg4 | 857 | 1146 | 1061 | 1148 | 884 | 1120 | 580 | 618 | 1364 | 1699 | 2777 | 836 | 873 | 706 | 767 | 1442 | 932 | 1451 | 906 | 491 | 623 | 1676 | 1353 | 1390 | 1173 | 802 | 737 | 650 | Cspg4 |
| Cspg4b | 765 | 614 | 437 | 397 | 420 | 641 | 292 | 1188 | 1670 | 1125 | 2290 | 880 | 1408 | 811 | 709 | 831 | 295 | 438 | 337 | 403 | 348 | 1041 | 1812 | 813 | 769 | 800 | 996 | 786 | Cspg4b |
| Cspg5 | 1315 | 1961 | 1800 | 2737 | 3917 | 3089 | 3178 | 3019 | 3001 | 3881 | 3255 | 2842 | 2247 | 2863 | 1581 | 1440 | 2478 | 3240 | 3305 | 2628 | 3518 | 4410 | 3053 | 3304 | 4193 | 2131 | 2556 | 2580 | Cspg5 |
| Cspp1 | 8946 | 6869 | 6463 | 8240 | 7674 | 8977 | 9440 | 8243 | 7409 | 7127 | 7376 | 7405 | 7302 | 8110 | 8694 | 8060 | 6617 | 7946 | 6990 | 9117 | 9355 | 6694 | 6867 | 7013 | 7502 | 5643 | 7495 | 7049 | Cspp1 |
| Csprs | 0 | 0 | 0 | 0 | 0 | 1 | 0 | 0 | 0 | 0 | 1 | 0 | 0 | 0 | 0 | 0 | 0 | 0 | 0 | 0 | 0 | 0 | 0 | 0 | 0 | 0 | 0 | 0 | Csprs |
| Csrnp1 | 504 | 577 | 516 | 607 | 412 | 423 | 358 | 117 | 215 | 153 | 184 | 124 | 201 | 146 | 368 | 588 | 589 | 547 | 388 | 348 | 451 | 117 | 223 | 151 | 112 | 154 | 144 | 126 | Csrnp1 |
| Csrnp2 | 2833 | 3293 | 2755 | 3809 | 3602 | 3277 | 3445 | 3799 | 3686 | 3904 | 3652 | 3762 | 3252 | 3683 | 3067 | 2908 | 3237 | 3990 | 3501 | 3320 | 3652 | 4485 | 3486 | 3741 | 4332 | 3579 | 3447 | 3342 | Csrnp2 |
| Csrnp3 | 6321 | 6156 | 4451 | 7847 | 11594 | 11380 | 11280 | 13268 | 9996 | 10751 | 9082 | 9128 | 7970 | 10176 | 8073 | 5917 | 6353 | 8195 | 9690 | 11402 | 12441 | 13102 | 9436 | 8487 | 12895 | 5197 | 8897 | 8434 | Csrnp3 |
| Csrp1 | 7059 | 6549 | 5829 | 4844 | 3293 | 4395 | 3368 | 3379 | 3813 | 3017 | 4245 | 3504 | 4043 | 3235 | 6523 | 7014 | 5205 | 5031 | 4073 | 3844 | 3665 | 2413 | 4586 | 3360 | 2668 | 3944 | 3640 | 3589 | Csrp1 |
| Csrp2 | 7081 | 7581 | 7590 | 6261 | 4786 | 5504 | 5112 | 4418 | 5487 | 4663 | 5245 | 5142 | 5568 | 5303 | 6763 | 8097 | 6579 | 5910 | 5303 | 5801 | 5307 | 3962 | 6183 | 5471 | 4875 | 6584 | 5557 | 6012 | Csrp2 |
| Csrp3 | 7 | 7 | 7 | 6 | 8 | 0 | 8 | 16 | 32 | 4 | 424 | 0 | 3 | 0 | 8 | 3 | 3 | 4 | 5 | 3 | 4 | 25 | 239 | 3 | 5 | 8 | 7 | 5 | Csrp3 |

| | | | | | | | | | | | | | | | | | | | | | | | | | | | | | |
|---|---|---|---|---|---|---|---|---|---|---|---|---|---|---|---|---|---|---|---|---|---|---|---|---|---|---|---|---|---|
| Cst12 | 0 | 0 | 0 | 0 | 0 | 0 | 0 | 2 | 0 | 0 | 0 | 3 | 0 | 0 | 0 | 0 | 0 | 2 | 0 | 0 | 0 | 2 | 0 | 0 | 1 | 0 | 0 | 1 | Cst12 |
| Cst3 | 4680 | 6334 | 5052 | 5550 | 4718 | 4592 | 4977 | 4684 | 5046 | 4504 | 4722 | 4407 | 4547 | 4803 | 5101 | 4957 | 5018 | 5268 | 5057 | 5416 | 5234 | 4119 | 5247 | 4793 | 4311 | 4721 | 4584 | 4886 | Cst3 |
| Cst6 | 2 | 15 | 4 | 10 | 7 | 6 | 3 | 13 | 23 | 30 | 17 | 14 | 18 | 26 | 8 | 14 | 2 | 2 | 3 | 18 | 14 | 12 | 19 | 13 | 22 | 8 | 20 | 20 | Cst6 |
| Cst7 | 0 | 1 | 0 | 0 | 2 | 0 | 0 | 2 | 3 | 0 | 0 | 0 | 2 | 0 | 3 | 0 | 2 | 0 | 5 | 0 | 0 | 7 | 2 | 2 | 2 | 0 | 1 | 5 | Cst7 |
| Csta1 | 0 | 0 | 0 | 0 | 0 | 0 | 0 | 0 | 3 | 0 | 2 | 0 | 0 | 0 | 0 | 0 | 0 | 0 | 0 | 0 | 0 | 0 | 6 | 0 | 1 | 0 | 0 | 0 | Csta1 |
| Csta2 | 1 | 0 | 0 | 4 | 2 | 0 | 2 | 1 | 2 | 2 | 9 | 3 | 1 | 2 | 0 | 0 | 0 | 0 | 1 | 0 | 2 | 0 | 1 | 3 | 2 | 6 | 1 | 2 | Csta2 |
| Csta3 | 1 | 0 | 0 | 4 | 1 | 0 | 0 | 0 | 0 | 0 | 2 | 0 | 0 | 0 | 0 | 0 | 0 | 0 | 0 | 0 | 0 | 3 | 3 | 0 | 0 | 0 | 0 | 0 | Csta3 |
| Cstad | 20 | 9 | 13 | 13 | 7 | 7 | 10 | 12 | 34 | 27 | 21 | 12 | 28 | 18 | 7 | 15 | 14 | 18 | 6 | 8 | 7 | 15 | 30 | 20 | 14 | 19 | 24 | 21 | Cstad |
| Cstb | 1452 | 1503 | 1459 | 1287 | 1044 | 1185 | 1354 | 1271 | 1238 | 1152 | 1259 | 1074 | 1233 | 1318 | 1345 | 1337 | 1378 | 1248 | 1146 | 1234 | 1277 | 871 | 1298 | 1272 | 1096 | 1334 | 1256 | 1372 | Cstb |
| Cstdc2 | 1 | 3 | 0 | 2 | 3 | 0 | 2 | 0 | 2 | 3 | 0 | 0 | 0 | 0 | 1 | 1 | 0 | 9 | 2 | 2 | 0 | 2 | 1 | 0 | 6 | 0 | 5 | 0 | Cstdc2 |
| Cstdc4 | 3 | 10 | 7 | 11 | 11 | 16 | 19 | 15 | 14 | 23 | 60 | 22 | 60 | 21 | 7 | 25 | 15 | 21 | 5 | 2 | 11 | 18 | 35 | 61 | 24 | 64 | 45 | 52 | Cstdc4 |
| Cstdc5 | 28 | 7 | 17 | 8 | 5 | 29 | 22 | 20 | 42 | 34 | 132 | 25 | 76 | 24 | 15 | 40 | 36 | 21 | 11 | 14 | 17 | 30 | 74 | 52 | 13 | 49 | 45 | 55 | Cstdc5 |
| Cstdc6 | 0 | 0 | 0 | 0 | 0 | 0 | 0 | 0 | 0 | 0 | 0 | 0 | 2 | 0 | 0 | 0 | 0 | 0 | 0 | 0 | 0 | 0 | 0 | 0 | 0 | 0 | 0 | 0 | Cstdc6 |
| Cstf1 | 2558 | 3019 | 3020 | 2570 | 2250 | 2387 | 2428 | 2740 | 3007 | 2917 | 2673 | 3614 | 3370 | 3294 | 2246 | 2530 | 2829 | 2397 | 2716 | 2282 | 2317 | 2465 | 2865 | 3168 | 3023 | 3889 | 3354 | 3355 | Cstf1 |
| Cstf2 | 7607 | 6721 | 6102 | 6865 | 8380 | 8171 | 8821 | 10906 | 9755 | 8915 | 8520 | 9982 | 8737 | 10042 | 7916 | 6551 | 6737 | 7118 | 8058 | 8971 | 8854 | 10088 | 8980 | 8979 | 9634 | 9116 | 9789 | 9625 | Cstf2 |
| Cstf2t | 8328 | 7819 | 8567 | 8007 | 8193 | 8279 | 8585 | 9905 | 9040 | 8793 | 8648 | 8982 | 8765 | 9408 | 8747 | 7127 | 8485 | 7702 | 7946 | 9148 | 8384 | 9027 | 8795 | 9032 | 9689 | 9600 | 9365 | 9585 | Cstf2t |
| Cstf3 | 11007 | 9929 | 10961 | 10916 | 9236 | 9205 | 11206 | 6995 | 5955 | 6542 | 5956 | 7130 | 6635 | 7850 | 11263 | 10435 | 10681 | 10377 | 9893 | 10417 | 10896 | 6577 | 5924 | 7034 | 6800 | 7594 | 7469 | 7697 | Cstf3 |
| Cstl1 | 1 | 0 | 0 | 0 | 2 | 3 | 2 | 2 | 5 | 4 | 7 | 3 | 8 | 3 | 0 | 1 | 0 | 0 | 0 | 0 | 0 | 3 | 1 | 4 | 0 | 4 | 1 | 1 | Cstl1 |
| Ct55 | 12 | 6 | 2 | 8 | 18 | 11 | 3 | 16 | 12 | 13 | 14 | 17 | 15 | 12 | 8 | 16 | 7 | 4 | 2 | 4 | 3 | 12 | 11 | 1 | 13 | 14 | 8 | 15 | Ct55 |
| Ctag2l1 | 1 | 2 | 6 | 1 | 1 | 5 | 0 | 1 | 4 | 0 | 5 | 0 | 8 | 3 | 3 | 1 | 7 | 10 | 2 | 10 | 2 | 0 | 1 | 2 | 1 | 1 | 0 | 9 | Ctag2l1 |
| Ctbp1 | 9930 | 12190 | 13529 | 13174 | 13644 | 12910 | 11505 | 15885 | 17364 | 20741 | 18584 | 23189 | 19509 | 20992 | 10030 | 10044 | 14132 | 12406 | 14808 | 11754 | 12646 | 18366 | 17196 | 22198 | 22013 | 24557 | 21090 | 21789 | Ctbp1 |
| Ctbp2 | 4546 | 4336 | 5162 | 4430 | 4898 | 4950 | 5028 | 6117 | 6105 | 6381 | 5548 | 7538 | 6520 | 7392 | 4469 | 4447 | 4972 | 4216 | 5034 | 4673 | 4725 | 6010 | 5771 | 7055 | 6512 | 8319 | 7354 | 7322 | Ctbp2 |
| Ctbs | 662 | 567 | 542 | 410 | 475 | 613 | 532 | 556 | 603 | 412 | 647 | 527 | 628 | 490 | 645 | 661 | 458 | 481 | 397 | 522 | 464 | 495 | 681 | 452 | 434 | 543 | 554 | 564 | Ctbs |
| Ctc1 | 4535 | 5363 | 4905 | 5080 | 5264 | 4793 | 5264 | 6337 | 7061 | 7324 | 6775 | 8524 | 7251 | 8165 | 4746 | 4727 | 5664 | 4994 | 5480 | 4462 | 5132 | 7267 | 6396 | 7875 | 7528 | 8347 | 7783 | 7744 | Ctc1 |
| Ctcf | 4985 | 4364 | 5870 | 5219 | 5749 | 5463 | 6055 | 11864 | 11040 | 12001 | 11315 | 14185 | 12895 | 14840 | 4834 | 4229 | 5718 | 5008 | 6138 | 5149 | 5529 | 12350 | 10675 | 13462 | 13335 | 15023 | 14339 | 14506 | Ctcf |
| Ctcfl | 0 | 0 | 0 | 0 | 0 | 0 | 0 | 0 | 0 | 0 | 0 | 0 | 0 | 1 | 3 | 0 | 0 | 2 | 0 | 0 | 0 | 0 | 3 | 4 | 0 | 0 | 0 | 2 | Ctcfl |
| Ctdnep1 | 2666 | 3018 | 3675 | 3166 | 3193 | 2830 | 2975 | 3552 | 3697 | 4191 | 4189 | 4600 | 3878 | 4071 | 2356 | 2576 | 3624 | 3133 | 3650 | 2662 | 3116 | 4124 | 3814 | 4692 | 4301 | 5285 | 4172 | 4284 | Ctdnep1 |
| Ctdp1 | 3179 | 3791 | 3704 | 3838 | 2812 | 3154 | 3058 | 1842 | 2129 | 2147 | 2076 | 2828 | 2471 | 2713 | 2962 | 3467 | 3656 | 3434 | 3368 | 2632 | 3140 | 2082 | 1948 | 2563 | 2330 | 3055 | 2653 | 2893 | Ctdp1 |
| Ctdsp1 | 3701 | 3808 | 3834 | 3435 | 2973 | 3061 | 2922 | 3548 | 5078 | 4383 | 5168 | 4040 | 4538 | 4370 | 3095 | 3672 | 3752 | 3242 | 3289 | 2690 | 2951 | 3375 | 5317 | 4544 | 3652 | 5664 | 4126 | 4313 | Ctdsp1 |
| Ctdsp2 | 10704 | 11008 | 11605 | 10238 | 9286 | 9576 | 9072 | 9739 | 12175 | 11832 | 13149 | 12295 | 12168 | 11097 | 8794 | 10685 | 11575 | 10128 | 10114 | 7887 | 9123 | 9972 | 12371 | 13082 | 10853 | 14446 | 11302 | 12035 | Ctdsp2 |
| Ctdspl | 1592 | 1539 | 1589 | 1487 | 1422 | 1485 | 1367 | 1498 | 2207 | 2168 | 2346 | 2053 | 2076 | 1800 | 1328 | 1717 | 1452 | 1563 | 1510 | 1203 | 1297 | 1960 | 2063 | 2112 | 1932 | 2527 | 1694 | 1799 | Ctdspl |
| Ctdspl2 | 8338 | 6373 | 5135 | 6249 | 5399 | 6519 | 6738 | 7146 | 6044 | 4777 | 5407 | 6248 | 5930 | 6129 | 7996 | 7211 | 5616 | 6205 | 5636 | 7490 | 6596 | 5319 | 5840 | 5241 | 5512 | 5619 | 6231 | 6621 | Ctdspl2 |
| Ctf1 | 212 | 127 | 133 | 192 | 132 | 150 | 150 | 322 | 353 | 297 | 384 | 253 | 293 | 237 | 169 | 165 | 134 | 132 | 108 | 159 | 146 | 234 | 333 | 320 | 213 | 311 | 210 | 210 | Ctf1 |
| Ctf2 | 1 | 2 | 0 | 0 | 0 | 0 | 1 | 5 | 1 | 1 | 7 | 0 | 0 | 0 | 1 | 1 | 0 | 0 | 0 | 0 | 1 | 0 | 5 | 20 | 0 | 3 | 1 | 0 | Ctf2 |
| Cth | 1452 | 1420 | 1258 | 1064 | 787 | 816 | 941 | 435 | 447 | 277 | 307 | 408 | 445 | 449 | 1427 | 1410 | 1129 | 971 | 924 | 1145 | 847 | 252 | 414 | 404 | 317 | 616 | 496 | 535 | Cth |
| Cthrc1 | 131 | 80 | 164 | 47 | 26 | 75 | 26 | 90 | 94 | 108 | 117 | 62 | 92 | 42 | 149 | 97 | 454 | 77 | 44 | 51 | 176 | 119 | 128 | 295 | 111 | 62 | 74 | 66 | Cthrc1 |
| Ctif | 5417 | 6269 | 7100 | 8128 | 8976 | 7785 | 7144 | 4938 | 5452 | 6919 | 5951 | 6339 | 5977 | 5911 | 5524 | 5577 | 8374 | 8289 | 9055 | 6711 | 7907 | 7503 | 4994 | 6541 | 7357 | 6004 | 5747 | 5886 | Ctif |
| Ctla2a | 60 | 29 | 86 | 32 | 47 | 32 | 56 | 65 | 45 | 74 | 50 | 52 | 68 | 82 | 53 | 55 | 30 | 35 | 37 | 43 | 40 | 50 | 82 | 22 | 47 | 79 | 38 | 58 | Ctla2a |
| Ctla2b | 1 | 0 | 1 | 0 | 2 | 1 | 1 | 4 | 4 | 0 | 4 | 1 | 3 | 6 | 0 | 2 | 0 | 0 | 1 | 4 | 3 | 2 | 3 | 5 | 2 | 4 | 1 | 7 | Ctla2b |
| Ctla4 | 0 | 0 | 0 | 0 | 0 | 3 | 0 | 0 | 0 | 0 | 3 | 0 | 0 | 1 | 0 | 0 | 0 | 0 | 0 | 0 | 1 | 0 | 0 | 0 | 0 | 0 | 1 | 2 | Ctla4 |
| Ctnna1 | 16198 | 15537 | 16582 | 12607 | 11104 | 12119 | 12433 | 14753 | 15077 | 14177 | 14310 | 17451 | 16429 | 17158 | 15369 | 15332 | 14180 | 12038 | 11823 | 12078 | 11911 | 11712 | 15386 | 15948 | 13270 | 19997 | 18273 | 18696 | Ctnna1 |
| Ctnna2 | 10367 | 12646 | 10192 | 14854 | 18781 | 16752 | 18552 | 16932 | 14950 | 15619 | 13017 | 13081 | 12609 | 13999 | 13839 | 10171 | 14124 | 16302 | 16984 | 18135 | 18843 | 18972 | 14446 | 13751 | 18409 | 10361 | 12482 | 12553 | Ctnna2 |
| Ctnna3 | 3 | 4 | 7 | 7 | 25 | 6 | 7 | 7 | 29 | 17 | 41 | 14 | 6 | 11 | 15 | 13 | 7 | 8 | 9 | 16 | 6 | 15 | 36 | 8 | 0 | 8 | 5 | 6 | Ctnna3 |
| Ctnnal1 | 5405 | 4529 | 4341 | 3602 | 3850 | 4119 | 3845 | 3374 | 3140 | 2540 | 2987 | 3227 | 3132 | 3101 | 4851 | 4954 | 3707 | 3390 | 3448 | 4289 | 3884 | 2635 | 3192 | 2536 | 2702 | 3164 | 3374 | 3323 | Ctnnal1 |
| Ctnnb1 | 132919 | 131302 | 119285 | 115170 | 96546 | 106830 | 110405 | 119827 | 109095 | 94064 | 91953 | 114695 | 103956 | 109996 | 130825 | 119780 | 116975 | 110915 | 112178 | 117360 | 110384 | 97596 | 107452 | 109152 | 101646 | 120223 | 121436 | 118283 | Ctnnb1 |
| Ctnnbip1 | 5549 | 5743 | 6364 | 6892 | 6489 | 5910 | 6178 | 3483 | 3765 | 4490 | 4159 | 4646 | 4092 | 4240 | 5321 | 5977 | 6770 | 6687 | 6782 | 5599 | 5967 | 4403 | 3878 | 4457 | 4760 | 4855 | 3856 | 4235 | Ctnnbip1 |
| Ctnnbl1 | 5714 | 7773 | 7407 | 6624 | 5812 | 5924 | 6172 | 4325 | 4317 | 4775 | 4183 | 5389 | 4682 | 5682 | 5995 | 6184 | 7265 | 6207 | 7092 | 6545 | 6604 | 3654 | 4089 | 5275 | 5135 | 5734 | 5650 | 5968 | Ctnnbl1 |
| Ctnnd1 | 2280 | 1849 | 2042 | 2181 | 1712 | 1755 | 2092 | 2049 | 1681 | 1758 | 1878 | 1638 | 1807 | 2354 | 2206 | 1630 | 2160 | 1926 | 1581 | 1778 | 2214 | 1786 | 1680 | 2154 | 1701 | 2019 | 1780 | 2235 | Ctnnd1 |
| Ctnnd2 | 3483 | 4227 | 4012 | 6991 | 8159 | 6628 | 7517 | 5277 | 5948 | 7022 | 5733 | 6753 | 4560 | 6442 | 4088 | 3349 | 6055 | 7399 | 7782 | 6193 | 7660 | 7713 | 4948 | 6761 | 7793 | 5710 | 4660 | 5242 | Ctnnd2 |
| Ctns | 2911 | 3419 | 2782 | 2894 | 2494 | 2605 | 2675 | 1795 | 1752 | 1697 | 1821 | 1690 | 1745 | 1695 | 2885 | 2975 | 2635 | 2838 | 2446 | 2649 | 2484 | 1945 | 1719 | 1671 | 1761 | 1473 | 1829 | 1670 | Ctns |
| Ctps | 6199 | 5827 | 5926 | 5363 | 5253 | 5413 | 6055 | 7262 | 6633 | 6309 | 6156 | 8160 | 6921 | 7799 | 6167 | 5434 | 5689 | 5429 | 5645 | 5957 | 5748 | 6221 | 6273 | 6896 | 6809 | 8774 | 8015 | 8124 | Ctps |
| Ctps2 | 4550 | 3778 | 3717 | 3335 | 3473 | 3932 | 3795 | 4763 | 4103 | 3993 | 3704 | 4031 | 3767 | 3924 | 4651 | 3826 | 3464 | 3586 | 3848 | 4315 | 3972 | 4350 | 4219 | 3697 | 3853 | 3842 | 4093 | 4033 | Ctps2 |
| Ctr9 | 17196 | 18216 | 14646 | 16150 | 14102 | 15257 | 15472 | 11228 | 10617 | 10189 | 9610 | 10277 | 9994 | 10338 | 17744 | 17231 | 15208 | 16214 | 14253 | 15786 | 15817 | 10709 | 10607 | 10127 | 10404 | 9808 | 10616 | 10790 | Ctr9 |
| Ctrb1 | 0 | 2 | 0 | 4 | 3 | 3 | 3 | 0 | 0 | 0 | 0 | 0 | 1 | 0 | 1 | 2 | 2 | 17 | 9 | 0 | 2 | 2 | 0 | 2 | 0 | 2 | 0 | 0 | Ctrb1 |
| Ctrl | 53 | 96 | 110 | 87 | 87 | 80 | 59 | 85 | 73 | 131 | 88 | 109 | 83 | 90 | 45 | 66 | 114 | 73 | 92 | 65 | 71 | 122 | 89 | 73 | 129 | 99 | 92 | 85 | Ctrl |
| Cts3 | 0 | 0 | 0 | 0 | 0 | 0 | 0 | 0 | 17 | 0 | 0 | 0 | 0 | 0 | 0 | 0 | 0 | 0 | 0 | 0 | 0 | 0 | 6 | 0 | 0 | 0 | 0 | 0 | Cts3 |
| Ctsa | 4339 | 5539 | 4266 | 4060 | 3834 | 4105 | 3550 | 3880 | 4996 | 4122 | 4666 | 4237 | 4528 | 4046 | 4300 | 5139 | 4120 | 4265 | 3925 | 3863 | 3567 | 4122 | 5249 | 4338 | 4075 | 4762 | 4183 | 4086 | Ctsa |
| Ctsb | 12647 | 14004 | 11783 | 11368 | 12093 | 12098 | 10827 | 11976 | 13701 | 10758 | 12356 | 9923 | 11069 | 9770 | 13115 | 14112 | 10653 | 12811 | 11381 | 12032 | 11735 | 11508 | 16364 | 10496 | 10455 | 11015 | 10578 | 10808 | Ctsb |
| Ctsc | 1338 | 1090 | 1254 | 752 | 852 | 1042 | 739 | 1153 | 1325 | 882 | 1330 | 1067 | 1292 | 1189 | 1365 | 1479 | 914 | 963 | 737 | 898 | 715 | 1253 | 1910 | 1160 | 973 | 1431 | 1180 | 1279 | Ctsc |
| Ctsd | 4272 | 5821 | 3876 | 4291 | 3929 | 3505 | 3951 | 3637 | 3893 | 3532 | 3828 | 3150 | 3400 | 3288 | 4498 | 4408 | 4031 | 4124 | 3770 | 3535 | 4346 | 3173 | 4526 | 3334 | 3265 | 3177 | 3013 | 3464 | Ctsd |
| Ctse | 10 | 6 | 1 | 2 | 9 | 1 | 1 | 2 | 5 | 13 | 7 | 5 | 25 | 17 | 4 | 3 | 5 | 3 | 2 | 10 | 4 | 7 | 5 | 2 | 7 | 2 | 6 | 23 | Ctse |
| Ctsf | 2237 | 2789 | 2332 | 2839 | 2768 | 2437 | 2246 | 1588 | 1661 | 1878 | 1914 | 1537 | 1695 | 1428 | 2529 | 2615 | 2662 | 2806 | 2675 | 2466 | 2616 | 1895 | 1783 | 1650 | 1688 | 1419 | 1651 | 1527 | Ctsf |
| Ctsg | 0 | 0 | 0 | 0 | 0 | 0 | 0 | 2 | 0 | 0 | 2 | 0 | 0 | 4 | 1 | 0 | 1 | 1 | 0 | 0 | 0 | 0 | 6 | 2 | 3 | 0 | 0 | 0 | Ctsg |
| Ctsh | 277 | 262 | 233 | 227 | 280 | 315 | 217 | 302 | 360 | 298 | 416 | 250 | 261 | 220 | 247 | 314 | 173 | 230 | 242 | 226 | 219 | 284 | 535 | 315 | 277 | 302 | 186 | 256 | Ctsh |
| Ctsj | 0 | 0 | 0 | 0 | 0 | 0 | 0 | 0 | 7 | 0 | 0 | 0 | 0 | 0 | 0 | 0 | 0 | 0 | 0 | 0 | 0 | 0 | 2 | 0 | 0 | 0 | 0 | 0 | Ctsj |
| Ctsk | 772 | 720 | 821 | 768 | 502 | 844 | 310 | 1038 | 1377 | 1056 | 1528 | 998 | 1471 | 1066 | 873 | 926 | 727 | 918 | 433 | 509 | 388 | 1133 | 1734 | 1133 | 1153 | 1011 | 1105 | 983 | Ctsk |
| Ctsl | 10381 | 10153 | 11042 | 8796 | 8024 | 8282 | 8755 | 5379 | 5876 | 4803 | 5282 | 4764 | 5659 | 5353 | 10196 | 10802 | 9640 | 9432 | 7827 | 9898 | 7771 | 4851 | 6444 | 4766 | 4212 | 5249 | 4890 | 5156 | Ctsl |
| Ctsm | 0 | 0 | 0 | 0 | 0 | 0 | 0 | 0 | 6 | 0 | 0 | 0 | 0 | 0 | 0 | 0 | 0 | 0 | 0 | 0 | 0 | 0 | 3 | 0 | 0 | 0 | 0 | 0 | Ctsm |
| Ctso | 359 | 308 | 320 | 247 | 347 | 340 | 343 | 607 | 643 | 567 | 616 | 567 | 706 | 540 | 405 | 319 | 261 | 255 | 264 | 400 | 290 | 580 | 731 | 496 | 477 | 685 | 581 | 523 | Ctso |
| Ctsq | 0 | 0 | 0 | 0 | 0 | 0 | 0 | 0 | 4 | 0 | 0 | 0 | 0 | 0 | 0 | 1 | 0 | 0 | 0 | 0 | 0 | 0 | 1 | 0 | 0 | 0 | 0 | 0 | Ctsq |
| Ctsr | 0 | 0 | 0 | 0 | 0 | 0 | 0 | 0 | 2 | 0 | 0 | 0 | 0 | 0 | 0 | 0 | 0 | 0 | 0 | 0 | 0 | 0 | 0 | 0 | 0 | 0 | 0 | 0 | Ctsr |
| Ctss | 404 | 376 | 363 | 318 | 390 | 360 | 364 | 438 | 489 | 389 | 581 | 390 | 394 | 382 | 417 | 455 | 276 | 375 | 293 | 437 | 377 | 500 | 613 | 360 | 370 | 472 | 361 | 383 | Ctss |
| Ctsw | 42 | 51 | 31 | 58 | 62 | 66 | 51 | 54 | 65 | 76 | 58 | 80 | 45 | 88 | 54 | 33 | 44 | 94 | 62 | 60 | 59 | 70 | 54 | 78 | 87 | 90 | 81 | 74 | Ctsw |
| Ctsz | 2826 | 3096 | 3161 | 2414 | 2167 | 2447 | 1867 | 2573 | 2932 | 2627 | 3098 | 2649 | 2744 | 2319 | 2635 | 3174 | 2892 | 2489 | 2410 | 1959 | 1983 | 2389 | 3378 | 2723 | 2108 | 2991 | 2716 | 2697 | Ctsz |
| Cttn | 14197 | 15774 | 17072 | 15231 | 13687 | 14374 | 12982 | 12995 | 14147 | 14298 | 13895 | 15032 | 14970 | 14319 | 14317 | 14882 | 15688 | 14477 | 14306 | 13477 | 12971 | 14282 | 14160 | 14904 | 14434 | 16050 | 15464 | 14747 | Cttn |
| Cttnbp2 | 2531 | 2357 | 1742 | 2682 | 3933 | 3803 | 4005 | 4311 | 4448 | 4472 | 4252 | 3076 | 3294 | 3473 | 2780 | 2354 | 2257 | 2777 | 3722 | 4121 | 4249 | 4147 | 4202 | 3599 | 5036 | 2101 | 3163 | 2649 | Cttnbp2 |
| Cttnbp2nl | 8334 | 8756 | 9058 | 10206 | 9924 | 9199 | 9741 | 9040 | 8198 | 9299 | 8323 | 9221 | 8599 | 9388 | 8690 | 9162 | 9689 | 9720 | 9481 | 8755 | 9542 | 9703 | 7887 | 9051 | 9743 | 8229 | 9017 | 9379 | Cttnbp2nl |
| Ctu1 | 1171 | 1274 | 1184 | 1321 | 1321 | 1269 | 1201 | 1076 | 1163 | 1226 | 1106 | 1297 | 1105 | 1075 | 1160 | 1041 | 1351 | 1183 | 1513 | 1303 | 1366 | 1298 | 1068 | 1253 | 1217 | 1340 | 1210 | 1190 | Ctu1 |
| Ctu2 | 1659 | 2090 | 1950 | 1893 | 1470 | 1569 | 1431 | 1031 | 1168 | 1185 | 1184 | 1366 | 1340 | 1202 | 1540 | 1731 | 1968 | 1802 | 1591 | 1468 | 1440 | 1008 | 1139 | 1365 | 1183 | 1514 | 1412 | 1327 | Ctu2 |

| | | | | | | | | | | | | | | | | | | | | | | | | | | | | | |
|---|---|---|---|---|---|---|---|---|---|---|---|---|---|---|---|---|---|---|---|---|---|---|---|---|---|---|---|---|---|
| Ctxn1 | 2757 | 3121 | 3039 | 3828 | 4300 | 3811 | 3817 | 4915 | 5119 | 6438 | 5462 | 6247 | 5181 | 6254 | 2724 | 2615 | 3869 | 3599 | 4493 | 3406 | 4077 | 6216 | 4764 | 6557 | 6795 | 6677 | 4781 | 5599 | Ctxn1 |
| Ctxn2 | 12 | 8 | 12 | 43 | 52 | 34 | 67 | 54 | 55 | 39 | 35 | 17 | 10 | 23 | 36 | 11 | 28 | 63 | 65 | 58 | 76 | 55 | 53 | 30 | 48 | 4 | 19 | 12 | Ctxn2 |
| Ctxn3 | 18 | 15 | 12 | 33 | 28 | 15 | 17 | 55 | 39 | 67 | 71 | 26 | 27 | 20 | 20 | 20 | 22 | 28 | 26 | 29 | 23 | 58 | 67 | 69 | 31 | 57 | 21 | 13 | Ctxn3 |
| Cubn | 194 | 219 | 163 | 106 | 128 | 155 | 88 | 258 | 248 | 201 | 177 | 224 | 242 | 219 | 281 | 171 | 146 | 110 | 141 | 230 | 131 | 207 | 249 | 154 | 148 | 148 | 270 | 215 | Cubn |
| Cuedc1 | 72 | 719 | 83 | 95 | 135 | 125 | 61 | 133 | 140 | 482 | 277 | 341 | 141 | 114 | 78 | 209 | 89 | 86 | 97 | 82 | 63 | 199 | 142 | 226 | 185 | 155 | 218 | 853 | Cuedc1 |
| Cuedc2 | 4193 | 5242 | 5012 | 5492 | 5222 | 4857 | 5287 | 4874 | 5504 | 5666 | 5327 | 5577 | 5316 | 6051 | 4390 | 4274 | 5183 | 4996 | 5678 | 5264 | 5537 | 5088 | 5491 | 5709 | 6084 | 6213 | 5693 | 5923 | Cuedc2 |
| Cul1 | 9796 | 8338 | 9033 | 7826 | 9115 | 9459 | 10673 | 17027 | 15139 | 13593 | 13551 | 16211 | 14521 | 16893 | 10048 | 8117 | 8771 | 7502 | 9442 | 10383 | 9262 | 14704 | 14490 | 14777 | 14783 | 17438 | 15932 | 16712 | Cul1 |
| Cul2 | 6104 | 4942 | 5336 | 4756 | 5065 | 5442 | 5858 | 6863 | 6027 | 5413 | 5294 | 5671 | 5754 | 6064 | 6295 | 5382 | 4926 | 4743 | 4878 | 5540 | 5645 | 6470 | 6261 | 5167 | 5676 | 5758 | 5758 | 6297 | Cul2 |
| Cul3 | 12144 | 8671 | 12787 | 12292 | 13576 | 13634 | 15345 | 15226 | 13017 | 14648 | 13727 | 16157 | 14804 | 16355 | 12233 | 10248 | 12369 | 12236 | 13356 | 12104 | 14271 | 16345 | 13185 | 14743 | 15260 | 16044 | 15669 | 16778 | Cul3 |
| Cul4a | 9574 | 9022 | 9235 | 8992 | 9136 | 9194 | 9182 | 8250 | 8175 | 7756 | 8604 | 8732 | 8532 | 8457 | 9126 | 8591 | 9084 | 8605 | 9345 | 8784 | 9206 | 7818 | 8327 | 8001 | 8044 | 8875 | 9028 | 8555 | Cul4a |
| Cul4b | 6615 | 4632 | 4824 | 4651 | 5079 | 5326 | 5551 | 7963 | 7179 | 6294 | 6978 | 6976 | 7634 | 7364 | 5980 | 5403 | 4578 | 4412 | 4885 | 5566 | 5259 | 6465 | 7353 | 6365 | 6507 | 7046 | 7411 | 7354 | Cul4b |
| Cul5 | 4320 | 3277 | 4221 | 3957 | 5240 | 4663 | 5177 | 7365 | 6470 | 6566 | 6515 | 7444 | 6857 | 7755 | 4361 | 3612 | 4234 | 4084 | 4577 | 4526 | 5108 | 7175 | 6311 | 6683 | 6953 | 6267 | 7342 | 7711 | Cul5 |
| Cul7 | 6978 | 7988 | 9119 | 7873 | 7415 | 7308 | 5954 | 9349 | 12803 | 12710 | 14032 | 13762 | 13888 | 12240 | 6220 | 7979 | 8440 | 7679 | 8152 | 5951 | 6375 | 11964 | 12872 | 13674 | 13132 | 13738 | 13179 | 13127 | Cul7 |
| Cul9 | 2276 | 3221 | 3095 | 4367 | 4639 | 4104 | 3571 | 2490 | 3197 | 4069 | 3695 | 3154 | 2971 | 2817 | 2409 | 2982 | 3957 | 4329 | 4401 | 3352 | 3832 | 4187 | 3050 | 3434 | 4039 | 1937 | 2782 | 2427 | Cul9 |
| Cuta | 2654 | 3572 | 2872 | 3387 | 2528 | 2439 | 3235 | 1331 | 1453 | 1567 | 1514 | 1697 | 1538 | 1905 | 2778 | 2742 | 3240 | 3009 | 2978 | 2670 | 3230 | 1287 | 1480 | 1580 | 1464 | 1889 | 1796 | 2123 | Cuta |
| Cutal | 6 | 4 | 9 | 3 | 4 | 7 | 1 | 10 | 13 | 5 | 8 | 13 | 8 | 6 | 15 | 6 | 5 | 3 | 9 | 12 | 3 | 15 | 3 | 16 | 20 | 8 | 13 | 11 | Cutal |
| Cutc | 200 | 206 | 248 | 222 | 214 | 166 | 188 | 399 | 363 | 366 | 349 | 426 | 473 | 466 | 286 | 213 | 287 | 195 | 184 | 193 | 162 | 289 | 350 | 411 | 298 | 560 | 504 | 438 | Cutc |
| Cux1 | 9839 | 11162 | 12096 | 12369 | 12051 | 11863 | 11980 | 11323 | 12092 | 13707 | 12005 | 16839 | 13217 | 16314 | 9526 | 9285 | 12811 | 11869 | 13087 | 10492 | 12142 | 12933 | 10390 | 14913 | 15669 | 15344 | 15616 | 16067 | Cux1 |
| Cux2 | 2258 | 3072 | 4253 | 4636 | 5998 | 4749 | 4292 | 3160 | 3676 | 5310 | 4093 | 4208 | 4701 | 3864 | 2731 | 2646 | 4741 | 4738 | 5375 | 3873 | 4912 | 5528 | 3225 | 4458 | 5360 | 3255 | 3685 | 3604 | Cux2 |
| Cuzd1 | 5 | 4 | 2 | 13 | 7 | 11 | 8 | 0 | 7 | 4 | 1 | 1 | 3 | 0 | 7 | 9 | 7 | 6 | 13 | 5 | 14 | 5 | 3 | 1 | 1 | 7 | 6 | 2 | Cuzd1 |
| Cwc15 | 5582 | 4964 | 5627 | 5241 | 4413 | 4289 | 5058 | 5840 | 4813 | 5300 | 4922 | 5712 | 5045 | 6606 | 5691 | 4623 | 5276 | 4737 | 4715 | 5007 | 4768 | 5230 | 4911 | 5833 | 5393 | 6263 | 5867 | 6359 | Cwc15 |
| Cwc22 | 2491 | 2103 | 2523 | 2067 | 2198 | 2265 | 2321 | 3777 | 3442 | 3221 | 3390 | 3998 | 3782 | 4179 | 2525 | 2080 | 2293 | 1888 | 2249 | 2299 | 2105 | 3220 | 3557 | 3510 | 3474 | 4531 | 4118 | 3966 | Cwc22 |
| Cwc25 | 2891 | 2790 | 2983 | 2847 | 2338 | 2499 | 2537 | 2188 | 2116 | 2318 | 2220 | 2383 | 2321 | 2434 | 2528 | 2781 | 2789 | 2659 | 2411 | 2404 | 2611 | 2052 | 2153 | 2456 | 2489 | 2581 | 2352 | 2457 | Cwc25 |
| Cwc27 | 1253 | 1140 | 1390 | 1355 | 1034 | 1153 | 1193 | 1838 | 1732 | 1710 | 1585 | 1946 | 1878 | 2262 | 1328 | 1395 | 1189 | 1121 | 1128 | 1449 | 1103 | 1469 | 1714 | 1767 | 1679 | 2517 | 2113 | 2377 | Cwc27 |
| Cwf19l1 | 2068 | 2098 | 1987 | 1734 | 1889 | 1782 | 2118 | 2369 | 2251 | 2153 | 1967 | 2331 | 2201 | 2449 | 2173 | 1724 | 2179 | 1818 | 1969 | 2105 | 1994 | 2274 | 2045 | 2256 | 2147 | 2110 | 2195 | 2270 | Cwf19l1 |
| Cwf19l2 | 2703 | 1748 | 2475 | 2073 | 2347 | 2339 | 2692 | 3306 | 2871 | 2421 | 2740 | 2994 | 3070 | 3272 | 2843 | 2313 | 2223 | 2027 | 2106 | 2616 | 2319 | 2607 | 2996 | 2935 | 2585 | 2961 | 3244 | 3235 | Cwf19l2 |
| Cwh43 | 1 | 4 | 2 | 3 | 6 | 0 | 9 | 10 | 7 | 15 | 5 | 1 | 5 | 1 | 5 | 1 | 1 | 7 | 0 | 0 | 4 | 8 | 4 | 2 | 1 | 8 | 8 | 4 | Cwh43 |
| Cx3cl1 | 690 | 880 | 1014 | 871 | 950 | 878 | 920 | 619 | 809 | 764 | 732 | 727 | 780 | 697 | 760 | 840 | 926 | 817 | 933 | 822 | 855 | 1041 | 812 | 811 | 758 | 702 | 696 | 738 | Cx3cl1 |
| Cx3cr1 | 401 | 374 | 534 | 350 | 590 | 483 | 424 | 807 | 887 | 868 | 1134 | 973 | 968 | 848 | 477 | 435 | 385 | 405 | 448 | 396 | 381 | 983 | 1314 | 854 | 798 | 947 | 873 | 892 | Cx3cr1 |
| Cxadr | 25256 | 22051 | 21278 | 25650 | 29139 | 29939 | 32020 | 29687 | 23633 | 22479 | 20077 | 23122 | 22167 | 24236 | 27948 | 23158 | 23854 | 27075 | 28545 | 33951 | 31968 | 26832 | 24390 | 22899 | 27096 | 17770 | 23812 | 23943 | Cxadr |
| Cxcl1 | 0 | 0 | 0 | 0 | 0 | 12 | 1 | 4 | 3 | 3 | 3 | 1 | 3 | 2 | 4 | 2 | 2 | 1 | 0 | 7 | 1 | 0 | 6 | 3 | 1 | 11 | 2 | 0 | Cxcl1 |
| Cxcl10 | 7 | 5 | 7 | 2 | 6 | 9 | 4 | 6 | 10 | 8 | 9 | 4 | 12 | 7 | 11 | 8 | 6 | 0 | 3 | 7 | 4 | 5 | 12 | 7 | 6 | 5 | 2 | 5 | Cxcl10 |
| Cxcl12 | 7005 | 6390 | 7050 | 4872 | 4214 | 4503 | 3822 | 8598 | 11139 | 8634 | 13369 | 8868 | 11219 | 8925 | 6355 | 9048 | 4928 | 5373 | 3641 | 3968 | 3965 | 7479 | 12565 | 9769 | 7142 | 11061 | 8639 | 9277 | Cxcl12 |
| Cxcl13 | 3 | 6 | 4 | 4 | 6 | 0 | 0 | 2 | 11 | 0 | 98 | 3 | 8 | 7 | 3 | 10 | 1 | 2 | 0 | 0 | 0 | 8 | 77 | 7 | 0 | 11 | 6 | 1 | Cxcl13 |
| Cxcl14 | 1542 | 952 | 781 | 631 | 227 | 1737 | 175 | 3731 | 6185 | 3153 | 3045 | 1945 | 3117 | 2040 | 1274 | 829 | 677 | 1399 | 628 | 295 | 252 | 1058 | 6656 | 3773 | 1525 | 1228 | 1779 | 1705 | Cxcl14 |
| Cxcl15 | 0 | 0 | 0 | 0 | 0 | 0 | 0 | 4 | 1 | 11 | 10 | 0 | 0 | 0 | 0 | 0 | 0 | 0 | 0 | 0 | 0 | 0 | 5 | 1 | 0 | 0 | 0 | 0 | Cxcl15 |
| Cxcl16 | 105 | 84 | 107 | 70 | 96 | 92 | 54 | 116 | 155 | 139 | 241 | 84 | 120 | 101 | 69 | 102 | 85 | 87 | 79 | 82 | 69 | 142 | 245 | 115 | 99 | 102 | 92 | 75 | Cxcl16 |
| Cxcl17 | 0 | 3 | 0 | 0 | 0 | 3 | 2 | 0 | 2 | 0 | 1 | 0 | 1 | 0 | 0 | 1 | 0 | 3 | 2 | 1 | 1 | 0 | 1 | 1 | 0 | 0 | 0 | 0 | Cxcl17 |
| Cxcl2 | 2 | 0 | 2 | 2 | 0 | 4 | 2 | 0 | 3 | 3 | 0 | 0 | 1 | 2 | 0 | 0 | 2 | 0 | 0 | 0 | 0 | 0 | 3 | 1 | 0 | 0 | 0 | 0 | Cxcl2 |
| Cxcl3 | 0 | 0 | 0 | 0 | 0 | 0 | 0 | 2 | 0 | 6 | 7 | 0 | 0 | 0 | 0 | 0 | 0 | 0 | 0 | 0 | 0 | 2 | 2 | 1 | 0 | 0 | 0 | 0 | Cxcl3 |
| Cxcl5 | 7 | 3 | 11 | 0 | 1 | 0 | 1 | 18 | 25 | 25 | 23 | 4 | 7 | 3 | 4 | 2 | 2 | 1 | 7 | 5 | 1 | 5 | 11 | 16 | 5 | 8 | 0 | 0 | Cxcl5 |
| Cxcl9 | 0 | 0 | 0 | 0 | 0 | 0 | 0 | 0 | 1 | 0 | 0 | 0 | 3 | 0 | 1 | 0 | 0 | 0 | 0 | 0 | 0 | 0 | 5 | 2 | 0 | 0 | 0 | 0 | Cxcl9 |
| Cxcr2 | 8 | 7 | 17 | 11 | 11 | 13 | 7 | 6 | 6 | 10 | 15 | 1 | 12 | 6 | 11 | 9 | 10 | 17 | 6 | 6 | 8 | 17 | 5 | 6 | 7 | 9 | 5 | 4 | Cxcr2 |
| Cxcr3 | 1 | 0 | 0 | 7 | 5 | 0 | 1 | 6 | 12 | 3 | 12 | 2 | 8 | 6 | 0 | 2 | 3 | 0 | 1 | 3 | 0 | 2 | 12 | 3 | 1 | 6 | 8 | 7 | Cxcr3 |
| Cxcr4 | 1749 | 2312 | 2673 | 1999 | 1658 | 1582 | 1422 | 2533 | 1884 | 1848 | 1684 | 1981 | 2797 | 2164 | 2255 | 2188 | 2299 | 1763 | 1711 | 1749 | 1644 | 2298 | 2143 | 1991 | 1769 | 2042 | 2654 | 2639 | Cxcr4 |
| Cxcr5 | 0 | 1 | 2 | 0 | 1 | 0 | 1 | 1 | 2 | 3 | 7 | 2 | 3 | 0 | 3 | 0 | 0 | 1 | 3 | 1 | 1 | 0 | 3 | 2 | 4 | 2 | 0 | 2 | Cxcr5 |
| Cxcr6 | 0 | 0 | 0 | 0 | 0 | 0 | 0 | 0 | 0 | 0 | 0 | 0 | 0 | 0 | 0 | 0 | 0 | 0 | 0 | 0 | 0 | 0 | 1 | 0 | 0 | 0 | 0 | 0 | Cxcr6 |
| Cxxc1 | 4439 | 5807 | 5689 | 6076 | 5225 | 4868 | 5213 | 4586 | 5225 | 5787 | 5000 | 6107 | 5340 | 5720 | 4386 | 4984 | 5909 | 5362 | 5923 | 5018 | 5288 | 4849 | 4806 | 6239 | 5437 | 6839 | 5632 | 5990 | Cxxc1 |
| Cxxc4 | 5728 | 4852 | 7232 | 8243 | 8363 | 8010 | 8656 | 9082 | 6464 | 8836 | 7626 | 10218 | 8641 | 10936 | 6336 | 4918 | 7927 | 7591 | 8496 | 7009 | 9002 | 11710 | 6277 | 9464 | 9965 | 8239 | 10909 | 11140 | Cxxc4 |
| Cxxc5 | 3519 | 3904 | 4998 | 5061 | 5834 | 4893 | 4955 | 3231 | 3203 | 4307 | 3758 | 4619 | 4058 | 4486 | 3505 | 3829 | 5312 | 5182 | 5940 | 4226 | 5453 | 4380 | 3500 | 4217 | 4421 | 4564 | 3830 | 4543 | Cxxc5 |
| Cyb561 | 1648 | 2058 | 1785 | 2098 | 2506 | 2082 | 2039 | 1230 | 1362 | 1395 | 1143 | 1046 | 1027 | 819 | 1912 | 2037 | 1962 | 2073 | 2475 | 2235 | 2083 | 1395 | 1493 | 1272 | 1069 | 1009 | 829 | 797 | Cyb561 |
| Cyb561a3 | 983 | 1091 | 1002 | 878 | 1024 | 1014 | 889 | 1106 | 1201 | 1104 | 1099 | 1082 | 1088 | 981 | 886 | 1030 | 925 | 899 | 985 | 995 | 905 | 1061 | 1223 | 1146 | 1024 | 1207 | 1088 | 1043 | Cyb561a3 |
| Cyb561d1 | 660 | 589 | 765 | 647 | 848 | 802 | 780 | 1263 | 1084 | 1290 | 1269 | 1484 | 1424 | 1517 | 655 | 518 | 782 | 633 | 804 | 715 | 759 | 1586 | 1014 | 1483 | 1580 | 1235 | 1536 | 1667 | Cyb561d1 |
| Cyb561d2 | 439 | 591 | 340 | 466 | 524 | 362 | 436 | 526 | 508 | 445 | 448 | 482 | 484 | 405 | 436 | 496 | 437 | 517 | 443 | 518 | 446 | 496 | 572 | 471 | 524 | 450 | 457 | 450 | Cyb561d2 |
| Cyb5a | 5642 | 5101 | 4696 | 4530 | 3910 | 3890 | 4237 | 3837 | 3985 | 3532 | 3341 | 3355 | 3975 | 3851 | 5451 | 4914 | 4515 | 3976 | 3810 | 4633 | 4077 | 3158 | 4045 | 3639 | 3150 | 4033 | 3438 | 4024 | Cyb5a |
| Cyb5b | 17950 | 17699 | 18863 | 16155 | 16143 | 16043 | 17454 | 14823 | 13158 | 13600 | 12615 | 16339 | 15190 | 16368 | 18306 | 17001 | 17588 | 15366 | 16323 | 17044 | 16467 | 13841 | 12870 | 14103 | 14264 | 16967 | 16940 | 17148 | Cyb5b |
| Cyb5d1 | 2049 | 2009 | 2395 | 1864 | 2009 | 2186 | 2038 | 1950 | 1989 | 1914 | 1868 | 2425 | 2398 | 2351 | 1917 | 1944 | 2019 | 2002 | 2071 | 2087 | 1935 | 1746 | 1929 | 2142 | 2227 | 2341 | 2666 | 2503 | Cyb5d1 |
| Cyb5d2 | 182 | 180 | 127 | 160 | 198 | 202 | 173 | 459 | 391 | 432 | 385 | 437 | 387 | 392 | 185 | 144 | 232 | 161 | 155 | 172 | 220 | 411 | 437 | 351 | 333 | 348 | 363 | 396 | Cyb5d2 |
| Cyb5r1 | 3310 | 3338 | 3220 | 2834 | 2432 | 2535 | 2596 | 1097 | 1344 | 1207 | 1336 | 1458 | 1503 | 1304 | 2875 | 3657 | 3133 | 2927 | 2528 | 2297 | 2460 | 1247 | 1346 | 1396 | 1256 | 1629 | 1399 | 1321 | Cyb5r1 |
| Cyb5r2 | 5 | 11 | 15 | 0 | 2 | 5 | 7 | 16 | 12 | 9 | 7 | 5 | 22 | 27 | 5 | 5 | 7 | 1 | 5 | 2 | 6 | 3 | 13 | 7 | 5 | 21 | 29 | 19 | Cyb5r2 |
| Cyb5r3 | 9223 | 11127 | 10931 | 9344 | 7967 | 8222 | 7758 | 6157 | 6993 | 6382 | 5841 | 7917 | 8040 | 7609 | 9312 | 11159 | 10019 | 8338 | 8511 | 9036 | 7511 | 6442 | 7176 | 7035 | 6681 | 9190 | 8383 | 8558 | Cyb5r3 |
| Cyb5r4 | 6938 | 6423 | 6948 | 6264 | 5616 | 5303 | 6534 | 6383 | 5107 | 5164 | 4278 | 6173 | 5328 | 6439 | 7787 | 6900 | 6733 | 5731 | 5827 | 6694 | 5947 | 5238 | 4905 | 5503 | 5050 | 6450 | 6224 | 6236 | Cyb5r4 |
| Cyb5rl | 1108 | 1386 | 1158 | 1131 | 843 | 927 | 813 | 350 | 415 | 376 | 475 | 510 | 516 | 419 | 938 | 1203 | 1179 | 958 | 786 | 760 | 756 | 369 | 459 | 398 | 398 | 479 | 458 | 462 | Cyb5rl |
| Cyba | 661 | 677 | 821 | 555 | 448 | 490 | 304 | 526 | 691 | 557 | 820 | 571 | 708 | 534 | 513 | 674 | 594 | 537 | 489 | 425 | 362 | 443 | 829 | 623 | 513 | 774 | 726 | 619 | Cyba |
| Cybb | 93 | 74 | 98 | 77 | 57 | 102 | 36 | 115 | 135 | 100 | 238 | 127 | 81 | 106 | 69 | 118 | 49 | 70 | 32 | 66 | 55 | 135 | 195 | 125 | 97 | 139 | 99 | 76 | Cybb |
| Cybc1 | 4188 | 4789 | 3781 | 4048 | 3965 | 4128 | 4088 | 3556 | 3462 | 2965 | 2901 | 3222 | 3178 | 3317 | 4133 | 4206 | 4151 | 4163 | 4168 | 4275 | 4191 | 3245 | 3342 | 3220 | 3312 | 3075 | 3276 | 3237 | Cybc1 |
| Cybrd1 | 600 | 438 | 398 | 420 | 682 | 558 | 330 | 811 | 1093 | 862 | 1024 | 245 | 420 | 291 | 460 | 498 | 371 | 639 | 368 | 301 | 385 | 912 | 1177 | 486 | 511 | 323 | 300 | 212 | Cybrd1 |
| Cyc1 | 7168 | 8488 | 8149 | 8374 | 7463 | 7267 | 7679 | 6224 | 6435 | 7029 | 6343 | 7137 | 6936 | 7009 | 6953 | 7134 | 8617 | 7726 | 7989 | 6845 | 7249 | 6031 | 6410 | 7481 | 6805 | 8413 | 7646 | 7107 | Cyc1 |
| Cycs | 5360 | 5163 | 5594 | 5263 | 5170 | 5246 | 6576 | 4714 | 4489 | 4370 | 4426 | 4781 | 4992 | 5580 | 5832 | 5537 | 5449 | 5166 | 5263 | 6485 | 5918 | 3974 | 4504 | 4799 | 4697 | 5749 | 5462 | 5890 | Cycs |
| Cyct | 1 | 0 | 2 | 0 | 2 | 5 | 7 | 1 | 4 | 0 | 9 | 1 | 11 | 6 | 5 | 2 | 0 | 1 | 0 | 4 | 0 | 0 | 1 | 2 | 2 | 0 | 5 | 0 | Cyct |
| Cyfip1 | 7109 | 6173 | 5582 | 4492 | 4678 | 5087 | 4492 | 7770 | 8608 | 6924 | 7501 | 7901 | 8210 | 7798 | 6204 | 6703 | 5072 | 4830 | 4677 | 4554 | 4579 | 6291 | 8550 | 7457 | 6924 | 8068 | 8008 | 7913 | Cyfip1 |
| Cyfip2 | 13643 | 14036 | 13216 | 18653 | 21935 | 18201 | 20315 | 10995 | 11107 | 14806 | 11620 | 13659 | 11954 | 12951 | 14586 | 13481 | 17399 | 18817 | 21442 | 14305 | 20862 | 17286 | 9490 | 13309 | 15586 | 9543 | 10422 | 11676 | Cyfip2 |
| Cygb | 374 | 431 | 434 | 496 | 822 | 646 | 507 | 436 | 641 | 586 | 637 | 329 | 380 | 331 | 361 | 555 | 413 | 635 | 717 | 557 | 617 | 643 | 625 | 532 | 611 | 352 | 279 | 288 | Cygb |
| Cyhr1 | 9020 | 11162 | 9631 | 10356 | 9328 | 9479 | 9366 | 7983 | 8326 | 8558 | 8083 | 9006 | 7969 | 8283 | 8483 | 9727 | 9910 | 10516 | 10342 | 9321 | 9634 | 8383 | 7814 | 8742 | 8651 | 8373 | 8112 | 8230 | Cyhr1 |
| Cyld | 6315 | 5636 | 5044 | 6418 | 6187 | 6211 | 6794 | 3640 | 3158 | 2863 | 3162 | 2849 | 2946 | 2932 | 6393 | 6295 | 5460 | 6489 | 5910 | 6362 | 6713 | 3439 | 3100 | 2835 | 3182 | 2460 | 2944 | 2701 | Cyld |
| Cym | 35 | 23 | 12 | 57 | 44 | 60 | 36 | 16 | 8 | 11 | 3 | 18 | 14 | 10 | 22 | 33 | 30 | 33 | 42 | 26 | 42 | 17 | 5 | 7 | 4 | 2 | 17 | 4 | Cym |
| Cyp11a1 | 59 | 77 | 47 | 166 | 137 | 79 | 123 | 12 | 27 | 23 | 31 | 14 | 17 | 15 | 58 | 41 | 101 | 120 | 112 | 99 | 106 | 30 | 31 | 10 | 25 | 8 | 16 | 15 | Cyp11a1 |
| Cyp17a1 | 0 | 0 | 0 | 0 | 0 | 0 | 3 | 2 | 1 | 2 | 0 | 2 | 1 | 4 | 3 | 2 | 0 | 0 | 1 | 0 | 3 | 2 | 1 | 2 | 0 | 4 | 0 | 2 | Cyp17a1 |
| Cyp19a1 | 2 | 3 | 2 | 12 | 5 | 0 | 4 | 4 | 2 | 9 | 0 | 10 | 0 | 1 | 1 | 0 | 7 | 9 | 4 | 0 | 6 | 2 | 1 | 4 | 12 | 9 | 8 | 1 | Cyp19a1 |

| | | | | | | | | | | | | | | | | | | | | | | | | | | | | | |
|---|---|---|---|---|---|---|---|---|---|---|---|---|---|---|---|---|---|---|---|---|---|---|---|---|---|---|---|---|---|
| Cyp1a1 | 1 | 2 | 14 | 4 | 11 | 6 | 3 | 1 | 8 | 4 | 5 | 3 | 1 | 2 | 7 | 4 | 5 | 18 | 4 | 16 | 21 | 5 | 1 | 1 | 6 | 0 | 0 | 7 | Cyp1a1 |
| Cyp1a2 | 0 | 1 | 0 | 5 | 0 | 0 | 0 | 0 | 0 | 2 | 0 | 2 | 0 | 0 | 0 | 0 | 0 | 0 | 3 | 1 | 0 | 0 | 0 | 0 | 0 | 0 | 0 | 0 | Cyp1a2 |
| Cyp1b1 | 767 | 485 | 589 | 565 | 1002 | 830 | 739 | 1209 | 1126 | 1204 | 1784 | 931 | 1202 | 905 | 835 | 813 | 502 | 721 | 775 | 861 | 878 | 1387 | 1881 | 1078 | 1272 | 1071 | 953 | 815 | Cyp1b1 |
| Cyp20a1 | 741 | 636 | 863 | 576 | 691 | 745 | 629 | 1231 | 1300 | 1275 | 1398 | 1440 | 1495 | 1427 | 873 | 793 | 767 | 702 | 684 | 731 | 651 | 1280 | 1458 | 1233 | 1165 | 1712 | 1577 | 1542 | Cyp20a1 |
| Cyp21a1 | 2 | 10 | 4 | 6 | 10 | 4 | 8 | 1 | 0 | 1 | 1 | 1 | 1 | 5 | 1 | 6 | 10 | 7 | 8 | 4 | 4 | 3 | 3 | 5 | 5 | 0 | 0 | 2 | Cyp21a1 |
| Cyp24a1 | 3 | 9 | 0 | 9 | 7 | 7 | 1 | 17 | 9 | 0 | 9 | 9 | 6 | 2 | 8 | 5 | 1 | 3 | 11 | 5 | 12 | 5 | 6 | 16 | 15 | 6 | 1 | 15 | Cyp24a1 |
| Cyp26a1 | 72 | 37 | 16 | 15 | 7 | 13 | 2 | 49 | 124 | 27 | 170 | 23 | 87 | 20 | 70 | 63 | 19 | 11 | 12 | 2 | 2 | 63 | 102 | 34 | 26 | 22 | 6 | 27 | Cyp26a1 |
| Cyp26b1 | 1746 | 2744 | 1234 | 3880 | 5471 | 3872 | 371 | 6687 | 7449 | 6937 | 4760 | 1487 | 9719 | 1187 | 3786 | 4208 | 3962 | 3430 | 2377 | 2991 | 1345 | 7650 | 6064 | 2181 | 4032 | 1659 | 1234 | 1270 | Cyp26b1 |
| Cyp26c1 | 32 | 41 | 2 | 2 | 0 | 12 | 0 | 20 | 37 | 12 | 35 | 0 | 29 | 2 | 34 | 30 | 7 | 6 | 0 | 0 | 0 | 12 | 30 | 1 | 3 | 1 | 3 | 2 | Cyp26c1 |
| Cyp27a1 | 375 | 267 | 351 | 213 | 151 | 180 | 124 | 158 | 190 | 173 | 158 | 137 | 227 | 107 | 287 | 419 | 222 | 188 | 156 | 159 | 102 | 112 | 255 | 176 | 100 | 205 | 189 | 132 | Cyp27a1 |
| Cyp27b1 | 61 | 76 | 85 | 87 | 94 | 112 | 135 | 31 | 49 | 32 | 42 | 54 | 35 | 36 | 76 | 82 | 104 | 114 | 111 | 83 | 113 | 28 | 45 | 37 | 37 | 50 | 16 | 28 | Cyp27b1 |
| Cyp2a5 | 0 | 1 | 0 | 1 | 7 | 6 | 0 | 13 | 16 | 23 | 17 | 4 | 16 | 4 | 0 | 1 | 0 | 0 | 2 | 1 | 1 | 12 | 14 | 11 | 5 | 4 | 7 | 1 | Cyp2a5 |
| Cyp2ab1 | 7 | 10 | 7 | 11 | 6 | 13 | 15 | 2 | 2 | 0 | 1 | 2 | 0 | 0 | 5 | 5 | 7 | 21 | 12 | 8 | 17 | 0 | 6 | 2 | 4 | 0 | 1 | 2 | Cyp2ab1 |
| Cyp2b10 | 0 | 1 | 0 | 0 | 0 | 0 | 0 | 0 | 2 | 0 | 0 | 0 | 0 | 0 | 0 | 0 | 0 | 0 | 0 | 0 | 0 | 2 | 0 | 0 | 0 | 0 | 0 | 0 | Cyp2b10 |
| Cyp2b23 | 0 | 0 | 0 | 0 | 5 | 0 | 0 | 2 | 2 | 0 | 0 | 0 | 1 | 0 | 0 | 1 | 0 | 0 | 0 | 0 | 0 | 2 | 0 | 1 | 2 | 0 | 0 | 0 | Cyp2b23 |
| Cyp2c23 | 1 | 1 | 0 | 4 | 0 | 2 | 1 | 0 | 5 | 0 | 1 | 2 | 3 | 1 | 3 | 2 | 0 | 0 | 3 | 0 | 5 | 3 | 5 | 2 | 1 | 0 | 1 | 1 | Cyp2c23 |
| Cyp2c29 | 0 | 0 | 0 | 0 | 0 | 0 | 0 | 0 | 0 | 0 | 1 | 0 | 0 | 0 | 0 | 0 | 0 | 0 | 0 | 0 | 0 | 0 | 0 | 0 | 0 | 0 | 0 | 0 | Cyp2c29 |
| Cyp2c40 | 0 | 0 | 0 | 0 | 0 | 0 | 0 | 0 | 0 | 1 | 0 | 0 | 0 | 0 | 0 | 0 | 0 | 0 | 0 | 0 | 0 | 0 | 0 | 0 | 0 | 0 | 0 | 0 | Cyp2c40 |
| Cyp2c55 | 21 | 33 | 39 | 46 | 41 | 26 | 40 | 6 | 4 | 13 | 10 | 3 | 8 | 3 | 16 | 28 | 53 | 54 | 33 | 21 | 27 | 10 | 3 | 7 | 10 | 0 | 9 | 5 | Cyp2c55 |
| Cyp2c65 | 2 | 7 | 1 | 20 | 7 | 5 | 9 | 1 | 2 | 10 | 4 | 0 | 3 | 0 | 5 | 16 | 6 | 29 | 5 | 15 | 22 | 7 | 1 | 6 | 0 | 2 | 2 | 2 | Cyp2c65 |
| Cyp2c68 | 0 | 0 | 0 | 0 | 0 | 0 | 0 | 0 | 0 | 0 | 0 | 0 | 0 | 0 | 0 | 0 | 0 | 0 | 0 | 0 | 0 | 0 | 1 | 0 | 0 | 0 | 0 | 0 | Cyp2c68 |
| Cyp2c69 | 0 | 0 | 0 | 0 | 0 | 0 | 0 | 0 | 0 | 0 | 0 | 0 | 0 | 0 | 0 | 0 | 0 | 0 | 0 | 0 | 0 | 0 | 1 | 0 | 0 | 0 | 0 | 0 | Cyp2c69 |
| Cyp2c70 | 0 | 0 | 0 | 0 | 0 | 0 | 0 | 0 | 0 | 0 | 0 | 2 | 0 | 0 | 0 | 0 | 0 | 0 | 0 | 0 | 0 | 0 | 1 | 0 | 0 | 1 | 0 | 1 | Cyp2c70 |
| Cyp2d10 | 0 | 0 | 0 | 0 | 0 | 0 | 0 | 0 | 0 | 0 | 0 | 0 | 0 | 1 | 0 | 0 | 0 | 0 | 0 | 0 | 0 | 0 | 0 | 0 | 0 | 0 | 0 | 0 | Cyp2d10 |
| Cyp2d11 | 0 | 0 | 0 | 0 | 0 | 0 | 0 | 0 | 0 | 0 | 0 | 0 | 0 | 0 | 0 | 0 | 0 | 0 | 0 | 0 | 0 | 0 | 0 | 0 | 0 | 1 | 0 | 0 | Cyp2d11 |
| Cyp2d12 | 0 | 0 | 0 | 0 | 0 | 0 | 0 | 0 | 0 | 0 | 0 | 0 | 1 | 0 | 0 | 0 | 0 | 0 | 0 | 0 | 0 | 0 | 0 | 0 | 0 | 0 | 0 | 0 | Cyp2d12 |
| Cyp2d22 | 496 | 531 | 628 | 514 | 457 | 531 | 459 | 347 | 441 | 434 | 586 | 371 | 443 | 330 | 458 | 590 | 572 | 545 | 498 | 361 | 461 | 317 | 528 | 377 | 309 | 446 | 289 | 280 | Cyp2d22 |
| Cyp2d26 | 24 | 34 | 32 | 34 | 26 | 25 | 30 | 9 | 16 | 25 | 6 | 20 | 28 | 10 | 11 | 20 | 28 | 46 | 39 | 31 | 32 | 15 | 12 | 7 | 17 | 13 | 6 | 6 | Cyp2d26 |
| Cyp2d40 | 8 | 11 | 6 | 9 | 11 | 6 | 15 | 0 | 2 | 16 | 1 | 3 | 3 | 3 | 8 | 6 | 10 | 26 | 12 | 27 | 21 | 3 | 3 | 4 | 9 | 3 | 0 | 0 | Cyp2d40 |
| Cyp2d9 | 1 | 1 | 1 | 6 | 1 | 0 | 0 | 0 | 0 | 0 | 4 | 0 | 0 | 0 | 5 | 0 | 3 | 3 | 0 | 0 | 0 | 0 | 0 | 0 | 0 | 0 | 3 | 0 | Cyp2d9 |
| Cyp2e1 | 0 | 0 | 0 | 0 | 0 | 0 | 0 | 0 | 0 | 0 | 0 | 0 | 1 | 0 | 0 | 0 | 0 | 0 | 0 | 0 | 0 | 0 | 0 | 1 | 0 | 0 | 0 | 0 | Cyp2e1 |
| Cyp2f2 | 0 | 0 | 0 | 0 | 0 | 0 | 0 | 17 | 44 | 39 | 12 | 0 | 0 | 0 | 0 | 0 | 0 | 0 | 0 | 0 | 0 | 3 | 83 | 18 | 0 | 0 | 0 | 0 | Cyp2f2 |
| Cyp2g1 | 0 | 0 | 0 | 4 | 0 | 0 | 0 | 82 | 51 | 82 | 50 | 0 | 7 | 4 | 0 | 0 | 0 | 0 | 0 | 0 | 0 | 0 | 104 | 113 | 0 | 19 | 0 | 0 | Cyp2g1 |
| Cyp2j11 | 0 | 0 | 0 | 0 | 0 | 0 | 0 | 0 | 0 | 2 | 0 | 0 | 0 | 0 | 0 | 0 | 0 | 0 | 0 | 0 | 0 | 2 | 0 | 0 | 0 | 0 | 0 | 0 | Cyp2j11 |
| Cyp2j12 | 0 | 0 | 0 | 0 | 0 | 0 | 0 | 2 | 1 | 0 | 0 | 0 | 0 | 0 | 0 | 0 | 0 | 0 | 0 | 0 | 0 | 0 | 0 | 0 | 0 | 0 | 0 | 0 | Cyp2j12 |
| Cyp2j13 | 2 | 1 | 3 | 4 | 7 | 1 | 1 | 11 | 0 | 9 | 4 | 9 | 9 | 3 | 1 | 3 | 1 | 0 | 3 | 2 | 4 | 0 | 1 | 11 | 1 | 5 | 2 | 9 | Cyp2j13 |
| Cyp2j5 | 0 | 0 | 0 | 0 | 0 | 0 | 0 | 0 | 2 | 0 | 4 | 0 | 0 | 0 | 0 | 0 | 0 | 0 | 0 | 0 | 1 | 0 | 0 | 0 | 0 | 0 | 0 | 0 | Cyp2j5 |
| Cyp2j6 | 870 | 631 | 590 | 561 | 770 | 629 | 618 | 877 | 892 | 813 | 842 | 665 | 761 | 591 | 973 | 886 | 530 | 635 | 575 | 661 | 564 | 825 | 873 | 614 | 644 | 687 | 542 | 606 | Cyp2j6 |
| Cyp2j7 | 0 | 0 | 0 | 0 | 0 | 0 | 0 | 0 | 0 | 0 | 0 | 0 | 0 | 0 | 0 | 0 | 0 | 0 | 0 | 0 | 0 | 0 | 0 | 1 | 0 | 0 | 0 | 0 | Cyp2j7 |
| Cyp2j8 | 8 | 6 | 7 | 0 | 2 | 9 | 7 | 9 | 2 | 2 | 8 | 3 | 6 | 1 | 21 | 7 | 2 | 6 | 19 | 6 | 6 | 7 | 1 | 0 | 0 | 0 | 3 | 0 | Cyp2j8 |
| Cyp2j9 | 57 | 72 | 44 | 55 | 117 | 81 | 101 | 161 | 196 | 133 | 161 | 115 | 132 | 137 | 73 | 61 | 56 | 69 | 80 | 90 | 90 | 127 | 154 | 99 | 102 | 152 | 116 | 101 | Cyp2j9 |
| Cyp2r1 | 36 | 48 | 18 | 20 | 26 | 37 | 16 | 125 | 103 | 52 | 102 | 73 | 87 | 66 | 59 | 60 | 20 | 17 | 18 | 37 | 29 | 63 | 119 | 59 | 60 | 32 | 72 | 50 | Cyp2r1 |
| Cyp2s1 | 205 | 102 | 66 | 48 | 76 | 54 | 30 | 78 | 146 | 106 | 180 | 74 | 75 | 34 | 108 | 206 | 44 | 87 | 52 | 28 | 30 | 119 | 117 | 60 | 56 | 52 | 27 | 42 | Cyp2s1 |
| Cyp2t4 | 0 | 0 | 0 | 0 | 4 | 0 | 0 | 0 | 0 | 1 | 0 | 0 | 0 | 0 | 0 | 0 | 0 | 0 | 0 | 0 | 0 | 0 | 3 | 0 | 0 | 0 | 0 | 0 | Cyp2t4 |
| Cyp2u1 | 374 | 409 | 604 | 615 | 576 | 548 | 596 | 136 | 141 | 155 | 129 | 127 | 142 | 151 | 401 | 465 | 606 | 626 | 596 | 386 | 582 | 182 | 149 | 173 | 225 | 182 | 80 | 136 | Cyp2u1 |
| Cyp2w1 | 43 | 18 | 6 | 0 | 0 | 18 | 0 | 186 | 302 | 128 | 91 | 45 | 64 | 27 | 21 | 14 | 1 | 10 | 4 | 0 | 0 | 12 | 353 | 102 | 4 | 25 | 35 | 28 | Cyp2w1 |
| Cyp39a1 | 3513 | 2988 | 3428 | 2829 | 2603 | 2681 | 3204 | 1853 | 1684 | 1516 | 1413 | 2284 | 1978 | 2186 | 3539 | 2921 | 3118 | 2689 | 3031 | 3235 | 2958 | 1425 | 1746 | 1859 | 1536 | 2355 | 2069 | 2388 | Cyp39a1 |
| Cyp3a13 | 0 | 0 | 0 | 0 | 0 | 0 | 0 | 1 | 0 | 0 | 1 | 0 | 1 | 0 | 0 | 0 | 0 | 0 | 0 | 0 | 1 | 3 | 4 | 2 | 0 | 0 | 3 | 0 | Cyp3a13 |
| Cyp3a57 | 3 | 0 | 0 | 0 | 0 | 1 | 2 | 1 | 0 | 3 | 0 | 3 | 0 | 0 | 1 | 0 | 2 | 2 | 0 | 2 | 0 | 0 | 0 | 0 | 0 | 1 | 0 | 0 | Cyp3a57 |
| Cyp46a1 | 2384 | 3081 | 3553 | 3931 | 2834 | 2906 | 3114 | 724 | 659 | 945 | 675 | 1181 | 1073 | 1057 | 2268 | 2808 | 3497 | 3594 | 3244 | 2723 | 3190 | 729 | 694 | 1113 | 1072 | 988 | 1062 | 1157 | Cyp46a1 |
| Cyp4a12a | 0 | 0 | 0 | 0 | 0 | 0 | 0 | 5 | 7 | 3 | 0 | 0 | 0 | 0 | 0 | 0 | 0 | 0 | 0 | 0 | 0 | 0 | 2 | 0 | 0 | 0 | 0 | 0 | Cyp4a12a |
| Cyp4a12b | 0 | 0 | 0 | 0 | 0 | 0 | 0 | 5 | 2 | 0 | 0 | 0 | 0 | 0 | 0 | 0 | 0 | 0 | 0 | 0 | 0 | 0 | 1 | 0 | 0 | 0 | 0 | 0 | Cyp4a12b |
| Cyp4a29 | 0 | 0 | 0 | 0 | 0 | 0 | 0 | 1 | 2 | 0 | 0 | 0 | 0 | 0 | 0 | 0 | 0 | 0 | 0 | 0 | 0 | 0 | 1 | 0 | 0 | 0 | 0 | 0 | Cyp4a29 |
| Cyp4a32 | 0 | 0 | 0 | 0 | 0 | 0 | 0 | 0 | 0 | 0 | 1 | 0 | 0 | 0 | 0 | 0 | 0 | 0 | 0 | 0 | 0 | 0 | 0 | 0 | 0 | 0 | 0 | 0 | Cyp4a32 |
| Cyp4b1 | 0 | 4 | 0 | 1 | 6 | 2 | 3 | 6 | 7 | 5 | 12 | 3 | 6 | 1 | 1 | 2 | 1 | 7 | 5 | 8 | 1 | 13 | 17 | 0 | 2 | 1 | 1 | 1 | Cyp4b1 |
| Cyp4f13 | 331 | 316 | 215 | 408 | 435 | 461 | 343 | 412 | 494 | 449 | 613 | 364 | 411 | 365 | 402 | 442 | 246 | 406 | 358 | 398 | 364 | 481 | 637 | 337 | 419 | 317 | 367 | 342 | Cyp4f13 |
| Cyp4f14 | 0 | 1 | 0 | 0 | 0 | 0 | 0 | 1 | 2 | 1 | 0 | 0 | 3 | 1 | 0 | 5 | 0 | 0 | 0 | 0 | 2 | 0 | 2 | 2 | 8 | 0 | 0 | 0 | Cyp4f14 |
| Cyp4f15 | 18 | 18 | 10 | 16 | 24 | 15 | 15 | 71 | 87 | 59 | 96 | 89 | 83 | 70 | 16 | 17 | 8 | 15 | 24 | 21 | 21 | 55 | 106 | 56 | 55 | 86 | 92 | 74 | Cyp4f15 |
| Cyp4f16 | 787 | 888 | 843 | 896 | 1005 | 953 | 885 | 1423 | 1582 | 1572 | 1463 | 1378 | 1405 | 1309 | 878 | 925 | 787 | 800 | 935 | 1025 | 985 | 1389 | 1528 | 1322 | 1383 | 1381 | 1384 | 1189 | Cyp4f16 |
| Cyp4f17 | 1322 | 1392 | 1657 | 1654 | 1622 | 1716 | 1525 | 839 | 989 | 1269 | 1267 | 1164 | 1181 | 1021 | 1234 | 1421 | 1785 | 1601 | 1742 | 1531 | 1536 | 1061 | 1007 | 1193 | 1129 | 998 | 1140 | 1144 | Cyp4f17 |
| Cyp4f18 | 8 | 4 | 4 | 5 | 6 | 2 | 1 | 4 | 2 | 5 | 9 | 1 | 3 | 3 | 5 | 7 | 6 | 4 | 3 | 1 | 6 | 12 | 10 | 8 | 1 | 8 | 3 | 6 | Cyp4f18 |
| Cyp4f37 | 0 | 1 | 0 | 0 | 0 | 1 | 0 | 0 | 2 | 1 | 0 | 0 | 0 | 1 | 0 | 0 | 0 | 0 | 1 | 0 | 0 | 0 | 1 | 0 | 0 | 0 | 0 | 0 | Cyp4f37 |
| Cyp4f39 | 38 | 36 | 57 | 27 | 40 | 31 | 19 | 17 | 42 | 9 | 36 | 13 | 17 | 9 | 45 | 54 | 35 | 33 | 13 | 15 | 26 | 23 | 37 | 7 | 13 | 6 | 12 | 8 | Cyp4f39 |
| Cyp4f40 | 0 | 0 | 0 | 0 | 7 | 0 | 1 | 2 | 1 | 4 | 0 | 0 | 2 | 0 | 0 | 0 | 0 | 0 | 0 | 0 | 1 | 3 | 1 | 4 | 2 | 6 | 0 | 0 | Cyp4f40 |
| Cyp4v3 | 236 | 185 | 231 | 202 | 172 | 169 | 158 | 225 | 179 | 218 | 191 | 111 | 184 | 170 | 200 | 171 | 156 | 124 | 151 | 140 | 173 | 159 | 229 | 264 | 138 | 229 | 180 | 189 | Cyp4v3 |
| Cyp4x1 | 118 | 99 | 125 | 180 | 344 | 204 | 251 | 475 | 461 | 506 | 489 | 304 | 329 | 275 | 186 | 138 | 167 | 200 | 188 | 170 | 210 | 739 | 417 | 290 | 442 | 129 | 243 | 201 | Cyp4x1 |
| Cyp51 | 18917 | 19790 | 17887 | 16871 | 15023 | 16529 | 16749 | 16150 | 12419 | 12806 | 11487 | 15008 | 14901 | 16544 | 21174 | 18654 | 16661 | 15921 | 16366 | 18520 | 17193 | 13856 | 12418 | 13225 | 14485 | 14861 | 17323 | 17893 | Cyp51 |
| Cyp7a1 | 1 | 0 | 0 | 0 | 0 | 0 | 0 | 0 | 0 | 0 | 0 | 0 | 0 | 0 | 0 | 0 | 0 | 0 | 0 | 0 | 0 | 0 | 1 | 0 | 0 | 0 | 0 | 1 | Cyp7a1 |
| Cyp7b1 | 77 | 70 | 78 | 43 | 92 | 124 | 106 | 358 | 373 | 239 | 382 | 262 | 270 | 216 | 94 | 98 | 56 | 78 | 106 | 120 | 118 | 219 | 404 | 282 | 247 | 266 | 282 | 217 | Cyp7b1 |
| Cypt12 | 0 | 0 | 0 | 0 | 0 | 0 | 0 | 0 | 0 | 0 | 0 | 0 | 0 | 0 | 0 | 0 | 0 | 0 | 0 | 0 | 0 | 0 | 0 | 0 | 0 | 0 | 0 | 0 | Cypt12 |
| Cypt3 | 2 | 2 | 0 | 2 | 2 | 0 | 1 | 1 | 0 | 0 | 0 | 3 | 0 | 2 | 3 | 2 | 0 | 0 | 0 | 3 | 1 | 2 | 1 | 0 | 1 | 2 | 1 | 5 | Cypt3 |
| Cyren | 764 | 860 | 964 | 905 | 887 | 819 | 797 | 797 | 957 | 1059 | 1034 | 964 | 1009 | 1033 | 719 | 934 | 986 | 919 | 990 | 797 | 810 | 862 | 891 | 1124 | 964 | 1135 | 1018 | 1027 | Cyren |
| Cyria | 5126 | 4904 | 4122 | 6455 | 8859 | 7960 | 9614 | 5407 | 5296 | 5172 | 4289 | 3222 | 2691 | 3581 | 6475 | 4717 | 5333 | 7585 | 8023 | 9254 | 9583 | 4588 | 4844 | 4703 | 6583 | 2364 | 2861 | 2725 | Cyria |
| Cyrib | 7501 | 5935 | 7273 | 7173 | 8067 | 8049 | 9231 | 8849 | 7860 | 7734 | 7438 | 8537 | 8082 | 9271 | 8359 | 6970 | 7049 | 6852 | 7307 | 8537 | 8206 | 8854 | 7743 | 7742 | 8751 | 8484 | 8526 | 8787 | Cyrib |
| Cys1 | 118 | 89 | 110 | 83 | 66 | 123 | 66 | 155 | 157 | 181 | 274 | 125 | 161 | 151 | 104 | 121 | 133 | 88 | 67 | 90 | 64 | 140 | 190 | 135 | 149 | 132 | 143 | 142 | Cys1 |
| Cysltr1 | 44 | 24 | 52 | 25 | 11 | 37 | 28 | 44 | 55 | 43 | 71 | 55 | 42 | 33 | 55 | 43 | 19 | 26 | 25 | 33 | 14 | 67 | 61 | 38 | 52 | 59 | 40 | 30 | Cysltr1 |
| Cysltr2 | 2 | 0 | 0 | 2 | 2 | 0 | 4 | 13 | 7 | 8 | 12 | 1 | 5 | 1 | 3 | 5 | 2 | 2 | 1 | 1 | 2 | 10 | 8 | 4 | 11 | 0 | 5 | 11 | Cysltr2 |
| Cysrt1 | 2 | 4 | 3 | 6 | 5 | 3 | 2 | 7 | 10 | 4 | 0 | 5 | 3 | 3 | 4 | 3 | 5 | 6 | 7 | 7 | 1 | 2 | 2 | 2 | 7 | 3 | 2 | 7 | Cysrt1 |
| Cystm1 | 318 | 438 | 322 | 498 | 489 | 399 | 424 | 241 | 294 | 330 | 259 | 182 | 257 | 156 | 398 | 400 | 385 | 426 | 422 | 487 | 449 | 262 | 357 | 302 | 201 | 201 | 195 | 164 | Cystm1 |

| | | | | | | | | | | | | | | | | | | | | | | | | | | | | | |
|---|---|---|---|---|---|---|---|---|---|---|---|---|---|---|---|---|---|---|---|---|---|---|---|---|---|---|---|---|---|
| Cyth1 | 3037 | 3525 | 3272 | 3802 | 3703 | 3777 | 3761 | 2807 | 2905 | 3364 | 3267 | 3288 | 3201 | 3325 | 2917 | 3157 | 3549 | 4044 | 4006 | 3283 | 3805 | 3260 | 2822 | 3306 | 3695 | 3013 | 3219 | 3109 | Cyth1 |
| Cyth2 | 3262 | 3898 | 4082 | 4618 | 4957 | 4480 | 4816 | 7780 | 7731 | 9303 | 7906 | 9488 | 7896 | 10240 | 3698 | 3262 | 4758 | 4273 | 5524 | 4298 | 5109 | 9157 | 7419 | 9943 | 9808 | 9651 | 8780 | 9693 | Cyth2 |
| Cyth3 | 5805 | 5741 | 4954 | 5386 | 5276 | 5131 | 4761 | 3307 | 3574 | 3323 | 3868 | 3206 | 3669 | 3175 | 5649 | 6381 | 5035 | 5369 | 4954 | 4717 | 4877 | 3601 | 4066 | 3139 | 3398 | 3210 | 3263 | 3163 | Cyth3 |
| Cyth4 | 121 | 165 | 171 | 148 | 162 | 198 | 129 | 357 | 449 | 349 | 487 | 482 | 441 | 396 | 117 | 195 | 162 | 138 | 128 | 149 | 130 | 404 | 640 | 379 | 322 | 413 | 397 | 348 | Cyth4 |
| Cytip | 9 | 10 | 11 | 4 | 8 | 24 | 7 | 7 | 33 | 41 | 41 | 10 | 6 | 9 | 13 | 16 | 10 | 14 | 7 | 20 | 5 | 22 | 27 | 3 | 7 | 12 | 13 | 21 | Cytip |
| Cytl1 | 20 | 28 | 10 | 8 | 8 | 60 | 3 | 96 | 406 | 263 | 640 | 11 | 42 | 13 | 18 | 23 | 10 | 25 | 10 | 3 | 4 | 239 | 287 | 190 | 140 | 24 | 0 | 5 | Cytl1 |
| Cyyr1 | 306 | 187 | 311 | 248 | 327 | 306 | 284 | 434 | 446 | 607 | 742 | 631 | 704 | 608 | 306 | 312 | 289 | 316 | 251 | 227 | 312 | 672 | 554 | 521 | 632 | 508 | 741 | 622 | Cyyr1 |
| Czib | 1790 | 1701 | 1865 | 1513 | 1461 | 1628 | 1793 | 3751 | 3882 | 3632 | 3734 | 3738 | 3430 | 3981 | 1814 | 1593 | 1727 | 1495 | 1764 | 1859 | 1648 | 3584 | 3938 | 3859 | 3717 | 3769 | 3671 | 3853 | Czib |
| D030056L22Rik | 4549 | 4311 | 4458 | 5169 | 4322 | 4521 | 5380 | 3654 | 2970 | 3083 | 2571 | 3860 | 3226 | 3775 | 4937 | 4194 | 4838 | 4640 | 5152 | 5494 | 4958 | 3225 | 2871 | 3697 | 3447 | 4244 | 3327 | 3811 | D030056L22Rik |
| D11Wsu47e | 2295 | 2829 | 1628 | 2361 | 1868 | 2145 | 2074 | 1597 | 1542 | 1499 | 1558 | 1718 | 1554 | 1571 | 2202 | 2287 | 1936 | 2194 | 1998 | 2172 | 2217 | 1522 | 1439 | 1473 | 1704 | 1248 | 1519 | 1548 | D11Wsu47e |
| D130040H23Rik | 633 | 532 | 441 | 389 | 480 | 663 | 449 | 1080 | 975 | 603 | 652 | 691 | 739 | 632 | 663 | 535 | 370 | 423 | 529 | 757 | 608 | 530 | 921 | 577 | 812 | 567 | 945 | 650 | D130040H23Rik |
| D130043K22Rik | 411 | 542 | 314 | 508 | 765 | 598 | 520 | 341 | 341 | 348 | 321 | 272 | 316 | 223 | 438 | 485 | 414 | 591 | 605 | 635 | 637 | 378 | 372 | 256 | 371 | 118 | 231 | 193 | D130043K22Rik |
| D13Ertd608e | 6 | 2 | 6 | 8 | 0 | 0 | 2 | 0 | 0 | 0 | 1 | 0 | 0 | 0 | 5 | 2 | 5 | 0 | 2 | 2 | 0 | 0 | 0 | 1 | 0 | 0 | 0 | 0 | D13Ertd608e |
| D16Ertd472e | 2979 | 2279 | 2178 | 2576 | 2031 | 2241 | 2514 | 1928 | 1865 | 1617 | 1770 | 1758 | 1915 | 1806 | 2418 | 2569 | 2304 | 2251 | 2023 | 2329 | 2313 | 1542 | 1938 | 1673 | 1637 | 1961 | 1922 | 1824 | D16Ertd472e |
| D17H6S53E | 606 | 669 | 842 | 697 | 642 | 657 | 606 | 547 | 585 | 548 | 683 | 710 | 687 | 645 | 554 | 580 | 730 | 653 | 718 | 619 | 599 | 511 | 611 | 646 | 726 | 790 | 662 | 660 | D17H6S53E |
| D1Pas1 | 43 | 20 | 49 | 32 | 18 | 28 | 15 | 1 | 12 | 20 | 24 | 12 | 35 | 7 | 32 | 43 | 36 | 13 | 2 | 15 | 10 | 3 | 3 | 2 | 12 | 3 | 6 | 14 | D1Pas1 |
| D2hgdh | 505 | 517 | 457 | 481 | 565 | 571 | 543 | 1166 | 1271 | 1155 | 1291 | 1366 | 1203 | 1079 | 525 | 526 | 548 | 592 | 583 | 550 | 523 | 1280 | 1196 | 1222 | 1210 | 1064 | 1147 | 1169 | D2hgdh |
| D3Ertd751e | 2553 | 1808 | 2158 | 1956 | 1573 | 2002 | 2069 | 1595 | 1457 | 1332 | 1341 | 1656 | 1557 | 1792 | 2532 | 2221 | 1938 | 1705 | 1506 | 2330 | 1731 | 1282 | 1427 | 1357 | 1281 | 1545 | 1633 | 1653 | D3Ertd751e |
| D430019H16Rik | 13848 | 14348 | 12559 | 16554 | 20435 | 18753 | 19270 | 15384 | 15556 | 18226 | 15593 | 16511 | 14635 | 17082 | 13876 | 13399 | 15410 | 17107 | 19610 | 16422 | 20190 | 21490 | 14101 | 17105 | 21457 | 12590 | 15342 | 15517 | D430019H16Rik |
| D430041D05Rik | 1405 | 1425 | 1044 | 2016 | 3728 | 2954 | 3219 | 3500 | 3916 | 4770 | 3668 | 3494 | 2929 | 3365 | 1888 | 1453 | 1598 | 2714 | 2982 | 2638 | 3348 | 4824 | 3405 | 3459 | 5236 | 2083 | 2998 | 2775 | D430041D05Rik |
| D5Ertd579e | 7442 | 7013 | 5190 | 6507 | 8112 | 8085 | 7983 | 10893 | 9683 | 9127 | 8415 | 9276 | 8921 | 9720 | 7953 | 6392 | 6058 | 6940 | 7289 | 8902 | 8118 | 9481 | 9144 | 8231 | 9984 | 5853 | 10103 | 8930 | D5Ertd579e |
| D630003M21Rik | 231 | 317 | 336 | 209 | 228 | 307 | 108 | 535 | 1006 | 748 | 926 | 635 | 934 | 534 | 274 | 369 | 292 | 238 | 217 | 161 | 170 | 827 | 852 | 672 | 587 | 666 | 695 | 612 | D630003M21Rik |
| D630023F18Rik | 13 | 10 | 34 | 30 | 24 | 24 | 30 | 4 | 23 | 16 | 21 | 17 | 10 | 16 | 17 | 15 | 19 | 39 | 27 | 24 | 23 | 13 | 22 | 25 | 13 | 18 | 9 | 11 | D630023F18Rik |
| D630039A03Rik | 22 | 7 | 23 | 10 | 13 | 11 | 4 | 16 | 25 | 10 | 24 | 11 | 15 | 5 | 17 | 17 | 0 | 11 | 5 | 5 | 5 | 7 | 25 | 23 | 11 | 8 | 2 | 5 | D630039A03Rik |
| D630045J12Rik | 3698 | 3664 | 4002 | 4923 | 6215 | 5560 | 5531 | 7573 | 8093 | 10512 | 8602 | 9799 | 8037 | 9139 | 3568 | 3623 | 4612 | 4830 | 6142 | 4178 | 5712 | 11090 | 7108 | 9448 | 11963 | 6383 | 8218 | 8162 | D630045J12Rik |
| D6Ertd527e | 57 | 77 | 78 | 91 | 102 | 87 | 78 | 12 | 22 | 14 | 14 | 24 | 22 | 23 | 47 | 102 | 102 | 107 | 57 | 42 | 57 | 17 | 14 | 12 | 8 | 21 | 10 | 15 | D6Ertd527e |
| D6Wsu163e | 2023 | 2051 | 1713 | 1835 | 1843 | 1773 | 1951 | 2175 | 2052 | 1916 | 1734 | 1764 | 1690 | 1991 | 2028 | 1793 | 1994 | 1776 | 1762 | 2196 | 2053 | 1903 | 2061 | 1874 | 1951 | 1654 | 1703 | 1879 | D6Wsu163e |
| D7Ertd443e | 2 | 4 | 4 | 7 | 13 | 14 | 6 | 20 | 47 | 27 | 43 | 8 | 28 | 3 | 13 | 4 | 6 | 15 | 3 | 4 | 3 | 25 | 39 | 8 | 21 | 14 | 10 | 5 | D7Ertd443e |
| D830030K20Rik | 105 | 90 | 82 | 80 | 110 | 121 | 110 | 99 | 105 | 89 | 81 | 83 | 84 | 101 | 113 | 108 | 88 | 70 | 115 | 149 | 103 | 117 | 92 | 68 | 88 | 64 | 84 | 94 | D830030K20Rik |
| D8Ertd738e | 3621 | 4685 | 4263 | 4142 | 3261 | 3148 | 3708 | 2542 | 2811 | 2832 | 3021 | 2914 | 2935 | 3557 | 3578 | 3547 | 4220 | 3639 | 3514 | 3946 | 3684 | 2117 | 2721 | 3121 | 2934 | 3606 | 3253 | 3364 | D8Ertd738e |
| D930020B18Rik | 225 | 116 | 87 | 102 | 72 | 85 | 83 | 125 | 69 | 135 | 77 | 8 | 28 | 24 | 133 | 157 | 96 | 78 | 40 | 75 | 53 | 17 | 102 | 119 | 13 | 174 | 2 | 5 | D930020B18Rik |
| D930048N14Rik | 256 | 168 | 209 | 124 | 173 | 149 | 146 | 115 | 168 | 135 | 293 | 125 | 180 | 116 | 160 | 245 | 214 | 228 | 148 | 145 | 142 | 132 | 215 | 105 | 133 | 154 | 131 | 138 | D930048N14Rik |
| Daam1 | 10308 | 10368 | 10099 | 12570 | 13994 | 13277 | 14613 | 9249 | 8421 | 9342 | 8359 | 8260 | 8022 | 8815 | 11570 | 10347 | 11346 | 13195 | 12752 | 13747 | 14447 | 9954 | 8134 | 8692 | 10193 | 6857 | 7861 | 7936 | Daam1 |
| Daam2 | 2179 | 1862 | 1537 | 1613 | 1992 | 1897 | 1101 | 3838 | 4108 | 4764 | 5036 | 2432 | 3813 | 2488 | 2000 | 2394 | 1683 | 2048 | 1400 | 1040 | 1249 | 3815 | 4113 | 3780 | 2996 | 2714 | 2411 | 2372 | Daam2 |
| Dab1 | 4159 | 4044 | 4026 | 5432 | 6998 | 7264 | 8122 | 6577 | 6613 | 7002 | 5690 | 7089 | 6358 | 7313 | 4923 | 3752 | 4997 | 6245 | 7196 | 7514 | 7745 | 7762 | 5818 | 6980 | 8695 | 5358 | 6202 | 6569 | Dab1 |
| Dab2 | 7453 | 6209 | 6838 | 3545 | 2864 | 4539 | 2482 | 6731 | 8869 | 7106 | 10184 | 7326 | 10010 | 7059 | 7425 | 9425 | 4582 | 4629 | 2320 | 3071 | 2468 | 7528 | 11204 | 7118 | 5748 | 7742 | 8486 | 7350 | Dab2 |
| Dab2ip | 6118 | 7178 | 7805 | 8533 | 9407 | 8143 | 7778 | 6483 | 8174 | 8852 | 8656 | 8755 | 7785 | 7778 | 5399 | 6777 | 8676 | 8873 | 9757 | 6619 | 8174 | 9095 | 7643 | 9037 | 9432 | 7652 | 7281 | 7436 | Dab2ip |
| Dach1 | 4736 | 3914 | 7819 | 6482 | 6567 | 6666 | 6563 | 6807 | 5685 | 6969 | 6582 | 9747 | 11297 | 10428 | 5304 | 4903 | 6999 | 5755 | 6656 | 5501 | 6148 | 7860 | 5711 | 7915 | 8177 | 9166 | 9630 | 11447 | Dach1 |
| Dach2 | 1039 | 1118 | 1049 | 1009 | 1065 | 1006 | 1047 | 1138 | 845 | 677 | 672 | 706 | 913 | 748 | 1503 | 1213 | 1210 | 946 | 851 | 1092 | 929 | 1078 | 900 | 641 | 756 | 737 | 851 | 818 | Dach2 |
| Dact1 | 4512 | 4044 | 4350 | 5026 | 6131 | 5461 | 4714 | 5939 | 6301 | 6841 | 6883 | 5036 | 5227 | 4696 | 4424 | 4854 | 4536 | 5382 | 5415 | 4359 | 5042 | 7095 | 5962 | 6349 | 6686 | 5574 | 4313 | 4494 | Dact1 |
| Dact2 | 219 | 282 | 269 | 279 | 254 | 267 | 189 | 169 | 276 | 219 | 262 | 154 | 185 | 130 | 187 | 250 | 200 | 246 | 263 | 206 | 187 | 122 | 257 | 192 | 132 | 168 | 128 | 132 | Dact2 |
| Dact3 | 4207 | 5878 | 4371 | 5939 | 4523 | 4573 | 4404 | 1711 | 1860 | 1966 | 2374 | 1777 | 2158 | 1669 | 4214 | 4777 | 5235 | 5899 | 4698 | 4053 | 4632 | 2184 | 2102 | 2004 | 1840 | 2187 | 1825 | 1673 | Dact3 |
| Dad1 | 4862 | 6276 | 4525 | 5125 | 4394 | 4490 | 4837 | 3644 | 3695 | 3278 | 3751 | 3161 | 3741 | 3688 | 4733 | 4898 | 4948 | 5071 | 4623 | 5342 | 4872 | 3227 | 3772 | 3538 | 3625 | 3980 | 3660 | 3603 | Dad1 |
| Dag1 | 14802 | 16707 | 18875 | 17069 | 15248 | 15012 | 13703 | 7850 | 9614 | 10733 | 11086 | 11556 | 11264 | 10249 | 14410 | 17913 | 17717 | 17785 | 15256 | 11745 | 14173 | 10828 | 10616 | 10912 | 10243 | 10525 | 10261 | 11186 | Dag1 |
| Dagla | 602 | 785 | 969 | 966 | 1424 | 1075 | 963 | 806 | 1126 | 1309 | 1316 | 1098 | 1170 | 895 | 626 | 855 | 969 | 1267 | 1218 | 860 | 1000 | 1549 | 1005 | 1285 | 1411 | 928 | 972 | 869 | Dagla |
| Daglb | 1108 | 1367 | 1205 | 1419 | 1233 | 1454 | 1226 | 1001 | 1065 | 1134 | 1134 | 1258 | 1120 | 1004 | 1117 | 1277 | 1238 | 1317 | 1347 | 1221 | 1205 | 1093 | 1070 | 1087 | 1211 | 1193 | 1189 | 1044 | Daglb |
| Dalrd3 | 2729 | 3572 | 3133 | 3254 | 3186 | 2846 | 3072 | 1937 | 2270 | 2559 | 2253 | 2482 | 2057 | 2071 | 2579 | 2890 | 3315 | 3333 | 3490 | 3031 | 3044 | 2264 | 2353 | 2491 | 2296 | 2580 | 2042 | 1917 | Dalrd3 |
| Dand5 | 197 | 201 | 211 | 252 | 271 | 291 | 235 | 220 | 298 | 325 | 353 | 310 | 278 | 300 | 211 | 174 | 255 | 270 | 277 | 276 | 298 | 307 | 322 | 369 | 373 | 332 | 296 | 300 | Dand5 |
| Dao | 0 | 2 | 2 | 1 | 1 | 0 | 2 | 0 | 0 | 3 | 0 | 0 | 0 | 1 | 0 | 1 | 0 | 1 | 0 | 0 | 2 | 0 | 1 | 0 | 0 | 2 | 3 | 1 | Dao |
| Dap | 1917 | 2005 | 1706 | 1595 | 1153 | 1471 | 1190 | 1293 | 1804 | 1607 | 1968 | 1329 | 1719 | 1259 | 1627 | 2171 | 1669 | 1556 | 1183 | 1185 | 1137 | 1424 | 2026 | 1486 | 1354 | 1835 | 1432 | 1392 | Dap |
| Dap3 | 8140 | 8241 | 6587 | 7882 | 7133 | 7015 | 8137 | 5377 | 5551 | 5311 | 5041 | 4866 | 4906 | 5395 | 7780 | 7497 | 6835 | 7515 | 7308 | 7657 | 8068 | 5158 | 5505 | 5119 | 5335 | 5395 | 5120 | 5600 | Dap3 |
| Dapk1 | 6252 | 6519 | 6210 | 8115 | 9870 | 8445 | 8851 | 8313 | 8767 | 10604 | 9255 | 9790 | 8267 | 9447 | 5912 | 6706 | 7435 | 8679 | 9705 | 7149 | 8961 | 11057 | 8025 | 10007 | 11589 | 7618 | 8179 | 8505 | Dapk1 |
| Dapk2 | 142 | 164 | 162 | 96 | 47 | 52 | 63 | 96 | 122 | 86 | 124 | 121 | 151 | 140 | 146 | 137 | 109 | 45 | 77 | 81 | 59 | 115 | 122 | 109 | 85 | 228 | 134 | 208 | Dapk2 |
| Dapk3 | 1197 | 1631 | 1569 | 1572 | 1310 | 1302 | 984 | 1523 | 1796 | 1894 | 1985 | 1922 | 1932 | 1756 | 1212 | 1321 | 1550 | 1301 | 1424 | 1100 | 1163 | 1880 | 1891 | 1927 | 1748 | 2147 | 1899 | 1698 | Dapk3 |
| Dapl1 | 36 | 27 | 19 | 9 | 28 | 16 | 12 | 34 | 28 | 15 | 39 | 10 | 17 | 12 | 18 | 23 | 16 | 13 | 7 | 29 | 16 | 7 | 43 | 35 | 10 | 17 | 6 | 0 | Dapl1 |
| Dapp1 | 441 | 357 | 420 | 351 | 308 | 410 | 446 | 518 | 482 | 542 | 509 | 489 | 406 | 415 | 385 | 436 | 326 | 359 | 379 | 383 | 371 | 286 | 546 | 639 | 382 | 742 | 392 | 379 | Dapp1 |
| Dars | 8618 | 8078 | 8676 | 7222 | 6822 | 7642 | 8146 | 10015 | 9234 | 7585 | 7895 | 8808 | 9649 | 9151 | 10108 | 8372 | 8234 | 7064 | 7355 | 9209 | 7545 | 8094 | 9527 | 8345 | 7844 | 9846 | 9584 | 10033 | Dars |
| Dars2 | 3344 | 3239 | 2905 | 2650 | 2259 | 2324 | 2765 | 2440 | 2092 | 1905 | 1764 | 2375 | 2195 | 2303 | 3512 | 2624 | 3074 | 2626 | 2621 | 2802 | 2726 | 1958 | 2036 | 2111 | 2077 | 2407 | 2607 | 2487 | Dars2 |
| Daw1 | 37 | 68 | 40 | 54 | 65 | 68 | 70 | 90 | 93 | 63 | 84 | 68 | 110 | 76 | 66 | 65 | 36 | 53 | 55 | 78 | 89 | 112 | 139 | 78 | 89 | 57 | 56 | 38 | Daw1 |
| Daxx | 3553 | 4386 | 4677 | 4718 | 3769 | 3805 | 4041 | 2407 | 2698 | 3237 | 3032 | 3662 | 3264 | 3474 | 3100 | 3675 | 4773 | 4384 | 4462 | 3608 | 4050 | 3024 | 2622 | 3531 | 3264 | 4168 | 3320 | 3823 | Daxx |
| Dazap1 | 6775 | 7845 | 9206 | 9799 | 9069 | 7992 | 8216 | 7425 | 8149 | 10132 | 9193 | 9322 | 8214 | 9387 | 6729 | 6664 | 9921 | 9164 | 9238 | 7747 | 8255 | 8485 | 7490 | 10671 | 9752 | 11100 | 8551 | 8803 | Dazap1 |
| Dazap2 | 8997 | 8717 | 8580 | 7126 | 6988 | 7574 | 8242 | 9330 | 8656 | 7693 | 7201 | 9935 | 9262 | 10078 | 8592 | 7765 | 8369 | 6570 | 7846 | 8817 | 7670 | 7678 | 8501 | 9109 | 8302 | 11165 | 10635 | 10447 | Dazap2 |
| Dazl | 17 | 4 | 14 | 8 | 14 | 26 | 22 | 17 | 21 | 4 | 18 | 11 | 7 | 18 | 9 | 15 | 8 | 11 | 18 | 27 | 23 | 20 | 20 | 5 | 12 | 14 | 3 | 14 | Dazl |
| Dbf4 | 5602 | 4360 | 4876 | 4403 | 3708 | 4033 | 4709 | 6038 | 4725 | 4460 | 4327 | 6417 | 5547 | 6676 | 5429 | 4615 | 4527 | 3818 | 4172 | 5010 | 4246 | 3888 | 4471 | 5392 | 4806 | 6983 | 6643 | 6992 | Dbf4 |
| Dbh | 2 | 57 | 8 | 49 | 65 | 79 | 9 | 57 | 69 | 64 | 223 | 5 | 91 | 1 | 44 | 51 | 66 | 101 | 23 | 66 | 21 | 77 | 226 | 1 | 68 | 8 | 8 | 0 | Dbh |
| Dbi | 5226 | 5664 | 5896 | 5676 | 6035 | 4727 | 8817 | 5300 | 4924 | 5056 | 5633 | 5709 | 4786 | 8023 | 5467 | 4231 | 5759 | 5592 | 6026 | 6435 | 7918 | 4179 | 4891 | 6340 | 5979 | 6815 | 5523 | 7230 | Dbi |
| Dbil5 | 7 | 10 | 19 | 5 | 5 | 10 | 7 | 9 | 6 | 1 | 9 | 11 | 2 | 9 | 11 | 11 | 7 | 8 | 12 | 7 | 13 | 7 | 16 | 15 | 10 | 8 | 5 | 11 | Dbil5 |
| Dbn1 | 16244 | 20003 | 21121 | 24122 | 25867 | 23165 | 22653 | 29263 | 33144 | 39939 | 33308 | 41429 | 33343 | 38928 | 16714 | 16768 | 25155 | 24070 | 28698 | 21585 | 24891 | 37095 | 30662 | 44168 | 43854 | 42230 | 36648 | 37048 | Dbn1 |
| Dbndd1 | 825 | 1060 | 1105 | 1144 | 1545 | 1233 | 1287 | 1603 | 1666 | 1843 | 1521 | 1410 | 1300 | 1457 | 989 | 833 | 1124 | 1161 | 1677 | 1442 | 1487 | 2002 | 1579 | 1727 | 1896 | 1214 | 1259 | 1415 | Dbndd1 |
| Dbndd2 | 623 | 655 | 901 | 1092 | 1231 | 1131 | 1187 | 708 | 488 | 734 | 591 | 565 | 606 | 672 | 717 | 545 | 1010 | 1062 | 1125 | 988 | 1266 | 825 | 435 | 677 | 769 | 440 | 574 | 587 | Dbndd2 |
| Dbnl | 5170 | 6697 | 6454 | 6439 | 6280 | 6019 | 5503 | 5326 | 6536 | 7124 | 6403 | 7432 | 6945 | 6956 | 5120 | 6146 | 6334 | 6406 | 6509 | 5653 | 5737 | 6079 | 6409 | 7096 | 6609 | 7918 | 7009 | 7035 | Dbnl |
| Dbp | 722 | 978 | 784 | 1083 | 784 | 750 | 802 | 398 | 480 | 463 | 473 | 318 | 390 | 343 | 651 | 713 | 979 | 969 | 834 | 760 | 835 | 341 | 518 | 464 | 412 | 356 | 302 | 375 | Dbp |
| Dbr1 | 2556 | 2310 | 2505 | 2312 | 1835 | 1870 | 2410 | 1735 | 1555 | 1481 | 1482 | 1786 | 1608 | 2132 | 2525 | 2123 | 1999 | 2176 | 1954 | 2069 | 2388 | 1499 | 1458 | 1664 | 1483 | 2007 | 1904 | 2220 | Dbr1 |
| Dbt | 3665 | 3172 | 3302 | 3266 | 2721 | 2897 | 3070 | 2081 | 1933 | 1784 | 1762 | 1776 | 1945 | 1894 | 3531 | 3534 | 3008 | 3218 | 2791 | 3234 | 2866 | 1763 | 1969 | 1901 | 1762 | 2121 | 1894 | 1919 | Dbt |
| Dbx1 | 1794 | 2401 | 6039 | 2450 | 1571 | 1909 | 987 | 1617 | 762 | 1393 | 1119 | 3593 | 3119 | 3088 | 1876 | 1636 | 3972 | 1486 | 2461 | 1641 | 1759 | 2139 | 1592 | 2294 | 1926 | 2202 | 4329 | 4306 | Dbx1 |
| Dbx2 | 23 | 25 | 20 | 22 | 31 | 17 | 19 | 22 | 23 | 54 | 30 | 22 | 37 | 19 | 36 | 24 | 41 | 26 | 22 | 15 | 11 | 37 | 23 | 17 | 39 | 21 | 26 | 30 | Dbx2 |
| Dcaf1 | 6251 | 5958 | 5690 | 5829 | 5938 | 6395 | 6254 | 6121 | 5839 | 5982 | 5214 | 7002 | 5941 | 6988 | 5753 | 5392 | 5813 | 5455 | 6387 | 5604 | 6381 | 5909 | 5436 | 6058 | 6380 | 5687 | 6832 | 6703 | Dcaf1 |
| Dcaf10 | 5196 | 4265 | 3760 | 4654 | 4395 | 5086 | 4911 | 3841 | 3295 | 3150 | 3322 | 3326 | 3260 | 3444 | 4859 | 4446 | 4005 | 4672 | 4285 | 4350 | 5167 | 3621 | 3350 | 2927 | 3449 | 2728 | 3346 | 3531 | Dcaf10 |
| Dcaf11 | 7034 | 8875 | 6646 | 8124 | 7212 | 7186 | 8096 | 5765 | 5568 | 5394 | 4913 | 5522 | 5254 | 5569 | 7683 | 7182 | 7201 | 8255 | 8012 | 7818 | 8250 | 5726 | 5310 | 5561 | 5952 | 5098 | 5642 | 5762 | Dcaf11 |

| | | | | | | | | | | | | | | | | | | | | | | | | | | | | | |
|---|---|---|---|---|---|---|---|---|---|---|---|---|---|---|---|---|---|---|---|---|---|---|---|---|---|---|---|---|---|
| Dcaf12 | 7171 | 7253 | 8126 | 8156 | 7666 | 7407 | 7976 | 6165 | 5820 | 5682 | 5655 | 6126 | 6028 | 6250 | 6928 | 6136 | 8051 | 8294 | 8419 | 7100 | 8123 | 6527 | 5767 | 6311 | 6950 | 6448 | 5808 | 6111 | Dcaf12 |
| Dcaf12l1 | 523 | 442 | 693 | 515 | 717 | 673 | 607 | 924 | 1071 | 910 | 1039 | 1090 | 1051 | 882 | 538 | 552 | 576 | 640 | 755 | 569 | 553 | 1140 | 1216 | 926 | 898 | 1194 | 968 | 933 | Dcaf12l1 |
| Dcaf12l2 | 53 | 36 | 62 | 31 | 56 | 42 | 49 | 82 | 126 | 91 | 99 | 88 | 112 | 73 | 46 | 61 | 59 | 40 | 48 | 30 | 26 | 95 | 139 | 76 | 112 | 80 | 92 | 96 | Dcaf12l2 |
| Dcaf13 | 2355 | 1761 | 2365 | 1596 | 1372 | 2275 | 2182 | 2521 | 1879 | 1330 | 1198 | 1537 | 1418 | 1622 | 2226 | 1826 | 1700 | 1525 | 1529 | 1815 | 1584 | 2141 | 1421 | 1423 | 1379 | 2042 | 1684 | 1702 | Dcaf13 |
| Dcaf15 | 2234 | 2616 | 2868 | 2869 | 3082 | 2680 | 2725 | 3515 | 4071 | 4377 | 3842 | 5227 | 4601 | 4635 | 2337 | 2275 | 2998 | 2746 | 3397 | 2725 | 2704 | 4612 | 3783 | 4533 | 4601 | 5514 | 4872 | 4615 | Dcaf15 |
| Dcaf17 | 2476 | 2051 | 1905 | 1916 | 1962 | 2330 | 2227 | 3464 | 3202 | 3055 | 2822 | 3407 | 3298 | 3143 | 2370 | 2106 | 2027 | 1840 | 1960 | 2301 | 2119 | 2866 | 3076 | 2827 | 3009 | 2573 | 3862 | 3259 | Dcaf17 |
| Dcaf4 | 2235 | 2375 | 1877 | 2183 | 2137 | 2339 | 2198 | 1961 | 2211 | 1914 | 1877 | 1711 | 1855 | 1878 | 2040 | 2069 | 2152 | 2376 | 2336 | 2156 | 2182 | 1856 | 2075 | 1994 | 1914 | 1819 | 1765 | 1792 | Dcaf4 |
| Dcaf5 | 5294 | 5175 | 6025 | 6202 | 7151 | 6531 | 6127 | 6883 | 6854 | 7681 | 7034 | 7772 | 7659 | 7255 | 5396 | 5065 | 6102 | 6191 | 7381 | 5763 | 6399 | 8263 | 6590 | 7135 | 8105 | 6988 | 7259 | 7471 | Dcaf5 |
| Dcaf6 | 4510 | 3898 | 3811 | 4349 | 4482 | 4777 | 4687 | 3076 | 2989 | 2972 | 3286 | 2673 | 2558 | 2394 | 4512 | 4059 | 3833 | 4562 | 4209 | 4223 | 4617 | 3377 | 3242 | 2735 | 2885 | 2261 | 2642 | 2388 | Dcaf6 |
| Dcaf7 | 25233 | 25179 | 25243 | 30790 | 31504 | 30536 | 31429 | 22427 | 23054 | 24732 | 23033 | 24522 | 22781 | 23586 | 26373 | 24593 | 28685 | 29653 | 32935 | 26380 | 32530 | 25887 | 21126 | 24390 | 26595 | 20697 | 23730 | 23326 | Dcaf7 |
| Dcaf8 | 8119 | 7971 | 7564 | 7265 | 7734 | 8116 | 7945 | 12355 | 12734 | 12157 | 13398 | 13025 | 13522 | 13387 | 7743 | 7068 | 7862 | 6895 | 8000 | 8067 | 7956 | 10875 | 13035 | 12687 | 12348 | 12940 | 14046 | 13492 | Dcaf8 |
| Dcakd | 7095 | 9618 | 9650 | 8077 | 7869 | 7366 | 6835 | 6999 | 9196 | 8885 | 8271 | 11267 | 9067 | 9464 | 6700 | 8535 | 9491 | 7754 | 8747 | 6544 | 7092 | 8128 | 8348 | 10015 | 9668 | 13354 | 9710 | 10109 | Dcakd |
| Dcbld1 | 2629 | 2406 | 2539 | 2182 | 2572 | 2440 | 2739 | 1501 | 1419 | 1424 | 1288 | 1552 | 1463 | 1580 | 2418 | 2307 | 2431 | 2265 | 2374 | 2153 | 2529 | 1253 | 1364 | 1396 | 1424 | 1761 | 1469 | 1601 | Dcbld1 |
| Dcbld2 | 12650 | 12570 | 12076 | 11724 | 10910 | 10870 | 11202 | 6708 | 5925 | 6256 | 6346 | 7275 | 7152 | 7358 | 13471 | 13196 | 11224 | 11438 | 10082 | 10221 | 10584 | 6924 | 5858 | 6273 | 6585 | 6993 | 7405 | 7665 | Dcbld2 |
| Dcc | 18475 | 19200 | 16649 | 24943 | 23653 | 28999 | 27412 | 23746 | 18709 | 21688 | 17619 | 24881 | 26244 | 26601 | 22801 | 19365 | 21261 | 21715 | 24582 | 26049 | 28310 | 22331 | 16589 | 19502 | 26568 | 14216 | 27902 | 24683 | Dcc |
| Dcdc2a | 264 | 237 | 345 | 236 | 337 | 278 | 242 | 326 | 278 | 299 | 298 | 428 | 456 | 382 | 332 | 354 | 234 | 277 | 287 | 396 | 263 | 374 | 474 | 287 | 327 | 334 | 292 | 375 | Dcdc2a |
| Dcdc2b | 249 | 323 | 286 | 384 | 452 | 373 | 330 | 308 | 279 | 295 | 282 | 263 | 234 | 312 | 257 | 288 | 442 | 446 | 432 | 315 | 385 | 424 | 300 | 303 | 322 | 244 | 224 | 224 | Dcdc2b |
| Dcdc2c | 0 | 0 | 0 | 0 | 0 | 0 | 2 | 0 | 2 | 0 | 0 | 0 | 2 | 0 | 0 | 0 | 0 | 3 | 0 | 0 | 0 | 2 | 3 | 5 | 0 | 0 | 0 | 0 | Dcdc2c |
| Dchs1 | 7814 | 9040 | 8763 | 8964 | 10731 | 10632 | 7659 | 15277 | 20163 | 22170 | 22052 | 23811 | 20720 | 20054 | 6749 | 9524 | 9525 | 9949 | 11123 | 6697 | 8668 | 23487 | 17404 | 22776 | 23456 | 14275 | 19107 | 18770 | Dchs1 |
| Dchs2 | 578 | 701 | 250 | 361 | 679 | 714 | 458 | 509 | 603 | 423 | 593 | 118 | 406 | 106 | 754 | 850 | 316 | 457 | 422 | 623 | 555 | 344 | 750 | 105 | 322 | 40 | 241 | 114 | Dchs2 |
| Dck | 7507 | 6220 | 7823 | 7081 | 6632 | 6938 | 8152 | 7184 | 5897 | 6292 | 5561 | 8616 | 7156 | 8689 | 7539 | 6488 | 7788 | 6682 | 7511 | 7347 | 7461 | 6315 | 5492 | 7491 | 7321 | 9483 | 8608 | 9252 | Dck |
| Dclk1 | 16222 | 14708 | 13806 | 22411 | 30354 | 28020 | 29376 | 29224 | 27870 | 31454 | 28149 | 26523 | 24930 | 28818 | 19386 | 15600 | 18212 | 24751 | 25988 | 24658 | 29775 | 37864 | 26088 | 26593 | 35960 | 18119 | 25048 | 25362 | Dclk1 |
| Dclk2 | 10388 | 11750 | 9210 | 14888 | 19676 | 17554 | 18355 | 16893 | 17959 | 20551 | 17291 | 21194 | 15536 | 20622 | 11122 | 9179 | 13420 | 16522 | 19518 | 15490 | 19606 | 21784 | 14087 | 19828 | 25244 | 12970 | 18152 | 17014 | Dclk2 |
| Dclk3 | 201 | 204 | 187 | 188 | 486 | 278 | 288 | 266 | 310 | 316 | 281 | 100 | 187 | 109 | 237 | 206 | 217 | 321 | 299 | 209 | 274 | 349 | 241 | 205 | 220 | 110 | 89 | 74 | Dclk3 |
| Dclre1a | 667 | 449 | 493 | 453 | 533 | 552 | 503 | 1272 | 1310 | 1201 | 1064 | 1212 | 1224 | 1189 | 683 | 526 | 469 | 436 | 568 | 635 | 512 | 1031 | 1269 | 1116 | 1048 | 1167 | 1291 | 1229 | Dclre1a |
| Dclre1b | 2143 | 1998 | 1774 | 1842 | 1683 | 1865 | 1989 | 1649 | 1655 | 1501 | 1498 | 2040 | 1724 | 2134 | 1882 | 2002 | 1869 | 1842 | 1841 | 1725 | 1936 | 1432 | 1570 | 1731 | 1647 | 2003 | 2097 | 2166 | Dclre1b |
| Dclre1c | 3095 | 2897 | 2021 | 2736 | 2172 | 2831 | 2778 | 1568 | 1521 | 1266 | 1393 | 1412 | 1491 | 1510 | 3112 | 3033 | 2341 | 2656 | 2173 | 2689 | 2638 | 1512 | 1498 | 1273 | 1340 | 1127 | 1403 | 1311 | Dclre1c |
| Dcn | 1944 | 1437 | 1554 | 1848 | 2388 | 2789 | 2014 | 3778 | 6058 | 3906 | 6694 | 2526 | 2911 | 2546 | 2193 | 1845 | 1497 | 3152 | 1947 | 2999 | 2122 | 4269 | 9378 | 3675 | 3850 | 2379 | 2493 | 2360 | Dcn |
| Dcp1a | 3758 | 3327 | 3482 | 3319 | 3181 | 3262 | 3386 | 3794 | 4128 | 3952 | 4240 | 4697 | 4710 | 4600 | 3047 | 3562 | 3479 | 3292 | 3338 | 3059 | 3267 | 3898 | 3704 | 4304 | 4247 | 4349 | 4502 | 4580 | Dcp1a |
| Dcp1b | 1481 | 1769 | 1919 | 2174 | 2360 | 2194 | 2416 | 2313 | 2348 | 2957 | 2546 | 2911 | 2675 | 2985 | 1655 | 1460 | 2274 | 2206 | 2392 | 1998 | 2183 | 2971 | 2261 | 2875 | 3062 | 2569 | 2881 | 2688 | Dcp1b |
| Dcp2 | 16879 | 13465 | 12074 | 12567 | 12925 | 14599 | 14124 | 12115 | 11136 | 10494 | 11901 | 13016 | 12277 | 13430 | 15093 | 16691 | 12672 | 13422 | 12211 | 13357 | 13691 | 9453 | 10452 | 10092 | 11053 | 9042 | 14061 | 12245 | Dcp2 |
| Dcps | 2092 | 2340 | 1971 | 1607 | 1398 | 1682 | 1662 | 2448 | 2378 | 2056 | 1959 | 2498 | 2198 | 2645 | 2189 | 1912 | 1920 | 1574 | 1800 | 1959 | 1658 | 2037 | 2379 | 2262 | 2225 | 3010 | 2700 | 2816 | Dcps |
| Dcst1 | 5 | 3 | 3 | 5 | 2 | 5 | 3 | 9 | 4 | 14 | 5 | 4 | 1 | 1 | 1 | 7 | 6 | 11 | 2 | 1 | 0 | 0 | 3 | 4 | 5 | 3 | 1 | 0 | Dcst1 |
| Dcst2 | 93 | 81 | 78 | 156 | 128 | 135 | 115 | 42 | 70 | 52 | 73 | 60 | 84 | 63 | 121 | 103 | 116 | 164 | 85 | 115 | 104 | 89 | 53 | 68 | 56 | 70 | 85 | 70 | Dcst2 |
| Dcstamp | 0 | 0 | 0 | 4 | 2 | 2 | 0 | 1 | 2 | 0 | 2 | 0 | 3 | 4 | 1 | 1 | 0 | 0 | 0 | 0 | 1 | 2 | 2 | 2 | 0 | 0 | 0 | 0 | Dcstamp |
| Dct | 233 | 155 | 152 | 167 | 204 | 317 | 396 | 402 | 699 | 493 | 465 | 469 | 414 | 502 | 199 | 149 | 126 | 234 | 265 | 271 | 360 | 82 | 310 | 510 | 469 | 472 | 329 | 422 | Dct |
| Dctd | 1414 | 1405 | 1542 | 1258 | 1021 | 1004 | 1075 | 1299 | 1521 | 1277 | 1593 | 1604 | 1591 | 1402 | 1350 | 1589 | 1325 | 1307 | 1099 | 1098 | 926 | 1384 | 1555 | 1649 | 1168 | 2147 | 1541 | 1620 | Dctd |
| Dctn1 | 13489 | 17652 | 18651 | 20944 | 19992 | 17109 | 16777 | 11617 | 14411 | 16513 | 15337 | 16252 | 15715 | 15382 | 13718 | 16733 | 20982 | 20881 | 20677 | 14672 | 17889 | 16289 | 13083 | 16766 | 16904 | 14897 | 13611 | 14709 | Dctn1 |
| Dctn2 | 11236 | 13040 | 12803 | 14270 | 13961 | 12762 | 13006 | 12634 | 13353 | 14681 | 13446 | 14192 | 13192 | 14742 | 11905 | 11695 | 14188 | 13553 | 14677 | 12600 | 14008 | 14375 | 13145 | 14424 | 15152 | 14259 | 13565 | 13905 | Dctn2 |
| Dctn3 | 3901 | 4768 | 4124 | 5031 | 4248 | 4096 | 4789 | 2820 | 2892 | 3194 | 2812 | 2880 | 2849 | 3626 | 3695 | 3750 | 4480 | 4592 | 4700 | 4561 | 4794 | 2775 | 3075 | 3259 | 3656 | 3113 | 3056 | 3322 | Dctn3 |
| Dctn4 | 7750 | 6862 | 6457 | 6238 | 6661 | 6737 | 7504 | 11363 | 9523 | 8516 | 8678 | 9689 | 9135 | 9810 | 7920 | 6653 | 6712 | 6079 | 6992 | 7933 | 7084 | 10059 | 9745 | 8917 | 9788 | 9308 | 10058 | 10047 | Dctn4 |
| Dctn5 | 6238 | 6278 | 6258 | 5814 | 5276 | 5306 | 5671 | 7330 | 6069 | 6180 | 6119 | 6562 | 6203 | 6673 | 6442 | 5822 | 5993 | 5447 | 5485 | 6149 | 5510 | 6727 | 6109 | 6458 | 6675 | 6858 | 6503 | 6958 | Dctn5 |
| Dctn6 | 2796 | 3085 | 2966 | 3111 | 3215 | 3131 | 3268 | 3335 | 3438 | 3681 | 3778 | 3631 | 3484 | 3865 | 3045 | 3001 | 3087 | 3034 | 3075 | 3506 | 3370 | 3317 | 3460 | 3720 | 3830 | 3794 | 3698 | 3648 | Dctn6 |
| Dctpp1 | 3552 | 4607 | 4128 | 4212 | 2955 | 3380 | 3498 | 2109 | 2315 | 2134 | 2130 | 2702 | 2469 | 2802 | 3294 | 3291 | 3898 | 3684 | 4030 | 4295 | 3780 | 1424 | 2128 | 2495 | 2506 | 3449 | 3181 | 3160 | Dctpp1 |
| Dcun1d1 | 5358 | 4038 | 5081 | 5025 | 4426 | 4643 | 5376 | 6361 | 5109 | 4368 | 4778 | 4993 | 5101 | 5764 | 5714 | 4878 | 4632 | 4625 | 4739 | 5789 | 5110 | 4979 | 5259 | 4930 | 4952 | 5684 | 5484 | 5801 | Dcun1d1 |
| Dcun1d2 | 3135 | 4101 | 2998 | 3596 | 3254 | 3238 | 3356 | 2860 | 2746 | 2646 | 2608 | 2366 | 2514 | 2527 | 3277 | 3477 | 3474 | 3607 | 3259 | 3621 | 3601 | 2520 | 2776 | 2529 | 2758 | 2558 | 2547 | 2481 | Dcun1d2 |
| Dcun1d3 | 2067 | 1963 | 2268 | 2333 | 2311 | 2302 | 2525 | 2204 | 1814 | 2017 | 1968 | 1832 | 1813 | 1961 | 2106 | 1997 | 2118 | 2415 | 2387 | 2200 | 2379 | 2269 | 1849 | 1831 | 2344 | 1609 | 1928 | 1969 | Dcun1d3 |
| Dcun1d4 | 10405 | 9502 | 9461 | 9407 | 9767 | 9714 | 10347 | 6690 | 5407 | 5474 | 5195 | 6597 | 6045 | 6786 | 11012 | 9392 | 9591 | 9341 | 9239 | 10227 | 9764 | 6542 | 5344 | 5583 | 6286 | 6010 | 6694 | 6781 | Dcun1d4 |
| Dcun1d5 | 7664 | 5970 | 6601 | 6527 | 5731 | 5971 | 7186 | 6944 | 5628 | 4928 | 5391 | 5471 | 5279 | 5863 | 8057 | 6865 | 6278 | 6005 | 6317 | 7646 | 6719 | 5533 | 5768 | 5862 | 5084 | 6411 | 5675 | 6087 | Dcun1d5 |
| Dcx | 31132 | 30083 | 22223 | 43250 | 68476 | 60987 | 64477 | 90966 | 78567 | 78125 | 70996 | 66549 | 64279 | 67724 | 46677 | 30951 | 35790 | 49737 | 53073 | 59566 | 65865 | 92779 | 70520 | 59506 | 92414 | 35894 | 61673 | 56926 | Dcx |
| Dcxr | 732 | 963 | 830 | 770 | 513 | 586 | 507 | 153 | 166 | 153 | 176 | 196 | 151 | 149 | 783 | 884 | 827 | 666 | 613 | 574 | 449 | 127 | 204 | 192 | 118 | 205 | 248 | 157 | Dcxr |
| Dda1 | 3698 | 5032 | 4732 | 5207 | 5331 | 4906 | 4539 | 3742 | 3962 | 4879 | 4382 | 4535 | 4135 | 4084 | 3845 | 3998 | 5421 | 4970 | 5449 | 4359 | 4793 | 4799 | 4042 | 4666 | 4890 | 4542 | 4014 | 4016 | Dda1 |
| Ddah1 | 4450 | 4438 | 3208 | 5141 | 5746 | 5831 | 7777 | 6815 | 7742 | 6973 | 6393 | 7303 | 3889 | 7403 | 5288 | 3358 | 4753 | 6096 | 6368 | 5593 | 6965 | 7117 | 6095 | 7394 | 7949 | 7965 | 5497 | 5879 | Ddah1 |
| Ddah2 | 11845 | 15457 | 15951 | 14615 | 12777 | 13194 | 11838 | 11645 | 12744 | 12758 | 12103 | 16202 | 15631 | 14674 | 11807 | 12153 | 16584 | 12813 | 16048 | 13910 | 12292 | 10382 | 12520 | 15465 | 14430 | 18432 | 17951 | 16470 | Ddah2 |
| Ddb1 | 70166 | 71011 | 59568 | 53284 | 48444 | 55190 | 50673 | 57338 | 57314 | 51641 | 52450 | 61087 | 60585 | 57595 | 65577 | 62481 | 58301 | 52683 | 55284 | 52884 | 53495 | 50542 | 57731 | 53411 | 52906 | 59663 | 64232 | 63502 | Ddb1 |
| Ddb2 | 1570 | 1475 | 1419 | 1456 | 1394 | 1459 | 1583 | 1116 | 1194 | 1088 | 1190 | 1030 | 1102 | 1081 | 1499 | 1431 | 1586 | 1525 | 1434 | 1424 | 1548 | 1277 | 1201 | 1126 | 1161 | 1029 | 972 | 1059 | Ddb2 |
| Ddc | 558 | 597 | 797 | 663 | 637 | 756 | 938 | 913 | 869 | 864 | 972 | 910 | 871 | 1017 | 620 | 483 | 640 | 638 | 782 | 986 | 821 | 889 | 1025 | 941 | 974 | 593 | 1178 | 1105 | Ddc |
| Ddhd1 | 5569 | 5207 | 5102 | 6509 | 7774 | 7465 | 8133 | 4722 | 4280 | 4524 | 4565 | 4134 | 3855 | 4423 | 5934 | 4939 | 5636 | 6943 | 7020 | 6638 | 8068 | 4929 | 3840 | 4104 | 4810 | 3191 | 3920 | 3948 | Ddhd1 |
| Ddhd2 | 3752 | 2856 | 2960 | 3351 | 3724 | 3804 | 3831 | 4247 | 4081 | 4019 | 4520 | 4035 | 4205 | 4030 | 3719 | 3393 | 3321 | 3706 | 3348 | 3401 | 3573 | 4629 | 3818 | 3789 | 4320 | 3211 | 4061 | 3687 | Ddhd2 |
| Ddi2 | 7998 | 8041 | 9440 | 9383 | 8030 | 7819 | 7829 | 5719 | 5841 | 6578 | 7006 | 6669 | 7214 | 7383 | 7734 | 9134 | 8810 | 8948 | 7476 | 6598 | 7901 | 6375 | 5567 | 6690 | 6752 | 6483 | 6964 | 7266 | Ddi2 |
| Ddias | 2263 | 1773 | 2187 | 1689 | 1087 | 1246 | 1634 | 1404 | 1228 | 1094 | 979 | 1494 | 1368 | 1495 | 2023 | 2079 | 1735 | 1425 | 1393 | 1692 | 1382 | 911 | 1144 | 1351 | 1192 | 1733 | 1539 | 1625 | Ddias |
| Ddit3 | 602 | 839 | 612 | 754 | 524 | 590 | 670 | 399 | 430 | 466 | 442 | 402 | 366 | 435 | 582 | 622 | 788 | 677 | 563 | 652 | 663 | 485 | 413 | 561 | 445 | 433 | 374 | 411 | Ddit3 |
| Ddit4 | 3893 | 6020 | 6862 | 8288 | 5827 | 4743 | 5405 | 3903 | 3891 | 5318 | 4586 | 3779 | 3742 | 4901 | 4000 | 4353 | 7788 | 6520 | 6251 | 4643 | 6463 | 3312 | 4119 | 5985 | 5046 | 5110 | 3540 | 4417 | Ddit4 |
| Ddit4l | 950 | 732 | 828 | 642 | 639 | 570 | 772 | 905 | 761 | 661 | 561 | 528 | 465 | 507 | 657 | 723 | 764 | 710 | 768 | 618 | 658 | 374 | 672 | 617 | 419 | 781 | 494 | 525 | Ddit4l |
| Ddn | 138 | 215 | 227 | 259 | 288 | 227 | 197 | 144 | 138 | 74 | 77 | 85 | 155 | 55 | 157 | 188 | 243 | 267 | 213 | 268 | 238 | 159 | 188 | 92 | 119 | 16 | 66 | 72 | Ddn |
| Ddo | 38 | 20 | 23 | 14 | 47 | 51 | 25 | 28 | 41 | 47 | 24 | 39 | 52 | 21 | 29 | 51 | 20 | 48 | 41 | 50 | 49 | 25 | 54 | 22 | 13 | 19 | 24 | 15 | Ddo |
| Ddost | 8574 | 9626 | 8457 | 7241 | 6372 | 7129 | 6244 | 7288 | 9035 | 7746 | 8590 | 8121 | 8338 | 7369 | 8247 | 9480 | 7358 | 7279 | 6742 | 6643 | 6380 | 7386 | 9689 | 7852 | 7470 | 9323 | 8094 | 8265 | Ddost |
| Ddr1 | 5195 | 7568 | 9584 | 7825 | 8318 | 7497 | 6299 | 11307 | 12013 | 13843 | 11455 | 17926 | 15289 | 15565 | 5794 | 6203 | 9810 | 7129 | 9483 | 6859 | 6676 | 13530 | 12290 | 16542 | 14831 | 17485 | 17168 | 16730 | Ddr1 |
| Ddr2 | 6850 | 3943 | 4011 | 2653 | 1502 | 3552 | 993 | 7394 | 9742 | 7142 | 12853 | 5748 | 9700 | 5572 | 5242 | 7628 | 2795 | 3301 | 1124 | 1456 | 1206 | 6423 | 11185 | 6755 | 5398 | 6130 | 6557 | 5607 | Ddr2 |
| Ddrgk1 | 1083 | 1207 | 1434 | 1453 | 1676 | 1359 | 1403 | 1942 | 2257 | 2494 | 2805 | 2454 | 2318 | 2415 | 1116 | 1149 | 1475 | 1333 | 1671 | 1432 | 1475 | 2381 | 2352 | 2842 | 2805 | 2606 | 2377 | 2572 | Ddrgk1 |
| Ddt | 129 | 164 | 123 | 96 | 102 | 107 | 133 | 51 | 48 | 102 | 51 | 61 | 50 | 70 | 119 | 105 | 156 | 75 | 105 | 121 | 120 | 58 | 56 | 69 | 68 | 73 | 77 | 93 | Ddt |
| Ddx1 | 13565 | 12133 | 13314 | 11178 | 10810 | 11205 | 12876 | 16637 | 14364 | 13814 | 12936 | 16204 | 14452 | 17153 | 13280 | 11689 | 12135 | 11189 | 11387 | 12715 | 11850 | 13670 | 14057 | 15045 | 14686 | 18251 | 16636 | 17971 | Ddx1 |
| Ddx10 | 6207 | 5233 | 6253 | 5368 | 4579 | 4748 | 5072 | 3441 | 3040 | 3003 | 2971 | 3366 | 3074 | 3292 | 6065 | 5717 | 5295 | 5157 | 4478 | 5328 | 4774 | 2777 | 3036 | 3181 | 3096 | 3398 | 3288 | 3474 | Ddx10 |
| Ddx11 | 5076 | 5206 | 4899 | 4760 | 3144 | 4017 | 4222 | 2379 | 2595 | 2507 | 2614 | 3574 | 3269 | 3430 | 4221 | 4697 | 4483 | 4196 | 4044 | 3680 | 3963 | 1910 | 2300 | 3179 | 2700 | 3532 | 3606 | 3509 | Ddx11 |
| Ddx17 | 88941 | 81731 | 79153 | 87104 | 87395 | 91893 | 92067 | 110315 | 99170 | 100122 | 105806 | 111927 | 107375 | 114883 | 87818 | 76184 | 84700 | 85580 | 91806 | 89558 | 95316 | 105976 | 95673 | 101752 | 113386 | 82616 | 116524 | 112101 | Ddx17 |
| Ddx18 | 3799 | 4248 | 4355 | 3507 | 2852 | 3135 | 3267 | 4228 | 4226 | 3771 | 3846 | 4747 | 4212 | 4677 | 3520 | 3903 | 3817 | 3267 | 3230 | 3622 | 3149 | 3187 | 4088 | 4461 | 4081 | 5528 | 4862 | 5039 | Ddx18 |
| Ddx19a | 5663 | 6365 | 6881 | 5730 | 5045 | 4805 | 5848 | 3865 | 3996 | 4626 | 4630 | 5995 | 5220 | 6313 | 5329 | 5412 | 6971 | 5786 | 5448 | 4545 | 5101 | 4147 | 3924 | 5293 | 4798 | 6458 | 5313 | 6649 | Ddx19a |
| Ddx19b | 3632 | 3389 | 3267 | 2956 | 3298 | 3211 | 3368 | 3635 | 3104 | 2956 | 2975 | 3952 | 3701 | 4043 | 3505 | 3123 | 3363 | 3113 | 3269 | 3252 | 3156 | 3322 | 3007 | 3587 | 3641 | 3103 | 4270 | 3947 | Ddx19b |
| Ddx20 | 4459 | 4005 | 3467 | 3708 | 3201 | 3413 | 3733 | 3876 | 3223 | 2996 | 2778 | 3747 | 3273 | 3644 | 4398 | 4101 | 3443 | 3499 | 3521 | 3717 | 3588 | 3275 | 3117 | 3185 | 3192 | 3940 | 3714 | 3852 | Ddx20 |

| | | | | | | | | | | | | | | | | | | | | | | | | | | | | | |
|---|---|---|---|---|---|---|---|---|---|---|---|---|---|---|---|---|---|---|---|---|---|---|---|---|---|---|---|---|---|
| Ddx21 | 16295 | 16308 | 14143 | 13036 | 10586 | 12025 | 12336 | 13345 | 11900 | 10946 | 11162 | 13344 | 12591 | 13839 | 15602 | 15021 | 12506 | 12265 | 11065 | 13298 | 11631 | 10691 | 11327 | 12055 | 12045 | 13547 | 14686 | 13992 | Ddx21 |
| Ddx23 | 7545 | 8598 | 10031 | 8527 | 7975 | 7759 | 7490 | 8264 | 8989 | 9259 | 8706 | 10526 | 9878 | 10116 | 7247 | 7623 | 9332 | 7776 | 8892 | 7861 | 7667 | 7912 | 8690 | 10421 | 9662 | 11479 | 10828 | 10839 | Ddx23 |
| Ddx24 | 8612 | 10095 | 8333 | 8512 | 8038 | 7953 | 8160 | 7215 | 7556 | 7813 | 7075 | 8072 | 7654 | 7733 | 8227 | 8273 | 8196 | 8115 | 8273 | 8208 | 8300 | 6894 | 7236 | 7770 | 7958 | 8070 | 8133 | 8275 | Ddx24 |
| Ddx25 | 2030 | 2082 | 1800 | 2348 | 3296 | 2594 | 3095 | 3123 | 2860 | 2852 | 2473 | 2262 | 2141 | 2358 | 2331 | 1858 | 1874 | 2615 | 2805 | 2882 | 3012 | 3434 | 2583 | 2315 | 2847 | 1826 | 2097 | 2074 | Ddx25 |
| Ddx27 | 4284 | 5030 | 5507 | 4914 | 3975 | 4054 | 4440 | 3042 | 3405 | 3589 | 3226 | 4403 | 3638 | 4310 | 4247 | 4415 | 5003 | 4408 | 4305 | 4001 | 4320 | 3190 | 3180 | 4120 | 3619 | 4743 | 4170 | 4457 | Ddx27 |
| Ddx28 | 786 | 1024 | 906 | 749 | 727 | 829 | 786 | 847 | 952 | 833 | 818 | 924 | 812 | 884 | 787 | 774 | 889 | 754 | 913 | 783 | 778 | 896 | 909 | 834 | 876 | 1127 | 877 | 878 | Ddx28 |
| Ddx31 | 1577 | 1748 | 1336 | 1499 | 1296 | 1367 | 1344 | 1265 | 1124 | 972 | 1038 | 1183 | 1139 | 1149 | 1481 | 1607 | 1374 | 1473 | 1428 | 1473 | 1347 | 1068 | 1116 | 1115 | 1143 | 1298 | 1158 | 1244 | Ddx31 |
| Ddx39a | 5269 | 5949 | 5883 | 5123 | 4124 | 4461 | 4456 | 4452 | 4367 | 4127 | 3862 | 6014 | 5256 | 5451 | 4803 | 5201 | 5532 | 4318 | 4960 | 4835 | 4391 | 3459 | 4071 | 5350 | 4628 | 7027 | 6358 | 6344 | Ddx39a |
| Ddx39b | 17154 | 20394 | 21185 | 18785 | 18538 | 18083 | 19600 | 22112 | 25828 | 26939 | 25215 | 33435 | 28846 | 33258 | 15662 | 16658 | 21554 | 17500 | 21589 | 17869 | 19091 | 25185 | 23692 | 30894 | 29530 | 36421 | 32115 | 34234 | Ddx39b |
| Ddx3x | 49189 | 35183 | 35853 | 32070 | 35886 | 40247 | 42039 | 68255 | 59926 | 50953 | 51179 | 66059 | 62438 | 69361 | 51907 | 41821 | 35371 | 34076 | 36246 | 48670 | 41409 | 60771 | 64668 | 61635 | 60520 | 70589 | 80580 | 74375 | Ddx3x |
| Ddx3y | 7459 | 4907 | 5753 | 6398 | 6215 | 7221 | 7940 | 5787 | 6751 | 5338 | 5806 | 6446 | 6445 | 6837 | 15 | 0 | 0 | 0 | 0 | 2 | 0 | 3 | 32 | 0 | 0 | 4 | 0 | 1 | Ddx3y |
| Ddx4 | 305 | 263 | 326 | 392 | 314 | 355 | 351 | 212 | 199 | 225 | 192 | 190 | 240 | 163 | 305 | 272 | 391 | 344 | 287 | 286 | 359 | 190 | 158 | 165 | 176 | 153 | 171 | 160 | Ddx4 |
| Ddx41 | 3429 | 3828 | 3293 | 3383 | 3093 | 3005 | 3248 | 3615 | 3814 | 3812 | 3499 | 4098 | 3565 | 3837 | 3059 | 3344 | 3376 | 3136 | 3313 | 3174 | 3191 | 3384 | 3592 | 3787 | 3793 | 4444 | 3934 | 4109 | Ddx41 |
| Ddx42 | 15476 | 13626 | 15190 | 15007 | 14205 | 14578 | 15193 | 14544 | 13222 | 13982 | 14012 | 15937 | 14466 | 16888 | 15616 | 14160 | 14915 | 14498 | 14039 | 14576 | 14273 | 14156 | 12860 | 15173 | 15386 | 15580 | 16014 | 15958 | Ddx42 |
| Ddx43 | 277 | 213 | 315 | 271 | 327 | 399 | 321 | 73 | 51 | 71 | 98 | 71 | 42 | 67 | 315 | 312 | 238 | 310 | 296 | 288 | 295 | 109 | 69 | 61 | 72 | 51 | 53 | 62 | Ddx43 |
| Ddx46 | 23836 | 21395 | 19653 | 20508 | 19367 | 21689 | 22693 | 17046 | 14885 | 14884 | 14378 | 17023 | 16147 | 17999 | 23911 | 22003 | 19883 | 20265 | 19368 | 22898 | 22518 | 13754 | 13893 | 14963 | 15552 | 15242 | 17235 | 16962 | Ddx46 |
| Ddx47 | 5045 | 4979 | 4716 | 4637 | 4123 | 4504 | 4523 | 5893 | 5699 | 5580 | 5505 | 6218 | 5924 | 6158 | 4634 | 4299 | 4715 | 4229 | 4562 | 4638 | 4322 | 5168 | 5363 | 5830 | 5445 | 6602 | 6157 | 6337 | Ddx47 |
| Ddx49 | 2864 | 3992 | 3218 | 3353 | 2738 | 2870 | 2662 | 2330 | 2616 | 2503 | 2461 | 2720 | 2508 | 2357 | 2852 | 3297 | 3314 | 3127 | 3039 | 2846 | 2847 | 2217 | 2614 | 2664 | 2704 | 3148 | 2731 | 2716 | Ddx49 |
| Ddx5 | 74058 | 63475 | 73504 | 70610 | 78983 | 75598 | 88357 | 128582 | 105177 | 108426 | 103415 | 123624 | 111468 | 131907 | 80926 | 64119 | 76166 | 72081 | 75768 | 84048 | 81584 | 121269 | 100349 | 116913 | 121632 | 121425 | 119463 | 123441 | Ddx5 |
| Ddx50 | 11575 | 8983 | 9742 | 9200 | 8314 | 9202 | 9971 | 9884 | 9410 | 8404 | 9843 | 8941 | 9045 | 9555 | 11402 | 10192 | 9136 | 9118 | 8389 | 10076 | 8786 | 8696 | 9417 | 9595 | 8747 | 9717 | 9174 | 9400 | Ddx50 |
| Ddx51 | 2140 | 2096 | 1938 | 2061 | 1777 | 1887 | 1963 | 2133 | 2122 | 2004 | 2167 | 2581 | 2326 | 2316 | 1942 | 1980 | 1801 | 1865 | 1825 | 1726 | 1885 | 2055 | 2042 | 2124 | 2113 | 2220 | 2420 | 2299 | Ddx51 |
| Ddx52 | 4474 | 3640 | 3881 | 3209 | 3187 | 3327 | 4027 | 5945 | 4789 | 4345 | 4322 | 5582 | 5034 | 5726 | 4675 | 3826 | 3501 | 2864 | 3422 | 4352 | 3556 | 4338 | 4842 | 4788 | 4482 | 5919 | 5944 | 5948 | Ddx52 |
| Ddx54 | 4818 | 6264 | 7611 | 6720 | 5773 | 5581 | 5377 | 3866 | 4697 | 5478 | 5184 | 6243 | 5451 | 5735 | 4555 | 5648 | 7544 | 6198 | 6371 | 4801 | 5617 | 4607 | 4734 | 6064 | 5474 | 6955 | 5670 | 5868 | Ddx54 |
| Ddx55 | 2606 | 2774 | 2869 | 3292 | 2846 | 2746 | 2971 | 2214 | 2210 | 2733 | 2524 | 2701 | 2677 | 3021 | 2505 | 2484 | 2965 | 3174 | 2865 | 2611 | 2936 | 2689 | 2163 | 2850 | 3080 | 2443 | 2487 | 2880 | Ddx55 |
| Ddx56 | 2707 | 3027 | 2851 | 2647 | 2583 | 2644 | 2690 | 2510 | 2949 | 2825 | 2886 | 3598 | 3197 | 3252 | 2454 | 2772 | 2707 | 2597 | 2766 | 2308 | 2630 | 2613 | 2868 | 3063 | 3059 | 3729 | 3510 | 3603 | Ddx56 |
| Ddx58 | 864 | 543 | 605 | 430 | 518 | 600 | 516 | 578 | 692 | 667 | 909 | 449 | 634 | 474 | 750 | 771 | 521 | 471 | 481 | 631 | 520 | 495 | 826 | 573 | 435 | 553 | 499 | 485 | Ddx58 |
| Ddx59 | 639 | 724 | 643 | 732 | 574 | 569 | 626 | 874 | 961 | 863 | 691 | 944 | 891 | 996 | 774 | 639 | 688 | 576 | 632 | 702 | 646 | 795 | 913 | 913 | 839 | 1041 | 1074 | 1010 | Ddx59 |
| Ddx6 | 23598 | 17797 | 22224 | 23552 | 25598 | 24042 | 28253 | 39787 | 34826 | 37316 | 38980 | 37866 | 38166 | 44878 | 22740 | 18235 | 23235 | 22734 | 25647 | 24508 | 28593 | 35511 | 34502 | 40603 | 42318 | 34811 | 40883 | 44794 | Ddx6 |
| Ddx60 | 39 | 26 | 12 | 24 | 24 | 44 | 17 | 50 | 44 | 78 | 75 | 64 | 60 | 61 | 29 | 40 | 22 | 41 | 33 | 58 | 31 | 62 | 101 | 37 | 45 | 75 | 80 | 65 | Ddx60 |
| Deaf1 | 1991 | 2475 | 2166 | 2744 | 3463 | 2731 | 2618 | 3533 | 3756 | 4175 | 4023 | 3796 | 3830 | 3604 | 2090 | 2106 | 2356 | 2542 | 3162 | 2741 | 2819 | 4415 | 3638 | 4138 | 4533 | 4069 | 3574 | 3628 | Deaf1 |
| Decr1 | 1644 | 1499 | 1755 | 1222 | 1172 | 1254 | 1143 | 1895 | 1676 | 1452 | 1802 | 1842 | 1983 | 1852 | 1631 | 1700 | 1322 | 1085 | 1116 | 1290 | 1188 | 1462 | 1972 | 1574 | 1525 | 1961 | 2186 | 1919 | Decr1 |
| Decr2 | 1142 | 1191 | 1307 | 1277 | 1244 | 1190 | 1241 | 1523 | 1643 | 1846 | 1797 | 1654 | 1761 | 1864 | 1077 | 1243 | 1265 | 1132 | 1216 | 1137 | 1146 | 1569 | 1709 | 1753 | 1695 | 1979 | 1668 | 1760 | Decr2 |
| Dedd | 2312 | 2463 | 2893 | 2711 | 2673 | 2340 | 2705 | 2742 | 2671 | 2990 | 2944 | 2902 | 2834 | 3285 | 2309 | 2313 | 3099 | 2530 | 2763 | 2444 | 2706 | 2836 | 2640 | 3145 | 3028 | 3164 | 3122 | 3125 | Dedd |
| Dedd2 | 3008 | 3416 | 3310 | 4214 | 3435 | 3314 | 3300 | 1362 | 1359 | 1795 | 1535 | 1309 | 1309 | 1343 | 2699 | 3004 | 3682 | 4004 | 3702 | 3008 | 3444 | 1579 | 1370 | 1730 | 1737 | 1430 | 1221 | 1321 | Dedd2 |
| Def6 | 98 | 85 | 133 | 96 | 97 | 111 | 75 | 187 | 263 | 256 | 274 | 164 | 235 | 179 | 92 | 122 | 105 | 77 | 102 | 96 | 104 | 232 | 330 | 242 | 207 | 267 | 180 | 198 | Def6 |
| Def8 | 2413 | 2540 | 2247 | 2046 | 2644 | 2608 | 2137 | 3861 | 4411 | 3839 | 3643 | 4043 | 3967 | 3780 | 2211 | 2111 | 2196 | 2115 | 2544 | 2372 | 2326 | 4002 | 4335 | 3701 | 4062 | 4236 | 3962 | 3779 | Def8 |
| Defa17 | 0 | 0 | 0 | 0 | 0 | 0 | 0 | 0 | 0 | 0 | 0 | 0 | 0 | 0 | 1 | 0 | 0 | 0 | 0 | 0 | 0 | 0 | 0 | 0 | 0 | 0 | 0 | 0 | Defa17 |
| Defa21 | 0 | 0 | 0 | 0 | 2 | 0 | 0 | 0 | 0 | 0 | 0 | 0 | 0 | 0 | 0 | 0 | 0 | 0 | 0 | 0 | 0 | 0 | 0 | 0 | 0 | 0 | 0 | 0 | Defa21 |
| Defa39 | 0 | 0 | 0 | 0 | 0 | 0 | 0 | 0 | 0 | 0 | 0 | 0 | 0 | 0 | 1 | 0 | 0 | 0 | 0 | 0 | 0 | 0 | 0 | 0 | 0 | 0 | 0 | 0 | Defa39 |
| Defb1 | 2 | 1 | 0 | 1 | 0 | 0 | 0 | 1 | 3 | 1 | 11 | 4 | 7 | 2 | 0 | 2 | 2 | 0 | 0 | 0 | 0 | 0 | 0 | 2 | 0 | 0 | 12 | 1 | Defb1 |
| Defb10 | 1 | 0 | 0 | 0 | 0 | 0 | 0 | 1 | 0 | 3 | 1 | 0 | 0 | 0 | 0 | 2 | 0 | 4 | 0 | 0 | 0 | 0 | 3 | 0 | 0 | 0 | 0 | 0 | Defb10 |
| Defb11 | 22 | 29 | 0 | 0 | 30 | 21 | 9 | 42 | 36 | 47 | 24 | 29 | 70 | 12 | 20 | 31 | 1 | 0 | 5 | 3 | 18 | 33 | 63 | 17 | 8 | 5 | 8 | 4 | Defb11 |
| Defb13 | 0 | 0 | 0 | 0 | 0 | 0 | 0 | 0 | 0 | 0 | 0 | 0 | 0 | 0 | 0 | 0 | 0 | 0 | 2 | 0 | 0 | 0 | 0 | 0 | 0 | 0 | 1 | 0 | Defb13 |
| Defb15 | 0 | 0 | 0 | 0 | 3 | 0 | 0 | 0 | 2 | 0 | 0 | 0 | 0 | 0 | 1 | 0 | 0 | 0 | 0 | 0 | 0 | 0 | 0 | 0 | 0 | 0 | 0 | 0 | Defb15 |
| Defb2 | 0 | 0 | 0 | 0 | 0 | 0 | 0 | 0 | 1 | 0 | 0 | 0 | 2 | 0 | 0 | 1 | 0 | 0 | 0 | 0 | 0 | 2 | 1 | 0 | 0 | 0 | 0 | 0 | Defb2 |
| Defb20 | 0 | 1 | 0 | 0 | 2 | 0 | 0 | 0 | 1 | 0 | 1 | 0 | 0 | 1 | 4 | 0 | 0 | 0 | 1 | 2 | 1 | 0 | 0 | 2 | 0 | 0 | 0 | 0 | Defb20 |
| Defb25 | 3 | 1 | 0 | 0 | 0 | 2 | 1 | 0 | 3 | 3 | 1 | 0 | 1 | 0 | 3 | 0 | 0 | 0 | 3 | 0 | 2 | 0 | 1 | 1 | 0 | 0 | 0 | 1 | Defb25 |
| Defb28 | 0 | 0 | 0 | 0 | 2 | 0 | 0 | 0 | 0 | 0 | 0 | 0 | 0 | 0 | 0 | 0 | 0 | 0 | 0 | 0 | 0 | 0 | 0 | 0 | 0 | 0 | 0 | 0 | Defb28 |
| Defb3 | 0 | 0 | 0 | 0 | 0 | 0 | 0 | 0 | 0 | 0 | 0 | 0 | 0 | 0 | 0 | 0 | 0 | 4 | 0 | 0 | 0 | 0 | 0 | 0 | 0 | 0 | 0 | 0 | Defb3 |
| Defb30 | 0 | 1 | 0 | 6 | 0 | 0 | 2 | 0 | 0 | 1 | 0 | 0 | 2 | 0 | 0 | 1 | 0 | 0 | 0 | 0 | 0 | 0 | 0 | 0 | 0 | 0 | 0 | 0 | Defb30 |
| Defb33 | 0 | 0 | 0 | 0 | 0 | 0 | 0 | 0 | 0 | 0 | 0 | 0 | 0 | 0 | 0 | 1 | 0 | 0 | 0 | 0 | 0 | 0 | 0 | 0 | 0 | 0 | 0 | 0 | Defb33 |
| Defb35 | 0 | 2 | 0 | 0 | 2 | 1 | 0 | 0 | 1 | 0 | 0 | 2 | 5 | 1 | 0 | 0 | 0 | 0 | 0 | 0 | 0 | 0 | 3 | 0 | 0 | 0 | 0 | 0 | Defb35 |
| Defb36 | 0 | 0 | 0 | 0 | 0 | 0 | 0 | 0 | 0 | 0 | 0 | 0 | 0 | 0 | 0 | 0 | 0 | 0 | 0 | 0 | 0 | 0 | 0 | 0 | 0 | 2 | 0 | 0 | Defb36 |
| Defb41 | 0 | 0 | 0 | 0 | 0 | 0 | 0 | 0 | 0 | 0 | 0 | 0 | 0 | 0 | 0 | 0 | 0 | 0 | 0 | 0 | 0 | 0 | 0 | 0 | 0 | 0 | 0 | 1 | Defb41 |
| Defb42 | 1 | 3 | 2 | 0 | 2 | 4 | 3 | 0 | 4 | 3 | 4 | 4 | 2 | 2 | 3 | 1 | 1 | 2 | 2 | 1 | 3 | 2 | 2 | 2 | 0 | 0 | 3 | 8 | Defb42 |
| Defb43 | 2 | 0 | 0 | 0 | 2 | 0 | 0 | 0 | 0 | 0 | 0 | 0 | 0 | 0 | 0 | 0 | 0 | 0 | 0 | 0 | 0 | 0 | 0 | 0 | 0 | 0 | 0 | 0 | Defb43 |
| Defb47 | 3 | 0 | 2 | 8 | 2 | 3 | 0 | 0 | 2 | 0 | 0 | 0 | 0 | 0 | 0 | 5 | 0 | 0 | 0 | 0 | 4 | 0 | 3 | 0 | 0 | 1 | 1 | 4 | Defb47 |
| Defb48 | 0 | 1 | 0 | 0 | 0 | 0 | 1 | 0 | 0 | 0 | 0 | 0 | 1 | 0 | 0 | 0 | 0 | 0 | 0 | 2 | 0 | 0 | 1 | 0 | 0 | 0 | 0 | 0 | Defb48 |
| Defb9 | 1 | 0 | 0 | 0 | 1 | 0 | 0 | 1 | 2 | 8 | 1 | 2 | 0 | 0 | 0 | 0 | 0 | 0 | 0 | 0 | 0 | 2 | 4 | 0 | 0 | 0 | 0 | 0 | Defb9 |
| Degs1 | 3313 | 2973 | 2548 | 2488 | 2287 | 2495 | 2364 | 2722 | 2831 | 2172 | 2810 | 2280 | 2541 | 2312 | 3335 | 3192 | 2372 | 2815 | 2216 | 2548 | 2419 | 2568 | 3172 | 2177 | 2242 | 2403 | 2244 | 2505 | Degs1 |
| Degs2 | 0 | 7 | 12 | 15 | 11 | 4 | 4 | 9 | 1 | 1 | 3 | 7 | 6 | 2 | 1 | 3 | 5 | 12 | 13 | 3 | 13 | 0 | 3 | 4 | 5 | 4 | 2 | 2 | Degs2 |
| Dek | 21554 | 15282 | 24813 | 18172 | 16175 | 18183 | 21773 | 19975 | 16738 | 16199 | 16220 | 24946 | 21227 | 25790 | 21522 | 18626 | 20668 | 17213 | 18729 | 19316 | 18453 | 16035 | 15812 | 21226 | 18379 | 31336 | 25598 | 27760 | Dek |
| Dele1 | 1977 | 2249 | 2218 | 2316 | 2570 | 2323 | 2405 | 3817 | 4330 | 4614 | 4358 | 5042 | 4843 | 5138 | 1873 | 2034 | 2275 | 2220 | 2614 | 2094 | 2325 | 4517 | 4009 | 4850 | 4863 | 4580 | 5025 | 4883 | Dele1 |
| Denn2b | 2428 | 2583 | 2797 | 2395 | 2240 | 2292 | 1888 | 2186 | 2422 | 2348 | 2735 | 2427 | 3049 | 2093 | 2405 | 2587 | 2488 | 2322 | 2260 | 1802 | 1970 | 2298 | 2740 | 2092 | 1955 | 2370 | 2528 | 2403 | Denn2b |
| Dennd10 | 2536 | 1970 | 2128 | 2073 | 1958 | 1960 | 2363 | 2246 | 2144 | 1947 | 1922 | 2002 | 1995 | 1959 | 2715 | 2407 | 2021 | 1961 | 1983 | 2563 | 2120 | 2017 | 2252 | 1846 | 1842 | 2023 | 2027 | 2102 | Dennd10 |
| Dennd11 | 4112 | 3432 | 3101 | 4366 | 5605 | 5748 | 6819 | 6858 | 6554 | 5985 | 5441 | 6369 | 5392 | 6909 | 3967 | 3167 | 3676 | 4824 | 5868 | 5838 | 6423 | 5312 | 5482 | 6767 | 8221 | 4670 | 6180 | 5944 | Dennd11 |
| Dennd1a | 2811 | 3216 | 2910 | 3362 | 3328 | 3450 | 3384 | 2845 | 3408 | 3449 | 3401 | 4139 | 3317 | 3865 | 2585 | 2660 | 3176 | 3497 | 3654 | 3329 | 3430 | 3506 | 3391 | 3902 | 3951 | 3970 | 3928 | 3538 | Dennd1a |
| Dennd1b | 1938 | 1533 | 1601 | 1633 | 1973 | 2030 | 2020 | 3263 | 2606 | 2362 | 2591 | 3166 | 3289 | 3280 | 2082 | 1600 | 1632 | 1539 | 1963 | 2133 | 2061 | 2650 | 2633 | 2476 | 3020 | 2276 | 3571 | 3426 | Dennd1b |
| Dennd1c | 429 | 495 | 537 | 530 | 619 | 571 | 609 | 601 | 626 | 647 | 515 | 746 | 467 | 775 | 427 | 347 | 661 | 513 | 584 | 593 | 645 | 693 | 480 | 712 | 699 | 818 | 594 | 713 | Dennd1c |
| Dennd2a | 3602 | 4586 | 6385 | 5122 | 5153 | 5072 | 4599 | 4065 | 4371 | 5091 | 5427 | 6742 | 6517 | 6236 | 3862 | 4485 | 5744 | 4831 | 5594 | 4166 | 4878 | 5140 | 4886 | 5993 | 5530 | 6185 | 6433 | 7243 | Dennd2a |
| Dennd2c | 1322 | 1081 | 945 | 831 | 682 | 820 | 672 | 600 | 714 | 609 | 623 | 399 | 650 | 462 | 1129 | 1363 | 852 | 839 | 704 | 694 | 610 | 394 | 825 | 509 | 353 | 465 | 531 | 589 | Dennd2c |
| Dennd2d | 136 | 91 | 82 | 36 | 63 | 60 | 34 | 154 | 119 | 109 | 65 | 49 | 70 | 56 | 108 | 105 | 59 | 67 | 46 | 47 | 46 | 23 | 118 | 102 | 33 | 104 | 52 | 46 | Dennd2d |
| Dennd3 | 1125 | 1220 | 1174 | 1306 | 1063 | 1129 | 952 | 281 | 377 | 328 | 462 | 329 | 371 | 301 | 1062 | 1284 | 1043 | 1279 | 1032 | 907 | 936 | 317 | 403 | 375 | 291 | 330 | 295 | 253 | Dennd3 |
| Dennd4a | 6090 | 5455 | 3477 | 4892 | 4776 | 5398 | 5591 | 3414 | 3154 | 2733 | 2987 | 2758 | 3014 | 2978 | 6252 | 5722 | 4234 | 4601 | 4565 | 5582 | 5482 | 2912 | 3076 | 2346 | 2941 | 1781 | 2926 | 2725 | Dennd4a |
| Dennd4b | 1120 | 1377 | 1916 | 1696 | 1910 | 1679 | 1492 | 1574 | 2063 | 2153 | 2278 | 2510 | 2219 | 2283 | 1096 | 1373 | 1931 | 1690 | 1784 | 1415 | 1465 | 2266 | 1961 | 2272 | 2209 | 2086 | 2280 | 2203 | Dennd4b |
| Dennd4c | 5464 | 4708 | 2850 | 3824 | 3850 | 4736 | 4336 | 3613 | 3496 | 2853 | 3206 | 3200 | 3081 | 3283 | 4952 | 5097 | 3435 | 3914 | 3824 | 4501 | 4480 | 2423 | 3441 | 2491 | 2892 | 2001 | 3347 | 3027 | Dennd4c |
| Dennd5a | 16436 | 18286 | 15232 | 18812 | 19639 | 18525 | 19829 | 15340 | 14926 | 15233 | 14654 | 15578 | 14276 | 15570 | 16991 | 15735 | 17291 | 19013 | 19861 | 17109 | 20344 | 15988 | 14240 | 15805 | 16559 | 13309 | 14407 | 15148 | Dennd5a |
| Dennd5b | 5497 | 4674 | 5155 | 6196 | 7987 | 8185 | 8448 | 7707 | 6283 | 7662 | 7291 | 8487 | 7957 | 8688 | 6111 | 4955 | 5407 | 6354 | 7098 | 6837 | 8151 | 8062 | 5957 | 6979 | 8908 | 5505 | 8577 | 8406 | Dennd5b |
| Dennd6a | 1747 | 1550 | 1594 | 1325 | 1311 | 1469 | 1428 | 1767 | 1682 | 1645 | 1655 | 1909 | 1889 | 1704 | 1689 | 1572 | 1511 | 1432 | 1351 | 1407 | 1432 | 1624 | 1739 | 1604 | 1591 | 1705 | 1937 | 1856 | Dennd6a |

| | | | | | | | | | | | | | | | | | | | | | | | | | | | | | |
|---|---|---|---|---|---|---|---|---|---|---|---|---|---|---|---|---|---|---|---|---|---|---|---|---|---|---|---|---|---|
| Dennd6b | 3164 | 4289 | 4412 | 5952 | 6499 | 5882 | 5519 | 4082 | 4191 | 5229 | 4391 | 5003 | 4787 | 4879 | 3684 | 3646 | 5093 | 5648 | 6287 | 5578 | 5654 | 6029 | 3491 | 4761 | 5878 | 4023 | 4821 | 4587 | Dennd6b |
| Denr | 5123 | 4230 | 5847 | 4660 | 3909 | 4023 | 4550 | 4559 | 3840 | 4625 | 4262 | 4830 | 4549 | 5537 | 5081 | 4186 | 4994 | 4160 | 3998 | 4460 | 4445 | 3962 | 3918 | 4850 | 4348 | 5910 | 4918 | 5569 | Denr |
| Depdc1a | 2231 | 1318 | 2009 | 1448 | 1198 | 1468 | 1663 | 2993 | 2622 | 1979 | 2001 | 2966 | 2721 | 2925 | 2074 | 1882 | 1561 | 1412 | 1496 | 2040 | 1432 | 1734 | 2556 | 2560 | 2253 | 3373 | 3151 | 3204 | Depdc1a |
| Depdc1b | 870 | 1008 | 1067 | 630 | 546 | 596 | 616 | 681 | 645 | 698 | 602 | 1142 | 890 | 1081 | 752 | 732 | 978 | 587 | 699 | 723 | 607 | 578 | 582 | 813 | 713 | 1256 | 1156 | 1098 | Depdc1b |
| Depdc5 | 8245 | 8727 | 6779 | 8128 | 7788 | 7646 | 8260 | 5040 | 5025 | 4983 | 4483 | 5176 | 4705 | 5220 | 8057 | 7795 | 7623 | 8065 | 8182 | 7327 | 8713 | 4751 | 4705 | 4796 | 5140 | 3839 | 5146 | 5258 | Depdc5 |
| Depdc7 | 644 | 573 | 553 | 553 | 394 | 498 | 543 | 158 | 145 | 115 | 147 | 194 | 99 | 162 | 510 | 474 | 562 | 483 | 518 | 487 | 544 | 157 | 132 | 203 | 157 | 234 | 170 | 154 | Depdc7 |
| Depp1 | 29 | 13 | 17 | 12 | 47 | 40 | 32 | 122 | 169 | 139 | 197 | 61 | 84 | 57 | 29 | 27 | 12 | 34 | 36 | 55 | 17 | 129 | 230 | 114 | 153 | 93 | 81 | 109 | Depp1 |
| Deptor | 1162 | 1012 | 922 | 949 | 956 | 880 | 801 | 299 | 397 | 269 | 766 | 184 | 248 | 209 | 1114 | 1368 | 958 | 1026 | 916 | 928 | 893 | 302 | 654 | 234 | 285 | 204 | 205 | 220 | Deptor |
| Dera | 1223 | 1172 | 1278 | 889 | 741 | 878 | 749 | 714 | 816 | 579 | 775 | 897 | 1015 | 941 | 1203 | 1322 | 1077 | 830 | 761 | 828 | 752 | 545 | 859 | 804 | 627 | 1159 | 1027 | 1081 | Dera |
| Derl1 | 5781 | 6220 | 5653 | 5515 | 5310 | 5179 | 5089 | 5100 | 5066 | 4702 | 5008 | 4540 | 4740 | 4271 | 5796 | 5768 | 5668 | 5732 | 5298 | 5655 | 5052 | 5135 | 5228 | 4789 | 4749 | 4706 | 4756 | 4543 | Derl1 |
| Derl2 | 4599 | 4285 | 4284 | 4630 | 4255 | 4720 | 4929 | 5083 | 4519 | 4428 | 4630 | 4667 | 4496 | 4832 | 4822 | 4312 | 4373 | 4677 | 4622 | 5270 | 4825 | 4400 | 4401 | 4664 | 4885 | 4428 | 4799 | 4726 | Derl2 |
| Derl3 | 165 | 191 | 112 | 144 | 49 | 72 | 87 | 32 | 38 | 60 | 51 | 37 | 44 | 28 | 149 | 162 | 145 | 141 | 82 | 109 | 99 | 32 | 47 | 40 | 40 | 51 | 34 | 30 | Derl3 |
| Des | 218 | 131 | 130 | 62 | 103 | 55 | 80 | 143 | 437 | 137 | 2175 | 67 | 146 | 82 | 102 | 288 | 96 | 79 | 63 | 85 | 71 | 287 | 1525 | 109 | 114 | 86 | 80 | 71 | Des |
| Desi1 | 5737 | 6084 | 7014 | 7428 | 6631 | 6300 | 6545 | 3083 | 2953 | 3885 | 3570 | 3905 | 3398 | 3709 | 5616 | 6232 | 6938 | 7288 | 7338 | 5224 | 6638 | 4147 | 3065 | 3740 | 4019 | 4238 | 3296 | 3857 | Desi1 |
| Desi2 | 236 | 174 | 147 | 308 | 293 | 186 | 143 | 96 | 77 | 56 | 235 | 83 | 77 | 70 | 218 | 178 | 159 | 391 | 141 | 120 | 145 | 132 | 181 | 74 | 82 | 78 | 208 | 65 | Desi2 |
| Det1 | 573 | 647 | 709 | 529 | 542 | 486 | 606 | 895 | 990 | 824 | 862 | 920 | 916 | 965 | 587 | 565 | 661 | 491 | 635 | 633 | 558 | 795 | 935 | 932 | 870 | 1032 | 1084 | 1062 | Det1 |
| Deup1 | 116 | 113 | 69 | 53 | 76 | 109 | 98 | 211 | 147 | 139 | 90 | 268 | 321 | 196 | 98 | 170 | 93 | 66 | 83 | 181 | 86 | 190 | 302 | 157 | 159 | 235 | 225 | 206 | Deup1 |
| Dexi | 1531 | 1851 | 1417 | 1575 | 1550 | 1507 | 1395 | 800 | 875 | 1011 | 942 | 947 | 830 | 841 | 1378 | 1396 | 1527 | 1558 | 1711 | 1444 | 1502 | 842 | 854 | 857 | 839 | 1017 | 836 | 849 | Dexi |
| Dffa | 3110 | 3595 | 3312 | 3067 | 2370 | 2479 | 2164 | 1856 | 1928 | 1899 | 1982 | 2164 | 2016 | 1932 | 3080 | 3291 | 3140 | 2639 | 2523 | 2487 | 2369 | 1840 | 1878 | 1968 | 2000 | 2274 | 2106 | 2009 | Dffa |
| Dffb | 666 | 612 | 597 | 589 | 717 | 728 | 670 | 692 | 763 | 852 | 741 | 855 | 856 | 815 | 603 | 513 | 717 | 545 | 669 | 656 | 588 | 829 | 748 | 831 | 822 | 869 | 865 | 780 | Dffb |
| Dgat1 | 2399 | 2479 | 2679 | 2730 | 2925 | 2942 | 2542 | 2648 | 2516 | 2803 | 2759 | 2917 | 2809 | 2711 | 2405 | 2427 | 2738 | 2937 | 2930 | 2588 | 2453 | 3253 | 2423 | 2866 | 3193 | 2933 | 2917 | 2573 | Dgat1 |
| Dgat2 | 806 | 958 | 1045 | 885 | 761 | 794 | 698 | 431 | 504 | 523 | 550 | 437 | 408 | 384 | 874 | 999 | 892 | 901 | 714 | 722 | 704 | 476 | 560 | 455 | 445 | 504 | 474 | 393 | Dgat2 |
| Dgat2l6 | 1 | 1 | 0 | 1 | 1 | 0 | 0 | 0 | 1 | 1 | 0 | 0 | 0 | 1 | 1 | 1 | 0 | 0 | 2 | 0 | 7 | 2 | 0 | 2 | 1 | 1 | 0 | 2 | Dgat2l6 |
| Dgcr2 | 6833 | 7502 | 8029 | 8299 | 7706 | 7241 | 7588 | 6064 | 6821 | 7631 | 7357 | 8131 | 7592 | 7615 | 6488 | 7127 | 8132 | 8386 | 8304 | 6576 | 8076 | 7738 | 6654 | 7981 | 7865 | 8680 | 7415 | 8119 | Dgcr2 |
| Dgcr6 | 0 | 0 | 0 | 0 | 0 | 3 | 0 | 0 | 1 | 0 | 0 | 0 | 0 | 1 | 0 | 0 | 0 | 0 | 0 | 0 | 0 | 0 | 0 | 0 | 0 | 0 | 0 | 0 | Dgcr6 |
| Dgcr8 | 0 | 0 | 0 | 0 | 0 | 0 | 0 | 0 | 1 | 0 | 0 | 0 | 0 | 0 | 0 | 0 | 0 | 0 | 0 | 0 | 0 | 0 | 0 | 0 | 0 | 0 | 0 | 0 | Dgcr8 |
| Dgka | 1078 | 904 | 969 | 797 | 928 | 873 | 648 | 862 | 995 | 948 | 914 | 623 | 785 | 664 | 897 | 1132 | 842 | 951 | 709 | 789 | 701 | 692 | 1000 | 831 | 588 | 628 | 595 | 621 | Dgka |
| Dgkb | 442 | 320 | 240 | 433 | 514 | 481 | 745 | 503 | 482 | 429 | 424 | 330 | 295 | 441 | 401 | 316 | 272 | 511 | 524 | 645 | 617 | 411 | 571 | 466 | 473 | 402 | 331 | 319 | Dgkb |
| Dgkd | 7360 | 7706 | 6789 | 8154 | 9047 | 8617 | 8895 | 7659 | 8001 | 8066 | 8050 | 8281 | 7421 | 7872 | 7361 | 7074 | 7589 | 8128 | 9334 | 8043 | 9063 | 8326 | 7421 | 8028 | 8871 | 6611 | 7786 | 7779 | Dgkd |
| Dgke | 5328 | 5067 | 4146 | 6502 | 6182 | 6505 | 7173 | 3543 | 3317 | 3716 | 3473 | 3304 | 3352 | 3284 | 5596 | 5139 | 4979 | 7068 | 5970 | 6358 | 7057 | 3638 | 3062 | 2971 | 3736 | 2408 | 3216 | 2789 | Dgke |
| Dgkg | 25 | 16 | 5 | 20 | 80 | 38 | 73 | 21 | 28 | 51 | 54 | 8 | 6 | 18 | 13 | 19 | 22 | 42 | 65 | 31 | 70 | 37 | 48 | 31 | 47 | 22 | 5 | 7 | Dgkg |
| Dgkh | 3256 | 2696 | 1975 | 2555 | 3608 | 4227 | 3994 | 2566 | 2728 | 2114 | 3010 | 1861 | 2100 | 2156 | 3285 | 3174 | 2444 | 2982 | 2926 | 3949 | 4376 | 2515 | 2418 | 1652 | 2428 | 1206 | 2055 | 1745 | Dgkh |
| Dgki | 735 | 854 | 143 | 517 | 898 | 1243 | 988 | 876 | 987 | 728 | 754 | 503 | 590 | 579 | 837 | 646 | 247 | 515 | 605 | 1158 | 1301 | 443 | 836 | 257 | 862 | 213 | 709 | 385 | Dgki |
| Dgkk | 188 | 60 | 110 | 49 | 128 | 54 | 69 | 59 | 83 | 56 | 111 | 14 | 65 | 60 | 75 | 91 | 93 | 233 | 105 | 124 | 128 | 85 | 28 | 16 | 22 | 12 | 52 | 18 | Dgkk |
| Dgkq | 4571 | 5271 | 5181 | 5760 | 6479 | 5675 | 5169 | 3451 | 3755 | 3901 | 3873 | 3684 | 3869 | 3343 | 4478 | 5014 | 5194 | 5908 | 5810 | 4958 | 5317 | 4480 | 3444 | 3635 | 4222 | 2995 | 3535 | 3317 | Dgkq |
| Dgkz | 5192 | 6551 | 6534 | 6831 | 6544 | 5894 | 5393 | 3844 | 5031 | 5144 | 4929 | 5084 | 4882 | 4473 | 4834 | 6250 | 6424 | 6980 | 6813 | 5183 | 5588 | 4690 | 5119 | 5069 | 5046 | 5772 | 4596 | 4575 | Dgkz |
| Dglucy | 80 | 92 | 67 | 86 | 94 | 147 | 102 | 249 | 279 | 188 | 227 | 144 | 164 | 144 | 103 | 112 | 67 | 94 | 93 | 93 | 87 | 192 | 254 | 141 | 152 | 130 | 143 | 134 | Dglucy |
| Dguok | 1190 | 1183 | 1122 | 1196 | 1084 | 1176 | 1203 | 1635 | 1697 | 1606 | 1807 | 1628 | 1526 | 1647 | 1266 | 1083 | 1170 | 995 | 1237 | 1115 | 1200 | 1619 | 1724 | 1671 | 1744 | 1523 | 1696 | 1544 | Dguok |
| Dhcr24 | 13449 | 17538 | 17407 | 15592 | 11720 | 11984 | 11051 | 6324 | 7075 | 7750 | 6925 | 10344 | 9846 | 9544 | 13156 | 15887 | 16309 | 14453 | 12832 | 10345 | 11802 | 7143 | 6468 | 8016 | 7979 | 11307 | 10197 | 10758 | Dhcr24 |
| Dhcr7 | 4625 | 6419 | 6171 | 5825 | 4715 | 4559 | 4558 | 2880 | 3011 | 3079 | 2967 | 3956 | 3882 | 3943 | 4800 | 5347 | 5556 | 5190 | 4926 | 4080 | 4539 | 3173 | 2871 | 3385 | 3211 | 4122 | 3908 | 4326 | Dhcr7 |
| Dhdds | 5615 | 6154 | 6544 | 6360 | 5175 | 4799 | 5566 | 3461 | 3819 | 4114 | 4031 | 4324 | 4016 | 4017 | 5284 | 5504 | 6339 | 6241 | 5852 | 4887 | 5488 | 3882 | 3706 | 4309 | 4205 | 4771 | 4159 | 4322 | Dhdds |
| Dhdh | 221 | 222 | 415 | 182 | 241 | 236 | 258 | 449 | 376 | 529 | 489 | 559 | 607 | 651 | 249 | 178 | 298 | 237 | 234 | 227 | 214 | 493 | 361 | 584 | 537 | 630 | 639 | 653 | Dhdh |
| Dhfr | 7276 | 6031 | 6584 | 5090 | 3864 | 4628 | 5721 | 3610 | 3081 | 2950 | 2730 | 4350 | 3734 | 4670 | 6425 | 5871 | 5794 | 4429 | 4645 | 5578 | 4933 | 2405 | 2781 | 3778 | 3195 | 5273 | 4735 | 5041 | Dhfr |
| Dhh | 131 | 191 | 182 | 135 | 113 | 68 | 84 | 57 | 51 | 46 | 79 | 27 | 32 | 32 | 115 | 160 | 118 | 96 | 71 | 61 | 83 | 69 | 72 | 24 | 18 | 30 | 30 | 30 | Dhh |
| Dhodh | 1338 | 1508 | 1645 | 1612 | 1272 | 1373 | 1203 | 1069 | 1371 | 1308 | 1256 | 1425 | 1353 | 1314 | 1269 | 1368 | 1401 | 1440 | 1383 | 1145 | 1142 | 1103 | 1321 | 1356 | 1278 | 1741 | 1368 | 1441 | Dhodh |
| Dhps | 1546 | 1725 | 1599 | 1993 | 1932 | 1839 | 1935 | 1661 | 1681 | 1798 | 1481 | 1809 | 1640 | 1906 | 1573 | 1399 | 1789 | 1854 | 2097 | 2008 | 2123 | 1820 | 1716 | 1845 | 1986 | 2023 | 1718 | 1877 | Dhps |
| Dhrs1 | 3869 | 4580 | 4048 | 3942 | 3534 | 3559 | 3393 | 2712 | 3129 | 2421 | 2675 | 2876 | 2812 | 2768 | 3906 | 4185 | 4051 | 3905 | 3809 | 4095 | 3346 | 2368 | 2895 | 2914 | 2721 | 3278 | 2911 | 2881 | Dhrs1 |
| Dhrs11 | 478 | 643 | 694 | 488 | 451 | 382 | 354 | 247 | 344 | 301 | 285 | 287 | 435 | 287 | 481 | 745 | 695 | 497 | 406 | 393 | 379 | 354 | 326 | 286 | 274 | 316 | 347 | 392 | Dhrs11 |
| Dhrs13 | 569 | 505 | 555 | 612 | 790 | 733 | 748 | 971 | 1016 | 1078 | 1008 | 918 | 844 | 871 | 587 | 460 | 621 | 738 | 820 | 671 | 729 | 1226 | 1040 | 1022 | 1148 | 996 | 905 | 895 | Dhrs13 |
| Dhrs2 | 2 | 5 | 0 | 5 | 7 | 16 | 2 | 16 | 24 | 13 | 27 | 7 | 6 | 6 | 7 | 2 | 1 | 1 | 9 | 8 | 4 | 27 | 27 | 13 | 7 | 9 | 5 | 1 | Dhrs2 |
| Dhrs3 | 280 | 174 | 200 | 202 | 250 | 214 | 123 | 623 | 823 | 748 | 972 | 321 | 472 | 250 | 190 | 229 | 149 | 219 | 182 | 156 | 153 | 603 | 961 | 516 | 470 | 477 | 248 | 228 | Dhrs3 |
| Dhrs4 | 1106 | 1056 | 1043 | 916 | 847 | 964 | 1100 | 1674 | 1837 | 1784 | 1781 | 1909 | 1704 | 1954 | 1087 | 995 | 1029 | 1007 | 1036 | 1201 | 969 | 1282 | 1868 | 2193 | 1736 | 2694 | 1923 | 1915 | Dhrs4 |
| Dhrs7 | 374 | 321 | 451 | 343 | 358 | 313 | 275 | 263 | 250 | 264 | 276 | 206 | 244 | 233 | 332 | 371 | 343 | 305 | 311 | 343 | 289 | 204 | 300 | 283 | 222 | 241 | 217 | 236 | Dhrs7 |
| Dhrs7b | 992 | 1150 | 989 | 986 | 934 | 952 | 1027 | 1194 | 1255 | 1175 | 1069 | 1180 | 1151 | 1099 | 994 | 1034 | 1088 | 1043 | 1161 | 1164 | 1051 | 1029 | 1354 | 1128 | 1108 | 1140 | 1149 | 1168 | Dhrs7b |
| Dhrs7c | 1 | 0 | 0 | 0 | 0 | 0 | 0 | 4 | 14 | 0 | 114 | 0 | 2 | 0 | 0 | 1 | 0 | 0 | 0 | 6 | 0 | 12 | 67 | 0 | 2 | 0 | 0 | 0 | Dhrs7c |
| Dhrs9 | 0 | 0 | 0 | 0 | 0 | 1 | 1 | 0 | 0 | 0 | 0 | 0 | 7 | 0 | 0 | 2 | 0 | 0 | 0 | 0 | 0 | 7 | 3 | 0 | 0 | 0 | 0 | 1 | Dhrs9 |
| Dhtkd1 | 2926 | 2841 | 2754 | 3632 | 3668 | 3363 | 3778 | 1375 | 1335 | 1585 | 1502 | 1386 | 1492 | 1538 | 3094 | 2963 | 3242 | 3832 | 3381 | 2874 | 3707 | 1681 | 1150 | 1421 | 1462 | 1148 | 1335 | 1234 | Dhtkd1 |
| Dhx15 | 26610 | 21113 | 23370 | 22168 | 21582 | 22949 | 23870 | 32205 | 26594 | 24939 | 24208 | 29572 | 27371 | 29792 | 26551 | 22374 | 23253 | 21879 | 22517 | 26273 | 23384 | 25693 | 25776 | 27311 | 26939 | 29583 | 30445 | 29719 | Dhx15 |
| Dhx16 | 5189 | 6306 | 6361 | 6619 | 4899 | 5017 | 5112 | 3367 | 3680 | 3859 | 3740 | 4369 | 3987 | 4204 | 4841 | 5603 | 6373 | 6070 | 5901 | 4584 | 5177 | 3539 | 3486 | 4479 | 4058 | 5213 | 4071 | 4498 | Dhx16 |
| Dhx29 | 9608 | 9265 | 8539 | 7722 | 7061 | 7448 | 8012 | 5451 | 5132 | 4808 | 4639 | 5324 | 5422 | 5520 | 9352 | 9097 | 7844 | 7722 | 7150 | 7965 | 7826 | 4654 | 4963 | 4715 | 5045 | 5157 | 5531 | 5593 | Dhx29 |
| Dhx30 | 6141 | 6943 | 6772 | 6987 | 7762 | 7173 | 7315 | 9976 | 11038 | 11237 | 9793 | 12359 | 10518 | 11161 | 6134 | 5960 | 7092 | 6910 | 7988 | 6965 | 7321 | 11207 | 9859 | 11509 | 11810 | 11818 | 11873 | 11629 | Dhx30 |
| Dhx32 | 4533 | 5058 | 4618 | 3989 | 3795 | 3927 | 4023 | 3152 | 2707 | 2470 | 2197 | 3574 | 3107 | 3349 | 4357 | 4206 | 4130 | 3773 | 4142 | 4451 | 4077 | 2548 | 2807 | 3000 | 2912 | 3775 | 3958 | 3933 | Dhx32 |
| Dhx33 | 5815 | 5573 | 3940 | 4801 | 4415 | 5206 | 5097 | 4319 | 4354 | 4113 | 4379 | 4013 | 4071 | 4094 | 5215 | 5423 | 4245 | 5092 | 4437 | 4763 | 4933 | 3957 | 4126 | 3835 | 3824 | 3594 | 3894 | 4094 | Dhx33 |
| Dhx34 | 743 | 1045 | 1283 | 1200 | 1240 | 986 | 951 | 1095 | 1330 | 1604 | 1710 | 1906 | 1755 | 1640 | 774 | 914 | 1224 | 1135 | 1331 | 989 | 1071 | 1502 | 1438 | 1788 | 1797 | 1792 | 1769 | 1663 | Dhx34 |
| Dhx35 | 1426 | 1551 | 1447 | 1238 | 1463 | 1501 | 1463 | 1695 | 2090 | 1911 | 1790 | 2358 | 2009 | 2202 | 1356 | 1370 | 1504 | 1235 | 1481 | 1452 | 1469 | 1619 | 1860 | 2018 | 2119 | 2334 | 2275 | 2251 | Dhx35 |
| Dhx36 | 14564 | 11625 | 11925 | 12589 | 13569 | 13801 | 15074 | 18061 | 14904 | 14145 | 13626 | 16209 | 15203 | 17014 | 15509 | 12546 | 12366 | 12646 | 12578 | 14323 | 14063 | 16192 | 15004 | 14271 | 15475 | 14392 | 16203 | 16086 | Dhx36 |
| Dhx37 | 2216 | 2725 | 2782 | 2997 | 2351 | 2316 | 2394 | 1866 | 2363 | 2354 | 2396 | 3041 | 2468 | 2734 | 2066 | 2487 | 2756 | 2787 | 2659 | 1778 | 2380 | 2511 | 2154 | 2837 | 2725 | 2956 | 2495 | 2839 | Dhx37 |
| Dhx38 | 3576 | 3992 | 4766 | 4409 | 3937 | 3893 | 3575 | 3828 | 4999 | 5218 | 5081 | 6218 | 5531 | 5999 | 3127 | 3470 | 4519 | 3771 | 4346 | 3415 | 3590 | 4766 | 4503 | 6008 | 5595 | 6206 | 5665 | 5955 | Dhx38 |
| Dhx40 | 10042 | 8657 | 9776 | 9100 | 7652 | 8400 | 9669 | 6256 | 5155 | 5375 | 4890 | 7261 | 6298 | 7147 | 9770 | 8754 | 9265 | 8439 | 8930 | 8396 | 9026 | 6144 | 5230 | 6208 | 6039 | 7192 | 7573 | 7784 | Dhx40 |
| Dhx57 | 8503 | 8404 | 6748 | 8066 | 9545 | 9163 | 9483 | 9183 | 8979 | 8294 | 8251 | 7546 | 7220 | 7685 | 8592 | 7559 | 7554 | 8675 | 8876 | 9199 | 9556 | 9580 | 8437 | 7758 | 9017 | 5887 | 7071 | 6925 | Dhx57 |
| Dhx58 | 197 | 238 | 386 | 329 | 326 | 301 | 244 | 116 | 138 | 186 | 192 | 193 | 191 | 184 | 228 | 318 | 298 | 366 | 288 | 278 | 244 | 219 | 152 | 168 | 140 | 173 | 171 | 179 | Dhx58 |
| Dhx8 | 5613 | 6202 | 5125 | 5208 | 4622 | 5149 | 4881 | 5560 | 6072 | 5665 | 5603 | 6396 | 6165 | 5827 | 5469 | 5460 | 5208 | 4651 | 5263 | 4895 | 5049 | 5424 | 5651 | 5740 | 5868 | 6200 | 6369 | 6160 | Dhx8 |
| Dhx9 | 54771 | 50096 | 46510 | 45327 | 44257 | 45039 | 52985 | 54324 | 46789 | 42213 | 41064 | 53293 | 48946 | 58315 | 54738 | 47430 | 46750 | 44649 | 45830 | 48912 | 49744 | 46951 | 43153 | 47542 | 46933 | 50072 | 55544 | 56957 | Dhx9 |
| Diablo | 944 | 984 | 566 | 902 | 846 | 775 | 933 | 805 | 632 | 697 | 760 | 659 | 756 | 804 | 870 | 842 | 740 | 814 | 728 | 785 | 867 | 645 | 631 | 637 | 740 | 375 | 696 | 732 | Diablo |
| Diaph1 | 5547 | 5334 | 6009 | 6385 | 6213 | 5818 | 6401 | 3705 | 3920 | 4441 | 4490 | 4834 | 4302 | 4705 | 4767 | 5638 | 6088 | 7039 | 6286 | 4688 | 6699 | 5130 | 3994 | 4916 | 4678 | 4579 | 4188 | 4555 | Diaph1 |
| Diaph2 | 1752 | 1406 | 1198 | 1494 | 1624 | 1781 | 1797 | 1641 | 1613 | 1593 | 1682 | 1867 | 1662 | 1720 | 1709 | 1847 | 1239 | 1613 | 1537 | 1652 | 1919 | 1509 | 1660 | 1508 | 1591 | 1163 | 1684 | 1491 | Diaph2 |
| Diaph3 | 1892 | 1363 | 1672 | 1263 | 934 | 1185 | 1248 | 1371 | 1388 | 1138 | 1357 | 1866 | 1526 | 1699 | 1563 | 1596 | 1344 | 1148 | 1069 | 1063 | 1026 | 956 | 1445 | 1543 | 1185 | 1821 | 1599 | 1642 | Diaph3 |
| Dido1 | 9750 | 10131 | 7558 | 9436 | 10204 | 10671 | 9876 | 11491 | 11786 | 11374 | 11478 | 13190 | 12526 | 12748 | 9336 | 9432 | 8790 | 8801 | 9894 | 10218 | 10185 | 11080 | 10747 | 11019 | 12144 | 8442 | 13174 | 12155 | Dido1 |
| Dimt1 | 2332 | 2440 | 2239 | 2517 | 2184 | 2227 | 2367 | 2269 | 2094 | 1810 | 2146 | 1876 | 1890 | 1900 | 2491 | 2521 | 2254 | 2400 | 2180 | 2655 | 2346 | 1873 | 2038 | 1991 | 1864 | 1985 | 1858 | 1938 | Dimt1 |

| | | | | | | | | | | | | | | | | | | | | | | | | | | | | | |
|---|---|---|---|---|---|---|---|---|---|---|---|---|---|---|---|---|---|---|---|---|---|---|---|---|---|---|---|---|---|
| Dio1 | 0 | 0 | 0 | 0 | 0 | 0 | 0 | 0 | 0 | 1 | 1 | 0 | 0 | 0 | 0 | 0 | 0 | 0 | 0 | 0 | 0 | 0 | 0 | 0 | 1 | 0 | 0 | 0 | Dio1 |
| Dio2 | 51 | 44 | 73 | 34 | 57 | 52 | 53 | 72 | 105 | 114 | 203 | 67 | 71 | 64 | 54 | 72 | 29 | 41 | 41 | 54 | 55 | 52 | 108 | 87 | 40 | 57 | 110 | 97 | Dio2 |
| Dio3 | 125 | 184 | 76 | 180 | 233 | 234 | 324 | 186 | 384 | 286 | 356 | 252 | 104 | 230 | 132 | 84 | 199 | 189 | 285 | 276 | 232 | 241 | 416 | 347 | 273 | 214 | 127 | 163 | Dio3 |
| Dip2a | 4289 | 4317 | 3887 | 4671 | 5432 | 5161 | 4695 | 4186 | 5186 | 5122 | 5236 | 4737 | 4763 | 4348 | 4178 | 4838 | 4252 | 5211 | 4903 | 4246 | 4946 | 5158 | 5145 | 4493 | 5155 | 3548 | 4449 | 4270 | Dip2a |
| Dip2b | 5518 | 5475 | 4762 | 6145 | 7193 | 7112 | 7017 | 6970 | 7111 | 7977 | 7607 | 8450 | 8022 | 8399 | 5222 | 5313 | 5483 | 6487 | 6938 | 5568 | 7036 | 8134 | 6471 | 7571 | 9090 | 6029 | 8486 | 8024 | Dip2b |
| Dip2c | 1819 | 1756 | 1431 | 2008 | 2941 | 2606 | 2652 | 4655 | 4588 | 4709 | 4559 | 4107 | 3639 | 4194 | 1925 | 1674 | 1852 | 2253 | 2318 | 2392 | 2569 | 5491 | 4152 | 3974 | 4916 | 2976 | 3674 | 3869 | Dip2c |
| Dipk1a | 2028 | 1593 | 2018 | 1942 | 2239 | 2235 | 2247 | 1713 | 1589 | 1646 | 1736 | 1474 | 1658 | 1644 | 2058 | 1737 | 1953 | 2125 | 1969 | 2066 | 2236 | 1977 | 1625 | 1527 | 1731 | 1446 | 1476 | 1663 | Dipk1a |
| Dipk1b | 549 | 605 | 582 | 609 | 844 | 709 | 715 | 1093 | 1182 | 1178 | 970 | 1098 | 1022 | 1014 | 517 | 487 | 703 | 594 | 888 | 706 | 654 | 1076 | 1155 | 1186 | 1252 | 1045 | 1126 | 1008 | Dipk1b |
| Dipk1c | 235 | 291 | 239 | 198 | 289 | 227 | 302 | 465 | 503 | 466 | 453 | 634 | 541 | 685 | 243 | 141 | 318 | 179 | 265 | 253 | 208 | 398 | 420 | 561 | 526 | 630 | 769 | 730 | Dipk1c |
| Dipk2a | 6635 | 5145 | 6876 | 5431 | 4556 | 4844 | 4756 | 6033 | 5482 | 4619 | 5688 | 4569 | 5680 | 4564 | 6671 | 7263 | 5173 | 5258 | 4193 | 4477 | 4243 | 5920 | 5305 | 4332 | 4223 | 4674 | 5648 | 5298 | Dipk2a |
| Dipk2b | 24 | 29 | 14 | 21 | 37 | 22 | 27 | 85 | 117 | 104 | 138 | 86 | 102 | 65 | 30 | 29 | 20 | 13 | 24 | 43 | 35 | 75 | 147 | 63 | 93 | 69 | 86 | 83 | Dipk2b |
| Diras1 | 534 | 676 | 561 | 1093 | 1640 | 1191 | 1119 | 1547 | 1846 | 1969 | 1776 | 1323 | 1200 | 1293 | 649 | 567 | 942 | 1172 | 1496 | 1222 | 1274 | 2194 | 1609 | 1642 | 2184 | 1010 | 1178 | 1137 | Diras1 |
| Diras2 | 385 | 352 | 238 | 623 | 1082 | 815 | 701 | 1005 | 1247 | 1184 | 932 | 532 | 962 | 545 | 452 | 343 | 419 | 698 | 811 | 779 | 791 | 1636 | 1138 | 615 | 1180 | 241 | 546 | 417 | Diras2 |
| Dis3 | 8711 | 7660 | 6729 | 6876 | 6122 | 7348 | 7698 | 6592 | 5983 | 5259 | 5883 | 6884 | 6324 | 6591 | 8213 | 7896 | 6612 | 6612 | 6125 | 7549 | 7349 | 5324 | 5655 | 5555 | 5895 | 6212 | 6936 | 6749 | Dis3 |
| Dis3l | 506 | 503 | 573 | 473 | 641 | 605 | 550 | 2613 | 2671 | 2506 | 2771 | 2658 | 2366 | 2856 | 570 | 427 | 497 | 462 | 605 | 519 | 536 | 2770 | 2609 | 2751 | 2764 | 2793 | 2680 | 2686 | Dis3l |
| Dis3l2 | 2098 | 2365 | 1967 | 2012 | 2141 | 1917 | 2140 | 2887 | 2844 | 2454 | 2469 | 2843 | 2649 | 2882 | 2090 | 1995 | 2002 | 1973 | 2173 | 2233 | 2120 | 2573 | 2707 | 2580 | 2800 | 2685 | 2836 | 2934 | Dis3l2 |
| Disc1 | 82 | 98 | 80 | 134 | 92 | 115 | 91 | 197 | 190 | 226 | 277 | 202 | 212 | 197 | 76 | 130 | 97 | 113 | 88 | 126 | 113 | 196 | 254 | 155 | 187 | 134 | 193 | 158 | Disc1 |
| Disp1 | 1572 | 1970 | 1623 | 1442 | 1452 | 1517 | 1142 | 1218 | 1634 | 1768 | 1876 | 1750 | 1796 | 1469 | 1319 | 1968 | 1741 | 1588 | 1413 | 1122 | 1213 | 1514 | 1839 | 1705 | 1548 | 1534 | 1538 | 1464 | Disp1 |
| Disp2 | 3122 | 4208 | 4186 | 6462 | 9344 | 6721 | 7232 | 5241 | 5405 | 7290 | 5866 | 5018 | 5078 | 4762 | 4284 | 3747 | 5846 | 7165 | 7889 | 6272 | 7651 | 9795 | 4606 | 5489 | 7404 | 2599 | 4559 | 4538 | Disp2 |
| Disp3 | 1089 | 2027 | 1498 | 3027 | 3369 | 2607 | 2614 | 1170 | 1499 | 1799 | 1507 | 1252 | 1504 | 1024 | 1561 | 1667 | 2328 | 3037 | 2943 | 2421 | 2875 | 1938 | 1396 | 1307 | 1840 | 557 | 1099 | 933 | Disp3 |
| Dixdc1 | 7689 | 7161 | 6470 | 7466 | 9289 | 8941 | 8287 | 6245 | 5177 | 5912 | 5524 | 5170 | 5891 | 5219 | 7950 | 7826 | 7283 | 8185 | 7782 | 8580 | 8714 | 6711 | 5537 | 5209 | 6056 | 3640 | 5344 | 5014 | Dixdc1 |
| Dkc1 | 6449 | 5181 | 6462 | 5210 | 3946 | 4624 | 5045 | 6149 | 5525 | 5269 | 5049 | 6762 | 6510 | 6970 | 5962 | 5431 | 5772 | 4475 | 4933 | 5150 | 4667 | 4588 | 5469 | 6119 | 5253 | 8396 | 7179 | 7891 | Dkc1 |
| Dkk1 | 555 | 267 | 300 | 148 | 106 | 156 | 74 | 506 | 442 | 380 | 211 | 239 | 320 | 211 | 252 | 349 | 177 | 131 | 94 | 137 | 116 | 119 | 337 | 327 | 154 | 431 | 282 | 317 | Dkk1 |
| Dkk2 | 2257 | 1919 | 3122 | 1372 | 723 | 1417 | 339 | 2102 | 2729 | 1903 | 3829 | 2657 | 3370 | 2654 | 2557 | 2775 | 2024 | 1632 | 390 | 632 | 386 | 2608 | 4017 | 2388 | 1967 | 2674 | 3658 | 3025 | Dkk2 |
| Dkk3 | 3422 | 3283 | 3520 | 3028 | 2729 | 3259 | 2277 | 2437 | 2951 | 2612 | 2796 | 2375 | 2700 | 2241 | 3483 | 4044 | 2855 | 3436 | 2481 | 2619 | 2318 | 2568 | 3785 | 2648 | 2573 | 2927 | 2487 | 2681 | Dkk3 |
| Dkk4 | 116 | 3 | 2 | 0 | 2 | 0 | 0 | 20 | 19 | 16 | 8 | 3 | 2 | 0 | 7 | 30 | 0 | 0 | 0 | 5 | 0 | 2 | 15 | 9 | 0 | 4 | 3 | 0 | Dkk4 |
| Dkkl1 | 102 | 83 | 102 | 89 | 103 | 62 | 50 | 99 | 78 | 89 | 70 | 73 | 91 | 95 | 125 | 107 | 90 | 77 | 62 | 61 | 58 | 105 | 76 | 68 | 75 | 103 | 98 | 65 | Dkkl1 |
| Dlat | 5279 | 4361 | 5458 | 5156 | 5141 | 4956 | 5530 | 3887 | 3640 | 3905 | 4166 | 4319 | 4043 | 4504 | 5039 | 4545 | 5135 | 5138 | 4997 | 4774 | 5368 | 3853 | 3729 | 4299 | 4235 | 4744 | 4029 | 4383 | Dlat |
| Dlc1 | 3047 | 3383 | 3322 | 3288 | 3382 | 2854 | 2613 | 2591 | 3010 | 3409 | 4421 | 2962 | 3703 | 2885 | 3028 | 4183 | 3095 | 3516 | 2619 | 2166 | 2716 | 3166 | 3315 | 3006 | 2989 | 2899 | 2760 | 2848 | Dlc1 |
| Dld | 13302 | 10726 | 12901 | 11107 | 10517 | 10906 | 12148 | 12245 | 10308 | 9539 | 10042 | 10936 | 10789 | 11239 | 14031 | 12789 | 11276 | 10904 | 10892 | 13514 | 11539 | 10188 | 10804 | 10234 | 10191 | 11850 | 11432 | 12114 | Dld |
| Dlec1 | 508 | 755 | 628 | 769 | 534 | 422 | 420 | 192 | 194 | 197 | 185 | 206 | 281 | 126 | 571 | 843 | 639 | 598 | 450 | 459 | 464 | 206 | 308 | 150 | 146 | 150 | 191 | 157 | Dlec1 |
| Dleu7 | 82 | 76 | 85 | 92 | 172 | 128 | 190 | 289 | 325 | 309 | 249 | 311 | 87 | 307 | 99 | 58 | 58 | 142 | 182 | 128 | 189 | 287 | 216 | 391 | 371 | 345 | 236 | 221 | Dleu7 |
| Dlg1 | 9133 | 7332 | 8177 | 7590 | 6910 | 7603 | 8444 | 7393 | 6938 | 6700 | 7327 | 8045 | 7925 | 8335 | 8116 | 8601 | 7486 | 8056 | 7307 | 7864 | 7838 | 5807 | 7671 | 7312 | 6608 | 8619 | 7861 | 8457 | Dlg1 |
| Dlg2 | 1644 | 1527 | 1139 | 2680 | 4675 | 3476 | 4109 | 3280 | 3241 | 3389 | 2866 | 1691 | 1049 | 1876 | 1896 | 1289 | 1662 | 3300 | 3695 | 3694 | 4770 | 3815 | 3286 | 2439 | 3521 | 1159 | 1341 | 1456 | Dlg2 |
| Dlg3 | 6569 | 6360 | 6650 | 7449 | 8084 | 7479 | 7886 | 6669 | 6124 | 6898 | 5989 | 6742 | 6152 | 6899 | 6650 | 6074 | 7057 | 7480 | 8036 | 7187 | 7881 | 7110 | 5666 | 6942 | 7621 | 6100 | 6507 | 6735 | Dlg3 |
| Dlg4 | 7139 | 9664 | 11038 | 14391 | 14938 | 12692 | 11999 | 9099 | 10153 | 13695 | 11594 | 11722 | 9678 | 10799 | 7346 | 8210 | 13217 | 14532 | 15325 | 11071 | 13434 | 13570 | 9932 | 13332 | 14347 | 11183 | 10110 | 10300 | Dlg4 |
| Dlg5 | 3332 | 3067 | 3359 | 3095 | 4147 | 3517 | 2909 | 7946 | 9737 | 10595 | 11151 | 10421 | 9460 | 9695 | 2756 | 3596 | 3541 | 3346 | 3730 | 2637 | 3162 | 10530 | 9085 | 10556 | 10750 | 8183 | 9022 | 9276 | Dlg5 |
| Dlgap1 | 431 | 498 | 254 | 949 | 1592 | 1119 | 1425 | 1014 | 1112 | 1300 | 1166 | 852 | 553 | 852 | 613 | 460 | 608 | 1173 | 1326 | 1138 | 1432 | 1384 | 1030 | 1048 | 1327 | 563 | 504 | 547 | Dlgap1 |
| Dlgap2 | 164 | 225 | 122 | 291 | 351 | 293 | 292 | 220 | 234 | 310 | 307 | 313 | 219 | 268 | 207 | 175 | 181 | 275 | 252 | 286 | 346 | 309 | 287 | 273 | 452 | 151 | 284 | 261 | Dlgap2 |
| Dlgap3 | 844 | 1357 | 1423 | 2065 | 2758 | 1887 | 1648 | 1185 | 1389 | 2115 | 1500 | 1389 | 1182 | 1183 | 1044 | 995 | 1822 | 2084 | 2452 | 1562 | 2095 | 2154 | 1282 | 1467 | 1884 | 1185 | 1027 | 889 | Dlgap3 |
| Dlgap4 | 7321 | 9339 | 9986 | 12100 | 13422 | 11573 | 11549 | 8691 | 10342 | 13966 | 11612 | 13544 | 10484 | 12448 | 7184 | 8320 | 12247 | 13058 | 13962 | 9144 | 11681 | 14825 | 9673 | 15027 | 15606 | 13258 | 10412 | 11522 | Dlgap4 |
| Dlgap5 | 3770 | 3527 | 4044 | 2785 | 1966 | 2511 | 2573 | 3609 | 3400 | 3277 | 2899 | 5031 | 4207 | 4950 | 3233 | 3341 | 3301 | 2505 | 2837 | 2684 | 2431 | 2453 | 3144 | 4323 | 3475 | 5682 | 5037 | 5098 | Dlgap5 |
| Dlk1 | 0 | 0 | 0 | 0 | 0 | 0 | 0 | 0 | 0 | 0 | 1 | 1 | 2 | 0 | 0 | 0 | 0 | 0 | 0 | 0 | 0 | 0 | 0 | 2 | 1 | 2 | 0 | 0 | Dlk1 |
| Dlk2 | 215 | 251 | 290 | 320 | 540 | 395 | 327 | 426 | 573 | 655 | 576 | 299 | 287 | 274 | 214 | 244 | 336 | 418 | 468 | 400 | 411 | 632 | 488 | 414 | 573 | 193 | 223 | 183 | Dlk2 |
| Dll1 | 2824 | 4202 | 5643 | 4408 | 4002 | 3708 | 3961 | 4347 | 3691 | 4272 | 3438 | 7758 | 5959 | 7175 | 3273 | 2962 | 5764 | 3795 | 4985 | 3495 | 3808 | 4329 | 3199 | 5724 | 5386 | 7561 | 7104 | 7495 | Dll1 |
| Dll3 | 570 | 771 | 1371 | 765 | 929 | 818 | 715 | 1308 | 1044 | 1345 | 1099 | 2436 | 2080 | 2311 | 698 | 463 | 1272 | 677 | 1043 | 674 | 771 | 1679 | 895 | 1964 | 1573 | 2322 | 2818 | 2619 | Dll3 |
| Dll4 | 340 | 386 | 400 | 389 | 522 | 453 | 380 | 491 | 492 | 572 | 530 | 500 | 506 | 431 | 284 | 412 | 415 | 446 | 503 | 426 | 417 | 568 | 533 | 611 | 483 | 558 | 561 | 460 | Dll4 |
| Dlst | 15268 | 14919 | 14007 | 13852 | 12661 | 13197 | 13475 | 11727 | 11605 | 10832 | 11130 | 11893 | 11494 | 12159 | 14838 | 14090 | 13908 | 13412 | 12800 | 12382 | 13276 | 11214 | 10829 | 11228 | 11759 | 12441 | 11870 | 12397 | Dlst |
| Dlx1 | 2697 | 2223 | 1953 | 4240 | 4734 | 4493 | 6784 | 4998 | 6349 | 7457 | 5695 | 10406 | 492 | 12543 | 2152 | 772 | 4566 | 4339 | 6119 | 3325 | 6176 | 8081 | 2930 | 11335 | 11021 | 14134 | 7039 | 8223 | Dlx1 |
| Dlx2 | 1563 | 1651 | 1271 | 2908 | 2351 | 1916 | 3061 | 1385 | 1678 | 1994 | 1741 | 3535 | 168 | 3374 | 1226 | 578 | 3176 | 2885 | 3215 | 1319 | 2840 | 2448 | 724 | 3255 | 2758 | 4769 | 1626 | 2137 | Dlx2 |
| Dlx3 | 298 | 205 | 167 | 96 | 61 | 105 | 30 | 250 | 299 | 232 | 289 | 45 | 236 | 54 | 125 | 218 | 121 | 149 | 74 | 45 | 39 | 65 | 258 | 189 | 59 | 127 | 46 | 55 | Dlx3 |
| Dlx4 | 76 | 41 | 30 | 12 | 6 | 32 | 2 | 13 | 28 | 12 | 39 | 1 | 29 | 7 | 21 | 36 | 23 | 21 | 8 | 3 | 9 | 3 | 32 | 25 | 0 | 11 | 6 | 1 | Dlx4 |
| Dlx5 | 691 | 672 | 532 | 1299 | 965 | 1013 | 1295 | 735 | 1067 | 1234 | 1066 | 1503 | 338 | 1295 | 492 | 289 | 1112 | 1096 | 1479 | 538 | 1184 | 1292 | 575 | 1637 | 1447 | 2211 | 911 | 1028 | Dlx5 |
| Dlx6 | 1309 | 974 | 768 | 1858 | 1216 | 1258 | 2143 | 783 | 862 | 1011 | 887 | 948 | 291 | 1256 | 1105 | 386 | 1369 | 1588 | 1615 | 1206 | 1939 | 951 | 512 | 1410 | 1067 | 1549 | 812 | 859 | Dlx6 |
| Dmac1 | 1599 | 1609 | 1798 | 1702 | 1348 | 1526 | 1544 | 1515 | 1526 | 1607 | 1660 | 1723 | 1668 | 1850 | 1535 | 1445 | 1713 | 1444 | 1670 | 1752 | 1605 | 1333 | 1573 | 1835 | 1766 | 1994 | 1833 | 1938 | Dmac1 |
| Dmac2 | 744 | 748 | 745 | 738 | 752 | 797 | 765 | 1093 | 1280 | 1227 | 1235 | 1237 | 1177 | 1136 | 574 | 713 | 702 | 662 | 817 | 747 | 743 | 1133 | 1237 | 1185 | 1190 | 1215 | 1178 | 1137 | Dmac2 |
| Dmac2l | 939 | 786 | 768 | 842 | 745 | 874 | 883 | 749 | 641 | 652 | 676 | 557 | 699 | 620 | 971 | 823 | 800 | 949 | 815 | 879 | 860 | 637 | 718 | 576 | 661 | 672 | 601 | 629 | Dmac2l |
| Dmap1 | 2172 | 2695 | 2206 | 2216 | 1879 | 1971 | 1995 | 2179 | 2376 | 2162 | 2080 | 2487 | 2339 | 2409 | 2239 | 2065 | 2133 | 1964 | 2173 | 2183 | 2011 | 2049 | 2346 | 2308 | 2379 | 2681 | 2454 | 2566 | Dmap1 |
| Dmbt1 | 0 | 1 | 0 | 0 | 0 | 1 | 0 | 1 | 1 | 0 | 0 | 0 | 1 | 0 | 0 | 2 | 0 | 0 | 0 | 0 | 0 | 0 | 0 | 0 | 0 | 0 | 0 | 0 | Dmbt1 |
| Dmbx1 | 694 | 922 | 1902 | 1332 | 737 | 786 | 196 | 795 | 704 | 942 | 544 | 781 | 1761 | 804 | 690 | 927 | 1374 | 792 | 637 | 606 | 502 | 1084 | 688 | 559 | 791 | 314 | 1010 | 1121 | Dmbx1 |
| Dmc1 | 652 | 485 | 493 | 575 | 448 | 584 | 625 | 369 | 334 | 354 | 446 | 326 | 383 | 324 | 589 | 628 | 465 | 525 | 441 | 569 | 496 | 478 | 296 | 283 | 338 | 267 | 408 | 375 | Dmc1 |
| Dmd | 2504 | 2033 | 1932 | 2493 | 2930 | 2973 | 3529 | 2341 | 2360 | 2221 | 2539 | 2181 | 2277 | 2220 | 2462 | 2364 | 2075 | 2820 | 2837 | 2936 | 3127 | 2394 | 2630 | 2116 | 2425 | 1673 | 1815 | 1920 | Dmd |
| Dmgdh | 12 | 16 | 3 | 11 | 13 | 14 | 9 | 12 | 18 | 12 | 28 | 8 | 16 | 14 | 7 | 16 | 0 | 9 | 5 | 18 | 10 | 30 | 29 | 6 | 10 | 8 | 3 | 5 | Dmgdh |
| Dmkn | 3 | 10 | 1 | 0 | 2 | 3 | 4 | 20 | 33 | 30 | 9 | 11 | 14 | 3 | 4 | 5 | 3 | 1 | 12 | 13 | 9 | 22 | 34 | 7 | 12 | 9 | 3 | 5 | Dmkn |
| Dmp1 | 23 | 31 | 37 | 55 | 20 | 25 | 29 | 49 | 47 | 56 | 42 | 70 | 57 | 82 | 41 | 12 | 44 | 35 | 39 | 28 | 29 | 63 | 33 | 65 | 38 | 80 | 60 | 90 | Dmp1 |
| Dmpk | 2331 | 3552 | 3523 | 4116 | 3394 | 3033 | 3083 | 2098 | 2202 | 2452 | 3294 | 2495 | 2345 | 2536 | 2155 | 2827 | 4022 | 3986 | 3645 | 2642 | 3087 | 2784 | 2399 | 2722 | 2634 | 2319 | 2298 | 2633 | Dmpk |
| Dmrt1 | 92 | 64 | 141 | 103 | 72 | 66 | 115 | 53 | 31 | 39 | 28 | 79 | 92 | 77 | 50 | 80 | 130 | 70 | 94 | 99 | 77 | 22 | 54 | 69 | 43 | 115 | 69 | 119 | Dmrt1 |
| Dmrt2 | 253 | 181 | 134 | 103 | 61 | 77 | 75 | 136 | 239 | 116 | 199 | 28 | 196 | 24 | 183 | 220 | 111 | 105 | 94 | 66 | 58 | 80 | 337 | 100 | 81 | 137 | 15 | 21 | Dmrt2 |
| Dmrt3 | 286 | 266 | 444 | 329 | 292 | 306 | 466 | 381 | 373 | 475 | 389 | 859 | 565 | 805 | 210 | 231 | 360 | 243 | 467 | 406 | 365 | 167 | 381 | 820 | 519 | 1192 | 528 | 838 | Dmrt3 |
| Dmrta1 | 1355 | 900 | 1455 | 1170 | 794 | 1133 | 1542 | 878 | 695 | 702 | 535 | 1615 | 1191 | 1738 | 1094 | 826 | 1139 | 844 | 1169 | 1362 | 995 | 241 | 432 | 1025 | 826 | 2007 | 1393 | 1860 | Dmrta1 |
| Dmrta2 | 1494 | 1617 | 2018 | 1722 | 1366 | 1383 | 1938 | 1307 | 1249 | 1524 | 1034 | 2414 | 1807 | 2451 | 1251 | 942 | 1753 | 1351 | 1921 | 1976 | 1628 | 754 | 1078 | 2326 | 1707 | 3355 | 2064 | 2597 | Dmrta2 |
| Dmrtb1 | 320 | 461 | 456 | 623 | 716 | 748 | 640 | 855 | 625 | 1060 | 753 | 1145 | 942 | 1189 | 433 | 333 | 623 | 609 | 789 | 690 | 756 | 1195 | 731 | 1244 | 1391 | 953 | 1221 | 1125 | Dmrtb1 |
| Dmrtc1a | 12 | 16 | 13 | 9 | 34 | 28 | 26 | 17 | 26 | 19 | 27 | 15 | 18 | 22 | 21 | 24 | 23 | 23 | 23 | 32 | 21 | 30 | 19 | 8 | 14 | 10 | 15 | 21 | Dmrtc1a |
| Dmrtc1b | 0 | 0 | 0 | 0 | 0 | 0 | 0 | 0 | 2 | 0 | 0 | 0 | 0 | 0 | 0 | 0 | 0 | 0 | 0 | 0 | 0 | 2 | 0 | 0 | 0 | 0 | 0 | 0 | Dmrtc1b |
| Dmrtc1c1 | 0 | 0 | 0 | 0 | 0 | 0 | 0 | 0 | 0 | 0 | 0 | 0 | 0 | 0 | 0 | 0 | 0 | 0 | 0 | 0 | 0 | 0 | 0 | 0 | 0 | 0 | 0 | 0 | Dmrtc1c1 |
| Dmrtc2 | 2 | 0 | 2 | 6 | 1 | 9 | 0 | 7 | 2 | 4 | 4 | 8 | 2 | 10 | 5 | 0 | 5 | 6 | 5 | 5 | 4 | 7 | 3 | 3 | 3 | 14 | 8 | 6 | Dmrtc2 |
| Dmtf1 | 10461 | 9333 | 9004 | 9457 | 9401 | 9649 | 10421 | 12665 | 10447 | 9470 | 9437 | 9928 | 9693 | 10165 | 10755 | 9302 | 8885 | 9480 | 9650 | 10548 | 10364 | 10890 | 10281 | 9373 | 9573 | 8912 | 10098 | 10303 | Dmtf1 |
| Dmtn | 723 | 1107 | 1080 | 1448 | 2123 | 1609 | 1598 | 1868 | 1979 | 2359 | 1913 | 1713 | 1827 | 1755 | 948 | 888 | 1504 | 1623 | 2136 | 1643 | 1773 | 2894 | 1937 | 2048 | 2528 | 1453 | 1617 | 1782 | Dmtn |
| Dmwd | 4469 | 6444 | 5515 | 6205 | 6060 | 5650 | 5185 | 4832 | 5762 | 6124 | 5323 | 6047 | 5383 | 5274 | 4592 | 4780 | 6312 | 6108 | 6657 | 5316 | 5611 | 5671 | 5398 | 6243 | 6142 | 6067 | 5575 | 5318 | Dmwd |

| | | | | | | | | | | | | | | | | | | | | | | | | | | | | | |
|---|---|---|---|---|---|---|---|---|---|---|---|---|---|---|---|---|---|---|---|---|---|---|---|---|---|---|---|---|---|
| Dmxl1 | 12299 | 9286 | 5191 | 7590 | 7318 | 10304 | 9683 | 9509 | 7823 | 5600 | 6084 | 6264 | 6595 | 6580 | 12212 | 10478 | 5990 | 7443 | 6742 | 11258 | 10059 | 5634 | 7515 | 4763 | 6161 | 3844 | 7182 | 5941 | Dmxl1 |
| Dmxl2 | 10175 | 10050 | 4815 | 9264 | 13533 | 14397 | 14275 | 12844 | 11631 | 9242 | 8956 | 7055 | 7370 | 7695 | 11707 | 9675 | 6937 | 9770 | 10751 | 15188 | 15189 | 9065 | 10460 | 6422 | 10315 | 3739 | 7168 | 6041 | Dmxl2 |
| Dna2 | 1611 | 1413 | 1471 | 1325 | 1357 | 1416 | 1530 | 2231 | 2285 | 2155 | 2149 | 3116 | 2628 | 2957 | 1395 | 1462 | 1392 | 1276 | 1580 | 1576 | 1417 | 1597 | 2073 | 2553 | 2414 | 3222 | 3139 | 3280 | Dna2 |
| Dnaaf1 | 15 | 35 | 13 | 26 | 28 | 11 | 20 | 28 | 16 | 15 | 12 | 22 | 33 | 31 | 12 | 33 | 9 | 24 | 25 | 17 | 29 | 25 | 40 | 29 | 20 | 31 | 19 | 21 | Dnaaf1 |
| Dnaaf2 | 570 | 471 | 490 | 508 | 281 | 367 | 515 | 222 | 184 | 230 | 179 | 332 | 272 | 285 | 513 | 460 | 468 | 445 | 346 | 384 | 461 | 197 | 195 | 218 | 243 | 341 | 267 | 339 | Dnaaf2 |
| Dnaaf3 | 1197 | 1857 | 2143 | 1534 | 1045 | 1001 | 980 | 211 | 297 | 216 | 220 | 343 | 380 | 251 | 1064 | 1670 | 1621 | 1179 | 1100 | 1011 | 819 | 244 | 252 | 239 | 228 | 343 | 349 | 297 | Dnaaf3 |
| Dnaaf4 | 39 | 27 | 22 | 27 | 35 | 23 | 57 | 56 | 51 | 52 | 51 | 58 | 82 | 53 | 67 | 30 | 36 | 21 | 37 | 37 | 29 | 70 | 41 | 56 | 46 | 51 | 51 | 56 | Dnaaf4 |
| Dnaaf5 | 1706 | 1876 | 1816 | 1814 | 1383 | 1306 | 1415 | 1033 | 1228 | 1233 | 1135 | 1529 | 1429 | 1310 | 1436 | 1668 | 1725 | 1438 | 1634 | 1244 | 1322 | 1115 | 1224 | 1319 | 1180 | 1654 | 1379 | 1333 | Dnaaf5 |
| Dnaaf6b | 12 | 3 | 11 | 0 | 4 | 4 | 3 | 11 | 4 | 0 | 7 | 7 | 2 | 1 | 0 | 4 | 0 | 4 | 0 | 10 | 4 | 5 | 8 | 6 | 3 | 4 | 2 | 2 | Dnaaf6b |
| Dnaaf9 | 10022 | 10246 | 9488 | 12314 | 14202 | 13194 | 12602 | 15218 | 13868 | 15946 | 14970 | 14723 | 14804 | 15107 | 11211 | 10161 | 11136 | 11905 | 13170 | 11325 | 13129 | 18083 | 12954 | 13922 | 16979 | 9926 | 14262 | 14001 | Dnaaf9 |
| Dnah1 | 1103 | 1071 | 1089 | 851 | 794 | 950 | 824 | 1014 | 1158 | 1230 | 1022 | 1239 | 1146 | 1104 | 939 | 1249 | 964 | 732 | 773 | 994 | 822 | 1307 | 1290 | 1177 | 998 | 1369 | 1390 | 1200 | Dnah1 |
| Dnah10 | 315 | 458 | 190 | 528 | 299 | 256 | 253 | 136 | 136 | 197 | 116 | 129 | 179 | 86 | 310 | 435 | 270 | 456 | 281 | 264 | 256 | 134 | 199 | 111 | 138 | 43 | 172 | 115 | Dnah10 |
| Dnah11 | 246 | 292 | 160 | 158 | 205 | 238 | 175 | 299 | 315 | 266 | 218 | 184 | 247 | 122 | 277 | 429 | 127 | 168 | 142 | 257 | 201 | 147 | 418 | 119 | 136 | 76 | 155 | 109 | Dnah11 |
| Dnah12 | 1 | 0 | 0 | 1 | 0 | 2 | 1 | 2 | 2 | 1 | 1 | 2 | 1 | 0 | 8 | 2 | 2 | 0 | 2 | 2 | 2 | 2 | 10 | 1 | 2 | 1 | 0 | 0 | Dnah12 |
| Dnah14 | 6 | 5 | 15 | 9 | 11 | 8 | 4 | 13 | 13 | 10 | 4 | 13 | 25 | 15 | 8 | 10 | 3 | 7 | 3 | 9 | 3 | 10 | 14 | 7 | 9 | 15 | 27 | 26 | Dnah14 |
| Dnah17 | 173 | 236 | 146 | 339 | 317 | 240 | 306 | 56 | 53 | 76 | 47 | 33 | 42 | 37 | 165 | 207 | 286 | 365 | 362 | 233 | 339 | 53 | 56 | 50 | 63 | 27 | 52 | 32 | Dnah17 |
| Dnah2 | 888 | 1018 | 597 | 1069 | 1328 | 1185 | 1192 | 700 | 869 | 862 | 749 | 610 | 767 | 541 | 843 | 1055 | 829 | 1191 | 1119 | 1182 | 1297 | 886 | 929 | 524 | 794 | 310 | 572 | 394 | Dnah2 |
| Dnah3 | 24 | 48 | 10 | 14 | 49 | 72 | 34 | 66 | 57 | 55 | 46 | 49 | 83 | 23 | 34 | 84 | 8 | 23 | 34 | 75 | 39 | 42 | 122 | 25 | 25 | 26 | 48 | 31 | Dnah3 |
| Dnah5 | 278 | 336 | 223 | 216 | 214 | 279 | 266 | 327 | 245 | 180 | 146 | 223 | 323 | 293 | 367 | 336 | 253 | 220 | 239 | 378 | 301 | 247 | 276 | 176 | 199 | 118 | 392 | 235 | Dnah5 |
| Dnah6 | 205 | 295 | 30 | 87 | 359 | 251 | 227 | 382 | 516 | 348 | 317 | 267 | 437 | 117 | 243 | 419 | 65 | 149 | 217 | 320 | 292 | 348 | 891 | 114 | 255 | 88 | 191 | 90 | Dnah6 |
| Dnah7a | 89 | 108 | 92 | 125 | 136 | 125 | 110 | 162 | 153 | 163 | 115 | 133 | 172 | 143 | 117 | 114 | 80 | 79 | 94 | 141 | 120 | 137 | 148 | 115 | 117 | 171 | 160 | 160 | Dnah7a |
| Dnah7b | 760 | 787 | 316 | 545 | 758 | 986 | 760 | 1473 | 1332 | 820 | 1023 | 1097 | 1042 | 1102 | 894 | 848 | 401 | 495 | 554 | 1156 | 857 | 1033 | 1418 | 632 | 888 | 551 | 1175 | 862 | Dnah7b |
| Dnah7c | 95 | 90 | 33 | 66 | 70 | 102 | 78 | 126 | 115 | 86 | 90 | 97 | 91 | 81 | 100 | 86 | 24 | 73 | 81 | 129 | 90 | 114 | 151 | 60 | 80 | 33 | 101 | 81 | Dnah7c |
| Dnah8 | 268 | 276 | 204 | 140 | 159 | 197 | 126 | 219 | 217 | 152 | 217 | 192 | 246 | 156 | 248 | 363 | 129 | 185 | 124 | 113 | 109 | 196 | 302 | 176 | 157 | 159 | 200 | 180 | Dnah8 |
| Dnah9 | 527 | 682 | 362 | 465 | 695 | 563 | 523 | 679 | 569 | 434 | 524 | 421 | 841 | 298 | 765 | 769 | 391 | 445 | 450 | 811 | 585 | 461 | 752 | 265 | 440 | 160 | 501 | 326 | Dnah9 |
| Dnai1 | 224 | 285 | 198 | 252 | 164 | 214 | 166 | 104 | 107 | 104 | 80 | 134 | 108 | 88 | 245 | 276 | 229 | 219 | 192 | 218 | 215 | 104 | 125 | 81 | 126 | 71 | 93 | 100 | Dnai1 |
| Dnai2 | 209 | 257 | 218 | 266 | 269 | 264 | 204 | 210 | 263 | 277 | 234 | 245 | 294 | 193 | 224 | 284 | 229 | 248 | 225 | 210 | 211 | 297 | 289 | 169 | 207 | 222 | 223 | 216 | Dnai2 |
| Dnai3 | 129 | 141 | 43 | 109 | 151 | 123 | 94 | 123 | 144 | 139 | 70 | 111 | 206 | 78 | 133 | 176 | 81 | 107 | 83 | 116 | 130 | 154 | 241 | 98 | 68 | 60 | 60 | 61 | Dnai3 |
| Dnai4 | 2666 | 2546 | 2104 | 2384 | 1956 | 2122 | 2318 | 1575 | 1366 | 1359 | 1273 | 1393 | 1602 | 1237 | 2606 | 2682 | 2401 | 2189 | 1959 | 2067 | 2260 | 1507 | 1720 | 1343 | 1070 | 1139 | 1304 | 1121 | Dnai4 |
| Dnaja1 | 24827 | 22956 | 28990 | 28159 | 29778 | 26533 | 32992 | 44133 | 35430 | 36676 | 35238 | 37694 | 36915 | 43570 | 30071 | 24464 | 28548 | 27413 | 28860 | 33982 | 29954 | 39883 | 35472 | 40081 | 39280 | 41431 | 39409 | 43747 | Dnaja1 |
| Dnaja2 | 16903 | 14937 | 16856 | 15748 | 14897 | 14937 | 16718 | 14256 | 12615 | 12613 | 12759 | 13579 | 13368 | 14200 | 17396 | 15800 | 16448 | 15408 | 15721 | 16412 | 16338 | 12838 | 12936 | 13692 | 13419 | 15232 | 14278 | 15234 | Dnaja2 |
| Dnaja3 | 2607 | 2823 | 2842 | 2951 | 2443 | 2321 | 2528 | 2535 | 2552 | 2751 | 2828 | 2678 | 2626 | 2726 | 2505 | 2544 | 2890 | 2852 | 2615 | 2401 | 2543 | 2552 | 2821 | 2857 | 2632 | 2894 | 2736 | 2874 | Dnaja3 |
| Dnaja4 | 617 | 595 | 606 | 548 | 738 | 638 | 502 | 546 | 636 | 556 | 740 | 367 | 523 | 402 | 630 | 716 | 535 | 668 | 556 | 537 | 575 | 573 | 843 | 499 | 448 | 348 | 403 | 403 | Dnaja4 |
| Dnajb1 | 4428 | 5372 | 4948 | 4681 | 3770 | 3782 | 3890 | 3814 | 3528 | 3507 | 3282 | 3713 | 3511 | 3571 | 4408 | 4342 | 4709 | 4305 | 4373 | 3950 | 4010 | 3775 | 3552 | 3889 | 3667 | 4546 | 3886 | 4009 | Dnajb1 |
| Dnajb11 | 6309 | 6263 | 7841 | 7066 | 5948 | 5728 | 6493 | 4732 | 4440 | 4869 | 5395 | 4687 | 4824 | 4996 | 6207 | 6284 | 6847 | 6891 | 6111 | 5448 | 6033 | 4931 | 4585 | 5265 | 5024 | 5505 | 4940 | 5239 | Dnajb11 |
| Dnajb12 | 741 | 751 | 1000 | 965 | 938 | 862 | 759 | 1503 | 1707 | 2036 | 1842 | 2056 | 1982 | 2065 | 661 | 651 | 876 | 814 | 1015 | 753 | 839 | 1853 | 1775 | 2006 | 1866 | 2347 | 1909 | 2198 | Dnajb12 |
| Dnajb13 | 37 | 86 | 29 | 48 | 97 | 109 | 51 | 129 | 95 | 85 | 72 | 114 | 167 | 49 | 83 | 83 | 28 | 35 | 52 | 87 | 48 | 112 | 179 | 56 | 103 | 65 | 56 | 69 | Dnajb13 |
| Dnajb14 | 2303 | 1932 | 1909 | 2295 | 2911 | 3047 | 2482 | 3949 | 3882 | 3874 | 4355 | 3726 | 3742 | 3521 | 2483 | 2467 | 1969 | 2702 | 2343 | 2464 | 2695 | 4055 | 3988 | 3348 | 4218 | 2641 | 3481 | 3278 | Dnajb14 |
| Dnajb2 | 1694 | 2013 | 2418 | 2298 | 2507 | 2272 | 2224 | 2168 | 2512 | 2806 | 2645 | 2457 | 2606 | 2317 | 1789 | 1840 | 2561 | 2397 | 2617 | 2205 | 2143 | 2615 | 2526 | 2879 | 2716 | 2501 | 2350 | 2402 | Dnajb2 |
| Dnajb3 | 170 | 191 | 280 | 169 | 253 | 234 | 242 | 201 | 225 | 312 | 330 | 224 | 295 | 339 | 181 | 147 | 189 | 143 | 208 | 200 | 223 | 212 | 254 | 259 | 244 | 258 | 281 | 265 | Dnajb3 |
| Dnajb4 | 3823 | 3334 | 4212 | 4006 | 4435 | 4062 | 4445 | 2793 | 2500 | 2682 | 2609 | 2615 | 2963 | 2837 | 4271 | 3974 | 4175 | 4471 | 3900 | 4374 | 4177 | 2846 | 2760 | 2668 | 2588 | 3019 | 2667 | 2937 | Dnajb4 |
| Dnajb5 | 3017 | 4664 | 4977 | 5319 | 6143 | 5520 | 5288 | 5076 | 5517 | 6372 | 5366 | 6323 | 5114 | 6224 | 3713 | 3529 | 5773 | 5523 | 6288 | 5436 | 5654 | 6876 | 5073 | 6445 | 6994 | 5210 | 5824 | 5941 | Dnajb5 |
| Dnajb6 | 10430 | 9493 | 10786 | 11398 | 10835 | 10349 | 12094 | 10978 | 9328 | 9890 | 8542 | 9744 | 8657 | 10272 | 11879 | 9569 | 10805 | 11074 | 11428 | 12658 | 11657 | 10169 | 9398 | 9852 | 10187 | 9751 | 9645 | 10001 | Dnajb6 |
| Dnajb7 | 2 | 2 | 0 | 0 | 1 | 0 | 0 | 0 | 0 | 1 | 0 | 2 | 2 | 4 | 0 | 1 | 0 | 0 | 0 | 0 | 4 | 0 | 3 | 0 | 0 | 0 | 2 | 7 | Dnajb7 |
| Dnajb8 | 0 | 0 | 0 | 2 | 0 | 0 | 2 | 0 | 0 | 0 | 0 | 0 | 0 | 0 | 0 | 0 | 0 | 0 | 0 | 0 | 0 | 0 | 0 | 0 | 0 | 0 | 0 | 0 | Dnajb8 |
| Dnajb9 | 2465 | 2025 | 2524 | 2809 | 2532 | 2388 | 2848 | 2462 | 2079 | 2009 | 2221 | 1892 | 2124 | 2120 | 2515 | 2448 | 2513 | 2854 | 2516 | 2469 | 2981 | 2647 | 2350 | 2161 | 2276 | 2090 | 1952 | 2191 | Dnajb9 |
| Dnajc1 | 6308 | 5821 | 4209 | 4706 | 4279 | 4874 | 5027 | 3881 | 3510 | 2881 | 3598 | 3509 | 3612 | 3866 | 6054 | 5374 | 3987 | 4559 | 4217 | 5263 | 5045 | 2837 | 3456 | 3134 | 3362 | 3350 | 3681 | 3756 | Dnajc1 |
| Dnajc10 | 13393 | 11245 | 11112 | 8767 | 8768 | 9731 | 9470 | 10153 | 9606 | 7864 | 8615 | 9086 | 9737 | 9250 | 13740 | 12728 | 9094 | 9185 | 8480 | 11210 | 8815 | 8363 | 9784 | 7710 | 7895 | 8945 | 9840 | 9133 | Dnajc10 |
| Dnajc11 | 6345 | 7192 | 7398 | 7055 | 6397 | 6256 | 6529 | 6128 | 6247 | 6694 | 6421 | 7459 | 6890 | 7824 | 6117 | 6200 | 7097 | 6092 | 6782 | 6701 | 6519 | 6137 | 5803 | 7380 | 7122 | 7584 | 7513 | 7973 | Dnajc11 |
| Dnajc12 | 2215 | 2563 | 2472 | 2309 | 2149 | 2133 | 2375 | 573 | 592 | 471 | 527 | 503 | 486 | 536 | 2629 | 2306 | 2622 | 2360 | 2389 | 2525 | 2389 | 590 | 609 | 499 | 431 | 447 | 579 | 545 | Dnajc12 |
| Dnajc13 | 10278 | 8847 | 6035 | 7221 | 7745 | 9249 | 8670 | 9816 | 9348 | 8198 | 8997 | 8685 | 8720 | 9022 | 9715 | 9164 | 6689 | 7565 | 7283 | 8829 | 9165 | 7698 | 8943 | 6816 | 8548 | 5553 | 9479 | 8389 | Dnajc13 |
| Dnajc14 | 5253 | 5339 | 6090 | 5481 | 5788 | 5090 | 5406 | 5324 | 5545 | 5679 | 6050 | 5889 | 5706 | 6423 | 5097 | 5061 | 5968 | 5699 | 5485 | 5127 | 5444 | 5631 | 5581 | 5929 | 6013 | 5977 | 5890 | 6088 | Dnajc14 |
| Dnajc15 | 3039 | 2959 | 2490 | 2801 | 2847 | 2882 | 3456 | 2302 | 2010 | 1744 | 1842 | 1925 | 1961 | 2325 | 3355 | 2727 | 2783 | 2673 | 2749 | 3645 | 3194 | 1985 | 1923 | 1817 | 1964 | 2012 | 2074 | 2461 | Dnajc15 |
| Dnajc16 | 3042 | 3758 | 3151 | 4099 | 3660 | 3488 | 3643 | 2375 | 2489 | 2863 | 2664 | 2785 | 2715 | 2600 | 3199 | 3400 | 3482 | 4002 | 3737 | 3130 | 3855 | 2979 | 2339 | 2601 | 2907 | 2232 | 2531 | 2620 | Dnajc16 |
| Dnajc17 | 353 | 465 | 563 | 530 | 585 | 517 | 542 | 832 | 838 | 908 | 905 | 991 | 856 | 1076 | 421 | 343 | 546 | 511 | 563 | 589 | 528 | 703 | 780 | 1082 | 975 | 1120 | 1038 | 997 | Dnajc17 |
| Dnajc18 | 3934 | 4209 | 3746 | 4237 | 4457 | 4111 | 4324 | 4045 | 3823 | 4266 | 4086 | 3999 | 3813 | 4042 | 4189 | 3847 | 4140 | 4510 | 4431 | 4041 | 4441 | 4662 | 3836 | 4083 | 4498 | 3541 | 3956 | 3877 | Dnajc18 |
| Dnajc19 | 997 | 1012 | 1126 | 940 | 931 | 773 | 1084 | 1458 | 1254 | 1190 | 1254 | 1337 | 1198 | 1607 | 1011 | 808 | 984 | 801 | 854 | 1095 | 1075 | 1203 | 1192 | 1376 | 1397 | 1452 | 1416 | 1629 | Dnajc19 |
| Dnajc2 | 7323 | 6328 | 7400 | 6373 | 5406 | 5781 | 6411 | 6270 | 5193 | 5212 | 4752 | 6140 | 5618 | 6371 | 7903 | 6537 | 6905 | 6218 | 5716 | 6885 | 6053 | 5198 | 4979 | 5754 | 5317 | 7066 | 6302 | 6990 | Dnajc2 |
| Dnajc21 | 3260 | 2737 | 3904 | 3294 | 2706 | 2830 | 2994 | 2583 | 2011 | 2402 | 2244 | 2497 | 2307 | 2839 | 3216 | 2967 | 3264 | 3132 | 2890 | 2999 | 2910 | 2296 | 2037 | 2488 | 2254 | 2909 | 2580 | 2732 | Dnajc21 |
| Dnajc22 | 31 | 66 | 100 | 79 | 111 | 105 | 44 | 110 | 68 | 85 | 64 | 97 | 172 | 70 | 76 | 75 | 137 | 66 | 107 | 98 | 57 | 105 | 52 | 58 | 82 | 45 | 76 | 82 | Dnajc22 |
| Dnajc24 | 1017 | 1070 | 1047 | 912 | 789 | 900 | 991 | 1016 | 947 | 791 | 805 | 882 | 874 | 1104 | 985 | 908 | 870 | 738 | 883 | 1095 | 925 | 648 | 924 | 852 | 851 | 961 | 918 | 1105 | Dnajc24 |
| Dnajc25 | 1061 | 828 | 876 | 952 | 690 | 796 | 736 | 725 | 611 | 760 | 760 | 628 | 642 | 647 | 901 | 1001 | 825 | 910 | 607 | 670 | 712 | 732 | 712 | 676 | 598 | 645 | 560 | 700 | Dnajc25 |
| Dnajc27 | 1536 | 1441 | 1536 | 1456 | 1652 | 1630 | 1633 | 2220 | 2283 | 2272 | 2174 | 2530 | 2328 | 2331 | 1382 | 1329 | 1428 | 1345 | 1588 | 1452 | 1607 | 2261 | 2086 | 1907 | 2212 | 2014 | 2439 | 2356 | Dnajc27 |
| Dnajc28 | 912 | 1063 | 1289 | 1043 | 868 | 865 | 945 | 716 | 580 | 693 | 585 | 657 | 629 | 635 | 1031 | 1023 | 1174 | 984 | 1018 | 1060 | 958 | 576 | 575 | 592 | 643 | 690 | 678 | 662 | Dnajc28 |
| Dnajc3 | 7201 | 6238 | 6905 | 6037 | 5114 | 5289 | 5539 | 4750 | 4356 | 4185 | 5317 | 3663 | 4199 | 3870 | 7234 | 7447 | 5953 | 6329 | 5034 | 6088 | 5146 | 4298 | 4969 | 4208 | 4111 | 3519 | 4011 | 3818 | Dnajc3 |
| Dnajc30 | 875 | 896 | 781 | 904 | 898 | 1033 | 835 | 923 | 943 | 1051 | 976 | 950 | 951 | 1007 | 849 | 780 | 932 | 833 | 880 | 792 | 955 | 1029 | 983 | 975 | 1039 | 1089 | 930 | 980 | Dnajc30 |
| Dnajc4 | 528 | 706 | 663 | 783 | 643 | 694 | 575 | 535 | 665 | 624 | 650 | 699 | 600 | 648 | 518 | 641 | 669 | 676 | 663 | 668 | 559 | 682 | 684 | 688 | 636 | 679 | 654 | 576 | Dnajc4 |
| Dnajc5 | 18758 | 19078 | 15828 | 20897 | 22951 | 21945 | 21994 | 18099 | 17279 | 17976 | 17256 | 17604 | 16278 | 16377 | 19494 | 16931 | 18373 | 21159 | 22068 | 21196 | 23158 | 20630 | 16824 | 17196 | 20116 | 14718 | 16867 | 16121 | Dnajc5 |
| Dnajc5b | 0 | 1 | 0 | 0 | 0 | 0 | 0 | 0 | 2 | 4 | 0 | 2 | 3 | 0 | 0 | 0 | 0 | 0 | 0 | 3 | 1 | 0 | 0 | 0 | 0 | 4 | 2 | 1 | Dnajc5b |
| Dnajc5g | 0 | 0 | 0 | 0 | 0 | 0 | 0 | 0 | 0 | 0 | 0 | 1 | 0 | 1 | 0 | 0 | 1 | 0 | 0 | 4 | 0 | 0 | 0 | 0 | 0 | 0 | 0 | 0 | Dnajc5g |
| Dnajc6 | 2335 | 2511 | 1767 | 2855 | 5428 | 4093 | 4310 | 3817 | 3957 | 4151 | 3767 | 2490 | 2804 | 2643 | 2935 | 2428 | 2700 | 3652 | 3949 | 4647 | 4296 | 5235 | 3841 | 3266 | 4630 | 1902 | 2155 | 2210 | Dnajc6 |
| Dnajc7 | 13776 | 13268 | 14277 | 13275 | 13938 | 13954 | 14180 | 21198 | 18004 | 18786 | 18241 | 20205 | 18274 | 21136 | 14209 | 11798 | 14317 | 12586 | 14647 | 15694 | 14040 | 19336 | 17876 | 19738 | 20203 | 21632 | 20571 | 20374 | Dnajc7 |
| Dnajc8 | 11580 | 10638 | 12502 | 11156 | 9223 | 9902 | 11126 | 9220 | 8320 | 8784 | 8089 | 9520 | 9140 | 10556 | 11735 | 10404 | 11434 | 10186 | 10196 | 11573 | 10771 | 7585 | 8090 | 9253 | 8837 | 11485 | 10447 | 10943 | Dnajc8 |
| Dnajc9 | 12530 | 10995 | 11970 | 10550 | 8125 | 8600 | 11619 | 7498 | 6568 | 7441 | 6248 | 9332 | 7231 | 10719 | 11530 | 9140 | 11419 | 9400 | 9976 | 9838 | 10440 | 5619 | 5853 | 8634 | 7492 | 11482 | 9207 | 10791 | Dnajc9 |
| Dnal1 | 4485 | 4292 | 3688 | 4768 | 4676 | 4936 | 5173 | 4027 | 3714 | 3694 | 3580 | 3624 | 3506 | 4006 | 4969 | 4255 | 3972 | 4735 | 4521 | 5418 | 5292 | 3913 | 3858 | 3376 | 4076 | 2880 | 3436 | 3483 | Dnal1 |
| Dnal4 | 2308 | 3006 | 2547 | 2742 | 2392 | 2502 | 2470 | 1824 | 2031 | 2141 | 2060 | 2325 | 2135 | 2123 | 2264 | 2479 | 2715 | 2481 | 2916 | 2351 | 2532 | 2050 | 2137 | 2460 | 2086 | 2430 | 2149 | 2239 | Dnal4 |
| Dnali1 | 23 | 43 | 25 | 39 | 49 | 45 | 46 | 67 | 75 | 63 | 50 | 77 | 98 | 32 | 44 | 51 | 23 | 34 | 51 | 67 | 51 | 57 | 124 | 59 | 40 | 31 | 44 | 44 | Dnali1 |
| Dnase1 | 0 | 0 | 0 | 0 | 0 | 2 | 2 | 4 | 2 | 3 | 0 | 1 | 5 | 3 | 0 | 0 | 0 | 0 | 1 | 1 | 2 | 2 | 1 | 2 | 0 | 1 | 2 | 2 | Dnase1 |
| Dnase1l1 | 254 | 211 | 195 | 169 | 155 | 165 | 165 | 158 | 201 | 208 | 230 | 188 | 248 | 192 | 243 | 259 | 146 | 200 | 143 | 200 | 130 | 140 | 252 | 203 | 178 | 218 | 199 | 224 | Dnase1l1 |

| | | | | | | | | | | | | | | | | | | | | | | | | | | | | | |
|---|---|---|---|---|---|---|---|---|---|---|---|---|---|---|---|---|---|---|---|---|---|---|---|---|---|---|---|---|---|
| Dnase1l2 | 261 | 574 | 563 | 682 | 699 | 585 | 577 | 424 | 466 | 561 | 624 | 463 | 488 | 437 | 336 | 479 | 696 | 726 | 793 | 498 | 654 | 650 | 437 | 583 | 661 | 413 | 471 | 471 | Dnase1l2 |
| Dnase1l3 | 7 | 6 | 0 | 1 | 2 | 5 | 2 | 6 | 9 | 17 | 6 | 10 | 6 | 4 | 9 | 10 | 2 | 8 | 4 | 2 | 1 | 10 | 19 | 10 | 15 | 8 | 12 | 6 | Dnase1l3 |
| Dnase2a | 2205 | 2331 | 2393 | 2139 | 1581 | 1680 | 1837 | 587 | 756 | 649 | 746 | 745 | 754 | 698 | 1701 | 2165 | 2261 | 2134 | 1880 | 1593 | 1784 | 518 | 723 | 878 | 553 | 1045 | 718 | 730 | Dnase2a |
| Dnase2b | 0 | 1 | 0 | 0 | 0 | 3 | 0 | 0 | 0 | 0 | 4 | 0 | 0 | 0 | 1 | 2 | 0 | 0 | 2 | 0 | 4 | 0 | 0 | 2 | 0 | 0 | 0 | 0 | Dnase2b |
| Dnd1 | 140 | 230 | 232 | 237 | 190 | 164 | 146 | 120 | 126 | 197 | 174 | 140 | 135 | 173 | 131 | 158 | 180 | 213 | 172 | 156 | 167 | 172 | 138 | 191 | 203 | 214 | 153 | 170 | Dnd1 |
| Dner | 3893 | 4565 | 4234 | 8803 | 13845 | 9536 | 10781 | 8120 | 8727 | 10648 | 9847 | 7449 | 7035 | 7503 | 5354 | 3903 | 7429 | 10222 | 10948 | 8478 | 11015 | 14253 | 8903 | 8054 | 11984 | 5722 | 5683 | 6590 | Dner |
| Dnhd1 | 1473 | 1932 | 982 | 1781 | 1573 | 1698 | 1656 | 666 | 686 | 660 | 653 | 550 | 582 | 521 | 1489 | 1769 | 1426 | 1966 | 1432 | 1489 | 1881 | 663 | 617 | 427 | 691 | 224 | 511 | 428 | Dnhd1 |
| Dnlz | 1437 | 1671 | 1571 | 1305 | 1457 | 1512 | 1361 | 2363 | 2477 | 2462 | 2184 | 2746 | 2663 | 2618 | 1410 | 1318 | 1562 | 1318 | 1602 | 1472 | 1408 | 2055 | 2460 | 2697 | 2527 | 3265 | 2960 | 2627 | Dnlz |
| Dnm1 | 4976 | 5168 | 5522 | 7120 | 8607 | 7105 | 5791 | 5635 | 7464 | 9457 | 9527 | 6543 | 6615 | 5584 | 4516 | 5831 | 6576 | 7801 | 7868 | 5172 | 6731 | 10190 | 6840 | 8027 | 8530 | 5389 | 5381 | 5105 | Dnm1 |
| Dnm1l | 14693 | 11710 | 12420 | 12910 | 16312 | 15876 | 16810 | 19267 | 16397 | 17012 | 16339 | 17136 | 16389 | 17536 | 15314 | 12636 | 13050 | 13230 | 14774 | 16172 | 16000 | 18198 | 16311 | 15416 | 17230 | 14734 | 16708 | 16813 | Dnm1l |
| Dnm2 | 9401 | 11606 | 10955 | 12564 | 10809 | 10611 | 10522 | 5083 | 5792 | 6481 | 6014 | 6429 | 5801 | 6074 | 9376 | 10344 | 11748 | 12404 | 11501 | 8452 | 10812 | 6450 | 5783 | 6432 | 6689 | 6352 | 5507 | 5791 | Dnm2 |
| Dnm3 | 2712 | 2834 | 1866 | 4458 | 8010 | 6447 | 6378 | 4863 | 5043 | 5702 | 5378 | 3252 | 4018 | 3260 | 3455 | 3100 | 3417 | 5650 | 5710 | 5299 | 6811 | 7229 | 4751 | 3551 | 5810 | 1646 | 2793 | 2639 | Dnm3 |
| Dnmbp | 2314 | 2771 | 2689 | 2362 | 1887 | 2074 | 1825 | 1778 | 2292 | 2175 | 2430 | 2463 | 2426 | 2357 | 2294 | 2769 | 2592 | 2384 | 2085 | 1781 | 1839 | 1988 | 2396 | 2317 | 1978 | 2522 | 2325 | 2322 | Dnmbp |
| Dnmt1 | 20788 | 21801 | 21711 | 20576 | 18681 | 20002 | 19801 | 15850 | 19207 | 20003 | 18594 | 25149 | 21244 | 22980 | 18296 | 20221 | 21797 | 20007 | 21446 | 18482 | 19538 | 16827 | 16508 | 21727 | 20726 | 22936 | 22896 | 23169 | Dnmt1 |
| Dnmt3a | 8589 | 8314 | 9868 | 9686 | 12317 | 10937 | 10184 | 18733 | 21136 | 23035 | 24500 | 22724 | 23181 | 22174 | 8225 | 8989 | 10754 | 10407 | 11985 | 9123 | 10460 | 20591 | 20300 | 23249 | 22411 | 18534 | 21066 | 21192 | Dnmt3a |
| Dnmt3b | 2271 | 2341 | 2348 | 1515 | 1294 | 1561 | 1482 | 768 | 873 | 725 | 848 | 971 | 1035 | 1017 | 1982 | 2579 | 1815 | 1624 | 1372 | 1406 | 1467 | 593 | 794 | 842 | 753 | 1108 | 1145 | 1060 | Dnmt3b |
| Dnmt3c | 7 | 4 | 2 | 1 | 1 | 1 | 3 | 2 | 5 | 14 | 5 | 11 | 9 | 11 | 7 | 4 | 0 | 1 | 4 | 1 | 0 | 13 | 6 | 7 | 6 | 13 | 13 | 6 | Dnmt3c |
| Dnmt3l | 0 | 0 | 3 | 0 | 0 | 0 | 0 | 1 | 0 | 0 | 3 | 0 | 0 | 1 | 0 | 0 | 0 | 0 | 1 | 0 | 0 | 0 | 1 | 2 | 3 | 2 | 0 | 0 | Dnmt3l |
| Dnpep | 6293 | 8274 | 6971 | 6475 | 6282 | 6052 | 5773 | 3916 | 4328 | 4296 | 3852 | 4369 | 4427 | 4388 | 6237 | 6804 | 7073 | 6000 | 6733 | 6246 | 5756 | 4142 | 4419 | 4207 | 4281 | 4927 | 4903 | 4587 | Dnpep |
| Dnph1 | 1467 | 1479 | 1552 | 1237 | 823 | 865 | 1083 | 951 | 852 | 851 | 741 | 1357 | 1124 | 1409 | 1278 | 1219 | 1494 | 1022 | 1064 | 1111 | 987 | 702 | 847 | 1186 | 1018 | 1612 | 1335 | 1576 | Dnph1 |
| Dntt | 0 | 3 | 0 | 7 | 5 | 2 | 2 | 1 | 2 | 3 | 2 | 8 | 8 | 1 | 3 | 7 | 0 | 2 | 0 | 9 | 2 | 7 | 8 | 3 | 5 | 0 | 3 | 9 | Dntt |
| Dnttip1 | 1844 | 2275 | 2276 | 2296 | 1906 | 1874 | 2190 | 1896 | 2138 | 2162 | 2084 | 2371 | 2188 | 2480 | 1791 | 1907 | 2313 | 2103 | 2046 | 1886 | 2069 | 2000 | 2040 | 2363 | 2266 | 2556 | 2461 | 2614 | Dnttip1 |
| Dnttip2 | 5906 | 5426 | 5595 | 4473 | 3921 | 4204 | 4706 | 5576 | 4779 | 4272 | 4405 | 5324 | 4955 | 5545 | 5987 | 5322 | 5161 | 4569 | 4210 | 5268 | 4276 | 4466 | 4827 | 5015 | 4521 | 5842 | 5298 | 5619 | Dnttip2 |
| Doc2a | 392 | 523 | 677 | 490 | 529 | 509 | 495 | 249 | 284 | 357 | 241 | 354 | 256 | 328 | 404 | 344 | 601 | 388 | 599 | 515 | 517 | 287 | 263 | 270 | 296 | 396 | 315 | 335 | Doc2a |
| Doc2b | 484 | 409 | 490 | 617 | 1064 | 679 | 845 | 708 | 903 | 956 | 838 | 398 | 373 | 353 | 398 | 425 | 622 | 774 | 860 | 656 | 852 | 1004 | 873 | 724 | 769 | 506 | 313 | 301 | Doc2b |
| Doc2g | 122 | 143 | 161 | 197 | 245 | 145 | 228 | 137 | 161 | 248 | 188 | 209 | 191 | 208 | 108 | 154 | 182 | 218 | 263 | 198 | 206 | 226 | 162 | 278 | 270 | 242 | 180 | 179 | Doc2g |
| Dock1 | 4609 | 3781 | 3683 | 2941 | 2563 | 3361 | 2713 | 4290 | 5306 | 4634 | 6069 | 4947 | 5558 | 4503 | 4063 | 5438 | 3007 | 3176 | 2561 | 2383 | 2404 | 4050 | 5727 | 4503 | 3827 | 4391 | 4686 | 4664 | Dock1 |
| Dock10 | 1502 | 1125 | 724 | 746 | 1126 | 1035 | 821 | 1413 | 1561 | 1072 | 1429 | 775 | 1044 | 699 | 1580 | 1629 | 746 | 892 | 736 | 951 | 820 | 1116 | 1463 | 809 | 831 | 657 | 893 | 687 | Dock10 |
| Dock11 | 5502 | 4196 | 3587 | 3611 | 4304 | 5007 | 4769 | 8109 | 6959 | 5866 | 6476 | 6827 | 5912 | 6806 | 5076 | 4692 | 3619 | 3557 | 4011 | 4826 | 4578 | 6007 | 6851 | 5620 | 6183 | 4907 | 6771 | 6252 | Dock11 |
| Dock2 | 168 | 187 | 143 | 149 | 128 | 196 | 108 | 194 | 203 | 146 | 270 | 136 | 174 | 149 | 215 | 210 | 94 | 163 | 122 | 161 | 130 | 179 | 280 | 120 | 145 | 99 | 146 | 132 | Dock2 |
| Dock3 | 1967 | 1968 | 1752 | 3270 | 5159 | 4608 | 4577 | 3968 | 4170 | 4715 | 3932 | 4236 | 3435 | 4239 | 2262 | 1699 | 2489 | 3650 | 4259 | 3863 | 4824 | 5309 | 3555 | 3867 | 5743 | 2392 | 3661 | 3559 | Dock3 |
| Dock4 | 4568 | 4050 | 2712 | 4304 | 5902 | 5826 | 5694 | 6449 | 5504 | 5367 | 5389 | 4474 | 4643 | 4911 | 4784 | 4541 | 3314 | 4929 | 4898 | 5169 | 5816 | 6126 | 5260 | 4538 | 6054 | 2717 | 4546 | 4330 | Dock4 |
| Dock5 | 1082 | 1063 | 1080 | 750 | 873 | 882 | 591 | 1231 | 1338 | 1316 | 1838 | 1437 | 1921 | 1391 | 1063 | 1401 | 831 | 902 | 577 | 663 | 659 | 1305 | 1646 | 1153 | 1116 | 905 | 2028 | 1529 | Dock5 |
| Dock6 | 4806 | 6439 | 7405 | 5649 | 4761 | 4339 | 3595 | 2583 | 3452 | 3704 | 4709 | 4300 | 4632 | 3526 | 4295 | 7391 | 6054 | 5798 | 4576 | 3594 | 3822 | 3795 | 4121 | 3863 | 3613 | 4104 | 4256 | 3710 | Dock6 |
| Dock7 | 23032 | 21114 | 15923 | 18653 | 20848 | 23304 | 22801 | 24569 | 22022 | 19675 | 19823 | 23284 | 21152 | 22356 | 23492 | 20724 | 18032 | 18351 | 19886 | 22559 | 22299 | 21497 | 21507 | 18623 | 22653 | 15843 | 22798 | 20874 | Dock7 |
| Dock8 | 223 | 207 | 159 | 118 | 105 | 182 | 68 | 335 | 470 | 325 | 507 | 288 | 279 | 196 | 191 | 251 | 95 | 118 | 117 | 114 | 97 | 199 | 605 | 362 | 198 | 139 | 289 | 252 | Dock8 |
| Dock9 | 3082 | 2999 | 1917 | 2298 | 3040 | 2767 | 2618 | 2676 | 3035 | 2650 | 2948 | 2156 | 2425 | 1874 | 2672 | 3305 | 2130 | 2580 | 2570 | 2751 | 2820 | 2695 | 3708 | 1772 | 2639 | 1259 | 2022 | 1620 | Dock9 |
| Dohh | 286 | 370 | 299 | 367 | 259 | 188 | 239 | 233 | 221 | 247 | 204 | 204 | 215 | 274 | 262 | 217 | 316 | 310 | 264 | 270 | 267 | 189 | 232 | 267 | 227 | 225 | 244 | 236 | Dohh |
| Dok1 | 520 | 688 | 711 | 529 | 443 | 499 | 416 | 208 | 324 | 284 | 343 | 300 | 361 | 283 | 501 | 628 | 664 | 536 | 481 | 462 | 473 | 206 | 397 | 288 | 268 | 473 | 361 | 309 | Dok1 |
| Dok2 | 152 | 133 | 143 | 129 | 131 | 116 | 123 | 236 | 308 | 232 | 388 | 263 | 315 | 229 | 117 | 159 | 184 | 123 | 127 | 123 | 123 | 264 | 337 | 311 | 247 | 289 | 281 | 202 | Dok2 |
| Dok3 | 769 | 1261 | 1249 | 1441 | 1532 | 1191 | 1155 | 536 | 588 | 772 | 645 | 667 | 624 | 627 | 764 | 1095 | 1415 | 1554 | 1380 | 1146 | 1227 | 862 | 531 | 746 | 765 | 687 | 585 | 579 | Dok3 |
| Dok4 | 2562 | 3363 | 2664 | 3712 | 4474 | 3859 | 3643 | 2949 | 3435 | 3372 | 3145 | 2500 | 2633 | 2336 | 2858 | 3197 | 3169 | 3927 | 4062 | 4104 | 3910 | 3773 | 3242 | 3115 | 3659 | 2423 | 2309 | 2191 | Dok4 |
| Dok5 | 622 | 745 | 840 | 1087 | 1614 | 1158 | 1314 | 999 | 920 | 1097 | 1208 | 830 | 981 | 880 | 795 | 701 | 957 | 1264 | 1407 | 1223 | 1490 | 1151 | 1016 | 953 | 1166 | 539 | 790 | 952 | Dok5 |
| Dok6 | 792 | 579 | 603 | 1067 | 2113 | 1697 | 1934 | 2513 | 2199 | 2203 | 2045 | 1604 | 1560 | 1790 | 986 | 732 | 744 | 1237 | 1415 | 1873 | 2000 | 2709 | 1957 | 1662 | 2639 | 1021 | 1485 | 1461 | Dok6 |
| Dok7 | 122 | 162 | 140 | 203 | 322 | 215 | 192 | 128 | 198 | 152 | 389 | 103 | 96 | 76 | 124 | 139 | 158 | 274 | 241 | 209 | 213 | 241 | 263 | 141 | 184 | 68 | 88 | 44 | Dok7 |
| Dolk | 1225 | 1513 | 1177 | 1353 | 1114 | 1023 | 938 | 801 | 961 | 879 | 1065 | 944 | 906 | 899 | 1121 | 1358 | 1114 | 1296 | 984 | 1073 | 1045 | 887 | 973 | 890 | 1019 | 895 | 927 | 978 | Dolk |
| Dolpp1 | 1888 | 2503 | 2227 | 2363 | 2008 | 2130 | 2034 | 1721 | 1955 | 1842 | 1852 | 2134 | 2068 | 2070 | 1809 | 1953 | 2380 | 2242 | 2297 | 2184 | 2065 | 1958 | 1820 | 2135 | 2191 | 2295 | 2388 | 2060 | Dolpp1 |
| Donson | 3178 | 2729 | 2887 | 2900 | 2749 | 2946 | 3242 | 2470 | 2015 | 2318 | 2177 | 2349 | 2346 | 2493 | 3109 | 2812 | 2968 | 3136 | 2895 | 2969 | 2940 | 2446 | 1874 | 2356 | 2296 | 2708 | 2520 | 2584 | Donson |
| Dop1a | 5031 | 4982 | 2316 | 4010 | 4943 | 5307 | 5351 | 7538 | 6641 | 5245 | 5083 | 4605 | 4264 | 4453 | 5557 | 4786 | 3264 | 4350 | 4416 | 6254 | 5609 | 5619 | 6238 | 3867 | 5753 | 2622 | 4402 | 3717 | Dop1a |
| Dop1b | 3560 | 4717 | 3521 | 5571 | 6014 | 5358 | 5166 | 2691 | 3288 | 4119 | 3802 | 3470 | 3388 | 3243 | 3762 | 4372 | 4829 | 6114 | 5690 | 3977 | 5486 | 4664 | 3202 | 3331 | 4427 | 1924 | 2785 | 2934 | Dop1b |
| Dot1l | 5745 | 7653 | 6973 | 8106 | 8176 | 7543 | 7250 | 6857 | 8401 | 9682 | 8824 | 11026 | 9361 | 9823 | 5478 | 6655 | 8284 | 8230 | 9022 | 6704 | 7473 | 9999 | 7448 | 10008 | 10772 | 7157 | 10072 | 9183 | Dot1l |
| Doxl1 | 0 | 3 | 0 | 0 | 0 | 0 | 0 | 0 | 0 | 0 | 1 | 0 | 0 | 1 | 0 | 2 | 0 | 0 | 0 | 0 | 1 | 0 | 2 | 0 | 0 | 3 | 0 | 0 | Doxl1 |
| Dpagt1 | 2139 | 2378 | 2192 | 1934 | 2059 | 1893 | 2235 | 2274 | 2208 | 2184 | 1959 | 2207 | 2376 | 2240 | 2194 | 2107 | 2106 | 2012 | 2039 | 2244 | 2017 | 2089 | 2202 | 2252 | 2094 | 2434 | 2457 | 2427 | Dpagt1 |
| DPCD | 1503 | 1697 | 2060 | 1826 | 1654 | 1562 | 1467 | 1272 | 1379 | 1696 | 1540 | 1670 | 1725 | 1427 | 1473 | 1667 | 2027 | 1624 | 1648 | 1828 | 1468 | 1248 | 1410 | 1671 | 1518 | 1916 | 1701 | 1641 | DPCD |
| Dpep1 | 33 | 22 | 24 | 8 | 1 | 16 | 1 | 51 | 44 | 14 | 104 | 29 | 25 | 19 | 29 | 34 | 10 | 23 | 0 | 1 | 0 | 23 | 87 | 39 | 19 | 39 | 33 | 59 | Dpep1 |
| Dpep2 | 0 | 0 | 0 | 0 | 0 | 0 | 0 | 0 | 0 | 0 | 0 | 0 | 0 | 0 | 0 | 0 | 0 | 0 | 0 | 0 | 0 | 2 | 1 | 0 | 0 | 0 | 0 | 1 | Dpep2 |
| Dpep2 | 0 | 0 | 0 | 0 | 0 | 0 | 0 | 0 | 0 | 0 | 0 | 0 | 0 | 0 | 0 | 0 | 0 | 0 | 0 | 0 | 0 | 0 | 1 | 0 | 0 | 0 | 0 | 0 | Dpep2 |
| Dpep2nb | 3 | 0 | 0 | 0 | 5 | 0 | 0 | 1 | 4 | 0 | 0 | 0 | 0 | 3 | 1 | 2 | 1 | 7 | 0 | 4 | 2 | 2 | 0 | 0 | 3 | 0 | 0 | 0 | Dpep2nb |
| Dpep3 | 0 | 0 | 0 | 0 | 0 | 0 | 0 | 0 | 0 | 0 | 0 | 0 | 0 | 0 | 0 | 0 | 0 | 0 | 0 | 0 | 1 | 0 | 0 | 0 | 0 | 0 | 0 | 0 | Dpep3 |
| Dpf1 | 4349 | 5805 | 6073 | 8209 | 7622 | 6974 | 7410 | 4564 | 4354 | 6094 | 4456 | 5921 | 4061 | 5709 | 4469 | 4448 | 7635 | 7630 | 8479 | 6340 | 7880 | 6151 | 3804 | 6602 | 6842 | 5452 | 4994 | 4913 | Dpf1 |
| Dpf2 | 9872 | 11562 | 11459 | 11521 | 10808 | 10095 | 10931 | 9381 | 9396 | 9589 | 9452 | 10757 | 10366 | 10794 | 9915 | 9910 | 11835 | 11191 | 11536 | 10317 | 10664 | 9700 | 8939 | 10771 | 10134 | 11985 | 10982 | 11256 | Dpf2 |
| Dpf3 | 2089 | 2773 | 1203 | 3580 | 4120 | 3710 | 3473 | 1142 | 1055 | 1476 | 1346 | 1014 | 1528 | 993 | 2691 | 2326 | 2462 | 3548 | 3729 | 3238 | 3826 | 1688 | 1125 | 921 | 1576 | 240 | 1155 | 943 | Dpf3 |
| Dph1 | 1020 | 1279 | 925 | 1086 | 798 | 933 | 788 | 547 | 698 | 609 | 657 | 679 | 616 | 553 | 976 | 1177 | 914 | 1006 | 834 | 921 | 742 | 587 | 656 | 528 | 592 | 608 | 694 | 586 | Dph1 |
| Dph2 | 1668 | 2312 | 2015 | 2106 | 1591 | 1671 | 1789 | 1065 | 1158 | 1247 | 1107 | 1383 | 1203 | 1294 | 1640 | 1684 | 2058 | 2047 | 1831 | 1653 | 1692 | 1093 | 1087 | 1270 | 1342 | 1690 | 1352 | 1375 | Dph2 |
| Dph3 | 1723 | 1638 | 1697 | 1577 | 1730 | 1681 | 1857 | 3347 | 3394 | 2737 | 2928 | 3358 | 2985 | 3399 | 1804 | 1414 | 1735 | 1546 | 1655 | 1808 | 1825 | 2923 | 3253 | 3107 | 3078 | 3553 | 3450 | 3710 | Dph3 |
| Dph5 | 677 | 643 | 679 | 530 | 592 | 629 | 625 | 874 | 766 | 783 | 844 | 895 | 815 | 950 | 709 | 551 | 666 | 554 | 533 | 690 | 580 | 710 | 852 | 914 | 767 | 911 | 859 | 929 | Dph5 |
| Dph6 | 1294 | 1080 | 1278 | 1334 | 1162 | 1352 | 1403 | 1878 | 1795 | 1728 | 1805 | 2060 | 2036 | 1928 | 1389 | 1084 | 1228 | 1249 | 1289 | 1246 | 1231 | 1706 | 1697 | 2019 | 1873 | 2111 | 2091 | 2077 | Dph6 |
| Dph7 | 1768 | 1758 | 1475 | 1683 | 1398 | 1468 | 1646 | 2050 | 1929 | 1918 | 1615 | 1833 | 1971 | 1803 | 1687 | 1420 | 1669 | 1510 | 1704 | 1711 | 1507 | 1840 | 1830 | 1836 | 1867 | 1993 | 1878 | 1919 | Dph7 |
| Dpm1 | 2008 | 2002 | 1675 | 2275 | 1489 | 1729 | 2016 | 1280 | 1055 | 1207 | 1164 | 1360 | 1192 | 1434 | 1961 | 1794 | 1671 | 1837 | 1710 | 1764 | 1907 | 1362 | 1042 | 1179 | 1232 | 1069 | 1262 | 1341 | Dpm1 |
| Dpm2 | 2445 | 2870 | 2610 | 2715 | 2258 | 2188 | 2630 | 2204 | 2110 | 2012 | 2087 | 2217 | 2172 | 2347 | 2318 | 2396 | 2577 | 2511 | 2364 | 2514 | 2590 | 2135 | 2204 | 2345 | 2242 | 2376 | 2238 | 2453 | Dpm2 |
| Dpm3 | 619 | 866 | 889 | 836 | 560 | 484 | 786 | 576 | 526 | 756 | 743 | 731 | 583 | 913 | 537 | 463 | 948 | 770 | 641 | 720 | 739 | 416 | 633 | 864 | 705 | 775 | 651 | 892 | Dpm3 |
| Dpp10 | 364 | 425 | 218 | 896 | 1965 | 1284 | 1339 | 1599 | 1568 | 1461 | 1445 | 507 | 651 | 484 | 873 | 477 | 474 | 1192 | 1026 | 1409 | 1549 | 2000 | 1588 | 827 | 1282 | 207 | 351 | 325 | Dpp10 |
| Dpp3 | 7559 | 9620 | 9585 | 7676 | 6724 | 7152 | 6434 | 7152 | 7807 | 7665 | 7503 | 9648 | 9406 | 9375 | 7456 | 8093 | 8990 | 7015 | 7870 | 6481 | 6729 | 7142 | 7742 | 8393 | 8025 | 10623 | 10068 | 9862 | Dpp3 |
| Dpp4 | 108 | 89 | 105 | 92 | 120 | 188 | 108 | 175 | 302 | 233 | 396 | 110 | 161 | 139 | 144 | 109 | 80 | 180 | 91 | 111 | 122 | 227 | 430 | 211 | 195 | 92 | 152 | 98 | Dpp4 |
| Dpp6 | 2510 | 3383 | 2658 | 4452 | 6748 | 5675 | 4816 | 5426 | 5518 | 6493 | 5348 | 4387 | 4738 | 4151 | 3498 | 2962 | 3936 | 5085 | 5167 | 5207 | 5378 | 8435 | 5018 | 4464 | 6278 | 3139 | 3936 | 3983 | Dpp6 |
| Dpp7 | 1325 | 1793 | 1591 | 1644 | 1527 | 1550 | 1554 | 1182 | 1266 | 1230 | 1095 | 1158 | 1139 | 1173 | 1385 | 1597 | 1671 | 1758 | 1588 | 1615 | 1563 | 1089 | 1245 | 1217 | 1172 | 1198 | 1008 | 1173 | Dpp7 |
| Dpp8 | 17834 | 15430 | 12089 | 14025 | 14297 | 16282 | 15930 | 16185 | 14141 | 13522 | 13730 | 14239 | 14079 | 14326 | 17856 | 15178 | 12860 | 13996 | 14361 | 15983 | 16461 | 13881 | 14187 | 12412 | 14637 | 10319 | 14777 | 13727 | Dpp8 |
| Dpp9 | 3309 | 4501 | 5317 | 4493 | 4007 | 3671 | 3347 | 3583 | 4593 | 5353 | 4927 | 5830 | 5090 | 5307 | 3138 | 4232 | 5247 | 4337 | 4690 | 3120 | 3549 | 5093 | 4336 | 5432 | 5379 | 6184 | 5202 | 5237 | Dpp9 |
| Dppa1 | 0 | 0 | 0 | 0 | 0 | 0 | 0 | 0 | 0 | 0 | 0 | 0 | 1 | 0 | 0 | 0 | 0 | 0 | 0 | 0 | 0 | 0 | 0 | 0 | 0 | 0 | 1 | 0 | Dppa1 |

| | | | | | | | | | | | | | | | | | | | | | | | | | | | | | |
|---|---|---|---|---|---|---|---|---|---|---|---|---|---|---|---|---|---|---|---|---|---|---|---|---|---|---|---|---|---|
| Dppa2 | 0 | 1 | 0 | 0 | 0 | 0 | 0 | 0 | 2 | 0 | 1 | 0 | 0 | 0 | 1 | 0 | 0 | 0 | 0 | 0 | 0 | 0 | 1 | 0 | 0 | 0 | 0 | 0 | Dppa2 |
| Dppa3 | 0 | 0 | 0 | 0 | 0 | 0 | 0 | 0 | 1 | 1 | 0 | 0 | 0 | 3 | 1 | 0 | 0 | 3 | 0 | 1 | 0 | 0 | 1 | 0 | 0 | 0 | 0 | 0 | Dppa3 |
| Dppa4 | 0 | 0 | 0 | 0 | 0 | 0 | 0 | 0 | 0 | 0 | 0 | 3 | 0 | 0 | 4 | 1 | 2 | 0 | 0 | 0 | 0 | 0 | 1 | 0 | 0 | 0 | 0 | 1 | Dppa4 |
| Dppa5a | 22 | 20 | 9 | 27 | 43 | 24 | 28 | 22 | 42 | 31 | 35 | 30 | 32 | 13 | 12 | 19 | 9 | 24 | 28 | 21 | 25 | 45 | 19 | 42 | 45 | 15 | 14 | 34 | Dppa5a |
| Dpt | 5 | 0 | 2 | 2 | 0 | 5 | 2 | 12 | 49 | 42 | 221 | 2 | 0 | 4 | 0 | 2 | 0 | 2 | 9 | 1 | 0 | 3 | 174 | 15 | 13 | 6 | 0 | 2 | Dpt |
| Dpy19l1 | 7011 | 5853 | 5914 | 6370 | 6852 | 7518 | 8046 | 6872 | 7887 | 6319 | 7243 | 6225 | 6171 | 6509 | 6794 | 6459 | 5556 | 6923 | 6780 | 7333 | 7471 | 6072 | 7431 | 6572 | 7657 | 6222 | 5978 | 5944 | Dpy19l1 |
| Dpy19l2 | 10 | 18 | 22 | 17 | 34 | 22 | 20 | 45 | 26 | 34 | 21 | 38 | 48 | 35 | 24 | 29 | 10 | 19 | 12 | 30 | 20 | 33 | 47 | 42 | 16 | 24 | 34 | 47 | Dpy19l2 |
| Dpy19l3 | 6097 | 6211 | 5206 | 6244 | 6532 | 6517 | 7090 | 3373 | 3686 | 3782 | 3579 | 3690 | 3667 | 3603 | 5644 | 5851 | 5386 | 6448 | 6272 | 5992 | 7264 | 3437 | 3476 | 3432 | 3646 | 2957 | 3275 | 3488 | Dpy19l3 |
| Dpy19l4 | 7491 | 5346 | 5640 | 6146 | 6057 | 7152 | 6710 | 5471 | 4963 | 4595 | 5398 | 4906 | 5616 | 5635 | 7479 | 6877 | 5349 | 6030 | 5570 | 6513 | 6439 | 4814 | 4882 | 4440 | 4762 | 3847 | 5388 | 5193 | Dpy19l4 |
| Dpy30 | 3053 | 2890 | 3316 | 2949 | 2466 | 2354 | 3301 | 2630 | 2322 | 2373 | 2232 | 2596 | 2527 | 3059 | 3235 | 2495 | 3095 | 2579 | 2685 | 3414 | 3012 | 2027 | 2341 | 2781 | 2431 | 3361 | 2841 | 3209 | Dpy30 |
| Dpyd | 730 | 501 | 333 | 394 | 380 | 410 | 479 | 546 | 640 | 381 | 488 | 442 | 201 | 416 | 551 | 483 | 473 | 474 | 450 | 466 | 443 | 448 | 553 | 410 | 333 | 462 | 316 | 282 | Dpyd |
| Dpys | 2 | 10 | 4 | 13 | 13 | 17 | 15 | 13 | 16 | 14 | 22 | 22 | 11 | 10 | 3 | 10 | 12 | 12 | 18 | 23 | 11 | 27 | 14 | 14 | 24 | 27 | 9 | 11 | Dpys |
| Dpysl2 | 5777 | 5868 | 10540 | 11178 | 11289 | 8895 | 11984 | 10998 | 10496 | 14776 | 12226 | 14213 | 12292 | 16534 | 6311 | 5492 | 11944 | 11016 | 11411 | 7929 | 11867 | 17106 | 9356 | 17487 | 18368 | 19445 | 12918 | 17526 | Dpysl2 |
| Dpysl3 | 46195 | 49185 | 45320 | 66662 | 90115 | 79570 | 80843 | 111014 | 99265 | 105832 | 92879 | 89112 | 79448 | 91313 | 58561 | 40074 | 59683 | 72971 | 80373 | 82218 | 85634 | 127084 | 96155 | 96507 | 123539 | 64492 | 87711 | 86427 | Dpysl3 |
| Dpysl4 | 313 | 418 | 371 | 644 | 675 | 629 | 419 | 399 | 377 | 887 | 289 | 576 | 534 | 291 | 605 | 413 | 447 | 360 | 438 | 316 | 301 | 565 | 344 | 315 | 364 | 304 | 390 | 1178 | Dpysl4 |
| Dpysl5 | 15061 | 19856 | 23638 | 30933 | 38731 | 31169 | 31255 | 23066 | 25700 | 34264 | 28481 | 32968 | 27862 | 30318 | 17268 | 16551 | 29870 | 32161 | 38556 | 25464 | 34538 | 39939 | 22784 | 34047 | 40622 | 25914 | 26598 | 29211 | Dpysl5 |
| Dqx1 | 315 | 468 | 532 | 494 | 335 | 302 | 324 | 111 | 166 | 122 | 155 | 134 | 146 | 185 | 357 | 455 | 517 | 506 | 349 | 266 | 315 | 122 | 126 | 159 | 132 | 165 | 143 | 127 | Dqx1 |
| Dr1 | 8055 | 5965 | 8509 | 7969 | 6402 | 7344 | 7437 | 5443 | 4434 | 4596 | 4817 | 5595 | 5347 | 5916 | 7912 | 7478 | 7312 | 7568 | 6728 | 6754 | 7316 | 5147 | 4817 | 5167 | 5275 | 6274 | 5829 | 6494 | Dr1 |
| Dram1 | 114 | 85 | 98 | 50 | 70 | 116 | 42 | 51 | 52 | 48 | 84 | 46 | 69 | 53 | 88 | 110 | 58 | 92 | 58 | 59 | 71 | 60 | 107 | 56 | 57 | 44 | 87 | 28 | Dram1 |
| Dram2 | 2968 | 2200 | 2559 | 2381 | 2731 | 2728 | 2898 | 2622 | 2306 | 2193 | 2336 | 2342 | 2542 | 2392 | 3169 | 2891 | 2493 | 2582 | 2544 | 3042 | 2624 | 2506 | 2582 | 2062 | 2170 | 2277 | 2342 | 2313 | Dram2 |
| Drap1 | 4268 | 5430 | 5464 | 6444 | 4939 | 4703 | 5182 | 3826 | 3887 | 5633 | 4755 | 4482 | 4234 | 5572 | 4242 | 3815 | 6100 | 5247 | 5554 | 4739 | 6137 | 3766 | 4175 | 5661 | 4919 | 4952 | 4415 | 5160 | Drap1 |
| Draxin | 3028 | 4078 | 6089 | 5600 | 8513 | 6610 | 5936 | 7133 | 7577 | 9745 | 7449 | 8738 | 9331 | 8109 | 4109 | 3667 | 5940 | 5822 | 8054 | 5977 | 7062 | 10652 | 8403 | 8767 | 10604 | 4524 | 8377 | 8710 | Draxin |
| Drc1 | 133 | 167 | 122 | 215 | 268 | 215 | 250 | 255 | 263 | 263 | 246 | 262 | 230 | 160 | 193 | 166 | 114 | 219 | 174 | 254 | 234 | 326 | 320 | 198 | 271 | 196 | 155 | 139 | Drc1 |
| Drc3 | 211 | 238 | 176 | 188 | 220 | 236 | 190 | 318 | 299 | 351 | 233 | 356 | 350 | 215 | 240 | 263 | 182 | 190 | 256 | 248 | 244 | 324 | 423 | 193 | 268 | 223 | 234 | 272 | Drc3 |
| Drc7 | 23 | 61 | 22 | 39 | 104 | 50 | 48 | 98 | 110 | 111 | 56 | 96 | 111 | 51 | 45 | 51 | 39 | 26 | 59 | 61 | 52 | 119 | 187 | 57 | 70 | 47 | 52 | 43 | Drc7 |
| Drd1 | 54 | 49 | 85 | 64 | 137 | 105 | 118 | 100 | 105 | 80 | 82 | 81 | 75 | 57 | 84 | 62 | 50 | 95 | 114 | 137 | 136 | 94 | 96 | 79 | 165 | 24 | 63 | 46 | Drd1 |
| Drd2 | 159 | 210 | 264 | 271 | 321 | 298 | 240 | 122 | 106 | 201 | 159 | 130 | 73 | 97 | 224 | 199 | 270 | 235 | 302 | 270 | 275 | 189 | 121 | 145 | 184 | 67 | 112 | 112 | Drd2 |
| Drd3 | 8 | 8 | 0 | 5 | 3 | 11 | 12 | 13 | 12 | 10 | 2 | 18 | 0 | 17 | 9 | 1 | 17 | 4 | 11 | 9 | 5 | 3 | 5 | 2 | 13 | 4 | 12 | 11 | Drd3 |
| Drd4 | 50 | 34 | 37 | 39 | 25 | 29 | 28 | 24 | 32 | 41 | 64 | 37 | 23 | 42 | 26 | 49 | 35 | 30 | 46 | 18 | 16 | 47 | 57 | 56 | 32 | 44 | 36 | 26 | Drd4 |
| Drd5 | 3 | 4 | 7 | 2 | 7 | 6 | 12 | 6 | 16 | 3 | 18 | 7 | 2 | 6 | 8 | 2 | 8 | 3 | 9 | 8 | 13 | 12 | 12 | 24 | 13 | 18 | 9 | 0 | Drd5 |
| Drg1 | 9173 | 8258 | 8523 | 8620 | 8174 | 8501 | 9370 | 7938 | 6912 | 6990 | 6090 | 8099 | 7717 | 8098 | 9654 | 7732 | 8786 | 8267 | 9323 | 9576 | 9542 | 6896 | 6866 | 7455 | 7504 | 8579 | 8879 | 8822 | Drg1 |
| Drg2 | 3994 | 4777 | 3984 | 3645 | 3166 | 3479 | 3302 | 4176 | 4461 | 3963 | 3783 | 4209 | 4151 | 4050 | 3897 | 4298 | 3631 | 3367 | 3561 | 3447 | 3464 | 3633 | 4310 | 3897 | 4018 | 4689 | 4421 | 4227 | Drg2 |
| Drosha | 10741 | 11005 | 13162 | 11691 | 11314 | 10572 | 10889 | 10831 | 11089 | 12875 | 11169 | 14229 | 12187 | 14203 | 10381 | 10289 | 12640 | 11421 | 11965 | 8890 | 11747 | 12978 | 10306 | 13238 | 13115 | 14669 | 12734 | 14158 | Drosha |
| Drp2 | 2257 | 2222 | 2174 | 2864 | 4165 | 3906 | 3759 | 5519 | 5135 | 5921 | 5135 | 4740 | 4430 | 4833 | 2422 | 1961 | 2560 | 3212 | 3744 | 3600 | 3924 | 7389 | 4682 | 4924 | 6331 | 3139 | 4737 | 4533 | Drp2 |
| Dsc1 | 3 | 0 | 0 | 0 | 2 | 5 | 1 | 24 | 21 | 3 | 5 | 2 | 2 | 2 | 1 | 1 | 1 | 2 | 2 | 4 | 2 | 3 | 20 | 3 | 2 | 0 | 0 | 0 | Dsc1 |
| Dsc2 | 322 | 81 | 57 | 74 | 46 | 78 | 18 | 697 | 640 | 436 | 382 | 184 | 246 | 178 | 153 | 164 | 49 | 48 | 19 | 28 | 20 | 124 | 787 | 396 | 42 | 293 | 137 | 199 | Dsc2 |
| Dsc3 | 97 | 4 | 6 | 4 | 5 | 11 | 1 | 302 | 438 | 269 | 177 | 10 | 85 | 13 | 9 | 12 | 0 | 13 | 1 | 0 | 1 | 38 | 548 | 136 | 2 | 107 | 10 | 7 | Dsc3 |
| Dscam | 1313 | 1662 | 1488 | 2904 | 4282 | 3846 | 3466 | 2807 | 3058 | 4162 | 3323 | 3110 | 3003 | 2767 | 1586 | 1582 | 2176 | 3355 | 3610 | 3380 | 4071 | 4864 | 2601 | 3149 | 4496 | 1050 | 2731 | 2505 | Dscam |
| Dscaml1 | 351 | 444 | 264 | 944 | 1376 | 1115 | 1298 | 909 | 1426 | 2021 | 1506 | 1366 | 508 | 1238 | 348 | 274 | 664 | 1131 | 1502 | 936 | 1360 | 2045 | 1181 | 1856 | 2209 | 1123 | 726 | 803 | Dscaml1 |
| Dscc1 | 833 | 583 | 668 | 513 | 454 | 510 | 562 | 708 | 777 | 671 | 663 | 984 | 807 | 920 | 740 | 583 | 571 | 408 | 517 | 673 | 494 | 518 | 667 | 719 | 687 | 1060 | 1044 | 1053 | Dscc1 |
| Dse | 2307 | 1586 | 2114 | 1190 | 1060 | 1319 | 777 | 2488 | 3002 | 2315 | 3537 | 2191 | 3052 | 2240 | 2048 | 2307 | 1446 | 1366 | 895 | 934 | 725 | 2321 | 3519 | 2335 | 1934 | 2689 | 2592 | 2575 | Dse |
| Dsel | 3252 | 2013 | 2298 | 2459 | 2185 | 2435 | 2280 | 2469 | 2435 | 2091 | 2559 | 1855 | 2033 | 1815 | 2881 | 2587 | 2123 | 2330 | 2011 | 2349 | 2204 | 2229 | 2716 | 1908 | 1793 | 1931 | 1893 | 2000 | Dsel |
| Dsg1a | 20 | 0 | 2 | 0 | 1 | 3 | 2 | 111 | 273 | 115 | 9 | 9 | 3 | 4 | 0 | 2 | 0 | 0 | 4 | 0 | 0 | 0 | 273 | 8 | 9 | 1 | 3 | 6 | Dsg1a |
| Dsg1b | 7 | 0 | 0 | 0 | 0 | 3 | 0 | 21 | 50 | 33 | 6 | 2 | 0 | 0 | 0 | 0 | 0 | 0 | 0 | 0 | 0 | 0 | 74 | 5 | 0 | 0 | 2 | 0 | Dsg1b |
| Dsg1c | 1 | 2 | 0 | 0 | 0 | 0 | 0 | 1 | 0 | 0 | 0 | 0 | 0 | 0 | 0 | 0 | 0 | 0 | 0 | 0 | 0 | 0 | 2 | 0 | 0 | 1 | 0 | 0 | Dsg1c |
| Dsg2 | 511 | 181 | 97 | 109 | 75 | 123 | 12 | 583 | 761 | 419 | 523 | 123 | 284 | 119 | 237 | 254 | 43 | 109 | 33 | 20 | 16 | 124 | 794 | 370 | 58 | 276 | 132 | 132 | Dsg2 |
| Dsg3 | 8 | 2 | 0 | 2 | 2 | 25 | 1 | 90 | 168 | 91 | 57 | 7 | 19 | 9 | 4 | 1 | 0 | 4 | 9 | 0 | 0 | 7 | 263 | 20 | 14 | 2 | 8 | 6 | Dsg3 |
| Dsg4 | 0 | 0 | 0 | 0 | 0 | 0 | 0 | 0 | 2 | 0 | 2 | 1 | 1 | 0 | 0 | 0 | 0 | 0 | 0 | 0 | 0 | 0 | 2 | 0 | 0 | 0 | 0 | 0 | Dsg4 |
| Dsn1 | 2365 | 1900 | 1993 | 1675 | 1359 | 1677 | 1652 | 2332 | 2197 | 1949 | 1945 | 2671 | 2445 | 2763 | 1971 | 1846 | 1811 | 1440 | 1562 | 1934 | 1637 | 1621 | 2076 | 2299 | 2056 | 2795 | 2728 | 2849 | Dsn1 |
| Dsp | 3232 | 1264 | 767 | 460 | 408 | 941 | 245 | 3579 | 5745 | 2872 | 2434 | 1059 | 1388 | 933 | 1365 | 2055 | 381 | 861 | 266 | 246 | 211 | 321 | 6058 | 1949 | 450 | 928 | 815 | 928 | Dsp |
| Dspp | 0 | 0 | 0 | 1 | 1 | 0 | 0 | 0 | 0 | 3 | 0 | 0 | 0 | 1 | 0 | 0 | 0 | 0 | 2 | 1 | 1 | 0 | 0 | 0 | 1 | 2 | 0 | 0 | Dspp |
| Dst | 20737 | 24190 | 9703 | 13137 | 18376 | 21279 | 15879 | 22827 | 25334 | 15231 | 19046 | 12492 | 16113 | 12341 | 24143 | 27577 | 11739 | 14367 | 13800 | 24847 | 20557 | 14547 | 22191 | 10645 | 15988 | 6376 | 16429 | 10842 | Dst |
| Dstn | 21821 | 18699 | 17569 | 16130 | 16598 | 17305 | 18519 | 20661 | 17657 | 15708 | 15191 | 16904 | 16564 | 17645 | 22833 | 20800 | 17194 | 17316 | 16767 | 20634 | 17444 | 17358 | 18568 | 16079 | 16142 | 18414 | 17585 | 18154 | Dstn |
| Dstyk | 3406 | 3222 | 2911 | 3285 | 3527 | 3761 | 3536 | 3483 | 3404 | 3278 | 3471 | 3469 | 3535 | 3356 | 3400 | 2931 | 3037 | 3419 | 3716 | 3524 | 3655 | 3292 | 3398 | 3080 | 3376 | 2695 | 3636 | 3433 | Dstyk |
| Dtd1 | 580 | 699 | 625 | 757 | 799 | 769 | 773 | 1083 | 1091 | 1149 | 1069 | 868 | 900 | 1037 | 614 | 632 | 693 | 736 | 848 | 975 | 744 | 1136 | 1131 | 1089 | 1197 | 1058 | 941 | 903 | Dtd1 |
| Dtd2 | 1152 | 1266 | 1209 | 1152 | 871 | 883 | 1107 | 628 | 660 | 614 | 660 | 775 | 706 | 849 | 1172 | 1059 | 1122 | 1095 | 940 | 1001 | 1057 | 535 | 638 | 684 | 696 | 818 | 812 | 854 | Dtd2 |
| Dthd1 | 0 | 0 | 0 | 0 | 0 | 1 | 0 | 0 | 0 | 0 | 0 | 0 | 0 | 0 | 0 | 0 | 0 | 0 | 0 | 0 | 0 | 0 | 0 | 0 | 0 | 0 | 1 | 0 | Dthd1 |
| Dtl | 7589 | 5458 | 6334 | 4927 | 3694 | 4670 | 5223 | 5151 | 4995 | 3993 | 4200 | 6147 | 5397 | 6515 | 6157 | 6031 | 5032 | 4200 | 4450 | 5120 | 4718 | 3088 | 4433 | 5123 | 4126 | 6979 | 6512 | 6552 | Dtl |
| Dtna | 4135 | 3976 | 3030 | 3857 | 4368 | 3862 | 4081 | 2446 | 2323 | 2150 | 2010 | 1564 | 1532 | 1589 | 4052 | 3720 | 3473 | 3668 | 3400 | 4381 | 3782 | 2794 | 2359 | 1742 | 1928 | 1797 | 1670 | 1585 | Dtna |
| Dtnb | 1694 | 1717 | 1385 | 1491 | 1724 | 1753 | 1640 | 1613 | 1919 | 1894 | 1883 | 1791 | 1818 | 1596 | 1526 | 1771 | 1480 | 1478 | 1599 | 1556 | 1675 | 1611 | 1845 | 1627 | 1846 | 1555 | 1536 | 1578 | Dtnb |
| Dtnbp1 | 2515 | 2917 | 2631 | 2981 | 2945 | 2713 | 2905 | 1898 | 1865 | 1996 | 1594 | 1807 | 1741 | 2014 | 2529 | 2647 | 3031 | 2965 | 3149 | 2797 | 3141 | 1873 | 1888 | 1996 | 1971 | 2142 | 1784 | 1924 | Dtnbp1 |
| Dtwd1 | 456 | 431 | 463 | 409 | 455 | 451 | 449 | 554 | 565 | 520 | 508 | 645 | 624 | 623 | 487 | 434 | 475 | 392 | 481 | 595 | 437 | 490 | 589 | 569 | 484 | 645 | 662 | 631 | Dtwd1 |
| Dtwd2 | 95 | 59 | 121 | 82 | 117 | 60 | 54 | 364 | 325 | 325 | 425 | 351 | 430 | 364 | 70 | 76 | 68 | 68 | 63 | 78 | 71 | 353 | 302 | 337 | 381 | 406 | 380 | 379 | Dtwd2 |
| Dtx1 | 1693 | 2268 | 1889 | 2877 | 3661 | 2980 | 2963 | 2026 | 2707 | 3404 | 2545 | 2206 | 1622 | 1950 | 1626 | 1723 | 2655 | 3110 | 3645 | 2609 | 3229 | 3599 | 2290 | 2796 | 3689 | 1932 | 1634 | 1577 | Dtx1 |
| Dtx2 | 1078 | 1392 | 1435 | 1332 | 1393 | 1246 | 1079 | 1182 | 1470 | 1371 | 1371 | 1442 | 1488 | 1357 | 1054 | 1119 | 1623 | 1151 | 1388 | 1064 | 1130 | 1253 | 1354 | 1561 | 1430 | 1725 | 1404 | 1527 | Dtx2 |
| Dtx3 | 10207 | 11854 | 13714 | 14227 | 14114 | 12981 | 13224 | 10723 | 11688 | 14082 | 12888 | 13454 | 12200 | 12752 | 9875 | 10576 | 14731 | 13724 | 15832 | 11466 | 13721 | 14145 | 12009 | 14673 | 13401 | 14498 | 13113 | 12822 | Dtx3 |
| Dtx3l | 238 | 142 | 214 | 167 | 218 | 172 | 240 | 523 | 493 | 582 | 536 | 582 | 621 | 662 | 169 | 193 | 152 | 184 | 219 | 308 | 208 | 394 | 492 | 534 | 509 | 754 | 544 | 652 | Dtx3l |
| Dtx4 | 5336 | 6375 | 7685 | 6652 | 7812 | 7023 | 6796 | 8644 | 7219 | 7957 | 6974 | 9716 | 10604 | 9293 | 6343 | 5835 | 7623 | 6234 | 7650 | 7000 | 7011 | 9957 | 7484 | 7473 | 8334 | 8012 | 10213 | 11253 | Dtx4 |
| Dtymk | 4617 | 4645 | 4889 | 4464 | 3879 | 3985 | 4674 | 5802 | 5293 | 5506 | 5557 | 6382 | 5486 | 6777 | 4572 | 4214 | 4639 | 4276 | 4368 | 4422 | 4262 | 4741 | 5438 | 6378 | 5664 | 7495 | 6329 | 7157 | Dtymk |
| Duox1 | 7 | 10 | 3 | 1 | 9 | 11 | 3 | 17 | 38 | 31 | 36 | 7 | 32 | 23 | 15 | 25 | 15 | 12 | 12 | 10 | 13 | 17 | 42 | 29 | 28 | 10 | 24 | 20 | Duox1 |
| Duox2 | 16 | 22 | 6 | 23 | 33 | 23 | 13 | 9 | 7 | 7 | 8 | 0 | 7 | 1 | 16 | 20 | 24 | 24 | 25 | 4 | 19 | 5 | 11 | 2 | 4 | 5 | 2 | 7 | Duox2 |
| Duoxa1 | 17 | 13 | 15 | 26 | 20 | 37 | 11 | 13 | 17 | 17 | 7 | 15 | 15 | 11 | 12 | 14 | 14 | 14 | 12 | 15 | 18 | 3 | 35 | 10 | 19 | 2 | 6 | 11 | Duoxa1 |
| Duoxa2 | 5 | 5 | 4 | 4 | 1 | 2 | 4 | 1 | 1 | 7 | 4 | 2 | 1 | 0 | 5 | 5 | 7 | 6 | 0 | 0 | 1 | 0 | 0 | 0 | 11 | 0 | 0 | 1 | Duoxa2 |
| Dupd1 | 1 | 2 | 4 | 3 | 2 | 0 | 0 | 2 | 2 | 1 | 0 | 0 | 0 | 2 | 0 | 7 | 5 | 0 | 2 | 1 | 1 | 0 | 5 | 6 | 0 | 5 | 0 | 0 | Dupd1 |
| Dus1l | 2983 | 3750 | 3199 | 3190 | 2590 | 2669 | 2639 | 2323 | 2488 | 2443 | 2368 | 2848 | 2669 | 2686 | 2727 | 3019 | 3210 | 2928 | 2843 | 2807 | 2586 | 2149 | 2397 | 2490 | 2540 | 3126 | 2944 | 2800 | Dus1l |
| Dus2 | 561 | 580 | 792 | 716 | 734 | 729 | 581 | 1025 | 1101 | 1227 | 1245 | 1164 | 1125 | 1092 | 597 | 554 | 734 | 713 | 745 | 577 | 554 | 1093 | 1038 | 1152 | 1151 | 1333 | 1139 | 1218 | Dus2 |
| Dus3l | 1330 | 1700 | 1775 | 2000 | 2180 | 1883 | 1633 | 2739 | 3426 | 3742 | 3408 | 3898 | 3392 | 3327 | 1366 | 1434 | 2107 | 1924 | 2311 | 1834 | 1886 | 3594 | 3245 | 3950 | 3703 | 3722 | 3404 | 3344 | Dus3l |
| Dus4l | 900 | 858 | 794 | 759 | 709 | 741 | 816 | 650 | 692 | 575 | 664 | 783 | 695 | 726 | 854 | 864 | 705 | 720 | 720 | 833 | 709 | 571 | 686 | 667 | 686 | 901 | 731 | 783 | Dus4l |
| Dusp1 | 814 | 832 | 762 | 1088 | 1084 | 963 | 1127 | 1441 | 1442 | 1747 | 1402 | 1762 | 1286 | 1835 | 897 | 694 | 1039 | 905 | 1152 | 1053 | 1046 | 1760 | 1377 | 1826 | 1703 | 2034 | 1563 | 1735 | Dusp1 |

| | | | | | | | | | | | | | | | | | | | | | | | | | | | | | |
|---|---|---|---|---|---|---|---|---|---|---|---|---|---|---|---|---|---|---|---|---|---|---|---|---|---|---|---|---|---|
| Dusp10 | 923 | 1012 | 1035 | 938 | 752 | 765 | 864 | 519 | 647 | 589 | 626 | 645 | 625 | 631 | 964 | 856 | 1078 | 869 | 715 | 815 | 807 | 419 | 646 | 712 | 501 | 708 | 721 | 657 | Dusp10 |
| Dusp11 | 12169 | 10826 | 10534 | 12176 | 13439 | 12588 | 13727 | 15115 | 13855 | 14444 | 15229 | 13012 | 13183 | 13106 | 11966 | 11189 | 11728 | 13417 | 12783 | 12764 | 13589 | 17049 | 13718 | 14183 | 15026 | 10434 | 12464 | 13416 | Dusp11 |
| Dusp12 | 1668 | 1522 | 1467 | 1659 | 1781 | 1775 | 1649 | 1811 | 1693 | 1747 | 1949 | 1725 | 1792 | 1675 | 1565 | 1558 | 1630 | 1812 | 1650 | 1664 | 1638 | 2060 | 1810 | 1909 | 1838 | 1798 | 1758 | 1646 | Dusp12 |
| Dusp13 | 6 | 5 | 2 | 0 | 1 | 3 | 2 | 1 | 5 | 2 | 7 | 3 | 2 | 1 | 3 | 3 | 3 | 3 | 2 | 0 | 6 | 2 | 10 | 2 | 0 | 2 | 0 | 1 | Dusp13 |
| Dusp14 | 683 | 731 | 508 | 707 | 709 | 657 | 799 | 372 | 444 | 322 | 300 | 340 | 287 | 366 | 613 | 646 | 744 | 686 | 711 | 726 | 860 | 231 | 400 | 453 | 357 | 407 | 286 | 325 | Dusp14 |
| Dusp15 | 384 | 539 | 478 | 621 | 988 | 666 | 761 | 302 | 337 | 420 | 372 | 237 | 276 | 221 | 462 | 549 | 599 | 845 | 771 | 645 | 794 | 510 | 315 | 266 | 413 | 160 | 208 | 220 | Dusp15 |
| Dusp16 | 4257 | 3728 | 4232 | 3647 | 3141 | 3338 | 3833 | 3268 | 3105 | 2959 | 3031 | 3887 | 3318 | 4031 | 3889 | 3439 | 3962 | 3625 | 3698 | 3152 | 3555 | 2294 | 2911 | 3292 | 3161 | 3949 | 4086 | 4005 | Dusp16 |
| Dusp18 | 2205 | 2141 | 2376 | 2667 | 2973 | 2843 | 2910 | 2425 | 2529 | 2715 | 2485 | 3313 | 2942 | 3051 | 2333 | 2188 | 2642 | 2541 | 2957 | 2596 | 2690 | 3282 | 2479 | 2819 | 3296 | 2892 | 2984 | 2968 | Dusp18 |
| Dusp19 | 606 | 541 | 649 | 641 | 731 | 678 | 749 | 775 | 750 | 770 | 835 | 808 | 763 | 775 | 687 | 670 | 651 | 683 | 645 | 696 | 761 | 747 | 779 | 771 | 764 | 608 | 753 | 787 | Dusp19 |
| Dusp2 | 111 | 101 | 104 | 95 | 118 | 98 | 98 | 89 | 79 | 131 | 119 | 92 | 121 | 102 | 95 | 129 | 127 | 72 | 113 | 111 | 102 | 119 | 110 | 110 | 96 | 121 | 112 | 81 | Dusp2 |
| Dusp22 | 923 | 818 | 956 | 891 | 1046 | 921 | 838 | 1189 | 1322 | 1374 | 1373 | 1208 | 1353 | 1141 | 1011 | 1022 | 934 | 993 | 946 | 827 | 867 | 1412 | 1432 | 1183 | 1165 | 1147 | 966 | 1208 | Dusp22 |
| Dusp23 | 400 | 514 | 504 | 509 | 538 | 498 | 430 | 184 | 292 | 258 | 267 | 191 | 243 | 212 | 425 | 500 | 583 | 518 | 535 | 436 | 444 | 197 | 285 | 224 | 190 | 270 | 253 | 209 | Dusp23 |
| Dusp26 | 1344 | 1944 | 1666 | 1968 | 2464 | 2085 | 2019 | 2282 | 1963 | 2390 | 1940 | 1748 | 1867 | 1564 | 1840 | 1408 | 2062 | 1794 | 2406 | 2259 | 2097 | 2951 | 1950 | 1970 | 2599 | 1415 | 1671 | 1545 | Dusp26 |
| Dusp27 | 167 | 199 | 258 | 106 | 104 | 103 | 64 | 199 | 247 | 258 | 593 | 278 | 376 | 276 | 189 | 231 | 137 | 105 | 119 | 79 | 79 | 349 | 471 | 261 | 264 | 347 | 403 | 286 | Dusp27 |
| Dusp28 | 529 | 696 | 470 | 530 | 484 | 574 | 577 | 414 | 383 | 280 | 295 | 391 | 387 | 399 | 518 | 533 | 535 | 472 | 563 | 621 | 550 | 297 | 366 | 342 | 388 | 373 | 423 | 425 | Dusp28 |
| Dusp3 | 956 | 988 | 1006 | 990 | 1105 | 1091 | 921 | 1235 | 1314 | 1237 | 1704 | 1239 | 1232 | 1157 | 980 | 1074 | 1007 | 906 | 1100 | 956 | 988 | 1327 | 1553 | 1173 | 1263 | 1243 | 1190 | 1243 | Dusp3 |
| Dusp4 | 3557 | 4064 | 3781 | 6865 | 8486 | 6962 | 7359 | 7835 | 7841 | 9126 | 7032 | 9355 | 7841 | 8573 | 4257 | 3227 | 5779 | 6771 | 8709 | 7376 | 7325 | 8864 | 6364 | 8908 | 10724 | 7844 | 7392 | 7781 | Dusp4 |
| Dusp5 | 408 | 434 | 347 | 316 | 223 | 245 | 254 | 101 | 88 | 115 | 122 | 87 | 103 | 89 | 327 | 453 | 311 | 265 | 228 | 248 | 233 | 110 | 122 | 102 | 85 | 177 | 113 | 75 | Dusp5 |
| Dusp6 | 3443 | 2153 | 2258 | 1939 | 1627 | 1590 | 1735 | 2387 | 2585 | 1870 | 2544 | 1975 | 1960 | 2058 | 2552 | 2820 | 1780 | 1644 | 1580 | 1813 | 1625 | 1913 | 2530 | 2003 | 1695 | 3108 | 1651 | 2150 | Dusp6 |
| Dusp7 | 2051 | 2560 | 2521 | 2243 | 1767 | 1927 | 1669 | 2109 | 2525 | 2449 | 2514 | 2027 | 2394 | 1931 | 1829 | 2575 | 2245 | 2281 | 1990 | 1582 | 1637 | 1970 | 2531 | 2314 | 1854 | 2988 | 1933 | 2115 | Dusp7 |
| Dusp8 | 3255 | 4306 | 4542 | 6979 | 8610 | 6536 | 6658 | 4998 | 5550 | 7298 | 5869 | 5991 | 4033 | 5760 | 3824 | 3266 | 6696 | 7339 | 7977 | 5781 | 7366 | 7787 | 4967 | 6803 | 7767 | 5764 | 4613 | 4953 | Dusp8 |
| Dusp9 | 896 | 758 | 844 | 441 | 326 | 458 | 303 | 299 | 435 | 223 | 364 | 295 | 449 | 291 | 636 | 784 | 546 | 401 | 407 | 265 | 280 | 147 | 406 | 364 | 178 | 591 | 309 | 345 | Dusp9 |
| Dut | 5777 | 5149 | 6463 | 5521 | 4599 | 4704 | 6046 | 4358 | 3724 | 4285 | 4112 | 5828 | 4927 | 6838 | 5091 | 4326 | 5718 | 5051 | 5274 | 5175 | 5342 | 3397 | 3403 | 5810 | 4612 | 7830 | 6250 | 6843 | Dut |
| Duxbl1 | 100 | 62 | 119 | 71 | 71 | 69 | 117 | 330 | 207 | 291 | 258 | 335 | 400 | 438 | 82 | 49 | 81 | 55 | 104 | 77 | 71 | 291 | 247 | 261 | 295 | 226 | 492 | 401 | Duxbl1 |
| Dvl1 | 5277 | 7093 | 6957 | 7466 | 7373 | 6397 | 6120 | 4491 | 4643 | 5298 | 4954 | 5256 | 4450 | 4368 | 5138 | 5825 | 7646 | 7495 | 7845 | 6226 | 6309 | 5406 | 4495 | 5446 | 5735 | 5121 | 4699 | 4357 | Dvl1 |
| Dvl2 | 4383 | 5136 | 5882 | 6056 | 4380 | 4561 | 4088 | 2775 | 3168 | 3551 | 3423 | 3751 | 3435 | 3529 | 3827 | 4876 | 5839 | 5675 | 4970 | 3676 | 4329 | 2984 | 3134 | 3769 | 3432 | 4481 | 3430 | 3356 | Dvl2 |
| Dvl3 | 7441 | 7965 | 9990 | 10900 | 11632 | 10113 | 9764 | 7834 | 8149 | 10797 | 10239 | 12744 | 10749 | 11850 | 7096 | 7844 | 11127 | 10496 | 11173 | 7904 | 10384 | 12195 | 7580 | 11408 | 11462 | 11160 | 10721 | 11155 | Dvl3 |
| Dxo | 1649 | 2343 | 1987 | 2073 | 1611 | 1541 | 1845 | 1207 | 1378 | 1564 | 1402 | 1632 | 1427 | 1673 | 1548 | 1787 | 1955 | 1867 | 1891 | 1351 | 1692 | 1350 | 1306 | 1593 | 1558 | 1847 | 1528 | 1771 | Dxo |
| Dydc1 | 1 | 3 | 7 | 5 | 2 | 18 | 8 | 0 | 7 | 1 | 3 | 10 | 9 | 1 | 5 | 10 | 2 | 2 | 6 | 4 | 3 | 2 | 13 | 1 | 4 | 0 | 7 | 7 | Dydc1 |
| Dydc2 | 7 | 7 | 12 | 3 | 16 | 10 | 7 | 26 | 11 | 18 | 10 | 10 | 23 | 10 | 9 | 12 | 10 | 9 | 8 | 8 | 9 | 22 | 23 | 10 | 22 | 8 | 16 | 8 | Dydc2 |
| Dym | 2264 | 2151 | 2380 | 2183 | 2118 | 1893 | 2048 | 1579 | 1610 | 1881 | 1549 | 1808 | 1631 | 1672 | 2076 | 2116 | 2218 | 2118 | 2196 | 1783 | 2046 | 1828 | 1614 | 1816 | 1796 | 1865 | 1610 | 1805 | Dym |
| Dynap | 0 | 0 | 0 | 0 | 0 | 0 | 0 | 11 | 20 | 27 | 0 | 0 | 0 | 0 | 0 | 0 | 0 | 0 | 0 | 0 | 0 | 0 | 16 | 0 | 0 | 0 | 0 | 0 | Dynap |
| Dync1h1 | 65891 | 74007 | 38100 | 63699 | 76399 | 88253 | 72032 | 58231 | 68392 | 69618 | 64663 | 64690 | 67375 | 63541 | 66756 | 72172 | 47298 | 64605 | 70381 | 69403 | 92290 | 53765 | 57961 | 45541 | 80511 | 33151 | 73849 | 54894 | Dync1h1 |
| Dync1i1 | 226 | 242 | 137 | 293 | 502 | 437 | 518 | 339 | 501 | 409 | 529 | 240 | 172 | 209 | 255 | 252 | 218 | 415 | 428 | 453 | 552 | 409 | 445 | 326 | 410 | 193 | 138 | 150 | Dync1i1 |
| Dync1i2 | 22283 | 21143 | 19518 | 20747 | 22803 | 21883 | 26176 | 27191 | 24070 | 23829 | 21937 | 23538 | 21691 | 25209 | 25168 | 20651 | 21903 | 21552 | 22559 | 25578 | 24592 | 26664 | 23049 | 23317 | 24761 | 23244 | 22910 | 23524 | Dync1i2 |
| Dync1li1 | 6689 | 6791 | 7538 | 8098 | 8430 | 7490 | 8796 | 8048 | 6839 | 7994 | 7678 | 8150 | 7778 | 9499 | 6957 | 6564 | 7859 | 7888 | 8314 | 7407 | 8316 | 9052 | 6928 | 8401 | 9239 | 8027 | 7603 | 9380 | Dync1li1 |
| Dync1li2 | 14173 | 13318 | 14647 | 15283 | 15934 | 15487 | 15356 | 18280 | 16094 | 17202 | 16780 | 18914 | 17438 | 19554 | 15072 | 13740 | 14916 | 15559 | 15252 | 16172 | 15169 | 18802 | 15742 | 17925 | 19784 | 17085 | 18923 | 18162 | Dync1li2 |
| Dync2h1 | 9370 | 8406 | 3723 | 4826 | 4631 | 6541 | 6415 | 8619 | 7172 | 4076 | 4643 | 5381 | 5410 | 5423 | 9124 | 8463 | 3950 | 4215 | 4318 | 9460 | 6680 | 3783 | 6649 | 3471 | 4834 | 3348 | 6746 | 4894 | Dync2h1 |
| Dync2i1 | 1941 | 1729 | 1671 | 1940 | 1912 | 2011 | 1947 | 2249 | 2250 | 2481 | 2489 | 2096 | 2150 | 2160 | 1891 | 1878 | 1860 | 1927 | 1800 | 1741 | 1801 | 2217 | 2236 | 2197 | 2033 | 1850 | 1970 | 2006 | Dync2i1 |
| Dync2i2 | 1392 | 1728 | 1322 | 1710 | 1836 | 1679 | 1614 | 1597 | 1898 | 1774 | 1855 | 1642 | 1669 | 1588 | 1226 | 1490 | 1611 | 1862 | 1816 | 1810 | 1726 | 1820 | 1989 | 1956 | 1983 | 1918 | 1574 | 1685 | Dync2i2 |
| Dync2li1 | 1112 | 1214 | 936 | 990 | 911 | 1064 | 1100 | 1276 | 1280 | 1123 | 1186 | 1250 | 1235 | 1338 | 1166 | 1220 | 998 | 941 | 977 | 1174 | 989 | 1091 | 1337 | 1231 | 1126 | 1362 | 1238 | 1345 | Dync2li1 |
| Dynll1 | 21389 | 24844 | 22469 | 22293 | 17840 | 16891 | 22598 | 15389 | 13560 | 13955 | 13911 | 13425 | 14326 | 16860 | 22158 | 19094 | 23409 | 20303 | 21094 | 22069 | 22550 | 13349 | 13761 | 15944 | 14859 | 16391 | 15325 | 17440 | Dynll1 |
| Dynll2 | 34537 | 36817 | 54137 | 66301 | 54645 | 42159 | 48912 | 27672 | 23236 | 38125 | 33490 | 34273 | 30531 | 37139 | 35711 | 31531 | 61754 | 60111 | 54727 | 40142 | 54593 | 36001 | 22703 | 41725 | 42936 | 37930 | 31571 | 36590 | Dynll2 |
| Dynlrb1 | 6191 | 6815 | 6042 | 6906 | 7022 | 6869 | 7329 | 7167 | 6998 | 7347 | 6437 | 6574 | 6501 | 7093 | 6733 | 6001 | 7006 | 6872 | 7915 | 8998 | 7655 | 6612 | 7172 | 7170 | 7262 | 7600 | 7204 | 7296 | Dynlrb1 |
| Dynlrb2 | 65 | 101 | 42 | 41 | 85 | 80 | 96 | 100 | 114 | 84 | 54 | 97 | 134 | 88 | 115 | 106 | 61 | 33 | 76 | 133 | 85 | 132 | 189 | 90 | 114 | 109 | 72 | 69 | Dynlrb2 |
| Dynlt1a | 1212 | 1220 | 944 | 1254 | 1673 | 1426 | 1430 | 1343 | 1316 | 1299 | 1141 | 1009 | 1264 | 1175 | 1298 | 1150 | 1170 | 1332 | 1498 | 1634 | 1385 | 1462 | 1289 | 1142 | 1390 | 1067 | 1223 | 1013 | Dynlt1a |
| Dynlt1b | 1442 | 937 | 871 | 978 | 1014 | 1395 | 1764 | 1826 | 2010 | 1135 | 1462 | 1554 | 1609 | 1225 | 1436 | 829 | 1103 | 1360 | 1061 | 1790 | 1148 | 1775 | 984 | 1636 | 1248 | 1503 | 1639 | 1150 | Dynlt1b |
| Dynlt1c | 745 | 733 | 572 | 751 | 847 | 1083 | 829 | 948 | 903 | 824 | 846 | 942 | 1128 | 930 | 749 | 856 | 858 | 919 | 908 | 1183 | 827 | 1128 | 854 | 789 | 942 | 845 | 1029 | 863 | Dynlt1c |
| Dynlt1f | 1675 | 1619 | 1748 | 1428 | 1495 | 1690 | 1458 | 2618 | 2432 | 2302 | 2452 | 2721 | 2787 | 2499 | 1817 | 1611 | 1571 | 1525 | 1626 | 1914 | 1369 | 2394 | 2561 | 2384 | 2555 | 3155 | 2932 | 2472 | Dynlt1f |
| Dynlt2a1 | 0 | 0 | 0 | 0 | 0 | 0 | 0 | 0 | 0 | 0 | 0 | 0 | 0 | 1 | 0 | 0 | 0 | 0 | 0 | 0 | 0 | 0 | 0 | 0 | 0 | 0 | 0 | 0 | Dynlt2a1 |
| Dynlt2a2 | 0 | 1 | 0 | 0 | 0 | 1 | 0 | 0 | 1 | 0 | 0 | 0 | 0 | 0 | 0 | 0 | 0 | 0 | 1 | 0 | 1 | 0 | 0 | 0 | 0 | 0 | 0 | 0 | Dynlt2a2 |
| Dynlt2a3 | 0 | 1 | 1 | 3 | 5 | 0 | 2 | 4 | 0 | 0 | 4 | 9 | 3 | 2 | 8 | 2 | 0 | 0 | 0 | 4 | 3 | 2 | 0 | 2 | 0 | 2 | 3 | 1 | Dynlt2a3 |
| Dynlt2b | 867 | 743 | 990 | 982 | 945 | 877 | 1073 | 817 | 782 | 758 | 830 | 958 | 792 | 1070 | 949 | 780 | 966 | 929 | 875 | 1017 | 980 | 897 | 808 | 912 | 763 | 1069 | 960 | 1019 | Dynlt2b |
| Dynlt3 | 2173 | 1615 | 1826 | 2027 | 2060 | 1892 | 2407 | 2423 | 2146 | 1940 | 1946 | 1965 | 1935 | 2090 | 2520 | 1883 | 2055 | 1982 | 1865 | 2352 | 2119 | 1903 | 2482 | 2089 | 1977 | 2247 | 2078 | 2042 | Dynlt3 |
| Dynlt4 | 0 | 3 | 0 | 0 | 2 | 1 | 0 | 0 | 3 | 1 | 4 | 3 | 14 | 1 | 3 | 5 | 0 | 0 | 1 | 0 | 0 | 5 | 8 | 2 | 1 | 0 | 1 | 2 | Dynlt4 |
| Dynlt5 | 27 | 32 | 29 | 13 | 29 | 16 | 12 | 44 | 30 | 38 | 28 | 27 | 53 | 9 | 34 | 49 | 25 | 15 | 18 | 13 | 10 | 48 | 63 | 24 | 14 | 31 | 20 | 21 | Dynlt5 |
| Dyrk1a | 5447 | 5009 | 6217 | 5871 | 6947 | 6447 | 6926 | 8573 | 7810 | 8617 | 8324 | 9609 | 8688 | 9900 | 5298 | 4618 | 6224 | 5746 | 6924 | 5663 | 6802 | 9683 | 7422 | 9087 | 9375 | 9260 | 9118 | 9609 | Dyrk1a |
| Dyrk1b | 595 | 944 | 479 | 833 | 1152 | 861 | 707 | 1658 | 1748 | 1920 | 1841 | 1796 | 1697 | 1637 | 555 | 709 | 643 | 837 | 1042 | 839 | 845 | 1856 | 1697 | 1437 | 2140 | 1233 | 1920 | 1343 | Dyrk1b |
| Dyrk2 | 5277 | 4673 | 5629 | 5673 | 6815 | 6371 | 6885 | 5965 | 5272 | 5921 | 5826 | 6094 | 5917 | 6245 | 5147 | 4893 | 5791 | 6129 | 6491 | 5711 | 6764 | 6234 | 5132 | 5915 | 6228 | 5133 | 6220 | 6220 | Dyrk2 |
| Dyrk3 | 1287 | 1662 | 1367 | 1507 | 1454 | 1352 | 1655 | 1099 | 972 | 1037 | 672 | 1247 | 1140 | 1265 | 1423 | 1273 | 1480 | 1633 | 1679 | 1542 | 1657 | 952 | 871 | 1111 | 1171 | 1326 | 1241 | 1336 | Dyrk3 |
| Dyrk4 | 2 | 3 | 3 | 9 | 7 | 4 | 3 | 17 | 16 | 28 | 19 | 10 | 8 | 10 | 8 | 9 | 5 | 4 | 8 | 3 | 1 | 13 | 16 | 14 | 2 | 17 | 8 | 9 | Dyrk4 |
| Dysf | 878 | 855 | 959 | 779 | 869 | 729 | 572 | 710 | 948 | 970 | 1392 | 772 | 947 | 793 | 816 | 1273 | 801 | 921 | 787 | 511 | 682 | 926 | 1111 | 851 | 767 | 740 | 722 | 742 | Dysf |
| Dytn | 2 | 6 | 4 | 0 | 6 | 7 | 6 | 11 | 21 | 10 | 11 | 1 | 10 | 1 | 15 | 15 | 6 | 11 | 4 | 7 | 3 | 8 | 12 | 2 | 13 | 0 | 9 | 1 | Dytn |
| Dzank1 | 2835 | 3124 | 2295 | 3377 | 4027 | 3405 | 3282 | 2154 | 1713 | 2012 | 1684 | 1369 | 1557 | 1481 | 3429 | 3113 | 2903 | 3564 | 3314 | 3312 | 3546 | 2558 | 1680 | 1531 | 2021 | 863 | 1455 | 1365 | Dzank1 |
| Dzip1 | 7379 | 7304 | 7077 | 8465 | 10203 | 8688 | 9280 | 8888 | 8295 | 9394 | 9037 | 7779 | 8348 | 8113 | 7820 | 7588 | 8268 | 9415 | 8962 | 8425 | 9781 | 10418 | 8038 | 8518 | 9292 | 6346 | 7875 | 7478 | Dzip1 |
| Dzip1l | 1796 | 1901 | 2015 | 1625 | 1748 | 1945 | 1665 | 2144 | 2104 | 2320 | 2149 | 2386 | 2455 | 2388 | 1714 | 1786 | 2027 | 1668 | 1933 | 1768 | 1650 | 1663 | 2049 | 2279 | 2055 | 2283 | 2558 | 2368 | Dzip1l |
| Dzip3 | 16394 | 11625 | 11447 | 14082 | 15174 | 16336 | 18163 | 15433 | 13532 | 12407 | 12357 | 13481 | 13269 | 13640 | 17508 | 13768 | 12501 | 13796 | 13574 | 17216 | 16332 | 13324 | 13323 | 11430 | 12804 | 9353 | 13768 | 12660 | Dzip3 |
| E030025P04Rik | 0 | 0 | 0 | 0 | 0 | 0 | 0 | 1 | 0 | 0 | 0 | 0 | 0 | 0 | 1 | 0 | 0 | 0 | 2 | 0 | 0 | 0 | 1 | 0 | 0 | 0 | 0 | 0 | E030025P04Rik |
| E030030I06Rik | 129 | 148 | 229 | 147 | 109 | 91 | 98 | 92 | 101 | 115 | 112 | 125 | 149 | 170 | 129 | 184 | 174 | 91 | 108 | 114 | 88 | 82 | 123 | 106 | 105 | 152 | 181 | 204 | E030030I06Rik |
| E130114P18Rik | 516 | 458 | 472 | 585 | 702 | 779 | 794 | 1947 | 1719 | 1564 | 1292 | 1950 | 1850 | 2014 | 680 | 402 | 508 | 524 | 715 | 883 | 764 | 1158 | 1386 | 1924 | 2125 | 1950 | 1691 | 1879 | E130114P18Rik |
| E130116L18Rik | 0 | 0 | 0 | 2 | 2 | 0 | 0 | 0 | 0 | 1 | 1 | 0 | 0 | 0 | 0 | 0 | 0 | 0 | 0 | 2 | 2 | 0 | 3 | 0 | 3 | 0 | 1 | 1 | E130116L18Rik |
| E130208F15Rik | 99 | 64 | 113 | 105 | 60 | 95 | 86 | 51 | 44 | 62 | 71 | 58 | 43 | 53 | 95 | 87 | 87 | 95 | 97 | 52 | 77 | 50 | 37 | 59 | 49 | 72 | 53 | 56 | E130208F15Rik |
| E130218I03Rik | 9 | 3 | 4 | 6 | 3 | 3 | 1 | 0 | 3 | 0 | 1 | 0 | 0 | 2 | 5 | 0 | 1 | 6 | 1 | 1 | 3 | 5 | 3 | 2 | 0 | 2 | 0 | 0 | E130218I03Rik |
| E130308A19Rik | 4380 | 4528 | 4018 | 5264 | 5374 | 5398 | 5018 | 4248 | 3731 | 4382 | 3821 | 4587 | 4940 | 4387 | 4497 | 4564 | 4450 | 4672 | 4835 | 4474 | 5451 | 4702 | 3633 | 3580 | 4666 | 3293 | 4780 | 4209 | E130308A19Rik |
| E130309D02Rik | 1581 | 1976 | 2227 | 1834 | 1789 | 1737 | 1635 | 1661 | 2013 | 2008 | 1977 | 2647 | 2182 | 2333 | 1535 | 1699 | 2191 | 1827 | 2233 | 1522 | 1717 | 2022 | 1908 | 2468 | 2307 | 3112 | 2502 | 2476 | E130309D02Rik |
| E130311K13Rik | 75 | 47 | 70 | 75 | 95 | 90 | 97 | 234 | 225 | 302 | 243 | 162 | 219 | 229 | 65 | 86 | 86 | 92 | 91 | 89 | 83 | 157 | 276 | 227 | 203 | 209 | 184 | 173 | E130311K13Rik |
| E230025N22Rik | 0 | 0 | 0 | 0 | 1 | 0 | 9 | 10 | 9 | 0 | 1 | 7 | 7 | 9 | 1 | 2 | 1 | 3 | 2 | 3 | 3 | 2 | 5 | 4 | 11 | 5 | 2 | 0 | E230025N22Rik |
| E2f1 | 2999 | 3883 | 4391 | 3584 | 2576 | 2841 | 2530 | 2148 | 2414 | 2542 | 2471 | 3881 | 3378 | 3421 | 2625 | 3103 | 3950 | 3134 | 3275 | 2429 | 2566 | 2032 | 2108 | 3389 | 2909 | 4427 | 4087 | 3853 | E2f1 |
| E2f2 | 2147 | 2491 | 3169 | 2829 | 2313 | 2331 | 2746 | 1194 | 1493 | 1505 | 1699 | 2777 | 1919 | 2511 | 1865 | 2220 | 3175 | 2687 | 3036 | 2230 | 2638 | 1168 | 1317 | 2214 | 1891 | 3059 | 2417 | 2600 | E2f2 |

| | | | | | | | | | | | | | | | | | | | | | | | | | | | | | |
|---|---|---|---|---|---|---|---|---|---|---|---|---|---|---|---|---|---|---|---|---|---|---|---|---|---|---|---|---|---|
| E2f3 | 3988 | 3561 | 4307 | 3987 | 4240 | 4255 | 4573 | 3876 | 3835 | 4124 | 3963 | 5749 | 5113 | 5593 | 3745 | 3459 | 4323 | 4001 | 4729 | 3735 | 4252 | 3397 | 3377 | 4602 | 4273 | 5657 | 5260 | 5610 | E2f3 |
| E2f4 | 4270 | 5012 | 5082 | 4556 | 3332 | 3566 | 3455 | 2729 | 3055 | 3086 | 3098 | 4113 | 3674 | 3734 | 3658 | 4653 | 4794 | 4391 | 3929 | 3211 | 3489 | 3267 | 2995 | 3697 | 3349 | 4448 | 3798 | 3904 | E2f4 |
| E2f5 | 2872 | 2393 | 2811 | 2446 | 1963 | 2197 | 2709 | 2737 | 2300 | 1932 | 1971 | 2856 | 2558 | 3044 | 3062 | 2655 | 2633 | 2224 | 2077 | 2496 | 2298 | 2353 | 2213 | 2319 | 2062 | 3265 | 2920 | 3260 | E2f5 |
| E2f6 | 7216 | 7457 | 7604 | 6181 | 5332 | 5563 | 6110 | 5804 | 5202 | 4867 | 4837 | 6179 | 5768 | 5842 | 6937 | 7233 | 7011 | 5795 | 5677 | 6214 | 5608 | 5110 | 5036 | 5343 | 5055 | 6645 | 6344 | 6611 | E2f6 |
| E2f7 | 3204 | 2923 | 3840 | 2834 | 2360 | 2541 | 2456 | 2192 | 2329 | 2344 | 2364 | 3982 | 3478 | 3896 | 2898 | 3467 | 3312 | 2697 | 2636 | 2287 | 2421 | 1962 | 2210 | 3121 | 2649 | 4021 | 3583 | 3939 | E2f7 |
| E2f8 | 1731 | 2046 | 2288 | 2005 | 1416 | 1442 | 1677 | 1417 | 1593 | 1614 | 1658 | 2625 | 1956 | 2428 | 1618 | 1712 | 2097 | 1811 | 1799 | 1420 | 1578 | 1198 | 1364 | 2153 | 1610 | 2882 | 2210 | 2248 | E2f8 |
| E330021D16Rik | 0 | 4 | 0 | 1 | 0 | 0 | 0 | 1 | 2 | 0 | 0 | 1 | 3 | 2 | 1 | 2 | 0 | 0 | 0 | 1 | 1 | 0 | 3 | 0 | 1 | 1 | 0 | 2 | E330021D16Rik |
| E430018J23Rik | 604 | 680 | 644 | 585 | 591 | 643 | 510 | 504 | 620 | 593 | 547 | 610 | 649 | 585 | 587 | 594 | 666 | 607 | 647 | 592 | 564 | 543 | 576 | 662 | 578 | 534 | 611 | 563 | E430018J23Rik |
| E4f1 | 1536 | 1887 | 2118 | 2565 | 2176 | 2180 | 1854 | 2109 | 2483 | 2791 | 2541 | 2738 | 2577 | 2562 | 1366 | 1632 | 2241 | 2403 | 2484 | 1919 | 1943 | 2535 | 2258 | 2848 | 2822 | 2698 | 2550 | 2415 | E4f1 |
| Eaf1 | 3766 | 3342 | 3522 | 3410 | 2752 | 3117 | 2938 | 3408 | 3110 | 3250 | 3442 | 3485 | 3402 | 3314 | 3458 | 3660 | 3416 | 3247 | 2857 | 2806 | 2881 | 3070 | 3136 | 3609 | 3006 | 3717 | 3399 | 3610 | Eaf1 |
| Eaf2 | 369 | 317 | 347 | 293 | 157 | 251 | 277 | 99 | 77 | 82 | 93 | 63 | 104 | 98 | 318 | 314 | 260 | 267 | 190 | 266 | 251 | 69 | 80 | 114 | 79 | 119 | 87 | 116 | Eaf2 |
| Eapp | 2152 | 1903 | 2450 | 2356 | 2413 | 2212 | 2720 | 3759 | 3049 | 3540 | 3070 | 3446 | 3288 | 3613 | 2385 | 1849 | 2573 | 2332 | 2365 | 2580 | 2491 | 3522 | 3172 | 3590 | 3730 | 3574 | 3307 | 3624 | Eapp |
| Ear1 | 0 | 0 | 0 | 0 | 2 | 0 | 0 | 0 | 0 | 0 | 5 | 7 | 0 | 2 | 0 | 0 | 1 | 1 | 0 | 0 | 1 | 2 | 1 | 2 | 0 | 3 | 0 | 1 | Ear1 |
| Ear10 | 0 | 0 | 0 | 0 | 0 | 0 | 0 | 0 | 0 | 0 | 0 | 0 | 2 | 0 | 0 | 0 | 0 | 0 | 0 | 0 | 0 | 0 | 0 | 0 | 0 | 0 | 0 | 0 | Ear10 |
| Ear2 | 0 | 1 | 2 | 4 | 0 | 0 | 0 | 0 | 0 | 0 | 0 | 0 | 1 | 1 | 0 | 2 | 0 | 0 | 0 | 0 | 0 | 0 | 1 | 2 | 0 | 4 | 0 | 4 | Ear2 |
| Ear6 | 2 | 0 | 4 | 0 | 0 | 0 | 0 | 0 | 0 | 0 | 5 | 0 | 0 | 3 | 0 | 2 | 0 | 0 | 0 | 0 | 1 | 0 | 0 | 2 | 0 | 5 | 0 | 0 | Ear6 |
| Ears2 | 1022 | 1153 | 782 | 902 | 900 | 925 | 959 | 1340 | 1333 | 1289 | 1249 | 1171 | 1376 | 1373 | 964 | 921 | 842 | 923 | 920 | 897 | 945 | 1051 | 1346 | 1231 | 1253 | 1046 | 1374 | 1428 | Ears2 |
| Ebag9 | 1857 | 1705 | 2500 | 1894 | 1906 | 1833 | 2001 | 1900 | 1714 | 1752 | 1716 | 2041 | 1990 | 2110 | 2099 | 1627 | 2085 | 1662 | 1867 | 2046 | 1939 | 1843 | 1741 | 1919 | 1743 | 2191 | 2200 | 2459 | Ebag9 |
| Ebf1 | 4065 | 4618 | 5290 | 6361 | 7482 | 6210 | 4644 | 8771 | 7860 | 9186 | 9651 | 6465 | 10602 | 6852 | 5958 | 6009 | 6414 | 6136 | 5743 | 4992 | 5372 | 11022 | 8459 | 6758 | 8647 | 5048 | 7208 | 7180 | Ebf1 |
| Ebf2 | 4024 | 4518 | 4262 | 5203 | 5033 | 4666 | 2697 | 5372 | 4687 | 5038 | 5666 | 4299 | 7655 | 3869 | 5515 | 5761 | 4955 | 4464 | 4003 | 4380 | 3414 | 6408 | 5583 | 4214 | 4568 | 3116 | 5058 | 4694 | Ebf2 |
| Ebf3 | 7211 | 7190 | 10543 | 11152 | 14292 | 11324 | 7043 | 15419 | 13388 | 19686 | 20498 | 13052 | 20637 | 13111 | 8910 | 9106 | 11276 | 9621 | 10442 | 8724 | 8883 | 23053 | 17429 | 13097 | 17680 | 6761 | 13990 | 15275 | Ebf3 |
| Ebf4 | 1458 | 1553 | 1495 | 1855 | 1767 | 1645 | 1481 | 1242 | 1352 | 1494 | 1372 | 1332 | 1176 | 1265 | 1221 | 1521 | 1705 | 1839 | 1723 | 1306 | 1524 | 1333 | 1152 | 1579 | 1384 | 1329 | 1260 | 1108 | Ebf4 |
| Ebi3 | 92 | 106 | 125 | 104 | 128 | 112 | 118 | 85 | 65 | 93 | 76 | 66 | 100 | 69 | 94 | 56 | 123 | 89 | 90 | 91 | 114 | 82 | 100 | 80 | 95 | 47 | 95 | 82 | Ebi3 |
| Ebna1bp2 | 5356 | 6195 | 5947 | 5463 | 5028 | 5264 | 5439 | 4891 | 4690 | 4603 | 4627 | 5462 | 4630 | 5426 | 5532 | 5281 | 5751 | 4971 | 5438 | 6028 | 5147 | 4157 | 4451 | 5447 | 5077 | 5783 | 5327 | 5371 | Ebna1bp2 |
| Ebp | 914 | 993 | 791 | 850 | 834 | 773 | 898 | 985 | 882 | 821 | 802 | 827 | 897 | 863 | 982 | 894 | 846 | 776 | 859 | 976 | 749 | 769 | 938 | 723 | 769 | 951 | 815 | 877 | Ebp |
| Ebpl | 630 | 663 | 678 | 607 | 663 | 716 | 696 | 1116 | 1184 | 1130 | 1174 | 1203 | 1207 | 1282 | 629 | 623 | 590 | 583 | 595 | 664 | 614 | 1068 | 1131 | 1294 | 1235 | 1169 | 1311 | 1209 | Ebpl |
| Ecd | 1976 | 1902 | 1962 | 1602 | 1906 | 1851 | 1845 | 4136 | 3984 | 3986 | 3846 | 4593 | 4238 | 4555 | 1887 | 1659 | 1780 | 1571 | 2018 | 1795 | 1737 | 3857 | 4103 | 4026 | 3991 | 4939 | 4551 | 4724 | Ecd |
| Ece1 | 5203 | 5709 | 6485 | 5446 | 5091 | 5026 | 4364 | 2971 | 3690 | 3886 | 3808 | 3543 | 4085 | 3299 | 4655 | 6097 | 5707 | 5873 | 5208 | 4028 | 4443 | 3544 | 4007 | 3881 | 3010 | 4336 | 3361 | 3369 | Ece1 |
| Ece2 | 125 | 111 | 178 | 190 | 286 | 236 | 276 | 247 | 233 | 359 | 239 | 228 | 206 | 319 | 119 | 114 | 218 | 175 | 235 | 174 | 306 | 369 | 199 | 319 | 320 | 203 | 267 | 318 | Ece2 |
| Ecel1 | 723 | 964 | 592 | 1415 | 2030 | 1452 | 1657 | 1356 | 1749 | 1747 | 1504 | 1553 | 878 | 1486 | 920 | 668 | 1181 | 1548 | 1901 | 1272 | 1526 | 2217 | 1215 | 1722 | 1909 | 1618 | 1059 | 976 | Ecel1 |
| Ech1 | 1309 | 1341 | 1113 | 1015 | 1051 | 1060 | 932 | 1788 | 2319 | 1845 | 2189 | 1776 | 2004 | 1611 | 1269 | 1402 | 1001 | 1056 | 890 | 1183 | 940 | 1584 | 2590 | 1835 | 1783 | 1973 | 1707 | 1797 | Ech1 |
| Echdc1 | 1891 | 1635 | 1957 | 1425 | 1391 | 1464 | 1553 | 1922 | 1728 | 1531 | 1646 | 1853 | 1664 | 2018 | 2054 | 1828 | 1793 | 1484 | 1349 | 1802 | 1524 | 1556 | 1759 | 1767 | 1590 | 1932 | 1972 | 2235 | Echdc1 |
| Echdc2 | 565 | 483 | 527 | 402 | 472 | 422 | 371 | 535 | 721 | 583 | 724 | 554 | 599 | 475 | 444 | 568 | 418 | 464 | 360 | 395 | 319 | 478 | 753 | 637 | 525 | 573 | 506 | 483 | Echdc2 |
| Echdc3 | 58 | 55 | 53 | 21 | 16 | 32 | 19 | 61 | 66 | 58 | 65 | 67 | 77 | 85 | 30 | 33 | 29 | 19 | 25 | 16 | 24 | 70 | 89 | 54 | 41 | 59 | 92 | 70 | Echdc3 |
| Echs1 | 4730 | 4796 | 4287 | 3773 | 3599 | 4041 | 3698 | 5053 | 5069 | 4345 | 4490 | 5109 | 5135 | 5156 | 4360 | 4460 | 3889 | 3648 | 3823 | 4472 | 3655 | 4393 | 5178 | 4893 | 4570 | 5418 | 5476 | 5540 | Echs1 |
| Eci1 | 569 | 606 | 796 | 533 | 525 | 529 | 463 | 832 | 889 | 971 | 907 | 1050 | 979 | 990 | 510 | 555 | 623 | 538 | 569 | 474 | 442 | 810 | 959 | 1000 | 883 | 1176 | 1062 | 1157 | Eci1 |
| Eci2 | 5269 | 4729 | 4429 | 4040 | 4141 | 4292 | 4623 | 5201 | 5080 | 4342 | 4714 | 4751 | 4934 | 4885 | 5037 | 4719 | 4029 | 4028 | 4288 | 5235 | 4383 | 4333 | 5510 | 4392 | 4410 | 5134 | 4997 | 5098 | Eci2 |
| Eci3 | 25 | 31 | 26 | 10 | 31 | 33 | 8 | 12 | 33 | 27 | 38 | 11 | 52 | 12 | 30 | 33 | 15 | 29 | 19 | 20 | 10 | 25 | 56 | 21 | 30 | 18 | 7 | 19 | Eci3 |
| Ecm1 | 361 | 480 | 628 | 511 | 524 | 433 | 454 | 416 | 470 | 620 | 545 | 673 | 672 | 659 | 359 | 401 | 639 | 535 | 586 | 392 | 507 | 732 | 482 | 721 | 618 | 833 | 608 | 720 | Ecm1 |
| Ecm2 | 215 | 242 | 180 | 214 | 373 | 302 | 325 | 360 | 338 | 326 | 399 | 299 | 335 | 304 | 210 | 220 | 199 | 341 | 175 | 276 | 368 | 386 | 479 | 239 | 264 | 229 | 336 | 303 | Ecm2 |
| Ecpas | 9164 | 7428 | 6422 | 6089 | 7692 | 8023 | 7843 | 14946 | 13017 | 10585 | 11838 | 11612 | 11751 | 11653 | 9403 | 8020 | 6193 | 6552 | 6871 | 8274 | 7586 | 11535 | 12890 | 10385 | 11816 | 8510 | 12035 | 11939 | Ecpas |
| Ecrg4 | 51 | 62 | 21 | 90 | 169 | 205 | 87 | 153 | 375 | 271 | 322 | 130 | 156 | 87 | 86 | 87 | 43 | 198 | 111 | 135 | 107 | 353 | 715 | 131 | 245 | 60 | 78 | 55 | Ecrg4 |
| Ecscr | 322 | 292 | 372 | 313 | 343 | 340 | 296 | 521 | 485 | 486 | 644 | 597 | 734 | 566 | 381 | 351 | 356 | 309 | 362 | 372 | 280 | 568 | 697 | 577 | 508 | 720 | 551 | 629 | Ecscr |
| Ecsit | 1554 | 2081 | 1736 | 1864 | 1745 | 1695 | 1587 | 1224 | 1509 | 1534 | 1410 | 1497 | 1436 | 1389 | 1441 | 1697 | 1750 | 1703 | 1887 | 1522 | 1772 | 1501 | 1525 | 1528 | 1580 | 1453 | 1465 | 1404 | Ecsit |
| Ect2 | 8111 | 6691 | 7775 | 5548 | 4565 | 5388 | 6502 | 7850 | 6291 | 5395 | 5137 | 9560 | 7626 | 9932 | 7489 | 6598 | 6591 | 5085 | 5574 | 6658 | 5563 | 5038 | 5655 | 7426 | 6624 | 9977 | 9692 | 10053 | Ect2 |
| Ect2l | 6 | 17 | 13 | 2 | 5 | 10 | 7 | 15 | 19 | 10 | 27 | 22 | 10 | 16 | 18 | 22 | 7 | 9 | 3 | 8 | 7 | 8 | 32 | 15 | 7 | 12 | 9 | 2 | Ect2l |
| Eda | 1495 | 1255 | 1237 | 1222 | 1028 | 989 | 1046 | 689 | 813 | 744 | 864 | 663 | 745 | 641 | 1246 | 1305 | 1197 | 1207 | 1022 | 853 | 910 | 735 | 843 | 660 | 670 | 676 | 654 | 774 | Eda |
| Eda2r | 581 | 530 | 517 | 416 | 479 | 465 | 536 | 280 | 253 | 188 | 249 | 270 | 232 | 270 | 596 | 505 | 533 | 439 | 526 | 565 | 483 | 226 | 293 | 233 | 276 | 165 | 279 | 286 | Eda2r |
| Edar | 144 | 125 | 159 | 104 | 65 | 139 | 118 | 110 | 135 | 120 | 65 | 66 | 43 | 73 | 98 | 111 | 110 | 142 | 127 | 101 | 111 | 55 | 111 | 108 | 95 | 110 | 70 | 78 | Edar |
| Edaradd | 230 | 143 | 300 | 157 | 173 | 154 | 132 | 299 | 456 | 260 | 335 | 340 | 338 | 268 | 150 | 161 | 187 | 100 | 177 | 165 | 137 | 206 | 390 | 239 | 199 | 258 | 375 | 343 | Edaradd |
| Edc3 | 5703 | 6062 | 4880 | 4995 | 4426 | 4874 | 4625 | 4631 | 4351 | 4111 | 3856 | 4497 | 4271 | 4313 | 5629 | 5199 | 4887 | 4636 | 4997 | 4803 | 5033 | 4209 | 4170 | 3933 | 4291 | 4433 | 4545 | 4561 | Edc3 |
| Edc4 | 7621 | 9168 | 8370 | 8224 | 7076 | 7530 | 6753 | 5614 | 6717 | 6551 | 6434 | 7517 | 6843 | 6656 | 7155 | 8509 | 7967 | 7789 | 7916 | 6529 | 6919 | 6602 | 6099 | 6821 | 7006 | 7030 | 7209 | 6929 | Edc4 |
| Eddm3b | 0 | 1 | 0 | 0 | 0 | 6 | 1 | 0 | 1 | 5 | 0 | 0 | 0 | 0 | 0 | 4 | 0 | 0 | 0 | 0 | 3 | 0 | 1 | 0 | 0 | 0 | 0 | 0 | Eddm3b |
| Edem1 | 4687 | 4019 | 4641 | 4570 | 4139 | 4313 | 4339 | 2964 | 2845 | 3017 | 3503 | 3591 | 3415 | 3394 | 4262 | 4547 | 4300 | 4490 | 4165 | 3441 | 4073 | 3110 | 2902 | 3234 | 3068 | 3377 | 3221 | 3369 | Edem1 |
| Edem2 | 1931 | 2204 | 2407 | 2300 | 2281 | 2130 | 2027 | 1614 | 1804 | 1858 | 1771 | 1754 | 1754 | 1804 | 1990 | 2083 | 2298 | 2388 | 2325 | 1778 | 2193 | 2124 | 1830 | 1981 | 2051 | 1702 | 1746 | 1768 | Edem2 |
| Edem3 | 15146 | 13137 | 10657 | 12619 | 11527 | 12697 | 12741 | 9351 | 7973 | 7562 | 8500 | 7407 | 7768 | 7879 | 15318 | 14939 | 10859 | 13358 | 10331 | 12881 | 12749 | 8542 | 8250 | 7169 | 8020 | 5650 | 7707 | 7167 | Edem3 |
| Edf1 | 3878 | 5707 | 5002 | 5052 | 4620 | 4334 | 4491 | 3146 | 3695 | 4171 | 3950 | 4130 | 4005 | 4393 | 3833 | 4172 | 5306 | 4869 | 5108 | 4775 | 4737 | 3384 | 3716 | 4658 | 4128 | 4859 | 4185 | 4228 | Edf1 |
| Edil3 | 5073 | 4244 | 4902 | 5177 | 5549 | 5973 | 5803 | 6669 | 5548 | 5491 | 5662 | 5449 | 5887 | 5565 | 6330 | 5514 | 5058 | 5438 | 5080 | 6005 | 5959 | 7212 | 6077 | 4970 | 6142 | 4247 | 5935 | 6258 | Edil3 |
| Edn1 | 104 | 94 | 210 | 108 | 94 | 107 | 107 | 144 | 163 | 144 | 141 | 121 | 183 | 165 | 149 | 117 | 126 | 140 | 83 | 103 | 112 | 206 | 198 | 115 | 121 | 129 | 162 | 135 | Edn1 |
| Edn2 | 10 | 3 | 7 | 1 | 4 | 8 | 10 | 16 | 12 | 19 | 10 | 1 | 2 | 2 | 0 | 1 | 5 | 4 | 13 | 9 | 13 | 2 | 38 | 7 | 4 | 3 | 6 | 4 | Edn2 |
| Edn3 | 339 | 342 | 476 | 259 | 355 | 361 | 257 | 609 | 886 | 717 | 1197 | 627 | 1024 | 581 | 330 | 493 | 326 | 280 | 243 | 384 | 282 | 673 | 1093 | 673 | 584 | 708 | 639 | 574 | Edn3 |
| Ednra | 1453 | 885 | 1024 | 457 | 417 | 660 | 269 | 1975 | 2406 | 1849 | 2381 | 1515 | 2222 | 1458 | 1092 | 1491 | 548 | 522 | 283 | 328 | 283 | 1253 | 2554 | 1817 | 1158 | 2045 | 1457 | 1519 | Ednra |
| Ednrb | 3821 | 4282 | 3781 | 3494 | 6536 | 5224 | 4406 | 12059 | 9448 | 8738 | 9097 | 8317 | 10689 | 8625 | 6279 | 4403 | 4621 | 4503 | 4475 | 4982 | 4826 | 11812 | 9982 | 7388 | 8815 | 6694 | 9613 | 9424 | Ednrb |
| Edrf1 | 5221 | 5607 | 4324 | 5150 | 4062 | 4476 | 4976 | 4274 | 3741 | 3226 | 3395 | 3884 | 3694 | 4011 | 5123 | 5498 | 4591 | 4910 | 4315 | 4906 | 4927 | 3604 | 3635 | 3445 | 3933 | 3038 | 3882 | 3934 | Edrf1 |
| Eea1 | 7660 | 6404 | 5168 | 6717 | 6604 | 7356 | 7459 | 5881 | 5618 | 4843 | 5721 | 4779 | 5142 | 5325 | 7775 | 7331 | 5621 | 6409 | 5851 | 7704 | 7090 | 4894 | 5692 | 4412 | 5005 | 3068 | 5081 | 4552 | Eea1 |
| Eed | 4600 | 3968 | 4739 | 4233 | 3540 | 3856 | 4435 | 4362 | 3560 | 3692 | 3572 | 4340 | 4085 | 4747 | 4785 | 4315 | 4354 | 3981 | 4086 | 4661 | 4108 | 3387 | 3552 | 4257 | 3725 | 5656 | 4649 | 4963 | Eed |
| Eef1a1 | 488756 | 496869 | 410652 | 382175 | 345919 | 368553 | 376321 | 479706 | 457035 | 384896 | 390270 | 398275 | 397336 | 427594 | 496438 | 429100 | 398801 | 388532 | 395001 | 422565 | 404994 | 397840 | 464276 | 401956 | 400820 | 471738 | 430568 | 456395 | Eef1a1 |
| Eef1a2 | 506 | 678 | 629 | 1843 | 3147 | 1951 | 1809 | 1495 | 2015 | 2869 | 2552 | 1489 | 1246 | 1477 | 707 | 579 | 1305 | 2223 | 2535 | 1497 | 2429 | 3292 | 1983 | 2074 | 2922 | 1254 | 999 | 1032 | Eef1a2 |
| Eef1akmt1 | 956 | 1046 | 970 | 839 | 893 | 835 | 1100 | 1314 | 1219 | 1006 | 1093 | 1069 | 1064 | 1177 | 1031 | 854 | 875 | 890 | 980 | 1066 | 1011 | 1071 | 1170 | 1146 | 1151 | 1216 | 1142 | 1156 | Eef1akmt1 |
| Eef1akmt2 | 1550 | 1252 | 1632 | 1579 | 1590 | 1497 | 2019 | 2440 | 1975 | 2223 | 1985 | 2406 | 2276 | 3004 | 1651 | 1255 | 1626 | 1453 | 1678 | 1997 | 1929 | 2197 | 2095 | 2326 | 2362 | 2639 | 2278 | 2795 | Eef1akmt2 |
| Eef1akmt3 | 31 | 54 | 81 | 47 | 28 | 28 | 32 | 17 | 27 | 25 | 51 | 12 | 23 | 34 | 28 | 36 | 51 | 33 | 41 | 48 | 57 | 50 | 34 | 36 | 20 | 22 | 16 | 19 | Eef1akmt3 |
| Eef1akmt4 | 384 | 403 | 286 | 283 | 214 | 255 | 331 | 120 | 155 | 148 | 188 | 186 | 191 | 210 | 342 | 280 | 334 | 336 | 244 | 269 | 332 | 127 | 156 | 203 | 170 | 257 | 205 | 212 | Eef1akmt4 |
| Eef1aknmt | 2932 | 2901 | 2514 | 2745 | 2159 | 2714 | 2527 | 2015 | 2130 | 2178 | 2058 | 2371 | 2201 | 2224 | 2734 | 2896 | 2240 | 2722 | 2289 | 2282 | 2364 | 1900 | 2071 | 2184 | 2168 | 2325 | 2411 | 2263 | Eef1aknmt |
| Eef1b2 | 30067 | 27709 | 26769 | 23815 | 20279 | 21004 | 25864 | 25733 | 24123 | 23550 | 25404 | 24116 | 24362 | 27472 | 28044 | 24353 | 24962 | 22232 | 22924 | 24172 | 25262 | 20326 | 25342 | 26346 | 22758 | 29408 | 25429 | 30766 | Eef1b2 |
| Eef1d | 14011 | 16929 | 14998 | 13847 | 10551 | 11807 | 11647 | 8574 | 9949 | 9531 | 9920 | 10637 | 10435 | 10271 | 12496 | 14597 | 14261 | 13411 | 11856 | 12397 | 11309 | 7379 | 9631 | 10413 | 9693 | 12519 | 10958 | 11447 | Eef1d |
| Eef1e1 | 2220 | 2093 | 2393 | 2488 | 2725 | 2449 | 2806 | 3893 | 3190 | 3186 | 2906 | 3092 | 3096 | 3634 | 2690 | 2207 | 2501 | 2359 | 2634 | 3265 | 2743 | 3344 | 3198 | 3186 | 3352 | 3472 | 3270 | 3516 | Eef1e1 |
| Eef1g | 51043 | 51290 | 50402 | 48998 | 41863 | 43395 | 46876 | 39128 | 40605 | 41109 | 43869 | 43784 | 45459 | 45459 | 46566 | 48430 | 49026 | 47998 | 48029 | 42062 | 47375 | 38046 | 43863 | 44171 | 41427 | 53655 | 46244 | 50534 | Eef1g |
| Eef2 | 159811 | 210353 | 169884 | 186157 | 159180 | 166847 | 146655 | 119084 | 150564 | 146419 | 139513 | 146505 | 141540 | 131874 | 151491 | 170701 | 177173 | 178473 | 190477 | 157564 | 169808 | 129729 | 150310 | 149141 | 150613 | 158772 | 148316 | 144236 | Eef2 |
| Eef2k | 2637 | 2454 | 2715 | 2267 | 2839 | 2717 | 1997 | 3148 | 3677 | 3980 | 4024 | 3320 | 3840 | 3202 | 2608 | 2826 | 2402 | 2289 | 2364 | 2104 | 2225 | 3623 | 3889 | 3598 | 3582 | 3471 | 3570 | 3440 | Eef2k |

| | | | | | | | | | | | | | | | | | | | | | | | | | | | | | |
|---|---|---|---|---|---|---|---|---|---|---|---|---|---|---|---|---|---|---|---|---|---|---|---|---|---|---|---|---|---|
| Eef2kmt | 475 | 528 | 483 | 390 | 376 | 378 | 358 | 445 | 520 | 452 | 534 | 525 | 507 | 412 | 398 | 471 | 375 | 396 | 382 | 389 | 325 | 393 | 565 | 512 | 401 | 610 | 582 | 509 | Eef2kmt |
| Eefsec | 1292 | 1404 | 1264 | 1033 | 871 | 1038 | 816 | 1192 | 1460 | 1237 | 1213 | 1514 | 1320 | 1212 | 1129 | 1249 | 1196 | 1012 | 1109 | 994 | 928 | 1084 | 1416 | 1340 | 1227 | 1591 | 1317 | 1303 | Eefsec |
| Eepd1 | 1170 | 1190 | 1174 | 961 | 929 | 830 | 827 | 557 | 669 | 530 | 650 | 433 | 473 | 388 | 1006 | 1229 | 959 | 1119 | 825 | 848 | 801 | 568 | 608 | 477 | 488 | 510 | 482 | 376 | Eepd1 |
| Efcab1 | 911 | 702 | 740 | 691 | 610 | 679 | 694 | 1136 | 1069 | 814 | 752 | 820 | 864 | 844 | 1106 | 938 | 643 | 530 | 639 | 923 | 668 | 1048 | 1067 | 658 | 694 | 761 | 724 | 858 | Efcab1 |
| Efcab10 | 24 | 25 | 14 | 24 | 16 | 27 | 38 | 22 | 43 | 50 | 30 | 28 | 37 | 27 | 25 | 34 | 22 | 22 | 22 | 30 | 45 | 30 | 34 | 30 | 23 | 20 | 10 | 33 | Efcab10 |
| Efcab11 | 974 | 993 | 1035 | 726 | 594 | 707 | 822 | 670 | 653 | 536 | 519 | 945 | 762 | 1023 | 1052 | 1026 | 984 | 617 | 668 | 823 | 684 | 443 | 627 | 742 | 647 | 955 | 1005 | 898 | Efcab11 |
| Efcab12 | 288 | 422 | 290 | 419 | 456 | 410 | 383 | 355 | 345 | 392 | 378 | 400 | 386 | 378 | 323 | 328 | 349 | 347 | 487 | 375 | 431 | 500 | 373 | 307 | 418 | 224 | 400 | 373 | Efcab12 |
| Efcab14 | 5717 | 5053 | 5050 | 4648 | 4028 | 4666 | 4634 | 3814 | 3573 | 3462 | 3709 | 3786 | 3721 | 3719 | 5036 | 5523 | 4815 | 4571 | 4209 | 4207 | 4757 | 3474 | 3453 | 3466 | 3294 | 3552 | 3916 | 3858 | Efcab14 |
| Efcab15 | 18 | 17 | 23 | 35 | 39 | 10 | 21 | 11 | 27 | 38 | 27 | 16 | 2 | 21 | 8 | 14 | 16 | 23 | 23 | 17 | 25 | 23 | 12 | 20 | 24 | 10 | 12 | 17 | Efcab15 |
| Efcab2 | 738 | 584 | 755 | 711 | 770 | 735 | 900 | 1237 | 1060 | 1200 | 1169 | 1311 | 1382 | 1337 | 840 | 807 | 677 | 713 | 775 | 956 | 798 | 1176 | 1192 | 1090 | 1279 | 1180 | 1306 | 1273 | Efcab2 |
| Efcab5 | 104 | 117 | 61 | 94 | 110 | 122 | 158 | 239 | 240 | 179 | 255 | 181 | 126 | 194 | 82 | 94 | 86 | 108 | 98 | 143 | 154 | 236 | 209 | 218 | 255 | 115 | 166 | 125 | Efcab5 |
| Efcab6 | 148 | 120 | 101 | 205 | 190 | 141 | 203 | 50 | 83 | 69 | 73 | 63 | 28 | 47 | 132 | 121 | 123 | 175 | 198 | 163 | 239 | 84 | 63 | 69 | 79 | 61 | 51 | 26 | Efcab6 |
| Efcab7 | 2 | 1 | 0 | 6 | 2 | 2 | 1 | 1 | 7 | 8 | 7 | 1 | 1 | 0 | 3 | 2 | 0 | 3 | 3 | 3 | 1 | 7 | 3 | 1 | 2 | 3 | 6 | 1 | Efcab7 |
| Efcab8 | 14 | 13 | 9 | 13 | 10 | 22 | 10 | 7 | 5 | 10 | 8 | 8 | 7 | 6 | 13 | 6 | 13 | 22 | 16 | 12 | 27 | 23 | 14 | 7 | 19 | 12 | 16 | 7 | Efcab8 |
| Efcab9 | 44 | 44 | 46 | 22 | 34 | 49 | 54 | 10 | 14 | 24 | 6 | 21 | 15 | 16 | 53 | 44 | 64 | 42 | 33 | 54 | 46 | 18 | 12 | 5 | 25 | 11 | 9 | 13 | Efcab9 |
| Efcc1 | 137 | 192 | 157 | 141 | 272 | 199 | 154 | 180 | 233 | 215 | 202 | 218 | 374 | 191 | 202 | 261 | 137 | 156 | 184 | 203 | 139 | 269 | 300 | 148 | 243 | 108 | 278 | 179 | Efcc1 |
| Efemp1 | 1210 | 800 | 922 | 526 | 277 | 721 | 173 | 1056 | 1645 | 899 | 1664 | 755 | 1077 | 785 | 1093 | 1331 | 747 | 743 | 258 | 265 | 210 | 859 | 1850 | 842 | 622 | 887 | 952 | 724 | Efemp1 |
| Efemp2 | 2028 | 2236 | 1965 | 1543 | 1420 | 1585 | 1043 | 2026 | 2754 | 2102 | 2914 | 2057 | 2549 | 1822 | 1908 | 2597 | 1808 | 1675 | 1323 | 1026 | 1075 | 2089 | 3097 | 2258 | 1676 | 2253 | 2116 | 2063 | Efemp2 |
| Efhb | 191 | 205 | 212 | 268 | 246 | 269 | 260 | 129 | 113 | 106 | 127 | 160 | 191 | 120 | 244 | 228 | 226 | 202 | 226 | 259 | 243 | 160 | 194 | 87 | 120 | 119 | 126 | 116 | Efhb |
| Efhc1 | 198 | 187 | 125 | 145 | 138 | 176 | 129 | 188 | 218 | 165 | 201 | 161 | 213 | 130 | 178 | 220 | 165 | 157 | 121 | 156 | 139 | 242 | 247 | 128 | 131 | 130 | 182 | 128 | Efhc1 |
| Efhc2 | 119 | 110 | 87 | 62 | 102 | 90 | 74 | 78 | 116 | 89 | 62 | 71 | 77 | 81 | 105 | 156 | 41 | 76 | 88 | 82 | 75 | 124 | 121 | 63 | 74 | 79 | 53 | 70 | Efhc2 |
| Efhd1 | 263 | 326 | 410 | 308 | 369 | 252 | 361 | 73 | 74 | 79 | 89 | 92 | 101 | 86 | 277 | 266 | 363 | 299 | 361 | 313 | 314 | 122 | 99 | 97 | 93 | 112 | 53 | 109 | Efhd1 |
| Efhd2 | 3079 | 3953 | 3920 | 4203 | 4308 | 3896 | 3892 | 2646 | 3112 | 3370 | 3270 | 3397 | 2852 | 3286 | 3072 | 3308 | 4589 | 4341 | 4846 | 3391 | 4246 | 3080 | 2944 | 3832 | 3520 | 3357 | 3024 | 3191 | Efhd2 |
| Efl1 | 2569 | 2645 | 2646 | 2360 | 1993 | 1985 | 2164 | 1856 | 1769 | 1619 | 1506 | 1748 | 1622 | 1754 | 2422 | 2562 | 2360 | 2206 | 2111 | 2153 | 2081 | 1357 | 1702 | 1687 | 1538 | 1748 | 1827 | 1720 | Efl1 |
| Efna1 | 610 | 638 | 709 | 556 | 448 | 531 | 400 | 365 | 675 | 554 | 612 | 468 | 568 | 484 | 576 | 798 | 541 | 551 | 462 | 369 | 407 | 409 | 726 | 577 | 410 | 625 | 457 | 498 | Efna1 |
| Efna2 | 1711 | 1993 | 3479 | 2315 | 2086 | 2122 | 1662 | 1795 | 1682 | 2509 | 2211 | 3018 | 2366 | 2819 | 1702 | 1895 | 2868 | 2083 | 2440 | 1574 | 2055 | 2376 | 1760 | 2844 | 2503 | 2687 | 2674 | 3031 | Efna2 |
| Efna3 | 2015 | 2630 | 1766 | 2504 | 2055 | 2321 | 1960 | 811 | 1019 | 1009 | 1045 | 607 | 730 | 615 | 1986 | 2441 | 2094 | 2698 | 2139 | 2218 | 1911 | 834 | 1002 | 869 | 948 | 656 | 539 | 570 | Efna3 |
| Efna4 | 1441 | 1658 | 1645 | 1524 | 1163 | 1171 | 1157 | 946 | 1202 | 1088 | 1027 | 1036 | 1253 | 1137 | 1201 | 1617 | 1455 | 1354 | 1260 | 1257 | 1144 | 677 | 1200 | 1232 | 980 | 1632 | 1135 | 1258 | Efna4 |
| Efna5 | 2985 | 2344 | 4686 | 3853 | 3209 | 3374 | 1998 | 4498 | 4239 | 5788 | 5546 | 5779 | 6343 | 6236 | 2711 | 3116 | 3962 | 3489 | 3469 | 2177 | 2973 | 5820 | 5029 | 6293 | 5447 | 5836 | 5935 | 7175 | Efna5 |
| Efnb1 | 7626 | 9873 | 11334 | 11181 | 8299 | 8786 | 8061 | 6935 | 8359 | 9133 | 8320 | 11979 | 9394 | 10381 | 7159 | 8486 | 10772 | 9841 | 10076 | 7974 | 7862 | 6199 | 7278 | 11070 | 9558 | 14327 | 9767 | 9692 | Efnb1 |
| Efnb2 | 8088 | 7986 | 8251 | 11711 | 10294 | 9933 | 11148 | 9108 | 10181 | 11240 | 11266 | 11788 | 10137 | 12493 | 7915 | 8731 | 10097 | 11669 | 10772 | 8114 | 10251 | 9015 | 8741 | 12828 | 13131 | 13920 | 9073 | 10652 | Efnb2 |
| Efnb3 | 8930 | 11564 | 10758 | 13322 | 13949 | 13578 | 12533 | 10087 | 10763 | 12008 | 10558 | 11246 | 10571 | 11626 | 9742 | 9948 | 12791 | 13442 | 14179 | 12382 | 13273 | 13964 | 10031 | 12402 | 13845 | 11283 | 10676 | 10867 | Efnb3 |
| Efr3a | 8929 | 7069 | 5936 | 6480 | 6824 | 7459 | 7564 | 4894 | 4514 | 3846 | 3997 | 3762 | 3936 | 3856 | 8687 | 8094 | 6147 | 7321 | 6138 | 7492 | 7276 | 3907 | 4783 | 3411 | 3455 | 3437 | 3977 | 3752 | Efr3a |
| Efr3b | 7200 | 7861 | 8082 | 11563 | 14708 | 12285 | 12524 | 8259 | 7874 | 10994 | 9266 | 9865 | 8917 | 9436 | 7752 | 7378 | 10348 | 12559 | 13390 | 10112 | 13101 | 12885 | 7416 | 9593 | 12610 | 6093 | 8638 | 8399 | Efr3b |
| Efs | 860 | 849 | 1005 | 731 | 1064 | 853 | 860 | 2186 | 3022 | 2952 | 3196 | 2928 | 2501 | 2650 | 702 | 794 | 1128 | 842 | 1139 | 823 | 785 | 2211 | 2999 | 3200 | 2530 | 3480 | 2704 | 2610 | Efs |
| Eftud2 | 15523 | 17179 | 15395 | 13804 | 11657 | 13138 | 12948 | 10699 | 11796 | 12038 | 10779 | 15127 | 13439 | 13750 | 13917 | 14620 | 14371 | 13085 | 14258 | 13124 | 13103 | 10432 | 10934 | 12870 | 12510 | 16481 | 14559 | 15455 | Eftud2 |
| Egf | 236 | 198 | 159 | 206 | 294 | 214 | 238 | 289 | 301 | 282 | 294 | 240 | 330 | 252 | 261 | 165 | 147 | 199 | 221 | 284 | 218 | 286 | 271 | 217 | 260 | 229 | 271 | 231 | Egf |
| Egfem1 | 339 | 404 | 568 | 379 | 263 | 313 | 268 | 337 | 187 | 218 | 163 | 399 | 312 | 362 | 382 | 286 | 353 | 263 | 286 | 366 | 261 | 338 | 229 | 261 | 268 | 327 | 425 | 515 | Egfem1 |
| Egfl6 | 1380 | 1061 | 1065 | 729 | 332 | 1315 | 142 | 1915 | 2871 | 2295 | 4106 | 1412 | 1876 | 1386 | 1298 | 1424 | 864 | 1468 | 502 | 271 | 159 | 1171 | 4424 | 2004 | 1272 | 1468 | 1453 | 1259 | Egfl6 |
| Egfl7 | 968 | 1185 | 1330 | 1011 | 1079 | 937 | 917 | 1043 | 1279 | 1485 | 1700 | 1470 | 1665 | 1304 | 968 | 1231 | 1114 | 971 | 957 | 1008 | 811 | 1176 | 1604 | 1346 | 1528 | 1605 | 1543 | 1310 | Egfl7 |
| Egfl8 | 88 | 136 | 114 | 131 | 94 | 81 | 115 | 94 | 117 | 112 | 123 | 98 | 75 | 74 | 62 | 110 | 148 | 118 | 127 | 114 | 109 | 129 | 88 | 107 | 104 | 95 | 88 | 83 | Egfl8 |
| Egflam | 1189 | 1493 | 1395 | 1178 | 1073 | 1157 | 554 | 1296 | 1812 | 1817 | 2456 | 1825 | 2467 | 1701 | 1265 | 1998 | 1070 | 1360 | 651 | 703 | 581 | 1803 | 2176 | 1707 | 1550 | 2015 | 1901 | 1818 | Egflam |
| Egfr | 2217 | 1317 | 1139 | 1015 | 808 | 1081 | 412 | 1956 | 2962 | 1973 | 3450 | 1322 | 2044 | 1414 | 1572 | 2296 | 906 | 1302 | 475 | 622 | 543 | 2094 | 3506 | 1742 | 1565 | 1463 | 1520 | 1171 | Egfr |
| Egln1 | 5236 | 5135 | 6350 | 5738 | 6383 | 6005 | 6201 | 5097 | 5501 | 6210 | 5904 | 6425 | 5640 | 6709 | 4892 | 4967 | 6471 | 5966 | 6708 | 5222 | 6091 | 5479 | 5258 | 6685 | 6466 | 7160 | 6015 | 6782 | Egln1 |
| Egln2 | 2526 | 3300 | 3064 | 3497 | 3423 | 3205 | 2672 | 4080 | 4584 | 5176 | 4475 | 5221 | 4617 | 4531 | 2467 | 2551 | 3497 | 3158 | 3607 | 2907 | 2979 | 4405 | 4202 | 5102 | 5098 | 5535 | 4852 | 4522 | Egln2 |
| Egln3 | 1316 | 1481 | 1354 | 1605 | 1405 | 1379 | 1256 | 670 | 755 | 707 | 674 | 491 | 670 | 518 | 1429 | 1427 | 1502 | 1615 | 1400 | 1322 | 1328 | 700 | 821 | 609 | 574 | 622 | 440 | 495 | Egln3 |
| Egr1 | 664 | 583 | 516 | 638 | 447 | 438 | 380 | 206 | 312 | 323 | 550 | 240 | 264 | 214 | 517 | 671 | 556 | 578 | 408 | 318 | 382 | 334 | 356 | 333 | 244 | 328 | 194 | 221 | Egr1 |
| Egr3 | 99 | 77 | 143 | 97 | 123 | 96 | 131 | 71 | 133 | 135 | 111 | 112 | 33 | 113 | 74 | 66 | 95 | 122 | 120 | 103 | 129 | 127 | 155 | 150 | 158 | 156 | 86 | 87 | Egr3 |
| Egr4 | 12 | 18 | 13 | 20 | 7 | 7 | 10 | 0 | 0 | 3 | 0 | 1 | 0 | 1 | 1 | 11 | 15 | 13 | 5 | 3 | 7 | 0 | 0 | 1 | 0 | 0 | 0 | 0 | Egr4 |
| Ehbp1 | 5733 | 5353 | 6647 | 7426 | 8846 | 7728 | 8760 | 8430 | 7453 | 8863 | 8423 | 8610 | 8805 | 9848 | 6381 | 5115 | 7234 | 7629 | 8319 | 7516 | 8728 | 9386 | 7612 | 8874 | 9626 | 7355 | 9337 | 9461 | Ehbp1 |
| Ehbp1l1 | 1204 | 1695 | 1734 | 1960 | 1744 | 1725 | 1469 | 1304 | 1537 | 1868 | 2966 | 1650 | 1873 | 1705 | 1266 | 1665 | 1671 | 1981 | 1788 | 1513 | 1569 | 1858 | 2116 | 1905 | 1976 | 1386 | 1873 | 1671 | Ehbp1l1 |
| Ehd2 | 1654 | 1945 | 1780 | 1374 | 1152 | 1490 | 830 | 1817 | 2895 | 2144 | 2976 | 1792 | 2171 | 1643 | 1667 | 2247 | 1367 | 1787 | 1021 | 863 | 898 | 1803 | 3504 | 2007 | 1629 | 1975 | 1934 | 1860 | Ehd2 |
| Ehd3 | 1739 | 1865 | 1888 | 1773 | 2477 | 1964 | 1947 | 1297 | 1538 | 1390 | 1412 | 1339 | 1396 | 1313 | 1779 | 1727 | 1889 | 1815 | 2138 | 1972 | 1982 | 1783 | 1266 | 1335 | 1460 | 1347 | 1156 | 1305 | Ehd3 |
| Ehd4 | 2836 | 2886 | 3159 | 2375 | 1822 | 2072 | 1684 | 985 | 1205 | 1125 | 1751 | 1251 | 1310 | 1005 | 2528 | 3357 | 2520 | 2458 | 1588 | 1722 | 1817 | 1063 | 1743 | 1184 | 958 | 1483 | 1309 | 1163 | Ehd4 |
| Ehf | 33 | 6 | 0 | 5 | 1 | 7 | 0 | 357 | 284 | 300 | 172 | 4 | 3 | 16 | 8 | 9 | 1 | 1 | 2 | 0 | 0 | 28 | 340 | 244 | 14 | 119 | 6 | 2 | Ehf |
| Ehhadh | 513 | 459 | 592 | 350 | 286 | 353 | 244 | 260 | 253 | 221 | 216 | 219 | 310 | 268 | 480 | 549 | 400 | 326 | 274 | 327 | 233 | 226 | 287 | 197 | 167 | 206 | 279 | 272 | Ehhadh |
| Ehmt1 | 8706 | 9277 | 9654 | 9329 | 9844 | 9593 | 9442 | 11195 | 11137 | 12630 | 10971 | 14618 | 12362 | 13805 | 8334 | 8030 | 9894 | 9259 | 10089 | 8915 | 9112 | 12607 | 10365 | 13759 | 13383 | 13852 | 13715 | 14142 | Ehmt1 |
| Ehmt2 | 11914 | 15227 | 16293 | 18190 | 17399 | 16226 | 15122 | 12977 | 15424 | 17567 | 16350 | 18977 | 15326 | 17235 | 10859 | 12214 | 17496 | 16976 | 19720 | 13623 | 17033 | 16807 | 14538 | 19708 | 19085 | 20381 | 16816 | 17169 | Ehmt2 |
| Ei24 | 13633 | 11888 | 11872 | 11780 | 11360 | 11005 | 12750 | 10128 | 8626 | 8525 | 8019 | 8992 | 8687 | 9950 | 13061 | 11576 | 12120 | 11775 | 11478 | 11130 | 12676 | 9663 | 8759 | 8943 | 8640 | 9758 | 8732 | 9932 | Ei24 |
| Eid1 | 28233 | 21865 | 23639 | 26541 | 28833 | 27929 | 33905 | 34775 | 29730 | 30363 | 30738 | 27766 | 28139 | 30850 | 29642 | 23855 | 24878 | 26875 | 27850 | 28479 | 32117 | 34808 | 31789 | 30040 | 31149 | 29589 | 27510 | 31393 | Eid1 |
| Eid2 | 1679 | 1682 | 1638 | 2073 | 2010 | 1692 | 2202 | 1728 | 1494 | 1817 | 1769 | 1674 | 1514 | 2026 | 1615 | 1325 | 2033 | 1913 | 1884 | 1677 | 2232 | 1888 | 1564 | 1892 | 1945 | 1747 | 1585 | 1886 | Eid2 |
| Eid2b | 995 | 879 | 896 | 1265 | 1629 | 1460 | 1496 | 1513 | 1451 | 1725 | 1351 | 1408 | 1210 | 1493 | 1029 | 761 | 1196 | 1226 | 1518 | 1313 | 1498 | 1713 | 1506 | 1537 | 1782 | 1416 | 1255 | 1351 | Eid2b |
| Eid3 | 47 | 35 | 24 | 45 | 21 | 48 | 28 | 28 | 31 | 20 | 30 | 27 | 20 | 13 | 45 | 37 | 41 | 18 | 26 | 31 | 41 | 27 | 33 | 23 | 25 | 23 | 33 | 19 | Eid3 |
| Eif1 | 14257 | 14699 | 16967 | 17049 | 16547 | 15925 | 17479 | 20206 | 18713 | 20028 | 20630 | 24422 | 22256 | 26739 | 14766 | 13221 | 17955 | 16351 | 18759 | 14909 | 18081 | 23051 | 19155 | 22249 | 23153 | 28851 | 23210 | 28154 | Eif1 |
| Eif1a | 9461 | 6940 | 7982 | 7039 | 5930 | 6393 | 6716 | 6128 | 5336 | 4476 | 5122 | 4643 | 5688 | 5206 | 9194 | 8937 | 6833 | 7228 | 5656 | 7535 | 6330 | 5153 | 5614 | 4989 | 4725 | 5615 | 5413 | 5830 | Eif1a |
| Eif1ad | 3930 | 4467 | 4736 | 3901 | 3350 | 3505 | 3879 | 3883 | 3482 | 3767 | 3430 | 4783 | 4301 | 4574 | 3774 | 3647 | 4483 | 3551 | 3784 | 3603 | 3548 | 3676 | 3237 | 4284 | 3750 | 5557 | 4786 | 5286 | Eif1ad |
| Eif1ad19 | 0 | 0 | 0 | 0 | 0 | 0 | 0 | 0 | 0 | 0 | 0 | 0 | 0 | 0 | 0 | 0 | 0 | 0 | 0 | 0 | 0 | 5 | 0 | 0 | 0 | 0 | 0 | 0 | Eif1ad19 |
| Eif1ad4 | 0 | 0 | 0 | 0 | 0 | 0 | 0 | 0 | 0 | 0 | 0 | 0 | 0 | 0 | 0 | 0 | 0 | 0 | 0 | 0 | 0 | 0 | 0 | 0 | 0 | 0 | 0 | 1 | Eif1ad4 |
| Eif1ax | 20086 | 17189 | 19118 | 17484 | 17437 | 17936 | 20642 | 16709 | 13869 | 13623 | 12703 | 17068 | 15282 | 18610 | 20634 | 16998 | 18535 | 17155 | 17687 | 21476 | 19350 | 13090 | 13521 | 15192 | 15102 | 18056 | 17540 | 18898 | Eif1ax |
| Eif1b | 4269 | 4356 | 4765 | 4958 | 4619 | 3920 | 5712 | 5814 | 5074 | 5632 | 5029 | 6357 | 4891 | 7771 | 4193 | 3153 | 5489 | 4784 | 5177 | 4331 | 5727 | 5638 | 4747 | 6636 | 6241 | 7522 | 5579 | 7239 | Eif1b |
| Eif2a | 5715 | 4658 | 4863 | 4280 | 3859 | 3953 | 4307 | 6741 | 5749 | 5045 | 5108 | 5640 | 5774 | 5844 | 5734 | 5373 | 4804 | 4181 | 4274 | 5032 | 4183 | 4983 | 5803 | 5458 | 5081 | 6247 | 6178 | 6220 | Eif2a |
| Eif2ak1 | 8823 | 8686 | 7437 | 7233 | 6822 | 6994 | 7805 | 5843 | 5498 | 5007 | 5151 | 5321 | 5833 | 5676 | 8591 | 7929 | 7221 | 7096 | 6996 | 7350 | 7542 | 5596 | 5273 | 5185 | 5387 | 5599 | 5663 | 5910 | Eif2ak1 |
| Eif2ak2 | 1849 | 1335 | 1313 | 1013 | 1051 | 1380 | 1040 | 1507 | 1537 | 1507 | 1689 | 1754 | 2157 | 1767 | 1680 | 1754 | 1092 | 989 | 900 | 1338 | 975 | 1252 | 1852 | 1402 | 1230 | 1572 | 2126 | 1711 | Eif2ak2 |
| Eif2ak3 | 3904 | 4453 | 3838 | 4437 | 4245 | 4336 | 4542 | 2446 | 2517 | 2641 | 2845 | 2525 | 2364 | 2528 | 3932 | 3885 | 4011 | 4362 | 4283 | 4080 | 4333 | 2702 | 2438 | 2589 | 2870 | 2262 | 2457 | 2267 | Eif2ak3 |
| Eif2ak4 | 5654 | 5768 | 4367 | 4891 | 4379 | 4930 | 5084 | 2919 | 3180 | 2974 | 2983 | 2860 | 3282 | 3005 | 5384 | 5599 | 4464 | 4773 | 4754 | 4879 | 5127 | 2811 | 2970 | 2694 | 2779 | 2620 | 2987 | 2962 | Eif2ak4 |
| Eif2b1 | 2843 | 3169 | 3035 | 2406 | 2047 | 2359 | 2421 | 3156 | 3013 | 2794 | 2756 | 3400 | 3030 | 2976 | 2971 | 2838 | 2668 | 2288 | 2426 | 2845 | 2196 | 2560 | 2987 | 3010 | 2971 | 3726 | 3517 | 3389 | Eif2b1 |
| Eif2b2 | 3290 | 4045 | 3762 | 3886 | 3152 | 3244 | 3246 | 2433 | 2630 | 2483 | 2370 | 2966 | 2676 | 2805 | 3229 | 3575 | 3590 | 3428 | 3583 | 3110 | 3281 | 2405 | 2617 | 2945 | 2723 | 3386 | 3040 | 2950 | Eif2b2 |
| Eif2b3 | 1967 | 1752 | 1739 | 1564 | 1384 | 1569 | 1738 | 1958 | 1671 | 1514 | 1526 | 1584 | 1517 | 1601 | 2078 | 1829 | 1652 | 1501 | 1414 | 1796 | 1531 | 1567 | 1672 | 1475 | 1518 | 1928 | 1641 | 1771 | Eif2b3 |
| Eif2b4 | 2045 | 2668 | 2418 | 2652 | 2269 | 2247 | 2160 | 2149 | 2503 | 2410 | 2427 | 2699 | 2505 | 2495 | 1960 | 2155 | 2294 | 2268 | 2491 | 2387 | 2131 | 2254 | 2393 | 2563 | 2546 | 2828 | 2640 | 2633 | Eif2b4 |

| | | | | | | | | | | | | | | | | | | | | | | | | | | | | | |
|---|---|---|---|---|---|---|---|---|---|---|---|---|---|---|---|---|---|---|---|---|---|---|---|---|---|---|---|---|---|
| Eif2b5 | 7160 | 7434 | 6457 | 6155 | 5496 | 6153 | 5998 | 5843 | 5906 | 5361 | 5225 | 6229 | 5926 | 5990 | 6690 | 6460 | 6185 | 5401 | 6214 | 5767 | 6288 | 5270 | 5793 | 5587 | 5733 | 6801 | 6269 | 6691 | Eif2b5 |
| Eif2d | 2573 | 2654 | 2026 | 2217 | 2075 | 2289 | 2109 | 2992 | 3087 | 2863 | 3030 | 3074 | 3061 | 2941 | 2268 | 2394 | 1971 | 2136 | 2164 | 2241 | 2328 | 2799 | 3205 | 2743 | 2954 | 3150 | 3126 | 3128 | Eif2d |
| Eif2s1 | 11100 | 8537 | 9994 | 8505 | 7265 | 8029 | 9829 | 10444 | 9030 | 8339 | 8297 | 9459 | 9334 | 10414 | 11045 | 9650 | 9073 | 8168 | 8021 | 9689 | 8523 | 7815 | 8907 | 9095 | 8736 | 10837 | 10513 | 11153 | Eif2s1 |
| Eif2s2 | 17788 | 13833 | 16844 | 14010 | 11889 | 12988 | 14855 | 14488 | 12560 | 11761 | 11593 | 13436 | 12735 | 14789 | 17359 | 16129 | 14768 | 13046 | 13196 | 16612 | 13938 | 10676 | 12902 | 12912 | 11742 | 15960 | 14205 | 16459 | Eif2s2 |
| Eif2s3x | 15060 | 13953 | 13697 | 12397 | 11142 | 12000 | 12896 | 12544 | 11221 | 9412 | 9401 | 10949 | 11071 | 11605 | 21074 | 19091 | 17717 | 16367 | 15054 | 18901 | 16929 | 14499 | 15912 | 15094 | 15611 | 16961 | 18131 | 18557 | Eif2s3x |
| Eif2s3y | 7981 | 6135 | 7438 | 7641 | 8112 | 7729 | 10240 | 7540 | 7236 | 7295 | 5487 | 8089 | 8238 | 9460 | 12 | 2 | 1 | 0 | 0 | 2 | 1 | 0 | 17 | 0 | 0 | 1 | 0 | 2 | Eif2s3y |
| Eif3a | 44240 | 37911 | 45395 | 42684 | 38647 | 40429 | 43067 | 47588 | 42207 | 42676 | 42504 | 46140 | 45409 | 51142 | 45039 | 41131 | 42767 | 40124 | 39200 | 44244 | 40031 | 38573 | 40944 | 44638 | 41791 | 45407 | 48693 | 48772 | Eif3a |
| Eif3b | 20765 | 22481 | 21588 | 20290 | 16904 | 17914 | 18393 | 14100 | 14997 | 14540 | 14132 | 16778 | 15610 | 16208 | 19060 | 19317 | 20578 | 19370 | 19999 | 16794 | 18941 | 13252 | 14354 | 15909 | 14461 | 19432 | 16899 | 17531 | Eif3b |
| Eif3c | 30349 | 32065 | 28959 | 25190 | 23860 | 25426 | 26333 | 25850 | 23933 | 21818 | 21179 | 24996 | 24116 | 25191 | 30793 | 27882 | 27081 | 24890 | 25976 | 27974 | 26656 | 21373 | 23680 | 23267 | 23127 | 26644 | 26714 | 26918 | Eif3c |
| Eif3d | 10171 | 11943 | 10194 | 9287 | 8336 | 9222 | 8752 | 9553 | 10401 | 9616 | 9434 | 11285 | 10593 | 10712 | 9595 | 9670 | 9978 | 8683 | 9988 | 9306 | 9123 | 8722 | 10396 | 10474 | 10243 | 13037 | 11979 | 11750 | Eif3d |
| Eif3e | 21609 | 16455 | 16606 | 15681 | 13677 | 15241 | 17108 | 20861 | 18761 | 15821 | 16151 | 16154 | 16780 | 18214 | 22045 | 19008 | 15723 | 14223 | 14882 | 19535 | 16566 | 15478 | 19442 | 16324 | 14924 | 19310 | 17767 | 19869 | Eif3e |
| Eif3f | 11220 | 11857 | 13684 | 13821 | 11872 | 11753 | 11015 | 10733 | 11515 | 12376 | 12436 | 13573 | 11636 | 12754 | 11079 | 11300 | 13555 | 13018 | 13858 | 9274 | 12687 | 12579 | 11990 | 13885 | 13058 | 16019 | 11920 | 13531 | Eif3f |
| Eif3g | 7303 | 8636 | 9100 | 8038 | 6357 | 6713 | 6840 | 5378 | 5812 | 6473 | 6061 | 7139 | 6913 | 7195 | 6926 | 7256 | 8208 | 6999 | 7697 | 6978 | 7242 | 4821 | 5662 | 6949 | 6537 | 8411 | 7470 | 7849 | Eif3g |
| Eif3h | 15915 | 16445 | 14784 | 14927 | 13111 | 14177 | 13662 | 14360 | 14138 | 13954 | 13312 | 13927 | 13729 | 13873 | 15306 | 14897 | 14865 | 13481 | 14690 | 15498 | 14541 | 12529 | 14903 | 14269 | 13519 | 15697 | 14419 | 14859 | Eif3h |
| Eif3i | 9596 | 11191 | 9110 | 8431 | 7106 | 6994 | 8212 | 7772 | 8210 | 6784 | 7198 | 8065 | 7905 | 7960 | 9537 | 9820 | 9006 | 7779 | 8270 | 9654 | 8589 | 6259 | 8167 | 7798 | 7159 | 8996 | 8559 | 8883 | Eif3i |
| Eif3j1 | 7294 | 5613 | 8223 | 6615 | 5659 | 6092 | 6748 | 5588 | 4988 | 5183 | 5496 | 5807 | 5709 | 6209 | 7130 | 6611 | 6843 | 6377 | 6278 | 6378 | 6377 | 5518 | 5136 | 5445 | 5523 | 7020 | 5853 | 6506 | Eif3j1 |
| Eif3j2 | 518 | 278 | 403 | 383 | 241 | 294 | 296 | 360 | 248 | 212 | 250 | 282 | 295 | 309 | 468 | 382 | 350 | 408 | 359 | 315 | 310 | 341 | 242 | 324 | 267 | 306 | 318 | 331 | Eif3j2 |
| Eif3k | 8221 | 9926 | 7654 | 7575 | 6554 | 7011 | 7125 | 7224 | 7901 | 7318 | 7112 | 7421 | 7592 | 7575 | 7917 | 8364 | 7596 | 6842 | 7875 | 8404 | 7606 | 6046 | 8225 | 7850 | 6967 | 8313 | 7967 | 8289 | Eif3k |
| Eif3l | 16257 | 17610 | 14082 | 12897 | 11362 | 11964 | 13194 | 13071 | 12778 | 10679 | 10691 | 12144 | 11731 | 13262 | 15387 | 15090 | 13243 | 12009 | 12738 | 14267 | 13483 | 10036 | 12695 | 11954 | 11200 | 14117 | 12796 | 14310 | Eif3l |
| Eif3m | 9165 | 7677 | 7580 | 6710 | 5904 | 6655 | 8170 | 9441 | 8740 | 7065 | 7128 | 8047 | 8010 | 8792 | 9454 | 8652 | 7004 | 6522 | 6831 | 8875 | 7309 | 6990 | 8827 | 7899 | 7097 | 9248 | 8788 | 9769 | Eif3m |
| Eif4a1 | 48734 | 46716 | 46586 | 41653 | 37372 | 38790 | 46736 | 70420 | 61801 | 55762 | 53842 | 67376 | 60214 | 77296 | 49265 | 41250 | 43343 | 39173 | 40297 | 47231 | 43617 | 54243 | 59101 | 65540 | 61830 | 78385 | 71290 | 78262 | Eif4a1 |
| Eif4a2 | 25155 | 20640 | 25579 | 28463 | 29622 | 26409 | 31176 | 50044 | 43063 | 40262 | 44608 | 36629 | 35434 | 38611 | 29488 | 25079 | 25644 | 29675 | 27938 | 31919 | 30280 | 46062 | 45140 | 40427 | 40499 | 38538 | 35307 | 36458 | Eif4a2 |
| Eif4a3 | 6899 | 8215 | 9087 | 8074 | 6905 | 7352 | 7735 | 5455 | 6331 | 7027 | 6655 | 8619 | 7538 | 8817 | 6287 | 6853 | 9144 | 7288 | 7928 | 6862 | 7714 | 5820 | 5811 | 8055 | 7227 | 10087 | 8766 | 9449 | Eif4a3 |
| Eif4a3l1 | 1 | 1 | 0 | 0 | 1 | 0 | 2 | 0 | 0 | 0 | 0 | 0 | 0 | 0 | 0 | 0 | 0 | 1 | 0 | 0 | 0 | 0 | 0 | 0 | 0 | 0 | 0 | 0 | Eif4a3l1 |
| Eif4a3l2 | 15 | 18 | 12 | 13 | 13 | 18 | 6 | 20 | 5 | 8 | 8 | 20 | 10 | 13 | 15 | 7 | 10 | 8 | 16 | 15 | 18 | 30 | 12 | 26 | 16 | 13 | 15 | 13 | Eif4a3l2 |
| Eif4b | 27644 | 25754 | 26125 | 24602 | 22863 | 24705 | 24453 | 31630 | 32023 | 30910 | 31217 | 32109 | 30709 | 32305 | 25709 | 24161 | 25362 | 24027 | 26221 | 24999 | 24903 | 27960 | 32134 | 30912 | 30456 | 34226 | 32511 | 34051 | Eif4b |
| Eif4e | 14504 | 11629 | 12966 | 11569 | 11302 | 11551 | 13943 | 15408 | 13282 | 11672 | 11319 | 13568 | 12744 | 14600 | 14778 | 11995 | 12250 | 12379 | 11972 | 14230 | 12720 | 13363 | 12516 | 12821 | 12780 | 15785 | 14274 | 15272 | Eif4e |
| Eif4e1b | 0 | 0 | 0 | 0 | 2 | 0 | 0 | 0 | 0 | 0 | 0 | 0 | 0 | 1 | 0 | 0 | 0 | 0 | 0 | 0 | 1 | 0 | 1 | 2 | 5 | 0 | 0 | 0 | Eif4e1b |
| Eif4e2 | 6676 | 6916 | 7645 | 7186 | 6469 | 6507 | 6419 | 7034 | 6756 | 6804 | 7381 | 7770 | 7402 | 7946 | 6421 | 6736 | 7128 | 6782 | 6919 | 6256 | 6644 | 7058 | 6886 | 7315 | 7409 | 8269 | 8157 | 8109 | Eif4e2 |
| Eif4e3 | 960 | 764 | 954 | 720 | 692 | 790 | 808 | 844 | 830 | 840 | 859 | 908 | 819 | 945 | 778 | 877 | 730 | 587 | 748 | 704 | 770 | 857 | 804 | 939 | 782 | 1174 | 948 | 1042 | Eif4e3 |
| Eif4ebp1 | 1264 | 1321 | 996 | 795 | 554 | 736 | 571 | 473 | 611 | 532 | 652 | 439 | 619 | 551 | 976 | 1291 | 899 | 750 | 591 | 708 | 511 | 369 | 742 | 503 | 397 | 695 | 589 | 638 | Eif4ebp1 |
| Eif4ebp2 | 28307 | 29014 | 36944 | 37292 | 30815 | 29917 | 34198 | 15707 | 16621 | 21179 | 18583 | 26295 | 19143 | 24609 | 25194 | 25797 | 37900 | 37824 | 36658 | 23485 | 35488 | 21896 | 14947 | 24572 | 23483 | 26831 | 20576 | 25023 | Eif4ebp2 |
| Eif4ebp3 | 60 | 81 | 85 | 73 | 75 | 40 | 82 | 39 | 44 | 51 | 67 | 81 | 77 | 69 | 50 | 71 | 86 | 86 | 65 | 74 | 63 | 62 | 42 | 74 | 46 | 71 | 59 | 64 | Eif4ebp3 |
| Eif4enif1 | 8082 | 7225 | 7428 | 7288 | 6602 | 6743 | 7297 | 7312 | 6917 | 7565 | 7286 | 8194 | 8032 | 8354 | 7472 | 6812 | 7502 | 6871 | 7037 | 6439 | 7083 | 7740 | 6754 | 7508 | 7489 | 8529 | 7946 | 8437 | Eif4enif1 |
| Eif4g1 | 40822 | 43738 | 45095 | 43475 | 40856 | 40875 | 39242 | 38521 | 42025 | 45032 | 44653 | 49945 | 48502 | 47691 | 38153 | 39822 | 44781 | 42238 | 43082 | 37344 | 39969 | 39921 | 40172 | 46233 | 44383 | 45809 | 49609 | 49422 | Eif4g1 |
| Eif4g2 | 120803 | 107417 | 105459 | 104975 | 121911 | 125529 | 129768 | 209889 | 172216 | 172981 | 158700 | 198371 | 181529 | 198947 | 129728 | 102102 | 111943 | 106627 | 121961 | 128599 | 129111 | 195080 | 169427 | 175090 | 187739 | 184555 | 202047 | 202790 | Eif4g2 |
| Eif4g3 | 13923 | 14731 | 18216 | 19951 | 21570 | 19771 | 19676 | 16330 | 17326 | 22861 | 20915 | 24453 | 23211 | 24011 | 13834 | 14576 | 19463 | 19818 | 21174 | 15392 | 20872 | 23208 | 16845 | 23104 | 23673 | 19552 | 21557 | 23182 | Eif4g3 |
| Eif4h | 21040 | 24000 | 26899 | 24962 | 22272 | 22333 | 22387 | 20549 | 22731 | 23989 | 25071 | 27604 | 26578 | 27513 | 19299 | 19857 | 25557 | 23366 | 25214 | 18929 | 22454 | 21253 | 22571 | 28097 | 26224 | 32739 | 27901 | 28776 | Eif4h |
| Eif5 | 22581 | 16325 | 21025 | 19036 | 21296 | 21799 | 24677 | 31365 | 27465 | 27320 | 27974 | 34362 | 31893 | 35484 | 22115 | 18493 | 20157 | 18913 | 21887 | 20981 | 23074 | 31052 | 27414 | 30394 | 30148 | 34647 | 35523 | 35881 | Eif5 |
| Eif5a | 32582 | 32529 | 34856 | 32347 | 27487 | 28425 | 31579 | 22416 | 26107 | 24499 | 23688 | 28477 | 26906 | 30817 | 28917 | 29588 | 33669 | 29067 | 32706 | 28260 | 30963 | 20493 | 23931 | 28455 | 24632 | 35744 | 29115 | 33429 | Eif5a |
| Eif5a2 | 6642 | 5262 | 4572 | 5215 | 4567 | 4996 | 5299 | 2842 | 2323 | 2089 | 2192 | 2380 | 2507 | 2317 | 6963 | 5847 | 4523 | 5142 | 4140 | 5671 | 4838 | 2354 | 2324 | 1802 | 2026 | 2149 | 2459 | 2300 | Eif5a2 |
| Eif5b | 23273 | 19206 | 26258 | 21982 | 18195 | 20349 | 20964 | 16399 | 14876 | 16384 | 15470 | 17019 | 16974 | 18264 | 23662 | 22039 | 23592 | 21202 | 18786 | 20717 | 19411 | 14967 | 14994 | 17153 | 15696 | 18610 | 17788 | 18409 | Eif5b |
| Eif6 | 5928 | 7226 | 6944 | 6817 | 5407 | 5401 | 5290 | 3644 | 3819 | 3757 | 3577 | 4451 | 4284 | 3885 | 5703 | 5765 | 6627 | 5914 | 6650 | 5945 | 5530 | 3106 | 3767 | 4291 | 4005 | 5214 | 4568 | 4417 | Eif6 |
| Eipr1 | 1916 | 2590 | 2208 | 2508 | 2405 | 2241 | 2107 | 1912 | 1989 | 2007 | 1758 | 1930 | 1738 | 1658 | 2156 | 2144 | 2444 | 2336 | 2448 | 2700 | 2347 | 1805 | 1962 | 1901 | 2060 | 1848 | 1928 | 1885 | Eipr1 |
| Elac1 | 1211 | 1063 | 1204 | 1125 | 1161 | 1061 | 1199 | 1329 | 1522 | 1429 | 1656 | 1436 | 1482 | 1471 | 1319 | 1181 | 1210 | 1070 | 943 | 1135 | 1081 | 1504 | 1440 | 1336 | 1495 | 1451 | 1402 | 1432 | Elac1 |
| Elac2 | 4223 | 4672 | 4399 | 4466 | 3633 | 3832 | 3785 | 2395 | 2803 | 2954 | 2651 | 3350 | 2860 | 3098 | 3798 | 4297 | 4179 | 3969 | 3941 | 3408 | 3880 | 2797 | 2676 | 2803 | 3070 | 3275 | 3031 | 3173 | Elac2 |
| Elane | 2 | 3 | 1 | 1 | 2 | 0 | 2 | 0 | 0 | 8 | 6 | 0 | 0 | 1 | 0 | 4 | 3 | 0 | 0 | 0 | 3 | 3 | 9 | 6 | 5 | 0 | 0 | 1 | Elane |
| Elapor1 | 382 | 486 | 433 | 717 | 868 | 828 | 796 | 506 | 535 | 695 | 592 | 502 | 578 | 583 | 442 | 408 | 588 | 763 | 767 | 769 | 786 | 802 | 486 | 650 | 799 | 575 | 733 | 517 | Elapor1 |
| Elapor2 | 991 | 1046 | 1058 | 1206 | 1924 | 1499 | 1383 | 1757 | 1497 | 1699 | 1922 | 1166 | 1438 | 1301 | 1328 | 1129 | 1168 | 1292 | 1345 | 1475 | 1408 | 1997 | 1661 | 1147 | 1632 | 907 | 1117 | 1272 | Elapor2 |
| Elavl1 | 9368 | 8458 | 12361 | 11242 | 9728 | 9283 | 10740 | 9062 | 7971 | 8946 | 8193 | 10551 | 9019 | 10390 | 9452 | 8761 | 11632 | 10103 | 10388 | 8830 | 10332 | 9842 | 7798 | 10263 | 9417 | 12247 | 9681 | 11284 | Elavl1 |
| Elavl2 | 12409 | 11717 | 12901 | 15975 | 18979 | 17053 | 20000 | 21534 | 16508 | 16228 | 14345 | 15782 | 15321 | 16637 | 17793 | 11754 | 15015 | 16287 | 17603 | 21032 | 19158 | 19569 | 15242 | 15939 | 19936 | 12900 | 16651 | 16391 | Elavl2 |
| Elavl3 | 12773 | 17427 | 22853 | 32064 | 34545 | 28899 | 29339 | 19877 | 20018 | 31033 | 24586 | 29541 | 24156 | 29447 | 15500 | 13464 | 28914 | 30637 | 35181 | 23922 | 32283 | 32950 | 18281 | 32838 | 36285 | 26715 | 24550 | 26628 | Elavl3 |
| Elavl4 | 7532 | 7111 | 8710 | 11102 | 14362 | 12126 | 13538 | 20469 | 14715 | 16700 | 14670 | 15874 | 17729 | 17335 | 11093 | 8439 | 10413 | 10953 | 12258 | 13637 | 12824 | 19831 | 13416 | 16709 | 18752 | 14251 | 15869 | 16643 | Elavl4 |
| Elf1 | 1264 | 898 | 1230 | 1043 | 972 | 998 | 949 | 1700 | 1711 | 1492 | 1624 | 1510 | 1681 | 1629 | 1065 | 1091 | 1056 | 919 | 980 | 816 | 915 | 1501 | 1847 | 1567 | 1420 | 1671 | 1592 | 1640 | Elf1 |
| Elf2 | 7534 | 7375 | 9021 | 8261 | 8407 | 8329 | 8723 | 7222 | 6553 | 7611 | 7145 | 8416 | 7669 | 8590 | 7272 | 6914 | 8858 | 8088 | 8951 | 7962 | 8808 | 7394 | 6735 | 8273 | 8268 | 8883 | 8555 | 8766 | Elf2 |
| Elf3 | 146 | 76 | 83 | 77 | 24 | 26 | 18 | 74 | 112 | 124 | 109 | 21 | 44 | 26 | 75 | 121 | 31 | 44 | 30 | 15 | 22 | 38 | 124 | 115 | 23 | 110 | 24 | 36 | Elf3 |
| Elf4 | 1197 | 1198 | 1640 | 1039 | 1038 | 1203 | 810 | 673 | 877 | 827 | 1354 | 764 | 788 | 639 | 1160 | 1403 | 1192 | 1235 | 1007 | 692 | 807 | 909 | 1160 | 732 | 698 | 709 | 845 | 648 | Elf4 |
| Elf5 | 82 | 12 | 3 | 41 | 0 | 1 | 0 | 491 | 285 | 416 | 183 | 7 | 11 | 16 | 32 | 30 | 5 | 4 | 2 | 4 | 4 | 5 | 391 | 446 | 1 | 323 | 3 | 19 | Elf5 |
| Elfn1 | 155 | 264 | 270 | 347 | 385 | 363 | 226 | 219 | 334 | 363 | 331 | 293 | 215 | 208 | 215 | 194 | 391 | 275 | 324 | 311 | 287 | 373 | 296 | 342 | 393 | 258 | 187 | 221 | Elfn1 |
| Elfn2 | 499 | 788 | 744 | 1294 | 2156 | 1524 | 1416 | 954 | 1158 | 1466 | 1218 | 628 | 708 | 632 | 661 | 618 | 1197 | 1439 | 1673 | 1263 | 1512 | 1698 | 992 | 986 | 1538 | 423 | 436 | 482 | Elfn2 |
| Elk1 | 4811 | 5311 | 3584 | 5565 | 5363 | 5657 | 5891 | 3634 | 3703 | 3787 | 3976 | 3662 | 3170 | 3835 | 4585 | 4738 | 4530 | 5828 | 5605 | 5082 | 6230 | 3738 | 3452 | 3819 | 4189 | 2629 | 3747 | 3315 | Elk1 |
| Elk3 | 2154 | 1636 | 1953 | 1420 | 1575 | 1677 | 1329 | 2384 | 3065 | 2906 | 3933 | 2199 | 2931 | 2318 | 1755 | 2179 | 1861 | 1549 | 1541 | 1327 | 1376 | 2525 | 3232 | 2688 | 2018 | 3411 | 2285 | 2364 | Elk3 |
| Elk4 | 2009 | 1830 | 1708 | 1878 | 1829 | 1753 | 1846 | 2113 | 1981 | 2043 | 2280 | 2156 | 2259 | 2207 | 1820 | 1822 | 1774 | 1844 | 1851 | 1710 | 1839 | 1858 | 2035 | 2089 | 2326 | 2223 | 2416 | 2383 | Elk4 |
| Ell | 1610 | 2208 | 2290 | 2463 | 1807 | 1778 | 1617 | 1279 | 1650 | 1791 | 1831 | 2055 | 1733 | 1734 | 1560 | 2129 | 2216 | 2213 | 1999 | 1648 | 1726 | 1758 | 1556 | 2142 | 1952 | 2268 | 1699 | 1883 | Ell |
| Ell2 | 1754 | 1570 | 1861 | 1685 | 1261 | 1288 | 1351 | 613 | 602 | 594 | 606 | 391 | 720 | 511 | 1816 | 1792 | 1453 | 1383 | 1217 | 1333 | 1208 | 426 | 717 | 491 | 360 | 614 | 494 | 708 | Ell2 |
| Ell3 | 2 | 3 | 4 | 3 | 0 | 3 | 1 | 9 | 0 | 13 | 4 | 2 | 2 | 0 | 1 | 2 | 3 | 1 | 1 | 2 | 0 | 3 | 1 | 8 | 1 | 8 | 3 | 2 | Ell3 |
| Elmo1 | 3279 | 3822 | 3489 | 4930 | 6334 | 5391 | 6015 | 4539 | 3854 | 4436 | 4213 | 4258 | 3662 | 4391 | 4348 | 3272 | 4702 | 5398 | 5538 | 5319 | 6500 | 5813 | 3918 | 4134 | 4905 | 3460 | 3720 | 4005 | Elmo1 |
| Elmo2 | 13739 | 17281 | 12627 | 15208 | 14163 | 13958 | 14454 | 6927 | 7979 | 7763 | 7650 | 8302 | 7269 | 7762 | 12879 | 14593 | 14300 | 16014 | 15116 | 14049 | 14690 | 7792 | 7365 | 7804 | 8488 | 7560 | 7482 | 7228 | Elmo2 |
| Elmo3 | 188 | 311 | 193 | 195 | 115 | 163 | 187 | 200 | 229 | 225 | 164 | 140 | 191 | 147 | 208 | 238 | 153 | 221 | 165 | 176 | 147 | 144 | 230 | 203 | 138 | 228 | 172 | 152 | Elmo3 |
| Elmod1 | 1024 | 1075 | 931 | 1587 | 2527 | 1777 | 2426 | 2102 | 1887 | 2167 | 1751 | 1416 | 1118 | 1431 | 1263 | 960 | 1376 | 1837 | 2219 | 2111 | 2660 | 2461 | 1682 | 1634 | 2471 | 929 | 1095 | 1120 | Elmod1 |
| Elmod2 | 6532 | 6025 | 5602 | 5848 | 5530 | 6308 | 6804 | 6740 | 5748 | 4786 | 5192 | 5202 | 4892 | 5519 | 6861 | 6526 | 5420 | 6365 | 5631 | 7101 | 6617 | 5548 | 5563 | 5022 | 5471 | 4444 | 5490 | 5140 | Elmod2 |
| Elmod3 | 1245 | 1592 | 1325 | 1467 | 1388 | 1321 | 1330 | 833 | 965 | 997 | 906 | 904 | 979 | 895 | 1269 | 1433 | 1452 | 1493 | 1375 | 1315 | 1303 | 795 | 983 | 954 | 811 | 971 | 875 | 774 | Elmod3 |
| Eln | 1035 | 812 | 699 | 1201 | 930 | 1538 | 497 | 2385 | 4072 | 4000 | 6533 | 1322 | 1420 | 992 | 744 | 1130 | 796 | 2122 | 867 | 657 | 573 | 3145 | 5998 | 3613 | 2848 | 1833 | 1212 | 846 | Eln |
| Eloa | 8273 | 8515 | 6939 | 6818 | 5557 | 6058 | 5791 | 5382 | 5065 | 4623 | 4926 | 4906 | 4678 | 4886 | 7697 | 7650 | 6738 | 6594 | 5786 | 6188 | 6086 | 4781 | 5077 | 4837 | 4622 | 5045 | 4904 | 5213 | Eloa |
| Elob | 5630 | 7695 | 7355 | 7579 | 6694 | 6016 | 7388 | 5858 | 6187 | 6918 | 6788 | 6689 | 6732 | 7826 | 5941 | 5470 | 7954 | 7100 | 7637 | 8229 | 7154 | 4854 | 6226 | 7575 | 7194 | 8033 | 7618 | 8049 | Elob |
| Elobl | 0 | 0 | 0 | 0 | 2 | 0 | 0 | 0 | 0 | 0 | 0 | 0 | 0 | 0 | 0 | 0 | 0 | 0 | 0 | 0 | 0 | 0 | 0 | 0 | 5 | 0 | 0 | 0 | Elobl |
| Eloc | 7832 | 6144 | 7808 | 7859 | 7630 | 7755 | 9316 | 9579 | 7500 | 7186 | 7257 | 7972 | 7485 | 9396 | 9030 | 7064 | 7400 | 7467 | 7814 | 9901 | 8707 | 8438 | 7760 | 7971 | 8317 | 9284 | 8668 | 9362 | Eloc |
| Elof1 | 3586 | 4171 | 4129 | 3532 | 3085 | 3208 | 3347 | 2694 | 2864 | 2509 | 2695 | 3030 | 3140 | 3241 | 3472 | 3653 | 3685 | 3360 | 3730 | 3802 | 3318 | 2425 | 2917 | 3032 | 2800 | 3697 | 3331 | 3374 | Elof1 |
| Elovl1 | 1377 | 1664 | 1546 | 1333 | 1235 | 1301 | 1000 | 1122 | 1333 | 1187 | 1444 | 1203 | 1302 | 1036 | 1325 | 1721 | 1358 | 1401 | 1267 | 1109 | 1023 | 1165 | 1495 | 1068 | 1063 | 1152 | 1173 | 1111 | Elovl1 |

| | | | | | | | | | | | | | | | | | | | | | | | | | | | | | |
|---|---|---|---|---|---|---|---|---|---|---|---|---|---|---|---|---|---|---|---|---|---|---|---|---|---|---|---|---|---|
| Elovl2 | 4174 | 4041 | 3796 | 4173 | 4791 | 4724 | 6040 | 4476 | 3853 | 4056 | 3160 | 4846 | 3978 | 5210 | 4327 | 3173 | 4567 | 4401 | 5259 | 5142 | 5689 | 4132 | 3366 | 4294 | 4671 | 4552 | 4701 | 5073 | Elovl2 |
| Elovl3 | 3 | 11 | 20 | 0 | 5 | 6 | 4 | 0 | 5 | 0 | 0 | 1 | 0 | 5 | 13 | 8 | 10 | 10 | 2 | 3 | 6 | 2 | 1 | 1 | 0 | 0 | 5 | 2 | Elovl3 |
| Elovl4 | 4404 | 3943 | 4509 | 4923 | 5048 | 4657 | 5729 | 3482 | 2927 | 2868 | 2810 | 2760 | 2650 | 3260 | 5003 | 4095 | 4516 | 5017 | 4683 | 5431 | 5512 | 3521 | 2795 | 2760 | 3413 | 2603 | 2657 | 3152 | Elovl4 |
| Elovl5 | 13147 | 13361 | 11761 | 11049 | 9442 | 10055 | 10863 | 5591 | 5071 | 4900 | 4643 | 5362 | 5319 | 5396 | 13676 | 13197 | 11266 | 11266 | 9746 | 11111 | 10366 | 4939 | 5215 | 5111 | 4841 | 6166 | 5722 | 5874 | Elovl5 |
| Elovl6 | 10054 | 8595 | 9230 | 8265 | 8364 | 9333 | 8998 | 7748 | 7382 | 7827 | 7429 | 8563 | 8211 | 9024 | 9480 | 9027 | 9021 | 8095 | 8416 | 8696 | 8770 | 7778 | 6102 | 7769 | 7960 | 8121 | 8997 | 8890 | Elovl6 |
| Elovl7 | 391 | 254 | 245 | 234 | 296 | 253 | 265 | 131 | 167 | 131 | 161 | 121 | 145 | 70 | 311 | 299 | 239 | 259 | 238 | 254 | 225 | 157 | 227 | 142 | 101 | 115 | 81 | 77 | Elovl7 |
| Elp1 | 17715 | 17011 | 14236 | 13179 | 13152 | 13997 | 14177 | 14875 | 14113 | 11924 | 12701 | 14307 | 14235 | 14564 | 17193 | 15568 | 13122 | 13287 | 13469 | 15055 | 14133 | 12554 | 13268 | 12110 | 13443 | 12576 | 15238 | 15167 | Elp1 |
| Elp2 | 14849 | 14349 | 11870 | 11935 | 11756 | 12748 | 12696 | 15717 | 14085 | 12027 | 11799 | 13070 | 12927 | 13246 | 15504 | 12945 | 12148 | 12184 | 12469 | 14441 | 13138 | 12942 | 13647 | 12773 | 13459 | 13989 | 13925 | 13315 | Elp2 |
| Elp3 | 7012 | 7242 | 6700 | 6451 | 6633 | 6705 | 6239 | 9191 | 8990 | 8709 | 7961 | 8875 | 8577 | 8887 | 7076 | 6413 | 6678 | 6037 | 6890 | 7089 | 6723 | 8363 | 8976 | 8778 | 8907 | 9589 | 9358 | 9371 | Elp3 |
| Elp4 | 2299 | 2103 | 2273 | 1857 | 1768 | 1884 | 2326 | 2273 | 1998 | 1957 | 1902 | 2331 | 2168 | 2494 | 2429 | 1899 | 2243 | 1652 | 1900 | 2211 | 2026 | 1768 | 1955 | 1863 | 2170 | 2273 | 2479 | 2614 | Elp4 |
| Elp5 | 6682 | 7270 | 6439 | 6500 | 5303 | 5556 | 6022 | 4305 | 4063 | 3830 | 3840 | 4464 | 4205 | 4249 | 7247 | 6368 | 6400 | 6567 | 6300 | 6819 | 5974 | 3878 | 4065 | 4249 | 4219 | 4971 | 4593 | 4554 | Elp5 |
| Elp6 | 468 | 488 | 511 | 465 | 515 | 590 | 505 | 1408 | 1481 | 1498 | 1424 | 1640 | 1545 | 1647 | 477 | 372 | 495 | 549 | 603 | 545 | 465 | 1497 | 1401 | 1597 | 1680 | 1711 | 1713 | 1677 | Elp6 |
| Emb | 1872 | 1157 | 1519 | 896 | 922 | 797 | 844 | 2010 | 1603 | 1476 | 1244 | 1335 | 1532 | 1503 | 1435 | 1557 | 1133 | 825 | 853 | 1002 | 821 | 1252 | 1710 | 1684 | 1016 | 2276 | 1675 | 1697 | Emb |
| Emc1 | 3387 | 3479 | 3718 | 3896 | 3642 | 3586 | 3532 | 3545 | 3789 | 4219 | 4256 | 4735 | 4427 | 4161 | 3170 | 3296 | 3892 | 3689 | 3667 | 3165 | 3397 | 4547 | 3766 | 4385 | 4542 | 4315 | 4181 | 4345 | Emc1 |
| Emc10 | 2683 | 3307 | 3255 | 3140 | 3543 | 3145 | 2665 | 4151 | 4720 | 4977 | 4572 | 4806 | 4566 | 4295 | 2532 | 2780 | 3318 | 3103 | 3735 | 2938 | 2936 | 4451 | 4739 | 4944 | 4898 | 5445 | 4761 | 4514 | Emc10 |
| Emc2 | 4150 | 3611 | 4882 | 3820 | 3177 | 3447 | 3948 | 3303 | 3090 | 2757 | 2884 | 2845 | 2911 | 3203 | 4759 | 4439 | 4015 | 3761 | 3444 | 4395 | 3583 | 3100 | 3204 | 2862 | 2821 | 3346 | 3220 | 3405 | Emc2 |
| Emc3 | 2691 | 3025 | 2893 | 2536 | 2607 | 2651 | 2575 | 3435 | 3498 | 3253 | 3355 | 3126 | 3030 | 3025 | 2676 | 2696 | 2612 | 2484 | 2657 | 2782 | 2607 | 3091 | 3657 | 3112 | 3092 | 3175 | 3025 | 3183 | Emc3 |
| Emc4 | 2305 | 2537 | 2443 | 2560 | 2390 | 2316 | 2377 | 2463 | 2315 | 2309 | 2382 | 2495 | 2359 | 2224 | 2574 | 2316 | 2591 | 2503 | 2494 | 2677 | 2298 | 2396 | 2402 | 2522 | 2611 | 2592 | 2596 | 2355 | Emc4 |
| Emc6 | 2201 | 2281 | 2397 | 2366 | 1981 | 1855 | 2128 | 2036 | 1782 | 1980 | 1799 | 2146 | 1813 | 2169 | 2359 | 2018 | 2408 | 2102 | 2032 | 2242 | 2173 | 1761 | 1805 | 2061 | 1850 | 2409 | 2172 | 2209 | Emc6 |
| Emc7 | 2728 | 2876 | 2640 | 2849 | 2864 | 2765 | 2776 | 2606 | 2445 | 2281 | 2546 | 2127 | 2080 | 2222 | 3011 | 2910 | 2553 | 2933 | 2544 | 3085 | 2859 | 2520 | 2619 | 2249 | 2531 | 2018 | 2120 | 2274 | Emc7 |
| Emc8 | 6743 | 6772 | 7422 | 7285 | 6443 | 6768 | 6771 | 3841 | 3630 | 4125 | 4175 | 4972 | 4483 | 4924 | 6352 | 6365 | 7507 | 7138 | 6932 | 5899 | 6983 | 3937 | 3670 | 4601 | 4589 | 4956 | 4916 | 4876 | Emc8 |
| Emc9 | 255 | 299 | 350 | 383 | 429 | 266 | 296 | 216 | 213 | 236 | 242 | 230 | 169 | 168 | 233 | 292 | 322 | 347 | 391 | 368 | 349 | 312 | 219 | 201 | 243 | 159 | 203 | 157 | Emc9 |
| Emcn | 731 | 584 | 607 | 442 | 343 | 419 | 334 | 999 | 1196 | 954 | 1108 | 812 | 1090 | 845 | 670 | 819 | 487 | 421 | 288 | 464 | 315 | 1026 | 1452 | 898 | 763 | 1102 | 949 | 819 | Emcn |
| Emd | 4717 | 5410 | 4838 | 4489 | 3472 | 3868 | 4804 | 3566 | 3625 | 3605 | 3433 | 4161 | 4046 | 5339 | 4320 | 4024 | 4673 | 3901 | 4186 | 4241 | 4571 | 2809 | 3625 | 4269 | 3776 | 5229 | 4671 | 5509 | Emd |
| Eme1 | 1523 | 1617 | 1723 | 1027 | 802 | 994 | 1102 | 1500 | 1571 | 1505 | 1137 | 2363 | 2180 | 2236 | 1332 | 1340 | 1432 | 975 | 1260 | 1207 | 965 | 969 | 1383 | 1924 | 1562 | 2915 | 2646 | 2502 | Eme1 |
| Eme2 | 1747 | 2116 | 1838 | 1777 | 1968 | 1908 | 1798 | 1430 | 1512 | 1684 | 1446 | 1706 | 1589 | 1542 | 1789 | 1776 | 1938 | 1952 | 1894 | 1693 | 1839 | 1688 | 1292 | 1526 | 1530 | 1488 | 1636 | 1780 | Eme2 |
| Emg1 | 4422 | 4756 | 5051 | 4706 | 3654 | 3686 | 4069 | 4825 | 4818 | 5216 | 4795 | 5722 | 5316 | 5421 | 4267 | 4164 | 4876 | 4046 | 4480 | 5192 | 3739 | 3603 | 4713 | 5787 | 5140 | 6966 | 6089 | 5693 | Emg1 |
| Emid1 | 520 | 515 | 592 | 463 | 484 | 517 | 455 | 504 | 719 | 622 | 599 | 517 | 589 | 512 | 446 | 554 | 474 | 524 | 500 | 322 | 411 | 426 | 606 | 554 | 542 | 614 | 476 | 529 | Emid1 |
| Emilin1 | 3722 | 4669 | 4484 | 2825 | 1441 | 3084 | 746 | 3693 | 6184 | 6068 | 9899 | 4826 | 7236 | 4349 | 3247 | 5791 | 3300 | 3165 | 1362 | 843 | 857 | 5324 | 7123 | 5927 | 4363 | 7223 | 5825 | 4981 | Emilin1 |
| Emilin2 | 1039 | 1170 | 1580 | 677 | 605 | 877 | 467 | 728 | 1003 | 772 | 955 | 941 | 1170 | 729 | 977 | 1282 | 1034 | 647 | 664 | 520 | 467 | 530 | 1118 | 715 | 520 | 1119 | 1219 | 1030 | Emilin2 |
| Emilin3 | 625 | 677 | 935 | 494 | 374 | 508 | 228 | 657 | 1315 | 822 | 2105 | 632 | 764 | 528 | 545 | 779 | 743 | 648 | 339 | 240 | 259 | 827 | 1378 | 847 | 582 | 917 | 837 | 656 | Emilin3 |
| Eml1 | 6101 | 7129 | 7241 | 6280 | 5730 | 5486 | 5661 | 5222 | 5121 | 5431 | 4982 | 6187 | 6421 | 5870 | 6397 | 6409 | 6962 | 5904 | 5941 | 5446 | 5611 | 5402 | 5022 | 5553 | 5342 | 6187 | 6638 | 6496 | Eml1 |
| Eml2 | 2414 | 3048 | 3386 | 4046 | 4101 | 3446 | 2999 | 1557 | 1742 | 2162 | 2139 | 2086 | 1953 | 1629 | 2256 | 2858 | 3980 | 3924 | 3934 | 3255 | 2960 | 2134 | 1860 | 2414 | 2179 | 1991 | 1947 | 1621 | Eml2 |
| Eml3 | 2212 | 2300 | 2655 | 2469 | 2269 | 2388 | 1816 | 1421 | 1826 | 1856 | 2329 | 1779 | 1965 | 1510 | 1953 | 2331 | 2444 | 2448 | 2360 | 1877 | 1861 | 1604 | 1819 | 1900 | 1740 | 1936 | 1854 | 1684 | Eml3 |
| Eml4 | 20594 | 17801 | 16622 | 17635 | 17507 | 18394 | 17609 | 12551 | 12600 | 12331 | 12753 | 13433 | 13935 | 12485 | 19561 | 20557 | 16435 | 18315 | 16443 | 17302 | 16776 | 11414 | 12389 | 12030 | 11756 | 11459 | 13249 | 12145 | Eml4 |
| Eml5 | 23636 | 20852 | 14905 | 20671 | 20196 | 27549 | 27487 | 13174 | 9032 | 9136 | 8331 | 13550 | 12140 | 14078 | 23997 | 19547 | 17208 | 19281 | 20762 | 27010 | 29402 | 8814 | 9023 | 8577 | 11302 | 7746 | 16540 | 12749 | Eml5 |
| Eml6 | 880 | 1052 | 687 | 1256 | 1850 | 1488 | 1691 | 1428 | 1269 | 1224 | 1253 | 789 | 880 | 817 | 996 | 839 | 976 | 1435 | 1454 | 1702 | 1660 | 1519 | 1289 | 843 | 1266 | 460 | 791 | 665 | Eml6 |
| Emp1 | 3205 | 2620 | 2729 | 1618 | 1574 | 3648 | 979 | 6572 | 9383 | 5970 | 10482 | 4909 | 7490 | 5425 | 3641 | 3433 | 1903 | 3299 | 1488 | 1648 | 1266 | 5106 | 12105 | 6436 | 4881 | 3983 | 5521 | 5804 | Emp1 |
| Emp2 | 617 | 434 | 445 | 226 | 328 | 426 | 194 | 607 | 851 | 588 | 668 | 399 | 489 | 372 | 425 | 610 | 319 | 357 | 248 | 203 | 192 | 476 | 936 | 561 | 405 | 463 | 480 | 484 | Emp2 |
| Emp3 | 471 | 570 | 497 | 394 | 290 | 517 | 192 | 700 | 975 | 740 | 1019 | 573 | 837 | 506 | 479 | 638 | 371 | 567 | 267 | 354 | 301 | 720 | 1147 | 659 | 664 | 656 | 744 | 597 | Emp3 |
| Emsy | 9312 | 8881 | 7945 | 8451 | 8750 | 9222 | 8944 | 10657 | 9642 | 8698 | 8524 | 10113 | 9019 | 9671 | 9283 | 8372 | 8229 | 8089 | 8914 | 9765 | 8973 | 9282 | 8936 | 8584 | 9885 | 8335 | 9827 | 9424 | Emsy |
| Emx1 | 621 | 758 | 766 | 1044 | 1001 | 1061 | 1387 | 829 | 1027 | 1212 | 839 | 1672 | 1220 | 1704 | 493 | 389 | 829 | 1020 | 1455 | 1183 | 1274 | 404 | 684 | 1765 | 1491 | 2051 | 1293 | 1602 | Emx1 |
| Emx2 | 3230 | 2339 | 4793 | 3739 | 3014 | 3284 | 4505 | 3210 | 2656 | 3786 | 2651 | 5327 | 4073 | 6348 | 2362 | 1652 | 4040 | 3186 | 4126 | 3270 | 4114 | 1977 | 2439 | 5593 | 4020 | 7180 | 4608 | 6886 | Emx2 |
| En1 | 415 | 533 | 887 | 694 | 665 | 733 | 272 | 733 | 627 | 870 | 656 | 930 | 1210 | 793 | 456 | 685 | 856 | 618 | 789 | 539 | 617 | 998 | 632 | 782 | 949 | 267 | 811 | 779 | En1 |
| En2 | 1303 | 1786 | 3147 | 2646 | 1792 | 2007 | 489 | 1403 | 1148 | 1449 | 1183 | 2149 | 3260 | 1933 | 1117 | 2234 | 3218 | 1853 | 2215 | 2005 | 1642 | 2171 | 1310 | 1300 | 2325 | 9 | 2372 | 2432 | En2 |
| Enah | 22287 | 19147 | 23137 | 24991 | 27534 | 26423 | 26983 | 29501 | 26555 | 32131 | 30868 | 33726 | 30804 | 33505 | 22234 | 21631 | 24308 | 25236 | 26105 | 22253 | 27536 | 33523 | 25770 | 32012 | 34113 | 31317 | 31860 | 32647 | Enah |
| Enam | 1 | 0 | 0 | 0 | 0 | 0 | 0 | 2 | 1 | 0 | 0 | 0 | 0 | 0 | 0 | 0 | 0 | 0 | 2 | 0 | 0 | 0 | 1 | 0 | 0 | 0 | 0 | 0 | Enam |
| Enc1 | 12576 | 15350 | 15242 | 23924 | 36878 | 26854 | 29987 | 30199 | 28381 | 35915 | 29170 | 25818 | 25188 | 27030 | 16885 | 12930 | 22045 | 26433 | 31823 | 28514 | 33089 | 39483 | 26227 | 30930 | 37861 | 20091 | 22263 | 23729 | Enc1 |
| Endod1 | 677 | 570 | 723 | 495 | 762 | 666 | 473 | 1188 | 1225 | 1298 | 1387 | 911 | 1331 | 1042 | 671 | 757 | 622 | 647 | 560 | 451 | 432 | 1340 | 1338 | 1009 | 1029 | 1081 | 1041 | 1093 | Endod1 |
| Endog | 131 | 149 | 119 | 144 | 152 | 169 | 120 | 184 | 236 | 183 | 210 | 207 | 223 | 158 | 127 | 148 | 139 | 122 | 150 | 182 | 113 | 192 | 211 | 197 | 184 | 232 | 220 | 235 | Endog |
| Endou | 7 | 5 | 16 | 3 | 11 | 13 | 10 | 13 | 42 | 27 | 33 | 17 | 20 | 21 | 8 | 18 | 15 | 10 | 7 | 12 | 14 | 10 | 40 | 16 | 14 | 18 | 12 | 23 | Endou |
| Endov | 2320 | 2829 | 2962 | 2843 | 2932 | 3006 | 2616 | 3000 | 3485 | 3854 | 3603 | 4473 | 4070 | 4038 | 2298 | 2416 | 3232 | 2853 | 2964 | 2630 | 2648 | 3300 | 3306 | 4059 | 3844 | 4042 | 4385 | 4296 | Endov |
| Eng | 1218 | 1661 | 1822 | 1464 | 1708 | 1532 | 1063 | 1373 | 2135 | 2279 | 2528 | 1867 | 2090 | 1487 | 1316 | 1785 | 1577 | 1774 | 1531 | 1091 | 1259 | 2321 | 2378 | 2299 | 1971 | 2239 | 1746 | 1527 | Eng |
| Engase | 1013 | 1388 | 1314 | 1419 | 1231 | 1135 | 1022 | 449 | 542 | 605 | 582 | 653 | 657 | 519 | 943 | 1325 | 1454 | 1502 | 1204 | 975 | 1020 | 553 | 574 | 543 | 597 | 705 | 626 | 598 | Engase |
| Enho | 873 | 1388 | 1030 | 1355 | 1773 | 1591 | 1417 | 2150 | 2299 | 2190 | 1933 | 1975 | 2045 | 1825 | 1266 | 1103 | 1523 | 1343 | 1742 | 1783 | 1524 | 2251 | 2201 | 2146 | 2187 | 1848 | 1938 | 1755 | Enho |
| Enkd1 | 1370 | 1666 | 1891 | 1785 | 1418 | 1482 | 1401 | 716 | 847 | 875 | 855 | 1190 | 1140 | 1236 | 1271 | 1342 | 1812 | 1591 | 1738 | 1386 | 1482 | 722 | 748 | 1110 | 860 | 1603 | 1255 | 1200 | Enkd1 |
| Enkur | 212 | 200 | 192 | 262 | 261 | 223 | 260 | 205 | 182 | 193 | 190 | 157 | 231 | 151 | 268 | 244 | 195 | 216 | 248 | 292 | 225 | 196 | 282 | 140 | 173 | 153 | 132 | 158 | Enkur |
| Eno1 | 51675 | 61222 | 48723 | 43985 | 37124 | 43965 | 37653 | 32960 | 42705 | 31993 | 35906 | 34553 | 38663 | 32049 | 46959 | 54578 | 46117 | 42310 | 45652 | 47956 | 39086 | 26923 | 42890 | 34577 | 33483 | 45304 | 40080 | 38394 | Eno1 |
| Eno1b | 1833 | 2151 | 1868 | 1616 | 1537 | 1772 | 1500 | 1115 | 1559 | 1064 | 1169 | 1322 | 1591 | 1403 | 1721 | 2103 | 1619 | 1718 | 1956 | 1946 | 1604 | 1008 | 1402 | 1148 | 1276 | 1699 | 1629 | 1562 | Eno1b |
| Eno2 | 2997 | 3225 | 3158 | 4094 | 5046 | 4024 | 4154 | 1254 | 1521 | 1746 | 1750 | 1155 | 1203 | 1044 | 3013 | 3172 | 3661 | 4410 | 4421 | 3529 | 4394 | 2054 | 1559 | 1333 | 1700 | 930 | 953 | 917 | Eno2 |
| Eno3 | 1121 | 1247 | 1371 | 1343 | 1338 | 1173 | 1098 | 933 | 1093 | 999 | 1932 | 1137 | 1245 | 1135 | 1166 | 1214 | 1424 | 1160 | 1414 | 1086 | 1147 | 1116 | 1589 | 1066 | 1169 | 1125 | 1260 | 1145 | Eno3 |
| Eno4 | 135 | 152 | 43 | 133 | 146 | 123 | 126 | 98 | 106 | 87 | 89 | 138 | 103 | 103 | 146 | 135 | 115 | 116 | 148 | 126 | 130 | 142 | 154 | 99 | 117 | 152 | 80 | 101 | Eno4 |
| Enoph1 | 2704 | 3115 | 3056 | 3152 | 2587 | 2596 | 2753 | 1496 | 1581 | 1725 | 1681 | 1842 | 1617 | 1619 | 2798 | 2628 | 2809 | 2899 | 3089 | 2852 | 2834 | 1464 | 1489 | 1754 | 1764 | 2026 | 1839 | 1800 | Enoph1 |
| Enox1 | 2028 | 2357 | 2077 | 2814 | 3206 | 2830 | 3337 | 2591 | 2553 | 2735 | 2398 | 2389 | 2104 | 2634 | 2434 | 2016 | 2735 | 3032 | 3261 | 2931 | 3395 | 2862 | 2388 | 2515 | 3141 | 2273 | 1893 | 2435 | Enox1 |
| Enox2 | 940 | 1121 | 926 | 980 | 1002 | 999 | 1014 | 1575 | 1322 | 1325 | 1053 | 1379 | 1423 | 1485 | 1274 | 973 | 1068 | 791 | 1001 | 1285 | 959 | 1405 | 1350 | 1049 | 1209 | 1529 | 1453 | 1365 | Enox2 |
| Enpep | 188 | 100 | 94 | 220 | 229 | 282 | 97 | 547 | 769 | 587 | 669 | 278 | 288 | 244 | 182 | 118 | 134 | 430 | 90 | 100 | 115 | 642 | 793 | 526 | 372 | 249 | 228 | 178 | Enpep |
| Enpp1 | 497 | 421 | 443 | 476 | 575 | 446 | 436 | 430 | 545 | 543 | 689 | 416 | 455 | 410 | 450 | 604 | 399 | 511 | 461 | 570 | 473 | 441 | 713 | 476 | 389 | 440 | 454 | 362 | Enpp1 |
| Enpp2 | 5833 | 4727 | 3389 | 3625 | 4662 | 3835 | 3674 | 3628 | 4002 | 3213 | 3556 | 2380 | 3946 | 1983 | 5828 | 6605 | 3451 | 3823 | 3574 | 4352 | 3975 | 3892 | 4908 | 2348 | 2331 | 2475 | 1993 | 2118 | Enpp2 |
| Enpp3 | 596 | 451 | 583 | 305 | 349 | 390 | 284 | 407 | 421 | 382 | 575 | 465 | 617 | 504 | 505 | 592 | 377 | 374 | 352 | 472 | 331 | 419 | 540 | 403 | 356 | 565 | 562 | 503 | Enpp3 |
| Enpp4 | 1107 | 1012 | 996 | 984 | 997 | 1025 | 968 | 778 | 737 | 758 | 713 | 834 | 746 | 762 | 1056 | 1279 | 911 | 909 | 962 | 1054 | 923 | 632 | 680 | 730 | 737 | 746 | 753 | 723 | Enpp4 |
| Enpp5 | 3748 | 4046 | 2993 | 4466 | 5688 | 4866 | 5289 | 5023 | 4612 | 4435 | 4058 | 3011 | 3040 | 3252 | 4697 | 3667 | 3980 | 4864 | 4909 | 5365 | 5372 | 5404 | 4548 | 3547 | 4655 | 2337 | 2678 | 2573 | Enpp5 |
| Enpp6 | 15 | 24 | 22 | 37 | 19 | 18 | 13 | 38 | 56 | 22 | 60 | 12 | 16 | 10 | 45 | 24 | 29 | 17 | 9 | 22 | 20 | 22 | 42 | 7 | 7 | 8 | 6 | 13 | Enpp6 |
| Enpp7 | 0 | 0 | 0 | 0 | 0 | 0 | 0 | 0 | 1 | 0 | 0 | 0 | 0 | 0 | 0 | 0 | 0 | 0 | 0 | 0 | 0 | 0 | 0 | 0 | 0 | 0 | 0 | 0 | Enpp7 |
| Ensa | 8756 | 8135 | 7912 | 7656 | 7969 | 8334 | 7908 | 13450 | 11643 | 11074 | 11095 | 12216 | 11060 | 12057 | 8565 | 7594 | 7538 | 7313 | 8399 | 8647 | 8461 | 11600 | 11850 | 11372 | 11812 | 12314 | 12673 | 12428 | Ensa |
| Enthd1 | 0 | 0 | 3 | 0 | 0 | 0 | 4 | 0 | 1 | 0 | 0 | 0 | 0 | 3 | 0 | 0 | 0 | 0 | 0 | 0 | 1 | 2 | 1 | 0 | 0 | 0 | 0 | 0 | Enthd1 |
| Entpd1 | 691 | 560 | 616 | 617 | 812 | 793 | 730 | 1349 | 1509 | 1375 | 1454 | 1360 | 1306 | 1288 | 775 | 678 | 647 | 601 | 684 | 719 | 718 | 1551 | 1467 | 1266 | 1195 | 1324 | 1223 | 1227 | Entpd1 |
| Entpd2 | 42 | 41 | 26 | 25 | 42 | 31 | 23 | 12 | 21 | 17 | 38 | 7 | 10 | 5 | 22 | 46 | 41 | 45 | 36 | 26 | 38 | 43 | 40 | 7 | 32 | 3 | 5 | 0 | Entpd2 |
| Entpd3 | 276 | 143 | 123 | 138 | 214 | 120 | 101 | 164 | 175 | 153 | 198 | 92 | 107 | 65 | 167 | 224 | 124 | 198 | 93 | 63 | 111 | 130 | 201 | 148 | 117 | 109 | 84 | 53 | Entpd3 |
| Entpd4 | 3 | 0 | 0 | 1 | 2 | 0 | 0 | 0 | 0 | 0 | 0 | 10 | 0 | 0 | 0 | 2 | 3 | 1 | 0 | 1 | 0 | 2 | 4 | 0 | 0 | 6 | 1 | 0 | Entpd4 |

| | | | | | | | | | | | | | | | | | | | | | | | | | | | | | |
|---|---|---|---|---|---|---|---|---|---|---|---|---|---|---|---|---|---|---|---|---|---|---|---|---|---|---|---|---|---|
| Entpd4b | 201 | 155 | 165 | 92 | 99 | 109 | 104 | 126 | 102 | 135 | 170 | 83 | 70 | 72 | 297 | 155 | 53 | 167 | 203 | 161 | 63 | 206 | 94 | 98 | 120 | 63 | 126 | 109 | Entpd4b |
| Entpd5 | 2581 | 2357 | 2103 | 2185 | 2476 | 2492 | 2505 | 2009 | 1762 | 1836 | 1725 | 1481 | 1593 | 1477 | 2670 | 2369 | 2179 | 2483 | 2385 | 2387 | 2380 | 1746 | 1955 | 1938 | 1492 | 1710 | 1439 | 1423 | Entpd5 |
| Entpd6 | 2298 | 2743 | 2334 | 3003 | 2768 | 2662 | 2586 | 1713 | 2159 | 2120 | 2325 | 1955 | 2046 | 1791 | 2498 | 2541 | 2719 | 3170 | 2731 | 2446 | 2775 | 2324 | 2026 | 1942 | 2312 | 1855 | 1920 | 1911 | Entpd6 |
| Entpd7 | 1288 | 1270 | 1691 | 1670 | 1443 | 1296 | 1367 | 807 | 812 | 1057 | 1068 | 1018 | 978 | 1023 | 1168 | 1283 | 1564 | 1592 | 1404 | 1066 | 1400 | 1146 | 708 | 1094 | 977 | 1128 | 941 | 1180 | Entpd7 |
| Entpd8 | 48 | 62 | 43 | 131 | 119 | 73 | 89 | 6 | 2 | 10 | 4 | 5 | 2 | 5 | 54 | 55 | 71 | 156 | 80 | 59 | 80 | 5 | 2 | 2 | 19 | 0 | 6 | 2 | Entpd8 |
| Entr1 | 5561 | 6506 | 6335 | 6882 | 6398 | 6112 | 6584 | 5779 | 6217 | 6866 | 6743 | 6453 | 6262 | 6574 | 5450 | 6052 | 6784 | 6882 | 6834 | 5752 | 6596 | 6851 | 5926 | 6957 | 6831 | 7021 | 6358 | 6516 | Entr1 |
| Eny2 | 9766 | 8293 | 10727 | 11099 | 10417 | 10703 | 12241 | 9605 | 8248 | 8859 | 9100 | 10392 | 9673 | 11997 | 10464 | 8418 | 10360 | 10335 | 10015 | 11706 | 11706 | 8748 | 7865 | 9946 | 9752 | 11086 | 10514 | 11932 | Eny2 |
| Eogt | 1638 | 1426 | 1661 | 1346 | 1668 | 1576 | 1410 | 1562 | 1593 | 1703 | 1886 | 1730 | 1780 | 1709 | 1639 | 1625 | 1417 | 1370 | 1400 | 1396 | 1420 | 1942 | 1734 | 1627 | 1658 | 1748 | 1732 | 1774 | Eogt |
| Eola1 | 1360 | 1451 | 1272 | 1397 | 1209 | 1377 | 1445 | 1248 | 1047 | 843 | 903 | 955 | 928 | 987 | 1614 | 1274 | 1298 | 1334 | 1308 | 1619 | 1415 | 904 | 1114 | 1022 | 987 | 980 | 1082 | 1011 | Eola1 |
| Eomes | 1736 | 1544 | 1558 | 2417 | 3614 | 3129 | 4923 | 4180 | 5047 | 5212 | 3657 | 6291 | 3002 | 6783 | 1652 | 775 | 2259 | 2686 | 4601 | 3621 | 4409 | 2413 | 3484 | 7561 | 7703 | 7234 | 3771 | 5001 | Eomes |
| Ep300 | 8641 | 8350 | 7033 | 8816 | 10708 | 10220 | 9600 | 11155 | 11685 | 11989 | 11793 | 14168 | 13146 | 14252 | 8205 | 8441 | 8429 | 8670 | 10217 | 8584 | 10115 | 12546 | 10418 | 11350 | 13555 | 8280 | 13584 | 12483 | Ep300 |
| Ep400 | 16625 | 17533 | 12826 | 18534 | 18358 | 18760 | 17724 | 15040 | 16139 | 18788 | 17410 | 21188 | 18625 | 19388 | 15539 | 16799 | 15175 | 17480 | 18060 | 15125 | 18922 | 16695 | 14655 | 15343 | 19712 | 12502 | 18750 | 17661 | Ep400 |
| Epas1 | 1789 | 2273 | 2910 | 2919 | 2901 | 2541 | 2455 | 1408 | 1854 | 2547 | 2215 | 2337 | 2444 | 2201 | 1791 | 2362 | 2497 | 3047 | 2677 | 1943 | 2582 | 2784 | 2172 | 2478 | 2662 | 2546 | 2173 | 1998 | Epas1 |
| Epb41 | 25945 | 29286 | 30700 | 25898 | 23325 | 23429 | 22005 | 22947 | 20477 | 22330 | 19095 | 27064 | 31161 | 26816 | 27805 | 27880 | 29787 | 23026 | 24031 | 22601 | 22338 | 21821 | 19537 | 22535 | 22629 | 22289 | 28653 | 30072 | Epb41 |
| Epb41l1 | 2967 | 2910 | 3165 | 3528 | 5462 | 4557 | 4074 | 6535 | 7929 | 9724 | 8880 | 6847 | 6557 | 6949 | 2981 | 3373 | 3483 | 4150 | 4419 | 3295 | 4544 | 11214 | 7583 | 8297 | 9968 | 5298 | 5905 | 6342 | Epb41l1 |
| Epb41l2 | 13723 | 11010 | 11602 | 10307 | 8915 | 10196 | 9346 | 7833 | 8536 | 7932 | 9428 | 6921 | 8695 | 6915 | 11744 | 13717 | 10077 | 10553 | 9204 | 9829 | 9194 | 6104 | 9598 | 7850 | 6383 | 8604 | 7313 | 7033 | Epb41l2 |
| Epb41l3 | 7916 | 6545 | 6526 | 6165 | 7492 | 6808 | 6232 | 9210 | 10503 | 8816 | 11129 | 7800 | 10049 | 7485 | 7402 | 8333 | 6494 | 6800 | 6397 | 6217 | 6164 | 9130 | 11011 | 8203 | 8179 | 8464 | 7645 | 7605 | Epb41l3 |
| Epb41l4a | 2060 | 2516 | 2985 | 2677 | 3108 | 2948 | 2283 | 3125 | 2726 | 3326 | 2538 | 3212 | 4146 | 2825 | 2602 | 2477 | 3085 | 2619 | 3137 | 2819 | 2683 | 4194 | 2932 | 2953 | 3509 | 2472 | 3395 | 3325 | Epb41l4a |
| Epb41l4b | 712 | 611 | 569 | 637 | 1146 | 788 | 664 | 878 | 908 | 1071 | 1010 | 529 | 835 | 502 | 790 | 834 | 625 | 746 | 747 | 616 | 733 | 1136 | 993 | 731 | 789 | 375 | 496 | 511 | Epb41l4b |
| Epb41l5 | 7587 | 7603 | 7117 | 7122 | 5886 | 6603 | 6889 | 3480 | 3057 | 3076 | 2881 | 4485 | 3734 | 4456 | 7146 | 6339 | 7187 | 6588 | 6752 | 6400 | 6653 | 2974 | 2774 | 3288 | 3694 | 3609 | 4800 | 4485 | Epb41l5 |
| Epb42 | 206 | 299 | 271 | 167 | 38 | 113 | 98 | 205 | 240 | 133 | 133 | 292 | 1077 | 502 | 294 | 294 | 111 | 55 | 91 | 191 | 47 | 109 | 223 | 124 | 144 | 336 | 425 | 972 | Epb42 |
| Epc1 | 2969 | 2965 | 3120 | 3948 | 4387 | 3899 | 4267 | 7537 | 6956 | 7211 | 6978 | 7065 | 6643 | 7880 | 3157 | 2602 | 3522 | 3540 | 4029 | 3783 | 4219 | 8126 | 6602 | 7494 | 8401 | 6738 | 7011 | 7589 | Epc1 |
| Epc2 | 2111 | 1572 | 1880 | 2127 | 2666 | 2360 | 2749 | 8716 | 7505 | 7435 | 6741 | 8336 | 7471 | 8462 | 2325 | 1562 | 2137 | 2003 | 2621 | 2584 | 2598 | 8260 | 7263 | 7638 | 8458 | 7299 | 7740 | 7987 | Epc2 |
| Epcam | 1445 | 714 | 422 | 487 | 275 | 259 | 114 | 1282 | 1475 | 1481 | 1244 | 283 | 612 | 386 | 856 | 850 | 256 | 304 | 175 | 154 | 118 | 418 | 1603 | 1451 | 212 | 1843 | 234 | 358 | Epcam |
| Epdr1 | 775 | 737 | 844 | 766 | 839 | 847 | 869 | 1263 | 1130 | 1136 | 1057 | 1194 | 1184 | 1278 | 964 | 579 | 757 | 777 | 828 | 925 | 870 | 1076 | 1273 | 1163 | 1214 | 1452 | 1232 | 1417 | Epdr1 |
| Epg5 | 2 | 0 | 1 | 0 | 0 | 1 | 0 | 0 | 0 | 0 | 0 | 1 | 0 | 0 | 1 | 0 | 1 | 0 | 0 | 0 | 2 | 0 | 0 | 1 | 0 | 0 | 0 | 1 | Epg5 |
| Epgn | 8 | 13 | 19 | 12 | 20 | 4 | 18 | 20 | 16 | 17 | 16 | 16 | 28 | 27 | 11 | 9 | 22 | 17 | 12 | 22 | 17 | 15 | 19 | 25 | 23 | 24 | 24 | 31 | Epgn |
| Epha1 | 730 | 402 | 372 | 200 | 95 | 213 | 61 | 288 | 480 | 306 | 284 | 146 | 247 | 159 | 397 | 544 | 176 | 204 | 73 | 51 | 30 | 105 | 431 | 319 | 103 | 267 | 131 | 150 | Epha1 |
| Epha10 | 2164 | 3245 | 4700 | 4287 | 4451 | 3920 | 3004 | 1258 | 1054 | 1357 | 1231 | 1715 | 2183 | 1514 | 2757 | 3219 | 4379 | 4258 | 4539 | 3640 | 3541 | 2099 | 862 | 1396 | 1613 | 422 | 2020 | 1792 | Epha10 |
| Epha2 | 1091 | 1566 | 1612 | 1391 | 810 | 1031 | 641 | 396 | 400 | 617 | 458 | 670 | 737 | 591 | 931 | 1636 | 1576 | 1169 | 1086 | 806 | 809 | 511 | 597 | 605 | 460 | 462 | 686 | 688 | Epha2 |
| Epha3 | 12001 | 9588 | 8031 | 8534 | 8687 | 10031 | 11500 | 11783 | 11755 | 8231 | 9556 | 9507 | 6352 | 10291 | 12932 | 9578 | 9075 | 8479 | 9047 | 12302 | 10238 | 9063 | 11518 | 8354 | 9597 | 8864 | 8745 | 9003 | Epha3 |
| Epha4 | 15201 | 14092 | 11128 | 15228 | 15338 | 16734 | 19215 | 16433 | 15205 | 13568 | 12270 | 15578 | 12119 | 16042 | 15195 | 12656 | 13261 | 15857 | 16984 | 17499 | 18603 | 12155 | 12549 | 15020 | 16269 | 12695 | 14064 | 13402 | Epha4 |
| Epha5 | 4459 | 4242 | 4743 | 7806 | 10444 | 10661 | 13404 | 10353 | 10284 | 10214 | 8406 | 9229 | 7904 | 10653 | 5803 | 3943 | 6231 | 8649 | 9728 | 10286 | 12230 | 9809 | 7892 | 10228 | 13738 | 5685 | 7908 | 7941 | Epha5 |
| Epha6 | 95 | 132 | 59 | 73 | 169 | 195 | 124 | 294 | 302 | 180 | 202 | 84 | 149 | 100 | 167 | 143 | 65 | 110 | 105 | 220 | 154 | 222 | 258 | 98 | 180 | 57 | 99 | 61 | Epha6 |
| Epha7 | 8051 | 7210 | 8583 | 6879 | 6817 | 7641 | 7478 | 8374 | 7403 | 7440 | 7857 | 7938 | 9877 | 9124 | 8380 | 8478 | 7007 | 6960 | 6138 | 7323 | 6727 | 9072 | 8383 | 6923 | 7612 | 6533 | 9581 | 9113 | Epha7 |
| Epha8 | 683 | 1080 | 1809 | 1583 | 1968 | 1379 | 1450 | 744 | 666 | 1034 | 931 | 982 | 818 | 726 | 786 | 945 | 1650 | 1777 | 2135 | 1277 | 1568 | 1465 | 780 | 1212 | 1250 | 415 | 961 | 877 | Epha8 |
| Ephb1 | 3128 | 3974 | 3802 | 5319 | 6545 | 6317 | 6254 | 5576 | 5239 | 6966 | 5358 | 6372 | 5732 | 6239 | 4018 | 3283 | 5149 | 5709 | 6698 | 5451 | 7145 | 8238 | 4706 | 6672 | 8040 | 5223 | 5470 | 5884 | Ephb1 |
| Ephb2 | 20288 | 27586 | 34978 | 35178 | 31108 | 29964 | 28649 | 15404 | 15895 | 22451 | 18118 | 26991 | 21535 | 24117 | 21903 | 23742 | 36848 | 34069 | 34284 | 23587 | 30202 | 24162 | 15723 | 23881 | 24069 | 18525 | 25530 | 26507 | Ephb2 |
| Ephb3 | 4295 | 4685 | 4956 | 5043 | 3902 | 3907 | 2909 | 2913 | 3957 | 3750 | 4221 | 3094 | 3615 | 2508 | 3239 | 5386 | 4770 | 4909 | 4031 | 2584 | 3406 | 2978 | 3734 | 3941 | 2593 | 4069 | 2837 | 2873 | Ephb3 |
| Ephb4 | 5529 | 5537 | 5939 | 3881 | 3026 | 3865 | 2489 | 4161 | 5649 | 4751 | 6442 | 4368 | 5574 | 4053 | 4398 | 6518 | 4717 | 4409 | 3415 | 2456 | 2588 | 3795 | 6302 | 4913 | 3671 | 6296 | 4781 | 4656 | Ephb4 |
| Ephb6 | 1151 | 1284 | 1373 | 1377 | 2107 | 1901 | 1505 | 2013 | 2542 | 2689 | 2364 | 2006 | 2065 | 1738 | 1122 | 1199 | 1130 | 1426 | 2001 | 1522 | 1585 | 2458 | 2391 | 2287 | 2692 | 1939 | 2002 | 1804 | Ephb6 |
| Ephx1 | 466 | 532 | 419 | 485 | 601 | 520 | 489 | 200 | 230 | 247 | 232 | 98 | 125 | 89 | 483 | 590 | 505 | 538 | 472 | 484 | 544 | 185 | 349 | 213 | 133 | 129 | 59 | 83 | Ephx1 |
| Ephx2 | 174 | 153 | 211 | 121 | 88 | 84 | 68 | 142 | 150 | 127 | 187 | 84 | 163 | 103 | 223 | 198 | 160 | 142 | 55 | 73 | 72 | 154 | 219 | 114 | 79 | 133 | 117 | 117 | Ephx2 |
| Ephx3 | 42 | 35 | 18 | 31 | 29 | 36 | 25 | 23 | 16 | 12 | 22 | 14 | 7 | 17 | 30 | 47 | 24 | 40 | 37 | 33 | 29 | 13 | 25 | 7 | 22 | 11 | 16 | 14 | Ephx3 |
| Ephx4 | 105 | 79 | 106 | 116 | 137 | 128 | 120 | 53 | 64 | 56 | 51 | 29 | 58 | 37 | 123 | 118 | 129 | 113 | 97 | 117 | 137 | 53 | 68 | 50 | 49 | 41 | 23 | 26 | Ephx4 |
| Epm2a | 120 | 102 | 131 | 121 | 187 | 170 | 132 | 546 | 513 | 573 | 576 | 470 | 513 | 466 | 141 | 81 | 108 | 152 | 130 | 202 | 139 | 667 | 595 | 431 | 554 | 474 | 549 | 519 | Epm2a |
| Epm2aip1 | 19478 | 16993 | 12239 | 18203 | 17500 | 18930 | 19937 | 20357 | 17692 | 14660 | 16005 | 13810 | 13133 | 13850 | 19969 | 18761 | 14207 | 19220 | 16472 | 20344 | 19673 | 17779 | 17248 | 13004 | 16184 | 9457 | 14473 | 13003 | Epm2aip1 |
| Epn1 | 7989 | 10599 | 11251 | 12454 | 11599 | 10367 | 9918 | 6567 | 8609 | 10110 | 9361 | 9929 | 8921 | 8395 | 7598 | 9402 | 11895 | 11530 | 12282 | 8740 | 10306 | 9545 | 8139 | 10373 | 10282 | 10518 | 8442 | 8378 | Epn1 |
| Epn2 | 5509 | 5776 | 5414 | 5388 | 5896 | 5728 | 5844 | 4501 | 4848 | 4450 | 4685 | 4462 | 4366 | 4458 | 5575 | 5484 | 5461 | 5554 | 5770 | 5867 | 5865 | 4812 | 4715 | 4600 | 5108 | 4308 | 4362 | 4313 | Epn2 |
| Epn3 | 135 | 137 | 136 | 79 | 94 | 73 | 58 | 122 | 148 | 131 | 152 | 76 | 130 | 47 | 96 | 147 | 61 | 109 | 79 | 64 | 66 | 74 | 150 | 123 | 32 | 169 | 72 | 55 | Epn3 |
| Epo | 6 | 11 | 11 | 3 | 5 | 6 | 3 | 4 | 1 | 8 | 4 | 1 | 0 | 3 | 3 | 16 | 15 | 1 | 3 | 2 | 0 | 2 | 3 | 2 | 3 | 10 | 10 | 12 | Epo |
| Epop | 2248 | 2967 | 3298 | 3071 | 2325 | 2198 | 2200 | 517 | 532 | 644 | 584 | 931 | 750 | 887 | 1875 | 2414 | 3102 | 2612 | 2696 | 1973 | 2331 | 518 | 497 | 887 | 719 | 1073 | 858 | 970 | Epop |
| Epor | 293 | 314 | 354 | 225 | 236 | 215 | 230 | 76 | 117 | 82 | 108 | 89 | 172 | 90 | 236 | 295 | 351 | 280 | 254 | 223 | 202 | 72 | 117 | 77 | 66 | 105 | 122 | 138 | Epor |
| Epp13 | 0 | 0 | 1 | 0 | 0 | 1 | 1 | 2 | 1 | 10 | 1 | 0 | 0 | 4 | 0 | 1 | 0 | 2 | 0 | 4 | 0 | 3 | 1 | 0 | 3 | 0 | 2 | 0 | Epp13 |
| Eppin | 3 | 0 | 4 | 2 | 0 | 0 | 0 | 1 | 1 | 0 | 0 | 0 | 2 | 0 | 0 | 0 | 6 | 3 | 0 | 0 | 1 | 7 | 0 | 0 | 0 | 2 | 1 | 0 | Eppin |
| Eprs | 30172 | 28325 | 24784 | 23224 | 22178 | 22820 | 24644 | 21633 | 19682 | 17449 | 17938 | 19214 | 18476 | 19208 | 30887 | 26715 | 24239 | 23468 | 22719 | 27384 | 23752 | 17067 | 19619 | 17295 | 17813 | 17535 | 19652 | 19253 | Eprs |
| Eps15 | 9912 | 8199 | 8564 | 10116 | 11931 | 11029 | 11135 | 11444 | 9568 | 9406 | 9522 | 7920 | 9429 | 8526 | 10920 | 9689 | 9062 | 10337 | 10039 | 11663 | 11212 | 11010 | 10346 | 8071 | 9407 | 6694 | 8485 | 8558 | Eps15 |
| Eps15l1 | 7005 | 8075 | 6427 | 7438 | 7401 | 7384 | 7415 | 7417 | 8176 | 8204 | 7944 | 8594 | 8082 | 8409 | 6802 | 6936 | 7035 | 7291 | 7407 | 6837 | 7217 | 8363 | 7384 | 8508 | 8591 | 7252 | 8211 | 7899 | Eps15l1 |
| Eps8 | 1651 | 1373 | 1501 | 1284 | 1267 | 1249 | 1111 | 1043 | 1149 | 1235 | 1334 | 791 | 1021 | 837 | 1672 | 1681 | 1272 | 1327 | 1151 | 1040 | 1147 | 996 | 1381 | 1059 | 742 | 1194 | 805 | 847 | Eps8 |
| Eps8l1 | 488 | 258 | 268 | 281 | 216 | 171 | 521 | 231 | 514 | 721 | 117 | 55 | 307 | 275 | 295 | 323 | 323 | 238 | 340 | 233 | 525 | 346 | 454 | 415 | 434 | 355 | 458 | 294 | Eps8l1 |
| Eps8l2 | 196 | 162 | 115 | 46 | 74 | 114 | 21 | 145 | 259 | 167 | 158 | 93 | 158 | 84 | 135 | 160 | 50 | 89 | 40 | 25 | 13 | 82 | 311 | 129 | 58 | 104 | 115 | 131 | Eps8l2 |
| Eps8l3 | 1 | 1 | 6 | 2 | 7 | 1 | 2 | 0 | 5 | 1 | 0 | 1 | 1 | 0 | 0 | 0 | 0 | 1 | 5 | 2 | 2 | 0 | 0 | 4 | 1 | 1 | 5 | 0 | Eps8l3 |
| Epsti1 | 22 | 28 | 22 | 8 | 18 | 11 | 20 | 21 | 16 | 37 | 39 | 16 | 23 | 28 | 30 | 41 | 5 | 14 | 3 | 18 | 29 | 22 | 35 | 32 | 14 | 29 | 37 | 18 | Epsti1 |
| Epx | 1 | 0 | 4 | 0 | 2 | 8 | 4 | 1 | 2 | 4 | 15 | 5 | 6 | 9 | 5 | 1 | 1 | 1 | 9 | 9 | 4 | 8 | 3 | 4 | 0 | 6 | 10 | 4 | Epx |
| Epyc | 300 | 238 | 306 | 164 | 33 | 228 | 32 | 326 | 673 | 339 | 950 | 390 | 511 | 449 | 407 | 343 | 254 | 210 | 65 | 47 | 43 | 511 | 691 | 418 | 362 | 400 | 657 | 539 | Epyc |
| Eqtn | 103 | 78 | 37 | 113 | 89 | 104 | 111 | 49 | 48 | 60 | 63 | 70 | 66 | 60 | 102 | 105 | 89 | 109 | 95 | 78 | 101 | 67 | 56 | 46 | 53 | 39 | 38 | 58 | Eqtn |
| Eral1 | 1215 | 1298 | 1119 | 1037 | 971 | 1059 | 1086 | 1537 | 1631 | 1474 | 1479 | 1622 | 1496 | 1585 | 1133 | 1167 | 1096 | 1080 | 1080 | 1156 | 1062 | 1529 | 1717 | 1673 | 1496 | 1780 | 1761 | 1632 | Eral1 |
| Erap1 | 384 | 292 | 259 | 183 | 329 | 335 | 235 | 590 | 672 | 512 | 612 | 529 | 558 | 534 | 360 | 382 | 222 | 240 | 197 | 254 | 239 | 480 | 672 | 536 | 462 | 649 | 553 | 633 | Erap1 |
| Eras | 7 | 14 | 10 | 9 | 10 | 10 | 12 | 4 | 2 | 2 | 3 | 2 | 8 | 1 | 3 | 12 | 10 | 19 | 12 | 10 | 9 | 2 | 3 | 3 | 3 | 8 | 6 | 1 | Eras |
| Erbb2 | 2438 | 2493 | 3031 | 2024 | 1751 | 2313 | 1465 | 2213 | 3118 | 2695 | 3536 | 2266 | 3030 | 1978 | 2064 | 2891 | 2737 | 2136 | 1999 | 1439 | 1500 | 2065 | 3176 | 2901 | 1945 | 3004 | 2501 | 2166 | Erbb2 |
| Erbb3 | 1174 | 739 | 464 | 556 | 589 | 388 | 213 | 1200 | 1565 | 1250 | 2384 | 260 | 981 | 272 | 795 | 1172 | 446 | 496 | 284 | 198 | 249 | 1023 | 1719 | 897 | 455 | 890 | 214 | 218 | Erbb3 |
| Erbb4 | 3044 | 2967 | 2398 | 3783 | 5134 | 5660 | 5816 | 4984 | 3976 | 4064 | 4067 | 3811 | 3450 | 4136 | 3794 | 2733 | 2840 | 3763 | 4315 | 5085 | 6117 | 5143 | 4306 | 3029 | 5102 | 1872 | 4396 | 3841 | Erbb4 |
| Erbin | 11941 | 7676 | 8878 | 8611 | 9432 | 11023 | 10584 | 21566 | 19758 | 16973 | 18794 | 19863 | 20373 | 20525 | 11909 | 10195 | 8569 | 8300 | 9192 | 10366 | 10215 | 16449 | 18993 | 17924 | 17432 | 16744 | 21122 | 20304 | Erbin |
| Erc1 | 11893 | 12692 | 9010 | 13302 | 15452 | 14589 | 14724 | 9208 | 9714 | 10528 | 10324 | 9135 | 9335 | 8952 | 12622 | 12159 | 11653 | 14105 | 14349 | 12485 | 15091 | 11312 | 8880 | 8764 | 10835 | 4670 | 8867 | 8362 | Erc1 |
| Erc2 | 3435 | 3689 | 2427 | 5286 | 7679 | 6496 | 7020 | 6555 | 5870 | 6180 | 5289 | 4259 | 4107 | 4448 | 4522 | 3256 | 3868 | 6085 | 6330 | 6871 | 7434 | 7738 | 5346 | 4545 | 6936 | 2578 | 3848 | 3579 | Erc2 |
| Ercc1 | 1316 | 1809 | 1739 | 1756 | 1762 | 1664 | 1505 | 1583 | 1807 | 1652 | 1805 | 2042 | 1902 | 2037 | 1370 | 1509 | 1866 | 1609 | 1951 | 1655 | 1620 | 1718 | 1696 | 2109 | 1971 | 2034 | 2201 | 2044 | Ercc1 |
| Ercc2 | 5357 | 7733 | 6898 | 7194 | 5435 | 6012 | 5474 | 3324 | 3725 | 4216 | 3699 | 4663 | 4208 | 3984 | 5281 | 6310 | 7091 | 6930 | 6460 | 5588 | 5695 | 3920 | 3595 | 4107 | 4349 | 4815 | 4336 | 4444 | Ercc2 |
| Ercc3 | 2613 | 2709 | 2330 | 2497 | 2059 | 2257 | 2192 | 2693 | 2961 | 2818 | 2819 | 2955 | 2947 | 2885 | 2467 | 2471 | 2213 | 2325 | 2105 | 2188 | 2129 | 2463 | 2952 | 2756 | 2679 | 3155 | 3075 | 3125 | Ercc3 |
| Ercc4 | 1554 | 1421 | 1435 | 1492 | 1912 | 1800 | 1633 | 1619 | 1797 | 2052 | 1893 | 1902 | 1853 | 1902 | 1505 | 1428 | 1506 | 1519 | 1728 | 1471 | 1666 | 1853 | 1917 | 1800 | 1892 | 1579 | 1956 | 1881 | Ercc4 |

| | | | | | | | | | | | | | | | | | | | | | | | | | | | | | |
|---|---|---|---|---|---|---|---|---|---|---|---|---|---|---|---|---|---|---|---|---|---|---|---|---|---|---|---|---|---|
| Ercc5 | 4756 | 4514 | 4560 | 4721 | 4498 | 4358 | 4706 | 5015 | 4343 | 4737 | 4419 | 5262 | 4682 | 5205 | 4930 | 4409 | 4904 | 4596 | 4643 | 4675 | 4582 | 4941 | 4286 | 4868 | 4937 | 4848 | 5020 | 5219 | Ercc5 |
| Ercc6 | 2175 | 2078 | 1724 | 1941 | 2238 | 2345 | 2137 | 4355 | 4351 | 4057 | 3934 | 4652 | 4368 | 4291 | 2153 | 2076 | 1966 | 2059 | 1965 | 2052 | 2209 | 3994 | 3869 | 3743 | 4491 | 3125 | 4523 | 4037 | Ercc6 |
| Ercc6l | 3795 | 3127 | 3097 | 2267 | 1725 | 2178 | 2444 | 3108 | 2567 | 2059 | 2071 | 3240 | 2744 | 3301 | 3497 | 3199 | 2823 | 2009 | 2123 | 2934 | 2246 | 1841 | 2327 | 2520 | 2336 | 3641 | 3362 | 3724 | Ercc6l |
| Ercc6l2 | 2873 | 2160 | 1962 | 1830 | 1972 | 2101 | 2086 | 4482 | 4212 | 3633 | 3704 | 3557 | 3694 | 3584 | 2643 | 2428 | 1861 | 1845 | 1729 | 2246 | 1949 | 3715 | 4054 | 3229 | 3167 | 3307 | 3892 | 3506 | Ercc6l2 |
| Ercc8 | 2318 | 2437 | 1709 | 1887 | 1647 | 2006 | 1939 | 1785 | 1543 | 1407 | 1297 | 1599 | 1472 | 1616 | 2156 | 2098 | 1692 | 1867 | 1833 | 2215 | 1992 | 1432 | 1401 | 1475 | 1590 | 1500 | 1575 | 1628 | Ercc8 |
| Ereg | 0 | 0 | 15 | 0 | 0 | 0 | 0 | 6 | 5 | 0 | 1 | 0 | 0 | 0 | 3 | 0 | 0 | 0 | 0 | 0 | 0 | 0 | 2 | 2 | 2 | 0 | 0 | 0 | Ereg |
| Erf | 2968 | 3831 | 3657 | 3779 | 2562 | 2685 | 2361 | 2616 | 3284 | 3123 | 3564 | 3009 | 3318 | 2942 | 2600 | 3242 | 3599 | 3425 | 2847 | 2684 | 2649 | 2212 | 3197 | 3411 | 2766 | 4276 | 2991 | 3115 | Erf |
| Erfe | 31 | 30 | 51 | 13 | 23 | 25 | 18 | 31 | 67 | 62 | 61 | 72 | 65 | 49 | 25 | 37 | 44 | 29 | 26 | 24 | 19 | 28 | 44 | 60 | 12 | 122 | 59 | 53 | Erfe |
| Erfl | 72 | 130 | 162 | 252 | 248 | 236 | 116 | 278 | 249 | 358 | 245 | 272 | 573 | 165 | 156 | 149 | 238 | 185 | 188 | 228 | 147 | 404 | 236 | 181 | 336 | 115 | 298 | 197 | Erfl |
| Erg | 696 | 845 | 816 | 651 | 444 | 762 | 411 | 1208 | 1664 | 1831 | 1974 | 1210 | 1910 | 1173 | 704 | 984 | 705 | 797 | 456 | 377 | 370 | 1439 | 1662 | 1622 | 1288 | 1876 | 1485 | 1345 | Erg |
| Erg28 | 2398 | 2252 | 2178 | 2009 | 1838 | 1969 | 2116 | 2328 | 2254 | 2168 | 2147 | 2147 | 2430 | 2539 | 2139 | 2354 | 2108 | 2107 | 1991 | 2166 | 1995 | 1992 | 2399 | 2249 | 2153 | 2627 | 2485 | 2643 | Erg28 |
| Ergic1 | 9727 | 10069 | 10815 | 10160 | 9493 | 9491 | 9379 | 7499 | 8625 | 9441 | 8881 | 9356 | 8275 | 8988 | 8992 | 9522 | 10706 | 9957 | 10037 | 8272 | 9554 | 8666 | 8787 | 9828 | 9047 | 10861 | 8451 | 9439 | Ergic1 |
| Ergic2 | 7078 | 5055 | 6415 | 4986 | 5375 | 5711 | 6102 | 9329 | 7807 | 7175 | 7601 | 8059 | 8190 | 8639 | 7065 | 6422 | 5674 | 5044 | 5125 | 6417 | 5768 | 8014 | 8130 | 7517 | 7713 | 8165 | 8232 | 8408 | Ergic2 |
| Ergic3 | 5090 | 6453 | 5800 | 6165 | 6040 | 6049 | 5437 | 5870 | 6849 | 6581 | 6900 | 7166 | 6713 | 6120 | 5095 | 5672 | 6092 | 5889 | 6427 | 5753 | 5563 | 6522 | 7374 | 6754 | 6992 | 6898 | 6892 | 6489 | Ergic3 |
| Erh | 12671 | 12962 | 14788 | 12903 | 10945 | 11419 | 13759 | 11692 | 10795 | 10864 | 10341 | 12991 | 12759 | 14382 | 13260 | 11720 | 13373 | 11567 | 12208 | 13708 | 12547 | 8993 | 10725 | 12222 | 11980 | 16227 | 13961 | 15640 | Erh |
| Eri1 | 6873 | 5842 | 5824 | 6358 | 5689 | 5980 | 6915 | 4606 | 3992 | 3833 | 3701 | 4190 | 3839 | 4155 | 7075 | 6115 | 5903 | 6280 | 6000 | 6750 | 6667 | 3589 | 4036 | 3882 | 3896 | 4515 | 4224 | 4410 | Eri1 |
| Eri2 | 2953 | 2394 | 2468 | 2305 | 2452 | 2606 | 2731 | 4449 | 3732 | 3250 | 3389 | 4215 | 3751 | 4256 | 2848 | 2459 | 2308 | 2213 | 2441 | 2948 | 2448 | 3247 | 3417 | 3479 | 3578 | 4090 | 4355 | 4367 | Eri2 |
| Eri3 | 3817 | 4674 | 4369 | 5341 | 5317 | 4823 | 4902 | 4112 | 5226 | 5148 | 5406 | 5189 | 4579 | 4783 | 3919 | 4111 | 4992 | 5096 | 5520 | 4799 | 5170 | 5020 | 5259 | 5324 | 5584 | 5605 | 4298 | 4885 | Eri3 |
| Erich1 | 934 | 1145 | 1229 | 1257 | 1136 | 1095 | 1037 | 658 | 596 | 710 | 597 | 743 | 706 | 699 | 1039 | 998 | 1179 | 1151 | 1087 | 1027 | 984 | 623 | 593 | 726 | 656 | 768 | 741 | 649 | Erich1 |
| Erich2 | 318 | 295 | 327 | 245 | 247 | 300 | 275 | 128 | 137 | 111 | 97 | 128 | 139 | 134 | 328 | 285 | 263 | 242 | 220 | 362 | 296 | 107 | 190 | 123 | 111 | 133 | 148 | 122 | Erich2 |
| Erich3 | 158 | 160 | 162 | 192 | 277 | 256 | 223 | 347 | 341 | 474 | 359 | 263 | 339 | 254 | 189 | 188 | 175 | 230 | 232 | 233 | 230 | 413 | 329 | 285 | 403 | 165 | 249 | 199 | Erich3 |
| Erich4 | 1 | 0 | 0 | 8 | 0 | 3 | 1 | 0 | 2 | 0 | 0 | 2 | 0 | 6 | 0 | 5 | 1 | 0 | 2 | 1 | 1 | 0 | 1 | 2 | 0 | 0 | 0 | 1 | Erich4 |
| Erich5 | 0 | 0 | 0 | 0 | 2 | 0 | 4 | 2 | 2 | 1 | 3 | 2 | 1 | 0 | 0 | 1 | 0 | 1 | 0 | 0 | 0 | 5 | 5 | 2 | 0 | 0 | 10 | 6 | Erich5 |
| Erich6 | 67 | 61 | 48 | 96 | 87 | 89 | 111 | 24 | 37 | 44 | 40 | 28 | 36 | 35 | 62 | 86 | 93 | 109 | 84 | 62 | 107 | 80 | 41 | 30 | 38 | 23 | 34 | 14 | Erich6 |
| Erich6b | 0 | 3 | 0 | 0 | 2 | 0 | 1 | 4 | 2 | 0 | 3 | 0 | 3 | 0 | 1 | 0 | 2 | 0 | 0 | 1 | 0 | 0 | 1 | 0 | 0 | 0 | 1 | 0 | Erich6b |
| Erlec1 | 2449 | 1765 | 1639 | 1782 | 2071 | 2171 | 1977 | 3429 | 2988 | 2768 | 2871 | 2817 | 3015 | 2852 | 2503 | 2101 | 1538 | 1632 | 1815 | 2070 | 2182 | 2739 | 3102 | 2294 | 2984 | 2499 | 2914 | 2702 | Erlec1 |
| Erlin1 | 3472 | 3220 | 3329 | 3052 | 2689 | 2786 | 2789 | 2491 | 2454 | 2322 | 2564 | 2356 | 2260 | 2318 | 3359 | 3465 | 3111 | 3306 | 2752 | 2945 | 2740 | 2540 | 2715 | 2311 | 2330 | 2520 | 2490 | 2462 | Erlin1 |
| Erlin2 | 7784 | 7849 | 7089 | 6775 | 6898 | 7088 | 7148 | 8391 | 7824 | 7551 | 7795 | 7860 | 8090 | 7932 | 7913 | 7756 | 7106 | 7026 | 6466 | 7059 | 7074 | 7991 | 8712 | 7290 | 7707 | 6999 | 8139 | 7937 | Erlin2 |
| Ermap | 303 | 382 | 389 | 176 | 77 | 160 | 154 | 355 | 275 | 336 | 269 | 605 | 1882 | 904 | 290 | 346 | 255 | 145 | 141 | 187 | 63 | 201 | 307 | 308 | 227 | 632 | 944 | 1786 | Ermap |
| Ermard | 431 | 401 | 386 | 400 | 400 | 442 | 443 | 937 | 861 | 757 | 797 | 855 | 788 | 867 | 517 | 447 | 477 | 335 | 421 | 478 | 443 | 817 | 819 | 737 | 834 | 788 | 974 | 789 | Ermard |
| Ermn | 271 | 160 | 80 | 195 | 165 | 190 | 285 | 299 | 236 | 265 | 228 | 143 | 86 | 242 | 214 | 101 | 232 | 221 | 194 | 205 | 250 | 159 | 219 | 358 | 179 | 453 | 153 | 161 | Ermn |
| Ermp1 | 3620 | 2557 | 2715 | 2148 | 2306 | 2380 | 1838 | 3386 | 3085 | 2921 | 2742 | 2810 | 3074 | 2611 | 2904 | 3365 | 2209 | 2269 | 2124 | 1924 | 1951 | 2568 | 3241 | 2720 | 2347 | 2846 | 2787 | 2853 | Ermp1 |
| Ern1 | 4395 | 4509 | 3968 | 5080 | 4522 | 4486 | 4710 | 1895 | 2090 | 2285 | 2434 | 2222 | 2080 | 2319 | 4102 | 4496 | 4285 | 5355 | 4582 | 3858 | 5014 | 2378 | 1942 | 2169 | 2489 | 1470 | 2166 | 2045 | Ern1 |
| Ern2 | 8 | 15 | 11 | 14 | 39 | 51 | 28 | 48 | 61 | 87 | 47 | 51 | 50 | 64 | 13 | 10 | 14 | 20 | 30 | 25 | 31 | 139 | 58 | 56 | 92 | 15 | 65 | 46 | Ern2 |
| Ero1a | 6241 | 4872 | 5226 | 4522 | 4596 | 4624 | 4860 | 3684 | 3262 | 3154 | 3354 | 3517 | 3442 | 3546 | 6008 | 5829 | 4653 | 4778 | 4222 | 4918 | 4520 | 3354 | 3312 | 2944 | 3226 | 3210 | 3569 | 3463 | Ero1a |
| Ero1b | 2256 | 2090 | 2091 | 2306 | 1901 | 1998 | 2411 | 1027 | 901 | 883 | 905 | 673 | 708 | 696 | 2206 | 2320 | 1856 | 2390 | 1976 | 2309 | 2323 | 1046 | 802 | 799 | 932 | 699 | 846 | 808 | Ero1b |
| Erp27 | 14 | 1 | 0 | 4 | 3 | 2 | 6 | 13 | 7 | 4 | 9 | 3 | 0 | 6 | 5 | 10 | 2 | 9 | 5 | 1 | 6 | 8 | 8 | 7 | 6 | 4 | 0 | 4 | Erp27 |
| Erp29 | 5045 | 5762 | 5506 | 4658 | 4638 | 4881 | 4546 | 4640 | 5446 | 5104 | 5460 | 4984 | 5502 | 5139 | 4721 | 5271 | 5180 | 4580 | 4905 | 4947 | 4564 | 4542 | 5559 | 5277 | 5034 | 5662 | 5456 | 5342 | Erp29 |
| Erp44 | 5540 | 4942 | 5734 | 4987 | 4539 | 4690 | 5433 | 3893 | 3680 | 3526 | 3966 | 3413 | 3645 | 3645 | 5722 | 5379 | 5359 | 4993 | 4871 | 4860 | 4994 | 3870 | 3998 | 3519 | 3437 | 3845 | 3643 | 3785 | Erp44 |
| Errfi1 | 2459 | 2299 | 2406 | 2328 | 1938 | 1981 | 1861 | 2088 | 1914 | 1807 | 2131 | 1842 | 2042 | 1719 | 2446 | 2793 | 2235 | 2411 | 1918 | 1914 | 1792 | 2152 | 2405 | 1743 | 1524 | 1926 | 1762 | 1790 | Errfi1 |
| Erv3 | 0 | 0 | 0 | 0 | 2 | 0 | 6 | 1 | 11 | 0 | 1 | 1 | 2 | 0 | 1 | 0 | 0 | 0 | 5 | 7 | 1 | 0 | 1 | 0 | 1 | 1 | 0 | 0 | Erv3 |
| Esam | 608 | 608 | 672 | 621 | 761 | 761 | 663 | 1018 | 1304 | 1373 | 1463 | 1325 | 1362 | 1130 | 525 | 677 | 547 | 693 | 722 | 628 | 659 | 1355 | 1419 | 1324 | 1386 | 1318 | 1245 | 1176 | Esam |
| Esco1 | 3185 | 2463 | 2961 | 2968 | 3002 | 2928 | 3290 | 3273 | 2502 | 2384 | 2326 | 2525 | 2428 | 2495 | 3413 | 2914 | 2790 | 2932 | 2904 | 3718 | 3080 | 2653 | 2769 | 2377 | 2395 | 2304 | 2345 | 2473 | Esco1 |
| Esco2 | 5321 | 3847 | 4552 | 3754 | 2943 | 3771 | 4437 | 3367 | 2825 | 2553 | 2582 | 3990 | 3187 | 3919 | 4895 | 4613 | 3950 | 3431 | 3657 | 4955 | 3878 | 2032 | 2566 | 3134 | 2784 | 4254 | 3939 | 3973 | Esco2 |
| Esd | 10968 | 11145 | 11519 | 9968 | 8611 | 9230 | 10741 | 7724 | 7118 | 6530 | 7094 | 7544 | 8473 | 8495 | 11653 | 10225 | 9999 | 9612 | 9264 | 11452 | 9687 | 5962 | 7184 | 7783 | 6885 | 9323 | 8505 | 9128 | Esd |
| Esf1 | 6536 | 4958 | 6885 | 5733 | 4525 | 5376 | 5582 | 4594 | 4004 | 3833 | 3789 | 4730 | 4605 | 5218 | 6546 | 6065 | 5960 | 5456 | 4993 | 5915 | 5084 | 3922 | 3882 | 4490 | 4103 | 5364 | 4929 | 5258 | Esf1 |
| Esm1 | 95 | 56 | 84 | 40 | 105 | 70 | 72 | 100 | 87 | 107 | 102 | 76 | 62 | 97 | 70 | 60 | 74 | 61 | 69 | 89 | 60 | 112 | 161 | 145 | 76 | 176 | 86 | 114 | Esm1 |
| Esp1 | 0 | 0 | 0 | 0 | 0 | 5 | 0 | 0 | 0 | 0 | 0 | 0 | 0 | 0 | 0 | 0 | 0 | 0 | 0 | 0 | 0 | 0 | 0 | 0 | 0 | 0 | 0 | 0 | Esp1 |
| Esp22 | 0 | 0 | 0 | 0 | 0 | 0 | 0 | 0 | 0 | 0 | 0 | 0 | 0 | 0 | 0 | 0 | 0 | 0 | 0 | 0 | 0 | 0 | 0 | 0 | 0 | 0 | 2 | 0 | Esp22 |
| Esp6 | 0 | 0 | 0 | 0 | 0 | 0 | 0 | 0 | 0 | 0 | 0 | 4 | 0 | 0 | 0 | 0 | 0 | 0 | 0 | 0 | 0 | 0 | 1 | 0 | 1 | 0 | 0 | 0 | Esp6 |
| Esp8 | 0 | 0 | 0 | 0 | 0 | 0 | 0 | 0 | 0 | 0 | 2 | 1 | 0 | 1 | 0 | 0 | 0 | 0 | 0 | 0 | 0 | 7 | 2 | 1 | 0 | 9 | 0 | 0 | Esp8 |
| Espl1 | 4790 | 5177 | 5829 | 4751 | 3211 | 3816 | 3917 | 3186 | 4034 | 3894 | 3714 | 6705 | 5704 | 6248 | 3761 | 4989 | 5668 | 4315 | 4514 | 3363 | 3623 | 2936 | 3463 | 5386 | 4297 | 5242 | 5937 | 6394 | Espl1 |
| Espn | 195 | 181 | 206 | 149 | 82 | 55 | 57 | 112 | 92 | 113 | 129 | 13 | 36 | 29 | 156 | 217 | 145 | 125 | 77 | 62 | 49 | 20 | 109 | 261 | 9 | 242 | 24 | 11 | Espn |
| Espnl | 61 | 77 | 42 | 76 | 66 | 57 | 68 | 20 | 30 | 21 | 39 | 16 | 27 | 30 | 49 | 90 | 65 | 92 | 76 | 50 | 55 | 27 | 25 | 33 | 30 | 22 | 15 | 23 | Espnl |
| Esr1 | 1 | 1 | 0 | 7 | 9 | 6 | 3 | 6 | 5 | 9 | 25 | 3 | 15 | 5 | 11 | 10 | 1 | 3 | 5 | 3 | 2 | 12 | 21 | 7 | 6 | 16 | 2 | 8 | Esr1 |
| Esr2 | 90 | 65 | 39 | 33 | 39 | 45 | 22 | 37 | 31 | 29 | 36 | 16 | 34 | 24 | 71 | 118 | 43 | 47 | 12 | 20 | 13 | 37 | 30 | 25 | 35 | 28 | 34 | 15 | Esr2 |
| Esrp1 | 1381 | 419 | 287 | 340 | 96 | 252 | 4 | 1060 | 1364 | 1030 | 971 | 357 | 720 | 378 | 575 | 639 | 127 | 241 | 68 | 53 | 29 | 211 | 1452 | 1008 | 139 | 925 | 359 | 450 | Esrp1 |
| Esrp2 | 234 | 106 | 78 | 76 | 41 | 67 | 21 | 374 | 498 | 322 | 330 | 90 | 142 | 90 | 112 | 139 | 41 | 58 | 19 | 14 | 19 | 37 | 649 | 347 | 34 | 330 | 81 | 97 | Esrp2 |
| Esrra | 1684 | 2269 | 2755 | 2687 | 2266 | 1949 | 2104 | 1373 | 1528 | 1881 | 1655 | 1893 | 1685 | 2003 | 1618 | 1743 | 2598 | 2464 | 2408 | 1994 | 2221 | 1815 | 1541 | 2192 | 2087 | 1666 | 2000 | 1767 | Esrra |
| Esrrb | 353 | 504 | 557 | 681 | 917 | 709 | 626 | 436 | 436 | 688 | 543 | 382 | 414 | 338 | 425 | 493 | 572 | 754 | 738 | 556 | 639 | 993 | 460 | 427 | 693 | 211 | 365 | 407 | Esrrb |
| Esrrg | 2496 | 2886 | 2619 | 4264 | 5154 | 4625 | 4745 | 3091 | 2882 | 2994 | 2591 | 1924 | 1964 | 1984 | 3922 | 2622 | 3410 | 4429 | 3987 | 3982 | 4516 | 4268 | 2339 | 2090 | 2866 | 1315 | 1584 | 1650 | Esrrg |
| Ess2 | 2373 | 2744 | 2554 | 2656 | 2443 | 2770 | 2566 | 2396 | 2818 | 3006 | 2567 | 3188 | 2938 | 3026 | 2268 | 2428 | 2603 | 2501 | 2904 | 2329 | 2719 | 2742 | 2614 | 3006 | 3127 | 3269 | 3037 | 3233 | Ess2 |
| Esx1 | 10 | 11 | 16 | 18 | 7 | 13 | 15 | 0 | 1 | 0 | 0 | 0 | 1 | 0 | 18 | 6 | 5 | 14 | 16 | 7 | 4 | 0 | 0 | 0 | 0 | 0 | 0 | 0 | Esx1 |
| Esyt1 | 5590 | 6025 | 6382 | 5558 | 4833 | 5338 | 3967 | 2340 | 3081 | 2814 | 4264 | 3020 | 4182 | 2765 | 5120 | 6671 | 5716 | 5264 | 4567 | 3992 | 3984 | 2961 | 3959 | 2841 | 2616 | 2826 | 3420 | 3045 | Esyt1 |
| Esyt2 | 7253 | 5677 | 5522 | 5595 | 5506 | 6418 | 6119 | 5234 | 5100 | 4470 | 5160 | 4419 | 5117 | 4685 | 7048 | 6625 | 5151 | 6001 | 5165 | 6003 | 5846 | 4739 | 5045 | 4249 | 4499 | 4155 | 4771 | 4555 | Esyt2 |
| Esyt3 | 202 | 200 | 196 | 243 | 185 | 192 | 153 | 34 | 54 | 54 | 56 | 28 | 20 | 18 | 187 | 242 | 147 | 211 | 154 | 189 | 142 | 30 | 49 | 34 | 25 | 20 | 10 | 19 | Esyt3 |
| Etaa1 | 3478 | 2644 | 2981 | 2670 | 2648 | 2772 | 3256 | 3112 | 2619 | 2458 | 2426 | 2955 | 2851 | 2975 | 3439 | 3056 | 2797 | 2815 | 2594 | 3102 | 2935 | 2246 | 2576 | 2714 | 2443 | 3444 | 3152 | 3250 | Etaa1 |
| Etd | 0 | 0 | 0 | 0 | 0 | 1 | 0 | 0 | 0 | 0 | 1 | 0 | 0 | 0 | 0 | 0 | 0 | 0 | 0 | 0 | 0 | 0 | 0 | 1 | 0 | 0 | 0 | 1 | Etd |
| Etf1 | 19117 | 16371 | 18479 | 16629 | 14180 | 15303 | 16670 | 13455 | 12120 | 11584 | 11653 | 14368 | 13537 | 14973 | 19155 | 17354 | 16840 | 15835 | 14675 | 16317 | 15951 | 11022 | 12145 | 12488 | 12297 | 15689 | 14923 | 15772 | Etf1 |
| Etfa | 5699 | 4714 | 5271 | 4304 | 3773 | 3746 | 4177 | 3883 | 3377 | 3200 | 3336 | 3748 | 3469 | 3760 | 5397 | 4586 | 4692 | 4136 | 3840 | 4097 | 3676 | 3076 | 3311 | 3546 | 3066 | 4457 | 3696 | 3977 | Etfa |
| Etfb | 129 | 136 | 131 | 91 | 90 | 92 | 139 | 104 | 94 | 122 | 82 | 115 | 116 | 163 | 113 | 72 | 111 | 97 | 89 | 73 | 124 | 85 | 90 | 147 | 85 | 134 | 84 | 170 | Etfb |
| Etfbkmt | 351 | 221 | 294 | 225 | 265 | 216 | 248 | 418 | 363 | 372 | 307 | 365 | 382 | 399 | 298 | 256 | 294 | 184 | 252 | 256 | 214 | 334 | 476 | 371 | 266 | 452 | 346 | 393 | Etfbkmt |
| Etfbl | 0 | 0 | 0 | 0 | 0 | 0 | 0 | 0 | 0 | 0 | 0 | 0 | 0 | 0 | 1 | 0 | 0 | 0 | 0 | 0 | 0 | 0 | 0 | 0 | 0 | 0 | 0 | 0 | Etfbl |
| Etfdh | 6373 | 6133 | 5918 | 5822 | 5218 | 5454 | 5579 | 3826 | 3589 | 3347 | 3554 | 3352 | 3382 | 3666 | 6682 | 6072 | 5405 | 5413 | 4850 | 6287 | 5402 | 3178 | 3709 | 3292 | 3282 | 3608 | 3447 | 3599 | Etfdh |
| Etfrf1 | 1171 | 872 | 1143 | 986 | 902 | 983 | 1026 | 1420 | 1168 | 1176 | 1262 | 1496 | 1537 | 1539 | 1222 | 1142 | 1146 | 1013 | 881 | 1023 | 945 | 1200 | 1364 | 1286 | 1139 | 1564 | 1486 | 1574 | Etfrf1 |
| Ethe1 | 1028 | 1300 | 1026 | 1031 | 884 | 830 | 910 | 365 | 427 | 420 | 357 | 324 | 372 | 377 | 985 | 1118 | 1079 | 980 | 923 | 927 | 858 | 324 | 450 | 350 | 323 | 415 | 485 | 382 | Ethe1 |
| Etl4 | 2271 | 1964 | 2033 | 1969 | 2003 | 2440 | 2311 | 2937 | 4071 | 3804 | 3814 | 4780 | 4217 | 4830 | 1925 | 2153 | 1871 | 2079 | 2257 | 1874 | 2200 | 2236 | 3524 | 4339 | 3913 | 3928 | 4038 | 4062 | Etl4 |
| Etnk1 | 26688 | 21782 | 18756 | 23911 | 25656 | 28218 | 25239 | 19045 | 18208 | 16624 | 18372 | 14710 | 16452 | 15255 | 27139 | 28436 | 20712 | 25741 | 22100 | 24777 | 27149 | 18282 | 17246 | 14153 | 17568 | 10538 | 15633 | 13833 | Etnk1 |
| Etnk2 | 678 | 875 | 1062 | 935 | 889 | 859 | 755 | 232 | 266 | 252 | 256 | 203 | 228 | 162 | 777 | 895 | 913 | 889 | 796 | 762 | 714 | 348 | 285 | 217 | 273 | 168 | 179 | 192 | Etnk2 |

| | | | | | | | | | | | | | | | | | | | | | | | | | | | | | |
|---|---|---|---|---|---|---|---|---|---|---|---|---|---|---|---|---|---|---|---|---|---|---|---|---|---|---|---|---|---|
| Etnppl | 3 | 1 | 0 | 0 | 1 | 0 | 0 | 0 | 2 | 5 | 1 | 2 | 0 | 0 | 4 | 4 | 0 | 0 | 0 | 0 | 0 | 2 | 3 | 2 | 0 | 2 | 0 | 0 | Etnppl |
| Ets1 | 3512 | 2584 | 3604 | 2668 | 2779 | 2627 | 2445 | 2976 | 3247 | 3308 | 4057 | 3257 | 3958 | 3256 | 3067 | 4054 | 2764 | 2992 | 2325 | 2552 | 2563 | 3542 | 3717 | 3315 | 3263 | 3571 | 3443 | 3210 | Ets1 |
| Ets2 | 2799 | 2682 | 2787 | 2498 | 2383 | 2571 | 1850 | 3131 | 4575 | 4222 | 5292 | 2154 | 3826 | 2348 | 2665 | 3155 | 2341 | 2605 | 1901 | 1710 | 1861 | 3838 | 4739 | 3603 | 2678 | 3446 | 2386 | 2337 | Ets2 |
| Etv1 | 2398 | 1760 | 1755 | 2115 | 2544 | 2352 | 2779 | 2315 | 2726 | 2219 | 2678 | 2708 | 1999 | 2792 | 2223 | 2053 | 2044 | 2165 | 2249 | 2300 | 2190 | 2480 | 2420 | 2344 | 2494 | 2714 | 2085 | 2047 | Etv1 |
| Etv2 | 1 | 2 | 5 | 6 | 2 | 4 | 0 | 0 | 0 | 0 | 2 | 0 | 2 | 0 | 4 | 5 | 13 | 0 | 3 | 0 | 4 | 2 | 0 | 0 | 0 | 0 | 0 | 0 | Etv2 |
| Etv3 | 2609 | 2792 | 2877 | 2914 | 2439 | 2480 | 2418 | 2082 | 2425 | 2475 | 2589 | 2563 | 2527 | 2570 | 2288 | 2977 | 2706 | 2847 | 2516 | 2384 | 2386 | 2152 | 2443 | 2495 | 2447 | 2385 | 2445 | 2537 | Etv3 |
| Etv3l | 6 | 6 | 29 | 5 | 18 | 8 | 4 | 17 | 15 | 23 | 16 | 15 | 64 | 1 | 18 | 23 | 30 | 11 | 13 | 20 | 8 | 35 | 16 | 2 | 26 | 7 | 15 | 2 | Etv3l |
| Etv4 | 556 | 470 | 480 | 334 | 362 | 332 | 272 | 470 | 696 | 557 | 744 | 454 | 590 | 437 | 352 | 572 | 422 | 277 | 327 | 321 | 231 | 381 | 514 | 392 | 368 | 799 | 463 | 490 | Etv4 |
| Etv5 | 2560 | 2048 | 2164 | 1867 | 1869 | 1629 | 1645 | 1881 | 2576 | 2210 | 2721 | 1858 | 2137 | 1844 | 2003 | 2576 | 1764 | 1888 | 1765 | 1393 | 1415 | 2064 | 2115 | 1716 | 1919 | 2312 | 1507 | 1718 | Etv5 |
| Etv6 | 2232 | 2008 | 2886 | 2655 | 2608 | 2366 | 2199 | 2326 | 2320 | 2632 | 2700 | 3483 | 2969 | 3004 | 2114 | 2121 | 2780 | 2355 | 2486 | 1784 | 2163 | 2386 | 2242 | 2860 | 2545 | 3393 | 3041 | 3252 | Etv6 |
| Eva1a | 164 | 109 | 88 | 71 | 68 | 82 | 47 | 123 | 140 | 135 | 135 | 117 | 121 | 88 | 112 | 129 | 81 | 67 | 67 | 53 | 47 | 67 | 147 | 135 | 47 | 185 | 78 | 70 | Eva1a |
| Eva1b | 353 | 450 | 384 | 254 | 217 | 280 | 135 | 423 | 653 | 510 | 752 | 362 | 562 | 358 | 328 | 484 | 313 | 291 | 194 | 180 | 161 | 436 | 744 | 470 | 413 | 571 | 481 | 430 | Eva1b |
| Eva1c | 96 | 122 | 85 | 131 | 86 | 123 | 113 | 54 | 115 | 146 | 95 | 122 | 65 | 116 | 96 | 81 | 93 | 108 | 122 | 117 | 116 | 80 | 105 | 131 | 107 | 128 | 101 | 96 | Eva1c |
| Evc | 3713 | 3439 | 2892 | 2567 | 2392 | 2936 | 2342 | 2639 | 3145 | 2748 | 2952 | 3022 | 3046 | 2872 | 2864 | 3434 | 2566 | 2474 | 2725 | 2219 | 2356 | 2094 | 2854 | 2884 | 2550 | 3029 | 3168 | 2976 | Evc |
| Evc2 | 1326 | 1245 | 1403 | 1062 | 936 | 984 | 792 | 1373 | 1794 | 1558 | 1958 | 1536 | 1674 | 1603 | 1011 | 1319 | 1096 | 914 | 928 | 943 | 795 | 1250 | 1817 | 1781 | 1443 | 1866 | 1684 | 1584 | Evc2 |
| Evi2 | 0 | 1 | 0 | 0 | 1 | 0 | 2 | 1 | 2 | 1 | 3 | 4 | 2 | 1 | 3 | 0 | 0 | 0 | 0 | 0 | 2 | 3 | 6 | 2 | 1 | 3 | 3 | 1 | Evi2 |
| Evi2a | 29 | 32 | 27 | 20 | 32 | 35 | 48 | 70 | 69 | 44 | 79 | 54 | 61 | 54 | 47 | 50 | 24 | 32 | 26 | 53 | 36 | 74 | 96 | 73 | 38 | 84 | 45 | 71 | Evi2a |
| Evi5 | 6202 | 4867 | 4630 | 4638 | 5240 | 5294 | 5396 | 6539 | 6112 | 5487 | 6242 | 5479 | 6424 | 5852 | 6129 | 5842 | 4624 | 4978 | 4521 | 5281 | 4979 | 5631 | 6090 | 5087 | 5502 | 4885 | 5671 | 5677 | Evi5 |
| Evi5l | 7288 | 8696 | 8352 | 10674 | 11241 | 10070 | 9997 | 5407 | 6114 | 7408 | 6713 | 6469 | 5955 | 6150 | 7117 | 7531 | 9608 | 10634 | 10926 | 8832 | 10324 | 8245 | 5685 | 6992 | 7676 | 5045 | 5459 | 5806 | Evi5l |
| Evl | 10142 | 11872 | 14742 | 17262 | 17652 | 15150 | 15036 | 12643 | 13716 | 19588 | 15537 | 17623 | 13375 | 17150 | 9945 | 9852 | 16472 | 16512 | 18561 | 13171 | 17567 | 17794 | 13652 | 19087 | 19184 | 18304 | 14162 | 15612 | Evl |
| Evpl | 208 | 244 | 220 | 140 | 111 | 222 | 89 | 325 | 517 | 313 | 262 | 176 | 239 | 165 | 167 | 239 | 117 | 198 | 98 | 76 | 75 | 125 | 562 | 354 | 89 | 142 | 216 | 225 | Evpl |
| Evx1 | 13 | 73 | 22 | 66 | 137 | 94 | 15 | 228 | 185 | 202 | 210 | 56 | 456 | 33 | 80 | 90 | 75 | 42 | 64 | 54 | 33 | 234 | 159 | 47 | 114 | 0 | 38 | 36 | Evx1 |
| Evx2 | 68 | 119 | 209 | 167 | 305 | 252 | 55 | 458 | 398 | 411 | 376 | 257 | 772 | 225 | 228 | 228 | 286 | 133 | 252 | 217 | 122 | 513 | 311 | 115 | 325 | 0 | 292 | 312 | Evx2 |
| Ewsr1 | 31910 | 33022 | 33519 | 31687 | 30540 | 31992 | 31947 | 39034 | 39962 | 40178 | 40641 | 45334 | 42710 | 46486 | 30956 | 30767 | 34128 | 31002 | 32028 | 32352 | 30555 | 36639 | 38341 | 45148 | 42495 | 48807 | 46502 | 44391 | Ewsr1 |
| Exd1 | 153 | 178 | 172 | 205 | 147 | 110 | 108 | 95 | 95 | 93 | 89 | 64 | 99 | 71 | 171 | 200 | 168 | 122 | 110 | 113 | 118 | 85 | 87 | 69 | 79 | 61 | 85 | 82 | Exd1 |
| Exd2 | 5488 | 5313 | 5129 | 5794 | 5881 | 5737 | 6586 | 5334 | 4780 | 4812 | 4702 | 5167 | 4657 | 5325 | 5350 | 4594 | 5588 | 5910 | 6114 | 5990 | 6331 | 5989 | 4538 | 5225 | 5189 | 4425 | 4912 | 5035 | Exd2 |
| Exo1 | 4788 | 4292 | 3777 | 3044 | 2318 | 2773 | 3349 | 2715 | 2750 | 2489 | 2268 | 4062 | 3598 | 3946 | 3964 | 3816 | 3413 | 2670 | 3068 | 3474 | 2901 | 1659 | 2211 | 2912 | 2627 | 4370 | 4092 | 4195 | Exo1 |
| Exo5 | 2217 | 2413 | 2648 | 2303 | 2187 | 2197 | 2413 | 2499 | 2281 | 2359 | 2232 | 2640 | 2406 | 2795 | 2202 | 2197 | 2542 | 2298 | 2220 | 2334 | 2332 | 2236 | 2250 | 2484 | 2409 | 2938 | 2581 | 2762 | Exo5 |
| Exoc1 | 8506 | 7958 | 6245 | 6804 | 7616 | 8419 | 8320 | 8617 | 7810 | 6930 | 6570 | 7579 | 7274 | 7740 | 8558 | 7394 | 6639 | 6519 | 6916 | 8745 | 7870 | 7072 | 7279 | 6587 | 7665 | 6352 | 7940 | 7340 | Exoc1 |
| Exoc2 | 5173 | 4560 | 4137 | 4300 | 4948 | 4755 | 4967 | 6132 | 5820 | 5567 | 5150 | 5614 | 5033 | 5534 | 5110 | 4995 | 4233 | 4365 | 4434 | 5233 | 4929 | 5893 | 5737 | 4962 | 5858 | 4960 | 5599 | 5478 | Exoc2 |
| Exoc3 | 7605 | 7779 | 6459 | 7054 | 7690 | 8116 | 8204 | 11372 | 9884 | 9128 | 8810 | 9305 | 8539 | 9456 | 8300 | 6574 | 6837 | 7531 | 7613 | 8409 | 8223 | 10099 | 9583 | 8929 | 9961 | 8676 | 9541 | 9172 | Exoc3 |
| Exoc3l | 298 | 328 | 400 | 321 | 344 | 319 | 273 | 265 | 322 | 401 | 394 | 421 | 479 | 328 | 290 | 329 | 378 | 353 | 415 | 319 | 317 | 344 | 384 | 462 | 467 | 498 | 492 | 434 | Exoc3l |
| Exoc3l2 | 116 | 133 | 146 | 77 | 89 | 61 | 61 | 153 | 166 | 144 | 190 | 164 | 187 | 163 | 104 | 144 | 127 | 99 | 76 | 77 | 63 | 179 | 151 | 176 | 109 | 223 | 180 | 157 | Exoc3l2 |
| Exoc3l4 | 104 | 145 | 132 | 89 | 57 | 123 | 44 | 122 | 208 | 139 | 206 | 123 | 174 | 98 | 116 | 181 | 112 | 145 | 54 | 51 | 45 | 109 | 274 | 135 | 90 | 127 | 142 | 90 | Exoc3l4 |
| Exoc4 | 9862 | 10138 | 7867 | 8406 | 8779 | 9107 | 9441 | 10148 | 9249 | 8819 | 8148 | 9652 | 8468 | 9279 | 10006 | 8379 | 8255 | 8644 | 8911 | 9651 | 9628 | 9102 | 9550 | 8811 | 9211 | 8006 | 9477 | 8816 | Exoc4 |
| Exoc5 | 13547 | 10253 | 14370 | 12952 | 12523 | 13905 | 14762 | 12684 | 9887 | 10126 | 10666 | 12211 | 11691 | 13244 | 14844 | 12182 | 12543 | 12513 | 12402 | 13813 | 14006 | 12830 | 10016 | 10920 | 11959 | 11282 | 12998 | 13540 | Exoc5 |
| Exoc6 | 834 | 676 | 650 | 717 | 848 | 752 | 935 | 1446 | 1458 | 1253 | 1330 | 1191 | 1175 | 1161 | 829 | 794 | 555 | 769 | 757 | 956 | 881 | 1435 | 1478 | 1152 | 1318 | 1207 | 1176 | 1127 | Exoc6 |
| Exoc6b | 6100 | 6001 | 4187 | 4984 | 6130 | 5937 | 5519 | 7746 | 7349 | 7055 | 6368 | 6143 | 6101 | 6144 | 6407 | 5523 | 4647 | 5138 | 5584 | 6112 | 6103 | 7426 | 7268 | 6207 | 7158 | 4774 | 6335 | 5946 | Exoc6b |
| Exoc7 | 5366 | 6811 | 5992 | 6663 | 6424 | 6168 | 6127 | 5113 | 5930 | 6121 | 5629 | 6298 | 5811 | 5786 | 5338 | 5608 | 6163 | 6415 | 7057 | 5839 | 6407 | 6106 | 5812 | 6105 | 6584 | 5740 | 5828 | 5619 | Exoc7 |
| Exoc8 | 2070 | 2122 | 2047 | 2372 | 2203 | 2253 | 2289 | 1860 | 1761 | 1962 | 1819 | 1831 | 1863 | 1892 | 2131 | 2029 | 2123 | 2422 | 2255 | 1997 | 2242 | 2074 | 1835 | 1910 | 2028 | 1686 | 2006 | 1876 | Exoc8 |
| Exog | 2555 | 3029 | 2410 | 2874 | 3090 | 2915 | 2970 | 2895 | 2767 | 2684 | 2565 | 2735 | 2590 | 2389 | 2734 | 2676 | 2896 | 2999 | 3023 | 2969 | 3176 | 2874 | 2471 | 2598 | 3031 | 2786 | 2624 | 2699 | Exog |
| Exosc1 | 1893 | 1923 | 1835 | 1778 | 1483 | 1722 | 1818 | 1666 | 1564 | 1492 | 1667 | 1819 | 1639 | 1861 | 1863 | 1798 | 1773 | 1672 | 1688 | 1945 | 1661 | 1335 | 1488 | 1686 | 1715 | 2012 | 1839 | 1917 | Exosc1 |
| Exosc10 | 2772 | 3140 | 2924 | 2599 | 3130 | 2910 | 2857 | 6859 | 7210 | 6517 | 6833 | 6963 | 6389 | 6874 | 2756 | 2657 | 2734 | 2683 | 3127 | 3095 | 2851 | 6196 | 7146 | 6753 | 6819 | 7422 | 7053 | 6897 | Exosc10 |
| Exosc2 | 3294 | 3286 | 3638 | 3241 | 2586 | 2640 | 2843 | 2606 | 2658 | 2660 | 2635 | 3086 | 2919 | 3120 | 2865 | 2891 | 3301 | 2889 | 2992 | 2747 | 2985 | 2314 | 2515 | 2956 | 2928 | 3856 | 3428 | 3652 | Exosc2 |
| Exosc3 | 1437 | 1458 | 1945 | 1582 | 1213 | 1367 | 1383 | 1330 | 1213 | 1441 | 1518 | 1950 | 1683 | 1999 | 1295 | 1378 | 1663 | 1362 | 1451 | 1411 | 1326 | 1084 | 1262 | 1867 | 1620 | 2299 | 1956 | 2090 | Exosc3 |
| Exosc4 | 1211 | 1275 | 1281 | 1164 | 1204 | 1191 | 1190 | 1108 | 1245 | 1268 | 1246 | 1395 | 1440 | 1422 | 1127 | 1053 | 1233 | 1272 | 1326 | 1102 | 1180 | 1153 | 1251 | 1426 | 1395 | 1672 | 1384 | 1441 | Exosc4 |
| Exosc5 | 1132 | 1276 | 1219 | 1327 | 1229 | 1225 | 1174 | 845 | 1009 | 991 | 1024 | 1072 | 1171 | 1063 | 1017 | 1111 | 1096 | 1354 | 1275 | 1133 | 1221 | 926 | 999 | 1140 | 1096 | 1260 | 1112 | 1114 | Exosc5 |
| Exosc7 | 1677 | 1639 | 1929 | 1703 | 1471 | 1366 | 1454 | 2132 | 1987 | 1909 | 1901 | 2337 | 2045 | 2371 | 1600 | 1503 | 1811 | 1417 | 1479 | 1736 | 1383 | 1529 | 1894 | 2197 | 2138 | 2702 | 2482 | 2381 | Exosc7 |
| Exosc8 | 3743 | 3207 | 3393 | 3019 | 2507 | 2749 | 3023 | 4567 | 3793 | 3370 | 3461 | 4329 | 3915 | 4304 | 3657 | 3231 | 3125 | 2849 | 2994 | 3499 | 3079 | 3546 | 3533 | 3835 | 3649 | 5153 | 4611 | 4516 | Exosc8 |
| Exosc9 | 6121 | 5457 | 6328 | 5217 | 4257 | 4372 | 5678 | 4804 | 3876 | 3804 | 3323 | 4634 | 4236 | 4736 | 6638 | 5608 | 5567 | 5033 | 4864 | 5889 | 4918 | 3867 | 3819 | 4101 | 3807 | 5029 | 4931 | 5200 | Exosc9 |
| Exph5 | 249 | 198 | 75 | 90 | 124 | 141 | 82 | 117 | 124 | 50 | 107 | 65 | 100 | 60 | 225 | 269 | 92 | 102 | 83 | 135 | 111 | 18 | 144 | 47 | 16 | 24 | 43 | 69 | Exph5 |
| Ext1 | 7524 | 7029 | 6847 | 5564 | 5154 | 5973 | 5645 | 4468 | 4868 | 4566 | 5153 | 4890 | 5439 | 4824 | 7184 | 7530 | 6136 | 6138 | 5243 | 5724 | 5547 | 4136 | 5270 | 4533 | 4270 | 5024 | 5137 | 5395 | Ext1 |
| Ext2 | 3637 | 4256 | 3778 | 3113 | 2898 | 3202 | 2521 | 3323 | 4305 | 3734 | 4082 | 3578 | 3765 | 3353 | 3504 | 4629 | 3321 | 3246 | 2883 | 2604 | 2538 | 3093 | 4319 | 3666 | 3361 | 4031 | 3638 | 3509 | Ext2 |
| Extl1 | 747 | 1052 | 786 | 984 | 654 | 585 | 570 | 77 | 190 | 203 | 254 | 80 | 96 | 50 | 632 | 1035 | 827 | 1050 | 640 | 502 | 639 | 177 | 166 | 183 | 107 | 150 | 70 | 62 | Extl1 |
| Extl2 | 3181 | 3055 | 2751 | 2876 | 3089 | 3051 | 3108 | 2071 | 1849 | 1642 | 1668 | 1850 | 1979 | 1739 | 3682 | 3157 | 2986 | 2993 | 2946 | 3603 | 2969 | 1785 | 2018 | 1655 | 1612 | 1893 | 1848 | 1907 | Extl2 |
| Extl3 | 5970 | 7069 | 8542 | 7766 | 8208 | 7137 | 6445 | 5879 | 6984 | 7870 | 7926 | 8105 | 7647 | 6889 | 5919 | 7429 | 8141 | 8139 | 8168 | 6067 | 7014 | 8081 | 6983 | 7782 | 8033 | 7884 | 7517 | 7282 | Extl3 |
| Eya1 | 1151 | 688 | 671 | 686 | 668 | 560 | 587 | 1724 | 1945 | 1945 | 2203 | 900 | 1650 | 1091 | 964 | 1157 | 542 | 549 | 568 | 574 | 513 | 1343 | 1862 | 1727 | 1014 | 2251 | 940 | 1075 | Eya1 |
| Eya2 | 976 | 851 | 813 | 713 | 511 | 491 | 292 | 1263 | 1243 | 1389 | 1109 | 671 | 1193 | 616 | 823 | 1230 | 754 | 616 | 420 | 382 | 339 | 869 | 1388 | 1424 | 627 | 1561 | 719 | 712 | Eya2 |
| Eya3 | 7759 | 7692 | 6300 | 6438 | 5411 | 5904 | 5898 | 4980 | 5285 | 5014 | 4889 | 5297 | 5243 | 5114 | 6661 | 7359 | 6438 | 6403 | 5652 | 5861 | 5737 | 4632 | 4907 | 4771 | 4742 | 5275 | 5348 | 5143 | Eya3 |
| Eya4 | 2059 | 1204 | 1169 | 1017 | 848 | 998 | 988 | 2707 | 2548 | 2695 | 3576 | 1615 | 2422 | 1727 | 1754 | 1883 | 955 | 941 | 898 | 1027 | 927 | 1434 | 2970 | 2846 | 1417 | 3688 | 1478 | 2015 | Eya4 |
| Ezh1 | 2635 | 2637 | 2220 | 2515 | 2630 | 2653 | 2417 | 2282 | 2276 | 2319 | 2430 | 1880 | 1991 | 1804 | 2432 | 2659 | 2300 | 2746 | 2456 | 2600 | 2494 | 2378 | 2471 | 2037 | 2358 | 1546 | 1877 | 1782 | Ezh1 |
| Ezh2 | 17349 | 14745 | 20017 | 17816 | 16377 | 16565 | 20398 | 19131 | 15690 | 18300 | 15880 | 24847 | 19449 | 26810 | 16780 | 13647 | 19232 | 16547 | 18749 | 17214 | 18842 | 16825 | 14436 | 21983 | 20976 | 27792 | 24054 | 26398 | Ezh2 |
| Ezhip | 2 | 0 | 0 | 3 | 5 | 3 | 4 | 4 | 3 | 1 | 4 | 2 | 0 | 3 | 4 | 0 | 2 | 3 | 3 | 9 | 4 | 8 | 6 | 3 | 6 | 0 | 7 | 7 | Ezhip |
| Ezr | 11369 | 12208 | 12883 | 11244 | 10600 | 11126 | 11103 | 6778 | 7682 | 7943 | 6122 | 9137 | 8997 | 8264 | 11182 | 12576 | 11800 | 11172 | 11633 | 10159 | 10904 | 5721 | 8636 | 8527 | 7350 | 10596 | 7800 | 8426 | Ezr |
| F10 | 20 | 18 | 30 | 23 | 29 | 11 | 22 | 13 | 23 | 5 | 28 | 49 | 40 | 18 | 25 | 27 | 25 | 19 | 15 | 20 | 32 | 15 | 31 | 22 | 13 | 57 | 28 | 14 | F10 |
| F11 | 5 | 3 | 1 | 10 | 4 | 0 | 0 | 0 | 2 | 5 | 0 | 1 | 3 | 0 | 4 | 2 | 5 | 2 | 3 | 1 | 3 | 0 | 2 | 0 | 1 | 2 | 2 | 0 | F11 |
| F11r | 1471 | 1112 | 898 | 914 | 862 | 801 | 556 | 811 | 841 | 766 | 839 | 674 | 816 | 546 | 1036 | 1426 | 807 | 858 | 664 | 784 | 566 | 667 | 1077 | 743 | 514 | 983 | 605 | 641 | F11r |
| F12 | 2 | 3 | 0 | 4 | 6 | 1 | 13 | 0 | 6 | 5 | 6 | 0 | 1 | 1 | 3 | 2 | 7 | 8 | 9 | 9 | 3 | 3 | 1 | 3 | 6 | 7 | 5 | 6 | F12 |
| F13a1 | 1685 | 1558 | 1679 | 1252 | 1131 | 1479 | 898 | 1746 | 2146 | 1310 | 2157 | 1940 | 2253 | 1908 | 1936 | 2026 | 1218 | 1379 | 933 | 1659 | 1154 | 1861 | 3131 | 1729 | 1619 | 1698 | 2260 | 2079 | F13a1 |
| F13b | 2 | 3 | 7 | 4 | 5 | 10 | 1 | 0 | 0 | 0 | 4 | 2 | 2 | 2 | 8 | 2 | 3 | 8 | 2 | 2 | 7 | 0 | 0 | 0 | 1 | 2 | 3 | 4 | F13b |
| F2 | 3 | 16 | 7 | 6 | 2 | 7 | 6 | 1 | 25 | 7 | 28 | 4 | 0 | 10 | 3 | 9 | 1 | 8 | 10 | 5 | 9 | 3 | 39 | 0 | 2 | 1 | 3 | 5 | F2 |
| F2r | 8545 | 7463 | 8062 | 6448 | 7575 | 7279 | 7458 | 8497 | 8173 | 7420 | 6747 | 7835 | 7951 | 8477 | 8168 | 7699 | 7457 | 6663 | 6995 | 8011 | 6887 | 7197 | 7598 | 7440 | 7293 | 8489 | 8307 | 8769 | F2r |
| F2rl1 | 407 | 239 | 243 | 231 | 115 | 143 | 112 | 120 | 141 | 99 | 197 | 21 | 40 | 26 | 282 | 326 | 160 | 136 | 108 | 109 | 93 | 43 | 215 | 85 | 17 | 124 | 36 | 36 | F2rl1 |
| F2rl2 | 36 | 26 | 33 | 9 | 11 | 31 | 13 | 50 | 26 | 20 | 35 | 37 | 66 | 40 | 34 | 26 | 28 | 21 | 12 | 26 | 15 | 37 | 36 | 34 | 26 | 104 | 89 | 32 | F2rl2 |
| F2rl3 | 91 | 80 | 47 | 85 | 52 | 66 | 69 | 39 | 41 | 47 | 53 | 46 | 44 | 38 | 54 | 61 | 58 | 64 | 52 | 49 | 64 | 63 | 45 | 40 | 35 | 53 | 43 | 37 | F2rl3 |
| F3 | 379 | 402 | 377 | 340 | 290 | 255 | 289 | 380 | 427 | 314 | 356 | 315 | 219 | 307 | 371 | 414 | 340 | 361 | 253 | 255 | 280 | 269 | 469 | 354 | 293 | 413 | 218 | 250 | F3 |
| F5 | 103 | 117 | 96 | 40 | 90 | 83 | 54 | 98 | 73 | 58 | 80 | 85 | 155 | 68 | 111 | 146 | 36 | 57 | 72 | 89 | 70 | 72 | 126 | 73 | 62 | 73 | 105 | 76 | F5 |
| F7 | 14 | 16 | 8 | 31 | 25 | 12 | 16 | 13 | 21 | 14 | 28 | 14 | 31 | 26 | 11 | 18 | 13 | 13 | 33 | 28 | 25 | 20 | 21 | 20 | 20 | 8 | 19 | 26 | F7 |
| F8 | 153 | 124 | 68 | 107 | 140 | 140 | 99 | 325 | 380 | 243 | 274 | 255 | 265 | 253 | 149 | 189 | 90 | 56 | 64 | 168 | 127 | 216 | 322 | 176 | 231 | 157 | 387 | 248 | F8 |

| | | | | | | | | | | | | | | | | | | | | | | | | | | | | | |
|---|---|---|---|---|---|---|---|---|---|---|---|---|---|---|---|---|---|---|---|---|---|---|---|---|---|---|---|---|---|
| F830016B08Rik | 14 | 3 | 0 | 0 | 4 | 4 | 1 | 9 | 12 | 11 | 21 | 9 | 1 | 4 | 8 | 13 | 0 | 3 | 5 | 9 | 6 | 2 | 26 | 11 | 2 | 1 | 2 | 6 | F830016B08Rik |
| F830045P16Rik | 0 | 0 | 0 | 0 | 0 | 0 | 0 | 2 | 2 | 0 | 1 | 0 | 2 | 0 | 0 | 0 | 0 | 0 | 0 | 1 | 0 | 0 | 5 | 2 | 0 | 1 | 0 | 0 | F830045P16Rik |
| F8a | 512 | 605 | 476 | 616 | 646 | 578 | 506 | 722 | 636 | 574 | 593 | 626 | 540 | 644 | 550 | 378 | 618 | 546 | 620 | 654 | 613 | 810 | 695 | 651 | 711 | 594 | 599 | 628 | F8a |
| F9 | 0 | 0 | 0 | 0 | 1 | 1 | 1 | 1 | 2 | 0 | 0 | 0 | 7 | 0 | 4 | 0 | 2 | 4 | 0 | 0 | 3 | 2 | 1 | 0 | 2 | 0 | 1 | 1 | F9 |
| Fa2h | 17 | 20 | 15 | 15 | 24 | 6 | 3 | 0 | 13 | 14 | 6 | 3 | 1 | 2 | 17 | 19 | 7 | 12 | 8 | 9 | 4 | 17 | 8 | 2 | 10 | 3 | 0 | 1 | Fa2h |
| Faah | 456 | 451 | 489 | 571 | 680 | 527 | 434 | 622 | 594 | 711 | 664 | 424 | 489 | 408 | 427 | 382 | 474 | 509 | 579 | 533 | 511 | 774 | 590 | 558 | 663 | 466 | 331 | 360 | Faah |
| Faap100 | 2096 | 2973 | 2412 | 3133 | 3349 | 2889 | 2746 | 2324 | 3015 | 3174 | 3062 | 3116 | 2807 | 2452 | 1990 | 2516 | 3153 | 3458 | 3419 | 2507 | 3028 | 3578 | 2926 | 3114 | 3330 | 2964 | 2652 | 2593 | Faap100 |
| Faap20 | 1047 | 1392 | 1236 | 1096 | 1203 | 1142 | 1079 | 1481 | 1812 | 1724 | 1723 | 2032 | 1947 | 1916 | 1172 | 1216 | 1329 | 1033 | 1247 | 1122 | 1002 | 1688 | 1779 | 2084 | 1773 | 2276 | 2206 | 1997 | Faap20 |
| Faap24 | 550 | 728 | 735 | 641 | 539 | 546 | 682 | 415 | 519 | 610 | 499 | 648 | 586 | 668 | 554 | 576 | 673 | 622 | 644 | 619 | 606 | 533 | 509 | 609 | 582 | 656 | 540 | 714 | Faap24 |
| Fabp1 | 1 | 0 | 0 | 0 | 0 | 0 | 0 | 0 | 1 | 0 | 0 | 0 | 0 | 0 | 0 | 0 | 0 | 0 | 0 | 0 | 0 | 0 | 0 | 0 | 0 | 0 | 0 | 0 | Fabp1 |
| Fabp12 | 0 | 1 | 0 | 0 | 0 | 0 | 0 | 0 | 0 | 0 | 1 | 0 | 0 | 2 | 7 | 1 | 0 | 0 | 2 | 0 | 0 | 0 | 0 | 0 | 0 | 0 | 0 | 0 | Fabp12 |
| Fabp2 | 2 | 0 | 0 | 0 | 0 | 0 | 0 | 1 | 0 | 0 | 0 | 0 | 0 | 0 | 3 | 0 | 0 | 0 | 0 | 0 | 0 | 0 | 1 | 0 | 0 | 0 | 0 | 0 | Fabp2 |
| Fabp3 | 315 | 448 | 376 | 557 | 830 | 611 | 686 | 813 | 880 | 893 | 935 | 468 | 606 | 598 | 497 | 409 | 520 | 627 | 662 | 798 | 740 | 916 | 949 | 690 | 923 | 408 | 572 | 517 | Fabp3 |
| Fabp4 | 12 | 6 | 6 | 4 | 18 | 23 | 0 | 16 | 30 | 20 | 89 | 8 | 10 | 22 | 11 | 12 | 16 | 10 | 3 | 6 | 3 | 42 | 104 | 14 | 58 | 8 | 7 | 7 | Fabp4 |
| Fabp5 | 8973 | 9902 | 9455 | 8921 | 9826 | 9448 | 11260 | 15382 | 13319 | 12679 | 11575 | 13739 | 13797 | 17981 | 10566 | 8413 | 9739 | 8382 | 10236 | 12608 | 10973 | 12617 | 13780 | 14101 | 15141 | 15409 | 14985 | 17576 | Fabp5 |
| Fabp6 | 0 | 0 | 0 | 1 | 0 | 0 | 1 | 0 | 0 | 0 | 0 | 0 | 0 | 0 | 0 | 0 | 2 | 0 | 0 | 1 | 1 | 0 | 0 | 0 | 0 | 0 | 0 | 0 | Fabp6 |
| Fabp7 | 13848 | 15034 | 18292 | 19145 | 26685 | 20151 | 24334 | 34377 | 24822 | 27402 | 22046 | 26079 | 27525 | 30060 | 23241 | 15990 | 20002 | 19979 | 22078 | 27037 | 23920 | 31885 | 27356 | 25759 | 29274 | 25078 | 26435 | 30787 | Fabp7 |
| Fadd | 1096 | 1102 | 1105 | 978 | 862 | 922 | 944 | 730 | 891 | 811 | 854 | 987 | 1000 | 1095 | 893 | 917 | 1079 | 869 | 904 | 795 | 913 | 744 | 801 | 980 | 837 | 1166 | 1025 | 993 | Fadd |
| Fads1 | 12547 | 13160 | 14639 | 13612 | 14280 | 13486 | 14549 | 12438 | 11973 | 13970 | 12221 | 16457 | 14982 | 16674 | 12531 | 12375 | 14366 | 13528 | 14860 | 11867 | 14386 | 15002 | 11622 | 14787 | 15040 | 17821 | 15706 | 18041 | Fads1 |
| Fads2 | 8966 | 10902 | 12595 | 10177 | 11154 | 10960 | 10150 | 14332 | 13876 | 15769 | 13102 | 19668 | 17585 | 18269 | 9361 | 9029 | 11375 | 9700 | 11853 | 9459 | 10284 | 16665 | 13023 | 16679 | 17485 | 20719 | 20296 | 19944 | Fads2 |
| Fads2b | 18 | 11 | 3 | 0 | 10 | 6 | 6 | 29 | 40 | 35 | 19 | 2 | 5 | 4 | 7 | 8 | 6 | 4 | 12 | 1 | 5 | 7 | 45 | 18 | 1 | 4 | 6 | 0 | Fads2b |
| Fads3 | 1127 | 1268 | 1164 | 1020 | 1301 | 1186 | 884 | 998 | 1298 | 1130 | 1270 | 784 | 1055 | 741 | 1292 | 1605 | 1002 | 1188 | 955 | 794 | 984 | 1365 | 1485 | 865 | 1074 | 789 | 745 | 752 | Fads3 |
| Fads6 | 125 | 150 | 103 | 115 | 228 | 199 | 139 | 182 | 225 | 188 | 226 | 153 | 193 | 159 | 125 | 159 | 176 | 205 | 188 | 143 | 136 | 251 | 181 | 164 | 218 | 219 | 144 | 170 | Fads6 |
| Faf1 | 6369 | 5897 | 5831 | 5089 | 5366 | 5517 | 6031 | 6451 | 5706 | 5071 | 4749 | 5495 | 4966 | 5380 | 6596 | 5713 | 5522 | 5330 | 5712 | 6019 | 5667 | 5743 | 5398 | 4913 | 5175 | 5413 | 5463 | 5974 | Faf1 |
| Faf2 | 5012 | 4964 | 5311 | 4979 | 4763 | 5111 | 4906 | 5854 | 5988 | 6220 | 6234 | 6570 | 6405 | 6635 | 4838 | 4662 | 5223 | 4875 | 5002 | 4428 | 5200 | 6156 | 6026 | 6416 | 6248 | 6476 | 6365 | 6679 | Faf2 |
| Fah | 326 | 385 | 349 | 326 | 383 | 441 | 273 | 286 | 370 | 315 | 257 | 293 | 319 | 292 | 334 | 390 | 289 | 344 | 303 | 343 | 368 | 351 | 349 | 254 | 283 | 200 | 325 | 274 | Fah |
| Fahd1 | 599 | 569 | 711 | 532 | 538 | 578 | 634 | 867 | 779 | 781 | 738 | 835 | 904 | 958 | 665 | 566 | 699 | 565 | 553 | 659 | 569 | 698 | 851 | 848 | 696 | 984 | 890 | 1034 | Fahd1 |
| Fahd2a | 653 | 607 | 552 | 570 | 699 | 672 | 653 | 728 | 859 | 730 | 733 | 887 | 847 | 744 | 734 | 621 | 567 | 622 | 609 | 675 | 584 | 799 | 777 | 758 | 871 | 849 | 820 | 813 | Fahd2a |
| Faim | 3325 | 3054 | 3069 | 3429 | 2963 | 2931 | 3197 | 2003 | 1945 | 2149 | 1992 | 2287 | 2214 | 2431 | 3144 | 3049 | 3212 | 3346 | 2834 | 3103 | 3373 | 1918 | 1865 | 2207 | 2314 | 2315 | 2359 | 2517 | Faim |
| Faim2 | 225 | 338 | 282 | 467 | 875 | 662 | 658 | 352 | 399 | 468 | 411 | 243 | 209 | 214 | 319 | 311 | 458 | 607 | 778 | 673 | 802 | 662 | 405 | 297 | 399 | 178 | 180 | 182 | Faim2 |
| Faiml | 0 | 0 | 0 | 0 | 0 | 0 | 2 | 0 | 0 | 0 | 0 | 0 | 0 | 0 | 0 | 0 | 0 | 0 | 0 | 0 | 0 | 0 | 0 | 0 | 0 | 0 | 0 | 0 | Faiml |
| Fam102a | 2630 | 3278 | 2927 | 3106 | 2665 | 2478 | 2282 | 1557 | 1907 | 1944 | 1985 | 1757 | 1957 | 1642 | 2223 | 3162 | 2931 | 3323 | 2670 | 2029 | 2332 | 1935 | 1750 | 1856 | 1619 | 2026 | 1550 | 1696 | Fam102a |
| Fam102b | 7127 | 6013 | 7257 | 7601 | 9697 | 8591 | 8872 | 5754 | 5494 | 5771 | 6370 | 4730 | 5159 | 4683 | 7913 | 6981 | 7409 | 8548 | 7848 | 7725 | 8737 | 6769 | 5741 | 4749 | 5650 | 3482 | 4915 | 4810 | Fam102b |
| Fam104a | 3351 | 3681 | 3620 | 3696 | 3014 | 3366 | 3462 | 2144 | 2159 | 2210 | 2205 | 2512 | 2275 | 2657 | 3377 | 3116 | 3546 | 3350 | 3511 | 3360 | 3398 | 2164 | 2235 | 2546 | 2391 | 2760 | 2388 | 2633 | Fam104a |
| Fam107a | 223 | 116 | 65 | 137 | 72 | 54 | 46 | 473 | 382 | 503 | 448 | 172 | 270 | 165 | 178 | 178 | 76 | 66 | 55 | 58 | 52 | 187 | 496 | 529 | 150 | 708 | 93 | 173 | Fam107a |
| Fam107b | 8033 | 6668 | 6646 | 6492 | 6454 | 6762 | 7082 | 5898 | 4876 | 4668 | 4615 | 5207 | 5049 | 5443 | 7834 | 7244 | 6306 | 6688 | 6604 | 6787 | 6774 | 5152 | 4829 | 5079 | 5280 | 5576 | 5173 | 5400 | Fam107b |
| Fam110a | 1701 | 2431 | 2873 | 2820 | 2375 | 2072 | 1973 | 1388 | 1637 | 1971 | 1630 | 2817 | 2226 | 2205 | 1896 | 2088 | 3202 | 2523 | 2839 | 1928 | 2169 | 1927 | 1529 | 2366 | 2178 | 2865 | 2494 | 2593 | Fam110a |
| Fam110b | 1696 | 1774 | 1630 | 2393 | 2844 | 2282 | 2511 | 2164 | 2374 | 2364 | 2345 | 2123 | 1801 | 1950 | 1762 | 1513 | 2055 | 2203 | 2685 | 2179 | 2634 | 2458 | 2287 | 2460 | 2495 | 2180 | 1833 | 1928 | Fam110b |
| Fam110c | 87 | 24 | 15 | 14 | 16 | 29 | 17 | 9 | 20 | 12 | 16 | 3 | 3 | 3 | 55 | 46 | 25 | 11 | 15 | 12 | 9 | 2 | 28 | 5 | 3 | 8 | 6 | 2 | Fam110c |
| Fam110d | 156 | 194 | 176 | 107 | 125 | 133 | 97 | 129 | 180 | 175 | 170 | 130 | 147 | 163 | 137 | 216 | 194 | 142 | 145 | 164 | 104 | 167 | 216 | 154 | 187 | 204 | 186 | 116 | Fam110d |
| Fam111a | 6774 | 5854 | 6453 | 4823 | 3880 | 4143 | 4636 | 3483 | 3470 | 2816 | 3168 | 3632 | 3661 | 3374 | 7338 | 6743 | 5569 | 4792 | 3858 | 5726 | 4131 | 2832 | 3505 | 3064 | 2848 | 4289 | 3905 | 3846 | Fam111a |
| Fam114a1 | 1574 | 1484 | 1453 | 954 | 721 | 1320 | 581 | 1642 | 2363 | 1716 | 2987 | 1455 | 2125 | 1474 | 1706 | 1988 | 1109 | 1165 | 592 | 701 | 525 | 1995 | 3141 | 1631 | 1606 | 1703 | 1879 | 1663 | Fam114a1 |
| Fam114a2 | 5230 | 5162 | 5005 | 5398 | 5434 | 5162 | 5742 | 6962 | 5909 | 6239 | 5876 | 5950 | 5971 | 6391 | 5640 | 4668 | 5532 | 5510 | 5487 | 5659 | 5611 | 6545 | 5998 | 6464 | 6235 | 6307 | 6236 | 6271 | Fam114a2 |
| Fam117a | 2577 | 2858 | 2217 | 2011 | 1289 | 1681 | 1431 | 568 | 770 | 620 | 776 | 707 | 1088 | 617 | 2080 | 2732 | 2062 | 1942 | 1554 | 1477 | 1483 | 481 | 849 | 672 | 483 | 891 | 700 | 881 | Fam117a |
| Fam117b | 7951 | 7092 | 9027 | 11828 | 13689 | 12159 | 13000 | 8348 | 7556 | 9643 | 8500 | 7708 | 7389 | 8052 | 8517 | 7782 | 10476 | 12661 | 12806 | 10346 | 12964 | 11658 | 7270 | 8342 | 10292 | 6501 | 7248 | 7361 | Fam117b |
| Fam118a | 2550 | 2362 | 2892 | 2320 | 2095 | 2029 | 2445 | 1369 | 1388 | 1507 | 1543 | 1637 | 1669 | 1442 | 2625 | 2438 | 2349 | 2432 | 2205 | 2336 | 2373 | 1464 | 1558 | 1550 | 1527 | 1780 | 1598 | 1722 | Fam118a |
| Fam118b | 4615 | 4218 | 3974 | 3895 | 3414 | 3879 | 4478 | 3460 | 2781 | 2693 | 2401 | 3078 | 2596 | 3168 | 4912 | 4058 | 3771 | 3916 | 3818 | 4527 | 4162 | 2769 | 2804 | 2910 | 2952 | 3332 | 2888 | 3294 | Fam118b |
| Fam120a | 14282 | 13205 | 15336 | 14273 | 14715 | 15027 | 15090 | 12324 | 13748 | 13369 | 15357 | 14438 | 14123 | 14352 | 13332 | 14620 | 14504 | 15207 | 14664 | 12608 | 14519 | 13059 | 13819 | 14468 | 14288 | 14856 | 13175 | 14182 | Fam120a |
| Fam120b | 7393 | 8000 | 5654 | 6631 | 6509 | 6776 | 6767 | 6324 | 6214 | 5297 | 5495 | 5197 | 5281 | 5216 | 8043 | 6873 | 6442 | 6528 | 6470 | 7511 | 7117 | 5760 | 6175 | 5033 | 5821 | 4983 | 5431 | 5283 | Fam120b |
| Fam120c | 2234 | 2086 | 1393 | 2408 | 2693 | 2879 | 2958 | 2002 | 1954 | 2053 | 1967 | 1689 | 1634 | 1762 | 2292 | 2003 | 1795 | 2609 | 2644 | 2437 | 3091 | 1932 | 1888 | 1497 | 2127 | 720 | 1576 | 1339 | Fam120c |
| Fam122a | 652 | 742 | 743 | 808 | 659 | 680 | 660 | 569 | 662 | 710 | 691 | 739 | 665 | 647 | 644 | 710 | 906 | 771 | 761 | 573 | 642 | 650 | 695 | 695 | 669 | 881 | 680 | 659 | Fam122a |
| Fam122b | 2591 | 1907 | 2335 | 1956 | 1734 | 1904 | 2078 | 1316 | 1361 | 1248 | 1507 | 1417 | 1353 | 1465 | 2380 | 2232 | 2084 | 2004 | 1813 | 2017 | 1851 | 1096 | 1386 | 1271 | 1097 | 1507 | 1514 | 1447 | Fam122b |
| Fam122c | 1 | 0 | 0 | 0 | 0 | 0 | 0 | 0 | 0 | 8 | 0 | 0 | 0 | 0 | 0 | 0 | 0 | 0 | 0 | 0 | 0 | 0 | 1 | 0 | 0 | 1 | 0 | 0 | Fam122c |
| Fam124a | 1016 | 1198 | 812 | 1640 | 2624 | 1999 | 2320 | 1363 | 1532 | 1707 | 1428 | 1466 | 1078 | 1457 | 1172 | 808 | 1423 | 1903 | 2312 | 2072 | 2318 | 1589 | 1404 | 1435 | 2026 | 1428 | 1105 | 1245 | Fam124a |
| Fam124b | 31 | 22 | 30 | 19 | 39 | 24 | 28 | 43 | 75 | 82 | 95 | 68 | 83 | 61 | 33 | 36 | 42 | 36 | 9 | 27 | 30 | 58 | 80 | 55 | 80 | 34 | 69 | 49 | Fam124b |
| Fam126a | 15156 | 14697 | 11374 | 15994 | 16531 | 16997 | 18035 | 10150 | 9198 | 8916 | 8518 | 9546 | 8679 | 9832 | 15862 | 13968 | 14073 | 16364 | 16572 | 17534 | 18068 | 8761 | 8635 | 8856 | 10156 | 7534 | 8956 | 8727 | Fam126a |
| Fam126b | 3834 | 2794 | 2175 | 3147 | 3144 | 3809 | 3933 | 3294 | 3019 | 2517 | 2776 | 2407 | 2376 | 2487 | 3941 | 3548 | 2538 | 3510 | 2972 | 3918 | 3705 | 2398 | 2933 | 2102 | 2753 | 1464 | 2434 | 2370 | Fam126b |
| Fam131a | 871 | 1386 | 1090 | 1387 | 1409 | 1338 | 1412 | 1087 | 1239 | 1371 | 1093 | 1149 | 951 | 1139 | 952 | 1012 | 1416 | 1539 | 1627 | 1390 | 1432 | 1190 | 1234 | 1363 | 1491 | 1162 | 1084 | 1048 | Fam131a |
| Fam131b | 3298 | 3645 | 4399 | 4531 | 5317 | 4524 | 4574 | 3149 | 3027 | 4014 | 3465 | 3307 | 3217 | 3655 | 3435 | 3302 | 4464 | 4273 | 5195 | 3879 | 4763 | 4067 | 2801 | 3568 | 4139 | 3037 | 3046 | 3401 | Fam131b |
| Fam131c | 187 | 262 | 282 | 247 | 316 | 251 | 177 | 99 | 139 | 194 | 152 | 89 | 91 | 72 | 166 | 245 | 257 | 219 | 268 | 208 | 239 | 256 | 169 | 133 | 206 | 72 | 59 | 80 | Fam131c |
| Fam133b | 4271 | 3604 | 4825 | 3923 | 3565 | 3922 | 4187 | 4929 | 3952 | 3914 | 3903 | 4397 | 4299 | 5250 | 4396 | 3854 | 4361 | 3568 | 3680 | 4397 | 4034 | 3778 | 3903 | 4738 | 4287 | 5537 | 5067 | 5276 | Fam133b |
| Fam135a | 13133 | 9843 | 8548 | 9710 | 9943 | 11356 | 12205 | 10032 | 8874 | 6989 | 7472 | 7835 | 8578 | 8574 | 13386 | 11411 | 8741 | 9230 | 9686 | 12397 | 11792 | 7623 | 8347 | 6818 | 7638 | 6538 | 8488 | 7875 | Fam135a |
| Fam135b | 115 | 114 | 48 | 186 | 466 | 386 | 445 | 517 | 605 | 494 | 481 | 217 | 215 | 353 | 161 | 103 | 111 | 274 | 301 | 451 | 502 | 550 | 528 | 291 | 547 | 174 | 203 | 179 | Fam135b |
| Fam136a | 4636 | 4028 | 4528 | 3650 | 3537 | 3535 | 4113 | 4253 | 3880 | 3642 | 3526 | 4145 | 3760 | 4093 | 4706 | 4069 | 4366 | 3802 | 4023 | 4555 | 3841 | 3506 | 4150 | 4037 | 3614 | 4743 | 4180 | 4634 | Fam136a |
| Fam13a | 1495 | 1386 | 1312 | 1477 | 1591 | 1532 | 1633 | 816 | 838 | 690 | 989 | 696 | 780 | 789 | 1514 | 1498 | 1260 | 1675 | 1496 | 1585 | 1612 | 941 | 909 | 758 | 934 | 640 | 744 | 662 | Fam13a |
| Fam13b | 6014 | 4988 | 4311 | 5617 | 6487 | 6291 | 7036 | 7573 | 6690 | 7016 | 6983 | 7194 | 6769 | 7625 | 5855 | 5108 | 5299 | 5749 | 5919 | 6016 | 6957 | 7630 | 6366 | 6493 | 7603 | 4768 | 7284 | 6886 | Fam13b |
| Fam13c | 1239 | 1176 | 1582 | 1305 | 1892 | 1572 | 1573 | 2470 | 2204 | 2471 | 2130 | 2518 | 2384 | 2481 | 1456 | 1206 | 1365 | 1238 | 1533 | 1813 | 1555 | 2677 | 2208 | 2274 | 2597 | 2055 | 2357 | 2339 | Fam13c |
| Fam149a | 773 | 728 | 707 | 555 | 635 | 645 | 507 | 1028 | 1190 | 989 | 1310 | 980 | 1089 | 804 | 762 | 809 | 590 | 611 | 549 | 600 | 483 | 981 | 1233 | 892 | 813 | 1095 | 932 | 924 | Fam149a |
| Fam149b | 2761 | 2858 | 2675 | 2537 | 2549 | 2784 | 2851 | 4501 | 4128 | 4322 | 3954 | 4438 | 4152 | 4617 | 2854 | 2552 | 2459 | 2363 | 2562 | 2854 | 2837 | 3974 | 3970 | 4207 | 4089 | 4210 | 4464 | 4628 | Fam149b |
| Fam151a | 1 | 0 | 0 | 0 | 0 | 5 | 0 | 0 | 3 | 0 | 0 | 0 | 3 | 0 | 0 | 0 | 0 | 0 | 0 | 0 | 0 | 0 | 1 | 1 | 3 | 1 | 3 | 0 | Fam151a |
| Fam151b | 48 | 58 | 77 | 85 | 57 | 73 | 104 | 172 | 146 | 135 | 181 | 113 | 128 | 188 | 84 | 62 | 72 | 92 | 58 | 62 | 80 | 114 | 138 | 211 | 190 | 138 | 156 | 170 | Fam151b |
| Fam155a | 737 | 774 | 868 | 1345 | 2010 | 1483 | 1656 | 1585 | 1463 | 1921 | 1520 | 1325 | 1374 | 1477 | 1036 | 710 | 1092 | 1416 | 1446 | 1616 | 1726 | 2343 | 1399 | 1565 | 2008 | 984 | 1310 | 1314 | Fam155a |
| Fam160a1 | 496 | 422 | 231 | 330 | 238 | 167 | 142 | 167 | 236 | 168 | 199 | 73 | 90 | 55 | 349 | 512 | 196 | 234 | 147 | 162 | 163 | 104 | 207 | 132 | 23 | 89 | 64 | 46 | Fam160a1 |
| Fam160a2 | 6861 | 7017 | 5902 | 7858 | 6906 | 7931 | 7867 | 4896 | 4966 | 5261 | 5338 | 5362 | 5138 | 5543 | 6357 | 7398 | 6599 | 7647 | 6719 | 6363 | 7935 | 5026 | 4609 | 4843 | 5708 | 4255 | 5460 | 5229 | Fam160a2 |
| Fam160b1 | 3862 | 3935 | 3787 | 4187 | 4327 | 4206 | 4291 | 4139 | 4273 | 4303 | 4384 | 4125 | 4160 | 4127 | 3719 | 3886 | 3978 | 4473 | 4204 | 3976 | 4318 | 4426 | 4039 | 3948 | 4261 | 3345 | 4228 | 3983 | Fam160b1 |
| Fam160b2 | 1857 | 2079 | 2700 | 2375 | 2734 | 2246 | 2118 | 1865 | 2184 | 2690 | 2354 | 2681 | 2758 | 2281 | 1655 | 2003 | 2958 | 2229 | 2695 | 1904 | 2085 | 2607 | 2010 | 2661 | 2449 | 2658 | 2494 | 2264 | Fam160b2 |
| Fam161a | 223 | 258 | 251 | 225 | 349 | 278 | 288 | 349 | 416 | 377 | 439 | 379 | 449 | 336 | 334 | 329 | 224 | 246 | 231 | 288 | 292 | 393 | 485 | 315 | 414 | 340 | 364 | 328 | Fam161a |
| Fam161b | 1602 | 1941 | 1639 | 1991 | 1942 | 1721 | 1991 | 1446 | 1289 | 1480 | 1368 | 1355 | 1222 | 1304 | 1656 | 1694 | 1839 | 1952 | 1778 | 1898 | 1943 | 1639 | 1238 | 1329 | 1603 | 1150 | 1154 | 1296 | Fam161b |
| Fam162a | 3110 | 3044 | 3049 | 3028 | 2147 | 2259 | 2984 | 2177 | 2241 | 2343 | 1960 | 2196 | 2127 | 2572 | 2988 | 2692 | 3019 | 2582 | 2536 | 2946 | 2698 | 1564 | 2430 | 2500 | 1924 | 2796 | 2452 | 2852 | Fam162a |

| | | | | | | | | | | | | | | | | | | | | | | | | | | | | | |
|---|---|---|---|---|---|---|---|---|---|---|---|---|---|---|---|---|---|---|---|---|---|---|---|---|---|---|---|---|---|
| Fam162b | 9 | 15 | 13 | 7 | 12 | 9 | 16 | 24 | 17 | 19 | 26 | 21 | 32 | 19 | 18 | 19 | 15 | 22 | 30 | 25 | 21 | 23 | 16 | 20 | 17 | 30 | 30 | 25 | Fam162b |
| Fam163a | 191 | 213 | 426 | 299 | 365 | 313 | 175 | 442 | 279 | 511 | 431 | 646 | 762 | 614 | 244 | 259 | 295 | 283 | 358 | 238 | 334 | 683 | 454 | 458 | 573 | 258 | 738 | 722 | Fam163a |
| Fam163b | 27 | 30 | 25 | 72 | 82 | 48 | 73 | 65 | 114 | 86 | 120 | 49 | 37 | 29 | 45 | 47 | 35 | 88 | 44 | 58 | 88 | 112 | 106 | 56 | 73 | 25 | 16 | 38 | Fam163b |
| Fam166a | 2 | 6 | 7 | 0 | 2 | 2 | 1 | 0 | 8 | 2 | 7 | 2 | 3 | 10 | 0 | 1 | 6 | 0 | 2 | 2 | 1 | 3 | 5 | 6 | 3 | 17 | 9 | 9 | Fam166a |
| Fam166b | 13 | 10 | 21 | 24 | 26 | 19 | 11 | 13 | 19 | 21 | 15 | 23 | 29 | 20 | 11 | 24 | 28 | 28 | 15 | 20 | 19 | 28 | 25 | 16 | 42 | 12 | 28 | 19 | Fam166b |
| Fam166c | 5 | 3 | 1 | 0 | 2 | 5 | 6 | 7 | 2 | 12 | 5 | 5 | 6 | 0 | 1 | 9 | 1 | 3 | 1 | 0 | 2 | 5 | 14 | 3 | 2 | 1 | 0 | 0 | Fam166c |
| Fam167a | 1430 | 1621 | 1580 | 1722 | 1910 | 1784 | 2123 | 1599 | 1396 | 1539 | 1295 | 2255 | 1531 | 2090 | 1593 | 1209 | 1882 | 1787 | 2058 | 1936 | 1925 | 1828 | 1447 | 1933 | 1968 | 2304 | 1829 | 2308 | Fam167a |
| Fam167b | 20 | 21 | 34 | 11 | 12 | 15 | 6 | 48 | 40 | 40 | 69 | 27 | 39 | 12 | 13 | 15 | 12 | 13 | 7 | 21 | 9 | 52 | 64 | 37 | 23 | 40 | 21 | 31 | Fam167b |
| Fam168a | 11871 | 11419 | 9778 | 13934 | 18090 | 16197 | 17308 | 17062 | 17254 | 20448 | 18272 | 18111 | 16519 | 18407 | 11786 | 10491 | 12843 | 15298 | 16745 | 14412 | 17660 | 20610 | 15771 | 18011 | 20936 | 13482 | 16207 | 17022 | Fam168a |
| Fam168b | 20515 | 19428 | 22475 | 24213 | 26549 | 24637 | 23688 | 30229 | 30447 | 34308 | 31597 | 39679 | 34033 | 36670 | 20505 | 19090 | 24156 | 24652 | 27041 | 22524 | 24279 | 33551 | 27934 | 37049 | 39229 | 37285 | 37770 | 36555 | Fam168b |
| Fam169a | 1821 | 1430 | 1689 | 1868 | 2528 | 2190 | 2306 | 3152 | 2730 | 2683 | 2504 | 2199 | 2606 | 2298 | 2176 | 1780 | 1615 | 1804 | 1972 | 2369 | 2141 | 3641 | 2564 | 2290 | 2918 | 1845 | 2179 | 2398 | Fam169a |
| Fam169b | 51 | 35 | 18 | 26 | 3 | 4 | 2 | 98 | 63 | 120 | 115 | 17 | 29 | 20 | 36 | 48 | 2 | 12 | 5 | 5 | 5 | 8 | 111 | 109 | 23 | 126 | 17 | 13 | Fam169b |
| Fam170a | 0 | 0 | 0 | 3 | 1 | 0 | 0 | 0 | 0 | 0 | 0 | 0 | 0 | 0 | 0 | 0 | 0 | 0 | 0 | 0 | 0 | 2 | 0 | 0 | 0 | 0 | 0 | 0 | Fam170a |
| Fam170b | 0 | 0 | 2 | 0 | 0 | 0 | 1 | 0 | 0 | 0 | 0 | 0 | 0 | 0 | 0 | 0 | 0 | 2 | 0 | 0 | 1 | 0 | 0 | 0 | 0 | 0 | 0 | 0 | Fam170b |
| Fam171a1 | 2620 | 2673 | 3241 | 2911 | 4082 | 3385 | 3075 | 5747 | 5590 | 6505 | 6264 | 6467 | 6402 | 5800 | 2730 | 2886 | 3086 | 2943 | 3488 | 2902 | 3184 | 7108 | 5562 | 5490 | 6210 | 5856 | 5832 | 6141 | Fam171a1 |
| Fam171a2 | 3418 | 4415 | 3524 | 4376 | 5239 | 4347 | 4098 | 5483 | 5742 | 6444 | 5609 | 5631 | 4851 | 4688 | 3214 | 3221 | 4743 | 4410 | 5646 | 4281 | 4211 | 6985 | 5429 | 6150 | 6703 | 5463 | 4951 | 4540 | Fam171a2 |
| Fam171b | 10694 | 9341 | 7889 | 11183 | 12780 | 12104 | 13647 | 12815 | 10061 | 10408 | 9168 | 11059 | 9970 | 11467 | 11813 | 7807 | 10131 | 11574 | 12109 | 12750 | 13391 | 11814 | 8859 | 9801 | 11806 | 8786 | 10885 | 10760 | Fam171b |
| Fam172a | 4979 | 4468 | 5299 | 5023 | 4945 | 5209 | 6429 | 5206 | 4549 | 4829 | 4552 | 5862 | 4543 | 6313 | 5595 | 4433 | 5443 | 5113 | 5176 | 4912 | 5768 | 5432 | 4896 | 5112 | 5352 | 5786 | 5480 | 6425 | Fam172a |
| Fam174a | 1232 | 963 | 1151 | 1078 | 902 | 918 | 984 | 964 | 945 | 683 | 852 | 685 | 856 | 861 | 1279 | 1097 | 999 | 969 | 948 | 1156 | 1055 | 835 | 1035 | 799 | 851 | 787 | 888 | 824 | Fam174a |
| Fam174b | 1827 | 1590 | 1700 | 1515 | 1419 | 1482 | 1060 | 2087 | 2300 | 2128 | 2150 | 2011 | 3354 | 1943 | 1789 | 2101 | 1595 | 1607 | 1213 | 1259 | 1076 | 1885 | 2531 | 2065 | 1894 | 2255 | 2289 | 2187 | Fam174b |
| Fam174c | 234 | 277 | 384 | 497 | 301 | 255 | 350 | 271 | 331 | 382 | 419 | 403 | 439 | 471 | 233 | 257 | 400 | 402 | 413 | 245 | 316 | 408 | 315 | 498 | 404 | 396 | 303 | 449 | Fam174c |
| Fam177a | 50 | 38 | 28 | 35 | 36 | 28 | 25 | 17 | 31 | 20 | 10 | 14 | 18 | 18 | 37 | 42 | 35 | 22 | 38 | 26 | 47 | 18 | 23 | 12 | 23 | 8 | 19 | 12 | Fam177a |
| Fam177a2 | 428 | 273 | 284 | 562 | 474 | 497 | 561 | 282 | 223 | 192 | 228 | 205 | 242 | 228 | 406 | 429 | 340 | 495 | 408 | 417 | 518 | 378 | 256 | 225 | 305 | 204 | 235 | 197 | Fam177a2 |
| Fam178b | 8 | 12 | 12 | 1 | 5 | 12 | 4 | 7 | 16 | 20 | 5 | 7 | 14 | 7 | 5 | 9 | 8 | 8 | 5 | 0 | 7 | 7 | 14 | 16 | 3 | 21 | 9 | 2 | Fam178b |
| Fam180a | 1 | 1 | 0 | 0 | 2 | 2 | 2 | 20 | 33 | 26 | 62 | 2 | 6 | 2 | 0 | 1 | 0 | 3 | 1 | 0 | 0 | 25 | 59 | 23 | 19 | 9 | 6 | 5 | Fam180a |
| Fam181a | 444 | 539 | 847 | 424 | 327 | 370 | 260 | 398 | 371 | 436 | 290 | 846 | 725 | 789 | 429 | 383 | 610 | 267 | 428 | 487 | 316 | 384 | 471 | 497 | 473 | 705 | 1067 | 989 | Fam181a |
| Fam181b | 1517 | 1940 | 2365 | 1827 | 1640 | 1422 | 1667 | 1083 | 1008 | 1304 | 1024 | 1685 | 1474 | 1886 | 1547 | 1233 | 2030 | 1610 | 1893 | 1646 | 1681 | 1101 | 1065 | 1248 | 1212 | 1575 | 1900 | 2147 | Fam181b |
| Fam183b | 38 | 60 | 63 | 20 | 43 | 30 | 40 | 72 | 42 | 56 | 52 | 46 | 113 | 55 | 63 | 57 | 37 | 61 | 48 | 90 | 61 | 75 | 65 | 47 | 69 | 37 | 44 | 44 | Fam183b |
| Fam184a | 3473 | 3275 | 2537 | 3217 | 3603 | 3669 | 3597 | 2277 | 2074 | 2094 | 1977 | 1771 | 1997 | 1822 | 3589 | 3656 | 2716 | 3371 | 2929 | 3559 | 3611 | 2097 | 2034 | 1487 | 1893 | 1362 | 1758 | 1766 | Fam184a |
| Fam184b | 264 | 241 | 221 | 297 | 471 | 419 | 514 | 307 | 416 | 350 | 391 | 390 | 367 | 423 | 222 | 181 | 267 | 361 | 489 | 521 | 466 | 247 | 388 | 431 | 436 | 462 | 350 | 390 | Fam184b |
| Fam185a | 1012 | 879 | 937 | 899 | 878 | 787 | 895 | 600 | 611 | 605 | 584 | 440 | 553 | 529 | 951 | 922 | 903 | 922 | 819 | 779 | 856 | 587 | 617 | 565 | 543 | 522 | 546 | 519 | Fam185a |
| Fam186a | 8 | 3 | 5 | 10 | 1 | 9 | 2 | 7 | 4 | 3 | 6 | 2 | 5 | 1 | 4 | 11 | 2 | 6 | 1 | 6 | 3 | 3 | 3 | 8 | 0 | 2 | 6 | 2 | Fam186a |
| Fam186b | 20 | 25 | 32 | 14 | 29 | 32 | 11 | 22 | 21 | 25 | 16 | 12 | 16 | 12 | 16 | 16 | 21 | 17 | 25 | 15 | 14 | 20 | 19 | 9 | 31 | 11 | 7 | 12 | Fam186b |
| Fam187a | 2 | 10 | 9 | 9 | 2 | 1 | 8 | 0 | 3 | 0 | 0 | 5 | 1 | 0 | 12 | 6 | 2 | 9 | 0 | 7 | 1 | 2 | 0 | 0 | 0 | 0 | 6 | 1 | Fam187a |
| Fam187b | 35 | 22 | 41 | 39 | 25 | 33 | 25 | 50 | 31 | 43 | 31 | 33 | 37 | 16 | 28 | 21 | 48 | 17 | 18 | 28 | 11 | 12 | 38 | 36 | 26 | 47 | 19 | 18 | Fam187b |
| Fam189a1 | 490 | 551 | 580 | 942 | 1265 | 936 | 887 | 913 | 1088 | 1258 | 1185 | 951 | 849 | 868 | 572 | 546 | 818 | 974 | 969 | 802 | 921 | 1786 | 1043 | 859 | 1300 | 641 | 718 | 839 | Fam189a1 |
| Fam189a2 | 601 | 860 | 1010 | 718 | 825 | 600 | 577 | 423 | 401 | 495 | 476 | 440 | 558 | 358 | 783 | 823 | 911 | 735 | 749 | 584 | 653 | 695 | 425 | 391 | 487 | 306 | 507 | 351 | Fam189a2 |
| Fam189b | 1382 | 1928 | 2071 | 1893 | 1991 | 1538 | 1454 | 1551 | 1969 | 2282 | 2310 | 2119 | 2193 | 1803 | 1331 | 1749 | 2121 | 1919 | 1964 | 1405 | 1558 | 2373 | 2001 | 2060 | 2147 | 2320 | 1766 | 1825 | Fam189b |
| Fam193a | 4983 | 5454 | 6395 | 6717 | 7112 | 6547 | 6154 | 6642 | 7331 | 8966 | 8075 | 9142 | 7879 | 8922 | 4814 | 5116 | 6538 | 6581 | 7390 | 5076 | 6820 | 8851 | 6987 | 8657 | 8923 | 8440 | 7990 | 7992 | Fam193a |
| Fam193b | 7186 | 8637 | 9832 | 10253 | 10247 | 9376 | 8838 | 9266 | 9873 | 11783 | 10696 | 11923 | 10611 | 11111 | 6994 | 8065 | 10607 | 10116 | 10616 | 8385 | 9301 | 11122 | 9135 | 11813 | 12315 | 10837 | 11384 | 10905 | Fam193b |
| Fam199x | 5444 | 3528 | 4465 | 4137 | 3959 | 4932 | 4529 | 2947 | 2531 | 2164 | 2530 | 2828 | 2865 | 2776 | 5097 | 4635 | 3946 | 4007 | 3898 | 4898 | 4421 | 2466 | 2247 | 2339 | 2523 | 2492 | 2923 | 2815 | Fam199x |
| Fam204a | 2353 | 1939 | 2415 | 2110 | 1915 | 2080 | 2220 | 3407 | 2745 | 2645 | 2783 | 3398 | 3656 | 3540 | 2557 | 2007 | 2173 | 1897 | 2188 | 2671 | 2168 | 2792 | 2844 | 2955 | 3050 | 2764 | 3674 | 3527 | Fam204a |
| Fam205a1 | 0 | 0 | 0 | 0 | 2 | 0 | 0 | 4 | 2 | 4 | 0 | 0 | 3 | 0 | 0 | 0 | 1 | 0 | 1 | 0 | 0 | 0 | 3 | 7 | 0 | 0 | 2 | 1 | Fam205a1 |
| Fam205a2 | 7 | 3 | 2 | 6 | 5 | 4 | 2 | 0 | 1 | 2 | 1 | 0 | 0 | 0 | 4 | 3 | 2 | 2 | 5 | 2 | 3 | 0 | 3 | 0 | 1 | 0 | 1 | 0 | Fam205a2 |
| Fam205a3 | 5 | 6 | 7 | 8 | 9 | 5 | 6 | 1 | 5 | 3 | 4 | 0 | 3 | 5 | 5 | 12 | 10 | 3 | 6 | 2 | 5 | 5 | 1 | 1 | 7 | 14 | 1 | 1 | Fam205a3 |
| Fam205c | 3 | 1 | 0 | 0 | 0 | 0 | 1 | 0 | 0 | 0 | 5 | 1 | 0 | 0 | 0 | 1 | 1 | 0 | 2 | 0 | 0 | 0 | 0 | 0 | 0 | 0 | 3 | 0 | Fam205c |
| Fam207a | 2155 | 2655 | 2554 | 2655 | 2323 | 2078 | 2081 | 2055 | 2265 | 2420 | 2243 | 2499 | 2279 | 2358 | 1985 | 2300 | 2691 | 2336 | 2340 | 2080 | 2063 | 2075 | 2157 | 2549 | 2336 | 2726 | 2335 | 2346 | Fam207a |
| Fam209 | 83 | 94 | 98 | 85 | 83 | 76 | 117 | 37 | 21 | 32 | 29 | 23 | 42 | 35 | 87 | 115 | 104 | 97 | 89 | 82 | 87 | 43 | 25 | 31 | 39 | 29 | 29 | 33 | Fam209 |
| Fam20a | 99 | 115 | 109 | 64 | 132 | 86 | 53 | 165 | 185 | 199 | 215 | 166 | 213 | 161 | 79 | 143 | 102 | 113 | 75 | 67 | 87 | 177 | 216 | 171 | 231 | 228 | 222 | 163 | Fam20a |
| Fam20b | 5278 | 5005 | 5282 | 5643 | 6242 | 6203 | 6102 | 6421 | 6307 | 6948 | 6999 | 6304 | 6408 | 5995 | 5548 | 5503 | 5490 | 5925 | 6224 | 5485 | 5970 | 7630 | 6462 | 6339 | 7140 | 6012 | 6199 | 6526 | Fam20b |
| Fam20c | 1288 | 1508 | 1543 | 2135 | 3216 | 2401 | 2194 | 2365 | 2581 | 3110 | 2669 | 2410 | 2042 | 2260 | 1495 | 1408 | 1942 | 2309 | 2732 | 2156 | 2298 | 3404 | 2669 | 2719 | 3323 | 2099 | 1961 | 1863 | Fam20c |
| Fam210a | 4926 | 4000 | 2921 | 3428 | 3121 | 3844 | 3557 | 4457 | 4182 | 3309 | 3766 | 3414 | 3635 | 3727 | 4522 | 4314 | 3029 | 3185 | 2723 | 3888 | 3565 | 3218 | 4198 | 3247 | 3545 | 3023 | 4065 | 3802 | Fam210a |
| Fam210b | 5817 | 5812 | 6211 | 7595 | 7382 | 7174 | 8471 | 2081 | 2315 | 2436 | 2076 | 2134 | 2848 | 2362 | 5759 | 5695 | 6541 | 7597 | 8044 | 6183 | 8497 | 1776 | 1915 | 2602 | 2842 | 1909 | 2076 | 2559 | Fam210b |
| Fam214a | 3495 | 3227 | 2347 | 3030 | 3324 | 3793 | 3656 | 4851 | 4286 | 4141 | 3604 | 4100 | 3631 | 4170 | 3392 | 2876 | 2692 | 2989 | 3062 | 3957 | 3694 | 4065 | 4036 | 3645 | 4361 | 3136 | 4135 | 3504 | Fam214a |
| Fam214b | 3535 | 5102 | 4935 | 5365 | 4807 | 4662 | 5050 | 3405 | 3363 | 3940 | 3476 | 4783 | 4182 | 4836 | 3951 | 3780 | 5510 | 5175 | 5657 | 4449 | 5124 | 4590 | 3313 | 4350 | 4666 | 4373 | 4464 | 4961 | Fam214b |
| Fam216a | 2882 | 3109 | 2950 | 3044 | 3133 | 2992 | 3066 | 3656 | 3129 | 3130 | 2925 | 3264 | 2967 | 3274 | 3171 | 2635 | 2929 | 2714 | 3373 | 3631 | 3316 | 3158 | 3144 | 3277 | 3252 | 3618 | 3056 | 3321 | Fam216a |
| Fam216b | 6 | 5 | 1 | 7 | 6 | 5 | 2 | 9 | 8 | 10 | 6 | 3 | 5 | 2 | 15 | 7 | 1 | 1 | 5 | 1 | 3 | 8 | 7 | 0 | 5 | 1 | 0 | 2 | Fam216b |
| Fam217a | 17 | 16 | 20 | 13 | 26 | 19 | 25 | 60 | 54 | 83 | 60 | 48 | 99 | 57 | 28 | 35 | 19 | 14 | 23 | 20 | 12 | 45 | 71 | 53 | 58 | 64 | 60 | 74 | Fam217a |
| Fam217b | 1023 | 969 | 1039 | 1307 | 1613 | 1254 | 1382 | 762 | 899 | 989 | 1071 | 496 | 666 | 470 | 1021 | 1043 | 1166 | 1428 | 1294 | 1203 | 1377 | 1061 | 871 | 689 | 771 | 493 | 470 | 491 | Fam217b |
| Fam219a | 4564 | 5883 | 7037 | 8471 | 8303 | 6737 | 6297 | 3031 | 3088 | 4279 | 3609 | 4059 | 3671 | 3494 | 4779 | 5241 | 7861 | 8582 | 8219 | 5426 | 6934 | 4869 | 2895 | 4171 | 4501 | 3781 | 3412 | 3248 | Fam219a |
| Fam219b | 1054 | 1069 | 1358 | 1187 | 1298 | 1268 | 1266 | 1806 | 1850 | 2090 | 2167 | 2166 | 1897 | 2101 | 1156 | 968 | 1326 | 1143 | 1422 | 1235 | 1228 | 2055 | 1961 | 2267 | 2111 | 2137 | 2027 | 2080 | Fam219b |
| Fam220a | 35 | 45 | 40 | 49 | 47 | 47 | 49 | 42 | 59 | 71 | 70 | 60 | 64 | 44 | 37 | 54 | 43 | 15 | 26 | 66 | 38 | 33 | 52 | 50 | 59 | 44 | 69 | 25 | Fam220a |
| Fam220a | 0 | 1 | 3 | 0 | 0 | 0 | 1 | 0 | 0 | 0 | 0 | 0 | 0 | 2 | 0 | 0 | 0 | 0 | 1 | 1 | 0 | 0 | 0 | 0 | 0 | 0 | 0 | 0 | Fam220a |
| Fam221a | 977 | 1009 | 1011 | 1175 | 1063 | 1121 | 1083 | 288 | 339 | 328 | 390 | 324 | 342 | 282 | 928 | 1043 | 1055 | 998 | 987 | 1068 | 1063 | 304 | 326 | 288 | 246 | 340 | 279 | 299 | Fam221a |
| Fam221b | 108 | 139 | 123 | 137 | 133 | 97 | 133 | 51 | 61 | 48 | 71 | 49 | 39 | 32 | 138 | 118 | 103 | 121 | 124 | 110 | 112 | 58 | 56 | 47 | 50 | 40 | 48 | 43 | Fam221b |
| Fam222a | 840 | 1402 | 1338 | 1675 | 1265 | 990 | 972 | 539 | 569 | 811 | 663 | 991 | 624 | 775 | 870 | 977 | 1541 | 1630 | 1195 | 801 | 1067 | 951 | 517 | 878 | 798 | 1055 | 712 | 803 | Fam222a |
| Fam222b | 2607 | 3080 | 3180 | 3534 | 3307 | 3160 | 3139 | 2280 | 2600 | 2860 | 2495 | 3202 | 2734 | 2897 | 2749 | 2993 | 3418 | 3592 | 3716 | 2638 | 3164 | 3654 | 2273 | 3036 | 3050 | 3387 | 2628 | 2909 | Fam222b |
| Fam227a | 2212 | 2149 | 1886 | 2479 | 2532 | 2619 | 2668 | 2775 | 2179 | 2854 | 2429 | 2826 | 2819 | 2907 | 2322 | 2012 | 2303 | 2455 | 2594 | 2572 | 2805 | 3175 | 2215 | 2614 | 3115 | 2072 | 2723 | 2683 | Fam227a |
| Fam227b | 2 | 3 | 9 | 3 | 1 | 4 | 1 | 2 | 1 | 3 | 0 | 1 | 0 | 1 | 5 | 1 | 6 | 4 | 0 | 1 | 4 | 5 | 2 | 2 | 2 | 1 | 3 | 0 | Fam227b |
| Fam228a | 57 | 62 | 52 | 46 | 61 | 98 | 66 | 166 | 194 | 182 | 250 | 174 | 164 | 163 | 54 | 72 | 52 | 67 | 77 | 105 | 108 | 187 | 190 | 175 | 206 | 126 | 192 | 154 | Fam228a |
| Fam228b | 171 | 95 | 56 | 111 | 138 | 197 | 129 | 487 | 484 | 368 | 424 | 433 | 366 | 377 | 162 | 123 | 81 | 95 | 114 | 182 | 146 | 351 | 460 | 293 | 410 | 273 | 388 | 313 | Fam228b |
| Fam229a | 0 | 1 | 0 | 0 | 1 | 2 | 3 | 0 | 0 | 1 | 0 | 1 | 2 | 4 | 0 | 11 | 1 | 7 | 0 | 4 | 3 | 0 | 0 | 4 | 1 | 0 | 0 | 1 | Fam229a |
| Fam229b | 272 | 294 | 251 | 341 | 370 | 451 | 410 | 731 | 629 | 752 | 760 | 664 | 612 | 788 | 359 | 320 | 336 | 317 | 372 | 346 | 432 | 866 | 655 | 766 | 752 | 719 | 701 | 642 | Fam229b |
| Fam234a | 2354 | 2737 | 2655 | 2771 | 2575 | 2294 | 2099 | 1434 | 1889 | 1738 | 1852 | 1730 | 1914 | 1486 | 2271 | 2811 | 2715 | 2700 | 2578 | 1949 | 2205 | 1992 | 1990 | 1689 | 1536 | 2040 | 1607 | 1526 | Fam234a |
| Fam234b | 3862 | 4422 | 3976 | 5744 | 7364 | 6029 | 6424 | 5250 | 5802 | 6347 | 5502 | 5361 | 4900 | 5303 | 4460 | 3744 | 5282 | 6349 | 6883 | 5619 | 6753 | 6776 | 4889 | 5788 | 6735 | 5024 | 4602 | 4916 | Fam234b |
| Fam240a | 0 | 0 | 0 | 0 | 0 | 0 | 0 | 0 | 0 | 0 | 0 | 0 | 0 | 0 | 0 | 0 | 0 | 0 | 0 | 0 | 0 | 0 | 1 | 0 | 0 | 0 | 0 | 0 | Fam240a |
| Fam240b | 0 | 0 | 0 | 0 | 0 | 0 | 1 | 0 | 0 | 0 | 1 | 0 | 0 | 0 | 0 | 0 | 0 | 0 | 0 | 0 | 1 | 2 | 0 | 0 | 0 | 0 | 0 | 0 | Fam240b |
| Fam241a | 814 | 737 | 848 | 1073 | 1059 | 963 | 1265 | 412 | 376 | 373 | 370 | 360 | 344 | 346 | 996 | 742 | 1007 | 1141 | 923 | 1050 | 1214 | 334 | 375 | 383 | 391 | 320 | 228 | 349 | Fam241a |
| Fam241b | 420 | 471 | 456 | 767 | 887 | 660 | 872 | 1261 | 1164 | 1448 | 1136 | 1135 | 811 | 1375 | 517 | 391 | 679 | 745 | 959 | 953 | 918 | 1313 | 1235 | 1422 | 1473 | 1178 | 992 | 1061 | Fam241b |

| | | | | | | | | | | | | | | | | | | | | | | | | | | | | | |
|---|---|---|---|---|---|---|---|---|---|---|---|---|---|---|---|---|---|---|---|---|---|---|---|---|---|---|---|---|---|
| Fam243 | 0 | 0 | 4 | 0 | 7 | 5 | 2 | 6 | 21 | 8 | 6 | 21 | 11 | 2 | 0 | 4 | 0 | 1 | 2 | 7 | 4 | 17 | 29 | 6 | 11 | 0 | 1 | 4 | Fam243 |
| Fam24a | 0 | 0 | 0 | 2 | 0 | 0 | 0 | 0 | 0 | 0 | 0 | 0 | 0 | 0 | 0 | 0 | 0 | 0 | 0 | 0 | 0 | 0 | 0 | 0 | 0 | 0 | 0 | 0 | Fam24a |
| Fam24b | 18 | 17 | 15 | 47 | 24 | 12 | 23 | 7 | 7 | 5 | 1 | 21 | 7 | 16 | 20 | 14 | 20 | 17 | 16 | 20 | 23 | 12 | 3 | 8 | 11 | 20 | 13 | 13 | Fam24b |
| Fam25c | 1 | 0 | 3 | 0 | 0 | 0 | 0 | 4 | 2 | 2 | 0 | 1 | 0 | 1 | 0 | 0 | 0 | 0 | 0 | 0 | 0 | 0 | 7 | 0 | 0 | 0 | 0 | 0 | Fam25c |
| Fam32a | 6001 | 6574 | 5504 | 5859 | 5999 | 6723 | 6634 | 9235 | 8545 | 8571 | 7440 | 7954 | 7510 | 8221 | 6988 | 5585 | 6415 | 6117 | 6636 | 7858 | 6975 | 7705 | 8224 | 8482 | 9558 | 8098 | 8939 | 8325 | Fam32a |
| Fam3a | 524 | 599 | 545 | 542 | 717 | 615 | 615 | 1221 | 1369 | 1404 | 1424 | 1430 | 1391 | 1442 | 570 | 575 | 645 | 508 | 616 | 621 | 605 | 1375 | 1372 | 1462 | 1432 | 1468 | 1334 | 1358 | Fam3a |
| Fam3b | 0 | 0 | 0 | 0 | 0 | 0 | 0 | 0 | 0 | 0 | 0 | 0 | 0 | 0 | 0 | 0 | 0 | 0 | 0 | 0 | 0 | 0 | 0 | 1 | 0 | 0 | 0 | 1 | Fam3b |
| Fam3c | 2532 | 1981 | 2776 | 2308 | 2320 | 2077 | 2221 | 2648 | 2516 | 2600 | 2975 | 2274 | 2352 | 2423 | 2529 | 2442 | 2490 | 2268 | 1976 | 2058 | 1999 | 2732 | 2937 | 2670 | 2570 | 2788 | 2377 | 2534 | Fam3c |
| Fam3d | 2 | 1 | 0 | 0 | 0 | 1 | 1 | 17 | 17 | 19 | 19 | 2 | 5 | 1 | 0 | 2 | 3 | 0 | 0 | 1 | 3 | 7 | 32 | 25 | 8 | 16 | 2 | 2 | Fam3d |
| Fam43a | 708 | 974 | 1175 | 1079 | 884 | 821 | 607 | 675 | 819 | 1037 | 1272 | 868 | 1149 | 934 | 736 | 1070 | 1048 | 972 | 778 | 685 | 717 | 894 | 1043 | 1059 | 811 | 1156 | 938 | 852 | Fam43a |
| Fam43b | 37 | 54 | 40 | 83 | 176 | 113 | 95 | 34 | 42 | 51 | 31 | 12 | 20 | 14 | 37 | 55 | 45 | 124 | 97 | 107 | 166 | 58 | 36 | 34 | 38 | 2 | 3 | 11 | Fam43b |
| Fam47e | 6 | 3 | 1 | 1 | 5 | 11 | 4 | 5 | 9 | 3 | 7 | 8 | 3 | 3 | 1 | 8 | 1 | 4 | 2 | 7 | 5 | 2 | 3 | 9 | 4 | 3 | 2 | 7 | Fam47e |
| Fam50a | 3934 | 3961 | 4841 | 4351 | 3474 | 3569 | 4074 | 2409 | 2357 | 2682 | 2470 | 2856 | 2565 | 3220 | 3978 | 3731 | 4661 | 4222 | 3708 | 3584 | 4075 | 2523 | 2381 | 3023 | 2753 | 3307 | 2901 | 3330 | Fam50a |
| Fam50b | 1 | 7 | 2 | 5 | 2 | 6 | 2 | 0 | 0 | 0 | 0 | 2 | 0 | 1 | 4 | 6 | 1 | 11 | 5 | 4 | 1 | 2 | 1 | 0 | 0 | 0 | 0 | 0 | Fam50b |
| Fam53a | 2158 | 2512 | 2658 | 2531 | 2084 | 2155 | 2116 | 1462 | 1720 | 1784 | 1842 | 2120 | 1888 | 1940 | 2090 | 2213 | 2767 | 2576 | 2271 | 1847 | 2019 | 1765 | 1605 | 2053 | 1825 | 2270 | 1877 | 1971 | Fam53a |
| Fam53b | 1848 | 2366 | 2307 | 2202 | 2412 | 2161 | 2340 | 1981 | 2520 | 2527 | 2396 | 3416 | 2355 | 2971 | 1870 | 1895 | 2734 | 2194 | 2678 | 2021 | 2329 | 2374 | 2076 | 3060 | 2792 | 3286 | 2854 | 2848 | Fam53b |
| Fam53c | 8343 | 8594 | 9345 | 9395 | 7692 | 7388 | 7242 | 5295 | 5218 | 5192 | 5401 | 5533 | 5091 | 4817 | 7718 | 7997 | 8788 | 8976 | 8119 | 7110 | 7116 | 5792 | 5147 | 6123 | 5657 | 6758 | 5441 | 5355 | Fam53c |
| Fam71a | 0 | 0 | 0 | 0 | 0 | 0 | 0 | 0 | 0 | 0 | 1 | 0 | 0 | 0 | 0 | 0 | 0 | 0 | 0 | 0 | 0 | 0 | 0 | 0 | 0 | 0 | 0 | 0 | Fam71a |
| Fam71b | 0 | 0 | 0 | 4 | 0 | 0 | 0 | 0 | 2 | 2 | 0 | 2 | 1 | 1 | 0 | 0 | 0 | 0 | 1 | 0 | 0 | 0 | 1 | 0 | 0 | 0 | 0 | 1 | Fam71b |
| Fam71d | 21 | 29 | 35 | 42 | 73 | 37 | 69 | 21 | 19 | 25 | 17 | 12 | 24 | 28 | 47 | 46 | 43 | 45 | 62 | 78 | 58 | 35 | 13 | 14 | 24 | 12 | 14 | 13 | Fam71d |
| Fam71e1 | 445 | 542 | 607 | 576 | 532 | 575 | 578 | 517 | 520 | 714 | 572 | 781 | 747 | 735 | 479 | 379 | 506 | 469 | 611 | 580 | 495 | 563 | 475 | 722 | 747 | 674 | 752 | 714 | Fam71e1 |
| Fam71e2 | 0 | 1 | 0 | 0 | 0 | 0 | 2 | 0 | 0 | 0 | 0 | 0 | 0 | 0 | 1 | 2 | 0 | 0 | 1 | 0 | 1 | 3 | 0 | 0 | 0 | 0 | 2 | 0 | Fam71e2 |
| Fam71f1 | 5 | 5 | 7 | 8 | 2 | 6 | 7 | 4 | 2 | 0 | 3 | 1 | 6 | 1 | 13 | 9 | 3 | 8 | 2 | 8 | 6 | 3 | 3 | 2 | 0 | 3 | 2 | 1 | Fam71f1 |
| Fam71f2 | 85 | 76 | 68 | 96 | 88 | 71 | 88 | 60 | 51 | 67 | 46 | 41 | 39 | 38 | 87 | 82 | 87 | 74 | 94 | 86 | 78 | 43 | 60 | 48 | 56 | 40 | 41 | 42 | Fam71f2 |
| Fam72a | 496 | 467 | 460 | 434 | 265 | 311 | 343 | 309 | 251 | 247 | 206 | 310 | 279 | 351 | 443 | 410 | 437 | 287 | 352 | 385 | 343 | 190 | 208 | 337 | 287 | 348 | 393 | 363 | Fam72a |
| Fam76a | 3537 | 3335 | 3579 | 3557 | 3110 | 3353 | 3366 | 2274 | 2254 | 2299 | 2433 | 2345 | 2488 | 2599 | 3576 | 3424 | 3480 | 3420 | 2845 | 3211 | 3403 | 2453 | 2467 | 2491 | 2633 | 2721 | 2408 | 2706 | Fam76a |
| Fam76b | 6873 | 4745 | 5292 | 5732 | 5992 | 6517 | 7304 | 7286 | 6071 | 5621 | 5908 | 7124 | 6674 | 7493 | 6719 | 5766 | 5388 | 5918 | 5744 | 6869 | 6591 | 6266 | 5733 | 6199 | 6398 | 6123 | 7488 | 6952 | Fam76b |
| Fam78a | 321 | 396 | 497 | 268 | 267 | 274 | 226 | 99 | 131 | 111 | 415 | 133 | 137 | 103 | 272 | 460 | 413 | 311 | 254 | 237 | 222 | 120 | 255 | 121 | 140 | 153 | 112 | 97 | Fam78a |
| Fam78b | 1846 | 2128 | 1982 | 2926 | 4232 | 3433 | 3747 | 3267 | 2862 | 3800 | 3028 | 3294 | 3369 | 3386 | 2326 | 2151 | 2677 | 3287 | 3792 | 3112 | 3708 | 5083 | 2888 | 3302 | 4106 | 2394 | 2814 | 2980 | Fam78b |
| Fam81a | 751 | 772 | 739 | 702 | 920 | 808 | 771 | 581 | 688 | 698 | 575 | 505 | 632 | 507 | 864 | 903 | 824 | 869 | 733 | 860 | 773 | 785 | 884 | 655 | 587 | 545 | 463 | 526 | Fam81a |
| Fam81b | 27 | 20 | 23 | 22 | 54 | 14 | 28 | 50 | 52 | 54 | 60 | 80 | 77 | 24 | 40 | 44 | 12 | 19 | 26 | 45 | 42 | 92 | 115 | 51 | 27 | 29 | 23 | 33 | Fam81b |
| Fam83b | 106 | 30 | 21 | 12 | 9 | 46 | 1 | 134 | 221 | 87 | 110 | 23 | 48 | 24 | 45 | 54 | 10 | 21 | 5 | 6 | 5 | 18 | 218 | 65 | 21 | 13 | 20 | 20 | Fam83b |
| Fam83c | 25 | 8 | 11 | 1 | 0 | 3 | 1 | 24 | 40 | 25 | 1 | 1 | 9 | 5 | 8 | 8 | 0 | 6 | 2 | 1 | 0 | 0 | 41 | 4 | 1 | 0 | 2 | 6 | Fam83c |
| Fam83d | 1836 | 2128 | 2133 | 1968 | 1217 | 1466 | 1579 | 1231 | 1432 | 1288 | 1431 | 2002 | 1736 | 1905 | 1619 | 1891 | 1774 | 1630 | 1562 | 1406 | 1413 | 1018 | 1302 | 1695 | 1256 | 2285 | 1886 | 1953 | Fam83d |
| Fam83e | 51 | 82 | 83 | 77 | 55 | 45 | 53 | 35 | 43 | 55 | 37 | 51 | 49 | 39 | 49 | 85 | 101 | 53 | 48 | 61 | 44 | 27 | 30 | 38 | 37 | 43 | 37 | 37 | Fam83e |
| Fam83f | 111 | 53 | 52 | 35 | 28 | 41 | 11 | 49 | 68 | 72 | 39 | 36 | 44 | 15 | 57 | 75 | 24 | 41 | 35 | 13 | 13 | 30 | 83 | 47 | 11 | 15 | 33 | 30 | Fam83f |
| Fam83g | 261 | 301 | 298 | 253 | 200 | 176 | 172 | 87 | 68 | 56 | 49 | 50 | 42 | 51 | 233 | 315 | 290 | 198 | 194 | 132 | 148 | 45 | 88 | 85 | 27 | 45 | 44 | 47 | Fam83g |
| Fam83h | 722 | 968 | 970 | 857 | 862 | 738 | 678 | 447 | 615 | 586 | 460 | 527 | 528 | 463 | 674 | 805 | 958 | 901 | 867 | 702 | 780 | 428 | 617 | 588 | 403 | 543 | 429 | 558 | Fam83h |
| Fam89a | 74 | 97 | 81 | 59 | 91 | 46 | 25 | 59 | 60 | 58 | 122 | 13 | 59 | 27 | 80 | 161 | 76 | 92 | 48 | 39 | 33 | 82 | 72 | 24 | 31 | 21 | 17 | 18 | Fam89a |
| Fam89b | 1 | 0 | 5 | 4 | 0 | 0 | 3 | 1 | 4 | 0 | 5 | 5 | 0 | 7 | 4 | 5 | 9 | 1 | 2 | 1 | 3 | 2 | 5 | 2 | 3 | 3 | 0 | 4 | Fam89b |
| Fam8a1 | 13567 | 10221 | 13649 | 14074 | 15325 | 15395 | 17588 | 14075 | 11520 | 11996 | 11857 | 12551 | 11443 | 13417 | 14395 | 11188 | 13997 | 14206 | 15606 | 15073 | 17039 | 14813 | 11378 | 12901 | 14538 | 12184 | 12362 | 13232 | Fam8a1 |
| Fam90a1a | 0 | 0 | 0 | 0 | 1 | 0 | 0 | 0 | 0 | 0 | 0 | 0 | 0 | 0 | 0 | 0 | 0 | 0 | 0 | 0 | 0 | 0 | 0 | 0 | 0 | 0 | 0 | 0 | Fam90a1a |
| Fam90a1b | 42 | 22 | 44 | 31 | 21 | 22 | 21 | 6 | 3 | 6 | 9 | 12 | 6 | 13 | 28 | 21 | 32 | 11 | 16 | 14 | 44 | 8 | 6 | 7 | 19 | 9 | 12 | 14 | Fam90a1b |
| Fam91a1 | 1 | 1 | 0 | 0 | 0 | 1 | 0 | 0 | 0 | 1 | 0 | 1 | 0 | 0 | 1 | 0 | 0 | 0 | 2 | 0 | 0 | 0 | 1 | 2 | 0 | 1 | 0 | 0 | Fam91a1 |
| Fam98a | 6811 | 6577 | 6378 | 6605 | 6466 | 6271 | 7117 | 6262 | 5987 | 5460 | 5983 | 5566 | 5668 | 5857 | 7415 | 6426 | 6342 | 6748 | 6648 | 7071 | 6834 | 5900 | 5874 | 5760 | 6375 | 5778 | 5966 | 6030 | Fam98a |
| Fam98b | 6599 | 5684 | 7871 | 6790 | 5646 | 6045 | 7083 | 7988 | 7000 | 7701 | 7347 | 10053 | 9006 | 11455 | 6504 | 5739 | 7166 | 5630 | 6348 | 6906 | 6591 | 7483 | 6757 | 9568 | 8819 | 11142 | 10542 | 11716 | Fam98b |
| Fam98c | 1142 | 1410 | 1490 | 1673 | 1475 | 1374 | 1288 | 488 | 540 | 595 | 565 | 617 | 518 | 481 | 1195 | 1277 | 1612 | 1470 | 1556 | 1272 | 1353 | 613 | 507 | 588 | 611 | 646 | 567 | 408 | Fam98c |
| Fan1 | 1367 | 1524 | 1163 | 1153 | 1321 | 1429 | 1288 | 1293 | 1445 | 1128 | 1267 | 1415 | 1420 | 1316 | 1374 | 1398 | 1176 | 1140 | 1319 | 1596 | 1331 | 1051 | 1369 | 1136 | 1296 | 991 | 1641 | 1275 | Fan1 |
| Fanca | 4726 | 4925 | 3540 | 4000 | 3175 | 3572 | 3700 | 1934 | 2176 | 1945 | 2029 | 3133 | 2722 | 2936 | 4032 | 4630 | 3931 | 3644 | 3615 | 3458 | 3502 | 1445 | 1836 | 2371 | 2044 | 2553 | 3124 | 3068 | Fanca |
| Fancb | 926 | 811 | 757 | 675 | 680 | 656 | 935 | 894 | 875 | 726 | 695 | 970 | 891 | 953 | 919 | 893 | 715 | 767 | 684 | 827 | 773 | 682 | 801 | 799 | 623 | 1001 | 984 | 943 | Fancb |
| Fancc | 749 | 788 | 692 | 561 | 657 | 749 | 659 | 1426 | 1365 | 1260 | 1191 | 1535 | 1466 | 1435 | 831 | 764 | 616 | 555 | 695 | 639 | 721 | 1315 | 1356 | 1199 | 1253 | 1384 | 1713 | 1580 | Fancc |
| Fancd2 | 1691 | 1488 | 1157 | 928 | 839 | 1051 | 1159 | 2365 | 2076 | 1708 | 1748 | 2620 | 2288 | 2593 | 1444 | 1192 | 998 | 917 | 1092 | 1428 | 1184 | 1293 | 1774 | 1986 | 1814 | 2443 | 2875 | 2543 | Fancd2 |
| Fancd2os | 1 | 2 | 5 | 1 | 2 | 0 | 2 | 4 | 5 | 2 | 3 | 4 | 7 | 4 | 0 | 1 | 3 | 0 | 3 | 3 | 4 | 3 | 6 | 3 | 5 | 4 | 1 | 2 | Fancd2os |
| Fance | 1876 | 2230 | 2043 | 1931 | 1573 | 1763 | 1557 | 1268 | 1515 | 1325 | 1392 | 1461 | 1360 | 1291 | 1733 | 1870 | 1928 | 1794 | 1710 | 1658 | 1574 | 1173 | 1397 | 1587 | 1309 | 1434 | 1453 | 1297 | Fance |
| Fancf | 411 | 421 | 421 | 374 | 340 | 413 | 415 | 580 | 476 | 475 | 461 | 456 | 370 | 532 | 459 | 332 | 381 | 314 | 387 | 453 | 408 | 421 | 469 | 472 | 424 | 477 | 457 | 535 | Fancf |
| Fancg | 1998 | 2300 | 2477 | 2278 | 2209 | 2020 | 2204 | 1285 | 1371 | 1400 | 1416 | 1494 | 1449 | 1420 | 1937 | 1931 | 2283 | 2138 | 2338 | 2120 | 2154 | 1357 | 1307 | 1517 | 1571 | 1608 | 1575 | 1517 | Fancg |
| Fanci | 2019 | 1818 | 1736 | 1259 | 953 | 1187 | 1456 | 1987 | 1952 | 1506 | 1518 | 2468 | 2084 | 2312 | 1908 | 1597 | 1564 | 1182 | 1156 | 1460 | 1158 | 1270 | 1604 | 1777 | 1650 | 2627 | 2673 | 2397 | Fanci |
| Fancl | 909 | 709 | 927 | 662 | 574 | 687 | 784 | 1150 | 1194 | 923 | 1047 | 1290 | 1219 | 1266 | 915 | 826 | 786 | 675 | 622 | 821 | 643 | 969 | 1028 | 1109 | 1035 | 1518 | 1369 | 1387 | Fancl |
| Fancm | 9994 | 8652 | 5427 | 7074 | 6008 | 7749 | 7688 | 4108 | 3546 | 3110 | 3197 | 3861 | 3436 | 3822 | 9038 | 8698 | 5371 | 6748 | 5845 | 7939 | 7738 | 2901 | 3223 | 2898 | 3363 | 2722 | 3946 | 3597 | Fancm |
| Fank1 | 245 | 275 | 227 | 320 | 251 | 279 | 295 | 175 | 131 | 184 | 171 | 173 | 151 | 121 | 284 | 252 | 280 | 331 | 267 | 338 | 296 | 179 | 178 | 154 | 176 | 122 | 98 | 100 | Fank1 |
| Fap | 403 | 147 | 226 | 129 | 86 | 116 | 47 | 460 | 653 | 490 | 1198 | 202 | 347 | 236 | 252 | 407 | 111 | 131 | 71 | 87 | 50 | 322 | 1097 | 379 | 205 | 476 | 228 | 218 | Fap |
| Far1 | 20199 | 13993 | 15110 | 15708 | 15161 | 16897 | 19027 | 16549 | 13556 | 11278 | 13475 | 12195 | 12951 | 13593 | 20596 | 18218 | 14753 | 16205 | 14449 | 19130 | 17463 | 12632 | 13331 | 11991 | 12078 | 11562 | 12766 | 13128 | Far1 |
| Far2 | 66 | 52 | 63 | 117 | 281 | 145 | 153 | 350 | 381 | 315 | 293 | 174 | 101 | 165 | 73 | 46 | 30 | 135 | 127 | 157 | 203 | 404 | 373 | 188 | 286 | 107 | 135 | 121 | Far2 |
| Farp1 | 5121 | 5766 | 5203 | 4719 | 4926 | 5426 | 3805 | 6691 | 9285 | 9326 | 9815 | 8020 | 9646 | 7453 | 4941 | 6887 | 4425 | 4928 | 4708 | 3657 | 4304 | 7850 | 9492 | 8868 | 8133 | 8729 | 7700 | 7321 | Farp1 |
| Farp2 | 2248 | 2334 | 1262 | 1475 | 1530 | 1888 | 1616 | 924 | 1021 | 673 | 809 | 458 | 631 | 535 | 2044 | 2301 | 1386 | 1773 | 1407 | 1745 | 1824 | 620 | 1046 | 504 | 550 | 441 | 579 | 431 | Farp2 |
| Fars2 | 556 | 652 | 469 | 471 | 450 | 525 | 468 | 916 | 844 | 785 | 757 | 912 | 786 | 776 | 529 | 562 | 563 | 474 | 548 | 554 | 532 | 834 | 961 | 760 | 800 | 962 | 984 | 927 | Fars2 |
| Farsa | 4215 | 4978 | 5039 | 5207 | 4489 | 4110 | 4137 | 4104 | 4481 | 4878 | 4438 | 5340 | 4796 | 5170 | 3864 | 4153 | 4983 | 4464 | 4969 | 4416 | 4452 | 4386 | 4161 | 5254 | 5157 | 5659 | 5194 | 5296 | Farsa |
| Farsb | 7975 | 7218 | 5864 | 5929 | 5249 | 5562 | 6269 | 6365 | 5936 | 4968 | 5055 | 5079 | 4957 | 5194 | 7867 | 7124 | 6129 | 5817 | 5879 | 6926 | 6258 | 5421 | 5860 | 4946 | 5190 | 5817 | 5385 | 5388 | Farsb |
| Fas | 142 | 105 | 91 | 74 | 47 | 69 | 55 | 72 | 123 | 68 | 96 | 23 | 39 | 26 | 117 | 142 | 80 | 86 | 36 | 67 | 46 | 65 | 100 | 56 | 24 | 37 | 36 | 26 | Fas |
| Fasl | 18 | 22 | 23 | 95 | 70 | 32 | 51 | 20 | 21 | 25 | 21 | 12 | 20 | 17 | 24 | 30 | 52 | 68 | 61 | 49 | 80 | 43 | 15 | 8 | 26 | 13 | 16 | 27 | Fasl |
| Fasn | 25110 | 35732 | 34412 | 36011 | 33480 | 32619 | 26856 | 25733 | 32360 | 35430 | 31320 | 47296 | 40676 | 39420 | 25655 | 32259 | 35740 | 35413 | 36987 | 24840 | 30853 | 38352 | 28131 | 37763 | 38784 | 26851 | 39739 | 42134 | Fasn |
| Fastk | 2721 | 3258 | 3281 | 2822 | 2734 | 2808 | 2617 | 3168 | 3804 | 3848 | 3689 | 3947 | 3720 | 3706 | 2616 | 2933 | 3224 | 2650 | 3185 | 2682 | 2615 | 3583 | 3763 | 3887 | 3530 | 4279 | 3970 | 3755 | Fastk |
| Fastkd1 | 2395 | 2137 | 2430 | 2271 | 2347 | 2358 | 2643 | 1970 | 1721 | 1885 | 1908 | 2001 | 1844 | 2027 | 2262 | 2224 | 2371 | 2200 | 2290 | 2231 | 2483 | 2079 | 1649 | 1835 | 1838 | 1928 | 2128 | 2060 | Fastkd1 |
| Fastkd2 | 2357 | 2122 | 1854 | 1744 | 1807 | 1993 | 2179 | 2880 | 2497 | 1907 | 2103 | 2230 | 2249 | 2419 | 2640 | 2330 | 1798 | 1832 | 1734 | 2211 | 1987 | 2120 | 2404 | 2010 | 1935 | 2298 | 2506 | 2538 | Fastkd2 |
| Fastkd3 | 1626 | 1646 | 1398 | 1553 | 1245 | 1247 | 1379 | 1542 | 1536 | 1409 | 1333 | 1715 | 1430 | 1700 | 1584 | 1369 | 1373 | 1343 | 1320 | 1409 | 1333 | 1385 | 1335 | 1430 | 1387 | 1730 | 1639 | 1653 | Fastkd3 |
| Fastkd5 | 401 | 396 | 426 | 261 | 317 | 274 | 332 | 514 | 388 | 337 | 382 | 443 | 353 | 419 | 436 | 365 | 312 | 292 | 336 | 377 | 346 | 338 | 351 | 371 | 359 | 461 | 407 | 521 | Fastkd5 |
| Fat1 | 22071 | 22308 | 13544 | 16839 | 18733 | 21715 | 17262 | 16308 | 21212 | 16478 | 20615 | 16833 | 20236 | 15875 | 19748 | 24569 | 14456 | 16900 | 16486 | 18169 | 20286 | 11911 | 18635 | 12696 | 15204 | 11585 | 20030 | 14313 | Fat1 |
| Fat2 | 77 | 33 | 8 | 39 | 146 | 103 | 37 | 349 | 639 | 396 | 584 | 93 | 124 | 70 | 36 | 66 | 42 | 138 | 31 | 48 | 64 | 261 | 809 | 221 | 279 | 43 | 105 | 45 | Fat2 |
| Fat3 | 9182 | 10071 | 6243 | 8307 | 9880 | 13459 | 11002 | 13363 | 11975 | 9591 | 9223 | 9586 | 9749 | 10126 | 10887 | 9305 | 7640 | 8143 | 9190 | 14382 | 12830 | 9097 | 11043 | 8000 | 11279 | 6285 | 12221 | 9645 | Fat3 |

| | | | | | | | | | | | | | | | | | | | | | | | | | | | | | |
|---|---|---|---|---|---|---|---|---|---|---|---|---|---|---|---|---|---|---|---|---|---|---|---|---|---|---|---|---|---|
| Fat4 | 11194 | 12093 | 7664 | 9403 | 11639 | 14699 | 10658 | 15740 | 17974 | 16277 | 18614 | 14698 | 16718 | 14366 | 11446 | 13061 | 8686 | 9992 | 10022 | 11093 | 12692 | 13216 | 16260 | 13321 | 15844 | 11354 | 18056 | 13398 | Fat4 |
| Fate1 | 5 | 13 | 0 | 5 | 2 | 3 | 6 | 4 | 5 | 0 | 5 | 0 | 1 | 2 | 15 | 7 | 0 | 4 | 4 | 8 | 9 | 2 | 2 | 2 | 1 | 0 | 5 | 4 | Fate1 |
| Fau | 15291 | 21622 | 18922 | 19373 | 16708 | 17314 | 17993 | 13420 | 15418 | 15628 | 18036 | 16537 | 18504 | 18763 | 14180 | 15657 | 18572 | 17142 | 18596 | 20649 | 17861 | 10550 | 15628 | 18834 | 17346 | 20381 | 19739 | 20445 | Fau |
| Faxc | 3700 | 3256 | 2297 | 3459 | 4943 | 5103 | 4164 | 9800 | 8709 | 8681 | 8369 | 8883 | 8500 | 8462 | 4056 | 3092 | 3174 | 3463 | 4200 | 4297 | 4505 | 9775 | 7508 | 7133 | 10455 | 3893 | 9089 | 7809 | Faxc |
| Fbf1 | 6569 | 8066 | 8924 | 9922 | 9882 | 9214 | 9275 | 7671 | 8963 | 10898 | 9386 | 11246 | 9527 | 10887 | 6435 | 7110 | 9598 | 9573 | 10002 | 7676 | 9269 | 10226 | 8209 | 11109 | 11437 | 9587 | 10119 | 9679 | Fbf1 |
| Fbh1 | 10299 | 10274 | 8518 | 8977 | 8773 | 8583 | 8394 | 11011 | 10270 | 8909 | 8998 | 9081 | 8972 | 8562 | 10747 | 9664 | 8721 | 9019 | 8817 | 9631 | 8934 | 9840 | 10398 | 8772 | 9094 | 8817 | 9343 | 8933 | Fbh1 |
| Fbl | 8113 | 10310 | 11140 | 10474 | 7125 | 7033 | 8751 | 5183 | 5800 | 6740 | 5946 | 7338 | 6474 | 8229 | 7303 | 7851 | 10791 | 8496 | 8313 | 7630 | 8739 | 4214 | 5505 | 8067 | 6321 | 9566 | 7864 | 9178 | Fbl |
| Fblim1 | 2475 | 2004 | 2038 | 1191 | 1762 | 1629 | 1082 | 1906 | 2583 | 2202 | 2737 | 1800 | 2638 | 1654 | 1975 | 2973 | 1426 | 1602 | 1281 | 1291 | 1063 | 2408 | 2805 | 1906 | 1753 | 2423 | 1984 | 1836 | Fblim1 |
| Fbll1 | 356 | 643 | 392 | 818 | 894 | 703 | 647 | 474 | 569 | 633 | 487 | 325 | 339 | 370 | 501 | 456 | 637 | 708 | 837 | 886 | 807 | 530 | 539 | 449 | 648 | 273 | 318 | 331 | Fbll1 |
| Fbln1 | 7434 | 8535 | 10061 | 5789 | 4156 | 4758 | 3286 | 5136 | 7101 | 5828 | 7696 | 7623 | 8571 | 6835 | 6485 | 10391 | 7389 | 6550 | 4386 | 3187 | 3514 | 5534 | 8139 | 7118 | 5288 | 9301 | 8972 | 7888 | Fbln1 |
| Fbln2 | 7279 | 8841 | 6121 | 5330 | 2324 | 5739 | 1107 | 6150 | 9092 | 8005 | 9583 | 6162 | 8631 | 5764 | 6068 | 10951 | 5204 | 6593 | 1795 | 1181 | 1117 | 6742 | 10128 | 7372 | 5942 | 7376 | 6516 | 6539 | Fbln2 |
| Fbln5 | 1381 | 1151 | 830 | 498 | 247 | 860 | 151 | 1439 | 2116 | 1288 | 2504 | 1076 | 1759 | 1022 | 1344 | 1559 | 717 | 952 | 226 | 175 | 159 | 1235 | 2629 | 1214 | 958 | 1119 | 1225 | 1086 | Fbln5 |
| Fbln7 | 98 | 117 | 143 | 90 | 90 | 107 | 60 | 125 | 194 | 220 | 317 | 124 | 163 | 121 | 108 | 150 | 80 | 101 | 78 | 54 | 61 | 224 | 247 | 193 | 192 | 162 | 178 | 174 | Fbln7 |
| Fbn1 | 11148 | 10607 | 6963 | 5990 | 4608 | 8152 | 2999 | 7562 | 10813 | 8905 | 16017 | 8816 | 13013 | 8343 | 9629 | 15190 | 5598 | 7701 | 3604 | 3625 | 3547 | 7907 | 14369 | 7643 | 7262 | 5427 | 10405 | 8495 | Fbn1 |
| Fbn2 | 17085 | 20849 | 15747 | 9951 | 5557 | 13368 | 3800 | 13406 | 22284 | 20252 | 30471 | 18225 | 30504 | 16467 | 16148 | 27495 | 12061 | 13197 | 5034 | 4381 | 4078 | 15527 | 24212 | 16709 | 14405 | 13641 | 22385 | 16916 | Fbn2 |
| Fbp1 | 1 | 1 | 0 | 1 | 3 | 0 | 0 | 6 | 12 | 4 | 18 | 3 | 3 | 2 | 3 | 1 | 6 | 9 | 0 | 0 | 1 | 22 | 9 | 6 | 5 | 2 | 1 | 4 | Fbp1 |
| Fbp2 | 0 | 7 | 0 | 7 | 9 | 1 | 0 | 4 | 9 | 4 | 24 | 1 | 1 | 0 | 0 | 5 | 2 | 4 | 3 | 0 | 0 | 2 | 17 | 4 | 6 | 2 | 6 | 0 | Fbp2 |
| Fbrs | 3049 | 4160 | 4942 | 4881 | 4468 | 3694 | 3578 | 2941 | 3368 | 4535 | 4055 | 4291 | 3796 | 3830 | 2949 | 3610 | 5021 | 4490 | 4753 | 3145 | 4078 | 4493 | 3271 | 4636 | 4115 | 4517 | 3478 | 3590 | Fbrs |
| Fbrsl1 | 4643 | 5922 | 6289 | 6709 | 6521 | 5579 | 5171 | 3709 | 4431 | 5282 | 4612 | 5868 | 4378 | 5033 | 4360 | 4897 | 6876 | 6724 | 6779 | 4836 | 5664 | 5419 | 3741 | 5619 | 5677 | 6536 | 4829 | 4467 | Fbrsl1 |
| Fbxl12 | 2185 | 2522 | 2426 | 2437 | 2032 | 1998 | 1986 | 1518 | 1608 | 1777 | 1738 | 1866 | 1899 | 1680 | 1920 | 2125 | 2343 | 2321 | 2109 | 2078 | 1837 | 1663 | 1685 | 1961 | 1737 | 1673 | 1812 | 1787 | Fbxl12 |
| Fbxl13 | 81 | 69 | 54 | 72 | 80 | 37 | 54 | 43 | 41 | 46 | 39 | 48 | 37 | 47 | 50 | 58 | 54 | 77 | 34 | 65 | 56 | 50 | 36 | 49 | 42 | 21 | 44 | 43 | Fbxl13 |
| Fbxl14 | 3301 | 3697 | 4324 | 4316 | 3310 | 3354 | 3213 | 2904 | 2679 | 3358 | 2886 | 3989 | 3301 | 3860 | 2951 | 3354 | 4111 | 3949 | 3490 | 3309 | 3380 | 2968 | 2441 | 3813 | 3533 | 4483 | 3579 | 3970 | Fbxl14 |
| Fbxl15 | 593 | 768 | 534 | 626 | 689 | 695 | 605 | 471 | 532 | 517 | 497 | 527 | 463 | 503 | 591 | 513 | 702 | 643 | 789 | 764 | 593 | 545 | 504 | 601 | 540 | 562 | 618 | 468 | Fbxl15 |
| Fbxl16 | 2449 | 3251 | 3296 | 3486 | 4089 | 3438 | 3549 | 2749 | 3019 | 3918 | 3091 | 3753 | 2989 | 3572 | 2438 | 2544 | 3498 | 3500 | 4076 | 3070 | 3559 | 4468 | 2739 | 3933 | 4343 | 3489 | 3285 | 3377 | Fbxl16 |
| Fbxl17 | 2431 | 2103 | 2009 | 2248 | 2716 | 3051 | 2131 | 3598 | 3186 | 3280 | 3378 | 3246 | 3297 | 3559 | 2321 | 2281 | 2190 | 2279 | 2501 | 2805 | 3130 | 3865 | 3740 | 3125 | 3837 | 2356 | 3588 | 3443 | Fbxl17 |
| Fbxl18 | 4086 | 5569 | 6320 | 7205 | 6437 | 6006 | 5663 | 3588 | 4051 | 5109 | 4789 | 5789 | 5278 | 5373 | 4188 | 4690 | 7047 | 6949 | 7077 | 5125 | 6068 | 5339 | 3880 | 5574 | 6119 | 4445 | 5385 | 5511 | Fbxl18 |
| Fbxl19 | 6885 | 8407 | 9158 | 10371 | 11709 | 9417 | 9047 | 11747 | 11725 | 15245 | 13352 | 14939 | 13674 | 13418 | 6906 | 6763 | 10880 | 10144 | 12507 | 9406 | 9956 | 14514 | 10956 | 15684 | 16236 | 15533 | 14106 | 14049 | Fbxl19 |
| Fbxl2 | 978 | 1111 | 798 | 1152 | 1289 | 1247 | 1259 | 1169 | 1225 | 1283 | 1158 | 975 | 1056 | 883 | 955 | 1110 | 1023 | 1260 | 1122 | 1275 | 1230 | 1390 | 1150 | 968 | 1269 | 754 | 927 | 819 | Fbxl2 |
| Fbxl20 | 7052 | 6471 | 5563 | 8181 | 8188 | 8342 | 8254 | 6218 | 5913 | 6509 | 6213 | 6498 | 6344 | 6762 | 7005 | 6681 | 6793 | 7771 | 7958 | 7079 | 8745 | 6706 | 5778 | 6089 | 7191 | 4113 | 6419 | 5873 | Fbxl20 |
| Fbxl21 | 621 | 559 | 506 | 583 | 575 | 608 | 527 | 171 | 176 | 161 | 189 | 176 | 183 | 199 | 603 | 714 | 495 | 573 | 503 | 543 | 536 | 226 | 223 | 139 | 155 | 150 | 175 | 166 | Fbxl21 |
| Fbxl22 | 325 | 428 | 429 | 504 | 479 | 432 | 507 | 275 | 286 | 308 | 402 | 394 | 347 | 400 | 352 | 360 | 599 | 447 | 537 | 434 | 515 | 371 | 275 | 388 | 367 | 399 | 378 | 407 | Fbxl22 |
| Fbxl3 | 3132 | 2676 | 2989 | 3048 | 3296 | 3321 | 3529 | 6172 | 5631 | 4881 | 5604 | 5436 | 5568 | 5591 | 3294 | 2959 | 2990 | 3050 | 3153 | 3454 | 3376 | 5693 | 5733 | 4998 | 5281 | 5125 | 5612 | 5446 | Fbxl3 |
| Fbxl4 | 940 | 894 | 808 | 751 | 870 | 831 | 932 | 996 | 988 | 882 | 874 | 884 | 848 | 821 | 1094 | 781 | 830 | 806 | 750 | 967 | 885 | 969 | 1043 | 753 | 765 | 771 | 919 | 897 | Fbxl4 |
| Fbxl5 | 7794 | 6124 | 5235 | 5712 | 6234 | 7129 | 7497 | 8989 | 7970 | 5830 | 6668 | 6387 | 6485 | 6957 | 7965 | 7039 | 5238 | 5873 | 5792 | 8491 | 6831 | 6665 | 7724 | 5821 | 6237 | 5986 | 6746 | 6543 | Fbxl5 |
| Fbxl6 | 2527 | 3412 | 2934 | 3189 | 2791 | 2669 | 2627 | 1824 | 2132 | 2128 | 2170 | 2339 | 2216 | 1966 | 2407 | 2867 | 3101 | 2869 | 3071 | 2702 | 2607 | 2017 | 2131 | 2418 | 2256 | 2727 | 2244 | 2074 | Fbxl6 |
| Fbxl7 | 1201 | 1069 | 1135 | 921 | 1118 | 1020 | 736 | 1864 | 2184 | 2258 | 2300 | 2070 | 2489 | 1808 | 1125 | 1281 | 998 | 983 | 855 | 684 | 760 | 2182 | 2303 | 2029 | 1733 | 2018 | 1962 | 2013 | Fbxl7 |
| Fbxl8 | 78 | 93 | 95 | 68 | 80 | 87 | 66 | 23 | 41 | 22 | 41 | 23 | 43 | 20 | 108 | 106 | 105 | 112 | 58 | 65 | 42 | 43 | 48 | 29 | 25 | 33 | 22 | 32 | Fbxl8 |
| Fbxo10 | 3881 | 4497 | 4198 | 4513 | 4325 | 4455 | 4000 | 2392 | 2858 | 2834 | 2935 | 2785 | 2406 | 2398 | 3570 | 3964 | 4313 | 4357 | 4219 | 3743 | 3836 | 2598 | 2810 | 2975 | 2799 | 2835 | 2658 | 2317 | Fbxo10 |
| Fbxo11 | 9491 | 7123 | 10786 | 11131 | 13029 | 11897 | 13615 | 15282 | 12484 | 15481 | 14724 | 16196 | 14749 | 17029 | 9854 | 8575 | 11097 | 11849 | 13280 | 10220 | 13147 | 18259 | 12745 | 15524 | 17085 | 14936 | 14494 | 16848 | Fbxo11 |
| Fbxo15 | 21 | 18 | 22 | 10 | 13 | 22 | 17 | 26 | 24 | 10 | 31 | 25 | 29 | 20 | 20 | 18 | 21 | 19 | 17 | 19 | 12 | 15 | 23 | 24 | 25 | 28 | 28 | 33 | Fbxo15 |
| Fbxo16 | 937 | 1019 | 1020 | 911 | 904 | 861 | 862 | 882 | 900 | 874 | 639 | 958 | 1006 | 995 | 1040 | 880 | 909 | 748 | 860 | 914 | 857 | 1053 | 844 | 882 | 879 | 943 | 951 | 1042 | Fbxo16 |
| Fbxo17 | 347 | 333 | 388 | 275 | 277 | 341 | 213 | 272 | 375 | 341 | 550 | 302 | 384 | 277 | 314 | 387 | 311 | 280 | 303 | 263 | 229 | 286 | 400 | 303 | 269 | 372 | 336 | 345 | Fbxo17 |
| Fbxo2 | 269 | 352 | 440 | 414 | 403 | 309 | 255 | 67 | 120 | 142 | 179 | 53 | 82 | 48 | 282 | 301 | 371 | 351 | 354 | 351 | 266 | 105 | 124 | 85 | 95 | 65 | 49 | 64 | Fbxo2 |
| Fbxo21 | 9401 | 10304 | 10656 | 11755 | 11959 | 10960 | 11616 | 10233 | 10635 | 11005 | 9963 | 11952 | 10317 | 11443 | 9434 | 8769 | 11454 | 11407 | 12638 | 10525 | 12151 | 12068 | 9493 | 11279 | 12526 | 11565 | 10461 | 10993 | Fbxo21 |
| Fbxo22 | 2628 | 2541 | 3232 | 3125 | 3340 | 2840 | 3224 | 4168 | 3827 | 4305 | 4418 | 4626 | 4197 | 5131 | 2538 | 2538 | 3242 | 2861 | 3274 | 2823 | 3125 | 4144 | 3729 | 4620 | 4693 | 5109 | 4668 | 4906 | Fbxo22 |
| Fbxo24 | 16 | 36 | 33 | 43 | 40 | 27 | 12 | 15 | 21 | 12 | 21 | 40 | 29 | 36 | 38 | 38 | 42 | 25 | 34 | 38 | 34 | 22 | 25 | 25 | 34 | 28 | 42 | 32 | Fbxo24 |
| Fbxo25 | 3233 | 3924 | 3092 | 4101 | 3974 | 3769 | 4179 | 3074 | 2952 | 3091 | 2730 | 2641 | 2467 | 2863 | 3494 | 3392 | 3342 | 4187 | 3770 | 4114 | 4386 | 3078 | 3055 | 2988 | 3384 | 2469 | 2506 | 2606 | Fbxo25 |
| Fbxo27 | 437 | 410 | 501 | 324 | 389 | 455 | 370 | 228 | 309 | 275 | 383 | 277 | 298 | 193 | 369 | 414 | 423 | 418 | 454 | 362 | 388 | 346 | 326 | 296 | 235 | 364 | 231 | 239 | Fbxo27 |
| Fbxo28 | 4969 | 4728 | 4625 | 5090 | 5307 | 4941 | 5432 | 5478 | 4668 | 4829 | 4692 | 5269 | 4863 | 4967 | 5002 | 4393 | 4720 | 4713 | 5286 | 5388 | 5378 | 5382 | 4625 | 4989 | 5520 | 4907 | 4852 | 5006 | Fbxo28 |
| Fbxo3 | 9168 | 7010 | 7952 | 7633 | 8134 | 8476 | 8639 | 8524 | 7129 | 6530 | 6729 | 7143 | 7121 | 7649 | 9357 | 7779 | 7649 | 7724 | 8221 | 8655 | 8345 | 8079 | 7315 | 7279 | 7016 | 7486 | 7491 | 7741 | Fbxo3 |
| Fbxo30 | 4815 | 3746 | 4646 | 3923 | 3978 | 4539 | 4472 | 5123 | 4333 | 3801 | 4046 | 4637 | 5065 | 5197 | 4814 | 4374 | 4538 | 4023 | 3977 | 4550 | 4302 | 4136 | 4330 | 4228 | 4023 | 4890 | 5519 | 5345 | Fbxo30 |
| Fbxo31 | 2874 | 3671 | 3502 | 4918 | 4195 | 3557 | 3570 | 1584 | 1808 | 2239 | 1944 | 1914 | 1602 | 1630 | 2680 | 3273 | 3946 | 4747 | 4399 | 3136 | 3898 | 2303 | 1728 | 2065 | 2140 | 1963 | 1642 | 1612 | Fbxo31 |
| Fbxo32 | 1 | 0 | 0 | 0 | 0 | 0 | 0 | 0 | 0 | 0 | 0 | 0 | 0 | 0 | 0 | 0 | 0 | 0 | 0 | 0 | 0 | 0 | 0 | 0 | 0 | 0 | 0 | 0 | Fbxo32 |
| Fbxo33 | 2268 | 2031 | 1840 | 2097 | 1642 | 1781 | 1964 | 1348 | 1153 | 1265 | 1104 | 1152 | 1217 | 1328 | 2102 | 2227 | 1752 | 1932 | 1682 | 1730 | 1894 | 1375 | 1223 | 1126 | 1283 | 1207 | 1311 | 1326 | Fbxo33 |
| Fbxo34 | 1884 | 2080 | 1921 | 1828 | 1938 | 2061 | 1795 | 1431 | 1551 | 1619 | 1407 | 1701 | 1710 | 1535 | 1921 | 1917 | 1917 | 1919 | 2111 | 1960 | 1853 | 1323 | 1561 | 1541 | 1557 | 1937 | 1622 | 1806 | Fbxo34 |
| Fbxo36 | 623 | 617 | 586 | 604 | 408 | 411 | 464 | 150 | 137 | 139 | 126 | 210 | 205 | 202 | 567 | 524 | 446 | 569 | 451 | 517 | 435 | 154 | 164 | 146 | 147 | 225 | 169 | 220 | Fbxo36 |
| Fbxo38 | 7504 | 7117 | 6033 | 6178 | 5991 | 6147 | 6393 | 7963 | 7137 | 5991 | 6362 | 7201 | 6818 | 7009 | 7377 | 7013 | 5771 | 5972 | 5929 | 6835 | 6342 | 6635 | 7271 | 6285 | 6847 | 5852 | 7308 | 6859 | Fbxo38 |
| Fbxo39 | 0 | 0 | 0 | 3 | 0 | 0 | 0 | 0 | 1 | 0 | 1 | 0 | 0 | 0 | 0 | 2 | 0 | 0 | 0 | 0 | 2 | 0 | 0 | 0 | 0 | 0 | 0 | 0 | Fbxo39 |
| Fbxo4 | 190 | 210 | 196 | 252 | 214 | 177 | 236 | 386 | 337 | 328 | 310 | 344 | 345 | 304 | 194 | 221 | 218 | 286 | 289 | 209 | 248 | 381 | 372 | 310 | 354 | 367 | 328 | 350 | Fbxo4 |
| Fbxo40 | 3 | 6 | 3 | 6 | 10 | 2 | 0 | 10 | 21 | 8 | 26 | 4 | 8 | 13 | 3 | 7 | 5 | 3 | 12 | 10 | 4 | 7 | 36 | 20 | 18 | 5 | 3 | 2 | Fbxo40 |
| Fbxo41 | 1269 | 1620 | 1547 | 2104 | 2978 | 2244 | 2357 | 1500 | 1703 | 1935 | 1571 | 1484 | 1295 | 1455 | 1328 | 1297 | 1905 | 2275 | 2751 | 2117 | 2534 | 2445 | 1488 | 1825 | 2267 | 1002 | 1316 | 1296 | Fbxo41 |
| Fbxo42 | 4733 | 5376 | 5527 | 6364 | 6012 | 5338 | 5488 | 3236 | 3742 | 4378 | 3942 | 4192 | 3657 | 3999 | 4685 | 5485 | 5670 | 6165 | 5953 | 4539 | 5652 | 4520 | 3574 | 3974 | 4292 | 3507 | 3632 | 3961 | Fbxo42 |
| Fbxo43 | 32 | 16 | 17 | 10 | 15 | 13 | 18 | 4 | 7 | 2 | 8 | 5 | 3 | 13 | 30 | 26 | 20 | 20 | 9 | 13 | 29 | 7 | 12 | 11 | 4 | 4 | 6 | 6 | Fbxo43 |
| Fbxo44 | 2096 | 2260 | 2171 | 2269 | 2701 | 2613 | 2546 | 2448 | 2385 | 2300 | 2077 | 2717 | 2588 | 2681 | 2269 | 1935 | 2606 | 2291 | 2851 | 2753 | 2663 | 2827 | 2275 | 2346 | 2924 | 2258 | 2660 | 2626 | Fbxo44 |
| Fbxo45 | 5779 | 4678 | 7400 | 7224 | 6800 | 6074 | 7216 | 5503 | 4745 | 5736 | 5082 | 6256 | 5549 | 6596 | 5719 | 5423 | 7392 | 6860 | 6762 | 5252 | 7000 | 6542 | 4346 | 6266 | 6542 | 7029 | 6082 | 6726 | Fbxo45 |
| Fbxo46 | 1232 | 1635 | 1594 | 1799 | 1549 | 1351 | 1456 | 975 | 1051 | 1297 | 1176 | 1249 | 1140 | 1214 | 1143 | 1544 | 1814 | 1778 | 1647 | 1199 | 1550 | 1221 | 1154 | 1224 | 1254 | 1265 | 1113 | 1131 | Fbxo46 |
| Fbxo47 | 106 | 89 | 68 | 88 | 102 | 111 | 142 | 82 | 76 | 82 | 80 | 106 | 82 | 85 | 152 | 71 | 86 | 100 | 117 | 145 | 138 | 90 | 87 | 93 | 81 | 68 | 103 | 81 | Fbxo47 |
| Fbxo48 | 624 | 470 | 327 | 456 | 349 | 504 | 572 | 482 | 422 | 399 | 445 | 509 | 431 | 656 | 483 | 474 | 400 | 409 | 428 | 488 | 564 | 292 | 383 | 440 | 438 | 219 | 642 | 586 | Fbxo48 |
| Fbxo5 | 2502 | 2162 | 2953 | 2187 | 1588 | 1898 | 2340 | 2727 | 2460 | 2389 | 2299 | 3556 | 2783 | 3674 | 2202 | 2233 | 2480 | 1736 | 2193 | 2143 | 2061 | 1798 | 2197 | 2880 | 2554 | 4535 | 3755 | 3938 | Fbxo5 |
| Fbxo6 | 364 | 460 | 407 | 349 | 466 | 456 | 445 | 403 | 381 | 414 | 417 | 475 | 519 | 390 | 426 | 386 | 423 | 369 | 447 | 447 | 399 | 451 | 438 | 427 | 486 | 455 | 440 | 492 | Fbxo6 |
| Fbxo7 | 553 | 451 | 446 | 390 | 394 | 339 | 350 | 904 | 937 | 850 | 961 | 690 | 872 | 735 | 530 | 553 | 387 | 304 | 298 | 383 | 259 | 866 | 1013 | 798 | 731 | 946 | 758 | 787 | Fbxo7 |
| Fbxo8 | 2031 | 1786 | 1790 | 1855 | 1676 | 1702 | 1988 | 1689 | 1355 | 1272 | 1320 | 1411 | 1247 | 1410 | 2127 | 1872 | 1729 | 1839 | 1723 | 2176 | 1883 | 1604 | 1455 | 1365 | 1389 | 1393 | 1374 | 1476 | Fbxo8 |
| Fbxo9 | 1914 | 2066 | 1733 | 2178 | 2413 | 1997 | 2145 | 2017 | 1977 | 1973 | 2119 | 1557 | 1910 | 1688 | 2045 | 2005 | 1874 | 2389 | 1926 | 2017 | 2105 | 2227 | 2014 | 1887 | 2115 | 1616 | 1578 | 1709 | Fbxo9 |
| Fbxw10 | 29 | 12 | 24 | 31 | 25 | 17 | 30 | 27 | 43 | 32 | 37 | 47 | 37 | 18 | 26 | 33 | 27 | 32 | 26 | 10 | 29 | 38 | 54 | 27 | 35 | 36 | 48 | 31 | Fbxw10 |
| Fbxw11 | 11252 | 11175 | 10553 | 11134 | 12030 | 11349 | 12151 | 10082 | 9231 | 9192 | 8869 | 9818 | 9073 | 9821 | 11341 | 10079 | 10699 | 10922 | 12100 | 11345 | 12079 | 10298 | 9161 | 9453 | 9812 | 9638 | 9659 | 9907 | Fbxw11 |
| Fbxw13 | 0 | 0 | 0 | 0 | 0 | 0 | 0 | 0 | 0 | 1 | 1 | 0 | 9 | 0 | 0 | 0 | 0 | 1 | 0 | 0 | 1 | 3 | 1 | 1 | 0 | 0 | 5 | 1 | Fbxw13 |
| Fbxw14 | 0 | 0 | 0 | 0 | 0 | 0 | 0 | 1 | 0 | 0 | 4 | 0 | 7 | 1 | 0 | 0 | 0 | 0 | 0 | 0 | 0 | 2 | 1 | 1 | 1 | 4 | 0 | 1 | Fbxw14 |
| Fbxw15 | 5 | 10 | 4 | 24 | 1 | 12 | 6 | 10 | 7 | 12 | 8 | 16 | 20 | 22 | 12 | 9 | 9 | 17 | 3 | 0 | 2 | 30 | 12 | 16 | 19 | 12 | 33 | 8 | Fbxw15 |
| Fbxw16 | 0 | 0 | 3 | 0 | 0 | 0 | 0 | 1 | 2 | 0 | 0 | 0 | 1 | 1 | 0 | 1 | 0 | 0 | 0 | 0 | 0 | 0 | 0 | 2 | 0 | 0 | 0 | 5 | Fbxw16 |

| | | | | | | | | | | | | | | | | | | | | | | | | | | | | | |
|---|---|---|---|---|---|---|---|---|---|---|---|---|---|---|---|---|---|---|---|---|---|---|---|---|---|---|---|---|---|
| Fbxw17 | 1276 | 1273 | 1043 | 1154 | 1196 | 1212 | 1101 | 1660 | 1780 | 1671 | 1728 | 1757 | 1684 | 1566 | 1187 | 1264 | 993 | 969 | 1075 | 1225 | 956 | 1718 | 1588 | 1494 | 1574 | 1437 | 1641 | 1539 | Fbxw17 |
| Fbxw18 | 1 | 3 | 0 | 0 | 3 | 0 | 0 | 1 | 8 | 0 | 10 | 5 | 3 | 11 | 1 | 8 | 1 | 1 | 1 | 1 | 1 | 5 | 16 | 1 | 5 | 6 | 21 | 7 | Fbxw18 |
| Fbxw19 | 0 | 0 | 0 | 0 | 2 | 1 | 0 | 0 | 1 | 0 | 2 | 1 | 0 | 0 | 0 | 0 | 5 | 0 | 0 | 0 | 0 | 0 | 0 | 2 | 0 | 1 | 0 | 0 | Fbxw19 |
| Fbxw2 | 4169 | 3767 | 3583 | 3746 | 3582 | 3630 | 4238 | 4129 | 3949 | 3666 | 3913 | 4211 | 3891 | 4109 | 4112 | 3608 | 3819 | 3807 | 3759 | 3865 | 3964 | 3922 | 3974 | 4108 | 3987 | 4448 | 3941 | 4314 | Fbxw2 |
| Fbxw20 | 0 | 0 | 8 | 0 | 0 | 2 | 0 | 0 | 0 | 0 | 0 | 0 | 1 | 3 | 1 | 0 | 0 | 0 | 0 | 0 | 0 | 7 | 3 | 2 | 2 | 0 | 1 | 4 | Fbxw20 |
| Fbxw21 | 0 | 0 | 0 | 0 | 1 | 0 | 0 | 0 | 1 | 0 | 1 | 0 | 1 | 0 | 0 | 0 | 0 | 0 | 0 | 0 | 0 | 3 | 0 | 0 | 2 | 0 | 0 | 0 | Fbxw21 |
| Fbxw22 | 0 | 1 | 0 | 0 | 0 | 0 | 0 | 1 | 0 | 0 | 0 | 0 | 2 | 0 | 0 | 1 | 0 | 0 | 0 | 0 | 0 | 2 | 1 | 0 | 0 | 0 | 1 | 0 | Fbxw22 |
| Fbxw24 | 15 | 16 | 7 | 4 | 0 | 7 | 1 | 4 | 12 | 4 | 16 | 7 | 7 | 2 | 7 | 10 | 7 | 1 | 0 | 6 | 7 | 13 | 17 | 5 | 1 | 0 | 12 | 5 | Fbxw24 |
| Fbxw25 | 6 | 4 | 0 | 0 | 2 | 5 | 0 | 5 | 4 | 1 | 7 | 10 | 1 | 2 | 0 | 5 | 1 | 0 | 1 | 1 | 2 | 2 | 8 | 1 | 3 | 6 | 2 | 5 | Fbxw25 |
| Fbxw26 | 5 | 5 | 0 | 0 | 0 | 0 | 2 | 2 | 0 | 0 | 0 | 0 | 8 | 2 | 4 | 2 | 0 | 0 | 1 | 0 | 1 | 0 | 6 | 2 | 1 | 0 | 5 | 1 | Fbxw26 |
| Fbxw27 | 2 | 0 | 4 | 2 | 0 | 0 | 4 | 0 | 3 | 0 | 5 | 0 | 7 | 6 | 1 | 0 | 2 | 2 | 0 | 1 | 0 | 8 | 6 | 10 | 2 | 3 | 2 | 5 | Fbxw27 |
| Fbxw28 | 0 | 0 | 0 | 0 | 0 | 0 | 0 | 0 | 2 | 0 | 0 | 0 | 0 | 0 | 0 | 0 | 0 | 0 | 0 | 0 | 0 | 2 | 1 | 0 | 0 | 0 | 0 | 0 | Fbxw28 |
| Fbxw4 | 439 | 454 | 488 | 432 | 594 | 479 | 454 | 769 | 838 | 1022 | 1012 | 888 | 820 | 863 | 431 | 430 | 499 | 556 | 508 | 361 | 469 | 892 | 942 | 1048 | 952 | 1070 | 944 | 879 | Fbxw4 |
| Fbxw5 | 3223 | 4254 | 3705 | 3899 | 3912 | 3643 | 3974 | 3650 | 4121 | 4090 | 3900 | 4531 | 3996 | 4304 | 3294 | 3327 | 4131 | 3842 | 4210 | 3532 | 3813 | 4502 | 3880 | 4796 | 4193 | 4319 | 4262 | 4079 | Fbxw5 |
| Fbxw7 | 5968 | 5769 | 6374 | 7079 | 7016 | 6763 | 7856 | 4191 | 3842 | 4044 | 3855 | 4545 | 3801 | 4599 | 6446 | 5667 | 6904 | 6818 | 6696 | 7040 | 7251 | 4024 | 3501 | 4339 | 4349 | 4679 | 4193 | 4423 | Fbxw7 |
| Fbxw8 | 7997 | 9580 | 8726 | 7461 | 6474 | 7146 | 6316 | 7474 | 8254 | 7354 | 7560 | 8619 | 8424 | 7362 | 7580 | 8328 | 7998 | 6929 | 7787 | 7314 | 6497 | 6904 | 8400 | 7862 | 7750 | 9503 | 8476 | 8076 | Fbxw8 |
| Fbxw9 | 1821 | 2202 | 2180 | 2470 | 2342 | 2232 | 2077 | 1633 | 1929 | 1877 | 1882 | 1912 | 1879 | 1616 | 1770 | 2103 | 2341 | 2556 | 2534 | 2093 | 2119 | 2224 | 1990 | 1979 | 2164 | 1825 | 1869 | 1709 | Fbxw9 |
| Fcamr | 1 | 1 | 3 | 2 | 1 | 0 | 0 | 7 | 4 | 2 | 2 | 2 | 3 | 4 | 3 | 2 | 1 | 0 | 2 | 0 | 0 | 2 | 5 | 1 | 2 | 5 | 9 | 5 | Fcamr |
| Fcer1a | 0 | 0 | 0 | 0 | 0 | 0 | 0 | 0 | 0 | 0 | 0 | 0 | 0 | 0 | 0 | 0 | 0 | 0 | 0 | 0 | 0 | 0 | 2 | 2 | 0 | 0 | 0 | 0 | Fcer1a |
| Fcer1g | 186 | 228 | 215 | 223 | 151 | 177 | 180 | 217 | 271 | 221 | 281 | 225 | 244 | 238 | 241 | 240 | 185 | 235 | 146 | 181 | 204 | 214 | 332 | 230 | 208 | 269 | 193 | 233 | Fcer1g |
| Fcer2a | 0 | 0 | 0 | 0 | 0 | 0 | 0 | 0 | 0 | 0 | 2 | 0 | 1 | 0 | 0 | 0 | 0 | 0 | 0 | 0 | 0 | 0 | 0 | 0 | 0 | 0 | 0 | 0 | Fcer2a |
| Fcf1 | 2877 | 2586 | 2861 | 2537 | 2162 | 2251 | 2504 | 2457 | 2155 | 2193 | 2033 | 2530 | 2420 | 2773 | 3108 | 2464 | 2675 | 2426 | 2495 | 2809 | 2358 | 1865 | 2141 | 2501 | 2206 | 3201 | 2815 | 2848 | Fcf1 |
| Fcgbp | 16 | 11 | 1 | 3 | 7 | 6 | 0 | 23 | 29 | 49 | 6 | 10 | 14 | 13 | 7 | 15 | 8 | 2 | 5 | 3 | 1 | 12 | 39 | 13 | 5 | 24 | 9 | 5 | Fcgbp |
| Fcgr1 | 59 | 59 | 67 | 64 | 42 | 62 | 32 | 116 | 171 | 131 | 165 | 115 | 127 | 116 | 69 | 76 | 38 | 54 | 41 | 62 | 71 | 127 | 237 | 131 | 107 | 175 | 123 | 134 | Fcgr1 |
| Fcgr2b | 6 | 45 | 22 | 46 | 30 | 53 | 29 | 26 | 40 | 31 | 47 | 25 | 29 | 32 | 25 | 44 | 15 | 52 | 19 | 39 | 30 | 63 | 85 | 31 | 58 | 38 | 34 | 39 | Fcgr2b |
| Fcgr3 | 231 | 194 | 221 | 200 | 174 | 216 | 180 | 364 | 361 | 256 | 419 | 278 | 315 | 253 | 260 | 282 | 160 | 235 | 103 | 251 | 206 | 291 | 530 | 279 | 292 | 344 | 278 | 298 | Fcgr3 |
| Fcgr4 | 9 | 10 | 6 | 12 | 2 | 7 | 1 | 2 | 15 | 3 | 25 | 15 | 12 | 11 | 7 | 12 | 8 | 6 | 6 | 14 | 9 | 18 | 12 | 7 | 3 | 6 | 12 | 13 | Fcgr4 |
| Fcgrt | 581 | 743 | 676 | 642 | 807 | 865 | 431 | 692 | 1210 | 950 | 1250 | 647 | 838 | 629 | 645 | 886 | 534 | 886 | 560 | 855 | 542 | 952 | 1799 | 867 | 853 | 746 | 816 | 622 | Fcgrt |
| Fcho1 | 3205 | 4962 | 5068 | 6933 | 6399 | 5815 | 5618 | 2519 | 2844 | 3491 | 2810 | 3151 | 2608 | 3061 | 3643 | 3801 | 6245 | 6720 | 7021 | 5309 | 5828 | 4152 | 2520 | 3493 | 4130 | 2937 | 2736 | 2692 | Fcho1 |
| Fcho2 | 5931 | 4188 | 6298 | 6402 | 7018 | 7419 | 6954 | 6889 | 6071 | 6550 | 7837 | 6563 | 7155 | 7297 | 5968 | 6203 | 6017 | 6495 | 6995 | 6199 | 7382 | 7554 | 6502 | 6788 | 7043 | 6252 | 6886 | 7204 | Fcho2 |
| Fchsd1 | 3430 | 4681 | 4058 | 5755 | 4858 | 4549 | 4508 | 1318 | 1426 | 1590 | 1350 | 1404 | 1412 | 1198 | 3455 | 4312 | 4909 | 5727 | 4848 | 4304 | 4690 | 1619 | 1362 | 1548 | 1639 | 1383 | 1259 | 1136 | Fchsd1 |
| Fchsd2 | 4223 | 3892 | 3957 | 5024 | 6391 | 5703 | 6290 | 6845 | 6375 | 7084 | 6536 | 6648 | 6144 | 6778 | 4511 | 3924 | 4383 | 5263 | 5913 | 5122 | 6361 | 7912 | 5988 | 6548 | 7705 | 5705 | 5672 | 6145 | Fchsd2 |
| Fcmr | 0 | 1 | 0 | 0 | 2 | 1 | 0 | 4 | 0 | 1 | 0 | 1 | 0 | 2 | 0 | 0 | 0 | 0 | 0 | 0 | 0 | 0 | 0 | 2 | 0 | 0 | 2 | 0 | Fcmr |
| Fcna | 32 | 40 | 34 | 64 | 61 | 46 | 38 | 61 | 68 | 62 | 87 | 59 | 37 | 61 | 37 | 46 | 61 | 51 | 34 | 71 | 34 | 50 | 95 | 56 | 36 | 41 | 30 | 68 | Fcna |
| Fcnb | 0 | 5 | 3 | 2 | 2 | 1 | 3 | 0 | 1 | 3 | 11 | 1 | 7 | 1 | 0 | 5 | 0 | 0 | 2 | 2 | 4 | 7 | 6 | 1 | 1 | 7 | 8 | 0 | Fcnb |
| Fcor | 37 | 23 | 25 | 13 | 23 | 11 | 35 | 60 | 44 | 76 | 100 | 43 | 62 | 87 | 22 | 8 | 23 | 9 | 16 | 25 | 29 | 57 | 35 | 53 | 68 | 54 | 55 | 78 | Fcor |
| Fcrl1 | 2 | 0 | 15 | 7 | 1 | 10 | 3 | 4 | 5 | 3 | 2 | 7 | 5 | 5 | 5 | 5 | 9 | 1 | 4 | 4 | 0 | 7 | 14 | 5 | 1 | 18 | 1 | 4 | Fcrl1 |
| Fcrl5 | 0 | 0 | 0 | 0 | 2 | 0 | 0 | 0 | 0 | 0 | 0 | 0 | 0 | 0 | 5 | 0 | 0 | 0 | 0 | 0 | 0 | 0 | 0 | 0 | 0 | 0 | 0 | 0 | Fcrl5 |
| Fcrl6 | 5 | 4 | 3 | 1 | 7 | 7 | 2 | 18 | 17 | 6 | 6 | 5 | 17 | 12 | 11 | 12 | 13 | 1 | 5 | 9 | 2 | 5 | 5 | 11 | 1 | 8 | 7 | 4 | Fcrl6 |
| Fcrla | 0 | 0 | 0 | 0 | 0 | 0 | 0 | 1 | 0 | 0 | 2 | 0 | 0 | 1 | 0 | 1 | 0 | 0 | 0 | 0 | 0 | 0 | 1 | 1 | 0 | 0 | 0 | 0 | Fcrla |
| Fcrlb | 6 | 22 | 21 | 15 | 11 | 9 | 10 | 2 | 2 | 3 | 0 | 2 | 0 | 0 | 17 | 14 | 24 | 13 | 11 | 18 | 5 | 2 | 1 | 3 | 0 | 8 | 3 | 0 | Fcrlb |
| Fcrls | 451 | 493 | 527 | 429 | 483 | 433 | 336 | 653 | 815 | 641 | 929 | 694 | 626 | 525 | 582 | 611 | 378 | 532 | 390 | 518 | 410 | 688 | 1148 | 699 | 612 | 688 | 723 | 574 | Fcrls |
| Fcsk | 1632 | 2007 | 1818 | 1885 | 1473 | 1491 | 1555 | 1122 | 1290 | 1160 | 1190 | 1282 | 1296 | 1177 | 1453 | 1619 | 1853 | 1738 | 1815 | 1442 | 1378 | 1068 | 1193 | 1436 | 1191 | 1529 | 1289 | 1266 | Fcsk |
| Fdft1 | 10062 | 12264 | 9390 | 9845 | 9456 | 9808 | 9919 | 9990 | 8869 | 8529 | 7338 | 8993 | 8902 | 9033 | 10962 | 9974 | 9754 | 9578 | 10677 | 11190 | 9805 | 9022 | 8046 | 8479 | 9452 | 9262 | 9817 | 9896 | Fdft1 |
| Fdps | 490 | 473 | 372 | 425 | 304 | 341 | 341 | 281 | 255 | 319 | 242 | 418 | 400 | 394 | 385 | 436 | 474 | 417 | 271 | 305 | 334 | 294 | 271 | 360 | 368 | 385 | 466 | 418 | Fdps |
| Fdx1 | 633 | 617 | 682 | 753 | 641 | 595 | 632 | 833 | 762 | 759 | 836 | 747 | 784 | 898 | 686 | 558 | 684 | 582 | 666 | 715 | 672 | 687 | 829 | 761 | 736 | 775 | 731 | 756 | Fdx1 |
| Fdx2 | 29 | 12 | 16 | 22 | 19 | 17 | 34 | 35 | 42 | 43 | 45 | 60 | 34 | 59 | 28 | 19 | 25 | 12 | 34 | 21 | 35 | 60 | 25 | 53 | 43 | 48 | 37 | 58 | Fdx2 |
| Fdxacb1 | 228 | 238 | 300 | 184 | 268 | 249 | 227 | 784 | 730 | 616 | 621 | 657 | 691 | 698 | 240 | 234 | 267 | 201 | 250 | 246 | 222 | 670 | 715 | 650 | 622 | 818 | 838 | 735 | Fdxacb1 |
| Fdxr | 950 | 1380 | 1493 | 1551 | 1350 | 1126 | 1245 | 898 | 950 | 1235 | 1019 | 1110 | 1058 | 1024 | 978 | 1219 | 1615 | 1538 | 1593 | 1172 | 1171 | 1150 | 849 | 1124 | 1134 | 1136 | 983 | 1030 | Fdxr |
| Fech | 2689 | 2685 | 2399 | 2062 | 2054 | 2362 | 2297 | 2379 | 3085 | 2186 | 2443 | 2197 | 3832 | 2702 | 2777 | 2516 | 2279 | 2198 | 2150 | 2327 | 2179 | 2171 | 2988 | 1904 | 2123 | 2306 | 2649 | 3673 | Fech |
| Fem1a | 5445 | 6012 | 4619 | 5926 | 5213 | 5648 | 5084 | 4606 | 4983 | 5007 | 4610 | 5070 | 4744 | 4784 | 5168 | 5386 | 5059 | 5710 | 5487 | 5031 | 5402 | 5055 | 4375 | 4808 | 5467 | 4306 | 4842 | 4703 | Fem1a |
| Fem1al | 10 | 9 | 7 | 7 | 3 | 0 | 10 | 5 | 6 | 8 | 9 | 7 | 10 | 9 | 5 | 2 | 1 | 12 | 12 | 9 | 5 | 5 | 6 | 9 | 16 | 6 | 2 | 4 | Fem1al |
| Fem1b | 28217 | 26857 | 25724 | 31041 | 33504 | 33329 | 34174 | 27500 | 21847 | 23287 | 21176 | 24085 | 21465 | 22525 | 30168 | 25031 | 29175 | 31996 | 33915 | 32301 | 34809 | 27532 | 21726 | 22716 | 26277 | 19091 | 24392 | 23295 | Fem1b |
| Fem1c | 7130 | 5762 | 6643 | 7540 | 7488 | 7570 | 7987 | 7333 | 6228 | 6323 | 6603 | 7051 | 6738 | 7217 | 7205 | 6462 | 7188 | 7081 | 7123 | 6944 | 7627 | 7568 | 6297 | 6808 | 7366 | 6771 | 7143 | 7250 | Fem1c |
| Fen1 | 5571 | 6296 | 5936 | 4556 | 3488 | 4220 | 4549 | 3605 | 3582 | 2984 | 3042 | 4636 | 4284 | 4573 | 5437 | 5111 | 5204 | 3952 | 4754 | 5458 | 4331 | 2289 | 3000 | 4245 | 3685 | 6341 | 5220 | 5426 | Fen1 |
| Fer | 3706 | 3407 | 2431 | 2709 | 2780 | 3085 | 3190 | 3352 | 2951 | 2355 | 2403 | 2292 | 2486 | 2617 | 3945 | 3443 | 2374 | 2699 | 2768 | 3863 | 3314 | 2279 | 3038 | 2117 | 2547 | 1847 | 2718 | 2472 | Fer |
| Fer1l4 | 10 | 6 | 1 | 11 | 11 | 3 | 2 | 4 | 5 | 2 | 1 | 0 | 3 | 3 | 9 | 3 | 12 | 15 | 13 | 3 | 7 | 2 | 3 | 3 | 5 | 1 | 8 | 7 | Fer1l4 |
| Fer1l5 | 1 | 1 | 4 | 2 | 4 | 5 | 3 | 4 | 3 | 12 | 2 | 10 | 1 | 4 | 0 | 0 | 5 | 2 | 3 | 0 | 4 | 7 | 3 | 0 | 7 | 5 | 3 | 7 | Fer1l5 |
| Fer1l6 | 18 | 19 | 4 | 10 | 21 | 6 | 8 | 24 | 24 | 14 | 22 | 26 | 31 | 18 | 29 | 20 | 15 | 8 | 5 | 8 | 4 | 17 | 17 | 3 | 7 | 4 | 21 | 15 | Fer1l6 |
| Ferd3l | 39 | 74 | 88 | 39 | 63 | 42 | 31 | 94 | 49 | 58 | 68 | 81 | 175 | 87 | 73 | 93 | 36 | 63 | 47 | 56 | 37 | 67 | 56 | 42 | 68 | 9 | 117 | 98 | Ferd3l |
| Fermt1 | 122 | 40 | 47 | 1 | 15 | 14 | 13 | 277 | 413 | 185 | 221 | 116 | 183 | 79 | 69 | 60 | 24 | 19 | 10 | 7 | 7 | 22 | 444 | 167 | 37 | 125 | 93 | 144 | Fermt1 |
| Fermt2 | 7448 | 6060 | 6400 | 5195 | 5096 | 5597 | 5021 | 6939 | 6662 | 5718 | 6981 | 5953 | 6493 | 5727 | 7072 | 7286 | 5731 | 5587 | 4802 | 5344 | 4660 | 5950 | 7296 | 5798 | 5379 | 6484 | 6172 | 6175 | Fermt2 |
| Fermt3 | 482 | 612 | 694 | 686 | 511 | 593 | 485 | 320 | 479 | 394 | 432 | 495 | 626 | 397 | 496 | 667 | 622 | 631 | 496 | 514 | 501 | 353 | 490 | 392 | 359 | 592 | 526 | 466 | Fermt3 |
| Fes | 223 | 266 | 282 | 268 | 244 | 297 | 180 | 509 | 589 | 539 | 772 | 680 | 863 | 597 | 270 | 283 | 261 | 295 | 259 | 204 | 174 | 590 | 788 | 668 | 562 | 730 | 741 | 749 | Fes |
| Fetub | 1 | 1 | 0 | 0 | 2 | 0 | 0 | 0 | 3 | 0 | 10 | 0 | 0 | 2 | 1 | 0 | 0 | 0 | 0 | 0 | 0 | 0 | 6 | 7 | 0 | 2 | 0 | 0 | Fetub |
| Fev | 8 | 22 | 16 | 16 | 35 | 15 | 13 | 74 | 44 | 63 | 37 | 23 | 90 | 16 | 34 | 28 | 29 | 19 | 16 | 19 | 32 | 62 | 65 | 29 | 29 | 1 | 21 | 6 | Fev |
| Fez1 | 10688 | 12442 | 10420 | 12266 | 14501 | 13704 | 16198 | 17165 | 14036 | 13900 | 11884 | 15496 | 12737 | 17299 | 12485 | 9277 | 12120 | 12521 | 14880 | 15213 | 16053 | 16193 | 13425 | 15032 | 16801 | 14300 | 15091 | 16434 | Fez1 |
| Fez2 | 1930 | 2110 | 2293 | 2027 | 2275 | 1948 | 1996 | 1542 | 1466 | 1437 | 1459 | 1494 | 1571 | 1488 | 2097 | 2047 | 2305 | 2155 | 2080 | 2073 | 1959 | 1571 | 1632 | 1390 | 1409 | 1587 | 1520 | 1726 | Fez2 |
| Fezf1 | 1000 | 847 | 907 | 801 | 524 | 590 | 840 | 1183 | 966 | 1075 | 569 | 1357 | 209 | 1243 | 849 | 470 | 1286 | 560 | 713 | 847 | 685 | 919 | 863 | 1465 | 876 | 2636 | 991 | 1317 | Fezf1 |
| Fezf2 | 705 | 694 | 801 | 913 | 1108 | 1003 | 1478 | 1205 | 1603 | 1677 | 1170 | 2292 | 1399 | 2230 | 618 | 410 | 943 | 873 | 1576 | 1352 | 1377 | 869 | 1419 | 2534 | 2375 | 2988 | 1807 | 2262 | Fezf2 |
| Ffar1 | 0 | 1 | 0 | 0 | 0 | 0 | 0 | 0 | 1 | 0 | 0 | 0 | 0 | 0 | 0 | 2 | 1 | 0 | 0 | 0 | 0 | 2 | 0 | 0 | 0 | 2 | 0 | 0 | Ffar1 |
| Ffar2 | 1 | 2 | 0 | 2 | 7 | 3 | 2 | 2 | 7 | 1 | 13 | 8 | 5 | 5 | 1 | 3 | 0 | 2 | 2 | 4 | 3 | 5 | 8 | 14 | 8 | 16 | 8 | 7 | Ffar2 |
| Ffar3 | 0 | 2 | 0 | 0 | 2 | 0 | 0 | 0 | 0 | 0 | 9 | 0 | 0 | 0 | 0 | 1 | 0 | 4 | 1 | 0 | 0 | 0 | 1 | 0 | 1 | 0 | 0 | 0 | Ffar3 |
| Ffar4 | 1 | 2 | 0 | 2 | 1 | 3 | 0 | 1 | 2 | 1 | 2 | 0 | 1 | 0 | 1 | 3 | 3 | 0 | 0 | 3 | 1 | 3 | 0 | 0 | 1 | 0 | 0 | 0 | Ffar4 |
| Fga | 0 | 1 | 0 | 0 | 8 | 0 | 0 | 0 | 20 | 0 | 0 | 0 | 0 | 0 | 0 | 7 | 0 | 0 | 0 | 0 | 1 | 0 | 1 | 0 | 0 | 0 | 0 | 0 | Fga |
| Fgb | 1 | 0 | 0 | 0 | 2 | 0 | 0 | 0 | 13 | 0 | 1 | 0 | 0 | 0 | 0 | 5 | 0 | 1 | 0 | 1 | 0 | 0 | 10 | 0 | 0 | 0 | 1 | 0 | Fgb |
| Fgd1 | 2459 | 2760 | 3187 | 3304 | 3633 | 3137 | 2867 | 2815 | 3284 | 3674 | 3630 | 3772 | 3219 | 3472 | 2371 | 2849 | 3369 | 3163 | 3685 | 2525 | 3221 | 3641 | 3220 | 3814 | 3695 | 4112 | 2961 | 3406 | Fgd1 |
| Fgd2 | 253 | 223 | 179 | 201 | 168 | 144 | 95 | 186 | 285 | 205 | 274 | 166 | 179 | 186 | 185 | 247 | 176 | 169 | 111 | 128 | 101 | 222 | 318 | 275 | 132 | 305 | 166 | 150 | Fgd2 |
| Fgd3 | 716 | 870 | 699 | 785 | 1341 | 885 | 959 | 1163 | 1369 | 1426 | 1116 | 1054 | 917 | 1247 | 865 | 781 | 898 | 859 | 1027 | 796 | 979 | 1678 | 880 | 1335 | 1592 | 1229 | 1048 | 1097 | Fgd3 |

| | | | | | | | | | | | | | | | | | | | | | | | | | | | | | |
|---|---|---|---|---|---|---|---|---|---|---|---|---|---|---|---|---|---|---|---|---|---|---|---|---|---|---|---|---|---|
| Fgd4 | 2300 | 1824 | 1611 | 2054 | 2950 | 2976 | 3174 | 7222 | 5693 | 4930 | 4634 | 5925 | 5214 | 6701 | 2745 | 2066 | 1909 | 2042 | 2667 | 3287 | 2995 | 5875 | 5073 | 4897 | 6265 | 4027 | 6043 | 6005 | Fgd4 |
| Fgd5 | 1103 | 1368 | 1372 | 1265 | 2014 | 1463 | 1380 | 1642 | 2021 | 2384 | 2629 | 1827 | 1810 | 1798 | 1237 | 1520 | 1460 | 1306 | 1564 | 1056 | 1373 | 2219 | 2017 | 1861 | 2151 | 1558 | 1722 | 1565 | Fgd5 |
| Fgd6 | 5096 | 5009 | 4914 | 5043 | 4922 | 5091 | 4947 | 1960 | 1976 | 2107 | 2269 | 2431 | 2837 | 2291 | 5035 | 5660 | 5188 | 5604 | 4604 | 4403 | 5132 | 2381 | 2142 | 2013 | 2042 | 1508 | 2502 | 2336 | Fgd6 |
| Fgf1 | 13 | 14 | 8 | 6 | 31 | 3 | 10 | 40 | 78 | 47 | 190 | 10 | 24 | 14 | 18 | 5 | 1 | 37 | 6 | 12 | 13 | 57 | 144 | 34 | 22 | 29 | 14 | 17 | Fgf1 |
| Fgf10 | 520 | 224 | 278 | 244 | 305 | 256 | 201 | 479 | 478 | 399 | 728 | 127 | 371 | 92 | 554 | 495 | 235 | 217 | 147 | 209 | 164 | 376 | 532 | 235 | 144 | 282 | 132 | 116 | Fgf10 |
| Fgf11 | 3630 | 3863 | 3938 | 5401 | 4969 | 4418 | 4811 | 1454 | 1535 | 1899 | 1793 | 1443 | 1435 | 1392 | 3609 | 3632 | 4536 | 5401 | 4793 | 3890 | 4950 | 2181 | 1585 | 1642 | 1789 | 1380 | 1115 | 1265 | Fgf11 |
| Fgf12 | 469 | 333 | 487 | 572 | 1057 | 761 | 745 | 967 | 1038 | 1062 | 1052 | 522 | 657 | 592 | 527 | 380 | 524 | 601 | 757 | 762 | 793 | 1313 | 993 | 767 | 1060 | 507 | 483 | 544 | Fgf12 |
| Fgf13 | 2413 | 2553 | 2339 | 3234 | 5933 | 3741 | 4433 | 4807 | 4846 | 4762 | 4991 | 3257 | 4261 | 3349 | 3492 | 3002 | 3091 | 4161 | 3909 | 3537 | 4400 | 7043 | 4243 | 3699 | 5248 | 3015 | 2973 | 3159 | Fgf13 |
| Fgf14 | 612 | 549 | 437 | 771 | 922 | 872 | 1064 | 1290 | 1205 | 905 | 851 | 907 | 689 | 923 | 866 | 624 | 618 | 670 | 829 | 1167 | 963 | 1362 | 962 | 804 | 1109 | 673 | 924 | 808 | Fgf14 |
| Fgf15 | 1465 | 1272 | 1727 | 1201 | 1391 | 1586 | 1493 | 2556 | 1879 | 1853 | 1376 | 3077 | 2105 | 2909 | 1530 | 1026 | 1837 | 945 | 1733 | 1562 | 1446 | 2540 | 1685 | 2330 | 2361 | 2948 | 3281 | 3032 | Fgf15 |
| Fgf16 | 28 | 14 | 16 | 20 | 32 | 3 | 18 | 17 | 21 | 20 | 43 | 2 | 15 | 3 | 18 | 12 | 13 | 11 | 10 | 14 | 15 | 17 | 35 | 12 | 6 | 21 | 0 | 2 | Fgf16 |
| Fgf17 | 185 | 262 | 285 | 332 | 201 | 221 | 227 | 298 | 330 | 278 | 219 | 589 | 601 | 488 | 214 | 244 | 258 | 296 | 302 | 229 | 243 | 189 | 146 | 302 | 322 | 591 | 488 | 544 | Fgf17 |
| Fgf18 | 52 | 97 | 95 | 38 | 113 | 85 | 54 | 150 | 201 | 184 | 299 | 139 | 155 | 103 | 65 | 109 | 81 | 97 | 83 | 54 | 52 | 201 | 283 | 129 | 205 | 127 | 109 | 89 | Fgf18 |
| Fgf2 | 210 | 119 | 153 | 139 | 142 | 202 | 146 | 471 | 503 | 644 | 587 | 324 | 357 | 374 | 170 | 189 | 195 | 165 | 96 | 108 | 109 | 424 | 601 | 450 | 368 | 382 | 311 | 333 | Fgf2 |
| Fgf20 | 37 | 12 | 3 | 36 | 13 | 9 | 8 | 64 | 46 | 74 | 60 | 16 | 25 | 27 | 13 | 12 | 14 | 23 | 10 | 20 | 13 | 25 | 33 | 54 | 35 | 68 | 21 | 25 | Fgf20 |
| Fgf21 | 1 | 0 | 0 | 2 | 0 | 4 | 0 | 10 | 6 | 5 | 4 | 0 | 0 | 0 | 0 | 4 | 0 | 4 | 0 | 0 | 0 | 2 | 8 | 1 | 0 | 2 | 1 | 4 | Fgf21 |
| Fgf22 | 39 | 55 | 82 | 54 | 29 | 66 | 49 | 12 | 12 | 8 | 11 | 4 | 16 | 9 | 42 | 51 | 46 | 73 | 63 | 63 | 39 | 0 | 12 | 5 | 3 | 8 | 22 | 17 | Fgf22 |
| Fgf23 | 14 | 9 | 8 | 7 | 13 | 7 | 8 | 18 | 7 | 9 | 18 | 12 | 8 | 20 | 9 | 9 | 13 | 15 | 26 | 16 | 13 | 20 | 10 | 7 | 9 | 8 | 15 | 9 | Fgf23 |
| Fgf3 | 87 | 77 | 112 | 81 | 160 | 118 | 108 | 221 | 367 | 289 | 221 | 118 | 73 | 118 | 57 | 62 | 88 | 90 | 122 | 100 | 80 | 426 | 299 | 129 | 364 | 159 | 76 | 82 | Fgf3 |
| Fgf4 | 10 | 6 | 22 | 6 | 6 | 5 | 9 | 1 | 2 | 0 | 22 | 0 | 0 | 0 | 5 | 7 | 2 | 6 | 8 | 2 | 5 | 2 | 13 | 0 | 0 | 0 | 0 | 0 | Fgf4 |
| Fgf5 | 29 | 27 | 19 | 20 | 44 | 42 | 35 | 26 | 28 | 56 | 55 | 15 | 17 | 16 | 25 | 29 | 27 | 33 | 35 | 22 | 29 | 45 | 54 | 45 | 39 | 23 | 16 | 30 | Fgf5 |
| Fgf6 | 3 | 1 | 3 | 4 | 3 | 0 | 1 | 5 | 22 | 0 | 146 | 4 | 3 | 0 | 5 | 11 | 0 | 3 | 2 | 2 | 0 | 10 | 144 | 2 | 5 | 0 | 0 | 2 | Fgf6 |
| Fgf7 | 136 | 96 | 171 | 243 | 188 | 196 | 223 | 147 | 233 | 172 | 398 | 105 | 125 | 111 | 120 | 118 | 224 | 278 | 250 | 138 | 196 | 204 | 260 | 125 | 170 | 125 | 122 | 114 | Fgf7 |
| Fgf8 | 92 | 78 | 96 | 68 | 29 | 48 | 38 | 59 | 54 | 65 | 29 | 124 | 103 | 89 | 54 | 104 | 112 | 59 | 65 | 50 | 33 | 43 | 42 | 56 | 55 | 150 | 92 | 88 | Fgf8 |
| Fgf9 | 363 | 319 | 284 | 458 | 479 | 444 | 377 | 403 | 451 | 469 | 485 | 205 | 372 | 194 | 353 | 312 | 304 | 464 | 397 | 394 | 422 | 413 | 482 | 344 | 374 | 275 | 229 | 201 | Fgf9 |
| Fgfbp1 | 21 | 23 | 10 | 11 | 21 | 32 | 17 | 53 | 121 | 47 | 101 | 23 | 59 | 33 | 37 | 29 | 8 | 31 | 23 | 16 | 31 | 40 | 129 | 54 | 43 | 27 | 20 | 38 | Fgfbp1 |
| Fgfbp3 | 1149 | 945 | 1534 | 1462 | 1907 | 1751 | 1956 | 2434 | 1709 | 2337 | 2224 | 2609 | 2601 | 3279 | 1456 | 749 | 1499 | 1439 | 1732 | 1237 | 2061 | 2379 | 1643 | 2469 | 2725 | 2426 | 2688 | 3387 | Fgfbp3 |
| Fgfr1 | 12618 | 13442 | 11830 | 12583 | 11733 | 11397 | 10238 | 9641 | 12031 | 11499 | 11676 | 10202 | 10406 | 9033 | 10995 | 13518 | 12596 | 12909 | 11621 | 9255 | 10507 | 9733 | 11535 | 11246 | 9692 | 12128 | 8825 | 9105 | Fgfr1 |
| Fgfr1op2 | 8554 | 7302 | 6889 | 7395 | 6117 | 6515 | 7507 | 6238 | 5535 | 4972 | 5702 | 5734 | 5709 | 5978 | 8587 | 8023 | 6993 | 7082 | 6283 | 7057 | 6958 | 5051 | 5693 | 5406 | 4764 | 6143 | 5965 | 6018 | Fgfr1op2 |
| Fgfr2 | 7190 | 6973 | 7891 | 6622 | 5292 | 6582 | 5939 | 6508 | 7664 | 7173 | 6690 | 7808 | 7478 | 7965 | 7011 | 7596 | 6928 | 6919 | 5739 | 5217 | 5672 | 5640 | 7348 | 7684 | 5919 | 7941 | 7460 | 8175 | Fgfr2 |
| Fgfr3 | 4109 | 6334 | 7367 | 5820 | 6069 | 6183 | 5939 | 6627 | 6816 | 8127 | 7324 | 12062 | 10150 | 11067 | 4849 | 4618 | 7243 | 5771 | 6742 | 6558 | 5588 | 7478 | 7681 | 10305 | 9031 | 11817 | 11512 | 12040 | Fgfr3 |
| Fgfr4 | 200 | 275 | 291 | 296 | 215 | 270 | 140 | 107 | 180 | 100 | 707 | 53 | 124 | 30 | 229 | 317 | 346 | 315 | 261 | 201 | 189 | 219 | 401 | 88 | 81 | 32 | 49 | 49 | Fgfr4 |
| Fgfrl1 | 2173 | 2756 | 2904 | 2586 | 1847 | 2021 | 1645 | 1198 | 1469 | 1446 | 2065 | 1388 | 1278 | 1190 | 2083 | 2521 | 2710 | 2571 | 2268 | 1489 | 1781 | 1335 | 1437 | 1577 | 1209 | 2193 | 1397 | 1351 | Fgfrl1 |
| Fgg | 1 | 3 | 1 | 4 | 0 | 0 | 0 | 2 | 10 | 0 | 4 | 0 | 0 | 2 | 1 | 5 | 3 | 1 | 4 | 2 | 0 | 0 | 6 | 0 | 0 | 0 | 0 | 0 | Fgg |
| Fggy | 412 | 352 | 286 | 309 | 275 | 347 | 322 | 436 | 369 | 288 | 359 | 378 | 369 | 362 | 386 | 361 | 285 | 310 | 291 | 360 | 293 | 391 | 364 | 311 | 390 | 369 | 425 | 360 | Fggy |
| Fgl1 | 57 | 56 | 53 | 78 | 58 | 43 | 55 | 34 | 32 | 27 | 36 | 12 | 34 | 38 | 51 | 62 | 64 | 50 | 47 | 51 | 70 | 40 | 39 | 41 | 34 | 43 | 36 | 25 | Fgl1 |
| Fgl2 | 251 | 347 | 354 | 334 | 392 | 302 | 192 | 404 | 326 | 223 | 378 | 179 | 382 | 206 | 550 | 428 | 322 | 288 | 221 | 384 | 227 | 336 | 386 | 184 | 248 | 222 | 242 | 227 | Fgl2 |
| Fgr | 6 | 12 | 8 | 6 | 8 | 7 | 13 | 6 | 14 | 23 | 16 | 15 | 8 | 11 | 3 | 9 | 3 | 8 | 4 | 9 | 5 | 13 | 13 | 12 | 2 | 21 | 5 | 5 | Fgr |
| Fh1 | 6240 | 5890 | 5532 | 4588 | 4342 | 4685 | 4434 | 5169 | 4752 | 3961 | 4270 | 4615 | 4604 | 4765 | 5970 | 5661 | 4800 | 4485 | 4648 | 5149 | 4303 | 4223 | 5036 | 4469 | 4272 | 5323 | 5226 | 5096 | Fh1 |
| Fhad1 | 127 | 145 | 118 | 145 | 236 | 195 | 123 | 238 | 210 | 274 | 283 | 244 | 395 | 161 | 210 | 235 | 161 | 131 | 136 | 172 | 168 | 262 | 337 | 168 | 206 | 125 | 203 | 208 | Fhad1 |
| Fhdc1 | 772 | 883 | 888 | 725 | 860 | 734 | 588 | 569 | 703 | 630 | 658 | 451 | 683 | 434 | 766 | 910 | 831 | 841 | 768 | 679 | 611 | 622 | 619 | 512 | 505 | 444 | 443 | 560 | Fhdc1 |
| Fhit | 83 | 89 | 85 | 93 | 104 | 83 | 102 | 116 | 128 | 75 | 104 | 92 | 94 | 66 | 108 | 83 | 71 | 99 | 88 | 97 | 114 | 107 | 110 | 78 | 94 | 66 | 89 | 87 | Fhit |
| Fhl1 | 6756 | 7007 | 6765 | 7751 | 9247 | 8694 | 10963 | 7851 | 7777 | 7844 | 7900 | 7603 | 6889 | 7904 | 7267 | 5999 | 7661 | 8832 | 9826 | 10478 | 10105 | 7930 | 8700 | 7952 | 8872 | 7294 | 7176 | 7533 | Fhl1 |
| Fhl2 | 235 | 220 | 208 | 138 | 187 | 163 | 102 | 271 | 361 | 304 | 353 | 167 | 266 | 170 | 196 | 253 | 147 | 143 | 125 | 109 | 99 | 296 | 438 | 284 | 211 | 222 | 237 | 178 | Fhl2 |
| Fhl3 | 811 | 1021 | 1142 | 835 | 554 | 616 | 432 | 495 | 646 | 592 | 839 | 542 | 862 | 541 | 719 | 970 | 936 | 763 | 588 | 450 | 534 | 369 | 729 | 692 | 441 | 864 | 715 | 572 | Fhl3 |
| Fhl4 | 135 | 67 | 73 | 88 | 66 | 81 | 102 | 115 | 124 | 83 | 92 | 91 | 86 | 115 | 105 | 86 | 92 | 77 | 74 | 94 | 105 | 104 | 133 | 96 | 116 | 94 | 110 | 80 | Fhl4 |
| Fhl5 | 1 | 0 | 0 | 0 | 1 | 0 | 0 | 0 | 0 | 8 | 14 | 0 | 0 | 0 | 0 | 1 | 1 | 0 | 0 | 0 | 0 | 8 | 4 | 3 | 0 | 0 | 0 | 0 | Fhl5 |
| Fhod1 | 2112 | 2656 | 3226 | 2651 | 1851 | 2034 | 1704 | 808 | 1174 | 1103 | 1142 | 1367 | 1424 | 1102 | 1750 | 2636 | 2826 | 2423 | 2125 | 1654 | 1737 | 929 | 1135 | 1277 | 1058 | 1803 | 1232 | 1272 | Fhod1 |
| Fhod3 | 2188 | 2315 | 3219 | 2562 | 3026 | 2670 | 2703 | 2503 | 2465 | 2712 | 2515 | 2927 | 2615 | 2815 | 2484 | 2203 | 2800 | 2464 | 3016 | 2619 | 2774 | 2867 | 2333 | 2798 | 2910 | 2471 | 2752 | 2964 | Fhod3 |
| Fibcd1 | 341 | 515 | 647 | 713 | 875 | 907 | 659 | 499 | 517 | 591 | 462 | 635 | 686 | 534 | 471 | 443 | 791 | 801 | 923 | 743 | 819 | 802 | 520 | 686 | 770 | 122 | 528 | 525 | Fibcd1 |
| Fibin | 526 | 436 | 475 | 307 | 128 | 541 | 55 | 891 | 1698 | 1281 | 4483 | 607 | 1546 | 694 | 568 | 714 | 262 | 453 | 274 | 71 | 37 | 1303 | 3581 | 900 | 1026 | 752 | 704 | 824 | Fibin |
| Fibp | 2722 | 3499 | 2648 | 3169 | 2892 | 3132 | 3333 | 2784 | 2998 | 2915 | 2755 | 2905 | 2970 | 3224 | 2888 | 2633 | 2702 | 2946 | 2982 | 3342 | 3336 | 2817 | 2808 | 2962 | 3133 | 3054 | 2975 | 3234 | Fibp |
| Ficd | 240 | 400 | 456 | 473 | 488 | 370 | 367 | 300 | 393 | 454 | 439 | 365 | 356 | 387 | 322 | 397 | 416 | 419 | 430 | 286 | 357 | 453 | 402 | 537 | 416 | 442 | 346 | 389 | Ficd |
| Fig4 | 4411 | 4872 | 3975 | 4393 | 4138 | 4002 | 4509 | 2489 | 2063 | 1929 | 1839 | 1955 | 1825 | 1994 | 5192 | 4224 | 4059 | 4380 | 4208 | 4600 | 4546 | 2344 | 2131 | 1851 | 2097 | 1766 | 2050 | 1880 | Fig4 |
| Figla | 1 | 1 | 1 | 3 | 5 | 2 | 2 | 0 | 2 | 4 | 0 | 0 | 0 | 0 | 0 | 0 | 3 | 0 | 2 | 3 | 4 | 3 | 1 | 0 | 0 | 3 | 1 | 0 | Figla |
| Fign | 1266 | 1261 | 2234 | 1968 | 2254 | 1950 | 2091 | 2119 | 1642 | 2528 | 2485 | 2735 | 2239 | 2886 | 1465 | 1350 | 2439 | 1874 | 2237 | 1506 | 2010 | 3526 | 1605 | 2558 | 2880 | 2824 | 2699 | 3023 | Fign |
| Fignl1 | 5001 | 3638 | 4681 | 3210 | 2596 | 2802 | 3389 | 4605 | 3981 | 3418 | 3183 | 4952 | 4064 | 5019 | 4606 | 3959 | 3863 | 2851 | 3116 | 3788 | 2998 | 2851 | 3480 | 4275 | 3214 | 6247 | 5260 | 5436 | Fignl1 |
| Fignl2 | 646 | 802 | 1353 | 642 | 603 | 548 | 446 | 432 | 361 | 437 | 474 | 593 | 880 | 580 | 596 | 900 | 918 | 489 | 506 | 426 | 380 | 431 | 400 | 462 | 365 | 732 | 870 | 818 | Fignl2 |
| Filip1 | 1359 | 1361 | 1827 | 1060 | 784 | 1011 | 977 | 804 | 666 | 708 | 855 | 1548 | 1245 | 1453 | 1321 | 1027 | 1467 | 797 | 981 | 1146 | 895 | 697 | 763 | 934 | 806 | 1284 | 1970 | 1650 | Filip1 |
| Filip1l | 1374 | 866 | 1000 | 527 | 365 | 646 | 293 | 998 | 1012 | 912 | 1535 | 841 | 1123 | 749 | 1014 | 1354 | 731 | 697 | 406 | 361 | 359 | 655 | 1410 | 968 | 575 | 1126 | 923 | 776 | Filip1l |
| Fip1l1 | 9538 | 8193 | 10519 | 10821 | 10276 | 10082 | 11419 | 9314 | 8588 | 10483 | 9932 | 11473 | 10227 | 12061 | 9244 | 8913 | 10353 | 10596 | 10491 | 8908 | 10843 | 11224 | 8620 | 11485 | 10756 | 11632 | 10419 | 11699 | Fip1l1 |
| Fis1 | 4553 | 5485 | 4782 | 5076 | 5063 | 5146 | 5500 | 5692 | 6060 | 5589 | 5782 | 5841 | 5903 | 6394 | 4969 | 4586 | 4941 | 4913 | 5327 | 5951 | 5543 | 5220 | 6041 | 6216 | 6325 | 6647 | 6542 | 6849 | Fis1 |
| Fitm1 | 5 | 3 | 0 | 7 | 1 | 0 | 3 | 4 | 28 | 1 | 174 | 5 | 14 | 2 | 7 | 2 | 3 | 1 | 0 | 2 | 2 | 8 | 84 | 0 | 0 | 1 | 0 | 0 | Fitm1 |
| Fitm2 | 1397 | 1489 | 1475 | 1568 | 1595 | 1500 | 1403 | 1137 | 1223 | 1220 | 1320 | 1260 | 1183 | 1229 | 1399 | 1541 | 1546 | 1818 | 1484 | 1448 | 1554 | 1272 | 1188 | 1218 | 1290 | 1250 | 1166 | 1221 | Fitm2 |
| Fiz1 | 1688 | 1864 | 1935 | 1734 | 1789 | 1788 | 1690 | 3736 | 3715 | 3628 | 2851 | 4375 | 3813 | 3981 | 1755 | 1550 | 1998 | 1414 | 2096 | 1848 | 1649 | 3357 | 3440 | 3959 | 3630 | 4831 | 4326 | 4174 | Fiz1 |
| Fjx1 | 1570 | 1830 | 1917 | 2244 | 2453 | 2087 | 2219 | 2331 | 2222 | 2731 | 2741 | 3107 | 2673 | 3128 | 1569 | 1429 | 2260 | 2344 | 2496 | 2005 | 2288 | 2714 | 2476 | 3028 | 3064 | 3205 | 2920 | 3078 | Fjx1 |
| Fkbp10 | 2900 | 3421 | 3284 | 2227 | 1539 | 2662 | 1066 | 2391 | 3870 | 3292 | 5202 | 2968 | 3841 | 2591 | 2711 | 4118 | 2359 | 2539 | 1409 | 1218 | 1128 | 3203 | 4691 | 3234 | 2882 | 3697 | 3349 | 3056 | Fkbp10 |
| Fkbp11 | 358 | 467 | 431 | 440 | 278 | 327 | 299 | 195 | 270 | 204 | 364 | 179 | 175 | 182 | 344 | 421 | 423 | 365 | 307 | 299 | 236 | 222 | 296 | 236 | 187 | 218 | 202 | 172 | Fkbp11 |
| Fkbp14 | 5154 | 5056 | 4597 | 5006 | 4613 | 4352 | 4782 | 2009 | 2039 | 1795 | 2603 | 1411 | 1826 | 1528 | 5620 | 5335 | 4839 | 5507 | 4479 | 4784 | 4742 | 2024 | 2497 | 1768 | 1765 | 1653 | 1677 | 1639 | Fkbp14 |
| Fkbp15 | 4457 | 4817 | 3604 | 3838 | 3630 | 4104 | 3704 | 3507 | 4046 | 3601 | 3547 | 3353 | 3661 | 3384 | 4275 | 4389 | 3494 | 3822 | 3539 | 4168 | 4009 | 3394 | 3943 | 3105 | 3478 | 2823 | 3507 | 3199 | Fkbp15 |
| Fkbp1a | 14821 | 13333 | 17183 | 15869 | 15352 | 13600 | 15492 | 12669 | 12475 | 13629 | 13692 | 13630 | 13959 | 14944 | 14629 | 13752 | 15541 | 14902 | 15534 | 13250 | 15748 | 12691 | 13017 | 14226 | 14159 | 15833 | 13294 | 15394 | Fkbp1a |
| Fkbp1b | 315 | 343 | 280 | 313 | 546 | 316 | 265 | 397 | 494 | 486 | 513 | 206 | 405 | 235 | 352 | 424 | 327 | 390 | 286 | 318 | 291 | 600 | 455 | 224 | 439 | 220 | 239 | 250 | Fkbp1b |
| Fkbp2 | 1145 | 1649 | 2076 | 2070 | 1838 | 1435 | 1592 | 1412 | 1645 | 2005 | 1881 | 1874 | 1776 | 1902 | 1135 | 1294 | 2107 | 1801 | 1855 | 1670 | 1604 | 1502 | 1545 | 2256 | 1958 | 2292 | 2017 | 1976 | Fkbp2 |
| Fkbp3 | 12457 | 10218 | 13270 | 11075 | 9345 | 9986 | 12800 | 12433 | 10055 | 10788 | 10448 | 13660 | 12775 | 15364 | 13177 | 11850 | 12118 | 10255 | 10391 | 13192 | 11229 | 10163 | 10552 | 12392 | 11294 | 15226 | 14264 | 16008 | Fkbp3 |
| Fkbp4 | 27148 | 29936 | 31668 | 26463 | 22299 | 23222 | 24818 | 23588 | 22079 | 22241 | 20037 | 25574 | 23056 | 26420 | 24878 | 24820 | 29744 | 23804 | 26095 | 25430 | 24436 | 18815 | 21065 | 25520 | 22564 | 30178 | 26624 | 27728 | Fkbp4 |
| Fkbp5 | 6714 | 6689 | 8551 | 7823 | 7355 | 7026 | 7259 | 3671 | 4033 | 4686 | 4315 | 5315 | 4568 | 5188 | 6128 | 7105 | 8774 | 7820 | 7782 | 5645 | 7398 | 4415 | 3667 | 5029 | 4983 | 5817 | 4486 | 5419 | Fkbp5 |
| Fkbp6 | 2 | 1 | 3 | 5 | 3 | 0 | 1 | 1 | 3 | 8 | 3 | 0 | 5 | 0 | 4 | 1 | 7 | 0 | 1 | 9 | 2 | 10 | 1 | 0 | 1 | 0 | 0 | 5 | Fkbp6 |
| Fkbp7 | 932 | 616 | 584 | 631 | 461 | 613 | 548 | 806 | 869 | 614 | 1134 | 540 | 666 | 587 | 881 | 908 | 499 | 638 | 370 | 465 | 521 | 710 | 1140 | 690 | 644 | 677 | 628 | 630 | Fkbp7 |
| Fkbp8 | 9397 | 12079 | 10081 | 10914 | 10216 | 10365 | 9312 | 10683 | 12710 | 12028 | 10763 | 12487 | 11330 | 11278 | 9381 | 9830 | 10802 | 10505 | 12059 | 10644 | 10383 | 10738 | 12323 | 12349 | 12309 | 13363 | 12154 | 11603 | Fkbp8 |

| | | | | | | | | | | | | | | | | | | | | | | | | | | | | | |
|---|---|---|---|---|---|---|---|---|---|---|---|---|---|---|---|---|---|---|---|---|---|---|---|---|---|---|---|---|---|
| Fkbp9 | 8522 | 7186 | 7493 | 4994 | 3660 | 5565 | 2890 | 7858 | 10031 | 8237 | 11706 | 6967 | 8486 | 6622 | 7812 | 9538 | 5398 | 5852 | 3232 | 3334 | 2966 | 7812 | 11378 | 7588 | 6810 | 8339 | 7699 | 7543 | Fkbp9 |
| Fkbpl | 563 | 783 | 706 | 647 | 489 | 604 | 573 | 497 | 559 | 434 | 464 | 489 | 497 | 584 | 634 | 759 | 709 | 609 | 574 | 644 | 673 | 428 | 489 | 503 | 452 | 574 | 592 | 557 | Fkbpl |
| Fkrp | 784 | 843 | 920 | 1007 | 1185 | 1070 | 889 | 2560 | 2250 | 2632 | 2570 | 2498 | 2371 | 2510 | 907 | 778 | 1055 | 964 | 1138 | 1047 | 960 | 2904 | 2531 | 2562 | 2505 | 2508 | 2597 | 2648 | Fkrp |
| Fktn | 3483 | 3097 | 3161 | 2886 | 2891 | 3318 | 3313 | 4710 | 4064 | 3913 | 3890 | 4294 | 3816 | 4576 | 3630 | 3135 | 2790 | 2610 | 3087 | 3435 | 3304 | 4248 | 3924 | 3755 | 3731 | 3390 | 4717 | 4565 | Fktn |
| Flacc1 | 0 | 1 | 0 | 0 | 0 | 0 | 1 | 1 | 1 | 0 | 0 | 2 | 1 | 1 | 3 | 0 | 0 | 0 | 2 | 0 | 0 | 0 | 0 | 3 | 0 | 1 | 2 | 0 | Flacc1 |
| Flad1 | 964 | 1164 | 1122 | 1102 | 1059 | 1118 | 1048 | 1502 | 1561 | 1669 | 1465 | 1795 | 1591 | 1686 | 901 | 900 | 1209 | 1081 | 1220 | 937 | 1098 | 1576 | 1425 | 1694 | 1771 | 1539 | 1581 | 1680 | Flad1 |
| Flcn | 2628 | 3028 | 2646 | 3003 | 2617 | 2530 | 2476 | 2364 | 2589 | 2645 | 2742 | 2509 | 2624 | 2448 | 2461 | 2929 | 2710 | 2995 | 2837 | 2523 | 2578 | 2787 | 2717 | 2599 | 2855 | 2726 | 2410 | 2490 | Flcn |
| Flg2 | 0 | 0 | 0 | 0 | 0 | 0 | 1 | 0 | 0 | 0 | 0 | 0 | 0 | 0 | 1 | 0 | 0 | 0 | 0 | 0 | 0 | 2 | 1 | 0 | 0 | 0 | 0 | 0 | Flg2 |
| Fli1 | 588 | 396 | 781 | 415 | 425 | 402 | 279 | 862 | 1025 | 971 | 1429 | 777 | 1107 | 751 | 493 | 630 | 388 | 391 | 345 | 409 | 328 | 1033 | 1182 | 1028 | 822 | 1316 | 737 | 853 | Fli1 |
| Flii | 5447 | 6765 | 6978 | 6161 | 6230 | 6025 | 5639 | 5901 | 6897 | 7076 | 6640 | 7493 | 7192 | 7201 | 5234 | 5969 | 6953 | 6122 | 6770 | 5577 | 5732 | 6916 | 6649 | 7483 | 7214 | 7330 | 7254 | 7220 | Flii |
| Flna | 76631 | 75685 | 48419 | 45525 | 40606 | 53312 | 35525 | 57968 | 72980 | 60436 | 68961 | 69483 | 81585 | 59190 | 66527 | 83224 | 45236 | 46857 | 42390 | 41013 | 40067 | 50319 | 74261 | 56719 | 58769 | 43208 | 77469 | 62386 | Flna |
| Flnb | 21058 | 22983 | 13453 | 16233 | 13398 | 18160 | 12635 | 10063 | 15796 | 13648 | 16638 | 12502 | 15660 | 11162 | 18327 | 24853 | 14202 | 17787 | 13858 | 12449 | 13928 | 12897 | 14856 | 12081 | 12717 | 8399 | 13709 | 10980 | Flnb |
| Flnc | 3646 | 4367 | 4644 | 3411 | 2860 | 4091 | 1753 | 3529 | 6504 | 4566 | 11184 | 3780 | 5271 | 3027 | 3513 | 5461 | 3752 | 4493 | 2876 | 2154 | 2073 | 4614 | 10245 | 4772 | 4163 | 2455 | 3765 | 3310 | Flnc |
| Flot1 | 2824 | 3429 | 3741 | 3915 | 3715 | 3406 | 3077 | 1713 | 2209 | 2377 | 2254 | 2419 | 2168 | 2191 | 2664 | 3076 | 3870 | 3581 | 3879 | 2987 | 3240 | 2239 | 2236 | 2434 | 2413 | 2606 | 2244 | 2195 | Flot1 |
| Flot2 | 4451 | 5233 | 5008 | 5182 | 5332 | 5032 | 4789 | 3623 | 4299 | 4136 | 3992 | 4428 | 4158 | 4205 | 4265 | 4689 | 5026 | 5230 | 5356 | 4597 | 5051 | 4288 | 4380 | 4108 | 4482 | 4806 | 4323 | 4376 | Flot2 |
| Flrt1 | 1069 | 1329 | 1201 | 1483 | 1746 | 1568 | 1302 | 810 | 1129 | 1294 | 1076 | 1196 | 1151 | 955 | 1006 | 1105 | 1627 | 1643 | 1771 | 1340 | 1540 | 1135 | 970 | 1274 | 1344 | 346 | 1120 | 889 | Flrt1 |
| Flrt2 | 6549 | 4942 | 5457 | 4896 | 6259 | 6782 | 5916 | 11163 | 11603 | 9541 | 13440 | 7786 | 9412 | 7751 | 6394 | 6939 | 4722 | 5657 | 5629 | 5944 | 5942 | 10136 | 13180 | 7896 | 8896 | 8038 | 7772 | 6773 | Flrt2 |
| Flrt3 | 8022 | 6300 | 6442 | 5530 | 4648 | 5369 | 5879 | 7544 | 6026 | 5724 | 5363 | 7168 | 7136 | 7453 | 7611 | 7103 | 5626 | 5011 | 5548 | 7047 | 5487 | 5964 | 6592 | 6083 | 5174 | 8173 | 8451 | 8738 | Flrt3 |
| Flt1 | 1774 | 1621 | 2228 | 2091 | 2044 | 2029 | 1779 | 1356 | 1475 | 1998 | 2083 | 1965 | 1895 | 2033 | 1631 | 2092 | 1998 | 2135 | 1813 | 1400 | 1984 | 2181 | 1690 | 2116 | 2078 | 1761 | 1837 | 2023 | Flt1 |
| Flt3 | 18 | 5 | 16 | 7 | 13 | 16 | 9 | 13 | 17 | 14 | 42 | 9 | 17 | 9 | 7 | 16 | 0 | 8 | 4 | 9 | 8 | 23 | 14 | 14 | 18 | 24 | 14 | 6 | Flt3 |
| Flt3l | 69 | 68 | 93 | 78 | 90 | 56 | 64 | 54 | 55 | 69 | 76 | 63 | 87 | 67 | 86 | 79 | 101 | 94 | 68 | 60 | 65 | 58 | 69 | 81 | 47 | 57 | 53 | 110 | Flt3l |
| Flt4 | 780 | 980 | 1058 | 645 | 703 | 692 | 564 | 535 | 690 | 736 | 1025 | 733 | 904 | 620 | 630 | 1045 | 870 | 731 | 706 | 452 | 612 | 747 | 764 | 854 | 628 | 833 | 652 | 602 | Flt4 |
| Flvcr1 | 3511 | 2991 | 3032 | 3268 | 3076 | 3075 | 3408 | 1599 | 1570 | 1463 | 1468 | 1827 | 1803 | 1964 | 3341 | 3312 | 3161 | 3317 | 3014 | 3272 | 3363 | 1997 | 1430 | 1609 | 1816 | 2064 | 1876 | 2172 | Flvcr1 |
| Flvcr2 | 295 | 294 | 358 | 321 | 311 | 236 | 256 | 195 | 185 | 232 | 185 | 174 | 205 | 180 | 249 | 316 | 202 | 370 | 257 | 255 | 290 | 271 | 249 | 164 | 253 | 157 | 163 | 163 | Flvcr2 |
| Flywch1 | 7157 | 8664 | 7941 | 8450 | 8917 | 8685 | 8079 | 7293 | 7654 | 8687 | 7287 | 9127 | 8540 | 8636 | 7199 | 7041 | 8952 | 8153 | 9606 | 8518 | 8603 | 8584 | 6830 | 8532 | 8631 | 8513 | 8638 | 8853 | Flywch1 |
| Flywch2 | 384 | 511 | 476 | 532 | 542 | 477 | 456 | 506 | 552 | 592 | 635 | 472 | 598 | 567 | 400 | 445 | 491 | 464 | 538 | 632 | 475 | 485 | 617 | 556 | 566 | 747 | 559 | 523 | Flywch2 |
| Fmc1 | 331 | 373 | 388 | 421 | 442 | 317 | 451 | 492 | 421 | 588 | 536 | 545 | 540 | 672 | 327 | 212 | 532 | 311 | 402 | 494 | 423 | 436 | 427 | 662 | 528 | 545 | 535 | 610 | Fmc1 |
| Fmn1 | 203 | 190 | 70 | 78 | 151 | 161 | 50 | 414 | 506 | 284 | 443 | 164 | 507 | 216 | 264 | 257 | 87 | 129 | 37 | 62 | 45 | 421 | 447 | 145 | 191 | 115 | 225 | 221 | Fmn1 |
| Fmn2 | 3941 | 3679 | 4992 | 5054 | 5469 | 4770 | 5140 | 3300 | 2808 | 3639 | 3118 | 4393 | 3999 | 4604 | 4118 | 3608 | 5261 | 4818 | 5227 | 3859 | 5087 | 4756 | 2624 | 3967 | 3916 | 3583 | 4379 | 4494 | Fmn2 |
| Fmnl1 | 343 | 461 | 601 | 882 | 1604 | 1028 | 1000 | 485 | 607 | 714 | 683 | 449 | 449 | 391 | 410 | 421 | 687 | 1073 | 1194 | 988 | 1186 | 989 | 592 | 681 | 779 | 370 | 442 | 312 | Fmnl1 |
| Fmnl2 | 14822 | 10978 | 12020 | 13419 | 14699 | 14673 | 16561 | 14337 | 12545 | 12891 | 12315 | 12796 | 12624 | 13253 | 14135 | 12603 | 12599 | 14038 | 14341 | 13665 | 16456 | 12673 | 11730 | 13328 | 13422 | 11195 | 11595 | 12487 | Fmnl2 |
| Fmnl3 | 3548 | 3373 | 3648 | 3294 | 2908 | 3087 | 2738 | 2926 | 3573 | 3518 | 3917 | 3565 | 3822 | 3101 | 2851 | 3808 | 3023 | 3220 | 3014 | 2350 | 2641 | 3227 | 3834 | 3603 | 3048 | 4147 | 3264 | 3444 | Fmnl3 |
| Fmo1 | 1025 | 1069 | 1059 | 839 | 732 | 1377 | 444 | 1873 | 2301 | 2113 | 2297 | 1627 | 2411 | 1868 | 1458 | 1377 | 705 | 1261 | 489 | 592 | 547 | 2904 | 3199 | 2080 | 2228 | 1102 | 1958 | 1741 | Fmo1 |
| Fmo2 | 43 | 35 | 6 | 26 | 27 | 26 | 11 | 105 | 103 | 113 | 129 | 20 | 43 | 18 | 53 | 44 | 32 | 44 | 20 | 20 | 7 | 52 | 194 | 64 | 26 | 24 | 5 | 7 | Fmo2 |
| Fmo3 | 2 | 1 | 0 | 4 | 0 | 1 | 0 | 0 | 3 | 0 | 0 | 0 | 1 | 1 | 0 | 0 | 0 | 0 | 0 | 0 | 0 | 0 | 3 | 1 | 1 | 0 | 2 | 0 | Fmo3 |
| Fmo4 | 12 | 21 | 47 | 23 | 17 | 36 | 8 | 51 | 39 | 46 | 50 | 23 | 56 | 32 | 37 | 35 | 12 | 29 | 5 | 18 | 12 | 60 | 60 | 24 | 17 | 18 | 36 | 31 | Fmo4 |
| Fmo5 | 106 | 79 | 60 | 41 | 64 | 71 | 47 | 114 | 145 | 103 | 133 | 71 | 119 | 85 | 83 | 131 | 60 | 102 | 45 | 35 | 33 | 95 | 170 | 105 | 74 | 131 | 102 | 84 | Fmo5 |
| Fmo6 | 21 | 3 | 0 | 20 | 0 | 0 | 0 | 93 | 50 | 126 | 54 | 0 | 16 | 13 | 8 | 12 | 2 | 0 | 0 | 0 | 0 | 2 | 126 | 128 | 0 | 62 | 0 | 1 | Fmo6 |
| Fmo9 | 0 | 1 | 0 | 0 | 0 | 0 | 0 | 0 | 0 | 0 | 1 | 0 | 0 | 0 | 0 | 2 | 0 | 0 | 0 | 5 | 0 | 3 | 1 | 0 | 0 | 1 | 0 | 1 | Fmo9 |
| Fmod | 772 | 1070 | 1059 | 718 | 497 | 957 | 313 | 664 | 1018 | 879 | 2052 | 1031 | 1427 | 951 | 1182 | 1276 | 984 | 1091 | 425 | 469 | 369 | 1377 | 1939 | 1017 | 1049 | 639 | 1347 | 999 | Fmod |
| Fmr1 | 11205 | 7942 | 10237 | 10385 | 9781 | 10925 | 11984 | 11019 | 9744 | 9230 | 11075 | 10620 | 10455 | 11707 | 10929 | 9927 | 9755 | 10310 | 9824 | 10102 | 11353 | 9672 | 10183 | 9896 | 10190 | 10430 | 10667 | 10837 | Fmr1 |
| Fmr1nb | 8 | 8 | 7 | 9 | 2 | 9 | 4 | 2 | 3 | 6 | 4 | 1 | 9 | 7 | 3 | 9 | 7 | 8 | 2 | 4 | 0 | 8 | 7 | 2 | 1 | 0 | 3 | 8 | Fmr1nb |
| Fn1 | 41956 | 45103 | 36707 | 37820 | 24754 | 43658 | 18471 | 31036 | 47705 | 53065 | 71870 | 51538 | 60510 | 47135 | 36644 | 55598 | 34078 | 46425 | 24285 | 18413 | 19461 | 47534 | 60609 | 50959 | 45031 | 36880 | 49015 | 42671 | Fn1 |
| Fn3k | 317 | 384 | 285 | 386 | 388 | 342 | 364 | 228 | 208 | 210 | 194 | 193 | 236 | 204 | 353 | 329 | 304 | 341 | 344 | 373 | 390 | 207 | 230 | 183 | 225 | 148 | 218 | 212 | Fn3k |
| Fn3krp | 2330 | 2505 | 2729 | 2691 | 2305 | 2161 | 2386 | 1337 | 1280 | 1272 | 1176 | 1429 | 1390 | 1464 | 2582 | 2263 | 2819 | 2437 | 2383 | 2370 | 2434 | 1390 | 1280 | 1330 | 1339 | 1575 | 1474 | 1589 | Fn3krp |
| Fnbp1 | 4941 | 5212 | 5156 | 5468 | 6041 | 5005 | 4584 | 4008 | 4963 | 5691 | 5882 | 4665 | 4981 | 4239 | 4804 | 5920 | 5372 | 5943 | 5033 | 4196 | 4704 | 5267 | 4853 | 4984 | 5086 | 4527 | 4038 | 4156 | Fnbp1 |
| Fnbp1l | 26813 | 21693 | 23326 | 30821 | 41919 | 39087 | 43409 | 66072 | 54651 | 57404 | 50971 | 51287 | 44719 | 57248 | 30387 | 22935 | 27983 | 33277 | 37477 | 42027 | 42906 | 66581 | 51486 | 55026 | 65499 | 43837 | 48162 | 48087 | Fnbp1l |
| Fnbp4 | 11808 | 10655 | 10367 | 10916 | 10780 | 11457 | 11462 | 11883 | 10828 | 11161 | 10800 | 12852 | 11624 | 12927 | 11214 | 10509 | 10624 | 10173 | 11065 | 10651 | 11448 | 12714 | 10092 | 11431 | 12164 | 11647 | 12857 | 12570 | Fnbp4 |
| Fnd3c2 | 2 | 1 | 3 | 0 | 2 | 0 | 0 | 0 | 2 | 0 | 0 | 1 | 0 | 0 | 0 | 1 | 0 | 0 | 0 | 0 | 0 | 0 | 3 | 0 | 0 | 0 | 1 | 0 | Fnd3c2 |
| Fndc1 | 381 | 251 | 74 | 184 | 268 | 322 | 171 | 419 | 774 | 594 | 1003 | 226 | 289 | 176 | 183 | 329 | 194 | 350 | 175 | 143 | 142 | 413 | 957 | 404 | 443 | 122 | 134 | 158 | Fndc1 |
| Fndc10 | 737 | 828 | 863 | 749 | 790 | 740 | 665 | 546 | 784 | 680 | 901 | 562 | 667 | 545 | 695 | 773 | 712 | 856 | 673 | 721 | 642 | 687 | 783 | 634 | 748 | 719 | 612 | 535 | Fndc10 |
| Fndc11 | 6 | 2 | 1 | 3 | 1 | 4 | 1 | 4 | 3 | 1 | 0 | 5 | 7 | 1 | 4 | 2 | 5 | 0 | 3 | 5 | 1 | 5 | 5 | 1 | 0 | 2 | 13 | 1 | Fndc11 |
| Fndc3a | 11724 | 9903 | 9974 | 9973 | 10730 | 11071 | 11086 | 13715 | 12057 | 12059 | 11975 | 13265 | 12597 | 13882 | 11836 | 10914 | 10036 | 9939 | 10406 | 11015 | 10882 | 13256 | 12102 | 12540 | 12911 | 11126 | 13539 | 13070 | Fndc3a |
| Fndc3b | 3357 | 2855 | 2765 | 2820 | 3086 | 3166 | 2644 | 3651 | 4386 | 3921 | 6153 | 3243 | 4136 | 3070 | 3034 | 3496 | 2669 | 3093 | 2892 | 2402 | 2737 | 4448 | 5178 | 3379 | 3256 | 2463 | 3532 | 3213 | Fndc3b |
| Fndc3c1 | 6571 | 4753 | 5173 | 2368 | 1333 | 1579 | 1646 | 1824 | 1720 | 969 | 2176 | 2300 | 2432 | 1864 | 4700 | 6044 | 3290 | 1779 | 1654 | 2284 | 1414 | 1268 | 1550 | 1375 | 992 | 3208 | 2832 | 2592 | Fndc3c1 |
| Fndc4 | 2741 | 2804 | 2988 | 2833 | 3250 | 3014 | 3042 | 3606 | 3934 | 3848 | 4312 | 3721 | 3485 | 3902 | 2581 | 2572 | 2877 | 2758 | 3423 | 2636 | 3086 | 3878 | 3990 | 3982 | 3932 | 4086 | 3612 | 3748 | Fndc4 |
| Fndc5 | 224 | 244 | 280 | 208 | 348 | 226 | 199 | 430 | 606 | 449 | 1717 | 230 | 305 | 280 | 244 | 301 | 243 | 293 | 232 | 252 | 229 | 495 | 1240 | 351 | 339 | 227 | 218 | 190 | Fndc5 |
| Fndc7 | 6 | 5 | 2 | 2 | 3 | 15 | 4 | 12 | 15 | 3 | 13 | 5 | 10 | 3 | 12 | 4 | 7 | 4 | 5 | 3 | 11 | 10 | 17 | 8 | 5 | 4 | 12 | 6 | Fndc7 |
| Fndc8 | 9 | 5 | 6 | 0 | 0 | 0 | 0 | 5 | 0 | 1 | 0 | 2 | 3 | 0 | 0 | 2 | 0 | 3 | 1 | 2 | 1 | 5 | 5 | 2 | 2 | 1 | 0 | 1 | Fndc8 |
| Fndc9 | 14 | 12 | 41 | 30 | 33 | 34 | 29 | 104 | 108 | 83 | 130 | 127 | 73 | 150 | 25 | 7 | 34 | 21 | 32 | 32 | 44 | 165 | 118 | 144 | 192 | 127 | 101 | 134 | Fndc9 |
| Fnip1 | 8273 | 6283 | 5311 | 7113 | 6394 | 7647 | 8153 | 5311 | 4460 | 4179 | 4635 | 4138 | 4188 | 4470 | 8048 | 7253 | 5704 | 7323 | 6242 | 7513 | 7521 | 4268 | 4420 | 3980 | 4529 | 3399 | 4434 | 4249 | Fnip1 |
| Fnip2 | 3696 | 3827 | 4234 | 4922 | 5415 | 5145 | 5485 | 2898 | 3026 | 3543 | 3414 | 3610 | 3508 | 3654 | 3811 | 4417 | 4520 | 5291 | 5273 | 4324 | 5276 | 3923 | 2842 | 3294 | 3838 | 2677 | 3254 | 3553 | Fnip2 |
| Fnta | 4440 | 4229 | 4062 | 3771 | 3555 | 3431 | 3778 | 5340 | 4622 | 4212 | 4368 | 4304 | 4393 | 4781 | 4572 | 4270 | 3773 | 3737 | 3508 | 3813 | 3777 | 4939 | 5113 | 4193 | 4275 | 4603 | 4319 | 4923 | Fnta |
| Fntb | 1164 | 1297 | 1426 | 1038 | 1164 | 1121 | 1106 | 1761 | 2233 | 2058 | 2076 | 2317 | 2131 | 2008 | 1139 | 1323 | 1207 | 1144 | 1296 | 1077 | 1119 | 2109 | 2183 | 2211 | 1983 | 2552 | 1930 | 2137 | Fntb |
| Focad | 3846 | 3841 | 2779 | 3101 | 3115 | 3237 | 3453 | 2378 | 2429 | 2117 | 2113 | 2229 | 2093 | 2173 | 3676 | 3621 | 2838 | 3147 | 3122 | 3598 | 3603 | 2022 | 2568 | 2036 | 2241 | 1740 | 2351 | 2263 | Focad |
| Folh1 | 8 | 5 | 7 | 4 | 18 | 4 | 11 | 15 | 26 | 27 | 18 | 5 | 7 | 6 | 15 | 11 | 0 | 6 | 14 | 3 | 13 | 23 | 27 | 16 | 26 | 11 | 13 | 9 | Folh1 |
| Folr1 | 91 | 192 | 119 | 78 | 250 | 135 | 146 | 198 | 264 | 231 | 171 | 310 | 302 | 146 | 141 | 164 | 76 | 116 | 178 | 176 | 164 | 294 | 441 | 149 | 235 | 170 | 220 | 215 | Folr1 |
| Folr2 | 51 | 70 | 77 | 56 | 63 | 84 | 42 | 85 | 87 | 113 | 124 | 137 | 126 | 86 | 58 | 90 | 79 | 77 | 68 | 82 | 68 | 120 | 138 | 109 | 116 | 112 | 108 | 94 | Folr2 |
| Fos | 127 | 197 | 130 | 183 | 94 | 140 | 117 | 73 | 93 | 81 | 100 | 58 | 66 | 44 | 140 | 266 | 129 | 220 | 129 | 112 | 85 | 40 | 107 | 80 | 64 | 65 | 34 | 39 | Fos |
| Fosb | 62 | 60 | 57 | 70 | 60 | 42 | 63 | 2 | 12 | 16 | 30 | 0 | 12 | 7 | 54 | 71 | 105 | 89 | 45 | 28 | 32 | 8 | 14 | 21 | 10 | 17 | 6 | 4 | Fosb |
| Fosl1 | 27 | 27 | 19 | 31 | 25 | 23 | 17 | 1 | 3 | 2 | 4 | 0 | 3 | 2 | 12 | 25 | 22 | 18 | 16 | 29 | 12 | 0 | 3 | 2 | 4 | 4 | 5 | 0 | Fosl1 |
| Fosl2 | 644 | 647 | 642 | 623 | 562 | 777 | 410 | 943 | 1243 | 1075 | 1145 | 677 | 799 | 742 | 703 | 791 | 622 | 908 | 515 | 392 | 526 | 615 | 1487 | 1050 | 854 | 631 | 652 | 738 | Fosl2 |
| Foxa1 | 378 | 295 | 904 | 339 | 346 | 327 | 467 | 843 | 530 | 555 | 526 | 769 | 536 | 721 | 388 | 276 | 563 | 266 | 463 | 513 | 379 | 703 | 575 | 690 | 582 | 598 | 938 | 896 | Foxa1 |
| Foxa2 | 196 | 211 | 567 | 220 | 200 | 190 | 254 | 238 | 145 | 188 | 173 | 404 | 400 | 369 | 243 | 217 | 321 | 167 | 240 | 206 | 200 | 274 | 118 | 254 | 188 | 97 | 468 | 501 | Foxa2 |
| Foxa3 | 2 | 3 | 3 | 6 | 7 | 0 | 3 | 2 | 7 | 8 | 36 | 0 | 0 | 0 | 1 | 5 | 6 | 2 | 9 | 0 | 6 | 3 | 3 | 4 | 1 | 1 | 0 | 0 | Foxa3 |
| Foxb1 | 263 | 314 | 646 | 391 | 477 | 368 | 361 | 360 | 302 | 469 | 374 | 465 | 318 | 441 | 245 | 248 | 477 | 324 | 478 | 389 | 428 | 533 | 471 | 373 | 475 | 376 | 513 | 498 | Foxb1 |
| Foxb2 | 21 | 27 | 59 | 29 | 81 | 46 | 22 | 59 | 111 | 81 | 65 | 34 | 62 | 43 | 47 | 24 | 64 | 55 | 19 | 27 | 15 | 102 | 101 | 27 | 67 | 43 | 51 | 36 | Foxb2 |
| Foxc1 | 1595 | 1234 | 2058 | 1234 | 952 | 1030 | 732 | 2028 | 1966 | 2142 | 2598 | 2161 | 2767 | 2192 | 1654 | 2130 | 1347 | 1193 | 628 | 808 | 659 | 2548 | 2533 | 2395 | 1773 | 3624 | 2600 | 2492 | Foxc1 |
| Foxc2 | 453 | 548 | 537 | 346 | 239 | 236 | 135 | 551 | 666 | 619 | 912 | 461 | 752 | 392 | 516 | 760 | 447 | 396 | 142 | 172 | 129 | 757 | 843 | 568 | 458 | 770 | 540 | 438 | Foxc2 |

| | | | | | | | | | | | | | | | | | | | | | | | | | | | | | |
|---|---|---|---|---|---|---|---|---|---|---|---|---|---|---|---|---|---|---|---|---|---|---|---|---|---|---|---|---|---|
| Foxd1 | 1351 | 1389 | 1892 | 982 | 692 | 714 | 435 | 1022 | 1136 | 915 | 1444 | 1108 | 1276 | 1206 | 1334 | 1537 | 1429 | 1022 | 364 | 489 | 408 | 1303 | 1312 | 981 | 889 | 1689 | 1630 | 1265 | Foxd1 |
| Foxd2 | 325 | 275 | 394 | 225 | 128 | 117 | 97 | 233 | 243 | 267 | 271 | 321 | 308 | 236 | 274 | 458 | 286 | 274 | 86 | 129 | 69 | 226 | 241 | 311 | 181 | 420 | 372 | 323 | Foxd2 |
| Foxd3 | 1 | 3 | 2 | 1 | 3 | 0 | 0 | 5 | 1 | 2 | 3 | 0 | 2 | 0 | 5 | 1 | 3 | 2 | 0 | 2 | 0 | 5 | 4 | 1 | 1 | 1 | 0 | 1 | Foxd3 |
| Foxd4 | 5 | 13 | 10 | 9 | 17 | 6 | 1 | 6 | 6 | 16 | 6 | 4 | 24 | 6 | 11 | 8 | 9 | 4 | 5 | 2 | 3 | 25 | 12 | 1 | 10 | 0 | 0 | 1 | Foxd4 |
| Foxe1 | 13 | 15 | 12 | 8 | 12 | 7 | 12 | 28 | 59 | 2 | 48 | 2 | 1 | 2 | 13 | 16 | 1 | 10 | 16 | 16 | 4 | 7 | 46 | 5 | 1 | 8 | 0 | 0 | Foxe1 |
| Foxe3 | 0 | 3 | 0 | 1 | 1 | 0 | 0 | 0 | 0 | 0 | 1 | 0 | 0 | 0 | 1 | 2 | 1 | 0 | 0 | 0 | 0 | 0 | 2 | 0 | 1 | 2 | 0 | 2 | Foxe3 |
| Foxf1 | 187 | 38 | 34 | 21 | 32 | 29 | 27 | 140 | 132 | 60 | 190 | 16 | 9 | 22 | 32 | 167 | 30 | 22 | 26 | 21 | 16 | 63 | 93 | 37 | 9 | 110 | 13 | 6 | Foxf1 |
| Foxf2 | 427 | 367 | 684 | 419 | 472 | 364 | 387 | 562 | 678 | 566 | 744 | 613 | 492 | 523 | 340 | 588 | 470 | 406 | 464 | 357 | 413 | 703 | 603 | 493 | 503 | 741 | 580 | 477 | Foxf2 |
| Foxg1 | 4721 | 3227 | 3298 | 7658 | 7234 | 6840 | 11615 | 5652 | 6212 | 8293 | 5817 | 11600 | 3536 | 12000 | 3728 | 1627 | 6504 | 7080 | 9288 | 5816 | 9919 | 5872 | 3351 | 12436 | 11225 | 14266 | 5950 | 8291 | Foxg1 |
| Foxh1 | 7 | 7 | 7 | 3 | 7 | 7 | 10 | 6 | 9 | 1 | 8 | 8 | 11 | 6 | 4 | 6 | 5 | 0 | 3 | 1 | 3 | 7 | 3 | 3 | 0 | 5 | 9 | 1 | Foxh1 |
| Foxi1 | 3 | 3 | 0 | 0 | 0 | 0 | 0 | 0 | 8 | 0 | 6 | 0 | 6 | 0 | 3 | 0 | 0 | 0 | 3 | 0 | 0 | 12 | 4 | 0 | 5 | 0 | 0 | 0 | Foxi1 |
| Foxi2 | 3 | 2 | 9 | 1 | 0 | 4 | 1 | 4 | 9 | 0 | 8 | 10 | 5 | 5 | 0 | 2 | 0 | 0 | 0 | 2 | 1 | 0 | 6 | 9 | 3 | 5 | 3 | 4 | Foxi2 |
| Foxi3 | 30 | 2 | 7 | 2 | 3 | 1 | 1 | 12 | 21 | 8 | 6 | 3 | 2 | 1 | 3 | 8 | 1 | 6 | 1 | 2 | 0 | 0 | 18 | 8 | 1 | 7 | 2 | 1 | Foxi3 |
| Foxj1 | 1095 | 1811 | 1709 | 1942 | 1863 | 1830 | 1525 | 1172 | 1148 | 1580 | 1070 | 1614 | 1822 | 1092 | 1313 | 1639 | 2022 | 1990 | 2051 | 1616 | 1634 | 1278 | 1665 | 1311 | 1326 | 1359 | 1181 | 1304 | Foxj1 |
| Foxj2 | 1571 | 1927 | 2108 | 2609 | 2661 | 2254 | 2224 | 1508 | 1619 | 2009 | 2075 | 1623 | 1594 | 1508 | 1564 | 1970 | 2333 | 2552 | 2569 | 1911 | 2325 | 2104 | 1690 | 1899 | 1896 | 1599 | 1378 | 1482 | Foxj2 |
| Foxj3 | 4854 | 3962 | 5004 | 5635 | 6114 | 5486 | 6055 | 5424 | 5169 | 5875 | 5438 | 5831 | 5066 | 5750 | 4615 | 4355 | 5339 | 5613 | 6003 | 4757 | 6061 | 6259 | 5143 | 5965 | 6121 | 5588 | 5080 | 5543 | Foxj3 |
| Foxk1 | 5678 | 7470 | 9407 | 8862 | 7557 | 7038 | 6143 | 3791 | 4797 | 5568 | 5962 | 5765 | 5722 | 5098 | 5310 | 6940 | 9051 | 8974 | 7677 | 5172 | 6848 | 5755 | 4822 | 5631 | 5884 | 4987 | 5190 | 5314 | Foxk1 |
| Foxk2 | 10672 | 11968 | 11577 | 12476 | 12156 | 12066 | 12100 | 9213 | 9300 | 10445 | 9030 | 10935 | 9347 | 10676 | 10010 | 9990 | 12557 | 11871 | 13025 | 10702 | 12283 | 9824 | 8765 | 10646 | 10791 | 10942 | 10062 | 10936 | Foxk2 |
| Foxl1 | 99 | 86 | 100 | 43 | 43 | 39 | 28 | 48 | 119 | 42 | 128 | 73 | 74 | 34 | 83 | 154 | 109 | 68 | 21 | 29 | 30 | 90 | 98 | 62 | 44 | 126 | 109 | 28 | Foxl1 |
| Foxl2 | 302 | 236 | 428 | 133 | 133 | 173 | 83 | 330 | 484 | 273 | 384 | 470 | 523 | 412 | 207 | 286 | 290 | 165 | 77 | 94 | 72 | 259 | 331 | 516 | 290 | 527 | 533 | 398 | Foxl2 |
| Foxl3 | 1 | 1 | 6 | 0 | 4 | 0 | 0 | 5 | 1 | 3 | 2 | 1 | 3 | 1 | 5 | 3 | 1 | 1 | 2 | 0 | 0 | 5 | 3 | 1 | 0 | 3 | 0 | 1 | Foxl3 |
| Foxm1 | 4505 | 4681 | 5465 | 4469 | 3353 | 3790 | 3933 | 4662 | 5178 | 5479 | 4513 | 8530 | 6339 | 8115 | 3512 | 3967 | 5014 | 3867 | 4369 | 3396 | 3866 | 4346 | 4158 | 6948 | 5909 | 8338 | 7415 | 8119 | Foxm1 |
| Foxn1 | 0 | 2 | 0 | 0 | 0 | 0 | 0 | 9 | 8 | 5 | 1 | 0 | 2 | 1 | 1 | 3 | 1 | 0 | 0 | 0 | 0 | 0 | 15 | 6 | 0 | 3 | 2 | 0 | Foxn1 |
| Foxn2 | 2420 | 1941 | 2430 | 2642 | 2465 | 2806 | 2761 | 1922 | 1846 | 1750 | 2084 | 2077 | 2224 | 2287 | 2421 | 2244 | 2323 | 2663 | 2491 | 2329 | 2630 | 1853 | 1835 | 1860 | 2169 | 2002 | 2048 | 2332 | Foxn2 |
| Foxn3 | 5713 | 5467 | 6865 | 7118 | 6717 | 6745 | 6738 | 5398 | 5058 | 5692 | 5602 | 6251 | 5642 | 6125 | 5844 | 5530 | 7004 | 7039 | 6663 | 5872 | 6786 | 5803 | 4893 | 6034 | 6286 | 6240 | 5970 | 6334 | Foxn3 |
| Foxn4 | 163 | 222 | 655 | 197 | 137 | 206 | 63 | 153 | 111 | 173 | 112 | 384 | 533 | 367 | 195 | 221 | 439 | 132 | 207 | 168 | 141 | 234 | 106 | 243 | 187 | 57 | 450 | 420 | Foxn4 |
| Foxo1 | 3505 | 3445 | 3914 | 3688 | 3095 | 3215 | 2883 | 1719 | 1926 | 1937 | 2112 | 1574 | 2110 | 1526 | 3413 | 4456 | 3363 | 3807 | 2754 | 2443 | 2938 | 2015 | 2059 | 1816 | 1784 | 2106 | 1525 | 1543 | Foxo1 |
| Foxo3 | 4991 | 5300 | 5818 | 7051 | 6712 | 5917 | 6217 | 3088 | 3685 | 4030 | 4347 | 3189 | 3540 | 3355 | 4735 | 5466 | 6153 | 7071 | 6606 | 4910 | 6543 | 3586 | 3809 | 3934 | 3691 | 2968 | 2800 | 3616 | Foxo3 |
| Foxo4 | 4492 | 4683 | 5008 | 4933 | 3915 | 3814 | 3758 | 2803 | 3144 | 3197 | 3919 | 2965 | 3289 | 3051 | 3972 | 4597 | 4976 | 4260 | 4422 | 3836 | 4329 | 2486 | 3407 | 3679 | 2719 | 3967 | 2903 | 3215 | Foxo4 |
| Foxo6 | 798 | 1097 | 737 | 1461 | 1528 | 1246 | 1274 | 692 | 946 | 942 | 813 | 719 | 518 | 575 | 713 | 743 | 1163 | 1558 | 1637 | 1335 | 1488 | 839 | 635 | 1168 | 1268 | 878 | 425 | 526 | Foxo6 |
| Foxp1 | 6583 | 5782 | 8056 | 7473 | 6222 | 6574 | 6488 | 4797 | 5225 | 6091 | 6762 | 5954 | 6492 | 5874 | 6253 | 7198 | 7409 | 7283 | 6159 | 5266 | 6596 | 5690 | 5040 | 6192 | 6024 | 6782 | 6292 | 5933 | Foxp1 |
| Foxp2 | 5546 | 4537 | 5866 | 5252 | 5136 | 6341 | 5350 | 9263 | 7874 | 8235 | 8471 | 7743 | 7872 | 7377 | 5751 | 6372 | 5262 | 5367 | 4988 | 5281 | 5095 | 11650 | 8124 | 7708 | 10094 | 6601 | 8559 | 7471 | Foxp2 |
| Foxp3 | 0 | 3 | 6 | 8 | 5 | 2 | 4 | 6 | 8 | 5 | 1 | 2 | 5 | 6 | 5 | 2 | 2 | 1 | 2 | 3 | 8 | 8 | 10 | 11 | 2 | 7 | 0 | 11 | Foxp3 |
| Foxp4 | 5758 | 7898 | 9386 | 7746 | 6222 | 6061 | 4759 | 4451 | 5899 | 6660 | 7018 | 7285 | 6634 | 6118 | 5413 | 7802 | 8269 | 7230 | 7253 | 4611 | 5469 | 5987 | 5022 | 7459 | 6819 | 9252 | 6738 | 6251 | Foxp4 |
| Foxq1 | 130 | 160 | 228 | 179 | 308 | 239 | 191 | 263 | 225 | 249 | 241 | 279 | 279 | 300 | 149 | 177 | 185 | 199 | 232 | 236 | 231 | 281 | 272 | 269 | 305 | 309 | 275 | 272 | Foxq1 |
| Foxr1 | 3 | 0 | 0 | 0 | 0 | 0 | 0 | 0 | 1 | 0 | 0 | 0 | 1 | 0 | 0 | 0 | 0 | 0 | 0 | 0 | 0 | 2 | 1 | 0 | 3 | 0 | 0 | 0 | Foxr1 |
| Foxr2 | 77 | 83 | 120 | 124 | 98 | 144 | 182 | 68 | 44 | 67 | 63 | 139 | 105 | 88 | 99 | 84 | 98 | 114 | 86 | 100 | 123 | 72 | 52 | 76 | 121 | 94 | 123 | 109 | Foxr2 |
| Foxred1 | 2933 | 3137 | 3130 | 3025 | 2824 | 2931 | 2892 | 3120 | 3065 | 3070 | 3148 | 3453 | 3348 | 3441 | 2908 | 2766 | 3179 | 2864 | 2877 | 2819 | 2965 | 3193 | 2954 | 3268 | 3282 | 3568 | 3509 | 3285 | Foxred1 |
| Foxred2 | 1494 | 1441 | 1441 | 1538 | 1748 | 1629 | 1791 | 1093 | 1399 | 1460 | 1163 | 1344 | 1235 | 1339 | 1370 | 1430 | 1494 | 1700 | 1655 | 1364 | 1762 | 1348 | 1106 | 1334 | 1145 | 1279 | 1240 | 1262 | Foxred2 |
| Foxs1 | 18 | 32 | 43 | 16 | 54 | 35 | 18 | 49 | 64 | 53 | 67 | 83 | 70 | 71 | 11 | 18 | 28 | 34 | 28 | 40 | 32 | 35 | 65 | 72 | 96 | 71 | 76 | 70 | Foxs1 |
| Fpgs | 1066 | 1195 | 1113 | 946 | 705 | 822 | 596 | 739 | 882 | 846 | 1024 | 899 | 864 | 756 | 924 | 1196 | 890 | 916 | 697 | 702 | 661 | 864 | 981 | 941 | 820 | 1089 | 803 | 767 | Fpgs |
| Fpgt | 1404 | 1189 | 1178 | 1353 | 1316 | 1225 | 1593 | 2246 | 1730 | 1526 | 1435 | 1677 | 1457 | 1733 | 1623 | 1230 | 1284 | 1323 | 1414 | 1625 | 1469 | 1821 | 1665 | 1572 | 1676 | 1548 | 1620 | 1797 | Fpgt |
| Fpr1 | 2 | 0 | 0 | 0 | 1 | 3 | 0 | 0 | 1 | 3 | 0 | 1 | 5 | 0 | 0 | 0 | 0 | 2 | 0 | 0 | 0 | 8 | 4 | 6 | 1 | 2 | 1 | 0 | Fpr1 |
| Fpr2 | 3 | 2 | 0 | 4 | 3 | 2 | 0 | 1 | 2 | 2 | 3 | 1 | 3 | 1 | 0 | 1 | 0 | 2 | 3 | 0 | 3 | 3 | 6 | 1 | 1 | 3 | 2 | 2 | Fpr2 |
| Fra10ac1 | 2079 | 1786 | 2450 | 2112 | 2074 | 2016 | 2421 | 2777 | 2347 | 2530 | 2272 | 2830 | 2470 | 3061 | 2120 | 1949 | 2254 | 2004 | 2276 | 2269 | 2271 | 2445 | 2333 | 2742 | 2548 | 2892 | 2759 | 2826 | Fra10ac1 |
| Fras1 | 5305 | 4940 | 2781 | 3035 | 3465 | 5028 | 3385 | 4308 | 6166 | 4117 | 5872 | 3871 | 4939 | 3508 | 4386 | 5875 | 2786 | 3316 | 3066 | 3882 | 3794 | 2881 | 6152 | 3327 | 3267 | 3040 | 4338 | 3312 | Fras1 |
| Frat1 | 1219 | 1427 | 1375 | 1995 | 1598 | 1494 | 1779 | 699 | 722 | 1088 | 795 | 893 | 653 | 1021 | 1276 | 1097 | 1738 | 1675 | 1803 | 1403 | 1836 | 1009 | 723 | 1159 | 1183 | 1012 | 797 | 1009 | Frat1 |
| Frat2 | 1676 | 2057 | 1665 | 2402 | 1554 | 1595 | 1809 | 1094 | 962 | 1127 | 970 | 1377 | 949 | 1237 | 1625 | 1399 | 1958 | 2229 | 1937 | 1842 | 1818 | 971 | 917 | 1388 | 1226 | 1662 | 1296 | 1136 | Frat2 |
| Frem1 | 6579 | 3169 | 2321 | 1750 | 1940 | 2305 | 1621 | 5002 | 4711 | 3493 | 3879 | 2540 | 3527 | 2286 | 3405 | 5597 | 2111 | 1848 | 1446 | 2239 | 1780 | 2266 | 4220 | 2921 | 1859 | 2633 | 2846 | 2357 | Frem1 |
| Frem2 | 5351 | 4239 | 2449 | 2370 | 2399 | 2930 | 2093 | 2900 | 3884 | 2316 | 3537 | 1651 | 2293 | 1340 | 3969 | 5485 | 2087 | 2527 | 2122 | 2359 | 2440 | 1586 | 4330 | 1402 | 1205 | 1202 | 1628 | 1227 | Frem2 |
| Frem3 | 1 | 3 | 2 | 5 | 2 | 4 | 6 | 4 | 5 | 5 | 7 | 5 | 2 | 1 | 0 | 4 | 0 | 0 | 3 | 7 | 1 | 10 | 4 | 4 | 5 | 7 | 3 | 6 | Frem3 |
| Frg1 | 3719 | 3503 | 4009 | 3465 | 3107 | 2954 | 3802 | 3149 | 2816 | 3281 | 3180 | 3421 | 3478 | 4139 | 3543 | 3208 | 3847 | 3526 | 3398 | 3280 | 3740 | 2984 | 2964 | 3752 | 3363 | 3749 | 3246 | 3947 | Frg1 |
| Frg2f1 | 210 | 210 | 175 | 290 | 286 | 248 | 261 | 331 | 343 | 441 | 321 | 287 | 251 | 259 | 237 | 157 | 224 | 272 | 216 | 213 | 243 | 428 | 298 | 353 | 357 | 290 | 209 | 182 | Frg2f1 |
| Frk | 291 | 174 | 180 | 141 | 89 | 191 | 65 | 680 | 695 | 585 | 702 | 313 | 578 | 336 | 256 | 303 | 132 | 120 | 67 | 90 | 40 | 366 | 786 | 537 | 240 | 780 | 344 | 386 | Frk |
| Frmd3 | 418 | 311 | 179 | 334 | 830 | 548 | 540 | 1522 | 1334 | 1147 | 1248 | 717 | 919 | 828 | 553 | 393 | 291 | 407 | 527 | 523 | 589 | 1758 | 1229 | 769 | 1205 | 479 | 647 | 642 | Frmd3 |
| Frmd4a | 6877 | 6464 | 6262 | 7355 | 10358 | 9305 | 8135 | 15242 | 15911 | 18306 | 16304 | 17182 | 18801 | 16909 | 6988 | 7045 | 7464 | 7667 | 8923 | 7764 | 8744 | 18797 | 15132 | 15987 | 18596 | 13155 | 16498 | 16321 | Frmd4a |
| Frmd4b | 4625 | 5096 | 4415 | 4583 | 4945 | 5230 | 4600 | 6749 | 5958 | 6107 | 5412 | 6727 | 6289 | 6427 | 5465 | 4549 | 4445 | 4110 | 4811 | 5073 | 4758 | 5848 | 6012 | 5955 | 6553 | 5944 | 6795 | 6045 | Frmd4b |
| Frmd5 | 3315 | 3733 | 3862 | 4904 | 5583 | 5204 | 5813 | 4373 | 4023 | 5169 | 3848 | 4901 | 4027 | 5034 | 3716 | 2757 | 4699 | 4561 | 5722 | 5016 | 5497 | 5604 | 3561 | 5011 | 5750 | 4263 | 4506 | 4501 | Frmd5 |
| Frmd6 | 5389 | 3911 | 3840 | 3096 | 2459 | 3023 | 2234 | 3080 | 3580 | 2911 | 3788 | 2311 | 3492 | 2266 | 4131 | 5782 | 3143 | 3848 | 2392 | 2165 | 2274 | 2257 | 3677 | 2781 | 2184 | 3260 | 2346 | 2471 | Frmd6 |
| Frmd7 | 15 | 15 | 17 | 13 | 26 | 21 | 13 | 20 | 19 | 8 | 20 | 5 | 18 | 13 | 12 | 24 | 17 | 18 | 15 | 16 | 16 | 3 | 16 | 17 | 6 | 4 | 12 | 12 | Frmd7 |
| Frmd8 | 3946 | 5086 | 3820 | 3970 | 3013 | 3223 | 3058 | 1788 | 2086 | 1764 | 2014 | 1913 | 2090 | 1809 | 3510 | 4833 | 3861 | 3912 | 3503 | 3342 | 3198 | 1591 | 2034 | 1940 | 1843 | 2301 | 2015 | 2060 | Frmd8 |
| Frmpd1 | 185 | 222 | 187 | 175 | 288 | 193 | 183 | 203 | 315 | 219 | 420 | 285 | 320 | 219 | 233 | 224 | 225 | 205 | 190 | 128 | 166 | 302 | 352 | 209 | 217 | 202 | 271 | 295 | Frmpd1 |
| Frmpd2 | 3 | 6 | 8 | 5 | 8 | 2 | 3 | 12 | 11 | 8 | 3 | 8 | 1 | 5 | 13 | 5 | 1 | 2 | 11 | 3 | 10 | 13 | 9 | 5 | 5 | 8 | 8 | 9 | Frmpd2 |
| Frmpd3 | 511 | 756 | 897 | 1102 | 1722 | 1260 | 943 | 891 | 983 | 1256 | 1147 | 893 | 1036 | 766 | 699 | 713 | 1130 | 1237 | 1399 | 950 | 1154 | 1596 | 838 | 873 | 1313 | 492 | 866 | 862 | Frmpd3 |
| Frmpd4 | 179 | 142 | 84 | 121 | 321 | 184 | 140 | 275 | 303 | 316 | 351 | 134 | 245 | 143 | 157 | 221 | 92 | 182 | 175 | 265 | 210 | 322 | 361 | 154 | 195 | 109 | 120 | 120 | Frmpd4 |
| Frrs1 | 536 | 336 | 596 | 314 | 421 | 374 | 392 | 583 | 668 | 649 | 665 | 734 | 962 | 719 | 402 | 406 | 387 | 383 | 448 | 320 | 336 | 553 | 706 | 636 | 546 | 811 | 673 | 770 | Frrs1 |
| Frrs1l | 1234 | 1237 | 1290 | 1651 | 2650 | 2179 | 2502 | 3181 | 2958 | 2833 | 2755 | 2402 | 2149 | 2819 | 1629 | 1334 | 1567 | 1935 | 2396 | 2481 | 2313 | 3218 | 2800 | 2701 | 3501 | 1671 | 2023 | 2207 | Frrs1l |
| Frs2 | 7304 | 7059 | 6967 | 7962 | 7321 | 7533 | 7502 | 6020 | 5986 | 6333 | 6130 | 6292 | 6225 | 6278 | 7163 | 7454 | 7142 | 8183 | 6995 | 6531 | 7597 | 6764 | 5743 | 5928 | 6563 | 5044 | 6235 | 6478 | Frs2 |
| Frs3 | 580 | 763 | 752 | 785 | 934 | 743 | 716 | 650 | 723 | 756 | 654 | 663 | 729 | 683 | 588 | 608 | 928 | 798 | 903 | 746 | 686 | 772 | 654 | 755 | 772 | 689 | 760 | 625 | Frs3 |
| Fry | 2359 | 2542 | 1580 | 3759 | 6769 | 5255 | 4951 | 4688 | 5563 | 6317 | 5791 | 3857 | 4557 | 3861 | 2863 | 2861 | 2758 | 4594 | 4547 | 4378 | 5292 | 7354 | 4601 | 3706 | 6700 | 1847 | 3349 | 3131 | Fry |
| Fryl | 11829 | 12730 | 10063 | 14321 | 15738 | 15636 | 16304 | 7272 | 7569 | 8766 | 7818 | 8963 | 8890 | 9148 | 11979 | 13181 | 11151 | 14232 | 14474 | 12932 | 17230 | 7842 | 6792 | 7275 | 9866 | 5913 | 8889 | 8094 | Fryl |
| Frzb | 1105 | 882 | 1107 | 510 | 652 | 496 | 454 | 961 | 1161 | 833 | 1106 | 682 | 1565 | 677 | 893 | 1114 | 702 | 552 | 430 | 561 | 416 | 764 | 1028 | 610 | 524 | 1104 | 766 | 876 | Frzb |
| Fsbp | 195 | 124 | 173 | 122 | 145 | 153 | 190 | 238 | 269 | 284 | 284 | 367 | 344 | 397 | 175 | 137 | 160 | 88 | 150 | 176 | 154 | 227 | 230 | 261 | 246 | 353 | 377 | 393 | Fsbp |
| Fscn1 | 23118 | 29005 | 32694 | 33182 | 30535 | 26962 | 24159 | 17493 | 21054 | 24646 | 23105 | 28313 | 23198 | 24828 | 21602 | 26531 | 34118 | 32502 | 33461 | 20990 | 28212 | 24231 | 20326 | 28174 | 26426 | 31950 | 22990 | 24369 | Fscn1 |
| Fscn2 | 7 | 10 | 5 | 7 | 12 | 10 | 15 | 17 | 15 | 36 | 14 | 21 | 15 | 12 | 5 | 9 | 21 | 14 | 9 | 5 | 13 | 25 | 16 | 16 | 23 | 12 | 8 | 21 | Fscn2 |
| Fscn3 | 8 | 16 | 22 | 19 | 15 | 8 | 13 | 5 | 12 | 8 | 0 | 10 | 11 | 14 | 8 | 14 | 6 | 40 | 15 | 19 | 14 | 8 | 6 | 7 | 3 | 2 | 13 | 6 | Fscn3 |
| Fsd1 | 2247 | 3393 | 3411 | 4018 | 4053 | 3402 | 3325 | 2066 | 2172 | 2965 | 2431 | 2548 | 2371 | 2258 | 2429 | 2869 | 4211 | 3710 | 4418 | 3202 | 3499 | 2734 | 2195 | 2730 | 2921 | 2566 | 2467 | 2288 | Fsd1 |
| Fsd1l | 8825 | 6791 | 7222 | 10178 | 11570 | 12453 | 14464 | 11061 | 9033 | 8867 | 8587 | 10554 | 8756 | 11400 | 9712 | 7781 | 8554 | 10277 | 11117 | 12800 | 13924 | 9939 | 8673 | 8987 | 11457 | 7181 | 10090 | 10104 | Fsd1l |
| Fsd2 | 13 | 10 | 10 | 16 | 20 | 14 | 15 | 22 | 42 | 41 | 60 | 36 | 46 | 34 | 36 | 19 | 14 | 9 | 16 | 9 | 10 | 50 | 50 | 37 | 41 | 23 | 29 | 46 | Fsd2 |

| | | | | | | | | | | | | | | | | | | | | | | | | | | | | | |
|---|---|---|---|---|---|---|---|---|---|---|---|---|---|---|---|---|---|---|---|---|---|---|---|---|---|---|---|---|---|
| Fshb | 0 | 0 | 0 | 0 | 0 | 0 | 0 | 1 | 1 | 0 | 0 | 0 | 0 | 0 | 0 | 0 | 0 | 0 | 0 | 0 | 0 | 0 | 0 | 0 | 0 | 0 | 0 | 0 | Fshb |
| Fshr | 0 | 0 | 0 | 0 | 0 | 0 | 0 | 0 | 0 | 0 | 1 | 0 | 0 | 0 | 0 | 0 | 0 | 0 | 0 | 0 | 0 | 2 | 1 | 0 | 0 | 0 | 0 | 0 | Fshr |
| Fsip1 | 142 | 120 | 106 | 166 | 175 | 237 | 218 | 153 | 179 | 186 | 143 | 189 | 144 | 142 | 136 | 141 | 141 | 166 | 170 | 202 | 215 | 137 | 163 | 132 | 193 | 133 | 145 | 129 | Fsip1 |
| Fsip2 | 0 | 0 | 0 | 0 | 1 | 0 | 1 | 6 | 1 | 0 | 0 | 0 | 0 | 0 | 0 | 0 | 0 | 0 | 0 | 1 | 0 | 0 | 3 | 0 | 0 | 0 | 0 | 0 | Fsip2 |
| Fsip2l | 1 | 4 | 1 | 6 | 1 | 0 | 1 | 0 | 0 | 0 | 3 | 1 | 0 | 0 | 4 | 4 | 1 | 4 | 2 | 3 | 0 | 0 | 0 | 0 | 0 | 0 | 0 | 0 | Fsip2l |
| Fst | 1052 | 727 | 1322 | 726 | 532 | 697 | 598 | 1136 | 950 | 876 | 1277 | 1290 | 1042 | 1388 | 836 | 818 | 898 | 647 | 704 | 662 | 583 | 789 | 1278 | 1276 | 840 | 1901 | 1572 | 1741 | Fst |
| Fstl1 | 34455 | 27240 | 26460 | 19032 | 14395 | 21540 | 11965 | 29063 | 36881 | 29123 | 41451 | 23261 | 32167 | 23752 | 31470 | 36670 | 20710 | 23350 | 12668 | 14711 | 11900 | 27664 | 45645 | 27117 | 22609 | 29268 | 24960 | 25072 | Fstl1 |
| Fstl3 | 305 | 348 | 289 | 252 | 163 | 184 | 127 | 50 | 84 | 81 | 125 | 65 | 64 | 32 | 219 | 365 | 212 | 222 | 155 | 130 | 138 | 50 | 109 | 72 | 47 | 76 | 62 | 64 | Fstl3 |
| Fstl4 | 102 | 84 | 62 | 122 | 291 | 197 | 131 | 215 | 280 | 267 | 288 | 182 | 215 | 96 | 136 | 173 | 111 | 208 | 115 | 173 | 154 | 267 | 197 | 140 | 236 | 50 | 100 | 110 | Fstl4 |
| Fstl5 | 2863 | 2457 | 1932 | 3438 | 4183 | 3197 | 3847 | 4529 | 3403 | 3757 | 2987 | 2420 | 2212 | 2571 | 3447 | 2476 | 2772 | 3204 | 3183 | 4450 | 3654 | 3217 | 3616 | 3266 | 3054 | 2607 | 2115 | 2207 | Fstl5 |
| Ftcd | 1 | 0 | 6 | 0 | 1 | 2 | 0 | 1 | 2 | 3 | 5 | 1 | 2 | 4 | 0 | 2 | 0 | 1 | 3 | 1 | 1 | 7 | 5 | 3 | 1 | 3 | 8 | 2 | Ftcd |
| Ftdc2 | 0 | 0 | 0 | 0 | 2 | 0 | 0 | 0 | 0 | 3 | 0 | 0 | 0 | 0 | 0 | 0 | 0 | 1 | 0 | 0 | 1 | 0 | 1 | 0 | 0 | 0 | 0 | 0 | Ftdc2 |
| Fth1 | 18636 | 20988 | 21030 | 21152 | 18755 | 17915 | 19754 | 15287 | 17183 | 17113 | 17221 | 16214 | 19979 | 18162 | 19256 | 18599 | 20018 | 20061 | 20160 | 18072 | 20214 | 16265 | 18964 | 16974 | 16604 | 19051 | 15281 | 21431 | Fth1 |
| Fthl17a | 0 | 0 | 0 | 0 | 0 | 0 | 0 | 0 | 2 | 0 | 1 | 0 | 0 | 0 | 0 | 0 | 0 | 1 | 0 | 0 | 0 | 0 | 1 | 0 | 0 | 0 | 0 | 1 | Fthl17a |
| Fthl17f | 0 | 0 | 0 | 0 | 0 | 0 | 0 | 0 | 0 | 0 | 0 | 2 | 0 | 0 | 0 | 0 | 0 | 0 | 0 | 0 | 0 | 0 | 1 | 0 | 0 | 0 | 0 | 0 | Fthl17f |
| Ftl1 | 39867 | 59947 | 47415 | 47948 | 45104 | 46246 | 51928 | 42424 | 52428 | 46384 | 48106 | 51829 | 53806 | 58464 | 41489 | 43828 | 50037 | 48441 | 51049 | 50456 | 51597 | 40153 | 52668 | 52351 | 51822 | 55851 | 53902 | 58986 | Ftl1 |
| Ftl1-ps1 | 1705 | 1754 | 1658 | 1678 | 1349 | 1364 | 1395 | 1600 | 1475 | 2111 | 1608 | 1983 | 1672 | 1833 | 1619 | 1352 | 2021 | 1488 | 1746 | 1589 | 1357 | 1521 | 1572 | 1812 | 1740 | 1913 | 1725 | 1911 | Ftl1-ps1 |
| Fto | 8865 | 9854 | 10681 | 11523 | 12080 | 11174 | 11296 | 9331 | 11332 | 12607 | 11748 | 12381 | 11367 | 11903 | 8826 | 9460 | 11720 | 11931 | 12232 | 9151 | 11861 | 13089 | 10917 | 12482 | 13292 | 12205 | 10606 | 11895 | Fto |
| Ftsj1 | 3313 | 3709 | 3139 | 3278 | 2874 | 3016 | 3251 | 2292 | 2346 | 2091 | 1961 | 2300 | 2197 | 2148 | 3235 | 3022 | 3120 | 2897 | 3089 | 3415 | 3049 | 1912 | 2162 | 2044 | 2158 | 2325 | 2492 | 2263 | Ftsj1 |
| Ftsj3 | 11423 | 10974 | 11294 | 9452 | 7655 | 8567 | 9036 | 6629 | 6239 | 5798 | 5767 | 7128 | 6884 | 7184 | 10763 | 10291 | 9683 | 8720 | 8286 | 10007 | 8455 | 5446 | 5946 | 6501 | 6118 | 8125 | 7281 | 7329 | Ftsj3 |
| Fubp1 | 37161 | 31752 | 33991 | 33231 | 33302 | 35908 | 39141 | 52817 | 43484 | 40187 | 42054 | 46936 | 45768 | 53138 | 37306 | 34001 | 34010 | 31261 | 34830 | 38777 | 38645 | 41664 | 41439 | 43749 | 44534 | 42908 | 50776 | 50849 | Fubp1 |
| Fubp3 | 7109 | 6874 | 7643 | 6854 | 6172 | 6689 | 6162 | 5479 | 5723 | 5881 | 6286 | 7101 | 6725 | 6891 | 6513 | 6822 | 7163 | 6510 | 6338 | 6491 | 5861 | 5533 | 5503 | 6490 | 6288 | 7834 | 7241 | 7028 | Fubp3 |
| Fuca1 | 3495 | 3943 | 3731 | 3775 | 3242 | 3601 | 3300 | 1773 | 1859 | 1799 | 1887 | 1779 | 1969 | 1718 | 3479 | 3706 | 3436 | 3406 | 3421 | 3485 | 3458 | 1968 | 2047 | 1874 | 1881 | 1876 | 2059 | 1829 | Fuca1 |
| Fuca2 | 1168 | 1087 | 1095 | 842 | 1145 | 1143 | 851 | 1475 | 1443 | 1259 | 1407 | 1048 | 1312 | 1149 | 1275 | 1237 | 1052 | 1094 | 934 | 1054 | 1031 | 1375 | 1634 | 1132 | 1153 | 1094 | 1119 | 1117 | Fuca2 |
| Fundc1 | 2112 | 1569 | 1714 | 1768 | 1971 | 2068 | 2218 | 3863 | 3544 | 2880 | 3090 | 3106 | 3109 | 3350 | 2325 | 1937 | 1809 | 1749 | 1815 | 2254 | 2015 | 3278 | 3609 | 3152 | 3050 | 3247 | 3148 | 3121 | Fundc1 |
| Fundc2 | 3329 | 2614 | 2704 | 2534 | 2555 | 2894 | 2861 | 4064 | 3862 | 3258 | 3721 | 3404 | 3645 | 3643 | 3401 | 2735 | 2435 | 2562 | 2661 | 2913 | 2923 | 3517 | 4102 | 3361 | 3267 | 3811 | 3588 | 4034 | Fundc2 |
| Fuom | 305 | 315 | 308 | 392 | 359 | 330 | 359 | 294 | 348 | 294 | 321 | 298 | 376 | 337 | 327 | 321 | 357 | 307 | 308 | 404 | 347 | 262 | 393 | 355 | 360 | 372 | 298 | 297 | Fuom |
| Furin | 1731 | 1850 | 2329 | 1895 | 1977 | 1800 | 1340 | 1461 | 2142 | 2301 | 2735 | 1804 | 2003 | 1609 | 1412 | 2153 | 1990 | 2053 | 1901 | 1228 | 1582 | 2079 | 2334 | 2156 | 1991 | 2424 | 1811 | 1658 | Furin |
| Fus | 39903 | 41887 | 60900 | 55332 | 49294 | 49492 | 50688 | 50557 | 50276 | 63858 | 67729 | 73675 | 70415 | 82387 | 36859 | 40757 | 57852 | 51926 | 50831 | 43526 | 51408 | 59411 | 47431 | 76975 | 71436 | 85613 | 75148 | 81083 | Fus |
| Fut1 | 21 | 6 | 3 | 20 | 2 | 11 | 1 | 45 | 44 | 73 | 44 | 9 | 22 | 7 | 11 | 11 | 6 | 0 | 4 | 1 | 4 | 2 | 72 | 85 | 3 | 57 | 2 | 1 | Fut1 |
| Fut10 | 5293 | 5211 | 4617 | 4132 | 3186 | 3776 | 3765 | 1895 | 1933 | 1714 | 1762 | 2182 | 2080 | 2153 | 4788 | 4991 | 3950 | 3864 | 3618 | 4049 | 3807 | 1257 | 1760 | 1881 | 1503 | 2396 | 2388 | 2257 | Fut10 |
| Fut11 | 2496 | 2357 | 2328 | 2836 | 3194 | 2928 | 2908 | 3242 | 2936 | 3045 | 3239 | 2785 | 3034 | 2780 | 2625 | 2427 | 2690 | 2990 | 2806 | 3011 | 3083 | 3586 | 3135 | 2828 | 3089 | 2533 | 2766 | 2713 | Fut11 |
| Fut2 | 3 | 3 | 3 | 3 | 0 | 1 | 0 | 10 | 4 | 6 | 11 | 0 | 6 | 0 | 5 | 4 | 1 | 3 | 5 | 3 | 1 | 5 | 14 | 12 | 2 | 5 | 6 | 6 | Fut2 |
| Fut4 | 120 | 98 | 182 | 68 | 140 | 145 | 88 | 209 | 201 | 150 | 217 | 137 | 163 | 115 | 113 | 128 | 136 | 70 | 111 | 101 | 87 | 159 | 193 | 169 | 132 | 224 | 122 | 122 | Fut4 |
| Fut7 | 24 | 57 | 52 | 41 | 47 | 43 | 39 | 5 | 9 | 1 | 13 | 9 | 6 | 7 | 21 | 25 | 41 | 42 | 57 | 32 | 48 | 15 | 5 | 10 | 16 | 8 | 3 | 6 | Fut7 |
| Fut8 | 3442 | 3200 | 3666 | 3442 | 3798 | 3582 | 3656 | 3987 | 3627 | 3626 | 3377 | 3768 | 3988 | 3816 | 4056 | 3436 | 3509 | 3429 | 3504 | 3644 | 3620 | 4500 | 3819 | 3791 | 3868 | 3773 | 3911 | 4084 | Fut8 |
| Fut9 | 3555 | 2146 | 1925 | 3557 | 4624 | 5477 | 6422 | 6216 | 5045 | 4142 | 3948 | 4891 | 4356 | 5458 | 3473 | 2446 | 2293 | 3367 | 4504 | 7039 | 6468 | 3960 | 5757 | 4254 | 5814 | 3529 | 5130 | 4934 | Fut9 |
| Fuz | 864 | 1113 | 1291 | 1118 | 1008 | 1022 | 1102 | 1071 | 1336 | 1351 | 1219 | 1450 | 1337 | 1491 | 905 | 888 | 1297 | 1022 | 1168 | 1134 | 1029 | 1186 | 1319 | 1459 | 1228 | 1652 | 1455 | 1532 | Fuz |
| Fv1 | 2202 | 2306 | 2136 | 2168 | 2072 | 2273 | 2182 | 1213 | 937 | 958 | 982 | 1098 | 1128 | 1164 | 2203 | 1896 | 2201 | 2105 | 2311 | 2436 | 2269 | 961 | 983 | 1037 | 1017 | 1125 | 1220 | 1167 | Fv1 |
| Fxn | 412 | 470 | 362 | 401 | 348 | 348 | 463 | 722 | 784 | 697 | 706 | 872 | 780 | 937 | 429 | 356 | 436 | 353 | 477 | 457 | 435 | 576 | 765 | 900 | 883 | 1020 | 909 | 928 | Fxn |
| Fxr1 | 12536 | 10122 | 12981 | 11291 | 10118 | 10883 | 12442 | 13714 | 11520 | 10753 | 12103 | 12225 | 11974 | 13757 | 12873 | 11955 | 11353 | 10910 | 11034 | 13303 | 11782 | 10911 | 12121 | 11905 | 11424 | 13313 | 12669 | 13389 | Fxr1 |
| Fxr2 | 7508 | 9062 | 9915 | 10368 | 9821 | 8988 | 9057 | 7889 | 7935 | 8867 | 8181 | 9427 | 8008 | 8837 | 7627 | 8168 | 10838 | 10456 | 10497 | 8957 | 9476 | 8372 | 7505 | 9374 | 9020 | 10640 | 8512 | 8660 | Fxr2 |
| Fxyd1 | 114 | 105 | 98 | 98 | 89 | 43 | 31 | 106 | 201 | 157 | 274 | 78 | 108 | 69 | 73 | 146 | 65 | 122 | 49 | 51 | 63 | 134 | 264 | 101 | 94 | 84 | 95 | 63 | Fxyd1 |
| Fxyd2 | 22 | 25 | 12 | 32 | 25 | 36 | 25 | 50 | 59 | 65 | 86 | 67 | 37 | 68 | 24 | 25 | 25 | 19 | 34 | 37 | 29 | 45 | 75 | 67 | 56 | 75 | 36 | 43 | Fxyd2 |
| Fxyd3 | 57 | 47 | 8 | 4 | 2 | 18 | 1 | 82 | 114 | 55 | 82 | 23 | 67 | 32 | 46 | 59 | 7 | 24 | 3 | 1 | 1 | 28 | 128 | 56 | 23 | 25 | 17 | 45 | Fxyd3 |
| Fxyd4 | 0 | 0 | 0 | 0 | 0 | 0 | 0 | 0 | 0 | 2 | 0 | 0 | 0 | 0 | 0 | 0 | 0 | 0 | 0 | 0 | 0 | 0 | 0 | 0 | 2 | 0 | 0 | 0 | Fxyd4 |
| Fxyd5 | 249 | 283 | 306 | 337 | 349 | 357 | 266 | 253 | 314 | 357 | 411 | 240 | 246 | 243 | 264 | 356 | 308 | 437 | 346 | 311 | 264 | 434 | 480 | 310 | 390 | 294 | 220 | 223 | Fxyd5 |
| Fxyd6 | 7082 | 8316 | 7123 | 7939 | 9378 | 8515 | 9022 | 9579 | 9389 | 8823 | 8129 | 9081 | 7400 | 9035 | 8007 | 7176 | 8223 | 8232 | 9603 | 10680 | 9467 | 9822 | 9838 | 9166 | 10473 | 9543 | 9276 | 8657 | Fxyd6 |
| Fxyd7 | 260 | 285 | 149 | 324 | 624 | 357 | 343 | 296 | 346 | 361 | 444 | 119 | 202 | 152 | 253 | 323 | 291 | 412 | 312 | 328 | 320 | 508 | 357 | 217 | 346 | 122 | 158 | 129 | Fxyd7 |
| Fyb | 113 | 112 | 124 | 113 | 93 | 130 | 78 | 198 | 292 | 245 | 344 | 225 | 236 | 212 | 128 | 177 | 89 | 105 | 58 | 90 | 73 | 209 | 359 | 230 | 185 | 204 | 248 | 177 | Fyb |
| Fyb2 | 69 | 45 | 19 | 64 | 41 | 36 | 13 | 57 | 40 | 40 | 57 | 27 | 34 | 27 | 46 | 47 | 21 | 45 | 14 | 33 | 31 | 38 | 42 | 38 | 30 | 41 | 28 | 19 | Fyb2 |
| Fyco1 | 3706 | 3956 | 2771 | 3458 | 2878 | 3344 | 3260 | 2097 | 2467 | 2155 | 2552 | 2269 | 2389 | 2054 | 3496 | 3791 | 2946 | 3239 | 2751 | 3387 | 3157 | 2264 | 2768 | 2047 | 2218 | 1508 | 2336 | 1943 | Fyco1 |
| Fyn | 8675 | 8782 | 8862 | 11422 | 13149 | 11961 | 13772 | 9912 | 9997 | 11431 | 10319 | 11086 | 9140 | 11540 | 9580 | 8115 | 10464 | 12047 | 13501 | 11747 | 13842 | 11688 | 9844 | 11314 | 12632 | 11001 | 9870 | 10363 | Fyn |
| Fyttd1 | 16082 | 11527 | 13537 | 13647 | 13388 | 14355 | 15626 | 13428 | 10500 | 9291 | 10471 | 10433 | 10629 | 10554 | 16313 | 14238 | 13151 | 13721 | 13275 | 15738 | 14595 | 11575 | 11346 | 9977 | 10324 | 10297 | 10569 | 10700 | Fyttd1 |
| Fzd1 | 4283 | 4406 | 5975 | 5428 | 4813 | 4906 | 4615 | 4628 | 4412 | 5855 | 5556 | 6730 | 5782 | 6540 | 3741 | 4148 | 5712 | 5214 | 5807 | 4065 | 5102 | 5927 | 5012 | 7117 | 6013 | 5948 | 6392 | 7039 | Fzd1 |
| Fzd10 | 3393 | 4669 | 7532 | 3719 | 2929 | 3364 | 1642 | 3008 | 2293 | 2497 | 2248 | 4108 | 4761 | 3805 | 3506 | 4033 | 5217 | 2818 | 3202 | 2882 | 2465 | 2699 | 3492 | 2959 | 2587 | 2228 | 5328 | 5477 | Fzd10 |
| Fzd2 | 6093 | 6026 | 6619 | 5530 | 4354 | 4630 | 4522 | 5282 | 6364 | 6011 | 6813 | 6642 | 6143 | 6287 | 5085 | 5858 | 6268 | 5012 | 4938 | 4123 | 4430 | 4339 | 5789 | 7209 | 4942 | 9044 | 6216 | 6518 | Fzd2 |
| Fzd3 | 32089 | 27067 | 25365 | 27963 | 30516 | 34492 | 35618 | 33452 | 24217 | 23698 | 21158 | 27775 | 27655 | 29176 | 33247 | 27878 | 26831 | 27088 | 30131 | 35828 | 35788 | 25913 | 24372 | 23310 | 27967 | 20578 | 31548 | 29673 | Fzd3 |
| Fzd4 | 3126 | 2516 | 1830 | 1778 | 1681 | 1655 | 1599 | 1484 | 1735 | 1254 | 2776 | 1114 | 1330 | 1099 | 2466 | 3281 | 1699 | 2053 | 1468 | 1813 | 1541 | 1440 | 2381 | 1217 | 1052 | 1276 | 1130 | 1002 | Fzd4 |
| Fzd5 | 960 | 1016 | 750 | 1095 | 689 | 602 | 730 | 369 | 446 | 336 | 455 | 337 | 184 | 353 | 810 | 951 | 934 | 982 | 623 | 584 | 719 | 592 | 362 | 466 | 437 | 591 | 323 | 343 | Fzd5 |
| Fzd6 | 52 | 54 | 12 | 13 | 14 | 21 | 12 | 45 | 32 | 29 | 22 | 29 | 34 | 15 | 33 | 26 | 9 | 17 | 12 | 21 | 8 | 89 | 42 | 37 | 23 | 19 | 38 | 88 | Fzd6 |
| Fzd7 | 4452 | 4462 | 4746 | 3782 | 2689 | 2930 | 2548 | 2638 | 3171 | 3037 | 3874 | 2886 | 3574 | 2612 | 4079 | 5639 | 4123 | 3714 | 2631 | 2378 | 2640 | 3165 | 3445 | 3144 | 2347 | 4157 | 2705 | 3249 | Fzd7 |
| Fzd8 | 3896 | 3844 | 3004 | 4437 | 3193 | 3560 | 4460 | 3407 | 3812 | 3593 | 3976 | 5236 | 2838 | 5160 | 3311 | 2584 | 3887 | 3945 | 4034 | 3660 | 4026 | 2732 | 2834 | 5067 | 3866 | 6604 | 4016 | 4085 | Fzd8 |
| Fzd9 | 562 | 873 | 804 | 997 | 781 | 895 | 791 | 535 | 769 | 819 | 873 | 837 | 743 | 753 | 593 | 696 | 875 | 955 | 918 | 870 | 797 | 580 | 841 | 933 | 910 | 902 | 730 | 779 | Fzd9 |
| Fzr1 | 4285 | 5771 | 5907 | 6624 | 5992 | 5637 | 4695 | 4218 | 5025 | 5437 | 4990 | 5933 | 5338 | 5209 | 4130 | 4908 | 6207 | 5848 | 6401 | 4760 | 5303 | 5103 | 4675 | 5935 | 5693 | 6802 | 5356 | 5464 | Fzr1 |
| G0s2 | 89 | 62 | 113 | 44 | 63 | 46 | 40 | 125 | 187 | 160 | 154 | 127 | 124 | 150 | 82 | 72 | 101 | 35 | 74 | 63 | 59 | 92 | 206 | 173 | 108 | 195 | 198 | 138 | G0s2 |
| G2e3 | 10185 | 6593 | 7499 | 7949 | 8764 | 10068 | 10965 | 10618 | 8924 | 7853 | 7994 | 10544 | 9131 | 11073 | 9916 | 8080 | 7853 | 7847 | 9019 | 10434 | 9980 | 7997 | 8295 | 8788 | 9495 | 8715 | 11086 | 10703 | G2e3 |
| G3bp1 | 26615 | 27147 | 27571 | 23848 | 19848 | 21110 | 22834 | 20643 | 21971 | 21974 | 21133 | 27233 | 24351 | 26237 | 24360 | 24836 | 25476 | 22514 | 21805 | 22269 | 21935 | 17970 | 20830 | 24579 | 22216 | 30475 | 27434 | 28391 | G3bp1 |
| G3bp2 | 31734 | 27621 | 27497 | 28041 | 31331 | 30070 | 32524 | 33890 | 28580 | 28039 | 26648 | 29878 | 30288 | 30651 | 34014 | 28330 | 28573 | 29335 | 30328 | 32947 | 31190 | 32049 | 27672 | 27071 | 30710 | 28327 | 30454 | 30858 | G3bp2 |
| G430095P16Rik | 17 | 16 | 13 | 12 | 7 | 5 | 16 | 46 | 51 | 31 | 43 | 43 | 41 | 52 | 17 | 23 | 13 | 17 | 16 | 19 | 9 | 20 | 42 | 39 | 29 | 29 | 40 | 49 | G430095P16Rik |
| G6pc | 0 | 0 | 0 | 0 | 0 | 0 | 0 | 0 | 0 | 3 | 4 | 0 | 0 | 0 | 0 | 0 | 0 | 0 | 0 | 0 | 0 | 0 | 1 | 1 | 0 | 0 | 1 | 0 | G6pc |
| G6pc2 | 15 | 18 | 2 | 45 | 31 | 28 | 29 | 7 | 10 | 11 | 4 | 5 | 9 | 3 | 15 | 16 | 20 | 37 | 13 | 22 | 34 | 2 | 3 | 2 | 4 | 4 | 0 | 7 | G6pc2 |
| G6pc3 | 1190 | 1569 | 1504 | 1432 | 1285 | 1306 | 1302 | 1044 | 1114 | 1257 | 1228 | 1460 | 1377 | 1396 | 1151 | 1357 | 1430 | 1260 | 1393 | 1292 | 1306 | 1348 | 1198 | 1331 | 1342 | 1484 | 1381 | 1442 | G6pc3 |
| G6pd2 | 2 | 3 | 0 | 0 | 1 | 0 | 1 | 4 | 1 | 1 | 0 | 0 | 0 | 1 | 7 | 1 | 0 | 0 | 1 | 0 | 0 | 2 | 0 | 1 | 1 | 2 | 1 | 0 | G6pd2 |
| G6pdx | 5384 | 5667 | 4471 | 4449 | 4331 | 4758 | 4530 | 4876 | 4706 | 4149 | 3435 | 4325 | 3851 | 4060 | 5177 | 4502 | 4613 | 4184 | 4856 | 5167 | 4771 | 4341 | 4477 | 4173 | 4360 | 4556 | 4448 | 4201 | G6pdx |
| Gaa | 4312 | 5897 | 5397 | 5988 | 5690 | 5307 | 4317 | 2980 | 4651 | 4612 | 5215 | 3867 | 3849 | 2949 | 4358 | 5946 | 5689 | 6270 | 5684 | 4244 | 4366 | 4859 | 4965 | 3957 | 4086 | 3517 | 3253 | 2961 | Gaa |
| Gab1 | 4248 | 3860 | 4097 | 4442 | 4714 | 4382 | 4204 | 3910 | 4844 | 4663 | 4974 | 4285 | 4574 | 3849 | 4112 | 4148 | 4393 | 4570 | 4674 | 3985 | 4200 | 4122 | 4948 | 4668 | 4354 | 4288 | 3763 | 3627 | Gab1 |
| Gab2 | 3061 | 2937 | 3149 | 3468 | 4875 | 4147 | 3749 | 3824 | 3766 | 3881 | 3595 | 3606 | 3866 | 3428 | 3483 | 2823 | 3483 | 3623 | 4347 | 3768 | 4131 | 4841 | 3364 | 3307 | 4215 | 2712 | 3490 | 3224 | Gab2 |
| Gab3 | 170 | 158 | 137 | 144 | 187 | 187 | 191 | 70 | 66 | 88 | 73 | 56 | 60 | 48 | 207 | 191 | 131 | 179 | 153 | 153 | 163 | 94 | 96 | 67 | 84 | 105 | 56 | 57 | Gab3 |

| | | | | | | | | | | | | | | | | | | | | | | | | | | | | | |
|---|---|---|---|---|---|---|---|---|---|---|---|---|---|---|---|---|---|---|---|---|---|---|---|---|---|---|---|---|---|
| Gabarap | 16057 | 16126 | 14421 | 20244 | 16553 | 15850 | 18391 | 17437 | 16579 | 17980 | 17312 | 15175 | 18003 | 16833 | 17971 | 14801 | 16219 | 16288 | 17276 | 17395 | 17220 | 15862 | 17054 | 16892 | 16682 | 17609 | 17187 | 17624 | Gabarap |
| Gabarapl1 | 8884 | 9421 | 9896 | 11031 | 10956 | 9277 | 10801 | 7399 | 6771 | 7930 | 6770 | 6278 | 6381 | 7045 | 9067 | 7893 | 9565 | 10146 | 10581 | 10501 | 10568 | 7347 | 6550 | 7437 | 7747 | 6455 | 6545 | 6700 | Gabarapl1 |
| Gabarapl2 | 5705 | 6684 | 6449 | 7190 | 7003 | 6391 | 8366 | 5559 | 4778 | 5388 | 4470 | 5377 | 4972 | 5786 | 6707 | 5690 | 6922 | 6949 | 7585 | 8264 | 7898 | 5250 | 4820 | 5329 | 5785 | 5325 | 5428 | 5905 | Gabarapl2 |
| Gabbr1 | 14519 | 15849 | 13835 | 23577 | 28648 | 25110 | 25490 | 19678 | 19128 | 24136 | 21370 | 19782 | 19984 | 19547 | 16210 | 13622 | 19045 | 23873 | 26767 | 21507 | 28133 | 27178 | 17125 | 19881 | 25122 | 13357 | 17980 | 17467 | Gabbr1 |
| Gabbr2 | 105 | 174 | 91 | 242 | 872 | 409 | 436 | 846 | 991 | 1255 | 1046 | 519 | 635 | 460 | 220 | 190 | 175 | 458 | 442 | 413 | 521 | 1711 | 931 | 697 | 1080 | 267 | 375 | 390 | Gabbr2 |
| Gabpa | 5264 | 3948 | 4997 | 4791 | 4540 | 4926 | 5614 | 7651 | 6524 | 6298 | 6733 | 7617 | 7312 | 8075 | 4999 | 4612 | 4832 | 4439 | 4931 | 5162 | 4967 | 6794 | 6941 | 7177 | 6946 | 7140 | 7299 | 7976 | Gabpa |
| Gabpb1 | 2784 | 2420 | 3400 | 3061 | 3067 | 2963 | 3394 | 4545 | 4043 | 4188 | 4095 | 5528 | 4570 | 5334 | 2683 | 2415 | 3019 | 3086 | 3296 | 2597 | 3283 | 4814 | 3962 | 4759 | 4539 | 5802 | 5263 | 5408 | Gabpb1 |
| Gabpb2 | 4567 | 3563 | 2806 | 3060 | 3197 | 3703 | 3481 | 8172 | 8412 | 7226 | 8082 | 8751 | 8641 | 8987 | 3789 | 3500 | 3049 | 3032 | 3242 | 3902 | 3624 | 5802 | 7774 | 6983 | 7446 | 6262 | 9678 | 9156 | Gabpb2 |
| Gabra1 | 321 | 197 | 363 | 368 | 547 | 568 | 560 | 1061 | 723 | 910 | 619 | 610 | 543 | 657 | 334 | 237 | 351 | 438 | 564 | 566 | 688 | 1110 | 772 | 847 | 969 | 359 | 669 | 612 | Gabra1 |
| Gabra2 | 557 | 450 | 662 | 1094 | 1573 | 1393 | 1728 | 1890 | 1569 | 2027 | 1623 | 1764 | 1337 | 2169 | 676 | 467 | 797 | 1142 | 1681 | 1493 | 1773 | 2480 | 1805 | 2352 | 2854 | 1369 | 1651 | 1734 | Gabra2 |
| Gabra3 | 1351 | 1175 | 1026 | 1681 | 2711 | 2366 | 2578 | 2455 | 2280 | 2341 | 2064 | 1527 | 1283 | 1576 | 1610 | 1178 | 1494 | 2245 | 2184 | 2478 | 2734 | 3217 | 2394 | 1848 | 2402 | 1317 | 1259 | 1304 | Gabra3 |
| Gabra4 | 1226 | 675 | 608 | 925 | 1332 | 1062 | 1474 | 492 | 567 | 456 | 528 | 288 | 308 | 325 | 905 | 935 | 781 | 1013 | 1240 | 1253 | 1415 | 508 | 649 | 394 | 487 | 317 | 222 | 293 | Gabra4 |
| Gabra5 | 107 | 102 | 86 | 152 | 475 | 292 | 417 | 398 | 405 | 388 | 305 | 151 | 141 | 140 | 129 | 86 | 123 | 245 | 344 | 324 | 390 | 416 | 376 | 261 | 416 | 108 | 129 | 128 | Gabra5 |
| Gabra6 | 0 | 0 | 0 | 0 | 0 | 0 | 0 | 0 | 0 | 0 | 0 | 0 | 0 | 1 | 7 | 0 | 0 | 0 | 0 | 0 | 0 | 0 | 3 | 0 | 0 | 0 | 0 | 0 | Gabra6 |
| Gabrb1 | 558 | 470 | 279 | 630 | 1087 | 950 | 1233 | 879 | 832 | 594 | 581 | 290 | 248 | 300 | 707 | 449 | 428 | 741 | 821 | 1317 | 1420 | 755 | 911 | 428 | 595 | 113 | 263 | 225 | Gabrb1 |
| Gabrb2 | 2150 | 1939 | 1073 | 1993 | 2630 | 2922 | 2692 | 3187 | 2031 | 1974 | 1636 | 1924 | 1733 | 1903 | 2430 | 1604 | 1342 | 1909 | 2352 | 3338 | 3131 | 2308 | 2019 | 1603 | 2527 | 303 | 2126 | 1831 | Gabrb2 |
| Gabrb3 | 3357 | 3372 | 2612 | 4823 | 7250 | 6090 | 7359 | 5860 | 5225 | 5534 | 4819 | 4024 | 3709 | 4302 | 4191 | 3242 | 3701 | 5465 | 5855 | 5925 | 7038 | 7451 | 5041 | 4578 | 6260 | 2575 | 3542 | 3620 | Gabrb3 |
| Gabrd | 2 | 3 | 0 | 19 | 40 | 27 | 18 | 1 | 4 | 2 | 4 | 3 | 7 | 0 | 1 | 9 | 2 | 32 | 17 | 16 | 31 | 10 | 6 | 7 | 15 | 2 | 6 | 0 | Gabrd |
| Gabre | 1216 | 1056 | 987 | 958 | 909 | 976 | 1063 | 446 | 455 | 396 | 470 | 403 | 399 | 365 | 1006 | 1091 | 1114 | 1050 | 1123 | 1052 | 1036 | 391 | 476 | 491 | 319 | 592 | 363 | 408 | Gabre |
| Gabrg1 | 639 | 574 | 646 | 785 | 1089 | 822 | 1027 | 291 | 291 | 226 | 227 | 167 | 169 | 267 | 715 | 567 | 672 | 839 | 946 | 1126 | 1138 | 439 | 331 | 212 | 357 | 147 | 225 | 227 | Gabrg1 |
| Gabrg2 | 1248 | 1158 | 1318 | 1826 | 3036 | 2474 | 3067 | 7376 | 5164 | 4992 | 4498 | 4432 | 3733 | 4633 | 2145 | 1086 | 1501 | 2060 | 2605 | 3239 | 2905 | 7596 | 5489 | 4437 | 5894 | 2761 | 4467 | 4367 | Gabrg2 |
| Gabrg3 | 57 | 43 | 22 | 74 | 181 | 168 | 184 | 151 | 165 | 140 | 177 | 112 | 41 | 104 | 87 | 48 | 52 | 154 | 134 | 149 | 187 | 185 | 147 | 77 | 163 | 86 | 59 | 82 | Gabrg3 |
| Gabrp | 661 | 662 | 562 | 1 | 1 | 613 | 1 | 1211 | 1481 | 593 | 1009 | 1377 | 1768 | 1510 | 1026 | 766 | 134 | 508 | 93 | 31 | 0 | 7 | 1558 | 1244 | 160 | 146 | 1237 | 1697 | Gabrp |
| Gabrq | 1433 | 1583 | 1110 | 1859 | 1589 | 1637 | 1971 | 586 | 460 | 433 | 382 | 495 | 348 | 478 | 1677 | 1192 | 1459 | 1618 | 1653 | 2024 | 1925 | 640 | 451 | 386 | 483 | 344 | 459 | 426 | Gabrq |
| Gabrr1 | 3 | 3 | 0 | 3 | 6 | 11 | 4 | 13 | 10 | 10 | 5 | 7 | 8 | 15 | 4 | 6 | 0 | 2 | 6 | 17 | 5 | 7 | 13 | 14 | 5 | 12 | 8 | 8 | Gabrr1 |
| Gabrr2 | 28 | 61 | 45 | 60 | 52 | 51 | 69 | 16 | 34 | 28 | 25 | 28 | 11 | 18 | 40 | 41 | 37 | 80 | 66 | 33 | 58 | 27 | 19 | 17 | 33 | 24 | 9 | 12 | Gabrr2 |
| Gabrr3 | 10 | 11 | 34 | 3 | 16 | 28 | 17 | 31 | 12 | 49 | 21 | 30 | 17 | 42 | 16 | 5 | 23 | 1 | 12 | 17 | 11 | 43 | 16 | 23 | 24 | 23 | 45 | 50 | Gabrr3 |
| Gad1 | 945 | 961 | 841 | 1721 | 3458 | 2698 | 3081 | 5709 | 4653 | 5464 | 4395 | 5046 | 3810 | 5163 | 1167 | 561 | 1424 | 1949 | 3066 | 2515 | 3143 | 7526 | 3614 | 5125 | 7001 | 3153 | 4610 | 4135 | Gad1 |
| Gad2 | 1562 | 1221 | 1518 | 2460 | 3984 | 3411 | 4019 | 8204 | 7188 | 9077 | 6835 | 10204 | 5192 | 11106 | 1742 | 769 | 2466 | 2400 | 3949 | 2818 | 3871 | 11221 | 4792 | 9469 | 11229 | 7576 | 8503 | 8109 | Gad2 |
| Gadd45a | 678 | 798 | 976 | 769 | 599 | 482 | 654 | 552 | 392 | 633 | 551 | 630 | 672 | 693 | 659 | 664 | 837 | 640 | 563 | 567 | 648 | 409 | 491 | 736 | 472 | 862 | 614 | 730 | Gadd45a |
| Gadd45b | 403 | 655 | 556 | 571 | 269 | 366 | 216 | 226 | 344 | 283 | 412 | 280 | 381 | 261 | 438 | 659 | 416 | 590 | 263 | 315 | 252 | 236 | 474 | 262 | 271 | 307 | 360 | 305 | Gadd45b |
| Gadd45g | 1340 | 2068 | 3268 | 2188 | 2044 | 1943 | 1559 | 1860 | 1671 | 2017 | 1793 | 3308 | 2537 | 2760 | 1530 | 1258 | 2883 | 1782 | 2503 | 1859 | 1649 | 1648 | 1581 | 2894 | 2512 | 2873 | 3309 | 3018 | Gadd45g |
| Gadd45gip1 | 858 | 964 | 1185 | 860 | 986 | 890 | 965 | 1445 | 1598 | 1666 | 1646 | 1707 | 1622 | 1852 | 769 | 878 | 1087 | 990 | 1180 | 895 | 930 | 1353 | 1638 | 1861 | 1585 | 1913 | 1587 | 1774 | Gadd45gip1 |
| Gadl1 | 8 | 4 | 2 | 19 | 11 | 3 | 1 | 33 | 45 | 30 | 53 | 22 | 14 | 22 | 9 | 8 | 1 | 7 | 2 | 5 | 10 | 13 | 46 | 51 | 42 | 16 | 2 | 7 | Gadl1 |
| Gak | 5247 | 5414 | 5419 | 5683 | 5643 | 5615 | 5260 | 4974 | 5852 | 6497 | 5936 | 6929 | 6032 | 6387 | 4987 | 5021 | 5175 | 5409 | 5870 | 4759 | 5672 | 6012 | 5664 | 6135 | 6561 | 6456 | 6108 | 6324 | Gak |
| Gal | 80 | 102 | 26 | 14 | 263 | 39 | 46 | 167 | 208 | 152 | 250 | 15 | 100 | 29 | 84 | 100 | 60 | 141 | 42 | 27 | 32 | 246 | 137 | 29 | 144 | 20 | 15 | 24 | Gal |
| Gal3st1 | 15 | 13 | 17 | 12 | 23 | 22 | 8 | 32 | 54 | 54 | 79 | 16 | 29 | 21 | 22 | 23 | 17 | 19 | 9 | 6 | 4 | 37 | 56 | 34 | 43 | 30 | 19 | 21 | Gal3st1 |
| Gal3st2 | 1 | 0 | 0 | 0 | 0 | 0 | 0 | 0 | 0 | 0 | 2 | 3 | 1 | 1 | 0 | 0 | 0 | 0 | 0 | 2 | 0 | 0 | 5 | 0 | 3 | 4 | 1 | 1 | Gal3st2 |
| Gal3st2b | 0 | 0 | 1 | 0 | 0 | 0 | 0 | 0 | 0 | 0 | 3 | 1 | 0 | 0 | 0 | 0 | 0 | 0 | 0 | 0 | 0 | 5 | 2 | 0 | 0 | 0 | 0 | 0 | Gal3st2b |
| Gal3st2c | 6 | 5 | 7 | 7 | 16 | 8 | 10 | 10 | 2 | 8 | 11 | 10 | 10 | 6 | 1 | 3 | 2 | 3 | 6 | 2 | 5 | 2 | 10 | 8 | 2 | 4 | 10 | 12 | Gal3st2c |
| Gal3st3 | 463 | 543 | 731 | 580 | 875 | 714 | 708 | 804 | 863 | 1086 | 853 | 959 | 755 | 818 | 476 | 496 | 772 | 666 | 860 | 623 | 705 | 1205 | 828 | 1063 | 1120 | 937 | 793 | 843 | Gal3st3 |
| Gal3st4 | 0 | 0 | 0 | 0 | 0 | 0 | 0 | 0 | 0 | 0 | 0 | 0 | 0 | 0 | 0 | 0 | 0 | 0 | 0 | 0 | 0 | 3 | 0 | 0 | 0 | 0 | 0 | 0 | Gal3st4 |
| Galc | 1788 | 1672 | 1942 | 2032 | 2124 | 2106 | 2216 | 2445 | 2295 | 2547 | 2387 | 2879 | 2555 | 2933 | 1945 | 1774 | 2005 | 2133 | 2136 | 1905 | 2070 | 2565 | 2380 | 2715 | 2656 | 2704 | 2554 | 2819 | Galc |
| Gale | 1448 | 1991 | 1690 | 1586 | 1314 | 1474 | 1260 | 694 | 943 | 833 | 941 | 812 | 785 | 680 | 1381 | 1755 | 1673 | 1465 | 1597 | 1407 | 1290 | 837 | 847 | 852 | 925 | 853 | 918 | 864 | Gale |
| Galk1 | 2834 | 3026 | 2648 | 2352 | 1616 | 2116 | 1638 | 1828 | 2716 | 2114 | 2457 | 2268 | 2497 | 1975 | 2462 | 2946 | 2387 | 2080 | 2018 | 2033 | 1591 | 1576 | 2978 | 2338 | 1851 | 2969 | 2586 | 2268 | Galk1 |
| Galk2 | 1689 | 1640 | 1577 | 1321 | 1236 | 1320 | 1226 | 1887 | 1911 | 1498 | 1693 | 1695 | 1839 | 1547 | 1691 | 1834 | 1396 | 1201 | 1270 | 1539 | 1189 | 1425 | 2081 | 1538 | 1444 | 1641 | 1629 | 1734 | Galk2 |
| Galm | 1606 | 1157 | 1122 | 1001 | 534 | 534 | 613 | 888 | 748 | 910 | 856 | 422 | 542 | 477 | 1199 | 1458 | 1010 | 654 | 581 | 666 | 525 | 297 | 994 | 1035 | 275 | 1379 | 446 | 443 | Galm |
| Galns | 753 | 915 | 777 | 732 | 669 | 608 | 449 | 357 | 486 | 508 | 577 | 430 | 479 | 427 | 671 | 896 | 684 | 706 | 589 | 482 | 542 | 386 | 623 | 342 | 376 | 508 | 500 | 437 | Galns |
| Galnt1 | 7745 | 6258 | 6989 | 5558 | 5827 | 6214 | 6187 | 9651 | 8932 | 7914 | 8737 | 9717 | 9768 | 9375 | 7764 | 7555 | 6152 | 5738 | 5849 | 6605 | 5746 | 8408 | 9291 | 8446 | 8140 | 10157 | 10287 | 10380 | Galnt1 |
| Galnt10 | 5577 | 6687 | 6759 | 6415 | 5155 | 5525 | 4255 | 2451 | 3171 | 2913 | 3469 | 2871 | 3148 | 2586 | 5189 | 6830 | 6420 | 6472 | 5591 | 3795 | 5112 | 2999 | 3432 | 2853 | 2773 | 2842 | 2907 | 2871 | Galnt10 |
| Galnt11 | 2233 | 2251 | 2162 | 2379 | 2374 | 2447 | 2337 | 2627 | 2896 | 2838 | 2988 | 2483 | 2815 | 2623 | 2285 | 2287 | 2443 | 2213 | 2285 | 2524 | 2406 | 2445 | 2906 | 2478 | 2828 | 2450 | 2889 | 2490 | Galnt11 |
| Galnt12 | 624 | 583 | 749 | 400 | 315 | 425 | 285 | 184 | 189 | 169 | 232 | 254 | 262 | 283 | 556 | 721 | 511 | 366 | 368 | 359 | 296 | 192 | 234 | 212 | 144 | 297 | 330 | 310 | Galnt12 |
| Galnt13 | 347 | 313 | 338 | 404 | 632 | 574 | 591 | 554 | 470 | 467 | 485 | 400 | 403 | 416 | 491 | 392 | 437 | 406 | 450 | 550 | 517 | 630 | 425 | 331 | 452 | 262 | 370 | 392 | Galnt13 |
| Galnt14 | 275 | 249 | 206 | 286 | 443 | 329 | 340 | 935 | 932 | 844 | 840 | 792 | 758 | 760 | 297 | 222 | 299 | 241 | 359 | 395 | 407 | 886 | 981 | 869 | 886 | 596 | 712 | 682 | Galnt14 |
| Galnt15 | 18 | 18 | 19 | 14 | 29 | 47 | 36 | 90 | 79 | 94 | 63 | 61 | 76 | 57 | 22 | 23 | 24 | 40 | 37 | 37 | 33 | 57 | 106 | 99 | 83 | 51 | 53 | 30 | Galnt15 |
| Galnt16 | 2939 | 3347 | 3233 | 3558 | 4119 | 3939 | 3778 | 3455 | 3691 | 3989 | 4070 | 4287 | 3880 | 4129 | 2988 | 2940 | 3519 | 3635 | 4330 | 3534 | 3925 | 4162 | 3842 | 3959 | 4167 | 3839 | 3836 | 4372 | Galnt16 |
| Galnt17 | 634 | 722 | 493 | 1042 | 2035 | 1512 | 1576 | 1341 | 1874 | 1994 | 2149 | 1118 | 1001 | 1127 | 804 | 690 | 804 | 1416 | 1629 | 1284 | 1769 | 2288 | 1959 | 1682 | 2203 | 1048 | 787 | 808 | Galnt17 |
| Galnt18 | 300 | 362 | 353 | 339 | 451 | 376 | 263 | 377 | 589 | 416 | 575 | 313 | 433 | 280 | 331 | 390 | 298 | 435 | 382 | 346 | 310 | 344 | 617 | 445 | 374 | 331 | 329 | 329 | Galnt18 |
| Galnt2 | 9 | 33 | 24 | 23 | 22 | 18 | 21 | 12 | 21 | 14 | 21 | 12 | 24 | 23 | 11 | 13 | 31 | 23 | 19 | 17 | 22 | 13 | 16 | 22 | 18 | 10 | 28 | 37 | Galnt2 |
| Galnt3 | 219 | 156 | 185 | 165 | 174 | 140 | 103 | 128 | 170 | 137 | 135 | 101 | 166 | 107 | 216 | 251 | 159 | 154 | 108 | 120 | 137 | 109 | 176 | 136 | 78 | 137 | 106 | 128 | Galnt3 |
| Galnt4 | 1212 | 783 | 1019 | 844 | 697 | 948 | 694 | 876 | 943 | 1043 | 1295 | 861 | 1035 | 799 | 1033 | 1095 | 820 | 846 | 733 | 652 | 668 | 906 | 1172 | 1000 | 938 | 895 | 761 | 913 | Galnt4 |
| Galnt5 | 158 | 73 | 59 | 31 | 16 | 75 | 1 | 223 | 271 | 227 | 366 | 101 | 229 | 106 | 91 | 108 | 30 | 63 | 20 | 6 | 3 | 184 | 284 | 196 | 165 | 118 | 145 | 108 | Galnt5 |
| Galnt6 | 68 | 65 | 53 | 55 | 71 | 33 | 29 | 12 | 29 | 48 | 48 | 10 | 19 | 9 | 70 | 113 | 57 | 61 | 49 | 25 | 41 | 20 | 60 | 21 | 12 | 19 | 8 | 8 | Galnt6 |
| Galnt7 | 1716 | 1347 | 1675 | 1374 | 1820 | 1556 | 1820 | 2440 | 2160 | 2095 | 2331 | 2570 | 2582 | 2452 | 1764 | 1632 | 1487 | 1530 | 1637 | 1863 | 1667 | 2003 | 2349 | 2290 | 2144 | 2768 | 2588 | 2616 | Galnt7 |
| Galnt9 | 307 | 441 | 512 | 626 | 938 | 631 | 511 | 480 | 582 | 719 | 533 | 539 | 490 | 416 | 404 | 408 | 716 | 683 | 738 | 583 | 648 | 1006 | 540 | 517 | 713 | 470 | 565 | 470 | Galnt9 |
| Galntl5 | 0 | 0 | 0 | 0 | 0 | 0 | 0 | 0 | 0 | 0 | 0 | 2 | 0 | 0 | 0 | 0 | 0 | 0 | 0 | 0 | 0 | 0 | 0 | 0 | 0 | 0 | 0 | 0 | Galntl5 |
| Galntl6 | 153 | 181 | 118 | 306 | 518 | 431 | 389 | 368 | 364 | 362 | 401 | 199 | 191 | 255 | 204 | 168 | 216 | 379 | 309 | 371 | 388 | 466 | 348 | 257 | 379 | 144 | 191 | 198 | Galntl6 |
| Galp | 0 | 0 | 0 | 0 | 0 | 0 | 0 | 0 | 0 | 0 | 0 | 0 | 1 | 0 | 0 | 1 | 0 | 2 | 0 | 0 | 0 | 0 | 0 | 0 | 0 | 0 | 0 | 0 | Galp |
| Galr1 | 2 | 17 | 3 | 13 | 47 | 23 | 32 | 43 | 42 | 69 | 64 | 24 | 61 | 15 | 37 | 8 | 10 | 13 | 14 | 22 | 13 | 48 | 54 | 26 | 29 | 14 | 30 | 9 | Galr1 |
| Galr2 | 12 | 7 | 3 | 0 | 11 | 4 | 6 | 7 | 7 | 3 | 9 | 12 | 7 | 1 | 8 | 2 | 2 | 7 | 2 | 3 | 0 | 17 | 9 | 2 | 9 | 11 | 1 | 1 | Galr2 |
| Galr3 | 10 | 6 | 2 | 2 | 9 | 7 | 4 | 0 | 0 | 6 | 0 | 0 | 3 | 0 | 8 | 12 | 9 | 1 | 3 | 3 | 6 | 0 | 3 | 1 | 5 | 0 | 0 | 5 | Galr3 |
| Gamt | 539 | 724 | 696 | 534 | 468 | 423 | 488 | 280 | 359 | 298 | 631 | 326 | 425 | 383 | 564 | 622 | 736 | 582 | 488 | 402 | 440 | 356 | 534 | 341 | 325 | 349 | 413 | 385 | Gamt |
| Gan | 3996 | 4127 | 3655 | 5079 | 5279 | 5313 | 4789 | 2839 | 3154 | 3466 | 3487 | 3458 | 3455 | 3402 | 3701 | 4259 | 4144 | 5253 | 5082 | 3994 | 5339 | 3267 | 2788 | 2925 | 4175 | 2665 | 3740 | 3217 | Gan |
| Ganab | 19288 | 20300 | 19505 | 18155 | 16708 | 18124 | 16668 | 20245 | 21517 | 21291 | 21954 | 22802 | 23092 | 21576 | 18228 | 19386 | 19459 | 17994 | 17824 | 16988 | 16764 | 21500 | 20877 | 21254 | 21316 | 23182 | 23145 | 22006 | Ganab |
| Ganc | 862 | 819 | 783 | 708 | 778 | 715 | 704 | 844 | 789 | 680 | 728 | 711 | 710 | 744 | 831 | 751 | 785 | 640 | 658 | 870 | 733 | 677 | 801 | 657 | 688 | 571 | 716 | 833 | Ganc |
| Gap43 | 9684 | 12476 | 11260 | 16912 | 23411 | 17471 | 19232 | 19853 | 18258 | 22120 | 18097 | 14185 | 16505 | 15483 | 13917 | 11092 | 14478 | 17982 | 19557 | 21644 | 18666 | 23509 | 18707 | 17732 | 22374 | 11660 | 14956 | 14523 | Gap43 |
| Gapdh | 845 | 1059 | 6213 | 2577 | 923 | 856 | 750 | 1070 | 1131 | 1765 | 896 | 1246 | 1190 | 1502 | 2181 | 2975 | 962 | 1907 | 775 | 769 | 1313 | 916 | 4306 | 1174 | 3153 | 2091 | 1235 | 1384 | Gapdh |
| Gapdhs | 1 | 3 | 5 | 1 | 0 | 2 | 2 | 2 | 4 | 9 | 5 | 8 | 7 | 13 | 0 | 1 | 0 | 0 | 2 | 2 | 6 | 8 | 9 | 2 | 9 | 3 | 6 | 4 | Gapdhs |
| Gapt | 1 | 2 | 8 | 0 | 1 | 3 | 0 | 2 | 7 | 4 | 4 | 0 | 0 | 2 | 1 | 8 | 2 | 3 | 2 | 7 | 2 | 0 | 5 | 2 | 1 | 3 | 1 | 0 | Gapt |

| | | | | | | | | | | | | | | | | | | | | | | | | | | | | | |
|---|---|---|---|---|---|---|---|---|---|---|---|---|---|---|---|---|---|---|---|---|---|---|---|---|---|---|---|---|---|
| Gapvd1 | 14962 | 14400 | 10874 | 13785 | 15026 | 15834 | 16134 | 15188 | 14075 | 14356 | 13714 | 14655 | 13514 | 14706 | 14668 | 13307 | 12560 | 14236 | 14724 | 15292 | 16601 | 14054 | 13282 | 12977 | 15346 | 10544 | 14164 | 13968 | Gapvd1 |
| Gar1 | 1593 | 1645 | 2983 | 2579 | 1736 | 1637 | 1971 | 1584 | 1688 | 2378 | 2384 | 2817 | 2508 | 3476 | 1431 | 1372 | 2664 | 1974 | 1975 | 1522 | 1844 | 1755 | 1629 | 3289 | 2466 | 3518 | 2826 | 3618 | Gar1 |
| Garem1 | 3233 | 3162 | 2070 | 2986 | 3355 | 3073 | 3023 | 1536 | 1722 | 1560 | 1629 | 1397 | 1712 | 1304 | 3157 | 3405 | 2491 | 3144 | 2818 | 2514 | 3051 | 1604 | 1609 | 1232 | 1528 | 754 | 1303 | 1333 | Garem1 |
| Garem2 | 2143 | 2978 | 2697 | 3144 | 3377 | 3148 | 3036 | 2842 | 2556 | 3089 | 2382 | 3994 | 3548 | 3649 | 2483 | 2039 | 3637 | 2731 | 3896 | 3179 | 3148 | 3160 | 2207 | 3344 | 3373 | 3326 | 3427 | 3701 | Garem2 |
| Garnl3 | 2939 | 3142 | 2476 | 3066 | 3675 | 3463 | 3296 | 1827 | 1852 | 1962 | 1883 | 1387 | 1698 | 1318 | 3063 | 3059 | 2671 | 3316 | 3247 | 3457 | 3481 | 2294 | 1909 | 1279 | 1673 | 1125 | 1410 | 1170 | Garnl3 |
| Garre1 | 5290 | 5498 | 4288 | 5474 | 5642 | 5554 | 5187 | 4853 | 4991 | 5014 | 5138 | 4989 | 4948 | 4895 | 5368 | 5374 | 5072 | 5585 | 5506 | 5168 | 5559 | 5095 | 5045 | 4515 | 4995 | 3485 | 4928 | 4735 | Garre1 |
| Gars | 14260 | 13635 | 13236 | 12429 | 12961 | 12823 | 13588 | 12208 | 11024 | 11048 | 10230 | 13259 | 11640 | 13256 | 14260 | 12167 | 12732 | 12444 | 13807 | 13502 | 13419 | 12629 | 10468 | 12268 | 12399 | 13462 | 12706 | 14156 | Gars |
| Gart | 15068 | 17094 | 16309 | 14672 | 11257 | 12518 | 12616 | 6260 | 7071 | 6819 | 6858 | 8256 | 7641 | 7990 | 13518 | 14396 | 14891 | 13661 | 13457 | 12866 | 12856 | 5335 | 6512 | 7820 | 7158 | 9801 | 8738 | 8759 | Gart |
| Gas1 | 5143 | 3978 | 5709 | 4099 | 3009 | 3903 | 3474 | 6293 | 6825 | 6856 | 7417 | 7879 | 8558 | 8226 | 4158 | 4150 | 4190 | 3542 | 3701 | 3276 | 3259 | 3155 | 6923 | 8055 | 5339 | 10588 | 7304 | 8980 | Gas1 |
| Gas2 | 1862 | 1425 | 1510 | 1229 | 1054 | 1162 | 1206 | 1910 | 2081 | 1605 | 3140 | 1908 | 2079 | 1884 | 1880 | 1787 | 1290 | 1209 | 1201 | 1191 | 1266 | 1755 | 2901 | 1561 | 1621 | 2426 | 1977 | 2016 | Gas2 |
| Gas2l1 | 3087 | 4391 | 4154 | 4078 | 3045 | 2922 | 2754 | 1021 | 1275 | 1431 | 1475 | 1459 | 1266 | 1188 | 2943 | 3620 | 4496 | 4013 | 3465 | 2614 | 2837 | 1348 | 1311 | 1468 | 1191 | 1583 | 1468 | 1178 | Gas2l1 |
| Gas2l2 | 5 | 4 | 2 | 0 | 15 | 17 | 1 | 7 | 28 | 21 | 19 | 26 | 31 | 7 | 3 | 13 | 5 | 0 | 5 | 12 | 6 | 23 | 47 | 9 | 11 | 8 | 0 | 2 | Gas2l2 |
| Gas2l3 | 3163 | 2681 | 2547 | 2907 | 2463 | 2842 | 3232 | 2182 | 2155 | 1796 | 1873 | 2509 | 2096 | 2684 | 2963 | 2628 | 2647 | 3090 | 2848 | 2809 | 3038 | 1679 | 1825 | 2269 | 2204 | 2244 | 2508 | 2409 | Gas2l3 |
| Gas6 | 1433 | 1921 | 1975 | 1560 | 1871 | 1607 | 1401 | 761 | 1232 | 1145 | 1508 | 1040 | 1153 | 972 | 1581 | 2034 | 1665 | 1794 | 1578 | 1460 | 1357 | 1312 | 1730 | 986 | 1117 | 1078 | 1009 | 903 | Gas6 |
| Gas7 | 872 | 849 | 795 | 1235 | 1664 | 1507 | 1530 | 1899 | 2568 | 2517 | 2555 | 1623 | 1008 | 1659 | 872 | 882 | 1036 | 1541 | 1504 | 1382 | 1858 | 2782 | 2804 | 2397 | 3061 | 1524 | 1254 | 1360 | Gas7 |
| Gas8 | 3881 | 4806 | 5273 | 5219 | 4495 | 4634 | 4619 | 2561 | 2829 | 3358 | 3269 | 3300 | 3241 | 3482 | 3874 | 4064 | 5174 | 5111 | 4811 | 4273 | 4740 | 2918 | 2650 | 3434 | 3361 | 3622 | 3441 | 3420 | Gas8 |
| Gask1a | 99 | 148 | 173 | 217 | 173 | 255 | 197 | 172 | 285 | 270 | 450 | 115 | 145 | 116 | 167 | 181 | 168 | 287 | 209 | 198 | 172 | 284 | 299 | 185 | 275 | 116 | 134 | 97 | Gask1a |
| Gask1b | 216 | 171 | 174 | 107 | 206 | 226 | 107 | 530 | 769 | 605 | 1189 | 337 | 628 | 358 | 240 | 213 | 169 | 215 | 115 | 150 | 76 | 592 | 1225 | 556 | 410 | 389 | 423 | 376 | Gask1b |
| Gast | 6 | 12 | 3 | 6 | 7 | 9 | 4 | 12 | 7 | 4 | 5 | 17 | 14 | 7 | 4 | 5 | 9 | 20 | 12 | 13 | 11 | 2 | 7 | 7 | 23 | 11 | 31 | 14 | Gast |
| Gata1 | 31 | 55 | 39 | 12 | 13 | 12 | 20 | 32 | 52 | 28 | 42 | 50 | 109 | 82 | 47 | 65 | 29 | 11 | 18 | 20 | 13 | 18 | 35 | 23 | 15 | 94 | 35 | 165 | Gata1 |
| Gata2 | 1021 | 812 | 1364 | 1003 | 754 | 674 | 402 | 1342 | 1082 | 1908 | 1124 | 1232 | 1668 | 1061 | 823 | 1001 | 1015 | 750 | 725 | 458 | 525 | 1133 | 1027 | 1698 | 991 | 1486 | 1455 | 1404 | Gata2 |
| Gata3 | 818 | 734 | 2076 | 1626 | 1763 | 1524 | 1153 | 1594 | 1101 | 2175 | 1670 | 2081 | 2613 | 2171 | 947 | 877 | 1727 | 1525 | 1809 | 1212 | 1591 | 2491 | 1230 | 2196 | 2199 | 404 | 2103 | 2298 | Gata3 |
| Gata4 | 5 | 16 | 7 | 4 | 0 | 2 | 0 | 0 | 7 | 0 | 1 | 0 | 0 | 0 | 7 | 9 | 2 | 13 | 4 | 0 | 2 | 0 | 1 | 2 | 0 | 0 | 0 | 0 | Gata4 |
| Gata5 | 7 | 10 | 6 | 2 | 2 | 5 | 6 | 0 | 2 | 2 | 2 | 3 | 11 | 0 | 1 | 5 | 3 | 3 | 0 | 1 | 7 | 2 | 5 | 0 | 1 | 2 | 1 | 4 | Gata5 |
| Gata6 | 72 | 97 | 78 | 66 | 71 | 72 | 45 | 37 | 78 | 42 | 139 | 50 | 93 | 47 | 103 | 100 | 64 | 47 | 30 | 47 | 38 | 70 | 213 | 43 | 38 | 33 | 34 | 44 | Gata6 |
| Gatad1 | 6010 | 5532 | 6346 | 5329 | 5169 | 5162 | 5654 | 5108 | 4491 | 4789 | 4468 | 5462 | 5290 | 6069 | 5676 | 5167 | 6057 | 4738 | 5610 | 5662 | 5798 | 4724 | 4385 | 5155 | 4737 | 6205 | 5951 | 6381 | Gatad1 |
| Gatad2a | 7524 | 8935 | 9438 | 9246 | 8472 | 8218 | 8210 | 4629 | 5581 | 5782 | 5705 | 6874 | 6180 | 6607 | 7371 | 8069 | 9853 | 8962 | 9380 | 6840 | 8566 | 5275 | 5410 | 6610 | 5931 | 8117 | 6415 | 6927 | Gatad2a |
| Gatad2b | 8646 | 7375 | 9912 | 11311 | 11010 | 10425 | 11356 | 9631 | 9474 | 11444 | 11223 | 11691 | 10947 | 12699 | 7907 | 7958 | 10214 | 11102 | 10938 | 8045 | 11738 | 11837 | 9198 | 12051 | 12572 | 10598 | 10570 | 12309 | Gatad2b |
| Gatb | 502 | 650 | 458 | 419 | 514 | 489 | 485 | 1112 | 930 | 937 | 845 | 1025 | 908 | 918 | 645 | 611 | 462 | 418 | 558 | 531 | 550 | 912 | 984 | 955 | 890 | 852 | 1092 | 1049 | Gatb |
| Gatc | 2163 | 2168 | 2160 | 1908 | 1850 | 1806 | 2026 | 2212 | 2244 | 2089 | 1837 | 2548 | 2171 | 2215 | 2172 | 1865 | 2230 | 1883 | 1967 | 2089 | 1893 | 2246 | 2145 | 2222 | 2250 | 2909 | 2294 | 2441 | Gatc |
| Gatd1 | 1443 | 1686 | 1796 | 1591 | 1456 | 1394 | 1347 | 1962 | 2217 | 2296 | 2313 | 2109 | 2301 | 2226 | 1383 | 1446 | 1655 | 1412 | 1511 | 1408 | 1316 | 2226 | 2309 | 2497 | 2278 | 2573 | 2027 | 2192 | Gatd1 |
| Gatd3a | 1865 | 2070 | 1871 | 1745 | 1488 | 1487 | 1517 | 1849 | 2043 | 1920 | 2031 | 1686 | 1825 | 2101 | 1717 | 1838 | 1892 | 1569 | 1645 | 1512 | 1653 | 1547 | 2188 | 2001 | 1846 | 2035 | 1941 | 2078 | Gatd3a |
| Gatm | 112 | 42 | 95 | 45 | 61 | 190 | 38 | 83 | 32 | 28 | 175 | 45 | 45 | 59 | 70 | 75 | 46 | 56 | 145 | 66 | 45 | 47 | 166 | 34 | 43 | 177 | 36 | 46 | Gatm |
| Gba | 2987 | 3626 | 2667 | 3143 | 2938 | 2990 | 2596 | 2075 | 2113 | 2045 | 1986 | 2056 | 2144 | 1872 | 3355 | 3237 | 2840 | 2927 | 2802 | 2979 | 2854 | 2214 | 2328 | 1778 | 2152 | 1850 | 2038 | 2015 | Gba |
| Gba2 | 6933 | 9410 | 7154 | 9037 | 8439 | 8374 | 7822 | 5046 | 5677 | 5932 | 5042 | 5858 | 5152 | 5213 | 7196 | 7139 | 7917 | 9031 | 9406 | 8329 | 8400 | 6014 | 5290 | 5771 | 6550 | 5417 | 5532 | 5116 | Gba2 |
| Gbe1 | 780 | 656 | 711 | 515 | 524 | 594 | 658 | 1133 | 1109 | 879 | 837 | 1019 | 1133 | 1077 | 868 | 714 | 626 | 452 | 538 | 783 | 584 | 964 | 1040 | 917 | 937 | 1313 | 1131 | 1328 | Gbe1 |
| Gbf1 | 10245 | 10994 | 9795 | 10545 | 10656 | 10807 | 9855 | 9165 | 10387 | 10066 | 10201 | 10143 | 10105 | 9490 | 9875 | 10739 | 10595 | 10699 | 10800 | 9194 | 10334 | 10955 | 9985 | 9666 | 10708 | 7712 | 9492 | 9202 | Gbf1 |
| Gbgt1 | 25 | 22 | 43 | 36 | 50 | 38 | 26 | 48 | 78 | 78 | 153 | 76 | 77 | 38 | 22 | 23 | 46 | 30 | 42 | 29 | 21 | 82 | 126 | 47 | 65 | 46 | 66 | 63 | Gbgt1 |
| Gbp10 | 3 | 3 | 0 | 6 | 16 | 3 | 4 | 13 | 21 | 21 | 26 | 15 | 5 | 12 | 9 | 0 | 0 | 12 | 5 | 4 | 9 | 27 | 40 | 29 | 16 | 6 | 14 | 6 | Gbp10 |
| Gbp2 | 15 | 17 | 10 | 20 | 23 | 37 | 18 | 35 | 40 | 36 | 31 | 40 | 46 | 45 | 22 | 23 | 10 | 20 | 11 | 31 | 21 | 43 | 109 | 28 | 36 | 35 | 42 | 49 | Gbp2 |
| Gbp2b | 0 | 0 | 0 | 0 | 0 | 0 | 0 | 0 | 0 | 0 | 1 | 0 | 0 | 0 | 0 | 0 | 0 | 0 | 0 | 0 | 0 | 0 | 0 | 0 | 0 | 1 | 0 | 0 | Gbp2b |
| Gbp3 | 17 | 18 | 25 | 17 | 18 | 24 | 25 | 48 | 71 | 87 | 109 | 71 | 81 | 68 | 30 | 24 | 17 | 24 | 19 | 63 | 19 | 104 | 92 | 75 | 110 | 90 | 73 | 69 | Gbp3 |
| Gbp4 | 35 | 38 | 60 | 48 | 125 | 71 | 75 | 101 | 167 | 165 | 222 | 60 | 81 | 89 | 42 | 47 | 37 | 83 | 66 | 89 | 100 | 177 | 299 | 101 | 116 | 71 | 46 | 53 | Gbp4 |
| Gbp5 | 8 | 8 | 15 | 9 | 9 | 4 | 2 | 12 | 21 | 10 | 12 | 12 | 7 | 10 | 5 | 13 | 2 | 11 | 1 | 5 | 8 | 22 | 25 | 17 | 18 | 17 | 13 | 18 | Gbp5 |
| Gbp6 | 12 | 4 | 16 | 12 | 38 | 37 | 10 | 89 | 76 | 58 | 82 | 55 | 60 | 49 | 16 | 12 | 12 | 48 | 12 | 10 | 17 | 145 | 161 | 76 | 85 | 40 | 53 | 58 | Gbp6 |
| Gbp7 | 87 | 66 | 85 | 67 | 109 | 82 | 88 | 170 | 190 | 170 | 287 | 171 | 197 | 155 | 90 | 80 | 74 | 88 | 78 | 133 | 80 | 201 | 255 | 183 | 135 | 200 | 184 | 237 | Gbp7 |
| Gbp8 | 28 | 26 | 21 | 31 | 54 | 49 | 27 | 73 | 78 | 63 | 131 | 35 | 49 | 46 | 38 | 25 | 15 | 51 | 29 | 62 | 29 | 82 | 109 | 74 | 76 | 56 | 20 | 36 | Gbp8 |
| Gbp9 | 100 | 86 | 126 | 98 | 159 | 119 | 122 | 199 | 274 | 318 | 450 | 211 | 246 | 220 | 117 | 125 | 79 | 140 | 72 | 189 | 113 | 349 | 472 | 259 | 306 | 226 | 318 | 233 | Gbp9 |
| Gbx1 | 65 | 88 | 74 | 156 | 557 | 225 | 232 | 231 | 434 | 644 | 834 | 428 | 159 | 471 | 136 | 104 | 192 | 232 | 208 | 133 | 207 | 456 | 610 | 369 | 443 | 498 | 213 | 305 | Gbx1 |
| Gbx2 | 463 | 524 | 666 | 773 | 970 | 677 | 858 | 895 | 499 | 1204 | 831 | 1023 | 350 | 1045 | 531 | 331 | 736 | 754 | 891 | 527 | 782 | 1365 | 727 | 1167 | 1016 | 1129 | 917 | 968 | Gbx2 |
| Gc | 0 | 4 | 0 | 0 | 3 | 4 | 1 | 1 | 10 | 0 | 2 | 2 | 7 | 2 | 7 | 4 | 3 | 0 | 4 | 8 | 4 | 2 | 5 | 1 | 0 | 0 | 1 | 1 | Gc |
| Gca | 2080 | 1791 | 1804 | 2555 | 2355 | 2383 | 3414 | 661 | 651 | 574 | 539 | 655 | 384 | 636 | 2168 | 1783 | 2094 | 2821 | 2514 | 2565 | 3056 | 550 | 545 | 775 | 700 | 668 | 503 | 580 | Gca |
| Gcat | 306 | 275 | 189 | 228 | 171 | 159 | 238 | 93 | 74 | 139 | 76 | 89 | 84 | 158 | 210 | 237 | 225 | 269 | 168 | 210 | 233 | 47 | 71 | 125 | 93 | 114 | 91 | 147 | Gcat |
| Gcat | 0 | 0 | 0 | 0 | 0 | 0 | 0 | 0 | 0 | 0 | 0 | 0 | 0 | 0 | 0 | 1 | 0 | 1 | 2 | 0 | 0 | 0 | 0 | 0 | 0 | 0 | 0 | 0 | Gcat |
| Gcc1 | 2346 | 2776 | 2494 | 2837 | 2315 | 2451 | 2358 | 2140 | 2089 | 2080 | 2011 | 2014 | 2200 | 2336 | 2368 | 2389 | 2502 | 2750 | 2387 | 2470 | 2388 | 2112 | 2008 | 2148 | 2222 | 1905 | 2117 | 2047 | Gcc1 |
| Gcc2 | 0 | 0 | 1 | 1 | 1 | 0 | 2 | 1 | 1 | 1 | 0 | 0 | 0 | 0 | 1 | 6 | 0 | 0 | 0 | 0 | 2 | 0 | 1 | 2 | 3 | 0 | 2 | 0 | Gcc2 |
| Gcdh | 4828 | 5139 | 4878 | 4728 | 3929 | 3925 | 4494 | 2258 | 2324 | 2035 | 2236 | 2260 | 2084 | 2152 | 4476 | 4762 | 4614 | 4551 | 4376 | 4433 | 4156 | 2115 | 2267 | 2329 | 2130 | 2522 | 2367 | 2363 | Gcdh |
| Gcfc2 | 2420 | 1772 | 1883 | 1787 | 1779 | 1986 | 2171 | 2318 | 2213 | 1849 | 2040 | 2303 | 2208 | 2316 | 2331 | 2041 | 1744 | 1634 | 1693 | 2146 | 1900 | 1646 | 2060 | 1964 | 1801 | 2132 | 2312 | 2229 | Gcfc2 |
| Gcg | 10 | 9 | 1 | 12 | 14 | 14 | 30 | 24 | 23 | 27 | 18 | 17 | 6 | 20 | 4 | 1 | 6 | 17 | 6 | 24 | 20 | 12 | 25 | 32 | 33 | 33 | 6 | 24 | Gcg |
| Gcgr | 21 | 36 | 38 | 37 | 19 | 14 | 9 | 13 | 23 | 26 | 45 | 8 | 20 | 4 | 21 | 39 | 30 | 19 | 30 | 13 | 6 | 8 | 33 | 16 | 11 | 22 | 16 | 14 | Gcgr |
| Gch1 | 809 | 876 | 880 | 945 | 637 | 664 | 572 | 112 | 128 | 158 | 159 | 136 | 147 | 121 | 711 | 954 | 980 | 795 | 677 | 686 | 685 | 172 | 181 | 102 | 158 | 113 | 179 | 145 | Gch1 |
| Gchfr | 100 | 135 | 131 | 144 | 74 | 70 | 58 | 40 | 60 | 60 | 41 | 35 | 52 | 28 | 65 | 124 | 88 | 95 | 72 | 117 | 59 | 45 | 77 | 41 | 27 | 35 | 38 | 20 | Gchfr |
| Gck | 114 | 151 | 82 | 143 | 164 | 153 | 143 | 194 | 227 | 230 | 246 | 219 | 116 | 196 | 119 | 112 | 185 | 151 | 112 | 121 | 147 | 264 | 261 | 285 | 266 | 269 | 173 | 183 | Gck |
| Gckr | 21 | 16 | 13 | 15 | 20 | 28 | 16 | 29 | 21 | 22 | 21 | 15 | 14 | 13 | 12 | 8 | 27 | 29 | 14 | 16 | 15 | 28 | 26 | 24 | 32 | 14 | 17 | 21 | Gckr |
| Gclc | 7587 | 7221 | 5713 | 7005 | 6001 | 6945 | 7718 | 4168 | 3678 | 3297 | 3489 | 3645 | 3383 | 3888 | 7544 | 6673 | 6163 | 6688 | 6498 | 7084 | 7326 | 3537 | 3541 | 3438 | 3431 | 3641 | 3510 | 3807 | Gclc |
| Gclm | 2226 | 2083 | 2475 | 2915 | 3094 | 2870 | 3572 | 2199 | 2309 | 2324 | 2257 | 2625 | 3068 | 2826 | 2354 | 2151 | 2458 | 2804 | 2910 | 2520 | 3132 | 2516 | 2379 | 2461 | 2515 | 2625 | 2398 | 3218 | Gclm |
| Gcm1 | 2 | 2 | 3 | 0 | 0 | 1 | 0 | 6 | 3 | 0 | 2 | 9 | 10 | 2 | 1 | 14 | 6 | 6 | 0 | 0 | 0 | 2 | 0 | 2 | 1 | 8 | 7 | 8 | Gcm1 |
| Gcm2 | 0 | 0 | 0 | 1 | 0 | 0 | 0 | 0 | 1 | 0 | 4 | 1 | 2 | 0 | 0 | 0 | 1 | 1 | 0 | 0 | 0 | 0 | 15 | 2 | 0 | 4 | 0 | 0 | Gcm2 |
| Gcn1 | 9709 | 12103 | 12629 | 12884 | 11669 | 11158 | 10288 | 9122 | 11800 | 12840 | 11878 | 14513 | 12254 | 11587 | 9082 | 11080 | 13023 | 12459 | 13111 | 9763 | 10911 | 11416 | 10778 | 13097 | 12492 | 8878 | 12293 | 12324 | Gcn1 |
| Gcnt1 | 724 | 631 | 584 | 349 | 228 | 340 | 225 | 319 | 529 | 299 | 550 | 299 | 687 | 348 | 687 | 951 | 370 | 350 | 170 | 272 | 173 | 341 | 470 | 359 | 285 | 463 | 396 | 553 | Gcnt1 |
| Gcnt2 | 689 | 654 | 636 | 675 | 815 | 694 | 656 | 701 | 700 | 763 | 686 | 562 | 642 | 603 | 804 | 671 | 549 | 623 | 683 | 745 | 637 | 749 | 764 | 730 | 648 | 772 | 609 | 534 | Gcnt2 |
| Gcnt3 | 2 | 0 | 0 | 0 | 2 | 0 | 0 | 2 | 2 | 0 | 2 | 1 | 0 | 1 | 0 | 1 | 3 | 1 | 0 | 0 | 0 | 0 | 1 | 1 | 0 | 4 | 3 | 0 | Gcnt3 |
| Gcnt4 | 166 | 133 | 148 | 104 | 90 | 146 | 63 | 55 | 73 | 65 | 79 | 41 | 53 | 49 | 182 | 185 | 140 | 121 | 107 | 92 | 82 | 69 | 47 | 49 | 33 | 29 | 60 | 45 | Gcnt4 |
| Gcnt7 | 0 | 0 | 0 | 0 | 0 | 0 | 0 | 2 | 2 | 2 | 2 | 0 | 0 | 0 | 0 | 0 | 0 | 0 | 0 | 0 | 0 | 0 | 5 | 4 | 6 | 0 | 0 | 2 | Gcnt7 |
| Gcsam | 6 | 9 | 2 | 2 | 2 | 2 | 3 | 7 | 8 | 9 | 8 | 3 | 5 | 3 | 4 | 9 | 2 | 4 | 5 | 1 | 5 | 8 | 4 | 6 | 2 | 7 | 7 | 12 | Gcsam |
| Gcsh | 6474 | 5340 | 5832 | 5124 | 4113 | 4419 | 5087 | 3532 | 3049 | 3125 | 3136 | 3811 | 3787 | 3996 | 5972 | 5442 | 5195 | 4620 | 4338 | 4998 | 4595 | 2687 | 2986 | 3392 | 3260 | 4496 | 3937 | 4180 | Gcsh |
| Gda | 85 | 60 | 17 | 103 | 188 | 106 | 163 | 73 | 106 | 45 | 120 | 30 | 19 | 32 | 96 | 108 | 53 | 136 | 141 | 126 | 200 | 104 | 166 | 69 | 101 | 45 | 28 | 13 | Gda |
| Gdap1 | 2581 | 2319 | 2182 | 3635 | 5681 | 4601 | 5341 | 7032 | 5877 | 6270 | 5174 | 5005 | 3800 | 5377 | 3407 | 2138 | 3170 | 4003 | 4770 | 5108 | 5285 | 7693 | 5348 | 5328 | 7155 | 3595 | 4481 | 4589 | Gdap1 |
| Gdap1l1 | 1557 | 2222 | 1811 | 2805 | 3941 | 3018 | 3140 | 3199 | 3361 | 4056 | 3289 | 3381 | 3144 | 3296 | 2066 | 1670 | 2644 | 3001 | 3628 | 3102 | 3512 | 4361 | 2947 | 3575 | 4390 | 2888 | 2933 | 3024 | Gdap1l1 |

| | | | | | | | | | | | | | | | | | | | | | | | | | | | | | |
|---|---|---|---|---|---|---|---|---|---|---|---|---|---|---|---|---|---|---|---|---|---|---|---|---|---|---|---|---|---|
| Gdap2 | 3391 | 3316 | 2704 | 2825 | 2707 | 2767 | 3226 | 2594 | 2144 | 1775 | 1754 | 2016 | 1853 | 2092 | 3564 | 3290 | 2735 | 2879 | 2763 | 3011 | 2872 | 2256 | 2117 | 1970 | 1971 | 1872 | 2073 | 2046 | Gdap2 |
| Gde1 | 5050 | 5376 | 4093 | 4627 | 4672 | 4508 | 4844 | 4024 | 3753 | 3409 | 3160 | 3214 | 3097 | 2944 | 5346 | 4469 | 4434 | 4577 | 4967 | 5172 | 4975 | 3868 | 3761 | 3236 | 3609 | 3258 | 3084 | 3091 | Gde1 |
| Gdf10 | 531 | 677 | 485 | 377 | 406 | 530 | 160 | 1036 | 2052 | 1578 | 2557 | 517 | 1094 | 485 | 688 | 742 | 413 | 628 | 293 | 295 | 211 | 1957 | 2197 | 1024 | 1024 | 753 | 568 | 430 | Gdf10 |
| Gdf11 | 7059 | 6974 | 9311 | 8928 | 7450 | 6999 | 7483 | 3889 | 3015 | 4391 | 4019 | 4170 | 4308 | 5044 | 6754 | 6687 | 8619 | 7876 | 7395 | 6203 | 8055 | 4124 | 3181 | 4750 | 4492 | 4583 | 4771 | 5510 | Gdf11 |
| Gdf15 | 3 | 2 | 2 | 6 | 0 | 2 | 0 | 2 | 3 | 4 | 3 | 1 | 2 | 0 | 0 | 2 | 1 | 1 | 1 | 0 | 0 | 0 | 6 | 3 | 6 | 3 | 2 | 0 | Gdf15 |
| Gdf2 | 2 | 2 | 0 | 0 | 4 | 0 | 1 | 0 | 2 | 3 | 6 | 0 | 2 | 1 | 3 | 2 | 6 | 3 | 0 | 2 | 2 | 3 | 1 | 0 | 4 | 0 | 0 | 0 | Gdf2 |
| Gdf3 | 1 | 3 | 2 | 0 | 2 | 3 | 0 | 4 | 1 | 3 | 5 | 2 | 0 | 4 | 4 | 7 | 3 | 0 | 2 | 0 | 1 | 0 | 3 | 1 | 0 | 1 | 7 | 4 | Gdf3 |
| Gdf5 | 61 | 45 | 57 | 44 | 74 | 47 | 41 | 143 | 192 | 139 | 240 | 177 | 150 | 98 | 57 | 37 | 37 | 57 | 81 | 59 | 50 | 132 | 252 | 113 | 102 | 170 | 126 | 106 | Gdf5 |
| Gdf6 | 181 | 124 | 188 | 84 | 44 | 54 | 35 | 151 | 134 | 139 | 246 | 54 | 81 | 53 | 116 | 188 | 126 | 84 | 62 | 30 | 44 | 65 | 226 | 95 | 44 | 150 | 27 | 56 | Gdf6 |
| Gdf7 | 57 | 67 | 48 | 20 | 38 | 9 | 9 | 116 | 97 | 96 | 50 | 89 | 108 | 52 | 55 | 69 | 79 | 24 | 12 | 20 | 9 | 62 | 116 | 83 | 33 | 10 | 81 | 69 | Gdf7 |
| Gdf9 | 50 | 53 | 48 | 46 | 56 | 34 | 40 | 96 | 72 | 120 | 81 | 87 | 83 | 94 | 37 | 43 | 54 | 46 | 66 | 40 | 34 | 129 | 101 | 79 | 99 | 58 | 92 | 113 | Gdf9 |
| Gdi1 | 29098 | 32864 | 31039 | 38481 | 40207 | 38402 | 40171 | 33987 | 31739 | 38931 | 32249 | 38814 | 33504 | 37789 | 31909 | 27649 | 36116 | 40091 | 41673 | 36811 | 41669 | 43565 | 31701 | 39392 | 42618 | 34869 | 36458 | 35858 | Gdi1 |
| Gdi2 | 45231 | 36449 | 39714 | 37677 | 36348 | 38696 | 42914 | 36278 | 32019 | 30672 | 30029 | 34269 | 32534 | 35157 | 45271 | 37668 | 38829 | 37640 | 39123 | 36691 | 42268 | 33713 | 32351 | 31726 | 32783 | 37142 | 35335 | 37205 | Gdi2 |
| Gdnf | 103 | 71 | 128 | 67 | 66 | 86 | 26 | 70 | 82 | 96 | 104 | 66 | 121 | 54 | 86 | 152 | 81 | 66 | 51 | 36 | 33 | 84 | 96 | 91 | 46 | 128 | 30 | 58 | Gdnf |
| Gdpd1 | 2731 | 2690 | 2788 | 3856 | 4772 | 4274 | 5023 | 7022 | 5611 | 6029 | 4899 | 5965 | 5530 | 6659 | 3748 | 2590 | 3356 | 3859 | 4623 | 4776 | 4859 | 7197 | 5487 | 6192 | 6923 | 5859 | 5865 | 6313 | Gdpd1 |
| Gdpd2 | 527 | 681 | 812 | 707 | 1220 | 892 | 951 | 1336 | 1218 | 1272 | 1018 | 939 | 1254 | 1036 | 749 | 615 | 805 | 793 | 888 | 941 | 995 | 1681 | 1361 | 1053 | 1317 | 856 | 1035 | 986 | Gdpd2 |
| Gdpd3 | 210 | 243 | 256 | 253 | 330 | 265 | 283 | 221 | 180 | 216 | 258 | 179 | 213 | 195 | 138 | 247 | 286 | 303 | 291 | 269 | 290 | 312 | 207 | 246 | 280 | 186 | 242 | 169 | Gdpd3 |
| Gdpd4 | 8 | 10 | 4 | 5 | 13 | 5 | 1 | 9 | 7 | 2 | 10 | 3 | 6 | 4 | 17 | 18 | 3 | 2 | 4 | 1 | 1 | 3 | 2 | 0 | 9 | 0 | 3 | 0 | Gdpd4 |
| Gdpd5 | 2765 | 3981 | 4384 | 4893 | 6005 | 4399 | 4290 | 2506 | 2898 | 3400 | 3209 | 3089 | 3019 | 2836 | 3162 | 3434 | 5014 | 5085 | 5446 | 3863 | 4808 | 3922 | 2673 | 3192 | 3679 | 2844 | 2893 | 2658 | Gdpd5 |
| Gdpgp1 | 1078 | 1271 | 1022 | 1072 | 893 | 964 | 1026 | 412 | 416 | 447 | 379 | 522 | 494 | 467 | 1000 | 1243 | 980 | 1205 | 956 | 956 | 1062 | 511 | 458 | 430 | 420 | 432 | 466 | 560 | Gdpgp1 |
| Gem | 390 | 369 | 397 | 359 | 307 | 354 | 334 | 153 | 219 | 144 | 241 | 173 | 150 | 186 | 375 | 353 | 374 | 384 | 369 | 368 | 320 | 130 | 209 | 212 | 182 | 242 | 185 | 166 | Gem |
| Gemin2 | 1375 | 1349 | 1607 | 1466 | 1088 | 1155 | 1245 | 1105 | 1027 | 1065 | 1222 | 1312 | 1267 | 1312 | 1435 | 1361 | 1563 | 1372 | 1227 | 1220 | 1198 | 983 | 1060 | 1137 | 1079 | 1557 | 1328 | 1462 | Gemin2 |
| Gemin4 | 18 | 14 | 9 | 9 | 11 | 9 | 12 | 7 | 21 | 6 | 21 | 9 | 29 | 21 | 11 | 14 | 21 | 11 | 16 | 12 | 12 | 8 | 11 | 14 | 7 | 8 | 16 | 21 | Gemin4 |
| Gemin5 | 9237 | 10039 | 7571 | 8102 | 6746 | 7935 | 7277 | 4146 | 4486 | 4090 | 4409 | 4766 | 4842 | 4571 | 8632 | 9326 | 7628 | 7707 | 7303 | 7280 | 7983 | 3728 | 4401 | 4114 | 4105 | 4277 | 5012 | 4930 | Gemin5 |
| Gemin6 | 693 | 582 | 808 | 541 | 536 | 623 | 617 | 954 | 812 | 736 | 738 | 845 | 925 | 1030 | 708 | 632 | 709 | 559 | 708 | 733 | 607 | 581 | 741 | 949 | 797 | 1200 | 1061 | 1142 | Gemin6 |
| Gemin7 | 1457 | 1750 | 1602 | 1570 | 1381 | 1282 | 1409 | 1197 | 1257 | 1204 | 1195 | 1378 | 1436 | 1418 | 1334 | 1379 | 1576 | 1466 | 1548 | 1348 | 1480 | 1105 | 1311 | 1405 | 1217 | 1577 | 1421 | 1454 | Gemin7 |
| Gemin8 | 1051 | 1133 | 1234 | 1114 | 930 | 910 | 1093 | 782 | 873 | 888 | 858 | 939 | 879 | 943 | 1000 | 1090 | 1082 | 1026 | 969 | 966 | 1001 | 673 | 832 | 810 | 832 | 1146 | 916 | 1027 | Gemin8 |
| Gen1 | 3173 | 2477 | 3311 | 2343 | 1478 | 2058 | 2368 | 2566 | 2108 | 1889 | 1949 | 3039 | 2834 | 3146 | 2826 | 2752 | 2742 | 2037 | 2046 | 2306 | 2111 | 1648 | 1926 | 2453 | 2338 | 3206 | 3393 | 3256 | Gen1 |
| Get1 | 4610 | 5099 | 5035 | 4729 | 4921 | 4563 | 4748 | 4727 | 4331 | 4222 | 3900 | 4373 | 4012 | 4289 | 4890 | 4558 | 4952 | 4754 | 5030 | 5174 | 4745 | 5108 | 4266 | 4418 | 4918 | 4333 | 4499 | 4441 | Get1 |
| Get3 | 5795 | 7288 | 6359 | 6859 | 5801 | 6261 | 5772 | 4903 | 5398 | 5464 | 4951 | 5536 | 5190 | 5217 | 6260 | 6222 | 6824 | 6113 | 6967 | 7350 | 6115 | 4284 | 5385 | 5279 | 5266 | 6280 | 5750 | 5543 | Get3 |
| Get4 | 1899 | 2325 | 2106 | 1828 | 1757 | 1764 | 1769 | 2216 | 2477 | 2315 | 2358 | 2528 | 2505 | 2428 | 1836 | 2026 | 1997 | 1852 | 2009 | 1864 | 1649 | 1917 | 2471 | 2397 | 2259 | 3150 | 2532 | 2533 | Get4 |
| Gfap | 7 | 10 | 14 | 4 | 7 | 3 | 8 | 7 | 13 | 8 | 21 | 8 | 24 | 11 | 9 | 5 | 13 | 7 | 9 | 6 | 7 | 7 | 15 | 4 | 8 | 10 | 20 | 9 | Gfap |
| Gfer | 1095 | 1161 | 1154 | 1119 | 1048 | 1071 | 1101 | 1504 | 1434 | 1472 | 1493 | 1566 | 1623 | 1771 | 1138 | 1088 | 1105 | 1052 | 1181 | 1101 | 1075 | 1317 | 1365 | 1665 | 1508 | 1940 | 1564 | 1829 | Gfer |
| Gfi1 | 18 | 17 | 7 | 26 | 11 | 16 | 8 | 7 | 5 | 4 | 16 | 7 | 12 | 10 | 11 | 15 | 8 | 25 | 16 | 7 | 15 | 7 | 9 | 8 | 3 | 8 | 7 | 6 | Gfi1 |
| Gfi1b | 17 | 29 | 51 | 9 | 11 | 4 | 3 | 15 | 17 | 27 | 15 | 17 | 61 | 28 | 13 | 30 | 22 | 4 | 8 | 14 | 10 | 8 | 12 | 12 | 21 | 49 | 44 | 68 | Gfi1b |
| Gfm1 | 3853 | 3730 | 3726 | 3375 | 3014 | 3317 | 3298 | 3149 | 3038 | 2866 | 3002 | 3460 | 3148 | 3450 | 3691 | 3610 | 3564 | 3125 | 3098 | 3282 | 3268 | 2954 | 3123 | 3047 | 3031 | 3539 | 3355 | 3366 | Gfm1 |
| Gfm2 | 3299 | 3344 | 3031 | 3040 | 2813 | 3175 | 3452 | 3701 | 3629 | 3532 | 3629 | 4179 | 4021 | 4247 | 3518 | 3167 | 3284 | 2925 | 3038 | 3330 | 3187 | 3536 | 3555 | 3657 | 3953 | 3923 | 4183 | 4146 | Gfm2 |
| Gfod1 | 1263 | 1367 | 1130 | 1861 | 2373 | 1922 | 1871 | 1134 | 1228 | 1409 | 1364 | 901 | 976 | 994 | 1361 | 1227 | 1577 | 2021 | 2083 | 1623 | 2138 | 1609 | 1199 | 1213 | 1520 | 653 | 655 | 984 | Gfod1 |
| Gfod2 | 3415 | 4111 | 3956 | 4617 | 5054 | 4275 | 4492 | 3425 | 3414 | 3809 | 3509 | 3850 | 3368 | 3663 | 3475 | 3420 | 4413 | 4738 | 5181 | 4197 | 4641 | 4311 | 3207 | 3902 | 4504 | 3652 | 3449 | 3661 | Gfod2 |
| Gfpt1 | 12380 | 10256 | 10495 | 13133 | 13679 | 13653 | 12905 | 10395 | 10852 | 11552 | 12707 | 9883 | 9909 | 9921 | 12538 | 12134 | 11208 | 13772 | 12602 | 12376 | 13412 | 12176 | 11042 | 10290 | 12095 | 7530 | 9069 | 8715 | Gfpt1 |
| Gfpt2 | 1774 | 1646 | 1581 | 1339 | 1275 | 1478 | 1373 | 699 | 1105 | 922 | 1185 | 722 | 938 | 691 | 1731 | 2003 | 1216 | 1626 | 1275 | 1220 | 1320 | 911 | 905 | 862 | 799 | 904 | 667 | 793 | Gfpt2 |
| Gfra1 | 2587 | 2998 | 2411 | 2573 | 4188 | 3179 | 2472 | 4851 | 4533 | 4526 | 5192 | 3643 | 5663 | 3253 | 3523 | 3344 | 2803 | 2941 | 3074 | 3051 | 2856 | 5812 | 4840 | 3002 | 4192 | 2266 | 3839 | 3792 | Gfra1 |
| Gfra2 | 388 | 408 | 626 | 526 | 928 | 688 | 610 | 674 | 925 | 967 | 1193 | 553 | 598 | 672 | 377 | 467 | 569 | 626 | 792 | 603 | 664 | 1178 | 1017 | 753 | 924 | 574 | 575 | 521 | Gfra2 |
| Gfra3 | 84 | 81 | 24 | 10 | 126 | 16 | 9 | 197 | 283 | 106 | 265 | 22 | 175 | 10 | 115 | 171 | 63 | 97 | 19 | 7 | 10 | 286 | 203 | 34 | 154 | 34 | 20 | 15 | Gfra3 |
| Gfra4 | 120 | 99 | 69 | 116 | 149 | 119 | 116 | 286 | 372 | 306 | 359 | 168 | 159 | 147 | 104 | 106 | 85 | 125 | 121 | 146 | 139 | 341 | 353 | 276 | 294 | 195 | 131 | 126 | Gfra4 |
| Gfral | 1 | 1 | 0 | 0 | 0 | 2 | 0 | 0 | 0 | 0 | 4 | 0 | 0 | 3 | 0 | 2 | 0 | 0 | 1 | 0 | 0 | 0 | 1 | 0 | 0 | 0 | 1 | 0 | Gfral |
| Gfus | 2723 | 3365 | 2897 | 3070 | 2575 | 2716 | 2626 | 2264 | 2652 | 2439 | 2195 | 2442 | 2470 | 2479 | 2616 | 3055 | 2782 | 2883 | 2922 | 2909 | 2608 | 2042 | 2491 | 2603 | 2616 | 2768 | 2615 | 2537 | Gfus |
| Gfy | 35 | 30 | 0 | 64 | 3 | 4 | 4 | 95 | 97 | 257 | 106 | 18 | 36 | 31 | 24 | 14 | 12 | 10 | 6 | 4 | 8 | 2 | 98 | 229 | 4 | 297 | 8 | 11 | Gfy |
| Gga1 | 2134 | 2581 | 3086 | 3254 | 3300 | 2839 | 2666 | 3091 | 3863 | 4608 | 4157 | 4929 | 4039 | 4228 | 1970 | 2257 | 3327 | 3111 | 3583 | 2433 | 2876 | 4101 | 3708 | 4688 | 4619 | 5370 | 4040 | 4001 | Gga1 |
| Gga2 | 10622 | 9980 | 10246 | 9575 | 8703 | 8816 | 8362 | 6517 | 7018 | 7111 | 8080 | 6985 | 7158 | 6921 | 8770 | 10664 | 9360 | 10193 | 9030 | 7623 | 8622 | 7050 | 7317 | 7322 | 7338 | 7765 | 7511 | 7083 | Gga2 |
| Gga3 | 4396 | 5616 | 5750 | 6379 | 6159 | 5467 | 5490 | 4396 | 4565 | 5492 | 5013 | 6012 | 5212 | 5227 | 4367 | 4767 | 6344 | 6104 | 6302 | 4691 | 5845 | 6176 | 4332 | 5426 | 5806 | 5374 | 5050 | 5406 | Gga3 |
| Ggact | 191 | 168 | 191 | 194 | 158 | 150 | 180 | 199 | 251 | 257 | 232 | 179 | 225 | 239 | 182 | 194 | 209 | 205 | 138 | 188 | 172 | 185 | 302 | 197 | 183 | 297 | 251 | 203 | Ggact |
| Ggct | 751 | 835 | 758 | 808 | 841 | 837 | 886 | 485 | 390 | 439 | 529 | 404 | 448 | 431 | 869 | 767 | 875 | 900 | 816 | 861 | 911 | 403 | 443 | 488 | 393 | 462 | 421 | 436 | Ggct |
| Ggcx | 231 | 168 | 75 | 100 | 280 | 251 | 179 | 110 | 56 | 70 | 89 | 62 | 54 | 44 | 157 | 87 | 78 | 138 | 56 | 71 | 136 | 339 | 84 | 193 | 233 | 94 | 196 | 74 | Ggcx |
| Ggh | 1216 | 951 | 1285 | 973 | 906 | 955 | 1086 | 1382 | 1239 | 1128 | 1111 | 1158 | 1202 | 1204 | 1386 | 1256 | 1159 | 921 | 988 | 1166 | 915 | 1168 | 1362 | 1156 | 1163 | 1249 | 1249 | 1358 | Ggh |
| Ggn | 167 | 215 | 231 | 201 | 190 | 185 | 182 | 74 | 80 | 115 | 100 | 84 | 99 | 66 | 191 | 206 | 249 | 209 | 225 | 153 | 172 | 92 | 95 | 102 | 102 | 116 | 86 | 95 | Ggn |
| Ggnbp1 | 127 | 143 | 159 | 208 | 164 | 154 | 173 | 131 | 132 | 152 | 138 | 156 | 128 | 107 | 129 | 140 | 144 | 169 | 171 | 191 | 165 | 159 | 135 | 151 | 170 | 87 | 102 | 128 | Ggnbp1 |
| Ggnbp2 | 7774 | 6683 | 7523 | 7907 | 7579 | 7320 | 8377 | 8950 | 7474 | 8176 | 7869 | 8238 | 8149 | 8418 | 8408 | 7628 | 7631 | 7989 | 7475 | 7669 | 7647 | 9276 | 7649 | 8118 | 8277 | 8168 | 8276 | 8186 | Ggnbp2 |
| Ggps1 | 5556 | 4840 | 5078 | 5163 | 4774 | 5085 | 5436 | 6977 | 5612 | 5397 | 5037 | 5654 | 5551 | 6116 | 6025 | 5302 | 5273 | 5170 | 4737 | 5753 | 5409 | 5994 | 5644 | 5416 | 5471 | 5554 | 6055 | 6035 | Ggps1 |
| Ggt1 | 36 | 47 | 84 | 37 | 74 | 53 | 50 | 56 | 53 | 63 | 62 | 89 | 96 | 80 | 50 | 70 | 59 | 62 | 60 | 44 | 63 | 99 | 101 | 92 | 68 | 52 | 80 | 70 | Ggt1 |
| Ggt5 | 62 | 82 | 108 | 107 | 128 | 163 | 123 | 156 | 162 | 180 | 235 | 169 | 205 | 156 | 78 | 68 | 80 | 125 | 130 | 94 | 122 | 221 | 207 | 225 | 246 | 178 | 186 | 174 | Ggt5 |
| Ggt6 | 1 | 1 | 0 | 0 | 1 | 0 | 0 | 2 | 9 | 7 | 15 | 0 | 0 | 0 | 0 | 0 | 0 | 0 | 0 | 0 | 0 | 0 | 12 | 1 | 0 | 0 | 0 | 5 | Ggt6 |
| Ggt7 | 1701 | 2423 | 2274 | 2903 | 3498 | 2919 | 2648 | 2385 | 2555 | 3120 | 2609 | 2775 | 2767 | 2554 | 1851 | 1959 | 2969 | 2762 | 3525 | 2821 | 2863 | 3324 | 2320 | 2798 | 3213 | 2310 | 2851 | 2594 | Ggt7 |
| Ggta1 | 866 | 720 | 842 | 615 | 821 | 716 | 571 | 866 | 1083 | 976 | 1415 | 770 | 978 | 790 | 767 | 869 | 637 | 648 | 598 | 715 | 584 | 871 | 1498 | 894 | 733 | 905 | 809 | 781 | Ggta1 |
| Gh | 0 | 0 | 0 | 0 | 0 | 0 | 0 | 0 | 0 | 0 | 0 | 0 | 0 | 0 | 0 | 0 | 0 | 0 | 0 | 0 | 0 | 0 | 1 | 0 | 0 | 0 | 0 | 0 | Gh |
| Ghdc | 540 | 696 | 766 | 635 | 499 | 535 | 305 | 316 | 447 | 515 | 498 | 355 | 398 | 304 | 555 | 802 | 608 | 611 | 398 | 409 | 356 | 418 | 482 | 355 | 353 | 462 | 363 | 361 | Ghdc |
| Ghitm | 13544 | 12021 | 13591 | 11976 | 10407 | 11033 | 11245 | 11260 | 9945 | 8924 | 9407 | 10012 | 10915 | 10179 | 13949 | 12715 | 12406 | 12746 | 11260 | 13001 | 10897 | 9185 | 10164 | 9666 | 9434 | 10681 | 10477 | 10535 | Ghitm |
| Ghr | 1908 | 1327 | 1582 | 1068 | 990 | 1321 | 662 | 2307 | 2745 | 2010 | 3466 | 1597 | 2518 | 1709 | 1764 | 2213 | 1139 | 1258 | 659 | 884 | 673 | 1888 | 3134 | 1856 | 1563 | 1789 | 1705 | 1778 | Ghr |
| Ghrh | 7 | 8 | 3 | 24 | 8 | 12 | 17 | 13 | 18 | 23 | 17 | 17 | 17 | 19 | 9 | 7 | 20 | 12 | 17 | 26 | 11 | 22 | 38 | 20 | 29 | 18 | 30 | 39 | Ghrh |
| Ghrhr | 0 | 0 | 0 | 0 | 0 | 0 | 0 | 1 | 1 | 0 | 0 | 0 | 0 | 1 | 0 | 0 | 1 | 0 | 0 | 0 | 0 | 0 | 3 | 0 | 3 | 0 | 0 | 0 | Ghrhr |
| Ghrl | 63 | 79 | 41 | 58 | 48 | 56 | 51 | 23 | 20 | 20 | 22 | 21 | 12 | 5 | 53 | 48 | 49 | 56 | 40 | 67 | 58 | 20 | 17 | 19 | 18 | 24 | 14 | 19 | Ghrl |
| Ghsr | 12 | 18 | 22 | 13 | 34 | 13 | 27 | 17 | 17 | 1 | 21 | 5 | 43 | 17 | 20 | 12 | 23 | 12 | 12 | 30 | 21 | 22 | 19 | 7 | 14 | 0 | 8 | 11 | Ghsr |
| Gid4 | 5954 | 6173 | 7690 | 7330 | 5582 | 5590 | 6816 | 3427 | 3489 | 4021 | 3733 | 4323 | 3887 | 4305 | 5794 | 5998 | 7502 | 7234 | 6539 | 5873 | 6411 | 3554 | 3579 | 4178 | 4023 | 4648 | 4050 | 4475 | Gid4 |
| Gid8 | 4398 | 4260 | 3833 | 3965 | 3647 | 3942 | 3928 | 4264 | 4158 | 4092 | 4110 | 4127 | 4287 | 4404 | 4330 | 3881 | 4058 | 3700 | 4062 | 3777 | 3913 | 4281 | 4106 | 4164 | 4172 | 4214 | 4140 | 4247 | Gid8 |
| Gigyf1 | 5989 | 7410 | 9519 | 10097 | 10073 | 8671 | 9302 | 6537 | 7335 | 9864 | 8883 | 10471 | 8981 | 9783 | 5600 | 6379 | 10159 | 9610 | 10757 | 7411 | 9400 | 9100 | 6746 | 10492 | 10051 | 10388 | 9308 | 9515 | Gigyf1 |
| Gigyf2 | 9609 | 9351 | 9045 | 9685 | 9453 | 9420 | 9747 | 10617 | 10115 | 9669 | 9815 | 10163 | 9941 | 10633 | 9392 | 9159 | 8955 | 9338 | 9360 | 9861 | 9534 | 9513 | 9901 | 10310 | 10345 | 8514 | 9999 | 10252 | Gigyf2 |
| Gimap1 | 60 | 74 | 117 | 74 | 66 | 71 | 59 | 75 | 82 | 85 | 116 | 72 | 120 | 101 | 62 | 65 | 63 | 74 | 92 | 79 | 56 | 72 | 102 | 103 | 95 | 110 | 101 | 81 | Gimap1 |
| Gimap3 | 0 | 0 | 0 | 0 | 0 | 1 | 1 | 0 | 0 | 0 | 1 | 0 | 0 | 0 | 0 | 1 | 0 | 1 | 0 | 0 | 0 | 0 | 1 | 0 | 0 | 0 | 0 | 0 | Gimap3 |
| Gimap4 | 172 | 135 | 189 | 96 | 116 | 154 | 104 | 300 | 305 | 271 | 340 | 220 | 266 | 243 | 158 | 167 | 137 | 140 | 122 | 155 | 132 | 261 | 390 | 300 | 221 | 323 | 270 | 252 | Gimap4 |

| | | | | | | | | | | | | | | | | | | | | | | | | | | | | | |
|---|---|---|---|---|---|---|---|---|---|---|---|---|---|---|---|---|---|---|---|---|---|---|---|---|---|---|---|---|---|
| Gimap5 | 51 | 42 | 53 | 41 | 29 | 45 | 32 | 29 | 42 | 56 | 87 | 51 | 43 | 44 | 40 | 35 | 36 | 40 | 37 | 38 | 36 | 28 | 56 | 42 | 34 | 37 | 36 | 28 | Gimap5 |
| Gimap6 | 430 | 286 | 380 | 226 | 319 | 300 | 275 | 487 | 552 | 501 | 720 | 440 | 511 | 414 | 421 | 357 | 305 | 314 | 330 | 408 | 290 | 460 | 731 | 463 | 369 | 447 | 435 | 377 | Gimap6 |
| Gimap7 | 0 | 0 | 0 | 0 | 0 | 0 | 0 | 0 | 0 | 0 | 0 | 0 | 0 | 0 | 0 | 0 | 0 | 0 | 0 | 0 | 0 | 0 | 1 | 0 | 0 | 2 | 0 | 0 | Gimap7 |
| Gimap8 | 83 | 48 | 100 | 74 | 66 | 89 | 59 | 204 | 168 | 189 | 213 | 166 | 214 | 191 | 74 | 85 | 65 | 59 | 87 | 64 | 79 | 197 | 230 | 151 | 157 | 184 | 237 | 155 | Gimap8 |
| Gimap9 | 176 | 122 | 171 | 72 | 88 | 116 | 97 | 184 | 225 | 114 | 240 | 98 | 202 | 137 | 157 | 169 | 82 | 101 | 92 | 107 | 93 | 125 | 256 | 163 | 114 | 219 | 162 | 183 | Gimap9 |
| Gimd1 | 0 | 0 | 0 | 0 | 0 | 0 | 0 | 0 | 0 | 0 | 0 | 0 | 0 | 0 | 0 | 0 | 0 | 0 | 0 | 0 | 0 | 0 | 0 | 0 | 0 | 2 | 0 | 0 | Gimd1 |
| Gin1 | 517 | 454 | 456 | 547 | 557 | 555 | 654 | 949 | 888 | 894 | 864 | 948 | 813 | 1043 | 662 | 484 | 528 | 516 | 567 | 603 | 600 | 739 | 821 | 1005 | 867 | 941 | 979 | 957 | Gin1 |
| Ginm1 | 671 | 561 | 629 | 588 | 481 | 616 | 591 | 597 | 512 | 472 | 597 | 584 | 524 | 549 | 633 | 640 | 537 | 587 | 504 | 641 | 581 | 463 | 534 | 484 | 554 | 512 | 623 | 572 | Ginm1 |
| Gins1 | 1396 | 1296 | 1603 | 1307 | 970 | 1109 | 1401 | 2003 | 1622 | 1648 | 1392 | 2359 | 2030 | 2511 | 1501 | 1181 | 1510 | 1176 | 1301 | 1437 | 1249 | 1427 | 1378 | 2239 | 1779 | 2981 | 2608 | 2817 | Gins1 |
| Gins2 | 2851 | 3304 | 3155 | 2570 | 1730 | 2071 | 1976 | 1307 | 1668 | 1330 | 1484 | 1992 | 1791 | 1821 | 2508 | 2801 | 2738 | 2216 | 2251 | 2434 | 2061 | 1016 | 1533 | 1605 | 1299 | 2507 | 2177 | 2059 | Gins2 |
| Gins3 | 2524 | 2529 | 2365 | 2000 | 1634 | 1679 | 2208 | 1247 | 1175 | 1129 | 1029 | 1516 | 1268 | 1523 | 2379 | 2350 | 2050 | 2053 | 1842 | 2096 | 1888 | 852 | 1031 | 1216 | 1084 | 1627 | 1521 | 1665 | Gins3 |
| Gins4 | 4167 | 3886 | 3478 | 3042 | 2195 | 2478 | 3064 | 2584 | 2267 | 1982 | 1920 | 2335 | 2326 | 2594 | 3687 | 3460 | 3209 | 2831 | 2650 | 2921 | 2753 | 1917 | 2190 | 2370 | 2037 | 2880 | 2568 | 2733 | Gins4 |
| Gip | 0 | 0 | 0 | 0 | 0 | 0 | 0 | 0 | 0 | 3 | 2 | 1 | 2 | 0 | 0 | 0 | 0 | 1 | 2 | 0 | 0 | 0 | 0 | 0 | 2 | 5 | 0 | 1 | Gip |
| Gipc1 | 5285 | 6542 | 6441 | 5691 | 4889 | 4610 | 4510 | 3303 | 3753 | 3872 | 3436 | 4504 | 3768 | 4103 | 5103 | 5490 | 5979 | 5332 | 5281 | 4547 | 4612 | 3233 | 3700 | 4120 | 3630 | 5102 | 4230 | 4269 | Gipc1 |
| Gipc2 | 141 | 136 | 129 | 130 | 180 | 136 | 93 | 98 | 89 | 80 | 88 | 66 | 107 | 55 | 157 | 140 | 139 | 127 | 129 | 162 | 136 | 125 | 89 | 63 | 89 | 59 | 67 | 75 | Gipc2 |
| Gipc3 | 17 | 14 | 11 | 38 | 17 | 8 | 10 | 22 | 30 | 48 | 34 | 7 | 26 | 19 | 12 | 19 | 14 | 9 | 7 | 8 | 13 | 10 | 20 | 71 | 9 | 60 | 5 | 9 | Gipc3 |
| Gipr | 489 | 602 | 782 | 864 | 928 | 989 | 869 | 260 | 349 | 472 | 351 | 406 | 369 | 376 | 603 | 645 | 889 | 952 | 989 | 779 | 928 | 476 | 259 | 509 | 650 | 230 | 323 | 345 | Gipr |
| Git1 | 8037 | 10750 | 10571 | 12508 | 11281 | 10271 | 9337 | 4941 | 5965 | 6799 | 6386 | 7489 | 6204 | 6339 | 7545 | 9502 | 11946 | 11931 | 12507 | 8644 | 10366 | 7249 | 5611 | 7136 | 6929 | 7784 | 6049 | 6102 | Git1 |
| Git2 | 4742 | 5255 | 4487 | 5084 | 4979 | 5074 | 4679 | 3485 | 4022 | 3925 | 4157 | 3631 | 3444 | 3423 | 4421 | 5252 | 4476 | 5157 | 4998 | 4667 | 4944 | 3818 | 3886 | 3542 | 3705 | 3245 | 3549 | 3183 | Git2 |
| Gja1 | 6153 | 4010 | 6429 | 3798 | 4452 | 5867 | 3553 | 12217 | 13150 | 10734 | 13602 | 12600 | 15667 | 13277 | 5658 | 5580 | 4802 | 4807 | 3726 | 4706 | 3283 | 11199 | 17996 | 13487 | 11451 | 14390 | 14609 | 13389 | Gja1 |
| Gja10 | 0 | 2 | 0 | 0 | 2 | 10 | 0 | 6 | 0 | 3 | 1 | 3 | 7 | 0 | 3 | 0 | 5 | 1 | 1 | 2 | 0 | 2 | 2 | 2 | 0 | 0 | 0 | 2 | Gja10 |
| Gja3 | 70 | 71 | 48 | 44 | 40 | 65 | 27 | 27 | 33 | 30 | 108 | 11 | 14 | 11 | 87 | 72 | 60 | 90 | 40 | 52 | 33 | 22 | 105 | 17 | 12 | 4 | 10 | 13 | Gja3 |
| Gja4 | 225 | 333 | 354 | 314 | 389 | 361 | 287 | 353 | 385 | 513 | 493 | 459 | 535 | 404 | 241 | 341 | 359 | 302 | 391 | 319 | 332 | 421 | 413 | 480 | 455 | 584 | 542 | 545 | Gja4 |
| Gja5 | 144 | 137 | 199 | 99 | 121 | 82 | 96 | 331 | 466 | 363 | 1033 | 379 | 355 | 364 | 152 | 219 | 173 | 76 | 97 | 109 | 103 | 441 | 797 | 480 | 257 | 754 | 401 | 523 | Gja5 |
| Gja6 | 2 | 3 | 4 | 3 | 2 | 4 | 4 | 7 | 10 | 10 | 1 | 17 | 7 | 9 | 11 | 9 | 3 | 4 | 10 | 6 | 3 | 10 | 14 | 13 | 1 | 15 | 3 | 5 | Gja6 |
| Gja8 | 3 | 0 | 3 | 0 | 0 | 0 | 0 | 7 | 9 | 4 | 5 | 0 | 0 | 0 | 0 | 0 | 0 | 0 | 0 | 0 | 0 | 0 | 7 | 7 | 0 | 16 | 0 | 0 | Gja8 |
| Gjb1 | 9 | 6 | 6 | 8 | 10 | 10 | 3 | 11 | 21 | 12 | 21 | 2 | 8 | 3 | 1 | 5 | 3 | 7 | 13 | 2 | 10 | 8 | 10 | 3 | 8 | 0 | 1 | 2 | Gjb1 |
| Gjb2 | 926 | 963 | 1011 | 192 | 510 | 1211 | 213 | 1324 | 2283 | 1175 | 1755 | 1375 | 1996 | 1322 | 1195 | 998 | 384 | 1284 | 358 | 305 | 253 | 797 | 2542 | 1618 | 752 | 478 | 1174 | 1313 | Gjb2 |
| Gjb3 | 29 | 36 | 34 | 36 | 25 | 48 | 18 | 23 | 41 | 24 | 22 | 13 | 26 | 20 | 32 | 55 | 36 | 23 | 39 | 22 | 21 | 3 | 45 | 26 | 8 | 9 | 24 | 12 | Gjb3 |
| Gjb4 | 7 | 5 | 1 | 1 | 2 | 0 | 3 | 1 | 7 | 7 | 2 | 7 | 9 | 1 | 4 | 5 | 3 | 0 | 3 | 4 | 4 | 3 | 2 | 5 | 1 | 2 | 2 | 4 | Gjb4 |
| Gjb5 | 25 | 33 | 29 | 8 | 12 | 12 | 0 | 5 | 8 | 10 | 9 | 2 | 10 | 3 | 41 | 45 | 27 | 21 | 8 | 20 | 13 | 7 | 16 | 4 | 2 | 1 | 3 | 2 | Gjb5 |
| Gjb6 | 149 | 130 | 142 | 103 | 232 | 196 | 104 | 833 | 1227 | 721 | 898 | 548 | 814 | 685 | 156 | 163 | 109 | 202 | 90 | 130 | 85 | 822 | 1608 | 800 | 628 | 409 | 547 | 491 | Gjb6 |
| Gjc1 | 4349 | 3836 | 5678 | 4962 | 5152 | 4552 | 4790 | 5029 | 4325 | 5313 | 5313 | 7078 | 6468 | 7094 | 4036 | 4113 | 5566 | 4680 | 5368 | 4035 | 4654 | 5294 | 4296 | 6236 | 5883 | 7783 | 6484 | 7213 | Gjc1 |
| Gjc2 | 52 | 87 | 54 | 116 | 118 | 89 | 80 | 37 | 37 | 37 | 38 | 20 | 18 | 20 | 58 | 70 | 67 | 123 | 112 | 91 | 82 | 40 | 29 | 33 | 34 | 25 | 23 | 11 | Gjc2 |
| Gjc3 | 13 | 5 | 2 | 4 | 23 | 6 | 1 | 42 | 80 | 51 | 119 | 18 | 56 | 13 | 11 | 8 | 1 | 2 | 2 | 2 | 2 | 84 | 79 | 23 | 47 | 18 | 12 | 13 | Gjc3 |
| Gjd2 | 363 | 327 | 377 | 555 | 849 | 674 | 778 | 1321 | 1129 | 1305 | 1078 | 1118 | 640 | 1160 | 382 | 213 | 636 | 621 | 858 | 882 | 745 | 1470 | 1091 | 1225 | 1561 | 978 | 1065 | 980 | Gjd2 |
| Gjd3 | 12 | 12 | 16 | 13 | 25 | 6 | 15 | 6 | 8 | 20 | 14 | 7 | 15 | 5 | 11 | 16 | 15 | 7 | 6 | 8 | 13 | 17 | 12 | 5 | 12 | 7 | 1 | 2 | Gjd3 |
| Gjd4 | 7 | 2 | 0 | 4 | 2 | 6 | 2 | 10 | 38 | 5 | 184 | 2 | 10 | 5 | 4 | 6 | 2 | 3 | 0 | 0 | 1 | 37 | 111 | 7 | 12 | 0 | 0 | 1 | Gjd4 |
| Gje1 | 0 | 0 | 0 | 0 | 0 | 1 | 1 | 0 | 3 | 0 | 2 | 0 | 0 | 1 | 0 | 0 | 0 | 0 | 0 | 3 | 0 | 0 | 0 | 0 | 0 | 0 | 0 | 0 | Gje1 |
| Gk | 2722 | 2137 | 2250 | 2122 | 1997 | 2393 | 2435 | 1270 | 1129 | 978 | 1041 | 1012 | 1187 | 1183 | 2768 | 2504 | 2113 | 2123 | 1977 | 2514 | 2255 | 1079 | 1223 | 989 | 1041 | 1040 | 1276 | 1225 | Gk |
| Gk5 | 1787 | 1332 | 1019 | 1108 | 1064 | 1343 | 1047 | 739 | 842 | 647 | 827 | 551 | 827 | 559 | 1589 | 1569 | 1086 | 1182 | 926 | 1089 | 968 | 665 | 804 | 535 | 486 | 436 | 579 | 477 | Gk5 |
| Gkap1 | 3155 | 2979 | 3716 | 3463 | 3107 | 3016 | 3471 | 2313 | 1880 | 2101 | 2302 | 2442 | 2209 | 2798 | 3312 | 2781 | 3536 | 3146 | 3467 | 3443 | 3390 | 2130 | 1995 | 2536 | 2270 | 2647 | 2621 | 2776 | Gkap1 |
| Gkn1 | 0 | 1 | 0 | 0 | 0 | 0 | 0 | 0 | 1 | 0 | 0 | 0 | 0 | 0 | 0 | 2 | 0 | 0 | 0 | 0 | 0 | 0 | 2 | 0 | 6 | 0 | 0 | 0 | Gkn1 |
| Gkn2 | 0 | 0 | 0 | 0 | 2 | 0 | 0 | 0 | 10 | 0 | 0 | 0 | 0 | 6 | 0 | 0 | 0 | 0 | 2 | 0 | 0 | 0 | 5 | 0 | 0 | 0 | 0 | 0 | Gkn2 |
| Gkn3 | 5 | 5 | 0 | 6 | 3 | 0 | 1 | 2 | 4 | 4 | 13 | 4 | 2 | 6 | 1 | 7 | 0 | 2 | 3 | 1 | 9 | 2 | 6 | 5 | 5 | 8 | 0 | 0 | Gkn3 |
| Gla | 1889 | 1938 | 1819 | 1777 | 1492 | 1549 | 1607 | 855 | 704 | 684 | 691 | 779 | 838 | 833 | 1851 | 1916 | 1757 | 1669 | 1345 | 1517 | 1531 | 750 | 732 | 713 | 670 | 950 | 838 | 930 | Gla |
| Glb1 | 1747 | 1919 | 1504 | 1540 | 1501 | 1512 | 1312 | 1321 | 1572 | 1327 | 1512 | 1347 | 1416 | 1237 | 1730 | 1933 | 1567 | 1502 | 1460 | 1524 | 1302 | 1506 | 1786 | 1274 | 1187 | 1460 | 1428 | 1250 | Glb1 |
| Glb1l | 1697 | 1614 | 1247 | 1372 | 1528 | 1758 | 1484 | 2201 | 2040 | 1905 | 2113 | 2068 | 2000 | 1953 | 1726 | 1634 | 1348 | 1567 | 1420 | 1507 | 1489 | 2364 | 2029 | 1889 | 2054 | 1692 | 2166 | 1926 | Glb1l |
| Glb1l2 | 687 | 504 | 575 | 487 | 422 | 522 | 457 | 413 | 377 | 389 | 337 | 444 | 423 | 350 | 571 | 538 | 447 | 394 | 415 | 418 | 340 | 291 | 350 | 333 | 291 | 426 | 403 | 370 | Glb1l2 |
| Glb1l3 | 0 | 5 | 0 | 0 | 2 | 3 | 0 | 0 | 1 | 0 | 0 | 0 | 1 | 0 | 0 | 4 | 1 | 1 | 0 | 1 | 0 | 3 | 3 | 2 | 1 | 0 | 1 | 0 | Glb1l3 |
| Glcci1 | 376 | 323 | 201 | 450 | 392 | 488 | 417 | 348 | 324 | 301 | 294 | 383 | 299 | 361 | 332 | 367 | 292 | 410 | 348 | 390 | 407 | 311 | 286 | 278 | 334 | 118 | 322 | 268 | Glcci1 |
| Glce | 6763 | 5046 | 4918 | 4365 | 3949 | 4298 | 4289 | 3864 | 3676 | 3057 | 3682 | 2776 | 3398 | 3101 | 6372 | 6122 | 4157 | 4609 | 3797 | 4604 | 4026 | 3120 | 3832 | 2877 | 2630 | 2894 | 3037 | 3071 | Glce |
| Gldc | 3953 | 4944 | 5324 | 4757 | 4052 | 4045 | 4859 | 2027 | 2230 | 2295 | 1741 | 4216 | 3087 | 3936 | 4050 | 3666 | 5467 | 4535 | 5114 | 4301 | 4479 | 1888 | 1650 | 3321 | 2904 | 5194 | 3922 | 4193 | Gldc |
| Gldn | 18 | 11 | 29 | 17 | 11 | 9 | 16 | 26 | 38 | 56 | 26 | 11 | 15 | 10 | 13 | 16 | 8 | 12 | 12 | 4 | 13 | 23 | 41 | 33 | 18 | 28 | 14 | 6 | Gldn |
| Gle1 | 5405 | 5985 | 4930 | 5329 | 4756 | 4775 | 5098 | 4717 | 4940 | 5108 | 4779 | 5333 | 4722 | 5120 | 5242 | 5358 | 5271 | 5209 | 5382 | 4878 | 5188 | 5157 | 4798 | 5151 | 5493 | 5678 | 5294 | 5211 | Gle1 |
| Glg1 | 19872 | 19567 | 17513 | 20134 | 21570 | 21743 | 20309 | 19154 | 19596 | 21174 | 22048 | 22540 | 21996 | 21335 | 19249 | 19710 | 19337 | 21375 | 20897 | 18014 | 21692 | 21225 | 18930 | 19550 | 22230 | 12802 | 21457 | 20178 | Glg1 |
| Gli1 | 1794 | 1748 | 1910 | 1204 | 972 | 937 | 838 | 1652 | 2396 | 1735 | 2257 | 1581 | 1354 | 1442 | 1217 | 1745 | 1823 | 1221 | 931 | 817 | 813 | 1566 | 2262 | 1554 | 1105 | 2227 | 1782 | 1677 | Gli1 |
| Gli2 | 2080 | 2126 | 2606 | 1922 | 1645 | 1931 | 1570 | 2511 | 3260 | 3492 | 3351 | 4613 | 3782 | 3874 | 1689 | 2074 | 2367 | 1869 | 2112 | 1348 | 1630 | 2374 | 2974 | 3895 | 2797 | 4578 | 4115 | 3991 | Gli2 |
| Gli3 | 6945 | 5979 | 7500 | 6572 | 5525 | 6576 | 6334 | 7266 | 7251 | 6929 | 6776 | 11922 | 10463 | 11048 | 6376 | 6115 | 6795 | 5567 | 6473 | 5824 | 5991 | 5838 | 6666 | 8371 | 7979 | 9896 | 11670 | 11774 | Gli3 |
| Glipr1 | 35 | 27 | 21 | 12 | 35 | 30 | 31 | 33 | 37 | 31 | 42 | 42 | 50 | 26 | 21 | 35 | 49 | 37 | 18 | 32 | 16 | 37 | 67 | 28 | 29 | 24 | 50 | 28 | Glipr1 |
| Glipr1l1 | 1 | 3 | 0 | 8 | 0 | 10 | 2 | 2 | 2 | 0 | 2 | 0 | 0 | 0 | 0 | 1 | 10 | 4 | 2 | 0 | 1 | 2 | 1 | 0 | 0 | 0 | 1 | 6 | Glipr1l1 |
| Glipr1l2 | 0 | 0 | 0 | 0 | 0 | 0 | 0 | 0 | 0 | 0 | 0 | 0 | 0 | 0 | 0 | 0 | 0 | 0 | 0 | 0 | 0 | 0 | 1 | 0 | 3 | 0 | 0 | 0 | Glipr1l2 |
| Glipr1l3 | 8 | 3 | 8 | 10 | 6 | 3 | 15 | 7 | 4 | 13 | 3 | 7 | 10 | 6 | 11 | 10 | 13 | 4 | 8 | 17 | 4 | 15 | 6 | 2 | 8 | 0 | 0 | 9 | Glipr1l3 |
| Glipr2 | 2874 | 2617 | 2404 | 1254 | 1281 | 1758 | 1091 | 2540 | 2909 | 1929 | 3131 | 2746 | 3222 | 2554 | 2888 | 3625 | 1884 | 1560 | 1221 | 1365 | 1060 | 2244 | 3653 | 2121 | 1772 | 2957 | 3038 | 2866 | Glipr2 |
| Glis1 | 221 | 225 | 200 | 143 | 82 | 74 | 64 | 169 | 234 | 138 | 159 | 89 | 177 | 64 | 165 | 238 | 145 | 112 | 111 | 62 | 71 | 60 | 216 | 130 | 75 | 149 | 45 | 64 | Glis1 |
| Glis2 | 3208 | 3992 | 3973 | 4044 | 3648 | 3797 | 3196 | 2591 | 2934 | 3140 | 3364 | 3555 | 3300 | 3222 | 2921 | 3932 | 4157 | 4063 | 3919 | 2861 | 3248 | 3165 | 3033 | 3365 | 3158 | 3645 | 3433 | 3328 | Glis2 |
| Glis3 | 773 | 465 | 543 | 475 | 620 | 484 | 482 | 1071 | 1036 | 914 | 589 | 722 | 797 | 584 | 551 | 646 | 484 | 483 | 515 | 510 | 491 | 810 | 1155 | 702 | 702 | 596 | 690 | 637 | Glis3 |
| Glmn | 2472 | 1722 | 1997 | 1857 | 1978 | 2220 | 2363 | 2749 | 2341 | 1898 | 2165 | 2271 | 2404 | 2229 | 2538 | 2034 | 1710 | 1936 | 1884 | 2455 | 2133 | 1957 | 2333 | 1888 | 2037 | 2302 | 2458 | 2402 | Glmn |
| Glmp | 3193 | 4325 | 3497 | 3457 | 2817 | 2827 | 2838 | 1802 | 2417 | 2139 | 2433 | 2135 | 2384 | 2201 | 3260 | 3863 | 3293 | 3410 | 2754 | 2929 | 2837 | 1898 | 2534 | 2086 | 2070 | 2611 | 2217 | 2084 | Glmp |
| Glo1 | 6377 | 6259 | 6675 | 5359 | 4780 | 5500 | 5709 | 5387 | 5125 | 4518 | 4703 | 5576 | 5389 | 5613 | 6494 | 6267 | 5697 | 5124 | 5361 | 6449 | 5274 | 4410 | 5322 | 5017 | 4723 | 6654 | 5971 | 6058 | Glo1 |
| Glod4 | 5186 | 4516 | 4879 | 4307 | 3832 | 4087 | 4491 | 5184 | 4537 | 4024 | 4248 | 4428 | 4447 | 5091 | 5114 | 4828 | 4771 | 4289 | 4027 | 5038 | 4356 | 4047 | 4669 | 4449 | 4470 | 5233 | 5002 | 5090 | Glod4 |
| Glod5 | 18 | 31 | 15 | 77 | 59 | 83 | 61 | 6 | 7 | 8 | 18 | 8 | 8 | 5 | 18 | 21 | 48 | 63 | 55 | 33 | 55 | 10 | 4 | 9 | 5 | 5 | 6 | 8 | Glod5 |
| Glp1r | 12 | 9 | 7 | 21 | 20 | 13 | 26 | 33 | 10 | 20 | 25 | 17 | 22 | 36 | 11 | 10 | 9 | 10 | 27 | 16 | 22 | 38 | 32 | 33 | 36 | 11 | 26 | 13 | Glp1r |
| Glp2r | 0 | 0 | 0 | 0 | 0 | 0 | 1 | 0 | 6 | 3 | 7 | 3 | 5 | 6 | 7 | 1 | 0 | 6 | 0 | 2 | 6 | 2 | 1 | 4 | 6 | 3 | 0 | 1 | Glp2r |
| Glra1 | 38 | 99 | 6 | 69 | 241 | 137 | 67 | 55 | 86 | 62 | 119 | 7 | 125 | 1 | 113 | 153 | 46 | 81 | 50 | 95 | 71 | 42 | 127 | 3 | 29 | 2 | 10 | 4 | Glra1 |
| Glra2 | 291 | 353 | 288 | 971 | 1991 | 1364 | 1759 | 1799 | 1795 | 2138 | 1757 | 899 | 761 | 1059 | 553 | 313 | 515 | 1497 | 1434 | 1524 | 1901 | 2555 | 1961 | 1580 | 2328 | 663 | 597 | 769 | Glra2 |
| Glra3 | 40 | 35 | 37 | 53 | 116 | 105 | 132 | 79 | 63 | 41 | 68 | 33 | 17 | 38 | 71 | 48 | 63 | 90 | 84 | 104 | 176 | 70 | 47 | 51 | 65 | 20 | 24 | 30 | Glra3 |
| Glra4 | 0 | 0 | 3 | 3 | 3 | 1 | 3 | 0 | 0 | 8 | 4 | 0 | 0 | 3 | 1 | 2 | 0 | 0 | 2 | 8 | 1 | 3 | 1 | 6 | 2 | 0 | 1 | 0 | Glra4 |
| Glrb | 975 | 777 | 818 | 972 | 1258 | 1239 | 1288 | 1313 | 1346 | 1095 | 1142 | 1079 | 1144 | 1167 | 1046 | 984 | 926 | 1152 | 1110 | 1206 | 1125 | 1160 | 1364 | 1093 | 1283 | 1015 | 1022 | 1043 | Glrb |

| | | | | | | | | | | | | | | | | | | | | | | | | | | | | | |
|---|---|---|---|---|---|---|---|---|---|---|---|---|---|---|---|---|---|---|---|---|---|---|---|---|---|---|---|---|---|
| Glrp1 | 2 | 3 | 0 | 9 | 2 | 8 | 4 | 5 | 0 | 1 | 0 | 7 | 1 | 1 | 5 | 3 | 0 | 3 | 0 | 3 | 4 | 2 | 1 | 0 | 0 | 0 | 0 | 0 | Glrp1 |
| Glrx | 1562 | 1607 | 1467 | 1328 | 1527 | 1472 | 1439 | 948 | 873 | 713 | 807 | 727 | 758 | 712 | 1809 | 1474 | 1318 | 1221 | 1432 | 1774 | 1468 | 846 | 874 | 677 | 675 | 646 | 747 | 765 | Glrx |
| Glrx2 | 4642 | 4209 | 4384 | 5713 | 5767 | 5835 | 6565 | 4673 | 3994 | 4782 | 4560 | 4417 | 3900 | 5125 | 4866 | 4161 | 5230 | 5861 | 5678 | 5545 | 6327 | 4811 | 3858 | 4465 | 4861 | 4273 | 4209 | 4351 | Glrx2 |
| Glrx3 | 12336 | 11436 | 10293 | 9569 | 9232 | 9491 | 10951 | 12684 | 11279 | 9955 | 8965 | 10478 | 10294 | 10989 | 13883 | 11400 | 10532 | 8982 | 10120 | 11491 | 10796 | 9957 | 11328 | 10146 | 9200 | 11210 | 10770 | 12056 | Glrx3 |
| Glrx5 | 3553 | 4524 | 4393 | 4455 | 3407 | 3231 | 3572 | 1842 | 1985 | 2306 | 2055 | 2223 | 2437 | 2500 | 3406 | 3461 | 4435 | 3882 | 3816 | 3208 | 3579 | 1791 | 2019 | 2469 | 2212 | 2656 | 2219 | 2766 | Glrx5 |
| Gls | 17320 | 12084 | 13311 | 16127 | 17950 | 19189 | 19135 | 16800 | 15858 | 13162 | 15400 | 13643 | 13856 | 13879 | 17842 | 15665 | 13874 | 17754 | 16210 | 18008 | 18308 | 16040 | 15373 | 12722 | 15386 | 10743 | 13877 | 12565 | Gls |
| Gls2 | 411 | 583 | 567 | 650 | 514 | 499 | 515 | 125 | 133 | 144 | 156 | 118 | 139 | 104 | 472 | 516 | 433 | 602 | 577 | 458 | 486 | 127 | 98 | 110 | 87 | 138 | 93 | 82 | Gls2 |
| Glt1d1 | 203 | 206 | 209 | 213 | 296 | 212 | 264 | 504 | 495 | 496 | 443 | 726 | 671 | 743 | 193 | 152 | 255 | 179 | 237 | 185 | 265 | 491 | 381 | 553 | 563 | 766 | 769 | 792 | Glt1d1 |
| Glt28d2 | 181 | 67 | 77 | 46 | 51 | 75 | 83 | 39 | 57 | 48 | 32 | 4 | 24 | 18 | 117 | 97 | 43 | 76 | 70 | 82 | 61 | 25 | 56 | 38 | 18 | 40 | 6 | 14 | Glt28d2 |
| Glt6d1 | 0 | 0 | 0 | 0 | 0 | 0 | 1 | 0 | 0 | 0 | 0 | 0 | 0 | 0 | 0 | 0 | 0 | 0 | 0 | 0 | 0 | 0 | 0 | 0 | 0 | 0 | 0 | 0 | Glt6d1 |
| Glt8d1 | 3445 | 3191 | 3652 | 3453 | 3393 | 3352 | 3752 | 5186 | 4485 | 4977 | 4816 | 4931 | 4796 | 5156 | 3618 | 3246 | 3781 | 3277 | 3582 | 3489 | 3620 | 5924 | 4700 | 4885 | 4916 | 5030 | 4857 | 5211 | Glt8d1 |
| Glt8d2 | 179 | 148 | 85 | 121 | 77 | 148 | 39 | 304 | 540 | 344 | 683 | 242 | 342 | 243 | 216 | 196 | 78 | 133 | 52 | 63 | 43 | 346 | 670 | 282 | 218 | 284 | 250 | 269 | Glt8d2 |
| Gltp | 2190 | 2463 | 2800 | 2219 | 1741 | 1889 | 1895 | 1215 | 1455 | 1552 | 1568 | 1997 | 1657 | 2017 | 1913 | 2255 | 2649 | 2038 | 2068 | 1629 | 1902 | 1288 | 1608 | 1758 | 1442 | 2216 | 1654 | 1915 | Gltp |
| Gltpd2 | 2 | 5 | 1 | 1 | 0 | 0 | 0 | 1 | 6 | 2 | 4 | 2 | 0 | 0 | 1 | 9 | 9 | 1 | 2 | 0 | 2 | 0 | 5 | 1 | 2 | 8 | 0 | 2 | Gltpd2 |
| Glud1 | 10635 | 9263 | 10902 | 10092 | 11776 | 10545 | 11358 | 10867 | 10168 | 10727 | 10263 | 10403 | 10023 | 10460 | 10603 | 9804 | 11068 | 10568 | 11091 | 9626 | 11231 | 11825 | 10558 | 10234 | 10507 | 11020 | 10004 | 10627 | Glud1 |
| Glul | 12716 | 13975 | 14445 | 13218 | 12070 | 11603 | 11769 | 11579 | 12229 | 13276 | 12277 | 13573 | 12377 | 13974 | 12936 | 12497 | 13847 | 12783 | 12056 | 11851 | 12388 | 12210 | 12751 | 13306 | 12938 | 14181 | 13386 | 14724 | Glul |
| Glyat | 0 | 1 | 0 | 7 | 0 | 6 | 4 | 0 | 2 | 0 | 0 | 0 | 0 | 1 | 4 | 2 | 0 | 3 | 0 | 2 | 4 | 0 | 3 | 1 | 0 | 2 | 0 | 2 | Glyat |
| Glyatl3 | 74 | 24 | 0 | 54 | 0 | 0 | 0 | 98 | 69 | 99 | 87 | 1 | 40 | 18 | 42 | 40 | 17 | 1 | 0 | 1 | 1 | 2 | 109 | 196 | 1 | 222 | 0 | 0 | Glyatl3 |
| Glyctk | 103 | 72 | 54 | 117 | 88 | 102 | 116 | 133 | 252 | 215 | 250 | 143 | 195 | 115 | 63 | 81 | 89 | 84 | 97 | 73 | 85 | 104 | 236 | 231 | 157 | 217 | 165 | 132 | Glyctk |
| Glyr1 | 22316 | 23482 | 22695 | 22129 | 21197 | 21216 | 25109 | 18636 | 17710 | 17940 | 17458 | 20141 | 19175 | 22717 | 22234 | 20065 | 22459 | 21818 | 22202 | 21656 | 24161 | 18205 | 16974 | 19214 | 20358 | 20382 | 20508 | 22375 | Glyr1 |
| Gm10033 | 2231 | 1477 | 1497 | 1592 | 1589 | 2257 | 1968 | 1931 | 1812 | 1699 | 1716 | 1866 | 2033 | 1939 | 2234 | 2160 | 1636 | 1465 | 1752 | 2281 | 2343 | 1674 | 1793 | 1557 | 1823 | 1661 | 2107 | 1978 | Gm10033 |
| Gm10037 | 171 | 96 | 207 | 201 | 250 | 206 | 185 | 292 | 246 | 317 | 293 | 258 | 221 | 228 | 137 | 140 | 182 | 271 | 235 | 167 | 170 | 359 | 245 | 250 | 237 | 276 | 213 | 240 | Gm10037 |
| Gm10053 | 386 | 343 | 389 | 336 | 276 | 232 | 365 | 360 | 220 | 247 | 266 | 291 | 296 | 336 | 388 | 300 | 364 | 326 | 281 | 357 | 381 | 282 | 263 | 303 | 297 | 313 | 325 | 386 | Gm10053 |
| Gm10100 | 0 | 0 | 0 | 0 | 0 | 0 | 0 | 0 | 0 | 0 | 0 | 0 | 0 | 0 | 0 | 0 | 0 | 0 | 0 | 0 | 0 | 0 | 1 | 0 | 0 | 0 | 0 | 0 | Gm10100 |
| Gm10128 | 8 | 7 | 1 | 1 | 12 | 5 | 3 | 23 | 33 | 3 | 18 | 5 | 17 | 3 | 7 | 6 | 1 | 2 | 7 | 0 | 4 | 17 | 8 | 15 | 15 | 7 | 9 | 0 | Gm10128 |
| Gm10130 | 376 | 318 | 280 | 282 | 302 | 306 | 231 | 330 | 401 | 341 | 425 | 282 | 392 | 318 | 315 | 333 | 340 | 357 | 281 | 339 | 247 | 344 | 409 | 371 | 331 | 307 | 408 | 375 | Gm10130 |
| Gm10131 | 2089 | 2060 | 2328 | 1730 | 1418 | 1594 | 2134 | 1773 | 1414 | 1380 | 1499 | 1893 | 1641 | 2335 | 2060 | 1578 | 1971 | 1620 | 1704 | 1868 | 2011 | 1395 | 1440 | 1801 | 1724 | 2211 | 1892 | 2414 | Gm10131 |
| Gm10135 | 0 | 0 | 0 | 0 | 0 | 0 | 0 | 0 | 0 | 0 | 0 | 0 | 0 | 1 | 0 | 0 | 0 | 2 | 0 | 0 | 0 | 0 | 0 | 0 | 0 | 1 | 1 | 0 | Gm10135 |
| Gm10142 | 0 | 0 | 0 | 0 | 0 | 0 | 0 | 0 | 0 | 0 | 0 | 0 | 0 | 0 | 0 | 0 | 0 | 0 | 0 | 0 | 1 | 0 | 0 | 0 | 0 | 0 | 0 | 0 | Gm10142 |
| Gm10153 | 0 | 0 | 0 | 0 | 0 | 0 | 0 | 0 | 0 | 0 | 0 | 0 | 0 | 0 | 0 | 0 | 0 | 0 | 0 | 0 | 0 | 0 | 0 | 0 | 0 | 0 | 0 | 2 | Gm10153 |
| Gm10160 | 5 | 7 | 6 | 3 | 7 | 17 | 3 | 23 | 3 | 7 | 10 | 3 | 6 | 1 | 1 | 9 | 1 | 10 | 4 | 10 | 12 | 10 | 7 | 1 | 10 | 6 | 17 | 7 | Gm10160 |
| Gm10184 | 691 | 689 | 599 | 499 | 369 | 378 | 469 | 536 | 389 | 325 | 346 | 608 | 417 | 609 | 649 | 552 | 589 | 416 | 513 | 638 | 415 | 343 | 397 | 419 | 380 | 655 | 607 | 631 | Gm10184 |
| Gm10188 | 0 | 0 | 0 | 0 | 0 | 0 | 1 | 0 | 0 | 0 | 0 | 0 | 0 | 0 | 0 | 0 | 0 | 2 | 0 | 0 | 0 | 2 | 0 | 0 | 0 | 0 | 0 | 0 | Gm10188 |
| Gm10190 | 1 | 3 | 1 | 4 | 5 | 0 | 3 | 4 | 2 | 9 | 6 | 5 | 8 | 3 | 1 | 4 | 0 | 2 | 2 | 1 | 4 | 5 | 8 | 5 | 8 | 9 | 8 | 0 | Gm10190 |
| Gm10197 | 0 | 0 | 4 | 4 | 0 | 0 | 0 | 0 | 0 | 3 | 0 | 5 | 6 | 1 | 0 | 0 | 2 | 4 | 0 | 0 | 2 | 0 | 0 | 5 | 5 | 3 | 0 | 1 | Gm10197 |
| Gm10220 | 0 | 0 | 0 | 0 | 0 | 0 | 0 | 0 | 0 | 0 | 1 | 0 | 0 | 0 | 0 | 0 | 0 | 0 | 0 | 0 | 0 | 0 | 1 | 0 | 0 | 0 | 0 | 0 | Gm10220 |
| Gm10226 | 301 | 328 | 370 | 344 | 358 | 307 | 339 | 381 | 278 | 203 | 218 | 280 | 264 | 214 | 355 | 338 | 311 | 372 | 333 | 402 | 284 | 304 | 263 | 218 | 251 | 284 | 307 | 217 | Gm10226 |
| Gm10267 | 0 | 0 | 0 | 0 | 0 | 0 | 0 | 1 | 0 | 0 | 0 | 0 | 0 | 0 | 0 | 0 | 0 | 0 | 0 | 0 | 0 | 2 | 0 | 0 | 0 | 0 | 0 | 0 | Gm10267 |
| Gm10271 | 0 | 0 | 0 | 0 | 0 | 0 | 0 | 0 | 0 | 0 | 0 | 0 | 2 | 0 | 0 | 0 | 1 | 0 | 0 | 2 | 0 | 2 | 0 | 0 | 0 | 0 | 0 | 0 | Gm10271 |
| Gm10272 | 0 | 0 | 0 | 0 | 0 | 0 | 0 | 0 | 0 | 0 | 0 | 0 | 0 | 0 | 0 | 0 | 2 | 0 | 0 | 0 | 0 | 0 | 0 | 0 | 0 | 0 | 0 | 0 | Gm10272 |
| Gm10277 | 0 | 1 | 0 | 0 | 0 | 0 | 0 | 0 | 0 | 0 | 0 | 0 | 1 | 1 | 0 | 0 | 0 | 0 | 0 | 0 | 0 | 0 | 0 | 1 | 0 | 0 | 0 | 0 | Gm10277 |
| Gm10282 | 3623 | 2638 | 3427 | 3551 | 1655 | 1307 | 2386 | 4034 | 1982 | 2472 | 2208 | 3496 | 2149 | 4726 | 2952 | 1419 | 3341 | 2091 | 2813 | 1630 | 2072 | 2363 | 1866 | 3119 | 3646 | 3234 | 3360 | 4614 | Gm10282 |
| Gm10309 | 0 | 0 | 0 | 0 | 0 | 0 | 0 | 0 | 1 | 0 | 4 | 0 | 5 | 0 | 0 | 0 | 0 | 0 | 0 | 1 | 0 | 0 | 1 | 2 | 0 | 0 | 0 | 0 | Gm10309 |
| Gm10318 | 0 | 0 | 0 | 0 | 0 | 0 | 2 | 0 | 2 | 1 | 0 | 0 | 0 | 4 | 0 | 0 | 0 | 0 | 0 | 0 | 0 | 0 | 0 | 0 | 0 | 0 | 0 | 1 | Gm10318 |
| Gm10320 | 40 | 39 | 25 | 27 | 20 | 8 | 22 | 24 | 23 | 19 | 23 | 15 | 11 | 22 | 36 | 30 | 29 | 22 | 15 | 18 | 34 | 35 | 21 | 24 | 22 | 24 | 27 | 28 | Gm10320 |
| Gm10334 | 1 | 3 | 3 | 0 | 2 | 0 | 2 | 4 | 3 | 10 | 1 | 7 | 9 | 2 | 8 | 7 | 8 | 7 | 0 | 0 | 0 | 12 | 5 | 0 | 2 | 8 | 3 | 0 | Gm10334 |
| Gm10337 | 0 | 2 | 2 | 0 | 2 | 1 | 3 | 1 | 1 | 8 | 3 | 5 | 5 | 1 | 1 | 2 | 1 | 2 | 2 | 1 | 0 | 7 | 1 | 2 | 10 | 0 | 5 | 1 | Gm10337 |
| Gm10338 | 3 | 0 | 1 | 5 | 1 | 8 | 1 | 5 | 3 | 1 | 3 | 0 | 1 | 2 | 0 | 2 | 0 | 0 | 1 | 4 | 2 | 2 | 1 | 0 | 7 | 1 | 1 | 2 | Gm10338 |
| Gm10340 | 0 | 1 | 0 | 4 | 1 | 2 | 0 | 1 | 5 | 3 | 5 | 1 | 2 | 1 | 3 | 0 | 5 | 3 | 5 | 4 | 0 | 7 | 3 | 8 | 5 | 8 | 3 | 8 | Gm10340 |
| Gm10358 | 0 | 0 | 0 | 0 | 0 | 0 | 1 | 0 | 0 | 0 | 0 | 0 | 0 | 0 | 0 | 1 | 0 | 0 | 1 | 0 | 0 | 0 | 0 | 0 | 0 | 0 | 0 | 0 | Gm10358 |
| Gm10375 | 0 | 0 | 0 | 4 | 0 | 0 | 0 | 0 | 0 | 0 | 0 | 0 | 0 | 0 | 0 | 0 | 0 | 0 | 0 | 0 | 0 | 0 | 0 | 0 | 0 | 0 | 0 | 0 | Gm10375 |
| Gm10401 | 2 | 3 | 14 | 14 | 11 | 11 | 2 | 1 | 3 | 0 | 0 | 3 | 3 | 3 | 9 | 11 | 7 | 4 | 8 | 4 | 5 | 2 | 2 | 5 | 0 | 6 | 3 | 4 | Gm10401 |
| Gm10406 | 96 | 47 | 81 | 83 | 118 | 121 | 113 | 346 | 346 | 344 | 297 | 291 | 310 | 304 | 83 | 55 | 52 | 50 | 77 | 120 | 84 | 329 | 269 | 337 | 357 | 315 | 288 | 233 | Gm10406 |
| Gm10408 | 0 | 0 | 1 | 0 | 0 | 0 | 0 | 0 | 0 | 1 | 0 | 2 | 0 | 0 | 0 | 0 | 0 | 0 | 0 | 0 | 0 | 0 | 0 | 0 | 0 | 3 | 0 | 0 | Gm10408 |
| Gm10409 | 2 | 0 | 1 | 0 | 7 | 3 | 3 | 10 | 3 | 16 | 4 | 1 | 3 | 3 | 0 | 0 | 6 | 0 | 2 | 1 | 3 | 7 | 10 | 6 | 16 | 1 | 1 | 2 | Gm10409 |
| Gm1043 | 165 | 229 | 49 | 344 | 696 | 608 | 610 | 371 | 481 | 501 | 430 | 185 | 191 | 162 | 274 | 181 | 177 | 473 | 488 | 523 | 762 | 465 | 385 | 185 | 557 | 46 | 159 | 96 | Gm1043 |
| Gm10439 | 0 | 0 | 0 | 0 | 0 | 0 | 0 | 0 | 0 | 0 | 0 | 0 | 0 | 0 | 0 | 0 | 0 | 0 | 2 | 0 | 0 | 0 | 0 | 0 | 0 | 0 | 0 | 0 | Gm10439 |
| Gm10471 | 0 | 0 | 0 | 0 | 0 | 0 | 0 | 1 | 1 | 0 | 0 | 1 | 0 | 0 | 0 | 0 | 0 | 0 | 1 | 0 | 0 | 0 | 0 | 0 | 0 | 0 | 0 | 0 | Gm10471 |
| Gm10490 | 2 | 8 | 3 | 1 | 1 | 0 | 6 | 2 | 2 | 0 | 0 | 3 | 0 | 1 | 0 | 1 | 3 | 7 | 9 | 7 | 4 | 3 | 1 | 0 | 1 | 0 | 0 | 2 | Gm10490 |
| Gm10491 | 0 | 0 | 0 | 0 | 0 | 0 | 0 | 0 | 0 | 0 | 0 | 0 | 1 | 0 | 0 | 0 | 0 | 0 | 0 | 0 | 0 | 0 | 0 | 0 | 0 | 1 | 0 | 0 | Gm10491 |
| Gm10521 | 6 | 3 | 9 | 3 | 2 | 5 | 1 | 9 | 2 | 12 | 24 | 7 | 10 | 6 | 1 | 5 | 7 | 4 | 9 | 8 | 3 | 7 | 12 | 3 | 3 | 2 | 8 | 12 | Gm10521 |
| Gm10563 | 13 | 8 | 38 | 13 | 13 | 26 | 13 | 9 | 12 | 23 | 10 | 16 | 25 | 11 | 17 | 6 | 28 | 13 | 22 | 25 | 13 | 13 | 8 | 27 | 16 | 32 | 14 | 23 | Gm10563 |
| Gm10591 | 16 | 45 | 23 | 26 | 29 | 33 | 42 | 31 | 40 | 14 | 42 | 20 | 20 | 36 | 40 | 13 | 21 | 15 | 17 | 42 | 40 | 28 | 8 | 53 | 20 | 65 | 53 | 8 | Gm10591 |
| Gm10643 | 186 | 174 | 194 | 185 | 137 | 114 | 153 | 158 | 188 | 233 | 141 | 175 | 184 | 204 | 144 | 181 | 198 | 147 | 167 | 168 | 173 | 152 | 157 | 194 | 193 | 176 | 182 | 158 | Gm10643 |
| Gm10663 | 0 | 0 | 0 | 0 | 0 | 0 | 0 | 0 | 0 | 0 | 1 | 0 | 0 | 0 | 0 | 0 | 0 | 0 | 0 | 0 | 0 | 0 | 0 | 0 | 0 | 0 | 2 | 0 | Gm10663 |
| Gm10718 | 0 | 0 | 0 | 0 | 0 | 2 | 0 | 0 | 0 | 0 | 0 | 0 | 0 | 0 | 0 | 0 | 0 | 0 | 1 | 0 | 0 | 0 | 0 | 0 | 0 | 0 | 0 | 0 | Gm10718 |
| Gm10719 | 5 | 0 | 0 | 0 | 0 | 0 | 0 | 0 | 0 | 1 | 0 | 0 | 0 | 0 | 0 | 1 | 0 | 0 | 0 | 0 | 1 | 0 | 0 | 0 | 0 | 0 | 0 | 2 | Gm10719 |
| Gm10722 | 1 | 0 | 0 | 0 | 2 | 0 | 0 | 0 | 0 | 0 | 0 | 0 | 0 | 0 | 0 | 0 | 0 | 2 | 1 | 1 | 3 | 0 | 0 | 1 | 0 | 3 | 0 | 0 | Gm10722 |
| Gm10767 | 39 | 21 | 27 | 41 | 95 | 74 | 66 | 187 | 111 | 125 | 135 | 128 | 110 | 119 | 47 | 26 | 31 | 37 | 51 | 60 | 62 | 201 | 140 | 123 | 141 | 149 | 121 | 98 | Gm10767 |
| Gm10775 | 3 | 3 | 2 | 5 | 0 | 4 | 6 | 0 | 2 | 4 | 2 | 2 | 2 | 0 | 4 | 3 | 1 | 0 | 5 | 3 | 4 | 0 | 3 | 0 | 1 | 2 | 0 | 0 | Gm10775 |
| Gm10778 | 138 | 83 | 132 | 111 | 82 | 108 | 113 | 122 | 85 | 123 | 77 | 85 | 74 | 89 | 109 | 120 | 88 | 103 | 74 | 112 | 97 | 65 | 76 | 95 | 117 | 111 | 96 | 84 | Gm10778 |
| Gm10800 | 9 | 3 | 4 | 7 | 15 | 18 | 16 | 10 | 7 | 9 | 3 | 5 | 7 | 15 | 12 | 10 | 8 | 12 | 9 | 5 | 17 | 32 | 6 | 14 | 17 | 2 | 14 | 14 | Gm10800 |
| Gm10801 | 1 | 0 | 0 | 0 | 0 | 0 | 2 | 1 | 0 | 0 | 0 | 0 | 0 | 6 | 0 | 0 | 0 | 0 | 0 | 0 | 0 | 2 | 0 | 1 | 0 | 0 | 0 | 0 | Gm10801 |
| Gm10945 | 0 | 0 | 0 | 0 | 0 | 0 | 0 | 0 | 0 | 0 | 0 | 0 | 0 | 0 | 0 | 0 | 5 | 0 | 0 | 0 | 0 | 0 | 0 | 0 | 0 | 0 | 0 | 0 | Gm10945 |
| Gm10959 | 3 | 1 | 3 | 6 | 2 | 0 | 1 | 4 | 2 | 0 | 3 | 0 | 1 | 2 | 4 | 1 | 1 | 0 | 0 | 5 | 2 | 0 | 4 | 1 | 9 | 2 | 5 | 8 | Gm10959 |
| Gm10961 | 6 | 2 | 7 | 0 | 2 | 3 | 2 | 2 | 2 | 6 | 6 | 8 | 1 | 5 | 0 | 4 | 1 | 2 | 2 | 0 | 4 | 2 | 3 | 2 | 1 | 5 | 9 | 6 | Gm10961 |
| Gm10964 | 0 | 0 | 3 | 0 | 0 | 0 | 0 | 1 | 0 | 0 | 0 | 0 | 2 | 1 | 0 | 0 | 6 | 0 | 0 | 0 | 0 | 2 | 1 | 0 | 0 | 2 | 1 | 6 | Gm10964 |
| Gm10974 | 0 | 0 | 0 | 0 | 0 | 0 | 0 | 0 | 0 | 0 | 0 | 0 | 0 | 0 | 1 | 0 | 0 | 0 | 0 | 0 | 0 | 0 | 0 | 0 | 0 | 0 | 0 | 0 | Gm10974 |
| Gm10999 | 1 | 0 | 0 | 0 | 2 | 0 | 1 | 0 | 0 | 0 | 0 | 0 | 0 | 0 | 0 | 0 | 2 | 0 | 1 | 0 | 0 | 0 | 1 | 3 | 0 | 0 | 0 | 1 | Gm10999 |

| | | | | | | | | | | | | | | | | | | | | | | | | | | | | | |
|---|---|---|---|---|---|---|---|---|---|---|---|---|---|---|---|---|---|---|---|---|---|---|---|---|---|---|---|---|---|
| Gm11007 | 98 | 44 | 19 | 51 | 53 | 447 | 13 | 67 | 390 | 48 | 34 | 40 | 158 | 28 | 480 | 62 | 435 | 22 | 37 | 47 | 23 | 79 | 82 | 20 | 58 | 23 | 41 | 27 | Gm11007 |
| Gm11009 | 0 | 0 | 0 | 0 | 0 | 0 | 0 | 0 | 0 | 0 | 0 | 0 | 0 | 0 | 1 | 0 | 1 | 0 | 0 | 0 | 0 | 0 | 0 | 0 | 0 | 0 | 0 | 0 | Gm11009 |
| Gm11020 | 0 | 0 | 0 | 0 | 0 | 0 | 0 | 0 | 0 | 0 | 0 | 1 | 0 | 0 | 0 | 2 | 2 | 0 | 0 | 0 | 0 | 0 | 0 | 0 | 0 | 0 | 0 | 0 | Gm11020 |
| Gm11037 | 5 | 1 | 0 | 2 | 9 | 0 | 1 | 1 | 9 | 4 | 14 | 16 | 2 | 12 | 3 | 5 | 2 | 0 | 2 | 0 | 4 | 5 | 8 | 3 | 19 | 1 | 12 | 11 | Gm11037 |
| Gm11042 | 17 | 12 | 13 | 7 | 21 | 12 | 10 | 33 | 40 | 48 | 37 | 45 | 41 | 64 | 9 | 12 | 22 | 14 | 21 | 19 | 18 | 38 | 19 | 55 | 44 | 35 | 50 | 37 | Gm11042 |
| Gm11077 | 0 | 0 | 0 | 4 | 0 | 0 | 2 | 1 | 0 | 1 | 3 | 0 | 3 | 0 | 0 | 1 | 0 | 0 | 0 | 5 | 5 | 2 | 1 | 0 | 2 | 0 | 2 | 0 | Gm11077 |
| Gm11084 | 3 | 0 | 0 | 6 | 0 | 2 | 0 | 1 | 1 | 3 | 4 | 2 | 3 | 1 | 1 | 0 | 8 | 2 | 2 | 1 | 2 | 3 | 3 | 2 | 0 | 0 | 3 | 1 | Gm11084 |
| Gm11096 | 0 | 0 | 0 | 0 | 0 | 0 | 0 | 1 | 1 | 0 | 0 | 0 | 0 | 0 | 0 | 0 | 0 | 0 | 0 | 0 | 0 | 0 | 0 | 0 | 0 | 0 | 2 | 0 | Gm11096 |
| Gm1110 | 8 | 4 | 7 | 4 | 19 | 13 | 21 | 23 | 26 | 11 | 9 | 12 | 26 | 16 | 7 | 2 | 8 | 7 | 13 | 14 | 13 | 22 | 41 | 7 | 16 | 21 | 7 | 7 | Gm1110 |
| Gm11100 | 2 | 2 | 2 | 3 | 0 | 3 | 0 | 2 | 0 | 0 | 3 | 7 | 0 | 2 | 1 | 2 | 1 | 0 | 1 | 0 | 4 | 2 | 3 | 7 | 0 | 0 | 1 | 0 | Gm11100 |
| Gm11111 | 0 | 0 | 0 | 0 | 0 | 0 | 0 | 0 | 0 | 0 | 0 | 0 | 0 | 0 | 0 | 0 | 0 | 0 | 0 | 0 | 0 | 0 | 1 | 0 | 0 | 0 | 0 | 0 | Gm11111 |
| Gm11127 | 0 | 1 | 0 | 0 | 0 | 0 | 0 | 2 | 0 | 0 | 0 | 0 | 1 | 0 | 0 | 0 | 0 | 0 | 1 | 0 | 0 | 0 | 0 | 0 | 0 | 0 | 0 | 2 | Gm11127 |
| Gm1113 | 0 | 0 | 0 | 0 | 0 | 0 | 0 | 0 | 0 | 0 | 0 | 0 | 0 | 0 | 0 | 0 | 0 | 0 | 0 | 0 | 0 | 0 | 0 | 0 | 0 | 0 | 0 | 1 | Gm1113 |
| Gm11146 | 1 | 0 | 0 | 0 | 0 | 0 | 0 | 5 | 0 | 0 | 0 | 0 | 1 | 0 | 0 | 0 | 1 | 1 | 1 | 0 | 1 | 0 | 0 | 3 | 1 | 0 | 0 | 0 | Gm11146 |
| Gm11168 | 0 | 0 | 0 | 0 | 0 | 0 | 0 | 0 | 1 | 0 | 0 | 0 | 0 | 0 | 0 | 0 | 0 | 1 | 0 | 0 | 0 | 0 | 0 | 0 | 0 | 0 | 0 | 0 | Gm11168 |
| Gm11213 | 0 | 0 | 0 | 0 | 0 | 2 | 0 | 0 | 0 | 0 | 0 | 0 | 0 | 0 | 0 | 0 | 0 | 0 | 0 | 0 | 0 | 0 | 0 | 0 | 0 | 0 | 0 | 0 | Gm11213 |
| Gm1123 | 6 | 16 | 12 | 32 | 6 | 10 | 21 | 13 | 2 | 13 | 9 | 1 | 3 | 3 | 7 | 10 | 7 | 11 | 12 | 19 | 16 | 7 | 5 | 3 | 1 | 0 | 0 | 5 | Gm1123 |
| Gm11232 | 0 | 2 | 2 | 0 | 1 | 0 | 1 | 5 | 1 | 3 | 2 | 0 | 0 | 4 | 0 | 3 | 5 | 2 | 0 | 0 | 0 | 2 | 5 | 2 | 4 | 4 | 0 | 0 | Gm11232 |
| Gm1140 | 0 | 0 | 0 | 0 | 0 | 0 | 0 | 0 | 2 | 0 | 0 | 1 | 1 | 0 | 0 | 0 | 0 | 0 | 0 | 0 | 0 | 0 | 1 | 0 | 0 | 0 | 0 | 0 | Gm1140 |
| Gm1141 | 180 | 172 | 166 | 271 | 223 | 193 | 284 | 117 | 88 | 106 | 104 | 104 | 79 | 129 | 164 | 206 | 219 | 317 | 242 | 222 | 252 | 127 | 107 | 137 | 141 | 98 | 101 | 112 | Gm1141 |
| Gm11437 | 1 | 3 | 4 | 0 | 0 | 1 | 0 | 2 | 0 | 0 | 0 | 0 | 0 | 0 | 1 | 0 | 0 | 0 | 0 | 1 | 1 | 0 | 0 | 0 | 0 | 2 | 0 | 0 | Gm11437 |
| Gm11541 | 3 | 2 | 0 | 0 | 2 | 0 | 0 | 15 | 21 | 19 | 9 | 3 | 6 | 1 | 5 | 2 | 0 | 1 | 0 | 0 | 3 | 5 | 17 | 3 | 4 | 2 | 0 | 1 | Gm11541 |
| Gm11569 | 0 | 0 | 0 | 0 | 0 | 0 | 1 | 0 | 0 | 0 | 0 | 0 | 0 | 0 | 0 | 0 | 0 | 0 | 0 | 0 | 0 | 0 | 0 | 0 | 0 | 0 | 0 | 0 | Gm11569 |
| Gm11596 | 0 | 0 | 0 | 0 | 0 | 0 | 0 | 0 | 0 | 0 | 0 | 0 | 0 | 0 | 0 | 0 | 0 | 0 | 0 | 3 | 0 | 0 | 0 | 0 | 0 | 0 | 0 | 0 | Gm11596 |
| Gm11627 | 113 | 111 | 179 | 92 | 70 | 107 | 120 | 205 | 241 | 271 | 264 | 310 | 263 | 375 | 84 | 85 | 155 | 99 | 117 | 112 | 110 | 134 | 186 | 356 | 273 | 380 | 306 | 325 | Gm11627 |
| Gm11639 | 2 | 7 | 0 | 0 | 7 | 1 | 4 | 5 | 12 | 4 | 5 | 4 | 1 | 1 | 8 | 6 | 0 | 4 | 9 | 6 | 8 | 10 | 3 | 1 | 8 | 1 | 0 | 1 | Gm11639 |
| Gm11733 | 9 | 7 | 5 | 2 | 10 | 8 | 18 | 2 | 14 | 15 | 4 | 5 | 14 | 4 | 13 | 5 | 5 | 9 | 5 | 5 | 8 | 25 | 6 | 2 | 19 | 10 | 13 | 18 | Gm11733 |
| Gm11756 | 0 | 0 | 0 | 0 | 0 | 0 | 0 | 0 | 0 | 0 | 1 | 0 | 0 | 0 | 0 | 0 | 0 | 0 | 0 | 0 | 0 | 0 | 0 | 0 | 0 | 0 | 0 | 0 | Gm11756 |
| Gm11808 | 3222 | 4426 | 4469 | 4368 | 3883 | 4326 | 2843 | 2937 | 3226 | 3369 | 3722 | 3526 | 4660 | 2913 | 3204 | 3784 | 3971 | 4092 | 4975 | 5350 | 3244 | 2562 | 3572 | 3656 | 3692 | 4719 | 5033 | 3258 | Gm11808 |
| Gm11992 | 159 | 204 | 154 | 142 | 97 | 162 | 124 | 45 | 44 | 55 | 31 | 52 | 51 | 21 | 187 | 175 | 136 | 167 | 109 | 134 | 97 | 33 | 67 | 35 | 44 | 57 | 34 | 37 | Gm11992 |
| Gm12166 | 137 | 118 | 110 | 100 | 79 | 123 | 102 | 85 | 82 | 83 | 71 | 72 | 66 | 80 | 157 | 146 | 109 | 139 | 108 | 126 | 108 | 84 | 89 | 76 | 73 | 88 | 80 | 77 | Gm12166 |
| Gm12169 | 1 | 0 | 1 | 0 | 0 | 0 | 0 | 0 | 0 | 0 | 0 | 0 | 0 | 0 | 0 | 0 | 0 | 0 | 0 | 0 | 0 | 0 | 0 | 0 | 0 | 0 | 0 | 0 | Gm12169 |
| Gm12184 | 761 | 807 | 644 | 543 | 478 | 517 | 466 | 1183 | 1076 | 1085 | 911 | 1404 | 1149 | 1249 | 587 | 557 | 628 | 405 | 569 | 590 | 481 | 877 | 934 | 1189 | 1112 | 1454 | 1239 | 1257 | Gm12184 |
| Gm12185 | 1 | 1 | 2 | 3 | 0 | 0 | 0 | 0 | 2 | 4 | 0 | 5 | 0 | 2 | 0 | 0 | 0 | 3 | 0 | 0 | 0 | 3 | 3 | 2 | 2 | 2 | 1 | 0 | Gm12185 |
| Gm12216 | 6 | 2 | 0 | 1 | 2 | 0 | 2 | 4 | 6 | 6 | 10 | 7 | 7 | 6 | 1 | 2 | 0 | 0 | 0 | 3 | 1 | 12 | 10 | 1 | 14 | 4 | 8 | 4 | Gm12216 |
| Gm12248 | 0 | 0 | 0 | 0 | 2 | 0 | 0 | 2 | 2 | 1 | 0 | 0 | 0 | 0 | 3 | 0 | 0 | 0 | 0 | 0 | 0 | 0 | 2 | 0 | 2 | 0 | 2 | 0 | Gm12248 |
| Gm12253 | 1 | 0 | 0 | 0 | 0 | 0 | 0 | 0 | 0 | 11 | 1 | 0 | 0 | 0 | 1 | 0 | 0 | 0 | 0 | 0 | 0 | 2 | 1 | 0 | 0 | 0 | 0 | 0 | Gm12253 |
| Gm12258 | 3278 | 3164 | 2337 | 2878 | 3018 | 3216 | 3033 | 3486 | 2923 | 2700 | 2581 | 2467 | 2283 | 2475 | 3575 | 3291 | 2593 | 2862 | 2718 | 3425 | 3075 | 3034 | 2935 | 2585 | 2931 | 2176 | 2372 | 2313 | Gm12258 |
| Gm12353 | 57 | 35 | 27 | 70 | 65 | 52 | 53 | 42 | 39 | 45 | 21 | 37 | 45 | 48 | 45 | 47 | 31 | 30 | 61 | 73 | 72 | 55 | 39 | 38 | 51 | 32 | 40 | 39 | Gm12353 |
| Gm12394 | 0 | 1 | 9 | 0 | 0 | 0 | 0 | 0 | 0 | 1 | 0 | 0 | 1 | 0 | 7 | 0 | 0 | 0 | 0 | 0 | 0 | 5 | 4 | 0 | 0 | 0 | 0 | 0 | Gm12394 |
| Gm12666 | 0 | 3 | 4 | 1 | 1 | 1 | 6 | 15 | 4 | 6 | 6 | 9 | 3 | 14 | 3 | 2 | 14 | 2 | 1 | 0 | 6 | 3 | 6 | 2 | 7 | 8 | 5 | 6 | Gm12666 |
| Gm12689 | 32 | 28 | 46 | 49 | 38 | 45 | 35 | 29 | 26 | 28 | 29 | 37 | 42 | 27 | 20 | 36 | 48 | 54 | 69 | 35 | 38 | 35 | 32 | 41 | 27 | 30 | 38 | 27 | Gm12689 |
| Gm12695 | 0 | 3 | 0 | 0 | 4 | 0 | 1 | 5 | 7 | 12 | 5 | 4 | 2 | 4 | 0 | 2 | 1 | 0 | 0 | 2 | 9 | 0 | 8 | 3 | 2 | 8 | 0 | 1 | Gm12695 |
| Gm128 | 13 | 12 | 17 | 11 | 11 | 6 | 7 | 10 | 19 | 15 | 15 | 5 | 8 | 3 | 1 | 16 | 13 | 17 | 8 | 6 | 13 | 13 | 19 | 14 | 7 | 9 | 2 | 6 | Gm128 |
| Gm12830 | 0 | 1 | 0 | 0 | 0 | 0 | 0 | 0 | 1 | 0 | 0 | 2 | 1 | 3 | 4 | 3 | 0 | 0 | 1 | 1 | 1 | 0 | 0 | 3 | 2 | 3 | 0 | 5 | Gm12830 |
| Gm12845 | 0 | 0 | 1 | 0 | 2 | 0 | 0 | 0 | 0 | 0 | 0 | 1 | 1 | 2 | 0 | 0 | 0 | 2 | 0 | 3 | 7 | 0 | 0 | 0 | 2 | 0 | 0 | 0 | Gm12845 |
| Gm13030 | 0 | 0 | 0 | 0 | 0 | 0 | 0 | 1 | 0 | 0 | 0 | 0 | 0 | 0 | 1 | 2 | 6 | 3 | 0 | 0 | 1 | 0 | 0 | 0 | 0 | 0 | 0 | 0 | Gm13030 |
| Gm13090 | 2 | 0 | 0 | 1 | 2 | 0 | 0 | 0 | 0 | 0 | 0 | 0 | 0 | 0 | 0 | 1 | 0 | 0 | 0 | 0 | 0 | 0 | 1 | 0 | 0 | 1 | 0 | 0 | Gm13090 |
| Gm13199 | 203 | 246 | 264 | 316 | 319 | 339 | 391 | 117 | 80 | 104 | 86 | 99 | 73 | 125 | 171 | 232 | 326 | 353 | 309 | 272 | 390 | 129 | 65 | 140 | 146 | 115 | 103 | 107 | Gm13199 |
| Gm13271 | 0 | 0 | 0 | 0 | 0 | 0 | 0 | 0 | 0 | 0 | 0 | 2 | 0 | 0 | 0 | 0 | 0 | 0 | 0 | 0 | 0 | 0 | 0 | 0 | 3 | 0 | 0 | 0 | Gm13271 |
| Gm13272 | 1 | 0 | 5 | 5 | 0 | 0 | 1 | 0 | 0 | 0 | 0 | 0 | 0 | 0 | 0 | 2 | 0 | 1 | 1 | 0 | 0 | 0 | 1 | 0 | 0 | 2 | 1 | 0 | Gm13272 |
| Gm13276 | 1 | 2 | 0 | 0 | 0 | 0 | 0 | 0 | 0 | 0 | 0 | 0 | 1 | 0 | 1 | 1 | 1 | 3 | 0 | 0 | 0 | 0 | 0 | 0 | 0 | 0 | 0 | 0 | Gm13276 |
| Gm13277 | 0 | 0 | 0 | 0 | 0 | 0 | 0 | 0 | 0 | 0 | 0 | 0 | 0 | 0 | 0 | 0 | 0 | 0 | 0 | 1 | 0 | 0 | 0 | 0 | 0 | 0 | 0 | 0 | Gm13277 |
| Gm13283 | 8 | 3 | 9 | 6 | 1 | 7 | 0 | 1 | 2 | 0 | 5 | 1 | 3 | 0 | 9 | 5 | 8 | 6 | 5 | 2 | 4 | 2 | 1 | 0 | 3 | 4 | 0 | 0 | Gm13283 |
| Gm13285 | 1 | 1 | 1 | 0 | 1 | 0 | 2 | 0 | 0 | 0 | 0 | 0 | 1 | 0 | 0 | 0 | 0 | 0 | 0 | 0 | 0 | 0 | 0 | 0 | 0 | 0 | 0 | 1 | Gm13285 |
| Gm13287 | 1 | 1 | 3 | 0 | 0 | 2 | 0 | 0 | 0 | 1 | 0 | 0 | 0 | 1 | 1 | 1 | 0 | 0 | 1 | 0 | 0 | 0 | 0 | 0 | 0 | 0 | 0 | 0 | Gm13287 |
| Gm13288 | 2 | 3 | 0 | 0 | 0 | 0 | 0 | 2 | 1 | 0 | 0 | 2 | 2 | 1 | 0 | 3 | 1 | 0 | 0 | 3 | 2 | 3 | 1 | 0 | 0 | 0 | 0 | 0 | Gm13288 |
| Gm13304 | 3 | 0 | 0 | 0 | 0 | 0 | 0 | 0 | 0 | 0 | 0 | 0 | 0 | 0 | 0 | 0 | 0 | 0 | 0 | 0 | 0 | 0 | 1 | 0 | 0 | 0 | 0 | 0 | Gm13304 |
| Gm13305 | 2 | 0 | 0 | 0 | 0 | 0 | 0 | 0 | 0 | 4 | 0 | 0 | 1 | 1 | 0 | 0 | 0 | 0 | 2 | 0 | 0 | 0 | 0 | 0 | 0 | 0 | 0 | 0 | Gm13305 |
| Gm13306 | 5 | 2 | 8 | 3 | 7 | 3 | 8 | 6 | 1 | 1 | 3 | 4 | 0 | 6 | 1 | 2 | 8 | 0 | 2 | 2 | 10 | 3 | 6 | 5 | 8 | 3 | 0 | 6 | Gm13306 |
| Gm13398 | 23 | 23 | 33 | 31 | 27 | 28 | 41 | 10 | 15 | 5 | 27 | 13 | 15 | 17 | 21 | 21 | 21 | 46 | 28 | 18 | 29 | 12 | 21 | 20 | 17 | 11 | 16 | 24 | Gm13398 |
| Gm13420 | 3 | 6 | 1 | 9 | 8 | 1 | 3 | 0 | 3 | 3 | 0 | 1 | 5 | 1 | 4 | 5 | 1 | 6 | 3 | 4 | 1 | 2 | 1 | 2 | 2 | 0 | 5 | 1 | Gm13420 |
| Gm13539 | 0 | 0 | 0 | 0 | 0 | 0 | 0 | 0 | 0 | 2 | 0 | 0 | 0 | 0 | 0 | 0 | 0 | 0 | 0 | 0 | 0 | 0 | 1 | 0 | 0 | 0 | 2 | 0 | Gm13539 |
| Gm13547 | 2 | 0 | 0 | 0 | 0 | 0 | 0 | 0 | 0 | 0 | 0 | 0 | 0 | 0 | 0 | 0 | 0 | 1 | 3 | 0 | 1 | 0 | 0 | 1 | 1 | 0 | 0 | 2 | Gm13547 |
| Gm136 | 1 | 1 | 6 | 1 | 1 | 4 | 7 | 0 | 2 | 0 | 5 | 2 | 0 | 2 | 0 | 3 | 1 | 2 | 7 | 1 | 2 | 2 | 3 | 2 | 0 | 1 | 2 | 4 | Gm136 |
| Gm13694 | 0 | 0 | 1 | 0 | 0 | 2 | 0 | 2 | 0 | 0 | 1 | 2 | 1 | 1 | 3 | 0 | 1 | 3 | 0 | 0 | 1 | 2 | 0 | 0 | 0 | 0 | 0 | 0 | Gm13694 |
| Gm13696 | 0 | 0 | 0 | 0 | 0 | 0 | 0 | 0 | 0 | 0 | 1 | 1 | 0 | 0 | 0 | 0 | 0 | 0 | 0 | 0 | 0 | 0 | 0 | 0 | 0 | 0 | 0 | 0 | Gm13696 |
| Gm13698 | 0 | 0 | 0 | 0 | 0 | 0 | 0 | 1 | 0 | 0 | 0 | 0 | 0 | 0 | 0 | 0 | 1 | 0 | 0 | 0 | 0 | 0 | 1 | 0 | 1 | 0 | 0 | 0 | Gm13698 |
| Gm13889 | 187 | 218 | 199 | 312 | 376 | 306 | 375 | 332 | 388 | 454 | 398 | 510 | 217 | 639 | 173 | 117 | 283 | 298 | 424 | 267 | 396 | 399 | 258 | 629 | 593 | 576 | 344 | 409 | Gm13889 |
| Gm13941 | 0 | 0 | 0 | 0 | 2 | 0 | 0 | 0 | 0 | 0 | 0 | 0 | 0 | 0 | 0 | 0 | 0 | 0 | 0 | 0 | 0 | 0 | 0 | 0 | 0 | 0 | 0 | 0 | Gm13941 |
| Gm14124 | 9 | 12 | 7 | 13 | 7 | 11 | 12 | 4 | 12 | 19 | 14 | 18 | 15 | 5 | 8 | 12 | 7 | 11 | 20 | 18 | 15 | 18 | 10 | 6 | 15 | 7 | 5 | 7 | Gm14124 |
| Gm14137 | 16 | 14 | 3 | 7 | 3 | 8 | 8 | 2 | 14 | 13 | 0 | 2 | 5 | 5 | 9 | 11 | 1 | 18 | 1 | 3 | 4 | 2 | 9 | 4 | 1 | 1 | 8 | 0 | Gm14137 |
| Gm14139 | 1 | 3 | 1 | 7 | 0 | 6 | 0 | 1 | 4 | 0 | 4 | 1 | 0 | 2 | 0 | 2 | 0 | 3 | 0 | 0 | 2 | 0 | 2 | 2 | 0 | 0 | 0 | 0 | Gm14139 |
| Gm14147 | 0 | 0 | 0 | 0 | 0 | 0 | 0 | 0 | 0 | 0 | 0 | 0 | 1 | 0 | 0 | 0 | 0 | 0 | 0 | 0 | 0 | 0 | 0 | 0 | 0 | 0 | 0 | 0 | Gm14147 |
| Gm14226 | 322 | 50 | 23 | 41 | 12 | 24 | 23 | 348 | 404 | 278 | 230 | 27 | 105 | 47 | 59 | 88 | 20 | 15 | 11 | 13 | 14 | 28 | 467 | 228 | 15 | 419 | 22 | 26 | Gm14226 |
| Gm14288 | 227 | 176 | 111 | 109 | 146 | 284 | 115 | 309 | 182 | 146 | 216 | 162 | 217 | 185 | 344 | 266 | 134 | 151 | 138 | 212 | 139 | 164 | 185 | 140 | 196 | 119 | 208 | 134 | Gm14288 |
| Gm14295 | 631 | 448 | 311 | 351 | 361 | 410 | 411 | 409 | 452 | 363 | 358 | 459 | 412 | 326 | 728 | 639 | 612 | 383 | 396 | 640 | 445 | 349 | 476 | 320 | 345 | 279 | 345 | 282 | Gm14295 |
| Gm14296 | 330 | 248 | 47 | 270 | 313 | 266 | 275 | 248 | 286 | 44 | 341 | 355 | 90 | 289 | 256 | 383 | 297 | 242 | 197 | 344 | 341 | 441 | 325 | 225 | 234 | 379 | 345 | 399 | Gm14296 |
| Gm14305 | 1250 | 730 | 735 | 696 | 813 | 962 | 883 | 911 | 854 | 664 | 699 | 749 | 750 | 695 | 1200 | 1013 | 788 | 900 | 712 | 969 | 875 | 700 | 832 | 643 | 679 | 510 | 686 | 723 | Gm14305 |

| | | | | | | | | | | | | | | | | | | | | | | | | | | | | | |
|---|---|---|---|---|---|---|---|---|---|---|---|---|---|---|---|---|---|---|---|---|---|---|---|---|---|---|---|---|---|
| Gm14308 | 39 | 20 | 17 | 13 | 11 | 25 | 18 | 51 | 16 | 7 | 11 | 42 | 22 | 17 | 30 | 58 | 13 | 33 | 28 | 43 | 29 | 17 | 38 | 19 | 20 | 8 | 24 | 15 | Gm14308 |
| Gm14322 | 494 | 343 | 475 | 411 | 430 | 392 | 447 | 358 | 512 | 380 | 324 | 312 | 499 | 316 | 376 | 475 | 599 | 551 | 356 | 455 | 302 | 302 | 446 | 266 | 361 | 245 | 457 | 452 | Gm14322 |
| Gm14325 | 1909 | 1378 | 1788 | 1593 | 1419 | 1693 | 1485 | 1633 | 1584 | 1390 | 1396 | 1480 | 1516 | 1453 | 1949 | 1930 | 1471 | 1477 | 1466 | 1896 | 1541 | 1221 | 1627 | 1279 | 1305 | 1462 | 1547 | 1584 | Gm14325 |
| Gm14326 | 1017 | 763 | 908 | 604 | 721 | 1077 | 723 | 1123 | 838 | 648 | 984 | 716 | 755 | 644 | 1030 | 994 | 932 | 579 | 704 | 1098 | 966 | 744 | 862 | 733 | 587 | 767 | 932 | 663 | Gm14326 |
| Gm14327 | 32 | 13 | 11 | 20 | 27 | 12 | 21 | 27 | 40 | 22 | 32 | 28 | 49 | 21 | 34 | 34 | 14 | 13 | 26 | 23 | 13 | 32 | 31 | 12 | 24 | 8 | 15 | 17 | Gm14327 |
| Gm14391 | 335 | 50 | 46 | 100 | 43 | 193 | 44 | 99 | 97 | 146 | 43 | 115 | 79 | 117 | 90 | 85 | 36 | 50 | 44 | 54 | 45 | 60 | 70 | 96 | 46 | 32 | 66 | 34 | Gm14391 |
| Gm14393 | 15 | 16 | 27 | 32 | 20 | 24 | 21 | 16 | 18 | 8 | 13 | 13 | 17 | 12 | 24 | 23 | 35 | 28 | 22 | 14 | 32 | 25 | 12 | 14 | 25 | 14 | 20 | 15 | Gm14393 |
| Gm14399 | 768 | 545 | 478 | 410 | 627 | 725 | 548 | 636 | 677 | 485 | 634 | 519 | 511 | 514 | 774 | 713 | 579 | 626 | 549 | 729 | 573 | 632 | 678 | 463 | 480 | 321 | 610 | 417 | Gm14399 |
| Gm14401 | 2 | 0 | 0 | 3 | 0 | 0 | 1 | 2 | 4 | 1 | 2 | 0 | 2 | 1 | 3 | 3 | 0 | 1 | 4 | 6 | 0 | 0 | 4 | 1 | 2 | 0 | 2 | 2 | Gm14401 |
| Gm14403 | 438 | 383 | 330 | 262 | 329 | 361 | 324 | 231 | 285 | 253 | 234 | 209 | 290 | 208 | 401 | 487 | 289 | 383 | 286 | 301 | 239 | 246 | 296 | 185 | 151 | 205 | 175 | 139 | Gm14403 |
| Gm14406 | 0 | 2 | 1 | 12 | 5 | 3 | 3 | 0 | 1 | 3 | 3 | 3 | 1 | 3 | 1 | 1 | 0 | 7 | 7 | 5 | 2 | 5 | 1 | 0 | 3 | 4 | 1 | 1 | Gm14406 |
| Gm14408 | 1 | 0 | 0 | 2 | 0 | 0 | 0 | 1 | 0 | 0 | 0 | 0 | 0 | 4 | 0 | 0 | 0 | 0 | 0 | 0 | 2 | 0 | 1 | 0 | 0 | 0 | 0 | 0 | Gm14408 |
| Gm14409 | 3 | 2 | 3 | 13 | 3 | 6 | 2 | 0 | 2 | 1 | 2 | 1 | 1 | 1 | 4 | 5 | 1 | 4 | 2 | 7 | 0 | 5 | 4 | 0 | 1 | 1 | 2 | 1 | Gm14409 |
| Gm14410 | 1498 | 1040 | 1333 | 1056 | 1103 | 1360 | 1253 | 1272 | 1213 | 1144 | 1267 | 1173 | 1319 | 1189 | 1435 | 1151 | 1305 | 953 | 1281 | 1464 | 1126 | 1081 | 1172 | 900 | 1044 | 1099 | 1330 | 1101 | Gm14410 |
| Gm14412 | 122 | 81 | 37 | 47 | 84 | 102 | 74 | 40 | 68 | 56 | 52 | 33 | 58 | 39 | 113 | 133 | 59 | 85 | 59 | 90 | 75 | 48 | 50 | 32 | 33 | 22 | 21 | 21 | Gm14412 |
| Gm14418 | 340 | 223 | 188 | 190 | 295 | 318 | 335 | 233 | 236 | 258 | 333 | 357 | 352 | 243 | 563 | 459 | 284 | 263 | 274 | 299 | 403 | 284 | 296 | 411 | 193 | 414 | 218 | 189 | Gm14418 |
| Gm14419 | 16 | 11 | 6 | 8 | 102 | 102 | 9 | 93 | 25 | 12 | 6 | 9 | 29 | 6 | 18 | 119 | 12 | 8 | 6 | 10 | 11 | 40 | 11 | 4 | 21 | 11 | 23 | 75 | Gm14419 |
| Gm14434 | 6 | 147 | 235 | 4 | 37 | 9 | 45 | 20 | 79 | 4 | 421 | 619 | 36 | 21 | 17 | 53 | 7 | 39 | 53 | 128 | 24 | 541 | 27 | 52 | 129 | 226 | 1232 | 28 | Gm14434 |
| Gm14440 | 18 | 18 | 10 | 21 | 11 | 36 | 19 | 27 | 30 | 32 | 25 | 24 | 26 | 14 | 11 | 24 | 10 | 6 | 38 | 23 | 14 | 7 | 21 | 17 | 19 | 7 | 31 | 9 | Gm14440 |
| Gm14443 | 340 | 287 | 322 | 276 | 373 | 406 | 459 | 264 | 233 | 205 | 291 | 218 | 253 | 239 | 355 | 368 | 268 | 284 | 327 | 380 | 391 | 274 | 244 | 209 | 207 | 197 | 192 | 243 | Gm14443 |
| Gm14444 | 0 | 2 | 1 | 0 | 2 | 2 | 0 | 0 | 2 | 1 | 3 | 1 | 0 | 0 | 0 | 5 | 1 | 3 | 4 | 1 | 1 | 2 | 1 | 2 | 6 | 0 | 0 | 1 | Gm14444 |
| Gm14569 | 6 | 12 | 0 | 3 | 13 | 6 | 4 | 2 | 7 | 6 | 1 | 4 | 0 | 2 | 3 | 4 | 2 | 3 | 2 | 4 | 4 | 5 | 3 | 0 | 10 | 0 | 1 | 0 | Gm14569 |
| Gm14692 | 0 | 0 | 0 | 0 | 0 | 0 | 0 | 0 | 0 | 0 | 0 | 0 | 0 | 0 | 0 | 0 | 0 | 0 | 0 | 0 | 0 | 3 | 0 | 0 | 0 | 0 | 0 | 0 | Gm14692 |
| Gm14698 | 428 | 242 | 246 | 277 | 318 | 275 | 334 | 297 | 252 | 180 | 264 | 207 | 243 | 217 | 456 | 323 | 238 | 229 | 245 | 352 | 291 | 274 | 250 | 250 | 217 | 205 | 201 | 224 | Gm14698 |
| Gm14862 | 5 | 1 | 1 | 2 | 0 | 1 | 3 | 2 | 4 | 3 | 1 | 9 | 31 | 13 | 4 | 3 | 0 | 0 | 1 | 6 | 2 | 0 | 3 | 0 | 4 | 16 | 3 | 34 | Gm14862 |
| Gm15023 | 0 | 3 | 0 | 0 | 0 | 0 | 0 | 0 | 0 | 0 | 1 | 0 | 0 | 0 | 0 | 0 | 0 | 0 | 0 | 0 | 0 | 0 | 0 | 0 | 0 | 0 | 0 | 0 | Gm15023 |
| Gm15097 | 0 | 0 | 0 | 0 | 0 | 0 | 0 | 0 | 0 | 0 | 0 | 0 | 0 | 0 | 0 | 0 | 0 | 0 | 0 | 0 | 0 | 0 | 0 | 2 | 0 | 0 | 0 | 0 | Gm15097 |
| Gm15128 | 0 | 0 | 0 | 0 | 0 | 0 | 0 | 0 | 0 | 0 | 1 | 0 | 0 | 0 | 0 | 0 | 0 | 0 | 0 | 0 | 0 | 0 | 0 | 0 | 0 | 0 | 0 | 0 | Gm15128 |
| Gm15155 | 3 | 2 | 1 | 0 | 4 | 0 | 8 | 1 | 7 | 2 | 0 | 1 | 2 | 0 | 1 | 5 | 1 | 0 | 1 | 5 | 3 | 2 | 5 | 3 | 6 | 0 | 2 | 2 | Gm15155 |
| Gm15217 | 0 | 0 | 0 | 0 | 0 | 0 | 0 | 1 | 0 | 4 | 0 | 0 | 0 | 0 | 0 | 1 | 0 | 0 | 0 | 0 | 0 | 0 | 0 | 0 | 0 | 0 | 0 | 0 | Gm15217 |
| Gm15262 | 0 | 0 | 0 | 0 | 0 | 0 | 0 | 0 | 0 | 0 | 0 | 0 | 0 | 0 | 0 | 0 | 0 | 0 | 0 | 0 | 0 | 0 | 2 | 0 | 0 | 0 | 0 | 0 | Gm15262 |
| Gm15319 | 35 | 60 | 8 | 17 | 32 | 57 | 37 | 23 | 18 | 11 | 18 | 8 | 18 | 14 | 74 | 44 | 12 | 42 | 39 | 68 | 52 | 28 | 8 | 6 | 12 | 2 | 16 | 15 | Gm15319 |
| Gm15433 | 15 | 15 | 10 | 5 | 16 | 17 | 12 | 1 | 5 | 0 | 0 | 11 | 1 | 6 | 8 | 4 | 9 | 19 | 5 | 12 | 11 | 0 | 2 | 4 | 7 | 4 | 1 | 2 | Gm15433 |
| Gm15446 | 2472 | 2068 | 2016 | 2363 | 2797 | 2991 | 3051 | 2328 | 2053 | 1865 | 1993 | 2070 | 1952 | 1910 | 2773 | 2327 | 2033 | 2281 | 2662 | 2878 | 3040 | 2187 | 2003 | 1627 | 2053 | 1577 | 1986 | 1842 | Gm15446 |
| Gm15557 | 0 | 0 | 0 | 0 | 0 | 0 | 0 | 0 | 0 | 0 | 0 | 0 | 1 | 0 | 0 | 0 | 0 | 0 | 0 | 0 | 0 | 0 | 2 | 0 | 0 | 0 | 0 | 0 | Gm15557 |
| Gm15737 | 0 | 0 | 0 | 0 | 0 | 1 | 1 | 0 | 1 | 0 | 5 | 3 | 0 | 0 | 0 | 1 | 0 | 0 | 0 | 0 | 0 | 0 | 3 | 0 | 0 | 0 | 0 | 0 | Gm15737 |
| Gm1604b | 0 | 0 | 0 | 0 | 2 | 0 | 0 | 0 | 0 | 1 | 0 | 1 | 0 | 1 | 0 | 0 | 0 | 2 | 0 | 0 | 1 | 0 | 0 | 0 | 0 | 0 | 0 | 0 | Gm1604b |
| Gm16181 | 1 | 0 | 0 | 3 | 0 | 2 | 0 | 0 | 0 | 0 | 0 | 0 | 0 | 0 | 0 | 0 | 0 | 0 | 0 | 0 | 1 | 0 | 0 | 0 | 0 | 5 | 0 | 2 | Gm16181 |
| Gm16223 | 1 | 3 | 5 | 1 | 3 | 3 | 1 | 1 | 2 | 5 | 7 | 5 | 5 | 3 | 1 | 0 | 3 | 0 | 5 | 1 | 1 | 10 | 7 | 2 | 7 | 0 | 5 | 1 | Gm16223 |
| Gm16253 | 6 | 0 | 10 | 0 | 8 | 4 | 1 | 4 | 6 | 2 | 11 | 1 | 9 | 4 | 4 | 2 | 0 | 4 | 2 | 3 | 0 | 10 | 3 | 7 | 3 | 7 | 3 | 0 | Gm16253 |
| Gm16434 | 0 | 0 | 6 | 0 | 0 | 0 | 0 | 0 | 0 | 0 | 0 | 0 | 0 | 0 | 0 | 1 | 0 | 1 | 0 | 0 | 0 | 0 | 0 | 0 | 0 | 0 | 0 | 0 | Gm16434 |
| Gm16440 | 2 | 3 | 11 | 4 | 14 | 1 | 1 | 6 | 10 | 10 | 4 | 1 | 15 | 6 | 0 | 3 | 16 | 3 | 6 | 4 | 5 | 7 | 3 | 5 | 4 | 4 | 7 | 2 | Gm16440 |
| Gm1673 | 1129 | 1493 | 1120 | 1501 | 1572 | 1224 | 1580 | 1508 | 1215 | 1576 | 1329 | 1369 | 1344 | 1580 | 1159 | 812 | 1588 | 1245 | 1511 | 1557 | 1539 | 1389 | 1210 | 1648 | 1740 | 1507 | 1511 | 1480 | Gm1673 |
| Gm16867 | 145 | 134 | 163 | 74 | 97 | 76 | 49 | 39 | 89 | 50 | 54 | 85 | 198 | 80 | 69 | 157 | 86 | 79 | 109 | 130 | 48 | 62 | 98 | 71 | 61 | 106 | 142 | 154 | Gm16867 |
| Gm17067 | 0 | 0 | 0 | 0 | 0 | 0 | 0 | 0 | 0 | 0 | 0 | 0 | 0 | 0 | 0 | 0 | 0 | 0 | 0 | 0 | 0 | 0 | 1 | 0 | 0 | 0 | 0 | 0 | Gm17067 |
| Gm17266 | 2 | 0 | 3 | 0 | 0 | 0 | 2 | 5 | 1 | 0 | 0 | 0 | 1 | 2 | 0 | 1 | 0 | 0 | 1 | 0 | 2 | 5 | 0 | 0 | 0 | 0 | 1 | 0 | Gm17266 |
| Gm17349 | 8 | 6 | 10 | 1 | 10 | 8 | 10 | 10 | 12 | 16 | 25 | 8 | 18 | 17 | 7 | 8 | 5 | 4 | 12 | 0 | 2 | 10 | 10 | 22 | 11 | 11 | 16 | 9 | Gm17349 |
| Gm17359 | 5 | 3 | 15 | 5 | 7 | 6 | 0 | 5 | 7 | 11 | 9 | 7 | 24 | 12 | 5 | 9 | 5 | 3 | 2 | 2 | 2 | 7 | 10 | 1 | 9 | 6 | 15 | 9 | Gm17359 |
| Gm17364 | 0 | 3 | 6 | 8 | 5 | 7 | 0 | 4 | 3 | 3 | 3 | 1 | 3 | 0 | 0 | 2 | 10 | 1 | 16 | 2 | 4 | 0 | 4 | 1 | 6 | 1 | 5 | 0 | Gm17364 |
| Gm17387 | 0 | 0 | 0 | 0 | 0 | 0 | 0 | 0 | 0 | 0 | 1 | 0 | 1 | 0 | 0 | 0 | 0 | 0 | 0 | 0 | 0 | 0 | 0 | 0 | 0 | 0 | 0 | 2 | Gm17387 |
| Gm17428 | 0 | 0 | 0 | 0 | 0 | 0 | 0 | 1 | 0 | 0 | 1 | 0 | 0 | 1 | 0 | 0 | 0 | 0 | 0 | 0 | 0 | 0 | 0 | 0 | 1 | 0 | 0 | 0 | Gm17428 |
| Gm17455 | 9 | 14 | 37 | 15 | 11 | 12 | 7 | 33 | 33 | 23 | 32 | 20 | 39 | 34 | 16 | 19 | 19 | 11 | 15 | 24 | 9 | 35 | 46 | 25 | 23 | 19 | 21 | 23 | Gm17455 |
| Gm17482 | 0 | 2 | 0 | 0 | 0 | 0 | 0 | 0 | 0 | 0 | 1 | 0 | 0 | 0 | 0 | 1 | 0 | 2 | 2 | 0 | 2 | 3 | 0 | 0 | 0 | 0 | 0 | 0 | Gm17482 |
| Gm17535 | 0 | 0 | 0 | 0 | 0 | 0 | 0 | 0 | 0 | 0 | 0 | 0 | 0 | 0 | 0 | 0 | 0 | 0 | 0 | 0 | 0 | 0 | 0 | 0 | 1 | 0 | 0 | 0 | Gm17535 |
| Gm17542 | 0 | 0 | 0 | 0 | 2 | 0 | 0 | 0 | 0 | 0 | 0 | 0 | 0 | 0 | 0 | 0 | 0 | 0 | 0 | 0 | 0 | 0 | 0 | 0 | 0 | 0 | 0 | 0 | Gm17542 |
| Gm17555 | 0 | 0 | 0 | 0 | 0 | 0 | 0 | 0 | 0 | 0 | 0 | 0 | 0 | 0 | 1 | 0 | 0 | 0 | 0 | 0 | 0 | 0 | 0 | 0 | 2 | 0 | 0 | 0 | Gm17555 |
| Gm17606 | 8 | 10 | 7 | 3 | 12 | 3 | 7 | 10 | 4 | 12 | 6 | 11 | 6 | 29 | 3 | 1 | 8 | 3 | 5 | 5 | 19 | 7 | 2 | 33 | 6 | 11 | 16 | 12 | Gm17606 |
| Gm17617 | 0 | 0 | 1 | 0 | 0 | 0 | 0 | 0 | 0 | 0 | 2 | 0 | 1 | 0 | 0 | 1 | 2 | 0 | 0 | 2 | 1 | 0 | 0 | 0 | 0 | 0 | 0 | 0 | Gm17617 |
| Gm17651 | 0 | 0 | 0 | 0 | 0 | 0 | 0 | 0 | 2 | 1 | 0 | 0 | 0 | 0 | 0 | 0 | 0 | 0 | 0 | 0 | 0 | 0 | 0 | 0 | 0 | 0 | 0 | 0 | Gm17651 |
| Gm17655 | 163 | 113 | 210 | 138 | 141 | 155 | 117 | 92 | 110 | 101 | 110 | 118 | 104 | 72 | 171 | 153 | 180 | 150 | 158 | 157 | 155 | 117 | 125 | 75 | 82 | 133 | 115 | 133 | Gm17655 |
| Gm17657 | 0 | 0 | 0 | 0 | 0 | 0 | 0 | 0 | 0 | 0 | 0 | 0 | 0 | 0 | 0 | 0 | 0 | 1 | 1 | 0 | 0 | 0 | 0 | 0 | 0 | 0 | 0 | 0 | Gm17657 |
| Gm17660 | 0 | 0 | 2 | 0 | 1 | 0 | 0 | 0 | 0 | 0 | 0 | 0 | 1 | 2 | 0 | 0 | 0 | 3 | 0 | 2 | 1 | 0 | 6 | 0 | 2 | 1 | 0 | 1 | Gm17660 |
| Gm17662 | 0 | 0 | 1 | 1 | 0 | 0 | 1 | 1 | 0 | 0 | 2 | 1 | 6 | 0 | 0 | 0 | 2 | 1 | 2 | 0 | 1 | 0 | 2 | 0 | 0 | 2 | 0 | 1 | Gm17662 |
| Gm17728 | 0 | 0 | 0 | 0 | 0 | 0 | 1 | 0 | 0 | 0 | 0 | 0 | 0 | 0 | 0 | 2 | 0 | 0 | 0 | 0 | 0 | 0 | 0 | 0 | 0 | 0 | 0 | 0 | Gm17728 |
| Gm17783 | 0 | 0 | 0 | 4 | 0 | 0 | 0 | 0 | 0 | 0 | 0 | 1 | 0 | 0 | 0 | 0 | 0 | 0 | 0 | 0 | 0 | 0 | 0 | 0 | 0 | 0 | 0 | 1 | Gm17783 |
| Gm17949 | 14 | 16 | 3 | 16 | 7 | 2 | 19 | 10 | 2 | 10 | 18 | 8 | 6 | 12 | 8 | 14 | 15 | 8 | 5 | 14 | 13 | 3 | 10 | 3 | 7 | 16 | 5 | 1 | Gm17949 |
| Gm18336 | 17 | 13 | 15 | 39 | 23 | 28 | 29 | 9 | 12 | 9 | 8 | 10 | 7 | 7 | 28 | 27 | 19 | 26 | 27 | 31 | 38 | 20 | 14 | 21 | 13 | 8 | 9 | 5 | Gm18336 |
| Gm19345 | 21 | 33 | 34 | 28 | 63 | 64 | 40 | 74 | 119 | 115 | 108 | 45 | 53 | 52 | 17 | 16 | 43 | 55 | 52 | 36 | 63 | 137 | 106 | 73 | 121 | 21 | 35 | 26 | Gm19345 |
| Gm19410 | 44 | 37 | 10 | 43 | 79 | 88 | 35 | 50 | 45 | 28 | 47 | 27 | 28 | 30 | 40 | 37 | 30 | 69 | 28 | 41 | 46 | 42 | 31 | 43 | 40 | 7 | 16 | 17 | Gm19410 |
| Gm19426 | 99 | 106 | 148 | 148 | 101 | 127 | 111 | 195 | 223 | 324 | 284 | 285 | 299 | 332 | 88 | 97 | 168 | 111 | 95 | 77 | 110 | 189 | 222 | 271 | 220 | 343 | 328 | 301 | Gm19426 |
| Gm19684 | 0 | 0 | 0 | 0 | 0 | 0 | 0 | 0 | 0 | 0 | 0 | 0 | 0 | 3 | 0 | 0 | 0 | 0 | 0 | 0 | 0 | 0 | 1 | 0 | 0 | 0 | 0 | 0 | Gm19684 |
| Gm1979 | 0 | 0 | 0 | 0 | 0 | 0 | 0 | 0 | 0 | 2 | 4 | 0 | 0 | 2 | 0 | 0 | 0 | 0 | 0 | 0 | 0 | 0 | 1 | 1 | 0 | 0 | 1 | 0 | Gm1979 |
| Gm19935 | 1 | 4 | 2 | 1 | 21 | 7 | 2 | 16 | 28 | 27 | 15 | 14 | 34 | 1 | 13 | 12 | 0 | 7 | 0 | 4 | 7 | 40 | 54 | 9 | 19 | 4 | 2 | 2 | Gm19935 |
| Gm19965 | 0 | 0 | 0 | 2 | 0 | 0 | 0 | 0 | 1 | 0 | 3 | 0 | 0 | 1 | 0 | 0 | 0 | 0 | 0 | 0 | 0 | 3 | 1 | 0 | 3 | 0 | 0 | 0 | Gm19965 |
| Gm2000 | 1418 | 1360 | 1219 | 1303 | 900 | 958 | 1306 | 1280 | 1084 | 1197 | 1151 | 1173 | 1161 | 1583 | 1423 | 1083 | 1132 | 1086 | 1153 | 1422 | 1403 | 792 | 1183 | 1340 | 1222 | 1251 | 1195 | 1542 | Gm2000 |
| Gm2004 | 68 | 231 | 400 | 239 | 17 | 50 | 146 | 59 | 255 | 31 | 214 | 198 | 335 | 234 | 80 | 64 | 163 | 22 | 237 | 150 | 149 | 69 | 80 | 235 | 79 | 48 | 33 | 45 | Gm2004 |
| Gm2007 | 1 | 0 | 1 | 0 | 0 | 2 | 7 | 0 | 0 | 0 | 0 | 1 | 2 | 0 | 0 | 0 | 0 | 0 | 2 | 18 | 1 | 2 | 0 | 1 | 0 | 0 | 0 | 0 | Gm2007 |
| Gm20075 | 1326 | 974 | 1031 | 844 | 901 | 862 | 1076 | 2229 | 1591 | 1422 | 1327 | 1880 | 1720 | 2014 | 1360 | 984 | 856 | 807 | 931 | 1162 | 987 | 1751 | 1693 | 1544 | 1619 | 2056 | 1733 | 2106 | Gm20075 |
| Gm2026 | 5 | 3 | 25 | 10 | 4 | 10 | 18 | 11 | 4 | 5 | 12 | 15 | 3 | 1 | 8 | 16 | 1 | 2 | 8 | 2 | 7 | 3 | 6 | 2 | 13 | 10 | 6 | 7 | Gm2026 |

| | | | | | | | | | | | | | | | | | | | | | | | | | | | | | |
|---|---|---|---|---|---|---|---|---|---|---|---|---|---|---|---|---|---|---|---|---|---|---|---|---|---|---|---|---|---|
| Gm20449 | 10 | 6 | 5 | 12 | 15 | 9 | 10 | 10 | 7 | 10 | 19 | 5 | 12 | 11 | 7 | 9 | 6 | 7 | 8 | 8 | 8 | 7 | 19 | 9 | 12 | 7 | 7 | 0 | Gm20449 |
| Gm20481 | 0 | 2 | 7 | 1 | 2 | 0 | 1 | 0 | 0 | 0 | 0 | 1 | 0 | 2 | 1 | 2 | 0 | 0 | 2 | 5 | 0 | 3 | 1 | 1 | 0 | 0 | 0 | 0 | Gm20481 |
| Gm20489 | 0 | 1 | 0 | 0 | 0 | 0 | 0 | 0 | 0 | 4 | 4 | 0 | 0 | 0 | 0 | 0 | 0 | 0 | 0 | 0 | 0 | 3 | 0 | 0 | 0 | 0 | 0 | 1 | Gm20489 |
| Gm20498 | 23 | 21 | 12 | 26 | 24 | 11 | 34 | 22 | 19 | 22 | 14 | 27 | 27 | 15 | 36 | 23 | 31 | 11 | 26 | 32 | 36 | 17 | 22 | 14 | 23 | 20 | 22 | 13 | Gm20498 |
| Gm20517 | 16 | 30 | 43 | 29 | 43 | 17 | 22 | 24 | 24 | 32 | 42 | 27 | 40 | 30 | 32 | 32 | 22 | 18 | 28 | 30 | 25 | 25 | 17 | 18 | 23 | 37 | 31 | 24 | Gm20517 |
| Gm20518 | 32 | 41 | 31 | 48 | 25 | 28 | 46 | 22 | 13 | 29 | 21 | 24 | 14 | 35 | 29 | 19 | 49 | 66 | 47 | 43 | 68 | 30 | 12 | 26 | 23 | 24 | 15 | 21 | Gm20518 |
| Gm20537 | 0 | 2 | 0 | 2 | 1 | 0 | 1 | 1 | 0 | 0 | 0 | 0 | 0 | 0 | 4 | 0 | 0 | 0 | 0 | 0 | 0 | 0 | 0 | 2 | 0 | 0 | 0 | 0 | Gm20537 |
| Gm20594 | 0 | 0 | 0 | 0 | 0 | 0 | 0 | 0 | 0 | 0 | 0 | 0 | 0 | 0 | 1 | 0 | 0 | 0 | 0 | 0 | 0 | 0 | 0 | 0 | 0 | 0 | 0 | 0 | Gm20594 |
| Gm20604 | 597 | 445 | 556 | 669 | 586 | 443 | 763 | 443 | 330 | 434 | 422 | 458 | 382 | 609 | 638 | 459 | 619 | 574 | 460 | 507 | 704 | 515 | 379 | 506 | 451 | 455 | 473 | 628 | Gm20604 |
| Gm20605 | 13 | 19 | 6 | 26 | 9 | 16 | 15 | 7 | 8 | 5 | 4 | 9 | 2 | 13 | 7 | 6 | 23 | 21 | 8 | 7 | 19 | 10 | 3 | 7 | 3 | 7 | 16 | 14 | Gm20605 |
| Gm20661 | 54 | 35 | 36 | 44 | 19 | 30 | 22 | 77 | 97 | 161 | 213 | 77 | 75 | 78 | 29 | 37 | 27 | 58 | 15 | 7 | 17 | 102 | 93 | 117 | 81 | 69 | 50 | 74 | Gm20661 |
| Gm20695 | 2 | 1 | 0 | 11 | 4 | 1 | 3 | 7 | 6 | 0 | 10 | 4 | 6 | 6 | 3 | 5 | 2 | 6 | 3 | 2 | 2 | 7 | 6 | 0 | 3 | 0 | 3 | 5 | Gm20695 |
| Gm20696 | 7 | 8 | 2 | 15 | 4 | 4 | 10 | 5 | 7 | 3 | 7 | 2 | 12 | 10 | 11 | 5 | 8 | 4 | 12 | 7 | 8 | 3 | 8 | 5 | 5 | 13 | 2 | 4 | Gm20696 |
| Gm20715 | 15 | 6 | 2 | 32 | 13 | 19 | 7 | 17 | 7 | 22 | 21 | 5 | 5 | 10 | 17 | 9 | 20 | 25 | 23 | 29 | 13 | 25 | 8 | 9 | 21 | 8 | 6 | 2 | Gm20715 |
| Gm20721 | 90 | 54 | 83 | 106 | 99 | 76 | 98 | 78 | 59 | 96 | 121 | 138 | 104 | 192 | 63 | 60 | 108 | 152 | 99 | 87 | 109 | 104 | 98 | 156 | 101 | 121 | 107 | 62 | Gm20721 |
| Gm20878 | 1 | 0 | 0 | 0 | 0 | 0 | 0 | 0 | 0 | 0 | 0 | 0 | 0 | 0 | 0 | 0 | 0 | 0 | 0 | 0 | 0 | 0 | 0 | 0 | 0 | 0 | 0 | 0 | Gm20878 |
| Gm20929 | 0 | 0 | 0 | 4 | 0 | 0 | 0 | 0 | 0 | 0 | 0 | 0 | 0 | 0 | 0 | 0 | 0 | 0 | 0 | 0 | 0 | 0 | 0 | 0 | 0 | 0 | 0 | 0 | Gm20929 |
| Gm21028 | 7 | 19 | 11 | 12 | 26 | 24 | 31 | 28 | 8 | 14 | 18 | 11 | 15 | 14 | 30 | 13 | 14 | 28 | 13 | 19 | 16 | 23 | 18 | 22 | 9 | 11 | 3 | 12 | Gm21028 |
| Gm21060 | 0 | 1 | 4 | 0 | 6 | 0 | 2 | 5 | 0 | 0 | 0 | 2 | 0 | 3 | 0 | 1 | 6 | 0 | 0 | 1 | 0 | 0 | 0 | 1 | 0 | 0 | 0 | 4 | Gm21060 |
| Gm21103 | 8 | 1 | 3 | 2 | 0 | 5 | 1 | 0 | 0 | 0 | 0 | 0 | 0 | 3 | 3 | 2 | 1 | 0 | 1 | 3 | 1 | 0 | 0 | 0 | 0 | 0 | 5 | 0 | Gm21103 |
| Gm21119 | 18 | 3 | 0 | 1 | 0 | 0 | 2 | 4 | 2 | 0 | 0 | 1 | 0 | 0 | 12 | 12 | 1 | 0 | 1 | 18 | 4 | 0 | 5 | 1 | 1 | 1 | 3 | 0 | Gm21119 |
| Gm2115 | 52 | 54 | 41 | 39 | 92 | 97 | 54 | 159 | 157 | 138 | 156 | 128 | 130 | 156 | 47 | 28 | 82 | 64 | 83 | 62 | 83 | 202 | 220 | 134 | 126 | 117 | 182 | 151 | Gm2115 |
| Gm21188 | 0 | 0 | 0 | 0 | 0 | 0 | 0 | 0 | 2 | 2 | 0 | 0 | 1 | 1 | 0 | 0 | 0 | 0 | 0 | 1 | 1 | 0 | 2 | 2 | 0 | 0 | 1 | 6 | Gm21188 |
| Gm21258 | 0 | 0 | 0 | 0 | 0 | 0 | 0 | 0 | 0 | 0 | 0 | 0 | 1 | 0 | 0 | 0 | 0 | 0 | 0 | 0 | 0 | 0 | 0 | 0 | 0 | 0 | 0 | 0 | Gm21258 |
| Gm21411 | 0 | 1 | 0 | 4 | 0 | 0 | 0 | 0 | 2 | 3 | 0 | 0 | 0 | 0 | 0 | 0 | 0 | 0 | 0 | 0 | 0 | 3 | 0 | 0 | 1 | 0 | 0 | 0 | Gm21411 |
| Gm21560 | 0 | 1 | 0 | 0 | 0 | 0 | 0 | 1 | 1 | 0 | 0 | 0 | 1 | 2 | 0 | 0 | 0 | 0 | 0 | 0 | 0 | 0 | 1 | 2 | 0 | 2 | 0 | 0 | Gm21560 |
| Gm21586 | 0 | 1 | 0 | 1 | 0 | 0 | 0 | 0 | 2 | 0 | 0 | 0 | 2 | 0 | 0 | 0 | 0 | 0 | 2 | 1 | 0 | 0 | 0 | 0 | 0 | 0 | 0 | 0 | Gm21586 |
| Gm21650 | 0 | 0 | 0 | 0 | 0 | 0 | 0 | 0 | 0 | 0 | 0 | 0 | 1 | 0 | 0 | 0 | 0 | 0 | 0 | 0 | 0 | 0 | 0 | 0 | 0 | 0 | 0 | 0 | Gm21650 |
| Gm21663 | 0 | 0 | 0 | 0 | 0 | 0 | 0 | 0 | 0 | 0 | 0 | 1 | 0 | 0 | 0 | 0 | 0 | 0 | 0 | 0 | 0 | 0 | 0 | 0 | 0 | 0 | 0 | 0 | Gm21663 |
| Gm21671 | 0 | 0 | 0 | 0 | 0 | 0 | 0 | 0 | 0 | 0 | 0 | 0 | 0 | 0 | 0 | 0 | 0 | 0 | 0 | 0 | 0 | 2 | 0 | 0 | 0 | 0 | 2 | 0 | Gm21671 |
| Gm21680 | 0 | 0 | 0 | 0 | 2 | 0 | 0 | 0 | 0 | 0 | 1 | 0 | 0 | 0 | 0 | 0 | 0 | 0 | 0 | 0 | 0 | 2 | 0 | 0 | 0 | 0 | 0 | 0 | Gm21680 |
| Gm21698 | 0 | 0 | 0 | 0 | 0 | 0 | 0 | 0 | 1 | 0 | 0 | 0 | 0 | 0 | 0 | 0 | 0 | 0 | 0 | 0 | 0 | 0 | 0 | 0 | 0 | 0 | 0 | 0 | Gm21698 |
| Gm21704 | 1 | 0 | 0 | 0 | 0 | 0 | 0 | 0 | 0 | 0 | 0 | 0 | 0 | 0 | 0 | 0 | 0 | 0 | 0 | 0 | 0 | 0 | 0 | 0 | 0 | 0 | 0 | 0 | Gm21704 |
| Gm2174 | 0 | 0 | 0 | 0 | 0 | 0 | 0 | 1 | 2 | 0 | 0 | 0 | 0 | 1 | 0 | 2 | 0 | 0 | 0 | 0 | 0 | 0 | 0 | 0 | 0 | 3 | 0 | 0 | Gm2174 |
| Gm21748 | 0 | 0 | 0 | 0 | 5 | 0 | 0 | 0 | 0 | 0 | 0 | 0 | 0 | 0 | 0 | 0 | 0 | 0 | 0 | 0 | 0 | 0 | 0 | 0 | 0 | 0 | 0 | 0 | Gm21748 |
| Gm21750 | 1 | 3 | 0 | 4 | 0 | 0 | 1 | 0 | 0 | 0 | 0 | 0 | 0 | 0 | 1 | 0 | 1 | 0 | 2 | 0 | 2 | 0 | 0 | 0 | 0 | 0 | 0 | 0 | Gm21750 |
| Gm21818 | 0 | 0 | 0 | 0 | 0 | 0 | 0 | 0 | 0 | 0 | 0 | 0 | 0 | 0 | 0 | 0 | 0 | 0 | 0 | 0 | 0 | 0 | 1 | 0 | 0 | 0 | 0 | 0 | Gm21818 |
| Gm21830 | 0 | 0 | 0 | 0 | 1 | 0 | 0 | 0 | 1 | 0 | 0 | 0 | 0 | 0 | 0 | 0 | 0 | 0 | 0 | 0 | 0 | 0 | 0 | 0 | 0 | 0 | 0 | 0 | Gm21830 |
| Gm21833 | 0 | 0 | 0 | 0 | 0 | 0 | 1 | 0 | 0 | 1 | 0 | 0 | 0 | 0 | 0 | 0 | 0 | 0 | 0 | 0 | 0 | 0 | 0 | 0 | 0 | 0 | 0 | 0 | Gm21833 |
| Gm21885 | 0 | 0 | 0 | 0 | 0 | 0 | 0 | 0 | 0 | 0 | 0 | 0 | 0 | 0 | 0 | 0 | 0 | 0 | 0 | 0 | 0 | 3 | 0 | 0 | 0 | 2 | 0 | 0 | Gm21885 |
| Gm21886 | 0 | 0 | 0 | 0 | 0 | 0 | 0 | 0 | 0 | 0 | 0 | 0 | 0 | 1 | 0 | 0 | 0 | 0 | 0 | 0 | 0 | 0 | 0 | 0 | 0 | 0 | 0 | 0 | Gm21886 |
| Gm21915 | 0 | 0 | 0 | 0 | 0 | 0 | 0 | 0 | 2 | 0 | 0 | 0 | 3 | 0 | 0 | 0 | 0 | 0 | 0 | 0 | 0 | 0 | 0 | 1 | 0 | 0 | 0 | 0 | Gm21915 |
| Gm21953 | 0 | 1 | 3 | 2 | 3 | 2 | 4 | 2 | 2 | 1 | 1 | 0 | 0 | 0 | 1 | 0 | 0 | 1 | 1 | 3 | 0 | 3 | 1 | 5 | 0 | 2 | 0 | 0 | Gm21953 |
| Gm21958 | 0 | 0 | 0 | 1 | 3 | 0 | 1 | 0 | 0 | 0 | 0 | 4 | 0 | 0 | 1 | 0 | 2 | 0 | 2 | 0 | 0 | 0 | 0 | 4 | 0 | 0 | 0 | 0 | Gm21958 |
| Gm21969 | 13 | 6 | 4 | 6 | 4 | 6 | 12 | 0 | 6 | 1 | 1 | 0 | 0 | 1 | 4 | 5 | 3 | 7 | 1 | 13 | 11 | 0 | 0 | 0 | 1 | 0 | 0 | 0 | Gm21969 |
| Gm21976 | 0 | 3 | 8 | 4 | 2 | 1 | 3 | 6 | 1 | 2 | 1 | 1 | 2 | 0 | 4 | 8 | 1 | 2 | 5 | 8 | 4 | 3 | 4 | 1 | 4 | 1 | 0 | 5 | Gm21976 |
| Gm21981 | 10 | 13 | 15 | 12 | 15 | 5 | 10 | 0 | 1 | 5 | 12 | 5 | 2 | 6 | 5 | 10 | 20 | 15 | 11 | 8 | 5 | 2 | 8 | 9 | 7 | 5 | 6 | 7 | Gm21981 |
| Gm21985 | 1 | 0 | 0 | 0 | 0 | 1 | 0 | 0 | 0 | 0 | 0 | 0 | 0 | 0 | 0 | 0 | 0 | 2 | 0 | 0 | 1 | 0 | 0 | 0 | 1 | 0 | 0 | 1 | Gm21985 |
| Gm21987 | 0 | 1 | 3 | 2 | 0 | 0 | 0 | 0 | 0 | 0 | 0 | 0 | 0 | 0 | 0 | 1 | 5 | 3 | 0 | 0 | 0 | 2 | 0 | 0 | 0 | 0 | 1 | 0 | Gm21987 |
| Gm21988 | 3 | 7 | 2 | 9 | 4 | 1 | 4 | 2 | 0 | 1 | 1 | 0 | 2 | 3 | 0 | 2 | 5 | 2 | 3 | 1 | 6 | 0 | 1 | 5 | 0 | 0 | 1 | 2 | Gm21988 |
| Gm21992 | 814 | 782 | 1244 | 1595 | 1046 | 1220 | 1582 | 277 | 289 | 475 | 483 | 538 | 562 | 630 | 663 | 719 | 1321 | 1563 | 1226 | 692 | 1459 | 506 | 237 | 558 | 459 | 656 | 524 | 570 | Gm21992 |
| Gm21994 | 6 | 4 | 0 | 8 | 1 | 3 | 1 | 6 | 4 | 1 | 3 | 5 | 6 | 1 | 3 | 2 | 1 | 7 | 3 | 3 | 3 | 0 | 3 | 0 | 4 | 1 | 1 | 8 | Gm21994 |
| Gm2237 | 345 | 234 | 431 | 460 | 488 | 405 | 408 | 713 | 632 | 647 | 645 | 730 | 610 | 736 | 389 | 328 | 337 | 358 | 388 | 446 | 436 | 662 | 555 | 669 | 722 | 742 | 623 | 654 | Gm2237 |
| Gm2244 | 0 | 0 | 3 | 2 | 2 | 0 | 6 | 0 | 2 | 0 | 3 | 2 | 0 | 2 | 4 | 2 | 8 | 0 | 2 | 6 | 3 | 0 | 1 | 1 | 3 | 8 | 3 | 1 | Gm2244 |
| Gm2296 | 403 | 366 | 332 | 300 | 244 | 305 | 217 | 286 | 207 | 201 | 204 | 307 | 254 | 220 | 413 | 354 | 297 | 273 | 300 | 333 | 199 | 257 | 195 | 273 | 265 | 349 | 294 | 247 | Gm2296 |
| Gm2381 | 0 | 0 | 1 | 0 | 0 | 0 | 0 | 1 | 0 | 0 | 1 | 1 | 1 | 0 | 0 | 0 | 0 | 1 | 0 | 0 | 0 | 2 | 0 | 1 | 0 | 0 | 0 | 0 | Gm2381 |
| Gm2411 | 0 | 3 | 0 | 1 | 2 | 6 | 0 | 0 | 0 | 0 | 0 | 3 | 0 | 2 | 4 | 3 | 0 | 6 | 0 | 0 | 0 | 0 | 5 | 0 | 5 | 0 | 1 | 1 | Gm2411 |
| Gm26566 | 2 | 3 | 9 | 3 | 7 | 1 | 7 | 5 | 15 | 5 | 6 | 9 | 12 | 9 | 3 | 1 | 3 | 6 | 9 | 5 | 9 | 17 | 18 | 13 | 13 | 6 | 12 | 8 | Gm26566 |
| Gm266 | 104 | 78 | 89 | 68 | 61 | 43 | 23 | 127 | 168 | 139 | 186 | 36 | 117 | 48 | 98 | 100 | 59 | 42 | 37 | 47 | 40 | 110 | 172 | 124 | 50 | 221 | 88 | 77 | Gm266 |
| Gm26602 | 0 | 0 | 0 | 2 | 1 | 0 | 0 | 0 | 2 | 0 | 2 | 0 | 0 | 1 | 1 | 1 | 0 | 1 | 1 | 0 | 1 | 3 | 1 | 0 | 3 | 1 | 1 | 2 | Gm26602 |
| Gm26620 | 12 | 5 | 8 | 7 | 6 | 9 | 16 | 13 | 7 | 3 | 6 | 3 | 7 | 12 | 3 | 4 | 25 | 11 | 17 | 8 | 5 | 7 | 11 | 9 | 12 | 8 | 2 | 8 | Gm26620 |
| Gm2663 | 0 | 0 | 0 | 0 | 0 | 0 | 0 | 0 | 0 | 0 | 0 | 0 | 0 | 0 | 0 | 0 | 0 | 0 | 0 | 0 | 0 | 0 | 0 | 0 | 0 | 0 | 1 | 0 | Gm2663 |
| Gm26657 | 0 | 0 | 0 | 0 | 0 | 0 | 0 | 0 | 0 | 0 | 0 | 0 | 0 | 1 | 0 | 0 | 0 | 0 | 0 | 0 | 0 | 0 | 0 | 0 | 0 | 0 | 0 | 0 | Gm26657 |
| Gm26741 | 0 | 0 | 0 | 0 | 0 | 0 | 0 | 0 | 0 | 0 | 0 | 0 | 1 | 2 | 0 | 0 | 2 | 0 | 0 | 0 | 0 | 0 | 0 | 0 | 0 | 0 | 1 | 0 | Gm26741 |
| Gm26897 | 0 | 0 | 0 | 0 | 1 | 0 | 1 | 0 | 0 | 0 | 0 | 0 | 0 | 2 | 0 | 0 | 0 | 0 | 0 | 0 | 0 | 0 | 0 | 0 | 3 | 0 | 1 | 0 | Gm26897 |
| Gm27027 | 3 | 7 | 10 | 12 | 8 | 10 | 13 | 17 | 14 | 9 | 11 | 8 | 8 | 12 | 3 | 2 | 17 | 14 | 16 | 7 | 8 | 8 | 5 | 25 | 19 | 30 | 17 | 18 | Gm27027 |
| Gm27029 | 31 | 20 | 30 | 38 | 30 | 32 | 20 | 22 | 23 | 19 | 24 | 18 | 24 | 23 | 24 | 22 | 35 | 41 | 31 | 40 | 21 | 28 | 16 | 18 | 25 | 20 | 14 | 30 | Gm27029 |
| Gm27179 | 0 | 1 | 0 | 0 | 0 | 0 | 0 | 0 | 0 | 0 | 0 | 0 | 0 | 0 | 0 | 0 | 0 | 0 | 0 | 0 | 0 | 2 | 0 | 0 | 0 | 0 | 0 | 0 | Gm27179 |
| Gm28035 | 0 | 0 | 0 | 4 | 0 | 0 | 1 | 0 | 0 | 0 | 0 | 0 | 0 | 1 | 0 | 0 | 0 | 2 | 0 | 3 | 2 | 0 | 0 | 2 | 0 | 0 | 1 | 0 | Gm28035 |
| Gm28040 | 2 | 0 | 2 | 2 | 2 | 0 | 4 | 0 | 0 | 0 | 1 | 4 | 1 | 0 | 0 | 0 | 0 | 0 | 2 | 1 | 3 | 0 | 0 | 0 | 0 | 0 | 0 | 0 | Gm28040 |
| Gm28041 | 0 | 1 | 0 | 0 | 0 | 0 | 0 | 0 | 0 | 0 | 0 | 0 | 0 | 0 | 0 | 0 | 0 | 0 | 0 | 0 | 0 | 0 | 0 | 0 | 0 | 0 | 0 | 0 | Gm28041 |
| Gm28043 | 145 | 167 | 85 | 209 | 210 | 162 | 181 | 154 | 130 | 126 | 126 | 187 | 134 | 171 | 182 | 121 | 124 | 169 | 136 | 180 | 194 | 117 | 119 | 119 | 138 | 68 | 169 | 136 | Gm28043 |
| Gm28044 | 0 | 0 | 0 | 0 | 1 | 0 | 0 | 2 | 1 | 0 | 0 | 0 | 0 | 2 | 0 | 0 | 0 | 0 | 0 | 1 | 0 | 0 | 2 | 2 | 0 | 1 | 0 | 0 | Gm28044 |
| Gm28053 | 0 | 0 | 0 | 0 | 0 | 3 | 2 | 0 | 0 | 0 | 0 | 2 | 1 | 0 | 0 | 0 | 0 | 3 | 0 | 0 | 0 | 0 | 0 | 0 | 0 | 1 | 0 | 0 | Gm28053 |
| Gm281 | 1 | 6 | 0 | 2 | 11 | 5 | 10 | 15 | 1 | 11 | 8 | 12 | 14 | 6 | 4 | 5 | 1 | 2 | 2 | 24 | 6 | 8 | 8 | 5 | 10 | 3 | 5 | 4 | Gm281 |
| Gm28230 | 2 | 0 | 0 | 0 | 8 | 5 | 0 | 1 | 5 | 4 | 23 | 1 | 0 | 0 | 0 | 0 | 0 | 2 | 1 | 0 | 0 | 2 | 10 | 2 | 1 | 0 | 0 | 0 | Gm28230 |
| Gm2832 | 29 | 9 | 17 | 12 | 21 | 6 | 11 | 10 | 16 | 14 | 14 | 9 | 20 | 14 | 13 | 18 | 12 | 21 | 10 | 17 | 6 | 22 | 19 | 18 | 22 | 28 | 12 | 6 | Gm2832 |
| Gm28372 | 0 | 0 | 0 | 0 | 0 | 0 | 1 | 1 | 0 | 0 | 0 | 0 | 0 | 0 | 0 | 0 | 0 | 0 | 0 | 0 | 0 | 0 | 0 | 0 | 0 | 0 | 0 | 0 | Gm28372 |
| Gm28374 | 7 | 3 | 1 | 4 | 2 | 2 | 4 | 5 | 6 | 0 | 0 | 4 | 2 | 4 | 4 | 0 | 7 | 0 | 2 | 3 | 4 | 12 | 0 | 7 | 3 | 4 | 2 | 0 | Gm28374 |

| | | | | | | | | | | | | | | | | | | | | | | | | | | | | | |
|---|---|---|---|---|---|---|---|---|---|---|---|---|---|---|---|---|---|---|---|---|---|---|---|---|---|---|---|---|---|
| Gm28510 | 0 | 0 | 0 | 0 | 0 | 0 | 0 | 0 | 0 | 0 | 0 | 0 | 0 | 0 | 0 | 0 | 0 | 0 | 0 | 0 | 0 | 0 | 1 | 0 | 0 | 0 | 0 | 0 | Gm28510 |
| Gm28551 | 7 | 6 | 15 | 8 | 20 | 26 | 30 | 7 | 5 | 8 | 3 | 3 | 12 | 4 | 18 | 16 | 9 | 18 | 13 | 12 | 15 | 3 | 12 | 2 | 7 | 5 | 6 | 1 | Gm28551 |
| Gm28557 | 69 | 73 | 81 | 56 | 45 | 51 | 44 | 71 | 67 | 36 | 74 | 101 | 69 | 57 | 62 | 68 | 48 | 45 | 30 | 44 | 39 | 62 | 72 | 69 | 64 | 85 | 73 | 65 | Gm28557 |
| Gm28710 | 1 | 1 | 1 | 0 | 4 | 0 | 0 | 2 | 7 | 2 | 1 | 0 | 2 | 0 | 1 | 2 | 0 | 0 | 0 | 1 | 1 | 0 | 8 | 4 | 0 | 0 | 0 | 0 | Gm28710 |
| Gm28729 | 8 | 10 | 0 | 4 | 8 | 10 | 7 | 16 | 21 | 21 | 17 | 16 | 25 | 9 | 4 | 9 | 2 | 15 | 4 | 6 | 4 | 17 | 43 | 9 | 20 | 11 | 5 | 4 | Gm28729 |
| Gm2888 | 2 | 0 | 0 | 0 | 0 | 0 | 0 | 0 | 0 | 0 | 0 | 0 | 0 | 1 | 0 | 0 | 0 | 0 | 0 | 0 | 0 | 0 | 0 | 0 | 0 | 1 | 0 | 0 | Gm2888 |
| Gm28919 | 0 | 0 | 0 | 0 | 0 | 0 | 0 | 0 | 0 | 0 | 0 | 0 | 0 | 2 | 0 | 0 | 0 | 0 | 0 | 0 | 0 | 0 | 0 | 0 | 0 | 0 | 0 | 0 | Gm28919 |
| Gm2897 | 234 | 201 | 250 | 206 | 403 | 370 | 306 | 462 | 456 | 416 | 470 | 500 | 391 | 361 | 260 | 223 | 206 | 253 | 331 | 338 | 292 | 444 | 355 | 419 | 502 | 389 | 380 | 371 | Gm2897 |
| Gm29106 | 0 | 2 | 0 | 0 | 0 | 2 | 0 | 0 | 0 | 0 | 0 | 0 | 2 | 0 | 0 | 0 | 0 | 2 | 0 | 3 | 0 | 0 | 0 | 0 | 0 | 0 | 0 | 0 | Gm29106 |
| Gm29133 | 0 | 0 | 0 | 0 | 1 | 0 | 1 | 0 | 0 | 0 | 0 | 0 | 0 | 0 | 0 | 0 | 0 | 0 | 0 | 0 | 1 | 0 | 2 | 0 | 0 | 0 | 0 | 0 | Gm29133 |
| Gm2916 | 0 | 1 | 0 | 0 | 0 | 0 | 0 | 0 | 0 | 0 | 0 | 0 | 0 | 0 | 0 | 0 | 0 | 0 | 0 | 0 | 0 | 0 | 0 | 0 | 0 | 0 | 0 | 0 | Gm2916 |
| Gm29289 | 1 | 0 | 0 | 0 | 0 | 0 | 0 | 0 | 0 | 0 | 0 | 0 | 0 | 0 | 0 | 0 | 0 | 0 | 0 | 0 | 0 | 0 | 0 | 0 | 0 | 0 | 0 | 0 | Gm29289 |
| Gm29427 | 6 | 22 | 23 | 54 | 31 | 31 | 45 | 7 | 2 | 2 | 12 | 13 | 6 | 7 | 7 | 6 | 30 | 46 | 37 | 14 | 29 | 22 | 8 | 0 | 9 | 5 | 6 | 5 | Gm29427 |
| Gm2956 | 2 | 1 | 1 | 0 | 11 | 0 | 1 | 13 | 7 | 5 | 6 | 4 | 2 | 5 | 3 | 3 | 0 | 3 | 2 | 2 | 2 | 8 | 6 | 14 | 11 | 5 | 2 | 6 | Gm2956 |
| Gm29609 | 61 | 85 | 82 | 78 | 62 | 70 | 106 | 72 | 70 | 77 | 72 | 91 | 78 | 98 | 58 | 65 | 93 | 81 | 69 | 64 | 64 | 110 | 74 | 121 | 51 | 60 | 56 | 107 | Gm29609 |
| Gm29666 | 429 | 365 | 329 | 441 | 409 | 390 | 483 | 326 | 339 | 400 | 320 | 412 | 312 | 402 | 398 | 342 | 461 | 450 | 404 | 434 | 430 | 306 | 271 | 300 | 351 | 413 | 371 | 366 | Gm29666 |
| Gm29695 | 422 | 564 | 657 | 485 | 740 | 526 | 590 | 1782 | 1342 | 1828 | 1442 | 2385 | 1956 | 2300 | 483 | 393 | 797 | 517 | 704 | 567 | 606 | 2326 | 1193 | 1919 | 2044 | 1709 | 2536 | 2459 | Gm29695 |
| Gm2974 | 47 | 22 | 32 | 57 | 97 | 88 | 87 | 281 | 217 | 218 | 233 | 187 | 121 | 175 | 45 | 33 | 37 | 72 | 75 | 123 | 105 | 277 | 250 | 181 | 322 | 226 | 202 | 169 | Gm2974 |
| Gm29776 | 3 | 0 | 1 | 2 | 4 | 7 | 0 | 0 | 4 | 1 | 3 | 1 | 9 | 1 | 1 | 2 | 2 | 2 | 2 | 1 | 3 | 2 | 6 | 5 | 2 | 4 | 5 | 4 | Gm29776 |
| Gm29797 | 25 | 40 | 28 | 30 | 43 | 40 | 42 | 15 | 24 | 25 | 30 | 12 | 41 | 32 | 24 | 38 | 46 | 47 | 51 | 42 | 50 | 20 | 30 | 18 | 39 | 25 | 17 | 26 | Gm29797 |
| Gm2a | 2121 | 2375 | 2065 | 2157 | 1921 | 1901 | 1983 | 1054 | 1219 | 1086 | 1218 | 955 | 1081 | 1050 | 2313 | 2087 | 2065 | 2080 | 1848 | 2172 | 2051 | 1113 | 1438 | 999 | 1109 | 953 | 1009 | 972 | Gm2a |
| Gm3002 | 12 | 6 | 27 | 13 | 25 | 25 | 31 | 114 | 102 | 105 | 103 | 76 | 81 | 57 | 24 | 16 | 23 | 22 | 30 | 27 | 37 | 104 | 106 | 83 | 94 | 77 | 56 | 71 | Gm3002 |
| Gm3005 | 14 | 13 | 17 | 15 | 27 | 33 | 17 | 98 | 94 | 109 | 69 | 66 | 65 | 53 | 17 | 13 | 28 | 15 | 20 | 36 | 20 | 67 | 69 | 65 | 72 | 75 | 80 | 89 | Gm3005 |
| Gm30083 | 0 | 0 | 0 | 1 | 0 | 0 | 0 | 0 | 0 | 0 | 0 | 0 | 0 | 0 | 0 | 0 | 0 | 0 | 0 | 0 | 0 | 0 | 0 | 0 | 2 | 0 | 0 | 4 | Gm30083 |
| Gm3012 | 0 | 0 | 0 | 0 | 0 | 0 | 1 | 0 | 1 | 0 | 0 | 0 | 0 | 0 | 0 | 0 | 0 | 0 | 2 | 0 | 0 | 0 | 0 | 0 | 1 | 0 | 0 | 0 | Gm3012 |
| Gm30191 | 1 | 5 | 10 | 1 | 2 | 0 | 0 | 4 | 4 | 7 | 4 | 8 | 9 | 0 | 5 | 2 | 1 | 0 | 2 | 1 | 1 | 3 | 9 | 0 | 3 | 0 | 1 | 0 | Gm30191 |
| Gm3020 | 1 | 0 | 1 | 2 | 2 | 0 | 0 | 4 | 2 | 1 | 3 | 7 | 9 | 1 | 0 | 5 | 0 | 0 | 2 | 1 | 3 | 2 | 18 | 0 | 2 | 3 | 13 | 4 | Gm3020 |
| Gm3029 | 0 | 0 | 0 | 0 | 0 | 1 | 0 | 0 | 0 | 0 | 0 | 0 | 1 | 0 | 0 | 0 | 0 | 0 | 0 | 0 | 0 | 0 | 0 | 0 | 0 | 0 | 0 | 0 | Gm3029 |
| Gm30302 | 0 | 0 | 0 | 0 | 0 | 0 | 0 | 0 | 0 | 0 | 0 | 0 | 0 | 0 | 0 | 0 | 0 | 0 | 0 | 0 | 0 | 0 | 0 | 0 | 0 | 0 | 1 | 0 | Gm30302 |
| Gm3043 | 16 | 16 | 17 | 27 | 34 | 11 | 12 | 17 | 21 | 14 | 14 | 17 | 0 | 15 | 3 | 16 | 23 | 8 | 16 | 8 | 21 | 18 | 12 | 2 | 22 | 14 | 10 | 26 | Gm3043 |
| Gm3047 | 2 | 0 | 1 | 0 | 0 | 0 | 0 | 0 | 1 | 1 | 0 | 1 | 1 | 0 | 0 | 0 | 0 | 0 | 0 | 0 | 0 | 2 | 0 | 0 | 0 | 0 | 0 | 0 | Gm3047 |
| Gm3055 | 149 | 101 | 118 | 137 | 131 | 169 | 156 | 278 | 221 | 189 | 261 | 229 | 171 | 235 | 179 | 128 | 109 | 113 | 125 | 198 | 154 | 237 | 219 | 203 | 231 | 222 | 170 | 240 | Gm3055 |
| Gm3095 | 5 | 4 | 10 | 2 | 6 | 4 | 15 | 15 | 16 | 30 | 16 | 14 | 19 | 21 | 5 | 5 | 17 | 3 | 12 | 35 | 4 | 30 | 8 | 29 | 23 | 11 | 3 | 7 | Gm3095 |
| Gm3099 | 0 | 0 | 0 | 0 | 0 | 0 | 0 | 0 | 0 | 1 | 0 | 0 | 0 | 1 | 0 | 0 | 0 | 0 | 0 | 0 | 0 | 0 | 0 | 0 | 0 | 0 | 1 | 0 | Gm3099 |
| Gm3115 | 0 | 0 | 0 | 0 | 0 | 0 | 0 | 0 | 0 | 1 | 0 | 0 | 0 | 0 | 0 | 0 | 0 | 0 | 2 | 0 | 0 | 0 | 0 | 2 | 0 | 0 | 0 | 2 | Gm3115 |
| Gm31160 | 0 | 9 | 0 | 7 | 5 | 22 | 8 | 11 | 12 | 8 | 12 | 7 | 12 | 12 | 0 | 0 | 5 | 1 | 9 | 8 | 1 | 32 | 19 | 5 | 19 | 19 | 22 | 6 | Gm31160 |
| Gm3127 | 1 | 0 | 0 | 0 | 0 | 0 | 0 | 0 | 1 | 1 | 1 | 3 | 0 | 1 | 0 | 0 | 0 | 0 | 1 | 1 | 3 | 0 | 0 | 0 | 0 | 0 | 3 | 5 | Gm3127 |
| Gm31371 | 13 | 12 | 3 | 17 | 5 | 13 | 19 | 0 | 1 | 3 | 1 | 0 | 6 | 2 | 9 | 14 | 0 | 4 | 3 | 13 | 7 | 0 | 5 | 0 | 1 | 0 | 2 | 0 | Gm31371 |
| Gm3138 | 0 | 0 | 0 | 1 | 1 | 0 | 0 | 1 | 0 | 0 | 0 | 2 | 2 | 0 | 0 | 0 | 0 | 0 | 0 | 0 | 0 | 2 | 0 | 0 | 0 | 0 | 0 | 0 | Gm3138 |
| Gm3149 | 0 | 2 | 0 | 0 | 0 | 1 | 0 | 0 | 1 | 0 | 0 | 0 | 0 | 1 | 0 | 0 | 0 | 0 | 0 | 0 | 0 | 0 | 0 | 0 | 0 | 0 | 0 | 1 | Gm3149 |
| Gm3159 | 0 | 0 | 1 | 0 | 2 | 3 | 0 | 1 | 1 | 0 | 0 | 0 | 0 | 0 | 0 | 0 | 1 | 0 | 1 | 0 | 1 | 2 | 1 | 0 | 1 | 2 | 1 | 1 | Gm3159 |
| Gm3164 | 40 | 32 | 58 | 37 | 80 | 50 | 78 | 175 | 130 | 148 | 142 | 111 | 100 | 129 | 37 | 40 | 35 | 45 | 46 | 69 | 46 | 189 | 135 | 124 | 95 | 150 | 116 | 100 | Gm3164 |
| Gm3173 | 44 | 16 | 14 | 9 | 34 | 44 | 30 | 115 | 68 | 94 | 78 | 146 | 85 | 78 | 16 | 35 | 43 | 21 | 48 | 33 | 14 | 74 | 78 | 79 | 108 | 91 | 89 | 161 | Gm3173 |
| Gm3182 | 0 | 0 | 0 | 0 | 0 | 0 | 0 | 1 | 2 | 0 | 1 | 1 | 2 | 1 | 0 | 0 | 0 | 0 | 0 | 0 | 0 | 0 | 2 | 1 | 4 | 1 | 0 | 0 | Gm3182 |
| Gm3194 | 0 | 0 | 0 | 0 | 0 | 0 | 0 | 1 | 1 | 0 | 4 | 0 | 10 | 1 | 0 | 0 | 0 | 0 | 0 | 0 | 0 | 0 | 1 | 1 | 0 | 1 | 0 | 2 | Gm3194 |
| Gm3233 | 0 | 0 | 0 | 0 | 0 | 0 | 0 | 0 | 0 | 0 | 0 | 0 | 0 | 0 | 0 | 0 | 0 | 0 | 0 | 0 | 0 | 0 | 0 | 2 | 0 | 0 | 0 | 0 | Gm3233 |
| Gm3239 | 5 | 3 | 4 | 21 | 19 | 4 | 7 | 22 | 7 | 21 | 29 | 14 | 44 | 15 | 7 | 2 | 1 | 6 | 9 | 0 | 8 | 40 | 17 | 13 | 17 | 9 | 40 | 8 | Gm3239 |
| Gm3248 | 63 | 39 | 65 | 31 | 77 | 108 | 104 | 76 | 99 | 102 | 80 | 70 | 61 | 70 | 42 | 43 | 61 | 41 | 56 | 110 | 75 | 109 | 78 | 74 | 90 | 60 | 62 | 58 | Gm3248 |
| Gm3250 | 0 | 0 | 0 | 0 | 0 | 0 | 0 | 0 | 0 | 0 | 0 | 0 | 0 | 1 | 0 | 0 | 0 | 0 | 0 | 0 | 0 | 0 | 1 | 0 | 0 | 0 | 0 | 0 | Gm3250 |
| Gm3252 | 50 | 18 | 63 | 30 | 69 | 84 | 66 | 204 | 161 | 168 | 151 | 163 | 151 | 139 | 47 | 18 | 44 | 39 | 52 | 81 | 71 | 206 | 138 | 176 | 189 | 190 | 121 | 151 | Gm3252 |
| Gm3264 | 24 | 29 | 21 | 19 | 24 | 38 | 42 | 115 | 112 | 118 | 132 | 116 | 137 | 117 | 21 | 13 | 52 | 17 | 19 | 36 | 41 | 159 | 98 | 87 | 130 | 94 | 145 | 72 | Gm3264 |
| Gm32687 | 135 | 122 | 111 | 137 | 135 | 223 | 191 | 388 | 335 | 306 | 290 | 285 | 350 | 319 | 161 | 141 | 112 | 106 | 154 | 174 | 140 | 356 | 311 | 284 | 258 | 287 | 332 | 333 | Gm32687 |
| Gm3269 | 0 | 1 | 1 | 0 | 2 | 0 | 1 | 9 | 6 | 8 | 8 | 1 | 2 | 4 | 0 | 3 | 0 | 1 | 1 | 4 | 0 | 8 | 4 | 6 | 10 | 0 | 6 | 0 | Gm3269 |
| Gm32717 | 5 | 8 | 4 | 6 | 5 | 4 | 1 | 7 | 4 | 2 | 3 | 10 | 1 | 5 | 9 | 4 | 5 | 2 | 4 | 5 | 2 | 3 | 5 | 4 | 2 | 4 | 2 | 1 | Gm32717 |
| Gm32742 | 6 | 4 | 0 | 3 | 10 | 1 | 11 | 13 | 9 | 14 | 18 | 11 | 15 | 2 | 15 | 8 | 6 | 3 | 7 | 7 | 5 | 28 | 20 | 22 | 5 | 9 | 16 | 6 | Gm32742 |
| Gm3278 | 5 | 1 | 1 | 2 | 9 | 5 | 9 | 5 | 8 | 12 | 7 | 10 | 7 | 10 | 7 | 3 | 8 | 14 | 6 | 5 | 11 | 12 | 5 | 13 | 8 | 10 | 2 | 11 | Gm3278 |
| Gm32802 | 29 | 24 | 25 | 20 | 16 | 23 | 28 | 38 | 26 | 16 | 12 | 26 | 15 | 32 | 24 | 19 | 25 | 15 | 21 | 14 | 22 | 23 | 16 | 16 | 15 | 21 | 34 | 33 | Gm32802 |
| Gm3285 | 1 | 0 | 0 | 0 | 0 | 0 | 0 | 0 | 1 | 0 | 0 | 0 | 0 | 2 | 1 | 0 | 5 | 0 | 0 | 0 | 1 | 0 | 2 | 0 | 0 | 0 | 0 | 0 | Gm3285 |
| Gm3298 | 6 | 4 | 6 | 0 | 3 | 3 | 2 | 15 | 10 | 12 | 2 | 8 | 12 | 21 | 17 | 3 | 2 | 0 | 3 | 3 | 0 | 45 | 23 | 20 | 14 | 7 | 1 | 13 | Gm3298 |
| Gm3317 | 12 | 7 | 42 | 4 | 19 | 51 | 20 | 22 | 26 | 68 | 12 | 16 | 34 | 39 | 1 | 19 | 12 | 32 | 21 | 24 | 25 | 57 | 117 | 38 | 96 | 65 | 45 | 97 | Gm3317 |
| Gm33543 | 0 | 1 | 0 | 0 | 0 | 0 | 0 | 0 | 2 | 1 | 0 | 0 | 0 | 1 | 3 | 0 | 0 | 3 | 2 | 0 | 2 | 0 | 2 | 0 | 0 | 0 | 0 | 0 | Gm33543 |
| Gm3373 | 0 | 0 | 4 | 0 | 2 | 0 | 1 | 0 | 2 | 2 | 2 | 0 | 2 | 0 | 0 | 2 | 1 | 1 | 1 | 5 | 0 | 0 | 4 | 1 | 5 | 4 | 7 | 1 | Gm3373 |
| Gm3376 | 0 | 2 | 0 | 0 | 0 | 0 | 0 | 0 | 0 | 0 | 0 | 0 | 0 | 0 | 0 | 0 | 0 | 1 | 0 | 0 | 0 | 0 | 0 | 0 | 0 | 0 | 0 | 0 | Gm3376 |
| Gm3383 | 27 | 22 | 37 | 28 | 18 | 6 | 25 | 31 | 30 | 65 | 21 | 110 | 40 | 60 | 33 | 15 | 28 | 28 | 27 | 20 | 25 | 48 | 21 | 48 | 62 | 23 | 22 | 56 | Gm3383 |
| Gm33933 | 20 | 9 | 16 | 35 | 20 | 23 | 22 | 68 | 49 | 65 | 31 | 33 | 34 | 68 | 17 | 16 | 43 | 18 | 9 | 27 | 11 | 105 | 32 | 38 | 25 | 46 | 37 | 61 | Gm33933 |
| Gm3402 | 0 | 0 | 0 | 0 | 1 | 0 | 0 | 0 | 0 | 0 | 0 | 0 | 0 | 0 | 0 | 0 | 0 | 0 | 0 | 0 | 0 | 0 | 0 | 0 | 0 | 0 | 0 | 0 | Gm3402 |
| Gm3404 | 0 | 0 | 0 | 0 | 0 | 0 | 1 | 0 | 0 | 0 | 0 | 0 | 0 | 0 | 0 | 0 | 0 | 0 | 0 | 0 | 0 | 0 | 0 | 0 | 0 | 0 | 0 | 0 | Gm3404 |
| Gm34066 | 37 | 29 | 20 | 73 | 44 | 40 | 59 | 6 | 1 | 4 | 4 | 0 | 7 | 6 | 40 | 23 | 15 | 51 | 54 | 36 | 55 | 3 | 6 | 2 | 0 | 7 | 3 | 1 | Gm34066 |
| Gm3411 | 54 | 35 | 74 | 16 | 55 | 38 | 40 | 192 | 127 | 133 | 119 | 103 | 142 | 125 | 73 | 49 | 53 | 17 | 79 | 54 | 44 | 162 | 151 | 109 | 179 | 96 | 110 | 134 | Gm3411 |
| Gm3415 | 0 | 0 | 0 | 1 | 0 | 0 | 0 | 0 | 0 | 0 | 0 | 0 | 0 | 0 | 0 | 0 | 0 | 0 | 0 | 0 | 0 | 0 | 0 | 0 | 0 | 0 | 0 | 0 | Gm3415 |
| Gm3424 | 0 | 0 | 0 | 0 | 1 | 1 | 0 | 0 | 0 | 0 | 0 | 0 | 0 | 1 | 0 | 3 | 1 | 0 | 1 | 0 | 4 | 0 | 1 | 0 | 0 | 0 | 0 | 0 | Gm3424 |
| Gm3435 | 40 | 91 | 60 | 79 | 74 | 71 | 53 | 136 | 80 | 71 | 95 | 91 | 91 | 111 | 88 | 42 | 41 | 81 | 42 | 54 | 80 | 70 | 113 | 64 | 114 | 127 | 115 | 151 | Gm3435 |
| Gm3453 | 2 | 0 | 1 | 0 | 0 | 0 | 0 | 1 | 0 | 0 | 0 | 0 | 0 | 0 | 0 | 1 | 5 | 0 | 0 | 1 | 0 | 2 | 3 | 2 | 0 | 0 | 1 | 1 | Gm3453 |
| Gm34653 | 12 | 4 | 7 | 6 | 4 | 10 | 12 | 6 | 2 | 7 | 1 | 1 | 6 | 5 | 4 | 10 | 9 | 6 | 16 | 9 | 11 | 5 | 4 | 4 | 6 | 4 | 1 | 8 | Gm34653 |
| Gm3468 | 137 | 78 | 137 | 119 | 181 | 171 | 143 | 424 | 288 | 383 | 341 | 378 | 367 | 355 | 140 | 126 | 108 | 88 | 188 | 175 | 163 | 349 | 293 | 352 | 326 | 417 | 346 | 358 | Gm3468 |
| Gm3476 | 8 | 10 | 0 | 2 | 0 | 1 | 2 | 21 | 12 | 5 | 3 | 8 | 6 | 1 | 3 | 4 | 1 | 0 | 0 | 1 | 2 | 3 | 10 | 0 | 11 | 0 | 5 | 6 | Gm3476 |
| Gm3486 | 0 | 0 | 0 | 0 | 0 | 0 | 0 | 0 | 0 | 0 | 0 | 0 | 1 | 0 | 1 | 0 | 0 | 0 | 0 | 0 | 0 | 0 | 1 | 0 | 0 | 0 | 0 | 0 | Gm3486 |
| Gm3488 | 3 | 2 | 0 | 3 | 13 | 7 | 10 | 35 | 10 | 19 | 42 | 15 | 9 | 19 | 3 | 1 | 5 | 6 | 18 | 5 | 5 | 27 | 11 | 25 | 23 | 54 | 36 | 15 | Gm3488 |
| Gm3500 | 10 | 10 | 15 | 7 | 25 | 27 | 17 | 49 | 30 | 21 | 18 | 28 | 32 | 49 | 12 | 6 | 27 | 10 | 13 | 18 | 20 | 38 | 15 | 41 | 28 | 36 | 34 | 55 | Gm3500 |

| | | | | | | | | | | | | | | | | | | | | | | | | | | | | | |
|---|---|---|---|---|---|---|---|---|---|---|---|---|---|---|---|---|---|---|---|---|---|---|---|---|---|---|---|---|---|
| Gm35060 | 0 | 0 | 0 | 0 | 0 | 0 | 0 | 2 | 3 | 0 | 4 | 0 | 1 | 0 | 1 | 0 | 0 | 0 | 0 | 0 | 1 | 0 | 0 | 0 | 0 | 0 | 0 | 1 | Gm35060 |
| Gm3512 | 7 | 11 | 17 | 10 | 25 | 15 | 15 | 66 | 51 | 74 | 63 | 41 | 82 | 23 | 3 | 33 | 13 | 15 | 42 | 10 | 17 | 75 | 25 | 15 | 106 | 64 | 20 | 20 | Gm3512 |
| Gm3526 | 0 | 0 | 1 | 0 | 0 | 0 | 1 | 0 | 0 | 0 | 0 | 0 | 0 | 2 | 0 | 0 | 0 | 0 | 0 | 0 | 0 | 0 | 0 | 1 | 0 | 0 | 0 | 0 | Gm3526 |
| Gm35315 | 225 | 145 | 306 | 243 | 210 | 220 | 277 | 170 | 163 | 190 | 151 | 220 | 230 | 189 | 309 | 231 | 270 | 221 | 236 | 209 | 209 | 149 | 164 | 193 | 134 | 256 | 244 | 284 | Gm35315 |
| Gm35339 | 879 | 1142 | 1244 | 1698 | 1438 | 1286 | 1301 | 934 | 798 | 1098 | 950 | 1026 | 920 | 972 | 814 | 945 | 1611 | 1684 | 1707 | 1249 | 1398 | 1250 | 746 | 1184 | 1174 | 759 | 946 | 846 | Gm35339 |
| Gm3542 | 0 | 0 | 0 | 0 | 0 | 0 | 0 | 0 | 0 | 1 | 0 | 0 | 0 | 0 | 0 | 0 | 0 | 0 | 0 | 0 | 0 | 0 | 0 | 0 | 0 | 0 | 0 | 0 | Gm3542 |
| Gm3543 | 0 | 1 | 0 | 0 | 0 | 0 | 0 | 4 | 0 | 0 | 0 | 0 | 0 | 0 | 0 | 0 | 0 | 0 | 0 | 0 | 0 | 0 | 0 | 0 | 0 | 0 | 2 | 0 | Gm3543 |
| Gm3550 | 2 | 2 | 3 | 2 | 0 | 0 | 1 | 0 | 0 | 1 | 0 | 0 | 1 | 1 | 0 | 1 | 5 | 2 | 5 | 1 | 3 | 2 | 1 | 0 | 4 | 4 | 5 | 2 | Gm3550 |
| Gm35549 | 1 | 0 | 0 | 0 | 1 | 0 | 0 | 0 | 0 | 1 | 0 | 0 | 0 | 0 | 0 | 0 | 0 | 0 | 0 | 4 | 0 | 0 | 1 | 0 | 0 | 0 | 1 | 0 | Gm35549 |
| Gm3558 | 128 | 79 | 100 | 133 | 213 | 171 | 161 | 133 | 149 | 108 | 161 | 109 | 83 | 97 | 137 | 138 | 117 | 105 | 132 | 147 | 160 | 152 | 100 | 94 | 135 | 110 | 119 | 78 | Gm3558 |
| Gm3573 | 0 | 1 | 0 | 0 | 0 | 0 | 0 | 0 | 0 | 0 | 0 | 0 | 0 | 0 | 0 | 0 | 0 | 0 | 0 | 0 | 0 | 0 | 0 | 0 | 0 | 0 | 0 | 0 | Gm3573 |
| Gm3594 | 0 | 0 | 0 | 0 | 0 | 0 | 0 | 0 | 0 | 2 | 0 | 0 | 0 | 0 | 0 | 0 | 0 | 0 | 0 | 0 | 0 | 0 | 0 | 0 | 0 | 0 | 0 | 0 | Gm3594 |
| Gm36028 | 7 | 14 | 10 | 13 | 13 | 5 | 9 | 16 | 18 | 24 | 11 | 21 | 24 | 29 | 11 | 5 | 17 | 0 | 5 | 4 | 6 | 18 | 10 | 34 | 10 | 37 | 27 | 8 | Gm36028 |
| Gm3604 | 233 | 133 | 239 | 125 | 151 | 151 | 187 | 216 | 178 | 138 | 160 | 153 | 214 | 132 | 256 | 210 | 189 | 161 | 171 | 164 | 163 | 155 | 207 | 144 | 104 | 200 | 159 | 158 | Gm3604 |
| Gm36210 | 15 | 12 | 15 | 16 | 12 | 13 | 28 | 26 | 44 | 16 | 23 | 24 | 12 | 23 | 20 | 5 | 20 | 37 | 33 | 37 | 17 | 17 | 28 | 29 | 33 | 35 | 20 | 24 | Gm36210 |
| Gm36255 | 0 | 0 | 0 | 0 | 0 | 0 | 0 | 1 | 0 | 1 | 0 | 0 | 0 | 0 | 0 | 0 | 0 | 0 | 0 | 0 | 0 | 2 | 0 | 0 | 0 | 0 | 0 | 0 | Gm36255 |
| Gm3629 | 1 | 2 | 0 | 3 | 7 | 2 | 1 | 0 | 0 | 0 | 0 | 0 | 1 | 0 | 0 | 0 | 1 | 0 | 4 | 0 | 1 | 2 | 0 | 0 | 0 | 3 | 3 | 0 | Gm3629 |
| Gm3633 | 0 | 0 | 0 | 0 | 0 | 0 | 0 | 0 | 0 | 0 | 2 | 0 | 0 | 0 | 0 | 0 | 0 | 0 | 0 | 0 | 0 | 0 | 0 | 0 | 0 | 0 | 0 | 0 | Gm3633 |
| Gm3636 | 456 | 381 | 544 | 485 | 639 | 538 | 567 | 865 | 777 | 940 | 834 | 757 | 786 | 953 | 527 | 400 | 549 | 470 | 605 | 501 | 509 | 897 | 773 | 962 | 921 | 882 | 819 | 854 | Gm3636 |
| Gm3642 | 0 | 0 | 4 | 0 | 0 | 0 | 0 | 0 | 0 | 3 | 0 | 3 | 2 | 0 | 0 | 0 | 0 | 0 | 0 | 0 | 0 | 0 | 0 | 0 | 0 | 0 | 0 | 1 | Gm3642 |
| Gm3646 | 0 | 0 | 0 | 0 | 0 | 0 | 1 | 0 | 0 | 0 | 0 | 2 | 0 | 0 | 0 | 0 | 0 | 3 | 3 | 0 | 0 | 0 | 0 | 0 | 0 | 2 | 0 | 0 | Gm3646 |
| Gm3667 | 74 | 44 | 40 | 38 | 126 | 71 | 80 | 281 | 241 | 301 | 253 | 300 | 172 | 251 | 51 | 44 | 51 | 48 | 79 | 93 | 69 | 299 | 243 | 201 | 251 | 209 | 227 | 138 | Gm3667 |
| Gm3676 | 0 | 0 | 0 | 2 | 2 | 0 | 0 | 0 | 0 | 0 | 4 | 0 | 5 | 0 | 0 | 0 | 1 | 0 | 2 | 0 | 0 | 0 | 0 | 0 | 0 | 0 | 0 | 0 | Gm3676 |
| Gm3685 | 3 | 2 | 1 | 0 | 1 | 3 | 3 | 1 | 3 | 4 | 4 | 3 | 1 | 0 | 0 | 3 | 0 | 4 | 2 | 2 | 8 | 0 | 1 | 2 | 0 | 1 | 2 | 5 | Gm3685 |
| Gm3696 | 63 | 68 | 70 | 63 | 88 | 93 | 115 | 320 | 209 | 226 | 250 | 240 | 298 | 316 | 71 | 63 | 67 | 69 | 126 | 84 | 78 | 291 | 225 | 265 | 230 | 233 | 298 | 203 | Gm3696 |
| Gm37240 | 7 | 1 | 0 | 3 | 0 | 1 | 0 | 0 | 1 | 2 | 0 | 1 | 6 | 0 | 3 | 6 | 3 | 0 | 2 | 1 | 1 | 0 | 1 | 0 | 3 | 0 | 0 | 1 | Gm37240 |
| Gm3727 | 0 | 5 | 0 | 3 | 0 | 0 | 0 | 0 | 2 | 8 | 3 | 1 | 0 | 4 | 0 | 0 | 5 | 3 | 0 | 0 | 2 | 2 | 1 | 5 | 0 | 13 | 0 | 0 | Gm3727 |
| Gm3739 | 270 | 190 | 228 | 223 | 343 | 317 | 321 | 492 | 431 | 416 | 436 | 507 | 430 | 511 | 278 | 239 | 250 | 206 | 329 | 302 | 301 | 496 | 397 | 384 | 522 | 444 | 444 | 508 | Gm3739 |
| Gm37419 | 0 | 0 | 0 | 0 | 0 | 0 | 0 | 0 | 2 | 0 | 0 | 0 | 0 | 2 | 0 | 0 | 2 | 0 | 0 | 0 | 0 | 0 | 0 | 1 | 0 | 1 | 0 | 0 | Gm37419 |
| Gm3752 | 50 | 29 | 82 | 21 | 128 | 42 | 104 | 181 | 181 | 150 | 201 | 272 | 163 | 217 | 55 | 51 | 61 | 58 | 57 | 69 | 69 | 311 | 240 | 208 | 313 | 197 | 209 | 177 | Gm3752 |
| Gm3755 | 13 | 9 | 10 | 9 | 8 | 13 | 35 | 17 | 9 | 5 | 17 | 37 | 20 | 11 | 34 | 5 | 10 | 15 | 4 | 12 | 13 | 15 | 7 | 16 | 16 | 13 | 13 | 7 | Gm3755 |
| Gm3776 | 1 | 0 | 0 | 0 | 0 | 0 | 0 | 0 | 0 | 0 | 0 | 0 | 0 | 0 | 0 | 0 | 0 | 0 | 0 | 0 | 0 | 0 | 0 | 1 | 0 | 0 | 0 | 0 | Gm3776 |
| Gm3839 | 12 | 4 | 6 | 4 | 7 | 5 | 3 | 4 | 7 | 1 | 1 | 8 | 7 | 2 | 7 | 12 | 5 | 11 | 5 | 10 | 4 | 3 | 8 | 4 | 7 | 6 | 3 | 5 | Gm3839 |
| Gm38392 | 8 | 6 | 0 | 3 | 5 | 8 | 6 | 2 | 1 | 10 | 1 | 7 | 5 | 6 | 7 | 5 | 3 | 4 | 4 | 4 | 4 | 2 | 8 | 7 | 3 | 0 | 1 | 5 | Gm38392 |
| Gm38393 | 4347 | 2507 | 1193 | 2859 | 2969 | 4476 | 3889 | 4107 | 3133 | 1958 | 2554 | 3216 | 3162 | 3654 | 4428 | 3624 | 2042 | 2608 | 2731 | 4628 | 4391 | 2306 | 2758 | 1749 | 2993 | 464 | 3872 | 3210 | Gm38393 |
| Gm38431 | 21 | 35 | 34 | 34 | 38 | 8 | 34 | 37 | 16 | 57 | 39 | 34 | 29 | 32 | 21 | 25 | 45 | 23 | 56 | 26 | 38 | 40 | 20 | 35 | 37 | 24 | 34 | 36 | Gm38431 |
| Gm3854 | 164 | 180 | 197 | 216 | 253 | 182 | 207 | 628 | 517 | 585 | 540 | 570 | 530 | 614 | 211 | 121 | 235 | 226 | 298 | 219 | 269 | 710 | 568 | 603 | 638 | 489 | 622 | 563 | Gm3854 |
| Gm3858 | 0 | 1 | 0 | 0 | 0 | 2 | 0 | 0 | 0 | 0 | 2 | 0 | 0 | 0 | 0 | 1 | 3 | 0 | 0 | 0 | 0 | 0 | 1 | 0 | 0 | 0 | 0 | 0 | Gm3858 |
| Gm3893 | 0 | 0 | 0 | 0 | 0 | 0 | 0 | 0 | 0 | 0 | 0 | 0 | 0 | 0 | 0 | 0 | 0 | 0 | 0 | 2 | 1 | 0 | 0 | 0 | 1 | 0 | 0 | 0 | Gm3893 |
| Gm39566 | 0 | 0 | 0 | 2 | 0 | 4 | 0 | 0 | 0 | 0 | 0 | 0 | 0 | 0 | 0 | 1 | 0 | 2 | 1 | 0 | 1 | 2 | 0 | 0 | 0 | 0 | 0 | 0 | Gm39566 |
| Gm39653 | 2 | 3 | 4 | 3 | 0 | 1 | 3 | 1 | 0 | 0 | 0 | 1 | 9 | 2 | 3 | 0 | 1 | 2 | 1 | 0 | 4 | 0 | 3 | 0 | 6 | 0 | 0 | 0 | Gm39653 |
| Gm4064 | 0 | 0 | 0 | 0 | 1 | 1 | 0 | 0 | 0 | 0 | 0 | 0 | 0 | 0 | 0 | 0 | 0 | 0 | 0 | 0 | 0 | 0 | 0 | 0 | 0 | 0 | 0 | 0 | Gm4064 |
| Gm4125 | 0 | 0 | 0 | 1 | 0 | 0 | 2 | 0 | 0 | 0 | 0 | 0 | 0 | 0 | 0 | 0 | 5 | 0 | 1 | 0 | 0 | 0 | 0 | 0 | 0 | 0 | 0 | 0 | Gm4125 |
| Gm4131 | 6 | 0 | 1 | 0 | 0 | 0 | 0 | 2 | 0 | 0 | 7 | 0 | 1 | 1 | 1 | 2 | 0 | 0 | 1 | 0 | 0 | 0 | 5 | 9 | 1 | 7 | 0 | 1 | Gm4131 |
| Gm4181 | 0 | 0 | 0 | 0 | 0 | 0 | 0 | 0 | 0 | 0 | 0 | 0 | 0 | 0 | 1 | 0 | 0 | 0 | 0 | 0 | 0 | 0 | 0 | 0 | 0 | 0 | 0 | 0 | Gm4181 |
| Gm42372 | 3527 | 4299 | 3519 | 5070 | 4534 | 4505 | 4280 | 1806 | 1681 | 1973 | 1634 | 1922 | 1472 | 1836 | 3994 | 3254 | 4155 | 4769 | 4507 | 4752 | 4705 | 2139 | 1580 | 1863 | 2046 | 1487 | 1855 | 1581 | Gm42372 |
| Gm42517 | 33 | 41 | 34 | 69 | 175 | 111 | 68 | 77 | 81 | 95 | 73 | 30 | 23 | 26 | 46 | 27 | 57 | 61 | 104 | 87 | 77 | 104 | 95 | 42 | 66 | 18 | 51 | 15 | Gm42517 |
| Gm42669 | 3 | 2 | 6 | 2 | 6 | 8 | 4 | 2 | 6 | 1 | 2 | 2 | 2 | 0 | 3 | 2 | 8 | 2 | 3 | 8 | 3 | 3 | 3 | 7 | 3 | 2 | 6 | 0 | Gm42669 |
| Gm42688 | 122 | 136 | 121 | 142 | 169 | 143 | 163 | 74 | 85 | 121 | 97 | 96 | 68 | 100 | 131 | 97 | 147 | 131 | 143 | 140 | 188 | 110 | 64 | 69 | 124 | 85 | 67 | 83 | Gm42688 |
| Gm42715 | 102 | 47 | 184 | 181 | 168 | 154 | 193 | 308 | 256 | 418 | 370 | 463 | 446 | 535 | 71 | 69 | 178 | 167 | 174 | 96 | 174 | 548 | 233 | 535 | 457 | 653 | 381 | 504 | Gm42715 |
| Gm42742 | 6 | 6 | 2 | 0 | 2 | 2 | 4 | 4 | 0 | 1 | 9 | 0 | 3 | 3 | 0 | 7 | 9 | 6 | 8 | 3 | 2 | 5 | 6 | 8 | 5 | 2 | 3 | 4 | Gm42742 |
| Gm42791 | 13 | 21 | 36 | 17 | 16 | 24 | 22 | 0 | 7 | 8 | 17 | 16 | 7 | 11 | 13 | 30 | 19 | 19 | 16 | 27 | 17 | 5 | 8 | 6 | 5 | 8 | 20 | 15 | Gm42791 |
| Gm42878 | 0 | 0 | 0 | 0 | 1 | 3 | 0 | 0 | 0 | 0 | 0 | 0 | 0 | 0 | 0 | 0 | 0 | 0 | 0 | 0 | 0 | 0 | 0 | 0 | 0 | 0 | 0 | 0 | Gm42878 |
| Gm42906 | 0 | 0 | 3 | 0 | 0 | 0 | 0 | 4 | 0 | 0 | 0 | 0 | 0 | 0 | 0 | 1 | 0 | 0 | 0 | 0 | 0 | 2 | 0 | 0 | 0 | 0 | 0 | 4 | Gm42906 |
| Gm42957 | 7 | 25 | 8 | 14 | 22 | 13 | 23 | 5 | 10 | 12 | 8 | 4 | 6 | 1 | 9 | 16 | 16 | 31 | 9 | 15 | 20 | 0 | 8 | 2 | 11 | 8 | 9 | 4 | Gm42957 |
| Gm43064 | 3 | 3 | 0 | 0 | 3 | 8 | 3 | 5 | 2 | 1 | 1 | 4 | 0 | 1 | 9 | 5 | 2 | 0 | 6 | 6 | 7 | 0 | 1 | 5 | 1 | 2 | 0 | 0 | Gm43064 |
| Gm43517 | 257 | 136 | 45 | 116 | 155 | 238 | 256 | 398 | 420 | 185 | 207 | 287 | 94 | 309 | 186 | 75 | 61 | 92 | 157 | 364 | 320 | 127 | 274 | 224 | 384 | 191 | 263 | 161 | Gm43517 |
| Gm43518 | 5 | 4 | 4 | 3 | 4 | 0 | 7 | 4 | 4 | 5 | 9 | 7 | 5 | 2 | 5 | 4 | 1 | 6 | 2 | 7 | 6 | 0 | 3 | 1 | 9 | 1 | 5 | 1 | Gm43518 |
| Gm4356 | 85 | 73 | 28 | 82 | 59 | 92 | 83 | 67 | 58 | 30 | 66 | 61 | 68 | 55 | 102 | 98 | 45 | 70 | 77 | 85 | 83 | 27 | 69 | 40 | 83 | 13 | 74 | 53 | Gm4356 |
| Gm43720 | 1 | 5 | 0 | 3 | 3 | 1 | 4 | 0 | 2 | 0 | 2 | 0 | 0 | 0 | 4 | 0 | 0 | 11 | 0 | 1 | 2 | 0 | 0 | 0 | 0 | 0 | 0 | 0 | Gm43720 |
| Gm43738 | 2 | 2 | 0 | 0 | 3 | 1 | 2 | 0 | 1 | 0 | 0 | 0 | 0 | 0 | 0 | 2 | 0 | 0 | 0 | 0 | 3 | 0 | 1 | 1 | 0 | 0 | 0 | 2 | Gm43738 |
| Gm44501 | 0 | 0 | 0 | 2 | 0 | 0 | 0 | 0 | 0 | 0 | 0 | 0 | 0 | 0 | 0 | 0 | 0 | 0 | 0 | 2 | 1 | 0 | 0 | 0 | 0 | 0 | 0 | 0 | Gm44501 |
| Gm4491 | 0 | 0 | 0 | 0 | 0 | 0 | 1 | 1 | 1 | 0 | 0 | 1 | 6 | 1 | 3 | 1 | 0 | 2 | 0 | 0 | 0 | 2 | 4 | 2 | 4 | 0 | 6 | 2 | Gm4491 |
| Gm45015 | 0 | 0 | 0 | 0 | 0 | 0 | 0 | 0 | 0 | 0 | 0 | 0 | 0 | 0 | 1 | 0 | 0 | 0 | 0 | 0 | 0 | 0 | 0 | 0 | 0 | 0 | 0 | 0 | Gm45015 |
| Gm45140 | 10 | 10 | 7 | 5 | 8 | 2 | 12 | 0 | 7 | 6 | 4 | 4 | 10 | 14 | 0 | 9 | 5 | 4 | 3 | 7 | 10 | 15 | 7 | 7 | 11 | 8 | 9 | 7 | Gm45140 |
| Gm45194 | 70 | 52 | 7 | 37 | 20 | 46 | 53 | 48 | 26 | 23 | 44 | 12 | 8 | 34 | 73 | 46 | 13 | 8 | 17 | 89 | 68 | 13 | 38 | 2 | 14 | 1 | 27 | 13 | Gm45194 |
| Gm45234 | 32 | 35 | 31 | 36 | 49 | 65 | 38 | 34 | 43 | 43 | 55 | 58 | 46 | 42 | 29 | 31 | 21 | 62 | 60 | 40 | 48 | 60 | 46 | 38 | 57 | 10 | 34 | 40 | Gm45234 |
| Gm45521 | 7 | 14 | 9 | 8 | 7 | 6 | 2 | 4 | 4 | 11 | 2 | 5 | 9 | 4 | 7 | 12 | 21 | 3 | 7 | 14 | 2 | 8 | 11 | 5 | 4 | 3 | 5 | 4 | Gm45521 |
| Gm45623 | 25 | 13 | 7 | 38 | 40 | 29 | 25 | 24 | 29 | 43 | 42 | 27 | 7 | 14 | 9 | 16 | 22 | 23 | 35 | 37 | 20 | 33 | 52 | 42 | 46 | 24 | 13 | 17 | Gm45623 |
| Gm45716 | 637 | 435 | 547 | 581 | 753 | 666 | 740 | 603 | 423 | 622 | 505 | 579 | 644 | 743 | 630 | 442 | 593 | 651 | 617 | 686 | 769 | 541 | 452 | 521 | 615 | 571 | 581 | 681 | Gm45716 |
| Gm45753 | 0 | 0 | 0 | 0 | 0 | 0 | 0 | 0 | 0 | 0 | 0 | 0 | 0 | 0 | 0 | 0 | 0 | 0 | 1 | 0 | 0 | 0 | 0 | 0 | 0 | 0 | 0 | 0 | Gm45753 |
| Gm45785 | 6 | 10 | 0 | 5 | 2 | 9 | 7 | 4 | 2 | 0 | 3 | 0 | 3 | 3 | 9 | 10 | 2 | 4 | 3 | 9 | 6 | 0 | 4 | 7 | 6 | 0 | 3 | 1 | Gm45785 |
| Gm45799 | 1 | 10 | 1 | 7 | 2 | 3 | 6 | 1 | 8 | 5 | 6 | 4 | 2 | 9 | 4 | 9 | 5 | 11 | 4 | 12 | 6 | 7 | 1 | 7 | 1 | 3 | 6 | 11 | Gm45799 |
| Gm45826 | 2 | 0 | 0 | 0 | 0 | 0 | 0 | 0 | 0 | 0 | 0 | 0 | 0 | 0 | 0 | 0 | 0 | 0 | 0 | 0 | 0 | 0 | 0 | 0 | 0 | 0 | 0 | 0 | Gm45826 |
| Gm45837 | 3 | 3 | 0 | 0 | 2 | 0 | 0 | 1 | 1 | 0 | 2 | 1 | 1 | 0 | 5 | 2 | 1 | 8 | 2 | 2 | 0 | 0 | 1 | 2 | 1 | 1 | 2 | 8 | Gm45837 |
| Gm45844 | 37 | 26 | 27 | 35 | 66 | 44 | 63 | 79 | 102 | 103 | 138 | 53 | 74 | 72 | 24 | 28 | 29 | 52 | 44 | 56 | 49 | 122 | 130 | 115 | 109 | 73 | 78 | 68 | Gm45844 |
| Gm45861 | 1 | 1 | 0 | 2 | 1 | 0 | 0 | 0 | 7 | 1 | 1 | 0 | 2 | 0 | 0 | 2 | 2 | 0 | 0 | 0 | 0 | 3 | 3 | 0 | 0 | 0 | 0 | 0 | Gm45861 |
| Gm45871 | 940 | 658 | 787 | 734 | 790 | 762 | 867 | 890 | 700 | 693 | 621 | 739 | 646 | 702 | 952 | 816 | 867 | 717 | 719 | 895 | 826 | 797 | 772 | 635 | 608 | 818 | 827 | 797 | Gm45871 |
| Gm45902 | 1781 | 1965 | 1462 | 1510 | 1614 | 1890 | 1658 | 1718 | 1453 | 1213 | 1250 | 1409 | 1331 | 1454 | 1899 | 1851 | 1586 | 1549 | 1460 | 1904 | 1750 | 1200 | 1349 | 1174 | 1314 | 1310 | 1482 | 1342 | Gm45902 |

| | | | | | | | | | | | | | | | | | | | | | | | | | | | | | |
|---|---|---|---|---|---|---|---|---|---|---|---|---|---|---|---|---|---|---|---|---|---|---|---|---|---|---|---|---|---|
| Gm4631 | 9 | 2 | 37 | 7 | 1 | 2 | 1 | 4 | 2 | 34 | 4 | 3 | 1 | 3 | 1 | 2 | 43 | 2 | 13 | 10 | 4 | 3 | 11 | 1 | 8 | 1 | 0 | 4 | Gm4631 |
| Gm4724 | 17 | 7 | 4 | 7 | 9 | 8 | 17 | 33 | 12 | 3 | 6 | 22 | 5 | 11 | 36 | 15 | 9 | 1 | 13 | 9 | 13 | 12 | 7 | 9 | 6 | 8 | 7 | 8 | Gm4724 |
| Gm4737 | 1497 | 1468 | 1391 | 953 | 738 | 889 | 683 | 1071 | 1234 | 836 | 1013 | 1312 | 1168 | 1092 | 1290 | 1365 | 1176 | 835 | 880 | 833 | 747 | 795 | 1097 | 1044 | 943 | 1463 | 1330 | 1275 | Gm4737 |
| Gm47429 | 0 | 0 | 0 | 0 | 0 | 0 | 0 | 0 | 1 | 0 | 1 | 0 | 0 | 0 | 0 | 0 | 0 | 0 | 0 | 0 | 0 | 0 | 1 | 0 | 0 | 0 | 1 | 0 | Gm47429 |
| Gm4756 | 0 | 0 | 0 | 0 | 0 | 0 | 1 | 0 | 0 | 0 | 0 | 0 | 0 | 0 | 0 | 0 | 0 | 0 | 0 | 0 | 0 | 3 | 0 | 0 | 0 | 0 | 0 | 1 | Gm4756 |
| Gm4767 | 8 | 9 | 1 | 6 | 7 | 5 | 2 | 7 | 3 | 3 | 3 | 3 | 41 | 14 | 53 | 8 | 5 | 12 | 8 | 9 | 14 | 7 | 14 | 17 | 7 | 8 | 7 | 13 | Gm4767 |
| Gm4779 | 5 | 7 | 1 | 0 | 4 | 3 | 2 | 15 | 3 | 5 | 21 | 8 | 20 | 19 | 0 | 2 | 23 | 0 | 0 | 3 | 0 | 35 | 14 | 0 | 2 | 1 | 0 | 1 | Gm4779 |
| Gm47791 | 1 | 0 | 0 | 0 | 0 | 0 | 0 | 0 | 0 | 0 | 0 | 0 | 0 | 0 | 0 | 0 | 0 | 0 | 2 | 1 | 0 | 0 | 0 | 0 | 0 | 1 | 0 | 0 | Gm47791 |
| Gm4787 | 61 | 86 | 91 | 73 | 79 | 109 | 72 | 111 | 96 | 41 | 68 | 46 | 81 | 77 | 90 | 97 | 71 | 72 | 60 | 104 | 62 | 107 | 63 | 74 | 94 | 60 | 108 | 84 | Gm4787 |
| Gm4791 | 5 | 0 | 0 | 1 | 0 | 5 | 0 | 11 | 9 | 3 | 7 | 0 | 0 | 2 | 1 | 0 | 0 | 0 | 0 | 0 | 0 | 0 | 10 | 17 | 4 | 16 | 0 | 0 | Gm4791 |
| Gm4792 | 1 | 2 | 5 | 2 | 4 | 1 | 3 | 4 | 5 | 5 | 3 | 10 | 6 | 11 | 3 | 1 | 2 | 0 | 5 | 0 | 0 | 3 | 4 | 5 | 3 | 3 | 14 | 8 | Gm4792 |
| Gm47955 | 0 | 0 | 0 | 0 | 0 | 0 | 0 | 0 | 0 | 0 | 0 | 0 | 0 | 0 | 0 | 0 | 0 | 0 | 0 | 0 | 0 | 0 | 0 | 1 | 0 | 0 | 0 | 1 | Gm47955 |
| Gm47959 | 0 | 0 | 0 | 0 | 0 | 0 | 0 | 0 | 0 | 0 | 0 | 0 | 2 | 0 | 0 | 0 | 0 | 1 | 0 | 0 | 0 | 0 | 0 | 0 | 0 | 0 | 0 | 2 | Gm47959 |
| Gm47969 | 0 | 0 | 0 | 0 | 0 | 0 | 1 | 1 | 0 | 2 | 0 | 0 | 0 | 0 | 0 | 0 | 0 | 0 | 0 | 0 | 2 | 0 | 0 | 2 | 0 | 0 | 0 | 0 | Gm47969 |
| Gm47985 | 2 | 8 | 10 | 1 | 10 | 3 | 7 | 23 | 18 | 8 | 1 | 18 | 70 | 24 | 12 | 2 | 6 | 9 | 3 | 13 | 0 | 8 | 10 | 7 | 5 | 12 | 20 | 72 | Gm47985 |
| Gm47995 | 0 | 0 | 0 | 0 | 0 | 0 | 0 | 0 | 0 | 0 | 0 | 0 | 0 | 1 | 0 | 0 | 2 | 0 | 0 | 0 | 0 | 0 | 2 | 2 | 0 | 5 | 2 | 4 | Gm47995 |
| Gm47996 | 0 | 0 | 0 | 0 | 0 | 0 | 0 | 0 | 0 | 0 | 0 | 0 | 0 | 0 | 0 | 0 | 0 | 0 | 0 | 0 | 0 | 0 | 1 | 0 | 0 | 1 | 0 | 1 | Gm47996 |
| Gm4841 | 0 | 0 | 0 | 0 | 1 | 0 | 0 | 5 | 3 | 3 | 2 | 1 | 2 | 0 | 0 | 0 | 0 | 0 | 0 | 1 | 0 | 0 | 3 | 16 | 1 | 18 | 0 | 1 | Gm4841 |
| Gm4846 | 0 | 0 | 0 | 0 | 0 | 0 | 0 | 0 | 2 | 0 | 0 | 0 | 0 | 0 | 0 | 0 | 0 | 0 | 0 | 0 | 0 | 3 | 1 | 1 | 0 | 9 | 3 | 0 | Gm4846 |
| Gm48551 | 16 | 22 | 4 | 15 | 16 | 19 | 27 | 22 | 7 | 8 | 7 | 15 | 9 | 16 | 22 | 13 | 21 | 32 | 9 | 24 | 22 | 13 | 5 | 15 | 11 | 1 | 5 | 9 | Gm48551 |
| Gm48552 | 108 | 135 | 136 | 186 | 102 | 116 | 107 | 89 | 69 | 80 | 61 | 74 | 71 | 49 | 119 | 143 | 134 | 164 | 129 | 104 | 84 | 94 | 69 | 67 | 68 | 71 | 49 | 53 | Gm48552 |
| Gm4871 | 0 | 0 | 0 | 0 | 0 | 0 | 0 | 0 | 0 | 0 | 0 | 0 | 1 | 0 | 0 | 0 | 0 | 0 | 0 | 0 | 0 | 0 | 1 | 0 | 0 | 0 | 0 | 0 | Gm4871 |
| Gm4894 | 0 | 0 | 0 | 0 | 2 | 1 | 0 | 0 | 1 | 0 | 0 | 0 | 3 | 0 | 0 | 0 | 0 | 0 | 0 | 0 | 0 | 0 | 1 | 0 | 1 | 0 | 1 | 0 | Gm4894 |
| Gm49027 | 93 | 103 | 72 | 110 | 165 | 111 | 111 | 83 | 133 | 126 | 132 | 136 | 115 | 110 | 98 | 114 | 96 | 122 | 108 | 92 | 103 | 165 | 93 | 91 | 158 | 73 | 115 | 117 | Gm49027 |
| Gm4922 | 0 | 0 | 0 | 0 | 0 | 0 | 0 | 1 | 0 | 0 | 0 | 0 | 0 | 0 | 0 | 0 | 0 | 0 | 0 | 0 | 0 | 0 | 0 | 0 | 0 | 0 | 0 | 0 | Gm4922 |
| Gm4924 | 384 | 351 | 679 | 453 | 576 | 511 | 611 | 965 | 1004 | 1056 | 892 | 1127 | 1112 | 1142 | 555 | 398 | 666 | 475 | 599 | 532 | 512 | 1200 | 942 | 1074 | 980 | 1157 | 1146 | 1101 | Gm4924 |
| Gm49322 | 95 | 143 | 178 | 155 | 143 | 134 | 120 | 147 | 151 | 194 | 217 | 293 | 220 | 222 | 74 | 93 | 145 | 154 | 177 | 134 | 123 | 196 | 170 | 265 | 225 | 199 | 237 | 225 | Gm49322 |
| Gm49336 | 10112 | 7432 | 8301 | 8658 | 8341 | 8744 | 9767 | 8327 | 6654 | 5633 | 5777 | 6347 | 6338 | 6760 | 10669 | 9371 | 7829 | 8307 | 7930 | 9856 | 9247 | 7265 | 6521 | 5937 | 6494 | 6419 | 6938 | 6868 | Gm49336 |
| Gm49355 | 5 | 3 | 1 | 1 | 5 | 8 | 4 | 7 | 3 | 6 | 2 | 5 | 1 | 1 | 1 | 9 | 1 | 0 | 2 | 2 | 10 | 0 | 5 | 1 | 4 | 2 | 9 | 1 | Gm49355 |
| Gm49373 | 3 | 2 | 9 | 9 | 5 | 1 | 6 | 10 | 3 | 4 | 8 | 9 | 6 | 10 | 5 | 1 | 7 | 0 | 2 | 3 | 8 | 8 | 6 | 9 | 8 | 2 | 1 | 7 | Gm49373 |
| Gm49380 | 15 | 2 | 23 | 22 | 22 | 12 | 17 | 2 | 6 | 3 | 12 | 8 | 8 | 13 | 16 | 11 | 6 | 26 | 12 | 5 | 4 | 7 | 6 | 12 | 12 | 15 | 5 | 17 | Gm49380 |
| Gm49383 | 29 | 10 | 11 | 36 | 33 | 35 | 28 | 13 | 5 | 12 | 8 | 5 | 5 | 10 | 15 | 28 | 31 | 46 | 19 | 22 | 27 | 7 | 4 | 3 | 11 | 7 | 14 | 11 | Gm49383 |
| Gm49394 | 0 | 0 | 2 | 0 | 2 | 0 | 0 | 2 | 7 | 10 | 28 | 0 | 2 | 0 | 0 | 3 | 5 | 2 | 0 | 0 | 1 | 5 | 21 | 1 | 5 | 0 | 2 | 2 | Gm49394 |
| Gm49395 | 7 | 6 | 7 | 17 | 20 | 9 | 11 | 10 | 3 | 5 | 19 | 13 | 10 | 17 | 11 | 5 | 5 | 6 | 14 | 7 | 4 | 10 | 6 | 7 | 18 | 5 | 1 | 5 | Gm49395 |
| Gm49396 | 104 | 73 | 116 | 104 | 137 | 118 | 159 | 289 | 199 | 313 | 214 | 264 | 244 | 330 | 99 | 83 | 103 | 114 | 118 | 109 | 141 | 324 | 206 | 304 | 272 | 283 | 231 | 311 | Gm49396 |
| Gm49405 | 180 | 233 | 242 | 295 | 317 | 230 | 350 | 263 | 291 | 403 | 307 | 311 | 253 | 368 | 231 | 151 | 309 | 352 | 353 | 219 | 356 | 359 | 284 | 404 | 359 | 310 | 252 | 332 | Gm49405 |
| Gm49425 | 0 | 0 | 0 | 0 | 0 | 1 | 0 | 0 | 0 | 0 | 0 | 0 | 0 | 0 | 0 | 0 | 0 | 0 | 0 | 0 | 0 | 0 | 0 | 0 | 0 | 0 | 0 | 0 | Gm49425 |
| Gm49486 | 69 | 47 | 75 | 116 | 88 | 72 | 78 | 38 | 39 | 41 | 49 | 27 | 41 | 48 | 67 | 57 | 97 | 64 | 69 | 61 | 93 | 50 | 43 | 56 | 37 | 46 | 26 | 51 | Gm49486 |
| Gm4951 | 2 | 1 | 1 | 6 | 3 | 11 | 4 | 13 | 6 | 5 | 17 | 8 | 12 | 13 | 7 | 6 | 0 | 0 | 0 | 24 | 4 | 32 | 20 | 15 | 19 | 5 | 21 | 28 | Gm4951 |
| Gm4952 | 14 | 16 | 13 | 10 | 12 | 19 | 17 | 2 | 4 | 5 | 6 | 0 | 1 | 2 | 24 | 20 | 7 | 22 | 7 | 9 | 16 | 5 | 2 | 2 | 1 | 5 | 1 | 2 | Gm4952 |
| Gm49601 | 257 | 359 | 486 | 453 | 540 | 462 | 435 | 414 | 452 | 506 | 500 | 532 | 417 | 622 | 289 | 291 | 421 | 517 | 509 | 363 | 556 | 642 | 372 | 601 | 633 | 426 | 431 | 446 | Gm49601 |
| Gm49673 | 7 | 5 | 3 | 5 | 2 | 7 | 0 | 0 | 3 | 0 | 0 | 0 | 9 | 4 | 7 | 2 | 2 | 2 | 2 | 1 | 4 | 0 | 5 | 0 | 2 | 0 | 1 | 0 | Gm49673 |
| Gm49711 | 21 | 20 | 17 | 21 | 40 | 18 | 34 | 2 | 2 | 10 | 8 | 1 | 8 | 12 | 44 | 23 | 36 | 34 | 42 | 24 | 50 | 3 | 10 | 7 | 5 | 13 | 3 | 1 | Gm49711 |
| Gm49721 | 7 | 7 | 0 | 7 | 7 | 7 | 10 | 2 | 5 | 4 | 3 | 3 | 12 | 5 | 7 | 9 | 9 | 3 | 9 | 9 | 4 | 3 | 9 | 0 | 8 | 0 | 8 | 0 | Gm49721 |
| Gm50092 | 0 | 1 | 0 | 3 | 1 | 1 | 1 | 6 | 1 | 3 | 1 | 1 | 3 | 2 | 7 | 0 | 3 | 3 | 1 | 3 | 1 | 0 | 3 | 4 | 6 | 2 | 5 | 0 | Gm50092 |
| Gm50253 | 18 | 20 | 20 | 21 | 18 | 15 | 11 | 13 | 9 | 14 | 10 | 23 | 15 | 15 | 12 | 10 | 29 | 13 | 13 | 7 | 43 | 23 | 9 | 11 | 19 | 20 | 8 | 27 | Gm50253 |
| Gm50364 | 9 | 9 | 0 | 11 | 6 | 8 | 8 | 18 | 14 | 6 | 8 | 14 | 9 | 4 | 13 | 8 | 8 | 6 | 7 | 15 | 4 | 8 | 8 | 6 | 12 | 13 | 7 | 9 | Gm50364 |
| Gm50387 | 265 | 282 | 275 | 333 | 270 | 227 | 365 | 223 | 189 | 184 | 203 | 167 | 124 | 223 | 276 | 254 | 286 | 362 | 253 | 256 | 371 | 269 | 184 | 214 | 209 | 184 | 160 | 202 | Gm50387 |
| Gm50388 | 1 | 0 | 0 | 0 | 0 | 0 | 1 | 0 | 0 | 0 | 1 | 1 | 0 | 0 | 5 | 2 | 0 | 0 | 0 | 1 | 4 | 0 | 0 | 1 | 0 | 0 | 0 | 0 | Gm50388 |
| Gm50457 | 2 | 1 | 6 | 3 | 4 | 1 | 3 | 7 | 5 | 1 | 2 | 1 | 1 | 3 | 3 | 1 | 1 | 0 | 0 | 2 | 2 | 3 | 3 | 2 | 0 | 1 | 0 | 0 | Gm50457 |
| Gm5065 | 0 | 0 | 0 | 0 | 1 | 0 | 0 | 0 | 0 | 0 | 0 | 0 | 0 | 0 | 0 | 0 | 0 | 7 | 0 | 3 | 1 | 0 | 0 | 0 | 0 | 0 | 0 | 0 | Gm5065 |
| Gm5096 | 1 | 2 | 0 | 0 | 3 | 2 | 1 | 2 | 6 | 8 | 3 | 3 | 5 | 0 | 1 | 5 | 0 | 0 | 2 | 2 | 3 | 8 | 10 | 0 | 0 | 0 | 0 | 0 | Gm5096 |
| Gm5105 | 8 | 5 | 15 | 16 | 16 | 6 | 9 | 2 | 2 | 8 | 17 | 4 | 8 | 1 | 8 | 9 | 13 | 8 | 8 | 14 | 6 | 10 | 9 | 4 | 6 | 2 | 0 | 5 | Gm5105 |
| Gm5108 | 0 | 1 | 0 | 0 | 2 | 0 | 0 | 0 | 0 | 0 | 0 | 0 | 0 | 0 | 0 | 0 | 2 | 0 | 0 | 0 | 1 | 0 | 0 | 0 | 0 | 0 | 0 | 0 | Gm5108 |
| Gm5127 | 47 | 37 | 67 | 13 | 21 | 14 | 4 | 29 | 33 | 37 | 33 | 34 | 48 | 15 | 49 | 96 | 31 | 9 | 4 | 4 | 9 | 35 | 16 | 28 | 17 | 51 | 52 | 21 | Gm5127 |
| Gm5134 | 1 | 3 | 2 | 1 | 2 | 3 | 7 | 2 | 11 | 1 | 4 | 1 | 8 | 7 | 7 | 3 | 9 | 3 | 2 | 2 | 4 | 13 | 6 | 2 | 7 | 12 | 3 | 4 | Gm5134 |
| Gm5136 | 5 | 1 | 9 | 18 | 10 | 12 | 19 | 4 | 2 | 14 | 6 | 16 | 15 | 11 | 1 | 6 | 15 | 15 | 9 | 3 | 20 | 17 | 1 | 9 | 13 | 16 | 3 | 12 | Gm5136 |
| Gm5141 | 901 | 645 | 932 | 787 | 938 | 940 | 949 | 1072 | 951 | 912 | 818 | 923 | 855 | 841 | 1068 | 942 | 862 | 813 | 1004 | 1078 | 919 | 937 | 884 | 903 | 905 | 1055 | 910 | 849 | Gm5141 |
| Gm5148 | 241 | 181 | 209 | 246 | 275 | 269 | 294 | 189 | 206 | 210 | 186 | 137 | 172 | 137 | 214 | 199 | 220 | 259 | 311 | 299 | 310 | 249 | 205 | 183 | 247 | 119 | 127 | 87 | Gm5148 |
| Gm5150 | 0 | 0 | 0 | 0 | 0 | 0 | 1 | 0 | 0 | 3 | 0 | 0 | 0 | 0 | 0 | 0 | 0 | 0 | 0 | 0 | 0 | 0 | 0 | 0 | 0 | 0 | 0 | 0 | Gm5150 |
| Gm5152 | 0 | 0 | 0 | 0 | 0 | 0 | 0 | 0 | 0 | 2 | 0 | 0 | 0 | 0 | 0 | 0 | 0 | 0 | 0 | 0 | 0 | 0 | 0 | 0 | 0 | 0 | 0 | 0 | Gm5152 |
| Gm5155 | 0 | 0 | 0 | 0 | 0 | 0 | 0 | 0 | 12 | 0 | 0 | 0 | 0 | 0 | 0 | 0 | 0 | 0 | 0 | 0 | 0 | 0 | 0 | 0 | 0 | 0 | 0 | 0 | Gm5155 |
| Gm5244 | 0 | 0 | 0 | 0 | 0 | 0 | 0 | 0 | 0 | 0 | 0 | 0 | 0 | 1 | 0 | 0 | 0 | 0 | 0 | 0 | 0 | 0 | 0 | 0 | 0 | 0 | 0 | 0 | Gm5244 |
| Gm525 | 2 | 0 | 1 | 2 | 7 | 2 | 6 | 2 | 8 | 1 | 9 | 1 | 3 | 1 | 5 | 0 | 2 | 2 | 2 | 2 | 1 | 2 | 10 | 5 | 10 | 0 | 0 | 0 | Gm525 |
| Gm527 | 518 | 408 | 353 | 533 | 466 | 439 | 466 | 506 | 418 | 380 | 501 | 403 | 438 | 454 | 608 | 493 | 450 | 636 | 432 | 466 | 510 | 556 | 449 | 374 | 447 | 316 | 418 | 438 | Gm527 |
| Gm52955 | 0 | 0 | 0 | 0 | 0 | 0 | 0 | 0 | 0 | 0 | 0 | 0 | 0 | 0 | 0 | 0 | 1 | 0 | 0 | 0 | 0 | 0 | 0 | 0 | 0 | 0 | 0 | 0 | Gm52955 |
| Gm52968 | 1 | 0 | 0 | 0 | 0 | 0 | 0 | 0 | 4 | 6 | 2 | 0 | 1 | 2 | 0 | 0 | 0 | 0 | 0 | 0 | 1 | 0 | 1 | 2 | 0 | 0 | 5 | 1 | Gm52968 |
| Gm53012 | 134 | 95 | 107 | 141 | 115 | 111 | 117 | 99 | 127 | 203 | 136 | 185 | 137 | 162 | 96 | 90 | 153 | 117 | 156 | 106 | 119 | 115 | 122 | 163 | 151 | 180 | 172 | 177 | Gm53012 |
| Gm5415 | 0 | 0 | 7 | 12 | 1 | 2 | 2 | 0 | 1 | 1 | 6 | 2 | 0 | 4 | 3 | 0 | 10 | 14 | 8 | 2 | 1 | 15 | 1 | 14 | 2 | 12 | 6 | 0 | Gm5415 |
| Gm5431 | 2 | 3 | 6 | 1 | 2 | 4 | 6 | 18 | 11 | 33 | 24 | 17 | 24 | 17 | 11 | 9 | 0 | 1 | 3 | 9 | 2 | 28 | 26 | 12 | 10 | 12 | 27 | 19 | Gm5431 |
| Gm5458 | 0 | 1 | 3 | 4 | 2 | 0 | 2 | 1 | 0 | 1 | 0 | 0 | 5 | 2 | 0 | 2 | 0 | 0 | 3 | 0 | 2 | 2 | 0 | 2 | 0 | 0 | 0 | 0 | Gm5458 |
| Gm5475 | 17 | 20 | 8 | 14 | 16 | 9 | 7 | 0 | 3 | 4 | 1 | 0 | 6 | 3 | 11 | 12 | 1 | 15 | 23 | 24 | 18 | 0 | 1 | 2 | 2 | 0 | 0 | 0 | Gm5475 |
| Gm5478 | 0 | 0 | 0 | 0 | 0 | 0 | 0 | 0 | 1 | 0 | 0 | 0 | 0 | 0 | 0 | 0 | 0 | 0 | 0 | 0 | 0 | 0 | 1 | 0 | 0 | 0 | 0 | 0 | Gm5478 |
| Gm5485 | 0 | 0 | 0 | 0 | 0 | 0 | 0 | 0 | 0 | 0 | 0 | 0 | 0 | 0 | 0 | 1 | 0 | 0 | 0 | 0 | 0 | 0 | 1 | 0 | 0 | 0 | 0 | 0 | Gm5485 |
| Gm550 | 257 | 238 | 110 | 165 | 160 | 152 | 151 | 134 | 149 | 106 | 85 | 126 | 150 | 138 | 207 | 181 | 140 | 135 | 175 | 182 | 164 | 69 | 102 | 83 | 96 | 123 | 241 | 115 | Gm550 |
| Gm5565 | 0 | 0 | 0 | 0 | 1 | 0 | 0 | 0 | 0 | 0 | 0 | 0 | 0 | 0 | 0 | 0 | 0 | 0 | 0 | 0 | 0 | 0 | 0 | 0 | 0 | 0 | 0 | 0 | Gm5565 |
| Gm5617 | 156 | 227 | 208 | 203 | 164 | 168 | 123 | 173 | 195 | 214 | 190 | 247 | 276 | 200 | 119 | 173 | 178 | 143 | 150 | 213 | 137 | 124 | 201 | 204 | 180 | 251 | 281 | 176 | Gm5617 |
| Gm5622 | 0 | 0 | 0 | 0 | 0 | 0 | 1 | 0 | 0 | 0 | 0 | 0 | 0 | 0 | 1 | 0 | 0 | 4 | 0 | 0 | 0 | 0 | 1 | 0 | 0 | 0 | 0 | 0 | Gm5622 |
| Gm5678 | 2 | 2 | 0 | 0 | 2 | 0 | 1 | 0 | 0 | 0 | 8 | 1 | 0 | 0 | 1 | 4 | 1 | 2 | 1 | 0 | 1 | 0 | 0 | 1 | 1 | 0 | 2 | 0 | Gm5678 |

| | | | | | | | | | | | | | | | | | | | | | | | | | | | | | |
|---|---|---|---|---|---|---|---|---|---|---|---|---|---|---|---|---|---|---|---|---|---|---|---|---|---|---|---|---|---|
| Gm572 | 1 | 0 | 0 | 0 | 1 | 2 | 0 | 1 | 5 | 0 | 10 | 0 | 0 | 1 | 0 | 1 | 0 | 0 | 2 | 6 | 0 | 5 | 12 | 2 | 0 | 9 | 1 | 0 | Gm572 |
| Gm5724 | 0 | 0 | 0 | 0 | 0 | 3 | 0 | 1 | 2 | 0 | 0 | 0 | 0 | 0 | 1 | 0 | 0 | 0 | 0 | 0 | 3 | 0 | 2 | 0 | 0 | 3 | 0 | 0 | Gm5724 |
| Gm5737 | 13 | 20 | 17 | 34 | 21 | 37 | 22 | 5 | 12 | 5 | 8 | 7 | 7 | 2 | 8 | 16 | 19 | 21 | 28 | 25 | 37 | 10 | 10 | 4 | 1 | 0 | 6 | 4 | Gm5737 |
| Gm5741 | 1 | 6 | 9 | 10 | 5 | 4 | 3 | 2 | 8 | 12 | 7 | 4 | 9 | 5 | 3 | 0 | 7 | 7 | 7 | 5 | 5 | 3 | 6 | 7 | 2 | 8 | 1 | 12 | Gm5741 |
| Gm5751 | 0 | 0 | 0 | 0 | 0 | 0 | 0 | 5 | 0 | 0 | 0 | 0 | 0 | 0 | 0 | 0 | 0 | 0 | 0 | 0 | 0 | 0 | 0 | 0 | 0 | 0 | 0 | 0 | Gm5751 |
| Gm5767 | 0 | 0 | 0 | 0 | 0 | 0 | 0 | 1 | 1 | 0 | 0 | 0 | 0 | 0 | 0 | 0 | 0 | 0 | 0 | 2 | 0 | 0 | 1 | 0 | 0 | 0 | 0 | 0 | Gm5767 |
| Gm5771 | 0 | 0 | 0 | 0 | 0 | 0 | 0 | 0 | 0 | 0 | 2 | 0 | 0 | 0 | 0 | 0 | 0 | 0 | 0 | 0 | 0 | 0 | 1 | 0 | 0 | 0 | 0 | 0 | Gm5771 |
| Gm5795 | 0 | 0 | 0 | 0 | 0 | 0 | 1 | 0 | 0 | 0 | 0 | 0 | 0 | 0 | 0 | 1 | 1 | 0 | 1 | 1 | 2 | 0 | 2 | 0 | 0 | 0 | 0 | 0 | Gm5795 |
| Gm5796 | 28 | 12 | 34 | 19 | 61 | 56 | 46 | 176 | 171 | 132 | 127 | 140 | 86 | 134 | 24 | 28 | 13 | 37 | 47 | 71 | 34 | 224 | 142 | 142 | 187 | 184 | 117 | 135 | Gm5796 |
| Gm5797 | 0 | 0 | 0 | 0 | 0 | 0 | 0 | 0 | 0 | 0 | 0 | 0 | 0 | 0 | 0 | 0 | 0 | 0 | 0 | 0 | 0 | 0 | 0 | 0 | 1 | 0 | 0 | 0 | Gm5797 |
| Gm5800 | 0 | 0 | 1 | 0 | 0 | 0 | 0 | 0 | 0 | 0 | 0 | 0 | 0 | 0 | 0 | 0 | 1 | 0 | 0 | 0 | 0 | 0 | 0 | 0 | 0 | 0 | 0 | 0 | Gm5800 |
| Gm5901 | 0 | 0 | 0 | 1 | 0 | 1 | 0 | 1 | 0 | 0 | 0 | 0 | 0 | 1 | 1 | 0 | 3 | 0 | 2 | 1 | 0 | 0 | 0 | 0 | 0 | 5 | 0 | 0 | Gm5901 |
| Gm5930 | 0 | 0 | 0 | 0 | 0 | 0 | 0 | 0 | 2 | 0 | 3 | 0 | 0 | 0 | 0 | 0 | 0 | 0 | 0 | 0 | 0 | 0 | 1 | 0 | 0 | 0 | 0 | 0 | Gm5930 |
| Gm5941 | 0 | 0 | 0 | 0 | 0 | 0 | 0 | 0 | 0 | 0 | 0 | 0 | 0 | 0 | 0 | 0 | 0 | 0 | 0 | 0 | 0 | 0 | 0 | 0 | 0 | 0 | 0 | 2 | Gm5941 |
| Gm5945 | 0 | 0 | 1 | 1 | 0 | 1 | 3 | 1 | 1 | 1 | 0 | 1 | 0 | 0 | 0 | 0 | 0 | 1 | 0 | 0 | 3 | 0 | 1 | 0 | 1 | 0 | 1 | 0 | Gm5945 |
| Gm6020 | 5 | 2 | 0 | 4 | 7 | 12 | 1 | 4 | 6 | 8 | 6 | 10 | 9 | 7 | 0 | 4 | 0 | 1 | 2 | 8 | 3 | 2 | 3 | 1 | 5 | 1 | 3 | 0 | Gm6020 |
| Gm6034 | 8 | 15 | 6 | 14 | 18 | 19 | 11 | 2 | 4 | 3 | 0 | 1 | 1 | 5 | 5 | 10 | 12 | 8 | 11 | 22 | 11 | 2 | 6 | 0 | 4 | 0 | 3 | 1 | Gm6034 |
| Gm609 | 0 | 0 | 0 | 0 | 0 | 0 | 0 | 0 | 0 | 6 | 0 | 0 | 0 | 0 | 0 | 0 | 0 | 0 | 0 | 0 | 0 | 0 | 0 | 0 | 0 | 0 | 0 | 1 | Gm609 |
| Gm6133 | 0 | 1 | 0 | 0 | 3 | 4 | 0 | 1 | 2 | 8 | 6 | 0 | 0 | 0 | 1 | 1 | 1 | 0 | 2 | 1 | 0 | 5 | 1 | 0 | 4 | 4 | 0 | 0 | Gm6133 |
| Gm614 | 0 | 1 | 0 | 0 | 0 | 0 | 0 | 0 | 3 | 0 | 0 | 1 | 2 | 0 | 1 | 0 | 0 | 0 | 2 | 5 | 0 | 0 | 1 | 1 | 0 | 4 | 0 | 0 | Gm614 |
| Gm6169 | 120 | 91 | 82 | 47 | 63 | 82 | 78 | 134 | 147 | 97 | 126 | 128 | 121 | 127 | 112 | 111 | 58 | 51 | 70 | 67 | 56 | 104 | 166 | 119 | 106 | 136 | 135 | 146 | Gm6169 |
| Gm6268 | 2 | 0 | 0 | 0 | 0 | 0 | 0 | 0 | 0 | 0 | 0 | 0 | 0 | 0 | 0 | 0 | 0 | 0 | 0 | 0 | 0 | 0 | 0 | 0 | 0 | 0 | 0 | 0 | Gm6268 |
| Gm6309 | 0 | 0 | 0 | 1 | 0 | 2 | 1 | 0 | 0 | 0 | 0 | 0 | 0 | 0 | 1 | 1 | 0 | 0 | 0 | 0 | 0 | 0 | 0 | 0 | 0 | 0 | 0 | 0 | Gm6309 |
| Gm6337 | 0 | 0 | 0 | 0 | 0 | 0 | 2 | 0 | 0 | 0 | 0 | 0 | 0 | 0 | 0 | 0 | 0 | 0 | 0 | 0 | 0 | 0 | 0 | 0 | 0 | 0 | 0 | 0 | Gm6337 |
| Gm6356 | 0 | 0 | 0 | 0 | 1 | 0 | 1 | 2 | 0 | 0 | 0 | 0 | 0 | 1 | 0 | 0 | 0 | 0 | 0 | 0 | 0 | 0 | 0 | 0 | 1 | 1 | 1 | 0 | Gm6356 |
| Gm6370 | 0 | 0 | 0 | 0 | 0 | 0 | 1 | 0 | 0 | 0 | 0 | 0 | 0 | 0 | 0 | 0 | 0 | 0 | 0 | 0 | 0 | 0 | 0 | 0 | 0 | 0 | 0 | 0 | Gm6370 |
| Gm6377 | 17 | 8 | 16 | 36 | 20 | 33 | 26 | 24 | 12 | 15 | 21 | 13 | 3 | 12 | 11 | 10 | 21 | 20 | 41 | 16 | 33 | 17 | 10 | 17 | 8 | 7 | 17 | 12 | Gm6377 |
| Gm6408 | 12 | 14 | 25 | 38 | 37 | 40 | 27 | 1 | 3 | 10 | 4 | 0 | 0 | 3 | 13 | 16 | 14 | 25 | 19 | 19 | 22 | 0 | 1 | 1 | 1 | 4 | 0 | 1 | Gm6408 |
| Gm648 | 3 | 14 | 16 | 10 | 7 | 10 | 8 | 9 | 7 | 14 | 9 | 4 | 3 | 6 | 15 | 7 | 14 | 26 | 16 | 5 | 12 | 3 | 6 | 11 | 19 | 7 | 3 | 8 | Gm648 |
| Gm6482 | 0 | 0 | 0 | 0 | 0 | 0 | 0 | 0 | 1 | 0 | 0 | 0 | 0 | 0 | 0 | 0 | 0 | 0 | 0 | 0 | 0 | 2 | 2 | 0 | 0 | 0 | 0 | 0 | Gm6482 |
| Gm6525 | 10 | 14 | 14 | 16 | 6 | 21 | 12 | 13 | 12 | 5 | 18 | 16 | 14 | 9 | 8 | 16 | 13 | 13 | 17 | 7 | 13 | 8 | 16 | 12 | 5 | 15 | 10 | 8 | Gm6525 |
| Gm6526 | 0 | 0 | 0 | 3 | 0 | 0 | 0 | 0 | 0 | 0 | 0 | 0 | 0 | 0 | 0 | 0 | 0 | 0 | 0 | 0 | 0 | 0 | 0 | 2 | 0 | 0 | 0 | 1 | Gm6526 |
| Gm6569 | 1 | 5 | 3 | 0 | 3 | 2 | 0 | 1 | 1 | 0 | 0 | 2 | 3 | 0 | 0 | 2 | 0 | 0 | 2 | 12 | 1 | 3 | 2 | 2 | 0 | 0 | 0 | 2 | Gm6569 |
| Gm6583 | 1 | 0 | 4 | 0 | 3 | 0 | 0 | 1 | 2 | 0 | 1 | 0 | 5 | 3 | 1 | 1 | 0 | 0 | 2 | 1 | 1 | 0 | 7 | 1 | 0 | 0 | 5 | 1 | Gm6583 |
| Gm6588 | 0 | 0 | 0 | 0 | 0 | 0 | 0 | 0 | 0 | 0 | 0 | 0 | 0 | 0 | 0 | 0 | 0 | 0 | 0 | 0 | 0 | 0 | 0 | 2 | 0 | 0 | 0 | 0 | Gm6588 |
| Gm6592 | 0 | 0 | 0 | 0 | 0 | 0 | 0 | 0 | 0 | 0 | 0 | 0 | 0 | 0 | 0 | 0 | 0 | 0 | 0 | 0 | 0 | 0 | 0 | 0 | 0 | 0 | 3 | 0 | Gm6592 |
| Gm6614 | 2 | 2 | 1 | 1 | 0 | 12 | 0 | 2 | 10 | 3 | 11 | 8 | 14 | 5 | 1 | 0 | 0 | 0 | 2 | 2 | 6 | 2 | 10 | 6 | 10 | 17 | 14 | 9 | Gm6614 |
| Gm6619 | 0 | 0 | 0 | 0 | 0 | 0 | 0 | 0 | 0 | 0 | 0 | 0 | 0 | 0 | 0 | 0 | 0 | 0 | 0 | 0 | 0 | 0 | 1 | 0 | 0 | 0 | 0 | 0 | Gm6619 |
| Gm6657 | 0 | 0 | 0 | 0 | 0 | 0 | 0 | 0 | 0 | 0 | 5 | 0 | 0 | 0 | 4 | 0 | 0 | 0 | 0 | 0 | 1 | 0 | 0 | 2 | 0 | 0 | 0 | 0 | Gm6657 |
| Gm6710 | 126 | 367 | 101 | 99 | 410 | 162 | 375 | 302 | 154 | 153 | 291 | 147 | 171 | 231 | 624 | 494 | 343 | 111 | 232 | 338 | 102 | 149 | 481 | 125 | 551 | 182 | 429 | 120 | Gm6710 |
| Gm6760 | 0 | 0 | 0 | 0 | 0 | 0 | 1 | 0 | 0 | 0 | 0 | 0 | 0 | 0 | 0 | 0 | 0 | 0 | 0 | 0 | 0 | 0 | 0 | 0 | 0 | 0 | 0 | 0 | Gm6760 |
| Gm6812 | 0 | 0 | 0 | 0 | 0 | 0 | 0 | 0 | 0 | 0 | 0 | 0 | 0 | 1 | 1 | 0 | 0 | 0 | 0 | 0 | 0 | 0 | 0 | 0 | 0 | 0 | 0 | 0 | Gm6812 |
| Gm6871 | 0 | 1 | 2 | 0 | 0 | 5 | 0 | 2 | 2 | 1 | 0 | 1 | 0 | 4 | 1 | 1 | 0 | 1 | 0 | 0 | 0 | 0 | 1 | 0 | 2 | 0 | 3 | 0 | Gm6871 |
| Gm7030 | 0 | 1 | 1 | 0 | 0 | 1 | 1 | 0 | 0 | 0 | 0 | 0 | 0 | 1 | 1 | 0 | 0 | 0 | 0 | 0 | 1 | 0 | 0 | 0 | 0 | 0 | 0 | 1 | Gm7030 |
| Gm7072 | 1961 | 1318 | 2016 | 1888 | 2026 | 2087 | 2148 | 2674 | 2556 | 2454 | 2266 | 2405 | 2384 | 2394 | 2106 | 1893 | 2073 | 1887 | 1907 | 2121 | 1841 | 2620 | 2374 | 2369 | 2272 | 2219 | 2322 | 2401 | Gm7072 |
| Gm7075 | 6 | 4 | 2 | 11 | 4 | 11 | 8 | 4 | 2 | 0 | 0 | 1 | 0 | 1 | 5 | 9 | 5 | 3 | 15 | 3 | 4 | 0 | 2 | 7 | 5 | 3 | 2 | 2 | Gm7075 |
| Gm7102 | 2 | 5 | 3 | 0 | 2 | 0 | 1 | 1 | 2 | 6 | 0 | 0 | 1 | 2 | 1 | 5 | 2 | 0 | 2 | 1 | 0 | 2 | 3 | 1 | 1 | 1 | 1 | 6 | Gm7102 |
| Gm7168 | 3 | 3 | 4 | 6 | 7 | 1 | 2 | 2 | 1 | 6 | 1 | 5 | 3 | 1 | 3 | 2 | 0 | 3 | 5 | 5 | 3 | 2 | 4 | 0 | 2 | 0 | 0 | 1 | Gm7168 |
| Gm7233 | 3 | 0 | 0 | 0 | 1 | 0 | 0 | 4 | 2 | 0 | 0 | 0 | 2 | 0 | 0 | 0 | 1 | 2 | 0 | 0 | 0 | 0 | 3 | 1 | 0 | 0 | 0 | 1 | Gm7233 |
| Gm7247 | 8 | 3 | 5 | 2 | 7 | 6 | 2 | 5 | 7 | 3 | 6 | 2 | 0 | 9 | 1 | 0 | 7 | 1 | 2 | 1 | 2 | 5 | 3 | 6 | 2 | 0 | 0 | 4 | Gm7247 |
| Gm7271 | 24 | 22 | 0 | 0 | 38 | 0 | 0 | 60 | 34 | 33 | 39 | 1 | 60 | 2 | 61 | 39 | 1 | 21 | 2 | 5 | 1 | 67 | 19 | 0 | 33 | 0 | 5 | 2 | Gm7271 |
| Gm7298 | 0 | 0 | 0 | 0 | 2 | 0 | 3 | 0 | 0 | 0 | 3 | 0 | 0 | 1 | 0 | 0 | 0 | 0 | 0 | 0 | 0 | 0 | 1 | 0 | 0 | 0 | 0 | 0 | Gm7298 |
| Gm7324 | 82 | 70 | 80 | 67 | 44 | 78 | 69 | 159 | 68 | 61 | 51 | 72 | 71 | 87 | 136 | 77 | 71 | 62 | 76 | 84 | 70 | 99 | 57 | 83 | 77 | 102 | 79 | 80 | Gm7324 |
| Gm7347 | 0 | 0 | 0 | 0 | 0 | 0 | 0 | 0 | 0 | 0 | 1 | 0 | 0 | 0 | 0 | 0 | 0 | 0 | 0 | 0 | 0 | 0 | 0 | 0 | 0 | 0 | 0 | 0 | Gm7347 |
| Gm7358 | 0 | 1 | 0 | 0 | 0 | 0 | 0 | 0 | 0 | 0 | 0 | 0 | 0 | 0 | 0 | 0 | 0 | 0 | 0 | 0 | 0 | 0 | 0 | 0 | 0 | 0 | 0 | 0 | Gm7358 |
| Gm7361 | 12 | 6 | 10 | 13 | 13 | 14 | 9 | 13 | 12 | 23 | 38 | 24 | 14 | 15 | 9 | 5 | 8 | 17 | 2 | 4 | 8 | 25 | 14 | 14 | 21 | 13 | 5 | 11 | Gm7361 |
| Gm7489 | 0 | 0 | 0 | 0 | 0 | 0 | 0 | 0 | 0 | 0 | 0 | 0 | 0 | 0 | 3 | 0 | 0 | 3 | 0 | 0 | 0 | 0 | 0 | 0 | 0 | 0 | 2 | 1 | Gm7489 |
| Gm7579 | 0 | 0 | 0 | 0 | 0 | 1 | 0 | 0 | 0 | 0 | 2 | 0 | 0 | 0 | 0 | 0 | 2 | 0 | 0 | 0 | 0 | 0 | 0 | 0 | 0 | 0 | 0 | 0 | Gm7579 |
| Gm7582 | 0 | 0 | 0 | 0 | 0 | 0 | 0 | 2 | 0 | 0 | 0 | 0 | 0 | 0 | 0 | 1 | 0 | 0 | 0 | 0 | 0 | 0 | 0 | 0 | 0 | 0 | 0 | 0 | Gm7582 |
| Gm7592 | 0 | 0 | 0 | 0 | 0 | 0 | 0 | 0 | 1 | 0 | 0 | 2 | 1 | 0 | 0 | 0 | 0 | 0 | 0 | 0 | 0 | 2 | 0 | 1 | 0 | 0 | 0 | 0 | Gm7592 |
| Gm7609 | 0 | 0 | 0 | 0 | 0 | 0 | 0 | 0 | 0 | 0 | 0 | 0 | 0 | 1 | 0 | 0 | 0 | 0 | 0 | 0 | 0 | 0 | 0 | 0 | 0 | 0 | 1 | 0 | Gm7609 |
| Gm7694 | 930 | 1164 | 1379 | 1333 | 1224 | 1199 | 1428 | 604 | 757 | 790 | 843 | 904 | 939 | 1013 | 1027 | 1089 | 1264 | 1357 | 1349 | 935 | 1286 | 832 | 660 | 1068 | 935 | 1063 | 776 | 1056 | Gm7694 |
| Gm7697 | 29 | 25 | 1 | 20 | 11 | 13 | 25 | 6 | 7 | 0 | 3 | 2 | 14 | 5 | 22 | 19 | 2 | 10 | 30 | 29 | 33 | 2 | 6 | 1 | 8 | 0 | 5 | 6 | Gm7697 |
| Gm773 | 0 | 0 | 0 | 0 | 0 | 0 | 0 | 0 | 0 | 0 | 0 | 0 | 0 | 0 | 0 | 0 | 1 | 0 | 0 | 0 | 0 | 2 | 0 | 0 | 0 | 0 | 0 | 0 | Gm773 |
| Gm7876 | 0 | 0 | 0 | 1 | 0 | 0 | 0 | 1 | 0 | 0 | 0 | 0 | 0 | 0 | 0 | 0 | 0 | 0 | 0 | 0 | 0 | 0 | 0 | 0 | 0 | 0 | 0 | 0 | Gm7876 |
| Gm7879 | 159 | 86 | 153 | 126 | 59 | 74 | 108 | 149 | 150 | 98 | 76 | 131 | 102 | 82 | 158 | 107 | 92 | 136 | 59 | 86 | 106 | 87 | 95 | 114 | 124 | 146 | 110 | 126 | Gm7879 |
| Gm7945 | 0 | 0 | 0 | 0 | 0 | 0 | 0 | 0 | 0 | 0 | 0 | 0 | 0 | 0 | 0 | 0 | 2 | 0 | 0 | 0 | 0 | 0 | 0 | 0 | 0 | 0 | 0 | 0 | Gm7945 |
| Gm7995 | 0 | 0 | 0 | 0 | 0 | 0 | 0 | 0 | 0 | 0 | 2 | 0 | 0 | 0 | 0 | 0 | 0 | 0 | 0 | 0 | 0 | 0 | 0 | 0 | 0 | 0 | 0 | 0 | Gm7995 |
| Gm8024 | 0 | 0 | 1 | 0 | 0 | 0 | 0 | 0 | 0 | 0 | 0 | 0 | 0 | 0 | 0 | 0 | 0 | 0 | 0 | 1 | 0 | 0 | 0 | 0 | 0 | 0 | 0 | 0 | Gm8024 |
| Gm8050 | 10 | 8 | 23 | 17 | 8 | 19 | 23 | 16 | 7 | 5 | 12 | 7 | 6 | 7 | 11 | 17 | 12 | 8 | 20 | 14 | 8 | 20 | 10 | 8 | 7 | 4 | 22 | 9 | Gm8050 |
| Gm8094 | 0 | 0 | 0 | 0 | 0 | 0 | 0 | 0 | 0 | 0 | 2 | 0 | 0 | 0 | 0 | 0 | 0 | 0 | 0 | 0 | 0 | 0 | 0 | 0 | 0 | 0 | 0 | 0 | Gm8094 |
| Gm8104 | 0 | 0 | 0 | 0 | 1 | 0 | 0 | 0 | 5 | 0 | 0 | 0 | 0 | 0 | 1 | 0 | 0 | 0 | 0 | 0 | 0 | 0 | 1 | 0 | 0 | 0 | 0 | 0 | Gm8104 |
| Gm8108 | 7 | 3 | 13 | 7 | 9 | 1 | 13 | 17 | 17 | 45 | 16 | 24 | 23 | 28 | 9 | 16 | 10 | 12 | 4 | 8 | 4 | 32 | 24 | 18 | 25 | 23 | 22 | 20 | Gm8108 |
| Gm8113 | 339 | 269 | 202 | 310 | 466 | 369 | 339 | 666 | 650 | 492 | 528 | 568 | 594 | 572 | 407 | 305 | 301 | 315 | 308 | 507 | 343 | 535 | 515 | 418 | 395 | 554 | 544 | 494 | Gm8113 |
| Gm8122 | 0 | 0 | 0 | 1 | 0 | 0 | 0 | 0 | 0 | 0 | 0 | 0 | 0 | 0 | 0 | 0 | 0 | 0 | 0 | 0 | 0 | 0 | 0 | 0 | 0 | 0 | 0 | 0 | Gm8122 |
| Gm8126 | 0 | 0 | 0 | 0 | 0 | 0 | 0 | 0 | 4 | 0 | 3 | 0 | 0 | 0 | 0 | 0 | 0 | 0 | 0 | 0 | 0 | 0 | 0 | 0 | 0 | 1 | 0 | 0 | Gm8126 |
| Gm815 | 3 | 0 | 0 | 0 | 0 | 0 | 0 | 0 | 0 | 2 | 0 | 0 | 0 | 0 | 0 | 0 | 0 | 0 | 0 | 0 | 0 | 0 | 0 | 2 | 0 | 0 | 0 | 0 | Gm815 |
| Gm8159 | 0 | 0 | 0 | 0 | 0 | 0 | 0 | 0 | 0 | 0 | 1 | 0 | 0 | 0 | 0 | 1 | 0 | 0 | 0 | 0 | 0 | 0 | 0 | 1 | 0 | 0 | 0 | 0 | Gm8159 |
| Gm8180 | 0 | 0 | 0 | 0 | 0 | 0 | 1 | 0 | 0 | 0 | 0 | 0 | 0 | 0 | 0 | 0 | 0 | 0 | 0 | 0 | 0 | 0 | 0 | 0 | 0 | 0 | 0 | 0 | Gm8180 |

| | | | | | | | | | | | | | | | | | | | | | | | | | | | | | |
|---|---|---|---|---|---|---|---|---|---|---|---|---|---|---|---|---|---|---|---|---|---|---|---|---|---|---|---|---|---|
| Gm8206 | 60 | 30 | 42 | 43 | 46 | 63 | 51 | 82 | 93 | 134 | 116 | 70 | 81 | 106 | 58 | 68 | 54 | 56 | 41 | 46 | 33 | 135 | 104 | 109 | 90 | 71 | 84 | 81 | Gm8206 |
| Gm8220 | 0 | 0 | 0 | 0 | 0 | 0 | 1 | 0 | 3 | 0 | 0 | 0 | 0 | 0 | 0 | 0 | 0 | 0 | 0 | 0 | 0 | 0 | 0 | 1 | 0 | 0 | 0 | 0 | Gm8220 |
| Gm8229 | 0 | 0 | 0 | 0 | 2 | 0 | 0 | 0 | 1 | 3 | 7 | 0 | 0 | 0 | 0 | 1 | 0 | 0 | 0 | 0 | 0 | 0 | 0 | 0 | 0 | 0 | 0 | 0 | Gm8229 |
| Gm8232 | 0 | 0 | 0 | 0 | 2 | 0 | 0 | 1 | 0 | 1 | 0 | 0 | 0 | 0 | 3 | 1 | 1 | 0 | 2 | 0 | 0 | 2 | 0 | 0 | 0 | 0 | 0 | 1 | Gm8232 |
| Gm8237 | 20 | 18 | 21 | 20 | 33 | 18 | 23 | 12 | 28 | 37 | 24 | 30 | 18 | 18 | 5 | 9 | 24 | 11 | 14 | 26 | 30 | 30 | 14 | 10 | 13 | 15 | 12 | 12 | Gm8237 |
| Gm8247 | 0 | 0 | 0 | 1 | 1 | 1 | 0 | 0 | 0 | 0 | 0 | 0 | 0 | 0 | 0 | 3 | 0 | 0 | 0 | 0 | 0 | 0 | 6 | 0 | 3 | 0 | 0 | 0 | Gm8247 |
| Gm8251 | 1220 | 995 | 355 | 654 | 530 | 676 | 731 | 260 | 183 | 128 | 101 | 150 | 166 | 147 | 1311 | 1089 | 553 | 722 | 596 | 1043 | 761 | 75 | 179 | 109 | 132 | 67 | 189 | 133 | Gm8251 |
| Gm826 | 3 | 5 | 6 | 1 | 2 | 2 | 2 | 9 | 4 | 5 | 6 | 3 | 3 | 6 | 4 | 8 | 19 | 9 | 5 | 4 | 6 | 3 | 13 | 6 | 6 | 0 | 6 | 5 | Gm826 |
| Gm8265 | 0 | 1 | 0 | 0 | 0 | 0 | 0 | 0 | 0 | 0 | 0 | 0 | 0 | 0 | 0 | 0 | 0 | 0 | 0 | 0 | 0 | 0 | 1 | 0 | 0 | 0 | 0 | 0 | Gm8265 |
| Gm8267 | 0 | 0 | 0 | 0 | 0 | 0 | 0 | 0 | 0 | 0 | 0 | 0 | 0 | 0 | 0 | 0 | 0 | 0 | 0 | 0 | 0 | 0 | 0 | 0 | 0 | 0 | 1 | 0 | Gm8267 |
| Gm8271 | 0 | 0 | 0 | 0 | 0 | 0 | 0 | 0 | 0 | 0 | 0 | 0 | 0 | 1 | 0 | 0 | 0 | 0 | 2 | 0 | 0 | 2 | 0 | 0 | 0 | 0 | 0 | 0 | Gm8271 |
| Gm8279 | 0 | 0 | 0 | 1 | 0 | 0 | 0 | 0 | 1 | 0 | 0 | 0 | 0 | 0 | 3 | 1 | 0 | 0 | 0 | 3 | 2 | 0 | 0 | 1 | 0 | 1 | 0 | 0 | Gm8279 |
| Gm8281 | 136 | 111 | 179 | 143 | 219 | 195 | 210 | 618 | 472 | 523 | 406 | 493 | 491 | 528 | 154 | 140 | 167 | 167 | 243 | 221 | 220 | 698 | 468 | 385 | 521 | 455 | 497 | 523 | Gm8281 |
| Gm8297 | 0 | 0 | 0 | 0 | 1 | 0 | 0 | 0 | 0 | 0 | 0 | 0 | 0 | 0 | 0 | 0 | 0 | 0 | 0 | 0 | 0 | 0 | 0 | 0 | 0 | 0 | 0 | 0 | Gm8297 |
| Gm8356 | 0 | 0 | 0 | 0 | 0 | 0 | 0 | 0 | 0 | 0 | 0 | 0 | 0 | 0 | 0 | 0 | 1 | 0 | 0 | 0 | 0 | 2 | 0 | 0 | 0 | 0 | 0 | 0 | Gm8356 |
| Gm8362 | 0 | 0 | 0 | 0 | 0 | 0 | 0 | 0 | 0 | 0 | 0 | 1 | 0 | 0 | 0 | 0 | 0 | 0 | 0 | 0 | 0 | 0 | 0 | 0 | 0 | 0 | 0 | 0 | Gm8362 |
| Gm8369 | 0 | 0 | 0 | 0 | 1 | 0 | 0 | 0 | 2 | 0 | 2 | 0 | 3 | 0 | 0 | 2 | 2 | 0 | 0 | 1 | 0 | 2 | 1 | 7 | 3 | 1 | 1 | 1 | Gm8369 |
| Gm8374 | 0 | 1 | 3 | 1 | 0 | 0 | 0 | 0 | 4 | 0 | 0 | 0 | 1 | 0 | 1 | 4 | 0 | 1 | 0 | 3 | 0 | 0 | 3 | 1 | 0 | 0 | 0 | 0 | Gm8374 |
| Gm8439 | 0 | 0 | 0 | 0 | 0 | 0 | 0 | 0 | 0 | 0 | 0 | 0 | 1 | 0 | 0 | 0 | 0 | 0 | 0 | 0 | 3 | 0 | 0 | 0 | 0 | 0 | 0 | 0 | Gm8439 |
| Gm867 | 22 | 37 | 44 | 21 | 41 | 32 | 7 | 34 | 36 | 27 | 35 | 27 | 83 | 32 | 42 | 44 | 38 | 13 | 9 | 14 | 12 | 47 | 56 | 18 | 23 | 26 | 57 | 77 | Gm867 |
| Gm8773 | 116 | 106 | 113 | 94 | 105 | 103 | 101 | 211 | 218 | 204 | 188 | 200 | 206 | 178 | 153 | 104 | 108 | 88 | 82 | 138 | 102 | 190 | 221 | 191 | 157 | 195 | 180 | 223 | Gm8773 |
| Gm8797 | 316 | 333 | 245 | 246 | 300 | 433 | 184 | 308 | 352 | 247 | 298 | 345 | 387 | 134 | 268 | 380 | 256 | 286 | 352 | 445 | 189 | 329 | 339 | 272 | 298 | 369 | 358 | 206 | Gm8797 |
| Gm884 | 2 | 0 | 0 | 0 | 0 | 1 | 0 | 1 | 0 | 0 | 1 | 0 | 5 | 0 | 1 | 0 | 0 | 3 | 0 | 0 | 8 | 0 | 0 | 1 | 0 | 0 | 1 | 0 | Gm884 |
| Gm8879 | 0 | 0 | 0 | 0 | 3 | 0 | 0 | 0 | 0 | 0 | 0 | 0 | 0 | 0 | 0 | 0 | 0 | 0 | 0 | 0 | 0 | 0 | 0 | 0 | 0 | 0 | 0 | 0 | Gm8879 |
| Gm8890 | 0 | 0 | 0 | 0 | 0 | 0 | 0 | 0 | 2 | 2 | 2 | 0 | 0 | 0 | 1 | 0 | 0 | 0 | 0 | 0 | 0 | 0 | 0 | 0 | 0 | 0 | 0 | 0 | Gm8890 |
| Gm8909 | 3 | 4 | 11 | 0 | 2 | 2 | 0 | 0 | 2 | 0 | 0 | 0 | 3 | 0 | 9 | 9 | 0 | 3 | 1 | 0 | 0 | 0 | 1 | 2 | 0 | 0 | 0 | 1 | Gm8909 |
| Gm8947 | 0 | 6 | 0 | 0 | 0 | 0 | 0 | 0 | 2 | 0 | 0 | 1 | 10 | 2 | 0 | 0 | 0 | 0 | 0 | 0 | 0 | 0 | 0 | 0 | 0 | 0 | 0 | 7 | Gm8947 |
| Gm9 | 0 | 0 | 0 | 0 | 0 | 0 | 0 | 0 | 0 | 0 | 0 | 1 | 0 | 0 | 0 | 2 | 0 | 0 | 0 | 0 | 0 | 0 | 0 | 2 | 0 | 0 | 0 | 0 | Gm9 |
| Gm9008 | 3 | 2 | 1 | 1 | 2 | 1 | 1 | 0 | 2 | 0 | 1 | 1 | 5 | 1 | 4 | 2 | 6 | 0 | 3 | 3 | 2 | 3 | 1 | 4 | 0 | 1 | 3 | 4 | Gm9008 |
| Gm9040 | 0 | 0 | 1 | 1 | 0 | 4 | 0 | 0 | 1 | 2 | 0 | 1 | 1 | 1 | 0 | 0 | 0 | 0 | 2 | 0 | 4 | 2 | 1 | 0 | 0 | 1 | 1 | 0 | Gm9040 |
| Gm9046 | 2 | 4 | 4 | 0 | 2 | 1 | 3 | 4 | 1 | 1 | 3 | 0 | 2 | 6 | 4 | 0 | 5 | 1 | 6 | 1 | 4 | 0 | 3 | 7 | 3 | 5 | 0 | 2 | Gm9046 |
| Gm9048 | 6 | 1 | 0 | 2 | 1 | 2 | 0 | 1 | 1 | 0 | 0 | 5 | 1 | 0 | 1 | 1 | 3 | 1 | 0 | 0 | 1 | 0 | 3 | 0 | 4 | 1 | 0 | 0 | Gm9048 |
| Gm9049 | 0 | 0 | 0 | 0 | 0 | 0 | 0 | 0 | 0 | 0 | 0 | 0 | 1 | 0 | 0 | 1 | 0 | 0 | 1 | 0 | 0 | 0 | 0 | 0 | 0 | 0 | 0 | 0 | Gm9049 |
| Gm9112 | 0 | 0 | 0 | 0 | 0 | 0 | 0 | 0 | 2 | 0 | 0 | 0 | 0 | 0 | 0 | 0 | 0 | 0 | 0 | 0 | 0 | 0 | 0 | 0 | 0 | 0 | 0 | 0 | Gm9112 |
| Gm9195 | 1 | 0 | 0 | 0 | 0 | 0 | 0 | 1 | 2 | 0 | 0 | 1 | 0 | 0 | 1 | 0 | 0 | 0 | 0 | 0 | 0 | 3 | 1 | 0 | 0 | 0 | 0 | 0 | Gm9195 |
| Gm9376 | 2 | 2 | 0 | 2 | 0 | 5 | 3 | 1 | 2 | 3 | 0 | 2 | 1 | 1 | 3 | 2 | 5 | 4 | 1 | 4 | 0 | 0 | 0 | 3 | 4 | 0 | 3 | 1 | Gm9376 |
| Gm9495 | 7 | 10 | 0 | 13 | 3 | 8 | 6 | 1 | 1 | 4 | 1 | 8 | 1 | 4 | 9 | 5 | 2 | 11 | 2 | 2 | 4 | 10 | 4 | 4 | 1 | 3 | 3 | 2 | Gm9495 |
| Gm9507 | 0 | 0 | 0 | 0 | 0 | 0 | 0 | 0 | 0 | 0 | 4 | 0 | 0 | 0 | 0 | 0 | 0 | 0 | 0 | 0 | 0 | 0 | 0 | 0 | 0 | 0 | 0 | 0 | Gm9507 |
| Gm960 | 344 | 344 | 196 | 462 | 416 | 430 | 505 | 134 | 118 | 133 | 145 | 99 | 81 | 115 | 313 | 362 | 285 | 464 | 371 | 430 | 448 | 170 | 135 | 119 | 217 | 96 | 62 | 55 | Gm960 |
| Gm9602 | 0 | 0 | 0 | 0 | 0 | 0 | 0 | 4 | 0 | 0 | 0 | 4 | 0 | 1 | 0 | 1 | 1 | 0 | 1 | 0 | 0 | 0 | 1 | 1 | 5 | 2 | 1 | 0 | Gm9602 |
| Gm9603 | 1 | 0 | 0 | 0 | 0 | 0 | 1 | 1 | 3 | 5 | 0 | 8 | 2 | 1 | 1 | 2 | 1 | 0 | 0 | 0 | 2 | 8 | 5 | 1 | 0 | 0 | 0 | 1 | Gm9603 |
| Gm973 | 65 | 112 | 83 | 69 | 101 | 83 | 65 | 212 | 191 | 170 | 157 | 211 | 237 | 142 | 136 | 136 | 94 | 101 | 82 | 117 | 71 | 256 | 256 | 116 | 183 | 93 | 189 | 177 | Gm973 |
| Gm9732 | 1 | 0 | 0 | 0 | 0 | 0 | 1 | 0 | 2 | 1 | 4 | 0 | 3 | 0 | 0 | 0 | 0 | 1 | 0 | 0 | 0 | 2 | 4 | 7 | 2 | 1 | 0 | 8 | Gm9732 |
| Gm9733 | 0 | 0 | 0 | 0 | 0 | 0 | 0 | 0 | 0 | 0 | 1 | 0 | 0 | 0 | 0 | 0 | 0 | 0 | 0 | 0 | 0 | 0 | 0 | 1 | 0 | 0 | 0 | 0 | Gm9733 |
| Gm9736 | 0 | 1 | 0 | 0 | 0 | 0 | 0 | 0 | 0 | 0 | 0 | 0 | 0 | 0 | 0 | 0 | 0 | 0 | 0 | 0 | 0 | 0 | 0 | 0 | 0 | 0 | 0 | 0 | Gm9736 |
| Gm9745 | 1 | 1 | 0 | 3 | 2 | 3 | 1 | 0 | 1 | 3 | 3 | 1 | 0 | 0 | 0 | 3 | 0 | 2 | 0 | 6 | 2 | 0 | 2 | 3 | 1 | 0 | 0 | 0 | Gm9745 |
| Gm9747 | 0 | 2 | 5 | 0 | 0 | 1 | 0 | 1 | 0 | 1 | 1 | 2 | 2 | 2 | 0 | 0 | 2 | 3 | 2 | 1 | 3 | 2 | 0 | 0 | 3 | 4 | 1 | 2 | Gm9747 |
| Gm9774 | 211 | 157 | 137 | 132 | 138 | 129 | 110 | 155 | 122 | 120 | 125 | 155 | 130 | 98 | 156 | 185 | 176 | 150 | 177 | 108 | 113 | 170 | 129 | 144 | 131 | 183 | 151 | 115 | Gm9774 |
| Gm9803 | 69 | 48 | 76 | 52 | 36 | 69 | 32 | 39 | 49 | 50 | 49 | 40 | 92 | 42 | 57 | 33 | 54 | 40 | 58 | 58 | 40 | 42 | 43 | 47 | 84 | 33 | 43 | 52 | Gm9803 |
| Gm9833 | 2278 | 1961 | 2315 | 2036 | 2316 | 2098 | 2580 | 4014 | 2960 | 2817 | 2653 | 3602 | 2982 | 4078 | 2586 | 2026 | 2208 | 2116 | 2314 | 2661 | 2463 | 3222 | 2845 | 3347 | 3389 | 3180 | 3711 | 3653 | Gm9833 |
| Gm9844 | 1451 | 1615 | 2295 | 2172 | 1845 | 1447 | 1352 | 1761 | 1424 | 3077 | 2077 | 2015 | 1588 | 1881 | 1457 | 1073 | 2226 | 1968 | 1887 | 1646 | 1809 | 1873 | 1553 | 3046 | 2466 | 1927 | 1689 | 1724 | Gm9844 |
| Gm9903 | 50 | 41 | 40 | 24 | 30 | 24 | 34 | 28 | 40 | 14 | 29 | 37 | 51 | 33 | 44 | 27 | 51 | 25 | 19 | 32 | 29 | 12 | 26 | 25 | 35 | 43 | 14 | 43 | Gm9903 |
| Gm9918 | 137 | 62 | 80 | 69 | 11 | 49 | 27 | 64 | 59 | 38 | 66 | 50 | 56 | 40 | 69 | 100 | 64 | 62 | 23 | 23 | 14 | 30 | 93 | 56 | 31 | 57 | 28 | 40 | Gm9918 |
| Gm9925 | 10 | 22 | 17 | 24 | 17 | 15 | 13 | 5 | 7 | 13 | 16 | 7 | 5 | 3 | 13 | 12 | 17 | 22 | 8 | 8 | 13 | 3 | 6 | 11 | 7 | 5 | 5 | 6 | Gm9925 |
| Gm9955 | 5 | 5 | 0 | 3 | 5 | 1 | 4 | 1 | 1 | 0 | 3 | 2 | 5 | 3 | 0 | 1 | 7 | 1 | 0 | 1 | 4 | 0 | 3 | 4 | 1 | 4 | 3 | 5 | Gm9955 |
| Gm9999 | 0 | 0 | 0 | 3 | 0 | 0 | 0 | 0 | 0 | 0 | 0 | 2 | 1 | 0 | 1 | 0 | 0 | 0 | 0 | 0 | 0 | 2 | 1 | 0 | 0 | 0 | 0 | 1 | Gm9999 |
| Gmcl1 | 5357 | 4119 | 4520 | 4939 | 4943 | 4873 | 5160 | 4942 | 4335 | 4520 | 4416 | 4573 | 4636 | 4677 | 4990 | 4741 | 4692 | 4908 | 4462 | 4341 | 4832 | 4752 | 4075 | 4516 | 4793 | 4911 | 4860 | 4555 | Gmcl1 |
| Gmds | 1336 | 1196 | 1300 | 930 | 969 | 914 | 893 | 1125 | 1263 | 1213 | 1164 | 1044 | 1182 | 1133 | 1284 | 1572 | 1102 | 1010 | 838 | 914 | 767 | 1228 | 1394 | 1228 | 1166 | 1431 | 1076 | 1159 | Gmds |
| Gmeb1 | 2668 | 2598 | 2866 | 2846 | 3035 | 3122 | 3087 | 5083 | 4416 | 5127 | 4529 | 5525 | 5204 | 5330 | 2619 | 2523 | 3038 | 2682 | 3225 | 2835 | 2918 | 5257 | 4391 | 5130 | 5126 | 5149 | 5516 | 5174 | Gmeb1 |
| Gmeb2 | 1105 | 969 | 1394 | 1291 | 1476 | 1329 | 1350 | 2849 | 2986 | 3303 | 3259 | 3796 | 3344 | 3782 | 1106 | 945 | 1442 | 1151 | 1612 | 1160 | 1387 | 3526 | 2755 | 3768 | 3305 | 3644 | 3828 | 3804 | Gmeb2 |
| Gmfb | 16916 | 13321 | 14373 | 14093 | 15218 | 15431 | 16337 | 18282 | 15108 | 13808 | 14110 | 14767 | 14540 | 15004 | 17823 | 15112 | 14054 | 14438 | 14555 | 17618 | 15616 | 15286 | 15601 | 13715 | 14877 | 13829 | 15307 | 15328 | Gmfb |
| Gmfg | 89 | 115 | 155 | 61 | 70 | 100 | 84 | 112 | 141 | 113 | 152 | 112 | 163 | 123 | 115 | 138 | 118 | 89 | 92 | 107 | 81 | 127 | 168 | 123 | 128 | 125 | 191 | 127 | Gmfg |
| Gmip | 2225 | 2744 | 2586 | 3589 | 4372 | 3527 | 3752 | 2842 | 3049 | 3579 | 2914 | 3146 | 3053 | 2987 | 2201 | 2523 | 3370 | 3746 | 4217 | 3248 | 3839 | 4029 | 2852 | 3389 | 4046 | 2408 | 2903 | 2720 | Gmip |
| Gml | 0 | 0 | 0 | 0 | 0 | 0 | 0 | 0 | 0 | 0 | 0 | 0 | 0 | 0 | 0 | 0 | 0 | 0 | 0 | 0 | 0 | 0 | 0 | 1 | 0 | 0 | 0 | 0 | Gml |
| Gml2 | 0 | 0 | 0 | 0 | 0 | 0 | 0 | 4 | 0 | 0 | 0 | 0 | 0 | 0 | 0 | 0 | 0 | 0 | 0 | 0 | 0 | 0 | 0 | 0 | 0 | 0 | 0 | 0 | Gml2 |
| Gmnc | 144 | 94 | 137 | 75 | 159 | 181 | 111 | 563 | 423 | 636 | 469 | 1111 | 1181 | 924 | 164 | 155 | 136 | 50 | 130 | 151 | 122 | 506 | 1087 | 583 | 479 | 984 | 857 | 1089 | Gmnc |
| Gmnn | 2382 | 2278 | 2855 | 2071 | 1703 | 1814 | 2207 | 2009 | 1797 | 1580 | 1550 | 2751 | 2183 | 2842 | 2496 | 2083 | 2516 | 1740 | 1966 | 2390 | 2013 | 1083 | 1572 | 2225 | 1847 | 3558 | 2800 | 3421 | Gmnn |
| Gmppa | 1777 | 2580 | 2671 | 2377 | 2216 | 2286 | 2134 | 1921 | 2176 | 2424 | 2467 | 2389 | 2322 | 2166 | 1928 | 1988 | 2605 | 2297 | 2341 | 1904 | 2019 | 2251 | 2181 | 2336 | 2246 | 2487 | 2424 | 2237 | Gmppa |
| Gmppb | 1376 | 1624 | 1344 | 1492 | 1142 | 1280 | 1071 | 746 | 999 | 903 | 988 | 834 | 1048 | 772 | 1451 | 1482 | 1387 | 1360 | 1226 | 1101 | 1154 | 744 | 896 | 896 | 851 | 900 | 923 | 832 | Gmppb |
| Gmpr | 925 | 1204 | 1003 | 965 | 968 | 1031 | 867 | 448 | 533 | 552 | 545 | 411 | 517 | 385 | 957 | 1085 | 1082 | 939 | 969 | 980 | 880 | 485 | 704 | 463 | 402 | 505 | 393 | 476 | Gmpr |
| Gmpr2 | 1851 | 1708 | 1897 | 1552 | 1348 | 1500 | 1516 | 2628 | 2638 | 2430 | 2410 | 2495 | 2577 | 2460 | 1783 | 1848 | 1719 | 1398 | 1568 | 1728 | 1423 | 2219 | 2491 | 2662 | 2371 | 2829 | 2594 | 2567 | Gmpr2 |
| Gmps | 17751 | 14154 | 17108 | 14391 | 12784 | 13315 | 14770 | 14345 | 11435 | 10391 | 10877 | 13351 | 12667 | 13692 | 17666 | 15209 | 14928 | 13558 | 13266 | 14929 | 13307 | 12342 | 10737 | 11769 | 11805 | 15157 | 14361 | 14639 | Gmps |
| Gna11 | 7369 | 8894 | 8132 | 9466 | 7784 | 8040 | 7285 | 4139 | 5333 | 5287 | 5302 | 5306 | 4746 | 5034 | 6487 | 8265 | 7988 | 9202 | 8391 | 6562 | 8048 | 5005 | 5102 | 5746 | 5545 | 5931 | 4926 | 5032 | Gna11 |
| Gna12 | 6959 | 9444 | 9110 | 9529 | 8795 | 8527 | 7519 | 5350 | 6898 | 7220 | 6805 | 8125 | 6708 | 6738 | 6891 | 8101 | 9428 | 9251 | 9400 | 7232 | 7925 | 6804 | 6785 | 7316 | 7393 | 8118 | 6960 | 6590 | Gna12 |
| Gna13 | 5736 | 5300 | 5686 | 5698 | 5830 | 5459 | 5927 | 6106 | 5764 | 5634 | 5378 | 6415 | 6131 | 6059 | 5496 | 5660 | 5777 | 5600 | 5598 | 5360 | 5754 | 5950 | 5639 | 5775 | 5950 | 5889 | 6109 | 6126 | Gna13 |
| Gna14 | 45 | 45 | 55 | 42 | 31 | 44 | 35 | 89 | 113 | 118 | 113 | 112 | 91 | 94 | 61 | 44 | 44 | 46 | 38 | 33 | 25 | 65 | 157 | 92 | 86 | 140 | 121 | 101 | Gna14 |
| Gna15 | 14 | 4 | 15 | 10 | 16 | 23 | 7 | 22 | 35 | 20 | 46 | 12 | 28 | 26 | 11 | 12 | 15 | 15 | 14 | 6 | 15 | 17 | 47 | 12 | 15 | 16 | 19 | 20 | Gna15 |
| Gnai1 | 4074 | 3016 | 3945 | 4792 | 6295 | 5672 | 6216 | 4219 | 3789 | 3552 | 3634 | 2813 | 2912 | 3075 | 4501 | 3644 | 3994 | 5197 | 5912 | 5236 | 6397 | 4729 | 3989 | 3402 | 3930 | 2539 | 2828 | 3033 | Gnai1 |
| Gnai2 | 30373 | 39301 | 41611 | 38213 | 28703 | 30428 | 27360 | 17520 | 19059 | 20324 | 18037 | 26913 | 22912 | 23257 | 30220 | 33982 | 40303 | 34094 | 34310 | 27664 | 29668 | 18561 | 19950 | 22678 | 20530 | 29291 | 26161 | 25106 | Gnai2 |

| | | | | | | | | | | | | | | | | | | | | | | | | | | | | | |
|---|---|---|---|---|---|---|---|---|---|---|---|---|---|---|---|---|---|---|---|---|---|---|---|---|---|---|---|---|---|
| Gnai3 | 27580 | 21648 | 23961 | 20754 | 19085 | 20611 | 21558 | 21002 | 17694 | 15491 | 16081 | 18317 | 18451 | 18194 | 28247 | 25351 | 22198 | 21529 | 19042 | 24143 | 19587 | 16954 | 17958 | 16512 | 16693 | 19955 | 19314 | 18924 | Gnai3 |
| Gnal | 1962 | 1837 | 1851 | 2418 | 4149 | 3017 | 2875 | 2836 | 2790 | 3295 | 2993 | 1954 | 2415 | 1877 | 2159 | 2063 | 2231 | 2874 | 2944 | 2542 | 3337 | 4761 | 2616 | 2137 | 3225 | 1033 | 1691 | 1640 | Gnal |
| Gnao1 | 18062 | 22811 | 24148 | 34980 | 40362 | 33315 | 34576 | 24748 | 26332 | 35370 | 26864 | 33406 | 25848 | 29345 | 19453 | 17383 | 32615 | 36783 | 40865 | 30723 | 37090 | 37034 | 22641 | 34397 | 38224 | 22714 | 26115 | 25560 | Gnao1 |
| Gnaq | 10763 | 9541 | 11162 | 14462 | 16975 | 15461 | 16201 | 18362 | 16222 | 18695 | 18330 | 17713 | 17243 | 18644 | 11699 | 9332 | 12880 | 14281 | 15489 | 13873 | 16491 | 23021 | 15731 | 19134 | 21824 | 14901 | 17249 | 17683 | Gnaq |
| Gnas | 52319 | 58703 | 60389 | 59368 | 62202 | 55561 | 53922 | 54694 | 63811 | 70014 | 79969 | 66206 | 63578 | 65319 | 50505 | 54869 | 61190 | 60445 | 58522 | 47764 | 56590 | 65079 | 72352 | 69154 | 64698 | 70226 | 60878 | 63973 | Gnas |
| Gnat1 | 40 | 68 | 63 | 99 | 63 | 49 | 51 | 11 | 13 | 12 | 13 | 11 | 14 | 6 | 36 | 75 | 52 | 95 | 70 | 50 | 69 | 7 | 9 | 12 | 9 | 12 | 14 | 5 | Gnat1 |
| Gnat2 | 27 | 29 | 17 | 20 | 14 | 14 | 11 | 11 | 14 | 24 | 11 | 17 | 16 | 26 | 24 | 30 | 25 | 20 | 12 | 24 | 12 | 28 | 23 | 10 | 6 | 13 | 22 | 17 | Gnat2 |
| Gnat3 | 25 | 28 | 27 | 15 | 5 | 5 | 8 | 26 | 12 | 8 | 12 | 18 | 69 | 13 | 82 | 30 | 25 | 6 | 6 | 12 | 8 | 8 | 10 | 6 | 9 | 11 | 14 | 28 | Gnat3 |
| Gnaz | 1524 | 2117 | 2071 | 2752 | 3062 | 2685 | 2554 | 1402 | 1408 | 1685 | 1618 | 1308 | 1353 | 1292 | 1857 | 1834 | 2520 | 2787 | 2995 | 2560 | 2822 | 1861 | 1393 | 1611 | 1978 | 1268 | 1247 | 1244 | Gnaz |
| Gnb1 | 38749 | 36695 | 41031 | 44188 | 48339 | 43793 | 50342 | 49272 | 46366 | 52403 | 46588 | 55236 | 48109 | 56392 | 40368 | 35181 | 44002 | 44721 | 47991 | 43181 | 49815 | 57383 | 45831 | 54022 | 56231 | 55648 | 51125 | 56703 | Gnb1 |
| Gnb1l | 632 | 651 | 514 | 684 | 598 | 531 | 542 | 377 | 402 | 415 | 379 | 386 | 342 | 346 | 533 | 607 | 555 | 602 | 602 | 570 | 668 | 363 | 400 | 335 | 395 | 412 | 310 | 370 | Gnb1l |
| Gnb2 | 8429 | 11501 | 11153 | 12165 | 11915 | 10598 | 9983 | 10690 | 12935 | 14372 | 12486 | 14403 | 12281 | 12306 | 8327 | 9502 | 12228 | 11592 | 13231 | 10158 | 10932 | 13085 | 13736 | 14551 | 14233 | 15279 | 12605 | 12251 | Gnb2 |
| Gnb3 | 81 | 98 | 104 | 113 | 133 | 106 | 70 | 85 | 93 | 82 | 96 | 127 | 162 | 134 | 87 | 120 | 88 | 113 | 140 | 93 | 108 | 139 | 125 | 127 | 74 | 40 | 194 | 148 | Gnb3 |
| Gnb4 | 5009 | 4438 | 4042 | 3829 | 4734 | 4750 | 4074 | 3991 | 4179 | 3569 | 4395 | 3057 | 3381 | 2957 | 5178 | 5256 | 3823 | 4418 | 3940 | 4512 | 4116 | 4132 | 4295 | 2932 | 3558 | 2769 | 3133 | 2892 | Gnb4 |
| Gnb5 | 3656 | 4101 | 3653 | 4147 | 3945 | 3716 | 4186 | 1865 | 2033 | 1841 | 1709 | 1700 | 1750 | 1736 | 3792 | 3851 | 3653 | 3736 | 3808 | 4024 | 4116 | 1998 | 1956 | 1749 | 1928 | 1758 | 1672 | 1831 | Gnb5 |
| Gne | 2125 | 2515 | 1685 | 2084 | 2446 | 2100 | 1968 | 1850 | 2001 | 1831 | 1934 | 1521 | 1682 | 1561 | 2289 | 2354 | 2082 | 2196 | 2212 | 2176 | 2166 | 1952 | 1994 | 1661 | 1854 | 1581 | 1482 | 1515 | Gne |
| Gng10 | 9 | 16 | 21 | 16 | 23 | 32 | 19 | 43 | 34 | 56 | 33 | 26 | 29 | 45 | 21 | 13 | 29 | 23 | 19 | 17 | 35 | 28 | 30 | 35 | 28 | 20 | 22 | 31 | Gng10 |
| Gng11 | 592 | 498 | 638 | 456 | 461 | 433 | 520 | 907 | 976 | 781 | 1021 | 754 | 922 | 918 | 550 | 528 | 461 | 421 | 391 | 572 | 477 | 775 | 1074 | 880 | 755 | 1006 | 780 | 981 | Gng11 |
| Gng12 | 6613 | 5174 | 5480 | 4306 | 4471 | 4635 | 4569 | 2898 | 3132 | 2655 | 3477 | 2573 | 2848 | 2753 | 6349 | 6277 | 4818 | 4710 | 4057 | 4205 | 4276 | 2856 | 3609 | 2468 | 2462 | 2647 | 2791 | 2848 | Gng12 |
| Gng13 | 10 | 13 | 7 | 9 | 10 | 1 | 3 | 34 | 24 | 23 | 40 | 3 | 9 | 18 | 4 | 6 | 2 | 1 | 3 | 0 | 3 | 7 | 21 | 65 | 5 | 24 | 5 | 11 | Gng13 |
| Gng2 | 16399 | 14524 | 15566 | 19944 | 26989 | 25519 | 27079 | 33729 | 29171 | 30401 | 26280 | 27542 | 25196 | 29793 | 20508 | 14850 | 17544 | 22341 | 24511 | 27260 | 27302 | 39495 | 28400 | 29813 | 36683 | 25195 | 26117 | 26892 | Gng2 |
| Gng3 | 3805 | 5496 | 4178 | 5641 | 7072 | 6044 | 6622 | 6703 | 6557 | 6810 | 6130 | 6246 | 6254 | 7167 | 4948 | 3908 | 5501 | 5504 | 6405 | 6802 | 7023 | 7498 | 5675 | 6539 | 7760 | 5128 | 6269 | 6680 | Gng3 |
| Gng4 | 908 | 759 | 525 | 1004 | 1654 | 1251 | 1398 | 821 | 966 | 891 | 748 | 484 | 474 | 436 | 835 | 653 | 1085 | 1265 | 1319 | 1180 | 1360 | 1243 | 845 | 632 | 937 | 328 | 310 | 330 | Gng4 |
| Gng5 | 4234 | 4616 | 4679 | 4134 | 3343 | 3400 | 5175 | 3638 | 3512 | 3426 | 3672 | 4050 | 4568 | 5776 | 4077 | 3785 | 4392 | 3629 | 3914 | 4411 | 4646 | 2801 | 3604 | 3809 | 3513 | 5329 | 4822 | 6093 | Gng5 |
| Gng7 | 1242 | 1508 | 1380 | 1669 | 1757 | 1465 | 1562 | 777 | 1037 | 1040 | 949 | 895 | 843 | 925 | 1338 | 1434 | 1690 | 2015 | 1873 | 1428 | 1646 | 1074 | 933 | 1058 | 1167 | 1126 | 721 | 894 | Gng7 |
| Gng8 | 610 | 688 | 590 | 727 | 416 | 580 | 491 | 492 | 399 | 507 | 542 | 422 | 644 | 501 | 549 | 743 | 691 | 643 | 520 | 445 | 458 | 374 | 439 | 660 | 465 | 677 | 449 | 456 | Gng8 |
| Gngt1 | 1 | 2 | 0 | 0 | 0 | 0 | 0 | 0 | 2 | 0 | 0 | 0 | 0 | 0 | 0 | 0 | 0 | 0 | 0 | 0 | 0 | 0 | 1 | 0 | 0 | 0 | 0 | 0 | Gngt1 |
| Gngt2 | 66 | 49 | 82 | 39 | 56 | 38 | 41 | 61 | 91 | 96 | 118 | 52 | 84 | 82 | 78 | 79 | 48 | 56 | 34 | 54 | 47 | 75 | 107 | 85 | 63 | 90 | 62 | 95 | Gngt2 |
| Gnl1 | 4909 | 5955 | 7275 | 7235 | 6407 | 5674 | 5481 | 2822 | 3677 | 4017 | 4284 | 4297 | 3652 | 3974 | 4416 | 5495 | 7470 | 7112 | 6876 | 4671 | 5920 | 3766 | 3600 | 4506 | 4128 | 5037 | 3842 | 4018 | Gnl1 |
| Gnl2 | 7560 | 7525 | 7046 | 6523 | 5919 | 6272 | 6609 | 6786 | 6021 | 6054 | 6001 | 6050 | 5865 | 6383 | 7369 | 7072 | 6678 | 6418 | 6092 | 7177 | 6462 | 6158 | 5897 | 5973 | 5976 | 6505 | 6331 | 6457 | Gnl2 |
| Gnl3 | 5117 | 4606 | 4849 | 4550 | 3466 | 3954 | 4356 | 5305 | 4586 | 4244 | 4708 | 5407 | 5095 | 5710 | 5097 | 4700 | 4175 | 4012 | 3530 | 4684 | 3887 | 4495 | 4581 | 5053 | 4479 | 6247 | 5297 | 6086 | Gnl3 |
| Gnl3l | 8809 | 9622 | 8690 | 10553 | 11810 | 11277 | 12170 | 12732 | 10977 | 11384 | 10662 | 10981 | 10819 | 11630 | 10183 | 7935 | 9815 | 10641 | 11366 | 11906 | 12112 | 12607 | 10947 | 11182 | 12741 | 9001 | 10793 | 11070 | Gnl3l |
| Gnmt | 182 | 254 | 192 | 303 | 200 | 261 | 220 | 50 | 55 | 76 | 44 | 67 | 44 | 44 | 175 | 206 | 211 | 285 | 259 | 207 | 232 | 75 | 35 | 59 | 48 | 59 | 56 | 50 | Gnmt |
| Gnpat | 7161 | 7570 | 6946 | 6003 | 5715 | 5805 | 6182 | 7636 | 7481 | 6901 | 6839 | 8486 | 8011 | 8318 | 7023 | 6238 | 6531 | 5706 | 6394 | 6435 | 6235 | 6891 | 7102 | 7319 | 7470 | 8565 | 8548 | 8812 | Gnpat |
| Gnpda1 | 1540 | 1573 | 1563 | 1462 | 1317 | 1312 | 1360 | 1159 | 1382 | 1256 | 1307 | 1456 | 1355 | 1411 | 1635 | 1595 | 1548 | 1354 | 1334 | 1480 | 1317 | 1285 | 1470 | 1244 | 1176 | 1605 | 1419 | 1318 | Gnpda1 |
| Gnpda2 | 1610 | 1183 | 1378 | 1569 | 1500 | 1535 | 1577 | 2159 | 1812 | 1607 | 1585 | 1502 | 1678 | 1625 | 1751 | 1544 | 1401 | 1479 | 1456 | 1819 | 1533 | 1683 | 1983 | 1557 | 1539 | 1755 | 1733 | 1650 | Gnpda2 |
| Gnpnat1 | 2307 | 1557 | 2044 | 1573 | 1432 | 1681 | 1748 | 1817 | 1742 | 1562 | 1607 | 1493 | 1691 | 1611 | 2244 | 2254 | 1654 | 1562 | 1397 | 1789 | 1569 | 1686 | 1909 | 1414 | 1437 | 1508 | 1614 | 1735 | Gnpnat1 |
| Gnptab | 4615 | 4454 | 3826 | 4890 | 6011 | 5574 | 5458 | 4103 | 4212 | 4530 | 4120 | 3541 | 3775 | 3574 | 4776 | 4586 | 4394 | 5080 | 5117 | 5094 | 5454 | 4672 | 4047 | 3819 | 4403 | 2914 | 3391 | 3509 | Gnptab |
| Gnptg | 2285 | 3002 | 2894 | 3237 | 2850 | 2860 | 2911 | 2326 | 2350 | 2922 | 2374 | 2843 | 2442 | 2819 | 2523 | 2444 | 3074 | 2979 | 3040 | 2887 | 2974 | 2558 | 2300 | 2694 | 2943 | 2575 | 2668 | 2763 | Gnptg |
| Gnrh1 | 248 | 335 | 255 | 381 | 259 | 357 | 285 | 123 | 92 | 156 | 123 | 141 | 170 | 125 | 225 | 328 | 346 | 418 | 312 | 247 | 288 | 124 | 108 | 181 | 141 | 184 | 142 | 154 | Gnrh1 |
| Gns | 10462 | 11545 | 10250 | 10488 | 8774 | 9367 | 8860 | 5245 | 6212 | 6180 | 6696 | 6076 | 6058 | 5464 | 9965 | 11993 | 10379 | 10549 | 8707 | 7725 | 8929 | 6358 | 6575 | 5704 | 5877 | 6199 | 5746 | 5574 | Gns |
| Golga1 | 3237 | 3236 | 2766 | 3022 | 3400 | 3478 | 3521 | 5708 | 4966 | 4724 | 4524 | 5091 | 4814 | 4932 | 3666 | 3116 | 2947 | 2864 | 3286 | 3927 | 3441 | 5048 | 5018 | 4605 | 5015 | 4075 | 5029 | 4926 | Golga1 |
| Golga2 | 5499 | 6280 | 6302 | 6658 | 6465 | 6269 | 6056 | 5223 | 5927 | 6468 | 6452 | 6043 | 5695 | 6098 | 5327 | 5647 | 6652 | 6743 | 6553 | 5360 | 6235 | 6711 | 6002 | 6400 | 6508 | 5772 | 5872 | 5733 | Golga2 |
| Golga3 | 6647 | 7039 | 5195 | 6436 | 6268 | 6384 | 6383 | 6748 | 6923 | 6481 | 6865 | 6467 | 6073 | 6299 | 6442 | 6952 | 5490 | 6655 | 5752 | 6704 | 6635 | 6363 | 6494 | 6108 | 6651 | 4690 | 6200 | 5929 | Golga3 |
| Golga4 | 16264 | 16980 | 12434 | 15234 | 15111 | 17046 | 16821 | 14801 | 14100 | 13022 | 13432 | 13399 | 12707 | 14250 | 16254 | 17263 | 14198 | 14860 | 14294 | 18044 | 16958 | 10881 | 13173 | 11810 | 13455 | 9211 | 13773 | 12155 | Golga4 |
| Golga5 | 3827 | 3730 | 3697 | 3450 | 3015 | 3126 | 3389 | 3230 | 2963 | 2808 | 3073 | 3028 | 2972 | 3307 | 3727 | 3475 | 3472 | 3361 | 3090 | 3244 | 3135 | 2906 | 3085 | 2988 | 3130 | 3214 | 3069 | 3063 | Golga5 |
| Golga7 | 8284 | 7914 | 9282 | 8473 | 7923 | 7872 | 8715 | 8841 | 8525 | 8125 | 7890 | 9146 | 8520 | 9327 | 8637 | 8681 | 8459 | 8156 | 8217 | 9215 | 8349 | 7511 | 8818 | 8536 | 8607 | 9628 | 9109 | 9202 | Golga7 |
| Golga7b | 315 | 439 | 366 | 836 | 1346 | 977 | 920 | 1215 | 1194 | 1441 | 1306 | 778 | 811 | 808 | 514 | 356 | 669 | 919 | 1177 | 842 | 1081 | 1534 | 1095 | 1077 | 1454 | 626 | 682 | 680 | Golga7b |
| Golgb1 | 21810 | 21243 | 11419 | 16065 | 15109 | 18754 | 16960 | 12581 | 12717 | 10673 | 12975 | 10010 | 10848 | 10277 | 20715 | 21577 | 12789 | 15451 | 13551 | 19110 | 17458 | 9205 | 12554 | 8105 | 10851 | 6057 | 11422 | 8915 | Golgb1 |
| Golim4 | 9124 | 7813 | 8706 | 7645 | 7351 | 7884 | 7888 | 6956 | 6536 | 6384 | 7392 | 7020 | 7313 | 7490 | 8627 | 8524 | 7587 | 8190 | 6997 | 7932 | 7612 | 6044 | 6494 | 6655 | 6220 | 7221 | 7290 | 7141 | Golim4 |
| Golm1 | 11191 | 11727 | 12533 | 8474 | 8231 | 8682 | 8555 | 9572 | 7809 | 7440 | 6722 | 11650 | 10872 | 11377 | 11661 | 11422 | 10581 | 8130 | 8556 | 9802 | 8163 | 7564 | 7578 | 9287 | 8811 | 12286 | 12984 | 12416 | Golm1 |
| Golm2 | 9461 | 7557 | 8160 | 10043 | 10407 | 10435 | 11691 | 7575 | 6713 | 6619 | 6568 | 6777 | 6336 | 6995 | 10419 | 8700 | 8389 | 9860 | 9391 | 9917 | 10914 | 8719 | 6922 | 6550 | 7339 | 7604 | 6381 | 6635 | Golm2 |
| Golph3 | 11252 | 8745 | 10100 | 8483 | 7772 | 8281 | 8780 | 8615 | 7617 | 6382 | 7182 | 7169 | 6940 | 7386 | 11468 | 9866 | 8671 | 9052 | 8242 | 8975 | 8507 | 8403 | 7732 | 6926 | 6641 | 7700 | 7079 | 7628 | Golph3 |
| Golph3l | 4645 | 4106 | 3750 | 3646 | 3740 | 3551 | 4297 | 3610 | 3221 | 2610 | 2656 | 3093 | 2857 | 3079 | 5236 | 4323 | 4115 | 3784 | 3838 | 4732 | 3959 | 2806 | 3200 | 2796 | 2828 | 3280 | 3002 | 3351 | Golph3l |
| Golt1a | 0 | 0 | 0 | 2 | 2 | 4 | 1 | 1 | 1 | 0 | 0 | 0 | 0 | 0 | 0 | 1 | 0 | 1 | 1 | 0 | 4 | 3 | 0 | 1 | 0 | 2 | 0 | 0 | Golt1a |
| Golt1b | 5875 | 5235 | 5086 | 4713 | 4309 | 4498 | 4907 | 4422 | 4213 | 3986 | 4250 | 4086 | 4045 | 4147 | 5736 | 5452 | 4862 | 4554 | 4407 | 4942 | 4558 | 4214 | 4418 | 4053 | 4060 | 4555 | 4316 | 4481 | Golt1b |
| Gon4l | 6650 | 7887 | 6885 | 8253 | 7940 | 7689 | 7768 | 6399 | 6854 | 7820 | 7257 | 8351 | 8019 | 8226 | 6562 | 7034 | 7926 | 8357 | 8127 | 6651 | 7370 | 7279 | 6236 | 7872 | 8153 | 6339 | 7997 | 7866 | Gon4l |
| Gon7 | 799 | 606 | 993 | 1025 | 953 | 944 | 1040 | 714 | 548 | 787 | 677 | 850 | 621 | 838 | 852 | 738 | 997 | 821 | 940 | 1006 | 978 | 734 | 570 | 878 | 902 | 802 | 783 | 954 | Gon7 |
| Gopc | 4084 | 3613 | 4056 | 4402 | 4375 | 4219 | 4508 | 3874 | 3412 | 3572 | 3948 | 3716 | 3648 | 3855 | 4121 | 4166 | 4084 | 4650 | 3750 | 3920 | 4405 | 3927 | 3477 | 3489 | 3865 | 3473 | 3447 | 3719 | Gopc |
| Gorab | 3064 | 2200 | 2188 | 2105 | 1559 | 2132 | 1852 | 2224 | 2229 | 1972 | 1980 | 2167 | 2173 | 1952 | 2781 | 2672 | 2050 | 2057 | 1683 | 2024 | 1581 | 1920 | 2199 | 1993 | 1793 | 2368 | 2008 | 1987 | Gorab |
| Gorasp1 | 1757 | 2188 | 1957 | 2076 | 2009 | 1891 | 1786 | 1418 | 1786 | 1796 | 1721 | 1679 | 1674 | 1479 | 1731 | 2056 | 2131 | 2047 | 2054 | 1800 | 1832 | 1644 | 1869 | 1711 | 1776 | 1830 | 1649 | 1531 | Gorasp1 |
| Gorasp2 | 5699 | 5945 | 5446 | 5009 | 4926 | 5000 | 5035 | 5025 | 5528 | 5173 | 5534 | 5572 | 5404 | 5592 | 5205 | 5445 | 5266 | 5264 | 4932 | 4922 | 5170 | 5335 | 5489 | 5521 | 5190 | 6072 | 5198 | 5686 | Gorasp2 |
| Gosr1 | 2049 | 1789 | 1926 | 1863 | 2276 | 2177 | 2194 | 3107 | 3265 | 3130 | 3257 | 3262 | 3028 | 3380 | 1921 | 1745 | 1764 | 1691 | 2111 | 2060 | 2024 | 3088 | 3207 | 3187 | 3300 | 2963 | 3158 | 3143 | Gosr1 |
| Gosr2 | 8142 | 8268 | 7433 | 6536 | 6902 | 7058 | 6900 | 8368 | 8433 | 7067 | 7217 | 7868 | 7819 | 7511 | 8215 | 7742 | 6773 | 6575 | 7208 | 7635 | 6493 | 7429 | 8621 | 7385 | 7818 | 7976 | 8123 | 7920 | Gosr2 |
| Got1 | 5305 | 7191 | 6056 | 6972 | 6254 | 5492 | 6468 | 2680 | 2860 | 2908 | 2739 | 3070 | 2704 | 3218 | 5523 | 5625 | 6030 | 7000 | 6478 | 6127 | 6547 | 3011 | 2570 | 3267 | 3036 | 3420 | 3089 | 3220 | Got1 |
| Got1l1 | 63 | 54 | 38 | 36 | 54 | 40 | 37 | 67 | 49 | 38 | 35 | 21 | 27 | 27 | 55 | 69 | 29 | 50 | 49 | 44 | 37 | 58 | 34 | 42 | 37 | 38 | 28 | 25 | Got1l1 |
| Got2 | 11394 | 13057 | 12870 | 11821 | 11293 | 11613 | 11049 | 8899 | 10542 | 11070 | 10599 | 12403 | 11895 | 11865 | 11037 | 11986 | 12601 | 11110 | 12765 | 10275 | 11614 | 10402 | 10215 | 11144 | 11036 | 13690 | 12112 | 13273 | Got2 |
| Gp1ba | 58 | 49 | 46 | 21 | 16 | 28 | 11 | 22 | 40 | 18 | 26 | 46 | 69 | 24 | 33 | 41 | 34 | 39 | 16 | 9 | 5 | 22 | 23 | 47 | 26 | 82 | 62 | 36 | Gp1ba |
| Gp1bb | 0 | 1 | 0 | 0 | 1 | 0 | 1 | 0 | 2 | 0 | 0 | 0 | 1 | 0 | 0 | 1 | 0 | 0 | 0 | 0 | 1 | 2 | 1 | 3 | 1 | 0 | 0 | 1 | Gp1bb |
| Gp2 | 0 | 0 | 0 | 0 | 2 | 0 | 0 | 0 | 0 | 0 | 0 | 3 | 0 | 0 | 0 | 0 | 0 | 0 | 0 | 0 | 0 | 0 | 0 | 0 | 0 | 0 | 0 | 2 | Gp2 |
| Gp5 | 40 | 29 | 58 | 12 | 16 | 27 | 19 | 26 | 35 | 36 | 23 | 39 | 58 | 34 | 15 | 39 | 39 | 17 | 24 | 12 | 21 | 40 | 23 | 52 | 27 | 79 | 53 | 46 | Gp5 |
| Gp6 | 33 | 17 | 28 | 20 | 12 | 7 | 15 | 24 | 22 | 24 | 15 | 41 | 59 | 30 | 20 | 37 | 15 | 8 | 5 | 16 | 12 | 18 | 18 | 22 | 9 | 41 | 48 | 27 | Gp6 |
| Gp9 | 30 | 46 | 43 | 14 | 13 | 17 | 13 | 27 | 44 | 24 | 14 | 31 | 81 | 24 | 26 | 39 | 29 | 10 | 11 | 37 | 13 | 18 | 30 | 24 | 14 | 69 | 65 | 26 | Gp9 |
| Gpa33 | 15 | 8 | 25 | 6 | 6 | 17 | 4 | 7 | 5 | 5 | 4 | 7 | 12 | 4 | 11 | 14 | 6 | 15 | 2 | 14 | 1 | 2 | 6 | 9 | 4 | 9 | 13 | 7 | Gpa33 |
| Gpaa1 | 3079 | 4101 | 3823 | 3883 | 3357 | 3348 | 3157 | 2180 | 2660 | 2776 | 2646 | 3138 | 2744 | 2731 | 2869 | 3730 | 3962 | 3871 | 3677 | 2862 | 3287 | 2677 | 2643 | 2723 | 2657 | 3030 | 2652 | 2528 | Gpaa1 |
| Gpalpp1 | 6429 | 5760 | 5615 | 5590 | 5001 | 5306 | 5542 | 5929 | 5040 | 4917 | 4627 | 5649 | 5651 | 5863 | 6388 | 5999 | 5168 | 5515 | 4980 | 5988 | 5465 | 5441 | 5447 | 5035 | 5681 | 4915 | 5762 | 5919 | Gpalpp1 |
| Gpam | 6123 | 6297 | 5761 | 5929 | 5840 | 6294 | 5753 | 4219 | 4261 | 4320 | 4337 | 4418 | 4823 | 4293 | 6005 | 6079 | 5756 | 6056 | 5972 | 5550 | 5974 | 4588 | 4133 | 3801 | 4358 | 3432 | 4846 | 4481 | Gpam |
| Gpank1 | 1095 | 1627 | 1410 | 1304 | 1250 | 1262 | 1247 | 878 | 891 | 866 | 848 | 967 | 1009 | 1093 | 1106 | 1126 | 1408 | 1254 | 1403 | 1337 | 1351 | 795 | 800 | 1086 | 910 | 1172 | 1095 | 1105 | Gpank1 |
| Gpat2 | 10 | 5 | 10 | 4 | 6 | 2 | 8 | 11 | 7 | 11 | 13 | 18 | 10 | 3 | 12 | 23 | 6 | 10 | 5 | 12 | 2 | 23 | 25 | 8 | 15 | 11 | 8 | 19 | Gpat2 |

| | | | | | | | | | | | | | | | | | | | | | | | | | | | | | |
|---|---|---|---|---|---|---|---|---|---|---|---|---|---|---|---|---|---|---|---|---|---|---|---|---|---|---|---|---|---|
| Gpat3 | 45 | 35 | 36 | 26 | 67 | 56 | 45 | 33 | 35 | 50 | 111 | 26 | 31 | 16 | 58 | 50 | 30 | 33 | 69 | 37 | 49 | 35 | 42 | 35 | 32 | 16 | 36 | 24 | Gpat3 |
| Gpat4 | 14239 | 14988 | 12949 | 13946 | 12493 | 13568 | 13134 | 10717 | 10560 | 10297 | 9955 | 11554 | 10685 | 10617 | 13519 | 13406 | 12964 | 13627 | 13512 | 12982 | 13229 | 10119 | 10082 | 10787 | 11234 | 11426 | 11622 | 11425 | Gpat4 |
| Gpatch1 | 2298 | 2429 | 2253 | 2309 | 2074 | 2095 | 2192 | 1628 | 1755 | 1911 | 1731 | 2021 | 1796 | 1881 | 2066 | 2228 | 2261 | 2114 | 2186 | 2134 | 2189 | 1726 | 1716 | 1923 | 1853 | 1883 | 1734 | 1761 | Gpatch1 |
| Gpatch11 | 2391 | 2579 | 2561 | 2714 | 2514 | 2363 | 2695 | 2282 | 2149 | 2233 | 2141 | 2144 | 2158 | 2266 | 2620 | 2464 | 2775 | 2852 | 2497 | 2922 | 2691 | 2020 | 2091 | 2283 | 2391 | 2150 | 2063 | 2231 | Gpatch11 |
| Gpatch2 | 3730 | 3657 | 3734 | 3675 | 3421 | 3450 | 3629 | 2917 | 2834 | 2856 | 2803 | 3413 | 3277 | 3477 | 3663 | 3473 | 3822 | 3449 | 3346 | 3319 | 3356 | 2804 | 2682 | 2905 | 3001 | 3049 | 3428 | 3450 | Gpatch2 |
| Gpatch2l | 3840 | 3866 | 3240 | 4135 | 3976 | 4206 | 4083 | 3604 | 3572 | 3642 | 3281 | 3368 | 3289 | 3419 | 4098 | 3503 | 3870 | 4156 | 4047 | 4013 | 4385 | 3790 | 3341 | 3282 | 3940 | 2841 | 3290 | 3312 | Gpatch2l |
| Gpatch3 | 339 | 414 | 510 | 329 | 503 | 420 | 407 | 410 | 462 | 489 | 480 | 638 | 545 | 566 | 336 | 324 | 423 | 401 | 441 | 374 | 393 | 446 | 471 | 543 | 520 | 607 | 603 | 555 | Gpatch3 |
| Gpatch4 | 1756 | 1837 | 2088 | 1830 | 1540 | 1619 | 1619 | 1873 | 1674 | 1911 | 1766 | 1897 | 1971 | 2257 | 1824 | 1703 | 1834 | 1632 | 1485 | 1699 | 1674 | 1601 | 1680 | 2019 | 1892 | 2638 | 1872 | 2381 | Gpatch4 |
| Gpatch8 | 9060 | 9087 | 9472 | 11173 | 11128 | 10567 | 10926 | 8905 | 9001 | 11395 | 10771 | 11516 | 10955 | 11441 | 8426 | 9054 | 10165 | 10914 | 10931 | 8434 | 11101 | 11670 | 8211 | 10910 | 11542 | 7963 | 10911 | 10890 | Gpatch8 |
| Gpbar1 | 0 | 0 | 0 | 0 | 1 | 0 | 1 | 0 | 5 | 0 | 0 | 0 | 2 | 1 | 0 | 2 | 0 | 0 | 1 | 3 | 5 | 0 | 3 | 0 | 1 | 2 | 2 | 1 | Gpbar1 |
| Gpbp1 | 10206 | 7664 | 9042 | 8935 | 8473 | 8359 | 10437 | 11313 | 9278 | 8716 | 8462 | 9633 | 8945 | 9815 | 11408 | 9168 | 8998 | 8702 | 8410 | 11078 | 9640 | 9254 | 9565 | 9340 | 9102 | 10239 | 9752 | 10268 | Gpbp1 |
| Gpbp1l1 | 6338 | 5347 | 5600 | 5380 | 4430 | 4843 | 5165 | 5683 | 4977 | 4656 | 5334 | 5387 | 5519 | 5862 | 5677 | 5666 | 5116 | 5432 | 4835 | 5071 | 5225 | 4906 | 4882 | 5121 | 5080 | 5707 | 5670 | 5616 | Gpbp1l1 |
| Gpc1 | 9164 | 10933 | 10639 | 10807 | 10126 | 9992 | 9020 | 10240 | 12084 | 13916 | 12340 | 16853 | 12328 | 14608 | 8036 | 8854 | 11466 | 10034 | 11252 | 8840 | 9096 | 11511 | 10222 | 15670 | 14361 | 19943 | 13544 | 14163 | Gpc1 |
| Gpc2 | 4905 | 5789 | 5782 | 6365 | 6176 | 5640 | 7353 | 5169 | 4520 | 5085 | 4547 | 4804 | 4526 | 5740 | 5028 | 4527 | 6731 | 6491 | 6624 | 4843 | 7211 | 6729 | 4057 | 5775 | 5643 | 4174 | 4718 | 5217 | Gpc2 |
| Gpc3 | 19075 | 14912 | 15247 | 8112 | 5105 | 8191 | 3736 | 10756 | 15242 | 10638 | 21582 | 9536 | 14341 | 9099 | 15347 | 21628 | 10138 | 9509 | 4114 | 5196 | 3699 | 11603 | 19572 | 10172 | 7892 | 11933 | 10793 | 10454 | Gpc3 |
| Gpc4 | 2419 | 2117 | 2205 | 2250 | 1829 | 2134 | 2302 | 2136 | 2739 | 2548 | 2586 | 3189 | 2232 | 3282 | 2019 | 1974 | 2167 | 2260 | 2251 | 1673 | 2250 | 1790 | 2361 | 3409 | 2490 | 3753 | 2511 | 2906 | Gpc4 |
| Gpc5 | 52 | 43 | 46 | 70 | 83 | 79 | 83 | 82 | 79 | 75 | 83 | 45 | 53 | 48 | 44 | 57 | 66 | 61 | 82 | 106 | 61 | 87 | 62 | 44 | 70 | 37 | 77 | 51 | Gpc5 |
| Gpc6 | 2985 | 2304 | 1413 | 1637 | 1368 | 1833 | 1279 | 1760 | 2338 | 1948 | 2781 | 1861 | 1953 | 1811 | 2012 | 2648 | 1415 | 1508 | 1260 | 1339 | 1272 | 1967 | 2355 | 1745 | 1677 | 1054 | 2066 | 1740 | Gpc6 |
| Gpcpd1 | 5157 | 3522 | 3551 | 4486 | 5177 | 5212 | 5830 | 7543 | 7374 | 7103 | 7221 | 6886 | 7395 | 7563 | 5631 | 4307 | 3969 | 4613 | 4437 | 5054 | 4844 | 7752 | 7308 | 6283 | 7092 | 5650 | 7004 | 7116 | Gpcpd1 |
| Gpd1 | 197 | 311 | 444 | 440 | 487 | 338 | 355 | 258 | 261 | 278 | 297 | 365 | 298 | 344 | 231 | 267 | 497 | 442 | 508 | 318 | 399 | 368 | 201 | 333 | 419 | 321 | 407 | 363 | Gpd1 |
| Gpd1l | 4594 | 5202 | 4401 | 5015 | 5131 | 4620 | 5161 | 2939 | 2908 | 2780 | 2485 | 2391 | 2239 | 2356 | 4799 | 4537 | 4475 | 4987 | 5108 | 5018 | 5322 | 2971 | 2850 | 2592 | 2866 | 2161 | 2365 | 2376 | Gpd1l |
| Gpd2 | 2936 | 2573 | 2482 | 2549 | 2646 | 2556 | 2921 | 2539 | 2577 | 2493 | 2262 | 2742 | 2563 | 2699 | 2931 | 2548 | 2540 | 2367 | 2706 | 2622 | 2600 | 2420 | 2357 | 2481 | 2302 | 2798 | 2511 | 2556 | Gpd2 |
| Gper1 | 68 | 93 | 69 | 65 | 167 | 96 | 118 | 162 | 225 | 228 | 219 | 146 | 163 | 180 | 54 | 76 | 46 | 109 | 145 | 142 | 125 | 224 | 258 | 194 | 277 | 165 | 163 | 133 | Gper1 |
| Gphb5 | 1 | 0 | 3 | 0 | 2 | 2 | 3 | 1 | 0 | 0 | 0 | 2 | 0 | 0 | 0 | 0 | 0 | 0 | 0 | 0 | 0 | 2 | 1 | 0 | 0 | 0 | 0 | 0 | Gphb5 |
| Gphn | 3517 | 3436 | 3333 | 3648 | 4096 | 3608 | 4270 | 3932 | 3213 | 3284 | 3046 | 3289 | 3026 | 3215 | 3798 | 3113 | 3410 | 3744 | 4102 | 4315 | 4280 | 3928 | 3429 | 3199 | 3667 | 2883 | 3201 | 3277 | Gphn |
| Gpi1 | 20608 | 23266 | 18787 | 19856 | 19983 | 19940 | 17438 | 15610 | 18398 | 17670 | 18035 | 16544 | 16833 | 15976 | 19312 | 20269 | 20183 | 19891 | 20229 | 19225 | 18525 | 16046 | 18332 | 16859 | 16860 | 18743 | 16867 | 16896 | Gpi1 |
| Gpihbp1 | 8 | 4 | 6 | 0 | 14 | 6 | 0 | 22 | 39 | 27 | 72 | 14 | 14 | 5 | 15 | 9 | 1 | 14 | 9 | 0 | 0 | 28 | 87 | 18 | 9 | 20 | 19 | 8 | Gpihbp1 |
| Gpkow | 6666 | 6739 | 5732 | 6430 | 5911 | 6125 | 6439 | 5327 | 5145 | 4561 | 4439 | 4906 | 4848 | 4810 | 7009 | 5918 | 5691 | 6229 | 6010 | 6583 | 6351 | 4602 | 4827 | 4461 | 4642 | 4721 | 5032 | 4809 | Gpkow |
| Gpld1 | 121 | 93 | 60 | 116 | 176 | 211 | 132 | 239 | 299 | 237 | 258 | 103 | 121 | 110 | 105 | 103 | 93 | 140 | 139 | 129 | 191 | 226 | 285 | 180 | 149 | 114 | 124 | 95 | Gpld1 |
| Gpm6a | 15362 | 13239 | 13868 | 21182 | 29490 | 27634 | 32959 | 43542 | 32722 | 32157 | 26282 | 29174 | 19664 | 31859 | 20961 | 11749 | 18707 | 24930 | 29476 | 36774 | 33541 | 41906 | 33379 | 34208 | 42958 | 24595 | 27870 | 28401 | Gpm6a |
| Gpm6b | 15777 | 16169 | 15863 | 21761 | 26041 | 26256 | 28879 | 29145 | 23777 | 24892 | 21004 | 26742 | 23651 | 28205 | 21943 | 14743 | 20678 | 23205 | 25694 | 27739 | 27920 | 28022 | 22565 | 26142 | 30851 | 23716 | 25100 | 25323 | Gpm6b |
| Gpn1 | 3720 | 3701 | 3101 | 3176 | 2953 | 2985 | 3408 | 3748 | 3690 | 3272 | 3298 | 3746 | 3522 | 3905 | 3518 | 3439 | 3257 | 3079 | 3222 | 3214 | 3193 | 3532 | 3492 | 3478 | 3535 | 3765 | 3439 | 4014 | Gpn1 |
| Gpn2 | 946 | 1216 | 980 | 953 | 732 | 800 | 767 | 735 | 725 | 628 | 630 | 800 | 792 | 756 | 927 | 1030 | 944 | 957 | 843 | 875 | 761 | 662 | 673 | 744 | 745 | 957 | 815 | 774 | Gpn2 |
| Gpn3 | 1787 | 1951 | 1973 | 2052 | 1564 | 1859 | 1623 | 1326 | 1371 | 1464 | 1457 | 1635 | 1516 | 1520 | 1623 | 1875 | 2034 | 1992 | 1855 | 1754 | 1605 | 1257 | 1382 | 1538 | 1437 | 1923 | 1691 | 1523 | Gpn3 |
| Gpnmb | 45 | 39 | 26 | 13 | 7 | 39 | 9 | 50 | 85 | 43 | 60 | 51 | 40 | 23 | 17 | 40 | 8 | 32 | 9 | 2 | 5 | 30 | 84 | 47 | 22 | 38 | 24 | 28 | Gpnmb |
| Gpr1 | 9 | 7 | 12 | 8 | 7 | 13 | 7 | 12 | 18 | 13 | 18 | 16 | 17 | 26 | 7 | 5 | 10 | 3 | 14 | 15 | 21 | 13 | 24 | 11 | 12 | 19 | 17 | 15 | Gpr1 |
| Gpr101 | 15 | 22 | 9 | 37 | 44 | 24 | 46 | 37 | 37 | 42 | 35 | 12 | 7 | 5 | 17 | 16 | 12 | 43 | 51 | 30 | 53 | 42 | 28 | 12 | 32 | 19 | 8 | 7 | Gpr101 |
| Gpr107 | 5063 | 5577 | 4887 | 5462 | 5314 | 5324 | 4766 | 5468 | 5564 | 5941 | 5788 | 6047 | 6026 | 5499 | 4932 | 5175 | 4920 | 5466 | 5635 | 4688 | 5198 | 5890 | 5602 | 5667 | 6169 | 5520 | 6057 | 5739 | Gpr107 |
| Gpr108 | 1183 | 1605 | 1597 | 1446 | 1296 | 1132 | 1093 | 1308 | 1439 | 1527 | 1540 | 1440 | 1356 | 1271 | 1241 | 1389 | 1542 | 1528 | 1312 | 1108 | 1189 | 1402 | 1564 | 1537 | 1446 | 1753 | 1432 | 1556 | Gpr108 |
| Gpr119 | 0 | 0 | 0 | 0 | 1 | 0 | 0 | 4 | 2 | 0 | 2 | 0 | 0 | 3 | 0 | 0 | 0 | 0 | 0 | 0 | 0 | 0 | 1 | 0 | 2 | 0 | 0 | 0 | Gpr119 |
| Gpr12 | 54 | 55 | 59 | 78 | 191 | 147 | 144 | 88 | 115 | 150 | 116 | 42 | 36 | 51 | 42 | 38 | 68 | 107 | 138 | 128 | 156 | 89 | 114 | 72 | 124 | 31 | 40 | 38 | Gpr12 |
| Gpr132 | 3 | 3 | 6 | 3 | 2 | 2 | 2 | 7 | 7 | 7 | 4 | 1 | 0 | 1 | 3 | 10 | 0 | 4 | 2 | 0 | 0 | 0 | 3 | 2 | 3 | 6 | 0 | 5 | Gpr132 |
| Gpr135 | 499 | 384 | 590 | 486 | 652 | 534 | 645 | 372 | 338 | 312 | 384 | 272 | 354 | 315 | 539 | 534 | 556 | 541 | 672 | 596 | 599 | 429 | 313 | 413 | 398 | 230 | 268 | 275 | Gpr135 |
| Gpr137 | 1752 | 2558 | 2035 | 2448 | 2341 | 2206 | 2099 | 1601 | 2028 | 1976 | 1929 | 1934 | 1837 | 1658 | 1849 | 2022 | 2217 | 2363 | 2632 | 2317 | 2229 | 2019 | 2043 | 1920 | 2016 | 1908 | 1696 | 1740 | Gpr137 |
| Gpr137b | 2019 | 1950 | 1697 | 2085 | 2227 | 2138 | 2135 | 1812 | 1648 | 1799 | 1590 | 1969 | 1777 | 2024 | 2132 | 1901 | 1838 | 1983 | 1866 | 2056 | 1944 | 1845 | 1427 | 1597 | 1919 | 1532 | 1909 | 1871 | Gpr137b |
| Gpr137c | 3157 | 2656 | 3119 | 4036 | 3654 | 3781 | 4390 | 2786 | 2309 | 2531 | 2570 | 2903 | 2358 | 2921 | 3241 | 2812 | 3472 | 3860 | 3594 | 3387 | 3949 | 3486 | 2131 | 2661 | 2799 | 2580 | 2530 | 2650 | Gpr137c |
| Gpr139 | 398 | 459 | 353 | 579 | 844 | 687 | 780 | 670 | 535 | 567 | 384 | 299 | 396 | 454 | 512 | 389 | 511 | 637 | 727 | 793 | 792 | 805 | 524 | 390 | 534 | 199 | 396 | 427 | Gpr139 |
| Gpr141 | 8 | 5 | 4 | 8 | 5 | 7 | 4 | 11 | 7 | 4 | 9 | 1 | 7 | 3 | 9 | 5 | 1 | 2 | 5 | 4 | 4 | 10 | 19 | 14 | 17 | 6 | 10 | 5 | Gpr141 |
| Gpr142 | 3 | 3 | 7 | 11 | 5 | 2 | 4 | 2 | 1 | 1 | 4 | 7 | 5 | 4 | 7 | 5 | 8 | 9 | 9 | 8 | 2 | 2 | 3 | 0 | 3 | 4 | 5 | 4 | Gpr142 |
| Gpr143 | 6 | 4 | 4 | 0 | 2 | 11 | 4 | 9 | 16 | 10 | 8 | 5 | 3 | 13 | 1 | 6 | 3 | 3 | 3 | 7 | 1 | 10 | 10 | 10 | 0 | 6 | 7 | 9 | Gpr143 |
| Gpr146 | 203 | 296 | 267 | 228 | 232 | 171 | 206 | 136 | 156 | 148 | 160 | 129 | 268 | 153 | 185 | 251 | 185 | 213 | 225 | 193 | 198 | 167 | 191 | 152 | 143 | 177 | 158 | 205 | Gpr146 |
| Gpr149 | 31 | 32 | 19 | 47 | 111 | 86 | 105 | 151 | 143 | 98 | 83 | 86 | 53 | 87 | 44 | 33 | 56 | 76 | 89 | 86 | 105 | 187 | 142 | 118 | 163 | 74 | 52 | 78 | Gpr149 |
| Gpr15 | 14 | 19 | 5 | 12 | 7 | 7 | 12 | 18 | 10 | 14 | 3 | 22 | 11 | 10 | 9 | 14 | 16 | 2 | 7 | 5 | 10 | 12 | 8 | 7 | 2 | 19 | 17 | 19 | Gpr15 |
| Gpr150 | 6 | 21 | 11 | 20 | 34 | 13 | 18 | 23 | 22 | 13 | 21 | 8 | 6 | 14 | 18 | 16 | 37 | 12 | 24 | 27 | 18 | 20 | 15 | 16 | 6 | 8 | 12 | 6 | Gpr150 |
| Gpr151 | 14 | 16 | 7 | 18 | 34 | 28 | 22 | 50 | 35 | 36 | 37 | 29 | 52 | 20 | 17 | 26 | 24 | 18 | 34 | 24 | 35 | 42 | 53 | 26 | 57 | 17 | 29 | 13 | Gpr151 |
| Gpr152 | 0 | 0 | 0 | 1 | 0 | 1 | 0 | 0 | 0 | 0 | 0 | 0 | 0 | 0 | 0 | 0 | 1 | 0 | 0 | 1 | 0 | 0 | 0 | 0 | 0 | 0 | 0 | 0 | Gpr152 |
| Gpr153 | 1606 | 1719 | 1906 | 1415 | 1753 | 1637 | 1060 | 1805 | 2096 | 2254 | 2621 | 1661 | 2074 | 1576 | 1338 | 1982 | 1732 | 1657 | 1431 | 1068 | 1129 | 2622 | 2258 | 2121 | 2082 | 1623 | 1785 | 1693 | Gpr153 |
| Gpr155 | 2629 | 2602 | 1744 | 2588 | 2617 | 2664 | 2944 | 1165 | 1082 | 1077 | 1090 | 847 | 915 | 814 | 2716 | 2690 | 2167 | 2891 | 2419 | 2620 | 2895 | 1275 | 1072 | 806 | 1098 | 634 | 839 | 739 | Gpr155 |
| Gpr156 | 814 | 840 | 971 | 1278 | 1530 | 1279 | 1291 | 822 | 769 | 976 | 840 | 1058 | 1046 | 1086 | 960 | 847 | 1307 | 1217 | 1488 | 1177 | 1387 | 1155 | 779 | 1034 | 1224 | 656 | 991 | 1061 | Gpr156 |
| Gpr157 | 544 | 582 | 602 | 464 | 418 | 391 | 273 | 143 | 201 | 170 | 217 | 164 | 177 | 140 | 472 | 642 | 542 | 451 | 335 | 298 | 320 | 180 | 234 | 182 | 150 | 145 | 121 | 182 | Gpr157 |
| Gpr158 | 300 | 309 | 121 | 471 | 755 | 632 | 620 | 543 | 473 | 462 | 415 | 324 | 354 | 312 | 427 | 266 | 283 | 561 | 581 | 665 | 660 | 683 | 421 | 347 | 522 | 148 | 293 | 262 | Gpr158 |
| Gpr160 | 115 | 69 | 53 | 53 | 20 | 34 | 20 | 57 | 59 | 43 | 69 | 45 | 56 | 52 | 71 | 104 | 20 | 65 | 30 | 27 | 28 | 42 | 65 | 50 | 34 | 60 | 36 | 34 | Gpr160 |
| Gpr161 | 4555 | 4275 | 4235 | 5395 | 7846 | 6717 | 6887 | 8718 | 8291 | 9590 | 8627 | 9688 | 8721 | 9765 | 4303 | 4041 | 5357 | 5702 | 7337 | 5907 | 7013 | 10753 | 7622 | 9580 | 11470 | 6341 | 9015 | 9143 | Gpr161 |
| Gpr162 | 1035 | 1659 | 1890 | 2199 | 3704 | 2633 | 2708 | 3362 | 3573 | 4904 | 3955 | 3505 | 3032 | 3439 | 1291 | 1132 | 2383 | 2660 | 3702 | 2512 | 2959 | 5008 | 3570 | 4469 | 5320 | 2891 | 3430 | 3242 | Gpr162 |
| Gpr165 | 131 | 130 | 102 | 155 | 262 | 265 | 241 | 325 | 344 | 253 | 276 | 297 | 260 | 297 | 138 | 93 | 144 | 212 | 249 | 277 | 264 | 434 | 311 | 222 | 358 | 154 | 304 | 268 | Gpr165 |
| Gpr17 | 46 | 27 | 59 | 8 | 41 | 17 | 18 | 225 | 304 | 258 | 314 | 168 | 193 | 153 | 26 | 55 | 39 | 22 | 27 | 22 | 24 | 231 | 329 | 206 | 133 | 227 | 169 | 177 | Gpr17 |
| Gpr171 | 0 | 2 | 0 | 0 | 0 | 0 | 0 | 4 | 3 | 1 | 0 | 1 | 0 | 1 | 0 | 1 | 0 | 0 | 1 | 3 | 0 | 3 | 1 | 2 | 0 | 11 | 2 | 0 | Gpr171 |
| Gpr173 | 2339 | 2355 | 2914 | 3308 | 3436 | 3129 | 3321 | 2931 | 2630 | 3495 | 3108 | 3476 | 3545 | 3811 | 2312 | 2145 | 3235 | 3300 | 3479 | 2612 | 3626 | 4226 | 2486 | 3544 | 3819 | 3075 | 3309 | 3707 | Gpr173 |
| Gpr174 | 1 | 0 | 0 | 0 | 2 | 1 | 2 | 1 | 7 | 10 | 12 | 1 | 5 | 6 | 5 | 3 | 0 | 0 | 0 | 0 | 1 | 2 | 7 | 6 | 10 | 9 | 5 | 1 | Gpr174 |
| Gpr176 | 118 | 117 | 84 | 159 | 228 | 160 | 190 | 171 | 218 | 271 | 194 | 101 | 82 | 136 | 125 | 105 | 116 | 184 | 212 | 176 | 212 | 251 | 203 | 169 | 231 | 93 | 108 | 72 | Gpr176 |
| Gpr179 | 35 | 20 | 29 | 44 | 65 | 61 | 63 | 44 | 75 | 93 | 88 | 74 | 66 | 68 | 50 | 19 | 60 | 65 | 87 | 43 | 73 | 134 | 78 | 80 | 120 | 54 | 63 | 71 | Gpr179 |
| Gpr18 | 0 | 0 | 0 | 0 | 0 | 0 | 1 | 0 | 2 | 1 | 0 | 0 | 0 | 0 | 0 | 0 | 0 | 0 | 0 | 0 | 0 | 0 | 1 | 2 | 0 | 0 | 0 | 0 | Gpr18 |
| Gpr180 | 1772 | 2074 | 1764 | 1803 | 1492 | 1686 | 1601 | 1224 | 1402 | 1187 | 1424 | 1142 | 1218 | 1090 | 1756 | 1835 | 1688 | 1610 | 1558 | 1700 | 1586 | 1111 | 1520 | 1242 | 1122 | 1135 | 1223 | 1139 | Gpr180 |
| Gpr182 | 53 | 70 | 123 | 89 | 69 | 79 | 53 | 42 | 60 | 52 | 85 | 64 | 61 | 51 | 71 | 73 | 73 | 78 | 44 | 42 | 55 | 65 | 92 | 69 | 64 | 61 | 83 | 71 | Gpr182 |
| Gpr183 | 47 | 15 | 12 | 6 | 19 | 22 | 35 | 55 | 92 | 73 | 89 | 65 | 95 | 51 | 15 | 29 | 7 | 6 | 12 | 12 | 12 | 72 | 109 | 76 | 37 | 105 | 87 | 52 | Gpr183 |
| Gpr19 | 1038 | 1120 | 1186 | 1202 | 1192 | 1145 | 1430 | 1239 | 1048 | 1156 | 989 | 1334 | 1122 | 1349 | 1172 | 879 | 1272 | 1197 | 1217 | 1295 | 1219 | 1218 | 998 | 1240 | 1201 | 1383 | 1406 | 1428 | Gpr19 |
| Gpr20 | 54 | 45 | 26 | 4 | 23 | 16 | 10 | 21 | 21 | 14 | 62 | 47 | 18 | 27 | 33 | 45 | 22 | 9 | 16 | 15 | 17 | 17 | 54 | 20 | 21 | 19 | 14 | 28 | Gpr20 |
| Gpr21 | 114 | 105 | 140 | 154 | 361 | 215 | 280 | 1143 | 1009 | 908 | 815 | 641 | 555 | 695 | 124 | 84 | 139 | 176 | 269 | 247 | 262 | 1211 | 1013 | 792 | 1166 | 411 | 506 | 587 | Gpr21 |
| Gpr22 | 439 | 473 | 467 | 679 | 981 | 964 | 849 | 1545 | 901 | 874 | 816 | 781 | 1058 | 852 | 857 | 546 | 610 | 665 | 798 | 1162 | 897 | 1338 | 1020 | 789 | 1034 | 487 | 877 | 946 | Gpr22 |

| | | | | | | | | | | | | | | | | | | | | | | | | | | | | | |
|---|---|---|---|---|---|---|---|---|---|---|---|---|---|---|---|---|---|---|---|---|---|---|---|---|---|---|---|---|---|
| Gpr25 | 15 | 3 | 13 | 22 | 20 | 15 | 12 | 9 | 7 | 20 | 20 | 0 | 3 | 0 | 15 | 7 | 2 | 14 | 31 | 8 | 30 | 18 | 12 | 14 | 8 | 7 | 1 | 1 | Gpr25 |
| Gpr26 | 419 | 403 | 232 | 660 | 1793 | 1510 | 1702 | 1788 | 2136 | 1993 | 1803 | 1302 | 717 | 1484 | 494 | 297 | 467 | 1008 | 1508 | 1554 | 1849 | 2212 | 1978 | 1439 | 2557 | 624 | 902 | 883 | Gpr26 |
| Gpr27 | 652 | 578 | 821 | 1139 | 1757 | 1261 | 1657 | 1920 | 1920 | 2324 | 2194 | 1605 | 1186 | 1702 | 745 | 551 | 1043 | 1322 | 1681 | 1475 | 1758 | 2455 | 1893 | 2329 | 2455 | 1664 | 1171 | 1422 | Gpr27 |
| Gpr3 | 186 | 263 | 178 | 247 | 199 | 152 | 221 | 24 | 17 | 34 | 32 | 10 | 32 | 17 | 177 | 228 | 233 | 284 | 171 | 221 | 246 | 33 | 14 | 21 | 13 | 18 | 6 | 27 | Gpr3 |
| Gpr31b | 0 | 0 | 0 | 0 | 0 | 0 | 0 | 0 | 3 | 0 | 3 | 2 | 0 | 0 | 0 | 1 | 0 | 0 | 0 | 0 | 0 | 2 | 1 | 0 | 0 | 0 | 0 | 0 | Gpr31b |
| Gpr34 | 25 | 11 | 26 | 6 | 38 | 16 | 15 | 187 | 163 | 93 | 144 | 119 | 127 | 111 | 33 | 12 | 24 | 17 | 39 | 16 | 17 | 145 | 190 | 135 | 102 | 128 | 117 | 120 | Gpr34 |
| Gpr35 | 160 | 185 | 184 | 384 | 323 | 293 | 308 | 79 | 84 | 87 | 79 | 81 | 76 | 72 | 181 | 181 | 276 | 453 | 350 | 201 | 277 | 140 | 89 | 79 | 108 | 87 | 81 | 80 | Gpr35 |
| Gpr37 | 235 | 243 | 218 | 321 | 515 | 286 | 331 | 438 | 428 | 332 | 507 | 159 | 287 | 195 | 297 | 255 | 256 | 255 | 351 | 278 | 306 | 493 | 382 | 202 | 303 | 216 | 194 | 167 | Gpr37 |
| Gpr37l1 | 2 | 0 | 0 | 0 | 0 | 1 | 2 | 2 | 1 | 7 | 11 | 3 | 5 | 0 | 0 | 0 | 0 | 2 | 0 | 0 | 1 | 0 | 10 | 2 | 3 | 0 | 0 | 0 | Gpr37l1 |
| Gpr39 | 1 | 3 | 1 | 1 | 7 | 7 | 4 | 7 | 11 | 3 | 12 | 8 | 2 | 2 | 3 | 2 | 0 | 0 | 5 | 2 | 2 | 23 | 5 | 6 | 9 | 5 | 15 | 4 | Gpr39 |
| Gpr4 | 539 | 637 | 505 | 437 | 434 | 365 | 341 | 111 | 144 | 163 | 174 | 117 | 117 | 127 | 476 | 614 | 535 | 478 | 420 | 386 | 414 | 147 | 159 | 135 | 98 | 159 | 122 | 113 | Gpr4 |
| Gpr45 | 451 | 416 | 455 | 450 | 652 | 611 | 646 | 1055 | 933 | 1079 | 902 | 1054 | 874 | 1106 | 500 | 386 | 484 | 447 | 676 | 678 | 652 | 981 | 832 | 1039 | 1120 | 957 | 1010 | 883 | Gpr45 |
| Gpr50 | 531 | 713 | 287 | 470 | 340 | 292 | 382 | 249 | 270 | 169 | 144 | 248 | 69 | 235 | 505 | 445 | 504 | 476 | 297 | 493 | 437 | 404 | 231 | 229 | 248 | 456 | 256 | 205 | Gpr50 |
| Gpr52 | 24 | 12 | 16 | 13 | 42 | 30 | 46 | 35 | 30 | 29 | 20 | 24 | 12 | 24 | 21 | 10 | 8 | 14 | 15 | 39 | 31 | 37 | 38 | 27 | 29 | 18 | 14 | 20 | Gpr52 |
| Gpr55 | 1 | 3 | 0 | 0 | 2 | 7 | 0 | 5 | 7 | 8 | 1 | 5 | 2 | 1 | 0 | 2 | 0 | 1 | 2 | 0 | 0 | 2 | 5 | 1 | 0 | 2 | 0 | 0 | Gpr55 |
| Gpr6 | 20 | 10 | 10 | 28 | 14 | 49 | 18 | 7 | 10 | 12 | 5 | 10 | 7 | 7 | 9 | 13 | 15 | 25 | 33 | 14 | 24 | 17 | 6 | 17 | 24 | 5 | 8 | 8 | Gpr6 |
| Gpr61 | 55 | 73 | 51 | 47 | 153 | 74 | 66 | 116 | 126 | 154 | 146 | 53 | 93 | 71 | 84 | 107 | 95 | 100 | 78 | 69 | 77 | 170 | 109 | 85 | 121 | 43 | 65 | 74 | Gpr61 |
| Gpr62 | 8 | 2 | 10 | 9 | 7 | 6 | 8 | 1 | 7 | 15 | 14 | 8 | 3 | 13 | 7 | 6 | 5 | 8 | 2 | 1 | 12 | 10 | 12 | 8 | 0 | 9 | 0 | 7 | Gpr62 |
| Gpr63 | 178 | 141 | 183 | 225 | 253 | 274 | 222 | 254 | 250 | 243 | 267 | 262 | 316 | 235 | 165 | 153 | 200 | 210 | 232 | 214 | 245 | 211 | 229 | 230 | 262 | 194 | 314 | 265 | Gpr63 |
| Gpr65 | 18 | 18 | 41 | 25 | 13 | 11 | 13 | 42 | 44 | 18 | 59 | 38 | 37 | 36 | 22 | 33 | 9 | 35 | 10 | 25 | 19 | 27 | 71 | 42 | 43 | 53 | 29 | 28 | Gpr65 |
| Gpr68 | 13 | 10 | 7 | 7 | 41 | 13 | 32 | 37 | 49 | 52 | 50 | 39 | 37 | 23 | 11 | 18 | 9 | 30 | 18 | 16 | 31 | 70 | 43 | 27 | 57 | 26 | 13 | 33 | Gpr68 |
| Gpr75 | 129 | 182 | 199 | 433 | 339 | 238 | 332 | 183 | 193 | 239 | 237 | 194 | 128 | 181 | 182 | 174 | 355 | 403 | 340 | 215 | 348 | 359 | 203 | 203 | 292 | 138 | 120 | 161 | Gpr75 |
| Gpr82 | 0 | 0 | 0 | 0 | 0 | 0 | 0 | 0 | 2 | 0 | 0 | 0 | 0 | 0 | 0 | 0 | 0 | 0 | 0 | 0 | 0 | 0 | 0 | 0 | 0 | 2 | 0 | 1 | Gpr82 |
| Gpr83 | 46 | 65 | 49 | 88 | 106 | 84 | 92 | 90 | 59 | 106 | 155 | 67 | 81 | 74 | 96 | 79 | 68 | 86 | 80 | 82 | 88 | 150 | 129 | 92 | 99 | 60 | 44 | 53 | Gpr83 |
| Gpr84 | 16 | 15 | 3 | 14 | 12 | 15 | 4 | 10 | 20 | 19 | 13 | 16 | 5 | 15 | 18 | 9 | 9 | 9 | 10 | 5 | 15 | 13 | 11 | 12 | 20 | 10 | 1 | 7 | Gpr84 |
| Gpr85 | 1353 | 1380 | 1485 | 2071 | 2912 | 2271 | 3056 | 3776 | 3058 | 3315 | 2713 | 2278 | 1834 | 2490 | 1859 | 1239 | 1816 | 2233 | 2703 | 2984 | 3021 | 3957 | 3014 | 2834 | 3826 | 1872 | 2223 | 2298 | Gpr85 |
| Gpr87 | 0 | 0 | 0 | 0 | 2 | 0 | 0 | 5 | 18 | 17 | 0 | 4 | 2 | 2 | 1 | 0 | 0 | 0 | 0 | 0 | 0 | 5 | 13 | 2 | 0 | 0 | 0 | 0 | Gpr87 |
| Gpr88 | 70 | 65 | 46 | 141 | 255 | 163 | 227 | 314 | 446 | 404 | 445 | 227 | 116 | 253 | 108 | 72 | 95 | 132 | 168 | 128 | 264 | 448 | 329 | 355 | 453 | 204 | 88 | 160 | Gpr88 |
| Gpr89 | 2999 | 3120 | 1969 | 1561 | 1730 | 2337 | 2053 | 2920 | 2832 | 2062 | 2386 | 2111 | 2156 | 2099 | 2728 | 2826 | 1627 | 1705 | 1725 | 2703 | 2292 | 1930 | 2660 | 1883 | 2324 | 1857 | 2453 | 2124 | Gpr89 |
| Gprasp1 | 37592 | 34973 | 32339 | 37915 | 45302 | 43069 | 45747 | 46394 | 38186 | 39777 | 37463 | 33787 | 31087 | 36388 | 41210 | 32630 | 36615 | 40689 | 41003 | 46889 | 45094 | 48408 | 37022 | 35972 | 42414 | 26834 | 32763 | 31811 | Gprasp1 |
| Gprasp2 | 9409 | 8848 | 8114 | 10138 | 11396 | 10899 | 10949 | 10204 | 8945 | 9453 | 8530 | 7992 | 7007 | 8280 | 9966 | 8173 | 9114 | 10698 | 10689 | 11589 | 11470 | 11683 | 8686 | 8797 | 10158 | 6767 | 7638 | 7522 | Gprasp2 |
| Gprc5a | 151 | 216 | 251 | 134 | 82 | 130 | 86 | 74 | 86 | 92 | 71 | 76 | 95 | 73 | 181 | 236 | 207 | 118 | 94 | 95 | 91 | 37 | 85 | 111 | 46 | 95 | 102 | 106 | Gprc5a |
| Gprc5b | 3715 | 4129 | 4561 | 4206 | 3814 | 3627 | 3510 | 3339 | 3431 | 3459 | 3113 | 3932 | 3848 | 3776 | 3542 | 3629 | 4451 | 3798 | 4122 | 3590 | 3758 | 2799 | 3390 | 3742 | 3317 | 4106 | 3720 | 4069 | Gprc5b |
| Gprc5c | 1347 | 1137 | 1205 | 760 | 808 | 922 | 583 | 964 | 1425 | 1176 | 1626 | 866 | 1461 | 734 | 1092 | 1441 | 924 | 855 | 662 | 553 | 496 | 1044 | 1557 | 1047 | 766 | 1258 | 1010 | 879 | Gprc5c |
| Gprc5d | 5 | 0 | 10 | 9 | 5 | 0 | 2 | 1 | 1 | 0 | 0 | 0 | 2 | 2 | 5 | 4 | 0 | 0 | 2 | 4 | 2 | 2 | 1 | 0 | 0 | 1 | 0 | 1 | Gprc5d |
| Gprc6a | 1 | 1 | 0 | 2 | 0 | 3 | 4 | 4 | 4 | 3 | 0 | 4 | 3 | 2 | 3 | 5 | 0 | 1 | 0 | 0 | 0 | 2 | 4 | 2 | 3 | 3 | 8 | 4 | Gprc6a |
| Gprin1 | 4530 | 5744 | 6067 | 9524 | 11586 | 9088 | 9182 | 7614 | 7563 | 10454 | 7951 | 8467 | 6599 | 8442 | 5458 | 4125 | 8100 | 9600 | 11737 | 7433 | 11069 | 12375 | 6885 | 9411 | 11458 | 6928 | 7075 | 7798 | Gprin1 |
| Gprin2 | 3530 | 3511 | 2513 | 4101 | 5411 | 4950 | 4241 | 2850 | 2811 | 3142 | 2922 | 1927 | 2249 | 2021 | 3534 | 3602 | 3414 | 4142 | 4635 | 4067 | 4689 | 3494 | 2610 | 2328 | 3419 | 1024 | 2109 | 1791 | Gprin2 |
| Gprin3 | 170 | 101 | 67 | 103 | 214 | 209 | 149 | 321 | 413 | 284 | 336 | 98 | 146 | 119 | 141 | 128 | 87 | 108 | 122 | 181 | 231 | 256 | 328 | 153 | 260 | 89 | 145 | 82 | Gprin3 |
| Gps1 | 6422 | 7891 | 6275 | 6573 | 6185 | 6412 | 6128 | 6188 | 6697 | 5689 | 5575 | 6201 | 5856 | 5879 | 6649 | 6489 | 6343 | 6384 | 6986 | 6868 | 6322 | 5939 | 6695 | 6006 | 6161 | 6831 | 6463 | 6355 | Gps1 |
| Gps2 | 2292 | 2594 | 2854 | 2816 | 2208 | 2181 | 2267 | 2240 | 2537 | 2855 | 2451 | 3304 | 2809 | 3493 | 2135 | 2193 | 2684 | 2338 | 2753 | 2067 | 2309 | 2657 | 2357 | 3331 | 3063 | 3679 | 2980 | 3303 | Gps2 |
| Gpsm1 | 4667 | 6235 | 6661 | 7342 | 8275 | 7234 | 6267 | 7436 | 8696 | 10187 | 9988 | 10486 | 9283 | 9285 | 4738 | 5608 | 7566 | 7175 | 8086 | 5894 | 7030 | 10804 | 8153 | 10565 | 11311 | 9210 | 9395 | 8853 | Gpsm1 |
| Gpsm2 | 5797 | 5872 | 5802 | 5300 | 4597 | 4587 | 5367 | 4754 | 4224 | 4486 | 4185 | 6037 | 5182 | 6216 | 5757 | 5434 | 5782 | 4878 | 5102 | 5144 | 5158 | 3818 | 4224 | 5168 | 4686 | 6773 | 6241 | 6953 | Gpsm2 |
| Gpsm3 | 72 | 103 | 140 | 95 | 93 | 98 | 59 | 74 | 105 | 115 | 106 | 96 | 81 | 83 | 79 | 103 | 101 | 92 | 101 | 63 | 81 | 90 | 96 | 92 | 92 | 98 | 110 | 89 | Gpsm3 |
| Gpt | 1102 | 1691 | 1800 | 1893 | 1809 | 1590 | 1771 | 855 | 836 | 1136 | 948 | 1144 | 873 | 1008 | 1023 | 1448 | 1951 | 1993 | 2280 | 1484 | 1721 | 1111 | 772 | 1124 | 1141 | 1041 | 1056 | 1040 | Gpt |
| Gpt2 | 5687 | 5661 | 6347 | 5523 | 4998 | 4798 | 5107 | 2677 | 3065 | 3131 | 3198 | 2640 | 2861 | 2753 | 5259 | 5514 | 5757 | 5340 | 5432 | 4732 | 5164 | 2994 | 3297 | 3047 | 2896 | 3019 | 2593 | 2855 | Gpt2 |
| Gpx1 | 6429 | 8674 | 7320 | 5425 | 4519 | 4981 | 5107 | 5907 | 7228 | 5233 | 5559 | 6375 | 9426 | 7035 | 6638 | 7357 | 6262 | 4824 | 5377 | 6387 | 4477 | 4487 | 7554 | 5880 | 5678 | 7927 | 7870 | 9127 | Gpx1 |
| Gpx2 | 3 | 13 | 12 | 13 | 18 | 10 | 8 | 6 | 7 | 28 | 41 | 1 | 11 | 3 | 9 | 19 | 10 | 22 | 3 | 8 | 3 | 12 | 14 | 28 | 8 | 26 | 0 | 0 | Gpx2 |
| Gpx3 | 1047 | 1320 | 1194 | 815 | 796 | 791 | 603 | 729 | 863 | 761 | 1131 | 794 | 1100 | 788 | 1212 | 1491 | 1042 | 787 | 676 | 700 | 597 | 715 | 1151 | 621 | 646 | 1000 | 796 | 904 | Gpx3 |
| Gpx4 | 5768 | 7261 | 6410 | 6309 | 5393 | 5170 | 5361 | 5068 | 5871 | 4523 | 4842 | 4571 | 5433 | 5195 | 5989 | 6777 | 6157 | 5505 | 6029 | 6346 | 5502 | 4151 | 5970 | 5151 | 5195 | 5885 | 5843 | 5762 | Gpx4 |
| Gpx5 | 0 | 0 | 0 | 0 | 0 | 0 | 0 | 0 | 0 | 0 | 0 | 0 | 0 | 0 | 1 | 0 | 0 | 0 | 0 | 0 | 0 | 0 | 0 | 0 | 0 | 0 | 0 | 0 | Gpx5 |
| Gpx6 | 14 | 3 | 4 | 0 | 2 | 0 | 3 | 0 | 2 | 0 | 0 | 0 | 0 | 0 | 12 | 10 | 3 | 1 | 1 | 0 | 0 | 2 | 1 | 0 | 0 | 0 | 0 | 1 | Gpx6 |
| Gpx7 | 654 | 756 | 720 | 392 | 347 | 509 | 256 | 931 | 1266 | 967 | 1326 | 918 | 1204 | 870 | 713 | 776 | 486 | 557 | 348 | 337 | 286 | 871 | 1594 | 972 | 896 | 1087 | 1192 | 1063 | Gpx7 |
| Gpx8 | 3639 | 2694 | 3298 | 2359 | 2144 | 2485 | 2386 | 3292 | 3268 | 2540 | 3216 | 2593 | 3103 | 2937 | 3501 | 3220 | 2785 | 2400 | 2328 | 2997 | 2249 | 2637 | 3842 | 2650 | 2263 | 3182 | 3016 | 3177 | Gpx8 |
| Gramd1a | 8023 | 10116 | 9967 | 11491 | 14662 | 11816 | 12295 | 16673 | 15145 | 17910 | 15574 | 16948 | 15280 | 16428 | 8538 | 9163 | 11786 | 12233 | 14641 | 10924 | 12627 | 20508 | 14007 | 17198 | 18854 | 14680 | 15558 | 15707 | Gramd1a |
| Gramd1b | 1902 | 1968 | 1690 | 2499 | 4200 | 3140 | 3195 | 3071 | 3679 | 3615 | 3626 | 2246 | 2513 | 2245 | 1974 | 1921 | 2225 | 3070 | 3641 | 2800 | 3455 | 4451 | 3377 | 2823 | 3900 | 1494 | 2087 | 1798 | Gramd1b |
| Gramd1c | 331 | 213 | 177 | 259 | 238 | 200 | 241 | 426 | 437 | 469 | 532 | 319 | 333 | 388 | 255 | 273 | 212 | 220 | 194 | 146 | 203 | 353 | 487 | 381 | 255 | 472 | 328 | 291 | Gramd1c |
| Gramd2 | 301 | 338 | 227 | 534 | 582 | 546 | 579 | 137 | 210 | 220 | 264 | 100 | 128 | 136 | 326 | 299 | 389 | 687 | 555 | 485 | 552 | 277 | 256 | 203 | 225 | 72 | 70 | 89 | Gramd2 |
| Gramd3 | 2335 | 2056 | 1990 | 1389 | 1254 | 1574 | 1219 | 1014 | 1141 | 914 | 1113 | 832 | 1150 | 947 | 2122 | 2290 | 1713 | 1550 | 1255 | 1611 | 1248 | 956 | 1389 | 827 | 704 | 1075 | 1090 | 924 | Gramd3 |
| Gramd4 | 2660 | 3140 | 3331 | 3922 | 4631 | 3828 | 4149 | 3960 | 4631 | 5254 | 4498 | 5071 | 3889 | 4732 | 2793 | 2805 | 3888 | 3998 | 4588 | 3615 | 4090 | 4817 | 4410 | 5377 | 5768 | 4263 | 4306 | 4400 | Gramd4 |
| Grap | 266 | 275 | 305 | 282 | 291 | 252 | 261 | 272 | 287 | 346 | 319 | 279 | 358 | 294 | 214 | 248 | 227 | 289 | 309 | 323 | 263 | 322 | 384 | 289 | 319 | 351 | 298 | 343 | Grap |
| Grap2 | 149 | 224 | 178 | 109 | 74 | 116 | 105 | 199 | 180 | 145 | 148 | 342 | 836 | 451 | 189 | 202 | 178 | 111 | 96 | 162 | 79 | 155 | 217 | 178 | 153 | 397 | 517 | 807 | Grap2 |
| Grb10 | 56371 | 47828 | 45460 | 30447 | 22930 | 31810 | 25241 | 53563 | 59145 | 46919 | 66491 | 52358 | 60287 | 53021 | 50759 | 56520 | 37994 | 31347 | 25757 | 27215 | 23837 | 45248 | 64498 | 49571 | 42083 | 59935 | 58486 | 59016 | Grb10 |
| Grb14 | 747 | 669 | 783 | 605 | 974 | 766 | 729 | 540 | 612 | 576 | 597 | 405 | 506 | 362 | 877 | 896 | 747 | 796 | 736 | 701 | 818 | 809 | 524 | 466 | 571 | 311 | 314 | 336 | Grb14 |
| Grb2 | 8905 | 9940 | 11410 | 12862 | 12347 | 10623 | 12271 | 6528 | 7322 | 8185 | 7936 | 8086 | 7601 | 8654 | 8883 | 9657 | 12026 | 13114 | 12140 | 9852 | 12652 | 7997 | 7394 | 8698 | 8723 | 8592 | 7453 | 8198 | Grb2 |
| Grb7 | 204 | 99 | 72 | 51 | 34 | 34 | 28 | 166 | 242 | 208 | 175 | 43 | 78 | 46 | 86 | 118 | 44 | 22 | 23 | 33 | 9 | 43 | 215 | 194 | 17 | 215 | 19 | 65 | Grb7 |
| Grcc10 | 236 | 219 | 221 | 260 | 272 | 209 | 321 | 459 | 400 | 446 | 412 | 516 | 477 | 452 | 222 | 186 | 254 | 281 | 307 | 290 | 306 | 485 | 416 | 547 | 433 | 498 | 416 | 534 | Grcc10 |
| Greb1 | 1223 | 1232 | 2010 | 969 | 954 | 1054 | 729 | 879 | 892 | 973 | 921 | 1476 | 1852 | 1270 | 1048 | 1590 | 1345 | 819 | 867 | 835 | 765 | 1021 | 805 | 805 | 879 | 941 | 1878 | 1670 | Greb1 |
| Greb1l | 1200 | 1281 | 1362 | 873 | 859 | 1107 | 666 | 1292 | 1194 | 1214 | 1225 | 1081 | 1902 | 1044 | 1605 | 1462 | 1174 | 802 | 715 | 1010 | 747 | 1333 | 1126 | 747 | 980 | 607 | 1536 | 1469 | Greb1l |
| Grem1 | 813 | 482 | 683 | 345 | 470 | 604 | 293 | 570 | 621 | 546 | 1054 | 327 | 794 | 305 | 708 | 726 | 379 | 518 | 298 | 306 | 331 | 461 | 830 | 492 | 436 | 356 | 301 | 381 | Grem1 |
| Grem2 | 428 | 399 | 384 | 182 | 236 | 436 | 95 | 652 | 779 | 552 | 823 | 558 | 964 | 446 | 550 | 526 | 348 | 359 | 205 | 178 | 109 | 470 | 937 | 553 | 378 | 492 | 585 | 579 | Grem2 |
| Grhl1 | 344 | 327 | 269 | 389 | 236 | 284 | 274 | 238 | 239 | 216 | 143 | 79 | 95 | 95 | 260 | 314 | 239 | 314 | 228 | 236 | 247 | 80 | 299 | 190 | 60 | 121 | 79 | 97 | Grhl1 |
| Grhl2 | 419 | 146 | 82 | 85 | 43 | 78 | 9 | 539 | 608 | 554 | 403 | 106 | 226 | 110 | 175 | 233 | 34 | 106 | 15 | 8 | 7 | 115 | 705 | 473 | 56 | 453 | 80 | 101 | Grhl2 |
| Grhl3 | 464 | 426 | 372 | 290 | 191 | 297 | 194 | 283 | 443 | 273 | 185 | 196 | 306 | 185 | 384 | 422 | 338 | 332 | 267 | 210 | 220 | 84 | 491 | 219 | 125 | 182 | 224 | 235 | Grhl3 |
| Grhpr | 1155 | 1359 | 1253 | 982 | 752 | 914 | 748 | 556 | 743 | 744 | 756 | 770 | 898 | 735 | 1042 | 1236 | 1102 | 980 | 885 | 852 | 832 | 503 | 799 | 719 | 703 | 980 | 716 | 890 | Grhpr |
| Gria1 | 1232 | 1234 | 791 | 2290 | 3789 | 3198 | 3948 | 2954 | 3374 | 3811 | 3158 | 2272 | 1529 | 2572 | 1444 | 986 | 1436 | 2870 | 3190 | 3224 | 4239 | 4751 | 2877 | 3167 | 4461 | 1571 | 1825 | 1746 | Gria1 |
| Gria2 | 15004 | 12196 | 7373 | 19146 | 27384 | 29323 | 31810 | 29962 | 26262 | 23223 | 21248 | 17716 | 16942 | 19597 | 18471 | 11292 | 12572 | 20690 | 24035 | 30945 | 33432 | 25340 | 21162 | 17992 | 30824 | 9160 | 16392 | 13924 | Gria2 |
| Gria3 | 1843 | 1454 | 1381 | 1911 | 2774 | 2658 | 2262 | 2992 | 3011 | 2775 | 3061 | 1757 | 2237 | 1819 | 1987 | 1822 | 1834 | 2012 | 2101 | 2583 | 2334 | 3101 | 2938 | 2119 | 2560 | 1360 | 1562 | 1581 | Gria3 |
| Gria4 | 1794 | 1842 | 1379 | 2690 | 4219 | 3205 | 3698 | 4439 | 3670 | 3945 | 3542 | 2929 | 3170 | 3103 | 2619 | 1896 | 2225 | 2955 | 3112 | 3582 | 3700 | 5462 | 3529 | 3003 | 4181 | 1766 | 3005 | 2598 | Gria4 |

| | | | | | | | | | | | | | | | | | | | | | | | | | | | | | |
|---|---|---|---|---|---|---|---|---|---|---|---|---|---|---|---|---|---|---|---|---|---|---|---|---|---|---|---|---|---|
| Grid1 | 420 | 521 | 395 | 854 | 1194 | 1035 | 1035 | 700 | 888 | 941 | 949 | 643 | 509 | 579 | 525 | 413 | 641 | 841 | 994 | 808 | 1021 | 1230 | 805 | 712 | 992 | 479 | 452 | 488 | Grid1 |
| Grid2 | 232 | 246 | 238 | 424 | 779 | 491 | 409 | 615 | 635 | 620 | 549 | 283 | 480 | 289 | 392 | 322 | 338 | 417 | 450 | 415 | 518 | 824 | 623 | 279 | 497 | 174 | 310 | 252 | Grid2 |
| Grid2ip | 215 | 308 | 253 | 219 | 282 | 204 | 229 | 77 | 91 | 87 | 84 | 124 | 79 | 92 | 262 | 288 | 308 | 286 | 241 | 217 | 230 | 80 | 60 | 79 | 98 | 52 | 79 | 95 | Grid2ip |
| Grifin | 0 | 1 | 0 | 0 | 2 | 9 | 4 | 2 | 2 | 0 | 0 | 1 | 5 | 0 | 0 | 0 | 0 | 2 | 0 | 2 | 0 | 2 | 4 | 2 | 2 | 2 | 1 | 1 | Grifin |
| Grik1 | 484 | 508 | 381 | 654 | 1196 | 959 | 1243 | 1112 | 1114 | 1085 | 915 | 534 | 576 | 649 | 717 | 507 | 585 | 985 | 1012 | 1115 | 1179 | 1225 | 994 | 790 | 1091 | 393 | 529 | 532 | Grik1 |
| Grik2 | 4804 | 4210 | 4401 | 6884 | 8841 | 8884 | 10650 | 6699 | 5437 | 6074 | 5711 | 6634 | 5012 | 6826 | 5141 | 3333 | 5456 | 7228 | 8047 | 8272 | 10003 | 7264 | 4941 | 6039 | 7731 | 4246 | 6149 | 6074 | Grik2 |
| Grik3 | 2159 | 2683 | 1865 | 4695 | 7413 | 5920 | 6062 | 5032 | 6106 | 8070 | 6298 | 6768 | 3818 | 6580 | 2606 | 2136 | 4332 | 5316 | 6481 | 4458 | 6398 | 9411 | 4409 | 7360 | 9693 | 4082 | 4874 | 4747 | Grik3 |
| Grik4 | 331 | 365 | 343 | 422 | 623 | 475 | 408 | 557 | 692 | 701 | 697 | 560 | 555 | 514 | 332 | 411 | 407 | 494 | 498 | 428 | 493 | 897 | 635 | 495 | 731 | 478 | 525 | 487 | Grik4 |
| Grik5 | 4188 | 5402 | 5856 | 7260 | 7940 | 6772 | 6277 | 6207 | 7712 | 9660 | 8178 | 9588 | 7549 | 8255 | 3970 | 4226 | 7116 | 7159 | 8316 | 5675 | 6810 | 10061 | 6895 | 9283 | 9984 | 8922 | 7835 | 7672 | Grik5 |
| Grin1 | 414 | 615 | 380 | 1060 | 2250 | 1424 | 1295 | 1130 | 1294 | 1548 | 1346 | 728 | 788 | 669 | 529 | 497 | 773 | 1304 | 1616 | 1340 | 1507 | 1855 | 1119 | 934 | 1641 | 385 | 561 | 542 | Grin1 |
| Grin2a | 123 | 128 | 137 | 141 | 260 | 209 | 182 | 440 | 400 | 425 | 537 | 318 | 306 | 263 | 146 | 132 | 121 | 146 | 199 | 201 | 181 | 515 | 422 | 283 | 430 | 206 | 364 | 342 | Grin2a |
| Grin2b | 929 | 892 | 542 | 2545 | 4967 | 4346 | 4993 | 5074 | 5331 | 6493 | 5637 | 2700 | 2113 | 3600 | 1220 | 762 | 1330 | 3387 | 3642 | 3942 | 5590 | 6672 | 4524 | 4152 | 7327 | 1139 | 2263 | 2002 | Grin2b |
| Grin2c | 66 | 114 | 121 | 95 | 97 | 46 | 42 | 31 | 39 | 21 | 48 | 24 | 54 | 17 | 73 | 112 | 103 | 83 | 98 | 52 | 76 | 35 | 44 | 29 | 64 | 24 | 21 | 31 | Grin2c |
| Grin2d | 159 | 313 | 346 | 534 | 913 | 659 | 518 | 738 | 763 | 1199 | 1043 | 590 | 798 | 442 | 310 | 366 | 452 | 713 | 695 | 511 | 582 | 1546 | 653 | 829 | 1131 | 413 | 513 | 407 | Grin2d |
| Grin3a | 431 | 458 | 254 | 705 | 1683 | 1067 | 1172 | 1695 | 1703 | 1730 | 1608 | 747 | 798 | 760 | 603 | 389 | 487 | 855 | 1033 | 1213 | 1203 | 2202 | 1518 | 958 | 1598 | 428 | 747 | 676 | Grin3a |
| Grin3b | 22 | 36 | 45 | 58 | 43 | 31 | 31 | 15 | 20 | 26 | 25 | 18 | 25 | 28 | 26 | 52 | 46 | 72 | 67 | 35 | 37 | 18 | 19 | 30 | 11 | 10 | 12 | 13 | Grin3b |
| Grina | 4928 | 6068 | 5358 | 6236 | 6441 | 6280 | 5584 | 4910 | 5585 | 6079 | 5225 | 5559 | 5747 | 5196 | 4847 | 5682 | 6169 | 6371 | 6917 | 5969 | 5571 | 6691 | 5389 | 5666 | 6395 | 4985 | 5109 | 5146 | Grina |
| Grip1 | 2562 | 3130 | 3125 | 3180 | 3296 | 3310 | 3123 | 2989 | 3405 | 3589 | 3328 | 3595 | 3405 | 3427 | 2910 | 2953 | 3208 | 3058 | 3006 | 2963 | 3313 | 3578 | 3128 | 3388 | 3674 | 3233 | 3517 | 3369 | Grip1 |
| Grip2 | 283 | 440 | 377 | 774 | 1294 | 951 | 872 | 634 | 859 | 1056 | 983 | 661 | 465 | 562 | 377 | 368 | 641 | 946 | 1085 | 855 | 945 | 1123 | 789 | 825 | 1065 | 377 | 503 | 464 | Grip2 |
| Gripap1 | 5120 | 5918 | 5655 | 6587 | 6812 | 6341 | 6602 | 6415 | 6405 | 7212 | 6396 | 7366 | 6747 | 7583 | 5595 | 4865 | 5988 | 6488 | 6551 | 6227 | 6714 | 7180 | 5825 | 7241 | 8053 | 6503 | 7330 | 7167 | Gripap1 |
| Grk1 | 246 | 228 | 139 | 281 | 201 | 264 | 248 | 27 | 32 | 30 | 37 | 39 | 35 | 45 | 241 | 210 | 187 | 261 | 178 | 185 | 245 | 30 | 19 | 38 | 20 | 25 | 37 | 28 | Grk1 |
| Grk2 | 3166 | 3523 | 4159 | 4138 | 4912 | 4059 | 3988 | 4842 | 5551 | 5854 | 5810 | 6096 | 5266 | 5488 | 3149 | 3159 | 4271 | 4030 | 4906 | 3445 | 4288 | 6127 | 5379 | 6077 | 6196 | 6493 | 5220 | 5387 | Grk2 |
| Grk3 | 2069 | 1997 | 1997 | 2497 | 3210 | 2955 | 2525 | 2262 | 2292 | 2610 | 2721 | 1947 | 2188 | 1957 | 2259 | 2177 | 2476 | 2747 | 2927 | 2630 | 2872 | 2872 | 2240 | 2254 | 2299 | 1601 | 2005 | 1777 | Grk3 |
| Grk4 | 298 | 229 | 241 | 241 | 303 | 247 | 204 | 285 | 284 | 262 | 338 | 315 | 354 | 255 | 256 | 301 | 239 | 278 | 284 | 235 | 249 | 336 | 317 | 329 | 318 | 291 | 301 | 292 | Grk4 |
| Grk5 | 4219 | 3822 | 3315 | 3880 | 3611 | 3755 | 3670 | 2235 | 2115 | 2004 | 2131 | 2065 | 2276 | 2140 | 3753 | 3699 | 3490 | 3351 | 3642 | 3680 | 3730 | 2112 | 1934 | 1968 | 2108 | 1749 | 2100 | 2190 | Grk5 |
| Grk6 | 6760 | 7784 | 8525 | 8471 | 7917 | 7883 | 7868 | 4634 | 5517 | 5641 | 5436 | 7154 | 6263 | 6628 | 5926 | 7012 | 8887 | 8283 | 9035 | 6632 | 8183 | 5210 | 4798 | 6364 | 6285 | 7875 | 6090 | 6597 | Grk6 |
| Grm1 | 196 | 185 | 126 | 401 | 607 | 544 | 684 | 573 | 665 | 523 | 470 | 449 | 128 | 480 | 218 | 142 | 284 | 509 | 598 | 610 | 634 | 707 | 521 | 467 | 743 | 293 | 300 | 343 | Grm1 |
| Grm2 | 123 | 230 | 377 | 273 | 345 | 230 | 220 | 275 | 330 | 344 | 289 | 445 | 389 | 428 | 186 | 210 | 351 | 249 | 346 | 280 | 261 | 438 | 258 | 441 | 497 | 338 | 382 | 447 | Grm2 |
| Grm3 | 239 | 291 | 208 | 150 | 342 | 386 | 298 | 648 | 689 | 455 | 522 | 382 | 464 | 363 | 376 | 238 | 176 | 185 | 284 | 470 | 434 | 546 | 931 | 300 | 494 | 201 | 530 | 427 | Grm3 |
| Grm4 | 242 | 298 | 461 | 514 | 743 | 515 | 474 | 485 | 534 | 930 | 731 | 804 | 669 | 673 | 293 | 307 | 598 | 591 | 734 | 412 | 551 | 1161 | 486 | 812 | 937 | 525 | 644 | 699 | Grm4 |
| Grm5 | 189 | 199 | 97 | 725 | 1878 | 1255 | 1340 | 1169 | 1242 | 1314 | 1295 | 355 | 396 | 348 | 360 | 192 | 262 | 1113 | 1112 | 1260 | 1608 | 1579 | 1592 | 729 | 1413 | 133 | 225 | 201 | Grm5 |
| Grm6 | 7 | 7 | 6 | 2 | 9 | 3 | 8 | 6 | 11 | 1 | 3 | 4 | 17 | 4 | 4 | 10 | 6 | 2 | 2 | 7 | 3 | 5 | 6 | 7 | 2 | 8 | 7 | 4 | Grm6 |
| Grm7 | 280 | 222 | 203 | 341 | 682 | 471 | 441 | 562 | 624 | 550 | 596 | 224 | 218 | 259 | 276 | 283 | 234 | 406 | 402 | 473 | 512 | 815 | 645 | 314 | 511 | 252 | 266 | 186 | Grm7 |
| Grm8 | 251 | 244 | 266 | 312 | 388 | 298 | 364 | 364 | 308 | 325 | 336 | 230 | 223 | 230 | 293 | 191 | 285 | 315 | 364 | 381 | 382 | 438 | 335 | 300 | 330 | 148 | 331 | 261 | Grm8 |
| Grn | 5959 | 6921 | 7375 | 6521 | 5294 | 5555 | 4947 | 2793 | 3697 | 3695 | 4366 | 3170 | 4123 | 3092 | 5124 | 7911 | 6521 | 6759 | 5568 | 4300 | 4979 | 3581 | 4562 | 3626 | 3171 | 4038 | 3393 | 3223 | Grn |
| Grp | 15 | 11 | 8 | 18 | 39 | 31 | 30 | 50 | 49 | 39 | 75 | 5 | 3 | 30 | 29 | 9 | 24 | 18 | 31 | 37 | 32 | 25 | 63 | 56 | 37 | 47 | 14 | 32 | Grp |
| Grpel1 | 5485 | 5881 | 4529 | 4388 | 3265 | 3730 | 4443 | 3544 | 3167 | 3026 | 2893 | 3117 | 2930 | 3670 | 5428 | 4953 | 4339 | 4052 | 3661 | 4633 | 4409 | 2777 | 3203 | 3103 | 3032 | 3463 | 3463 | 3585 | Grpel1 |
| Grpel2 | 5004 | 4798 | 4000 | 4678 | 4395 | 4656 | 5045 | 4940 | 4051 | 3810 | 3635 | 4096 | 3823 | 4242 | 4974 | 4590 | 4428 | 4514 | 4228 | 5312 | 4755 | 4341 | 4010 | 3818 | 4133 | 3881 | 3957 | 4082 | Grpel2 |
| Grpr | 12 | 22 | 0 | 15 | 34 | 25 | 27 | 43 | 40 | 14 | 55 | 5 | 3 | 14 | 9 | 8 | 9 | 52 | 22 | 9 | 28 | 25 | 60 | 11 | 29 | 1 | 13 | 8 | Grpr |
| Grsf1 | 9890 | 8665 | 10754 | 9573 | 9234 | 9004 | 10647 | 10186 | 8429 | 8544 | 8312 | 10671 | 9400 | 11310 | 10406 | 8592 | 9868 | 9196 | 10036 | 10002 | 9924 | 9327 | 8381 | 9787 | 9630 | 11667 | 10842 | 11954 | Grsf1 |
| Grtp1 | 1685 | 2032 | 1535 | 1925 | 1656 | 1811 | 1906 | 738 | 745 | 796 | 707 | 821 | 727 | 837 | 1582 | 1706 | 1783 | 1924 | 1918 | 1931 | 1853 | 605 | 780 | 817 | 827 | 916 | 791 | 812 | Grtp1 |
| Grwd1 | 1624 | 1985 | 1984 | 1698 | 1397 | 1547 | 1395 | 1475 | 1709 | 1602 | 1717 | 1899 | 1979 | 1829 | 1419 | 1673 | 1810 | 1574 | 1513 | 1447 | 1386 | 1402 | 1741 | 1882 | 1695 | 2543 | 2107 | 2115 | Grwd1 |
| Grxcr1 | 3 | 2 | 0 | 0 | 4 | 1 | 1 | 0 | 12 | 0 | 5 | 1 | 10 | 0 | 1 | 8 | 0 | 0 | 0 | 0 | 3 | 17 | 5 | 0 | 9 | 0 | 0 | 7 | Grxcr1 |
| Grxcr2 | 1 | 0 | 0 | 0 | 0 | 0 | 0 | 0 | 1 | 5 | 8 | 0 | 0 | 0 | 0 | 0 | 0 | 0 | 0 | 0 | 0 | 2 | 1 | 0 | 1 | 0 | 0 | 0 | Grxcr2 |
| Gsap | 212 | 147 | 226 | 220 | 413 | 325 | 292 | 669 | 681 | 629 | 616 | 432 | 388 | 430 | 262 | 219 | 247 | 307 | 342 | 332 | 343 | 844 | 776 | 472 | 677 | 341 | 341 | 325 | Gsap |
| Gsc | 77 | 15 | 19 | 21 | 11 | 8 | 4 | 107 | 98 | 103 | 116 | 8 | 25 | 12 | 22 | 51 | 16 | 12 | 3 | 5 | 4 | 45 | 71 | 154 | 15 | 180 | 2 | 17 | Gsc |
| Gsc2 | 8 | 5 | 14 | 2 | 6 | 19 | 3 | 9 | 9 | 13 | 6 | 1 | 17 | 6 | 7 | 3 | 9 | 7 | 12 | 2 | 5 | 28 | 10 | 13 | 16 | 6 | 8 | 9 | Gsc2 |
| Gsdma | 3 | 0 | 0 | 0 | 0 | 0 | 0 | 6 | 4 | 3 | 0 | 0 | 5 | 0 | 0 | 0 | 0 | 1 | 1 | 0 | 0 | 0 | 3 | 2 | 3 | 3 | 3 | 0 | Gsdma |
| Gsdma2 | 0 | 2 | 0 | 0 | 2 | 0 | 0 | 1 | 2 | 3 | 0 | 0 | 3 | 2 | 0 | 0 | 0 | 0 | 0 | 0 | 0 | 0 | 2 | 0 | 4 | 0 | 0 | 0 | Gsdma2 |
| Gsdma3 | 0 | 0 | 0 | 0 | 0 | 0 | 0 | 0 | 1 | 0 | 0 | 0 | 0 | 0 | 0 | 0 | 0 | 0 | 0 | 0 | 0 | 0 | 0 | 0 | 0 | 0 | 0 | 0 | Gsdma3 |
| Gsdmc | 2 | 0 | 0 | 0 | 0 | 0 | 0 | 0 | 14 | 4 | 4 | 0 | 2 | 1 | 0 | 2 | 0 | 0 | 0 | 0 | 0 | 0 | 15 | 0 | 0 | 1 | 0 | 0 | Gsdmc |
| Gsdmc2 | 0 | 0 | 0 | 0 | 0 | 0 | 0 | 0 | 0 | 1 | 0 | 0 | 0 | 0 | 0 | 0 | 0 | 0 | 0 | 0 | 0 | 0 | 0 | 0 | 0 | 0 | 1 | 0 | Gsdmc2 |
| Gsdmc3 | 0 | 0 | 0 | 0 | 0 | 0 | 0 | 0 | 0 | 0 | 0 | 0 | 0 | 0 | 0 | 0 | 0 | 0 | 0 | 0 | 0 | 2 | 0 | 0 | 0 | 0 | 0 | 0 | Gsdmc3 |
| Gsdmc4 | 9 | 10 | 4 | 16 | 18 | 25 | 8 | 2 | 9 | 8 | 9 | 1 | 6 | 3 | 5 | 9 | 10 | 20 | 16 | 2 | 5 | 5 | 9 | 7 | 4 | 0 | 5 | 2 | Gsdmc4 |
| Gsdmd | 120 | 105 | 130 | 116 | 134 | 163 | 106 | 335 | 388 | 342 | 478 | 325 | 335 | 260 | 157 | 112 | 111 | 140 | 113 | 147 | 108 | 264 | 458 | 351 | 306 | 336 | 294 | 254 | Gsdmd |
| Gsdme | 315 | 366 | 370 | 342 | 502 | 428 | 375 | 1009 | 882 | 956 | 826 | 831 | 863 | 777 | 393 | 292 | 393 | 341 | 471 | 469 | 410 | 1054 | 837 | 818 | 1041 | 706 | 872 | 863 | Gsdme |
| Gse1 | 4797 | 5887 | 6985 | 7283 | 7922 | 6599 | 5525 | 5873 | 7007 | 8903 | 7646 | 10836 | 7542 | 9007 | 4263 | 5234 | 8128 | 7199 | 8286 | 5344 | 6457 | 9474 | 6065 | 10066 | 9530 | 8376 | 8793 | 8731 | Gse1 |
| Gsg1 | 8 | 14 | 5 | 2 | 7 | 7 | 9 | 9 | 11 | 8 | 18 | 16 | 10 | 27 | 8 | 6 | 0 | 2 | 5 | 2 | 6 | 5 | 14 | 11 | 17 | 18 | 6 | 8 | Gsg1 |
| Gsg1l | 424 | 696 | 343 | 808 | 1186 | 1035 | 654 | 1248 | 1308 | 1280 | 1030 | 838 | 1327 | 694 | 720 | 795 | 740 | 874 | 987 | 1128 | 902 | 1516 | 1361 | 900 | 1516 | 354 | 780 | 727 | Gsg1l |
| Gsg1l2 | 0 | 0 | 4 | 3 | 2 | 4 | 2 | 0 | 4 | 0 | 1 | 0 | 0 | 2 | 4 | 0 | 0 | 0 | 0 | 0 | 1 | 0 | 0 | 2 | 4 | 2 | 0 | 1 | Gsg1l2 |
| Gsk3a | 7521 | 9388 | 8788 | 9505 | 9340 | 8231 | 8600 | 6991 | 8017 | 8686 | 8126 | 9074 | 7735 | 8760 | 7251 | 7888 | 9887 | 9239 | 9870 | 7502 | 9245 | 8306 | 7606 | 9200 | 8837 | 9533 | 8042 | 8216 | Gsk3a |
| Gsk3b | 18972 | 16936 | 18534 | 24644 | 26061 | 25301 | 27291 | 21068 | 19228 | 23331 | 20973 | 26726 | 24343 | 26786 | 20274 | 17365 | 21774 | 24689 | 24919 | 21509 | 27827 | 26212 | 17785 | 23701 | 27031 | 18924 | 25794 | 26464 | Gsk3b |
| Gskip | 1885 | 1798 | 1744 | 1633 | 1336 | 1458 | 1498 | 1793 | 1423 | 1239 | 1318 | 1596 | 1645 | 1517 | 1879 | 1823 | 1567 | 1581 | 1401 | 1646 | 1509 | 1554 | 1541 | 1349 | 1436 | 1511 | 1763 | 1664 | Gskip |
| Gsn | 4354 | 4633 | 3217 | 2246 | 1730 | 2831 | 912 | 2364 | 3769 | 2627 | 4377 | 2025 | 3306 | 1917 | 3984 | 5956 | 2666 | 3043 | 1479 | 1246 | 959 | 2433 | 4864 | 2406 | 1994 | 2519 | 2327 | 2140 | Gsn |
| Gspt1 | 21299 | 16890 | 16661 | 15565 | 15866 | 18629 | 19600 | 15699 | 13958 | 13263 | 12987 | 16035 | 15278 | 16670 | 20168 | 16785 | 16708 | 15737 | 16959 | 17922 | 18834 | 13147 | 13599 | 13870 | 14222 | 13912 | 16668 | 16736 | Gspt1 |
| Gspt2 | 1908 | 1642 | 2020 | 2277 | 2356 | 2096 | 2122 | 1508 | 1281 | 1522 | 1459 | 1511 | 1506 | 1627 | 1934 | 1686 | 2060 | 2042 | 2269 | 1867 | 2211 | 1786 | 1309 | 1606 | 1542 | 1256 | 1370 | 1519 | Gspt2 |
| Gsr | 1325 | 1170 | 1703 | 1314 | 1295 | 1186 | 1259 | 1787 | 1957 | 1784 | 1963 | 2321 | 2195 | 2320 | 1315 | 1343 | 1451 | 1295 | 1220 | 1312 | 1231 | 1501 | 1891 | 2226 | 1929 | 2974 | 2138 | 2325 | Gsr |
| Gss | 773 | 763 | 666 | 619 | 641 | 672 | 773 | 811 | 954 | 737 | 774 | 790 | 807 | 799 | 690 | 688 | 607 | 588 | 773 | 722 | 749 | 618 | 922 | 890 | 691 | 1013 | 755 | 767 | Gss |
| Gsta1 | 0 | 0 | 0 | 0 | 0 | 0 | 0 | 0 | 0 | 0 | 0 | 0 | 0 | 1 | 3 | 0 | 0 | 0 | 0 | 0 | 0 | 0 | 0 | 0 | 0 | 0 | 0 | 0 | Gsta1 |
| Gsta2 | 0 | 0 | 0 | 0 | 0 | 0 | 1 | 2 | 1 | 4 | 0 | 0 | 0 | 1 | 0 | 0 | 0 | 2 | 0 | 0 | 1 | 2 | 3 | 0 | 0 | 0 | 0 | 1 | Gsta2 |
| Gsta3 | 28 | 10 | 24 | 13 | 25 | 42 | 28 | 49 | 73 | 41 | 34 | 36 | 43 | 28 | 22 | 21 | 16 | 24 | 16 | 17 | 30 | 32 | 56 | 42 | 38 | 48 | 24 | 27 | Gsta3 |
| Gsta4 | 4181 | 3660 | 3518 | 2902 | 1862 | 2170 | 2363 | 1325 | 1275 | 1228 | 1478 | 982 | 1344 | 948 | 3444 | 3713 | 3216 | 2458 | 2180 | 3027 | 2001 | 1019 | 1464 | 1320 | 876 | 2133 | 1110 | 1205 | Gsta4 |
| Gsta5 | 0 | 0 | 0 | 0 | 0 | 0 | 0 | 0 | 2 | 0 | 0 | 0 | 0 | 0 | 0 | 0 | 0 | 0 | 0 | 0 | 0 | 0 | 1 | 0 | 0 | 0 | 0 | 0 | Gsta5 |
| Gstcd | 2782 | 2433 | 2813 | 2311 | 2124 | 2226 | 2788 | 2806 | 2478 | 2321 | 2460 | 3321 | 3017 | 3794 | 2640 | 2262 | 2467 | 2170 | 2406 | 2419 | 2585 | 2199 | 2397 | 2701 | 2830 | 3491 | 3359 | 3883 | Gstcd |
| Gstk1 | 670 | 738 | 567 | 649 | 493 | 535 | 428 | 403 | 426 | 387 | 385 | 399 | 464 | 341 | 666 | 791 | 672 | 512 | 483 | 593 | 441 | 287 | 399 | 380 | 342 | 429 | 331 | 401 | Gstk1 |
| Gstm1 | 1335 | 1205 | 1143 | 1280 | 1133 | 1134 | 1002 | 1674 | 1814 | 1940 | 1671 | 684 | 695 | 653 | 1023 | 1195 | 1159 | 1017 | 1025 | 1062 | 946 | 846 | 2089 | 1957 | 772 | 1214 | 751 | 681 | Gstm1 |
| Gstm2 | 469 | 411 | 340 | 219 | 156 | 206 | 83 | 410 | 540 | 410 | 558 | 332 | 485 | 327 | 456 | 566 | 279 | 261 | 109 | 172 | 118 | 341 | 680 | 388 | 233 | 437 | 392 | 351 | Gstm2 |
| Gstm3 | 2 | 0 | 0 | 0 | 2 | 0 | 0 | 0 | 0 | 3 | 0 | 0 | 0 | 0 | 0 | 0 | 0 | 0 | 1 | 0 | 0 | 0 | 1 | 3 | 0 | 0 | 0 | 0 | Gstm3 |
| Gstm4 | 512 | 441 | 460 | 441 | 456 | 419 | 392 | 504 | 493 | 448 | 489 | 467 | 452 | 475 | 473 | 499 | 458 | 461 | 368 | 406 | 413 | 470 | 562 | 422 | 458 | 555 | 439 | 450 | Gstm4 |

| | | | | | | | | | | | | | | | | | | | | | | | | | | | | | |
|---|---|---|---|---|---|---|---|---|---|---|---|---|---|---|---|---|---|---|---|---|---|---|---|---|---|---|---|---|---|
| Gstm5 | 4218 | 4313 | 3965 | 3718 | 2939 | 3483 | 3329 | 5230 | 4739 | 4138 | 4188 | 4027 | 4197 | 4517 | 4516 | 3704 | 3567 | 3178 | 3370 | 4060 | 3442 | 3873 | 4785 | 4265 | 3803 | 4998 | 4456 | 4841 | Gstm5 |
| Gstm6 | 368 | 428 | 468 | 296 | 329 | 264 | 370 | 376 | 303 | 348 | 268 | 344 | 335 | 400 | 340 | 403 | 335 | 330 | 320 | 363 | 312 | 438 | 305 | 333 | 337 | 472 | 395 | 470 | Gstm6 |
| Gstm7 | 1699 | 2122 | 1923 | 2381 | 2418 | 2246 | 2471 | 2330 | 2093 | 2654 | 2344 | 3028 | 2539 | 3270 | 1949 | 1500 | 2515 | 2079 | 2581 | 2108 | 2632 | 2617 | 1964 | 2839 | 2925 | 2867 | 2923 | 3052 | Gstm7 |
| Gsto1 | 2692 | 2655 | 2424 | 2335 | 2266 | 2244 | 2333 | 1362 | 1625 | 1679 | 1603 | 1385 | 1566 | 1340 | 2393 | 2797 | 2366 | 2202 | 2239 | 2369 | 2303 | 1469 | 1756 | 1497 | 1441 | 1607 | 1343 | 1518 | Gsto1 |
| Gsto2 | 31 | 39 | 24 | 20 | 8 | 36 | 20 | 42 | 55 | 30 | 52 | 42 | 57 | 60 | 24 | 35 | 22 | 34 | 34 | 18 | 27 | 42 | 57 | 29 | 45 | 57 | 65 | 42 | Gsto2 |
| Gstp1 | 6 | 10 | 5 | 8 | 10 | 6 | 4 | 9 | 7 | 10 | 7 | 7 | 5 | 4 | 5 | 9 | 10 | 2 | 4 | 7 | 8 | 3 | 52 | 6 | 3 | 11 | 13 | 6 | Gstp1 |
| Gstp2 | 0 | 0 | 0 | 0 | 0 | 0 | 0 | 0 | 0 | 0 | 0 | 0 | 0 | 0 | 0 | 0 | 0 | 0 | 0 | 0 | 0 | 0 | 3 | 0 | 0 | 0 | 1 | 0 | Gstp2 |
| Gstt1 | 426 | 546 | 460 | 393 | 337 | 393 | 329 | 219 | 324 | 252 | 257 | 164 | 205 | 182 | 393 | 471 | 457 | 417 | 315 | 371 | 286 | 251 | 399 | 225 | 206 | 168 | 199 | 216 | Gstt1 |
| Gstt2 | 397 | 432 | 494 | 366 | 326 | 306 | 297 | 159 | 185 | 183 | 250 | 182 | 249 | 186 | 407 | 474 | 377 | 335 | 311 | 258 | 292 | 180 | 254 | 175 | 185 | 254 | 279 | 183 | Gstt2 |
| Gstt3 | 361 | 259 | 208 | 224 | 104 | 131 | 83 | 61 | 79 | 67 | 145 | 29 | 57 | 37 | 264 | 380 | 173 | 204 | 80 | 104 | 89 | 70 | 182 | 60 | 54 | 89 | 17 | 21 | Gstt3 |
| Gstt4 | 2 | 2 | 7 | 3 | 2 | 4 | 2 | 0 | 2 | 1 | 4 | 1 | 1 | 1 | 5 | 2 | 7 | 0 | 2 | 4 | 3 | 2 | 0 | 3 | 0 | 0 | 0 | 0 | Gstt4 |
| Gstz1 | 1887 | 1887 | 1574 | 1627 | 1856 | 1977 | 1944 | 2728 | 2726 | 2386 | 2691 | 2521 | 2525 | 2677 | 1896 | 1751 | 1576 | 1647 | 1824 | 2072 | 1924 | 2366 | 2872 | 2261 | 2446 | 2361 | 2739 | 2606 | Gstz1 |
| Gsx1 | 84 | 131 | 205 | 136 | 204 | 147 | 146 | 232 | 198 | 298 | 236 | 405 | 306 | 435 | 108 | 53 | 249 | 90 | 243 | 117 | 162 | 354 | 183 | 361 | 299 | 377 | 423 | 503 | Gsx1 |
| Gsx2 | 191 | 283 | 197 | 264 | 231 | 202 | 317 | 281 | 218 | 290 | 185 | 492 | 60 | 479 | 241 | 72 | 403 | 240 | 378 | 243 | 298 | 229 | 93 | 373 | 243 | 639 | 259 | 386 | Gsx2 |
| Gtdc1 | 1731 | 1730 | 1468 | 1848 | 1848 | 1966 | 2143 | 2426 | 2209 | 2028 | 2188 | 2160 | 1810 | 2279 | 1996 | 1611 | 1534 | 1918 | 1857 | 2162 | 2176 | 2109 | 2379 | 1966 | 2377 | 2269 | 2017 | 2124 | Gtdc1 |
| Gtf2a1 | 3148 | 2293 | 4820 | 4070 | 4178 | 3719 | 4694 | 4866 | 4374 | 5854 | 5354 | 5928 | 5729 | 6182 | 3310 | 2541 | 4464 | 3962 | 4379 | 3663 | 3967 | 6112 | 4314 | 6258 | 6177 | 7557 | 6070 | 6575 | Gtf2a1 |
| Gtf2a1l | 12 | 11 | 5 | 7 | 2 | 3 | 8 | 7 | 6 | 3 | 7 | 9 | 10 | 4 | 7 | 13 | 9 | 1 | 5 | 8 | 10 | 2 | 5 | 5 | 5 | 1 | 13 | 0 | Gtf2a1l |
| Gtf2a2 | 6165 | 5055 | 5625 | 6092 | 5578 | 6028 | 7002 | 6591 | 5582 | 6283 | 6004 | 6857 | 6555 | 7696 | 6030 | 4979 | 5660 | 5653 | 5947 | 6651 | 6972 | 5770 | 5289 | 6511 | 6524 | 6823 | 7055 | 7340 | Gtf2a2 |
| Gtf2b | 4171 | 4326 | 3804 | 3824 | 3341 | 3411 | 4061 | 3694 | 3271 | 2939 | 2694 | 3148 | 3164 | 3673 | 4676 | 4338 | 3737 | 3524 | 3451 | 4551 | 3639 | 3001 | 3197 | 3257 | 3005 | 3729 | 3435 | 3573 | Gtf2b |
| Gtf2e1 | 5568 | 5296 | 5650 | 4895 | 4004 | 4361 | 4708 | 4759 | 4472 | 3909 | 3878 | 4625 | 4377 | 4895 | 5892 | 5004 | 5355 | 4774 | 4315 | 5057 | 4347 | 3999 | 4272 | 4410 | 4434 | 5549 | 4873 | 5256 | Gtf2e1 |
| Gtf2e2 | 6750 | 5907 | 6615 | 6318 | 5277 | 5300 | 6060 | 4361 | 3623 | 3492 | 3639 | 3844 | 3923 | 3962 | 7267 | 6548 | 6610 | 6168 | 5644 | 6671 | 5588 | 3609 | 3772 | 3932 | 3776 | 4735 | 4071 | 4215 | Gtf2e2 |
| Gtf2f1 | 5072 | 6404 | 6284 | 5755 | 5197 | 5193 | 5536 | 4739 | 5046 | 5089 | 4731 | 5305 | 4936 | 5893 | 5238 | 5095 | 6049 | 5162 | 5908 | 6053 | 5230 | 4114 | 4685 | 5485 | 5648 | 6106 | 5796 | 5807 | Gtf2f1 |
| Gtf2f2 | 2991 | 2910 | 2713 | 2776 | 2737 | 2740 | 3338 | 1906 | 1817 | 1821 | 1525 | 2219 | 1805 | 2177 | 3191 | 2929 | 2754 | 2812 | 2819 | 3674 | 3188 | 1440 | 1843 | 1894 | 1926 | 2274 | 2059 | 2398 | Gtf2f2 |
| Gtf2h1 | 5952 | 5253 | 5368 | 4536 | 3987 | 4473 | 4985 | 4421 | 3910 | 3669 | 3789 | 4127 | 3788 | 4221 | 5913 | 5615 | 4738 | 4424 | 4356 | 5177 | 4592 | 3639 | 3924 | 3901 | 3776 | 4304 | 3893 | 4463 | Gtf2h1 |
| Gtf2h2 | 2536 | 2066 | 2924 | 2349 | 2420 | 2418 | 2882 | 3276 | 2942 | 3066 | 3100 | 3401 | 3432 | 3391 | 2854 | 2576 | 2175 | 2343 | 2289 | 2736 | 2619 | 3018 | 3022 | 2992 | 3111 | 3513 | 3578 | 3705 | Gtf2h2 |
| Gtf2h3 | 3887 | 4093 | 4234 | 3612 | 2700 | 2757 | 3125 | 3054 | 3074 | 3001 | 3021 | 3486 | 3210 | 3432 | 3517 | 3539 | 3586 | 3286 | 3089 | 3160 | 2958 | 2542 | 2988 | 3160 | 2914 | 3914 | 3529 | 3681 | Gtf2h3 |
| Gtf2h4 | 1076 | 1238 | 1249 | 1103 | 947 | 945 | 917 | 1092 | 1228 | 1120 | 1091 | 1297 | 1219 | 1187 | 952 | 1143 | 1286 | 1009 | 1117 | 1062 | 893 | 924 | 1185 | 1315 | 1173 | 1517 | 1407 | 1385 | Gtf2h4 |
| Gtf2h5 | 5647 | 4895 | 5796 | 5732 | 5560 | 5189 | 5996 | 5882 | 5085 | 5266 | 5147 | 5183 | 5048 | 5601 | 5846 | 4611 | 5591 | 5493 | 5598 | 6040 | 5630 | 5095 | 5024 | 5581 | 5682 | 5878 | 5398 | 5772 | Gtf2h5 |
| Gtf2i | 12704 | 12471 | 17091 | 14435 | 14882 | 14265 | 12942 | 26796 | 29042 | 35421 | 33256 | 39753 | 33886 | 34686 | 12092 | 12329 | 16236 | 13370 | 16315 | 11283 | 14161 | 32871 | 29170 | 37034 | 35002 | 40386 | 32418 | 36519 | Gtf2i |
| Gtf2ird1 | 3246 | 3131 | 3955 | 3247 | 3578 | 3443 | 3057 | 4461 | 5085 | 5177 | 5348 | 5865 | 5141 | 5386 | 2867 | 2904 | 3582 | 3115 | 3754 | 2962 | 3253 | 5203 | 4871 | 5453 | 5176 | 5652 | 5571 | 5515 | Gtf2ird1 |
| Gtf2ird2 | 520 | 650 | 696 | 636 | 895 | 786 | 726 | 1544 | 1675 | 1933 | 1716 | 2026 | 1691 | 1793 | 492 | 539 | 809 | 684 | 868 | 739 | 760 | 2010 | 1628 | 1961 | 1964 | 1693 | 1791 | 1850 | Gtf2ird2 |
| Gtf3a | 1806 | 2065 | 2067 | 1812 | 1793 | 1729 | 1749 | 1778 | 2089 | 2308 | 2077 | 2338 | 2097 | 2109 | 1636 | 1822 | 1853 | 1708 | 1888 | 1577 | 1782 | 1935 | 2073 | 2245 | 2025 | 2675 | 2069 | 2050 | Gtf3a |
| Gtf3c1 | 11386 | 12289 | 11758 | 14216 | 14496 | 13986 | 12514 | 12395 | 14538 | 15740 | 14035 | 17567 | 15136 | 15044 | 10490 | 11576 | 13332 | 14103 | 15195 | 11200 | 13817 | 14715 | 13384 | 15374 | 16052 | 11605 | 14647 | 15196 | Gtf3c1 |
| Gtf3c2 | 4557 | 3858 | 4722 | 4582 | 3844 | 4241 | 4875 | 4181 | 3859 | 4843 | 4356 | 5168 | 4983 | 5677 | 4191 | 3787 | 4753 | 4318 | 4336 | 3916 | 4403 | 4274 | 3585 | 5132 | 4900 | 5515 | 4957 | 5756 | Gtf3c2 |
| Gtf3c3 | 2803 | 2298 | 2503 | 2218 | 2408 | 2532 | 2514 | 5486 | 4674 | 4204 | 4174 | 4608 | 4782 | 4937 | 2835 | 2277 | 2391 | 2070 | 2314 | 2950 | 2557 | 4451 | 4553 | 4288 | 4263 | 4503 | 5103 | 5041 | Gtf3c3 |
| Gtf3c4 | 16122 | 15897 | 11976 | 14170 | 13394 | 15033 | 14076 | 12167 | 10825 | 10499 | 10232 | 11308 | 10895 | 10852 | 15289 | 14035 | 13316 | 14705 | 14023 | 13935 | 15508 | 10733 | 10483 | 9775 | 11518 | 7972 | 11803 | 11096 | Gtf3c4 |
| Gtf3c5 | 1854 | 2019 | 2266 | 2060 | 2117 | 2072 | 1943 | 3309 | 3959 | 4008 | 3959 | 4923 | 4613 | 4591 | 1795 | 1921 | 2296 | 1920 | 2388 | 1940 | 1892 | 3962 | 3698 | 4390 | 4234 | 4911 | 4700 | 4787 | Gtf3c5 |
| Gtf3c6 | 2052 | 1811 | 2328 | 1688 | 1956 | 2028 | 2056 | 6074 | 4833 | 4807 | 4489 | 5979 | 5767 | 6562 | 2334 | 1704 | 2042 | 1484 | 1919 | 2224 | 1885 | 5000 | 4735 | 5704 | 5174 | 6688 | 6078 | 6903 | Gtf3c6 |
| Gtpbp1 | 7223 | 8638 | 7088 | 8054 | 7327 | 7844 | 7255 | 5453 | 5875 | 5912 | 5197 | 6424 | 5452 | 5919 | 6800 | 6655 | 7280 | 8084 | 8464 | 7477 | 7800 | 5556 | 5428 | 6306 | 6344 | 6801 | 6380 | 6184 | Gtpbp1 |
| Gtpbp10 | 2279 | 1797 | 2031 | 1679 | 1794 | 1807 | 2153 | 2745 | 2522 | 2011 | 2070 | 2755 | 2506 | 2710 | 2217 | 2011 | 1742 | 1784 | 1783 | 2212 | 1951 | 2117 | 2445 | 2259 | 2242 | 2864 | 2966 | 2903 | Gtpbp10 |
| Gtpbp2 | 5282 | 6382 | 7032 | 8140 | 7675 | 6669 | 6764 | 4860 | 5608 | 6732 | 6343 | 6427 | 6208 | 6296 | 5105 | 6235 | 8129 | 7897 | 7826 | 6057 | 6995 | 6801 | 5225 | 6863 | 6515 | 6025 | 6093 | 6365 | Gtpbp2 |
| Gtpbp3 | 3002 | 3803 | 3632 | 3670 | 2441 | 2677 | 2624 | 1239 | 1407 | 1384 | 1580 | 1587 | 1592 | 1484 | 2645 | 3553 | 3427 | 3334 | 2662 | 2657 | 2457 | 1292 | 1447 | 1639 | 1378 | 1871 | 1712 | 1467 | Gtpbp3 |
| Gtpbp4 | 7623 | 6847 | 7807 | 6568 | 6005 | 6570 | 6955 | 8089 | 7055 | 6593 | 6609 | 8193 | 7496 | 8439 | 8181 | 7155 | 7071 | 6069 | 6512 | 7809 | 6352 | 6381 | 6985 | 7343 | 7146 | 8957 | 8923 | 8808 | Gtpbp4 |
| Gtpbp6 | 1119 | 1727 | 1703 | 1781 | 1717 | 1437 | 1376 | 1364 | 1496 | 1590 | 1544 | 1560 | 1435 | 1406 | 1158 | 1235 | 1855 | 1678 | 1813 | 1532 | 1529 | 1557 | 1425 | 1789 | 1682 | 1707 | 1576 | 1349 | Gtpbp6 |
| Gtpbp8 | 1889 | 1503 | 1619 | 1536 | 1771 | 1893 | 2105 | 2699 | 2383 | 2415 | 2307 | 2472 | 2364 | 2489 | 2103 | 1560 | 1643 | 1448 | 1745 | 1978 | 1809 | 2704 | 2305 | 2080 | 2612 | 2137 | 2404 | 2529 | Gtpbp8 |
| Gtse1 | 1801 | 2158 | 2673 | 1692 | 1363 | 1624 | 1516 | 1497 | 1752 | 1599 | 1482 | 2835 | 2342 | 2647 | 1456 | 1879 | 2153 | 1550 | 1785 | 1601 | 1406 | 1156 | 1447 | 2202 | 1855 | 3504 | 2664 | 2741 | Gtse1 |
| Gtsf1 | 0 | 0 | 0 | 0 | 0 | 0 | 0 | 0 | 2 | 0 | 1 | 0 | 1 | 0 | 0 | 0 | 0 | 0 | 0 | 0 | 0 | 0 | 0 | 0 | 0 | 1 | 0 | 0 | Gtsf1 |
| Gtsf1l | 0 | 1 | 0 | 0 | 0 | 0 | 0 | 0 | 2 | 3 | 0 | 1 | 3 | 0 | 1 | 2 | 0 | 0 | 2 | 0 | 0 | 5 | 5 | 0 | 0 | 1 | 1 | 0 | Gtsf1l |
| Gtsf2 | 0 | 0 | 1 | 0 | 3 | 1 | 0 | 2 | 2 | 3 | 1 | 2 | 0 | 0 | 5 | 3 | 2 | 0 | 0 | 1 | 3 | 0 | 1 | 2 | 4 | 8 | 0 | 4 | Gtsf2 |
| Guca2a | 3 | 1 | 1 | 5 | 3 | 1 | 1 | 1 | 0 | 4 | 0 | 3 | 1 | 4 | 4 | 5 | 1 | 2 | 5 | 1 | 4 | 0 | 3 | 2 | 5 | 10 | 6 | 2 | Guca2a |
| Guca2b | 0 | 0 | 0 | 0 | 0 | 2 | 2 | 0 | 16 | 0 | 0 | 0 | 0 | 0 | 0 | 1 | 0 | 2 | 0 | 0 | 0 | 0 | 3 | 0 | 0 | 0 | 0 | 0 | Guca2b |
| Gucd1 | 2993 | 3252 | 3141 | 2707 | 2639 | 2888 | 2542 | 2302 | 2637 | 2374 | 2505 | 2784 | 2869 | 2451 | 2716 | 3035 | 2826 | 2580 | 2696 | 2635 | 2490 | 2080 | 2561 | 2465 | 2339 | 2955 | 2988 | 2694 | Gucd1 |
| Gucy1a1 | 1159 | 957 | 942 | 1300 | 2091 | 1830 | 1849 | 1779 | 1875 | 1906 | 1815 | 1263 | 1086 | 1203 | 1251 | 1097 | 1074 | 1588 | 1583 | 1648 | 1784 | 2174 | 1737 | 1522 | 2022 | 1052 | 926 | 920 | Gucy1a1 |
| Gucy1a2 | 448 | 410 | 106 | 168 | 147 | 436 | 315 | 1016 | 861 | 311 | 388 | 405 | 388 | 457 | 431 | 323 | 104 | 117 | 175 | 588 | 461 | 302 | 749 | 218 | 375 | 242 | 629 | 348 | Gucy1a2 |
| Gucy1b1 | 3201 | 2959 | 2747 | 3622 | 4379 | 3851 | 4474 | 2058 | 1902 | 1694 | 1614 | 1322 | 1261 | 1269 | 3589 | 3073 | 3006 | 3857 | 3861 | 4715 | 4255 | 2024 | 1889 | 1508 | 1958 | 1191 | 1159 | 1094 | Gucy1b1 |
| Gucy1b2 | 76 | 62 | 60 | 52 | 88 | 119 | 116 | 253 | 244 | 224 | 185 | 318 | 122 | 287 | 46 | 23 | 60 | 69 | 102 | 94 | 130 | 159 | 140 | 221 | 278 | 210 | 212 | 172 | Gucy1b2 |
| Gucy2c | 29 | 18 | 34 | 41 | 38 | 21 | 58 | 53 | 42 | 43 | 29 | 27 | 25 | 33 | 21 | 18 | 16 | 29 | 39 | 55 | 45 | 20 | 37 | 52 | 33 | 30 | 35 | 27 | Gucy2c |
| Gucy2d | 0 | 3 | 0 | 0 | 3 | 1 | 0 | 5 | 2 | 5 | 9 | 0 | 0 | 0 | 1 | 0 | 0 | 0 | 0 | 0 | 0 | 2 | 5 | 2 | 0 | 2 | 0 | 0 | Gucy2d |
| Gucy2e | 22 | 42 | 49 | 28 | 35 | 48 | 23 | 28 | 52 | 52 | 71 | 27 | 41 | 19 | 37 | 19 | 24 | 26 | 34 | 54 | 38 | 33 | 57 | 50 | 36 | 31 | 29 | 26 | Gucy2e |
| Gucy2f | 9 | 9 | 2 | 12 | 19 | 33 | 13 | 46 | 72 | 49 | 81 | 11 | 48 | 18 | 29 | 20 | 2 | 9 | 14 | 29 | 14 | 53 | 61 | 16 | 37 | 4 | 29 | 33 | Gucy2f |
| Gucy2g | 0 | 0 | 0 | 0 | 8 | 0 | 0 | 5 | 4 | 9 | 2 | 0 | 8 | 2 | 1 | 0 | 0 | 1 | 2 | 1 | 1 | 3 | 7 | 2 | 3 | 0 | 1 | 0 | Gucy2g |
| Guf1 | 7153 | 5301 | 5463 | 6023 | 6209 | 6898 | 7109 | 9476 | 8282 | 8233 | 8982 | 8517 | 8750 | 9043 | 7395 | 6534 | 5590 | 6140 | 5599 | 7126 | 6654 | 8970 | 8317 | 8077 | 7963 | 7360 | 8268 | 8160 | Guf1 |
| Guk1 | 1684 | 2031 | 1633 | 1896 | 1834 | 1661 | 1894 | 1722 | 1786 | 1494 | 1611 | 1430 | 1506 | 1473 | 1706 | 1636 | 1761 | 1853 | 1921 | 2081 | 1985 | 1661 | 1769 | 1601 | 1612 | 1610 | 1467 | 1539 | Guk1 |
| Gulo | 21 | 5 | 5 | 9 | 12 | 10 | 6 | 35 | 14 | 53 | 21 | 1 | 10 | 3 | 4 | 7 | 2 | 9 | 3 | 4 | 4 | 17 | 30 | 32 | 0 | 25 | 3 | 9 | Gulo |
| Gulp1 | 2598 | 1618 | 2111 | 1407 | 1106 | 1581 | 1155 | 2542 | 2512 | 1800 | 2281 | 2292 | 2900 | 2407 | 2349 | 2569 | 1650 | 1346 | 1071 | 1421 | 1084 | 2237 | 2873 | 2067 | 1666 | 2707 | 2637 | 2601 | Gulp1 |
| Gusb | 1455 | 1499 | 1714 | 1120 | 1324 | 1161 | 891 | 1483 | 1828 | 1886 | 2081 | 1706 | 2147 | 1517 | 1469 | 1632 | 1258 | 1250 | 1124 | 1111 | 995 | 1744 | 2110 | 1641 | 1559 | 1971 | 1915 | 1730 | Gusb |
| Gvin1 | 10 | 4 | 1 | 0 | 7 | 0 | 0 | 5 | 11 | 3 | 14 | 5 | 8 | 4 | 0 | 4 | 0 | 9 | 7 | 10 | 5 | 7 | 8 | 4 | 2 | 3 | 8 | 6 | Gvin1 |
| Gvin2 | 5 | 2 | 0 | 4 | 6 | 3 | 6 | 2 | 7 | 4 | 10 | 1 | 14 | 2 | 3 | 7 | 1 | 8 | 2 | 6 | 3 | 12 | 16 | 7 | 4 | 1 | 8 | 8 | Gvin2 |
| Gxylt1 | 10217 | 8485 | 9125 | 9137 | 7117 | 8798 | 8490 | 5183 | 5227 | 5258 | 6198 | 6397 | 6440 | 6651 | 9126 | 12200 | 8413 | 9193 | 7137 | 7670 | 8047 | 4742 | 5066 | 5808 | 5705 | 5720 | 6482 | 6424 | Gxylt1 |
| Gxylt2 | 2496 | 1913 | 1709 | 1360 | 861 | 1782 | 546 | 1984 | 2857 | 2665 | 3904 | 1265 | 2243 | 1302 | 2093 | 2869 | 1380 | 1916 | 662 | 599 | 527 | 2196 | 4072 | 2061 | 1674 | 1240 | 1446 | 1336 | Gxylt2 |
| Gyg | 2000 | 1687 | 1597 | 1241 | 1188 | 1143 | 1241 | 1605 | 1596 | 1392 | 1510 | 1344 | 1511 | 1479 | 1965 | 2005 | 1508 | 1335 | 1016 | 1091 | 1106 | 1470 | 1753 | 1479 | 1327 | 1826 | 1507 | 1585 | Gyg |
| Gykl1 | 2 | 2 | 0 | 0 | 2 | 0 | 0 | 0 | 5 | 3 | 2 | 4 | 0 | 1 | 0 | 5 | 0 | 0 | 0 | 2 | 0 | 0 | 1 | 1 | 0 | 0 | 1 | 1 | Gykl1 |
| Gypa | 173 | 186 | 262 | 103 | 72 | 72 | 169 | 188 | 316 | 164 | 138 | 167 | 380 | 271 | 189 | 218 | 107 | 103 | 117 | 202 | 125 | 130 | 282 | 107 | 123 | 212 | 195 | 777 | Gypa |
| Gypc | 788 | 707 | 847 | 444 | 432 | 577 | 358 | 1108 | 1328 | 1135 | 1441 | 1194 | 1535 | 1218 | 766 | 931 | 550 | 486 | 423 | 442 | 365 | 1073 | 1434 | 1097 | 889 | 1467 | 1336 | 1453 | Gypc |
| Gys1 | 3787 | 4296 | 4136 | 3600 | 3364 | 3624 | 2809 | 2562 | 3183 | 3361 | 3348 | 3495 | 3698 | 2886 | 3638 | 4498 | 4117 | 3494 | 3688 | 2842 | 3076 | 2841 | 3346 | 3365 | 2815 | 3686 | 3471 | 3419 | Gys1 |
| Gys2 | 3 | 10 | 2 | 7 | 3 | 0 | 1 | 2 | 7 | 12 | 1 | 0 | 2 | 2 | 3 | 7 | 2 | 9 | 6 | 3 | 4 | 10 | 1 | 3 | 4 | 0 | 2 | 2 | Gys2 |
| Gzf1 | 6529 | 6714 | 5600 | 5726 | 4557 | 5030 | 5523 | 4162 | 3819 | 3174 | 3240 | 3654 | 3646 | 3715 | 6306 | 6332 | 5311 | 5565 | 4930 | 5620 | 5445 | 3243 | 3610 | 3504 | 3539 | 3532 | 4097 | 3863 | Gzf1 |
| Gzma | 0 | 1 | 0 | 0 | 0 | 0 | 0 | 0 | 0 | 0 | 0 | 3 | 0 | 0 | 0 | 0 | 2 | 0 | 0 | 0 | 0 | 0 | 1 | 0 | 0 | 0 | 0 | 0 | Gzma |
| Gzmb | 1 | 0 | 0 | 0 | 0 | 0 | 0 | 5 | 10 | 4 | 0 | 1 | 2 | 7 | 0 | 0 | 0 | 1 | 0 | 0 | 0 | 0 | 8 | 5 | 3 | 2 | 0 | 0 | Gzmb |

| | | | | | | | | | | | | | | | | | | | | | | | | | | | | | |
|---|---|---|---|---|---|---|---|---|---|---|---|---|---|---|---|---|---|---|---|---|---|---|---|---|---|---|---|---|---|
| Gzmc | 5 | 0 | 0 | 0 | 0 | 0 | 0 | 2 | 4 | 0 | 3 | 1 | 5 | 2 | 0 | 0 | 0 | 0 | 0 | 0 | 0 | 0 | 5 | 2 | 0 | 0 | 0 | 0 | Gzmc |
| Gzmd | 0 | 1 | 0 | 0 | 0 | 0 | 0 | 0 | 6 | 0 | 0 | 0 | 0 | 0 | 0 | 0 | 0 | 0 | 0 | 0 | 0 | 2 | 2 | 0 | 0 | 0 | 0 | 0 | Gzmd |
| Gzme | 0 | 0 | 0 | 0 | 0 | 0 | 0 | 0 | 2 | 0 | 0 | 0 | 0 | 0 | 0 | 0 | 0 | 0 | 0 | 0 | 0 | 3 | 1 | 0 | 0 | 0 | 0 | 0 | Gzme |
| Gzmf | 0 | 1 | 0 | 0 | 0 | 0 | 0 | 0 | 10 | 0 | 0 | 0 | 0 | 0 | 0 | 0 | 0 | 0 | 0 | 0 | 0 | 0 | 6 | 0 | 0 | 0 | 0 | 0 | Gzmf |
| Gzmg | 0 | 0 | 0 | 0 | 0 | 0 | 0 | 0 | 6 | 0 | 0 | 0 | 0 | 0 | 1 | 0 | 0 | 0 | 0 | 0 | 0 | 0 | 2 | 0 | 0 | 0 | 0 | 0 | Gzmg |
| Gzmk | 2 | 3 | 0 | 2 | 1 | 0 | 1 | 2 | 2 | 3 | 2 | 0 | 0 | 1 | 5 | 0 | 3 | 0 | 7 | 8 | 0 | 0 | 2 | 1 | 8 | 4 | 6 | 0 | Gzmk |
| Gzmm | 159 | 206 | 202 | 231 | 245 | 211 | 238 | 87 | 103 | 106 | 121 | 124 | 79 | 98 | 162 | 172 | 227 | 238 | 249 | 199 | 249 | 69 | 112 | 103 | 142 | 119 | 95 | 104 | Gzmm |
| H13 | 7742 | 8706 | 7289 | 8436 | 8997 | 9141 | 8419 | 10110 | 9650 | 10569 | 10078 | 9637 | 9372 | 9636 | 8057 | 7742 | 8112 | 8292 | 9012 | 8405 | 9061 | 11045 | 9306 | 9780 | 11176 | 8771 | 9380 | 9352 | H13 |
| H1f0 | 97290 | 93247 | 103697 | 102212 | 62258 | 70967 | 80068 | 23100 | 23314 | 27837 | 23083 | 32680 | 26484 | 33752 | 93663 | 76621 | 99659 | 85365 | 78084 | 74852 | 80360 | 17687 | 22485 | 32661 | 27323 | 39949 | 33073 | 34684 | H1f0 |
| H1f1 | 33 | 24 | 6 | 29 | 8 | 31 | 16 | 7 | 16 | 16 | 20 | 16 | 5 | 17 | 13 | 19 | 21 | 22 | 26 | 17 | 5 | 3 | 7 | 27 | 16 | 25 | 27 | 7 | H1f1 |
| H1f10 | 6225 | 7389 | 7324 | 8074 | 5267 | 4536 | 5020 | 2981 | 3048 | 3934 | 3187 | 5027 | 3229 | 4362 | 5498 | 4825 | 8144 | 6455 | 6199 | 4433 | 5655 | 3116 | 2696 | 4595 | 3772 | 6002 | 4030 | 4258 | H1f10 |
| H1f2 | 1339 | 1637 | 1139 | 2231 | 1348 | 1226 | 1766 | 307 | 466 | 385 | 453 | 420 | 460 | 342 | 1531 | 1354 | 1627 | 2192 | 1678 | 1529 | 1539 | 272 | 504 | 523 | 312 | 509 | 396 | 323 | H1f2 |
| H1f3 | 88 | 80 | 62 | 139 | 56 | 69 | 124 | 20 | 36 | 27 | 38 | 46 | 28 | 29 | 94 | 91 | 89 | 121 | 76 | 76 | 110 | 10 | 27 | 42 | 23 | 26 | 38 | 30 | H1f3 |
| H1f4 | 75 | 99 | 54 | 134 | 57 | 55 | 80 | 31 | 18 | 43 | 17 | 29 | 19 | 27 | 76 | 87 | 97 | 102 | 72 | 62 | 66 | 30 | 21 | 34 | 23 | 28 | 34 | 19 | H1f4 |
| H1f5 | 46 | 50 | 18 | 51 | 15 | 28 | 32 | 17 | 10 | 10 | 8 | 33 | 8 | 15 | 25 | 26 | 22 | 55 | 23 | 14 | 35 | 5 | 6 | 16 | 16 | 15 | 22 | 14 | H1f5 |
| H1f6 | 2 | 3 | 5 | 6 | 0 | 0 | 2 | 0 | 2 | 1 | 6 | 0 | 0 | 2 | 1 | 2 | 5 | 3 | 3 | 2 | 0 | 0 | 0 | 2 | 1 | 4 | 0 | 0 | H1f6 |
| H1f8 | 7 | 7 | 5 | 8 | 1 | 3 | 2 | 1 | 0 | 0 | 4 | 0 | 1 | 0 | 3 | 7 | 1 | 0 | 3 | 4 | 8 | 3 | 1 | 2 | 0 | 2 | 3 | 0 | H1f8 |
| H1f9 | 0 | 1 | 0 | 0 | 0 | 0 | 1 | 0 | 2 | 2 | 0 | 0 | 7 | 0 | 1 | 0 | 0 | 0 | 0 | 0 | 0 | 0 | 2 | 0 | 0 | 0 | 0 | 0 | H1f9 |
| H2-Aa | 3 | 1 | 0 | 2 | 7 | 1 | 1 | 13 | 18 | 11 | 15 | 1 | 1 | 2 | 3 | 5 | 2 | 7 | 2 | 2 | 2 | 13 | 23 | 7 | 4 | 7 | 3 | 0 | H2-Aa |
| H2-Ab1 | 526 | 592 | 737 | 474 | 299 | 389 | 351 | 32 | 26 | 19 | 39 | 29 | 34 | 20 | 402 | 459 | 676 | 369 | 443 | 341 | 386 | 18 | 39 | 27 | 23 | 45 | 35 | 38 | H2-Ab1 |
| H2ac1 | 0 | 0 | 2 | 0 | 0 | 0 | 0 | 0 | 0 | 0 | 0 | 0 | 0 | 0 | 0 | 0 | 0 | 0 | 0 | 0 | 0 | 0 | 0 | 0 | 0 | 0 | 0 | 0 | H2ac1 |
| H2ac10 | 149 | 154 | 54 | 90 | 89 | 119 | 178 | 223 | 264 | 131 | 153 | 160 | 203 | 173 | 142 | 134 | 53 | 94 | 135 | 157 | 94 | 139 | 168 | 206 | 173 | 219 | 246 | 201 | H2ac10 |
| H2ac11 | 95 | 79 | 91 | 76 | 61 | 90 | 59 | 68 | 45 | 56 | 53 | 91 | 92 | 86 | 99 | 85 | 70 | 69 | 75 | 83 | 49 | 48 | 64 | 74 | 67 | 147 | 95 | 57 | H2ac11 |
| H2ac12 | 9 | 13 | 1 | 22 | 8 | 7 | 10 | 10 | 7 | 10 | 12 | 8 | 12 | 17 | 16 | 20 | 3 | 24 | 6 | 8 | 1 | 7 | 6 | 11 | 2 | 9 | 14 | 1 | H2ac12 |
| H2ac13 | 51 | 45 | 35 | 59 | 29 | 40 | 25 | 27 | 38 | 36 | 50 | 33 | 44 | 29 | 37 | 61 | 41 | 26 | 43 | 70 | 39 | 47 | 36 | 48 | 34 | 73 | 55 | 28 | H2ac13 |
| H2ac15 | 33 | 27 | 27 | 21 | 14 | 32 | 19 | 10 | 12 | 20 | 24 | 37 | 18 | 31 | 20 | 23 | 22 | 14 | 18 | 24 | 10 | 12 | 19 | 9 | 19 | 41 | 36 | 14 | H2ac15 |
| H2ac19 | 10 | 8 | 4 | 17 | 19 | 21 | 6 | 12 | 26 | 16 | 17 | 15 | 12 | 6 | 7 | 4 | 23 | 11 | 24 | 8 | 12 | 35 | 17 | 22 | 20 | 19 | 16 | 9 | H2ac19 |
| H2ac20 | 9 | 6 | 2 | 10 | 3 | 8 | 3 | 1 | 2 | 0 | 3 | 2 | 0 | 3 | 4 | 5 | 0 | 1 | 2 | 4 | 2 | 2 | 2 | 5 | 0 | 2 | 8 | 0 | H2ac20 |
| H2ac21 | 12 | 22 | 10 | 24 | 14 | 46 | 31 | 24 | 23 | 34 | 11 | 33 | 20 | 48 | 15 | 12 | 22 | 25 | 18 | 30 | 26 | 20 | 23 | 25 | 43 | 21 | 23 | 24 | H2ac21 |
| H2ac22 | 3 | 2 | 1 | 8 | 7 | 4 | 2 | 1 | 1 | 4 | 5 | 7 | 6 | 3 | 4 | 2 | 6 | 3 | 5 | 9 | 0 | 0 | 1 | 2 | 2 | 4 | 10 | 1 | H2ac22 |
| H2ac23 | 662 | 415 | 533 | 496 | 276 | 380 | 202 | 466 | 335 | 315 | 257 | 475 | 382 | 226 | 647 | 340 | 460 | 366 | 535 | 503 | 248 | 292 | 256 | 525 | 390 | 1061 | 631 | 333 | H2ac23 |
| H2ac24 | 368 | 181 | 194 | 168 | 101 | 218 | 68 | 233 | 138 | 111 | 95 | 238 | 197 | 72 | 306 | 180 | 166 | 154 | 252 | 253 | 71 | 139 | 106 | 162 | 152 | 296 | 302 | 114 | H2ac24 |
| H2ac4 | 29 | 18 | 11 | 21 | 8 | 19 | 4 | 13 | 12 | 8 | 11 | 9 | 3 | 11 | 18 | 5 | 21 | 12 | 19 | 12 | 8 | 12 | 16 | 17 | 13 | 16 | 26 | 6 | H2ac4 |
| H2ac6 | 82 | 73 | 66 | 68 | 74 | 64 | 80 | 68 | 52 | 76 | 39 | 59 | 51 | 36 | 75 | 66 | 68 | 98 | 79 | 103 | 68 | 48 | 60 | 59 | 29 | 63 | 56 | 28 | H2ac6 |
| H2ac7 | 18 | 9 | 24 | 9 | 8 | 9 | 12 | 21 | 12 | 10 | 4 | 12 | 7 | 12 | 26 | 18 | 8 | 9 | 9 | 5 | 6 | 7 | 9 | 9 | 8 | 13 | 6 | 7 | H2ac7 |
| H2ac8 | 149 | 143 | 134 | 113 | 72 | 119 | 78 | 92 | 111 | 79 | 101 | 103 | 174 | 74 | 132 | 149 | 104 | 156 | 108 | 167 | 97 | 72 | 101 | 101 | 95 | 138 | 164 | 85 | H2ac8 |
| H2aj | 2791 | 3322 | 3296 | 4393 | 3148 | 3127 | 3431 | 1608 | 1497 | 1854 | 1650 | 1905 | 1804 | 1886 | 2800 | 2710 | 3739 | 3900 | 3491 | 3377 | 3237 | 1719 | 1392 | 2102 | 2094 | 2229 | 1966 | 1778 | H2aj |
| H2al1a | 0 | 0 | 0 | 0 | 0 | 0 | 0 | 0 | 0 | 0 | 0 | 0 | 0 | 0 | 0 | 0 | 0 | 0 | 0 | 0 | 0 | 0 | 0 | 5 | 0 | 0 | 0 | 0 | H2al1a |
| H2al1b | 0 | 0 | 0 | 1 | 0 | 0 | 0 | 0 | 0 | 0 | 0 | 0 | 0 | 0 | 0 | 0 | 0 | 0 | 0 | 0 | 0 | 0 | 0 | 0 | 0 | 0 | 0 | 0 | H2al1b |
| H2al1f | 0 | 0 | 0 | 0 | 0 | 0 | 0 | 0 | 0 | 0 | 0 | 0 | 0 | 0 | 0 | 0 | 0 | 0 | 0 | 0 | 0 | 0 | 2 | 0 | 0 | 0 | 0 | 0 | H2al1f |
| H2al1g | 0 | 0 | 0 | 0 | 1 | 0 | 0 | 0 | 0 | 0 | 0 | 0 | 0 | 0 | 0 | 0 | 0 | 0 | 0 | 0 | 0 | 0 | 0 | 0 | 0 | 0 | 0 | 0 | H2al1g |
| H2al1i | 0 | 0 | 0 | 1 | 0 | 0 | 0 | 0 | 0 | 0 | 0 | 0 | 0 | 0 | 0 | 0 | 0 | 0 | 0 | 0 | 0 | 0 | 0 | 0 | 0 | 0 | 0 | 0 | H2al1i |
| H2al1m | 3 | 2 | 0 | 0 | 1 | 0 | 0 | 4 | 3 | 12 | 2 | 3 | 6 | 6 | 4 | 2 | 0 | 4 | 3 | 0 | 1 | 0 | 0 | 0 | 3 | 6 | 7 | 1 | H2al1m |
| H2al1o | 0 | 0 | 0 | 0 | 0 | 0 | 0 | 0 | 0 | 1 | 0 | 0 | 0 | 0 | 0 | 0 | 0 | 0 | 1 | 0 | 0 | 0 | 0 | 0 | 0 | 0 | 0 | 1 | H2al1o |
| H2al2a | 1 | 0 | 2 | 3 | 2 | 4 | 2 | 2 | 0 | 3 | 0 | 1 | 0 | 0 | 1 | 2 | 6 | 0 | 0 | 1 | 2 | 2 | 2 | 0 | 0 | 3 | 1 | 1 | H2al2a |
| H2al3 | 0 | 0 | 2 | 0 | 0 | 0 | 0 | 0 | 0 | 0 | 0 | 0 | 0 | 0 | 0 | 0 | 0 | 7 | 2 | 4 | 0 | 0 | 0 | 0 | 0 | 0 | 0 | 0 | H2al3 |
| H2ap | 0 | 0 | 0 | 0 | 0 | 0 | 2 | 0 | 1 | 2 | 0 | 0 | 0 | 2 | 0 | 0 | 0 | 0 | 0 | 0 | 3 | 0 | 0 | 0 | 0 | 0 | 0 | 0 | H2ap |
| H2aw | 1099 | 1011 | 1036 | 1749 | 1852 | 1508 | 2168 | 1485 | 1175 | 1657 | 1325 | 1426 | 1002 | 1571 | 1510 | 829 | 1600 | 1858 | 1792 | 1798 | 2264 | 1542 | 1271 | 1805 | 1635 | 1111 | 1189 | 1322 | H2aw |
| H2ax | 6962 | 7863 | 8978 | 7818 | 6122 | 5856 | 6159 | 4689 | 5333 | 5978 | 5411 | 8661 | 6386 | 7995 | 5902 | 6257 | 8762 | 7132 | 7222 | 5786 | 6460 | 4318 | 4689 | 7923 | 6168 | 11061 | 7205 | 7711 | H2ax |
| H2az1 | 27925 | 31452 | 33210 | 27435 | 21897 | 24915 | 25787 | 25756 | 25862 | 23094 | 20826 | 33032 | 29203 | 32186 | 27534 | 28147 | 29415 | 23293 | 29050 | 34596 | 25368 | 15869 | 22132 | 29253 | 25412 | 43082 | 37927 | 35348 | H2az1 |
| H2az2 | 8656 | 10360 | 11929 | 10432 | 7700 | 7944 | 9821 | 6588 | 6841 | 7755 | 6959 | 10547 | 8915 | 11681 | 7985 | 7995 | 11777 | 9034 | 9828 | 9300 | 9527 | 5310 | 6320 | 10138 | 8856 | 12839 | 10708 | 11661 | H2az2 |
| H2bc1 | 0 | 0 | 0 | 0 | 0 | 0 | 0 | 0 | 0 | 0 | 0 | 0 | 0 | 0 | 0 | 0 | 0 | 0 | 0 | 0 | 1 | 0 | 0 | 0 | 0 | 0 | 0 | 0 | H2bc1 |
| H2bc11 | 55 | 54 | 44 | 60 | 24 | 31 | 32 | 33 | 21 | 34 | 43 | 46 | 39 | 34 | 50 | 47 | 60 | 62 | 43 | 50 | 49 | 17 | 26 | 29 | 21 | 41 | 62 | 89 | H2bc11 |
| H2bc12 | 50 | 54 | 84 | 49 | 20 | 55 | 31 | 34 | 24 | 30 | 32 | 42 | 48 | 71 | 40 | 62 | 36 | 32 | 48 | 40 | 36 | 13 | 24 | 41 | 37 | 66 | 80 | 71 | H2bc12 |
| H2bc13 | 18 | 17 | 12 | 16 | 9 | 13 | 19 | 10 | 17 | 5 | 12 | 25 | 16 | 20 | 13 | 10 | 3 | 15 | 9 | 22 | 15 | 20 | 13 | 22 | 14 | 36 | 27 | 24 | H2bc13 |
| H2bc14 | 32 | 24 | 15 | 20 | 16 | 22 | 10 | 12 | 4 | 15 | 1 | 29 | 16 | 16 | 17 | 8 | 8 | 29 | 17 | 24 | 15 | 5 | 13 | 32 | 13 | 15 | 17 | 30 | H2bc14 |
| H2bc15 | 18 | 19 | 11 | 20 | 12 | 16 | 11 | 15 | 10 | 9 | 14 | 8 | 16 | 15 | 7 | 5 | 20 | 22 | 9 | 10 | 6 | 5 | 6 | 7 | 6 | 7 | 31 | 17 | H2bc15 |
| H2bc18 | 0 | 0 | 2 | 2 | 0 | 0 | 0 | 0 | 1 | 0 | 0 | 0 | 0 | 0 | 0 | 0 | 1 | 1 | 0 | 0 | 1 | 0 | 0 | 0 | 0 | 0 | 2 | 0 | H2bc18 |
| H2bc21 | 1751 | 1641 | 1516 | 1890 | 1544 | 1624 | 1769 | 1756 | 1519 | 1565 | 1221 | 1493 | 1425 | 1523 | 1668 | 1518 | 1761 | 1580 | 1738 | 2008 | 1667 | 1357 | 1447 | 1553 | 1516 | 1904 | 1590 | 1477 | H2bc21 |
| H2bc22 | 13 | 22 | 24 | 10 | 11 | 13 | 8 | 6 | 15 | 19 | 25 | 15 | 14 | 18 | 12 | 15 | 8 | 13 | 12 | 9 | 4 | 8 | 8 | 29 | 13 | 35 | 42 | 9 | H2bc22 |
| H2bc23 | 8 | 2 | 1 | 0 | 1 | 2 | 1 | 7 | 4 | 2 | 0 | 1 | 2 | 2 | 0 | 3 | 0 | 0 | 12 | 6 | 1 | 3 | 5 | 8 | 12 | 2 | 3 | 4 | H2bc23 |
| H2bc24 | 3 | 7 | 9 | 6 | 1 | 1 | 4 | 1 | 2 | 1 | 0 | 1 | 8 | 1 | 11 | 3 | 2 | 7 | 5 | 9 | 3 | 2 | 3 | 2 | 2 | 0 | 3 | 1 | H2bc24 |
| H2bc3 | 8 | 13 | 7 | 15 | 14 | 21 | 9 | 15 | 7 | 16 | 11 | 25 | 18 | 20 | 13 | 13 | 14 | 11 | 14 | 14 | 8 | 8 | 11 | 18 | 14 | 22 | 38 | 17 | H2bc3 |
| H2bc4 | 294 | 330 | 222 | 382 | 329 | 281 | 398 | 263 | 280 | 357 | 343 | 291 | 280 | 358 | 297 | 232 | 360 | 406 | 400 | 388 | 481 | 219 | 324 | 445 | 330 | 375 | 309 | 300 | H2bc4 |
| H2bc6 | 153 | 168 | 185 | 217 | 125 | 141 | 222 | 96 | 82 | 82 | 67 | 94 | 83 | 77 | 190 | 176 | 189 | 207 | 168 | 159 | 172 | 58 | 88 | 116 | 71 | 119 | 78 | 95 | H2bc6 |
| H2bc7 | 12 | 28 | 13 | 6 | 2 | 10 | 11 | 7 | 5 | 10 | 3 | 14 | 11 | 18 | 13 | 13 | 20 | 9 | 8 | 10 | 9 | 10 | 2 | 8 | 16 | 10 | 10 | 7 | H2bc7 |
| H2bc8 | 15 | 24 | 21 | 15 | 20 | 17 | 23 | 31 | 34 | 31 | 33 | 27 | 26 | 18 | 21 | 18 | 14 | 30 | 17 | 31 | 9 | 18 | 30 | 20 | 15 | 13 | 65 | 15 | H2bc8 |
| H2bc9 | 85 | 47 | 57 | 38 | 43 | 30 | 45 | 27 | 28 | 23 | 13 | 38 | 24 | 28 | 54 | 45 | 38 | 44 | 47 | 43 | 44 | 15 | 17 | 39 | 19 | 30 | 40 | 33 | H2bc9 |
| H2bu1-ps | 9 | 22 | 15 | 29 | 15 | 13 | 12 | 15 | 22 | 32 | 28 | 41 | 33 | 30 | 12 | 12 | 10 | 21 | 23 | 10 | 21 | 22 | 31 | 29 | 29 | 24 | 27 | 25 | H2bu1-ps |
| H2bu2 | 2016 | 2914 | 1938 | 3055 | 3084 | 3168 | 2849 | 3617 | 3625 | 3656 | 3366 | 3960 | 3399 | 4076 | 2414 | 2021 | 3146 | 2930 | 3537 | 3819 | 3010 | 3638 | 3540 | 4307 | 4444 | 4126 | 4242 | 3410 | H2bu2 |
| H2-D1 | 2485 | 3476 | 4149 | 3775 | 3244 | 3014 | 3024 | 915 | 938 | 1090 | 1039 | 1250 | 1174 | 1181 | 2611 | 2893 | 3799 | 3415 | 3625 | 2990 | 2960 | 1103 | 1059 | 1199 | 1109 | 1311 | 1182 | 1270 | H2-D1 |
| H2-DMa | 396 | 458 | 464 | 500 | 484 | 459 | 523 | 208 | 234 | 207 | 261 | 258 | 209 | 252 | 393 | 435 | 496 | 530 | 464 | 383 | 412 | 241 | 223 | 262 | 217 | 234 | 234 | 215 | H2-DMa |
| H2-DMb1 | 7 | 27 | 8 | 5 | 19 | 10 | 6 | 22 | 40 | 34 | 27 | 21 | 39 | 9 | 11 | 16 | 2 | 33 | 12 | 6 | 16 | 25 | 64 | 40 | 35 | 24 | 10 | 24 | H2-DMb1 |
| H2-DMb2 | 6 | 3 | 2 | 3 | 8 | 0 | 9 | 1 | 11 | 4 | 3 | 5 | 3 | 6 | 4 | 8 | 1 | 2 | 9 | 0 | 6 | 10 | 10 | 5 | 1 | 0 | 7 | 4 | H2-DMb2 |
| H2-Eb1 | 22 | 11 | 17 | 9 | 7 | 7 | 6 | 11 | 26 | 38 | 35 | 11 | 12 | 17 | 7 | 9 | 6 | 10 | 9 | 7 | 4 | 22 | 35 | 14 | 8 | 16 | 12 | 14 | H2-Eb1 |
| H2-Eb2 | 3 | 1 | 10 | 11 | 11 | 12 | 11 | 9 | 13 | 9 | 27 | 5 | 5 | 6 | 7 | 5 | 6 | 2 | 2 | 14 | 10 | 35 | 11 | 3 | 6 | 5 | 14 | 6 | H2-Eb2 |
| H2-K1 | 2821 | 4280 | 5094 | 3484 | 2203 | 2441 | 2489 | 568 | 431 | 531 | 383 | 747 | 672 | 797 | 2972 | 3134 | 4047 | 2839 | 2912 | 2942 | 2442 | 441 | 509 | 556 | 490 | 1011 | 847 | 1028 | H2-K1 |
| H2-Ke6 | 2 | 9 | 2 | 9 | 2 | 10 | 7 | 2 | 0 | 9 | 0 | 1 | 0 | 3 | 3 | 5 | 7 | 3 | 4 | 5 | 9 | 5 | 4 | 2 | 7 | 5 | 5 | 5 | H2-Ke6 |
| H2-M1 | 0 | 0 | 0 | 0 | 0 | 0 | 0 | 0 | 2 | 0 | 0 | 0 | 0 | 0 | 4 | 1 | 0 | 0 | 0 | 0 | 0 | 0 | 0 | 0 | 0 | 0 | 0 | 0 | H2-M1 |

| | | | | | | | | | | | | | | | | | | | | | | | | | | | | | |
|---|---|---|---|---|---|---|---|---|---|---|---|---|---|---|---|---|---|---|---|---|---|---|---|---|---|---|---|---|---|
| H2-M10.3 | 0 | 0 | 0 | 0 | 0 | 0 | 0 | 0 | 0 | 1 | 0 | 0 | 0 | 0 | 0 | 0 | 0 | 0 | 0 | 0 | 0 | 0 | 0 | 0 | 0 | 0 | 0 | 0 | H2-M10.3 |
| H2-M10.5 | 0 | 0 | 0 | 0 | 0 | 0 | 0 | 0 | 0 | 0 | 0 | 0 | 0 | 0 | 0 | 1 | 0 | 0 | 0 | 0 | 0 | 0 | 0 | 0 | 0 | 0 | 0 | 0 | H2-M10.5 |
| H2-M11 | 3 | 4 | 2 | 0 | 2 | 0 | 0 | 4 | 3 | 3 | 3 | 0 | 1 | 0 | 11 | 2 | 0 | 3 | 1 | 1 | 4 | 5 | 1 | 1 | 1 | 0 | 1 | 0 | H2-M11 |
| H2-M2 | 0 | 0 | 0 | 0 | 0 | 1 | 0 | 0 | 0 | 0 | 5 | 0 | 0 | 0 | 0 | 0 | 0 | 0 | 0 | 0 | 0 | 0 | 0 | 0 | 0 | 5 | 0 | 0 | H2-M2 |
| H2-M3 | 78 | 81 | 109 | 119 | 83 | 79 | 66 | 57 | 50 | 58 | 35 | 61 | 57 | 52 | 78 | 68 | 74 | 37 | 85 | 97 | 67 | 47 | 43 | 45 | 82 | 69 | 69 | 63 | H2-M3 |
| H2-M5 | 414 | 458 | 575 | 728 | 745 | 802 | 799 | 913 | 798 | 1163 | 1008 | 1068 | 1298 | 1288 | 467 | 448 | 611 | 744 | 831 | 690 | 830 | 1440 | 731 | 1176 | 1412 | 820 | 1152 | 1104 | H2-M5 |
| H2-Oa | 18 | 16 | 9 | 11 | 5 | 15 | 3 | 6 | 7 | 3 | 15 | 3 | 5 | 11 | 12 | 9 | 5 | 8 | 9 | 9 | 6 | 2 | 3 | 5 | 5 | 1 | 0 | 1 | H2-Oa |
| H2-Ob | 7 | 0 | 2 | 1 | 0 | 0 | 2 | 0 | 2 | 0 | 0 | 1 | 0 | 0 | 5 | 2 | 1 | 2 | 0 | 3 | 1 | 0 | 2 | 0 | 10 | 1 | 0 | 1 | H2-Ob |
| H2-Q1 | 3 | 3 | 3 | 6 | 0 | 1 | 1 | 10 | 2 | 10 | 13 | 13 | 3 | 4 | 1 | 1 | 2 | 1 | 9 | 1 | 5 | 5 | 10 | 2 | 8 | 4 | 6 | 11 | H2-Q1 |
| H2-Q10 | 25 | 19 | 37 | 20 | 25 | 23 | 21 | 0 | 2 | 0 | 1 | 0 | 0 | 0 | 18 | 40 | 25 | 28 | 13 | 7 | 13 | 3 | 6 | 1 | 1 | 0 | 5 | 4 | H2-Q10 |
| H2-Q2 | 20 | 30 | 16 | 13 | 29 | 4 | 7 | 10 | 17 | 14 | 14 | 9 | 15 | 10 | 20 | 23 | 13 | 20 | 14 | 10 | 9 | 15 | 21 | 9 | 7 | 9 | 5 | 7 | H2-Q2 |
| H2-Q4 | 1125 | 1101 | 1860 | 1119 | 1047 | 1080 | 955 | 267 | 307 | 254 | 306 | 427 | 434 | 434 | 1005 | 1111 | 1505 | 952 | 1166 | 941 | 875 | 296 | 324 | 363 | 223 | 618 | 533 | 540 | H2-Q4 |
| H2-Q6 | 213 | 201 | 288 | 173 | 152 | 137 | 122 | 6 | 20 | 12 | 11 | 20 | 23 | 13 | 178 | 283 | 219 | 147 | 151 | 156 | 139 | 22 | 15 | 20 | 33 | 28 | 30 | 31 | H2-Q6 |
| H2-Q7 | 376 | 489 | 739 | 387 | 436 | 355 | 337 | 116 | 103 | 117 | 129 | 174 | 162 | 156 | 347 | 410 | 601 | 386 | 377 | 360 | 306 | 92 | 84 | 118 | 106 | 196 | 213 | 252 | H2-Q7 |
| H2-T22 | 1122 | 1228 | 1109 | 811 | 864 | 974 | 763 | 744 | 990 | 820 | 948 | 734 | 839 | 734 | 1164 | 1294 | 968 | 1073 | 860 | 954 | 764 | 724 | 1124 | 779 | 638 | 896 | 724 | 751 | H2-T22 |
| H2-T23 | 218 | 211 | 251 | 198 | 275 | 247 | 251 | 243 | 365 | 298 | 313 | 272 | 361 | 275 | 161 | 155 | 277 | 178 | 310 | 238 | 230 | 197 | 326 | 309 | 224 | 347 | 300 | 331 | H2-T23 |
| H2-T24 | 28 | 46 | 29 | 16 | 6 | 14 | 13 | 39 | 39 | 49 | 30 | 68 | 221 | 135 | 26 | 23 | 36 | 12 | 10 | 14 | 14 | 27 | 52 | 34 | 32 | 101 | 145 | 225 | H2-T24 |
| H3c1 | 36 | 41 | 9 | 26 | 16 | 24 | 16 | 18 | 17 | 24 | 14 | 15 | 27 | 17 | 37 | 33 | 20 | 17 | 22 | 30 | 13 | 13 | 8 | 25 | 24 | 28 | 46 | 23 | H3c1 |
| H3c10 | 54 | 68 | 44 | 21 | 34 | 34 | 41 | 40 | 41 | 14 | 41 | 55 | 36 | 43 | 62 | 51 | 39 | 40 | 50 | 55 | 21 | 22 | 38 | 38 | 47 | 53 | 51 | 52 | H3c10 |
| H3c11 | 30 | 38 | 34 | 25 | 24 | 32 | 16 | 13 | 13 | 14 | 20 | 12 | 19 | 9 | 17 | 26 | 32 | 26 | 27 | 20 | 13 | 10 | 19 | 16 | 9 | 23 | 34 | 27 | H3c11 |
| H3c13 | 5 | 13 | 14 | 13 | 8 | 14 | 7 | 6 | 8 | 4 | 3 | 11 | 10 | 2 | 7 | 5 | 2 | 2 | 1 | 0 | 4 | 3 | 6 | 2 | 4 | 10 | 15 | 6 | H3c13 |
| H3c15 | 32 | 35 | 26 | 39 | 38 | 73 | 57 | 43 | 46 | 77 | 49 | 45 | 46 | 56 | 45 | 29 | 37 | 50 | 76 | 25 | 57 | 63 | 64 | 74 | 36 | 59 | 51 | 31 | H3c15 |
| H3c2 | 21 | 11 | 8 | 2 | 7 | 25 | 4 | 4 | 3 | 1 | 5 | 13 | 9 | 4 | 12 | 9 | 6 | 10 | 6 | 17 | 3 | 2 | 10 | 8 | 2 | 6 | 5 | 1 | H3c2 |
| H3c3 | 211 | 208 | 184 | 176 | 94 | 136 | 89 | 110 | 109 | 76 | 128 | 153 | 132 | 136 | 195 | 167 | 131 | 102 | 163 | 150 | 91 | 63 | 109 | 175 | 139 | 296 | 187 | 135 | H3c3 |
| H3c4 | 23 | 30 | 22 | 33 | 25 | 30 | 22 | 21 | 28 | 27 | 34 | 16 | 15 | 16 | 32 | 28 | 34 | 33 | 34 | 27 | 26 | 15 | 21 | 27 | 21 | 22 | 37 | 28 | H3c4 |
| H3c6 | 37 | 48 | 16 | 36 | 36 | 32 | 18 | 21 | 16 | 17 | 35 | 37 | 29 | 24 | 34 | 37 | 17 | 34 | 25 | 39 | 21 | 12 | 25 | 40 | 33 | 37 | 49 | 23 | H3c6 |
| H3c7 | 12 | 18 | 10 | 10 | 6 | 13 | 4 | 7 | 9 | 0 | 7 | 5 | 6 | 11 | 5 | 5 | 13 | 2 | 5 | 13 | 8 | 8 | 8 | 7 | 11 | 14 | 15 | 13 | H3c7 |
| H3c8 | 35 | 23 | 16 | 15 | 16 | 25 | 29 | 16 | 20 | 4 | 21 | 28 | 16 | 19 | 18 | 18 | 23 | 9 | 14 | 21 | 13 | 10 | 16 | 18 | 2 | 17 | 20 | 9 | H3c8 |
| H3f3a | 67509 | 60205 | 60083 | 65367 | 60298 | 60752 | 81282 | 73832 | 62860 | 57855 | 56080 | 61620 | 60920 | 81538 | 72120 | 61610 | 61428 | 59740 | 65401 | 77211 | 75844 | 60337 | 64440 | 65008 | 64660 | 69080 | 65941 | 78146 | H3f3a |
| H3f3b | 123614 | 105153 | 115396 | 133176 | 111208 | 118062 | 131166 | 84306 | 79664 | 81039 | 76409 | 88270 | 81339 | 89851 | 132609 | 107376 | 121425 | 125332 | 117191 | 140807 | 125713 | 73576 | 79715 | 90328 | 85727 | 102159 | 89772 | 88945 | H3f3b |
| H3f4 | 5 | 7 | 3 | 12 | 2 | 14 | 4 | 1 | 4 | 3 | 1 | 0 | 2 | 2 | 5 | 5 | 5 | 8 | 3 | 2 | 4 | 2 | 3 | 5 | 4 | 3 | 5 | 0 | H3f4 |
| H4c1 | 25 | 17 | 17 | 15 | 9 | 15 | 29 | 11 | 9 | 13 | 10 | 39 | 18 | 31 | 12 | 9 | 12 | 17 | 8 | 9 | 25 | 5 | 8 | 27 | 12 | 12 | 16 | 28 | H4c1 |
| H4c11 | 39 | 10 | 17 | 35 | 35 | 17 | 23 | 31 | 36 | 24 | 20 | 18 | 7 | 35 | 18 | 19 | 17 | 24 | 18 | 5 | 18 | 12 | 26 | 28 | 23 | 33 | 35 | 8 | H4c11 |
| H4c12 | 48 | 60 | 63 | 76 | 37 | 66 | 57 | 31 | 30 | 29 | 53 | 41 | 32 | 37 | 34 | 47 | 65 | 64 | 53 | 35 | 56 | 22 | 21 | 42 | 16 | 31 | 46 | 18 | H4c12 |
| H4c14 | 44 | 33 | 12 | 49 | 14 | 31 | 22 | 43 | 31 | 50 | 29 | 35 | 39 | 26 | 25 | 26 | 24 | 40 | 38 | 21 | 40 | 13 | 28 | 56 | 23 | 12 | 21 | 18 | H4c14 |
| H4c17 | 27 | 11 | 8 | 31 | 11 | 13 | 16 | 9 | 7 | 10 | 12 | 13 | 17 | 9 | 4 | 21 | 9 | 12 | 30 | 10 | 17 | 8 | 12 | 18 | 17 | 15 | 10 | 14 | H4c17 |
| H4c18 | 9 | 15 | 3 | 14 | 8 | 10 | 7 | 4 | 12 | 10 | 5 | 14 | 7 | 7 | 9 | 10 | 12 | 11 | 9 | 12 | 6 | 2 | 12 | 3 | 3 | 13 | 27 | 9 | H4c18 |
| H4c2 | 50 | 46 | 62 | 79 | 19 | 33 | 36 | 39 | 30 | 55 | 51 | 42 | 57 | 86 | 37 | 40 | 66 | 47 | 47 | 39 | 42 | 15 | 16 | 60 | 24 | 67 | 43 | 83 | H4c2 |
| H4c3 | 25 | 19 | 12 | 24 | 16 | 19 | 6 | 32 | 20 | 17 | 20 | 15 | 16 | 19 | 16 | 11 | 3 | 19 | 18 | 18 | 12 | 12 | 14 | 16 | 23 | 24 | 27 | 18 | H4c3 |
| H4c4 | 31 | 46 | 27 | 43 | 38 | 47 | 38 | 29 | 24 | 28 | 47 | 30 | 28 | 39 | 34 | 33 | 35 | 41 | 28 | 25 | 36 | 23 | 23 | 46 | 40 | 51 | 57 | 39 | H4c4 |
| H4c6 | 2 | 15 | 8 | 8 | 2 | 5 | 9 | 1 | 4 | 5 | 5 | 9 | 2 | 6 | 4 | 5 | 5 | 7 | 0 | 7 | 6 | 0 | 0 | 9 | 4 | 2 | 10 | 8 | H4c6 |
| H4c8 | 16 | 22 | 11 | 39 | 19 | 29 | 26 | 13 | 19 | 34 | 25 | 20 | 15 | 28 | 16 | 23 | 19 | 18 | 16 | 29 | 12 | 22 | 16 | 26 | 13 | 11 | 29 | 15 | H4c8 |
| H4c9 | 727 | 905 | 1011 | 879 | 556 | 723 | 744 | 633 | 560 | 580 | 517 | 898 | 706 | 863 | 691 | 658 | 990 | 620 | 813 | 952 | 606 | 344 | 476 | 830 | 654 | 1244 | 1055 | 924 | H4c9 |
| H4f16 | 27 | 29 | 14 | 27 | 25 | 29 | 21 | 35 | 21 | 32 | 38 | 26 | 22 | 50 | 25 | 23 | 20 | 45 | 44 | 24 | 36 | 33 | 24 | 18 | 42 | 30 | 30 | 24 | H4f16 |
| H60b | 298 | 278 | 389 | 223 | 242 | 273 | 189 | 132 | 91 | 139 | 70 | 167 | 197 | 150 | 322 | 343 | 382 | 241 | 222 | 247 | 211 | 137 | 133 | 112 | 90 | 89 | 216 | 166 | H60b |
| H60c | 0 | 3 | 0 | 0 | 0 | 1 | 0 | 11 | 42 | 20 | 14 | 3 | 5 | 4 | 0 | 1 | 0 | 3 | 0 | 0 | 0 | 3 | 57 | 20 | 6 | 0 | 0 | 5 | H60c |
| H6pd | 761 | 867 | 850 | 573 | 529 | 689 | 365 | 794 | 1253 | 1102 | 1551 | 866 | 1088 | 769 | 667 | 994 | 689 | 687 | 442 | 403 | 374 | 1106 | 1552 | 956 | 912 | 856 | 996 | 774 | H6pd |
| Haao | 10 | 12 | 0 | 2 | 2 | 4 | 6 | 6 | 7 | 5 | 8 | 10 | 3 | 6 | 1 | 5 | 0 | 3 | 4 | 2 | 4 | 3 | 17 | 2 | 4 | 6 | 0 | 4 | Haao |
| Habp2 | 0 | 0 | 0 | 0 | 0 | 0 | 0 | 0 | 2 | 0 | 0 | 0 | 0 | 0 | 0 | 2 | 0 | 0 | 0 | 0 | 0 | 0 | 0 | 0 | 0 | 0 | 0 | 1 | Habp2 |
| Habp4 | 4456 | 5237 | 5625 | 5904 | 5400 | 4927 | 4983 | 1755 | 1911 | 2161 | 2656 | 1807 | 2144 | 1971 | 4383 | 5306 | 5861 | 6308 | 5065 | 4165 | 5059 | 2351 | 2315 | 2077 | 2130 | 1828 | 1892 | 1825 | Habp4 |
| Hacd1 | 241 | 225 | 275 | 249 | 269 | 233 | 223 | 761 | 551 | 464 | 651 | 432 | 422 | 499 | 266 | 210 | 277 | 256 | 295 | 312 | 266 | 617 | 751 | 445 | 529 | 446 | 464 | 539 | Hacd1 |
| Hacd2 | 1843 | 1641 | 1693 | 1629 | 1361 | 1432 | 1636 | 857 | 931 | 941 | 838 | 976 | 906 | 971 | 1727 | 1677 | 1497 | 1443 | 1418 | 1404 | 1443 | 941 | 978 | 977 | 923 | 1095 | 1027 | 1032 | Hacd2 |
| Hacd3 | 12230 | 10726 | 11059 | 10752 | 11473 | 11091 | 11696 | 11398 | 9025 | 9345 | 8373 | 9115 | 8885 | 9412 | 12897 | 11134 | 10703 | 11306 | 11539 | 11931 | 11766 | 11185 | 9210 | 8928 | 10381 | 8784 | 9350 | 9659 | Hacd3 |
| Hacd4 | 938 | 626 | 665 | 467 | 414 | 627 | 391 | 879 | 803 | 659 | 795 | 565 | 699 | 492 | 906 | 1008 | 524 | 664 | 316 | 541 | 405 | 652 | 1040 | 538 | 456 | 567 | 636 | 534 | Hacd4 |
| Hace1 | 3202 | 2420 | 2867 | 2618 | 2395 | 2677 | 2657 | 3303 | 3054 | 2860 | 2990 | 3151 | 3127 | 3318 | 3028 | 2699 | 2597 | 2346 | 2442 | 2723 | 2490 | 3083 | 3003 | 2601 | 3024 | 2928 | 3131 | 3184 | Hace1 |
| Hacl1 | 80 | 65 | 70 | 47 | 62 | 72 | 73 | 34 | 26 | 52 | 30 | 25 | 45 | 42 | 86 | 53 | 39 | 43 | 68 | 51 | 70 | 13 | 39 | 47 | 46 | 23 | 64 | 47 | Hacl1 |
| Hadh | 6996 | 7224 | 6937 | 5676 | 3744 | 4369 | 4416 | 2600 | 3136 | 2563 | 3063 | 2937 | 3392 | 3214 | 6253 | 6960 | 5954 | 5328 | 4552 | 4679 | 4378 | 2049 | 3356 | 2914 | 2303 | 3799 | 3380 | 3411 | Hadh |
| Hadha | 14613 | 16084 | 13484 | 12568 | 10595 | 11362 | 12003 | 9104 | 9424 | 7893 | 8781 | 8649 | 9166 | 8848 | 14286 | 15036 | 12653 | 12414 | 11031 | 12434 | 11712 | 7451 | 9970 | 8199 | 8211 | 9742 | 9186 | 9018 | Hadha |
| Hadhb | 6612 | 6555 | 5418 | 5257 | 3877 | 4589 | 5206 | 3348 | 3615 | 2579 | 3183 | 3363 | 3361 | 3358 | 6418 | 6929 | 5032 | 5031 | 4203 | 5437 | 4598 | 2602 | 3643 | 2911 | 2884 | 3842 | 3381 | 3550 | Hadhb |
| Hagh | 1867 | 2637 | 1974 | 2241 | 1937 | 1855 | 1978 | 804 | 826 | 772 | 709 | 741 | 751 | 697 | 2082 | 2218 | 2114 | 2338 | 2022 | 2373 | 1953 | 795 | 825 | 773 | 855 | 776 | 812 | 720 | Hagh |
| Haghl | 3066 | 3769 | 4096 | 4188 | 4230 | 4451 | 3646 | 2748 | 3405 | 3456 | 2940 | 4371 | 3798 | 3703 | 3161 | 3226 | 4142 | 3789 | 4430 | 4278 | 3726 | 3088 | 2964 | 3636 | 3934 | 4532 | 4306 | 3787 | Haghl |
| Hal | 12 | 19 | 16 | 25 | 12 | 17 | 10 | 9 | 4 | 8 | 8 | 13 | 18 | 1 | 7 | 21 | 17 | 26 | 10 | 7 | 12 | 5 | 11 | 8 | 4 | 3 | 9 | 11 | Hal |
| Hamp | 5 | 3 | 6 | 4 | 1 | 2 | 1 | 0 | 4 | 1 | 4 | 5 | 2 | 0 | 3 | 2 | 5 | 0 | 3 | 5 | 1 | 0 | 0 | 0 | 0 | 1 | 0 | 1 | Hamp |
| Hamp2 | 2 | 0 | 0 | 0 | 2 | 1 | 0 | 1 | 5 | 7 | 5 | 2 | 2 | 2 | 0 | 0 | 0 | 0 | 0 | 0 | 0 | 0 | 6 | 5 | 1 | 4 | 3 | 4 | Hamp2 |
| Hand1 | 2 | 6 | 0 | 0 | 0 | 0 | 0 | 0 | 7 | 0 | 49 | 0 | 3 | 0 | 0 | 1 | 1 | 0 | 0 | 0 | 0 | 2 | 41 | 0 | 1 | 2 | 0 | 0 | Hand1 |
| Hand2 | 31 | 10 | 8 | 2 | 12 | 5 | 4 | 50 | 35 | 26 | 391 | 5 | 5 | 3 | 15 | 21 | 3 | 2 | 5 | 8 | 1 | 17 | 183 | 2 | 7 | 0 | 1 | 1 | Hand2 |
| Hao1 | 2 | 0 | 0 | 0 | 4 | 4 | 8 | 5 | 2 | 4 | 2 | 0 | 6 | 4 | 5 | 0 | 2 | 7 | 2 | 0 | 1 | 0 | 1 | 2 | 2 | 1 | 1 | 1 | Hao1 |
| Hao2 | 3 | 3 | 1 | 0 | 0 | 0 | 0 | 2 | 5 | 0 | 2 | 0 | 3 | 0 | 0 | 3 | 0 | 0 | 0 | 0 | 3 | 5 | 0 | 2 | 0 | 1 | 7 | 9 | Hao2 |
| Hap1 | 10434 | 13757 | 13937 | 18492 | 18960 | 17039 | 18903 | 6996 | 7539 | 9836 | 7259 | 9447 | 6074 | 9652 | 10580 | 10360 | 18428 | 18471 | 21250 | 16585 | 19374 | 10813 | 7163 | 11008 | 11777 | 9769 | 8182 | 8304 | Hap1 |
| Hapln1 | 1125 | 942 | 1637 | 642 | 315 | 880 | 227 | 2168 | 4478 | 4305 | 8754 | 1572 | 2062 | 1495 | 1319 | 1544 | 764 | 895 | 336 | 366 | 251 | 3917 | 2922 | 3800 | 2545 | 1883 | 1941 | 1895 | Hapln1 |
| Hapln2 | 27 | 40 | 11 | 33 | 30 | 37 | 31 | 15 | 12 | 5 | 9 | 10 | 7 | 9 | 18 | 23 | 42 | 33 | 30 | 44 | 29 | 15 | 6 | 9 | 15 | 4 | 6 | 11 | Hapln2 |
| Hapln3 | 165 | 172 | 204 | 139 | 204 | 134 | 153 | 142 | 157 | 98 | 119 | 124 | 151 | 92 | 121 | 192 | 140 | 176 | 154 | 157 | 134 | 104 | 170 | 123 | 114 | 148 | 120 | 100 | Hapln3 |
| Hapln4 | 403 | 580 | 673 | 578 | 559 | 513 | 429 | 16 | 21 | 29 | 19 | 20 | 14 | 15 | 509 | 559 | 672 | 583 | 492 | 408 | 453 | 47 | 19 | 24 | 27 | 16 | 14 | 14 | Hapln4 |
| Harbi1 | 1061 | 1035 | 1125 | 971 | 973 | 970 | 1014 | 1082 | 981 | 1108 | 924 | 1005 | 1027 | 948 | 1094 | 1055 | 1169 | 951 | 1072 | 1003 | 965 | 924 | 1042 | 1052 | 998 | 1034 | 1072 | 1057 | Harbi1 |
| Hars | 6259 | 6851 | 6198 | 5909 | 5807 | 5891 | 6006 | 5928 | 6297 | 6071 | 5862 | 6329 | 6123 | 6673 | 6190 | 6002 | 6269 | 5529 | 5975 | 6376 | 6117 | 5427 | 6043 | 6331 | 6319 | 7305 | 6622 | 6697 | Hars |
| Hars2 | 4270 | 4049 | 4303 | 4657 | 4194 | 4292 | 4442 | 4849 | 4337 | 4611 | 4870 | 4716 | 4661 | 4999 | 4175 | 4016 | 4028 | 4527 | 4387 | 4151 | 4304 | 5155 | 4272 | 4783 | 4902 | 5092 | 4914 | 4945 | Hars2 |
| Has1 | 1 | 0 | 0 | 2 | 2 | 0 | 0 | 0 | 1 | 3 | 0 | 2 | 0 | 0 | 0 | 2 | 0 | 1 | 2 | 2 | 1 | 0 | 1 | 0 | 10 | 0 | 1 | 0 | Has1 |
| Has2 | 653 | 367 | 412 | 295 | 339 | 376 | 282 | 1093 | 1494 | 946 | 1519 | 554 | 1235 | 531 | 616 | 650 | 417 | 350 | 261 | 304 | 276 | 956 | 1295 | 751 | 591 | 1052 | 542 | 598 | Has2 |
| Has3 | 676 | 511 | 384 | 654 | 1192 | 1145 | 1340 | 1341 | 1362 | 1261 | 1019 | 1145 | 458 | 1199 | 489 | 351 | 717 | 835 | 1253 | 941 | 1215 | 1437 | 1076 | 1296 | 1512 | 927 | 703 | 777 | Has3 |

| | | | | | | | | | | | | | | | | | | | | | | | | | | | | | |
|---|---|---|---|---|---|---|---|---|---|---|---|---|---|---|---|---|---|---|---|---|---|---|---|---|---|---|---|---|---|
| Haspin | 1435 | 1249 | 1340 | 1289 | 808 | 975 | 1171 | 712 | 691 | 705 | 702 | 915 | 913 | 1043 | 1245 | 1363 | 1278 | 1027 | 1077 | 905 | 1032 | 570 | 604 | 730 | 743 | 1197 | 919 | 992 | Haspin |
| Hat1 | 11220 | 9112 | 10554 | 8680 | 6397 | 7684 | 10081 | 7224 | 5988 | 5622 | 5011 | 7689 | 6149 | 8394 | 11247 | 9009 | 9732 | 8547 | 8493 | 10087 | 8624 | 4938 | 5384 | 6646 | 5857 | 9991 | 8218 | 8838 | Hat1 |
| Haus1 | 518 | 89 | 97 | 393 | 100 | 110 | 95 | 78 | 59 | 514 | 137 | 272 | 591 | 115 | 480 | 131 | 771 | 348 | 493 | 158 | 209 | 488 | 80 | 83 | 54 | 513 | 306 | 424 | Haus1 |
| Haus2 | 3579 | 2940 | 3166 | 2842 | 2610 | 2994 | 2920 | 2994 | 2647 | 2366 | 2410 | 3352 | 3040 | 3236 | 3422 | 2912 | 2818 | 2662 | 2749 | 3005 | 2855 | 2513 | 2681 | 2786 | 2648 | 3395 | 3307 | 3402 | Haus2 |
| Haus3 | 3383 | 2419 | 3157 | 2675 | 2117 | 2339 | 2776 | 2292 | 1801 | 1794 | 1660 | 2442 | 1987 | 2529 | 3231 | 3132 | 2899 | 2616 | 2300 | 2879 | 2621 | 1760 | 1927 | 2131 | 1821 | 2607 | 2496 | 2726 | Haus3 |
| Haus4 | 1411 | 1426 | 1289 | 1258 | 1216 | 1190 | 1452 | 1034 | 1168 | 982 | 1091 | 1489 | 1251 | 1297 | 1226 | 1216 | 1407 | 1205 | 1233 | 1335 | 1191 | 839 | 1082 | 1232 | 1046 | 1689 | 1470 | 1481 | Haus4 |
| Haus5 | 239 | 204 | 234 | 242 | 142 | 196 | 268 | 211 | 188 | 216 | 189 | 207 | 169 | 289 | 227 | 148 | 254 | 158 | 218 | 188 | 242 | 139 | 114 | 201 | 201 | 199 | 253 | 267 | Haus5 |
| Haus6 | 9465 | 7216 | 7719 | 7303 | 6029 | 7511 | 8295 | 5135 | 4618 | 4498 | 4470 | 6643 | 5939 | 6729 | 8542 | 7959 | 7157 | 6934 | 6322 | 7581 | 7128 | 4219 | 4095 | 5151 | 4971 | 6181 | 7096 | 6939 | Haus6 |
| Haus7 | 2706 | 2543 | 2527 | 2194 | 2247 | 2241 | 2265 | 2501 | 2182 | 2188 | 2042 | 2531 | 2327 | 2636 | 2527 | 2401 | 2299 | 2123 | 2303 | 2500 | 2112 | 2206 | 2061 | 2346 | 2383 | 2795 | 2499 | 2779 | Haus7 |
| Haus8 | 1813 | 1989 | 1732 | 1720 | 1661 | 1673 | 1720 | 1804 | 1889 | 1761 | 1730 | 1890 | 1812 | 1916 | 1659 | 1638 | 1880 | 1688 | 1960 | 1699 | 1651 | 1693 | 1764 | 1888 | 1955 | 2098 | 2024 | 1911 | Haus8 |
| Havcr1 | 0 | 0 | 5 | 0 | 0 | 0 | 3 | 2 | 5 | 0 | 19 | 1 | 2 | 1 | 0 | 1 | 0 | 0 | 0 | 3 | 0 | 0 | 8 | 2 | 0 | 1 | 2 | 0 | Havcr1 |
| Havcr2 | 54 | 38 | 66 | 90 | 43 | 66 | 57 | 40 | 36 | 35 | 41 | 49 | 39 | 48 | 98 | 55 | 61 | 58 | 61 | 56 | 60 | 55 | 32 | 29 | 40 | 56 | 35 | 33 | Havcr2 |
| Hax1 | 2855 | 2989 | 3162 | 3099 | 2620 | 2876 | 2920 | 3623 | 3477 | 3280 | 3315 | 3753 | 3569 | 4093 | 2695 | 2630 | 2994 | 2718 | 3151 | 3255 | 2862 | 2956 | 3368 | 3884 | 3368 | 4230 | 3975 | 4391 | Hax1 |
| Hba-a1 | 21548 | 38861 | 24379 | 30206 | 20882 | 36170 | 30623 | 25794 | 61007 | 27031 | 30807 | 23502 | 96020 | 24391 | 28396 | 35423 | 20272 | 36246 | 44593 | 58980 | 19475 | 21744 | 64180 | 22695 | 37705 | 28864 | 39341 | 43913 | Hba-a1 |
| Hba-a2 | 1623 | 2841 | 1279 | 1248 | 1155 | 1416 | 3427 | 1334 | 3966 | 2476 | 1697 | 1390 | 4555 | 3447 | 2291 | 1065 | 1417 | 1695 | 1502 | 3009 | 2725 | 1669 | 4337 | 2170 | 2305 | 737 | 1703 | 4937 | Hba-a2 |
| Hba-x | 20307 | 46426 | 38562 | 15859 | 6568 | 10481 | 11453 | 8673 | 16500 | 11541 | 9768 | 16892 | 66921 | 36133 | 22219 | 36913 | 22275 | 10519 | 10970 | 16553 | 7135 | 5452 | 17267 | 8974 | 9980 | 26256 | 30100 | 85468 | Hba-x |
| Hbb-bh1 | 10355 | 13663 | 16522 | 3214 | 897 | 2101 | 1685 | 2379 | 2359 | 1918 | 1254 | 5168 | 20418 | 14831 | 9351 | 15114 | 7565 | 1362 | 1422 | 3180 | 973 | 916 | 2240 | 1399 | 1267 | 8951 | 9324 | 39491 | Hbb-bh1 |
| Hbb-bh2 | 0 | 5 | 10 | 0 | 2 | 2 | 6 | 1 | 4 | 3 | 2 | 4 | 6 | 5 | 3 | 2 | 0 | 2 | 2 | 6 | 2 | 0 | 4 | 4 | 0 | 1 | 5 | 7 | Hbb-bh2 |
| Hbb-bs | 2653 | 4545 | 1970 | 7662 | 9552 | 12230 | 18810 | 11130 | 33642 | 12368 | 13265 | 3855 | 12410 | 4399 | 4460 | 3606 | 2086 | 12997 | 16657 | 23429 | 12240 | 9274 | 34685 | 8959 | 16988 | 3262 | 4359 | 5708 | Hbb-bs |
| Hbb-bt | 3353 | 4355 | 3815 | 5115 | 4915 | 6320 | 10413 | 6709 | 13042 | 7542 | 6375 | 4109 | 12225 | 5836 | 4495 | 4477 | 3089 | 7142 | 8684 | 12457 | 7702 | 4610 | 15687 | 5569 | 8957 | 4265 | 5143 | 11166 | Hbb-bt |
| Hbb-y | 120673 | 182802 | 140702 | 107429 | 52658 | 77832 | 107296 | 98152 | 134913 | 103672 | 80826 | 99017 | 353904 | 161345 | 150138 | 135259 | 108038 | 82915 | 95332 | 148225 | 64596 | 53773 | 153808 | 74427 | 89905 | 124072 | 146702 | 331227 | Hbb-y |
| Hbegf | 526 | 453 | 483 | 539 | 707 | 591 | 580 | 413 | 378 | 428 | 515 | 348 | 288 | 310 | 588 | 517 | 503 | 618 | 660 | 547 | 609 | 508 | 530 | 414 | 484 | 418 | 285 | 306 | Hbegf |
| Hbp1 | 6760 | 5377 | 5278 | 4793 | 4878 | 5302 | 5586 | 9086 | 8330 | 6869 | 7440 | 7021 | 7557 | 7233 | 6597 | 5964 | 5045 | 4897 | 5189 | 6076 | 5369 | 7157 | 8644 | 7473 | 6772 | 7420 | 7340 | 7456 | Hbp1 |
| Hbq1a | 2 | 16 | 5 | 7 | 7 | 0 | 12 | 9 | 6 | 15 | 5 | 14 | 16 | 13 | 9 | 2 | 2 | 7 | 4 | 15 | 10 | 10 | 19 | 2 | 2 | 2 | 6 | 17 | Hbq1a |
| Hbq1b | 29 | 30 | 39 | 17 | 13 | 23 | 36 | 21 | 42 | 20 | 39 | 36 | 145 | 60 | 36 | 39 | 27 | 35 | 12 | 27 | 13 | 20 | 44 | 23 | 15 | 56 | 65 | 96 | Hbq1b |
| Hbs1l | 8346 | 7935 | 7976 | 7152 | 7041 | 7188 | 8094 | 7377 | 6882 | 7054 | 6858 | 8092 | 7469 | 8338 | 8333 | 7685 | 7723 | 7175 | 7238 | 7274 | 7857 | 7220 | 6765 | 7389 | 7438 | 7762 | 8092 | 8139 | Hbs1l |
| Hc | 22 | 11 | 9 | 16 | 11 | 25 | 28 | 21 | 21 | 20 | 12 | 29 | 37 | 37 | 26 | 12 | 13 | 25 | 10 | 10 | 16 | 23 | 21 | 35 | 38 | 39 | 20 | 38 | Hc |
| Hcar1 | 8 | 11 | 21 | 6 | 11 | 5 | 9 | 20 | 46 | 49 | 35 | 25 | 27 | 19 | 4 | 5 | 7 | 4 | 7 | 5 | 4 | 47 | 58 | 26 | 22 | 17 | 15 | 14 | Hcar1 |
| Hcar2 | 0 | 1 | 0 | 0 | 0 | 0 | 0 | 1 | 10 | 4 | 2 | 0 | 7 | 0 | 0 | 2 | 0 | 1 | 0 | 0 | 0 | 0 | 9 | 1 | 1 | 0 | 1 | 1 | Hcar2 |
| Hccs | 3692 | 2936 | 3487 | 3274 | 2998 | 3104 | 3530 | 2994 | 2394 | 1988 | 2240 | 2757 | 2623 | 2663 | 3911 | 3419 | 3436 | 3310 | 3203 | 3590 | 3187 | 2440 | 2432 | 2562 | 2357 | 2844 | 2839 | 2976 | Hccs |
| Hcfc1 | 9672 | 10347 | 11478 | 10673 | 12104 | 11362 | 10275 | 19098 | 22385 | 24706 | 22338 | 32108 | 26333 | 27654 | 9141 | 9838 | 12140 | 10599 | 12994 | 9276 | 10629 | 24638 | 19540 | 26833 | 25495 | 20767 | 26040 | 27823 | Hcfc1 |
| Hcfc1r1 | 1486 | 2126 | 1613 | 1686 | 1958 | 1959 | 1456 | 1335 | 1763 | 1649 | 1689 | 1158 | 1555 | 1142 | 1528 | 1953 | 1716 | 1973 | 1947 | 1961 | 1528 | 1636 | 1745 | 1521 | 1636 | 1427 | 1228 | 1168 | Hcfc1r1 |
| Hcfc2 | 3328 | 2988 | 2665 | 3035 | 2855 | 2904 | 3122 | 3380 | 3034 | 2840 | 3243 | 2790 | 2768 | 3088 | 3361 | 3036 | 2815 | 2928 | 2825 | 3013 | 2917 | 3579 | 3108 | 3058 | 3165 | 2698 | 2817 | 2989 | Hcfc2 |
| Hck | 110 | 88 | 117 | 44 | 73 | 74 | 58 | 134 | 143 | 81 | 139 | 85 | 98 | 77 | 83 | 136 | 96 | 76 | 53 | 85 | 64 | 82 | 152 | 94 | 76 | 98 | 100 | 77 | Hck |
| Hcls1 | 136 | 150 | 206 | 181 | 121 | 159 | 113 | 167 | 214 | 186 | 268 | 210 | 209 | 234 | 157 | 185 | 167 | 149 | 107 | 132 | 124 | 159 | 327 | 225 | 193 | 290 | 206 | 274 | Hcls1 |
| Hcn1 | 134 | 68 | 63 | 162 | 300 | 120 | 129 | 254 | 279 | 289 | 297 | 70 | 135 | 95 | 169 | 124 | 72 | 176 | 110 | 152 | 167 | 329 | 256 | 142 | 170 | 136 | 83 | 81 | Hcn1 |
| Hcn2 | 476 | 793 | 813 | 1027 | 1116 | 813 | 796 | 147 | 194 | 265 | 205 | 137 | 120 | 111 | 497 | 665 | 934 | 1158 | 1065 | 720 | 906 | 307 | 192 | 226 | 213 | 154 | 78 | 127 | Hcn2 |
| Hcn3 | 1968 | 3163 | 3048 | 4798 | 4857 | 3629 | 3718 | 1479 | 1604 | 2273 | 1926 | 1793 | 1550 | 1621 | 2203 | 2459 | 4101 | 4581 | 4666 | 3374 | 4009 | 2542 | 1550 | 1977 | 2292 | 1425 | 1599 | 1435 | Hcn3 |
| Hcn4 | 1377 | 1482 | 2318 | 2671 | 3355 | 2824 | 2429 | 2052 | 2075 | 2970 | 2237 | 2535 | 2197 | 2543 | 1536 | 1485 | 2555 | 2605 | 3402 | 2310 | 2839 | 3477 | 2012 | 2730 | 3345 | 2089 | 2164 | 2458 | Hcn4 |
| Hcrt | 0 | 0 | 2 | 0 | 0 | 0 | 1 | 0 | 2 | 3 | 1 | 0 | 0 | 0 | 0 | 1 | 2 | 0 | 6 | 0 | 3 | 0 | 1 | 2 | 3 | 0 | 0 | 0 | Hcrt |
| Hcrtr1 | 10 | 22 | 13 | 12 | 20 | 14 | 11 | 23 | 28 | 21 | 18 | 8 | 51 | 22 | 25 | 30 | 25 | 21 | 17 | 28 | 14 | 18 | 17 | 15 | 12 | 11 | 19 | 13 | Hcrtr1 |
| Hcrtr2 | 80 | 70 | 82 | 85 | 165 | 106 | 146 | 192 | 161 | 110 | 105 | 78 | 105 | 96 | 96 | 88 | 104 | 96 | 132 | 146 | 141 | 206 | 146 | 96 | 131 | 73 | 86 | 122 | Hcrtr2 |
| Hcst | 6 | 11 | 23 | 39 | 13 | 38 | 17 | 18 | 11 | 17 | 18 | 18 | 8 | 14 | 22 | 17 | 25 | 22 | 12 | 25 | 23 | 25 | 11 | 16 | 14 | 26 | 15 | 13 | Hcst |
| Hdac1 | 16523 | 17237 | 18104 | 15783 | 12994 | 14043 | 14549 | 8821 | 8861 | 8066 | 8348 | 10460 | 10057 | 10619 | 15923 | 16173 | 17051 | 14901 | 15061 | 15164 | 14121 | 7346 | 8914 | 9868 | 8272 | 13321 | 11269 | 11794 | Hdac1 |
| Hdac10 | 912 | 1094 | 1043 | 1148 | 1241 | 1173 | 1136 | 1255 | 1505 | 1498 | 1379 | 1597 | 1306 | 1458 | 1055 | 1018 | 1203 | 1212 | 1290 | 1188 | 1115 | 1432 | 1424 | 1610 | 1615 | 1402 | 1479 | 1364 | Hdac10 |
| Hdac11 | 3934 | 5275 | 4403 | 5768 | 5648 | 5156 | 4988 | 3331 | 3475 | 3881 | 3733 | 3277 | 3206 | 3259 | 4122 | 4508 | 5552 | 6015 | 6104 | 5170 | 5232 | 4187 | 3769 | 3488 | 3747 | 3229 | 3174 | 3108 | Hdac11 |
| Hdac2 | 25962 | 22019 | 24828 | 24348 | 24700 | 23455 | 28994 | 35418 | 28719 | 28712 | 25268 | 29856 | 27025 | 33355 | 28300 | 21973 | 25678 | 23618 | 25591 | 27221 | 27262 | 30798 | 28017 | 30332 | 29638 | 32547 | 28651 | 32401 | Hdac2 |
| Hdac3 | 12064 | 12048 | 10374 | 11186 | 9755 | 9851 | 10598 | 9221 | 8426 | 7930 | 7721 | 9127 | 8570 | 8958 | 11782 | 11081 | 10467 | 10620 | 10459 | 10372 | 10346 | 8763 | 8418 | 8532 | 9105 | 9972 | 9216 | 9142 | Hdac3 |
| Hdac4 | 2385 | 2661 | 2641 | 3008 | 3175 | 2691 | 2753 | 2213 | 2505 | 2861 | 2845 | 3012 | 2826 | 2994 | 2155 | 2560 | 2772 | 2945 | 3216 | 2210 | 2906 | 2806 | 2537 | 2852 | 3008 | 2521 | 2973 | 2968 | Hdac4 |
| Hdac5 | 7665 | 11130 | 10908 | 12939 | 11191 | 10558 | 9344 | 5759 | 6642 | 7699 | 6410 | 7198 | 6451 | 6170 | 8147 | 9252 | 12407 | 12537 | 12514 | 9235 | 10595 | 7123 | 6297 | 8130 | 7787 | 6928 | 6553 | 6271 | Hdac5 |
| Hdac6 | 11978 | 14137 | 12518 | 13547 | 12508 | 12848 | 12979 | 7286 | 7598 | 8104 | 7234 | 9754 | 8565 | 9350 | 11213 | 10944 | 13318 | 12735 | 13637 | 11846 | 12744 | 8522 | 6957 | 8175 | 8397 | 9163 | 9713 | 9390 | Hdac6 |
| Hdac7 | 2209 | 2702 | 3592 | 2743 | 3287 | 2904 | 2402 | 4421 | 5606 | 6631 | 6340 | 6357 | 6199 | 5518 | 1993 | 2882 | 3432 | 2836 | 3451 | 2514 | 2524 | 6507 | 5342 | 6336 | 6243 | 6125 | 5655 | 5165 | Hdac7 |
| Hdac8 | 1482 | 1360 | 1342 | 1350 | 1347 | 1262 | 1292 | 1354 | 1199 | 1240 | 1022 | 1214 | 1162 | 1260 | 1420 | 1322 | 1401 | 1251 | 1333 | 1427 | 1310 | 1105 | 1255 | 1203 | 1211 | 1490 | 1177 | 1290 | Hdac8 |
| Hdac9 | 1294 | 1113 | 1431 | 1117 | 1253 | 1324 | 1245 | 1807 | 1681 | 1607 | 1600 | 1745 | 1854 | 1742 | 1365 | 1398 | 1247 | 1270 | 1441 | 1265 | 1331 | 1719 | 1759 | 1527 | 1659 | 1509 | 2001 | 1790 | Hdac9 |
| Hdc | 57 | 49 | 24 | 22 | 26 | 64 | 17 | 89 | 136 | 103 | 140 | 102 | 82 | 40 | 50 | 48 | 12 | 47 | 22 | 29 | 27 | 94 | 188 | 95 | 87 | 72 | 52 | 45 | Hdc |
| Hddc2 | 1375 | 1117 | 1067 | 1025 | 911 | 1050 | 1113 | 1282 | 1458 | 1075 | 1164 | 1239 | 1438 | 1329 | 1331 | 1262 | 1058 | 939 | 1016 | 1301 | 1007 | 984 | 1219 | 1226 | 1134 | 1516 | 1307 | 1473 | Hddc2 |
| Hddc3 | 779 | 757 | 737 | 596 | 645 | 671 | 740 | 1031 | 1149 | 1035 | 934 | 1054 | 1041 | 1044 | 799 | 692 | 683 | 576 | 717 | 716 | 679 | 904 | 1113 | 992 | 860 | 1132 | 984 | 1158 | Hddc3 |
| Hdgf | 20724 | 19990 | 22708 | 19606 | 17493 | 18118 | 19769 | 17700 | 18161 | 19382 | 17180 | 21768 | 19351 | 24027 | 18924 | 17782 | 21709 | 18107 | 19984 | 18195 | 19543 | 16660 | 17442 | 22400 | 19928 | 27312 | 20948 | 24194 | Hdgf |
| Hdgfl1 | 0 | 0 | 0 | 0 | 0 | 0 | 0 | 0 | 1 | 0 | 0 | 0 | 0 | 0 | 0 | 0 | 0 | 0 | 0 | 0 | 3 | 0 | 1 | 0 | 0 | 2 | 0 | 0 | Hdgfl1 |
| Hdgfl2 | 10920 | 13611 | 15222 | 15426 | 11713 | 11812 | 12006 | 5881 | 6744 | 8042 | 7399 | 9359 | 7952 | 8747 | 10235 | 12714 | 15317 | 14519 | 13596 | 10068 | 11919 | 7090 | 6499 | 8925 | 8506 | 10255 | 8279 | 9054 | Hdgfl2 |
| Hdgfl3 | 11651 | 8452 | 10185 | 12617 | 16908 | 15925 | 17417 | 26768 | 21609 | 22900 | 20228 | 22733 | 19707 | 24665 | 13028 | 8826 | 11483 | 13152 | 15037 | 16844 | 16948 | 26034 | 20370 | 21723 | 25738 | 18465 | 21651 | 22219 | Hdgfl3 |
| Hdhd2 | 768 | 860 | 568 | 693 | 738 | 792 | 878 | 655 | 597 | 650 | 570 | 690 | 607 | 678 | 822 | 736 | 801 | 769 | 747 | 828 | 877 | 670 | 638 | 624 | 689 | 658 | 679 | 700 | Hdhd2 |
| Hdhd3 | 433 | 519 | 538 | 487 | 413 | 506 | 422 | 254 | 251 | 216 | 284 | 223 | 277 | 242 | 434 | 514 | 480 | 484 | 467 | 498 | 354 | 189 | 324 | 226 | 205 | 291 | 335 | 227 | Hdhd3 |
| Hdhd5 | 1892 | 2262 | 1974 | 2226 | 1978 | 1998 | 1941 | 1434 | 1525 | 1435 | 1521 | 1529 | 1565 | 1307 | 1865 | 2069 | 1941 | 2096 | 2065 | 1992 | 1871 | 1419 | 1454 | 1382 | 1650 | 1418 | 1452 | 1386 | Hdhd5 |
| Hdlbp | 31231 | 29504 | 22321 | 19268 | 17931 | 21899 | 17907 | 31530 | 32740 | 27838 | 32197 | 27440 | 29518 | 25789 | 28506 | 29080 | 20941 | 20375 | 18826 | 20641 | 18140 | 26580 | 34662 | 26937 | 25730 | 24479 | 29936 | 27537 | Hdlbp |
| Hdx | 1067 | 950 | 743 | 1074 | 1245 | 1196 | 1297 | 2168 | 1821 | 1552 | 1508 | 1647 | 1761 | 1729 | 1286 | 1002 | 935 | 1094 | 1211 | 1289 | 1295 | 1821 | 1718 | 1201 | 1736 | 1194 | 1826 | 1619 | Hdx |
| Heatr1 | 13005 | 13369 | 8048 | 8817 | 7271 | 9055 | 9388 | 7654 | 6956 | 5702 | 5856 | 7037 | 6599 | 6704 | 12088 | 11372 | 8369 | 8305 | 8065 | 10210 | 9418 | 5045 | 6445 | 5496 | 5897 | 5222 | 7422 | 7030 | Heatr1 |
| Heatr3 | 5004 | 4340 | 4373 | 3866 | 3706 | 4037 | 4194 | 4106 | 3976 | 3240 | 3897 | 3921 | 3995 | 3971 | 4762 | 4379 | 3727 | 4010 | 3829 | 4064 | 4014 | 3853 | 4112 | 3402 | 3686 | 4039 | 4071 | 4287 | Heatr3 |
| Heatr4 | 1 | 0 | 0 | 0 | 1 | 3 | 0 | 0 | 2 | 4 | 3 | 4 | 3 | 0 | 1 | 0 | 0 | 2 | 0 | 5 | 0 | 5 | 0 | 2 | 0 | 0 | 0 | 2 | Heatr4 |
| Heatr5a | 7013 | 7094 | 5070 | 5047 | 4523 | 5634 | 5195 | 4097 | 4641 | 3910 | 4649 | 4452 | 4284 | 4240 | 6576 | 6976 | 5066 | 5620 | 4736 | 5201 | 5109 | 3798 | 4860 | 3742 | 3989 | 3258 | 4509 | 4314 | Heatr5a |
| Heatr5b | 6108 | 7002 | 3975 | 5471 | 5761 | 6169 | 5803 | 4963 | 5118 | 4689 | 4871 | 4322 | 4265 | 4451 | 5900 | 5900 | 4617 | 5560 | 5380 | 5857 | 6075 | 4490 | 4725 | 4005 | 5161 | 2601 | 4621 | 3970 | Heatr5b |
| Heatr6 | 2445 | 2599 | 2787 | 2996 | 3310 | 2927 | 3129 | 4186 | 4393 | 4731 | 4256 | 5008 | 4647 | 4909 | 2488 | 2321 | 2965 | 2684 | 3192 | 2856 | 3178 | 4725 | 3879 | 4714 | 4943 | 4067 | 4799 | 4685 | Heatr6 |
| Hebp1 | 513 | 608 | 748 | 501 | 367 | 422 | 358 | 247 | 400 | 272 | 373 | 277 | 389 | 266 | 492 | 739 | 512 | 505 | 359 | 449 | 329 | 244 | 506 | 284 | 257 | 285 | 309 | 408 | Hebp1 |
| Hebp2 | 428 | 487 | 593 | 323 | 348 | 378 | 291 | 363 | 459 | 344 | 409 | 392 | 450 | 321 | 480 | 494 | 450 | 293 | 346 | 357 | 243 | 334 | 518 | 305 | 237 | 342 | 330 | 319 | Hebp2 |
| Heca | 5352 | 4230 | 6058 | 6698 | 6709 | 6528 | 7169 | 3751 | 3537 | 4415 | 4239 | 3700 | 3740 | 3776 | 5407 | 5233 | 5948 | 6888 | 6639 | 5600 | 6852 | 4311 | 3756 | 3945 | 4562 | 3570 | 3539 | 3823 | Heca |
| Hectd1 | 11247 | 9577 | 6387 | 7445 | 9588 | 10267 | 9676 | 21837 | 20144 | 16275 | 17870 | 17530 | 17892 | 17523 | 10969 | 10111 | 6747 | 7703 | 8273 | 10667 | 9388 | 15796 | 19024 | 13628 | 17374 | 10792 | 18576 | 16140 | Hectd1 |
| Hectd2 | 2000 | 1469 | 1324 | 2142 | 2648 | 2762 | 3256 | 2933 | 2824 | 2106 | 2180 | 2319 | 2402 | 2449 | 2246 | 1577 | 1530 | 2458 | 2507 | 3076 | 3037 | 2379 | 2262 | 2036 | 2535 | 1647 | 2037 | 1989 | Hectd2 |
| Hectd3 | 5971 | 7246 | 6539 | 8041 | 7103 | 6944 | 6830 | 3519 | 3883 | 3892 | 3589 | 3763 | 3447 | 3380 | 5636 | 6378 | 7230 | 7869 | 8015 | 6250 | 7493 | 4338 | 3624 | 3898 | 4117 | 3382 | 3288 | 3425 | Hectd3 |
| Hectd4 | 12019 | 15484 | 9854 | 17736 | 22522 | 20360 | 19133 | 13216 | 16198 | 18959 | 16461 | 15862 | 14488 | 15270 | 11718 | 13511 | 13320 | 19623 | 20098 | 16668 | 21654 | 17523 | 13663 | 12624 | 21936 | 7513 | 15190 | 12597 | Hectd4 |

| | | | | | | | | | | | | | | | | | | | | | | | | | | | | | |
|---|---|---|---|---|---|---|---|---|---|---|---|---|---|---|---|---|---|---|---|---|---|---|---|---|---|---|---|---|---|
| Hecw1 | 1889 | 1789 | 963 | 2753 | 4937 | 4586 | 4679 | 7767 | 7523 | 7136 | 6076 | 4744 | 4714 | 5273 | 2422 | 1650 | 1874 | 3256 | 3998 | 4399 | 4886 | 7780 | 6376 | 4834 | 8109 | 2648 | 4394 | 4043 | Hecw1 |
| Hecw2 | 648 | 695 | 682 | 684 | 899 | 768 | 608 | 1955 | 1813 | 1796 | 1844 | 1366 | 1865 | 1388 | 808 | 916 | 625 | 642 | 692 | 725 | 665 | 1800 | 1922 | 1331 | 1599 | 980 | 1554 | 1346 | Hecw2 |
| Heg1 | 3844 | 3768 | 5094 | 4351 | 4359 | 4883 | 4235 | 5061 | 5122 | 6222 | 6154 | 9517 | 8039 | 8496 | 3895 | 3712 | 4824 | 4199 | 4880 | 3691 | 4410 | 5589 | 4946 | 7017 | 7034 | 7071 | 10101 | 9184 | Heg1 |
| Helb | 3136 | 3328 | 2918 | 2929 | 2470 | 2726 | 2600 | 902 | 1101 | 950 | 1006 | 996 | 1010 | 1026 | 2901 | 3124 | 2800 | 2923 | 2553 | 2683 | 2627 | 807 | 1113 | 1058 | 916 | 1044 | 1065 | 998 | Helb |
| Hells | 10615 | 7008 | 9799 | 7563 | 6815 | 8420 | 9269 | 8465 | 7259 | 6675 | 6443 | 10465 | 8908 | 10651 | 10138 | 8620 | 8442 | 7155 | 7871 | 9424 | 7998 | 5355 | 6064 | 8065 | 6859 | 11677 | 11252 | 10765 | Hells |
| Helq | 2746 | 2844 | 2232 | 2686 | 2610 | 2754 | 2912 | 1950 | 1945 | 1937 | 1938 | 2208 | 1988 | 2144 | 2633 | 2653 | 2432 | 2681 | 2406 | 2819 | 2646 | 1841 | 1808 | 1829 | 2152 | 1748 | 2292 | 2004 | Helq |
| Helt | 121 | 170 | 300 | 169 | 126 | 169 | 106 | 220 | 80 | 190 | 158 | 443 | 356 | 428 | 123 | 75 | 234 | 157 | 214 | 128 | 155 | 241 | 87 | 306 | 255 | 207 | 522 | 517 | Helt |
| Helz | 6915 | 6398 | 7177 | 7866 | 7987 | 7893 | 8351 | 4946 | 5409 | 6583 | 6571 | 6328 | 6098 | 6210 | 6192 | 7319 | 7394 | 8337 | 8034 | 6288 | 8225 | 6139 | 5088 | 5730 | 6833 | 4830 | 6004 | 5637 | Helz |
| Helz2 | 674 | 780 | 572 | 523 | 549 | 580 | 324 | 388 | 555 | 563 | 771 | 464 | 678 | 410 | 566 | 1013 | 589 | 618 | 439 | 475 | 445 | 578 | 610 | 580 | 456 | 330 | 557 | 440 | Helz2 |
| Hemgn | 39 | 46 | 112 | 3 | 1 | 11 | 4 | 24 | 33 | 23 | 37 | 75 | 117 | 142 | 36 | 83 | 21 | 12 | 6 | 17 | 4 | 12 | 27 | 24 | 7 | 93 | 79 | 460 | Hemgn |
| Hemk1 | 867 | 960 | 1065 | 1151 | 1191 | 1246 | 1131 | 1418 | 1538 | 1791 | 1721 | 1667 | 1648 | 1484 | 984 | 953 | 1158 | 1155 | 1194 | 1118 | 1000 | 1734 | 1493 | 1621 | 1708 | 1042 | 1635 | 1441 | Hemk1 |
| Henmt1 | 23 | 26 | 27 | 28 | 37 | 23 | 21 | 24 | 23 | 36 | 65 | 23 | 39 | 39 | 9 | 34 | 29 | 26 | 26 | 25 | 21 | 37 | 40 | 26 | 27 | 24 | 33 | 42 | Henmt1 |
| Hepacam | 16 | 28 | 5 | 39 | 120 | 62 | 34 | 82 | 147 | 157 | 150 | 24 | 37 | 26 | 18 | 26 | 3 | 52 | 55 | 40 | 43 | 190 | 182 | 42 | 93 | 19 | 26 | 18 | Hepacam |
| Hepacam2 | 107 | 92 | 142 | 116 | 123 | 89 | 118 | 156 | 123 | 94 | 105 | 139 | 128 | 159 | 164 | 90 | 148 | 132 | 164 | 177 | 127 | 165 | 95 | 157 | 153 | 92 | 220 | 199 | Hepacam2 |
| Heph | 3069 | 2367 | 2279 | 1269 | 1042 | 1543 | 948 | 2782 | 2811 | 2019 | 2714 | 2103 | 2973 | 2047 | 2624 | 2948 | 1633 | 1412 | 956 | 1298 | 906 | 1873 | 3330 | 2101 | 1579 | 2473 | 2487 | 2314 | Heph |
| Hephl1 | 2 | 0 | 1 | 2 | 0 | 0 | 2 | 0 | 0 | 3 | 8 | 0 | 0 | 0 | 3 | 0 | 0 | 0 | 0 | 0 | 0 | 0 | 6 | 0 | 0 | 0 | 0 | 0 | Hephl1 |
| Herc1 | 33625 | 32586 | 14790 | 21076 | 24826 | 33130 | 28168 | 31142 | 27971 | 18703 | 19517 | 19049 | 19159 | 20181 | 35313 | 30985 | 17399 | 20915 | 22222 | 36933 | 32726 | 17246 | 25150 | 15078 | 22229 | 11580 | 23625 | 17279 | Herc1 |
| Herc2 | 23023 | 23866 | 11013 | 18842 | 22805 | 27685 | 23646 | 19583 | 21012 | 17158 | 20245 | 14709 | 16187 | 15619 | 22879 | 23126 | 13983 | 19140 | 19231 | 25032 | 29319 | 14608 | 17960 | 12358 | 20680 | 8322 | 18827 | 13092 | Herc2 |
| Herc3 | 4074 | 3941 | 3212 | 4557 | 5526 | 5025 | 5487 | 2645 | 2689 | 2713 | 2446 | 2348 | 2412 | 2346 | 4332 | 3805 | 3877 | 4956 | 4795 | 5067 | 5674 | 2949 | 2469 | 2266 | 2941 | 2003 | 2337 | 2225 | Herc3 |
| Herc4 | 7829 | 5739 | 6406 | 5780 | 5323 | 6134 | 6257 | 6166 | 5118 | 4690 | 4805 | 5414 | 5412 | 5643 | 6807 | 6389 | 6067 | 5639 | 5570 | 6329 | 5769 | 5048 | 5209 | 5103 | 4875 | 5388 | 5678 | 5633 | Herc4 |
| Herc6 | 175 | 124 | 178 | 132 | 131 | 181 | 175 | 158 | 190 | 160 | 202 | 213 | 245 | 235 | 171 | 156 | 160 | 127 | 150 | 157 | 134 | 152 | 181 | 216 | 141 | 218 | 261 | 204 | Herc6 |
| Herpud1 | 1494 | 1851 | 1624 | 1708 | 1559 | 1652 | 1465 | 1537 | 1696 | 1621 | 1763 | 1485 | 1477 | 1478 | 1542 | 1563 | 1775 | 1587 | 1699 | 1689 | 1616 | 1559 | 1783 | 1651 | 1582 | 1612 | 1546 | 1481 | Herpud1 |
| Herpud2 | 3760 | 3129 | 3934 | 4261 | 3700 | 3819 | 3965 | 3915 | 3374 | 3767 | 3645 | 4480 | 3842 | 4297 | 3513 | 3303 | 4070 | 3925 | 3783 | 3391 | 4028 | 4060 | 3498 | 4050 | 4022 | 4182 | 3904 | 4352 | Herpud2 |
| Hes1 | 1270 | 1212 | 1293 | 1388 | 1105 | 989 | 1258 | 1573 | 1657 | 1637 | 1864 | 2148 | 1826 | 2235 | 1130 | 1145 | 1150 | 1252 | 1162 | 1201 | 1205 | 1312 | 1816 | 2303 | 1647 | 2827 | 1772 | 2332 | Hes1 |
| Hes2 | 20 | 10 | 24 | 22 | 7 | 28 | 10 | 23 | 20 | 16 | 6 | 36 | 69 | 28 | 16 | 26 | 37 | 13 | 12 | 13 | 3 | 10 | 3 | 15 | 11 | 5 | 48 | 18 | Hes2 |
| Hes3 | 8 | 22 | 43 | 4 | 7 | 12 | 8 | 13 | 12 | 14 | 17 | 63 | 67 | 24 | 4 | 26 | 36 | 4 | 7 | 4 | 6 | 27 | 19 | 11 | 24 | 19 | 78 | 70 | Hes3 |
| Hes5 | 1653 | 2803 | 4519 | 2579 | 2295 | 2100 | 1676 | 2060 | 1760 | 2550 | 1786 | 4825 | 3880 | 4732 | 1904 | 1489 | 4719 | 1810 | 3027 | 1895 | 2227 | 2498 | 1758 | 3827 | 3253 | 4446 | 5051 | 5458 | Hes5 |
| Hes6 | 3587 | 5517 | 5237 | 4730 | 4210 | 4212 | 4133 | 4865 | 4531 | 4787 | 4580 | 6650 | 6175 | 6398 | 4133 | 3724 | 5862 | 3949 | 5114 | 4792 | 4194 | 4461 | 4598 | 6317 | 5654 | 6697 | 7283 | 6878 | Hes6 |
| Hes7 | 269 | 284 | 269 | 290 | 213 | 244 | 187 | 20 | 12 | 23 | 26 | 10 | 11 | 9 | 193 | 287 | 362 | 263 | 225 | 174 | 217 | 25 | 27 | 19 | 6 | 16 | 24 | 18 | Hes7 |
| Hesx1 | 14 | 3 | 3 | 1 | 4 | 1 | 0 | 0 | 7 | 0 | 4 | 4 | 1 | 3 | 8 | 8 | 2 | 2 | 0 | 0 | 0 | 7 | 5 | 2 | 1 | 10 | 5 | 4 | Hesx1 |
| Hexa | 2575 | 2978 | 2745 | 2974 | 2757 | 2726 | 2424 | 2339 | 2843 | 2888 | 3516 | 2651 | 2885 | 2583 | 2666 | 3204 | 2851 | 3178 | 2755 | 2352 | 2578 | 3098 | 3350 | 2871 | 2750 | 2923 | 2672 | 2574 | Hexa |
| Hexb | 1989 | 1864 | 1866 | 1790 | 1905 | 1904 | 1589 | 1297 | 1549 | 1395 | 1666 | 1164 | 1194 | 1224 | 1985 | 2151 | 1633 | 2118 | 1688 | 1710 | 1742 | 1470 | 1814 | 1158 | 1237 | 1206 | 1210 | 1190 | Hexb |
| Hexdc | 1611 | 1843 | 1601 | 1948 | 2035 | 1951 | 1906 | 1541 | 1771 | 1929 | 1761 | 1840 | 1708 | 1924 | 1480 | 1616 | 1765 | 2032 | 1860 | 1846 | 1902 | 1811 | 1652 | 1902 | 2010 | 1786 | 1669 | 1712 | Hexdc |
| Hexim1 | 3205 | 4001 | 3905 | 4202 | 3674 | 3335 | 3381 | 2805 | 3372 | 3540 | 3259 | 3126 | 3607 | 3131 | 2942 | 3403 | 4117 | 3921 | 3929 | 3731 | 3299 | 2854 | 3242 | 3698 | 3246 | 3764 | 3297 | 3201 | Hexim1 |
| Hexim2 | 236 | 231 | 235 | 278 | 345 | 331 | 354 | 459 | 469 | 530 | 427 | 365 | 371 | 404 | 243 | 188 | 246 | 307 | 343 | 337 | 365 | 444 | 477 | 401 | 521 | 419 | 385 | 430 | Hexim2 |
| Hey1 | 2587 | 2336 | 2393 | 2306 | 2258 | 2034 | 2367 | 2911 | 2283 | 2649 | 2162 | 2965 | 3112 | 3185 | 2238 | 2237 | 2225 | 2058 | 2491 | 2334 | 2166 | 1963 | 2207 | 3099 | 2533 | 4233 | 2936 | 3336 | Hey1 |
| Hey2 | 1288 | 879 | 900 | 745 | 893 | 792 | 702 | 1498 | 1405 | 1079 | 1041 | 867 | 1048 | 842 | 1006 | 1293 | 712 | 903 | 531 | 689 | 628 | 907 | 1504 | 1001 | 775 | 1052 | 782 | 774 | Hey2 |
| Heyl | 550 | 721 | 784 | 545 | 787 | 473 | 525 | 818 | 892 | 892 | 1229 | 1002 | 1199 | 926 | 625 | 959 | 735 | 671 | 524 | 379 | 539 | 1158 | 1146 | 937 | 862 | 1124 | 937 | 965 | Heyl |
| Hfe | 211 | 230 | 233 | 158 | 199 | 164 | 124 | 197 | 249 | 170 | 225 | 241 | 262 | 167 | 249 | 280 | 141 | 155 | 133 | 159 | 155 | 266 | 310 | 127 | 163 | 189 | 156 | 151 | Hfe |
| Hfm1 | 738 | 637 | 612 | 632 | 839 | 812 | 817 | 1552 | 1281 | 1198 | 1156 | 1248 | 1281 | 1430 | 897 | 725 | 588 | 715 | 755 | 1053 | 899 | 1357 | 1225 | 1003 | 1406 | 915 | 1379 | 1167 | Hfm1 |
| Hgd | 3 | 13 | 9 | 6 | 1 | 8 | 7 | 9 | 12 | 1 | 11 | 10 | 15 | 12 | 11 | 6 | 2 | 10 | 15 | 9 | 6 | 5 | 5 | 19 | 5 | 7 | 22 | 13 | Hgd |
| Hgf | 144 | 59 | 56 | 51 | 34 | 18 | 12 | 161 | 212 | 129 | 198 | 80 | 113 | 50 | 80 | 92 | 34 | 8 | 19 | 13 | 28 | 80 | 194 | 129 | 76 | 122 | 37 | 66 | Hgf |
| Hgfac | 90 | 163 | 147 | 173 | 124 | 148 | 103 | 77 | 91 | 119 | 93 | 79 | 95 | 51 | 131 | 179 | 158 | 182 | 146 | 153 | 122 | 85 | 95 | 96 | 79 | 66 | 89 | 64 | Hgfac |
| Hgh1 | 1053 | 1268 | 1213 | 1477 | 1196 | 1080 | 959 | 896 | 894 | 967 | 1035 | 1038 | 949 | 881 | 1019 | 1125 | 1234 | 1335 | 1303 | 1055 | 1237 | 897 | 927 | 1053 | 930 | 1170 | 1037 | 1031 | Hgh1 |
| Hgs | 4705 | 6228 | 6318 | 6963 | 5919 | 5408 | 5254 | 4256 | 5100 | 6105 | 5245 | 6100 | 5129 | 5286 | 4770 | 5474 | 6587 | 6776 | 6604 | 4742 | 5811 | 5997 | 5001 | 6106 | 6200 | 6111 | 5072 | 5270 | Hgs |
| Hgsnat | 2579 | 2567 | 1993 | 2024 | 1816 | 2102 | 1885 | 1704 | 1989 | 1685 | 1718 | 1636 | 2014 | 1748 | 2607 | 2569 | 1812 | 1990 | 1941 | 1963 | 1841 | 1663 | 2073 | 1618 | 1721 | 1748 | 1764 | 1870 | Hgsnat |
| Hhat | 91 | 78 | 85 | 54 | 43 | 76 | 42 | 311 | 279 | 270 | 368 | 316 | 366 | 351 | 86 | 115 | 81 | 50 | 57 | 46 | 43 | 294 | 322 | 290 | 297 | 394 | 427 | 322 | Hhat |
| Hhatl | 45 | 62 | 35 | 98 | 45 | 43 | 42 | 6 | 15 | 34 | 35 | 8 | 15 | 1 | 24 | 43 | 73 | 81 | 56 | 39 | 51 | 17 | 27 | 11 | 3 | 7 | 6 | 9 | Hhatl |
| Hhex | 12 | 7 | 14 | 20 | 23 | 17 | 8 | 17 | 5 | 4 | 8 | 4 | 14 | 6 | 8 | 16 | 8 | 12 | 8 | 7 | 10 | 13 | 16 | 5 | 7 | 16 | 8 | 14 | Hhex |
| Hhip | 766 | 668 | 540 | 454 | 542 | 475 | 560 | 667 | 840 | 470 | 884 | 563 | 454 | 570 | 741 | 728 | 447 | 532 | 401 | 501 | 468 | 530 | 680 | 408 | 390 | 463 | 551 | 540 | Hhip |
| Hhipl1 | 264 | 320 | 543 | 280 | 376 | 386 | 310 | 490 | 521 | 575 | 517 | 994 | 872 | 868 | 276 | 299 | 435 | 320 | 424 | 300 | 327 | 580 | 520 | 808 | 769 | 784 | 972 | 996 | Hhipl1 |
| Hhipl2 | 152 | 275 | 188 | 343 | 239 | 236 | 234 | 82 | 71 | 100 | 63 | 70 | 45 | 81 | 210 | 236 | 220 | 364 | 282 | 211 | 270 | 70 | 60 | 71 | 68 | 73 | 67 | 61 | Hhipl2 |
| Hhla1 | 7 | 0 | 0 | 0 | 1 | 4 | 3 | 0 | 8 | 1 | 2 | 0 | 3 | 3 | 4 | 6 | 0 | 0 | 1 | 3 | 3 | 0 | 2 | 0 | 6 | 0 | 0 | 0 | Hhla1 |
| Hibadh | 3706 | 3208 | 3278 | 3317 | 2879 | 3063 | 3144 | 2646 | 2548 | 2286 | 2490 | 2312 | 2353 | 2380 | 3592 | 3267 | 3237 | 3381 | 2900 | 2785 | 3083 | 2237 | 2679 | 2236 | 2197 | 2467 | 2473 | 2478 | Hibadh |
| Hibch | 2712 | 2602 | 2765 | 2497 | 2380 | 2446 | 2615 | 2099 | 1857 | 1806 | 1796 | 1702 | 2053 | 1921 | 3013 | 2608 | 2646 | 2590 | 2524 | 2900 | 2405 | 1861 | 1838 | 1751 | 1745 | 1816 | 1937 | 1975 | Hibch |
| Hic1 | 354 | 335 | 244 | 251 | 391 | 282 | 135 | 710 | 1321 | 945 | 2726 | 534 | 828 | 503 | 302 | 389 | 261 | 331 | 271 | 189 | 180 | 800 | 1926 | 974 | 653 | 915 | 519 | 481 | Hic1 |
| Hic2 | 2847 | 3066 | 3640 | 3234 | 2690 | 2685 | 2463 | 2284 | 2421 | 2498 | 2570 | 3383 | 3216 | 3237 | 2450 | 3124 | 3493 | 2593 | 2922 | 2230 | 2472 | 2801 | 2230 | 2862 | 2966 | 2755 | 3793 | 3696 | Hic2 |
| Hid1 | 4613 | 6447 | 5870 | 7022 | 6827 | 6269 | 6156 | 2262 | 2450 | 2910 | 2324 | 2808 | 2413 | 2530 | 4989 | 5058 | 6806 | 6737 | 7555 | 5763 | 6421 | 3399 | 2319 | 2972 | 3051 | 2693 | 2604 | 2605 | Hid1 |
| Hif1a | 11671 | 8532 | 9992 | 9518 | 10429 | 10551 | 11044 | 17144 | 14700 | 13358 | 13865 | 15077 | 15111 | 16087 | 12059 | 10220 | 9889 | 9595 | 9910 | 9864 | 10808 | 14917 | 14573 | 14755 | 14046 | 16051 | 15682 | 16010 | Hif1a |
| Hif1an | 8394 | 8391 | 7013 | 7776 | 6237 | 6901 | 6631 | 7706 | 7439 | 7609 | 7728 | 8359 | 7969 | 8281 | 7779 | 7625 | 7198 | 7495 | 7077 | 6353 | 7150 | 7528 | 7502 | 7609 | 8132 | 6604 | 8849 | 8655 | Hif1an |
| Hif3a | 3636 | 3545 | 3230 | 3577 | 3221 | 3310 | 3366 | 2226 | 2760 | 2839 | 2939 | 2402 | 2707 | 2340 | 3182 | 3791 | 3123 | 3821 | 2995 | 2656 | 3062 | 2074 | 2802 | 2696 | 1834 | 2622 | 2054 | 2174 | Hif3a |
| Higd1a | 4170 | 4486 | 4945 | 4659 | 4002 | 3707 | 4834 | 2959 | 2636 | 2830 | 3115 | 2934 | 2973 | 3380 | 4418 | 3827 | 4891 | 4328 | 4285 | 4818 | 4135 | 2488 | 2590 | 3166 | 2839 | 3384 | 3368 | 3607 | Higd1a |
| Higd1b | 3 | 1 | 0 | 5 | 19 | 1 | 10 | 12 | 16 | 18 | 13 | 2 | 0 | 2 | 0 | 2 | 0 | 4 | 23 | 8 | 26 | 17 | 19 | 16 | 9 | 4 | 2 | 2 | Higd1b |
| Higd2a | 1444 | 1869 | 1854 | 1919 | 1720 | 1474 | 1916 | 1519 | 1657 | 1750 | 1632 | 1757 | 1677 | 1870 | 1524 | 1471 | 1958 | 1776 | 1715 | 1871 | 1861 | 1385 | 1597 | 1916 | 1784 | 1971 | 1689 | 1873 | Higd2a |
| Hikeshi | 2718 | 2534 | 2999 | 2862 | 2785 | 2898 | 3187 | 2637 | 2307 | 2430 | 2436 | 2569 | 2519 | 2703 | 3125 | 2651 | 3065 | 2799 | 3052 | 3158 | 2869 | 2364 | 2358 | 2665 | 2583 | 2890 | 2530 | 2782 | Hikeshi |
| Hilpda | 712 | 682 | 694 | 605 | 646 | 649 | 630 | 708 | 635 | 609 | 682 | 661 | 596 | 597 | 673 | 660 | 744 | 655 | 739 | 712 | 534 | 550 | 669 | 665 | 626 | 708 | 674 | 598 | Hilpda |
| Hinfp | 6525 | 7154 | 4543 | 6581 | 5107 | 5673 | 5719 | 4450 | 3621 | 3541 | 3769 | 4085 | 3920 | 4022 | 6686 | 6606 | 5394 | 6597 | 4968 | 5697 | 5898 | 3989 | 3502 | 3426 | 4223 | 2718 | 4288 | 3828 | Hinfp |
| Hint1 | 6871 | 7550 | 7123 | 7511 | 7553 | 7086 | 8618 | 7559 | 7412 | 7629 | 7182 | 8020 | 7344 | 9108 | 6906 | 6057 | 7675 | 6609 | 7967 | 8713 | 8416 | 6204 | 7436 | 8523 | 8021 | 9549 | 8484 | 9768 | Hint1 |
| Hint2 | 794 | 806 | 813 | 844 | 670 | 627 | 666 | 545 | 604 | 680 | 603 | 641 | 697 | 764 | 687 | 650 | 714 | 681 | 701 | 647 | 724 | 555 | 642 | 637 | 610 | 754 | 660 | 770 | Hint2 |
| Hint3 | 359 | 292 | 292 | 261 | 321 | 411 | 350 | 568 | 558 | 501 | 625 | 458 | 481 | 519 | 320 | 257 | 187 | 294 | 286 | 405 | 398 | 461 | 568 | 452 | 596 | 315 | 514 | 470 | Hint3 |
| Hip1 | 13432 | 13842 | 13762 | 12379 | 14655 | 12518 | 12579 | 13163 | 14183 | 15057 | 16690 | 16367 | 17542 | 16086 | 13991 | 14166 | 14004 | 13317 | 13523 | 10927 | 12790 | 17114 | 13553 | 13873 | 15016 | 11957 | 16069 | 16515 | Hip1 |
| Hip1r | 3460 | 4481 | 4238 | 5721 | 5830 | 5218 | 4982 | 1818 | 2378 | 2721 | 2134 | 2035 | 1992 | 1904 | 3423 | 4118 | 5328 | 5656 | 6316 | 4767 | 5512 | 2349 | 2232 | 2359 | 2645 | 1935 | 1744 | 1932 | Hip1r |
| Hipk1 | 13040 | 12447 | 11698 | 13865 | 13769 | 14194 | 14685 | 10273 | 9708 | 9717 | 9981 | 11155 | 11374 | 11456 | 12722 | 12744 | 12675 | 14039 | 13889 | 12279 | 15166 | 10835 | 9313 | 9451 | 11350 | 8347 | 11240 | 11439 | Hipk1 |
| Hipk2 | 3849 | 4034 | 867 | 3446 | 4959 | 5208 | 3739 | 5293 | 5562 | 5645 | 5749 | 5440 | 6141 | 5071 | 3798 | 3853 | 1635 | 3341 | 4057 | 3839 | 4659 | 3721 | 4930 | 2053 | 7206 | 770 | 7346 | 3936 | Hipk2 |
| Hipk3 | 2498 | 1787 | 2234 | 1973 | 2756 | 2533 | 2162 | 4571 | 4833 | 4175 | 5974 | 4073 | 4566 | 3858 | 2389 | 2579 | 2011 | 2469 | 2218 | 2069 | 2263 | 4796 | 5649 | 3884 | 4146 | 3447 | 4112 | 3979 | Hipk3 |
| Hipk4 | 146 | 147 | 161 | 181 | 128 | 180 | 152 | 120 | 156 | 193 | 156 | 186 | 181 | 170 | 127 | 165 | 178 | 222 | 181 | 131 | 130 | 147 | 146 | 209 | 131 | 214 | 202 | 153 | Hipk4 |
| Hira | 1516 | 1475 | 2277 | 2271 | 2051 | 1923 | 2048 | 1412 | 1319 | 1701 | 1627 | 1929 | 1862 | 2071 | 1333 | 1276 | 2159 | 2060 | 2159 | 1596 | 2054 | 1569 | 1190 | 2147 | 2002 | 2271 | 1922 | 2085 | Hira |
| Hirip3 | 3403 | 3311 | 4501 | 3408 | 3336 | 3400 | 3819 | 5868 | 6067 | 7030 | 6450 | 9364 | 7638 | 10173 | 3104 | 2602 | 4196 | 3148 | 4111 | 3800 | 3634 | 5157 | 5387 | 9243 | 7787 | 12407 | 10091 | 9960 | Hirip3 |
| Hivep1 | 7727 | 8443 | 7414 | 10051 | 10411 | 9532 | 9340 | 5699 | 5869 | 7644 | 6863 | 8063 | 7100 | 7465 | 7783 | 8058 | 8903 | 9873 | 10429 | 7507 | 9768 | 7979 | 5647 | 6916 | 7774 | 4607 | 6847 | 6435 | Hivep1 |

| | | | | | | | | | | | | | | | | | | | | | | | | | | | | | |
|---|---|---|---|---|---|---|---|---|---|---|---|---|---|---|---|---|---|---|---|---|---|---|---|---|---|---|---|---|---|
| Hivep2 | 1869 | 1981 | 1464 | 2224 | 3395 | 3095 | 2981 | 3215 | 3776 | 3873 | 4790 | 2930 | 3512 | 2799 | 2024 | 2012 | 1881 | 2588 | 2845 | 2372 | 3187 | 4171 | 3933 | 2643 | 4397 | 1524 | 3140 | 2530 | Hivep2 |
| Hivep3 | 496 | 669 | 519 | 920 | 1372 | 735 | 840 | 673 | 880 | 1044 | 1055 | 510 | 686 | 493 | 549 | 585 | 693 | 1072 | 1121 | 657 | 979 | 1440 | 668 | 695 | 1205 | 233 | 367 | 368 | Hivep3 |
| Hjurp | 20633 | 19758 | 16866 | 17067 | 16318 | 18384 | 16936 | 26216 | 26632 | 24628 | 27085 | 29932 | 28738 | 28525 | 17951 | 20590 | 16439 | 16537 | 17362 | 17903 | 18130 | 20954 | 25388 | 24814 | 25758 | 25481 | 31606 | 28660 | Hjurp |
| Hjv | 77 | 42 | 68 | 51 | 41 | 29 | 25 | 39 | 67 | 27 | 585 | 26 | 45 | 18 | 45 | 58 | 57 | 25 | 52 | 42 | 41 | 75 | 361 | 25 | 30 | 31 | 9 | 23 | Hjv |
| Hk1 | 10269 | 11274 | 11430 | 12450 | 12603 | 11185 | 10747 | 5409 | 6657 | 7286 | 6730 | 5346 | 5700 | 5253 | 10123 | 11099 | 11953 | 13137 | 12571 | 9606 | 11749 | 8101 | 6331 | 6293 | 7026 | 5418 | 4951 | 5187 | Hk1 |
| Hk2 | 5169 | 5513 | 5995 | 5721 | 3855 | 4729 | 4622 | 1884 | 2071 | 2269 | 2304 | 2876 | 2355 | 2935 | 4685 | 4932 | 5646 | 5291 | 4810 | 4036 | 4649 | 1395 | 1910 | 2710 | 2174 | 3197 | 2922 | 3343 | Hk2 |
| Hk3 | 97 | 112 | 108 | 56 | 83 | 84 | 74 | 67 | 128 | 133 | 180 | 110 | 111 | 93 | 95 | 129 | 73 | 117 | 81 | 89 | 108 | 104 | 148 | 106 | 139 | 98 | 122 | 91 | Hk3 |
| Hkdc1 | 30 | 22 | 28 | 17 | 42 | 28 | 10 | 37 | 62 | 63 | 94 | 52 | 101 | 28 | 17 | 55 | 21 | 22 | 12 | 21 | 19 | 72 | 128 | 28 | 36 | 33 | 59 | 46 | Hkdc1 |
| Hlcs | 1195 | 1177 | 1578 | 1198 | 1087 | 1055 | 1134 | 1025 | 1155 | 1253 | 1189 | 1565 | 1592 | 1459 | 1017 | 1132 | 1421 | 1311 | 1166 | 920 | 1014 | 1101 | 1065 | 1508 | 1170 | 1578 | 1471 | 1613 | Hlcs |
| Hlf | 1521 | 1475 | 1696 | 1865 | 1568 | 1541 | 1682 | 821 | 1018 | 882 | 1110 | 942 | 988 | 958 | 1551 | 1749 | 1604 | 2076 | 1397 | 1402 | 1771 | 1198 | 1115 | 1092 | 1174 | 849 | 941 | 816 | Hlf |
| Hltf | 5567 | 4523 | 4933 | 4967 | 4786 | 5201 | 5673 | 4835 | 3927 | 3705 | 3543 | 4038 | 3755 | 4059 | 5914 | 5256 | 4755 | 5042 | 4729 | 5677 | 5354 | 4151 | 3846 | 4041 | 3829 | 3481 | 3814 | 3888 | Hltf |
| Hlx | 63 | 108 | 87 | 97 | 74 | 43 | 47 | 71 | 133 | 101 | 204 | 108 | 99 | 78 | 86 | 102 | 107 | 79 | 73 | 59 | 63 | 110 | 228 | 113 | 111 | 106 | 77 | 78 | Hlx |
| Hmbox1 | 5456 | 5687 | 4347 | 5211 | 5970 | 6058 | 5442 | 7233 | 7404 | 6779 | 7444 | 7761 | 7649 | 7211 | 5203 | 5393 | 4857 | 5192 | 5764 | 5493 | 5728 | 6624 | 6684 | 6077 | 7921 | 5879 | 8455 | 6760 | Hmbox1 |
| Hmbs | 1284 | 1475 | 1416 | 850 | 695 | 764 | 810 | 1860 | 2022 | 1626 | 1633 | 2152 | 2461 | 2363 | 1246 | 1446 | 1020 | 824 | 880 | 851 | 784 | 1592 | 2027 | 1814 | 1569 | 2561 | 2350 | 3037 | Hmbs |
| Hmces | 2070 | 1841 | 2027 | 1979 | 1653 | 1765 | 1874 | 1502 | 1373 | 1466 | 1332 | 1526 | 1331 | 1573 | 1742 | 1768 | 1955 | 1783 | 1975 | 1689 | 1857 | 1353 | 1377 | 1609 | 1469 | 1953 | 1557 | 1670 | Hmces |
| Hmcn1 | 8538 | 6245 | 2276 | 1988 | 1028 | 2925 | 565 | 6014 | 7364 | 3771 | 6097 | 2688 | 4656 | 2808 | 6152 | 11108 | 1736 | 2430 | 723 | 1155 | 811 | 2558 | 7218 | 3425 | 2283 | 3214 | 3857 | 2808 | Hmcn1 |
| Hmcn2 | 20 | 35 | 38 | 55 | 85 | 36 | 54 | 33 | 27 | 54 | 45 | 43 | 40 | 28 | 30 | 58 | 22 | 63 | 31 | 44 | 59 | 67 | 48 | 31 | 40 | 28 | 48 | 18 | Hmcn2 |
| Hmg20a | 4885 | 4644 | 4411 | 4538 | 5468 | 5119 | 5111 | 9064 | 8829 | 8839 | 9196 | 8780 | 8386 | 9207 | 4917 | 4707 | 4551 | 4788 | 5032 | 4997 | 5511 | 9633 | 8447 | 8644 | 9072 | 7898 | 8910 | 8651 | Hmg20a |
| Hmg20b | 4334 | 5809 | 6521 | 5835 | 4246 | 4162 | 3913 | 2275 | 2648 | 2939 | 2880 | 3687 | 3371 | 3232 | 3998 | 5037 | 6283 | 5119 | 5068 | 3734 | 4061 | 2341 | 2549 | 3505 | 2691 | 4137 | 3383 | 3329 | Hmg20b |
| Hmga1 | 6685 | 8066 | 9242 | 5982 | 4858 | 5362 | 4958 | 4418 | 5311 | 4966 | 4746 | 6683 | 5998 | 6112 | 5519 | 7197 | 8039 | 5357 | 6200 | 5150 | 4950 | 3855 | 4997 | 5946 | 4551 | 8759 | 6817 | 6966 | Hmga1 |
| Hmga1b | 865 | 1029 | 1404 | 778 | 623 | 518 | 594 | 567 | 549 | 717 | 602 | 896 | 696 | 1021 | 696 | 706 | 1207 | 683 | 782 | 485 | 693 | 418 | 481 | 895 | 666 | 1110 | 784 | 940 | Hmga1b |
| Hmga2 | 11267 | 6431 | 8824 | 3765 | 2289 | 3204 | 3024 | 7723 | 8259 | 6303 | 8133 | 10183 | 12559 | 10511 | 7796 | 9576 | 5913 | 3153 | 2800 | 2989 | 2239 | 4453 | 6223 | 7430 | 4265 | 19064 | 10478 | 12384 | Hmga2 |
| Hmgb1 | 26150 | 17244 | 27746 | 21482 | 17392 | 20961 | 23191 | 29990 | 25738 | 25111 | 24941 | 33006 | 33140 | 35326 | 24586 | 22506 | 20531 | 19175 | 20427 | 28438 | 18964 | 22586 | 26433 | 28683 | 28246 | 44125 | 36678 | 35339 | Hmgb1 |
| Hmgb2 | 20159 | 19232 | 26155 | 19295 | 14223 | 17056 | 19762 | 24730 | 22874 | 22341 | 20874 | 34427 | 29130 | 40192 | 20892 | 17243 | 21529 | 15417 | 18711 | 22893 | 17982 | 16115 | 19533 | 30824 | 26066 | 46248 | 37809 | 41314 | Hmgb2 |
| Hmgb3 | 14572 | 13755 | 16810 | 15116 | 14207 | 15556 | 16477 | 20250 | 17159 | 19350 | 16649 | 26070 | 22769 | 28332 | 14984 | 12079 | 15662 | 13291 | 15401 | 16755 | 15708 | 18700 | 15488 | 23517 | 22869 | 30494 | 26884 | 28227 | Hmgb3 |
| Hmgb4 | 1 | 0 | 0 | 0 | 0 | 0 | 0 | 1 | 1 | 0 | 0 | 0 | 0 | 1 | 0 | 0 | 0 | 0 | 0 | 0 | 0 | 2 | 0 | 0 | 1 | 0 | 0 | 2 | Hmgb4 |
| Hmgcl | 2611 | 3265 | 3036 | 2813 | 2317 | 2620 | 2615 | 1165 | 1472 | 1338 | 1458 | 1439 | 1279 | 1490 | 2378 | 3016 | 2866 | 2915 | 2506 | 2330 | 2592 | 1345 | 1423 | 1518 | 1382 | 1483 | 1427 | 1486 | Hmgcl |
| Hmgcll1 | 569 | 532 | 593 | 580 | 908 | 768 | 907 | 1933 | 1626 | 1823 | 1489 | 1721 | 1598 | 1703 | 762 | 583 | 621 | 666 | 742 | 883 | 778 | 2358 | 1531 | 1640 | 1932 | 1652 | 1670 | 1691 | Hmgcll1 |
| Hmgcr | 25100 | 28218 | 23000 | 23227 | 20706 | 22025 | 24055 | 22280 | 18402 | 18194 | 16476 | 21590 | 21383 | 24145 | 26703 | 24017 | 23008 | 23071 | 20591 | 23976 | 22563 | 19502 | 17129 | 18917 | 21762 | 19010 | 24634 | 24404 | Hmgcr |
| Hmgcs1 | 48992 | 54488 | 57148 | 51671 | 49139 | 48707 | 51419 | 44781 | 35608 | 37607 | 33524 | 45963 | 47175 | 49537 | 54268 | 51072 | 54001 | 49164 | 45586 | 52602 | 47740 | 40503 | 33778 | 39339 | 42640 | 44211 | 52801 | 52677 | Hmgcs1 |
| Hmgcs2 | 257 | 241 | 220 | 192 | 465 | 311 | 239 | 556 | 808 | 538 | 723 | 387 | 634 | 393 | 293 | 308 | 148 | 316 | 260 | 420 | 292 | 770 | 1036 | 333 | 465 | 243 | 431 | 389 | Hmgcs2 |
| Hmgn1 | 31661 | 25780 | 31088 | 26340 | 22808 | 23860 | 25786 | 29703 | 25870 | 27814 | 27434 | 33394 | 30550 | 33985 | 29715 | 26003 | 29251 | 25211 | 25476 | 25984 | 23815 | 25217 | 25347 | 33064 | 29125 | 39730 | 33865 | 34803 | Hmgn1 |
| Hmgn2 | 30238 | 25703 | 30186 | 23172 | 20675 | 24012 | 28455 | 33895 | 32145 | 31097 | 30050 | 42381 | 36855 | 45215 | 28710 | 25220 | 26480 | 21421 | 24157 | 26518 | 25857 | 27718 | 31739 | 37444 | 33265 | 54905 | 44510 | 47841 | Hmgn2 |
| Hmgn3 | 5511 | 4441 | 5537 | 5298 | 5105 | 4919 | 6389 | 3640 | 3148 | 3451 | 3038 | 3401 | 3151 | 4092 | 5504 | 4547 | 5057 | 5283 | 5444 | 6073 | 5919 | 2836 | 3185 | 3759 | 3483 | 4678 | 3493 | 3814 | Hmgn3 |
| Hmgn5 | 6374 | 5009 | 9370 | 7297 | 5854 | 6734 | 7897 | 4748 | 3695 | 4972 | 3532 | 6057 | 4899 | 7126 | 7097 | 5629 | 8456 | 6764 | 7208 | 7886 | 6807 | 3606 | 3586 | 5985 | 5194 | 7698 | 6255 | 6640 | Hmgn5 |
| Hmgxb3 | 3805 | 4159 | 4338 | 4507 | 4275 | 3986 | 3816 | 3269 | 3591 | 3911 | 3832 | 4023 | 3828 | 3730 | 3465 | 3994 | 4614 | 4387 | 4563 | 3505 | 4190 | 4040 | 3469 | 3832 | 4002 | 3691 | 3870 | 4010 | Hmgxb3 |
| Hmgxb4 | 3549 | 2683 | 3040 | 2813 | 2933 | 3008 | 3245 | 5995 | 5436 | 4781 | 5121 | 5587 | 5376 | 5883 | 3268 | 2644 | 2796 | 2586 | 2998 | 3371 | 2929 | 5041 | 5032 | 5176 | 5029 | 6097 | 5836 | 5809 | Hmgxb4 |
| Hmmr | 5976 | 4741 | 5270 | 3635 | 2906 | 3715 | 4041 | 5517 | 4662 | 4093 | 3980 | 6126 | 5110 | 6424 | 5496 | 4857 | 4595 | 3355 | 4039 | 4753 | 3685 | 3245 | 4369 | 5275 | 4093 | 6868 | 6623 | 6363 | Hmmr |
| Hmox1 | 1539 | 1694 | 1627 | 1119 | 1122 | 1226 | 1052 | 850 | 1183 | 815 | 1133 | 841 | 1052 | 822 | 1483 | 1685 | 1263 | 1226 | 1095 | 1230 | 1000 | 817 | 1573 | 849 | 775 | 946 | 803 | 770 | Hmox1 |
| Hmox2 | 4263 | 4181 | 3731 | 3768 | 3579 | 3877 | 3786 | 3844 | 3586 | 3260 | 3528 | 3219 | 3379 | 3380 | 4639 | 3964 | 3680 | 3735 | 3755 | 4087 | 3873 | 3283 | 3943 | 3455 | 3389 | 3891 | 3402 | 3501 | Hmox2 |
| Hmx1 | 123 | 172 | 69 | 58 | 88 | 84 | 19 | 167 | 206 | 124 | 320 | 26 | 249 | 21 | 178 | 160 | 88 | 133 | 30 | 23 | 13 | 206 | 219 | 87 | 113 | 73 | 41 | 34 | Hmx1 |
| Hmx2 | 63 | 77 | 80 | 79 | 125 | 91 | 80 | 111 | 161 | 185 | 157 | 110 | 98 | 148 | 127 | 60 | 66 | 87 | 97 | 76 | 89 | 194 | 121 | 123 | 159 | 128 | 110 | 89 | Hmx2 |
| Hmx3 | 42 | 35 | 21 | 22 | 65 | 39 | 41 | 67 | 79 | 111 | 91 | 64 | 37 | 74 | 75 | 30 | 67 | 41 | 28 | 26 | 47 | 82 | 81 | 83 | 74 | 114 | 40 | 26 | Hmx3 |
| Hnf1a | 1 | 0 | 0 | 1 | 0 | 0 | 0 | 6 | 1 | 0 | 8 | 0 | 1 | 0 | 0 | 0 | 0 | 0 | 1 | 0 | 0 | 2 | 0 | 0 | 0 | 0 | 0 | 1 | Hnf1a |
| Hnf1b | 2 | 4 | 10 | 8 | 2 | 0 | 1 | 4 | 7 | 5 | 1 | 5 | 2 | 4 | 0 | 4 | 2 | 0 | 3 | 0 | 3 | 3 | 8 | 1 | 5 | 3 | 0 | 0 | Hnf1b |
| Hnf4a | 0 | 2 | 0 | 0 | 0 | 1 | 0 | 0 | 6 | 0 | 1 | 0 | 0 | 0 | 0 | 2 | 0 | 0 | 0 | 0 | 0 | 0 | 4 | 0 | 0 | 4 | 0 | 1 | Hnf4a |
| Hnf4g | 7 | 3 | 0 | 1 | 2 | 4 | 4 | 5 | 6 | 3 | 0 | 0 | 1 | 7 | 1 | 4 | 6 | 3 | 11 | 2 | 2 | 5 | 4 | 2 | 0 | 3 | 1 | 9 | Hnf4g |
| Hnmt | 51 | 35 | 50 | 39 | 71 | 73 | 77 | 92 | 97 | 115 | 112 | 94 | 104 | 69 | 46 | 51 | 74 | 53 | 44 | 61 | 73 | 145 | 123 | 87 | 90 | 99 | 91 | 81 | Hnmt |
| Hnrnpa0 | 34238 | 28233 | 45479 | 42884 | 38152 | 34521 | 40971 | 35795 | 31642 | 39965 | 38465 | 52100 | 41809 | 57068 | 31813 | 27545 | 45497 | 38800 | 40064 | 31680 | 41492 | 39664 | 30156 | 51288 | 46057 | 59640 | 45732 | 53330 | Hnrnpa0 |
| Hnrnpa1 | 128502 | 119630 | 158769 | 129255 | 111577 | 114896 | 131419 | 141880 | 124095 | 130202 | 127744 | 160674 | 150207 | 185624 | 131855 | 115834 | 140179 | 115951 | 124016 | 136448 | 123199 | 115901 | 119549 | 155963 | 149430 | 193096 | 172232 | 181273 | Hnrnpa1 |
| Hnrnpa2b1 | 140104 | 117670 | 162938 | 142421 | 134023 | 136128 | 155719 | 175886 | 141584 | 162604 | 158424 | 199603 | 182225 | 224029 | 140610 | 115573 | 155649 | 134284 | 140319 | 148524 | 141893 | 160889 | 136902 | 199212 | 175882 | 230668 | 207156 | 219310 | Hnrnpa2b1 |
| Hnrnpa3 | 21882 | 17354 | 20726 | 19606 | 20846 | 21650 | 24917 | 26826 | 22802 | 21517 | 20683 | 26032 | 23327 | 27570 | 23591 | 20767 | 20028 | 20557 | 20346 | 23523 | 22052 | 24610 | 22151 | 23077 | 24384 | 25360 | 25840 | 26552 | Hnrnpa3 |
| Hnrnpab | 32845 | 30499 | 44120 | 37605 | 36194 | 34734 | 39112 | 46231 | 42758 | 54567 | 49628 | 68235 | 56433 | 71571 | 30882 | 28045 | 41250 | 34459 | 39311 | 32760 | 37613 | 46727 | 40112 | 65832 | 61505 | 80422 | 63022 | 70895 | Hnrnpab |
| Hnrnpc | 43467 | 39849 | 42754 | 38105 | 32195 | 34453 | 42762 | 42812 | 36535 | 35499 | 34201 | 39784 | 36731 | 44538 | 44423 | 37760 | 41360 | 36943 | 36129 | 40802 | 38826 | 36225 | 36555 | 40723 | 36025 | 46080 | 41597 | 45797 | Hnrnpc |
| Hnrnpd | 31680 | 24027 | 45894 | 41534 | 37225 | 34646 | 40746 | 29502 | 26590 | 37106 | 35637 | 48265 | 36782 | 52561 | 29940 | 25654 | 44447 | 38158 | 38867 | 28554 | 40871 | 35227 | 25040 | 45787 | 40262 | 56989 | 40399 | 50262 | Hnrnpd |
| Hnrnpdl | 44394 | 37060 | 42728 | 43463 | 42245 | 45432 | 49691 | 47180 | 40541 | 43438 | 41877 | 56983 | 50613 | 60312 | 45195 | 39635 | 41058 | 42716 | 42850 | 48785 | 47719 | 43042 | 38525 | 47755 | 48867 | 58835 | 57308 | 57002 | Hnrnpdl |
| Hnrnpf | 26069 | 25235 | 31485 | 24131 | 21597 | 24761 | 24284 | 20897 | 22396 | 18665 | 21467 | 26653 | 26541 | 23987 | 25870 | 27053 | 27182 | 24168 | 25549 | 28884 | 21008 | 16121 | 21650 | 23024 | 21689 | 33762 | 29491 | 26801 | Hnrnpf |
| Hnrnph1 | 73623 | 58433 | 78251 | 69139 | 69422 | 72926 | 79353 | 119876 | 100389 | 98114 | 102803 | 127099 | 114710 | 137848 | 75490 | 66354 | 73364 | 64194 | 73131 | 83287 | 74019 | 104407 | 96341 | 118444 | 112842 | 125819 | 129187 | 131335 | Hnrnph1 |
| Hnrnph2 | 14626 | 12653 | 17152 | 15347 | 14094 | 14628 | 16798 | 15516 | 13752 | 13463 | 13642 | 16671 | 15489 | 17966 | 15447 | 13146 | 15705 | 14486 | 14838 | 16549 | 14860 | 14080 | 13738 | 15670 | 15297 | 18944 | 17472 | 17833 | Hnrnph2 |
| Hnrnph3 | 10278 | 8762 | 15835 | 13852 | 12199 | 12213 | 12150 | 14931 | 12310 | 16961 | 14880 | 18735 | 17010 | 19402 | 10772 | 8699 | 14191 | 11565 | 13190 | 12307 | 11630 | 16639 | 11525 | 20029 | 18523 | 21917 | 19098 | 18677 | Hnrnph3 |
| Hnrnpk | 61772 | 57619 | 64862 | 56657 | 54073 | 55902 | 63998 | 79877 | 69576 | 66373 | 65939 | 86512 | 78200 | 92741 | 64092 | 56537 | 62453 | 53023 | 57084 | 64918 | 59627 | 66523 | 67976 | 78910 | 75873 | 97672 | 89299 | 92429 | Hnrnpk |
| Hnrnpl | 18905 | 19788 | 24341 | 21775 | 19362 | 19388 | 20083 | 26813 | 26601 | 29209 | 27406 | 34848 | 30518 | 35854 | 18220 | 18160 | 23079 | 19279 | 21409 | 18682 | 20431 | 26777 | 26161 | 32896 | 29750 | 39004 | 33154 | 35826 | Hnrnpl |
| Hnrnpll | 9398 | 7405 | 9135 | 8646 | 8907 | 9079 | 9945 | 14080 | 11440 | 11357 | 11134 | 14556 | 13101 | 15058 | 9347 | 7989 | 8805 | 8001 | 9556 | 9280 | 9775 | 11949 | 11093 | 13057 | 12851 | 15307 | 14466 | 15622 | Hnrnpll |
| Hnrnpm | 28168 | 25596 | 36336 | 26210 | 22539 | 23621 | 24834 | 36276 | 33064 | 35009 | 33839 | 44786 | 40241 | 44867 | 26843 | 25131 | 31690 | 22423 | 25693 | 25701 | 22637 | 31874 | 31435 | 45321 | 37320 | 53699 | 46314 | 45425 | Hnrnpm |
| Hnrnpr | 24891 | 21007 | 26350 | 25765 | 25995 | 26560 | 29492 | 31671 | 27855 | 29672 | 28800 | 34360 | 31216 | 37167 | 24840 | 22560 | 25683 | 23843 | 26095 | 28030 | 28360 | 27106 | 26532 | 32043 | 32007 | 37238 | 35391 | 36330 | Hnrnpr |
| Hnrnpu | 3666 | 1824 | 2573 | 1955 | 1748 | 2484 | 2209 | 5236 | 2833 | 2791 | 3205 | 4115 | 4316 | 3655 | 3488 | 2120 | 2143 | 1761 | 2143 | 2687 | 1842 | 4431 | 2664 | 3272 | 3430 | 3967 | 4052 | 3902 | Hnrnpu |
| Hnrnpul1 | 13689 | 13771 | 15894 | 14800 | 14345 | 14879 | 14524 | 14996 | 17045 | 17901 | 17525 | 21192 | 18276 | 20307 | 13473 | 12548 | 15783 | 13679 | 15141 | 13833 | 14058 | 15550 | 15533 | 20067 | 18319 | 23224 | 19295 | 20066 | Hnrnpul1 |
| Hnrnpul2 | 23444 | 21310 | 28661 | 29970 | 26912 | 25256 | 28945 | 19554 | 17881 | 22630 | 21485 | 23649 | 21188 | 26327 | 22536 | 20703 | 29143 | 28816 | 26776 | 21816 | 28175 | 22421 | 17109 | 25392 | 24010 | 24204 | 22625 | 24722 | Hnrnpul2 |
| Hoatz | 7 | 18 | 10 | 5 | 14 | 22 | 22 | 28 | 40 | 47 | 59 | 29 | 49 | 22 | 33 | 22 | 6 | 7 | 14 | 27 | 17 | 55 | 87 | 21 | 32 | 18 | 10 | 13 | Hoatz |
| Hoga1 | 67 | 104 | 92 | 52 | 82 | 91 | 54 | 74 | 72 | 99 | 138 | 104 | 84 | 87 | 75 | 109 | 65 | 72 | 90 | 74 | 63 | 109 | 100 | 68 | 52 | 105 | 85 | 93 | Hoga1 |
| Homer1 | 6834 | 5548 | 5807 | 6897 | 6217 | 6994 | 7430 | 4854 | 3780 | 3613 | 3966 | 3937 | 4077 | 4201 | 7230 | 6094 | 6249 | 6160 | 6022 | 7090 | 7178 | 4303 | 3976 | 3703 | 4382 | 3751 | 4000 | 4093 | Homer1 |
| Homer2 | 3571 | 3063 | 2135 | 3369 | 4282 | 4424 | 3997 | 5095 | 4964 | 4968 | 4714 | 4275 | 4458 | 4492 | 3865 | 3317 | 2793 | 3623 | 3702 | 3918 | 3813 | 3740 | 4624 | 4497 | 4825 | 3802 | 4195 | 3940 | Homer2 |
| Homer3 | 2841 | 3302 | 3949 | 3347 | 3481 | 3338 | 2999 | 3413 | 3723 | 3945 | 3714 | 5251 | 4762 | 4767 | 2561 | 2987 | 3950 | 3102 | 4015 | 3121 | 3188 | 3552 | 3687 | 4740 | 4353 | 5977 | 5289 | 5120 | Homer3 |
| Homez | 2299 | 2362 | 2472 | 2528 | 2335 | 2463 | 2349 | 1331 | 1188 | 1230 | 1102 | 1459 | 1271 | 1316 | 2244 | 2129 | 2429 | 2349 | 2247 | 2181 | 2386 | 1358 | 1166 | 1293 | 1251 | 1541 | 1402 | 1392 | Homez |
| Hook1 | 10059 | 7775 | 7085 | 9217 | 9925 | 10259 | 11241 | 9279 | 7583 | 6471 | 6237 | 7623 | 6685 | 7915 | 10693 | 8608 | 7736 | 9185 | 9142 | 12419 | 10690 | 7122 | 6831 | 6231 | 7603 | 5375 | 7426 | 7295 | Hook1 |
| Hook2 | 1512 | 1618 | 1907 | 2186 | 2004 | 1798 | 1882 | 1254 | 1478 | 1713 | 1533 | 1502 | 1382 | 1432 | 1333 | 1431 | 1976 | 2138 | 2068 | 1646 | 1885 | 1628 | 1323 | 1705 | 1580 | 1423 | 1349 | 1449 | Hook2 |
| Hook3 | 19627 | 14447 | 10052 | 12276 | 13694 | 17728 | 16166 | 18802 | 15999 | 11976 | 14352 | 12848 | 14269 | 13922 | 18606 | 16882 | 10721 | 11912 | 12639 | 19559 | 17080 | 12711 | 15404 | 10842 | 14198 | 8970 | 14908 | 12073 | Hook3 |
| Hopx | 322 | 396 | 393 | 500 | 582 | 407 | 560 | 412 | 499 | 504 | 544 | 364 | 446 | 417 | 375 | 295 | 346 | 454 | 429 | 560 | 496 | 338 | 722 | 364 | 383 | 462 | 255 | 320 | Hopx |
| Hormad1 | 2 | 0 | 0 | 0 | 0 | 0 | 0 | 0 | 1 | 2 | 0 | 0 | 0 | 0 | 3 | 0 | 0 | 0 | 0 | 0 | 0 | 0 | 0 | 0 | 0 | 0 | 0 | 0 | Hormad1 |

| | | | | | | | | | | | | | | | | | | | | | | | | | | | | | |
|---|---|---|---|---|---|---|---|---|---|---|---|---|---|---|---|---|---|---|---|---|---|---|---|---|---|---|---|---|---|
| Hormad2 | 8 | 8 | 7 | 13 | 4 | 10 | 10 | 2 | 8 | 8 | 3 | 11 | 7 | 10 | 16 | 16 | 8 | 13 | 7 | 5 | 10 | 13 | 11 | 2 | 17 | 0 | 9 | 11 | Hormad2 |
| Hoxa1 | 48 | 79 | 68 | 13 | 70 | 39 | 7 | 57 | 64 | 41 | 112 | 60 | 95 | 36 | 98 | 93 | 19 | 48 | 14 | 7 | 2 | 27 | 115 | 35 | 20 | 7 | 51 | 43 | Hoxa1 |
| Hoxa10 | 1 | 0 | 1 | 0 | 0 | 0 | 0 | 0 | 0 | 0 | 8 | 0 | 0 | 0 | 0 | 0 | 0 | 0 | 0 | 0 | 0 | 0 | 3 | 0 | 0 | 0 | 0 | 0 | Hoxa10 |
| Hoxa11 | 0 | 1 | 0 | 0 | 0 | 0 | 0 | 0 | 0 | 0 | 0 | 0 | 0 | 0 | 0 | 0 | 0 | 0 | 0 | 0 | 0 | 0 | 1 | 0 | 0 | 0 | 0 | 0 | Hoxa11 |
| Hoxa13 | 14 | 7 | 10 | 9 | 1 | 5 | 4 | 1 | 0 | 0 | 2 | 3 | 3 | 3 | 9 | 6 | 0 | 4 | 6 | 10 | 1 | 0 | 1 | 4 | 0 | 0 | 0 | 0 | Hoxa13 |
| Hoxa2 | 8 | 46 | 9 | 3 | 251 | 57 | 0 | 308 | 396 | 445 | 754 | 12 | 634 | 9 | 162 | 74 | 14 | 28 | 13 | 1 | 0 | 366 | 722 | 11 | 38 | 0 | 7 | 5 | Hoxa2 |
| Hoxa3 | 2 | 8 | 6 | 0 | 457 | 18 | 1 | 241 | 361 | 797 | 1214 | 25 | 316 | 17 | 96 | 30 | 2 | 4 | 4 | 4 | 0 | 404 | 819 | 28 | 10 | 0 | 22 | 24 | Hoxa3 |
| Hoxa4 | 0 | 2 | 6 | 0 | 318 | 6 | 0 | 60 | 178 | 495 | 512 | 0 | 26 | 2 | 13 | 0 | 3 | 1 | 1 | 0 | 4 | 149 | 438 | 0 | 1 | 0 | 0 | 4 | Hoxa4 |
| Hoxa5 | 2 | 3 | 2 | 0 | 1630 | 6 | 1 | 35 | 180 | 1195 | 1892 | 1 | 3 | 3 | 4 | 2 | 10 | 2 | 6 | 2 | 0 | 53 | 1591 | 9 | 2 | 0 | 0 | 1 | Hoxa5 |
| Hoxa6 | 0 | 0 | 0 | 0 | 281 | 2 | 0 | 0 | 2 | 102 | 316 | 0 | 0 | 0 | 0 | 0 | 0 | 0 | 0 | 0 | 0 | 2 | 200 | 0 | 3 | 0 | 0 | 0 | Hoxa6 |
| Hoxa7 | 6 | 5 | 7 | 13 | 38 | 4 | 6 | 1 | 5 | 13 | 440 | 0 | 3 | 0 | 3 | 8 | 5 | 8 | 5 | 2 | 1 | 2 | 21 | 4 | 0 | 1 | 0 | 0 | Hoxa7 |
| Hoxa9 | 0 | 1 | 1 | 2 | 0 | 4 | 3 | 0 | 2 | 1 | 95 | 2 | 0 | 0 | 0 | 2 | 0 | 0 | 0 | 0 | 0 | 0 | 10 | 0 | 0 | 2 | 0 | 0 | Hoxa9 |
| Hoxb1 | 1 | 1 | 2 | 0 | 24 | 2 | 0 | 1 | 1 | 1 | 25 | 0 | 11 | 0 | 12 | 23 | 0 | 1 | 0 | 0 | 0 | 3 | 9 | 0 | 0 | 0 | 0 | 6 | Hoxb1 |
| Hoxb13 | 0 | 0 | 1 | 0 | 0 | 0 | 0 | 0 | 0 | 0 | 0 | 2 | 0 | 0 | 0 | 0 | 0 | 0 | 0 | 0 | 0 | 2 | 1 | 0 | 0 | 0 | 0 | 0 | Hoxb13 |
| Hoxb2 | 2 | 16 | 5 | 6 | 325 | 28 | 0 | 293 | 283 | 415 | 826 | 15 | 532 | 7 | 88 | 55 | 8 | 20 | 6 | 0 | 0 | 359 | 590 | 10 | 50 | 2 | 8 | 5 | Hoxb2 |
| Hoxb3 | 0 | 10 | 4 | 3 | 1845 | 21 | 1 | 1007 | 1259 | 2369 | 3394 | 5 | 1025 | 1 | 253 | 29 | 12 | 19 | 9 | 2 | 0 | 1561 | 2403 | 4 | 44 | 2 | 1 | 2 | Hoxb3 |
| Hoxb4 | 6 | 10 | 20 | 2 | 742 | 7 | 2 | 56 | 209 | 734 | 1051 | 3 | 43 | 2 | 62 | 7 | 9 | 10 | 10 | 12 | 11 | 137 | 656 | 1 | 0 | 4 | 3 | 6 | Hoxb4 |
| Hoxb5 | 0 | 1 | 1 | 0 | 1293 | 0 | 0 | 18 | 147 | 1549 | 1737 | 3 | 0 | 0 | 0 | 0 | 1 | 0 | 0 | 0 | 0 | 57 | 1106 | 0 | 5 | 0 | 0 | 0 | Hoxb5 |
| Hoxb6 | 0 | 0 | 0 | 0 | 497 | 0 | 0 | 0 | 2 | 245 | 1043 | 0 | 0 | 0 | 0 | 0 | 0 | 0 | 0 | 0 | 0 | 0 | 612 | 0 | 0 | 2 | 0 | 0 | Hoxb6 |
| Hoxb7 | 3 | 3 | 0 | 1 | 191 | 0 | 0 | 0 | 1 | 23 | 425 | 0 | 0 | 0 | 0 | 1 | 3 | 9 | 2 | 0 | 2 | 0 | 150 | 0 | 0 | 0 | 0 | 1 | Hoxb7 |
| Hoxb8 | 1 | 2 | 0 | 1 | 764 | 2 | 1 | 1 | 2 | 287 | 2837 | 1 | 0 | 0 | 0 | 3 | 1 | 2 | 2 | 1 | 1 | 0 | 847 | 0 | 0 | 0 | 1 | 4 | Hoxb8 |
| Hoxb9 | 2 | 3 | 1 | 3 | 98 | 4 | 3 | 4 | 8 | 8 | 1892 | 3 | 7 | 6 | 1 | 2 | 0 | 1 | 4 | 0 | 3 | 0 | 32 | 5 | 1 | 15 | 0 | 12 | Hoxb9 |
| Hoxc10 | 0 | 0 | 0 | 0 | 0 | 0 | 1 | 0 | 0 | 0 | 0 | 1 | 0 | 1 | 0 | 0 | 0 | 0 | 0 | 0 | 0 | 0 | 0 | 0 | 0 | 0 | 0 | 0 | Hoxc10 |
| Hoxc12 | 1 | 1 | 0 | 0 | 0 | 1 | 0 | 0 | 2 | 0 | 0 | 0 | 2 | 1 | 1 | 2 | 0 | 2 | 0 | 0 | 0 | 0 | 1 | 0 | 0 | 0 | 0 | 0 | Hoxc12 |
| Hoxc13 | 17 | 8 | 11 | 5 | 9 | 16 | 6 | 29 | 31 | 11 | 17 | 13 | 19 | 13 | 9 | 9 | 12 | 15 | 10 | 5 | 2 | 0 | 36 | 15 | 5 | 0 | 15 | 12 | Hoxc13 |
| Hoxc4 | 0 | 6 | 5 | 0 | 918 | 10 | 1 | 76 | 218 | 814 | 1251 | 3 | 44 | 4 | 7 | 2 | 2 | 11 | 2 | 1 | 2 | 202 | 731 | 3 | 7 | 0 | 2 | 2 | Hoxc4 |
| Hoxc5 | 1 | 6 | 0 | 0 | 358 | 0 | 0 | 57 | 95 | 506 | 796 | 4 | 24 | 13 | 1 | 2 | 0 | 9 | 2 | 1 | 1 | 37 | 600 | 21 | 5 | 0 | 7 | 11 | Hoxc5 |
| Hoxc6 | 1 | 2 | 0 | 0 | 74 | 2 | 2 | 0 | 1 | 18 | 456 | 0 | 0 | 0 | 0 | 0 | 0 | 0 | 0 | 0 | 0 | 0 | 50 | 2 | 1 | 0 | 0 | 0 | Hoxc6 |
| Hoxc8 | 0 | 1 | 0 | 4 | 0 | 0 | 2 | 0 | 3 | 0 | 533 | 3 | 0 | 2 | 0 | 3 | 0 | 0 | 0 | 0 | 0 | 0 | 7 | 2 | 3 | 1 | 0 | 0 | Hoxc8 |
| Hoxc9 | 0 | 0 | 0 | 0 | 1 | 1 | 0 | 0 | 0 | 0 | 21 | 0 | 0 | 0 | 0 | 0 | 0 | 0 | 0 | 0 | 0 | 0 | 1 | 0 | 0 | 0 | 0 | 0 | Hoxc9 |
| Hoxd1 | 141 | 65 | 29 | 67 | 122 | 107 | 6 | 126 | 256 | 100 | 197 | 53 | 94 | 67 | 79 | 65 | 42 | 186 | 9 | 20 | 4 | 134 | 159 | 150 | 133 | 8 | 36 | 34 | Hoxd1 |
| Hoxd10 | 0 | 1 | 0 | 0 | 0 | 2 | 0 | 1 | 1 | 0 | 0 | 0 | 1 | 0 | 0 | 0 | 0 | 0 | 0 | 0 | 3 | 0 | 3 | 0 | 0 | 0 | 0 | 0 | Hoxd10 |
| Hoxd11 | 0 | 0 | 0 | 0 | 0 | 1 | 0 | 0 | 0 | 0 | 0 | 0 | 0 | 0 | 0 | 0 | 0 | 0 | 0 | 0 | 0 | 0 | 0 | 0 | 0 | 0 | 0 | 0 | Hoxd11 |
| Hoxd13 | 0 | 6 | 16 | 14 | 14 | 8 | 3 | 12 | 10 | 12 | 10 | 8 | 16 | 4 | 4 | 13 | 17 | 22 | 7 | 12 | 6 | 13 | 6 | 5 | 9 | 0 | 1 | 0 | Hoxd13 |
| Hoxd3 | 0 | 3 | 3 | 1 | 492 | 11 | 0 | 305 | 360 | 547 | 767 | 2 | 282 | 5 | 116 | 12 | 0 | 13 | 2 | 1 | 0 | 428 | 543 | 25 | 12 | 0 | 0 | 0 | Hoxd3 |
| Hoxd4 | 5 | 3 | 2 | 3 | 155 | 15 | 0 | 26 | 54 | 90 | 295 | 7 | 26 | 5 | 12 | 3 | 3 | 9 | 2 | 6 | 0 | 23 | 155 | 5 | 4 | 3 | 2 | 1 | Hoxd4 |
| Hoxd8 | 65 | 18 | 34 | 14 | 13 | 81 | 9 | 118 | 225 | 137 | 355 | 53 | 60 | 45 | 30 | 34 | 25 | 145 | 30 | 26 | 13 | 38 | 315 | 166 | 81 | 15 | 35 | 28 | Hoxd8 |
| Hoxd9 | 5 | 1 | 3 | 0 | 2 | 15 | 0 | 5 | 24 | 2 | 29 | 7 | 9 | 2 | 4 | 2 | 0 | 21 | 5 | 3 | 1 | 0 | 46 | 15 | 13 | 0 | 0 | 4 | Hoxd9 |
| Hp | 5 | 3 | 1 | 5 | 2 | 9 | 8 | 1 | 19 | 25 | 32 | 11 | 27 | 1 | 9 | 12 | 7 | 7 | 6 | 0 | 1 | 5 | 40 | 8 | 6 | 27 | 8 | 6 | Hp |
| Hp1bp3 | 14981 | 13503 | 14113 | 13900 | 15657 | 15752 | 15759 | 27553 | 25239 | 26427 | 24939 | 28480 | 25722 | 30380 | 14652 | 12738 | 14422 | 14098 | 15176 | 15337 | 15966 | 27222 | 24078 | 28668 | 28408 | 27913 | 27485 | 28766 | Hp1bp3 |
| Hpca | 353 | 550 | 474 | 809 | 1445 | 1012 | 1148 | 609 | 868 | 846 | 846 | 554 | 476 | 461 | 500 | 371 | 634 | 1094 | 1246 | 1066 | 1180 | 962 | 805 | 775 | 1058 | 431 | 378 | 405 | Hpca |
| Hpcal1 | 2571 | 3065 | 2433 | 2729 | 2626 | 2226 | 2258 | 1462 | 1638 | 1554 | 1447 | 1389 | 1477 | 1330 | 2525 | 2796 | 2593 | 2633 | 2382 | 2450 | 2329 | 1389 | 1647 | 1385 | 1318 | 1442 | 1305 | 1423 | Hpcal1 |
| Hpcal4 | 338 | 396 | 379 | 622 | 1093 | 802 | 919 | 435 | 538 | 592 | 478 | 245 | 206 | 279 | 447 | 286 | 502 | 683 | 1008 | 853 | 942 | 810 | 544 | 399 | 608 | 169 | 170 | 234 | Hpcal4 |
| Hpd | 0 | 0 | 0 | 0 | 0 | 0 | 0 | 0 | 2 | 0 | 0 | 0 | 0 | 1 | 0 | 0 | 0 | 0 | 0 | 0 | 0 | 0 | 0 | 0 | 0 | 0 | 0 | 0 | Hpd |
| Hpdl | 89 | 131 | 89 | 70 | 67 | 69 | 58 | 20 | 43 | 43 | 29 | 25 | 45 | 35 | 69 | 78 | 89 | 53 | 65 | 86 | 61 | 13 | 27 | 27 | 23 | 41 | 27 | 23 | Hpdl |
| Hpf1 | 5104 | 3889 | 5464 | 4661 | 4056 | 4326 | 5197 | 4715 | 4138 | 4065 | 4373 | 5149 | 4689 | 5600 | 4954 | 4315 | 4589 | 4240 | 4108 | 5033 | 4653 | 4005 | 3880 | 4790 | 4723 | 5551 | 5246 | 5793 | Hpf1 |
| Hpgd | 717 | 360 | 321 | 242 | 268 | 402 | 203 | 712 | 756 | 554 | 525 | 383 | 641 | 289 | 479 | 525 | 224 | 251 | 242 | 234 | 195 | 520 | 1023 | 443 | 285 | 560 | 395 | 337 | Hpgd |
| Hpgds | 483 | 544 | 577 | 522 | 452 | 507 | 350 | 706 | 630 | 524 | 600 | 782 | 1012 | 690 | 554 | 702 | 662 | 459 | 402 | 620 | 350 | 734 | 823 | 604 | 637 | 588 | 839 | 749 | Hpgds |
| Hpn | 63 | 44 | 27 | 32 | 29 | 53 | 10 | 37 | 82 | 82 | 78 | 49 | 67 | 14 | 58 | 60 | 34 | 35 | 23 | 21 | 11 | 32 | 85 | 58 | 30 | 77 | 37 | 28 | Hpn |
| Hprt | 3015 | 2354 | 3058 | 2885 | 3009 | 2807 | 3363 | 2491 | 2314 | 2268 | 2587 | 2427 | 2431 | 2574 | 3132 | 2921 | 2803 | 2963 | 2862 | 3412 | 3037 | 2413 | 2501 | 2374 | 2361 | 3004 | 2396 | 2598 | Hprt |
| Hps1 | 955 | 1292 | 1218 | 1279 | 1337 | 1393 | 1294 | 1327 | 1657 | 1896 | 1647 | 2014 | 1683 | 1739 | 907 | 1135 | 1420 | 1289 | 1327 | 1031 | 1205 | 1897 | 1480 | 1822 | 1710 | 1751 | 1640 | 1698 | Hps1 |
| Hps3 | 2821 | 2686 | 2347 | 2472 | 2313 | 2406 | 2571 | 2158 | 1705 | 1607 | 1659 | 1756 | 1924 | 1861 | 2968 | 2769 | 2202 | 2573 | 2159 | 2579 | 2560 | 2080 | 1799 | 1616 | 1794 | 1692 | 1749 | 1651 | Hps3 |
| Hps4 | 2623 | 3524 | 3151 | 3562 | 2777 | 2679 | 2759 | 2039 | 2304 | 2390 | 2311 | 2309 | 2293 | 2039 | 2669 | 3328 | 3705 | 3339 | 3181 | 2508 | 2824 | 2483 | 2193 | 2321 | 2454 | 2180 | 2264 | 2149 | Hps4 |
| Hps5 | 4669 | 4702 | 4232 | 4685 | 4172 | 4298 | 4412 | 3584 | 3633 | 3390 | 3694 | 3645 | 3664 | 3589 | 4627 | 4777 | 4291 | 4787 | 4197 | 4022 | 4294 | 3664 | 3430 | 3755 | 3595 | 3176 | 3627 | 3786 | Hps5 |
| Hps6 | 415 | 523 | 566 | 494 | 433 | 416 | 288 | 128 | 211 | 218 | 214 | 152 | 195 | 122 | 347 | 560 | 492 | 443 | 476 | 324 | 341 | 179 | 227 | 173 | 159 | 204 | 156 | 138 | Hps6 |
| Hpse | 150 | 100 | 94 | 88 | 59 | 113 | 47 | 140 | 163 | 150 | 175 | 123 | 137 | 100 | 119 | 125 | 97 | 128 | 57 | 65 | 24 | 134 | 167 | 155 | 107 | 98 | 119 | 138 | Hpse |
| Hpse2 | 932 | 358 | 177 | 294 | 20 | 85 | 6 | 1073 | 639 | 1096 | 383 | 313 | 501 | 244 | 390 | 925 | 198 | 110 | 16 | 10 | 1 | 69 | 579 | 998 | 106 | 1376 | 157 | 352 | Hpse2 |
| Hpx | 74 | 94 | 78 | 109 | 65 | 77 | 91 | 20 | 21 | 23 | 34 | 12 | 16 | 15 | 79 | 101 | 115 | 109 | 83 | 61 | 67 | 23 | 15 | 23 | 10 | 31 | 8 | 25 | Hpx |
| Hr | 224 | 236 | 227 | 269 | 203 | 234 | 215 | 249 | 386 | 354 | 315 | 134 | 143 | 137 | 158 | 266 | 251 | 221 | 228 | 156 | 181 | 211 | 359 | 315 | 248 | 170 | 96 | 120 | Hr |
| Hras | 2658 | 3301 | 2845 | 3472 | 3277 | 2964 | 2942 | 2467 | 2637 | 2620 | 2423 | 2549 | 2368 | 2540 | 2649 | 2888 | 3274 | 2998 | 3441 | 2862 | 3282 | 2379 | 2660 | 2844 | 2545 | 2912 | 2460 | 2553 | Hras |
| Hrc | 60 | 68 | 56 | 48 | 51 | 43 | 38 | 66 | 138 | 53 | 656 | 36 | 45 | 18 | 63 | 82 | 88 | 91 | 34 | 63 | 57 | 122 | 424 | 62 | 93 | 47 | 45 | 44 | Hrc |
| Hrct1 | 2 | 10 | 5 | 5 | 1 | 9 | 12 | 12 | 7 | 8 | 26 | 16 | 6 | 10 | 7 | 5 | 1 | 3 | 3 | 4 | 10 | 10 | 12 | 2 | 6 | 10 | 6 | 5 | Hrct1 |
| Hrg | 1 | 3 | 2 | 7 | 3 | 6 | 2 | 11 | 15 | 12 | 17 | 15 | 5 | 14 | 1 | 5 | 5 | 3 | 4 | 3 | 3 | 10 | 8 | 3 | 3 | 15 | 13 | 17 | Hrg |
| Hrh1 | 201 | 267 | 219 | 280 | 227 | 245 | 234 | 18 | 15 | 32 | 18 | 16 | 16 | 20 | 270 | 179 | 298 | 216 | 218 | 180 | 214 | 32 | 19 | 10 | 17 | 26 | 7 | 12 | Hrh1 |
| Hrh2 | 13 | 27 | 13 | 21 | 26 | 42 | 26 | 16 | 30 | 31 | 26 | 16 | 17 | 28 | 16 | 19 | 27 | 32 | 34 | 28 | 43 | 35 | 28 | 22 | 32 | 8 | 14 | 9 | Hrh2 |
| Hrh3 | 746 | 1243 | 1222 | 1381 | 1274 | 1114 | 1117 | 402 | 322 | 444 | 325 | 528 | 572 | 550 | 808 | 950 | 1538 | 1293 | 1468 | 1107 | 1104 | 525 | 293 | 529 | 493 | 598 | 553 | 546 | Hrh3 |
| Hrh4 | 3 | 0 | 0 | 0 | 1 | 1 | 1 | 1 | 2 | 0 | 1 | 2 | 1 | 1 | 3 | 0 | 0 | 1 | 1 | 0 | 0 | 2 | 2 | 0 | 0 | 0 | 0 | 1 | Hrh4 |
| Hrk | 278 | 290 | 214 | 385 | 795 | 444 | 413 | 545 | 573 | 720 | 661 | 307 | 559 | 334 | 409 | 358 | 295 | 512 | 438 | 480 | 395 | 725 | 561 | 484 | 542 | 316 | 206 | 266 | Hrk |
| Hrnr | 0 | 0 | 0 | 0 | 0 | 0 | 0 | 0 | 5 | 3 | 0 | 0 | 0 | 0 | 0 | 0 | 0 | 0 | 0 | 0 | 0 | 0 | 5 | 0 | 0 | 0 | 0 | 0 | Hrnr |
| Hrob | 1162 | 1373 | 1482 | 1160 | 649 | 805 | 865 | 541 | 653 | 688 | 553 | 976 | 814 | 922 | 952 | 1273 | 1234 | 1106 | 983 | 707 | 882 | 409 | 559 | 860 | 635 | 1215 | 845 | 909 | Hrob |
| Hs1bp3 | 1544 | 1850 | 1870 | 1582 | 1383 | 1382 | 1268 | 851 | 1233 | 1033 | 1071 | 1061 | 1175 | 962 | 1480 | 1872 | 1760 | 1495 | 1562 | 1389 | 1266 | 1031 | 1251 | 1133 | 889 | 1309 | 1083 | 997 | Hs1bp3 |
| Hs2st1 | 15102 | 13487 | 13068 | 14292 | 13636 | 15067 | 16820 | 9863 | 8829 | 8449 | 8112 | 10406 | 9140 | 10423 | 14848 | 13085 | 13016 | 14516 | 14364 | 16309 | 16051 | 8654 | 9333 | 9582 | 10236 | 10725 | 10313 | 10690 | Hs2st1 |
| Hs3st1 | 332 | 415 | 356 | 453 | 512 | 542 | 469 | 495 | 568 | 497 | 565 | 409 | 513 | 554 | 367 | 384 | 404 | 508 | 503 | 438 | 481 | 481 | 597 | 601 | 584 | 475 | 436 | 465 | Hs3st1 |
| Hs3st2 | 98 | 119 | 63 | 125 | 318 | 161 | 124 | 122 | 142 | 153 | 195 | 77 | 137 | 27 | 138 | 120 | 66 | 146 | 175 | 167 | 173 | 172 | 138 | 78 | 122 | 11 | 48 | 34 | Hs3st2 |
| Hs3st3a1 | 291 | 253 | 388 | 229 | 189 | 147 | 122 | 363 | 275 | 428 | 401 | 395 | 525 | 338 | 262 | 420 | 245 | 184 | 168 | 82 | 130 | 316 | 274 | 368 | 264 | 541 | 351 | 415 | Hs3st3a1 |
| Hs3st3b1 | 1497 | 1459 | 1919 | 1574 | 914 | 951 | 720 | 1154 | 1023 | 1204 | 1234 | 1413 | 1706 | 1396 | 1391 | 1843 | 1450 | 1440 | 1059 | 629 | 884 | 1201 | 1015 | 1450 | 1115 | 1273 | 1381 | 1762 | Hs3st3b1 |
| Hs3st4 | 255 | 327 | 180 | 730 | 1240 | 1043 | 1009 | 922 | 992 | 965 | 800 | 508 | 497 | 388 | 444 | 325 | 509 | 907 | 1086 | 1117 | 1381 | 1380 | 852 | 817 | 1173 | 367 | 323 | 341 | Hs3st4 |
| Hs3st5 | 318 | 306 | 248 | 363 | 614 | 427 | 342 | 603 | 564 | 566 | 558 | 333 | 566 | 329 | 409 | 435 | 320 | 420 | 501 | 455 | 395 | 675 | 626 | 482 | 576 | 315 | 301 | 323 | Hs3st5 |
| Hs3st6 | 105 | 108 | 79 | 43 | 27 | 55 | 28 | 65 | 56 | 75 | 29 | 8 | 9 | 6 | 37 | 109 | 54 | 50 | 31 | 27 | 26 | 22 | 47 | 87 | 5 | 60 | 22 | 9 | Hs3st6 |

| | | | | | | | | | | | | | | | | | | | | | | | | | | | | | |
|---|---|---|---|---|---|---|---|---|---|---|---|---|---|---|---|---|---|---|---|---|---|---|---|---|---|---|---|---|---|
| Hs6st1 | 7011 | 9512 | 10772 | 11717 | 11233 | 9671 | 8499 | 5056 | 5235 | 6883 | 6389 | 6635 | 6400 | 6110 | 7460 | 8604 | 11305 | 11611 | 10518 | 7260 | 9502 | 7972 | 5482 | 6684 | 6807 | 5722 | 5779 | 6461 | Hs6st1 |
| Hs6st2 | 2553 | 1884 | 1660 | 2304 | 3016 | 2857 | 3121 | 3392 | 3477 | 3290 | 3900 | 3107 | 2875 | 3061 | 2316 | 2156 | 1810 | 2466 | 2703 | 2855 | 2955 | 3551 | 3869 | 3322 | 3629 | 3541 | 2802 | 2781 | Hs6st2 |
| Hs6st3 | 54 | 71 | 33 | 146 | 229 | 194 | 139 | 162 | 158 | 202 | 177 | 101 | 113 | 81 | 69 | 58 | 79 | 164 | 149 | 195 | 212 | 199 | 184 | 117 | 216 | 4 | 56 | 58 | Hs6st3 |
| Hsbp1 | 8338 | 9088 | 9157 | 9227 | 9429 | 8630 | 9759 | 9395 | 8409 | 9495 | 8416 | 8376 | 8282 | 9596 | 9121 | 8062 | 9387 | 9264 | 9372 | 10569 | 9861 | 9336 | 8738 | 9141 | 9162 | 8711 | 8365 | 9016 | Hsbp1 |
| Hsbp1l1 | 13 | 13 | 29 | 11 | 16 | 4 | 13 | 26 | 23 | 30 | 20 | 15 | 20 | 18 | 13 | 34 | 20 | 19 | 17 | 15 | 22 | 20 | 16 | 23 | 35 | 17 | 17 | 15 | Hsbp1l1 |
| Hscb | 546 | 539 | 617 | 518 | 394 | 394 | 398 | 269 | 284 | 295 | 273 | 298 | 318 | 339 | 473 | 492 | 545 | 481 | 417 | 462 | 437 | 206 | 328 | 305 | 274 | 385 | 344 | 377 | Hscb |
| Hsd11b1 | 9 | 8 | 2 | 6 | 2 | 3 | 6 | 4 | 3 | 3 | 11 | 5 | 3 | 1 | 5 | 13 | 2 | 14 | 5 | 2 | 8 | 10 | 13 | 5 | 1 | 4 | 1 | 0 | Hsd11b1 |
| Hsd11b2 | 414 | 592 | 869 | 417 | 400 | 393 | 341 | 541 | 433 | 549 | 438 | 954 | 916 | 943 | 485 | 505 | 765 | 330 | 429 | 367 | 294 | 658 | 548 | 653 | 592 | 992 | 1058 | 1065 | Hsd11b2 |
| Hsd17b1 | 17 | 73 | 47 | 87 | 78 | 56 | 49 | 23 | 23 | 38 | 8 | 16 | 16 | 15 | 59 | 57 | 82 | 133 | 63 | 60 | 57 | 30 | 19 | 16 | 30 | 16 | 16 | 9 | Hsd17b1 |
| Hsd17b10 | 4617 | 4273 | 4233 | 3559 | 3269 | 3721 | 3295 | 3969 | 4209 | 3782 | 3999 | 3811 | 4188 | 3957 | 4102 | 4175 | 4025 | 3462 | 3709 | 4243 | 3480 | 2988 | 4370 | 3930 | 3579 | 4980 | 4221 | 4328 | Hsd17b10 |
| Hsd17b11 | 693 | 577 | 869 | 688 | 708 | 653 | 643 | 845 | 1022 | 959 | 1171 | 681 | 824 | 753 | 708 | 781 | 581 | 678 | 596 | 571 | 602 | 902 | 1187 | 963 | 760 | 971 | 694 | 676 | Hsd17b11 |
| Hsd17b12 | 7444 | 6489 | 7018 | 6463 | 5893 | 5728 | 6631 | 7166 | 6258 | 5614 | 5469 | 6030 | 5947 | 6956 | 7387 | 6585 | 6557 | 6104 | 6093 | 6979 | 6391 | 6025 | 6354 | 6178 | 5905 | 6921 | 6501 | 6834 | Hsd17b12 |
| Hsd17b13 | 9 | 2 | 3 | 2 | 0 | 2 | 0 | 2 | 2 | 1 | 3 | 0 | 3 | 0 | 0 | 5 | 1 | 0 | 1 | 1 | 0 | 0 | 1 | 2 | 0 | 0 | 2 | 1 | Hsd17b13 |
| Hsd17b14 | 17 | 7 | 10 | 6 | 7 | 2 | 2 | 6 | 24 | 13 | 34 | 4 | 14 | 2 | 5 | 25 | 9 | 7 | 16 | 0 | 5 | 12 | 11 | 8 | 13 | 17 | 7 | 6 | Hsd17b14 |
| Hsd17b2 | 0 | 0 | 0 | 0 | 1 | 1 | 0 | 1 | 7 | 12 | 2 | 0 | 0 | 0 | 0 | 1 | 1 | 0 | 0 | 1 | 0 | 0 | 1 | 0 | 0 | 7 | 3 | 2 | Hsd17b2 |
| Hsd17b3 | 17 | 18 | 18 | 4 | 3 | 13 | 13 | 20 | 12 | 12 | 13 | 38 | 29 | 30 | 9 | 13 | 14 | 9 | 21 | 20 | 12 | 23 | 5 | 22 | 14 | 41 | 36 | 44 | Hsd17b3 |
| Hsd17b4 | 9490 | 9114 | 9373 | 8297 | 7927 | 7760 | 7968 | 7266 | 6975 | 7284 | 6686 | 7819 | 7673 | 7877 | 9152 | 9120 | 9025 | 8299 | 8292 | 7794 | 7968 | 6676 | 7435 | 7648 | 7037 | 8579 | 8106 | 8184 | Hsd17b4 |
| Hsd17b6 | 0 | 0 | 0 | 0 | 0 | 0 | 2 | 2 | 2 | 0 | 8 | 0 | 0 | 0 | 0 | 0 | 0 | 0 | 0 | 0 | 0 | 2 | 6 | 4 | 0 | 5 | 0 | 0 | Hsd17b6 |
| Hsd17b7 | 4774 | 4570 | 4285 | 4272 | 3404 | 3937 | 3543 | 2235 | 2011 | 1983 | 1916 | 2135 | 2546 | 2334 | 4824 | 5077 | 3996 | 4210 | 3137 | 3544 | 3550 | 1898 | 1952 | 1763 | 1927 | 1997 | 2650 | 2428 | Hsd17b7 |
| Hsd3b1 | 23 | 22 | 7 | 0 | 0 | 3 | 6 | 6 | 7 | 3 | 3 | 25 | 66 | 18 | 17 | 3 | 2 | 7 | 2 | 12 | 2 | 0 | 14 | 5 | 10 | 24 | 30 | 49 | Hsd3b1 |
| Hsd3b2 | 0 | 0 | 0 | 0 | 0 | 0 | 0 | 0 | 0 | 0 | 0 | 0 | 0 | 0 | 0 | 0 | 0 | 0 | 0 | 0 | 0 | 0 | 1 | 0 | 0 | 0 | 0 | 0 | Hsd3b2 |
| Hsd3b3 | 0 | 0 | 0 | 0 | 0 | 1 | 0 | 2 | 0 | 0 | 0 | 0 | 0 | 0 | 0 | 0 | 0 | 0 | 0 | 0 | 0 | 0 | 0 | 0 | 0 | 0 | 1 | 0 | Hsd3b3 |
| Hsd3b6 | 870 | 1304 | 861 | 681 | 245 | 476 | 495 | 672 | 854 | 493 | 440 | 613 | 2591 | 1012 | 1080 | 1119 | 661 | 430 | 394 | 716 | 206 | 297 | 834 | 391 | 439 | 937 | 1052 | 2255 | Hsd3b6 |
| Hsd3b7 | 69 | 71 | 82 | 101 | 68 | 57 | 107 | 45 | 70 | 45 | 44 | 65 | 44 | 85 | 54 | 61 | 97 | 99 | 79 | 63 | 99 | 72 | 53 | 76 | 92 | 73 | 69 | 69 | Hsd3b7 |
| Hsdl1 | 6138 | 7400 | 7134 | 8643 | 9515 | 8438 | 8543 | 5952 | 5516 | 6607 | 5415 | 6089 | 5236 | 5704 | 6879 | 6020 | 8129 | 9172 | 9939 | 8426 | 8880 | 7229 | 5569 | 6381 | 7382 | 5575 | 5684 | 5554 | Hsdl1 |
| Hsdl2 | 5087 | 4594 | 5005 | 4906 | 4357 | 4161 | 4510 | 2477 | 2188 | 2067 | 2085 | 2004 | 2147 | 2153 | 5466 | 4781 | 4879 | 4787 | 4156 | 4950 | 4597 | 2065 | 2262 | 1938 | 1987 | 2031 | 2035 | 2201 | Hsdl2 |
| Hsf1 | 2279 | 3131 | 2961 | 2899 | 2822 | 2553 | 2551 | 2699 | 3080 | 3350 | 3218 | 3396 | 3057 | 3044 | 2190 | 2673 | 3264 | 3086 | 3215 | 2444 | 2744 | 3166 | 2982 | 3449 | 3191 | 3828 | 2880 | 3077 | Hsf1 |
| Hsf2 | 9569 | 8201 | 8908 | 9935 | 9637 | 9593 | 11451 | 11086 | 7751 | 7885 | 7442 | 9929 | 9643 | 10969 | 9981 | 8452 | 8920 | 9400 | 9637 | 10285 | 11147 | 9421 | 7625 | 8331 | 9296 | 8527 | 10402 | 11275 | Hsf2 |
| Hsf2bp | 17 | 26 | 25 | 10 | 11 | 12 | 11 | 16 | 26 | 26 | 19 | 47 | 32 | 32 | 20 | 19 | 24 | 23 | 12 | 17 | 19 | 25 | 32 | 16 | 23 | 30 | 30 | 31 | Hsf2bp |
| Hsf3 | 20 | 6 | 11 | 3 | 2 | 6 | 0 | 16 | 6 | 7 | 12 | 4 | 18 | 3 | 5 | 12 | 0 | 1 | 7 | 5 | 4 | 12 | 10 | 9 | 2 | 9 | 5 | 9 | Hsf3 |
| Hsf4 | 12 | 13 | 21 | 21 | 34 | 29 | 12 | 10 | 7 | 8 | 13 | 21 | 8 | 7 | 18 | 16 | 19 | 24 | 26 | 3 | 16 | 23 | 12 | 5 | 10 | 10 | 6 | 5 | Hsf4 |
| Hsf5 | 40 | 32 | 47 | 59 | 39 | 39 | 56 | 39 | 21 | 23 | 36 | 46 | 16 | 34 | 50 | 27 | 54 | 46 | 50 | 48 | 50 | 42 | 24 | 31 | 24 | 31 | 36 | 23 | Hsf5 |
| Hsh2d | 36 | 31 | 42 | 47 | 29 | 31 | 15 | 1 | 2 | 10 | 5 | 5 | 5 | 6 | 29 | 37 | 37 | 29 | 37 | 5 | 19 | 7 | 3 | 7 | 5 | 15 | 6 | 4 | Hsh2d |
| Hsp90aa1 | 96005 | 88377 | 106246 | 85128 | 80048 | 79025 | 98694 | 111856 | 89032 | 82009 | 78422 | 98944 | 90917 | 105642 | 111188 | 93946 | 96437 | 83520 | 83970 | 104939 | 88816 | 91001 | 89799 | 90626 | 87677 | 106991 | 105930 | 108956 | Hsp90aa1 |
| Hsp90ab1 | 171157 | 173241 | 206219 | 175467 | 157062 | 158678 | 170765 | 152109 | 167473 | 174680 | 170818 | 204833 | 188495 | 212739 | 161258 | 172237 | 189938 | 166388 | 176894 | 147066 | 167261 | 169952 | 166389 | 199281 | 183284 | 242929 | 195973 | 223492 | Hsp90ab1 |
| Hsp90b1 | 61696 | 49831 | 57075 | 44895 | 34946 | 36557 | 45207 | 48586 | 43255 | 41354 | 45592 | 38750 | 42796 | 45178 | 59597 | 55499 | 50832 | 45090 | 38982 | 41594 | 40213 | 42870 | 45888 | 44431 | 36826 | 45045 | 41916 | 46512 | Hsp90b1 |
| Hspa12a | 6346 | 6878 | 5364 | 7566 | 11373 | 8214 | 8522 | 6804 | 6414 | 6663 | 6571 | 4856 | 6860 | 4879 | 8225 | 7555 | 7078 | 8525 | 8179 | 8215 | 9150 | 8794 | 5779 | 4844 | 6821 | 2885 | 4644 | 4524 | Hspa12a |
| Hspa12b | 962 | 1090 | 1144 | 716 | 651 | 923 | 478 | 1470 | 1859 | 1650 | 2272 | 1519 | 2167 | 1298 | 961 | 1287 | 957 | 759 | 573 | 487 | 462 | 1425 | 2210 | 1725 | 1337 | 1740 | 1768 | 1616 | Hspa12b |
| Hspa13 | 9105 | 7749 | 7409 | 7646 | 7627 | 7782 | 8834 | 8684 | 7000 | 5970 | 6716 | 6783 | 6397 | 7066 | 9860 | 8639 | 7260 | 8216 | 7426 | 9538 | 8400 | 7352 | 7210 | 6276 | 6680 | 6425 | 7093 | 7017 | Hspa13 |
| Hspa14 | 6057 | 5808 | 5178 | 5012 | 3670 | 4235 | 5075 | 3076 | 2900 | 2728 | 2741 | 3256 | 3044 | 3544 | 5466 | 5153 | 5078 | 4836 | 4046 | 4654 | 4362 | 2553 | 2764 | 3057 | 2612 | 3931 | 3309 | 3592 | Hspa14 |
| Hspa1a | 406 | 553 | 447 | 511 | 387 | 309 | 286 | 56 | 81 | 77 | 73 | 45 | 82 | 48 | 310 | 438 | 475 | 526 | 381 | 319 | 287 | 85 | 113 | 95 | 63 | 49 | 36 | 45 | Hspa1a |
| Hspa1b | 618 | 852 | 930 | 857 | 650 | 515 | 489 | 101 | 176 | 111 | 144 | 94 | 124 | 77 | 603 | 724 | 844 | 938 | 697 | 559 | 493 | 127 | 228 | 143 | 107 | 112 | 98 | 115 | Hspa1b |
| Hspa1l | 29 | 36 | 42 | 52 | 39 | 27 | 29 | 10 | 11 | 12 | 22 | 11 | 7 | 7 | 28 | 35 | 41 | 36 | 40 | 30 | 55 | 10 | 16 | 16 | 9 | 5 | 13 | 15 | Hspa1l |
| Hspa2 | 1407 | 2141 | 2246 | 1949 | 1429 | 1435 | 1471 | 232 | 272 | 254 | 293 | 285 | 325 | 246 | 1427 | 1854 | 2176 | 1907 | 1678 | 1370 | 1541 | 277 | 334 | 259 | 285 | 276 | 280 | 248 | Hspa2 |
| Hspa4 | 40997 | 37094 | 43804 | 39203 | 36889 | 37253 | 42693 | 31414 | 27493 | 29554 | 28082 | 33520 | 31208 | 36259 | 40923 | 36377 | 42198 | 39752 | 38072 | 36918 | 40510 | 29086 | 27119 | 31846 | 30036 | 33658 | 33655 | 35835 | Hspa4 |
| Hspa4l | 799 | 327 | 300 | 290 | 299 | 352 | 302 | 440 | 224 | 161 | 351 | 226 | 231 | 347 | 559 | 440 | 283 | 792 | 361 | 741 | 301 | 371 | 287 | 395 | 182 | 267 | 310 | 170 | Hspa4l |
| Hspa5 | 53318 | 52609 | 54958 | 43728 | 32028 | 32875 | 35197 | 47809 | 44199 | 41709 | 50588 | 36855 | 41764 | 38420 | 52909 | 54252 | 47503 | 44120 | 36423 | 41150 | 32643 | 41883 | 46016 | 45405 | 38184 | 45401 | 41097 | 42058 | Hspa5 |
| Hspa8 | 114426 | 132005 | 128970 | 111950 | 107713 | 98773 | 117895 | 156864 | 141174 | 125518 | 127078 | 116459 | 117598 | 128027 | 118158 | 113366 | 127547 | 111802 | 133105 | 121506 | 122102 | 131017 | 143144 | 135992 | 133383 | 137604 | 128610 | 136669 | Hspa8 |
| Hspa9 | 28005 | 24258 | 27531 | 24237 | 21561 | 23403 | 25349 | 22810 | 19861 | 20258 | 20672 | 24013 | 21897 | 25823 | 26791 | 23018 | 24671 | 23320 | 23643 | 23449 | 24742 | 20299 | 19680 | 22538 | 21539 | 27043 | 24818 | 25897 | Hspa9 |
| Hspb1 | 309 | 384 | 348 | 190 | 137 | 300 | 122 | 289 | 480 | 258 | 738 | 209 | 319 | 172 | 301 | 398 | 169 | 281 | 144 | 111 | 89 | 147 | 792 | 252 | 129 | 129 | 202 | 229 | Hspb1 |
| Hspb11 | 715 | 574 | 799 | 681 | 704 | 631 | 807 | 607 | 535 | 601 | 569 | 532 | 562 | 644 | 688 | 656 | 790 | 588 | 614 | 692 | 760 | 526 | 568 | 654 | 618 | 657 | 616 | 627 | Hspb11 |
| Hspb2 | 14 | 0 | 4 | 1 | 10 | 4 | 0 | 18 | 49 | 21 | 217 | 4 | 15 | 6 | 9 | 11 | 1 | 2 | 1 | 2 | 2 | 13 | 132 | 9 | 9 | 1 | 3 | 4 | Hspb2 |
| Hspb3 | 3 | 0 | 4 | 0 | 3 | 1 | 2 | 5 | 8 | 7 | 104 | 5 | 3 | 2 | 4 | 2 | 0 | 0 | 0 | 2 | 4 | 3 | 58 | 2 | 3 | 3 | 0 | 1 | Hspb3 |
| Hspb6 | 197 | 251 | 246 | 241 | 173 | 208 | 241 | 200 | 258 | 258 | 225 | 268 | 187 | 275 | 223 | 160 | 234 | 160 | 213 | 222 | 213 | 177 | 253 | 279 | 240 | 267 | 222 | 246 | Hspb6 |
| Hspb7 | 0 | 0 | 0 | 0 | 1 | 0 | 0 | 5 | 1 | 1 | 25 | 0 | 3 | 3 | 1 | 1 | 0 | 2 | 0 | 0 | 0 | 3 | 38 | 1 | 2 | 1 | 0 | 2 | Hspb7 |
| Hspb8 | 946 | 749 | 693 | 472 | 454 | 535 | 466 | 701 | 803 | 686 | 681 | 606 | 784 | 669 | 777 | 847 | 563 | 547 | 515 | 570 | 442 | 505 | 1111 | 720 | 564 | 786 | 682 | 813 | Hspb8 |
| Hspb9 | 0 | 0 | 0 | 0 | 0 | 0 | 1 | 0 | 1 | 1 | 0 | 0 | 5 | 4 | 0 | 0 | 0 | 0 | 0 | 0 | 0 | 0 | 1 | 0 | 0 | 3 | 0 | 5 | Hspb9 |
| Hspbap1 | 2017 | 2274 | 1950 | 2171 | 1713 | 1799 | 1912 | 1061 | 968 | 944 | 1027 | 1032 | 1135 | 1096 | 2078 | 2069 | 1866 | 1955 | 1809 | 1980 | 1891 | 1033 | 1027 | 970 | 1191 | 1208 | 1075 | 1063 | Hspbap1 |
| Hspbp1 | 1405 | 2007 | 1728 | 1740 | 1780 | 1794 | 1618 | 1906 | 2241 | 1938 | 1896 | 2182 | 2146 | 2041 | 1557 | 1640 | 1983 | 1577 | 1936 | 1854 | 1607 | 1856 | 2177 | 2168 | 2211 | 2571 | 2317 | 2146 | Hspbp1 |
| Hspd1 | 36267 | 32405 | 41285 | 31572 | 27458 | 28398 | 33224 | 29388 | 26383 | 27153 | 28455 | 32140 | 31013 | 35853 | 35948 | 34177 | 36784 | 30915 | 29945 | 32504 | 30810 | 24595 | 26956 | 30889 | 28060 | 38892 | 34300 | 36413 | Hspd1 |
| Hspe1 | 7401 | 6677 | 8632 | 7426 | 5437 | 5230 | 7706 | 6544 | 5829 | 7507 | 7058 | 7029 | 7180 | 9508 | 6940 | 6065 | 7816 | 6426 | 5936 | 7068 | 7225 | 4847 | 6135 | 7873 | 6702 | 9133 | 7714 | 9957 | Hspe1 |
| Hspe1-rs1 | 330 | 251 | 343 | 337 | 181 | 142 | 253 | 309 | 188 | 357 | 242 | 298 | 168 | 381 | 313 | 200 | 356 | 201 | 191 | 217 | 301 | 184 | 227 | 388 | 232 | 318 | 261 | 363 | Hspe1-rs1 |
| Hspg2 | 12653 | 18215 | 19514 | 14440 | 10316 | 12800 | 7287 | 6483 | 11146 | 11649 | 15315 | 12662 | 13939 | 9441 | 10350 | 19877 | 15927 | 15860 | 11281 | 7557 | 8241 | 12013 | 12659 | 13694 | 10257 | 9549 | 11622 | 9795 | Hspg2 |
| Hsph1 | 19057 | 19918 | 23268 | 21311 | 20166 | 18569 | 22998 | 14458 | 12388 | 15033 | 13807 | 14790 | 12923 | 15080 | 19235 | 17637 | 22928 | 22278 | 22329 | 19200 | 21730 | 14275 | 12481 | 16557 | 15310 | 14766 | 14092 | 15531 | Hsph1 |
| Htatip2 | 272 | 241 | 304 | 219 | 122 | 148 | 156 | 195 | 177 | 181 | 164 | 197 | 315 | 201 | 247 | 247 | 228 | 154 | 112 | 168 | 126 | 137 | 227 | 185 | 142 | 187 | 274 | 335 | Htatip2 |
| Htatsf1 | 10320 | 9179 | 11378 | 10182 | 9463 | 9641 | 11094 | 10805 | 9843 | 10542 | 9746 | 11335 | 10324 | 12218 | 11082 | 10149 | 10917 | 9863 | 9576 | 10899 | 9994 | 10576 | 9551 | 11418 | 10286 | 11611 | 11072 | 11389 | Htatsf1 |
| Htr1a | 17 | 23 | 9 | 33 | 84 | 37 | 47 | 66 | 87 | 76 | 82 | 20 | 68 | 32 | 26 | 32 | 34 | 33 | 34 | 31 | 38 | 110 | 98 | 42 | 81 | 35 | 22 | 23 | Htr1a |
| Htr1b | 209 | 182 | 250 | 218 | 308 | 303 | 184 | 332 | 347 | 317 | 458 | 127 | 330 | 170 | 216 | 246 | 246 | 276 | 214 | 202 | 248 | 461 | 378 | 221 | 270 | 197 | 180 | 217 | Htr1b |
| Htr1d | 24 | 20 | 24 | 9 | 37 | 7 | 15 | 11 | 12 | 23 | 13 | 9 | 3 | 12 | 29 | 27 | 19 | 28 | 12 | 9 | 13 | 22 | 16 | 11 | 2 | 21 | 16 | 5 | Htr1d |
| Htr1f | 12 | 3 | 4 | 6 | 0 | 3 | 7 | 12 | 18 | 3 | 19 | 7 | 6 | 6 | 8 | 4 | 7 | 1 | 1 | 5 | 11 | 7 | 10 | 10 | 7 | 7 | 16 | 8 | Htr1f |
| Htr2a | 107 | 81 | 104 | 124 | 209 | 149 | 114 | 283 | 236 | 206 | 209 | 142 | 211 | 162 | 142 | 83 | 103 | 140 | 144 | 159 | 128 | 267 | 259 | 167 | 246 | 144 | 179 | 170 | Htr2a |
| Htr2b | 2 | 0 | 0 | 0 | 0 | 3 | 0 | 2 | 1 | 0 | 8 | 1 | 0 | 1 | 0 | 0 | 1 | 1 | 0 | 3 | 0 | 0 | 6 | 0 | 0 | 0 | 0 | 0 | Htr2b |
| Htr2c | 280 | 267 | 155 | 210 | 771 | 442 | 365 | 696 | 719 | 703 | 645 | 704 | 810 | 317 | 326 | 403 | 213 | 339 | 436 | 385 | 567 | 1039 | 1407 | 406 | 519 | 281 | 307 | 297 | Htr2c |
| Htr3a | 105 | 105 | 48 | 104 | 128 | 113 | 124 | 114 | 142 | 100 | 173 | 75 | 92 | 74 | 105 | 81 | 100 | 130 | 121 | 165 | 133 | 182 | 194 | 103 | 161 | 75 | 78 | 68 | Htr3a |
| Htr3b | 13 | 2 | 1 | 5 | 9 | 3 | 6 | 23 | 23 | 10 | 24 | 2 | 5 | 2 | 16 | 9 | 0 | 8 | 8 | 2 | 5 | 37 | 36 | 8 | 22 | 25 | 0 | 2 | Htr3b |
| Htr4 | 7 | 6 | 3 | 7 | 16 | 19 | 18 | 34 | 30 | 49 | 31 | 2 | 10 | 20 | 8 | 9 | 6 | 8 | 23 | 20 | 18 | 48 | 35 | 25 | 28 | 2 | 9 | 8 | Htr4 |
| Htr5a | 35 | 36 | 27 | 63 | 169 | 130 | 130 | 136 | 151 | 188 | 110 | 97 | 76 | 76 | 44 | 30 | 59 | 100 | 129 | 128 | 145 | 179 | 118 | 110 | 175 | 42 | 43 | 64 | Htr5a |
| Htr5b | 48 | 44 | 48 | 41 | 50 | 38 | 50 | 6 | 12 | 10 | 10 | 5 | 15 | 6 | 55 | 37 | 37 | 25 | 43 | 33 | 32 | 5 | 17 | 13 | 7 | 8 | 3 | 4 | Htr5b |
| Htr6 | 42 | 48 | 47 | 50 | 58 | 46 | 44 | 21 | 30 | 49 | 24 | 18 | 7 | 14 | 26 | 44 | 28 | 55 | 36 | 47 | 48 | 23 | 28 | 22 | 12 | 14 | 8 | 5 | Htr6 |

| | | | | | | | | | | | | | | | | | | | | | | | | | | | | | |
|---|---|---|---|---|---|---|---|---|---|---|---|---|---|---|---|---|---|---|---|---|---|---|---|---|---|---|---|---|---|
| Htr7 | 42 | 61 | 38 | 76 | 180 | 71 | 111 | 129 | 157 | 192 | 312 | 88 | 94 | 99 | 50 | 46 | 87 | 103 | 101 | 63 | 99 | 217 | 234 | 98 | 163 | 69 | 69 | 62 | Htr7 |
| Htra1 | 911 | 1020 | 774 | 744 | 772 | 1235 | 591 | 526 | 943 | 683 | 1263 | 420 | 665 | 458 | 944 | 1113 | 599 | 1169 | 664 | 532 | 681 | 682 | 1580 | 717 | 789 | 192 | 411 | 443 | Htra1 |
| Htra2 | 2856 | 3710 | 3640 | 3522 | 3014 | 2884 | 2984 | 1583 | 1856 | 2078 | 2037 | 2081 | 1986 | 2142 | 2662 | 3035 | 3713 | 3162 | 3192 | 2746 | 3012 | 1693 | 1915 | 2261 | 2053 | 2425 | 2062 | 2103 | Htra2 |
| Htra3 | 254 | 343 | 306 | 326 | 464 | 400 | 265 | 277 | 435 | 446 | 633 | 495 | 371 | 411 | 194 | 235 | 306 | 392 | 407 | 233 | 310 | 558 | 584 | 443 | 564 | 336 | 351 | 392 | Htra3 |
| Htra4 | 8 | 4 | 1 | 9 | 7 | 1 | 9 | 16 | 16 | 8 | 22 | 1 | 0 | 1 | 1 | 11 | 2 | 3 | 2 | 9 | 17 | 13 | 8 | 9 | 13 | 9 | 0 | 1 | Htra4 |
| Htt | 9827 | 9858 | 5485 | 7690 | 8909 | 9954 | 9302 | 8841 | 8976 | 8196 | 9107 | 8516 | 8749 | 8742 | 9155 | 9655 | 6264 | 7859 | 7763 | 9050 | 10285 | 7279 | 8540 | 6191 | 9362 | 5026 | 9704 | 7809 | Htt |
| Hunk | 2989 | 2585 | 3423 | 2873 | 3540 | 3228 | 3232 | 4091 | 5035 | 5117 | 4922 | 5396 | 5040 | 5343 | 2525 | 2823 | 3177 | 2979 | 3677 | 2545 | 3055 | 4089 | 4594 | 5609 | 5147 | 6090 | 4896 | 5302 | Hunk |
| Hus1 | 2539 | 2436 | 2455 | 2122 | 1864 | 2076 | 2160 | 2507 | 2265 | 2078 | 2171 | 2760 | 2566 | 2758 | 2403 | 2189 | 2394 | 1888 | 1972 | 2173 | 2006 | 1895 | 2142 | 2277 | 2102 | 2918 | 2915 | 2820 | Hus1 |
| Hus1b | 2 | 3 | 4 | 1 | 0 | 3 | 6 | 4 | 7 | 3 | 2 | 4 | 9 | 4 | 5 | 0 | 0 | 9 | 5 | 2 | 4 | 2 | 4 | 5 | 2 | 2 | 3 | 1 | Hus1b |
| Huwe1 | 50712 | 47487 | 29161 | 38227 | 45015 | 53687 | 48496 | 60037 | 62706 | 52893 | 56758 | 53410 | 53577 | 55297 | 49402 | 49043 | 33011 | 36740 | 39876 | 47167 | 54440 | 42737 | 54587 | 40227 | 58525 | 31850 | 64979 | 45560 | Huwe1 |
| Hvcn1 | 189 | 167 | 220 | 115 | 118 | 144 | 77 | 388 | 444 | 346 | 399 | 551 | 556 | 516 | 157 | 179 | 158 | 105 | 161 | 140 | 101 | 296 | 464 | 399 | 275 | 626 | 733 | 615 | Hvcn1 |
| Hyal1 | 516 | 623 | 549 | 466 | 382 | 404 | 363 | 274 | 293 | 293 | 341 | 346 | 349 | 311 | 422 | 607 | 547 | 420 | 432 | 332 | 384 | 256 | 351 | 390 | 310 | 387 | 411 | 338 | Hyal1 |
| Hyal2 | 2538 | 3450 | 2957 | 2495 | 2362 | 2346 | 1912 | 2970 | 3286 | 2930 | 2781 | 3364 | 3174 | 2972 | 2531 | 2820 | 2812 | 2484 | 2894 | 2466 | 2151 | 2764 | 3441 | 2919 | 2827 | 3847 | 3619 | 2957 | Hyal2 |
| Hyal3 | 46 | 79 | 43 | 79 | 74 | 45 | 47 | 45 | 47 | 53 | 55 | 47 | 44 | 48 | 58 | 51 | 43 | 52 | 46 | 54 | 55 | 50 | 60 | 39 | 69 | 62 | 37 | 37 | Hyal3 |
| Hyal5 | 0 | 0 | 0 | 0 | 0 | 0 | 0 | 0 | 0 | 0 | 0 | 0 | 0 | 0 | 0 | 0 | 0 | 0 | 0 | 0 | 0 | 0 | 0 | 1 | 0 | 0 | 0 | 0 | Hyal5 |
| Hyal6 | 1 | 2 | 0 | 0 | 2 | 0 | 0 | 0 | 0 | 0 | 0 | 0 | 0 | 2 | 1 | 0 | 0 | 0 | 0 | 0 | 0 | 0 | 0 | 0 | 3 | 0 | 0 | 0 | Hyal6 |
| Hydin | 134 | 140 | 42 | 110 | 154 | 203 | 106 | 167 | 216 | 96 | 152 | 141 | 194 | 92 | 120 | 205 | 57 | 88 | 105 | 181 | 133 | 107 | 319 | 86 | 145 | 66 | 114 | 68 | Hydin |
| Hykk | 1706 | 1628 | 1633 | 1525 | 1345 | 1368 | 1263 | 744 | 778 | 831 | 1009 | 696 | 923 | 751 | 1601 | 1997 | 1533 | 1418 | 1182 | 1308 | 1172 | 834 | 856 | 694 | 698 | 796 | 876 | 849 | Hykk |
| Hyls1 | 3092 | 2688 | 2739 | 2895 | 2577 | 2698 | 2893 | 2722 | 2231 | 2029 | 1797 | 2326 | 2258 | 2623 | 3018 | 2665 | 2530 | 2676 | 2495 | 3037 | 2749 | 2065 | 2065 | 2107 | 2133 | 2396 | 2603 | 2460 | Hyls1 |
| Hyou1 | 16050 | 18347 | 20400 | 18843 | 14681 | 14808 | 14393 | 9785 | 10342 | 11467 | 12063 | 11174 | 11546 | 10968 | 15207 | 17569 | 20312 | 18525 | 16781 | 13114 | 14932 | 11418 | 9901 | 12522 | 11203 | 11839 | 11427 | 12067 | Hyou1 |
| Hypk | 624 | 676 | 411 | 626 | 477 | 587 | 581 | 336 | 318 | 305 | 297 | 331 | 330 | 402 | 506 | 559 | 523 | 584 | 451 | 537 | 581 | 311 | 293 | 311 | 328 | 273 | 320 | 352 | Hypk |
| I0C0044D17Rik | 0 | 0 | 0 | 2 | 1 | 1 | 1 | 0 | 0 | 0 | 0 | 0 | 2 | 2 | 0 | 0 | 0 | 0 | 1 | 0 | 1 | 0 | 0 | 0 | 0 | 0 | 0 | 0 | I0C0044D17Rik |
| I830077J02Rik | 35 | 25 | 66 | 27 | 45 | 36 | 37 | 42 | 40 | 71 | 73 | 52 | 73 | 69 | 29 | 48 | 25 | 22 | 28 | 48 | 32 | 67 | 76 | 61 | 36 | 82 | 38 | 56 | I830077J02Rik |
| Iah1 | 1664 | 1722 | 1680 | 1590 | 1467 | 1419 | 1533 | 1325 | 1293 | 1182 | 1297 | 1415 | 1338 | 1378 | 1549 | 1650 | 1553 | 1481 | 1488 | 1491 | 1471 | 1180 | 1328 | 1435 | 1273 | 1387 | 1449 | 1434 | Iah1 |
| Iapp | 8 | 12 | 18 | 0 | 9 | 11 | 7 | 7 | 2 | 8 | 5 | 3 | 22 | 19 | 9 | 9 | 25 | 6 | 2 | 18 | 4 | 3 | 3 | 9 | 5 | 3 | 16 | 28 | Iapp |
| Iars | 16168 | 16726 | 14519 | 12671 | 11051 | 12295 | 12226 | 10535 | 10174 | 9405 | 9658 | 12825 | 11420 | 13103 | 15191 | 14308 | 13822 | 12145 | 11790 | 11589 | 11856 | 9379 | 9314 | 10794 | 11027 | 12180 | 12821 | 14158 | Iars |
| Iars2 | 4421 | 4026 | 3706 | 3349 | 3069 | 3355 | 3543 | 4377 | 4364 | 3769 | 3748 | 4387 | 4143 | 4163 | 4090 | 3727 | 3321 | 3147 | 3021 | 3510 | 3304 | 3790 | 4117 | 3908 | 3931 | 4372 | 4406 | 4323 | Iars2 |
| Iba57 | 198 | 254 | 289 | 372 | 376 | 321 | 297 | 625 | 771 | 819 | 746 | 886 | 823 | 748 | 222 | 238 | 349 | 261 | 411 | 313 | 318 | 810 | 723 | 780 | 983 | 748 | 862 | 758 | Iba57 |
| Ibsp | 3 | 3 | 7 | 6 | 11 | 87 | 20 | 131 | 302 | 376 | 279 | 8 | 7 | 12 | 5 | 9 | 0 | 232 | 3 | 5 | 7 | 515 | 553 | 150 | 623 | 5 | 1 | 1 | Ibsp |
| Ibtk | 5410 | 4276 | 4810 | 4404 | 5332 | 5348 | 5194 | 8248 | 7889 | 7550 | 7900 | 8348 | 8490 | 8403 | 5147 | 4892 | 4663 | 4528 | 5032 | 5212 | 4918 | 7879 | 7821 | 7670 | 7982 | 6817 | 8542 | 8175 | Ibtk |
| Ica1 | 646 | 633 | 614 | 665 | 975 | 729 | 705 | 1112 | 1098 | 1242 | 1052 | 749 | 874 | 856 | 717 | 758 | 633 | 690 | 767 | 802 | 754 | 1206 | 1144 | 945 | 1025 | 830 | 774 | 757 | Ica1 |
| Ica1l | 130 | 159 | 120 | 270 | 695 | 490 | 542 | 888 | 979 | 1132 | 939 | 570 | 560 | 634 | 215 | 169 | 226 | 403 | 427 | 503 | 541 | 1348 | 971 | 687 | 1027 | 383 | 497 | 527 | Ica1l |
| Icam1 | 642 | 660 | 542 | 416 | 313 | 481 | 298 | 274 | 419 | 304 | 508 | 290 | 392 | 295 | 584 | 812 | 447 | 624 | 296 | 283 | 296 | 286 | 542 | 297 | 296 | 321 | 274 | 301 | Icam1 |
| Icam2 | 205 | 174 | 135 | 144 | 187 | 170 | 113 | 319 | 407 | 421 | 412 | 278 | 369 | 298 | 116 | 207 | 129 | 147 | 168 | 193 | 162 | 281 | 527 | 378 | 246 | 455 | 287 | 336 | Icam2 |
| Icam4 | 9 | 14 | 41 | 1 | 2 | 1 | 4 | 2 | 6 | 1 | 0 | 9 | 29 | 18 | 8 | 17 | 10 | 6 | 3 | 2 | 4 | 0 | 4 | 16 | 5 | 13 | 15 | 66 | Icam4 |
| Icam5 | 240 | 295 | 245 | 248 | 164 | 166 | 123 | 15 | 14 | 25 | 18 | 4 | 5 | 7 | 152 | 271 | 170 | 246 | 139 | 164 | 131 | 10 | 16 | 12 | 11 | 16 | 5 | 5 | Icam5 |
| Ice1 | 15043 | 16203 | 10891 | 14573 | 16535 | 16617 | 16075 | 15961 | 15815 | 15349 | 15790 | 14127 | 14072 | 13732 | 14968 | 14985 | 12758 | 15042 | 15101 | 15411 | 16373 | 15308 | 15166 | 12353 | 16464 | 8076 | 14145 | 12589 | Ice1 |
| Ice2 | 1977 | 1621 | 1750 | 1428 | 1351 | 1433 | 1672 | 3220 | 3083 | 2318 | 2448 | 2690 | 2637 | 2595 | 2064 | 1872 | 1728 | 1221 | 1512 | 1824 | 1536 | 2129 | 2966 | 2365 | 2250 | 2406 | 2639 | 2572 | Ice2 |
| Icmt | 5114 | 5529 | 5676 | 5401 | 4829 | 4798 | 4126 | 3579 | 3814 | 3846 | 3940 | 4108 | 3967 | 3406 | 4588 | 5605 | 5373 | 5406 | 4827 | 4484 | 4364 | 3588 | 3662 | 3957 | 4008 | 3733 | 3871 | 3764 | Icmt |
| Icos | 1 | 4 | 5 | 0 | 4 | 0 | 0 | 7 | 7 | 10 | 16 | 8 | 7 | 3 | 8 | 3 | 0 | 2 | 1 | 3 | 1 | 7 | 5 | 7 | 0 | 8 | 7 | 2 | Icos |
| Icosl | 270 | 255 | 357 | 204 | 190 | 195 | 177 | 140 | 179 | 119 | 176 | 173 | 166 | 162 | 270 | 360 | 220 | 215 | 163 | 180 | 138 | 110 | 186 | 134 | 100 | 197 | 166 | 166 | Icosl |
| Id1 | 1666 | 1980 | 1665 | 1273 | 982 | 1026 | 529 | 1843 | 2208 | 1852 | 2020 | 1184 | 2244 | 1097 | 1694 | 2231 | 1509 | 1152 | 858 | 867 | 598 | 1457 | 2673 | 1516 | 1108 | 1676 | 1323 | 1380 | Id1 |
| Id2 | 4685 | 3936 | 3541 | 2922 | 2912 | 2958 | 2704 | 5699 | 5613 | 4744 | 4709 | 3811 | 4948 | 4097 | 4349 | 4878 | 2780 | 3116 | 2442 | 2901 | 2926 | 4189 | 5934 | 4349 | 3925 | 4076 | 4005 | 4690 | Id2 |
| Id3 | 3262 | 3537 | 4080 | 2269 | 1671 | 1821 | 1084 | 3573 | 4033 | 3349 | 4335 | 3447 | 4524 | 3288 | 2668 | 3861 | 2810 | 2028 | 1421 | 1622 | 994 | 2921 | 4762 | 3774 | 2494 | 4662 | 4184 | 4355 | Id3 |
| Id4 | 11701 | 9987 | 11464 | 20191 | 22058 | 19702 | 24465 | 22791 | 17115 | 26027 | 19860 | 30158 | 21798 | 33433 | 12864 | 8527 | 16507 | 20778 | 22415 | 20187 | 24570 | 23841 | 16601 | 32038 | 31055 | 32520 | 24076 | 27966 | Id4 |
| Ide | 2 | 1 | 0 | 0 | 0 | 0 | 0 | 0 | 0 | 0 | 0 | 0 | 0 | 0 | 4 | 0 | 0 | 0 | 0 | 0 | 1 | 0 | 0 | 0 | 0 | 0 | 0 | 0 | Ide |
| Idh1 | 18590 | 17602 | 17941 | 18802 | 20918 | 20552 | 22897 | 28482 | 21550 | 21537 | 18957 | 23381 | 22093 | 25740 | 22411 | 17775 | 19520 | 19444 | 21052 | 25243 | 21913 | 25950 | 21854 | 22750 | 24981 | 22984 | 25747 | 26182 | Idh1 |
| Idh2 | 9567 | 9986 | 9148 | 8193 | 7333 | 8102 | 7248 | 9492 | 11089 | 9543 | 10388 | 10914 | 10404 | 10182 | 8315 | 9396 | 8321 | 7672 | 8369 | 8523 | 7543 | 8214 | 11328 | 10730 | 9996 | 12690 | 11341 | 10822 | Idh2 |
| Idh3a | 10811 | 10236 | 9624 | 9787 | 8843 | 9085 | 10658 | 5737 | 5167 | 4756 | 4557 | 5118 | 4612 | 5438 | 10205 | 9091 | 9488 | 9993 | 9436 | 10020 | 10395 | 5125 | 5141 | 5159 | 5671 | 5311 | 4882 | 5431 | Idh3a |
| Idh3b | 7398 | 8187 | 8199 | 7485 | 6862 | 6921 | 7357 | 8004 | 7669 | 7302 | 7507 | 7585 | 7440 | 8150 | 7440 | 7114 | 7755 | 7268 | 7484 | 7590 | 7183 | 7503 | 7720 | 7926 | 7820 | 8890 | 7915 | 8071 | Idh3b |
| Idh3g | 4556 | 5513 | 4375 | 4507 | 4631 | 5010 | 4537 | 6050 | 6347 | 5548 | 5313 | 6339 | 5907 | 5776 | 4861 | 4435 | 4444 | 4401 | 5018 | 5401 | 4791 | 5476 | 6174 | 6058 | 5883 | 6808 | 6369 | 6068 | Idh3g |
| Idi1 | 7578 | 6979 | 7953 | 6267 | 4975 | 5664 | 6534 | 4509 | 3278 | 3317 | 3134 | 5405 | 5281 | 6231 | 7962 | 7396 | 6828 | 6304 | 5227 | 6417 | 5689 | 3514 | 3304 | 4015 | 4005 | 6189 | 6213 | 6992 | Idi1 |
| Idi2 | 0 | 0 | 0 | 0 | 0 | 0 | 0 | 5 | 7 | 0 | 5 | 0 | 1 | 1 | 0 | 0 | 0 | 0 | 0 | 0 | 0 | 0 | 8 | 0 | 0 | 1 | 0 | 2 | Idi2 |
| Idnk | 3045 | 3414 | 3846 | 4037 | 3617 | 3442 | 3965 | 1611 | 1507 | 1773 | 1658 | 1816 | 1759 | 1909 | 3165 | 3109 | 3910 | 3822 | 4119 | 3752 | 3951 | 1826 | 1495 | 1784 | 1962 | 1714 | 1906 | 2093 | Idnk |
| Ido1 | 3 | 8 | 3 | 2 | 7 | 1 | 4 | 4 | 5 | 1 | 3 | 4 | 7 | 2 | 4 | 5 | 10 | 3 | 5 | 2 | 4 | 13 | 1 | 0 | 2 | 4 | 0 | 0 | Ido1 |
| Ido2 | 16 | 11 | 10 | 8 | 25 | 22 | 10 | 48 | 28 | 43 | 40 | 34 | 39 | 32 | 11 | 10 | 16 | 20 | 9 | 18 | 12 | 35 | 29 | 52 | 59 | 31 | 35 | 36 | Ido2 |
| Ids | 7845 | 6398 | 6038 | 9969 | 14177 | 12375 | 14023 | 11836 | 11535 | 11056 | 11250 | 9066 | 9483 | 9865 | 9206 | 7416 | 7868 | 11475 | 11798 | 12044 | 13848 | 16439 | 10381 | 10100 | 13291 | 7038 | 8297 | 8810 | Ids |
| Idua | 429 | 533 | 578 | 574 | 652 | 576 | 530 | 658 | 859 | 991 | 931 | 853 | 995 | 884 | 500 | 603 | 581 | 517 | 585 | 427 | 467 | 912 | 867 | 1027 | 940 | 854 | 924 | 898 | Idua |
| Ier2 | 1578 | 2145 | 1837 | 2100 | 1222 | 1325 | 1330 | 833 | 983 | 956 | 972 | 1043 | 1088 | 933 | 1598 | 1931 | 1874 | 1777 | 1556 | 1248 | 1351 | 675 | 937 | 1021 | 864 | 1491 | 1012 | 1016 | Ier2 |
| Ier3 | 384 | 394 | 304 | 300 | 272 | 371 | 268 | 125 | 180 | 137 | 172 | 117 | 154 | 125 | 347 | 418 | 408 | 354 | 305 | 352 | 268 | 117 | 195 | 161 | 140 | 145 | 152 | 121 | Ier3 |
| Ier3ip1 | 3599 | 2669 | 3343 | 3565 | 3024 | 3089 | 3613 | 3297 | 2550 | 2274 | 2378 | 2551 | 2682 | 2698 | 4031 | 3432 | 3118 | 3156 | 3187 | 4105 | 3253 | 2978 | 2644 | 2532 | 2674 | 2621 | 2580 | 2840 | Ier3ip1 |
| Ier5 | 3286 | 3692 | 3705 | 4549 | 3477 | 3682 | 4307 | 2560 | 2684 | 3106 | 2811 | 3350 | 2730 | 3530 | 3082 | 3089 | 4310 | 4461 | 4078 | 3595 | 4412 | 2483 | 2577 | 3767 | 3674 | 4007 | 2625 | 3165 | Ier5 |
| Ier5l | 837 | 1059 | 1061 | 981 | 767 | 714 | 664 | 1055 | 979 | 1147 | 1022 | 1019 | 1034 | 908 | 749 | 863 | 1034 | 865 | 897 | 773 | 701 | 957 | 978 | 1113 | 978 | 1321 | 933 | 938 | Ier5l |
| Iffo1 | 1922 | 2174 | 2790 | 2904 | 2510 | 2331 | 2202 | 1249 | 1589 | 1668 | 2444 | 1806 | 1653 | 1706 | 1851 | 2196 | 2574 | 2632 | 2498 | 1873 | 2223 | 1739 | 1772 | 1791 | 1793 | 1779 | 1685 | 1746 | Iffo1 |
| Iffo2 | 2141 | 2121 | 2607 | 3025 | 2767 | 2529 | 2291 | 1075 | 1077 | 1293 | 1599 | 1198 | 1456 | 1214 | 2023 | 2284 | 2780 | 3005 | 2606 | 1917 | 2397 | 1370 | 1058 | 1412 | 1283 | 1337 | 1160 | 1386 | Iffo2 |
| Ifi202b | 1 | 0 | 3 | 0 | 2 | 1 | 0 | 10 | 16 | 5 | 9 | 0 | 0 | 4 | 1 | 2 | 0 | 2 | 0 | 0 | 0 | 0 | 17 | 1 | 3 | 2 | 0 | 0 | Ifi202b |
| Ifi203 | 8 | 4 | 10 | 15 | 56 | 29 | 38 | 59 | 58 | 90 | 98 | 35 | 25 | 36 | 9 | 6 | 12 | 23 | 31 | 62 | 45 | 62 | 126 | 102 | 108 | 19 | 28 | 14 | Ifi203 |
| Ifi204 | 54 | 60 | 52 | 28 | 36 | 52 | 41 | 65 | 59 | 100 | 122 | 78 | 65 | 63 | 76 | 74 | 27 | 43 | 38 | 105 | 55 | 95 | 102 | 65 | 71 | 80 | 102 | 87 | Ifi204 |
| Ifi205 | 3 | 0 | 0 | 3 | 0 | 0 | 0 | 2 | 5 | 0 | 0 | 0 | 0 | 2 | 1 | 0 | 0 | 0 | 1 | 2 | 9 | 5 | 6 | 3 | 3 | 3 | 0 | 2 | Ifi205 |
| Ifi206 | 2 | 0 | 1 | 0 | 0 | 1 | 2 | 2 | 5 | 5 | 9 | 1 | 5 | 2 | 0 | 0 | 0 | 0 | 0 | 4 | 4 | 2 | 4 | 3 | 1 | 10 | 6 | 6 | Ifi206 |
| Ifi207 | 54 | 30 | 62 | 15 | 31 | 31 | 32 | 109 | 95 | 110 | 142 | 111 | 168 | 103 | 59 | 41 | 43 | 45 | 22 | 58 | 31 | 109 | 181 | 118 | 115 | 114 | 120 | 85 | Ifi207 |
| Ifi208 | 10 | 8 | 0 | 10 | 4 | 1 | 4 | 4 | 2 | 30 | 18 | 10 | 7 | 7 | 7 | 25 | 20 | 7 | 2 | 28 | 8 | 5 | 14 | 5 | 18 | 9 | 28 | 11 | Ifi208 |
| Ifi209 | 8 | 11 | 4 | 21 | 6 | 8 | 1 | 24 | 7 | 10 | 16 | 15 | 11 | 9 | 9 | 11 | 12 | 15 | 8 | 19 | 16 | 15 | 22 | 13 | 9 | 5 | 15 | 12 | Ifi209 |
| Ifi211 | 10 | 12 | 1 | 9 | 4 | 4 | 11 | 10 | 13 | 21 | 19 | 3 | 12 | 13 | 9 | 15 | 5 | 12 | 0 | 10 | 7 | 13 | 17 | 17 | 12 | 14 | 23 | 17 | Ifi211 |
| Ifi213 | 3 | 1 | 0 | 3 | 2 | 4 | 0 | 1 | 0 | 14 | 6 | 4 | 3 | 3 | 1 | 0 | 1 | 3 | 0 | 0 | 0 | 7 | 5 | 2 | 8 | 1 | 6 | 5 | Ifi213 |
| Ifi214 | 0 | 0 | 0 | 0 | 0 | 0 | 0 | 0 | 0 | 0 | 3 | 0 | 0 | 0 | 0 | 0 | 0 | 0 | 0 | 0 | 0 | 0 | 1 | 0 | 0 | 0 | 0 | 0 | Ifi214 |
| Ifi27 | 172 | 140 | 125 | 148 | 146 | 198 | 75 | 228 | 367 | 363 | 525 | 210 | 282 | 209 | 156 | 179 | 117 | 288 | 89 | 172 | 105 | 485 | 561 | 374 | 329 | 217 | 243 | 227 | Ifi27 |
| Ifi27l2a | 24 | 28 | 15 | 35 | 25 | 16 | 34 | 26 | 41 | 63 | 68 | 40 | 26 | 47 | 15 | 21 | 10 | 45 | 18 | 26 | 50 | 32 | 37 | 65 | 66 | 19 | 22 | 49 | Ifi27l2a |
| Ifi30 | 6 | 14 | 24 | 18 | 5 | 12 | 8 | 6 | 20 | 6 | 14 | 7 | 6 | 1 | 18 | 19 | 7 | 17 | 15 | 14 | 16 | 13 | 3 | 3 | 4 | 8 | 3 | 6 | Ifi30 |

| | | | | | | | | | | | | | | | | | | | | | | | | | | | | | |
|---|---|---|---|---|---|---|---|---|---|---|---|---|---|---|---|---|---|---|---|---|---|---|---|---|---|---|---|---|---|
| Ifi35 | 108 | 122 | 123 | 90 | 91 | 92 | 60 | 96 | 119 | 126 | 152 | 84 | 115 | 98 | 96 | 160 | 80 | 97 | 95 | 128 | 85 | 97 | 165 | 114 | 85 | 124 | 107 | 109 | Ifi35 |
| Ifi44 | 3 | 3 | 2 | 3 | 2 | 6 | 0 | 55 | 21 | 45 | 57 | 26 | 19 | 21 | 3 | 5 | 7 | 9 | 2 | 28 | 4 | 62 | 63 | 48 | 23 | 40 | 28 | 21 | Ifi44 |
| Ifi44l | 0 | 0 | 0 | 0 | 0 | 0 | 0 | 0 | 2 | 0 | 1 | 0 | 0 | 0 | 0 | 0 | 0 | 0 | 0 | 4 | 0 | 0 | 0 | 0 | 0 | 0 | 0 | 0 | Ifi44l |
| Ifi47 | 0 | 0 | 0 | 0 | 0 | 0 | 0 | 0 | 0 | 0 | 0 | 0 | 0 | 0 | 0 | 0 | 0 | 0 | 0 | 1 | 0 | 0 | 1 | 1 | 0 | 0 | 0 | 0 | Ifi47 |
| Ifih1 | 204 | 155 | 163 | 150 | 182 | 200 | 187 | 255 | 234 | 221 | 234 | 229 | 218 | 246 | 208 | 238 | 137 | 158 | 135 | 281 | 196 | 227 | 299 | 202 | 166 | 220 | 231 | 193 | Ifih1 |
| Ifit1 | 6 | 4 | 1 | 4 | 2 | 4 | 3 | 16 | 12 | 55 | 22 | 22 | 25 | 43 | 4 | 12 | 9 | 2 | 1 | 25 | 7 | 18 | 34 | 25 | 21 | 25 | 36 | 20 | Ifit1 |
| Ifit1bl1 | 20 | 29 | 28 | 13 | 39 | 13 | 19 | 18 | 35 | 52 | 41 | 10 | 24 | 29 | 21 | 26 | 17 | 12 | 16 | 15 | 12 | 35 | 38 | 17 | 20 | 16 | 15 | 20 | Ifit1bl1 |
| Ifit1bl2 | 18 | 11 | 19 | 6 | 7 | 5 | 9 | 42 | 29 | 14 | 28 | 31 | 36 | 11 | 21 | 25 | 10 | 18 | 5 | 5 | 4 | 40 | 32 | 27 | 17 | 17 | 16 | 14 | Ifit1bl2 |
| Ifit2 | 662 | 364 | 468 | 381 | 437 | 462 | 484 | 896 | 974 | 803 | 914 | 909 | 829 | 873 | 578 | 474 | 381 | 353 | 478 | 582 | 407 | 732 | 896 | 932 | 816 | 1038 | 803 | 904 | Ifit2 |
| Ifit3 | 15 | 18 | 7 | 8 | 15 | 17 | 12 | 20 | 24 | 39 | 70 | 34 | 37 | 30 | 24 | 8 | 10 | 13 | 11 | 15 | 21 | 52 | 54 | 49 | 36 | 57 | 33 | 28 | Ifit3 |
| Ifit3b | 3 | 3 | 0 | 5 | 1 | 1 | 1 | 5 | 11 | 5 | 15 | 5 | 10 | 2 | 4 | 2 | 0 | 1 | 5 | 5 | 4 | 8 | 6 | 6 | 4 | 3 | 2 | 5 | Ifit3b |
| Ifitm1 | 166 | 143 | 179 | 115 | 122 | 133 | 63 | 231 | 321 | 227 | 246 | 135 | 220 | 131 | 100 | 200 | 98 | 151 | 80 | 147 | 111 | 155 | 379 | 230 | 88 | 201 | 149 | 166 | Ifitm1 |
| Ifitm10 | 13 | 14 | 1 | 8 | 13 | 16 | 13 | 6 | 15 | 12 | 20 | 18 | 5 | 12 | 13 | 11 | 13 | 36 | 27 | 18 | 25 | 30 | 21 | 19 | 18 | 12 | 2 | 8 | Ifitm10 |
| Ifitm2 | 475 | 358 | 232 | 235 | 146 | 136 | 251 | 419 | 451 | 337 | 457 | 185 | 278 | 460 | 386 | 296 | 211 | 160 | 116 | 136 | 212 | 254 | 519 | 387 | 196 | 270 | 195 | 471 | Ifitm2 |
| Ifitm3 | 231 | 187 | 148 | 187 | 165 | 227 | 142 | 199 | 277 | 304 | 395 | 206 | 264 | 182 | 164 | 263 | 177 | 191 | 157 | 294 | 259 | 251 | 377 | 317 | 223 | 262 | 255 | 197 | Ifitm3 |
| Ifitm5 | 22 | 19 | 25 | 17 | 7 | 23 | 4 | 7 | 18 | 27 | 37 | 17 | 15 | 10 | 13 | 31 | 20 | 19 | 7 | 10 | 10 | 5 | 15 | 24 | 16 | 8 | 17 | 17 | Ifitm5 |
| Ifitm6 | 1 | 1 | 0 | 5 | 3 | 0 | 0 | 0 | 2 | 10 | 18 | 3 | 10 | 1 | 0 | 2 | 1 | 0 | 2 | 1 | 5 | 0 | 5 | 4 | 12 | 4 | 8 | 8 | Ifitm6 |
| Ifitm7 | 25 | 32 | 19 | 6 | 36 | 34 | 21 | 104 | 76 | 74 | 107 | 130 | 143 | 118 | 25 | 21 | 34 | 18 | 39 | 47 | 35 | 85 | 90 | 51 | 76 | 75 | 141 | 109 | Ifitm7 |
| Ifna11 | 0 | 0 | 0 | 0 | 0 | 0 | 0 | 0 | 0 | 0 | 0 | 0 | 0 | 0 | 0 | 0 | 0 | 0 | 0 | 0 | 0 | 0 | 1 | 0 | 0 | 0 | 0 | 0 | Ifna11 |
| Ifna12 | 0 | 0 | 0 | 0 | 0 | 0 | 0 | 0 | 0 | 0 | 0 | 0 | 0 | 0 | 0 | 2 | 0 | 0 | 0 | 0 | 0 | 0 | 0 | 0 | 0 | 0 | 0 | 0 | Ifna12 |
| Ifna16 | 0 | 0 | 0 | 0 | 1 | 0 | 0 | 0 | 0 | 0 | 0 | 0 | 0 | 0 | 0 | 0 | 0 | 0 | 0 | 0 | 0 | 0 | 0 | 0 | 0 | 0 | 0 | 0 | Ifna16 |
| Ifna2 | 0 | 0 | 2 | 0 | 0 | 0 | 1 | 2 | 0 | 0 | 3 | 0 | 1 | 0 | 0 | 0 | 0 | 0 | 0 | 0 | 0 | 0 | 0 | 0 | 0 | 0 | 0 | 0 | Ifna2 |
| Ifna7 | 0 | 0 | 0 | 3 | 0 | 0 | 0 | 0 | 0 | 0 | 0 | 0 | 0 | 0 | 0 | 1 | 0 | 0 | 0 | 0 | 0 | 0 | 0 | 0 | 0 | 0 | 0 | 0 | Ifna7 |
| Ifnab | 1 | 0 | 0 | 1 | 3 | 0 | 0 | 0 | 0 | 0 | 0 | 0 | 0 | 1 | 0 | 2 | 1 | 0 | 1 | 0 | 2 | 0 | 1 | 1 | 1 | 0 | 1 | 0 | Ifnab |
| Ifnar1 | 7359 | 7696 | 5762 | 7052 | 6690 | 7046 | 6683 | 7173 | 7601 | 7083 | 7666 | 7297 | 7514 | 7327 | 7129 | 7256 | 6298 | 6940 | 6443 | 6579 | 6693 | 7061 | 7299 | 6879 | 7646 | 6100 | 7598 | 7122 | Ifnar1 |
| Ifnar2 | 443 | 381 | 488 | 344 | 439 | 388 | 355 | 854 | 887 | 973 | 1019 | 891 | 913 | 851 | 352 | 384 | 372 | 343 | 413 | 393 | 393 | 904 | 933 | 963 | 900 | 1061 | 883 | 870 | Ifnar2 |
| Ifng | 0 | 0 | 0 | 0 | 0 | 0 | 0 | 0 | 0 | 0 | 1 | 0 | 0 | 0 | 0 | 0 | 0 | 0 | 0 | 0 | 0 | 0 | 0 | 0 | 0 | 0 | 0 | 0 | Ifng |
| Ifngr1 | 2022 | 1996 | 1712 | 1534 | 1368 | 1386 | 1369 | 1036 | 1135 | 956 | 1193 | 1046 | 1229 | 1046 | 1904 | 2051 | 1737 | 1549 | 1322 | 1518 | 1276 | 1135 | 1340 | 1076 | 903 | 1181 | 1173 | 1156 | Ifngr1 |
| Ifngr2 | 2594 | 2695 | 2647 | 2925 | 3129 | 2911 | 2979 | 1934 | 2387 | 2203 | 2282 | 1735 | 1909 | 1759 | 2734 | 2974 | 2539 | 2996 | 2949 | 3066 | 3141 | 2115 | 2453 | 2127 | 2422 | 1855 | 1847 | 1879 | Ifngr2 |
| Ifnlr1 | 287 | 205 | 258 | 179 | 148 | 201 | 171 | 164 | 173 | 226 | 163 | 97 | 127 | 85 | 187 | 234 | 210 | 196 | 196 | 163 | 139 | 100 | 206 | 181 | 63 | 172 | 89 | 88 | Ifnlr1 |
| Ifnz | 0 | 3 | 0 | 3 | 0 | 2 | 1 | 0 | 2 | 0 | 0 | 0 | 1 | 0 | 3 | 6 | 0 | 2 | 0 | 0 | 2 | 0 | 1 | 0 | 0 | 0 | 0 | 0 | Ifnz |
| Ifrd1 | 4573 | 4166 | 4240 | 4859 | 4327 | 4705 | 5965 | 3972 | 3260 | 3167 | 2962 | 3973 | 3822 | 4674 | 4812 | 3585 | 4561 | 4587 | 4716 | 5065 | 5354 | 3631 | 3225 | 3561 | 3627 | 4400 | 4247 | 4518 | Ifrd1 |
| Ifrd2 | 1024 | 1164 | 1249 | 1073 | 691 | 741 | 770 | 856 | 1056 | 1002 | 1166 | 1021 | 1038 | 981 | 860 | 1113 | 1129 | 1008 | 809 | 749 | 809 | 714 | 1111 | 1076 | 956 | 1419 | 972 | 1135 | Ifrd2 |
| Ift122 | 1817 | 1965 | 1722 | 1783 | 1775 | 1851 | 1682 | 2876 | 3339 | 3352 | 2928 | 3145 | 3046 | 3001 | 1892 | 1884 | 1861 | 1744 | 1862 | 1700 | 1763 | 3009 | 3149 | 3148 | 3058 | 2884 | 3253 | 2986 | Ift122 |
| Ift140 | 7844 | 8383 | 5706 | 7769 | 7460 | 7810 | 7283 | 4013 | 4761 | 4382 | 4018 | 4561 | 4786 | 4385 | 6902 | 7076 | 6537 | 7594 | 7779 | 7114 | 7805 | 4040 | 4266 | 4202 | 4450 | 3715 | 4664 | 4347 | Ift140 |
| Ift172 | 5034 | 5801 | 4461 | 5864 | 5515 | 5701 | 5703 | 5051 | 6294 | 6417 | 6410 | 5886 | 6023 | 5497 | 4971 | 5590 | 5203 | 5748 | 5670 | 5075 | 5653 | 5474 | 6100 | 5936 | 6058 | 4413 | 5957 | 5652 | Ift172 |
| Ift20 | 3656 | 3992 | 3842 | 3953 | 3346 | 3607 | 3701 | 2599 | 2529 | 2306 | 2640 | 2202 | 2324 | 2490 | 3825 | 3829 | 3845 | 4121 | 3471 | 4098 | 3674 | 2336 | 2710 | 2757 | 2678 | 2614 | 2418 | 2544 | Ift20 |
| Ift22 | 2772 | 3514 | 3151 | 3101 | 2789 | 2925 | 3027 | 2137 | 2174 | 1925 | 1890 | 2346 | 2194 | 2439 | 2937 | 3046 | 3180 | 2881 | 3038 | 3231 | 2975 | 1997 | 2223 | 2236 | 2349 | 2553 | 2464 | 2511 | Ift22 |
| Ift27 | 2776 | 3844 | 3109 | 3271 | 2995 | 3017 | 3287 | 1699 | 1834 | 1845 | 1840 | 2368 | 2092 | 2253 | 2719 | 2678 | 3401 | 3254 | 3370 | 2952 | 3010 | 1589 | 1710 | 2054 | 2165 | 2667 | 2443 | 2490 | Ift27 |
| Ift43 | 1314 | 1476 | 1561 | 1479 | 1208 | 1336 | 1321 | 1077 | 1172 | 1292 | 1100 | 1453 | 1505 | 1484 | 1341 | 1163 | 1640 | 1394 | 1433 | 1409 | 1337 | 959 | 1189 | 1393 | 1321 | 1723 | 1507 | 1512 | Ift43 |
| Ift46 | 5324 | 5795 | 4385 | 4995 | 4793 | 5018 | 5481 | 4631 | 4458 | 4221 | 3959 | 4265 | 3977 | 4443 | 5438 | 4846 | 4876 | 5093 | 5072 | 5342 | 5487 | 4368 | 4450 | 4316 | 4351 | 4330 | 4503 | 4549 | Ift46 |
| Ift52 | 2504 | 2432 | 2472 | 2033 | 1852 | 2164 | 2162 | 2869 | 2859 | 2911 | 2747 | 3257 | 3178 | 3399 | 2445 | 2211 | 2211 | 1841 | 1931 | 2080 | 2045 | 2637 | 2929 | 2924 | 2759 | 3220 | 3381 | 3538 | Ift52 |
| Ift57 | 2742 | 2452 | 2675 | 2930 | 2654 | 2483 | 3100 | 1583 | 1503 | 1480 | 1362 | 1487 | 1466 | 1587 | 2807 | 2498 | 2921 | 2834 | 2629 | 2894 | 2759 | 1626 | 1525 | 1496 | 1564 | 1537 | 1396 | 1559 | Ift57 |
| Ift74 | 6281 | 5254 | 6756 | 5383 | 4619 | 5214 | 5292 | 4799 | 3847 | 3743 | 3465 | 5184 | 4901 | 5227 | 6629 | 5748 | 5991 | 4581 | 5073 | 6267 | 4789 | 3820 | 3893 | 4479 | 3799 | 6016 | 5320 | 5682 | Ift74 |
| Ift80 | 7101 | 5734 | 5986 | 5432 | 5119 | 5475 | 5843 | 4317 | 3896 | 3366 | 3597 | 3845 | 3809 | 4189 | 6965 | 6316 | 5601 | 5509 | 5344 | 6378 | 5952 | 3233 | 4016 | 3658 | 3236 | 3610 | 4161 | 3926 | Ift80 |
| Ift81 | 4321 | 3890 | 4436 | 3849 | 4060 | 3848 | 4441 | 5117 | 4264 | 4177 | 3920 | 4217 | 4124 | 4716 | 4704 | 4105 | 4284 | 3855 | 4185 | 4907 | 4086 | 4697 | 4322 | 4260 | 4208 | 4351 | 4610 | 4513 | Ift81 |
| Ift88 | 3740 | 3333 | 3275 | 3310 | 3301 | 3157 | 3477 | 3186 | 2821 | 2835 | 2998 | 2740 | 3044 | 3113 | 3798 | 3519 | 3174 | 3146 | 3068 | 3597 | 3421 | 2822 | 3077 | 2747 | 2716 | 2823 | 2993 | 3214 | Ift88 |
| Iftap | 1742 | 1310 | 1751 | 1384 | 1316 | 1519 | 1616 | 1822 | 1523 | 1330 | 1505 | 1603 | 1609 | 1804 | 2029 | 1701 | 1484 | 1437 | 1320 | 1916 | 1433 | 1504 | 1612 | 1503 | 1364 | 1929 | 1739 | 1804 | Iftap |
| Igbp1 | 4384 | 4085 | 3259 | 3192 | 2965 | 3269 | 3306 | 4316 | 3902 | 3302 | 3250 | 3524 | 3504 | 3650 | 4510 | 3807 | 3262 | 3043 | 3051 | 3807 | 3296 | 3395 | 3967 | 3285 | 3270 | 3643 | 3818 | 3830 | Igbp1 |
| Igbp1b | 6 | 2 | 0 | 8 | 6 | 5 | 9 | 7 | 6 | 10 | 2 | 10 | 1 | 17 | 0 | 1 | 17 | 7 | 0 | 4 | 6 | 18 | 5 | 9 | 17 | 6 | 9 | 14 | Igbp1b |
| Igdcc3 | 1179 | 1569 | 3289 | 1537 | 1424 | 1356 | 1210 | 1541 | 1320 | 1967 | 1518 | 3829 | 2918 | 3295 | 1170 | 1304 | 2476 | 1317 | 1781 | 1087 | 1259 | 2079 | 1249 | 2382 | 2186 | 3592 | 4043 | 4211 | Igdcc3 |
| Igdcc4 | 2953 | 2975 | 3513 | 3066 | 3268 | 3858 | 2813 | 4726 | 5699 | 6221 | 7795 | 6218 | 5782 | 5557 | 2930 | 3368 | 3559 | 3526 | 3636 | 2572 | 2792 | 6116 | 6295 | 6561 | 5860 | 4219 | 5826 | 5613 | Igdcc4 |
| Igf1 | 3026 | 1407 | 1943 | 956 | 669 | 1172 | 434 | 3631 | 5430 | 3273 | 10378 | 2433 | 4227 | 2287 | 2051 | 3031 | 1136 | 1159 | 399 | 539 | 441 | 4079 | 10464 | 2617 | 2533 | 2550 | 2817 | 2784 | Igf1 |
| Igf1r | 13239 | 12935 | 18214 | 19223 | 18483 | 17528 | 17488 | 11370 | 12147 | 15960 | 15735 | 15038 | 15777 | 15090 | 12871 | 14610 | 18489 | 18026 | 18637 | 12760 | 17140 | 16238 | 11607 | 16011 | 16723 | 13970 | 15218 | 15717 | Igf1r |
| Igf2 | 1369 | 1917 | 1336 | 1029 | 546 | 932 | 247 | 1822 | 3505 | 2619 | 5351 | 1729 | 2601 | 1479 | 1195 | 1796 | 985 | 1205 | 422 | 316 | 295 | 2879 | 4574 | 2596 | 1971 | 2067 | 1799 | 1629 | Igf2 |
| Igf2bp1 | 8200 | 8225 | 16362 | 9008 | 7392 | 7673 | 7949 | 6727 | 6478 | 8112 | 8102 | 13301 | 11677 | 13448 | 7292 | 9129 | 13545 | 8692 | 9207 | 6118 | 7590 | 7484 | 6234 | 10769 | 8965 | 16424 | 12987 | 16327 | Igf2bp1 |
| Igf2bp2 | 6526 | 6097 | 8708 | 5461 | 5213 | 5108 | 4875 | 7165 | 7783 | 7732 | 8509 | 9670 | 9208 | 9637 | 5902 | 6801 | 7132 | 5301 | 5459 | 4417 | 4514 | 7090 | 7361 | 9166 | 7514 | 12205 | 9067 | 10433 | Igf2bp2 |
| Igf2bp3 | 13325 | 10432 | 15617 | 9831 | 8336 | 9493 | 9788 | 8899 | 8459 | 8608 | 8995 | 12676 | 11995 | 13054 | 12188 | 12342 | 12391 | 9186 | 9303 | 7640 | 8981 | 8350 | 8303 | 9930 | 9058 | 13813 | 13309 | 15016 | Igf2bp3 |
| Igf2r | 26812 | 27902 | 17817 | 22116 | 20371 | 23333 | 19824 | 12505 | 16554 | 15227 | 25078 | 13467 | 14894 | 13330 | 23343 | 28148 | 19570 | 22895 | 18984 | 19038 | 19384 | 13334 | 19014 | 11683 | 13601 | 8661 | 14149 | 11730 | Igf2r |
| Igfals | 7 | 16 | 0 | 23 | 8 | 15 | 12 | 6 | 7 | 0 | 5 | 2 | 6 | 0 | 9 | 16 | 9 | 25 | 19 | 19 | 24 | 8 | 3 | 2 | 7 | 0 | 1 | 2 | Igfals |
| Igfbp1 | 13 | 27 | 20 | 28 | 32 | 39 | 25 | 16 | 6 | 7 | 30 | 4 | 19 | 9 | 18 | 17 | 16 | 50 | 25 | 21 | 38 | 18 | 13 | 7 | 3 | 8 | 10 | 5 | Igfbp1 |
| Igfbp2 | 14302 | 17899 | 17453 | 13033 | 11306 | 13882 | 8690 | 11091 | 12889 | 11581 | 10559 | 13404 | 15039 | 11835 | 14400 | 17171 | 14552 | 13291 | 12472 | 12588 | 9558 | 8245 | 15599 | 13489 | 10067 | 15058 | 14747 | 13734 | Igfbp2 |
| Igfbp3 | 1239 | 797 | 685 | 668 | 724 | 694 | 582 | 1806 | 2883 | 2039 | 5270 | 919 | 1232 | 872 | 820 | 1118 | 601 | 921 | 609 | 549 | 622 | 1766 | 4406 | 1740 | 1086 | 1547 | 682 | 1055 | Igfbp3 |
| Igfbp4 | 15482 | 14498 | 14759 | 10645 | 7166 | 9899 | 5131 | 14381 | 19111 | 17443 | 22684 | 13812 | 16788 | 12534 | 13336 | 19303 | 11288 | 11753 | 6306 | 5237 | 5418 | 14101 | 22381 | 17298 | 12093 | 19506 | 13331 | 12703 | Igfbp4 |
| Igfbp5 | 17195 | 12038 | 12493 | 12295 | 13405 | 11245 | 10327 | 29593 | 39809 | 35261 | 60197 | 20459 | 24004 | 19565 | 13764 | 16934 | 13085 | 12181 | 10430 | 11007 | 9434 | 30941 | 48590 | 33537 | 18075 | 34072 | 18632 | 21669 | Igfbp5 |
| Igfbp6 | 27 | 12 | 16 | 33 | 26 | 15 | 17 | 33 | 42 | 43 | 65 | 3 | 17 | 11 | 16 | 14 | 10 | 47 | 4 | 14 | 13 | 38 | 57 | 34 | 36 | 12 | 16 | 14 | Igfbp6 |
| Igfbp7 | 450 | 508 | 387 | 523 | 705 | 643 | 630 | 598 | 595 | 664 | 641 | 614 | 515 | 637 | 469 | 347 | 535 | 612 | 697 | 762 | 631 | 703 | 688 | 697 | 782 | 587 | 581 | 555 | Igfbp7 |
| Igfbpl1 | 8732 | 12358 | 11496 | 15553 | 15658 | 15316 | 16239 | 13247 | 12094 | 16437 | 12380 | 18167 | 13345 | 18545 | 10532 | 7555 | 14599 | 14134 | 17849 | 16537 | 17975 | 14708 | 13593 | 18284 | 20108 | 17390 | 17335 | 17881 | Igfbpl1 |
| Igfn1 | 3 | 0 | 1 | 0 | 6 | 1 | 0 | 4 | 17 | 10 | 20 | 8 | 15 | 1 | 0 | 2 | 6 | 3 | 2 | 0 | 1 | 17 | 14 | 3 | 2 | 1 | 2 | 0 | Igfn1 |
| Ighmbp2 | 648 | 708 | 836 | 744 | 920 | 802 | 703 | 979 | 1291 | 1321 | 1340 | 1547 | 1412 | 1224 | 663 | 625 | 689 | 667 | 825 | 690 | 663 | 1024 | 1210 | 1200 | 1427 | 1224 | 1331 | 1286 | Ighmbp2 |
| Igip | 580 | 494 | 909 | 1101 | 1165 | 980 | 1319 | 713 | 714 | 908 | 1008 | 753 | 711 | 842 | 665 | 582 | 1035 | 1325 | 1059 | 853 | 1109 | 1223 | 651 | 915 | 1021 | 752 | 679 | 749 | Igip |
| Igll1 | 0 | 0 | 0 | 4 | 0 | 4 | 0 | 4 | 7 | 5 | 11 | 21 | 6 | 6 | 0 | 1 | 5 | 1 | 3 | 2 | 0 | 10 | 6 | 9 | 8 | 3 | 1 | 4 | Igll1 |
| Iglon5 | 2782 | 3649 | 4378 | 4755 | 5312 | 4375 | 3436 | 3733 | 3915 | 5210 | 4348 | 4958 | 4407 | 4119 | 2863 | 2822 | 4561 | 4495 | 5241 | 3611 | 3942 | 5135 | 3654 | 4814 | 5148 | 4063 | 4092 | 3900 | Iglon5 |
| Igsf1 | 314 | 376 | 220 | 214 | 436 | 293 | 299 | 288 | 365 | 282 | 318 | 239 | 290 | 197 | 444 | 276 | 226 | 309 | 320 | 276 | 284 | 453 | 396 | 203 | 276 | 154 | 171 | 158 | Igsf1 |
| Igsf10 | 4892 | 3606 | 1925 | 2255 | 2366 | 3128 | 1867 | 4675 | 5749 | 4024 | 5558 | 2561 | 3873 | 2621 | 4121 | 4733 | 1961 | 2641 | 1935 | 2213 | 1742 | 3026 | 6615 | 2748 | 2875 | 2151 | 2888 | 2330 | Igsf10 |
| Igsf11 | 584 | 571 | 660 | 834 | 833 | 708 | 889 | 680 | 824 | 1027 | 749 | 1051 | 875 | 1066 | 682 | 437 | 839 | 777 | 813 | 677 | 813 | 744 | 627 | 1026 | 830 | 1175 | 823 | 965 | Igsf11 |
| Igsf21 | 295 | 529 | 632 | 773 | 902 | 819 | 751 | 589 | 610 | 962 | 722 | 804 | 654 | 721 | 359 | 357 | 779 | 829 | 1110 | 727 | 885 | 1044 | 718 | 908 | 1037 | 702 | 794 | 693 | Igsf21 |
| Igsf23 | 1 | 9 | 18 | 0 | 2 | 0 | 1 | 1 | 2 | 3 | 1 | 0 | 1 | 1 | 0 | 3 | 7 | 4 | 2 | 2 | 1 | 0 | 6 | 0 | 4 | 1 | 0 | 0 | Igsf23 |
| Igsf3 | 12829 | 13531 | 12667 | 13896 | 14392 | 13971 | 11342 | 7696 | 9753 | 10829 | 11327 | 7686 | 9834 | 7327 | 11355 | 15306 | 12362 | 14083 | 13409 | 11179 | 13036 | 10726 | 9490 | 9072 | 9619 | 6275 | 7304 | 7528 | Igsf3 |

| | | | | | | | | | | | | | | | | | | | | | | | | | | | | | |
|---|---|---|---|---|---|---|---|---|---|---|---|---|---|---|---|---|---|---|---|---|---|---|---|---|---|---|---|---|---|
| Igsf5 | 2 | 3 | 0 | 3 | 3 | 1 | 0 | 11 | 2 | 0 | 2 | 2 | 1 | 0 | 0 | 0 | 2 | 0 | 1 | 0 | 2 | 7 | 10 | 2 | 0 | 1 | 1 | 2 | Igsf5 |
| Igsf6 | 27 | 38 | 14 | 21 | 11 | 26 | 4 | 53 | 68 | 60 | 85 | 54 | 46 | 51 | 34 | 40 | 13 | 23 | 14 | 21 | 28 | 45 | 88 | 61 | 59 | 50 | 67 | 50 | Igsf6 |
| Igsf8 | 6770 | 10555 | 10120 | 11704 | 9908 | 9458 | 8563 | 4448 | 5019 | 5931 | 5178 | 7288 | 6255 | 6009 | 7561 | 7801 | 12406 | 11137 | 11696 | 8382 | 9167 | 5259 | 4555 | 7309 | 6755 | 7702 | 5836 | 5534 | Igsf8 |
| Igsf9 | 8634 | 11411 | 11767 | 12987 | 12006 | 11478 | 11734 | 7413 | 8309 | 9173 | 7064 | 11373 | 9595 | 9633 | 8020 | 9573 | 12655 | 11954 | 13722 | 10566 | 11228 | 8285 | 7100 | 10782 | 9763 | 10691 | 9681 | 9452 | Igsf9 |
| Igsf9b | 7609 | 8634 | 7043 | 11329 | 12586 | 11789 | 11931 | 4529 | 4608 | 5414 | 5704 | 5374 | 5670 | 5717 | 7353 | 7872 | 8859 | 11905 | 10899 | 9772 | 12077 | 5466 | 4451 | 4664 | 6562 | 3250 | 5478 | 5085 | Igsf9b |
| Igtp | 61 | 60 | 73 | 63 | 40 | 60 | 34 | 126 | 114 | 93 | 141 | 100 | 94 | 110 | 73 | 85 | 51 | 57 | 38 | 93 | 39 | 104 | 124 | 103 | 76 | 175 | 132 | 145 | Igtp |
| Ihh | 8 | 15 | 17 | 8 | 17 | 10 | 8 | 17 | 17 | 30 | 60 | 21 | 18 | 21 | 5 | 7 | 16 | 7 | 11 | 2 | 11 | 43 | 25 | 41 | 25 | 28 | 23 | 19 | Ihh |
| Iho1 | 228 | 190 | 271 | 293 | 197 | 276 | 234 | 132 | 152 | 207 | 170 | 187 | 177 | 175 | 175 | 220 | 205 | 212 | 201 | 229 | 225 | 164 | 170 | 149 | 224 | 147 | 171 | 177 | Iho1 |
| Iigp1 | 25 | 5 | 7 | 19 | 5 | 9 | 3 | 99 | 40 | 78 | 105 | 31 | 32 | 32 | 21 | 18 | 9 | 9 | 2 | 7 | 1 | 22 | 79 | 93 | 8 | 72 | 22 | 32 | Iigp1 |
| Ik | 17507 | 16806 | 18787 | 17032 | 13830 | 14355 | 16213 | 15886 | 13283 | 13712 | 13096 | 13987 | 13311 | 16550 | 18386 | 16163 | 17803 | 15550 | 14821 | 18087 | 15908 | 12995 | 13212 | 14698 | 14066 | 16579 | 14765 | 15902 | Ik |
| Ikbip | 2909 | 2594 | 2775 | 2361 | 1963 | 2299 | 2014 | 1541 | 1821 | 1492 | 2355 | 1339 | 1701 | 1439 | 2966 | 3185 | 2664 | 2480 | 1964 | 2171 | 2040 | 1472 | 2225 | 1443 | 1386 | 1522 | 1447 | 1533 | Ikbip |
| Ikbkb | 8235 | 9944 | 7032 | 8828 | 8434 | 8600 | 8631 | 5882 | 5993 | 6378 | 6000 | 6227 | 5880 | 6325 | 8296 | 8562 | 7617 | 8822 | 8385 | 8519 | 9230 | 5828 | 5595 | 5768 | 6274 | 5063 | 6539 | 5952 | Ikbkb |
| Ikbke | 285 | 324 | 194 | 313 | 286 | 413 | 250 | 260 | 384 | 342 | 549 | 212 | 363 | 244 | 264 | 329 | 299 | 372 | 290 | 267 | 288 | 411 | 520 | 308 | 260 | 194 | 261 | 208 | Ikbke |
| Ikbkg | 3549 | 3200 | 2880 | 2765 | 2720 | 3097 | 3000 | 4222 | 4002 | 3222 | 3584 | 4029 | 4367 | 3904 | 3850 | 3256 | 2721 | 2749 | 2815 | 3298 | 2997 | 3487 | 4030 | 3300 | 3556 | 3629 | 4369 | 4078 | Ikbkg |
| Ikzf1 | 159 | 152 | 314 | 206 | 249 | 211 | 185 | 278 | 290 | 340 | 363 | 339 | 395 | 333 | 187 | 207 | 189 | 232 | 219 | 187 | 225 | 394 | 286 | 242 | 280 | 361 | 322 | 428 | Ikzf1 |
| Ikzf2 | 1529 | 1088 | 1426 | 1469 | 1253 | 1230 | 1248 | 1539 | 1306 | 1031 | 1089 | 917 | 998 | 864 | 1390 | 1280 | 1141 | 1251 | 1257 | 1320 | 1290 | 846 | 1290 | 948 | 833 | 787 | 1017 | 1006 | Ikzf2 |
| Ikzf3 | 9 | 1 | 0 | 4 | 18 | 4 | 1 | 11 | 30 | 14 | 14 | 1 | 5 | 5 | 9 | 2 | 0 | 2 | 1 | 1 | 6 | 15 | 28 | 12 | 3 | 4 | 3 | 4 | Ikzf3 |
| Ikzf4 | 1008 | 1016 | 1294 | 1297 | 2181 | 1639 | 1714 | 2096 | 2121 | 2649 | 2391 | 2304 | 2165 | 2358 | 965 | 929 | 1486 | 1509 | 1852 | 1402 | 1731 | 3334 | 1890 | 2506 | 2903 | 1905 | 1890 | 2206 | Ikzf4 |
| Ikzf5 | 5578 | 4710 | 4478 | 4869 | 4025 | 4728 | 5080 | 4382 | 3849 | 3117 | 3453 | 4157 | 4113 | 4451 | 5542 | 5179 | 4394 | 4533 | 4332 | 4941 | 5024 | 3596 | 3562 | 3731 | 3938 | 3838 | 4371 | 4421 | Ikzf5 |
| Il10 | 0 | 0 | 0 | 0 | 0 | 0 | 0 | 0 | 2 | 0 | 2 | 0 | 5 | 1 | 0 | 0 | 0 | 0 | 1 | 1 | 0 | 0 | 1 | 0 | 0 | 3 | 0 | 5 | Il10 |
| Il10ra | 129 | 136 | 149 | 71 | 96 | 96 | 80 | 89 | 99 | 60 | 86 | 78 | 88 | 72 | 96 | 132 | 88 | 127 | 70 | 85 | 69 | 84 | 142 | 74 | 61 | 89 | 80 | 89 | Il10ra |
| Il10rb | 137 | 119 | 147 | 104 | 150 | 127 | 135 | 247 | 246 | 254 | 373 | 250 | 240 | 220 | 164 | 132 | 123 | 102 | 128 | 106 | 105 | 239 | 335 | 254 | 198 | 219 | 250 | 235 | Il10rb |
| Il11 | 28 | 48 | 54 | 44 | 53 | 52 | 51 | 24 | 12 | 27 | 18 | 12 | 11 | 14 | 34 | 37 | 48 | 57 | 35 | 61 | 65 | 18 | 21 | 22 | 37 | 29 | 21 | 9 | Il11 |
| Il11ra1 | 4482 | 4366 | 3677 | 3293 | 2244 | 2929 | 1809 | 2970 | 4087 | 3343 | 4277 | 2112 | 3133 | 1779 | 3775 | 5579 | 3170 | 3660 | 2271 | 2293 | 1863 | 2615 | 4522 | 3084 | 2166 | 3326 | 2069 | 1997 | Il11ra1 |
| Il11ra2 | 12 | 6 | 7 | 24 | 4 | 7 | 3 | 7 | 21 | 7 | 6 | 4 | 2 | 3 | 16 | 16 | 6 | 4 | 3 | 2 | 1 | 10 | 8 | 2 | 4 | 2 | 1 | 6 | Il11ra2 |
| Il12a | 0 | 1 | 0 | 1 | 0 | 3 | 2 | 0 | 0 | 2 | 4 | 1 | 1 | 0 | 1 | 0 | 2 | 1 | 0 | 4 | 3 | 0 | 3 | 2 | 0 | 0 | 0 | 1 | Il12a |
| Il12b | 0 | 3 | 0 | 1 | 4 | 0 | 2 | 0 | 3 | 0 | 0 | 1 | 3 | 0 | 0 | 1 | 0 | 1 | 0 | 0 | 0 | 2 | 1 | 2 | 0 | 0 | 1 | 0 | Il12b |
| Il12rb1 | 0 | 0 | 3 | 0 | 2 | 3 | 0 | 0 | 0 | 0 | 0 | 0 | 1 | 0 | 1 | 1 | 2 | 0 | 0 | 4 | 0 | 0 | 3 | 1 | 0 | 0 | 0 | 0 | Il12rb1 |
| Il12rb2 | 14 | 12 | 34 | 7 | 4 | 14 | 7 | 24 | 20 | 2 | 15 | 25 | 16 | 12 | 12 | 14 | 19 | 11 | 8 | 10 | 7 | 22 | 17 | 24 | 28 | 17 | 12 | 9 | Il12rb2 |
| Il13 | 1 | 3 | 0 | 0 | 0 | 0 | 0 | 0 | 0 | 0 | 0 | 0 | 1 | 1 | 0 | 1 | 0 | 0 | 0 | 0 | 2 | 0 | 2 | 0 | 0 | 0 | 0 | 0 | Il13 |
| Il13ra1 | 644 | 448 | 452 | 379 | 407 | 377 | 356 | 641 | 775 | 592 | 731 | 432 | 556 | 490 | 642 | 659 | 353 | 380 | 363 | 475 | 332 | 708 | 990 | 624 | 445 | 706 | 474 | 474 | Il13ra1 |
| Il13ra2 | 1 | 1 | 0 | 1 | 6 | 2 | 0 | 5 | 3 | 1 | 3 | 0 | 2 | 3 | 1 | 7 | 2 | 0 | 1 | 3 | 3 | 15 | 8 | 1 | 8 | 0 | 0 | 1 | Il13ra2 |
| Il15 | 78 | 59 | 51 | 37 | 29 | 43 | 40 | 56 | 52 | 22 | 80 | 50 | 42 | 43 | 69 | 67 | 41 | 31 | 31 | 39 | 38 | 53 | 67 | 25 | 18 | 42 | 27 | 43 | Il15 |
| Il15ra | 60 | 54 | 38 | 45 | 42 | 50 | 40 | 17 | 18 | 17 | 20 | 14 | 20 | 19 | 32 | 58 | 35 | 42 | 63 | 43 | 37 | 42 | 34 | 14 | 23 | 8 | 26 | 12 | Il15ra |
| Il16 | 80 | 75 | 99 | 39 | 50 | 87 | 46 | 173 | 259 | 173 | 321 | 125 | 179 | 89 | 76 | 97 | 56 | 73 | 62 | 60 | 46 | 135 | 318 | 146 | 136 | 138 | 134 | 119 | Il16 |
| Il17b | 0 | 0 | 2 | 0 | 8 | 7 | 0 | 4 | 29 | 2 | 210 | 3 | 7 | 0 | 1 | 3 | 3 | 2 | 1 | 5 | 0 | 13 | 113 | 2 | 3 | 2 | 5 | 2 | Il17b |
| Il17c | 2 | 10 | 11 | 14 | 7 | 6 | 6 | 5 | 3 | 3 | 4 | 3 | 2 | 6 | 11 | 8 | 10 | 10 | 12 | 21 | 9 | 7 | 3 | 5 | 5 | 10 | 5 | 5 | Il17c |
| Il17d | 146 | 139 | 120 | 108 | 149 | 160 | 153 | 226 | 268 | 273 | 337 | 198 | 195 | 220 | 152 | 101 | 124 | 149 | 136 | 111 | 161 | 282 | 321 | 261 | 276 | 203 | 187 | 223 | Il17d |
| Il17f | 80 | 79 | 43 | 178 | 66 | 94 | 115 | 20 | 20 | 12 | 17 | 21 | 23 | 38 | 91 | 52 | 82 | 95 | 79 | 78 | 78 | 20 | 19 | 29 | 23 | 34 | 33 | 30 | Il17f |
| Il17ra | 2365 | 3149 | 2603 | 3716 | 3233 | 3075 | 2927 | 696 | 862 | 937 | 858 | 873 | 1142 | 834 | 2618 | 3058 | 3121 | 3876 | 2960 | 2498 | 2903 | 787 | 942 | 762 | 807 | 755 | 681 | 751 | Il17ra |
| Il17rb | 9 | 4 | 1 | 5 | 12 | 10 | 6 | 12 | 25 | 26 | 12 | 10 | 7 | 0 | 5 | 1 | 2 | 3 | 6 | 5 | 6 | 13 | 10 | 4 | 9 | 5 | 0 | 12 | Il17rb |
| Il17rc | 48 | 79 | 61 | 49 | 37 | 41 | 16 | 87 | 140 | 137 | 143 | 83 | 133 | 87 | 69 | 84 | 49 | 36 | 38 | 28 | 32 | 122 | 203 | 103 | 97 | 82 | 101 | 83 | Il17rc |
| Il17rd | 9564 | 7896 | 5951 | 7611 | 5761 | 6342 | 5513 | 4081 | 4210 | 4195 | 4925 | 4312 | 5592 | 4191 | 7067 | 9094 | 6830 | 6814 | 5974 | 5784 | 6063 | 3263 | 4188 | 3826 | 3298 | 3820 | 4855 | 4710 | Il17rd |
| Il17re | 38 | 21 | 18 | 9 | 11 | 26 | 1 | 101 | 115 | 106 | 64 | 48 | 45 | 28 | 28 | 21 | 3 | 25 | 2 | 6 | 3 | 20 | 139 | 69 | 15 | 41 | 35 | 47 | Il17re |
| Il18 | 212 | 194 | 295 | 220 | 256 | 339 | 277 | 427 | 324 | 401 | 356 | 582 | 445 | 610 | 252 | 168 | 242 | 217 | 250 | 325 | 239 | 451 | 372 | 369 | 472 | 394 | 601 | 541 | Il18 |
| Il18bp | 1138 | 1210 | 684 | 949 | 1024 | 990 | 990 | 962 | 896 | 881 | 986 | 832 | 872 | 811 | 1109 | 971 | 880 | 891 | 920 | 1094 | 1068 | 986 | 849 | 805 | 1016 | 753 | 927 | 814 | Il18bp |
| Il18r1 | 0 | 1 | 0 | 0 | 0 | 0 | 0 | 6 | 13 | 1 | 8 | 0 | 0 | 1 | 1 | 0 | 0 | 2 | 1 | 1 | 0 | 18 | 13 | 3 | 1 | 2 | 0 | 1 | Il18r1 |
| Il18rap | 38 | 17 | 23 | 7 | 18 | 15 | 23 | 34 | 30 | 35 | 63 | 65 | 48 | 55 | 17 | 23 | 9 | 17 | 13 | 16 | 17 | 28 | 50 | 42 | 13 | 90 | 63 | 59 | Il18rap |
| Il19 | 2 | 6 | 6 | 4 | 10 | 1 | 11 | 4 | 9 | 3 | 4 | 0 | 9 | 1 | 4 | 7 | 2 | 14 | 5 | 1 | 5 | 2 | 1 | 5 | 7 | 1 | 0 | 5 | Il19 |
| Il1a | 3 | 3 | 2 | 5 | 3 | 7 | 6 | 6 | 7 | 9 | 5 | 4 | 7 | 4 | 9 | 3 | 7 | 3 | 1 | 6 | 6 | 10 | 12 | 13 | 7 | 5 | 9 | 8 | Il1a |
| Il1b | 0 | 1 | 0 | 0 | 0 | 3 | 1 | 1 | 2 | 3 | 3 | 1 | 1 | 2 | 1 | 0 | 0 | 0 | 2 | 0 | 0 | 2 | 4 | 0 | 3 | 7 | 2 | 7 | Il1b |
| Il1r1 | 386 | 234 | 228 | 176 | 180 | 292 | 153 | 366 | 461 | 313 | 523 | 240 | 282 | 221 | 318 | 381 | 174 | 265 | 115 | 219 | 165 | 277 | 491 | 331 | 221 | 241 | 292 | 243 | Il1r1 |
| Il1r2 | 14 | 8 | 29 | 11 | 10 | 7 | 6 | 22 | 10 | 13 | 6 | 3 | 18 | 5 | 20 | 18 | 12 | 10 | 4 | 8 | 13 | 10 | 23 | 4 | 7 | 7 | 3 | 5 | Il1r2 |
| Il1rap | 1762 | 1372 | 1388 | 1080 | 1631 | 1502 | 1352 | 1203 | 1037 | 789 | 1076 | 803 | 1065 | 735 | 1960 | 1880 | 1137 | 1213 | 1248 | 1433 | 1358 | 904 | 1210 | 636 | 736 | 569 | 956 | 905 | Il1rap |
| Il1rapl1 | 152 | 136 | 68 | 297 | 362 | 389 | 345 | 388 | 339 | 302 | 316 | 151 | 201 | 184 | 174 | 161 | 181 | 296 | 288 | 329 | 325 | 328 | 340 | 263 | 394 | 101 | 203 | 176 | Il1rapl1 |
| Il1rapl2 | 52 | 44 | 24 | 48 | 34 | 87 | 16 | 87 | 102 | 87 | 90 | 42 | 56 | 63 | 49 | 62 | 39 | 75 | 30 | 63 | 55 | 85 | 144 | 87 | 103 | 30 | 41 | 44 | Il1rapl2 |
| Il1rl1 | 8 | 7 | 19 | 3 | 3 | 19 | 7 | 42 | 74 | 60 | 56 | 30 | 29 | 31 | 11 | 16 | 6 | 6 | 9 | 2 | 3 | 20 | 102 | 41 | 27 | 14 | 24 | 17 | Il1rl1 |
| Il1rl2 | 278 | 169 | 164 | 116 | 123 | 206 | 101 | 379 | 439 | 365 | 403 | 252 | 318 | 216 | 215 | 220 | 148 | 134 | 105 | 123 | 122 | 239 | 500 | 335 | 188 | 327 | 212 | 183 | Il1rl2 |
| Il1rn | 3 | 0 | 2 | 2 | 2 | 1 | 0 | 2 | 10 | 6 | 1 | 2 | 7 | 3 | 12 | 3 | 0 | 3 | 0 | 2 | 4 | 7 | 9 | 5 | 4 | 7 | 6 | 9 | Il1rn |
| Il2 | 2 | 1 | 0 | 5 | 3 | 0 | 9 | 4 | 2 | 4 | 1 | 0 | 3 | 12 | 8 | 2 | 9 | 1 | 3 | 2 | 3 | 3 | 3 | 0 | 5 | 0 | 6 | 2 | Il2 |
| Il20 | 0 | 1 | 0 | 1 | 1 | 0 | 1 | 0 | 0 | 1 | 0 | 0 | 0 | 0 | 0 | 0 | 10 | 0 | 0 | 0 | 1 | 0 | 0 | 0 | 4 | 0 | 0 | 0 | Il20 |
| Il20ra | 17 | 4 | 6 | 10 | 7 | 8 | 9 | 4 | 13 | 8 | 7 | 0 | 1 | 1 | 5 | 9 | 6 | 3 | 9 | 7 | 5 | 10 | 15 | 7 | 7 | 2 | 0 | 8 | Il20ra |
| Il20rb | 351 | 312 | 280 | 344 | 266 | 361 | 344 | 134 | 147 | 150 | 167 | 245 | 236 | 193 | 335 | 310 | 272 | 281 | 309 | 325 | 277 | 164 | 139 | 178 | 170 | 234 | 257 | 179 | Il20rb |
| Il21 | 1 | 5 | 0 | 6 | 0 | 4 | 4 | 1 | 0 | 1 | 1 | 0 | 0 | 0 | 3 | 2 | 1 | 8 | 1 | 0 | 3 | 0 | 3 | 1 | 0 | 0 | 6 | 0 | Il21 |
| Il21r | 25 | 17 | 27 | 17 | 29 | 43 | 20 | 55 | 62 | 73 | 90 | 76 | 118 | 51 | 18 | 20 | 15 | 18 | 16 | 23 | 27 | 47 | 82 | 54 | 84 | 81 | 89 | 62 | Il21r |
| Il22 | 31 | 95 | 86 | 93 | 38 | 52 | 53 | 4 | 9 | 12 | 2 | 7 | 7 | 12 | 55 | 51 | 67 | 92 | 64 | 42 | 72 | 17 | 1 | 5 | 3 | 9 | 1 | 14 | Il22 |
| Il22b | 0 | 0 | 0 | 0 | 0 | 0 | 0 | 0 | 0 | 0 | 0 | 0 | 0 | 0 | 0 | 2 | 0 | 0 | 0 | 0 | 0 | 0 | 0 | 0 | 1 | 0 | 0 | 0 | Il22b |
| Il22ra1 | 5 | 2 | 0 | 2 | 3 | 2 | 1 | 9 | 7 | 21 | 11 | 5 | 12 | 4 | 0 | 1 | 1 | 1 | 2 | 3 | 3 | 7 | 23 | 18 | 8 | 1 | 7 | 8 | Il22ra1 |
| Il22ra2 | 0 | 0 | 0 | 0 | 1 | 0 | 0 | 0 | 0 | 0 | 0 | 1 | 3 | 0 | 0 | 2 | 0 | 1 | 0 | 0 | 0 | 0 | 1 | 0 | 2 | 0 | 0 | 0 | Il22ra2 |
| Il23a | 1 | 3 | 3 | 2 | 2 | 12 | 2 | 4 | 8 | 10 | 1 | 5 | 2 | 10 | 1 | 2 | 1 | 0 | 3 | 3 | 1 | 13 | 8 | 7 | 10 | 0 | 6 | 4 | Il23a |
| Il23r | 0 | 0 | 2 | 0 | 0 | 0 | 0 | 2 | 0 | 0 | 6 | 0 | 0 | 0 | 1 | 0 | 0 | 0 | 0 | 0 | 0 | 2 | 3 | 4 | 1 | 3 | 0 | 1 | Il23r |
| Il24 | 0 | 0 | 0 | 1 | 0 | 0 | 0 | 0 | 0 | 0 | 0 | 0 | 0 | 0 | 0 | 0 | 0 | 0 | 0 | 0 | 0 | 0 | 1 | 0 | 0 | 0 | 0 | 0 | Il24 |
| Il25 | 5 | 20 | 14 | 3 | 6 | 0 | 4 | 4 | 9 | 19 | 17 | 4 | 3 | 1 | 8 | 9 | 2 | 11 | 4 | 3 | 5 | 8 | 12 | 7 | 7 | 5 | 2 | 5 | Il25 |
| Il27 | 0 | 0 | 0 | 1 | 0 | 0 | 0 | 0 | 0 | 0 | 0 | 0 | 1 | 0 | 0 | 0 | 0 | 0 | 0 | 0 | 0 | 0 | 1 | 1 | 0 | 0 | 0 | 0 | Il27 |
| Il27ra | 9 | 14 | 5 | 1 | 9 | 10 | 12 | 17 | 33 | 34 | 61 | 26 | 23 | 17 | 11 | 17 | 5 | 21 | 9 | 10 | 18 | 37 | 45 | 33 | 19 | 13 | 10 | 13 | Il27ra |
| Il2ra | 0 | 0 | 0 | 1 | 0 | 0 | 0 | 1 | 2 | 3 | 5 | 1 | 5 | 2 | 1 | 0 | 2 | 0 | 2 | 6 | 1 | 10 | 5 | 5 | 0 | 2 | 6 | 1 | Il2ra |
| Il2rb | 20 | 15 | 0 | 4 | 7 | 5 | 1 | 9 | 11 | 17 | 15 | 1 | 7 | 2 | 12 | 21 | 3 | 22 | 4 | 4 | 4 | 8 | 8 | 7 | 8 | 2 | 2 | 0 | Il2rb |
| Il2rg | 38 | 35 | 35 | 20 | 35 | 26 | 21 | 39 | 43 | 54 | 41 | 21 | 26 | 36 | 33 | 40 | 19 | 44 | 12 | 31 | 33 | 50 | 49 | 39 | 26 | 31 | 41 | 45 | Il2rg |
| Il3 | 0 | 0 | 0 | 0 | 1 | 0 | 0 | 0 | 0 | 0 | 0 | 0 | 0 | 0 | 0 | 0 | 0 | 0 | 0 | 0 | 0 | 0 | 0 | 0 | 0 | 0 | 0 | 0 | Il3 |

| | | | | | | | | | | | | | | | | | | | | | | | | | | | | | |
|---|---|---|---|---|---|---|---|---|---|---|---|---|---|---|---|---|---|---|---|---|---|---|---|---|---|---|---|---|---|
| Il31 | 0 | 0 | 0 | 0 | 0 | 3 | 0 | 0 | 0 | 0 | 0 | 0 | 0 | 1 | 4 | 0 | 1 | 1 | 2 | 0 | 0 | 0 | 0 | 0 | 0 | 0 | 0 | 2 | Il31 |
| Il31ra | 10 | 1 | 0 | 18 | 18 | 6 | 11 | 17 | 25 | 25 | 22 | 11 | 3 | 16 | 4 | 5 | 6 | 20 | 13 | 28 | 30 | 5 | 38 | 20 | 33 | 3 | 2 | 13 | Il31ra |
| Il33 | 88 | 82 | 53 | 48 | 55 | 86 | 61 | 134 | 209 | 93 | 182 | 47 | 90 | 45 | 128 | 101 | 73 | 95 | 48 | 58 | 39 | 170 | 201 | 93 | 101 | 70 | 67 | 84 | Il33 |
| Il34 | 7 | 19 | 14 | 12 | 16 | 9 | 12 | 44 | 54 | 67 | 42 | 29 | 58 | 16 | 11 | 10 | 16 | 14 | 23 | 17 | 27 | 28 | 88 | 41 | 41 | 30 | 46 | 24 | Il34 |
| Il36g | 0 | 0 | 0 | 0 | 0 | 0 | 0 | 0 | 1 | 0 | 0 | 3 | 1 | 0 | 0 | 0 | 0 | 0 | 0 | 0 | 0 | 0 | 2 | 2 | 0 | 0 | 0 | 0 | Il36g |
| Il36rn | 1 | 0 | 0 | 0 | 0 | 1 | 1 | 4 | 19 | 11 | 18 | 2 | 0 | 7 | 1 | 0 | 0 | 0 | 0 | 1 | 0 | 2 | 15 | 9 | 0 | 4 | 0 | 0 | Il36rn |
| Il3ra | 257 | 327 | 438 | 357 | 290 | 290 | 183 | 232 | 291 | 282 | 277 | 362 | 380 | 254 | 252 | 322 | 351 | 336 | 337 | 263 | 256 | 328 | 282 | 385 | 297 | 335 | 336 | 304 | Il3ra |
| Il4 | 1 | 0 | 2 | 2 | 5 | 2 | 0 | 5 | 2 | 3 | 2 | 0 | 1 | 0 | 0 | 1 | 6 | 0 | 4 | 0 | 1 | 0 | 1 | 2 | 2 | 0 | 1 | 4 | Il4 |
| Il4i1 | 31 | 15 | 31 | 18 | 30 | 18 | 28 | 35 | 23 | 30 | 29 | 29 | 46 | 39 | 29 | 18 | 27 | 37 | 30 | 32 | 25 | 32 | 45 | 34 | 48 | 53 | 31 | 56 | Il4i1 |
| Il4ra | 701 | 777 | 836 | 662 | 619 | 605 | 389 | 419 | 587 | 435 | 583 | 446 | 501 | 346 | 711 | 910 | 635 | 636 | 545 | 498 | 423 | 404 | 661 | 449 | 436 | 563 | 585 | 415 | Il4ra |
| Il5 | 1 | 2 | 4 | 0 | 4 | 1 | 2 | 5 | 2 | 2 | 2 | 0 | 1 | 1 | 1 | 1 | 0 | 2 | 5 | 4 | 2 | 3 | 1 | 0 | 0 | 0 | 6 | 1 | Il5 |
| Il5ra | 0 | 0 | 0 | 1 | 2 | 2 | 1 | 0 | 0 | 7 | 0 | 0 | 0 | 2 | 1 | 0 | 0 | 0 | 0 | 0 | 0 | 2 | 4 | 1 | 0 | 0 | 1 | 2 | Il5ra |
| Il6 | 8 | 3 | 1 | 8 | 10 | 8 | 13 | 5 | 6 | 5 | 5 | 0 | 1 | 0 | 9 | 5 | 0 | 13 | 12 | 5 | 13 | 5 | 2 | 2 | 0 | 1 | 2 | 0 | Il6 |
| Il6ra | 80 | 111 | 198 | 113 | 145 | 188 | 129 | 122 | 152 | 139 | 182 | 162 | 149 | 155 | 119 | 139 | 127 | 146 | 140 | 136 | 129 | 164 | 227 | 139 | 183 | 140 | 149 | 115 | Il6ra |
| Il6st | 15402 | 14891 | 12311 | 11876 | 10269 | 11815 | 8752 | 8988 | 10550 | 10036 | 13202 | 9613 | 11872 | 9039 | 14679 | 17531 | 10999 | 12864 | 8886 | 9151 | 9742 | 9832 | 12591 | 9278 | 8169 | 7530 | 10270 | 9428 | Il6st |
| Il7 | 7 | 3 | 7 | 3 | 11 | 6 | 6 | 11 | 7 | 0 | 11 | 0 | 3 | 4 | 9 | 2 | 7 | 3 | 8 | 10 | 2 | 3 | 11 | 5 | 3 | 27 | 7 | 1 | Il7 |
| Il7r | 7 | 2 | 6 | 1 | 6 | 0 | 4 | 10 | 11 | 5 | 30 | 4 | 7 | 3 | 1 | 2 | 0 | 3 | 1 | 0 | 0 | 17 | 26 | 11 | 2 | 11 | 5 | 5 | Il7r |
| Il9r | 6 | 7 | 17 | 5 | 6 | 6 | 1 | 6 | 7 | 5 | 2 | 1 | 9 | 4 | 1 | 5 | 2 | 1 | 7 | 1 | 1 | 7 | 8 | 4 | 1 | 5 | 5 | 5 | Il9r |
| Ildr1 | 70 | 20 | 5 | 26 | 9 | 10 | 4 | 137 | 153 | 175 | 125 | 3 | 31 | 5 | 28 | 37 | 9 | 9 | 0 | 5 | 1 | 30 | 131 | 123 | 19 | 151 | 2 | 2 | Ildr1 |
| Ildr2 | 7081 | 6933 | 9395 | 7782 | 7335 | 8179 | 8303 | 5345 | 4291 | 5073 | 4487 | 9182 | 7483 | 9329 | 7572 | 5882 | 9364 | 7327 | 7924 | 7401 | 8275 | 5945 | 4697 | 5928 | 6532 | 6448 | 10537 | 9862 | Ildr2 |
| Ilf2 | 20054 | 18916 | 19907 | 19983 | 18920 | 18908 | 21225 | 22699 | 19366 | 19866 | 18327 | 22537 | 21106 | 24379 | 20054 | 17203 | 19812 | 18735 | 20274 | 20606 | 20922 | 19836 | 18579 | 21692 | 21366 | 24313 | 23140 | 24275 | Ilf2 |
| Ilf3 | 13243 | 14239 | 15210 | 14626 | 15653 | 14915 | 15137 | 21415 | 23049 | 23443 | 21933 | 26711 | 24446 | 27560 | 13006 | 11851 | 15579 | 13991 | 16338 | 14540 | 15472 | 21631 | 21010 | 26213 | 26106 | 25293 | 26690 | 27333 | Ilf3 |
| Ilk | 8573 | 9389 | 8608 | 8106 | 7263 | 8049 | 7545 | 6959 | 7552 | 6988 | 7605 | 7053 | 7328 | 7405 | 8412 | 8783 | 8402 | 7931 | 8158 | 7042 | 7973 | 6918 | 8262 | 7199 | 6802 | 8203 | 7299 | 7785 | Ilk |
| Ilkap | 6033 | 5813 | 5843 | 5848 | 5262 | 5494 | 5928 | 5891 | 5381 | 5451 | 5617 | 6137 | 5632 | 6058 | 5826 | 5605 | 5822 | 5838 | 5804 | 5666 | 5758 | 5449 | 5488 | 5695 | 5944 | 6175 | 6090 | 6261 | Ilkap |
| Ilrun | 8067 | 8517 | 9395 | 9119 | 7916 | 7660 | 6817 | 5368 | 6175 | 6734 | 6953 | 6636 | 6783 | 6153 | 7030 | 9091 | 9063 | 8998 | 8567 | 6188 | 7558 | 6191 | 6422 | 7329 | 6965 | 7405 | 6408 | 6424 | Ilrun |
| Ilvbl | 2308 | 2648 | 2615 | 2805 | 2282 | 2643 | 1973 | 1662 | 1910 | 2006 | 1932 | 1972 | 2137 | 1680 | 2313 | 2443 | 2630 | 2737 | 2602 | 2487 | 2114 | 1865 | 2033 | 1749 | 1892 | 1964 | 2007 | 1692 | Ilvbl |
| Immp1l | 2707 | 2334 | 2800 | 2475 | 1681 | 1938 | 2727 | 1986 | 1799 | 1716 | 1756 | 1687 | 1661 | 1855 | 2952 | 2453 | 2422 | 2260 | 2502 | 2936 | 2496 | 1480 | 1646 | 1879 | 1685 | 2098 | 1876 | 2135 | Immp1l |
| Immp2l | 100 | 104 | 135 | 98 | 106 | 122 | 99 | 131 | 111 | 101 | 95 | 89 | 127 | 94 | 91 | 110 | 90 | 68 | 86 | 107 | 64 | 62 | 107 | 103 | 100 | 111 | 109 | 122 | Immp2l |
| Immt | 11623 | 10799 | 10522 | 9093 | 8962 | 9375 | 9417 | 12128 | 10925 | 10448 | 10229 | 11205 | 11272 | 11907 | 11815 | 10118 | 9953 | 8868 | 9380 | 10115 | 9698 | 11202 | 10853 | 10579 | 10330 | 11865 | 11617 | 12589 | Immt |
| Imp3 | 2520 | 2572 | 2030 | 2277 | 1816 | 1821 | 2195 | 1867 | 1683 | 1738 | 1613 | 1630 | 1682 | 1805 | 2305 | 2093 | 2221 | 1913 | 2046 | 2466 | 2139 | 1539 | 1783 | 1910 | 1672 | 1937 | 1741 | 1752 | Imp3 |
| Imp4 | 8156 | 8208 | 7546 | 6910 | 5814 | 6349 | 6518 | 4676 | 4492 | 4022 | 4077 | 4719 | 4525 | 4534 | 7878 | 7402 | 6920 | 6913 | 6372 | 6521 | 6379 | 4239 | 4408 | 4332 | 4401 | 4674 | 4638 | 4525 | Imp4 |
| Impa1 | 2802 | 2751 | 3056 | 2843 | 2972 | 2683 | 3072 | 4035 | 3339 | 3044 | 3117 | 3279 | 3379 | 3508 | 3301 | 2783 | 3132 | 2881 | 2773 | 3290 | 2942 | 3253 | 3457 | 3401 | 3306 | 3791 | 3477 | 3843 | Impa1 |
| Impa2 | 1652 | 1551 | 1669 | 1548 | 1305 | 1324 | 1169 | 451 | 599 | 596 | 768 | 648 | 752 | 655 | 1501 | 1879 | 1528 | 1566 | 1304 | 1026 | 1247 | 461 | 708 | 651 | 543 | 848 | 630 | 646 | Impa2 |
| Impact | 5426 | 4113 | 5228 | 4387 | 4376 | 4358 | 4212 | 5145 | 4507 | 4667 | 5324 | 5299 | 5367 | 2600 | 5215 | 5322 | 4781 | 4560 | 4074 | 3956 | 3985 | 5559 | 4904 | 4932 | 5063 | 5259 | 5430 | 5413 | Impact |
| Impdh1 | 3038 | 3911 | 4982 | 4436 | 4017 | 3921 | 3513 | 1824 | 2321 | 2623 | 2901 | 2616 | 2503 | 2417 | 3090 | 3537 | 4803 | 4390 | 4394 | 3323 | 3611 | 2750 | 2435 | 2751 | 2607 | 2815 | 2421 | 2528 | Impdh1 |
| Impdh2 | 10659 | 10932 | 10603 | 10209 | 7842 | 8842 | 9505 | 6066 | 7667 | 7129 | 7476 | 8169 | 7725 | 7928 | 8898 | 10530 | 10024 | 10150 | 9297 | 8695 | 9221 | 5543 | 7236 | 8112 | 7106 | 10431 | 7977 | 8974 | Impdh2 |
| Impg1 | 0 | 0 | 0 | 0 | 0 | 0 | 0 | 0 | 1 | 0 | 2 | 1 | 0 | 0 | 0 | 0 | 0 | 0 | 0 | 1 | 0 | 0 | 0 | 0 | 0 | 0 | 0 | 0 | Impg1 |
| Impg2 | 456 | 429 | 351 | 571 | 648 | 586 | 719 | 270 | 253 | 320 | 276 | 193 | 255 | 225 | 462 | 513 | 498 | 677 | 629 | 549 | 651 | 271 | 239 | 265 | 296 | 170 | 228 | 193 | Impg2 |
| Ina | 21659 | 27414 | 25503 | 44207 | 57355 | 43618 | 42918 | 36032 | 33492 | 45678 | 36126 | 33383 | 39253 | 32215 | 30565 | 25550 | 39456 | 46845 | 47528 | 40823 | 46808 | 58372 | 32312 | 33401 | 43840 | 24335 | 28344 | 30572 | Ina |
| Inafm1 | 1092 | 1202 | 1165 | 1326 | 1343 | 1254 | 1167 | 869 | 940 | 1080 | 1154 | 957 | 937 | 917 | 1125 | 1064 | 1515 | 1332 | 1341 | 1086 | 1260 | 1140 | 962 | 1155 | 1061 | 1042 | 932 | 971 | Inafm1 |
| Inafm2 | 2909 | 3086 | 3663 | 4012 | 3089 | 3204 | 3769 | 1559 | 1336 | 1604 | 1546 | 1804 | 1711 | 2091 | 2851 | 2697 | 3737 | 3805 | 3258 | 3048 | 3732 | 1646 | 1325 | 1849 | 1860 | 1973 | 1722 | 2063 | Inafm2 |
| Inava | 1339 | 1341 | 1532 | 1287 | 1352 | 1236 | 1143 | 1003 | 1185 | 1057 | 997 | 1358 | 1202 | 1172 | 1108 | 1271 | 1560 | 1311 | 1513 | 1108 | 1277 | 876 | 1020 | 1285 | 867 | 1759 | 1345 | 1258 | Inava |
| Inca1 | 72 | 73 | 93 | 48 | 41 | 49 | 59 | 43 | 43 | 28 | 42 | 47 | 36 | 45 | 78 | 55 | 89 | 68 | 49 | 48 | 67 | 58 | 32 | 25 | 29 | 25 | 56 | 33 | Inca1 |
| Incenp | 7455 | 8656 | 10979 | 9375 | 6640 | 6993 | 7645 | 5655 | 6765 | 7979 | 7329 | 11508 | 9318 | 11509 | 6665 | 8001 | 9740 | 7888 | 8542 | 6518 | 7416 | 5718 | 5919 | 10150 | 8493 | 13621 | 10458 | 11841 | Incenp |
| Inf2 | 1063 | 1292 | 1365 | 1267 | 1379 | 1330 | 1120 | 700 | 1197 | 1271 | 1534 | 1052 | 1208 | 822 | 943 | 1439 | 1166 | 1782 | 1316 | 966 | 1141 | 1036 | 1114 | 1196 | 1267 | 1062 | 932 | 841 | Inf2 |
| Ing1 | 5917 | 6293 | 6441 | 6687 | 5875 | 6210 | 6455 | 4846 | 4328 | 4839 | 4431 | 5513 | 4630 | 5647 | 6046 | 5727 | 6426 | 6454 | 6198 | 6027 | 6184 | 4792 | 4311 | 5084 | 5153 | 6315 | 5352 | 5878 | Ing1 |
| Ing2 | 1601 | 1570 | 1853 | 1832 | 1522 | 1482 | 1428 | 966 | 839 | 900 | 958 | 991 | 1034 | 1030 | 1576 | 1618 | 1561 | 1577 | 1480 | 1567 | 1440 | 956 | 962 | 936 | 876 | 1093 | 1062 | 1097 | Ing2 |
| Ing3 | 2144 | 1999 | 2186 | 2136 | 1557 | 1667 | 2189 | 2694 | 2439 | 2277 | 2229 | 2778 | 2345 | 2775 | 2172 | 1848 | 1982 | 1930 | 1903 | 2104 | 1965 | 2406 | 2251 | 2574 | 2436 | 3060 | 2496 | 2858 | Ing3 |
| Ing4 | 163 | 98 | 100 | 166 | 139 | 165 | 179 | 275 | 293 | 352 | 264 | 301 | 308 | 369 | 148 | 155 | 174 | 120 | 154 | 174 | 118 | 286 | 241 | 346 | 336 | 172 | 314 | 292 | Ing4 |
| Ing5 | 1163 | 1092 | 1175 | 963 | 985 | 1083 | 1020 | 2341 | 2312 | 2363 | 2399 | 2989 | 2740 | 2944 | 1091 | 1051 | 1274 | 979 | 1086 | 1009 | 1015 | 2129 | 2244 | 2578 | 2408 | 2733 | 3076 | 2894 | Ing5 |
| Inha | 480 | 533 | 598 | 865 | 1016 | 814 | 825 | 775 | 983 | 1154 | 1119 | 784 | 856 | 752 | 509 | 573 | 657 | 832 | 990 | 669 | 854 | 1158 | 1056 | 1042 | 1273 | 754 | 752 | 764 | Inha |
| Inhba | 848 | 421 | 466 | 263 | 111 | 497 | 131 | 893 | 948 | 717 | 907 | 407 | 483 | 395 | 609 | 775 | 331 | 396 | 154 | 114 | 124 | 302 | 946 | 595 | 289 | 385 | 422 | 375 | Inhba |
| Inhbb | 272 | 303 | 374 | 481 | 705 | 586 | 610 | 553 | 655 | 781 | 694 | 550 | 668 | 549 | 225 | 252 | 458 | 562 | 758 | 637 | 644 | 495 | 669 | 854 | 1014 | 620 | 442 | 497 | Inhbb |
| Inhbc | 0 | 1 | 0 | 0 | 0 | 1 | 0 | 0 | 2 | 0 | 0 | 1 | 0 | 1 | 0 | 1 | 1 | 0 | 0 | 0 | 0 | 0 | 1 | 0 | 0 | 0 | 2 | 2 | Inhbc |
| Inhbe | 5 | 0 | 0 | 0 | 1 | 0 | 2 | 0 | 1 | 0 | 1 | 0 | 0 | 0 | 0 | 0 | 0 | 2 | 0 | 0 | 0 | 0 | 4 | 1 | 0 | 0 | 2 | 0 | Inhbe |
| Inip | 6095 | 5775 | 5849 | 6131 | 5375 | 5723 | 6548 | 4087 | 3484 | 3952 | 3290 | 4933 | 4308 | 5024 | 5858 | 5521 | 6039 | 6022 | 6004 | 5650 | 6422 | 3648 | 3255 | 4175 | 4381 | 5184 | 4926 | 5199 | Inip |
| Inka1 | 379 | 432 | 346 | 329 | 216 | 301 | 191 | 347 | 493 | 365 | 633 | 314 | 473 | 250 | 335 | 497 | 327 | 321 | 231 | 282 | 210 | 329 | 575 | 333 | 308 | 475 | 343 | 352 | Inka1 |
| Inka2 | 1197 | 1478 | 1164 | 1233 | 1294 | 1211 | 1303 | 591 | 590 | 662 | 778 | 676 | 640 | 714 | 1311 | 1320 | 1396 | 1337 | 1351 | 1211 | 1194 | 680 | 685 | 636 | 658 | 893 | 522 | 710 | Inka2 |
| Inmt | 35 | 10 | 4 | 20 | 3 | 0 | 0 | 35 | 44 | 52 | 38 | 2 | 19 | 10 | 8 | 15 | 0 | 8 | 0 | 0 | 0 | 0 | 39 | 85 | 9 | 89 | 5 | 1 | Inmt |
| Ino80 | 2247 | 2100 | 1879 | 1977 | 2026 | 2108 | 2120 | 3462 | 3547 | 3417 | 3473 | 4032 | 3750 | 3805 | 2053 | 2009 | 1797 | 1866 | 2102 | 1929 | 2103 | 3593 | 3320 | 3279 | 3608 | 2761 | 3846 | 3682 | Ino80 |
| Ino80b | 1587 | 1982 | 1993 | 2144 | 1888 | 1812 | 1692 | 1014 | 1168 | 1136 | 1217 | 1348 | 1195 | 1272 | 1673 | 1664 | 2097 | 2005 | 1924 | 1739 | 1881 | 962 | 1080 | 1382 | 1247 | 1401 | 1271 | 1262 | Ino80b |
| Ino80c | 603 | 655 | 613 | 510 | 607 | 675 | 637 | 1829 | 1781 | 1734 | 1627 | 1833 | 1903 | 2098 | 704 | 562 | 618 | 529 | 609 | 742 | 651 | 1683 | 1708 | 1745 | 1884 | 1919 | 2065 | 2163 | Ino80c |
| Ino80d | 2370 | 1816 | 2653 | 2899 | 3905 | 3511 | 3457 | 4722 | 4593 | 5452 | 5500 | 5683 | 5272 | 5682 | 2345 | 2159 | 2931 | 2830 | 3582 | 2777 | 3443 | 5546 | 4209 | 5220 | 5975 | 4475 | 5554 | 5448 | Ino80d |
| Ino80e | 3030 | 3702 | 4732 | 4249 | 4278 | 3946 | 3962 | 3516 | 3867 | 4114 | 4329 | 4629 | 4268 | 4623 | 2776 | 3082 | 4520 | 4064 | 4472 | 3324 | 4111 | 3838 | 3728 | 4952 | 4467 | 5261 | 4134 | 4722 | Ino80e |
| Inpp1 | 728 | 578 | 750 | 685 | 917 | 898 | 750 | 1518 | 1595 | 1468 | 1699 | 1417 | 1548 | 1400 | 767 | 780 | 721 | 723 | 750 | 712 | 752 | 1761 | 1585 | 1363 | 1599 | 1364 | 1440 | 1347 | Inpp1 |
| Inpp4a | 5494 | 5596 | 5747 | 8258 | 9220 | 7749 | 8470 | 3987 | 4203 | 4902 | 4864 | 4392 | 4259 | 4341 | 5662 | 5575 | 7041 | 8641 | 8267 | 6811 | 8490 | 5850 | 4103 | 4548 | 5581 | 3363 | 3914 | 4014 | Inpp4a |
| Inpp4b | 694 | 416 | 364 | 325 | 371 | 437 | 307 | 799 | 1041 | 662 | 1516 | 542 | 768 | 553 | 703 | 861 | 271 | 358 | 274 | 440 | 361 | 780 | 1211 | 502 | 550 | 532 | 601 | 602 | Inpp4b |
| Inpp5a | 3234 | 3374 | 3081 | 2779 | 2572 | 2743 | 2546 | 1784 | 1997 | 1819 | 2091 | 1819 | 1996 | 1665 | 2925 | 3447 | 2929 | 2807 | 2572 | 2397 | 2508 | 1763 | 2056 | 1759 | 1529 | 2078 | 1833 | 1813 | Inpp5a |
| Inpp5b | 4779 | 5056 | 3950 | 4190 | 3933 | 4104 | 4155 | 2594 | 2444 | 2634 | 2447 | 2624 | 2273 | 2401 | 4898 | 4379 | 4209 | 4214 | 4171 | 4040 | 4100 | 2882 | 2529 | 2287 | 2532 | 2248 | 2379 | 2340 | Inpp5b |
| Inpp5d | 258 | 259 | 347 | 258 | 255 | 233 | 199 | 371 | 551 | 503 | 527 | 428 | 466 | 366 | 268 | 339 | 236 | 276 | 271 | 245 | 219 | 483 | 656 | 411 | 426 | 462 | 523 | 405 | Inpp5d |
| Inpp5e | 2247 | 2434 | 2340 | 2376 | 2608 | 2480 | 2484 | 2574 | 2747 | 3025 | 2786 | 3149 | 3038 | 2876 | 2142 | 2298 | 2475 | 2432 | 2679 | 2215 | 2359 | 3309 | 2709 | 2893 | 2945 | 3122 | 3009 | 3019 | Inpp5e |
| Inpp5f | 12129 | 12288 | 10897 | 15223 | 18683 | 17501 | 19916 | 17122 | 14233 | 15264 | 12398 | 13387 | 11363 | 14571 | 14861 | 10671 | 13433 | 16445 | 17211 | 19926 | 19401 | 19141 | 13237 | 13844 | 17259 | 11054 | 13240 | 12847 | Inpp5f |
| Inpp5j | 45 | 33 | 12 | 61 | 104 | 48 | 38 | 82 | 162 | 227 | 201 | 78 | 124 | 65 | 41 | 33 | 31 | 54 | 56 | 60 | 42 | 180 | 140 | 150 | 131 | 77 | 31 | 46 | Inpp5j |
| Inpp5k | 3520 | 4216 | 2856 | 3950 | 3787 | 3931 | 3905 | 2458 | 2431 | 2441 | 2315 | 2366 | 2371 | 2212 | 3634 | 3962 | 3409 | 4258 | 3857 | 3721 | 4052 | 2632 | 2408 | 2215 | 2917 | 2161 | 2384 | 2118 | Inpp5k |
| Inppl1 | 4779 | 5938 | 7636 | 6314 | 5427 | 5076 | 5147 | 4788 | 5644 | 5670 | 6967 | 7243 | 6750 | 6623 | 4440 | 4960 | 7389 | 5589 | 6579 | 4517 | 5464 | 5070 | 5775 | 7038 | 5286 | 7986 | 6552 | 7155 | Inppl1 |
| Ins2 | 0 | 3 | 0 | 0 | 7 | 5 | 0 | 5 | 8 | 10 | 6 | 3 | 7 | 0 | 0 | 0 | 0 | 4 | 2 | 1 | 0 | 7 | 17 | 1 | 4 | 4 | 0 | 0 | Ins2 |
| Insc | 31 | 41 | 24 | 12 | 56 | 12 | 15 | 66 | 123 | 120 | 154 | 41 | 176 | 47 | 54 | 90 | 28 | 39 | 13 | 3 | 4 | 132 | 133 | 42 | 58 | 45 | 85 | 50 | Insc |
| Insig1 | 6247 | 5932 | 6510 | 5680 | 3858 | 5001 | 4765 | 6271 | 4850 | 5457 | 4880 | 7163 | 7327 | 7687 | 6268 | 6274 | 5799 | 4724 | 4344 | 5086 | 4293 | 5145 | 4900 | 6111 | 6184 | 7117 | 8513 | 8413 | Insig1 |
| Insig2 | 2993 | 2836 | 2704 | 3058 | 3006 | 3007 | 3108 | 2505 | 2193 | 1978 | 1838 | 2096 | 1994 | 1984 | 3163 | 2837 | 2880 | 2951 | 2966 | 3231 | 2943 | 2453 | 2057 | 1976 | 2084 | 2073 | 2034 | 2050 | Insig2 |

| | | | | | | | | | | | | | | | | | | | | | | | | | | | | | |
|---|---|---|---|---|---|---|---|---|---|---|---|---|---|---|---|---|---|---|---|---|---|---|---|---|---|---|---|---|---|
| Insl3 | 6 | 9 | 17 | 13 | 11 | 17 | 7 | 2 | 4 | 4 | 9 | 2 | 1 | 7 | 3 | 11 | 3 | 15 | 9 | 4 | 10 | 3 | 11 | 3 | 9 | 3 | 3 | 12 | Insl3 |
| Insl5 | 141 | 154 | 193 | 232 | 204 | 180 | 245 | 21 | 16 | 25 | 17 | 24 | 18 | 32 | 170 | 154 | 312 | 297 | 242 | 136 | 219 | 30 | 19 | 25 | 23 | 35 | 13 | 21 | Insl5 |
| Insl6 | 373 | 374 | 289 | 241 | 211 | 173 | 255 | 53 | 55 | 34 | 44 | 67 | 66 | 57 | 377 | 404 | 355 | 296 | 217 | 264 | 210 | 40 | 53 | 44 | 21 | 72 | 36 | 80 | Insl6 |
| Insm1 | 2669 | 2627 | 4376 | 4148 | 4067 | 3476 | 3783 | 4104 | 3507 | 5409 | 3939 | 6760 | 4458 | 6580 | 2699 | 2211 | 4979 | 3825 | 4752 | 2774 | 4337 | 4976 | 2755 | 6780 | 6290 | 6689 | 5287 | 6270 | Insm1 |
| Insm2 | 65 | 79 | 102 | 101 | 124 | 91 | 126 | 78 | 74 | 68 | 62 | 87 | 64 | 92 | 86 | 56 | 97 | 113 | 96 | 89 | 106 | 120 | 88 | 91 | 113 | 108 | 44 | 80 | Insm2 |
| Insr | 9897 | 8892 | 9093 | 11291 | 12354 | 12692 | 13531 | 8433 | 8174 | 9539 | 9173 | 9400 | 9280 | 10330 | 9866 | 9465 | 10275 | 11241 | 11533 | 10120 | 12953 | 9722 | 7326 | 9084 | 10692 | 7219 | 9186 | 9117 | Insr |
| Insrr | 385 | 444 | 394 | 421 | 553 | 289 | 272 | 342 | 518 | 470 | 780 | 329 | 552 | 327 | 348 | 699 | 416 | 472 | 322 | 305 | 267 | 745 | 465 | 330 | 459 | 397 | 265 | 294 | Insrr |
| Insyn1 | 609 | 711 | 720 | 1004 | 1402 | 1087 | 1096 | 1182 | 1391 | 1625 | 1688 | 1284 | 1203 | 1064 | 558 | 631 | 927 | 1148 | 1356 | 1103 | 1155 | 1671 | 1443 | 1580 | 1735 | 1382 | 1080 | 1042 | Insyn1 |
| Insyn2a | 161 | 152 | 202 | 301 | 496 | 374 | 436 | 442 | 508 | 655 | 452 | 401 | 272 | 456 | 171 | 170 | 250 | 355 | 526 | 359 | 450 | 725 | 381 | 524 | 672 | 343 | 217 | 292 | Insyn2a |
| Insyn2b | 361 | 348 | 270 | 390 | 649 | 549 | 597 | 962 | 886 | 783 | 687 | 535 | 606 | 553 | 426 | 344 | 389 | 426 | 561 | 588 | 517 | 1071 | 868 | 610 | 854 | 369 | 470 | 490 | Insyn2b |
| Ints1 | 7645 | 10194 | 11292 | 10572 | 8979 | 8484 | 7495 | 5523 | 7340 | 8077 | 7517 | 9375 | 7574 | 7538 | 7083 | 9604 | 11206 | 10418 | 10097 | 6865 | 8148 | 8255 | 6576 | 8853 | 8281 | 7684 | 7259 | 7139 | Ints1 |
| Ints10 | 6450 | 6516 | 5737 | 6124 | 6531 | 6048 | 7215 | 6103 | 5556 | 5299 | 5000 | 5219 | 4663 | 5359 | 6463 | 5829 | 5656 | 6012 | 6658 | 7222 | 6955 | 5162 | 5330 | 5081 | 5761 | 4539 | 4832 | 5009 | Ints10 |
| Ints11 | 4204 | 4600 | 4526 | 4461 | 3649 | 3875 | 4188 | 3281 | 3388 | 3407 | 3105 | 3936 | 3352 | 3819 | 3852 | 4078 | 4304 | 4150 | 4422 | 3978 | 4131 | 3230 | 3489 | 3974 | 3795 | 4406 | 4051 | 3952 | Ints11 |
| Ints12 | 1832 | 1589 | 1775 | 1583 | 1518 | 1677 | 1626 | 2980 | 2437 | 2212 | 2046 | 2430 | 2421 | 2525 | 1899 | 1602 | 1738 | 1541 | 1627 | 2018 | 1517 | 2562 | 2467 | 2323 | 2325 | 2795 | 2707 | 2671 | Ints12 |
| Ints13 | 6262 | 6093 | 5663 | 5775 | 4328 | 4938 | 5370 | 6135 | 5062 | 4693 | 4666 | 5560 | 5237 | 5855 | 6136 | 5816 | 5762 | 5326 | 4697 | 5544 | 5143 | 4931 | 4966 | 4921 | 5018 | 5856 | 5560 | 6054 | Ints13 |
| Ints14 | 1746 | 1697 | 1680 | 1227 | 1564 | 1569 | 1519 | 4153 | 4120 | 3341 | 3351 | 4277 | 4042 | 4153 | 1902 | 1551 | 1591 | 1301 | 1589 | 1773 | 1429 | 3330 | 4072 | 3737 | 3822 | 4614 | 4625 | 4587 | Ints14 |
| Ints2 | 5966 | 4634 | 4450 | 4254 | 4383 | 4708 | 4835 | 7806 | 6879 | 6125 | 6454 | 7577 | 7303 | 7412 | 5512 | 4981 | 4429 | 4407 | 4525 | 4967 | 4797 | 6378 | 6272 | 6728 | 6510 | 6238 | 7704 | 7435 | Ints2 |
| Ints3 | 6567 | 7184 | 7844 | 7693 | 7650 | 6837 | 7245 | 7581 | 8275 | 8987 | 8218 | 9904 | 8558 | 9156 | 6153 | 6423 | 7844 | 7232 | 8382 | 5969 | 7192 | 9022 | 7892 | 9077 | 8931 | 9477 | 9073 | 9365 | Ints3 |
| Ints4 | 5917 | 6216 | 4624 | 4556 | 4423 | 4764 | 5188 | 5973 | 5691 | 4763 | 4439 | 5403 | 4998 | 5377 | 6101 | 5579 | 4647 | 4811 | 4781 | 5768 | 5078 | 4593 | 5277 | 4842 | 4653 | 5471 | 5319 | 5447 | Ints4 |
| Ints5 | 3013 | 3498 | 3592 | 3726 | 2874 | 2974 | 3049 | 2104 | 2298 | 2532 | 2322 | 2805 | 2368 | 2633 | 2585 | 3337 | 3522 | 3436 | 3169 | 2512 | 3133 | 2204 | 2203 | 2741 | 2475 | 3025 | 2392 | 2918 | Ints5 |
| Ints6 | 2895 | 2343 | 2568 | 2543 | 2723 | 2753 | 2997 | 4516 | 3953 | 3494 | 4089 | 4341 | 4043 | 4437 | 2929 | 2454 | 2605 | 2461 | 2874 | 2885 | 3039 | 3873 | 3880 | 3897 | 3932 | 4064 | 4217 | 4351 | Ints6 |
| Ints6l | 6909 | 5218 | 4723 | 5142 | 6110 | 6773 | 6533 | 9932 | 8164 | 7434 | 8450 | 8660 | 8748 | 9092 | 6773 | 5656 | 4832 | 5313 | 5515 | 6620 | 6596 | 8504 | 8106 | 6879 | 7861 | 5360 | 8814 | 8380 | Ints6l |
| Ints7 | 4432 | 4345 | 3924 | 4206 | 3818 | 3930 | 4442 | 6878 | 7517 | 7789 | 7232 | 8591 | 8109 | 8976 | 3965 | 3984 | 3945 | 3871 | 4371 | 3898 | 4303 | 6724 | 7213 | 7701 | 7571 | 8170 | 8849 | 8968 | Ints7 |
| Ints8 | 7414 | 6982 | 7168 | 6898 | 6260 | 6032 | 7306 | 9003 | 6863 | 6273 | 6124 | 7865 | 7012 | 7936 | 7832 | 7185 | 7085 | 6775 | 6837 | 7705 | 6856 | 7097 | 6592 | 7149 | 6956 | 8112 | 7697 | 7973 | Ints8 |
| Ints9 | 4366 | 5008 | 4188 | 3812 | 3026 | 3286 | 3677 | 2691 | 2663 | 2457 | 2181 | 2878 | 2740 | 2980 | 4143 | 3853 | 3968 | 3738 | 3578 | 3478 | 3548 | 2396 | 2638 | 2727 | 2543 | 3472 | 3086 | 3055 | Ints9 |
| Intu | 212 | 89 | 194 | 76 | 106 | 84 | 211 | 84 | 59 | 35 | 145 | 93 | 78 | 45 | 161 | 70 | 133 | 123 | 56 | 146 | 66 | 62 | 46 | 35 | 144 | 39 | 193 | 20 | Intu |
| Invs | 1299 | 1417 | 1236 | 1352 | 1209 | 1312 | 1332 | 1719 | 1817 | 1752 | 1970 | 1654 | 1862 | 1750 | 1363 | 1262 | 1442 | 1402 | 1314 | 1212 | 1418 | 1975 | 1904 | 1619 | 1532 | 1578 | 1632 | 1728 | Invs |
| Ip6k1 | 8774 | 10342 | 10425 | 11999 | 12559 | 11621 | 11126 | 9669 | 10046 | 11228 | 11054 | 10823 | 10892 | 10467 | 8658 | 9189 | 11651 | 11898 | 12834 | 10324 | 12150 | 11214 | 9836 | 11441 | 12295 | 11082 | 10119 | 10900 | Ip6k1 |
| Ip6k2 | 566 | 718 | 939 | 1069 | 1298 | 1005 | 758 | 1249 | 1325 | 1670 | 1595 | 1514 | 1624 | 1375 | 527 | 590 | 1049 | 969 | 1206 | 1094 | 878 | 1125 | 1292 | 1775 | 2088 | 1823 | 1871 | 1278 | Ip6k2 |
| Ip6k3 | 6 | 5 | 3 | 0 | 2 | 8 | 4 | 11 | 5 | 5 | 14 | 11 | 17 | 14 | 4 | 6 | 2 | 0 | 2 | 1 | 6 | 13 | 14 | 7 | 2 | 19 | 17 | 13 | Ip6k3 |
| Ipcef1 | 48 | 30 | 48 | 50 | 116 | 67 | 110 | 281 | 270 | 218 | 224 | 264 | 186 | 246 | 41 | 37 | 49 | 55 | 52 | 117 | 80 | 241 | 258 | 248 | 267 | 222 | 230 | 223 | Ipcef1 |
| Ipmk | 5512 | 5218 | 5487 | 6093 | 5785 | 6167 | 6693 | 3495 | 3128 | 3459 | 3236 | 3893 | 3370 | 3866 | 5434 | 5079 | 5667 | 6006 | 5697 | 5445 | 6231 | 3502 | 3036 | 3561 | 3819 | 3556 | 3586 | 3949 | Ipmk |
| Ipo11 | 7374 | 6678 | 6552 | 5504 | 5410 | 6067 | 6521 | 6155 | 5348 | 4612 | 4544 | 6064 | 5700 | 6027 | 7667 | 6510 | 5921 | 5567 | 5697 | 7003 | 6129 | 4533 | 5291 | 4972 | 5050 | 5936 | 6523 | 6354 | Ipo11 |
| Ipo13 | 8169 | 10860 | 8414 | 10536 | 9353 | 9376 | 8983 | 4615 | 5240 | 5301 | 4858 | 5660 | 5058 | 5227 | 8039 | 9067 | 9447 | 10423 | 10068 | 8539 | 9584 | 5274 | 4897 | 5084 | 6005 | 5087 | 5480 | 5035 | Ipo13 |
| Ipo4 | 1712 | 2100 | 2214 | 2184 | 2124 | 1876 | 1925 | 1512 | 1914 | 2346 | 2392 | 2287 | 2152 | 1984 | 1643 | 2079 | 2199 | 2210 | 2055 | 1779 | 2096 | 2151 | 2058 | 2442 | 2111 | 2477 | 1864 | 2044 | Ipo4 |
| Ipo5 | 41666 | 40506 | 36993 | 30601 | 26649 | 29870 | 33053 | 27114 | 24767 | 21777 | 20871 | 30173 | 26008 | 30203 | 38733 | 36007 | 33428 | 29591 | 31018 | 34141 | 31709 | 20277 | 23753 | 25021 | 23430 | 31218 | 31568 | 32261 | Ipo5 |
| Ipo7 | 33561 | 28234 | 28482 | 27999 | 25737 | 27648 | 32830 | 28482 | 23524 | 20693 | 21381 | 24510 | 23602 | 25891 | 34661 | 30313 | 28385 | 28225 | 27231 | 32537 | 30534 | 21960 | 23231 | 22106 | 22763 | 24041 | 25816 | 26819 | Ipo7 |
| Ipo8 | 5472 | 4622 | 5385 | 5022 | 4901 | 4786 | 5168 | 4731 | 4578 | 4717 | 4682 | 5296 | 5379 | 5281 | 5105 | 5083 | 4905 | 4829 | 4701 | 4559 | 4783 | 4461 | 4484 | 4744 | 4812 | 5201 | 5691 | 5319 | Ipo8 |
| Ipo9 | 21311 | 20797 | 19595 | 20157 | 20630 | 20861 | 22154 | 20252 | 20673 | 21370 | 19422 | 24203 | 21965 | 23977 | 19978 | 18922 | 20334 | 20010 | 21503 | 19904 | 22096 | 20286 | 18911 | 22006 | 22611 | 22377 | 23580 | 24192 | Ipo9 |
| Ipp | 1061 | 1019 | 836 | 1061 | 970 | 959 | 1056 | 1263 | 1130 | 984 | 975 | 1300 | 1047 | 1156 | 1197 | 868 | 968 | 951 | 992 | 1269 | 1055 | 759 | 1052 | 1097 | 1015 | 1077 | 1235 | 1135 | Ipp |
| Ippk | 1335 | 1367 | 1147 | 1057 | 884 | 927 | 921 | 1133 | 1008 | 838 | 928 | 999 | 937 | 915 | 1283 | 1271 | 1065 | 943 | 921 | 1118 | 906 | 974 | 995 | 919 | 954 | 1021 | 926 | 1044 | Ippk |
| Iqank1 | 141 | 146 | 198 | 166 | 146 | 219 | 204 | 154 | 156 | 203 | 157 | 136 | 186 | 140 | 124 | 111 | 175 | 150 | 168 | 166 | 143 | 170 | 161 | 185 | 195 | 168 | 153 | 165 | Iqank1 |
| Iqca | 21 | 26 | 12 | 22 | 45 | 16 | 28 | 42 | 45 | 28 | 23 | 43 | 45 | 17 | 17 | 30 | 14 | 18 | 25 | 14 | 26 | 32 | 67 | 25 | 20 | 26 | 28 | 23 | Iqca |
| Iqca1l | 30 | 33 | 20 | 50 | 36 | 46 | 60 | 53 | 52 | 64 | 51 | 77 | 36 | 92 | 21 | 17 | 48 | 43 | 39 | 53 | 54 | 37 | 53 | 96 | 76 | 62 | 69 | 51 | Iqca1l |
| Iqcb1 | 4975 | 4186 | 5222 | 4425 | 3848 | 4220 | 4769 | 4714 | 3609 | 3567 | 3470 | 3893 | 3647 | 4392 | 5356 | 4808 | 4735 | 4066 | 4052 | 4732 | 4352 | 4189 | 3696 | 3913 | 3621 | 4507 | 4335 | 4405 | Iqcb1 |
| Iqcc | 1057 | 1243 | 1296 | 1293 | 1348 | 1218 | 1225 | 1467 | 1495 | 1529 | 1577 | 1574 | 1580 | 1761 | 1182 | 1099 | 1234 | 1218 | 1340 | 1188 | 1383 | 1429 | 1518 | 1606 | 1649 | 1573 | 1590 | 1571 | Iqcc |
| Iqcd | 130 | 202 | 226 | 140 | 160 | 152 | 149 | 118 | 138 | 155 | 117 | 202 | 191 | 114 | 167 | 199 | 194 | 153 | 163 | 199 | 151 | 137 | 188 | 136 | 138 | 133 | 122 | 110 | Iqcd |
| Iqce | 2947 | 3455 | 3936 | 4175 | 4271 | 3996 | 3785 | 2400 | 3097 | 3532 | 3454 | 3712 | 3588 | 3454 | 2832 | 3217 | 4040 | 4365 | 4143 | 3286 | 3991 | 3574 | 3157 | 3730 | 3650 | 3131 | 3256 | 3456 | Iqce |
| Iqcf1 | 0 | 1 | 0 | 0 | 0 | 0 | 0 | 0 | 0 | 0 | 0 | 0 | 0 | 0 | 0 | 0 | 0 | 0 | 2 | 1 | 0 | 0 | 0 | 0 | 0 | 0 | 0 | 0 | Iqcf1 |
| Iqcf4 | 0 | 0 | 0 | 0 | 0 | 0 | 0 | 0 | 0 | 0 | 0 | 0 | 0 | 0 | 0 | 0 | 0 | 0 | 2 | 0 | 0 | 0 | 0 | 0 | 0 | 0 | 0 | 0 | Iqcf4 |
| Iqcf5 | 1 | 4 | 0 | 0 | 2 | 7 | 2 | 0 | 1 | 0 | 0 | 0 | 0 | 1 | 4 | 8 | 8 | 1 | 2 | 1 | 0 | 0 | 1 | 0 | 0 | 2 | 5 | 1 | Iqcf5 |
| Iqcf6 | 0 | 0 | 0 | 0 | 0 | 0 | 0 | 0 | 0 | 0 | 2 | 0 | 0 | 0 | 0 | 0 | 0 | 0 | 0 | 0 | 0 | 0 | 0 | 0 | 0 | 0 | 0 | 0 | Iqcf6 |
| Iqcg | 993 | 1233 | 1099 | 1081 | 969 | 980 | 933 | 535 | 461 | 379 | 414 | 471 | 586 | 474 | 1216 | 1239 | 907 | 986 | 937 | 1164 | 945 | 536 | 640 | 414 | 395 | 459 | 474 | 470 | Iqcg |
| Iqch | 3 | 5 | 28 | 5 | 13 | 6 | 2 | 9 | 15 | 18 | 17 | 12 | 12 | 6 | 7 | 8 | 5 | 10 | 0 | 18 | 6 | 13 | 17 | 5 | 19 | 11 | 3 | 6 | Iqch |
| Iqcj | 5 | 2 | 10 | 12 | 4 | 6 | 13 | 7 | 10 | 2 | 3 | 5 | 9 | 6 | 13 | 2 | 2 | 0 | 5 | 7 | 7 | 17 | 5 | 6 | 3 | 11 | 12 | 18 | Iqcj |
| Iqck | 231 | 336 | 224 | 349 | 271 | 262 | 302 | 278 | 260 | 247 | 196 | 231 | 239 | 249 | 324 | 259 | 309 | 296 | 281 | 319 | 257 | 261 | 267 | 306 | 194 | 287 | 255 | 252 | Iqck |
| Iqcm | 1 | 4 | 5 | 1 | 1 | 23 | 0 | 1 | 3 | 8 | 2 | 0 | 11 | 3 | 5 | 0 | 6 | 1 | 3 | 1 | 0 | 7 | 16 | 2 | 5 | 3 | 5 | 1 | Iqcm |
| Iqcn | 0 | 1 | 0 | 0 | 0 | 0 | 0 | 0 | 0 | 0 | 0 | 0 | 0 | 0 | 0 | 1 | 0 | 0 | 0 | 0 | 0 | 2 | 0 | 0 | 0 | 0 | 0 | 0 | Iqcn |
| Iqgap1 | 11690 | 10366 | 7173 | 7201 | 6617 | 8303 | 6779 | 8085 | 9779 | 8065 | 9508 | 7310 | 8808 | 7399 | 10395 | 11528 | 6778 | 7737 | 6582 | 7702 | 6803 | 6580 | 10877 | 7048 | 6917 | 5626 | 7592 | 7067 | Iqgap1 |
| Iqgap2 | 4869 | 4399 | 4189 | 3338 | 2914 | 3723 | 3492 | 4248 | 4220 | 3515 | 4012 | 5313 | 4725 | 5140 | 4795 | 4341 | 3935 | 2914 | 3199 | 3558 | 3193 | 3639 | 4232 | 3866 | 3287 | 4625 | 5281 | 5272 | Iqgap2 |
| Iqgap3 | 6240 | 6592 | 4910 | 4738 | 3476 | 4298 | 4404 | 2941 | 3491 | 3071 | 3009 | 4428 | 3869 | 3949 | 5045 | 5381 | 4881 | 4229 | 4696 | 4351 | 4319 | 2104 | 2941 | 3428 | 3010 | 3716 | 4478 | 4017 | Iqgap3 |
| Iqsec1 | 4584 | 5771 | 6355 | 6588 | 7355 | 6321 | 5315 | 4949 | 5701 | 6869 | 6383 | 7222 | 6424 | 6075 | 4879 | 5453 | 7168 | 6771 | 7009 | 4919 | 6058 | 7361 | 5467 | 6391 | 6682 | 5891 | 5996 | 5916 | Iqsec1 |
| Iqsec2 | 1503 | 1660 | 1823 | 2070 | 2516 | 2159 | 1916 | 1150 | 1443 | 1564 | 1506 | 1247 | 1208 | 1132 | 1436 | 1685 | 1647 | 2265 | 2353 | 1733 | 2112 | 1579 | 1480 | 1585 | 1497 | 1228 | 1052 | 1009 | Iqsec2 |
| Iqsec3 | 1173 | 1515 | 1512 | 3178 | 5746 | 3638 | 3586 | 2023 | 2197 | 3366 | 2962 | 1836 | 1694 | 1640 | 1692 | 1252 | 2446 | 3524 | 3938 | 3212 | 3757 | 3928 | 2204 | 2051 | 2945 | 1075 | 1373 | 1410 | Iqsec3 |
| Iqub | 77 | 105 | 84 | 99 | 167 | 129 | 112 | 221 | 182 | 166 | 133 | 188 | 297 | 116 | 138 | 148 | 94 | 123 | 110 | 139 | 178 | 244 | 327 | 96 | 139 | 133 | 114 | 112 | Iqub |
| Irag1 | 83 | 98 | 77 | 57 | 104 | 77 | 60 | 160 | 178 | 198 | 317 | 253 | 346 | 279 | 63 | 109 | 81 | 88 | 95 | 64 | 79 | 234 | 229 | 206 | 232 | 238 | 291 | 173 | Irag1 |
| Irak1 | 2358 | 2269 | 2593 | 2240 | 2329 | 2398 | 2116 | 3892 | 4258 | 4668 | 4568 | 5116 | 4831 | 4982 | 2111 | 2320 | 2308 | 2201 | 2229 | 1950 | 2053 | 4580 | 4198 | 4743 | 4633 | 5291 | 4671 | 5002 | Irak1 |
| Irak1bp1 | 1248 | 965 | 1140 | 1181 | 1402 | 1209 | 1379 | 1301 | 1143 | 1125 | 1056 | 1207 | 1114 | 1307 | 1284 | 995 | 1176 | 1140 | 1237 | 1393 | 1301 | 1252 | 1186 | 1222 | 1375 | 1362 | 1171 | 1232 | Irak1bp1 |
| Irak2 | 78 | 130 | 127 | 97 | 175 | 87 | 77 | 192 | 213 | 219 | 263 | 178 | 197 | 151 | 117 | 137 | 104 | 113 | 101 | 61 | 60 | 184 | 276 | 195 | 178 | 206 | 229 | 188 | Irak2 |
| Irak3 | 213 | 195 | 227 | 100 | 156 | 171 | 92 | 112 | 184 | 142 | 218 | 52 | 145 | 51 | 210 | 241 | 124 | 121 | 72 | 94 | 111 | 142 | 199 | 108 | 53 | 98 | 65 | 62 | Irak3 |
| Irak4 | 463 | 480 | 559 | 456 | 374 | 327 | 370 | 222 | 280 | 286 | 294 | 243 | 295 | 243 | 380 | 497 | 534 | 454 | 372 | 373 | 345 | 216 | 347 | 242 | 218 | 272 | 309 | 218 | Irak4 |
| Ireb2 | 24694 | 18565 | 18885 | 21044 | 21813 | 22724 | 25155 | 26357 | 21554 | 20202 | 20779 | 22929 | 21962 | 23838 | 24704 | 21981 | 20929 | 21437 | 21577 | 22889 | 24315 | 23515 | 20867 | 20046 | 22369 | 18842 | 23632 | 23533 | Ireb2 |
| Irf1 | 556 | 564 | 523 | 447 | 331 | 381 | 317 | 266 | 270 | 255 | 270 | 248 | 225 | 193 | 429 | 624 | 399 | 478 | 308 | 418 | 300 | 229 | 337 | 297 | 235 | 330 | 249 | 215 | Irf1 |
| Irf2 | 737 | 810 | 1138 | 1031 | 932 | 807 | 926 | 659 | 743 | 794 | 798 | 762 | 796 | 780 | 750 | 721 | 1017 | 933 | 1023 | 697 | 877 | 700 | 784 | 958 | 767 | 1007 | 651 | 807 | Irf2 |
| Irf2bp1 | 2872 | 3816 | 3614 | 3802 | 3382 | 2977 | 2904 | 2790 | 3175 | 3257 | 3289 | 3540 | 2740 | 3099 | 2603 | 3028 | 3806 | 3702 | 3819 | 2704 | 3287 | 3293 | 3102 | 3588 | 3513 | 4013 | 3059 | 3210 | Irf2bp1 |
| Irf2bp2 | 1835 | 1787 | 2331 | 2613 | 2150 | 1820 | 1887 | 2408 | 2365 | 2821 | 2867 | 2363 | 2130 | 2175 | 1617 | 1881 | 2300 | 2502 | 2170 | 1689 | 1991 | 3021 | 2577 | 3205 | 2583 | 3304 | 2092 | 2225 | Irf2bp2 |
| Irf2bpl | 3257 | 3437 | 4838 | 5232 | 5434 | 4389 | 4418 | 4881 | 4838 | 6749 | 5773 | 6982 | 4758 | 6378 | 2946 | 3100 | 4872 | 5169 | 5230 | 3502 | 4813 | 7671 | 4718 | 7456 | 7139 | 7306 | 5127 | 5498 | Irf2bpl |
| Irf3 | 3793 | 5007 | 4827 | 4523 | 3654 | 3651 | 3747 | 2495 | 2728 | 2848 | 2630 | 2688 | 2733 | 2569 | 3948 | 4245 | 4921 | 4351 | 4247 | 4002 | 3802 | 2261 | 2804 | 2953 | 2555 | 3015 | 2768 | 2687 | Irf3 |
| Irf4 | 51 | 56 | 76 | 23 | 26 | 15 | 19 | 4 | 8 | 10 | 6 | 0 | 3 | 9 | 46 | 81 | 44 | 44 | 9 | 42 | 23 | 13 | 14 | 2 | 2 | 12 | 0 | 1 | Irf4 |

| | | | | | | | | | | | | | | | | | | | | | | | | | | | | | |
|---|---|---|---|---|---|---|---|---|---|---|---|---|---|---|---|---|---|---|---|---|---|---|---|---|---|---|---|---|---|
| Irf5 | 92 | 136 | 148 | 95 | 133 | 98 | 80 | 90 | 112 | 133 | 184 | 123 | 128 | 79 | 113 | 142 | 180 | 123 | 129 | 103 | 84 | 102 | 206 | 121 | 89 | 109 | 103 | 83 | Irf5 |
| Irf6 | 1371 | 523 | 431 | 327 | 156 | 341 | 98 | 705 | 1023 | 547 | 720 | 181 | 324 | 191 | 698 | 1031 | 228 | 428 | 100 | 126 | 86 | 169 | 1243 | 498 | 113 | 351 | 199 | 228 | Irf6 |
| Irf7 | 35 | 45 | 40 | 31 | 36 | 30 | 17 | 38 | 31 | 56 | 63 | 27 | 25 | 36 | 37 | 53 | 37 | 29 | 29 | 35 | 29 | 33 | 52 | 38 | 38 | 67 | 52 | 42 | Irf7 |
| Irf8 | 333 | 432 | 560 | 409 | 273 | 284 | 276 | 147 | 165 | 147 | 259 | 192 | 169 | 166 | 334 | 452 | 440 | 360 | 303 | 280 | 284 | 206 | 221 | 163 | 169 | 220 | 170 | 165 | Irf8 |
| Irf9 | 1837 | 2177 | 2127 | 2667 | 2423 | 2337 | 2451 | 1515 | 1412 | 1745 | 1796 | 1607 | 1752 | 1672 | 1718 | 2232 | 2427 | 2611 | 2431 | 2075 | 2561 | 1977 | 1430 | 1879 | 1821 | 1687 | 1841 | 1830 | Irf9 |
| Irgc1 | 0 | 0 | 0 | 0 | 0 | 0 | 0 | 0 | 1 | 0 | 1 | 0 | 0 | 0 | 0 | 0 | 0 | 0 | 2 | 0 | 0 | 0 | 0 | 0 | 0 | 0 | 1 | 0 | Irgc1 |
| Irgm1 | 40 | 24 | 59 | 30 | 51 | 50 | 38 | 71 | 67 | 73 | 98 | 59 | 67 | 54 | 42 | 51 | 42 | 26 | 51 | 66 | 35 | 85 | 104 | 36 | 57 | 132 | 60 | 71 | Irgm1 |
| Irgm2 | 102 | 91 | 110 | 76 | 97 | 123 | 57 | 115 | 130 | 183 | 237 | 150 | 138 | 105 | 121 | 124 | 80 | 94 | 49 | 133 | 69 | 174 | 200 | 150 | 96 | 139 | 134 | 154 | Irgm2 |
| Irgq | 7974 | 9685 | 9707 | 11508 | 12201 | 10768 | 10330 | 7409 | 7963 | 9491 | 8375 | 9518 | 8325 | 8690 | 7759 | 8268 | 10980 | 11465 | 12543 | 8667 | 11395 | 11102 | 7175 | 9436 | 10555 | 7293 | 7927 | 8193 | Irgq |
| Irs1 | 5084 | 4224 | 6441 | 5544 | 5831 | 5908 | 5702 | 5208 | 5155 | 5861 | 6649 | 6378 | 6411 | 6144 | 5093 | 4708 | 5439 | 5585 | 6026 | 4702 | 5511 | 5803 | 5584 | 6225 | 6180 | 8465 | 6061 | 6659 | Irs1 |
| Irs2 | 3823 | 4333 | 4847 | 4813 | 3919 | 3434 | 3546 | 2738 | 2968 | 3378 | 3043 | 3255 | 3376 | 3345 | 3473 | 4279 | 5018 | 4648 | 4204 | 3020 | 3827 | 3375 | 2766 | 3554 | 3137 | 3793 | 3119 | 3560 | Irs2 |
| Irs3 | 30 | 28 | 54 | 26 | 12 | 15 | 10 | 4 | 7 | 2 | 12 | 4 | 8 | 10 | 20 | 41 | 31 | 24 | 6 | 20 | 20 | 3 | 3 | 2 | 0 | 10 | 3 | 6 | Irs3 |
| Irs4 | 1220 | 1136 | 828 | 1181 | 1220 | 1033 | 1019 | 432 | 615 | 463 | 549 | 544 | 263 | 528 | 1031 | 967 | 1255 | 1432 | 944 | 872 | 1165 | 1026 | 539 | 543 | 618 | 819 | 514 | 523 | Irs4 |
| Irx1 | 684 | 1283 | 1810 | 1421 | 1060 | 1093 | 635 | 1139 | 1148 | 1433 | 1287 | 1244 | 1827 | 1114 | 1015 | 1090 | 1685 | 1102 | 1189 | 924 | 904 | 1459 | 1320 | 1288 | 1370 | 505 | 1264 | 1329 | Irx1 |
| Irx2 | 1375 | 2499 | 3558 | 2685 | 2286 | 2379 | 1266 | 2738 | 2028 | 2728 | 2218 | 2353 | 4431 | 2118 | 2450 | 2824 | 3428 | 2317 | 2360 | 2028 | 1913 | 3380 | 2522 | 1897 | 2584 | 767 | 2502 | 2359 | Irx2 |
| Irx3 | 1647 | 2116 | 3678 | 1992 | 1244 | 1377 | 518 | 1685 | 1489 | 1837 | 1582 | 2141 | 2866 | 2311 | 1708 | 1776 | 2670 | 1437 | 1538 | 1000 | 1271 | 1835 | 1526 | 2104 | 1777 | 1423 | 2537 | 2841 | Irx3 |
| Irx4 | 59 | 113 | 76 | 93 | 135 | 113 | 4 | 237 | 452 | 322 | 262 | 114 | 491 | 117 | 120 | 150 | 145 | 112 | 65 | 96 | 21 | 227 | 441 | 167 | 101 | 55 | 112 | 117 | Irx4 |
| Irx5 | 985 | 1217 | 1901 | 1125 | 878 | 1096 | 526 | 1292 | 1032 | 1145 | 1145 | 1376 | 1951 | 1299 | 1114 | 1143 | 1465 | 982 | 1065 | 762 | 877 | 1288 | 1133 | 1189 | 1193 | 676 | 1546 | 1648 | Irx5 |
| Irx6 | 91 | 109 | 172 | 120 | 114 | 112 | 54 | 140 | 150 | 137 | 185 | 137 | 153 | 149 | 65 | 110 | 123 | 90 | 175 | 116 | 109 | 211 | 187 | 163 | 200 | 79 | 146 | 171 | Irx6 |
| Isca1 | 6853 | 7578 | 6361 | 6942 | 6389 | 6590 | 7610 | 4528 | 4233 | 3741 | 3968 | 3958 | 4124 | 3998 | 7709 | 7258 | 6540 | 6859 | 6400 | 7731 | 7234 | 4064 | 4493 | 3878 | 4235 | 3895 | 4036 | 4330 | Isca1 |
| Isca2 | 2255 | 2243 | 2466 | 2166 | 1877 | 1833 | 2360 | 2429 | 1841 | 1977 | 1897 | 2171 | 2168 | 2490 | 2368 | 2135 | 2146 | 1976 | 1988 | 2082 | 2121 | 2102 | 2008 | 2263 | 2016 | 2463 | 2403 | 2616 | Isca2 |
| Iscu | 1852 | 2275 | 1918 | 1961 | 1403 | 1640 | 1662 | 1286 | 1280 | 1320 | 1198 | 1464 | 1370 | 1442 | 1871 | 1923 | 1841 | 1754 | 1670 | 1971 | 1674 | 1135 | 1298 | 1336 | 1254 | 1565 | 1528 | 1524 | Iscu |
| Isg15 | 21 | 16 | 23 | 29 | 25 | 16 | 22 | 23 | 35 | 38 | 50 | 40 | 32 | 52 | 13 | 53 | 23 | 22 | 12 | 132 | 60 | 38 | 34 | 51 | 36 | 56 | 63 | 63 | Isg15 |
| Isg20 | 130 | 193 | 199 | 220 | 123 | 145 | 159 | 12 | 47 | 25 | 43 | 33 | 27 | 33 | 109 | 169 | 195 | 194 | 148 | 144 | 140 | 18 | 24 | 32 | 45 | 20 | 44 | 45 | Isg20 |
| Isg20l2 | 4760 | 4660 | 5178 | 4507 | 3768 | 3907 | 4261 | 4460 | 4061 | 4345 | 4327 | 5264 | 4782 | 5401 | 4619 | 4123 | 4651 | 4048 | 4290 | 3961 | 4196 | 4107 | 3928 | 4997 | 4533 | 6387 | 5211 | 5620 | Isg20l2 |
| Isl1 | 2756 | 2280 | 1447 | 2559 | 3548 | 2487 | 3760 | 5202 | 4832 | 4094 | 4186 | 3955 | 2724 | 3977 | 3192 | 2132 | 2481 | 2626 | 2985 | 2591 | 3227 | 6122 | 2840 | 3871 | 4835 | 3771 | 3228 | 3189 | Isl1 |
| Isl2 | 215 | 276 | 182 | 174 | 320 | 105 | 65 | 399 | 371 | 325 | 424 | 70 | 513 | 65 | 265 | 333 | 197 | 228 | 114 | 98 | 102 | 585 | 183 | 57 | 272 | 35 | 109 | 108 | Isl2 |
| Islr | 1660 | 1869 | 1773 | 1624 | 1582 | 1720 | 921 | 2451 | 3846 | 2899 | 5775 | 2034 | 2671 | 1856 | 1667 | 2284 | 1513 | 1942 | 1094 | 1404 | 1018 | 2984 | 5421 | 2631 | 2211 | 2912 | 2273 | 1784 | Islr |
| Islr2 | 10766 | 13611 | 11281 | 21078 | 23740 | 22582 | 25019 | 22311 | 26592 | 27851 | 20792 | 32487 | 12651 | 29794 | 10615 | 7805 | 18607 | 21810 | 29603 | 21682 | 25419 | 28649 | 18489 | 36280 | 39359 | 32990 | 25003 | 23949 | Islr2 |
| Ism1 | 151 | 98 | 92 | 73 | 56 | 101 | 23 | 145 | 224 | 202 | 271 | 46 | 136 | 88 | 99 | 157 | 44 | 112 | 42 | 42 | 31 | 99 | 238 | 102 | 81 | 64 | 87 | 84 | Ism1 |
| Ism2 | 148 | 134 | 152 | 125 | 156 | 135 | 95 | 118 | 108 | 141 | 100 | 184 | 204 | 129 | 124 | 139 | 154 | 138 | 110 | 124 | 113 | 95 | 166 | 143 | 105 | 187 | 124 | 163 | Ism2 |
| Isoc1 | 10810 | 8064 | 9167 | 8117 | 8065 | 8364 | 8726 | 7844 | 8064 | 7225 | 7825 | 8460 | 8543 | 8442 | 9724 | 9745 | 8189 | 8464 | 8108 | 8033 | 8267 | 7204 | 8309 | 7979 | 6961 | 9475 | 8496 | 9012 | Isoc1 |
| Isoc2a | 1908 | 2682 | 2548 | 2636 | 2006 | 2038 | 1912 | 894 | 1212 | 1140 | 1182 | 1188 | 1024 | 1156 | 1921 | 2291 | 2555 | 2264 | 2104 | 1960 | 1900 | 876 | 1220 | 1175 | 1118 | 1436 | 1152 | 1241 | Isoc2a |
| Isoc2b | 80 | 65 | 56 | 71 | 65 | 54 | 57 | 68 | 96 | 94 | 62 | 58 | 66 | 56 | 63 | 56 | 50 | 55 | 57 | 42 | 55 | 79 | 98 | 62 | 67 | 68 | 71 | 77 | Isoc2b |
| Ist1 | 15479 | 15299 | 12877 | 14215 | 14173 | 13717 | 14397 | 17954 | 14862 | 14394 | 13900 | 14828 | 14212 | 14844 | 15335 | 14435 | 13626 | 14038 | 14276 | 15457 | 14263 | 15720 | 14532 | 14407 | 15379 | 14475 | 14760 | 14455 | Ist1 |
| Isx | 0 | 0 | 0 | 0 | 0 | 0 | 0 | 0 | 0 | 0 | 0 | 0 | 2 | 0 | 0 | 0 | 0 | 0 | 0 | 0 | 0 | 0 | 1 | 0 | 0 | 0 | 0 | 0 | Isx |
| Isy1 | 964 | 729 | 819 | 732 | 736 | 775 | 858 | 1381 | 1320 | 1382 | 1287 | 1586 | 1387 | 1683 | 828 | 735 | 758 | 661 | 644 | 806 | 733 | 1163 | 1180 | 1387 | 1517 | 1551 | 1598 | 1730 | Isy1 |
| Isyna1 | 5375 | 7121 | 6639 | 5908 | 4807 | 4779 | 4169 | 3268 | 4164 | 3475 | 4046 | 3401 | 3963 | 2760 | 4824 | 6538 | 6387 | 5420 | 5586 | 4560 | 4206 | 3456 | 4162 | 3860 | 3271 | 4838 | 3433 | 3151 | Isyna1 |
| Itch | 8556 | 7197 | 6569 | 6104 | 6828 | 7349 | 7569 | 10190 | 8897 | 7634 | 7859 | 8774 | 8128 | 8913 | 8465 | 7462 | 6675 | 6381 | 6764 | 7958 | 7269 | 8193 | 8920 | 7702 | 8334 | 7455 | 9022 | 8871 | Itch |
| Itfg1 | 8327 | 7351 | 6889 | 7717 | 9875 | 9581 | 10087 | 13087 | 11098 | 10751 | 9716 | 9698 | 9662 | 10004 | 9853 | 7791 | 7314 | 8734 | 8635 | 10045 | 9862 | 13139 | 11804 | 9647 | 10948 | 9072 | 9917 | 9862 | Itfg1 |
| Itfg2 | 911 | 886 | 1015 | 968 | 1126 | 1054 | 940 | 1788 | 1934 | 2059 | 2146 | 2373 | 2181 | 2293 | 793 | 803 | 969 | 963 | 1174 | 962 | 1014 | 2239 | 1878 | 2302 | 2393 | 2342 | 2322 | 2217 | Itfg2 |
| Itga1 | 1555 | 1433 | 1161 | 1257 | 1624 | 1633 | 1661 | 1504 | 1766 | 1515 | 1976 | 1640 | 1830 | 1833 | 1638 | 1594 | 1095 | 1263 | 1345 | 1677 | 1715 | 1831 | 2016 | 1428 | 1835 | 1325 | 1753 | 1591 | Itga1 |
| Itga10 | 604 | 781 | 732 | 810 | 765 | 655 | 713 | 442 | 472 | 568 | 823 | 348 | 484 | 441 | 556 | 724 | 736 | 894 | 869 | 571 | 752 | 700 | 437 | 574 | 535 | 372 | 395 | 409 | Itga10 |
| Itga11 | 912 | 1419 | 1411 | 1194 | 569 | 1354 | 437 | 872 | 1074 | 1409 | 1541 | 1718 | 1917 | 1561 | 1015 | 1640 | 1064 | 1448 | 599 | 416 | 487 | 1196 | 1447 | 1672 | 1406 | 1362 | 1705 | 1569 | Itga11 |
| Itga2 | 551 | 435 | 287 | 347 | 490 | 482 | 383 | 972 | 1059 | 905 | 1063 | 821 | 976 | 832 | 630 | 437 | 350 | 395 | 312 | 562 | 417 | 906 | 1127 | 791 | 845 | 608 | 912 | 831 | Itga2 |
| Itga2b | 498 | 598 | 624 | 442 | 378 | 387 | 277 | 302 | 372 | 351 | 336 | 517 | 790 | 391 | 508 | 714 | 556 | 475 | 422 | 386 | 369 | 287 | 368 | 398 | 305 | 635 | 625 | 497 | Itga2b |
| Itga3 | 1803 | 1973 | 2225 | 1949 | 2575 | 1864 | 1629 | 1249 | 1742 | 1947 | 1495 | 1138 | 1372 | 955 | 1532 | 2359 | 2187 | 2220 | 2204 | 1641 | 1854 | 1726 | 1589 | 1349 | 1381 | 1097 | 1025 | 955 | Itga3 |
| Itga4 | 3037 | 2344 | 2264 | 1572 | 2184 | 2284 | 1919 | 2955 | 2993 | 2377 | 4332 | 2860 | 3894 | 2843 | 3233 | 3523 | 1738 | 1758 | 1600 | 2158 | 1908 | 2577 | 3573 | 1946 | 2360 | 1983 | 3412 | 2932 | Itga4 |
| Itga5 | 2765 | 2973 | 3732 | 2314 | 2023 | 2781 | 1570 | 2199 | 3087 | 2700 | 4686 | 3129 | 3729 | 2530 | 2450 | 3478 | 2956 | 2523 | 2288 | 1502 | 1633 | 2525 | 3893 | 3043 | 2358 | 3450 | 3218 | 3132 | Itga5 |
| Itga6 | 10668 | 8585 | 7921 | 8603 | 8395 | 8672 | 9878 | 6003 | 6217 | 5838 | 6357 | 6429 | 5635 | 6282 | 9244 | 8753 | 7907 | 8507 | 8293 | 8841 | 9235 | 5158 | 6419 | 5848 | 5677 | 6199 | 5955 | 5722 | Itga6 |
| Itga7 | 444 | 715 | 728 | 661 | 956 | 479 | 373 | 278 | 427 | 359 | 669 | 129 | 439 | 129 | 593 | 913 | 707 | 724 | 547 | 435 | 479 | 587 | 456 | 143 | 401 | 154 | 146 | 152 | Itga7 |
| Itga8 | 1427 | 891 | 737 | 556 | 456 | 861 | 449 | 1307 | 1359 | 970 | 1367 | 1070 | 1239 | 961 | 1110 | 1326 | 483 | 810 | 440 | 505 | 412 | 680 | 1428 | 1079 | 705 | 821 | 902 | 863 | Itga8 |
| Itga9 | 2503 | 2152 | 2133 | 1322 | 931 | 1485 | 586 | 2023 | 2638 | 2530 | 3353 | 2301 | 3274 | 2101 | 2134 | 3353 | 1431 | 1615 | 709 | 669 | 611 | 2294 | 2651 | 2485 | 1844 | 2520 | 2628 | 2279 | Itga9 |
| Itgad | 0 | 1 | 3 | 0 | 0 | 2 | 0 | 0 | 0 | 0 | 1 | 1 | 6 | 0 | 1 | 0 | 0 | 0 | 0 | 0 | 2 | 3 | 2 | 0 | 0 | 1 | 0 | 0 | Itgad |
| Itgae | 20 | 23 | 30 | 36 | 25 | 43 | 21 | 28 | 30 | 16 | 21 | 28 | 42 | 24 | 21 | 29 | 24 | 24 | 13 | 18 | 23 | 32 | 21 | 20 | 37 | 30 | 21 | 26 | Itgae |
| Itgal | 91 | 105 | 62 | 52 | 70 | 89 | 53 | 89 | 108 | 78 | 131 | 80 | 135 | 86 | 78 | 126 | 93 | 79 | 55 | 60 | 67 | 102 | 131 | 98 | 91 | 93 | 99 | 109 | Itgal |
| Itgam | 14 | 27 | 53 | 36 | 33 | 31 | 44 | 37 | 35 | 34 | 56 | 18 | 28 | 40 | 8 | 35 | 27 | 53 | 32 | 38 | 27 | 63 | 59 | 40 | 18 | 50 | 36 | 43 | Itgam |
| Itgav | 4244 | 3005 | 3184 | 2889 | 2892 | 3461 | 2674 | 4026 | 4103 | 3429 | 4909 | 3972 | 4509 | 3973 | 3926 | 4294 | 2983 | 3149 | 2606 | 2546 | 2676 | 3643 | 4445 | 3618 | 3360 | 3420 | 4131 | 4176 | Itgav |
| Itgax | 6 | 4 | 6 | 1 | 4 | 8 | 8 | 10 | 9 | 5 | 20 | 4 | 7 | 2 | 8 | 3 | 10 | 13 | 2 | 6 | 10 | 10 | 21 | 13 | 6 | 3 | 3 | 13 | Itgax |
| Itgb1 | 21820 | 17121 | 18558 | 13952 | 13115 | 15216 | 12122 | 23786 | 23871 | 19732 | 24857 | 19979 | 23009 | 20400 | 22140 | 22878 | 14968 | 16104 | 11825 | 14633 | 11867 | 20082 | 27033 | 20069 | 18495 | 21482 | 21171 | 20751 | Itgb1 |
| Itgb1bp1 | 1859 | 1697 | 1617 | 1494 | 1497 | 1449 | 1519 | 1458 | 1347 | 1121 | 1220 | 1391 | 1376 | 1308 | 1857 | 1554 | 1488 | 1447 | 1502 | 1717 | 1483 | 1103 | 1364 | 1361 | 1217 | 1237 | 1389 | 1348 | Itgb1bp1 |
| Itgb1bp2 | 188 | 174 | 173 | 169 | 239 | 209 | 230 | 304 | 330 | 148 | 1098 | 215 | 221 | 217 | 194 | 127 | 133 | 167 | 185 | 198 | 212 | 247 | 795 | 240 | 228 | 188 | 245 | 183 | Itgb1bp2 |
| Itgb2 | 62 | 60 | 76 | 42 | 61 | 63 | 26 | 46 | 65 | 66 | 68 | 55 | 51 | 53 | 54 | 94 | 58 | 53 | 41 | 78 | 53 | 43 | 91 | 75 | 61 | 80 | 76 | 52 | Itgb2 |
| Itgb2l | 1 | 1 | 0 | 0 | 0 | 0 | 0 | 0 | 0 | 0 | 0 | 4 | 7 | 0 | 1 | 1 | 1 | 0 | 1 | 1 | 2 | 0 | 4 | 2 | 1 | 1 | 7 | 0 | Itgb2l |
| Itgb3 | 634 | 599 | 719 | 358 | 380 | 357 | 273 | 504 | 619 | 533 | 707 | 635 | 954 | 463 | 526 | 858 | 430 | 468 | 298 | 322 | 314 | 588 | 656 | 530 | 373 | 757 | 829 | 577 | Itgb3 |
| Itgb3bp | 804 | 524 | 770 | 373 | 173 | 442 | 605 | 454 | 305 | 122 | 321 | 178 | 472 | 259 | 671 | 315 | 576 | 276 | 489 | 627 | 405 | 480 | 165 | 592 | 550 | 829 | 809 | 453 | Itgb3bp |
| Itgb4 | 251 | 136 | 147 | 52 | 94 | 123 | 22 | 369 | 814 | 421 | 475 | 146 | 271 | 175 | 124 | 203 | 72 | 143 | 50 | 36 | 27 | 139 | 815 | 344 | 143 | 165 | 135 | 154 | Itgb4 |
| Itgb5 | 4828 | 4767 | 4336 | 2818 | 1975 | 2936 | 1269 | 3659 | 5220 | 4345 | 4930 | 3161 | 3984 | 2506 | 3960 | 6095 | 3148 | 3739 | 1919 | 1376 | 1444 | 3258 | 5929 | 4099 | 2868 | 4007 | 3153 | 2931 | Itgb5 |
| Itgb6 | 66 | 27 | 18 | 13 | 7 | 12 | 1 | 110 | 98 | 85 | 209 | 27 | 58 | 26 | 36 | 41 | 0 | 18 | 5 | 1 | 4 | 37 | 164 | 60 | 10 | 81 | 19 | 26 | Itgb6 |
| Itgb7 | 43 | 49 | 43 | 28 | 41 | 15 | 21 | 23 | 40 | 53 | 69 | 33 | 27 | 15 | 44 | 49 | 46 | 36 | 28 | 26 | 28 | 20 | 54 | 41 | 21 | 41 | 13 | 8 | Itgb7 |
| Itgb8 | 5877 | 5252 | 5162 | 4259 | 5231 | 6375 | 5278 | 9591 | 6841 | 5474 | 5158 | 8736 | 9284 | 8960 | 7286 | 5219 | 5201 | 4094 | 5514 | 6010 | 5844 | 7105 | 5844 | 5745 | 6613 | 5073 | 11590 | 10223 | Itgb8 |
| Itgbl1 | 120 | 91 | 50 | 146 | 83 | 161 | 67 | 142 | 245 | 235 | 335 | 134 | 144 | 182 | 164 | 105 | 88 | 206 | 81 | 75 | 62 | 343 | 377 | 189 | 258 | 126 | 217 | 131 | Itgbl1 |
| Itih1 | 0 | 0 | 0 | 0 | 4 | 0 | 1 | 0 | 0 | 0 | 0 | 0 | 0 | 0 | 1 | 0 | 0 | 0 | 0 | 0 | 0 | 0 | 3 | 0 | 0 | 1 | 0 | 1 | Itih1 |
| Itih2 | 29 | 27 | 34 | 73 | 108 | 79 | 55 | 78 | 124 | 135 | 180 | 56 | 67 | 73 | 41 | 45 | 38 | 78 | 49 | 101 | 71 | 147 | 196 | 105 | 118 | 56 | 44 | 40 | Itih2 |
| Itih3 | 8 | 7 | 7 | 5 | 16 | 0 | 0 | 35 | 37 | 29 | 28 | 16 | 16 | 12 | 17 | 15 | 1 | 23 | 5 | 8 | 8 | 43 | 39 | 15 | 13 | 10 | 35 | 12 | Itih3 |
| Itih4 | 48 | 70 | 59 | 87 | 53 | 67 | 74 | 6 | 4 | 6 | 7 | 1 | 8 | 10 | 38 | 65 | 45 | 77 | 64 | 55 | 87 | 7 | 9 | 11 | 5 | 4 | 9 | 7 | Itih4 |
| Itih5 | 2126 | 2064 | 2253 | 1810 | 2293 | 2064 | 1567 | 1430 | 1547 | 1872 | 2550 | 1943 | 2303 | 1816 | 2123 | 2779 | 1672 | 2072 | 1539 | 1985 | 1766 | 1955 | 2297 | 1573 | 1706 | 1544 | 2056 | 1667 | Itih5 |
| Itk | 1 | 3 | 0 | 4 | 2 | 1 | 2 | 4 | 3 | 2 | 9 | 0 | 1 | 0 | 1 | 2 | 0 | 0 | 2 | 2 | 2 | 0 | 4 | 16 | 0 | 1 | 0 | 0 | Itk |
| Itm2a | 8823 | 6167 | 7215 | 5877 | 5293 | 6087 | 4851 | 7562 | 10082 | 6704 | 19947 | 5074 | 6564 | 5505 | 7781 | 8897 | 5788 | 6457 | 4998 | 6069 | 4917 | 7643 | 17290 | 6628 | 6079 | 7033 | 5554 | 5351 | Itm2a |

| | | | | | | | | | | | | | | | | | | | | | | | | | | | | | |
|---|---|---|---|---|---|---|---|---|---|---|---|---|---|---|---|---|---|---|---|---|---|---|---|---|---|---|---|---|---|
| Itm2b | 12777 | 11537 | 11668 | 12232 | 13436 | 12604 | 13168 | 12181 | 13050 | 12480 | 13797 | 10219 | 11484 | 10492 | 13338 | 12899 | 11070 | 13338 | 12034 | 13876 | 12868 | 12442 | 15190 | 11610 | 12028 | 11500 | 9861 | 10341 | Itm2b |
| Itm2c | 7094 | 8005 | 7055 | 7547 | 6861 | 7128 | 6041 | 6482 | 8013 | 7416 | 7737 | 6053 | 6673 | 5647 | 6626 | 7627 | 6744 | 7071 | 7065 | 7405 | 6478 | 6181 | 9068 | 6821 | 6415 | 6742 | 6177 | 5999 | Itm2c |
| Itpa | 5264 | 6248 | 6303 | 5348 | 4789 | 4947 | 5334 | 4838 | 4874 | 4228 | 4173 | 5204 | 4931 | 5185 | 5432 | 6068 | 5742 | 4876 | 5353 | 6812 | 5067 | 3925 | 5130 | 4857 | 4626 | 5760 | 5771 | 5478 | Itpa |
| Itpk1 | 2798 | 3756 | 3159 | 3970 | 4118 | 3531 | 3977 | 1175 | 1542 | 1499 | 1325 | 1524 | 1295 | 1400 | 2831 | 2848 | 3611 | 4185 | 4430 | 3275 | 4050 | 1502 | 1231 | 1635 | 1641 | 1529 | 1094 | 1267 | Itpk1 |
| Itpka | 328 | 411 | 459 | 424 | 251 | 246 | 248 | 18 | 22 | 14 | 11 | 23 | 18 | 32 | 272 | 368 | 399 | 308 | 321 | 250 | 264 | 13 | 16 | 27 | 23 | 55 | 24 | 42 | Itpka |
| Itpkb | 2497 | 2293 | 2798 | 2509 | 2416 | 2584 | 1821 | 2177 | 2912 | 2755 | 3498 | 2380 | 2816 | 2206 | 2143 | 2519 | 2684 | 2542 | 2757 | 1776 | 2128 | 2470 | 3234 | 2806 | 2260 | 2663 | 2368 | 2584 | Itpkb |
| Itpkc | 1423 | 1735 | 1472 | 1736 | 1420 | 1497 | 1224 | 1010 | 1132 | 1075 | 1137 | 1213 | 1145 | 1075 | 1304 | 1703 | 1537 | 1817 | 1578 | 1169 | 1406 | 1173 | 1220 | 1176 | 1220 | 1100 | 1113 | 1127 | Itpkc |
| Itpr1 | 7816 | 7029 | 4348 | 4851 | 5002 | 5538 | 5335 | 3058 | 3407 | 2949 | 3769 | 2931 | 3495 | 2960 | 6592 | 8214 | 4028 | 5327 | 4788 | 6140 | 6001 | 2374 | 4339 | 2566 | 2921 | 2084 | 3397 | 2718 | Itpr1 |
| Itpr2 | 607 | 724 | 582 | 527 | 606 | 644 | 459 | 558 | 695 | 694 | 917 | 465 | 632 | 514 | 608 | 825 | 481 | 618 | 497 | 577 | 582 | 657 | 791 | 460 | 620 | 384 | 494 | 446 | Itpr2 |
| Itpr3 | 2193 | 2273 | 1755 | 1807 | 1573 | 1734 | 1052 | 838 | 1265 | 1244 | 1331 | 964 | 1298 | 818 | 1652 | 2960 | 1581 | 2154 | 1218 | 908 | 1078 | 1011 | 1417 | 988 | 850 | 794 | 967 | 856 | Itpr3 |
| Itprid1 | 2 | 3 | 0 | 13 | 19 | 18 | 29 | 26 | 38 | 60 | 25 | 28 | 19 | 21 | 1 | 5 | 6 | 17 | 6 | 13 | 29 | 12 | 38 | 33 | 45 | 17 | 12 | 14 | Itprid1 |
| Itprid2 | 4099 | 3965 | 3279 | 3270 | 3632 | 3521 | 3549 | 2275 | 2524 | 2092 | 2590 | 1821 | 2257 | 1845 | 4067 | 4569 | 3286 | 3822 | 3070 | 3414 | 3418 | 2276 | 2840 | 1940 | 2078 | 1656 | 2012 | 1812 | Itprid2 |
| Itprip | 2173 | 2635 | 2383 | 2418 | 1987 | 2185 | 2056 | 1065 | 1350 | 1264 | 1206 | 1784 | 1731 | 1622 | 1913 | 2573 | 2420 | 2079 | 2291 | 1890 | 1970 | 1079 | 1223 | 1454 | 1194 | 1934 | 1771 | 1792 | Itprip |
| Itpripl1 | 1262 | 1278 | 1283 | 1182 | 1178 | 1174 | 1179 | 738 | 912 | 904 | 907 | 961 | 1033 | 1007 | 1073 | 1146 | 1287 | 1170 | 1268 | 1122 | 1107 | 695 | 962 | 993 | 871 | 1192 | 975 | 1106 | Itpripl1 |
| Itpripl2 | 1388 | 1072 | 1619 | 891 | 1141 | 1158 | 725 | 1854 | 2305 | 2173 | 3565 | 1742 | 2612 | 1798 | 1216 | 1350 | 1181 | 1146 | 873 | 794 | 778 | 1963 | 2742 | 2072 | 1778 | 1943 | 2026 | 1936 | Itpripl2 |
| Itsn1 | 17642 | 19960 | 20777 | 24962 | 27576 | 24965 | 26942 | 16767 | 17213 | 21699 | 18751 | 19733 | 18770 | 20694 | 18762 | 20150 | 22482 | 25409 | 25306 | 20475 | 28364 | 20301 | 15872 | 19619 | 23938 | 15324 | 18389 | 18924 | Itsn1 |
| Itsn2 | 2076 | 1622 | 1329 | 1503 | 1919 | 1940 | 1673 | 3336 | 3369 | 3107 | 3702 | 2468 | 2844 | 2561 | 1967 | 2014 | 1395 | 1742 | 1679 | 1713 | 1768 | 3014 | 3580 | 2523 | 2702 | 1986 | 2474 | 2432 | Itsn2 |
| Ivd | 6368 | 6657 | 6159 | 6548 | 6193 | 5905 | 6338 | 4882 | 5745 | 5644 | 5975 | 5781 | 5745 | 5794 | 5943 | 6261 | 6624 | 6542 | 6563 | 5996 | 6471 | 5013 | 5583 | 5947 | 5755 | 6309 | 5913 | 5859 | Ivd |
| Ivl | 1 | 0 | 0 | 0 | 0 | 0 | 0 | 0 | 2 | 0 | 1 | 0 | 0 | 0 | 0 | 1 | 0 | 0 | 0 | 0 | 0 | 0 | 2 | 0 | 0 | 0 | 0 | 0 | Ivl |
| Ivns1abp | 41720 | 31947 | 34401 | 33208 | 33513 | 35264 | 37632 | 51296 | 41982 | 38349 | 35043 | 44184 | 39344 | 46699 | 40643 | 35621 | 33925 | 32325 | 34279 | 40318 | 36572 | 37580 | 38574 | 42072 | 40413 | 45907 | 45233 | 44404 | Ivns1abp |
| Iws1 | 7442 | 5846 | 5540 | 5992 | 6551 | 7388 | 6946 | 11177 | 9534 | 8868 | 9724 | 10755 | 10024 | 11352 | 7214 | 6076 | 6077 | 5611 | 6427 | 6999 | 6940 | 9556 | 9071 | 9145 | 9680 | 8850 | 11369 | 10205 | Iws1 |
| Iyd | 5 | 11 | 9 | 18 | 30 | 15 | 44 | 18 | 26 | 16 | 37 | 5 | 7 | 9 | 8 | 9 | 7 | 13 | 20 | 7 | 13 | 18 | 21 | 16 | 23 | 10 | 1 | 7 | Iyd |
| Izumo1 | 16 | 25 | 27 | 20 | 23 | 16 | 18 | 26 | 21 | 38 | 34 | 9 | 25 | 19 | 22 | 26 | 28 | 14 | 15 | 13 | 14 | 15 | 30 | 49 | 7 | 20 | 16 | 4 | Izumo1 |
| Izumo1r | 0 | 0 | 0 | 0 | 0 | 2 | 2 | 4 | 2 | 0 | 1 | 0 | 0 | 2 | 0 | 0 | 0 | 0 | 2 | 0 | 0 | 0 | 8 | 7 | 2 | 6 | 0 | 0 | Izumo1r |
| Izumo4 | 850 | 1067 | 1061 | 1189 | 1217 | 1224 | 1142 | 811 | 971 | 1133 | 1159 | 942 | 1082 | 1102 | 858 | 893 | 1179 | 1199 | 1196 | 1024 | 1139 | 1138 | 849 | 1142 | 1240 | 986 | 1078 | 1000 | Izumo4 |
| Jade1 | 1810 | 1752 | 2087 | 1793 | 2117 | 2081 | 2025 | 2939 | 2918 | 2825 | 2821 | 3587 | 2882 | 3748 | 1746 | 1606 | 1978 | 1724 | 2208 | 2034 | 2061 | 2374 | 2693 | 3099 | 3078 | 3321 | 3484 | 3488 | Jade1 |
| Jade2 | 241 | 197 | 293 | 245 | 290 | 246 | 200 | 220 | 402 | 287 | 382 | 244 | 260 | 199 | 185 | 178 | 264 | 235 | 263 | 212 | 215 | 204 | 370 | 288 | 233 | 238 | 172 | 203 | Jade2 |
| Jade3 | 2330 | 1867 | 1967 | 2257 | 2644 | 2469 | 2626 | 2836 | 2315 | 2477 | 2283 | 2633 | 2430 | 2663 | 2362 | 2048 | 2256 | 2122 | 2430 | 2439 | 2335 | 2854 | 2150 | 2453 | 2759 | 2399 | 2398 | 2593 | Jade3 |
| Jag1 | 3223 | 3445 | 4730 | 4204 | 3800 | 3454 | 2883 | 3556 | 3300 | 3916 | 3722 | 4246 | 5125 | 4224 | 3200 | 3958 | 4109 | 4031 | 3565 | 3140 | 3363 | 3850 | 3698 | 3836 | 3542 | 3587 | 4788 | 4679 | Jag1 |
| Jag2 | 3052 | 3798 | 4350 | 4606 | 4453 | 3947 | 3045 | 2334 | 2919 | 3244 | 3065 | 2342 | 2647 | 1895 | 2682 | 3810 | 4390 | 4152 | 4390 | 3385 | 3311 | 3146 | 2754 | 3027 | 2882 | 2346 | 2017 | 1804 | Jag2 |
| Jagn1 | 1564 | 1664 | 1655 | 1418 | 1546 | 1409 | 1535 | 1864 | 1764 | 1517 | 1507 | 1670 | 1548 | 1669 | 1691 | 1453 | 1627 | 1392 | 1499 | 1619 | 1453 | 1686 | 1604 | 1570 | 1546 | 1845 | 1764 | 1716 | Jagn1 |
| Jak1 | 12115 | 10811 | 9456 | 9543 | 10179 | 9725 | 8993 | 8877 | 9095 | 8335 | 9433 | 6698 | 7743 | 6537 | 11737 | 12016 | 9371 | 10289 | 8701 | 8867 | 9408 | 8494 | 10153 | 7158 | 7456 | 6518 | 7161 | 6874 | Jak1 |
| Jak2 | 7841 | 6151 | 6141 | 6553 | 8040 | 8217 | 9475 | 7984 | 6716 | 6366 | 5722 | 5787 | 5159 | 6311 | 7920 | 6442 | 6449 | 6984 | 7838 | 8493 | 8614 | 7807 | 6436 | 6072 | 6920 | 4889 | 5406 | 5629 | Jak2 |
| Jak3 | 278 | 405 | 466 | 326 | 408 | 265 | 246 | 223 | 328 | 357 | 411 | 326 | 354 | 270 | 298 | 386 | 309 | 359 | 317 | 277 | 284 | 398 | 357 | 360 | 347 | 372 | 335 | 263 | Jak3 |
| Jakmip1 | 51 | 74 | 61 | 118 | 113 | 68 | 105 | 72 | 60 | 113 | 74 | 41 | 40 | 50 | 86 | 51 | 92 | 94 | 122 | 86 | 131 | 92 | 54 | 74 | 113 | 41 | 35 | 42 | Jakmip1 |
| Jakmip2 | 4392 | 5443 | 3886 | 5241 | 6198 | 6017 | 6220 | 6653 | 5281 | 5047 | 4387 | 5703 | 5044 | 5586 | 5851 | 4358 | 5055 | 5051 | 6007 | 7318 | 6324 | 5556 | 4944 | 4799 | 5967 | 4274 | 5407 | 4917 | Jakmip2 |
| Jakmip3 | 5 | 2 | 2 | 13 | 35 | 2 | 6 | 6 | 6 | 19 | 4 | 4 | 15 | 3 | 1 | 5 | 2 | 12 | 3 | 1 | 11 | 35 | 12 | 4 | 13 | 13 | 3 | 6 | Jakmip3 |
| Jam2 | 5973 | 6369 | 6797 | 5273 | 5934 | 6105 | 6391 | 7477 | 5116 | 5392 | 4601 | 8429 | 8108 | 8496 | 7482 | 5828 | 6594 | 5209 | 6053 | 6933 | 6044 | 6577 | 5340 | 5963 | 5985 | 6361 | 9687 | 9320 | Jam2 |
| Jam3 | 7835 | 8631 | 8240 | 7698 | 6384 | 6938 | 7679 | 6260 | 5500 | 5437 | 5927 | 7424 | 7160 | 7397 | 8389 | 7957 | 8017 | 7065 | 7003 | 8164 | 6861 | 5491 | 5730 | 6056 | 5807 | 7933 | 8220 | 7987 | Jam3 |
| Jaml | 1 | 1 | 0 | 2 | 2 | 0 | 2 | 4 | 12 | 15 | 12 | 8 | 14 | 5 | 1 | 3 | 2 | 3 | 5 | 2 | 1 | 15 | 10 | 3 | 6 | 2 | 8 | 13 | Jaml |
| Jarid2 | 12843 | 12929 | 14771 | 14526 | 12297 | 12497 | 12569 | 10179 | 8960 | 10986 | 10515 | 11535 | 11834 | 11663 | 12591 | 12723 | 14857 | 13201 | 12432 | 11168 | 12088 | 11334 | 8922 | 10569 | 11035 | 10173 | 11537 | 11345 | Jarid2 |
| Jazf1 | 446 | 334 | 387 | 421 | 703 | 537 | 536 | 1012 | 1093 | 1071 | 1118 | 782 | 697 | 808 | 431 | 444 | 413 | 492 | 655 | 549 | 593 | 1440 | 991 | 891 | 1205 | 618 | 638 | 726 | Jazf1 |
| Jcad | 4038 | 3960 | 3761 | 3083 | 3613 | 3683 | 2894 | 2628 | 2950 | 2759 | 3865 | 2758 | 3337 | 2624 | 3763 | 4629 | 3013 | 3739 | 3120 | 3349 | 3205 | 3292 | 3921 | 2530 | 3262 | 2090 | 2914 | 2725 | Jcad |
| Jchain | 21 | 24 | 42 | 30 | 45 | 49 | 54 | 29 | 33 | 54 | 52 | 54 | 40 | 57 | 40 | 19 | 75 | 63 | 29 | 30 | 41 | 57 | 16 | 56 | 42 | 53 | 34 | 44 | Jchain |
| Jdp2 | 290 | 371 | 424 | 357 | 384 | 366 | 210 | 318 | 503 | 539 | 730 | 332 | 422 | 384 | 324 | 408 | 330 | 371 | 318 | 279 | 234 | 475 | 555 | 496 | 431 | 398 | 368 | 360 | Jdp2 |
| Jhy | 36 | 43 | 54 | 42 | 72 | 56 | 39 | 110 | 92 | 95 | 83 | 96 | 174 | 101 | 63 | 63 | 59 | 26 | 44 | 50 | 46 | 109 | 102 | 48 | 92 | 51 | 108 | 107 | Jhy |
| Jkamp | 3846 | 3558 | 3581 | 4165 | 4058 | 4024 | 4552 | 4412 | 3720 | 3789 | 3532 | 3746 | 3895 | 4061 | 4243 | 3878 | 3961 | 4175 | 4087 | 4687 | 4370 | 4134 | 3816 | 3755 | 3889 | 3803 | 4054 | 4133 | Jkamp |
| Jmjd1c | 7770 | 6531 | 1566 | 4255 | 5131 | 6898 | 6122 | 7746 | 7448 | 4816 | 5200 | 4845 | 5049 | 4891 | 7528 | 6382 | 2792 | 4274 | 4630 | 7718 | 6768 | 3664 | 6717 | 2881 | 5262 | 963 | 5606 | 3814 | Jmjd1c |
| Jmjd4 | 4131 | 4439 | 2542 | 3380 | 3426 | 3763 | 3755 | 2963 | 2815 | 2473 | 2513 | 2767 | 2539 | 2877 | 4023 | 3724 | 2896 | 3461 | 3575 | 4170 | 3867 | 2532 | 2808 | 2448 | 2583 | 2143 | 2898 | 2651 | Jmjd4 |
| Jmjd6 | 3512 | 4180 | 3839 | 3917 | 3752 | 3604 | 3482 | 3504 | 3259 | 3222 | 3354 | 3449 | 3265 | 3708 | 3402 | 3646 | 4129 | 3782 | 3847 | 3820 | 3611 | 3357 | 3480 | 3718 | 3587 | 3669 | 3545 | 3966 | Jmjd6 |
| Jmjd7 | 51 | 69 | 49 | 69 | 93 | 97 | 82 | 79 | 98 | 92 | 105 | 98 | 98 | 70 | 49 | 59 | 89 | 88 | 88 | 53 | 73 | 94 | 63 | 109 | 104 | 24 | 116 | 114 | Jmjd7 |
| Jmjd8 | 2081 | 2565 | 2362 | 2193 | 2153 | 2276 | 2095 | 3438 | 3620 | 4024 | 3396 | 4037 | 3597 | 3729 | 2174 | 2150 | 2308 | 2122 | 2397 | 2102 | 2131 | 3477 | 3656 | 3797 | 3670 | 4518 | 3856 | 3728 | Jmjd8 |
| Jmy | 4151 | 3499 | 3530 | 5035 | 5242 | 4946 | 5226 | 4325 | 4058 | 4619 | 4882 | 4031 | 4020 | 4377 | 3829 | 3766 | 4057 | 5303 | 4684 | 4006 | 5385 | 4886 | 3866 | 4219 | 4902 | 3254 | 4051 | 4194 | Jmy |
| Josd1 | 4926 | 5216 | 5023 | 5467 | 4546 | 4651 | 4417 | 3300 | 3210 | 3132 | 3612 | 3038 | 3130 | 2922 | 4833 | 5371 | 5162 | 5534 | 5020 | 4114 | 5110 | 3467 | 3432 | 3416 | 3355 | 3327 | 3024 | 3114 | Josd1 |
| Josd2 | 1459 | 1918 | 2106 | 1909 | 1640 | 1639 | 1635 | 1581 | 1979 | 2185 | 2263 | 1989 | 2011 | 1909 | 1522 | 1549 | 2123 | 1819 | 1753 | 1457 | 1654 | 1795 | 2001 | 2266 | 1967 | 2257 | 1875 | 2091 | Josd2 |
| Jph1 | 709 | 588 | 661 | 702 | 641 | 689 | 692 | 424 | 506 | 511 | 621 | 632 | 486 | 691 | 654 | 587 | 657 | 682 | 693 | 636 | 621 | 312 | 502 | 663 | 556 | 655 | 471 | 546 | Jph1 |
| Jph2 | 88 | 58 | 91 | 67 | 67 | 35 | 32 | 98 | 183 | 100 | 698 | 131 | 128 | 79 | 69 | 93 | 76 | 72 | 62 | 58 | 31 | 159 | 395 | 85 | 97 | 120 | 106 | 120 | Jph2 |
| Jph3 | 661 | 1020 | 1209 | 1607 | 2502 | 1585 | 1552 | 1451 | 1775 | 2369 | 1841 | 1626 | 1541 | 1496 | 839 | 819 | 1552 | 1624 | 2097 | 1562 | 1713 | 2441 | 1441 | 1683 | 2350 | 1271 | 1302 | 1196 | Jph3 |
| Jph4 | 2635 | 3314 | 3495 | 5854 | 8567 | 6169 | 7515 | 5495 | 6579 | 8259 | 6515 | 5258 | 4069 | 5512 | 2796 | 2492 | 5158 | 7159 | 8336 | 6039 | 7944 | 9633 | 5525 | 7755 | 9560 | 4192 | 3873 | 4125 | Jph4 |
| Jpt1 | 23930 | 29491 | 27598 | 30147 | 28937 | 28455 | 32465 | 24200 | 24281 | 24526 | 21137 | 27065 | 22424 | 28248 | 24902 | 23672 | 29595 | 29597 | 32466 | 32782 | 31705 | 22749 | 22951 | 27820 | 29110 | 29003 | 26893 | 27670 | Jpt1 |
| Jpt2 | 16236 | 15330 | 16793 | 10765 | 9299 | 11005 | 10854 | 13652 | 12680 | 11155 | 11563 | 16865 | 15856 | 17784 | 14584 | 14076 | 14161 | 9832 | 10910 | 11467 | 10575 | 10684 | 12411 | 13861 | 12572 | 20395 | 18706 | 20216 | Jpt2 |
| Jrk | 1173 | 1303 | 1246 | 1091 | 1075 | 1092 | 1132 | 1158 | 1333 | 1176 | 1285 | 1334 | 1421 | 1340 | 1118 | 1069 | 1225 | 1089 | 1161 | 1201 | 1157 | 1074 | 1188 | 1409 | 1250 | 1520 | 1305 | 1404 | Jrk |
| Jrkl | 2384 | 1913 | 2266 | 2379 | 1864 | 2074 | 2333 | 2221 | 1731 | 1454 | 1586 | 1694 | 1686 | 1811 | 2683 | 2470 | 2037 | 2304 | 1991 | 2489 | 2274 | 1887 | 1893 | 1752 | 1670 | 1794 | 1690 | 1787 | Jrkl |
| Jsrp1 | 12 | 21 | 23 | 4 | 11 | 9 | 11 | 20 | 26 | 8 | 82 | 22 | 25 | 20 | 11 | 20 | 15 | 20 | 12 | 15 | 6 | 35 | 52 | 11 | 12 | 15 | 8 | 12 | Jsrp1 |
| Jtb | 1644 | 1445 | 1606 | 1603 | 1549 | 1704 | 1829 | 1951 | 1684 | 1767 | 1825 | 2004 | 1954 | 2132 | 1680 | 1537 | 1577 | 1651 | 1689 | 1991 | 1546 | 1830 | 1886 | 2008 | 1896 | 2242 | 2076 | 2019 | Jtb |
| Jun | 3440 | 4084 | 4725 | 5327 | 4345 | 3639 | 4102 | 3406 | 3965 | 4495 | 4773 | 4881 | 3920 | 4973 | 3160 | 3417 | 5120 | 4913 | 4786 | 3584 | 4318 | 3056 | 3732 | 5577 | 4540 | 6328 | 3871 | 4403 | Jun |
| Junb | 248 | 358 | 313 | 390 | 242 | 242 | 223 | 127 | 161 | 150 | 162 | 105 | 120 | 92 | 240 | 312 | 315 | 329 | 280 | 255 | 226 | 125 | 192 | 134 | 125 | 137 | 94 | 75 | Junb |
| Jund | 291 | 292 | 323 | 325 | 254 | 289 | 174 | 280 | 209 | 247 | 226 | 306 | 240 | 193 | 302 | 229 | 294 | 275 | 310 | 254 | 190 | 349 | 208 | 265 | 239 | 290 | 261 | 215 | Jund |
| Jup | 3405 | 3946 | 3409 | 3421 | 4043 | 3351 | 2434 | 2977 | 4127 | 4024 | 3866 | 2956 | 3402 | 2464 | 3055 | 4433 | 3669 | 4048 | 3414 | 2515 | 2998 | 4137 | 4469 | 3790 | 3468 | 3596 | 2517 | 2711 | Jup |
| Kalrn | 8750 | 9701 | 7613 | 12147 | 14810 | 16419 | 16040 | 12314 | 12000 | 13553 | 13320 | 12854 | 12816 | 13779 | 9970 | 9583 | 9931 | 12192 | 13024 | 13064 | 17686 | 14763 | 10799 | 11889 | 16176 | 7550 | 13520 | 11970 | Kalrn |
| Kank1 | 2494 | 2357 | 2500 | 1513 | 1694 | 1856 | 1133 | 1392 | 2120 | 1749 | 2554 | 1371 | 1931 | 1272 | 2090 | 2817 | 1808 | 1708 | 1580 | 1428 | 1228 | 1449 | 2397 | 1560 | 1215 | 1303 | 1663 | 1641 | Kank1 |
| Kank2 | 2491 | 2475 | 3554 | 2340 | 2013 | 2040 | 1657 | 2424 | 2909 | 2891 | 3896 | 2714 | 3415 | 2672 | 2082 | 2813 | 2664 | 2384 | 2028 | 1516 | 1626 | 2406 | 3432 | 3097 | 2322 | 3313 | 2879 | 2828 | Kank2 |
| Kank3 | 1521 | 1666 | 1768 | 1770 | 1670 | 1578 | 1409 | 942 | 1093 | 1242 | 1250 | 1185 | 1275 | 1061 | 1375 | 1746 | 1757 | 1906 | 1634 | 1406 | 1443 | 1287 | 1342 | 1177 | 1318 | 1384 | 1112 | 1129 | Kank3 |
| Kank4 | 1576 | 1321 | 1343 | 930 | 598 | 866 | 415 | 1100 | 1278 | 1379 | 1625 | 821 | 1313 | 807 | 1232 | 1926 | 948 | 900 | 533 | 387 | 440 | 973 | 1636 | 1263 | 820 | 1495 | 934 | 1002 | Kank4 |
| Kansl1 | 6879 | 6725 | 7194 | 7164 | 8265 | 7315 | 8208 | 10052 | 10256 | 10470 | 10251 | 11353 | 10261 | 11201 | 6836 | 6160 | 7234 | 6747 | 8344 | 7197 | 7843 | 11028 | 9359 | 10770 | 11096 | 9858 | 10848 | 11178 | Kansl1 |
| Kansl1l | 1514 | 1274 | 1386 | 1476 | 1641 | 1503 | 1706 | 3171 | 2944 | 2723 | 2737 | 2333 | 2523 | 2444 | 1621 | 1331 | 1710 | 1630 | 1624 | 1553 | 1698 | 3238 | 2987 | 2343 | 2576 | 2287 | 2269 | 2254 | Kansl1l |
| Kansl2 | 4310 | 4923 | 5171 | 5143 | 4812 | 4756 | 4902 | 6063 | 6063 | 6403 | 6264 | 7247 | 6481 | 7360 | 4458 | 4388 | 5056 | 4889 | 5202 | 4750 | 4895 | 6005 | 5902 | 6850 | 6979 | 7540 | 7166 | 7595 | Kansl2 |
| Kansl3 | 7552 | 8190 | 7611 | 8126 | 7625 | 7900 | 7685 | 6274 | 6683 | 6612 | 6681 | 7229 | 6733 | 6827 | 7135 | 7741 | 8170 | 7828 | 7974 | 7352 | 7635 | 7285 | 6352 | 6606 | 6803 | 6650 | 6835 | 6808 | Kansl3 |
| Kantr | 1826 | 1476 | 2489 | 2843 | 2461 | 2308 | 2678 | 1475 | 1343 | 1204 | 1431 | 1434 | 1580 | 1526 | 1953 | 1821 | 2486 | 2650 | 2809 | 2082 | 2856 | 2060 | 1287 | 1649 | 1588 | 1589 | 1446 | 1539 | Kantr |

| | | | | | | | | | | | | | | | | | | | | | | | | | | | | | |
|---|---|---|---|---|---|---|---|---|---|---|---|---|---|---|---|---|---|---|---|---|---|---|---|---|---|---|---|---|---|
| Kars | 14589 | 16330 | 15397 | 13560 | 11729 | 12646 | 12275 | 9789 | 10499 | 9853 | 9660 | 11337 | 10680 | 11573 | 13979 | 15182 | 14296 | 13288 | 13503 | 13920 | 12649 | 8589 | 10172 | 10776 | 9991 | 14001 | 12149 | 12067 | Kars |
| Kash5 | 63 | 70 | 122 | 85 | 51 | 62 | 48 | 67 | 95 | 64 | 105 | 97 | 78 | 98 | 71 | 80 | 131 | 45 | 72 | 59 | 48 | 85 | 85 | 99 | 86 | 110 | 110 | 121 | Kash5 |
| Kat14 | 1757 | 1802 | 2237 | 1701 | 1726 | 1682 | 1642 | 2499 | 2502 | 2619 | 2627 | 3222 | 2880 | 3158 | 1617 | 1795 | 1983 | 1766 | 1795 | 1409 | 1642 | 2446 | 2410 | 2968 | 2663 | 3346 | 2984 | 3297 | Kat14 |
| Kat2a | 2124 | 2771 | 3358 | 3207 | 3307 | 2844 | 2684 | 3491 | 3968 | 4546 | 4256 | 4845 | 4223 | 4251 | 2230 | 2545 | 3486 | 3182 | 3437 | 2489 | 2865 | 4182 | 3941 | 4551 | 4247 | 5618 | 4014 | 4322 | Kat2a |
| Kat2b | 1338 | 1040 | 1219 | 1298 | 1794 | 1670 | 1643 | 2859 | 2469 | 2342 | 2183 | 2586 | 2174 | 2504 | 1369 | 1005 | 1203 | 1210 | 1785 | 1576 | 1629 | 2724 | 2255 | 2373 | 2722 | 2152 | 2335 | 2387 | Kat2b |
| Kat5 | 1776 | 1937 | 2103 | 1987 | 2165 | 2176 | 2143 | 3896 | 4182 | 4119 | 3690 | 4524 | 4149 | 4336 | 1960 | 1791 | 2160 | 2040 | 2378 | 2269 | 2103 | 3917 | 3903 | 4287 | 4333 | 4800 | 4558 | 4138 | Kat5 |
| Kat6a | 15331 | 15247 | 11391 | 17489 | 17763 | 17972 | 18418 | 17531 | 16576 | 18287 | 17330 | 19683 | 18185 | 20665 | 14781 | 14094 | 14302 | 17792 | 16892 | 15146 | 19305 | 17179 | 14934 | 16263 | 20460 | 10778 | 19253 | 18830 | Kat6a |
| Kat6b | 3788 | 3726 | 2977 | 4657 | 5046 | 4988 | 5740 | 9979 | 11493 | 12047 | 12267 | 12166 | 11687 | 13206 | 3533 | 3214 | 3905 | 4784 | 5365 | 4932 | 5484 | 9613 | 9916 | 12272 | 13316 | 7868 | 11436 | 11331 | Kat6b |
| Kat7 | 4954 | 4434 | 4810 | 4018 | 4392 | 4304 | 4805 | 5816 | 5779 | 5444 | 5193 | 6244 | 6018 | 6326 | 4834 | 3916 | 4755 | 4006 | 4499 | 4436 | 4629 | 5872 | 5538 | 5990 | 5749 | 6841 | 6185 | 6556 | Kat7 |
| Kat8 | 1640 | 2014 | 1886 | 1864 | 1783 | 1666 | 1786 | 2067 | 2108 | 2062 | 1895 | 2120 | 1886 | 2098 | 1820 | 1727 | 1990 | 1651 | 1944 | 1895 | 1724 | 1953 | 1996 | 2286 | 2127 | 2383 | 2206 | 2194 | Kat8 |
| Katna1 | 0 | 3 | 2 | 0 | 2 | 1 | 2 | 2 | 0 | 3 | 1 | 0 | 0 | 1 | 4 | 2 | 6 | 3 | 1 | 1 | 1 | 0 | 2 | 1 | 8 | 3 | 7 | 0 | Katna1 |
| Katnal1 | 1602 | 1513 | 1635 | 2035 | 2029 | 1810 | 2017 | 3232 | 3161 | 3812 | 3183 | 3874 | 3459 | 4158 | 1597 | 1698 | 2042 | 1881 | 1886 | 1686 | 2014 | 4072 | 2895 | 3936 | 4146 | 3266 | 3840 | 3953 | Katnal1 |
| Katnal2 | 880 | 1018 | 1135 | 1054 | 945 | 818 | 984 | 928 | 812 | 890 | 780 | 1287 | 1077 | 1245 | 984 | 893 | 1161 | 919 | 899 | 918 | 948 | 862 | 823 | 956 | 974 | 1321 | 1432 | 1272 | Katnal2 |
| Katnb1 | 3851 | 5326 | 4244 | 5055 | 4364 | 4339 | 4078 | 2031 | 2497 | 2618 | 2354 | 2834 | 2490 | 2463 | 3752 | 4345 | 4687 | 4555 | 5133 | 4132 | 4114 | 2491 | 2242 | 2624 | 2904 | 2904 | 2622 | 2439 | Katnb1 |
| Katnbl1 | 2996 | 1905 | 2769 | 2779 | 2708 | 2722 | 3387 | 3079 | 2274 | 2200 | 2562 | 2739 | 2692 | 3147 | 3022 | 2558 | 2545 | 2529 | 2612 | 3146 | 2979 | 2780 | 2539 | 2611 | 2682 | 2959 | 2798 | 3072 | Katnbl1 |
| Katnip | 4630 | 5188 | 3577 | 4624 | 3956 | 4125 | 4275 | 3181 | 3447 | 3090 | 2936 | 3688 | 3385 | 3264 | 4283 | 4887 | 3868 | 4449 | 4190 | 3962 | 4312 | 3038 | 3461 | 2993 | 3256 | 2619 | 3570 | 3222 | Katnip |
| Kazald1 | 110 | 114 | 90 | 102 | 93 | 107 | 75 | 131 | 213 | 116 | 232 | 30 | 100 | 37 | 141 | 104 | 103 | 121 | 99 | 72 | 55 | 170 | 197 | 105 | 114 | 55 | 34 | 32 | Kazald1 |
| Kazn | 126 | 154 | 88 | 536 | 218 | 331 | 102 | 110 | 105 | 223 | 334 | 226 | 92 | 330 | 390 | 132 | 246 | 92 | 436 | 133 | 204 | 363 | 174 | 257 | 118 | 121 | 181 | 103 | Kazn |
| Kbtbd11 | 2136 | 2476 | 2953 | 2992 | 3989 | 3520 | 3429 | 4522 | 4280 | 4979 | 4130 | 5048 | 5380 | 5122 | 2718 | 2358 | 3307 | 3223 | 3910 | 3150 | 3673 | 4518 | 3864 | 5000 | 5488 | 4026 | 4500 | 5005 | Kbtbd11 |
| Kbtbd12 | 2 | 1 | 0 | 1 | 1 | 12 | 4 | 2 | 14 | 14 | 19 | 8 | 11 | 6 | 3 | 1 | 0 | 0 | 1 | 3 | 5 | 3 | 10 | 7 | 7 | 12 | 2 | 4 | Kbtbd12 |
| Kbtbd13 | 32 | 22 | 30 | 21 | 25 | 28 | 21 | 44 | 67 | 40 | 78 | 36 | 59 | 29 | 38 | 39 | 19 | 32 | 16 | 19 | 26 | 53 | 60 | 46 | 53 | 61 | 34 | 49 | Kbtbd13 |
| Kbtbd2 | 3651 | 3197 | 3689 | 3727 | 3977 | 3663 | 3862 | 5145 | 4294 | 4079 | 4369 | 4757 | 4488 | 4840 | 3688 | 2996 | 3531 | 3815 | 4057 | 3790 | 3828 | 5000 | 4112 | 4762 | 5035 | 5145 | 4767 | 4968 | Kbtbd2 |
| Kbtbd3 | 744 | 557 | 553 | 563 | 557 | 568 | 689 | 656 | 504 | 422 | 533 | 534 | 551 | 524 | 847 | 704 | 546 | 559 | 507 | 782 | 684 | 478 | 557 | 494 | 511 | 489 | 635 | 553 | Kbtbd3 |
| Kbtbd6 | 415 | 373 | 518 | 459 | 389 | 438 | 440 | 315 | 299 | 335 | 230 | 430 | 330 | 429 | 447 | 399 | 460 | 474 | 481 | 384 | 527 | 254 | 265 | 412 | 342 | 552 | 399 | 479 | Kbtbd6 |
| Kbtbd7 | 3216 | 2352 | 3606 | 3865 | 4187 | 3946 | 4594 | 4085 | 3488 | 3660 | 3682 | 3712 | 3687 | 4015 | 3434 | 3027 | 3854 | 4006 | 4359 | 3536 | 4479 | 4375 | 3366 | 3900 | 4136 | 3975 | 3498 | 3796 | Kbtbd7 |
| Kbtbd8 | 1694 | 1665 | 1789 | 1888 | 1472 | 1565 | 1605 | 1542 | 1093 | 937 | 932 | 1340 | 1108 | 1323 | 2036 | 1854 | 1703 | 1654 | 1564 | 2019 | 1522 | 1110 | 1014 | 1102 | 1146 | 1401 | 1396 | 1333 | Kbtbd8 |
| Kcmf1 | 3610 | 3429 | 3925 | 3732 | 3885 | 3750 | 4289 | 4940 | 4538 | 4558 | 4621 | 4986 | 4557 | 5318 | 3683 | 3269 | 3710 | 3712 | 4158 | 3865 | 4176 | 4669 | 4539 | 4725 | 4560 | 5249 | 4853 | 5542 | Kcmf1 |
| Kcna1 | 520 | 417 | 392 | 471 | 628 | 526 | 472 | 247 | 298 | 257 | 359 | 149 | 273 | 160 | 516 | 495 | 391 | 486 | 498 | 505 | 513 | 324 | 282 | 151 | 307 | 29 | 173 | 192 | Kcna1 |
| Kcna10 | 0 | 1 | 0 | 0 | 1 | 0 | 0 | 0 | 0 | 4 | 0 | 0 | 0 | 0 | 1 | 1 | 0 | 0 | 2 | 1 | 1 | 0 | 0 | 0 | 0 | 0 | 0 | 0 | Kcna10 |
| Kcna2 | 1020 | 1167 | 709 | 1472 | 2555 | 2045 | 1895 | 1653 | 1446 | 1571 | 1625 | 874 | 1236 | 994 | 1477 | 1026 | 1079 | 1478 | 1697 | 1935 | 2003 | 1711 | 1396 | 847 | 1588 | 441 | 1026 | 781 | Kcna2 |
| Kcna3 | 487 | 578 | 91 | 488 | 568 | 765 | 688 | 553 | 565 | 383 | 394 | 488 | 302 | 483 | 440 | 359 | 260 | 447 | 535 | 848 | 799 | 311 | 392 | 308 | 614 | 40 | 467 | 273 | Kcna3 |
| Kcna4 | 337 | 336 | 328 | 390 | 442 | 373 | 260 | 354 | 355 | 379 | 359 | 279 | 384 | 329 | 364 | 308 | 353 | 386 | 385 | 436 | 376 | 446 | 384 | 245 | 335 | 102 | 309 | 368 | Kcna4 |
| Kcna5 | 174 | 129 | 185 | 133 | 186 | 122 | 116 | 245 | 258 | 245 | 365 | 196 | 198 | 204 | 149 | 121 | 137 | 107 | 147 | 96 | 127 | 211 | 328 | 286 | 175 | 259 | 207 | 231 | Kcna5 |
| Kcna6 | 2759 | 2910 | 3195 | 2971 | 4268 | 3578 | 3096 | 1833 | 2085 | 2269 | 2480 | 1372 | 1982 | 1422 | 2873 | 2998 | 3259 | 3693 | 3563 | 2555 | 3545 | 2914 | 1721 | 1476 | 2140 | 1436 | 1538 | 1380 | Kcna6 |
| Kcna7 | 5 | 3 | 5 | 9 | 4 | 6 | 1 | 0 | 0 | 0 | 0 | 0 | 1 | 0 | 3 | 5 | 0 | 4 | 1 | 4 | 0 | 0 | 0 | 0 | 4 | 0 | 2 | 2 | Kcna7 |
| Kcnab1 | 260 | 250 | 288 | 143 | 314 | 261 | 143 | 368 | 411 | 449 | 497 | 341 | 489 | 349 | 392 | 419 | 251 | 254 | 164 | 196 | 148 | 478 | 562 | 275 | 362 | 235 | 558 | 479 | Kcnab1 |
| Kcnab2 | 2440 | 3206 | 3535 | 3906 | 3715 | 3356 | 2973 | 1272 | 1563 | 1598 | 1866 | 1578 | 1755 | 1325 | 2293 | 2602 | 3756 | 3688 | 3991 | 2859 | 3271 | 1980 | 1479 | 1481 | 1941 | 1603 | 1475 | 1518 | Kcnab2 |
| Kcnab3 | 1485 | 1649 | 1894 | 2059 | 1777 | 1584 | 1942 | 372 | 384 | 469 | 419 | 504 | 403 | 521 | 1352 | 1389 | 2052 | 2002 | 1894 | 1500 | 1865 | 531 | 334 | 517 | 544 | 520 | 423 | 458 | Kcnab3 |
| Kcnb1 | 1727 | 1748 | 1557 | 2349 | 3023 | 2545 | 2704 | 1551 | 1657 | 1805 | 1826 | 1346 | 1419 | 1292 | 1644 | 1650 | 2150 | 2540 | 2836 | 2317 | 2873 | 2100 | 1436 | 1664 | 1917 | 1070 | 1293 | 1254 | Kcnb1 |
| Kcnb2 | 383 | 275 | 192 | 360 | 708 | 692 | 616 | 1791 | 1681 | 1211 | 1281 | 876 | 899 | 994 | 514 | 450 | 279 | 384 | 408 | 705 | 625 | 1462 | 1495 | 997 | 1438 | 528 | 940 | 851 | Kcnb2 |
| Kcnc1 | 1416 | 2160 | 2067 | 3880 | 5375 | 4213 | 4361 | 1983 | 2178 | 3117 | 2568 | 2347 | 1858 | 2183 | 1931 | 1772 | 3668 | 4504 | 4923 | 3371 | 4694 | 3599 | 1880 | 2491 | 3586 | 1214 | 1749 | 1765 | Kcnc1 |
| Kcnc2 | 261 | 179 | 267 | 337 | 539 | 380 | 468 | 455 | 425 | 517 | 478 | 321 | 281 | 354 | 299 | 220 | 345 | 363 | 414 | 384 | 446 | 622 | 441 | 373 | 464 | 221 | 367 | 343 | Kcnc2 |
| Kcnc3 | 5728 | 8424 | 8808 | 11660 | 9442 | 7996 | 7519 | 1305 | 1328 | 2288 | 1686 | 1718 | 1609 | 1657 | 5683 | 6797 | 9853 | 10837 | 9664 | 6543 | 8444 | 2440 | 1158 | 1937 | 2039 | 1375 | 1578 | 1686 | Kcnc3 |
| Kcnc4 | 191 | 248 | 233 | 358 | 597 | 410 | 327 | 178 | 231 | 320 | 243 | 113 | 125 | 88 | 171 | 261 | 238 | 447 | 457 | 352 | 454 | 386 | 221 | 182 | 287 | 75 | 59 | 71 | Kcnc4 |
| Kcnd1 | 736 | 962 | 858 | 1049 | 935 | 824 | 884 | 480 | 552 | 629 | 570 | 553 | 612 | 536 | 783 | 829 | 969 | 1076 | 917 | 710 | 819 | 775 | 499 | 525 | 584 | 497 | 443 | 515 | Kcnd1 |
| Kcnd2 | 647 | 568 | 570 | 1047 | 1885 | 1555 | 1577 | 1573 | 1475 | 1573 | 1550 | 1249 | 1098 | 1343 | 829 | 539 | 856 | 1149 | 1404 | 1477 | 1625 | 2207 | 1524 | 1380 | 1858 | 1122 | 1109 | 1314 | Kcnd2 |
| Kcnd3 | 1389 | 1477 | 1230 | 2496 | 3854 | 3742 | 3629 | 4653 | 4387 | 4792 | 3986 | 3814 | 3957 | 4067 | 1970 | 1502 | 2128 | 2836 | 3399 | 3411 | 3969 | 5972 | 3658 | 3860 | 5704 | 1915 | 3267 | 3048 | Kcnd3 |
| Kcne1 | 30 | 20 | 24 | 6 | 2 | 4 | 10 | 7 | 7 | 4 | 5 | 7 | 7 | 0 | 16 | 19 | 21 | 12 | 5 | 6 | 8 | 7 | 5 | 1 | 2 | 2 | 5 | 6 | Kcne1 |
| Kcne1l | 316 | 421 | 372 | 283 | 398 | 363 | 324 | 458 | 598 | 536 | 830 | 349 | 496 | 448 | 380 | 386 | 324 | 296 | 335 | 393 | 373 | 439 | 805 | 407 | 486 | 416 | 478 | 383 | Kcne1l |
| Kcne2 | 142 | 211 | 55 | 202 | 554 | 333 | 267 | 591 | 849 | 810 | 565 | 605 | 824 | 244 | 165 | 269 | 87 | 224 | 230 | 305 | 332 | 936 | 1524 | 395 | 509 | 208 | 191 | 259 | Kcne2 |
| Kcne3 | 264 | 117 | 94 | 68 | 45 | 102 | 63 | 150 | 172 | 145 | 155 | 45 | 81 | 65 | 144 | 171 | 87 | 80 | 72 | 89 | 58 | 97 | 181 | 125 | 74 | 103 | 96 | 88 | Kcne3 |
| Kcne4 | 152 | 86 | 104 | 153 | 59 | 164 | 73 | 259 | 349 | 240 | 542 | 153 | 163 | 152 | 109 | 147 | 60 | 189 | 97 | 66 | 55 | 221 | 585 | 268 | 217 | 154 | 102 | 135 | Kcne4 |
| Kcnf1 | 290 | 213 | 252 | 390 | 608 | 436 | 546 | 447 | 494 | 539 | 418 | 398 | 274 | 416 | 287 | 188 | 375 | 434 | 514 | 415 | 470 | 536 | 429 | 593 | 494 | 491 | 378 | 390 | Kcnf1 |
| Kcng1 | 411 | 601 | 626 | 795 | 791 | 744 | 839 | 325 | 499 | 439 | 339 | 308 | 149 | 331 | 392 | 484 | 823 | 788 | 966 | 713 | 979 | 317 | 278 | 507 | 673 | 284 | 206 | 211 | Kcng1 |
| Kcng2 | 15 | 18 | 9 | 13 | 30 | 25 | 22 | 35 | 44 | 43 | 36 | 20 | 18 | 26 | 8 | 21 | 13 | 24 | 4 | 15 | 12 | 32 | 55 | 27 | 27 | 10 | 29 | 27 | Kcng2 |
| Kcng3 | 38 | 32 | 0 | 1 | 17 | 25 | 11 | 6 | 11 | 10 | 4 | 12 | 8 | 5 | 25 | 40 | 2 | 4 | 10 | 35 | 27 | 5 | 14 | 2 | 12 | 3 | 2 | 6 | Kcng3 |
| Kcng4 | 3 | 8 | 6 | 5 | 6 | 2 | 2 | 13 | 7 | 18 | 12 | 9 | 11 | 13 | 5 | 5 | 2 | 1 | 5 | 10 | 7 | 13 | 20 | 14 | 2 | 12 | 6 | 13 | Kcng4 |
| Kcnh1 | 120 | 118 | 77 | 147 | 395 | 220 | 192 | 368 | 464 | 549 | 424 | 247 | 307 | 238 | 135 | 161 | 104 | 259 | 234 | 169 | 230 | 627 | 431 | 295 | 479 | 154 | 221 | 184 | Kcnh1 |
| Kcnh2 | 1716 | 2467 | 2714 | 2936 | 3034 | 2709 | 2321 | 1784 | 1605 | 2241 | 1859 | 2435 | 2056 | 1957 | 1954 | 2041 | 3079 | 2659 | 2942 | 2354 | 2356 | 2550 | 1686 | 2176 | 2254 | 2155 | 2363 | 2163 | Kcnh2 |
| Kcnh3 | 343 | 331 | 284 | 492 | 557 | 410 | 549 | 186 | 306 | 310 | 221 | 133 | 103 | 134 | 237 | 304 | 298 | 589 | 535 | 425 | 572 | 224 | 346 | 340 | 329 | 152 | 66 | 56 | Kcnh3 |
| Kcnh4 | 473 | 787 | 693 | 1081 | 1581 | 1251 | 1215 | 1155 | 1208 | 1661 | 1221 | 1411 | 1254 | 1468 | 582 | 633 | 1077 | 1141 | 1566 | 1146 | 1222 | 2169 | 1060 | 1649 | 1688 | 1158 | 1325 | 1390 | Kcnh4 |
| Kcnh5 | 104 | 86 | 22 | 51 | 98 | 121 | 93 | 125 | 176 | 105 | 98 | 53 | 53 | 64 | 88 | 100 | 44 | 52 | 72 | 153 | 145 | 53 | 128 | 37 | 113 | 36 | 88 | 27 | Kcnh5 |
| Kcnh6 | 72 | 95 | 49 | 67 | 185 | 55 | 57 | 160 | 281 | 196 | 295 | 46 | 175 | 47 | 92 | 133 | 88 | 102 | 83 | 66 | 46 | 264 | 135 | 98 | 177 | 59 | 52 | 33 | Kcnh6 |
| Kcnh7 | 864 | 730 | 311 | 1119 | 1909 | 2208 | 2230 | 3057 | 2910 | 2272 | 2183 | 1845 | 1601 | 2221 | 1170 | 779 | 621 | 1227 | 1424 | 2117 | 2431 | 2232 | 2207 | 1911 | 2774 | 1278 | 1682 | 1298 | Kcnh7 |
| Kcnh8 | 170 | 187 | 136 | 275 | 319 | 259 | 240 | 710 | 638 | 601 | 568 | 537 | 441 | 607 | 257 | 162 | 161 | 213 | 199 | 320 | 233 | 678 | 519 | 511 | 578 | 425 | 533 | 458 | Kcnh8 |
| Kcnip1 | 466 | 608 | 323 | 789 | 1397 | 1260 | 1316 | 1365 | 1327 | 1457 | 1258 | 770 | 963 | 944 | 854 | 527 | 590 | 1044 | 1057 | 1268 | 1329 | 1628 | 1353 | 1022 | 1422 | 724 | 682 | 888 | Kcnip1 |
| Kcnip2 | 258 | 334 | 294 | 495 | 616 | 351 | 430 | 238 | 348 | 298 | 264 | 175 | 147 | 212 | 289 | 294 | 404 | 427 | 542 | 462 | 457 | 328 | 315 | 257 | 300 | 243 | 152 | 166 | Kcnip2 |
| Kcnip3 | 484 | 538 | 584 | 723 | 888 | 806 | 864 | 819 | 868 | 994 | 801 | 1090 | 714 | 906 | 462 | 441 | 695 | 717 | 1005 | 851 | 910 | 1043 | 860 | 1130 | 1113 | 891 | 881 | 882 | Kcnip3 |
| Kcnip4 | 171 | 107 | 130 | 135 | 310 | 188 | 207 | 300 | 281 | 287 | 226 | 138 | 191 | 159 | 153 | 106 | 96 | 151 | 157 | 219 | 255 | 393 | 341 | 208 | 262 | 127 | 141 | 151 | Kcnip4 |
| Kcnj1 | 0 | 0 | 2 | 0 | 1 | 0 | 0 | 1 | 0 | 5 | 0 | 1 | 1 | 1 | 1 | 0 | 0 | 2 | 2 | 0 | 1 | 0 | 3 | 3 | 0 | 0 | 0 | 1 | Kcnj1 |
| Kcnj10 | 1418 | 1808 | 2059 | 2300 | 1999 | 1842 | 1969 | 712 | 674 | 818 | 707 | 665 | 623 | 732 | 1501 | 1466 | 2269 | 2149 | 1942 | 1494 | 1834 | 815 | 533 | 717 | 929 | 525 | 618 | 634 | Kcnj10 |
| Kcnj11 | 165 | 187 | 233 | 279 | 506 | 403 | 403 | 230 | 327 | 386 | 349 | 218 | 194 | 160 | 171 | 150 | 293 | 410 | 529 | 385 | 428 | 311 | 271 | 315 | 436 | 217 | 155 | 120 | Kcnj11 |
| Kcnj12 | 61 | 63 | 33 | 68 | 184 | 66 | 83 | 177 | 179 | 171 | 220 | 31 | 203 | 26 | 98 | 105 | 67 | 89 | 95 | 81 | 64 | 339 | 139 | 65 | 158 | 22 | 36 | 36 | Kcnj12 |
| Kcnj13 | 82 | 84 | 15 | 31 | 160 | 103 | 77 | 142 | 181 | 107 | 138 | 142 | 138 | 54 | 90 | 126 | 37 | 44 | 73 | 83 | 104 | 197 | 357 | 65 | 73 | 32 | 46 | 30 | Kcnj13 |
| Kcnj14 | 66 | 89 | 116 | 97 | 92 | 86 | 89 | 79 | 70 | 74 | 94 | 67 | 85 | 71 | 49 | 82 | 94 | 116 | 83 | 65 | 77 | 94 | 62 | 68 | 68 | 69 | 50 | 76 | Kcnj14 |
| Kcnj15 | 78 | 28 | 48 | 20 | 36 | 28 | 25 | 73 | 101 | 88 | 93 | 28 | 42 | 44 | 26 | 38 | 37 | 34 | 29 | 10 | 17 | 35 | 140 | 47 | 48 | 43 | 62 | 38 | Kcnj15 |
| Kcnj16 | 5 | 3 | 5 | 4 | 20 | 3 | 1 | 38 | 26 | 34 | 42 | 5 | 28 | 15 | 4 | 5 | 6 | 10 | 12 | 12 | 2 | 35 | 45 | 7 | 23 | 8 | 20 | 15 | Kcnj16 |
| Kcnj2 | 933 | 680 | 659 | 553 | 494 | 479 | 412 | 1005 | 1032 | 792 | 973 | 415 | 665 | 478 | 786 | 951 | 564 | 548 | 396 | 503 | 487 | 687 | 1055 | 681 | 540 | 728 | 420 | 527 | Kcnj2 |
| Kcnj3 | 277 | 371 | 375 | 658 | 1196 | 874 | 792 | 762 | 744 | 647 | 617 | 412 | 513 | 420 | 464 | 379 | 452 | 815 | 936 | 869 | 977 | 852 | 674 | 516 | 865 | 332 | 352 | 371 | Kcnj3 |

| | | | | | | | | | | | | | | | | | | | | | | | | | | | | | |
|---|---|---|---|---|---|---|---|---|---|---|---|---|---|---|---|---|---|---|---|---|---|---|---|---|---|---|---|---|---|
| Kcnj4 | 141 | 187 | 182 | 161 | 165 | 122 | 105 | 2 | 30 | 25 | 4 | 18 | 3 | 5 | 133 | 182 | 172 | 196 | 163 | 131 | 134 | 23 | 11 | 11 | 14 | 10 | 12 | 11 | Kcnj4 |
| Kcnj5 | 67 | 54 | 59 | 64 | 153 | 131 | 124 | 219 | 184 | 229 | 208 | 215 | 120 | 200 | 69 | 72 | 87 | 97 | 118 | 165 | 124 | 177 | 194 | 279 | 267 | 249 | 195 | 189 | Kcnj5 |
| Kcnj6 | 23 | 41 | 20 | 57 | 140 | 102 | 63 | 115 | 111 | 98 | 112 | 45 | 60 | 52 | 41 | 26 | 20 | 52 | 93 | 115 | 113 | 124 | 124 | 62 | 178 | 8 | 46 | 38 | Kcnj6 |
| Kcnj8 | 58 | 61 | 112 | 55 | 182 | 120 | 102 | 342 | 410 | 468 | 522 | 436 | 395 | 412 | 45 | 51 | 108 | 74 | 154 | 108 | 130 | 418 | 551 | 503 | 506 | 422 | 370 | 388 | Kcnj8 |
| Kcnj9 | 233 | 342 | 388 | 539 | 502 | 567 | 526 | 238 | 247 | 315 | 289 | 422 | 305 | 386 | 302 | 227 | 505 | 522 | 507 | 527 | 587 | 319 | 245 | 381 | 418 | 308 | 380 | 320 | Kcnj9 |
| Kcnk1 | 535 | 383 | 314 | 290 | 361 | 313 | 245 | 361 | 537 | 395 | 510 | 237 | 299 | 250 | 434 | 468 | 249 | 317 | 289 | 266 | 291 | 369 | 533 | 373 | 272 | 378 | 249 | 274 | Kcnk1 |
| Kcnk10 | 1722 | 1732 | 2205 | 2171 | 2410 | 2405 | 2501 | 3328 | 2588 | 2979 | 2448 | 4026 | 2799 | 4041 | 1934 | 1239 | 2363 | 1892 | 2646 | 2265 | 2668 | 3832 | 2262 | 3517 | 3973 | 3365 | 4102 | 4124 | Kcnk10 |
| Kcnk12 | 16 | 16 | 36 | 37 | 37 | 29 | 21 | 39 | 21 | 8 | 23 | 17 | 11 | 10 | 18 | 9 | 20 | 22 | 48 | 22 | 36 | 23 | 25 | 28 | 34 | 10 | 6 | 15 | Kcnk12 |
| Kcnk13 | 62 | 34 | 116 | 73 | 91 | 47 | 110 | 32 | 36 | 25 | 105 | 46 | 46 | 40 | 78 | 67 | 72 | 92 | 64 | 84 | 82 | 28 | 85 | 36 | 23 | 37 | 49 | 32 | Kcnk13 |
| Kcnk15 | 1 | 0 | 3 | 1 | 3 | 3 | 1 | 0 | 0 | 0 | 0 | 1 | 0 | 0 | 7 | 3 | 0 | 4 | 2 | 4 | 1 | 0 | 2 | 2 | 4 | 0 | 2 | 0 | Kcnk15 |
| Kcnk16 | 0 | 2 | 0 | 0 | 0 | 0 | 0 | 0 | 1 | 0 | 6 | 0 | 0 | 0 | 0 | 0 | 0 | 0 | 0 | 0 | 0 | 2 | 2 | 0 | 0 | 0 | 0 | 0 | Kcnk16 |
| Kcnk18 | 0 | 2 | 0 | 0 | 2 | 1 | 0 | 0 | 1 | 0 | 0 | 0 | 0 | 0 | 0 | 1 | 2 | 0 | 0 | 0 | 0 | 0 | 1 | 1 | 0 | 0 | 0 | 0 | Kcnk18 |
| Kcnk2 | 486 | 444 | 673 | 587 | 1055 | 872 | 789 | 1955 | 2066 | 2272 | 2011 | 1863 | 1860 | 1910 | 496 | 445 | 613 | 637 | 951 | 835 | 775 | 2186 | 2360 | 2309 | 2297 | 2005 | 1902 | 1761 | Kcnk2 |
| Kcnk3 | 53 | 100 | 59 | 144 | 388 | 176 | 161 | 180 | 223 | 246 | 270 | 96 | 144 | 89 | 116 | 79 | 89 | 212 | 251 | 201 | 225 | 304 | 214 | 118 | 251 | 73 | 71 | 69 | Kcnk3 |
| Kcnk4 | 42 | 50 | 77 | 76 | 107 | 83 | 65 | 22 | 22 | 34 | 36 | 12 | 11 | 24 | 62 | 54 | 71 | 98 | 83 | 71 | 79 | 33 | 20 | 23 | 26 | 16 | 15 | 12 | Kcnk4 |
| Kcnk5 | 1283 | 1698 | 1518 | 1398 | 809 | 1019 | 1002 | 474 | 508 | 550 | 441 | 760 | 798 | 740 | 1122 | 1445 | 1393 | 1148 | 1068 | 1112 | 877 | 349 | 491 | 619 | 471 | 1076 | 908 | 827 | Kcnk5 |
| Kcnk6 | 217 | 184 | 246 | 138 | 126 | 196 | 107 | 233 | 316 | 220 | 592 | 220 | 236 | 184 | 183 | 237 | 192 | 211 | 106 | 134 | 124 | 302 | 364 | 235 | 216 | 231 | 231 | 173 | Kcnk6 |
| Kcnk7 | 1 | 3 | 2 | 5 | 7 | 0 | 0 | 0 | 2 | 2 | 3 | 2 | 2 | 2 | 4 | 8 | 1 | 7 | 1 | 3 | 0 | 0 | 3 | 0 | 1 | 2 | 2 | 0 | Kcnk7 |
| Kcnk9 | 333 | 366 | 291 | 833 | 2249 | 1198 | 950 | 1784 | 1862 | 2248 | 2105 | 781 | 1621 | 744 | 600 | 511 | 571 | 1169 | 1026 | 1124 | 1193 | 2602 | 1889 | 829 | 1562 | 211 | 775 | 694 | Kcnk9 |
| Kcnma1 | 1203 | 1331 | 740 | 2240 | 3938 | 3362 | 3386 | 2978 | 2728 | 2800 | 3089 | 1527 | 1787 | 1541 | 1854 | 1615 | 1254 | 2648 | 2546 | 3324 | 3747 | 3562 | 2630 | 1554 | 2982 | 690 | 1355 | 1161 | Kcnma1 |
| Kcnmb1 | 7 | 11 | 27 | 5 | 7 | 11 | 9 | 27 | 16 | 13 | 15 | 13 | 34 | 4 | 7 | 13 | 12 | 3 | 7 | 15 | 9 | 15 | 16 | 9 | 6 | 18 | 23 | 12 | Kcnmb1 |
| Kcnmb2 | 242 | 228 | 168 | 166 | 367 | 203 | 187 | 635 | 556 | 502 | 563 | 254 | 553 | 226 | 405 | 381 | 204 | 201 | 161 | 242 | 194 | 734 | 469 | 278 | 442 | 176 | 265 | 221 | Kcnmb2 |
| Kcnmb3 | 2 | 2 | 6 | 0 | 4 | 0 | 2 | 0 | 1 | 1 | 3 | 0 | 0 | 0 | 4 | 2 | 0 | 0 | 0 | 2 | 8 | 2 | 0 | 1 | 3 | 2 | 0 | 0 | Kcnmb3 |
| Kcnmb4 | 549 | 669 | 585 | 655 | 814 | 606 | 593 | 1018 | 1046 | 1147 | 1168 | 1077 | 1131 | 923 | 633 | 678 | 621 | 680 | 595 | 493 | 585 | 1151 | 1043 | 1034 | 1133 | 1071 | 953 | 940 | Kcnmb4 |
| Kcnn1 | 704 | 1011 | 820 | 904 | 758 | 746 | 667 | 812 | 752 | 853 | 720 | 1150 | 940 | 874 | 719 | 741 | 976 | 880 | 921 | 804 | 671 | 897 | 695 | 1026 | 1077 | 883 | 1109 | 992 | Kcnn1 |
| Kcnn2 | 585 | 651 | 497 | 634 | 749 | 727 | 872 | 658 | 687 | 691 | 609 | 570 | 387 | 559 | 699 | 600 | 683 | 735 | 776 | 721 | 936 | 553 | 512 | 628 | 740 | 486 | 389 | 399 | Kcnn2 |
| Kcnn3 | 2820 | 2870 | 1639 | 3116 | 3649 | 3686 | 3884 | 1396 | 1367 | 1245 | 1498 | 786 | 971 | 800 | 3001 | 2881 | 2135 | 3548 | 2990 | 3437 | 4211 | 1394 | 1530 | 842 | 1314 | 451 | 705 | 571 | Kcnn3 |
| Kcnn4 | 14 | 22 | 20 | 13 | 16 | 12 | 10 | 22 | 58 | 27 | 33 | 36 | 64 | 33 | 15 | 24 | 10 | 9 | 20 | 15 | 8 | 22 | 36 | 15 | 18 | 46 | 52 | 107 | Kcnn4 |
| Kcnq1 | 68 | 55 | 72 | 53 | 79 | 44 | 38 | 55 | 73 | 69 | 48 | 68 | 70 | 48 | 47 | 65 | 34 | 46 | 64 | 40 | 55 | 72 | 71 | 60 | 76 | 73 | 88 | 64 | Kcnq1 |
| Kcnq2 | 4858 | 6179 | 4059 | 8184 | 13565 | 11163 | 10515 | 13814 | 13475 | 15532 | 13164 | 12200 | 11746 | 11729 | 6253 | 5114 | 7214 | 9384 | 11676 | 10157 | 11721 | 19041 | 11441 | 11466 | 18037 | 5919 | 11405 | 10325 | Kcnq2 |
| Kcnq3 | 1227 | 1330 | 756 | 1777 | 2943 | 2818 | 2977 | 3626 | 3681 | 3415 | 2823 | 2583 | 2690 | 2617 | 1688 | 1391 | 1137 | 1929 | 2392 | 2726 | 3455 | 3390 | 3178 | 2396 | 4081 | 1341 | 2418 | 1988 | Kcnq3 |
| Kcnq4 | 498 | 615 | 681 | 505 | 642 | 648 | 432 | 530 | 760 | 799 | 755 | 570 | 687 | 537 | 508 | 654 | 594 | 592 | 520 | 341 | 461 | 762 | 839 | 668 | 600 | 520 | 608 | 506 | Kcnq4 |
| Kcnq5 | 215 | 159 | 139 | 112 | 230 | 214 | 168 | 390 | 516 | 403 | 530 | 213 | 324 | 255 | 223 | 280 | 140 | 188 | 169 | 115 | 196 | 429 | 378 | 323 | 326 | 251 | 286 | 239 | Kcnq5 |
| Kcnrg | 261 | 249 | 180 | 296 | 347 | 335 | 360 | 227 | 167 | 175 | 155 | 164 | 142 | 161 | 301 | 189 | 191 | 329 | 305 | 370 | 424 | 224 | 166 | 161 | 223 | 133 | 145 | 146 | Kcnrg |
| Kcns1 | 60 | 84 | 81 | 64 | 56 | 117 | 29 | 48 | 101 | 101 | 153 | 37 | 76 | 37 | 51 | 41 | 65 | 80 | 51 | 26 | 33 | 109 | 96 | 66 | 78 | 50 | 53 | 26 | Kcns1 |
| Kcns2 | 60 | 17 | 14 | 31 | 115 | 47 | 54 | 125 | 124 | 192 | 140 | 43 | 26 | 26 | 36 | 43 | 27 | 30 | 84 | 65 | 71 | 149 | 113 | 105 | 151 | 64 | 24 | 38 | Kcns2 |
| Kcns3 | 39 | 24 | 33 | 29 | 48 | 28 | 25 | 37 | 73 | 53 | 105 | 20 | 45 | 37 | 44 | 49 | 34 | 23 | 23 | 49 | 34 | 74 | 73 | 44 | 32 | 35 | 21 | 17 | Kcns3 |
| Kcnt1 | 431 | 669 | 378 | 1039 | 2110 | 1330 | 1355 | 1056 | 1316 | 1619 | 1334 | 781 | 821 | 834 | 688 | 557 | 791 | 1241 | 1361 | 1184 | 1356 | 1988 | 1141 | 1043 | 1605 | 462 | 751 | 592 | Kcnt1 |
| Kcnt2 | 464 | 355 | 491 | 529 | 611 | 662 | 753 | 474 | 503 | 371 | 437 | 516 | 464 | 545 | 471 | 486 | 413 | 532 | 529 | 554 | 708 | 622 | 496 | 481 | 550 | 375 | 453 | 482 | Kcnt2 |
| Kcnu1 | 47 | 72 | 15 | 82 | 121 | 111 | 94 | 162 | 190 | 175 | 138 | 125 | 110 | 118 | 95 | 50 | 74 | 79 | 97 | 119 | 100 | 184 | 174 | 138 | 174 | 86 | 112 | 75 | Kcnu1 |
| Kcnv1 | 83 | 74 | 86 | 115 | 246 | 141 | 208 | 305 | 301 | 234 | 243 | 127 | 160 | 154 | 78 | 69 | 104 | 140 | 196 | 243 | 238 | 234 | 261 | 179 | 325 | 108 | 119 | 132 | Kcnv1 |
| Kcnv2 | 3 | 19 | 0 | 6 | 13 | 6 | 7 | 45 | 35 | 33 | 22 | 22 | 40 | 15 | 9 | 16 | 1 | 7 | 7 | 5 | 0 | 33 | 45 | 11 | 3 | 12 | 26 | 17 | Kcnv2 |
| Kcp | 2065 | 1909 | 2119 | 1736 | 2204 | 1953 | 1214 | 1794 | 2210 | 2378 | 2827 | 2076 | 2707 | 1722 | 1462 | 2703 | 1949 | 1968 | 1634 | 1160 | 1271 | 2520 | 2391 | 2181 | 1897 | 1839 | 2078 | 1824 | Kcp |
| Kctd1 | 1222 | 1284 | 1373 | 1448 | 1614 | 1284 | 1313 | 1595 | 1604 | 1885 | 1482 | 1489 | 1539 | 1398 | 1185 | 1221 | 1562 | 1356 | 1680 | 1350 | 1484 | 1599 | 1675 | 1689 | 1670 | 1630 | 1389 | 1403 | Kctd1 |
| Kctd10 | 6827 | 7471 | 7182 | 6600 | 6221 | 6735 | 5424 | 6206 | 6648 | 6304 | 6525 | 7123 | 7350 | 6532 | 6347 | 7534 | 6996 | 6506 | 6464 | 5801 | 5624 | 6403 | 6599 | 6627 | 6836 | 7645 | 7180 | 7149 | Kctd10 |
| Kctd11 | 399 | 555 | 560 | 332 | 402 | 352 | 227 | 260 | 311 | 304 | 311 | 310 | 367 | 243 | 400 | 480 | 538 | 368 | 327 | 370 | 264 | 282 | 401 | 269 | 199 | 397 | 318 | 301 | Kctd11 |
| Kctd12 | 14723 | 8630 | 11352 | 9424 | 10166 | 10840 | 6681 | 14680 | 17666 | 16284 | 26934 | 11458 | 15580 | 12023 | 12389 | 15905 | 9678 | 12567 | 6950 | 6187 | 7570 | 20516 | 20316 | 15801 | 14697 | 11795 | 12996 | 11532 | Kctd12 |
| Kctd12b | 1202 | 726 | 1115 | 694 | 930 | 1011 | 812 | 1583 | 1573 | 1347 | 1589 | 1148 | 1528 | 1139 | 1042 | 967 | 867 | 810 | 813 | 1068 | 877 | 1365 | 1825 | 1127 | 1148 | 1153 | 1281 | 1248 | Kctd12b |
| Kctd13 | 845 | 1146 | 972 | 1535 | 1645 | 1225 | 1422 | 998 | 1023 | 1312 | 1021 | 1023 | 824 | 1074 | 893 | 879 | 1298 | 1556 | 1475 | 1247 | 1488 | 1253 | 968 | 1272 | 1214 | 1004 | 897 | 943 | Kctd13 |
| Kctd14 | 96 | 70 | 56 | 69 | 110 | 101 | 80 | 140 | 189 | 206 | 207 | 236 | 232 | 186 | 65 | 93 | 72 | 87 | 83 | 91 | 80 | 157 | 197 | 259 | 227 | 156 | 171 | 185 | Kctd14 |
| Kctd15 | 1693 | 1883 | 2366 | 2222 | 2888 | 2587 | 2252 | 4042 | 4436 | 5360 | 4730 | 5352 | 5037 | 4952 | 1569 | 1675 | 2497 | 2140 | 3184 | 2346 | 2236 | 4314 | 4210 | 5653 | 5509 | 5675 | 4993 | 4970 | Kctd15 |
| Kctd16 | 40 | 72 | 11 | 36 | 61 | 36 | 37 | 195 | 260 | 80 | 134 | 54 | 79 | 19 | 59 | 75 | 7 | 36 | 18 | 90 | 62 | 154 | 201 | 53 | 89 | 24 | 62 | 30 | Kctd16 |
| Kctd17 | 4586 | 6233 | 6222 | 6187 | 5986 | 5526 | 5420 | 3721 | 4508 | 5005 | 4525 | 4936 | 4502 | 4626 | 4527 | 5057 | 6514 | 5891 | 6057 | 4963 | 5513 | 4450 | 4283 | 5370 | 5164 | 5573 | 4449 | 4446 | Kctd17 |
| Kctd18 | 2209 | 1741 | 1664 | 2167 | 2309 | 2457 | 2272 | 2665 | 2611 | 2502 | 2747 | 2719 | 2655 | 2699 | 2273 | 1961 | 1854 | 1995 | 2051 | 2305 | 2231 | 2159 | 2590 | 2396 | 2615 | 2253 | 2629 | 2286 | Kctd18 |
| Kctd19 | 44 | 36 | 38 | 42 | 18 | 15 | 22 | 4 | 2 | 1 | 3 | 10 | 5 | 3 | 38 | 40 | 37 | 25 | 15 | 20 | 15 | 2 | 1 | 0 | 7 | 2 | 1 | 1 | Kctd19 |
| Kctd2 | 2501 | 2654 | 2410 | 2778 | 2675 | 2719 | 2619 | 2595 | 2705 | 2694 | 2475 | 3049 | 2767 | 2849 | 2351 | 2266 | 2556 | 2436 | 2794 | 2658 | 2607 | 2685 | 2622 | 2856 | 3001 | 3243 | 3086 | 2991 | Kctd2 |
| Kctd20 | 11772 | 11453 | 11760 | 11983 | 11406 | 11714 | 11250 | 10485 | 10291 | 10386 | 10186 | 11535 | 10765 | 11009 | 11693 | 11073 | 11683 | 11850 | 12103 | 10964 | 11461 | 11376 | 9846 | 10966 | 11654 | 11739 | 10756 | 11356 | Kctd20 |
| Kctd21 | 734 | 892 | 836 | 881 | 768 | 712 | 820 | 697 | 559 | 545 | 590 | 569 | 634 | 533 | 849 | 774 | 783 | 728 | 803 | 709 | 807 | 765 | 630 | 612 | 685 | 564 | 552 | 573 | Kctd21 |
| Kctd3 | 4523 | 4783 | 4627 | 4909 | 4780 | 4528 | 4460 | 5471 | 5558 | 5639 | 5274 | 5868 | 5407 | 5575 | 4193 | 4643 | 4779 | 4632 | 4872 | 4573 | 4531 | 5564 | 5321 | 5865 | 6144 | 6001 | 5655 | 5574 | Kctd3 |
| Kctd4 | 57 | 31 | 33 | 117 | 259 | 235 | 257 | 413 | 405 | 312 | 340 | 299 | 183 | 274 | 73 | 30 | 81 | 163 | 207 | 295 | 272 | 384 | 425 | 336 | 375 | 148 | 250 | 178 | Kctd4 |
| Kctd5 | 4901 | 5290 | 4829 | 4954 | 4705 | 4705 | 4857 | 4935 | 4940 | 4526 | 4555 | 5168 | 4974 | 5234 | 4836 | 4659 | 5070 | 4555 | 4998 | 4789 | 4843 | 4645 | 4755 | 4987 | 4931 | 5903 | 5276 | 5496 | Kctd5 |
| Kctd6 | 1068 | 860 | 807 | 1165 | 1459 | 1276 | 1630 | 1097 | 1126 | 1141 | 1122 | 1222 | 1009 | 1229 | 1017 | 998 | 1044 | 1327 | 1296 | 1312 | 1518 | 1248 | 1066 | 1195 | 1229 | 1138 | 1053 | 1034 | Kctd6 |
| Kctd7 | 699 | 811 | 784 | 858 | 918 | 803 | 891 | 807 | 789 | 824 | 870 | 773 | 705 | 808 | 752 | 664 | 761 | 974 | 899 | 818 | 917 | 971 | 856 | 764 | 906 | 627 | 657 | 784 | Kctd7 |
| Kctd8 | 186 | 207 | 181 | 270 | 465 | 337 | 295 | 308 | 310 | 287 | 338 | 188 | 218 | 194 | 286 | 186 | 212 | 330 | 330 | 404 | 358 | 403 | 369 | 229 | 365 | 136 | 234 | 202 | Kctd8 |
| Kctd9 | 3636 | 3011 | 3116 | 3100 | 2449 | 2957 | 3047 | 2148 | 2015 | 1902 | 2052 | 2260 | 2362 | 2538 | 3388 | 3603 | 2770 | 3028 | 2272 | 2735 | 2968 | 1775 | 1994 | 2142 | 1908 | 2369 | 2313 | 2509 | Kctd9 |
| Kdelr1 | 4467 | 4632 | 4842 | 4257 | 4169 | 4164 | 4181 | 5151 | 5790 | 5886 | 6415 | 5351 | 5725 | 5928 | 4344 | 4597 | 4671 | 4135 | 4306 | 3919 | 4352 | 5815 | 6380 | 5937 | 5217 | 6328 | 5631 | 5948 | Kdelr1 |
| Kdelr2 | 6472 | 6042 | 6105 | 5532 | 4051 | 4854 | 4644 | 4369 | 5176 | 4322 | 5791 | 4271 | 4850 | 4301 | 5996 | 6900 | 5638 | 5710 | 4269 | 5151 | 4632 | 4338 | 5706 | 4677 | 4105 | 5267 | 4322 | 4789 | Kdelr2 |
| Kdelr3 | 1305 | 1299 | 1119 | 875 | 645 | 954 | 510 | 772 | 1218 | 866 | 1393 | 647 | 814 | 573 | 1396 | 1574 | 915 | 1125 | 679 | 667 | 543 | 1111 | 1469 | 681 | 887 | 635 | 680 | 765 | Kdelr3 |
| Kdf1 | 87 | 38 | 6 | 29 | 4 | 21 | 0 | 61 | 97 | 76 | 74 | 16 | 15 | 19 | 36 | 56 | 20 | 18 | 5 | 3 | 0 | 10 | 108 | 83 | 23 | 78 | 9 | 13 | Kdf1 |
| Kdm1a | 12245 | 11166 | 14187 | 13460 | 14429 | 13027 | 14565 | 17054 | 15348 | 16899 | 15018 | 18710 | 16032 | 18023 | 11876 | 10784 | 13891 | 12698 | 14993 | 12421 | 13516 | 18782 | 14666 | 18237 | 19092 | 19082 | 17224 | 17940 | Kdm1a |
| Kdm1b | 8183 | 8111 | 7693 | 7880 | 7117 | 7782 | 7959 | 3809 | 3410 | 3404 | 3090 | 3583 | 3366 | 3530 | 8231 | 7543 | 7577 | 8396 | 7611 | 8468 | 7771 | 3634 | 3534 | 3389 | 3694 | 3081 | 3511 | 3494 | Kdm1b |
| Kdm2a | 9874 | 9347 | 10107 | 10587 | 10840 | 10144 | 10411 | 11118 | 10597 | 12023 | 12398 | 12441 | 12305 | 12689 | 9037 | 9230 | 10436 | 10738 | 10910 | 8140 | 10836 | 12415 | 10686 | 11840 | 12197 | 9975 | 11675 | 12493 | Kdm2a |
| Kdm2b | 6592 | 6984 | 8404 | 8631 | 9750 | 8724 | 8765 | 6834 | 7190 | 8844 | 8052 | 9232 | 7607 | 8594 | 6465 | 6646 | 8759 | 8579 | 10361 | 6980 | 9307 | 9451 | 6797 | 8973 | 9714 | 8914 | 7519 | 8653 | Kdm2b |
| Kdm3a | 13989 | 12842 | 10172 | 11575 | 11638 | 12748 | 12723 | 14222 | 11506 | 10861 | 10136 | 11847 | 11712 | 12700 | 14231 | 11980 | 11808 | 11118 | 12011 | 13676 | 12534 | 10848 | 11734 | 10868 | 11493 | 10802 | 12127 | 12458 | Kdm3a |
| Kdm3b | 8877 | 8642 | 9337 | 9791 | 10492 | 10281 | 10528 | 9546 | 9887 | 10624 | 10987 | 12718 | 11550 | 12145 | 8263 | 8269 | 9875 | 9743 | 10941 | 8305 | 10525 | 10649 | 9387 | 11159 | 11228 | 9729 | 11687 | 11754 | Kdm3b |
| Kdm4a | 7217 | 8365 | 8993 | 9301 | 9803 | 8848 | 9081 | 11372 | 13174 | 14681 | 13413 | 16140 | 14758 | 15827 | 7213 | 7460 | 9651 | 9022 | 10242 | 7762 | 9268 | 14745 | 12099 | 15456 | 15882 | 14671 | 14508 | 15891 | Kdm4a |
| Kdm4b | 6629 | 8699 | 7458 | 9688 | 9823 | 8875 | 8154 | 5606 | 6873 | 7596 | 6886 | 7693 | 6727 | 6396 | 6765 | 7853 | 8893 | 9756 | 10336 | 7652 | 8879 | 7670 | 6543 | 7527 | 8406 | 6590 | 6649 | 6548 | Kdm4b |
| Kdm4c | 3599 | 3268 | 3089 | 3246 | 3507 | 3690 | 3862 | 3463 | 3054 | 2787 | 2978 | 2800 | 2650 | 2931 | 3865 | 3203 | 2996 | 3217 | 3214 | 3877 | 3762 | 3011 | 2920 | 2587 | 3013 | 2663 | 2765 | 2622 | Kdm4c |
| Kdm4d | 43 | 25 | 21 | 36 | 42 | 48 | 41 | 17 | 32 | 19 | 24 | 13 | 35 | 14 | 25 | 46 | 20 | 22 | 32 | 31 | 41 | 25 | 30 | 17 | 11 | 10 | 21 | 13 | Kdm4d |

| | | | | | | | | | | | | | | | | | | | | | | | | | | | | | |
|---|---|---|---|---|---|---|---|---|---|---|---|---|---|---|---|---|---|---|---|---|---|---|---|---|---|---|---|---|---|
| Kdm5a | 16019 | 13455 | 10311 | 12463 | 10903 | 13027 | 12925 | 13877 | 11453 | 9875 | 10453 | 11318 | 11349 | 12100 | 16166 | 13448 | 10780 | 11681 | 10621 | 13501 | 12848 | 9894 | 10952 | 9575 | 10641 | 7652 | 12450 | 11055 | Kdm5a |
| Kdm5b | 19622 | 17694 | 19055 | 20826 | 23014 | 22710 | 22343 | 20263 | 19831 | 22233 | 20631 | 21758 | 21063 | 22362 | 19019 | 18510 | 21075 | 21798 | 23098 | 17801 | 22770 | 23758 | 19297 | 21630 | 23315 | 17789 | 20994 | 21967 | Kdm5b |
| Kdm5c | 8996 | 10028 | 7991 | 9060 | 9281 | 9356 | 8963 | 11696 | 12770 | 12618 | 12228 | 13414 | 13040 | 13071 | 10338 | 10304 | 10159 | 10557 | 11145 | 10576 | 11054 | 14875 | 15684 | 15947 | 17629 | 13313 | 18512 | 17267 | Kdm5c |
| Kdm5d | 5173 | 4934 | 3868 | 4318 | 4666 | 5139 | 4998 | 6592 | 7813 | 6821 | 5951 | 7517 | 7267 | 8003 | 5 | 0 | 0 | 1 | 1 | 7 | 0 | 0 | 14 | 0 | 0 | 1 | 0 | 0 | Kdm5d |
| Kdm6a | 5800 | 4769 | 4391 | 4950 | 5226 | 5920 | 5877 | 6008 | 5353 | 4950 | 4957 | 5538 | 5642 | 5532 | 8681 | 7298 | 6944 | 6858 | 7057 | 8199 | 8051 | 7414 | 7768 | 7155 | 7382 | 7105 | 8702 | 8267 | Kdm6a |
| Kdm6b | 4713 | 6431 | 8816 | 8306 | 7873 | 6410 | 5896 | 6471 | 7460 | 10265 | 9575 | 11308 | 8408 | 9552 | 4117 | 5339 | 9320 | 7723 | 8667 | 4941 | 6687 | 10945 | 7064 | 11962 | 11119 | 11395 | 8894 | 9280 | Kdm6b |
| Kdm7a | 6192 | 4765 | 5093 | 5411 | 5921 | 6750 | 6382 | 7441 | 5777 | 6182 | 5984 | 6616 | 7050 | 7223 | 6435 | 5485 | 5911 | 5027 | 5940 | 6265 | 6288 | 6017 | 5908 | 5854 | 6891 | 5166 | 7222 | 7111 | Kdm7a |
| Kdm8 | 171 | 183 | 256 | 122 | 133 | 180 | 130 | 529 | 600 | 518 | 499 | 701 | 713 | 732 | 129 | 144 | 229 | 133 | 168 | 182 | 174 | 556 | 557 | 593 | 579 | 729 | 732 | 732 | Kdm8 |
| Kdr | 4358 | 4343 | 4963 | 4290 | 5186 | 4910 | 4203 | 4797 | 5177 | 6366 | 6711 | 6617 | 7448 | 6363 | 4349 | 4844 | 4393 | 4478 | 4422 | 4499 | 4429 | 6256 | 6187 | 6039 | 6302 | 5501 | 6689 | 6128 | Kdr |
| Kdsr | 2639 | 2211 | 2321 | 2986 | 2779 | 2918 | 2925 | 1807 | 1673 | 1773 | 2029 | 1727 | 1889 | 1788 | 2629 | 2555 | 2391 | 2863 | 2642 | 2756 | 2990 | 1917 | 1746 | 1768 | 1920 | 1578 | 1725 | 1870 | Kdsr |
| Keap1 | 2225 | 2357 | 2576 | 2425 | 2416 | 2486 | 2428 | 2878 | 3541 | 3758 | 3838 | 3418 | 3256 | 3580 | 2014 | 2076 | 2566 | 2259 | 2588 | 2238 | 2446 | 3235 | 3489 | 3851 | 3734 | 3881 | 3395 | 3483 | Keap1 |
| Keg1 | 0 | 0 | 0 | 0 | 0 | 0 | 0 | 0 | 2 | 0 | 0 | 0 | 0 | 0 | 0 | 0 | 0 | 0 | 0 | 0 | 0 | 0 | 0 | 0 | 0 | 0 | 0 | 0 | Keg1 |
| Kel | 325 | 478 | 365 | 194 | 78 | 167 | 156 | 319 | 328 | 258 | 131 | 622 | 1864 | 942 | 384 | 405 | 212 | 108 | 76 | 311 | 90 | 139 | 329 | 194 | 128 | 642 | 893 | 1860 | Kel |
| Kera | 68 | 14 | 29 | 10 | 4 | 42 | 9 | 106 | 244 | 67 | 161 | 37 | 59 | 23 | 38 | 53 | 10 | 56 | 38 | 9 | 4 | 90 | 326 | 55 | 23 | 49 | 51 | 31 | Kera |
| Khdc1a | 0 | 0 | 0 | 0 | 0 | 0 | 0 | 0 | 1 | 0 | 0 | 0 | 3 | 0 | 0 | 0 | 0 | 0 | 0 | 0 | 0 | 0 | 1 | 2 | 0 | 1 | 0 | 0 | Khdc1a |
| Khdc1b | 0 | 0 | 0 | 0 | 0 | 0 | 0 | 0 | 0 | 0 | 0 | 0 | 0 | 0 | 0 | 0 | 0 | 0 | 0 | 0 | 0 | 0 | 0 | 0 | 1 | 0 | 0 | 0 | Khdc1b |
| Khdc1c | 0 | 0 | 0 | 0 | 0 | 0 | 0 | 1 | 0 | 0 | 1 | 0 | 0 | 0 | 0 | 1 | 0 | 0 | 0 | 0 | 0 | 0 | 0 | 2 | 0 | 2 | 0 | 0 | Khdc1c |
| Khdc3 | 0 | 1 | 0 | 0 | 1 | 0 | 2 | 5 | 0 | 3 | 2 | 0 | 1 | 2 | 5 | 0 | 0 | 6 | 0 | 3 | 1 | 2 | 3 | 2 | 2 | 0 | 0 | 1 | Khdc3 |
| Khdc4 | 8824 | 7960 | 8289 | 8828 | 8588 | 8635 | 9170 | 11876 | 9981 | 10325 | 10124 | 11239 | 10768 | 11718 | 8814 | 7754 | 8041 | 8298 | 8890 | 8743 | 9053 | 11000 | 9916 | 10621 | 10594 | 11204 | 10727 | 11450 | Khdc4 |
| Khdrbs1 | 37957 | 30469 | 42308 | 38629 | 38494 | 38528 | 43590 | 34878 | 31411 | 34933 | 35696 | 43814 | 40216 | 47861 | 36979 | 32009 | 40132 | 37527 | 38877 | 35515 | 42568 | 36469 | 30277 | 40027 | 40988 | 47552 | 41976 | 47786 | Khdrbs1 |
| Khdrbs2 | 788 | 809 | 975 | 1213 | 1494 | 1274 | 1433 | 1501 | 1341 | 1292 | 1188 | 1185 | 1047 | 1280 | 994 | 814 | 998 | 1118 | 1320 | 1512 | 1415 | 1547 | 1346 | 1323 | 1472 | 1078 | 1178 | 1183 | Khdrbs2 |
| Khdrbs3 | 6908 | 6796 | 7685 | 8402 | 8516 | 7435 | 10391 | 6897 | 6125 | 7363 | 5617 | 7153 | 5200 | 7903 | 7564 | 5907 | 8438 | 8045 | 9230 | 8555 | 9683 | 6904 | 5699 | 7421 | 7893 | 8086 | 6529 | 7522 | Khdrbs3 |
| Khk | 352 | 419 | 448 | 350 | 313 | 306 | 254 | 199 | 286 | 282 | 357 | 229 | 235 | 186 | 318 | 379 | 357 | 336 | 320 | 365 | 239 | 227 | 336 | 269 | 206 | 267 | 277 | 192 | Khk |
| Khnyn | 5144 | 5482 | 6990 | 6098 | 5174 | 4963 | 4713 | 3401 | 3568 | 4329 | 4619 | 4488 | 4630 | 4162 | 4636 | 6017 | 6507 | 5692 | 5566 | 4209 | 4887 | 3843 | 3862 | 4240 | 3606 | 4481 | 4391 | 4300 | Khnyn |
| Khsrp | 7741 | 9164 | 13467 | 11939 | 10433 | 8722 | 9317 | 9788 | 11688 | 14021 | 13401 | 17024 | 13632 | 15671 | 7201 | 8541 | 13385 | 10853 | 11536 | 7789 | 9322 | 12318 | 10777 | 16944 | 14310 | 20067 | 14573 | 16117 | Khsrp |
| Kidins220 | 30613 | 32669 | 22969 | 35000 | 45382 | 43320 | 42972 | 42393 | 40153 | 40767 | 35692 | 33052 | 31905 | 33476 | 34585 | 29354 | 28912 | 38919 | 39537 | 42573 | 45271 | 45101 | 37475 | 31971 | 45096 | 17854 | 31052 | 27855 | Kidins220 |
| Kif11 | 927 | 755 | 390 | 458 | 277 | 337 | 923 | 1192 | 637 | 399 | 767 | 697 | 1084 | 508 | 767 | 558 | 1929 | 478 | 458 | 502 | 307 | 488 | 365 | 437 | 561 | 694 | 648 | 998 | Kif11 |
| Kif12 | 1 | 2 | 0 | 7 | 0 | 2 | 0 | 0 | 1 | 1 | 3 | 1 | 1 | 0 | 0 | 3 | 0 | 7 | 0 | 0 | 4 | 3 | 4 | 1 | 5 | 5 | 0 | 0 | Kif12 |
| Kif13a | 2679 | 2697 | 2593 | 2487 | 2917 | 2927 | 2189 | 4482 | 5001 | 5119 | 5518 | 5710 | 5697 | 5259 | 2591 | 2779 | 2484 | 2426 | 2498 | 2252 | 2364 | 5040 | 5103 | 4913 | 4847 | 4041 | 5765 | 5451 | Kif13a |
| Kif13b | 3741 | 3788 | 2716 | 4083 | 4463 | 4345 | 4114 | 2873 | 3239 | 3157 | 3353 | 2864 | 2971 | 2732 | 3756 | 4115 | 3473 | 4624 | 4197 | 3719 | 4189 | 3076 | 2992 | 2698 | 3295 | 1654 | 2705 | 2575 | Kif13b |
| Kif14 | 2453 | 1928 | 1752 | 1632 | 1254 | 1528 | 1667 | 2251 | 2351 | 2344 | 2071 | 3018 | 2578 | 3182 | 1919 | 1885 | 1591 | 1448 | 1541 | 1419 | 1490 | 1601 | 1920 | 2486 | 2249 | 2244 | 3232 | 2961 | Kif14 |
| Kif15 | 9439 | 7449 | 7457 | 5945 | 4345 | 6189 | 6554 | 7952 | 7143 | 5970 | 5486 | 9977 | 8255 | 9777 | 8294 | 7289 | 6468 | 4925 | 5587 | 7081 | 5679 | 4767 | 5950 | 7279 | 6769 | 9773 | 10689 | 9882 | Kif15 |
| Kif16b | 1703 | 1412 | 1418 | 1119 | 975 | 1187 | 1076 | 1545 | 1669 | 1494 | 1702 | 1427 | 1678 | 1563 | 1515 | 1777 | 1243 | 1137 | 950 | 1116 | 945 | 1404 | 1789 | 1364 | 1309 | 1413 | 1618 | 1650 | Kif16b |
| Kif17 | 473 | 583 | 484 | 811 | 797 | 656 | 565 | 134 | 179 | 188 | 160 | 90 | 170 | 123 | 522 | 632 | 649 | 769 | 708 | 587 | 617 | 226 | 166 | 134 | 132 | 88 | 102 | 85 | Kif17 |
| Kif18a | 3592 | 2932 | 2797 | 2526 | 1939 | 2220 | 2699 | 3132 | 2548 | 1956 | 1977 | 3388 | 2866 | 3572 | 3349 | 3135 | 2522 | 2214 | 2383 | 3134 | 2507 | 1983 | 2291 | 2668 | 2231 | 3725 | 3843 | 3628 | Kif18a |
| Kif18b | 2684 | 3143 | 3441 | 2802 | 1886 | 2175 | 2202 | 2004 | 2310 | 2152 | 1898 | 3151 | 2535 | 2931 | 2281 | 2629 | 3275 | 2422 | 2580 | 2225 | 2070 | 1462 | 1929 | 2742 | 2317 | 3874 | 2952 | 3010 | Kif18b |
| Kif19a | 571 | 566 | 420 | 450 | 429 | 212 | 174 | 365 | 434 | 567 | 412 | 71 | 403 | 95 | 613 | 808 | 474 | 371 | 252 | 180 | 200 | 276 | 373 | 420 | 220 | 644 | 81 | 80 | Kif19a |
| Kif19b | 59 | 93 | 75 | 105 | 83 | 80 | 77 | 11 | 16 | 5 | 10 | 11 | 22 | 20 | 59 | 93 | 78 | 106 | 73 | 59 | 83 | 23 | 25 | 10 | 21 | 17 | 16 | 8 | Kif19b |
| Kif1a | 23218 | 29704 | 25209 | 39441 | 47284 | 40813 | 39267 | 24504 | 29245 | 33622 | 27065 | 28874 | 26409 | 27037 | 25477 | 25996 | 33943 | 41247 | 45796 | 34951 | 43328 | 38345 | 24629 | 29977 | 39367 | 15424 | 23301 | 24417 | Kif1a |
| Kif1b | 46036 | 46497 | 30166 | 48993 | 59828 | 57330 | 57203 | 54063 | 49029 | 51445 | 45728 | 48137 | 46233 | 49520 | 50858 | 43354 | 39468 | 51103 | 51152 | 55591 | 60969 | 52647 | 44839 | 39259 | 58317 | 26996 | 50138 | 44362 | Kif1b |
| Kif1c | 7542 | 8920 | 10431 | 8659 | 7086 | 7242 | 5989 | 3991 | 5456 | 5471 | 6363 | 6024 | 6348 | 5446 | 6858 | 9668 | 9244 | 8262 | 7492 | 6100 | 6280 | 4542 | 5847 | 6365 | 5430 | 6650 | 5788 | 5860 | Kif1c |
| Kif20a | 7055 | 6217 | 6291 | 4149 | 3323 | 4094 | 4014 | 7216 | 6803 | 5609 | 5465 | 8399 | 7541 | 8483 | 5765 | 5643 | 5247 | 3504 | 4344 | 4985 | 3810 | 4408 | 6227 | 7196 | 5565 | 10247 | 8793 | 8539 | Kif20a |
| Kif20b | 6304 | 4983 | 5409 | 4444 | 3412 | 4237 | 4842 | 5697 | 4492 | 3694 | 3837 | 5802 | 5011 | 6079 | 6050 | 5828 | 4727 | 4006 | 4112 | 5542 | 4488 | 3275 | 4146 | 4593 | 4039 | 4998 | 6064 | 5734 | Kif20b |
| Kif21a | 15648 | 16916 | 16488 | 17905 | 18359 | 18525 | 18565 | 19844 | 15781 | 17726 | 14716 | 20992 | 18046 | 21020 | 17977 | 14422 | 17321 | 16657 | 18672 | 19225 | 18788 | 18586 | 14947 | 18346 | 21016 | 13943 | 22167 | 20704 | Kif21a |
| Kif21b | 16308 | 21208 | 18470 | 33062 | 39739 | 33393 | 34820 | 22379 | 27656 | 37205 | 28533 | 32246 | 23362 | 30372 | 16635 | 18364 | 27692 | 35249 | 39675 | 26833 | 37957 | 37552 | 22227 | 34550 | 44888 | 14852 | 22779 | 24961 | Kif21b |
| Kif22 | 6749 | 6678 | 7130 | 5250 | 3671 | 4634 | 4807 | 4899 | 4889 | 4309 | 3948 | 6958 | 5417 | 6790 | 5796 | 5535 | 6047 | 4559 | 5256 | 5402 | 4486 | 3352 | 4031 | 5663 | 4963 | 8750 | 6958 | 6842 | Kif22 |
| Kif23 | 7922 | 7284 | 7168 | 6484 | 5035 | 6047 | 6873 | 6453 | 6083 | 5659 | 5464 | 8407 | 7438 | 8986 | 7249 | 6816 | 6339 | 6012 | 5786 | 6542 | 6395 | 4355 | 5339 | 6636 | 6121 | 7894 | 9001 | 8432 | Kif23 |
| Kif24 | 3601 | 3575 | 2251 | 2045 | 1777 | 2472 | 2481 | 1461 | 1468 | 1174 | 1283 | 1843 | 1812 | 1718 | 3167 | 3183 | 2386 | 1804 | 1903 | 2748 | 2523 | 926 | 1317 | 1273 | 1199 | 1336 | 2210 | 1822 | Kif24 |
| Kif26a | 4935 | 6094 | 6795 | 8252 | 8055 | 7705 | 7772 | 4491 | 5880 | 6452 | 5554 | 8276 | 5012 | 6787 | 4369 | 4400 | 7999 | 8208 | 10160 | 6899 | 7743 | 6236 | 4766 | 8288 | 8148 | 8222 | 6671 | 6372 | Kif26a |
| Kif26b | 4404 | 3444 | 2534 | 3089 | 4281 | 4127 | 2160 | 8372 | 9227 | 11490 | 10249 | 5848 | 9327 | 5668 | 3142 | 5022 | 2307 | 3725 | 2669 | 2310 | 2508 | 8585 | 7555 | 7378 | 7597 | 5864 | 6236 | 5832 | Kif26b |
| Kif27 | 1069 | 1018 | 925 | 1064 | 1037 | 1074 | 997 | 445 | 469 | 515 | 533 | 545 | 481 | 443 | 1130 | 1094 | 940 | 1086 | 902 | 1108 | 1032 | 510 | 641 | 394 | 452 | 383 | 408 | 445 | Kif27 |
| Kif28 | 10 | 2 | 6 | 0 | 5 | 5 | 3 | 7 | 4 | 14 | 2 | 9 | 14 | 7 | 3 | 2 | 3 | 2 | 3 | 2 | 4 | 13 | 14 | 3 | 7 | 8 | 10 | 6 | Kif28 |
| Kif2a | 9135 | 7207 | 9641 | 10539 | 12188 | 11830 | 13052 | 11264 | 9898 | 10988 | 10863 | 11564 | 11090 | 12651 | 9483 | 8242 | 10384 | 11145 | 11368 | 10316 | 13213 | 13072 | 9449 | 11031 | 12311 | 10979 | 11386 | 12657 | Kif2a |
| Kif2c | 4698 | 4425 | 4517 | 3325 | 2611 | 2945 | 2981 | 4844 | 4484 | 3653 | 3409 | 6054 | 4656 | 5661 | 3836 | 3886 | 3858 | 2727 | 3469 | 3479 | 2881 | 3006 | 3926 | 4949 | 4136 | 7468 | 6138 | 6300 | Kif2c |
| Kif3a | 10606 | 10827 | 8157 | 10487 | 11888 | 11714 | 11855 | 11571 | 10513 | 9973 | 9791 | 8755 | 8574 | 9401 | 11188 | 10231 | 9180 | 10730 | 11233 | 12495 | 12270 | 10569 | 9805 | 9021 | 11164 | 6428 | 8797 | 8562 | Kif3a |
| Kif3b | 9040 | 9508 | 7344 | 9414 | 9272 | 9336 | 9785 | 7768 | 6615 | 6244 | 6148 | 6361 | 6048 | 6479 | 9621 | 8133 | 8150 | 9426 | 9028 | 9699 | 9940 | 7503 | 6486 | 6013 | 7274 | 4881 | 6197 | 6252 | Kif3b |
| Kif3c | 7564 | 9060 | 7178 | 10760 | 13397 | 11557 | 11686 | 10437 | 10802 | 12354 | 10457 | 10351 | 8969 | 10441 | 8403 | 8029 | 9675 | 11569 | 12456 | 10741 | 13175 | 13867 | 9967 | 11172 | 14252 | 8120 | 9096 | 9551 | Kif3c |
| Kif4 | 6691 | 6091 | 5906 | 4216 | 3388 | 4308 | 4842 | 6759 | 5995 | 5200 | 4902 | 8995 | 7074 | 8773 | 6157 | 5040 | 5071 | 3913 | 4429 | 5174 | 4393 | 4273 | 5152 | 6635 | 5975 | 8802 | 9058 | 8748 | Kif4 |
| Kif5a | 12038 | 14860 | 11835 | 17918 | 24125 | 18700 | 18676 | 17018 | 17372 | 19411 | 16673 | 15135 | 15688 | 15122 | 14184 | 12857 | 15983 | 19487 | 20755 | 18457 | 20309 | 24262 | 14711 | 15929 | 21466 | 9007 | 13835 | 13681 | Kif5a |
| Kif5b | 18850 | 15007 | 15337 | 17187 | 18098 | 18921 | 19883 | 20668 | 17943 | 16711 | 17532 | 19156 | 18432 | 19840 | 18863 | 17654 | 15577 | 16845 | 17309 | 18747 | 19157 | 16915 | 18376 | 16938 | 18124 | 15086 | 19100 | 19553 | Kif5b |
| Kif5c | 35384 | 35206 | 30235 | 45574 | 57286 | 51414 | 55075 | 52831 | 43994 | 49159 | 41305 | 46074 | 40946 | 46375 | 42581 | 33561 | 41215 | 48972 | 50416 | 54337 | 55202 | 57423 | 38395 | 43345 | 56145 | 30175 | 43022 | 41416 | Kif5c |
| Kif6 | 358 | 403 | 221 | 345 | 316 | 342 | 282 | 200 | 197 | 199 | 156 | 207 | 221 | 181 | 324 | 382 | 290 | 273 | 317 | 295 | 289 | 196 | 224 | 130 | 183 | 160 | 152 | 132 | Kif6 |
| Kif7 | 3160 | 3972 | 4624 | 4163 | 3183 | 3317 | 3348 | 1215 | 1622 | 1579 | 1668 | 1943 | 1854 | 1729 | 2851 | 3820 | 4289 | 4123 | 3631 | 3210 | 3286 | 1305 | 1518 | 1818 | 1558 | 2113 | 1800 | 1920 | Kif7 |
| Kif9 | 235 | 277 | 161 | 204 | 344 | 229 | 203 | 319 | 301 | 278 | 261 | 255 | 313 | 238 | 252 | 310 | 174 | 250 | 284 | 294 | 246 | 321 | 432 | 188 | 220 | 194 | 209 | 227 | Kif9 |
| Kifap3 | 10986 | 8539 | 8570 | 9445 | 12457 | 11128 | 11754 | 15397 | 14774 | 14216 | 13997 | 11729 | 12238 | 12182 | 11383 | 10909 | 8759 | 11014 | 10522 | 10589 | 11740 | 17346 | 15003 | 12291 | 14852 | 10600 | 11065 | 11718 | Kifap3 |
| Kifbp | 4152 | 4715 | 4303 | 4819 | 4572 | 4625 | 4548 | 4505 | 4029 | 4322 | 3893 | 4129 | 4054 | 4249 | 4584 | 4197 | 4825 | 4574 | 4769 | 4274 | 4666 | 4943 | 4126 | 4344 | 4507 | 4286 | 4084 | 4276 | Kifbp |
| Kifc1 | 4111 | 4611 | 4914 | 4293 | 3179 | 3525 | 3620 | 2753 | 3388 | 3093 | 3114 | 5078 | 4132 | 4707 | 3444 | 3984 | 4606 | 3706 | 4104 | 3334 | 3354 | 2199 | 2904 | 4238 | 3568 | 5815 | 4771 | 4874 | Kifc1 |
| Kifc2 | 2894 | 3930 | 3804 | 5165 | 5249 | 4350 | 4417 | 1473 | 1677 | 2044 | 1900 | 1257 | 1313 | 1145 | 3051 | 3369 | 4582 | 4903 | 4893 | 4052 | 4555 | 2296 | 1671 | 1683 | 1795 | 1017 | 1202 | 1031 | Kifc2 |
| Kifc3 | 1361 | 1664 | 1430 | 1744 | 2155 | 1726 | 1551 | 982 | 1224 | 1249 | 1283 | 920 | 998 | 827 | 1452 | 1522 | 1749 | 1692 | 1833 | 1655 | 1608 | 1260 | 1180 | 1066 | 1267 | 936 | 777 | 749 | Kifc3 |
| Kifc5b | 2234 | 2818 | 2791 | 2724 | 1746 | 2042 | 2034 | 1109 | 1341 | 1324 | 1244 | 1932 | 1618 | 1911 | 1816 | 2247 | 2584 | 2450 | 2265 | 2019 | 2129 | 936 | 1054 | 1709 | 1527 | 2230 | 2005 | 1844 | Kifc5b |
| Kin | 2475 | 2670 | 3112 | 2964 | 2390 | 2304 | 2582 | 2390 | 2042 | 2247 | 2042 | 2483 | 2269 | 2763 | 2491 | 2578 | 2848 | 2724 | 2349 | 2829 | 2438 | 2274 | 1895 | 2661 | 2496 | 3001 | 2631 | 2690 | Kin |
| Kir3dl1 | 0 | 1 | 0 | 0 | 0 | 0 | 0 | 0 | 0 | 0 | 0 | 0 | 0 | 0 | 3 | 0 | 0 | 0 | 0 | 0 | 0 | 0 | 0 | 0 | 0 | 0 | 0 | 0 | Kir3dl1 |
| Kir3dl2 | 1 | 0 | 0 | 0 | 5 | 0 | 2 | 1 | 5 | 0 | 8 | 1 | 0 | 2 | 0 | 0 | 0 | 7 | 5 | 3 | 3 | 3 | 8 | 2 | 10 | 6 | 3 | 2 | Kir3dl2 |
| Kirrel | 5945 | 5922 | 6681 | 5564 | 6015 | 5911 | 5065 | 5510 | 6377 | 6790 | 8312 | 7225 | 8107 | 6795 | 5000 | 6457 | 6380 | 6003 | 6198 | 4496 | 5286 | 6607 | 6581 | 7108 | 6085 | 6914 | 7060 | 7182 | Kirrel |
| Kirrel2 | 63 | 418 | 92 | 422 | 325 | 298 | 61 | 462 | 499 | 429 | 375 | 105 | 1360 | 95 | 339 | 660 | 722 | 172 | 236 | 431 | 71 | 498 | 554 | 136 | 375 | 126 | 100 | 109 | Kirrel2 |
| Kirrel3 | 745 | 1014 | 650 | 1301 | 2121 | 1812 | 1677 | 1370 | 1910 | 1815 | 1657 | 1174 | 988 | 992 | 942 | 884 | 924 | 1564 | 1759 | 1713 | 1937 | 1947 | 1608 | 1312 | 2012 | 754 | 939 | 813 | Kirrel3 |
| Kiss1r | 189 | 224 | 241 | 268 | 210 | 164 | 175 | 81 | 115 | 147 | 109 | 108 | 103 | 96 | 166 | 243 | 238 | 295 | 212 | 130 | 139 | 130 | 88 | 116 | 121 | 73 | 114 | 70 | Kiss1r |
| Kit | 2863 | 3064 | 3040 | 3680 | 5766 | 4333 | 3824 | 4407 | 4970 | 5422 | 4959 | 3775 | 4784 | 3853 | 3522 | 3455 | 3559 | 4138 | 4369 | 4190 | 4184 | 5634 | 5376 | 4430 | 5453 | 3520 | 3534 | 3398 | Kit |
| Kitl | 4410 | 3138 | 2598 | 4169 | 5237 | 5165 | 7304 | 4813 | 4576 | 4165 | 4320 | 3542 | 2765 | 3598 | 4447 | 3614 | 3307 | 4910 | 5442 | 5211 | 6439 | 4879 | 4789 | 4328 | 4764 | 3382 | 2443 | 2807 | Kitl |

| | | | | | | | | | | | | | | | | | | | | | | | | | | | | | |
|---|---|---|---|---|---|---|---|---|---|---|---|---|---|---|---|---|---|---|---|---|---|---|---|---|---|---|---|---|---|
| Kiz | 2268 | 2051 | 1997 | 1665 | 1657 | 1603 | 1937 | 1844 | 1633 | 1498 | 1343 | 1617 | 1488 | 1606 | 2248 | 1805 | 1878 | 1552 | 1548 | 1953 | 1661 | 1509 | 1658 | 1423 | 1492 | 1641 | 1710 | 1803 | Kiz |
| Kl | 21 | 35 | 38 | 52 | 52 | 49 | 41 | 62 | 54 | 80 | 82 | 89 | 99 | 63 | 18 | 33 | 36 | 26 | 41 | 50 | 68 | 87 | 69 | 78 | 107 | 47 | 71 | 74 | Kl |
| Klb | 14 | 19 | 39 | 26 | 13 | 18 | 20 | 9 | 12 | 10 | 9 | 9 | 10 | 14 | 8 | 26 | 14 | 28 | 16 | 29 | 24 | 17 | 6 | 6 | 17 | 10 | 23 | 15 | Klb |
| Klc1 | 20172 | 26531 | 25183 | 30702 | 31397 | 27677 | 27450 | 23007 | 22251 | 27099 | 22769 | 25350 | 24145 | 24834 | 23179 | 22119 | 29488 | 29729 | 31977 | 28059 | 29756 | 26963 | 21462 | 25355 | 28246 | 19633 | 24535 | 24309 | Klc1 |
| Klc2 | 3838 | 5200 | 5639 | 6240 | 5960 | 5457 | 4965 | 4658 | 5007 | 5860 | 5156 | 5423 | 5383 | 4979 | 4163 | 4476 | 6095 | 5601 | 6448 | 5326 | 5168 | 5930 | 4757 | 5297 | 5942 | 5453 | 5289 | 4926 | Klc2 |
| Klc3 | 281 | 237 | 286 | 199 | 188 | 126 | 95 | 122 | 166 | 215 | 158 | 78 | 132 | 70 | 225 | 249 | 158 | 144 | 153 | 130 | 97 | 67 | 213 | 217 | 32 | 263 | 72 | 69 | Klc3 |
| Klc4 | 3129 | 4428 | 4334 | 4848 | 4148 | 3994 | 3864 | 2630 | 2898 | 3184 | 2929 | 3358 | 3095 | 3318 | 3386 | 3514 | 4663 | 4478 | 4789 | 4008 | 4025 | 3003 | 2790 | 3521 | 3616 | 3177 | 3280 | 3262 | Klc4 |
| Klf1 | 53 | 82 | 67 | 44 | 31 | 44 | 30 | 26 | 35 | 51 | 61 | 58 | 93 | 60 | 61 | 77 | 74 | 45 | 65 | 48 | 33 | 20 | 38 | 39 | 22 | 78 | 62 | 128 | Klf1 |
| Klf10 | 1667 | 1661 | 1785 | 1588 | 1470 | 1497 | 1402 | 1566 | 1566 | 1575 | 1559 | 1350 | 1484 | 1426 | 1584 | 1729 | 1586 | 1669 | 1438 | 1433 | 1410 | 1491 | 1805 | 1480 | 1342 | 1688 | 1484 | 1508 | Klf10 |
| Klf11 | 1083 | 922 | 1150 | 1119 | 1163 | 1141 | 1045 | 1642 | 1874 | 1738 | 1687 | 1897 | 1861 | 1662 | 969 | 1014 | 1095 | 1174 | 1079 | 1086 | 998 | 1848 | 1818 | 1763 | 1843 | 1717 | 1743 | 1684 | Klf11 |
| Klf12 | 3682 | 3399 | 1284 | 3041 | 3781 | 4811 | 4051 | 6191 | 6012 | 4793 | 5807 | 5301 | 4883 | 5770 | 3649 | 3122 | 1845 | 3002 | 3129 | 4067 | 4571 | 4445 | 5535 | 3373 | 6211 | 1498 | 5733 | 4321 | Klf12 |
| Klf13 | 3598 | 4222 | 7039 | 6021 | 5799 | 4834 | 4852 | 3522 | 3794 | 4917 | 4674 | 5383 | 5302 | 5513 | 3497 | 4150 | 6431 | 5621 | 6241 | 4094 | 5149 | 4525 | 3508 | 5652 | 5579 | 5932 | 5039 | 5622 | Klf13 |
| Klf14 | 42 | 38 | 53 | 37 | 49 | 43 | 20 | 29 | 79 | 37 | 315 | 29 | 18 | 23 | 40 | 51 | 31 | 45 | 30 | 36 | 25 | 62 | 173 | 30 | 26 | 37 | 45 | 24 | Klf14 |
| Klf15 | 471 | 566 | 426 | 622 | 557 | 508 | 467 | 261 | 324 | 322 | 289 | 178 | 245 | 187 | 434 | 477 | 417 | 588 | 507 | 551 | 524 | 296 | 300 | 283 | 346 | 156 | 184 | 161 | Klf15 |
| Klf16 | 1691 | 1981 | 2064 | 2428 | 1818 | 1704 | 1630 | 783 | 924 | 961 | 1044 | 1033 | 911 | 1000 | 1439 | 1812 | 2085 | 2254 | 1943 | 1499 | 1742 | 866 | 892 | 1114 | 1081 | 1396 | 884 | 926 | Klf16 |
| Klf17 | 0 | 1 | 0 | 3 | 0 | 0 | 1 | 0 | 2 | 0 | 0 | 0 | 0 | 0 | 0 | 0 | 0 | 0 | 2 | 0 | 1 | 2 | 1 | 1 | 0 | 1 | 0 | 1 | Klf17 |
| Klf2 | 218 | 272 | 248 | 373 | 201 | 234 | 166 | 245 | 293 | 300 | 372 | 360 | 259 | 301 | 248 | 306 | 257 | 358 | 190 | 124 | 151 | 333 | 339 | 346 | 380 | 332 | 267 | 311 | Klf2 |
| Klf3 | 3007 | 2421 | 3285 | 2804 | 3003 | 3113 | 2841 | 3751 | 4057 | 4166 | 4929 | 4447 | 5319 | 4338 | 2888 | 3030 | 2992 | 3081 | 3019 | 2520 | 2878 | 3776 | 4301 | 4361 | 4126 | 4384 | 4250 | 4593 | Klf3 |
| Klf4 | 893 | 791 | 792 | 818 | 458 | 800 | 367 | 877 | 1351 | 1290 | 1584 | 784 | 1071 | 707 | 746 | 923 | 750 | 976 | 475 | 419 | 395 | 1074 | 1610 | 1114 | 1052 | 1063 | 795 | 718 | Klf4 |
| Klf5 | 475 | 344 | 548 | 499 | 381 | 398 | 359 | 369 | 540 | 396 | 576 | 232 | 238 | 212 | 335 | 407 | 428 | 568 | 408 | 302 | 375 | 317 | 681 | 385 | 230 | 287 | 245 | 201 | Klf5 |
| Klf6 | 5133 | 4933 | 5734 | 5248 | 4570 | 5150 | 4661 | 4951 | 4665 | 4525 | 5565 | 5229 | 6131 | 5417 | 5657 | 5852 | 4908 | 5681 | 4299 | 4610 | 4186 | 5467 | 5436 | 4938 | 4952 | 4735 | 5389 | 5510 | Klf6 |
| Klf7 | 5013 | 4761 | 4797 | 6595 | 7967 | 6628 | 7423 | 8096 | 7792 | 8311 | 7810 | 7417 | 7172 | 7878 | 5780 | 4408 | 5788 | 6549 | 6783 | 6674 | 6997 | 10696 | 7009 | 7903 | 9515 | 5507 | 6793 | 7037 | Klf7 |
| Klf8 | 1574 | 1211 | 1138 | 1325 | 1637 | 1637 | 1573 | 1425 | 1463 | 1362 | 1151 | 1200 | 1082 | 1299 | 1596 | 1328 | 1249 | 1473 | 1410 | 1632 | 1591 | 1604 | 1331 | 1118 | 1400 | 807 | 1288 | 1252 | Klf8 |
| Klf9 | 1376 | 1302 | 1387 | 1537 | 1004 | 1147 | 1181 | 335 | 401 | 355 | 347 | 238 | 254 | 215 | 1278 | 1392 | 1262 | 1457 | 1026 | 980 | 1168 | 322 | 442 | 305 | 272 | 303 | 272 | 236 | Klf9 |
| Klhdc1 | 181 | 139 | 134 | 241 | 319 | 285 | 292 | 366 | 445 | 351 | 348 | 273 | 243 | 318 | 222 | 168 | 165 | 215 | 263 | 284 | 307 | 403 | 444 | 378 | 355 | 256 | 261 | 261 | Klhdc1 |
| Klhdc10 | 19738 | 17062 | 17766 | 21839 | 24004 | 24258 | 23643 | 15687 | 15010 | 17158 | 19299 | 17373 | 14907 | 16820 | 18858 | 16678 | 19870 | 22518 | 23332 | 19142 | 24497 | 20229 | 15213 | 16114 | 19228 | 13527 | 16727 | 15702 | Klhdc10 |
| Klhdc2 | 10501 | 9599 | 10771 | 10951 | 12390 | 11962 | 11854 | 16967 | 13648 | 14018 | 12954 | 14963 | 14519 | 15580 | 11540 | 9442 | 10874 | 10638 | 12494 | 13519 | 12243 | 15421 | 13619 | 14996 | 16546 | 14895 | 15739 | 16008 | Klhdc2 |
| Klhdc3 | 8063 | 9562 | 9589 | 10564 | 9861 | 9667 | 9081 | 6359 | 7246 | 7458 | 7587 | 7716 | 7270 | 7337 | 7615 | 8294 | 9781 | 10210 | 10816 | 8314 | 9748 | 7454 | 7150 | 7996 | 8181 | 8321 | 6952 | 7151 | Klhdc3 |
| Klhdc4 | 2886 | 3664 | 4355 | 3940 | 3373 | 3369 | 3334 | 2352 | 2717 | 3177 | 2988 | 3681 | 3185 | 3343 | 3082 | 3248 | 4298 | 3650 | 4016 | 2937 | 3555 | 2637 | 2584 | 3310 | 3351 | 4139 | 3439 | 3674 | Klhdc4 |
| Klhdc7a | 58 | 41 | 53 | 26 | 99 | 45 | 57 | 200 | 261 | 317 | 217 | 307 | 266 | 213 | 44 | 40 | 60 | 42 | 84 | 84 | 70 | 266 | 446 | 219 | 240 | 187 | 242 | 197 | Klhdc7a |
| Klhdc7b | 0 | 0 | 4 | 1 | 2 | 2 | 0 | 0 | 0 | 0 | 5 | 1 | 5 | 0 | 0 | 1 | 3 | 2 | 7 | 1 | 2 | 2 | 1 | 0 | 9 | 2 | 1 | 2 | Klhdc7b |
| Klhdc8a | 307 | 364 | 575 | 459 | 525 | 435 | 463 | 580 | 500 | 625 | 445 | 466 | 606 | 477 | 360 | 324 | 548 | 383 | 574 | 440 | 465 | 714 | 417 | 431 | 599 | 363 | 655 | 600 | Klhdc8a |
| Klhdc8b | 3125 | 3821 | 3836 | 3944 | 4033 | 3893 | 3401 | 2567 | 2892 | 2858 | 2834 | 3511 | 2947 | 2886 | 2955 | 3302 | 4325 | 4034 | 4282 | 3876 | 3632 | 3237 | 3020 | 3263 | 3312 | 3554 | 3246 | 3030 | Klhdc8b |
| Klhdc9 | 116 | 117 | 153 | 119 | 131 | 147 | 123 | 57 | 130 | 136 | 61 | 151 | 104 | 123 | 117 | 83 | 147 | 145 | 136 | 185 | 127 | 85 | 113 | 88 | 101 | 124 | 98 | 100 | Klhdc9 |
| Klhl1 | 271 | 340 | 511 | 445 | 700 | 497 | 494 | 850 | 713 | 713 | 582 | 560 | 637 | 547 | 423 | 321 | 385 | 502 | 559 | 735 | 577 | 1066 | 717 | 553 | 781 | 206 | 693 | 637 | Klhl1 |
| Klhl10 | 37 | 27 | 35 | 48 | 42 | 23 | 26 | 64 | 70 | 90 | 85 | 55 | 65 | 61 | 24 | 38 | 30 | 34 | 30 | 33 | 28 | 62 | 70 | 54 | 41 | 63 | 78 | 62 | Klhl10 |
| Klhl11 | 2068 | 2232 | 482 | 1551 | 1511 | 1955 | 1635 | 2074 | 1844 | 1376 | 1395 | 1481 | 1267 | 1432 | 1900 | 1845 | 913 | 1473 | 1283 | 2002 | 1780 | 1105 | 1707 | 1035 | 1441 | 200 | 1577 | 1176 | Klhl11 |
| Klhl12 | 4587 | 4103 | 4231 | 4270 | 4139 | 4286 | 4330 | 3690 | 3522 | 3622 | 3617 | 3809 | 3841 | 3976 | 4498 | 4172 | 4194 | 4402 | 4087 | 3973 | 4254 | 3798 | 3432 | 3771 | 3960 | 3718 | 3624 | 3915 | Klhl12 |
| Klhl13 | 7751 | 6761 | 6955 | 6211 | 7419 | 7161 | 8210 | 12611 | 9313 | 8826 | 9751 | 9606 | 8828 | 10355 | 8550 | 6218 | 6672 | 6333 | 7455 | 8432 | 7879 | 11976 | 10223 | 9511 | 9725 | 8969 | 10717 | 10631 | Klhl13 |
| Klhl14 | 189 | 170 | 344 | 201 | 476 | 180 | 168 | 266 | 267 | 399 | 364 | 220 | 348 | 147 | 273 | 248 | 308 | 193 | 272 | 197 | 217 | 309 | 250 | 203 | 214 | 164 | 221 | 184 | Klhl14 |
| Klhl15 | 1895 | 1367 | 1968 | 2361 | 1920 | 1940 | 2062 | 1401 | 1048 | 1081 | 1229 | 1348 | 1213 | 1321 | 1886 | 1720 | 2070 | 2102 | 1816 | 1640 | 1941 | 1399 | 1082 | 1174 | 1200 | 1258 | 1403 | 1538 | Klhl15 |
| Klhl18 | 1615 | 1868 | 1785 | 1900 | 2522 | 2281 | 2042 | 1928 | 2162 | 2597 | 2422 | 2184 | 2161 | 2081 | 1647 | 1854 | 1943 | 2283 | 2242 | 1895 | 2134 | 2765 | 2245 | 2069 | 2581 | 1836 | 2020 | 2035 | Klhl18 |
| Klhl2 | 3031 | 3076 | 3387 | 3858 | 4364 | 4094 | 4320 | 3348 | 2984 | 3191 | 2986 | 3163 | 2565 | 3195 | 3268 | 2746 | 3748 | 3858 | 4146 | 4273 | 4387 | 3695 | 2969 | 3319 | 3637 | 3243 | 2901 | 2967 | Klhl2 |
| Klhl20 | 2955 | 2601 | 2560 | 2666 | 2767 | 2702 | 2825 | 2977 | 2876 | 2730 | 2971 | 3095 | 2905 | 3073 | 2879 | 2555 | 2700 | 2779 | 2558 | 2620 | 2807 | 3023 | 2739 | 2848 | 3023 | 2681 | 3133 | 3019 | Klhl20 |
| Klhl21 | 1269 | 1697 | 1853 | 1531 | 1087 | 1162 | 911 | 480 | 597 | 599 | 979 | 600 | 747 | 511 | 1130 | 1710 | 1461 | 1644 | 1212 | 988 | 1034 | 555 | 776 | 668 | 511 | 829 | 751 | 532 | Klhl21 |
| Klhl22 | 4932 | 5053 | 4529 | 4747 | 4703 | 5006 | 4853 | 5256 | 5787 | 5163 | 5927 | 4604 | 4835 | 4625 | 4898 | 5045 | 4493 | 4626 | 4918 | 4799 | 4889 | 4769 | 5827 | 5108 | 5231 | 4640 | 4871 | 4899 | Klhl22 |
| Klhl23 | 3924 | 3107 | 3389 | 3496 | 5518 | 4883 | 5377 | 8462 | 7225 | 7615 | 7012 | 7140 | 6290 | 7647 | 4094 | 3284 | 3552 | 4056 | 4711 | 4938 | 5012 | 9307 | 7451 | 7162 | 8278 | 6638 | 6957 | 7355 | Klhl23 |
| Klhl24 | 8992 | 7710 | 5852 | 8281 | 9524 | 9536 | 9833 | 18997 | 14455 | 13805 | 13144 | 13891 | 13053 | 15032 | 9570 | 7657 | 7176 | 8319 | 9149 | 10057 | 9991 | 14953 | 14408 | 12720 | 15632 | 9594 | 14141 | 13513 | Klhl24 |
| Klhl25 | 4302 | 5642 | 4123 | 5121 | 4585 | 4944 | 4469 | 2480 | 2553 | 2252 | 2165 | 2353 | 2259 | 2192 | 4399 | 4530 | 4561 | 5205 | 5527 | 4700 | 4986 | 2356 | 2776 | 2445 | 2538 | 2498 | 2303 | 2267 | Klhl25 |
| Klhl26 | 2207 | 2372 | 2368 | 2063 | 2108 | 2069 | 1836 | 1218 | 1453 | 1346 | 1449 | 1303 | 1272 | 1161 | 1890 | 2280 | 2189 | 2122 | 2130 | 1781 | 1818 | 1285 | 1342 | 1392 | 1222 | 1483 | 1127 | 1120 | Klhl26 |
| Klhl28 | 7314 | 6837 | 5285 | 8000 | 6129 | 7480 | 7240 | 6135 | 4636 | 3789 | 4424 | 4353 | 4628 | 4576 | 8466 | 8020 | 5835 | 8030 | 6212 | 8234 | 7422 | 4786 | 4798 | 4333 | 4629 | 3026 | 4618 | 4476 | Klhl28 |
| Klhl29 | 2640 | 3085 | 3073 | 4022 | 4892 | 4174 | 4158 | 2678 | 3275 | 4065 | 3634 | 3369 | 3148 | 3497 | 2860 | 2843 | 3612 | 4377 | 4532 | 3590 | 4552 | 4236 | 3071 | 3667 | 4425 | 2764 | 2875 | 2910 | Klhl29 |
| Klhl3 | 316 | 406 | 451 | 537 | 992 | 635 | 662 | 1039 | 1104 | 1394 | 1480 | 1240 | 1212 | 1243 | 480 | 375 | 542 | 574 | 697 | 607 | 662 | 1614 | 1139 | 1320 | 1485 | 883 | 991 | 1069 | Klhl3 |
| Klhl30 | 12 | 13 | 10 | 15 | 11 | 7 | 6 | 16 | 39 | 21 | 90 | 14 | 19 | 7 | 7 | 9 | 7 | 23 | 4 | 7 | 4 | 23 | 80 | 11 | 16 | 9 | 9 | 9 | Klhl30 |
| Klhl31 | 12 | 5 | 15 | 2 | 4 | 1 | 2 | 31 | 54 | 31 | 339 | 5 | 39 | 15 | 20 | 18 | 1 | 1 | 3 | 4 | 2 | 50 | 201 | 36 | 11 | 20 | 5 | 4 | Klhl31 |
| Klhl32 | 1313 | 1393 | 1022 | 1782 | 1956 | 1660 | 1893 | 921 | 814 | 942 | 723 | 779 | 737 | 773 | 1538 | 1216 | 1477 | 1806 | 1705 | 1920 | 1935 | 979 | 878 | 704 | 966 | 612 | 704 | 710 | Klhl32 |
| Klhl33 | 23 | 30 | 49 | 46 | 124 | 52 | 47 | 77 | 60 | 76 | 102 | 43 | 68 | 66 | 22 | 21 | 51 | 58 | 44 | 65 | 52 | 117 | 64 | 68 | 69 | 40 | 70 | 49 | Klhl33 |
| Klhl34 | 3 | 0 | 6 | 4 | 1 | 1 | 0 | 2 | 5 | 1 | 5 | 5 | 12 | 7 | 4 | 5 | 0 | 0 | 3 | 0 | 0 | 2 | 5 | 3 | 4 | 8 | 0 | 6 | Klhl34 |
| Klhl35 | 50 | 106 | 99 | 57 | 92 | 103 | 57 | 429 | 311 | 511 | 393 | 638 | 912 | 619 | 65 | 70 | 144 | 70 | 79 | 68 | 59 | 645 | 394 | 441 | 569 | 416 | 889 | 782 | Klhl35 |
| Klhl36 | 847 | 1342 | 1065 | 1176 | 1071 | 1071 | 938 | 331 | 432 | 416 | 401 | 441 | 396 | 358 | 938 | 992 | 1348 | 1166 | 1188 | 988 | 962 | 403 | 467 | 433 | 426 | 411 | 417 | 405 | Klhl36 |
| Klhl4 | 662 | 418 | 425 | 300 | 526 | 545 | 408 | 1014 | 1102 | 660 | 1112 | 607 | 763 | 548 | 645 | 551 | 315 | 419 | 431 | 481 | 485 | 902 | 1409 | 678 | 617 | 583 | 661 | 506 | Klhl4 |
| Klhl40 | 16 | 11 | 8 | 9 | 28 | 19 | 15 | 13 | 50 | 18 | 334 | 17 | 20 | 11 | 7 | 17 | 13 | 9 | 16 | 10 | 19 | 40 | 245 | 14 | 14 | 8 | 5 | 11 | Klhl40 |
| Klhl41 | 120 | 77 | 67 | 87 | 77 | 95 | 78 | 118 | 252 | 36 | 1549 | 35 | 79 | 53 | 74 | 126 | 74 | 87 | 71 | 60 | 100 | 222 | 928 | 51 | 51 | 38 | 62 | 49 | Klhl41 |
| Klhl42 | 1804 | 1486 | 1752 | 1505 | 1777 | 1806 | 1587 | 2394 | 2272 | 2131 | 2331 | 1997 | 2095 | 2019 | 1614 | 1710 | 1708 | 1388 | 1714 | 1554 | 1572 | 2393 | 2271 | 1985 | 2128 | 1872 | 2000 | 1996 | Klhl42 |
| Klhl5 | 5422 | 5534 | 5254 | 5821 | 6843 | 6693 | 7119 | 5081 | 4813 | 5034 | 4497 | 5090 | 4454 | 5019 | 5556 | 5354 | 5852 | 6855 | 6864 | 6627 | 6748 | 5167 | 5031 | 4903 | 5833 | 5310 | 4573 | 4593 | Klhl5 |
| Klhl6 | 375 | 374 | 437 | 244 | 331 | 291 | 255 | 459 | 412 | 410 | 480 | 431 | 506 | 459 | 356 | 328 | 301 | 300 | 254 | 346 | 256 | 393 | 516 | 379 | 345 | 559 | 487 | 433 | Klhl6 |
| Klhl7 | 16031 | 13653 | 13144 | 15281 | 16908 | 17212 | 19227 | 18678 | 14865 | 13472 | 12149 | 14688 | 12250 | 16040 | 18076 | 13858 | 14484 | 15254 | 16670 | 20164 | 18445 | 15563 | 14397 | 14831 | 16228 | 13936 | 14627 | 15116 | Klhl7 |
| Klhl8 | 1564 | 1678 | 1480 | 2007 | 2110 | 1856 | 1981 | 1518 | 1753 | 1714 | 1384 | 1322 | 1132 | 1284 | 1720 | 1515 | 1671 | 1951 | 2126 | 2053 | 2001 | 1733 | 1660 | 1469 | 1704 | 1254 | 1145 | 1050 | Klhl8 |
| Klhl9 | 16253 | 13996 | 14473 | 14403 | 13990 | 14107 | 15552 | 19342 | 16876 | 13935 | 14846 | 15146 | 15222 | 16239 | 17318 | 14971 | 14936 | 14544 | 14314 | 17194 | 15332 | 16516 | 17118 | 15128 | 15113 | 16103 | 16343 | 17068 | Klhl9 |
| Klk1 | 0 | 0 | 0 | 0 | 0 | 0 | 0 | 0 | 4 | 0 | 0 | 0 | 0 | 2 | 0 | 0 | 0 | 0 | 0 | 0 | 0 | 0 | 1 | 0 | 0 | 0 | 0 | 0 | Klk1 |
| Klk10 | 1 | 3 | 6 | 3 | 2 | 0 | 0 | 4 | 18 | 5 | 14 | 4 | 2 | 6 | 0 | 3 | 0 | 0 | 4 | 4 | 0 | 7 | 8 | 3 | 8 | 3 | 8 | 5 | Klk10 |
| Klk11 | 1 | 0 | 0 | 0 | 0 | 0 | 0 | 15 | 8 | 8 | 5 | 0 | 0 | 3 | 0 | 1 | 0 | 0 | 4 | 0 | 0 | 0 | 12 | 1 | 0 | 0 | 3 | 0 | Klk11 |
| Klk12 | 0 | 0 | 0 | 0 | 0 | 0 | 0 | 4 | 1 | 0 | 0 | 0 | 0 | 1 | 0 | 0 | 0 | 0 | 0 | 0 | 0 | 0 | 1 | 5 | 0 | 0 | 0 | 0 | Klk12 |
| Klk13 | 12 | 8 | 6 | 3 | 11 | 7 | 2 | 11 | 24 | 11 | 3 | 8 | 0 | 2 | 8 | 9 | 7 | 7 | 2 | 2 | 3 | 2 | 15 | 3 | 5 | 1 | 7 | 4 | Klk13 |
| Klk14 | 7 | 15 | 9 | 4 | 11 | 11 | 11 | 2 | 6 | 5 | 2 | 4 | 5 | 7 | 8 | 4 | 10 | 4 | 9 | 7 | 5 | 3 | 5 | 4 | 0 | 2 | 3 | 5 | Klk14 |
| Klk1b24 | 0 | 6 | 3 | 1 | 0 | 2 | 0 | 0 | 1 | 0 | 3 | 5 | 1 | 1 | 1 | 2 | 5 | 3 | 2 | 1 | 1 | 0 | 1 | 1 | 4 | 7 | 0 | 1 | Klk1b24 |
| Klk1b26 | 0 | 0 | 0 | 0 | 0 | 0 | 0 | 0 | 1 | 0 | 4 | 0 | 0 | 0 | 0 | 0 | 0 | 0 | 0 | 0 | 0 | 0 | 3 | 0 | 0 | 0 | 0 | 0 | Klk1b26 |
| Klk1b27 | 0 | 0 | 0 | 0 | 0 | 0 | 0 | 0 | 1 | 0 | 0 | 0 | 0 | 0 | 0 | 0 | 0 | 0 | 0 | 0 | 0 | 2 | 2 | 0 | 0 | 0 | 0 | 0 | Klk1b27 |
| Klk1b3 | 0 | 0 | 0 | 0 | 0 | 0 | 0 | 0 | 0 | 1 | 0 | 0 | 0 | 0 | 0 | 0 | 0 | 0 | 0 | 0 | 0 | 2 | 0 | 0 | 3 | 0 | 0 | 1 | Klk1b3 |

| | | | | | | | | | | | | | | | | | | | | | | | | | | | | | |
|---|---|---|---|---|---|---|---|---|---|---|---|---|---|---|---|---|---|---|---|---|---|---|---|---|---|---|---|---|---|
| Klk1b5 | 0 | 0 | 0 | 0 | 0 | 0 | 0 | 0 | 0 | 0 | 0 | 0 | 0 | 0 | 0 | 0 | 0 | 3 | 2 | 0 | 0 | 0 | 0 | 0 | 0 | 0 | 0 | 0 | Klk1b5 |
| Klk1b8 | 0 | 0 | 0 | 0 | 0 | 0 | 0 | 0 | 0 | 0 | 0 | 0 | 0 | 0 | 0 | 0 | 0 | 0 | 0 | 0 | 0 | 0 | 0 | 0 | 0 | 3 | 0 | 0 | Klk1b8 |
| Klk4 | 3 | 12 | 0 | 0 | 6 | 8 | 0 | 9 | 7 | 5 | 3 | 14 | 9 | 0 | 8 | 4 | 2 | 0 | 0 | 0 | 0 | 17 | 30 | 0 | 2 | 2 | 0 | 1 | Klk4 |
| Klk5 | 5 | 13 | 1 | 0 | 0 | 7 | 0 | 5 | 12 | 4 | 0 | 4 | 12 | 6 | 3 | 7 | 0 | 4 | 0 | 0 | 0 | 0 | 8 | 5 | 0 | 0 | 10 | 12 | Klk5 |
| Klk6 | 0 | 0 | 0 | 0 | 0 | 0 | 0 | 0 | 0 | 0 | 0 | 0 | 0 | 0 | 0 | 0 | 0 | 0 | 0 | 0 | 0 | 0 | 1 | 2 | 0 | 0 | 0 | 0 | Klk6 |
| Klk7 | 9 | 7 | 19 | 2 | 9 | 12 | 2 | 0 | 8 | 7 | 3 | 4 | 2 | 10 | 4 | 9 | 1 | 11 | 12 | 0 | 8 | 2 | 8 | 1 | 7 | 8 | 5 | 0 | Klk7 |
| Klk8 | 233 | 182 | 191 | 131 | 154 | 148 | 123 | 217 | 227 | 192 | 207 | 198 | 226 | 158 | 140 | 222 | 146 | 118 | 118 | 101 | 95 | 135 | 297 | 149 | 99 | 224 | 170 | 170 | Klk8 |
| Klk9 | 1 | 0 | 0 | 0 | 0 | 1 | 0 | 0 | 2 | 2 | 0 | 0 | 0 | 0 | 0 | 2 | 0 | 0 | 0 | 0 | 2 | 0 | 0 | 0 | 0 | 0 | 0 | 0 | Klk9 |
| Klkb1 | 13 | 11 | 16 | 15 | 9 | 10 | 8 | 6 | 6 | 25 | 2 | 4 | 9 | 11 | 8 | 12 | 28 | 19 | 5 | 8 | 7 | 3 | 9 | 11 | 13 | 8 | 6 | 4 | Klkb1 |
| Klra17 | 0 | 0 | 0 | 0 | 0 | 0 | 0 | 0 | 0 | 0 | 0 | 0 | 2 | 0 | 0 | 0 | 0 | 0 | 0 | 0 | 0 | 0 | 0 | 0 | 0 | 0 | 0 | 0 | Klra17 |
| Klra2 | 12 | 3 | 1 | 9 | 20 | 16 | 16 | 11 | 10 | 12 | 21 | 5 | 5 | 2 | 5 | 6 | 10 | 13 | 19 | 15 | 12 | 17 | 6 | 3 | 16 | 6 | 2 | 11 | Klra2 |
| Klra9 | 0 | 0 | 0 | 0 | 0 | 0 | 0 | 0 | 0 | 0 | 0 | 0 | 0 | 0 | 0 | 0 | 0 | 0 | 0 | 0 | 0 | 0 | 1 | 0 | 0 | 0 | 0 | 0 | Klra9 |
| Klrb1 | 0 | 1 | 0 | 0 | 0 | 0 | 0 | 4 | 1 | 4 | 2 | 0 | 0 | 0 | 0 | 0 | 0 | 0 | 5 | 0 | 0 | 0 | 1 | 0 | 3 | 0 | 1 | 8 | Klrb1 |
| Klrb1a | 0 | 1 | 2 | 0 | 0 | 0 | 0 | 1 | 1 | 0 | 0 | 0 | 0 | 0 | 0 | 0 | 0 | 0 | 0 | 0 | 0 | 0 | 0 | 0 | 0 | 0 | 0 | 0 | Klrb1a |
| Klrb1b | 0 | 0 | 0 | 0 | 0 | 0 | 0 | 0 | 2 | 0 | 5 | 0 | 0 | 0 | 0 | 2 | 2 | 2 | 0 | 0 | 0 | 0 | 1 | 2 | 0 | 1 | 0 | 1 | Klrb1b |
| Klrb1c | 0 | 0 | 0 | 0 | 2 | 0 | 0 | 1 | 2 | 1 | 5 | 0 | 2 | 4 | 1 | 0 | 0 | 3 | 1 | 0 | 0 | 3 | 1 | 3 | 0 | 0 | 0 | 1 | Klrb1c |
| Klrb1f | 7 | 5 | 6 | 4 | 15 | 2 | 2 | 11 | 18 | 2 | 5 | 7 | 8 | 5 | 13 | 8 | 0 | 8 | 5 | 12 | 6 | 8 | 12 | 3 | 13 | 9 | 5 | 9 | Klrb1f |
| Klrc1 | 0 | 0 | 0 | 0 | 0 | 0 | 0 | 0 | 1 | 0 | 0 | 0 | 0 | 0 | 0 | 0 | 0 | 0 | 0 | 0 | 0 | 0 | 1 | 0 | 0 | 0 | 0 | 0 | Klrc1 |
| Klrd1 | 10 | 9 | 14 | 12 | 7 | 16 | 19 | 5 | 3 | 9 | 15 | 2 | 1 | 0 | 8 | 5 | 8 | 10 | 16 | 4 | 8 | 3 | 7 | 6 | 0 | 3 | 2 | 6 | Klrd1 |
| Klre1 | 43 | 35 | 57 | 71 | 39 | 66 | 72 | 13 | 8 | 12 | 21 | 18 | 10 | 7 | 40 | 68 | 45 | 55 | 47 | 45 | 57 | 3 | 8 | 16 | 10 | 14 | 12 | 7 | Klre1 |
| Klrg1 | 7 | 6 | 1 | 0 | 3 | 12 | 8 | 21 | 37 | 9 | 10 | 15 | 24 | 20 | 12 | 7 | 6 | 7 | 13 | 19 | 10 | 13 | 25 | 15 | 32 | 12 | 14 | 7 | Klrg1 |
| Klrg2 | 126 | 108 | 40 | 65 | 70 | 58 | 49 | 13 | 16 | 16 | 13 | 4 | 15 | 5 | 99 | 115 | 81 | 116 | 51 | 53 | 53 | 5 | 27 | 11 | 8 | 3 | 12 | 6 | Klrg2 |
| Klri1 | 0 | 0 | 0 | 0 | 0 | 0 | 0 | 2 | 1 | 0 | 2 | 1 | 2 | 1 | 3 | 2 | 0 | 1 | 0 | 2 | 1 | 7 | 3 | 3 | 0 | 2 | 0 | 2 | Klri1 |
| Klri2 | 0 | 0 | 0 | 0 | 0 | 0 | 0 | 1 | 0 | 0 | 3 | 0 | 0 | 0 | 0 | 0 | 0 | 0 | 0 | 0 | 0 | 0 | 1 | 0 | 0 | 2 | 0 | 0 | Klri2 |
| Klrk1 | 0 | 0 | 0 | 0 | 0 | 0 | 0 | 0 | 1 | 0 | 0 | 0 | 0 | 0 | 0 | 0 | 0 | 2 | 0 | 0 | 0 | 0 | 1 | 1 | 0 | 0 | 0 | 0 | Klrk1 |
| Kmo | 3 | 1 | 10 | 9 | 0 | 1 | 0 | 9 | 18 | 15 | 9 | 7 | 5 | 9 | 3 | 3 | 3 | 3 | 1 | 2 | 0 | 10 | 17 | 6 | 7 | 11 | 9 | 2 | Kmo |
| Kmt2a | 16970 | 18537 | 11930 | 17580 | 21140 | 23692 | 21730 | 17804 | 19106 | 19297 | 20837 | 18970 | 18356 | 19821 | 15728 | 17763 | 14371 | 17750 | 18800 | 19218 | 25062 | 16330 | 16378 | 14823 | 23550 | 11219 | 21910 | 16636 | Kmt2a |
| Kmt2b | 7109 | 8542 | 9856 | 10885 | 10725 | 10056 | 9664 | 8475 | 9977 | 11751 | 11114 | 13912 | 12026 | 12759 | 6327 | 7556 | 10754 | 10762 | 11809 | 7775 | 9837 | 12171 | 8712 | 13089 | 12935 | 9814 | 11723 | 12087 | Kmt2b |
| Kmt2c | 13767 | 13118 | 7080 | 10977 | 12188 | 16220 | 14214 | 14520 | 14760 | 11068 | 12198 | 10865 | 10841 | 12005 | 13439 | 12758 | 8778 | 10989 | 11686 | 15892 | 16169 | 10581 | 12578 | 9519 | 13273 | 7385 | 13323 | 10026 | Kmt2c |
| Kmt2d | 8588 | 11579 | 15472 | 17315 | 16737 | 14457 | 13396 | 9739 | 11726 | 16079 | 14887 | 17552 | 14503 | 15323 | 7675 | 9968 | 17117 | 17032 | 17836 | 10968 | 14536 | 16068 | 10363 | 18146 | 17587 | 12501 | 14794 | 15278 | Kmt2d |
| Kmt2e | 13268 | 11689 | 14761 | 16595 | 18843 | 17843 | 16920 | 20353 | 18845 | 21536 | 20588 | 22458 | 20632 | 21707 | 13500 | 11562 | 15966 | 15685 | 17944 | 14833 | 17832 | 24922 | 17513 | 21632 | 24111 | 18190 | 19926 | 20055 | Kmt2e |
| Kmt5a | 6061 | 6286 | 8978 | 7253 | 6120 | 6044 | 6461 | 5097 | 4837 | 6097 | 5423 | 7436 | 6217 | 7084 | 5770 | 6200 | 8124 | 7043 | 7049 | 5140 | 6285 | 5569 | 4747 | 7437 | 6062 | 8578 | 6364 | 7665 | Kmt5a |
| Kmt5b | 1260 | 1189 | 1518 | 1534 | 2706 | 2091 | 2062 | 11616 | 10796 | 11040 | 11501 | 11989 | 10817 | 12034 | 1440 | 1091 | 1644 | 1405 | 2138 | 2212 | 2057 | 12791 | 10306 | 11231 | 12847 | 8871 | 11681 | 11333 | Kmt5b |
| Kmt5c | 4773 | 6229 | 6872 | 7222 | 6343 | 5687 | 5391 | 4300 | 4703 | 5941 | 5326 | 6422 | 5517 | 5968 | 4548 | 5597 | 7434 | 6512 | 6791 | 5036 | 5476 | 5629 | 4394 | 6478 | 5779 | 6863 | 6062 | 5790 | Kmt5c |
| Kncn | 27 | 54 | 44 | 60 | 43 | 44 | 42 | 4 | 2 | 8 | 1 | 2 | 6 | 7 | 29 | 39 | 53 | 78 | 72 | 35 | 86 | 8 | 6 | 6 | 1 | 2 | 1 | 5 | Kncn |
| Kndc1 | 358 | 564 | 341 | 1155 | 2609 | 1506 | 1513 | 1233 | 1661 | 2244 | 1668 | 880 | 852 | 789 | 537 | 533 | 760 | 1538 | 1859 | 1427 | 1793 | 2839 | 1519 | 1302 | 1985 | 377 | 531 | 559 | Kndc1 |
| Kng1 | 0 | 0 | 0 | 0 | 1 | 0 | 0 | 0 | 2 | 2 | 2 | 0 | 0 | 0 | 0 | 2 | 0 | 1 | 0 | 0 | 0 | 0 | 7 | 0 | 0 | 0 | 0 | 0 | Kng1 |
| Kng2 | 0 | 1 | 0 | 0 | 0 | 0 | 0 | 1 | 2 | 5 | 1 | 0 | 0 | 0 | 0 | 2 | 0 | 0 | 0 | 0 | 0 | 0 | 3 | 1 | 0 | 0 | 0 | 0 | Kng2 |
| Knl1 | 6667 | 4352 | 3699 | 3840 | 3144 | 4516 | 4714 | 6587 | 5629 | 4349 | 4142 | 6969 | 5916 | 7276 | 5789 | 5024 | 3661 | 3291 | 3899 | 5307 | 4578 | 3227 | 4699 | 4980 | 4952 | 4588 | 7525 | 6670 | Knl1 |
| Knop1 | 21980 | 22615 | 20197 | 18910 | 15632 | 17640 | 18672 | 12901 | 11382 | 11398 | 10584 | 13416 | 12780 | 13747 | 22101 | 20187 | 20182 | 18493 | 16325 | 18078 | 17432 | 10938 | 11344 | 12243 | 11285 | 13356 | 13702 | 14250 | Knop1 |
| Knstrn | 6713 | 6279 | 6071 | 4794 | 3571 | 4413 | 5011 | 5558 | 4915 | 4777 | 4080 | 7500 | 5907 | 7714 | 6483 | 5650 | 5573 | 4081 | 4488 | 5472 | 4592 | 3601 | 4268 | 6137 | 5035 | 8605 | 7910 | 7838 | Knstrn |
| Kntc1 | 8463 | 7548 | 4865 | 4952 | 4002 | 5490 | 6049 | 5539 | 5257 | 3835 | 4000 | 6175 | 5211 | 6168 | 7108 | 6760 | 5092 | 4518 | 5033 | 6459 | 5684 | 2984 | 4467 | 4131 | 4130 | 4049 | 6529 | 5823 | Kntc1 |
| Kpna1 | 11592 | 9979 | 9109 | 9519 | 10481 | 11344 | 12364 | 13814 | 10908 | 9682 | 9257 | 10982 | 10198 | 11239 | 12498 | 9550 | 9301 | 10095 | 10347 | 12330 | 11483 | 11707 | 10656 | 9660 | 10953 | 9814 | 11301 | 11445 | Kpna1 |
| Kpna2 | 26843 | 26388 | 25858 | 21190 | 16450 | 17570 | 21893 | 21670 | 17378 | 15559 | 14249 | 24518 | 20449 | 24955 | 27221 | 24260 | 23255 | 18742 | 19529 | 25147 | 19200 | 14369 | 16481 | 19874 | 16977 | 29953 | 25946 | 26520 | Kpna2 |
| Kpna3 | 18191 | 14602 | 15276 | 15012 | 14406 | 15163 | 17159 | 15234 | 12080 | 10999 | 10867 | 12854 | 11764 | 12930 | 18937 | 16406 | 14275 | 14861 | 13866 | 17765 | 15885 | 12554 | 12090 | 11471 | 12298 | 12816 | 13058 | 13325 | Kpna3 |
| Kpna4 | 16295 | 13470 | 15918 | 16810 | 16138 | 16248 | 18261 | 14660 | 12124 | 12325 | 12034 | 14411 | 13614 | 15027 | 16384 | 14143 | 15842 | 16315 | 16370 | 15398 | 17933 | 13760 | 12295 | 12988 | 13316 | 13965 | 14238 | 15349 | Kpna4 |
| Kpna6 | 8498 | 8593 | 7368 | 7970 | 7977 | 7921 | 8243 | 7343 | 6628 | 6203 | 6296 | 6200 | 6081 | 6920 | 8511 | 7784 | 7669 | 8135 | 7663 | 7588 | 8565 | 7279 | 6741 | 6475 | 6788 | 5718 | 6307 | 6764 | Kpna6 |
| Kpna7 | 3 | 0 | 0 | 0 | 0 | 0 | 1 | 1 | 1 | 0 | 1 | 2 | 5 | 4 | 0 | 0 | 0 | 0 | 0 | 3 | 0 | 0 | 0 | 3 | 0 | 0 | 3 | 0 | Kpna7 |
| Kpnb1 | 37850 | 33468 | 43681 | 33653 | 32100 | 33779 | 34306 | 49010 | 44801 | 45822 | 45031 | 61035 | 57671 | 62532 | 36076 | 32376 | 39273 | 31711 | 35051 | 30022 | 33530 | 46170 | 42887 | 52103 | 48640 | 64019 | 64632 | 66351 | Kpnb1 |
| Kprp | 0 | 0 | 0 | 0 | 0 | 0 | 0 | 0 | 0 | 0 | 2 | 0 | 0 | 0 | 0 | 0 | 0 | 0 | 0 | 0 | 0 | 0 | 1 | 0 | 0 | 0 | 0 | 0 | Kprp |
| Kptn | 536 | 662 | 621 | 639 | 641 | 672 | 646 | 1088 | 1332 | 1380 | 1350 | 1263 | 1205 | 1191 | 593 | 544 | 575 | 612 | 696 | 491 | 635 | 1310 | 1199 | 1267 | 1321 | 1378 | 1228 | 1270 | Kptn |
| Kras | 18728 | 13439 | 17801 | 18463 | 19338 | 20356 | 22070 | 14652 | 12086 | 12735 | 12571 | 14329 | 13237 | 15031 | 18581 | 15990 | 18208 | 18352 | 18936 | 18816 | 21173 | 15107 | 11835 | 13626 | 14867 | 14475 | 13927 | 14452 | Kras |
| Krba1 | 5277 | 5307 | 3792 | 5369 | 6536 | 6157 | 5566 | 6344 | 6935 | 6797 | 6949 | 6345 | 6520 | 5930 | 4990 | 4833 | 4398 | 5240 | 5807 | 5701 | 5932 | 6293 | 6414 | 5512 | 7335 | 4324 | 6319 | 5375 | Krba1 |
| Krcc1 | 2015 | 1467 | 2041 | 1472 | 1406 | 1378 | 1463 | 2010 | 1729 | 1817 | 1918 | 1946 | 2110 | 1959 | 2007 | 1745 | 1766 | 1499 | 1428 | 1538 | 1335 | 1739 | 2072 | 2021 | 1592 | 2332 | 2083 | 2186 | Krcc1 |
| Kremen1 | 5980 | 6709 | 6797 | 6372 | 5696 | 6154 | 5297 | 2085 | 2612 | 2358 | 3141 | 2512 | 2892 | 2243 | 5434 | 6974 | 6495 | 6825 | 5956 | 4872 | 5345 | 2104 | 3007 | 2516 | 2017 | 2578 | 2460 | 2355 | Kremen1 |
| Kremen2 | 253 | 178 | 240 | 157 | 151 | 186 | 89 | 259 | 538 | 218 | 490 | 109 | 215 | 98 | 117 | 222 | 210 | 135 | 150 | 124 | 87 | 70 | 603 | 210 | 91 | 89 | 112 | 108 | Kremen2 |
| Kri1 | 3257 | 4009 | 3952 | 3978 | 3157 | 3418 | 3539 | 2594 | 2753 | 3201 | 3034 | 3299 | 3105 | 3402 | 3235 | 3379 | 3800 | 3685 | 3573 | 3258 | 3390 | 2881 | 2706 | 3362 | 3276 | 3600 | 3535 | 3376 | Kri1 |
| Krit1 | 10250 | 8532 | 8106 | 8761 | 9280 | 10103 | 11165 | 10083 | 8165 | 7601 | 8023 | 8380 | 8437 | 9208 | 10557 | 9340 | 8844 | 9347 | 9193 | 10946 | 10891 | 8682 | 7747 | 7885 | 8640 | 7053 | 8627 | 8967 | Krit1 |
| Krr1 | 6292 | 5507 | 5828 | 5634 | 4760 | 4907 | 5614 | 5751 | 5047 | 4491 | 4673 | 5428 | 5036 | 5679 | 6418 | 5713 | 5454 | 5208 | 4959 | 6390 | 5444 | 4742 | 4860 | 5211 | 5049 | 5749 | 5453 | 5813 | Krr1 |
| Krt1 | 30 | 45 | 47 | 56 | 85 | 85 | 69 | 172 | 499 | 375 | 125 | 126 | 81 | 98 | 42 | 37 | 46 | 92 | 72 | 66 | 91 | 137 | 652 | 143 | 145 | 136 | 87 | 101 | Krt1 |
| Krt10 | 133 | 120 | 173 | 165 | 174 | 162 | 209 | 331 | 533 | 686 | 342 | 375 | 353 | 494 | 144 | 76 | 148 | 167 | 139 | 206 | 184 | 244 | 527 | 397 | 371 | 435 | 341 | 456 | Krt10 |
| Krt12 | 0 | 1 | 0 | 1 | 0 | 0 | 1 | 0 | 1 | 0 | 1 | 0 | 0 | 0 | 1 | 0 | 0 | 0 | 0 | 0 | 1 | 2 | 3 | 0 | 2 | 0 | 0 | 0 | Krt12 |
| Krt13 | 0 | 0 | 0 | 0 | 0 | 0 | 0 | 1 | 2 | 0 | 2 | 0 | 0 | 0 | 0 | 0 | 0 | 0 | 0 | 1 | 0 | 2 | 8 | 1 | 1 | 0 | 1 | 0 | Krt13 |
| Krt14 | 427 | 158 | 66 | 11 | 7 | 376 | 4 | 1695 | 3404 | 1549 | 1245 | 338 | 625 | 464 | 149 | 109 | 45 | 271 | 76 | 18 | 0 | 33 | 3862 | 1325 | 263 | 284 | 358 | 438 | Krt14 |
| Krt15 | 322 | 259 | 177 | 10 | 19 | 405 | 3 | 713 | 2174 | 875 | 1671 | 462 | 610 | 445 | 302 | 199 | 76 | 446 | 57 | 22 | 2 | 48 | 2811 | 832 | 235 | 94 | 282 | 475 | Krt15 |
| Krt16 | 0 | 0 | 0 | 0 | 0 | 0 | 0 | 0 | 4 | 3 | 0 | 0 | 0 | 1 | 0 | 0 | 0 | 0 | 0 | 0 | 0 | 0 | 4 | 0 | 0 | 0 | 0 | 0 | Krt16 |
| Krt17 | 54 | 18 | 2 | 8 | 6 | 100 | 6 | 283 | 756 | 399 | 294 | 47 | 120 | 61 | 26 | 24 | 12 | 72 | 30 | 4 | 5 | 47 | 800 | 192 | 43 | 37 | 74 | 44 | Krt17 |
| Krt18 | 1102 | 723 | 335 | 426 | 317 | 264 | 113 | 717 | 880 | 844 | 826 | 218 | 408 | 175 | 645 | 773 | 236 | 234 | 151 | 164 | 165 | 324 | 1023 | 818 | 135 | 1022 | 156 | 188 | Krt18 |
| Krt19 | 111 | 148 | 127 | 63 | 22 | 108 | 11 | 164 | 360 | 172 | 267 | 143 | 174 | 123 | 125 | 180 | 51 | 64 | 26 | 28 | 23 | 22 | 408 | 174 | 38 | 82 | 132 | 192 | Krt19 |
| Krt2 | 0 | 0 | 0 | 0 | 0 | 0 | 0 | 1 | 0 | 5 | 3 | 0 | 0 | 0 | 0 | 0 | 0 | 2 | 0 | 1 | 0 | 2 | 0 | 0 | 0 | 0 | 0 | 0 | Krt2 |
| Krt20 | 5 | 8 | 14 | 0 | 6 | 5 | 1 | 5 | 12 | 12 | 6 | 12 | 7 | 12 | 20 | 23 | 8 | 1 | 2 | 2 | 3 | 7 | 9 | 5 | 0 | 10 | 7 | 15 | Krt20 |
| Krt222 | 84 | 60 | 69 | 86 | 170 | 105 | 159 | 158 | 161 | 179 | 159 | 96 | 103 | 85 | 116 | 87 | 87 | 128 | 104 | 143 | 130 | 204 | 158 | 122 | 161 | 84 | 77 | 97 | Krt222 |
| Krt23 | 20 | 7 | 0 | 23 | 0 | 0 | 0 | 71 | 66 | 129 | 52 | 2 | 17 | 3 | 7 | 20 | 1 | 2 | 1 | 4 | 0 | 2 | 74 | 118 | 0 | 80 | 0 | 1 | Krt23 |
| Krt24 | 2 | 0 | 7 | 1 | 2 | 6 | 0 | 0 | 6 | 0 | 16 | 2 | 0 | 2 | 1 | 2 | 0 | 0 | 1 | 4 | 0 | 5 | 10 | 0 | 0 | 0 | 0 | 0 | Krt24 |
| Krt25 | 1 | 0 | 3 | 0 | 2 | 0 | 2 | 6 | 3 | 7 | 1 | 2 | 1 | 4 | 4 | 3 | 1 | 0 | 0 | 2 | 0 | 0 | 1 | 7 | 0 | 2 | 6 | 6 | Krt25 |
| Krt26 | 8 | 14 | 18 | 12 | 11 | 4 | 4 | 17 | 8 | 5 | 8 | 21 | 16 | 22 | 15 | 16 | 8 | 6 | 12 | 3 | 5 | 27 | 13 | 17 | 20 | 8 | 36 | 7 | Krt26 |
| Krt27 | 1 | 2 | 0 | 0 | 3 | 0 | 0 | 2 | 2 | 1 | 7 | 0 | 2 | 0 | 0 | 1 | 0 | 0 | 0 | 1 | 2 | 3 | 2 | 0 | 1 | 0 | 0 | 1 | Krt27 |
| Krt28 | 9 | 10 | 5 | 2 | 7 | 5 | 4 | 15 | 9 | 19 | 14 | 7 | 16 | 14 | 1 | 9 | 10 | 14 | 6 | 5 | 2 | 7 | 18 | 7 | 7 | 14 | 20 | 9 | Krt28 |
| Krt31 | 0 | 0 | 0 | 0 | 0 | 0 | 0 | 0 | 4 | 0 | 1 | 0 | 0 | 0 | 0 | 0 | 0 | 0 | 0 | 0 | 0 | 0 | 4 | 0 | 0 | 0 | 0 | 0 | Krt31 |

| | | | | | | | | | | | | | | | | | | | | | | | | | | | | | |
|---|---|---|---|---|---|---|---|---|---|---|---|---|---|---|---|---|---|---|---|---|---|---|---|---|---|---|---|---|---|
| Krt33a | 0 | 0 | 0 | 0 | 0 | 0 | 0 | 0 | 0 | 0 | 0 | 0 | 0 | 0 | 0 | 1 | 0 | 0 | 0 | 0 | 0 | 0 | 0 | 2 | 1 | 0 | 0 | 2 | Krt33a |
| Krt33b | 0 | 1 | 0 | 0 | 0 | 0 | 0 | 0 | 3 | 0 | 2 | 1 | 0 | 0 | 0 | 0 | 0 | 0 | 0 | 1 | 0 | 2 | 5 | 2 | 1 | 0 | 0 | 1 | Krt33b |
| Krt34 | 1 | 0 | 0 | 0 | 0 | 0 | 0 | 0 | 0 | 0 | 0 | 0 | 0 | 0 | 0 | 0 | 0 | 0 | 0 | 0 | 0 | 0 | 0 | 0 | 0 | 0 | 0 | 0 | Krt34 |
| Krt35 | 0 | 0 | 0 | 1 | 0 | 0 | 0 | 5 | 1 | 0 | 0 | 0 | 0 | 0 | 0 | 0 | 0 | 0 | 0 | 0 | 0 | 0 | 1 | 0 | 0 | 0 | 0 | 0 | Krt35 |
| Krt36 | 0 | 0 | 0 | 0 | 0 | 0 | 0 | 1 | 0 | 0 | 0 | 0 | 0 | 0 | 0 | 0 | 0 | 0 | 0 | 0 | 0 | 0 | 1 | 0 | 0 | 0 | 0 | 0 | Krt36 |
| Krt4 | 0 | 1 | 0 | 1 | 0 | 6 | 0 | 9 | 24 | 4 | 40 | 0 | 0 | 1 | 7 | 0 | 0 | 1 | 4 | 0 | 0 | 0 | 41 | 16 | 0 | 0 | 2 | 2 | Krt4 |
| Krt40 | 0 | 0 | 0 | 0 | 0 | 0 | 0 | 0 | 0 | 0 | 0 | 0 | 0 | 0 | 0 | 2 | 0 | 0 | 0 | 0 | 0 | 0 | 0 | 0 | 0 | 0 | 0 | 0 | Krt40 |
| Krt42 | 0 | 7 | 4 | 0 | 0 | 0 | 0 | 0 | 0 | 0 | 0 | 0 | 2 | 0 | 3 | 2 | 8 | 0 | 4 | 0 | 0 | 0 | 2 | 0 | 0 | 0 | 0 | 0 | Krt42 |
| Krt5 | 2030 | 1070 | 511 | 55 | 33 | 1764 | 7 | 3500 | 8093 | 3688 | 3501 | 1388 | 2174 | 1531 | 1283 | 895 | 206 | 1617 | 265 | 51 | 0 | 23 | 9525 | 3224 | 623 | 478 | 1076 | 1533 | Krt5 |
| Krt6a | 16 | 0 | 3 | 0 | 1 | 2 | 0 | 204 | 264 | 188 | 32 | 0 | 10 | 5 | 1 | 0 | 0 | 0 | 0 | 0 | 0 | 0 | 217 | 18 | 2 | 5 | 1 | 2 | Krt6a |
| Krt6b | 0 | 0 | 0 | 0 | 0 | 0 | 0 | 0 | 1 | 0 | 0 | 0 | 0 | 0 | 0 | 0 | 0 | 0 | 0 | 0 | 0 | 0 | 0 | 0 | 0 | 0 | 0 | 0 | Krt6b |
| Krt7 | 277 | 280 | 251 | 20 | 25 | 239 | 15 | 387 | 658 | 312 | 414 | 381 | 523 | 300 | 340 | 306 | 75 | 172 | 26 | 25 | 14 | 77 | 682 | 489 | 82 | 195 | 385 | 470 | Krt7 |
| Krt71 | 0 | 0 | 0 | 0 | 4 | 0 | 0 | 2 | 12 | 2 | 2 | 0 | 0 | 0 | 0 | 0 | 0 | 0 | 0 | 0 | 0 | 0 | 11 | 0 | 1 | 0 | 0 | 1 | Krt71 |
| Krt72 | 1 | 0 | 0 | 0 | 0 | 0 | 0 | 0 | 11 | 4 | 1 | 0 | 1 | 0 | 0 | 1 | 0 | 3 | 0 | 0 | 0 | 2 | 3 | 4 | 0 | 2 | 1 | 0 | Krt72 |
| Krt73 | 2 | 4 | 2 | 4 | 8 | 1 | 3 | 22 | 38 | 43 | 32 | 28 | 18 | 23 | 0 | 2 | 0 | 8 | 4 | 7 | 9 | 55 | 52 | 10 | 46 | 23 | 26 | 23 | Krt73 |
| Krt75 | 8 | 0 | 0 | 1 | 7 | 0 | 0 | 22 | 33 | 16 | 32 | 4 | 16 | 2 | 4 | 13 | 2 | 0 | 1 | 0 | 0 | 12 | 44 | 17 | 9 | 16 | 3 | 8 | Krt75 |
| Krt76 | 0 | 0 | 0 | 0 | 0 | 3 | 0 | 2 | 5 | 4 | 0 | 0 | 0 | 0 | 0 | 0 | 0 | 0 | 0 | 0 | 0 | 0 | 12 | 2 | 0 | 0 | 0 | 0 | Krt76 |
| Krt77 | 3 | 0 | 7 | 7 | 13 | 8 | 4 | 1 | 6 | 13 | 5 | 4 | 3 | 3 | 1 | 0 | 0 | 0 | 5 | 8 | 7 | 5 | 12 | 5 | 4 | 0 | 0 | 0 | Krt77 |
| Krt78 | 0 | 0 | 1 | 0 | 4 | 1 | 2 | 5 | 3 | 5 | 0 | 4 | 6 | 0 | 1 | 2 | 0 | 1 | 0 | 0 | 0 | 2 | 5 | 1 | 0 | 5 | 5 | 0 | Krt78 |
| Krt79 | 1 | 7 | 11 | 5 | 4 | 4 | 7 | 6 | 35 | 24 | 7 | 4 | 2 | 0 | 5 | 3 | 5 | 3 | 2 | 5 | 1 | 3 | 43 | 12 | 2 | 9 | 14 | 1 | Krt79 |
| Krt8 | 1181 | 731 | 339 | 472 | 206 | 280 | 55 | 866 | 981 | 904 | 719 | 243 | 542 | 242 | 814 | 836 | 183 | 224 | 91 | 93 | 68 | 294 | 1038 | 832 | 94 | 1130 | 248 | 333 | Krt8 |
| Krt80 | 8 | 6 | 2 | 0 | 2 | 9 | 0 | 22 | 49 | 25 | 36 | 11 | 7 | 10 | 4 | 9 | 0 | 7 | 0 | 0 | 2 | 12 | 80 | 7 | 9 | 19 | 5 | 9 | Krt80 |
| Krt81 | 0 | 0 | 0 | 0 | 0 | 5 | 0 | 1 | 1 | 2 | 1 | 0 | 0 | 0 | 0 | 0 | 0 | 0 | 0 | 2 | 0 | 0 | 1 | 0 | 0 | 0 | 0 | 0 | Krt81 |
| Krt83 | 0 | 0 | 3 | 0 | 0 | 0 | 0 | 0 | 0 | 0 | 0 | 0 | 1 | 0 | 0 | 0 | 0 | 0 | 0 | 0 | 0 | 0 | 1 | 0 | 0 | 0 | 0 | 0 | Krt83 |
| Krt84 | 0 | 0 | 3 | 0 | 0 | 0 | 0 | 0 | 0 | 0 | 1 | 0 | 1 | 0 | 1 | 0 | 0 | 0 | 0 | 0 | 0 | 0 | 2 | 0 | 0 | 0 | 2 | 0 | Krt84 |
| Krt86 | 0 | 0 | 8 | 1 | 0 | 0 | 0 | 0 | 1 | 2 | 1 | 0 | 0 | 0 | 0 | 0 | 0 | 1 | 2 | 3 | 0 | 3 | 3 | 0 | 0 | 1 | 0 | 1 | Krt86 |
| Krt87 | 2 | 2 | 0 | 0 | 2 | 8 | 3 | 17 | 15 | 29 | 24 | 5 | 10 | 2 | 3 | 1 | 1 | 0 | 2 | 0 | 1 | 10 | 17 | 8 | 9 | 4 | 3 | 12 | Krt87 |
| Krt88 | 2 | 0 | 0 | 12 | 0 | 0 | 1 | 0 | 2 | 0 | 3 | 1 | 0 | 0 | 0 | 4 | 0 | 1 | 1 | 0 | 2 | 0 | 0 | 0 | 0 | 0 | 0 | 1 | Krt88 |
| Krt9 | 0 | 0 | 0 | 0 | 3 | 2 | 2 | 5 | 12 | 2 | 11 | 0 | 6 | 0 | 3 | 1 | 0 | 0 | 5 | 0 | 1 | 7 | 12 | 5 | 1 | 9 | 0 | 1 | Krt9 |
| Krt90 | 13 | 7 | 6 | 7 | 5 | 3 | 2 | 20 | 4 | 13 | 5 | 10 | 8 | 5 | 3 | 19 | 6 | 6 | 4 | 4 | 7 | 3 | 4 | 3 | 1 | 12 | 7 | 9 | Krt90 |
| Krtap10-10 | 0 | 0 | 0 | 0 | 0 | 0 | 0 | 0 | 2 | 0 | 0 | 0 | 0 | 0 | 0 | 0 | 0 | 0 | 0 | 0 | 0 | 0 | 1 | 0 | 2 | 0 | 0 | 0 | Krtap10-10 |
| Krtap12-1 | 0 | 0 | 0 | 0 | 0 | 0 | 0 | 1 | 0 | 0 | 0 | 0 | 0 | 0 | 0 | 0 | 0 | 0 | 0 | 0 | 0 | 0 | 0 | 0 | 0 | 0 | 0 | 0 | Krtap12-1 |
| Krtap15 | 0 | 0 | 0 | 0 | 0 | 0 | 0 | 0 | 0 | 0 | 2 | 0 | 0 | 0 | 0 | 0 | 0 | 0 | 0 | 0 | 0 | 0 | 3 | 0 | 0 | 0 | 0 | 0 | Krtap15 |
| Krtap16-1 | 0 | 1 | 0 | 0 | 0 | 0 | 0 | 0 | 0 | 0 | 0 | 0 | 0 | 0 | 0 | 0 | 0 | 0 | 0 | 0 | 0 | 0 | 4 | 1 | 3 | 0 | 0 | 0 | Krtap16-1 |
| Krtap17-1 | 0 | 0 | 0 | 0 | 0 | 0 | 0 | 0 | 0 | 0 | 1 | 2 | 0 | 0 | 3 | 0 | 0 | 0 | 0 | 0 | 2 | 2 | 3 | 2 | 6 | 0 | 2 | 1 | Krtap17-1 |
| Krtap21-1 | 0 | 0 | 0 | 0 | 0 | 0 | 0 | 0 | 0 | 0 | 0 | 0 | 0 | 0 | 0 | 1 | 0 | 0 | 0 | 0 | 0 | 0 | 0 | 0 | 0 | 0 | 0 | 0 | Krtap21-1 |
| Krtap24-1 | 0 | 0 | 0 | 0 | 0 | 0 | 0 | 0 | 0 | 0 | 0 | 1 | 0 | 0 | 0 | 0 | 0 | 0 | 0 | 0 | 0 | 0 | 0 | 0 | 0 | 0 | 0 | 0 | Krtap24-1 |
| Krtap26-1 | 0 | 0 | 0 | 0 | 0 | 0 | 4 | 0 | 0 | 0 | 0 | 0 | 0 | 0 | 0 | 0 | 0 | 0 | 0 | 0 | 0 | 0 | 0 | 0 | 0 | 0 | 0 | 0 | Krtap26-1 |
| Krtap28-13 | 0 | 0 | 0 | 0 | 0 | 0 | 0 | 0 | 0 | 0 | 0 | 0 | 0 | 1 | 0 | 0 | 0 | 0 | 0 | 0 | 0 | 0 | 1 | 0 | 0 | 0 | 0 | 0 | Krtap28-13 |
| Krtap29-1 | 0 | 0 | 0 | 0 | 0 | 0 | 0 | 0 | 0 | 0 | 0 | 0 | 0 | 0 | 0 | 0 | 0 | 0 | 0 | 0 | 0 | 2 | 0 | 0 | 0 | 0 | 0 | 0 | Krtap29-1 |
| Krtap3-3 | 0 | 0 | 0 | 0 | 0 | 0 | 0 | 0 | 0 | 0 | 0 | 0 | 0 | 0 | 0 | 0 | 0 | 0 | 0 | 0 | 0 | 0 | 0 | 2 | 0 | 0 | 0 | 0 | Krtap3-3 |
| Krtap4-16 | 0 | 0 | 0 | 0 | 0 | 0 | 0 | 0 | 0 | 0 | 1 | 0 | 0 | 0 | 0 | 0 | 0 | 0 | 0 | 0 | 0 | 0 | 0 | 0 | 0 | 0 | 0 | 0 | Krtap4-16 |
| Krtap5-3 | 0 | 3 | 0 | 0 | 4 | 0 | 0 | 0 | 0 | 0 | 0 | 0 | 0 | 0 | 0 | 0 | 0 | 0 | 0 | 0 | 1 | 0 | 0 | 1 | 0 | 0 | 0 | 0 | Krtap5-3 |
| Krtap5-4 | 0 | 0 | 0 | 0 | 0 | 0 | 0 | 0 | 0 | 0 | 0 | 0 | 0 | 0 | 0 | 0 | 0 | 0 | 0 | 0 | 0 | 0 | 1 | 0 | 0 | 0 | 0 | 0 | Krtap5-4 |
| Krtap6-2 | 0 | 0 | 0 | 0 | 0 | 0 | 0 | 0 | 0 | 0 | 0 | 0 | 0 | 0 | 0 | 0 | 0 | 0 | 0 | 0 | 0 | 0 | 0 | 1 | 0 | 0 | 0 | 0 | Krtap6-2 |
| Krtap7-1 | 3 | 2 | 0 | 5 | 1 | 3 | 8 | 9 | 2 | 10 | 3 | 7 | 7 | 7 | 1 | 2 | 21 | 1 | 9 | 0 | 6 | 3 | 3 | 3 | 16 | 10 | 10 | 12 | Krtap7-1 |
| Krtap8-1 | 1 | 0 | 0 | 0 | 0 | 0 | 1 | 0 | 2 | 0 | 0 | 0 | 0 | 2 | 0 | 1 | 0 | 0 | 0 | 0 | 0 | 2 | 2 | 0 | 0 | 3 | 5 | 0 | Krtap8-1 |
| Krtap9-1 | 0 | 0 | 0 | 0 | 0 | 0 | 0 | 0 | 0 | 0 | 0 | 0 | 0 | 0 | 0 | 0 | 0 | 0 | 0 | 0 | 0 | 0 | 0 | 2 | 0 | 0 | 0 | 0 | Krtap9-1 |
| Krtcap2 | 2970 | 3912 | 4011 | 3762 | 3139 | 3179 | 3589 | 2098 | 2524 | 2530 | 2764 | 2599 | 2814 | 2931 | 2993 | 3077 | 3684 | 3455 | 3857 | 3868 | 3408 | 2047 | 2561 | 2753 | 2595 | 3433 | 2998 | 3102 | Krtcap2 |
| Krtcap3 | 87 | 78 | 87 | 68 | 69 | 58 | 57 | 105 | 89 | 101 | 97 | 79 | 105 | 77 | 45 | 79 | 81 | 69 | 78 | 63 | 53 | 55 | 128 | 145 | 43 | 156 | 86 | 71 | Krtcap3 |
| Krtdap | 1 | 0 | 0 | 0 | 1 | 3 | 0 | 7 | 57 | 25 | 8 | 2 | 0 | 0 | 0 | 0 | 0 | 3 | 0 | 0 | 0 | 0 | 67 | 9 | 0 | 0 | 2 | 0 | Krtdap |
| Ksr1 | 1425 | 1816 | 1807 | 2114 | 2561 | 1959 | 2037 | 1399 | 1773 | 1806 | 2066 | 1879 | 1486 | 1755 | 1404 | 1593 | 2286 | 2163 | 2269 | 1645 | 2081 | 1930 | 1801 | 1914 | 2014 | 1704 | 1526 | 1576 | Ksr1 |
| Ksr2 | 812 | 891 | 814 | 1635 | 2434 | 1996 | 1681 | 1929 | 1926 | 2843 | 2353 | 1920 | 1910 | 1810 | 924 | 921 | 1415 | 1640 | 1907 | 1495 | 2003 | 3153 | 1615 | 2094 | 2806 | 1154 | 1555 | 1529 | Ksr2 |
| Kti12 | 1345 | 1557 | 1359 | 1177 | 1040 | 962 | 1031 | 1142 | 1151 | 1027 | 1046 | 1286 | 1244 | 1124 | 1205 | 1324 | 1236 | 1192 | 1050 | 1152 | 1028 | 1036 | 1125 | 1133 | 1091 | 1347 | 1220 | 1309 | Kti12 |
| Ktn1 | 17272 | 15068 | 18309 | 16886 | 18187 | 18123 | 19691 | 18058 | 15125 | 15712 | 15085 | 17422 | 16450 | 18173 | 18478 | 16120 | 17328 | 16677 | 17117 | 19302 | 18102 | 16140 | 14804 | 15809 | 16360 | 15316 | 17317 | 17340 | Ktn1 |
| Kxd1 | 1198 | 1417 | 1332 | 1112 | 1148 | 1273 | 1063 | 1407 | 1701 | 1533 | 1434 | 1764 | 1594 | 1665 | 1209 | 1245 | 1260 | 1125 | 1248 | 1194 | 1088 | 1358 | 1714 | 1668 | 1585 | 1877 | 1733 | 1728 | Kxd1 |
| Ky | 37 | 50 | 38 | 33 | 75 | 41 | 41 | 54 | 96 | 91 | 246 | 81 | 120 | 47 | 34 | 40 | 49 | 51 | 59 | 42 | 46 | 134 | 172 | 39 | 89 | 54 | 49 | 56 | Ky |
| Kyat1 | 1002 | 1278 | 1187 | 1220 | 1443 | 1253 | 1222 | 1051 | 1195 | 1101 | 1075 | 1127 | 1091 | 1020 | 1087 | 1116 | 1366 | 1362 | 1467 | 1265 | 1281 | 1190 | 1061 | 1189 | 1284 | 964 | 1125 | 993 | Kyat1 |
| Kyat3 | 1924 | 1733 | 1335 | 1472 | 1252 | 1189 | 1340 | 497 | 643 | 476 | 587 | 496 | 570 | 469 | 2004 | 1719 | 1421 | 1461 | 1196 | 1303 | 1322 | 373 | 580 | 474 | 384 | 632 | 539 | 502 | Kyat3 |
| Kynu | 59 | 42 | 63 | 31 | 14 | 18 | 16 | 49 | 54 | 29 | 63 | 61 | 110 | 83 | 46 | 60 | 39 | 30 | 22 | 32 | 26 | 60 | 80 | 38 | 31 | 64 | 87 | 85 | Kynu |
| L1cam | 6266 | 7573 | 6195 | 12947 | 19152 | 14913 | 15826 | 13438 | 14604 | 18568 | 14368 | 13989 | 11662 | 13023 | 8081 | 7243 | 10335 | 14550 | 16819 | 13600 | 16360 | 21950 | 12407 | 15440 | 21355 | 9249 | 11208 | 10843 | L1cam |
| L1td1 | 322 | 187 | 254 | 146 | 88 | 190 | 56 | 83 | 111 | 120 | 120 | 66 | 124 | 73 | 274 | 319 | 155 | 161 | 101 | 90 | 63 | 100 | 168 | 78 | 66 | 103 | 50 | 107 | L1td1 |
| L2hgdh | 4237 | 3525 | 3529 | 3937 | 3657 | 3818 | 4210 | 1533 | 1506 | 1417 | 1640 | 1485 | 1484 | 1521 | 3774 | 4079 | 3654 | 4200 | 3668 | 4226 | 3961 | 1250 | 1628 | 1585 | 1470 | 1759 | 1520 | 1552 | L2hgdh |
| L3hypdh | 767 | 590 | 573 | 704 | 744 | 772 | 625 | 449 | 558 | 473 | 595 | 350 | 458 | 408 | 678 | 752 | 639 | 577 | 645 | 682 | 695 | 404 | 548 | 450 | 458 | 359 | 430 | 336 | L3hypdh |
| L3mbtl1 | 231 | 258 | 303 | 362 | 479 | 362 | 253 | 123 | 153 | 213 | 167 | 101 | 151 | 100 | 210 | 289 | 304 | 465 | 426 | 301 | 381 | 244 | 162 | 127 | 190 | 55 | 121 | 85 | L3mbtl1 |
| L3mbtl2 | 2829 | 3298 | 3775 | 3403 | 3059 | 3114 | 2914 | 1947 | 2218 | 2135 | 2119 | 2967 | 2535 | 2668 | 2648 | 3037 | 3408 | 3261 | 3269 | 3096 | 2924 | 2303 | 2123 | 2540 | 2637 | 3300 | 2945 | 2921 | L3mbtl2 |
| L3mbtl3 | 4191 | 3952 | 4453 | 4723 | 6091 | 5373 | 5768 | 5228 | 5056 | 5992 | 5811 | 5448 | 5147 | 5545 | 4611 | 3907 | 4694 | 5048 | 5561 | 5043 | 5928 | 6505 | 5082 | 5887 | 5713 | 4902 | 5211 | 5635 | L3mbtl3 |
| L3mbtl4 | 15 | 14 | 2 | 38 | 54 | 26 | 37 | 31 | 50 | 46 | 42 | 18 | 20 | 21 | 16 | 10 | 13 | 25 | 35 | 71 | 37 | 50 | 50 | 53 | 37 | 10 | 16 | 17 | L3mbtl4 |
| Lacc1 | 849 | 654 | 623 | 488 | 503 | 495 | 439 | 468 | 501 | 436 | 590 | 405 | 510 | 403 | 754 | 833 | 461 | 495 | 440 | 472 | 397 | 451 | 584 | 368 | 344 | 405 | 407 | 349 | Lacc1 |
| Lactb | 2287 | 2380 | 2485 | 2181 | 1861 | 1866 | 2046 | 869 | 759 | 849 | 835 | 947 | 889 | 905 | 2395 | 2350 | 2319 | 2298 | 1915 | 2090 | 1944 | 1008 | 766 | 784 | 853 | 880 | 841 | 946 | Lactb |
| Lactb2 | 1982 | 1672 | 1419 | 1374 | 1391 | 1740 | 1777 | 1045 | 961 | 886 | 909 | 1040 | 1066 | 944 | 2003 | 1928 | 1378 | 1420 | 1265 | 1919 | 1641 | 652 | 1073 | 840 | 642 | 1046 | 1031 | 1024 | Lactb2 |
| Lactbl1 | 3 | 0 | 0 | 6 | 1 | 2 | 2 | 0 | 7 | 1 | 3 | 4 | 0 | 3 | 1 | 3 | 0 | 1 | 0 | 2 | 4 | 2 | 4 | 7 | 1 | 4 | 0 | 1 | Lactbl1 |
| Lad1 | 146 | 48 | 30 | 62 | 26 | 30 | 18 | 147 | 203 | 178 | 186 | 15 | 45 | 23 | 63 | 45 | 28 | 21 | 12 | 13 | 4 | 35 | 250 | 150 | 23 | 215 | 27 | 34 | Lad1 |
| Lag3 | 155 | 190 | 185 | 279 | 290 | 236 | 258 | 255 | 197 | 265 | 245 | 286 | 248 | 226 | 136 | 185 | 209 | 259 | 342 | 253 | 261 | 309 | 222 | 220 | 310 | 185 | 187 | 249 | Lag3 |
| Lage3 | 1427 | 1462 | 1610 | 1273 | 1119 | 1193 | 1199 | 1526 | 1585 | 1452 | 1466 | 1748 | 1693 | 1939 | 1255 | 1238 | 1444 | 1032 | 1223 | 1341 | 1098 | 1292 | 1594 | 1596 | 1550 | 2174 | 1820 | 2006 | Lage3 |
| Lair1 | 112 | 121 | 73 | 91 | 178 | 157 | 148 | 252 | 218 | 227 | 239 | 240 | 222 | 289 | 125 | 89 | 88 | 108 | 155 | 169 | 147 | 271 | 191 | 237 | 216 | 209 | 198 | 228 | Lair1 |
| Lalba | 1 | 0 | 0 | 0 | 0 | 0 | 0 | 0 | 3 | 0 | 1 | 0 | 3 | 1 | 0 | 1 | 0 | 0 | 0 | 0 | 1 | 0 | 1 | 0 | 0 | 1 | 5 | 0 | Lalba |
| Lama1 | 6215 | 7540 | 7084 | 4980 | 5056 | 5523 | 4646 | 3231 | 2745 | 3740 | 3515 | 6090 | 6332 | 5292 | 5639 | 8089 | 5682 | 5201 | 4901 | 4827 | 5100 | 3156 | 3460 | 3611 | 4076 | 4384 | 7498 | 5841 | Lama1 |
| Lama2 | 3904 | 3532 | 2286 | 2347 | 2346 | 2947 | 1560 | 2705 | 2886 | 3052 | 4303 | 2873 | 4009 | 2845 | 3575 | 4786 | 1830 | 2568 | 1531 | 2374 | 2064 | 2727 | 4301 | 2453 | 2455 | 2100 | 3291 | 2689 | Lama2 |

| | | | | | | | | | | | | | | | | | | | | | | | | | | | | | |
|---|---|---|---|---|---|---|---|---|---|---|---|---|---|---|---|---|---|---|---|---|---|---|---|---|---|---|---|---|---|
| Lama3 | 188 | 160 | 73 | 125 | 176 | 204 | 98 | 537 | 856 | 469 | 552 | 223 | 418 | 204 | 204 | 223 | 109 | 186 | 122 | 112 | 116 | 262 | 907 | 291 | 274 | 104 | 221 | 182 | Lama3 |
| Lama4 | 4577 | 4042 | 3463 | 3730 | 4178 | 4118 | 2834 | 4949 | 6512 | 6103 | 8404 | 4376 | 5641 | 4114 | 3972 | 5091 | 3393 | 4901 | 3102 | 2920 | 3397 | 5905 | 7879 | 5487 | 5208 | 4378 | 4408 | 3693 | Lama4 |
| Lama5 | 5609 | 5458 | 5994 | 5168 | 4159 | 4438 | 3728 | 1870 | 3194 | 2014 | 2127 | 1900 | 1971 | 1374 | 3883 | 6292 | 5284 | 5217 | 4795 | 3672 | 3620 | 1103 | 3118 | 2335 | 1163 | 1493 | 1367 | 1435 | Lama5 |
| Lamb1 | 9009 | 8455 | 8209 | 6103 | 6411 | 6870 | 4616 | 8294 | 11142 | 10341 | 14780 | 8768 | 10754 | 8104 | 7570 | 10875 | 6219 | 8152 | 5408 | 5305 | 5219 | 8883 | 13762 | 9962 | 8739 | 9602 | 8814 | 8064 | Lamb1 |
| Lamb2 | 6574 | 8221 | 8086 | 7162 | 5616 | 6661 | 4545 | 4268 | 6203 | 6011 | 6924 | 6810 | 6648 | 5329 | 5788 | 7868 | 7722 | 7130 | 6351 | 5049 | 4867 | 5723 | 6492 | 6317 | 5490 | 6598 | 6694 | 5737 | Lamb2 |
| Lamb3 | 61 | 31 | 9 | 24 | 26 | 43 | 30 | 95 | 212 | 103 | 94 | 60 | 37 | 49 | 22 | 42 | 30 | 31 | 15 | 17 | 21 | 52 | 178 | 75 | 50 | 56 | 43 | 55 | Lamb3 |
| Lamc1 | 15177 | 15784 | 13148 | 13115 | 12612 | 13392 | 10321 | 7847 | 10708 | 10766 | 13522 | 10025 | 12242 | 9805 | 13175 | 18210 | 12245 | 15152 | 11120 | 9883 | 11304 | 9618 | 11768 | 10110 | 9657 | 7885 | 10206 | 9252 | Lamc1 |
| Lamc2 | 509 | 209 | 109 | 86 | 141 | 167 | 48 | 421 | 677 | 378 | 430 | 100 | 177 | 95 | 154 | 306 | 57 | 173 | 79 | 77 | 59 | 167 | 623 | 221 | 128 | 179 | 94 | 96 | Lamc2 |
| Lamc3 | 1410 | 2087 | 2692 | 1633 | 1664 | 1684 | 1268 | 916 | 946 | 1359 | 1153 | 2184 | 2465 | 1788 | 1444 | 2147 | 2101 | 1571 | 1583 | 1539 | 1464 | 1465 | 1288 | 1334 | 1476 | 1696 | 2617 | 1968 | Lamc3 |
| Lamp1 | 14645 | 15964 | 15273 | 13955 | 13270 | 13868 | 12398 | 13213 | 14032 | 13322 | 13570 | 12686 | 13782 | 12309 | 15217 | 16240 | 14326 | 14198 | 13061 | 12718 | 12948 | 13356 | 15655 | 12469 | 12174 | 13705 | 12511 | 13099 | Lamp1 |
| Lamp2 | 15628 | 11693 | 12346 | 10883 | 9667 | 10570 | 10017 | 10461 | 10165 | 7912 | 10234 | 8232 | 9162 | 8113 | 15136 | 14952 | 10720 | 10962 | 9294 | 11530 | 9377 | 8915 | 11908 | 7911 | 7147 | 8285 | 8451 | 8274 | Lamp2 |
| Lamp3 | 21 | 16 | 32 | 12 | 11 | 9 | 13 | 16 | 12 | 10 | 15 | 22 | 12 | 22 | 12 | 9 | 25 | 21 | 7 | 16 | 14 | 12 | 13 | 24 | 8 | 10 | 16 | 12 | Lamp3 |
| Lamp5 | 61 | 123 | 62 | 79 | 939 | 211 | 68 | 1570 | 1653 | 1816 | 2203 | 202 | 1750 | 140 | 347 | 213 | 129 | 114 | 76 | 98 | 57 | 1684 | 1913 | 149 | 271 | 128 | 194 | 172 | Lamp5 |
| Lamtor1 | 1623 | 1660 | 1369 | 1237 | 1322 | 1454 | 1403 | 3160 | 3454 | 3013 | 2826 | 3186 | 3030 | 3423 | 1524 | 1554 | 1280 | 1268 | 1425 | 1430 | 1367 | 2690 | 3416 | 3152 | 2930 | 3662 | 3340 | 3376 | Lamtor1 |
| Lamtor2 | 3751 | 4905 | 6354 | 6308 | 4898 | 4838 | 4495 | 2622 | 2952 | 3527 | 3332 | 3704 | 3839 | 3437 | 4120 | 4483 | 6071 | 5490 | 5715 | 5928 | 4245 | 2391 | 3036 | 3771 | 3819 | 4622 | 4314 | 3779 | Lamtor2 |
| Lamtor3 | 3752 | 3913 | 3764 | 4007 | 3435 | 3570 | 4193 | 2967 | 2726 | 2856 | 2608 | 2836 | 2759 | 3125 | 3767 | 3573 | 3722 | 3695 | 3722 | 4274 | 4073 | 2650 | 2807 | 2862 | 2934 | 3210 | 2815 | 3169 | Lamtor3 |
| Lamtor3-ps | 45 | 12 | 34 | 16 | 65 | 45 | 19 | 140 | 132 | 154 | 130 | 113 | 116 | 126 | 24 | 27 | 21 | 12 | 54 | 53 | 29 | 159 | 137 | 147 | 144 | 151 | 156 | 112 | Lamtor3-ps |
| Lamtor4 | 1010 | 1370 | 1282 | 1361 | 1332 | 1284 | 1205 | 1435 | 1461 | 1584 | 1545 | 1417 | 1456 | 1420 | 1167 | 1048 | 1469 | 1318 | 1303 | 1479 | 1353 | 1211 | 1496 | 1683 | 1625 | 1391 | 1449 | 1499 | Lamtor4 |
| Lamtor5 | 2692 | 2587 | 2951 | 2701 | 2520 | 2525 | 3009 | 2403 | 2258 | 2339 | 2373 | 2407 | 2388 | 2621 | 2846 | 2470 | 2775 | 2634 | 2729 | 3261 | 2719 | 2052 | 2269 | 2611 | 2439 | 2883 | 2422 | 2651 | Lamtor5 |
| Lancl1 | 5252 | 5884 | 4031 | 4987 | 5174 | 5156 | 5223 | 4820 | 4668 | 4120 | 4062 | 3917 | 3758 | 3903 | 5557 | 5224 | 4612 | 5152 | 4993 | 5489 | 5411 | 4518 | 4847 | 3875 | 4359 | 3704 | 3903 | 3858 | Lancl1 |
| Lancl2 | 2417 | 2160 | 2281 | 2249 | 2894 | 2799 | 2758 | 6883 | 5867 | 6604 | 5680 | 6652 | 5991 | 6842 | 2622 | 1912 | 2351 | 2234 | 2780 | 2888 | 2703 | 7691 | 5749 | 6388 | 7095 | 6658 | 6577 | 7031 | Lancl2 |
| Lancl3 | 364 | 296 | 176 | 381 | 619 | 658 | 625 | 573 | 477 | 572 | 446 | 463 | 480 | 423 | 340 | 250 | 302 | 464 | 556 | 516 | 536 | 449 | 334 | 378 | 607 | 99 | 463 | 310 | Lancl3 |
| Lao1 | 0 | 0 | 0 | 0 | 0 | 0 | 0 | 0 | 0 | 0 | 0 | 0 | 0 | 0 | 0 | 0 | 0 | 0 | 0 | 0 | 0 | 0 | 1 | 0 | 0 | 0 | 0 | 0 | Lao1 |
| Lap3 | 4169 | 4475 | 4856 | 3688 | 3170 | 3586 | 3675 | 2959 | 3160 | 2811 | 3492 | 3280 | 3461 | 3586 | 3918 | 4179 | 3942 | 3554 | 3698 | 3662 | 3365 | 2386 | 3405 | 3349 | 2876 | 4490 | 3860 | 3796 | Lap3 |
| Laptm4a | 22571 | 18944 | 21041 | 20348 | 18425 | 19462 | 20619 | 21090 | 20337 | 19554 | 21539 | 18169 | 19046 | 20266 | 22524 | 20814 | 19169 | 20323 | 18686 | 21908 | 19951 | 17817 | 23488 | 20466 | 18624 | 21833 | 18999 | 20749 | Laptm4a |
| Laptm4b | 4274 | 4206 | 4087 | 3505 | 3599 | 3402 | 3378 | 2533 | 2622 | 2345 | 2684 | 2223 | 2609 | 2221 | 4199 | 4399 | 3640 | 3458 | 3497 | 3693 | 3521 | 2226 | 3109 | 2065 | 1994 | 2443 | 2224 | 2509 | Laptm4b |
| Laptm5 | 648 | 846 | 1040 | 693 | 788 | 812 | 584 | 639 | 972 | 1026 | 1123 | 972 | 1004 | 721 | 630 | 917 | 720 | 916 | 824 | 669 | 764 | 1046 | 1265 | 879 | 946 | 1077 | 953 | 802 | Laptm5 |
| Large1 | 4014 | 4106 | 4040 | 4965 | 5359 | 4652 | 5317 | 3863 | 4205 | 4621 | 4133 | 3999 | 3816 | 3843 | 4002 | 3724 | 4638 | 5102 | 5532 | 3840 | 5519 | 4684 | 4049 | 4073 | 4496 | 3703 | 3546 | 3783 | Large1 |
| Large2 | 464 | 371 | 575 | 248 | 172 | 198 | 118 | 250 | 391 | 261 | 308 | 335 | 348 | 277 | 352 | 499 | 356 | 179 | 186 | 143 | 119 | 227 | 399 | 266 | 156 | 429 | 379 | 421 | Large2 |
| Larp1 | 11590 | 12323 | 15957 | 15475 | 15219 | 13911 | 13132 | 9515 | 10704 | 13082 | 13263 | 13799 | 12447 | 12526 | 11006 | 12626 | 15921 | 15699 | 15127 | 10999 | 13950 | 13256 | 10290 | 13166 | 13715 | 12542 | 11431 | 12773 | Larp1 |
| Larp1b | 1175 | 1058 | 1393 | 1505 | 1386 | 1390 | 1581 | 426 | 431 | 478 | 568 | 441 | 468 | 488 | 1139 | 1113 | 1376 | 1531 | 1340 | 1119 | 1465 | 578 | 399 | 476 | 538 | 413 | 451 | 502 | Larp1b |
| Larp4 | 9003 | 7477 | 7973 | 8558 | 7586 | 8764 | 8533 | 8081 | 7042 | 5987 | 6937 | 7193 | 7007 | 7641 | 9356 | 8383 | 7756 | 7899 | 7786 | 9411 | 8334 | 6547 | 7159 | 6584 | 7073 | 6452 | 7632 | 7510 | Larp4 |
| Larp4b | 14021 | 12120 | 11652 | 13560 | 12249 | 12656 | 14190 | 9664 | 9154 | 9135 | 8836 | 9764 | 9003 | 10040 | 13471 | 12350 | 12150 | 12536 | 12066 | 13697 | 13942 | 8731 | 8700 | 9075 | 9554 | 8941 | 9861 | 9855 | Larp4b |
| Larp6 | 99 | 105 | 98 | 107 | 241 | 133 | 123 | 269 | 353 | 359 | 363 | 223 | 255 | 237 | 100 | 90 | 143 | 147 | 171 | 113 | 163 | 429 | 343 | 324 | 383 | 189 | 242 | 220 | Larp6 |
| Larp7 | 2786 | 2558 | 3629 | 2242 | 1888 | 2103 | 2278 | 5797 | 4952 | 4784 | 4896 | 6600 | 5865 | 7152 | 2846 | 2316 | 2802 | 1955 | 2323 | 2691 | 2189 | 4346 | 4885 | 5924 | 5051 | 8040 | 7599 | 7552 | Larp7 |
| Lars | 10657 | 9758 | 9782 | 8351 | 8330 | 8447 | 9451 | 7988 | 7731 | 6698 | 6705 | 8393 | 7834 | 8509 | 10411 | 9794 | 8717 | 8212 | 8301 | 10228 | 8636 | 6198 | 7337 | 7359 | 6941 | 8415 | 8610 | 8552 | Lars |
| Lars2 | 14493 | 24509 | 13629 | 21050 | 13820 | 28120 | 17465 | 17267 | 13058 | 21891 | 11971 | 38706 | 12528 | 22297 | 9047 | 9868 | 12465 | 21665 | 18463 | 10960 | 12908 | 12651 | 11128 | 15707 | 11626 | 13237 | 34586 | 14432 | Lars2 |
| Las1l | 7754 | 7915 | 7881 | 8248 | 8603 | 8199 | 8532 | 6007 | 5629 | 5988 | 5861 | 5681 | 5662 | 5988 | 8093 | 7080 | 7985 | 8445 | 7774 | 8593 | 8268 | 6107 | 5231 | 5800 | 6557 | 5627 | 5768 | 5834 | Las1l |
| Lasp1 | 10530 | 11222 | 12557 | 13064 | 13651 | 13012 | 11962 | 10182 | 12039 | 13370 | 13269 | 13091 | 12086 | 12476 | 9939 | 10790 | 13770 | 13727 | 14125 | 10613 | 12784 | 14130 | 11814 | 13691 | 13653 | 14082 | 11640 | 11746 | Lasp1 |
| Lat | 105 | 152 | 190 | 206 | 142 | 137 | 165 | 120 | 114 | 114 | 121 | 133 | 146 | 74 | 111 | 139 | 220 | 202 | 195 | 129 | 126 | 160 | 117 | 98 | 131 | 124 | 145 | 116 | Lat |
| Lat2 | 559 | 797 | 985 | 1189 | 904 | 846 | 920 | 232 | 247 | 322 | 229 | 349 | 349 | 319 | 601 | 648 | 1167 | 1221 | 1011 | 777 | 976 | 373 | 234 | 393 | 342 | 434 | 280 | 366 | Lat2 |
| Lats1 | 0 | 0 | 0 | 0 | 0 | 0 | 0 | 0 | 0 | 1 | 0 | 0 | 0 | 0 | 0 | 0 | 0 | 0 | 2 | 0 | 0 | 0 | 0 | 0 | 0 | 0 | 0 | 1 | Lats1 |
| Lats2 | 1067 | 893 | 1398 | 1189 | 1293 | 1220 | 1033 | 1152 | 990 | 1248 | 1417 | 1228 | 1387 | 1228 | 940 | 1137 | 1402 | 1262 | 1149 | 895 | 1139 | 1290 | 1188 | 1273 | 1146 | 1464 | 1177 | 1260 | Lats2 |
| Lax1 | 240 | 284 | 238 | 270 | 326 | 253 | 222 | 56 | 62 | 52 | 92 | 36 | 60 | 30 | 269 | 401 | 283 | 344 | 238 | 225 | 245 | 37 | 91 | 38 | 37 | 31 | 42 | 44 | Lax1 |
| Layn | 393 | 258 | 272 | 236 | 273 | 303 | 216 | 92 | 124 | 79 | 129 | 75 | 85 | 60 | 282 | 329 | 270 | 224 | 237 | 226 | 230 | 97 | 142 | 83 | 129 | 92 | 56 | 69 | Layn |
| Lbh | 6448 | 7283 | 7956 | 8776 | 9696 | 8051 | 8119 | 10352 | 10358 | 10944 | 10327 | 10469 | 10269 | 10981 | 7238 | 6149 | 8576 | 8228 | 8659 | 8612 | 7797 | 12392 | 9638 | 10490 | 12041 | 10497 | 10629 | 10905 | Lbh |
| Lbhd1 | 1254 | 1029 | 1052 | 998 | 947 | 968 | 1141 | 819 | 742 | 720 | 664 | 822 | 759 | 862 | 1228 | 1078 | 951 | 884 | 997 | 977 | 1107 | 692 | 680 | 773 | 701 | 894 | 834 | 1041 | Lbhd1 |
| Lbhd2 | 0 | 1 | 0 | 0 | 1 | 4 | 2 | 2 | 8 | 7 | 21 | 1 | 1 | 0 | 3 | 3 | 1 | 4 | 2 | 6 | 0 | 12 | 10 | 7 | 7 | 2 | 8 | 0 | Lbhd2 |
| Lbp | 107 | 98 | 178 | 63 | 92 | 103 | 28 | 114 | 163 | 159 | 179 | 148 | 184 | 137 | 86 | 173 | 74 | 98 | 47 | 61 | 49 | 145 | 246 | 130 | 104 | 165 | 158 | 135 | Lbp |
| Lbr | 10924 | 10260 | 12049 | 9808 | 6872 | 8458 | 8860 | 6543 | 5916 | 5304 | 6108 | 8690 | 8367 | 9226 | 10508 | 10866 | 10166 | 8501 | 8386 | 8935 | 7864 | 4979 | 5746 | 7544 | 6139 | 10169 | 10010 | 10224 | Lbr |
| Lbx1 | 3 | 42 | 0 | 15 | 314 | 67 | 2 | 177 | 158 | 322 | 508 | 1 | 366 | 3 | 112 | 83 | 48 | 46 | 1 | 2 | 2 | 261 | 317 | 1 | 56 | 2 | 3 | 6 | Lbx1 |
| Lbx2 | 10 | 23 | 25 | 16 | 15 | 10 | 11 | 7 | 2 | 15 | 6 | 10 | 5 | 6 | 16 | 17 | 15 | 19 | 20 | 24 | 11 | 7 | 9 | 11 | 6 | 21 | 6 | 2 | Lbx2 |
| Lca5 | 2526 | 2272 | 2471 | 2711 | 3181 | 2961 | 3085 | 2621 | 2534 | 2482 | 2387 | 2805 | 2448 | 2820 | 2578 | 2234 | 2617 | 2835 | 2952 | 3021 | 3157 | 2757 | 2214 | 2623 | 2809 | 2616 | 2840 | 2592 | Lca5 |
| Lca5l | 279 | 288 | 293 | 308 | 316 | 302 | 276 | 242 | 240 | 306 | 217 | 249 | 324 | 225 | 319 | 318 | 268 | 272 | 258 | 289 | 264 | 262 | 276 | 229 | 213 | 243 | 249 | 211 | Lca5l |
| Lcat | 115 | 159 | 159 | 116 | 85 | 116 | 97 | 88 | 128 | 154 | 195 | 83 | 100 | 68 | 102 | 203 | 136 | 146 | 100 | 55 | 75 | 92 | 159 | 118 | 103 | 126 | 102 | 89 | Lcat |
| Lce1g | 0 | 0 | 0 | 0 | 0 | 0 | 0 | 0 | 0 | 0 | 1 | 0 | 0 | 0 | 0 | 0 | 0 | 0 | 0 | 0 | 0 | 0 | 0 | 0 | 0 | 0 | 0 | 0 | Lce1g |
| Lce1m | 0 | 0 | 0 | 0 | 0 | 0 | 0 | 0 | 0 | 0 | 0 | 0 | 0 | 0 | 1 | 0 | 0 | 0 | 0 | 0 | 0 | 0 | 0 | 0 | 0 | 0 | 0 | 0 | Lce1m |
| Lce3a | 0 | 0 | 0 | 0 | 0 | 0 | 0 | 0 | 0 | 0 | 2 | 0 | 0 | 0 | 0 | 0 | 0 | 0 | 0 | 0 | 0 | 0 | 1 | 0 | 0 | 0 | 0 | 0 | Lce3a |
| Lce3f | 0 | 1 | 0 | 0 | 0 | 0 | 0 | 0 | 0 | 0 | 0 | 0 | 0 | 0 | 0 | 0 | 0 | 0 | 0 | 0 | 0 | 0 | 0 | 0 | 3 | 0 | 0 | 0 | Lce3f |
| Lce6a | 0 | 0 | 0 | 0 | 0 | 0 | 1 | 2 | 0 | 0 | 0 | 0 | 0 | 0 | 0 | 1 | 0 | 0 | 0 | 0 | 0 | 0 | 0 | 1 | 0 | 0 | 0 | 0 | Lce6a |
| Lck | 223 | 239 | 198 | 193 | 210 | 243 | 229 | 276 | 325 | 287 | 263 | 298 | 318 | 268 | 203 | 225 | 219 | 160 | 262 | 213 | 250 | 222 | 354 | 267 | 258 | 367 | 352 | 298 | Lck |
| Lclat1 | 3896 | 3558 | 3336 | 3241 | 3238 | 3340 | 3456 | 2379 | 2297 | 2298 | 2270 | 2086 | 2249 | 2158 | 3700 | 3505 | 3130 | 3595 | 3125 | 3094 | 3238 | 2465 | 2350 | 2063 | 2127 | 1967 | 2042 | 2112 | Lclat1 |
| Lcmt1 | 4412 | 4964 | 4086 | 4325 | 3870 | 3944 | 4849 | 2866 | 2712 | 2449 | 2297 | 2447 | 2468 | 2623 | 4830 | 4379 | 4333 | 4415 | 4293 | 4924 | 4570 | 2480 | 2548 | 2421 | 2693 | 2523 | 2602 | 2744 | Lcmt1 |
| Lcmt2 | 920 | 895 | 938 | 854 | 836 | 907 | 1034 | 710 | 935 | 846 | 926 | 925 | 873 | 941 | 955 | 800 | 999 | 949 | 1027 | 881 | 1061 | 832 | 860 | 1069 | 906 | 1094 | 883 | 948 | Lcmt2 |
| Lcn11 | 1 | 0 | 0 | 2 | 0 | 0 | 0 | 0 | 0 | 0 | 1 | 0 | 0 | 0 | 0 | 0 | 0 | 0 | 0 | 0 | 2 | 0 | 0 | 0 | 0 | 0 | 0 | 0 | Lcn11 |
| Lcn12 | 0 | 1 | 0 | 0 | 0 | 0 | 0 | 0 | 0 | 1 | 1 | 0 | 0 | 2 | 0 | 1 | 0 | 0 | 0 | 0 | 0 | 3 | 3 | 2 | 0 | 9 | 2 | 1 | Lcn12 |
| Lcn2 | 2 | 3 | 5 | 9 | 0 | 7 | 2 | 4 | 15 | 9 | 38 | 9 | 26 | 6 | 1 | 19 | 8 | 6 | 3 | 6 | 2 | 13 | 23 | 19 | 4 | 10 | 19 | 19 | Lcn2 |
| Lcn3 | 0 | 0 | 0 | 0 | 0 | 0 | 2 | 1 | 0 | 0 | 0 | 0 | 0 | 0 | 3 | 1 | 0 | 2 | 0 | 0 | 0 | 0 | 0 | 0 | 0 | 0 | 0 | 0 | Lcn3 |
| Lcn5 | 0 | 0 | 0 | 0 | 0 | 0 | 0 | 0 | 0 | 0 | 0 | 0 | 0 | 0 | 0 | 0 | 0 | 0 | 0 | 0 | 0 | 0 | 1 | 0 | 0 | 0 | 0 | 0 | Lcn5 |
| Lcn6 | 0 | 0 | 0 | 0 | 0 | 3 | 0 | 0 | 0 | 0 | 0 | 0 | 0 | 0 | 0 | 0 | 0 | 1 | 0 | 0 | 0 | 0 | 0 | 0 | 0 | 0 | 0 | 0 | Lcn6 |
| Lcn8 | 2 | 0 | 0 | 2 | 1 | 3 | 2 | 4 | 3 | 1 | 1 | 0 | 7 | 1 | 0 | 1 | 3 | 0 | 0 | 0 | 4 | 0 | 3 | 2 | 4 | 1 | 3 | 0 | Lcn8 |
| Lcor | 5214 | 4495 | 576 | 2734 | 3634 | 5285 | 4357 | 7260 | 5441 | 3503 | 4189 | 4216 | 4585 | 4503 | 5352 | 4255 | 1267 | 2552 | 2865 | 6094 | 5038 | 2426 | 4890 | 1920 | 4572 | 461 | 5905 | 3503 | Lcor |
| Lcorl | 5374 | 3827 | 3617 | 4977 | 5917 | 6528 | 7032 | 9220 | 7222 | 6047 | 5978 | 6794 | 5960 | 7355 | 6397 | 4974 | 4077 | 5388 | 5594 | 7752 | 6579 | 7135 | 7272 | 6217 | 7202 | 5382 | 7065 | 6636 | Lcorl |
| Lcp1 | 2798 | 2390 | 2466 | 2075 | 1861 | 2090 | 1943 | 2291 | 2591 | 2379 | 2516 | 2456 | 2559 | 2534 | 2511 | 2571 | 1953 | 2195 | 1800 | 2005 | 1963 | 2503 | 2940 | 2346 | 2201 | 2976 | 2485 | 2465 | Lcp1 |
| Lcp2 | 95 | 78 | 74 | 60 | 70 | 61 | 50 | 151 | 198 | 174 | 234 | 167 | 160 | 154 | 103 | 115 | 49 | 79 | 90 | 84 | 76 | 211 | 261 | 166 | 117 | 211 | 189 | 150 | Lcp2 |
| Lct | 205 | 249 | 225 | 180 | 106 | 198 | 150 | 72 | 86 | 78 | 69 | 110 | 84 | 99 | 171 | 239 | 199 | 160 | 159 | 153 | 144 | 65 | 60 | 66 | 57 | 80 | 63 | 78 | Lct |
| Lctl | 1 | 0 | 0 | 1 | 2 | 0 | 2 | 1 | 1 | 0 | 0 | 1 | 0 | 2 | 1 | 2 | 1 | 1 | 0 | 0 | 0 | 3 | 2 | 1 | 1 | 9 | 6 | 4 | Lctl |
| Ldah | 2678 | 2672 | 2652 | 2214 | 2340 | 2241 | 2312 | 3853 | 3274 | 3284 | 3201 | 3379 | 3514 | 3412 | 2806 | 2572 | 2526 | 2085 | 2462 | 2464 | 2162 | 3352 | 3396 | 3109 | 3234 | 3507 | 3555 | 3527 | Ldah |
| Ldb1 | 11539 | 13623 | 13860 | 15177 | 14545 | 13698 | 12890 | 15975 | 17077 | 18809 | 16499 | 20433 | 17386 | 18173 | 10535 | 11623 | 15333 | 14582 | 15888 | 12678 | 13687 | 17355 | 15612 | 20771 | 19875 | 22064 | 18288 | 18199 | Ldb1 |

| | | | | | | | | | | | | | | | | | | | | | | | | | | | | | |
|---|---|---|---|---|---|---|---|---|---|---|---|---|---|---|---|---|---|---|---|---|---|---|---|---|---|---|---|---|---|
| Ldb2 | 1397 | 1263 | 1256 | 1237 | 1342 | 1324 | 1256 | 1474 | 1647 | 1668 | 2055 | 1196 | 1650 | 1415 | 1356 | 1672 | 1041 | 1434 | 1360 | 1280 | 1500 | 1828 | 2069 | 1622 | 1718 | 1499 | 1374 | 1329 | Ldb2 |
| Ldb3 | 38 | 41 | 46 | 33 | 89 | 70 | 42 | 103 | 180 | 89 | 991 | 36 | 79 | 60 | 47 | 48 | 34 | 44 | 57 | 35 | 42 | 175 | 658 | 58 | 53 | 51 | 60 | 56 | Ldb3 |
| Ldc1 | 25 | 23 | 0 | 18 | 29 | 19 | 13 | 23 | 35 | 34 | 11 | 40 | 40 | 2 | 13 | 43 | 2 | 14 | 29 | 20 | 18 | 22 | 99 | 7 | 19 | 1 | 5 | 8 | Ldc1 |
| Ldha | 31661 | 28917 | 26723 | 21318 | 17201 | 22658 | 18134 | 22297 | 26592 | 21885 | 23027 | 23315 | 25484 | 23379 | 27606 | 26827 | 24839 | 20520 | 21850 | 23190 | 19025 | 17326 | 28405 | 24246 | 20451 | 32416 | 27041 | 28769 | Ldha |
| Ldhal6b | 42 | 46 | 25 | 59 | 35 | 43 | 48 | 53 | 67 | 91 | 55 | 66 | 50 | 51 | 42 | 50 | 22 | 39 | 52 | 55 | 42 | 43 | 63 | 48 | 57 | 10 | 41 | 68 | Ldhal6b |
| Ldhb | 10912 | 12958 | 10810 | 11374 | 10566 | 10076 | 11706 | 10833 | 10821 | 10085 | 9510 | 10820 | 9443 | 10904 | 11236 | 11139 | 11502 | 9798 | 11956 | 12518 | 11701 | 8758 | 10174 | 10722 | 10486 | 12612 | 10412 | 11056 | Ldhb |
| Ldhc | 0 | 0 | 0 | 0 | 2 | 0 | 1 | 1 | 2 | 3 | 2 | 0 | 0 | 0 | 3 | 0 | 0 | 0 | 0 | 4 | 0 | 0 | 1 | 2 | 0 | 0 | 6 | 0 | Ldhc |
| Ldhd | 14 | 21 | 14 | 19 | 25 | 16 | 12 | 20 | 39 | 32 | 22 | 12 | 34 | 16 | 9 | 23 | 16 | 17 | 24 | 19 | 15 | 50 | 30 | 29 | 25 | 25 | 21 | 18 | Ldhd |
| Ldlr | 2337 | 2728 | 3257 | 2237 | 1919 | 2060 | 1805 | 2252 | 2528 | 2685 | 2671 | 3950 | 4082 | 4023 | 2197 | 2626 | 2843 | 2204 | 1928 | 1570 | 1770 | 2369 | 2236 | 3180 | 2792 | 3954 | 4645 | 4653 | Ldlr |
| Ldlrad1 | 6 | 7 | 0 | 0 | 4 | 3 | 1 | 1 | 0 | 0 | 1 | 4 | 5 | 0 | 1 | 5 | 3 | 2 | 3 | 3 | 5 | 2 | 2 | 3 | 0 | 2 | 1 | 2 | Ldlrad1 |
| Ldlrad2 | 17 | 10 | 7 | 3 | 8 | 19 | 7 | 9 | 22 | 10 | 13 | 26 | 25 | 4 | 5 | 10 | 2 | 8 | 5 | 1 | 9 | 42 | 32 | 4 | 10 | 8 | 7 | 2 | Ldlrad2 |
| Ldlrad3 | 4841 | 4304 | 4660 | 4431 | 4345 | 4441 | 4293 | 4330 | 4186 | 4454 | 4903 | 4907 | 4897 | 4781 | 4874 | 4279 | 4256 | 4329 | 4147 | 4332 | 4372 | 4472 | 4503 | 4363 | 4434 | 4938 | 4865 | 5124 | Ldlrad3 |
| Ldlrad4 | 450 | 453 | 227 | 389 | 551 | 577 | 381 | 532 | 637 | 631 | 772 | 457 | 578 | 424 | 419 | 575 | 311 | 501 | 444 | 361 | 474 | 478 | 751 | 447 | 580 | 204 | 553 | 349 | Ldlrad4 |
| Ldlrap1 | 1780 | 1866 | 1589 | 1378 | 898 | 1177 | 746 | 700 | 930 | 808 | 999 | 556 | 756 | 500 | 1255 | 2167 | 1352 | 1405 | 1051 | 747 | 730 | 525 | 1095 | 891 | 525 | 892 | 660 | 616 | Ldlrap1 |
| Ldoc1 | 30 | 34 | 35 | 20 | 59 | 36 | 38 | 33 | 43 | 28 | 41 | 15 | 31 | 21 | 26 | 45 | 24 | 22 | 48 | 36 | 28 | 72 | 24 | 20 | 21 | 13 | 20 | 12 | Ldoc1 |
| Leap2 | 3 | 6 | 0 | 2 | 2 | 6 | 7 | 4 | 2 | 7 | 0 | 2 | 3 | 2 | 3 | 5 | 6 | 0 | 4 | 4 | 2 | 7 | 0 | 5 | 4 | 2 | 7 | 11 | Leap2 |
| Lect2 | 0 | 0 | 0 | 2 | 0 | 0 | 0 | 0 | 0 | 0 | 0 | 0 | 0 | 0 | 0 | 1 | 2 | 0 | 0 | 0 | 0 | 0 | 0 | 0 | 1 | 0 | 0 | 0 | Lect2 |
| Lef1 | 2450 | 1810 | 3169 | 2188 | 1856 | 1862 | 1793 | 1932 | 1977 | 2233 | 1687 | 3132 | 2879 | 2983 | 1863 | 2091 | 2460 | 2092 | 2427 | 1646 | 1963 | 2065 | 1960 | 2997 | 2236 | 2875 | 2989 | 3551 | Lef1 |
| Lefty1 | 44 | 87 | 68 | 68 | 21 | 37 | 23 | 7 | 27 | 12 | 24 | 10 | 20 | 14 | 78 | 88 | 42 | 59 | 30 | 20 | 25 | 13 | 36 | 20 | 18 | 20 | 26 | 19 | Lefty1 |
| Lefty2 | 8 | 6 | 0 | 3 | 15 | 8 | 7 | 4 | 9 | 4 | 19 | 4 | 1 | 3 | 4 | 5 | 3 | 17 | 4 | 15 | 15 | 10 | 9 | 4 | 22 | 3 | 0 | 2 | Lefty2 |
| Lekr1 | 679 | 523 | 604 | 566 | 650 | 599 | 637 | 512 | 487 | 447 | 442 | 458 | 451 | 498 | 675 | 657 | 581 | 653 | 485 | 621 | 620 | 508 | 553 | 476 | 472 | 467 | 458 | 496 | Lekr1 |
| Lemd1 | 200 | 219 | 172 | 313 | 327 | 319 | 344 | 482 | 447 | 564 | 535 | 642 | 541 | 731 | 227 | 174 | 350 | 335 | 348 | 264 | 382 | 622 | 432 | 546 | 659 | 365 | 733 | 654 | Lemd1 |
| Lemd2 | 2844 | 3504 | 3694 | 3916 | 3805 | 3647 | 3081 | 3171 | 3660 | 3986 | 3986 | 4264 | 4102 | 3673 | 2651 | 3016 | 3833 | 3761 | 4094 | 3073 | 3296 | 4067 | 3620 | 4380 | 4498 | 4327 | 4106 | 4190 | Lemd2 |
| Lemd3 | 3279 | 2389 | 3241 | 3568 | 3530 | 3490 | 3572 | 3022 | 2771 | 3429 | 3260 | 3382 | 3302 | 3455 | 3144 | 2928 | 3008 | 3425 | 3444 | 3062 | 3417 | 3369 | 2683 | 3394 | 3533 | 3380 | 3134 | 3474 | Lemd3 |
| Lenep | 1 | 1 | 0 | 8 | 0 | 0 | 2 | 0 | 1 | 3 | 0 | 0 | 0 | 1 | 0 | 0 | 0 | 1 | 0 | 2 | 1 | 3 | 0 | 0 | 0 | 5 | 0 | 2 | Lenep |
| Leng1 | 1336 | 1757 | 2061 | 2033 | 1577 | 1471 | 1493 | 771 | 778 | 1012 | 945 | 976 | 892 | 1007 | 1316 | 1527 | 2129 | 1826 | 1635 | 1431 | 1599 | 867 | 764 | 987 | 936 | 1117 | 949 | 1025 | Leng1 |
| Leng8 | 15246 | 18565 | 20099 | 21803 | 21419 | 20924 | 22218 | 19615 | 20818 | 25813 | 23339 | 29515 | 26528 | 29711 | 14893 | 14968 | 22130 | 20588 | 22696 | 17914 | 21510 | 27294 | 18396 | 28089 | 27357 | 26326 | 26164 | 26631 | Leng8 |
| Leng9 | 257 | 242 | 249 | 256 | 275 | 254 | 222 | 219 | 239 | 220 | 300 | 172 | 191 | 170 | 214 | 240 | 258 | 288 | 264 | 207 | 187 | 172 | 237 | 266 | 206 | 226 | 192 | 217 | Leng9 |
| Leo1 | 3466 | 3848 | 4572 | 3953 | 3471 | 3392 | 3476 | 3274 | 3241 | 3754 | 3659 | 4096 | 3775 | 4552 | 3224 | 3227 | 4183 | 3441 | 3496 | 3378 | 3437 | 3527 | 3154 | 4162 | 3819 | 4756 | 4331 | 4376 | Leo1 |
| Lep | 0 | 0 | 0 | 0 | 0 | 0 | 0 | 0 | 0 | 0 | 0 | 0 | 0 | 0 | 0 | 0 | 0 | 0 | 0 | 0 | 0 | 0 | 1 | 0 | 0 | 0 | 0 | 0 | Lep |
| Lepr | 1215 | 609 | 571 | 554 | 810 | 1033 | 660 | 1341 | 1629 | 1198 | 2076 | 732 | 1103 | 772 | 1092 | 1212 | 561 | 999 | 450 | 646 | 632 | 1579 | 2289 | 1115 | 1031 | 699 | 790 | 642 | Lepr |
| Leprot | 4624 | 3904 | 4357 | 3928 | 4104 | 4084 | 3918 | 4473 | 4394 | 3868 | 3902 | 3738 | 3936 | 3880 | 4623 | 4125 | 3797 | 4040 | 4035 | 4501 | 4156 | 4229 | 4779 | 3898 | 3734 | 4257 | 3993 | 4010 | Leprot |
| Leprotl1 | 3937 | 3778 | 4205 | 5207 | 5443 | 5153 | 5461 | 5318 | 4597 | 5606 | 4775 | 5419 | 5202 | 5617 | 4377 | 3534 | 4895 | 5133 | 5073 | 5351 | 5171 | 5558 | 4368 | 5760 | 5921 | 5372 | 5761 | 5416 | Leprotl1 |
| Letm1 | 2915 | 2738 | 3002 | 3184 | 3427 | 3483 | 3001 | 4162 | 4287 | 5211 | 4353 | 4779 | 4348 | 4649 | 2554 | 2692 | 3298 | 3159 | 3564 | 3046 | 3181 | 4502 | 4171 | 4981 | 5041 | 5137 | 4565 | 4594 | Letm1 |
| Letm2 | 765 | 827 | 942 | 1105 | 1207 | 1084 | 999 | 1161 | 1141 | 1324 | 1382 | 1165 | 1263 | 1206 | 869 | 725 | 1160 | 1054 | 1205 | 1045 | 1040 | 1335 | 1169 | 1280 | 1254 | 955 | 1281 | 1085 | Letm2 |
| Letmd1 | 2788 | 3256 | 3037 | 3183 | 3308 | 2859 | 3329 | 3326 | 3540 | 3551 | 3406 | 3729 | 3509 | 3828 | 2909 | 2846 | 3318 | 3223 | 3313 | 2969 | 3309 | 3606 | 3387 | 3662 | 3649 | 3719 | 3542 | 3796 | Letmd1 |
| Lexm | 2 | 13 | 3 | 5 | 0 | 6 | 3 | 5 | 12 | 6 | 8 | 8 | 34 | 24 | 4 | 2 | 9 | 0 | 5 | 4 | 2 | 7 | 16 | 8 | 17 | 16 | 17 | 34 | Lexm |
| Lfng | 1044 | 1475 | 1382 | 1143 | 1152 | 1066 | 1131 | 1704 | 1786 | 1777 | 1655 | 2467 | 2146 | 2271 | 1076 | 1048 | 1549 | 1124 | 1441 | 1168 | 1144 | 1394 | 1772 | 2155 | 1942 | 2560 | 2328 | 2336 | Lfng |
| Lgals1 | 13205 | 11501 | 9888 | 6187 | 3365 | 7686 | 2546 | 8954 | 12994 | 7622 | 15478 | 8060 | 11551 | 7493 | 10802 | 12365 | 7310 | 9102 | 3585 | 4062 | 2968 | 6086 | 17018 | 9130 | 6595 | 8889 | 9360 | 8568 | Lgals1 |
| Lgals12 | 74 | 34 | 44 | 60 | 67 | 38 | 44 | 73 | 123 | 81 | 70 | 72 | 43 | 51 | 41 | 57 | 39 | 39 | 80 | 48 | 60 | 69 | 94 | 101 | 66 | 58 | 42 | 44 | Lgals12 |
| Lgals2 | 2 | 0 | 0 | 1 | 1 | 0 | 0 | 0 | 6 | 0 | 1 | 0 | 1 | 0 | 0 | 2 | 0 | 6 | 0 | 1 | 0 | 2 | 1 | 2 | 0 | 0 | 0 | 0 | Lgals2 |
| Lgals3 | 30 | 61 | 16 | 63 | 58 | 55 | 74 | 42 | 74 | 51 | 100 | 23 | 22 | 24 | 42 | 51 | 57 | 89 | 59 | 48 | 70 | 20 | 58 | 38 | 44 | 10 | 31 | 28 | Lgals3 |
| Lgals3bp | 213 | 280 | 240 | 270 | 301 | 238 | 228 | 153 | 245 | 326 | 294 | 180 | 237 | 160 | 187 | 266 | 227 | 326 | 242 | 286 | 299 | 272 | 294 | 237 | 245 | 155 | 193 | 193 | Lgals3bp |
| Lgals4 | 98 | 126 | 102 | 127 | 145 | 114 | 124 | 160 | 183 | 213 | 235 | 125 | 181 | 186 | 140 | 131 | 166 | 142 | 149 | 104 | 125 | 224 | 211 | 173 | 136 | 164 | 137 | 142 | Lgals4 |
| Lgals7 | 89 | 142 | 85 | 140 | 247 | 240 | 92 | 267 | 398 | 328 | 248 | 152 | 335 | 166 | 127 | 156 | 126 | 217 | 169 | 210 | 139 | 259 | 412 | 288 | 251 | 129 | 209 | 190 | Lgals7 |
| Lgals8 | 2431 | 2110 | 2146 | 2105 | 2194 | 2315 | 2168 | 1314 | 1354 | 1344 | 1571 | 1148 | 1368 | 1227 | 2414 | 2317 | 2059 | 2298 | 1979 | 2259 | 2151 | 1507 | 1504 | 1111 | 1360 | 1132 | 1231 | 1162 | Lgals8 |
| Lgals9 | 299 | 330 | 283 | 219 | 209 | 343 | 143 | 408 | 516 | 418 | 582 | 356 | 424 | 332 | 380 | 381 | 198 | 342 | 178 | 292 | 231 | 530 | 789 | 486 | 425 | 385 | 350 | 324 | Lgals9 |
| Lgalsl | 3252 | 2255 | 2986 | 2716 | 3637 | 3290 | 3588 | 4364 | 4412 | 3955 | 4009 | 3221 | 3351 | 3381 | 3451 | 2672 | 2726 | 3090 | 3165 | 3559 | 3593 | 4258 | 4412 | 3716 | 4012 | 3300 | 3192 | 3414 | Lgalsl |
| Lgi1 | 133 | 81 | 59 | 134 | 336 | 216 | 219 | 546 | 637 | 810 | 729 | 467 | 420 | 361 | 140 | 79 | 108 | 142 | 243 | 261 | 273 | 879 | 840 | 456 | 697 | 223 | 328 | 295 | Lgi1 |
| Lgi2 | 1414 | 966 | 1279 | 1297 | 1497 | 1504 | 1164 | 2034 | 1906 | 1910 | 2172 | 996 | 1310 | 957 | 1158 | 1301 | 1198 | 1422 | 1244 | 1024 | 1124 | 1875 | 2208 | 1654 | 1451 | 988 | 898 | 885 | Lgi2 |
| Lgi3 | 97 | 211 | 123 | 176 | 397 | 197 | 144 | 214 | 247 | 225 | 302 | 76 | 294 | 79 | 245 | 220 | 165 | 223 | 203 | 215 | 167 | 276 | 254 | 102 | 139 | 63 | 59 | 57 | Lgi3 |
| Lgi4 | 114 | 124 | 117 | 66 | 91 | 57 | 38 | 159 | 209 | 208 | 249 | 106 | 294 | 104 | 117 | 195 | 122 | 101 | 44 | 47 | 30 | 232 | 201 | 128 | 192 | 123 | 155 | 155 | Lgi4 |
| Lgmn | 3567 | 4071 | 3579 | 3607 | 3504 | 3350 | 2816 | 2707 | 3124 | 2753 | 3049 | 2349 | 2581 | 2197 | 3584 | 4224 | 3490 | 3515 | 3057 | 3373 | 2963 | 2648 | 3607 | 2377 | 2254 | 2751 | 2303 | 2376 | Lgmn |
| Lgr4 | 7235 | 6263 | 6608 | 6224 | 5613 | 5581 | 4727 | 4402 | 4107 | 3758 | 5041 | 3823 | 5559 | 3844 | 7299 | 8020 | 6230 | 5854 | 4532 | 4395 | 4980 | 4406 | 4416 | 3818 | 3186 | 4011 | 4328 | 4074 | Lgr4 |
| Lgr5 | 1411 | 1024 | 843 | 697 | 553 | 785 | 358 | 1130 | 1585 | 977 | 1552 | 630 | 930 | 490 | 1225 | 1504 | 793 | 781 | 398 | 485 | 407 | 581 | 1244 | 834 | 459 | 450 | 589 | 487 | Lgr5 |
| Lgr6 | 276 | 66 | 79 | 60 | 66 | 103 | 29 | 332 | 438 | 277 | 273 | 109 | 120 | 90 | 76 | 132 | 22 | 105 | 48 | 36 | 40 | 154 | 484 | 251 | 203 | 131 | 110 | 82 | Lgr6 |
| Lgsn | 0 | 1 | 0 | 0 | 0 | 0 | 0 | 0 | 1 | 1 | 0 | 5 | 0 | 0 | 0 | 0 | 0 | 0 | 2 | 0 | 0 | 0 | 0 | 0 | 0 | 0 | 0 | 0 | Lgsn |
| Lhb | 0 | 0 | 0 | 0 | 0 | 0 | 0 | 0 | 0 | 0 | 0 | 0 | 0 | 0 | 0 | 0 | 0 | 1 | 0 | 0 | 0 | 0 | 0 | 0 | 0 | 2 | 0 | 0 | Lhb |
| Lhb | 23 | 49 | 12 | 18 | 56 | 35 | 23 | 38 | 39 | 56 | 27 | 33 | 62 | 16 | 32 | 45 | 15 | 36 | 20 | 35 | 29 | 47 | 97 | 34 | 39 | 16 | 26 | 19 | Lhb |
| Lhcgr | 14 | 14 | 9 | 5 | 52 | 18 | 8 | 85 | 85 | 71 | 44 | 31 | 64 | 24 | 24 | 18 | 5 | 10 | 13 | 18 | 16 | 84 | 109 | 42 | 26 | 10 | 21 | 26 | Lhcgr |
| Lhfp | 2331 | 2184 | 2524 | 2066 | 2179 | 2198 | 1590 | 2216 | 2281 | 2244 | 2531 | 2033 | 2710 | 2000 | 2575 | 2530 | 2143 | 2260 | 1746 | 1820 | 1615 | 2618 | 2975 | 2158 | 2253 | 2283 | 2398 | 2255 | Lhfp |
| Lhfpl1 | 28 | 13 | 10 | 12 | 16 | 7 | 4 | 44 | 62 | 21 | 115 | 12 | 15 | 7 | 15 | 34 | 0 | 19 | 0 | 8 | 4 | 50 | 74 | 21 | 17 | 32 | 5 | 12 | Lhfpl1 |
| Lhfpl2 | 3564 | 3137 | 3188 | 2449 | 2257 | 2659 | 1854 | 3762 | 4818 | 4006 | 7282 | 3879 | 4811 | 3648 | 3104 | 4167 | 2573 | 2890 | 1972 | 2101 | 1983 | 4176 | 7074 | 4278 | 3658 | 4753 | 3898 | 4260 | Lhfpl2 |
| Lhfpl3 | 165 | 143 | 112 | 241 | 525 | 348 | 477 | 413 | 394 | 375 | 297 | 180 | 147 | 219 | 241 | 136 | 232 | 372 | 333 | 427 | 472 | 475 | 399 | 292 | 442 | 132 | 149 | 160 | Lhfpl3 |
| Lhfpl4 | 5095 | 6532 | 9089 | 11052 | 12838 | 11284 | 9941 | 7383 | 7321 | 9700 | 7605 | 9526 | 7996 | 8594 | 6485 | 5710 | 10423 | 10806 | 12558 | 10326 | 10613 | 11533 | 6530 | 9769 | 11544 | 8087 | 8836 | 8419 | Lhfpl4 |
| Lhfpl5 | 21 | 20 | 22 | 25 | 65 | 38 | 30 | 33 | 49 | 58 | 37 | 37 | 34 | 19 | 21 | 19 | 9 | 23 | 34 | 28 | 29 | 42 | 50 | 27 | 42 | 22 | 29 | 32 | Lhfpl5 |
| Lhpp | 735 | 923 | 816 | 658 | 707 | 670 | 559 | 722 | 901 | 768 | 668 | 668 | 663 | 559 | 781 | 906 | 677 | 836 | 771 | 746 | 634 | 675 | 910 | 630 | 698 | 656 | 700 | 662 | Lhpp |
| Lhx1 | 1889 | 2703 | 4338 | 4725 | 4745 | 4112 | 2887 | 4340 | 3112 | 5336 | 4637 | 5653 | 7626 | 5184 | 2545 | 2598 | 4584 | 3934 | 4090 | 3517 | 3173 | 6617 | 3793 | 4510 | 5379 | 2550 | 5878 | 6394 | Lhx1 |
| Lhx2 | 6347 | 6938 | 8096 | 10109 | 9255 | 8756 | 11644 | 8131 | 7003 | 10771 | 7432 | 16411 | 11231 | 16100 | 5599 | 4402 | 9547 | 9127 | 12292 | 9897 | 11451 | 7668 | 6890 | 15729 | 12588 | 19402 | 12944 | 16141 | Lhx2 |
| Lhx3 | 35 | 52 | 21 | 31 | 111 | 13 | 8 | 67 | 129 | 74 | 114 | 8 | 142 | 9 | 66 | 28 | 9 | 66 | 10 | 18 | 18 | 162 | 67 | 4 | 7 | 8 | 15 | 5 | Lhx3 |
| Lhx4 | 120 | 182 | 202 | 210 | 385 | 229 | 144 | 319 | 264 | 369 | 333 | 217 | 696 | 140 | 268 | 209 | 224 | 210 | 171 | 121 | 170 | 446 | 157 | 148 | 249 | 5 | 232 | 206 | Lhx4 |
| Lhx5 | 1055 | 1692 | 2470 | 2132 | 1806 | 1718 | 1210 | 1840 | 1499 | 2414 | 1884 | 2643 | 3158 | 2424 | 1250 | 1193 | 2651 | 1860 | 1864 | 1541 | 1515 | 2692 | 1556 | 2209 | 2649 | 1990 | 2508 | 2754 | Lhx5 |
| Lhx6 | 1046 | 652 | 468 | 1330 | 1411 | 1311 | 1884 | 774 | 1846 | 1896 | 1505 | 2489 | 175 | 2464 | 454 | 491 | 1336 | 1163 | 1869 | 724 | 1768 | 2769 | 852 | 2452 | 2503 | 2946 | 1327 | 1543 | Lhx6 |
| Lhx8 | 444 | 214 | 52 | 317 | 390 | 330 | 537 | 733 | 1225 | 762 | 701 | 1128 | 65 | 1288 | 148 | 129 | 421 | 322 | 588 | 203 | 522 | 1031 | 394 | 1168 | 968 | 1506 | 747 | 749 | Lhx8 |
| Lhx9 | 2759 | 2553 | 4848 | 5114 | 6587 | 6676 | 6053 | 9233 | 4660 | 8984 | 6192 | 11687 | 10153 | 13243 | 3394 | 2392 | 5775 | 4631 | 8023 | 6593 | 7365 | 11988 | 7002 | 11510 | 12471 | 7205 | 12667 | 13620 | Lhx9 |
| Lias | 2359 | 2194 | 2176 | 2168 | 2313 | 2228 | 2472 | 2746 | 2626 | 2564 | 2623 | 2994 | 2767 | 3157 | 2499 | 2082 | 2103 | 2249 | 2334 | 2387 | 2507 | 2532 | 2659 | 2915 | 2712 | 3309 | 3127 | 3285 | Lias |
| Lif | 0 | 3 | 5 | 0 | 2 | 3 | 0 | 7 | 7 | 5 | 12 | 8 | 3 | 4 | 11 | 11 | 6 | 3 | 0 | 0 | 2 | 3 | 8 | 5 | 2 | 2 | 5 | 1 | Lif |
| Lifr | 3421 | 3200 | 3309 | 3199 | 3662 | 2817 | 2559 | 2462 | 2958 | 2535 | 3627 | 1757 | 2899 | 1657 | 3880 | 4517 | 2766 | 3861 | 2493 | 2330 | 2613 | 3420 | 3291 | 1761 | 2404 | 1842 | 1685 | 1721 | Lifr |
| Lig1 | 10676 | 10460 | 11396 | 9080 | 7755 | 9108 | 8810 | 7388 | 8609 | 8387 | 8063 | 12205 | 10034 | 11191 | 9087 | 9245 | 9797 | 8389 | 9277 | 8516 | 8333 | 6229 | 7275 | 10019 | 8633 | 14764 | 11886 | 12055 | Lig1 |

| | | | | | | | | | | | | | | | | | | | | | | | | | | | | | |
|---|---|---|---|---|---|---|---|---|---|---|---|---|---|---|---|---|---|---|---|---|---|---|---|---|---|---|---|---|---|
| Lig3 | 7248 | 7178 | 5889 | 6125 | 6098 | 6924 | 7017 | 9662 | 9141 | 8472 | 7819 | 9350 | 8970 | 9302 | 7232 | 5977 | 5904 | 5442 | 6496 | 6998 | 6603 | 8241 | 8761 | 8349 | 8603 | 8020 | 9518 | 9145 | Lig3 |
| Lig4 | 2766 | 3029 | 2837 | 3288 | 2825 | 2778 | 3101 | 2816 | 2168 | 2299 | 2063 | 2285 | 2130 | 2274 | 2970 | 2742 | 3042 | 3270 | 2748 | 3166 | 2943 | 2632 | 2137 | 2258 | 2506 | 2230 | 2434 | 2403 | Lig4 |
| Lilr4b | 3 | 1 | 0 | 1 | 2 | 0 | 3 | 4 | 12 | 0 | 23 | 7 | 16 | 10 | 4 | 2 | 1 | 0 | 0 | 7 | 5 | 7 | 16 | 5 | 7 | 7 | 3 | 4 | Lilr4b |
| Lilra5 | 3 | 0 | 0 | 7 | 1 | 1 | 3 | 2 | 2 | 8 | 6 | 4 | 1 | 1 | 0 | 0 | 0 | 3 | 0 | 1 | 2 | 3 | 4 | 2 | 3 | 2 | 1 | 6 | Lilra5 |
| Lilra6 | 0 | 0 | 0 | 0 | 0 | 0 | 0 | 1 | 0 | 0 | 0 | 0 | 5 | 1 | 0 | 1 | 0 | 0 | 0 | 0 | 0 | 0 | 3 | 0 | 5 | 0 | 0 | 0 | Lilra6 |
| Lilrb4a | 12 | 8 | 9 | 1 | 11 | 13 | 15 | 31 | 8 | 16 | 38 | 5 | 15 | 14 | 9 | 13 | 10 | 20 | 3 | 12 | 6 | 18 | 42 | 15 | 23 | 22 | 6 | 25 | Lilrb4a |
| Lima1 | 8056 | 8200 | 7365 | 7672 | 6388 | 7249 | 6754 | 5598 | 6703 | 5996 | 7509 | 5434 | 5675 | 5430 | 7354 | 7795 | 7493 | 7275 | 7237 | 6207 | 6346 | 4562 | 7009 | 6579 | 5042 | 7452 | 4898 | 5003 | Lima1 |
| Limch1 | 5725 | 5036 | 4914 | 3803 | 4673 | 5536 | 4063 | 7468 | 8268 | 7516 | 7942 | 7090 | 8921 | 7313 | 5906 | 6257 | 4285 | 4801 | 4321 | 4329 | 4261 | 6269 | 8201 | 7174 | 7001 | 5858 | 7211 | 7283 | Limch1 |
| Limd1 | 6810 | 7097 | 6448 | 6215 | 5678 | 5809 | 5757 | 4325 | 5044 | 4445 | 4674 | 5483 | 5382 | 4851 | 6211 | 6805 | 6261 | 6083 | 6502 | 5464 | 5658 | 3721 | 4827 | 4885 | 4191 | 5732 | 5930 | 5245 | Limd1 |
| Limd2 | 2997 | 3600 | 3624 | 2765 | 2729 | 3092 | 2678 | 5062 | 5984 | 5886 | 5553 | 6539 | 6747 | 6820 | 2892 | 3221 | 3233 | 2591 | 2976 | 2732 | 2747 | 4296 | 5960 | 6481 | 5528 | 8108 | 6763 | 7131 | Limd2 |
| Lime1 | 3 | 7 | 0 | 1 | 0 | 0 | 7 | 1 | 0 | 2 | 2 | 2 | 0 | 1 | 1 | 2 | 0 | 0 | 1 | 0 | 2 | 3 | 0 | 0 | 0 | 0 | 1 | 1 | Lime1 |
| Limk1 | 6747 | 8718 | 9047 | 10167 | 9984 | 8669 | 8076 | 4297 | 5050 | 5706 | 5589 | 5982 | 5051 | 5348 | 6707 | 7418 | 10343 | 10324 | 10387 | 6930 | 9071 | 6004 | 4846 | 5618 | 6202 | 5708 | 5055 | 5345 | Limk1 |
| Limk2 | 9878 | 11485 | 10011 | 11778 | 13493 | 12414 | 11579 | 8701 | 9074 | 9791 | 8837 | 9018 | 9161 | 8240 | 10309 | 10722 | 11631 | 12765 | 13150 | 11121 | 12291 | 10333 | 8791 | 9121 | 9962 | 7258 | 8684 | 8496 | Limk2 |
| Lims1 | 15741 | 11486 | 11997 | 10640 | 8832 | 10699 | 11268 | 9062 | 8230 | 6483 | 8162 | 7999 | 9039 | 8541 | 15191 | 14495 | 10119 | 11115 | 9192 | 11725 | 10002 | 7240 | 8758 | 7138 | 7123 | 8438 | 8666 | 9062 | Lims1 |
| Lims2 | 67 | 109 | 116 | 85 | 106 | 89 | 74 | 84 | 101 | 96 | 150 | 121 | 151 | 117 | 78 | 114 | 100 | 80 | 96 | 47 | 67 | 112 | 102 | 98 | 94 | 87 | 182 | 127 | Lims2 |
| Lin28a | 359 | 292 | 586 | 256 | 177 | 220 | 181 | 94 | 63 | 111 | 119 | 304 | 338 | 346 | 306 | 298 | 392 | 174 | 223 | 199 | 155 | 157 | 69 | 203 | 125 | 267 | 507 | 495 | Lin28a |
| Lin28b | 2620 | 2035 | 2879 | 2169 | 2346 | 2413 | 2699 | 2301 | 1771 | 2059 | 1835 | 2575 | 2129 | 2839 | 2781 | 2163 | 2553 | 2068 | 2394 | 2342 | 2490 | 2582 | 1630 | 2020 | 2233 | 2457 | 3004 | 2948 | Lin28b |
| Lin37 | 1100 | 1486 | 1206 | 1310 | 975 | 1097 | 1069 | 830 | 742 | 845 | 796 | 863 | 736 | 852 | 1073 | 1012 | 1250 | 1032 | 1116 | 1055 | 1087 | 744 | 757 | 881 | 828 | 843 | 787 | 949 | Lin37 |
| Lin52 | 1857 | 2011 | 1763 | 1762 | 1713 | 1748 | 2200 | 1596 | 1388 | 1286 | 1233 | 1505 | 1448 | 1610 | 1987 | 1837 | 1757 | 1864 | 1766 | 2017 | 2113 | 1439 | 1316 | 1485 | 1576 | 1702 | 1548 | 1631 | Lin52 |
| Lin7a | 563 | 673 | 438 | 684 | 1168 | 945 | 860 | 1520 | 1308 | 1139 | 1148 | 771 | 1146 | 805 | 930 | 727 | 643 | 859 | 919 | 1050 | 902 | 1581 | 1197 | 687 | 1155 | 295 | 782 | 744 | Lin7a |
| Lin7b | 386 | 626 | 575 | 610 | 475 | 527 | 421 | 71 | 84 | 100 | 85 | 83 | 66 | 80 | 398 | 594 | 492 | 706 | 500 | 483 | 424 | 92 | 98 | 64 | 86 | 88 | 57 | 72 | Lin7b |
| Lin7c | 16470 | 13432 | 15637 | 17223 | 18920 | 18170 | 20775 | 23338 | 17664 | 17020 | 16346 | 18409 | 17430 | 19839 | 20181 | 15108 | 16989 | 17216 | 17218 | 21965 | 19418 | 21221 | 17877 | 17417 | 20939 | 16139 | 18986 | 18778 | Lin7c |
| Lin9 | 1210 | 917 | 1102 | 1044 | 1193 | 1066 | 1187 | 2331 | 1888 | 1732 | 1733 | 2165 | 1792 | 2354 | 1246 | 951 | 1089 | 865 | 1329 | 1388 | 1171 | 1815 | 1768 | 2065 | 1979 | 2402 | 2243 | 2249 | Lin9 |
| Lingo1 | 614 | 954 | 920 | 1368 | 2450 | 1602 | 1341 | 1934 | 2380 | 3021 | 2398 | 1568 | 1852 | 1462 | 828 | 879 | 1189 | 1487 | 1895 | 1407 | 1533 | 3385 | 2263 | 1987 | 2673 | 1120 | 1291 | 1327 | Lingo1 |
| Lingo2 | 206 | 194 | 69 | 233 | 461 | 463 | 374 | 650 | 591 | 510 | 527 | 281 | 227 | 289 | 235 | 158 | 151 | 274 | 248 | 422 | 453 | 465 | 672 | 422 | 481 | 358 | 217 | 215 | Lingo2 |
| Lingo3 | 246 | 202 | 215 | 200 | 281 | 197 | 203 | 92 | 95 | 128 | 105 | 63 | 52 | 65 | 164 | 217 | 217 | 219 | 225 | 172 | 223 | 110 | 97 | 93 | 82 | 106 | 57 | 69 | Lingo3 |
| Lingo4 | 54 | 60 | 23 | 47 | 115 | 72 | 102 | 46 | 60 | 32 | 49 | 43 | 52 | 49 | 47 | 32 | 48 | 58 | 70 | 86 | 70 | 47 | 67 | 37 | 68 | 11 | 79 | 51 | Lingo4 |
| Lins1 | 1347 | 1234 | 1120 | 1169 | 1176 | 1189 | 1312 | 2019 | 1851 | 1763 | 1832 | 1869 | 1858 | 1958 | 1349 | 1205 | 1167 | 1176 | 1230 | 1211 | 1213 | 1947 | 1820 | 1711 | 1948 | 1686 | 1887 | 1889 | Lins1 |
| Lipa | 5188 | 4548 | 4877 | 3803 | 3325 | 3933 | 3148 | 2475 | 2501 | 2428 | 2773 | 2207 | 2541 | 2105 | 5023 | 5339 | 3768 | 4044 | 3368 | 3502 | 3196 | 2618 | 3275 | 2411 | 2134 | 2251 | 2306 | 2270 | Lipa |
| Lipc | 0 | 0 | 0 | 0 | 2 | 1 | 1 | 2 | 7 | 3 | 4 | 2 | 5 | 0 | 0 | 0 | 0 | 0 | 0 | 2 | 3 | 3 | 8 | 2 | 1 | 1 | 3 | 9 | Lipc |
| Lipe | 1035 | 1452 | 1387 | 1443 | 1293 | 1261 | 1190 | 508 | 741 | 796 | 604 | 690 | 640 | 607 | 974 | 1261 | 1504 | 1371 | 1455 | 1194 | 1183 | 640 | 633 | 770 | 703 | 738 | 645 | 584 | Lipe |
| Lipf | 0 | 0 | 0 | 0 | 0 | 0 | 0 | 0 | 0 | 0 | 2 | 0 | 0 | 0 | 0 | 0 | 0 | 0 | 0 | 0 | 0 | 0 | 1 | 0 | 0 | 0 | 0 | 0 | Lipf |
| Lipg | 863 | 1040 | 783 | 759 | 781 | 775 | 882 | 805 | 840 | 725 | 576 | 709 | 531 | 728 | 957 | 914 | 972 | 746 | 890 | 799 | 818 | 630 | 484 | 835 | 909 | 730 | 560 | 468 | Lipg |
| Liph | 29 | 27 | 36 | 7 | 21 | 11 | 18 | 28 | 38 | 18 | 31 | 23 | 16 | 21 | 17 | 36 | 15 | 10 | 19 | 7 | 18 | 15 | 36 | 16 | 17 | 17 | 26 | 13 | Liph |
| Lipi | 0 | 0 | 0 | 4 | 1 | 6 | 2 | 4 | 1 | 5 | 1 | 0 | 0 | 0 | 0 | 1 | 0 | 0 | 0 | 1 | 2 | 12 | 4 | 1 | 5 | 5 | 7 | 6 | Lipi |
| Lipk | 0 | 0 | 0 | 0 | 2 | 0 | 0 | 1 | 2 | 0 | 2 | 0 | 0 | 0 | 0 | 0 | 0 | 0 | 0 | 0 | 0 | 0 | 2 | 0 | 0 | 0 | 0 | 0 | Lipk |
| Lipm | 8 | 0 | 11 | 0 | 0 | 1 | 0 | 72 | 45 | 65 | 7 | 0 | 3 | 3 | 4 | 1 | 0 | 0 | 0 | 0 | 0 | 3 | 56 | 15 | 0 | 6 | 1 | 0 | Lipm |
| Lipo2 | 0 | 4 | 15 | 4 | 9 | 10 | 8 | 23 | 21 | 38 | 25 | 42 | 14 | 37 | 5 | 7 | 9 | 0 | 3 | 3 | 10 | 27 | 22 | 17 | 46 | 26 | 20 | 18 | Lipo2 |
| Lipo3 | 349 | 256 | 401 | 322 | 447 | 387 | 314 | 1003 | 922 | 922 | 885 | 1102 | 1072 | 1109 | 407 | 336 | 299 | 281 | 327 | 373 | 348 | 1308 | 1019 | 999 | 1012 | 937 | 1074 | 1069 | Lipo3 |
| Lipo4 | 0 | 1 | 0 | 0 | 0 | 0 | 0 | 0 | 0 | 0 | 0 | 0 | 0 | 0 | 0 | 0 | 0 | 0 | 0 | 0 | 0 | 2 | 0 | 2 | 0 | 2 | 0 | 0 | Lipo4 |
| Lipt1 | 316 | 347 | 358 | 249 | 236 | 286 | 279 | 308 | 291 | 223 | 276 | 261 | 247 | 246 | 351 | 368 | 355 | 300 | 293 | 341 | 252 | 266 | 308 | 270 | 230 | 281 | 324 | 277 | Lipt1 |
| Lipt2 | 465 | 381 | 318 | 296 | 275 | 249 | 341 | 506 | 492 | 483 | 428 | 450 | 522 | 471 | 454 | 351 | 363 | 310 | 341 | 290 | 298 | 453 | 542 | 489 | 394 | 514 | 475 | 549 | Lipt2 |
| Litaf | 3910 | 3587 | 3585 | 3013 | 2699 | 2812 | 2826 | 1988 | 2278 | 2165 | 2442 | 2095 | 2352 | 2066 | 3588 | 3921 | 3262 | 2883 | 2802 | 2810 | 2655 | 1716 | 2158 | 2061 | 1867 | 2655 | 1978 | 2149 | Litaf |
| Lix1 | 3438 | 3149 | 3734 | 2560 | 2316 | 2909 | 3148 | 3197 | 2865 | 2476 | 2380 | 4379 | 3584 | 4376 | 3287 | 2650 | 3270 | 2306 | 2724 | 3456 | 2502 | 1962 | 2308 | 3140 | 3005 | 5536 | 3983 | 4797 | Lix1 |
| Lix1l | 12701 | 11512 | 14433 | 15217 | 14395 | 13659 | 14777 | 9673 | 8873 | 10519 | 10325 | 10317 | 10104 | 11544 | 12083 | 11078 | 15425 | 15162 | 14994 | 11656 | 15073 | 10972 | 9304 | 11261 | 11089 | 11055 | 9716 | 11208 | Lix1l |
| Llcfc1 | 3 | 0 | 0 | 0 | 1 | 0 | 0 | 1 | 0 | 0 | 0 | 0 | 0 | 0 | 0 | 0 | 0 | 2 | 0 | 0 | 1 | 0 | 0 | 1 | 0 | 0 | 0 | 0 | Llcfc1 |
| Llgl1 | 9118 | 12139 | 12710 | 11512 | 10196 | 10177 | 9595 | 8398 | 10205 | 10466 | 9445 | 14356 | 11639 | 11602 | 8805 | 10121 | 12863 | 10836 | 11871 | 9015 | 9572 | 10151 | 9027 | 12293 | 10835 | 14699 | 12191 | 12447 | Llgl1 |
| Llgl2 | 674 | 809 | 701 | 622 | 378 | 378 | 295 | 193 | 288 | 218 | 205 | 136 | 236 | 103 | 562 | 946 | 616 | 621 | 401 | 325 | 279 | 139 | 305 | 252 | 94 | 251 | 134 | 148 | Llgl2 |
| Llph | 3028 | 3303 | 4797 | 4142 | 3031 | 3313 | 3733 | 2303 | 2084 | 2683 | 2620 | 3004 | 2794 | 3325 | 3158 | 3115 | 4220 | 3623 | 3376 | 3945 | 3270 | 1686 | 2204 | 3067 | 2859 | 3557 | 3068 | 3398 | Llph |
| Lman1 | 10135 | 8899 | 8724 | 6985 | 6081 | 6621 | 5912 | 8680 | 8860 | 7686 | 9953 | 7679 | 8597 | 7588 | 9759 | 10449 | 7676 | 7289 | 5532 | 6811 | 5990 | 8166 | 9820 | 7663 | 6912 | 7823 | 8364 | 8003 | Lman1 |
| Lman1l | 5 | 42 | 37 | 61 | 50 | 48 | 44 | 16 | 2 | 17 | 17 | 14 | 14 | 19 | 24 | 43 | 37 | 62 | 60 | 35 | 38 | 12 | 8 | 11 | 9 | 10 | 8 | 11 | Lman1l |
| Lman2 | 7538 | 8389 | 7234 | 6290 | 5516 | 6205 | 5398 | 6830 | 7641 | 6404 | 7511 | 7319 | 7395 | 6813 | 7203 | 7902 | 7050 | 6277 | 6352 | 6189 | 5720 | 6744 | 7609 | 7013 | 6593 | 7690 | 7722 | 7150 | Lman2 |
| Lman2l | 3518 | 3781 | 3483 | 4225 | 4474 | 4365 | 4188 | 4019 | 3952 | 4508 | 4425 | 4602 | 4420 | 4225 | 3603 | 3649 | 3755 | 4362 | 4442 | 3340 | 4373 | 4837 | 3937 | 4285 | 4556 | 3830 | 4235 | 4092 | Lman2l |
| Lmbr1 | 1961 | 1842 | 1918 | 1956 | 2106 | 1823 | 1989 | 2202 | 1928 | 1857 | 1761 | 1889 | 1797 | 1817 | 2209 | 1999 | 1991 | 1984 | 1904 | 2151 | 1916 | 2400 | 1979 | 1789 | 2035 | 1623 | 1800 | 1750 | Lmbr1 |
| Lmbr1l | 4564 | 5595 | 4994 | 6672 | 5938 | 5678 | 5439 | 3245 | 3345 | 3642 | 3247 | 4078 | 4021 | 3646 | 4785 | 4876 | 5562 | 6346 | 6124 | 5259 | 5625 | 3888 | 3007 | 3906 | 4287 | 3364 | 4212 | 3609 | Lmbr1l |
| Lmbrd1 | 2601 | 2463 | 2584 | 2818 | 3340 | 3284 | 3365 | 3315 | 3074 | 3117 | 3325 | 2804 | 3080 | 2970 | 2885 | 2584 | 2753 | 2704 | 3217 | 3116 | 3323 | 3897 | 3287 | 2733 | 3310 | 2603 | 2887 | 3072 | Lmbrd1 |
| Lmbrd2 | 182 | 140 | 213 | 205 | 154 | 236 | 241 | 153 | 111 | 110 | 141 | 129 | 142 | 136 | 207 | 210 | 147 | 198 | 190 | 212 | 182 | 182 | 143 | 109 | 168 | 122 | 113 | 125 | Lmbrd2 |
| Lmcd1 | 332 | 302 | 277 | 396 | 398 | 390 | 331 | 415 | 629 | 572 | 782 | 318 | 372 | 288 | 265 | 308 | 305 | 368 | 363 | 332 | 349 | 391 | 658 | 535 | 439 | 306 | 275 | 227 | Lmcd1 |
| Lmf1 | 570 | 644 | 511 | 530 | 649 | 602 | 450 | 794 | 1094 | 975 | 1133 | 1016 | 1092 | 827 | 514 | 706 | 571 | 581 | 558 | 439 | 461 | 894 | 1029 | 980 | 959 | 918 | 1103 | 920 | Lmf1 |
| Lmf2 | 2358 | 2992 | 3246 | 3064 | 2730 | 2662 | 2221 | 1906 | 2551 | 2636 | 2792 | 3301 | 2933 | 2789 | 2184 | 2582 | 3102 | 2806 | 2991 | 2191 | 2312 | 2465 | 2572 | 2942 | 2769 | 3775 | 3103 | 2846 | Lmf2 |
| Lmln | 1517 | 1537 | 1453 | 1760 | 1647 | 1745 | 1759 | 1163 | 1172 | 1266 | 1135 | 1615 | 1465 | 1458 | 1441 | 1381 | 1627 | 1755 | 1644 | 1608 | 1681 | 1290 | 1094 | 1262 | 1347 | 1298 | 1540 | 1504 | Lmln |
| Lmna | 4404 | 4396 | 3920 | 3665 | 2770 | 3794 | 1976 | 2413 | 3621 | 2891 | 3807 | 1843 | 2292 | 1762 | 3665 | 5039 | 3675 | 4508 | 2724 | 2141 | 2311 | 2206 | 4383 | 2901 | 2116 | 1893 | 2099 | 1907 | Lmna |
| Lmnb1 | 23125 | 21281 | 34098 | 26086 | 21950 | 21161 | 24717 | 24940 | 23333 | 28629 | 26094 | 39793 | 30823 | 42962 | 21366 | 19411 | 30960 | 22948 | 25118 | 20716 | 24649 | 23541 | 21381 | 37786 | 32004 | 48364 | 36631 | 43353 | Lmnb1 |
| Lmnb2 | 5943 | 7557 | 7635 | 7072 | 5737 | 5822 | 5623 | 4526 | 5057 | 4999 | 5112 | 6007 | 5434 | 5421 | 5590 | 6472 | 7511 | 6422 | 6675 | 5875 | 5686 | 4319 | 4778 | 5862 | 5541 | 6993 | 5882 | 5548 | Lmnb2 |
| Lmntd1 | 16 | 19 | 13 | 36 | 16 | 50 | 30 | 72 | 72 | 57 | 88 | 47 | 66 | 66 | 33 | 22 | 22 | 32 | 26 | 46 | 33 | 70 | 89 | 65 | 74 | 53 | 59 | 52 | Lmntd1 |
| Lmntd2 | 404 | 465 | 410 | 395 | 342 | 394 | 394 | 388 | 373 | 320 | 343 | 540 | 594 | 519 | 385 | 460 | 417 | 432 | 385 | 392 | 323 | 383 | 364 | 473 | 401 | 524 | 651 | 530 | Lmntd2 |
| Lmo1 | 716 | 917 | 1016 | 1074 | 1134 | 1071 | 1053 | 1345 | 1093 | 1442 | 991 | 1833 | 1125 | 1859 | 766 | 473 | 1246 | 889 | 1578 | 950 | 1264 | 1532 | 929 | 1839 | 1671 | 1673 | 1582 | 1816 | Lmo1 |
| Lmo2 | 726 | 698 | 973 | 682 | 943 | 815 | 810 | 1503 | 1150 | 1307 | 1382 | 1607 | 1841 | 1730 | 760 | 674 | 699 | 732 | 856 | 1002 | 777 | 1529 | 1597 | 1405 | 1514 | 1958 | 1826 | 1943 | Lmo2 |
| Lmo3 | 2743 | 2550 | 3106 | 3801 | 4592 | 4505 | 4843 | 5382 | 4135 | 4996 | 3610 | 5173 | 3720 | 5492 | 3142 | 1851 | 3489 | 3833 | 5040 | 5306 | 5364 | 6241 | 4398 | 5258 | 6358 | 3762 | 4939 | 5165 | Lmo3 |
| Lmo4 | 5206 | 5197 | 5055 | 6463 | 6982 | 6259 | 7000 | 6399 | 6589 | 6404 | 6458 | 4941 | 4740 | 5340 | 6076 | 5139 | 6071 | 6278 | 6258 | 6050 | 7024 | 5893 | 6512 | 6040 | 6118 | 6347 | 4616 | 5228 | Lmo4 |
| Lmo7 | 1557 | 731 | 707 | 590 | 472 | 693 | 463 | 1062 | 1253 | 1018 | 1379 | 487 | 634 | 552 | 852 | 1010 | 604 | 739 | 478 | 475 | 483 | 483 | 1557 | 1025 | 576 | 821 | 538 | 557 | Lmo7 |
| Lmod1 | 150 | 165 | 162 | 94 | 45 | 71 | 22 | 77 | 80 | 78 | 122 | 125 | 160 | 126 | 182 | 206 | 125 | 72 | 9 | 21 | 29 | 72 | 119 | 86 | 45 | 127 | 178 | 119 | Lmod1 |
| Lmod2 | 0 | 0 | 0 | 0 | 0 | 0 | 0 | 4 | 11 | 0 | 57 | 0 | 0 | 0 | 4 | 0 | 0 | 0 | 0 | 0 | 0 | 2 | 52 | 0 | 0 | 0 | 0 | 2 | Lmod2 |
| Lmod3 | 7 | 0 | 0 | 8 | 2 | 3 | 2 | 28 | 65 | 8 | 444 | 7 | 19 | 0 | 3 | 8 | 5 | 0 | 3 | 2 | 0 | 77 | 286 | 7 | 6 | 0 | 2 | 1 | Lmod3 |
| Lmtk2 | 3690 | 4117 | 4200 | 5810 | 5579 | 5300 | 5113 | 3103 | 3353 | 3947 | 3738 | 4349 | 3751 | 3905 | 3498 | 4058 | 4864 | 5757 | 5580 | 4093 | 5124 | 4166 | 2942 | 3957 | 4597 | 2923 | 3865 | 3774 | Lmtk2 |
| Lmtk3 | 7740 | 10537 | 11934 | 17050 | 18049 | 14084 | 13927 | 9156 | 9616 | 15318 | 11748 | 12928 | 10265 | 11407 | 7903 | 8252 | 15534 | 16469 | 18519 | 11997 | 15197 | 17782 | 8185 | 14767 | 16706 | 11255 | 10517 | 9975 | Lmtk3 |
| Lmx1a | 660 | 802 | 1177 | 649 | 659 | 661 | 631 | 967 | 892 | 1007 | 825 | 1505 | 1594 | 1432 | 773 | 823 | 986 | 541 | 704 | 893 | 560 | 1196 | 1273 | 1016 | 1130 | 1147 | 1802 | 2007 | Lmx1a |
| Lmx1b | 1114 | 1864 | 2764 | 1651 | 1704 | 1680 | 648 | 1001 | 1313 | 1703 | 1893 | 1479 | 2681 | 1272 | 1444 | 2097 | 1942 | 1689 | 1050 | 1004 | 747 | 1686 | 1732 | 1271 | 1268 | 956 | 1617 | 1608 | Lmx1b |
| Lnp1 | 0 | 0 | 0 | 0 | 3 | 2 | 1 | 2 | 7 | 0 | 4 | 0 | 1 | 1 | 3 | 0 | 0 | 0 | 0 | 0 | 1 | 5 | 5 | 2 | 3 | 0 | 1 | 2 | Lnp1 |

| | | | | | | | | | | | | | | | | | | | | | | | | | | | | | |
|---|---|---|---|---|---|---|---|---|---|---|---|---|---|---|---|---|---|---|---|---|---|---|---|---|---|---|---|---|---|
| Lnpep | 8421 | 6267 | 6208 | 7212 | 7508 | 8670 | 7786 | 6398 | 6159 | 5690 | 6582 | 5854 | 6035 | 6160 | 7559 | 7067 | 6149 | 7103 | 6685 | 7191 | 7635 | 6071 | 5809 | 5172 | 6360 | 4701 | 5922 | 5417 | Lnpep |
| Lnpk | 7614 | 5573 | 5079 | 5895 | 6155 | 7417 | 7347 | 5680 | 4778 | 4305 | 4592 | 4964 | 5346 | 5623 | 8446 | 6903 | 5468 | 6209 | 5692 | 7853 | 7242 | 4650 | 4642 | 4051 | 4858 | 3755 | 5542 | 5178 | Lnpk |
| Lnx1 | 119 | 83 | 36 | 74 | 185 | 94 | 78 | 105 | 136 | 146 | 116 | 59 | 51 | 68 | 49 | 79 | 61 | 59 | 104 | 104 | 100 | 135 | 136 | 101 | 137 | 71 | 83 | 51 | Lnx1 |
| Lnx2 | 1301 | 1360 | 1444 | 1282 | 1245 | 1225 | 1108 | 835 | 964 | 846 | 887 | 779 | 852 | 783 | 1188 | 1404 | 1319 | 1275 | 1237 | 1108 | 1101 | 695 | 974 | 877 | 702 | 1067 | 739 | 771 | Lnx2 |
| Lonp1 | 8996 | 10706 | 11713 | 11320 | 8642 | 8850 | 8739 | 5263 | 6251 | 6823 | 6487 | 7297 | 6899 | 7119 | 8443 | 9864 | 11288 | 10270 | 9663 | 8072 | 8980 | 5922 | 5834 | 7418 | 6985 | 8381 | 7331 | 7411 | Lonp1 |
| Lonp2 | 9703 | 10255 | 8495 | 9010 | 9070 | 8750 | 9531 | 6735 | 6157 | 5431 | 5688 | 5924 | 5598 | 5846 | 10533 | 9252 | 9135 | 9098 | 8961 | 10150 | 9519 | 6061 | 6395 | 5393 | 5903 | 5452 | 5871 | 5694 | Lonp2 |
| Lonrf1 | 1696 | 1773 | 1786 | 2261 | 2437 | 2086 | 1999 | 1647 | 1747 | 1707 | 1737 | 1634 | 1632 | 1655 | 1800 | 1648 | 2153 | 2225 | 2177 | 2024 | 2123 | 2124 | 1733 | 1617 | 1814 | 1593 | 1600 | 1589 | Lonrf1 |
| Lonrf2 | 3429 | 3760 | 3204 | 6134 | 9507 | 7115 | 7968 | 6702 | 6299 | 7502 | 6787 | 5027 | 4339 | 4953 | 4225 | 3401 | 5132 | 6759 | 7657 | 6462 | 8461 | 9929 | 5974 | 5828 | 8401 | 3003 | 4198 | 4330 | Lonrf2 |
| Lonrf3 | 1172 | 1266 | 577 | 979 | 1136 | 1286 | 1335 | 393 | 339 | 225 | 232 | 414 | 268 | 391 | 1212 | 1010 | 817 | 1004 | 1141 | 1585 | 1339 | 169 | 279 | 215 | 326 | 130 | 390 | 357 | Lonrf3 |
| Lor | 14 | 25 | 53 | 54 | 61 | 49 | 23 | 40 | 51 | 96 | 116 | 73 | 70 | 54 | 36 | 20 | 37 | 47 | 45 | 30 | 23 | 100 | 76 | 74 | 60 | 70 | 60 | 53 | Lor |
| Lox | 3665 | 2581 | 3250 | 2945 | 1707 | 4995 | 1275 | 6662 | 10764 | 7727 | 13335 | 5436 | 6489 | 5947 | 3948 | 3804 | 2807 | 5820 | 1354 | 1616 | 1260 | 6667 | 14444 | 8560 | 7239 | 5110 | 5831 | 5250 | Lox |
| Loxhd1 | 2 | 0 | 0 | 6 | 0 | 0 | 1 | 0 | 6 | 5 | 0 | 0 | 0 | 0 | 3 | 2 | 8 | 0 | 5 | 1 | 4 | 10 | 3 | 0 | 0 | 0 | 0 | 1 | Loxhd1 |
| Loxl1 | 1059 | 1402 | 1312 | 897 | 616 | 912 | 435 | 717 | 1185 | 1086 | 1254 | 797 | 1090 | 777 | 1013 | 1616 | 1085 | 1091 | 591 | 472 | 463 | 1039 | 1482 | 999 | 877 | 1065 | 966 | 745 | Loxl1 |
| Loxl2 | 3609 | 3097 | 3273 | 1797 | 1252 | 2246 | 1048 | 2644 | 2961 | 2679 | 4891 | 2628 | 3200 | 2666 | 3306 | 3931 | 2336 | 2341 | 1252 | 964 | 1103 | 2488 | 4383 | 2857 | 2110 | 3052 | 2786 | 3012 | Loxl2 |
| Loxl3 | 1082 | 1348 | 1343 | 901 | 837 | 1033 | 594 | 878 | 1455 | 1365 | 1997 | 1143 | 1449 | 774 | 1122 | 1571 | 1053 | 930 | 872 | 789 | 655 | 1268 | 1540 | 1106 | 1053 | 1244 | 1115 | 1029 | Loxl3 |
| Loxl4 | 20 | 15 | 26 | 4 | 6 | 11 | 4 | 17 | 35 | 40 | 114 | 18 | 43 | 15 | 16 | 23 | 27 | 14 | 12 | 2 | 4 | 32 | 49 | 33 | 15 | 21 | 23 | 9 | Loxl4 |
| Lpar1 | 2833 | 1833 | 2414 | 1377 | 1192 | 1650 | 1269 | 3013 | 3825 | 2648 | 4509 | 3094 | 4420 | 3244 | 2232 | 2707 | 1647 | 1526 | 1323 | 1405 | 1113 | 2481 | 4313 | 3155 | 2490 | 4174 | 3560 | 3588 | Lpar1 |
| Lpar2 | 1797 | 1858 | 1827 | 1790 | 1777 | 1740 | 1762 | 1315 | 1391 | 1290 | 1313 | 1546 | 1506 | 1534 | 1582 | 1819 | 1878 | 1727 | 1965 | 1704 | 1791 | 1417 | 1312 | 1376 | 1429 | 1768 | 1370 | 1549 | Lpar2 |
| Lpar3 | 53 | 41 | 93 | 103 | 44 | 31 | 41 | 27 | 33 | 31 | 32 | 18 | 24 | 33 | 59 | 100 | 73 | 46 | 30 | 27 | 29 | 30 | 26 | 29 | 21 | 25 | 33 | 21 | Lpar3 |
| Lpar4 | 2503 | 1595 | 2244 | 1763 | 1609 | 1848 | 1680 | 2198 | 2210 | 1654 | 3133 | 1698 | 2338 | 1825 | 2467 | 2331 | 1824 | 1858 | 1640 | 2132 | 1614 | 1915 | 2698 | 1778 | 1591 | 2160 | 2183 | 2086 | Lpar4 |
| Lpar5 | 9 | 10 | 7 | 11 | 8 | 7 | 12 | 6 | 12 | 12 | 13 | 11 | 16 | 6 | 5 | 4 | 16 | 18 | 8 | 13 | 17 | 23 | 14 | 19 | 17 | 13 | 9 | 17 | Lpar5 |
| Lpar6 | 1057 | 674 | 801 | 657 | 738 | 807 | 765 | 1457 | 1339 | 1221 | 1264 | 1455 | 1809 | 1511 | 989 | 805 | 735 | 547 | 773 | 903 | 807 | 1043 | 1621 | 1392 | 1203 | 1562 | 1751 | 1778 | Lpar6 |
| Lpcat1 | 6473 | 7197 | 7707 | 6972 | 6514 | 6475 | 6583 | 5534 | 5535 | 5910 | 5638 | 6714 | 6749 | 6754 | 6628 | 6942 | 7238 | 6464 | 6782 | 6073 | 6485 | 5937 | 5520 | 6147 | 6396 | 7097 | 6598 | 7124 | Lpcat1 |
| Lpcat2 | 654 | 562 | 524 | 583 | 614 | 552 | 513 | 435 | 508 | 480 | 536 | 345 | 375 | 301 | 686 | 759 | 462 | 577 | 506 | 637 | 491 | 493 | 582 | 403 | 309 | 379 | 225 | 254 | Lpcat2 |
| Lpcat2b | 0 | 1 | 6 | 0 | 0 | 2 | 1 | 0 | 0 | 0 | 0 | 0 | 0 | 0 | 0 | 0 | 0 | 3 | 1 | 0 | 0 | 0 | 0 | 0 | 0 | 0 | 0 | 0 | Lpcat2b |
| Lpcat3 | 1821 | 2172 | 1907 | 1644 | 1661 | 1597 | 1604 | 1275 | 1508 | 1345 | 1386 | 1294 | 1437 | 1253 | 1795 | 2230 | 1734 | 1799 | 1438 | 1625 | 1555 | 1280 | 1566 | 1179 | 1242 | 1415 | 1323 | 1317 | Lpcat3 |
| Lpcat4 | 851 | 898 | 1091 | 1001 | 1109 | 955 | 824 | 757 | 891 | 1022 | 954 | 949 | 902 | 833 | 692 | 967 | 1118 | 910 | 1162 | 857 | 821 | 1071 | 877 | 969 | 946 | 937 | 872 | 805 | Lpcat4 |
| Lpgat1 | 7827 | 6028 | 6319 | 6842 | 9007 | 8799 | 8685 | 12174 | 11161 | 11418 | 11941 | 11334 | 11429 | 11718 | 7793 | 7094 | 6995 | 7434 | 8187 | 7803 | 8773 | 11939 | 10788 | 10962 | 12620 | 8613 | 11347 | 11013 | Lpgat1 |
| Lpin1 | 2700 | 3221 | 2765 | 3653 | 3814 | 3369 | 3356 | 2179 | 2042 | 2333 | 2105 | 2192 | 2370 | 2117 | 2880 | 3152 | 3274 | 3550 | 3595 | 3280 | 3622 | 2774 | 1994 | 2254 | 2491 | 1566 | 2077 | 2134 | Lpin1 |
| Lpin2 | 7919 | 7796 | 5967 | 6608 | 7159 | 6988 | 6963 | 5964 | 6264 | 5846 | 5904 | 5944 | 5836 | 6023 | 7207 | 8170 | 6383 | 6973 | 6857 | 7199 | 7057 | 5459 | 5834 | 5501 | 5675 | 5245 | 5817 | 6045 | Lpin2 |
| Lpin3 | 156 | 84 | 93 | 90 | 64 | 60 | 50 | 60 | 105 | 67 | 81 | 59 | 79 | 20 | 105 | 132 | 79 | 65 | 43 | 66 | 42 | 62 | 126 | 83 | 18 | 60 | 43 | 46 | Lpin3 |
| Lpl | 1624 | 1351 | 1718 | 1129 | 1140 | 1303 | 1353 | 1678 | 1469 | 1457 | 1722 | 2133 | 1875 | 2100 | 1834 | 1744 | 1410 | 1209 | 1214 | 1749 | 1273 | 1290 | 1696 | 1554 | 1639 | 2294 | 2251 | 2080 | Lpl |
| Lpo | 9 | 10 | 8 | 6 | 4 | 10 | 17 | 18 | 19 | 25 | 22 | 29 | 48 | 33 | 11 | 9 | 6 | 1 | 16 | 5 | 14 | 35 | 21 | 23 | 39 | 24 | 14 | 23 | Lpo |
| Lpp | 4469 | 3125 | 3389 | 2446 | 2174 | 3335 | 2061 | 3717 | 4112 | 3818 | 5066 | 3630 | 4834 | 3488 | 3765 | 4988 | 2728 | 3119 | 1911 | 2094 | 1881 | 3628 | 4866 | 3544 | 3119 | 3408 | 4088 | 3877 | Lpp |
| Lpxn | 116 | 125 | 93 | 81 | 98 | 108 | 97 | 87 | 77 | 57 | 118 | 88 | 81 | 88 | 117 | 132 | 98 | 79 | 119 | 135 | 80 | 63 | 100 | 80 | 71 | 98 | 110 | 91 | Lpxn |
| Lrat | 20 | 30 | 16 | 28 | 32 | 12 | 21 | 7 | 7 | 10 | 8 | 7 | 5 | 6 | 18 | 31 | 13 | 19 | 19 | 24 | 18 | 23 | 11 | 7 | 5 | 8 | 16 | 4 | Lrat |
| Lratd1 | 2524 | 2637 | 3329 | 3464 | 3850 | 3355 | 3604 | 3556 | 3647 | 3740 | 3582 | 3152 | 3276 | 3361 | 2674 | 2632 | 3516 | 4051 | 3684 | 2846 | 3521 | 4625 | 3435 | 3604 | 4373 | 2639 | 2847 | 2923 | Lratd1 |
| Lratd2 | 2157 | 1849 | 2570 | 2034 | 2027 | 1787 | 1862 | 1973 | 2183 | 2135 | 1869 | 2412 | 2759 | 1923 | 1961 | 2307 | 2166 | 1944 | 1743 | 1917 | 1850 | 2373 | 3100 | 1901 | 1729 | 2508 | 2086 | 2540 | Lratd2 |
| Lrba | 4493 | 3992 | 3392 | 3312 | 3185 | 3778 | 2890 | 4428 | 4343 | 3703 | 4148 | 3978 | 5262 | 3937 | 4276 | 3848 | 3062 | 2838 | 2957 | 3660 | 3308 | 3686 | 4406 | 3095 | 3663 | 2135 | 5168 | 4470 | Lrba |
| Lrch1 | 1057 | 917 | 1120 | 1201 | 1472 | 1417 | 1351 | 1324 | 1494 | 1620 | 1628 | 1200 | 1357 | 1290 | 1048 | 1146 | 1124 | 1180 | 1456 | 1094 | 1219 | 1576 | 1493 | 1388 | 1488 | 1318 | 1008 | 1195 | Lrch1 |
| Lrch2 | 6668 | 4659 | 5335 | 6995 | 8306 | 8974 | 9662 | 11085 | 8719 | 7925 | 7968 | 9179 | 8283 | 9822 | 7317 | 5718 | 6112 | 6930 | 8041 | 8884 | 9222 | 10198 | 8486 | 8693 | 9627 | 7627 | 9108 | 9248 | Lrch2 |
| Lrch3 | 9513 | 9462 | 7341 | 8528 | 9022 | 9459 | 9300 | 7999 | 7860 | 7585 | 8006 | 7993 | 7889 | 8340 | 8780 | 8541 | 7920 | 8800 | 8596 | 8907 | 9542 | 7762 | 7158 | 7249 | 8207 | 6602 | 8178 | 7471 | Lrch3 |
| Lrch4 | 17 | 22 | 10 | 27 | 20 | 16 | 13 | 16 | 14 | 12 | 13 | 15 | 11 | 13 | 20 | 22 | 38 | 33 | 16 | 22 | 17 | 20 | 11 | 19 | 9 | 8 | 7 | 11 | Lrch4 |
| Lrcol1 | 1 | 0 | 3 | 1 | 0 | 0 | 1 | 1 | 0 | 0 | 1 | 0 | 0 | 1 | 0 | 0 | 1 | 3 | 0 | 0 | 2 | 2 | 0 | 0 | 1 | 0 | 0 | 0 | Lrcol1 |
| Lrfn1 | 1891 | 2567 | 2434 | 2602 | 2723 | 2533 | 2113 | 1570 | 1761 | 2252 | 1862 | 1897 | 1675 | 1729 | 1642 | 2083 | 2808 | 2551 | 2866 | 2037 | 2359 | 2471 | 1676 | 1916 | 2145 | 1910 | 1591 | 1513 | Lrfn1 |
| Lrfn2 | 72 | 87 | 106 | 164 | 335 | 202 | 241 | 239 | 366 | 357 | 397 | 232 | 144 | 234 | 121 | 108 | 109 | 231 | 225 | 215 | 241 | 428 | 301 | 346 | 424 | 237 | 160 | 196 | Lrfn2 |
| Lrfn3 | 1314 | 1704 | 1877 | 1790 | 2116 | 1760 | 1526 | 960 | 1180 | 1389 | 1286 | 1240 | 1125 | 1025 | 1228 | 1452 | 1871 | 1909 | 1869 | 1333 | 1632 | 1526 | 1063 | 1267 | 1243 | 1245 | 1048 | 1002 | Lrfn3 |
| Lrfn4 | 1491 | 1681 | 1864 | 1764 | 2396 | 1979 | 1790 | 3684 | 4158 | 4349 | 3980 | 4512 | 3918 | 3692 | 1333 | 1335 | 2104 | 1752 | 2626 | 1928 | 1805 | 4084 | 3802 | 4484 | 4617 | 4751 | 4288 | 3793 | Lrfn4 |
| Lrfn5 | 451 | 406 | 253 | 677 | 1387 | 1036 | 1133 | 2069 | 1918 | 1784 | 1461 | 960 | 909 | 977 | 670 | 434 | 476 | 923 | 1072 | 1106 | 1197 | 2124 | 1564 | 1207 | 1918 | 684 | 710 | 706 | Lrfn5 |
| Lrg1 | 0 | 2 | 0 | 5 | 0 | 5 | 0 | 0 | 2 | 0 | 4 | 3 | 1 | 2 | 0 | 4 | 0 | 2 | 2 | 1 | 1 | 0 | 5 | 0 | 1 | 0 | 0 | 4 | Lrg1 |
| Lrguk | 852 | 761 | 627 | 799 | 824 | 779 | 961 | 418 | 442 | 439 | 537 | 379 | 449 | 410 | 910 | 884 | 758 | 835 | 778 | 759 | 886 | 344 | 432 | 309 | 348 | 267 | 503 | 382 | Lrguk |
| Lrif1 | 2812 | 2110 | 2699 | 3081 | 2583 | 2893 | 3144 | 2361 | 2150 | 2187 | 2066 | 2216 | 2159 | 2158 | 3116 | 2657 | 2760 | 2649 | 2726 | 3166 | 2858 | 2413 | 2222 | 2104 | 2123 | 2163 | 2181 | 2276 | Lrif1 |
| Lrig1 | 5046 | 6217 | 5130 | 5812 | 4709 | 5062 | 4126 | 4136 | 5015 | 4075 | 5130 | 4506 | 5046 | 4047 | 5073 | 5953 | 5191 | 5848 | 4899 | 4844 | 4460 | 3658 | 5249 | 4068 | 3722 | 4625 | 4356 | 4374 | Lrig1 |
| Lrig2 | 6537 | 6036 | 4679 | 6424 | 6601 | 6547 | 6641 | 6532 | 5347 | 5143 | 5394 | 5403 | 5270 | 5561 | 6749 | 5872 | 5578 | 6344 | 5819 | 6425 | 6757 | 6498 | 5200 | 4743 | 5700 | 3924 | 5269 | 5230 | Lrig2 |
| Lrig3 | 2548 | 2451 | 2011 | 1751 | 1084 | 1334 | 941 | 1621 | 2165 | 1567 | 2489 | 1265 | 1940 | 1107 | 2169 | 2875 | 1664 | 1664 | 1078 | 1018 | 1073 | 1425 | 2155 | 1457 | 973 | 1720 | 1395 | 1345 | Lrig3 |
| Lrit1 | 0 | 0 | 0 | 2 | 0 | 0 | 0 | 1 | 0 | 0 | 0 | 2 | 2 | 0 | 0 | 0 | 0 | 0 | 0 | 0 | 0 | 0 | 1 | 0 | 0 | 0 | 0 | 2 | Lrit1 |
| Lrit2 | 0 | 0 | 0 | 0 | 1 | 0 | 0 | 0 | 0 | 0 | 0 | 0 | 0 | 0 | 0 | 0 | 0 | 0 | 0 | 0 | 1 | 0 | 0 | 0 | 0 | 0 | 0 | 0 | Lrit2 |
| Lrit3 | 22 | 14 | 12 | 28 | 43 | 52 | 44 | 15 | 11 | 26 | 17 | 10 | 6 | 10 | 20 | 23 | 42 | 62 | 32 | 38 | 46 | 15 | 6 | 19 | 23 | 13 | 9 | 12 | Lrit3 |
| Lrmda | 27 | 15 | 18 | 15 | 16 | 4 | 15 | 18 | 10 | 19 | 8 | 11 | 18 | 34 | 21 | 11 | 9 | 19 | 7 | 20 | 6 | 27 | 29 | 16 | 21 | 24 | 23 | 19 | Lrmda |
| Lrmp | 57 | 74 | 76 | 39 | 42 | 70 | 32 | 59 | 66 | 69 | 97 | 98 | 120 | 81 | 75 | 101 | 56 | 53 | 55 | 62 | 44 | 57 | 105 | 79 | 92 | 126 | 93 | 84 | Lrmp |
| Lrp1 | 18653 | 24540 | 23488 | 29072 | 30728 | 29337 | 21018 | 21151 | 31347 | 37492 | 36068 | 36546 | 33932 | 30103 | 17186 | 26272 | 27146 | 32991 | 30807 | 19518 | 25023 | 38320 | 29582 | 34204 | 36155 | 17698 | 29551 | 27461 | Lrp1 |
| Lrp10 | 3388 | 4527 | 4357 | 4571 | 4239 | 4025 | 3555 | 1579 | 2286 | 2331 | 2400 | 2164 | 2379 | 1877 | 3280 | 4013 | 4217 | 4428 | 4369 | 3226 | 3761 | 2242 | 2304 | 2308 | 2357 | 2586 | 2033 | 2139 | Lrp10 |
| Lrp11 | 2 | 0 | 0 | 5 | 1 | 0 | 0 | 1 | 0 | 0 | 0 | 1 | 2 | 0 | 0 | 0 | 2 | 0 | 0 | 0 | 1 | 0 | 0 | 0 | 0 | 1 | 0 | 0 | Lrp11 |
| Lrp12 | 5035 | 4908 | 4357 | 5528 | 6624 | 6057 | 6092 | 6963 | 6288 | 6000 | 6053 | 6041 | 5717 | 6026 | 5632 | 4497 | 5551 | 5558 | 6410 | 5993 | 6116 | 7018 | 5871 | 5989 | 6652 | 5378 | 6082 | 5671 | Lrp12 |
| Lrp1b | 347 | 428 | 226 | 390 | 647 | 670 | 596 | 1099 | 920 | 559 | 631 | 389 | 511 | 338 | 702 | 444 | 211 | 501 | 477 | 974 | 714 | 620 | 871 | 292 | 542 | 110 | 350 | 306 | Lrp1b |
| Lrp2 | 4345 | 5024 | 3055 | 2549 | 2725 | 3613 | 2518 | 3449 | 2356 | 1814 | 1665 | 2022 | 3009 | 2293 | 4817 | 4804 | 3270 | 2409 | 2476 | 3471 | 3280 | 2154 | 2011 | 1628 | 1886 | 1154 | 3575 | 2587 | Lrp2 |
| Lrp2bp | 40 | 36 | 37 | 51 | 35 | 26 | 30 | 40 | 35 | 21 | 21 | 28 | 25 | 22 | 38 | 35 | 66 | 19 | 50 | 21 | 30 | 43 | 20 | 35 | 13 | 34 | 27 | 23 | Lrp2bp |
| Lrp3 | 3375 | 4468 | 4843 | 4840 | 5406 | 4672 | 4120 | 5217 | 5658 | 6771 | 6171 | 7219 | 6172 | 6243 | 3444 | 3973 | 4970 | 4679 | 5594 | 3884 | 4519 | 7444 | 5112 | 7100 | 7616 | 6447 | 6386 | 6440 | Lrp3 |
| Lrp4 | 3475 | 3622 | 3845 | 3482 | 3702 | 4088 | 3636 | 4590 | 4635 | 5824 | 5153 | 6626 | 5460 | 6561 | 3220 | 3523 | 3797 | 3584 | 4286 | 3620 | 3742 | 4577 | 5088 | 6398 | 5919 | 5184 | 6419 | 6329 | Lrp4 |
| Lrp5 | 3237 | 3582 | 3923 | 3098 | 2883 | 3098 | 2006 | 2253 | 3292 | 3208 | 4087 | 2953 | 3231 | 2524 | 2507 | 4125 | 3483 | 3447 | 3126 | 1988 | 2397 | 2578 | 3328 | 3477 | 2666 | 3573 | 2818 | 2584 | Lrp5 |
| Lrp6 | 11215 | 9608 | 7867 | 9639 | 9265 | 10631 | 9918 | 12097 | 12241 | 11647 | 13751 | 11630 | 12432 | 11921 | 10508 | 11357 | 8231 | 9764 | 8751 | 8944 | 9939 | 11015 | 11486 | 10399 | 11395 | 8994 | 12490 | 11177 | Lrp6 |
| Lrp8 | 7423 | 7447 | 7062 | 10467 | 15042 | 13543 | 14634 | 14051 | 12850 | 16495 | 13734 | 15096 | 12414 | 16156 | 8091 | 6706 | 9111 | 11828 | 13532 | 12089 | 15169 | 17644 | 11822 | 15132 | 19888 | 9684 | 13953 | 13363 | Lrp8 |
| Lrpap1 | 7256 | 8158 | 6532 | 7218 | 7155 | 7320 | 6994 | 4942 | 5601 | 4893 | 5122 | 4849 | 4875 | 4863 | 7332 | 7703 | 6898 | 7657 | 7026 | 7415 | 7421 | 5160 | 5891 | 4975 | 5288 | 5012 | 5025 | 4588 | Lrpap1 |
| Lrpprc | 13055 | 12140 | 10598 | 10428 | 9220 | 9525 | 10891 | 7400 | 6859 | 6037 | 6264 | 6749 | 6371 | 6837 | 12406 | 11716 | 10968 | 10349 | 9252 | 10384 | 10383 | 6721 | 6640 | 6286 | 6581 | 6749 | 6959 | 6961 | Lrpprc |
| Lrr1 | 980 | 878 | 861 | 700 | 466 | 536 | 731 | 482 | 385 | 356 | 282 | 564 | 400 | 566 | 858 | 847 | 776 | 603 | 609 | 767 | 650 | 349 | 368 | 455 | 405 | 660 | 559 | 625 | Lrr1 |
| Lrrc1 | 3392 | 3206 | 2989 | 3079 | 2583 | 2818 | 3303 | 1862 | 1831 | 1742 | 1824 | 1995 | 1726 | 2089 | 2992 | 3024 | 3002 | 2939 | 2945 | 3077 | 2945 | 1312 | 1671 | 1922 | 1791 | 2720 | 1737 | 1975 | Lrrc1 |
| Lrrc10 | 0 | 3 | 0 | 3 | 0 | 7 | 2 | 2 | 2 | 3 | 0 | 1 | 0 | 0 | 5 | 4 | 3 | 1 | 4 | 5 | 1 | 0 | 3 | 0 | 0 | 0 | 0 | 8 | Lrrc10 |
| Lrrc10b | 172 | 185 | 210 | 200 | 142 | 151 | 136 | 53 | 66 | 79 | 61 | 87 | 100 | 73 | 162 | 184 | 139 | 138 | 119 | 142 | 122 | 79 | 89 | 76 | 70 | 125 | 78 | 61 | Lrrc10b |
| Lrrc14 | 1908 | 2224 | 2079 | 2039 | 1755 | 1715 | 1685 | 1793 | 1949 | 1840 | 1747 | 2057 | 1842 | 1763 | 1813 | 1900 | 2052 | 1939 | 1903 | 1789 | 1750 | 1813 | 1867 | 1901 | 1817 | 2236 | 1807 | 1836 | Lrrc14 |

| | | | | | | | | | | | | | | | | | | | | | | | | | | | | | |
|---|---|---|---|---|---|---|---|---|---|---|---|---|---|---|---|---|---|---|---|---|---|---|---|---|---|---|---|---|---|
| Lrrc14b | 0 | 2 | 4 | 0 | 1 | 0 | 0 | 0 | 0 | 0 | 7 | 0 | 1 | 0 | 1 | 1 | 1 | 1 | 0 | 4 | 4 | 5 | 3 | 2 | 0 | 0 | 2 | 1 | Lrrc14b |
| Lrrc15 | 349 | 318 | 318 | 169 | 79 | 201 | 66 | 142 | 152 | 186 | 177 | 203 | 305 | 188 | 266 | 497 | 155 | 193 | 93 | 59 | 72 | 199 | 256 | 181 | 165 | 180 | 246 | 285 | Lrrc15 |
| Lrrc17 | 1789 | 1562 | 1314 | 844 | 397 | 1335 | 275 | 2125 | 3195 | 2452 | 3692 | 1826 | 2529 | 1668 | 1685 | 2132 | 833 | 1174 | 474 | 516 | 270 | 1973 | 4333 | 2065 | 1593 | 2111 | 1768 | 1914 | Lrrc17 |
| Lrrc18 | 100 | 83 | 117 | 64 | 128 | 121 | 98 | 37 | 48 | 40 | 46 | 39 | 45 | 24 | 95 | 138 | 82 | 87 | 85 | 92 | 77 | 42 | 93 | 46 | 44 | 61 | 31 | 30 | Lrrc18 |
| Lrrc19 | 0 | 0 | 0 | 0 | 2 | 0 | 0 | 0 | 0 | 0 | 0 | 0 | 0 | 0 | 0 | 2 | 0 | 0 | 2 | 0 | 0 | 0 | 0 | 0 | 0 | 0 | 0 | 0 | Lrrc19 |
| Lrrc2 | 5 | 7 | 1 | 7 | 5 | 1 | 6 | 10 | 24 | 28 | 31 | 8 | 6 | 16 | 11 | 5 | 0 | 2 | 1 | 5 | 7 | 17 | 17 | 14 | 18 | 9 | 8 | 8 | Lrrc2 |
| Lrrc20 | 2263 | 2525 | 2389 | 2738 | 2666 | 2611 | 2497 | 1664 | 1917 | 2077 | 1946 | 1980 | 1748 | 2148 | 2023 | 2246 | 2664 | 2572 | 2839 | 2262 | 2641 | 1778 | 1779 | 2178 | 2237 | 2255 | 2083 | 2066 | Lrrc20 |
| Lrrc23 | 153 | 264 | 261 | 191 | 282 | 207 | 215 | 223 | 278 | 280 | 266 | 349 | 429 | 278 | 239 | 260 | 200 | 183 | 217 | 234 | 215 | 301 | 444 | 270 | 237 | 315 | 275 | 288 | Lrrc23 |
| Lrrc24 | 143 | 244 | 215 | 322 | 398 | 324 | 283 | 231 | 267 | 273 | 253 | 210 | 201 | 192 | 216 | 213 | 287 | 349 | 353 | 317 | 328 | 336 | 253 | 185 | 306 | 119 | 156 | 176 | Lrrc24 |
| Lrrc25 | 22 | 29 | 31 | 25 | 25 | 23 | 13 | 45 | 64 | 60 | 82 | 56 | 49 | 44 | 22 | 35 | 24 | 15 | 14 | 19 | 22 | 42 | 88 | 52 | 39 | 49 | 45 | 50 | Lrrc25 |
| Lrrc26 | 1 | 0 | 0 | 0 | 1 | 0 | 0 | 0 | 2 | 0 | 0 | 1 | 0 | 0 | 0 | 0 | 1 | 0 | 0 | 0 | 1 | 0 | 4 | 0 | 0 | 0 | 0 | 0 | Lrrc26 |
| Lrrc27 | 22 | 14 | 6 | 8 | 6 | 7 | 8 | 38 | 16 | 10 | 10 | 13 | 15 | 4 | 12 | 6 | 9 | 4 | 17 | 13 | 4 | 20 | 8 | 17 | 17 | 10 | 73 | 11 | Lrrc27 |
| Lrrc28 | 2646 | 2907 | 2345 | 2710 | 2404 | 2206 | 2949 | 1564 | 1471 | 1355 | 1250 | 1347 | 1234 | 1465 | 2718 | 2585 | 2727 | 2669 | 2466 | 2587 | 2639 | 1382 | 1378 | 1391 | 1400 | 1409 | 1290 | 1445 | Lrrc28 |
| Lrrc29 | 182 | 148 | 95 | 110 | 138 | 119 | 114 | 137 | 162 | 161 | 169 | 178 | 204 | 166 | 157 | 122 | 129 | 105 | 117 | 143 | 128 | 159 | 152 | 157 | 113 | 203 | 152 | 145 | Lrrc29 |
| Lrrc3 | 1801 | 2211 | 1927 | 2007 | 2161 | 1827 | 1772 | 712 | 909 | 978 | 894 | 717 | 878 | 666 | 1907 | 2120 | 2055 | 1937 | 2095 | 1931 | 1975 | 998 | 923 | 753 | 883 | 630 | 607 | 608 | Lrrc3 |
| Lrrc30 | 2 | 1 | 0 | 1 | 0 | 0 | 0 | 1 | 4 | 1 | 8 | 0 | 2 | 0 | 0 | 0 | 0 | 0 | 0 | 0 | 0 | 0 | 6 | 0 | 0 | 2 | 0 | 0 | Lrrc30 |
| Lrrc31 | 2 | 2 | 0 | 1 | 6 | 0 | 1 | 5 | 9 | 7 | 3 | 0 | 2 | 0 | 1 | 5 | 0 | 3 | 0 | 3 | 4 | 8 | 5 | 0 | 5 | 0 | 2 | 2 | Lrrc31 |
| Lrrc32 | 624 | 886 | 839 | 589 | 615 | 620 | 438 | 338 | 567 | 426 | 691 | 569 | 724 | 499 | 530 | 958 | 620 | 898 | 472 | 430 | 482 | 511 | 622 | 521 | 455 | 508 | 522 | 470 | Lrrc32 |
| Lrrc34 | 108 | 116 | 140 | 85 | 79 | 84 | 63 | 83 | 108 | 71 | 72 | 108 | 175 | 79 | 174 | 181 | 87 | 79 | 71 | 111 | 57 | 99 | 139 | 56 | 49 | 89 | 58 | 83 | Lrrc34 |
| Lrrc36 | 52 | 66 | 54 | 58 | 59 | 60 | 53 | 164 | 97 | 101 | 79 | 160 | 196 | 145 | 70 | 76 | 93 | 50 | 59 | 101 | 57 | 107 | 99 | 88 | 88 | 96 | 203 | 154 | Lrrc36 |
| Lrrc37a | 0 | 0 | 0 | 0 | 0 | 0 | 0 | 0 | 0 | 0 | 0 | 0 | 0 | 0 | 0 | 0 | 0 | 0 | 0 | 0 | 0 | 0 | 0 | 2 | 0 | 0 | 0 | 0 | Lrrc37a |
| Lrrc39 | 15 | 10 | 18 | 7 | 10 | 6 | 7 | 13 | 12 | 18 | 43 | 21 | 46 | 17 | 7 | 10 | 12 | 4 | 6 | 15 | 10 | 7 | 41 | 16 | 15 | 30 | 15 | 37 | Lrrc39 |
| Lrrc3b | 51 | 25 | 47 | 53 | 107 | 68 | 72 | 76 | 84 | 139 | 98 | 34 | 91 | 61 | 59 | 62 | 57 | 35 | 52 | 59 | 93 | 134 | 141 | 78 | 113 | 57 | 31 | 65 | Lrrc3b |
| Lrrc3c | 0 | 0 | 0 | 1 | 0 | 0 | 0 | 0 | 0 | 0 | 0 | 0 | 0 | 0 | 1 | 0 | 0 | 0 | 0 | 1 | 0 | 0 | 0 | 0 | 0 | 0 | 0 | 0 | Lrrc3c |
| Lrrc4 | 204 | 241 | 334 | 313 | 716 | 503 | 495 | 1319 | 1147 | 1534 | 1293 | 1336 | 1198 | 1364 | 305 | 167 | 357 | 326 | 595 | 499 | 529 | 1726 | 1334 | 1475 | 1731 | 1166 | 1263 | 1406 | Lrrc4 |
| Lrrc40 | 8529 | 6658 | 6384 | 6312 | 5308 | 5811 | 6677 | 5893 | 4970 | 4149 | 4202 | 5246 | 4977 | 5544 | 8451 | 7011 | 6230 | 5816 | 5289 | 7172 | 6250 | 4503 | 4877 | 4789 | 4533 | 5446 | 5528 | 5572 | Lrrc40 |
| Lrrc41 | 2918 | 3486 | 3120 | 2864 | 3224 | 3017 | 2959 | 4512 | 4982 | 4256 | 4302 | 4281 | 4216 | 3895 | 2935 | 2956 | 3137 | 2792 | 3532 | 3308 | 3161 | 4139 | 4980 | 4015 | 4112 | 4590 | 4314 | 4450 | Lrrc41 |
| Lrrc42 | 6501 | 6412 | 7021 | 6171 | 5402 | 5557 | 5959 | 3395 | 3288 | 3262 | 3024 | 3385 | 3496 | 3671 | 6367 | 5833 | 6392 | 5751 | 6094 | 5737 | 5973 | 2904 | 3445 | 3395 | 3191 | 4404 | 3660 | 4018 | Lrrc42 |
| Lrrc43 | 14 | 24 | 4 | 17 | 34 | 26 | 15 | 15 | 13 | 34 | 5 | 27 | 35 | 10 | 17 | 17 | 2 | 13 | 22 | 19 | 21 | 20 | 34 | 14 | 23 | 9 | 26 | 18 | Lrrc43 |
| Lrrc45 | 5432 | 7173 | 6983 | 8020 | 7518 | 7136 | 6776 | 4410 | 4944 | 5710 | 5360 | 5727 | 5281 | 5096 | 5020 | 6361 | 7715 | 7649 | 7929 | 6272 | 6923 | 5561 | 4423 | 5938 | 5920 | 5134 | 5233 | 4853 | Lrrc45 |
| Lrrc46 | 206 | 270 | 216 | 227 | 187 | 172 | 145 | 138 | 135 | 176 | 133 | 155 | 167 | 128 | 220 | 188 | 220 | 152 | 147 | 147 | 159 | 150 | 146 | 145 | 144 | 184 | 135 | 151 | Lrrc46 |
| Lrrc47 | 6249 | 6565 | 7643 | 7959 | 6759 | 6484 | 5651 | 4822 | 4965 | 6054 | 5611 | 6528 | 5770 | 5446 | 5831 | 5515 | 7992 | 7242 | 7606 | 5866 | 6142 | 5658 | 4754 | 6682 | 6290 | 6949 | 6099 | 5765 | Lrrc47 |
| Lrrc49 | 3011 | 3035 | 2545 | 2564 | 3159 | 3075 | 3446 | 7299 | 6016 | 5543 | 4527 | 5829 | 4939 | 5925 | 3423 | 2699 | 2716 | 2607 | 3243 | 4030 | 3267 | 6554 | 5594 | 5095 | 6179 | 5423 | 5758 | 5795 | Lrrc49 |
| Lrrc4b | 1167 | 1749 | 1824 | 2319 | 3310 | 2542 | 2357 | 5468 | 5530 | 7287 | 5605 | 6924 | 4974 | 6077 | 1315 | 1090 | 2352 | 2073 | 3352 | 2508 | 2616 | 7944 | 5050 | 7054 | 7991 | 6099 | 5960 | 5542 | Lrrc4b |
| Lrrc4c | 787 | 688 | 655 | 946 | 1424 | 1260 | 1557 | 1404 | 1310 | 1413 | 1241 | 1249 | 1181 | 1256 | 994 | 636 | 833 | 1058 | 1230 | 1363 | 1372 | 1601 | 1296 | 1093 | 1392 | 972 | 1177 | 1151 | Lrrc4c |
| Lrrc51 | 219 | 260 | 160 | 238 | 187 | 173 | 172 | 167 | 181 | 160 | 135 | 142 | 156 | 149 | 237 | 260 | 231 | 204 | 167 | 212 | 201 | 149 | 236 | 127 | 161 | 191 | 143 | 134 | Lrrc51 |
| Lrrc52 | 0 | 0 | 0 | 0 | 0 | 3 | 0 | 0 | 0 | 0 | 4 | 0 | 0 | 0 | 0 | 0 | 0 | 0 | 0 | 0 | 0 | 0 | 0 | 0 | 1 | 0 | 0 | 0 | Lrrc52 |
| Lrrc55 | 356 | 283 | 301 | 536 | 1113 | 785 | 916 | 1053 | 1140 | 1084 | 894 | 668 | 533 | 765 | 426 | 264 | 458 | 590 | 953 | 822 | 935 | 1348 | 917 | 794 | 1109 | 478 | 638 | 497 | Lrrc55 |
| Lrrc56 | 1054 | 1201 | 961 | 955 | 965 | 915 | 981 | 1069 | 1149 | 1175 | 951 | 1317 | 1278 | 1177 | 928 | 908 | 1095 | 809 | 939 | 954 | 982 | 988 | 1122 | 1096 | 1114 | 1267 | 1277 | 1260 | Lrrc56 |
| Lrrc57 | 1490 | 1321 | 1350 | 1271 | 1373 | 1266 | 1529 | 1638 | 1503 | 1393 | 1347 | 1606 | 1388 | 1573 | 1556 | 1350 | 1344 | 1247 | 1495 | 1528 | 1368 | 1643 | 1530 | 1548 | 1574 | 1625 | 1557 | 1631 | Lrrc57 |
| Lrrc58 | 5832 | 3809 | 7247 | 6806 | 7234 | 7432 | 7364 | 6861 | 6732 | 8053 | 8764 | 8410 | 9121 | 8644 | 5764 | 5603 | 6691 | 7178 | 7021 | 5769 | 6948 | 8590 | 6954 | 8803 | 8464 | 10918 | 8748 | 8746 | Lrrc58 |
| Lrrc59 | 17117 | 18354 | 18436 | 16193 | 13995 | 14796 | 14251 | 11510 | 12162 | 12086 | 12297 | 13179 | 12998 | 12105 | 16220 | 15432 | 17260 | 15249 | 16250 | 14244 | 14654 | 11498 | 11989 | 12544 | 12153 | 15085 | 13204 | 13884 | Lrrc59 |
| Lrrc6 | 16 | 15 | 10 | 11 | 32 | 26 | 16 | 40 | 40 | 53 | 31 | 56 | 64 | 12 | 40 | 17 | 14 | 20 | 14 | 28 | 25 | 15 | 43 | 19 | 33 | 17 | 20 | 17 | Lrrc6 |
| Lrrc61 | 591 | 718 | 827 | 727 | 855 | 737 | 717 | 1599 | 1582 | 1807 | 1661 | 2034 | 1917 | 1963 | 703 | 551 | 772 | 664 | 821 | 679 | 675 | 1925 | 1535 | 2042 | 1962 | 2052 | 2136 | 2198 | Lrrc61 |
| Lrrc63 | 33 | 17 | 28 | 15 | 25 | 28 | 32 | 12 | 8 | 12 | 7 | 21 | 15 | 24 | 29 | 24 | 39 | 14 | 26 | 25 | 20 | 7 | 10 | 17 | 9 | 27 | 17 | 20 | Lrrc63 |
| Lrrc66 | 15 | 24 | 28 | 13 | 13 | 6 | 6 | 0 | 2 | 1 | 2 | 1 | 9 | 7 | 22 | 26 | 15 | 10 | 16 | 14 | 18 | 5 | 1 | 1 | 0 | 0 | 7 | 6 | Lrrc66 |
| Lrrc69 | 3 | 5 | 1 | 6 | 6 | 12 | 6 | 2 | 5 | 1 | 9 | 5 | 0 | 3 | 8 | 5 | 16 | 9 | 12 | 6 | 14 | 2 | 6 | 1 | 6 | 2 | 1 | 2 | Lrrc69 |
| Lrrc7 | 682 | 590 | 273 | 648 | 862 | 960 | 997 | 1851 | 2057 | 1149 | 917 | 892 | 582 | 898 | 752 | 498 | 457 | 580 | 795 | 1398 | 1050 | 1108 | 1613 | 1076 | 1361 | 575 | 702 | 665 | Lrrc7 |
| Lrrc71 | 22 | 42 | 25 | 13 | 31 | 21 | 6 | 20 | 29 | 31 | 34 | 33 | 52 | 17 | 17 | 45 | 20 | 17 | 17 | 10 | 14 | 60 | 46 | 11 | 28 | 24 | 31 | 11 | Lrrc71 |
| Lrrc72 | 1 | 0 | 0 | 0 | 0 | 0 | 0 | 0 | 0 | 0 | 4 | 4 | 0 | 0 | 0 | 1 | 0 | 0 | 0 | 0 | 0 | 0 | 3 | 1 | 0 | 0 | 0 | 0 | Lrrc72 |
| Lrrc73 | 576 | 881 | 713 | 1026 | 995 | 811 | 781 | 256 | 385 | 437 | 339 | 341 | 269 | 270 | 621 | 694 | 867 | 1070 | 926 | 756 | 819 | 495 | 335 | 337 | 393 | 337 | 303 | 248 | Lrrc73 |
| Lrrc74a | 0 | 0 | 0 | 3 | 0 | 3 | 0 | 0 | 1 | 0 | 0 | 0 | 1 | 1 | 1 | 1 | 0 | 0 | 0 | 1 | 2 | 5 | 2 | 0 | 2 | 0 | 0 | 4 | Lrrc74a |
| Lrrc74b | 37 | 30 | 18 | 21 | 48 | 34 | 32 | 37 | 44 | 40 | 38 | 46 | 86 | 22 | 37 | 59 | 16 | 26 | 37 | 55 | 52 | 57 | 69 | 31 | 35 | 24 | 29 | 36 | Lrrc74b |
| Lrrc75a | 2375 | 2492 | 2098 | 2074 | 1933 | 1993 | 1899 | 1066 | 1073 | 1127 | 1035 | 1058 | 1019 | 1020 | 2218 | 2364 | 1997 | 2169 | 2112 | 1816 | 2101 | 1086 | 1170 | 1066 | 950 | 1245 | 952 | 1073 | Lrrc75a |
| Lrrc75b | 359 | 432 | 509 | 585 | 1321 | 934 | 847 | 2274 | 2628 | 3413 | 2982 | 2130 | 2212 | 1957 | 419 | 384 | 549 | 742 | 973 | 768 | 917 | 3985 | 2518 | 2708 | 3617 | 1634 | 1730 | 1877 | Lrrc75b |
| Lrrc8a | 499 | 461 | 753 | 868 | 716 | 564 | 744 | 653 | 618 | 891 | 789 | 868 | 822 | 900 | 489 | 495 | 721 | 721 | 613 | 521 | 698 | 851 | 616 | 981 | 897 | 1037 | 857 | 1035 | Lrrc8a |
| Lrrc8b | 3181 | 2478 | 2457 | 2637 | 3625 | 3576 | 3576 | 4877 | 4801 | 4831 | 4366 | 4602 | 4596 | 4482 | 3256 | 3032 | 2518 | 2780 | 2980 | 3096 | 3335 | 5155 | 4307 | 4138 | 4952 | 3320 | 4392 | 4180 | Lrrc8b |
| Lrrc8c | 2061 | 1879 | 2155 | 1957 | 2334 | 2074 | 1840 | 1756 | 1996 | 2467 | 2273 | 2073 | 2037 | 2009 | 1845 | 2341 | 2004 | 1961 | 1968 | 1752 | 1937 | 2386 | 2163 | 1941 | 1882 | 1955 | 1928 | 1871 | Lrrc8c |
| Lrrc8d | 1810 | 1949 | 2296 | 2585 | 2933 | 2362 | 2532 | 2645 | 2970 | 3462 | 3020 | 3477 | 2933 | 3325 | 1983 | 1812 | 2457 | 2391 | 2611 | 2136 | 2548 | 3651 | 2660 | 3284 | 3378 | 3612 | 2963 | 3296 | Lrrc8d |
| Lrrc8e | 82 | 50 | 38 | 18 | 29 | 16 | 15 | 45 | 23 | 36 | 27 | 7 | 7 | 15 | 57 | 48 | 20 | 11 | 10 | 10 | 13 | 13 | 53 | 25 | 13 | 30 | 8 | 4 | Lrrc8e |
| Lrrc9 | 911 | 976 | 310 | 616 | 740 | 818 | 753 | 677 | 606 | 435 | 390 | 482 | 482 | 392 | 988 | 960 | 447 | 563 | 590 | 962 | 879 | 495 | 475 | 290 | 508 | 263 | 406 | 300 | Lrrc9 |
| Lrrcc1 | 5292 | 3237 | 3544 | 3551 | 4234 | 4813 | 4442 | 5892 | 5460 | 4858 | 5018 | 5787 | 5604 | 5746 | 5418 | 4549 | 3635 | 3530 | 3642 | 4957 | 4380 | 5038 | 5345 | 4548 | 4770 | 4287 | 6051 | 5657 | Lrrcc1 |
| Lrrd1 | 9 | 73 | 34 | 51 | 25 | 31 | 22 | 23 | 15 | 19 | 29 | 26 | 26 | 27 | 13 | 32 | 43 | 51 | 42 | 37 | 34 | 22 | 17 | 16 | 25 | 24 | 24 | 27 | Lrrd1 |
| Lrrfip1 | 3334 | 2872 | 2781 | 2723 | 2801 | 2805 | 2558 | 2275 | 2339 | 2217 | 2817 | 1837 | 2160 | 1970 | 3215 | 3485 | 2685 | 2857 | 2558 | 2698 | 2597 | 2197 | 2524 | 2142 | 2114 | 2099 | 1866 | 1773 | Lrrfip1 |
| Lrrfip2 | 3217 | 3302 | 3576 | 3486 | 3359 | 3158 | 3453 | 3188 | 2926 | 3241 | 3255 | 3474 | 3164 | 3697 | 3320 | 3206 | 3468 | 3294 | 3296 | 3238 | 3329 | 3202 | 2845 | 3331 | 3522 | 3322 | 3460 | 3566 | Lrrfip2 |
| Lrriq1 | 467 | 350 | 168 | 277 | 283 | 370 | 259 | 626 | 384 | 346 | 216 | 239 | 418 | 226 | 436 | 497 | 218 | 208 | 193 | 385 | 303 | 267 | 517 | 190 | 191 | 220 | 206 | 186 | Lrriq1 |
| Lrriq3 | 62 | 50 | 58 | 63 | 64 | 56 | 50 | 100 | 75 | 70 | 77 | 111 | 52 | 71 | 74 | 64 | 63 | 56 | 72 | 43 | 64 | 102 | 78 | 60 | 57 | 73 | 94 | 80 | Lrriq3 |
| Lrriq4 | 5 | 3 | 0 | 0 | 1 | 0 | 0 | 0 | 1 | 2 | 0 | 0 | 1 | 2 | 5 | 2 | 0 | 2 | 0 | 4 | 0 | 0 | 0 | 1 | 1 | 2 | 0 | 0 | Lrriq4 |
| Lrrk1 | 422 | 507 | 466 | 284 | 311 | 403 | 190 | 943 | 1406 | 1081 | 1346 | 918 | 1188 | 838 | 456 | 536 | 392 | 368 | 257 | 245 | 227 | 892 | 1546 | 1066 | 882 | 786 | 1053 | 841 | Lrrk1 |
| Lrrk2 | 622 | 443 | 345 | 276 | 344 | 525 | 278 | 1025 | 1122 | 793 | 1001 | 857 | 1102 | 897 | 501 | 643 | 307 | 340 | 211 | 336 | 315 | 708 | 1282 | 693 | 724 | 602 | 966 | 868 | Lrrk2 |
| Lrrn1 | 11125 | 11825 | 11300 | 8754 | 9250 | 8727 | 8932 | 12090 | 9723 | 9085 | 8940 | 13076 | 15933 | 12775 | 12895 | 11727 | 10682 | 8134 | 9168 | 10551 | 8368 | 9408 | 9600 | 8501 | 8569 | 12846 | 14241 | 15022 | Lrrn1 |
| Lrrn2 | 764 | 1071 | 1148 | 1304 | 1913 | 1542 | 1464 | 1126 | 1461 | 1773 | 1425 | 1549 | 946 | 1358 | 981 | 954 | 1512 | 1358 | 1809 | 1352 | 1549 | 1990 | 1449 | 1646 | 1894 | 1346 | 1082 | 1225 | Lrrn2 |
| Lrrn3 | 4008 | 3864 | 3345 | 4475 | 5969 | 5450 | 5935 | 9575 | 7308 | 6832 | 5839 | 5069 | 5615 | 5228 | 5607 | 3467 | 4117 | 4456 | 5497 | 6923 | 5715 | 8473 | 6935 | 5757 | 6973 | 4603 | 5031 | 4934 | Lrrn3 |
| Lrrn4 | 73 | 67 | 67 | 25 | 36 | 63 | 44 | 11 | 12 | 1 | 3 | 7 | 10 | 5 | 76 | 97 | 41 | 31 | 17 | 47 | 24 | 5 | 9 | 7 | 9 | 14 | 10 | 9 | Lrrn4 |
| Lrrn4cl | 111 | 184 | 156 | 206 | 191 | 152 | 184 | 77 | 79 | 82 | 105 | 72 | 130 | 100 | 146 | 131 | 177 | 209 | 182 | 195 | 175 | 117 | 108 | 111 | 77 | 73 | 85 | 63 | Lrrn4cl |
| Lrrtm1 | 216 | 243 | 279 | 207 | 396 | 343 | 320 | 847 | 1009 | 974 | 886 | 757 | 711 | 661 | 266 | 191 | 290 | 250 | 390 | 304 | 332 | 849 | 853 | 869 | 988 | 661 | 642 | 704 | Lrrtm1 |
| Lrrtm2 | 1047 | 978 | 686 | 1710 | 2050 | 2166 | 2225 | 1840 | 1256 | 1153 | 1056 | 1089 | 683 | 1038 | 1588 | 833 | 1181 | 1861 | 1923 | 2427 | 2529 | 1512 | 1278 | 1118 | 1447 | 694 | 977 | 831 | Lrrtm2 |
| Lrrtm3 | 266 | 273 | 229 | 365 | 573 | 424 | 474 | 558 | 635 | 507 | 417 | 230 | 336 | 251 | 406 | 238 | 331 | 441 | 476 | 503 | 434 | 637 | 605 | 364 | 455 | 217 | 155 | 224 | Lrrtm3 |
| Lrrtm4 | 201 | 234 | 280 | 297 | 476 | 316 | 402 | 670 | 659 | 575 | 528 | 458 | 345 | 424 | 319 | 210 | 245 | 292 | 398 | 476 | 407 | 764 | 608 | 464 | 626 | 362 | 370 | 398 | Lrrtm4 |

| | | | | | | | | | | | | | | | | | | | | | | | | | | | | | |
|---|---|---|---|---|---|---|---|---|---|---|---|---|---|---|---|---|---|---|---|---|---|---|---|---|---|---|---|---|---|
| Lrsam1 | 1558 | 1756 | 1676 | 1826 | 1878 | 1539 | 1549 | 1938 | 2077 | 2301 | 2178 | 2066 | 2033 | 2039 | 1435 | 1565 | 1871 | 1760 | 1803 | 1577 | 1641 | 2346 | 2091 | 2083 | 2299 | 1725 | 2064 | 1836 | Lrsam1 |
| Lrtm1 | 40 | 18 | 39 | 6 | 3 | 13 | 4 | 32 | 39 | 40 | 77 | 78 | 99 | 38 | 22 | 30 | 3 | 0 | 0 | 5 | 4 | 42 | 75 | 14 | 10 | 28 | 73 | 68 | Lrtm1 |
| Lrtm2 | 186 | 201 | 235 | 377 | 679 | 557 | 542 | 228 | 253 | 361 | 254 | 161 | 88 | 128 | 229 | 193 | 319 | 504 | 647 | 429 | 548 | 384 | 285 | 219 | 314 | 72 | 76 | 72 | Lrtm2 |
| Lrwd1 | 2094 | 2459 | 2243 | 2135 | 2092 | 1988 | 1981 | 2451 | 2802 | 2827 | 2603 | 3151 | 2965 | 2936 | 1869 | 1998 | 2356 | 1872 | 2383 | 2031 | 1838 | 2408 | 2651 | 3117 | 2905 | 3773 | 3319 | 2959 | Lrwd1 |
| Lsamp | 3699 | 3459 | 2686 | 4815 | 6559 | 6131 | 6090 | 4818 | 4415 | 4928 | 4688 | 3726 | 3725 | 4200 | 4191 | 3248 | 3716 | 5366 | 5262 | 5927 | 7028 | 6176 | 4192 | 3854 | 5508 | 2417 | 4005 | 3588 | Lsamp |
| Lsg1 | 4196 | 4628 | 4487 | 3889 | 3511 | 3761 | 3528 | 3556 | 3654 | 3429 | 3215 | 4073 | 3861 | 3955 | 4146 | 4425 | 4016 | 3947 | 3680 | 4047 | 3382 | 3031 | 3509 | 3832 | 3560 | 4335 | 4250 | 4122 | Lsg1 |
| Lsm1 | 2225 | 2166 | 2056 | 2014 | 1726 | 1766 | 2144 | 1735 | 1481 | 1492 | 1304 | 1500 | 1405 | 1647 | 2203 | 2023 | 1893 | 2001 | 1746 | 2110 | 2102 | 1512 | 1468 | 1517 | 1497 | 1676 | 1593 | 1756 | Lsm1 |
| Lsm10 | 367 | 501 | 458 | 367 | 455 | 409 | 406 | 629 | 643 | 653 | 594 | 596 | 619 | 629 | 429 | 421 | 445 | 385 | 451 | 402 | 390 | 592 | 679 | 673 | 582 | 727 | 632 | 733 | Lsm10 |
| Lsm11 | 3496 | 3736 | 3606 | 5148 | 6276 | 5420 | 5463 | 3765 | 3395 | 4030 | 3866 | 3610 | 3602 | 3444 | 3529 | 3499 | 4591 | 5695 | 5866 | 4732 | 5944 | 4704 | 3402 | 3551 | 4683 | 2548 | 3489 | 3569 | Lsm11 |
| Lsm12 | 3164 | 3140 | 3663 | 3338 | 2847 | 2886 | 3368 | 3775 | 3582 | 3729 | 3412 | 4110 | 3758 | 4148 | 3076 | 2855 | 3385 | 3224 | 3061 | 3156 | 3155 | 3715 | 3592 | 3925 | 3920 | 4946 | 3900 | 4595 | Lsm12 |
| Lsm14a | 10080 | 9048 | 12046 | 10367 | 9513 | 9668 | 10102 | 11839 | 10249 | 10374 | 10456 | 12756 | 11621 | 13074 | 10001 | 9523 | 11184 | 9672 | 9891 | 9614 | 9773 | 10646 | 10471 | 12281 | 11554 | 14139 | 13034 | 13943 | Lsm14a |
| Lsm14b | 12339 | 13723 | 15866 | 17437 | 17117 | 14586 | 18519 | 12485 | 11908 | 16132 | 12981 | 15825 | 13359 | 19778 | 12585 | 11714 | 17263 | 16321 | 17725 | 13681 | 19448 | 15154 | 11207 | 17240 | 17685 | 16004 | 15347 | 18493 | Lsm14b |
| Lsm2 | 2152 | 2155 | 2557 | 2156 | 1625 | 1702 | 2198 | 1757 | 1828 | 2046 | 2110 | 2791 | 2419 | 3101 | 1779 | 1878 | 2417 | 1697 | 1932 | 1691 | 1872 | 1856 | 1706 | 2603 | 2105 | 3482 | 2618 | 3268 | Lsm2 |
| Lsm3 | 1491 | 1303 | 1378 | 1226 | 879 | 804 | 1241 | 1566 | 1301 | 1360 | 1054 | 1508 | 1128 | 1948 | 1532 | 830 | 1401 | 896 | 980 | 1053 | 1265 | 1108 | 1197 | 1674 | 1298 | 1759 | 1511 | 2040 | Lsm3 |
| Lsm4 | 879 | 981 | 1586 | 1155 | 929 | 998 | 867 | 990 | 1208 | 1422 | 1315 | 1455 | 1477 | 1609 | 872 | 840 | 1227 | 911 | 1146 | 960 | 777 | 844 | 970 | 1550 | 1365 | 1979 | 1640 | 1588 | Lsm4 |
| Lsm5 | 2024 | 1557 | 2530 | 2163 | 1494 | 1291 | 2322 | 1592 | 1194 | 1527 | 1390 | 1767 | 1602 | 2497 | 2181 | 1448 | 2162 | 1565 | 1768 | 1893 | 2335 | 892 | 1116 | 1824 | 1495 | 2233 | 1987 | 2624 | Lsm5 |
| Lsm6 | 3609 | 3404 | 4719 | 3419 | 2983 | 3192 | 3376 | 5202 | 4426 | 4635 | 4888 | 5635 | 5283 | 6377 | 3870 | 3074 | 3872 | 2945 | 3208 | 3692 | 3235 | 4294 | 4444 | 5465 | 5229 | 6756 | 5961 | 6681 | Lsm6 |
| Lsm7 | 1825 | 2121 | 2291 | 2240 | 1811 | 1687 | 1942 | 1838 | 2106 | 2095 | 2492 | 2447 | 2533 | 2444 | 1767 | 1703 | 2194 | 1909 | 1922 | 2322 | 1979 | 1526 | 2030 | 2564 | 2501 | 2908 | 2689 | 2730 | Lsm7 |
| Lsm8 | 3633 | 3321 | 3786 | 3788 | 3256 | 3094 | 4111 | 3951 | 3142 | 3209 | 3031 | 4069 | 3504 | 5232 | 3814 | 3224 | 3746 | 3398 | 3476 | 4100 | 3967 | 2953 | 3047 | 4157 | 3810 | 4576 | 4305 | 4771 | Lsm8 |
| Lsmem1 | 9 | 12 | 12 | 28 | 17 | 11 | 27 | 6 | 7 | 6 | 37 | 4 | 9 | 1 | 11 | 11 | 20 | 12 | 18 | 22 | 22 | 8 | 25 | 10 | 8 | 5 | 3 | 7 | Lsmem1 |
| Lsmem2 | 10 | 18 | 21 | 22 | 20 | 24 | 21 | 9 | 5 | 16 | 21 | 12 | 18 | 4 | 20 | 27 | 22 | 6 | 25 | 12 | 18 | 3 | 16 | 10 | 9 | 2 | 15 | 6 | Lsmem2 |
| Lsp1 | 427 | 428 | 441 | 269 | 200 | 276 | 144 | 642 | 844 | 639 | 1134 | 655 | 855 | 559 | 402 | 516 | 336 | 218 | 210 | 170 | 161 | 608 | 1174 | 609 | 546 | 715 | 759 | 707 | Lsp1 |
| Lsr | 1051 | 1058 | 1236 | 1090 | 884 | 1011 | 960 | 723 | 844 | 916 | 733 | 906 | 855 | 959 | 876 | 1032 | 878 | 863 | 987 | 892 | 820 | 551 | 968 | 1025 | 778 | 1428 | 837 | 1030 | Lsr |
| Lss | 8772 | 10648 | 7062 | 8148 | 6789 | 7847 | 6987 | 6211 | 5827 | 5743 | 5255 | 7277 | 7473 | 7367 | 8793 | 9583 | 7426 | 7593 | 6984 | 7237 | 6975 | 5833 | 5023 | 5547 | 6591 | 5556 | 8229 | 7649 | Lss |
| Lst1 | 46 | 50 | 43 | 38 | 49 | 33 | 60 | 61 | 78 | 66 | 92 | 39 | 69 | 82 | 30 | 48 | 31 | 61 | 14 | 29 | 46 | 52 | 92 | 63 | 82 | 86 | 58 | 72 | Lst1 |
| Lta | 5 | 3 | 2 | 2 | 9 | 9 | 8 | 2 | 4 | 9 | 2 | 2 | 9 | 5 | 5 | 1 | 3 | 3 | 4 | 1 | 8 | 10 | 6 | 8 | 1 | 7 | 5 | 1 | Lta |
| Lta4h | 10010 | 9218 | 8458 | 7997 | 7244 | 7564 | 7926 | 5503 | 5599 | 4986 | 5590 | 5236 | 5303 | 5287 | 9517 | 9347 | 8265 | 8048 | 7526 | 7768 | 7794 | 5023 | 6009 | 5058 | 4698 | 6216 | 5453 | 5650 | Lta4h |
| Ltb | 5 | 7 | 5 | 5 | 5 | 1 | 1 | 6 | 19 | 10 | 25 | 26 | 10 | 7 | 4 | 5 | 8 | 2 | 2 | 3 | 2 | 17 | 30 | 11 | 7 | 9 | 10 | 11 | Ltb |
| Ltb4r1 | 7 | 4 | 19 | 16 | 7 | 25 | 6 | 7 | 14 | 10 | 23 | 21 | 11 | 3 | 11 | 9 | 16 | 11 | 9 | 10 | 4 | 12 | 15 | 20 | 16 | 32 | 12 | 19 | Ltb4r1 |
| Ltb4r2 | 18 | 19 | 30 | 3 | 29 | 16 | 12 | 15 | 22 | 35 | 34 | 10 | 20 | 26 | 17 | 15 | 16 | 15 | 16 | 10 | 10 | 20 | 30 | 34 | 26 | 33 | 27 | 27 | Ltb4r2 |
| Ltbp1 | 3224 | 2070 | 1782 | 1500 | 1358 | 2163 | 1123 | 3224 | 5104 | 4098 | 6084 | 2108 | 2848 | 1888 | 2275 | 2974 | 1539 | 2436 | 1406 | 1113 | 1190 | 2732 | 5500 | 3755 | 2743 | 2487 | 2005 | 1950 | Ltbp1 |
| Ltbp2 | 47 | 50 | 30 | 118 | 80 | 98 | 13 | 172 | 386 | 284 | 485 | 66 | 128 | 60 | 37 | 85 | 37 | 231 | 12 | 23 | 7 | 234 | 408 | 285 | 264 | 56 | 77 | 42 | Ltbp2 |
| Ltbp3 | 1994 | 2128 | 2079 | 2578 | 2657 | 2686 | 1998 | 3121 | 4452 | 4794 | 5928 | 2852 | 3082 | 2598 | 1812 | 2299 | 2289 | 3112 | 2632 | 1883 | 2151 | 4111 | 4818 | 4551 | 3965 | 2995 | 2658 | 2357 | Ltbp3 |
| Ltbp4 | 3576 | 4388 | 5608 | 5161 | 3870 | 4039 | 3493 | 1768 | 2704 | 2606 | 3369 | 2687 | 2029 | 2107 | 2884 | 4021 | 5359 | 5211 | 4969 | 3252 | 3619 | 2316 | 2977 | 3106 | 2492 | 3182 | 2097 | 2019 | Ltbp4 |
| Ltbr | 404 | 448 | 398 | 321 | 384 | 383 | 242 | 835 | 1046 | 1079 | 1094 | 971 | 997 | 772 | 339 | 458 | 374 | 370 | 345 | 261 | 289 | 926 | 1249 | 874 | 739 | 1076 | 965 | 825 | Ltbr |
| Ltc4s | 12 | 11 | 12 | 13 | 10 | 29 | 10 | 9 | 11 | 10 | 25 | 3 | 10 | 18 | 13 | 14 | 7 | 14 | 13 | 14 | 13 | 18 | 23 | 18 | 23 | 8 | 21 | 20 | Ltc4s |
| Ltf | 14 | 0 | 6 | 31 | 7 | 22 | 17 | 21 | 27 | 27 | 86 | 37 | 102 | 26 | 28 | 35 | 20 | 50 | 13 | 20 | 10 | 47 | 82 | 65 | 14 | 61 | 57 | 64 | Ltf |
| Ltk | 20 | 31 | 17 | 39 | 31 | 17 | 29 | 38 | 37 | 47 | 42 | 35 | 36 | 21 | 28 | 20 | 35 | 28 | 26 | 27 | 38 | 70 | 47 | 44 | 28 | 31 | 19 | 28 | Ltk |
| Ltn1 | 10207 | 9253 | 6508 | 7610 | 7395 | 8059 | 8766 | 8862 | 7989 | 6645 | 7193 | 7342 | 7341 | 7622 | 10006 | 9314 | 6816 | 7658 | 7197 | 9050 | 8259 | 6679 | 7798 | 6179 | 7071 | 5295 | 7551 | 7068 | Ltn1 |
| LTO1 | 2403 | 2798 | 2885 | 2698 | 2233 | 2283 | 2334 | 2026 | 1884 | 2286 | 1866 | 2298 | 2002 | 2269 | 2285 | 2439 | 2772 | 2466 | 2638 | 2649 | 2222 | 1968 | 1912 | 2259 | 2261 | 2367 | 2312 | 2391 | LTO1 |
| Ltv1 | 3511 | 3135 | 3409 | 2656 | 2224 | 2273 | 2680 | 2942 | 2744 | 2722 | 3056 | 3058 | 2821 | 3216 | 3278 | 2947 | 2880 | 2426 | 2332 | 2830 | 2395 | 2445 | 2898 | 2685 | 2537 | 3476 | 3339 | 3377 | Ltv1 |
| Luc7l | 11951 | 11623 | 11390 | 13171 | 14307 | 14256 | 13604 | 19411 | 15999 | 16944 | 16794 | 17610 | 16629 | 17765 | 12686 | 10629 | 12573 | 13592 | 14324 | 14528 | 14480 | 18828 | 15172 | 17125 | 19071 | 15377 | 17462 | 17061 | Luc7l |
| Luc7l2 | 15939 | 12615 | 18753 | 19512 | 21341 | 22696 | 22256 | 26374 | 24568 | 29725 | 33335 | 33049 | 31433 | 37029 | 16804 | 14369 | 18884 | 19065 | 19734 | 19604 | 21287 | 30537 | 23498 | 31188 | 32157 | 30458 | 32584 | 33108 | Luc7l2 |
| Luc7l3 | 28813 | 23844 | 29120 | 31328 | 31560 | 32343 | 34564 | 40273 | 31494 | 33288 | 31424 | 34629 | 31439 | 36077 | 31898 | 26258 | 29030 | 30101 | 32028 | 37590 | 33410 | 36008 | 31127 | 34568 | 35251 | 33565 | 32947 | 34069 | Luc7l3 |
| Lum | 7917 | 5657 | 7524 | 4103 | 2702 | 6069 | 2525 | 8852 | 12888 | 7510 | 11144 | 9145 | 11853 | 9272 | 8482 | 8530 | 4899 | 5804 | 2554 | 3499 | 2319 | 8671 | 15572 | 8264 | 7863 | 9305 | 10000 | 9772 | Lum |
| Lurap1 | 1252 | 1027 | 1026 | 962 | 1042 | 1002 | 1102 | 359 | 325 | 278 | 284 | 248 | 235 | 253 | 1096 | 1083 | 1007 | 1011 | 1006 | 976 | 1018 | 284 | 330 | 253 | 227 | 322 | 235 | 239 | Lurap1 |
| Lurap1l | 371 | 358 | 358 | 399 | 435 | 341 | 394 | 348 | 355 | 398 | 365 | 236 | 346 | 277 | 376 | 339 | 396 | 417 | 458 | 543 | 421 | 384 | 389 | 301 | 340 | 246 | 306 | 267 | Lurap1l |
| Luzp1 | 9867 | 9306 | 6376 | 7573 | 7357 | 7907 | 7456 | 7380 | 6857 | 6156 | 6639 | 7357 | 7101 | 6886 | 9174 | 9224 | 6946 | 7679 | 7293 | 8057 | 7817 | 5773 | 6674 | 6105 | 6601 | 5289 | 7682 | 7363 | Luzp1 |
| Luzp2 | 672 | 550 | 634 | 766 | 1754 | 1527 | 1050 | 2441 | 2425 | 1933 | 1741 | 1155 | 1397 | 1139 | 876 | 706 | 709 | 1224 | 1188 | 1604 | 1384 | 2737 | 2202 | 1250 | 2154 | 666 | 1190 | 1186 | Luzp2 |
| Luzp4 | 0 | 0 | 0 | 0 | 0 | 0 | 0 | 0 | 2 | 0 | 0 | 0 | 0 | 0 | 0 | 0 | 0 | 0 | 0 | 0 | 0 | 0 | 1 | 0 | 0 | 0 | 0 | 0 | Luzp4 |
| Lvrn | 143 | 123 | 90 | 122 | 84 | 117 | 143 | 98 | 142 | 87 | 181 | 84 | 151 | 90 | 170 | 139 | 88 | 112 | 62 | 99 | 119 | 67 | 190 | 98 | 77 | 93 | 139 | 126 | Lvrn |
| Lxn | 799 | 901 | 814 | 753 | 1316 | 984 | 730 | 1806 | 1697 | 1761 | 1925 | 1460 | 1562 | 1380 | 1058 | 789 | 804 | 930 | 1117 | 977 | 1010 | 1975 | 2060 | 1542 | 1730 | 1293 | 1381 | 1246 | Lxn |
| Ly6a | 5 | 2 | 0 | 1 | 3 | 5 | 1 | 5 | 14 | 1 | 1 | 3 | 7 | 0 | 1 | 1 | 0 | 7 | 0 | 0 | 2 | 8 | 6 | 0 | 0 | 10 | 0 | 6 | Ly6a |
| Ly6c1 | 17 | 13 | 16 | 2 | 9 | 8 | 11 | 34 | 19 | 19 | 27 | 0 | 16 | 20 | 20 | 15 | 27 | 10 | 14 | 26 | 19 | 20 | 25 | 23 | 16 | 8 | 28 | 12 | Ly6c1 |
| Ly6c2 | 1 | 0 | 2 | 0 | 0 | 0 | 0 | 4 | 2 | 6 | 6 | 0 | 6 | 0 | 0 | 2 | 1 | 7 | 0 | 1 | 1 | 2 | 5 | 1 | 0 | 1 | 0 | 2 | Ly6c2 |
| Ly6d | 0 | 0 | 0 | 0 | 0 | 0 | 0 | 1 | 8 | 7 | 0 | 0 | 0 | 0 | 0 | 0 | 0 | 0 | 0 | 0 | 0 | 0 | 12 | 7 | 0 | 0 | 0 | 0 | Ly6d |
| Ly6e | 2162 | 2017 | 2142 | 1899 | 1972 | 1936 | 1964 | 2209 | 2628 | 2545 | 2697 | 2547 | 2538 | 2655 | 1917 | 2045 | 1938 | 1721 | 2067 | 1915 | 1879 | 2294 | 2406 | 2549 | 2492 | 3098 | 2364 | 2552 | Ly6e |
| Ly6f | 1 | 0 | 0 | 0 | 0 | 0 | 0 | 0 | 1 | 0 | 0 | 0 | 0 | 0 | 0 | 0 | 0 | 0 | 0 | 0 | 0 | 0 | 0 | 0 | 0 | 0 | 0 | 0 | Ly6f |
| Ly6g | 0 | 0 | 0 | 0 | 0 | 0 | 0 | 0 | 0 | 0 | 0 | 1 | 1 | 0 | 0 | 0 | 0 | 0 | 0 | 0 | 0 | 0 | 4 | 0 | 0 | 0 | 0 | 0 | Ly6g |
| Ly6g2 | 0 | 0 | 0 | 0 | 0 | 0 | 0 | 0 | 0 | 0 | 0 | 0 | 0 | 0 | 0 | 0 | 0 | 0 | 0 | 0 | 0 | 0 | 1 | 0 | 0 | 0 | 0 | 0 | Ly6g2 |
| Ly6g5b | 110 | 131 | 94 | 115 | 116 | 132 | 127 | 56 | 73 | 78 | 63 | 59 | 54 | 65 | 76 | 101 | 138 | 106 | 112 | 116 | 106 | 77 | 63 | 50 | 73 | 55 | 63 | 63 | Ly6g5b |
| Ly6g5c | 0 | 0 | 0 | 0 | 0 | 0 | 0 | 1 | 0 | 0 | 0 | 0 | 2 | 0 | 0 | 1 | 0 | 0 | 0 | 1 | 0 | 0 | 0 | 0 | 0 | 0 | 0 | 0 | Ly6g5c |
| Ly6g6c | 10 | 9 | 6 | 5 | 10 | 12 | 15 | 6 | 2 | 4 | 10 | 1 | 0 | 5 | 12 | 9 | 21 | 19 | 6 | 9 | 6 | 3 | 3 | 9 | 0 | 14 | 8 | 6 | Ly6g6c |
| Ly6g6d | 7 | 21 | 24 | 13 | 11 | 13 | 11 | 29 | 18 | 21 | 11 | 14 | 32 | 33 | 16 | 25 | 13 | 9 | 12 | 24 | 9 | 18 | 18 | 22 | 34 | 24 | 29 | 18 | Ly6g6d |
| Ly6g6e | 6 | 3 | 2 | 13 | 0 | 8 | 3 | 12 | 21 | 27 | 6 | 17 | 10 | 13 | 1 | 0 | 3 | 7 | 4 | 5 | 8 | 5 | 15 | 15 | 18 | 7 | 7 | 6 | Ly6g6e |
| Ly6g6f | 21 | 17 | 17 | 9 | 5 | 4 | 8 | 24 | 10 | 14 | 21 | 10 | 25 | 23 | 13 | 23 | 19 | 14 | 11 | 22 | 6 | 17 | 16 | 13 | 6 | 24 | 23 | 8 | Ly6g6f |
| Ly6g6g | 1 | 0 | 0 | 0 | 0 | 0 | 0 | 2 | 8 | 0 | 3 | 0 | 0 | 0 | 1 | 1 | 0 | 0 | 0 | 0 | 0 | 0 | 3 | 1 | 0 | 2 | 0 | 0 | Ly6g6g |
| Ly6h | 1749 | 2427 | 1876 | 3101 | 4479 | 3593 | 3337 | 4372 | 4796 | 5151 | 4414 | 3699 | 3238 | 3295 | 2110 | 1759 | 2720 | 3177 | 4189 | 4186 | 3754 | 5392 | 4835 | 4523 | 5463 | 3491 | 3055 | 2811 | Ly6h |
| Ly6i | 0 | 0 | 0 | 0 | 0 | 0 | 0 | 0 | 0 | 0 | 0 | 0 | 0 | 0 | 0 | 0 | 0 | 0 | 1 | 0 | 0 | 0 | 0 | 1 | 0 | 0 | 0 | 0 | Ly6i |
| Ly6k | 1 | 0 | 0 | 2 | 0 | 0 | 0 | 0 | 0 | 0 | 0 | 2 | 1 | 1 | 0 | 1 | 0 | 0 | 0 | 0 | 0 | 2 | 1 | 2 | 0 | 0 | 3 | 1 | Ly6k |
| Ly6m | 0 | 0 | 0 | 0 | 0 | 0 | 0 | 0 | 0 | 3 | 0 | 0 | 0 | 0 | 0 | 1 | 0 | 0 | 0 | 0 | 0 | 0 | 0 | 0 | 0 | 0 | 0 | 0 | Ly6m |
| Ly75 | 1020 | 689 | 763 | 625 | 628 | 648 | 473 | 390 | 405 | 368 | 442 | 344 | 413 | 282 | 822 | 1153 | 647 | 612 | 457 | 465 | 510 | 237 | 363 | 335 | 238 | 376 | 394 | 333 | Ly75 |
| Ly86 | 134 | 124 | 144 | 110 | 131 | 141 | 120 | 192 | 240 | 202 | 236 | 167 | 188 | 149 | 178 | 167 | 88 | 150 | 90 | 146 | 128 | 194 | 313 | 179 | 163 | 169 | 158 | 179 | Ly86 |
| Ly9 | 16 | 6 | 2 | 3 | 4 | 7 | 6 | 16 | 16 | 5 | 27 | 11 | 9 | 11 | 12 | 9 | 6 | 11 | 10 | 7 | 13 | 17 | 23 | 30 | 8 | 5 | 9 | 6 | Ly9 |
| Ly96 | 28 | 16 | 26 | 26 | 20 | 23 | 30 | 88 | 66 | 120 | 105 | 80 | 79 | 76 | 29 | 28 | 29 | 30 | 34 | 22 | 29 | 60 | 86 | 100 | 41 | 87 | 66 | 103 | Ly96 |
| Lyar | 2836 | 2622 | 3205 | 2478 | 1852 | 1888 | 2302 | 2191 | 2057 | 2385 | 2101 | 2905 | 2713 | 3135 | 2614 | 2681 | 2964 | 2130 | 2083 | 2324 | 2013 | 1798 | 2029 | 2799 | 2258 | 3777 | 2864 | 3167 | Lyar |
| Lyg1 | 0 | 0 | 5 | 0 | 0 | 0 | 0 | 0 | 0 | 4 | 1 | 0 | 0 | 0 | 0 | 0 | 0 | 0 | 0 | 0 | 0 | 0 | 0 | 0 | 0 | 0 | 0 | 0 | Lyg1 |
| Lyg2 | 0 | 0 | 0 | 0 | 0 | 0 | 0 | 0 | 2 | 0 | 1 | 1 | 1 | 2 | 0 | 3 | 0 | 0 | 0 | 1 | 0 | 0 | 2 | 3 | 1 | 0 | 0 | 2 | Lyg2 |
| Lyl1 | 105 | 164 | 156 | 148 | 151 | 148 | 107 | 162 | 258 | 203 | 265 | 256 | 380 | 211 | 129 | 173 | 154 | 128 | 168 | 116 | 120 | 196 | 326 | 222 | 178 | 299 | 202 | 271 | Lyl1 |

| | | | | | | | | | | | | | | | | | | | | | | | | | | | | | |
|---|---|---|---|---|---|---|---|---|---|---|---|---|---|---|---|---|---|---|---|---|---|---|---|---|---|---|---|---|---|
| Lyn | 669 | 547 | 680 | 460 | 469 | 520 | 384 | 738 | 691 | 574 | 764 | 610 | 677 | 661 | 646 | 723 | 531 | 481 | 393 | 545 | 474 | 760 | 895 | 627 | 671 | 698 | 751 | 676 | Lyn |
| Lynx1 | 768 | 860 | 1084 | 837 | 1003 | 846 | 769 | 194 | 291 | 234 | 273 | 169 | 186 | 189 | 741 | 777 | 985 | 767 | 947 | 801 | 752 | 266 | 370 | 199 | 163 | 136 | 245 | 153 | Lynx1 |
| Lypd1 | 775 | 931 | 776 | 1164 | 1976 | 1580 | 1329 | 1573 | 1851 | 2087 | 2011 | 1011 | 1210 | 1032 | 1038 | 922 | 1049 | 1328 | 1419 | 1450 | 1527 | 2176 | 1834 | 1356 | 1951 | 691 | 926 | 815 | Lypd1 |
| Lypd10 | 0 | 0 | 0 | 0 | 0 | 0 | 0 | 0 | 0 | 0 | 0 | 0 | 0 | 0 | 0 | 0 | 0 | 0 | 0 | 1 | 0 | 0 | 0 | 0 | 0 | 0 | 0 | 0 | Lypd10 |
| Lypd11 | 0 | 0 | 0 | 0 | 0 | 0 | 0 | 0 | 0 | 0 | 0 | 0 | 0 | 0 | 0 | 1 | 0 | 0 | 2 | 0 | 0 | 0 | 0 | 2 | 0 | 0 | 0 | 0 | Lypd11 |
| Lypd2 | 5 | 5 | 0 | 0 | 4 | 3 | 1 | 21 | 39 | 20 | 30 | 0 | 1 | 3 | 5 | 6 | 0 | 2 | 2 | 0 | 1 | 20 | 36 | 24 | 2 | 13 | 0 | 0 | Lypd2 |
| Lypd3 | 116 | 85 | 77 | 55 | 44 | 54 | 23 | 55 | 94 | 65 | 27 | 40 | 44 | 31 | 83 | 80 | 60 | 48 | 53 | 44 | 24 | 2 | 128 | 38 | 10 | 10 | 28 | 53 | Lypd3 |
| Lypd4 | 1 | 0 | 5 | 0 | 3 | 0 | 2 | 0 | 2 | 0 | 0 | 0 | 0 | 0 | 0 | 0 | 3 | 0 | 0 | 0 | 2 | 2 | 0 | 0 | 4 | 3 | 1 | 0 | Lypd4 |
| Lypd5 | 0 | 0 | 0 | 6 | 4 | 0 | 0 | 6 | 3 | 10 | 0 | 0 | 0 | 1 | 0 | 0 | 0 | 1 | 0 | 0 | 1 | 3 | 6 | 0 | 0 | 2 | 0 | 0 | Lypd5 |
| Lypd6 | 2222 | 1715 | 1667 | 1569 | 1743 | 1745 | 2281 | 1033 | 1135 | 1000 | 1004 | 1059 | 1067 | 1020 | 1994 | 1970 | 1579 | 1786 | 1725 | 1928 | 2122 | 1066 | 1241 | 1043 | 1093 | 1402 | 923 | 960 | Lypd6 |
| Lypd6b | 140 | 94 | 60 | 82 | 126 | 161 | 95 | 265 | 352 | 205 | 252 | 140 | 138 | 195 | 116 | 104 | 75 | 76 | 97 | 124 | 147 | 249 | 390 | 254 | 229 | 156 | 109 | 153 | Lypd6b |
| Lypd8 | 0 | 0 | 0 | 0 | 0 | 0 | 1 | 0 | 0 | 0 | 0 | 0 | 0 | 0 | 0 | 0 | 0 | 0 | 0 | 0 | 0 | 0 | 0 | 1 | 0 | 0 | 0 | 0 | Lypd8 |
| Lypd8l | 0 | 0 | 0 | 0 | 0 | 0 | 0 | 0 | 0 | 0 | 0 | 0 | 0 | 0 | 0 | 0 | 0 | 0 | 0 | 0 | 0 | 0 | 0 | 2 | 0 | 0 | 0 | 0 | Lypd8l |
| Lypd9 | 0 | 0 | 0 | 0 | 0 | 0 | 0 | 0 | 2 | 0 | 0 | 0 | 0 | 0 | 0 | 0 | 0 | 0 | 0 | 0 | 0 | 0 | 0 | 0 | 0 | 0 | 0 | 0 | Lypd9 |
| Lypla1 | 3652 | 2180 | 3353 | 2878 | 2743 | 3009 | 3179 | 3296 | 2874 | 2431 | 2865 | 3290 | 3500 | 3836 | 3493 | 3197 | 2938 | 2979 | 2709 | 2882 | 2997 | 2872 | 3014 | 2956 | 2845 | 3637 | 3466 | 4062 | Lypla1 |
| Lypla2 | 4418 | 5961 | 6258 | 6014 | 5919 | 4999 | 4986 | 3660 | 4491 | 4972 | 4258 | 4730 | 4645 | 4845 | 4618 | 5154 | 6411 | 5690 | 6482 | 5184 | 5222 | 4388 | 4212 | 5013 | 4983 | 5647 | 5075 | 4675 | Lypla2 |
| Lyplal1 | 482 | 376 | 533 | 391 | 348 | 383 | 417 | 795 | 718 | 611 | 629 | 638 | 743 | 718 | 501 | 483 | 415 | 314 | 355 | 418 | 363 | 642 | 671 | 604 | 661 | 705 | 600 | 694 | Lyplal1 |
| Lyrm1 | 281 | 242 | 253 | 285 | 322 | 312 | 291 | 388 | 335 | 354 | 312 | 306 | 273 | 330 | 305 | 240 | 245 | 289 | 302 | 389 | 319 | 343 | 343 | 289 | 392 | 248 | 331 | 301 | Lyrm1 |
| Lyrm2 | 1476 | 1341 | 1632 | 1337 | 1359 | 1418 | 1543 | 2286 | 2196 | 2268 | 2173 | 2662 | 2639 | 2880 | 1669 | 1301 | 1508 | 1301 | 1544 | 1637 | 1541 | 2169 | 2222 | 2380 | 2201 | 2989 | 3086 | 3053 | Lyrm2 |
| Lyrm4 | 1028 | 937 | 926 | 1120 | 964 | 923 | 1252 | 992 | 748 | 895 | 804 | 849 | 755 | 1111 | 1100 | 854 | 979 | 1030 | 990 | 1260 | 1196 | 851 | 851 | 914 | 885 | 967 | 900 | 1023 | Lyrm4 |
| Lyrm7 | 592 | 445 | 466 | 511 | 509 | 489 | 563 | 503 | 459 | 445 | 528 | 478 | 379 | 528 | 539 | 442 | 451 | 484 | 488 | 564 | 513 | 515 | 455 | 445 | 451 | 415 | 380 | 466 | Lyrm7 |
| Lyrm9 | 2907 | 3189 | 2300 | 3350 | 3695 | 3407 | 3696 | 1793 | 1814 | 1964 | 1665 | 1679 | 1416 | 1791 | 3112 | 2743 | 2928 | 3307 | 3178 | 3443 | 3674 | 2161 | 1654 | 1630 | 2149 | 1174 | 1613 | 1519 | Lyrm9 |
| Lysmd1 | 529 | 556 | 546 | 684 | 813 | 810 | 832 | 548 | 602 | 530 | 539 | 530 | 521 | 566 | 580 | 569 | 687 | 660 | 841 | 730 | 844 | 610 | 577 | 644 | 680 | 548 | 475 | 533 | Lysmd1 |
| Lysmd2 | 107 | 101 | 150 | 136 | 246 | 156 | 160 | 375 | 497 | 467 | 545 | 277 | 332 | 292 | 121 | 129 | 137 | 132 | 190 | 124 | 149 | 363 | 462 | 378 | 377 | 385 | 351 | 354 | Lysmd2 |
| Lysmd3 | 5399 | 3911 | 4067 | 3948 | 3161 | 3872 | 3966 | 3485 | 2910 | 2306 | 2730 | 2773 | 2901 | 2722 | 5304 | 5362 | 3739 | 4036 | 3242 | 4530 | 3858 | 2822 | 3170 | 2435 | 2522 | 2621 | 2817 | 3021 | Lysmd3 |
| Lysmd4 | 1713 | 1913 | 1823 | 2070 | 2264 | 1988 | 2171 | 2251 | 2397 | 2328 | 2230 | 2086 | 1898 | 2162 | 1878 | 1763 | 1991 | 2099 | 2162 | 2075 | 2110 | 2261 | 2399 | 2215 | 2489 | 1793 | 2072 | 1983 | Lysmd4 |
| Lyst | 4464 | 4546 | 1843 | 3165 | 3995 | 4659 | 4656 | 3593 | 3573 | 2481 | 2773 | 2225 | 2322 | 2349 | 4425 | 4180 | 2381 | 3344 | 2988 | 5399 | 4794 | 1970 | 3315 | 1800 | 2528 | 1318 | 2509 | 1818 | Lyst |
| Lyve1 | 295 | 275 | 421 | 261 | 340 | 359 | 248 | 468 | 596 | 535 | 684 | 477 | 761 | 550 | 359 | 315 | 315 | 306 | 267 | 436 | 285 | 560 | 1056 | 610 | 593 | 483 | 624 | 470 | Lyve1 |
| Lyz1 | 3 | 5 | 3 | 1 | 3 | 2 | 6 | 0 | 2 | 0 | 5 | 0 | 1 | 5 | 7 | 5 | 9 | 7 | 5 | 7 | 4 | 3 | 1 | 0 | 5 | 1 | 5 | 5 | Lyz1 |
| Lyz2 | 270 | 280 | 264 | 265 | 284 | 323 | 257 | 375 | 465 | 322 | 655 | 320 | 359 | 328 | 295 | 426 | 178 | 340 | 197 | 288 | 276 | 379 | 909 | 377 | 436 | 370 | 338 | 392 | Lyz2 |
| Lyzl4 | 3 | 3 | 2 | 1 | 4 | 2 | 1 | 0 | 2 | 5 | 4 | 1 | 1 | 0 | 1 | 0 | 0 | 4 | 0 | 1 | 3 | 5 | 5 | 7 | 0 | 1 | 0 | 1 | Lyzl4 |
| Lyzl6 | 0 | 0 | 0 | 0 | 0 | 0 | 1 | 0 | 1 | 0 | 0 | 0 | 0 | 0 | 0 | 0 | 0 | 0 | 1 | 0 | 1 | 2 | 0 | 0 | 0 | 0 | 0 | 0 | Lyzl6 |
| Lzic | 2201 | 2284 | 2340 | 2174 | 2009 | 1854 | 2107 | 2491 | 2507 | 2546 | 2468 | 2647 | 2537 | 2705 | 2040 | 1930 | 2161 | 2031 | 2122 | 2056 | 2025 | 2351 | 2537 | 2729 | 2549 | 2645 | 2585 | 2777 | Lzic |
| Lztfl1 | 7266 | 5807 | 4688 | 5577 | 5015 | 6069 | 6354 | 5969 | 5414 | 4379 | 5054 | 4485 | 4567 | 4630 | 7647 | 6822 | 5093 | 5567 | 4828 | 6896 | 6049 | 5116 | 5302 | 4195 | 4691 | 4136 | 4656 | 4574 | Lztfl1 |
| Lztr1 | 5538 | 6386 | 5796 | 6275 | 6964 | 6568 | 6150 | 7668 | 8586 | 8577 | 8386 | 8276 | 8247 | 8071 | 5811 | 5841 | 6255 | 6268 | 6958 | 6382 | 6575 | 8590 | 8427 | 8526 | 8889 | 7730 | 8373 | 8218 | Lztr1 |
| Lzts1 | 2806 | 3544 | 3404 | 4154 | 4111 | 4039 | 4021 | 3577 | 3672 | 4136 | 3425 | 4651 | 3851 | 5036 | 3264 | 2730 | 3885 | 3933 | 4331 | 3521 | 4025 | 3992 | 3324 | 4707 | 5521 | 3738 | 4385 | 5029 | Lzts1 |
| Lzts2 | 2977 | 3680 | 4246 | 3225 | 3058 | 3126 | 2476 | 3263 | 3822 | 3856 | 4007 | 4503 | 4186 | 3795 | 2620 | 3307 | 3837 | 2952 | 3513 | 2713 | 2561 | 3489 | 3713 | 4264 | 3627 | 5019 | 4499 | 4087 | Lzts2 |
| Lzts3 | 2376 | 3144 | 3238 | 3748 | 3393 | 2934 | 3120 | 1353 | 1545 | 1881 | 1751 | 1546 | 1371 | 1505 | 2136 | 2646 | 3686 | 3859 | 3497 | 2765 | 3362 | 1975 | 1478 | 1816 | 1972 | 1681 | 1525 | 1510 | Lzts3 |
| M1ap | 1 | 8 | 4 | 2 | 2 | 4 | 6 | 7 | 11 | 3 | 11 | 2 | 14 | 4 | 12 | 7 | 0 | 3 | 5 | 3 | 1 | 13 | 8 | 4 | 3 | 8 | 5 | 7 | M1ap |
| M6pr | 9401 | 9350 | 8902 | 7443 | 5897 | 6971 | 7015 | 5526 | 5436 | 4753 | 5116 | 5207 | 5289 | 5807 | 9312 | 9407 | 8006 | 7384 | 5919 | 7360 | 6506 | 4832 | 5572 | 5275 | 4674 | 6078 | 5426 | 6027 | M6pr |
| Mab21l1 | 1837 | 2492 | 3199 | 2895 | 3483 | 3251 | 1649 | 4979 | 4128 | 4619 | 3775 | 3905 | 5722 | 3841 | 2740 | 2419 | 3175 | 2386 | 3235 | 3130 | 2681 | 5817 | 4938 | 3903 | 4550 | 1552 | 4534 | 4550 | Mab21l1 |
| Mab21l2 | 2075 | 2063 | 3605 | 2638 | 5137 | 2600 | 1606 | 4146 | 4606 | 5923 | 5933 | 2699 | 6664 | 2782 | 2606 | 2146 | 3145 | 2345 | 2629 | 2105 | 2238 | 5778 | 4932 | 2676 | 2769 | 779 | 3271 | 3965 | Mab21l2 |
| Mab21l3 | 456 | 380 | 337 | 283 | 249 | 234 | 217 | 1127 | 211 | 113 | 156 | 179 | 166 | 164 | 373 | 389 | 337 | 277 | 293 | 286 | 245 | 85 | 197 | 168 | 135 | 185 | 194 | 180 | Mab21l3 |
| Mab21l4 | 8 | 4 | 6 | 1 | 11 | 12 | 1 | 21 | 34 | 10 | 10 | 15 | 22 | 12 | 1 | 8 | 6 | 3 | 2 | 2 | 2 | 5 | 36 | 10 | 12 | 3 | 10 | 17 | Mab21l4 |
| Macc1 | 212 | 55 | 31 | 35 | 36 | 41 | 19 | 241 | 233 | 190 | 163 | 62 | 125 | 70 | 123 | 113 | 16 | 51 | 12 | 20 | 21 | 67 | 242 | 141 | 26 | 146 | 73 | 81 | Macc1 |
| Macf1 | 33778 | 32302 | 17322 | 23459 | 29318 | 37941 | 32229 | 54362 | 52476 | 37981 | 42575 | 38572 | 38299 | 40854 | 34596 | 31962 | 19483 | 24922 | 25489 | 37701 | 38297 | 33652 | 48259 | 31022 | 42651 | 23050 | 47259 | 35198 | Macf1 |
| Macir | 2603 | 2154 | 2708 | 3010 | 3165 | 2960 | 3741 | 4295 | 3337 | 3277 | 3143 | 3795 | 3009 | 4119 | 2794 | 1950 | 2876 | 2926 | 3388 | 3370 | 3499 | 3984 | 3254 | 3682 | 3931 | 4189 | 3948 | 4118 | Macir |
| Maco1 | 8290 | 7233 | 7954 | 9302 | 10130 | 9677 | 10123 | 11623 | 10086 | 12056 | 11105 | 11593 | 10915 | 12108 | 8438 | 7449 | 8427 | 8899 | 9853 | 8550 | 9939 | 13774 | 10213 | 12015 | 13151 | 10584 | 10915 | 11357 | Maco1 |
| Macrod1 | 221 | 229 | 195 | 232 | 211 | 200 | 173 | 93 | 146 | 159 | 195 | 99 | 119 | 117 | 189 | 220 | 238 | 231 | 215 | 195 | 199 | 139 | 201 | 136 | 146 | 176 | 100 | 102 | Macrod1 |
| Macrod2 | 2127 | 1945 | 2462 | 2588 | 2960 | 2806 | 3488 | 3835 | 2295 | 3042 | 2573 | 2880 | 2224 | 3465 | 2536 | 1878 | 2508 | 2691 | 2940 | 3602 | 3263 | 3982 | 3355 | 3097 | 3387 | 2951 | 3382 | 3650 | Macrod2 |
| Macroh2a1 | 16909 | 19937 | 21537 | 18406 | 16442 | 15902 | 18948 | 14537 | 15473 | 16437 | 14956 | 20693 | 17932 | 21450 | 16072 | 16824 | 20286 | 16830 | 18898 | 17742 | 17832 | 13426 | 14402 | 18949 | 17965 | 23409 | 20428 | 22009 | Macroh2a1 |
| Macroh2a2 | 7880 | 7934 | 8232 | 6154 | 6806 | 6942 | 7115 | 13636 | 14133 | 14766 | 14001 | 17078 | 16217 | 18237 | 7411 | 7262 | 7757 | 6090 | 6858 | 7245 | 6606 | 14766 | 13675 | 16896 | 16599 | 19961 | 17453 | 18364 | Macroh2a2 |
| Mad1l1 | 2891 | 3090 | 3303 | 2760 | 2377 | 2370 | 2367 | 2409 | 2675 | 2579 | 2385 | 3149 | 2624 | 3038 | 2587 | 2657 | 3069 | 2534 | 2696 | 2422 | 2434 | 2296 | 2367 | 2970 | 2674 | 3662 | 2975 | 3255 | Mad1l1 |
| Mad2l1 | 9912 | 7990 | 9291 | 7031 | 4922 | 6290 | 7016 | 6482 | 5515 | 4808 | 4945 | 7211 | 6270 | 7788 | 9351 | 8659 | 7516 | 6015 | 6130 | 7878 | 6467 | 3902 | 5167 | 6262 | 5688 | 9117 | 7884 | 8103 | Mad2l1 |
| Mad2l1bp | 910 | 1283 | 1004 | 991 | 712 | 777 | 754 | 700 | 718 | 751 | 784 | 821 | 696 | 751 | 865 | 1116 | 970 | 902 | 860 | 1000 | 765 | 630 | 760 | 846 | 742 | 959 | 894 | 965 | Mad2l1bp |
| Mad2l2 | 2365 | 2846 | 2624 | 3009 | 2782 | 2493 | 2691 | 1860 | 2071 | 2276 | 2055 | 2237 | 2208 | 2032 | 2480 | 2424 | 2772 | 2935 | 2991 | 2707 | 2913 | 2296 | 1964 | 2335 | 2313 | 2393 | 2189 | 2013 | Mad2l2 |
| Madcam1 | 52 | 36 | 24 | 21 | 15 | 28 | 8 | 32 | 89 | 54 | 89 | 38 | 65 | 30 | 44 | 46 | 28 | 35 | 14 | 13 | 7 | 67 | 87 | 56 | 26 | 64 | 72 | 42 | Madcam1 |
| Madd | 8762 | 11118 | 10689 | 15276 | 16720 | 14489 | 14462 | 6797 | 7353 | 9717 | 7923 | 8516 | 7860 | 7873 | 9792 | 9759 | 13584 | 15391 | 16274 | 12750 | 15661 | 10482 | 6709 | 8875 | 10917 | 5512 | 7538 | 7304 | Madd |
| Maea | 9875 | 12016 | 11278 | 9739 | 8536 | 8914 | 8353 | 7311 | 7604 | 7148 | 7365 | 7948 | 7647 | 7076 | 9740 | 10587 | 10021 | 9300 | 9190 | 9682 | 8417 | 6856 | 7908 | 7513 | 7207 | 8474 | 8436 | 7976 | Maea |
| Mael | 2 | 3 | 7 | 0 | 2 | 11 | 0 | 2 | 5 | 6 | 5 | 12 | 0 | 5 | 4 | 2 | 3 | 2 | 0 | 2 | 2 | 5 | 8 | 2 | 0 | 0 | 6 | 0 | Mael |
| Maf | 1945 | 2118 | 2805 | 2606 | 3051 | 2641 | 1771 | 3709 | 3787 | 4377 | 4997 | 3230 | 4179 | 3319 | 2707 | 2897 | 2541 | 3061 | 1873 | 2024 | 2089 | 5424 | 4829 | 4113 | 3670 | 3623 | 3137 | 3420 | Maf |
| Maf1 | 4947 | 5470 | 5739 | 5623 | 5507 | 5384 | 5129 | 5630 | 5627 | 5817 | 5293 | 6417 | 5527 | 6216 | 4605 | 4351 | 5607 | 4888 | 6781 | 5181 | 5729 | 5768 | 5462 | 6655 | 6557 | 6788 | 6245 | 6324 | Maf1 |
| Mafa | 175 | 234 | 237 | 386 | 319 | 221 | 216 | 100 | 136 | 161 | 302 | 71 | 82 | 62 | 191 | 214 | 275 | 375 | 261 | 236 | 234 | 139 | 157 | 67 | 105 | 48 | 37 | 47 | Mafa |
| Mafb | 734 | 714 | 809 | 759 | 1125 | 787 | 695 | 1243 | 1528 | 1514 | 1544 | 961 | 1031 | 997 | 692 | 831 | 735 | 776 | 710 | 543 | 696 | 1554 | 1596 | 1396 | 1175 | 1142 | 837 | 835 | Mafb |
| Maff | 310 | 308 | 415 | 400 | 283 | 249 | 236 | 184 | 189 | 195 | 268 | 156 | 203 | 171 | 299 | 345 | 348 | 350 | 327 | 290 | 281 | 174 | 239 | 212 | 233 | 215 | 169 | 198 | Maff |
| Mafg | 4645 | 5890 | 6379 | 7069 | 6927 | 6431 | 6170 | 4841 | 5614 | 6702 | 6546 | 5951 | 5947 | 5817 | 4424 | 5281 | 6937 | 7744 | 7419 | 5681 | 6401 | 6365 | 5325 | 6481 | 7147 | 4974 | 5652 | 5826 | Mafg |
| Mafk | 842 | 1202 | 985 | 1275 | 898 | 803 | 798 | 697 | 756 | 829 | 703 | 775 | 724 | 801 | 729 | 1044 | 1097 | 1360 | 898 | 787 | 911 | 812 | 767 | 833 | 943 | 827 | 705 | 809 | Mafk |
| Mag | 38 | 67 | 61 | 62 | 106 | 39 | 51 | 37 | 46 | 70 | 67 | 37 | 44 | 29 | 62 | 76 | 58 | 64 | 48 | 54 | 56 | 63 | 52 | 33 | 54 | 30 | 37 | 28 | Mag |
| Magea10 | 1 | 0 | 0 | 0 | 0 | 0 | 1 | 0 | 0 | 1 | 0 | 2 | 0 | 0 | 0 | 1 | 0 | 0 | 0 | 0 | 1 | 0 | 0 | 0 | 0 | 0 | 1 | 0 | Magea10 |
| Magea13 | 1 | 6 | 0 | 0 | 3 | 4 | 1 | 2 | 5 | 2 | 0 | 3 | 3 | 6 | 5 | 2 | 0 | 3 | 2 | 7 | 2 | 2 | 2 | 7 | 8 | 3 | 7 | 4 | Magea13 |
| Magea14 | 0 | 0 | 0 | 0 | 0 | 0 | 0 | 0 | 0 | 0 | 0 | 0 | 0 | 0 | 0 | 0 | 0 | 0 | 0 | 0 | 0 | 0 | 1 | 0 | 0 | 0 | 0 | 0 | Magea14 |
| Magea9 | 20 | 23 | 11 | 0 | 0 | 14 | 0 | 23 | 26 | 1 | 18 | 31 | 40 | 21 | 29 | 36 | 1 | 9 | 0 | 0 | 0 | 2 | 32 | 20 | 3 | 4 | 30 | 38 | Magea9 |
| Mageb16 | 1 | 0 | 0 | 0 | 1 | 0 | 0 | 0 | 5 | 0 | 1 | 0 | 0 | 0 | 0 | 1 | 0 | 0 | 1 | 0 | 0 | 2 | 0 | 0 | 0 | 0 | 0 | 0 | Mageb16 |
| Mageb18 | 6 | 0 | 0 | 0 | 7 | 5 | 3 | 1 | 2 | 0 | 0 | 0 | 3 | 2 | 3 | 2 | 2 | 4 | 4 | 4 | 1 | 0 | 1 | 3 | 0 | 4 | 1 | 4 | Mageb18 |
| Mageb2 | 1 | 0 | 0 | 0 | 0 | 0 | 0 | 0 | 0 | 0 | 0 | 0 | 0 | 0 | 0 | 0 | 0 | 0 | 0 | 0 | 0 | 0 | 0 | 0 | 0 | 0 | 0 | 0 | Mageb2 |
| Mageb3 | 16 | 25 | 14 | 15 | 33 | 14 | 31 | 32 | 25 | 38 | 20 | 28 | 28 | 29 | 21 | 11 | 30 | 11 | 19 | 15 | 15 | 32 | 34 | 22 | 27 | 29 | 30 | 21 | Mageb3 |
| Mageb4 | 0 | 0 | 0 | 0 | 0 | 0 | 0 | 0 | 0 | 0 | 0 | 0 | 0 | 0 | 0 | 0 | 0 | 2 | 0 | 0 | 0 | 0 | 0 | 0 | 0 | 0 | 0 | 0 | Mageb4 |
| Magec2 | 0 | 0 | 0 | 0 | 0 | 0 | 0 | 1 | 11 | 0 | 0 | 0 | 0 | 0 | 0 | 6 | 0 | 0 | 0 | 0 | 0 | 2 | 3 | 0 | 0 | 0 | 0 | 0 | Magec2 |
| Maged1 | 47137 | 60572 | 44061 | 50254 | 52130 | 53187 | 46930 | 58438 | 65508 | 61839 | 57981 | 60160 | 54797 | 54234 | 50209 | 49212 | 49343 | 49444 | 54760 | 55869 | 49649 | 61102 | 64966 | 58816 | 61467 | 60374 | 57711 | 52678 | Maged1 |

| | | | | | | | | | | | | | | | | | | | | | | | | | | | | | |
|---|---|---|---|---|---|---|---|---|---|---|---|---|---|---|---|---|---|---|---|---|---|---|---|---|---|---|---|---|---|
| Maged2 | 12712 | 14027 | 11375 | 10609 | 10951 | 12048 | 9917 | 14399 | 17859 | 13793 | 18571 | 13156 | 14743 | 12506 | 12444 | 13946 | 10843 | 11206 | 10516 | 11677 | 10043 | 13588 | 20510 | 13364 | 13268 | 14450 | 13643 | 12507 | Maged2 |
| Magee1 | 6081 | 6420 | 7733 | 8185 | 10679 | 8904 | 9341 | 7491 | 7504 | 8691 | 8021 | 7387 | 6857 | 7543 | 6419 | 5738 | 8876 | 8672 | 9802 | 8295 | 9427 | 11463 | 6963 | 8321 | 9798 | 6902 | 6248 | 7026 | Magee1 |
| Magee2 | 519 | 522 | 574 | 551 | 719 | 577 | 706 | 614 | 618 | 598 | 624 | 477 | 459 | 527 | 537 | 431 | 597 | 543 | 722 | 664 | 690 | 824 | 681 | 612 | 674 | 490 | 497 | 477 | Magee2 |
| Mageh1 | 3154 | 3642 | 3778 | 3818 | 4129 | 3631 | 4128 | 4643 | 4002 | 4445 | 3947 | 4733 | 4194 | 5024 | 3587 | 2862 | 4022 | 3715 | 4176 | 4437 | 4068 | 4246 | 4097 | 4586 | 4643 | 4576 | 4897 | 4843 | Mageh1 |
| Magel2 | 3601 | 5218 | 3907 | 6624 | 5451 | 4583 | 6167 | 4600 | 4687 | 5655 | 5161 | 6188 | 3356 | 6189 | 3790 | 3468 | 5964 | 6490 | 5406 | 4640 | 5390 | 7768 | 4110 | 7142 | 6847 | 7258 | 5251 | 4875 | Magel2 |
| Magi1 | 5451 | 5751 | 5773 | 6687 | 7860 | 7695 | 7244 | 12401 | 11962 | 13521 | 11496 | 14302 | 12611 | 14920 | 5658 | 5298 | 6230 | 6311 | 7236 | 7142 | 7607 | 13363 | 11240 | 13094 | 15323 | 10497 | 14299 | 13492 | Magi1 |
| Magi2 | 2232 | 2300 | 2316 | 3683 | 4416 | 3625 | 4237 | 3104 | 3029 | 4079 | 3261 | 3609 | 2693 | 3660 | 2618 | 2117 | 3186 | 3921 | 4121 | 3073 | 4313 | 4485 | 3034 | 3650 | 4216 | 2665 | 3095 | 3214 | Magi2 |
| Magi3 | 5801 | 5490 | 4710 | 6212 | 6437 | 6374 | 6555 | 4360 | 4116 | 4249 | 4442 | 3993 | 3862 | 4105 | 5896 | 5379 | 5390 | 6024 | 5811 | 6361 | 6767 | 4228 | 4286 | 3949 | 4202 | 3018 | 3693 | 3917 | Magi3 |
| Magix | 2 | 2 | 10 | 1 | 8 | 4 | 6 | 6 | 10 | 6 | 10 | 10 | 9 | 11 | 5 | 8 | 5 | 4 | 12 | 6 | 4 | 5 | 5 | 5 | 9 | 5 | 5 | 9 | Magix |
| Magoh | 5146 | 5598 | 6563 | 5497 | 4223 | 4643 | 5716 | 4183 | 4116 | 4189 | 3891 | 5131 | 4549 | 5625 | 5246 | 5062 | 6849 | 4892 | 5543 | 6502 | 5717 | 3024 | 3901 | 4894 | 4126 | 6424 | 5505 | 6008 | Magoh |
| Magohb | 562 | 475 | 648 | 682 | 550 | 589 | 696 | 818 | 714 | 857 | 845 | 971 | 797 | 1145 | 583 | 464 | 656 | 548 | 584 | 691 | 676 | 645 | 720 | 810 | 838 | 978 | 938 | 1050 | Magohb |
| Magt1 | 8359 | 6135 | 6854 | 5622 | 4492 | 5199 | 4974 | 5762 | 5901 | 5217 | 6552 | 4974 | 5850 | 5343 | 7640 | 8921 | 6178 | 5861 | 4453 | 4922 | 4790 | 5080 | 6637 | 5401 | 4490 | 5413 | 5333 | 5463 | Magt1 |
| Maip1 | 1528 | 1320 | 1546 | 1517 | 1479 | 1456 | 1659 | 1767 | 1624 | 1683 | 1568 | 1702 | 1714 | 1888 | 1568 | 1270 | 1539 | 1349 | 1448 | 1536 | 1624 | 1823 | 1706 | 1655 | 1717 | 1953 | 1730 | 1930 | Maip1 |
| Mak | 1115 | 1108 | 770 | 930 | 907 | 975 | 1025 | 220 | 212 | 196 | 148 | 153 | 156 | 101 | 955 | 1002 | 913 | 908 | 868 | 1054 | 1150 | 175 | 240 | 139 | 117 | 98 | 124 | 93 | Mak |
| Mak16 | 5259 | 5069 | 5082 | 4591 | 4093 | 4439 | 4941 | 4444 | 3835 | 3718 | 3480 | 4070 | 3855 | 4213 | 5470 | 4804 | 4682 | 4402 | 4228 | 5299 | 4548 | 3544 | 3593 | 3980 | 3739 | 4692 | 4094 | 4402 | Mak16 |
| Mal | 55 | 44 | 42 | 46 | 79 | 68 | 47 | 151 | 126 | 161 | 118 | 88 | 86 | 96 | 67 | 54 | 42 | 80 | 48 | 60 | 75 | 137 | 150 | 87 | 135 | 62 | 73 | 102 | Mal |
| Mal2 | 323 | 125 | 72 | 125 | 156 | 103 | 74 | 283 | 265 | 319 | 196 | 47 | 70 | 36 | 199 | 194 | 48 | 109 | 76 | 44 | 83 | 187 | 318 | 176 | 96 | 182 | 41 | 28 | Mal2 |
| Mall | 57 | 38 | 62 | 66 | 35 | 60 | 41 | 93 | 125 | 100 | 122 | 73 | 91 | 68 | 54 | 63 | 43 | 42 | 46 | 33 | 31 | 75 | 169 | 109 | 61 | 100 | 71 | 68 | Mall |
| Malrd1 | 10 | 9 | 10 | 7 | 11 | 9 | 7 | 5 | 15 | 8 | 14 | 4 | 7 | 10 | 17 | 19 | 0 | 11 | 8 | 1 | 13 | 8 | 13 | 10 | 7 | 5 | 10 | 13 | Malrd1 |
| Malsu1 | 1088 | 1056 | 962 | 1027 | 934 | 964 | 1104 | 1098 | 943 | 955 | 776 | 968 | 826 | 1022 | 1280 | 960 | 964 | 820 | 1007 | 1139 | 1087 | 854 | 865 | 958 | 910 | 1080 | 999 | 979 | Malsu1 |
| Malt1 | 2998 | 2194 | 1839 | 2004 | 1659 | 1850 | 1808 | 1004 | 1120 | 915 | 1216 | 731 | 961 | 778 | 2462 | 2736 | 1907 | 1913 | 1659 | 1814 | 1958 | 825 | 1095 | 749 | 666 | 843 | 891 | 746 | Malt1 |
| Mamdc2 | 114 | 124 | 165 | 80 | 121 | 104 | 87 | 231 | 182 | 140 | 282 | 247 | 248 | 202 | 148 | 122 | 112 | 86 | 84 | 118 | 95 | 239 | 339 | 218 | 197 | 210 | 223 | 178 | Mamdc2 |
| Mamdc4 | 458 | 646 | 778 | 843 | 677 | 696 | 611 | 876 | 873 | 1094 | 1139 | 1235 | 1098 | 967 | 541 | 649 | 876 | 647 | 764 | 619 | 612 | 1039 | 823 | 1131 | 1080 | 1033 | 1135 | 1104 | Mamdc4 |
| Maml1 | 2480 | 2407 | 2870 | 2453 | 2459 | 2442 | 1991 | 2884 | 3321 | 3479 | 3525 | 4361 | 4063 | 3827 | 2308 | 2439 | 2856 | 2489 | 2627 | 1854 | 2132 | 3603 | 3246 | 3956 | 3569 | 4345 | 3624 | 3882 | Maml1 |
| Maml2 | 1712 | 1413 | 1525 | 1620 | 1537 | 1737 | 1402 | 1890 | 1850 | 1877 | 2002 | 1925 | 2353 | 2006 | 1464 | 1515 | 1528 | 1469 | 1464 | 1467 | 1480 | 1793 | 2018 | 1832 | 1747 | 1881 | 2116 | 2034 | Maml2 |
| Maml3 | 2545 | 2939 | 2976 | 3060 | 3023 | 2812 | 2547 | 2131 | 2082 | 2675 | 2246 | 2749 | 2777 | 2651 | 2408 | 2955 | 2975 | 2807 | 2993 | 2578 | 2802 | 2807 | 2015 | 2545 | 2488 | 2318 | 2703 | 2696 | Maml3 |
| Mamld1 | 1564 | 1724 | 1759 | 1848 | 2292 | 2172 | 2131 | 1432 | 1441 | 1718 | 1508 | 2155 | 1676 | 2006 | 1619 | 1509 | 1788 | 1844 | 2280 | 1822 | 2063 | 1801 | 1309 | 1803 | 1887 | 2311 | 1775 | 1914 | Mamld1 |
| Mamstr | 170 | 164 | 183 | 223 | 186 | 157 | 178 | 145 | 225 | 216 | 465 | 159 | 205 | 193 | 127 | 181 | 229 | 206 | 136 | 129 | 174 | 216 | 358 | 239 | 230 | 185 | 141 | 173 | Mamstr |
| Man1a | 1828 | 1509 | 1359 | 1473 | 1520 | 1609 | 1391 | 1078 | 1509 | 1173 | 1808 | 711 | 1216 | 656 | 1811 | 2174 | 1307 | 1826 | 1253 | 1352 | 1496 | 1325 | 1713 | 939 | 996 | 835 | 682 | 727 | Man1a |
| Man1a2 | 10156 | 8332 | 7346 | 7454 | 7715 | 8758 | 8094 | 8658 | 8513 | 7557 | 8315 | 7896 | 8344 | 7870 | 9803 | 9657 | 6840 | 8037 | 6876 | 8260 | 8091 | 8111 | 8835 | 6980 | 7309 | 6539 | 8034 | 7980 | Man1a2 |
| Man1b1 | 9574 | 11073 | 10069 | 9886 | 8552 | 9396 | 8775 | 8620 | 9181 | 9230 | 9297 | 10411 | 9780 | 9702 | 9256 | 10125 | 10026 | 10031 | 9619 | 8515 | 8971 | 8918 | 8939 | 9872 | 9817 | 10557 | 10105 | 10313 | Man1b1 |
| Man1c1 | 2284 | 2823 | 3516 | 2906 | 2746 | 2398 | 2249 | 891 | 1184 | 1262 | 1389 | 1340 | 1330 | 1194 | 2413 | 2950 | 3457 | 3073 | 2657 | 1979 | 2459 | 1384 | 1458 | 1297 | 1232 | 1501 | 1319 | 1220 | Man1c1 |
| Man2a1 | 5116 | 3748 | 4233 | 4116 | 4239 | 5132 | 3898 | 3980 | 4445 | 4475 | 5703 | 3905 | 4732 | 4024 | 4680 | 4981 | 3772 | 5081 | 3525 | 3624 | 3994 | 4284 | 5392 | 4159 | 4144 | 2988 | 4143 | 3739 | Man2a1 |
| Man2a2 | 7472 | 7797 | 7819 | 9052 | 10158 | 9357 | 8819 | 6537 | 7772 | 9291 | 8819 | 8526 | 8121 | 7797 | 6748 | 7518 | 8719 | 9452 | 9709 | 7312 | 9124 | 8607 | 7067 | 8550 | 8833 | 6325 | 7709 | 7620 | Man2a2 |
| Man2b1 | 3956 | 4575 | 4605 | 4521 | 3883 | 3768 | 3576 | 1984 | 2534 | 2443 | 2788 | 2383 | 2730 | 2206 | 3608 | 4790 | 4281 | 4552 | 3741 | 3218 | 3595 | 2597 | 2652 | 2526 | 2371 | 2615 | 2503 | 2353 | Man2b1 |
| Man2b2 | 711 | 985 | 850 | 648 | 736 | 724 | 419 | 823 | 1066 | 986 | 945 | 1023 | 1117 | 880 | 687 | 954 | 807 | 739 | 659 | 589 | 471 | 1044 | 1212 | 863 | 823 | 1078 | 1140 | 866 | Man2b2 |
| Man2c1 | 2850 | 3491 | 3084 | 3632 | 3634 | 3340 | 3073 | 3826 | 4594 | 4783 | 4732 | 4659 | 4495 | 4437 | 2848 | 3028 | 3139 | 3628 | 3525 | 3195 | 3357 | 4861 | 4402 | 4654 | 4889 | 4474 | 4422 | 4400 | Man2c1 |
| Manba | 1103 | 1170 | 862 | 857 | 946 | 729 | 873 | 462 | 513 | 571 | 540 | 434 | 500 | 380 | 1092 | 1167 | 840 | 979 | 958 | 992 | 911 | 483 | 664 | 391 | 388 | 484 | 485 | 409 | Manba |
| Manbal | 1160 | 1332 | 1391 | 1297 | 1268 | 1366 | 1279 | 1688 | 1810 | 2062 | 1948 | 2054 | 1839 | 2055 | 1155 | 1122 | 1358 | 1246 | 1431 | 1209 | 1448 | 1639 | 1935 | 2116 | 2178 | 2379 | 1995 | 2040 | Manbal |
| Manea | 3989 | 2773 | 3378 | 3454 | 3714 | 3664 | 3749 | 3613 | 2962 | 2734 | 2891 | 3521 | 3453 | 3202 | 4026 | 3513 | 3452 | 3129 | 3433 | 3592 | 3579 | 3474 | 3313 | 2663 | 2995 | 3098 | 3511 | 3398 | Manea |
| Maneal | 1340 | 1574 | 1732 | 1658 | 2231 | 1912 | 2073 | 1905 | 1792 | 2142 | 1776 | 2179 | 1821 | 2131 | 1424 | 1185 | 1825 | 1692 | 2208 | 1801 | 2052 | 2453 | 1628 | 2103 | 2253 | 1848 | 1951 | 2072 | Maneal |
| Manf | 7609 | 8192 | 9335 | 8557 | 6345 | 6078 | 7538 | 4993 | 4593 | 5249 | 5440 | 4566 | 4873 | 5403 | 7953 | 7346 | 8489 | 7908 | 7268 | 7331 | 7253 | 4360 | 4671 | 5880 | 5164 | 5429 | 5435 | 5645 | Manf |
| Mansc1 | 51 | 53 | 57 | 45 | 74 | 69 | 73 | 79 | 113 | 79 | 105 | 105 | 86 | 67 | 62 | 80 | 65 | 79 | 65 | 49 | 61 | 85 | 106 | 76 | 78 | 89 | 83 | 81 | Mansc1 |
| Mansc4 | 15 | 10 | 9 | 14 | 11 | 5 | 3 | 37 | 27 | 15 | 38 | 23 | 14 | 15 | 7 | 11 | 3 | 8 | 12 | 0 | 4 | 8 | 27 | 14 | 16 | 12 | 13 | 14 | Mansc4 |
| Maoa | 5822 | 4177 | 4404 | 4010 | 4290 | 4015 | 4364 | 3960 | 3686 | 3073 | 3265 | 2910 | 3035 | 3003 | 5641 | 4866 | 3911 | 4371 | 3596 | 4710 | 4051 | 3190 | 4100 | 2988 | 2842 | 3201 | 2897 | 3128 | Maoa |
| Maob | 608 | 362 | 217 | 337 | 95 | 171 | 77 | 518 | 499 | 571 | 381 | 177 | 354 | 284 | 448 | 482 | 214 | 161 | 124 | 123 | 99 | 175 | 537 | 556 | 195 | 739 | 228 | 250 | Maob |
| Map10 | 1110 | 1071 | 1249 | 1118 | 1117 | 1123 | 1244 | 717 | 878 | 728 | 839 | 897 | 886 | 840 | 1213 | 1012 | 1093 | 1128 | 1137 | 1087 | 1134 | 769 | 819 | 856 | 778 | 812 | 865 | 819 | Map10 |
| Map1a | 8630 | 11594 | 12230 | 16338 | 16332 | 14777 | 13415 | 5565 | 6629 | 8769 | 8409 | 10665 | 7997 | 9623 | 9531 | 10690 | 15688 | 16620 | 15911 | 10791 | 15530 | 10312 | 6085 | 9464 | 10013 | 5258 | 8157 | 9476 | Map1a |
| Map1b | 81904 | 80030 | 68428 | 107528 | 149839 | 146972 | 141545 | 142886 | 146175 | 168344 | 151018 | 154072 | 142516 | 165873 | 97990 | 87628 | 84070 | 107546 | 120107 | 123732 | 152054 | 152830 | 121819 | 120538 | 195267 | 89194 | 158394 | 138597 | Map1b |
| Map1lc3a | 2324 | 2947 | 2526 | 3053 | 3305 | 2630 | 2520 | 2542 | 3161 | 3357 | 3193 | 2500 | 2486 | 2547 | 2134 | 2332 | 2827 | 2878 | 3181 | 2484 | 3155 | 2963 | 3294 | 3018 | 3082 | 2523 | 2238 | 2550 | Map1lc3a |
| Map1lc3b | 14243 | 18075 | 13690 | 15920 | 14937 | 13949 | 15834 | 10610 | 10447 | 9480 | 9076 | 9299 | 9764 | 9562 | 15539 | 16098 | 14796 | 16174 | 15392 | 16144 | 16341 | 10313 | 10379 | 9877 | 10721 | 9229 | 9620 | 9848 | Map1lc3b |
| Map1s | 1744 | 2208 | 2602 | 2166 | 2117 | 1935 | 1644 | 2411 | 2882 | 3243 | 3060 | 3249 | 3002 | 2605 | 1473 | 1768 | 2579 | 2079 | 2466 | 1899 | 1725 | 3156 | 2664 | 3189 | 3133 | 3625 | 2988 | 2740 | Map1s |
| Map2 | 42667 | 42319 | 33942 | 51498 | 68001 | 67123 | 70265 | 108321 | 85847 | 87013 | 73289 | 81011 | 75618 | 85912 | 55809 | 39697 | 44839 | 53669 | 59038 | 73510 | 72001 | 97407 | 80255 | 72200 | 101077 | 49410 | 81954 | 78279 | Map2 |
| Map2k1 | 5975 | 5823 | 6098 | 5598 | 4408 | 4818 | 4512 | 3102 | 3022 | 3030 | 3457 | 2918 | 3407 | 2957 | 5756 | 5833 | 5745 | 5183 | 4972 | 5247 | 4556 | 2765 | 3585 | 3202 | 2709 | 3581 | 3223 | 3258 | Map2k1 |
| Map2k2 | 4126 | 5063 | 5045 | 4920 | 4896 | 4203 | 4294 | 4404 | 5408 | 5572 | 5483 | 6443 | 5857 | 5513 | 3824 | 4562 | 5296 | 4606 | 5261 | 3999 | 4260 | 5514 | 5092 | 5962 | 5881 | 6989 | 6252 | 5923 | Map2k2 |
| Map2k3 | 3210 | 3261 | 3011 | 2936 | 1932 | 2246 | 1933 | 1418 | 1681 | 1749 | 1853 | 1567 | 1820 | 1513 | 2715 | 3740 | 2700 | 2854 | 2020 | 1909 | 1932 | 1409 | 1838 | 1772 | 1538 | 2059 | 1461 | 1568 | Map2k3 |
| Map2k4 | 8485 | 7398 | 7005 | 6524 | 7168 | 6944 | 7708 | 7806 | 6875 | 6400 | 6233 | 6877 | 6701 | 6874 | 8818 | 7248 | 6623 | 6348 | 6614 | 7437 | 6965 | 7658 | 6658 | 6307 | 6666 | 6879 | 6694 | 6959 | Map2k4 |
| Map2k5 | 1616 | 1640 | 1891 | 1745 | 1771 | 1778 | 1753 | 1859 | 2080 | 2216 | 2158 | 2355 | 2273 | 2311 | 1534 | 1635 | 1778 | 1573 | 1804 | 1593 | 1706 | 2120 | 1876 | 2248 | 2223 | 2507 | 2307 | 2415 | Map2k5 |
| Map2k6 | 4301 | 3822 | 3489 | 3691 | 3839 | 3877 | 4146 | 2472 | 2458 | 2550 | 2057 | 2322 | 2347 | 2533 | 3840 | 3657 | 3615 | 3346 | 4029 | 4024 | 4120 | 1917 | 2266 | 2377 | 2455 | 2605 | 2567 | 2504 | Map2k6 |
| Map2k7 | 1248 | 1376 | 1807 | 1824 | 1998 | 1651 | 1700 | 2733 | 2952 | 3776 | 3501 | 3825 | 3484 | 3803 | 1251 | 1187 | 1874 | 1687 | 1896 | 1471 | 1847 | 3290 | 2950 | 3812 | 3688 | 3836 | 3410 | 3894 | Map2k7 |
| Map3k1 | 2863 | 2636 | 3037 | 3300 | 3367 | 3210 | 3282 | 2621 | 2734 | 2829 | 3049 | 2800 | 2808 | 2907 | 2834 | 3157 | 3314 | 3272 | 3135 | 2658 | 3190 | 3110 | 2735 | 2808 | 2937 | 2586 | 2715 | 2839 | Map3k1 |
| Map3k10 | 3856 | 5692 | 5236 | 6355 | 6612 | 5483 | 4998 | 3176 | 3556 | 4079 | 3880 | 3600 | 3259 | 3214 | 3877 | 4308 | 6052 | 6238 | 6846 | 5121 | 5623 | 4214 | 3430 | 3791 | 4438 | 3444 | 3357 | 3081 | Map3k10 |
| Map3k11 | 2207 | 2699 | 2775 | 2400 | 2228 | 2255 | 1810 | 1849 | 2098 | 2053 | 2221 | 2344 | 2428 | 1950 | 1838 | 2349 | 2757 | 2181 | 2579 | 2025 | 1888 | 2037 | 2137 | 2328 | 1963 | 2613 | 2248 | 2338 | Map3k11 |
| Map3k12 | 5377 | 6321 | 5871 | 6871 | 7862 | 7079 | 7194 | 6202 | 6200 | 7101 | 6281 | 6588 | 6229 | 6514 | 5769 | 5537 | 6704 | 7453 | 7872 | 6398 | 7699 | 8096 | 5858 | 6913 | 7683 | 5959 | 6383 | 6636 | Map3k12 |
| Map3k13 | 2987 | 3704 | 2676 | 4336 | 5109 | 4554 | 4599 | 2820 | 2551 | 3446 | 3138 | 3221 | 3259 | 3181 | 3438 | 3224 | 3581 | 4678 | 4692 | 3970 | 5027 | 3867 | 2629 | 2665 | 3831 | 1593 | 3532 | 3266 | Map3k13 |
| Map3k14 | 690 | 1015 | 1003 | 1292 | 798 | 796 | 780 | 440 | 447 | 500 | 531 | 547 | 531 | 510 | 582 | 953 | 1038 | 1139 | 962 | 735 | 784 | 470 | 516 | 549 | 502 | 549 | 589 | 598 | Map3k14 |
| Map3k15 | 323 | 346 | 234 | 345 | 412 | 412 | 426 | 203 | 153 | 124 | 178 | 203 | 175 | 170 | 389 | 329 | 302 | 388 | 327 | 397 | 425 | 194 | 177 | 166 | 163 | 137 | 265 | 196 | Map3k15 |
| Map3k19 | 10 | 14 | 8 | 14 | 50 | 18 | 16 | 93 | 77 | 98 | 55 | 71 | 118 | 31 | 25 | 36 | 9 | 15 | 21 | 36 | 20 | 99 | 148 | 38 | 33 | 19 | 37 | 38 | Map3k19 |
| Map3k2 | 6418 | 5200 | 3844 | 5366 | 5838 | 6944 | 6661 | 6620 | 5444 | 4746 | 5372 | 4697 | 4586 | 5205 | 6243 | 6076 | 4113 | 5789 | 5111 | 6599 | 6882 | 5115 | 5313 | 3888 | 5375 | 3052 | 5163 | 4476 | Map3k2 |
| Map3k20 | 1776 | 1564 | 1949 | 1773 | 1878 | 1912 | 1888 | 1237 | 1481 | 1497 | 1750 | 1466 | 1208 | 1348 | 1788 | 1674 | 1878 | 1887 | 2074 | 1454 | 1879 | 1353 | 1695 | 1502 | 1335 | 1244 | 1369 | 1320 | Map3k20 |
| Map3k21 | 141 | 124 | 105 | 161 | 173 | 146 | 115 | 48 | 40 | 58 | 60 | 40 | 42 | 29 | 137 | 200 | 116 | 167 | 94 | 94 | 103 | 62 | 50 | 45 | 67 | 22 | 19 | 28 | Map3k21 |
| Map3k3 | 4737 | 4971 | 4358 | 4367 | 4281 | 4712 | 3883 | 3215 | 3540 | 3319 | 3810 | 3669 | 3434 | 3310 | 4317 | 5062 | 4060 | 4283 | 4428 | 3789 | 4061 | 3526 | 3596 | 3383 | 3488 | 3233 | 3787 | 3386 | Map3k3 |
| Map3k4 | 4439 | 4964 | 4704 | 4447 | 4226 | 4225 | 4101 | 3864 | 4423 | 4597 | 4326 | 5314 | 4790 | 5028 | 4257 | 4827 | 4533 | 4624 | 4447 | 4155 | 4249 | 4072 | 3983 | 4779 | 4499 | 4737 | 4889 | 4968 | Map3k4 |
| Map3k5 | 420 | 301 | 279 | 214 | 329 | 239 | 230 | 581 | 662 | 581 | 784 | 286 | 401 | 270 | 306 | 318 | 174 | 297 | 288 | 233 | 227 | 613 | 627 | 524 | 433 | 425 | 308 | 292 | Map3k5 |
| Map3k6 | 96 | 81 | 129 | 56 | 45 | 46 | 49 | 62 | 89 | 91 | 105 | 89 | 103 | 82 | 86 | 107 | 118 | 67 | 69 | 84 | 59 | 67 | 129 | 100 | 100 | 101 | 85 | 87 | Map3k6 |
| Map3k7 | 18264 | 15648 | 13795 | 14576 | 13442 | 15543 | 15430 | 12185 | 11142 | 10635 | 10648 | 11920 | 11295 | 11863 | 17450 | 15805 | 13752 | 14060 | 13495 | 14765 | 15253 | 10908 | 10784 | 10310 | 11191 | 10638 | 11850 | 11719 | Map3k7 |
| Map3k7cl | 10 | 3 | 8 | 5 | 7 | 11 | 17 | 9 | 20 | 13 | 105 | 10 | 12 | 11 | 3 | 8 | 20 | 10 | 9 | 5 | 8 | 22 | 98 | 16 | 17 | 15 | 26 | 17 | Map3k7cl |
| Map3k8 | 89 | 58 | 55 | 35 | 81 | 95 | 48 | 71 | 87 | 51 | 66 | 79 | 65 | 55 | 86 | 107 | 43 | 63 | 46 | 67 | 64 | 70 | 115 | 65 | 40 | 55 | 65 | 69 | Map3k8 |
| Map3k9 | 1969 | 2456 | 1498 | 3158 | 4419 | 3828 | 3223 | 2332 | 2648 | 3170 | 2647 | 2391 | 2028 | 2126 | 1969 | 1953 | 2468 | 3345 | 3907 | 3174 | 3883 | 3287 | 2172 | 2131 | 3649 | 457 | 2012 | 1627 | Map3k9 |

| | | | | | | | | | | | | | | | | | | | | | | | | | | | | | |
|---|---|---|---|---|---|---|---|---|---|---|---|---|---|---|---|---|---|---|---|---|---|---|---|---|---|---|---|---|---|
| Map4 | 22942 | 25004 | 24775 | 27199 | 30035 | 27918 | 26021 | 21159 | 23487 | 26222 | 26190 | 24637 | 24619 | 23258 | 24100 | 24649 | 26501 | 28773 | 29184 | 21412 | 26504 | 28642 | 23407 | 23623 | 26457 | 20551 | 22887 | 22528 | Map4 |
| Map4k1 | 228 | 285 | 346 | 270 | 277 | 269 | 277 | 283 | 312 | 319 | 377 | 413 | 387 | 394 | 225 | 277 | 327 | 248 | 252 | 251 | 223 | 259 | 332 | 336 | 325 | 456 | 381 | 368 | Map4k1 |
| Map4k2 | 0 | 1 | 0 | 0 | 1 | 2 | 0 | 0 | 1 | 3 | 0 | 0 | 0 | 0 | 0 | 1 | 0 | 0 | 0 | 1 | 0 | 0 | 0 | 0 | 0 | 0 | 0 | 0 | Map4k2 |
| Map4k3 | 6688 | 5180 | 5140 | 6001 | 6828 | 7152 | 7139 | 8639 | 8115 | 7708 | 7445 | 7656 | 7143 | 7845 | 6744 | 5831 | 5423 | 6545 | 6803 | 6332 | 6917 | 8169 | 8042 | 7670 | 7996 | 7219 | 7184 | 7514 | Map4k3 |
| Map4k4 | 25906 | 26673 | 27868 | 29325 | 32499 | 30431 | 30855 | 45250 | 43797 | 49493 | 45114 | 52384 | 45914 | 54171 | 25582 | 25020 | 29713 | 28563 | 32521 | 28464 | 31603 | 48127 | 39667 | 51023 | 55330 | 41993 | 49023 | 50409 | Map4k4 |
| Map4k5 | 5777 | 4435 | 4431 | 4427 | 6024 | 5611 | 5858 | 7102 | 6733 | 5795 | 6783 | 5647 | 5838 | 5807 | 5884 | 5344 | 4501 | 4949 | 4994 | 5898 | 5673 | 6388 | 7000 | 5903 | 6136 | 5591 | 5653 | 5473 | Map4k5 |
| Map6 | 7500 | 9253 | 10871 | 15649 | 19871 | 15201 | 14977 | 11430 | 10485 | 15863 | 12758 | 13071 | 11796 | 11889 | 9366 | 8203 | 14473 | 16353 | 18181 | 12980 | 16655 | 19704 | 9688 | 14083 | 17376 | 9348 | 11212 | 11405 | Map6 |
| Map6d1 | 125 | 133 | 114 | 81 | 192 | 146 | 115 | 173 | 126 | 144 | 148 | 99 | 116 | 100 | 119 | 93 | 115 | 143 | 205 | 122 | 136 | 149 | 115 | 106 | 149 | 78 | 120 | 81 | Map6d1 |
| Map7 | 830 | 684 | 928 | 701 | 976 | 717 | 757 | 861 | 1121 | 844 | 801 | 739 | 836 | 661 | 847 | 833 | 869 | 752 | 718 | 785 | 692 | 861 | 956 | 721 | 695 | 847 | 707 | 724 | Map7 |
| Map7d1 | 9568 | 13295 | 16486 | 16367 | 17286 | 14764 | 11432 | 10555 | 11680 | 16441 | 14598 | 15864 | 13613 | 13606 | 9939 | 11588 | 18068 | 16143 | 17615 | 11812 | 13403 | 16157 | 11451 | 16232 | 16780 | 15996 | 13508 | 13287 | Map7d1 |
| Map7d2 | 2705 | 2705 | 3693 | 3920 | 4672 | 4058 | 4280 | 2451 | 1812 | 2371 | 2004 | 2027 | 2184 | 2143 | 3443 | 2622 | 3780 | 3873 | 4263 | 4089 | 4096 | 2871 | 1790 | 2168 | 2428 | 1629 | 2192 | 2231 | Map7d2 |
| Map7d3 | 742 | 671 | 728 | 443 | 403 | 539 | 420 | 234 | 240 | 173 | 260 | 249 | 362 | 283 | 856 | 694 | 542 | 442 | 401 | 541 | 413 | 199 | 272 | 176 | 214 | 248 | 315 | 282 | Map7d3 |
| Map9 | 5655 | 5265 | 3577 | 6730 | 7708 | 7539 | 8995 | 7331 | 6471 | 6411 | 5267 | 5123 | 3703 | 5493 | 6436 | 4718 | 5016 | 7035 | 7136 | 8760 | 8577 | 7334 | 6094 | 5570 | 7722 | 3524 | 4499 | 4516 | Map9 |
| Mapk1 | 25620 | 22715 | 22081 | 23897 | 22949 | 23454 | 25871 | 19712 | 17252 | 16951 | 16635 | 18634 | 17775 | 18601 | 25938 | 22123 | 23244 | 24375 | 23170 | 22525 | 24965 | 19727 | 17259 | 16878 | 18590 | 18379 | 18503 | 19130 | Mapk1 |
| Mapk10 | 4252 | 4989 | 4344 | 6286 | 8004 | 7310 | 6809 | 7619 | 5879 | 7423 | 5645 | 6521 | 6872 | 6455 | 5955 | 4375 | 5826 | 6546 | 7226 | 7607 | 7669 | 9125 | 5811 | 6565 | 8387 | 4442 | 6603 | 6234 | Mapk10 |
| Mapk11 | 2450 | 2717 | 3696 | 3306 | 3535 | 3136 | 3366 | 3283 | 3244 | 4048 | 3307 | 5082 | 4233 | 4816 | 2532 | 2281 | 3846 | 3112 | 3854 | 2933 | 3329 | 4343 | 2836 | 4747 | 4558 | 4727 | 4947 | 4640 | Mapk11 |
| Mapk12 | 2021 | 2179 | 2710 | 1963 | 1714 | 1778 | 1759 | 1082 | 851 | 967 | 797 | 1460 | 1276 | 1382 | 1965 | 1906 | 2415 | 1683 | 1998 | 1873 | 1628 | 882 | 810 | 1144 | 1106 | 1490 | 1535 | 1575 | Mapk12 |
| Mapk13 | 414 | 187 | 113 | 176 | 72 | 115 | 30 | 219 | 310 | 288 | 278 | 62 | 127 | 80 | 219 | 293 | 65 | 119 | 62 | 43 | 36 | 52 | 285 | 258 | 38 | 253 | 38 | 62 | Mapk13 |
| Mapk14 | 9330 | 9264 | 8690 | 8498 | 7869 | 8257 | 8252 | 5560 | 5523 | 5209 | 5514 | 5888 | 5571 | 5436 | 8337 | 9404 | 8856 | 8357 | 8191 | 7598 | 8166 | 5243 | 5317 | 5561 | 5419 | 5884 | 5764 | 5655 | Mapk14 |
| Mapk15 | 566 | 680 | 805 | 696 | 506 | 488 | 491 | 148 | 200 | 218 | 166 | 216 | 243 | 165 | 498 | 684 | 708 | 548 | 588 | 507 | 420 | 177 | 240 | 217 | 158 | 247 | 188 | 186 | Mapk15 |
| Mapk1ip1 | 1008 | 1398 | 1123 | 1457 | 1615 | 1352 | 1406 | 1545 | 1631 | 1885 | 1774 | 1483 | 1524 | 1562 | 1063 | 1191 | 1424 | 1549 | 1432 | 1301 | 1432 | 1818 | 1603 | 1781 | 1844 | 1407 | 1485 | 1385 | Mapk1ip1 |
| Mapk1ip1l | 8779 | 7525 | 11315 | 9761 | 9809 | 9607 | 10107 | 9355 | 10059 | 11551 | 11434 | 13805 | 12920 | 13382 | 7928 | 8911 | 10306 | 9630 | 10524 | 8446 | 9288 | 10596 | 9917 | 12394 | 12056 | 14318 | 13316 | 13513 | Mapk1ip1l |
| Mapk3 | 3637 | 4533 | 3824 | 4071 | 4664 | 4140 | 4248 | 4236 | 4785 | 4751 | 4252 | 4313 | 4098 | 4156 | 3620 | 3694 | 4164 | 4034 | 4625 | 4027 | 4214 | 4846 | 5064 | 4484 | 4704 | 4232 | 3940 | 3823 | Mapk3 |
| Mapk4 | 1014 | 1146 | 1409 | 1462 | 1765 | 1477 | 1396 | 1178 | 1257 | 1458 | 1210 | 1669 | 1461 | 1604 | 1143 | 1206 | 1501 | 1433 | 1548 | 1277 | 1407 | 1693 | 1296 | 1421 | 1555 | 1544 | 1352 | 1410 | Mapk4 |
| Mapk6 | 14164 | 13879 | 13480 | 13181 | 14613 | 14344 | 14475 | 20727 | 17495 | 17739 | 15619 | 21794 | 20222 | 21715 | 15096 | 12774 | 14054 | 12835 | 14027 | 15010 | 14126 | 21016 | 16543 | 18156 | 19776 | 20318 | 23007 | 21975 | Mapk6 |
| Mapk7 | 4402 | 5553 | 5868 | 6312 | 5698 | 5289 | 5006 | 3953 | 4668 | 5429 | 5010 | 6461 | 5131 | 5349 | 4096 | 4910 | 6371 | 6117 | 6223 | 4299 | 4909 | 5289 | 4457 | 5857 | 5464 | 6569 | 5166 | 5222 | Mapk7 |
| Mapk8 | 10716 | 9709 | 7844 | 10612 | 13146 | 12625 | 14171 | 18447 | 14292 | 12735 | 11948 | 12230 | 11650 | 12709 | 12808 | 9898 | 9505 | 11285 | 12460 | 15527 | 13861 | 15984 | 14013 | 11941 | 14559 | 10251 | 11739 | 11725 | Mapk8 |
| Mapk8ip1 | 10493 | 14255 | 14323 | 16101 | 15586 | 13321 | 13860 | 8361 | 8885 | 10732 | 10019 | 10187 | 9754 | 9590 | 11044 | 11562 | 16005 | 15106 | 17121 | 12936 | 15451 | 11010 | 9156 | 10623 | 11492 | 10251 | 9408 | 10240 | Mapk8ip1 |
| Mapk8ip2 | 4450 | 5630 | 6969 | 7526 | 9468 | 7244 | 7508 | 7083 | 6970 | 9706 | 7558 | 8967 | 7562 | 8820 | 4688 | 4273 | 7900 | 7684 | 9742 | 6451 | 8386 | 10769 | 6336 | 9211 | 10587 | 7650 | 7831 | 8388 | Mapk8ip2 |
| Mapk8ip3 | 14229 | 18652 | 16716 | 21141 | 20163 | 18529 | 18239 | 12469 | 15109 | 16481 | 15744 | 16058 | 14382 | 14655 | 14458 | 17612 | 19491 | 21578 | 20878 | 15913 | 19066 | 16753 | 13790 | 15685 | 17662 | 11977 | 14120 | 13699 | Mapk8ip3 |
| Mapk9 | 13029 | 12207 | 10002 | 13812 | 14676 | 14920 | 14507 | 8856 | 8030 | 8373 | 8269 | 7545 | 7787 | 7647 | 13557 | 12985 | 11564 | 14311 | 13097 | 14346 | 15179 | 9095 | 7956 | 6753 | 8428 | 5858 | 7509 | 7209 | Mapk9 |
| Mapkap1 | 6102 | 6810 | 6533 | 6067 | 5563 | 5426 | 5594 | 5326 | 5380 | 5396 | 4917 | 5946 | 5548 | 5660 | 5989 | 5787 | 6488 | 5931 | 5672 | 5665 | 5550 | 5110 | 5262 | 5560 | 5448 | 6076 | 5914 | 5886 | Mapkap1 |
| Mapkapk2 | 10809 | 11756 | 12350 | 10843 | 8523 | 9670 | 8536 | 6288 | 6470 | 6073 | 6145 | 8638 | 7629 | 7289 | 10214 | 10547 | 11300 | 9894 | 10149 | 8715 | 9217 | 6117 | 6524 | 7342 | 6574 | 8950 | 8671 | 8651 | Mapkapk2 |
| Mapkapk3 | 1466 | 1410 | 1303 | 1036 | 666 | 805 | 759 | 231 | 280 | 266 | 292 | 252 | 365 | 238 | 1185 | 1413 | 1060 | 1029 | 765 | 748 | 686 | 212 | 368 | 265 | 189 | 291 | 259 | 233 | Mapkapk3 |
| Mapkapk5 | 130 | 97 | 119 | 198 | 142 | 124 | 112 | 153 | 132 | 153 | 118 | 156 | 118 | 144 | 96 | 94 | 159 | 127 | 138 | 103 | 105 | 137 | 122 | 150 | 166 | 145 | 120 | 154 | Mapkapk5 |
| Mapkbp1 | 4563 | 5471 | 5019 | 6194 | 6677 | 6296 | 6032 | 3841 | 4565 | 5058 | 4757 | 5194 | 4913 | 4769 | 4485 | 5057 | 5894 | 6385 | 6617 | 5281 | 6338 | 5496 | 4443 | 4994 | 5413 | 3395 | 4645 | 4593 | Mapkbp1 |
| Mapre1 | 20179 | 17517 | 19335 | 19548 | 21624 | 21230 | 23525 | 28722 | 24597 | 25250 | 24654 | 28379 | 26578 | 29959 | 21664 | 17178 | 19335 | 19026 | 20821 | 22726 | 22515 | 27315 | 24699 | 25647 | 28031 | 27388 | 28217 | 29110 | Mapre1 |
| Mapre2 | 14782 | 15505 | 14910 | 17935 | 20423 | 18019 | 18154 | 12075 | 12153 | 14347 | 12949 | 12041 | 13354 | 11947 | 17037 | 16080 | 16822 | 18843 | 17238 | 17180 | 18506 | 17029 | 12209 | 12245 | 14457 | 9923 | 11102 | 11763 | Mapre2 |
| Mapre3 | 3744 | 4571 | 4447 | 6876 | 7193 | 6410 | 6293 | 3422 | 3680 | 5063 | 4541 | 4224 | 3787 | 4361 | 3853 | 3991 | 5924 | 6971 | 7015 | 5384 | 6808 | 5207 | 3531 | 4923 | 5516 | 3491 | 3688 | 4081 | Mapre3 |
| Mapt | 14829 | 18236 | 17096 | 35550 | 56637 | 41259 | 41607 | 35367 | 38257 | 49991 | 40951 | 28268 | 29011 | 27951 | 19810 | 17552 | 27403 | 42247 | 46331 | 37410 | 46151 | 62337 | 34995 | 37087 | 54611 | 18397 | 22791 | 23468 | Mapt |
| Marchf1 | 668 | 790 | 703 | 1413 | 2016 | 1698 | 1522 | 1893 | 1598 | 1628 | 1283 | 1123 | 1412 | 1087 | 1139 | 872 | 1014 | 1465 | 1531 | 2024 | 1634 | 2400 | 1647 | 1093 | 1737 | 524 | 1026 | 1100 | Marchf1 |
| Marchf10 | 17 | 17 | 2 | 16 | 13 | 18 | 15 | 7 | 16 | 14 | 11 | 15 | 15 | 6 | 7 | 17 | 19 | 17 | 22 | 12 | 17 | 5 | 17 | 11 | 20 | 12 | 23 | 2 | Marchf10 |
| Marchf11 | 50 | 66 | 61 | 98 | 187 | 104 | 108 | 164 | 161 | 175 | 149 | 88 | 100 | 78 | 74 | 71 | 90 | 98 | 147 | 133 | 108 | 269 | 166 | 123 | 151 | 98 | 74 | 85 | Marchf11 |
| Marchf2 | 4026 | 4547 | 3147 | 3904 | 3922 | 4089 | 4022 | 1896 | 2102 | 1738 | 1913 | 1826 | 2110 | 1833 | 4158 | 3914 | 3482 | 3973 | 3719 | 4299 | 4236 | 1781 | 2108 | 1687 | 1995 | 1490 | 1951 | 1925 | Marchf2 |
| Marchf3 | 308 | 494 | 333 | 429 | 408 | 338 | 296 | 161 | 217 | 174 | 166 | 147 | 229 | 153 | 411 | 467 | 362 | 375 | 298 | 415 | 353 | 187 | 291 | 118 | 164 | 135 | 180 | 246 | Marchf3 |
| Marchf4 | 187 | 247 | 257 | 565 | 1325 | 793 | 658 | 949 | 1179 | 1396 | 1192 | 502 | 641 | 537 | 353 | 199 | 424 | 646 | 922 | 805 | 870 | 1591 | 1214 | 924 | 1325 | 335 | 437 | 430 | Marchf4 |
| Marchf5 | 2 | 3 | 0 | 0 | 1 | 3 | 0 | 0 | 0 | 0 | 0 | 1 | 0 | 1 | 1 | 1 | 0 | 0 | 2 | 0 | 0 | 0 | 0 | 2 | 0 | 0 | 1 | 0 | Marchf5 |
| Marchf6 | 20262 | 18170 | 18887 | 25475 | 28629 | 28275 | 30422 | 22884 | 21261 | 23137 | 21910 | 24035 | 20516 | 23899 | 21220 | 18693 | 22305 | 26767 | 27977 | 24670 | 30912 | 27165 | 20250 | 23091 | 27426 | 18338 | 23538 | 22527 | Marchf6 |
| Marchf7 | 11395 | 7859 | 7994 | 9711 | 10543 | 11028 | 11942 | 15398 | 13098 | 11090 | 13246 | 11815 | 12415 | 13650 | 11432 | 10078 | 7779 | 9531 | 9777 | 11712 | 11213 | 13007 | 13561 | 11162 | 12599 | 10635 | 12808 | 12781 | Marchf7 |
| Marchf8 | 9035 | 9875 | 8546 | 9288 | 10870 | 9970 | 9959 | 10580 | 9220 | 9453 | 8945 | 8840 | 9115 | 9000 | 10216 | 8784 | 9788 | 9617 | 10078 | 10027 | 10376 | 11394 | 9103 | 8599 | 10038 | 6921 | 8920 | 8725 | Marchf8 |
| Marchf9 | 1829 | 2105 | 2386 | 2746 | 3267 | 2663 | 2768 | 2376 | 2630 | 3295 | 2860 | 2912 | 2276 | 2548 | 1822 | 1738 | 2808 | 2804 | 3404 | 2527 | 3027 | 3556 | 2480 | 3118 | 3526 | 2662 | 2415 | 2448 | Marchf9 |
| Marcks | 38936 | 28141 | 46991 | 39018 | 39620 | 41556 | 45503 | 83140 | 71986 | 76242 | 74642 | 93545 | 89738 | 98478 | 42722 | 33975 | 43624 | 38228 | 41627 | 41635 | 42038 | 86934 | 72027 | 89814 | 88473 | 105012 | 95934 | 103998 | Marcks |
| Marcksl1 | 35221 | 39768 | 39969 | 39222 | 41756 | 40497 | 41814 | 53224 | 53221 | 54609 | 51558 | 62972 | 56845 | 68933 | 37194 | 34362 | 41315 | 37442 | 44191 | 40032 | 43594 | 56581 | 50644 | 62153 | 62385 | 71064 | 59951 | 67513 | Marcksl1 |
| Marco | 0 | 4 | 7 | 6 | 16 | 5 | 0 | 7 | 7 | 7 | 8 | 2 | 2 | 19 | 0 | 13 | 0 | 12 | 2 | 8 | 1 | 7 | 11 | 19 | 6 | 6 | 6 | 31 | Marco |
| Marf1 | 10448 | 9799 | 8883 | 9835 | 9707 | 10140 | 10352 | 11342 | 11209 | 11660 | 11746 | 11255 | 11247 | 11404 | 9872 | 10235 | 9676 | 10197 | 9773 | 9571 | 10535 | 10609 | 10413 | 10192 | 11745 | 8427 | 11757 | 10753 | Marf1 |
| Mark1 | 3380 | 3511 | 4262 | 4828 | 6621 | 5315 | 5621 | 6139 | 6452 | 7480 | 6572 | 7358 | 5763 | 6923 | 3574 | 3238 | 5246 | 4990 | 5959 | 4684 | 5782 | 8519 | 5660 | 6932 | 8253 | 6004 | 6079 | 6060 | Mark1 |
| Mark2 | 4230 | 5186 | 6689 | 6870 | 6334 | 5811 | 5661 | 3850 | 4646 | 5679 | 5317 | 6269 | 5230 | 5736 | 3916 | 4945 | 6915 | 7096 | 6771 | 4465 | 5851 | 6183 | 4194 | 6386 | 6116 | 6210 | 4931 | 5630 | Mark2 |
| Mark3 | 12695 | 12132 | 12721 | 13436 | 13160 | 12995 | 13995 | 10767 | 9907 | 11012 | 9327 | 11472 | 10284 | 11376 | 12535 | 11574 | 13331 | 12855 | 13662 | 12556 | 13979 | 10713 | 9669 | 10953 | 11274 | 11250 | 11280 | 11736 | Mark3 |
| Mark4 | 4811 | 6423 | 7080 | 8544 | 8072 | 6995 | 6722 | 3549 | 4292 | 5280 | 4883 | 5072 | 4352 | 4606 | 4530 | 5479 | 8285 | 8361 | 8444 | 5312 | 7666 | 5736 | 4103 | 5293 | 5261 | 5186 | 4322 | 4329 | Mark4 |
| Mars1 | 7591 | 8278 | 7886 | 7846 | 7020 | 7133 | 7009 | 5383 | 5892 | 5716 | 5536 | 6900 | 6228 | 6635 | 7251 | 6943 | 8433 | 7202 | 8133 | 7024 | 7422 | 5287 | 5510 | 6106 | 6120 | 7314 | 6892 | 7135 | Mars1 |
| Mars2 | 6008 | 4965 | 4147 | 5988 | 4674 | 5914 | 6072 | 2833 | 2459 | 2342 | 2361 | 2559 | 2360 | 2568 | 5685 | 5843 | 4233 | 5840 | 4488 | 5606 | 6318 | 2550 | 2358 | 2134 | 2601 | 1907 | 2661 | 2369 | Mars2 |
| Marveld1 | 1870 | 1792 | 1997 | 1065 | 1117 | 1336 | 896 | 1758 | 2572 | 2177 | 3022 | 2156 | 2503 | 1777 | 1720 | 1994 | 1555 | 1390 | 1111 | 963 | 863 | 2034 | 2840 | 2129 | 1661 | 2369 | 2008 | 1958 | Marveld1 |
| Marveld2 | 180 | 86 | 132 | 63 | 92 | 63 | 55 | 177 | 232 | 225 | 252 | 137 | 210 | 138 | 132 | 131 | 48 | 50 | 52 | 56 | 44 | 175 | 278 | 239 | 128 | 236 | 124 | 138 | Marveld2 |
| Marveld3 | 205 | 199 | 265 | 260 | 236 | 199 | 268 | 72 | 83 | 83 | 59 | 34 | 37 | 70 | 208 | 206 | 221 | 248 | 221 | 232 | 190 | 43 | 95 | 100 | 55 | 78 | 50 | 51 | Marveld3 |
| Mas1 | 31 | 23 | 18 | 0 | 0 | 6 | 0 | 24 | 48 | 8 | 32 | 8 | 29 | 11 | 12 | 33 | 6 | 7 | 0 | 2 | 1 | 10 | 27 | 19 | 8 | 17 | 15 | 13 | Mas1 |
| Maskbp3 | 105 | 86 | 59 | 113 | 153 | 179 | 158 | 98 | 77 | 74 | 83 | 80 | 101 | 81 | 123 | 93 | 81 | 123 | 150 | 163 | 158 | 87 | 62 | 75 | 65 | 57 | 102 | 70 | Maskbp3 |
| Masp1 | 1344 | 1602 | 1550 | 1625 | 2539 | 2844 | 2392 | 5404 | 5470 | 5962 | 4847 | 6081 | 6146 | 6048 | 1726 | 1093 | 1814 | 1897 | 2793 | 2463 | 2458 | 4804 | 5742 | 6188 | 6899 | 4879 | 5582 | 5889 | Masp1 |
| Masp2 | 65 | 106 | 71 | 103 | 104 | 103 | 140 | 33 | 35 | 81 | 38 | 39 | 31 | 38 | 80 | 83 | 93 | 116 | 109 | 78 | 145 | 53 | 44 | 53 | 43 | 31 | 57 | 23 | Masp2 |
| Mast1 | 2347 | 3258 | 3602 | 5081 | 6774 | 4990 | 5041 | 4483 | 5216 | 7550 | 5758 | 5915 | 4544 | 5221 | 2664 | 2311 | 4942 | 5497 | 6727 | 4611 | 5395 | 9108 | 4447 | 6552 | 7919 | 4785 | 4312 | 4498 | Mast1 |
| Mast2 | 9908 | 10994 | 10245 | 12357 | 13322 | 12673 | 12239 | 10387 | 11581 | 13318 | 11674 | 12592 | 11111 | 11855 | 9298 | 10738 | 11422 | 12431 | 13472 | 10504 | 12853 | 14155 | 10448 | 12188 | 14340 | 9840 | 11080 | 11130 | Mast2 |
| Mast3 | 582 | 264 | 271 | 656 | 732 | 1001 | 779 | 260 | 163 | 80 | 409 | 205 | 177 | 187 | 335 | 748 | 739 | 346 | 732 | 565 | 888 | 246 | 307 | 485 | 228 | 220 | 106 | 399 | Mast3 |
| Mast4 | 4731 | 5719 | 5267 | 7753 | 8708 | 7691 | 6375 | 4152 | 4373 | 5381 | 5335 | 4535 | 6191 | 4261 | 5542 | 6447 | 6707 | 7793 | 7545 | 6035 | 6993 | 6273 | 4217 | 4032 | 5360 | 2855 | 4338 | 4119 | Mast4 |
| Mastl | 2794 | 2059 | 1977 | 1662 | 1366 | 1755 | 1858 | 1799 | 1749 | 1447 | 1440 | 2109 | 1784 | 2268 | 2665 | 2364 | 1637 | 1504 | 1439 | 2119 | 1665 | 1221 | 1526 | 1629 | 1490 | 2147 | 2150 | 2148 | Mastl |
| Mat1a | 2 | 1 | 0 | 0 | 2 | 3 | 2 | 5 | 9 | 0 | 0 | 0 | 0 | 1 | 4 | 2 | 0 | 1 | 2 | 0 | 3 | 2 | 1 | 3 | 1 | 1 | 1 | 0 | Mat1a |
| Mat2a | 1489 | 676 | 697 | 668 | 1778 | 1281 | 602 | 1700 | 994 | 821 | 885 | 1117 | 980 | 1435 | 1481 | 734 | 1779 | 1011 | 1106 | 1826 | 701 | 1703 | 1972 | 896 | 1295 | 1212 | 1484 | 1362 | Mat2a |
| Mat2b | 6425 | 5690 | 5927 | 5624 | 4420 | 4983 | 5382 | 4883 | 4249 | 3885 | 4011 | 4839 | 4748 | 4786 | 6262 | 6057 | 5393 | 5302 | 4953 | 5841 | 5028 | 3989 | 4493 | 4451 | 3975 | 5150 | 5089 | 5149 | Mat2b |
| Matk | 393 | 572 | 548 | 671 | 875 | 657 | 620 | 228 | 368 | 416 | 430 | 224 | 268 | 199 | 384 | 520 | 743 | 786 | 795 | 548 | 583 | 389 | 350 | 331 | 347 | 250 | 182 | 202 | Matk |
| Matn1 | 14 | 21 | 0 | 60 | 28 | 213 | 0 | 79 | 1307 | 1082 | 2581 | 50 | 136 | 40 | 15 | 8 | 9 | 222 | 44 | 0 | 2 | 1412 | 386 | 616 | 879 | 38 | 60 | 32 | Matn1 |
| Matn2 | 621 | 531 | 464 | 452 | 557 | 547 | 188 | 979 | 1508 | 1021 | 1719 | 594 | 849 | 506 | 574 | 763 | 290 | 748 | 192 | 287 | 246 | 1106 | 1925 | 863 | 858 | 545 | 553 | 458 | Matn2 |

| | | | | | | | | | | | | | | | | | | | | | | | | | | | | | |
|---|---|---|---|---|---|---|---|---|---|---|---|---|---|---|---|---|---|---|---|---|---|---|---|---|---|---|---|---|---|
| Matn3 | 110 | 104 | 109 | 71 | 33 | 65 | 40 | 50 | 215 | 197 | 801 | 23 | 58 | 33 | 109 | 147 | 61 | 65 | 56 | 41 | 26 | 252 | 190 | 82 | 142 | 56 | 43 | 63 | Matn3 |
| Matn4 | 316 | 402 | 141 | 744 | 280 | 1268 | 4 | 1122 | 3930 | 4586 | 6898 | 722 | 1227 | 575 | 407 | 534 | 363 | 1532 | 394 | 7 | 3 | 5681 | 3164 | 2696 | 3215 | 870 | 933 | 508 | Matn4 |
| Matr3 | 21607 | 14169 | 19500 | 20001 | 21413 | 20749 | 27064 | 32145 | 24713 | 23851 | 22695 | 26373 | 23626 | 29896 | 24976 | 18010 | 20179 | 19815 | 21121 | 24780 | 25673 | 29085 | 24755 | 25680 | 26592 | 27585 | 25293 | 28736 | Matr3 |
| Mau2 | 9109 | 9497 | 5456 | 8560 | 8859 | 9572 | 8765 | 11352 | 11127 | 10465 | 10270 | 10657 | 10218 | 10284 | 8962 | 8346 | 6675 | 8470 | 8952 | 9212 | 9587 | 10723 | 10187 | 9256 | 11570 | 7054 | 10779 | 9930 | Mau2 |
| Mavs | 1954 | 2244 | 1625 | 1579 | 1122 | 1344 | 1088 | 982 | 1107 | 864 | 980 | 956 | 1005 | 847 | 1751 | 2035 | 1556 | 1438 | 1180 | 1460 | 1046 | 780 | 1113 | 935 | 713 | 1004 | 1088 | 979 | Mavs |
| Max | 5873 | 6886 | 7236 | 7530 | 6473 | 6488 | 7042 | 3754 | 3704 | 4023 | 3868 | 4069 | 3708 | 4303 | 6028 | 6153 | 7105 | 6999 | 6759 | 6238 | 6977 | 3877 | 3721 | 4310 | 4374 | 4641 | 4038 | 4379 | Max |
| Maz | 9982 | 11213 | 13530 | 13380 | 13463 | 12038 | 12207 | 17397 | 17932 | 22852 | 19721 | 25632 | 19689 | 23572 | 9473 | 8572 | 14568 | 12043 | 14480 | 10754 | 13247 | 20317 | 16111 | 24995 | 22429 | 28238 | 21487 | 23985 | Maz |
| Mb | 13 | 14 | 3 | 14 | 8 | 5 | 4 | 11 | 30 | 10 | 97 | 3 | 2 | 3 | 12 | 17 | 13 | 14 | 8 | 7 | 4 | 23 | 97 | 20 | 1 | 8 | 5 | 2 | Mb |
| Mb21d2 | 1474 | 1445 | 1664 | 1739 | 2513 | 1831 | 1882 | 2553 | 2425 | 2515 | 2215 | 1993 | 2146 | 1860 | 1663 | 1494 | 1859 | 1738 | 2279 | 1961 | 1989 | 3193 | 2466 | 1998 | 2393 | 2065 | 1970 | 1930 | Mb21d2 |
| Mbd1 | 6683 | 6893 | 6755 | 7111 | 6352 | 6531 | 6415 | 5543 | 5962 | 6017 | 5820 | 6156 | 6254 | 6122 | 6000 | 6263 | 6573 | 6803 | 6597 | 6193 | 6284 | 5822 | 5352 | 6190 | 6482 | 5710 | 6365 | 6080 | Mbd1 |
| Mbd2 | 3109 | 3049 | 3292 | 3042 | 2632 | 2496 | 2603 | 2276 | 2200 | 2170 | 2264 | 2131 | 2228 | 2166 | 2917 | 3041 | 3065 | 2497 | 2573 | 2394 | 2693 | 2211 | 2409 | 2312 | 1911 | 2565 | 2258 | 2483 | Mbd2 |
| Mbd3 | 3528 | 4448 | 5075 | 4257 | 4178 | 4021 | 3475 | 6160 | 7442 | 8426 | 8270 | 10354 | 8489 | 9060 | 3058 | 3565 | 5038 | 4021 | 5178 | 3485 | 3918 | 7005 | 7368 | 9765 | 8766 | 12283 | 9645 | 9301 | Mbd3 |
| Mbd3l1 | 0 | 0 | 0 | 0 | 0 | 0 | 0 | 0 | 0 | 0 | 0 | 2 | 0 | 2 | 0 | 0 | 0 | 0 | 0 | 0 | 0 | 0 | 0 | 0 | 0 | 0 | 0 | 1 | Mbd3l1 |
| Mbd3l2 | 0 | 0 | 0 | 2 | 0 | 0 | 0 | 0 | 0 | 0 | 0 | 0 | 0 | 0 | 0 | 0 | 0 | 0 | 0 | 0 | 1 | 0 | 0 | 0 | 0 | 0 | 0 | 0 | Mbd3l2 |
| Mbd4 | 3494 | 3239 | 2842 | 2938 | 2535 | 2825 | 2813 | 2307 | 2127 | 2124 | 2172 | 2434 | 2237 | 2564 | 3186 | 3142 | 2688 | 2747 | 2627 | 3035 | 2799 | 1860 | 1959 | 2134 | 2028 | 2128 | 2656 | 2480 | Mbd4 |
| Mbd5 | 3767 | 3343 | 2315 | 3333 | 3692 | 3771 | 3723 | 4107 | 3707 | 3396 | 3482 | 3414 | 3246 | 3396 | 3746 | 3395 | 2917 | 3371 | 3543 | 3772 | 4084 | 3907 | 3335 | 2920 | 3524 | 2210 | 3460 | 3101 | Mbd5 |
| Mbd6 | 3727 | 4945 | 6410 | 6696 | 6649 | 5566 | 5656 | 4246 | 4934 | 6573 | 5848 | 7089 | 5940 | 6519 | 3698 | 4311 | 6898 | 6128 | 7003 | 4472 | 5805 | 6413 | 4497 | 7229 | 6796 | 7954 | 6415 | 6149 | Mbd6 |
| Mbip | 2645 | 2021 | 1778 | 2044 | 2059 | 2308 | 2758 | 2762 | 2551 | 2285 | 2242 | 2858 | 1936 | 2962 | 2459 | 1856 | 2188 | 1991 | 2128 | 2711 | 2597 | 2511 | 2136 | 2462 | 2482 | 3249 | 2578 | 2541 | Mbip |
| Mbl1 | 0 | 0 | 0 | 0 | 0 | 0 | 0 | 1 | 0 | 0 | 0 | 0 | 0 | 0 | 0 | 0 | 0 | 0 | 0 | 0 | 0 | 0 | 0 | 0 | 0 | 0 | 0 | 0 | Mbl1 |
| Mbl2 | 0 | 3 | 0 | 0 | 0 | 2 | 0 | 4 | 2 | 1 | 0 | 1 | 2 | 1 | 0 | 0 | 0 | 0 | 0 | 0 | 0 | 0 | 1 | 3 | 0 | 0 | 0 | 0 | Mbl2 |
| Mblac1 | 284 | 271 | 323 | 285 | 320 | 269 | 253 | 270 | 278 | 342 | 236 | 300 | 282 | 271 | 257 | 207 | 343 | 232 | 286 | 298 | 250 | 206 | 272 | 297 | 235 | 332 | 299 | 212 | Mblac1 |
| Mblac2 | 1746 | 1464 | 1978 | 2573 | 2542 | 2508 | 2865 | 1032 | 946 | 934 | 1107 | 852 | 883 | 874 | 1841 | 1703 | 2033 | 2600 | 2272 | 2193 | 2738 | 1544 | 1003 | 819 | 1132 | 706 | 739 | 891 | Mblac2 |
| Mbnl1 | 4245 | 3047 | 2766 | 3300 | 3683 | 3847 | 3903 | 5490 | 5795 | 4208 | 6026 | 3290 | 4524 | 3824 | 4399 | 4274 | 2438 | 3708 | 3167 | 3789 | 3766 | 4371 | 5531 | 4288 | 3810 | 3893 | 3914 | 3885 | Mbnl1 |
| Mbnl2 | 10905 | 7001 | 6794 | 6599 | 6414 | 7913 | 7434 | 6530 | 7247 | 5670 | 7782 | 4456 | 5830 | 4852 | 9349 | 9787 | 6509 | 7362 | 5725 | 6804 | 6997 | 5900 | 7575 | 5576 | 4420 | 5247 | 5037 | 4859 | Mbnl2 |
| Mbnl3 | 1119 | 636 | 346 | 379 | 369 | 678 | 541 | 1087 | 1251 | 642 | 1149 | 768 | 1023 | 790 | 823 | 917 | 371 | 432 | 408 | 617 | 501 | 451 | 1273 | 568 | 562 | 355 | 990 | 767 | Mbnl3 |
| Mboat1 | 514 | 431 | 488 | 294 | 229 | 287 | 229 | 280 | 345 | 273 | 328 | 307 | 302 | 311 | 411 | 583 | 375 | 327 | 205 | 200 | 189 | 249 | 287 | 271 | 230 | 402 | 336 | 344 | Mboat1 |
| Mboat2 | 7722 | 6612 | 5920 | 6998 | 7274 | 7503 | 7386 | 6161 | 5235 | 5561 | 5359 | 5733 | 5557 | 5812 | 7861 | 6979 | 5997 | 6974 | 6868 | 7651 | 7411 | 6084 | 5387 | 5078 | 6094 | 4615 | 5819 | 5692 | Mboat2 |
| Mboat4 | 2 | 9 | 4 | 0 | 2 | 1 | 1 | 2 | 4 | 1 | 6 | 2 | 6 | 0 | 4 | 2 | 6 | 2 | 12 | 0 | 3 | 2 | 4 | 9 | 8 | 2 | 1 | 0 | Mboat4 |
| Mboat7 | 6210 | 6732 | 6543 | 7550 | 7910 | 7006 | 7330 | 3379 | 3704 | 4027 | 4109 | 3512 | 3367 | 3376 | 5748 | 6266 | 7113 | 7745 | 7638 | 5822 | 7376 | 4455 | 3678 | 3680 | 3631 | 3325 | 3005 | 3188 | Mboat7 |
| Mbp | 2002 | 2175 | 2587 | 2229 | 2585 | 2118 | 2114 | 1542 | 1635 | 1751 | 1719 | 2157 | 1842 | 1945 | 1838 | 2201 | 2574 | 2068 | 2570 | 2037 | 2063 | 1928 | 1598 | 1916 | 1883 | 1790 | 1905 | 1851 | Mbp |
| Mbtd1 | 18430 | 13924 | 15480 | 16114 | 16025 | 17090 | 19542 | 20229 | 16245 | 16652 | 16345 | 19894 | 18329 | 21468 | 17934 | 15066 | 15723 | 15191 | 16491 | 17642 | 18112 | 17704 | 15655 | 17580 | 19252 | 18042 | 21016 | 21318 | Mbtd1 |
| Mbtps1 | 9777 | 9833 | 8588 | 7714 | 7273 | 8447 | 7038 | 10209 | 12275 | 10879 | 12634 | 12194 | 12396 | 11487 | 8865 | 10435 | 8066 | 7996 | 7671 | 6994 | 7262 | 10517 | 12013 | 11254 | 10685 | 11331 | 12117 | 11628 | Mbtps1 |
| Mbtps2 | 3538 | 3036 | 2726 | 3441 | 4285 | 3885 | 4510 | 4893 | 4181 | 3957 | 4062 | 3543 | 3706 | 3515 | 3918 | 3273 | 3266 | 3777 | 3847 | 4288 | 4272 | 4689 | 4067 | 3741 | 4051 | 3337 | 3452 | 3455 | Mbtps2 |
| Mc1r | 23 | 29 | 20 | 13 | 31 | 29 | 18 | 10 | 8 | 5 | 6 | 5 | 16 | 16 | 16 | 12 | 13 | 23 | 30 | 23 | 21 | 2 | 15 | 7 | 7 | 7 | 8 | 5 | Mc1r |
| Mc2r | 0 | 0 | 0 | 0 | 1 | 0 | 0 | 0 | 2 | 2 | 7 | 0 | 0 | 1 | 0 | 0 | 0 | 0 | 0 | 0 | 0 | 0 | 6 | 0 | 0 | 0 | 0 | 0 | Mc2r |
| Mc3r | 1 | 1 | 2 | 4 | 7 | 5 | 6 | 10 | 8 | 22 | 13 | 11 | 0 | 4 | 9 | 4 | 2 | 13 | 7 | 3 | 10 | 10 | 6 | 4 | 6 | 4 | 0 | 1 | Mc3r |
| Mc4r | 238 | 214 | 210 | 159 | 249 | 211 | 160 | 162 | 174 | 216 | 132 | 73 | 130 | 99 | 223 | 185 | 239 | 169 | 168 | 183 | 190 | 110 | 161 | 123 | 95 | 146 | 116 | 94 | Mc4r |
| Mc5r | 125 | 92 | 118 | 134 | 224 | 153 | 130 | 125 | 112 | 108 | 154 | 58 | 79 | 53 | 121 | 129 | 116 | 178 | 136 | 155 | 193 | 147 | 183 | 105 | 67 | 53 | 50 | 40 | Mc5r |
| Mcam | 6687 | 6626 | 5969 | 5106 | 4650 | 4849 | 4872 | 2927 | 3270 | 3082 | 3727 | 2888 | 2982 | 2818 | 5720 | 6146 | 5696 | 5195 | 4753 | 5242 | 4508 | 3502 | 3575 | 3364 | 3006 | 3534 | 2865 | 2708 | Mcam |
| Mcat | 1490 | 1487 | 1464 | 1385 | 1226 | 1371 | 1257 | 1788 | 1865 | 1740 | 1515 | 1845 | 1838 | 1807 | 1346 | 1266 | 1509 | 1227 | 1452 | 1284 | 1255 | 1566 | 1781 | 1779 | 1646 | 2298 | 1949 | 1851 | Mcat |
| Mcc | 904 | 809 | 864 | 824 | 910 | 816 | 628 | 818 | 952 | 946 | 1136 | 649 | 1159 | 639 | 793 | 1007 | 912 | 950 | 764 | 655 | 667 | 1051 | 1173 | 699 | 783 | 626 | 796 | 752 | Mcc |
| Mccc1 | 2850 | 2649 | 1971 | 2602 | 2748 | 2860 | 2845 | 3413 | 3343 | 2768 | 3118 | 2966 | 3087 | 3081 | 2869 | 2511 | 2186 | 2500 | 2564 | 2958 | 2926 | 2719 | 3109 | 2736 | 3181 | 2418 | 3103 | 2948 | Mccc1 |
| Mccc2 | 1899 | 1956 | 1915 | 1641 | 1483 | 1789 | 1616 | 1490 | 1666 | 1418 | 1622 | 1644 | 1528 | 1605 | 1950 | 1963 | 1875 | 1725 | 1643 | 1759 | 1672 | 1362 | 1759 | 1584 | 1595 | 1758 | 1620 | 1570 | Mccc2 |
| Mcee | 776 | 726 | 714 | 719 | 604 | 617 | 767 | 558 | 552 | 532 | 490 | 445 | 510 | 563 | 761 | 610 | 705 | 676 | 627 | 796 | 693 | 429 | 548 | 588 | 449 | 517 | 438 | 548 | Mcee |
| Mcemp1 | 3 | 8 | 9 | 16 | 13 | 7 | 9 | 5 | 7 | 6 | 2 | 10 | 14 | 3 | 28 | 16 | 9 | 13 | 6 | 7 | 7 | 8 | 14 | 5 | 10 | 8 | 6 | 13 | Mcemp1 |
| Mcf2 | 66 | 71 | 56 | 49 | 90 | 53 | 79 | 144 | 102 | 95 | 122 | 75 | 76 | 120 | 69 | 90 | 86 | 46 | 78 | 77 | 99 | 154 | 148 | 95 | 73 | 112 | 121 | 136 | Mcf2 |
| Mcf2l | 2719 | 3485 | 3561 | 4975 | 7305 | 5599 | 5842 | 8927 | 9363 | 11627 | 10051 | 10413 | 8968 | 11049 | 3189 | 2929 | 4606 | 5468 | 6726 | 5071 | 6309 | 13774 | 8908 | 11237 | 13767 | 7454 | 9391 | 9609 | Mcf2l |
| Mcfd2 | 4316 | 4184 | 3872 | 3485 | 2891 | 3285 | 3157 | 2947 | 2988 | 2513 | 3042 | 2654 | 2697 | 2632 | 4114 | 4132 | 3787 | 3372 | 3343 | 3511 | 3145 | 2707 | 3268 | 2720 | 2469 | 3108 | 2843 | 2760 | Mcfd2 |
| Mchr1 | 67 | 73 | 73 | 75 | 122 | 102 | 108 | 44 | 54 | 60 | 66 | 38 | 17 | 34 | 65 | 58 | 79 | 96 | 115 | 104 | 94 | 65 | 56 | 55 | 34 | 31 | 29 | 44 | Mchr1 |
| Mcidas | 51 | 34 | 50 | 50 | 52 | 47 | 54 | 138 | 125 | 143 | 112 | 325 | 301 | 239 | 40 | 55 | 89 | 40 | 62 | 81 | 50 | 130 | 231 | 183 | 140 | 357 | 267 | 268 | Mcidas |
| Mcl1 | 12545 | 10537 | 13516 | 12377 | 13227 | 12309 | 13011 | 16361 | 13577 | 14670 | 14017 | 15499 | 14775 | 16173 | 12177 | 11158 | 13139 | 12378 | 13801 | 12084 | 13271 | 16494 | 13693 | 15377 | 16169 | 16504 | 15583 | 17255 | Mcl1 |
| Mcm10 | 3736 | 3688 | 3591 | 2612 | 2058 | 2487 | 2713 | 2716 | 2745 | 2291 | 2042 | 3631 | 2752 | 3433 | 3204 | 3009 | 3045 | 2290 | 2616 | 2697 | 2383 | 1654 | 2162 | 2808 | 2301 | 4131 | 3754 | 3639 | Mcm10 |
| Mcm2 | 11626 | 13420 | 13105 | 10387 | 7725 | 8986 | 9157 | 8734 | 10011 | 8492 | 7752 | 13408 | 10993 | 12438 | 10526 | 10869 | 12216 | 9202 | 9838 | 9934 | 8890 | 6363 | 8365 | 10914 | 9109 | 15816 | 13770 | 13804 | Mcm2 |
| Mcm3 | 10800 | 10928 | 11154 | 8237 | 6371 | 7625 | 7938 | 8695 | 9352 | 7752 | 7397 | 12459 | 10295 | 11529 | 9111 | 9048 | 9319 | 6949 | 8486 | 7947 | 7533 | 5718 | 8176 | 10150 | 8196 | 15664 | 12755 | 12797 | Mcm3 |
| Mcm3ap | 2230 | 2229 | 2283 | 2081 | 2828 | 2640 | 2227 | 7470 | 8633 | 8343 | 8060 | 8993 | 8388 | 8663 | 2102 | 2130 | 2356 | 1965 | 2786 | 2238 | 2326 | 7907 | 7721 | 8395 | 8836 | 6763 | 8761 | 8791 | Mcm3ap |
| Mcm4 | 11506 | 10216 | 11145 | 7643 | 5789 | 7074 | 7658 | 8573 | 8645 | 7467 | 7490 | 10408 | 9377 | 10271 | 9509 | 9756 | 8633 | 6945 | 6603 | 7624 | 6478 | 6086 | 7863 | 8613 | 7068 | 12369 | 10369 | 10745 | Mcm4 |
| Mcm5 | 11221 | 11822 | 10908 | 8151 | 5701 | 7377 | 7311 | 8338 | 9611 | 7770 | 7929 | 11958 | 10182 | 10602 | 9068 | 9521 | 9042 | 6875 | 8194 | 7963 | 7068 | 5633 | 8220 | 9529 | 8133 | 15003 | 12454 | 12219 | Mcm5 |
| Mcm6 | 17264 | 16897 | 17426 | 13392 | 10978 | 12911 | 13698 | 12830 | 13107 | 11377 | 11346 | 15345 | 13902 | 15193 | 16644 | 15890 | 15285 | 12115 | 12732 | 13605 | 12627 | 9474 | 12782 | 12861 | 11266 | 19440 | 16444 | 17061 | Mcm6 |
| Mcm7 | 13568 | 14962 | 14516 | 12170 | 10648 | 12193 | 13027 | 14073 | 14802 | 13439 | 12288 | 20411 | 16268 | 20010 | 12526 | 11727 | 13517 | 11669 | 13691 | 14384 | 12754 | 10019 | 12927 | 16966 | 15428 | 23542 | 21810 | 21236 | Mcm7 |
| Mcm8 | 1805 | 1590 | 1751 | 1240 | 922 | 1050 | 1376 | 1779 | 1722 | 1485 | 1297 | 2126 | 1869 | 2183 | 1630 | 1570 | 1639 | 1092 | 1194 | 1414 | 1145 | 1093 | 1494 | 1785 | 1419 | 2412 | 2206 | 2288 | Mcm8 |
| Mcm9 | 952 | 828 | 980 | 953 | 839 | 959 | 875 | 1188 | 1188 | 1029 | 1226 | 1370 | 1203 | 1194 | 1052 | 900 | 946 | 870 | 942 | 917 | 863 | 1034 | 1177 | 1149 | 937 | 1446 | 1316 | 1297 | Mcm9 |
| Mcmbp | 7974 | 7492 | 7682 | 7754 | 7450 | 7966 | 8490 | 6831 | 6697 | 7140 | 6631 | 8155 | 7194 | 8350 | 7564 | 7355 | 7740 | 7634 | 8200 | 7566 | 8210 | 6742 | 6326 | 7437 | 7318 | 7968 | 8261 | 8632 | Mcmbp |
| Mcmdc2 | 161 | 130 | 147 | 110 | 133 | 173 | 194 | 166 | 128 | 188 | 105 | 152 | 169 | 160 | 214 | 132 | 146 | 125 | 204 | 206 | 186 | 184 | 127 | 136 | 182 | 160 | 141 | 170 | Mcmdc2 |
| Mcoln1 | 4448 | 4774 | 4590 | 5739 | 5829 | 5513 | 5430 | 3389 | 3515 | 4449 | 3695 | 4326 | 3799 | 3782 | 4245 | 4224 | 5463 | 5936 | 5824 | 4732 | 5303 | 4525 | 3447 | 4189 | 4341 | 3625 | 3799 | 3678 | Mcoln1 |
| Mcoln2 | 40 | 32 | 20 | 28 | 23 | 26 | 15 | 20 | 40 | 22 | 57 | 25 | 44 | 14 | 26 | 43 | 37 | 14 | 20 | 24 | 13 | 65 | 49 | 30 | 19 | 27 | 41 | 21 | Mcoln2 |
| Mcoln3 | 354 | 275 | 169 | 188 | 92 | 151 | 56 | 328 | 313 | 283 | 297 | 230 | 329 | 197 | 352 | 368 | 146 | 147 | 96 | 79 | 74 | 150 | 356 | 318 | 147 | 397 | 255 | 215 | Mcoln3 |
| Mcph1 | 1465 | 1294 | 1194 | 1214 | 1162 | 1232 | 1206 | 1420 | 1400 | 1292 | 1325 | 1569 | 1581 | 1497 | 1316 | 1248 | 1152 | 1094 | 1154 | 1101 | 1140 | 1282 | 1383 | 1318 | 1499 | 1598 | 1565 | 1504 | Mcph1 |
| Mcpt1 | 0 | 0 | 0 | 0 | 0 | 0 | 0 | 0 | 0 | 0 | 1 | 0 | 0 | 0 | 0 | 0 | 0 | 0 | 1 | 0 | 0 | 0 | 0 | 0 | 0 | 0 | 0 | 0 | Mcpt1 |
| Mcpt2 | 0 | 0 | 0 | 0 | 0 | 0 | 0 | 0 | 2 | 0 | 0 | 0 | 3 | 0 | 1 | 0 | 0 | 0 | 5 | 0 | 1 | 0 | 0 | 2 | 0 | 0 | 0 | 0 | Mcpt2 |
| Mcpt4 | 0 | 0 | 0 | 0 | 0 | 0 | 0 | 0 | 4 | 1 | 0 | 0 | 0 | 0 | 0 | 0 | 0 | 0 | 3 | 0 | 0 | 0 | 5 | 0 | 0 | 0 | 0 | 1 | Mcpt4 |
| Mcpt8 | 1 | 0 | 0 | 0 | 0 | 1 | 0 | 1 | 3 | 0 | 3 | 0 | 0 | 0 | 0 | 0 | 0 | 0 | 0 | 0 | 2 | 0 | 3 | 0 | 0 | 0 | 0 | 0 | Mcpt8 |
| Mcrip1 | 4185 | 5078 | 4342 | 4649 | 4020 | 4279 | 4331 | 3653 | 4219 | 4152 | 3923 | 4029 | 3958 | 4192 | 4002 | 3894 | 4746 | 4309 | 4663 | 4215 | 4378 | 3646 | 4243 | 4186 | 4130 | 4932 | 4114 | 4169 | Mcrip1 |
| Mcrip2 | 299 | 406 | 300 | 452 | 268 | 280 | 293 | 72 | 130 | 121 | 98 | 134 | 77 | 76 | 248 | 304 | 329 | 366 | 286 | 321 | 321 | 94 | 116 | 128 | 118 | 91 | 99 | 82 | Mcrip2 |
| Mcrs1 | 2354 | 3381 | 2636 | 2546 | 2890 | 2928 | 2733 | 4733 | 5675 | 4991 | 4814 | 5645 | 5254 | 5736 | 2499 | 2460 | 2615 | 2384 | 2895 | 2921 | 2744 | 4792 | 5409 | 5413 | 5313 | 6264 | 5975 | 5966 | Mcrs1 |
| Mctp1 | 374 | 295 | 356 | 298 | 340 | 380 | 440 | 363 | 375 | 315 | 277 | 363 | 370 | 470 | 476 | 315 | 342 | 326 | 291 | 487 | 362 | 480 | 337 | 273 | 390 | 388 | 506 | 458 | Mctp1 |
| Mctp2 | 260 | 149 | 96 | 131 | 90 | 95 | 65 | 87 | 108 | 101 | 117 | 75 | 66 | 66 | 210 | 224 | 76 | 124 | 55 | 63 | 55 | 109 | 114 | 52 | 53 | 37 | 123 | 97 | Mctp2 |
| Mcts1 | 3032 | 2460 | 2851 | 3167 | 3616 | 3471 | 4013 | 4643 | 3952 | 3978 | 4270 | 3915 | 4206 | 4603 | 3559 | 2850 | 2958 | 3045 | 3285 | 3881 | 3817 | 4465 | 4102 | 4083 | 4412 | 3648 | 4131 | 4285 | Mcts1 |
| Mcts2 | 991 | 1013 | 808 | 855 | 926 | 986 | 1096 | 1155 | 985 | 947 | 929 | 936 | 989 | 1116 | 1138 | 947 | 939 | 895 | 853 | 1109 | 971 | 784 | 948 | 922 | 990 | 958 | 1026 | 1138 | Mcts2 |
| Mcu | 1976 | 1875 | 1958 | 2041 | 2072 | 2021 | 1942 | 1529 | 1756 | 1555 | 2342 | 1398 | 1670 | 1509 | 1814 | 2077 | 2041 | 2141 | 1867 | 1679 | 1907 | 1693 | 2060 | 1562 | 1569 | 1614 | 1515 | 1531 | Mcu |

| | | | | | | | | | | | | | | | | | | | | | | | | | | | | | |
|---|---|---|---|---|---|---|---|---|---|---|---|---|---|---|---|---|---|---|---|---|---|---|---|---|---|---|---|---|---|
| Mcub | 178 | 185 | 149 | 194 | 190 | 187 | 245 | 87 | 92 | 99 | 74 | 94 | 64 | 82 | 165 | 174 | 147 | 169 | 199 | 191 | 210 | 94 | 86 | 90 | 98 | 92 | 101 | 83 | Mcub |
| Mcur1 | 1519 | 1418 | 1244 | 1404 | 1176 | 1240 | 1477 | 1218 | 1248 | 1213 | 1398 | 1374 | 1386 | 1458 | 1381 | 1382 | 1364 | 1267 | 1205 | 1199 | 1281 | 1245 | 1342 | 1350 | 1272 | 1380 | 1469 | 1431 | Mcur1 |
| Mdc1 | 13480 | 13460 | 13097 | 14418 | 12016 | 12171 | 13162 | 8763 | 9508 | 10489 | 9842 | 13605 | 11332 | 13474 | 12183 | 13006 | 14183 | 13906 | 13526 | 11098 | 12883 | 8696 | 8259 | 11376 | 10373 | 9870 | 12570 | 12392 | Mdc1 |
| Mdfi | 2194 | 2640 | 2298 | 1597 | 1073 | 1466 | 739 | 1209 | 1779 | 1475 | 2060 | 1134 | 1621 | 845 | 1909 | 2960 | 2044 | 1546 | 1180 | 1010 | 804 | 1146 | 1985 | 1449 | 930 | 1978 | 1166 | 1035 | Mdfi |
| Mdfic | 1025 | 655 | 941 | 682 | 626 | 881 | 568 | 1656 | 1874 | 1582 | 1993 | 1718 | 2071 | 1632 | 935 | 878 | 835 | 817 | 690 | 519 | 544 | 1945 | 2331 | 1735 | 1463 | 1897 | 1858 | 1809 | Mdfic |
| Mdfic2 | 0 | 3 | 0 | 4 | 1 | 0 | 0 | 5 | 2 | 12 | 6 | 0 | 5 | 1 | 1 | 2 | 0 | 3 | 0 | 0 | 0 | 7 | 4 | 0 | 4 | 2 | 8 | 0 | Mdfic2 |
| Mdga1 | 3147 | 4062 | 5611 | 7075 | 10186 | 7527 | 6813 | 5632 | 6171 | 8619 | 6794 | 7585 | 6294 | 6277 | 3574 | 3559 | 7130 | 7845 | 9861 | 6391 | 7872 | 10281 | 6319 | 8043 | 9941 | 4479 | 6505 | 6509 | Mdga1 |
| Mdga2 | 1497 | 1108 | 1032 | 1221 | 1645 | 1667 | 1443 | 1848 | 2023 | 1671 | 1678 | 1358 | 1534 | 1394 | 1659 | 1482 | 990 | 1512 | 1230 | 1579 | 1582 | 2142 | 1775 | 1298 | 1605 | 829 | 1390 | 1229 | Mdga2 |
| Mdh1 | 26579 | 25543 | 21647 | 24003 | 19851 | 21593 | 23006 | 18421 | 16022 | 14143 | 13386 | 14698 | 14446 | 14791 | 28193 | 24540 | 23235 | 22455 | 22788 | 29041 | 22144 | 13904 | 15738 | 14917 | 13919 | 16153 | 15966 | 15281 | Mdh1 |
| Mdh1b | 18 | 10 | 23 | 13 | 32 | 23 | 23 | 27 | 43 | 24 | 43 | 45 | 35 | 18 | 40 | 26 | 21 | 31 | 12 | 29 | 19 | 38 | 41 | 16 | 26 | 34 | 14 | 25 | Mdh1b |
| Mdh2 | 14113 | 17493 | 14251 | 14616 | 13075 | 12989 | 13739 | 10989 | 11839 | 11572 | 10836 | 12313 | 12018 | 12457 | 13818 | 14599 | 14698 | 13542 | 13878 | 13983 | 13844 | 10586 | 11259 | 11772 | 12147 | 13008 | 12832 | 13208 | Mdh2 |
| Mdk | 10024 | 9497 | 10508 | 6886 | 4755 | 6368 | 4600 | 9574 | 12008 | 11003 | 13325 | 11468 | 13577 | 11606 | 7617 | 10413 | 8112 | 6167 | 5539 | 6038 | 4345 | 6721 | 12158 | 12914 | 8814 | 19033 | 13237 | 12525 | Mdk |
| Mdm1 | 11400 | 12065 | 9610 | 10669 | 8754 | 9697 | 10363 | 4362 | 3820 | 3791 | 3520 | 4650 | 4440 | 4654 | 11127 | 11135 | 9564 | 10365 | 9163 | 10664 | 10138 | 3559 | 3740 | 4144 | 4075 | 4627 | 5075 | 4705 | Mdm1 |
| Mdm2 | 4127 | 4256 | 4855 | 5488 | 4979 | 4654 | 5167 | 5935 | 4859 | 6159 | 5656 | 7128 | 6150 | 7403 | 4127 | 3967 | 5241 | 5218 | 4863 | 4419 | 5320 | 6597 | 5020 | 6976 | 6749 | 7331 | 6783 | 7189 | Mdm2 |
| Mdm4 | 17548 | 14455 | 11635 | 14877 | 15590 | 17344 | 17298 | 24845 | 20555 | 19448 | 21324 | 20600 | 20053 | 20934 | 17271 | 15608 | 12983 | 15220 | 14830 | 16908 | 17368 | 19976 | 19787 | 17768 | 20364 | 13517 | 21264 | 18646 | Mdm4 |
| Mdn1 | 19777 | 20638 | 8701 | 12178 | 11598 | 16932 | 13977 | 12320 | 12152 | 7682 | 8916 | 7811 | 8035 | 8398 | 19020 | 18600 | 9786 | 11946 | 11033 | 19072 | 16964 | 6542 | 10532 | 6858 | 8782 | 5245 | 9857 | 7379 | Mdn1 |
| Mdp1 | 1979 | 2027 | 2165 | 1919 | 2104 | 1983 | 2133 | 2583 | 2488 | 2452 | 2367 | 2837 | 2726 | 2892 | 2109 | 1932 | 2289 | 1860 | 2110 | 2479 | 2007 | 2348 | 2497 | 2656 | 2584 | 2993 | 3001 | 2909 | Mdp1 |
| Me1 | 1059 | 1030 | 994 | 926 | 1365 | 1149 | 938 | 1537 | 1589 | 1563 | 1937 | 1278 | 1762 | 1256 | 1134 | 1319 | 962 | 1139 | 1084 | 929 | 1087 | 2052 | 1937 | 1332 | 1649 | 1164 | 1187 | 1289 | Me1 |
| Me2 | 3226 | 3004 | 2787 | 2284 | 2270 | 2523 | 2592 | 2915 | 2954 | 2544 | 2620 | 2914 | 3071 | 3044 | 3190 | 3013 | 2540 | 2163 | 2241 | 2761 | 2385 | 2453 | 2834 | 2592 | 2532 | 2741 | 3295 | 3315 | Me2 |
| Me3 | 66 | 48 | 46 | 46 | 70 | 60 | 59 | 99 | 105 | 89 | 100 | 58 | 44 | 61 | 49 | 65 | 53 | 42 | 51 | 60 | 63 | 109 | 105 | 85 | 90 | 85 | 71 | 53 | Me3 |
| Mea1 | 2654 | 3722 | 3150 | 3211 | 2964 | 2896 | 3123 | 2989 | 3273 | 3248 | 3002 | 2904 | 3004 | 3769 | 3086 | 2896 | 3358 | 2974 | 3218 | 3602 | 3345 | 2593 | 3319 | 3291 | 3232 | 3407 | 3253 | 3533 | Mea1 |
| Meaf6 | 2129 | 2166 | 2451 | 3006 | 3513 | 3259 | 3289 | 4968 | 5061 | 6210 | 5371 | 6461 | 5505 | 6209 | 2093 | 1798 | 2898 | 3198 | 3647 | 3149 | 3368 | 6452 | 4923 | 6544 | 7106 | 6798 | 5799 | 5985 | Meaf6 |
| Meak7 | 937 | 1023 | 1011 | 973 | 999 | 952 | 1064 | 942 | 1086 | 1024 | 1037 | 1105 | 1082 | 1049 | 829 | 892 | 1109 | 988 | 1106 | 905 | 1037 | 976 | 1043 | 1205 | 1110 | 1284 | 1152 | 1144 | Meak7 |
| Mecom | 1671 | 1444 | 2025 | 868 | 936 | 983 | 637 | 2014 | 2348 | 1788 | 2636 | 1423 | 2765 | 1492 | 1851 | 1653 | 1307 | 716 | 805 | 929 | 757 | 1972 | 2449 | 1178 | 1202 | 1571 | 2086 | 1773 | Mecom |
| Mecp2 | 7608 | 7305 | 5736 | 8623 | 11225 | 10955 | 10869 | 14428 | 13228 | 14101 | 14051 | 14674 | 13442 | 14895 | 7745 | 6456 | 7139 | 8819 | 9760 | 9017 | 11667 | 15672 | 12213 | 12131 | 16603 | 8640 | 14752 | 13460 | Mecp2 |
| Mecr | 2149 | 2760 | 2146 | 2417 | 1730 | 1999 | 1983 | 989 | 1102 | 1062 | 1105 | 1214 | 1145 | 1182 | 2054 | 2428 | 2216 | 2418 | 2015 | 1871 | 2004 | 936 | 1080 | 1038 | 1103 | 1322 | 1293 | 1275 | Mecr |
| Med1 | 9059 | 8608 | 7745 | 9605 | 9178 | 9093 | 9683 | 9108 | 8538 | 8940 | 8788 | 9854 | 9703 | 10010 | 8881 | 8224 | 8770 | 8904 | 9030 | 8813 | 9554 | 8779 | 8195 | 9237 | 9424 | 7459 | 10162 | 10026 | Med1 |
| Med10 | 3644 | 4440 | 4247 | 4542 | 3654 | 4004 | 4067 | 2638 | 2731 | 2610 | 2569 | 2967 | 2896 | 3050 | 3889 | 4198 | 4161 | 4225 | 3892 | 4153 | 3938 | 2480 | 2642 | 2965 | 2793 | 3282 | 3153 | 3266 | Med10 |
| Med11 | 452 | 525 | 555 | 420 | 411 | 428 | 440 | 849 | 952 | 830 | 877 | 879 | 967 | 961 | 414 | 449 | 512 | 379 | 468 | 631 | 415 | 782 | 957 | 853 | 761 | 1030 | 888 | 898 | Med11 |
| Med12 | 5395 | 5392 | 4539 | 5462 | 6430 | 6367 | 5754 | 7458 | 8427 | 8700 | 9032 | 9841 | 9440 | 9012 | 5229 | 5664 | 5187 | 5822 | 6028 | 5498 | 6067 | 8946 | 7568 | 8241 | 9083 | 5376 | 9097 | 8816 | Med12 |
| Med12l | 2024 | 1744 | 1683 | 2834 | 3569 | 3082 | 3443 | 2847 | 2623 | 3556 | 2984 | 2866 | 2742 | 3043 | 1990 | 2011 | 1992 | 2571 | 2915 | 2558 | 3519 | 3171 | 2301 | 2799 | 3171 | 2283 | 2535 | 2194 | Med12l |
| Med13 | 12966 | 9878 | 10551 | 12429 | 13045 | 13966 | 14427 | 15917 | 13935 | 13270 | 13469 | 14657 | 14537 | 15895 | 12585 | 11812 | 11420 | 12041 | 12553 | 12296 | 14120 | 13812 | 12439 | 12924 | 15662 | 11903 | 15322 | 14853 | Med13 |
| Med13l | 6170 | 6956 | 6524 | 9192 | 9205 | 8366 | 9018 | 5906 | 6762 | 7904 | 8111 | 7865 | 7302 | 7661 | 5933 | 6555 | 7559 | 9245 | 9077 | 7274 | 9100 | 7546 | 6427 | 7572 | 8744 | 5311 | 6760 | 6741 | Med13l |
| Med14 | 12132 | 10322 | 7643 | 9094 | 9136 | 10832 | 11177 | 9744 | 8892 | 7052 | 7377 | 8698 | 7370 | 8575 | 11900 | 9609 | 8633 | 8718 | 9132 | 11251 | 11100 | 7294 | 7915 | 7116 | 8038 | 6713 | 8901 | 8051 | Med14 |
| Med15 | 2646 | 3042 | 2931 | 3112 | 3080 | 2972 | 2728 | 2705 | 3234 | 3444 | 3404 | 3685 | 3198 | 3260 | 2534 | 2860 | 3048 | 3159 | 3263 | 2407 | 2638 | 3501 | 3095 | 3556 | 3653 | 3953 | 3086 | 2989 | Med15 |
| Med16 | 1370 | 1672 | 1770 | 1685 | 1685 | 1636 | 1406 | 2700 | 3239 | 3297 | 3167 | 3735 | 3587 | 3061 | 1480 | 1604 | 1686 | 1584 | 1799 | 1464 | 1449 | 3240 | 3204 | 3515 | 3307 | 3971 | 3526 | 3363 | Med16 |
| Med17 | 3811 | 3831 | 3629 | 3306 | 3049 | 3261 | 3540 | 3655 | 3117 | 2998 | 2966 | 3556 | 3164 | 3671 | 3740 | 3440 | 3252 | 3121 | 3088 | 3694 | 3287 | 3050 | 3055 | 3183 | 3277 | 3537 | 3485 | 3604 | Med17 |
| Med18 | 623 | 667 | 744 | 765 | 450 | 527 | 574 | 447 | 451 | 403 | 405 | 597 | 526 | 510 | 589 | 676 | 703 | 651 | 547 | 547 | 493 | 331 | 467 | 536 | 509 | 627 | 678 | 579 | Med18 |
| Med19 | 3402 | 3642 | 3655 | 3360 | 3388 | 3270 | 3846 | 3009 | 2748 | 2849 | 2381 | 3367 | 2892 | 3306 | 3622 | 3410 | 3614 | 3355 | 3561 | 3431 | 3553 | 3008 | 2801 | 3179 | 3049 | 3769 | 3151 | 3501 | Med19 |
| Med20 | 3166 | 3216 | 2662 | 2816 | 2399 | 2502 | 2462 | 3088 | 2893 | 2614 | 2807 | 2629 | 2614 | 2712 | 2943 | 3244 | 2745 | 2698 | 2636 | 2835 | 2423 | 2780 | 2759 | 2729 | 2555 | 2539 | 2712 | 2721 | Med20 |
| Med21 | 2603 | 2205 | 2526 | 2372 | 2171 | 2189 | 2732 | 2577 | 2242 | 1803 | 2184 | 2136 | 2285 | 2622 | 2670 | 2279 | 2360 | 2060 | 2364 | 2734 | 2498 | 2396 | 2236 | 2242 | 1990 | 2599 | 2471 | 2684 | Med21 |
| Med22 | 7968 | 8522 | 6989 | 7691 | 6464 | 6854 | 6962 | 5103 | 5553 | 6099 | 5378 | 5527 | 5466 | 5481 | 7038 | 7388 | 7990 | 7263 | 7442 | 6853 | 6655 | 5088 | 5358 | 5973 | 5565 | 6661 | 6021 | 5517 | Med22 |
| Med23 | 6382 | 6613 | 5152 | 5313 | 5445 | 5531 | 6117 | 7482 | 6908 | 6645 | 6325 | 7205 | 6666 | 7316 | 6206 | 5794 | 5054 | 5195 | 5584 | 5975 | 5909 | 6898 | 6653 | 6459 | 6908 | 6219 | 7059 | 6894 | Med23 |
| Med24 | 4405 | 5171 | 4856 | 4576 | 4951 | 4682 | 4676 | 6953 | 7719 | 7760 | 6815 | 8771 | 7813 | 8019 | 4350 | 4188 | 4850 | 4615 | 5609 | 4889 | 4964 | 7823 | 6983 | 7965 | 8392 | 8037 | 8003 | 8078 | Med24 |
| Med25 | 7958 | 11159 | 11905 | 11717 | 10110 | 9557 | 9198 | 6056 | 6895 | 7902 | 7309 | 10170 | 7931 | 8520 | 7899 | 9294 | 12381 | 11221 | 11524 | 8435 | 9489 | 7919 | 6768 | 8685 | 8389 | 10444 | 8546 | 8638 | Med25 |
| Med26 | 858 | 1209 | 1038 | 1486 | 980 | 882 | 919 | 679 | 819 | 780 | 823 | 969 | 811 | 819 | 785 | 1052 | 1179 | 1139 | 1020 | 797 | 948 | 819 | 777 | 891 | 906 | 1070 | 795 | 870 | Med26 |
| Med27 | 1593 | 2116 | 1598 | 1907 | 1807 | 1713 | 1649 | 1645 | 1673 | 1511 | 1405 | 1665 | 1500 | 1724 | 1687 | 1569 | 1785 | 1736 | 1821 | 1896 | 1829 | 1517 | 1498 | 1744 | 1718 | 1817 | 1861 | 1750 | Med27 |
| Med28 | 3607 | 3812 | 3540 | 3362 | 3360 | 3048 | 3923 | 3453 | 3306 | 3042 | 3096 | 3383 | 3040 | 3604 | 3454 | 3437 | 3741 | 3350 | 3473 | 4032 | 3752 | 3073 | 3333 | 3461 | 3381 | 3898 | 3483 | 3761 | Med28 |
| Med29 | 881 | 1180 | 972 | 886 | 771 | 769 | 883 | 728 | 744 | 807 | 850 | 908 | 839 | 893 | 882 | 836 | 859 | 817 | 846 | 874 | 787 | 715 | 869 | 896 | 846 | 983 | 836 | 933 | Med29 |
| Med30 | 1534 | 1386 | 1366 | 1437 | 1195 | 1044 | 1433 | 1666 | 1257 | 1324 | 1223 | 1392 | 1347 | 1821 | 1643 | 1349 | 1414 | 1206 | 1252 | 1564 | 1486 | 1247 | 1404 | 1433 | 1373 | 1613 | 1533 | 1765 | Med30 |
| Med31 | 1280 | 1208 | 1499 | 1442 | 1357 | 1350 | 1472 | 1336 | 1231 | 1157 | 1019 | 1402 | 1348 | 1395 | 1428 | 1168 | 1402 | 1357 | 1537 | 1614 | 1357 | 1171 | 1289 | 1362 | 1506 | 1564 | 1479 | 1410 | Med31 |
| Med4 | 3979 | 3987 | 3811 | 3597 | 3016 | 3183 | 3934 | 3314 | 2787 | 2779 | 2707 | 3170 | 2858 | 3878 | 3782 | 3589 | 3715 | 3462 | 3292 | 3852 | 3670 | 2826 | 2806 | 3344 | 3165 | 3733 | 3213 | 3603 | Med4 |
| Med6 | 1804 | 1550 | 1882 | 1401 | 1334 | 1546 | 1589 | 2698 | 2371 | 2159 | 2193 | 2547 | 2459 | 2992 | 2022 | 1611 | 1589 | 1349 | 1541 | 1873 | 1502 | 2252 | 2414 | 2598 | 2329 | 2990 | 2781 | 3229 | Med6 |
| Med7 | 3149 | 2664 | 2791 | 2678 | 2307 | 2292 | 3028 | 2617 | 2183 | 1995 | 1943 | 2673 | 2453 | 2814 | 3236 | 2785 | 2787 | 2497 | 2362 | 2868 | 2679 | 2207 | 2260 | 2275 | 2260 | 2767 | 2574 | 2927 | Med7 |
| Med8 | 3701 | 3978 | 4393 | 3647 | 3196 | 3367 | 3581 | 3543 | 3380 | 3723 | 3458 | 4441 | 4025 | 4413 | 3785 | 3400 | 4004 | 3344 | 3624 | 3850 | 3462 | 3420 | 3494 | 3981 | 3950 | 4999 | 4592 | 4799 | Med8 |
| Med9 | 1262 | 1354 | 1294 | 1085 | 1074 | 1185 | 1019 | 1294 | 1336 | 1279 | 1290 | 1575 | 1514 | 1560 | 1121 | 1148 | 1381 | 1177 | 1243 | 1133 | 1034 | 1230 | 1251 | 1573 | 1350 | 1942 | 1542 | 1481 | Med9 |
| Medag | 45 | 35 | 27 | 63 | 82 | 81 | 93 | 81 | 77 | 79 | 149 | 86 | 100 | 90 | 21 | 32 | 56 | 52 | 86 | 70 | 81 | 84 | 143 | 84 | 115 | 87 | 88 | 81 | Medag |
| Mef2a | 4959 | 3749 | 4439 | 5209 | 6406 | 6211 | 6614 | 7006 | 6840 | 7571 | 8374 | 6607 | 6930 | 7088 | 5308 | 4731 | 4814 | 5905 | 5650 | 5533 | 6302 | 8388 | 7298 | 7089 | 7688 | 5752 | 6298 | 6278 | Mef2a |
| Mef2b | 63 | 100 | 60 | 110 | 65 | 51 | 66 | 39 | 36 | 91 | 59 | 26 | 29 | 26 | 76 | 86 | 110 | 97 | 50 | 30 | 54 | 22 | 52 | 135 | 19 | 79 | 29 | 27 | Mef2b |
| Mef2c | 3483 | 2386 | 1916 | 3591 | 4232 | 4123 | 5323 | 3341 | 4294 | 3264 | 6022 | 2131 | 1752 | 2376 | 3200 | 2773 | 2829 | 4222 | 4018 | 4481 | 5008 | 3228 | 5271 | 3087 | 3392 | 2217 | 1842 | 1926 | Mef2c |
| Mef2d | 5495 | 6253 | 6546 | 9085 | 8241 | 7917 | 7812 | 3926 | 4357 | 5286 | 5728 | 5093 | 4347 | 4688 | 4982 | 6025 | 7801 | 9209 | 8761 | 6135 | 8420 | 5792 | 4394 | 5525 | 5640 | 5115 | 4085 | 4378 | Mef2d |
| Mefv | 0 | 0 | 0 | 0 | 0 | 0 | 2 | 0 | 1 | 0 | 0 | 0 | 0 | 0 | 0 | 0 | 3 | 0 | 0 | 0 | 0 | 0 | 0 | 3 | 0 | 0 | 0 | 0 | Mefv |
| Megf10 | 2100 | 2139 | 2708 | 2354 | 2719 | 2905 | 2477 | 4443 | 4462 | 4368 | 4379 | 5710 | 5280 | 5536 | 2392 | 2158 | 2749 | 2321 | 2802 | 2751 | 2644 | 4408 | 3902 | 4242 | 4865 | 3881 | 5710 | 4991 | Megf10 |
| Megf11 | 634 | 723 | 617 | 857 | 1694 | 1229 | 1305 | 1410 | 1115 | 1279 | 1147 | 960 | 981 | 917 | 949 | 647 | 818 | 1054 | 1311 | 1164 | 1331 | 1679 | 1239 | 983 | 1204 | 488 | 986 | 881 | Megf11 |
| Megf6 | 6484 | 8096 | 7696 | 5772 | 3958 | 5909 | 2856 | 4142 | 6252 | 5338 | 6336 | 4940 | 6771 | 3919 | 6108 | 8833 | 6587 | 6399 | 4690 | 3135 | 2916 | 4279 | 6698 | 5437 | 3987 | 4758 | 5122 | 4584 | Megf6 |
| Megf8 | 9033 | 11815 | 13480 | 13850 | 13917 | 12392 | 10627 | 8145 | 10364 | 11636 | 11611 | 11697 | 10993 | 9290 | 8508 | 10624 | 14551 | 14442 | 14918 | 9874 | 11621 | 12228 | 9655 | 12020 | 11989 | 9431 | 10461 | 9672 | Megf8 |
| Megf9 | 14616 | 12186 | 12856 | 15738 | 16222 | 16550 | 17796 | 11881 | 9750 | 10796 | 9823 | 10597 | 10475 | 12002 | 16230 | 13138 | 13761 | 15431 | 14651 | 15411 | 17379 | 11939 | 9124 | 10233 | 12513 | 8467 | 10758 | 11005 | Megf9 |
| Mei1 | 8 | 14 | 5 | 2 | 8 | 11 | 6 | 0 | 8 | 10 | 11 | 15 | 12 | 11 | 9 | 11 | 10 | 15 | 5 | 1 | 2 | 7 | 8 | 5 | 12 | 7 | 7 | 5 | Mei1 |
| Mei4 | 37 | 33 | 35 | 20 | 43 | 16 | 27 | 64 | 49 | 21 | 34 | 41 | 48 | 55 | 24 | 41 | 13 | 51 | 37 | 36 | 27 | 40 | 56 | 39 | 32 | 43 | 48 | 43 | Mei4 |
| Meig1 | 29 | 46 | 38 | 37 | 33 | 39 | 38 | 21 | 23 | 43 | 17 | 50 | 43 | 31 | 33 | 37 | 48 | 33 | 26 | 72 | 42 | 27 | 45 | 23 | 23 | 21 | 14 | 23 | Meig1 |
| Meikin | 0 | 0 | 2 | 4 | 2 | 0 | 1 | 0 | 0 | 0 | 0 | 0 | 0 | 0 | 1 | 0 | 0 | 1 | 1 | 0 | 1 | 0 | 2 | 0 | 0 | 2 | 0 | 0 | Meikin |
| Meiob | 16 | 7 | 3 | 8 | 0 | 1 | 0 | 2 | 0 | 0 | 1 | 0 | 1 | 4 | 9 | 9 | 3 | 1 | 2 | 0 | 1 | 0 | 3 | 1 | 3 | 8 | 0 | 0 | Meiob |
| Meioc | 324 | 213 | 275 | 177 | 83 | 139 | 65 | 67 | 59 | 42 | 92 | 52 | 65 | 70 | 302 | 478 | 160 | 154 | 66 | 68 | 53 | 70 | 95 | 25 | 22 | 17 | 58 | 19 | Meioc |
| Meiosin | 150 | 108 | 116 | 134 | 233 | 208 | 170 | 203 | 306 | 238 | 295 | 133 | 197 | 177 | 115 | 118 | 110 | 165 | 132 | 153 | 171 | 232 | 267 | 160 | 237 | 206 | 114 | 138 | Meiosin |
| Meis1 | 2637 | 3192 | 3044 | 3816 | 3906 | 3096 | 2381 | 4203 | 4027 | 4228 | 5603 | 2507 | 4817 | 2517 | 3617 | 3225 | 3494 | 2934 | 2470 | 2592 | 2478 | 4284 | 4805 | 2330 | 2641 | 2645 | 2169 | 2265 | Meis1 |
| Meis2 | 8433 | 8635 | 9055 | 13193 | 18559 | 14244 | 15489 | 16963 | 17863 | 20474 | 17411 | 18038 | 18849 | 20515 | 10921 | 8154 | 12327 | 13184 | 15389 | 11349 | 16037 | 19081 | 14445 | 19187 | 20727 | 17647 | 13953 | 16351 | Meis2 |
| Meis3 | 2906 | 3591 | 3759 | 4383 | 4935 | 4027 | 4036 | 3280 | 3376 | 4022 | 3517 | 4298 | 4216 | 4143 | 3212 | 3314 | 4498 | 4324 | 4795 | 3887 | 4237 | 4433 | 3109 | 4329 | 4154 | 3972 | 3973 | 4155 | Meis3 |
| Melk | 4644 | 4204 | 4404 | 3143 | 2219 | 2976 | 3292 | 3634 | 3280 | 2704 | 2611 | 4485 | 3655 | 4531 | 4149 | 3593 | 3883 | 2736 | 3088 | 3773 | 3031 | 2156 | 2831 | 3551 | 2956 | 5318 | 4969 | 4781 | Melk |
| Meltf | 16 | 22 | 22 | 10 | 16 | 29 | 12 | 28 | 151 | 122 | 197 | 18 | 24 | 11 | 20 | 12 | 9 | 24 | 15 | 6 | 10 | 107 | 49 | 62 | 83 | 8 | 8 | 9 | Meltf |

| | | | | | | | | | | | | | | | | | | | | | | | | | | | | | |
|---|---|---|---|---|---|---|---|---|---|---|---|---|---|---|---|---|---|---|---|---|---|---|---|---|---|---|---|---|---|
| Memo1 | 4054 | 3048 | 4132 | 3476 | 3634 | 3780 | 4016 | 4943 | 4048 | 4104 | 4282 | 4694 | 4643 | 5204 | 3972 | 3234 | 3629 | 3206 | 3494 | 3742 | 3971 | 3913 | 4073 | 4581 | 4316 | 5050 | 4974 | 5467 | Memo1 |
| Meox1 | 75 | 43 | 25 | 21 | 25 | 23 | 13 | 121 | 168 | 91 | 809 | 42 | 100 | 39 | 61 | 86 | 19 | 20 | 19 | 14 | 21 | 175 | 419 | 105 | 43 | 71 | 36 | 39 | Meox1 |
| Meox2 | 200 | 101 | 136 | 75 | 88 | 183 | 63 | 231 | 712 | 298 | 1223 | 93 | 344 | 54 | 183 | 252 | 82 | 134 | 122 | 95 | 47 | 541 | 1325 | 154 | 251 | 118 | 63 | 81 | Meox2 |
| Mep1a | 1 | 1 | 2 | 1 | 2 | 0 | 2 | 7 | 6 | 3 | 18 | 5 | 0 | 4 | 3 | 5 | 2 | 0 | 3 | 8 | 3 | 5 | 10 | 2 | 2 | 8 | 2 | 5 | Mep1a |
| Mep1b | 0 | 0 | 0 | 0 | 0 | 0 | 0 | 1 | 1 | 2 | 1 | 0 | 1 | 1 | 0 | 0 | 0 | 0 | 0 | 0 | 0 | 0 | 3 | 0 | 0 | 0 | 1 | 0 | Mep1b |
| Mepce | 1802 | 2275 | 2636 | 2622 | 2568 | 2194 | 2319 | 3072 | 3142 | 3593 | 3382 | 3688 | 3276 | 3640 | 1725 | 1916 | 2626 | 2619 | 2862 | 2120 | 2406 | 3061 | 3216 | 3825 | 3489 | 4303 | 3590 | 3650 | Mepce |
| Mertk | 1481 | 1210 | 1281 | 872 | 749 | 905 | 640 | 920 | 1065 | 1039 | 1370 | 741 | 1103 | 690 | 1180 | 1712 | 927 | 939 | 720 | 609 | 624 | 847 | 1127 | 902 | 616 | 1142 | 766 | 749 | Mertk |
| Mesd | 4049 | 3981 | 3954 | 3327 | 2981 | 2961 | 3279 | 3301 | 3075 | 2939 | 3144 | 3329 | 3339 | 3150 | 3981 | 3965 | 3337 | 3211 | 3183 | 3217 | 3129 | 2924 | 3298 | 3063 | 2883 | 3613 | 3498 | 3437 | Mesd |
| Mesp1 | 0 | 1 | 0 | 6 | 0 | 0 | 0 | 0 | 0 | 1 | 1 | 0 | 2 | 2 | 1 | 0 | 1 | 0 | 2 | 3 | 0 | 2 | 0 | 0 | 0 | 2 | 2 | 0 | Mesp1 |
| Mesp2 | 12 | 13 | 22 | 8 | 17 | 25 | 13 | 32 | 24 | 13 | 21 | 24 | 17 | 16 | 8 | 11 | 24 | 10 | 20 | 15 | 14 | 15 | 33 | 26 | 23 | 28 | 20 | 27 | Mesp2 |
| Mest | 44731 | 38352 | 39360 | 23165 | 20663 | 24268 | 24274 | 37758 | 43055 | 31409 | 58759 | 42525 | 41012 | 42737 | 42660 | 43919 | 31333 | 23895 | 20768 | 24579 | 20182 | 36755 | 52323 | 34602 | 30694 | 57390 | 42467 | 45623 | Mest |
| Met | 912 | 622 | 533 | 426 | 637 | 542 | 453 | 625 | 756 | 565 | 914 | 338 | 502 | 349 | 789 | 741 | 457 | 474 | 500 | 504 | 518 | 388 | 884 | 420 | 262 | 232 | 396 | 362 | Met |
| Metap1 | 7029 | 6664 | 5762 | 6255 | 5760 | 6036 | 6067 | 5324 | 4501 | 4252 | 4433 | 4316 | 4368 | 4240 | 6910 | 6798 | 5759 | 6209 | 5791 | 6689 | 6139 | 4395 | 4482 | 4144 | 4313 | 4294 | 4564 | 4474 | Metap1 |
| Metap1d | 2530 | 2097 | 1701 | 1838 | 1891 | 2184 | 2338 | 3482 | 3178 | 2820 | 2838 | 2774 | 2530 | 3195 | 2458 | 2074 | 1971 | 2101 | 2067 | 2445 | 2344 | 2640 | 3029 | 2567 | 2669 | 2480 | 2818 | 2777 | Metap1d |
| Metap2 | 17928 | 15132 | 17761 | 14929 | 14058 | 14713 | 16054 | 17514 | 15451 | 15823 | 14328 | 16937 | 16071 | 17263 | 18609 | 15595 | 16120 | 14175 | 14908 | 17089 | 15166 | 14646 | 15297 | 15923 | 15455 | 18772 | 17242 | 18111 | Metap2 |
| Methig1 | 0 | 0 | 0 | 0 | 0 | 0 | 0 | 2 | 4 | 0 | 3 | 0 | 3 | 0 | 0 | 1 | 0 | 3 | 0 | 0 | 0 | 2 | 0 | 0 | 0 | 0 | 0 | 0 | Methig1 |
| Metrn | 6245 | 8231 | 7571 | 6507 | 3845 | 4173 | 4646 | 2191 | 2168 | 2064 | 1845 | 3276 | 2536 | 3181 | 5798 | 5828 | 7550 | 4787 | 4705 | 4663 | 4494 | 1815 | 1723 | 2301 | 2117 | 3580 | 3277 | 3449 | Metrn |
| Metrnl | 245 | 371 | 308 | 232 | 251 | 247 | 156 | 221 | 282 | 232 | 290 | 156 | 353 | 98 | 310 | 308 | 292 | 224 | 205 | 242 | 166 | 184 | 317 | 191 | 140 | 156 | 205 | 141 | Metrnl |
| Mettl1 | 1218 | 1476 | 1609 | 1310 | 1130 | 1168 | 1225 | 1015 | 1346 | 1179 | 1399 | 1366 | 1249 | 1361 | 1124 | 1423 | 1483 | 1294 | 1128 | 1142 | 1067 | 941 | 1274 | 1309 | 1157 | 1531 | 1383 | 1451 | Mettl1 |
| Mettl11b | 48 | 16 | 31 | 5 | 14 | 14 | 19 | 42 | 65 | 18 | 29 | 55 | 76 | 62 | 54 | 19 | 10 | 9 | 10 | 17 | 10 | 27 | 54 | 25 | 54 | 38 | 42 | 55 | Mettl11b |
| Mettl14 | 3178 | 2881 | 3580 | 3031 | 2785 | 2633 | 3108 | 3440 | 3204 | 3229 | 3066 | 3976 | 3677 | 4519 | 2937 | 2829 | 3420 | 2788 | 3026 | 3092 | 2985 | 3130 | 3181 | 3779 | 3616 | 4495 | 4093 | 4308 | Mettl14 |
| Mettl15 | 257 | 211 | 227 | 183 | 156 | 177 | 202 | 360 | 343 | 251 | 300 | 329 | 290 | 365 | 194 | 207 | 197 | 175 | 207 | 251 | 150 | 234 | 310 | 337 | 308 | 343 | 337 | 347 | Mettl15 |
| Mettl16 | 3517 | 3606 | 3517 | 3266 | 3534 | 3742 | 3546 | 5797 | 5206 | 4678 | 5006 | 5330 | 4782 | 5355 | 3538 | 3565 | 3501 | 3404 | 3390 | 3806 | 3639 | 5095 | 4944 | 4850 | 5301 | 4964 | 5294 | 5353 | Mettl16 |
| Mettl17 | 3081 | 3337 | 2867 | 3584 | 3313 | 3003 | 3208 | 2506 | 2384 | 2430 | 2336 | 2447 | 2184 | 2485 | 2774 | 2824 | 2967 | 3375 | 3296 | 3345 | 3197 | 2669 | 2320 | 2537 | 2635 | 2196 | 2243 | 2158 | Mettl17 |
| Mettl18 | 475 | 375 | 388 | 438 | 444 | 432 | 503 | 491 | 381 | 363 | 428 | 401 | 386 | 421 | 479 | 406 | 435 | 368 | 469 | 578 | 441 | 322 | 430 | 412 | 440 | 476 | 393 | 425 | Mettl18 |
| Mettl2 | 2584 | 2393 | 2631 | 2315 | 1830 | 2147 | 2107 | 2140 | 2146 | 2117 | 2310 | 2304 | 2383 | 2456 | 2329 | 2410 | 2380 | 2188 | 2067 | 1997 | 2104 | 2102 | 2135 | 2302 | 2157 | 2794 | 2700 | 2451 | Mettl2 |
| Mettl21a | 652 | 612 | 919 | 691 | 747 | 633 | 650 | 998 | 1035 | 1075 | 1023 | 1083 | 1034 | 1153 | 799 | 728 | 628 | 749 | 695 | 697 | 661 | 993 | 1087 | 1119 | 982 | 1242 | 1052 | 1123 | Mettl21a |
| Mettl21c | 27 | 16 | 25 | 20 | 22 | 39 | 44 | 131 | 90 | 65 | 83 | 91 | 75 | 79 | 38 | 25 | 15 | 24 | 23 | 31 | 40 | 95 | 72 | 65 | 68 | 63 | 62 | 78 | Mettl21c |
| Mettl21e | 0 | 0 | 0 | 0 | 3 | 0 | 1 | 0 | 1 | 3 | 3 | 0 | 1 | 0 | 0 | 2 | 0 | 0 | 1 | 0 | 0 | 2 | 2 | 1 | 0 | 0 | 0 | 1 | Mettl21e |
| Mettl22 | 254 | 339 | 280 | 247 | 417 | 403 | 306 | 590 | 758 | 672 | 671 | 678 | 640 | 650 | 282 | 275 | 275 | 296 | 333 | 466 | 331 | 693 | 761 | 659 | 844 | 617 | 753 | 619 | Mettl22 |
| Mettl23 | 2283 | 2194 | 1804 | 1966 | 1758 | 2083 | 2297 | 2081 | 1865 | 1764 | 1546 | 1782 | 1782 | 1950 | 2350 | 1819 | 2100 | 1931 | 2287 | 2296 | 2380 | 1865 | 1878 | 1870 | 1828 | 1909 | 1855 | 2013 | Mettl23 |
| Mettl24 | 3 | 8 | 11 | 6 | 5 | 5 | 6 | 20 | 17 | 23 | 37 | 17 | 22 | 20 | 4 | 9 | 6 | 4 | 2 | 1 | 7 | 15 | 19 | 17 | 16 | 12 | 3 | 14 | Mettl24 |
| Mettl25 | 218 | 233 | 229 | 308 | 298 | 255 | 310 | 760 | 741 | 702 | 657 | 860 | 637 | 847 | 265 | 184 | 292 | 244 | 284 | 286 | 288 | 774 | 688 | 760 | 739 | 831 | 722 | 725 | Mettl25 |
| Mettl26 | 916 | 1308 | 1199 | 1051 | 950 | 935 | 964 | 939 | 966 | 917 | 830 | 1120 | 1082 | 1122 | 1021 | 1079 | 1218 | 910 | 1044 | 1240 | 980 | 932 | 962 | 1058 | 982 | 1173 | 1119 | 1054 | Mettl26 |
| Mettl27 | 1129 | 1215 | 1234 | 1180 | 806 | 1171 | 882 | 410 | 479 | 517 | 487 | 426 | 539 | 490 | 1077 | 1208 | 1394 | 1106 | 895 | 892 | 803 | 356 | 493 | 471 | 344 | 634 | 470 | 460 | Mettl27 |
| Mettl3 | 3756 | 3924 | 4101 | 3974 | 3334 | 3207 | 3613 | 3616 | 3262 | 3423 | 3145 | 3647 | 3200 | 3625 | 3744 | 3465 | 3955 | 3561 | 3772 | 3613 | 3531 | 3546 | 3221 | 3637 | 3397 | 3768 | 3822 | 3789 | Mettl3 |
| Mettl4 | 2084 | 1895 | 2015 | 1958 | 2077 | 2076 | 2377 | 5127 | 4390 | 4118 | 3769 | 4355 | 4008 | 4443 | 2382 | 1927 | 2001 | 1914 | 2198 | 2534 | 2233 | 4029 | 4138 | 3983 | 4249 | 3667 | 4341 | 4498 | Mettl4 |
| Mettl5 | 859 | 724 | 861 | 845 | 884 | 827 | 825 | 724 | 613 | 683 | 644 | 857 | 805 | 895 | 773 | 674 | 926 | 732 | 780 | 780 | 862 | 708 | 671 | 771 | 610 | 792 | 891 | 902 | Mettl5 |
| Mettl6 | 5394 | 4919 | 4844 | 4720 | 3895 | 4319 | 4598 | 4019 | 3459 | 3137 | 3315 | 3709 | 3708 | 3898 | 5351 | 4992 | 4495 | 4258 | 4215 | 4804 | 4593 | 3255 | 3348 | 3575 | 3480 | 4029 | 3747 | 4297 | Mettl6 |
| Mettl7a1 | 721 | 798 | 979 | 771 | 731 | 629 | 656 | 723 | 679 | 741 | 786 | 720 | 692 | 810 | 676 | 766 | 953 | 786 | 803 | 756 | 726 | 643 | 737 | 876 | 722 | 894 | 838 | 885 | Mettl7a1 |
| Mettl7a2 | 0 | 1 | 0 | 0 | 0 | 0 | 0 | 0 | 0 | 0 | 0 | 0 | 0 | 0 | 0 | 0 | 0 | 0 | 0 | 0 | 0 | 0 | 0 | 0 | 0 | 0 | 0 | 0 | Mettl7a2 |
| Mettl7a3 | 5 | 13 | 4 | 9 | 20 | 0 | 13 | 15 | 9 | 8 | 7 | 2 | 10 | 1 | 9 | 11 | 6 | 19 | 7 | 5 | 5 | 28 | 12 | 2 | 11 | 3 | 3 | 1 | Mettl7a3 |
| Mettl7b | 0 | 0 | 0 | 1 | 0 | 0 | 1 | 0 | 2 | 2 | 0 | 0 | 0 | 0 | 0 | 2 | 0 | 0 | 0 | 0 | 0 | 0 | 0 | 1 | 0 | 0 | 0 | 0 | Mettl7b |
| Mettl8 | 2412 | 2363 | 2115 | 2231 | 1828 | 2037 | 2358 | 1616 | 1462 | 1384 | 1329 | 1555 | 1552 | 1431 | 2614 | 2163 | 2249 | 2025 | 2145 | 2169 | 2211 | 1322 | 1320 | 1302 | 1319 | 1233 | 1633 | 1470 | Mettl8 |
| Mettl9 | 7041 | 5984 | 7421 | 6664 | 7055 | 6435 | 8507 | 9733 | 8363 | 9426 | 8729 | 10510 | 9237 | 11933 | 7373 | 6083 | 7226 | 6531 | 7357 | 6425 | 7866 | 9845 | 8226 | 10321 | 9377 | 11850 | 10046 | 11743 | Mettl9 |
| Mex3a | 13004 | 12645 | 19171 | 17319 | 19945 | 17883 | 18020 | 25350 | 23628 | 29211 | 27663 | 34989 | 29659 | 36638 | 12398 | 11270 | 19586 | 16505 | 21404 | 14506 | 18587 | 31697 | 21575 | 35789 | 36351 | 34217 | 32065 | 36566 | Mex3a |
| Mex3b | 4455 | 3982 | 5098 | 4331 | 5710 | 5368 | 5674 | 10481 | 9887 | 9396 | 9186 | 11119 | 10155 | 11912 | 4134 | 3734 | 5004 | 4401 | 6198 | 5021 | 5605 | 11070 | 9193 | 11664 | 11513 | 11919 | 10315 | 11552 | Mex3b |
| Mex3c | 8187 | 6754 | 7744 | 7035 | 7171 | 7201 | 7818 | 8971 | 7314 | 6964 | 6527 | 8025 | 7449 | 8275 | 7956 | 7211 | 7646 | 6837 | 7594 | 7661 | 7662 | 7957 | 6987 | 7405 | 7268 | 8868 | 7977 | 8535 | Mex3c |
| Mex3d | 3432 | 4473 | 5026 | 4959 | 4588 | 3884 | 4121 | 4082 | 4082 | 5155 | 4636 | 5440 | 4487 | 5950 | 3353 | 3445 | 5872 | 4543 | 4988 | 3569 | 4590 | 5138 | 3983 | 5815 | 5737 | 5709 | 5291 | 5765 | Mex3d |
| Mfap1a | 4056 | 3558 | 3974 | 3917 | 3301 | 3589 | 3902 | 3145 | 2730 | 2935 | 2800 | 2899 | 2893 | 3193 | 4147 | 3930 | 4081 | 3919 | 3506 | 4138 | 3603 | 3026 | 2807 | 2854 | 2908 | 3282 | 3073 | 3280 | Mfap1a |
| Mfap1b | 2299 | 2168 | 2122 | 2157 | 1763 | 1942 | 2225 | 1828 | 1490 | 1762 | 1705 | 1538 | 1533 | 1833 | 2405 | 2174 | 2147 | 2139 | 1883 | 2366 | 2154 | 1609 | 1604 | 1779 | 1808 | 1709 | 1611 | 1987 | Mfap1b |
| Mfap2 | 4878 | 5950 | 5157 | 4271 | 3784 | 4450 | 2997 | 5996 | 8304 | 6495 | 8126 | 6800 | 7935 | 5660 | 4543 | 6160 | 4553 | 4636 | 4044 | 4030 | 2967 | 5335 | 8480 | 7016 | 6363 | 8613 | 7019 | 6012 | Mfap2 |
| Mfap3 | 11858 | 11463 | 8952 | 9654 | 9406 | 10568 | 10661 | 9573 | 8244 | 7398 | 7339 | 7844 | 7753 | 8444 | 12306 | 10711 | 9463 | 9872 | 9718 | 11453 | 10554 | 8298 | 7827 | 7072 | 7998 | 6718 | 8417 | 8330 | Mfap3 |
| Mfap3l | 1448 | 1270 | 1089 | 1471 | 1553 | 1393 | 1503 | 780 | 689 | 503 | 747 | 489 | 662 | 516 | 1519 | 1579 | 1359 | 1536 | 1279 | 1532 | 1392 | 737 | 686 | 418 | 544 | 428 | 539 | 464 | Mfap3l |
| Mfap4 | 4437 | 4495 | 2853 | 2300 | 1712 | 2841 | 1234 | 5628 | 7456 | 5498 | 8082 | 4441 | 6442 | 4274 | 4013 | 5555 | 2687 | 3125 | 1668 | 1687 | 1489 | 4588 | 8732 | 5794 | 4343 | 6546 | 5116 | 4756 | Mfap4 |
| Mfap5 | 22 | 36 | 51 | 27 | 23 | 29 | 13 | 56 | 73 | 49 | 216 | 42 | 59 | 43 | 26 | 37 | 10 | 50 | 20 | 23 | 25 | 65 | 208 | 71 | 59 | 33 | 44 | 31 | Mfap5 |
| Mff | 11303 | 9671 | 9192 | 9916 | 10792 | 10919 | 12643 | 14175 | 12640 | 10701 | 10603 | 10595 | 10404 | 11506 | 12573 | 9963 | 9619 | 9717 | 10472 | 13211 | 11966 | 11911 | 11981 | 10573 | 11582 | 10056 | 10836 | 11177 | Mff |
| Mfge8 | 3850 | 4642 | 4056 | 3451 | 3210 | 3264 | 2820 | 2102 | 3306 | 2482 | 2655 | 2060 | 2335 | 2093 | 3494 | 4343 | 3569 | 3412 | 3249 | 2930 | 3073 | 2241 | 3600 | 2388 | 2310 | 2681 | 2401 | 2162 | Mfge8 |
| Mfhas1 | 2876 | 3119 | 3861 | 3778 | 4516 | 3833 | 3109 | 2426 | 2615 | 3006 | 3264 | 2687 | 3153 | 2598 | 2926 | 3670 | 3756 | 3911 | 3809 | 2785 | 3507 | 3613 | 2730 | 2760 | 3146 | 2201 | 2582 | 2918 | Mfhas1 |
| Mfn1 | 7270 | 6400 | 7611 | 7682 | 7888 | 7362 | 8106 | 6084 | 5501 | 6129 | 6031 | 6120 | 6178 | 6373 | 6873 | 7038 | 7435 | 7748 | 7398 | 6796 | 7686 | 6734 | 5460 | 5880 | 6532 | 6134 | 6080 | 6019 | Mfn1 |
| Mfn2 | 5373 | 6025 | 6537 | 6703 | 7400 | 6729 | 6514 | 6765 | 7375 | 8239 | 7530 | 8042 | 7643 | 7691 | 5482 | 5132 | 6408 | 6749 | 7374 | 6008 | 7116 | 9043 | 6851 | 8040 | 8851 | 7000 | 7541 | 7452 | Mfn2 |
| Mfng | 1536 | 2132 | 2090 | 1841 | 1754 | 1738 | 1899 | 3416 | 3014 | 3624 | 2698 | 6084 | 4374 | 5658 | 1603 | 1284 | 2454 | 1443 | 2358 | 1904 | 1764 | 2911 | 2721 | 4944 | 4376 | 6681 | 6178 | 6043 | Mfng |
| Mfrp | 39 | 45 | 25 | 35 | 101 | 50 | 21 | 59 | 101 | 88 | 47 | 105 | 78 | 27 | 22 | 64 | 6 | 40 | 42 | 32 | 35 | 140 | 205 | 29 | 45 | 19 | 29 | 25 | Mfrp |
| Mfsd1 | 7726 | 7264 | 8470 | 6719 | 5640 | 6149 | 5686 | 4269 | 4088 | 3789 | 3967 | 4186 | 4661 | 4146 | 7751 | 7739 | 7243 | 6804 | 5970 | 6297 | 5929 | 3905 | 4084 | 4106 | 3791 | 4723 | 4503 | 4689 | Mfsd1 |
| Mfsd10 | 910 | 1086 | 1184 | 867 | 861 | 900 | 742 | 1478 | 1862 | 1700 | 2015 | 1883 | 1954 | 1785 | 793 | 1033 | 912 | 810 | 958 | 817 | 779 | 1587 | 1903 | 1964 | 1823 | 2293 | 1927 | 1807 | Mfsd10 |
| Mfsd11 | 2767 | 3182 | 2505 | 3078 | 2887 | 2781 | 2738 | 1980 | 1963 | 1981 | 2078 | 1951 | 1930 | 1888 | 2888 | 3157 | 2837 | 3045 | 2773 | 2669 | 2894 | 2152 | 1999 | 1867 | 2068 | 1773 | 1766 | 1829 | Mfsd11 |
| Mfsd12 | 3585 | 5095 | 5434 | 5176 | 3804 | 3741 | 2889 | 922 | 1046 | 1043 | 1251 | 954 | 964 | 868 | 3384 | 4357 | 4880 | 4930 | 4018 | 3310 | 3387 | 1006 | 1198 | 1155 | 1064 | 1067 | 997 | 863 | Mfsd12 |
| Mfsd13a | 456 | 666 | 743 | 651 | 664 | 519 | 572 | 230 | 305 | 369 | 336 | 336 | 304 | 351 | 489 | 582 | 712 | 741 | 676 | 543 | 584 | 301 | 233 | 365 | 360 | 469 | 320 | 336 | Mfsd13a |
| Mfsd13b | 14 | 27 | 26 | 35 | 18 | 22 | 27 | 18 | 33 | 15 | 21 | 23 | 15 | 15 | 50 | 38 | 21 | 24 | 19 | 30 | 45 | 25 | 27 | 9 | 18 | 24 | 26 | 18 | Mfsd13b |
| Mfsd14a | 3895 | 3233 | 3211 | 3447 | 3080 | 3291 | 3660 | 4357 | 3662 | 3484 | 3353 | 3750 | 3436 | 3942 | 3997 | 3421 | 3082 | 3259 | 3091 | 3680 | 3489 | 3676 | 3522 | 3235 | 3514 | 4000 | 3631 | 4133 | Mfsd14a |
| Mfsd14b | 5952 | 4785 | 4967 | 4815 | 4250 | 4731 | 4966 | 4113 | 3754 | 3300 | 3892 | 3760 | 4020 | 3651 | 5889 | 5322 | 4661 | 4755 | 4116 | 5258 | 4405 | 3452 | 3904 | 3535 | 3256 | 4062 | 3801 | 3718 | Mfsd14b |
| Mfsd2a | 1134 | 1659 | 1166 | 1377 | 1546 | 1434 | 1463 | 1242 | 1148 | 1218 | 911 | 1149 | 1272 | 1247 | 1360 | 1294 | 1359 | 1393 | 1493 | 1508 | 1394 | 1233 | 1065 | 1085 | 1335 | 1098 | 1187 | 1327 | Mfsd2a |
| Mfsd2b | 427 | 518 | 474 | 671 | 747 | 664 | 674 | 321 | 384 | 416 | 344 | 389 | 552 | 384 | 468 | 558 | 557 | 820 | 685 | 589 | 642 | 458 | 315 | 360 | 396 | 365 | 457 | 504 | Mfsd2b |
| Mfsd3 | 242 | 346 | 363 | 347 | 346 | 273 | 320 | 193 | 244 | 212 | 244 | 204 | 263 | 205 | 216 | 285 | 370 | 329 | 339 | 311 | 299 | 239 | 244 | 226 | 204 | 222 | 200 | 225 | Mfsd3 |
| Mfsd4a | 21 | 10 | 26 | 21 | 18 | 10 | 26 | 59 | 56 | 67 | 69 | 21 | 27 | 23 | 9 | 12 | 9 | 21 | 28 | 10 | 11 | 53 | 57 | 38 | 35 | 45 | 28 | 9 | Mfsd4a |
| Mfsd4b1 | 28 | 16 | 27 | 27 | 23 | 11 | 19 | 22 | 19 | 22 | 11 | 15 | 8 | 23 | 36 | 24 | 22 | 29 | 31 | 29 | 16 | 15 | 39 | 5 | 20 | 8 | 12 | 2 | Mfsd4b1 |
| Mfsd4b4 | 1103 | 891 | 956 | 967 | 1006 | 926 | 955 | 1319 | 1434 | 1268 | 1306 | 1352 | 1479 | 1301 | 996 | 1001 | 1096 | 825 | 938 | 955 | 909 | 1205 | 1218 | 1226 | 1224 | 1019 | 1362 | 1201 | Mfsd4b4 |
| Mfsd4b5 | 7 | 8 | 1 | 6 | 5 | 6 | 9 | 5 | 0 | 3 | 3 | 5 | 1 | 4 | 4 | 9 | 10 | 6 | 9 | 7 | 4 | 3 | 0 | 2 | 0 | 0 | 0 | 1 | Mfsd4b5 |
| Mfsd5 | 1223 | 1456 | 1409 | 1145 | 1158 | 1205 | 1001 | 1130 | 1326 | 1210 | 1244 | 1252 | 1220 | 1170 | 1094 | 1403 | 1224 | 1279 | 1198 | 1151 | 1036 | 1131 | 1357 | 1220 | 1166 | 1364 | 1225 | 1284 | Mfsd5 |

| | | | | | | | | | | | | | | | | | | | | | | | | | | | | | |
|---|---|---|---|---|---|---|---|---|---|---|---|---|---|---|---|---|---|---|---|---|---|---|---|---|---|---|---|---|---|
| Mfsd6 | 399 | 466 | 415 | 531 | 893 | 570 | 592 | 1843 | 1576 | 1856 | 1605 | 1438 | 1477 | 1683 | 481 | 421 | 445 | 465 | 708 | 532 | 593 | 2242 | 1489 | 1567 | 1926 | 1071 | 1363 | 1424 | Mfsd6 |
| Mfsd6l | 9 | 12 | 5 | 2 | 11 | 10 | 7 | 5 | 13 | 10 | 22 | 5 | 9 | 7 | 7 | 14 | 10 | 12 | 4 | 2 | 4 | 7 | 7 | 17 | 18 | 14 | 5 | 1 | Mfsd6l |
| Mfsd7a | 51 | 54 | 54 | 64 | 47 | 43 | 42 | 78 | 82 | 93 | 151 | 67 | 93 | 93 | 80 | 66 | 60 | 53 | 44 | 51 | 31 | 95 | 117 | 75 | 74 | 71 | 106 | 81 | Mfsd7a |
| Mfsd8 | 24 | 11 | 27 | 48 | 23 | 106 | 67 | 142 | 62 | 78 | 126 | 47 | 174 | 143 | 32 | 44 | 20 | 17 | 29 | 83 | 38 | 254 | 124 | 91 | 128 | 45 | 232 | 108 | Mfsd8 |
| Mfsd9 | 1029 | 1127 | 786 | 881 | 994 | 943 | 962 | 335 | 363 | 310 | 381 | 262 | 315 | 248 | 1063 | 948 | 918 | 1016 | 818 | 1074 | 891 | 343 | 445 | 284 | 320 | 273 | 249 | 280 | Mfsd9 |
| Mga | 18457 | 15682 | 9040 | 10959 | 12195 | 15864 | 14871 | 18809 | 16396 | 12169 | 13918 | 12133 | 12989 | 13227 | 18515 | 16047 | 9441 | 10668 | 11161 | 17414 | 15296 | 10923 | 15659 | 9854 | 13530 | 7683 | 14435 | 11811 | Mga |
| Mgam | 5 | 20 | 10 | 11 | 11 | 9 | 10 | 17 | 18 | 18 | 35 | 12 | 2 | 9 | 20 | 15 | 1 | 11 | 11 | 10 | 14 | 5 | 38 | 2 | 17 | 2 | 17 | 13 | Mgam |
| Mgam2-ps | 0 | 1 | 0 | 0 | 0 | 0 | 2 | 1 | 4 | 0 | 0 | 0 | 0 | 0 | 0 | 0 | 0 | 0 | 0 | 0 | 2 | 0 | 2 | 0 | 0 | 0 | 0 | 4 | Mgam2-ps |
| Mgarp | 460 | 435 | 651 | 326 | 180 | 320 | 140 | 125 | 129 | 102 | 150 | 267 | 262 | 250 | 340 | 576 | 429 | 350 | 239 | 263 | 204 | 94 | 140 | 167 | 139 | 102 | 344 | 325 | Mgarp |
| Mgat1 | 2285 | 2917 | 2877 | 2819 | 2517 | 2723 | 2187 | 2800 | 3642 | 3473 | 3954 | 3720 | 3758 | 3475 | 2288 | 2673 | 2701 | 2824 | 2624 | 2219 | 2238 | 3369 | 3677 | 3634 | 3572 | 3898 | 3664 | 3415 | Mgat1 |
| Mgat2 | 3887 | 3266 | 3023 | 2946 | 2588 | 3034 | 2842 | 3146 | 2808 | 2437 | 2794 | 2633 | 2756 | 2540 | 3655 | 3340 | 2640 | 3051 | 2598 | 2923 | 2731 | 2951 | 3228 | 2468 | 2618 | 2717 | 2755 | 2788 | Mgat2 |
| Mgat3 | 3071 | 2946 | 3937 | 3647 | 5072 | 3959 | 3674 | 5451 | 5330 | 6766 | 6762 | 6130 | 6052 | 5912 | 3033 | 3093 | 4203 | 3746 | 4458 | 3316 | 3816 | 7770 | 5198 | 6320 | 6864 | 5667 | 5328 | 5811 | Mgat3 |
| Mgat4a | 2255 | 1844 | 1446 | 1979 | 2740 | 2417 | 2494 | 1893 | 1894 | 1831 | 2034 | 1254 | 1305 | 1377 | 2443 | 2048 | 1603 | 2217 | 2216 | 2280 | 2308 | 1962 | 1988 | 1447 | 1770 | 998 | 1288 | 1194 | Mgat4a |
| Mgat4b | 3305 | 3634 | 3932 | 4230 | 4928 | 3921 | 3772 | 3178 | 3425 | 4116 | 3884 | 3617 | 3141 | 3258 | 3189 | 3323 | 4457 | 4278 | 4542 | 3204 | 4222 | 4396 | 3370 | 3912 | 4080 | 4268 | 3148 | 3222 | Mgat4b |
| Mgat4c | 280 | 257 | 216 | 466 | 779 | 561 | 541 | 677 | 567 | 579 | 608 | 299 | 482 | 374 | 518 | 350 | 393 | 527 | 486 | 664 | 574 | 832 | 623 | 375 | 662 | 235 | 315 | 354 | Mgat4c |
| Mgat4d | 0 | 1 | 0 | 0 | 0 | 0 | 0 | 4 | 0 | 0 | 0 | 0 | 0 | 0 | 0 | 0 | 0 | 0 | 0 | 0 | 0 | 0 | 0 | 0 | 0 | 0 | 0 | 0 | Mgat4d |
| Mgat4e | 0 | 1 | 0 | 0 | 3 | 0 | 2 | 5 | 2 | 10 | 2 | 2 | 0 | 1 | 3 | 2 | 7 | 2 | 0 | 2 | 4 | 5 | 4 | 4 | 3 | 3 | 2 | 2 | Mgat4e |
| Mgat4f | 1 | 0 | 0 | 2 | 0 | 0 | 1 | 1 | 0 | 1 | 0 | 0 | 0 | 0 | 0 | 1 | 0 | 0 | 0 | 0 | 0 | 0 | 0 | 2 | 0 | 1 | 0 | 1 | Mgat4f |
| Mgat5 | 5413 | 5444 | 5471 | 7278 | 8852 | 7968 | 7672 | 5506 | 5782 | 6774 | 6130 | 5839 | 5781 | 5684 | 5740 | 5503 | 6601 | 7789 | 8286 | 7013 | 8018 | 7481 | 5203 | 5911 | 7526 | 4562 | 5691 | 5715 | Mgat5 |
| Mgat5b | 1177 | 1764 | 1982 | 2753 | 4206 | 3057 | 2811 | 3495 | 4500 | 5763 | 4573 | 5229 | 3882 | 4342 | 1403 | 1279 | 2729 | 3098 | 3952 | 2568 | 3095 | 5765 | 3720 | 5216 | 6401 | 4438 | 4005 | 3993 | Mgat5b |
| Mgl2 | 8 | 10 | 8 | 15 | 7 | 26 | 11 | 20 | 14 | 14 | 27 | 22 | 8 | 7 | 12 | 14 | 10 | 22 | 16 | 12 | 8 | 27 | 56 | 19 | 13 | 14 | 15 | 12 | Mgl2 |
| Mgll | 305 | 329 | 240 | 358 | 539 | 392 | 477 | 331 | 615 | 542 | 692 | 237 | 269 | 268 | 282 | 379 | 316 | 533 | 593 | 411 | 493 | 488 | 653 | 425 | 520 | 289 | 216 | 204 | Mgll |
| Mgme1 | 443 | 478 | 603 | 496 | 394 | 364 | 428 | 790 | 896 | 886 | 840 | 971 | 934 | 990 | 451 | 497 | 553 | 419 | 458 | 450 | 406 | 660 | 876 | 891 | 857 | 1086 | 1004 | 1015 | Mgme1 |
| Mgmt | 189 | 213 | 142 | 139 | 97 | 127 | 69 | 220 | 227 | 181 | 200 | 178 | 203 | 138 | 164 | 213 | 123 | 157 | 98 | 116 | 73 | 140 | 269 | 163 | 157 | 238 | 192 | 150 | Mgmt |
| Mgp | 81 | 85 | 36 | 96 | 38 | 124 | 18 | 291 | 538 | 543 | 676 | 148 | 179 | 151 | 104 | 147 | 66 | 164 | 25 | 21 | 13 | 364 | 646 | 391 | 304 | 181 | 98 | 122 | Mgp |
| Mgrn1 | 8586 | 10237 | 7968 | 8999 | 9607 | 9676 | 8480 | 6299 | 7795 | 7614 | 7241 | 7723 | 7324 | 6542 | 8068 | 9511 | 8785 | 9192 | 9552 | 8551 | 8759 | 8039 | 7522 | 6956 | 8030 | 6730 | 7234 | 6591 | Mgrn1 |
| Mgst1 | 991 | 777 | 859 | 644 | 812 | 752 | 800 | 1025 | 924 | 800 | 947 | 582 | 788 | 648 | 1135 | 1000 | 720 | 699 | 713 | 931 | 733 | 854 | 1186 | 624 | 615 | 755 | 572 | 685 | Mgst1 |
| Mgst2 | 12 | 6 | 11 | 35 | 12 | 5 | 7 | 12 | 35 | 6 | 68 | 9 | 12 | 15 | 7 | 19 | 13 | 24 | 6 | 8 | 16 | 23 | 20 | 17 | 4 | 11 | 17 | 9 | Mgst2 |
| Mgst3 | 1523 | 2249 | 1711 | 2114 | 1973 | 1762 | 1780 | 1252 | 1441 | 1277 | 1160 | 912 | 1215 | 1006 | 1816 | 2093 | 1861 | 2051 | 2017 | 2358 | 1866 | 1123 | 1425 | 1109 | 1239 | 1131 | 1137 | 1199 | Mgst3 |
| Mia | 10 | 13 | 10 | 13 | 20 | 12 | 16 | 24 | 11 | 19 | 36 | 14 | 10 | 16 | 12 | 10 | 23 | 21 | 9 | 7 | 18 | 43 | 12 | 25 | 18 | 19 | 12 | 5 | Mia |
| Mia2 | 5463 | 4958 | 4666 | 5279 | 4532 | 5040 | 5417 | 4822 | 4444 | 4156 | 4641 | 4737 | 4534 | 4821 | 5480 | 5054 | 4866 | 5163 | 4743 | 5162 | 5051 | 4014 | 4267 | 4322 | 4017 | 4491 | 4743 | 4671 | Mia2 |
| Mia3 | 11585 | 11048 | 11089 | 11975 | 11940 | 12252 | 12675 | 10147 | 10289 | 10566 | 11942 | 10642 | 10856 | 11276 | 11534 | 11552 | 11235 | 12328 | 11189 | 11166 | 12270 | 10450 | 10443 | 10717 | 10665 | 8826 | 10841 | 10291 | Mia3 |
| Mib1 | 19482 | 15513 | 21487 | 20069 | 18687 | 19619 | 20013 | 17443 | 14403 | 15308 | 16014 | 19287 | 19756 | 19918 | 19252 | 17644 | 20160 | 18284 | 18280 | 17700 | 19291 | 16599 | 14629 | 16476 | 17275 | 17172 | 20704 | 20924 | Mib1 |
| Mib2 | 3419 | 4349 | 4627 | 5169 | 4854 | 4248 | 3774 | 2689 | 3283 | 3582 | 3552 | 3145 | 2897 | 2744 | 3071 | 4028 | 5109 | 5060 | 5294 | 3925 | 4093 | 3568 | 3261 | 3929 | 3667 | 3240 | 3000 | 2595 | Mib2 |
| Mical1 | 1673 | 1767 | 2024 | 1535 | 1723 | 1788 | 1457 | 2495 | 3174 | 3220 | 2886 | 3484 | 3424 | 3336 | 1584 | 1688 | 1924 | 1436 | 1728 | 1519 | 1464 | 2727 | 3043 | 3105 | 2999 | 4016 | 3890 | 3640 | Mical1 |
| Mical2 | 1252 | 1465 | 1215 | 1294 | 1217 | 1144 | 1055 | 441 | 555 | 584 | 737 | 451 | 400 | 405 | 1214 | 1618 | 1242 | 1416 | 1094 | 1015 | 1186 | 481 | 794 | 689 | 440 | 610 | 389 | 389 | Mical2 |
| Mical3 | 5762 | 7115 | 7703 | 9015 | 9370 | 7905 | 7049 | 4400 | 5359 | 6457 | 5949 | 5133 | 4442 | 4683 | 5569 | 6962 | 8377 | 9235 | 8398 | 6123 | 7994 | 7199 | 4812 | 5495 | 6758 | 3613 | 4309 | 4265 | Mical3 |
| Micall1 | 3136 | 3173 | 3969 | 3391 | 4815 | 3967 | 3419 | 4169 | 4660 | 5508 | 5397 | 5063 | 5169 | 4508 | 2967 | 3505 | 4066 | 3960 | 4347 | 3300 | 3600 | 5713 | 4855 | 5167 | 5144 | 3833 | 4578 | 4570 | Micall1 |
| Micall2 | 1479 | 1962 | 1903 | 2026 | 1631 | 1632 | 1196 | 670 | 860 | 842 | 807 | 817 | 949 | 683 | 1611 | 2012 | 1883 | 1971 | 1581 | 1376 | 1421 | 937 | 806 | 847 | 847 | 936 | 736 | 806 | Micall2 |
| Micos10 | 5700 | 5379 | 7017 | 6616 | 5974 | 5539 | 6698 | 5040 | 4666 | 5110 | 5331 | 5070 | 5184 | 5629 | 5662 | 5185 | 6285 | 6121 | 5908 | 6902 | 6244 | 4326 | 4799 | 5741 | 5363 | 6379 | 5700 | 5982 | Micos10 |
| Micos13 | 2429 | 2747 | 2771 | 2722 | 2480 | 2666 | 2563 | 2334 | 2576 | 2391 | 2815 | 2509 | 2894 | 2860 | 2407 | 2436 | 2843 | 2652 | 2766 | 3316 | 2565 | 1947 | 2791 | 2989 | 2793 | 3142 | 2861 | 2872 | Micos13 |
| Micu1 | 6880 | 7763 | 7161 | 7447 | 6472 | 6566 | 7062 | 3886 | 3929 | 3593 | 3923 | 3610 | 3535 | 3911 | 7221 | 6953 | 7106 | 6974 | 7208 | 6852 | 7357 | 3559 | 4347 | 3528 | 3544 | 4002 | 3871 | 3875 | Micu1 |
| Micu2 | 2834 | 2262 | 2612 | 2149 | 2170 | 2343 | 2342 | 5482 | 5029 | 4857 | 4998 | 4975 | 4925 | 5020 | 2782 | 2602 | 2218 | 2035 | 2124 | 2551 | 2203 | 4779 | 5410 | 4805 | 4436 | 5664 | 5016 | 5077 | Micu2 |
| Micu3 | 1735 | 1319 | 1606 | 2127 | 2389 | 2232 | 2466 | 2787 | 2090 | 2159 | 2197 | 2071 | 2210 | 2415 | 2094 | 1587 | 1805 | 2211 | 2006 | 2355 | 2431 | 2923 | 2357 | 2093 | 2543 | 1926 | 2097 | 2307 | Micu3 |
| Mid1 | 2474 | 3446 | 3749 | 3723 | 2213 | 3586 | 4697 | 2296 | 2565 | 3429 | 2074 | 6442 | 4132 | 4457 | 3132 | 2780 | 5699 | 3323 | 3782 | 3262 | 4417 | 4926 | 3476 | 5011 | 4808 | 4577 | 5626 | 5872 | Mid1 |
| Mid1ip1 | 1600 | 2204 | 2268 | 2395 | 2709 | 2091 | 2121 | 1867 | 2344 | 2346 | 2200 | 2451 | 2258 | 2447 | 1751 | 1818 | 2597 | 2362 | 2579 | 2253 | 2257 | 2226 | 2210 | 2525 | 2476 | 2829 | 2305 | 2218 | Mid1ip1 |
| Mid2 | 1166 | 953 | 1007 | 1044 | 1555 | 1216 | 1083 | 1404 | 1523 | 1448 | 1633 | 925 | 1329 | 939 | 1271 | 1172 | 1145 | 1053 | 1160 | 1323 | 1176 | 1703 | 1572 | 1065 | 1241 | 812 | 1031 | 1046 | Mid2 |
| Mideas | 3091 | 3006 | 3742 | 3915 | 3790 | 3566 | 3513 | 2216 | 2497 | 2722 | 2974 | 2913 | 2734 | 2991 | 2598 | 3052 | 3854 | 3879 | 3769 | 2973 | 3764 | 2379 | 2436 | 2978 | 2641 | 2666 | 2611 | 2957 | Mideas |
| Midn | 7237 | 9626 | 12673 | 11639 | 11736 | 9618 | 9089 | 14831 | 16360 | 21375 | 18522 | 24186 | 18861 | 22488 | 6939 | 7994 | 12808 | 10881 | 12709 | 8163 | 9818 | 20128 | 14843 | 24511 | 22569 | 26724 | 20578 | 21739 | Midn |
| Mief1 | 3224 | 3262 | 3588 | 3589 | 3164 | 3009 | 3361 | 3263 | 3181 | 3391 | 3481 | 3948 | 3659 | 3803 | 3096 | 3101 | 3511 | 3607 | 3460 | 2966 | 3203 | 3882 | 3166 | 3700 | 3743 | 4068 | 3731 | 3964 | Mief1 |
| Mief2 | 744 | 937 | 920 | 868 | 947 | 942 | 784 | 817 | 1037 | 979 | 1059 | 1101 | 1026 | 946 | 756 | 809 | 1021 | 934 | 888 | 813 | 793 | 921 | 1085 | 1055 | 1019 | 1099 | 981 | 961 | Mief2 |
| Mien1 | 1504 | 1960 | 1692 | 2176 | 2152 | 1907 | 2390 | 1991 | 1864 | 2290 | 1873 | 1846 | 1753 | 2202 | 1750 | 1517 | 1977 | 1902 | 2167 | 2153 | 2330 | 1875 | 1904 | 2051 | 2196 | 1794 | 1820 | 2188 | Mien1 |
| Mier1 | 7740 | 5387 | 6684 | 6516 | 6274 | 6814 | 7261 | 7906 | 6993 | 7126 | 7750 | 7693 | 7902 | 8154 | 7088 | 7167 | 6560 | 6692 | 6564 | 6509 | 6991 | 7346 | 7274 | 7638 | 7264 | 7659 | 7955 | 8313 | Mier1 |
| Mier2 | 2895 | 3459 | 3849 | 4126 | 3957 | 3693 | 3200 | 2387 | 2867 | 3213 | 2929 | 3616 | 3163 | 3050 | 2643 | 3148 | 4193 | 4111 | 4410 | 3255 | 3363 | 2949 | 2675 | 3429 | 3107 | 3613 | 3004 | 3078 | Mier2 |
| Mier3 | 3851 | 2929 | 3258 | 3354 | 3482 | 3583 | 3907 | 5108 | 4469 | 3963 | 4821 | 4054 | 4529 | 4307 | 3812 | 3185 | 3288 | 3334 | 3561 | 3940 | 3872 | 4567 | 4455 | 4262 | 4434 | 3830 | 4298 | 4531 | Mier3 |
| Mif | 8436 | 11902 | 8589 | 8568 | 8201 | 8064 | 7861 | 7491 | 8970 | 7380 | 8391 | 8760 | 10034 | 9091 | 7777 | 8214 | 9063 | 8057 | 9609 | 10226 | 7839 | 6345 | 8143 | 9659 | 9043 | 11432 | 10366 | 9558 | Mif |
| Mif4gd | 647 | 714 | 931 | 812 | 667 | 675 | 676 | 480 | 495 | 605 | 577 | 585 | 682 | 586 | 617 | 656 | 793 | 650 | 777 | 723 | 610 | 409 | 542 | 704 | 467 | 816 | 623 | 610 | Mif4gd |
| Miga1 | 5291 | 4024 | 4493 | 4926 | 4988 | 5078 | 5658 | 4892 | 4361 | 4584 | 4725 | 4910 | 5007 | 5104 | 5072 | 4843 | 5001 | 5014 | 4581 | 4950 | 5136 | 4482 | 4369 | 4598 | 4760 | 4541 | 5147 | 4873 | Miga1 |
| Miga2 | 3497 | 4407 | 3389 | 3890 | 3463 | 3749 | 3363 | 1867 | 2227 | 2162 | 2203 | 2208 | 2188 | 2147 | 3488 | 4160 | 3668 | 4017 | 3720 | 3437 | 3516 | 2042 | 2163 | 2209 | 2192 | 2186 | 2287 | 2178 | Miga2 |
| Miip | 4131 | 4972 | 4719 | 4599 | 3849 | 4192 | 4331 | 2801 | 3028 | 3364 | 2870 | 3808 | 3416 | 3764 | 3770 | 3854 | 4609 | 4571 | 4107 | 4188 | 4175 | 2809 | 2747 | 3680 | 3608 | 4103 | 3792 | 3744 | Miip |
| Mill1 | 0 | 0 | 0 | 0 | 0 | 0 | 0 | 2 | 2 | 0 | 2 | 0 | 0 | 0 | 0 | 0 | 0 | 0 | 0 | 0 | 0 | 0 | 4 | 0 | 0 | 0 | 0 | 0 | Mill1 |
| Mill2 | 48 | 49 | 82 | 55 | 73 | 58 | 49 | 96 | 98 | 109 | 164 | 139 | 155 | 118 | 46 | 75 | 63 | 73 | 19 | 68 | 55 | 172 | 157 | 125 | 160 | 82 | 188 | 94 | Mill2 |
| Milr1 | 15 | 16 | 11 | 7 | 7 | 14 | 6 | 38 | 51 | 24 | 60 | 39 | 29 | 35 | 18 | 24 | 6 | 12 | 12 | 13 | 10 | 22 | 65 | 34 | 45 | 45 | 44 | 38 | Milr1 |
| Minar1 | 467 | 409 | 407 | 432 | 1012 | 653 | 676 | 344 | 379 | 453 | 371 | 154 | 345 | 134 | 593 | 424 | 484 | 467 | 633 | 585 | 659 | 408 | 325 | 139 | 203 | 93 | 120 | 136 | Minar1 |
| Minar2 | 2476 | 2209 | 1903 | 2703 | 3792 | 3163 | 3723 | 5584 | 4295 | 4641 | 3840 | 3668 | 3926 | 3831 | 3045 | 2090 | 2289 | 2834 | 3318 | 3670 | 3520 | 6092 | 4464 | 3670 | 4612 | 3200 | 3791 | 3679 | Minar2 |
| Mindy1 | 1818 | 1762 | 1622 | 1773 | 2040 | 1835 | 1769 | 2934 | 3070 | 2987 | 2909 | 3026 | 2919 | 2915 | 1862 | 1617 | 1598 | 1612 | 1829 | 1988 | 1808 | 3090 | 2958 | 3069 | 2958 | 2985 | 3015 | 2792 | Mindy1 |
| Mindy2 | 7002 | 5725 | 4907 | 6655 | 7781 | 7610 | 7525 | 5327 | 4604 | 4758 | 4932 | 4154 | 4536 | 4256 | 7125 | 6603 | 5882 | 6778 | 6720 | 6771 | 7526 | 5725 | 4664 | 3740 | 5230 | 2568 | 4248 | 4289 | Mindy2 |
| Mindy3 | 4254 | 3613 | 4091 | 4182 | 4289 | 4067 | 4851 | 3500 | 3096 | 2691 | 2930 | 2905 | 3174 | 3246 | 4978 | 4221 | 3872 | 4217 | 4050 | 4403 | 4538 | 3669 | 2775 | 2822 | 3148 | 2923 | 3221 | 3256 | Mindy3 |
| Mindy4 | 250 | 193 | 169 | 189 | 177 | 158 | 136 | 944 | 1023 | 900 | 833 | 995 | 907 | 907 | 229 | 251 | 188 | 171 | 177 | 178 | 177 | 747 | 973 | 894 | 832 | 1099 | 1019 | 974 | Mindy4 |
| Mindy4b-ps | 3 | 5 | 0 | 0 | 2 | 1 | 2 | 4 | 8 | 5 | 3 | 1 | 1 | 3 | 5 | 9 | 0 | 3 | 0 | 1 | 1 | 3 | 7 | 2 | 2 | 1 | 0 | 1 | Mindy4b-ps |
| Mink1 | 5647 | 7036 | 8008 | 8995 | 8320 | 7660 | 7718 | 4450 | 5302 | 6259 | 6241 | 6278 | 5687 | 5928 | 5590 | 6526 | 8219 | 8934 | 8458 | 6171 | 7907 | 6144 | 5057 | 6754 | 6341 | 6494 | 5197 | 5501 | Mink1 |
| Minpp1 | 2971 | 2865 | 3286 | 2795 | 2484 | 2333 | 2833 | 2347 | 2256 | 2440 | 2362 | 2258 | 2315 | 2502 | 2745 | 3166 | 2975 | 2594 | 2446 | 2270 | 2552 | 2383 | 2342 | 2378 | 2074 | 2787 | 2303 | 2557 | Minpp1 |
| Mios | 2825 | 2342 | 2178 | 2298 | 2503 | 2738 | 2762 | 3649 | 3334 | 3178 | 3042 | 3224 | 3108 | 3358 | 3037 | 2446 | 2183 | 2412 | 2523 | 2844 | 2829 | 2852 | 3144 | 2901 | 2976 | 3183 | 3596 | 3105 | Mios |
| Miox | 0 | 1 | 0 | 0 | 0 | 0 | 0 | 0 | 0 | 0 | 4 | 0 | 0 | 0 | 0 | 0 | 0 | 0 | 0 | 0 | 0 | 0 | 1 | 0 | 0 | 0 | 0 | 2 | Miox |
| Mip | 44 | 19 | 14 | 20 | 24 | 21 | 17 | 29 | 30 | 27 | 35 | 39 | 40 | 32 | 18 | 27 | 9 | 4 | 12 | 27 | 22 | 35 | 43 | 51 | 14 | 44 | 27 | 30 | Mip |
| Mipep | 2070 | 1892 | 1950 | 1688 | 1419 | 1534 | 1724 | 1749 | 1634 | 1519 | 1442 | 1588 | 1664 | 1702 | 1971 | 1866 | 1579 | 1670 | 1553 | 1568 | 1621 | 1607 | 1807 | 1565 | 1466 | 1849 | 1723 | 1909 | Mipep |
| Mipol1 | 474 | 360 | 382 | 357 | 341 | 388 | 389 | 448 | 472 | 412 | 393 | 428 | 423 | 442 | 450 | 431 | 386 | 339 | 356 | 449 | 389 | 349 | 442 | 417 | 376 | 402 | 447 | 434 | Mipol1 |
| Mis12 | 5973 | 5822 | 6460 | 5530 | 4849 | 5123 | 5538 | 5619 | 4478 | 4553 | 4387 | 5660 | 5126 | 5706 | 5881 | 5643 | 5974 | 5286 | 5066 | 5595 | 5147 | 4685 | 4267 | 5017 | 4868 | 6509 | 5884 | 6183 | Mis12 |
| Mis18a | 2827 | 2681 | 2659 | 2404 | 1494 | 1854 | 2315 | 1474 | 1399 | 1147 | 1222 | 1724 | 1569 | 1831 | 2606 | 2699 | 2608 | 1987 | 2153 | 2414 | 2076 | 1063 | 1312 | 1598 | 1437 | 2193 | 1844 | 1909 | Mis18a |

| | | | | | | | | | | | | | | | | | | | | | | | | | | | | | |
|---|---|---|---|---|---|---|---|---|---|---|---|---|---|---|---|---|---|---|---|---|---|---|---|---|---|---|---|---|---|
| Mis18bp1 | 5914 | 3947 | 4975 | 3542 | 3026 | 3931 | 4323 | 5820 | 5144 | 4454 | 3858 | 6518 | 5704 | 6803 | 5600 | 4609 | 4252 | 3290 | 3744 | 4768 | 3968 | 3452 | 4282 | 4957 | 4628 | 6925 | 7028 | 6748 | Mis18bp1 |
| Misp | 174 | 306 | 333 | 452 | 510 | 337 | 322 | 140 | 130 | 176 | 149 | 65 | 95 | 62 | 182 | 341 | 431 | 569 | 507 | 331 | 366 | 164 | 159 | 176 | 100 | 149 | 52 | 69 | Misp |
| Misp3 | 28 | 31 | 31 | 22 | 23 | 31 | 20 | 2 | 4 | 3 | 3 | 1 | 1 | 0 | 28 | 25 | 46 | 10 | 23 | 35 | 20 | 0 | 6 | 2 | 3 | 0 | 3 | 2 | Misp3 |
| Mitd1 | 1073 | 874 | 1027 | 1067 | 907 | 995 | 1076 | 1117 | 977 | 1036 | 977 | 1046 | 1082 | 1113 | 1112 | 1093 | 1080 | 894 | 940 | 1253 | 965 | 914 | 992 | 1008 | 927 | 1175 | 990 | 1076 | Mitd1 |
| Mitf | 759 | 617 | 700 | 808 | 646 | 655 | 735 | 348 | 388 | 254 | 414 | 256 | 290 | 258 | 831 | 761 | 749 | 876 | 622 | 633 | 715 | 304 | 495 | 313 | 265 | 267 | 218 | 262 | Mitf |
| Mixl1 | 3 | 4 | 8 | 1 | 6 | 3 | 4 | 1 | 1 | 0 | 6 | 0 | 2 | 0 | 8 | 7 | 0 | 4 | 3 | 0 | 1 | 5 | 1 | 1 | 1 | 0 | 6 | 1 | Mixl1 |
| Mki67 | 47566 | 37961 | 29125 | 26433 | 22669 | 30608 | 29536 | 50570 | 46035 | 37954 | 34774 | 52982 | 45329 | 50005 | 41190 | 35829 | 27409 | 22663 | 26651 | 34887 | 27907 | 26533 | 38520 | 37750 | 38235 | 42084 | 58804 | 46520 | Mki67 |
| Mkks | 762 | 790 | 692 | 795 | 707 | 603 | 720 | 902 | 807 | 693 | 666 | 634 | 677 | 656 | 822 | 842 | 677 | 743 | 700 | 801 | 701 | 805 | 814 | 634 | 714 | 682 | 661 | 653 | Mkks |
| Mkln1 | 10967 | 9072 | 8201 | 9310 | 9655 | 10467 | 10969 | 11241 | 9657 | 8273 | 8519 | 9130 | 8960 | 9545 | 10897 | 9976 | 8485 | 8890 | 9204 | 11223 | 10745 | 9249 | 9176 | 7899 | 8791 | 8108 | 9706 | 9304 | Mkln1 |
| Mknk1 | 4120 | 5003 | 4611 | 4695 | 3878 | 4194 | 4161 | 1882 | 1922 | 2140 | 2257 | 2026 | 2015 | 1970 | 4209 | 4619 | 4616 | 4809 | 4320 | 4004 | 4467 | 1967 | 2106 | 1992 | 1950 | 1721 | 2128 | 2066 | Mknk1 |
| Mknk2 | 10770 | 14158 | 11039 | 11646 | 8067 | 8888 | 7868 | 5098 | 6015 | 5715 | 6134 | 6430 | 6095 | 5627 | 9816 | 13126 | 10987 | 11606 | 8425 | 7783 | 8197 | 5738 | 5978 | 6293 | 5867 | 7238 | 5917 | 5708 | Mknk2 |
| Mkrn1 | 6503 | 7471 | 7914 | 9586 | 11864 | 9750 | 10384 | 11617 | 10528 | 13039 | 10187 | 11473 | 10459 | 11217 | 7791 | 5992 | 9500 | 9481 | 11642 | 10358 | 10786 | 14058 | 10383 | 12477 | 13936 | 9935 | 10867 | 10846 | Mkrn1 |
| Mkrn2 | 2712 | 2709 | 2209 | 2464 | 2836 | 2769 | 3109 | 2883 | 2974 | 2612 | 2564 | 2474 | 2533 | 2746 | 2891 | 2435 | 2487 | 2749 | 2731 | 2979 | 3044 | 2801 | 2910 | 2612 | 2665 | 2634 | 2558 | 2567 | Mkrn2 |
| Mkrn2os | 10 | 2 | 1 | 2 | 3 | 3 | 0 | 11 | 7 | 9 | 16 | 0 | 1 | 4 | 5 | 2 | 2 | 4 | 0 | 2 | 1 | 0 | 8 | 19 | 0 | 7 | 0 | 1 | Mkrn2os |
| Mkrn3 | 1222 | 1190 | 1465 | 1509 | 1276 | 1240 | 1380 | 1342 | 1057 | 1103 | 970 | 1502 | 1176 | 1502 | 1352 | 1045 | 1371 | 1312 | 1410 | 1442 | 1246 | 1340 | 1120 | 1137 | 1419 | 1399 | 1535 | 1460 | Mkrn3 |
| Mks1 | 592 | 604 | 778 | 604 | 727 | 717 | 724 | 1139 | 1287 | 1323 | 1373 | 1510 | 1479 | 1531 | 559 | 608 | 864 | 559 | 759 | 628 | 649 | 1310 | 1251 | 1405 | 1238 | 1452 | 1481 | 1495 | Mks1 |
| Mkx | 963 | 813 | 1113 | 751 | 515 | 643 | 401 | 449 | 520 | 480 | 580 | 431 | 634 | 490 | 878 | 1068 | 694 | 733 | 444 | 481 | 374 | 491 | 734 | 409 | 428 | 691 | 466 | 529 | Mkx |
| Mlana | 1 | 2 | 1 | 0 | 0 | 0 | 0 | 7 | 9 | 8 | 11 | 7 | 3 | 6 | 7 | 4 | 3 | 0 | 2 | 0 | 0 | 3 | 13 | 7 | 0 | 3 | 6 | 5 | Mlana |
| Mlc1 | 180 | 219 | 215 | 283 | 415 | 346 | 350 | 818 | 967 | 858 | 674 | 1023 | 586 | 1037 | 231 | 160 | 263 | 276 | 408 | 412 | 364 | 708 | 902 | 997 | 1133 | 1190 | 890 | 890 | Mlc1 |
| Mlec | 32766 | 32920 | 32564 | 27943 | 22826 | 25276 | 23648 | 17088 | 20018 | 19185 | 23329 | 19020 | 20742 | 17689 | 28887 | 33597 | 29722 | 29915 | 24849 | 21426 | 23213 | 17157 | 21333 | 19269 | 17121 | 17608 | 19758 | 19170 | Mlec |
| Mlf1 | 1450 | 1373 | 1213 | 1120 | 1016 | 919 | 1000 | 858 | 734 | 588 | 512 | 755 | 890 | 620 | 1551 | 1558 | 1194 | 1015 | 974 | 1297 | 977 | 715 | 898 | 598 | 591 | 713 | 585 | 687 | Mlf1 |
| Mlf2 | 12964 | 16505 | 15798 | 17465 | 15852 | 14664 | 14984 | 10343 | 11623 | 12962 | 12528 | 13044 | 11997 | 13178 | 12547 | 13618 | 18004 | 16687 | 17927 | 14460 | 16232 | 11991 | 11809 | 13924 | 12959 | 14283 | 12745 | 12720 | Mlf2 |
| Mlh1 | 2110 | 1978 | 1929 | 1632 | 1537 | 1823 | 1790 | 2820 | 2605 | 2470 | 2301 | 2942 | 2764 | 2943 | 2047 | 1987 | 1767 | 1484 | 1575 | 1927 | 1585 | 2351 | 2611 | 2523 | 2476 | 3053 | 3199 | 3145 | Mlh1 |
| Mlh3 | 1390 | 1213 | 1104 | 1191 | 1327 | 1393 | 1377 | 2540 | 2469 | 2271 | 2115 | 2102 | 2224 | 2156 | 1452 | 1261 | 1233 | 1287 | 1261 | 1412 | 1395 | 1983 | 2355 | 2051 | 2167 | 1659 | 2230 | 2158 | Mlh3 |
| Mlip | 14 | 6 | 9 | 16 | 20 | 26 | 22 | 18 | 15 | 32 | 32 | 14 | 11 | 7 | 5 | 11 | 9 | 7 | 24 | 10 | 17 | 17 | 32 | 28 | 24 | 21 | 3 | 5 | Mlip |
| Mlkl | 51 | 28 | 61 | 49 | 33 | 60 | 21 | 28 | 24 | 33 | 21 | 7 | 14 | 15 | 30 | 34 | 30 | 36 | 30 | 41 | 39 | 18 | 32 | 14 | 21 | 26 | 20 | 23 | Mlkl |
| Mllt1 | 7892 | 10496 | 10346 | 10836 | 9839 | 9092 | 8697 | 6988 | 8973 | 9539 | 8904 | 10453 | 8528 | 9390 | 7229 | 8663 | 10819 | 10225 | 11610 | 7828 | 9731 | 8976 | 8220 | 9897 | 9608 | 11040 | 8668 | 9374 | Mllt1 |
| Mllt10 | 4803 | 4342 | 4416 | 4322 | 4517 | 4673 | 4920 | 6909 | 5996 | 6410 | 6032 | 7315 | 7011 | 7342 | 4766 | 4029 | 4269 | 4439 | 4698 | 4423 | 4673 | 6390 | 5861 | 6643 | 6566 | 6592 | 7653 | 7328 | Mllt10 |
| Mllt11 | 8387 | 8596 | 7202 | 11872 | 17965 | 14400 | 17150 | 24380 | 19263 | 20964 | 17466 | 16778 | 15547 | 19505 | 11780 | 7572 | 10440 | 13116 | 14668 | 16955 | 17671 | 26520 | 18281 | 19331 | 23880 | 15338 | 16063 | 17789 | Mllt11 |
| Mllt3 | 6132 | 5047 | 5517 | 5816 | 6944 | 6454 | 7011 | 10601 | 9773 | 9689 | 8260 | 9090 | 8761 | 9142 | 6326 | 5103 | 5625 | 5853 | 6844 | 7507 | 7121 | 9962 | 9401 | 9562 | 10506 | 7947 | 9139 | 8866 | Mllt3 |
| Mllt6 | 6487 | 7589 | 10637 | 10745 | 11515 | 9971 | 9434 | 8425 | 8843 | 11516 | 10595 | 10821 | 10656 | 10152 | 6248 | 6731 | 11192 | 11088 | 12405 | 8779 | 10162 | 10881 | 8060 | 12361 | 12020 | 10389 | 9797 | 10301 | Mllt6 |
| Mlph | 53 | 50 | 56 | 49 | 16 | 47 | 28 | 60 | 69 | 70 | 87 | 72 | 79 | 57 | 46 | 81 | 44 | 65 | 26 | 28 | 19 | 45 | 80 | 75 | 41 | 35 | 48 | 55 | Mlph |
| Mlst8 | 1118 | 1209 | 1281 | 1147 | 1454 | 1408 | 1211 | 3043 | 3479 | 3240 | 3126 | 3712 | 3474 | 3516 | 1150 | 1018 | 1292 | 1075 | 1426 | 1283 | 1205 | 3322 | 3130 | 3474 | 3608 | 3869 | 3825 | 3824 | Mlst8 |
| Mlx | 2062 | 2513 | 1996 | 2313 | 2142 | 1914 | 2302 | 2253 | 2008 | 1861 | 1812 | 1816 | 1803 | 1870 | 2300 | 2083 | 2067 | 2287 | 2067 | 2242 | 2237 | 2104 | 2114 | 2001 | 2049 | 1918 | 1743 | 1834 | Mlx |
| Mlxip | 11844 | 12836 | 12450 | 16009 | 13574 | 13130 | 13614 | 5620 | 5668 | 6813 | 6421 | 7151 | 6549 | 6564 | 11058 | 12907 | 14303 | 15472 | 14303 | 10581 | 13563 | 7285 | 5378 | 7048 | 7660 | 5497 | 6694 | 6566 | Mlxip |
| Mlxipl | 910 | 971 | 1189 | 972 | 602 | 686 | 685 | 149 | 178 | 229 | 178 | 202 | 187 | 198 | 707 | 1038 | 1030 | 858 | 743 | 601 | 603 | 147 | 160 | 256 | 160 | 351 | 192 | 227 | Mlxipl |
| Mlycd | 866 | 1191 | 961 | 1014 | 1040 | 882 | 868 | 723 | 882 | 739 | 788 | 828 | 790 | 759 | 907 | 956 | 1198 | 946 | 946 | 930 | 928 | 715 | 935 | 858 | 831 | 916 | 761 | 757 | Mlycd |
| Mmaa | 1250 | 1283 | 1082 | 1736 | 1592 | 1591 | 1888 | 1955 | 1914 | 1898 | 1747 | 1755 | 1640 | 1775 | 1336 | 1162 | 1402 | 1700 | 1527 | 1652 | 1690 | 2064 | 1930 | 1942 | 1893 | 1704 | 1667 | 1544 | Mmaa |
| Mmab | 962 | 1158 | 1152 | 1039 | 992 | 1060 | 884 | 1559 | 1965 | 1956 | 2010 | 2044 | 1968 | 1868 | 955 | 1114 | 1270 | 1061 | 1057 | 874 | 869 | 1768 | 2003 | 1959 | 1832 | 2125 | 2076 | 1909 | Mmab |
| Mmachc | 1204 | 1127 | 1323 | 1061 | 957 | 1017 | 991 | 1827 | 1619 | 1547 | 1480 | 2110 | 1816 | 2059 | 1213 | 1079 | 1274 | 865 | 1054 | 1220 | 973 | 1544 | 1703 | 1734 | 1756 | 2360 | 1994 | 2250 | Mmachc |
| Mmadhc | 4665 | 4686 | 4468 | 3914 | 3537 | 3607 | 3924 | 4680 | 4247 | 3940 | 3875 | 4419 | 4383 | 4749 | 5077 | 4350 | 4562 | 3865 | 3735 | 4116 | 3815 | 3984 | 4285 | 4330 | 4026 | 4971 | 4817 | 4911 | Mmadhc |
| Mmd | 6819 | 5223 | 6584 | 7873 | 8498 | 7370 | 8917 | 5461 | 4342 | 5680 | 4684 | 4708 | 4611 | 5106 | 7089 | 5856 | 7436 | 8029 | 8182 | 7311 | 9228 | 6630 | 4798 | 5079 | 5601 | 4568 | 4131 | 5071 | Mmd |
| Mmd2 | 929 | 989 | 639 | 1386 | 2277 | 1892 | 2293 | 2234 | 2286 | 1987 | 1808 | 1404 | 1457 | 1693 | 1269 | 868 | 1081 | 1788 | 2175 | 2282 | 2346 | 2296 | 2180 | 2031 | 2558 | 1531 | 1364 | 1542 | Mmd2 |
| Mme | 4334 | 2684 | 2759 | 1533 | 797 | 2080 | 523 | 4898 | 7321 | 5051 | 6681 | 3279 | 5213 | 3164 | 3869 | 4703 | 1744 | 1607 | 718 | 867 | 479 | 3628 | 6805 | 3876 | 2675 | 3275 | 3475 | 3146 | Mme |
| Mmel1 | 17 | 30 | 28 | 25 | 21 | 22 | 19 | 23 | 18 | 5 | 33 | 26 | 31 | 12 | 32 | 29 | 22 | 25 | 20 | 22 | 13 | 12 | 16 | 20 | 23 | 34 | 27 | 27 | Mmel1 |
| Mmgt1 | 7882 | 6626 | 6980 | 7376 | 7773 | 7651 | 8786 | 9365 | 7456 | 7055 | 6589 | 7658 | 7184 | 8151 | 8418 | 7044 | 6906 | 7264 | 7439 | 8589 | 8549 | 7807 | 7219 | 6995 | 7628 | 7170 | 7483 | 8246 | Mmgt1 |
| Mmgt2 | 905 | 866 | 944 | 877 | 935 | 760 | 831 | 1186 | 1204 | 1086 | 1186 | 1107 | 1160 | 1228 | 944 | 899 | 829 | 747 | 835 | 929 | 849 | 966 | 1271 | 1269 | 940 | 1642 | 1328 | 1304 | Mmgt2 |
| Mmp10 | 0 | 0 | 0 | 0 | 2 | 0 | 0 | 0 | 0 | 0 | 1 | 0 | 0 | 0 | 0 | 0 | 0 | 0 | 0 | 0 | 0 | 0 | 0 | 0 | 0 | 1 | 0 | 0 | Mmp10 |
| Mmp11 | 2209 | 2142 | 2007 | 1487 | 1230 | 1690 | 1141 | 2022 | 2896 | 2065 | 2594 | 1981 | 2410 | 1882 | 2052 | 2534 | 1658 | 1665 | 1187 | 1107 | 1046 | 1552 | 2907 | 2219 | 1584 | 2352 | 2067 | 1877 | Mmp11 |
| Mmp12 | 1 | 0 | 0 | 0 | 2 | 0 | 1 | 0 | 2 | 2 | 0 | 0 | 0 | 1 | 0 | 1 | 0 | 4 | 2 | 1 | 0 | 2 | 4 | 0 | 0 | 1 | 0 | 0 | Mmp12 |
| Mmp13 | 10 | 6 | 3 | 6 | 7 | 5 | 20 | 0 | 2 | 5 | 1 | 5 | 9 | 5 | 13 | 8 | 13 | 9 | 15 | 19 | 11 | 8 | 6 | 2 | 8 | 10 | 5 | 2 | Mmp13 |
| Mmp14 | 3237 | 3407 | 3515 | 2777 | 2474 | 3708 | 1720 | 7024 | 9189 | 8806 | 10578 | 8210 | 9096 | 7054 | 3301 | 4263 | 2907 | 3414 | 2505 | 1904 | 1872 | 8042 | 10259 | 8684 | 7452 | 8822 | 7669 | 7315 | Mmp14 |
| Mmp15 | 2354 | 2909 | 3750 | 3882 | 4816 | 3544 | 3167 | 3307 | 3607 | 4389 | 4408 | 4487 | 4082 | 4250 | 2389 | 2674 | 3846 | 3794 | 4028 | 2711 | 3428 | 4455 | 3548 | 4632 | 4346 | 4866 | 3689 | 4118 | Mmp15 |
| Mmp16 | 2609 | 2052 | 799 | 1176 | 1814 | 2786 | 2019 | 3441 | 3701 | 2342 | 3180 | 2038 | 2482 | 2218 | 2607 | 2543 | 883 | 1402 | 1205 | 2735 | 2300 | 1912 | 3736 | 1403 | 2501 | 1029 | 2324 | 1711 | Mmp16 |
| Mmp17 | 2234 | 3403 | 3645 | 3759 | 3542 | 3073 | 3063 | 949 | 1126 | 1256 | 937 | 1358 | 1038 | 1233 | 2450 | 2527 | 4088 | 3627 | 3953 | 3017 | 3498 | 1285 | 945 | 1423 | 1481 | 1376 | 1142 | 1351 | Mmp17 |
| Mmp19 | 29 | 35 | 22 | 29 | 43 | 46 | 34 | 43 | 47 | 38 | 57 | 36 | 61 | 50 | 26 | 31 | 34 | 42 | 37 | 37 | 45 | 74 | 66 | 63 | 31 | 49 | 70 | 43 | Mmp19 |
| Mmp2 | 4600 | 4710 | 4124 | 2412 | 1543 | 3034 | 807 | 4457 | 7697 | 6114 | 8781 | 3770 | 6027 | 3329 | 4064 | 6514 | 2786 | 3464 | 1045 | 1186 | 847 | 4879 | 8492 | 5619 | 4009 | 5783 | 4278 | 3909 | Mmp2 |
| Mmp20 | 0 | 0 | 0 | 0 | 0 | 0 | 0 | 1 | 0 | 0 | 0 | 0 | 0 | 0 | 0 | 1 | 0 | 0 | 0 | 0 | 0 | 0 | 0 | 0 | 0 | 0 | 0 | 0 | Mmp20 |
| Mmp21 | 7 | 12 | 2 | 11 | 10 | 6 | 6 | 1 | 0 | 18 | 0 | 7 | 9 | 3 | 13 | 5 | 3 | 12 | 11 | 18 | 4 | 8 | 3 | 5 | 4 | 3 | 6 | 6 | Mmp21 |
| Mmp23 | 188 | 364 | 351 | 436 | 364 | 316 | 361 | 172 | 225 | 207 | 202 | 127 | 150 | 125 | 203 | 300 | 380 | 511 | 428 | 309 | 337 | 165 | 231 | 185 | 200 | 135 | 134 | 136 | Mmp23 |
| Mmp24 | 922 | 1252 | 1152 | 2062 | 3629 | 2652 | 2461 | 2904 | 3546 | 4109 | 3647 | 2884 | 2672 | 2655 | 1323 | 1083 | 1954 | 2588 | 2949 | 2273 | 2809 | 5287 | 3206 | 3309 | 4515 | 2040 | 2321 | 2366 | Mmp24 |
| Mmp25 | 119 | 129 | 207 | 112 | 209 | 208 | 124 | 167 | 258 | 227 | 254 | 203 | 271 | 228 | 162 | 190 | 154 | 180 | 146 | 107 | 158 | 333 | 183 | 223 | 240 | 209 | 157 | 188 | Mmp25 |
| Mmp28 | 373 | 538 | 559 | 526 | 543 | 536 | 380 | 346 | 507 | 592 | 709 | 643 | 608 | 560 | 439 | 558 | 571 | 573 | 481 | 368 | 425 | 724 | 604 | 632 | 710 | 644 | 517 | 580 | Mmp28 |
| Mmp3 | 0 | 0 | 0 | 0 | 0 | 0 | 0 | 0 | 1 | 0 | 0 | 0 | 0 | 0 | 1 | 0 | 0 | 0 | 0 | 0 | 0 | 0 | 0 | 0 | 0 | 0 | 0 | 0 | Mmp3 |
| Mmp7 | 1 | 0 | 0 | 0 | 0 | 0 | 0 | 0 | 0 | 0 | 0 | 0 | 0 | 0 | 0 | 0 | 0 | 0 | 0 | 0 | 0 | 0 | 1 | 0 | 0 | 0 | 0 | 0 | Mmp7 |
| Mmp8 | 3 | 3 | 2 | 0 | 1 | 2 | 1 | 1 | 5 | 1 | 15 | 2 | 17 | 2 | 4 | 9 | 0 | 4 | 0 | 0 | 3 | 7 | 14 | 17 | 12 | 6 | 23 | 8 | Mmp8 |
| Mmp9 | 217 | 291 | 319 | 619 | 963 | 757 | 543 | 443 | 707 | 997 | 823 | 470 | 464 | 428 | 328 | 255 | 454 | 790 | 749 | 405 | 653 | 1041 | 652 | 757 | 837 | 277 | 285 | 274 | Mmp9 |
| Mmrn1 | 1062 | 841 | 996 | 650 | 674 | 858 | 662 | 909 | 894 | 741 | 1078 | 1113 | 1270 | 976 | 959 | 1086 | 771 | 654 | 591 | 973 | 684 | 846 | 1132 | 909 | 919 | 1071 | 1214 | 957 | Mmrn1 |
| Mmrn2 | 646 | 707 | 803 | 670 | 691 | 723 | 627 | 716 | 864 | 980 | 989 | 969 | 1014 | 933 | 659 | 762 | 771 | 647 | 710 | 609 | 615 | 792 | 884 | 879 | 792 | 1176 | 970 | 913 | Mmrn2 |
| Mms19 | 9123 | 10258 | 6559 | 7715 | 6995 | 7602 | 8234 | 4996 | 5245 | 4564 | 4754 | 5166 | 4899 | 4856 | 8472 | 9045 | 7291 | 8047 | 7254 | 8096 | 7906 | 4644 | 4902 | 4415 | 4567 | 4728 | 4969 | 4777 | Mms19 |
| Mms22l | 4794 | 4161 | 4183 | 3856 | 3022 | 3729 | 4387 | 2937 | 2802 | 2540 | 2137 | 3836 | 3034 | 3742 | 4245 | 3960 | 3875 | 3648 | 3749 | 3940 | 4067 | 1938 | 2048 | 2857 | 2870 | 3787 | 3611 | 3593 | Mms22l |
| Mmut | 3021 | 2383 | 2437 | 2211 | 1921 | 2138 | 2341 | 2983 | 3038 | 2489 | 2856 | 2627 | 2850 | 2818 | 2865 | 2773 | 2296 | 1986 | 1985 | 2290 | 1922 | 2473 | 2983 | 2615 | 2422 | 2615 | 2832 | 2720 | Mmut |
| Mn1 | 4638 | 6678 | 7630 | 9180 | 7798 | 7462 | 6577 | 5950 | 7448 | 8877 | 8129 | 9957 | 8114 | 8767 | 4705 | 6067 | 8967 | 9400 | 8690 | 5530 | 7421 | 9229 | 5610 | 10888 | 10323 | 11114 | 7547 | 8032 | Mn1 |
| Mnat1 | 2635 | 1967 | 2973 | 1959 | 1724 | 1698 | 1767 | 2027 | 1959 | 2146 | 2048 | 2309 | 2236 | 2350 | 2495 | 2446 | 2355 | 1811 | 1699 | 2012 | 1610 | 1646 | 2126 | 2245 | 1896 | 2634 | 2396 | 2624 | Mnat1 |
| Mnd1 | 246 | 205 | 300 | 156 | 124 | 164 | 178 | 245 | 234 | 190 | 234 | 275 | 257 | 305 | 229 | 214 | 219 | 128 | 175 | 186 | 151 | 147 | 229 | 221 | 171 | 320 | 296 | 291 | Mnd1 |
| Mndal | 25 | 10 | 25 | 19 | 45 | 28 | 32 | 57 | 72 | 85 | 112 | 43 | 28 | 31 | 29 | 28 | 14 | 43 | 15 | 48 | 32 | 82 | 91 | 77 | 59 | 64 | 26 | 30 | Mndal |
| Mns1 | 5059 | 4770 | 5599 | 4053 | 2997 | 3604 | 3738 | 2928 | 2486 | 2441 | 2206 | 3781 | 3195 | 4010 | 4803 | 4337 | 4810 | 3544 | 3719 | 4251 | 3590 | 1957 | 2340 | 3225 | 2693 | 4057 | 4178 | 4086 | Mns1 |
| Mnt | 2808 | 3595 | 4223 | 5006 | 4552 | 3882 | 3492 | 2665 | 3063 | 4092 | 3756 | 4003 | 3289 | 3664 | 2620 | 3110 | 4514 | 4968 | 4661 | 3030 | 3967 | 4263 | 2832 | 4202 | 4259 | 4224 | 3206 | 3478 | Mnt |
| Mnx1 | 9 | 17 | 7 | 11 | 16 | 5 | 9 | 0 | 2 | 7 | 23 | 0 | 2 | 3 | 7 | 21 | 12 | 11 | 7 | 9 | 10 | 0 | 9 | 1 | 0 | 5 | 1 | 6 | Mnx1 |

| | | | | | | | | | | | | | | | | | | | | | | | | | | | | | |
|---|---|---|---|---|---|---|---|---|---|---|---|---|---|---|---|---|---|---|---|---|---|---|---|---|---|---|---|---|---|
| Moap1 | 1 | 3 | 5 | 7 | 2 | 1 | 2 | 0 | 0 | 2 | 3 | 5 | 5 | 0 | 4 | 4 | 3 | 0 | 2 | 1 | 1 | 8 | 3 | 1 | 1 | 5 | 0 | 9 | Moap1 |
| Mob1a | 13432 | 11238 | 11043 | 8975 | 7206 | 8733 | 8990 | 9716 | 8967 | 7352 | 8523 | 9067 | 8981 | 8995 | 12092 | 12213 | 9798 | 9239 | 7930 | 9658 | 8476 | 7855 | 9455 | 8274 | 7625 | 9519 | 9460 | 9608 | Mob1a |
| Mob1b | 6422 | 4400 | 6029 | 5918 | 5767 | 6586 | 6832 | 4949 | 4508 | 4465 | 5011 | 5564 | 5361 | 5739 | 5983 | 5909 | 6000 | 6070 | 5626 | 5787 | 6573 | 5280 | 4365 | 5018 | 5268 | 5454 | 5545 | 5835 | Mob1b |
| Mob2 | 2173 | 2452 | 1929 | 2228 | 1913 | 2011 | 1986 | 1533 | 1695 | 1431 | 1486 | 1554 | 1397 | 1481 | 2205 | 2301 | 2045 | 2144 | 1975 | 2273 | 2015 | 1425 | 1614 | 1487 | 1488 | 1633 | 1576 | 1448 | Mob2 |
| Mob3a | 4651 | 6207 | 4494 | 4471 | 3624 | 4267 | 3406 | 2616 | 2871 | 2818 | 2864 | 3060 | 3071 | 2748 | 4531 | 5568 | 4444 | 4512 | 4150 | 4036 | 3653 | 2842 | 3021 | 2817 | 2655 | 3306 | 3109 | 3119 | Mob3a |
| Mob3b | 896 | 879 | 1353 | 975 | 868 | 835 | 865 | 840 | 811 | 954 | 715 | 1388 | 1489 | 1431 | 1005 | 857 | 1276 | 809 | 923 | 856 | 875 | 785 | 737 | 1150 | 839 | 1542 | 1603 | 1641 | Mob3b |
| Mob3c | 329 | 287 | 383 | 366 | 358 | 398 | 294 | 429 | 474 | 541 | 559 | 496 | 595 | 469 | 262 | 294 | 346 | 377 | 302 | 309 | 280 | 578 | 503 | 532 | 533 | 567 | 467 | 523 | Mob3c |
| Mob4 | 6687 | 4884 | 6729 | 6833 | 6562 | 6686 | 7517 | 8448 | 6908 | 6913 | 6961 | 7484 | 7094 | 8087 | 7239 | 6234 | 6363 | 6655 | 6505 | 7172 | 6886 | 8024 | 7420 | 7453 | 7555 | 8072 | 7262 | 8043 | Mob4 |
| Mobp | 20 | 12 | 12 | 9 | 6 | 13 | 8 | 17 | 23 | 10 | 13 | 43 | 0 | 36 | 21 | 14 | 16 | 8 | 8 | 5 | 11 | 12 | 8 | 20 | 25 | 65 | 29 | 42 | Mobp |
| Mocos | 32 | 18 | 33 | 7 | 17 | 30 | 8 | 28 | 31 | 59 | 41 | 2 | 11 | 6 | 29 | 34 | 10 | 40 | 14 | 26 | 19 | 5 | 55 | 32 | 10 | 24 | 1 | 5 | Mocos |
| Mocs1 | 662 | 775 | 681 | 656 | 793 | 802 | 612 | 678 | 801 | 800 | 834 | 881 | 906 | 786 | 591 | 688 | 739 | 666 | 654 | 723 | 634 | 702 | 837 | 911 | 793 | 981 | 809 | 764 | Mocs1 |
| Mocs2 | 5685 | 5217 | 5588 | 5741 | 4852 | 4747 | 5566 | 3922 | 3385 | 3253 | 3396 | 3335 | 3035 | 3356 | 5972 | 5553 | 5399 | 5317 | 5186 | 6385 | 5239 | 3180 | 3361 | 3510 | 3384 | 3741 | 3424 | 3563 | Mocs2 |
| Mocs3 | 615 | 619 | 578 | 682 | 652 | 613 | 582 | 462 | 485 | 465 | 520 | 522 | 536 | 484 | 550 | 640 | 681 | 747 | 645 | 552 | 636 | 449 | 484 | 549 | 470 | 479 | 506 | 522 | Mocs3 |
| Mog | 3 | 3 | 2 | 4 | 11 | 0 | 0 | 0 | 0 | 2 | 0 | 3 | 2 | 0 | 1 | 1 | 8 | 6 | 4 | 0 | 3 | 0 | 0 | 6 | 2 | 0 | 2 | 0 | Mog |
| Mogat1 | 9 | 6 | 0 | 0 | 0 | 0 | 1 | 2 | 7 | 16 | 11 | 7 | 3 | 5 | 7 | 7 | 0 | 0 | 1 | 1 | 3 | 2 | 10 | 7 | 4 | 6 | 0 | 12 | Mogat1 |
| Mogat2 | 311 | 402 | 480 | 255 | 208 | 197 | 196 | 101 | 127 | 119 | 88 | 147 | 198 | 115 | 351 | 502 | 308 | 217 | 149 | 230 | 182 | 95 | 146 | 128 | 120 | 183 | 209 | 206 | Mogat2 |
| Mogs | 2974 | 3807 | 3505 | 3203 | 2626 | 2486 | 2486 | 2235 | 2764 | 2612 | 2774 | 2757 | 2654 | 2413 | 2722 | 3544 | 3472 | 3026 | 2987 | 2588 | 2389 | 2308 | 2815 | 2689 | 2393 | 3314 | 2452 | 2511 | Mogs |
| Mok | 873 | 1176 | 792 | 1019 | 976 | 911 | 929 | 524 | 565 | 592 | 542 | 677 | 515 | 557 | 936 | 977 | 1080 | 958 | 1031 | 1051 | 976 | 617 | 564 | 527 | 617 | 540 | 538 | 498 | Mok |
| Mon1a | 549 | 790 | 758 | 759 | 945 | 845 | 710 | 981 | 976 | 1115 | 979 | 982 | 923 | 1011 | 667 | 684 | 811 | 724 | 1015 | 714 | 844 | 1081 | 971 | 1072 | 1246 | 1076 | 1117 | 1062 | Mon1a |
| Mon1b | 2533 | 3073 | 3388 | 3976 | 3562 | 3028 | 3581 | 2180 | 2198 | 2660 | 2172 | 2684 | 2470 | 2694 | 2649 | 2714 | 3585 | 3836 | 3673 | 2754 | 3750 | 2565 | 2135 | 2740 | 2877 | 2512 | 2352 | 2706 | Mon1b |
| Mon2 | 11610 | 11302 | 8226 | 10634 | 10424 | 11151 | 11107 | 8348 | 8730 | 8032 | 9024 | 7895 | 7952 | 8185 | 11305 | 11247 | 9079 | 10796 | 9831 | 10016 | 11657 | 8457 | 8501 | 7277 | 8682 | 5999 | 7961 | 7692 | Mon2 |
| Morc1 | 1 | 3 | 0 | 6 | 3 | 1 | 2 | 6 | 7 | 9 | 3 | 1 | 3 | 3 | 3 | 9 | 3 | 9 | 5 | 1 | 1 | 5 | 6 | 1 | 0 | 0 | 6 | 11 | Morc1 |
| Morc2a | 12129 | 11747 | 11968 | 14015 | 13255 | 13439 | 13641 | 11295 | 10785 | 11937 | 11651 | 12509 | 11456 | 12153 | 11719 | 11279 | 12311 | 13682 | 13333 | 11151 | 13974 | 12644 | 10255 | 11745 | 12893 | 10205 | 11900 | 11614 | Morc2a |
| Morc2b | 33 | 42 | 15 | 62 | 56 | 30 | 55 | 12 | 16 | 13 | 25 | 9 | 7 | 7 | 49 | 35 | 27 | 52 | 60 | 32 | 49 | 25 | 17 | 20 | 13 | 7 | 6 | 8 | Morc2b |
| Morc3 | 3459 | 3017 | 3025 | 3254 | 3340 | 3343 | 3386 | 4689 | 4210 | 3960 | 4206 | 4278 | 3821 | 4217 | 3491 | 3127 | 3330 | 3130 | 3225 | 3698 | 3250 | 4370 | 4212 | 3935 | 4222 | 3846 | 4194 | 4111 | Morc3 |
| Morc4 | 2290 | 1788 | 1892 | 1619 | 1490 | 1574 | 1421 | 1154 | 1266 | 993 | 1546 | 991 | 1133 | 1022 | 2069 | 2200 | 1603 | 1545 | 1307 | 1393 | 1333 | 1108 | 1276 | 1105 | 961 | 1022 | 1033 | 1123 | Morc4 |
| Morf4l1 | 15441 | 16035 | 18597 | 15786 | 15552 | 15517 | 16689 | 23395 | 22883 | 21774 | 21450 | 23500 | 23098 | 25357 | 16634 | 15776 | 17216 | 14835 | 16056 | 18936 | 15826 | 21012 | 24390 | 23684 | 22388 | 26231 | 25763 | 26798 | Morf4l1 |
| Morf4l2 | 35696 | 33640 | 35513 | 33567 | 30633 | 32178 | 34797 | 36009 | 34595 | 30162 | 32228 | 32583 | 30938 | 34629 | 38133 | 33364 | 34762 | 32424 | 32433 | 39147 | 33797 | 30878 | 35485 | 33315 | 31781 | 37271 | 33792 | 35081 | Morf4l2 |
| Morn1 | 870 | 963 | 683 | 1028 | 880 | 838 | 924 | 289 | 395 | 359 | 381 | 375 | 416 | 391 | 831 | 860 | 844 | 925 | 864 | 880 | 886 | 359 | 402 | 328 | 450 | 384 | 431 | 377 | Morn1 |
| Morn2 | 405 | 438 | 475 | 468 | 470 | 392 | 591 | 504 | 461 | 470 | 494 | 554 | 468 | 691 | 400 | 373 | 454 | 406 | 460 | 575 | 584 | 399 | 475 | 546 | 536 | 527 | 531 | 612 | Morn2 |
| Morn3 | 0 | 5 | 6 | 5 | 9 | 11 | 9 | 18 | 23 | 32 | 8 | 25 | 22 | 5 | 5 | 18 | 8 | 1 | 14 | 6 | 10 | 22 | 17 | 19 | 5 | 15 | 24 | 11 | Morn3 |
| Morn4 | 3081 | 3834 | 3991 | 4852 | 4969 | 4541 | 4522 | 3510 | 3808 | 4371 | 3742 | 4800 | 3933 | 4724 | 3219 | 3121 | 4341 | 4731 | 4881 | 4307 | 4740 | 4634 | 3489 | 4498 | 5107 | 4454 | 4172 | 4230 | Morn4 |
| Morn5 | 18 | 18 | 13 | 20 | 21 | 13 | 17 | 16 | 21 | 12 | 11 | 12 | 27 | 18 | 25 | 23 | 27 | 12 | 19 | 23 | 40 | 23 | 35 | 12 | 23 | 18 | 9 | 13 | Morn5 |
| Mos | 21 | 16 | 17 | 27 | 25 | 19 | 9 | 22 | 21 | 20 | 19 | 29 | 32 | 24 | 26 | 18 | 16 | 17 | 26 | 23 | 25 | 18 | 16 | 18 | 9 | 30 | 27 | 13 | Mos |
| Mosmo | 5472 | 4210 | 5181 | 5752 | 5102 | 5583 | 5906 | 5311 | 4367 | 4433 | 5046 | 5153 | 4945 | 5302 | 5404 | 4784 | 5109 | 5543 | 5293 | 5158 | 5691 | 5182 | 4370 | 4799 | 5397 | 4733 | 5488 | 5255 | Mosmo |
| Mospd1 | 3102 | 2602 | 2941 | 2734 | 2831 | 2699 | 3226 | 3497 | 2684 | 2228 | 2977 | 2590 | 2879 | 3059 | 3690 | 3044 | 2730 | 2903 | 2696 | 3574 | 2883 | 3056 | 2910 | 2410 | 2555 | 2791 | 2941 | 3201 | Mospd1 |
| Mospd2 | 3485 | 2729 | 2622 | 2801 | 2761 | 2958 | 3181 | 2759 | 2388 | 1891 | 2252 | 2230 | 2337 | 2347 | 3775 | 3417 | 2561 | 2956 | 2554 | 3336 | 2926 | 2431 | 2359 | 2139 | 2135 | 2058 | 2458 | 2365 | Mospd2 |
| Mospd3 | 1137 | 1432 | 1247 | 1683 | 1508 | 1330 | 1294 | 934 | 1121 | 1285 | 1210 | 1218 | 1038 | 1047 | 1030 | 1308 | 1549 | 1710 | 1584 | 1243 | 1405 | 1303 | 1014 | 1222 | 1321 | 1220 | 874 | 1057 | Mospd3 |
| Mov10 | 964 | 1038 | 1201 | 781 | 766 | 850 | 621 | 1191 | 1665 | 1609 | 1995 | 1302 | 1695 | 1138 | 836 | 1192 | 1002 | 796 | 749 | 520 | 587 | 1377 | 1885 | 1660 | 1071 | 1947 | 1307 | 1208 | Mov10 |
| Mov10l1 | 0 | 0 | 0 | 0 | 1 | 0 | 2 | 0 | 0 | 0 | 6 | 0 | 3 | 0 | 0 | 0 | 0 | 0 | 0 | 0 | 1 | 0 | 4 | 0 | 0 | 0 | 1 | 0 | Mov10l1 |
| Moxd1 | 1067 | 709 | 538 | 354 | 495 | 376 | 152 | 662 | 966 | 855 | 1355 | 269 | 555 | 155 | 981 | 1458 | 338 | 456 | 219 | 202 | 201 | 690 | 1176 | 645 | 286 | 695 | 252 | 171 | Moxd1 |
| Mpc1 | 1456 | 1466 | 1235 | 1513 | 1567 | 1765 | 1796 | 1326 | 1215 | 1154 | 1271 | 1108 | 1055 | 1053 | 1542 | 1530 | 1322 | 1808 | 1626 | 1851 | 1849 | 1313 | 1315 | 1135 | 1455 | 987 | 1138 | 894 | Mpc1 |
| Mpc2 | 2136 | 2050 | 2097 | 2497 | 2302 | 2054 | 2924 | 1909 | 1606 | 2103 | 1883 | 1660 | 1791 | 2008 | 2310 | 1872 | 2421 | 2444 | 2274 | 2691 | 2691 | 1876 | 1763 | 1995 | 2006 | 1661 | 1644 | 1915 | Mpc2 |
| Mpdu1 | 1388 | 1454 | 1664 | 1509 | 1631 | 1737 | 1599 | 3031 | 3138 | 3121 | 3067 | 3673 | 3504 | 3537 | 1383 | 1302 | 1604 | 1479 | 1776 | 1724 | 1642 | 3019 | 3082 | 3389 | 3400 | 4122 | 4011 | 3865 | Mpdu1 |
| Mpdz | 12702 | 11577 | 9009 | 9052 | 8818 | 10556 | 10676 | 11358 | 11742 | 9924 | 11031 | 14066 | 13267 | 14119 | 12221 | 12444 | 9523 | 9333 | 9171 | 11193 | 10299 | 9105 | 10965 | 10598 | 10793 | 10113 | 14878 | 13324 | Mpdz |
| Mpeg1 | 617 | 546 | 447 | 520 | 624 | 552 | 476 | 834 | 830 | 660 | 1213 | 709 | 582 | 577 | 694 | 684 | 499 | 614 | 511 | 551 | 575 | 939 | 1153 | 730 | 697 | 732 | 597 | 610 | Mpeg1 |
| Mpg | 1720 | 1792 | 1780 | 1605 | 1780 | 1688 | 1638 | 1332 | 1591 | 1591 | 1655 | 1607 | 1733 | 1479 | 1701 | 1597 | 1924 | 1679 | 1880 | 1760 | 1608 | 1609 | 1564 | 1637 | 1578 | 1720 | 1646 | 1610 | Mpg |
| Mphosph10 | 4104 | 3714 | 4080 | 3511 | 2890 | 3036 | 3412 | 3527 | 3062 | 2786 | 2884 | 3631 | 3353 | 3863 | 4304 | 3773 | 3650 | 3179 | 3068 | 4089 | 3032 | 2767 | 3202 | 3329 | 3028 | 4259 | 3794 | 3938 | Mphosph10 |
| Mphosph6 | 1566 | 1379 | 1829 | 1479 | 1492 | 1500 | 1600 | 1537 | 1366 | 1687 | 1557 | 1742 | 1577 | 1966 | 1716 | 1336 | 1647 | 1533 | 1502 | 1668 | 1564 | 1485 | 1544 | 1749 | 1478 | 2043 | 1708 | 1976 | Mphosph6 |
| Mphosph8 | 8867 | 7694 | 9719 | 8965 | 8198 | 8849 | 9302 | 7113 | 5773 | 6250 | 5822 | 7204 | 6944 | 7445 | 9723 | 8270 | 9181 | 8771 | 8218 | 9755 | 8645 | 6126 | 5858 | 6787 | 6697 | 7816 | 7355 | 7436 | Mphosph8 |
| Mphosph9 | 2788 | 2455 | 2058 | 2759 | 3167 | 3454 | 3672 | 4082 | 3877 | 4128 | 4145 | 4884 | 4425 | 4693 | 2685 | 2402 | 2373 | 2896 | 3225 | 3205 | 3490 | 3952 | 3409 | 3907 | 4418 | 3476 | 4929 | 4479 | Mphosph9 |
| Mpi | 4158 | 4842 | 4134 | 4147 | 3723 | 4097 | 3873 | 3256 | 2813 | 2790 | 2594 | 3087 | 3032 | 3047 | 4464 | 4081 | 3958 | 3945 | 3950 | 4255 | 3720 | 2928 | 2759 | 2796 | 2811 | 3172 | 3209 | 3341 | Mpi |
| Mpig6b | 51 | 60 | 53 | 44 | 31 | 34 | 35 | 53 | 37 | 48 | 67 | 52 | 49 | 40 | 51 | 62 | 42 | 39 | 43 | 48 | 22 | 32 | 43 | 60 | 19 | 83 | 49 | 43 | Mpig6b |
| Mpl | 40 | 35 | 34 | 7 | 14 | 11 | 13 | 17 | 16 | 11 | 11 | 40 | 59 | 19 | 25 | 37 | 19 | 14 | 12 | 6 | 4 | 17 | 19 | 18 | 5 | 54 | 42 | 36 | Mpl |
| Mplkip | 2067 | 1424 | 1635 | 1682 | 1496 | 1683 | 1739 | 2124 | 1884 | 1714 | 1753 | 1813 | 1657 | 2056 | 2073 | 1688 | 1621 | 1395 | 1528 | 2007 | 1669 | 1656 | 1940 | 1704 | 1672 | 2050 | 1818 | 1920 | Mplkip |
| Mpnd | 5916 | 7515 | 6682 | 7421 | 6879 | 6794 | 6429 | 5567 | 6547 | 6657 | 5850 | 7367 | 6402 | 6835 | 5648 | 6396 | 7552 | 6820 | 8023 | 6314 | 6759 | 6269 | 5941 | 7129 | 7009 | 8052 | 7273 | 6931 | Mpnd |
| Mpo | 35 | 54 | 56 | 22 | 10 | 16 | 12 | 15 | 17 | 27 | 73 | 20 | 37 | 11 | 16 | 74 | 25 | 26 | 6 | 13 | 20 | 23 | 69 | 26 | 4 | 64 | 35 | 50 | Mpo |
| Mpp1 | 2564 | 2208 | 2259 | 1959 | 1967 | 1931 | 1858 | 1911 | 2187 | 1738 | 2120 | 1829 | 1932 | 1775 | 2378 | 2398 | 1962 | 2047 | 1771 | 2003 | 1944 | 1688 | 2435 | 1726 | 1536 | 1888 | 1914 | 1831 | Mpp1 |
| Mpp2 | 4269 | 5537 | 6576 | 6882 | 6442 | 5876 | 5460 | 2538 | 2676 | 3211 | 3227 | 3698 | 3079 | 3450 | 4234 | 5044 | 7197 | 6756 | 7047 | 5019 | 6047 | 3208 | 2560 | 3855 | 3616 | 3986 | 3112 | 3267 | Mpp2 |
| Mpp3 | 2505 | 3265 | 2947 | 4013 | 4040 | 3661 | 4060 | 2385 | 2282 | 2985 | 2554 | 3156 | 2587 | 3175 | 2803 | 2805 | 3694 | 3921 | 4307 | 3520 | 3799 | 2744 | 2022 | 3011 | 3192 | 2453 | 2952 | 2935 | Mpp3 |
| Mpp4 | 223 | 252 | 185 | 402 | 422 | 294 | 393 | 140 | 138 | 142 | 120 | 74 | 103 | 96 | 268 | 287 | 280 | 463 | 357 | 294 | 364 | 172 | 124 | 99 | 131 | 61 | 99 | 70 | Mpp4 |
| Mpp7 | 3748 | 3096 | 3347 | 3573 | 3782 | 3743 | 4583 | 2209 | 1650 | 2044 | 1832 | 2294 | 1955 | 2570 | 4069 | 3406 | 3859 | 3811 | 4003 | 3591 | 3935 | 2146 | 1876 | 1973 | 2194 | 2290 | 2497 | 2349 | Mpp7 |
| Mppe1 | 250 | 162 | 327 | 149 | 192 | 190 | 126 | 451 | 508 | 557 | 589 | 468 | 718 | 513 | 219 | 293 | 225 | 178 | 161 | 176 | 136 | 456 | 576 | 496 | 396 | 607 | 572 | 489 | Mppe1 |
| Mpped1 | 1765 | 1951 | 1408 | 2624 | 3444 | 3173 | 4317 | 2171 | 3526 | 3332 | 2432 | 2996 | 1339 | 2767 | 1502 | 1181 | 2299 | 2917 | 4148 | 2818 | 4437 | 2105 | 2257 | 3735 | 4783 | 3373 | 1941 | 2161 | Mpped1 |
| Mpped2 | 2265 | 1488 | 1716 | 2677 | 3605 | 3072 | 3567 | 5804 | 6016 | 6998 | 6334 | 7798 | 6081 | 8393 | 1952 | 1616 | 2181 | 2633 | 3713 | 2735 | 3615 | 5536 | 5012 | 7853 | 7929 | 8193 | 6057 | 7164 | Mpped2 |
| Mprip | 26946 | 30485 | 27553 | 31285 | 32136 | 31672 | 29533 | 22659 | 25906 | 27650 | 26965 | 27317 | 24495 | 25143 | 25976 | 28810 | 29848 | 32926 | 32558 | 26681 | 32211 | 25044 | 25392 | 26279 | 28802 | 24512 | 24478 | 24343 | Mprip |
| Mpst | 2155 | 2691 | 2369 | 2606 | 2244 | 2177 | 2148 | 1421 | 1673 | 1695 | 1395 | 1745 | 1529 | 1469 | 1946 | 2184 | 2608 | 2406 | 2630 | 2343 | 2152 | 1454 | 1754 | 1772 | 1648 | 2182 | 1582 | 1645 | Mpst |
| Mptx2 | 0 | 0 | 0 | 0 | 0 | 0 | 0 | 0 | 0 | 0 | 0 | 0 | 0 | 0 | 0 | 0 | 0 | 0 | 0 | 0 | 1 | 0 | 0 | 0 | 0 | 0 | 0 | 0 | Mptx2 |
| Mpv17 | 1133 | 1139 | 1179 | 1146 | 895 | 960 | 1106 | 1330 | 1111 | 1284 | 1155 | 1304 | 1162 | 1554 | 1218 | 1069 | 1078 | 1054 | 971 | 886 | 1107 | 1407 | 1341 | 1352 | 1176 | 1472 | 1297 | 1572 | Mpv17 |
| Mpv17l | 982 | 745 | 835 | 1055 | 1337 | 1037 | 1217 | 564 | 552 | 531 | 552 | 373 | 519 | 449 | 1060 | 906 | 740 | 1030 | 1017 | 1212 | 1269 | 623 | 645 | 462 | 482 | 333 | 438 | 351 | Mpv17l |
| Mpz | 87 | 84 | 18 | 41 | 350 | 28 | 17 | 260 | 385 | 280 | 599 | 18 | 184 | 38 | 102 | 149 | 44 | 166 | 17 | 19 | 5 | 453 | 359 | 118 | 264 | 70 | 35 | 40 | Mpz |
| Mpzl1 | 11655 | 11921 | 12493 | 11888 | 10823 | 10656 | 10064 | 7299 | 6953 | 7464 | 7425 | 7111 | 6534 | 6579 | 11613 | 12325 | 11811 | 11855 | 11160 | 11341 | 10645 | 6776 | 7592 | 7473 | 7281 | 7649 | 6929 | 6687 | Mpzl1 |
| Mpzl2 | 420 | 174 | 160 | 75 | 65 | 167 | 22 | 457 | 588 | 371 | 436 | 146 | 276 | 180 | 233 | 242 | 87 | 122 | 39 | 40 | 26 | 174 | 641 | 400 | 75 | 328 | 115 | 142 | Mpzl2 |
| Mpzl3 | 62 | 55 | 50 | 27 | 39 | 58 | 42 | 245 | 239 | 182 | 172 | 231 | 265 | 226 | 70 | 40 | 49 | 41 | 36 | 37 | 48 | 209 | 236 | 222 | 198 | 171 | 265 | 234 | Mpzl3 |
| Mr1 | 546 | 325 | 196 | 284 | 262 | 335 | 182 | 276 | 420 | 313 | 367 | 118 | 229 | 110 | 451 | 562 | 217 | 277 | 161 | 225 | 205 | 237 | 485 | 268 | 179 | 190 | 127 | 135 | Mr1 |
| Mrap | 9 | 10 | 8 | 4 | 2 | 4 | 3 | 0 | 2 | 3 | 0 | 3 | 1 | 4 | 5 | 16 | 2 | 4 | 3 | 8 | 1 | 12 | 3 | 3 | 1 | 2 | 5 | 6 | Mrap |
| Mrap2 | 33 | 30 | 46 | 43 | 77 | 53 | 49 | 22 | 23 | 21 | 22 | 11 | 23 | 17 | 54 | 47 | 46 | 52 | 54 | 40 | 47 | 35 | 32 | 25 | 28 | 8 | 13 | 7 | Mrap2 |
| Mras | 2986 | 3183 | 2955 | 4167 | 5228 | 4812 | 4497 | 3817 | 4028 | 4479 | 3829 | 3582 | 3167 | 3539 | 3136 | 3131 | 3817 | 4656 | 5156 | 4181 | 4763 | 4852 | 3812 | 4171 | 4759 | 3398 | 2955 | 3164 | Mras |
| Mrc1 | 1901 | 1682 | 2117 | 1428 | 1707 | 1931 | 1538 | 1840 | 2382 | 1887 | 2787 | 2456 | 2766 | 2121 | 1902 | 2120 | 1556 | 2028 | 1195 | 2314 | 1607 | 2425 | 3717 | 2129 | 2206 | 1859 | 2642 | 2046 | Mrc1 |

| | | | | | | | | | | | | | | | | | | | | | | | | | | | | | |
|---|---|---|---|---|---|---|---|---|---|---|---|---|---|---|---|---|---|---|---|---|---|---|---|---|---|---|---|---|---|
| Mrc2 | 3665 | 4418 | 5515 | 3541 | 2855 | 3943 | 2196 | 4195 | 6387 | 5991 | 7588 | 6358 | 7386 | 5437 | 3330 | 4718 | 4118 | 4265 | 2904 | 2354 | 2221 | 5310 | 7022 | 6615 | 5038 | 7041 | 6465 | 5814 | Mrc2 |
| Mre11a | 3720 | 3586 | 2931 | 2894 | 2522 | 3133 | 3149 | 3323 | 3221 | 2950 | 3150 | 3839 | 3540 | 4133 | 3580 | 3262 | 2647 | 2542 | 2623 | 3463 | 3296 | 2540 | 3014 | 3185 | 2938 | 3825 | 3936 | 4078 | Mre11a |
| Mreg | 670 | 670 | 540 | 585 | 641 | 376 | 445 | 371 | 269 | 258 | 293 | 126 | 494 | 125 | 874 | 945 | 782 | 589 | 404 | 374 | 396 | 297 | 195 | 200 | 221 | 242 | 131 | 155 | Mreg |
| Mrfap1 | 18481 | 14785 | 19728 | 14954 | 13853 | 14400 | 14191 | 20262 | 18208 | 17786 | 20980 | 21411 | 21673 | 19674 | 17033 | 15798 | 16606 | 14801 | 14869 | 15825 | 12296 | 17645 | 18528 | 21194 | 18887 | 26704 | 22724 | 21748 | Mrfap1 |
| Mrgbp | 2384 | 2598 | 2668 | 2745 | 2609 | 2564 | 2769 | 2870 | 2661 | 2793 | 2627 | 3023 | 2768 | 3126 | 2469 | 2201 | 2840 | 2509 | 2740 | 2821 | 2738 | 2630 | 2564 | 3164 | 3132 | 3464 | 2915 | 3043 | Mrgbp |
| Mrgpra2a | 0 | 0 | 0 | 0 | 0 | 0 | 0 | 0 | 0 | 0 | 0 | 0 | 0 | 0 | 0 | 0 | 0 | 0 | 0 | 0 | 0 | 0 | 0 | 0 | 0 | 1 | 0 | 0 | Mrgpra2a |
| Mrgpra2b | 0 | 0 | 0 | 0 | 0 | 0 | 0 | 0 | 0 | 0 | 1 | 0 | 1 | 0 | 0 | 0 | 0 | 0 | 0 | 0 | 0 | 0 | 0 | 0 | 0 | 0 | 0 | 0 | Mrgpra2b |
| Mrgprd | 0 | 0 | 0 | 0 | 0 | 0 | 0 | 0 | 2 | 0 | 0 | 0 | 0 | 0 | 0 | 0 | 0 | 0 | 0 | 0 | 0 | 0 | 0 | 0 | 0 | 0 | 0 | 0 | Mrgprd |
| Mrgpre | 255 | 297 | 291 | 305 | 479 | 386 | 335 | 452 | 474 | 554 | 484 | 493 | 573 | 504 | 344 | 282 | 379 | 280 | 394 | 322 | 349 | 702 | 497 | 498 | 536 | 443 | 470 | 510 | Mrgpre |
| Mrgprf | 12 | 13 | 7 | 25 | 48 | 37 | 35 | 81 | 91 | 86 | 99 | 96 | 61 | 38 | 12 | 4 | 19 | 40 | 31 | 31 | 28 | 102 | 205 | 101 | 76 | 63 | 45 | 52 | Mrgprf |
| Mrgprh | 10 | 1 | 5 | 2 | 1 | 7 | 2 | 2 | 7 | 0 | 8 | 1 | 8 | 2 | 5 | 16 | 0 | 2 | 0 | 0 | 2 | 12 | 5 | 2 | 5 | 0 | 1 | 1 | Mrgprh |
| Mrgprx1 | 0 | 0 | 0 | 0 | 0 | 0 | 0 | 0 | 1 | 0 | 0 | 0 | 0 | 0 | 0 | 0 | 0 | 0 | 0 | 0 | 0 | 0 | 1 | 1 | 0 | 0 | 0 | 0 | Mrgprx1 |
| Mri1 | 1384 | 1470 | 1431 | 1505 | 1422 | 1423 | 1445 | 1327 | 1346 | 1401 | 1388 | 1565 | 1470 | 1594 | 1226 | 1260 | 1628 | 1459 | 1645 | 1365 | 1410 | 1477 | 1232 | 1470 | 1667 | 1586 | 1599 | 1603 | Mri1 |
| Mrln | 30 | 16 | 25 | 38 | 62 | 35 | 66 | 24 | 37 | 34 | 136 | 34 | 37 | 29 | 34 | 40 | 28 | 66 | 42 | 20 | 57 | 79 | 114 | 22 | 16 | 30 | 19 | 26 | Mrln |
| Mrm1 | 964 | 1278 | 1254 | 1205 | 1019 | 1039 | 953 | 539 | 593 | 583 | 595 | 665 | 552 | 583 | 1051 | 1017 | 1320 | 1165 | 1179 | 1124 | 1034 | 473 | 579 | 530 | 500 | 721 | 657 | 686 | Mrm1 |
| Mrm2 | 601 | 783 | 677 | 710 | 641 | 657 | 583 | 568 | 612 | 575 | 507 | 529 | 532 | 511 | 637 | 725 | 692 | 664 | 668 | 654 | 632 | 583 | 612 | 594 | 637 | 632 | 604 | 587 | Mrm2 |
| Mrm3 | 386 | 502 | 416 | 477 | 395 | 433 | 373 | 503 | 602 | 522 | 458 | 512 | 493 | 495 | 433 | 412 | 442 | 344 | 355 | 448 | 347 | 535 | 561 | 493 | 518 | 623 | 539 | 476 | Mrm3 |
| Mrnip | 874 | 935 | 901 | 973 | 843 | 868 | 926 | 772 | 885 | 803 | 793 | 1013 | 770 | 912 | 916 | 819 | 972 | 801 | 934 | 1025 | 953 | 812 | 869 | 925 | 877 | 909 | 906 | 845 | Mrnip |
| Mro | 332 | 372 | 366 | 488 | 500 | 443 | 558 | 217 | 280 | 304 | 290 | 300 | 271 | 261 | 316 | 308 | 445 | 429 | 533 | 416 | 488 | 256 | 245 | 284 | 279 | 258 | 275 | 260 | Mro |
| Mroh1 | 5266 | 7082 | 6482 | 7994 | 7251 | 6969 | 6090 | 2992 | 3353 | 3951 | 3562 | 3463 | 3317 | 2864 | 5413 | 6883 | 7579 | 7746 | 7412 | 5709 | 6290 | 4132 | 3295 | 3507 | 3717 | 2604 | 3194 | 2928 | Mroh1 |
| Mroh2a | 1 | 5 | 7 | 7 | 5 | 7 | 9 | 6 | 6 | 10 | 6 | 7 | 6 | 6 | 1 | 2 | 5 | 10 | 2 | 15 | 3 | 3 | 8 | 12 | 6 | 3 | 7 | 2 | Mroh2a |
| Mroh2b | 0 | 0 | 0 | 0 | 0 | 0 | 0 | 0 | 0 | 0 | 3 | 0 | 0 | 0 | 0 | 0 | 0 | 0 | 0 | 0 | 0 | 0 | 0 | 0 | 0 | 0 | 0 | 0 | Mroh2b |
| Mroh3 | 141 | 130 | 86 | 124 | 116 | 102 | 158 | 61 | 43 | 34 | 41 | 68 | 71 | 48 | 94 | 132 | 85 | 86 | 139 | 133 | 126 | 38 | 55 | 49 | 75 | 76 | 69 | 69 | Mroh3 |
| Mroh4 | 0 | 0 | 0 | 0 | 2 | 0 | 0 | 0 | 0 | 0 | 0 | 0 | 0 | 0 | 0 | 0 | 0 | 0 | 0 | 0 | 1 | 0 | 1 | 0 | 0 | 0 | 0 | 0 | Mroh4 |
| Mroh5 | 1 | 2 | 0 | 0 | 0 | 7 | 2 | 4 | 3 | 1 | 0 | 3 | 2 | 0 | 4 | 0 | 0 | 1 | 3 | 1 | 2 | 2 | 3 | 1 | 1 | 1 | 0 | 4 | Mroh5 |
| Mroh6 | 48 | 82 | 62 | 103 | 61 | 57 | 83 | 9 | 11 | 10 | 9 | 10 | 6 | 7 | 59 | 101 | 73 | 142 | 69 | 31 | 70 | 8 | 12 | 19 | 14 | 8 | 16 | 19 | Mroh6 |
| Mroh7 | 38 | 67 | 24 | 72 | 77 | 88 | 53 | 56 | 74 | 85 | 99 | 52 | 57 | 36 | 45 | 65 | 44 | 85 | 65 | 74 | 46 | 84 | 89 | 64 | 81 | 59 | 37 | 31 | Mroh7 |
| Mroh8 | 43 | 60 | 37 | 39 | 56 | 33 | 67 | 28 | 31 | 30 | 52 | 28 | 32 | 40 | 24 | 28 | 42 | 61 | 48 | 39 | 49 | 32 | 27 | 36 | 15 | 37 | 34 | 39 | Mroh8 |
| Mroh9 | 2 | 6 | 0 | 0 | 0 | 0 | 2 | 2 | 0 | 0 | 1 | 0 | 1 | 3 | 3 | 9 | 0 | 2 | 0 | 0 | 1 | 0 | 0 | 0 | 2 | 0 | 0 | 1 | Mroh9 |
| Mrpl1 | 1651 | 1410 | 1485 | 1445 | 1346 | 1538 | 1460 | 1833 | 1513 | 1455 | 1475 | 1677 | 1659 | 1789 | 1568 | 1557 | 1488 | 1358 | 1400 | 1512 | 1397 | 1663 | 1581 | 1620 | 1477 | 1833 | 1751 | 1858 | Mrpl1 |
| Mrpl10 | 2534 | 2747 | 2634 | 2305 | 2018 | 2136 | 2215 | 2026 | 2506 | 2516 | 2271 | 2461 | 2545 | 2491 | 2283 | 2523 | 2600 | 2149 | 2122 | 2157 | 2010 | 2141 | 2544 | 2431 | 2155 | 2887 | 2646 | 2549 | Mrpl10 |
| Mrpl11 | 2471 | 2587 | 2679 | 2285 | 2288 | 2352 | 2458 | 3178 | 3073 | 3096 | 3155 | 3413 | 3299 | 3865 | 2502 | 2220 | 2420 | 2127 | 2446 | 2840 | 2458 | 2562 | 2969 | 3389 | 3192 | 3946 | 3703 | 4017 | Mrpl11 |
| Mrpl12 | 3585 | 4380 | 3284 | 3438 | 2869 | 3126 | 3178 | 2928 | 3411 | 2823 | 2864 | 3106 | 3043 | 3185 | 3287 | 3443 | 3218 | 2960 | 3580 | 3436 | 3420 | 2383 | 3350 | 3101 | 2884 | 3796 | 3317 | 3419 | Mrpl12 |
| Mrpl13 | 4706 | 4025 | 3832 | 3743 | 3399 | 3602 | 4282 | 3872 | 3439 | 3227 | 3275 | 3910 | 3621 | 4500 | 4437 | 3797 | 3812 | 3320 | 3633 | 4222 | 3855 | 2943 | 3594 | 3614 | 3360 | 4711 | 4115 | 4374 | Mrpl13 |
| Mrpl14 | 1213 | 1468 | 1262 | 1312 | 1037 | 1122 | 1145 | 862 | 1016 | 980 | 988 | 828 | 971 | 961 | 1113 | 1100 | 1355 | 1292 | 1246 | 1289 | 1138 | 759 | 1086 | 987 | 951 | 1100 | 998 | 1024 | Mrpl14 |
| Mrpl15 | 3321 | 3146 | 2976 | 2792 | 2533 | 2538 | 3285 | 2985 | 2614 | 2721 | 2582 | 2914 | 2801 | 3080 | 3510 | 2788 | 2906 | 2598 | 2825 | 3136 | 3051 | 2675 | 2754 | 2880 | 2722 | 3256 | 2948 | 3432 | Mrpl15 |
| Mrpl16 | 1413 | 1519 | 1623 | 1370 | 1334 | 1370 | 1464 | 1461 | 1491 | 1375 | 1406 | 1517 | 1591 | 1713 | 1424 | 1305 | 1490 | 1387 | 1437 | 1449 | 1407 | 1250 | 1520 | 1499 | 1506 | 1881 | 1678 | 1791 | Mrpl16 |
| Mrpl17 | 2819 | 2819 | 2709 | 3088 | 2691 | 2764 | 3075 | 2165 | 2084 | 1933 | 2241 | 2189 | 2106 | 2471 | 2749 | 2447 | 2783 | 2714 | 2774 | 2687 | 2942 | 1875 | 2316 | 2294 | 1991 | 2475 | 2215 | 2294 | Mrpl17 |
| Mrpl18 | 2607 | 2531 | 2859 | 2501 | 2175 | 2190 | 2623 | 2855 | 2653 | 2888 | 2605 | 3261 | 2993 | 3599 | 2304 | 2308 | 2699 | 2250 | 2497 | 2700 | 2434 | 2182 | 2577 | 3339 | 2984 | 3717 | 3370 | 3769 | Mrpl18 |
| Mrpl19 | 2476 | 2524 | 2430 | 2308 | 2186 | 2424 | 2283 | 2839 | 2525 | 2644 | 2566 | 2721 | 2827 | 2968 | 2472 | 2262 | 2294 | 2112 | 2304 | 2359 | 2382 | 2319 | 2749 | 2586 | 2627 | 2887 | 3216 | 3110 | Mrpl19 |
| Mrpl2 | 1614 | 2151 | 1953 | 1691 | 1312 | 1512 | 1471 | 1348 | 1545 | 1560 | 1514 | 1725 | 1566 | 1851 | 1596 | 1625 | 1738 | 1562 | 1644 | 1505 | 1549 | 1135 | 1459 | 1789 | 1521 | 2180 | 1847 | 1775 | Mrpl2 |
| Mrpl20 | 3886 | 4156 | 3621 | 4045 | 3379 | 3401 | 4136 | 3259 | 3273 | 3055 | 2941 | 3125 | 3156 | 3931 | 4096 | 3581 | 3772 | 3401 | 3540 | 4471 | 4254 | 2627 | 3192 | 3286 | 3216 | 3529 | 3546 | 3982 | Mrpl20 |
| Mrpl21 | 1231 | 1367 | 1643 | 1258 | 1167 | 1172 | 1265 | 1769 | 1691 | 1736 | 1756 | 1831 | 1855 | 2274 | 1300 | 1241 | 1394 | 1045 | 1167 | 1471 | 1136 | 1379 | 1730 | 1961 | 1813 | 2229 | 2028 | 2127 | Mrpl21 |
| Mrpl22 | 1308 | 1447 | 1410 | 1281 | 1111 | 1070 | 1388 | 1114 | 1236 | 1115 | 1082 | 1266 | 1247 | 1450 | 1258 | 1334 | 1341 | 1121 | 1144 | 1295 | 1273 | 932 | 1173 | 1241 | 1137 | 1513 | 1363 | 1591 | Mrpl22 |
| Mrpl23 | 1140 | 1373 | 1398 | 1172 | 1141 | 1396 | 1323 | 1623 | 1886 | 1754 | 1700 | 1861 | 1913 | 2068 | 1031 | 1088 | 1337 | 1216 | 1277 | 1502 | 1252 | 1379 | 1687 | 1869 | 1930 | 2217 | 2060 | 2193 | Mrpl23 |
| Mrpl24 | 6809 | 7551 | 7545 | 7450 | 6166 | 6665 | 7342 | 5488 | 4941 | 5229 | 5239 | 5555 | 5593 | 6051 | 7304 | 6894 | 7519 | 7244 | 6895 | 7559 | 7069 | 5212 | 4963 | 5824 | 5350 | 6246 | 6015 | 6163 | Mrpl24 |
| Mrpl27 | 704 | 774 | 808 | 609 | 663 | 657 | 736 | 1428 | 1487 | 1505 | 1481 | 1672 | 1526 | 1700 | 766 | 675 | 801 | 595 | 655 | 746 | 650 | 1166 | 1494 | 1637 | 1391 | 1832 | 1687 | 1639 | Mrpl27 |
| Mrpl28 | 4092 | 5081 | 3905 | 3895 | 2923 | 3219 | 3461 | 2595 | 2815 | 2531 | 2306 | 2931 | 2749 | 2894 | 3816 | 3692 | 3801 | 3499 | 3808 | 3704 | 3363 | 2283 | 2619 | 2737 | 2784 | 3336 | 3008 | 3357 | Mrpl28 |
| Mrpl3 | 3027 | 2812 | 2858 | 2893 | 3019 | 3264 | 3148 | 4590 | 4348 | 4180 | 4083 | 4642 | 4591 | 4841 | 3045 | 2685 | 2745 | 2719 | 2998 | 3054 | 2990 | 4000 | 4313 | 4256 | 4383 | 5003 | 5017 | 4999 | Mrpl3 |
| Mrpl30 | 3793 | 3999 | 3405 | 3717 | 3245 | 3329 | 3844 | 3270 | 3431 | 2976 | 2911 | 2999 | 3097 | 3510 | 3944 | 3858 | 3686 | 3384 | 3671 | 4297 | 3651 | 2951 | 3270 | 3192 | 3123 | 3522 | 3318 | 3652 | Mrpl30 |
| Mrpl32 | 1987 | 1826 | 2132 | 2253 | 2334 | 2027 | 2538 | 2862 | 2693 | 2397 | 2328 | 2169 | 2380 | 2670 | 2347 | 1863 | 1900 | 2045 | 2349 | 2761 | 2447 | 2376 | 2716 | 2580 | 2514 | 2524 | 2574 | 2693 | Mrpl32 |
| Mrpl33 | 1208 | 941 | 794 | 829 | 736 | 890 | 993 | 1106 | 865 | 840 | 902 | 873 | 937 | 998 | 1216 | 921 | 773 | 765 | 735 | 910 | 882 | 829 | 881 | 876 | 876 | 801 | 841 | 924 | Mrpl33 |
| Mrpl34 | 2023 | 2701 | 2407 | 2384 | 2147 | 2024 | 2086 | 1550 | 1462 | 1544 | 1479 | 1929 | 1777 | 1798 | 1937 | 1825 | 2363 | 1964 | 2402 | 2352 | 2074 | 1402 | 1419 | 1959 | 1811 | 1942 | 1988 | 1901 | Mrpl34 |
| Mrpl35 | 1273 | 1245 | 1185 | 974 | 1019 | 1021 | 1181 | 1988 | 1648 | 1676 | 1427 | 1575 | 1675 | 1911 | 1352 | 1209 | 1109 | 1049 | 1054 | 1356 | 1124 | 1634 | 1643 | 1661 | 1596 | 1884 | 1694 | 2067 | Mrpl35 |
| Mrpl36 | 1466 | 1766 | 1477 | 1258 | 1183 | 1188 | 1336 | 1335 | 1383 | 1248 | 1320 | 1418 | 1542 | 1500 | 1485 | 1474 | 1577 | 1165 | 1270 | 1402 | 1260 | 1188 | 1390 | 1347 | 1215 | 1782 | 1468 | 1533 | Mrpl36 |
| Mrpl37 | 3325 | 3600 | 3140 | 2906 | 2806 | 2914 | 2723 | 3748 | 3985 | 3901 | 3589 | 4230 | 4253 | 4045 | 3088 | 3294 | 3034 | 2670 | 3094 | 2994 | 2751 | 3666 | 3797 | 3909 | 3950 | 4204 | 4479 | 4370 | Mrpl37 |
| Mrpl38 | 2321 | 2947 | 2555 | 2483 | 2036 | 1995 | 2005 | 1702 | 1914 | 1824 | 1663 | 2073 | 2025 | 1989 | 2107 | 2231 | 2649 | 2262 | 2279 | 2026 | 1926 | 1612 | 1812 | 2034 | 1874 | 2388 | 2143 | 2131 | Mrpl38 |
| Mrpl39 | 3015 | 2789 | 2880 | 2650 | 2294 | 2440 | 2750 | 2956 | 2489 | 2258 | 2354 | 2899 | 2631 | 2954 | 3049 | 2447 | 2651 | 2550 | 2526 | 3006 | 2766 | 2455 | 2473 | 2607 | 2413 | 3219 | 2963 | 3024 | Mrpl39 |
| Mrpl4 | 2362 | 2948 | 2439 | 2528 | 2041 | 2076 | 1845 | 1810 | 2062 | 1910 | 1956 | 2112 | 2081 | 1940 | 2119 | 2421 | 2474 | 2105 | 2254 | 2124 | 1909 | 1806 | 1878 | 2071 | 1914 | 2427 | 1987 | 2108 | Mrpl4 |
| Mrpl40 | 1359 | 1278 | 1425 | 972 | 1057 | 1012 | 1206 | 1584 | 1542 | 1591 | 1450 | 1732 | 1626 | 1932 | 1365 | 1162 | 1162 | 988 | 1019 | 1113 | 1083 | 1216 | 1521 | 1579 | 1477 | 1921 | 1693 | 1966 | Mrpl40 |
| Mrpl41 | 2598 | 2897 | 2268 | 2628 | 2271 | 2267 | 2729 | 1871 | 1947 | 1740 | 1634 | 1770 | 1871 | 2126 | 2774 | 2383 | 2642 | 2300 | 2212 | 3128 | 2737 | 1661 | 1908 | 1897 | 1647 | 2058 | 1849 | 2066 | Mrpl41 |
| Mrpl42 | 2738 | 2826 | 3101 | 2460 | 2231 | 2148 | 2471 | 2746 | 2574 | 2388 | 2406 | 2792 | 2951 | 2936 | 2846 | 2664 | 2808 | 2410 | 2525 | 3012 | 2309 | 2316 | 2708 | 3006 | 2561 | 3552 | 3108 | 3298 | Mrpl42 |
| Mrpl43 | 2647 | 3159 | 2867 | 2845 | 2468 | 2421 | 2797 | 1489 | 1681 | 1905 | 1665 | 2056 | 1842 | 1945 | 2567 | 2453 | 3120 | 2776 | 2712 | 2596 | 2574 | 1604 | 1689 | 1991 | 1954 | 2262 | 2012 | 1906 | Mrpl43 |
| Mrpl44 | 1051 | 1137 | 1310 | 1119 | 1059 | 1093 | 1201 | 1769 | 1546 | 1569 | 1482 | 1846 | 1651 | 1845 | 1242 | 977 | 1253 | 1020 | 1254 | 1302 | 1138 | 1322 | 1578 | 1871 | 1676 | 2044 | 2011 | 1901 | Mrpl44 |
| Mrpl45 | 1253 | 1389 | 1476 | 1150 | 1163 | 1200 | 1162 | 2400 | 2404 | 2307 | 2215 | 2537 | 2354 | 2474 | 1309 | 1307 | 1236 | 1161 | 1248 | 1250 | 1101 | 2127 | 2438 | 2428 | 2257 | 2887 | 2551 | 2759 | Mrpl45 |
| Mrpl46 | 1002 | 1160 | 1148 | 1000 | 943 | 799 | 1052 | 1225 | 1187 | 1244 | 1245 | 1355 | 1302 | 1387 | 997 | 970 | 1187 | 968 | 1035 | 1139 | 954 | 901 | 1188 | 1311 | 1275 | 1630 | 1539 | 1570 | Mrpl46 |
| Mrpl47 | 2396 | 2092 | 2187 | 1957 | 1642 | 1906 | 2257 | 2332 | 1966 | 1804 | 1723 | 2119 | 2074 | 2533 | 2505 | 2069 | 2231 | 1946 | 1969 | 2347 | 2141 | 1905 | 1945 | 2185 | 2018 | 2325 | 2360 | 2389 | Mrpl47 |
| Mrpl48 | 1125 | 1226 | 1356 | 1175 | 1171 | 1214 | 1340 | 2467 | 2374 | 2399 | 2234 | 2258 | 2210 | 2427 | 1327 | 1115 | 1258 | 1103 | 1248 | 1484 | 1293 | 2141 | 2395 | 2268 | 2361 | 2357 | 2337 | 2460 | Mrpl48 |
| Mrpl49 | 4791 | 5332 | 4421 | 4055 | 3203 | 3589 | 3921 | 2980 | 2726 | 2498 | 2365 | 2760 | 2740 | 3081 | 4770 | 4569 | 3898 | 4004 | 3659 | 4079 | 3608 | 2597 | 2702 | 2800 | 2715 | 3427 | 2993 | 3143 | Mrpl49 |
| Mrpl50 | 6007 | 4799 | 5618 | 5028 | 4674 | 4852 | 5668 | 5785 | 4631 | 4397 | 4581 | 4889 | 4835 | 6123 | 6589 | 5329 | 5317 | 4907 | 4606 | 5908 | 5167 | 4836 | 4932 | 5162 | 4761 | 6136 | 5156 | 5995 | Mrpl50 |
| Mrpl51 | 3045 | 3329 | 3350 | 3125 | 2783 | 2794 | 2946 | 2908 | 2894 | 2835 | 2938 | 3360 | 3309 | 3866 | 2879 | 2783 | 3275 | 2697 | 2787 | 3207 | 2836 | 2400 | 2975 | 3528 | 3185 | 4082 | 3567 | 3776 | Mrpl51 |
| Mrpl52 | 1632 | 1937 | 1800 | 1900 | 1666 | 1444 | 2190 | 1603 | 1589 | 1558 | 1694 | 1585 | 1685 | 2206 | 1547 | 1232 | 2173 | 1639 | 1591 | 1622 | 2057 | 1390 | 1504 | 2082 | 1912 | 2038 | 1862 | 2241 | Mrpl52 |
| Mrpl54 | 497 | 603 | 456 | 424 | 300 | 355 | 326 | 995 | 1047 | 1086 | 1035 | 1081 | 1004 | 1243 | 546 | 436 | 448 | 380 | 327 | 327 | 382 | 705 | 1090 | 1077 | 934 | 1200 | 1120 | 1309 | Mrpl54 |
| Mrpl55 | 1753 | 2240 | 1776 | 1754 | 1528 | 1632 | 1662 | 1233 | 1342 | 1259 | 1270 | 1325 | 1376 | 1278 | 1553 | 1798 | 1866 | 1679 | 1838 | 2023 | 1676 | 1021 | 1260 | 1344 | 1277 | 1689 | 1561 | 1500 | Mrpl55 |
| Mrpl57 | 873 | 972 | 922 | 876 | 864 | 734 | 920 | 1033 | 986 | 978 | 1022 | 973 | 1024 | 1125 | 810 | 705 | 1067 | 831 | 929 | 889 | 886 | 1018 | 1047 | 1057 | 1156 | 1206 | 1033 | 1173 | Mrpl57 |
| Mrpl58 | 1727 | 2112 | 2048 | 1701 | 1728 | 1891 | 1996 | 2668 | 2386 | 2566 | 2482 | 3228 | 3029 | 3402 | 1788 | 1424 | 1948 | 1591 | 2062 | 1977 | 1868 | 2547 | 2427 | 3014 | 2936 | 3377 | 3466 | 3634 | Mrpl58 |
| Mrpl9 | 3305 | 3819 | 3799 | 3501 | 3174 | 3175 | 3362 | 3646 | 3350 | 3279 | 3184 | 3667 | 3537 | 3955 | 3422 | 3530 | 3874 | 3352 | 3239 | 3403 | 3301 | 3252 | 3414 | 3621 | 3368 | 4345 | 3987 | 4087 | Mrpl9 |
| Mrps11 | 2016 | 2555 | 2157 | 2375 | 1999 | 2075 | 1921 | 1628 | 1833 | 1660 | 1813 | 1913 | 2022 | 1827 | 1861 | 2120 | 2245 | 2346 | 2288 | 2229 | 1906 | 1536 | 1777 | 2035 | 1786 | 2273 | 2236 | 1919 | Mrps11 |
| Mrps12 | 1242 | 1557 | 1502 | 1421 | 1089 | 1095 | 1212 | 869 | 1000 | 962 | 1026 | 1023 | 1009 | 1043 | 1124 | 1214 | 1506 | 1282 | 1171 | 1253 | 1049 | 678 | 1006 | 1115 | 959 | 1199 | 1016 | 1004 | Mrps12 |

| | | | | | | | | | | | | | | | | | | | | | | | | | | | | | |
|---|---|---|---|---|---|---|---|---|---|---|---|---|---|---|---|---|---|---|---|---|---|---|---|---|---|---|---|---|---|
| Mrps14 | 2040 | 1773 | 2052 | 1844 | 1619 | 1541 | 2044 | 2205 | 1952 | 1881 | 1883 | 2353 | 2176 | 2552 | 2144 | 1671 | 2038 | 1699 | 1674 | 2143 | 1902 | 1634 | 1994 | 2344 | 2105 | 2737 | 2692 | 2705 | Mrps14 |
| Mrps15 | 1705 | 1756 | 1525 | 1317 | 1441 | 1335 | 1549 | 2397 | 2184 | 2140 | 2081 | 2173 | 2294 | 2421 | 1702 | 1611 | 1581 | 1465 | 1476 | 1594 | 1481 | 2309 | 2139 | 2211 | 2132 | 2440 | 2337 | 2464 | Mrps15 |
| Mrps16 | 919 | 1137 | 1118 | 1009 | 993 | 870 | 964 | 1055 | 1392 | 1198 | 1443 | 1227 | 1397 | 1400 | 1005 | 1016 | 1110 | 797 | 984 | 1094 | 953 | 988 | 1478 | 1490 | 1169 | 1442 | 1477 | 1466 | Mrps16 |
| Mrps17 | 5852 | 5919 | 5285 | 5573 | 5210 | 5262 | 6327 | 4938 | 4879 | 4746 | 4539 | 4508 | 4578 | 5193 | 5917 | 5478 | 5743 | 5741 | 5534 | 6114 | 6002 | 4535 | 4887 | 4881 | 4813 | 4821 | 4728 | 5064 | Mrps17 |
| Mrps18a | 1046 | 1131 | 1245 | 1107 | 1135 | 1116 | 1054 | 1491 | 1788 | 1718 | 1731 | 1682 | 1738 | 1855 | 1005 | 1076 | 1205 | 1052 | 1199 | 1103 | 1092 | 1516 | 1720 | 1830 | 1670 | 1907 | 1804 | 1902 | Mrps18a |
| Mrps18b | 961 | 1128 | 945 | 875 | 867 | 826 | 811 | 863 | 1128 | 938 | 973 | 1111 | 1097 | 978 | 959 | 956 | 1045 | 856 | 1004 | 988 | 785 | 809 | 1137 | 990 | 1061 | 1288 | 1303 | 1044 | Mrps18b |
| Mrps18c | 588 | 438 | 393 | 422 | 352 | 265 | 574 | 536 | 369 | 458 | 461 | 444 | 424 | 738 | 494 | 282 | 435 | 333 | 309 | 404 | 639 | 500 | 347 | 560 | 461 | 463 | 447 | 762 | Mrps18c |
| Mrps2 | 2951 | 2886 | 2887 | 2560 | 2535 | 2648 | 2830 | 2787 | 2824 | 2880 | 2623 | 3133 | 2852 | 3242 | 2867 | 2761 | 2958 | 2714 | 2846 | 2701 | 2737 | 2653 | 2713 | 3103 | 2676 | 3673 | 2933 | 3217 | Mrps2 |
| Mrps21 | 1556 | 1831 | 2189 | 1963 | 1760 | 1562 | 2202 | 1835 | 1859 | 2276 | 2238 | 2411 | 2224 | 2831 | 1560 | 1401 | 2145 | 1811 | 2000 | 2001 | 2078 | 1419 | 1925 | 2675 | 2452 | 2752 | 2458 | 3014 | Mrps21 |
| Mrps22 | 2886 | 2663 | 2562 | 2498 | 2167 | 2316 | 2570 | 2235 | 1939 | 1725 | 1671 | 2155 | 2059 | 2205 | 3096 | 2670 | 2553 | 2261 | 2359 | 2954 | 2398 | 1744 | 1950 | 1960 | 1847 | 2320 | 2080 | 2124 | Mrps22 |
| Mrps23 | 145 | 71 | 83 | 216 | 77 | 77 | 77 | 110 | 48 | 248 | 59 | 80 | 268 | 54 | 132 | 178 | 227 | 136 | 83 | 101 | 87 | 122 | 60 | 168 | 75 | 66 | 91 | 74 | Mrps23 |
| Mrps24 | 1183 | 1286 | 1052 | 1233 | 1068 | 988 | 1268 | 1242 | 1242 | 1189 | 1218 | 1315 | 1173 | 1323 | 1091 | 1017 | 1258 | 1082 | 1206 | 1411 | 1253 | 1016 | 1276 | 1441 | 1393 | 1419 | 1352 | 1426 | Mrps24 |
| Mrps25 | 2835 | 2769 | 2813 | 2506 | 2060 | 2311 | 2542 | 2678 | 2464 | 2460 | 2341 | 2438 | 2660 | 2726 | 2694 | 2671 | 2760 | 2249 | 2412 | 2556 | 2481 | 2059 | 2481 | 2647 | 2574 | 2820 | 2767 | 3007 | Mrps25 |
| Mrps26 | 2074 | 2256 | 2211 | 2322 | 2120 | 2032 | 2134 | 1579 | 1543 | 1806 | 1767 | 1782 | 1610 | 1802 | 1805 | 1962 | 2257 | 2222 | 2238 | 1771 | 2117 | 1791 | 1557 | 1775 | 1733 | 1961 | 1809 | 1718 | Mrps26 |
| Mrps27 | 2130 | 2371 | 2119 | 1876 | 1552 | 1680 | 1783 | 2104 | 2202 | 1848 | 1908 | 2364 | 2357 | 2218 | 2024 | 2122 | 1827 | 1601 | 1792 | 1734 | 1736 | 1718 | 2198 | 2118 | 2159 | 2698 | 2248 | 2612 | Mrps27 |
| Mrps28 | 325 | 413 | 404 | 337 | 299 | 326 | 345 | 457 | 515 | 554 | 504 | 641 | 578 | 641 | 326 | 303 | 415 | 284 | 402 | 432 | 319 | 299 | 505 | 618 | 529 | 693 | 652 | 675 | Mrps28 |
| Mrps30 | 2008 | 1751 | 1569 | 1644 | 1553 | 1437 | 1771 | 2395 | 2233 | 1977 | 2082 | 2194 | 2265 | 2374 | 1924 | 1659 | 1677 | 1443 | 1741 | 1558 | 1800 | 2202 | 2308 | 2223 | 2031 | 2668 | 2184 | 2572 | Mrps30 |
| Mrps31 | 1737 | 1721 | 1708 | 1554 | 1489 | 1516 | 1735 | 1658 | 1695 | 1505 | 1563 | 1642 | 1569 | 1840 | 1957 | 1635 | 1773 | 1498 | 1544 | 1908 | 1615 | 1424 | 1636 | 1597 | 1504 | 1784 | 1671 | 1860 | Mrps31 |
| Mrps33 | 2315 | 2298 | 2358 | 2258 | 1843 | 1674 | 2615 | 2677 | 2174 | 2266 | 2017 | 2194 | 1923 | 2946 | 2583 | 1884 | 2250 | 1881 | 2050 | 2278 | 2611 | 2002 | 2158 | 2240 | 2257 | 2549 | 2363 | 3021 | Mrps33 |
| Mrps34 | 1353 | 1469 | 1741 | 1622 | 1458 | 1383 | 1351 | 1213 | 1278 | 1547 | 1475 | 1651 | 1504 | 1535 | 1280 | 1270 | 1763 | 1465 | 1577 | 1330 | 1331 | 1285 | 1363 | 1687 | 1551 | 1932 | 1553 | 1476 | Mrps34 |
| Mrps35 | 1081 | 952 | 1035 | 1041 | 1069 | 1012 | 1154 | 1745 | 1809 | 1701 | 1792 | 1935 | 1776 | 1980 | 984 | 884 | 999 | 861 | 1181 | 1010 | 1048 | 1586 | 1692 | 1934 | 1884 | 2221 | 1898 | 2034 | Mrps35 |
| Mrps36 | 681 | 745 | 916 | 828 | 821 | 792 | 898 | 807 | 856 | 884 | 1001 | 876 | 1108 | 1165 | 686 | 663 | 867 | 734 | 906 | 979 | 931 | 708 | 856 | 1018 | 955 | 1015 | 1063 | 1174 | Mrps36 |
| Mrps5 | 3749 | 4150 | 3916 | 3903 | 4081 | 4099 | 4157 | 5620 | 5370 | 4895 | 4978 | 5272 | 5006 | 5753 | 3831 | 3593 | 3919 | 3735 | 4198 | 4631 | 4088 | 4881 | 5133 | 5329 | 5689 | 5389 | 5698 | 5603 | Mrps5 |
| Mrps6 | 1753 | 1675 | 1571 | 1848 | 1462 | 1301 | 2080 | 1147 | 921 | 1140 | 1062 | 1017 | 1012 | 1232 | 1780 | 1151 | 1633 | 1613 | 1556 | 1568 | 2003 | 1208 | 1009 | 1213 | 1078 | 1048 | 815 | 1276 | Mrps6 |
| Mrps7 | 2135 | 2562 | 2307 | 2037 | 1905 | 1873 | 1922 | 2130 | 2240 | 2336 | 2386 | 2610 | 2498 | 2527 | 1993 | 2157 | 2213 | 2012 | 2079 | 2245 | 2037 | 1820 | 2368 | 2378 | 2166 | 2933 | 2619 | 2735 | Mrps7 |
| Mrps9 | 3841 | 3842 | 3924 | 3536 | 3224 | 3559 | 3589 | 2999 | 2981 | 2528 | 2673 | 3372 | 3275 | 3432 | 3927 | 3716 | 3473 | 3230 | 3527 | 4072 | 3475 | 2351 | 3104 | 2979 | 2823 | 3446 | 3565 | 3612 | Mrps9 |
| Mrrf | 1039 | 1073 | 1013 | 1132 | 1118 | 1071 | 1188 | 1986 | 1963 | 1745 | 1695 | 1727 | 1727 | 1711 | 1121 | 1024 | 1077 | 1001 | 1206 | 1180 | 1125 | 1895 | 1961 | 1876 | 1875 | 1712 | 1804 | 1786 | Mrrf |
| Mrs2 | 9720 | 9653 | 7514 | 9670 | 9709 | 10472 | 10594 | 7720 | 7297 | 7877 | 7041 | 8326 | 8031 | 8357 | 10213 | 9332 | 8905 | 9206 | 9056 | 9677 | 10389 | 7352 | 6750 | 6716 | 8158 | 5649 | 8280 | 7834 | Mrs2 |
| Mrtfa | 2971 | 3841 | 3934 | 4454 | 4300 | 3695 | 3247 | 2212 | 2653 | 2895 | 3101 | 3169 | 2465 | 2707 | 2807 | 3666 | 4195 | 4478 | 4157 | 2769 | 3598 | 3349 | 2658 | 3099 | 3126 | 3249 | 2618 | 2759 | Mrtfa |
| Mrtfb | 2982 | 2711 | 2189 | 2482 | 2671 | 2727 | 2448 | 3309 | 4005 | 3668 | 3895 | 3755 | 4140 | 3663 | 2676 | 3125 | 2283 | 2442 | 2311 | 2194 | 2279 | 3370 | 3848 | 3276 | 3696 | 2804 | 3710 | 3401 | Mrtfb |
| Mrto4 | 1897 | 2020 | 2001 | 1883 | 1664 | 1828 | 1914 | 2081 | 2132 | 2193 | 2147 | 2578 | 2490 | 2498 | 1833 | 1783 | 1984 | 1719 | 1838 | 1779 | 1896 | 1923 | 1953 | 2537 | 2400 | 3185 | 2662 | 2820 | Mrto4 |
| Ms4a1 | 0 | 1 | 0 | 0 | 0 | 0 | 0 | 0 | 0 | 0 | 0 | 0 | 0 | 0 | 0 | 0 | 1 | 0 | 0 | 0 | 0 | 0 | 0 | 0 | 0 | 4 | 0 | 0 | Ms4a1 |
| Ms4a10 | 0 | 0 | 0 | 0 | 0 | 0 | 0 | 2 | 0 | 0 | 3 | 0 | 0 | 3 | 1 | 1 | 0 | 0 | 0 | 0 | 0 | 0 | 0 | 0 | 0 | 0 | 0 | 4 | Ms4a10 |
| Ms4a13 | 0 | 0 | 0 | 0 | 0 | 0 | 0 | 0 | 0 | 0 | 0 | 0 | 0 | 0 | 0 | 0 | 2 | 0 | 0 | 0 | 0 | 0 | 0 | 0 | 0 | 0 | 0 | 0 | Ms4a13 |
| Ms4a14 | 17 | 13 | 15 | 22 | 31 | 18 | 11 | 10 | 21 | 24 | 38 | 23 | 18 | 26 | 20 | 23 | 6 | 24 | 16 | 8 | 15 | 32 | 52 | 28 | 8 | 10 | 24 | 17 | Ms4a14 |
| Ms4a15 | 0 | 0 | 0 | 0 | 0 | 0 | 0 | 1 | 0 | 0 | 0 | 0 | 0 | 0 | 0 | 0 | 0 | 0 | 0 | 0 | 0 | 0 | 0 | 0 | 0 | 0 | 0 | 0 | Ms4a15 |
| Ms4a2 | 2 | 0 | 10 | 2 | 3 | 10 | 0 | 12 | 8 | 10 | 27 | 4 | 3 | 0 | 9 | 2 | 5 | 0 | 4 | 4 | 1 | 13 | 9 | 0 | 4 | 1 | 3 | 2 | Ms4a2 |
| Ms4a20 | 0 | 0 | 0 | 0 | 0 | 2 | 0 | 0 | 0 | 0 | 0 | 0 | 0 | 0 | 0 | 0 | 0 | 0 | 0 | 0 | 0 | 0 | 0 | 0 | 0 | 0 | 0 | 0 | Ms4a20 |
| Ms4a3 | 1 | 1 | 0 | 1 | 0 | 0 | 0 | 0 | 0 | 0 | 6 | 1 | 0 | 0 | 0 | 2 | 0 | 0 | 0 | 0 | 4 | 0 | 2 | 3 | 0 | 0 | 0 | 0 | Ms4a3 |
| Ms4a4a | 32 | 29 | 52 | 26 | 38 | 56 | 27 | 60 | 69 | 59 | 126 | 61 | 70 | 65 | 49 | 61 | 35 | 25 | 26 | 35 | 28 | 77 | 112 | 83 | 70 | 93 | 67 | 74 | Ms4a4a |
| Ms4a4b | 0 | 0 | 0 | 1 | 0 | 0 | 0 | 0 | 0 | 1 | 0 | 0 | 2 | 1 | 0 | 0 | 0 | 0 | 0 | 0 | 0 | 2 | 1 | 0 | 0 | 0 | 2 | 5 | Ms4a4b |
| Ms4a4c | 3 | 2 | 0 | 5 | 2 | 2 | 9 | 2 | 5 | 11 | 6 | 3 | 5 | 6 | 3 | 10 | 0 | 1 | 0 | 2 | 5 | 3 | 5 | 2 | 4 | 0 | 0 | 1 | Ms4a4c |
| Ms4a4d | 3 | 8 | 4 | 0 | 0 | 0 | 0 | 4 | 7 | 3 | 10 | 10 | 2 | 1 | 3 | 4 | 6 | 0 | 0 | 1 | 0 | 3 | 6 | 1 | 9 | 4 | 6 | 7 | Ms4a4d |
| Ms4a5 | 0 | 0 | 0 | 0 | 0 | 0 | 0 | 0 | 0 | 0 | 0 | 0 | 0 | 0 | 0 | 0 | 0 | 0 | 0 | 1 | 0 | 0 | 0 | 0 | 0 | 0 | 0 | 0 | Ms4a5 |
| Ms4a6b | 122 | 86 | 146 | 65 | 125 | 122 | 85 | 186 | 250 | 181 | 238 | 242 | 226 | 176 | 144 | 137 | 92 | 68 | 73 | 133 | 79 | 221 | 381 | 167 | 202 | 214 | 216 | 198 | Ms4a6b |
| Ms4a6c | 144 | 113 | 150 | 99 | 121 | 102 | 68 | 186 | 222 | 221 | 304 | 240 | 257 | 218 | 149 | 165 | 116 | 123 | 74 | 120 | 103 | 249 | 423 | 197 | 206 | 253 | 221 | 237 | Ms4a6c |
| Ms4a6d | 96 | 105 | 97 | 78 | 70 | 72 | 74 | 121 | 140 | 89 | 154 | 75 | 100 | 77 | 121 | 162 | 95 | 77 | 55 | 116 | 69 | 110 | 181 | 103 | 73 | 126 | 115 | 116 | Ms4a6d |
| Ms4a7 | 63 | 53 | 44 | 58 | 59 | 65 | 49 | 81 | 103 | 60 | 142 | 59 | 75 | 63 | 47 | 63 | 28 | 78 | 50 | 59 | 36 | 97 | 188 | 99 | 100 | 53 | 89 | 93 | Ms4a7 |
| Ms4a8a | 0 | 0 | 0 | 1 | 0 | 3 | 0 | 0 | 0 | 0 | 0 | 0 | 0 | 0 | 0 | 0 | 0 | 0 | 0 | 0 | 0 | 0 | 0 | 0 | 0 | 0 | 0 | 4 | Ms4a8a |
| Msantd1 | 202 | 337 | 249 | 329 | 277 | 305 | 294 | 201 | 160 | 188 | 217 | 203 | 187 | 198 | 202 | 290 | 279 | 347 | 278 | 278 | 268 | 231 | 157 | 210 | 215 | 208 | 180 | 147 | Msantd1 |
| Msantd2 | 3598 | 3081 | 4342 | 4303 | 4394 | 4367 | 4929 | 4169 | 3639 | 4509 | 4708 | 5073 | 4935 | 5313 | 3670 | 3331 | 4726 | 4233 | 4564 | 3477 | 4669 | 5360 | 3488 | 4555 | 4713 | 5018 | 4977 | 5231 | Msantd2 |
| Msantd3 | 1769 | 1811 | 2374 | 1956 | 2048 | 1842 | 2093 | 3185 | 2862 | 2992 | 2700 | 3619 | 3323 | 3891 | 1797 | 1546 | 2245 | 1711 | 2248 | 2113 | 1912 | 2775 | 2633 | 3550 | 3552 | 3876 | 3828 | 4106 | Msantd3 |
| Msantd4 | 9811 | 8734 | 9729 | 8529 | 8046 | 8351 | 9014 | 9745 | 7944 | 7172 | 6576 | 8962 | 7785 | 9627 | 10226 | 8866 | 9048 | 7846 | 8387 | 10184 | 8524 | 8174 | 7531 | 8237 | 7858 | 9491 | 9828 | 9765 | Msantd4 |
| Msc | 43 | 27 | 37 | 6 | 41 | 24 | 15 | 90 | 129 | 38 | 523 | 14 | 134 | 4 | 34 | 51 | 42 | 15 | 16 | 20 | 11 | 105 | 326 | 24 | 51 | 15 | 7 | 11 | Msc |
| Msgn1 | 0 | 0 | 0 | 0 | 0 | 0 | 0 | 0 | 0 | 0 | 0 | 0 | 0 | 0 | 0 | 0 | 1 | 1 | 0 | 0 | 1 | 0 | 0 | 0 | 0 | 0 | 0 | 0 | Msgn1 |
| Msh2 | 12555 | 10797 | 12497 | 11039 | 10381 | 11141 | 11504 | 8255 | 7703 | 7451 | 7065 | 9644 | 8837 | 9703 | 11884 | 11073 | 12307 | 10763 | 11263 | 11054 | 11261 | 7436 | 7120 | 8594 | 8035 | 10723 | 9976 | 10017 | Msh2 |
| Msh3 | 2936 | 2621 | 2379 | 2332 | 2392 | 2409 | 2910 | 3932 | 3696 | 3176 | 3103 | 3547 | 3339 | 3689 | 2988 | 2590 | 2199 | 2286 | 2384 | 3097 | 2668 | 3334 | 3553 | 3434 | 3413 | 3836 | 3678 | 3723 | Msh3 |
| Msh4 | 42 | 20 | 5 | 27 | 21 | 61 | 20 | 21 | 25 | 8 | 25 | 11 | 7 | 27 | 28 | 33 | 8 | 23 | 21 | 29 | 51 | 20 | 21 | 10 | 14 | 21 | 19 | 18 | Msh4 |
| Msh5 | 983 | 1051 | 993 | 1271 | 844 | 1013 | 1097 | 294 | 343 | 433 | 329 | 591 | 451 | 572 | 938 | 939 | 1161 | 1161 | 1099 | 897 | 1066 | 391 | 327 | 425 | 422 | 430 | 652 | 528 | Msh5 |
| Msh6 | 13478 | 12740 | 12515 | 11203 | 9043 | 10090 | 11165 | 9544 | 8483 | 7756 | 7454 | 10454 | 9560 | 10681 | 12511 | 12144 | 11471 | 11190 | 10403 | 10978 | 10450 | 7327 | 7513 | 9072 | 7958 | 11918 | 11169 | 11244 | Msh6 |
| Msi1 | 3774 | 4612 | 9596 | 6417 | 7074 | 5792 | 5660 | 9695 | 9127 | 13898 | 10894 | 20661 | 15423 | 20506 | 3952 | 3646 | 8842 | 5567 | 8008 | 5058 | 6200 | 13353 | 8600 | 18646 | 16349 | 20984 | 18384 | 20952 | Msi1 |
| Msi2 | 12973 | 10824 | 18140 | 18378 | 17346 | 17719 | 19109 | 14736 | 12779 | 17089 | 16014 | 19548 | 19463 | 21645 | 13486 | 11642 | 18250 | 17201 | 17146 | 15040 | 18586 | 18554 | 12286 | 19741 | 19029 | 19045 | 20087 | 22574 | Msi2 |
| Msl1 | 8315 | 8258 | 10907 | 10934 | 12040 | 11179 | 10829 | 18387 | 15856 | 19342 | 18818 | 20897 | 19666 | 21577 | 8213 | 7379 | 11121 | 10274 | 12866 | 10014 | 11130 | 21118 | 15153 | 21223 | 21372 | 21737 | 20692 | 21942 | Msl1 |
| Msl2 | 6029 | 5145 | 6201 | 6596 | 6713 | 6752 | 7295 | 7157 | 6249 | 6443 | 6388 | 7063 | 6746 | 7204 | 6074 | 5248 | 6260 | 6537 | 6847 | 6120 | 7006 | 7590 | 6021 | 6723 | 7337 | 7484 | 6976 | 7342 | Msl2 |
| Msl3 | 2975 | 2394 | 2657 | 2448 | 2228 | 2574 | 2586 | 3191 | 3159 | 2669 | 2700 | 2987 | 3040 | 3057 | 2855 | 2654 | 2415 | 2367 | 2620 | 2798 | 2499 | 2379 | 3059 | 2915 | 2746 | 2901 | 3149 | 3214 | Msl3 |
| Msl3l2 | 269 | 221 | 293 | 214 | 299 | 291 | 287 | 519 | 532 | 533 | 494 | 495 | 538 | 520 | 293 | 232 | 246 | 231 | 262 | 259 | 226 | 511 | 431 | 564 | 498 | 524 | 513 | 503 | Msl3l2 |
| Msln | 2 | 3 | 0 | 0 | 2 | 7 | 4 | 4 | 14 | 5 | 10 | 5 | 0 | 11 | 0 | 3 | 1 | 9 | 2 | 1 | 1 | 3 | 12 | 19 | 6 | 2 | 6 | 6 | Msln |
| Mslnl | 18 | 21 | 17 | 24 | 6 | 9 | 11 | 62 | 43 | 80 | 33 | 1 | 17 | 6 | 12 | 6 | 10 | 18 | 15 | 9 | 3 | 0 | 50 | 109 | 1 | 73 | 5 | 7 | Mslnl |
| Msmb | 0 | 0 | 0 | 0 | 0 | 0 | 1 | 0 | 0 | 0 | 0 | 0 | 0 | 1 | 0 | 0 | 0 | 0 | 0 | 0 | 0 | 0 | 0 | 0 | 0 | 0 | 0 | 2 | Msmb |
| Msmo1 | 10629 | 10969 | 10286 | 8799 | 6171 | 7251 | 8598 | 7066 | 5058 | 5251 | 4286 | 6667 | 6885 | 7701 | 11606 | 10530 | 9173 | 8469 | 6777 | 9359 | 7670 | 5213 | 5291 | 5337 | 5778 | 6981 | 7975 | 8619 | Msmo1 |
| Msmp | 1 | 0 | 5 | 0 | 1 | 0 | 0 | 0 | 2 | 3 | 2 | 0 | 5 | 2 | 0 | 1 | 5 | 0 | 3 | 0 | 2 | 0 | 0 | 2 | 0 | 2 | 0 | 1 | Msmp |
| Msn | 12177 | 11828 | 13148 | 8519 | 8401 | 9641 | 9048 | 9205 | 9426 | 8193 | 9375 | 11391 | 11361 | 11382 | 11838 | 11523 | 11223 | 8911 | 9000 | 9455 | 8459 | 8278 | 9804 | 9282 | 8450 | 11850 | 11569 | 11981 | Msn |
| Msr1 | 51 | 34 | 44 | 33 | 35 | 42 | 39 | 65 | 101 | 82 | 91 | 70 | 107 | 70 | 41 | 68 | 30 | 51 | 28 | 62 | 36 | 77 | 139 | 65 | 88 | 86 | 72 | 83 | Msr1 |
| Msra | 685 | 811 | 754 | 702 | 740 | 799 | 849 | 899 | 929 | 846 | 790 | 905 | 818 | 980 | 799 | 678 | 714 | 609 | 937 | 996 | 852 | 777 | 851 | 856 | 953 | 983 | 1077 | 1054 | Msra |
| Msrb1 | 1848 | 2360 | 1964 | 1935 | 1640 | 1545 | 1716 | 906 | 961 | 901 | 898 | 861 | 922 | 986 | 1746 | 2012 | 1951 | 1813 | 1730 | 1904 | 1737 | 657 | 974 | 968 | 896 | 1091 | 958 | 1095 | Msrb1 |
| Msrb2 | 136 | 110 | 116 | 63 | 92 | 78 | 91 | 371 | 419 | 322 | 359 | 353 | 415 | 337 | 127 | 87 | 98 | 57 | 93 | 123 | 83 | 269 | 439 | 374 | 270 | 473 | 386 | 329 | Msrb2 |
| Msrb3 | 2754 | 1918 | 2241 | 1684 | 1102 | 1631 | 1329 | 1622 | 1792 | 1657 | 1973 | 2156 | 2719 | 2282 | 2213 | 2569 | 1672 | 1585 | 1228 | 1331 | 1157 | 1255 | 1948 | 2279 | 1568 | 2697 | 2386 | 2312 | Msrb3 |
| Mss51 | 424 | 425 | 575 | 543 | 498 | 502 | 560 | 431 | 349 | 473 | 548 | 634 | 570 | 618 | 392 | 498 | 606 | 524 | 597 | 468 | 487 | 548 | 356 | 526 | 519 | 683 | 675 | 605 | Mss51 |
| Mst1 | 9 | 12 | 3 | 3 | 0 | 1 | 4 | 4 | 7 | 8 | 9 | 2 | 10 | 11 | 4 | 12 | 8 | 7 | 5 | 1 | 2 | 0 | 5 | 5 | 1 | 10 | 20 | 18 | Mst1 |

| | | | | | | | | | | | | | | | | | | | | | | | | | | | | | |
|---|---|---|---|---|---|---|---|---|---|---|---|---|---|---|---|---|---|---|---|---|---|---|---|---|---|---|---|---|---|
| Mst1r | 407 | 441 | 508 | 413 | 416 | 394 | 220 | 238 | 308 | 347 | 453 | 264 | 347 | 231 | 415 | 549 | 428 | 450 | 379 | 300 | 281 | 329 | 383 | 303 | 272 | 206 | 278 | 266 | Mst1r |
| Mstn | 134 | 92 | 43 | 80 | 80 | 76 | 64 | 42 | 119 | 49 | 470 | 11 | 64 | 17 | 123 | 202 | 59 | 77 | 45 | 59 | 57 | 104 | 340 | 25 | 27 | 14 | 7 | 13 | Mstn |
| Msto1 | 1499 | 1809 | 1667 | 1602 | 1537 | 1639 | 1462 | 1461 | 1668 | 1661 | 1675 | 2086 | 1754 | 1940 | 1487 | 1730 | 1550 | 1472 | 1713 | 1404 | 1550 | 1491 | 1748 | 1701 | 1684 | 2180 | 1933 | 1971 | Msto1 |
| Msx1 | 3725 | 2884 | 3298 | 1282 | 920 | 1284 | 687 | 2727 | 1926 | 1838 | 982 | 1614 | 2273 | 1506 | 2271 | 3549 | 1683 | 1188 | 716 | 1034 | 671 | 1046 | 2265 | 1717 | 934 | 2244 | 1888 | 2051 | Msx1 |
| Msx2 | 1399 | 1557 | 1722 | 717 | 246 | 897 | 104 | 793 | 962 | 886 | 706 | 1158 | 2028 | 1072 | 1568 | 2079 | 948 | 810 | 249 | 213 | 113 | 1161 | 1069 | 720 | 751 | 995 | 1383 | 1488 | Msx2 |
| Msx3 | 281 | 793 | 279 | 578 | 716 | 610 | 352 | 308 | 346 | 403 | 339 | 159 | 955 | 93 | 774 | 1069 | 769 | 436 | 410 | 686 | 288 | 368 | 495 | 70 | 255 | 105 | 83 | 87 | Msx3 |
| Mt1 | 714 | 853 | 782 | 734 | 606 | 736 | 799 | 315 | 481 | 385 | 497 | 447 | 494 | 583 | 705 | 661 | 721 | 705 | 863 | 917 | 749 | 207 | 526 | 503 | 462 | 474 | 415 | 705 | Mt1 |
| Mt2 | 497 | 599 | 648 | 441 | 456 | 527 | 725 | 355 | 547 | 442 | 547 | 541 | 574 | 758 | 600 | 472 | 495 | 507 | 499 | 617 | 550 | 122 | 463 | 657 | 522 | 542 | 411 | 921 | Mt2 |
| Mt3 | 460 | 758 | 415 | 1232 | 1298 | 862 | 1766 | 954 | 817 | 1179 | 824 | 799 | 386 | 1448 | 571 | 239 | 954 | 1119 | 1184 | 1021 | 1803 | 779 | 709 | 1533 | 1345 | 932 | 501 | 813 | Mt3 |
| Mt4 | 0 | 0 | 4 | 0 | 0 | 0 | 0 | 0 | 0 | 0 | 0 | 0 | 1 | 1 | 0 | 0 | 0 | 0 | 0 | 0 | 0 | 0 | 0 | 0 | 0 | 0 | 0 | 0 | Mt4 |
| Mta1 | 5287 | 6173 | 8451 | 7516 | 6667 | 6037 | 5541 | 6006 | 6906 | 8782 | 7904 | 10153 | 7562 | 9326 | 4787 | 5597 | 8069 | 6769 | 7634 | 4924 | 6543 | 7971 | 6694 | 10055 | 8725 | 11687 | 8105 | 8981 | Mta1 |
| Mta2 | 13867 | 14076 | 15254 | 15479 | 14117 | 14527 | 14328 | 13122 | 13631 | 13975 | 13803 | 16376 | 14983 | 16429 | 13152 | 12690 | 15383 | 13980 | 16295 | 13460 | 15040 | 13100 | 12852 | 15886 | 15194 | 18617 | 15903 | 16773 | Mta2 |
| Mta3 | 1027 | 1088 | 1160 | 1053 | 1532 | 1430 | 1272 | 1776 | 2037 | 2137 | 2166 | 1972 | 2131 | 1906 | 1114 | 1105 | 1043 | 1104 | 1366 | 1157 | 1281 | 1953 | 2028 | 2027 | 2026 | 1988 | 1954 | 1943 | Mta3 |
| Mtap | 8412 | 6825 | 7529 | 5559 | 4525 | 5054 | 5674 | 3821 | 3756 | 3487 | 3535 | 4237 | 4196 | 4499 | 7283 | 7009 | 6304 | 5362 | 5142 | 5343 | 4994 | 2989 | 3746 | 3840 | 3197 | 5845 | 4369 | 4965 | Mtap |
| Mtarc1 | 23 | 29 | 29 | 32 | 45 | 29 | 27 | 7 | 11 | 9 | 11 | 8 | 5 | 6 | 29 | 30 | 17 | 47 | 31 | 24 | 17 | 17 | 5 | 4 | 4 | 4 | 8 | 6 | Mtarc1 |
| Mtarc2 | 3609 | 3507 | 2898 | 2933 | 2876 | 2800 | 2542 | 2164 | 2330 | 2065 | 2192 | 1919 | 2291 | 1755 | 3538 | 3571 | 2950 | 2972 | 2849 | 3058 | 2632 | 1868 | 2452 | 2014 | 1872 | 2090 | 1948 | 1958 | Mtarc2 |
| mt-Atp6 | 3358 | 693 | 928 | 1017 | 957 | 1196 | 901 | 3309 | 919 | 921 | 943 | 854 | 1035 | 1003 | 3939 | 1058 | 816 | 6116 | 1509 | 1379 | 1217 | 3716 | 1058 | 961 | 1131 | 1047 | 1233 | 900 | mt-Atp6 |
| mt-Atp8 | 5 | 5 | 4 | 7 | 7 | 6 | 6 | 12 | 10 | 5 | 5 | 4 | 5 | 5 | 7 | 3 | 5 | 6 | 2 | 5 | 7 | 3 | 4 | 5 | 2 | 8 | 3 | 8 | mt-Atp8 |
| Mtbp | 2342 | 1887 | 1945 | 1597 | 1416 | 1556 | 1666 | 2143 | 2076 | 1793 | 2012 | 2465 | 2218 | 2458 | 1924 | 2028 | 1690 | 1563 | 1543 | 1846 | 1509 | 1494 | 1881 | 1924 | 1827 | 2436 | 2560 | 2430 | Mtbp |
| Mtch1 | 20965 | 27227 | 22638 | 24899 | 22477 | 22340 | 20966 | 18315 | 20795 | 19568 | 18769 | 19987 | 19002 | 18632 | 21001 | 22286 | 24270 | 23776 | 24622 | 22040 | 22673 | 18859 | 20650 | 20447 | 20997 | 21382 | 19601 | 19378 | Mtch1 |
| Mtch2 | 4571 | 4563 | 4103 | 4094 | 4256 | 4271 | 4666 | 3854 | 3527 | 2842 | 3162 | 3492 | 3405 | 3363 | 4737 | 4597 | 4247 | 4215 | 4324 | 5099 | 4239 | 3400 | 3562 | 2993 | 3348 | 3503 | 4007 | 3579 | Mtch2 |
| Mtcl1 | 5259 | 7724 | 8851 | 9430 | 9543 | 8117 | 7373 | 4664 | 4476 | 5812 | 5429 | 7417 | 7392 | 7275 | 6173 | 6669 | 10032 | 8911 | 9075 | 6642 | 8191 | 6492 | 4598 | 6219 | 6580 | 4967 | 7749 | 7882 | Mtcl1 |
| mt-Co1 | 612083 | 482357 | 491775 | 474052 | 510775 | 542890 | 555449 | 646653 | 570404 | 481101 | 473496 | 494983 | 501397 | 505962 | 684471 | 563998 | 479373 | 524917 | 520070 | 642841 | 577214 | 611214 | 576603 | 483962 | 500296 | 552232 | 514897 | 543478 | mt-Co1 |
| mt-Co2 | 4485 | 1975 | 2296 | 2602 | 2317 | 2824 | 2451 | 4135 | 2401 | 1949 | 2280 | 2115 | 2854 | 2567 | 4623 | 2534 | 2020 | 2456 | 2686 | 2934 | 2311 | 4712 | 2542 | 2414 | 2304 | 2557 | 2517 | 2591 | mt-Co2 |
| mt-Co3 | 4209 | 1437 | 1398 | 1835 | 1787 | 2251 | 1923 | 4881 | 1962 | 1514 | 1628 | 1669 | 1932 | 1738 | 4825 | 2064 | 1482 | 1847 | 1905 | 2660 | 2019 | 4463 | 2232 | 1753 | 1731 | 1872 | 1827 | 1911 | mt-Co3 |
| Mtcp1 | 1362 | 945 | 700 | 972 | 1019 | 1326 | 1209 | 1669 | 1435 | 1121 | 1317 | 1337 | 1374 | 1364 | 1387 | 1143 | 743 | 935 | 895 | 1395 | 1301 | 931 | 1452 | 1044 | 1191 | 724 | 1544 | 1262 | Mtcp1 |
| mt-Cytb | 298285 | 238366 | 255487 | 266796 | 287271 | 287412 | 288098 | 313755 | 276978 | 256119 | 274960 | 258308 | 276587 | 279458 | 325859 | 255688 | 260438 | 280112 | 293635 | 316974 | 308867 | 332210 | 281899 | 285025 | 276917 | 304655 | 275721 | 287895 | mt-Cytb |
| Mtdh | 13704 | 9784 | 12381 | 12541 | 11999 | 13428 | 13131 | 13455 | 12039 | 12186 | 12679 | 13084 | 12921 | 13381 | 12576 | 11719 | 11281 | 12253 | 12274 | 11801 | 12388 | 12509 | 12200 | 12678 | 12835 | 13825 | 13343 | 13828 | Mtdh |
| Mterf1a | 505 | 357 | 566 | 476 | 496 | 518 | 501 | 519 | 381 | 410 | 416 | 508 | 407 | 513 | 563 | 448 | 495 | 548 | 483 | 574 | 492 | 446 | 441 | 486 | 432 | 569 | 483 | 544 | Mterf1a |
| Mterf1b | 226 | 191 | 341 | 246 | 265 | 293 | 305 | 231 | 246 | 247 | 259 | 212 | 212 | 278 | 228 | 202 | 261 | 265 | 266 | 214 | 262 | 259 | 219 | 292 | 268 | 296 | 225 | 241 | Mterf1b |
| Mterf2 | 1361 | 1299 | 1329 | 1231 | 1219 | 1245 | 1332 | 1712 | 1393 | 1247 | 1208 | 1489 | 1429 | 1514 | 1697 | 1454 | 1311 | 1082 | 1177 | 1531 | 1289 | 1397 | 1456 | 1324 | 1219 | 1369 | 1464 | 1431 | Mterf2 |
| Mterf3 | 2664 | 2179 | 2023 | 2148 | 2299 | 2598 | 3067 | 4062 | 3381 | 3106 | 3220 | 3292 | 3107 | 3447 | 2666 | 2266 | 2058 | 2319 | 2315 | 2753 | 2806 | 3521 | 3407 | 3244 | 3167 | 2932 | 3359 | 3404 | Mterf3 |
| Mterf4 | 1465 | 1433 | 1515 | 1350 | 1220 | 1226 | 1424 | 1619 | 1430 | 1283 | 1226 | 1402 | 1351 | 1536 | 1494 | 1319 | 1384 | 1281 | 1392 | 1590 | 1364 | 1404 | 1476 | 1393 | 1336 | 1539 | 1536 | 1690 | Mterf4 |
| Mtf1 | 1478 | 1496 | 1775 | 1767 | 1859 | 1645 | 1734 | 1724 | 1678 | 2007 | 1901 | 2191 | 2022 | 2094 | 1333 | 1447 | 1685 | 1771 | 1845 | 1380 | 1706 | 2092 | 1691 | 2017 | 2093 | 1754 | 2105 | 2191 | Mtf1 |
| Mtf2 | 11788 | 8761 | 10305 | 10110 | 10260 | 11127 | 11591 | 15746 | 12131 | 11043 | 11392 | 13667 | 13388 | 14427 | 12217 | 10734 | 10265 | 9652 | 10378 | 12530 | 10739 | 13383 | 11130 | 12428 | 12960 | 13131 | 14423 | 14150 | Mtf2 |
| Mtfmt | 2209 | 2314 | 2007 | 2274 | 2042 | 2034 | 2215 | 1384 | 1427 | 1422 | 1512 | 1326 | 1433 | 1405 | 2074 | 2139 | 2132 | 2201 | 2018 | 1961 | 2275 | 1519 | 1372 | 1354 | 1428 | 1289 | 1388 | 1352 | Mtfmt |
| Mtfp1 | 430 | 536 | 551 | 470 | 496 | 448 | 447 | 154 | 219 | 199 | 190 | 270 | 217 | 214 | 411 | 463 | 584 | 434 | 513 | 556 | 509 | 159 | 188 | 250 | 188 | 280 | 212 | 248 | Mtfp1 |
| Mtfr1 | 6406 | 6062 | 5769 | 5704 | 4751 | 5428 | 5582 | 4399 | 3563 | 3602 | 3318 | 4467 | 4016 | 4328 | 6392 | 5649 | 5706 | 5359 | 5216 | 5668 | 5386 | 3307 | 3746 | 3813 | 3759 | 4579 | 4530 | 4686 | Mtfr1 |
| Mtfr1l | 8561 | 9133 | 8076 | 8116 | 7351 | 7065 | 8753 | 5514 | 5257 | 4939 | 4804 | 5139 | 4970 | 5552 | 8614 | 7884 | 8218 | 8098 | 7794 | 7669 | 8457 | 5152 | 5145 | 5132 | 5162 | 5242 | 4931 | 5402 | Mtfr1l |
| Mtfr2 | 1302 | 1195 | 1339 | 1093 | 787 | 911 | 1132 | 628 | 527 | 447 | 460 | 726 | 661 | 777 | 1118 | 1205 | 1183 | 983 | 1115 | 1066 | 948 | 389 | 473 | 646 | 532 | 917 | 793 | 797 | Mtfr2 |
| Mtg1 | 499 | 602 | 490 | 573 | 705 | 627 | 644 | 1189 | 1155 | 1107 | 1116 | 1158 | 1171 | 1213 | 531 | 504 | 549 | 628 | 687 | 706 | 598 | 1091 | 1195 | 1095 | 1229 | 1096 | 1314 | 1102 | Mtg1 |
| Mtg2 | 1136 | 1310 | 1313 | 1529 | 1401 | 1385 | 1450 | 1406 | 1582 | 1701 | 1487 | 1887 | 1718 | 1810 | 1118 | 1178 | 1433 | 1444 | 1572 | 1327 | 1427 | 1684 | 1488 | 1856 | 1812 | 1736 | 1733 | 1774 | Mtg2 |
| Mthfd1 | 5685 | 5999 | 5214 | 4633 | 3742 | 4321 | 4278 | 3418 | 4128 | 3486 | 3367 | 4148 | 3725 | 4174 | 4594 | 5449 | 4723 | 4539 | 4690 | 4570 | 4392 | 2588 | 3700 | 3921 | 3584 | 5070 | 4324 | 4393 | Mthfd1 |
| Mthfd1l | 849 | 730 | 847 | 489 | 430 | 596 | 429 | 900 | 1144 | 915 | 1030 | 1372 | 1335 | 1308 | 721 | 877 | 597 | 517 | 458 | 488 | 470 | 672 | 1016 | 945 | 873 | 1534 | 1499 | 1622 | Mthfd1l |
| Mthfd2 | 2788 | 3312 | 3282 | 2874 | 2072 | 2141 | 2481 | 1467 | 1449 | 1522 | 1522 | 1820 | 1662 | 2119 | 2413 | 3173 | 2982 | 2905 | 2316 | 2521 | 2478 | 1303 | 1377 | 1655 | 1570 | 2192 | 1928 | 2277 | Mthfd2 |
| Mthfd2l | 2197 | 1823 | 1993 | 1745 | 2149 | 2276 | 2454 | 4681 | 3461 | 3004 | 2937 | 3698 | 3390 | 3980 | 2484 | 1893 | 2000 | 1739 | 2167 | 2625 | 2318 | 3723 | 3281 | 3308 | 3403 | 3599 | 3940 | 3995 | Mthfd2l |
| Mthfr | 4013 | 5053 | 4554 | 4267 | 4183 | 3963 | 3774 | 3712 | 4138 | 4483 | 4186 | 5196 | 5153 | 4886 | 3836 | 4724 | 4400 | 4118 | 4078 | 3919 | 3747 | 4353 | 4143 | 4344 | 4279 | 3685 | 5387 | 4974 | Mthfr |
| Mthfs | 43 | 30 | 31 | 35 | 43 | 34 | 38 | 65 | 64 | 78 | 94 | 62 | 79 | 100 | 29 | 27 | 39 | 29 | 48 | 35 | 37 | 87 | 82 | 60 | 82 | 44 | 60 | 71 | Mthfs |
| Mthfsd | 3286 | 3937 | 3628 | 3789 | 3525 | 3204 | 3270 | 2856 | 3111 | 3213 | 3247 | 3125 | 3031 | 3149 | 3092 | 3794 | 3714 | 3685 | 3456 | 3206 | 3268 | 3242 | 3008 | 3016 | 3131 | 2714 | 3182 | 3017 | Mthfsd |
| Mthfsl | 161 | 163 | 164 | 168 | 196 | 199 | 238 | 493 | 460 | 372 | 408 | 406 | 353 | 521 | 175 | 193 | 172 | 136 | 128 | 267 | 219 | 403 | 420 | 369 | 325 | 385 | 389 | 369 | Mthfsl |
| Mtif2 | 3644 | 3176 | 4227 | 3930 | 3950 | 3881 | 4342 | 4615 | 4154 | 5015 | 4698 | 5233 | 5014 | 5638 | 3816 | 3387 | 4063 | 3816 | 4213 | 4035 | 4268 | 4741 | 4250 | 5120 | 5058 | 5201 | 5141 | 5180 | Mtif2 |
| Mtif3 | 1043 | 1183 | 1092 | 1063 | 1128 | 1077 | 1083 | 1172 | 1170 | 1116 | 998 | 1027 | 1038 | 1084 | 1124 | 1048 | 1087 | 1012 | 1234 | 1181 | 1111 | 1106 | 1192 | 1111 | 1190 | 1189 | 1115 | 1081 | Mtif3 |
| Mtln | 246 | 238 | 346 | 270 | 304 | 310 | 418 | 289 | 275 | 312 | 336 | 342 | 278 | 358 | 266 | 241 | 362 | 263 | 298 | 389 | 382 | 231 | 316 | 455 | 292 | 456 | 352 | 395 | Mtln |
| Mtm1 | 1896 | 1525 | 1395 | 1550 | 1604 | 1761 | 2071 | 657 | 603 | 480 | 484 | 569 | 644 | 659 | 2169 | 1638 | 1374 | 1578 | 1442 | 2031 | 1814 | 585 | 559 | 483 | 530 | 598 | 507 | 611 | Mtm1 |
| Mtmr1 | 2008 | 1678 | 1436 | 1990 | 2658 | 2716 | 2614 | 4075 | 3795 | 3352 | 3353 | 3112 | 2852 | 3274 | 2284 | 1771 | 1768 | 1951 | 2330 | 2595 | 2586 | 3703 | 3408 | 3059 | 3493 | 2528 | 3069 | 2880 | Mtmr1 |
| Mtmr10 | 3467 | 3511 | 2975 | 2994 | 2822 | 3069 | 3134 | 2075 | 2111 | 2110 | 1856 | 2077 | 2028 | 1942 | 3556 | 3446 | 2752 | 3032 | 2699 | 3156 | 3151 | 1845 | 2170 | 1722 | 1806 | 1717 | 2228 | 2076 | Mtmr10 |
| Mtmr11 | 1622 | 1310 | 1120 | 1238 | 1244 | 1361 | 1206 | 909 | 1181 | 1037 | 1256 | 798 | 1215 | 878 | 1197 | 1498 | 1185 | 1217 | 1240 | 1168 | 1180 | 800 | 1176 | 1003 | 778 | 1087 | 865 | 900 | Mtmr11 |
| Mtmr12 | 4101 | 3791 | 3636 | 3592 | 3141 | 3527 | 3391 | 2270 | 2377 | 2192 | 2358 | 2518 | 2582 | 2565 | 3798 | 4066 | 3618 | 3416 | 3257 | 3118 | 3363 | 2252 | 2246 | 2303 | 2172 | 2514 | 2518 | 2517 | Mtmr12 |
| Mtmr14 | 4305 | 4771 | 3724 | 3988 | 3103 | 3407 | 3685 | 2530 | 2503 | 2227 | 2227 | 2692 | 2464 | 2624 | 4027 | 4084 | 3676 | 3662 | 3585 | 3514 | 3635 | 2232 | 2400 | 2432 | 2527 | 2864 | 2866 | 2770 | Mtmr14 |
| Mtmr2 | 4527 | 3677 | 3843 | 3217 | 3547 | 3923 | 4123 | 6638 | 5952 | 5306 | 5622 | 6145 | 6009 | 6698 | 4454 | 3861 | 3693 | 3314 | 3546 | 4151 | 3740 | 5641 | 5774 | 5617 | 5507 | 6445 | 6287 | 6840 | Mtmr2 |
| Mtmr3 | 9190 | 9662 | 7200 | 8874 | 8972 | 8934 | 9100 | 9441 | 9331 | 9152 | 8680 | 9623 | 8748 | 9318 | 8922 | 8026 | 7907 | 8601 | 9263 | 8675 | 9227 | 9910 | 8943 | 8887 | 9651 | 7604 | 9331 | 9314 | Mtmr3 |
| Mtmr4 | 13098 | 13405 | 12896 | 13678 | 12697 | 12642 | 12382 | 12247 | 10918 | 11726 | 11518 | 12466 | 12056 | 12046 | 12795 | 12809 | 13140 | 13049 | 13811 | 11694 | 12569 | 13983 | 10051 | 12325 | 13537 | 10477 | 11902 | 12620 | Mtmr4 |
| Mtmr6 | 9649 | 8830 | 8187 | 9284 | 9556 | 9654 | 10328 | 8532 | 7097 | 6791 | 6412 | 6713 | 6129 | 6821 | 10777 | 8996 | 8652 | 9950 | 9331 | 11122 | 9862 | 7865 | 7039 | 6922 | 7602 | 6164 | 6651 | 6473 | Mtmr6 |
| Mtmr7 | 2028 | 1938 | 1522 | 2122 | 2648 | 2407 | 2822 | 2501 | 2202 | 1963 | 1804 | 1794 | 1708 | 1846 | 2205 | 1717 | 1927 | 2251 | 2476 | 2931 | 2687 | 2289 | 1972 | 1790 | 2184 | 1713 | 1718 | 1660 | Mtmr7 |
| Mtmr9 | 4684 | 4875 | 4146 | 5029 | 6186 | 6174 | 5725 | 6644 | 6653 | 6753 | 6129 | 7115 | 6564 | 6397 | 5306 | 4684 | 5146 | 5207 | 5815 | 5340 | 5810 | 7369 | 6207 | 6110 | 7490 | 4853 | 6732 | 6618 | Mtmr9 |
| mt-Nd1 | 198278 | 147656 | 176389 | 182572 | 181947 | 202904 | 191151 | 199635 | 183288 | 178387 | 181379 | 189655 | 193391 | 201455 | 221807 | 174241 | 171832 | 177529 | 194885 | 216695 | 214164 | 196367 | 188863 | 186299 | 178434 | 224755 | 192510 | 213486 | mt-Nd1 |
| mt-Nd2 | 117669 | 91748 | 97053 | 95570 | 102593 | 107082 | 89237 | 114512 | 100926 | 95702 | 100235 | 98938 | 103719 | 95182 | 122874 | 92793 | 97683 | 102248 | 110084 | 111030 | 110267 | 124168 | 104549 | 110110 | 101438 | 116079 | 103358 | 100578 | mt-Nd2 |
| mt-Nd3 | 610 | 173 | 239 | 430 | 272 | 353 | 562 | 785 | 201 | 305 | 435 | 276 | 310 | 496 | 662 | 246 | 275 | 294 | 281 | 348 | 514 | 536 | 220 | 1057 | 363 | 270 | 252 | 452 | mt-Nd3 |
| mt-Nd4 | 98243 | 78746 | 84048 | 93318 | 97258 | 103631 | 97216 | 106245 | 97291 | 90059 | 97439 | 88942 | 98501 | 92509 | 108910 | 90900 | 85723 | 98880 | 100999 | 107701 | 107835 | 117882 | 99783 | 98118 | 98797 | 108482 | 92350 | 97935 | mt-Nd4 |
| mt-Nd4l | 126 | 69 | 47 | 63 | 42 | 35 | 54 | 147 | 44 | 32 | 53 | 39 | 48 | 97 | 133 | 30 | 54 | 51 | 46 | 38 | 107 | 145 | 41 | 68 | 49 | 42 | 41 | 63 | mt-Nd4l |
| mt-Nd5 | 195125 | 135466 | 160549 | 118835 | 150405 | 173034 | 127474 | 159196 | 157881 | 131827 | 162427 | 158088 | 176029 | 138669 | 203623 | 161459 | 149587 | 146268 | 150567 | 169656 | 145744 | 168273 | 157859 | 152967 | 148183 | 176890 | 167113 | 148575 | mt-Nd5 |
| mt-Nd6 | 6536 | 3053 | 2854 | 3642 | 3256 | 4747 | 3641 | 4936 | 4233 | 3688 | 4691 | 3930 | 4936 | 3908 | 5828 | 4478 | 3576 | 3588 | 3629 | 5088 | 4021 | 3454 | 4644 | 4213 | 3869 | 2765 | 5417 | 3987 | mt-Nd6 |
| Mtnr1a | 21 | 23 | 6 | 7 | 33 | 17 | 10 | 23 | 18 | 16 | 29 | 22 | 24 | 10 | 26 | 26 | 27 | 4 | 16 | 24 | 13 | 35 | 44 | 7 | 13 | 13 | 26 | 19 | Mtnr1a |
| Mtnr1b | 1 | 3 | 0 | 0 | 2 | 0 | 0 | 1 | 0 | 0 | 0 | 3 | 0 | 0 | 0 | 2 | 0 | 0 | 1 | 0 | 1 | 0 | 0 | 0 | 0 | 0 | 0 | 1 | Mtnr1b |
| Mto1 | 1254 | 1229 | 1119 | 1096 | 1082 | 1170 | 1284 | 1613 | 1197 | 1188 | 1188 | 1360 | 1194 | 1368 | 1197 | 1115 | 1036 | 1055 | 1221 | 1266 | 1160 | 1327 | 1251 | 1354 | 1396 | 1433 | 1290 | 1432 | Mto1 |
| Mtor | 11426 | 13698 | 10228 | 13607 | 13261 | 13450 | 12997 | 7603 | 8426 | 8943 | 7950 | 9081 | 8737 | 9056 | 11425 | 12453 | 11899 | 13554 | 13994 | 12137 | 13895 | 8554 | 7518 | 7750 | 9228 | 6155 | 9140 | 8797 | Mtor |
| Mtpap | 2173 | 1752 | 1628 | 1751 | 1762 | 1792 | 2159 | 2696 | 2207 | 2118 | 2105 | 2153 | 2205 | 2432 | 2135 | 1763 | 1600 | 1598 | 1573 | 2003 | 1933 | 2343 | 2291 | 2002 | 2073 | 2326 | 2404 | 2340 | Mtpap |
| Mtpn | 26125 | 19460 | 24041 | 22274 | 22834 | 24369 | 26540 | 22304 | 19611 | 19808 | 19960 | 21976 | 21391 | 23156 | 26057 | 22677 | 22957 | 22793 | 22863 | 21099 | 25743 | 23469 | 20450 | 21270 | 22366 | 22682 | 22025 | 24231 | Mtpn |
| Mtr | 5501 | 4182 | 4387 | 4365 | 3924 | 4425 | 4584 | 3615 | 3798 | 3541 | 4115 | 3897 | 4173 | 4175 | 4751 | 4696 | 4378 | 4116 | 3712 | 3762 | 4148 | 3524 | 3843 | 3835 | 3428 | 3493 | 4169 | 3909 | Mtr |

| | | | | | | | | | | | | | | | | | | | | | | | | | | | | | |
|---|---|---|---|---|---|---|---|---|---|---|---|---|---|---|---|---|---|---|---|---|---|---|---|---|---|---|---|---|---|
| Mtres1 | 1523 | 1571 | 1619 | 1707 | 1492 | 1446 | 1617 | 1364 | 1250 | 1237 | 1259 | 1248 | 1377 | 1508 | 1582 | 1421 | 1576 | 1535 | 1372 | 1684 | 1617 | 1216 | 1207 | 1455 | 1235 | 1369 | 1406 | 1556 | Mtres1 |
| Mtrex | 12913 | 11518 | 11082 | 10628 | 10334 | 10725 | 11707 | 12893 | 10936 | 9731 | 9742 | 11579 | 10743 | 11885 | 13585 | 11946 | 11232 | 10287 | 10689 | 12747 | 11214 | 9778 | 10173 | 9984 | 10407 | 11432 | 11576 | 11824 | Mtrex |
| Mtrf1 | 253 | 244 | 331 | 266 | 287 | 261 | 273 | 526 | 572 | 487 | 489 | 527 | 475 | 550 | 306 | 283 | 324 | 263 | 316 | 355 | 290 | 448 | 570 | 492 | 510 | 513 | 580 | 513 | Mtrf1 |
| Mtrf1l | 1193 | 1254 | 1387 | 1375 | 1180 | 1236 | 1301 | 918 | 797 | 872 | 921 | 976 | 792 | 859 | 1166 | 1184 | 1388 | 1425 | 1233 | 1213 | 1244 | 809 | 799 | 956 | 870 | 816 | 820 | 840 | Mtrf1l |
| Mtrfr | 1231 | 1389 | 1388 | 1331 | 1149 | 1162 | 1273 | 1247 | 1186 | 1176 | 1237 | 1362 | 1384 | 1323 | 1279 | 1213 | 1498 | 1258 | 1297 | 1349 | 1251 | 951 | 1239 | 1396 | 1378 | 1579 | 1431 | 1442 | Mtrfr |
| Mtrr | 866 | 929 | 1049 | 950 | 953 | 871 | 870 | 1321 | 1396 | 1461 | 1457 | 1458 | 1436 | 1445 | 849 | 884 | 1009 | 868 | 861 | 850 | 802 | 1283 | 1350 | 1392 | 1488 | 1559 | 1538 | 1584 | Mtrr |
| Mtss1 | 5021 | 5283 | 7262 | 8881 | 11277 | 8976 | 10537 | 8230 | 8491 | 11219 | 9275 | 9838 | 8470 | 11207 | 5618 | 5177 | 8408 | 9212 | 10392 | 8527 | 11037 | 11007 | 8230 | 11324 | 13245 | 8678 | 8756 | 9766 | Mtss1 |
| Mtss2 | 2985 | 3901 | 5243 | 4805 | 5938 | 4579 | 3824 | 4993 | 6247 | 7875 | 7314 | 6274 | 5746 | 5295 | 3034 | 3782 | 5420 | 5246 | 5459 | 3795 | 4502 | 8288 | 6361 | 7269 | 7675 | 6940 | 4600 | 5032 | Mtss2 |
| Mttp | 732 | 796 | 767 | 774 | 705 | 747 | 711 | 862 | 849 | 748 | 748 | 1046 | 951 | 911 | 804 | 684 | 684 | 660 | 683 | 772 | 752 | 857 | 806 | 865 | 751 | 923 | 1033 | 1053 | Mttp |
| Mturn | 12469 | 14416 | 11903 | 16727 | 19426 | 17714 | 18529 | 11385 | 10710 | 11477 | 10100 | 9078 | 8934 | 9488 | 14024 | 12495 | 13891 | 17708 | 18727 | 18491 | 20745 | 12673 | 10928 | 10001 | 13540 | 5813 | 8846 | 8626 | Mturn |
| Mtus1 | 2902 | 2465 | 2607 | 2620 | 3341 | 2875 | 2794 | 5874 | 5676 | 6198 | 5820 | 5786 | 5514 | 6123 | 2846 | 2717 | 2556 | 2803 | 2791 | 2463 | 2831 | 7645 | 5488 | 5878 | 6756 | 5128 | 5492 | 5895 | Mtus1 |
| Mtus2 | 1305 | 2081 | 1604 | 2280 | 3329 | 2599 | 1972 | 3119 | 2838 | 3314 | 3085 | 2344 | 3596 | 2157 | 2301 | 2063 | 2272 | 2476 | 2520 | 2555 | 2374 | 4234 | 2784 | 2320 | 3213 | 1380 | 2280 | 2290 | Mtus2 |
| Mtx1 | 1551 | 1881 | 1905 | 1400 | 1413 | 1335 | 1572 | 2028 | 2146 | 2055 | 1903 | 2388 | 2371 | 2362 | 1429 | 1573 | 1595 | 1389 | 1575 | 1579 | 1411 | 1783 | 2038 | 2268 | 2116 | 2683 | 2563 | 2575 | Mtx1 |
| Mtx2 | 3962 | 3028 | 3342 | 3074 | 3395 | 3102 | 3633 | 5250 | 4447 | 3727 | 3885 | 4192 | 3977 | 4617 | 4221 | 3433 | 3201 | 3166 | 3272 | 3869 | 3470 | 4435 | 4592 | 4191 | 3911 | 4373 | 4113 | 4723 | Mtx2 |
| Mtx3 | 2374 | 1939 | 1695 | 1763 | 2459 | 2376 | 2293 | 6631 | 6138 | 5412 | 6064 | 5843 | 5790 | 6075 | 2480 | 2098 | 1840 | 1872 | 2071 | 2527 | 2277 | 5177 | 5823 | 5112 | 5741 | 4271 | 6450 | 5555 | Mtx3 |
| Muc1 | 532 | 627 | 718 | 648 | 554 | 602 | 536 | 601 | 507 | 699 | 545 | 996 | 798 | 878 | 485 | 461 | 798 | 581 | 697 | 591 | 528 | 690 | 520 | 924 | 824 | 904 | 1004 | 958 | Muc1 |
| Muc13 | 22 | 15 | 34 | 13 | 7 | 10 | 18 | 9 | 19 | 8 | 0 | 20 | 18 | 13 | 16 | 23 | 25 | 31 | 29 | 15 | 24 | 12 | 9 | 12 | 14 | 25 | 7 | 11 | Muc13 |
| Muc15 | 5 | 1 | 4 | 0 | 5 | 2 | 0 | 23 | 37 | 21 | 38 | 0 | 0 | 2 | 1 | 4 | 0 | 0 | 0 | 0 | 0 | 3 | 44 | 9 | 4 | 1 | 0 | 0 | Muc15 |
| Muc16 | 0 | 0 | 0 | 0 | 0 | 0 | 2 | 10 | 7 | 1 | 2 | 0 | 5 | 1 | 4 | 0 | 0 | 1 | 0 | 0 | 0 | 3 | 14 | 4 | 0 | 0 | 0 | 0 | Muc16 |
| Muc2 | 7 | 10 | 3 | 5 | 5 | 5 | 0 | 7 | 2 | 0 | 1 | 0 | 5 | 0 | 4 | 3 | 0 | 2 | 5 | 3 | 1 | 3 | 1 | 0 | 1 | 0 | 0 | 0 | Muc2 |
| Muc20 | 1 | 1 | 0 | 0 | 0 | 3 | 2 | 0 | 7 | 9 | 15 | 1 | 0 | 0 | 5 | 3 | 2 | 0 | 5 | 5 | 3 | 3 | 7 | 3 | 6 | 7 | 2 | 1 | Muc20 |
| Muc21 | 0 | 0 | 0 | 0 | 0 | 0 | 0 | 0 | 1 | 2 | 0 | 0 | 0 | 0 | 0 | 0 | 0 | 0 | 0 | 0 | 0 | 0 | 0 | 0 | 0 | 0 | 0 | 0 | Muc21 |
| Muc3 | 0 | 0 | 0 | 0 | 0 | 0 | 0 | 0 | 0 | 0 | 0 | 1 | 0 | 1 | 0 | 1 | 0 | 2 | 0 | 0 | 0 | 0 | 0 | 4 | 0 | 0 | 0 | 0 | Muc3 |
| Muc3a | 14 | 17 | 12 | 25 | 54 | 54 | 35 | 15 | 18 | 6 | 20 | 3 | 23 | 12 | 20 | 15 | 23 | 26 | 49 | 33 | 39 | 38 | 19 | 21 | 23 | 12 | 21 | 9 | Muc3a |
| Muc5ac | 0 | 1 | 0 | 0 | 0 | 1 | 0 | 0 | 0 | 0 | 2 | 0 | 0 | 0 | 1 | 0 | 0 | 1 | 0 | 0 | 2 | 0 | 0 | 2 | 0 | 0 | 0 | 0 | Muc5ac |
| Muc5b | 0 | 0 | 0 | 0 | 0 | 0 | 1 | 0 | 0 | 0 | 0 | 0 | 0 | 0 | 0 | 0 | 0 | 0 | 0 | 0 | 1 | 2 | 1 | 0 | 0 | 0 | 0 | 0 | Muc5b |
| Mucl3 | 27 | 39 | 50 | 36 | 15 | 37 | 18 | 16 | 28 | 7 | 8 | 20 | 35 | 14 | 32 | 40 | 39 | 28 | 22 | 25 | 17 | 20 | 30 | 14 | 18 | 15 | 20 | 20 | Mucl3 |
| Mug1 | 0 | 0 | 0 | 0 | 0 | 0 | 0 | 0 | 2 | 0 | 0 | 0 | 1 | 1 | 3 | 2 | 3 | 0 | 0 | 0 | 0 | 0 | 0 | 0 | 0 | 0 | 0 | 0 | Mug1 |
| Mug2 | 0 | 0 | 0 | 0 | 0 | 0 | 0 | 1 | 0 | 0 | 0 | 0 | 0 | 0 | 0 | 0 | 0 | 0 | 0 | 0 | 0 | 0 | 0 | 0 | 0 | 0 | 0 | 0 | Mug2 |
| Mul1 | 3054 | 3886 | 3546 | 3601 | 2971 | 3215 | 2808 | 1917 | 2053 | 2206 | 2193 | 2430 | 2276 | 2201 | 2769 | 3292 | 3610 | 3681 | 3600 | 2705 | 3095 | 2080 | 2208 | 2416 | 2393 | 2409 | 2374 | 2504 | Mul1 |
| Mup12 | 0 | 0 | 0 | 0 | 0 | 0 | 0 | 0 | 1 | 0 | 0 | 0 | 0 | 0 | 0 | 0 | 0 | 0 | 0 | 0 | 0 | 0 | 0 | 0 | 0 | 0 | 0 | 0 | Mup12 |
| Mup15 | 0 | 0 | 5 | 0 | 0 | 0 | 0 | 0 | 0 | 0 | 0 | 0 | 0 | 0 | 0 | 0 | 0 | 0 | 0 | 0 | 0 | 0 | 2 | 0 | 0 | 0 | 0 | 0 | Mup15 |
| Mup18 | 0 | 0 | 1 | 0 | 0 | 0 | 0 | 0 | 0 | 0 | 0 | 0 | 0 | 0 | 0 | 0 | 0 | 0 | 0 | 0 | 0 | 0 | 0 | 0 | 0 | 0 | 0 | 0 | Mup18 |
| Mup21 | 0 | 0 | 0 | 0 | 0 | 0 | 2 | 0 | 0 | 0 | 1 | 1 | 0 | 0 | 0 | 0 | 0 | 1 | 1 | 0 | 2 | 5 | 0 | 0 | 0 | 0 | 0 | 1 | Mup21 |
| Mup4 | 0 | 0 | 0 | 0 | 0 | 0 | 0 | 0 | 0 | 0 | 0 | 0 | 0 | 0 | 0 | 0 | 0 | 0 | 0 | 0 | 0 | 0 | 0 | 1 | 0 | 0 | 0 | 0 | Mup4 |
| Mup5 | 0 | 0 | 0 | 0 | 0 | 0 | 0 | 0 | 2 | 0 | 0 | 0 | 0 | 0 | 0 | 0 | 0 | 0 | 0 | 0 | 0 | 0 | 2 | 0 | 0 | 0 | 0 | 0 | Mup5 |
| Mup6 | 1 | 0 | 0 | 0 | 2 | 0 | 2 | 0 | 0 | 0 | 0 | 2 | 0 | 1 | 1 | 0 | 0 | 3 | 0 | 0 | 2 | 0 | 1 | 1 | 0 | 0 | 6 | 0 | Mup6 |
| Mup8 | 0 | 0 | 0 | 0 | 0 | 0 | 0 | 0 | 0 | 0 | 0 | 0 | 2 | 0 | 0 | 0 | 0 | 0 | 0 | 0 | 0 | 2 | 0 | 0 | 0 | 0 | 0 | 0 | Mup8 |
| Mus81 | 1096 | 1387 | 1372 | 1263 | 1289 | 1185 | 1154 | 1225 | 1550 | 1448 | 1381 | 1547 | 1585 | 1407 | 1030 | 1137 | 1365 | 1129 | 1364 | 1143 | 1128 | 1260 | 1496 | 1565 | 1506 | 1570 | 1730 | 1543 | Mus81 |
| Musk | 5 | 1 | 0 | 1 | 3 | 0 | 1 | 21 | 30 | 6 | 196 | 4 | 18 | 0 | 7 | 8 | 0 | 4 | 0 | 0 | 1 | 18 | 145 | 9 | 2 | 0 | 0 | 4 | Musk |
| Mustn1 | 67 | 72 | 75 | 83 | 69 | 88 | 82 | 37 | 73 | 87 | 218 | 77 | 73 | 73 | 42 | 66 | 114 | 85 | 94 | 66 | 78 | 70 | 148 | 39 | 64 | 78 | 51 | 63 | Mustn1 |
| Mutyh | 798 | 1088 | 1147 | 941 | 696 | 788 | 893 | 567 | 582 | 615 | 628 | 843 | 841 | 822 | 798 | 951 | 1160 | 987 | 879 | 852 | 836 | 526 | 553 | 797 | 720 | 1043 | 912 | 859 | Mutyh |
| Mvb12a | 1863 | 2265 | 2048 | 2174 | 1767 | 1848 | 1781 | 1097 | 1234 | 1288 | 1294 | 1105 | 1148 | 1109 | 1733 | 1995 | 1962 | 1898 | 1858 | 1950 | 1793 | 1105 | 1258 | 1226 | 1128 | 1260 | 1264 | 1133 | Mvb12a |
| Mvb12b | 3878 | 4763 | 5316 | 6524 | 8313 | 7113 | 6700 | 8921 | 11081 | 13337 | 11143 | 13889 | 10989 | 12435 | 4187 | 4072 | 6574 | 7017 | 8361 | 5816 | 7296 | 14165 | 9910 | 13615 | 15024 | 11890 | 11201 | 11759 | Mvb12b |
| Mvd | 3180 | 4790 | 3506 | 3628 | 2977 | 2961 | 2779 | 1595 | 1864 | 1767 | 1589 | 2043 | 1975 | 1924 | 3098 | 3798 | 3683 | 3422 | 3065 | 3205 | 3004 | 1492 | 1739 | 1824 | 1840 | 2086 | 2167 | 1977 | Mvd |
| Mvk | 3517 | 4753 | 3897 | 4336 | 3175 | 3389 | 3284 | 1697 | 1986 | 2087 | 1981 | 2332 | 2169 | 2196 | 3368 | 3851 | 4224 | 3988 | 3921 | 3053 | 3354 | 1778 | 1895 | 2030 | 2153 | 2646 | 2401 | 2432 | Mvk |
| Mvp | 771 | 1044 | 920 | 900 | 900 | 846 | 698 | 425 | 502 | 570 | 583 | 526 | 607 | 475 | 775 | 1063 | 956 | 907 | 782 | 777 | 711 | 535 | 606 | 541 | 587 | 640 | 538 | 589 | Mvp |
| Mxd1 | 3486 | 4534 | 4489 | 5854 | 4350 | 4296 | 4347 | 2202 | 2148 | 2520 | 2427 | 2565 | 2554 | 2754 | 3600 | 4414 | 5124 | 5698 | 4601 | 3694 | 4790 | 2954 | 2043 | 2756 | 2809 | 2414 | 2477 | 2599 | Mxd1 |
| Mxd3 | 1331 | 1697 | 1394 | 1459 | 1018 | 1194 | 1206 | 849 | 1104 | 922 | 933 | 1423 | 1065 | 1376 | 1058 | 1241 | 1468 | 1150 | 1387 | 1189 | 1199 | 729 | 940 | 1300 | 1138 | 1694 | 1463 | 1486 | Mxd3 |
| Mxd4 | 6938 | 7571 | 6175 | 7067 | 7646 | 7598 | 6602 | 7989 | 8688 | 8281 | 8741 | 7256 | 7515 | 6813 | 6699 | 7330 | 6234 | 7341 | 7455 | 7033 | 7258 | 8560 | 9100 | 7696 | 8232 | 6481 | 7201 | 6899 | Mxd4 |
| Mxi1 | 5482 | 5816 | 6163 | 6913 | 6953 | 6015 | 7013 | 4321 | 3845 | 4671 | 4080 | 4612 | 4216 | 5084 | 5743 | 5475 | 6889 | 6763 | 7066 | 5901 | 7133 | 4909 | 3832 | 4837 | 4918 | 4798 | 4040 | 4905 | Mxi1 |
| Mxra7 | 3029 | 3736 | 4102 | 3786 | 3244 | 3036 | 2617 | 1596 | 1595 | 1751 | 1902 | 1717 | 2210 | 1619 | 3223 | 3541 | 4115 | 3394 | 3274 | 2733 | 2792 | 1863 | 1821 | 1649 | 1539 | 1632 | 1902 | 1886 | Mxra7 |
| Mxra8 | 1546 | 1911 | 1779 | 1299 | 1121 | 1136 | 921 | 954 | 1201 | 1193 | 1703 | 1180 | 1401 | 1110 | 1535 | 1996 | 1510 | 1209 | 986 | 1078 | 861 | 1275 | 1642 | 1071 | 990 | 1288 | 1261 | 1243 | Mxra8 |
| Myadm | 0 | 0 | 6 | 0 | 0 | 2 | 1 | 1 | 0 | 1 | 0 | 0 | 0 | 1 | 0 | 2 | 1 | 0 | 2 | 1 | 0 | 2 | 0 | 0 | 1 | 0 | 2 | 0 | Myadm |
| Myadml2 | 10 | 11 | 15 | 2 | 11 | 8 | 6 | 9 | 16 | 16 | 25 | 13 | 15 | 16 | 12 | 10 | 2 | 4 | 5 | 12 | 9 | 18 | 30 | 16 | 18 | 13 | 20 | 27 | Myadml2 |
| Myb | 812 | 760 | 891 | 781 | 668 | 885 | 806 | 474 | 732 | 716 | 706 | 1039 | 878 | 1089 | 752 | 673 | 919 | 725 | 899 | 899 | 812 | 590 | 672 | 1066 | 898 | 1257 | 1098 | 993 | Myb |
| Mybbp1a | 14710 | 15366 | 14911 | 12855 | 10807 | 12330 | 11025 | 11618 | 14153 | 14239 | 13287 | 16698 | 15529 | 15704 | 12455 | 13691 | 13825 | 11653 | 13250 | 11011 | 11597 | 11769 | 12985 | 15475 | 13951 | 18497 | 16147 | 16959 | Mybbp1a |
| Mybl1 | 3570 | 2756 | 3327 | 3451 | 2396 | 3118 | 3590 | 2218 | 1691 | 1742 | 1689 | 2231 | 2463 | 2299 | 3715 | 3576 | 3160 | 3191 | 2782 | 2958 | 3211 | 1323 | 1482 | 2164 | 1667 | 2191 | 2192 | 2232 | Mybl1 |
| Mybl2 | 2798 | 3050 | 3720 | 2585 | 1843 | 2402 | 2425 | 1684 | 2150 | 2135 | 1932 | 3471 | 2886 | 3279 | 2258 | 2729 | 3228 | 2178 | 2616 | 2111 | 2188 | 1507 | 1750 | 2736 | 2331 | 4230 | 3205 | 3580 | Mybl2 |
| Mybpc1 | 356 | 371 | 209 | 133 | 375 | 620 | 182 | 615 | 1280 | 699 | 3090 | 214 | 945 | 264 | 663 | 437 | 189 | 315 | 253 | 295 | 223 | 1058 | 2328 | 237 | 525 | 99 | 345 | 306 | Mybpc1 |
| Mybpc2 | 2 | 4 | 12 | 9 | 2 | 3 | 3 | 4 | 2 | 10 | 1 | 2 | 5 | 2 | 9 | 9 | 2 | 0 | 3 | 1 | 0 | 2 | 4 | 1 | 3 | 1 | 2 | 1 | Mybpc2 |
| Mybpc3 | 0 | 1 | 0 | 3 | 4 | 0 | 1 | 0 | 0 | 0 | 8 | 2 | 3 | 0 | 0 | 2 | 0 | 2 | 0 | 0 | 0 | 3 | 5 | 1 | 0 | 2 | 0 | 0 | Mybpc3 |
| Mybph | 57 | 9 | 0 | 3 | 16 | 2 | 9 | 117 | 363 | 44 | 2515 | 29 | 100 | 11 | 15 | 72 | 12 | 12 | 5 | 1 | 1 | 237 | 1423 | 47 | 48 | 12 | 14 | 7 | Mybph |
| Mybphl | 36 | 35 | 19 | 43 | 25 | 42 | 37 | 2 | 3 | 20 | 11 | 5 | 12 | 7 | 17 | 23 | 34 | 24 | 38 | 27 | 30 | 12 | 8 | 3 | 9 | 9 | 3 | 7 | Mybphl |
| Myc | 484 | 422 | 472 | 630 | 778 | 594 | 555 | 1044 | 1257 | 1521 | 1704 | 903 | 847 | 898 | 401 | 456 | 526 | 626 | 764 | 431 | 550 | 1325 | 1306 | 1348 | 1120 | 1157 | 700 | 711 | Myc |
| Mycbp | 1985 | 1545 | 1441 | 1337 | 1019 | 1181 | 1250 | 885 | 817 | 809 | 867 | 769 | 769 | 846 | 1843 | 1600 | 1451 | 1188 | 1004 | 1386 | 1262 | 727 | 857 | 720 | 693 | 956 | 820 | 888 | Mycbp |
| Mycbp2 | 27996 | 27938 | 12081 | 19664 | 25852 | 30473 | 27230 | 28620 | 27329 | 19168 | 21313 | 15790 | 18547 | 16746 | 30734 | 27722 | 15643 | 20496 | 20736 | 33055 | 31609 | 19305 | 24372 | 13541 | 21258 | 8889 | 18835 | 13934 | Mycbp2 |
| Mycbpap | 352 | 510 | 490 | 325 | 328 | 343 | 284 | 211 | 259 | 240 | 200 | 298 | 321 | 213 | 361 | 488 | 468 | 309 | 372 | 360 | 324 | 259 | 296 | 244 | 200 | 210 | 322 | 269 | Mycbpap |
| Mycl | 6412 | 6675 | 6447 | 6306 | 7060 | 6624 | 6998 | 8046 | 6664 | 6968 | 5470 | 7978 | 7562 | 7794 | 6546 | 5328 | 6807 | 5972 | 7501 | 7032 | 7183 | 7539 | 6741 | 7445 | 7344 | 7416 | 8457 | 8518 | Mycl |
| Mycn | 2202 | 2262 | 3279 | 2602 | 2899 | 2435 | 2904 | 3283 | 3204 | 3307 | 3049 | 4907 | 3557 | 4648 | 1981 | 2213 | 3120 | 2629 | 3261 | 2871 | 2886 | 3190 | 2781 | 4518 | 4137 | 5193 | 3833 | 4889 | Mycn |
| Mycs | 0 | 0 | 0 | 0 | 0 | 0 | 0 | 1 | 0 | 3 | 0 | 1 | 0 | 0 | 0 | 0 | 0 | 0 | 0 | 0 | 1 | 2 | 0 | 0 | 3 | 0 | 0 | 0 | Mycs |
| Myct1 | 141 | 81 | 103 | 95 | 101 | 112 | 94 | 205 | 284 | 154 | 306 | 160 | 222 | 166 | 127 | 114 | 101 | 90 | 59 | 83 | 71 | 182 | 284 | 199 | 204 | 239 | 187 | 159 | Myct1 |
| Myd88 | 291 | 282 | 271 | 224 | 255 | 262 | 211 | 304 | 438 | 375 | 421 | 324 | 341 | 293 | 280 | 286 | 267 | 283 | 220 | 225 | 201 | 373 | 444 | 392 | 377 | 374 | 302 | 285 | Myd88 |
| Mydgf | 2125 | 2351 | 2072 | 1832 | 1641 | 1642 | 1668 | 2096 | 2408 | 2079 | 1975 | 1959 | 2117 | 1987 | 2164 | 2328 | 1866 | 1809 | 1700 | 2033 | 1608 | 1863 | 2626 | 2128 | 1986 | 2457 | 2141 | 2216 | Mydgf |
| Myef2 | 18182 | 14744 | 18134 | 18045 | 20687 | 20877 | 22236 | 30619 | 25626 | 25157 | 24440 | 29250 | 26236 | 30946 | 20740 | 16743 | 18389 | 18015 | 19978 | 23453 | 20148 | 26553 | 23780 | 27161 | 29634 | 27350 | 30654 | 28542 | Myef2 |
| Myf5 | 23 | 3 | 8 | 0 | 0 | 1 | 1 | 26 | 32 | 4 | 210 | 10 | 8 | 3 | 8 | 30 | 0 | 0 | 0 | 0 | 0 | 27 | 140 | 2 | 11 | 7 | 8 | 6 | Myf5 |
| Myf6 | 5 | 0 | 0 | 0 | 1 | 1 | 0 | 7 | 2 | 2 | 49 | 1 | 0 | 2 | 1 | 1 | 6 | 0 | 5 | 1 | 1 | 0 | 37 | 2 | 0 | 1 | 0 | 2 | Myf6 |
| Myg1 | 2834 | 3268 | 2780 | 2756 | 2212 | 2227 | 2608 | 2216 | 2239 | 2159 | 1929 | 2424 | 2245 | 2407 | 2567 | 2826 | 2684 | 2675 | 2508 | 2533 | 2524 | 2100 | 2110 | 2345 | 2142 | 2482 | 2407 | 2435 | Myg1 |
| Myh1 | 10 | 3 | 0 | 3 | 5 | 2 | 0 | 20 | 47 | 15 | 247 | 4 | 5 | 0 | 5 | 0 | 0 | 0 | 0 | 0 | 1 | 30 | 188 | 13 | 14 | 9 | 0 | 4 | Myh1 |

| | | | | | | | | | | | | | | | | | | | | | | | | | | | | | |
|---|---|---|---|---|---|---|---|---|---|---|---|---|---|---|---|---|---|---|---|---|---|---|---|---|---|---|---|---|---|
| Myh10 | 46332 | 47225 | 40582 | 39225 | 39310 | 43114 | 40629 | 40961 | 46326 | 46788 | 46522 | 51260 | 50018 | 52672 | 44179 | 50785 | 40026 | 39078 | 38071 | 38788 | 40682 | 37578 | 43851 | 45543 | 47604 | 34156 | 53461 | 49434 | Myh10 |
| Myh11 | 120 | 133 | 112 | 90 | 133 | 112 | 65 | 107 | 174 | 150 | 435 | 168 | 178 | 166 | 166 | 147 | 83 | 138 | 47 | 66 | 77 | 149 | 261 | 112 | 113 | 67 | 138 | 125 | Myh11 |
| Myh13 | 1 | 5 | 0 | 4 | 3 | 7 | 0 | 13 | 7 | 12 | 18 | 8 | 10 | 9 | 8 | 5 | 5 | 6 | 4 | 1 | 2 | 15 | 17 | 2 | 7 | 6 | 1 | 2 | Myh13 |
| Myh14 | 855 | 724 | 697 | 810 | 558 | 513 | 362 | 821 | 889 | 985 | 582 | 342 | 421 | 346 | 582 | 747 | 641 | 664 | 458 | 331 | 398 | 374 | 967 | 1096 | 247 | 773 | 272 | 350 | Myh14 |
| Myh15 | 2 | 5 | 7 | 11 | 12 | 3 | 8 | 13 | 15 | 14 | 21 | 29 | 20 | 20 | 7 | 5 | 5 | 7 | 9 | 7 | 14 | 30 | 16 | 11 | 23 | 10 | 2 | 12 | Myh15 |
| Myh2 | 9 | 7 | 0 | 2 | 1 | 15 | 1 | 16 | 12 | 10 | 5 | 26 | 9 | 31 | 8 | 6 | 7 | 8 | 0 | 0 | 1 | 7 | 27 | 18 | 10 | 8 | 22 | 19 | Myh2 |
| Myh3 | 782 | 126 | 103 | 62 | 160 | 92 | 61 | 1387 | 3374 | 444 | 24153 | 168 | 718 | 112 | 343 | 769 | 123 | 103 | 51 | 72 | 47 | 2069 | 15444 | 462 | 342 | 81 | 152 | 112 | Myh3 |
| Myh4 | 0 | 1 | 0 | 0 | 0 | 2 | 1 | 1 | 12 | 0 | 24 | 3 | 10 | 11 | 5 | 1 | 0 | 6 | 3 | 0 | 0 | 0 | 12 | 1 | 1 | 3 | 3 | 5 | Myh4 |
| Myh6 | 7 | 15 | 9 | 13 | 7 | 9 | 6 | 15 | 22 | 28 | 86 | 16 | 23 | 9 | 11 | 9 | 6 | 12 | 16 | 4 | 5 | 22 | 84 | 7 | 7 | 10 | 7 | 20 | Myh6 |
| Myh7 | 100 | 79 | 65 | 65 | 119 | 126 | 130 | 249 | 545 | 203 | 3396 | 151 | 170 | 117 | 86 | 86 | 56 | 109 | 148 | 111 | 112 | 388 | 2113 | 192 | 258 | 90 | 102 | 97 | Myh7 |
| Myh7b | 51 | 58 | 50 | 103 | 55 | 50 | 54 | 46 | 111 | 32 | 669 | 40 | 57 | 50 | 62 | 64 | 81 | 96 | 52 | 50 | 67 | 112 | 449 | 52 | 38 | 41 | 34 | 75 | Myh7b |
| Myh8 | 172 | 93 | 8 | 245 | 184 | 354 | 23 | 625 | 1280 | 446 | 7321 | 162 | 413 | 219 | 138 | 150 | 43 | 285 | 38 | 265 | 43 | 1096 | 6019 | 304 | 446 | 30 | 145 | 84 | Myh8 |
| Myh9 | 21465 | 23766 | 26425 | 19199 | 15230 | 16738 | 13133 | 10828 | 15677 | 14698 | 15460 | 15940 | 18012 | 14262 | 18980 | 28264 | 20240 | 20051 | 15543 | 12477 | 13897 | 11640 | 16106 | 15874 | 12618 | 13846 | 15651 | 14987 | Myh9 |
| Myl1 | 228 | 127 | 60 | 151 | 106 | 116 | 141 | 172 | 522 | 61 | 4025 | 46 | 78 | 20 | 135 | 213 | 127 | 182 | 102 | 182 | 128 | 361 | 2614 | 55 | 46 | 17 | 16 | 17 | Myl1 |
| Myl10 | 0 | 1 | 0 | 0 | 0 | 0 | 0 | 0 | 0 | 0 | 0 | 2 | 0 | 0 | 0 | 0 | 0 | 0 | 0 | 0 | 0 | 0 | 0 | 0 | 0 | 0 | 0 | 0 | Myl10 |
| Myl12a | 2962 | 2553 | 2673 | 2247 | 1247 | 1541 | 1960 | 1893 | 1668 | 2018 | 2347 | 1811 | 2068 | 2822 | 2658 | 2109 | 2292 | 1906 | 1439 | 1347 | 1854 | 1288 | 2043 | 2331 | 1532 | 2333 | 1963 | 2839 | Myl12a |
| Myl12b | 414 | 391 | 428 | 455 | 407 | 381 | 445 | 282 | 246 | 305 | 272 | 300 | 228 | 258 | 394 | 371 | 414 | 448 | 381 | 341 | 449 | 314 | 217 | 286 | 287 | 293 | 246 | 284 | Myl12b |
| Myl2 | 6 | 4 | 0 | 13 | 6 | 1 | 6 | 5 | 15 | 14 | 38 | 3 | 8 | 4 | 1 | 2 | 0 | 3 | 8 | 3 | 4 | 3 | 34 | 3 | 9 | 8 | 6 | 4 | Myl2 |
| Myl3 | 12 | 8 | 10 | 13 | 4 | 19 | 8 | 2 | 12 | 12 | 114 | 4 | 3 | 4 | 11 | 5 | 12 | 4 | 9 | 8 | 3 | 17 | 63 | 12 | 5 | 10 | 17 | 9 | Myl3 |
| Myl4 | 84 | 19 | 16 | 21 | 49 | 25 | 30 | 171 | 427 | 91 | 2963 | 51 | 115 | 42 | 42 | 108 | 19 | 36 | 34 | 44 | 24 | 284 | 1794 | 93 | 114 | 38 | 29 | 40 | Myl4 |
| Myl6 | 15873 | 19274 | 19677 | 17791 | 15190 | 15840 | 14098 | 14880 | 17993 | 16274 | 18622 | 15854 | 18081 | 17483 | 17392 | 18580 | 17891 | 16995 | 16096 | 18343 | 15305 | 13017 | 19821 | 17416 | 16471 | 18280 | 18914 | 17836 | Myl6 |
| Myl6b | 679 | 963 | 1206 | 1273 | 1410 | 1118 | 1057 | 579 | 687 | 729 | 1440 | 752 | 539 | 645 | 769 | 768 | 1176 | 1326 | 1467 | 956 | 1230 | 824 | 1043 | 754 | 768 | 692 | 624 | 660 | Myl6b |
| Myl7 | 8 | 8 | 12 | 9 | 4 | 11 | 3 | 11 | 4 | 7 | 7 | 2 | 6 | 6 | 3 | 6 | 15 | 10 | 5 | 6 | 9 | 8 | 9 | 7 | 7 | 12 | 6 | 12 | Myl7 |
| Myl9 | 3159 | 4185 | 3073 | 1558 | 716 | 1689 | 541 | 1875 | 2399 | 1776 | 2440 | 2717 | 4017 | 2651 | 3579 | 4480 | 2322 | 1755 | 658 | 739 | 560 | 1355 | 3012 | 2105 | 1500 | 2956 | 3488 | 2899 | Myl9 |
| Mylip | 2425 | 2129 | 1848 | 1627 | 1040 | 1219 | 905 | 1627 | 1819 | 1547 | 1760 | 1048 | 1707 | 1054 | 1902 | 2751 | 1446 | 1782 | 830 | 961 | 834 | 1419 | 2001 | 1478 | 1037 | 1622 | 1175 | 1117 | Mylip |
| Mylk | 1955 | 1574 | 2157 | 1562 | 1485 | 1652 | 1189 | 1784 | 2002 | 2048 | 2223 | 1891 | 2067 | 1915 | 1617 | 2082 | 1659 | 1763 | 1236 | 1256 | 1265 | 1973 | 2159 | 2012 | 1693 | 1766 | 2158 | 1837 | Mylk |
| Mylk2 | 0 | 6 | 1 | 3 | 3 | 1 | 0 | 1 | 2 | 0 | 3 | 0 | 7 | 2 | 4 | 8 | 1 | 0 | 4 | 1 | 1 | 2 | 3 | 2 | 4 | 3 | 2 | 2 | Mylk2 |
| Mylk3 | 72 | 91 | 71 | 46 | 21 | 26 | 42 | 61 | 102 | 46 | 62 | 62 | 260 | 118 | 65 | 90 | 53 | 50 | 57 | 44 | 39 | 32 | 98 | 42 | 56 | 120 | 130 | 253 | Mylk3 |
| Mylk4 | 59 | 7 | 9 | 19 | 34 | 34 | 20 | 107 | 246 | 45 | 1354 | 25 | 76 | 35 | 30 | 49 | 23 | 17 | 21 | 16 | 16 | 167 | 769 | 27 | 41 | 18 | 29 | 24 | Mylk4 |
| Mylpf | 337 | 317 | 327 | 340 | 181 | 222 | 245 | 372 | 739 | 155 | 5109 | 138 | 237 | 170 | 306 | 343 | 401 | 215 | 242 | 264 | 264 | 480 | 3163 | 203 | 187 | 151 | 157 | 151 | Mylpf |
| Mymk | 15 | 3 | 0 | 0 | 0 | 2 | 1 | 44 | 110 | 16 | 558 | 1 | 18 | 1 | 5 | 14 | 0 | 0 | 4 | 0 | 0 | 55 | 326 | 7 | 4 | 2 | 0 | 0 | Mymk |
| Mymx | 9 | 1 | 1 | 0 | 2 | 2 | 0 | 15 | 42 | 7 | 245 | 3 | 14 | 2 | 5 | 4 | 0 | 4 | 4 | 3 | 0 | 28 | 104 | 2 | 5 | 0 | 0 | 2 | Mymx |
| Mynn | 4076 | 3682 | 3624 | 3974 | 3415 | 3635 | 3853 | 4183 | 3510 | 3025 | 3165 | 3140 | 3336 | 3412 | 4337 | 3840 | 3612 | 3700 | 3418 | 4249 | 3738 | 3272 | 3538 | 3097 | 3354 | 3126 | 3546 | 3400 | Mynn |
| Myo10 | 6125 | 6312 | 7037 | 6371 | 5938 | 5928 | 5152 | 5516 | 6082 | 6573 | 6116 | 8939 | 7235 | 8495 | 5300 | 6506 | 7121 | 6254 | 6255 | 4514 | 5306 | 6201 | 5124 | 7881 | 7027 | 6662 | 8623 | 8183 | Myo10 |
| Myo15 | 33 | 56 | 44 | 31 | 36 | 43 | 26 | 22 | 72 | 63 | 69 | 28 | 42 | 36 | 51 | 43 | 36 | 57 | 30 | 47 | 23 | 79 | 65 | 56 | 30 | 31 | 38 | 58 | Myo15 |
| Myo15b | 105 | 113 | 64 | 55 | 23 | 27 | 22 | 10 | 40 | 35 | 29 | 26 | 36 | 21 | 83 | 171 | 52 | 77 | 17 | 12 | 13 | 10 | 34 | 16 | 5 | 24 | 12 | 27 | Myo15b |
| Myo16 | 1246 | 1395 | 1962 | 2406 | 3274 | 2736 | 2514 | 1856 | 2038 | 2763 | 2204 | 2175 | 2101 | 1951 | 1509 | 1541 | 2301 | 2516 | 2953 | 2087 | 2927 | 3472 | 1858 | 2174 | 2847 | 1213 | 1935 | 1924 | Myo16 |
| Myo18a | 7250 | 9211 | 9783 | 12464 | 13918 | 12786 | 11624 | 9928 | 12133 | 15492 | 13548 | 14885 | 12957 | 13792 | 8060 | 8360 | 11323 | 12327 | 14555 | 9920 | 13222 | 15047 | 11034 | 15111 | 16166 | 10059 | 13919 | 13610 | Myo18a |
| Myo18b | 150 | 163 | 234 | 269 | 202 | 181 | 206 | 231 | 249 | 284 | 844 | 276 | 243 | 202 | 189 | 150 | 221 | 265 | 243 | 221 | 217 | 311 | 524 | 233 | 280 | 137 | 199 | 241 | Myo18b |
| Myo19 | 1993 | 2135 | 2081 | 2121 | 2621 | 2455 | 2171 | 2172 | 2088 | 2380 | 2381 | 2502 | 2431 | 2333 | 1911 | 2046 | 2248 | 2437 | 2533 | 2158 | 2151 | 2373 | 2026 | 2223 | 2380 | 2088 | 2487 | 2451 | Myo19 |
| Myo1a | 742 | 1179 | 1120 | 1508 | 962 | 1001 | 1059 | 142 | 171 | 201 | 200 | 196 | 161 | 212 | 717 | 1081 | 1235 | 1544 | 1078 | 852 | 1129 | 187 | 150 | 235 | 201 | 190 | 196 | 234 | Myo1a |
| Myo1b | 8304 | 6956 | 7440 | 6549 | 7635 | 8258 | 7813 | 16908 | 14514 | 14249 | 14011 | 17643 | 16577 | 18142 | 8207 | 7520 | 7181 | 6983 | 7511 | 8221 | 7612 | 13714 | 13512 | 15522 | 15932 | 16252 | 18300 | 17631 | Myo1b |
| Myo1c | 2895 | 3085 | 3371 | 2589 | 2226 | 2649 | 1769 | 2149 | 3214 | 2726 | 3594 | 2360 | 2855 | 2172 | 2651 | 3396 | 2650 | 2838 | 2230 | 1761 | 1896 | 2172 | 3625 | 2867 | 2275 | 2496 | 2561 | 2530 | Myo1c |
| Myo1d | 746 | 621 | 567 | 489 | 453 | 442 | 372 | 390 | 524 | 558 | 528 | 362 | 573 | 351 | 625 | 859 | 477 | 573 | 397 | 300 | 352 | 321 | 539 | 471 | 304 | 478 | 342 | 414 | Myo1d |
| Myo1e | 5488 | 5433 | 5608 | 4869 | 4493 | 4500 | 4174 | 1456 | 1528 | 1561 | 1596 | 1290 | 1613 | 1285 | 5417 | 5733 | 5110 | 4908 | 4244 | 3741 | 4353 | 1273 | 1840 | 1421 | 1282 | 1580 | 1302 | 1519 | Myo1e |
| Myo1f | 148 | 157 | 149 | 148 | 114 | 157 | 104 | 167 | 262 | 232 | 305 | 193 | 210 | 185 | 164 | 204 | 122 | 158 | 94 | 111 | 122 | 194 | 326 | 192 | 180 | 240 | 187 | 189 | Myo1f |
| Myo1g | 30 | 52 | 53 | 30 | 34 | 25 | 42 | 45 | 91 | 65 | 87 | 39 | 78 | 46 | 51 | 48 | 29 | 32 | 27 | 36 | 44 | 20 | 100 | 48 | 41 | 65 | 70 | 52 | Myo1g |
| Myo1h | 146 | 74 | 31 | 87 | 29 | 22 | 12 | 107 | 107 | 147 | 145 | 48 | 74 | 65 | 57 | 136 | 50 | 25 | 29 | 38 | 21 | 77 | 138 | 196 | 44 | 195 | 56 | 44 | Myo1h |
| Myo3a | 10 | 14 | 6 | 18 | 16 | 17 | 22 | 13 | 10 | 17 | 9 | 11 | 7 | 19 | 8 | 5 | 20 | 10 | 12 | 16 | 14 | 25 | 13 | 10 | 18 | 8 | 7 | 5 | Myo3a |
| Myo3b | 48 | 52 | 23 | 65 | 93 | 71 | 104 | 71 | 77 | 72 | 48 | 71 | 34 | 50 | 63 | 28 | 49 | 72 | 62 | 78 | 91 | 48 | 60 | 52 | 68 | 39 | 26 | 27 | Myo3b |
| Myo5a | 14239 | 13084 | 9025 | 13499 | 19052 | 20052 | 19427 | 22815 | 21388 | 20236 | 19734 | 20670 | 20420 | 21613 | 16016 | 13478 | 11603 | 14051 | 15195 | 18044 | 19762 | 20501 | 19004 | 15864 | 23553 | 12800 | 21599 | 18260 | Myo5a |
| Myo5b | 538 | 301 | 189 | 235 | 275 | 221 | 196 | 484 | 549 | 549 | 853 | 408 | 458 | 387 | 352 | 377 | 173 | 207 | 223 | 237 | 190 | 401 | 726 | 476 | 308 | 621 | 451 | 400 | Myo5b |
| Myo5c | 461 | 309 | 124 | 237 | 162 | 177 | 111 | 303 | 245 | 287 | 252 | 155 | 204 | 133 | 331 | 382 | 176 | 143 | 150 | 237 | 161 | 104 | 329 | 279 | 101 | 178 | 177 | 166 | Myo5c |
| Myo6 | 7887 | 6848 | 5868 | 7268 | 7212 | 7624 | 7567 | 4054 | 3845 | 4154 | 4052 | 3406 | 3404 | 3601 | 8453 | 8008 | 6596 | 7785 | 6205 | 6725 | 6945 | 3933 | 3808 | 3708 | 3793 | 3068 | 3361 | 3077 | Myo6 |
| Myo7a | 1209 | 1544 | 1492 | 1633 | 1959 | 1947 | 1612 | 2207 | 2773 | 3338 | 2942 | 3068 | 2878 | 2795 | 1282 | 1596 | 1576 | 1712 | 1918 | 1531 | 1786 | 3177 | 2806 | 3038 | 3286 | 2136 | 2862 | 2664 | Myo7a |
| Myo7b | 2 | 0 | 0 | 0 | 0 | 0 | 1 | 0 | 3 | 0 | 0 | 1 | 2 | 0 | 1 | 1 | 2 | 1 | 0 | 5 | 1 | 2 | 1 | 0 | 0 | 1 | 1 | 1 | Myo7b |
| Myo9a | 14885 | 12837 | 8078 | 11364 | 13447 | 15837 | 15809 | 15966 | 13144 | 10883 | 11577 | 10292 | 11085 | 10485 | 16427 | 14104 | 9089 | 11870 | 11768 | 16704 | 15698 | 11548 | 12956 | 8111 | 11930 | 5991 | 11288 | 9541 | Myo9a |
| Myo9b | 14027 | 16923 | 11373 | 15205 | 15609 | 16419 | 14796 | 9225 | 11197 | 10617 | 11019 | 11008 | 10278 | 9710 | 13243 | 14919 | 12987 | 15784 | 15815 | 14004 | 15847 | 10017 | 10160 | 9292 | 11009 | 7203 | 10708 | 8897 | Myo9b |
| Myoc | 35 | 28 | 34 | 38 | 35 | 31 | 36 | 0 | 2 | 11 | 4 | 4 | 5 | 2 | 21 | 26 | 24 | 26 | 69 | 25 | 36 | 2 | 5 | 4 | 1 | 10 | 1 | 4 | Myoc |
| Myocd | 72 | 40 | 101 | 12 | 20 | 25 | 21 | 50 | 30 | 49 | 71 | 67 | 105 | 43 | 44 | 70 | 70 | 50 | 14 | 37 | 15 | 42 | 44 | 38 | 18 | 56 | 138 | 44 | Myocd |
| Myocos | 2 | 0 | 0 | 0 | 2 | 0 | 1 | 0 | 2 | 0 | 0 | 0 | 1 | 1 | 1 | 0 | 0 | 0 | 3 | 0 | 0 | 0 | 0 | 0 | 0 | 0 | 0 | 0 | Myocos |
| Myod1 | 45 | 24 | 14 | 23 | 22 | 16 | 26 | 64 | 158 | 23 | 769 | 10 | 42 | 0 | 26 | 42 | 43 | 39 | 15 | 5 | 11 | 127 | 390 | 7 | 30 | 3 | 0 | 4 | Myod1 |
| Myof | 2110 | 1575 | 1129 | 871 | 659 | 1106 | 488 | 1469 | 2190 | 1553 | 2880 | 1121 | 1661 | 1098 | 1697 | 2345 | 769 | 1071 | 484 | 574 | 448 | 1347 | 2666 | 1377 | 1206 | 957 | 1263 | 1100 | Myof |
| Myog | 76 | 11 | 0 | 1 | 18 | 3 | 4 | 143 | 531 | 41 | 2555 | 14 | 118 | 6 | 24 | 67 | 0 | 9 | 0 | 2 | 4 | 276 | 1424 | 25 | 47 | 2 | 5 | 0 | Myog |
| Myom1 | 335 | 271 | 230 | 251 | 234 | 275 | 244 | 293 | 599 | 356 | 2267 | 328 | 347 | 297 | 303 | 336 | 227 | 210 | 242 | 251 | 231 | 401 | 1546 | 221 | 285 | 297 | 345 | 323 | Myom1 |
| Myom2 | 15 | 10 | 5 | 10 | 23 | 0 | 2 | 76 | 156 | 32 | 1381 | 9 | 18 | 6 | 7 | 16 | 9 | 11 | 0 | 0 | 9 | 112 | 758 | 29 | 23 | 5 | 2 | 7 | Myom2 |
| Myom3 | 61 | 35 | 2 | 14 | 22 | 23 | 17 | 45 | 107 | 29 | 889 | 11 | 22 | 7 | 38 | 61 | 34 | 34 | 8 | 30 | 14 | 147 | 567 | 26 | 25 | 19 | 9 | 12 | Myom3 |
| Myorg | 3802 | 4495 | 3949 | 4044 | 3056 | 3406 | 3003 | 935 | 1053 | 1118 | 1049 | 1081 | 1303 | 999 | 3705 | 4408 | 4220 | 3571 | 3409 | 2874 | 3203 | 1029 | 1259 | 1063 | 997 | 1072 | 1135 | 1140 | Myorg |
| Myot | 13 | 6 | 4 | 1 | 0 | 1 | 0 | 24 | 44 | 1 | 362 | 0 | 18 | 4 | 12 | 12 | 0 | 0 | 0 | 0 | 0 | 45 | 258 | 13 | 0 | 3 | 7 | 1 | Myot |
| Myoz1 | 5 | 5 | 5 | 4 | 0 | 1 | 0 | 4 | 10 | 0 | 50 | 7 | 10 | 3 | 3 | 5 | 1 | 7 | 5 | 4 | 6 | 15 | 36 | 4 | 2 | 4 | 7 | 2 | Myoz1 |
| Myoz2 | 12 | 3 | 0 | 0 | 2 | 4 | 1 | 13 | 38 | 18 | 376 | 0 | 6 | 2 | 3 | 4 | 5 | 0 | 0 | 0 | 0 | 38 | 240 | 1 | 1 | 0 | 0 | 0 | Myoz2 |
| Myoz3 | 3 | 7 | 14 | 0 | 5 | 5 | 2 | 5 | 0 | 3 | 3 | 13 | 5 | 3 | 7 | 19 | 6 | 11 | 0 | 0 | 0 | 8 | 2 | 4 | 8 | 10 | 3 | 1 | Myoz3 |
| Mypn | 1 | 5 | 9 | 1 | 11 | 3 | 8 | 26 | 52 | 8 | 398 | 3 | 12 | 3 | 7 | 15 | 6 | 7 | 13 | 2 | 4 | 43 | 212 | 14 | 9 | 1 | 6 | 9 | Mypn |
| Mypop | 855 | 1066 | 1131 | 1182 | 1132 | 1065 | 905 | 488 | 499 | 621 | 658 | 535 | 506 | 504 | 728 | 820 | 1166 | 1097 | 1050 | 761 | 920 | 536 | 525 | 593 | 583 | 531 | 459 | 488 | Mypop |
| Myrf | 487 | 616 | 744 | 398 | 382 | 448 | 232 | 278 | 327 | 335 | 380 | 409 | 561 | 386 | 471 | 866 | 555 | 374 | 299 | 296 | 245 | 316 | 399 | 362 | 286 | 405 | 490 | 444 | Myrf |
| Myrfl | 67 | 103 | 66 | 113 | 119 | 111 | 99 | 21 | 25 | 39 | 24 | 12 | 27 | 29 | 75 | 85 | 125 | 160 | 99 | 75 | 136 | 12 | 25 | 26 | 19 | 21 | 33 | 24 | Myrfl |
| Myrip | 317 | 413 | 347 | 647 | 840 | 608 | 575 | 315 | 336 | 433 | 431 | 324 | 315 | 295 | 385 | 347 | 556 | 755 | 701 | 599 | 659 | 496 | 383 | 282 | 444 | 174 | 258 | 307 | Myrip |
| Mysm1 | 5840 | 4438 | 3604 | 4305 | 4144 | 5080 | 4966 | 6232 | 5684 | 4893 | 5686 | 5308 | 5399 | 5753 | 5387 | 5198 | 3688 | 4324 | 4066 | 5180 | 4842 | 4687 | 5561 | 4457 | 5223 | 3428 | 5722 | 5177 | Mysm1 |
| Myt1 | 5021 | 5955 | 4603 | 6060 | 7616 | 7832 | 7557 | 11434 | 9247 | 10251 | 8898 | 13368 | 11428 | 13762 | 6410 | 4327 | 5846 | 5809 | 7416 | 7607 | 7531 | 11244 | 8321 | 11453 | 13638 | 9216 | 13852 | 13536 | Myt1 |

| | | | | | | | | | | | | | | | | | | | | | | | | | | | | | |
|---|---|---|---|---|---|---|---|---|---|---|---|---|---|---|---|---|---|---|---|---|---|---|---|---|---|---|---|---|---|
| Myt1l | 2795 | 2972 | 1795 | 6468 | 11869 | 9439 | 9626 | 10273 | 10756 | 12207 | 9584 | 6695 | 5816 | 7309 | 3745 | 2583 | 3970 | 7850 | 8743 | 9072 | 10687 | 13665 | 9182 | 8618 | 13245 | 3496 | 5187 | 5038 | Myt1l |
| Myzap | 824 | 657 | 751 | 478 | 467 | 412 | 343 | 377 | 527 | 330 | 1316 | 358 | 460 | 293 | 767 | 969 | 553 | 515 | 341 | 466 | 355 | 506 | 1017 | 345 | 328 | 318 | 365 | 267 | Myzap |
| Mzb1 | 37 | 57 | 59 | 62 | 55 | 40 | 40 | 13 | 26 | 25 | 15 | 14 | 16 | 9 | 34 | 58 | 57 | 62 | 53 | 67 | 55 | 20 | 28 | 13 | 19 | 24 | 23 | 23 | Mzb1 |
| Mzf1 | 150 | 149 | 128 | 148 | 201 | 160 | 197 | 663 | 712 | 761 | 655 | 764 | 703 | 846 | 115 | 121 | 172 | 191 | 235 | 182 | 230 | 866 | 663 | 789 | 878 | 624 | 752 | 706 | Mzf1 |
| Mzt1 | 9244 | 6714 | 9494 | 9245 | 9001 | 8731 | 11035 | 8160 | 6448 | 6182 | 6574 | 8275 | 7667 | 8730 | 9911 | 8233 | 9503 | 8871 | 8970 | 9784 | 9463 | 7543 | 6745 | 7865 | 7679 | 8825 | 8212 | 8984 | Mzt1 |
| Mzt2 | 1022 | 1155 | 1174 | 1177 | 1001 | 1037 | 1048 | 863 | 881 | 844 | 802 | 925 | 917 | 925 | 1015 | 913 | 1270 | 1030 | 1169 | 1210 | 1097 | 734 | 817 | 942 | 984 | 1195 | 1051 | 974 | Mzt2 |
| N4bp1 | 5890 | 6372 | 6591 | 6984 | 6632 | 6677 | 6453 | 5503 | 5279 | 5923 | 5735 | 6202 | 5757 | 6213 | 5459 | 6400 | 6880 | 7064 | 6830 | 5926 | 6741 | 6328 | 5066 | 5765 | 6212 | 5293 | 5625 | 6172 | N4bp1 |
| N4bp2 | 9184 | 6851 | 6707 | 7152 | 8395 | 8961 | 8843 | 13080 | 11839 | 11345 | 12777 | 12475 | 12295 | 13678 | 8790 | 7881 | 7098 | 6728 | 7816 | 8324 | 8723 | 11003 | 11202 | 10856 | 12250 | 9466 | 13345 | 12448 | N4bp2 |
| N4bp2l1 | 774 | 841 | 673 | 934 | 1195 | 1205 | 1214 | 488 | 456 | 499 | 469 | 406 | 354 | 409 | 829 | 868 | 781 | 1081 | 1010 | 1057 | 1262 | 566 | 482 | 416 | 498 | 318 | 384 | 291 | N4bp2l1 |
| N4bp2l2 | 6496 | 4900 | 4975 | 5548 | 4982 | 5891 | 5618 | 8563 | 7412 | 6278 | 6743 | 6774 | 6813 | 7083 | 6381 | 5583 | 5079 | 5140 | 5093 | 6225 | 5728 | 6916 | 7426 | 6326 | 6934 | 6469 | 7145 | 7202 | N4bp2l2 |
| N4bp3 | 704 | 797 | 828 | 733 | 604 | 602 | 393 | 442 | 707 | 562 | 809 | 472 | 577 | 421 | 626 | 787 | 695 | 603 | 705 | 636 | 520 | 546 | 845 | 545 | 422 | 510 | 463 | 465 | N4bp3 |
| N6amt1 | 400 | 278 | 405 | 375 | 427 | 411 | 360 | 1391 | 1416 | 1544 | 1609 | 1564 | 1593 | 1581 | 392 | 357 | 391 | 288 | 393 | 362 | 399 | 1288 | 1533 | 1439 | 1460 | 1606 | 1557 | 1669 | N6amt1 |
| Naa10 | 1015 | 1124 | 1181 | 1077 | 907 | 841 | 1123 | 1058 | 998 | 1275 | 1109 | 1233 | 1258 | 1527 | 885 | 849 | 1148 | 913 | 948 | 894 | 989 | 1034 | 880 | 1325 | 1198 | 1384 | 1174 | 1520 | Naa10 |
| Naa11 | 0 | 2 | 0 | 2 | 0 | 0 | 0 | 0 | 0 | 0 | 0 | 0 | 0 | 0 | 0 | 0 | 0 | 2 | 1 | 0 | 1 | 0 | 0 | 0 | 0 | 0 | 0 | 1 | Naa11 |
| Naa12 | 4988 | 5712 | 5210 | 4594 | 4444 | 4748 | 4609 | 5296 | 5609 | 5161 | 4854 | 5670 | 5402 | 5531 | 5061 | 4865 | 4968 | 4554 | 4695 | 4859 | 4645 | 4645 | 5237 | 5244 | 5144 | 5991 | 5771 | 5906 | Naa12 |
| Naa15 | 16067 | 12265 | 16571 | 15538 | 14550 | 16028 | 17267 | 15997 | 13867 | 14124 | 13922 | 17167 | 16113 | 17599 | 15875 | 14792 | 15445 | 14876 | 15322 | 15316 | 16038 | 14962 | 13297 | 14833 | 15289 | 17093 | 17648 | 17709 | Naa15 |
| Naa16 | 1878 | 1488 | 1653 | 1760 | 1595 | 1763 | 1797 | 2396 | 1973 | 2021 | 2066 | 2465 | 2269 | 2483 | 1770 | 1883 | 1417 | 1799 | 1569 | 1761 | 1841 | 2042 | 1963 | 2229 | 2173 | 2440 | 2446 | 2302 | Naa16 |
| Naa20 | 4315 | 3568 | 3935 | 3628 | 3190 | 3592 | 3660 | 3644 | 3154 | 2960 | 2949 | 2876 | 2944 | 3216 | 4163 | 3833 | 3657 | 3518 | 3546 | 3937 | 3597 | 2924 | 3219 | 3064 | 3001 | 3386 | 3303 | 3418 | Naa20 |
| Naa25 | 7128 | 7283 | 5849 | 6535 | 6356 | 6657 | 6805 | 7100 | 6214 | 5818 | 5459 | 6205 | 5746 | 5957 | 7483 | 6542 | 6098 | 6162 | 6266 | 7293 | 6664 | 5892 | 6055 | 5632 | 6059 | 5300 | 6295 | 5972 | Naa25 |
| Naa30 | 6085 | 4130 | 7033 | 7164 | 6383 | 6225 | 6735 | 5267 | 4331 | 4818 | 5078 | 5533 | 5071 | 5666 | 5946 | 5650 | 6545 | 6792 | 5842 | 5647 | 6524 | 6244 | 4447 | 5069 | 5538 | 5319 | 5619 | 5660 | Naa30 |
| Naa35 | 12441 | 11796 | 10724 | 11020 | 10226 | 10456 | 11736 | 10698 | 8979 | 8594 | 8018 | 9799 | 8843 | 10064 | 12552 | 10758 | 11426 | 10652 | 10841 | 11531 | 11689 | 9311 | 8714 | 9323 | 9451 | 9965 | 9798 | 9894 | Naa35 |
| Naa38 | 664 | 880 | 862 | 672 | 588 | 501 | 723 | 686 | 799 | 835 | 897 | 805 | 905 | 949 | 608 | 584 | 767 | 601 | 617 | 690 | 692 | 561 | 827 | 881 | 828 | 1002 | 1006 | 1041 | Naa38 |
| Naa40 | 5986 | 6633 | 6823 | 6418 | 6047 | 5901 | 6373 | 5152 | 5267 | 5418 | 5179 | 6429 | 5917 | 6407 | 5988 | 6162 | 6714 | 6241 | 6438 | 5761 | 6212 | 5566 | 5014 | 5806 | 5811 | 6572 | 6422 | 6501 | Naa40 |
| Naa50 | 17347 | 11593 | 18627 | 16916 | 14022 | 15573 | 17577 | 11031 | 9501 | 9262 | 11208 | 11870 | 11329 | 12965 | 16736 | 15421 | 16314 | 16437 | 15047 | 15158 | 16437 | 10358 | 10248 | 11212 | 10567 | 14374 | 12411 | 14010 | Naa50 |
| Naa60 | 2833 | 3105 | 2869 | 2896 | 3236 | 2903 | 3180 | 2558 | 2759 | 2910 | 2997 | 3072 | 2694 | 2995 | 2701 | 2921 | 3002 | 3083 | 3074 | 2593 | 3040 | 3133 | 2790 | 2939 | 2905 | 3124 | 2725 | 2912 | Naa60 |
| Naa80 | 546 | 726 | 667 | 521 | 520 | 541 | 425 | 613 | 732 | 663 | 616 | 649 | 638 | 545 | 443 | 718 | 685 | 530 | 587 | 418 | 448 | 647 | 751 | 666 | 635 | 791 | 565 | 592 | Naa80 |
| Naaa | 352 | 357 | 350 | 359 | 518 | 450 | 279 | 379 | 517 | 452 | 453 | 339 | 479 | 292 | 413 | 453 | 370 | 336 | 363 | 349 | 264 | 498 | 627 | 347 | 396 | 271 | 317 | 318 | Naaa |
| Naalad2 | 272 | 174 | 192 | 167 | 183 | 194 | 194 | 401 | 434 | 250 | 527 | 303 | 375 | 260 | 264 | 328 | 173 | 162 | 160 | 212 | 168 | 538 | 465 | 365 | 346 | 419 | 259 | 329 | Naalad2 |
| Naaladl1 | 458 | 538 | 623 | 485 | 300 | 393 | 497 | 166 | 262 | 290 | 227 | 445 | 374 | 424 | 368 | 468 | 504 | 416 | 446 | 370 | 383 | 187 | 226 | 364 | 291 | 478 | 478 | 317 | Naaladl1 |
| Naaladl2 | 189 | 86 | 66 | 71 | 78 | 107 | 46 | 139 | 145 | 131 | 279 | 88 | 136 | 138 | 129 | 136 | 94 | 123 | 56 | 76 | 56 | 157 | 218 | 110 | 96 | 55 | 126 | 112 | Naaladl2 |
| Nab1 | 1678 | 1523 | 1645 | 1995 | 2442 | 2236 | 2514 | 2096 | 2256 | 2079 | 2055 | 1887 | 1973 | 1986 | 1924 | 1729 | 1798 | 2128 | 2377 | 2211 | 2389 | 2181 | 2375 | 2114 | 2116 | 2231 | 1711 | 1741 | Nab1 |
| Nab2 | 1113 | 1227 | 1492 | 1191 | 1027 | 1032 | 774 | 1039 | 1422 | 1302 | 1677 | 896 | 1224 | 840 | 981 | 1319 | 1308 | 1052 | 1172 | 848 | 819 | 951 | 1557 | 1214 | 820 | 1506 | 1004 | 922 | Nab2 |
| Nabp1 | 1626 | 1312 | 1475 | 1644 | 1438 | 1499 | 1643 | 767 | 562 | 550 | 612 | 641 | 579 | 655 | 1710 | 1456 | 1544 | 1732 | 1411 | 1555 | 1584 | 677 | 693 | 623 | 507 | 633 | 568 | 503 | Nabp1 |
| Nabp2 | 2343 | 3089 | 3269 | 3272 | 2981 | 2852 | 2908 | 2606 | 2850 | 3423 | 2949 | 3918 | 3274 | 3688 | 2276 | 2209 | 3306 | 2885 | 3607 | 3074 | 3104 | 2837 | 2743 | 3832 | 3654 | 4362 | 3597 | 3814 | Nabp2 |
| Naca | 30417 | 29940 | 31160 | 28448 | 23448 | 24475 | 30716 | 30963 | 28796 | 27428 | 30693 | 30344 | 30181 | 37955 | 30623 | 27001 | 30178 | 25874 | 26745 | 31754 | 29309 | 24495 | 29152 | 33186 | 29140 | 37122 | 32895 | 38887 | Naca |
| Nacad | 1390 | 2046 | 2410 | 2891 | 4410 | 3337 | 2881 | 4704 | 5278 | 7343 | 6059 | 7117 | 5846 | 5751 | 1659 | 1624 | 3029 | 2963 | 4348 | 2689 | 3270 | 8863 | 4754 | 7065 | 8314 | 5453 | 5439 | 5478 | Nacad |
| Nacc1 | 6312 | 7411 | 9088 | 7911 | 7729 | 7571 | 6708 | 6571 | 8232 | 8593 | 8083 | 10780 | 8913 | 9290 | 5908 | 6547 | 8876 | 7291 | 8870 | 6225 | 7312 | 7568 | 7976 | 10126 | 9041 | 12142 | 9389 | 9817 | Nacc1 |
| Nacc2 | 1004 | 1136 | 1290 | 1363 | 1715 | 1458 | 1305 | 1221 | 1428 | 1729 | 1769 | 1620 | 1533 | 1458 | 1122 | 1178 | 1442 | 1545 | 1417 | 1138 | 1179 | 1883 | 1492 | 1733 | 1823 | 1457 | 1427 | 1400 | Nacc2 |
| Nadk | 7194 | 7819 | 6659 | 6992 | 6559 | 6596 | 7021 | 5998 | 6115 | 5455 | 5521 | 5994 | 5722 | 5875 | 7137 | 7061 | 6665 | 7346 | 7135 | 6767 | 7167 | 5817 | 6091 | 6152 | 5797 | 6168 | 6019 | 6158 | Nadk |
| Nadk2 | 5 | 2 | 9 | 1 | 3 | 0 | 2 | 1 | 3 | 0 | 1 | 1 | 0 | 2 | 0 | 2 | 2 | 1 | 3 | 1 | 1 | 2 | 1 | 1 | 3 | 2 | 0 | 1 | Nadk2 |
| Nadsyn1 | 630 | 929 | 826 | 719 | 823 | 784 | 695 | 681 | 695 | 612 | 682 | 705 | 778 | 620 | 728 | 829 | 796 | 754 | 794 | 722 | 644 | 597 | 698 | 672 | 710 | 735 | 862 | 662 | Nadsyn1 |
| Nae1 | 5683 | 4417 | 5121 | 4607 | 4911 | 5236 | 5511 | 6891 | 5871 | 5192 | 5646 | 6091 | 6055 | 6157 | 5851 | 5204 | 4821 | 4590 | 4867 | 5942 | 5084 | 5553 | 5729 | 5670 | 5876 | 6318 | 6207 | 6395 | Nae1 |
| Naf1 | 1888 | 1521 | 2410 | 2108 | 1910 | 1917 | 2038 | 1437 | 1437 | 1459 | 1541 | 1806 | 1641 | 1806 | 1859 | 1826 | 2022 | 1951 | 1887 | 1673 | 1999 | 1430 | 1459 | 1531 | 1356 | 1949 | 1452 | 1885 | Naf1 |
| Naga | 2261 | 2757 | 2783 | 3034 | 2459 | 2299 | 2101 | 1134 | 1500 | 1383 | 1575 | 1357 | 1414 | 1197 | 2244 | 2822 | 2756 | 3023 | 2402 | 2063 | 2080 | 1370 | 1570 | 1450 | 1217 | 1419 | 1285 | 1353 | Naga |
| Nagk | 1138 | 1369 | 1363 | 1396 | 1625 | 1453 | 1359 | 1734 | 1999 | 2182 | 2130 | 1945 | 1979 | 1985 | 1145 | 1178 | 1418 | 1472 | 1644 | 1477 | 1391 | 2301 | 2148 | 2173 | 2332 | 2014 | 1813 | 1887 | Nagk |
| Naglu | 715 | 988 | 893 | 911 | 776 | 777 | 618 | 443 | 558 | 587 | 579 | 645 | 686 | 487 | 734 | 967 | 988 | 801 | 828 | 660 | 688 | 622 | 656 | 482 | 563 | 600 | 595 | 487 | Naglu |
| Nagpa | 369 | 498 | 423 | 296 | 374 | 314 | 298 | 355 | 436 | 446 | 397 | 441 | 447 | 371 | 378 | 418 | 422 | 420 | 429 | 308 | 305 | 379 | 454 | 387 | 362 | 454 | 402 | 422 | Nagpa |
| Nags | 10 | 22 | 17 | 9 | 3 | 2 | 4 | 4 | 4 | 0 | 0 | 0 | 2 | 4 | 7 | 19 | 14 | 9 | 16 | 3 | 8 | 0 | 3 | 0 | 0 | 0 | 0 | 1 | Nags |
| Naif1 | 170 | 197 | 269 | 210 | 188 | 147 | 163 | 458 | 510 | 500 | 503 | 553 | 440 | 475 | 177 | 148 | 184 | 150 | 217 | 142 | 178 | 491 | 484 | 543 | 472 | 585 | 513 | 484 | Naif1 |
| Naip1 | 8 | 12 | 7 | 4 | 12 | 12 | 23 | 9 | 9 | 9 | 14 | 1 | 10 | 3 | 5 | 3 | 8 | 14 | 12 | 7 | 13 | 7 | 10 | 1 | 3 | 4 | 0 | 4 | Naip1 |
| Naip2 | 88 | 86 | 88 | 59 | 40 | 50 | 44 | 62 | 64 | 41 | 46 | 56 | 70 | 51 | 88 | 115 | 70 | 59 | 34 | 72 | 40 | 75 | 103 | 51 | 62 | 70 | 50 | 68 | Naip2 |
| Naip5 | 21 | 42 | 44 | 41 | 30 | 37 | 35 | 45 | 38 | 32 | 44 | 50 | 66 | 52 | 30 | 64 | 32 | 32 | 36 | 25 | 25 | 47 | 59 | 41 | 62 | 31 | 41 | 52 | Naip5 |
| Naip6 | 36 | 17 | 15 | 17 | 33 | 11 | 11 | 45 | 51 | 34 | 54 | 45 | 39 | 32 | 16 | 40 | 22 | 18 | 22 | 26 | 24 | 67 | 63 | 18 | 22 | 44 | 42 | 26 | Naip6 |
| Nalcn | 1023 | 983 | 702 | 1066 | 1647 | 1475 | 1498 | 2192 | 2314 | 2014 | 1800 | 1634 | 1269 | 1535 | 1138 | 941 | 972 | 1338 | 1264 | 1708 | 1570 | 2440 | 1906 | 1398 | 2285 | 978 | 1564 | 1413 | Nalcn |
| Nampt | 6734 | 5394 | 5690 | 5389 | 5128 | 5434 | 6039 | 5947 | 4924 | 4326 | 4359 | 5133 | 4807 | 5485 | 6697 | 6104 | 5366 | 5373 | 5460 | 6460 | 5849 | 4951 | 4877 | 4611 | 4817 | 5075 | 5070 | 5508 | Nampt |
| Nanog | 13 | 4 | 5 | 9 | 11 | 4 | 7 | 21 | 24 | 28 | 24 | 15 | 29 | 29 | 12 | 4 | 9 | 8 | 14 | 8 | 11 | 12 | 23 | 20 | 23 | 47 | 17 | 25 | Nanog |
| Nanos1 | 1751 | 1665 | 1545 | 1878 | 2184 | 2391 | 2436 | 2315 | 1773 | 2109 | 1536 | 1546 | 1342 | 1644 | 2124 | 1699 | 1782 | 2189 | 2272 | 2506 | 2359 | 2473 | 1831 | 1556 | 2219 | 1227 | 1379 | 1447 | Nanos1 |
| Nanos2 | 163 | 207 | 244 | 476 | 491 | 382 | 406 | 39 | 76 | 88 | 74 | 103 | 116 | 126 | 177 | 167 | 404 | 516 | 407 | 292 | 429 | 135 | 45 | 108 | 142 | 129 | 92 | 101 | Nanos2 |
| Nanos3 | 50 | 44 | 67 | 29 | 56 | 58 | 48 | 111 | 138 | 150 | 129 | 180 | 172 | 148 | 41 | 51 | 35 | 21 | 41 | 51 | 70 | 109 | 107 | 152 | 168 | 166 | 179 | 164 | Nanos3 |
| Nanp | 65 | 46 | 67 | 74 | 44 | 89 | 83 | 60 | 55 | 36 | 41 | 62 | 35 | 48 | 108 | 62 | 79 | 67 | 41 | 64 | 62 | 37 | 43 | 38 | 51 | 66 | 62 | 51 | Nanp |
| Nans | 6100 | 7320 | 6786 | 6490 | 5300 | 5480 | 5801 | 3276 | 3827 | 3400 | 3647 | 3357 | 3317 | 3738 | 6145 | 6319 | 6397 | 6025 | 5823 | 5974 | 6067 | 3056 | 3917 | 3805 | 3526 | 3989 | 3527 | 4002 | Nans |
| Nap1l1 | 49532 | 39505 | 56121 | 42068 | 37837 | 40577 | 49061 | 46099 | 38681 | 40282 | 40146 | 55041 | 49142 | 57967 | 48450 | 43623 | 47130 | 40439 | 42109 | 45078 | 44238 | 38185 | 39702 | 47394 | 42378 | 63347 | 57282 | 61441 | Nap1l1 |
| Nap1l2 | 1743 | 1679 | 2210 | 2592 | 3980 | 3309 | 3443 | 4023 | 3222 | 3669 | 3155 | 2917 | 3300 | 3092 | 2397 | 1663 | 2500 | 2815 | 3250 | 3611 | 3337 | 4358 | 3124 | 3021 | 3671 | 2338 | 2981 | 3067 | Nap1l2 |
| Nap1l3 | 1374 | 1368 | 1499 | 1937 | 2414 | 2223 | 2144 | 2135 | 1707 | 1916 | 1695 | 1472 | 1441 | 1385 | 1560 | 1333 | 1751 | 2063 | 2100 | 2064 | 2354 | 2717 | 1887 | 1528 | 1937 | 1206 | 1362 | 1422 | Nap1l3 |
| Nap1l4 | 17574 | 18905 | 19491 | 18292 | 16963 | 16553 | 17305 | 17453 | 16361 | 17621 | 15977 | 19678 | 17994 | 19317 | 17359 | 16214 | 19102 | 17085 | 17963 | 18617 | 17161 | 15909 | 15957 | 18730 | 18637 | 21045 | 20219 | 20403 | Nap1l4 |
| Nap1l5 | 3453 | 4240 | 3382 | 5751 | 5746 | 5293 | 6001 | 2991 | 2349 | 2968 | 2587 | 2286 | 2278 | 2348 | 4928 | 3913 | 4881 | 5880 | 5168 | 6080 | 5717 | 3783 | 2398 | 2386 | 3038 | 1757 | 2278 | 2263 | Nap1l5 |
| Napa | 7673 | 9808 | 7140 | 8268 | 7349 | 7294 | 7460 | 5277 | 6246 | 6044 | 6204 | 5835 | 5989 | 5811 | 7512 | 8407 | 7541 | 7586 | 7077 | 6798 | 7245 | 5994 | 6125 | 5708 | 5992 | 6251 | 5871 | 6026 | Napa |
| Napb | 4359 | 4382 | 4123 | 5953 | 6789 | 6090 | 6671 | 5689 | 4252 | 4450 | 4215 | 3872 | 3762 | 4008 | 5470 | 4244 | 5064 | 5849 | 5778 | 6741 | 6805 | 6234 | 4188 | 3843 | 5135 | 2555 | 3715 | 3401 | Napb |
| Napepld | 2415 | 2721 | 3051 | 2647 | 1974 | 2219 | 2459 | 1034 | 864 | 932 | 698 | 1738 | 1041 | 1718 | 2541 | 2246 | 3007 | 2341 | 2263 | 2388 | 2197 | 916 | 827 | 1397 | 1112 | 1800 | 1906 | 1839 | Napepld |
| Napg | 4725 | 4781 | 4131 | 4822 | 5046 | 5037 | 5396 | 4860 | 4360 | 4034 | 4226 | 4015 | 3929 | 4469 | 4974 | 5161 | 4394 | 5062 | 4398 | 5376 | 5151 | 4553 | 4525 | 3943 | 4660 | 3347 | 4051 | 3997 | Napg |
| Naprt | 157 | 276 | 204 | 362 | 186 | 215 | 192 | 145 | 191 | 181 | 225 | 168 | 174 | 122 | 153 | 289 | 301 | 370 | 186 | 221 | 203 | 184 | 212 | 196 | 192 | 176 | 149 | 125 | Naprt |
| Napsa | 7 | 3 | 4 | 2 | 2 | 4 | 1 | 5 | 29 | 22 | 9 | 2 | 5 | 3 | 5 | 5 | 3 | 1 | 3 | 6 | 1 | 3 | 25 | 15 | 1 | 10 | 7 | 2 | Napsa |
| Narf | 9896 | 10336 | 9437 | 9561 | 9492 | 9237 | 8843 | 10569 | 9316 | 9331 | 8927 | 8770 | 8796 | 9304 | 10392 | 9503 | 9284 | 9770 | 9826 | 10046 | 10094 | 9783 | 9681 | 9494 | 10307 | 8052 | 8737 | 9588 | Narf |
| Nars | 15309 | 15222 | 16425 | 14927 | 12612 | 13427 | 13940 | 12308 | 11398 | 11485 | 11074 | 12339 | 11367 | 14128 | 15361 | 14147 | 15616 | 13684 | 13719 | 15087 | 13789 | 10460 | 11071 | 12423 | 11818 | 14118 | 13289 | 14601 | Nars |
| Nars2 | 2321 | 2065 | 1943 | 1936 | 1660 | 1714 | 1964 | 1170 | 1127 | 862 | 976 | 984 | 1014 | 1069 | 2428 | 2223 | 1964 | 1810 | 1663 | 1950 | 1784 | 864 | 1148 | 891 | 898 | 1103 | 1137 | 1075 | Nars2 |
| Nasp | 35803 | 28652 | 40593 | 28105 | 24821 | 28890 | 32232 | 41471 | 34719 | 33179 | 32326 | 48362 | 42845 | 52812 | 34094 | 30343 | 34040 | 24955 | 28901 | 34075 | 28896 | 29875 | 31241 | 41155 | 36655 | 57258 | 51408 | 51837 | Nasp |
| Nat1 | 8 | 6 | 13 | 4 | 2 | 5 | 6 | 12 | 7 | 0 | 9 | 7 | 12 | 4 | 16 | 14 | 2 | 13 | 2 | 6 | 3 | 8 | 10 | 2 | 4 | 17 | 0 | 6 | Nat1 |
| Nat10 | 5602 | 5975 | 5373 | 5911 | 5041 | 5591 | 5630 | 3605 | 4196 | 4175 | 4089 | 4912 | 4361 | 4790 | 5012 | 5364 | 5554 | 5574 | 5720 | 5133 | 5700 | 3847 | 3782 | 4376 | 4425 | 4472 | 4965 | 4883 | Nat10 |
| Nat14 | 412 | 563 | 669 | 579 | 698 | 530 | 648 | 699 | 707 | 914 | 720 | 831 | 646 | 876 | 404 | 420 | 629 | 578 | 693 | 669 | 592 | 941 | 702 | 954 | 934 | 876 | 761 | 835 | Nat14 |

| | | | | | | | | | | | | | | | | | | | | | | | | | | | | | |
|---|---|---|---|---|---|---|---|---|---|---|---|---|---|---|---|---|---|---|---|---|---|---|---|---|---|---|---|---|---|
| Nat2 | 288 | 231 | 355 | 178 | 230 | 200 | 203 | 232 | 242 | 191 | 184 | 210 | 320 | 194 | 282 | 293 | 261 | 172 | 252 | 245 | 172 | 150 | 238 | 146 | 110 | 207 | 202 | 211 | Nat2 |
| Nat3 | 0 | 1 | 0 | 0 | 1 | 0 | 0 | 0 | 0 | 0 | 1 | 0 | 0 | 0 | 3 | 0 | 3 | 0 | 0 | 0 | 1 | 3 | 1 | 0 | 0 | 0 | 0 | 0 | Nat3 |
| Nat8 | 0 | 1 | 2 | 0 | 0 | 0 | 0 | 0 | 1 | 2 | 0 | 0 | 0 | 0 | 0 | 1 | 0 | 0 | 0 | 0 | 0 | 0 | 0 | 0 | 0 | 2 | 1 | 0 | Nat8 |
| Nat8f1 | 194 | 198 | 191 | 165 | 245 | 168 | 248 | 216 | 246 | 184 | 218 | 244 | 178 | 228 | 183 | 146 | 207 | 167 | 196 | 249 | 220 | 237 | 249 | 199 | 218 | 195 | 264 | 209 | Nat8f1 |
| Nat8f2 | 6 | 5 | 0 | 13 | 3 | 4 | 9 | 1 | 5 | 3 | 7 | 9 | 6 | 1 | 3 | 4 | 6 | 9 | 1 | 8 | 6 | 2 | 3 | 3 | 5 | 4 | 0 | 1 | Nat8f2 |
| Nat8f3 | 0 | 0 | 0 | 0 | 1 | 0 | 0 | 0 | 0 | 1 | 0 | 0 | 0 | 0 | 0 | 0 | 0 | 0 | 0 | 0 | 0 | 0 | 0 | 0 | 0 | 0 | 0 | 0 | Nat8f3 |
| Nat8f4 | 497 | 448 | 479 | 445 | 465 | 501 | 497 | 793 | 662 | 666 | 517 | 768 | 636 | 732 | 538 | 359 | 506 | 506 | 520 | 531 | 477 | 663 | 619 | 680 | 692 | 821 | 784 | 821 | Nat8f4 |
| Nat8f5 | 0 | 0 | 0 | 0 | 0 | 0 | 0 | 1 | 0 | 0 | 1 | 0 | 0 | 0 | 0 | 0 | 0 | 0 | 0 | 0 | 0 | 0 | 0 | 0 | 1 | 0 | 0 | 0 | Nat8f5 |
| Nat8f6 | 0 | 0 | 0 | 0 | 0 | 0 | 0 | 0 | 0 | 0 | 0 | 0 | 0 | 0 | 0 | 0 | 0 | 0 | 0 | 0 | 0 | 0 | 1 | 0 | 0 | 0 | 0 | 0 | Nat8f6 |
| Nat8l | 1972 | 2683 | 3521 | 3897 | 4196 | 3368 | 3320 | 2099 | 2464 | 3001 | 2552 | 2996 | 2383 | 2624 | 2037 | 2155 | 3782 | 4077 | 4199 | 3134 | 3425 | 3068 | 2218 | 3281 | 3730 | 2729 | 2482 | 2687 | Nat8l |
| Nat9 | 516 | 662 | 696 | 679 | 547 | 560 | 493 | 613 | 676 | 694 | 748 | 716 | 711 | 641 | 568 | 575 | 643 | 495 | 613 | 571 | 580 | 545 | 715 | 752 | 808 | 757 | 793 | 792 | Nat9 |
| Natd1 | 5328 | 6719 | 4817 | 5897 | 5435 | 5325 | 5465 | 4319 | 4468 | 4656 | 4239 | 4647 | 4356 | 4344 | 5363 | 5791 | 5311 | 5673 | 5636 | 4982 | 5363 | 5060 | 4406 | 4631 | 4888 | 4649 | 4018 | 4222 | Natd1 |
| Nav1 | 13836 | 13744 | 10833 | 19437 | 26871 | 23870 | 24582 | 17437 | 19556 | 25672 | 22125 | 21301 | 17859 | 21144 | 13828 | 14360 | 15051 | 21737 | 23185 | 18606 | 25659 | 25927 | 16476 | 20199 | 27246 | 13849 | 17282 | 17329 | Nav1 |
| Nav2 | 3101 | 4202 | 5906 | 5868 | 6944 | 4832 | 4188 | 3604 | 3779 | 5190 | 5287 | 4765 | 5866 | 4328 | 3783 | 5061 | 6006 | 5913 | 5531 | 3506 | 4868 | 5655 | 4167 | 4622 | 4733 | 2427 | 4652 | 4844 | Nav2 |
| Nav3 | 2410 | 2632 | 1778 | 4148 | 6069 | 5185 | 5280 | 4454 | 4735 | 5047 | 5073 | 3669 | 3850 | 3667 | 3369 | 3038 | 2792 | 4351 | 4796 | 4480 | 5861 | 5740 | 4675 | 3501 | 5567 | 1747 | 3524 | 3004 | Nav3 |
| Naxd | 2132 | 2607 | 2327 | 2681 | 2507 | 2648 | 2429 | 2814 | 3321 | 3238 | 3189 | 3492 | 3219 | 3419 | 2119 | 2323 | 2471 | 2360 | 2884 | 2536 | 2585 | 2946 | 3100 | 3429 | 3389 | 3352 | 3605 | 3445 | Naxd |
| Naxe | 2457 | 2610 | 2375 | 2385 | 2019 | 2185 | 2309 | 2467 | 2499 | 2490 | 2295 | 2248 | 2480 | 2432 | 2466 | 2311 | 2328 | 2273 | 2293 | 2194 | 2383 | 2354 | 2753 | 2643 | 2463 | 2917 | 2367 | 2669 | Naxe |
| Nbas | 4022 | 4361 | 2365 | 2843 | 3209 | 3464 | 3095 | 4509 | 4642 | 3879 | 4095 | 4018 | 4177 | 3736 | 4131 | 3777 | 2887 | 3066 | 3144 | 3531 | 3465 | 3987 | 4321 | 3179 | 4047 | 2460 | 4398 | 3819 | Nbas |
| Nbdy | 1428 | 1456 | 1330 | 1458 | 1447 | 1479 | 1645 | 1896 | 1769 | 1689 | 1704 | 1554 | 1588 | 1708 | 1481 | 1406 | 1415 | 1416 | 1494 | 1550 | 1590 | 1648 | 1728 | 1619 | 1853 | 1855 | 1614 | 1651 | Nbdy |
| Nbea | 14650 | 13563 | 9142 | 17954 | 23581 | 24917 | 25836 | 20743 | 19655 | 21430 | 18964 | 17439 | 15973 | 18435 | 15387 | 12931 | 12389 | 18324 | 20326 | 22253 | 28607 | 19763 | 18970 | 15606 | 24307 | 9120 | 17809 | 15270 | Nbea |
| Nbeal1 | 9360 | 7173 | 4462 | 6715 | 6939 | 8893 | 9371 | 7267 | 6292 | 3784 | 4718 | 3932 | 4533 | 4395 | 9904 | 8287 | 5302 | 6684 | 6092 | 10540 | 9421 | 4538 | 5965 | 3595 | 4448 | 2715 | 4578 | 3889 | Nbeal1 |
| Nbeal2 | 1299 | 1825 | 2016 | 2174 | 2094 | 1720 | 1595 | 677 | 826 | 910 | 725 | 959 | 713 | 783 | 1208 | 1674 | 1953 | 1874 | 2025 | 1700 | 1742 | 911 | 721 | 908 | 950 | 593 | 863 | 818 | Nbeal2 |
| Nbl1 | 1483 | 1504 | 1325 | 1040 | 1128 | 1330 | 606 | 2208 | 2897 | 2322 | 2973 | 1563 | 1961 | 1288 | 1415 | 1928 | 1039 | 1476 | 705 | 799 | 669 | 1623 | 3256 | 2307 | 1544 | 2249 | 1535 | 1333 | Nbl1 |
| Nbn | 782 | 588 | 809 | 537 | 598 | 606 | 641 | 2792 | 2536 | 2086 | 2076 | 2623 | 2369 | 2578 | 773 | 618 | 664 | 508 | 645 | 747 | 619 | 2082 | 2387 | 2268 | 2106 | 3026 | 2671 | 2856 | Nbn |
| Nbr1 | 17292 | 17443 | 13867 | 17123 | 16179 | 16097 | 18256 | 21174 | 18080 | 17693 | 16740 | 17781 | 16595 | 19171 | 18626 | 15575 | 15527 | 17090 | 16289 | 17629 | 17972 | 18307 | 17764 | 18064 | 19257 | 16012 | 17959 | 17920 | Nbr1 |
| Ncald | 3344 | 3608 | 4039 | 4067 | 5006 | 4541 | 4555 | 2469 | 2506 | 2691 | 2528 | 1748 | 1711 | 1951 | 4073 | 3259 | 3722 | 4344 | 4255 | 4911 | 4782 | 3235 | 3139 | 2017 | 2787 | 1658 | 1983 | 2013 | Ncald |
| Ncam1 | 15859 | 18139 | 22167 | 31045 | 35902 | 27718 | 28972 | 22904 | 23693 | 33563 | 27973 | 24852 | 25810 | 25763 | 19010 | 17973 | 28239 | 31559 | 31959 | 24984 | 31704 | 39766 | 22376 | 30591 | 36410 | 18190 | 21062 | 23883 | Ncam1 |
| Ncam2 | 1114 | 898 | 687 | 1320 | 2050 | 2032 | 1963 | 1907 | 1814 | 1661 | 1609 | 1259 | 1233 | 1396 | 1367 | 846 | 1050 | 1395 | 1792 | 2026 | 2171 | 2029 | 1865 | 1258 | 1709 | 671 | 1288 | 1121 | Ncam2 |
| Ncan | 7999 | 10948 | 14112 | 18159 | 22264 | 18366 | 18832 | 17170 | 18389 | 25757 | 19065 | 25861 | 19578 | 23425 | 10337 | 8061 | 18099 | 18403 | 23300 | 15295 | 19966 | 28990 | 15822 | 26081 | 30633 | 18042 | 21043 | 22335 | Ncan |
| Ncapd2 | 22228 | 23470 | 25399 | 21475 | 17463 | 18784 | 20091 | 17407 | 19630 | 20453 | 18093 | 29212 | 23689 | 28359 | 19417 | 20023 | 24094 | 19585 | 22039 | 18183 | 19882 | 16225 | 16176 | 24914 | 21989 | 32228 | 28229 | 28591 | Ncapd2 |
| Ncapd3 | 8731 | 8738 | 5435 | 5854 | 5194 | 6148 | 6671 | 5482 | 4790 | 4082 | 3855 | 5106 | 4217 | 5144 | 7965 | 7477 | 5575 | 6078 | 5577 | 6966 | 6757 | 3639 | 4361 | 3924 | 4384 | 4166 | 5179 | 4836 | Ncapd3 |
| Ncapg | 9063 | 6440 | 6518 | 5362 | 3908 | 5288 | 6176 | 7248 | 6073 | 4675 | 4532 | 7898 | 6844 | 8116 | 7987 | 7182 | 5371 | 4786 | 5032 | 7088 | 5408 | 3721 | 5367 | 5873 | 5361 | 8184 | 8400 | 8261 | Ncapg |
| Ncapg2 | 8135 | 6391 | 5893 | 5670 | 4320 | 5722 | 6505 | 6281 | 5553 | 4953 | 4754 | 6924 | 6004 | 7206 | 7203 | 6872 | 5676 | 4927 | 5277 | 6677 | 5626 | 3801 | 5005 | 5743 | 4706 | 6025 | 7076 | 6912 | Ncapg2 |
| Ncaph | 7789 | 7850 | 6994 | 6411 | 4852 | 5645 | 6594 | 4670 | 4787 | 4618 | 3986 | 6920 | 5579 | 6645 | 6825 | 6587 | 6966 | 5615 | 5938 | 6136 | 6094 | 3228 | 4295 | 5538 | 4531 | 7974 | 6754 | 7036 | Ncaph |
| Ncaph2 | 10738 | 13676 | 11507 | 10738 | 8740 | 9619 | 9681 | 8922 | 9985 | 8723 | 8431 | 10434 | 9727 | 10470 | 10266 | 10830 | 10931 | 9588 | 10919 | 10458 | 10165 | 7429 | 9519 | 10043 | 9206 | 11870 | 11059 | 10903 | Ncaph2 |
| Ncbp1 | 15597 | 14922 | 13384 | 13266 | 11736 | 12623 | 13876 | 12224 | 10401 | 9420 | 8644 | 11169 | 9762 | 11390 | 15927 | 13567 | 13174 | 11913 | 12996 | 14238 | 13609 | 9180 | 9825 | 10205 | 10148 | 11449 | 11355 | 11510 | Ncbp1 |
| Ncbp2 | 10259 | 9481 | 11744 | 9396 | 7706 | 8346 | 8825 | 9404 | 8291 | 8167 | 8419 | 9676 | 9302 | 9594 | 10203 | 10276 | 9864 | 8621 | 8433 | 9780 | 8270 | 7738 | 8515 | 9166 | 8537 | 11128 | 10135 | 10251 | Ncbp2 |
| Ncbp3 | 4967 | 4444 | 4426 | 4843 | 4714 | 4667 | 4870 | 5968 | 5083 | 5180 | 5151 | 5515 | 5281 | 5807 | 4957 | 4338 | 4507 | 4590 | 4623 | 4854 | 5059 | 5372 | 5182 | 5181 | 5088 | 5521 | 5649 | 5734 | Ncbp3 |
| Nccrp1 | 35 | 22 | 9 | 13 | 15 | 13 | 16 | 4 | 6 | 12 | 1 | 1 | 3 | 1 | 5 | 26 | 14 | 10 | 9 | 2 | 9 | 2 | 2 | 8 | 7 | 6 | 7 | 2 | Nccrp1 |
| Ncdn | 3891 | 5073 | 5065 | 6173 | 6715 | 5624 | 5630 | 3225 | 4217 | 4768 | 4404 | 4309 | 3708 | 3862 | 3833 | 4314 | 5842 | 6641 | 6748 | 4859 | 6050 | 4918 | 3972 | 4584 | 4966 | 4464 | 3435 | 3612 | Ncdn |
| Nceh1 | 1634 | 1552 | 1418 | 1602 | 1620 | 1531 | 1749 | 586 | 507 | 431 | 489 | 521 | 511 | 443 | 1631 | 1429 | 1389 | 1640 | 1470 | 1696 | 1773 | 623 | 534 | 445 | 428 | 479 | 495 | 434 | Nceh1 |
| Ncf1 | 110 | 117 | 149 | 159 | 97 | 92 | 94 | 205 | 257 | 262 | 311 | 226 | 247 | 193 | 91 | 132 | 97 | 119 | 155 | 97 | 101 | 214 | 355 | 232 | 251 | 265 | 244 | 201 | Ncf1 |
| Ncf2 | 185 | 209 | 244 | 151 | 124 | 184 | 110 | 234 | 255 | 216 | 269 | 236 | 217 | 182 | 189 | 220 | 143 | 164 | 102 | 175 | 144 | 142 | 325 | 258 | 160 | 236 | 201 | 227 | Ncf2 |
| Ncf4 | 46 | 49 | 69 | 28 | 29 | 22 | 15 | 22 | 56 | 24 | 42 | 38 | 95 | 68 | 30 | 42 | 30 | 21 | 22 | 42 | 16 | 33 | 45 | 35 | 20 | 31 | 58 | 93 | Ncf4 |
| Nck1 | 3182 | 2848 | 3476 | 3198 | 2686 | 2763 | 3132 | 2992 | 2434 | 2191 | 2394 | 2301 | 2647 | 2870 | 3443 | 3132 | 3101 | 2958 | 2938 | 3390 | 2920 | 2587 | 2750 | 2454 | 2230 | 2686 | 2607 | 2797 | Nck1 |
| Nck2 | 2147 | 2108 | 2573 | 2393 | 2585 | 2414 | 2305 | 2567 | 2838 | 2942 | 3134 | 2802 | 2954 | 2751 | 1990 | 2337 | 2239 | 2216 | 2439 | 1971 | 2108 | 2812 | 2899 | 3096 | 3068 | 3423 | 2592 | 2656 | Nck2 |
| Nckap1 | 17467 | 14774 | 15711 | 15558 | 17651 | 17954 | 18796 | 22278 | 20337 | 20390 | 18918 | 21767 | 20634 | 21288 | 17991 | 15896 | 15939 | 15944 | 17252 | 17346 | 17991 | 22368 | 20740 | 19505 | 21073 | 20184 | 21474 | 21021 | Nckap1 |
| Nckap1l | 438 | 405 | 437 | 291 | 329 | 344 | 265 | 459 | 562 | 405 | 541 | 461 | 416 | 398 | 538 | 591 | 254 | 343 | 273 | 440 | 292 | 486 | 683 | 346 | 374 | 423 | 373 | 373 | Nckap1l |
| Nckap5 | 1419 | 1534 | 1375 | 1968 | 2439 | 2255 | 2398 | 2085 | 2362 | 2267 | 1985 | 2229 | 2093 | 2107 | 1667 | 1354 | 1596 | 1932 | 2149 | 2285 | 2413 | 2204 | 1889 | 2209 | 2655 | 1420 | 1888 | 1737 | Nckap5 |
| Nckap5l | 1693 | 2504 | 3158 | 3141 | 3080 | 2668 | 2104 | 2107 | 2656 | 3258 | 3310 | 3148 | 2781 | 2455 | 1601 | 2173 | 3312 | 2969 | 3508 | 1927 | 2421 | 3606 | 2629 | 3312 | 3336 | 3566 | 2708 | 2420 | Nckap5l |
| Nckipsd | 0 | 0 | 1 | 0 | 0 | 0 | 1 | 0 | 0 | 0 | 0 | 0 | 0 | 0 | 0 | 0 | 0 | 0 | 0 | 0 | 0 | 0 | 0 | 0 | 0 | 0 | 0 | 0 | Nckipsd |
| Ncl | 54253 | 44068 | 60922 | 46140 | 39673 | 45627 | 47630 | 59248 | 53968 | 55309 | 56171 | 67705 | 65787 | 74163 | 52445 | 50048 | 51024 | 41928 | 42853 | 49334 | 42158 | 50672 | 51870 | 63040 | 56561 | 82770 | 73436 | 75364 | Ncl |
| Ncln | 4847 | 6662 | 6602 | 6307 | 5027 | 5091 | 4159 | 3043 | 3971 | 3959 | 4153 | 4331 | 4103 | 3338 | 4567 | 6336 | 6360 | 6016 | 5725 | 4080 | 4455 | 4077 | 4018 | 4283 | 3887 | 4640 | 3977 | 3576 | Ncln |
| Ncmap | 82 | 117 | 87 | 119 | 106 | 103 | 108 | 51 | 171 | 113 | 176 | 33 | 60 | 39 | 71 | 81 | 74 | 114 | 86 | 63 | 95 | 100 | 104 | 69 | 59 | 36 | 41 | 32 | Ncmap |
| Ncoa1 | 3688 | 3630 | 3640 | 4831 | 6169 | 5628 | 5776 | 4341 | 4894 | 5714 | 5900 | 5267 | 4295 | 5220 | 3607 | 3653 | 4155 | 5216 | 5750 | 4420 | 5745 | 5304 | 4912 | 5750 | 5717 | 3757 | 4178 | 4237 | Ncoa1 |
| Ncoa2 | 5045 | 4729 | 4399 | 5305 | 6572 | 6334 | 5813 | 6995 | 7865 | 8202 | 8240 | 8118 | 7919 | 7911 | 5115 | 5565 | 4908 | 5406 | 5693 | 5297 | 5973 | 8011 | 7442 | 7205 | 8447 | 5602 | 7489 | 7334 | Ncoa2 |
| Ncoa3 | 5842 | 6440 | 6480 | 6706 | 6343 | 6065 | 5559 | 3913 | 4345 | 4859 | 4983 | 4612 | 4851 | 4605 | 5313 | 6879 | 6496 | 7130 | 5898 | 4788 | 6065 | 5046 | 4291 | 4487 | 4941 | 3463 | 4259 | 4429 | Ncoa3 |
| Ncoa4 | 4394 | 4521 | 3402 | 3576 | 3901 | 3779 | 3967 | 5238 | 4805 | 3828 | 3830 | 3695 | 4711 | 4085 | 4940 | 4193 | 3600 | 3821 | 3928 | 4776 | 3979 | 4022 | 5035 | 3766 | 4121 | 3864 | 4267 | 5084 | Ncoa4 |
| Ncoa5 | 6294 | 7015 | 8394 | 7909 | 8501 | 7745 | 7529 | 8995 | 9648 | 10685 | 9359 | 11596 | 9800 | 10698 | 5848 | 6114 | 8707 | 7826 | 9420 | 7152 | 7670 | 10505 | 9030 | 11842 | 11630 | 11409 | 10780 | 10828 | Ncoa5 |
| Ncoa6 | 7499 | 7068 | 6585 | 7372 | 8885 | 8671 | 7943 | 10005 | 10187 | 11055 | 10114 | 11674 | 10550 | 10739 | 7606 | 6618 | 7356 | 7132 | 8680 | 7713 | 8454 | 11348 | 8925 | 10333 | 11367 | 8775 | 10508 | 9800 | Ncoa6 |
| Ncoa7 | 980 | 702 | 842 | 928 | 1315 | 940 | 1206 | 2175 | 2130 | 2028 | 2227 | 1784 | 2011 | 2063 | 942 | 917 | 811 | 987 | 998 | 886 | 1062 | 2338 | 2078 | 1865 | 2112 | 1691 | 1680 | 1767 | Ncoa7 |
| Ncor1 | 18880 | 17639 | 12113 | 17276 | 19625 | 21361 | 21070 | 22462 | 22634 | 22470 | 21841 | 23015 | 21526 | 23034 | 18485 | 17263 | 14659 | 17563 | 19010 | 20472 | 21270 | 19963 | 21399 | 19712 | 23350 | 13895 | 22822 | 20788 | Ncor1 |
| Ncor2 | 9517 | 13129 | 15663 | 16980 | 15871 | 13396 | 10372 | 7365 | 10159 | 12769 | 12369 | 13148 | 10250 | 9657 | 8578 | 11562 | 16169 | 16796 | 16255 | 10116 | 12717 | 12061 | 9341 | 13602 | 11939 | 11488 | 9907 | 8880 | Ncor2 |
| Ncs1 | 3151 | 3118 | 3776 | 4296 | 6586 | 4922 | 4581 | 7697 | 8600 | 10160 | 9057 | 8880 | 8316 | 8511 | 3273 | 3322 | 4486 | 5041 | 5655 | 4106 | 4972 | 11139 | 7778 | 9335 | 11366 | 8575 | 7853 | 8567 | Ncs1 |
| Ncstn | 4542 | 4912 | 4998 | 4742 | 5228 | 4972 | 4651 | 6503 | 7154 | 7660 | 7151 | 8503 | 8316 | 8338 | 4866 | 4868 | 5074 | 4702 | 4953 | 4497 | 4732 | 7379 | 7304 | 7511 | 7637 | 8095 | 8097 | 8495 | Ncstn |
| Ndc1 | 6658 | 5161 | 5580 | 4372 | 3949 | 4554 | 4635 | 5560 | 5061 | 4557 | 4661 | 5940 | 5251 | 5628 | 5727 | 5663 | 4646 | 4127 | 4235 | 4932 | 4333 | 4346 | 4866 | 5182 | 4763 | 6642 | 6123 | 6241 | Ndc1 |
| Ndc80 | 3894 | 3410 | 3960 | 2816 | 2081 | 2550 | 2956 | 3261 | 2832 | 2489 | 2449 | 3994 | 3387 | 4221 | 3696 | 3380 | 3591 | 2450 | 2753 | 3462 | 2693 | 1965 | 2527 | 3496 | 2834 | 4891 | 4328 | 4209 | Ndc80 |
| Nde1 | 7435 | 7852 | 7963 | 6175 | 4864 | 5494 | 5712 | 5951 | 5967 | 5580 | 5012 | 8387 | 7199 | 8605 | 6758 | 6572 | 6997 | 5609 | 5783 | 5768 | 5379 | 4184 | 5269 | 7173 | 5987 | 9996 | 9043 | 9156 | Nde1 |
| Ndel1 | 4606 | 4559 | 3981 | 4434 | 4204 | 4221 | 4309 | 3670 | 3672 | 3681 | 3691 | 3261 | 3300 | 3172 | 4655 | 4551 | 4205 | 4447 | 4044 | 4233 | 4286 | 3820 | 3851 | 3268 | 3634 | 3201 | 3134 | 3183 | Ndel1 |
| Ndfip1 | 17212 | 13669 | 17175 | 19585 | 21486 | 19107 | 21120 | 16822 | 13870 | 15632 | 14884 | 14339 | 13571 | 14624 | 19781 | 15617 | 17971 | 19688 | 19590 | 19961 | 20979 | 19798 | 15296 | 14548 | 16565 | 13302 | 13345 | 13986 | Ndfip1 |
| Ndfip2 | 4733 | 4253 | 4397 | 4809 | 5524 | 5065 | 5453 | 5105 | 4271 | 4211 | 4111 | 3892 | 3950 | 4103 | 5603 | 4646 | 4669 | 4901 | 4895 | 5579 | 5295 | 4741 | 4412 | 3885 | 4200 | 3651 | 4048 | 4133 | Ndfip2 |
| Ndn | 9738 | 11504 | 10492 | 16043 | 19046 | 15068 | 16650 | 13437 | 15128 | 14500 | 14469 | 11769 | 11230 | 11138 | 11408 | 9873 | 12020 | 14385 | 16520 | 16734 | 17924 | 16083 | 14701 | 13210 | 16480 | 10426 | 11033 | 10510 | Ndn |
| Ndnf | 2501 | 1922 | 1688 | 1615 | 1908 | 1918 | 1381 | 2154 | 2104 | 1687 | 1982 | 1385 | 1973 | 1401 | 2341 | 2588 | 1869 | 1581 | 1476 | 1787 | 1495 | 2107 | 2597 | 1147 | 1436 | 1383 | 1416 | 1361 | Ndnf |
| Ndor1 | 884 | 1082 | 795 | 989 | 821 | 857 | 977 | 547 | 672 | 794 | 616 | 671 | 648 | 712 | 767 | 899 | 928 | 957 | 837 | 711 | 908 | 695 | 500 | 664 | 677 | 529 | 644 | 687 | Ndor1 |
| Ndp | 113 | 130 | 131 | 127 | 215 | 135 | 204 | 366 | 314 | 291 | 282 | 328 | 338 | 330 | 148 | 86 | 172 | 105 | 227 | 184 | 161 | 289 | 308 | 346 | 391 | 315 | 322 | 326 | Ndp |
| Ndrg1 | 1595 | 1481 | 1270 | 1383 | 1628 | 1490 | 1229 | 1171 | 1613 | 1384 | 1495 | 495 | 623 | 567 | 1465 | 1566 | 1279 | 1513 | 1267 | 1152 | 1297 | 1287 | 1859 | 1395 | 1294 | 484 | 440 | 373 | Ndrg1 |
| Ndrg2 | 2515 | 2338 | 1865 | 3159 | 4858 | 3954 | 4138 | 3468 | 4294 | 4328 | 3787 | 2249 | 2427 | 2159 | 2532 | 2294 | 2260 | 3831 | 4252 | 3810 | 4384 | 3497 | 4471 | 3866 | 4034 | 2034 | 1668 | 1722 | Ndrg2 |
| Ndrg3 | 7838 | 8320 | 6719 | 7824 | 7803 | 7902 | 8916 | 5015 | 4889 | 4904 | 4482 | 4187 | 4206 | 4271 | 7940 | 7379 | 7626 | 8511 | 8249 | 8014 | 9017 | 5437 | 4823 | 4784 | 5167 | 4379 | 4265 | 4146 | Ndrg3 |
| Ndrg4 | 6824 | 8479 | 7264 | 9812 | 12053 | 9694 | 9722 | 6443 | 7000 | 7571 | 7019 | 5293 | 5689 | 5321 | 7884 | 7704 | 8983 | 10554 | 10668 | 9717 | 9947 | 9367 | 6613 | 6077 | 7911 | 4413 | 4964 | 4873 | Ndrg4 |

| | | | | | | | | | | | | | | | | | | | | | | | | | | | | | |
|---|---|---|---|---|---|---|---|---|---|---|---|---|---|---|---|---|---|---|---|---|---|---|---|---|---|---|---|---|---|
| Ndst1 | 8209 | 8573 | 8347 | 8832 | 10722 | 9827 | 8418 | 9583 | 11831 | 12276 | 12458 | 12381 | 11878 | 11259 | 7448 | 8586 | 9020 | 9652 | 10310 | 7757 | 8798 | 10835 | 10773 | 11604 | 12011 | 9381 | 11338 | 11089 | Ndst1 |
| Ndst2 | 1742 | 1995 | 1920 | 1810 | 1463 | 1707 | 1618 | 1945 | 2363 | 2333 | 2440 | 2414 | 2557 | 2230 | 1710 | 1911 | 1736 | 1738 | 1655 | 1449 | 1435 | 2212 | 2328 | 2420 | 2134 | 2450 | 2280 | 2279 | Ndst2 |
| Ndst3 | 1589 | 1566 | 1524 | 1653 | 2095 | 2090 | 1949 | 3646 | 2778 | 2726 | 2167 | 3036 | 2976 | 3142 | 2041 | 1266 | 1545 | 1689 | 2061 | 2179 | 2413 | 2907 | 2566 | 2648 | 3376 | 1193 | 3495 | 3194 | Ndst3 |
| Ndst4 | 427 | 407 | 245 | 542 | 854 | 682 | 792 | 741 | 755 | 571 | 561 | 354 | 379 | 385 | 690 | 516 | 455 | 602 | 633 | 869 | 777 | 794 | 641 | 425 | 700 | 236 | 328 | 344 | Ndst4 |
| Ndufa1 | 1630 | 1983 | 2083 | 2045 | 1846 | 1210 | 2319 | 1691 | 1459 | 2023 | 1675 | 1738 | 1800 | 2630 | 1708 | 1191 | 2022 | 1630 | 1623 | 1907 | 2218 | 1459 | 1579 | 2244 | 1980 | 1711 | 1716 | 2656 | Ndufa1 |
| Ndufa10 | 5555 | 5925 | 5201 | 4804 | 4578 | 4825 | 4657 | 5522 | 5544 | 5058 | 4929 | 5227 | 5134 | 5681 | 5410 | 5153 | 5188 | 4492 | 4722 | 4913 | 4830 | 4771 | 5358 | 5359 | 5060 | 6085 | 5425 | 5769 | Ndufa10 |
| Ndufa11 | 2118 | 2943 | 2255 | 2497 | 2407 | 2514 | 2470 | 1969 | 2656 | 2269 | 2465 | 2221 | 2501 | 2353 | 2124 | 2245 | 2466 | 2376 | 2801 | 2937 | 2355 | 1942 | 2578 | 2234 | 2430 | 2547 | 2558 | 2409 | Ndufa11 |
| Ndufa12 | 1429 | 2822 | 1858 | 2997 | 1937 | 1872 | 2497 | 1397 | 2063 | 2111 | 2172 | 2238 | 2392 | 1940 | 2639 | 2064 | 2296 | 2687 | 2243 | 3190 | 3038 | 2117 | 2409 | 1864 | 2684 | 3330 | 1697 | 3567 | Ndufa12 |
| Ndufa13 | 2001 | 2520 | 2805 | 2615 | 2497 | 2250 | 2506 | 2452 | 2656 | 2805 | 3046 | 2930 | 3047 | 3411 | 1908 | 2036 | 2486 | 2461 | 2734 | 2828 | 2472 | 2097 | 2480 | 3325 | 2857 | 3203 | 3230 | 3236 | Ndufa13 |
| Ndufa2 | 1718 | 2107 | 2272 | 2192 | 1762 | 1571 | 2187 | 2127 | 2155 | 2773 | 2536 | 2638 | 2429 | 3160 | 1698 | 1583 | 2275 | 1721 | 1813 | 2317 | 2066 | 1614 | 2241 | 2917 | 2564 | 2985 | 2659 | 3330 | Ndufa2 |
| Ndufa3 | 3044 | 3581 | 3560 | 3869 | 3090 | 2541 | 4761 | 3013 | 2715 | 3527 | 3693 | 2999 | 3221 | 5593 | 3119 | 1963 | 4063 | 3187 | 3021 | 3652 | 4849 | 2336 | 2673 | 4253 | 3682 | 3559 | 3398 | 5153 | Ndufa3 |
| Ndufa4 | 8679 | 8853 | 9611 | 9171 | 8344 | 7503 | 10747 | 8600 | 7708 | 8030 | 8710 | 7604 | 8251 | 9963 | 9800 | 7245 | 9831 | 8055 | 8505 | 10346 | 10354 | 6946 | 7935 | 9041 | 8335 | 9067 | 8087 | 10504 | Ndufa4 |
| Ndufa4l2 | 76 | 98 | 76 | 108 | 148 | 109 | 139 | 100 | 143 | 133 | 179 | 89 | 138 | 131 | 74 | 74 | 98 | 130 | 99 | 115 | 117 | 192 | 151 | 141 | 150 | 155 | 134 | 138 | Ndufa4l2 |
| Ndufa5 | 1831 | 1992 | 1904 | 2239 | 1920 | 1593 | 2847 | 1903 | 1570 | 1786 | 1869 | 1760 | 1777 | 2355 | 1925 | 1379 | 2027 | 1870 | 1778 | 2055 | 2381 | 1614 | 1607 | 2140 | 1890 | 1952 | 1840 | 2565 | Ndufa5 |
| Ndufa6 | 2029 | 2230 | 2209 | 2134 | 2036 | 1743 | 2417 | 2160 | 1981 | 2338 | 2309 | 2208 | 2201 | 3015 | 2006 | 1804 | 2449 | 2045 | 2016 | 2213 | 2729 | 2099 | 2089 | 2653 | 2295 | 2203 | 2163 | 2889 | Ndufa6 |
| Ndufa7 | 2865 | 3472 | 3272 | 3095 | 2923 | 2908 | 2959 | 3598 | 3617 | 4035 | 3760 | 3716 | 3891 | 4244 | 2894 | 2646 | 3395 | 2737 | 3081 | 3317 | 3179 | 2979 | 3695 | 4116 | 3923 | 4058 | 4058 | 4392 | Ndufa7 |
| Ndufa8 | 2790 | 3527 | 3366 | 3607 | 3174 | 2998 | 3303 | 2286 | 2709 | 2855 | 2692 | 2689 | 2595 | 2879 | 2765 | 2959 | 3525 | 3292 | 3420 | 3358 | 3497 | 2187 | 2773 | 2895 | 2859 | 3171 | 2897 | 2994 | Ndufa8 |
| Ndufa9 | 5522 | 5207 | 5370 | 4538 | 4336 | 4627 | 4237 | 6886 | 6058 | 5610 | 5719 | 6292 | 6083 | 5830 | 5641 | 5020 | 5099 | 4414 | 4643 | 5407 | 4060 | 5755 | 6047 | 6127 | 6148 | 7148 | 6892 | 6330 | Ndufa9 |
| Ndufab1 | 3778 | 3733 | 4170 | 3953 | 3422 | 3521 | 4324 | 3976 | 3657 | 3736 | 4050 | 4231 | 4068 | 4792 | 3716 | 3366 | 3982 | 3554 | 3723 | 3567 | 3939 | 3566 | 3479 | 4530 | 4101 | 4776 | 4558 | 5079 | Ndufab1 |
| Ndufaf1 | 917 | 676 | 1048 | 702 | 808 | 767 | 888 | 1561 | 1337 | 1268 | 1485 | 1420 | 1439 | 1564 | 919 | 725 | 762 | 743 | 729 | 777 | 732 | 1352 | 1447 | 1354 | 1276 | 1589 | 1436 | 1646 | Ndufaf1 |
| Ndufaf2 | 1861 | 1878 | 2198 | 2114 | 1753 | 1759 | 2472 | 1272 | 1116 | 1372 | 1244 | 1350 | 1302 | 1856 | 1841 | 1688 | 2224 | 2157 | 1979 | 2349 | 2419 | 1131 | 1163 | 1626 | 1489 | 1594 | 1488 | 1686 | Ndufaf2 |
| Ndufaf3 | 752 | 904 | 751 | 763 | 841 | 792 | 892 | 976 | 924 | 902 | 846 | 883 | 789 | 923 | 807 | 678 | 800 | 706 | 925 | 1020 | 793 | 874 | 962 | 907 | 891 | 1045 | 924 | 853 | Ndufaf3 |
| Ndufaf4 | 3811 | 3471 | 3613 | 3655 | 3406 | 3345 | 4265 | 3132 | 2625 | 2742 | 2494 | 2682 | 2659 | 2853 | 4412 | 3809 | 3582 | 3793 | 3453 | 4360 | 3851 | 3003 | 2697 | 2627 | 2614 | 3103 | 2772 | 2885 | Ndufaf4 |
| Ndufaf5 | 1232 | 1221 | 1051 | 1204 | 1313 | 1129 | 1414 | 1246 | 1087 | 1055 | 1014 | 976 | 934 | 1182 | 1432 | 1131 | 1145 | 1207 | 1196 | 1513 | 1386 | 1006 | 1219 | 1100 | 1040 | 1155 | 1033 | 1101 | Ndufaf5 |
| Ndufaf6 | 601 | 514 | 573 | 525 | 488 | 461 | 463 | 383 | 372 | 337 | 376 | 413 | 413 | 376 | 539 | 515 | 457 | 450 | 514 | 432 | 521 | 361 | 402 | 415 | 371 | 452 | 417 | 533 | Ndufaf6 |
| Ndufaf7 | 1257 | 1266 | 1382 | 1404 | 1515 | 1339 | 1385 | 2249 | 1972 | 2273 | 2201 | 2034 | 2070 | 2267 | 1313 | 1320 | 1452 | 1502 | 1365 | 1479 | 1539 | 2019 | 2144 | 2251 | 2231 | 2255 | 1965 | 2248 | Ndufaf7 |
| Ndufaf8 | 193 | 274 | 289 | 188 | 190 | 180 | 191 | 348 | 404 | 414 | 450 | 475 | 464 | 437 | 216 | 147 | 299 | 190 | 203 | 257 | 231 | 236 | 377 | 459 | 422 | 617 | 481 | 501 | Ndufaf8 |
| Ndufb1 | 1637 | 1506 | 1467 | 1871 | 1531 | 1107 | 3175 | 1871 | 1389 | 2182 | 1813 | 1636 | 1605 | 4172 | 1511 | 791 | 1732 | 1278 | 1269 | 1733 | 3219 | 1352 | 1194 | 2680 | 1944 | 1666 | 1691 | 3287 | Ndufb1 |
| Ndufb10 | 4032 | 4502 | 4754 | 5098 | 4322 | 4256 | 4457 | 3335 | 3330 | 4060 | 3820 | 3596 | 3699 | 3878 | 3818 | 3379 | 4684 | 4415 | 4734 | 4482 | 4535 | 3183 | 3307 | 4223 | 3968 | 4248 | 4037 | 3974 | Ndufb10 |
| Ndufb11 | 3452 | 4421 | 3598 | 3742 | 3526 | 3486 | 3865 | 4184 | 4110 | 3909 | 3912 | 3982 | 4102 | 4508 | 3696 | 3445 | 3814 | 3465 | 3801 | 4437 | 4029 | 3405 | 4264 | 4394 | 4083 | 4583 | 4470 | 4459 | Ndufb11 |
| Ndufb11b | 0 | 0 | 0 | 0 | 0 | 0 | 1 | 0 | 0 | 0 | 0 | 0 | 0 | 0 | 0 | 0 | 0 | 0 | 0 | 0 | 0 | 0 | 1 | 0 | 0 | 0 | 0 | 0 | Ndufb11b |
| Ndufb2 | 880 | 860 | 862 | 986 | 1116 | 930 | 1101 | 1270 | 1071 | 1093 | 1213 | 978 | 962 | 1059 | 897 | 829 | 1083 | 922 | 1016 | 1056 | 1030 | 1402 | 1118 | 1159 | 1282 | 969 | 1068 | 1012 | Ndufb2 |
| Ndufb3 | 2105 | 1992 | 2875 | 2710 | 2125 | 1960 | 2647 | 2770 | 2481 | 3073 | 2889 | 3041 | 2834 | 3727 | 2293 | 1727 | 2924 | 2327 | 2250 | 2765 | 2647 | 2192 | 2498 | 3450 | 2982 | 3426 | 2980 | 3635 | Ndufb3 |
| Ndufb4 | 1806 | 2268 | 2122 | 2439 | 2286 | 2369 | 2659 | 2047 | 2251 | 2184 | 2542 | 2152 | 2503 | 2866 | 1974 | 1843 | 2201 | 2183 | 2500 | 2737 | 2692 | 1713 | 2319 | 2616 | 2445 | 2584 | 2617 | 2908 | Ndufb4 |
| Ndufb5 | 4196 | 4483 | 3747 | 3842 | 3780 | 3698 | 4483 | 4935 | 4439 | 4221 | 4278 | 4207 | 4146 | 4857 | 4315 | 3941 | 3721 | 3660 | 3871 | 4910 | 4337 | 3880 | 4390 | 4177 | 4179 | 4456 | 4495 | 4803 | Ndufb5 |
| Ndufb6 | 2990 | 2988 | 3176 | 3254 | 2818 | 2570 | 3562 | 2232 | 2054 | 2355 | 2291 | 2191 | 2401 | 2862 | 2987 | 2446 | 3326 | 3017 | 3057 | 3462 | 3481 | 1673 | 2025 | 2675 | 2156 | 2884 | 2471 | 2964 | Ndufb6 |
| Ndufb7 | 2859 | 3844 | 3670 | 3298 | 2932 | 2997 | 3035 | 3243 | 3356 | 3407 | 3232 | 3652 | 3606 | 3931 | 3022 | 2563 | 3763 | 3144 | 3476 | 3468 | 3181 | 2872 | 3311 | 4033 | 3722 | 4362 | 3948 | 4292 | Ndufb7 |
| Ndufb8 | 1081 | 1184 | 1513 | 1217 | 1052 | 796 | 1151 | 1136 | 1004 | 1495 | 1269 | 1266 | 1127 | 1663 | 1117 | 886 | 1447 | 998 | 954 | 980 | 1289 | 986 | 1115 | 1576 | 1291 | 1409 | 1199 | 1667 | Ndufb8 |
| Ndufb9 | 745 | 829 | 563 | 745 | 642 | 483 | 1077 | 900 | 697 | 915 | 803 | 819 | 689 | 1512 | 764 | 366 | 695 | 610 | 532 | 627 | 1116 | 650 | 677 | 1192 | 804 | 773 | 791 | 1334 | Ndufb9 |
| Ndufc1 | 1706 | 1794 | 1815 | 1842 | 1736 | 1547 | 2571 | 1901 | 1556 | 1866 | 1900 | 1934 | 1905 | 2734 | 1712 | 1262 | 1896 | 1564 | 1715 | 1887 | 2475 | 1596 | 1658 | 2231 | 1849 | 2130 | 2098 | 2648 | Ndufc1 |
| Ndufc2 | 3009 | 3297 | 3307 | 3853 | 3648 | 3412 | 3843 | 3351 | 3292 | 3749 | 3344 | 3395 | 3349 | 3790 | 3152 | 2821 | 3403 | 3445 | 4020 | 4666 | 4244 | 2510 | 3389 | 3994 | 3700 | 4096 | 3553 | 3941 | Ndufc2 |
| Ndufs1 | 9703 | 8194 | 8587 | 7653 | 7736 | 7447 | 8035 | 11021 | 9462 | 8197 | 8758 | 9354 | 9046 | 9623 | 10598 | 8841 | 7795 | 7342 | 7642 | 8857 | 7411 | 9443 | 9375 | 8612 | 8711 | 9973 | 9596 | 9714 | Ndufs1 |
| Ndufs2 | 8870 | 10141 | 8979 | 8650 | 8025 | 8226 | 8018 | 9678 | 10324 | 9495 | 8690 | 9994 | 9755 | 9773 | 9154 | 9034 | 9101 | 8523 | 9032 | 8748 | 8469 | 8742 | 10354 | 9886 | 9442 | 11032 | 10560 | 10822 | Ndufs2 |
| Ndufs3 | 0 | 1 | 0 | 0 | 0 | 0 | 2 | 0 | 0 | 0 | 0 | 0 | 0 | 0 | 0 | 0 | 0 | 1 | 0 | 0 | 1 | 10 | 0 | 0 | 0 | 1 | 0 | 0 | Ndufs3 |
| Ndufs4 | 4139 | 3792 | 4287 | 3762 | 3580 | 3551 | 4312 | 4691 | 3921 | 4125 | 3520 | 4176 | 4121 | 4853 | 4477 | 3715 | 4025 | 3560 | 3717 | 4881 | 4262 | 3646 | 4113 | 4304 | 4160 | 4630 | 4462 | 5235 | Ndufs4 |
| Ndufs5 | 1865 | 2515 | 2447 | 2664 | 2442 | 2187 | 2930 | 1911 | 1874 | 2574 | 2448 | 2386 | 2464 | 3179 | 1900 | 1734 | 2739 | 2356 | 2413 | 2689 | 2789 | 1634 | 1909 | 2683 | 2400 | 2754 | 2439 | 2909 | Ndufs5 |
| Ndufs6 | 1684 | 2172 | 2436 | 2128 | 1820 | 1627 | 2251 | 1727 | 1823 | 2141 | 2134 | 2252 | 2250 | 2843 | 1709 | 1571 | 2416 | 1722 | 1902 | 2234 | 2192 | 1380 | 1763 | 2631 | 2189 | 2712 | 2391 | 2993 | Ndufs6 |
| Ndufs7 | 1132 | 1572 | 1354 | 1285 | 1118 | 1176 | 1124 | 1508 | 1673 | 1527 | 1561 | 1644 | 1539 | 1556 | 1088 | 1271 | 1357 | 1110 | 1347 | 1339 | 1096 | 1330 | 1731 | 1830 | 1600 | 2000 | 1772 | 1530 | Ndufs7 |
| Ndufs8 | 1664 | 2162 | 2063 | 1773 | 1648 | 1781 | 1523 | 2542 | 3015 | 2371 | 2578 | 2506 | 2922 | 2535 | 1783 | 1764 | 1795 | 1440 | 1774 | 2110 | 1581 | 2015 | 2924 | 2617 | 2464 | 3059 | 2955 | 2730 | Ndufs8 |
| Ndufv1 | 6612 | 8570 | 7476 | 7241 | 6542 | 6814 | 6175 | 5627 | 6180 | 6441 | 5974 | 6656 | 6317 | 5910 | 6504 | 7126 | 7346 | 7027 | 7473 | 7332 | 6267 | 5198 | 6153 | 6492 | 6302 | 7334 | 6711 | 6281 | Ndufv1 |
| Ndufv2 | 7394 | 6594 | 6577 | 6637 | 6332 | 6126 | 8082 | 7925 | 6577 | 6409 | 6262 | 6592 | 6403 | 8215 | 7760 | 6361 | 7119 | 5971 | 6724 | 7972 | 7918 | 6592 | 6776 | 7349 | 6367 | 7563 | 6973 | 8454 | Ndufv2 |
| Ndufv3 | 4332 | 5696 | 4226 | 5060 | 4382 | 4450 | 4827 | 2908 | 2982 | 3014 | 3300 | 2898 | 3137 | 3067 | 4473 | 4244 | 4735 | 4474 | 4573 | 5272 | 4743 | 2613 | 2996 | 3483 | 3200 | 3054 | 2939 | 3146 | Ndufv3 |
| Neb | 324 | 52 | 11 | 44 | 92 | 102 | 108 | 468 | 1284 | 240 | 4168 | 99 | 142 | 109 | 138 | 261 | 36 | 84 | 62 | 126 | 122 | 578 | 3997 | 172 | 304 | 66 | 76 | 53 | Neb |
| Nebl | 1418 | 1012 | 737 | 828 | 1022 | 1533 | 885 | 1381 | 1724 | 1219 | 1395 | 1026 | 1201 | 1091 | 1515 | 1481 | 870 | 1161 | 895 | 940 | 911 | 1061 | 1777 | 1091 | 937 | 588 | 858 | 955 | Nebl |
| Necab1 | 542 | 462 | 381 | 589 | 819 | 754 | 620 | 670 | 592 | 602 | 586 | 492 | 623 | 453 | 713 | 518 | 491 | 758 | 572 | 719 | 648 | 627 | 669 | 539 | 540 | 358 | 556 | 522 | Necab1 |
| Necab2 | 1039 | 1320 | 1320 | 1160 | 1574 | 1275 | 1284 | 1271 | 1285 | 1513 | 1311 | 1609 | 1355 | 1531 | 1114 | 1100 | 1378 | 1281 | 1533 | 1333 | 1381 | 1796 | 1295 | 1323 | 1491 | 1463 | 1706 | 1720 | Necab2 |
| Necab3 | 452 | 715 | 652 | 786 | 822 | 791 | 801 | 437 | 510 | 539 | 488 | 702 | 579 | 615 | 492 | 591 | 773 | 870 | 815 | 701 | 877 | 516 | 449 | 590 | 715 | 441 | 582 | 497 | Necab3 |
| Necap1 | 4385 | 4347 | 3937 | 4755 | 4951 | 4694 | 5270 | 5166 | 4557 | 4353 | 4296 | 3899 | 4065 | 4353 | 4630 | 4036 | 4413 | 4708 | 4874 | 4831 | 5098 | 5367 | 4462 | 4268 | 4589 | 4016 | 4172 | 4179 | Necap1 |
| Necap2 | 3008 | 3576 | 3696 | 3712 | 3089 | 3111 | 2785 | 1936 | 2408 | 2401 | 2554 | 2546 | 2537 | 2334 | 2718 | 3756 | 3445 | 3734 | 3087 | 2852 | 2708 | 2338 | 2504 | 2584 | 2603 | 2823 | 2745 | 2415 | Necap2 |
| Nectin1 | 2406 | 3156 | 4667 | 5171 | 5307 | 3991 | 3550 | 1983 | 2365 | 3178 | 2958 | 3214 | 2480 | 2638 | 2531 | 2663 | 5113 | 5058 | 5058 | 3072 | 3962 | 3532 | 2291 | 3324 | 3351 | 4193 | 2614 | 2643 | Nectin1 |
| Nectin2 | 4692 | 5912 | 5198 | 5251 | 4597 | 4452 | 3760 | 2287 | 2941 | 2636 | 2837 | 2455 | 3029 | 2195 | 4431 | 5611 | 5330 | 4920 | 4731 | 3907 | 3968 | 2289 | 2902 | 2533 | 2212 | 3189 | 2472 | 2375 | Nectin2 |
| Nectin3 | 12125 | 10982 | 9978 | 9038 | 8306 | 9036 | 10188 | 8924 | 7272 | 6549 | 6574 | 9795 | 9567 | 10268 | 12881 | 10208 | 8795 | 8128 | 8161 | 9969 | 9111 | 6490 | 7123 | 6575 | 7056 | 7998 | 10670 | 10554 | Nectin3 |
| Nectin4 | 218 | 182 | 214 | 165 | 160 | 180 | 136 | 304 | 503 | 251 | 249 | 175 | 232 | 159 | 215 | 217 | 181 | 190 | 167 | 160 | 125 | 119 | 498 | 222 | 85 | 130 | 146 | 176 | Nectin4 |
| Nedd1 | 4594 | 3899 | 4153 | 3378 | 2963 | 3339 | 3952 | 3659 | 3433 | 2726 | 2940 | 3899 | 3762 | 3865 | 4284 | 4181 | 3778 | 3487 | 3483 | 4275 | 3376 | 2653 | 3400 | 3108 | 2976 | 4139 | 3892 | 4052 | Nedd1 |
| Nedd4 | 153986 | 138526 | 118895 | 106666 | 105922 | 119223 | 111977 | 171889 | 165213 | 148079 | 155384 | 173619 | 170244 | 166693 | 147886 | 138176 | 116075 | 109102 | 107023 | 120159 | 109839 | 148375 | 162852 | 151623 | 148084 | 153645 | 183364 | 172393 | Nedd4 |
| Nedd4l | 5391 | 5500 | 5825 | 7638 | 9748 | 8374 | 8766 | 8048 | 8827 | 11533 | 9919 | 9843 | 8005 | 9524 | 5627 | 5199 | 7507 | 8308 | 9598 | 6845 | 9106 | 12915 | 7981 | 10619 | 13043 | 8290 | 7784 | 8488 | Nedd4l |
| Nedd8 | 7433 | 7856 | 8842 | 7911 | 6455 | 6614 | 7811 | 6980 | 6460 | 6099 | 6168 | 6915 | 6716 | 8560 | 8012 | 7256 | 8094 | 6892 | 7439 | 8329 | 7437 | 5469 | 6604 | 7774 | 6760 | 8694 | 7754 | 8796 | Nedd8 |
| Nedd9 | 4282 | 3940 | 3135 | 3710 | 3341 | 3786 | 3730 | 2783 | 2911 | 2415 | 2419 | 2732 | 2631 | 2547 | 3796 | 3840 | 3333 | 3803 | 3625 | 3814 | 3653 | 2209 | 3094 | 2626 | 2676 | 2650 | 2795 | 2720 | Nedd9 |
| Nefh | 119 | 135 | 98 | 92 | 311 | 123 | 77 | 269 | 280 | 277 | 363 | 111 | 215 | 109 | 156 | 154 | 115 | 158 | 122 | 147 | 94 | 378 | 274 | 136 | 254 | 107 | 139 | 107 | Nefh |
| Nefl | 9501 | 14239 | 11499 | 12111 | 24118 | 12624 | 8934 | 20011 | 18416 | 17179 | 19523 | 6811 | 22096 | 6123 | 20692 | 17897 | 13859 | 14486 | 11181 | 12986 | 10238 | 24662 | 17630 | 6016 | 12903 | 3366 | 8409 | 7945 | Nefl |
| Nefm | 10927 | 14696 | 20070 | 15821 | 26349 | 13169 | 10365 | 16927 | 16900 | 22037 | 23673 | 9009 | 28779 | 9771 | 21021 | 22784 | 21273 | 19027 | 12337 | 10701 | 10781 | 28604 | 13615 | 8112 | 17104 | 5834 | 11217 | 11744 | Nefm |
| Negr1 | 3584 | 3509 | 2203 | 6015 | 7814 | 7215 | 8133 | 7182 | 6682 | 6400 | 5664 | 4468 | 3670 | 4701 | 4787 | 3427 | 4203 | 6797 | 7438 | 8339 | 8606 | 8482 | 6797 | 5309 | 7488 | 3212 | 3461 | 3623 | Negr1 |
| Neil1 | 200 | 285 | 285 | 253 | 374 | 367 | 306 | 595 | 684 | 752 | 699 | 696 | 708 | 646 | 269 | 240 | 322 | 298 | 334 | 310 | 299 | 759 | 662 | 828 | 694 | 628 | 654 | 614 | Neil1 |
| Neil2 | 241 | 226 | 168 | 203 | 258 | 275 | 269 | 250 | 259 | 267 | 245 | 326 | 328 | 353 | 210 | 191 | 214 | 256 | 243 | 263 | 225 | 189 | 260 | 247 | 268 | 207 | 389 | 316 | Neil2 |
| Neil3 | 1895 | 1304 | 1552 | 1040 | 1113 | 1415 | 1482 | 2087 | 2146 | 1732 | 1819 | 2838 | 2300 | 2827 | 1515 | 1386 | 1298 | 909 | 1344 | 1813 | 1354 | 1589 | 1733 | 2125 | 2252 | 2511 | 3068 | 2659 | Neil3 |
| Nek1 | 8116 | 6612 | 6294 | 7083 | 6589 | 6732 | 7613 | 4932 | 4450 | 4454 | 4625 | 3905 | 4154 | 4292 | 8553 | 7639 | 6696 | 7322 | 6271 | 7172 | 7015 | 4802 | 4565 | 3775 | 4141 | 3451 | 3942 | 3999 | Nek1 |
| Nek10 | 29 | 22 | 21 | 32 | 47 | 50 | 34 | 50 | 43 | 28 | 49 | 45 | 59 | 43 | 54 | 42 | 15 | 32 | 50 | 48 | 55 | 25 | 71 | 30 | 28 | 31 | 15 | 24 | Nek10 |
| Nek11 | 98 | 118 | 53 | 156 | 144 | 128 | 114 | 162 | 143 | 128 | 85 | 125 | 159 | 99 | 129 | 132 | 124 | 133 | 108 | 141 | 118 | 135 | 183 | 124 | 149 | 90 | 91 | 80 | Nek11 |
| Nek2 | 396 | 61 | 56 | 60 | 46 | 57 | 38 | 87 | 55 | 43 | 55 | 186 | 64 | 189 | 112 | 65 | 71 | 47 | 61 | 73 | 46 | 42 | 45 | 67 | 59 | 83 | 88 | 504 | Nek2 |

| | | | | | | | | | | | | | | | | | | | | | | | | | | | | | |
|---|---|---|---|---|---|---|---|---|---|---|---|---|---|---|---|---|---|---|---|---|---|---|---|---|---|---|---|---|---|
| Nek3 | 265 | 358 | 437 | 402 | 302 | 329 | 296 | 521 | 477 | 552 | 462 | 640 | 593 | 749 | 331 | 277 | 416 | 344 | 356 | 291 | 321 | 647 | 491 | 584 | 638 | 705 | 553 | 691 | Nek3 |
| Nek4 | 2012 | 1756 | 2076 | 1814 | 1743 | 1751 | 1817 | 2783 | 2344 | 2409 | 2371 | 2897 | 2630 | 2793 | 2090 | 1646 | 1818 | 1716 | 1712 | 1749 | 1751 | 2610 | 2311 | 2631 | 2462 | 2682 | 3082 | 3017 | Nek4 |
| Nek5 | 20 | 35 | 19 | 24 | 38 | 48 | 26 | 56 | 68 | 49 | 41 | 47 | 92 | 29 | 41 | 38 | 20 | 14 | 25 | 35 | 41 | 74 | 96 | 30 | 42 | 31 | 12 | 38 | Nek5 |
| Nek6 | 4561 | 4617 | 3760 | 3863 | 3534 | 4143 | 3823 | 4035 | 4528 | 3790 | 4165 | 4740 | 4194 | 4494 | 4382 | 4145 | 3843 | 3674 | 3875 | 3752 | 3896 | 3379 | 4417 | 4282 | 4127 | 5083 | 4286 | 4454 | Nek6 |
| Nek7 | 2912 | 1978 | 2551 | 2383 | 2227 | 2632 | 2724 | 3135 | 2748 | 2546 | 3044 | 3091 | 2948 | 3195 | 2920 | 2466 | 2538 | 2553 | 2512 | 2770 | 2540 | 2697 | 3281 | 2984 | 2471 | 3140 | 3211 | 3191 | Nek7 |
| Nek8 | 642 | 709 | 716 | 787 | 620 | 676 | 565 | 590 | 751 | 701 | 878 | 717 | 852 | 637 | 644 | 758 | 778 | 766 | 634 | 617 | 587 | 570 | 779 | 735 | 617 | 921 | 818 | 720 | Nek8 |
| Nek9 | 9896 | 8822 | 8337 | 7900 | 8119 | 8737 | 7816 | 10832 | 11519 | 10982 | 11985 | 11712 | 12440 | 11355 | 8790 | 9111 | 7755 | 8358 | 8045 | 7700 | 7962 | 9892 | 11514 | 11181 | 10383 | 10778 | 11489 | 11678 | Nek9 |
| Nelfa | 1974 | 2184 | 2380 | 2130 | 1902 | 1878 | 1880 | 2502 | 2782 | 2513 | 2642 | 3297 | 2939 | 2972 | 1869 | 1992 | 2255 | 1873 | 2125 | 1951 | 1852 | 2632 | 2591 | 3289 | 2778 | 3744 | 2912 | 3258 | Nelfa |
| Nelfb | 4645 | 4895 | 5616 | 4619 | 4472 | 3998 | 4075 | 5127 | 5379 | 5897 | 5835 | 6307 | 6187 | 6154 | 4259 | 4583 | 5055 | 4519 | 4791 | 3893 | 4166 | 6126 | 5502 | 6283 | 5731 | 6846 | 6365 | 6573 | Nelfb |
| Nelfcd | 5000 | 6154 | 5735 | 4858 | 4656 | 5057 | 4322 | 5296 | 5665 | 5574 | 5413 | 6896 | 6263 | 6355 | 4743 | 5048 | 5577 | 4427 | 5312 | 5140 | 4531 | 4904 | 5602 | 6285 | 6020 | 8018 | 7186 | 6792 | Nelfcd |
| Nelfe | 3878 | 4301 | 4100 | 3903 | 2925 | 3091 | 3394 | 1796 | 2150 | 2481 | 2207 | 2776 | 2564 | 2747 | 3305 | 3713 | 3901 | 3474 | 3382 | 2993 | 3309 | 1801 | 2060 | 2541 | 2325 | 3289 | 2705 | 2887 | Nelfe |
| Nell1 | 253 | 263 | 191 | 366 | 641 | 417 | 378 | 512 | 591 | 488 | 693 | 270 | 381 | 246 | 306 | 287 | 245 | 370 | 401 | 321 | 516 | 708 | 645 | 310 | 463 | 178 | 235 | 328 | Nell1 |
| Nell2 | 5668 | 7121 | 6685 | 8344 | 10289 | 9039 | 9233 | 9272 | 9168 | 11027 | 9014 | 10824 | 8758 | 10554 | 6994 | 5044 | 8562 | 8566 | 10011 | 8451 | 9940 | 12489 | 9046 | 10652 | 11710 | 9935 | 10142 | 10427 | Nell2 |
| Nemf | 10490 | 8467 | 9913 | 9731 | 8929 | 9389 | 9973 | 8668 | 7175 | 7403 | 7176 | 7842 | 7595 | 8183 | 10664 | 9728 | 9270 | 9425 | 9065 | 10844 | 9561 | 7315 | 7092 | 7415 | 7404 | 7479 | 8148 | 8039 | Nemf |
| Nemp1 | 5975 | 6036 | 5012 | 5639 | 4286 | 4939 | 5602 | 2049 | 1864 | 1948 | 1930 | 2608 | 2589 | 2723 | 5664 | 5611 | 5101 | 5324 | 4945 | 5220 | 5508 | 1741 | 1707 | 2268 | 2199 | 2227 | 2867 | 2669 | Nemp1 |
| Nemp2 | 2746 | 2198 | 2210 | 2516 | 2520 | 2682 | 3061 | 2446 | 2428 | 2296 | 2412 | 2638 | 2454 | 2570 | 2951 | 2326 | 2268 | 2650 | 2526 | 2827 | 2590 | 2105 | 2168 | 2269 | 2389 | 2271 | 2632 | 2158 | Nemp2 |
| Nenf | 644 | 661 | 583 | 596 | 641 | 609 | 598 | 1011 | 1029 | 997 | 1073 | 896 | 941 | 1136 | 609 | 568 | 607 | 633 | 588 | 654 | 711 | 959 | 1142 | 1005 | 1051 | 852 | 943 | 1162 | Nenf |
| Neo1 | 16120 | 14749 | 12379 | 14752 | 17457 | 17619 | 16791 | 17565 | 18680 | 19040 | 18447 | 16227 | 17097 | 16783 | 15848 | 15601 | 12837 | 16413 | 16066 | 15513 | 17832 | 19794 | 17712 | 16923 | 19044 | 13253 | 16275 | 16107 | Neo1 |
| Nepn | 2 | 2 | 0 | 0 | 5 | 4 | 7 | 1 | 1 | 5 | 0 | 0 | 1 | 2 | 1 | 9 | 7 | 3 | 3 | 4 | 3 | 8 | 3 | 1 | 4 | 6 | 3 | 1 | Nepn |
| Nepro | 1932 | 1844 | 1578 | 1663 | 1369 | 1562 | 1494 | 1962 | 1710 | 1620 | 1698 | 1887 | 1893 | 1744 | 1709 | 1726 | 1334 | 1372 | 1460 | 1604 | 1511 | 1369 | 1680 | 1611 | 1671 | 1485 | 2185 | 1985 | Nepro |
| Nes | 68751 | 81869 | 115280 | 81071 | 61102 | 70587 | 67799 | 43399 | 37952 | 43599 | 41667 | 80556 | 63788 | 74002 | 76324 | 75226 | 101487 | 70098 | 74370 | 71567 | 64725 | 39567 | 34797 | 56170 | 52419 | 74096 | 84858 | 80408 | Nes |
| Net1 | 4444 | 2744 | 3200 | 2265 | 1631 | 2122 | 1843 | 3565 | 4195 | 3473 | 4904 | 3269 | 4017 | 3305 | 2723 | 3825 | 2557 | 2088 | 1818 | 1699 | 1705 | 2303 | 4303 | 3908 | 2513 | 4910 | 2984 | 3394 | Net1 |
| Neto1 | 495 | 362 | 516 | 669 | 875 | 858 | 859 | 725 | 694 | 707 | 718 | 649 | 462 | 679 | 530 | 333 | 550 | 500 | 895 | 621 | 813 | 1033 | 586 | 681 | 903 | 464 | 622 | 580 | Neto1 |
| Neto2 | 3714 | 3108 | 3086 | 4687 | 7539 | 6520 | 7441 | 10984 | 8914 | 8231 | 7511 | 8294 | 7102 | 8814 | 4814 | 3110 | 3945 | 5050 | 6547 | 7165 | 7539 | 10846 | 8952 | 7918 | 10209 | 5902 | 7901 | 7978 | Neto2 |
| Neu1 | 2271 | 2884 | 2791 | 3004 | 2445 | 2372 | 2132 | 1189 | 1170 | 1322 | 1286 | 1134 | 1073 | 1123 | 2267 | 2959 | 2479 | 2725 | 2321 | 2293 | 2246 | 1308 | 1233 | 1268 | 1246 | 1011 | 1099 | 1121 | Neu1 |
| Neu2 | 0 | 3 | 0 | 1 | 1 | 2 | 0 | 4 | 0 | 0 | 0 | 0 | 3 | 5 | 0 | 2 | 0 | 0 | 1 | 1 | 1 | 2 | 1 | 0 | 0 | 0 | 0 | 0 | Neu2 |
| Neu3 | 519 | 462 | 399 | 525 | 553 | 461 | 468 | 313 | 278 | 267 | 337 | 239 | 337 | 251 | 514 | 530 | 399 | 554 | 487 | 525 | 531 | 266 | 284 | 253 | 246 | 177 | 258 | 254 | Neu3 |
| Neu4 | 0 | 0 | 0 | 0 | 0 | 0 | 0 | 0 | 0 | 3 | 3 | 0 | 0 | 0 | 0 | 0 | 0 | 0 | 0 | 0 | 0 | 0 | 1 | 1 | 0 | 0 | 0 | 0 | Neu4 |
| Neurl1a | 789 | 922 | 893 | 943 | 1168 | 990 | 715 | 696 | 747 | 896 | 1038 | 545 | 584 | 495 | 728 | 1019 | 968 | 984 | 913 | 817 | 849 | 994 | 834 | 633 | 820 | 575 | 474 | 481 | Neurl1a |
| Neurl1b | 582 | 722 | 773 | 952 | 1114 | 898 | 977 | 576 | 770 | 972 | 795 | 604 | 454 | 589 | 566 | 681 | 1022 | 1107 | 1153 | 843 | 1024 | 971 | 693 | 920 | 997 | 732 | 482 | 509 | Neurl1b |
| Neurl2 | 90 | 92 | 119 | 76 | 97 | 76 | 73 | 89 | 147 | 129 | 132 | 135 | 203 | 87 | 66 | 114 | 95 | 145 | 104 | 86 | 77 | 152 | 144 | 114 | 110 | 181 | 152 | 123 | Neurl2 |
| Neurl3 | 193 | 181 | 205 | 336 | 209 | 240 | 204 | 65 | 71 | 79 | 125 | 89 | 85 | 61 | 219 | 230 | 176 | 341 | 187 | 150 | 220 | 92 | 111 | 92 | 79 | 76 | 51 | 50 | Neurl3 |
| Neurl4 | 7260 | 9804 | 11391 | 12633 | 11742 | 10636 | 10353 | 7772 | 9600 | 11597 | 10387 | 12159 | 10143 | 10815 | 7457 | 8413 | 12107 | 12110 | 12619 | 8744 | 10946 | 11373 | 8662 | 12118 | 12538 | 10433 | 10330 | 10607 | Neurl4 |
| Neurod1 | 1366 | 1239 | 1599 | 2023 | 2473 | 2149 | 2428 | 2925 | 2375 | 2600 | 2162 | 2936 | 2750 | 3246 | 1833 | 1204 | 1803 | 1928 | 2318 | 2125 | 2372 | 3009 | 2521 | 3130 | 3154 | 3280 | 2888 | 3446 | Neurod1 |
| Neurod2 | 453 | 528 | 248 | 1506 | 2608 | 1932 | 2714 | 2616 | 3597 | 3879 | 2821 | 2503 | 1327 | 2885 | 425 | 160 | 833 | 1870 | 2810 | 1761 | 2864 | 2521 | 2445 | 4609 | 5412 | 2557 | 1297 | 1440 | Neurod2 |
| Neurod4 | 559 | 594 | 885 | 384 | 373 | 372 | 367 | 890 | 494 | 496 | 406 | 988 | 1051 | 956 | 872 | 739 | 614 | 361 | 322 | 486 | 411 | 645 | 463 | 627 | 565 | 693 | 1256 | 1326 | Neurod4 |
| Neurod6 | 795 | 718 | 307 | 2081 | 5220 | 4134 | 6472 | 7551 | 9051 | 8097 | 6280 | 4978 | 2514 | 6118 | 1217 | 384 | 1063 | 3067 | 5537 | 5070 | 6119 | 4271 | 5818 | 9311 | 12413 | 4534 | 2718 | 3040 | Neurod6 |
| Neurog1 | 225 | 297 | 737 | 215 | 260 | 252 | 149 | 423 | 323 | 607 | 338 | 848 | 841 | 884 | 270 | 241 | 547 | 193 | 316 | 195 | 215 | 376 | 342 | 816 | 579 | 1029 | 807 | 977 | Neurog1 |
| Neurog2 | 2972 | 3235 | 6312 | 4345 | 5402 | 4709 | 6479 | 6031 | 4643 | 6285 | 4658 | 9519 | 5000 | 9683 | 3448 | 1488 | 6063 | 4095 | 7379 | 5901 | 5651 | 4944 | 4265 | 9839 | 8698 | 11160 | 8334 | 9317 | Neurog2 |
| Neurog3 | 5 | 12 | 9 | 14 | 7 | 2 | 9 | 17 | 25 | 15 | 14 | 17 | 10 | 23 | 13 | 13 | 6 | 6 | 3 | 7 | 3 | 12 | 14 | 32 | 28 | 28 | 28 | 7 | Neurog3 |
| Nexmif | 1588 | 1045 | 779 | 1487 | 3107 | 3173 | 3261 | 6184 | 5650 | 4486 | 4646 | 3225 | 2828 | 3618 | 1774 | 1008 | 1134 | 1906 | 2544 | 3585 | 3488 | 5230 | 4992 | 3322 | 5920 | 1707 | 3196 | 2734 | Nexmif |
| Nexn | 311 | 234 | 468 | 219 | 208 | 344 | 171 | 474 | 524 | 438 | 1662 | 569 | 712 | 578 | 475 | 407 | 432 | 245 | 207 | 177 | 172 | 351 | 980 | 507 | 366 | 626 | 746 | 648 | Nexn |
| Nf1 | 7399 | 6584 | 5614 | 7209 | 8041 | 8859 | 8695 | 9911 | 9690 | 9685 | 10246 | 10191 | 10003 | 10555 | 6936 | 6403 | 6218 | 6715 | 7518 | 7496 | 9349 | 9491 | 9032 | 7942 | 10899 | 7116 | 10404 | 9575 | Nf1 |
| Nf2 | 13627 | 14259 | 14560 | 14907 | 15387 | 14669 | 15064 | 12603 | 12833 | 13101 | 12920 | 13144 | 12414 | 13595 | 13630 | 13199 | 15366 | 14765 | 15932 | 13723 | 15389 | 13882 | 12515 | 13500 | 14301 | 13246 | 13381 | 13468 | Nf2 |
| Nfam1 | 230 | 241 | 339 | 170 | 184 | 215 | 161 | 263 | 318 | 262 | 381 | 325 | 460 | 335 | 194 | 295 | 224 | 235 | 175 | 210 | 200 | 304 | 377 | 319 | 308 | 347 | 432 | 421 | Nfam1 |
| Nfasc | 8822 | 12255 | 11927 | 20096 | 24112 | 20315 | 20702 | 11779 | 12478 | 16438 | 12589 | 14404 | 14292 | 13320 | 11512 | 11664 | 17389 | 20673 | 21964 | 17046 | 21508 | 17829 | 10648 | 14427 | 18317 | 7697 | 12217 | 12092 | Nfasc |
| Nfat5 | 20528 | 14867 | 11300 | 15193 | 17593 | 20439 | 18803 | 16391 | 15975 | 14332 | 17885 | 12353 | 14698 | 13225 | 18282 | 19048 | 12238 | 15225 | 14728 | 17137 | 19763 | 14105 | 15519 | 10978 | 13638 | 8286 | 13441 | 11160 | Nfat5 |
| Nfatc1 | 594 | 689 | 679 | 512 | 405 | 524 | 327 | 639 | 929 | 1036 | 1410 | 622 | 948 | 569 | 596 | 867 | 570 | 643 | 349 | 254 | 336 | 911 | 1145 | 863 | 707 | 845 | 686 | 637 | Nfatc1 |
| Nfatc2 | 640 | 740 | 660 | 871 | 893 | 705 | 792 | 316 | 484 | 440 | 506 | 289 | 278 | 280 | 527 | 669 | 642 | 1053 | 874 | 619 | 847 | 416 | 421 | 435 | 406 | 253 | 208 | 192 | Nfatc2 |
| Nfatc2ip | 3120 | 2871 | 3571 | 3677 | 2978 | 3387 | 3507 | 1480 | 1317 | 1499 | 1308 | 1806 | 1667 | 1899 | 2921 | 2747 | 3424 | 3704 | 3258 | 3003 | 3376 | 1447 | 1131 | 1632 | 1732 | 2123 | 1870 | 1864 | Nfatc2ip |
| Nfatc3 | 9206 | 6987 | 6262 | 5615 | 4825 | 5844 | 5556 | 5714 | 5481 | 4785 | 5421 | 5359 | 5550 | 5289 | 7760 | 8027 | 5763 | 5564 | 5058 | 5247 | 5303 | 4508 | 5481 | 4788 | 4220 | 4877 | 5560 | 5604 | Nfatc3 |
| Nfatc4 | 2998 | 2955 | 3512 | 2414 | 1976 | 2357 | 1846 | 4382 | 5992 | 5813 | 6526 | 4524 | 5914 | 3726 | 2334 | 3224 | 2709 | 2486 | 2339 | 1818 | 1785 | 3551 | 6117 | 6251 | 3851 | 6650 | 4217 | 4457 | Nfatc4 |
| Nfe2 | 60 | 105 | 120 | 53 | 23 | 60 | 35 | 78 | 96 | 87 | 91 | 162 | 301 | 164 | 55 | 121 | 59 | 40 | 37 | 63 | 35 | 70 | 114 | 81 | 49 | 195 | 192 | 331 | Nfe2 |
| Nfe2l1 | 10415 | 12747 | 14465 | 13930 | 12430 | 11955 | 10861 | 8806 | 10343 | 11270 | 12496 | 11892 | 11702 | 11488 | 10581 | 13059 | 14373 | 14192 | 11982 | 10071 | 11176 | 11658 | 10321 | 12034 | 11973 | 11328 | 11984 | 11624 | Nfe2l1 |
| Nfe2l2 | 2751 | 2714 | 2750 | 2521 | 2095 | 2108 | 1998 | 1780 | 1737 | 1529 | 2052 | 1416 | 1692 | 1607 | 2561 | 2961 | 2569 | 2290 | 2352 | 2409 | 2098 | 1434 | 1988 | 1739 | 1395 | 1922 | 1574 | 1619 | Nfe2l2 |
| Nfe2l3 | 1449 | 1245 | 1232 | 1222 | 1308 | 1129 | 1230 | 432 | 327 | 302 | 262 | 198 | 320 | 264 | 1493 | 1342 | 1201 | 1225 | 1271 | 1382 | 1324 | 376 | 368 | 397 | 354 | 227 | 277 | 325 | Nfe2l3 |
| Nfia | 2799 | 2376 | 3921 | 6022 | 6350 | 5377 | 6950 | 6436 | 7188 | 9796 | 9454 | 8526 | 7790 | 8734 | 2978 | 2696 | 4759 | 6194 | 6688 | 4400 | 6606 | 8031 | 7111 | 11677 | 10952 | 10995 | 6410 | 7774 | Nfia |
| Nfib | 7130 | 4975 | 6285 | 12887 | 17731 | 15820 | 19964 | 21389 | 25902 | 30479 | 29126 | 21963 | 20238 | 23833 | 6956 | 5557 | 9175 | 14932 | 17644 | 13140 | 19096 | 21614 | 24840 | 32484 | 36533 | 23052 | 17816 | 19950 | Nfib |
| Nfic | 1696 | 2091 | 2334 | 2228 | 2160 | 2181 | 1468 | 2104 | 3283 | 3563 | 4480 | 2663 | 3142 | 2167 | 1622 | 2335 | 2227 | 2406 | 2086 | 1431 | 1583 | 3369 | 3594 | 3639 | 3257 | 3139 | 2293 | 2155 | Nfic |
| Nfil3 | 1159 | 1304 | 1361 | 1532 | 1614 | 1281 | 1350 | 1220 | 1327 | 1490 | 1267 | 1183 | 1125 | 1193 | 1244 | 1124 | 1349 | 1652 | 1576 | 1199 | 1574 | 1592 | 1293 | 1370 | 1436 | 1155 | 895 | 1012 | Nfil3 |
| Nfix | 1628 | 1698 | 2245 | 4180 | 6133 | 4967 | 5084 | 4974 | 7672 | 9843 | 10632 | 7287 | 5433 | 6715 | 1549 | 1481 | 2941 | 5443 | 6410 | 3709 | 5988 | 6789 | 7895 | 12566 | 12594 | 7796 | 4894 | 5521 | Nfix |
| Nfkb1 | 1412 | 1358 | 1559 | 1260 | 1575 | 1476 | 1130 | 2334 | 2996 | 2633 | 3104 | 2796 | 2762 | 2525 | 1290 | 1491 | 1484 | 1238 | 1301 | 1215 | 1171 | 2159 | 3067 | 2759 | 2328 | 2914 | 2632 | 2552 | Nfkb1 |
| Nfkb2 | 1744 | 2443 | 2488 | 2196 | 1302 | 1540 | 1152 | 462 | 708 | 568 | 840 | 477 | 612 | 417 | 1619 | 2424 | 2311 | 2145 | 1566 | 1129 | 1364 | 503 | 793 | 577 | 505 | 540 | 494 | 516 | Nfkb2 |
| Nfkbia | 580 | 567 | 506 | 482 | 585 | 493 | 473 | 989 | 1056 | 1112 | 1118 | 865 | 856 | 891 | 487 | 465 | 469 | 472 | 553 | 534 | 466 | 964 | 1242 | 1088 | 851 | 959 | 932 | 934 | Nfkbia |
| Nfkbib | 953 | 1202 | 1037 | 1143 | 1018 | 954 | 787 | 675 | 761 | 755 | 729 | 717 | 730 | 656 | 928 | 970 | 1136 | 1119 | 1086 | 895 | 861 | 668 | 749 | 762 | 700 | 801 | 811 | 678 | Nfkbib |
| Nfkbid | 113 | 133 | 149 | 130 | 138 | 115 | 134 | 107 | 113 | 100 | 134 | 87 | 111 | 102 | 125 | 109 | 170 | 118 | 147 | 96 | 124 | 129 | 138 | 87 | 113 | 124 | 88 | 102 | Nfkbid |
| Nfkbie | 1054 | 1125 | 1343 | 1136 | 1123 | 938 | 984 | 248 | 246 | 260 | 287 | 305 | 289 | 342 | 940 | 1228 | 1267 | 1213 | 1008 | 847 | 985 | 252 | 274 | 289 | 253 | 358 | 313 | 358 | Nfkbie |
| Nfkbil1 | 436 | 429 | 556 | 483 | 518 | 527 | 466 | 482 | 673 | 652 | 610 | 669 | 649 | 611 | 396 | 389 | 561 | 491 | 610 | 526 | 438 | 546 | 654 | 684 | 645 | 860 | 687 | 560 | Nfkbil1 |
| Nfkbiz | 731 | 530 | 499 | 690 | 520 | 615 | 569 | 383 | 519 | 466 | 526 | 339 | 463 | 367 | 624 | 683 | 560 | 636 | 478 | 456 | 541 | 314 | 556 | 485 | 322 | 468 | 344 | 310 | Nfkbiz |
| Nfrkb | 5507 | 6334 | 4807 | 6372 | 5658 | 5911 | 5806 | 4661 | 5162 | 4895 | 5046 | 5319 | 5154 | 5228 | 5322 | 5699 | 5333 | 6097 | 6010 | 5397 | 5871 | 4933 | 4632 | 4808 | 5594 | 4425 | 5534 | 4780 | Nfrkb |
| Nfs1 | 1030 | 1238 | 1051 | 1062 | 955 | 915 | 945 | 935 | 993 | 864 | 885 | 1072 | 1027 | 1046 | 1062 | 1015 | 1250 | 951 | 1047 | 967 | 1024 | 837 | 1032 | 1095 | 1061 | 1203 | 967 | 1105 | Nfs1 |
| Nfu1 | 1115 | 1418 | 1418 | 1312 | 1331 | 1202 | 1391 | 1780 | 1902 | 1791 | 1828 | 1700 | 1848 | 1987 | 1415 | 1290 | 1378 | 1319 | 1353 | 1490 | 1406 | 1614 | 1699 | 1887 | 1792 | 1926 | 1888 | 2114 | Nfu1 |
| Nfx1 | 12126 | 11759 | 9887 | 11049 | 11234 | 11407 | 11853 | 11307 | 11089 | 10012 | 10434 | 10199 | 9605 | 9961 | 11774 | 10745 | 10057 | 11330 | 10790 | 12441 | 11851 | 10449 | 10554 | 9539 | 10937 | 9231 | 10039 | 9818 | Nfx1 |
| Nfxl1 | 5285 | 4018 | 4720 | 4576 | 4720 | 4880 | 5192 | 4205 | 3608 | 3597 | 3955 | 3884 | 4146 | 4037 | 5255 | 4539 | 4648 | 4570 | 4453 | 4953 | 4822 | 3683 | 3743 | 3591 | 3433 | 3903 | 4399 | 4101 | Nfxl1 |
| Nfya | 294 | 213 | 585 | 231 | 222 | 258 | 196 | 242 | 434 | 139 | 324 | 220 | 225 | 170 | 601 | 203 | 182 | 223 | 165 | 163 | 147 | 374 | 168 | 201 | 206 | 219 | 212 | 173 | Nfya |
| Nfyb | 3853 | 2906 | 3729 | 3328 | 3428 | 3199 | 3940 | 6081 | 5138 | 4673 | 4477 | 5587 | 4969 | 5792 | 3967 | 3250 | 3709 | 3189 | 3594 | 4006 | 3513 | 4906 | 4908 | 5814 | 5507 | 5946 | 5767 | 5830 | Nfyb |
| Nfyc | 1936 | 1906 | 1853 | 1877 | 2054 | 1921 | 1843 | 4005 | 4061 | 3826 | 4110 | 4312 | 4322 | 4274 | 1874 | 1808 | 1920 | 1780 | 2265 | 2185 | 1815 | 3761 | 3948 | 4092 | 4176 | 5068 | 4639 | 4168 | Nfyc |
| Ngb | 42 | 71 | 65 | 90 | 66 | 53 | 74 | 48 | 31 | 40 | 34 | 25 | 33 | 38 | 18 | 48 | 81 | 108 | 73 | 70 | 65 | 55 | 31 | 40 | 49 | 42 | 23 | 39 | Ngb |
| Ngdn | 2714 | 3147 | 3094 | 3162 | 2465 | 2599 | 2775 | 2196 | 2207 | 2070 | 1981 | 2364 | 2174 | 2397 | 2742 | 2838 | 3116 | 2971 | 3032 | 3476 | 2727 | 1663 | 2047 | 2171 | 2076 | 2904 | 2484 | 2430 | Ngdn |

| | | | | | | | | | | | | | | | | | | | | | | | | | | | | | |
|---|---|---|---|---|---|---|---|---|---|---|---|---|---|---|---|---|---|---|---|---|---|---|---|---|---|---|---|---|---|
| Ngef | 332 | 434 | 627 | 475 | 767 | 541 | 559 | 204 | 232 | 335 | 275 | 241 | 268 | 234 | 400 | 410 | 521 | 522 | 673 | 513 | 654 | 374 | 294 | 303 | 348 | 176 | 182 | 220 | Ngef |
| Ngf | 96 | 54 | 73 | 35 | 9 | 45 | 18 | 72 | 132 | 87 | 149 | 61 | 120 | 89 | 51 | 107 | 46 | 70 | 17 | 12 | 13 | 75 | 168 | 126 | 47 | 45 | 101 | 78 | Ngf |
| Ngfr | 2175 | 2315 | 2196 | 1500 | 2603 | 1198 | 971 | 2306 | 3032 | 2642 | 3678 | 1765 | 3098 | 1572 | 2139 | 3419 | 1673 | 1816 | 1233 | 1054 | 1041 | 3252 | 3333 | 1632 | 1989 | 1685 | 1973 | 1870 | Ngfr |
| Ngly1 | 1543 | 1205 | 1284 | 1305 | 1375 | 1393 | 1502 | 2391 | 2158 | 1768 | 2064 | 1924 | 1813 | 1968 | 1680 | 1334 | 1155 | 1361 | 1276 | 1545 | 1318 | 1858 | 2275 | 1914 | 1808 | 2060 | 1898 | 1919 | Ngly1 |
| Ngp | 20 | 3 | 11 | 10 | 15 | 10 | 15 | 16 | 45 | 23 | 75 | 55 | 85 | 19 | 24 | 39 | 15 | 23 | 16 | 7 | 7 | 43 | 83 | 74 | 9 | 65 | 66 | 51 | Ngp |
| Ngrn | 1957 | 2034 | 2042 | 2143 | 1965 | 1981 | 2152 | 2027 | 2016 | 1852 | 1862 | 1987 | 2081 | 2030 | 1895 | 1710 | 2064 | 1917 | 2179 | 2296 | 2145 | 1796 | 1946 | 2031 | 2025 | 2097 | 1972 | 2029 | Ngrn |
| Nhej1 | 145 | 153 | 137 | 88 | 108 | 113 | 123 | 269 | 275 | 236 | 274 | 256 | 294 | 260 | 153 | 125 | 112 | 123 | 89 | 100 | 97 | 227 | 305 | 223 | 229 | 283 | 314 | 324 | Nhej1 |
| Nhlh1 | 1613 | 2467 | 2306 | 2843 | 2135 | 2151 | 1961 | 3472 | 2432 | 3464 | 2837 | 3783 | 5297 | 3991 | 2198 | 2070 | 2634 | 2401 | 2181 | 1934 | 2190 | 2949 | 2393 | 3778 | 3726 | 4305 | 3569 | 4628 | Nhlh1 |
| Nhlh2 | 1836 | 2199 | 3213 | 2514 | 3629 | 2971 | 2556 | 8405 | 5085 | 6878 | 5505 | 8218 | 11917 | 8613 | 3181 | 2354 | 3539 | 2103 | 3003 | 3652 | 2636 | 8159 | 5994 | 6518 | 7011 | 7496 | 10168 | 10790 | Nhlh2 |
| Nhlrc1 | 648 | 663 | 769 | 854 | 907 | 740 | 761 | 380 | 407 | 393 | 401 | 355 | 353 | 456 | 642 | 586 | 820 | 919 | 839 | 722 | 886 | 543 | 447 | 429 | 453 | 339 | 334 | 350 | Nhlrc1 |
| Nhlrc2 | 3125 | 2725 | 2574 | 2637 | 2862 | 3172 | 3035 | 4931 | 4942 | 4446 | 4308 | 5130 | 4815 | 5223 | 2835 | 2581 | 2444 | 2346 | 2982 | 3183 | 3154 | 3878 | 4522 | 4202 | 4738 | 4013 | 5267 | 5020 | Nhlrc2 |
| Nhlrc3 | 2502 | 2369 | 2035 | 2176 | 1814 | 2016 | 1906 | 2332 | 2380 | 2241 | 2371 | 2440 | 2270 | 2302 | 2404 | 2315 | 1912 | 1791 | 1646 | 2093 | 1855 | 2115 | 2334 | 2163 | 2113 | 2150 | 2580 | 2375 | Nhlrc3 |
| Nhlrc4 | 0 | 3 | 4 | 2 | 4 | 0 | 0 | 0 | 2 | 2 | 3 | 4 | 9 | 1 | 4 | 3 | 0 | 10 | 2 | 0 | 2 | 5 | 3 | 0 | 1 | 5 | 0 | 12 | Nhlrc4 |
| Nhp2 | 3785 | 4625 | 4393 | 3906 | 2881 | 3341 | 3311 | 2419 | 2824 | 2550 | 2446 | 2821 | 3006 | 3130 | 3645 | 3657 | 4096 | 3644 | 3718 | 4229 | 3522 | 1860 | 2728 | 2851 | 2650 | 3841 | 3704 | 3364 | Nhp2 |
| Nhs | 3632 | 2521 | 3216 | 3109 | 2937 | 3294 | 2899 | 3033 | 2699 | 3017 | 3243 | 3711 | 3918 | 3663 | 3475 | 3111 | 2699 | 2693 | 2746 | 2748 | 2969 | 2816 | 3004 | 3016 | 2883 | 2857 | 3977 | 3957 | Nhs |
| Nhsl1 | 3527 | 3850 | 4273 | 5394 | 6449 | 5419 | 4964 | 4427 | 4932 | 5796 | 5649 | 5428 | 5584 | 5041 | 3967 | 4295 | 5106 | 5026 | 5509 | 4547 | 5240 | 5573 | 4774 | 5574 | 6148 | 4175 | 4786 | 4721 | Nhsl1 |
| Nhsl2 | 2095 | 1715 | 1507 | 1458 | 1710 | 1814 | 1338 | 1903 | 2244 | 2213 | 2921 | 2030 | 2588 | 2104 | 1756 | 2217 | 1455 | 1516 | 1425 | 1388 | 1448 | 2052 | 2530 | 1761 | 1926 | 1690 | 2358 | 1869 | Nhsl2 |
| Niban1 | 329 | 269 | 204 | 201 | 295 | 297 | 218 | 424 | 536 | 384 | 1083 | 408 | 480 | 325 | 287 | 327 | 185 | 308 | 231 | 284 | 216 | 364 | 973 | 381 | 303 | 322 | 422 | 324 | Niban1 |
| Niban2 | 5029 | 5723 | 5731 | 5406 | 5470 | 5207 | 3796 | 3600 | 4813 | 5006 | 5029 | 3704 | 4175 | 3328 | 4821 | 6555 | 5277 | 5548 | 5097 | 4057 | 4033 | 4236 | 5202 | 4516 | 4298 | 4257 | 3557 | 3463 | Niban2 |
| Niban3 | 287 | 411 | 472 | 468 | 532 | 392 | 386 | 321 | 354 | 465 | 440 | 369 | 371 | 352 | 265 | 411 | 568 | 494 | 504 | 343 | 442 | 471 | 352 | 378 | 405 | 321 | 339 | 331 | Niban3 |
| Nicn1 | 4315 | 4970 | 4282 | 4509 | 5063 | 4662 | 5190 | 4991 | 4838 | 4707 | 4328 | 4104 | 4297 | 4379 | 4977 | 4397 | 4588 | 4884 | 4997 | 5368 | 5176 | 5399 | 4799 | 4293 | 4925 | 3508 | 4221 | 4317 | Nicn1 |
| Nid1 | 15363 | 14638 | 14601 | 11025 | 11381 | 12331 | 7511 | 9795 | 12284 | 12437 | 16090 | 12671 | 16500 | 11959 | 14310 | 19413 | 12271 | 13231 | 7631 | 8741 | 7992 | 14400 | 15927 | 11347 | 11736 | 11199 | 13350 | 11932 | Nid1 |
| Nid2 | 7686 | 9644 | 10001 | 8529 | 7436 | 7824 | 5646 | 5255 | 7615 | 7774 | 11186 | 7156 | 9131 | 6242 | 7755 | 10921 | 8416 | 9535 | 6685 | 5950 | 6036 | 8194 | 10642 | 7474 | 6940 | 7304 | 6940 | 6157 | Nid2 |
| Nif3l1 | 2804 | 2714 | 2591 | 2305 | 2419 | 2371 | 2699 | 3765 | 3353 | 3012 | 3119 | 3598 | 3404 | 3835 | 2970 | 2480 | 2517 | 2183 | 2563 | 2663 | 2506 | 3148 | 3389 | 3343 | 3234 | 3929 | 3620 | 3614 | Nif3l1 |
| Nifk | 4731 | 4302 | 4652 | 4016 | 3338 | 3686 | 3981 | 4703 | 4005 | 3510 | 3712 | 4329 | 4077 | 4259 | 4967 | 4403 | 4153 | 3609 | 3505 | 4540 | 3833 | 3811 | 3985 | 4008 | 3663 | 5071 | 4264 | 4732 | Nifk |
| Nim1k | 633 | 718 | 682 | 521 | 709 | 701 | 732 | 492 | 462 | 337 | 401 | 449 | 260 | 487 | 641 | 551 | 600 | 566 | 754 | 870 | 721 | 485 | 393 | 463 | 448 | 507 | 442 | 414 | Nim1k |
| Nin | 4760 | 4029 | 2667 | 3390 | 4054 | 4496 | 3886 | 5348 | 5277 | 4938 | 5671 | 4687 | 5074 | 4919 | 4410 | 4831 | 3217 | 3514 | 3479 | 4054 | 4118 | 4709 | 5529 | 4491 | 5230 | 3478 | 5273 | 4383 | Nin |
| Ninj1 | 1185 | 1380 | 938 | 973 | 875 | 1030 | 875 | 308 | 510 | 427 | 456 | 302 | 452 | 301 | 988 | 1277 | 892 | 1084 | 966 | 1126 | 822 | 307 | 566 | 354 | 345 | 377 | 303 | 298 | Ninj1 |
| Ninj2 | 0 | 1 | 0 | 0 | 0 | 3 | 0 | 5 | 5 | 2 | 8 | 0 | 3 | 0 | 1 | 0 | 0 | 0 | 1 | 0 | 1 | 8 | 1 | 1 | 0 | 0 | 0 | 2 | Ninj2 |
| Ninl | 2675 | 2656 | 2702 | 2421 | 2237 | 2416 | 2692 | 2363 | 2382 | 2507 | 2088 | 3477 | 2870 | 3437 | 2242 | 2263 | 2478 | 2319 | 2454 | 2636 | 2368 | 1668 | 1967 | 2858 | 2585 | 3289 | 3627 | 3469 | Ninl |
| Nip7 | 3885 | 3788 | 3236 | 3155 | 2738 | 2808 | 3013 | 3225 | 3049 | 2646 | 2748 | 3170 | 3098 | 2842 | 3775 | 3714 | 3225 | 2928 | 3060 | 3578 | 2794 | 2493 | 3183 | 3020 | 2923 | 3684 | 3226 | 3213 | Nip7 |
| Nipa1 | 1354 | 1261 | 1211 | 1467 | 1991 | 1726 | 1814 | 2122 | 2043 | 1813 | 1877 | 1822 | 2020 | 1953 | 1489 | 1282 | 1308 | 1499 | 1695 | 1855 | 1819 | 2189 | 1922 | 1696 | 2097 | 1147 | 1878 | 1862 | Nipa1 |
| Nipa2 | 5888 | 4736 | 5103 | 5187 | 4765 | 5167 | 5539 | 4583 | 3631 | 3387 | 3572 | 4228 | 3961 | 4158 | 6136 | 5675 | 5139 | 5294 | 4733 | 5401 | 5198 | 3893 | 3698 | 3505 | 3837 | 4190 | 4210 | 4316 | Nipa2 |
| Nipal1 | 275 | 231 | 273 | 172 | 114 | 219 | 120 | 93 | 98 | 79 | 64 | 90 | 58 | 71 | 278 | 262 | 147 | 228 | 120 | 174 | 128 | 45 | 129 | 89 | 39 | 95 | 46 | 74 | Nipal1 |
| Nipal2 | 102 | 53 | 60 | 55 | 89 | 118 | 91 | 324 | 330 | 277 | 330 | 168 | 197 | 181 | 73 | 59 | 48 | 94 | 55 | 85 | 62 | 331 | 443 | 273 | 228 | 161 | 144 | 172 | Nipal2 |
| Nipal3 | 3094 | 3883 | 3841 | 4220 | 4721 | 3673 | 4014 | 1646 | 1767 | 2056 | 1811 | 1719 | 1559 | 1545 | 3266 | 3806 | 4198 | 4316 | 4623 | 3705 | 4278 | 2246 | 1719 | 1762 | 2219 | 1343 | 1374 | 1556 | Nipal3 |
| Nipal4 | 142 | 172 | 134 | 87 | 52 | 116 | 63 | 90 | 87 | 66 | 70 | 98 | 127 | 109 | 161 | 194 | 96 | 111 | 51 | 67 | 68 | 37 | 124 | 118 | 76 | 102 | 141 | 161 | Nipal4 |
| Nipbl | 18121 | 14753 | 11369 | 14586 | 15226 | 18677 | 18471 | 20359 | 17923 | 17259 | 16831 | 18397 | 18397 | 19974 | 18509 | 16140 | 13063 | 14077 | 14911 | 18605 | 18375 | 14801 | 16942 | 14315 | 17086 | 11888 | 20711 | 17858 | Nipbl |
| Nipsnap1 | 5044 | 4974 | 4478 | 4912 | 5276 | 4945 | 5339 | 6149 | 5506 | 5719 | 5099 | 5677 | 5754 | 6107 | 5144 | 4552 | 4682 | 4525 | 5567 | 5827 | 5186 | 6158 | 5305 | 5940 | 6437 | 6011 | 5793 | 6169 | Nipsnap1 |
| Nipsnap2 | 2728 | 2560 | 2541 | 2399 | 2372 | 2705 | 2380 | 3815 | 4079 | 4392 | 4304 | 4607 | 4314 | 4880 | 2563 | 2509 | 2450 | 2240 | 2146 | 2612 | 2434 | 3663 | 4159 | 4149 | 4296 | 5213 | 4487 | 4686 | Nipsnap2 |
| Nipsnap3a | 0 | 0 | 0 | 0 | 0 | 0 | 0 | 1 | 3 | 0 | 0 | 0 | 2 | 0 | 0 | 0 | 0 | 0 | 0 | 0 | 0 | 0 | 1 | 0 | 3 | 0 | 0 | 0 | Nipsnap3a |
| Nipsnap3b | 2457 | 2527 | 2446 | 2456 | 2480 | 2590 | 2996 | 3154 | 2633 | 2242 | 2128 | 2503 | 2586 | 2888 | 3053 | 2344 | 2385 | 2450 | 2692 | 3056 | 2921 | 3106 | 2735 | 2176 | 2519 | 2293 | 2705 | 3004 | Nipsnap3b |
| Nisch | 30179 | 33151 | 30058 | 31587 | 35699 | 34610 | 30486 | 41573 | 50323 | 51319 | 49003 | 54164 | 50240 | 47560 | 28972 | 31563 | 32244 | 32429 | 36629 | 28952 | 32634 | 51315 | 47962 | 48164 | 50104 | 43160 | 48972 | 46687 | Nisch |
| Nit1 | 2952 | 3285 | 3189 | 3294 | 3065 | 2779 | 3305 | 2409 | 2257 | 2378 | 2454 | 2640 | 2459 | 2882 | 3022 | 2679 | 3445 | 3300 | 3290 | 2877 | 3394 | 2486 | 2388 | 2712 | 2849 | 2826 | 2833 | 2709 | Nit1 |
| Nit2 | 1788 | 1498 | 1498 | 1281 | 1191 | 1429 | 1483 | 1937 | 1915 | 1492 | 1849 | 1549 | 1593 | 1823 | 1600 | 1467 | 1350 | 1348 | 1204 | 1513 | 1416 | 1557 | 1939 | 1739 | 1427 | 1845 | 1725 | 1841 | Nit2 |
| Nkain1 | 332 | 300 | 242 | 264 | 287 | 292 | 226 | 393 | 278 | 258 | 336 | 343 | 363 | 336 | 339 | 325 | 293 | 237 | 303 | 294 | 281 | 414 | 278 | 315 | 311 | 306 | 356 | 299 | Nkain1 |
| Nkain2 | 385 | 329 | 172 | 607 | 1116 | 864 | 1123 | 879 | 724 | 676 | 710 | 458 | 367 | 544 | 435 | 266 | 355 | 836 | 967 | 921 | 1074 | 941 | 652 | 768 | 932 | 394 | 323 | 319 | Nkain2 |
| Nkain3 | 916 | 844 | 168 | 393 | 582 | 1381 | 1006 | 1413 | 1214 | 633 | 780 | 530 | 724 | 664 | 1064 | 861 | 216 | 380 | 478 | 1566 | 1361 | 348 | 1022 | 273 | 867 | 160 | 913 | 472 | Nkain3 |
| Nkain4 | 720 | 986 | 964 | 1049 | 1213 | 1064 | 866 | 1305 | 1121 | 1090 | 1343 | 1528 | 1690 | 1281 | 968 | 861 | 1060 | 903 | 1096 | 1142 | 949 | 1285 | 1318 | 1174 | 1418 | 1136 | 1610 | 1373 | Nkain4 |
| Nkap | 3082 | 2825 | 3635 | 3562 | 3614 | 3627 | 3802 | 4344 | 3540 | 3795 | 3459 | 4342 | 3957 | 4255 | 3746 | 2800 | 3465 | 3399 | 3584 | 3784 | 3784 | 4430 | 3504 | 3923 | 4219 | 4042 | 4190 | 4387 | Nkap |
| Nkapd1 | 3609 | 2830 | 3228 | 2902 | 3018 | 3401 | 3637 | 3898 | 3453 | 3680 | 3551 | 3802 | 3802 | 4188 | 3731 | 2950 | 3089 | 3126 | 2873 | 3255 | 3371 | 3598 | 3574 | 3777 | 3580 | 3770 | 3747 | 4069 | Nkapd1 |
| Nkapl | 155 | 158 | 182 | 116 | 134 | 158 | 150 | 95 | 131 | 125 | 91 | 130 | 126 | 116 | 199 | 151 | 125 | 129 | 183 | 201 | 143 | 142 | 101 | 147 | 101 | 121 | 112 | 145 | Nkapl |
| Nkd1 | 6298 | 7535 | 8386 | 7008 | 5552 | 5799 | 4855 | 3496 | 3717 | 3690 | 3193 | 4261 | 4508 | 3772 | 6146 | 8114 | 6887 | 6363 | 6209 | 4945 | 5454 | 3883 | 4357 | 3994 | 3694 | 3888 | 3947 | 4161 | Nkd1 |
| Nkd2 | 1963 | 1570 | 1674 | 771 | 843 | 866 | 568 | 1705 | 1706 | 1344 | 1353 | 1155 | 1850 | 1044 | 1499 | 2123 | 962 | 841 | 768 | 772 | 617 | 964 | 1862 | 1266 | 790 | 1231 | 1399 | 1248 | Nkd2 |
| Nkg7 | 0 | 0 | 2 | 0 | 1 | 0 | 0 | 1 | 2 | 5 | 2 | 0 | 2 | 1 | 0 | 1 | 0 | 3 | 0 | 0 | 0 | 0 | 9 | 6 | 0 | 0 | 2 | 0 | Nkg7 |
| Nkiras1 | 1969 | 2157 | 2020 | 2553 | 2368 | 2263 | 2560 | 1279 | 1098 | 1229 | 1263 | 1095 | 1084 | 1029 | 2243 | 1908 | 2330 | 2638 | 2291 | 2363 | 2455 | 1275 | 1153 | 1173 | 1326 | 1083 | 1020 | 1006 | Nkiras1 |
| Nkiras2 | 4931 | 6420 | 5615 | 5681 | 5056 | 4760 | 4881 | 3694 | 4149 | 4335 | 4001 | 4716 | 4432 | 4174 | 4833 | 5486 | 5591 | 5224 | 5638 | 4797 | 4924 | 4216 | 4328 | 4204 | 4432 | 4903 | 4521 | 4682 | Nkiras2 |
| Nkpd1 | 17 | 18 | 32 | 28 | 29 | 26 | 21 | 6 | 6 | 17 | 11 | 13 | 12 | 3 | 21 | 22 | 39 | 42 | 19 | 14 | 21 | 7 | 12 | 7 | 2 | 6 | 3 | 8 | Nkpd1 |
| Nkrf | 1501 | 1260 | 1632 | 1697 | 1927 | 1687 | 2066 | 2144 | 1710 | 2201 | 1731 | 2033 | 1882 | 2111 | 1633 | 1417 | 1713 | 1751 | 2010 | 1672 | 2051 | 2191 | 1683 | 1975 | 2022 | 1655 | 1958 | 2008 | Nkrf |
| Nktr | 32131 | 26968 | 16416 | 23258 | 25126 | 30642 | 27975 | 27077 | 25367 | 20582 | 23961 | 19931 | 21698 | 21349 | 31204 | 28283 | 19006 | 23929 | 22096 | 30241 | 29730 | 19597 | 23649 | 17584 | 21333 | 11417 | 22636 | 18440 | Nktr |
| Nkx1-1 | 1 | 5 | 6 | 6 | 31 | 8 | 4 | 51 | 40 | 38 | 35 | 8 | 88 | 3 | 13 | 11 | 14 | 6 | 10 | 8 | 4 | 58 | 32 | 1 | 12 | 6 | 10 | 8 | Nkx1-1 |
| Nkx2-1 | 1177 | 1055 | 756 | 1436 | 1545 | 1279 | 2123 | 1749 | 1864 | 2313 | 1767 | 4110 | 145 | 4378 | 840 | 374 | 1965 | 1308 | 1980 | 1130 | 1860 | 3559 | 1034 | 3729 | 3177 | 6092 | 2916 | 2734 | Nkx2-1 |
| Nkx2-2 | 264 | 198 | 513 | 317 | 263 | 271 | 318 | 276 | 157 | 300 | 234 | 651 | 256 | 559 | 200 | 113 | 438 | 217 | 375 | 294 | 296 | 485 | 159 | 453 | 412 | 618 | 610 | 684 | Nkx2-2 |
| Nkx2-3 | 189 | 222 | 118 | 378 | 184 | 129 | 272 | 109 | 132 | 154 | 94 | 147 | 27 | 230 | 144 | 97 | 270 | 281 | 231 | 136 | 258 | 144 | 36 | 209 | 138 | 268 | 129 | 119 | Nkx2-3 |
| Nkx2-4 | 70 | 88 | 130 | 107 | 73 | 71 | 94 | 128 | 68 | 124 | 107 | 152 | 23 | 221 | 51 | 45 | 94 | 69 | 74 | 108 | 113 | 180 | 101 | 179 | 186 | 284 | 191 | 225 | Nkx2-4 |
| Nkx2-5 | 0 | 3 | 1 | 1 | 1 | 0 | 0 | 0 | 1 | 0 | 27 | 0 | 3 | 0 | 0 | 2 | 0 | 3 | 2 | 0 | 0 | 0 | 3 | 0 | 0 | 0 | 0 | 0 | Nkx2-5 |
| Nkx2-6 | 1 | 1 | 0 | 0 | 0 | 0 | 0 | 0 | 0 | 0 | 0 | 0 | 0 | 0 | 1 | 1 | 0 | 0 | 0 | 0 | 0 | 0 | 0 | 0 | 0 | 0 | 0 | 0 | Nkx2-6 |
| Nkx2-9 | 22 | 16 | 49 | 10 | 8 | 17 | 4 | 7 | 7 | 21 | 16 | 37 | 39 | 49 | 22 | 18 | 31 | 4 | 16 | 6 | 7 | 18 | 5 | 18 | 12 | 39 | 56 | 65 | Nkx2-9 |
| Nkx3-1 | 171 | 128 | 163 | 88 | 67 | 57 | 59 | 6 | 11 | 12 | 18 | 15 | 20 | 19 | 145 | 142 | 116 | 109 | 83 | 87 | 73 | 15 | 19 | 16 | 19 | 9 | 13 | 17 | Nkx3-1 |
| Nkx3-2 | 17 | 12 | 6 | 6 | 9 | 8 | 1 | 2 | 29 | 11 | 127 | 1 | 8 | 1 | 13 | 30 | 9 | 12 | 3 | 3 | 0 | 27 | 28 | 11 | 9 | 3 | 10 | 0 | Nkx3-2 |
| Nkx6-1 | 333 | 530 | 1129 | 514 | 686 | 424 | 246 | 514 | 434 | 697 | 620 | 513 | 1413 | 466 | 655 | 741 | 878 | 473 | 334 | 237 | 293 | 847 | 350 | 258 | 309 | 257 | 723 | 600 | Nkx6-1 |
| Nkx6-2 | 218 | 227 | 479 | 215 | 149 | 128 | 78 | 162 | 162 | 197 | 146 | 303 | 333 | 266 | 196 | 266 | 330 | 189 | 178 | 103 | 105 | 224 | 107 | 199 | 176 | 170 | 371 | 366 | Nkx6-2 |
| Nkx6-3 | 0 | 3 | 0 | 1 | 83 | 3 | 1 | 18 | 49 | 63 | 69 | 2 | 6 | 1 | 3 | 0 | 0 | 3 | 2 | 2 | 2 | 22 | 33 | 2 | 2 | 1 | 1 | 1 | Nkx6-3 |
| Nle1 | 1180 | 1572 | 1366 | 1569 | 1207 | 1319 | 1063 | 1122 | 1312 | 1171 | 1296 | 1395 | 1209 | 1261 | 1046 | 1358 | 1380 | 1346 | 1422 | 1210 | 1152 | 1158 | 1217 | 1290 | 1321 | 1454 | 1392 | 1171 | Nle1 |
| Nlgn1 | 1538 | 1551 | 964 | 1942 | 2428 | 2675 | 2829 | 2711 | 2350 | 2105 | 1784 | 1626 | 1473 | 1585 | 2102 | 1349 | 1467 | 2125 | 2295 | 2990 | 2779 | 2433 | 2222 | 1535 | 2448 | 1332 | 1641 | 1392 | Nlgn1 |
| Nlgn2 | 11711 | 14581 | 19030 | 19144 | 19880 | 16509 | 16896 | 11865 | 13176 | 18111 | 16087 | 16523 | 14192 | 15430 | 11028 | 12450 | 21105 | 19785 | 20832 | 14145 | 17608 | 19148 | 12130 | 18884 | 17984 | 16142 | 14563 | 14919 | Nlgn2 |
| Nlgn3 | 1039 | 908 | 1671 | 1993 | 2483 | 2007 | 2249 | 1519 | 1519 | 2295 | 1666 | 1895 | 1615 | 1912 | 1146 | 1053 | 1982 | 2335 | 2311 | 1688 | 2264 | 3016 | 1252 | 2268 | 2452 | 2268 | 1477 | 1721 | Nlgn3 |
| Nlk | 6913 | 6641 | 7202 | 8318 | 7765 | 7257 | 9366 | 5519 | 4626 | 5101 | 4707 | 5860 | 4719 | 6440 | 7457 | 6101 | 8069 | 8053 | 7871 | 7357 | 8843 | 5914 | 4116 | 6083 | 5813 | 6227 | 5439 | 6192 | Nlk |
| Nln | 5388 | 5048 | 4630 | 4489 | 5042 | 5031 | 4910 | 3405 | 3127 | 3242 | 2944 | 2788 | 3113 | 2811 | 5723 | 5010 | 4772 | 4732 | 4676 | 5462 | 5036 | 3501 | 3253 | 2951 | 3072 | 2431 | 2778 | 2732 | Nln |

| | | | | | | | | | | | | | | | | | | | | | | | | | | | | | |
|---|---|---|---|---|---|---|---|---|---|---|---|---|---|---|---|---|---|---|---|---|---|---|---|---|---|---|---|---|---|
| Nlrc3 | 134 | 200 | 159 | 182 | 214 | 191 | 202 | 156 | 157 | 172 | 222 | 212 | 198 | 177 | 173 | 157 | 165 | 179 | 153 | 189 | 247 | 216 | 180 | 175 | 232 | 138 | 212 | 147 | Nlrc3 |
| Nlrc4 | 9 | 13 | 7 | 4 | 13 | 29 | 9 | 13 | 21 | 17 | 24 | 16 | 16 | 22 | 15 | 15 | 13 | 7 | 9 | 10 | 9 | 22 | 36 | 12 | 21 | 24 | 23 | 14 | Nlrc4 |
| Nlrc5 | 13 | 21 | 7 | 7 | 53 | 26 | 19 | 32 | 49 | 34 | 36 | 34 | 22 | 21 | 42 | 38 | 16 | 25 | 19 | 23 | 38 | 62 | 43 | 29 | 44 | 29 | 24 | 31 | Nlrc5 |
| Nlrp10 | 3 | 0 | 0 | 1 | 2 | 0 | 3 | 9 | 2 | 1 | 2 | 0 | 1 | 1 | 1 | 1 | 0 | 0 | 0 | 3 | 0 | 5 | 3 | 3 | 1 | 4 | 1 | 1 | Nlrp10 |
| Nlrp14 | 0 | 0 | 0 | 0 | 0 | 0 | 0 | 0 | 2 | 0 | 0 | 0 | 1 | 0 | 0 | 0 | 0 | 0 | 0 | 0 | 0 | 0 | 0 | 0 | 0 | 0 | 0 | 0 | Nlrp14 |
| Nlrp1a | 1 | 0 | 0 | 0 | 0 | 0 | 0 | 1 | 1 | 0 | 0 | 0 | 1 | 0 | 1 | 0 | 0 | 0 | 0 | 0 | 0 | 0 | 2 | 0 | 0 | 0 | 6 | 1 | Nlrp1a |
| Nlrp1b | 1 | 9 | 0 | 1 | 5 | 2 | 0 | 7 | 5 | 7 | 7 | 2 | 2 | 5 | 0 | 9 | 2 | 2 | 5 | 6 | 4 | 5 | 14 | 8 | 3 | 1 | 0 | 13 | Nlrp1b |
| Nlrp3 | 12 | 16 | 16 | 8 | 15 | 7 | 10 | 22 | 43 | 26 | 39 | 29 | 52 | 51 | 7 | 7 | 12 | 28 | 15 | 15 | 13 | 37 | 50 | 36 | 22 | 17 | 19 | 25 | Nlrp3 |
| Nlrp4a | 0 | 0 | 0 | 0 | 0 | 0 | 0 | 0 | 1 | 0 | 0 | 0 | 0 | 1 | 0 | 0 | 0 | 0 | 0 | 0 | 0 | 0 | 1 | 0 | 1 | 0 | 0 | 0 | Nlrp4a |
| Nlrp4c | 1 | 0 | 0 | 0 | 0 | 0 | 0 | 0 | 0 | 0 | 0 | 1 | 0 | 0 | 1 | 2 | 7 | 0 | 0 | 1 | 0 | 0 | 0 | 0 | 0 | 4 | 0 | 0 | Nlrp4c |
| Nlrp4f | 7 | 5 | 3 | 4 | 5 | 5 | 16 | 2 | 10 | 8 | 13 | 1 | 6 | 10 | 20 | 8 | 5 | 6 | 5 | 7 | 5 | 5 | 3 | 2 | 6 | 9 | 10 | 5 | Nlrp4f |
| Nlrp5 | 0 | 0 | 0 | 0 | 0 | 0 | 0 | 0 | 0 | 0 | 0 | 0 | 0 | 0 | 0 | 2 | 0 | 1 | 0 | 0 | 0 | 0 | 1 | 0 | 0 | 0 | 0 | 0 | Nlrp5 |
| Nlrp6 | 28 | 18 | 15 | 16 | 16 | 17 | 28 | 16 | 13 | 20 | 14 | 24 | 10 | 12 | 18 | 16 | 21 | 19 | 19 | 24 | 25 | 20 | 17 | 16 | 9 | 17 | 6 | 12 | Nlrp6 |
| Nlrp9b | 0 | 0 | 0 | 0 | 0 | 0 | 0 | 0 | 2 | 0 | 0 | 0 | 0 | 0 | 0 | 0 | 0 | 0 | 0 | 0 | 0 | 0 | 1 | 0 | 0 | 0 | 0 | 0 | Nlrp9b |
| Nlrx1 | 131 | 122 | 161 | 169 | 191 | 191 | 143 | 379 | 461 | 411 | 527 | 425 | 422 | 332 | 116 | 149 | 184 | 219 | 167 | 157 | 182 | 354 | 559 | 449 | 411 | 510 | 415 | 363 | Nlrx1 |
| Nmb | 150 | 230 | 162 | 222 | 218 | 225 | 273 | 114 | 134 | 170 | 150 | 165 | 159 | 199 | 167 | 148 | 189 | 204 | 233 | 169 | 267 | 217 | 120 | 195 | 170 | 173 | 151 | 191 | Nmb |
| Nmbr | 30 | 41 | 20 | 27 | 24 | 35 | 22 | 38 | 55 | 36 | 37 | 40 | 39 | 40 | 21 | 29 | 27 | 21 | 30 | 45 | 46 | 77 | 52 | 33 | 43 | 33 | 38 | 43 | Nmbr |
| Nmd3 | 4383 | 3612 | 3669 | 3840 | 3247 | 3478 | 3941 | 3226 | 2901 | 2593 | 2723 | 3014 | 2967 | 3025 | 4469 | 3987 | 3600 | 3516 | 3397 | 4050 | 3486 | 2769 | 2924 | 2865 | 2744 | 3305 | 3118 | 3418 | Nmd3 |
| Nme1 | 2005 | 1883 | 1486 | 1866 | 1939 | 1738 | 2390 | 2668 | 1896 | 2111 | 1796 | 1892 | 1714 | 2587 | 2254 | 1494 | 1899 | 1812 | 1842 | 2183 | 2290 | 2386 | 1805 | 2142 | 2074 | 2062 | 1929 | 2680 | Nme1 |
| Nme2 | 85 | 59 | 99 | 105 | 120 | 100 | 122 | 49 | 49 | 49 | 62 | 64 | 56 | 60 | 94 | 51 | 89 | 87 | 100 | 105 | 88 | 89 | 45 | 64 | 97 | 42 | 65 | 82 | Nme2 |
| Nme4 | 1065 | 1037 | 973 | 693 | 570 | 694 | 624 | 1089 | 1269 | 1103 | 1199 | 1113 | 1245 | 1166 | 947 | 1049 | 875 | 695 | 623 | 710 | 634 | 762 | 1326 | 1235 | 812 | 1327 | 1188 | 1253 | Nme4 |
| Nme5 | 291 | 289 | 249 | 350 | 240 | 245 | 273 | 167 | 162 | 153 | 157 | 191 | 248 | 125 | 316 | 298 | 256 | 316 | 273 | 292 | 293 | 247 | 234 | 160 | 166 | 127 | 134 | 165 | Nme5 |
| Nme6 | 863 | 942 | 1011 | 804 | 757 | 674 | 808 | 1202 | 1191 | 980 | 1053 | 1341 | 1275 | 1361 | 878 | 757 | 917 | 745 | 832 | 927 | 760 | 1111 | 1192 | 1325 | 1095 | 1695 | 1603 | 1466 | Nme6 |
| Nme7 | 2208 | 1991 | 2001 | 1841 | 1537 | 1584 | 2008 | 1834 | 1755 | 1488 | 1365 | 1706 | 1462 | 1842 | 2072 | 2004 | 1793 | 1717 | 1850 | 2105 | 1827 | 1322 | 1621 | 1705 | 1496 | 2070 | 1655 | 1966 | Nme7 |
| Nme8 | 2 | 6 | 2 | 1 | 7 | 2 | 7 | 4 | 7 | 3 | 4 | 1 | 0 | 1 | 13 | 8 | 8 | 6 | 6 | 8 | 2 | 5 | 10 | 2 | 3 | 3 | 8 | 0 | Nme8 |
| Nme9 | 13 | 27 | 2 | 15 | 45 | 30 | 13 | 15 | 30 | 36 | 26 | 25 | 27 | 17 | 26 | 51 | 24 | 42 | 24 | 30 | 25 | 13 | 43 | 11 | 16 | 14 | 7 | 12 | Nme9 |
| Nmi | 136 | 95 | 121 | 51 | 77 | 92 | 65 | 244 | 261 | 265 | 258 | 234 | 266 | 233 | 141 | 138 | 56 | 68 | 129 | 100 | 90 | 197 | 330 | 239 | 180 | 293 | 277 | 252 | Nmi |
| Nmnat1 | 158 | 200 | 203 | 197 | 200 | 179 | 178 | 629 | 677 | 616 | 610 | 733 | 649 | 868 | 189 | 139 | 261 | 171 | 207 | 211 | 190 | 617 | 680 | 820 | 740 | 940 | 875 | 800 | Nmnat1 |
| Nmnat2 | 2249 | 3087 | 2693 | 4412 | 6374 | 4932 | 5022 | 3927 | 4066 | 4920 | 4121 | 3882 | 3633 | 3764 | 3237 | 2997 | 3955 | 5197 | 5191 | 4395 | 5316 | 6400 | 3896 | 4502 | 5882 | 3092 | 3199 | 3503 | Nmnat2 |
| Nmnat3 | 296 | 247 | 224 | 334 | 186 | 200 | 196 | 94 | 116 | 109 | 101 | 108 | 110 | 122 | 219 | 206 | 246 | 211 | 251 | 168 | 206 | 67 | 106 | 127 | 87 | 119 | 120 | 122 | Nmnat3 |
| Nmral1 | 4451 | 5294 | 5186 | 4211 | 3471 | 4138 | 3756 | 4496 | 4486 | 4225 | 3982 | 5861 | 5314 | 5667 | 3683 | 4267 | 4825 | 3785 | 4868 | 4406 | 4035 | 3278 | 3855 | 5026 | 4646 | 6892 | 6143 | 5445 | Nmral1 |
| Nmrk1 | 743 | 599 | 636 | 520 | 514 | 542 | 394 | 383 | 481 | 410 | 469 | 311 | 378 | 268 | 661 | 785 | 470 | 551 | 422 | 407 | 388 | 339 | 419 | 322 | 272 | 279 | 279 | 298 | Nmrk1 |
| Nmrk2 | 17 | 32 | 23 | 48 | 45 | 16 | 27 | 5 | 1 | 2 | 16 | 9 | 0 | 1 | 17 | 20 | 36 | 46 | 50 | 45 | 42 | 12 | 7 | 9 | 3 | 11 | 2 | 2 | Nmrk2 |
| Nms | 15 | 7 | 14 | 2 | 5 | 3 | 12 | 9 | 5 | 21 | 8 | 4 | 14 | 3 | 15 | 17 | 9 | 3 | 6 | 6 | 4 | 3 | 10 | 3 | 11 | 10 | 13 | 8 | Nms |
| Nmt1 | 7827 | 8324 | 8082 | 7138 | 6793 | 6585 | 6881 | 6791 | 6522 | 6724 | 6212 | 7114 | 6823 | 7216 | 7184 | 7499 | 7509 | 7477 | 7305 | 6778 | 7098 | 6605 | 6502 | 7104 | 6753 | 7628 | 7296 | 7332 | Nmt1 |
| Nmt2 | 13506 | 11759 | 11814 | 12863 | 13209 | 13581 | 14902 | 9432 | 8062 | 8482 | 7884 | 8568 | 8471 | 9297 | 13508 | 11831 | 11514 | 12744 | 12447 | 13798 | 14551 | 8736 | 7445 | 7598 | 9220 | 7177 | 8796 | 8710 | Nmt2 |
| Nmu | 7 | 10 | 5 | 5 | 3 | 0 | 1 | 1 | 1 | 0 | 10 | 1 | 2 | 0 | 5 | 4 | 7 | 4 | 2 | 1 | 5 | 3 | 4 | 0 | 0 | 0 | 7 | 1 | Nmu |
| Nmur1 | 7 | 3 | 0 | 2 | 2 | 1 | 0 | 4 | 8 | 1 | 8 | 0 | 1 | 3 | 4 | 6 | 7 | 0 | 2 | 1 | 0 | 2 | 5 | 4 | 0 | 2 | 3 | 0 | Nmur1 |
| Nmur2 | 5 | 9 | 3 | 13 | 16 | 17 | 17 | 4 | 9 | 4 | 13 | 2 | 3 | 6 | 5 | 6 | 5 | 20 | 3 | 16 | 9 | 5 | 4 | 1 | 10 | 4 | 5 | 0 | Nmur2 |
| Nnat | 77766 | 101529 | 70497 | 109486 | 108659 | 101148 | 131245 | 110235 | 101034 | 109208 | 82871 | 105318 | 72583 | 128902 | 88481 | 62529 | 101136 | 104321 | 121363 | 122403 | 141850 | 125550 | 91715 | 120591 | 126504 | 113771 | 98583 | 113703 | Nnat |
| Nnmt | 24 | 34 | 36 | 30 | 15 | 58 | 11 | 38 | 75 | 70 | 98 | 70 | 74 | 60 | 50 | 48 | 25 | 26 | 15 | 17 | 13 | 40 | 95 | 57 | 46 | 66 | 71 | 76 | Nnmt |
| Nnt | 182 | 105 | 81 | 92 | 78 | 117 | 107 | 150 | 109 | 65 | 90 | 127 | 111 | 129 | 173 | 104 | 60 | 58 | 65 | 146 | 126 | 74 | 92 | 97 | 94 | 71 | 157 | 121 | Nnt |
| Noa1 | 2528 | 2784 | 2695 | 3160 | 2430 | 2328 | 2566 | 1360 | 1352 | 1518 | 1416 | 1582 | 1518 | 1549 | 2429 | 2557 | 2802 | 2932 | 2682 | 2180 | 2639 | 1546 | 1254 | 1531 | 1580 | 1770 | 1553 | 1613 | Noa1 |
| Nob1 | 2553 | 3391 | 2824 | 2925 | 2518 | 2461 | 2396 | 2831 | 3142 | 3128 | 2862 | 2994 | 2918 | 3185 | 2322 | 2709 | 2758 | 2796 | 2745 | 2766 | 2496 | 2677 | 2871 | 3117 | 2995 | 3455 | 3256 | 3197 | Nob1 |
| Nobox | 5 | 1 | 0 | 4 | 0 | 1 | 3 | 4 | 3 | 4 | 3 | 0 | 0 | 6 | 1 | 1 | 0 | 2 | 2 | 0 | 2 | 2 | 3 | 5 | 5 | 0 | 0 | 1 | Nobox |
| Noc2l | 8436 | 9606 | 10335 | 9742 | 8106 | 8615 | 8801 | 7284 | 7813 | 7821 | 8089 | 9072 | 8469 | 8773 | 7801 | 8165 | 9938 | 9198 | 9503 | 7716 | 8703 | 7541 | 7646 | 9031 | 8376 | 10457 | 8883 | 9174 | Noc2l |
| Noc3l | 4265 | 4222 | 3993 | 4053 | 3121 | 3262 | 3784 | 3687 | 3151 | 3117 | 3128 | 3636 | 3299 | 3792 | 4296 | 4268 | 3905 | 3899 | 3324 | 3955 | 3673 | 3071 | 3086 | 3203 | 3404 | 3498 | 3468 | 3957 | Noc3l |
| Noc4l | 2524 | 3289 | 2625 | 2443 | 2036 | 2006 | 2105 | 1506 | 1893 | 1591 | 1593 | 1928 | 1897 | 1818 | 2478 | 2685 | 2457 | 2170 | 2146 | 2144 | 1994 | 1355 | 1532 | 1709 | 1702 | 2249 | 1826 | 1963 | Noc4l |
| Noct | 1979 | 2099 | 3059 | 3578 | 3621 | 3038 | 3352 | 1649 | 1656 | 2296 | 2245 | 1939 | 1937 | 1862 | 2057 | 2171 | 3284 | 3935 | 3516 | 2304 | 3372 | 2836 | 1718 | 2037 | 2243 | 1982 | 1619 | 1918 | Noct |
| Nod1 | 257 | 355 | 307 | 264 | 260 | 325 | 144 | 307 | 451 | 396 | 564 | 341 | 488 | 339 | 270 | 371 | 293 | 285 | 203 | 198 | 201 | 434 | 479 | 378 | 400 | 365 | 342 | 275 | Nod1 |
| Nod2 | 268 | 242 | 245 | 296 | 216 | 220 | 177 | 44 | 46 | 59 | 35 | 18 | 43 | 26 | 218 | 277 | 228 | 235 | 179 | 200 | 226 | 25 | 53 | 56 | 31 | 53 | 23 | 27 | Nod2 |
| Nodal | 9 | 20 | 18 | 2 | 16 | 4 | 7 | 1 | 2 | 7 | 4 | 4 | 5 | 2 | 13 | 21 | 3 | 14 | 15 | 8 | 2 | 0 | 3 | 2 | 3 | 1 | 2 | 4 | Nodal |
| Nog | 366 | 305 | 326 | 375 | 403 | 369 | 344 | 484 | 433 | 631 | 591 | 635 | 431 | 639 | 310 | 232 | 394 | 331 | 394 | 386 | 354 | 465 | 432 | 625 | 482 | 659 | 485 | 533 | Nog |
| Nol10 | 2982 | 2776 | 3526 | 2936 | 2639 | 2582 | 2875 | 2689 | 2664 | 2723 | 2710 | 3305 | 3003 | 3295 | 2900 | 2692 | 3008 | 2761 | 2855 | 2410 | 2552 | 2573 | 2540 | 2813 | 2917 | 3736 | 3178 | 3431 | Nol10 |
| Nol11 | 5420 | 4749 | 4866 | 4635 | 3976 | 4261 | 4737 | 5189 | 4804 | 4557 | 4330 | 5407 | 5277 | 5848 | 5374 | 5123 | 4695 | 4342 | 4136 | 4889 | 4580 | 4446 | 4849 | 4976 | 4740 | 6034 | 5523 | 5648 | Nol11 |
| Nol12 | 1235 | 1748 | 1690 | 1508 | 1376 | 1338 | 1373 | 1479 | 1778 | 1775 | 1667 | 1691 | 1655 | 1777 | 1292 | 1322 | 1574 | 1372 | 1555 | 1400 | 1358 | 1544 | 1656 | 1883 | 1774 | 2003 | 1721 | 1881 | Nol12 |
| Nol3 | 125 | 138 | 200 | 151 | 157 | 139 | 127 | 133 | 169 | 142 | 185 | 179 | 186 | 156 | 183 | 164 | 175 | 163 | 133 | 205 | 142 | 209 | 217 | 191 | 179 | 163 | 264 | 223 | Nol3 |
| Nol4 | 2737 | 2737 | 2186 | 4385 | 5608 | 5084 | 6354 | 4164 | 3546 | 3674 | 2955 | 3391 | 2012 | 3776 | 3185 | 1866 | 3500 | 4686 | 5579 | 5794 | 6205 | 4313 | 3604 | 4094 | 4493 | 3448 | 2983 | 3151 | Nol4 |
| Nol4l | 4601 | 5008 | 5540 | 6951 | 8718 | 6866 | 7546 | 6704 | 6910 | 9038 | 7462 | 8093 | 6700 | 7793 | 4813 | 4961 | 6671 | 7134 | 8173 | 6041 | 7766 | 10330 | 6246 | 8199 | 9383 | 7318 | 6267 | 7303 | Nol4l |
| Nol6 | 5144 | 6210 | 6167 | 6431 | 5858 | 5527 | 5669 | 3662 | 4926 | 4828 | 5112 | 5264 | 4784 | 4880 | 4669 | 5957 | 6187 | 6852 | 6156 | 4586 | 5597 | 4974 | 4423 | 5355 | 4993 | 5390 | 4305 | 4690 | Nol6 |
| Nol7 | 3821 | 3456 | 4267 | 3952 | 3453 | 3613 | 4342 | 5249 | 4139 | 4589 | 4493 | 5855 | 5080 | 6708 | 3880 | 3259 | 4076 | 3519 | 3677 | 4313 | 4026 | 4319 | 4154 | 5547 | 5123 | 6698 | 5915 | 6500 | Nol7 |
| Nol8 | 3565 | 3251 | 3502 | 3297 | 2867 | 3165 | 3464 | 4604 | 3798 | 3635 | 3353 | 4236 | 3950 | 4181 | 3825 | 3678 | 3232 | 3320 | 2992 | 3755 | 3100 | 3596 | 3765 | 4000 | 3901 | 4020 | 4350 | 4146 | Nol8 |
| Nol9 | 3301 | 3560 | 3778 | 3765 | 3406 | 3405 | 3315 | 2860 | 3193 | 3416 | 3395 | 3837 | 3525 | 3527 | 3107 | 3545 | 3727 | 3674 | 3552 | 2735 | 3382 | 3192 | 3146 | 3630 | 3441 | 4135 | 3521 | 3787 | Nol9 |
| Nolc1 | 13823 | 15844 | 14557 | 13458 | 10554 | 11961 | 11447 | 8193 | 8238 | 8503 | 7856 | 9963 | 9461 | 9992 | 13002 | 13285 | 13428 | 12386 | 11588 | 12253 | 11287 | 7384 | 7849 | 9198 | 8823 | 11110 | 10343 | 9760 | Nolc1 |
| Nom1 | 3655 | 3586 | 3581 | 3552 | 2943 | 3496 | 3385 | 3059 | 2827 | 2743 | 2770 | 3189 | 3248 | 3529 | 3415 | 3321 | 3311 | 3262 | 3129 | 3135 | 3268 | 2841 | 2772 | 2993 | 2936 | 3447 | 3228 | 3345 | Nom1 |
| Nomo1 | 6812 | 8370 | 8980 | 9794 | 8269 | 8162 | 7483 | 3714 | 4617 | 5696 | 5040 | 5391 | 4908 | 4758 | 6380 | 7997 | 8960 | 9783 | 9149 | 6277 | 8209 | 5444 | 4442 | 5496 | 5763 | 5372 | 4676 | 5042 | Nomo1 |
| Nono | 51661 | 48387 | 48664 | 42420 | 39798 | 42928 | 47158 | 59353 | 50761 | 48988 | 45997 | 60175 | 54079 | 65154 | 51651 | 42988 | 46554 | 39568 | 44807 | 48742 | 44992 | 46630 | 49084 | 57686 | 52960 | 67204 | 63837 | 66219 | Nono |
| Nop10 | 3848 | 4108 | 4116 | 3848 | 2942 | 3202 | 4520 | 2623 | 2414 | 2351 | 2497 | 2357 | 2300 | 3261 | 3955 | 3048 | 3931 | 3389 | 3384 | 4112 | 4265 | 1988 | 2414 | 2716 | 2565 | 3027 | 2760 | 3475 | Nop10 |
| Nop14 | 5339 | 5105 | 5702 | 4849 | 4349 | 4526 | 4840 | 3799 | 3455 | 3627 | 3484 | 3963 | 3635 | 4197 | 5269 | 4932 | 5000 | 4858 | 4606 | 4821 | 4629 | 3215 | 3333 | 3814 | 3560 | 4144 | 4278 | 4049 | Nop14 |
| Nop16 | 2971 | 3121 | 3416 | 3090 | 2587 | 2617 | 2527 | 1896 | 2104 | 2036 | 2237 | 2180 | 2201 | 2245 | 2709 | 3065 | 3212 | 2977 | 2798 | 2920 | 2607 | 1676 | 2152 | 2338 | 2052 | 2878 | 2281 | 2295 | Nop16 |
| Nop2 | 4350 | 5638 | 4659 | 4751 | 3778 | 4257 | 3863 | 3357 | 3674 | 3568 | 3617 | 4013 | 3674 | 3814 | 3973 | 4735 | 4459 | 4596 | 4286 | 4015 | 4046 | 3039 | 3493 | 3845 | 3611 | 4490 | 3908 | 4291 | Nop2 |
| Nop53 | 3620 | 4039 | 4198 | 3979 | 3453 | 3381 | 3447 | 4544 | 5439 | 5990 | 5557 | 6176 | 5812 | 6504 | 3386 | 3703 | 4084 | 3690 | 3892 | 3126 | 3477 | 4824 | 5511 | 6426 | 5908 | 7153 | 6068 | 6412 | Nop53 |
| Nop56 | 11345 | 11674 | 11830 | 10784 | 10169 | 10079 | 11199 | 13577 | 12603 | 11965 | 11580 | 14388 | 12528 | 14577 | 11579 | 10761 | 11288 | 10568 | 10748 | 12313 | 10702 | 11327 | 11923 | 13586 | 12079 | 16847 | 14126 | 15145 | Nop56 |
| Nop58 | 14425 | 11756 | 13843 | 11751 | 9816 | 11168 | 12231 | 12688 | 10903 | 10549 | 10769 | 12545 | 12027 | 13887 | 14400 | 12316 | 12305 | 11054 | 10334 | 13440 | 11312 | 9962 | 10424 | 12146 | 11048 | 15315 | 13567 | 14589 | Nop58 |
| Nop9 | 4730 | 5083 | 4816 | 4639 | 3785 | 4083 | 4244 | 2823 | 2844 | 2724 | 2822 | 3304 | 3289 | 3396 | 4121 | 4306 | 4705 | 4394 | 4274 | 3823 | 4317 | 2476 | 2676 | 3262 | 3172 | 3964 | 3348 | 3713 | Nop9 |
| Nopchap1 | 13329 | 11936 | 10660 | 12989 | 11392 | 13751 | 14377 | 5847 | 5447 | 5536 | 5555 | 6351 | 6214 | 6757 | 11880 | 12101 | 10792 | 11944 | 11270 | 11662 | 14500 | 5237 | 4733 | 5379 | 6144 | 6054 | 6878 | 6299 | Nopchap1 |
| Nos1 | 155 | 199 | 124 | 272 | 776 | 504 | 462 | 539 | 889 | 941 | 649 | 316 | 592 | 316 | 306 | 348 | 289 | 454 | 507 | 370 | 554 | 1126 | 689 | 458 | 953 | 163 | 259 | 249 | Nos1 |
| Nos1ap | 480 | 536 | 419 | 603 | 746 | 711 | 698 | 578 | 720 | 748 | 696 | 729 | 515 | 733 | 440 | 467 | 513 | 671 | 784 | 744 | 757 | 722 | 633 | 686 | 968 | 461 | 607 | 592 | Nos1ap |
| Nos2 | 57 | 41 | 68 | 24 | 77 | 65 | 36 | 81 | 90 | 164 | 142 | 118 | 178 | 102 | 44 | 46 | 42 | 55 | 37 | 35 | 52 | 109 | 123 | 158 | 109 | 127 | 115 | 94 | Nos2 |

| | | | | | | | | | | | | | | | | | | | | | | | | | | | | | |
|---|---|---|---|---|---|---|---|---|---|---|---|---|---|---|---|---|---|---|---|---|---|---|---|---|---|---|---|---|---|
| Nos3 | 338 | 405 | 449 | 385 | 443 | 379 | 298 | 482 | 704 | 864 | 778 | 841 | 974 | 681 | 361 | 447 | 385 | 370 | 392 | 325 | 388 | 795 | 768 | 874 | 860 | 1045 | 895 | 716 | Nos3 |
| Nosip | 1345 | 1479 | 1394 | 1296 | 1440 | 1401 | 1199 | 1994 | 2329 | 2254 | 2182 | 2382 | 2080 | 2037 | 1290 | 1299 | 1413 | 1294 | 1516 | 1471 | 1321 | 1866 | 2295 | 2315 | 2276 | 2600 | 2221 | 2224 | Nosip |
| Nostrin | 118 | 83 | 82 | 70 | 83 | 90 | 96 | 64 | 59 | 74 | 55 | 58 | 53 | 77 | 115 | 77 | 105 | 81 | 90 | 105 | 88 | 60 | 74 | 56 | 57 | 53 | 59 | 51 | Nostrin |
| Notch1 | 8010 | 12653 | 13790 | 12973 | 10735 | 10270 | 9162 | 8983 | 10631 | 12550 | 10057 | 19348 | 13347 | 16117 | 8596 | 10608 | 15433 | 12309 | 12757 | 8637 | 9902 | 11617 | 9207 | 15656 | 13163 | 12547 | 15297 | 15618 | Notch1 |
| Notch2 | 10947 | 10020 | 8623 | 10411 | 9130 | 10244 | 10169 | 7820 | 10931 | 11131 | 11674 | 10786 | 11118 | 10191 | 9034 | 11083 | 9062 | 11624 | 8937 | 8514 | 9738 | 9307 | 10910 | 10561 | 9189 | 8614 | 9434 | 9254 | Notch2 |
| Notch3 | 4511 | 5708 | 6855 | 5164 | 4489 | 4628 | 3365 | 6974 | 8178 | 8725 | 8895 | 12658 | 11991 | 10252 | 4269 | 5789 | 6256 | 5050 | 5047 | 3379 | 3643 | 8280 | 7917 | 10493 | 8059 | 10693 | 12196 | 11515 | Notch3 |
| Notch4 | 287 | 358 | 459 | 348 | 508 | 381 | 344 | 773 | 892 | 1245 | 1121 | 1271 | 1208 | 1227 | 351 | 318 | 445 | 380 | 428 | 308 | 351 | 1158 | 931 | 1245 | 1122 | 1314 | 1278 | 1163 | Notch4 |
| Noto | 0 | 0 | 0 | 0 | 0 | 0 | 2 | 0 | 0 | 0 | 0 | 0 | 0 | 0 | 0 | 0 | 0 | 0 | 2 | 0 | 1 | 0 | 0 | 0 | 0 | 0 | 0 | 0 | Noto |
| Notum | 95 | 96 | 95 | 77 | 84 | 73 | 61 | 112 | 135 | 96 | 189 | 61 | 70 | 46 | 86 | 119 | 133 | 81 | 85 | 68 | 51 | 53 | 150 | 80 | 91 | 80 | 56 | 58 | Notum |
| Nova1 | 4451 | 4162 | 5329 | 7385 | 10353 | 8846 | 8472 | 16212 | 11543 | 14401 | 12302 | 12277 | 13058 | 13201 | 5975 | 4091 | 6238 | 7383 | 8969 | 8563 | 9492 | 19682 | 10895 | 11802 | 15432 | 8461 | 13457 | 13200 | Nova1 |
| Nova2 | 5072 | 7229 | 11202 | 12925 | 12047 | 9857 | 10105 | 6567 | 6818 | 11014 | 8032 | 12611 | 9197 | 11359 | 6074 | 5640 | 12683 | 12049 | 12699 | 7941 | 12094 | 12841 | 6093 | 12079 | 13201 | 12053 | 9668 | 11818 | Nova2 |
| Nox1 | 16 | 10 | 8 | 12 | 10 | 9 | 13 | 6 | 2 | 3 | 0 | 2 | 3 | 0 | 12 | 6 | 9 | 4 | 23 | 2 | 8 | 0 | 1 | 1 | 0 | 8 | 8 | 0 | Nox1 |
| Nox3 | 2 | 5 | 0 | 0 | 0 | 0 | 0 | 1 | 49 | 8 | 8 | 2 | 14 | 0 | 1 | 0 | 0 | 0 | 16 | 0 | 2 | 22 | 11 | 7 | 19 | 0 | 0 | 1 | Nox3 |
| Nox4 | 239 | 154 | 177 | 166 | 178 | 203 | 134 | 499 | 554 | 445 | 454 | 529 | 603 | 445 | 240 | 238 | 163 | 141 | 140 | 156 | 142 | 466 | 563 | 401 | 441 | 455 | 493 | 478 | Nox4 |
| Noxa1 | 10 | 10 | 7 | 2 | 11 | 7 | 7 | 5 | 13 | 16 | 8 | 1 | 7 | 1 | 4 | 13 | 0 | 1 | 4 | 4 | 2 | 2 | 6 | 17 | 2 | 8 | 6 | 0 | Noxa1 |
| Noxo1 | 175 | 181 | 247 | 258 | 259 | 253 | 206 | 276 | 316 | 304 | 278 | 249 | 337 | 258 | 210 | 210 | 247 | 245 | 250 | 232 | 252 | 326 | 275 | 312 | 309 | 340 | 257 | 275 | Noxo1 |
| Noxred1 | 360 | 427 | 272 | 401 | 353 | 384 | 359 | 238 | 195 | 210 | 202 | 201 | 177 | 195 | 334 | 430 | 313 | 406 | 369 | 389 | 426 | 302 | 197 | 196 | 224 | 128 | 229 | 165 | Noxred1 |
| Npas1 | 171 | 226 | 208 | 402 | 335 | 342 | 377 | 53 | 95 | 136 | 64 | 88 | 33 | 79 | 179 | 191 | 351 | 460 | 378 | 308 | 343 | 122 | 74 | 118 | 152 | 87 | 40 | 63 | Npas1 |
| Npas2 | 32 | 13 | 5 | 30 | 34 | 36 | 34 | 74 | 63 | 82 | 101 | 21 | 17 | 27 | 13 | 16 | 9 | 23 | 25 | 24 | 25 | 57 | 96 | 112 | 31 | 89 | 8 | 19 | Npas2 |
| Npas3 | 2028 | 2046 | 3141 | 3479 | 3378 | 3363 | 3434 | 2458 | 2090 | 2636 | 2406 | 3091 | 2699 | 3175 | 2368 | 1940 | 3335 | 3368 | 3416 | 2675 | 3238 | 3751 | 2047 | 2944 | 3418 | 2917 | 2811 | 3459 | Npas3 |
| Npas4 | 113 | 153 | 234 | 232 | 309 | 192 | 239 | 188 | 180 | 258 | 227 | 229 | 164 | 264 | 117 | 118 | 234 | 241 | 239 | 154 | 251 | 389 | 158 | 283 | 315 | 195 | 192 | 165 | Npas4 |
| Npat | 8394 | 7054 | 5149 | 6147 | 5241 | 6515 | 6423 | 5062 | 4372 | 3693 | 4104 | 4583 | 4469 | 4323 | 7854 | 7636 | 5543 | 5989 | 5248 | 6859 | 6122 | 3638 | 4336 | 3666 | 3792 | 3881 | 4659 | 4441 | Npat |
| Npb | 15 | 22 | 13 | 34 | 20 | 16 | 17 | 10 | 12 | 11 | 9 | 23 | 16 | 14 | 8 | 15 | 10 | 10 | 8 | 17 | 8 | 12 | 15 | 14 | 7 | 8 | 12 | 26 | Npb |
| Npbwr1 | 9 | 14 | 8 | 19 | 38 | 17 | 21 | 2 | 16 | 8 | 3 | 2 | 12 | 9 | 26 | 6 | 7 | 10 | 18 | 21 | 31 | 27 | 23 | 2 | 9 | 1 | 3 | 6 | Npbwr1 |
| Npc1 | 5974 | 6350 | 6176 | 6970 | 7103 | 7064 | 6496 | 3403 | 3611 | 4144 | 4077 | 3629 | 4187 | 3454 | 5811 | 6343 | 6452 | 6994 | 7097 | 5674 | 7047 | 4361 | 3639 | 3482 | 3956 | 3065 | 3485 | 3513 | Npc1 |
| Npc1l1 | 42 | 55 | 42 | 75 | 65 | 57 | 76 | 5 | 12 | 9 | 11 | 20 | 7 | 5 | 40 | 32 | 79 | 84 | 42 | 28 | 51 | 12 | 12 | 21 | 20 | 1 | 12 | 9 | Npc1l1 |
| Npc2 | 12696 | 13686 | 14144 | 12366 | 11305 | 12447 | 11451 | 9699 | 9905 | 9394 | 9622 | 10873 | 10990 | 10646 | 12753 | 12930 | 13364 | 12425 | 12275 | 12607 | 11496 | 8627 | 10392 | 10447 | 10158 | 12211 | 12167 | 11776 | Npc2 |
| Npdc1 | 4202 | 6308 | 5506 | 7778 | 9431 | 8054 | 7936 | 6837 | 7524 | 8833 | 7145 | 8214 | 7176 | 7562 | 4770 | 4751 | 7204 | 8111 | 9909 | 7496 | 8342 | 9618 | 7083 | 9063 | 9997 | 7075 | 7017 | 6896 | Npdc1 |
| Npepl1 | 1141 | 1382 | 1258 | 1077 | 1242 | 1213 | 924 | 1870 | 2292 | 1945 | 2372 | 2259 | 2163 | 1966 | 1068 | 1274 | 1170 | 1010 | 1084 | 862 | 1045 | 1688 | 2190 | 2151 | 1909 | 2643 | 2407 | 2095 | Npepl1 |
| Npepps | 15282 | 14098 | 18969 | 19517 | 20684 | 18561 | 21101 | 15736 | 14595 | 18103 | 16240 | 18394 | 15854 | 18488 | 15141 | 13841 | 20163 | 19908 | 21766 | 15622 | 21156 | 20307 | 14457 | 18941 | 19611 | 17417 | 16080 | 19039 | Npepps |
| Npff | 107 | 128 | 160 | 175 | 138 | 112 | 145 | 85 | 93 | 101 | 166 | 131 | 127 | 134 | 99 | 125 | 155 | 158 | 132 | 120 | 179 | 190 | 114 | 145 | 127 | 170 | 113 | 147 | Npff |
| Npffr1 | 33 | 31 | 33 | 21 | 49 | 36 | 39 | 51 | 36 | 42 | 29 | 48 | 24 | 19 | 30 | 31 | 23 | 33 | 52 | 51 | 49 | 57 | 39 | 22 | 57 | 41 | 34 | 28 | Npffr1 |
| Npffr2 | 16 | 14 | 5 | 7 | 7 | 8 | 9 | 5 | 12 | 6 | 1 | 2 | 15 | 5 | 12 | 15 | 16 | 7 | 7 | 18 | 5 | 12 | 10 | 1 | 0 | 3 | 2 | 2 | Npffr2 |
| Nphp1 | 2024 | 1972 | 2040 | 1484 | 1432 | 1516 | 1526 | 2318 | 2152 | 1999 | 1953 | 2462 | 2275 | 2620 | 2041 | 1802 | 1742 | 1369 | 1486 | 1766 | 1445 | 1840 | 2104 | 2147 | 1986 | 2724 | 2753 | 2782 | Nphp1 |
| Nphp3 | 2701 | 2869 | 2859 | 2592 | 2342 | 2488 | 2496 | 1126 | 1325 | 1259 | 1562 | 1493 | 1503 | 1421 | 2418 | 2921 | 2606 | 2595 | 2258 | 2139 | 2245 | 1073 | 1322 | 1213 | 1162 | 1246 | 1484 | 1399 | Nphp3 |
| Nphp4 | 1006 | 1426 | 1688 | 1657 | 1465 | 1269 | 1101 | 669 | 846 | 978 | 834 | 1222 | 1167 | 978 | 1054 | 1292 | 1663 | 1435 | 1444 | 1069 | 1214 | 1024 | 790 | 994 | 993 | 990 | 1005 | 994 | Nphp4 |
| Nphs1 | 33 | 400 | 45 | 342 | 223 | 327 | 13 | 248 | 331 | 331 | 255 | 49 | 1058 | 38 | 236 | 522 | 346 | 162 | 107 | 233 | 38 | 343 | 307 | 47 | 279 | 41 | 26 | 55 | Nphs1 |
| Nphs2 | 8 | 6 | 5 | 27 | 13 | 17 | 12 | 17 | 14 | 15 | 14 | 26 | 42 | 13 | 24 | 17 | 12 | 9 | 6 | 8 | 12 | 23 | 17 | 25 | 27 | 23 | 23 | 32 | Nphs2 |
| Npl | 1032 | 1183 | 1226 | 920 | 690 | 806 | 824 | 463 | 498 | 393 | 407 | 508 | 540 | 488 | 1296 | 1217 | 1045 | 891 | 813 | 1002 | 784 | 461 | 596 | 409 | 360 | 621 | 503 | 635 | Npl |
| Nploc4 | 7304 | 7740 | 6851 | 7536 | 6689 | 6981 | 6786 | 5431 | 5689 | 6165 | 5412 | 7177 | 6273 | 6281 | 6980 | 6977 | 7132 | 7158 | 7251 | 5661 | 6677 | 5930 | 5526 | 6187 | 6179 | 6765 | 6313 | 6820 | Nploc4 |
| Npm1 | 69279 | 53165 | 73106 | 54169 | 43359 | 48261 | 55673 | 62457 | 55853 | 54774 | 56967 | 65482 | 62920 | 76469 | 67517 | 63963 | 62625 | 49974 | 49809 | 63618 | 50411 | 47046 | 58330 | 64031 | 53275 | 87768 | 73156 | 79150 | Npm1 |
| Npm2 | 242 | 208 | 191 | 196 | 169 | 233 | 192 | 117 | 97 | 164 | 145 | 124 | 134 | 133 | 175 | 202 | 192 | 229 | 228 | 197 | 221 | 149 | 82 | 119 | 94 | 146 | 129 | 155 | Npm2 |
| Npm3 | 2593 | 2802 | 2730 | 2253 | 1739 | 1784 | 2451 | 1691 | 1793 | 1765 | 1843 | 1702 | 1660 | 2253 | 2217 | 2192 | 2542 | 2300 | 2206 | 2059 | 2423 | 1263 | 1848 | 2237 | 1697 | 2614 | 1997 | 2403 | Npm3 |
| Npnt | 3117 | 2909 | 2427 | 2130 | 2950 | 2591 | 1535 | 3667 | 4306 | 3576 | 4145 | 2860 | 3680 | 1506 | 2556 | 3925 | 2042 | 2451 | 2381 | 2549 | 2214 | 3796 | 5981 | 2323 | 2661 | 1526 | 1686 | 1520 | Npnt |
| Nppa | 2 | 2 | 2 | 12 | 2 | 4 | 10 | 12 | 11 | 18 | 21 | 5 | 6 | 31 | 1 | 4 | 3 | 3 | 9 | 8 | 16 | 17 | 16 | 18 | 30 | 8 | 27 | 14 | Nppa |
| Nppb | 5 | 0 | 0 | 1 | 0 | 0 | 2 | 5 | 3 | 3 | 5 | 1 | 0 | 3 | 5 | 5 | 1 | 0 | 1 | 0 | 0 | 2 | 3 | 0 | 5 | 3 | 1 | 1 | Nppb |
| Nppc | 28 | 37 | 27 | 39 | 70 | 32 | 37 | 43 | 49 | 71 | 84 | 14 | 33 | 14 | 40 | 21 | 37 | 25 | 25 | 35 | 33 | 40 | 107 | 34 | 42 | 26 | 10 | 13 | Nppc |
| Npr1 | 466 | 595 | 736 | 536 | 676 | 620 | 406 | 291 | 410 | 460 | 689 | 384 | 429 | 388 | 529 | 620 | 691 | 548 | 504 | 406 | 411 | 515 | 617 | 400 | 469 | 445 | 504 | 341 | Npr1 |
| Npr2 | 2060 | 2485 | 2310 | 1949 | 2567 | 2111 | 1752 | 3467 | 3892 | 4032 | 4366 | 3442 | 4473 | 3251 | 2210 | 3026 | 2432 | 2122 | 2155 | 1743 | 1728 | 4451 | 3677 | 3581 | 3933 | 3150 | 3426 | 3534 | Npr2 |
| Npr3 | 1158 | 864 | 1251 | 553 | 841 | 945 | 459 | 2005 | 2322 | 1594 | 2309 | 1702 | 2437 | 1392 | 1357 | 1341 | 848 | 741 | 595 | 673 | 543 | 1567 | 3051 | 1442 | 1176 | 1340 | 1536 | 1540 | Npr3 |
| Nprl2 | 2748 | 3841 | 2703 | 3416 | 2673 | 2561 | 2879 | 1642 | 1662 | 1503 | 1436 | 1496 | 1264 | 1533 | 2834 | 3040 | 2765 | 3098 | 2992 | 3012 | 2890 | 1509 | 1598 | 1577 | 1520 | 1452 | 1456 | 1386 | Nprl2 |
| Nprl3 | 1485 | 1528 | 1263 | 1368 | 1528 | 1525 | 1403 | 1701 | 2134 | 1885 | 1939 | 2162 | 2076 | 1927 | 1544 | 1383 | 1320 | 1352 | 1443 | 1572 | 1432 | 1572 | 1911 | 1810 | 1990 | 1741 | 2072 | 1841 | Nprl3 |
| Nps | 13 | 5 | 4 | 3 | 7 | 18 | 6 | 4 | 1 | 0 | 9 | 7 | 7 | 3 | 7 | 11 | 2 | 11 | 11 | 1 | 4 | 0 | 3 | 2 | 6 | 8 | 1 | 4 | Nps |
| Npsr1 | 0 | 3 | 0 | 4 | 2 | 1 | 10 | 4 | 12 | 1 | 7 | 9 | 7 | 3 | 8 | 1 | 1 | 4 | 0 | 0 | 4 | 8 | 5 | 12 | 16 | 2 | 0 | 4 | Npsr1 |
| Nptn | 8611 | 7515 | 7165 | 8149 | 9239 | 8496 | 9623 | 7166 | 6752 | 6090 | 6474 | 4836 | 5322 | 5498 | 9837 | 8178 | 7419 | 9063 | 8338 | 9444 | 9452 | 7391 | 7219 | 5395 | 6285 | 4871 | 4635 | 5193 | Nptn |
| Nptx1 | 731 | 1031 | 1056 | 1817 | 2077 | 1326 | 1476 | 933 | 1029 | 1340 | 1331 | 942 | 1232 | 962 | 940 | 1015 | 1464 | 1825 | 1605 | 1278 | 1556 | 1544 | 1074 | 1126 | 1377 | 789 | 841 | 930 | Nptx1 |
| Nptx2 | 726 | 961 | 979 | 1093 | 878 | 808 | 988 | 302 | 330 | 383 | 323 | 374 | 271 | 374 | 684 | 780 | 1061 | 1013 | 923 | 881 | 894 | 396 | 324 | 400 | 441 | 515 | 285 | 342 | Nptx2 |
| Nptxr | 1745 | 1911 | 2012 | 3443 | 5297 | 3912 | 3692 | 2329 | 2685 | 3591 | 3067 | 2080 | 1906 | 2154 | 2054 | 1928 | 2924 | 4150 | 4082 | 3002 | 4138 | 4508 | 2841 | 3101 | 3672 | 1693 | 1593 | 1915 | Nptxr |
| Npvf | 1 | 0 | 0 | 3 | 0 | 0 | 2 | 5 | 21 | 3 | 17 | 5 | 9 | 2 | 0 | 2 | 0 | 0 | 1 | 2 | 3 | 12 | 14 | 6 | 1 | 0 | 2 | 0 | Npvf |
| Npw | 2 | 4 | 2 | 1 | 2 | 0 | 0 | 5 | 2 | 0 | 0 | 0 | 1 | 2 | 5 | 4 | 2 | 0 | 1 | 0 | 0 | 0 | 1 | 0 | 1 | 0 | 0 | 0 | Npw |
| Npy | 48 | 71 | 35 | 125 | 533 | 164 | 126 | 183 | 216 | 467 | 649 | 177 | 99 | 211 | 44 | 35 | 97 | 117 | 199 | 184 | 216 | 256 | 548 | 301 | 333 | 118 | 129 | 133 | Npy |
| Npy1r | 242 | 230 | 333 | 254 | 454 | 276 | 280 | 322 | 374 | 369 | 399 | 173 | 308 | 223 | 313 | 312 | 248 | 262 | 271 | 259 | 272 | 428 | 386 | 230 | 311 | 164 | 178 | 255 | Npy1r |
| Npy2r | 9 | 8 | 11 | 11 | 35 | 29 | 36 | 71 | 80 | 78 | 80 | 26 | 39 | 42 | 17 | 11 | 22 | 15 | 29 | 28 | 45 | 77 | 93 | 60 | 69 | 40 | 35 | 32 | Npy2r |
| Npy4r | 30 | 11 | 17 | 27 | 35 | 21 | 18 | 23 | 7 | 12 | 11 | 4 | 5 | 3 | 18 | 23 | 6 | 22 | 26 | 21 | 30 | 7 | 10 | 13 | 4 | 7 | 0 | 1 | Npy4r |
| Npy5r | 24 | 24 | 41 | 28 | 47 | 35 | 32 | 22 | 20 | 18 | 34 | 2 | 28 | 9 | 40 | 23 | 27 | 30 | 27 | 29 | 19 | 35 | 27 | 12 | 11 | 6 | 14 | 14 | Npy5r |
| Npy6r | 0 | 0 | 0 | 0 | 0 | 0 | 0 | 0 | 0 | 0 | 0 | 0 | 0 | 0 | 0 | 0 | 0 | 0 | 0 | 0 | 0 | 0 | 0 | 0 | 0 | 1 | 0 | 0 | Npy6r |
| Nqo1 | 233 | 181 | 193 | 102 | 84 | 135 | 105 | 242 | 291 | 235 | 301 | 180 | 253 | 195 | 225 | 222 | 108 | 138 | 104 | 142 | 101 | 229 | 349 | 202 | 167 | 253 | 191 | 259 | Nqo1 |
| Nqo2 | 828 | 728 | 900 | 461 | 595 | 616 | 559 | 1050 | 948 | 796 | 990 | 1037 | 1154 | 1097 | 881 | 845 | 686 | 526 | 574 | 600 | 513 | 964 | 1014 | 916 | 780 | 1055 | 1081 | 1099 | Nqo2 |
| Nr0b1 | 30 | 57 | 57 | 14 | 13 | 12 | 16 | 20 | 21 | 17 | 17 | 8 | 7 | 17 | 26 | 38 | 32 | 13 | 19 | 24 | 22 | 42 | 17 | 18 | 16 | 45 | 15 | 21 | Nr0b1 |
| Nr0b2 | 0 | 0 | 0 | 0 | 0 | 0 | 0 | 2 | 1 | 0 | 1 | 3 | 0 | 0 | 0 | 0 | 0 | 1 | 0 | 0 | 0 | 0 | 0 | 0 | 0 | 3 | 0 | 0 | Nr0b2 |
| Nr1d1 | 205 | 398 | 240 | 540 | 286 | 275 | 279 | 89 | 137 | 142 | 176 | 137 | 103 | 134 | 196 | 278 | 294 | 526 | 371 | 272 | 369 | 174 | 123 | 164 | 198 | 202 | 94 | 151 | Nr1d1 |
| Nr1d2 | 1881 | 1422 | 1397 | 1710 | 1249 | 1478 | 1633 | 1452 | 1406 | 1255 | 1368 | 1322 | 1075 | 1420 | 1539 | 1635 | 1298 | 1622 | 1379 | 1468 | 1583 | 1196 | 1274 | 1396 | 1406 | 1348 | 1306 | 1221 | Nr1d2 |
| Nr1h2 | 2817 | 3798 | 3554 | 3563 | 3315 | 3159 | 2980 | 2491 | 3054 | 2855 | 2744 | 3338 | 2999 | 3055 | 2871 | 2889 | 3724 | 3453 | 3527 | 3146 | 3154 | 2777 | 2752 | 3133 | 3041 | 3656 | 3195 | 3088 | Nr1h2 |
| Nr1h3 | 42 | 54 | 59 | 58 | 53 | 34 | 47 | 77 | 91 | 77 | 94 | 123 | 107 | 115 | 38 | 48 | 52 | 55 | 56 | 46 | 34 | 85 | 117 | 90 | 104 | 99 | 103 | 121 | Nr1h3 |
| Nr1h4 | 5 | 0 | 4 | 0 | 1 | 3 | 2 | 0 | 2 | 1 | 2 | 1 | 3 | 0 | 0 | 3 | 0 | 0 | 0 | 0 | 1 | 2 | 2 | 1 | 1 | 0 | 6 | 5 | Nr1h4 |
| Nr1h5 | 5 | 11 | 6 | 5 | 9 | 11 | 7 | 7 | 10 | 8 | 21 | 18 | 7 | 16 | 5 | 3 | 3 | 3 | 1 | 5 | 7 | 13 | 9 | 4 | 10 | 4 | 26 | 9 | Nr1h5 |
| Nr1i2 | 2 | 3 | 0 | 7 | 3 | 3 | 2 | 0 | 2 | 0 | 1 | 3 | 0 | 0 | 0 | 4 | 0 | 3 | 1 | 0 | 0 | 7 | 3 | 3 | 0 | 3 | 0 | 6 | Nr1i2 |
| Nr1i3 | 0 | 4 | 0 | 0 | 0 | 1 | 1 | 7 | 2 | 0 | 0 | 2 | 3 | 14 | 0 | 3 | 1 | 0 | 8 | 2 | 5 | 8 | 5 | 6 | 4 | 0 | 3 | 1 | Nr1i3 |
| Nr2c1 | 1466 | 1209 | 1642 | 1497 | 1305 | 1347 | 1332 | 1793 | 1823 | 1856 | 1798 | 1896 | 2004 | 2082 | 1476 | 1368 | 1488 | 1228 | 1350 | 1234 | 1339 | 2044 | 1785 | 1971 | 1768 | 2075 | 2024 | 2061 | Nr2c1 |

| | | | | | | | | | | | | | | | | | | | | | | | | | | | | | |
|---|---|---|---|---|---|---|---|---|---|---|---|---|---|---|---|---|---|---|---|---|---|---|---|---|---|---|---|---|---|
| Nr2c2 | 9274 | 8396 | 7095 | 9639 | 12070 | 11747 | 12178 | 14741 | 13463 | 13594 | 13635 | 14205 | 12528 | 14389 | 9315 | 7330 | 8696 | 9838 | 11671 | 10320 | 12237 | 14546 | 12550 | 13078 | 15964 | 10150 | 13726 | 13410 | Nr2c2 |
| Nr2c2ap | 1277 | 1733 | 1452 | 1735 | 1420 | 1478 | 1466 | 821 | 797 | 783 | 750 | 904 | 796 | 764 | 1244 | 1378 | 1564 | 1720 | 1668 | 1609 | 1587 | 705 | 830 | 874 | 886 | 1073 | 827 | 878 | Nr2c2ap |
| Nr2e1 | 3746 | 3771 | 4738 | 4035 | 3467 | 3663 | 5140 | 3378 | 3307 | 3479 | 2460 | 7059 | 3191 | 6916 | 3268 | 1787 | 4811 | 3491 | 4770 | 4278 | 4590 | 2562 | 2144 | 5771 | 4734 | 8865 | 5605 | 6772 | Nr2e1 |
| Nr2e3 | 8 | 3 | 0 | 3 | 0 | 8 | 0 | 17 | 40 | 19 | 15 | 12 | 27 | 6 | 13 | 5 | 0 | 6 | 3 | 0 | 1 | 0 | 38 | 18 | 13 | 10 | 7 | 4 | Nr2e3 |
| Nr2f1 | 8116 | 8954 | 13294 | 11736 | 12257 | 11095 | 13330 | 16185 | 14093 | 18246 | 14708 | 23292 | 16755 | 25014 | 9792 | 7400 | 13243 | 10161 | 13023 | 11433 | 12499 | 20927 | 15359 | 21234 | 20377 | 25751 | 20983 | 25249 | Nr2f1 |
| Nr2f2 | 7510 | 6981 | 15170 | 12564 | 12791 | 10844 | 14591 | 13115 | 10870 | 14331 | 13716 | 16887 | 13304 | 19295 | 9268 | 7708 | 13997 | 11704 | 11873 | 10395 | 12607 | 20110 | 12220 | 17155 | 16898 | 22477 | 15171 | 20766 | Nr2f2 |
| Nr2f6 | 2599 | 3048 | 3208 | 2396 | 2162 | 1890 | 1741 | 1715 | 1682 | 1707 | 1759 | 1775 | 1878 | 1520 | 2349 | 2796 | 2855 | 2248 | 2308 | 1972 | 1847 | 1556 | 1874 | 1799 | 1340 | 2038 | 1880 | 1766 | Nr2f6 |
| Nr3c1 | 2141 | 1617 | 1591 | 1354 | 1502 | 1654 | 1530 | 2429 | 2701 | 2008 | 3012 | 1509 | 1856 | 1677 | 1795 | 1878 | 1518 | 1615 | 1160 | 1549 | 1516 | 1850 | 3243 | 1813 | 1611 | 1238 | 1472 | 1549 | Nr3c1 |
| Nr3c2 | 341 | 379 | 340 | 352 | 425 | 301 | 302 | 189 | 237 | 161 | 315 | 98 | 109 | 87 | 368 | 374 | 313 | 381 | 287 | 301 | 324 | 212 | 337 | 147 | 170 | 58 | 101 | 91 | Nr3c2 |
| Nr4a1 | 411 | 479 | 447 | 471 | 392 | 306 | 334 | 347 | 493 | 440 | 577 | 404 | 415 | 348 | 352 | 489 | 443 | 408 | 388 | 325 | 333 | 389 | 532 | 458 | 380 | 387 | 392 | 318 | Nr4a1 |
| Nr4a2 | 2090 | 2494 | 3483 | 4148 | 3218 | 2974 | 4041 | 2097 | 1915 | 2514 | 1966 | 2813 | 2829 | 2546 | 1973 | 2459 | 3851 | 3978 | 3750 | 3219 | 3572 | 3187 | 1821 | 2701 | 2895 | 2414 | 2339 | 2790 | Nr4a2 |
| Nr4a3 | 611 | 739 | 812 | 1012 | 1060 | 984 | 1003 | 630 | 620 | 794 | 625 | 378 | 667 | 368 | 754 | 776 | 909 | 1215 | 1043 | 784 | 1186 | 919 | 641 | 498 | 925 | 280 | 375 | 370 | Nr4a3 |
| Nr5a1 | 55 | 51 | 35 | 56 | 54 | 37 | 51 | 123 | 105 | 126 | 75 | 74 | 9 | 64 | 40 | 35 | 64 | 51 | 41 | 46 | 62 | 167 | 144 | 149 | 158 | 152 | 37 | 25 | Nr5a1 |
| Nr5a2 | 187 | 139 | 187 | 175 | 290 | 237 | 263 | 429 | 287 | 333 | 612 | 302 | 259 | 300 | 193 | 182 | 207 | 262 | 327 | 188 | 325 | 456 | 326 | 411 | 408 | 171 | 351 | 360 | Nr5a2 |
| Nr6a1 | 3417 | 3323 | 2611 | 2732 | 2720 | 2725 | 2985 | 909 | 736 | 860 | 787 | 693 | 810 | 751 | 3556 | 3290 | 2547 | 2710 | 2291 | 2579 | 2834 | 842 | 806 | 689 | 676 | 552 | 793 | 765 | Nr6a1 |
| Nradd | 631 | 721 | 857 | 616 | 569 | 608 | 515 | 933 | 1071 | 880 | 864 | 959 | 1124 | 894 | 612 | 643 | 745 | 594 | 727 | 700 | 587 | 744 | 1072 | 990 | 826 | 1137 | 1130 | 1046 | Nradd |
| Nrap | 50 | 75 | 48 | 73 | 85 | 92 | 73 | 77 | 87 | 46 | 295 | 79 | 73 | 69 | 70 | 72 | 46 | 76 | 47 | 75 | 58 | 105 | 199 | 85 | 98 | 72 | 67 | 71 | Nrap |
| Nrarp | 3002 | 3972 | 4414 | 3608 | 2949 | 3079 | 3195 | 4145 | 3460 | 3936 | 3023 | 5549 | 4951 | 6237 | 3620 | 3140 | 4385 | 3078 | 3455 | 3139 | 3390 | 3659 | 3374 | 5103 | 4169 | 6244 | 6047 | 6843 | Nrarp |
| Nras | 15045 | 13094 | 13937 | 12702 | 11968 | 12834 | 13527 | 19867 | 16628 | 15987 | 15701 | 19578 | 18758 | 20089 | 14666 | 13164 | 13005 | 11798 | 12742 | 13273 | 12977 | 17306 | 16241 | 17562 | 17792 | 19930 | 20583 | 21010 | Nras |
| Nrbf2 | 0 | 0 | 0 | 0 | 0 | 0 | 0 | 0 | 0 | 0 | 0 | 0 | 0 | 0 | 0 | 0 | 0 | 0 | 0 | 1 | 0 | 0 | 0 | 0 | 0 | 0 | 0 | 0 | Nrbf2 |
| Nrbp1 | 6305 | 6715 | 6415 | 6825 | 6320 | 6687 | 6312 | 8330 | 7969 | 7580 | 7460 | 7707 | 7530 | 7697 | 6279 | 5812 | 6603 | 6268 | 6860 | 6837 | 6621 | 8136 | 8279 | 7531 | 7856 | 8048 | 7937 | 8332 | Nrbp1 |
| Nrbp2 | 4106 | 5454 | 5807 | 5873 | 7040 | 6171 | 5266 | 4914 | 5008 | 5928 | 5404 | 4566 | 5824 | 4478 | 5093 | 5604 | 6122 | 5948 | 6060 | 5176 | 5635 | 7431 | 5000 | 4830 | 5857 | 3245 | 4501 | 4342 | Nrbp2 |
| Nrcam | 9067 | 10713 | 5575 | 9330 | 12936 | 13212 | 12785 | 17363 | 14228 | 12758 | 11113 | 11313 | 11110 | 12472 | 12008 | 8526 | 8020 | 9434 | 10953 | 15072 | 14091 | 13849 | 13573 | 9702 | 14331 | 5983 | 12666 | 10763 | Nrcam |
| Nrd1 | 25077 | 23211 | 21709 | 23190 | 21972 | 22936 | 25530 | 20314 | 18475 | 18995 | 18248 | 20309 | 19224 | 21120 | 24684 | 22831 | 21398 | 22958 | 21722 | 22126 | 25128 | 19392 | 18109 | 18519 | 19968 | 18077 | 20737 | 20507 | Nrd1 |
| Nrde2 | 561 | 454 | 449 | 455 | 576 | 523 | 514 | 2058 | 2078 | 2065 | 2055 | 2288 | 2074 | 2272 | 513 | 414 | 528 | 408 | 613 | 545 | 531 | 1771 | 2013 | 2018 | 2295 | 1996 | 2128 | 2073 | Nrde2 |
| Nrep | 44494 | 38932 | 43343 | 52195 | 62623 | 58005 | 65748 | 77396 | 67518 | 68888 | 68754 | 64019 | 66395 | 72771 | 53920 | 43370 | 47482 | 54542 | 56961 | 64489 | 63569 | 77605 | 71323 | 71179 | 78487 | 64936 | 62742 | 65348 | Nrep |
| Nrf1 | 3746 | 3966 | 4247 | 4349 | 4010 | 4143 | 3961 | 4224 | 4322 | 4484 | 4172 | 5432 | 4923 | 5212 | 3460 | 3464 | 4149 | 3875 | 4571 | 3866 | 4147 | 4079 | 4029 | 5006 | 5052 | 6072 | 5605 | 5457 | Nrf1 |
| Nrg1 | 1694 | 1492 | 1065 | 1695 | 2275 | 2133 | 2376 | 2107 | 2013 | 1862 | 1761 | 1728 | 1456 | 1872 | 1916 | 1551 | 1469 | 1596 | 1975 | 2019 | 2352 | 1696 | 1547 | 1535 | 1838 | 1489 | 1575 | 1358 | Nrg1 |
| Nrg2 | 128 | 161 | 156 | 220 | 327 | 222 | 197 | 112 | 124 | 148 | 140 | 93 | 124 | 99 | 137 | 160 | 172 | 198 | 298 | 214 | 241 | 177 | 156 | 159 | 165 | 62 | 114 | 84 | Nrg2 |
| Nrg3 | 314 | 347 | 248 | 581 | 1001 | 772 | 821 | 659 | 644 | 803 | 661 | 513 | 502 | 458 | 402 | 349 | 437 | 770 | 789 | 691 | 918 | 947 | 632 | 554 | 895 | 253 | 366 | 379 | Nrg3 |
| Nrg4 | 38 | 32 | 5 | 20 | 24 | 15 | 10 | 7 | 27 | 10 | 11 | 11 | 11 | 12 | 22 | 33 | 19 | 32 | 9 | 32 | 19 | 17 | 17 | 9 | 6 | 17 | 9 | 7 | Nrg4 |
| Nrgn | 305 | 488 | 420 | 502 | 502 | 479 | 336 | 247 | 235 | 314 | 299 | 139 | 332 | 128 | 367 | 415 | 470 | 471 | 351 | 344 | 292 | 244 | 261 | 169 | 226 | 134 | 132 | 131 | Nrgn |
| Nrip1 | 3068 | 2238 | 2357 | 2852 | 3390 | 3468 | 3120 | 3323 | 3189 | 3265 | 3737 | 3118 | 2994 | 3353 | 2726 | 2697 | 2509 | 2724 | 3012 | 2657 | 3294 | 3342 | 3136 | 3212 | 3197 | 2603 | 3045 | 2807 | Nrip1 |
| Nrip2 | 20 | 37 | 15 | 42 | 70 | 54 | 54 | 117 | 144 | 130 | 139 | 92 | 92 | 106 | 49 | 38 | 29 | 34 | 53 | 69 | 80 | 170 | 143 | 96 | 132 | 99 | 100 | 82 | Nrip2 |
| Nrip3 | 1411 | 1507 | 1258 | 1741 | 1843 | 1732 | 2275 | 320 | 299 | 272 | 337 | 200 | 153 | 269 | 1741 | 1354 | 1369 | 1933 | 1661 | 1928 | 2129 | 274 | 311 | 273 | 285 | 192 | 173 | 165 | Nrip3 |
| Nrk | 2384 | 1384 | 1098 | 992 | 637 | 1454 | 546 | 1925 | 3703 | 2289 | 10898 | 1614 | 2339 | 1612 | 1996 | 2663 | 852 | 1595 | 598 | 566 | 564 | 2249 | 7440 | 1865 | 1689 | 1274 | 1245 | 1374 | Nrk |
| Nrl | 14 | 21 | 10 | 20 | 20 | 4 | 26 | 15 | 12 | 8 | 11 | 23 | 18 | 12 | 26 | 18 | 10 | 18 | 20 | 17 | 28 | 10 | 14 | 8 | 8 | 11 | 1 | 17 | Nrl |
| Nrm | 1377 | 1557 | 1580 | 1376 | 1159 | 1200 | 1519 | 1051 | 1232 | 1156 | 1199 | 1809 | 1690 | 1788 | 1369 | 1289 | 1658 | 1367 | 1316 | 1338 | 1254 | 1089 | 1179 | 1527 | 1234 | 2205 | 1816 | 1836 | Nrm |
| Nrn1 | 1078 | 1139 | 1798 | 1681 | 2734 | 2158 | 2733 | 3850 | 2809 | 3420 | 2881 | 3999 | 3656 | 5047 | 1493 | 980 | 1978 | 1788 | 2684 | 2619 | 2684 | 3678 | 2919 | 4094 | 4354 | 3213 | 4349 | 5045 | Nrn1 |
| Nrn1l | 0 | 3 | 0 | 1 | 0 | 0 | 0 | 0 | 0 | 0 | 0 | 0 | 0 | 0 | 0 | 0 | 0 | 0 | 0 | 0 | 0 | 0 | 0 | 0 | 0 | 0 | 0 | 0 | Nrn1l |
| Nrp1 | 2220 | 2233 | 1457 | 1694 | 3296 | 2493 | 2081 | 4799 | 5769 | 4551 | 6078 | 3221 | 3631 | 2959 | 2490 | 2359 | 1635 | 2316 | 2272 | 2551 | 2298 | 5071 | 5922 | 3533 | 4518 | 2175 | 3054 | 2659 | Nrp1 |
| Nrp2 | 11129 | 11326 | 10774 | 12451 | 13345 | 12609 | 11763 | 11611 | 14283 | 16343 | 16147 | 13636 | 13840 | 13448 | 11701 | 13926 | 11556 | 13705 | 11513 | 10501 | 11324 | 16681 | 13138 | 15379 | 15399 | 12003 | 11846 | 12056 | Nrp2 |
| Nrros | 318 | 251 | 217 | 189 | 277 | 233 | 187 | 353 | 465 | 403 | 399 | 339 | 417 | 282 | 272 | 273 | 258 | 252 | 261 | 247 | 221 | 341 | 552 | 400 | 361 | 416 | 403 | 363 | Nrros |
| Nrsn1 | 1001 | 1104 | 835 | 1557 | 2410 | 1696 | 2029 | 1037 | 1001 | 1275 | 1127 | 817 | 716 | 819 | 1295 | 1021 | 1161 | 1869 | 1898 | 1792 | 2127 | 1686 | 1033 | 1022 | 1231 | 692 | 795 | 856 | Nrsn1 |
| Nrsn2 | 428 | 460 | 484 | 761 | 981 | 679 | 744 | 482 | 528 | 726 | 616 | 618 | 481 | 675 | 493 | 507 | 753 | 763 | 842 | 664 | 819 | 792 | 457 | 600 | 800 | 475 | 500 | 596 | Nrsn2 |
| Nrtn | 333 | 347 | 271 | 276 | 151 | 176 | 152 | 88 | 91 | 80 | 67 | 62 | 86 | 42 | 264 | 256 | 270 | 260 | 224 | 186 | 142 | 43 | 81 | 56 | 47 | 56 | 83 | 55 | Nrtn |
| Nrxn1 | 3541 | 3728 | 3611 | 7199 | 10597 | 9192 | 9184 | 8024 | 7196 | 9097 | 7895 | 6003 | 5964 | 6704 | 4958 | 3883 | 5435 | 7663 | 8312 | 7985 | 9548 | 11802 | 7682 | 6933 | 10347 | 4059 | 5493 | 5190 | Nrxn1 |
| Nrxn2 | 0 | 1 | 0 | 0 | 1 | 0 | 1 | 0 | 0 | 0 | 0 | 0 | 0 | 0 | 0 | 0 | 0 | 0 | 0 | 0 | 0 | 0 | 0 | 0 | 0 | 0 | 2 | 0 | Nrxn2 |
| Nrxn3 | 2869 | 3308 | 2880 | 6212 | 8065 | 7055 | 7453 | 4643 | 4918 | 6070 | 4939 | 5003 | 4118 | 5157 | 3579 | 2507 | 4766 | 6584 | 7079 | 5466 | 7339 | 8109 | 3585 | 4939 | 7404 | 3316 | 4118 | 3788 | Nrxn3 |
| Nsa2 | 10035 | 8509 | 9527 | 8318 | 7031 | 7931 | 8933 | 10279 | 9391 | 8251 | 8827 | 8939 | 9190 | 10575 | 10512 | 9111 | 8722 | 7604 | 7397 | 10092 | 8278 | 7910 | 9629 | 9183 | 8423 | 10634 | 9640 | 10850 | Nsa2 |
| Nscme3l | 137 | 142 | 67 | 160 | 155 | 177 | 133 | 116 | 146 | 139 | 143 | 135 | 130 | 126 | 95 | 135 | 111 | 177 | 168 | 115 | 126 | 140 | 95 | 74 | 151 | 23 | 138 | 81 | Nscme3l |
| Nsd1 | 16842 | 14559 | 10888 | 13086 | 17177 | 18913 | 16815 | 29366 | 28056 | 24750 | 27163 | 26092 | 26589 | 26428 | 16498 | 13974 | 12432 | 12829 | 15321 | 17918 | 18173 | 21524 | 25765 | 21440 | 27952 | 18724 | 29398 | 24953 | Nsd1 |
| Nsd2 | 29433 | 26849 | 24189 | 26500 | 29757 | 32261 | 32710 | 41570 | 35670 | 34658 | 33071 | 42239 | 37245 | 43463 | 29993 | 24952 | 24976 | 26201 | 29483 | 32443 | 32693 | 34116 | 31826 | 33971 | 40751 | 29590 | 45725 | 40502 | Nsd2 |
| Nsd3 | 16251 | 14976 | 12992 | 17188 | 16661 | 17383 | 17597 | 11351 | 10827 | 10929 | 11266 | 10820 | 10260 | 11123 | 15749 | 15229 | 14028 | 16942 | 16121 | 15175 | 18180 | 11219 | 9926 | 9967 | 11428 | 8488 | 11006 | 10385 | Nsd3 |
| Nsdhl | 4456 | 4912 | 4378 | 3559 | 3253 | 3232 | 3654 | 2981 | 2456 | 2609 | 2141 | 2961 | 3091 | 3428 | 4704 | 4050 | 3962 | 3408 | 3332 | 3565 | 3667 | 2785 | 2592 | 2571 | 2489 | 3405 | 3333 | 3794 | Nsdhl |
| Nsf | 6943 | 8152 | 7567 | 8482 | 10351 | 9039 | 10058 | 5680 | 6100 | 6305 | 5804 | 5604 | 5459 | 5546 | 7776 | 6733 | 8057 | 9376 | 9548 | 9723 | 9899 | 6948 | 5853 | 5638 | 6509 | 4947 | 5713 | 5594 | Nsf |
| Nsfl1c | 9499 | 11194 | 10952 | 9933 | 8669 | 9039 | 9088 | 9780 | 9848 | 8971 | 8627 | 9546 | 9470 | 10138 | 9844 | 10129 | 10361 | 9205 | 9616 | 10275 | 8995 | 8565 | 9536 | 9769 | 9481 | 11347 | 10437 | 10644 | Nsfl1c |
| Nsg1 | 5212 | 5958 | 5114 | 8452 | 13086 | 10515 | 10912 | 19913 | 19017 | 20625 | 17941 | 16336 | 14322 | 18484 | 6530 | 4780 | 7122 | 9568 | 10932 | 11296 | 12492 | 21768 | 18999 | 19675 | 23940 | 15595 | 15206 | 16643 | Nsg1 |
| Nsg2 | 12680 | 16262 | 13319 | 26566 | 36705 | 29989 | 35215 | 34474 | 33708 | 39212 | 30514 | 30919 | 23323 | 33191 | 16994 | 10352 | 20325 | 28804 | 34547 | 33404 | 38662 | 40213 | 31922 | 38051 | 46932 | 25093 | 26989 | 27652 | Nsg2 |
| Nsl1 | 1885 | 1688 | 2102 | 1568 | 1181 | 1424 | 1332 | 1547 | 1514 | 1466 | 1347 | 2341 | 1850 | 2115 | 1548 | 1675 | 1745 | 1240 | 1350 | 1322 | 1217 | 1188 | 1363 | 1911 | 1516 | 2883 | 2431 | 2218 | Nsl1 |
| Nsmaf | 2219 | 1849 | 2095 | 1480 | 1909 | 1805 | 1604 | 4477 | 4107 | 3400 | 3732 | 4276 | 4080 | 3809 | 2081 | 2023 | 1893 | 1641 | 1802 | 1888 | 1717 | 3730 | 4110 | 3486 | 3470 | 3859 | 4371 | 4041 | Nsmaf |
| Nsmce1 | 1384 | 1323 | 1208 | 1067 | 1082 | 1097 | 1267 | 2133 | 2113 | 1711 | 1781 | 2091 | 1911 | 2379 | 1396 | 1205 | 1129 | 1031 | 1206 | 1397 | 1247 | 1795 | 2057 | 2112 | 1915 | 2357 | 2216 | 2370 | Nsmce1 |
| Nsmce2 | 3140 | 2785 | 3309 | 2379 | 2016 | 2048 | 2505 | 2589 | 2188 | 2135 | 2055 | 2620 | 2336 | 2637 | 3059 | 2864 | 2641 | 2319 | 2197 | 2688 | 2409 | 2204 | 2224 | 2513 | 2093 | 3058 | 2623 | 2809 | Nsmce2 |
| Nsmce3 | 1816 | 1838 | 2014 | 1988 | 1766 | 1713 | 1939 | 1304 | 1208 | 1225 | 1363 | 1321 | 1219 | 1448 | 1727 | 1722 | 1792 | 2060 | 1746 | 1726 | 2027 | 1397 | 1208 | 1357 | 1329 | 1428 | 1316 | 1393 | Nsmce3 |
| Nsmce4a | 5892 | 5293 | 5493 | 4811 | 4148 | 4647 | 5386 | 5619 | 5227 | 4796 | 4484 | 5751 | 4916 | 5779 | 5856 | 5295 | 4989 | 4239 | 4773 | 5611 | 4935 | 3984 | 4632 | 5486 | 4735 | 6468 | 5777 | 6380 | Nsmce4a |
| Nsmf | 5202 | 6994 | 5444 | 7407 | 8070 | 7421 | 7575 | 4968 | 5867 | 5807 | 5350 | 4968 | 4772 | 4660 | 5401 | 5686 | 6576 | 7793 | 8546 | 7269 | 7937 | 5391 | 5475 | 5850 | 6347 | 4877 | 4423 | 4287 | Nsmf |
| Nsrp1 | 2602 | 2255 | 3499 | 2877 | 2623 | 2578 | 2623 | 3118 | 2881 | 3185 | 2921 | 3329 | 3188 | 3550 | 2962 | 2412 | 3278 | 2743 | 2821 | 3081 | 2559 | 2744 | 2857 | 3644 | 3158 | 4103 | 3510 | 3565 | Nsrp1 |
| Nsun2 | 8931 | 8320 | 7931 | 6685 | 5919 | 6804 | 6569 | 7454 | 7457 | 6844 | 7208 | 7652 | 7456 | 7733 | 8048 | 8408 | 6978 | 6764 | 6178 | 6789 | 6498 | 6268 | 7482 | 7298 | 6957 | 8534 | 7689 | 7863 | Nsun2 |
| Nsun3 | 757 | 628 | 767 | 818 | 758 | 822 | 978 | 1144 | 999 | 1042 | 946 | 1326 | 1190 | 1178 | 877 | 600 | 790 | 750 | 767 | 807 | 849 | 1061 | 1076 | 1096 | 1051 | 1178 | 1270 | 1196 | Nsun3 |
| Nsun4 | 980 | 864 | 1169 | 862 | 848 | 933 | 859 | 1288 | 1195 | 1202 | 1246 | 1371 | 1521 | 1373 | 951 | 873 | 1086 | 704 | 893 | 908 | 841 | 1255 | 1309 | 1226 | 1211 | 1368 | 1557 | 1406 | Nsun4 |
| Nsun5 | 1182 | 1489 | 1313 | 1496 | 1341 | 1346 | 1402 | 1321 | 1454 | 1549 | 1309 | 1529 | 1526 | 1577 | 1171 | 1267 | 1344 | 1479 | 1515 | 1164 | 1275 | 1571 | 1343 | 1637 | 1641 | 1741 | 1489 | 1506 | Nsun5 |
| Nsun6 | 1765 | 1399 | 1223 | 1335 | 1215 | 1536 | 1634 | 1336 | 1233 | 1075 | 1079 | 1059 | 1119 | 1349 | 1869 | 1506 | 1189 | 1352 | 1209 | 1619 | 1490 | 1081 | 1170 | 1061 | 1228 | 823 | 1168 | 1124 | Nsun6 |
| Nsun7 | 465 | 396 | 376 | 471 | 567 | 593 | 568 | 402 | 502 | 473 | 527 | 426 | 500 | 436 | 520 | 492 | 467 | 543 | 482 | 538 | 533 | 563 | 569 | 407 | 452 | 387 | 464 | 307 | Nsun7 |
| Nt5c | 967 | 1234 | 1065 | 1060 | 1205 | 1276 | 1117 | 1619 | 1823 | 1735 | 1676 | 1751 | 1863 | 1858 | 960 | 1064 | 1236 | 1011 | 1199 | 1377 | 1146 | 1475 | 1805 | 1899 | 1821 | 2011 | 1972 | 1807 | Nt5c |
| Nt5c1a | 1 | 4 | 7 | 7 | 9 | 4 | 4 | 4 | 9 | 5 | 4 | 3 | 2 | 4 | 3 | 9 | 7 | 14 | 12 | 10 | 7 | 2 | 5 | 2 | 1 | 0 | 5 | 1 | Nt5c1a |
| Nt5c1b | 0 | 0 | 0 | 0 | 0 | 0 | 0 | 2 | 2 | 3 | 0 | 0 | 1 | 0 | 0 | 0 | 1 | 0 | 0 | 1 | 0 | 0 | 1 | 0 | 0 | 0 | 0 | 0 | Nt5c1b |
| Nt5c2 | 6336 | 6574 | 5682 | 6100 | 6370 | 6379 | 6300 | 6768 | 6035 | 6105 | 5892 | 6397 | 6208 | 6502 | 6701 | 6270 | 5861 | 5983 | 5822 | 6769 | 6321 | 6432 | 6169 | 5788 | 6626 | 5972 | 6406 | 6616 | Nt5c2 |
| Nt5c3 | 3165 | 2947 | 2932 | 3100 | 3285 | 3142 | 3690 | 3802 | 3098 | 3106 | 2669 | 2885 | 3015 | 3099 | 3674 | 3129 | 2896 | 2868 | 3133 | 3924 | 3388 | 3708 | 3111 | 2906 | 3050 | 2801 | 2828 | 3303 | Nt5c3 |
| Nt5c3b | 2871 | 2750 | 2553 | 2534 | 2989 | 2962 | 2695 | 3750 | 4196 | 3945 | 4254 | 3557 | 3521 | 3866 | 2677 | 2891 | 2533 | 2625 | 2964 | 2990 | 2754 | 3738 | 4335 | 3997 | 3868 | 4139 | 3973 | 3614 | Nt5c3b |

| | | | | | | | | | | | | | | | | | | | | | | | | | | | | | |
|---|---|---|---|---|---|---|---|---|---|---|---|---|---|---|---|---|---|---|---|---|---|---|---|---|---|---|---|---|---|
| Nt5dc1 | 463 | 282 | 416 | 302 | 281 | 299 | 294 | 396 | 441 | 306 | 421 | 405 | 485 | 444 | 456 | 356 | 248 | 277 | 281 | 345 | 239 | 286 | 443 | 381 | 310 | 515 | 487 | 441 | Nt5dc1 |
| Nt5dc2 | 14132 | 18484 | 16458 | 14560 | 12007 | 12666 | 11999 | 8512 | 9769 | 9263 | 9019 | 11715 | 11460 | 10673 | 13291 | 15850 | 16157 | 13419 | 13757 | 12121 | 12207 | 8017 | 9849 | 10643 | 9561 | 13263 | 12126 | 11966 | Nt5dc2 |
| Nt5dc3 | 3270 | 3672 | 3399 | 5820 | 6667 | 5747 | 5730 | 2325 | 2492 | 2963 | 2694 | 1490 | 1714 | 1480 | 3460 | 3400 | 4216 | 6203 | 6351 | 5452 | 6207 | 3307 | 2407 | 2174 | 3094 | 1342 | 1383 | 1252 | Nt5dc3 |
| Nt5e | 236 | 155 | 197 | 94 | 97 | 237 | 63 | 175 | 372 | 166 | 373 | 133 | 215 | 150 | 244 | 248 | 111 | 223 | 115 | 83 | 78 | 107 | 370 | 185 | 125 | 89 | 127 | 117 | Nt5e |
| Nt5el | 1 | 1 | 0 | 0 | 2 | 3 | 0 | 0 | 0 | 0 | 0 | 0 | 0 | 2 | 1 | 2 | 1 | 1 | 1 | 0 | 1 | 2 | 2 | 0 | 0 | 0 | 0 | 0 | Nt5el |
| Nt5m | 495 | 658 | 658 | 549 | 737 | 689 | 615 | 1109 | 1221 | 1354 | 1340 | 1284 | 1191 | 1223 | 550 | 497 | 656 | 598 | 804 | 633 | 670 | 1340 | 1141 | 1314 | 1433 | 1292 | 1246 | 1296 | Nt5m |
| Ntan1 | 3120 | 2782 | 2662 | 2539 | 2952 | 2789 | 3414 | 4085 | 3728 | 3738 | 3435 | 3542 | 3319 | 4010 | 3223 | 2572 | 2897 | 2544 | 2866 | 3219 | 3224 | 3581 | 3625 | 3676 | 3769 | 4065 | 3581 | 3963 | Ntan1 |
| Ntf3 | 331 | 190 | 214 | 143 | 132 | 114 | 148 | 176 | 173 | 153 | 177 | 68 | 104 | 59 | 233 | 239 | 197 | 155 | 149 | 147 | 129 | 67 | 178 | 144 | 39 | 186 | 62 | 81 | Ntf3 |
| Ntf5 | 68 | 26 | 42 | 37 | 17 | 19 | 12 | 103 | 105 | 57 | 90 | 39 | 58 | 42 | 45 | 38 | 23 | 24 | 11 | 16 | 6 | 25 | 107 | 65 | 16 | 57 | 38 | 20 | Ntf5 |
| Nthl1 | 401 | 428 | 637 | 419 | 434 | 337 | 365 | 496 | 551 | 540 | 574 | 617 | 621 | 541 | 345 | 447 | 451 | 403 | 406 | 371 | 332 | 446 | 579 | 647 | 543 | 828 | 621 | 656 | Nthl1 |
| Ntm | 2139 | 2388 | 1945 | 3867 | 6127 | 4540 | 4939 | 4664 | 4643 | 5400 | 5299 | 3274 | 3222 | 3490 | 3075 | 2203 | 3045 | 4747 | 4523 | 5166 | 5310 | 6378 | 5138 | 3958 | 5509 | 2675 | 2847 | 3084 | Ntm |
| Ntmt1 | 3211 | 4391 | 3336 | 4279 | 3224 | 3234 | 3206 | 2082 | 2128 | 2262 | 2060 | 2494 | 2446 | 2230 | 3189 | 3573 | 3714 | 3826 | 3811 | 3630 | 3413 | 1912 | 2222 | 2356 | 2439 | 2632 | 2650 | 2435 | Ntmt1 |
| Ntn1 | 4863 | 7317 | 7604 | 6959 | 7969 | 7319 | 5322 | 6701 | 6983 | 7721 | 8179 | 6999 | 9191 | 6116 | 7232 | 6991 | 8781 | 7707 | 6903 | 5473 | 5910 | 8278 | 6625 | 6367 | 6944 | 5597 | 7066 | 6457 | Ntn1 |
| Ntn3 | 1166 | 1254 | 917 | 693 | 580 | 759 | 397 | 1017 | 1561 | 1162 | 1501 | 1222 | 1557 | 1050 | 938 | 1373 | 762 | 790 | 461 | 424 | 432 | 1218 | 1431 | 1183 | 1088 | 1312 | 1220 | 1105 | Ntn3 |
| Ntn4 | 642 | 746 | 1016 | 826 | 909 | 890 | 855 | 1377 | 1341 | 1384 | 1099 | 2104 | 1660 | 2098 | 748 | 502 | 984 | 723 | 995 | 697 | 938 | 1370 | 1042 | 1620 | 1723 | 2020 | 1951 | 1940 | Ntn4 |
| Ntn5 | 14 | 26 | 12 | 12 | 40 | 17 | 13 | 55 | 53 | 95 | 59 | 23 | 59 | 24 | 17 | 24 | 37 | 42 | 11 | 10 | 12 | 58 | 70 | 77 | 39 | 22 | 26 | 19 | Ntn5 |
| Ntng1 | 571 | 520 | 577 | 815 | 1427 | 1054 | 1331 | 1258 | 1024 | 1752 | 1418 | 963 | 871 | 977 | 691 | 600 | 691 | 1063 | 1142 | 950 | 1239 | 2196 | 1213 | 1416 | 1689 | 612 | 837 | 792 | Ntng1 |
| Ntng2 | 762 | 853 | 1169 | 1395 | 1906 | 1619 | 1796 | 1213 | 1440 | 1852 | 1436 | 2011 | 1164 | 1852 | 754 | 776 | 1502 | 1512 | 2141 | 1389 | 1908 | 2172 | 1351 | 2155 | 2649 | 1131 | 1685 | 1625 | Ntng2 |
| Ntpcr | 731 | 800 | 615 | 651 | 619 | 644 | 601 | 615 | 690 | 700 | 656 | 595 | 575 | 558 | 611 | 665 | 734 | 666 | 561 | 608 | 576 | 503 | 695 | 574 | 542 | 655 | 580 | 565 | Ntpcr |
| Ntrk1 | 466 | 764 | 160 | 166 | 1550 | 157 | 55 | 1288 | 1572 | 1086 | 2010 | 99 | 1516 | 88 | 827 | 1283 | 433 | 955 | 111 | 130 | 69 | 2381 | 880 | 85 | 1185 | 59 | 131 | 97 | Ntrk1 |
| Ntrk2 | 3797 | 4455 | 4439 | 5755 | 9399 | 7433 | 7548 | 7566 | 7939 | 8885 | 9169 | 7314 | 7201 | 7009 | 5025 | 4005 | 5561 | 7023 | 7540 | 6444 | 7878 | 10920 | 9042 | 7133 | 9492 | 4921 | 6194 | 6222 | Ntrk2 |
| Ntrk3 | 2987 | 2896 | 2062 | 4602 | 7055 | 6767 | 6683 | 7584 | 7735 | 7613 | 8018 | 5786 | 5576 | 6250 | 4013 | 3181 | 3362 | 5203 | 5483 | 6623 | 7392 | 8769 | 7148 | 6504 | 9190 | 3757 | 5480 | 4719 | Ntrk3 |
| Nts | 328 | 47 | 32 | 157 | 237 | 209 | 400 | 473 | 386 | 285 | 205 | 149 | 104 | 195 | 91 | 77 | 63 | 260 | 261 | 221 | 337 | 175 | 403 | 289 | 424 | 96 | 86 | 123 | Nts |
| Ntsr1 | 46 | 47 | 46 | 104 | 199 | 140 | 160 | 150 | 199 | 196 | 202 | 134 | 44 | 107 | 44 | 25 | 54 | 111 | 199 | 143 | 187 | 216 | 190 | 171 | 250 | 92 | 69 | 76 | Ntsr1 |
| Ntsr2 | 6 | 12 | 6 | 13 | 65 | 25 | 34 | 26 | 30 | 36 | 21 | 12 | 17 | 22 | 22 | 10 | 3 | 28 | 28 | 26 | 47 | 23 | 17 | 26 | 17 | 14 | 15 | 13 | Ntsr2 |
| Nuak1 | 4901 | 5793 | 6463 | 7419 | 7531 | 6990 | 6752 | 6024 | 6751 | 7316 | 6842 | 6668 | 6225 | 6646 | 5317 | 5094 | 6816 | 7516 | 7577 | 5985 | 7496 | 6600 | 5872 | 7728 | 9070 | 5990 | 5885 | 6227 | Nuak1 |
| Nuak2 | 3028 | 4377 | 4899 | 5032 | 3715 | 3972 | 3619 | 2215 | 1725 | 2145 | 1537 | 3747 | 2905 | 3593 | 3390 | 3222 | 4543 | 4409 | 4456 | 3475 | 3875 | 2428 | 1662 | 2638 | 2442 | 3248 | 4077 | 4239 | Nuak2 |
| Nub1 | 5338 | 5953 | 4955 | 5760 | 5690 | 5482 | 6051 | 4925 | 5182 | 5015 | 5064 | 5059 | 4613 | 5103 | 5461 | 5403 | 5385 | 6133 | 5690 | 5703 | 6134 | 5574 | 5223 | 5257 | 5726 | 4823 | 4682 | 4826 | Nub1 |
| Nubp1 | 1211 | 1225 | 1139 | 982 | 1137 | 1081 | 882 | 2027 | 2217 | 2075 | 2020 | 2603 | 2667 | 2475 | 1005 | 1050 | 1072 | 861 | 1093 | 982 | 882 | 2079 | 1956 | 2360 | 2163 | 2921 | 2658 | 2600 | Nubp1 |
| Nubp2 | 3477 | 4548 | 3443 | 3886 | 2682 | 2929 | 2982 | 1713 | 1823 | 1745 | 1616 | 1897 | 1917 | 1802 | 3364 | 3457 | 3189 | 3497 | 3406 | 3307 | 3062 | 1475 | 1798 | 1834 | 1879 | 2302 | 2193 | 2015 | Nubp2 |
| Nubpl | 888 | 650 | 683 | 518 | 494 | 647 | 636 | 865 | 692 | 548 | 637 | 778 | 746 | 764 | 881 | 701 | 605 | 508 | 553 | 795 | 600 | 643 | 717 | 634 | 610 | 939 | 817 | 862 | Nubpl |
| Nucb1 | 7035 | 8712 | 9140 | 8681 | 7192 | 7338 | 6014 | 4583 | 5645 | 5794 | 6714 | 5189 | 5389 | 4779 | 6737 | 8301 | 8502 | 8602 | 7552 | 5603 | 6700 | 5265 | 6554 | 5695 | 5121 | 5966 | 4818 | 5243 | Nucb1 |
| Nucb2 | 1874 | 1474 | 1712 | 1752 | 1954 | 2129 | 1852 | 1726 | 1476 | 1415 | 1687 | 1352 | 1407 | 1442 | 1925 | 1640 | 1781 | 1807 | 1643 | 2055 | 1855 | 1517 | 1601 | 1439 | 1437 | 1256 | 1324 | 1314 | Nucb2 |
| Nucks1 | 14867 | 10044 | 20033 | 13870 | 12566 | 12940 | 14182 | 18169 | 16140 | 17538 | 17698 | 23020 | 20781 | 23191 | 13924 | 12012 | 16523 | 12940 | 13263 | 12456 | 12994 | 17483 | 15502 | 21717 | 19199 | 32387 | 23699 | 26067 | Nucks1 |
| Nudc | 8679 | 10388 | 12059 | 9736 | 8174 | 7721 | 8880 | 7063 | 7617 | 8758 | 8032 | 9873 | 9260 | 10548 | 8159 | 8638 | 10998 | 8844 | 9354 | 8740 | 8300 | 6714 | 7484 | 10069 | 8640 | 12729 | 10437 | 10977 | Nudc |
| Nudcd1 | 5868 | 4672 | 5155 | 4817 | 4003 | 4570 | 5235 | 3990 | 3288 | 2775 | 2913 | 3649 | 3375 | 3898 | 5686 | 5085 | 4819 | 4616 | 4559 | 5406 | 4883 | 2989 | 3205 | 3251 | 3157 | 4004 | 3907 | 4090 | Nudcd1 |
| Nudcd2 | 4670 | 3753 | 5234 | 3951 | 3328 | 3719 | 4245 | 4158 | 3492 | 3251 | 3465 | 4607 | 4389 | 5029 | 4622 | 4213 | 4297 | 3628 | 3711 | 4505 | 3699 | 2872 | 3511 | 3991 | 3465 | 5863 | 5063 | 5435 | Nudcd2 |
| Nudcd3 | 7769 | 7855 | 6591 | 6776 | 7037 | 7565 | 6949 | 7819 | 7874 | 7704 | 7410 | 8210 | 7805 | 8078 | 7435 | 6947 | 6696 | 6947 | 7248 | 7184 | 7394 | 7758 | 7477 | 7757 | 8836 | 8397 | 8457 | 8053 | Nudcd3 |
| Nudt1 | 431 | 497 | 514 | 394 | 394 | 424 | 379 | 797 | 834 | 781 | 769 | 1001 | 1080 | 1038 | 404 | 399 | 504 | 402 | 454 | 404 | 341 | 668 | 928 | 938 | 733 | 1177 | 1151 | 1130 | Nudt1 |
| Nudt10 | 1392 | 1286 | 1517 | 1966 | 2208 | 1991 | 2199 | 2088 | 1631 | 1847 | 1588 | 1685 | 1529 | 1917 | 1557 | 1253 | 1666 | 1696 | 2040 | 1857 | 2103 | 2197 | 1550 | 1777 | 2081 | 1633 | 1417 | 1772 | Nudt10 |
| Nudt11 | 2986 | 2686 | 3091 | 3797 | 4139 | 3925 | 4383 | 3051 | 2429 | 2739 | 2301 | 2608 | 2222 | 2525 | 3257 | 2525 | 3518 | 3843 | 4171 | 4022 | 4226 | 3345 | 2355 | 2545 | 2955 | 2358 | 2295 | 2387 | Nudt11 |
| Nudt12 | 1205 | 1137 | 1293 | 986 | 1097 | 971 | 1044 | 703 | 699 | 722 | 683 | 780 | 743 | 750 | 1266 | 1139 | 1264 | 1095 | 1027 | 1142 | 977 | 658 | 661 | 634 | 604 | 781 | 906 | 748 | Nudt12 |
| Nudt13 | 955 | 964 | 1214 | 1098 | 1109 | 1040 | 1054 | 808 | 872 | 926 | 970 | 934 | 849 | 992 | 982 | 886 | 1059 | 1115 | 1016 | 1045 | 1025 | 830 | 970 | 949 | 895 | 959 | 960 | 884 | Nudt13 |
| Nudt14 | 586 | 672 | 669 | 615 | 576 | 549 | 515 | 679 | 822 | 766 | 805 | 680 | 712 | 637 | 558 | 594 | 637 | 601 | 566 | 627 | 500 | 615 | 935 | 812 | 677 | 954 | 781 | 724 | Nudt14 |
| Nudt15 | 103 | 81 | 134 | 88 | 100 | 109 | 94 | 501 | 468 | 521 | 446 | 655 | 616 | 574 | 108 | 75 | 96 | 70 | 125 | 134 | 101 | 438 | 487 | 506 | 393 | 613 | 625 | 676 | Nudt15 |
| Nudt16 | 424 | 437 | 501 | 579 | 586 | 505 | 580 | 620 | 579 | 694 | 557 | 723 | 615 | 653 | 462 | 355 | 481 | 473 | 543 | 480 | 549 | 645 | 574 | 699 | 690 | 655 | 607 | 610 | Nudt16 |
| Nudt16l1 | 2119 | 2854 | 2401 | 2435 | 2216 | 2121 | 2125 | 2943 | 3179 | 3020 | 2922 | 2965 | 3070 | 2721 | 2113 | 2287 | 2395 | 2272 | 2361 | 2340 | 2188 | 2582 | 3117 | 2953 | 2935 | 3379 | 3020 | 2894 | Nudt16l1 |
| Nudt16l2 | 0 | 0 | 0 | 3 | 0 | 0 | 2 | 0 | 0 | 0 | 0 | 0 | 0 | 0 | 0 | 0 | 0 | 0 | 1 | 0 | 0 | 0 | 0 | 0 | 0 | 0 | 2 | 0 | Nudt16l2 |
| Nudt17 | 135 | 198 | 201 | 140 | 161 | 105 | 158 | 166 | 214 | 201 | 158 | 176 | 210 | 180 | 144 | 192 | 177 | 155 | 160 | 152 | 155 | 204 | 225 | 153 | 170 | 224 | 205 | 193 | Nudt17 |
| Nudt18 | 1009 | 1191 | 1262 | 1229 | 1136 | 996 | 1039 | 468 | 420 | 532 | 518 | 469 | 485 | 469 | 1029 | 1037 | 1297 | 1238 | 1190 | 1067 | 1074 | 451 | 507 | 599 | 583 | 480 | 543 | 507 | Nudt18 |
| Nudt19 | 1591 | 1373 | 1267 | 1287 | 1389 | 1510 | 1571 | 1625 | 1520 | 1384 | 1135 | 1302 | 1210 | 1257 | 1601 | 1333 | 1125 | 1237 | 1400 | 1595 | 1400 | 1322 | 1342 | 1288 | 1311 | 1281 | 1205 | 1237 | Nudt19 |
| Nudt2 | 1227 | 1273 | 1180 | 1153 | 1141 | 1034 | 1367 | 953 | 982 | 877 | 885 | 1035 | 990 | 1178 | 1295 | 1260 | 1195 | 1013 | 1131 | 1460 | 1259 | 744 | 1134 | 991 | 883 | 1213 | 1176 | 1305 | Nudt2 |
| Nudt21 | 18709 | 16341 | 17645 | 18228 | 14546 | 16530 | 18733 | 14933 | 12411 | 12135 | 11459 | 14795 | 13301 | 16080 | 19246 | 16568 | 17685 | 16976 | 17042 | 19133 | 18481 | 12001 | 11874 | 13532 | 13935 | 14981 | 15514 | 17106 | Nudt21 |
| Nudt22 | 1320 | 1816 | 1395 | 1538 | 1128 | 1196 | 1012 | 393 | 519 | 542 | 591 | 461 | 486 | 485 | 1155 | 1472 | 1221 | 1449 | 1132 | 1210 | 1037 | 396 | 546 | 486 | 504 | 534 | 535 | 431 | Nudt22 |
| Nudt3 | 3000 | 3422 | 4202 | 4884 | 5749 | 5132 | 5137 | 6704 | 7486 | 9107 | 8429 | 8290 | 7593 | 9325 | 3117 | 2884 | 4930 | 4762 | 5930 | 4850 | 5970 | 7798 | 7378 | 9547 | 9974 | 8613 | 7675 | 8701 | Nudt3 |
| Nudt4 | 18859 | 16539 | 18656 | 17325 | 16874 | 16349 | 19528 | 14621 | 13046 | 12707 | 12914 | 14404 | 19289 | 17052 | 20653 | 19000 | 17565 | 17185 | 16352 | 16921 | 18368 | 13044 | 13595 | 13601 | 13051 | 16512 | 15163 | 20661 | Nudt4 |
| Nudt5 | 4239 | 3740 | 4094 | 2984 | 2792 | 3071 | 3140 | 4099 | 3701 | 3462 | 3365 | 4201 | 4175 | 4453 | 3836 | 3779 | 3445 | 2880 | 2985 | 3401 | 2974 | 3242 | 3779 | 3937 | 3663 | 4927 | 4819 | 4931 | Nudt5 |
| Nudt6 | 173 | 168 | 113 | 132 | 115 | 159 | 155 | 189 | 173 | 165 | 161 | 176 | 142 | 178 | 181 | 144 | 123 | 135 | 121 | 178 | 135 | 150 | 188 | 141 | 166 | 155 | 128 | 142 | Nudt6 |
| Nudt7 | 619 | 591 | 683 | 489 | 475 | 438 | 443 | 358 | 339 | 260 | 334 | 329 | 383 | 308 | 646 | 681 | 511 | 470 | 428 | 548 | 426 | 274 | 380 | 314 | 231 | 345 | 311 | 382 | Nudt7 |
| Nudt8 | 9 | 4 | 7 | 13 | 9 | 3 | 11 | 9 | 16 | 10 | 2 | 1 | 3 | 11 | 3 | 5 | 8 | 12 | 5 | 4 | 11 | 5 | 5 | 6 | 6 | 11 | 13 | 11 | Nudt8 |
| Nudt9 | 2006 | 2300 | 2208 | 1946 | 1843 | 1652 | 1705 | 1912 | 2098 | 1893 | 1842 | 1904 | 2000 | 1906 | 1903 | 2291 | 2064 | 1823 | 1745 | 1875 | 1672 | 1716 | 2033 | 1949 | 1788 | 2246 | 1923 | 1923 | Nudt9 |
| Nuf2 | 4301 | 3461 | 4593 | 2939 | 2473 | 2957 | 3771 | 5308 | 4473 | 4133 | 3742 | 6192 | 5675 | 6797 | 4134 | 3439 | 3940 | 2575 | 3171 | 3842 | 3052 | 3380 | 3963 | 5267 | 4845 | 8174 | 6796 | 7436 | Nuf2 |
| Nufip1 | 2290 | 2198 | 2404 | 2346 | 1933 | 2015 | 2191 | 1851 | 1797 | 1806 | 1900 | 2056 | 1910 | 2232 | 2297 | 2185 | 2247 | 2102 | 2093 | 1907 | 2127 | 1903 | 1792 | 1887 | 1874 | 2383 | 2029 | 2321 | Nufip1 |
| Nufip2 | 4172 | 3429 | 3204 | 3830 | 3566 | 4022 | 3753 | 4676 | 3862 | 3694 | 4358 | 3886 | 4034 | 4081 | 4071 | 3741 | 3290 | 3671 | 3341 | 4227 | 3885 | 3755 | 4047 | 3448 | 4140 | 3319 | 4349 | 4014 | Nufip2 |
| Nuggc | 14 | 3 | 8 | 5 | 7 | 7 | 6 | 0 | 8 | 3 | 11 | 14 | 15 | 18 | 1 | 9 | 12 | 19 | 4 | 6 | 3 | 10 | 12 | 11 | 15 | 3 | 14 | 2 | Nuggc |
| Numa1 | 12064 | 13318 | 12971 | 13761 | 12720 | 12733 | 12336 | 18966 | 21685 | 22527 | 21540 | 26512 | 24482 | 25238 | 11144 | 12391 | 13842 | 12898 | 14024 | 12035 | 12599 | 19448 | 19601 | 24044 | 23440 | 17971 | 24894 | 24239 | Numa1 |
| Numb | 5067 | 4818 | 6246 | 5525 | 5078 | 5266 | 5377 | 2450 | 2386 | 2857 | 3130 | 3306 | 3510 | 3332 | 4507 | 4994 | 5923 | 5689 | 5368 | 4287 | 4992 | 3009 | 2436 | 3067 | 2821 | 3440 | 3111 | 3304 | Numb |
| Numbl | 3788 | 5185 | 4890 | 4971 | 4910 | 4641 | 4005 | 4031 | 4439 | 5103 | 4779 | 5086 | 4447 | 4315 | 3815 | 3980 | 5107 | 4899 | 4874 | 3928 | 4303 | 5319 | 4456 | 4951 | 5062 | 4416 | 4800 | 4152 | Numbl |
| Nup107 | 5847 | 5840 | 5364 | 4758 | 3943 | 4303 | 4782 | 4940 | 4873 | 4270 | 4250 | 5223 | 4821 | 5420 | 5488 | 5481 | 5029 | 4456 | 4447 | 5230 | 4505 | 3584 | 4544 | 4868 | 4321 | 6004 | 5356 | 5540 | Nup107 |
| Nup133 | 6282 | 6169 | 6387 | 5694 | 4847 | 5303 | 5601 | 4507 | 4700 | 4533 | 4436 | 6020 | 5388 | 5813 | 5629 | 5703 | 6088 | 5403 | 5520 | 4640 | 5541 | 4303 | 4372 | 4989 | 4767 | 6268 | 5739 | 6075 | Nup133 |
| Nup153 | 16073 | 14824 | 11399 | 12388 | 11966 | 13141 | 13984 | 13826 | 12240 | 10381 | 10342 | 12761 | 11835 | 12310 | 15816 | 14347 | 12330 | 12270 | 12423 | 13887 | 13196 | 11404 | 11690 | 10668 | 11246 | 10176 | 13116 | 12933 | Nup153 |
| Nup155 | 11206 | 10022 | 8236 | 8567 | 7211 | 8439 | 8983 | 9413 | 8155 | 7320 | 7658 | 9294 | 8668 | 9345 | 10673 | 10009 | 8452 | 7997 | 7480 | 9322 | 8494 | 7287 | 7776 | 7920 | 7594 | 8149 | 9802 | 9660 | Nup155 |
| Nup160 | 6451 | 5518 | 3978 | 4049 | 3854 | 4410 | 4587 | 6007 | 5205 | 4254 | 4614 | 5551 | 5390 | 5514 | 5970 | 5474 | 3958 | 3954 | 3928 | 4976 | 4623 | 4428 | 5002 | 4558 | 4689 | 4820 | 6007 | 5569 | Nup160 |
| Nup188 | 10247 | 11262 | 9263 | 8856 | 7693 | 8414 | 8015 | 6346 | 7702 | 7251 | 6617 | 8950 | 7633 | 7816 | 9246 | 10348 | 9502 | 8904 | 8830 | 7753 | 8269 | 6380 | 7194 | 7592 | 7190 | 8156 | 8470 | 8261 | Nup188 |
| Nup205 | 10243 | 9833 | 6609 | 6975 | 7128 | 8031 | 8155 | 14586 | 13743 | 11230 | 10519 | 13102 | 11690 | 12935 | 9878 | 9040 | 7306 | 6969 | 7543 | 9198 | 8388 | 10484 | 12459 | 10730 | 11861 | 10714 | 13877 | 13379 | Nup205 |
| Nup210 | 2831 | 3214 | 3426 | 3628 | 3565 | 3766 | 3479 | 3245 | 4050 | 4241 | 3858 | 6118 | 4636 | 5516 | 2635 | 2752 | 3657 | 3200 | 3843 | 3195 | 3435 | 4458 | 3559 | 4755 | 5024 | 4628 | 5073 | 5670 | Nup210 |
| Nup210l | 1194 | 1374 | 887 | 1202 | 795 | 920 | 1082 | 435 | 365 | 350 | 409 | 384 | 375 | 433 | 1191 | 1203 | 1085 | 1196 | 987 | 851 | 1015 | 391 | 373 | 351 | 351 | 304 | 425 | 376 | Nup210l |
| Nup214 | 5332 | 5563 | 4791 | 4516 | 4180 | 4405 | 4172 | 4285 | 5503 | 4848 | 4901 | 6113 | 5560 | 5423 | 4700 | 5433 | 4594 | 4510 | 4390 | 4172 | 4208 | 4914 | 4754 | 5228 | 5121 | 4448 | 5462 | 5343 | Nup214 |
| Nup35 | 3049 | 3125 | 2913 | 2840 | 2401 | 2601 | 2856 | 2965 | 2591 | 2377 | 2094 | 2808 | 2447 | 2726 | 3264 | 3046 | 2803 | 2667 | 2582 | 3073 | 2534 | 2453 | 2537 | 2380 | 2283 | 3055 | 2788 | 2887 | Nup35 |

| | | | | | | | | | | | | | | | | | | | | | | | | | | | | | |
|---|---|---|---|---|---|---|---|---|---|---|---|---|---|---|---|---|---|---|---|---|---|---|---|---|---|---|---|---|---|
| Nup37 | 1698 | 1293 | 1701 | 1151 | 905 | 1008 | 1237 | 1884 | 1757 | 1496 | 1284 | 1970 | 1850 | 1876 | 1639 | 1466 | 1350 | 927 | 1058 | 1319 | 1070 | 1325 | 1608 | 1618 | 1371 | 2390 | 1998 | 2172 | Nup37 |
| Nup43 | 1 | 0 | 1 | 1 | 0 | 0 | 0 | 0 | 0 | 0 | 0 | 0 | 0 | 0 | 0 | 1 | 0 | 0 | 0 | 0 | 0 | 0 | 0 | 0 | 0 | 1 | 0 | 0 | Nup43 |
| Nup50 | 9059 | 9627 | 11571 | 10312 | 9061 | 9197 | 9261 | 7684 | 7175 | 8231 | 7754 | 10516 | 9377 | 9845 | 8768 | 9290 | 11097 | 9750 | 10075 | 8086 | 9232 | 8012 | 7150 | 9073 | 8645 | 10411 | 10089 | 10834 | Nup50 |
| Nup54 | 4618 | 4023 | 4194 | 3820 | 3560 | 3844 | 4276 | 4251 | 3593 | 3321 | 3401 | 3910 | 3627 | 4320 | 4762 | 4144 | 3894 | 3745 | 3781 | 4330 | 4010 | 3611 | 3660 | 3503 | 3551 | 4358 | 4107 | 4196 | Nup54 |
| Nup62 | 5479 | 5869 | 5643 | 4946 | 4626 | 4644 | 4612 | 5175 | 5112 | 4590 | 4487 | 5800 | 5186 | 5331 | 4992 | 4817 | 5337 | 4605 | 5307 | 5191 | 4818 | 4231 | 4803 | 5426 | 4861 | 7210 | 5977 | 6157 | Nup62 |
| Nup62cl | 657 | 555 | 597 | 566 | 392 | 412 | 465 | 199 | 157 | 102 | 84 | 113 | 59 | 133 | 686 | 719 | 501 | 565 | 445 | 616 | 436 | 142 | 164 | 117 | 110 | 213 | 128 | 109 | Nup62cl |
| Nup85 | 7782 | 8325 | 7819 | 7275 | 5599 | 5919 | 6686 | 5594 | 5632 | 5185 | 5006 | 6780 | 5761 | 6404 | 7038 | 6968 | 7430 | 6929 | 6743 | 6710 | 6366 | 4941 | 5467 | 6044 | 5512 | 7721 | 6782 | 6838 | Nup85 |
| Nup88 | 9852 | 9615 | 8615 | 8388 | 8182 | 8091 | 9227 | 10683 | 9194 | 8503 | 8266 | 10405 | 9074 | 10513 | 10042 | 8256 | 8574 | 8051 | 8712 | 9644 | 9128 | 8864 | 8958 | 9584 | 9290 | 10849 | 9952 | 10453 | Nup88 |
| Nup93 | 10934 | 11685 | 10587 | 9048 | 7243 | 8177 | 9020 | 5808 | 5618 | 4985 | 4871 | 6808 | 6099 | 6879 | 10085 | 9544 | 9929 | 8692 | 8770 | 8789 | 8842 | 4772 | 5242 | 5760 | 5268 | 7766 | 7127 | 7021 | Nup93 |
| Nup98 | 10785 | 10682 | 9185 | 9083 | 7565 | 8896 | 8933 | 8182 | 8175 | 7589 | 7717 | 8788 | 8345 | 9082 | 9969 | 10596 | 8951 | 8910 | 8504 | 7940 | 8807 | 7641 | 7780 | 7514 | 8082 | 7241 | 9128 | 9175 | Nup98 |
| Nupl1 | 3 | 2 | 10 | 8 | 4 | 11 | 2 | 0 | 0 | 0 | 0 | 0 | 0 | 0 | 1 | 2 | 0 | 11 | 1 | 2 | 4 | 2 | 2 | 3 | 0 | 3 | 0 | 1 | Nupl1 |
| Nupl2 | 670 | 628 | 688 | 578 | 646 | 640 | 677 | 981 | 924 | 921 | 858 | 1037 | 972 | 1008 | 720 | 620 | 644 | 603 | 581 | 610 | 653 | 881 | 927 | 910 | 1023 | 1113 | 992 | 970 | Nupl2 |
| Nupr1 | 37 | 34 | 73 | 44 | 55 | 51 | 19 | 101 | 153 | 101 | 180 | 88 | 102 | 95 | 28 | 55 | 32 | 51 | 25 | 43 | 38 | 134 | 203 | 117 | 104 | 177 | 134 | 89 | Nupr1 |
| Nupr1l | 133 | 123 | 137 | 143 | 121 | 130 | 142 | 73 | 89 | 99 | 93 | 119 | 96 | 110 | 112 | 123 | 140 | 127 | 112 | 96 | 125 | 92 | 90 | 118 | 108 | 106 | 84 | 116 | Nupr1l |
| Nus1 | 8176 | 6424 | 8498 | 8141 | 7531 | 8000 | 8496 | 7457 | 6576 | 7129 | 7408 | 7651 | 7439 | 7549 | 8044 | 7139 | 7954 | 7955 | 8231 | 7594 | 8169 | 8189 | 6701 | 7462 | 7492 | 7874 | 7524 | 7834 | Nus1 |
| Nusap1 | 9024 | 7311 | 9655 | 6704 | 5499 | 6811 | 7996 | 8745 | 7066 | 6322 | 6235 | 10115 | 8217 | 10989 | 8997 | 7357 | 8274 | 5969 | 7221 | 8596 | 7055 | 5715 | 6679 | 8541 | 7367 | 12222 | 10339 | 11089 | Nusap1 |
| Nutf2 | 4035 | 4774 | 4974 | 4088 | 3317 | 3256 | 3768 | 3275 | 3368 | 3160 | 3065 | 3898 | 3823 | 4415 | 3774 | 4161 | 4292 | 3813 | 3821 | 3845 | 3593 | 2906 | 3565 | 3709 | 3420 | 5076 | 4773 | 4652 | Nutf2 |
| Nutf2-ps1 | 501 | 422 | 562 | 397 | 357 | 421 | 399 | 337 | 393 | 337 | 277 | 449 | 549 | 408 | 525 | 465 | 400 | 387 | 498 | 424 | 348 | 312 | 355 | 382 | 375 | 511 | 508 | 476 | Nutf2-ps1 |
| Nutm1 | 0 | 1 | 0 | 1 | 0 | 3 | 0 | 2 | 1 | 0 | 0 | 0 | 2 | 0 | 3 | 3 | 0 | 7 | 0 | 1 | 3 | 0 | 0 | 2 | 0 | 0 | 0 | 0 | Nutm1 |
| Nutm2 | 0 | 3 | 0 | 0 | 0 | 0 | 1 | 0 | 1 | 0 | 0 | 0 | 0 | 0 | 0 | 0 | 0 | 0 | 0 | 0 | 0 | 0 | 0 | 0 | 0 | 0 | 0 | 0 | Nutm2 |
| Nvl | 11389 | 10477 | 8416 | 9395 | 9226 | 10205 | 10830 | 12233 | 11471 | 10646 | 10900 | 11643 | 11208 | 12398 | 10714 | 10103 | 8914 | 9085 | 9245 | 10609 | 10797 | 10594 | 10804 | 10323 | 11296 | 9481 | 11517 | 11585 | Nvl |
| Nwd1 | 728 | 862 | 449 | 1229 | 1633 | 1421 | 1305 | 686 | 854 | 693 | 654 | 440 | 392 | 460 | 890 | 770 | 734 | 1438 | 1194 | 1108 | 1480 | 734 | 628 | 466 | 835 | 233 | 347 | 260 | Nwd1 |
| Nwd2 | 157 | 195 | 197 | 286 | 760 | 436 | 421 | 1172 | 1210 | 1325 | 1325 | 500 | 871 | 550 | 322 | 210 | 196 | 353 | 386 | 450 | 442 | 1527 | 1017 | 566 | 867 | 261 | 629 | 579 | Nwd2 |
| Nxf1 | 21917 | 23590 | 19498 | 19621 | 16827 | 17818 | 19986 | 17068 | 15888 | 15059 | 14168 | 15541 | 14995 | 16594 | 21606 | 20654 | 19986 | 19202 | 18770 | 19646 | 18812 | 15542 | 14870 | 15634 | 15198 | 16256 | 15933 | 16369 | Nxf1 |
| Nxf2 | 52 | 46 | 61 | 27 | 22 | 15 | 21 | 26 | 64 | 56 | 40 | 38 | 56 | 34 | 45 | 54 | 21 | 26 | 24 | 37 | 22 | 35 | 52 | 44 | 31 | 43 | 42 | 31 | Nxf2 |
| Nxf3 | 3 | 2 | 0 | 1 | 6 | 1 | 2 | 2 | 5 | 8 | 2 | 1 | 5 | 2 | 3 | 4 | 2 | 7 | 3 | 2 | 2 | 5 | 3 | 1 | 3 | 1 | 1 | 1 | Nxf3 |
| Nxf7 | 2 | 4 | 3 | 1 | 3 | 4 | 1 | 1 | 9 | 3 | 5 | 5 | 9 | 0 | 1 | 2 | 0 | 2 | 5 | 1 | 1 | 0 | 6 | 0 | 1 | 0 | 5 | 2 | Nxf7 |
| Nxn | 8155 | 9297 | 8920 | 6983 | 5905 | 6451 | 5707 | 4962 | 5912 | 4985 | 5224 | 5906 | 5994 | 5223 | 7727 | 8375 | 7674 | 6972 | 6551 | 6607 | 5830 | 3952 | 6609 | 5322 | 4243 | 6568 | 6465 | 6518 | Nxn |
| Nxnl1 | 2 | 4 | 10 | 9 | 6 | 1 | 11 | 1 | 5 | 1 | 7 | 0 | 3 | 3 | 1 | 2 | 7 | 0 | 0 | 10 | 2 | 0 | 5 | 7 | 1 | 2 | 0 | 1 | Nxnl1 |
| Nxnl2 | 183 | 176 | 122 | 131 | 101 | 93 | 105 | 33 | 69 | 56 | 71 | 21 | 76 | 23 | 141 | 147 | 131 | 111 | 110 | 60 | 120 | 35 | 63 | 45 | 29 | 73 | 22 | 21 | Nxnl2 |
| Nxpe2 | 37 | 23 | 24 | 31 | 24 | 25 | 37 | 22 | 39 | 21 | 26 | 27 | 39 | 27 | 26 | 37 | 31 | 62 | 35 | 37 | 37 | 28 | 32 | 16 | 29 | 41 | 36 | 43 | Nxpe2 |
| Nxpe3 | 1449 | 1283 | 1601 | 1473 | 1622 | 1497 | 1578 | 882 | 707 | 745 | 710 | 789 | 780 | 827 | 1493 | 1504 | 1424 | 1536 | 1515 | 1488 | 1487 | 866 | 659 | 830 | 856 | 843 | 780 | 853 | Nxpe3 |
| Nxpe4 | 1183 | 1057 | 941 | 971 | 945 | 1044 | 874 | 978 | 1051 | 917 | 935 | 1153 | 1193 | 1045 | 1305 | 1148 | 988 | 857 | 812 | 953 | 880 | 1084 | 1060 | 804 | 913 | 1094 | 1270 | 1123 | Nxpe4 |
| Nxpe5 | 1 | 0 | 0 | 0 | 1 | 0 | 0 | 0 | 2 | 0 | 0 | 0 | 0 | 0 | 0 | 0 | 0 | 0 | 1 | 0 | 0 | 0 | 1 | 0 | 4 | 0 | 0 | 0 | Nxpe5 |
| Nxph1 | 296 | 322 | 304 | 462 | 877 | 646 | 636 | 1688 | 1548 | 1619 | 1387 | 958 | 1024 | 1008 | 458 | 287 | 396 | 416 | 608 | 797 | 697 | 2164 | 1713 | 1066 | 1475 | 659 | 1030 | 958 | Nxph1 |
| Nxph2 | 2 | 5 | 0 | 10 | 16 | 9 | 35 | 53 | 78 | 102 | 74 | 18 | 18 | 28 | 8 | 3 | 3 | 12 | 18 | 30 | 27 | 74 | 74 | 81 | 105 | 16 | 10 | 17 | Nxph2 |
| Nxph3 | 625 | 677 | 754 | 664 | 696 | 693 | 562 | 253 | 339 | 416 | 268 | 181 | 202 | 180 | 462 | 603 | 728 | 595 | 772 | 618 | 644 | 244 | 332 | 403 | 371 | 250 | 172 | 153 | Nxph3 |
| Nxph4 | 614 | 902 | 744 | 1488 | 1960 | 1371 | 1192 | 1274 | 1267 | 1879 | 1521 | 1596 | 1430 | 1505 | 612 | 700 | 1138 | 1383 | 1568 | 1302 | 1381 | 1810 | 1374 | 1541 | 1721 | 1542 | 1191 | 1301 | Nxph4 |
| Nxt1 | 2013 | 2273 | 2086 | 1943 | 1150 | 1443 | 1715 | 1285 | 1236 | 1129 | 1055 | 1460 | 1245 | 1706 | 1814 | 1890 | 1910 | 1448 | 1538 | 1837 | 1535 | 882 | 1178 | 1493 | 1281 | 1891 | 1680 | 1856 | Nxt1 |
| Nxt2 | 1431 | 948 | 1243 | 1150 | 1126 | 1371 | 1506 | 1360 | 1219 | 1031 | 1021 | 1111 | 959 | 1221 | 1490 | 1262 | 1159 | 1113 | 1338 | 1704 | 1398 | 1091 | 1229 | 1066 | 1135 | 1343 | 1163 | 1227 | Nxt2 |
| Nyap1 | 5230 | 7768 | 6797 | 10202 | 10759 | 9523 | 9332 | 6977 | 7030 | 9236 | 7053 | 7752 | 6210 | 7535 | 6001 | 5439 | 9043 | 9678 | 11187 | 9248 | 10735 | 9000 | 6748 | 9334 | 10104 | 6872 | 6923 | 6961 | Nyap1 |
| Nyap2 | 284 | 265 | 273 | 666 | 1469 | 1022 | 1012 | 1340 | 1351 | 1408 | 1470 | 960 | 984 | 1040 | 494 | 342 | 495 | 815 | 831 | 765 | 944 | 2132 | 1272 | 974 | 1671 | 525 | 827 | 875 | Nyap2 |
| Nynrin | 10150 | 11295 | 14566 | 10353 | 9914 | 10248 | 8027 | 9163 | 12395 | 12093 | 13797 | 13523 | 14731 | 11897 | 8817 | 11728 | 12694 | 10469 | 10952 | 7361 | 8643 | 11757 | 12693 | 13197 | 10861 | 13299 | 13618 | 12936 | Nynrin |
| Nyx | 14 | 20 | 19 | 23 | 46 | 29 | 37 | 57 | 53 | 50 | 63 | 35 | 56 | 37 | 25 | 21 | 36 | 23 | 26 | 38 | 25 | 80 | 39 | 41 | 54 | 44 | 56 | 58 | Nyx |
| Oacyl | 60 | 63 | 30 | 48 | 79 | 58 | 60 | 90 | 73 | 80 | 105 | 91 | 155 | 116 | 45 | 41 | 54 | 55 | 49 | 42 | 71 | 99 | 89 | 69 | 69 | 64 | 123 | 87 | Oacyl |
| Oaf | 1938 | 2179 | 1735 | 1269 | 734 | 1188 | 761 | 1166 | 1638 | 1142 | 2046 | 1065 | 1500 | 953 | 1681 | 2549 | 1199 | 1346 | 850 | 907 | 760 | 1089 | 2008 | 1223 | 955 | 1475 | 1203 | 1116 | Oaf |
| Oard1 | 495 | 450 | 803 | 456 | 98 | 127 | 422 | 551 | 587 | 548 | 135 | 291 | 389 | 188 | 695 | 136 | 486 | 364 | 157 | 278 | 104 | 906 | 867 | 588 | 491 | 1265 | 809 | 222 | Oard1 |
| Oas1a | 22 | 30 | 13 | 22 | 11 | 18 | 4 | 20 | 25 | 29 | 45 | 22 | 14 | 20 | 20 | 23 | 43 | 17 | 9 | 41 | 21 | 32 | 39 | 34 | 30 | 37 | 36 | 49 | Oas1a |
| Oas1c | 68 | 60 | 52 | 55 | 61 | 55 | 40 | 42 | 50 | 34 | 50 | 39 | 49 | 57 | 75 | 54 | 92 | 44 | 47 | 66 | 55 | 40 | 47 | 43 | 46 | 83 | 48 | 33 | Oas1c |
| Oas1d | 0 | 0 | 0 | 0 | 0 | 0 | 0 | 0 | 1 | 0 | 1 | 2 | 1 | 1 | 4 | 0 | 0 | 0 | 2 | 0 | 0 | 0 | 0 | 0 | 0 | 0 | 0 | 0 | Oas1d |
| Oas1e | 0 | 0 | 0 | 3 | 2 | 2 | 3 | 5 | 3 | 0 | 7 | 8 | 6 | 5 | 4 | 2 | 3 | 0 | 2 | 4 | 3 | 5 | 5 | 1 | 0 | 4 | 2 | 7 | Oas1e |
| Oas1f | 1 | 0 | 0 | 0 | 0 | 0 | 1 | 0 | 2 | 3 | 1 | 2 | 0 | 0 | 0 | 1 | 0 | 0 | 1 | 0 | 0 | 0 | 1 | 0 | 0 | 0 | 0 | 0 | Oas1f |
| Oas1g | 3 | 3 | 5 | 4 | 5 | 2 | 3 | 2 | 3 | 0 | 6 | 10 | 7 | 6 | 1 | 3 | 3 | 1 | 2 | 4 | 5 | 10 | 10 | 9 | 5 | 8 | 5 | 8 | Oas1g |
| Oas2 | 15 | 16 | 8 | 38 | 37 | 14 | 22 | 13 | 33 | 69 | 74 | 72 | 45 | 46 | 20 | 20 | 25 | 21 | 12 | 54 | 37 | 80 | 66 | 49 | 25 | 62 | 107 | 55 | Oas2 |
| Oas3 | 10 | 14 | 1 | 11 | 25 | 19 | 23 | 27 | 33 | 23 | 27 | 21 | 9 | 15 | 5 | 18 | 22 | 23 | 25 | 21 | 18 | 57 | 31 | 39 | 46 | 12 | 26 | 26 | Oas3 |
| Oasl1 | 8 | 3 | 9 | 6 | 3 | 5 | 7 | 2 | 8 | 36 | 20 | 8 | 12 | 5 | 13 | 11 | 6 | 3 | 5 | 21 | 7 | 0 | 15 | 6 | 7 | 21 | 15 | 17 | Oasl1 |
| Oasl2 | 68 | 39 | 53 | 33 | 62 | 45 | 70 | 182 | 145 | 218 | 227 | 148 | 137 | 143 | 67 | 80 | 58 | 45 | 45 | 142 | 95 | 149 | 166 | 163 | 175 | 203 | 159 | 161 | Oasl2 |
| Oat | 9523 | 9607 | 8441 | 7341 | 5986 | 6969 | 6836 | 7607 | 8768 | 6501 | 7577 | 6820 | 6870 | 6672 | 9758 | 9141 | 7842 | 7191 | 7338 | 7548 | 6970 | 6351 | 8371 | 6703 | 6249 | 7508 | 7304 | 6906 | Oat |
| Oaz1 | 15047 | 19635 | 20089 | 18543 | 14042 | 14287 | 14434 | 12752 | 14169 | 14530 | 14456 | 17531 | 17126 | 16388 | 13878 | 15673 | 19229 | 17314 | 17197 | 15163 | 14102 | 12078 | 13698 | 17254 | 15389 | 22077 | 18371 | 18247 | Oaz1 |
| Oaz2 | 11553 | 11549 | 12197 | 12591 | 11144 | 11283 | 10752 | 8913 | 8558 | 8660 | 9203 | 8687 | 8768 | 8375 | 11395 | 11670 | 12165 | 12440 | 12372 | 10833 | 11234 | 9125 | 8965 | 8832 | 9080 | 8937 | 8801 | 8294 | Oaz2 |
| Oaz3 | 43 | 54 | 40 | 80 | 55 | 25 | 67 | 2 | 10 | 12 | 4 | 11 | 9 | 12 | 28 | 47 | 80 | 85 | 58 | 37 | 62 | 7 | 9 | 13 | 2 | 8 | 3 | 9 | Oaz3 |
| Obi1 | 4241 | 3956 | 5317 | 4272 | 4299 | 3961 | 3904 | 3885 | 3374 | 3545 | 3315 | 4007 | 4093 | 4085 | 4362 | 4178 | 4870 | 4164 | 4419 | 4098 | 3782 | 3725 | 3327 | 3720 | 3592 | 4636 | 4341 | 4371 | Obi1 |
| Obox6 | 0 | 0 | 0 | 0 | 0 | 0 | 0 | 0 | 0 | 0 | 0 | 0 | 0 | 0 | 0 | 0 | 0 | 0 | 0 | 0 | 0 | 0 | 0 | 0 | 0 | 4 | 0 | 0 | Obox6 |
| Obox7 | 0 | 0 | 0 | 0 | 0 | 0 | 0 | 0 | 0 | 0 | 1 | 0 | 0 | 3 | 0 | 0 | 0 | 0 | 0 | 0 | 0 | 0 | 0 | 0 | 0 | 0 | 0 | 0 | Obox7 |
| Obp1a | 0 | 0 | 0 | 0 | 1 | 0 | 0 | 1 | 0 | 5 | 0 | 2 | 0 | 1 | 0 | 0 | 0 | 0 | 0 | 0 | 1 | 0 | 0 | 0 | 2 | 8 | 0 | 4 | Obp1a |
| Obp2a | 0 | 3 | 0 | 0 | 0 | 0 | 0 | 0 | 0 | 0 | 0 | 0 | 0 | 0 | 0 | 1 | 0 | 0 | 0 | 0 | 1 | 0 | 0 | 0 | 0 | 0 | 0 | 0 | Obp2a |
| Obp2b | 3 | 3 | 12 | 3 | 4 | 1 | 6 | 0 | 0 | 2 | 0 | 1 | 0 | 3 | 8 | 2 | 12 | 11 | 4 | 6 | 4 | 3 | 0 | 0 | 0 | 0 | 2 | 0 | Obp2b |
| Obscn | 45 | 44 | 42 | 43 | 163 | 83 | 60 | 122 | 202 | 128 | 985 | 53 | 85 | 56 | 51 | 60 | 31 | 54 | 62 | 39 | 62 | 227 | 548 | 52 | 125 | 8 | 44 | 34 | Obscn |
| Obsl1 | 2738 | 2777 | 3371 | 2279 | 2313 | 2348 | 1501 | 3816 | 5756 | 5485 | 8913 | 4591 | 5316 | 3965 | 2076 | 3292 | 2707 | 2348 | 2426 | 1571 | 1615 | 5103 | 6978 | 5449 | 4383 | 5242 | 4829 | 4796 | Obsl1 |
| Oc90 | 559 | 1168 | 379 | 5 | 61 | 456 | 7 | 595 | 3248 | 1354 | 2851 | 9 | 1958 | 4 | 2710 | 519 | 12 | 4 | 49 | 5 | 14 | 2172 | 1480 | 11 | 651 | 6 | 7 | 0 | Oc90 |
| Oca2 | 14 | 13 | 6 | 18 | 43 | 21 | 19 | 32 | 44 | 45 | 25 | 39 | 52 | 15 | 9 | 22 | 8 | 23 | 22 | 15 | 17 | 43 | 120 | 10 | 23 | 13 | 9 | 5 | Oca2 |
| Ocel1 | 115 | 131 | 136 | 133 | 232 | 188 | 204 | 264 | 333 | 401 | 373 | 232 | 252 | 252 | 141 | 112 | 166 | 229 | 224 | 188 | 212 | 411 | 367 | 385 | 444 | 212 | 199 | 248 | Ocel1 |
| Ociad1 | 9789 | 9493 | 9224 | 9471 | 9499 | 9151 | 10208 | 13161 | 12163 | 11275 | 11779 | 10844 | 11303 | 11698 | 10249 | 9233 | 8951 | 9523 | 9828 | 10367 | 9979 | 11952 | 12629 | 10755 | 11497 | 11465 | 11647 | 11822 | Ociad1 |
| Ociad2 | 516 | 458 | 673 | 573 | 704 | 649 | 650 | 888 | 791 | 930 | 827 | 916 | 855 | 1025 | 596 | 502 | 670 | 593 | 796 | 679 | 685 | 891 | 889 | 933 | 1046 | 904 | 913 | 1056 | Ociad2 |
| Ocln | 571 | 329 | 467 | 325 | 470 | 392 | 411 | 386 | 386 | 368 | 393 | 249 | 322 | 227 | 512 | 486 | 273 | 316 | 295 | 390 | 389 | 399 | 599 | 321 | 225 | 316 | 196 | 212 | Ocln |
| Ocm | 0 | 0 | 0 | 0 | 0 | 0 | 0 | 0 | 1 | 0 | 0 | 2 | 0 | 0 | 0 | 0 | 0 | 0 | 0 | 0 | 0 | 3 | 0 | 0 | 0 | 4 | 0 | 0 | Ocm |
| Ocrl | 5527 | 4885 | 4513 | 4815 | 5640 | 5388 | 5884 | 6466 | 6110 | 5686 | 5816 | 5791 | 5390 | 5799 | 5705 | 4933 | 4904 | 5132 | 5301 | 5531 | 5694 | 6488 | 5930 | 5190 | 5909 | 4812 | 5588 | 5521 | Ocrl |
| Ocstamp | 18 | 12 | 27 | 13 | 8 | 7 | 10 | 6 | 4 | 2 | 3 | 4 | 3 | 2 | 5 | 18 | 20 | 17 | 5 | 10 | 14 | 2 | 1 | 3 | 4 | 7 | 7 | 5 | Ocstamp |
| Odad1 | 416 | 555 | 644 | 519 | 626 | 514 | 438 | 339 | 401 | 405 | 435 | 433 | 476 | 324 | 477 | 662 | 565 | 573 | 488 | 509 | 441 | 428 | 515 | 311 | 347 | 363 | 420 | 376 | Odad1 |

| | | | | | | | | | | | | | | | | | | | | | | | | | | | | | |
|---|---|---|---|---|---|---|---|---|---|---|---|---|---|---|---|---|---|---|---|---|---|---|---|---|---|---|---|---|---|
| Odad2 | 40 | 56 | 20 | 33 | 41 | 61 | 49 | 34 | 32 | 34 | 34 | 68 | 53 | 38 | 54 | 58 | 43 | 52 | 48 | 41 | 42 | 84 | 72 | 39 | 26 | 16 | 28 | 30 | Odad2 |
| Odad3 | 474 | 698 | 751 | 680 | 521 | 492 | 520 | 380 | 358 | 491 | 277 | 478 | 460 | 466 | 512 | 523 | 643 | 643 | 641 | 603 | 530 | 416 | 440 | 433 | 437 | 447 | 431 | 436 | Odad3 |
| Odad4 | 170 | 222 | 235 | 192 | 244 | 201 | 228 | 171 | 180 | 207 | 133 | 237 | 225 | 217 | 204 | 194 | 191 | 212 | 204 | 248 | 194 | 222 | 274 | 182 | 214 | 194 | 217 | 189 | Odad4 |
| Odam | 0 | 0 | 0 | 0 | 0 | 0 | 0 | 2 | 0 | 0 | 1 | 0 | 0 | 0 | 0 | 0 | 0 | 0 | 0 | 0 | 0 | 0 | 2 | 0 | 0 | 0 | 0 | 0 | Odam |
| Odaph | 8 | 4 | 2 | 2 | 2 | 1 | 2 | 2 | 6 | 0 | 4 | 4 | 9 | 4 | 1 | 6 | 9 | 3 | 4 | 4 | 10 | 0 | 1 | 2 | 0 | 2 | 1 | 4 | Odaph |
| Odc1 | 34783 | 34089 | 30395 | 30007 | 21669 | 25326 | 26585 | 18070 | 18728 | 16491 | 16711 | 22518 | 20924 | 22311 | 34474 | 31755 | 28875 | 28080 | 24188 | 27006 | 25556 | 14878 | 17697 | 18336 | 18070 | 25274 | 21896 | 23518 | Odc1 |
| Odf1 | 3 | 2 | 2 | 3 | 3 | 2 | 2 | 1 | 2 | 1 | 1 | 0 | 0 | 0 | 5 | 10 | 2 | 3 | 0 | 3 | 1 | 0 | 0 | 0 | 0 | 0 | 0 | 0 | Odf1 |
| Odf2 | 16504 | 17855 | 15772 | 16979 | 15419 | 16390 | 16508 | 9343 | 9527 | 10382 | 9518 | 11983 | 11455 | 12456 | 15523 | 14789 | 16220 | 16011 | 16673 | 16482 | 17130 | 8913 | 8709 | 10145 | 10901 | 11571 | 12459 | 11921 | Odf2 |
| Odf2l | 2321 | 1853 | 1960 | 1932 | 1874 | 2129 | 2274 | 4131 | 3466 | 3527 | 3408 | 4449 | 3957 | 4413 | 2301 | 2011 | 2060 | 1749 | 2068 | 2276 | 2215 | 3355 | 3345 | 3703 | 3605 | 3611 | 4482 | 4296 | Odf2l |
| Odf3 | 0 | 0 | 0 | 0 | 1 | 0 | 0 | 0 | 1 | 2 | 2 | 0 | 0 | 0 | 0 | 0 | 0 | 0 | 0 | 0 | 0 | 2 | 2 | 2 | 0 | 1 | 0 | 0 | Odf3 |
| Odf3b | 3 | 16 | 8 | 11 | 16 | 13 | 28 | 13 | 26 | 18 | 21 | 24 | 32 | 13 | 9 | 24 | 16 | 7 | 19 | 17 | 12 | 12 | 33 | 5 | 31 | 13 | 6 | 17 | Odf3b |
| Odf3l1 | 1 | 0 | 0 | 0 | 1 | 0 | 2 | 2 | 13 | 10 | 18 | 2 | 12 | 1 | 0 | 0 | 0 | 0 | 2 | 0 | 2 | 10 | 24 | 4 | 0 | 6 | 0 | 0 | Odf3l1 |
| Odf3l2 | 0 | 0 | 0 | 0 | 0 | 0 | 0 | 1 | 1 | 5 | 2 | 1 | 0 | 2 | 0 | 0 | 0 | 0 | 0 | 0 | 0 | 0 | 1 | 0 | 7 | 0 | 0 | 0 | Odf3l2 |
| Odf4 | 24 | 29 | 22 | 103 | 118 | 91 | 61 | 57 | 44 | 74 | 58 | 63 | 68 | 50 | 33 | 28 | 54 | 78 | 84 | 82 | 82 | 102 | 59 | 75 | 98 | 36 | 45 | 36 | Odf4 |
| Odr4 | 1939 | 1581 | 1900 | 1736 | 1871 | 1991 | 1953 | 3745 | 3540 | 3144 | 3679 | 3113 | 3319 | 3350 | 2197 | 1896 | 1773 | 1827 | 1755 | 2080 | 1885 | 3430 | 3724 | 3322 | 3367 | 3240 | 3197 | 3336 | Odr4 |
| Ofcc1 | 2 | 0 | 3 | 2 | 4 | 1 | 1 | 1 | 4 | 0 | 8 | 1 | 3 | 1 | 0 | 1 | 0 | 4 | 3 | 0 | 4 | 8 | 2 | 1 | 6 | 0 | 1 | 0 | Ofcc1 |
| Ofd1 | 1885 | 1476 | 1455 | 1531 | 1818 | 1852 | 1804 | 2214 | 2071 | 2252 | 2095 | 2437 | 2394 | 2529 | 1975 | 1656 | 1727 | 1585 | 1785 | 1629 | 1726 | 2383 | 2053 | 2128 | 2243 | 2513 | 2474 | 2425 | Ofd1 |
| Oga | 25622 | 19587 | 22909 | 25126 | 26124 | 25303 | 28641 | 30879 | 25818 | 26951 | 25440 | 28241 | 27372 | 30404 | 25913 | 19805 | 23549 | 25312 | 25860 | 24713 | 27142 | 32174 | 25509 | 27400 | 29593 | 24686 | 28469 | 28324 | Oga |
| Ogdh | 9912 | 10940 | 12073 | 11950 | 11582 | 10786 | 11039 | 7222 | 8713 | 8955 | 9728 | 10461 | 9964 | 10068 | 9646 | 11130 | 11932 | 12143 | 12446 | 8174 | 11563 | 10029 | 8393 | 9459 | 9717 | 9749 | 9009 | 10610 | Ogdh |
| Ogdhl | 2489 | 2984 | 3040 | 3023 | 3126 | 2930 | 2854 | 2380 | 2756 | 3080 | 2647 | 3077 | 2718 | 3012 | 2457 | 2842 | 2955 | 2840 | 3096 | 2492 | 2918 | 2694 | 2359 | 3103 | 2976 | 2779 | 3004 | 2847 | Ogdhl |
| Ogfod1 | 6195 | 5581 | 5057 | 5234 | 4565 | 5147 | 5357 | 2072 | 2174 | 2124 | 2168 | 1988 | 2139 | 2186 | 5408 | 5942 | 4926 | 5714 | 4710 | 4649 | 5126 | 2034 | 2092 | 2175 | 2081 | 1937 | 2037 | 2034 | Ogfod1 |
| Ogfod2 | 1559 | 1957 | 1807 | 2334 | 1789 | 1651 | 1757 | 767 | 839 | 1015 | 995 | 825 | 886 | 811 | 1501 | 1606 | 1976 | 2042 | 1912 | 1554 | 1833 | 967 | 813 | 1056 | 981 | 918 | 786 | 838 | Ogfod2 |
| Ogfod3 | 200 | 221 | 270 | 251 | 243 | 210 | 208 | 379 | 419 | 425 | 472 | 428 | 407 | 501 | 231 | 243 | 226 | 286 | 251 | 214 | 233 | 501 | 478 | 469 | 463 | 561 | 496 | 478 | Ogfod3 |
| Ogfr | 5066 | 6183 | 6047 | 5336 | 5099 | 5395 | 4828 | 5915 | 6297 | 6481 | 6200 | 7014 | 6319 | 6976 | 4837 | 5145 | 6128 | 4934 | 5424 | 5309 | 4866 | 5870 | 6090 | 6785 | 6657 | 7555 | 7277 | 7321 | Ogfr |
| Ogfrl1 | 3895 | 2888 | 3435 | 3633 | 3871 | 3839 | 4405 | 2186 | 2183 | 2164 | 2280 | 2048 | 1972 | 2074 | 3944 | 3389 | 3264 | 3779 | 3532 | 3685 | 3948 | 2216 | 2059 | 1931 | 2306 | 2209 | 2011 | 2007 | Ogfrl1 |
| Ogg1 | 827 | 917 | 1016 | 982 | 745 | 788 | 853 | 618 | 702 | 742 | 657 | 764 | 744 | 765 | 845 | 904 | 914 | 872 | 847 | 752 | 827 | 483 | 691 | 690 | 585 | 1024 | 873 | 809 | Ogg1 |
| Ogn | 1912 | 1320 | 1292 | 1264 | 883 | 1749 | 419 | 3907 | 5570 | 3541 | 6732 | 2318 | 3139 | 2263 | 2339 | 2163 | 1455 | 2272 | 532 | 607 | 567 | 4318 | 9123 | 3501 | 2873 | 2661 | 2593 | 2136 | Ogn |
| Ogt | 35634 | 29627 | 22961 | 29332 | 35346 | 36886 | 35933 | 60137 | 50507 | 45685 | 47719 | 45581 | 45245 | 46249 | 36636 | 31291 | 26103 | 30463 | 32515 | 39933 | 38873 | 49504 | 49458 | 39795 | 48392 | 30511 | 46586 | 42925 | Ogt |
| Oip5 | 952 | 822 | 972 | 780 | 524 | 695 | 810 | 790 | 641 | 741 | 654 | 1108 | 885 | 1041 | 985 | 768 | 846 | 646 | 726 | 851 | 755 | 550 | 734 | 894 | 687 | 1360 | 1148 | 1164 | Oip5 |
| Oit3 | 22 | 43 | 15 | 30 | 24 | 17 | 15 | 21 | 37 | 23 | 23 | 25 | 28 | 29 | 21 | 36 | 20 | 31 | 26 | 17 | 7 | 40 | 52 | 25 | 23 | 18 | 33 | 20 | Oit3 |
| Ola1 | 9358 | 7372 | 8434 | 8587 | 7494 | 7307 | 9952 | 5924 | 5241 | 5502 | 5673 | 5764 | 5586 | 6829 | 8930 | 7812 | 8538 | 8538 | 7783 | 7483 | 9099 | 5299 | 5640 | 5929 | 5685 | 7171 | 5711 | 6812 | Ola1 |
| Olah | 0 | 0 | 0 | 0 | 0 | 3 | 1 | 0 | 2 | 1 | 1 | 1 | 0 | 4 | 0 | 1 | 0 | 0 | 1 | 0 | 2 | 0 | 3 | 2 | 1 | 2 | 2 | 1 | Olah |
| Olfm1 | 4791 | 5958 | 6607 | 8297 | 13394 | 8995 | 8599 | 9962 | 10214 | 12964 | 10983 | 9036 | 10103 | 8527 | 7292 | 5766 | 8553 | 9564 | 9845 | 7627 | 9671 | 16001 | 9985 | 10003 | 11878 | 7028 | 7031 | 7659 | Olfm1 |
| Olfm2 | 1466 | 2207 | 1855 | 2691 | 3503 | 2608 | 2855 | 1928 | 2347 | 2853 | 2105 | 1767 | 1852 | 1713 | 1799 | 1567 | 2504 | 2751 | 3250 | 2546 | 3153 | 3105 | 2221 | 2111 | 2715 | 1434 | 1411 | 1447 | Olfm2 |
| Olfm3 | 221 | 244 | 325 | 442 | 776 | 525 | 588 | 570 | 492 | 480 | 460 | 260 | 435 | 229 | 414 | 329 | 337 | 471 | 471 | 694 | 589 | 802 | 557 | 282 | 519 | 104 | 225 | 279 | Olfm3 |
| Olfm4 | 38 | 36 | 50 | 84 | 54 | 54 | 66 | 65 | 53 | 65 | 130 | 77 | 91 | 56 | 71 | 60 | 67 | 86 | 43 | 61 | 58 | 82 | 110 | 95 | 52 | 77 | 70 | 69 | Olfm4 |
| Olfm5 | 8 | 0 | 0 | 0 | 0 | 0 | 0 | 277 | 190 | 357 | 149 | 2 | 2 | 0 | 7 | 8 | 0 | 0 | 0 | 0 | 0 | 2 | 366 | 358 | 0 | 56 | 0 | 0 | Olfm5 |
| Olfml1 | 638 | 447 | 358 | 288 | 149 | 281 | 86 | 646 | 751 | 561 | 715 | 401 | 645 | 392 | 512 | 642 | 283 | 329 | 119 | 126 | 115 | 460 | 889 | 556 | 448 | 619 | 444 | 417 | Olfml1 |
| Olfml2a | 762 | 772 | 760 | 630 | 893 | 753 | 324 | 1159 | 1866 | 1477 | 2025 | 993 | 1302 | 1001 | 880 | 970 | 611 | 942 | 389 | 336 | 348 | 1927 | 2318 | 1104 | 1204 | 837 | 896 | 926 | Olfml2a |
| Olfml2b | 423 | 482 | 436 | 477 | 551 | 461 | 349 | 496 | 775 | 629 | 1345 | 533 | 533 | 486 | 430 | 452 | 433 | 475 | 536 | 416 | 369 | 590 | 1019 | 599 | 559 | 571 | 453 | 468 | Olfml2b |
| Olfml3 | 971 | 961 | 770 | 635 | 458 | 859 | 268 | 1495 | 2208 | 1793 | 2785 | 1149 | 1890 | 1230 | 956 | 1206 | 671 | 940 | 348 | 320 | 325 | 1653 | 2589 | 1750 | 1280 | 1807 | 1363 | 1347 | Olfml3 |
| Olfr1 | 0 | 0 | 0 | 0 | 0 | 0 | 0 | 0 | 0 | 1 | 2 | 0 | 0 | 0 | 0 | 0 | 0 | 0 | 0 | 0 | 0 | 0 | 0 | 0 | 0 | 0 | 0 | 0 | Olfr1 |
| Olfr10 | 0 | 0 | 0 | 4 | 4 | 0 | 0 | 0 | 0 | 0 | 0 | 4 | 1 | 0 | 0 | 0 | 5 | 6 | 0 | 0 | 4 | 0 | 0 | 2 | 3 | 1 | 0 | 0 | Olfr10 |
| Olfr1000 | 0 | 0 | 0 | 0 | 0 | 0 | 0 | 0 | 0 | 0 | 0 | 0 | 0 | 0 | 0 | 0 | 0 | 0 | 0 | 0 | 0 | 0 | 0 | 0 | 0 | 1 | 0 | 0 | Olfr1000 |
| Olfr1002 | 0 | 0 | 0 | 0 | 0 | 0 | 0 | 0 | 0 | 0 | 2 | 0 | 0 | 0 | 0 | 0 | 0 | 0 | 0 | 0 | 0 | 0 | 0 | 0 | 0 | 0 | 0 | 0 | Olfr1002 |
| Olfr1006 | 0 | 0 | 0 | 1 | 0 | 0 | 0 | 2 | 3 | 6 | 6 | 0 | 0 | 0 | 0 | 1 | 0 | 0 | 0 | 0 | 0 | 0 | 5 | 4 | 0 | 5 | 0 | 1 | Olfr1006 |
| Olfr101 | 0 | 0 | 0 | 0 | 0 | 0 | 0 | 2 | 2 | 0 | 0 | 0 | 0 | 0 | 0 | 0 | 0 | 0 | 0 | 0 | 0 | 0 | 0 | 0 | 0 | 0 | 0 | 0 | Olfr101 |
| Olfr1012 | 0 | 0 | 0 | 0 | 0 | 0 | 0 | 0 | 0 | 0 | 0 | 0 | 0 | 0 | 0 | 0 | 0 | 0 | 0 | 0 | 0 | 0 | 1 | 0 | 0 | 0 | 0 | 0 | Olfr1012 |
| Olfr1013 | 0 | 0 | 0 | 0 | 0 | 0 | 0 | 0 | 0 | 1 | 0 | 0 | 0 | 0 | 0 | 0 | 0 | 0 | 0 | 0 | 0 | 0 | 1 | 0 | 0 | 2 | 0 | 0 | Olfr1013 |
| Olfr1014 | 0 | 0 | 0 | 0 | 0 | 0 | 0 | 0 | 0 | 0 | 1 | 0 | 0 | 0 | 0 | 0 | 0 | 0 | 0 | 0 | 0 | 0 | 1 | 0 | 0 | 0 | 0 | 0 | Olfr1014 |
| Olfr1015 | 0 | 0 | 0 | 0 | 0 | 0 | 0 | 0 | 2 | 0 | 0 | 0 | 0 | 0 | 0 | 0 | 0 | 0 | 0 | 0 | 0 | 0 | 0 | 0 | 0 | 0 | 0 | 0 | Olfr1015 |
| Olfr1016 | 0 | 0 | 0 | 0 | 0 | 0 | 0 | 1 | 2 | 0 | 1 | 0 | 0 | 0 | 0 | 0 | 0 | 0 | 0 | 0 | 0 | 0 | 1 | 2 | 0 | 3 | 0 | 0 | Olfr1016 |
| Olfr1018 | 0 | 0 | 0 | 0 | 0 | 0 | 0 | 0 | 0 | 0 | 1 | 0 | 1 | 0 | 0 | 0 | 0 | 0 | 0 | 0 | 0 | 0 | 2 | 0 | 0 | 0 | 0 | 0 | Olfr1018 |
| Olfr1019 | 1 | 0 | 0 | 0 | 0 | 0 | 0 | 0 | 0 | 0 | 0 | 0 | 1 | 0 | 0 | 0 | 0 | 0 | 0 | 0 | 0 | 0 | 0 | 0 | 0 | 0 | 0 | 0 | Olfr1019 |
| Olfr102 | 0 | 0 | 0 | 2 | 0 | 0 | 0 | 0 | 0 | 0 | 1 | 0 | 0 | 0 | 0 | 0 | 0 | 0 | 0 | 0 | 0 | 0 | 0 | 0 | 0 | 0 | 0 | 0 | Olfr102 |
| Olfr1020 | 0 | 0 | 0 | 0 | 0 | 0 | 0 | 0 | 0 | 0 | 3 | 0 | 0 | 0 | 0 | 0 | 0 | 0 | 0 | 0 | 0 | 0 | 1 | 2 | 0 | 1 | 0 | 0 | Olfr1020 |
| Olfr1023 | 0 | 0 | 0 | 0 | 0 | 0 | 0 | 2 | 0 | 0 | 3 | 0 | 0 | 0 | 0 | 0 | 0 | 0 | 0 | 0 | 0 | 0 | 0 | 0 | 0 | 0 | 0 | 0 | Olfr1023 |
| Olfr1024 | 0 | 0 | 0 | 0 | 0 | 0 | 0 | 0 | 0 | 0 | 0 | 0 | 0 | 0 | 0 | 0 | 0 | 0 | 0 | 0 | 0 | 0 | 0 | 4 | 0 | 0 | 0 | 0 | Olfr1024 |
| Olfr1025-ps1 | 0 | 0 | 0 | 1 | 0 | 0 | 0 | 4 | 9 | 3 | 1 | 0 | 2 | 0 | 0 | 0 | 0 | 0 | 0 | 0 | 0 | 0 | 14 | 8 | 0 | 0 | 0 | 0 | Olfr1025-ps1 |
| Olfr1026 | 0 | 0 | 0 | 0 | 0 | 0 | 0 | 0 | 0 | 0 | 7 | 0 | 0 | 0 | 0 | 0 | 0 | 0 | 0 | 0 | 0 | 0 | 1 | 5 | 0 | 7 | 0 | 0 | Olfr1026 |
| Olfr1028 | 0 | 0 | 0 | 0 | 0 | 0 | 0 | 0 | 0 | 1 | 2 | 0 | 0 | 0 | 0 | 0 | 0 | 0 | 0 | 0 | 0 | 0 | 2 | 1 | 0 | 0 | 0 | 0 | Olfr1028 |
| Olfr1029 | 0 | 0 | 0 | 0 | 0 | 0 | 0 | 1 | 0 | 0 | 0 | 0 | 0 | 0 | 0 | 0 | 0 | 0 | 0 | 0 | 0 | 0 | 0 | 0 | 0 | 0 | 0 | 2 | Olfr1029 |
| Olfr103 | 0 | 0 | 0 | 0 | 0 | 0 | 0 | 0 | 0 | 0 | 0 | 0 | 0 | 0 | 0 | 0 | 0 | 0 | 0 | 0 | 0 | 0 | 0 | 3 | 0 | 0 | 0 | 0 | Olfr103 |
| Olfr1030 | 0 | 0 | 0 | 0 | 0 | 0 | 0 | 0 | 2 | 4 | 0 | 0 | 2 | 0 | 0 | 0 | 0 | 0 | 0 | 0 | 0 | 0 | 2 | 3 | 0 | 4 | 0 | 0 | Olfr1030 |
| Olfr1031 | 0 | 0 | 0 | 0 | 0 | 0 | 0 | 5 | 2 | 1 | 1 | 0 | 0 | 0 | 0 | 0 | 0 | 0 | 0 | 0 | 0 | 0 | 1 | 5 | 0 | 2 | 0 | 0 | Olfr1031 |
| Olfr1032 | 0 | 0 | 0 | 0 | 0 | 0 | 0 | 0 | 1 | 6 | 2 | 0 | 0 | 0 | 0 | 0 | 0 | 0 | 0 | 0 | 0 | 0 | 1 | 0 | 0 | 1 | 0 | 0 | Olfr1032 |
| Olfr1033 | 6 | 6 | 14 | 1 | 13 | 0 | 3 | 22 | 12 | 19 | 76 | 10 | 10 | 10 | 7 | 8 | 0 | 1 | 6 | 1 | 4 | 25 | 62 | 14 | 12 | 17 | 3 | 7 | Olfr1033 |
| Olfr1034 | 0 | 0 | 0 | 5 | 0 | 0 | 1 | 6 | 3 | 9 | 7 | 0 | 2 | 2 | 0 | 0 | 0 | 0 | 0 | 0 | 1 | 0 | 8 | 13 | 0 | 1 | 0 | 0 | Olfr1034 |
| Olfr1036 | 0 | 0 | 0 | 0 | 0 | 0 | 0 | 1 | 0 | 0 | 0 | 0 | 0 | 0 | 0 | 0 | 0 | 0 | 0 | 0 | 0 | 0 | 1 | 2 | 0 | 0 | 0 | 0 | Olfr1036 |
| Olfr1037 | 0 | 0 | 0 | 1 | 0 | 0 | 0 | 0 | 0 | 0 | 0 | 0 | 0 | 0 | 0 | 0 | 0 | 0 | 0 | 0 | 0 | 0 | 0 | 0 | 0 | 1 | 0 | 0 | Olfr1037 |
| Olfr1039 | 0 | 0 | 0 | 0 | 0 | 0 | 0 | 2 | 0 | 0 | 0 | 0 | 0 | 0 | 0 | 0 | 0 | 0 | 0 | 0 | 0 | 0 | 0 | 0 | 0 | 0 | 0 | 0 | Olfr1039 |
| Olfr1040 | 0 | 0 | 0 | 0 | 0 | 0 | 0 | 0 | 0 | 0 | 0 | 0 | 0 | 0 | 0 | 0 | 0 | 0 | 0 | 0 | 0 | 0 | 0 | 0 | 0 | 3 | 0 | 0 | Olfr1040 |
| Olfr1042 | 0 | 0 | 0 | 0 | 0 | 0 | 0 | 2 | 3 | 0 | 0 | 0 | 0 | 0 | 0 | 0 | 0 | 0 | 0 | 0 | 0 | 0 | 1 | 0 | 0 | 0 | 0 | 0 | Olfr1042 |
| Olfr1043 | 0 | 0 | 0 | 0 | 0 | 0 | 0 | 1 | 2 | 0 | 0 | 0 | 0 | 0 | 0 | 0 | 0 | 0 | 0 | 0 | 0 | 0 | 1 | 0 | 0 | 1 | 0 | 1 | Olfr1043 |
| Olfr1044 | 0 | 0 | 0 | 0 | 0 | 0 | 0 | 1 | 2 | 1 | 1 | 0 | 0 | 2 | 0 | 0 | 0 | 0 | 0 | 0 | 0 | 0 | 0 | 6 | 0 | 0 | 0 | 0 | Olfr1044 |
| Olfr1045 | 0 | 0 | 0 | 0 | 0 | 0 | 0 | 1 | 1 | 0 | 0 | 0 | 0 | 0 | 0 | 0 | 0 | 0 | 0 | 0 | 0 | 0 | 1 | 0 | 0 | 0 | 0 | 0 | Olfr1045 |
| Olfr1046 | 0 | 0 | 0 | 0 | 0 | 0 | 0 | 0 | 0 | 1 | 0 | 0 | 0 | 0 | 0 | 0 | 0 | 0 | 0 | 0 | 0 | 0 | 1 | 0 | 0 | 1 | 0 | 0 | Olfr1046 |
| Olfr1047 | 0 | 0 | 0 | 0 | 0 | 0 | 0 | 0 | 1 | 0 | 0 | 0 | 0 | 0 | 0 | 0 | 0 | 0 | 0 | 0 | 0 | 0 | 1 | 0 | 0 | 0 | 0 | 0 | Olfr1047 |
| Olfr1048 | 0 | 0 | 0 | 0 | 0 | 0 | 0 | 1 | 0 | 1 | 0 | 0 | 0 | 0 | 0 | 0 | 0 | 0 | 0 | 0 | 0 | 0 | 0 | 1 | 0 | 0 | 0 | 0 | Olfr1048 |

| | | | | | | | | | | | | | | | | | | | | | | | | | | | | | |
|---|---|---|---|---|---|---|---|---|---|---|---|---|---|---|---|---|---|---|---|---|---|---|---|---|---|---|---|---|---|
| Olfr104-ps | 0 | 0 | 0 | 0 | 0 | 0 | 0 | 0 | 0 | 0 | 0 | 0 | 0 | 0 | 0 | 0 | 0 | 0 | 0 | 0 | 0 | 0 | 3 | 2 | 0 | 3 | 0 | 0 | Olfr104-ps |
| Olfr1051 | 0 | 0 | 0 | 0 | 0 | 0 | 0 | 0 | 0 | 0 | 0 | 0 | 0 | 0 | 0 | 0 | 0 | 0 | 0 | 0 | 0 | 0 | 1 | 1 | 0 | 0 | 0 | 0 | Olfr1051 |
| Olfr1052 | 0 | 0 | 0 | 0 | 0 | 0 | 0 | 0 | 1 | 0 | 0 | 0 | 0 | 0 | 0 | 0 | 0 | 0 | 0 | 0 | 0 | 0 | 0 | 0 | 0 | 0 | 0 | 0 | Olfr1052 |
| Olfr1054 | 0 | 0 | 0 | 0 | 0 | 0 | 0 | 0 | 0 | 0 | 0 | 0 | 0 | 0 | 0 | 0 | 0 | 0 | 0 | 0 | 0 | 0 | 0 | 2 | 0 | 0 | 0 | 0 | Olfr1054 |
| Olfr1056 | 1 | 0 | 0 | 0 | 0 | 0 | 0 | 4 | 2 | 1 | 2 | 0 | 0 | 0 | 0 | 0 | 0 | 0 | 0 | 0 | 0 | 0 | 4 | 2 | 0 | 3 | 0 | 0 | Olfr1056 |
| Olfr1057 | 0 | 0 | 0 | 0 | 0 | 0 | 0 | 2 | 1 | 1 | 0 | 0 | 0 | 0 | 0 | 0 | 0 | 0 | 0 | 0 | 0 | 0 | 2 | 2 | 0 | 0 | 0 | 0 | Olfr1057 |
| Olfr1058 | 0 | 0 | 0 | 0 | 0 | 0 | 0 | 0 | 0 | 0 | 3 | 0 | 0 | 0 | 0 | 0 | 0 | 0 | 0 | 0 | 0 | 0 | 1 | 0 | 0 | 0 | 0 | 0 | Olfr1058 |
| Olfr105-ps | 0 | 0 | 0 | 0 | 0 | 0 | 0 | 4 | 2 | 0 | 1 | 0 | 0 | 0 | 0 | 0 | 0 | 0 | 0 | 0 | 0 | 0 | 3 | 2 | 0 | 0 | 0 | 0 | Olfr105-ps |
| Olfr1062 | 0 | 0 | 0 | 0 | 0 | 0 | 0 | 0 | 1 | 0 | 0 | 0 | 0 | 0 | 0 | 0 | 0 | 0 | 0 | 0 | 0 | 0 | 0 | 1 | 0 | 0 | 0 | 0 | Olfr1062 |
| Olfr1065 | 0 | 0 | 0 | 0 | 0 | 0 | 0 | 0 | 0 | 0 | 1 | 0 | 1 | 0 | 0 | 0 | 0 | 0 | 0 | 0 | 0 | 0 | 0 | 2 | 0 | 0 | 0 | 0 | Olfr1065 |
| Olfr1066 | 0 | 0 | 0 | 0 | 0 | 0 | 0 | 1 | 2 | 0 | 0 | 0 | 0 | 0 | 0 | 0 | 0 | 0 | 0 | 0 | 0 | 0 | 2 | 4 | 0 | 0 | 0 | 0 | Olfr1066 |
| Olfr106-ps | 0 | 0 | 0 | 0 | 0 | 0 | 0 | 0 | 0 | 0 | 0 | 0 | 0 | 0 | 0 | 0 | 0 | 0 | 0 | 0 | 0 | 0 | 1 | 2 | 0 | 0 | 0 | 0 | Olfr106-ps |
| Olfr107 | 0 | 0 | 0 | 0 | 0 | 0 | 0 | 2 | 2 | 0 | 0 | 0 | 3 | 0 | 0 | 0 | 0 | 0 | 0 | 0 | 0 | 0 | 5 | 6 | 0 | 0 | 0 | 0 | Olfr107 |
| Olfr1079 | 0 | 0 | 0 | 0 | 0 | 0 | 0 | 1 | 5 | 1 | 4 | 0 | 0 | 0 | 0 | 0 | 0 | 0 | 0 | 0 | 0 | 0 | 1 | 0 | 0 | 0 | 0 | 0 | Olfr1079 |
| Olfr1082 | 0 | 0 | 0 | 0 | 0 | 0 | 0 | 0 | 0 | 0 | 0 | 0 | 0 | 2 | 0 | 0 | 0 | 0 | 0 | 0 | 0 | 0 | 0 | 0 | 0 | 0 | 0 | 0 | Olfr1082 |
| Olfr1085 | 0 | 0 | 0 | 0 | 0 | 0 | 0 | 0 | 0 | 0 | 0 | 0 | 0 | 0 | 0 | 0 | 0 | 0 | 0 | 0 | 0 | 0 | 0 | 2 | 0 | 0 | 0 | 0 | Olfr1085 |
| Olfr1086 | 0 | 0 | 0 | 0 | 0 | 0 | 0 | 1 | 0 | 0 | 0 | 0 | 0 | 0 | 0 | 0 | 0 | 0 | 0 | 0 | 0 | 0 | 0 | 0 | 0 | 0 | 0 | 0 | Olfr1086 |
| Olfr1089 | 0 | 0 | 0 | 0 | 0 | 0 | 0 | 0 | 0 | 8 | 0 | 0 | 0 | 1 | 0 | 0 | 0 | 0 | 0 | 0 | 0 | 0 | 1 | 0 | 0 | 0 | 0 | 0 | Olfr1089 |
| Olfr109 | 0 | 0 | 0 | 0 | 0 | 0 | 0 | 0 | 0 | 0 | 0 | 0 | 0 | 0 | 0 | 0 | 0 | 0 | 0 | 0 | 0 | 0 | 2 | 1 | 0 | 0 | 0 | 0 | Olfr109 |
| Olfr1090 | 0 | 0 | 0 | 0 | 0 | 0 | 0 | 1 | 0 | 0 | 1 | 0 | 0 | 0 | 0 | 0 | 0 | 0 | 0 | 0 | 0 | 0 | 0 | 0 | 0 | 0 | 0 | 0 | Olfr1090 |
| Olfr1094 | 0 | 0 | 0 | 0 | 0 | 0 | 0 | 0 | 0 | 0 | 0 | 0 | 0 | 0 | 0 | 0 | 0 | 0 | 0 | 0 | 0 | 0 | 0 | 1 | 0 | 0 | 0 | 0 | Olfr1094 |
| Olfr1099 | 0 | 0 | 0 | 0 | 0 | 0 | 0 | 0 | 0 | 1 | 0 | 0 | 0 | 0 | 0 | 0 | 0 | 0 | 0 | 0 | 0 | 0 | 0 | 0 | 0 | 0 | 0 | 0 | Olfr1099 |
| Olfr11 | 0 | 0 | 0 | 0 | 0 | 0 | 0 | 1 | 2 | 2 | 1 | 0 | 0 | 0 | 0 | 0 | 0 | 0 | 0 | 0 | 0 | 0 | 1 | 0 | 0 | 0 | 0 | 0 | Olfr11 |
| Olfr110 | 0 | 0 | 0 | 0 | 0 | 0 | 0 | 1 | 7 | 0 | 2 | 0 | 0 | 0 | 0 | 0 | 0 | 0 | 0 | 0 | 0 | 2 | 0 | 2 | 0 | 0 | 0 | 1 | Olfr110 |
| Olfr1101 | 0 | 0 | 0 | 0 | 0 | 0 | 0 | 0 | 0 | 0 | 0 | 0 | 0 | 0 | 0 | 0 | 0 | 0 | 0 | 0 | 0 | 0 | 1 | 0 | 0 | 0 | 0 | 0 | Olfr1101 |
| Olfr1102 | 1 | 0 | 0 | 0 | 0 | 0 | 0 | 1 | 0 | 0 | 0 | 0 | 0 | 0 | 0 | 0 | 0 | 0 | 0 | 0 | 0 | 0 | 0 | 0 | 0 | 0 | 0 | 0 | Olfr1102 |
| Olfr1104 | 0 | 0 | 0 | 0 | 0 | 0 | 0 | 2 | 2 | 5 | 3 | 0 | 0 | 0 | 0 | 0 | 0 | 0 | 0 | 0 | 0 | 0 | 2 | 0 | 0 | 0 | 0 | 0 | Olfr1104 |
| Olfr1105 | 1 | 0 | 0 | 1 | 0 | 0 | 0 | 1 | 2 | 10 | 0 | 0 | 0 | 0 | 0 | 0 | 0 | 0 | 0 | 0 | 0 | 0 | 1 | 2 | 0 | 1 | 0 | 0 | Olfr1105 |
| Olfr1106 | 0 | 0 | 0 | 0 | 0 | 0 | 0 | 0 | 0 | 3 | 0 | 0 | 0 | 0 | 0 | 0 | 0 | 0 | 0 | 0 | 0 | 0 | 0 | 0 | 0 | 0 | 0 | 0 | Olfr1106 |
| Olfr1107 | 0 | 0 | 0 | 0 | 0 | 0 | 0 | 2 | 0 | 0 | 0 | 0 | 0 | 0 | 0 | 0 | 0 | 0 | 0 | 0 | 0 | 0 | 0 | 0 | 0 | 0 | 0 | 0 | Olfr1107 |
| Olfr1109 | 0 | 0 | 0 | 0 | 0 | 0 | 0 | 1 | 0 | 0 | 0 | 0 | 0 | 0 | 0 | 0 | 0 | 0 | 0 | 0 | 0 | 0 | 0 | 0 | 0 | 0 | 0 | 0 | Olfr1109 |
| Olfr111 | 0 | 2 | 0 | 0 | 1 | 5 | 1 | 6 | 5 | 8 | 6 | 10 | 2 | 4 | 1 | 0 | 0 | 0 | 0 | 1 | 2 | 5 | 15 | 0 | 5 | 0 | 10 | 6 | Olfr111 |
| Olfr1110 | 0 | 0 | 0 | 0 | 0 | 0 | 0 | 2 | 5 | 0 | 0 | 0 | 0 | 0 | 0 | 0 | 0 | 0 | 0 | 0 | 0 | 0 | 1 | 3 | 0 | 0 | 0 | 0 | Olfr1110 |
| Olfr1111 | 0 | 0 | 0 | 0 | 0 | 0 | 0 | 0 | 0 | 0 | 0 | 0 | 0 | 2 | 0 | 0 | 0 | 0 | 0 | 0 | 0 | 0 | 0 | 0 | 0 | 1 | 0 | 0 | Olfr1111 |
| Olfr1115 | 0 | 0 | 0 | 0 | 0 | 0 | 0 | 0 | 1 | 0 | 0 | 0 | 0 | 0 | 0 | 0 | 0 | 0 | 0 | 0 | 0 | 0 | 0 | 0 | 0 | 0 | 0 | 0 | Olfr1115 |
| Olfr1116 | 0 | 0 | 0 | 0 | 0 | 0 | 0 | 0 | 2 | 0 | 0 | 0 | 0 | 0 | 0 | 0 | 0 | 0 | 0 | 0 | 0 | 0 | 0 | 0 | 0 | 0 | 0 | 0 | Olfr1116 |
| Olfr112 | 0 | 0 | 0 | 1 | 0 | 0 | 0 | 1 | 3 | 1 | 0 | 0 | 0 | 0 | 0 | 0 | 0 | 0 | 0 | 0 | 0 | 0 | 3 | 2 | 0 | 0 | 0 | 0 | Olfr112 |
| Olfr1121 | 0 | 0 | 0 | 0 | 0 | 0 | 0 | 0 | 0 | 0 | 0 | 0 | 0 | 0 | 0 | 0 | 0 | 0 | 0 | 0 | 0 | 0 | 1 | 0 | 0 | 0 | 0 | 0 | Olfr1121 |
| Olfr1122 | 5 | 1 | 0 | 0 | 0 | 0 | 0 | 0 | 0 | 9 | 5 | 0 | 0 | 0 | 0 | 0 | 0 | 0 | 0 | 0 | 0 | 0 | 3 | 5 | 0 | 8 | 0 | 0 | Olfr1122 |
| Olfr1123 | 0 | 0 | 0 | 0 | 0 | 0 | 0 | 0 | 2 | 2 | 0 | 0 | 0 | 0 | 0 | 0 | 0 | 0 | 0 | 0 | 0 | 0 | 0 | 2 | 0 | 0 | 0 | 0 | Olfr1123 |
| Olfr1124 | 0 | 0 | 0 | 0 | 0 | 0 | 0 | 0 | 2 | 0 | 0 | 0 | 0 | 0 | 0 | 0 | 0 | 0 | 0 | 0 | 0 | 0 | 0 | 0 | 0 | 0 | 0 | 0 | Olfr1124 |
| Olfr1129 | 0 | 0 | 0 | 0 | 0 | 0 | 0 | 0 | 1 | 0 | 0 | 0 | 0 | 0 | 0 | 0 | 0 | 0 | 0 | 0 | 0 | 0 | 0 | 0 | 0 | 0 | 0 | 0 | Olfr1129 |
| Olfr1130 | 0 | 0 | 0 | 0 | 0 | 0 | 0 | 0 | 0 | 0 | 0 | 0 | 0 | 0 | 0 | 0 | 0 | 0 | 0 | 0 | 0 | 0 | 0 | 0 | 0 | 1 | 0 | 0 | Olfr1130 |
| Olfr1133 | 0 | 0 | 0 | 0 | 0 | 0 | 0 | 4 | 0 | 0 | 0 | 0 | 0 | 0 | 0 | 0 | 0 | 0 | 0 | 0 | 0 | 0 | 0 | 0 | 0 | 0 | 0 | 0 | Olfr1133 |
| Olfr1136 | 0 | 0 | 0 | 0 | 0 | 0 | 0 | 0 | 0 | 0 | 0 | 0 | 0 | 0 | 0 | 0 | 0 | 0 | 0 | 0 | 0 | 0 | 1 | 0 | 0 | 4 | 0 | 0 | Olfr1136 |
| Olfr1137 | 0 | 0 | 0 | 0 | 0 | 0 | 0 | 1 | 0 | 1 | 0 | 0 | 0 | 0 | 0 | 0 | 0 | 0 | 0 | 0 | 0 | 0 | 0 | 2 | 0 | 2 | 0 | 0 | Olfr1137 |
| Olfr1138 | 0 | 0 | 0 | 0 | 0 | 0 | 0 | 0 | 0 | 0 | 1 | 0 | 0 | 0 | 0 | 0 | 0 | 0 | 0 | 0 | 0 | 0 | 0 | 0 | 0 | 0 | 0 | 0 | Olfr1138 |
| Olfr114 | 0 | 0 | 0 | 0 | 0 | 0 | 0 | 0 | 0 | 0 | 1 | 0 | 0 | 0 | 0 | 0 | 0 | 0 | 0 | 0 | 0 | 0 | 0 | 0 | 0 | 0 | 0 | 0 | Olfr114 |
| Olfr1140 | 0 | 0 | 0 | 0 | 0 | 0 | 0 | 0 | 0 | 0 | 0 | 0 | 1 | 0 | 0 | 0 | 0 | 0 | 0 | 0 | 0 | 0 | 1 | 0 | 0 | 0 | 0 | 0 | Olfr1140 |
| Olfr1143 | 0 | 0 | 0 | 0 | 0 | 0 | 0 | 0 | 2 | 0 | 0 | 0 | 0 | 0 | 0 | 0 | 0 | 0 | 0 | 0 | 0 | 0 | 3 | 1 | 0 | 0 | 0 | 0 | Olfr1143 |
| Olfr1145 | 0 | 0 | 0 | 0 | 0 | 0 | 0 | 0 | 0 | 0 | 2 | 0 | 0 | 0 | 0 | 0 | 0 | 0 | 0 | 0 | 0 | 0 | 0 | 2 | 0 | 0 | 0 | 0 | Olfr1145 |
| Olfr115 | 1 | 0 | 0 | 0 | 0 | 0 | 0 | 0 | 0 | 0 | 1 | 0 | 0 | 0 | 0 | 2 | 0 | 0 | 0 | 0 | 0 | 0 | 2 | 2 | 0 | 0 | 0 | 0 | Olfr115 |
| Olfr1151 | 0 | 0 | 0 | 0 | 0 | 0 | 0 | 0 | 0 | 0 | 0 | 0 | 0 | 0 | 0 | 0 | 0 | 0 | 0 | 0 | 0 | 0 | 0 | 1 | 0 | 0 | 0 | 0 | Olfr1151 |
| Olfr1152 | 0 | 0 | 0 | 0 | 0 | 0 | 0 | 0 | 0 | 1 | 0 | 0 | 0 | 0 | 0 | 0 | 0 | 0 | 0 | 0 | 0 | 0 | 0 | 1 | 0 | 0 | 0 | 0 | Olfr1152 |
| Olfr1154 | 0 | 0 | 0 | 0 | 0 | 0 | 0 | 0 | 0 | 0 | 0 | 0 | 0 | 0 | 0 | 0 | 0 | 0 | 0 | 0 | 0 | 0 | 0 | 0 | 0 | 2 | 0 | 0 | Olfr1154 |
| Olfr1155 | 0 | 0 | 0 | 0 | 0 | 0 | 0 | 1 | 0 | 0 | 0 | 0 | 0 | 0 | 0 | 0 | 0 | 0 | 0 | 0 | 0 | 0 | 0 | 0 | 0 | 0 | 0 | 0 | Olfr1155 |
| Olfr1156 | 0 | 0 | 0 | 0 | 0 | 0 | 0 | 1 | 0 | 0 | 0 | 0 | 0 | 0 | 0 | 0 | 0 | 0 | 0 | 0 | 0 | 0 | 0 | 2 | 0 | 0 | 0 | 0 | Olfr1156 |
| Olfr116 | 0 | 0 | 0 | 0 | 0 | 0 | 0 | 0 | 2 | 3 | 0 | 0 | 0 | 0 | 0 | 0 | 0 | 0 | 0 | 0 | 0 | 0 | 1 | 2 | 0 | 0 | 0 | 0 | Olfr116 |
| Olfr1160 | 0 | 0 | 0 | 5 | 0 | 0 | 0 | 0 | 1 | 2 | 0 | 0 | 0 | 0 | 0 | 0 | 0 | 0 | 0 | 0 | 0 | 0 | 1 | 3 | 0 | 2 | 0 | 0 | Olfr1160 |
| Olfr1161 | 0 | 0 | 0 | 0 | 0 | 0 | 0 | 0 | 0 | 0 | 0 | 0 | 0 | 0 | 0 | 0 | 0 | 0 | 0 | 0 | 0 | 0 | 0 | 0 | 0 | 2 | 0 | 0 | Olfr1161 |
| Olfr1162 | 0 | 0 | 0 | 0 | 0 | 0 | 0 | 1 | 1 | 0 | 0 | 0 | 0 | 0 | 0 | 0 | 0 | 0 | 0 | 0 | 0 | 0 | 1 | 3 | 0 | 0 | 0 | 0 | Olfr1162 |
| Olfr1166 | 0 | 0 | 0 | 0 | 0 | 0 | 0 | 0 | 1 | 1 | 0 | 0 | 0 | 0 | 0 | 0 | 0 | 0 | 0 | 0 | 0 | 0 | 0 | 1 | 0 | 0 | 0 | 0 | Olfr1166 |
| Olfr1170 | 0 | 0 | 0 | 0 | 0 | 0 | 0 | 1 | 0 | 0 | 0 | 0 | 0 | 0 | 0 | 0 | 0 | 0 | 0 | 0 | 0 | 0 | 0 | 0 | 0 | 0 | 0 | 0 | Olfr1170 |
| Olfr1173 | 0 | 0 | 2 | 0 | 0 | 0 | 0 | 0 | 0 | 1 | 0 | 0 | 0 | 0 | 0 | 1 | 0 | 0 | 0 | 0 | 1 | 0 | 1 | 3 | 0 | 0 | 0 | 0 | Olfr1173 |
| Olfr1174 | 0 | 0 | 0 | 0 | 0 | 0 | 0 | 0 | 1 | 0 | 0 | 0 | 0 | 0 | 0 | 0 | 0 | 0 | 0 | 0 | 0 | 0 | 2 | 0 | 0 | 0 | 0 | 0 | Olfr1174 |
| Olfr1176 | 0 | 0 | 0 | 0 | 0 | 0 | 0 | 0 | 0 | 2 | 0 | 0 | 0 | 0 | 0 | 0 | 0 | 0 | 0 | 0 | 0 | 0 | 1 | 0 | 0 | 0 | 0 | 0 | Olfr1176 |
| Olfr1177-ps | 0 | 0 | 0 | 0 | 0 | 0 | 0 | 2 | 0 | 0 | 0 | 0 | 0 | 0 | 0 | 0 | 0 | 0 | 0 | 0 | 0 | 0 | 0 | 1 | 0 | 0 | 0 | 0 | Olfr1177-ps |
| Olfr1178 | 0 | 0 | 0 | 0 | 0 | 0 | 0 | 0 | 0 | 4 | 0 | 0 | 0 | 0 | 0 | 0 | 0 | 0 | 0 | 0 | 0 | 0 | 5 | 0 | 0 | 0 | 0 | 0 | Olfr1178 |
| Olfr1179 | 0 | 0 | 0 | 0 | 0 | 0 | 0 | 0 | 0 | 1 | 0 | 0 | 0 | 0 | 0 | 0 | 0 | 0 | 0 | 0 | 0 | 0 | 0 | 0 | 0 | 0 | 0 | 0 | Olfr1179 |
| Olfr1182 | 0 | 0 | 0 | 0 | 0 | 0 | 0 | 0 | 0 | 3 | 0 | 0 | 0 | 0 | 0 | 0 | 0 | 0 | 0 | 0 | 0 | 0 | 0 | 0 | 0 | 0 | 0 | 0 | Olfr1182 |
| Olfr1184 | 0 | 0 | 0 | 0 | 0 | 0 | 0 | 0 | 0 | 3 | 0 | 0 | 0 | 0 | 0 | 0 | 0 | 0 | 0 | 0 | 0 | 0 | 0 | 0 | 0 | 0 | 0 | 0 | Olfr1184 |
| Olfr1189 | 0 | 0 | 0 | 0 | 0 | 0 | 0 | 0 | 0 | 0 | 0 | 0 | 0 | 0 | 0 | 0 | 0 | 0 | 0 | 0 | 0 | 0 | 0 | 2 | 0 | 0 | 0 | 0 | Olfr1189 |
| Olfr1191-ps1 | 0 | 0 | 0 | 0 | 0 | 0 | 0 | 0 | 0 | 4 | 0 | 0 | 0 | 0 | 0 | 0 | 0 | 0 | 0 | 0 | 0 | 0 | 0 | 0 | 0 | 0 | 0 | 0 | Olfr1191-ps1 |
| Olfr1192-ps1 | 0 | 0 | 0 | 0 | 0 | 0 | 0 | 0 | 0 | 1 | 1 | 0 | 0 | 0 | 0 | 0 | 0 | 0 | 0 | 0 | 0 | 2 | 1 | 0 | 0 | 0 | 0 | 0 | Olfr1192-ps1 |
| Olfr1196 | 0 | 0 | 0 | 0 | 0 | 0 | 0 | 0 | 0 | 0 | 2 | 0 | 0 | 0 | 0 | 0 | 0 | 0 | 0 | 0 | 0 | 0 | 0 | 2 | 0 | 0 | 0 | 0 | Olfr1196 |
| Olfr1197 | 0 | 0 | 0 | 0 | 0 | 0 | 0 | 0 | 0 | 0 | 0 | 0 | 0 | 0 | 0 | 0 | 0 | 0 | 0 | 0 | 0 | 0 | 1 | 0 | 0 | 0 | 0 | 0 | Olfr1197 |
| Olfr1198 | 0 | 0 | 0 | 0 | 0 | 0 | 0 | 0 | 1 | 0 | 0 | 0 | 0 | 0 | 0 | 0 | 0 | 0 | 0 | 0 | 0 | 0 | 0 | 0 | 0 | 0 | 0 | 0 | Olfr1198 |
| Olfr12 | 0 | 2 | 1 | 1 | 0 | 3 | 0 | 4 | 1 | 5 | 0 | 1 | 6 | 2 | 1 | 0 | 0 | 2 | 0 | 0 | 0 | 2 | 3 | 1 | 0 | 5 | 0 | 6 | Olfr12 |
| Olfr120 | 0 | 0 | 0 | 0 | 0 | 0 | 0 | 0 | 0 | 0 | 1 | 0 | 0 | 0 | 0 | 0 | 0 | 0 | 0 | 0 | 0 | 0 | 1 | 0 | 0 | 0 | 0 | 0 | Olfr120 |

| | | | | | | | | | | | | | | | | | | | | | | | | | | | | | |
|---|---|---|---|---|---|---|---|---|---|---|---|---|---|---|---|---|---|---|---|---|---|---|---|---|---|---|---|---|---|
| Olfr1200 | 0 | 0 | 0 | 0 | 0 | 0 | 0 | 0 | 0 | 0 | 0 | 0 | 0 | 0 | 0 | 0 | 0 | 0 | 0 | 0 | 0 | 0 | 0 | 0 | 0 | 4 | 0 | 0 | Olfr1200 |
| Olfr1208 | 0 | 0 | 0 | 0 | 0 | 0 | 0 | 0 | 0 | 0 | 1 | 0 | 0 | 0 | 0 | 0 | 0 | 0 | 0 | 0 | 0 | 0 | 1 | 0 | 0 | 0 | 0 | 0 | Olfr1208 |
| Olfr1211 | 0 | 0 | 0 | 0 | 0 | 0 | 0 | 0 | 0 | 0 | 0 | 0 | 1 | 0 | 0 | 0 | 0 | 0 | 0 | 0 | 0 | 0 | 0 | 0 | 0 | 0 | 0 | 0 | Olfr1211 |
| Olfr1213 | 0 | 0 | 0 | 0 | 0 | 0 | 0 | 0 | 0 | 1 | 0 | 0 | 0 | 0 | 0 | 0 | 0 | 0 | 0 | 0 | 0 | 0 | 0 | 0 | 0 | 0 | 0 | 0 | Olfr1213 |
| Olfr1214 | 0 | 0 | 0 | 0 | 0 | 0 | 0 | 1 | 0 | 0 | 0 | 0 | 0 | 0 | 0 | 0 | 0 | 0 | 0 | 0 | 0 | 0 | 0 | 0 | 0 | 0 | 0 | 0 | Olfr1214 |
| Olfr1216 | 0 | 0 | 0 | 0 | 0 | 0 | 0 | 2 | 0 | 1 | 6 | 0 | 0 | 0 | 0 | 0 | 0 | 0 | 0 | 0 | 0 | 0 | 4 | 1 | 0 | 2 | 0 | 0 | Olfr1216 |
| Olfr1219 | 0 | 0 | 0 | 0 | 0 | 0 | 0 | 0 | 1 | 0 | 0 | 0 | 0 | 0 | 0 | 0 | 0 | 0 | 0 | 0 | 0 | 0 | 0 | 0 | 0 | 0 | 0 | 0 | Olfr1219 |
| Olfr122 | 0 | 0 | 0 | 0 | 0 | 0 | 0 | 0 | 0 | 3 | 0 | 0 | 0 | 0 | 0 | 0 | 0 | 0 | 0 | 0 | 0 | 0 | 0 | 0 | 0 | 0 | 0 | 0 | Olfr122 |
| Olfr1220 | 0 | 0 | 0 | 0 | 0 | 0 | 0 | 1 | 0 | 0 | 0 | 0 | 0 | 0 | 0 | 0 | 0 | 0 | 0 | 0 | 0 | 0 | 0 | 0 | 0 | 0 | 0 | 0 | Olfr1220 |
| Olfr1221 | 0 | 0 | 0 | 0 | 0 | 0 | 0 | 1 | 1 | 0 | 0 | 0 | 0 | 0 | 0 | 0 | 0 | 0 | 0 | 0 | 0 | 0 | 5 | 0 | 0 | 0 | 0 | 0 | Olfr1221 |
| Olfr1222 | 0 | 0 | 0 | 2 | 0 | 0 | 0 | 0 | 0 | 0 | 0 | 0 | 0 | 0 | 0 | 0 | 0 | 0 | 0 | 0 | 0 | 0 | 0 | 3 | 0 | 0 | 0 | 0 | Olfr1222 |
| Olfr1225 | 0 | 0 | 0 | 0 | 0 | 0 | 0 | 0 | 0 | 0 | 1 | 0 | 0 | 0 | 0 | 0 | 0 | 0 | 0 | 0 | 0 | 0 | 1 | 0 | 0 | 0 | 0 | 0 | Olfr1225 |
| Olfr1226 | 0 | 0 | 0 | 0 | 0 | 0 | 0 | 2 | 1 | 2 | 1 | 0 | 0 | 0 | 0 | 0 | 0 | 0 | 0 | 0 | 0 | 0 | 1 | 0 | 0 | 0 | 0 | 0 | Olfr1226 |
| Olfr1231 | 0 | 0 | 0 | 0 | 0 | 0 | 0 | 1 | 0 | 0 | 0 | 0 | 0 | 0 | 0 | 0 | 0 | 0 | 0 | 0 | 0 | 0 | 0 | 0 | 0 | 0 | 0 | 0 | Olfr1231 |
| Olfr1234 | 0 | 0 | 0 | 0 | 0 | 0 | 0 | 0 | 0 | 0 | 0 | 0 | 0 | 0 | 0 | 0 | 0 | 0 | 0 | 0 | 0 | 0 | 1 | 2 | 0 | 0 | 0 | 0 | Olfr1234 |
| Olfr1238 | 0 | 0 | 0 | 0 | 0 | 0 | 0 | 0 | 0 | 0 | 0 | 0 | 0 | 0 | 0 | 0 | 0 | 0 | 0 | 0 | 0 | 0 | 3 | 0 | 0 | 0 | 0 | 0 | Olfr1238 |
| Olfr124 | 0 | 0 | 0 | 0 | 0 | 0 | 0 | 5 | 2 | 1 | 8 | 0 | 0 | 0 | 0 | 0 | 0 | 0 | 0 | 0 | 0 | 0 | 8 | 3 | 0 | 0 | 0 | 0 | Olfr124 |
| Olfr1240 | 0 | 0 | 0 | 0 | 0 | 0 | 0 | 1 | 2 | 0 | 0 | 0 | 0 | 0 | 0 | 0 | 0 | 0 | 0 | 0 | 0 | 0 | 3 | 4 | 0 | 0 | 0 | 0 | Olfr1240 |
| Olfr1246 | 0 | 0 | 0 | 0 | 0 | 0 | 0 | 1 | 1 | 0 | 0 | 0 | 0 | 0 | 0 | 0 | 0 | 0 | 0 | 0 | 0 | 0 | 1 | 0 | 0 | 0 | 0 | 0 | Olfr1246 |
| Olfr1247 | 0 | 0 | 0 | 0 | 0 | 0 | 1 | 0 | 0 | 0 | 0 | 0 | 0 | 0 | 0 | 0 | 0 | 0 | 0 | 0 | 0 | 0 | 3 | 0 | 0 | 0 | 0 | 0 | Olfr1247 |
| Olfr1250 | 0 | 0 | 0 | 0 | 0 | 0 | 0 | 0 | 0 | 0 | 0 | 0 | 0 | 0 | 0 | 0 | 0 | 0 | 0 | 1 | 0 | 0 | 3 | 0 | 0 | 0 | 0 | 0 | Olfr1250 |
| Olfr1251 | 0 | 0 | 0 | 0 | 0 | 0 | 0 | 2 | 1 | 0 | 0 | 0 | 0 | 0 | 0 | 0 | 0 | 0 | 0 | 0 | 0 | 0 | 1 | 0 | 0 | 0 | 0 | 0 | Olfr1251 |
| Olfr1252 | 0 | 0 | 0 | 0 | 0 | 0 | 0 | 0 | 0 | 0 | 0 | 0 | 0 | 0 | 0 | 0 | 0 | 0 | 0 | 0 | 0 | 0 | 0 | 3 | 0 | 0 | 0 | 0 | Olfr1252 |
| Olfr1253 | 0 | 0 | 0 | 0 | 0 | 0 | 0 | 2 | 0 | 0 | 0 | 0 | 0 | 0 | 0 | 0 | 0 | 0 | 0 | 0 | 0 | 0 | 1 | 0 | 0 | 2 | 0 | 0 | Olfr1253 |
| Olfr1255 | 0 | 0 | 0 | 0 | 0 | 0 | 0 | 0 | 0 | 0 | 0 | 0 | 0 | 0 | 0 | 0 | 0 | 0 | 0 | 0 | 0 | 0 | 0 | 1 | 0 | 0 | 0 | 0 | Olfr1255 |
| Olfr1256 | 0 | 0 | 0 | 0 | 0 | 0 | 0 | 0 | 2 | 0 | 0 | 0 | 0 | 0 | 0 | 0 | 0 | 0 | 0 | 0 | 0 | 0 | 0 | 3 | 0 | 0 | 0 | 0 | Olfr1256 |
| Olfr1257 | 0 | 0 | 0 | 0 | 0 | 0 | 0 | 0 | 0 | 0 | 0 | 0 | 0 | 0 | 0 | 0 | 0 | 0 | 0 | 0 | 1 | 0 | 0 | 0 | 0 | 0 | 2 | 1 | Olfr1257 |
| Olfr1258 | 0 | 0 | 0 | 0 | 0 | 0 | 0 | 1 | 3 | 5 | 3 | 0 | 0 | 0 | 0 | 0 | 0 | 0 | 0 | 0 | 0 | 0 | 8 | 2 | 0 | 5 | 0 | 0 | Olfr1258 |
| Olfr1259 | 0 | 0 | 0 | 0 | 0 | 0 | 0 | 1 | 3 | 2 | 1 | 0 | 0 | 0 | 0 | 0 | 0 | 0 | 0 | 0 | 0 | 0 | 5 | 8 | 0 | 1 | 0 | 0 | Olfr1259 |
| Olfr1260 | 0 | 0 | 0 | 0 | 0 | 0 | 0 | 0 | 2 | 2 | 0 | 0 | 0 | 0 | 0 | 0 | 0 | 0 | 0 | 0 | 0 | 2 | 2 | 0 | 0 | 0 | 0 | 0 | Olfr1260 |
| Olfr1262 | 0 | 0 | 0 | 0 | 0 | 0 | 0 | 0 | 0 | 0 | 0 | 0 | 0 | 0 | 0 | 0 | 0 | 0 | 0 | 0 | 0 | 0 | 1 | 1 | 0 | 0 | 0 | 0 | Olfr1262 |
| Olfr1264 | 1 | 0 | 0 | 0 | 0 | 0 | 0 | 6 | 9 | 23 | 4 | 0 | 0 | 0 | 0 | 0 | 0 | 0 | 0 | 0 | 0 | 0 | 9 | 7 | 0 | 7 | 0 | 0 | Olfr1264 |
| Olfr1265 | 0 | 0 | 0 | 0 | 0 | 0 | 0 | 0 | 0 | 0 | 2 | 0 | 0 | 0 | 0 | 0 | 0 | 0 | 0 | 0 | 0 | 0 | 0 | 0 | 0 | 0 | 0 | 0 | Olfr1265 |
| Olfr1269 | 0 | 0 | 0 | 0 | 0 | 0 | 0 | 1 | 0 | 1 | 0 | 0 | 0 | 0 | 0 | 0 | 0 | 0 | 0 | 0 | 0 | 0 | 0 | 0 | 0 | 0 | 0 | 0 | Olfr1269 |
| Olfr1270 | 0 | 0 | 0 | 0 | 0 | 0 | 0 | 0 | 0 | 0 | 0 | 0 | 0 | 0 | 0 | 0 | 0 | 2 | 0 | 0 | 0 | 0 | 1 | 1 | 0 | 0 | 0 | 1 | Olfr1270 |
| Olfr1271 | 0 | 0 | 0 | 0 | 0 | 0 | 0 | 0 | 3 | 8 | 4 | 0 | 0 | 0 | 0 | 0 | 0 | 0 | 0 | 0 | 0 | 0 | 1 | 2 | 0 | 1 | 0 | 0 | Olfr1271 |
| Olfr1274 | 2 | 0 | 0 | 2 | 0 | 0 | 0 | 0 | 2 | 2 | 2 | 0 | 1 | 1 | 4 | 0 | 0 | 0 | 0 | 0 | 0 | 0 | 4 | 2 | 0 | 1 | 0 | 0 | Olfr1274 |
| Olfr1276 | 0 | 0 | 0 | 0 | 0 | 0 | 0 | 0 | 0 | 0 | 2 | 0 | 0 | 0 | 0 | 0 | 0 | 0 | 0 | 0 | 0 | 0 | 0 | 0 | 0 | 0 | 0 | 0 | Olfr1276 |
| Olfr128 | 0 | 0 | 0 | 0 | 0 | 0 | 0 | 1 | 0 | 0 | 0 | 0 | 0 | 0 | 0 | 0 | 0 | 0 | 0 | 0 | 0 | 0 | 1 | 0 | 0 | 0 | 0 | 0 | Olfr128 |
| Olfr1280 | 0 | 0 | 0 | 0 | 0 | 0 | 0 | 0 | 1 | 1 | 2 | 0 | 0 | 0 | 0 | 0 | 0 | 0 | 0 | 3 | 0 | 0 | 1 | 0 | 0 | 0 | 0 | 0 | Olfr1280 |
| Olfr1281 | 0 | 0 | 0 | 0 | 0 | 0 | 0 | 0 | 1 | 0 | 0 | 0 | 0 | 0 | 0 | 0 | 0 | 0 | 0 | 0 | 0 | 0 | 0 | 0 | 0 | 0 | 0 | 0 | Olfr1281 |
| Olfr129 | 0 | 0 | 0 | 0 | 0 | 0 | 0 | 1 | 1 | 0 | 0 | 0 | 0 | 0 | 0 | 0 | 0 | 0 | 0 | 0 | 0 | 0 | 1 | 3 | 0 | 0 | 0 | 0 | Olfr129 |
| Olfr1290 | 0 | 0 | 0 | 0 | 0 | 1 | 0 | 0 | 0 | 0 | 0 | 0 | 0 | 0 | 0 | 1 | 0 | 0 | 0 | 0 | 0 | 0 | 0 | 0 | 0 | 0 | 0 | 0 | Olfr1290 |
| Olfr1297 | 0 | 0 | 0 | 0 | 0 | 0 | 0 | 1 | 0 | 0 | 0 | 0 | 0 | 0 | 3 | 0 | 0 | 0 | 0 | 0 | 0 | 0 | 0 | 1 | 0 | 0 | 0 | 0 | Olfr1297 |
| Olfr1299 | 0 | 0 | 0 | 0 | 0 | 0 | 0 | 0 | 0 | 0 | 0 | 0 | 0 | 0 | 0 | 0 | 0 | 0 | 0 | 0 | 0 | 0 | 0 | 0 | 0 | 0 | 2 | 0 | Olfr1299 |
| Olfr13 | 0 | 0 | 0 | 0 | 0 | 0 | 0 | 1 | 1 | 0 | 2 | 1 | 0 | 0 | 0 | 0 | 0 | 0 | 0 | 0 | 0 | 0 | 1 | 1 | 0 | 1 | 0 | 0 | Olfr13 |
| Olfr130 | 0 | 0 | 0 | 0 | 0 | 0 | 0 | 1 | 0 | 0 | 0 | 0 | 0 | 0 | 0 | 1 | 0 | 1 | 0 | 0 | 0 | 0 | 0 | 0 | 0 | 0 | 0 | 0 | Olfr130 |
| Olfr1301 | 0 | 0 | 0 | 0 | 0 | 0 | 0 | 0 | 0 | 0 | 0 | 2 | 0 | 0 | 0 | 0 | 0 | 0 | 0 | 0 | 0 | 0 | 1 | 2 | 0 | 0 | 0 | 0 | Olfr1301 |
| Olfr1302 | 0 | 0 | 0 | 0 | 0 | 0 | 0 | 0 | 1 | 0 | 0 | 0 | 1 | 0 | 0 | 0 | 0 | 0 | 0 | 0 | 0 | 0 | 1 | 0 | 0 | 0 | 0 | 0 | Olfr1302 |
| Olfr1303 | 0 | 0 | 0 | 0 | 0 | 0 | 0 | 0 | 0 | 0 | 0 | 0 | 0 | 0 | 0 | 0 | 0 | 0 | 0 | 0 | 0 | 0 | 0 | 0 | 0 | 1 | 0 | 0 | Olfr1303 |
| Olfr1307 | 0 | 0 | 0 | 0 | 0 | 0 | 0 | 0 | 0 | 0 | 1 | 0 | 0 | 0 | 0 | 0 | 0 | 0 | 0 | 0 | 0 | 0 | 0 | 0 | 0 | 0 | 0 | 0 | Olfr1307 |
| Olfr1309 | 0 | 0 | 0 | 0 | 5 | 0 | 0 | 0 | 0 | 0 | 0 | 0 | 0 | 0 | 3 | 0 | 0 | 2 | 0 | 0 | 4 | 0 | 0 | 0 | 0 | 0 | 0 | 0 | Olfr1309 |
| Olfr131 | 0 | 0 | 0 | 0 | 0 | 0 | 0 | 2 | 0 | 3 | 1 | 0 | 0 | 0 | 0 | 0 | 0 | 0 | 0 | 0 | 0 | 0 | 0 | 2 | 0 | 0 | 0 | 0 | Olfr131 |
| Olfr1310 | 0 | 0 | 0 | 3 | 0 | 0 | 0 | 0 | 0 | 0 | 0 | 0 | 0 | 0 | 0 | 0 | 0 | 0 | 0 | 0 | 0 | 3 | 0 | 0 | 0 | 0 | 0 | 0 | Olfr1310 |
| Olfr1312 | 0 | 0 | 0 | 0 | 0 | 0 | 0 | 0 | 0 | 0 | 0 | 0 | 0 | 0 | 0 | 0 | 0 | 0 | 0 | 0 | 3 | 0 | 1 | 0 | 0 | 0 | 0 | 0 | Olfr1312 |
| Olfr1314 | 1 | 0 | 0 | 0 | 0 | 3 | 1 | 0 | 2 | 0 | 1 | 0 | 0 | 0 | 3 | 0 | 0 | 0 | 0 | 0 | 0 | 5 | 1 | 0 | 0 | 2 | 0 | 0 | Olfr1314 |
| Olfr1316 | 0 | 0 | 0 | 0 | 0 | 1 | 0 | 0 | 0 | 0 | 0 | 2 | 0 | 0 | 0 | 0 | 0 | 0 | 0 | 0 | 0 | 0 | 0 | 0 | 0 | 0 | 0 | 0 | Olfr1316 |
| Olfr132 | 0 | 0 | 0 | 0 | 0 | 0 | 0 | 0 | 0 | 0 | 1 | 0 | 0 | 0 | 0 | 0 | 0 | 0 | 0 | 0 | 0 | 0 | 0 | 0 | 0 | 0 | 0 | 0 | Olfr132 |
| Olfr1320 | 0 | 0 | 0 | 0 | 0 | 0 | 0 | 0 | 0 | 0 | 2 | 0 | 0 | 0 | 0 | 0 | 0 | 0 | 0 | 0 | 0 | 0 | 2 | 0 | 0 | 0 | 0 | 0 | Olfr1320 |
| Olfr1321 | 0 | 0 | 0 | 0 | 0 | 0 | 0 | 0 | 0 | 0 | 0 | 0 | 0 | 0 | 0 | 0 | 0 | 0 | 0 | 0 | 0 | 0 | 1 | 0 | 0 | 0 | 0 | 0 | Olfr1321 |
| Olfr1324 | 0 | 0 | 0 | 0 | 0 | 0 | 0 | 0 | 1 | 0 | 5 | 0 | 0 | 0 | 3 | 1 | 0 | 0 | 0 | 0 | 0 | 0 | 1 | 4 | 0 | 5 | 0 | 0 | Olfr1324 |
| Olfr1325 | 0 | 0 | 0 | 0 | 0 | 0 | 0 | 1 | 2 | 3 | 1 | 0 | 0 | 0 | 0 | 0 | 0 | 0 | 0 | 0 | 0 | 0 | 0 | 2 | 0 | 2 | 0 | 0 | Olfr1325 |
| Olfr1326-ps1 | 0 | 0 | 0 | 0 | 0 | 0 | 0 | 4 | 1 | 10 | 0 | 0 | 0 | 0 | 0 | 0 | 0 | 0 | 0 | 0 | 0 | 0 | 0 | 0 | 0 | 0 | 0 | 0 | Olfr1326-ps1 |
| Olfr1328 | 2 | 0 | 0 | 1 | 0 | 1 | 0 | 0 | 0 | 2 | 2 | 0 | 0 | 0 | 0 | 0 | 0 | 0 | 0 | 0 | 0 | 0 | 0 | 5 | 0 | 0 | 0 | 0 | Olfr1328 |
| Olfr1329 | 0 | 0 | 0 | 0 | 0 | 0 | 0 | 0 | 1 | 0 | 0 | 0 | 0 | 0 | 0 | 0 | 0 | 0 | 0 | 0 | 0 | 0 | 0 | 2 | 0 | 0 | 0 | 0 | Olfr1329 |
| Olfr133 | 0 | 0 | 0 | 0 | 0 | 0 | 0 | 0 | 0 | 1 | 3 | 0 | 0 | 0 | 0 | 0 | 0 | 0 | 0 | 0 | 0 | 0 | 0 | 0 | 0 | 0 | 0 | 0 | Olfr133 |
| Olfr1330 | 0 | 0 | 0 | 0 | 0 | 0 | 0 | 2 | 0 | 5 | 1 | 0 | 0 | 0 | 0 | 0 | 0 | 0 | 0 | 0 | 0 | 0 | 1 | 7 | 0 | 5 | 0 | 0 | Olfr1330 |
| Olfr1331 | 1 | 0 | 0 | 0 | 0 | 0 | 0 | 0 | 2 | 0 | 2 | 0 | 0 | 0 | 0 | 0 | 0 | 0 | 0 | 0 | 0 | 0 | 0 | 2 | 0 | 0 | 0 | 0 | Olfr1331 |
| Olfr1333 | 0 | 0 | 0 | 0 | 0 | 0 | 0 | 1 | 0 | 0 | 0 | 0 | 0 | 0 | 0 | 1 | 0 | 0 | 0 | 0 | 0 | 0 | 0 | 0 | 0 | 2 | 0 | 0 | Olfr1333 |
| Olfr1336 | 1 | 1 | 0 | 0 | 2 | 1 | 0 | 0 | 0 | 2 | 2 | 0 | 0 | 0 | 0 | 2 | 0 | 0 | 0 | 0 | 2 | 0 | 1 | 3 | 2 | 0 | 0 | 0 | Olfr1336 |
| Olfr1337 | 0 | 0 | 0 | 0 | 0 | 0 | 0 | 1 | 0 | 0 | 0 | 0 | 0 | 0 | 0 | 0 | 0 | 0 | 0 | 0 | 0 | 0 | 0 | 0 | 0 | 0 | 0 | 0 | Olfr1337 |
| Olfr1338 | 0 | 0 | 0 | 0 | 0 | 0 | 0 | 1 | 0 | 3 | 5 | 0 | 0 | 0 | 1 | 0 | 0 | 0 | 0 | 0 | 0 | 0 | 1 | 1 | 0 | 0 | 0 | 0 | Olfr1338 |
| Olfr1339 | 0 | 0 | 0 | 1 | 0 | 0 | 0 | 1 | 2 | 12 | 7 | 0 | 0 | 0 | 3 | 0 | 0 | 0 | 0 | 0 | 0 | 0 | 6 | 2 | 0 | 7 | 0 | 1 | Olfr1339 |
| Olfr134 | 0 | 0 | 0 | 0 | 0 | 0 | 0 | 0 | 1 | 0 | 0 | 0 | 0 | 0 | 0 | 0 | 0 | 0 | 0 | 0 | 0 | 0 | 0 | 1 | 0 | 0 | 0 | 0 | Olfr134 |
| Olfr1340 | 0 | 0 | 0 | 0 | 0 | 0 | 0 | 4 | 0 | 2 | 0 | 0 | 1 | 0 | 0 | 0 | 0 | 0 | 0 | 0 | 0 | 0 | 5 | 0 | 0 | 1 | 0 | 0 | Olfr1340 |
| Olfr1341 | 3 | 0 | 0 | 0 | 0 | 0 | 0 | 7 | 5 | 12 | 1 | 0 | 5 | 0 | 1 | 0 | 0 | 0 | 1 | 0 | 0 | 0 | 6 | 6 | 4 | 10 | 0 | 0 | Olfr1341 |
| Olfr1342 | 0 | 0 | 0 | 0 | 0 | 0 | 0 | 5 | 3 | 16 | 0 | 5 | 5 | 5 | 3 | 2 | 0 | 0 | 0 | 0 | 3 | 2 | 5 | 5 | 1 | 7 | 0 | 1 | Olfr1342 |
| Olfr1344 | 1 | 0 | 0 | 0 | 3 | 3 | 2 | 0 | 0 | 0 | 0 | 0 | 0 | 2 | 0 | 0 | 0 | 2 | 1 | 2 | 2 | 0 | 0 | 0 | 0 | 0 | 0 | 0 | Olfr1344 |
| Olfr1346 | 0 | 0 | 0 | 0 | 0 | 0 | 0 | 0 | 0 | 0 | 0 | 0 | 0 | 0 | 1 | 2 | 0 | 0 | 0 | 0 | 0 | 0 | 1 | 0 | 0 | 0 | 0 | 0 | Olfr1346 |

| | | | | | | | | | | | | | | | | | | | | | | | | | | | | | |
|---|---|---|---|---|---|---|---|---|---|---|---|---|---|---|---|---|---|---|---|---|---|---|---|---|---|---|---|---|---|
| Olfr1348 | 0 | 0 | 0 | 0 | 0 | 0 | 0 | 0 | 0 | 0 | 0 | 0 | 0 | 0 | 0 | 0 | 0 | 0 | 0 | 0 | 0 | 0 | 0 | 2 | 0 | 0 | 0 | 0 | Olfr1348 |
| Olfr1349 | 1 | 0 | 2 | 0 | 0 | 0 | 0 | 0 | 2 | 0 | 2 | 2 | 0 | 0 | 0 | 0 | 0 | 1 | 0 | 0 | 0 | 2 | 0 | 0 | 2 | 8 | 0 | 0 | Olfr1349 |
| Olfr1350 | 0 | 0 | 0 | 0 | 0 | 3 | 0 | 0 | 0 | 0 | 0 | 0 | 0 | 0 | 0 | 0 | 0 | 0 | 0 | 0 | 0 | 0 | 2 | 2 | 0 | 0 | 0 | 0 | Olfr1350 |
| Olfr1351 | 0 | 0 | 0 | 0 | 0 | 0 | 0 | 1 | 0 | 1 | 0 | 0 | 0 | 1 | 0 | 0 | 0 | 0 | 0 | 0 | 0 | 0 | 0 | 0 | 0 | 0 | 0 | 0 | Olfr1351 |
| Olfr1352 | 0 | 0 | 0 | 2 | 0 | 0 | 0 | 1 | 0 | 3 | 4 | 0 | 0 | 0 | 0 | 0 | 0 | 0 | 0 | 0 | 0 | 0 | 0 | 0 | 0 | 0 | 0 | 0 | Olfr1352 |
| Olfr1353 | 0 | 0 | 0 | 0 | 0 | 0 | 0 | 0 | 0 | 0 | 2 | 0 | 0 | 0 | 0 | 0 | 0 | 0 | 0 | 0 | 0 | 0 | 0 | 0 | 0 | 0 | 0 | 0 | Olfr1353 |
| Olfr1354 | 0 | 0 | 0 | 0 | 0 | 0 | 0 | 2 | 0 | 3 | 0 | 0 | 0 | 0 | 0 | 0 | 0 | 0 | 0 | 0 | 0 | 0 | 0 | 0 | 0 | 0 | 0 | 0 | Olfr1354 |
| Olfr1355 | 0 | 0 | 0 | 0 | 0 | 0 | 0 | 0 | 0 | 0 | 1 | 0 | 0 | 0 | 0 | 0 | 0 | 0 | 0 | 0 | 0 | 0 | 0 | 0 | 0 | 0 | 0 | 0 | Olfr1355 |
| Olfr1356 | 0 | 0 | 0 | 0 | 0 | 0 | 0 | 0 | 0 | 0 | 0 | 0 | 0 | 0 | 0 | 0 | 0 | 0 | 0 | 0 | 0 | 0 | 0 | 1 | 0 | 0 | 0 | 0 | Olfr1356 |
| Olfr1357 | 0 | 0 | 0 | 0 | 0 | 0 | 0 | 0 | 1 | 0 | 1 | 0 | 0 | 0 | 0 | 0 | 0 | 0 | 0 | 0 | 0 | 0 | 1 | 0 | 0 | 0 | 0 | 0 | Olfr1357 |
| Olfr1358 | 0 | 0 | 0 | 0 | 0 | 0 | 0 | 4 | 0 | 1 | 0 | 0 | 1 | 0 | 0 | 0 | 0 | 0 | 0 | 0 | 0 | 0 | 3 | 3 | 0 | 1 | 0 | 0 | Olfr1358 |
| Olfr1359 | 0 | 0 | 0 | 0 | 0 | 0 | 0 | 0 | 2 | 0 | 0 | 0 | 0 | 0 | 0 | 0 | 0 | 0 | 0 | 0 | 0 | 0 | 1 | 6 | 0 | 0 | 0 | 0 | Olfr1359 |
| Olfr136 | 0 | 0 | 0 | 1 | 0 | 0 | 0 | 2 | 1 | 0 | 2 | 0 | 0 | 0 | 0 | 0 | 0 | 0 | 0 | 0 | 0 | 0 | 5 | 2 | 0 | 3 | 0 | 0 | Olfr136 |
| Olfr1360 | 0 | 0 | 0 | 0 | 0 | 0 | 0 | 0 | 1 | 0 | 0 | 0 | 0 | 0 | 0 | 0 | 0 | 0 | 0 | 0 | 0 | 0 | 0 | 0 | 0 | 0 | 0 | 0 | Olfr1360 |
| Olfr1361 | 0 | 0 | 0 | 0 | 0 | 0 | 0 | 6 | 1 | 1 | 1 | 0 | 0 | 0 | 0 | 1 | 0 | 0 | 0 | 0 | 0 | 0 | 3 | 2 | 0 | 0 | 0 | 0 | Olfr1361 |
| Olfr1364 | 0 | 0 | 0 | 0 | 0 | 0 | 0 | 1 | 0 | 5 | 0 | 0 | 1 | 0 | 0 | 0 | 0 | 0 | 0 | 0 | 0 | 0 | 1 | 4 | 0 | 0 | 0 | 0 | Olfr1364 |
| Olfr1366 | 0 | 0 | 0 | 0 | 0 | 0 | 0 | 0 | 0 | 0 | 0 | 0 | 0 | 0 | 0 | 0 | 0 | 0 | 0 | 0 | 0 | 0 | 1 | 4 | 0 | 2 | 0 | 0 | Olfr1366 |
| Olfr1368 | 0 | 0 | 0 | 0 | 0 | 0 | 0 | 0 | 0 | 2 | 0 | 0 | 0 | 0 | 0 | 0 | 0 | 0 | 0 | 0 | 0 | 0 | 0 | 0 | 0 | 1 | 0 | 0 | Olfr1368 |
| Olfr1369-ps1 | 0 | 0 | 5 | 7 | 0 | 2 | 0 | 0 | 1 | 1 | 2 | 1 | 1 | 2 | 1 | 0 | 1 | 1 | 5 | 0 | 0 | 0 | 1 | 7 | 0 | 1 | 1 | 1 | Olfr1369-ps1 |
| Olfr1370 | 0 | 1 | 0 | 0 | 0 | 0 | 0 | 7 | 4 | 3 | 5 | 0 | 1 | 0 | 3 | 0 | 0 | 0 | 0 | 0 | 0 | 0 | 6 | 8 | 0 | 6 | 0 | 0 | Olfr1370 |
| Olfr1371 | 0 | 0 | 0 | 0 | 0 | 0 | 0 | 1 | 0 | 0 | 0 | 0 | 0 | 0 | 0 | 0 | 0 | 0 | 0 | 0 | 3 | 0 | 0 | 0 | 0 | 0 | 0 | 0 | Olfr1371 |
| Olfr1375 | 0 | 0 | 0 | 0 | 0 | 0 | 0 | 2 | 2 | 4 | 0 | 0 | 0 | 0 | 0 | 0 | 0 | 0 | 0 | 0 | 0 | 0 | 1 | 3 | 0 | 0 | 0 | 0 | Olfr1375 |
| Olfr1377 | 1 | 0 | 0 | 0 | 0 | 0 | 0 | 1 | 2 | 0 | 4 | 0 | 1 | 1 | 0 | 0 | 0 | 0 | 0 | 0 | 0 | 0 | 0 | 1 | 0 | 2 | 0 | 0 | Olfr1377 |
| Olfr1378 | 2 | 1 | 0 | 0 | 0 | 0 | 0 | 0 | 0 | 0 | 0 | 0 | 0 | 0 | 0 | 0 | 0 | 0 | 0 | 2 | 0 | 0 | 0 | 0 | 0 | 2 | 0 | 0 | Olfr1378 |
| Olfr138 | 0 | 0 | 0 | 0 | 0 | 0 | 0 | 0 | 0 | 0 | 0 | 0 | 0 | 0 | 0 | 0 | 0 | 0 | 0 | 0 | 0 | 0 | 1 | 0 | 0 | 0 | 0 | 0 | Olfr138 |
| Olfr1380 | 0 | 0 | 0 | 0 | 0 | 0 | 0 | 0 | 0 | 4 | 0 | 0 | 0 | 0 | 0 | 0 | 0 | 0 | 0 | 0 | 0 | 0 | 0 | 2 | 0 | 0 | 0 | 0 | Olfr1380 |
| Olfr1381 | 0 | 0 | 0 | 0 | 0 | 0 | 0 | 0 | 0 | 0 | 1 | 0 | 0 | 0 | 0 | 0 | 0 | 0 | 0 | 0 | 0 | 0 | 1 | 2 | 0 | 0 | 0 | 0 | Olfr1381 |
| Olfr1382 | 0 | 0 | 0 | 0 | 0 | 0 | 0 | 0 | 0 | 0 | 0 | 0 | 0 | 0 | 0 | 0 | 0 | 0 | 0 | 0 | 0 | 0 | 0 | 0 | 0 | 1 | 0 | 0 | Olfr1382 |
| Olfr1383 | 0 | 0 | 0 | 0 | 0 | 0 | 0 | 0 | 2 | 0 | 0 | 0 | 1 | 0 | 0 | 0 | 1 | 0 | 0 | 0 | 0 | 0 | 0 | 3 | 0 | 0 | 0 | 0 | Olfr1383 |
| Olfr1384 | 0 | 0 | 0 | 0 | 0 | 0 | 0 | 0 | 1 | 0 | 0 | 0 | 0 | 0 | 1 | 0 | 0 | 0 | 0 | 0 | 0 | 0 | 3 | 0 | 0 | 0 | 0 | 0 | Olfr1384 |
| Olfr1385 | 0 | 0 | 0 | 0 | 0 | 0 | 0 | 0 | 0 | 0 | 3 | 0 | 0 | 0 | 0 | 0 | 0 | 0 | 0 | 1 | 0 | 0 | 0 | 0 | 0 | 1 | 0 | 0 | Olfr1385 |
| Olfr1386 | 0 | 0 | 0 | 0 | 0 | 0 | 0 | 4 | 1 | 0 | 0 | 0 | 0 | 0 | 0 | 0 | 0 | 0 | 0 | 0 | 0 | 0 | 1 | 2 | 0 | 0 | 0 | 0 | Olfr1386 |
| Olfr1387 | 0 | 0 | 0 | 0 | 0 | 0 | 0 | 0 | 2 | 0 | 0 | 0 | 0 | 0 | 0 | 0 | 0 | 0 | 0 | 0 | 0 | 0 | 2 | 1 | 0 | 2 | 0 | 0 | Olfr1387 |
| Olfr1388 | 3 | 0 | 0 | 0 | 0 | 0 | 0 | 0 | 0 | 5 | 2 | 0 | 0 | 1 | 0 | 0 | 0 | 0 | 0 | 0 | 2 | 0 | 0 | 3 | 0 | 0 | 0 | 0 | Olfr1388 |
| Olfr1389 | 0 | 1 | 0 | 1 | 0 | 0 | 0 | 1 | 2 | 0 | 3 | 0 | 0 | 0 | 1 | 2 | 0 | 0 | 0 | 0 | 1 | 0 | 1 | 0 | 0 | 0 | 0 | 1 | Olfr1389 |
| Olfr139 | 0 | 0 | 0 | 0 | 0 | 0 | 0 | 2 | 0 | 2 | 1 | 0 | 1 | 0 | 0 | 0 | 0 | 0 | 0 | 0 | 0 | 0 | 0 | 2 | 0 | 0 | 0 | 0 | Olfr139 |
| Olfr1390 | 1 | 2 | 0 | 2 | 0 | 0 | 0 | 0 | 0 | 0 | 0 | 0 | 0 | 0 | 0 | 2 | 0 | 0 | 0 | 0 | 1 | 0 | 0 | 0 | 0 | 0 | 0 | 1 | Olfr1390 |
| Olfr1391 | 0 | 1 | 0 | 0 | 0 | 0 | 2 | 0 | 0 | 0 | 0 | 0 | 0 | 0 | 0 | 1 | 0 | 0 | 2 | 0 | 0 | 0 | 0 | 0 | 0 | 0 | 0 | 0 | Olfr1391 |
| Olfr1392 | 25 | 19 | 18 | 30 | 21 | 29 | 40 | 15 | 16 | 12 | 11 | 14 | 12 | 15 | 29 | 27 | 13 | 35 | 17 | 22 | 34 | 13 | 9 | 8 | 8 | 6 | 6 | 18 | Olfr1392 |
| Olfr1393 | 33 | 29 | 35 | 37 | 29 | 37 | 30 | 16 | 11 | 20 | 16 | 14 | 25 | 18 | 34 | 25 | 17 | 39 | 18 | 24 | 26 | 20 | 14 | 28 | 28 | 19 | 12 | 18 | Olfr1393 |
| Olfr1394 | 0 | 0 | 0 | 0 | 0 | 0 | 0 | 0 | 0 | 0 | 0 | 0 | 0 | 0 | 3 | 0 | 0 | 0 | 0 | 0 | 0 | 0 | 1 | 1 | 0 | 0 | 0 | 0 | Olfr1394 |
| Olfr1395 | 0 | 0 | 0 | 2 | 0 | 0 | 0 | 0 | 1 | 2 | 1 | 0 | 1 | 4 | 0 | 0 | 0 | 0 | 0 | 0 | 0 | 0 | 3 | 9 | 0 | 3 | 0 | 0 | Olfr1395 |
| Olfr1396 | 0 | 0 | 0 | 0 | 0 | 0 | 0 | 0 | 0 | 0 | 0 | 0 | 0 | 0 | 0 | 0 | 0 | 0 | 0 | 0 | 0 | 0 | 1 | 0 | 0 | 0 | 0 | 0 | Olfr1396 |
| Olfr140 | 3 | 0 | 0 | 0 | 0 | 0 | 0 | 0 | 2 | 3 | 0 | 0 | 0 | 0 | 0 | 0 | 0 | 0 | 0 | 0 | 0 | 0 | 3 | 0 | 0 | 0 | 0 | 0 | Olfr140 |
| Olfr1402 | 1 | 0 | 0 | 0 | 0 | 0 | 0 | 0 | 0 | 4 | 0 | 0 | 0 | 0 | 0 | 0 | 0 | 0 | 0 | 0 | 0 | 0 | 0 | 0 | 0 | 0 | 0 | 0 | Olfr1402 |
| Olfr1404 | 1 | 0 | 0 | 2 | 0 | 0 | 0 | 6 | 2 | 0 | 10 | 0 | 5 | 0 | 3 | 0 | 0 | 0 | 0 | 0 | 0 | 0 | 12 | 7 | 0 | 8 | 0 | 0 | Olfr1404 |
| Olfr1406 | 0 | 0 | 0 | 0 | 0 | 0 | 0 | 2 | 4 | 0 | 0 | 0 | 0 | 0 | 0 | 0 | 0 | 0 | 0 | 0 | 0 | 0 | 1 | 7 | 0 | 4 | 0 | 0 | Olfr1406 |
| Olfr1408 | 0 | 0 | 0 | 2 | 0 | 0 | 0 | 0 | 1 | 0 | 0 | 0 | 0 | 0 | 0 | 0 | 0 | 0 | 0 | 0 | 0 | 0 | 5 | 7 | 0 | 6 | 0 | 0 | Olfr1408 |
| Olfr1410 | 0 | 2 | 0 | 0 | 0 | 0 | 0 | 0 | 1 | 0 | 0 | 0 | 0 | 0 | 0 | 0 | 0 | 0 | 0 | 0 | 0 | 0 | 0 | 0 | 0 | 1 | 0 | 0 | Olfr1410 |
| Olfr1411 | 0 | 0 | 0 | 0 | 0 | 0 | 0 | 0 | 0 | 0 | 0 | 0 | 0 | 0 | 0 | 0 | 0 | 0 | 0 | 0 | 0 | 0 | 1 | 0 | 0 | 0 | 0 | 0 | Olfr1411 |
| Olfr1413 | 0 | 0 | 0 | 0 | 0 | 0 | 0 | 0 | 0 | 0 | 0 | 0 | 0 | 0 | 0 | 0 | 0 | 0 | 0 | 0 | 0 | 0 | 0 | 1 | 0 | 0 | 0 | 0 | Olfr1413 |
| Olfr1414 | 0 | 0 | 0 | 0 | 1 | 0 | 0 | 2 | 2 | 5 | 0 | 0 | 2 | 0 | 0 | 0 | 0 | 0 | 0 | 0 | 0 | 0 | 1 | 2 | 0 | 1 | 0 | 2 | Olfr1414 |
| Olfr1415 | 0 | 0 | 0 | 1 | 0 | 0 | 0 | 0 | 0 | 0 | 0 | 0 | 0 | 0 | 0 | 0 | 0 | 0 | 0 | 0 | 0 | 0 | 1 | 0 | 0 | 2 | 0 | 0 | Olfr1415 |
| Olfr1416 | 0 | 2 | 0 | 0 | 0 | 0 | 0 | 0 | 0 | 0 | 0 | 0 | 0 | 0 | 0 | 0 | 0 | 0 | 0 | 0 | 0 | 0 | 0 | 1 | 0 | 0 | 0 | 0 | Olfr1416 |
| Olfr1418 | 0 | 0 | 0 | 0 | 0 | 0 | 0 | 0 | 0 | 0 | 0 | 0 | 0 | 0 | 0 | 0 | 0 | 0 | 0 | 0 | 0 | 0 | 0 | 1 | 0 | 0 | 0 | 0 | Olfr1418 |
| Olfr1419 | 0 | 0 | 0 | 0 | 0 | 2 | 2 | 1 | 0 | 1 | 4 | 0 | 0 | 0 | 0 | 0 | 0 | 2 | 1 | 6 | 2 | 3 | 1 | 2 | 2 | 3 | 1 | 0 | Olfr1419 |
| Olfr142 | 0 | 0 | 0 | 0 | 0 | 0 | 0 | 1 | 0 | 0 | 0 | 0 | 0 | 0 | 0 | 0 | 0 | 0 | 0 | 0 | 0 | 0 | 1 | 0 | 0 | 1 | 0 | 0 | Olfr142 |
| Olfr1420 | 0 | 0 | 0 | 0 | 0 | 0 | 1 | 2 | 2 | 10 | 0 | 0 | 6 | 0 | 0 | 0 | 0 | 0 | 0 | 0 | 0 | 0 | 0 | 0 | 0 | 1 | 0 | 0 | Olfr1420 |
| Olfr1423 | 0 | 0 | 0 | 0 | 0 | 0 | 0 | 2 | 0 | 0 | 0 | 0 | 0 | 0 | 0 | 1 | 0 | 0 | 2 | 0 | 0 | 0 | 2 | 0 | 0 | 3 | 0 | 4 | Olfr1423 |
| Olfr1426 | 0 | 0 | 0 | 0 | 0 | 0 | 0 | 0 | 0 | 0 | 0 | 0 | 0 | 0 | 0 | 0 | 0 | 0 | 0 | 0 | 0 | 0 | 3 | 0 | 0 | 0 | 0 | 1 | Olfr1426 |
| Olfr1427 | 0 | 0 | 0 | 0 | 0 | 0 | 0 | 0 | 0 | 0 | 0 | 0 | 0 | 0 | 0 | 0 | 0 | 0 | 0 | 0 | 0 | 0 | 1 | 0 | 0 | 0 | 0 | 0 | Olfr1427 |
| Olfr1428 | 0 | 0 | 0 | 0 | 2 | 0 | 0 | 0 | 0 | 4 | 3 | 0 | 0 | 0 | 0 | 0 | 0 | 0 | 0 | 0 | 0 | 0 | 1 | 0 | 0 | 0 | 0 | 0 | Olfr1428 |
| Olfr143 | 0 | 0 | 0 | 0 | 0 | 0 | 0 | 0 | 0 | 0 | 0 | 0 | 0 | 0 | 0 | 0 | 0 | 0 | 0 | 0 | 0 | 0 | 0 | 1 | 0 | 0 | 0 | 0 | Olfr143 |
| Olfr1431 | 0 | 0 | 0 | 0 | 0 | 0 | 0 | 0 | 2 | 0 | 0 | 0 | 0 | 0 | 1 | 0 | 0 | 0 | 1 | 0 | 0 | 0 | 0 | 1 | 0 | 1 | 0 | 0 | Olfr1431 |
| Olfr1432 | 0 | 0 | 0 | 0 | 0 | 0 | 0 | 1 | 0 | 5 | 1 | 0 | 0 | 0 | 0 | 0 | 0 | 0 | 0 | 0 | 0 | 0 | 0 | 0 | 0 | 0 | 0 | 0 | Olfr1432 |
| Olfr1434 | 0 | 0 | 0 | 0 | 0 | 0 | 0 | 6 | 11 | 19 | 3 | 0 | 1 | 3 | 0 | 0 | 0 | 0 | 0 | 0 | 0 | 0 | 10 | 11 | 0 | 2 | 0 | 0 | Olfr1434 |
| Olfr1436 | 0 | 0 | 0 | 0 | 0 | 0 | 0 | 0 | 0 | 0 | 3 | 0 | 0 | 0 | 0 | 0 | 0 | 0 | 0 | 0 | 0 | 0 | 0 | 0 | 0 | 0 | 0 | 0 | Olfr1436 |
| Olfr1437 | 0 | 0 | 0 | 0 | 0 | 0 | 0 | 6 | 2 | 1 | 2 | 0 | 0 | 0 | 0 | 0 | 0 | 0 | 0 | 0 | 0 | 0 | 6 | 4 | 0 | 4 | 0 | 0 | Olfr1437 |
| Olfr1438-ps1 | 0 | 0 | 0 | 0 | 0 | 0 | 0 | 0 | 1 | 0 | 0 | 0 | 0 | 0 | 0 | 0 | 0 | 0 | 0 | 0 | 0 | 0 | 0 | 0 | 0 | 5 | 0 | 0 | Olfr1438-ps1 |
| Olfr1440 | 3 | 1 | 0 | 8 | 0 | 0 | 0 | 2 | 3 | 5 | 1 | 0 | 0 | 0 | 3 | 5 | 0 | 1 | 0 | 0 | 0 | 0 | 8 | 22 | 0 | 12 | 0 | 0 | Olfr1440 |
| Olfr1441 | 0 | 0 | 0 | 0 | 0 | 0 | 0 | 4 | 0 | 2 | 3 | 0 | 0 | 0 | 0 | 0 | 0 | 0 | 0 | 0 | 0 | 3 | 0 | 2 | 0 | 0 | 0 | 0 | Olfr1441 |
| Olfr1443 | 0 | 0 | 0 | 0 | 0 | 0 | 0 | 0 | 0 | 0 | 0 | 0 | 0 | 0 | 0 | 0 | 0 | 0 | 0 | 0 | 0 | 0 | 0 | 1 | 0 | 0 | 0 | 0 | Olfr1443 |
| Olfr1444 | 0 | 0 | 0 | 0 | 0 | 0 | 0 | 0 | 2 | 5 | 0 | 1 | 0 | 0 | 0 | 0 | 0 | 0 | 0 | 0 | 0 | 0 | 0 | 0 | 0 | 0 | 0 | 0 | Olfr1444 |
| Olfr1445 | 0 | 0 | 0 | 1 | 0 | 0 | 0 | 0 | 0 | 0 | 0 | 0 | 0 | 0 | 0 | 0 | 0 | 0 | 0 | 0 | 0 | 0 | 0 | 0 | 0 | 0 | 0 | 0 | Olfr1445 |
| Olfr1446 | 0 | 0 | 0 | 0 | 0 | 0 | 0 | 0 | 0 | 0 | 5 | 0 | 0 | 0 | 0 | 0 | 0 | 0 | 0 | 0 | 0 | 0 | 3 | 1 | 0 | 1 | 0 | 0 | Olfr1446 |
| Olfr1447 | 0 | 0 | 0 | 0 | 0 | 0 | 0 | 0 | 0 | 0 | 0 | 0 | 0 | 0 | 0 | 0 | 0 | 0 | 0 | 0 | 0 | 0 | 1 | 0 | 0 | 0 | 0 | 0 | Olfr1447 |
| Olfr1448 | 0 | 0 | 0 | 0 | 0 | 0 | 0 | 2 | 2 | 0 | 0 | 0 | 0 | 0 | 0 | 0 | 0 | 0 | 0 | 0 | 0 | 0 | 1 | 5 | 0 | 0 | 0 | 0 | Olfr1448 |
| Olfr1449 | 0 | 0 | 0 | 0 | 0 | 0 | 0 | 0 | 0 | 0 | 1 | 0 | 0 | 0 | 0 | 0 | 0 | 0 | 0 | 0 | 0 | 0 | 1 | 4 | 0 | 0 | 0 | 0 | Olfr1449 |
| Olfr145 | 2 | 0 | 0 | 0 | 0 | 0 | 0 | 6 | 4 | 4 | 1 | 0 | 0 | 0 | 0 | 0 | 0 | 0 | 0 | 0 | 0 | 0 | 8 | 7 | 0 | 6 | 0 | 0 | Olfr145 |

| | | | | | | | | | | | | | | | | | | | | | | | | | | | | | |
|---|---|---|---|---|---|---|---|---|---|---|---|---|---|---|---|---|---|---|---|---|---|---|---|---|---|---|---|---|---|
| Olfr1450 | 0 | 0 | 0 | 0 | 0 | 0 | 0 | 0 | 0 | 1 | 1 | 0 | 0 | 0 | 0 | 0 | 0 | 0 | 0 | 0 | 0 | 0 | 1 | 2 | 0 | 0 | 0 | 0 | Olfr1450 |
| Olfr1453 | 0 | 0 | 0 | 0 | 0 | 0 | 0 | 0 | 0 | 0 | 0 | 0 | 0 | 0 | 0 | 0 | 0 | 0 | 0 | 0 | 0 | 0 | 1 | 0 | 0 | 0 | 0 | 0 | Olfr1453 |
| Olfr1457 | 0 | 0 | 0 | 0 | 0 | 0 | 0 | 0 | 0 | 0 | 0 | 0 | 0 | 0 | 0 | 0 | 0 | 0 | 0 | 0 | 0 | 0 | 1 | 0 | 0 | 1 | 0 | 0 | Olfr1457 |
| Olfr1459 | 0 | 0 | 0 | 0 | 0 | 0 | 0 | 0 | 1 | 0 | 0 | 0 | 0 | 0 | 0 | 0 | 0 | 0 | 1 | 0 | 0 | 0 | 2 | 0 | 0 | 0 | 0 | 1 | Olfr1459 |
| Olfr146 | 0 | 0 | 0 | 0 | 0 | 0 | 0 | 0 | 0 | 0 | 0 | 0 | 1 | 0 | 0 | 0 | 0 | 1 | 0 | 0 | 0 | 0 | 0 | 3 | 0 | 0 | 0 | 0 | Olfr146 |
| Olfr1462 | 0 | 0 | 0 | 0 | 0 | 0 | 0 | 0 | 2 | 3 | 0 | 0 | 0 | 3 | 0 | 0 | 0 | 0 | 0 | 0 | 0 | 0 | 6 | 7 | 0 | 1 | 0 | 0 | Olfr1462 |
| Olfr1465 | 0 | 0 | 0 | 0 | 0 | 0 | 0 | 4 | 0 | 3 | 2 | 0 | 0 | 0 | 0 | 0 | 0 | 0 | 0 | 0 | 0 | 0 | 3 | 5 | 0 | 0 | 0 | 0 | Olfr1465 |
| Olfr1466 | 0 | 0 | 0 | 0 | 0 | 0 | 0 | 1 | 0 | 0 | 0 | 0 | 0 | 0 | 3 | 0 | 0 | 0 | 0 | 0 | 0 | 0 | 0 | 2 | 0 | 0 | 0 | 0 | Olfr1466 |
| Olfr1469 | 0 | 0 | 0 | 0 | 0 | 0 | 0 | 0 | 0 | 0 | 1 | 0 | 0 | 0 | 0 | 0 | 0 | 0 | 0 | 0 | 0 | 0 | 0 | 2 | 0 | 0 | 0 | 0 | Olfr1469 |
| Olfr147 | 0 | 0 | 0 | 0 | 0 | 0 | 0 | 0 | 0 | 0 | 0 | 0 | 0 | 0 | 0 | 0 | 0 | 0 | 0 | 0 | 0 | 0 | 1 | 0 | 0 | 1 | 0 | 0 | Olfr147 |
| Olfr1471 | 0 | 0 | 0 | 0 | 0 | 0 | 0 | 0 | 0 | 0 | 0 | 0 | 0 | 1 | 0 | 0 | 0 | 0 | 0 | 0 | 0 | 0 | 0 | 0 | 0 | 0 | 0 | 0 | Olfr1471 |
| Olfr1472 | 0 | 0 | 0 | 0 | 0 | 0 | 0 | 0 | 0 | 1 | 0 | 0 | 0 | 0 | 0 | 0 | 0 | 0 | 0 | 0 | 0 | 0 | 0 | 0 | 0 | 0 | 0 | 0 | Olfr1472 |
| Olfr1474 | 0 | 0 | 0 | 0 | 0 | 0 | 0 | 0 | 0 | 0 | 0 | 0 | 0 | 0 | 0 | 0 | 0 | 0 | 0 | 0 | 0 | 0 | 1 | 0 | 0 | 0 | 0 | 0 | Olfr1474 |
| Olfr148 | 0 | 0 | 0 | 0 | 1 | 3 | 1 | 0 | 1 | 0 | 0 | 0 | 0 | 0 | 1 | 0 | 3 | 1 | 0 | 0 | 3 | 0 | 1 | 0 | 0 | 0 | 0 | 0 | Olfr148 |
| Olfr1484 | 0 | 0 | 0 | 0 | 0 | 0 | 0 | 0 | 0 | 0 | 0 | 0 | 0 | 0 | 0 | 0 | 0 | 0 | 0 | 0 | 0 | 0 | 0 | 2 | 0 | 0 | 0 | 0 | Olfr1484 |
| Olfr1489 | 0 | 0 | 0 | 0 | 0 | 0 | 0 | 0 | 1 | 0 | 0 | 0 | 0 | 0 | 0 | 0 | 0 | 0 | 0 | 0 | 0 | 0 | 0 | 0 | 0 | 0 | 0 | 0 | Olfr1489 |
| Olfr149 | 0 | 0 | 0 | 0 | 0 | 1 | 0 | 0 | 1 | 2 | 1 | 0 | 0 | 0 | 0 | 0 | 0 | 0 | 0 | 0 | 0 | 2 | 4 | 2 | 0 | 0 | 0 | 1 | Olfr149 |
| Olfr1490 | 0 | 0 | 0 | 0 | 0 | 0 | 0 | 0 | 1 | 0 | 0 | 0 | 0 | 0 | 0 | 0 | 0 | 0 | 0 | 0 | 0 | 0 | 0 | 2 | 0 | 0 | 0 | 0 | Olfr1490 |
| Olfr1495 | 0 | 0 | 0 | 0 | 0 | 0 | 0 | 0 | 0 | 0 | 2 | 0 | 0 | 0 | 0 | 0 | 0 | 0 | 0 | 0 | 0 | 0 | 0 | 0 | 0 | 0 | 0 | 0 | Olfr1495 |
| Olfr1496 | 0 | 0 | 0 | 0 | 0 | 0 | 0 | 0 | 1 | 0 | 0 | 0 | 0 | 0 | 0 | 0 | 0 | 0 | 0 | 0 | 0 | 0 | 0 | 0 | 0 | 0 | 0 | 0 | Olfr1496 |
| Olfr1497 | 0 | 0 | 0 | 0 | 0 | 0 | 0 | 0 | 0 | 0 | 0 | 0 | 0 | 0 | 0 | 0 | 0 | 0 | 0 | 0 | 0 | 0 | 0 | 2 | 0 | 0 | 0 | 0 | Olfr1497 |
| Olfr1499 | 1 | 0 | 0 | 0 | 0 | 0 | 0 | 0 | 0 | 0 | 1 | 0 | 0 | 0 | 0 | 0 | 0 | 0 | 0 | 0 | 0 | 0 | 0 | 0 | 0 | 0 | 0 | 0 | Olfr1499 |
| Olfr15 | 48 | 24 | 0 | 47 | 0 | 5 | 0 | 109 | 72 | 145 | 64 | 54 | 101 | 31 | 50 | 32 | 12 | 8 | 0 | 0 | 14 | 3 | 89 | 142 | 6 | 175 | 19 | 31 | Olfr15 |
| Olfr150 | 0 | 0 | 0 | 0 | 0 | 0 | 0 | 1 | 0 | 0 | 0 | 0 | 0 | 0 | 0 | 0 | 0 | 0 | 0 | 0 | 0 | 0 | 1 | 0 | 0 | 2 | 0 | 0 | Olfr150 |
| Olfr1500 | 1 | 0 | 0 | 0 | 0 | 0 | 0 | 0 | 1 | 1 | 3 | 0 | 0 | 0 | 0 | 0 | 0 | 0 | 2 | 0 | 0 | 0 | 5 | 4 | 0 | 0 | 0 | 0 | Olfr1500 |
| Olfr1501 | 1 | 0 | 0 | 0 | 0 | 0 | 0 | 2 | 2 | 3 | 1 | 0 | 0 | 0 | 0 | 3 | 0 | 0 | 0 | 0 | 0 | 0 | 0 | 1 | 0 | 4 | 0 | 0 | Olfr1501 |
| Olfr1504 | 0 | 0 | 0 | 0 | 0 | 0 | 0 | 0 | 0 | 0 | 0 | 0 | 0 | 0 | 0 | 0 | 0 | 0 | 0 | 0 | 0 | 0 | 1 | 0 | 0 | 2 | 0 | 0 | Olfr1504 |
| Olfr1505 | 0 | 0 | 0 | 0 | 0 | 0 | 0 | 0 | 0 | 1 | 0 | 0 | 0 | 0 | 0 | 0 | 0 | 0 | 0 | 0 | 0 | 0 | 0 | 0 | 0 | 0 | 0 | 0 | Olfr1505 |
| Olfr1506 | 1 | 0 | 0 | 0 | 0 | 0 | 1 | 1 | 0 | 1 | 1 | 0 | 0 | 0 | 0 | 0 | 0 | 0 | 0 | 0 | 0 | 0 | 0 | 0 | 0 | 0 | 0 | 0 | Olfr1506 |
| Olfr1507 | 0 | 1 | 0 | 0 | 0 | 0 | 0 | 2 | 1 | 0 | 3 | 0 | 2 | 0 | 0 | 0 | 0 | 0 | 1 | 0 | 3 | 2 | 5 | 0 | 1 | 0 | 0 | 0 | Olfr1507 |
| Olfr1508 | 0 | 0 | 0 | 0 | 0 | 0 | 0 | 0 | 1 | 0 | 1 | 0 | 0 | 0 | 0 | 0 | 0 | 0 | 0 | 0 | 0 | 3 | 3 | 2 | 0 | 0 | 0 | 0 | Olfr1508 |
| Olfr1510 | 0 | 0 | 0 | 0 | 0 | 0 | 0 | 0 | 0 | 4 | 0 | 0 | 0 | 0 | 0 | 0 | 0 | 0 | 0 | 0 | 0 | 0 | 0 | 2 | 0 | 0 | 0 | 0 | Olfr1510 |
| Olfr1511 | 0 | 0 | 0 | 0 | 0 | 0 | 0 | 4 | 1 | 1 | 0 | 0 | 0 | 0 | 0 | 0 | 0 | 0 | 0 | 0 | 0 | 0 | 0 | 5 | 0 | 0 | 0 | 0 | Olfr1511 |
| Olfr1512 | 0 | 0 | 0 | 0 | 0 | 0 | 0 | 1 | 1 | 2 | 5 | 0 | 0 | 0 | 0 | 0 | 0 | 0 | 0 | 0 | 0 | 0 | 1 | 3 | 0 | 0 | 0 | 0 | Olfr1512 |
| Olfr1513 | 0 | 0 | 0 | 0 | 0 | 0 | 0 | 1 | 1 | 0 | 3 | 0 | 0 | 0 | 0 | 0 | 0 | 0 | 0 | 0 | 0 | 0 | 5 | 0 | 0 | 0 | 0 | 0 | Olfr1513 |
| Olfr152 | 0 | 0 | 0 | 0 | 0 | 0 | 0 | 0 | 1 | 0 | 0 | 0 | 0 | 0 | 0 | 0 | 0 | 0 | 0 | 0 | 0 | 0 | 1 | 0 | 0 | 0 | 0 | 0 | Olfr152 |
| Olfr153 | 0 | 0 | 0 | 0 | 2 | 0 | 0 | 0 | 0 | 0 | 3 | 0 | 0 | 2 | 1 | 0 | 0 | 0 | 0 | 0 | 0 | 0 | 0 | 3 | 0 | 0 | 0 | 0 | Olfr153 |
| Olfr1532-ps1 | 0 | 0 | 0 | 0 | 0 | 0 | 0 | 0 | 1 | 0 | 0 | 0 | 0 | 0 | 0 | 0 | 0 | 2 | 0 | 0 | 0 | 0 | 0 | 0 | 0 | 0 | 0 | 0 | Olfr1532-ps1 |
| Olfr1535 | 0 | 0 | 0 | 0 | 0 | 0 | 0 | 0 | 1 | 0 | 0 | 0 | 3 | 0 | 0 | 0 | 0 | 0 | 0 | 0 | 0 | 0 | 1 | 0 | 0 | 0 | 0 | 0 | Olfr1535 |
| Olfr1537 | 0 | 0 | 0 | 0 | 0 | 0 | 0 | 1 | 0 | 0 | 0 | 0 | 1 | 2 | 0 | 0 | 0 | 0 | 0 | 0 | 0 | 0 | 1 | 2 | 0 | 6 | 0 | 0 | Olfr1537 |
| Olfr155 | 0 | 0 | 0 | 1 | 0 | 0 | 0 | 0 | 3 | 1 | 0 | 0 | 1 | 0 | 0 | 0 | 0 | 0 | 0 | 0 | 0 | 0 | 1 | 2 | 0 | 1 | 0 | 0 | Olfr155 |
| Olfr1555-ps1 | 0 | 0 | 0 | 0 | 0 | 0 | 0 | 0 | 1 | 3 | 0 | 0 | 0 | 0 | 0 | 0 | 0 | 0 | 0 | 0 | 0 | 0 | 5 | 2 | 0 | 0 | 0 | 0 | Olfr1555-ps1 |
| Olfr156 | 1 | 0 | 0 | 0 | 0 | 0 | 0 | 0 | 2 | 0 | 1 | 0 | 0 | 0 | 0 | 0 | 0 | 0 | 0 | 0 | 0 | 0 | 0 | 0 | 0 | 0 | 0 | 0 | Olfr156 |
| Olfr1564 | 53 | 38 | 20 | 86 | 56 | 72 | 80 | 22 | 16 | 20 | 21 | 23 | 46 | 27 | 51 | 47 | 42 | 56 | 62 | 55 | 64 | 38 | 27 | 18 | 24 | 17 | 27 | 20 | Olfr1564 |
| Olfr157 | 0 | 0 | 0 | 0 | 0 | 0 | 0 | 1 | 1 | 3 | 2 | 0 | 0 | 0 | 1 | 0 | 0 | 2 | 0 | 2 | 0 | 0 | 3 | 1 | 0 | 3 | 0 | 0 | Olfr157 |
| Olfr159 | 0 | 0 | 0 | 0 | 0 | 0 | 0 | 0 | 0 | 0 | 0 | 0 | 0 | 0 | 0 | 0 | 0 | 0 | 0 | 0 | 0 | 0 | 0 | 2 | 0 | 0 | 0 | 0 | Olfr159 |
| Olfr16 | 0 | 0 | 0 | 0 | 0 | 0 | 0 | 0 | 0 | 5 | 0 | 0 | 0 | 0 | 0 | 0 | 0 | 0 | 0 | 0 | 0 | 0 | 1 | 2 | 0 | 0 | 0 | 0 | Olfr16 |
| Olfr160 | 0 | 0 | 0 | 0 | 0 | 0 | 0 | 0 | 0 | 0 | 0 | 0 | 0 | 0 | 0 | 0 | 0 | 0 | 0 | 0 | 0 | 2 | 1 | 1 | 0 | 0 | 0 | 0 | Olfr160 |
| Olfr161 | 0 | 0 | 0 | 0 | 0 | 0 | 0 | 2 | 0 | 3 | 0 | 0 | 0 | 0 | 1 | 0 | 0 | 0 | 0 | 0 | 0 | 0 | 0 | 0 | 0 | 0 | 0 | 0 | Olfr161 |
| Olfr164 | 0 | 0 | 0 | 0 | 0 | 0 | 0 | 0 | 0 | 0 | 0 | 0 | 0 | 2 | 0 | 0 | 0 | 0 | 0 | 0 | 0 | 0 | 1 | 1 | 0 | 0 | 0 | 0 | Olfr164 |
| Olfr165 | 0 | 0 | 0 | 0 | 0 | 0 | 0 | 0 | 0 | 0 | 0 | 0 | 0 | 0 | 0 | 0 | 0 | 0 | 0 | 0 | 0 | 0 | 0 | 4 | 0 | 0 | 0 | 0 | Olfr165 |
| Olfr166 | 0 | 0 | 0 | 0 | 0 | 0 | 0 | 6 | 0 | 0 | 6 | 0 | 0 | 0 | 0 | 0 | 0 | 0 | 0 | 0 | 0 | 0 | 0 | 0 | 0 | 0 | 0 | 0 | Olfr166 |
| Olfr17 | 0 | 0 | 0 | 0 | 0 | 0 | 0 | 0 | 0 | 1 | 1 | 0 | 0 | 0 | 0 | 0 | 0 | 0 | 0 | 0 | 0 | 0 | 1 | 2 | 0 | 0 | 0 | 0 | Olfr17 |
| Olfr170 | 0 | 0 | 0 | 0 | 0 | 0 | 0 | 0 | 2 | 0 | 0 | 0 | 0 | 0 | 0 | 0 | 0 | 0 | 0 | 0 | 0 | 0 | 0 | 2 | 0 | 0 | 0 | 0 | Olfr170 |
| Olfr172 | 0 | 0 | 0 | 0 | 0 | 0 | 0 | 0 | 1 | 4 | 0 | 0 | 0 | 0 | 0 | 0 | 0 | 0 | 0 | 0 | 0 | 0 | 2 | 0 | 0 | 0 | 0 | 0 | Olfr172 |
| Olfr173 | 0 | 0 | 0 | 0 | 0 | 0 | 0 | 2 | 1 | 0 | 5 | 0 | 0 | 0 | 0 | 0 | 0 | 0 | 0 | 0 | 0 | 0 | 3 | 2 | 0 | 1 | 0 | 0 | Olfr173 |
| Olfr175 | 0 | 0 | 0 | 0 | 0 | 0 | 0 | 0 | 1 | 0 | 0 | 0 | 0 | 0 | 0 | 0 | 0 | 0 | 0 | 0 | 0 | 0 | 0 | 0 | 0 | 0 | 0 | 0 | Olfr175 |
| Olfr178 | 0 | 0 | 0 | 0 | 0 | 0 | 0 | 0 | 0 | 0 | 0 | 0 | 0 | 0 | 0 | 0 | 0 | 0 | 0 | 0 | 0 | 0 | 0 | 1 | 0 | 0 | 0 | 0 | Olfr178 |
| Olfr18 | 9 | 1 | 0 | 5 | 8 | 8 | 6 | 5 | 2 | 11 | 3 | 4 | 5 | 4 | 0 | 2 | 9 | 0 | 3 | 5 | 4 | 15 | 0 | 7 | 1 | 4 | 3 | 7 | Olfr18 |
| Olfr180 | 0 | 0 | 0 | 0 | 0 | 0 | 0 | 2 | 2 | 0 | 0 | 0 | 0 | 0 | 0 | 0 | 0 | 0 | 0 | 0 | 0 | 0 | 0 | 0 | 0 | 0 | 0 | 0 | Olfr180 |
| Olfr183 | 0 | 0 | 0 | 0 | 0 | 0 | 0 | 0 | 0 | 0 | 0 | 0 | 0 | 0 | 0 | 0 | 2 | 0 | 0 | 0 | 0 | 0 | 0 | 0 | 0 | 0 | 1 | 0 | Olfr183 |
| Olfr187 | 0 | 0 | 0 | 0 | 0 | 0 | 0 | 0 | 0 | 0 | 0 | 0 | 0 | 0 | 0 | 0 | 0 | 0 | 0 | 0 | 0 | 0 | 1 | 2 | 0 | 0 | 0 | 0 | Olfr187 |
| Olfr19 | 1 | 0 | 0 | 0 | 0 | 0 | 1 | 5 | 0 | 2 | 4 | 0 | 0 | 2 | 0 | 0 | 3 | 0 | 5 | 2 | 0 | 0 | 1 | 2 | 0 | 0 | 0 | 1 | Olfr19 |
| Olfr190 | 0 | 0 | 0 | 0 | 0 | 0 | 0 | 0 | 1 | 0 | 1 | 0 | 0 | 0 | 0 | 0 | 0 | 0 | 0 | 0 | 0 | 0 | 1 | 0 | 0 | 0 | 0 | 0 | Olfr190 |
| Olfr191 | 0 | 0 | 0 | 0 | 0 | 0 | 0 | 0 | 0 | 3 | 0 | 0 | 0 | 0 | 0 | 0 | 0 | 0 | 0 | 0 | 0 | 0 | 0 | 0 | 0 | 0 | 0 | 0 | Olfr191 |
| Olfr194 | 0 | 0 | 0 | 0 | 0 | 0 | 0 | 2 | 0 | 0 | 0 | 0 | 0 | 0 | 0 | 0 | 0 | 0 | 0 | 0 | 0 | 0 | 0 | 0 | 0 | 0 | 0 | 0 | Olfr194 |
| Olfr201 | 0 | 0 | 0 | 0 | 0 | 0 | 0 | 0 | 0 | 0 | 0 | 0 | 0 | 0 | 0 | 0 | 0 | 0 | 0 | 0 | 0 | 0 | 0 | 0 | 0 | 0 | 0 | 2 | Olfr201 |
| Olfr202 | 0 | 0 | 0 | 0 | 0 | 0 | 0 | 0 | 2 | 0 | 0 | 0 | 0 | 0 | 0 | 0 | 0 | 0 | 0 | 0 | 0 | 0 | 0 | 0 | 0 | 1 | 0 | 0 | Olfr202 |
| Olfr203 | 0 | 0 | 0 | 0 | 0 | 0 | 0 | 0 | 0 | 0 | 0 | 0 | 0 | 0 | 0 | 0 | 0 | 0 | 0 | 0 | 0 | 0 | 1 | 0 | 0 | 0 | 0 | 0 | Olfr203 |
| Olfr204 | 0 | 0 | 0 | 0 | 0 | 0 | 0 | 0 | 0 | 0 | 0 | 0 | 0 | 0 | 0 | 0 | 0 | 0 | 0 | 3 | 0 | 0 | 0 | 2 | 0 | 0 | 0 | 0 | Olfr204 |
| Olfr205 | 0 | 0 | 0 | 0 | 0 | 0 | 0 | 0 | 0 | 0 | 0 | 0 | 1 | 0 | 0 | 0 | 0 | 0 | 0 | 0 | 0 | 0 | 0 | 0 | 0 | 0 | 0 | 0 | Olfr205 |
| Olfr206 | 0 | 0 | 0 | 0 | 0 | 0 | 0 | 0 | 1 | 1 | 0 | 0 | 0 | 0 | 0 | 0 | 0 | 0 | 0 | 0 | 0 | 0 | 0 | 0 | 0 | 0 | 0 | 0 | Olfr206 |
| Olfr209 | 0 | 0 | 0 | 0 | 0 | 0 | 0 | 0 | 0 | 0 | 0 | 0 | 0 | 0 | 0 | 0 | 0 | 0 | 0 | 0 | 0 | 0 | 0 | 2 | 0 | 0 | 0 | 0 | Olfr209 |
| Olfr211 | 8 | 0 | 0 | 7 | 5 | 0 | 4 | 4 | 3 | 0 | 0 | 3 | 0 | 0 | 3 | 4 | 1 | 3 | 4 | 0 | 2 | 0 | 5 | 8 | 0 | 5 | 1 | 0 | Olfr211 |
| Olfr212 | 1 | 10 | 0 | 12 | 10 | 4 | 12 | 4 | 0 | 1 | 0 | 0 | 2 | 0 | 4 | 4 | 3 | 12 | 3 | 3 | 7 | 0 | 1 | 0 | 0 | 2 | 0 | 1 | Olfr212 |
| Olfr213 | 0 | 0 | 0 | 0 | 0 | 0 | 0 | 0 | 0 | 0 | 0 | 0 | 0 | 0 | 0 | 0 | 0 | 0 | 0 | 5 | 2 | 0 | 0 | 0 | 0 | 0 | 0 | 0 | Olfr213 |
| Olfr214 | 0 | 0 | 0 | 3 | 2 | 0 | 1 | 0 | 0 | 0 | 0 | 0 | 0 | 0 | 0 | 2 | 1 | 2 | 0 | 0 | 1 | 0 | 0 | 0 | 0 | 0 | 0 | 0 | Olfr214 |
| Olfr215 | 0 | 0 | 0 | 2 | 0 | 6 | 0 | 2 | 2 | 0 | 0 | 2 | 0 | 0 | 3 | 0 | 0 | 0 | 0 | 0 | 0 | 0 | 1 | 0 | 0 | 0 | 0 | 0 | Olfr215 |
| Olfr218 | 0 | 0 | 0 | 0 | 0 | 0 | 0 | 0 | 2 | 0 | 0 | 0 | 0 | 0 | 0 | 0 | 0 | 0 | 0 | 0 | 0 | 0 | 0 | 1 | 0 | 0 | 0 | 0 | Olfr218 |

| | | | | | | | | | | | | | | | | | | | | | | | | | | | | | |
|---|---|---|---|---|---|---|---|---|---|---|---|---|---|---|---|---|---|---|---|---|---|---|---|---|---|---|---|---|---|
| Olfr220 | 0 | 0 | 0 | 1 | 0 | 0 | 0 | 0 | 1 | 0 | 0 | 0 | 0 | 0 | 0 | 0 | 0 | 0 | 0 | 0 | 0 | 0 | 3 | 1 | 0 | 0 | 0 | 0 | Olfr220 |
| Olfr221 | 0 | 0 | 0 | 7 | 0 | 0 | 0 | 0 | 1 | 12 | 0 | 0 | 0 | 0 | 3 | 0 | 0 | 0 | 0 | 0 | 2 | 0 | 2 | 3 | 0 | 0 | 0 | 1 | Olfr221 |
| Olfr222 | 0 | 0 | 0 | 1 | 0 | 0 | 0 | 2 | 4 | 3 | 11 | 0 | 6 | 1 | 0 | 0 | 0 | 0 | 0 | 0 | 0 | 0 | 8 | 8 | 0 | 13 | 0 | 0 | Olfr222 |
| Olfr223 | 0 | 0 | 0 | 0 | 0 | 0 | 0 | 0 | 0 | 0 | 0 | 0 | 2 | 1 | 1 | 0 | 0 | 0 | 0 | 0 | 0 | 0 | 0 | 2 | 0 | 2 | 0 | 0 | Olfr223 |
| Olfr224 | 2 | 1 | 0 | 0 | 0 | 0 | 0 | 1 | 2 | 0 | 0 | 0 | 0 | 0 | 3 | 0 | 0 | 0 | 0 | 0 | 0 | 0 | 1 | 0 | 1 | 0 | 0 | 0 | Olfr224 |
| Olfr228 | 0 | 0 | 0 | 0 | 0 | 0 | 0 | 2 | 2 | 0 | 0 | 0 | 0 | 0 | 0 | 0 | 0 | 0 | 0 | 0 | 0 | 0 | 1 | 5 | 0 | 1 | 0 | 0 | Olfr228 |
| Olfr23 | 0 | 0 | 0 | 0 | 0 | 0 | 0 | 1 | 0 | 0 | 0 | 0 | 0 | 0 | 0 | 0 | 0 | 0 | 0 | 0 | 0 | 0 | 0 | 0 | 0 | 2 | 0 | 0 | Olfr23 |
| Olfr231 | 24 | 16 | 0 | 13 | 0 | 0 | 0 | 21 | 10 | 25 | 27 | 22 | 56 | 29 | 28 | 19 | 9 | 0 | 0 | 1 | 4 | 0 | 19 | 46 | 6 | 60 | 15 | 24 | Olfr231 |
| Olfr235 | 3 | 0 | 0 | 0 | 0 | 0 | 0 | 7 | 3 | 4 | 3 | 0 | 0 | 0 | 0 | 0 | 0 | 0 | 0 | 0 | 0 | 0 | 15 | 5 | 0 | 3 | 0 | 0 | Olfr235 |
| Olfr237 | 1 | 0 | 0 | 0 | 1 | 0 | 2 | 2 | 2 | 0 | 1 | 1 | 0 | 1 | 0 | 0 | 0 | 0 | 0 | 2 | 0 | 0 | 2 | 1 | 0 | 0 | 0 | 1 | Olfr237 |
| Olfr239 | 0 | 0 | 0 | 2 | 0 | 0 | 0 | 4 | 2 | 2 | 4 | 0 | 0 | 0 | 0 | 0 | 0 | 0 | 0 | 0 | 0 | 0 | 1 | 3 | 0 | 0 | 0 | 0 | Olfr239 |
| Olfr24 | 2 | 1 | 0 | 8 | 1 | 0 | 0 | 0 | 4 | 1 | 5 | 1 | 3 | 3 | 3 | 0 | 0 | 0 | 0 | 0 | 0 | 0 | 7 | 13 | 0 | 12 | 0 | 0 | Olfr24 |
| Olfr25 | 1 | 0 | 0 | 0 | 0 | 0 | 0 | 2 | 12 | 1 | 13 | 0 | 1 | 0 | 0 | 1 | 0 | 0 | 0 | 0 | 0 | 0 | 8 | 12 | 0 | 3 | 0 | 0 | Olfr25 |
| Olfr250 | 0 | 0 | 0 | 0 | 0 | 0 | 0 | 0 | 1 | 0 | 0 | 0 | 0 | 0 | 0 | 0 | 0 | 0 | 0 | 0 | 0 | 0 | 1 | 0 | 0 | 0 | 0 | 0 | Olfr250 |
| Olfr251 | 0 | 0 | 0 | 0 | 0 | 0 | 0 | 0 | 0 | 0 | 0 | 0 | 0 | 0 | 0 | 0 | 0 | 0 | 0 | 0 | 0 | 0 | 0 | 2 | 0 | 0 | 0 | 0 | Olfr251 |
| Olfr263 | 0 | 0 | 0 | 0 | 0 | 0 | 0 | 0 | 0 | 0 | 0 | 0 | 0 | 0 | 0 | 0 | 0 | 0 | 0 | 0 | 0 | 0 | 4 | 1 | 0 | 0 | 0 | 0 | Olfr263 |
| Olfr266 | 1 | 0 | 0 | 0 | 0 | 0 | 1 | 0 | 2 | 0 | 0 | 0 | 2 | 0 | 1 | 1 | 0 | 0 | 2 | 0 | 1 | 2 | 1 | 0 | 3 | 0 | 0 | 1 | Olfr266 |
| Olfr267 | 21 | 10 | 6 | 21 | 27 | 16 | 15 | 23 | 9 | 20 | 16 | 17 | 10 | 13 | 11 | 12 | 23 | 9 | 8 | 19 | 20 | 18 | 10 | 18 | 10 | 16 | 23 | 17 | Olfr267 |
| Olfr270 | 0 | 0 | 0 | 2 | 0 | 0 | 0 | 0 | 0 | 0 | 0 | 0 | 0 | 0 | 0 | 0 | 0 | 0 | 0 | 0 | 0 | 0 | 0 | 2 | 0 | 0 | 0 | 0 | Olfr270 |
| Olfr272 | 0 | 0 | 0 | 3 | 0 | 0 | 0 | 0 | 0 | 0 | 0 | 0 | 0 | 0 | 1 | 1 | 0 | 0 | 0 | 0 | 0 | 0 | 0 | 0 | 0 | 0 | 0 | 0 | Olfr272 |
| Olfr273 | 0 | 0 | 0 | 0 | 0 | 0 | 0 | 0 | 1 | 0 | 0 | 0 | 0 | 1 | 0 | 0 | 0 | 0 | 0 | 0 | 0 | 0 | 3 | 0 | 0 | 0 | 0 | 0 | Olfr273 |
| Olfr275 | 0 | 0 | 0 | 0 | 0 | 0 | 0 | 0 | 0 | 0 | 0 | 0 | 0 | 0 | 0 | 0 | 0 | 0 | 0 | 0 | 0 | 0 | 0 | 1 | 0 | 0 | 0 | 0 | Olfr275 |
| Olfr279 | 0 | 0 | 0 | 0 | 0 | 0 | 0 | 0 | 2 | 0 | 0 | 0 | 0 | 0 | 0 | 0 | 0 | 0 | 0 | 0 | 0 | 0 | 1 | 6 | 0 | 0 | 0 | 0 | Olfr279 |
| Olfr281 | 1 | 0 | 0 | 0 | 0 | 0 | 0 | 9 | 5 | 5 | 6 | 0 | 1 | 0 | 0 | 0 | 0 | 0 | 1 | 0 | 0 | 0 | 5 | 0 | 0 | 4 | 0 | 0 | Olfr281 |
| Olfr282 | 0 | 0 | 0 | 0 | 0 | 0 | 0 | 0 | 0 | 0 | 2 | 0 | 2 | 0 | 0 | 0 | 0 | 0 | 0 | 0 | 0 | 0 | 0 | 3 | 0 | 0 | 0 | 0 | Olfr282 |
| Olfr283 | 0 | 0 | 0 | 0 | 0 | 0 | 0 | 1 | 0 | 0 | 0 | 0 | 0 | 0 | 3 | 0 | 0 | 0 | 3 | 0 | 0 | 0 | 0 | 0 | 1 | 0 | 0 | 1 | Olfr283 |
| Olfr284 | 0 | 0 | 0 | 0 | 0 | 0 | 0 | 2 | 1 | 0 | 1 | 0 | 0 | 0 | 4 | 0 | 0 | 0 | 0 | 0 | 0 | 0 | 1 | 0 | 0 | 1 | 0 | 0 | Olfr284 |
| Olfr285 | 0 | 0 | 0 | 0 | 0 | 0 | 0 | 0 | 1 | 0 | 0 | 0 | 0 | 0 | 0 | 0 | 0 | 0 | 0 | 0 | 0 | 0 | 0 | 0 | 0 | 0 | 0 | 0 | Olfr285 |
| Olfr286 | 0 | 0 | 0 | 0 | 0 | 0 | 0 | 0 | 0 | 5 | 2 | 0 | 0 | 0 | 0 | 0 | 0 | 0 | 0 | 0 | 0 | 0 | 0 | 0 | 0 | 0 | 0 | 0 | Olfr286 |
| Olfr287 | 0 | 0 | 0 | 0 | 0 | 0 | 0 | 0 | 0 | 0 | 0 | 0 | 0 | 0 | 3 | 0 | 0 | 0 | 0 | 0 | 0 | 0 | 0 | 2 | 0 | 0 | 0 | 0 | Olfr287 |
| Olfr290 | 0 | 0 | 0 | 0 | 1 | 0 | 0 | 2 | 2 | 0 | 0 | 0 | 1 | 0 | 0 | 0 | 0 | 0 | 0 | 0 | 0 | 0 | 1 | 0 | 0 | 0 | 0 | 0 | Olfr290 |
| Olfr291 | 0 | 0 | 0 | 0 | 0 | 0 | 0 | 0 | 1 | 0 | 1 | 0 | 0 | 0 | 0 | 0 | 0 | 0 | 0 | 0 | 0 | 0 | 0 | 3 | 0 | 0 | 7 | 0 | Olfr291 |
| Olfr292 | 0 | 0 | 0 | 0 | 0 | 0 | 0 | 1 | 4 | 3 | 5 | 0 | 0 | 0 | 0 | 0 | 0 | 0 | 0 | 0 | 0 | 0 | 1 | 4 | 0 | 0 | 0 | 0 | Olfr292 |
| Olfr293 | 0 | 0 | 0 | 0 | 0 | 0 | 0 | 2 | 2 | 0 | 4 | 0 | 0 | 0 | 0 | 0 | 0 | 0 | 0 | 0 | 0 | 0 | 3 | 7 | 0 | 0 | 0 | 0 | Olfr293 |
| Olfr295 | 0 | 0 | 0 | 0 | 0 | 0 | 0 | 0 | 3 | 2 | 1 | 0 | 0 | 1 | 0 | 0 | 0 | 0 | 0 | 0 | 0 | 0 | 2 | 0 | 0 | 2 | 0 | 0 | Olfr295 |
| Olfr297 | 0 | 0 | 0 | 0 | 0 | 0 | 0 | 1 | 0 | 0 | 2 | 0 | 0 | 0 | 0 | 0 | 0 | 0 | 0 | 0 | 0 | 0 | 2 | 7 | 0 | 0 | 0 | 0 | Olfr297 |
| Olfr298 | 0 | 0 | 0 | 0 | 0 | 0 | 0 | 1 | 3 | 0 | 2 | 0 | 0 | 0 | 0 | 0 | 0 | 0 | 0 | 0 | 0 | 0 | 1 | 8 | 0 | 4 | 0 | 0 | Olfr298 |
| Olfr299 | 0 | 0 | 0 | 0 | 0 | 0 | 0 | 1 | 4 | 8 | 1 | 2 | 0 | 0 | 0 | 0 | 0 | 0 | 0 | 0 | 0 | 0 | 2 | 1 | 0 | 3 | 0 | 0 | Olfr299 |
| Olfr30 | 0 | 0 | 0 | 0 | 0 | 0 | 0 | 0 | 0 | 0 | 0 | 0 | 0 | 0 | 0 | 1 | 0 | 0 | 0 | 0 | 0 | 0 | 1 | 1 | 0 | 0 | 0 | 0 | Olfr30 |
| Olfr301 | 0 | 0 | 0 | 0 | 0 | 0 | 0 | 1 | 0 | 1 | 1 | 0 | 0 | 0 | 0 | 0 | 0 | 0 | 0 | 0 | 0 | 0 | 1 | 0 | 0 | 1 | 0 | 0 | Olfr301 |
| Olfr303 | 0 | 0 | 0 | 0 | 0 | 0 | 0 | 1 | 4 | 10 | 5 | 0 | 0 | 0 | 0 | 0 | 0 | 0 | 0 | 0 | 0 | 0 | 5 | 3 | 0 | 0 | 0 | 0 | Olfr303 |
| Olfr304 | 0 | 0 | 0 | 0 | 0 | 0 | 0 | 0 | 0 | 0 | 0 | 0 | 0 | 0 | 0 | 0 | 0 | 0 | 0 | 0 | 0 | 0 | 1 | 0 | 0 | 0 | 0 | 0 | Olfr304 |
| Olfr305 | 0 | 0 | 0 | 0 | 0 | 0 | 0 | 0 | 0 | 3 | 0 | 0 | 0 | 0 | 0 | 0 | 0 | 0 | 0 | 0 | 0 | 0 | 0 | 0 | 0 | 0 | 0 | 0 | Olfr305 |
| Olfr307 | 0 | 0 | 0 | 0 | 0 | 0 | 0 | 0 | 0 | 0 | 0 | 0 | 0 | 0 | 0 | 0 | 0 | 0 | 0 | 0 | 0 | 0 | 1 | 0 | 0 | 0 | 0 | 0 | Olfr307 |
| Olfr308 | 0 | 0 | 0 | 0 | 0 | 0 | 0 | 0 | 2 | 4 | 0 | 0 | 0 | 0 | 0 | 0 | 0 | 0 | 0 | 0 | 0 | 0 | 0 | 1 | 0 | 0 | 0 | 0 | Olfr308 |
| Olfr309 | 0 | 0 | 0 | 0 | 0 | 0 | 0 | 1 | 2 | 0 | 4 | 0 | 0 | 0 | 0 | 1 | 0 | 0 | 0 | 0 | 0 | 0 | 5 | 7 | 0 | 0 | 0 | 0 | Olfr309 |
| Olfr31 | 0 | 0 | 0 | 0 | 0 | 0 | 0 | 2 | 2 | 1 | 2 | 0 | 0 | 0 | 0 | 0 | 0 | 0 | 0 | 0 | 1 | 0 | 1 | 1 | 0 | 0 | 0 | 0 | Olfr31 |
| Olfr311 | 0 | 0 | 0 | 0 | 0 | 0 | 0 | 1 | 1 | 0 | 0 | 0 | 0 | 0 | 3 | 0 | 0 | 0 | 0 | 0 | 0 | 0 | 4 | 4 | 0 | 1 | 0 | 0 | Olfr311 |
| Olfr312 | 1 | 0 | 0 | 0 | 0 | 0 | 0 | 0 | 3 | 2 | 0 | 0 | 0 | 0 | 0 | 0 | 0 | 0 | 1 | 0 | 0 | 2 | 2 | 3 | 0 | 4 | 0 | 0 | Olfr312 |
| Olfr313 | 0 | 1 | 0 | 0 | 0 | 0 | 0 | 1 | 0 | 0 | 1 | 0 | 0 | 0 | 0 | 0 | 0 | 0 | 0 | 0 | 0 | 2 | 1 | 3 | 0 | 5 | 0 | 2 | Olfr313 |
| Olfr315 | 0 | 0 | 0 | 0 | 0 | 0 | 0 | 1 | 1 | 0 | 0 | 0 | 0 | 0 | 0 | 0 | 0 | 0 | 0 | 0 | 0 | 0 | 2 | 0 | 2 | 0 | 0 | 0 | Olfr315 |
| Olfr316 | 0 | 0 | 0 | 0 | 0 | 0 | 0 | 1 | 0 | 0 | 1 | 0 | 0 | 0 | 0 | 0 | 0 | 0 | 0 | 0 | 0 | 0 | 3 | 0 | 0 | 0 | 0 | 0 | Olfr316 |
| Olfr317 | 0 | 1 | 0 | 0 | 7 | 5 | 6 | 6 | 12 | 2 | 6 | 3 | 14 | 0 | 1 | 2 | 3 | 2 | 2 | 2 | 1 | 5 | 12 | 9 | 3 | 2 | 2 | 0 | Olfr317 |
| Olfr318 | 0 | 0 | 0 | 0 | 1 | 0 | 0 | 0 | 0 | 0 | 0 | 0 | 0 | 0 | 0 | 0 | 0 | 0 | 0 | 0 | 0 | 0 | 0 | 0 | 0 | 0 | 0 | 0 | Olfr318 |
| Olfr319 | 0 | 0 | 0 | 0 | 0 | 0 | 0 | 0 | 2 | 3 | 0 | 0 | 0 | 0 | 0 | 0 | 0 | 0 | 0 | 0 | 0 | 0 | 1 | 0 | 0 | 0 | 0 | 0 | Olfr319 |
| Olfr32 | 7 | 0 | 0 | 13 | 0 | 0 | 0 | 23 | 7 | 25 | 11 | 2 | 2 | 0 | 1 | 0 | 0 | 0 | 0 | 0 | 0 | 2 | 30 | 29 | 2 | 18 | 0 | 0 | Olfr32 |
| Olfr322 | 2 | 0 | 0 | 0 | 0 | 0 | 1 | 1 | 6 | 0 | 4 | 1 | 7 | 2 | 0 | 1 | 0 | 0 | 0 | 0 | 0 | 2 | 8 | 7 | 2 | 6 | 3 | 5 | Olfr322 |
| Olfr323 | 0 | 0 | 0 | 0 | 0 | 0 | 0 | 1 | 1 | 3 | 0 | 0 | 0 | 0 | 0 | 0 | 0 | 0 | 0 | 0 | 0 | 0 | 2 | 5 | 0 | 0 | 0 | 0 | Olfr323 |
| Olfr324 | 0 | 0 | 0 | 0 | 0 | 0 | 1 | 0 | 6 | 0 | 1 | 0 | 1 | 0 | 0 | 0 | 0 | 0 | 0 | 0 | 0 | 2 | 0 | 1 | 0 | 0 | 1 | 0 | Olfr324 |
| Olfr325 | 0 | 0 | 0 | 0 | 0 | 0 | 0 | 2 | 0 | 0 | 2 | 0 | 0 | 0 | 0 | 0 | 0 | 0 | 0 | 0 | 0 | 0 | 0 | 0 | 0 | 0 | 0 | 0 | Olfr325 |
| Olfr328 | 0 | 0 | 0 | 0 | 0 | 0 | 0 | 0 | 0 | 0 | 0 | 0 | 0 | 0 | 0 | 1 | 0 | 0 | 0 | 0 | 0 | 0 | 2 | 0 | 0 | 0 | 0 | 0 | Olfr328 |
| Olfr329 | 0 | 0 | 0 | 0 | 0 | 0 | 0 | 0 | 0 | 0 | 0 | 0 | 0 | 0 | 0 | 0 | 0 | 0 | 0 | 0 | 0 | 0 | 2 | 2 | 4 | 0 | 0 | 0 | Olfr329 |
| Olfr33 | 0 | 0 | 0 | 0 | 0 | 0 | 0 | 0 | 0 | 0 | 0 | 0 | 0 | 0 | 0 | 0 | 0 | 0 | 0 | 0 | 0 | 0 | 2 | 0 | 0 | 0 | 0 | 0 | Olfr33 |
| Olfr330 | 0 | 0 | 0 | 0 | 0 | 0 | 0 | 6 | 0 | 2 | 5 | 0 | 0 | 0 | 0 | 0 | 0 | 0 | 0 | 0 | 0 | 0 | 3 | 1 | 0 | 0 | 0 | 0 | Olfr330 |
| Olfr332 | 0 | 0 | 0 | 0 | 0 | 0 | 0 | 0 | 0 | 0 | 0 | 0 | 3 | 0 | 0 | 0 | 0 | 0 | 0 | 0 | 0 | 0 | 0 | 0 | 0 | 0 | 0 | 0 | Olfr332 |
| Olfr338 | 0 | 0 | 0 | 0 | 0 | 0 | 0 | 0 | 0 | 0 | 0 | 0 | 0 | 0 | 0 | 0 | 0 | 0 | 0 | 0 | 0 | 0 | 0 | 3 | 0 | 0 | 0 | 0 | Olfr338 |
| Olfr341 | 0 | 0 | 0 | 4 | 0 | 0 | 0 | 0 | 1 | 3 | 1 | 0 | 0 | 0 | 1 | 0 | 0 | 0 | 0 | 0 | 0 | 0 | 2 | 2 | 0 | 1 | 0 | 0 | Olfr341 |
| Olfr344 | 0 | 0 | 0 | 0 | 0 | 0 | 0 | 0 | 1 | 4 | 0 | 0 | 0 | 0 | 0 | 2 | 0 | 0 | 0 | 0 | 0 | 0 | 1 | 3 | 0 | 0 | 0 | 1 | Olfr344 |
| Olfr347 | 0 | 0 | 0 | 0 | 0 | 0 | 0 | 0 | 2 | 0 | 0 | 0 | 0 | 0 | 0 | 0 | 0 | 2 | 0 | 0 | 0 | 0 | 0 | 0 | 0 | 0 | 0 | 0 | Olfr347 |
| Olfr348 | 0 | 0 | 0 | 0 | 0 | 0 | 0 | 0 | 0 | 0 | 0 | 0 | 0 | 0 | 0 | 0 | 0 | 0 | 0 | 0 | 0 | 0 | 0 | 0 | 0 | 1 | 0 | 0 | Olfr348 |
| Olfr350 | 0 | 0 | 0 | 0 | 0 | 0 | 0 | 0 | 0 | 3 | 0 | 0 | 0 | 0 | 0 | 0 | 0 | 0 | 0 | 0 | 0 | 0 | 0 | 0 | 0 | 0 | 0 | 0 | Olfr350 |
| Olfr351 | 0 | 0 | 0 | 0 | 0 | 0 | 0 | 0 | 1 | 0 | 0 | 0 | 0 | 0 | 0 | 0 | 0 | 0 | 0 | 0 | 0 | 0 | 1 | 0 | 0 | 0 | 0 | 0 | Olfr351 |
| Olfr352 | 0 | 0 | 0 | 0 | 0 | 0 | 0 | 0 | 0 | 0 | 6 | 0 | 1 | 0 | 0 | 0 | 0 | 0 | 0 | 0 | 0 | 0 | 0 | 0 | 0 | 0 | 0 | 0 | Olfr352 |
| Olfr354 | 0 | 0 | 0 | 0 | 0 | 0 | 0 | 0 | 0 | 0 | 0 | 0 | 0 | 1 | 0 | 0 | 0 | 0 | 0 | 0 | 0 | 0 | 1 | 0 | 0 | 2 | 0 | 0 | Olfr354 |
| Olfr355 | 0 | 0 | 0 | 0 | 0 | 0 | 0 | 0 | 1 | 0 | 0 | 0 | 0 | 0 | 0 | 0 | 0 | 0 | 0 | 0 | 0 | 0 | 0 | 0 | 0 | 2 | 0 | 0 | Olfr355 |
| Olfr357 | 1 | 0 | 0 | 0 | 0 | 0 | 0 | 0 | 0 | 0 | 3 | 0 | 0 | 0 | 0 | 0 | 0 | 0 | 0 | 0 | 0 | 0 | 0 | 0 | 0 | 0 | 0 | 0 | Olfr357 |
| Olfr358 | 0 | 0 | 0 | 0 | 0 | 0 | 0 | 0 | 0 | 0 | 0 | 0 | 0 | 0 | 0 | 0 | 0 | 0 | 0 | 0 | 0 | 0 | 0 | 2 | 0 | 0 | 0 | 0 | Olfr358 |
| Olfr360 | 2 | 0 | 0 | 0 | 0 | 0 | 0 | 0 | 0 | 0 | 0 | 0 | 0 | 0 | 0 | 0 | 0 | 0 | 0 | 0 | 0 | 0 | 0 | 0 | 0 | 0 | 0 | 0 | Olfr360 |
| Olfr364-ps1 | 0 | 0 | 0 | 0 | 0 | 0 | 0 | 0 | 0 | 0 | 0 | 0 | 0 | 0 | 0 | 0 | 0 | 0 | 0 | 0 | 0 | 0 | 1 | 2 | 0 | 0 | 0 | 0 | Olfr364-ps1 |

| | | | | | | | | | | | | | | | | | | | | | | | | | | | | | |
|---|---|---|---|---|---|---|---|---|---|---|---|---|---|---|---|---|---|---|---|---|---|---|---|---|---|---|---|---|---|
| Olfr365 | 0 | 0 | 0 | 0 | 0 | 0 | 0 | 0 | 0 | 0 | 0 | 0 | 0 | 0 | 0 | 0 | 0 | 0 | 0 | 0 | 0 | 0 | 1 | 0 | 0 | 0 | 0 | 0 | Olfr365 |
| Olfr368 | 0 | 0 | 0 | 3 | 0 | 0 | 0 | 2 | 1 | 0 | 1 | 0 | 0 | 0 | 1 | 0 | 0 | 0 | 0 | 0 | 0 | 0 | 3 | 0 | 0 | 2 | 0 | 0 | Olfr368 |
| Olfr370 | 0 | 7 | 7 | 3 | 2 | 0 | 1 | 31 | 8 | 3 | 3 | 3 | 7 | 1 | 3 | 3 | 3 | 0 | 2 | 10 | 1 | 2 | 19 | 1 | 0 | 4 | 3 | 2 | Olfr370 |
| Olfr371 | 0 | 0 | 0 | 0 | 0 | 0 | 0 | 0 | 0 | 1 | 0 | 0 | 1 | 0 | 0 | 0 | 0 | 0 | 0 | 0 | 0 | 0 | 0 | 2 | 0 | 1 | 0 | 0 | Olfr371 |
| Olfr372 | 2 | 0 | 0 | 1 | 5 | 0 | 0 | 0 | 1 | 0 | 0 | 0 | 0 | 0 | 0 | 0 | 0 | 0 | 0 | 3 | 0 | 0 | 0 | 0 | 0 | 0 | 0 | 0 | Olfr372 |
| Olfr373 | 0 | 0 | 0 | 0 | 2 | 0 | 0 | 0 | 0 | 0 | 0 | 0 | 0 | 0 | 0 | 0 | 0 | 0 | 0 | 0 | 0 | 0 | 0 | 1 | 0 | 0 | 0 | 1 | Olfr373 |
| Olfr374 | 1 | 1 | 0 | 4 | 1 | 0 | 1 | 2 | 2 | 12 | 14 | 0 | 0 | 0 | 0 | 0 | 1 | 0 | 0 | 0 | 4 | 0 | 3 | 5 | 0 | 7 | 0 | 0 | Olfr374 |
| Olfr376 | 0 | 1 | 0 | 0 | 0 | 0 | 0 | 1 | 0 | 1 | 0 | 0 | 0 | 0 | 0 | 0 | 0 | 0 | 0 | 0 | 0 | 0 | 1 | 2 | 0 | 5 | 0 | 0 | Olfr376 |
| Olfr378 | 0 | 0 | 0 | 0 | 0 | 0 | 0 | 0 | 0 | 0 | 0 | 0 | 0 | 0 | 0 | 0 | 0 | 0 | 0 | 0 | 0 | 0 | 0 | 0 | 0 | 1 | 0 | 0 | Olfr378 |
| Olfr380 | 0 | 0 | 0 | 0 | 0 | 0 | 0 | 0 | 0 | 0 | 1 | 0 | 0 | 0 | 0 | 0 | 0 | 0 | 0 | 0 | 0 | 0 | 0 | 0 | 0 | 0 | 0 | 0 | Olfr380 |
| Olfr381 | 0 | 0 | 0 | 0 | 0 | 0 | 0 | 0 | 3 | 0 | 0 | 0 | 0 | 0 | 0 | 0 | 0 | 0 | 0 | 0 | 0 | 0 | 1 | 2 | 0 | 0 | 0 | 0 | Olfr381 |
| Olfr384 | 0 | 0 | 0 | 0 | 0 | 0 | 0 | 0 | 0 | 0 | 1 | 0 | 0 | 0 | 0 | 0 | 0 | 0 | 0 | 0 | 0 | 0 | 0 | 0 | 0 | 0 | 0 | 0 | Olfr384 |
| Olfr385 | 0 | 0 | 0 | 0 | 0 | 0 | 0 | 1 | 0 | 0 | 0 | 0 | 0 | 0 | 0 | 0 | 0 | 0 | 0 | 0 | 0 | 0 | 0 | 0 | 0 | 0 | 0 | 0 | Olfr385 |
| Olfr39 | 0 | 0 | 0 | 0 | 0 | 0 | 0 | 1 | 0 | 0 | 0 | 0 | 0 | 0 | 0 | 0 | 0 | 0 | 0 | 0 | 0 | 0 | 1 | 2 | 0 | 0 | 0 | 0 | Olfr39 |
| Olfr390 | 0 | 0 | 0 | 0 | 0 | 0 | 0 | 0 | 0 | 0 | 0 | 0 | 0 | 0 | 0 | 0 | 0 | 0 | 0 | 0 | 0 | 0 | 0 | 2 | 0 | 0 | 0 | 0 | Olfr390 |
| Olfr391 | 0 | 0 | 0 | 0 | 0 | 0 | 0 | 2 | 0 | 3 | 0 | 0 | 0 | 0 | 0 | 0 | 0 | 0 | 0 | 0 | 0 | 0 | 0 | 0 | 0 | 0 | 0 | 0 | Olfr391 |
| Olfr392 | 0 | 0 | 0 | 0 | 0 | 0 | 0 | 0 | 0 | 0 | 0 | 0 | 0 | 0 | 0 | 0 | 0 | 0 | 0 | 0 | 0 | 0 | 1 | 0 | 0 | 0 | 0 | 0 | Olfr392 |
| Olfr393 | 3 | 1 | 0 | 0 | 0 | 0 | 0 | 1 | 2 | 8 | 7 | 1 | 0 | 0 | 0 | 1 | 0 | 0 | 0 | 0 | 0 | 0 | 5 | 12 | 0 | 10 | 0 | 0 | Olfr393 |
| Olfr394 | 0 | 0 | 0 | 0 | 0 | 0 | 0 | 1 | 1 | 0 | 1 | 0 | 0 | 0 | 0 | 0 | 0 | 0 | 0 | 0 | 1 | 0 | 2 | 0 | 0 | 0 | 0 | 0 | Olfr394 |
| Olfr395 | 0 | 0 | 0 | 0 | 0 | 0 | 0 | 0 | 2 | 0 | 7 | 0 | 0 | 0 | 0 | 0 | 0 | 0 | 0 | 0 | 0 | 0 | 1 | 1 | 0 | 1 | 0 | 0 | Olfr395 |
| Olfr399 | 0 | 0 | 0 | 0 | 0 | 0 | 0 | 0 | 2 | 0 | 0 | 0 | 0 | 0 | 0 | 0 | 0 | 0 | 0 | 0 | 0 | 0 | 0 | 0 | 0 | 2 | 0 | 0 | Olfr399 |
| Olfr402 | 0 | 0 | 0 | 0 | 0 | 0 | 0 | 0 | 0 | 0 | 0 | 0 | 0 | 0 | 0 | 0 | 0 | 0 | 0 | 0 | 0 | 0 | 0 | 4 | 0 | 0 | 0 | 0 | Olfr402 |
| Olfr403 | 0 | 0 | 0 | 0 | 0 | 0 | 0 | 1 | 0 | 1 | 0 | 0 | 0 | 0 | 0 | 0 | 0 | 0 | 0 | 0 | 0 | 0 | 0 | 2 | 0 | 0 | 0 | 0 | Olfr403 |
| Olfr406 | 0 | 0 | 0 | 0 | 0 | 0 | 0 | 0 | 0 | 0 | 0 | 0 | 0 | 0 | 0 | 0 | 0 | 0 | 0 | 0 | 0 | 0 | 1 | 0 | 0 | 0 | 0 | 0 | Olfr406 |
| Olfr410 | 0 | 0 | 0 | 0 | 0 | 0 | 0 | 2 | 0 | 5 | 2 | 0 | 0 | 0 | 0 | 0 | 1 | 0 | 0 | 0 | 0 | 0 | 2 | 7 | 0 | 5 | 0 | 0 | Olfr410 |
| Olfr411 | 0 | 0 | 0 | 0 | 0 | 0 | 0 | 0 | 0 | 0 | 1 | 0 | 0 | 0 | 0 | 0 | 0 | 0 | 0 | 0 | 0 | 0 | 4 | 2 | 0 | 1 | 0 | 0 | Olfr411 |
| Olfr412 | 0 | 0 | 0 | 0 | 0 | 0 | 0 | 0 | 2 | 0 | 0 | 0 | 0 | 0 | 0 | 0 | 0 | 0 | 0 | 0 | 0 | 0 | 0 | 1 | 0 | 1 | 0 | 0 | Olfr412 |
| Olfr414 | 0 | 0 | 0 | 0 | 0 | 0 | 0 | 0 | 0 | 0 | 0 | 0 | 0 | 0 | 0 | 0 | 0 | 0 | 0 | 0 | 0 | 0 | 0 | 0 | 0 | 1 | 0 | 0 | Olfr414 |
| Olfr417 | 0 | 0 | 0 | 0 | 0 | 0 | 0 | 0 | 0 | 1 | 0 | 0 | 0 | 0 | 0 | 0 | 0 | 0 | 0 | 0 | 0 | 0 | 2 | 1 | 0 | 0 | 0 | 0 | Olfr417 |
| Olfr418 | 0 | 0 | 0 | 0 | 0 | 0 | 0 | 1 | 0 | 2 | 0 | 0 | 0 | 0 | 0 | 0 | 0 | 0 | 0 | 0 | 0 | 0 | 1 | 2 | 0 | 0 | 0 | 0 | Olfr418 |
| Olfr419 | 0 | 0 | 0 | 0 | 0 | 0 | 0 | 1 | 2 | 0 | 1 | 0 | 0 | 1 | 0 | 0 | 2 | 0 | 0 | 0 | 0 | 0 | 0 | 2 | 0 | 0 | 0 | 0 | Olfr419 |
| Olfr420 | 1 | 0 | 0 | 0 | 0 | 0 | 0 | 1 | 2 | 5 | 3 | 0 | 0 | 0 | 0 | 0 | 0 | 0 | 0 | 0 | 0 | 0 | 0 | 2 | 0 | 0 | 0 | 0 | Olfr420 |
| Olfr424 | 0 | 0 | 0 | 0 | 0 | 0 | 0 | 0 | 0 | 2 | 4 | 0 | 0 | 0 | 3 | 0 | 0 | 0 | 0 | 0 | 0 | 0 | 2 | 3 | 0 | 2 | 0 | 1 | Olfr424 |
| Olfr427 | 0 | 0 | 0 | 0 | 0 | 0 | 0 | 1 | 0 | 0 | 0 | 0 | 0 | 0 | 0 | 0 | 0 | 0 | 0 | 0 | 0 | 0 | 3 | 1 | 0 | 3 | 0 | 0 | Olfr427 |
| Olfr429 | 0 | 0 | 0 | 0 | 0 | 0 | 0 | 1 | 1 | 1 | 1 | 0 | 0 | 0 | 0 | 0 | 0 | 0 | 0 | 0 | 0 | 0 | 3 | 2 | 0 | 0 | 0 | 0 | Olfr429 |
| Olfr430 | 3 | 0 | 0 | 0 | 0 | 0 | 0 | 10 | 5 | 18 | 0 | 0 | 0 | 0 | 0 | 0 | 0 | 0 | 0 | 0 | 0 | 0 | 7 | 30 | 0 | 14 | 0 | 1 | Olfr430 |
| Olfr432 | 0 | 0 | 0 | 0 | 1 | 0 | 0 | 0 | 0 | 1 | 0 | 0 | 0 | 0 | 0 | 4 | 0 | 0 | 0 | 0 | 0 | 0 | 4 | 1 | 0 | 0 | 0 | 0 | Olfr432 |
| Olfr433 | 0 | 0 | 0 | 0 | 0 | 0 | 0 | 0 | 2 | 4 | 0 | 1 | 0 | 0 | 0 | 0 | 2 | 0 | 0 | 0 | 0 | 0 | 1 | 3 | 0 | 4 | 6 | 1 | Olfr433 |
| Olfr434 | 0 | 0 | 0 | 1 | 0 | 0 | 0 | 2 | 0 | 1 | 0 | 0 | 0 | 0 | 0 | 0 | 0 | 0 | 0 | 0 | 0 | 0 | 1 | 0 | 0 | 3 | 0 | 0 | Olfr434 |
| Olfr435 | 0 | 0 | 0 | 0 | 0 | 0 | 0 | 2 | 0 | 3 | 0 | 0 | 1 | 0 | 0 | 0 | 0 | 0 | 0 | 0 | 0 | 0 | 3 | 2 | 0 | 1 | 0 | 0 | Olfr435 |
| Olfr441 | 0 | 0 | 2 | 0 | 0 | 0 | 0 | 1 | 0 | 0 | 0 | 0 | 0 | 0 | 0 | 0 | 0 | 0 | 0 | 0 | 1 | 0 | 0 | 1 | 0 | 0 | 0 | 0 | Olfr441 |
| Olfr444 | 0 | 0 | 0 | 0 | 0 | 0 | 0 | 0 | 0 | 0 | 0 | 0 | 0 | 0 | 1 | 0 | 0 | 0 | 0 | 0 | 0 | 0 | 0 | 0 | 0 | 0 | 0 | 0 | Olfr444 |
| Olfr446 | 0 | 0 | 0 | 0 | 1 | 0 | 0 | 6 | 2 | 0 | 2 | 0 | 1 | 0 | 0 | 0 | 0 | 0 | 0 | 0 | 0 | 0 | 6 | 2 | 1 | 2 | 0 | 0 | Olfr446 |
| Olfr447 | 0 | 0 | 0 | 0 | 0 | 0 | 0 | 0 | 0 | 0 | 0 | 0 | 0 | 0 | 0 | 0 | 0 | 0 | 0 | 0 | 0 | 0 | 0 | 2 | 0 | 0 | 0 | 0 | Olfr447 |
| Olfr448 | 0 | 0 | 0 | 0 | 0 | 0 | 0 | 0 | 1 | 2 | 3 | 0 | 0 | 0 | 0 | 0 | 0 | 0 | 0 | 0 | 0 | 0 | 1 | 2 | 0 | 0 | 0 | 0 | Olfr448 |
| Olfr449 | 0 | 0 | 0 | 0 | 0 | 0 | 0 | 0 | 0 | 0 | 0 | 0 | 0 | 0 | 0 | 0 | 0 | 0 | 0 | 0 | 0 | 0 | 1 | 0 | 0 | 0 | 0 | 0 | Olfr449 |
| Olfr45 | 1 | 1 | 0 | 0 | 0 | 0 | 0 | 0 | 2 | 3 | 0 | 0 | 0 | 0 | 0 | 0 | 0 | 0 | 0 | 0 | 0 | 0 | 0 | 0 | 0 | 0 | 1 | 0 | Olfr45 |
| Olfr450 | 0 | 0 | 0 | 0 | 0 | 0 | 0 | 0 | 0 | 0 | 0 | 0 | 0 | 0 | 0 | 0 | 0 | 0 | 0 | 0 | 0 | 0 | 1 | 0 | 0 | 0 | 0 | 0 | Olfr450 |
| Olfr453 | 0 | 0 | 0 | 0 | 0 | 0 | 0 | 0 | 0 | 1 | 0 | 0 | 0 | 0 | 0 | 0 | 0 | 0 | 0 | 0 | 0 | 0 | 0 | 0 | 0 | 0 | 0 | 0 | Olfr453 |
| Olfr455 | 0 | 0 | 0 | 2 | 0 | 0 | 0 | 1 | 2 | 0 | 0 | 0 | 0 | 0 | 0 | 0 | 0 | 0 | 0 | 0 | 0 | 0 | 2 | 0 | 0 | 0 | 0 | 1 | Olfr455 |
| Olfr457 | 0 | 0 | 0 | 0 | 0 | 0 | 0 | 0 | 1 | 0 | 0 | 0 | 0 | 0 | 0 | 0 | 0 | 0 | 0 | 0 | 0 | 0 | 0 | 0 | 0 | 1 | 0 | 0 | Olfr457 |
| Olfr458 | 0 | 0 | 0 | 0 | 0 | 0 | 0 | 0 | 0 | 0 | 0 | 0 | 0 | 0 | 0 | 0 | 0 | 0 | 0 | 0 | 0 | 0 | 0 | 0 | 0 | 8 | 0 | 0 | Olfr458 |
| Olfr459 | 3 | 0 | 0 | 0 | 0 | 0 | 0 | 1 | 0 | 2 | 0 | 0 | 0 | 0 | 0 | 0 | 0 | 0 | 0 | 0 | 0 | 0 | 0 | 2 | 0 | 7 | 0 | 0 | Olfr459 |
| Olfr46 | 0 | 0 | 0 | 1 | 1 | 0 | 0 | 5 | 2 | 1 | 1 | 1 | 0 | 0 | 0 | 0 | 1 | 0 | 0 | 0 | 0 | 0 | 0 | 0 | 0 | 1 | 1 | 0 | Olfr46 |
| Olfr460 | 24 | 29 | 27 | 82 | 61 | 37 | 49 | 4 | 3 | 1 | 5 | 8 | 7 | 11 | 33 | 29 | 45 | 83 | 37 | 21 | 58 | 10 | 8 | 8 | 11 | 16 | 8 | 6 | Olfr460 |
| Olfr461 | 3 | 0 | 2 | 0 | 0 | 1 | 3 | 0 | 4 | 0 | 1 | 0 | 0 | 0 | 4 | 2 | 1 | 0 | 0 | 0 | 0 | 0 | 1 | 2 | 0 | 0 | 0 | 0 | Olfr461 |
| Olfr462 | 5 | 0 | 0 | 0 | 0 | 0 | 0 | 1 | 0 | 0 | 0 | 0 | 0 | 0 | 0 | 0 | 0 | 0 | 0 | 0 | 0 | 0 | 0 | 0 | 0 | 0 | 0 | 0 | Olfr462 |
| Olfr463 | 2 | 0 | 0 | 0 | 3 | 1 | 0 | 0 | 1 | 1 | 6 | 0 | 0 | 0 | 0 | 0 | 0 | 0 | 0 | 0 | 1 | 0 | 1 | 1 | 5 | 3 | 0 | 0 | Olfr463 |
| Olfr464 | 3 | 2 | 0 | 0 | 2 | 0 | 1 | 6 | 4 | 3 | 5 | 1 | 5 | 1 | 1 | 0 | 0 | 0 | 1 | 1 | 0 | 7 | 6 | 12 | 2 | 3 | 1 | 1 | Olfr464 |
| Olfr465-ps1 | 0 | 0 | 0 | 0 | 0 | 0 | 0 | 0 | 0 | 0 | 0 | 0 | 0 | 0 | 0 | 0 | 0 | 0 | 0 | 0 | 0 | 0 | 0 | 0 | 7 | 0 | 0 | 0 | Olfr465-ps1 |
| Olfr467 | 0 | 0 | 0 | 0 | 0 | 0 | 0 | 0 | 1 | 0 | 0 | 0 | 0 | 0 | 0 | 0 | 0 | 0 | 0 | 0 | 0 | 0 | 0 | 0 | 0 | 0 | 0 | 0 | Olfr467 |
| Olfr469 | 0 | 0 | 0 | 0 | 0 | 0 | 0 | 0 | 0 | 4 | 0 | 0 | 0 | 0 | 0 | 0 | 0 | 0 | 0 | 0 | 0 | 0 | 0 | 0 | 0 | 0 | 0 | 0 | Olfr469 |
| Olfr47 | 2 | 0 | 0 | 0 | 0 | 0 | 0 | 0 | 0 | 2 | 0 | 0 | 0 | 0 | 0 | 0 | 0 | 0 | 0 | 0 | 0 | 0 | 1 | 1 | 0 | 2 | 0 | 0 | Olfr47 |
| Olfr477 | 0 | 0 | 0 | 0 | 0 | 0 | 0 | 0 | 0 | 0 | 0 | 0 | 0 | 0 | 0 | 0 | 0 | 0 | 4 | 0 | 0 | 0 | 0 | 0 | 0 | 0 | 0 | 0 | Olfr477 |
| Olfr478 | 0 | 0 | 0 | 0 | 0 | 0 | 0 | 0 | 0 | 0 | 0 | 0 | 0 | 0 | 0 | 0 | 0 | 0 | 0 | 0 | 0 | 0 | 0 | 4 | 0 | 0 | 0 | 0 | Olfr478 |
| Olfr48 | 0 | 0 | 0 | 0 | 0 | 0 | 0 | 1 | 2 | 0 | 2 | 0 | 2 | 0 | 0 | 0 | 0 | 0 | 0 | 0 | 0 | 3 | 3 | 1 | 0 | 0 | 0 | 0 | Olfr48 |
| Olfr480 | 0 | 0 | 0 | 0 | 0 | 0 | 0 | 0 | 0 | 0 | 0 | 0 | 0 | 0 | 0 | 0 | 0 | 0 | 0 | 0 | 0 | 0 | 0 | 1 | 0 | 0 | 0 | 0 | Olfr480 |
| Olfr481 | 0 | 0 | 0 | 0 | 0 | 0 | 0 | 0 | 2 | 3 | 0 | 0 | 0 | 0 | 0 | 0 | 0 | 0 | 0 | 0 | 0 | 0 | 4 | 2 | 0 | 1 | 0 | 0 | Olfr481 |
| Olfr482 | 10 | 6 | 8 | 9 | 7 | 11 | 10 | 12 | 6 | 8 | 9 | 12 | 8 | 12 | 12 | 11 | 8 | 10 | 8 | 7 | 16 | 15 | 12 | 5 | 17 | 18 | 16 | 6 | Olfr482 |
| Olfr483 | 0 | 0 | 0 | 0 | 0 | 0 | 0 | 0 | 2 | 1 | 0 | 0 | 0 | 0 | 0 | 0 | 0 | 0 | 0 | 0 | 0 | 0 | 0 | 0 | 0 | 0 | 0 | 0 | Olfr483 |
| Olfr485 | 0 | 0 | 0 | 0 | 0 | 0 | 0 | 0 | 1 | 8 | 1 | 0 | 0 | 0 | 0 | 0 | 0 | 0 | 0 | 0 | 0 | 0 | 3 | 0 | 0 | 2 | 0 | 0 | Olfr485 |
| Olfr486 | 0 | 0 | 0 | 0 | 0 | 0 | 0 | 2 | 0 | 0 | 0 | 0 | 0 | 0 | 0 | 0 | 0 | 0 | 0 | 0 | 0 | 0 | 0 | 0 | 0 | 0 | 0 | 0 | Olfr486 |
| Olfr487 | 0 | 0 | 0 | 0 | 0 | 0 | 0 | 0 | 0 | 3 | 2 | 0 | 0 | 0 | 0 | 0 | 0 | 0 | 0 | 0 | 0 | 0 | 0 | 0 | 0 | 3 | 0 | 0 | Olfr487 |
| Olfr488 | 0 | 0 | 0 | 0 | 0 | 0 | 0 | 1 | 0 | 0 | 0 | 0 | 1 | 0 | 0 | 0 | 0 | 0 | 0 | 0 | 0 | 0 | 0 | 0 | 0 | 0 | 0 | 0 | Olfr488 |
| Olfr49 | 0 | 0 | 0 | 4 | 0 | 2 | 0 | 0 | 4 | 3 | 5 | 2 | 0 | 0 | 3 | 0 | 0 | 0 | 0 | 0 | 0 | 0 | 6 | 2 | 0 | 0 | 1 | 0 | Olfr49 |
| Olfr490 | 0 | 0 | 0 | 0 | 0 | 0 | 0 | 0 | 0 | 0 | 1 | 0 | 0 | 0 | 0 | 0 | 0 | 0 | 0 | 0 | 0 | 0 | 0 | 0 | 0 | 0 | 0 | 0 | Olfr490 |
| Olfr492 | 0 | 0 | 0 | 0 | 0 | 0 | 0 | 4 | 6 | 0 | 4 | 0 | 0 | 0 | 0 | 0 | 0 | 0 | 0 | 0 | 0 | 0 | 1 | 0 | 0 | 0 | 0 | 0 | Olfr492 |
| Olfr495 | 0 | 0 | 0 | 0 | 0 | 0 | 0 | 0 | 4 | 1 | 1 | 0 | 0 | 0 | 0 | 0 | 0 | 0 | 0 | 0 | 0 | 0 | 3 | 2 | 0 | 5 | 0 | 0 | Olfr495 |
| Olfr497 | 0 | 0 | 0 | 0 | 0 | 0 | 0 | 0 | 2 | 0 | 0 | 0 | 0 | 0 | 0 | 0 | 0 | 0 | 0 | 0 | 0 | 0 | 7 | 0 | 0 | 0 | 0 | 0 | Olfr497 |

| | | | | | | | | | | | | | | | | | | | | | | | | | | | | | |
|---|---|---|---|---|---|---|---|---|---|---|---|---|---|---|---|---|---|---|---|---|---|---|---|---|---|---|---|---|---|
| Olfr498 | 0 | 0 | 0 | 0 | 0 | 0 | 0 | 2 | 2 | 4 | 2 | 0 | 0 | 0 | 0 | 0 | 0 | 0 | 0 | 0 | 0 | 0 | 3 | 7 | 0 | 2 | 0 | 0 | Olfr498 |
| Olfr5 | 0 | 0 | 0 | 0 | 0 | 0 | 0 | 0 | 0 | 0 | 0 | 1 | 0 | 0 | 0 | 0 | 0 | 0 | 0 | 0 | 0 | 2 | 1 | 0 | 0 | 0 | 0 | 0 | Olfr5 |
| Olfr50 | 0 | 0 | 0 | 0 | 0 | 0 | 0 | 4 | 2 | 0 | 0 | 0 | 0 | 0 | 0 | 0 | 0 | 0 | 0 | 0 | 0 | 0 | 3 | 2 | 0 | 0 | 0 | 0 | Olfr50 |
| Olfr502 | 0 | 0 | 0 | 0 | 0 | 0 | 0 | 0 | 2 | 0 | 0 | 0 | 0 | 0 | 0 | 0 | 0 | 0 | 0 | 0 | 0 | 0 | 0 | 0 | 0 | 0 | 0 | 0 | Olfr502 |
| Olfr506 | 0 | 0 | 0 | 1 | 0 | 0 | 0 | 0 | 0 | 9 | 0 | 0 | 0 | 0 | 1 | 0 | 0 | 0 | 2 | 0 | 1 | 0 | 3 | 2 | 0 | 0 | 0 | 0 | Olfr506 |
| Olfr508 | 0 | 0 | 0 | 0 | 0 | 0 | 0 | 0 | 0 | 0 | 0 | 0 | 0 | 0 | 0 | 0 | 0 | 0 | 0 | 0 | 0 | 0 | 0 | 1 | 0 | 0 | 0 | 0 | Olfr508 |
| Olfr509 | 1 | 0 | 0 | 2 | 0 | 0 | 0 | 11 | 0 | 7 | 3 | 0 | 1 | 5 | 0 | 2 | 0 | 0 | 0 | 0 | 0 | 0 | 7 | 8 | 0 | 7 | 0 | 0 | Olfr509 |
| Olfr510 | 0 | 2 | 0 | 0 | 0 | 0 | 0 | 0 | 1 | 3 | 0 | 0 | 0 | 0 | 0 | 0 | 0 | 0 | 0 | 0 | 0 | 0 | 0 | 0 | 0 | 0 | 0 | 0 | Olfr510 |
| Olfr512 | 0 | 0 | 0 | 0 | 0 | 0 | 0 | 0 | 0 | 0 | 1 | 0 | 0 | 0 | 0 | 0 | 0 | 0 | 0 | 0 | 0 | 0 | 1 | 0 | 0 | 0 | 0 | 0 | Olfr512 |
| Olfr513 | 0 | 0 | 0 | 0 | 0 | 0 | 0 | 0 | 0 | 0 | 0 | 0 | 0 | 0 | 0 | 0 | 0 | 0 | 0 | 0 | 0 | 0 | 0 | 2 | 0 | 0 | 0 | 0 | Olfr513 |
| Olfr516 | 0 | 0 | 0 | 0 | 0 | 0 | 0 | 0 | 0 | 0 | 0 | 0 | 0 | 0 | 0 | 0 | 0 | 0 | 0 | 0 | 0 | 0 | 0 | 1 | 0 | 0 | 0 | 0 | Olfr516 |
| Olfr518 | 0 | 0 | 0 | 0 | 0 | 0 | 0 | 0 | 1 | 0 | 0 | 0 | 0 | 0 | 0 | 0 | 0 | 0 | 0 | 0 | 0 | 0 | 1 | 0 | 0 | 0 | 0 | 0 | Olfr518 |
| Olfr519 | 0 | 0 | 0 | 0 | 0 | 0 | 0 | 2 | 0 | 0 | 0 | 0 | 0 | 0 | 0 | 0 | 0 | 0 | 0 | 0 | 0 | 0 | 0 | 0 | 0 | 0 | 0 | 0 | Olfr519 |
| Olfr52 | 0 | 0 | 0 | 0 | 0 | 0 | 0 | 0 | 0 | 4 | 0 | 0 | 0 | 0 | 0 | 0 | 0 | 1 | 0 | 0 | 0 | 0 | 0 | 0 | 0 | 0 | 0 | 0 | Olfr52 |
| Olfr520 | 0 | 0 | 0 | 0 | 0 | 0 | 2 | 1 | 0 | 7 | 2 | 0 | 0 | 0 | 0 | 0 | 0 | 0 | 0 | 0 | 0 | 0 | 0 | 0 | 0 | 0 | 1 | 0 | Olfr520 |
| Olfr521 | 0 | 0 | 0 | 0 | 0 | 0 | 0 | 2 | 0 | 0 | 0 | 0 | 0 | 0 | 0 | 0 | 0 | 0 | 0 | 0 | 0 | 0 | 2 | 0 | 0 | 0 | 0 | 0 | Olfr521 |
| Olfr522 | 0 | 2 | 0 | 0 | 0 | 0 | 0 | 0 | 1 | 0 | 5 | 0 | 0 | 1 | 0 | 0 | 0 | 0 | 0 | 0 | 0 | 0 | 1 | 2 | 0 | 0 | 0 | 0 | Olfr522 |
| Olfr523 | 0 | 0 | 0 | 0 | 0 | 0 | 0 | 1 | 5 | 8 | 5 | 0 | 0 | 0 | 0 | 0 | 0 | 0 | 0 | 0 | 0 | 0 | 11 | 15 | 0 | 4 | 0 | 0 | Olfr523 |
| Olfr525 | 0 | 0 | 0 | 0 | 0 | 0 | 0 | 0 | 0 | 0 | 0 | 0 | 0 | 0 | 0 | 0 | 0 | 0 | 0 | 0 | 0 | 0 | 0 | 2 | 0 | 0 | 0 | 0 | Olfr525 |
| Olfr527 | 0 | 1 | 0 | 0 | 0 | 0 | 0 | 0 | 0 | 0 | 0 | 0 | 0 | 0 | 0 | 0 | 0 | 0 | 0 | 0 | 0 | 0 | 0 | 0 | 0 | 0 | 0 | 0 | Olfr527 |
| Olfr53 | 0 | 0 | 0 | 0 | 0 | 0 | 0 | 0 | 0 | 0 | 0 | 0 | 0 | 0 | 0 | 0 | 0 | 0 | 0 | 0 | 0 | 0 | 1 | 0 | 0 | 0 | 0 | 0 | Olfr53 |
| Olfr530 | 0 | 0 | 0 | 0 | 0 | 0 | 0 | 0 | 4 | 3 | 5 | 0 | 0 | 0 | 0 | 0 | 0 | 0 | 0 | 0 | 0 | 0 | 1 | 1 | 0 | 0 | 0 | 0 | Olfr530 |
| Olfr531 | 0 | 0 | 0 | 0 | 0 | 0 | 0 | 0 | 0 | 0 | 4 | 0 | 0 | 0 | 0 | 0 | 0 | 0 | 0 | 0 | 0 | 0 | 1 | 0 | 0 | 1 | 0 | 0 | Olfr531 |
| Olfr533 | 0 | 0 | 0 | 0 | 0 | 0 | 1 | 1 | 0 | 1 | 1 | 0 | 0 | 0 | 0 | 0 | 0 | 0 | 0 | 0 | 0 | 0 | 0 | 3 | 0 | 0 | 0 | 0 | Olfr533 |
| Olfr535 | 0 | 0 | 0 | 0 | 0 | 0 | 0 | 2 | 0 | 0 | 0 | 0 | 0 | 0 | 0 | 0 | 0 | 0 | 0 | 0 | 0 | 0 | 0 | 0 | 0 | 0 | 0 | 0 | Olfr535 |
| Olfr536 | 0 | 0 | 0 | 0 | 0 | 0 | 0 | 4 | 3 | 0 | 1 | 0 | 0 | 0 | 0 | 0 | 0 | 0 | 0 | 0 | 0 | 0 | 1 | 2 | 0 | 0 | 0 | 0 | Olfr536 |
| Olfr538 | 0 | 0 | 0 | 0 | 0 | 0 | 0 | 0 | 0 | 0 | 0 | 0 | 0 | 0 | 1 | 0 | 0 | 0 | 0 | 0 | 0 | 0 | 0 | 0 | 0 | 2 | 0 | 0 | Olfr538 |
| Olfr539 | 3 | 2 | 0 | 0 | 1 | 4 | 1 | 17 | 28 | 10 | 12 | 2 | 9 | 2 | 1 | 2 | 0 | 0 | 0 | 1 | 0 | 0 | 39 | 7 | 2 | 8 | 0 | 1 | Olfr539 |
| Olfr54 | 0 | 0 | 0 | 0 | 0 | 0 | 0 | 2 | 4 | 6 | 2 | 0 | 0 | 1 | 0 | 0 | 0 | 0 | 0 | 0 | 0 | 0 | 1 | 2 | 0 | 2 | 0 | 0 | Olfr54 |
| Olfr541 | 0 | 0 | 0 | 0 | 0 | 0 | 0 | 0 | 0 | 5 | 0 | 0 | 0 | 0 | 0 | 0 | 0 | 0 | 0 | 0 | 0 | 0 | 0 | 0 | 0 | 0 | 0 | 0 | Olfr541 |
| Olfr543 | 0 | 1 | 0 | 0 | 0 | 0 | 0 | 0 | 0 | 0 | 0 | 0 | 0 | 0 | 0 | 0 | 0 | 0 | 0 | 0 | 0 | 0 | 0 | 1 | 0 | 2 | 0 | 0 | Olfr543 |
| Olfr544 | 2 | 0 | 0 | 0 | 1 | 1 | 0 | 0 | 0 | 10 | 11 | 0 | 0 | 1 | 0 | 0 | 1 | 0 | 0 | 0 | 0 | 0 | 0 | 1 | 0 | 1 | 0 | 0 | Olfr544 |
| Olfr545 | 0 | 1 | 0 | 0 | 0 | 0 | 0 | 1 | 1 | 4 | 0 | 0 | 0 | 0 | 0 | 0 | 6 | 0 | 1 | 1 | 0 | 0 | 0 | 8 | 0 | 0 | 3 | 0 | Olfr545 |
| Olfr547 | 2 | 1 | 0 | 0 | 0 | 0 | 0 | 0 | 0 | 0 | 0 | 0 | 0 | 0 | 0 | 0 | 0 | 0 | 0 | 0 | 0 | 0 | 0 | 0 | 0 | 0 | 0 | 0 | Olfr547 |
| Olfr548-ps1 | 3 | 0 | 0 | 1 | 2 | 0 | 2 | 0 | 1 | 0 | 0 | 0 | 0 | 0 | 1 | 2 | 0 | 0 | 1 | 0 | 1 | 2 | 1 | 0 | 0 | 0 | 2 | 0 | Olfr548-ps1 |
| Olfr549 | 0 | 0 | 0 | 2 | 0 | 0 | 0 | 2 | 2 | 5 | 0 | 0 | 0 | 0 | 0 | 0 | 0 | 0 | 0 | 0 | 0 | 0 | 0 | 0 | 0 | 1 | 0 | 1 | Olfr549 |
| Olfr55 | 1 | 0 | 0 | 0 | 0 | 2 | 0 | 4 | 3 | 8 | 2 | 0 | 0 | 0 | 0 | 0 | 0 | 0 | 2 | 1 | 0 | 0 | 6 | 0 | 0 | 3 | 0 | 0 | Olfr55 |
| Olfr550 | 0 | 0 | 0 | 12 | 3 | 0 | 1 | 11 | 5 | 21 | 7 | 3 | 0 | 4 | 3 | 0 | 1 | 0 | 0 | 1 | 0 | 0 | 6 | 10 | 0 | 5 | 2 | 1 | Olfr550 |
| Olfr551 | 0 | 0 | 0 | 0 | 0 | 0 | 0 | 2 | 1 | 1 | 0 | 0 | 0 | 0 | 0 | 0 | 0 | 0 | 0 | 0 | 0 | 0 | 0 | 0 | 0 | 0 | 0 | 0 | Olfr551 |
| Olfr552 | 0 | 0 | 1 | 0 | 2 | 0 | 0 | 1 | 7 | 1 | 0 | 0 | 0 | 2 | 0 | 1 | 0 | 0 | 0 | 0 | 0 | 0 | 1 | 6 | 0 | 1 | 0 | 0 | Olfr552 |
| Olfr553 | 0 | 0 | 0 | 0 | 0 | 0 | 0 | 9 | 2 | 8 | 3 | 0 | 0 | 0 | 0 | 0 | 0 | 0 | 0 | 0 | 0 | 0 | 5 | 1 | 0 | 0 | 0 | 0 | Olfr553 |
| Olfr554 | 0 | 0 | 0 | 0 | 0 | 0 | 0 | 0 | 0 | 0 | 0 | 0 | 0 | 0 | 0 | 0 | 0 | 0 | 0 | 0 | 0 | 0 | 2 | 0 | 0 | 0 | 0 | 0 | Olfr554 |
| Olfr557 | 0 | 0 | 0 | 3 | 0 | 0 | 0 | 0 | 0 | 1 | 1 | 0 | 0 | 1 | 0 | 0 | 0 | 0 | 0 | 0 | 0 | 0 | 0 | 0 | 0 | 0 | 0 | 0 | Olfr557 |
| Olfr558 | 9 | 8 | 0 | 0 | 36 | 0 | 1 | 94 | 83 | 56 | 52 | 0 | 58 | 2 | 17 | 14 | 6 | 7 | 2 | 0 | 0 | 97 | 39 | 2 | 39 | 4 | 0 | 1 | Olfr558 |
| Olfr56 | 0 | 3 | 2 | 0 | 1 | 2 | 2 | 2 | 5 | 15 | 3 | 7 | 6 | 3 | 0 | 0 | 2 | 1 | 2 | 2 | 7 | 7 | 3 | 9 | 0 | 8 | 1 | 0 | Olfr56 |
| Olfr560 | 0 | 0 | 0 | 0 | 3 | 0 | 0 | 4 | 1 | 0 | 1 | 0 | 0 | 0 | 1 | 2 | 0 | 2 | 0 | 0 | 0 | 0 | 0 | 0 | 0 | 0 | 0 | 0 | Olfr560 |
| Olfr561 | 0 | 0 | 0 | 0 | 2 | 0 | 0 | 0 | 2 | 1 | 1 | 0 | 0 | 0 | 0 | 0 | 0 | 3 | 0 | 0 | 0 | 3 | 2 | 0 | 0 | 0 | 0 | 0 | Olfr561 |
| Olfr566 | 0 | 0 | 0 | 0 | 0 | 0 | 0 | 0 | 0 | 0 | 1 | 0 | 0 | 0 | 0 | 0 | 0 | 0 | 0 | 0 | 0 | 0 | 1 | 2 | 4 | 0 | 2 | 0 | Olfr566 |
| Olfr568 | 0 | 0 | 0 | 0 | 0 | 0 | 0 | 0 | 0 | 1 | 0 | 0 | 0 | 0 | 0 | 0 | 0 | 0 | 0 | 0 | 0 | 0 | 0 | 2 | 0 | 0 | 0 | 0 | Olfr568 |
| Olfr569 | 0 | 2 | 0 | 0 | 0 | 0 | 0 | 1 | 0 | 0 | 0 | 0 | 2 | 0 | 0 | 0 | 0 | 0 | 0 | 0 | 0 | 0 | 2 | 1 | 0 | 0 | 0 | 0 | Olfr569 |
| Olfr57 | 0 | 0 | 0 | 0 | 0 | 0 | 0 | 0 | 3 | 0 | 0 | 0 | 0 | 0 | 0 | 0 | 0 | 0 | 0 | 0 | 0 | 0 | 0 | 0 | 0 | 0 | 0 | 0 | Olfr57 |
| Olfr570 | 0 | 0 | 0 | 0 | 3 | 0 | 0 | 0 | 4 | 1 | 6 | 0 | 0 | 0 | 1 | 0 | 0 | 0 | 0 | 3 | 0 | 3 | 3 | 2 | 0 | 0 | 0 | 0 | Olfr570 |
| Olfr571 | 13 | 4 | 7 | 11 | 2 | 1 | 0 | 43 | 20 | 30 | 17 | 11 | 24 | 12 | 16 | 11 | 5 | 0 | 2 | 3 | 1 | 8 | 13 | 21 | 6 | 47 | 10 | 19 | Olfr571 |
| Olfr574 | 0 | 0 | 0 | 0 | 0 | 0 | 0 | 0 | 1 | 0 | 0 | 0 | 0 | 0 | 0 | 0 | 0 | 0 | 0 | 0 | 1 | 2 | 1 | 0 | 0 | 0 | 0 | 0 | Olfr574 |
| Olfr576 | 0 | 0 | 0 | 0 | 0 | 0 | 0 | 0 | 0 | 0 | 3 | 0 | 0 | 0 | 0 | 0 | 0 | 0 | 0 | 0 | 0 | 0 | 0 | 0 | 0 | 0 | 0 | 0 | Olfr576 |
| Olfr577 | 0 | 0 | 0 | 0 | 0 | 0 | 0 | 0 | 1 | 0 | 0 | 0 | 0 | 0 | 0 | 0 | 0 | 0 | 0 | 0 | 0 | 0 | 3 | 1 | 0 | 0 | 0 | 0 | Olfr577 |
| Olfr578 | 2 | 1 | 0 | 0 | 0 | 0 | 0 | 6 | 1 | 8 | 0 | 0 | 3 | 1 | 0 | 1 | 0 | 0 | 0 | 0 | 0 | 0 | 5 | 14 | 0 | 2 | 0 | 0 | Olfr578 |
| Olfr58 | 0 | 1 | 0 | 0 | 0 | 4 | 1 | 0 | 0 | 0 | 0 | 0 | 2 | 0 | 5 | 0 | 0 | 0 | 1 | 0 | 0 | 0 | 0 | 1 | 0 | 0 | 1 | 0 | Olfr58 |
| Olfr583 | 0 | 0 | 0 | 0 | 0 | 0 | 0 | 0 | 0 | 0 | 0 | 0 | 0 | 0 | 0 | 0 | 0 | 0 | 0 | 0 | 0 | 0 | 0 | 0 | 0 | 2 | 0 | 0 | Olfr583 |
| Olfr584 | 0 | 0 | 0 | 1 | 0 | 0 | 0 | 0 | 0 | 0 | 1 | 0 | 0 | 0 | 0 | 0 | 0 | 0 | 0 | 0 | 0 | 0 | 1 | 0 | 0 | 0 | 0 | 0 | Olfr584 |
| Olfr585 | 0 | 0 | 0 | 0 | 0 | 0 | 0 | 0 | 0 | 0 | 1 | 0 | 0 | 0 | 0 | 0 | 0 | 0 | 0 | 0 | 0 | 0 | 1 | 0 | 0 | 0 | 0 | 0 | Olfr585 |
| Olfr588-ps1 | 0 | 0 | 0 | 0 | 0 | 0 | 0 | 1 | 0 | 0 | 1 | 0 | 0 | 0 | 0 | 0 | 0 | 0 | 0 | 0 | 0 | 0 | 0 | 0 | 0 | 0 | 0 | 0 | Olfr588-ps1 |
| Olfr591 | 0 | 0 | 0 | 0 | 0 | 0 | 0 | 2 | 5 | 1 | 2 | 0 | 0 | 1 | 0 | 0 | 0 | 0 | 0 | 0 | 0 | 0 | 3 | 0 | 0 | 0 | 0 | 0 | Olfr591 |
| Olfr592 | 0 | 0 | 0 | 0 | 0 | 0 | 0 | 0 | 0 | 0 | 0 | 0 | 0 | 0 | 0 | 0 | 0 | 0 | 0 | 0 | 0 | 0 | 2 | 2 | 0 | 0 | 0 | 0 | Olfr592 |
| Olfr594 | 1 | 0 | 0 | 0 | 0 | 0 | 0 | 1 | 0 | 0 | 0 | 0 | 0 | 0 | 0 | 0 | 0 | 0 | 0 | 0 | 0 | 0 | 0 | 0 | 0 | 2 | 0 | 0 | Olfr594 |
| Olfr596 | 0 | 0 | 0 | 0 | 0 | 0 | 0 | 0 | 0 | 0 | 0 | 0 | 0 | 0 | 0 | 2 | 0 | 0 | 0 | 0 | 0 | 0 | 0 | 2 | 0 | 0 | 0 | 0 | Olfr596 |
| Olfr597 | 1 | 0 | 0 | 0 | 0 | 0 | 0 | 0 | 0 | 0 | 0 | 0 | 0 | 0 | 0 | 0 | 0 | 0 | 0 | 0 | 0 | 0 | 0 | 0 | 0 | 0 | 0 | 0 | Olfr597 |
| Olfr598 | 0 | 0 | 0 | 0 | 0 | 0 | 0 | 0 | 0 | 3 | 1 | 0 | 0 | 0 | 0 | 0 | 0 | 0 | 0 | 0 | 0 | 0 | 2 | 0 | 0 | 0 | 0 | 0 | Olfr598 |
| Olfr599 | 0 | 0 | 0 | 0 | 0 | 0 | 0 | 0 | 0 | 0 | 0 | 0 | 0 | 0 | 0 | 0 | 0 | 0 | 0 | 0 | 0 | 0 | 0 | 1 | 0 | 0 | 0 | 0 | Olfr599 |
| Olfr60 | 0 | 0 | 0 | 3 | 0 | 0 | 0 | 0 | 0 | 0 | 0 | 0 | 0 | 0 | 0 | 0 | 0 | 0 | 0 | 0 | 0 | 0 | 1 | 1 | 0 | 0 | 0 | 0 | Olfr60 |
| Olfr606 | 0 | 0 | 0 | 0 | 0 | 0 | 0 | 0 | 0 | 0 | 3 | 0 | 0 | 0 | 0 | 0 | 0 | 0 | 0 | 0 | 0 | 0 | 0 | 0 | 0 | 0 | 0 | 0 | Olfr606 |
| Olfr61 | 0 | 0 | 0 | 0 | 0 | 0 | 0 | 0 | 2 | 5 | 0 | 0 | 1 | 0 | 0 | 0 | 0 | 0 | 0 | 0 | 0 | 0 | 0 | 0 | 0 | 3 | 0 | 0 | Olfr61 |
| Olfr610 | 0 | 0 | 0 | 3 | 0 | 0 | 0 | 0 | 0 | 0 | 0 | 0 | 0 | 0 | 0 | 0 | 0 | 0 | 0 | 0 | 0 | 0 | 0 | 0 | 0 | 0 | 0 | 0 | Olfr610 |
| Olfr613 | 1 | 0 | 0 | 0 | 0 | 0 | 0 | 5 | 0 | 12 | 3 | 0 | 0 | 0 | 1 | 0 | 0 | 0 | 0 | 0 | 0 | 0 | 5 | 0 | 0 | 3 | 0 | 0 | Olfr613 |
| Olfr615 | 0 | 0 | 0 | 1 | 0 | 0 | 0 | 0 | 1 | 0 | 1 | 0 | 0 | 0 | 0 | 0 | 0 | 0 | 0 | 0 | 0 | 0 | 3 | 2 | 0 | 4 | 0 | 0 | Olfr615 |
| Olfr616 | 0 | 0 | 0 | 0 | 0 | 0 | 0 | 2 | 0 | 2 | 1 | 0 | 0 | 0 | 0 | 0 | 0 | 0 | 0 | 0 | 0 | 0 | 0 | 7 | 0 | 0 | 0 | 0 | Olfr616 |
| Olfr617 | 0 | 0 | 0 | 0 | 0 | 0 | 0 | 0 | 0 | 0 | 0 | 0 | 0 | 0 | 0 | 0 | 0 | 0 | 0 | 0 | 0 | 0 | 0 | 0 | 0 | 4 | 0 | 0 | Olfr617 |
| Olfr618 | 0 | 0 | 0 | 0 | 0 | 0 | 0 | 2 | 2 | 0 | 0 | 0 | 3 | 2 | 0 | 0 | 0 | 0 | 0 | 0 | 0 | 0 | 1 | 0 | 0 | 1 | 1 | 0 | Olfr618 |
| Olfr619 | 0 | 0 | 0 | 0 | 0 | 0 | 0 | 0 | 0 | 0 | 0 | 0 | 0 | 0 | 0 | 0 | 0 | 0 | 0 | 0 | 0 | 3 | 0 | 0 | 0 | 0 | 0 | 0 | Olfr619 |

| | | | | | | | | | | | | | | | | | | | | | | | | | | | | |
|---|---|---|---|---|---|---|---|---|---|---|---|---|---|---|---|---|---|---|---|---|---|---|---|---|---|---|---|---|
| Olfr62 | 0 | 3 | 0 | 2 | 2 | 0 | 1 | 1 | 2 | 0 | 2 | 0 | 1 | 0 | 0 | 0 | 0 | 0 | 0 | 0 | 1 | 0 | 6 | 2 | 0 | 1 | 0 | 0 | Olfr62 |
| Olfr620 | 0 | 0 | 0 | 0 | 0 | 0 | 0 | 1 | 0 | 0 | 3 | 0 | 0 | 0 | 0 | 0 | 0 | 0 | 0 | 0 | 0 | 0 | 3 | 2 | 0 | 2 | 0 | 0 | Olfr620 |
| Olfr622 | 1 | 0 | 0 | 0 | 2 | 0 | 0 | 2 | 0 | 7 | 3 | 0 | 0 | 0 | 0 | 0 | 0 | 0 | 0 | 0 | 0 | 0 | 1 | 2 | 0 | 3 | 0 | 0 | Olfr622 |
| Olfr624 | 0 | 0 | 0 | 1 | 0 | 0 | 1 | 0 | 0 | 0 | 0 | 1 | 0 | 0 | 0 | 0 | 0 | 0 | 0 | 1 | 0 | 0 | 1 | 0 | 0 | 0 | 1 | 0 | Olfr624 |
| Olfr63 | 0 | 0 | 0 | 0 | 0 | 0 | 0 | 0 | 0 | 1 | 0 | 0 | 0 | 0 | 0 | 0 | 0 | 0 | 0 | 0 | 0 | 0 | 3 | 0 | 0 | 0 | 0 | 0 | Olfr63 |
| Olfr630 | 0 | 0 | 0 | 0 | 0 | 0 | 0 | 0 | 0 | 0 | 0 | 0 | 0 | 0 | 0 | 0 | 0 | 0 | 0 | 0 | 0 | 0 | 0 | 0 | 0 | 0 | 0 | 1 | Olfr630 |
| Olfr631 | 0 | 0 | 0 | 0 | 0 | 0 | 0 | 0 | 0 | 0 | 0 | 2 | 1 | 3 | 0 | 0 | 0 | 0 | 1 | 0 | 0 | 0 | 0 | 0 | 0 | 0 | 0 | 0 | Olfr631 |
| Olfr632 | 0 | 0 | 0 | 0 | 0 | 2 | 0 | 2 | 5 | 0 | 0 | 5 | 3 | 14 | 0 | 0 | 0 | 0 | 0 | 0 | 0 | 0 | 0 | 0 | 0 | 5 | 3 | 13 | Olfr632 |
| Olfr633 | 0 | 0 | 0 | 0 | 0 | 0 | 0 | 1 | 0 | 3 | 0 | 1 | 5 | 0 | 0 | 0 | 0 | 0 | 0 | 0 | 0 | 0 | 0 | 0 | 0 | 0 | 1 | 0 | Olfr633 |
| Olfr635 | 0 | 0 | 0 | 0 | 0 | 0 | 0 | 0 | 0 | 1 | 0 | 0 | 0 | 2 | 0 | 0 | 0 | 0 | 0 | 0 | 0 | 0 | 2 | 1 | 0 | 0 | 5 | 0 | Olfr635 |
| Olfr638 | 1 | 0 | 0 | 0 | 0 | 0 | 0 | 0 | 0 | 0 | 0 | 0 | 0 | 0 | 0 | 0 | 0 | 0 | 0 | 0 | 0 | 0 | 0 | 0 | 0 | 0 | 0 | 0 | Olfr638 |
| Olfr639 | 0 | 0 | 0 | 0 | 0 | 0 | 0 | 2 | 0 | 2 | 0 | 0 | 0 | 0 | 0 | 0 | 0 | 0 | 0 | 0 | 0 | 0 | 0 | 1 | 0 | 0 | 0 | 0 | Olfr639 |
| Olfr64 | 0 | 0 | 0 | 0 | 0 | 0 | 0 | 0 | 0 | 0 | 0 | 0 | 0 | 0 | 0 | 0 | 0 | 0 | 0 | 0 | 0 | 0 | 0 | 5 | 0 | 0 | 0 | 0 | Olfr64 |
| Olfr640 | 0 | 0 | 0 | 0 | 0 | 0 | 0 | 1 | 0 | 0 | 0 | 0 | 0 | 0 | 0 | 0 | 0 | 0 | 0 | 0 | 0 | 0 | 3 | 2 | 0 | 0 | 0 | 0 | Olfr640 |
| Olfr641 | 0 | 0 | 0 | 2 | 0 | 0 | 0 | 2 | 1 | 0 | 0 | 0 | 0 | 0 | 0 | 0 | 0 | 0 | 0 | 0 | 0 | 0 | 0 | 1 | 0 | 0 | 0 | 0 | Olfr641 |
| Olfr642 | 0 | 0 | 0 | 0 | 0 | 0 | 0 | 1 | 0 | 8 | 1 | 0 | 0 | 0 | 0 | 0 | 0 | 0 | 0 | 0 | 0 | 0 | 4 | 3 | 0 | 0 | 0 | 0 | Olfr642 |
| Olfr643 | 0 | 0 | 0 | 0 | 0 | 0 | 0 | 1 | 1 | 4 | 3 | 0 | 0 | 0 | 0 | 0 | 0 | 0 | 0 | 0 | 0 | 0 | 2 | 2 | 0 | 0 | 0 | 0 | Olfr643 |
| Olfr644 | 0 | 0 | 0 | 0 | 0 | 0 | 0 | 1 | 2 | 4 | 2 | 0 | 0 | 0 | 0 | 0 | 0 | 0 | 0 | 0 | 0 | 0 | 4 | 9 | 0 | 0 | 0 | 0 | Olfr644 |
| Olfr645 | 0 | 0 | 0 | 0 | 0 | 0 | 0 | 1 | 0 | 0 | 0 | 0 | 0 | 0 | 0 | 0 | 0 | 0 | 0 | 0 | 0 | 0 | 0 | 0 | 0 | 0 | 0 | 0 | Olfr645 |
| Olfr646 | 0 | 0 | 0 | 0 | 0 | 0 | 0 | 0 | 0 | 0 | 0 | 0 | 0 | 0 | 0 | 0 | 0 | 0 | 0 | 0 | 0 | 0 | 3 | 0 | 0 | 0 | 0 | 0 | Olfr646 |
| Olfr648 | 0 | 0 | 0 | 0 | 0 | 2 | 0 | 4 | 3 | 5 | 0 | 0 | 1 | 0 | 0 | 0 | 0 | 0 | 0 | 0 | 0 | 0 | 0 | 3 | 0 | 1 | 0 | 0 | Olfr648 |
| Olfr649 | 0 | 0 | 0 | 0 | 0 | 0 | 0 | 0 | 0 | 0 | 0 | 0 | 0 | 0 | 0 | 0 | 0 | 0 | 0 | 0 | 0 | 0 | 3 | 0 | 0 | 6 | 0 | 0 | Olfr649 |
| Olfr65 | 2 | 1 | 0 | 0 | 0 | 0 | 0 | 2 | 2 | 6 | 4 | 0 | 0 | 1 | 0 | 0 | 0 | 0 | 0 | 0 | 0 | 0 | 3 | 2 | 0 | 7 | 0 | 0 | Olfr65 |
| Olfr651 | 0 | 0 | 0 | 3 | 0 | 0 | 0 | 0 | 1 | 0 | 0 | 0 | 0 | 0 | 0 | 0 | 0 | 0 | 0 | 0 | 0 | 0 | 0 | 0 | 0 | 0 | 0 | 0 | Olfr651 |
| Olfr652 | 0 | 0 | 0 | 0 | 0 | 0 | 0 | 1 | 0 | 0 | 0 | 0 | 0 | 0 | 0 | 0 | 0 | 0 | 1 | 0 | 0 | 0 | 3 | 0 | 0 | 0 | 0 | 0 | Olfr652 |
| Olfr653 | 1 | 1 | 0 | 6 | 0 | 0 | 0 | 15 | 3 | 13 | 7 | 1 | 10 | 1 | 4 | 2 | 0 | 0 | 0 | 0 | 0 | 0 | 12 | 11 | 0 | 8 | 1 | 0 | Olfr653 |
| Olfr654 | 0 | 0 | 0 | 0 | 0 | 0 | 0 | 1 | 0 | 0 | 0 | 0 | 0 | 0 | 0 | 0 | 0 | 0 | 0 | 0 | 0 | 0 | 1 | 0 | 0 | 0 | 0 | 0 | Olfr654 |
| Olfr655 | 0 | 0 | 0 | 0 | 0 | 0 | 0 | 0 | 0 | 1 | 2 | 0 | 0 | 0 | 0 | 0 | 0 | 0 | 0 | 0 | 0 | 0 | 1 | 0 | 3 | 0 | 0 | 0 | Olfr655 |
| Olfr656 | 0 | 0 | 0 | 0 | 0 | 0 | 0 | 0 | 0 | 0 | 4 | 0 | 0 | 0 | 0 | 0 | 0 | 0 | 0 | 0 | 0 | 0 | 1 | 0 | 0 | 0 | 0 | 0 | Olfr656 |
| Olfr657 | 0 | 0 | 0 | 0 | 0 | 0 | 0 | 0 | 0 | 0 | 1 | 0 | 0 | 0 | 0 | 0 | 0 | 0 | 0 | 0 | 0 | 0 | 1 | 0 | 0 | 0 | 0 | 0 | Olfr657 |
| Olfr658 | 0 | 2 | 1 | 0 | 2 | 0 | 0 | 18 | 2 | 16 | 5 | 9 | 8 | 4 | 3 | 3 | 0 | 0 | 1 | 5 | 0 | 3 | 2 | 1 | 6 | 7 | 8 | 14 | Olfr658 |
| Olfr659 | 0 | 0 | 0 | 0 | 0 | 0 | 0 | 0 | 1 | 4 | 2 | 0 | 0 | 0 | 0 | 0 | 0 | 0 | 0 | 0 | 0 | 0 | 3 | 1 | 0 | 0 | 0 | 0 | Olfr659 |
| Olfr661 | 0 | 0 | 0 | 0 | 0 | 0 | 0 | 0 | 0 | 1 | 1 | 0 | 0 | 0 | 0 | 0 | 0 | 0 | 0 | 0 | 0 | 0 | 1 | 0 | 0 | 0 | 0 | 0 | Olfr661 |
| Olfr666 | 0 | 0 | 0 | 0 | 0 | 0 | 0 | 0 | 0 | 2 | 1 | 0 | 0 | 0 | 0 | 0 | 0 | 0 | 0 | 0 | 0 | 0 | 2 | 0 | 0 | 0 | 0 | 0 | Olfr666 |
| Olfr667 | 1 | 0 | 0 | 0 | 0 | 0 | 0 | 0 | 0 | 0 | 0 | 0 | 0 | 0 | 0 | 0 | 0 | 0 | 0 | 0 | 0 | 0 | 0 | 0 | 0 | 0 | 0 | 0 | Olfr667 |
| Olfr668 | 0 | 0 | 0 | 0 | 0 | 0 | 0 | 0 | 1 | 0 | 0 | 0 | 0 | 0 | 0 | 0 | 0 | 0 | 0 | 0 | 0 | 0 | 0 | 2 | 0 | 3 | 0 | 0 | Olfr668 |
| Olfr669 | 0 | 0 | 0 | 0 | 0 | 0 | 0 | 1 | 0 | 0 | 0 | 0 | 0 | 0 | 0 | 0 | 0 | 0 | 0 | 0 | 0 | 0 | 0 | 0 | 0 | 0 | 0 | 0 | Olfr669 |
| Olfr67 | 0 | 0 | 0 | 0 | 0 | 0 | 0 | 0 | 0 | 1 | 0 | 0 | 0 | 0 | 0 | 0 | 0 | 1 | 0 | 0 | 0 | 0 | 1 | 0 | 0 | 0 | 0 | 0 | Olfr67 |
| Olfr670 | 0 | 0 | 0 | 0 | 0 | 0 | 0 | 0 | 2 | 0 | 0 | 0 | 0 | 0 | 0 | 0 | 0 | 0 | 0 | 0 | 0 | 0 | 1 | 2 | 0 | 0 | 0 | 0 | Olfr670 |
| Olfr671 | 0 | 0 | 0 | 0 | 0 | 0 | 0 | 4 | 2 | 0 | 3 | 0 | 0 | 0 | 0 | 0 | 0 | 0 | 0 | 0 | 0 | 0 | 1 | 2 | 0 | 0 | 0 | 0 | Olfr671 |
| Olfr672 | 0 | 0 | 0 | 0 | 0 | 0 | 0 | 0 | 1 | 6 | 0 | 0 | 0 | 0 | 0 | 0 | 0 | 0 | 0 | 0 | 0 | 0 | 1 | 0 | 0 | 0 | 0 | 0 | Olfr672 |
| Olfr675 | 0 | 0 | 0 | 0 | 0 | 0 | 0 | 0 | 0 | 0 | 0 | 0 | 0 | 0 | 0 | 0 | 0 | 0 | 0 | 0 | 0 | 0 | 1 | 0 | 0 | 1 | 0 | 0 | Olfr675 |
| Olfr677 | 0 | 0 | 0 | 0 | 0 | 0 | 0 | 0 | 0 | 2 | 0 | 0 | 0 | 0 | 0 | 0 | 0 | 0 | 0 | 0 | 0 | 0 | 3 | 2 | 0 | 0 | 0 | 0 | Olfr677 |
| Olfr679 | 0 | 0 | 0 | 0 | 0 | 0 | 0 | 0 | 0 | 5 | 0 | 0 | 0 | 0 | 0 | 0 | 0 | 0 | 0 | 0 | 0 | 0 | 1 | 4 | 0 | 0 | 0 | 0 | Olfr679 |
| Olfr68 | 0 | 0 | 0 | 0 | 0 | 0 | 0 | 0 | 3 | 0 | 0 | 0 | 0 | 0 | 0 | 0 | 0 | 0 | 0 | 0 | 0 | 0 | 4 | 2 | 0 | 2 | 0 | 0 | Olfr68 |
| Olfr681 | 0 | 0 | 0 | 0 | 0 | 0 | 0 | 0 | 2 | 3 | 1 | 0 | 0 | 0 | 0 | 0 | 0 | 0 | 0 | 0 | 0 | 0 | 6 | 8 | 0 | 0 | 0 | 0 | Olfr681 |
| Olfr683 | 0 | 0 | 0 | 2 | 2 | 0 | 0 | 0 | 2 | 0 | 0 | 0 | 0 | 0 | 0 | 0 | 0 | 0 | 0 | 0 | 0 | 0 | 0 | 0 | 0 | 0 | 0 | 0 | Olfr683 |
| Olfr684 | 0 | 0 | 0 | 0 | 0 | 0 | 0 | 0 | 0 | 0 | 0 | 0 | 0 | 0 | 0 | 0 | 0 | 0 | 0 | 0 | 0 | 0 | 0 | 2 | 0 | 0 | 0 | 0 | Olfr684 |
| Olfr685 | 2 | 0 | 0 | 0 | 0 | 0 | 0 | 1 | 1 | 3 | 2 | 0 | 1 | 0 | 0 | 0 | 0 | 0 | 0 | 0 | 1 | 0 | 3 | 8 | 0 | 1 | 0 | 0 | Olfr685 |
| Olfr686 | 0 | 0 | 0 | 0 | 1 | 1 | 0 | 0 | 0 | 0 | 0 | 0 | 0 | 0 | 0 | 0 | 0 | 6 | 0 | 0 | 0 | 0 | 0 | 0 | 0 | 3 | 0 | 0 | Olfr686 |
| Olfr689 | 1 | 0 | 0 | 0 | 0 | 0 | 0 | 6 | 2 | 2 | 5 | 0 | 0 | 0 | 0 | 0 | 0 | 0 | 0 | 0 | 0 | 0 | 8 | 5 | 0 | 0 | 0 | 0 | Olfr689 |
| Olfr69 | 0 | 0 | 0 | 0 | 0 | 0 | 0 | 1 | 2 | 0 | 3 | 0 | 0 | 0 | 0 | 0 | 0 | 0 | 0 | 0 | 0 | 3 | 1 | 4 | 0 | 0 | 0 | 0 | Olfr69 |
| Olfr690 | 33 | 42 | 22 | 48 | 52 | 53 | 46 | 4 | 8 | 5 | 8 | 8 | 1 | 5 | 44 | 41 | 36 | 40 | 43 | 29 | 44 | 12 | 4 | 2 | 3 | 0 | 2 | 6 | Olfr690 |
| Olfr691 | 9 | 29 | 30 | 49 | 20 | 27 | 21 | 17 | 9 | 6 | 6 | 2 | 2 | 7 | 33 | 27 | 31 | 25 | 16 | 45 | 28 | 3 | 8 | 9 | 7 | 0 | 3 | 0 | Olfr691 |
| Olfr692 | 0 | 0 | 0 | 0 | 0 | 0 | 0 | 2 | 1 | 1 | 0 | 0 | 0 | 0 | 0 | 0 | 0 | 0 | 0 | 0 | 0 | 0 | 1 | 2 | 0 | 5 | 0 | 0 | Olfr692 |
| Olfr693 | 0 | 0 | 0 | 0 | 0 | 0 | 0 | 2 | 0 | 0 | 2 | 0 | 0 | 2 | 0 | 0 | 0 | 0 | 0 | 0 | 0 | 0 | 1 | 0 | 0 | 0 | 0 | 1 | Olfr693 |
| Olfr694 | 0 | 0 | 0 | 0 | 0 | 0 | 0 | 0 | 0 | 1 | 0 | 0 | 0 | 0 | 0 | 0 | 0 | 0 | 0 | 0 | 0 | 0 | 1 | 1 | 0 | 0 | 0 | 0 | Olfr694 |
| Olfr695 | 0 | 0 | 0 | 0 | 0 | 0 | 0 | 0 | 0 | 0 | 1 | 0 | 0 | 0 | 0 | 0 | 0 | 0 | 0 | 0 | 0 | 0 | 0 | 0 | 0 | 0 | 0 | 0 | Olfr695 |
| Olfr698 | 0 | 0 | 0 | 0 | 0 | 0 | 0 | 1 | 0 | 0 | 0 | 0 | 0 | 0 | 0 | 0 | 0 | 0 | 0 | 0 | 0 | 0 | 0 | 3 | 0 | 3 | 0 | 0 | Olfr698 |
| Olfr70 | 0 | 0 | 0 | 0 | 0 | 0 | 2 | 5 | 1 | 5 | 0 | 0 | 0 | 0 | 0 | 0 | 0 | 0 | 0 | 0 | 1 | 0 | 1 | 0 | 0 | 3 | 0 | 0 | Olfr70 |
| Olfr701 | 0 | 2 | 0 | 0 | 0 | 0 | 0 | 2 | 0 | 4 | 0 | 0 | 1 | 0 | 0 | 1 | 0 | 0 | 0 | 0 | 0 | 0 | 3 | 0 | 0 | 0 | 0 | 0 | Olfr701 |
| Olfr703 | 0 | 1 | 0 | 0 | 0 | 0 | 0 | 0 | 0 | 0 | 0 | 0 | 0 | 0 | 0 | 0 | 0 | 0 | 0 | 0 | 0 | 0 | 0 | 0 | 0 | 0 | 0 | 0 | Olfr703 |
| Olfr704 | 0 | 0 | 0 | 0 | 0 | 0 | 0 | 0 | 1 | 0 | 0 | 0 | 0 | 0 | 0 | 0 | 0 | 0 | 0 | 0 | 0 | 0 | 0 | 0 | 0 | 0 | 0 | 0 | Olfr704 |
| Olfr705 | 0 | 0 | 0 | 0 | 0 | 0 | 0 | 0 | 0 | 2 | 0 | 0 | 0 | 0 | 0 | 0 | 0 | 0 | 0 | 0 | 0 | 0 | 1 | 0 | 0 | 0 | 0 | 0 | Olfr705 |
| Olfr706 | 0 | 0 | 0 | 0 | 0 | 0 | 0 | 0 | 0 | 1 | 4 | 0 | 0 | 0 | 0 | 0 | 0 | 0 | 0 | 0 | 0 | 0 | 3 | 5 | 0 | 1 | 0 | 0 | Olfr706 |
| Olfr707 | 0 | 0 | 0 | 0 | 0 | 0 | 0 | 0 | 0 | 0 | 0 | 0 | 0 | 0 | 0 | 0 | 0 | 0 | 0 | 0 | 0 | 0 | 0 | 2 | 0 | 0 | 0 | 0 | Olfr707 |
| Olfr709-ps1 | 0 | 0 | 0 | 0 | 0 | 0 | 0 | 0 | 0 | 0 | 0 | 0 | 0 | 0 | 0 | 0 | 0 | 0 | 0 | 0 | 0 | 0 | 0 | 2 | 0 | 0 | 0 | 0 | Olfr709-ps1 |
| Olfr71 | 0 | 0 | 0 | 2 | 0 | 0 | 0 | 2 | 3 | 5 | 1 | 0 | 0 | 0 | 0 | 0 | 0 | 0 | 1 | 0 | 0 | 0 | 0 | 13 | 0 | 3 | 0 | 0 | Olfr71 |
| Olfr713 | 0 | 0 | 0 | 0 | 0 | 0 | 0 | 0 | 2 | 1 | 0 | 0 | 0 | 0 | 0 | 0 | 0 | 0 | 0 | 0 | 0 | 0 | 0 | 0 | 0 | 0 | 0 | 0 | Olfr713 |
| Olfr714 | 0 | 0 | 0 | 0 | 0 | 0 | 0 | 1 | 0 | 0 | 0 | 0 | 0 | 0 | 0 | 0 | 0 | 0 | 0 | 0 | 0 | 0 | 0 | 0 | 0 | 0 | 0 | 0 | Olfr714 |
| Olfr715 | 0 | 0 | 0 | 2 | 0 | 0 | 0 | 1 | 0 | 0 | 0 | 0 | 0 | 0 | 0 | 0 | 0 | 0 | 0 | 0 | 0 | 0 | 0 | 0 | 0 | 0 | 0 | 0 | Olfr715 |
| Olfr715b | 0 | 0 | 0 | 0 | 0 | 0 | 0 | 0 | 0 | 1 | 0 | 0 | 0 | 0 | 0 | 0 | 0 | 0 | 0 | 0 | 0 | 0 | 0 | 0 | 0 | 0 | 0 | 0 | Olfr715b |
| Olfr716 | 0 | 0 | 0 | 0 | 0 | 0 | 0 | 0 | 1 | 0 | 0 | 0 | 0 | 0 | 0 | 0 | 0 | 0 | 0 | 0 | 0 | 0 | 2 | 3 | 0 | 0 | 0 | 0 | Olfr716 |
| Olfr718-ps1 | 1 | 4 | 0 | 8 | 0 | 0 | 0 | 15 | 7 | 13 | 12 | 0 | 3 | 4 | 3 | 3 | 3 | 0 | 0 | 0 | 0 | 0 | 17 | 29 | 0 | 21 | 0 | 0 | Olfr718-ps1 |
| Olfr720 | 5 | 0 | 0 | 2 | 0 | 0 | 0 | 11 | 2 | 9 | 3 | 0 | 0 | 0 | 4 | 1 | 5 | 0 | 0 | 0 | 0 | 0 | 7 | 11 | 0 | 11 | 0 | 0 | Olfr720 |
| Olfr721-ps1 | 0 | 0 | 0 | 0 | 0 | 0 | 0 | 6 | 4 | 19 | 4 | 0 | 0 | 1 | 0 | 0 | 6 | 0 | 0 | 0 | 0 | 0 | 5 | 5 | 0 | 8 | 0 | 0 | Olfr721-ps1 |
| Olfr722 | 0 | 0 | 0 | 0 | 0 | 0 | 0 | 0 | 0 | 0 | 0 | 0 | 0 | 0 | 0 | 0 | 0 | 0 | 0 | 0 | 0 | 0 | 1 | 0 | 0 | 0 | 0 | 0 | Olfr722 |
| Olfr724 | 0 | 0 | 0 | 0 | 0 | 0 | 0 | 1 | 0 | 0 | 0 | 0 | 0 | 0 | 0 | 0 | 0 | 0 | 0 | 0 | 0 | 0 | 0 | 0 | 0 | 0 | 0 | 0 | Olfr724 |
| Olfr725 | 2 | 0 | 0 | 0 | 0 | 0 | 0 | 0 | 0 | 0 | 0 | 0 | 1 | 0 | 0 | 0 | 0 | 0 | 0 | 0 | 0 | 0 | 0 | 0 | 0 | 0 | 1 | 0 | Olfr725 |

| | | | | | | | | | | | | | | | | | | | | | | | | | | | | | |
|---|---|---|---|---|---|---|---|---|---|---|---|---|---|---|---|---|---|---|---|---|---|---|---|---|---|---|---|---|---|
| Olfr726 | 0 | 0 | 0 | 0 | 0 | 0 | 0 | 0 | 0 | 0 | 0 | 0 | 0 | 0 | 0 | 0 | 0 | 0 | 0 | 0 | 0 | 0 | 1 | 0 | 0 | 0 | 0 | 0 | Olfr726 |
| Olfr727 | 0 | 0 | 0 | 0 | 1 | 0 | 0 | 0 | 1 | 3 | 1 | 0 | 0 | 0 | 0 | 0 | 0 | 0 | 2 | 0 | 1 | 0 | 1 | 8 | 0 | 0 | 0 | 1 | Olfr727 |
| Olfr728 | 0 | 0 | 0 | 0 | 0 | 0 | 0 | 0 | 0 | 4 | 0 | 0 | 0 | 0 | 0 | 0 | 0 | 0 | 0 | 0 | 0 | 0 | 1 | 0 | 0 | 0 | 0 | 0 | Olfr728 |
| Olfr73 | 0 | 0 | 0 | 0 | 0 | 0 | 0 | 2 | 0 | 0 | 0 | 0 | 2 | 0 | 0 | 0 | 0 | 0 | 0 | 0 | 0 | 0 | 0 | 0 | 0 | 0 | 0 | 0 | Olfr73 |
| Olfr730 | 0 | 0 | 0 | 0 | 0 | 0 | 0 | 0 | 0 | 0 | 1 | 0 | 0 | 0 | 0 | 0 | 0 | 0 | 0 | 0 | 0 | 0 | 3 | 0 | 0 | 0 | 0 | 0 | Olfr730 |
| Olfr731 | 0 | 0 | 0 | 0 | 0 | 0 | 0 | 2 | 0 | 0 | 0 | 0 | 0 | 0 | 0 | 0 | 0 | 0 | 0 | 0 | 0 | 0 | 0 | 0 | 0 | 0 | 0 | 0 | Olfr731 |
| Olfr733 | 0 | 0 | 0 | 0 | 0 | 0 | 0 | 0 | 0 | 0 | 0 | 0 | 0 | 0 | 0 | 0 | 0 | 0 | 0 | 0 | 0 | 0 | 0 | 1 | 0 | 0 | 0 | 0 | Olfr733 |
| Olfr734 | 0 | 0 | 0 | 0 | 0 | 0 | 0 | 0 | 0 | 1 | 8 | 0 | 0 | 0 | 0 | 0 | 0 | 0 | 0 | 0 | 0 | 0 | 0 | 4 | 0 | 0 | 0 | 0 | Olfr734 |
| Olfr735 | 0 | 0 | 0 | 0 | 0 | 0 | 0 | 0 | 0 | 0 | 0 | 0 | 0 | 0 | 0 | 0 | 0 | 0 | 0 | 0 | 0 | 0 | 0 | 3 | 0 | 0 | 0 | 0 | Olfr735 |
| Olfr736 | 0 | 0 | 0 | 6 | 0 | 0 | 0 | 1 | 0 | 8 | 1 | 0 | 3 | 0 | 0 | 0 | 0 | 0 | 0 | 0 | 0 | 0 | 3 | 7 | 0 | 2 | 0 | 0 | Olfr736 |
| Olfr738 | 0 | 0 | 0 | 0 | 0 | 0 | 0 | 0 | 0 | 8 | 0 | 0 | 0 | 0 | 0 | 0 | 0 | 0 | 0 | 0 | 0 | 0 | 0 | 5 | 0 | 0 | 0 | 0 | Olfr738 |
| Olfr739 | 0 | 0 | 0 | 0 | 0 | 0 | 0 | 0 | 2 | 5 | 0 | 0 | 0 | 0 | 0 | 0 | 0 | 0 | 0 | 0 | 0 | 0 | 5 | 2 | 0 | 0 | 0 | 0 | Olfr739 |
| Olfr74 | 0 | 0 | 0 | 0 | 0 | 0 | 0 | 1 | 0 | 0 | 0 | 0 | 0 | 0 | 0 | 0 | 0 | 0 | 0 | 0 | 0 | 0 | 0 | 0 | 0 | 0 | 0 | 0 | Olfr74 |
| Olfr740 | 0 | 0 | 0 | 0 | 0 | 0 | 0 | 1 | 1 | 0 | 0 | 0 | 0 | 0 | 0 | 0 | 0 | 0 | 0 | 0 | 0 | 0 | 3 | 0 | 0 | 0 | 0 | 0 | Olfr740 |
| Olfr741 | 0 | 0 | 0 | 0 | 0 | 0 | 0 | 0 | 2 | 0 | 1 | 0 | 0 | 0 | 0 | 0 | 0 | 0 | 0 | 0 | 0 | 0 | 0 | 0 | 0 | 0 | 0 | 0 | Olfr741 |
| Olfr743 | 0 | 0 | 0 | 0 | 0 | 0 | 0 | 0 | 0 | 0 | 0 | 0 | 0 | 0 | 0 | 0 | 0 | 0 | 0 | 0 | 0 | 0 | 1 | 0 | 0 | 0 | 0 | 0 | Olfr743 |
| Olfr744 | 0 | 0 | 0 | 0 | 0 | 0 | 0 | 0 | 0 | 1 | 0 | 0 | 0 | 0 | 0 | 0 | 0 | 0 | 0 | 0 | 0 | 0 | 1 | 2 | 0 | 0 | 0 | 0 | Olfr744 |
| Olfr745 | 0 | 3 | 0 | 11 | 0 | 0 | 0 | 11 | 13 | 19 | 11 | 0 | 7 | 0 | 0 | 0 | 5 | 0 | 0 | 0 | 0 | 0 | 12 | 13 | 0 | 8 | 0 | 0 | Olfr745 |
| Olfr746 | 0 | 0 | 0 | 0 | 0 | 0 | 0 | 0 | 0 | 0 | 1 | 0 | 0 | 0 | 0 | 0 | 0 | 0 | 0 | 0 | 0 | 0 | 0 | 0 | 0 | 0 | 0 | 0 | Olfr746 |
| Olfr748 | 0 | 0 | 0 | 1 | 0 | 0 | 0 | 1 | 0 | 0 | 2 | 0 | 0 | 0 | 0 | 0 | 0 | 0 | 0 | 0 | 0 | 0 | 1 | 1 | 0 | 2 | 0 | 0 | Olfr748 |
| Olfr749 | 0 | 0 | 0 | 0 | 2 | 3 | 0 | 2 | 0 | 0 | 1 | 3 | 0 | 0 | 1 | 0 | 0 | 2 | 0 | 3 | 0 | 0 | 1 | 2 | 0 | 0 | 0 | 0 | Olfr749 |
| Olfr750 | 0 | 0 | 0 | 0 | 1 | 0 | 0 | 0 | 1 | 0 | 1 | 0 | 0 | 0 | 0 | 0 | 0 | 0 | 2 | 0 | 0 | 0 | 1 | 0 | 0 | 0 | 0 | 0 | Olfr750 |
| Olfr76 | 0 | 0 | 0 | 0 | 0 | 0 | 0 | 9 | 1 | 0 | 0 | 0 | 0 | 0 | 0 | 0 | 0 | 0 | 0 | 0 | 0 | 0 | 3 | 0 | 0 | 2 | 0 | 0 | Olfr76 |
| Olfr763 | 0 | 0 | 0 | 0 | 0 | 0 | 0 | 0 | 0 | 0 | 0 | 0 | 0 | 0 | 0 | 0 | 0 | 0 | 0 | 0 | 0 | 0 | 0 | 0 | 0 | 1 | 0 | 0 | Olfr763 |
| Olfr764-ps1 | 0 | 0 | 0 | 1 | 0 | 0 | 0 | 0 | 0 | 0 | 0 | 0 | 0 | 0 | 0 | 0 | 0 | 0 | 0 | 0 | 0 | 0 | 1 | 0 | 0 | 0 | 0 | 0 | Olfr764-ps1 |
| Olfr765 | 0 | 0 | 0 | 0 | 0 | 0 | 0 | 0 | 0 | 0 | 0 | 0 | 0 | 0 | 0 | 0 | 0 | 0 | 0 | 0 | 0 | 0 | 0 | 1 | 0 | 0 | 0 | 0 | Olfr765 |
| Olfr766-ps1 | 0 | 0 | 0 | 0 | 0 | 0 | 0 | 0 | 0 | 0 | 3 | 0 | 0 | 0 | 0 | 0 | 0 | 0 | 0 | 0 | 0 | 0 | 1 | 0 | 0 | 0 | 1 | 0 | Olfr766-ps1 |
| Olfr767 | 0 | 0 | 0 | 0 | 0 | 0 | 0 | 1 | 0 | 0 | 0 | 0 | 0 | 0 | 0 | 0 | 0 | 0 | 0 | 0 | 1 | 0 | 0 | 0 | 0 | 0 | 0 | 0 | Olfr767 |
| Olfr768 | 0 | 0 | 0 | 0 | 0 | 0 | 0 | 0 | 0 | 0 | 0 | 0 | 0 | 0 | 0 | 0 | 0 | 0 | 0 | 0 | 0 | 0 | 1 | 1 | 0 | 4 | 0 | 0 | Olfr768 |
| Olfr769 | 1 | 0 | 0 | 0 | 0 | 0 | 0 | 2 | 2 | 8 | 2 | 0 | 0 | 2 | 0 | 0 | 0 | 0 | 0 | 0 | 0 | 0 | 5 | 2 | 0 | 2 | 0 | 0 | Olfr769 |
| Olfr77 | 0 | 0 | 0 | 0 | 0 | 0 | 0 | 0 | 0 | 0 | 0 | 0 | 0 | 0 | 0 | 0 | 0 | 0 | 1 | 0 | 0 | 0 | 3 | 0 | 0 | 0 | 0 | 0 | Olfr77 |
| Olfr771 | 0 | 0 | 0 | 0 | 0 | 0 | 0 | 0 | 0 | 0 | 0 | 0 | 0 | 0 | 0 | 0 | 0 | 0 | 0 | 0 | 0 | 0 | 2 | 2 | 0 | 0 | 0 | 0 | Olfr771 |
| Olfr772 | 0 | 0 | 0 | 0 | 0 | 0 | 0 | 0 | 0 | 0 | 0 | 0 | 0 | 0 | 0 | 1 | 0 | 0 | 0 | 0 | 0 | 0 | 0 | 1 | 0 | 0 | 0 | 0 | Olfr772 |
| Olfr775 | 0 | 0 | 0 | 1 | 0 | 0 | 0 | 2 | 0 | 7 | 1 | 0 | 1 | 2 | 0 | 0 | 0 | 0 | 0 | 0 | 0 | 0 | 2 | 2 | 0 | 4 | 0 | 0 | Olfr775 |
| Olfr776 | 0 | 0 | 0 | 0 | 0 | 0 | 0 | 1 | 0 | 0 | 0 | 0 | 0 | 0 | 0 | 0 | 0 | 0 | 0 | 0 | 0 | 0 | 0 | 0 | 0 | 0 | 0 | 0 | Olfr776 |
| Olfr777 | 0 | 0 | 0 | 0 | 0 | 0 | 0 | 0 | 1 | 0 | 0 | 0 | 0 | 0 | 0 | 0 | 0 | 0 | 0 | 0 | 0 | 0 | 1 | 4 | 0 | 2 | 0 | 0 | Olfr777 |
| Olfr78 | 24 | 9 | 0 | 3 | 11 | 0 | 0 | 33 | 30 | 36 | 30 | 5 | 42 | 1 | 7 | 16 | 10 | 4 | 0 | 0 | 4 | 38 | 25 | 9 | 12 | 14 | 1 | 6 | Olfr78 |
| Olfr781 | 0 | 0 | 0 | 0 | 0 | 0 | 0 | 0 | 0 | 0 | 2 | 0 | 0 | 0 | 0 | 0 | 0 | 0 | 0 | 0 | 0 | 0 | 0 | 0 | 0 | 0 | 0 | 0 | Olfr781 |
| Olfr782 | 0 | 0 | 0 | 0 | 0 | 0 | 0 | 1 | 1 | 3 | 0 | 0 | 0 | 0 | 0 | 0 | 0 | 0 | 0 | 0 | 0 | 0 | 0 | 1 | 0 | 1 | 0 | 0 | Olfr782 |
| Olfr784 | 0 | 0 | 0 | 0 | 0 | 0 | 0 | 0 | 1 | 2 | 0 | 0 | 0 | 0 | 0 | 0 | 0 | 0 | 2 | 0 | 0 | 0 | 0 | 0 | 0 | 0 | 0 | 0 | Olfr784 |
| Olfr788 | 0 | 0 | 0 | 0 | 0 | 0 | 0 | 0 | 1 | 4 | 0 | 0 | 2 | 0 | 0 | 0 | 0 | 0 | 0 | 0 | 0 | 0 | 0 | 0 | 0 | 0 | 0 | 0 | Olfr788 |
| Olfr790 | 0 | 0 | 0 | 0 | 0 | 0 | 0 | 0 | 0 | 0 | 0 | 0 | 0 | 0 | 0 | 0 | 0 | 0 | 0 | 0 | 0 | 0 | 0 | 2 | 0 | 0 | 0 | 0 | Olfr790 |
| Olfr791 | 0 | 0 | 0 | 0 | 0 | 0 | 0 | 0 | 2 | 0 | 0 | 0 | 0 | 0 | 0 | 0 | 0 | 0 | 0 | 0 | 0 | 0 | 3 | 0 | 0 | 0 | 0 | 0 | Olfr791 |
| Olfr792 | 0 | 0 | 0 | 0 | 0 | 0 | 0 | 2 | 1 | 0 | 0 | 0 | 0 | 0 | 0 | 0 | 0 | 0 | 0 | 0 | 0 | 0 | 0 | 3 | 0 | 0 | 0 | 0 | Olfr792 |
| Olfr796 | 0 | 0 | 0 | 0 | 0 | 0 | 0 | 2 | 0 | 0 | 0 | 0 | 0 | 0 | 0 | 0 | 0 | 0 | 0 | 0 | 0 | 0 | 0 | 1 | 0 | 0 | 0 | 0 | Olfr796 |
| Olfr798 | 0 | 0 | 0 | 0 | 0 | 0 | 0 | 0 | 0 | 0 | 0 | 0 | 0 | 0 | 0 | 0 | 0 | 0 | 0 | 0 | 0 | 0 | 1 | 1 | 0 | 1 | 0 | 0 | Olfr798 |
| Olfr8 | 0 | 0 | 0 | 0 | 0 | 0 | 0 | 0 | 0 | 0 | 1 | 0 | 0 | 0 | 0 | 0 | 0 | 0 | 0 | 0 | 0 | 0 | 1 | 3 | 0 | 0 | 0 | 0 | Olfr8 |
| Olfr800 | 0 | 0 | 0 | 1 | 0 | 0 | 0 | 0 | 1 | 0 | 1 | 0 | 0 | 0 | 0 | 0 | 0 | 0 | 0 | 0 | 0 | 0 | 0 | 0 | 0 | 0 | 0 | 0 | Olfr800 |
| Olfr801 | 0 | 0 | 0 | 0 | 0 | 0 | 0 | 0 | 0 | 0 | 1 | 0 | 0 | 0 | 0 | 0 | 0 | 0 | 0 | 0 | 0 | 0 | 0 | 0 | 0 | 0 | 0 | 0 | Olfr801 |
| Olfr802 | 0 | 0 | 0 | 0 | 0 | 0 | 0 | 0 | 0 | 0 | 0 | 0 | 0 | 0 | 0 | 0 | 0 | 0 | 0 | 0 | 0 | 0 | 1 | 0 | 0 | 0 | 0 | 0 | Olfr802 |
| Olfr803 | 0 | 0 | 0 | 0 | 0 | 0 | 0 | 0 | 0 | 0 | 0 | 0 | 0 | 0 | 0 | 0 | 0 | 0 | 0 | 0 | 0 | 0 | 0 | 1 | 0 | 0 | 0 | 0 | Olfr803 |
| Olfr804 | 0 | 0 | 0 | 0 | 0 | 0 | 0 | 0 | 0 | 0 | 0 | 0 | 0 | 0 | 0 | 0 | 0 | 0 | 0 | 0 | 0 | 0 | 2 | 2 | 0 | 0 | 0 | 0 | Olfr804 |
| Olfr805 | 0 | 0 | 0 | 0 | 0 | 0 | 0 | 0 | 0 | 0 | 0 | 0 | 0 | 0 | 0 | 0 | 0 | 0 | 0 | 0 | 0 | 0 | 0 | 1 | 0 | 0 | 0 | 0 | Olfr805 |
| Olfr806 | 0 | 0 | 0 | 0 | 0 | 0 | 0 | 0 | 0 | 0 | 0 | 0 | 0 | 0 | 0 | 0 | 0 | 0 | 0 | 0 | 0 | 0 | 0 | 0 | 0 | 2 | 0 | 0 | Olfr806 |
| Olfr808 | 0 | 0 | 0 | 0 | 0 | 0 | 0 | 0 | 1 | 0 | 0 | 0 | 0 | 0 | 0 | 0 | 0 | 0 | 0 | 0 | 0 | 0 | 0 | 0 | 0 | 0 | 0 | 0 | Olfr808 |
| Olfr809 | 0 | 0 | 0 | 0 | 0 | 0 | 0 | 0 | 2 | 0 | 0 | 0 | 0 | 0 | 0 | 0 | 0 | 0 | 0 | 0 | 0 | 0 | 0 | 2 | 0 | 0 | 0 | 0 | Olfr809 |
| Olfr810 | 1 | 3 | 0 | 2 | 0 | 0 | 0 | 1 | 7 | 2 | 10 | 0 | 0 | 0 | 0 | 1 | 0 | 0 | 0 | 0 | 0 | 0 | 3 | 12 | 0 | 1 | 0 | 0 | Olfr810 |
| Olfr812 | 0 | 0 | 0 | 0 | 0 | 0 | 0 | 0 | 0 | 0 | 0 | 0 | 0 | 0 | 0 | 0 | 0 | 0 | 0 | 0 | 0 | 0 | 0 | 1 | 0 | 0 | 0 | 0 | Olfr812 |
| Olfr813 | 0 | 0 | 0 | 3 | 0 | 0 | 0 | 4 | 1 | 0 | 5 | 0 | 0 | 0 | 0 | 0 | 0 | 0 | 0 | 0 | 0 | 0 | 7 | 10 | 0 | 2 | 0 | 0 | Olfr813 |
| Olfr814 | 0 | 0 | 0 | 0 | 0 | 0 | 0 | 0 | 0 | 0 | 0 | 0 | 0 | 0 | 0 | 0 | 0 | 0 | 0 | 0 | 0 | 0 | 1 | 0 | 0 | 0 | 1 | 0 | Olfr814 |
| Olfr816 | 0 | 0 | 0 | 0 | 0 | 0 | 0 | 0 | 1 | 0 | 0 | 0 | 0 | 0 | 0 | 0 | 0 | 0 | 0 | 0 | 0 | 0 | 0 | 0 | 0 | 0 | 0 | 0 | Olfr816 |
| Olfr819 | 0 | 0 | 0 | 0 | 0 | 0 | 0 | 0 | 0 | 0 | 0 | 0 | 0 | 0 | 0 | 0 | 0 | 0 | 0 | 0 | 0 | 0 | 2 | 0 | 0 | 0 | 0 | 0 | Olfr819 |
| Olfr820 | 0 | 0 | 0 | 0 | 0 | 0 | 0 | 0 | 2 | 1 | 0 | 0 | 0 | 0 | 0 | 0 | 0 | 0 | 0 | 0 | 0 | 0 | 1 | 0 | 0 | 0 | 0 | 0 | Olfr820 |
| Olfr821 | 0 | 0 | 0 | 0 | 0 | 0 | 0 | 2 | 0 | 0 | 5 | 0 | 0 | 0 | 0 | 0 | 0 | 0 | 0 | 0 | 0 | 0 | 0 | 0 | 0 | 0 | 0 | 0 | Olfr821 |
| Olfr822 | 0 | 0 | 0 | 0 | 0 | 0 | 0 | 0 | 0 | 0 | 3 | 0 | 0 | 0 | 0 | 0 | 0 | 0 | 0 | 0 | 0 | 0 | 0 | 0 | 0 | 0 | 0 | 0 | Olfr822 |
| Olfr824 | 0 | 0 | 0 | 0 | 0 | 0 | 0 | 0 | 0 | 0 | 0 | 0 | 0 | 0 | 0 | 0 | 0 | 0 | 0 | 0 | 0 | 0 | 3 | 0 | 0 | 0 | 0 | 0 | Olfr824 |
| Olfr825 | 0 | 0 | 0 | 0 | 0 | 0 | 0 | 0 | 1 | 0 | 0 | 0 | 2 | 0 | 0 | 0 | 0 | 0 | 0 | 0 | 0 | 0 | 0 | 1 | 0 | 2 | 0 | 1 | Olfr825 |
| Olfr826 | 0 | 0 | 2 | 0 | 0 | 0 | 2 | 0 | 2 | 0 | 0 | 0 | 5 | 0 | 0 | 0 | 0 | 3 | 0 | 0 | 0 | 0 | 1 | 4 | 0 | 0 | 1 | 0 | Olfr826 |
| Olfr827 | 0 | 0 | 0 | 0 | 0 | 0 | 0 | 1 | 6 | 0 | 0 | 1 | 2 | 2 | 0 | 0 | 0 | 0 | 1 | 0 | 0 | 0 | 3 | 2 | 3 | 0 | 0 | 0 | Olfr827 |
| Olfr828 | 0 | 0 | 0 | 0 | 0 | 0 | 0 | 0 | 1 | 0 | 0 | 0 | 0 | 0 | 0 | 0 | 0 | 0 | 0 | 0 | 0 | 0 | 0 | 2 | 0 | 0 | 0 | 0 | Olfr828 |
| Olfr829 | 0 | 0 | 0 | 3 | 0 | 0 | 0 | 0 | 0 | 3 | 1 | 0 | 0 | 1 | 0 | 0 | 0 | 0 | 0 | 0 | 1 | 0 | 0 | 1 | 0 | 0 | 0 | 1 | Olfr829 |
| Olfr832 | 0 | 0 | 0 | 0 | 0 | 0 | 0 | 0 | 2 | 0 | 0 | 0 | 0 | 0 | 0 | 0 | 0 | 0 | 0 | 0 | 1 | 3 | 0 | 1 | 0 | 2 | 0 | 0 | Olfr832 |
| Olfr835 | 0 | 0 | 0 | 3 | 0 | 2 | 0 | 1 | 0 | 2 | 1 | 0 | 0 | 0 | 0 | 0 | 0 | 0 | 0 | 0 | 0 | 0 | 5 | 4 | 0 | 0 | 0 | 0 | Olfr835 |
| Olfr836 | 0 | 0 | 0 | 0 | 0 | 0 | 0 | 0 | 0 | 0 | 4 | 0 | 0 | 0 | 0 | 0 | 0 | 0 | 0 | 0 | 0 | 0 | 0 | 0 | 0 | 0 | 0 | 0 | Olfr836 |
| Olfr837 | 0 | 0 | 0 | 0 | 0 | 0 | 0 | 1 | 0 | 0 | 0 | 0 | 0 | 0 | 0 | 0 | 0 | 0 | 0 | 0 | 0 | 0 | 0 | 2 | 0 | 0 | 0 | 0 | Olfr837 |
| Olfr839-ps1 | 0 | 0 | 0 | 0 | 0 | 0 | 0 | 0 | 1 | 0 | 4 | 0 | 0 | 0 | 0 | 0 | 0 | 0 | 0 | 0 | 0 | 0 | 1 | 0 | 0 | 0 | 0 | 0 | Olfr839-ps1 |
| Olfr843 | 0 | 0 | 0 | 0 | 0 | 0 | 0 | 0 | 0 | 0 | 4 | 0 | 0 | 0 | 0 | 0 | 0 | 0 | 0 | 0 | 0 | 0 | 0 | 0 | 0 | 0 | 0 | 0 | Olfr843 |
| Olfr845 | 0 | 0 | 0 | 0 | 0 | 0 | 0 | 2 | 2 | 5 | 0 | 0 | 0 | 0 | 0 | 0 | 0 | 0 | 0 | 0 | 0 | 0 | 1 | 0 | 0 | 0 | 0 | 0 | Olfr845 |
| Olfr846 | 1 | 0 | 0 | 0 | 0 | 0 | 0 | 0 | 0 | 0 | 0 | 0 | 0 | 1 | 0 | 0 | 0 | 0 | 0 | 0 | 0 | 0 | 0 | 0 | 0 | 0 | 0 | 0 | Olfr846 |

| | | | | | | | | | | | | | | | | | | | | | | | | | | | | | |
|---|---|---|---|---|---|---|---|---|---|---|---|---|---|---|---|---|---|---|---|---|---|---|---|---|---|---|---|---|---|
| Olfr849 | 0 | 0 | 0 | 0 | 0 | 0 | 0 | 0 | 1 | 1 | 2 | 0 | 0 | 0 | 0 | 0 | 0 | 0 | 0 | 0 | 0 | 0 | 2 | 0 | 0 | 2 | 0 | 0 | Olfr849 |
| Olfr851 | 0 | 0 | 0 | 0 | 0 | 0 | 0 | 0 | 0 | 0 | 1 | 0 | 0 | 0 | 0 | 0 | 0 | 0 | 0 | 0 | 0 | 0 | 1 | 0 | 0 | 0 | 0 | 0 | Olfr851 |
| Olfr854 | 0 | 0 | 0 | 0 | 0 | 0 | 0 | 0 | 0 | 1 | 0 | 0 | 0 | 0 | 0 | 0 | 0 | 0 | 0 | 0 | 0 | 0 | 0 | 0 | 0 | 0 | 0 | 0 | Olfr854 |
| Olfr855 | 0 | 0 | 0 | 0 | 0 | 0 | 0 | 2 | 2 | 0 | 3 | 0 | 0 | 0 | 1 | 0 | 0 | 0 | 0 | 0 | 0 | 0 | 3 | 6 | 0 | 2 | 0 | 0 | Olfr855 |
| Olfr857 | 1 | 2 | 7 | 7 | 5 | 4 | 3 | 2 | 2 | 10 | 4 | 3 | 2 | 7 | 4 | 0 | 2 | 2 | 2 | 0 | 3 | 3 | 0 | 7 | 0 | 1 | 3 | 5 | Olfr857 |
| Olfr859 | 0 | 0 | 0 | 5 | 0 | 0 | 1 | 1 | 2 | 0 | 1 | 0 | 0 | 1 | 1 | 0 | 0 | 4 | 3 | 0 | 0 | 3 | 3 | 4 | 0 | 1 | 1 | 1 | Olfr859 |
| Olfr860 | 0 | 0 | 0 | 0 | 0 | 0 | 0 | 5 | 0 | 0 | 0 | 0 | 0 | 0 | 0 | 0 | 0 | 0 | 0 | 0 | 0 | 0 | 1 | 0 | 0 | 0 | 0 | 0 | Olfr860 |
| Olfr867 | 0 | 0 | 0 | 8 | 0 | 0 | 0 | 0 | 1 | 4 | 0 | 0 | 0 | 0 | 0 | 0 | 0 | 0 | 0 | 0 | 0 | 0 | 3 | 2 | 0 | 0 | 0 | 0 | Olfr867 |
| Olfr868 | 0 | 0 | 0 | 0 | 0 | 0 | 0 | 0 | 0 | 0 | 0 | 0 | 0 | 0 | 0 | 0 | 0 | 0 | 0 | 0 | 0 | 0 | 0 | 2 | 0 | 0 | 0 | 0 | Olfr868 |
| Olfr870 | 0 | 0 | 0 | 1 | 0 | 0 | 0 | 0 | 1 | 3 | 0 | 0 | 1 | 0 | 0 | 0 | 0 | 0 | 0 | 0 | 0 | 0 | 1 | 1 | 0 | 0 | 0 | 0 | Olfr870 |
| Olfr872 | 0 | 0 | 0 | 0 | 0 | 0 | 0 | 1 | 0 | 0 | 0 | 0 | 0 | 0 | 0 | 1 | 0 | 0 | 0 | 0 | 0 | 0 | 0 | 0 | 0 | 1 | 0 | 0 | Olfr872 |
| Olfr873 | 0 | 1 | 0 | 0 | 0 | 0 | 0 | 0 | 2 | 3 | 0 | 0 | 1 | 4 | 0 | 0 | 0 | 0 | 0 | 0 | 0 | 0 | 3 | 8 | 0 | 5 | 0 | 0 | Olfr873 |
| Olfr874 | 0 | 1 | 0 | 0 | 0 | 0 | 0 | 1 | 1 | 1 | 6 | 0 | 1 | 0 | 0 | 0 | 0 | 0 | 1 | 0 | 0 | 0 | 1 | 2 | 0 | 0 | 0 | 0 | Olfr874 |
| Olfr875 | 0 | 0 | 0 | 0 | 0 | 1 | 1 | 1 | 0 | 1 | 0 | 0 | 0 | 0 | 0 | 2 | 0 | 0 | 0 | 0 | 1 | 0 | 2 | 2 | 5 | 0 | 0 | 0 | Olfr875 |
| Olfr876 | 0 | 0 | 0 | 7 | 0 | 0 | 6 | 2 | 0 | 4 | 2 | 1 | 2 | 0 | 1 | 1 | 0 | 0 | 0 | 1 | 1 | 0 | 4 | 2 | 0 | 2 | 1 | 2 | Olfr876 |
| Olfr877 | 0 | 0 | 0 | 0 | 0 | 0 | 0 | 0 | 1 | 6 | 0 | 0 | 0 | 0 | 0 | 1 | 0 | 0 | 0 | 0 | 0 | 0 | 2 | 2 | 0 | 6 | 0 | 0 | Olfr877 |
| Olfr878 | 0 | 0 | 0 | 0 | 0 | 0 | 0 | 1 | 2 | 0 | 0 | 0 | 0 | 0 | 0 | 0 | 0 | 0 | 0 | 0 | 0 | 0 | 0 | 2 | 0 | 1 | 0 | 0 | Olfr878 |
| Olfr884 | 0 | 0 | 0 | 0 | 0 | 0 | 0 | 1 | 2 | 0 | 0 | 0 | 0 | 0 | 0 | 0 | 0 | 0 | 0 | 0 | 0 | 0 | 1 | 0 | 0 | 0 | 0 | 0 | Olfr884 |
| Olfr885 | 0 | 0 | 0 | 0 | 0 | 0 | 0 | 1 | 0 | 0 | 0 | 0 | 0 | 0 | 0 | 0 | 0 | 0 | 0 | 0 | 0 | 0 | 0 | 0 | 0 | 3 | 0 | 0 | Olfr885 |
| Olfr888 | 0 | 0 | 0 | 0 | 0 | 0 | 0 | 0 | 0 | 0 | 0 | 0 | 0 | 0 | 0 | 0 | 0 | 0 | 0 | 0 | 0 | 0 | 0 | 0 | 0 | 2 | 0 | 0 | Olfr888 |
| Olfr889 | 0 | 2 | 0 | 0 | 0 | 0 | 0 | 0 | 0 | 0 | 0 | 2 | 1 | 0 | 0 | 0 | 0 | 0 | 0 | 0 | 0 | 0 | 0 | 0 | 0 | 0 | 0 | 0 | Olfr889 |
| Olfr890 | 0 | 0 | 0 | 0 | 0 | 0 | 0 | 4 | 0 | 0 | 0 | 0 | 0 | 0 | 0 | 0 | 0 | 0 | 0 | 0 | 0 | 0 | 0 | 1 | 0 | 0 | 0 | 0 | Olfr890 |
| Olfr893 | 2 | 0 | 0 | 0 | 0 | 0 | 0 | 0 | 2 | 1 | 1 | 0 | 0 | 1 | 0 | 0 | 0 | 0 | 0 | 0 | 0 | 0 | 0 | 0 | 0 | 0 | 0 | 0 | Olfr893 |
| Olfr895 | 0 | 0 | 0 | 0 | 0 | 0 | 0 | 0 | 0 | 1 | 0 | 0 | 0 | 0 | 0 | 0 | 0 | 1 | 0 | 0 | 0 | 0 | 0 | 0 | 0 | 0 | 0 | 0 | Olfr895 |
| Olfr896-ps1 | 0 | 0 | 0 | 0 | 0 | 0 | 0 | 0 | 0 | 3 | 0 | 0 | 0 | 0 | 0 | 0 | 0 | 0 | 0 | 0 | 0 | 0 | 1 | 0 | 0 | 0 | 0 | 0 | Olfr896-ps1 |
| Olfr898 | 2 | 0 | 0 | 0 | 0 | 0 | 0 | 0 | 1 | 0 | 0 | 0 | 0 | 0 | 0 | 0 | 0 | 0 | 0 | 0 | 0 | 0 | 1 | 0 | 0 | 6 | 0 | 0 | Olfr898 |
| Olfr9 | 0 | 0 | 0 | 0 | 0 | 0 | 0 | 0 | 0 | 0 | 0 | 0 | 0 | 0 | 0 | 0 | 0 | 0 | 0 | 0 | 0 | 0 | 0 | 2 | 0 | 0 | 0 | 0 | Olfr9 |
| Olfr90 | 0 | 0 | 0 | 0 | 0 | 0 | 0 | 1 | 0 | 0 | 4 | 0 | 0 | 1 | 0 | 1 | 0 | 0 | 0 | 0 | 0 | 0 | 0 | 4 | 0 | 0 | 0 | 0 | Olfr90 |
| Olfr901 | 0 | 0 | 0 | 0 | 0 | 0 | 0 | 0 | 0 | 0 | 0 | 0 | 0 | 0 | 0 | 0 | 0 | 0 | 0 | 0 | 0 | 0 | 0 | 4 | 0 | 0 | 0 | 0 | Olfr901 |
| Olfr904 | 0 | 0 | 0 | 0 | 0 | 0 | 0 | 0 | 0 | 0 | 0 | 0 | 0 | 0 | 0 | 0 | 0 | 0 | 0 | 0 | 0 | 0 | 0 | 1 | 0 | 0 | 0 | 0 | Olfr904 |
| Olfr91 | 0 | 1 | 0 | 0 | 0 | 0 | 0 | 0 | 0 | 0 | 0 | 0 | 0 | 0 | 0 | 0 | 0 | 0 | 0 | 0 | 0 | 0 | 2 | 1 | 0 | 0 | 0 | 0 | Olfr91 |
| Olfr911-ps1 | 0 | 0 | 0 | 0 | 0 | 0 | 0 | 0 | 0 | 3 | 0 | 0 | 0 | 0 | 0 | 0 | 0 | 0 | 0 | 0 | 0 | 0 | 0 | 0 | 0 | 0 | 0 | 0 | Olfr911-ps1 |
| Olfr912 | 0 | 0 | 0 | 0 | 0 | 0 | 0 | 0 | 1 | 0 | 0 | 0 | 0 | 0 | 0 | 0 | 0 | 0 | 0 | 0 | 0 | 0 | 0 | 0 | 0 | 0 | 0 | 0 | Olfr912 |
| Olfr913 | 0 | 0 | 0 | 0 | 0 | 0 | 0 | 2 | 0 | 0 | 0 | 0 | 0 | 0 | 0 | 0 | 0 | 0 | 0 | 0 | 0 | 0 | 0 | 0 | 0 | 3 | 0 | 0 | Olfr913 |
| Olfr917 | 0 | 0 | 0 | 0 | 0 | 0 | 0 | 0 | 0 | 0 | 0 | 0 | 0 | 0 | 0 | 0 | 0 | 0 | 0 | 0 | 0 | 0 | 0 | 0 | 0 | 2 | 0 | 0 | Olfr917 |
| Olfr918 | 0 | 0 | 0 | 0 | 0 | 0 | 0 | 2 | 0 | 4 | 1 | 0 | 0 | 0 | 0 | 0 | 0 | 0 | 0 | 0 | 0 | 0 | 0 | 1 | 0 | 0 | 0 | 0 | Olfr918 |
| Olfr92 | 0 | 0 | 0 | 0 | 0 | 0 | 0 | 0 | 1 | 0 | 0 | 0 | 0 | 0 | 0 | 0 | 0 | 0 | 0 | 0 | 0 | 0 | 0 | 0 | 0 | 0 | 0 | 0 | Olfr92 |
| Olfr920 | 35 | 17 | 18 | 37 | 20 | 19 | 15 | 10 | 11 | 5 | 15 | 10 | 19 | 5 | 45 | 40 | 16 | 33 | 16 | 23 | 13 | 10 | 14 | 16 | 4 | 5 | 9 | 7 | Olfr920 |
| Olfr921 | 1 | 0 | 4 | 2 | 0 | 0 | 1 | 0 | 1 | 0 | 0 | 0 | 0 | 1 | 0 | 1 | 5 | 0 | 2 | 0 | 1 | 0 | 0 | 3 | 0 | 0 | 0 | 0 | Olfr921 |
| Olfr922 | 0 | 1 | 0 | 0 | 0 | 0 | 0 | 0 | 0 | 0 | 0 | 0 | 0 | 0 | 0 | 0 | 0 | 3 | 0 | 0 | 0 | 0 | 0 | 0 | 0 | 0 | 0 | 0 | Olfr922 |
| Olfr923 | 0 | 0 | 0 | 0 | 0 | 0 | 0 | 1 | 0 | 0 | 4 | 0 | 0 | 0 | 0 | 2 | 0 | 0 | 0 | 1 | 0 | 0 | 1 | 3 | 0 | 1 | 0 | 0 | Olfr923 |
| Olfr93 | 0 | 0 | 0 | 0 | 0 | 0 | 0 | 2 | 1 | 0 | 3 | 0 | 0 | 0 | 0 | 0 | 0 | 0 | 0 | 0 | 0 | 0 | 3 | 0 | 0 | 0 | 0 | 0 | Olfr93 |
| Olfr930 | 0 | 0 | 0 | 0 | 0 | 0 | 0 | 0 | 0 | 0 | 1 | 0 | 0 | 0 | 0 | 0 | 0 | 0 | 0 | 0 | 0 | 0 | 0 | 1 | 0 | 0 | 0 | 0 | Olfr930 |
| Olfr933 | 0 | 0 | 0 | 0 | 0 | 0 | 0 | 0 | 0 | 0 | 2 | 0 | 0 | 0 | 0 | 0 | 0 | 0 | 0 | 0 | 0 | 0 | 1 | 1 | 0 | 0 | 0 | 0 | Olfr933 |
| Olfr935 | 0 | 0 | 0 | 0 | 0 | 0 | 0 | 0 | 0 | 0 | 0 | 0 | 0 | 0 | 0 | 0 | 0 | 0 | 0 | 0 | 0 | 0 | 3 | 0 | 0 | 0 | 0 | 0 | Olfr935 |
| Olfr937 | 0 | 0 | 0 | 0 | 0 | 0 | 0 | 0 | 0 | 0 | 0 | 0 | 0 | 0 | 0 | 0 | 0 | 0 | 0 | 0 | 0 | 0 | 2 | 0 | 0 | 5 | 0 | 0 | Olfr937 |
| Olfr938 | 0 | 0 | 0 | 0 | 0 | 0 | 0 | 0 | 0 | 0 | 0 | 0 | 0 | 0 | 0 | 0 | 0 | 0 | 0 | 0 | 0 | 0 | 1 | 0 | 0 | 0 | 0 | 0 | Olfr938 |
| Olfr94 | 0 | 0 | 0 | 0 | 0 | 0 | 0 | 4 | 8 | 2 | 2 | 0 | 0 | 0 | 0 | 0 | 0 | 0 | 0 | 0 | 0 | 0 | 2 | 3 | 0 | 4 | 0 | 0 | Olfr94 |
| Olfr943 | 0 | 0 | 0 | 0 | 0 | 0 | 0 | 0 | 0 | 0 | 0 | 0 | 0 | 0 | 0 | 0 | 0 | 0 | 0 | 0 | 0 | 0 | 1 | 0 | 0 | 0 | 0 | 0 | Olfr943 |
| Olfr944 | 0 | 0 | 0 | 3 | 0 | 0 | 0 | 0 | 0 | 0 | 0 | 0 | 0 | 0 | 0 | 0 | 0 | 0 | 0 | 0 | 0 | 0 | 0 | 0 | 0 | 0 | 0 | 0 | Olfr944 |
| Olfr945 | 0 | 0 | 0 | 0 | 0 | 0 | 0 | 1 | 1 | 0 | 3 | 0 | 0 | 0 | 0 | 0 | 0 | 0 | 0 | 0 | 0 | 0 | 0 | 0 | 0 | 0 | 0 | 0 | Olfr945 |
| Olfr95 | 0 | 0 | 0 | 0 | 0 | 2 | 0 | 0 | 2 | 3 | 6 | 1 | 0 | 0 | 1 | 2 | 2 | 0 | 0 | 0 | 0 | 0 | 5 | 5 | 0 | 0 | 0 | 1 | Olfr95 |
| Olfr951 | 0 | 0 | 0 | 0 | 0 | 0 | 0 | 2 | 2 | 1 | 0 | 0 | 0 | 0 | 0 | 0 | 0 | 0 | 0 | 0 | 0 | 0 | 0 | 0 | 0 | 0 | 0 | 0 | Olfr951 |
| Olfr952 | 0 | 0 | 0 | 0 | 0 | 0 | 0 | 0 | 0 | 0 | 2 | 0 | 0 | 1 | 0 | 0 | 0 | 0 | 0 | 0 | 0 | 0 | 0 | 0 | 0 | 0 | 0 | 0 | Olfr952 |
| Olfr954 | 0 | 0 | 0 | 0 | 0 | 0 | 0 | 0 | 0 | 0 | 1 | 0 | 0 | 0 | 0 | 0 | 0 | 0 | 0 | 0 | 0 | 0 | 0 | 0 | 0 | 0 | 0 | 0 | Olfr954 |
| Olfr955 | 0 | 0 | 0 | 0 | 0 | 0 | 0 | 0 | 0 | 0 | 0 | 0 | 0 | 0 | 0 | 0 | 0 | 0 | 0 | 0 | 0 | 0 | 1 | 0 | 0 | 0 | 0 | 0 | Olfr955 |
| Olfr957 | 0 | 0 | 0 | 0 | 0 | 0 | 0 | 1 | 6 | 0 | 1 | 2 | 0 | 0 | 0 | 0 | 0 | 0 | 0 | 0 | 1 | 0 | 4 | 0 | 0 | 2 | 0 | 0 | Olfr957 |
| Olfr958 | 1 | 0 | 0 | 0 | 0 | 0 | 0 | 5 | 3 | 0 | 1 | 0 | 0 | 1 | 0 | 0 | 0 | 0 | 0 | 0 | 0 | 0 | 5 | 1 | 0 | 0 | 0 | 0 | Olfr958 |
| Olfr959 | 1 | 0 | 0 | 0 | 1 | 0 | 3 | 7 | 2 | 8 | 1 | 0 | 0 | 0 | 0 | 1 | 0 | 2 | 3 | 1 | 3 | 3 | 3 | 8 | 0 | 0 | 0 | 1 | Olfr959 |
| Olfr96 | 3 | 0 | 0 | 0 | 1 | 0 | 0 | 1 | 2 | 1 | 4 | 0 | 0 | 0 | 0 | 0 | 0 | 0 | 0 | 0 | 0 | 0 | 6 | 4 | 0 | 2 | 0 | 0 | Olfr96 |
| Olfr960 | 0 | 0 | 3 | 0 | 0 | 0 | 0 | 0 | 0 | 0 | 0 | 0 | 0 | 0 | 0 | 0 | 0 | 0 | 0 | 1 | 0 | 0 | 0 | 0 | 0 | 2 | 0 | 0 | Olfr960 |
| Olfr961 | 0 | 0 | 0 | 0 | 0 | 0 | 0 | 6 | 3 | 0 | 2 | 5 | 0 | 0 | 0 | 0 | 0 | 0 | 0 | 0 | 0 | 0 | 2 | 1 | 0 | 3 | 0 | 0 | Olfr961 |
| Olfr963 | 0 | 0 | 0 | 0 | 0 | 0 | 0 | 0 | 0 | 2 | 0 | 0 | 0 | 0 | 0 | 0 | 0 | 0 | 0 | 0 | 0 | 0 | 0 | 3 | 0 | 0 | 0 | 0 | Olfr963 |
| Olfr965 | 0 | 0 | 0 | 0 | 0 | 0 | 0 | 0 | 0 | 0 | 0 | 0 | 0 | 0 | 0 | 0 | 0 | 0 | 0 | 0 | 0 | 0 | 1 | 0 | 0 | 0 | 0 | 0 | Olfr965 |
| Olfr968 | 0 | 0 | 0 | 0 | 0 | 0 | 0 | 0 | 0 | 1 | 1 | 0 | 0 | 0 | 0 | 0 | 0 | 0 | 0 | 0 | 0 | 0 | 2 | 0 | 3 | 0 | 0 | 0 | Olfr968 |
| Olfr969 | 0 | 0 | 0 | 0 | 0 | 0 | 0 | 2 | 2 | 0 | 0 | 0 | 0 | 0 | 0 | 0 | 0 | 0 | 0 | 0 | 0 | 0 | 1 | 0 | 0 | 0 | 0 | 0 | Olfr969 |
| Olfr970 | 0 | 0 | 0 | 0 | 0 | 0 | 0 | 1 | 0 | 0 | 0 | 0 | 1 | 1 | 0 | 0 | 0 | 0 | 0 | 0 | 0 | 0 | 0 | 0 | 0 | 0 | 0 | 0 | Olfr970 |
| Olfr972 | 0 | 0 | 0 | 0 | 0 | 0 | 0 | 1 | 0 | 0 | 0 | 0 | 0 | 0 | 0 | 0 | 0 | 0 | 0 | 0 | 0 | 0 | 2 | 8 | 0 | 1 | 0 | 0 | Olfr972 |
| Olfr974 | 0 | 0 | 0 | 0 | 0 | 0 | 0 | 2 | 1 | 0 | 0 | 0 | 0 | 0 | 0 | 0 | 0 | 0 | 0 | 0 | 0 | 0 | 0 | 2 | 0 | 0 | 0 | 0 | Olfr974 |
| Olfr975 | 0 | 0 | 0 | 0 | 0 | 0 | 0 | 0 | 0 | 0 | 0 | 0 | 0 | 0 | 0 | 0 | 0 | 0 | 0 | 0 | 0 | 0 | 0 | 0 | 0 | 1 | 0 | 0 | Olfr975 |
| Olfr976 | 0 | 1 | 0 | 1 | 2 | 0 | 0 | 0 | 0 | 0 | 0 | 0 | 0 | 0 | 0 | 0 | 0 | 0 | 0 | 0 | 0 | 0 | 0 | 0 | 0 | 0 | 0 | 0 | Olfr976 |
| Olfr979 | 0 | 0 | 0 | 0 | 0 | 0 | 0 | 0 | 0 | 0 | 0 | 0 | 1 | 0 | 0 | 0 | 2 | 0 | 0 | 0 | 0 | 0 | 1 | 0 | 0 | 0 | 0 | 0 | Olfr979 |
| Olfr980 | 0 | 0 | 0 | 0 | 0 | 0 | 0 | 1 | 0 | 0 | 0 | 0 | 0 | 0 | 0 | 0 | 0 | 0 | 0 | 0 | 0 | 0 | 1 | 2 | 0 | 0 | 0 | 0 | Olfr980 |
| Olfr981 | 5 | 1 | 0 | 4 | 0 | 0 | 0 | 6 | 6 | 3 | 0 | 0 | 0 | 0 | 1 | 0 | 0 | 0 | 0 | 0 | 0 | 0 | 16 | 13 | 0 | 8 | 3 | 0 | Olfr981 |
| Olfr982 | 0 | 0 | 0 | 0 | 0 | 0 | 0 | 1 | 1 | 0 | 0 | 0 | 5 | 0 | 0 | 0 | 0 | 0 | 0 | 0 | 0 | 0 | 0 | 0 | 0 | 3 | 0 | 0 | Olfr982 |
| Olfr984 | 0 | 0 | 0 | 0 | 2 | 0 | 0 | 0 | 0 | 0 | 0 | 0 | 0 | 0 | 0 | 0 | 0 | 0 | 0 | 3 | 1 | 0 | 1 | 0 | 0 | 0 | 0 | 0 | Olfr984 |
| Olfr985 | 0 | 0 | 0 | 0 | 1 | 0 | 0 | 0 | 1 | 0 | 8 | 0 | 1 | 0 | 0 | 0 | 0 | 0 | 0 | 3 | 0 | 0 | 0 | 3 | 0 | 0 | 0 | 0 | Olfr985 |
| Olfr986 | 0 | 0 | 0 | 1 | 0 | 0 | 0 | 0 | 0 | 0 | 0 | 0 | 0 | 0 | 1 | 0 | 0 | 0 | 0 | 0 | 0 | 0 | 0 | 0 | 0 | 0 | 0 | 0 | Olfr986 |
| Olfr988 | 0 | 0 | 0 | 0 | 0 | 0 | 0 | 0 | 0 | 0 | 0 | 0 | 0 | 0 | 0 | 0 | 0 | 0 | 0 | 0 | 0 | 0 | 0 | 2 | 0 | 0 | 0 | 0 | Olfr988 |

| | | | | | | | | | | | | | | | | | | | | | | | | | | | | | |
|---|---|---|---|---|---|---|---|---|---|---|---|---|---|---|---|---|---|---|---|---|---|---|---|---|---|---|---|---|---|
| Olfr99 | 0 | 0 | 0 | 0 | 0 | 0 | 0 | 0 | 0 | 0 | 0 | 0 | 0 | 0 | 0 | 0 | 0 | 0 | 0 | 0 | 0 | 0 | 1 | 0 | 0 | 0 | 0 | 0 | Olfr99 |
| Olfr992 | 0 | 0 | 0 | 0 | 0 | 0 | 0 | 1 | 0 | 0 | 1 | 0 | 0 | 0 | 0 | 0 | 0 | 0 | 0 | 0 | 0 | 0 | 0 | 2 | 0 | 0 | 0 | 0 | Olfr992 |
| Olfr994 | 0 | 0 | 0 | 0 | 0 | 0 | 0 | 0 | 1 | 0 | 0 | 0 | 0 | 0 | 0 | 0 | 0 | 0 | 0 | 0 | 0 | 0 | 0 | 0 | 0 | 2 | 0 | 0 | Olfr994 |
| Olfr998 | 0 | 0 | 0 | 0 | 0 | 0 | 0 | 0 | 0 | 0 | 0 | 0 | 0 | 1 | 0 | 0 | 3 | 0 | 0 | 0 | 0 | 0 | 0 | 0 | 0 | 0 | 0 | 0 | Olfr998 |
| Olig1 | 465 | 588 | 552 | 631 | 417 | 347 | 424 | 201 | 267 | 425 | 249 | 184 | 124 | 170 | 426 | 224 | 577 | 414 | 435 | 292 | 334 | 286 | 256 | 324 | 167 | 462 | 158 | 158 | Olig1 |
| Olig2 | 1077 | 1310 | 1497 | 1416 | 1003 | 1018 | 1389 | 1114 | 1030 | 1277 | 861 | 1865 | 415 | 2072 | 1065 | 516 | 1787 | 1083 | 1187 | 934 | 1173 | 1046 | 630 | 1741 | 1203 | 3198 | 1549 | 1774 | Olig2 |
| Olig3 | 589 | 873 | 1680 | 774 | 618 | 599 | 678 | 766 | 334 | 434 | 496 | 1139 | 645 | 1159 | 1014 | 467 | 1276 | 552 | 686 | 501 | 523 | 859 | 542 | 699 | 621 | 1499 | 1404 | 1456 | Olig3 |
| Olr1 | 1 | 2 | 0 | 0 | 1 | 2 | 1 | 4 | 7 | 0 | 2 | 3 | 2 | 2 | 0 | 1 | 6 | 0 | 0 | 0 | 0 | 2 | 2 | 3 | 0 | 1 | 0 | 4 | Olr1 |
| Oma1 | 415 | 463 | 447 | 410 | 396 | 361 | 422 | 551 | 594 | 513 | 552 | 496 | 549 | 586 | 440 | 463 | 481 | 439 | 390 | 453 | 345 | 468 | 655 | 628 | 541 | 749 | 610 | 598 | Oma1 |
| Omd | 5 | 10 | 2 | 0 | 2 | 8 | 7 | 76 | 80 | 32 | 94 | 5 | 14 | 9 | 11 | 1 | 2 | 1 | 5 | 6 | 0 | 27 | 82 | 17 | 15 | 18 | 26 | 15 | Omd |
| Omg | 9 | 13 | 13 | 7 | 14 | 13 | 17 | 78 | 68 | 67 | 60 | 92 | 45 | 77 | 28 | 5 | 9 | 11 | 34 | 19 | 21 | 117 | 76 | 72 | 70 | 73 | 72 | 77 | Omg |
| Omp | 13 | 10 | 15 | 16 | 12 | 8 | 10 | 24 | 30 | 30 | 36 | 15 | 8 | 13 | 11 | 14 | 16 | 15 | 16 | 14 | 14 | 12 | 33 | 71 | 28 | 24 | 19 | 18 | Omp |
| Onecut1 | 265 | 302 | 174 | 308 | 306 | 339 | 188 | 233 | 172 | 216 | 209 | 294 | 324 | 236 | 295 | 308 | 263 | 326 | 252 | 300 | 266 | 284 | 162 | 144 | 262 | 20 | 352 | 262 | Onecut1 |
| Onecut2 | 7401 | 6966 | 7541 | 8022 | 7846 | 9014 | 8573 | 6676 | 4586 | 4754 | 4368 | 5621 | 7690 | 5893 | 8785 | 9170 | 7747 | 7156 | 6773 | 8606 | 9609 | 6475 | 3577 | 4014 | 5637 | 2656 | 7861 | 6896 | Onecut2 |
| Onecut3 | 508 | 651 | 1035 | 763 | 705 | 667 | 611 | 691 | 492 | 805 | 472 | 926 | 1046 | 846 | 683 | 723 | 1037 | 797 | 704 | 666 | 549 | 1023 | 381 | 755 | 733 | 486 | 948 | 1015 | Onecut3 |
| Ooep | 232 | 251 | 205 | 222 | 272 | 321 | 249 | 382 | 340 | 384 | 276 | 330 | 296 | 320 | 289 | 226 | 210 | 167 | 264 | 318 | 283 | 346 | 342 | 336 | 371 | 301 | 307 | 341 | Ooep |
| Oosp1 | 0 | 0 | 0 | 0 | 0 | 0 | 0 | 0 | 0 | 0 | 0 | 0 | 0 | 0 | 0 | 0 | 0 | 0 | 0 | 0 | 0 | 0 | 0 | 0 | 0 | 0 | 1 | 0 | Oosp1 |
| Oosp2 | 0 | 0 | 0 | 0 | 0 | 0 | 0 | 0 | 0 | 0 | 0 | 0 | 1 | 0 | 0 | 2 | 0 | 0 | 0 | 0 | 0 | 0 | 0 | 0 | 0 | 0 | 0 | 0 | Oosp2 |
| Opa1 | 10154 | 8745 | 7591 | 8063 | 9263 | 9444 | 9769 | 11955 | 10576 | 10722 | 9779 | 11102 | 10176 | 11058 | 10323 | 8688 | 7953 | 8207 | 8695 | 10044 | 9742 | 10920 | 9811 | 9638 | 10978 | 8541 | 10691 | 10426 | Opa1 |
| Opa3 | 2182 | 2251 | 2388 | 2229 | 2193 | 2146 | 2025 | 3457 | 3568 | 3927 | 3565 | 4472 | 4373 | 4328 | 2053 | 1792 | 2336 | 1878 | 2578 | 1920 | 2134 | 3609 | 3548 | 4117 | 3935 | 5161 | 4307 | 4671 | Opa3 |
| Opalin | 3 | 3 | 0 | 0 | 2 | 1 | 6 | 0 | 0 | 2 | 4 | 3 | 3 | 6 | 1 | 2 | 0 | 1 | 7 | 2 | 4 | 0 | 3 | 6 | 2 | 4 | 2 | 1 | Opalin |
| Opcml | 520 | 491 | 547 | 903 | 1636 | 1308 | 1416 | 1316 | 1444 | 1343 | 1417 | 818 | 743 | 830 | 675 | 618 | 736 | 1357 | 1217 | 1266 | 1568 | 2167 | 1397 | 1118 | 1644 | 657 | 743 | 742 | Opcml |
| Ophn1 | 2862 | 2837 | 2430 | 3314 | 4120 | 3923 | 4034 | 4763 | 4242 | 4579 | 4194 | 3987 | 3953 | 4316 | 3357 | 2542 | 2954 | 3618 | 3948 | 4067 | 4274 | 5165 | 4284 | 4112 | 4714 | 3079 | 3929 | 3837 | Ophn1 |
| Oplah | 610 | 893 | 660 | 961 | 887 | 852 | 648 | 701 | 1034 | 884 | 1040 | 820 | 811 | 682 | 651 | 855 | 813 | 1009 | 941 | 793 | 693 | 936 | 1067 | 853 | 875 | 803 | 865 | 748 | Oplah |
| Opn1mw | 0 | 0 | 3 | 3 | 4 | 0 | 0 | 4 | 2 | 0 | 0 | 0 | 1 | 0 | 1 | 2 | 0 | 3 | 0 | 0 | 0 | 0 | 1 | 0 | 0 | 0 | 0 | 0 | Opn1mw |
| Opn1sw | 1 | 1 | 3 | 1 | 4 | 8 | 1 | 4 | 6 | 12 | 12 | 16 | 0 | 13 | 4 | 2 | 7 | 4 | 10 | 4 | 4 | 15 | 23 | 3 | 12 | 6 | 3 | 9 | Opn1sw |
| Opn3 | 714 | 726 | 673 | 770 | 490 | 570 | 549 | 215 | 185 | 204 | 216 | 313 | 282 | 301 | 662 | 652 | 590 | 675 | 491 | 592 | 633 | 211 | 193 | 199 | 205 | 313 | 336 | 318 | Opn3 |
| Opn4 | 0 | 2 | 3 | 0 | 0 | 0 | 1 | 6 | 2 | 0 | 5 | 10 | 10 | 11 | 4 | 5 | 0 | 0 | 2 | 2 | 3 | 0 | 8 | 9 | 2 | 16 | 3 | 11 | Opn4 |
| Opn5 | 3 | 2 | 8 | 21 | 10 | 9 | 10 | 6 | 6 | 10 | 11 | 4 | 2 | 2 | 7 | 6 | 7 | 19 | 10 | 4 | 12 | 7 | 5 | 8 | 2 | 2 | 0 | 5 | Opn5 |
| Oprd1 | 22 | 9 | 1 | 23 | 33 | 21 | 15 | 33 | 63 | 36 | 57 | 24 | 42 | 19 | 22 | 32 | 17 | 29 | 18 | 20 | 21 | 62 | 50 | 30 | 26 | 16 | 15 | 17 | Oprd1 |
| Oprk1 | 294 | 282 | 295 | 209 | 346 | 236 | 250 | 140 | 83 | 101 | 108 | 90 | 82 | 73 | 314 | 341 | 277 | 209 | 209 | 305 | 277 | 119 | 102 | 56 | 98 | 106 | 119 | 84 | Oprk1 |
| Oprl1 | 1624 | 1928 | 1236 | 2243 | 3732 | 3117 | 2881 | 6064 | 4810 | 4685 | 4077 | 4275 | 3941 | 4228 | 2234 | 1407 | 1914 | 2333 | 3170 | 3527 | 3104 | 6286 | 4559 | 3982 | 5605 | 2841 | 4188 | 3696 | Oprl1 |
| Oprm1 | 17 | 13 | 0 | 2 | 25 | 23 | 22 | 33 | 46 | 16 | 34 | 21 | 3 | 31 | 17 | 9 | 2 | 12 | 20 | 31 | 29 | 18 | 53 | 8 | 46 | 8 | 27 | 19 | Oprm1 |
| Optc | 121 | 206 | 176 | 124 | 205 | 215 | 169 | 333 | 265 | 300 | 241 | 317 | 454 | 303 | 186 | 186 | 195 | 178 | 192 | 215 | 190 | 279 | 215 | 304 | 258 | 341 | 349 | 337 | Optc |
| Optn | 1517 | 1378 | 1338 | 1464 | 1320 | 1343 | 1359 | 427 | 437 | 441 | 453 | 378 | 473 | 364 | 1470 | 1624 | 1378 | 1515 | 1197 | 1335 | 1247 | 607 | 535 | 375 | 422 | 361 | 389 | 338 | Optn |
| OR5BS1P | 0 | 0 | 0 | 0 | 0 | 0 | 0 | 5 | 1 | 0 | 2 | 0 | 0 | 0 | 0 | 0 | 0 | 0 | 0 | 0 | 0 | 0 | 1 | 5 | 0 | 2 | 0 | 0 | OR5BS1P |
| Orai1 | 819 | 1216 | 786 | 1041 | 851 | 833 | 877 | 371 | 415 | 372 | 424 | 429 | 434 | 436 | 757 | 896 | 961 | 1088 | 958 | 948 | 950 | 408 | 463 | 455 | 435 | 451 | 374 | 418 | Orai1 |
| Orai2 | 2040 | 2398 | 1929 | 2789 | 3436 | 2908 | 2722 | 1766 | 2089 | 2063 | 1821 | 1658 | 1633 | 1677 | 2157 | 2244 | 2590 | 3061 | 3106 | 2997 | 3106 | 2264 | 1935 | 1951 | 2394 | 1532 | 1518 | 1551 | Orai2 |
| Orai3 | 1095 | 1353 | 1120 | 1080 | 914 | 864 | 873 | 537 | 640 | 585 | 840 | 567 | 557 | 507 | 1031 | 1261 | 1075 | 1192 | 1014 | 842 | 877 | 615 | 723 | 570 | 525 | 670 | 530 | 570 | Orai3 |
| Orc1 | 1580 | 1424 | 1327 | 822 | 581 | 842 | 846 | 580 | 682 | 460 | 525 | 806 | 770 | 767 | 1396 | 1488 | 1002 | 818 | 856 | 956 | 701 | 361 | 628 | 570 | 507 | 933 | 904 | 853 | Orc1 |
| Orc2 | 7000 | 5966 | 6061 | 5860 | 5042 | 5689 | 5989 | 5109 | 4471 | 4283 | 4113 | 5340 | 5036 | 5469 | 6854 | 6263 | 5609 | 5443 | 5295 | 6034 | 5646 | 4055 | 4165 | 4729 | 4502 | 5786 | 5330 | 5566 | Orc2 |
| Orc3 | 5377 | 4345 | 4242 | 3789 | 3806 | 4524 | 4461 | 7577 | 6796 | 5539 | 5802 | 6772 | 6538 | 6395 | 5562 | 4560 | 4006 | 3623 | 3758 | 5050 | 4129 | 5914 | 6766 | 5787 | 5614 | 6149 | 6801 | 6689 | Orc3 |
| Orc4 | 8559 | 6670 | 7574 | 8714 | 7642 | 8721 | 9675 | 6950 | 5497 | 5655 | 5650 | 6294 | 5824 | 6774 | 9049 | 8277 | 7990 | 8364 | 7768 | 8771 | 9055 | 6376 | 5559 | 6005 | 6052 | 6042 | 6082 | 6350 | Orc4 |
| Orc5 | 1613 | 1392 | 1326 | 1237 | 1251 | 1292 | 1386 | 1859 | 1698 | 1445 | 1287 | 1706 | 1464 | 1632 | 1718 | 1411 | 1268 | 1295 | 1250 | 1596 | 1381 | 1648 | 1556 | 1545 | 1439 | 1867 | 1642 | 1782 | Orc5 |
| Orc6 | 6324 | 5603 | 5442 | 5072 | 4076 | 4628 | 5273 | 4764 | 4057 | 3763 | 3721 | 4854 | 4373 | 4996 | 6220 | 5819 | 5362 | 4636 | 4333 | 5311 | 4835 | 4079 | 3970 | 4142 | 4038 | 5409 | 4898 | 5288 | Orc6 |
| Orm1 | 0 | 2 | 9 | 0 | 0 | 0 | 1 | 0 | 0 | 2 | 5 | 0 | 0 | 0 | 0 | 0 | 1 | 0 | 0 | 0 | 0 | 0 | 1 | 0 | 0 | 0 | 0 | 0 | Orm1 |
| Orm3 | 0 | 0 | 0 | 0 | 1 | 0 | 0 | 0 | 0 | 0 | 2 | 0 | 0 | 0 | 3 | 0 | 0 | 0 | 0 | 0 | 0 | 0 | 0 | 1 | 2 | 0 | 0 | 0 | Orm3 |
| Ormdl1 | 3035 | 2524 | 2634 | 2306 | 2317 | 2216 | 2975 | 2717 | 2128 | 2123 | 1638 | 2609 | 2168 | 2599 | 3455 | 2703 | 2726 | 2356 | 2482 | 2991 | 2443 | 2169 | 2141 | 2236 | 2079 | 2645 | 2625 | 2873 | Ormdl1 |
| Ormdl2 | 2290 | 2596 | 2206 | 2016 | 1468 | 1512 | 2135 | 910 | 759 | 719 | 625 | 863 | 768 | 1008 | 2661 | 2213 | 2044 | 1914 | 1762 | 2257 | 1903 | 583 | 823 | 814 | 709 | 1016 | 953 | 1076 | Ormdl2 |
| Ormdl3 | 6288 | 8146 | 8169 | 8333 | 7197 | 7267 | 6755 | 3538 | 4168 | 4112 | 4473 | 4237 | 4280 | 3965 | 6161 | 7193 | 8049 | 8336 | 7530 | 7470 | 6719 | 3917 | 4132 | 4598 | 4517 | 4932 | 4291 | 4041 | Ormdl3 |
| Os9 | 10621 | 13985 | 14055 | 15350 | 13628 | 13174 | 12799 | 7260 | 7941 | 9214 | 8742 | 8962 | 8667 | 8859 | 11351 | 12042 | 14947 | 14823 | 14217 | 12560 | 13752 | 8179 | 7816 | 9031 | 9507 | 8759 | 8263 | 8623 | Os9 |
| Osbp | 7815 | 7754 | 8731 | 8673 | 7789 | 7862 | 7691 | 6036 | 6006 | 7014 | 7081 | 7071 | 6653 | 6757 | 7369 | 7917 | 8628 | 8649 | 8172 | 6516 | 7856 | 7112 | 6093 | 6900 | 7047 | 6715 | 6463 | 6732 | Osbp |
| Osbp2 | 223 | 244 | 163 | 371 | 607 | 440 | 466 | 509 | 608 | 649 | 508 | 404 | 378 | 403 | 252 | 235 | 315 | 545 | 508 | 404 | 567 | 737 | 557 | 552 | 689 | 284 | 361 | 401 | Osbp2 |
| Osbpl10 | 189 | 241 | 284 | 191 | 225 | 251 | 141 | 413 | 535 | 425 | 521 | 384 | 420 | 377 | 212 | 273 | 188 | 197 | 171 | 203 | 162 | 556 | 512 | 354 | 417 | 435 | 415 | 354 | Osbpl10 |
| Osbpl11 | 4773 | 4970 | 4564 | 4469 | 4129 | 4214 | 4118 | 3693 | 3399 | 3195 | 3173 | 3598 | 3662 | 3551 | 4776 | 4865 | 4581 | 4617 | 3967 | 4104 | 4203 | 3484 | 3407 | 3234 | 3588 | 3534 | 3775 | 3833 | Osbpl11 |
| Osbpl1a | 2906 | 2688 | 2405 | 2314 | 2490 | 2417 | 2515 | 2875 | 2741 | 2403 | 2687 | 2342 | 2742 | 2356 | 3014 | 2762 | 2184 | 2198 | 2208 | 2470 | 2349 | 2702 | 2746 | 2247 | 2504 | 2509 | 2568 | 2569 | Osbpl1a |
| Osbpl2 | 4758 | 5912 | 4932 | 6367 | 5767 | 5397 | 6019 | 3461 | 3483 | 3543 | 3500 | 3416 | 3442 | 3672 | 5069 | 4991 | 5516 | 6316 | 5535 | 5241 | 6419 | 3969 | 3472 | 3465 | 3940 | 3517 | 3548 | 3493 | Osbpl2 |
| Osbpl3 | 3106 | 2734 | 2700 | 2603 | 2354 | 2544 | 2563 | 1415 | 1420 | 1447 | 1389 | 1531 | 1499 | 1523 | 2843 | 3092 | 2350 | 2634 | 2512 | 2331 | 2605 | 1474 | 1631 | 1430 | 1267 | 1376 | 1664 | 1601 | Osbpl3 |
| Osbpl5 | 1494 | 1874 | 1499 | 1736 | 2174 | 2232 | 1654 | 2220 | 3019 | 2922 | 3126 | 2205 | 2624 | 2082 | 1629 | 2082 | 1636 | 2107 | 1910 | 1668 | 1750 | 2814 | 3072 | 2527 | 2702 | 2193 | 2117 | 1983 | Osbpl5 |
| Osbpl6 | 2378 | 2124 | 1875 | 2605 | 3492 | 2908 | 2845 | 2713 | 2662 | 2899 | 2936 | 2144 | 2597 | 2326 | 2455 | 2463 | 2141 | 2687 | 2702 | 2682 | 3017 | 3457 | 2400 | 2223 | 2862 | 1648 | 2109 | 2083 | Osbpl6 |
| Osbpl7 | 1156 | 1511 | 1543 | 1597 | 1472 | 1416 | 1157 | 833 | 1113 | 1213 | 1095 | 1162 | 1092 | 1144 | 1188 | 1598 | 1520 | 1573 | 1383 | 1267 | 1292 | 1148 | 1035 | 1164 | 1219 | 1138 | 1153 | 1136 | Osbpl7 |
| Osbpl8 | 8290 | 6334 | 6225 | 7853 | 9811 | 10078 | 10248 | 8723 | 7977 | 7078 | 7614 | 6693 | 6840 | 7056 | 9152 | 8023 | 6977 | 8496 | 8313 | 9624 | 9998 | 8779 | 7981 | 6646 | 8157 | 4594 | 6697 | 6735 | Osbpl8 |
| Osbpl9 | 6674 | 5891 | 5969 | 6177 | 6024 | 5876 | 6499 | 7544 | 6431 | 6305 | 5978 | 6192 | 6358 | 6558 | 7185 | 6308 | 5821 | 6524 | 5882 | 6907 | 6272 | 7269 | 6697 | 6099 | 6783 | 6433 | 6258 | 6444 | Osbpl9 |
| Oscar | 133 | 171 | 127 | 240 | 205 | 192 | 185 | 62 | 60 | 100 | 57 | 52 | 57 | 66 | 127 | 148 | 181 | 199 | 148 | 155 | 201 | 77 | 50 | 56 | 59 | 54 | 55 | 64 | Oscar |
| Oscp1 | 1745 | 2044 | 1892 | 1714 | 1874 | 1930 | 1934 | 1795 | 1596 | 1641 | 1450 | 1728 | 1685 | 1699 | 1931 | 1703 | 1944 | 1877 | 2152 | 2071 | 2002 | 1773 | 1660 | 1638 | 1724 | 1609 | 1698 | 1768 | Oscp1 |
| Oser1 | 2053 | 2144 | 1890 | 2021 | 1488 | 1738 | 1522 | 1614 | 1449 | 1266 | 1287 | 1329 | 1457 | 1435 | 2045 | 2071 | 1774 | 1926 | 1663 | 1818 | 1593 | 1429 | 1522 | 1422 | 1406 | 1658 | 1612 | 1603 | Oser1 |
| Osgep | 3320 | 3556 | 2853 | 3200 | 2613 | 2927 | 3079 | 2926 | 3023 | 2537 | 2667 | 2947 | 2615 | 2985 | 3265 | 3100 | 3026 | 3026 | 2918 | 3171 | 2961 | 2445 | 2796 | 2889 | 2598 | 3228 | 3053 | 3045 | Osgep |
| Osgepl1 | 1400 | 1113 | 758 | 1011 | 1043 | 1112 | 1382 | 1331 | 1195 | 989 | 1029 | 1177 | 1057 | 1265 | 1369 | 1143 | 966 | 972 | 934 | 1341 | 1205 | 916 | 1054 | 1021 | 1035 | 964 | 1377 | 1245 | Osgepl1 |
| Osgin1 | 13 | 10 | 11 | 6 | 7 | 9 | 7 | 7 | 21 | 9 | 21 | 13 | 12 | 9 | 7 | 12 | 8 | 12 | 7 | 7 | 11 | 13 | 28 | 17 | 8 | 19 | 6 | 7 | Osgin1 |
| Osgin2 | 2278 | 2044 | 2355 | 2494 | 2485 | 2550 | 2862 | 2163 | 1917 | 1711 | 1935 | 2020 | 1986 | 2252 | 2466 | 2174 | 2226 | 2390 | 2243 | 2487 | 2386 | 2124 | 1857 | 1949 | 2029 | 2036 | 2045 | 2210 | Osgin2 |
| Osm | 0 | 1 | 0 | 0 | 2 | 0 | 0 | 0 | 0 | 2 | 5 | 0 | 3 | 1 | 1 | 1 | 0 | 0 | 0 | 0 | 2 | 2 | 1 | 2 | 0 | 4 | 1 | 0 | Osm |
| Osmr | 180 | 136 | 157 | 107 | 201 | 186 | 143 | 228 | 205 | 227 | 254 | 207 | 219 | 158 | 174 | 201 | 124 | 189 | 160 | 149 | 163 | 296 | 263 | 137 | 210 | 109 | 148 | 153 | Osmr |
| Osr1 | 502 | 363 | 536 | 259 | 259 | 319 | 166 | 1080 | 1609 | 835 | 1554 | 604 | 937 | 523 | 377 | 409 | 377 | 281 | 222 | 257 | 143 | 645 | 2002 | 917 | 515 | 674 | 818 | 567 | Osr1 |
| Osr2 | 649 | 265 | 140 | 133 | 25 | 184 | 6 | 731 | 1285 | 722 | 925 | 419 | 593 | 289 | 298 | 483 | 119 | 232 | 84 | 15 | 16 | 251 | 1154 | 758 | 311 | 632 | 288 | 299 | Osr2 |
| Ost4 | 3541 | 4264 | 3915 | 3702 | 2458 | 2499 | 3542 | 2833 | 2741 | 2696 | 3018 | 2772 | 2782 | 3833 | 3427 | 2931 | 3854 | 3046 | 3013 | 3344 | 3497 | 2214 | 3049 | 3330 | 2753 | 3382 | 3352 | 4200 | Ost4 |
| Ostc | 8959 | 7788 | 7758 | 6735 | 5303 | 5422 | 6902 | 6450 | 5885 | 4994 | 5570 | 5201 | 5249 | 6233 | 8968 | 7716 | 7243 | 6628 | 5713 | 6850 | 6416 | 5233 | 6340 | 5614 | 4843 | 6101 | 5606 | 6508 | Ostc |
| Ostf1 | 2173 | 2407 | 2264 | 2121 | 1789 | 1812 | 1607 | 1069 | 1277 | 1063 | 1265 | 978 | 1171 | 954 | 2342 | 2514 | 1935 | 2189 | 1547 | 1851 | 1700 | 1016 | 1491 | 945 | 916 | 1100 | 1112 | 1008 | Ostf1 |
| Ostm1 | 6124 | 5607 | 5080 | 5803 | 5295 | 5550 | 6383 | 2551 | 2167 | 2274 | 1931 | 2669 | 2002 | 2654 | 6719 | 5023 | 5234 | 5728 | 5651 | 6568 | 6186 | 2480 | 2156 | 2469 | 2549 | 2775 | 2406 | 2599 | Ostm1 |
| Ostn | 0 | 0 | 0 | 3 | 0 | 0 | 0 | 17 | 18 | 5 | 33 | 0 | 0 | 0 | 0 | 0 | 0 | 0 | 0 | 0 | 0 | 17 | 25 | 7 | 3 | 4 | 0 | 0 | Ostn |

| | | | | | | | | | | | | | | | | | | | | | | | | | | | | | |
|---|---|---|---|---|---|---|---|---|---|---|---|---|---|---|---|---|---|---|---|---|---|---|---|---|---|---|---|---|---|
| Otc | 0 | 0 | 0 | 0 | 2 | 4 | 0 | 4 | 6 | 3 | 0 | 0 | 0 | 1 | 0 | 0 | 0 | 0 | 1 | 3 | 1 | 7 | 1 | 0 | 0 | 0 | 3 | 4 | Otc |
| Otoa | 43 | 54 | 50 | 64 | 47 | 62 | 64 | 54 | 56 | 62 | 87 | 40 | 79 | 33 | 63 | 48 | 30 | 73 | 48 | 51 | 63 | 77 | 38 | 42 | 32 | 27 | 52 | 47 | Otoa |
| Otof | 78 | 155 | 113 | 276 | 407 | 293 | 265 | 264 | 353 | 374 | 322 | 147 | 350 | 158 | 140 | 218 | 175 | 365 | 275 | 302 | 298 | 443 | 320 | 168 | 444 | 87 | 174 | 152 | Otof |
| Otog | 70 | 85 | 54 | 6 | 14 | 35 | 6 | 121 | 195 | 159 | 307 | 21 | 109 | 4 | 144 | 65 | 7 | 11 | 13 | 8 | 5 | 207 | 117 | 8 | 65 | 10 | 16 | 11 | Otog |
| Otogl | 31 | 20 | 12 | 6 | 14 | 25 | 6 | 109 | 172 | 96 | 271 | 5 | 66 | 10 | 57 | 40 | 1 | 9 | 1 | 7 | 1 | 97 | 109 | 21 | 66 | 19 | 9 | 8 | Otogl |
| Otol1 | 10 | 10 | 4 | 7 | 15 | 25 | 2 | 27 | 66 | 39 | 81 | 2 | 41 | 4 | 42 | 23 | 0 | 8 | 3 | 0 | 4 | 52 | 21 | 0 | 19 | 11 | 0 | 2 | Otol1 |
| Otop1 | 8 | 13 | 14 | 5 | 4 | 15 | 6 | 6 | 3 | 3 | 3 | 2 | 2 | 1 | 5 | 10 | 6 | 3 | 8 | 4 | 9 | 12 | 8 | 3 | 3 | 3 | 3 | 0 | Otop1 |
| Otop2 | 1 | 4 | 2 | 2 | 8 | 2 | 6 | 6 | 4 | 10 | 0 | 2 | 1 | 1 | 7 | 2 | 0 | 3 | 5 | 8 | 0 | 3 | 3 | 4 | 2 | 4 | 1 | 1 | Otop2 |
| Otop3 | 12 | 6 | 25 | 11 | 16 | 7 | 9 | 5 | 7 | 14 | 26 | 8 | 3 | 10 | 12 | 9 | 6 | 4 | 10 | 12 | 9 | 8 | 14 | 8 | 7 | 17 | 12 | 1 | Otop3 |
| Otor | 265 | 126 | 87 | 215 | 47 | 137 | 13 | 287 | 563 | 464 | 576 | 204 | 295 | 172 | 170 | 114 | 132 | 149 | 76 | 98 | 47 | 438 | 476 | 433 | 308 | 387 | 175 | 252 | Otor |
| Otos | 5 | 5 | 10 | 10 | 6 | 6 | 9 | 5 | 7 | 3 | 5 | 2 | 1 | 7 | 7 | 4 | 8 | 6 | 12 | 3 | 5 | 3 | 5 | 3 | 10 | 4 | 9 | 2 | Otos |
| Otp | 450 | 497 | 366 | 830 | 1108 | 1019 | 657 | 1766 | 1496 | 2219 | 1658 | 1880 | 2567 | 1442 | 711 | 525 | 1268 | 760 | 897 | 871 | 698 | 2949 | 1220 | 1633 | 2102 | 2620 | 1211 | 1498 | Otp |
| Otub1 | 5389 | 6096 | 5831 | 5444 | 5386 | 5342 | 5475 | 7452 | 7617 | 7532 | 7014 | 8655 | 8055 | 8384 | 5515 | 5101 | 5949 | 5631 | 6082 | 6143 | 5526 | 7329 | 7172 | 7888 | 7680 | 9493 | 8832 | 8839 | Otub1 |
| Otub2 | 801 | 831 | 861 | 728 | 938 | 807 | 854 | 1246 | 1149 | 1291 | 1057 | 1117 | 1099 | 1055 | 818 | 808 | 853 | 825 | 953 | 842 | 827 | 1404 | 1289 | 1050 | 1255 | 1061 | 1054 | 1168 | Otub2 |
| Otud1 | 329 | 344 | 415 | 468 | 379 | 408 | 427 | 232 | 232 | 220 | 248 | 189 | 211 | 214 | 338 | 390 | 384 | 513 | 380 | 332 | 377 | 289 | 270 | 228 | 240 | 263 | 149 | 183 | Otud1 |
| Otud3 | 1123 | 1371 | 1035 | 1378 | 1125 | 1180 | 1019 | 703 | 747 | 723 | 883 | 780 | 853 | 775 | 1000 | 1231 | 1129 | 1418 | 1079 | 1061 | 1118 | 702 | 683 | 802 | 819 | 471 | 804 | 754 | Otud3 |
| Otud4 | 9983 | 7912 | 8360 | 9882 | 10234 | 10992 | 11307 | 10269 | 9114 | 9289 | 9636 | 11239 | 10525 | 10999 | 9835 | 9414 | 9441 | 10394 | 9644 | 9619 | 10610 | 9763 | 8613 | 9207 | 9917 | 8954 | 10854 | 10657 | Otud4 |
| Otud5 | 7246 | 7990 | 8037 | 7735 | 7073 | 7043 | 7108 | 7080 | 7238 | 7591 | 6837 | 7915 | 7124 | 7641 | 7151 | 6875 | 7876 | 7331 | 7767 | 6718 | 7467 | 7192 | 6983 | 7756 | 7569 | 7756 | 7160 | 7525 | Otud5 |
| Otud6a | 0 | 2 | 0 | 4 | 0 | 2 | 2 | 0 | 0 | 0 | 0 | 1 | 0 | 1 | 1 | 1 | 0 | 0 | 6 | 0 | 0 | 0 | 0 | 0 | 0 | 0 | 0 | 0 | Otud6a |
| Otud6b | 7139 | 6123 | 6537 | 5950 | 5222 | 5246 | 6319 | 5970 | 5063 | 4250 | 4525 | 4580 | 4632 | 5177 | 7516 | 6643 | 5777 | 5872 | 5192 | 6656 | 5798 | 4859 | 5160 | 4654 | 4765 | 5286 | 5062 | 5394 | Otud6b |
| Otud7a | 211 | 340 | 355 | 527 | 650 | 436 | 417 | 326 | 358 | 448 | 399 | 374 | 320 | 355 | 284 | 260 | 445 | 461 | 550 | 411 | 441 | 461 | 316 | 357 | 477 | 302 | 291 | 269 | Otud7a |
| Otud7b | 6159 | 5699 | 6044 | 6165 | 5083 | 5441 | 5283 | 3794 | 4056 | 4390 | 5242 | 4760 | 4942 | 4784 | 5462 | 5903 | 5829 | 5754 | 4885 | 4507 | 5095 | 3878 | 3943 | 4557 | 4282 | 4251 | 4950 | 4919 | Otud7b |
| Otulin | 1732 | 1971 | 1703 | 1761 | 1357 | 1341 | 1548 | 1276 | 1224 | 1090 | 1084 | 1190 | 1116 | 1324 | 1692 | 1803 | 1651 | 1538 | 1477 | 1429 | 1477 | 1004 | 1191 | 1222 | 1126 | 1587 | 1321 | 1348 | Otulin |
| Otulinl | 942 | 811 | 905 | 755 | 628 | 694 | 686 | 487 | 470 | 383 | 436 | 370 | 336 | 365 | 960 | 989 | 710 | 687 | 655 | 671 | 605 | 421 | 551 | 445 | 349 | 465 | 343 | 329 | Otulinl |
| Otx1 | 2457 | 2731 | 5054 | 3321 | 2276 | 2452 | 3194 | 1981 | 1603 | 2323 | 1656 | 4769 | 3301 | 4710 | 2036 | 1716 | 3660 | 2636 | 3193 | 2476 | 2860 | 1933 | 1558 | 3553 | 2886 | 4457 | 4627 | 5289 | Otx1 |
| Otx2 | 5262 | 5229 | 10722 | 5962 | 4672 | 5453 | 4292 | 6464 | 3427 | 5626 | 3826 | 9643 | 8379 | 10031 | 5349 | 3455 | 7459 | 4282 | 6423 | 5483 | 5785 | 6691 | 5155 | 7236 | 6498 | 5823 | 12290 | 12728 | Otx2 |
| Ovca2 | 2103 | 1794 | 1721 | 1608 | 1269 | 1410 | 1465 | 1508 | 1376 | 1235 | 1225 | 1275 | 1241 | 1331 | 2029 | 1943 | 1669 | 1484 | 1372 | 1590 | 1510 | 1362 | 1437 | 1271 | 1153 | 1438 | 1362 | 1512 | Ovca2 |
| Ovgp1 | 922 | 1030 | 886 | 1455 | 1164 | 1045 | 1309 | 488 | 371 | 458 | 442 | 398 | 440 | 433 | 997 | 1004 | 1160 | 1436 | 1167 | 1151 | 1409 | 587 | 358 | 422 | 502 | 370 | 451 | 376 | Ovgp1 |
| Ovol1 | 27 | 16 | 19 | 2 | 6 | 8 | 2 | 32 | 35 | 20 | 11 | 14 | 26 | 4 | 36 | 20 | 5 | 3 | 9 | 4 | 6 | 2 | 72 | 31 | 0 | 18 | 6 | 14 | Ovol1 |
| Ovol2 | 42 | 22 | 27 | 19 | 7 | 14 | 12 | 20 | 18 | 25 | 21 | 9 | 7 | 7 | 20 | 33 | 29 | 12 | 9 | 23 | 8 | 8 | 34 | 8 | 6 | 15 | 5 | 5 | Ovol2 |
| Ovol3 | 27 | 56 | 42 | 54 | 29 | 26 | 34 | 9 | 1 | 2 | 5 | 2 | 3 | 6 | 28 | 40 | 38 | 41 | 30 | 24 | 32 | 3 | 6 | 2 | 5 | 8 | 2 | 5 | Ovol3 |
| Oxa1l | 3624 | 4005 | 3355 | 3372 | 3565 | 3557 | 3603 | 3955 | 4205 | 3930 | 3625 | 3981 | 3858 | 3892 | 3605 | 3289 | 3692 | 3267 | 3832 | 3809 | 3726 | 3750 | 4025 | 3863 | 3937 | 4194 | 3900 | 3902 | Oxa1l |
| Oxct1 | 17604 | 15051 | 15054 | 12910 | 14511 | 15038 | 15234 | 18795 | 17721 | 15649 | 15646 | 16944 | 15984 | 17005 | 17703 | 15264 | 14279 | 13570 | 13916 | 16486 | 14421 | 15959 | 17274 | 16029 | 15821 | 17558 | 17236 | 17302 | Oxct1 |
| Oxct2a | 8 | 25 | 1 | 2 | 14 | 10 | 7 | 1 | 0 | 1 | 0 | 0 | 2 | 0 | 4 | 19 | 14 | 25 | 9 | 3 | 11 | 2 | 0 | 2 | 1 | 0 | 3 | 1 | Oxct2a |
| Oxct2b | 9 | 9 | 12 | 6 | 5 | 2 | 4 | 5 | 14 | 13 | 5 | 1 | 5 | 5 | 7 | 8 | 0 | 6 | 4 | 5 | 11 | 8 | 6 | 4 | 13 | 0 | 7 | 5 | Oxct2b |
| Oxgr1 | 1 | 1 | 2 | 0 | 1 | 3 | 1 | 0 | 2 | 1 | 0 | 1 | 1 | 1 | 1 | 0 | 0 | 9 | 0 | 1 | 2 | 2 | 3 | 7 | 0 | 4 | 2 | 2 | Oxgr1 |
| Oxld1 | 138 | 130 | 123 | 114 | 107 | 113 | 93 | 159 | 219 | 136 | 174 | 165 | 152 | 166 | 140 | 153 | 161 | 87 | 131 | 133 | 111 | 135 | 181 | 155 | 172 | 198 | 164 | 171 | Oxld1 |
| Oxnad1 | 1254 | 1334 | 1414 | 1256 | 1186 | 1280 | 1288 | 1330 | 1322 | 1385 | 1279 | 1493 | 1308 | 1603 | 1279 | 1087 | 1305 | 1159 | 1277 | 1286 | 1338 | 1270 | 1212 | 1442 | 1367 | 1463 | 1542 | 1580 | Oxnad1 |
| Oxr1 | 6635 | 4880 | 5668 | 5106 | 5033 | 5415 | 5683 | 6268 | 5338 | 4846 | 5133 | 5172 | 4778 | 5163 | 6229 | 6061 | 5036 | 5381 | 4690 | 5633 | 4701 | 5345 | 5642 | 4886 | 4995 | 5592 | 5248 | 5172 | Oxr1 |
| Oxsm | 579 | 466 | 624 | 516 | 574 | 523 | 543 | 563 | 609 | 641 | 682 | 684 | 705 | 844 | 541 | 478 | 565 | 544 | 572 | 541 | 527 | 605 | 662 | 713 | 639 | 745 | 709 | 732 | Oxsm |
| Oxsr1 | 13302 | 11479 | 12073 | 12168 | 11117 | 11437 | 12037 | 7675 | 6899 | 6761 | 6597 | 7615 | 7060 | 7503 | 12676 | 11210 | 11458 | 11293 | 10841 | 11715 | 11433 | 7082 | 6403 | 7135 | 7485 | 7598 | 7671 | 7954 | Oxsr1 |
| Oxt | 2 | 9 | 10 | 32 | 4 | 7 | 3 | 0 | 4 | 8 | 1 | 1 | 0 | 0 | 4 | 9 | 12 | 8 | 13 | 5 | 4 | 0 | 0 | 3 | 6 | 2 | 0 | 1 | Oxt |
| Oxtr | 104 | 83 | 111 | 87 | 142 | 82 | 74 | 83 | 80 | 48 | 49 | 41 | 54 | 35 | 105 | 143 | 102 | 116 | 97 | 76 | 117 | 79 | 74 | 48 | 48 | 26 | 43 | 27 | Oxtr |
| P2rx1 | 52 | 37 | 68 | 28 | 24 | 43 | 16 | 53 | 46 | 27 | 62 | 39 | 68 | 39 | 45 | 69 | 27 | 34 | 19 | 41 | 19 | 33 | 49 | 56 | 29 | 67 | 65 | 52 | P2rx1 |
| P2rx2 | 67 | 58 | 41 | 70 | 81 | 56 | 66 | 162 | 133 | 167 | 145 | 64 | 87 | 76 | 67 | 47 | 30 | 43 | 86 | 67 | 63 | 179 | 186 | 104 | 109 | 134 | 37 | 71 | P2rx2 |
| P2rx3 | 498 | 689 | 169 | 133 | 1221 | 192 | 106 | 1192 | 1272 | 793 | 1360 | 148 | 1090 | 193 | 1040 | 1078 | 356 | 728 | 139 | 152 | 110 | 1623 | 834 | 111 | 806 | 151 | 229 | 225 | P2rx3 |
| P2rx4 | 115 | 159 | 191 | 124 | 144 | 150 | 104 | 299 | 436 | 431 | 458 | 300 | 457 | 291 | 125 | 157 | 134 | 119 | 136 | 140 | 107 | 401 | 481 | 351 | 381 | 293 | 294 | 292 | P2rx4 |
| P2rx5 | 8 | 10 | 23 | 11 | 20 | 16 | 21 | 13 | 31 | 41 | 72 | 13 | 16 | 4 | 15 | 14 | 16 | 35 | 14 | 12 | 15 | 22 | 61 | 20 | 25 | 19 | 14 | 11 | P2rx5 |
| P2rx6 | 35 | 32 | 66 | 34 | 43 | 45 | 42 | 84 | 96 | 71 | 102 | 64 | 133 | 66 | 34 | 48 | 52 | 44 | 40 | 63 | 37 | 79 | 134 | 47 | 82 | 71 | 95 | 76 | P2rx6 |
| P2rx7 | 81 | 130 | 91 | 131 | 242 | 158 | 94 | 352 | 355 | 308 | 350 | 169 | 379 | 184 | 161 | 165 | 190 | 132 | 138 | 157 | 129 | 413 | 339 | 188 | 304 | 137 | 231 | 179 | P2rx7 |
| P2ry1 | 708 | 560 | 713 | 572 | 551 | 567 | 510 | 404 | 454 | 421 | 522 | 499 | 472 | 417 | 571 | 795 | 645 | 643 | 493 | 606 | 553 | 538 | 507 | 422 | 428 | 475 | 475 | 412 | P2ry1 |
| P2ry10 | 6 | 0 | 0 | 0 | 0 | 0 | 1 | 1 | 3 | 4 | 18 | 5 | 7 | 0 | 0 | 2 | 0 | 0 | 0 | 0 | 0 | 5 | 19 | 8 | 0 | 10 | 3 | 5 | P2ry10 |
| P2ry10b | 28 | 3 | 13 | 30 | 47 | 32 | 18 | 76 | 82 | 41 | 180 | 53 | 60 | 38 | 22 | 26 | 29 | 13 | 29 | 25 | 32 | 58 | 140 | 34 | 56 | 43 | 67 | 47 | P2ry10b |
| P2ry12 | 90 | 89 | 123 | 103 | 177 | 142 | 146 | 321 | 256 | 229 | 246 | 223 | 229 | 233 | 92 | 95 | 73 | 94 | 143 | 114 | 114 | 294 | 313 | 217 | 234 | 226 | 210 | 190 | P2ry12 |
| P2ry13 | 46 | 44 | 41 | 42 | 59 | 39 | 56 | 92 | 117 | 80 | 107 | 83 | 91 | 86 | 41 | 48 | 22 | 58 | 34 | 58 | 51 | 119 | 106 | 87 | 68 | 119 | 48 | 84 | P2ry13 |
| P2ry14 | 8 | 11 | 0 | 5 | 13 | 15 | 8 | 22 | 38 | 45 | 38 | 14 | 36 | 14 | 9 | 11 | 1 | 11 | 11 | 13 | 10 | 53 | 38 | 37 | 53 | 28 | 12 | 24 | P2ry14 |
| P2ry2 | 70 | 60 | 31 | 35 | 38 | 37 | 17 | 23 | 42 | 52 | 48 | 5 | 19 | 12 | 74 | 84 | 38 | 73 | 14 | 16 | 13 | 30 | 49 | 25 | 10 | 10 | 15 | 20 | P2ry2 |
| P2ry4 | 10 | 16 | 9 | 31 | 17 | 15 | 16 | 4 | 2 | 0 | 6 | 0 | 1 | 10 | 11 | 16 | 12 | 35 | 8 | 12 | 12 | 2 | 6 | 3 | 8 | 0 | 6 | 2 | P2ry4 |
| P2ry6 | 45 | 36 | 57 | 31 | 67 | 31 | 36 | 68 | 137 | 124 | 132 | 63 | 108 | 94 | 55 | 49 | 54 | 33 | 34 | 31 | 32 | 109 | 146 | 116 | 95 | 122 | 114 | 87 | P2ry6 |
| P3h1 | 4186 | 4816 | 4014 | 3633 | 2750 | 3468 | 2341 | 2036 | 2960 | 2601 | 3764 | 2103 | 2617 | 1854 | 3679 | 4917 | 3620 | 3776 | 2650 | 2457 | 2362 | 2523 | 3246 | 2569 | 2165 | 2398 | 2413 | 2059 | P3h1 |
| P3h2 | 332 | 249 | 336 | 229 | 198 | 233 | 211 | 463 | 529 | 555 | 982 | 505 | 692 | 451 | 218 | 393 | 199 | 184 | 220 | 186 | 174 | 389 | 800 | 521 | 371 | 678 | 374 | 457 | P3h2 |
| P3h3 | 2400 | 2810 | 2466 | 2818 | 2625 | 2867 | 2095 | 3543 | 5058 | 5361 | 6433 | 4210 | 4839 | 3976 | 2248 | 3023 | 2569 | 3136 | 2570 | 1918 | 1995 | 4809 | 5434 | 5340 | 4687 | 5201 | 3932 | 3836 | P3h3 |
| P3h4 | 1159 | 1339 | 1190 | 912 | 895 | 1044 | 651 | 1020 | 1507 | 1292 | 1819 | 1008 | 1200 | 899 | 1189 | 1497 | 1048 | 938 | 775 | 835 | 644 | 1180 | 1838 | 1177 | 1090 | 1238 | 1108 | 1049 | P3h4 |
| P3r3urf | 0 | 2 | 0 | 0 | 0 | 0 | 0 | 0 | 0 | 0 | 0 | 0 | 0 | 1 | 0 | 0 | 0 | 4 | 2 | 0 | 0 | 0 | 0 | 0 | 0 | 0 | 0 | 0 | P3r3urf |
| P4ha1 | 5059 | 4736 | 4542 | 4085 | 3272 | 3706 | 3287 | 4285 | 4577 | 4446 | 5760 | 2898 | 3432 | 3380 | 5095 | 5393 | 4502 | 4648 | 3264 | 3632 | 3457 | 4306 | 5502 | 4189 | 3591 | 3645 | 3196 | 3288 | P4ha1 |
| P4ha2 | 2076 | 2140 | 1885 | 1616 | 1417 | 1740 | 1123 | 1102 | 1762 | 1481 | 2430 | 1071 | 1414 | 890 | 1816 | 2730 | 1826 | 2273 | 1283 | 1126 | 1170 | 1360 | 1786 | 1382 | 1377 | 1256 | 1280 | 999 | P4ha2 |
| P4ha3 | 597 | 715 | 606 | 542 | 592 | 554 | 277 | 1388 | 1864 | 1887 | 1796 | 908 | 1425 | 809 | 745 | 906 | 547 | 692 | 348 | 319 | 275 | 1679 | 1780 | 1581 | 1128 | 1527 | 918 | 784 | P4ha3 |
| P4hb | 35174 | 34260 | 30502 | 25309 | 20997 | 24259 | 21507 | 25866 | 28206 | 24381 | 30268 | 24239 | 27053 | 23276 | 33280 | 35908 | 26445 | 26662 | 22775 | 21499 | 22204 | 25795 | 31448 | 24999 | 23173 | 28166 | 26598 | 26046 | P4hb |
| P4htm | 303 | 401 | 344 | 445 | 601 | 471 | 484 | 509 | 669 | 779 | 610 | 698 | 675 | 690 | 349 | 316 | 410 | 454 | 592 | 448 | 474 | 787 | 597 | 694 | 707 | 780 | 608 | 674 | P4htm |
| Pa2g4 | 18363 | 17645 | 18689 | 16838 | 14315 | 14204 | 16918 | 12410 | 11375 | 11443 | 10981 | 13283 | 12487 | 14712 | 17578 | 15507 | 17781 | 15096 | 15633 | 14932 | 16533 | 10322 | 11040 | 13215 | 11916 | 16005 | 13894 | 16060 | Pa2g4 |
| Pabpc1 | 56338 | 53208 | 68515 | 57818 | 48010 | 46288 | 52299 | 38412 | 38904 | 41630 | 43504 | 50268 | 48889 | 51058 | 51764 | 54194 | 60841 | 57388 | 53946 | 46168 | 53297 | 40155 | 40926 | 47439 | 41423 | 60486 | 46365 | 55723 | Pabpc1 |
| Pabpc1l | 324 | 452 | 336 | 623 | 445 | 455 | 456 | 98 | 101 | 117 | 93 | 81 | 101 | 97 | 348 | 365 | 416 | 572 | 423 | 400 | 475 | 132 | 118 | 135 | 145 | 128 | 103 | 120 | Pabpc1l |
| Pabpc4 | 9689 | 9515 | 12763 | 10279 | 8582 | 8408 | 8679 | 7489 | 8586 | 9369 | 10329 | 10819 | 10119 | 10944 | 9141 | 10185 | 10950 | 10085 | 9719 | 7830 | 8732 | 8233 | 8739 | 10069 | 9209 | 12142 | 9828 | 11731 | Pabpc4 |
| Pabpc4l | 1047 | 648 | 825 | 538 | 529 | 570 | 536 | 1110 | 1232 | 778 | 1297 | 733 | 1101 | 797 | 870 | 903 | 705 | 580 | 515 | 642 | 486 | 729 | 1338 | 926 | 687 | 1098 | 859 | 828 | Pabpc4l |
| Pabpc5 | 55 | 36 | 32 | 58 | 71 | 106 | 108 | 138 | 144 | 111 | 110 | 76 | 76 | 78 | 75 | 30 | 30 | 25 | 78 | 103 | 122 | 92 | 153 | 66 | 100 | 15 | 119 | 65 | Pabpc5 |
| Pabpc6 | 1 | 3 | 0 | 0 | 0 | 0 | 2 | 1 | 1 | 0 | 4 | 0 | 7 | 0 | 1 | 2 | 0 | 2 | 0 | 4 | 0 | 2 | 0 | 1 | 1 | 2 | 7 | 0 | Pabpc6 |
| Pabpn1 | 2848 | 3201 | 3913 | 4346 | 4118 | 3817 | 3998 | 4404 | 4053 | 5331 | 4634 | 5564 | 4535 | 5515 | 2668 | 2562 | 4507 | 3982 | 4249 | 3551 | 4152 | 5043 | 3809 | 5990 | 5510 | 5530 | 5168 | 5174 | Pabpn1 |
| Pabpn1l | 2 | 5 | 21 | 11 | 17 | 11 | 15 | 7 | 6 | 9 | 22 | 8 | 2 | 4 | 5 | 14 | 5 | 23 | 13 | 12 | 14 | 15 | 8 | 8 | 18 | 8 | 6 | 6 | Pabpn1l |
| Pacc1 | 1564 | 1894 | 1522 | 1572 | 1535 | 1683 | 1656 | 1831 | 1838 | 1604 | 1578 | 1943 | 1715 | 1867 | 1672 | 1616 | 1458 | 1536 | 1642 | 1850 | 1695 | 1696 | 1789 | 1914 | 1879 | 1854 | 2015 | 1830 | Pacc1 |

| | | | | | | | | | | | | | | | | | | | | | | | | | | | | | |
|---|---|---|---|---|---|---|---|---|---|---|---|---|---|---|---|---|---|---|---|---|---|---|---|---|---|---|---|---|---|
| Pacrg | 208 | 317 | 223 | 281 | 355 | 302 | 283 | 236 | 250 | 207 | 260 | 203 | 276 | 206 | 320 | 242 | 229 | 235 | 308 | 331 | 299 | 252 | 284 | 191 | 204 | 213 | 178 | 235 | Pacrg |
| Pacs1 | 6037 | 6264 | 6275 | 7033 | 7706 | 6639 | 6669 | 5057 | 5735 | 5970 | 5830 | 5623 | 5218 | 5248 | 5790 | 6136 | 6720 | 7113 | 7416 | 5784 | 6957 | 7090 | 5615 | 5873 | 5898 | 5453 | 5081 | 5056 | Pacs1 |
| Pacs2 | 11274 | 12758 | 14254 | 15404 | 15403 | 14854 | 13987 | 9867 | 11416 | 13403 | 12367 | 14148 | 13177 | 12803 | 10883 | 12108 | 15238 | 15789 | 15897 | 11771 | 14754 | 14352 | 10463 | 13540 | 14592 | 12437 | 12246 | 12712 | Pacs2 |
| Pacsin1 | 805 | 901 | 956 | 1331 | 2023 | 1539 | 1409 | 860 | 1013 | 1376 | 1152 | 907 | 679 | 852 | 682 | 872 | 1367 | 1415 | 1858 | 1223 | 1541 | 1539 | 930 | 1267 | 1336 | 946 | 611 | 644 | Pacsin1 |
| Pacsin2 | 5023 | 5161 | 6724 | 5412 | 4714 | 4595 | 4403 | 2942 | 3398 | 3317 | 4058 | 3778 | 3928 | 3541 | 4758 | 5785 | 5743 | 5366 | 4736 | 3911 | 4217 | 3171 | 3667 | 3758 | 3285 | 4119 | 3764 | 3944 | Pacsin2 |
| Pacsin3 | 1414 | 1504 | 1852 | 1516 | 1510 | 1482 | 1412 | 923 | 1210 | 1169 | 1270 | 1317 | 1203 | 1331 | 1201 | 1274 | 1753 | 1531 | 1844 | 1362 | 1414 | 772 | 1168 | 1436 | 1073 | 1781 | 1377 | 1356 | Pacsin3 |
| Padi1 | 10 | 13 | 15 | 18 | 15 | 4 | 3 | 11 | 16 | 25 | 13 | 9 | 32 | 11 | 12 | 21 | 13 | 10 | 23 | 10 | 9 | 13 | 8 | 16 | 6 | 12 | 9 | 8 | Padi1 |
| Padi2 | 301 | 387 | 268 | 244 | 291 | 191 | 112 | 184 | 186 | 240 | 285 | 37 | 244 | 54 | 458 | 638 | 297 | 338 | 178 | 126 | 153 | 145 | 174 | 126 | 110 | 139 | 59 | 72 | Padi2 |
| Padi3 | 8 | 6 | 14 | 2 | 1 | 3 | 3 | 1 | 8 | 5 | 5 | 5 | 7 | 7 | 8 | 8 | 14 | 8 | 0 | 0 | 4 | 2 | 16 | 1 | 5 | 7 | 0 | 9 | Padi3 |
| Padi4 | 30 | 33 | 35 | 54 | 15 | 24 | 15 | 9 | 10 | 6 | 10 | 58 | 34 | 29 | 30 | 24 | 31 | 14 | 18 | 17 | 11 | 13 | 8 | 15 | 27 | 53 | 55 | 47 | Padi4 |
| Padi6 | 0 | 1 | 0 | 0 | 0 | 0 | 0 | 2 | 1 | 1 | 0 | 0 | 0 | 0 | 0 | 0 | 1 | 0 | 0 | 3 | 1 | 0 | 0 | 0 | 0 | 0 | 0 | 0 | Padi6 |
| Paf1 | 10322 | 11119 | 11732 | 12211 | 9356 | 9625 | 10326 | 4949 | 4637 | 5345 | 4753 | 5488 | 4932 | 5800 | 10493 | 10013 | 11145 | 11052 | 10124 | 9257 | 10060 | 5115 | 4574 | 5753 | 5337 | 5971 | 5196 | 5525 | Paf1 |
| Pafah1b1 | 24751 | 20411 | 22075 | 24728 | 29935 | 27783 | 31112 | 39449 | 31978 | 32043 | 32073 | 32350 | 32484 | 35074 | 27257 | 21390 | 24189 | 26319 | 27720 | 28980 | 29875 | 37368 | 31936 | 31331 | 36118 | 27661 | 33539 | 33495 | Pafah1b1 |
| Pafah1b2 | 20584 | 17314 | 19699 | 19589 | 21503 | 20896 | 22060 | 27196 | 21835 | 21315 | 19063 | 23232 | 20862 | 22668 | 22396 | 16992 | 20678 | 19597 | 21516 | 22715 | 21817 | 25298 | 21252 | 22238 | 23814 | 23052 | 23573 | 23830 | Pafah1b2 |
| Pafah1b3 | 4983 | 6489 | 5802 | 6708 | 6974 | 6464 | 6305 | 6996 | 6873 | 7480 | 6695 | 6894 | 6510 | 7397 | 5325 | 4645 | 6640 | 6640 | 7420 | 7228 | 7132 | 6861 | 6893 | 7959 | 8205 | 7687 | 7693 | 7312 | Pafah1b3 |
| Pafah2 | 1619 | 1878 | 1765 | 1684 | 1545 | 1523 | 1538 | 949 | 1257 | 972 | 1110 | 989 | 1238 | 904 | 1654 | 1769 | 1759 | 1641 | 1460 | 1322 | 1376 | 1078 | 1184 | 1052 | 1017 | 1080 | 996 | 967 | Pafah2 |
| Pag1 | 5018 | 4712 | 4281 | 7069 | 7743 | 7559 | 7633 | 3123 | 3351 | 3560 | 3855 | 3060 | 3144 | 3012 | 5370 | 5480 | 5167 | 7799 | 6629 | 5855 | 7828 | 3700 | 3212 | 3285 | 4100 | 1968 | 2869 | 2566 | Pag1 |
| Pagr1a | 0 | 1 | 5 | 0 | 0 | 0 | 0 | 0 | 0 | 1 | 0 | 0 | 2 | 2 | 0 | 2 | 0 | 0 | 1 | 0 | 1 | 2 | 1 | 0 | 2 | 1 | 0 | 0 | Pagr1a |
| Pah | 3 | 7 | 13 | 3 | 0 | 4 | 2 | 4 | 0 | 3 | 0 | 2 | 8 | 11 | 8 | 6 | 2 | 3 | 4 | 1 | 3 | 3 | 1 | 1 | 0 | 0 | 7 | 9 | Pah |
| Paics | 32491 | 28322 | 28839 | 23762 | 19734 | 21981 | 26463 | 22150 | 19639 | 15889 | 16264 | 22752 | 19992 | 24053 | 32127 | 28931 | 26675 | 22966 | 23529 | 28791 | 23584 | 14552 | 17865 | 19398 | 17986 | 27617 | 24080 | 25497 | Paics |
| Paip1 | 9716 | 7401 | 10394 | 10609 | 11597 | 11515 | 12729 | 15618 | 12149 | 12091 | 13668 | 14316 | 15093 | 17882 | 10334 | 7712 | 10512 | 10094 | 11817 | 5893 | 11397 | 14831 | 11958 | 15972 | 15739 | 16753 | 16014 | 18058 | Paip1 |
| Paip2 | 12453 | 10726 | 12946 | 11712 | 11273 | 11219 | 12996 | 18075 | 14243 | 14998 | 14237 | 15640 | 14785 | 16998 | 13172 | 11158 | 12896 | 11364 | 12473 | 13591 | 12970 | 16300 | 14950 | 16116 | 15039 | 15928 | 16593 | 17888 | Paip2 |
| Paip2b | 16699 | 15095 | 14662 | 15305 | 14645 | 14866 | 16359 | 11093 | 9919 | 9157 | 8589 | 10328 | 9631 | 10423 | 16966 | 14427 | 14841 | 15649 | 16190 | 16636 | 15851 | 9277 | 9323 | 9707 | 10480 | 10995 | 10438 | 10593 | Paip2b |
| Pak1 | 8746 | 8961 | 11159 | 9795 | 11444 | 10277 | 10575 | 13273 | 10231 | 12206 | 9501 | 12527 | 10997 | 12673 | 10152 | 7507 | 11104 | 9937 | 11357 | 10093 | 10709 | 15742 | 9872 | 11937 | 13459 | 11079 | 12305 | 13168 | Pak1 |
| Pak1ip1 | 4624 | 5349 | 4979 | 4726 | 3823 | 3553 | 4625 | 3555 | 3033 | 3038 | 2626 | 3598 | 3360 | 4041 | 4646 | 4567 | 4954 | 4328 | 4215 | 5172 | 4270 | 2777 | 2988 | 3358 | 3320 | 3915 | 3829 | 3821 | Pak1ip1 |
| Pak2 | 11297 | 9284 | 11713 | 12021 | 11669 | 11354 | 12754 | 12242 | 10706 | 11373 | 11002 | 13810 | 11923 | 14140 | 11035 | 9848 | 11798 | 11675 | 11596 | 10209 | 12574 | 12795 | 10467 | 11964 | 12951 | 12947 | 13498 | 14175 | Pak2 |
| Pak3 | 15789 | 14618 | 9654 | 17179 | 19124 | 20189 | 22502 | 19604 | 16295 | 13823 | 12558 | 15597 | 11995 | 16559 | 19507 | 13471 | 14081 | 17510 | 17993 | 23313 | 22254 | 16282 | 13050 | 12839 | 17437 | 10215 | 15186 | 14497 | Pak3 |
| Pak4 | 848 | 1088 | 1430 | 1146 | 1091 | 964 | 955 | 1315 | 1711 | 1888 | 1657 | 2219 | 1885 | 2093 | 785 | 951 | 1396 | 1108 | 1262 | 904 | 978 | 1594 | 1651 | 2309 | 1747 | 2907 | 1993 | 2169 | Pak4 |
| Pak5 | 1660 | 1790 | 1628 | 2431 | 3272 | 2675 | 2690 | 4484 | 3526 | 3720 | 3131 | 3261 | 3341 | 3325 | 2207 | 1572 | 2086 | 2380 | 2915 | 3157 | 2752 | 4764 | 3387 | 3373 | 4570 | 2555 | 2972 | 3051 | Pak5 |
| Pak6 | 346 | 432 | 288 | 852 | 1570 | 1234 | 1274 | 971 | 1321 | 1616 | 1218 | 895 | 702 | 827 | 381 | 331 | 714 | 1104 | 1519 | 1047 | 1427 | 1942 | 1137 | 1286 | 1846 | 782 | 644 | 688 | Pak6 |
| Pakap | 7 | 7 | 6 | 9 | 9 | 5 | 7 | 1 | 4 | 12 | 4 | 7 | 16 | 6 | 11 | 11 | 14 | 9 | 8 | 2 | 5 | 10 | 3 | 13 | 12 | 24 | 12 | 14 | Pakap |
| Pakap | 0 | 0 | 0 | 0 | 2 | 0 | 1 | 0 | 1 | 3 | 0 | 0 | 0 | 0 | 0 | 0 | 0 | 1 | 0 | 3 | 2 | 3 | 1 | 0 | 0 | 0 | 0 | 0 | Pakap |
| Pakap | 59 | 41 | 16 | 56 | 178 | 115 | 84 | 259 | 314 | 271 | 296 | 104 | 164 | 117 | 74 | 60 | 36 | 81 | 95 | 85 | 111 | 358 | 277 | 157 | 250 | 18 | 130 | 84 | Pakap |
| Palb2 | 1472 | 1301 | 1287 | 1270 | 889 | 1183 | 1134 | 1042 | 1011 | 822 | 817 | 1133 | 1034 | 1130 | 1329 | 1274 | 1250 | 1020 | 921 | 1231 | 1086 | 688 | 852 | 936 | 884 | 1142 | 1233 | 1171 | Palb2 |
| Pald1 | 3916 | 4388 | 4653 | 3812 | 3802 | 3439 | 2894 | 2319 | 2706 | 2320 | 2619 | 2492 | 3039 | 2074 | 4089 | 4956 | 4145 | 3571 | 3609 | 3354 | 2911 | 2481 | 2975 | 2221 | 2200 | 2847 | 2757 | 2295 | Pald1 |
| Palld | 6927 | 5856 | 7280 | 5277 | 4381 | 5276 | 3846 | 5935 | 6950 | 6402 | 8598 | 6820 | 7592 | 6557 | 5806 | 6992 | 6510 | 5268 | 4796 | 3412 | 4103 | 5482 | 7843 | 7134 | 5353 | 7555 | 6826 | 6900 | Palld |
| Palm | 6679 | 8447 | 10331 | 11236 | 12570 | 10468 | 8595 | 7730 | 8833 | 12407 | 10327 | 12353 | 9410 | 9353 | 6401 | 7047 | 11408 | 11000 | 13303 | 8161 | 10246 | 11921 | 8748 | 12463 | 13276 | 12085 | 9761 | 9382 | Palm |
| Palm3 | 990 | 1158 | 1613 | 1103 | 808 | 790 | 543 | 426 | 498 | 609 | 639 | 388 | 549 | 412 | 1093 | 1178 | 1216 | 1024 | 616 | 618 | 545 | 369 | 536 | 480 | 339 | 577 | 503 | 526 | Palm3 |
| Palmd | 328 | 282 | 305 | 426 | 575 | 557 | 578 | 882 | 932 | 814 | 1317 | 811 | 803 | 839 | 371 | 282 | 404 | 449 | 584 | 656 | 669 | 874 | 1214 | 918 | 1117 | 788 | 776 | 701 | Palmd |
| Pals1 | 6995 | 6243 | 6271 | 6400 | 5556 | 5954 | 6712 | 5069 | 4338 | 3756 | 3895 | 4537 | 4349 | 4531 | 7743 | 7063 | 6293 | 6413 | 5591 | 7059 | 5957 | 3888 | 4394 | 3947 | 3686 | 3911 | 4472 | 4475 | Pals1 |
| Pals2 | 4908 | 4214 | 5253 | 4327 | 3986 | 4343 | 4749 | 3683 | 3511 | 3429 | 3909 | 4301 | 4344 | 4265 | 4836 | 4678 | 4636 | 4406 | 4316 | 4186 | 4452 | 3479 | 3860 | 3833 | 3758 | 4586 | 4463 | 4691 | Pals2 |
| Pam | 5253 | 4228 | 4575 | 4540 | 4896 | 4921 | 5494 | 6697 | 6842 | 6662 | 6193 | 5813 | 5264 | 6551 | 4743 | 4104 | 4465 | 4775 | 4849 | 5075 | 5323 | 6171 | 6814 | 6722 | 7099 | 5751 | 5549 | 5655 | Pam |
| Pam16 | 884 | 1053 | 990 | 1196 | 1180 | 1065 | 1160 | 1084 | 1195 | 1155 | 1145 | 997 | 1114 | 1207 | 897 | 917 | 1057 | 929 | 1234 | 1324 | 1068 | 946 | 1082 | 1226 | 1269 | 1153 | 1232 | 1114 | Pam16 |
| Pamr1 | 174 | 188 | 201 | 186 | 225 | 305 | 112 | 244 | 395 | 338 | 1570 | 167 | 325 | 162 | 249 | 354 | 195 | 256 | 161 | 178 | 149 | 376 | 1122 | 194 | 256 | 143 | 170 | 146 | Pamr1 |
| Pan2 | 3938 | 3943 | 3756 | 3350 | 3310 | 3434 | 3550 | 6179 | 6355 | 6628 | 6234 | 8315 | 6949 | 7751 | 3578 | 3453 | 3706 | 3230 | 3671 | 3296 | 3335 | 6244 | 5824 | 7021 | 6692 | 7409 | 8283 | 7990 | Pan2 |
| Pan3 | 4050 | 3667 | 4744 | 4812 | 5016 | 4905 | 5464 | 5682 | 4814 | 5944 | 6039 | 6314 | 5757 | 6611 | 3850 | 3685 | 4636 | 4840 | 5130 | 4318 | 5197 | 6450 | 4818 | 6132 | 6283 | 5853 | 5997 | 6236 | Pan3 |
| Pank1 | 1293 | 1148 | 1484 | 1306 | 1442 | 1227 | 1446 | 1831 | 1726 | 1808 | 1667 | 2013 | 1905 | 2229 | 1274 | 1119 | 1415 | 1326 | 1506 | 1184 | 1457 | 2166 | 1741 | 2008 | 2035 | 2205 | 1912 | 2281 | Pank1 |
| Pank2 | 5592 | 4992 | 6199 | 5408 | 5038 | 5411 | 5506 | 5598 | 4583 | 5061 | 4974 | 6392 | 5446 | 6342 | 5347 | 4906 | 5882 | 5328 | 5451 | 4773 | 5380 | 5332 | 4583 | 5559 | 5441 | 6573 | 5796 | 6261 | Pank2 |
| Pank3 | 17342 | 14033 | 11196 | 14655 | 15919 | 17111 | 17311 | 21594 | 16808 | 14349 | 15767 | 16634 | 16223 | 17455 | 18180 | 16002 | 12944 | 15340 | 14808 | 17553 | 17744 | 17649 | 16421 | 13884 | 17806 | 10979 | 18009 | 16802 | Pank3 |
| Pank4 | 2682 | 3113 | 2615 | 2833 | 2825 | 2980 | 2769 | 3473 | 3849 | 3661 | 3511 | 4035 | 3751 | 3702 | 2622 | 2754 | 2934 | 2843 | 3129 | 2990 | 2837 | 3429 | 3387 | 3681 | 3573 | 3614 | 4019 | 3690 | Pank4 |
| Panx1 | 3926 | 4061 | 4167 | 5661 | 7549 | 6769 | 6753 | 8629 | 7448 | 8798 | 7271 | 7868 | 7066 | 8299 | 4480 | 3376 | 5030 | 5743 | 7423 | 6189 | 7309 | 10218 | 7023 | 8013 | 9764 | 6248 | 7308 | 7783 | Panx1 |
| Panx2 | 111 | 157 | 191 | 235 | 382 | 250 | 263 | 364 | 441 | 538 | 430 | 417 | 342 | 395 | 131 | 120 | 275 | 226 | 352 | 283 | 315 | 637 | 353 | 474 | 657 | 329 | 323 | 309 | Panx2 |
| Panx3 | 3 | 5 | 5 | 9 | 2 | 53 | 0 | 103 | 132 | 182 | 431 | 34 | 35 | 40 | 8 | 5 | 6 | 41 | 2 | 0 | 0 | 185 | 162 | 109 | 211 | 49 | 62 | 43 | Panx3 |
| Paox | 1731 | 1637 | 1568 | 1595 | 1538 | 1621 | 1623 | 865 | 961 | 778 | 1033 | 882 | 1066 | 794 | 1568 | 1673 | 1504 | 1586 | 1354 | 1491 | 1383 | 810 | 1020 | 856 | 725 | 960 | 1031 | 872 | Paox |
| Papln | 62 | 85 | 39 | 47 | 88 | 90 | 45 | 123 | 258 | 133 | 172 | 41 | 66 | 62 | 45 | 52 | 57 | 125 | 62 | 43 | 49 | 50 | 223 | 95 | 57 | 36 | 43 | 27 | Papln |
| Papola | 22843 | 18465 | 20133 | 19320 | 18663 | 19249 | 21926 | 26050 | 21583 | 21043 | 21143 | 25362 | 22992 | 25344 | 22173 | 18679 | 19030 | 18180 | 19805 | 20407 | 21005 | 22378 | 21052 | 23118 | 22866 | 23957 | 26293 | 25751 | Papola |
| Papolb | 122 | 63 | 101 | 100 | 85 | 88 | 82 | 44 | 46 | 52 | 43 | 58 | 81 | 48 | 103 | 81 | 76 | 58 | 56 | 70 | 56 | 57 | 67 | 53 | 78 | 41 | 45 | 49 | Papolb |
| Papolg | 4097 | 3382 | 3521 | 3794 | 3444 | 3584 | 3923 | 4897 | 3803 | 3430 | 3303 | 3818 | 3517 | 4143 | 4425 | 3886 | 3459 | 3724 | 3628 | 4280 | 4059 | 3882 | 3689 | 3414 | 4065 | 3322 | 3755 | 3775 | Papolg |
| Pappa | 859 | 598 | 609 | 470 | 453 | 672 | 522 | 1506 | 1601 | 1021 | 1599 | 1544 | 1312 | 1668 | 795 | 555 | 663 | 379 | 533 | 513 | 491 | 1034 | 1873 | 1010 | 1054 | 1479 | 1725 | 1642 | Pappa |
| Pappa2 | 1649 | 1921 | 954 | 1212 | 1133 | 1577 | 943 | 1671 | 2267 | 2069 | 1976 | 1807 | 2427 | 1939 | 1866 | 2991 | 1175 | 1531 | 860 | 901 | 1017 | 1932 | 2067 | 1624 | 2232 | 1137 | 1718 | 1753 | Pappa2 |
| Papss1 | 10199 | 11010 | 9954 | 10517 | 10233 | 9782 | 10782 | 6467 | 7472 | 7314 | 7312 | 6484 | 6465 | 6856 | 9635 | 9447 | 10396 | 10702 | 11039 | 9847 | 10948 | 6505 | 7583 | 7406 | 7180 | 7607 | 6708 | 6972 | Papss1 |
| Papss2 | 392 | 279 | 202 | 335 | 80 | 350 | 38 | 1056 | 1772 | 1694 | 2517 | 545 | 724 | 483 | 384 | 485 | 168 | 343 | 124 | 66 | 52 | 1646 | 1590 | 1803 | 886 | 1462 | 521 | 494 | Papss2 |
| Paqr3 | 2569 | 3040 | 2207 | 3133 | 2696 | 2777 | 2822 | 1479 | 1439 | 1473 | 1249 | 1392 | 1246 | 1187 | 2537 | 2519 | 2560 | 2980 | 2842 | 2770 | 3034 | 1502 | 1171 | 1245 | 1699 | 999 | 1419 | 1063 | Paqr3 |
| Paqr4 | 2485 | 3151 | 2912 | 3172 | 3293 | 3026 | 3029 | 1550 | 1575 | 1709 | 1461 | 1667 | 1482 | 1558 | 2661 | 2466 | 3122 | 2907 | 3441 | 3083 | 3209 | 2055 | 1563 | 1502 | 1802 | 1377 | 1679 | 1720 | Paqr4 |
| Paqr5 | 69 | 84 | 45 | 39 | 38 | 53 | 42 | 33 | 50 | 23 | 26 | 17 | 16 | 14 | 88 | 85 | 36 | 61 | 33 | 39 | 55 | 15 | 71 | 22 | 13 | 9 | 3 | 19 | Paqr5 |
| Paqr6 | 90 | 109 | 98 | 72 | 76 | 81 | 44 | 33 | 67 | 65 | 93 | 38 | 39 | 24 | 74 | 148 | 122 | 98 | 37 | 58 | 45 | 58 | 73 | 34 | 39 | 49 | 38 | 37 | Paqr6 |
| Paqr7 | 373 | 392 | 465 | 356 | 575 | 450 | 410 | 499 | 583 | 671 | 594 | 433 | 592 | 392 | 363 | 417 | 483 | 419 | 542 | 446 | 479 | 688 | 686 | 466 | 543 | 332 | 418 | 414 | Paqr7 |
| Paqr8 | 1453 | 1450 | 1761 | 1723 | 2592 | 2084 | 2082 | 2649 | 2873 | 2826 | 2710 | 2142 | 2552 | 2317 | 1696 | 1528 | 1583 | 1838 | 2011 | 1774 | 1942 | 2938 | 2912 | 2274 | 2677 | 1917 | 1908 | 2099 | Paqr8 |
| Paqr9 | 337 | 331 | 214 | 563 | 1435 | 858 | 988 | 832 | 830 | 897 | 880 | 411 | 533 | 430 | 460 | 308 | 404 | 755 | 894 | 724 | 996 | 1263 | 703 | 594 | 892 | 235 | 350 | 343 | Paqr9 |
| Pard3 | 5187 | 4508 | 5144 | 4750 | 4940 | 4956 | 4740 | 5918 | 6379 | 7188 | 7128 | 7308 | 7070 | 7326 | 4739 | 5139 | 4906 | 4680 | 4899 | 3979 | 4808 | 6415 | 6467 | 7477 | 6379 | 7250 | 6966 | 7403 | Pard3 |
| Pard3b | 1791 | 1738 | 1555 | 1706 | 1522 | 1606 | 1468 | 1531 | 1877 | 1626 | 2118 | 1944 | 1948 | 1829 | 1668 | 1891 | 1582 | 1760 | 1286 | 1570 | 1524 | 1571 | 1843 | 1556 | 1699 | 1307 | 1907 | 1920 | Pard3b |
| Pard6a | 571 | 681 | 527 | 700 | 723 | 693 | 730 | 586 | 729 | 676 | 566 | 600 | 585 | 577 | 553 | 550 | 661 | 734 | 788 | 770 | 659 | 622 | 646 | 628 | 661 | 624 | 673 | 641 | Pard6a |
| Pard6b | 1412 | 1841 | 2114 | 1937 | 1523 | 1332 | 1374 | 680 | 728 | 911 | 692 | 1004 | 915 | 946 | 1389 | 1619 | 1961 | 1848 | 1569 | 1338 | 1451 | 835 | 705 | 1057 | 891 | 1150 | 960 | 1010 | Pard6b |
| Pard6g | 5705 | 5331 | 5699 | 4878 | 4593 | 5404 | 4790 | 3679 | 4008 | 3978 | 3751 | 4543 | 4588 | 4572 | 5138 | 5445 | 5139 | 4758 | 4972 | 4311 | 4644 | 3584 | 3939 | 4061 | 3869 | 4942 | 4496 | 4791 | Pard6g |
| Parg | 7063 | 6087 | 6406 | 5792 | 5105 | 5622 | 5669 | 5456 | 4801 | 4270 | 4407 | 5155 | 4880 | 4974 | 7047 | 6534 | 5623 | 5479 | 5093 | 6079 | 5321 | 4543 | 4589 | 4804 | 4833 | 5199 | 5302 | 5139 | Parg |
| Park7 | 6237 | 7386 | 8327 | 7579 | 6585 | 6675 | 6463 | 5785 | 6535 | 7898 | 7451 | 8034 | 7734 | 8162 | 5657 | 6072 | 7985 | 6807 | 7321 | 7155 | 6879 | 5324 | 6384 | 8670 | 7841 | 9793 | 8700 | 8850 | Park7 |
| Parl | 2651 | 2349 | 2399 | 2237 | 2167 | 2049 | 2243 | 2784 | 2562 | 2368 | 2404 | 3078 | 3065 | 3268 | 2524 | 2164 | 2298 | 1917 | 2258 | 2526 | 2064 | 2393 | 2392 | 2697 | 2675 | 3255 | 3407 | 3233 | Parl |
| Parm1 | 2970 | 2429 | 3175 | 1577 | 1434 | 1721 | 1382 | 1712 | 2218 | 1517 | 3180 | 1832 | 2375 | 1854 | 2567 | 3734 | 1900 | 1769 | 1281 | 1502 | 1321 | 1549 | 2741 | 1776 | 1181 | 2057 | 2228 | 2176 | Parm1 |
| Parn | 1945 | 1892 | 1759 | 1550 | 1636 | 1788 | 1609 | 2959 | 2876 | 2618 | 2634 | 2931 | 2926 | 2892 | 1905 | 1646 | 1644 | 1438 | 1633 | 1813 | 1625 | 2672 | 2838 | 2736 | 2669 | 3244 | 3233 | 3131 | Parn |

| | | | | | | | | | | | | | | | | | | | | | | | | | | | | | |
|---|---|---|---|---|---|---|---|---|---|---|---|---|---|---|---|---|---|---|---|---|---|---|---|---|---|---|---|---|---|
| Parp1 | 21665 | 24082 | 27493 | 23803 | 20248 | 21227 | 22254 | 14566 | 15823 | 17371 | 16377 | 23792 | 19560 | 23125 | 19777 | 22312 | 26259 | 21740 | 22948 | 18298 | 21340 | 14783 | 14255 | 20566 | 19153 | 26075 | 22079 | 24208 | Parp1 |
| Parp10 | 58 | 48 | 42 | 31 | 51 | 71 | 46 | 74 | 84 | 120 | 143 | 126 | 119 | 97 | 70 | 51 | 42 | 66 | 51 | 63 | 59 | 142 | 138 | 105 | 108 | 113 | 129 | 112 | Parp10 |
| Parp11 | 2753 | 2820 | 2796 | 2848 | 3326 | 3204 | 3576 | 6618 | 5652 | 5489 | 4778 | 5868 | 4994 | 5905 | 3187 | 2333 | 3195 | 2862 | 3486 | 3679 | 3589 | 6417 | 5415 | 5654 | 6129 | 5219 | 5626 | 5777 | Parp11 |
| Parp12 | 218 | 183 | 222 | 151 | 196 | 251 | 136 | 479 | 601 | 532 | 721 | 477 | 559 | 446 | 198 | 243 | 187 | 166 | 171 | 217 | 147 | 506 | 740 | 488 | 465 | 540 | 489 | 411 | Parp12 |
| Parp14 | 97 | 54 | 95 | 63 | 89 | 74 | 46 | 184 | 207 | 255 | 297 | 134 | 181 | 131 | 82 | 121 | 64 | 57 | 79 | 144 | 97 | 201 | 294 | 163 | 142 | 133 | 216 | 163 | Parp14 |
| Parp16 | 554 | 500 | 810 | 585 | 497 | 526 | 507 | 369 | 389 | 435 | 499 | 482 | 511 | 477 | 489 | 614 | 550 | 571 | 525 | 464 | 469 | 386 | 370 | 510 | 391 | 558 | 438 | 483 | Parp16 |
| Parp2 | 2162 | 1802 | 2418 | 1893 | 1943 | 1973 | 2175 | 2989 | 3211 | 3139 | 2904 | 3729 | 3214 | 3772 | 2186 | 1807 | 2067 | 1855 | 2029 | 2065 | 2027 | 2699 | 2825 | 3382 | 3006 | 4150 | 3485 | 3687 | Parp2 |
| Parp3 | 232 | 299 | 243 | 190 | 105 | 225 | 85 | 111 | 179 | 203 | 204 | 148 | 214 | 123 | 228 | 329 | 178 | 185 | 86 | 107 | 78 | 154 | 264 | 195 | 192 | 186 | 159 | 153 | Parp3 |
| Parp4 | 478 | 432 | 578 | 270 | 452 | 368 | 307 | 364 | 370 | 499 | 496 | 444 | 509 | 435 | 455 | 556 | 388 | 421 | 326 | 275 | 360 | 466 | 513 | 375 | 422 | 376 | 380 | 411 | Parp4 |
| Parp6 | 7074 | 7789 | 6513 | 9902 | 11508 | 10069 | 11676 | 9682 | 8473 | 9951 | 8301 | 9306 | 8096 | 9259 | 8292 | 6458 | 8772 | 10399 | 11318 | 9552 | 11970 | 12225 | 7636 | 9340 | 11098 | 8086 | 8142 | 8622 | Parp6 |
| Parp8 | 993 | 946 | 921 | 958 | 1483 | 1306 | 1291 | 2682 | 2583 | 2579 | 2766 | 2131 | 2076 | 2126 | 1013 | 966 | 919 | 1160 | 1319 | 1133 | 1356 | 2876 | 2468 | 2218 | 2483 | 1730 | 1909 | 1987 | Parp8 |
| Parp9 | 172 | 129 | 127 | 149 | 137 | 168 | 140 | 363 | 377 | 363 | 471 | 416 | 375 | 385 | 153 | 132 | 114 | 130 | 107 | 184 | 126 | 279 | 395 | 458 | 353 | 564 | 388 | 390 | Parp9 |
| Parpbp | 1714 | 1321 | 1367 | 931 | 641 | 929 | 940 | 1420 | 1244 | 956 | 1034 | 1563 | 1348 | 1545 | 1366 | 1565 | 973 | 739 | 839 | 1121 | 846 | 774 | 1183 | 1160 | 1007 | 1565 | 1680 | 1607 | Parpbp |
| Pars2 | 572 | 603 | 601 | 524 | 461 | 574 | 537 | 574 | 668 | 649 | 561 | 712 | 655 | 666 | 632 | 486 | 590 | 450 | 543 | 549 | 563 | 506 | 674 | 649 | 643 | 790 | 760 | 689 | Pars2 |
| Parva | 9054 | 7763 | 7502 | 5679 | 4583 | 5936 | 4437 | 9110 | 10415 | 9365 | 10535 | 9581 | 10350 | 8986 | 8450 | 9170 | 6412 | 6008 | 4453 | 4068 | 4398 | 8585 | 11672 | 10031 | 7174 | 11453 | 9393 | 10011 | Parva |
| Parvb | 1481 | 1876 | 2794 | 1732 | 2135 | 1629 | 1210 | 1618 | 1778 | 2177 | 2244 | 2160 | 2642 | 2091 | 1652 | 2345 | 2154 | 1819 | 1581 | 1178 | 1563 | 2777 | 1724 | 1869 | 2218 | 2102 | 2328 | 2533 | Parvb |
| Parvg | 106 | 130 | 163 | 123 | 105 | 95 | 61 | 79 | 93 | 96 | 119 | 68 | 95 | 66 | 107 | 185 | 129 | 119 | 92 | 75 | 75 | 67 | 129 | 97 | 93 | 64 | 101 | 89 | Parvg |
| Pask | 4209 | 3978 | 3304 | 3253 | 2748 | 3401 | 3396 | 2523 | 2705 | 2514 | 2351 | 3447 | 3149 | 3621 | 3518 | 3653 | 3423 | 2983 | 3285 | 3266 | 3130 | 2025 | 2465 | 2602 | 2637 | 3145 | 3729 | 3646 | Pask |
| Pate1 | 0 | 0 | 0 | 0 | 0 | 0 | 0 | 0 | 0 | 2 | 0 | 0 | 0 | 0 | 0 | 0 | 0 | 0 | 0 | 0 | 0 | 0 | 0 | 0 | 0 | 0 | 0 | 0 | Pate1 |
| Pate10 | 0 | 1 | 0 | 1 | 3 | 3 | 3 | 4 | 1 | 0 | 1 | 0 | 2 | 1 | 0 | 1 | 1 | 8 | 6 | 0 | 2 | 0 | 1 | 0 | 1 | 0 | 0 | 0 | Pate10 |
| Pate13 | 0 | 0 | 0 | 0 | 1 | 3 | 0 | 1 | 0 | 0 | 1 | 0 | 0 | 3 | 4 | 0 | 0 | 0 | 0 | 2 | 2 | 0 | 3 | 0 | 0 | 4 | 0 | 0 | Pate13 |
| Pate14 | 0 | 0 | 0 | 0 | 0 | 0 | 0 | 0 | 0 | 0 | 0 | 0 | 0 | 0 | 0 | 0 | 0 | 0 | 0 | 0 | 0 | 0 | 0 | 0 | 0 | 0 | 1 | 0 | Pate14 |
| Pate2 | 296 | 282 | 263 | 208 | 220 | 256 | 277 | 210 | 207 | 172 | 146 | 203 | 215 | 229 | 302 | 245 | 277 | 196 | 259 | 272 | 234 | 152 | 181 | 203 | 135 | 189 | 246 | 221 | Pate2 |
| Pate3 | 0 | 0 | 0 | 0 | 0 | 0 | 0 | 0 | 0 | 3 | 0 | 0 | 0 | 0 | 0 | 0 | 0 | 0 | 0 | 0 | 0 | 0 | 0 | 0 | 0 | 0 | 1 | 0 | Pate3 |
| Patj | 1057 | 949 | 780 | 691 | 755 | 857 | 723 | 1813 | 2174 | 1849 | 2007 | 1678 | 1980 | 1800 | 947 | 980 | 754 | 735 | 715 | 767 | 733 | 1594 | 2214 | 1785 | 1586 | 1770 | 1684 | 1756 | Patj |
| Patl1 | 4944 | 5186 | 5371 | 5980 | 4989 | 4793 | 5439 | 4745 | 4481 | 4891 | 4506 | 5579 | 5029 | 5776 | 4970 | 5120 | 5358 | 5810 | 4999 | 4471 | 5373 | 5412 | 4396 | 5071 | 5084 | 5538 | 5089 | 5695 | Patl1 |
| Patl2 | 1 | 1 | 2 | 0 | 0 | 1 | 0 | 0 | 2 | 0 | 1 | 3 | 1 | 1 | 3 | 1 | 1 | 0 | 0 | 0 | 0 | 2 | 1 | 0 | 4 | 0 | 2 | 2 | Patl2 |
| Patz1 | 3196 | 3264 | 3911 | 3552 | 4193 | 3880 | 3655 | 6735 | 6561 | 7599 | 6772 | 9269 | 8046 | 8913 | 3140 | 2979 | 4091 | 3488 | 4408 | 3532 | 3902 | 7765 | 6220 | 8311 | 8250 | 9994 | 8861 | 9646 | Patz1 |
| Pawr | 1691 | 1469 | 1982 | 1721 | 1315 | 1568 | 1469 | 720 | 596 | 603 | 509 | 781 | 743 | 779 | 1667 | 1558 | 1842 | 1576 | 1517 | 1499 | 1398 | 436 | 586 | 838 | 610 | 1167 | 738 | 854 | Pawr |
| Pax1 | 1016 | 334 | 104 | 121 | 93 | 241 | 11 | 724 | 1489 | 776 | 1061 | 330 | 671 | 309 | 425 | 538 | 206 | 259 | 37 | 31 | 12 | 319 | 1078 | 589 | 287 | 463 | 237 | 262 | Pax1 |
| Pax2 | 77 | 155 | 94 | 119 | 1088 | 158 | 29 | 471 | 630 | 1051 | 1036 | 88 | 785 | 38 | 294 | 252 | 180 | 176 | 78 | 73 | 33 | 976 | 632 | 54 | 143 | 93 | 42 | 27 | Pax2 |
| Pax3 | 2064 | 1909 | 3436 | 2308 | 1667 | 1663 | 685 | 2765 | 1583 | 2484 | 1440 | 2068 | 3679 | 2266 | 2010 | 2718 | 3201 | 1492 | 1584 | 1302 | 1366 | 2087 | 2172 | 2389 | 1727 | 1293 | 2836 | 3010 | Pax3 |
| Pax4 | 0 | 0 | 0 | 0 | 0 | 5 | 0 | 4 | 1 | 0 | 0 | 1 | 0 | 3 | 0 | 0 | 0 | 0 | 0 | 0 | 0 | 0 | 6 | 2 | 0 | 0 | 0 | 0 | Pax4 |
| Pax5 | 1523 | 1792 | 2774 | 2837 | 2463 | 2844 | 777 | 1793 | 1411 | 2004 | 1596 | 3129 | 4240 | 1961 | 1458 | 2501 | 3121 | 2374 | 3239 | 1907 | 2295 | 2966 | 805 | 1994 | 2893 | 63 | 3258 | 2219 | Pax5 |
| Pax6 | 6384 | 6683 | 8301 | 7404 | 5974 | 7069 | 9268 | 5964 | 5183 | 5841 | 4324 | 8795 | 5235 | 9470 | 6331 | 3789 | 7205 | 6401 | 8055 | 7830 | 7492 | 4223 | 4066 | 7793 | 7134 | 10717 | 8477 | 9620 | Pax6 |
| Pax7 | 646 | 558 | 1427 | 1094 | 1028 | 1009 | 443 | 1463 | 899 | 1836 | 1339 | 1721 | 2122 | 1441 | 487 | 871 | 1279 | 809 | 1367 | 790 | 1004 | 2015 | 1101 | 2084 | 1990 | 770 | 1478 | 1651 | Pax7 |
| Pax8 | 98 | 111 | 128 | 94 | 518 | 129 | 28 | 335 | 404 | 440 | 475 | 152 | 557 | 65 | 272 | 224 | 117 | 162 | 132 | 87 | 80 | 403 | 333 | 85 | 128 | 9 | 146 | 80 | Pax8 |
| Pax9 | 574 | 130 | 52 | 186 | 56 | 29 | 18 | 915 | 728 | 897 | 722 | 7 | 60 | 23 | 117 | 516 | 75 | 73 | 10 | 12 | 8 | 234 | 678 | 713 | 36 | 1099 | 34 | 7 | Pax9 |
| Paxbp1 | 14141 | 11574 | 10625 | 12361 | 12952 | 14169 | 14941 | 15663 | 12817 | 12942 | 13447 | 14202 | 13199 | 14751 | 13992 | 12364 | 11282 | 12553 | 12272 | 14101 | 14761 | 14302 | 12628 | 12654 | 13690 | 11710 | 13974 | 12978 | Paxbp1 |
| Paxip1 | 5509 | 5859 | 5364 | 5749 | 5302 | 5448 | 5387 | 4484 | 4639 | 4690 | 4508 | 5856 | 5173 | 5460 | 4944 | 5423 | 5836 | 5478 | 5857 | 4970 | 5397 | 4336 | 4286 | 5247 | 4827 | 5930 | 5703 | 5550 | Paxip1 |
| Paxx | 533 | 625 | 565 | 715 | 1049 | 853 | 831 | 927 | 1047 | 1062 | 1121 | 1070 | 902 | 942 | 555 | 519 | 656 | 757 | 831 | 784 | 826 | 1240 | 1056 | 1042 | 1236 | 872 | 953 | 828 | Paxx |
| Pbdc1 | 3434 | 3257 | 3725 | 3009 | 2344 | 2315 | 2920 | 2678 | 2273 | 2116 | 2162 | 2673 | 2685 | 3002 | 4158 | 3794 | 3972 | 2922 | 2833 | 3772 | 3092 | 2593 | 3016 | 3348 | 2792 | 4683 | 4104 | 4486 | Pbdc1 |
| Pbk | 4570 | 3715 | 4962 | 3638 | 2928 | 3717 | 4529 | 5078 | 4397 | 4048 | 3577 | 6857 | 5305 | 7280 | 4556 | 3626 | 4357 | 3349 | 4126 | 4365 | 3904 | 3322 | 3580 | 5701 | 5150 | 7854 | 6933 | 7444 | Pbk |
| Pbld1 | 24 | 38 | 12 | 33 | 45 | 41 | 22 | 12 | 21 | 5 | 28 | 13 | 11 | 13 | 26 | 25 | 35 | 30 | 34 | 27 | 43 | 27 | 19 | 18 | 25 | 22 | 31 | 8 | Pbld1 |
| Pbld2 | 98 | 99 | 89 | 163 | 99 | 79 | 93 | 106 | 109 | 184 | 139 | 127 | 127 | 139 | 104 | 118 | 107 | 113 | 124 | 106 | 105 | 132 | 139 | 146 | 127 | 177 | 148 | 123 | Pbld2 |
| Pbp2 | 1 | 0 | 0 | 0 | 0 | 0 | 0 | 0 | 2 | 3 | 2 | 0 | 0 | 1 | 0 | 0 | 0 | 1 | 0 | 0 | 2 | 0 | 1 | 0 | 0 | 0 | 0 | 1 | Pbp2 |
| Pbrm1 | 11726 | 9631 | 8641 | 9232 | 10637 | 12290 | 11364 | 22666 | 20534 | 19163 | 20197 | 23661 | 22416 | 23812 | 11780 | 10489 | 9312 | 9250 | 10227 | 11474 | 11449 | 18783 | 20035 | 18309 | 21422 | 15759 | 25400 | 23260 | Pbrm1 |
| Pbx1 | 22758 | 19593 | 18363 | 23284 | 23805 | 25314 | 27336 | 30337 | 27414 | 26350 | 25839 | 28760 | 26525 | 29766 | 22400 | 19156 | 21489 | 22394 | 25593 | 24145 | 27554 | 29604 | 24654 | 28714 | 31848 | 22884 | 29230 | 29265 | Pbx1 |
| Pbx2 | 5852 | 6374 | 7851 | 5502 | 5083 | 5229 | 5108 | 7552 | 7629 | 8231 | 7883 | 10627 | 9331 | 10702 | 5326 | 5552 | 6757 | 5043 | 5686 | 4777 | 5252 | 7454 | 7288 | 9707 | 8317 | 12630 | 11087 | 11703 | Pbx2 |
| Pbx3 | 5577 | 6805 | 6674 | 8992 | 10028 | 8990 | 7518 | 10367 | 8745 | 11008 | 9071 | 8767 | 10204 | 8813 | 7692 | 6315 | 9065 | 8685 | 8533 | 8251 | 8194 | 12183 | 8541 | 8529 | 10295 | 7372 | 7971 | 8777 | Pbx3 |
| Pbx4 | 1087 | 1428 | 1332 | 1918 | 1667 | 1484 | 1557 | 526 | 541 | 737 | 529 | 853 | 735 | 804 | 1042 | 1217 | 1694 | 1796 | 1933 | 1485 | 1526 | 703 | 509 | 815 | 856 | 747 | 815 | 784 | Pbx4 |
| Pbxip1 | 3252 | 4080 | 4553 | 3603 | 3175 | 3380 | 2532 | 1832 | 2419 | 2354 | 3145 | 2384 | 3014 | 1966 | 3303 | 4377 | 4222 | 3723 | 2914 | 2455 | 2673 | 2394 | 3029 | 2125 | 2081 | 2592 | 2373 | 2155 | Pbxip1 |
| Pcare | 0 | 0 | 0 | 0 | 0 | 0 | 0 | 0 | 2 | 0 | 0 | 0 | 1 | 0 | 1 | 0 | 0 | 0 | 2 | 0 | 0 | 2 | 6 | 0 | 1 | 1 | 0 | 0 | Pcare |
| Pcbd1 | 213 | 231 | 233 | 179 | 220 | 144 | 247 | 398 | 270 | 301 | 298 | 351 | 358 | 414 | 243 | 209 | 263 | 184 | 215 | 230 | 257 | 426 | 287 | 301 | 273 | 342 | 361 | 438 | Pcbd1 |
| Pcbd2 | 172 | 178 | 250 | 212 | 178 | 136 | 159 | 210 | 216 | 236 | 248 | 206 | 251 | 262 | 167 | 132 | 229 | 142 | 157 | 175 | 181 | 115 | 192 | 277 | 213 | 271 | 217 | 255 | Pcbd2 |
| Pcbp1 | 9006 | 10186 | 12462 | 10475 | 10119 | 8693 | 9404 | 13681 | 14664 | 15887 | 15365 | 18065 | 15188 | 18535 | 8283 | 8213 | 12090 | 9009 | 11543 | 8189 | 10094 | 15020 | 14557 | 19002 | 15985 | 21117 | 16210 | 18314 | Pcbp1 |
| Pcbp2 | 21856 | 19431 | 24932 | 22838 | 20197 | 18920 | 23026 | 24082 | 21777 | 24953 | 24098 | 27999 | 24384 | 31544 | 20226 | 18291 | 24649 | 21478 | 22589 | 17014 | 24843 | 27818 | 21711 | 28079 | 25328 | 33333 | 25134 | 32660 | Pcbp2 |
| Pcbp3 | 3500 | 4701 | 5511 | 4629 | 4666 | 3844 | 3664 | 1726 | 2054 | 2213 | 2396 | 2315 | 2223 | 2107 | 3918 | 4365 | 5416 | 4562 | 4546 | 3204 | 3948 | 2453 | 1936 | 2105 | 2291 | 2235 | 2029 | 2218 | Pcbp3 |
| Pcbp4 | 12032 | 15236 | 14969 | 13318 | 12974 | 12841 | 11134 | 13998 | 13899 | 15852 | 13194 | 17932 | 15490 | 16001 | 12101 | 12026 | 14748 | 11334 | 14727 | 13104 | 11853 | 15319 | 12984 | 16814 | 16526 | 18617 | 17325 | 16702 | Pcbp4 |
| Pcca | 3400 | 3356 | 3142 | 2809 | 2216 | 2384 | 2476 | 2113 | 1789 | 1594 | 1626 | 2039 | 1987 | 2007 | 3616 | 3401 | 2790 | 2725 | 2413 | 2804 | 2455 | 1689 | 1912 | 1693 | 1744 | 2202 | 2026 | 2149 | Pcca |
| Pccb | 1816 | 1829 | 1824 | 1836 | 1807 | 2109 | 1662 | 2803 | 3180 | 3171 | 3449 | 3113 | 2899 | 2947 | 1693 | 1812 | 1827 | 1699 | 1894 | 1764 | 1791 | 3148 | 3290 | 3398 | 3269 | 3375 | 3113 | 3049 | Pccb |
| Pcdh1 | 1992 | 2636 | 2864 | 3545 | 5034 | 3760 | 3385 | 2342 | 2957 | 4360 | 3629 | 3038 | 2689 | 2950 | 1730 | 2540 | 3789 | 4307 | 4789 | 2574 | 3753 | 4630 | 2856 | 4184 | 4438 | 3210 | 2488 | 2351 | Pcdh1 |
| Pcdh10 | 8569 | 8193 | 6560 | 11614 | 12629 | 12413 | 13218 | 7845 | 7796 | 8253 | 7412 | 6393 | 5489 | 6692 | 8919 | 6797 | 9108 | 12178 | 11881 | 11025 | 14148 | 9738 | 6513 | 7026 | 9262 | 4590 | 5521 | 5516 | Pcdh10 |
| Pcdh11x | 1111 | 841 | 608 | 1231 | 1568 | 1548 | 1730 | 1638 | 1352 | 1034 | 1083 | 741 | 667 | 808 | 1407 | 910 | 815 | 1268 | 1286 | 1702 | 1700 | 1511 | 1236 | 870 | 1231 | 591 | 841 | 708 | Pcdh11x |
| Pcdh12 | 822 | 891 | 756 | 734 | 735 | 827 | 704 | 760 | 853 | 888 | 955 | 737 | 878 | 774 | 750 | 848 | 654 | 671 | 815 | 739 | 817 | 772 | 919 | 905 | 858 | 741 | 857 | 745 | Pcdh12 |
| Pcdh15 | 551 | 583 | 184 | 637 | 1009 | 835 | 821 | 1083 | 1096 | 882 | 847 | 548 | 836 | 487 | 760 | 823 | 477 | 727 | 702 | 976 | 883 | 934 | 889 | 398 | 813 | 425 | 397 | 415 | Pcdh15 |
| Pcdh17 | 4737 | 4526 | 5018 | 7892 | 8739 | 7901 | 8406 | 4030 | 3909 | 4999 | 4457 | 3217 | 3497 | 3150 | 5347 | 5102 | 6313 | 7937 | 7825 | 7105 | 8539 | 5638 | 3808 | 3608 | 5002 | 2477 | 2839 | 2807 | Pcdh17 |
| Pcdh18 | 8240 | 6221 | 6586 | 4662 | 4502 | 5341 | 4168 | 10232 | 9795 | 8413 | 10670 | 8416 | 11208 | 8764 | 7119 | 7764 | 5295 | 5055 | 4517 | 5053 | 4561 | 6945 | 10389 | 8545 | 6450 | 9068 | 10318 | 10239 | Pcdh18 |
| Pcdh19 | 7818 | 6479 | 7033 | 11072 | 10438 | 11811 | 13642 | 5917 | 6091 | 7528 | 6900 | 9036 | 7313 | 9556 | 7514 | 6763 | 8820 | 10524 | 11536 | 9658 | 12719 | 7356 | 6229 | 8062 | 9179 | 8707 | 8125 | 9274 | Pcdh19 |
| Pcdh20 | 736 | 670 | 678 | 614 | 695 | 689 | 721 | 476 | 489 | 385 | 310 | 238 | 184 | 222 | 833 | 769 | 620 | 612 | 648 | 890 | 766 | 376 | 374 | 343 | 400 | 268 | 222 | 190 | Pcdh20 |
| Pcdh7 | 3265 | 2268 | 2421 | 2815 | 4260 | 3702 | 3620 | 4511 | 4937 | 4636 | 4545 | 2722 | 3797 | 2974 | 3182 | 2974 | 2319 | 3035 | 3131 | 3196 | 3504 | 4585 | 4887 | 3564 | 3900 | 2497 | 3130 | 2900 | Pcdh7 |
| Pcdh8 | 1555 | 2328 | 3184 | 3560 | 3305 | 2658 | 2790 | 1884 | 1828 | 2541 | 1950 | 2712 | 2174 | 2342 | 1921 | 1970 | 3438 | 3750 | 3473 | 2397 | 2979 | 3136 | 1651 | 2714 | 2658 | 1965 | 2501 | 2586 | Pcdh8 |
| Pcdh9 | 2881 | 2713 | 1959 | 2715 | 3764 | 3399 | 3418 | 4446 | 4120 | 3582 | 3301 | 2677 | 2971 | 2681 | 3498 | 2781 | 2103 | 3119 | 2898 | 3716 | 3245 | 4229 | 3796 | 2451 | 3693 | 1730 | 2487 | 2299 | Pcdh9 |
| Pcdha1 | 3 | 7 | 6 | 6 | 42 | 7 | 17 | 11 | 13 | 18 | 20 | 8 | 17 | 6 | 4 | 7 | 5 | 24 | 15 | 22 | 19 | 7 | 20 | 7 | 18 | 4 | 7 | 6 | Pcdha1 |
| Pcdha10 | 23 | 41 | 23 | 54 | 46 | 40 | 26 | 92 | 87 | 85 | 105 | 84 | 51 | 63 | 42 | 35 | 21 | 47 | 52 | 56 | 80 | 169 | 93 | 65 | 123 | 27 | 67 | 72 | Pcdha10 |
| Pcdha11 | 53 | 32 | 31 | 61 | 129 | 57 | 86 | 194 | 171 | 169 | 163 | 126 | 84 | 128 | 80 | 42 | 51 | 74 | 90 | 104 | 130 | 251 | 189 | 150 | 201 | 59 | 136 | 125 | Pcdha11 |
| Pcdha12 | 66 | 66 | 55 | 143 | 154 | 126 | 122 | 197 | 223 | 190 | 188 | 148 | 144 | 180 | 115 | 99 | 123 | 133 | 188 | 222 | 223 | 256 | 212 | 174 | 294 | 106 | 117 | 171 | Pcdha12 |
| Pcdha2 | 30 | 49 | 23 | 68 | 113 | 95 | 56 | 154 | 142 | 195 | 136 | 71 | 100 | 92 | 66 | 45 | 61 | 91 | 69 | 101 | 112 | 241 | 168 | 95 | 179 | 27 | 93 | 84 | Pcdha2 |
| Pcdha3 | 35 | 49 | 57 | 64 | 151 | 98 | 88 | 176 | 181 | 186 | 164 | 115 | 133 | 130 | 51 | 50 | 53 | 78 | 95 | 99 | 94 | 224 | 204 | 118 | 212 | 69 | 100 | 101 | Pcdha3 |

| | | | | | | | | | | | | | | | | | | | | | | | | | | | | | |
|---|---|---|---|---|---|---|---|---|---|---|---|---|---|---|---|---|---|---|---|---|---|---|---|---|---|---|---|---|---|
| Pcdha4 | 0 | 0 | 0 | 1 | 0 | 0 | 0 | 2 | 1 | 0 | 0 | 1 | 0 | 2 | 0 | 0 | 0 | 0 | 3 | 0 | 0 | 0 | 0 | 0 | 3 | 0 | 0 | 2 | Pcdha4 |
| Pcdha5 | 20 | 29 | 25 | 29 | 121 | 82 | 84 | 126 | 182 | 164 | 160 | 89 | 104 | 106 | 62 | 39 | 54 | 75 | 75 | 76 | 83 | 224 | 164 | 123 | 206 | 71 | 101 | 87 | Pcdha5 |
| Pcdha6 | 24 | 27 | 25 | 38 | 110 | 89 | 76 | 145 | 143 | 147 | 101 | 73 | 74 | 84 | 36 | 37 | 59 | 57 | 61 | 114 | 74 | 175 | 153 | 67 | 138 | 33 | 62 | 55 | Pcdha6 |
| Pcdha7 | 38 | 59 | 27 | 64 | 124 | 100 | 114 | 128 | 192 | 148 | 189 | 122 | 110 | 111 | 59 | 43 | 48 | 92 | 82 | 100 | 108 | 289 | 188 | 161 | 217 | 57 | 99 | 63 | Pcdha7 |
| Pcdha8 | 2 | 4 | 0 | 6 | 12 | 12 | 8 | 24 | 11 | 13 | 27 | 23 | 5 | 11 | 7 | 5 | 13 | 3 | 9 | 12 | 20 | 20 | 14 | 11 | 19 | 13 | 13 | 7 | Pcdha8 |
| Pcdha9 | 50 | 61 | 32 | 93 | 160 | 175 | 123 | 204 | 226 | 181 | 213 | 117 | 150 | 143 | 73 | 46 | 78 | 96 | 135 | 140 | 128 | 257 | 177 | 133 | 253 | 56 | 117 | 96 | Pcdha9 |
| Pcdhac1 | 83 | 113 | 60 | 122 | 174 | 166 | 162 | 155 | 155 | 146 | 108 | 111 | 70 | 134 | 120 | 107 | 74 | 129 | 159 | 152 | 191 | 102 | 96 | 162 | 138 | 96 | 72 | 72 | Pcdhac1 |
| Pcdhac2 | 468 | 442 | 319 | 547 | 790 | 705 | 681 | 541 | 672 | 622 | 493 | 452 | 353 | 440 | 349 | 307 | 370 | 605 | 739 | 627 | 695 | 480 | 572 | 518 | 649 | 316 | 306 | 332 | Pcdhac2 |
| Pcdhb1 | 5 | 0 | 0 | 5 | 2 | 0 | 0 | 15 | 7 | 5 | 18 | 0 | 0 | 0 | 1 | 2 | 1 | 0 | 0 | 0 | 0 | 0 | 9 | 13 | 0 | 5 | 0 | 1 | Pcdhb1 |
| Pcdhb10 | 42 | 30 | 60 | 42 | 81 | 60 | 42 | 158 | 183 | 180 | 162 | 125 | 141 | 113 | 46 | 39 | 50 | 56 | 54 | 43 | 68 | 172 | 219 | 146 | 214 | 107 | 117 | 125 | Pcdhb10 |
| Pcdhb11 | 43 | 48 | 36 | 51 | 139 | 86 | 88 | 201 | 213 | 183 | 221 | 129 | 130 | 142 | 57 | 40 | 54 | 85 | 75 | 146 | 70 | 328 | 333 | 151 | 189 | 118 | 85 | 155 | Pcdhb11 |
| Pcdhb12 | 18 | 21 | 23 | 42 | 32 | 33 | 37 | 118 | 120 | 83 | 95 | 43 | 88 | 77 | 25 | 20 | 22 | 37 | 36 | 63 | 46 | 139 | 121 | 104 | 142 | 75 | 72 | 52 | Pcdhb12 |
| Pcdhb13 | 10 | 14 | 15 | 4 | 61 | 45 | 19 | 70 | 85 | 78 | 105 | 34 | 41 | 59 | 20 | 18 | 21 | 34 | 30 | 32 | 26 | 122 | 84 | 75 | 89 | 41 | 62 | 28 | Pcdhb13 |
| Pcdhb14 | 43 | 40 | 43 | 77 | 93 | 79 | 48 | 200 | 213 | 172 | 212 | 101 | 137 | 100 | 73 | 47 | 54 | 57 | 61 | 90 | 70 | 244 | 267 | 136 | 178 | 123 | 91 | 112 | Pcdhb14 |
| Pcdhb15 | 28 | 16 | 13 | 22 | 32 | 30 | 27 | 115 | 156 | 113 | 112 | 76 | 76 | 100 | 41 | 26 | 28 | 28 | 45 | 49 | 51 | 99 | 178 | 69 | 98 | 52 | 83 | 46 | Pcdhb15 |
| Pcdhb16 | 89 | 79 | 45 | 68 | 125 | 170 | 143 | 531 | 577 | 310 | 374 | 202 | 234 | 245 | 145 | 99 | 73 | 107 | 91 | 197 | 152 | 409 | 568 | 204 | 372 | 105 | 282 | 199 | Pcdhb16 |
| Pcdhb17 | 138 | 151 | 146 | 221 | 409 | 270 | 245 | 832 | 849 | 679 | 798 | 487 | 595 | 592 | 243 | 157 | 196 | 295 | 288 | 404 | 318 | 1014 | 890 | 657 | 797 | 426 | 479 | 551 | Pcdhb17 |
| Pcdhb18 | 90 | 75 | 47 | 77 | 208 | 193 | 124 | 496 | 425 | 314 | 332 | 191 | 245 | 186 | 121 | 86 | 124 | 87 | 131 | 175 | 175 | 473 | 446 | 249 | 301 | 198 | 220 | 191 | Pcdhb18 |
| Pcdhb19 | 157 | 162 | 59 | 178 | 371 | 325 | 230 | 891 | 945 | 705 | 774 | 411 | 567 | 473 | 245 | 203 | 114 | 213 | 247 | 334 | 329 | 825 | 891 | 449 | 851 | 198 | 487 | 420 | Pcdhb19 |
| Pcdhb2 | 38 | 25 | 20 | 53 | 70 | 41 | 51 | 182 | 176 | 226 | 151 | 86 | 100 | 144 | 47 | 30 | 22 | 48 | 48 | 54 | 67 | 120 | 155 | 99 | 188 | 33 | 114 | 91 | Pcdhb2 |
| Pcdhb20 | 105 | 109 | 89 | 203 | 366 | 239 | 219 | 623 | 629 | 688 | 618 | 368 | 443 | 411 | 182 | 141 | 151 | 295 | 290 | 258 | 290 | 906 | 646 | 585 | 740 | 401 | 342 | 463 | Pcdhb20 |
| Pcdhb21 | 70 | 75 | 24 | 71 | 115 | 142 | 75 | 341 | 311 | 265 | 236 | 128 | 222 | 147 | 102 | 86 | 45 | 69 | 60 | 122 | 128 | 227 | 359 | 132 | 301 | 51 | 157 | 139 | Pcdhb21 |
| Pcdhb22 | 216 | 190 | 226 | 311 | 496 | 422 | 288 | 677 | 712 | 697 | 822 | 427 | 562 | 490 | 281 | 287 | 234 | 390 | 335 | 408 | 358 | 1041 | 821 | 649 | 705 | 469 | 545 | 457 | Pcdhb22 |
| Pcdhb3 | 60 | 72 | 63 | 77 | 132 | 110 | 104 | 315 | 306 | 311 | 340 | 200 | 240 | 222 | 113 | 69 | 93 | 113 | 130 | 109 | 116 | 421 | 366 | 264 | 319 | 219 | 246 | 235 | Pcdhb3 |
| Pcdhb4 | 103 | 93 | 52 | 95 | 122 | 90 | 110 | 319 | 346 | 306 | 187 | 204 | 256 | 213 | 103 | 50 | 65 | 99 | 77 | 169 | 112 | 354 | 276 | 302 | 259 | 161 | 235 | 236 | Pcdhb4 |
| Pcdhb5 | 93 | 84 | 61 | 96 | 198 | 145 | 145 | 358 | 410 | 323 | 281 | 222 | 362 | 305 | 107 | 93 | 67 | 123 | 147 | 119 | 177 | 443 | 340 | 309 | 352 | 222 | 268 | 249 | Pcdhb5 |
| Pcdhb6 | 53 | 57 | 37 | 48 | 79 | 61 | 68 | 195 | 267 | 183 | 190 | 90 | 133 | 133 | 45 | 62 | 61 | 55 | 61 | 94 | 57 | 219 | 254 | 146 | 195 | 116 | 162 | 117 | Pcdhb6 |
| Pcdhb7 | 170 | 129 | 132 | 195 | 363 | 299 | 275 | 808 | 732 | 598 | 590 | 506 | 409 | 515 | 210 | 145 | 188 | 266 | 270 | 340 | 320 | 715 | 647 | 575 | 640 | 429 | 403 | 462 | Pcdhb7 |
| Pcdhb8 | 89 | 54 | 62 | 70 | 106 | 112 | 101 | 380 | 298 | 249 | 282 | 200 | 181 | 198 | 92 | 70 | 81 | 74 | 103 | 138 | 136 | 346 | 334 | 311 | 296 | 202 | 223 | 201 | Pcdhb8 |
| Pcdhb9 | 70 | 59 | 53 | 86 | 154 | 102 | 99 | 348 | 409 | 306 | 312 | 193 | 242 | 206 | 99 | 53 | 63 | 102 | 135 | 131 | 130 | 434 | 394 | 278 | 340 | 191 | 214 | 178 | Pcdhb9 |
| Pcdhga1 | 150 | 129 | 138 | 160 | 253 | 161 | 170 | 297 | 367 | 361 | 325 | 247 | 243 | 248 | 174 | 149 | 108 | 156 | 165 | 189 | 177 | 326 | 326 | 256 | 228 | 151 | 238 | 196 | Pcdhga1 |
| Pcdhga10 | 314 | 390 | 381 | 385 | 488 | 397 | 339 | 491 | 616 | 673 | 623 | 613 | 695 | 634 | 380 | 357 | 424 | 468 | 499 | 365 | 383 | 749 | 679 | 652 | 705 | 539 | 605 | 638 | Pcdhga10 |
| Pcdhga11 | 480 | 530 | 449 | 590 | 819 | 607 | 731 | 1045 | 1118 | 1361 | 1219 | 1295 | 1245 | 1359 | 509 | 478 | 571 | 657 | 711 | 640 | 823 | 1612 | 1117 | 1352 | 1603 | 1117 | 1191 | 1272 | Pcdhga11 |
| Pcdhga12 | 322 | 360 | 563 | 608 | 704 | 535 | 580 | 548 | 702 | 975 | 842 | 1075 | 818 | 841 | 324 | 394 | 685 | 642 | 778 | 464 | 674 | 1009 | 590 | 1133 | 1095 | 980 | 779 | 845 | Pcdhga12 |
| Pcdhga2 | 366 | 408 | 474 | 466 | 711 | 615 | 486 | 573 | 714 | 741 | 760 | 633 | 827 | 627 | 385 | 456 | 431 | 612 | 526 | 442 | 470 | 924 | 685 | 561 | 755 | 408 | 580 | 593 | Pcdhga2 |
| Pcdhga3 | 438 | 469 | 458 | 528 | 731 | 800 | 573 | 708 | 822 | 853 | 857 | 722 | 857 | 685 | 527 | 503 | 430 | 514 | 553 | 469 | 592 | 1063 | 833 | 688 | 904 | 561 | 758 | 682 | Pcdhga3 |
| Pcdhga4 | 164 | 203 | 180 | 248 | 266 | 273 | 253 | 335 | 368 | 374 | 394 | 428 | 440 | 367 | 187 | 185 | 172 | 242 | 341 | 211 | 285 | 493 | 349 | 385 | 424 | 251 | 354 | 295 | Pcdhga4 |
| Pcdhga5 | 388 | 411 | 347 | 419 | 520 | 456 | 415 | 473 | 540 | 628 | 583 | 590 | 505 | 552 | 330 | 347 | 355 | 375 | 474 | 364 | 461 | 645 | 511 | 555 | 558 | 415 | 540 | 500 | Pcdhga5 |
| Pcdhga6 | 226 | 253 | 176 | 268 | 377 | 311 | 292 | 518 | 498 | 482 | 569 | 544 | 399 | 536 | 258 | 216 | 240 | 315 | 365 | 329 | 345 | 602 | 618 | 574 | 696 | 397 | 453 | 483 | Pcdhga6 |
| Pcdhga7 | 354 | 419 | 328 | 456 | 542 | 505 | 491 | 669 | 640 | 756 | 821 | 620 | 776 | 678 | 406 | 403 | 402 | 468 | 440 | 369 | 545 | 956 | 778 | 778 | 787 | 588 | 730 | 646 | Pcdhga7 |
| Pcdhga8 | 212 | 215 | 249 | 293 | 330 | 291 | 296 | 347 | 370 | 434 | 369 | 399 | 432 | 396 | 208 | 203 | 290 | 341 | 335 | 295 | 365 | 543 | 375 | 383 | 458 | 247 | 381 | 320 | Pcdhga8 |
| Pcdhga9 | 180 | 215 | 199 | 227 | 301 | 320 | 201 | 308 | 370 | 448 | 397 | 374 | 361 | 343 | 220 | 194 | 248 | 291 | 274 | 236 | 275 | 556 | 362 | 455 | 498 | 373 | 409 | 387 | Pcdhga9 |
| Pcdhgb1 | 171 | 174 | 147 | 216 | 277 | 360 | 245 | 303 | 410 | 438 | 447 | 416 | 388 | 319 | 183 | 138 | 190 | 223 | 217 | 220 | 265 | 495 | 364 | 320 | 389 | 209 | 332 | 292 | Pcdhgb1 |
| Pcdhgb2 | 308 | 300 | 242 | 350 | 421 | 389 | 332 | 477 | 505 | 641 | 511 | 592 | 577 | 461 | 285 | 266 | 290 | 337 | 372 | 291 | 372 | 642 | 432 | 555 | 572 | 455 | 452 | 457 | Pcdhgb2 |
| Pcdhgb4 | 333 | 277 | 265 | 272 | 255 | 344 | 272 | 299 | 434 | 517 | 496 | 449 | 466 | 456 | 295 | 217 | 256 | 283 | 282 | 258 | 277 | 444 | 424 | 459 | 472 | 317 | 400 | 318 | Pcdhgb4 |
| Pcdhgb5 | 268 | 303 | 278 | 322 | 430 | 437 | 345 | 462 | 526 | 611 | 582 | 547 | 524 | 603 | 305 | 236 | 362 | 420 | 432 | 328 | 387 | 715 | 560 | 626 | 599 | 494 | 644 | 598 | Pcdhgb5 |
| Pcdhgb6 | 607 | 655 | 678 | 862 | 1186 | 914 | 866 | 1088 | 1230 | 1489 | 1513 | 1410 | 1338 | 1198 | 628 | 694 | 797 | 924 | 1057 | 718 | 939 | 1693 | 1274 | 1509 | 1478 | 1189 | 1039 | 1243 | Pcdhgb6 |
| Pcdhgb7 | 374 | 338 | 367 | 412 | 538 | 502 | 421 | 646 | 769 | 793 | 766 | 841 | 618 | 679 | 404 | 396 | 397 | 478 | 474 | 344 | 444 | 844 | 750 | 721 | 830 | 657 | 614 | 553 | Pcdhgb7 |
| Pcdhgc3 | 5505 | 6344 | 7139 | 8502 | 9218 | 7424 | 7348 | 4498 | 5365 | 6861 | 6316 | 5828 | 4986 | 4921 | 5054 | 5952 | 7599 | 8615 | 9028 | 5749 | 8023 | 6184 | 5429 | 6299 | 6223 | 5475 | 4392 | 4405 | Pcdhgc3 |
| Pcdhgc4 | 781 | 942 | 781 | 1538 | 1803 | 1389 | 1373 | 860 | 976 | 1130 | 1025 | 770 | 764 | 767 | 778 | 859 | 1036 | 1467 | 1523 | 1178 | 1544 | 1337 | 887 | 947 | 1293 | 583 | 664 | 675 | Pcdhgc4 |
| Pcdhgc5 | 158 | 194 | 179 | 292 | 376 | 310 | 286 | 258 | 302 | 333 | 372 | 419 | 348 | 313 | 166 | 191 | 316 | 328 | 358 | 218 | 286 | 439 | 284 | 363 | 503 | 202 | 329 | 277 | Pcdhgc5 |
| Pced1a | 707 | 871 | 832 | 828 | 947 | 843 | 865 | 1627 | 1912 | 2011 | 1931 | 1778 | 1845 | 1853 | 802 | 807 | 907 | 879 | 952 | 862 | 861 | 1900 | 1996 | 1977 | 1967 | 1966 | 1875 | 1926 | Pced1a |
| Pced1b | 102 | 126 | 210 | 135 | 171 | 93 | 121 | 151 | 173 | 281 | 187 | 219 | 320 | 230 | 113 | 127 | 166 | 177 | 186 | 128 | 132 | 261 | 210 | 230 | 249 | 323 | 258 | 284 | Pced1b |
| Pcf11 | 6628 | 5968 | 5841 | 6231 | 6091 | 6094 | 6365 | 8241 | 6715 | 6331 | 6929 | 7140 | 7182 | 7451 | 6538 | 6397 | 5742 | 6245 | 5978 | 6624 | 6105 | 6766 | 6735 | 6684 | 7015 | 6382 | 7217 | 6857 | Pcf11 |
| Pcgf1 | 592 | 672 | 634 | 694 | 743 | 649 | 632 | 910 | 870 | 899 | 841 | 936 | 921 | 871 | 658 | 669 | 718 | 632 | 743 | 738 | 626 | 866 | 927 | 935 | 873 | 993 | 844 | 919 | Pcgf1 |
| Pcgf2 | 2632 | 3111 | 3866 | 4174 | 5303 | 4271 | 3928 | 5117 | 6072 | 7384 | 6503 | 7400 | 6693 | 7056 | 2517 | 2785 | 4230 | 4064 | 5135 | 3581 | 4181 | 7305 | 5398 | 7544 | 7563 | 7302 | 6349 | 6553 | Pcgf2 |
| Pcgf3 | 8612 | 7787 | 8715 | 9421 | 9684 | 9189 | 9382 | 11144 | 10210 | 11333 | 10541 | 11943 | 10783 | 11697 | 8312 | 8141 | 9649 | 8771 | 9589 | 7979 | 9521 | 12337 | 9878 | 11573 | 12259 | 10338 | 11376 | 11516 | Pcgf3 |
| Pcgf5 | 1804 | 1634 | 1628 | 2253 | 2219 | 2354 | 2995 | 1603 | 1438 | 1470 | 1242 | 1816 | 1484 | 1911 | 1878 | 1738 | 1888 | 2388 | 2494 | 2509 | 2647 | 1111 | 1235 | 1660 | 1791 | 1573 | 1627 | 1773 | Pcgf5 |
| Pcgf6 | 4121 | 3626 | 3790 | 4762 | 5293 | 4858 | 5417 | 4086 | 3515 | 3883 | 3826 | 4120 | 3783 | 4189 | 4174 | 3790 | 4218 | 5317 | 5168 | 4868 | 5470 | 4513 | 3364 | 3891 | 4292 | 3696 | 3927 | 4103 | Pcgf6 |
| Pcid2 | 2623 | 2896 | 3078 | 2756 | 2468 | 2439 | 2581 | 2676 | 3035 | 3220 | 3112 | 3716 | 3223 | 3444 | 2488 | 2589 | 2897 | 2571 | 2818 | 2257 | 2715 | 2996 | 2748 | 3311 | 3394 | 3919 | 3467 | 3728 | Pcid2 |
| Pcif1 | 2936 | 3456 | 3708 | 3654 | 3951 | 3764 | 3362 | 4226 | 5041 | 5310 | 4925 | 6000 | 5007 | 5568 | 2686 | 2838 | 3690 | 3536 | 4454 | 3184 | 3593 | 5223 | 4564 | 5942 | 5743 | 6065 | 5160 | 5245 | Pcif1 |
| Pck1 | 0 | 0 | 0 | 1 | 6 | 0 | 2 | 2 | 0 | 2 | 1 | 0 | 0 | 0 | 0 | 1 | 1 | 3 | 2 | 0 | 2 | 0 | 1 | 1 | 0 | 1 | 2 | 0 | Pck1 |
| Pck2 | 889 | 1015 | 1228 | 866 | 880 | 970 | 707 | 1093 | 1500 | 1391 | 1718 | 1360 | 1396 | 1349 | 800 | 1115 | 951 | 927 | 902 | 762 | 821 | 1203 | 1612 | 1517 | 1400 | 1569 | 1378 | 1337 | Pck2 |
| Pclaf | 14244 | 9728 | 14586 | 9042 | 6293 | 7919 | 10393 | 9653 | 8864 | 8112 | 7701 | 12687 | 10260 | 13154 | 12150 | 10893 | 11641 | 7673 | 8736 | 9977 | 8401 | 6193 | 7937 | 10848 | 8469 | 17000 | 13330 | 14325 | Pclaf |
| Pclo | 3513 | 3533 | 1956 | 5153 | 7006 | 7305 | 6862 | 6018 | 5949 | 5873 | 5580 | 3358 | 4012 | 3882 | 4152 | 3766 | 3216 | 5374 | 5434 | 6552 | 7969 | 5842 | 5007 | 3635 | 6236 | 1972 | 3833 | 2908 | Pclo |
| Pcm1 | 18718 | 13694 | 11176 | 13491 | 15863 | 18368 | 18213 | 24976 | 21368 | 19417 | 19340 | 22024 | 20747 | 22225 | 18842 | 15157 | 12969 | 13671 | 15256 | 18396 | 17823 | 18284 | 20179 | 17079 | 19680 | 14049 | 23121 | 20842 | Pcm1 |
| Pcmt1 | 6 | 5 | 6 | 4 | 2 | 8 | 2 | 5 | 5 | 3 | 1 | 2 | 6 | 2 | 3 | 7 | 5 | 2 | 2 | 8 | 4 | 5 | 0 | 2 | 1 | 2 | 1 | 2 | Pcmt1 |
| Pcmtd1 | 7366 | 4901 | 5909 | 7115 | 7671 | 8515 | 9536 | 6868 | 6224 | 5815 | 6121 | 6107 | 6264 | 6501 | 8112 | 6402 | 6306 | 7242 | 7119 | 8199 | 8618 | 6462 | 5826 | 5799 | 6561 | 5040 | 5983 | 6100 | Pcmtd1 |
| Pcmtd2 | 6634 | 5129 | 6769 | 7804 | 7832 | 7914 | 8967 | 6708 | 5800 | 5991 | 6524 | 6159 | 6591 | 6778 | 6999 | 6308 | 7342 | 7898 | 7957 | 7258 | 8711 | 7058 | 5829 | 6224 | 6546 | 5824 | 6107 | 6461 | Pcmtd2 |
| Pcna | 17280 | 15993 | 17514 | 14383 | 11441 | 13720 | 16452 | 17540 | 16120 | 13751 | 13194 | 20600 | 16786 | 22071 | 17041 | 15007 | 15602 | 12842 | 14582 | 18213 | 14880 | 10886 | 14361 | 17552 | 15692 | 26738 | 21530 | 24206 | Pcna |
| Pcnp | 11485 | 9001 | 12857 | 10343 | 9287 | 9734 | 10655 | 11590 | 9565 | 9785 | 9860 | 11177 | 10827 | 11938 | 11584 | 10192 | 11064 | 9783 | 10279 | 11372 | 9972 | 10357 | 9614 | 11011 | 10205 | 13086 | 12484 | 12566 | Pcnp |
| Pcnt | 1960 | 2158 | 1906 | 2027 | 2247 | 2127 | 2149 | 3501 | 4065 | 4562 | 4282 | 5879 | 5211 | 5809 | 1853 | 2059 | 2216 | 1861 | 2406 | 1973 | 2102 | 3823 | 3602 | 4788 | 4790 | 3777 | 5915 | 5393 | Pcnt |
| Pcnx | 6494 | 6121 | 4763 | 6376 | 8332 | 8582 | 7970 | 7630 | 8283 | 8448 | 9199 | 7677 | 7727 | 7452 | 6072 | 6674 | 5527 | 6896 | 7325 | 7100 | 8492 | 7827 | 7550 | 6761 | 8830 | 5143 | 7595 | 6848 | Pcnx |
| Pcnx2 | 302 | 427 | 329 | 643 | 1021 | 718 | 708 | 595 | 789 | 965 | 767 | 638 | 475 | 620 | 394 | 340 | 474 | 650 | 780 | 658 | 749 | 1056 | 652 | 722 | 1002 | 390 | 521 | 469 | Pcnx2 |
| Pcnx3 | 5382 | 7377 | 8195 | 7841 | 7409 | 6413 | 6295 | 5681 | 7486 | 8004 | 7961 | 9093 | 7940 | 7583 | 4953 | 6473 | 8310 | 7845 | 7742 | 5753 | 6279 | 8111 | 6704 | 8699 | 8205 | 7293 | 7768 | 7755 | Pcnx3 |
| Pcnx4 | 5409 | 5928 | 4193 | 5639 | 5389 | 5541 | 5612 | 2657 | 2548 | 2356 | 2172 | 2408 | 2307 | 2291 | 5537 | 5506 | 4774 | 5639 | 5263 | 5957 | 5519 | 2445 | 2458 | 2203 | 2264 | 2062 | 2391 | 2118 | Pcnx4 |
| Pcolce | 1523 | 2062 | 1758 | 1125 | 487 | 1369 | 279 | 1489 | 2080 | 1895 | 2606 | 2245 | 3006 | 2048 | 1544 | 2571 | 1265 | 1373 | 391 | 343 | 256 | 1780 | 2621 | 2211 | 1744 | 2787 | 2644 | 2413 | Pcolce |
| Pcolce2 | 75 | 42 | 34 | 36 | 95 | 57 | 38 | 230 | 272 | 212 | 371 | 177 | 240 | 153 | 66 | 76 | 72 | 62 | 54 | 43 | 40 | 219 | 263 | 220 | 189 | 270 | 164 | 154 | Pcolce2 |
| Pcp2 | 1 | 0 | 0 | 0 | 4 | 3 | 1 | 2 | 2 | 0 | 8 | 3 | 0 | 1 | 0 | 0 | 1 | 1 | 2 | 5 | 1 | 2 | 2 | 2 | 0 | 7 | 1 | 0 | Pcp2 |

| | | | | | | | | | | | | | | | | | | | | | | | | | | | | | |
|---|---|---|---|---|---|---|---|---|---|---|---|---|---|---|---|---|---|---|---|---|---|---|---|---|---|---|---|---|---|
| Pcp4 | 17 | 21 | 2 | 30 | 80 | 39 | 69 | 56 | 33 | 80 | 33 | 23 | 25 | 39 | 40 | 8 | 24 | 29 | 16 | 44 | 104 | 52 | 46 | 69 | 43 | 22 | 8 | 33 | Pcp4 |
| Pcp4l1 | 1637 | 1926 | 2366 | 1651 | 1482 | 1401 | 1081 | 614 | 655 | 724 | 524 | 756 | 1074 | 456 | 1598 | 2174 | 1807 | 1396 | 1386 | 1387 | 1151 | 787 | 972 | 522 | 419 | 522 | 592 | 674 | Pcp4l1 |
| Pcsk1 | 281 | 291 | 259 | 358 | 565 | 389 | 446 | 332 | 306 | 308 | 224 | 209 | 193 | 187 | 410 | 291 | 322 | 307 | 359 | 517 | 387 | 404 | 265 | 155 | 232 | 174 | 187 | 190 | Pcsk1 |
| Pcsk1n | 1046 | 1653 | 1583 | 2441 | 2860 | 1785 | 1688 | 1542 | 1441 | 2280 | 1941 | 1637 | 1446 | 1391 | 1280 | 992 | 2095 | 2222 | 2649 | 1874 | 1942 | 2791 | 1588 | 1832 | 2294 | 1265 | 1273 | 1264 | Pcsk1n |
| Pcsk2 | 1655 | 2205 | 1858 | 2642 | 3523 | 2972 | 2820 | 2179 | 1638 | 2103 | 1783 | 1781 | 1762 | 1707 | 2300 | 1717 | 2392 | 2612 | 3337 | 3500 | 3425 | 2938 | 2007 | 1735 | 2512 | 970 | 1847 | 1754 | Pcsk2 |
| Pcsk4 | 228 | 333 | 429 | 361 | 281 | 221 | 235 | 99 | 115 | 144 | 142 | 134 | 153 | 100 | 223 | 273 | 380 | 352 | 295 | 242 | 173 | 125 | 123 | 131 | 112 | 186 | 102 | 95 | Pcsk4 |
| Pcsk5 | 1927 | 1600 | 1603 | 1206 | 1010 | 1490 | 943 | 1664 | 1574 | 1639 | 2167 | 1605 | 1877 | 1592 | 1713 | 2131 | 1296 | 1369 | 923 | 891 | 859 | 1619 | 2041 | 1579 | 1467 | 1880 | 1766 | 1787 | Pcsk5 |
| Pcsk6 | 555 | 555 | 512 | 449 | 397 | 541 | 374 | 669 | 924 | 893 | 1191 | 770 | 854 | 741 | 571 | 639 | 385 | 519 | 442 | 443 | 422 | 493 | 1001 | 933 | 603 | 980 | 623 | 873 | Pcsk6 |
| Pcsk7 | 3280 | 3861 | 2668 | 3190 | 2950 | 3199 | 2876 | 2518 | 2642 | 2565 | 2501 | 2518 | 2652 | 2566 | 3212 | 3523 | 2877 | 3147 | 2869 | 3085 | 2857 | 2533 | 2422 | 2449 | 2827 | 2197 | 2797 | 2461 | Pcsk7 |
| Pcsk9 | 209 | 231 | 269 | 155 | 182 | 170 | 166 | 100 | 120 | 96 | 130 | 207 | 229 | 203 | 191 | 208 | 217 | 145 | 184 | 209 | 126 | 90 | 155 | 153 | 143 | 381 | 256 | 282 | Pcsk9 |
| Pctp | 836 | 892 | 1055 | 602 | 465 | 449 | 487 | 358 | 344 | 269 | 292 | 388 | 454 | 448 | 742 | 854 | 823 | 490 | 473 | 515 | 426 | 206 | 297 | 356 | 225 | 615 | 517 | 538 | Pctp |
| Pcx | 3513 | 4257 | 3630 | 3413 | 3103 | 3115 | 2549 | 1722 | 2296 | 2057 | 2155 | 2170 | 2337 | 1983 | 3123 | 4266 | 3355 | 3498 | 3085 | 2462 | 2940 | 1893 | 2307 | 2130 | 2115 | 2386 | 2148 | 1990 | Pcx |
| Pcyox1 | 12412 | 12608 | 13174 | 11794 | 11073 | 11911 | 11283 | 8994 | 8574 | 9115 | 8891 | 10151 | 9601 | 9724 | 12676 | 12361 | 11874 | 11766 | 11200 | 11139 | 11008 | 8950 | 8833 | 8785 | 9111 | 9915 | 10667 | 10227 | Pcyox1 |
| Pcyox1l | 940 | 1251 | 1714 | 1471 | 1583 | 1300 | 1200 | 1348 | 1448 | 1661 | 1470 | 1882 | 1647 | 1522 | 1033 | 1186 | 1753 | 1366 | 1604 | 1232 | 1296 | 1698 | 1383 | 2020 | 1803 | 1993 | 1699 | 1567 | Pcyox1l |
| Pcyt1a | 9476 | 9048 | 8633 | 7719 | 7180 | 7730 | 7744 | 6553 | 6347 | 6020 | 6029 | 6384 | 6424 | 6512 | 9337 | 9006 | 8316 | 7990 | 7398 | 8114 | 7340 | 6223 | 6216 | 5966 | 6412 | 6060 | 6951 | 6515 | Pcyt1a |
| Pcyt1b | 1831 | 2044 | 1577 | 2113 | 2788 | 2527 | 2649 | 5179 | 4698 | 5232 | 4143 | 4557 | 4131 | 4633 | 2213 | 1746 | 2073 | 2188 | 2416 | 2536 | 2657 | 5910 | 4251 | 4255 | 5662 | 3670 | 4350 | 4025 | Pcyt1b |
| Pcyt2 | 4642 | 6549 | 5870 | 6225 | 5139 | 5049 | 4797 | 3047 | 3474 | 3879 | 3518 | 3529 | 3730 | 3653 | 4487 | 6012 | 5945 | 6352 | 5500 | 4628 | 4766 | 3681 | 3435 | 3750 | 3917 | 3990 | 3987 | 3725 | Pcyt2 |
| Pdap1 | 11418 | 12836 | 17147 | 15172 | 12003 | 11468 | 14005 | 9944 | 10515 | 13872 | 12002 | 13743 | 12287 | 17133 | 11074 | 10520 | 15755 | 12995 | 13552 | 11744 | 13744 | 9519 | 10103 | 15039 | 13200 | 17579 | 13655 | 16576 | Pdap1 |
| Pdc | 6 | 5 | 4 | 0 | 1 | 3 | 1 | 5 | 8 | 3 | 5 | 18 | 17 | 0 | 7 | 7 | 1 | 1 | 0 | 2 | 0 | 2 | 3 | 1 | 2 | 5 | 6 | 7 | Pdc |
| Pdcd1 | 0 | 1 | 1 | 0 | 0 | 0 | 0 | 5 | 6 | 9 | 4 | 3 | 1 | 0 | 0 | 2 | 0 | 0 | 0 | 1 | 0 | 2 | 4 | 1 | 1 | 7 | 0 | 1 | Pdcd1 |
| Pdcd10 | 4790 | 3444 | 4085 | 4093 | 3669 | 3846 | 4641 | 4172 | 3363 | 3004 | 2974 | 3114 | 3307 | 3544 | 5156 | 4406 | 4014 | 3879 | 4048 | 4765 | 4424 | 2909 | 3555 | 2987 | 3210 | 3480 | 3282 | 3665 | Pdcd10 |
| Pdcd11 | 7972 | 9286 | 7393 | 7914 | 6035 | 7002 | 6510 | 5631 | 6290 | 6071 | 6082 | 6928 | 6541 | 6786 | 7159 | 8276 | 7615 | 7334 | 6723 | 6666 | 6669 | 5021 | 5787 | 6101 | 6539 | 5899 | 7445 | 6696 | Pdcd11 |
| Pdcd1lg2 | 0 | 0 | 2 | 0 | 2 | 0 | 0 | 0 | 0 | 1 | 0 | 0 | 0 | 0 | 1 | 5 | 0 | 1 | 0 | 1 | 0 | 0 | 0 | 0 | 1 | 0 | 0 | 0 | Pdcd1lg2 |
| Pdcd2 | 1711 | 1592 | 1645 | 1597 | 1562 | 1519 | 1804 | 1599 | 1285 | 1431 | 1343 | 1486 | 1386 | 1574 | 1685 | 1468 | 1559 | 1558 | 1552 | 1670 | 1600 | 1504 | 1325 | 1472 | 1443 | 1624 | 1438 | 1575 | Pdcd2 |
| Pdcd2l | 1429 | 1540 | 1375 | 1404 | 1230 | 1237 | 1288 | 1216 | 1187 | 1227 | 1222 | 1446 | 1309 | 1400 | 1396 | 1186 | 1462 | 1414 | 1405 | 1399 | 1341 | 1230 | 1110 | 1276 | 1313 | 1370 | 1291 | 1412 | Pdcd2l |
| Pdcd4 | 7199 | 5809 | 7839 | 7711 | 6419 | 6580 | 8561 | 6681 | 6049 | 6912 | 6788 | 7248 | 7750 | 7648 | 6682 | 6892 | 7369 | 7773 | 7198 | 7119 | 7586 | 6629 | 6552 | 8238 | 7209 | 8711 | 7096 | 7971 | Pdcd4 |
| Pdcd5 | 2200 | 2024 | 2497 | 2330 | 2071 | 1907 | 2360 | 2591 | 2164 | 2603 | 2489 | 2582 | 2510 | 3259 | 2135 | 1791 | 2416 | 1991 | 2036 | 2580 | 2374 | 2247 | 2208 | 2741 | 2564 | 3058 | 2638 | 3133 | Pdcd5 |
| Pdcd6 | 4549 | 4084 | 3225 | 3265 | 3053 | 3145 | 3695 | 3640 | 3159 | 2713 | 3016 | 3001 | 3055 | 3350 | 4406 | 3811 | 3310 | 3185 | 3157 | 3736 | 3568 | 2916 | 3236 | 2798 | 2756 | 3086 | 2925 | 3165 | Pdcd6 |
| Pdcd6ip | 7494 | 6998 | 8551 | 7589 | 6884 | 7129 | 7222 | 5875 | 6143 | 6975 | 7061 | 7425 | 7307 | 7760 | 6943 | 7902 | 7787 | 7902 | 6919 | 5414 | 7129 | 7225 | 6685 | 7044 | 6538 | 7050 | 6594 | 7523 | Pdcd6ip |
| Pdcd7 | 1812 | 1784 | 2045 | 2114 | 1595 | 1330 | 1674 | 1704 | 1588 | 1571 | 1655 | 1640 | 1648 | 1734 | 1705 | 1576 | 1931 | 1673 | 1826 | 1683 | 1674 | 1551 | 1552 | 1777 | 1769 | 2003 | 1648 | 1799 | Pdcd7 |
| Pdcl | 5357 | 4773 | 5926 | 5072 | 5479 | 5246 | 6032 | 7955 | 6358 | 5625 | 5557 | 6894 | 6590 | 7637 | 6177 | 5013 | 5665 | 5080 | 5843 | 6673 | 5756 | 6350 | 6501 | 6636 | 6614 | 7077 | 7369 | 7440 | Pdcl |
| Pdcl2 | 0 | 0 | 0 | 0 | 1 | 0 | 0 | 0 | 0 | 0 | 0 | 0 | 0 | 0 | 0 | 1 | 0 | 0 | 0 | 0 | 0 | 0 | 0 | 0 | 0 | 0 | 0 | 0 | Pdcl2 |
| Pdcl3 | 5016 | 4922 | 5266 | 5015 | 4101 | 4041 | 5011 | 4115 | 3502 | 3330 | 3393 | 3616 | 3585 | 4205 | 5178 | 4693 | 5055 | 4863 | 4560 | 5267 | 4780 | 3461 | 3634 | 4087 | 3826 | 4281 | 3713 | 4195 | Pdcl3 |
| Pde10a | 1405 | 1559 | 1213 | 1718 | 2986 | 2391 | 2738 | 3212 | 3238 | 3800 | 3101 | 2864 | 2498 | 2984 | 1573 | 1348 | 1659 | 1897 | 2607 | 2147 | 2829 | 4139 | 3012 | 3154 | 4323 | 1839 | 2393 | 2566 | Pde10a |
| Pde11a | 18 | 23 | 19 | 17 | 70 | 51 | 47 | 44 | 97 | 52 | 105 | 20 | 40 | 38 | 34 | 27 | 41 | 35 | 23 | 66 | 42 | 69 | 91 | 62 | 61 | 8 | 48 | 49 | Pde11a |
| Pde1a | 258 | 149 | 159 | 221 | 338 | 255 | 389 | 383 | 486 | 346 | 589 | 307 | 202 | 350 | 231 | 161 | 147 | 269 | 294 | 367 | 383 | 403 | 499 | 333 | 450 | 300 | 251 | 250 | Pde1a |
| Pde1b | 2682 | 3359 | 3276 | 4135 | 5636 | 4831 | 4773 | 5145 | 5135 | 5418 | 4553 | 5473 | 5389 | 5187 | 3190 | 2695 | 4085 | 4616 | 5791 | 4713 | 5292 | 5461 | 4612 | 5480 | 6072 | 3897 | 4762 | 4824 | Pde1b |
| Pde1c | 1366 | 1144 | 819 | 1145 | 1847 | 1843 | 1555 | 2818 | 2558 | 1909 | 2108 | 1486 | 1643 | 1622 | 1524 | 1353 | 907 | 1152 | 1280 | 1899 | 1561 | 1980 | 2548 | 1390 | 2175 | 1231 | 1680 | 1329 | Pde1c |
| Pde2a | 17 | 22 | 6 | 17 | 30 | 23 | 28 | 10 | 8 | 20 | 14 | 8 | 6 | 19 | 20 | 6 | 17 | 17 | 18 | 13 | 29 | 22 | 26 | 19 | 17 | 4 | 3 | 17 | Pde2a |
| Pde3a | 1444 | 980 | 1090 | 982 | 946 | 1394 | 830 | 1590 | 2124 | 1641 | 2621 | 1416 | 1719 | 1469 | 1273 | 1542 | 802 | 1092 | 926 | 825 | 840 | 2087 | 2452 | 1527 | 1628 | 1111 | 1684 | 1450 | Pde3a |
| Pde3b | 3275 | 2293 | 3144 | 2819 | 2598 | 2636 | 2950 | 2900 | 2619 | 2618 | 2721 | 3331 | 3345 | 3480 | 3069 | 2894 | 2843 | 2608 | 2724 | 2389 | 2549 | 2876 | 2372 | 2953 | 3022 | 3423 | 3410 | 3706 | Pde3b |
| Pde4a | 1159 | 1476 | 1385 | 2257 | 2601 | 2052 | 2276 | 894 | 1134 | 1527 | 1213 | 1344 | 972 | 1272 | 1162 | 1275 | 1990 | 2281 | 2507 | 1869 | 2175 | 1684 | 994 | 1450 | 1597 | 1190 | 1084 | 1180 | Pde4a |
| Pde4b | 1031 | 822 | 801 | 1067 | 1629 | 1410 | 1415 | 2003 | 2001 | 1861 | 1999 | 1130 | 1330 | 1322 | 1113 | 786 | 907 | 1078 | 1441 | 1438 | 1500 | 2396 | 2117 | 1553 | 2123 | 964 | 1263 | 1081 | Pde4b |
| Pde4c | 15 | 25 | 36 | 13 | 5 | 1 | 3 | 5 | 0 | 3 | 6 | 2 | 2 | 0 | 7 | 5 | 6 | 4 | 5 | 1 | 2 | 8 | 3 | 0 | 5 | 3 | 2 | 2 | Pde4c |
| Pde4d | 1736 | 1906 | 1591 | 2460 | 2876 | 2774 | 2537 | 2199 | 2290 | 2368 | 2083 | 1891 | 1964 | 1710 | 2035 | 1883 | 2154 | 2702 | 2661 | 2532 | 2707 | 2772 | 2102 | 1979 | 2539 | 1235 | 1898 | 1810 | Pde4d |
| Pde4dip | 9797 | 10818 | 7796 | 10940 | 13357 | 11562 | 11693 | 10187 | 10854 | 10679 | 10696 | 7478 | 8160 | 7933 | 10599 | 10069 | 9250 | 11898 | 11531 | 11716 | 12566 | 11931 | 9483 | 8292 | 11275 | 5426 | 7316 | 7098 | Pde4dip |
| Pde5a | 2656 | 1915 | 2153 | 1925 | 2437 | 2343 | 2617 | 3301 | 2909 | 2782 | 2883 | 3102 | 2880 | 3135 | 2703 | 2034 | 2265 | 1955 | 2471 | 2508 | 2439 | 2540 | 2966 | 2793 | 2680 | 2777 | 3290 | 3287 | Pde5a |
| Pde6a | 0 | 4 | 2 | 8 | 1 | 1 | 3 | 11 | 10 | 10 | 10 | 4 | 9 | 2 | 4 | 9 | 0 | 0 | 6 | 0 | 1 | 7 | 11 | 11 | 8 | 8 | 7 | 2 | Pde6a |
| Pde6b | 0 | 0 | 1 | 1 | 0 | 8 | 3 | 0 | 0 | 0 | 0 | 0 | 0 | 0 | 0 | 0 | 0 | 2 | 1 | 1 | 3 | 0 | 0 | 0 | 0 | 0 | 0 | 0 | Pde6b |
| Pde6c | 6 | 14 | 5 | 11 | 22 | 15 | 7 | 28 | 36 | 42 | 26 | 31 | 64 | 16 | 20 | 30 | 2 | 9 | 19 | 7 | 12 | 50 | 56 | 11 | 23 | 2 | 9 | 12 | Pde6c |
| Pde6d | 2683 | 2994 | 3138 | 2882 | 2834 | 2395 | 2849 | 2120 | 2056 | 2407 | 1916 | 2293 | 2047 | 2206 | 2592 | 2674 | 3124 | 2448 | 2990 | 2839 | 2630 | 2014 | 2179 | 2221 | 2072 | 2537 | 2366 | 2277 | Pde6d |
| Pde6g | 14 | 14 | 38 | 36 | 33 | 10 | 21 | 13 | 9 | 16 | 26 | 22 | 24 | 12 | 21 | 26 | 21 | 23 | 20 | 28 | 26 | 23 | 14 | 21 | 15 | 17 | 13 | 14 | Pde6g |
| Pde6h | 0 | 3 | 0 | 0 | 6 | 1 | 3 | 9 | 3 | 0 | 3 | 0 | 14 | 1 | 1 | 2 | 1 | 0 | 2 | 0 | 0 | 5 | 6 | 1 | 2 | 1 | 0 | 0 | Pde6h |
| Pde7a | 10138 | 8895 | 9389 | 10489 | 11164 | 10871 | 12010 | 13103 | 10752 | 10953 | 10735 | 12433 | 11633 | 12911 | 10493 | 9240 | 9873 | 10294 | 10963 | 11026 | 11748 | 12358 | 10029 | 11624 | 12493 | 11241 | 12433 | 12644 | Pde7a |
| Pde7b | 499 | 369 | 349 | 312 | 383 | 447 | 327 | 741 | 1046 | 816 | 953 | 499 | 785 | 521 | 404 | 484 | 346 | 421 | 343 | 256 | 304 | 931 | 1128 | 732 | 728 | 642 | 516 | 425 | Pde7b |
| Pde8a | 472 | 439 | 629 | 391 | 520 | 580 | 390 | 556 | 603 | 620 | 708 | 477 | 594 | 502 | 622 | 563 | 424 | 464 | 387 | 312 | 427 | 566 | 696 | 628 | 583 | 540 | 480 | 432 | Pde8a |
| Pde8b | 106 | 110 | 100 | 151 | 292 | 200 | 196 | 223 | 284 | 329 | 359 | 241 | 229 | 248 | 164 | 112 | 148 | 155 | 217 | 152 | 181 | 414 | 330 | 249 | 287 | 203 | 238 | 225 | Pde8b |
| Pde9a | 3026 | 3349 | 3023 | 3457 | 3339 | 3203 | 3462 | 2142 | 2061 | 2029 | 1847 | 2002 | 1658 | 2079 | 3157 | 2821 | 3093 | 3202 | 3316 | 3512 | 3472 | 2204 | 1950 | 2189 | 2508 | 2235 | 1907 | 1942 | Pde9a |
| Pdf | 110 | 129 | 83 | 104 | 129 | 201 | 155 | 94 | 96 | 110 | 84 | 115 | 143 | 114 | 125 | 108 | 79 | 130 | 143 | 129 | 130 | 99 | 102 | 100 | 155 | 116 | 101 | 113 | Pdf |
| Pdgfa | 999 | 878 | 765 | 804 | 919 | 785 | 743 | 858 | 1362 | 1131 | 1287 | 625 | 710 | 566 | 725 | 822 | 893 | 945 | 855 | 849 | 712 | 637 | 1461 | 1020 | 705 | 1033 | 543 | 503 | Pdgfa |
| Pdgfb | 467 | 546 | 643 | 543 | 592 | 483 | 453 | 448 | 667 | 759 | 815 | 672 | 653 | 545 | 450 | 479 | 606 | 546 | 508 | 451 | 501 | 685 | 763 | 744 | 775 | 835 | 600 | 595 | Pdgfb |
| Pdgfc | 1684 | 919 | 770 | 700 | 701 | 909 | 650 | 2076 | 2312 | 1730 | 2404 | 1240 | 1807 | 1397 | 1092 | 1386 | 789 | 691 | 838 | 667 | 651 | 1338 | 2428 | 1713 | 1294 | 2244 | 1257 | 1437 | Pdgfc |
| Pdgfd | 187 | 142 | 198 | 102 | 146 | 113 | 69 | 243 | 266 | 227 | 364 | 148 | 210 | 156 | 183 | 171 | 144 | 87 | 84 | 113 | 78 | 212 | 327 | 186 | 193 | 270 | 192 | 220 | Pdgfd |
| Pdgfra | 7988 | 4868 | 4475 | 2983 | 2488 | 4169 | 1659 | 9068 | 12366 | 9525 | 17457 | 6106 | 10521 | 6123 | 5500 | 7720 | 3430 | 3814 | 1994 | 2048 | 1768 | 7305 | 14751 | 8120 | 5411 | 7103 | 6438 | 5942 | Pdgfra |
| Pdgfrb | 4982 | 4918 | 4414 | 3360 | 3004 | 3661 | 2227 | 3621 | 5262 | 5488 | 5964 | 4442 | 6178 | 3901 | 3848 | 6816 | 3617 | 3847 | 2660 | 2128 | 2305 | 4448 | 5464 | 5032 | 4016 | 5079 | 4672 | 4037 | Pdgfrb |
| Pdgfrl | 247 | 214 | 188 | 203 | 134 | 307 | 80 | 216 | 336 | 233 | 367 | 234 | 244 | 246 | 245 | 245 | 172 | 262 | 103 | 121 | 93 | 281 | 474 | 305 | 253 | 193 | 261 | 229 | Pdgfrl |
| Pdha1 | 15303 | 14196 | 16264 | 13242 | 12246 | 12441 | 13829 | 11900 | 10416 | 9471 | 9331 | 10812 | 10376 | 11500 | 16311 | 13685 | 14171 | 12619 | 12809 | 14666 | 13413 | 9951 | 11154 | 10165 | 9857 | 11560 | 12058 | 12536 | Pdha1 |
| Pdha2 | 0 | 0 | 3 | 0 | 0 | 0 | 2 | 1 | 0 | 0 | 0 | 0 | 0 | 0 | 5 | 0 | 0 | 0 | 0 | 0 | 0 | 0 | 1 | 0 | 0 | 0 | 0 | 0 | Pdha2 |
| Pdhb | 9186 | 8878 | 8932 | 9177 | 8328 | 8423 | 9569 | 8965 | 8156 | 7979 | 7387 | 8415 | 7850 | 8715 | 9490 | 8109 | 8895 | 8197 | 9220 | 9843 | 9381 | 8134 | 8092 | 8218 | 8217 | 8989 | 8501 | 8858 | Pdhb |
| Pdhx | 3524 | 3575 | 3146 | 3617 | 3252 | 3358 | 3707 | 3113 | 2881 | 2478 | 2374 | 2833 | 2671 | 2937 | 3754 | 3380 | 3355 | 3486 | 3436 | 3938 | 3613 | 2682 | 2708 | 2653 | 2825 | 2789 | 2695 | 2872 | Pdhx |
| Pdia2 | 23 | 13 | 5 | 8 | 22 | 0 | 6 | 37 | 30 | 37 | 33 | 11 | 14 | 4 | 15 | 20 | 17 | 6 | 27 | 4 | 11 | 7 | 37 | 60 | 6 | 61 | 6 | 12 | Pdia2 |
| Pdia3 | 42135 | 35633 | 42601 | 32196 | 27777 | 29526 | 30248 | 33994 | 31948 | 30074 | 36145 | 29668 | 33654 | 29757 | 42300 | 41310 | 36506 | 32917 | 30511 | 29747 | 28003 | 33272 | 35525 | 31620 | 27714 | 35863 | 32256 | 32888 | Pdia3 |
| Pdia4 | 15346 | 15040 | 14844 | 12124 | 8905 | 9669 | 9751 | 12313 | 11578 | 10268 | 11553 | 9472 | 10291 | 10136 | 15598 | 15410 | 13230 | 11881 | 10225 | 10837 | 9665 | 10617 | 11879 | 11173 | 9280 | 11447 | 10167 | 10779 | Pdia4 |
| Pdia5 | 1192 | 1549 | 1406 | 1211 | 1064 | 989 | 917 | 761 | 1009 | 851 | 1139 | 850 | 988 | 736 | 1193 | 1576 | 1279 | 1282 | 1026 | 1096 | 906 | 824 | 1113 | 885 | 718 | 925 | 874 | 865 | Pdia5 |
| Pdia6 | 25793 | 28210 | 30316 | 23227 | 17915 | 19304 | 20245 | 19814 | 21620 | 18776 | 21750 | 19220 | 19294 | 19776 | 25983 | 28853 | 25942 | 23009 | 20393 | 22988 | 18231 | 16802 | 21773 | 21002 | 18374 | 23382 | 21208 | 20912 | Pdia6 |
| Pdik1l | 3058 | 2723 | 2646 | 3196 | 3449 | 3270 | 3406 | 4068 | 3410 | 3461 | 3062 | 3388 | 2956 | 3399 | 3418 | 2820 | 3000 | 3224 | 3219 | 3941 | 3500 | 3681 | 3172 | 3410 | 3917 | 3127 | 3330 | 3451 | Pdik1l |
| Pdilt | 2 | 2 | 0 | 1 | 4 | 8 | 13 | 11 | 10 | 8 | 6 | 9 | 16 | 4 | 3 | 0 | 0 | 13 | 5 | 4 | 3 | 2 | 9 | 7 | 8 | 1 | 5 | 5 | Pdilt |
| Pdk1 | 5669 | 4477 | 3759 | 4533 | 3853 | 4765 | 4528 | 3342 | 4249 | 3931 | 4398 | 3058 | 3328 | 3191 | 4929 | 4836 | 4343 | 5044 | 4087 | 4790 | 4313 | 2941 | 4757 | 3806 | 3129 | 3616 | 3166 | 2947 | Pdk1 |
| Pdk2 | 1956 | 2490 | 2305 | 2816 | 2788 | 2374 | 2417 | 1001 | 1408 | 1334 | 1392 | 1184 | 1260 | 1004 | 2064 | 2493 | 2737 | 2923 | 2740 | 2161 | 2507 | 1333 | 1533 | 1225 | 1312 | 1250 | 961 | 1095 | Pdk2 |

| | | | | | | | | | | | | | | | | | | | | | | | | | | | | | |
|---|---|---|---|---|---|---|---|---|---|---|---|---|---|---|---|---|---|---|---|---|---|---|---|---|---|---|---|---|---|
| Pdk3 | 3690 | 3156 | 2316 | 2110 | 1829 | 2364 | 2482 | 2394 | 2346 | 1848 | 1846 | 1981 | 2147 | 2060 | 3397 | 3113 | 2140 | 2121 | 2186 | 2574 | 2401 | 1636 | 2405 | 1754 | 1683 | 2378 | 2138 | 2160 | Pdk3 |
| Pdk4 | 366 | 308 | 230 | 296 | 241 | 351 | 292 | 128 | 186 | 155 | 134 | 118 | 107 | 93 | 336 | 320 | 234 | 306 | 238 | 257 | 271 | 104 | 169 | 127 | 83 | 106 | 78 | 101 | Pdk4 |
| Pdlim1 | 1329 | 1342 | 1382 | 822 | 664 | 795 | 748 | 902 | 1071 | 668 | 708 | 905 | 909 | 782 | 1192 | 1400 | 1038 | 822 | 865 | 769 | 636 | 423 | 996 | 766 | 401 | 985 | 830 | 1003 | Pdlim1 |
| Pdlim2 | 311 | 319 | 348 | 160 | 140 | 197 | 93 | 285 | 351 | 258 | 256 | 241 | 396 | 261 | 282 | 378 | 190 | 151 | 129 | 184 | 134 | 211 | 409 | 273 | 222 | 253 | 372 | 343 | Pdlim2 |
| Pdlim3 | 599 | 566 | 925 | 448 | 458 | 557 | 457 | 766 | 550 | 574 | 1098 | 701 | 458 | 791 | 550 | 397 | 649 | 424 | 563 | 432 | 491 | 632 | 886 | 625 | 606 | 688 | 1067 | 891 | Pdlim3 |
| Pdlim4 | 1734 | 1887 | 1551 | 1542 | 1202 | 1251 | 1009 | 847 | 1069 | 828 | 1020 | 755 | 941 | 661 | 1441 | 2021 | 1442 | 1415 | 1308 | 1039 | 1057 | 627 | 1065 | 900 | 532 | 1025 | 804 | 758 | Pdlim4 |
| Pdlim5 | 6144 | 5578 | 6763 | 5646 | 4875 | 4910 | 5031 | 2379 | 2536 | 2277 | 2687 | 2526 | 2903 | 2409 | 6581 | 6511 | 5991 | 5907 | 4740 | 4824 | 4862 | 2817 | 3209 | 2281 | 2258 | 2495 | 2807 | 2428 | Pdlim5 |
| Pdlim7 | 3676 | 5279 | 4536 | 4736 | 3625 | 3706 | 3596 | 2201 | 2639 | 2868 | 3519 | 2870 | 3098 | 2783 | 3560 | 4365 | 4530 | 4813 | 3956 | 3199 | 3852 | 3038 | 2999 | 3324 | 2918 | 3170 | 2959 | 2713 | Pdlim7 |
| Pdp1 | 5293 | 4953 | 4637 | 5629 | 5839 | 6014 | 6178 | 5390 | 4296 | 3907 | 3953 | 3847 | 3683 | 3907 | 6078 | 5136 | 5034 | 5675 | 5671 | 6466 | 6179 | 5534 | 4297 | 3858 | 4684 | 3107 | 3799 | 3841 | Pdp1 |
| Pdp2 | 2336 | 2660 | 2652 | 2714 | 2716 | 2577 | 2682 | 2013 | 1768 | 2004 | 1891 | 2199 | 2080 | 2201 | 2308 | 2290 | 2789 | 2787 | 2978 | 2535 | 2725 | 2099 | 1726 | 2022 | 2120 | 1882 | 2152 | 2230 | Pdp2 |
| Pdpk1 | 11845 | 11062 | 8605 | 10954 | 10568 | 11241 | 11171 | 9111 | 8033 | 7595 | 7432 | 7782 | 7525 | 7734 | 11856 | 10697 | 9379 | 10741 | 10361 | 10832 | 11582 | 9478 | 8061 | 7034 | 8730 | 5662 | 7843 | 7644 | Pdpk1 |
| Pdpr | 3319 | 3177 | 1434 | 2366 | 2562 | 2877 | 2744 | 3142 | 2976 | 2379 | 2688 | 2588 | 2727 | 2758 | 3298 | 2845 | 1847 | 2340 | 2388 | 2843 | 2979 | 2463 | 2847 | 1927 | 2826 | 871 | 3115 | 2470 | Pdpr |
| Pdrg1 | 4657 | 5309 | 4655 | 5729 | 5096 | 5199 | 5281 | 3947 | 3723 | 3760 | 3843 | 3393 | 3627 | 3681 | 5163 | 4914 | 5012 | 5734 | 5329 | 5755 | 5542 | 4080 | 3667 | 3790 | 3924 | 3792 | 3689 | 3826 | Pdrg1 |
| Pds5a | 19568 | 14892 | 15189 | 14939 | 16368 | 17704 | 18464 | 22022 | 18342 | 16517 | 16858 | 20503 | 19619 | 21688 | 19055 | 16049 | 14731 | 14070 | 15844 | 18492 | 17210 | 16078 | 17416 | 17492 | 17974 | 17605 | 21895 | 21289 | Pds5a |
| Pds5b | 14542 | 11629 | 9775 | 10531 | 11852 | 13099 | 12552 | 19833 | 17291 | 14885 | 16286 | 15462 | 15636 | 15669 | 14546 | 12974 | 10398 | 10995 | 11278 | 13564 | 12365 | 15575 | 16570 | 13800 | 15363 | 12408 | 16324 | 15060 | Pds5b |
| Pdss1 | 1876 | 1437 | 1408 | 1689 | 1690 | 1781 | 2097 | 1595 | 1247 | 1257 | 1225 | 1473 | 1371 | 1567 | 1913 | 1546 | 1479 | 1653 | 1670 | 1818 | 2062 | 1465 | 1238 | 1155 | 1487 | 1203 | 1390 | 1455 | Pdss1 |
| Pdss2 | 452 | 495 | 555 | 533 | 460 | 442 | 531 | 702 | 650 | 666 | 622 | 698 | 649 | 786 | 448 | 476 | 453 | 492 | 456 | 582 | 517 | 677 | 622 | 696 | 649 | 752 | 745 | 801 | Pdss2 |
| Pdx1 | 1 | 0 | 0 | 0 | 0 | 0 | 1 | 0 | 0 | 0 | 0 | 0 | 0 | 0 | 0 | 0 | 0 | 0 | 0 | 0 | 1 | 0 | 0 | 0 | 2 | 0 | 0 | 0 | Pdx1 |
| Pdxdc1 | 10526 | 9634 | 8068 | 7917 | 7477 | 8395 | 7842 | 9424 | 9083 | 7724 | 8691 | 7719 | 7977 | 7794 | 10127 | 9256 | 7776 | 7614 | 7557 | 8361 | 8135 | 8163 | 9622 | 7764 | 7897 | 7852 | 8278 | 8065 | Pdxdc1 |
| Pdxk | 3555 | 4391 | 4075 | 4556 | 4227 | 3855 | 3503 | 1506 | 1875 | 1962 | 2067 | 1655 | 1838 | 1599 | 3500 | 4472 | 3745 | 4537 | 4090 | 3279 | 3559 | 2259 | 2094 | 1790 | 2053 | 1809 | 1764 | 1595 | Pdxk |
| Pdxp | 636 | 733 | 650 | 984 | 973 | 795 | 1054 | 615 | 554 | 864 | 648 | 589 | 411 | 675 | 580 | 435 | 877 | 934 | 1050 | 777 | 1166 | 754 | 485 | 824 | 858 | 674 | 499 | 616 | Pdxp |
| Pdyn | 81 | 161 | 78 | 215 | 373 | 256 | 237 | 272 | 260 | 247 | 228 | 93 | 237 | 99 | 322 | 172 | 145 | 265 | 202 | 242 | 242 | 334 | 258 | 114 | 160 | 49 | 126 | 106 | Pdyn |
| Pdzd11 | 2644 | 2761 | 2767 | 2334 | 2219 | 2296 | 2596 | 3209 | 2863 | 2623 | 2517 | 3036 | 2964 | 3512 | 2946 | 2482 | 2650 | 2363 | 2447 | 2761 | 2510 | 2488 | 2832 | 2962 | 2612 | 3311 | 3177 | 3529 | Pdzd11 |
| Pdzd2 | 791 | 863 | 880 | 718 | 1038 | 858 | 628 | 1108 | 1521 | 1865 | 2489 | 1302 | 1419 | 1222 | 715 | 1054 | 766 | 934 | 803 | 554 | 682 | 1602 | 2012 | 1352 | 1395 | 761 | 1418 | 1295 | Pdzd2 |
| Pdzd3 | 7 | 23 | 8 | 34 | 39 | 21 | 26 | 40 | 55 | 58 | 40 | 40 | 34 | 42 | 12 | 12 | 7 | 19 | 42 | 15 | 35 | 47 | 63 | 53 | 38 | 69 | 38 | 89 | Pdzd3 |
| Pdzd4 | 5691 | 6476 | 6529 | 11828 | 13502 | 11073 | 12585 | 8157 | 8665 | 12304 | 9657 | 9531 | 7379 | 10103 | 5979 | 4581 | 9693 | 11826 | 13965 | 9712 | 14334 | 12741 | 8244 | 11804 | 13329 | 8705 | 7898 | 9163 | Pdzd4 |
| Pdzd7 | 314 | 383 | 365 | 604 | 799 | 582 | 584 | 225 | 278 | 366 | 265 | 198 | 172 | 193 | 347 | 391 | 521 | 661 | 661 | 453 | 569 | 408 | 269 | 313 | 358 | 182 | 166 | 179 | Pdzd7 |
| Pdzd8 | 10795 | 8952 | 8825 | 10133 | 9561 | 10131 | 10123 | 6825 | 6597 | 6387 | 7019 | 6721 | 7263 | 6790 | 10461 | 10011 | 8903 | 10489 | 9001 | 9126 | 9631 | 6577 | 6209 | 5966 | 6507 | 6126 | 7055 | 6737 | Pdzd8 |
| Pdzd9 | 22 | 15 | 47 | 17 | 13 | 27 | 18 | 16 | 16 | 10 | 16 | 7 | 16 | 18 | 21 | 26 | 19 | 7 | 10 | 14 | 12 | 12 | 7 | 16 | 24 | 10 | 20 | 19 | Pdzd9 |
| Pdzk1 | 7 | 16 | 15 | 28 | 19 | 9 | 13 | 2 | 12 | 9 | 15 | 7 | 11 | 6 | 16 | 18 | 19 | 19 | 11 | 16 | 20 | 5 | 4 | 7 | 8 | 3 | 5 | 1 | Pdzk1 |
| Pdzk1ip1 | 116 | 184 | 141 | 124 | 82 | 122 | 111 | 158 | 236 | 180 | 148 | 178 | 435 | 229 | 136 | 158 | 105 | 111 | 130 | 157 | 102 | 120 | 217 | 105 | 129 | 193 | 248 | 485 | Pdzk1ip1 |
| Pdzph1 | 148 | 149 | 243 | 104 | 85 | 132 | 149 | 269 | 185 | 159 | 165 | 381 | 352 | 446 | 146 | 121 | 183 | 90 | 143 | 175 | 133 | 175 | 156 | 217 | 257 | 402 | 439 | 576 | Pdzph1 |
| Pdzrn3 | 3714 | 4438 | 3628 | 3570 | 3660 | 3537 | 2890 | 3621 | 3547 | 4000 | 4305 | 3087 | 3845 | 2845 | 4189 | 4817 | 3582 | 3791 | 3114 | 3362 | 3060 | 4137 | 4401 | 3459 | 3674 | 2925 | 3094 | 2797 | Pdzrn3 |
| Pdzrn4 | 519 | 781 | 686 | 803 | 1297 | 922 | 981 | 1442 | 1350 | 1723 | 1400 | 878 | 1452 | 948 | 790 | 600 | 829 | 876 | 959 | 1050 | 1059 | 1793 | 1555 | 999 | 1303 | 699 | 1002 | 946 | Pdzrn4 |
| Pea15a | 27508 | 33071 | 26658 | 29244 | 28526 | 30228 | 32742 | 26710 | 23206 | 22872 | 18909 | 26066 | 21127 | 27327 | 32690 | 25297 | 30057 | 28159 | 31948 | 31747 | 32877 | 24182 | 22319 | 23222 | 25372 | 26870 | 25544 | 26748 | Pea15a |
| Peak1 | 4218 | 4043 | 2429 | 3086 | 3768 | 3981 | 3108 | 2133 | 2639 | 2217 | 2789 | 1596 | 2163 | 1527 | 4048 | 4582 | 2422 | 3469 | 3042 | 3439 | 3731 | 1995 | 2691 | 1490 | 1883 | 1042 | 1696 | 1456 | Peak1 |
| Pear1 | 791 | 1063 | 1124 | 788 | 758 | 701 | 462 | 701 | 1132 | 1178 | 1649 | 1051 | 1382 | 865 | 778 | 1276 | 890 | 954 | 556 | 484 | 433 | 1258 | 1452 | 1119 | 1008 | 1262 | 1209 | 879 | Pear1 |
| Pebp1 | 19887 | 23570 | 18609 | 21476 | 19372 | 19963 | 20956 | 18005 | 18938 | 17808 | 16607 | 18323 | 18787 | 18996 | 19551 | 19390 | 20561 | 19657 | 22875 | 22018 | 21356 | 16619 | 18921 | 18994 | 19372 | 20694 | 19113 | 20103 | Pebp1 |
| Pebp4 | 2 | 2 | 1 | 3 | 1 | 0 | 0 | 1 | 0 | 0 | 1 | 0 | 0 | 0 | 0 | 2 | 0 | 0 | 0 | 2 | 0 | 3 | 2 | 2 | 0 | 0 | 0 | 0 | Pebp4 |
| Pecam1 | 2906 | 2700 | 2441 | 2310 | 2271 | 2487 | 2005 | 3035 | 3193 | 3045 | 3547 | 2678 | 2895 | 2445 | 2753 | 2798 | 2062 | 2532 | 2343 | 2575 | 2200 | 3128 | 3903 | 2978 | 2659 | 3552 | 2851 | 2665 | Pecam1 |
| Pecr | 576 | 596 | 627 | 598 | 455 | 540 | 543 | 418 | 443 | 378 | 436 | 379 | 510 | 360 | 567 | 645 | 538 | 634 | 460 | 599 | 561 | 292 | 491 | 385 | 337 | 496 | 431 | 430 | Pecr |
| Pef1 | 1369 | 1716 | 1413 | 1192 | 1273 | 1313 | 1247 | 2414 | 2505 | 2138 | 1975 | 2270 | 2333 | 2272 | 1396 | 1467 | 1460 | 1246 | 1455 | 1459 | 1180 | 1973 | 2557 | 2253 | 2361 | 2688 | 2581 | 2515 | Pef1 |
| Peg10 | 26994 | 22340 | 29279 | 32683 | 30315 | 29514 | 33839 | 17239 | 20516 | 23350 | 29417 | 24359 | 17964 | 25450 | 25500 | 25160 | 37454 | 32611 | 30877 | 21228 | 32213 | 34499 | 18898 | 23367 | 22208 | 27372 | 21601 | 22783 | Peg10 |
| Peg12 | 1213 | 1453 | 2019 | 1249 | 1222 | 1218 | 1437 | 1783 | 1645 | 1818 | 1747 | 3207 | 2801 | 3588 | 1340 | 1086 | 1870 | 1038 | 1454 | 1226 | 1301 | 1485 | 1543 | 2531 | 2191 | 3711 | 3519 | 3710 | Peg12 |
| Peg3 | 64735 | 58680 | 35775 | 46169 | 40679 | 50576 | 45561 | 42985 | 54437 | 42038 | 71535 | 37247 | 41891 | 37439 | 61430 | 64224 | 40806 | 48730 | 38641 | 45742 | 44537 | 46404 | 54839 | 34587 | 39334 | 28581 | 38822 | 32144 | Peg3 |
| Peli1 | 8173 | 6808 | 7932 | 8605 | 9051 | 8882 | 10731 | 10640 | 8819 | 8962 | 8005 | 10467 | 9228 | 11230 | 8607 | 6988 | 8177 | 8670 | 9180 | 9373 | 10221 | 10368 | 8331 | 9789 | 10671 | 9851 | 10036 | 11017 | Peli1 |
| Peli2 | 2624 | 2540 | 4048 | 3595 | 3693 | 3168 | 3363 | 1953 | 2167 | 2481 | 2473 | 2836 | 2767 | 2864 | 2806 | 2875 | 3650 | 3386 | 3564 | 2674 | 3377 | 2764 | 2023 | 2553 | 2511 | 2221 | 2506 | 2855 | Peli2 |
| Peli3 | 1151 | 1172 | 1297 | 1408 | 1526 | 1495 | 1364 | 647 | 690 | 828 | 676 | 609 | 543 | 498 | 1027 | 1110 | 1395 | 1371 | 1424 | 1211 | 1358 | 864 | 598 | 701 | 829 | 517 | 475 | 449 | Peli3 |
| Pelo | 300 | 517 | 474 | 512 | 380 | 308 | 310 | 227 | 297 | 320 | 266 | 330 | 269 | 293 | 290 | 388 | 452 | 463 | 419 | 337 | 328 | 311 | 313 | 318 | 303 | 379 | 253 | 293 | Pelo |
| Pelp1 | 3820 | 4316 | 5201 | 4811 | 3976 | 3862 | 3856 | 3594 | 4330 | 4666 | 4755 | 5685 | 4849 | 5752 | 3439 | 4083 | 4960 | 4661 | 4371 | 2910 | 4124 | 4453 | 4186 | 5622 | 4930 | 7205 | 4831 | 5781 | Pelp1 |
| Pemt | 61 | 66 | 52 | 44 | 48 | 32 | 45 | 125 | 103 | 76 | 74 | 86 | 109 | 131 | 51 | 51 | 44 | 32 | 39 | 49 | 46 | 112 | 102 | 86 | 90 | 107 | 130 | 159 | Pemt |
| Penk | 1374 | 1311 | 991 | 582 | 846 | 644 | 222 | 2414 | 3236 | 1995 | 2821 | 1410 | 1955 | 1138 | 1363 | 1915 | 1064 | 953 | 197 | 317 | 248 | 1917 | 3253 | 1518 | 984 | 1402 | 1722 | 1174 | Penk |
| Pepd | 2351 | 3294 | 2596 | 2549 | 2050 | 2017 | 2014 | 1583 | 1585 | 1281 | 1477 | 1527 | 1655 | 1444 | 2523 | 2943 | 2629 | 2266 | 2334 | 2486 | 2037 | 1320 | 1766 | 1461 | 1431 | 1656 | 1701 | 1676 | Pepd |
| Per1 | 3201 | 4981 | 4559 | 5799 | 3706 | 3781 | 3434 | 1777 | 2112 | 2457 | 2311 | 2363 | 2036 | 2027 | 3069 | 4423 | 4953 | 5641 | 4314 | 2992 | 3878 | 2438 | 2229 | 2670 | 2462 | 2587 | 2094 | 2033 | Per1 |
| Per2 | 1084 | 1403 | 1192 | 1761 | 1424 | 1328 | 1434 | 731 | 837 | 980 | 1027 | 741 | 637 | 676 | 1017 | 1461 | 1287 | 1909 | 1404 | 1103 | 1418 | 1138 | 785 | 985 | 1048 | 543 | 581 | 647 | Per2 |
| Per3 | 1742 | 1690 | 1485 | 1668 | 1342 | 1442 | 1364 | 772 | 906 | 761 | 956 | 775 | 904 | 711 | 1600 | 1784 | 1581 | 1644 | 1325 | 1429 | 1375 | 835 | 931 | 766 | 768 | 738 | 908 | 640 | Per3 |
| Percc1 | 0 | 0 | 0 | 4 | 1 | 0 | 0 | 1 | 0 | 2 | 1 | 0 | 0 | 1 | 3 | 0 | 0 | 2 | 0 | 0 | 0 | 0 | 1 | 0 | 0 | 0 | 1 | 0 | Percc1 |
| Perm1 | 285 | 524 | 453 | 556 | 362 | 324 | 364 | 117 | 146 | 148 | 111 | 190 | 205 | 165 | 402 | 437 | 556 | 537 | 418 | 401 | 422 | 144 | 135 | 154 | 127 | 162 | 160 | 189 | Perm1 |
| Perp | 561 | 256 | 267 | 58 | 128 | 260 | 88 | 1025 | 1511 | 797 | 830 | 344 | 494 | 397 | 384 | 391 | 97 | 239 | 140 | 99 | 83 | 189 | 1902 | 524 | 174 | 218 | 339 | 460 | Perp |
| Pes1 | 7968 | 8773 | 7832 | 6930 | 5775 | 6926 | 6555 | 6902 | 6993 | 6257 | 6124 | 6908 | 6989 | 7608 | 7601 | 7227 | 7429 | 6585 | 6999 | 7051 | 6834 | 5471 | 6956 | 6825 | 6584 | 8131 | 7827 | 8105 | Pes1 |
| Pet100 | 807 | 741 | 827 | 781 | 764 | 776 | 978 | 894 | 681 | 927 | 779 | 872 | 865 | 1256 | 785 | 600 | 765 | 756 | 696 | 708 | 978 | 745 | 701 | 1109 | 802 | 887 | 893 | 1137 | Pet100 |
| Pex1 | 1961 | 1900 | 1513 | 1905 | 1949 | 1993 | 1979 | 2663 | 2559 | 2376 | 2227 | 2258 | 2266 | 2230 | 1927 | 1884 | 1701 | 1881 | 1944 | 2073 | 2068 | 2383 | 2346 | 2161 | 2531 | 1900 | 2529 | 2103 | Pex1 |
| Pex10 | 711 | 799 | 779 | 779 | 615 | 601 | 617 | 587 | 724 | 705 | 706 | 695 | 703 | 786 | 576 | 615 | 776 | 754 | 711 | 614 | 683 | 618 | 631 | 797 | 695 | 825 | 783 | 745 | Pex10 |
| Pex11a | 1068 | 1235 | 1074 | 1138 | 1002 | 951 | 1216 | 501 | 559 | 594 | 569 | 652 | 594 | 576 | 996 | 1063 | 1177 | 1164 | 1086 | 1128 | 1079 | 560 | 526 | 675 | 547 | 709 | 603 | 649 | Pex11a |
| Pex11b | 907 | 871 | 759 | 785 | 700 | 716 | 864 | 891 | 827 | 722 | 848 | 806 | 730 | 946 | 945 | 643 | 809 | 655 | 766 | 775 | 864 | 822 | 821 | 868 | 853 | 971 | 784 | 921 | Pex11b |
| Pex11g | 130 | 147 | 149 | 144 | 147 | 136 | 150 | 220 | 297 | 271 | 315 | 285 | 299 | 330 | 104 | 165 | 138 | 146 | 147 | 132 | 114 | 237 | 276 | 306 | 254 | 300 | 330 | 260 | Pex11g |
| Pex12 | 398 | 393 | 382 | 357 | 409 | 433 | 394 | 604 | 545 | 519 | 487 | 585 | 601 | 503 | 439 | 367 | 400 | 340 | 422 | 464 | 343 | 526 | 553 | 586 | 652 | 597 | 579 | 574 | Pex12 |
| Pex13 | 7005 | 5724 | 6171 | 6204 | 5184 | 5386 | 6254 | 4451 | 3649 | 3335 | 3321 | 3648 | 3605 | 3797 | 7354 | 6482 | 6117 | 6020 | 5413 | 6650 | 5960 | 3775 | 3721 | 3501 | 3639 | 4050 | 3660 | 4034 | Pex13 |
| Pex14 | 1467 | 1704 | 1844 | 1566 | 1510 | 1651 | 1454 | 1348 | 1702 | 1782 | 1666 | 1753 | 1655 | 1703 | 1379 | 1572 | 1769 | 1523 | 1643 | 1413 | 1466 | 1684 | 1741 | 1856 | 1758 | 2041 | 1777 | 1616 | Pex14 |
| Pex16 | 1053 | 1313 | 1032 | 1200 | 965 | 1068 | 951 | 784 | 910 | 857 | 813 | 884 | 845 | 700 | 940 | 1033 | 1213 | 1127 | 1070 | 1112 | 994 | 710 | 859 | 856 | 882 | 1007 | 883 | 859 | Pex16 |
| Pex19 | 10265 | 10793 | 9915 | 10026 | 9162 | 9776 | 9659 | 8215 | 7861 | 7948 | 8148 | 8863 | 8608 | 8744 | 10435 | 9962 | 9865 | 9625 | 9512 | 9763 | 9862 | 7730 | 8272 | 8388 | 7895 | 9060 | 9121 | 9228 | Pex19 |
| Pex2 | 3828 | 3061 | 3379 | 2945 | 2880 | 2974 | 3522 | 3819 | 3329 | 2943 | 3132 | 3715 | 3539 | 3959 | 4036 | 3206 | 3357 | 2958 | 3218 | 3816 | 3283 | 3248 | 3454 | 3312 | 3240 | 4091 | 4025 | 4035 | Pex2 |
| Pex26 | 2343 | 3047 | 3299 | 3609 | 3321 | 3129 | 3045 | 1561 | 1704 | 1723 | 1912 | 1648 | 1627 | 1608 | 2263 | 2694 | 3398 | 3565 | 3411 | 2805 | 3188 | 1768 | 1744 | 1800 | 2032 | 1751 | 1534 | 1577 | Pex26 |
| Pex3 | 2035 | 1583 | 1446 | 1449 | 1630 | 1643 | 1711 | 2067 | 2024 | 1784 | 1848 | 1770 | 1797 | 1702 | 2152 | 1699 | 1345 | 1478 | 1498 | 1878 | 1658 | 2059 | 2143 | 1523 | 1725 | 1749 | 1800 | 1842 | Pex3 |
| Pex5 | 6568 | 8040 | 8896 | 9274 | 8799 | 8075 | 8149 | 4449 | 5071 | 5422 | 5363 | 5843 | 5142 | 5328 | 6356 | 7415 | 9292 | 9022 | 9618 | 7014 | 8350 | 5621 | 4804 | 6028 | 6192 | 5735 | 5170 | 5414 | Pex5 |
| Pex5l | 652 | 652 | 455 | 639 | 824 | 656 | 577 | 815 | 769 | 673 | 668 | 553 | 881 | 624 | 872 | 791 | 523 | 616 | 606 | 687 | 603 | 732 | 731 | 589 | 608 | 468 | 587 | 584 | Pex5l |
| Pex6 | 4032 | 5266 | 5109 | 5287 | 4129 | 3818 | 3825 | 2651 | 3556 | 3541 | 3488 | 3368 | 3353 | 2665 | 3688 | 4889 | 5127 | 5281 | 5058 | 3498 | 3920 | 3153 | 3694 | 3585 | 3256 | 3621 | 3052 | 3010 | Pex6 |
| Pex7 | 388 | 408 | 485 | 539 | 553 | 521 | 534 | 1188 | 1218 | 1093 | 1061 | 1161 | 1258 | 1270 | 530 | 400 | 518 | 480 | 507 | 520 | 498 | 1195 | 1111 | 1209 | 1117 | 1237 | 1239 | 1161 | Pex7 |
| Pf4 | 408 | 421 | 658 | 353 | 310 | 313 | 303 | 336 | 399 | 357 | 446 | 488 | 780 | 483 | 367 | 500 | 385 | 338 | 290 | 395 | 267 | 328 | 533 | 414 | 353 | 735 | 676 | 525 | Pf4 |

| | | | | | | | | | | | | | | | | | | | | | | | | | | | | | |
|---|---|---|---|---|---|---|---|---|---|---|---|---|---|---|---|---|---|---|---|---|---|---|---|---|---|---|---|---|---|
| Pfas | 6151 | 6829 | 7024 | 7075 | 5507 | 5707 | 5602 | 4671 | 5238 | 5462 | 5445 | 7402 | 6321 | 6359 | 5540 | 6487 | 7017 | 6454 | 6315 | 5115 | 5384 | 4674 | 4841 | 6357 | 5770 | 6545 | 6890 | 6796 | Pfas |
| Pfdn1 | 2870 | 2737 | 3199 | 2435 | 2151 | 2169 | 2770 | 4284 | 3597 | 3675 | 3846 | 3920 | 3875 | 5647 | 2914 | 2526 | 2858 | 2324 | 2162 | 2866 | 2630 | 3096 | 3886 | 4332 | 3784 | 4493 | 4171 | 5327 | Pfdn1 |
| Pfdn2 | 5242 | 5346 | 7415 | 6884 | 5702 | 5546 | 6968 | 5203 | 4403 | 5602 | 5200 | 6192 | 5391 | 7715 | 5401 | 4407 | 7310 | 6001 | 6317 | 6005 | 7189 | 4991 | 4434 | 6838 | 6235 | 7152 | 6307 | 7562 | Pfdn2 |
| Pfdn4 | 2309 | 2138 | 2315 | 2628 | 2360 | 2300 | 2909 | 2424 | 1984 | 2238 | 1662 | 1987 | 2066 | 2505 | 2457 | 2120 | 2293 | 2374 | 2226 | 2987 | 2828 | 1763 | 1956 | 2173 | 2083 | 2129 | 2183 | 2529 | Pfdn4 |
| Pfdn5 | 7916 | 7749 | 7661 | 7965 | 7135 | 7060 | 8072 | 8144 | 7383 | 7075 | 7150 | 6923 | 7159 | 8288 | 8064 | 6508 | 7296 | 7090 | 7690 | 8653 | 8284 | 6958 | 7558 | 7797 | 7537 | 8320 | 7891 | 8505 | Pfdn5 |
| Pfdn6 | 4828 | 5941 | 8189 | 7520 | 5743 | 5697 | 6023 | 3110 | 3509 | 4303 | 4015 | 4898 | 4501 | 4860 | 5047 | 5074 | 7738 | 6534 | 7063 | 6424 | 6012 | 2879 | 3123 | 5308 | 4905 | 6242 | 5316 | 5381 | Pfdn6 |
| Pfkfb1 | 251 | 181 | 199 | 181 | 148 | 203 | 202 | 281 | 321 | 270 | 298 | 320 | 327 | 331 | 227 | 188 | 163 | 138 | 158 | 173 | 144 | 246 | 324 | 203 | 265 | 322 | 311 | 323 | Pfkfb1 |
| Pfkfb2 | 2885 | 2580 | 2093 | 2954 | 3052 | 3441 | 3194 | 2260 | 2191 | 2032 | 2166 | 2094 | 2034 | 2091 | 3169 | 2680 | 2405 | 3003 | 2935 | 3070 | 3244 | 2413 | 1990 | 1950 | 2312 | 1613 | 2099 | 1803 | Pfkfb2 |
| Pfkfb3 | 1651 | 1614 | 1434 | 1764 | 2715 | 2222 | 2321 | 4263 | 4238 | 4678 | 4494 | 3919 | 3770 | 4492 | 1783 | 1562 | 1763 | 1955 | 2377 | 2274 | 2318 | 4944 | 4030 | 4241 | 5142 | 3721 | 3628 | 3823 | Pfkfb3 |
| Pfkfb4 | 1407 | 1823 | 1509 | 1861 | 1865 | 1655 | 1585 | 1304 | 1525 | 1508 | 1475 | 1389 | 1360 | 1303 | 1495 | 1716 | 1618 | 1805 | 1894 | 1662 | 1618 | 1559 | 1484 | 1349 | 1566 | 1260 | 1370 | 1219 | Pfkfb4 |
| Pfkl | 7491 | 8492 | 7862 | 7638 | 7041 | 7274 | 6126 | 7119 | 9334 | 8927 | 8494 | 8823 | 8557 | 7650 | 6915 | 7871 | 8190 | 7653 | 7608 | 6648 | 6464 | 7434 | 9397 | 9628 | 8481 | 10890 | 8832 | 8705 | Pfkl |
| Pfkm | 5869 | 6830 | 6354 | 7407 | 7616 | 7191 | 6795 | 6378 | 7130 | 7744 | 7248 | 7262 | 6561 | 6519 | 5991 | 6054 | 6873 | 7558 | 8121 | 7065 | 7311 | 7177 | 7282 | 7387 | 7949 | 7439 | 6937 | 6594 | Pfkm |
| Pfkp | 17466 | 19196 | 10467 | 16703 | 26365 | 24239 | 22248 | 14001 | 13505 | 12295 | 12724 | 5740 | 8448 | 6002 | 21097 | 18890 | 12396 | 18757 | 18809 | 25689 | 24972 | 12579 | 11524 | 5671 | 12644 | 2283 | 5921 | 4480 | Pfkp |
| Pfn1 | 10282 | 11118 | 12066 | 8742 | 7769 | 7508 | 8118 | 10243 | 12128 | 10464 | 11872 | 12530 | 12694 | 13100 | 9260 | 10465 | 10711 | 7894 | 8871 | 7385 | 8073 | 9712 | 12317 | 13041 | 10384 | 15633 | 12729 | 14223 | Pfn1 |
| Pfn2 | 19910 | 16628 | 18226 | 21125 | 24296 | 22731 | 24132 | 26555 | 22171 | 24069 | 20930 | 23381 | 20158 | 24028 | 21841 | 16401 | 20619 | 21860 | 23053 | 22934 | 24377 | 29066 | 22427 | 23265 | 26150 | 23311 | 21673 | 23356 | Pfn2 |
| Pfn3 | 2 | 3 | 3 | 1 | 6 | 0 | 3 | 0 | 0 | 0 | 1 | 0 | 0 | 1 | 1 | 5 | 0 | 1 | 4 | 12 | 2 | 0 | 3 | 2 | 1 | 0 | 2 | 2 | Pfn3 |
| Pfn4 | 54 | 47 | 74 | 78 | 142 | 85 | 104 | 365 | 311 | 399 | 327 | 332 | 295 | 303 | 91 | 51 | 83 | 69 | 127 | 121 | 97 | 353 | 347 | 313 | 385 | 318 | 323 | 295 | Pfn4 |
| Pfpl | 0 | 0 | 0 | 0 | 0 | 0 | 0 | 2 | 0 | 0 | 0 | 0 | 0 | 0 | 0 | 2 | 0 | 0 | 1 | 0 | 0 | 0 | 0 | 0 | 0 | 0 | 0 | 0 | Pfpl |
| Pga5 | 1 | 2 | 0 | 1 | 5 | 0 | 1 | 0 | 9 | 0 | 1 | 0 | 0 | 2 | 1 | 0 | 1 | 2 | 5 | 0 | 3 | 0 | 8 | 6 | 2 | 0 | 0 | 1 | Pga5 |
| Pgam1 | 14690 | 15812 | 15547 | 13230 | 11898 | 12700 | 12583 | 8926 | 10332 | 9921 | 9403 | 11376 | 11323 | 10434 | 13676 | 14988 | 14638 | 13265 | 13897 | 12432 | 11881 | 8813 | 10861 | 10685 | 9599 | 13984 | 11207 | 11993 | Pgam1 |
| Pgam2 | 37 | 79 | 75 | 6 | 3 | 25 | 0 | 27 | 91 | 25 | 366 | 23 | 107 | 32 | 57 | 100 | 29 | 44 | 2 | 6 | 4 | 62 | 261 | 47 | 24 | 47 | 60 | 46 | Pgam2 |
| Pgam5 | 5756 | 5461 | 4449 | 4525 | 4147 | 4558 | 4902 | 4756 | 4211 | 3784 | 3595 | 4313 | 3798 | 4192 | 5933 | 4880 | 4602 | 4220 | 4552 | 5388 | 4696 | 3887 | 4145 | 3853 | 3796 | 4523 | 4317 | 4263 | Pgam5 |
| Pgap1 | 19229 | 16306 | 11878 | 18078 | 20805 | 22972 | 25251 | 16919 | 13500 | 11296 | 10320 | 11604 | 9787 | 12140 | 22504 | 16080 | 14278 | 18832 | 19187 | 28970 | 24771 | 11787 | 12691 | 9822 | 13953 | 7910 | 11489 | 10703 | Pgap1 |
| Pgap2 | 1151 | 957 | 993 | 1048 | 943 | 1003 | 1076 | 1263 | 1110 | 1351 | 1301 | 1165 | 1205 | 1227 | 1153 | 1120 | 962 | 1020 | 923 | 886 | 1134 | 1094 | 1221 | 1253 | 1214 | 1409 | 1097 | 1231 | Pgap2 |
| Pgap3 | 127 | 158 | 165 | 80 | 144 | 142 | 137 | 518 | 551 | 552 | 567 | 571 | 606 | 552 | 146 | 121 | 147 | 102 | 134 | 151 | 118 | 495 | 594 | 588 | 564 | 712 | 589 | 592 | Pgap3 |
| Pgap4 | 950 | 1161 | 1010 | 1090 | 1532 | 1360 | 1131 | 1533 | 1790 | 1617 | 1575 | 1135 | 1264 | 1121 | 1054 | 928 | 999 | 1231 | 1413 | 1381 | 1225 | 1811 | 1694 | 1289 | 1607 | 1076 | 1184 | 1150 | Pgap4 |
| Pgap6 | 322 | 420 | 496 | 374 | 416 | 391 | 382 | 346 | 339 | 455 | 352 | 339 | 386 | 298 | 361 | 371 | 509 | 414 | 421 | 391 | 387 | 396 | 400 | 383 | 404 | 373 | 330 | 361 | Pgap6 |
| Pgbd1 | 449 | 461 | 679 | 462 | 443 | 434 | 446 | 683 | 600 | 730 | 618 | 774 | 743 | 824 | 483 | 426 | 451 | 376 | 457 | 461 | 437 | 568 | 522 | 780 | 650 | 932 | 824 | 794 | Pgbd1 |
| Pgbd5 | 1430 | 2338 | 2122 | 2914 | 2838 | 2423 | 2446 | 946 | 966 | 1196 | 1053 | 1065 | 801 | 909 | 1735 | 1950 | 2473 | 2915 | 2725 | 2220 | 2511 | 1180 | 988 | 1039 | 1135 | 1089 | 915 | 999 | Pgbd5 |
| Pgc | 0 | 0 | 1 | 0 | 0 | 0 | 1 | 1 | 2 | 10 | 2 | 0 | 9 | 0 | 0 | 0 | 0 | 0 | 1 | 0 | 0 | 2 | 4 | 1 | 2 | 1 | 5 | 0 | Pgc |
| Pgd | 10477 | 13499 | 13354 | 11851 | 10064 | 10820 | 9899 | 7228 | 8150 | 7759 | 8247 | 8842 | 9461 | 9197 | 10303 | 11559 | 12120 | 11017 | 11198 | 10717 | 10608 | 6811 | 8287 | 8579 | 8323 | 10433 | 10361 | 10338 | Pgd |
| Pgf | 356 | 355 | 249 | 266 | 264 | 384 | 235 | 374 | 449 | 339 | 453 | 331 | 384 | 355 | 332 | 404 | 267 | 328 | 312 | 309 | 262 | 414 | 498 | 371 | 406 | 413 | 303 | 324 | Pgf |
| Pgghg | 978 | 1051 | 1061 | 739 | 746 | 853 | 591 | 1230 | 1578 | 1427 | 1396 | 1521 | 1631 | 1260 | 931 | 1136 | 822 | 738 | 764 | 654 | 608 | 1088 | 1676 | 1354 | 1014 | 1739 | 1529 | 1491 | Pgghg |
| Pggt1b | 5696 | 4321 | 5237 | 5566 | 4959 | 5434 | 5401 | 5947 | 5188 | 5195 | 5522 | 5861 | 5671 | 5814 | 5898 | 5227 | 4877 | 4880 | 4867 | 5783 | 5362 | 4868 | 5242 | 4857 | 5259 | 4778 | 5798 | 5587 | Pggt1b |
| Pgk1 | 34702 | 34446 | 31617 | 26016 | 23404 | 25589 | 26156 | 22556 | 23152 | 18591 | 19610 | 22619 | 23126 | 23302 | 32591 | 32126 | 28963 | 26133 | 26634 | 30624 | 26059 | 17276 | 23607 | 21533 | 19931 | 29621 | 25221 | 25686 | Pgk1 |
| Pgk2 | 0 | 0 | 0 | 0 | 0 | 0 | 0 | 0 | 1 | 0 | 0 | 0 | 0 | 0 | 0 | 0 | 0 | 0 | 0 | 0 | 0 | 0 | 0 | 0 | 0 | 0 | 0 | 0 | Pgk2 |
| Pgls | 2705 | 3216 | 2527 | 2519 | 2295 | 2322 | 2130 | 2810 | 2864 | 2819 | 2647 | 2850 | 2961 | 2624 | 2442 | 2319 | 2462 | 2157 | 2674 | 2437 | 2249 | 2513 | 2753 | 2855 | 2626 | 3237 | 2887 | 2784 | Pgls |
| Pglyrp1 | 27 | 49 | 44 | 51 | 85 | 87 | 85 | 67 | 87 | 113 | 63 | 61 | 65 | 69 | 28 | 19 | 36 | 69 | 79 | 72 | 95 | 75 | 88 | 87 | 100 | 53 | 56 | 65 | Pglyrp1 |
| Pglyrp2 | 1 | 0 | 0 | 0 | 0 | 0 | 0 | 0 | 2 | 0 | 0 | 1 | 5 | 1 | 3 | 1 | 0 | 2 | 0 | 0 | 0 | 0 | 3 | 1 | 0 | 6 | 1 | 0 | Pglyrp2 |
| Pglyrp3 | 0 | 0 | 0 | 0 | 0 | 0 | 0 | 6 | 9 | 9 | 1 | 0 | 0 | 0 | 0 | 0 | 0 | 0 | 0 | 0 | 0 | 0 | 12 | 0 | 0 | 0 | 0 | 0 | Pglyrp3 |
| Pglyrp4 | 0 | 0 | 0 | 0 | 0 | 0 | 0 | 0 | 0 | 0 | 0 | 0 | 0 | 0 | 0 | 0 | 0 | 1 | 0 | 0 | 0 | 0 | 0 | 0 | 0 | 0 | 0 | 0 | Pglyrp4 |
| Pgm1 | 0 | 0 | 0 | 0 | 1 | 0 | 0 | 0 | 1 | 0 | 0 | 0 | 1 | 0 | 0 | 0 | 0 | 0 | 1 | 0 | 0 | 0 | 1 | 1 | 1 | 0 | 0 | 0 | Pgm1 |
| Pgm2 | 4384 | 4274 | 3814 | 3957 | 3738 | 3924 | 4073 | 2119 | 2167 | 2292 | 2185 | 2111 | 2141 | 2313 | 3952 | 4203 | 4074 | 4001 | 4083 | 4138 | 4161 | 1823 | 2255 | 2339 | 2068 | 2562 | 2219 | 2254 | Pgm2 |
| Pgm2l1 | 12093 | 9887 | 9854 | 13020 | 16336 | 16037 | 15320 | 14696 | 11059 | 12822 | 11400 | 10663 | 12580 | 11424 | 13260 | 11751 | 11561 | 13063 | 13606 | 14784 | 16303 | 14639 | 11035 | 10089 | 13337 | 6538 | 11718 | 11783 | Pgm2l1 |
| Pgm3 | 4105 | 3539 | 2907 | 3250 | 3156 | 3395 | 3405 | 2523 | 2302 | 2042 | 2091 | 2027 | 2101 | 2164 | 4139 | 3708 | 2866 | 3169 | 3174 | 3728 | 3336 | 2135 | 2351 | 2059 | 2211 | 1792 | 2271 | 2169 | Pgm3 |
| Pgm5 | 890 | 668 | 363 | 668 | 583 | 1158 | 220 | 1494 | 2366 | 1465 | 2665 | 727 | 1339 | 802 | 897 | 1082 | 520 | 1035 | 281 | 510 | 317 | 1230 | 3030 | 1154 | 1198 | 592 | 661 | 699 | Pgm5 |
| Pgp | 3580 | 4399 | 4004 | 4565 | 3510 | 3586 | 3690 | 1876 | 1808 | 1944 | 1972 | 2323 | 2048 | 2101 | 3471 | 3777 | 4132 | 4325 | 3670 | 3234 | 3808 | 1953 | 1879 | 2117 | 2162 | 2486 | 2158 | 2245 | Pgp |
| Pgpep1 | 6234 | 8978 | 7745 | 7938 | 5830 | 6086 | 5596 | 4257 | 3854 | 4166 | 3926 | 4628 | 5441 | 4558 | 6613 | 7675 | 8306 | 7154 | 6547 | 5900 | 5987 | 3940 | 4259 | 4327 | 4234 | 4442 | 5396 | 5207 | Pgpep1 |
| Pgpep1l | 29 | 28 | 21 | 29 | 27 | 22 | 22 | 6 | 7 | 19 | 31 | 10 | 17 | 6 | 21 | 45 | 21 | 34 | 12 | 18 | 13 | 15 | 19 | 11 | 2 | 7 | 9 | 11 | Pgpep1l |
| Pgr | 260 | 232 | 194 | 167 | 180 | 292 | 188 | 161 | 210 | 155 | 109 | 89 | 136 | 84 | 214 | 226 | 169 | 133 | 182 | 264 | 243 | 164 | 199 | 94 | 157 | 62 | 89 | 88 | Pgr |
| Pgr15l | 78 | 65 | 35 | 38 | 24 | 32 | 30 | 1 | 8 | 8 | 2 | 4 | 0 | 2 | 59 | 67 | 32 | 39 | 16 | 43 | 19 | 7 | 3 | 2 | 2 | 4 | 2 | 1 | Pgr15l |
| Pgrmc1 | 20948 | 18645 | 19555 | 20102 | 20810 | 20609 | 22264 | 24076 | 20472 | 20946 | 18838 | 21603 | 20693 | 21212 | 22871 | 19439 | 20594 | 19461 | 21122 | 23121 | 21895 | 24134 | 22453 | 21013 | 21936 | 21823 | 21214 | 21421 | Pgrmc1 |
| Pgrmc2 | 6942 | 6284 | 7702 | 7462 | 5779 | 5827 | 6250 | 3631 | 3451 | 3901 | 4153 | 4238 | 3977 | 4228 | 6502 | 6671 | 7051 | 7037 | 5915 | 4898 | 6279 | 4431 | 3670 | 4403 | 3896 | 4794 | 3677 | 4540 | Pgrmc2 |
| Pgs1 | 2581 | 3191 | 2771 | 3048 | 2828 | 2520 | 2516 | 2274 | 2706 | 2871 | 2606 | 2766 | 2776 | 2848 | 2546 | 2940 | 2876 | 3028 | 2818 | 2302 | 2606 | 2542 | 2515 | 2696 | 3158 | 2857 | 2909 | 2705 | Pgs1 |
| Phactr1 | 1445 | 1634 | 1638 | 2712 | 3908 | 3410 | 3328 | 3326 | 3668 | 4764 | 3934 | 3548 | 3058 | 3323 | 1735 | 1445 | 2194 | 2954 | 3561 | 2741 | 3752 | 4796 | 3328 | 4305 | 5201 | 2829 | 2841 | 3031 | Phactr1 |
| Phactr2 | 3699 | 3074 | 2936 | 2620 | 2623 | 2993 | 2514 | 3948 | 4701 | 4171 | 6545 | 4323 | 4737 | 4289 | 3572 | 3826 | 2736 | 2543 | 2242 | 2629 | 2449 | 4234 | 5450 | 3754 | 3648 | 3714 | 4661 | 4129 | Phactr2 |
| Phactr3 | 744 | 864 | 1019 | 1602 | 2437 | 1886 | 1994 | 2194 | 2234 | 3140 | 2434 | 2502 | 1808 | 2607 | 1062 | 752 | 1352 | 1699 | 2067 | 1786 | 2153 | 3298 | 2093 | 2880 | 3358 | 2055 | 2043 | 2101 | Phactr3 |
| Phactr4 | 3388 | 3592 | 4687 | 4463 | 3965 | 3841 | 3800 | 3777 | 4256 | 4991 | 4804 | 6532 | 5330 | 6297 | 3046 | 3581 | 4642 | 4237 | 4535 | 3066 | 3941 | 4461 | 3829 | 5898 | 5394 | 6744 | 5174 | 6437 | Phactr4 |
| Phaf1 | 3637 | 3936 | 3370 | 3614 | 3189 | 3345 | 3365 | 2341 | 2233 | 2243 | 2049 | 2260 | 2288 | 2156 | 3659 | 3708 | 3271 | 3587 | 3158 | 3384 | 3247 | 2314 | 2255 | 2110 | 2415 | 2135 | 2241 | 2308 | Phaf1 |
| Phax | 3067 | 3266 | 3608 | 3305 | 3298 | 3312 | 3509 | 4148 | 4001 | 3920 | 3677 | 4015 | 3963 | 4104 | 3494 | 3028 | 3452 | 3170 | 3439 | 4040 | 3150 | 3741 | 3871 | 4192 | 4248 | 4791 | 4202 | 4207 | Phax |
| Phb | 7841 | 9316 | 8278 | 7675 | 6545 | 7038 | 6696 | 5835 | 6700 | 5700 | 5861 | 5663 | 6014 | 5484 | 7842 | 8629 | 8070 | 7636 | 7779 | 7880 | 6973 | 4734 | 6686 | 5956 | 5883 | 7243 | 6446 | 6270 | Phb |
| Phb2 | 10064 | 11711 | 11331 | 10510 | 8800 | 9201 | 9247 | 7245 | 8071 | 8175 | 8227 | 9117 | 8932 | 9151 | 9198 | 9933 | 11019 | 9708 | 10302 | 9270 | 9414 | 6853 | 8260 | 9127 | 8526 | 11015 | 9395 | 10049 | Phb2 |
| Phc1 | 8168 | 8980 | 8753 | 9751 | 10287 | 9853 | 9192 | 8838 | 9535 | 10528 | 9417 | 11966 | 10557 | 10977 | 7692 | 8023 | 9457 | 9413 | 10344 | 8531 | 9240 | 11339 | 8473 | 10919 | 11308 | 10952 | 10280 | 10332 | Phc1 |
| Phc2 | 9185 | 11994 | 12375 | 12394 | 11081 | 10795 | 9988 | 8421 | 9783 | 11079 | 10123 | 14861 | 11132 | 13293 | 8616 | 10305 | 12526 | 12027 | 12441 | 9065 | 10844 | 10535 | 8870 | 13349 | 12482 | 15459 | 11935 | 13013 | Phc2 |
| Phc3 | 7425 | 7111 | 4791 | 7542 | 8225 | 9289 | 8994 | 6837 | 6891 | 6115 | 7278 | 5686 | 6001 | 6187 | 7626 | 7580 | 5500 | 8151 | 7187 | 8658 | 9688 | 5910 | 6331 | 5054 | 7051 | 3613 | 5963 | 5146 | Phc3 |
| Pheta1 | 2969 | 4047 | 3741 | 4264 | 3734 | 3624 | 3248 | 2158 | 2357 | 2571 | 2394 | 2671 | 2501 | 2313 | 2871 | 3516 | 3975 | 4248 | 4338 | 3500 | 3503 | 2640 | 2319 | 2899 | 2897 | 2974 | 2581 | 2391 | Pheta1 |
| Pheta2 | 135 | 86 | 101 | 74 | 91 | 87 | 74 | 276 | 377 | 335 | 418 | 255 | 379 | 268 | 137 | 105 | 82 | 81 | 73 | 70 | 46 | 227 | 432 | 288 | 213 | 281 | 298 | 356 | Pheta2 |
| Phex | 115 | 71 | 75 | 155 | 174 | 158 | 174 | 136 | 127 | 149 | 509 | 104 | 109 | 103 | 150 | 142 | 144 | 183 | 135 | 124 | 189 | 134 | 480 | 101 | 113 | 60 | 110 | 88 | Phex |
| Phf1 | 1588 | 1961 | 1979 | 2516 | 2213 | 2046 | 1963 | 1568 | 1845 | 2173 | 2150 | 1833 | 1796 | 1789 | 1589 | 1991 | 2094 | 2467 | 2340 | 1814 | 2018 | 2191 | 1868 | 2093 | 2096 | 1809 | 1772 | 1664 | Phf1 |
| Phf10 | 10837 | 8444 | 9659 | 9380 | 8260 | 8853 | 9232 | 8501 | 7806 | 8176 | 8529 | 8590 | 8129 | 8622 | 9415 | 10076 | 9021 | 9327 | 8278 | 7893 | 8597 | 8224 | 7769 | 8466 | 7952 | 8620 | 8149 | 8874 | Phf10 |
| Phf11a | 1 | 0 | 0 | 2 | 2 | 1 | 1 | 1 | 5 | 0 | 0 | 0 | 0 | 0 | 1 | 0 | 0 | 1 | 0 | 1 | 1 | 0 | 2 | 2 | 0 | 0 | 0 | 2 | Phf11a |
| Phf11b | 14 | 13 | 4 | 9 | 8 | 6 | 17 | 7 | 8 | 14 | 10 | 9 | 7 | 4 | 20 | 11 | 13 | 13 | 10 | 7 | 21 | 10 | 7 | 9 | 13 | 9 | 6 | 8 | Phf11b |
| Phf11c | 691 | 646 | 628 | 783 | 574 | 635 | 654 | 239 | 220 | 258 | 282 | 241 | 208 | 252 | 612 | 794 | 689 | 741 | 531 | 570 | 609 | 216 | 218 | 244 | 198 | 243 | 246 | 237 | Phf11c |
| Phf11d | 42 | 57 | 39 | 78 | 30 | 41 | 50 | 11 | 17 | 28 | 28 | 20 | 7 | 9 | 67 | 74 | 78 | 58 | 34 | 47 | 65 | 5 | 23 | 15 | 14 | 14 | 20 | 11 | Phf11d |
| Phf12 | 7194 | 8612 | 9622 | 10219 | 9898 | 8958 | 9075 | 6161 | 7055 | 7999 | 7448 | 9392 | 7765 | 8608 | 6699 | 7750 | 10179 | 9799 | 10077 | 7363 | 9297 | 7870 | 6260 | 8274 | 8184 | 8874 | 8291 | 8196 | Phf12 |
| Phf13 | 3730 | 3957 | 4649 | 4503 | 3879 | 3683 | 3486 | 2375 | 2496 | 2331 | 2763 | 2459 | 2431 | 2168 | 3335 | 3897 | 4590 | 4390 | 4127 | 3254 | 3600 | 2431 | 2575 | 2287 | 2466 | 2935 | 2616 | 2334 | Phf13 |
| Phf14 | 7423 | 6197 | 7824 | 7708 | 9133 | 9466 | 9390 | 14884 | 12726 | 13585 | 12943 | 14777 | 14174 | 16244 | 8252 | 6636 | 7538 | 7690 | 8654 | 8963 | 9173 | 14979 | 12389 | 13798 | 14698 | 14488 | 15339 | 15044 | Phf14 |
| Phf19 | 232 | 268 | 313 | 266 | 275 | 271 | 236 | 305 | 466 | 397 | 401 | 532 | 307 | 459 | 208 | 203 | 285 | 205 | 299 | 253 | 215 | 326 | 395 | 410 | 410 | 554 | 457 | 422 | Phf19 |
| Phf2 | 3192 | 3851 | 5633 | 4848 | 5919 | 4741 | 4394 | 6171 | 7472 | 9688 | 8807 | 10341 | 8063 | 9600 | 3153 | 3469 | 5539 | 4712 | 6302 | 3830 | 4935 | 9112 | 6658 | 10791 | 9899 | 10373 | 8147 | 8953 | Phf2 |

| | | | | | | | | | | | | | | | | | | | | | | | | | | | | | |
|---|---|---|---|---|---|---|---|---|---|---|---|---|---|---|---|---|---|---|---|---|---|---|---|---|---|---|---|---|---|
| Phf20 | 9326 | 9070 | 7653 | 9146 | 9659 | 9674 | 10254 | 7995 | 7392 | 7811 | 6980 | 7714 | 7038 | 7496 | 9808 | 8449 | 8620 | 9487 | 9104 | 10089 | 9886 | 7133 | 7091 | 7303 | 7812 | 6397 | 7764 | 7334 | Phf20 |
| Phf20l1 | 12901 | 9044 | 8624 | 11511 | 12066 | 14017 | 15046 | 15560 | 13632 | 12650 | 12657 | 12734 | 12037 | 13729 | 13121 | 11292 | 9684 | 11642 | 11208 | 14045 | 14799 | 12469 | 12558 | 12374 | 14354 | 9638 | 12605 | 12057 | Phf20l1 |
| Phf21a | 7575 | 7552 | 8638 | 9596 | 10933 | 9760 | 10632 | 7999 | 7977 | 9370 | 8818 | 9713 | 8977 | 9771 | 7452 | 6827 | 9868 | 9695 | 11010 | 9180 | 11138 | 8748 | 7536 | 9997 | 9635 | 8399 | 9238 | 9079 | Phf21a |
| Phf21b | 4854 | 6631 | 8481 | 8664 | 9056 | 8076 | 7222 | 6090 | 6008 | 9052 | 6865 | 12767 | 9046 | 11000 | 5164 | 5021 | 9962 | 7711 | 9745 | 6455 | 7649 | 8779 | 5080 | 10763 | 10505 | 11594 | 10931 | 11165 | Phf21b |
| Phf23 | 4420 | 5447 | 5926 | 5598 | 5254 | 5164 | 5363 | 4730 | 4686 | 4871 | 5160 | 5648 | 5245 | 6009 | 4210 | 4399 | 5726 | 5487 | 5871 | 4636 | 5470 | 4604 | 4916 | 5727 | 5224 | 6668 | 5497 | 6404 | Phf23 |
| Phf24 | 2212 | 2718 | 1806 | 3847 | 5507 | 4755 | 4241 | 2599 | 2946 | 3300 | 2888 | 1876 | 2097 | 1955 | 2781 | 2720 | 2788 | 4503 | 4216 | 3579 | 4782 | 4361 | 2421 | 2214 | 3627 | 981 | 1664 | 1540 | Phf24 |
| Phf3 | 10720 | 9896 | 8728 | 10645 | 10971 | 11147 | 11336 | 13643 | 11947 | 11901 | 12231 | 12029 | 12014 | 12837 | 10445 | 9890 | 9507 | 10528 | 10644 | 11329 | 11526 | 11921 | 11753 | 11450 | 12864 | 8734 | 12414 | 11769 | Phf3 |
| Phf5a | 4390 | 4996 | 4919 | 4211 | 3724 | 3991 | 4255 | 3845 | 3945 | 3971 | 3797 | 4605 | 4568 | 5029 | 4247 | 4509 | 4395 | 4140 | 4408 | 4696 | 4127 | 3394 | 3926 | 4545 | 4072 | 5841 | 4969 | 5292 | Phf5a |
| Phf6 | 12952 | 9448 | 10440 | 9782 | 11083 | 10747 | 12501 | 16321 | 12947 | 12005 | 10955 | 14139 | 13061 | 15520 | 12539 | 10552 | 10362 | 9556 | 11113 | 12595 | 12071 | 13226 | 12079 | 13473 | 13814 | 14244 | 14704 | 14663 | Phf6 |
| Phf7 | 1054 | 833 | 900 | 910 | 839 | 822 | 1009 | 805 | 807 | 863 | 782 | 907 | 908 | 945 | 915 | 879 | 845 | 815 | 747 | 948 | 936 | 752 | 836 | 734 | 842 | 959 | 873 | 904 | Phf7 |
| Phf8 | 5879 | 5408 | 5325 | 5156 | 5455 | 5438 | 5188 | 5302 | 4815 | 5088 | 4492 | 5910 | 5218 | 6102 | 5559 | 4780 | 5683 | 4872 | 5602 | 5011 | 5501 | 5868 | 4324 | 5689 | 5685 | 4519 | 5536 | 5895 | Phf8 |
| Phgdh | 11195 | 14700 | 12576 | 10254 | 8446 | 9437 | 8765 | 8072 | 9532 | 7287 | 6951 | 9816 | 8960 | 9427 | 10772 | 11533 | 11330 | 8945 | 10584 | 10336 | 9075 | 5518 | 8813 | 8338 | 7467 | 11708 | 10644 | 10671 | Phgdh |
| Phgr1 | 2 | 0 | 0 | 4 | 0 | 1 | 1 | 0 | 1 | 0 | 0 | 0 | 0 | 0 | 0 | 0 | 3 | 0 | 0 | 0 | 1 | 0 | 0 | 0 | 0 | 0 | 0 | 0 | Phgr1 |
| Phip | 21780 | 15038 | 12006 | 16096 | 17893 | 22248 | 21245 | 34854 | 27465 | 21457 | 22044 | 25151 | 23414 | 27412 | 21618 | 18303 | 13004 | 15256 | 17379 | 24305 | 21823 | 20633 | 25475 | 19632 | 25574 | 17394 | 27502 | 24733 | Phip |
| Phka1 | 1716 | 1255 | 1216 | 1132 | 1384 | 1459 | 1310 | 1556 | 1652 | 1586 | 1899 | 1537 | 1486 | 1413 | 1472 | 1502 | 1325 | 1339 | 1313 | 1212 | 1320 | 1507 | 1600 | 1460 | 1410 | 1569 | 1485 | 1340 | Phka1 |
| Phka2 | 4104 | 3885 | 2829 | 2873 | 2765 | 3469 | 3213 | 2742 | 2683 | 2455 | 2487 | 2989 | 2936 | 2916 | 3749 | 3578 | 2706 | 2849 | 2953 | 3218 | 3441 | 2191 | 2629 | 2307 | 2521 | 2334 | 3138 | 2727 | Phka2 |
| Phkb | 4314 | 3688 | 3282 | 3012 | 3264 | 3365 | 3074 | 5178 | 4752 | 4267 | 4543 | 4288 | 4661 | 4461 | 4263 | 3996 | 2977 | 2987 | 2894 | 3739 | 3233 | 4169 | 4810 | 3684 | 3768 | 3808 | 4652 | 4364 | Phkb |
| Phkg1 | 14 | 20 | 12 | 14 | 15 | 28 | 19 | 11 | 11 | 14 | 8 | 13 | 8 | 10 | 11 | 13 | 12 | 18 | 15 | 17 | 13 | 15 | 8 | 9 | 10 | 15 | 10 | 13 | Phkg1 |
| Phkg2 | 3542 | 4267 | 3850 | 4420 | 3938 | 4008 | 4121 | 2999 | 2960 | 3421 | 3021 | 3320 | 3239 | 3338 | 3504 | 3428 | 4204 | 3974 | 4304 | 3986 | 4125 | 3337 | 2963 | 3441 | 3369 | 3363 | 3242 | 3369 | Phkg2 |
| Phlda1 | 794 | 843 | 1069 | 1196 | 1492 | 933 | 870 | 1377 | 1472 | 1833 | 2052 | 1072 | 1471 | 1014 | 808 | 1001 | 1082 | 1069 | 1177 | 944 | 978 | 1604 | 1664 | 1447 | 1370 | 1313 | 886 | 869 | Phlda1 |
| Phlda2 | 42 | 29 | 19 | 18 | 11 | 14 | 6 | 13 | 25 | 14 | 62 | 9 | 23 | 6 | 17 | 30 | 28 | 31 | 23 | 7 | 1 | 17 | 43 | 8 | 23 | 15 | 17 | 7 | Phlda2 |
| Phlda3 | 449 | 386 | 453 | 297 | 418 | 429 | 260 | 470 | 568 | 486 | 584 | 398 | 438 | 337 | 316 | 424 | 448 | 352 | 426 | 324 | 282 | 338 | 711 | 554 | 357 | 494 | 482 | 360 | Phlda3 |
| Phldb1 | 7171 | 9503 | 10468 | 10124 | 10538 | 10064 | 8995 | 7527 | 9354 | 10387 | 10111 | 11280 | 10578 | 10167 | 7119 | 8904 | 10454 | 9858 | 10956 | 8975 | 9246 | 10686 | 8730 | 10835 | 11312 | 9757 | 10524 | 9990 | Phldb1 |
| Phldb2 | 8059 | 6258 | 7134 | 4304 | 4271 | 5910 | 3950 | 7238 | 7969 | 6720 | 9437 | 8617 | 10816 | 8713 | 7470 | 8835 | 5439 | 5001 | 3854 | 4357 | 3858 | 5250 | 8896 | 7276 | 5639 | 8367 | 9747 | 9404 | Phldb2 |
| Phldb3 | 242 | 328 | 268 | 300 | 150 | 156 | 137 | 53 | 95 | 78 | 35 | 26 | 24 | 16 | 195 | 273 | 338 | 254 | 156 | 129 | 155 | 13 | 121 | 52 | 7 | 68 | 20 | 21 | Phldb3 |
| Phlpp1 | 4009 | 4196 | 4156 | 4886 | 5339 | 4712 | 5301 | 3678 | 3423 | 3989 | 3351 | 4502 | 3724 | 4379 | 4050 | 3597 | 4854 | 5082 | 5291 | 4255 | 5117 | 3967 | 3193 | 3843 | 4299 | 3866 | 3918 | 4342 | Phlpp1 |
| Phlpp2 | 3128 | 2822 | 2467 | 2990 | 3502 | 3663 | 3360 | 3742 | 3727 | 3532 | 4015 | 3652 | 3711 | 3812 | 2800 | 2634 | 2658 | 2982 | 3434 | 3199 | 3404 | 3541 | 3418 | 3335 | 3905 | 2678 | 3810 | 3824 | Phlpp2 |
| Phospho1 | 5 | 6 | 4 | 0 | 2 | 2 | 0 | 0 | 0 | 12 | 3 | 0 | 0 | 2 | 1 | 4 | 0 | 6 | 1 | 1 | 3 | 0 | 7 | 1 | 0 | 0 | 3 | 0 | Phospho1 |
| Phospho2 | 2649 | 2524 | 2465 | 2587 | 2472 | 2300 | 2946 | 2639 | 1996 | 1833 | 1668 | 1972 | 1953 | 2107 | 3098 | 2527 | 2575 | 2424 | 2731 | 3082 | 2780 | 2102 | 2067 | 1849 | 1983 | 2082 | 1945 | 2159 | Phospho2 |
| Phox2a | 38 | 44 | 76 | 39 | 84 | 35 | 22 | 50 | 44 | 54 | 103 | 11 | 65 | 1 | 82 | 44 | 70 | 43 | 26 | 33 | 23 | 77 | 83 | 8 | 20 | 0 | 7 | 6 | Phox2a |
| Phox2b | 52 | 244 | 98 | 172 | 548 | 231 | 19 | 843 | 700 | 1025 | 724 | 55 | 1218 | 34 | 525 | 377 | 300 | 180 | 51 | 115 | 32 | 1054 | 632 | 32 | 204 | 2 | 43 | 59 | Phox2b |
| Phpt1 | 62 | 68 | 95 | 86 | 128 | 68 | 111 | 95 | 82 | 159 | 126 | 130 | 115 | 118 | 50 | 38 | 125 | 67 | 126 | 90 | 119 | 155 | 89 | 151 | 122 | 98 | 95 | 138 | Phpt1 |
| Phrf1 | 7004 | 8330 | 9751 | 9816 | 8404 | 8219 | 7638 | 6737 | 8122 | 9391 | 8615 | 10987 | 9254 | 9832 | 6643 | 8174 | 9763 | 8902 | 9245 | 6578 | 7669 | 9331 | 7438 | 10276 | 9716 | 9687 | 9399 | 9946 | Phrf1 |
| Phtf1 | 4180 | 3622 | 3674 | 2998 | 3656 | 3577 | 3501 | 6013 | 5397 | 5200 | 5074 | 5827 | 5503 | 6157 | 4142 | 3582 | 3334 | 3103 | 3324 | 3756 | 3502 | 5243 | 5205 | 5276 | 5277 | 5665 | 5721 | 6159 | Phtf1 |
| Phtf2 | 4898 | 3960 | 4132 | 4620 | 4637 | 4827 | 5339 | 3596 | 3034 | 2865 | 2956 | 3130 | 2823 | 3364 | 4876 | 4277 | 4429 | 4790 | 4541 | 5064 | 5111 | 3364 | 2849 | 3109 | 3344 | 2887 | 3011 | 3043 | Phtf2 |
| Phxr2 | 0 | 0 | 0 | 0 | 0 | 0 | 0 | 0 | 0 | 0 | 0 | 0 | 0 | 0 | 0 | 0 | 0 | 0 | 0 | 3 | 0 | 0 | 0 | 0 | 2 | 0 | 0 | 0 | Phxr2 |
| Phyh | 1164 | 1056 | 1152 | 1081 | 1208 | 1045 | 1209 | 1623 | 1599 | 1604 | 1632 | 1553 | 1674 | 1610 | 1284 | 1134 | 1137 | 1073 | 1036 | 1055 | 1030 | 1681 | 1658 | 1423 | 1395 | 1597 | 1462 | 1530 | Phyh |
| Phyhd1 | 81 | 114 | 112 | 143 | 154 | 127 | 183 | 140 | 115 | 131 | 100 | 93 | 82 | 144 | 74 | 96 | 140 | 201 | 170 | 115 | 176 | 130 | 82 | 136 | 128 | 86 | 80 | 144 | Phyhd1 |
| Phyhip | 240 | 244 | 188 | 236 | 301 | 245 | 278 | 123 | 157 | 142 | 134 | 117 | 134 | 121 | 196 | 239 | 210 | 255 | 288 | 253 | 294 | 94 | 160 | 189 | 114 | 236 | 134 | 88 | Phyhip |
| Phyhipl | 2102 | 2445 | 2430 | 3930 | 5657 | 3936 | 4550 | 3783 | 3269 | 3816 | 3100 | 2763 | 3023 | 2842 | 2991 | 2079 | 3396 | 4193 | 4480 | 4063 | 4824 | 5048 | 3106 | 3001 | 3686 | 2168 | 2269 | 2582 | Phyhipl |
| Phykpl | 426 | 407 | 448 | 430 | 401 | 438 | 393 | 502 | 473 | 467 | 570 | 513 | 569 | 479 | 485 | 449 | 469 | 465 | 399 | 356 | 397 | 455 | 537 | 507 | 515 | 459 | 608 | 496 | Phykpl |
| Pi15 | 20 | 10 | 14 | 12 | 16 | 15 | 12 | 46 | 59 | 69 | 86 | 27 | 41 | 15 | 15 | 17 | 7 | 28 | 9 | 17 | 20 | 70 | 166 | 42 | 34 | 25 | 27 | 13 | Pi15 |
| Pi16 | 45 | 60 | 93 | 97 | 70 | 81 | 76 | 136 | 154 | 125 | 125 | 162 | 147 | 152 | 71 | 65 | 54 | 62 | 85 | 60 | 85 | 160 | 162 | 146 | 178 | 222 | 157 | 177 | Pi16 |
| Pi4k2a | 7414 | 9014 | 7451 | 9033 | 7252 | 7573 | 7399 | 3367 | 3560 | 3914 | 3600 | 4043 | 3669 | 3522 | 7279 | 7983 | 8362 | 8851 | 7827 | 7086 | 7345 | 3673 | 3520 | 3672 | 3739 | 4114 | 3791 | 3785 | Pi4k2a |
| Pi4k2b | 4744 | 3534 | 4511 | 3777 | 2536 | 3206 | 3376 | 1594 | 1460 | 1537 | 1752 | 1694 | 1850 | 1807 | 4336 | 4315 | 3734 | 3417 | 2854 | 3395 | 2966 | 1275 | 1752 | 1481 | 1324 | 1919 | 1888 | 1835 | Pi4k2b |
| Pi4ka | 9844 | 10551 | 8848 | 11444 | 13136 | 12197 | 12350 | 10232 | 10491 | 11573 | 10175 | 11416 | 10218 | 11062 | 9882 | 9783 | 10185 | 12128 | 12695 | 10681 | 13201 | 12277 | 9846 | 10440 | 12836 | 7586 | 10544 | 10473 | Pi4ka |
| Pi4kb | 3981 | 4545 | 3664 | 4014 | 3667 | 3934 | 3631 | 4252 | 4412 | 4077 | 4112 | 4238 | 3993 | 3930 | 4209 | 3921 | 3823 | 3795 | 3811 | 3748 | 3674 | 4055 | 4441 | 3730 | 4089 | 4156 | 4063 | 4012 | Pi4kb |
| Pianp | 659 | 767 | 807 | 1277 | 2144 | 1463 | 1588 | 977 | 1237 | 1575 | 1407 | 893 | 644 | 915 | 608 | 653 | 1161 | 1604 | 1871 | 1362 | 1657 | 1741 | 975 | 1417 | 1719 | 772 | 611 | 681 | Pianp |
| Pias1 | 6322 | 5672 | 5392 | 5504 | 5313 | 5382 | 5935 | 5802 | 4926 | 4327 | 4276 | 4914 | 4498 | 5305 | 6570 | 5564 | 5428 | 5192 | 5329 | 6224 | 5470 | 4939 | 4963 | 4646 | 4652 | 4659 | 5055 | 5073 | Pias1 |
| Pias2 | 8795 | 7260 | 8206 | 8798 | 9245 | 9627 | 10254 | 8255 | 7163 | 7181 | 7579 | 8504 | 8013 | 8629 | 8743 | 7952 | 8304 | 8357 | 9230 | 8896 | 10086 | 8438 | 7042 | 8114 | 8545 | 7880 | 7989 | 8751 | Pias2 |
| Pias3 | 1751 | 1668 | 2138 | 2001 | 1753 | 1756 | 1736 | 2183 | 2804 | 2779 | 2819 | 2839 | 2972 | 2552 | 1465 | 1759 | 2155 | 1777 | 1946 | 1574 | 1633 | 2625 | 2910 | 2718 | 2349 | 3145 | 2773 | 2673 | Pias3 |
| Pias4 | 2413 | 3373 | 3446 | 3418 | 2582 | 2642 | 2217 | 1912 | 2228 | 2438 | 2355 | 2675 | 2463 | 2546 | 2337 | 2840 | 3298 | 2957 | 2854 | 2453 | 2428 | 2448 | 2147 | 2776 | 2523 | 3169 | 2310 | 2541 | Pias4 |
| Pibf1 | 1671 | 1385 | 1648 | 1561 | 1200 | 1395 | 1482 | 1937 | 1793 | 1360 | 1521 | 1703 | 1552 | 1750 | 1729 | 1511 | 1517 | 1253 | 1374 | 1662 | 1466 | 1678 | 1777 | 1558 | 1549 | 1778 | 1722 | 1771 | Pibf1 |
| Picalm | 22704 | 18962 | 20447 | 21819 | 21792 | 22088 | 24534 | 22777 | 20059 | 20424 | 20662 | 19984 | 19591 | 21177 | 24166 | 21078 | 21072 | 22226 | 20603 | 21940 | 23374 | 22207 | 20127 | 19860 | 20544 | 19803 | 19473 | 20480 | Picalm |
| Pick1 | 931 | 993 | 1072 | 1299 | 1246 | 974 | 1030 | 1119 | 1135 | 1802 | 1522 | 1480 | 1418 | 1483 | 836 | 842 | 1365 | 1141 | 1195 | 1000 | 1068 | 1377 | 1116 | 1662 | 1646 | 1600 | 1389 | 1453 | Pick1 |
| Pid1 | 2117 | 2364 | 1777 | 2810 | 3858 | 3256 | 2724 | 4822 | 4660 | 4516 | 4249 | 3255 | 4715 | 3234 | 2819 | 2232 | 2615 | 2987 | 2904 | 3084 | 2805 | 5003 | 4474 | 3478 | 4401 | 3337 | 2898 | 2839 | Pid1 |
| Pidd1 | 1475 | 1696 | 1790 | 1551 | 1416 | 1444 | 1510 | 946 | 1148 | 1203 | 1093 | 1824 | 1459 | 1570 | 1218 | 1334 | 1796 | 1417 | 1568 | 1313 | 1304 | 947 | 1069 | 1510 | 1307 | 2150 | 1618 | 1752 | Pidd1 |
| Piezo1 | 2074 | 2693 | 2800 | 1972 | 1461 | 2130 | 1073 | 1918 | 3001 | 2831 | 3868 | 2539 | 3255 | 2113 | 1924 | 3221 | 1865 | 2351 | 1360 | 1023 | 1068 | 2705 | 3599 | 2910 | 2377 | 2202 | 2723 | 2436 | Piezo1 |
| Piezo2 | 1570 | 1221 | 747 | 767 | 1393 | 1032 | 544 | 1894 | 3158 | 2171 | 4678 | 1047 | 1999 | 1218 | 1216 | 1736 | 626 | 1237 | 534 | 502 | 504 | 2638 | 3275 | 1393 | 1862 | 992 | 1325 | 1127 | Piezo2 |
| Pif1 | 2286 | 2553 | 2872 | 2326 | 1679 | 1910 | 1949 | 1768 | 1769 | 1808 | 1813 | 2967 | 2443 | 2901 | 1954 | 2257 | 2657 | 2049 | 2368 | 1801 | 1790 | 1552 | 1476 | 2580 | 2206 | 3318 | 3156 | 3025 | Pif1 |
| Pifo | 39 | 43 | 41 | 33 | 79 | 48 | 67 | 111 | 117 | 87 | 80 | 84 | 119 | 57 | 58 | 72 | 57 | 37 | 59 | 60 | 80 | 87 | 175 | 63 | 76 | 57 | 48 | 37 | Pifo |
| Piga | 4212 | 3556 | 3412 | 3163 | 2407 | 2763 | 3026 | 2120 | 1836 | 1619 | 1899 | 2051 | 1912 | 2104 | 3699 | 3742 | 3088 | 3051 | 2421 | 2789 | 2681 | 1546 | 1814 | 1965 | 1721 | 2037 | 2069 | 2105 | Piga |
| Pigb | 1133 | 1085 | 893 | 982 | 1096 | 1172 | 1017 | 1386 | 1445 | 1494 | 1392 | 1587 | 1453 | 1390 | 1125 | 999 | 1004 | 1020 | 1121 | 1111 | 1098 | 1432 | 1388 | 1426 | 1405 | 1263 | 1589 | 1519 | Pigb |
| Pigbos1 | 529 | 580 | 604 | 681 | 565 | 534 | 721 | 569 | 532 | 638 | 595 | 628 | 649 | 802 | 653 | 505 | 613 | 512 | 512 | 667 | 644 | 476 | 603 | 726 | 661 | 773 | 545 | 777 | Pigbos1 |
| Pigc | 1351 | 1120 | 1254 | 1132 | 1211 | 1199 | 1309 | 2262 | 1986 | 2085 | 2001 | 1990 | 2105 | 2181 | 1432 | 1237 | 1261 | 1206 | 1241 | 1252 | 1180 | 2069 | 2081 | 2024 | 1968 | 2030 | 2047 | 2207 | Pigc |
| Pigf | 2225 | 2397 | 2247 | 2150 | 1536 | 1918 | 2065 | 988 | 868 | 690 | 715 | 897 | 890 | 907 | 2490 | 2264 | 2176 | 1949 | 1848 | 2327 | 2041 | 725 | 825 | 845 | 752 | 1084 | 965 | 1019 | Pigf |
| Pigg | 1486 | 1428 | 1317 | 1261 | 1289 | 1438 | 1385 | 1428 | 1442 | 1248 | 1337 | 1420 | 1428 | 1402 | 1580 | 1355 | 1338 | 1115 | 1218 | 1585 | 1359 | 1267 | 1337 | 1222 | 1377 | 1230 | 1504 | 1435 | Pigg |
| Pigh | 1040 | 822 | 1023 | 762 | 817 | 812 | 908 | 1261 | 1099 | 1163 | 1150 | 1315 | 1296 | 1288 | 1036 | 877 | 891 | 717 | 822 | 915 | 827 | 1036 | 1162 | 1200 | 1011 | 1444 | 1228 | 1305 | Pigh |
| Pigk | 2866 | 2792 | 2757 | 2075 | 2530 | 2475 | 2016 | 3797 | 3781 | 3385 | 3456 | 3321 | 3644 | 3190 | 2935 | 3245 | 2395 | 2410 | 1966 | 2442 | 1953 | 3623 | 3996 | 3061 | 3496 | 3120 | 3626 | 3331 | Pigk |
| Pigl | 964 | 1032 | 1217 | 908 | 938 | 962 | 908 | 739 | 760 | 723 | 753 | 862 | 955 | 878 | 1004 | 1037 | 969 | 947 | 923 | 995 | 955 | 702 | 793 | 718 | 685 | 934 | 949 | 809 | Pigl |
| Pigm | 6983 | 5572 | 7243 | 6352 | 4673 | 5483 | 5535 | 3580 | 3215 | 3257 | 3464 | 4243 | 4278 | 4304 | 6347 | 6699 | 6457 | 6196 | 4739 | 4797 | 5217 | 3776 | 3126 | 3675 | 3439 | 4960 | 4388 | 4619 | Pigm |
| Pign | 7711 | 6112 | 4974 | 5617 | 5331 | 5785 | 6136 | 4572 | 3948 | 3708 | 3722 | 3968 | 3962 | 3980 | 7663 | 7094 | 5310 | 5794 | 4854 | 6400 | 5802 | 3653 | 4088 | 3158 | 3546 | 2768 | 4142 | 3762 | Pign |
| Pigo | 1390 | 1678 | 1579 | 1711 | 1780 | 1733 | 1526 | 1998 | 2446 | 2533 | 2415 | 2721 | 2777 | 2351 | 1270 | 1555 | 1702 | 1723 | 1774 | 1389 | 1579 | 2513 | 2342 | 2639 | 2771 | 2650 | 2667 | 2554 | Pigo |
| Pigp | 1585 | 1478 | 1456 | 1521 | 1458 | 1665 | 1871 | 2678 | 2516 | 2314 | 2551 | 2465 | 2520 | 2795 | 1767 | 1445 | 1540 | 1350 | 1501 | 1834 | 1709 | 2288 | 2715 | 2497 | 2356 | 2542 | 2547 | 2904 | Pigp |
| Pigq | 3419 | 3523 | 3197 | 3202 | 3220 | 3351 | 2963 | 4162 | 4544 | 4760 | 4475 | 5232 | 5090 | 4791 | 3136 | 3212 | 3413 | 3091 | 3411 | 3112 | 3091 | 5035 | 4713 | 4634 | 4642 | 4684 | 5126 | 5369 | Pigq |
| Pigr | 0 | 0 | 2 | 0 | 0 | 4 | 1 | 1 | 1 | 0 | 0 | 2 | 0 | 0 | 0 | 3 | 0 | 2 | 1 | 0 | 2 | 2 | 2 | 0 | 0 | 0 | 0 | 0 | Pigr |
| Pigs | 5940 | 7189 | 6795 | 6778 | 5900 | 5516 | 6297 | 2654 | 3347 | 3517 | 3516 | 3802 | 3698 | 3339 | 5627 | 6966 | 6875 | 6911 | 6218 | 4820 | 5722 | 3624 | 3599 | 3489 | 3349 | 3920 | 3268 | 3418 | Pigs |
| Pigt | 3074 | 3936 | 4321 | 4589 | 4350 | 3939 | 3342 | 3416 | 4266 | 4979 | 4796 | 4843 | 4157 | 4267 | 3026 | 3604 | 4182 | 4580 | 4676 | 3171 | 4052 | 5497 | 4432 | 4899 | 4807 | 5215 | 4028 | 4028 | Pigt |
| Pigu | 1962 | 2095 | 1830 | 1663 | 1976 | 2030 | 2100 | 3069 | 2692 | 2341 | 2577 | 2619 | 2650 | 2582 | 2060 | 1979 | 1867 | 1782 | 1944 | 2157 | 2120 | 2872 | 2696 | 2577 | 2658 | 2903 | 2651 | 2865 | Pigu |

| | | | | | | | | | | | | | | | | | | | | | | | | | | | | | |
|---|---|---|---|---|---|---|---|---|---|---|---|---|---|---|---|---|---|---|---|---|---|---|---|---|---|---|---|---|---|
| Pigv | 1429 | 1748 | 1728 | 1715 | 1505 | 1502 | 1698 | 965 | 1093 | 1173 | 1198 | 1326 | 1306 | 1169 | 1333 | 1446 | 1964 | 1829 | 1817 | 1501 | 1593 | 1287 | 1025 | 1245 | 1287 | 1206 | 1121 | 1205 | Pigv |
| Pigw | 311 | 260 | 427 | 383 | 330 | 352 | 367 | 724 | 648 | 609 | 725 | 737 | 839 | 763 | 298 | 326 | 384 | 317 | 355 | 371 | 333 | 760 | 658 | 694 | 683 | 777 | 824 | 773 | Pigw |
| Pigx | 3402 | 3156 | 2901 | 3165 | 2670 | 2604 | 3262 | 2848 | 2334 | 2319 | 2000 | 2586 | 2358 | 2819 | 3501 | 2939 | 3159 | 2850 | 2879 | 3281 | 3262 | 2426 | 2355 | 2498 | 2333 | 2646 | 2496 | 2921 | Pigx |
| Pigyl | 814 | 1101 | 908 | 1138 | 963 | 995 | 1135 | 642 | 853 | 920 | 747 | 762 | 712 | 750 | 940 | 835 | 1004 | 966 | 1079 | 1295 | 1004 | 580 | 783 | 779 | 689 | 780 | 761 | 706 | Pigyl |
| Pigz | 6 | 8 | 13 | 3 | 16 | 7 | 10 | 29 | 32 | 34 | 29 | 20 | 27 | 29 | 8 | 7 | 8 | 3 | 3 | 14 | 4 | 18 | 14 | 13 | 35 | 15 | 30 | 19 | Pigz |
| Pih1d1 | 1691 | 2103 | 1815 | 1707 | 1933 | 1920 | 1762 | 3030 | 3163 | 2993 | 2980 | 3451 | 3332 | 3325 | 1731 | 1655 | 1913 | 1625 | 2018 | 2067 | 1874 | 3023 | 3149 | 3349 | 3280 | 3457 | 3756 | 3387 | Pih1d1 |
| Pih1d2 | 183 | 223 | 153 | 195 | 183 | 191 | 197 | 296 | 232 | 325 | 210 | 288 | 310 | 221 | 264 | 209 | 158 | 154 | 142 | 183 | 192 | 227 | 284 | 228 | 247 | 192 | 307 | 295 | Pih1d2 |
| Pik3ap1 | 423 | 420 | 454 | 319 | 236 | 254 | 158 | 276 | 313 | 288 | 472 | 422 | 438 | 330 | 438 | 556 | 312 | 337 | 145 | 257 | 200 | 421 | 476 | 331 | 298 | 494 | 490 | 395 | Pik3ap1 |
| Pik3c2a | 11733 | 7901 | 6337 | 7368 | 6941 | 8885 | 8504 | 7723 | 7273 | 5431 | 6138 | 6478 | 6468 | 6564 | 11195 | 10054 | 6466 | 7264 | 6555 | 8976 | 8241 | 5073 | 7203 | 4813 | 5568 | 4305 | 6424 | 6081 | Pik3c2a |
| Pik3c2b | 2125 | 2377 | 3398 | 3192 | 3627 | 3156 | 3256 | 4015 | 4047 | 5200 | 4550 | 5941 | 5197 | 5963 | 2145 | 2051 | 3639 | 3061 | 3761 | 2749 | 3328 | 5304 | 3502 | 5833 | 6111 | 4828 | 5535 | 6435 | Pik3c2b |
| Pik3c2g | 9 | 8 | 5 | 2 | 15 | 2 | 11 | 23 | 32 | 8 | 13 | 11 | 8 | 6 | 13 | 22 | 2 | 4 | 11 | 9 | 8 | 22 | 37 | 12 | 13 | 1 | 19 | 6 | Pik3c2g |
| Pik3c3 | 4576 | 4169 | 3873 | 4019 | 4044 | 4088 | 4507 | 3808 | 3690 | 4215 | 3509 | 3720 | 3672 | 3738 | 4820 | 4089 | 3745 | 3508 | 4101 | 4381 | 4652 | 3847 | 3747 | 3683 | 3827 | 3616 | 3602 | 3726 | Pik3c3 |
| Pik3ca | 10670 | 9834 | 8585 | 10282 | 11601 | 11834 | 12585 | 12002 | 9824 | 9370 | 9355 | 9422 | 9279 | 9600 | 11267 | 9717 | 9668 | 10566 | 11245 | 12099 | 12378 | 10637 | 9560 | 9207 | 10108 | 7722 | 9304 | 9126 | Pik3ca |
| Pik3cb | 2568 | 2465 | 2351 | 2990 | 3035 | 2724 | 3158 | 1591 | 1659 | 1677 | 1504 | 1519 | 1574 | 1617 | 2516 | 2382 | 2454 | 3053 | 2802 | 2709 | 3092 | 1596 | 1467 | 1656 | 1811 | 1385 | 1314 | 1537 | Pik3cb |
| Pik3cd | 1301 | 1689 | 1819 | 1721 | 2211 | 1645 | 1547 | 1453 | 1806 | 2261 | 1975 | 1750 | 1951 | 1674 | 1348 | 1632 | 2086 | 1799 | 2067 | 1400 | 1517 | 2371 | 1604 | 1784 | 2075 | 1297 | 1591 | 1622 | Pik3cd |
| Pik3cg | 157 | 127 | 124 | 97 | 115 | 151 | 111 | 277 | 327 | 190 | 343 | 270 | 316 | 214 | 115 | 156 | 87 | 87 | 94 | 132 | 94 | 259 | 442 | 180 | 225 | 218 | 257 | 256 | Pik3cg |
| Pik3ip1 | 604 | 533 | 608 | 573 | 864 | 666 | 757 | 944 | 948 | 1065 | 928 | 861 | 844 | 918 | 654 | 557 | 590 | 594 | 819 | 681 | 698 | 1011 | 968 | 999 | 1021 | 818 | 812 | 777 | Pik3ip1 |
| Pik3r1 | 6110 | 5380 | 6096 | 7096 | 8642 | 7422 | 7612 | 7010 | 7590 | 7984 | 9309 | 6074 | 6444 | 6251 | 6223 | 6578 | 6480 | 8358 | 7116 | 6281 | 7602 | 8507 | 7818 | 6886 | 7265 | 5008 | 6046 | 5775 | Pik3r1 |
| Pik3r2 | 525 | 506 | 215 | 603 | 261 | 268 | 683 | 248 | 177 | 567 | 155 | 199 | 127 | 130 | 280 | 545 | 242 | 508 | 305 | 464 | 630 | 491 | 570 | 270 | 200 | 154 | 246 | 396 | Pik3r2 |
| Pik3r3 | 8406 | 7546 | 8327 | 9188 | 11835 | 10140 | 11398 | 16773 | 12152 | 13193 | 10956 | 13993 | 12505 | 14332 | 9695 | 7143 | 8908 | 9353 | 10970 | 11202 | 11266 | 16648 | 12114 | 13767 | 15802 | 12306 | 13923 | 13728 | Pik3r3 |
| Pik3r4 | 4847 | 5375 | 4355 | 4166 | 3603 | 3892 | 3786 | 3232 | 3597 | 3372 | 3182 | 3552 | 3394 | 3425 | 4472 | 4848 | 4048 | 4267 | 3891 | 3758 | 3822 | 3237 | 3474 | 3289 | 3208 | 3156 | 3622 | 3519 | Pik3r4 |
| Pik3r5 | 326 | 383 | 543 | 232 | 205 | 226 | 166 | 92 | 124 | 120 | 134 | 110 | 115 | 115 | 386 | 500 | 338 | 297 | 172 | 185 | 101 | 77 | 135 | 97 | 64 | 143 | 107 | 103 | Pik3r5 |
| Pik3r6 | 175 | 217 | 199 | 199 | 206 | 182 | 171 | 115 | 165 | 139 | 153 | 153 | 170 | 156 | 202 | 235 | 174 | 183 | 192 | 199 | 229 | 135 | 182 | 141 | 153 | 166 | 185 | 173 | Pik3r6 |
| Pikfyve | 14596 | 13562 | 8670 | 13188 | 12714 | 14639 | 15499 | 7849 | 6461 | 5411 | 5654 | 5845 | 5689 | 6039 | 15559 | 13935 | 10266 | 13763 | 11403 | 15505 | 15357 | 5883 | 6106 | 4929 | 6312 | 4060 | 6270 | 5721 | Pikfyve |
| Pilra | 3 | 3 | 0 | 9 | 4 | 1 | 9 | 9 | 11 | 14 | 16 | 8 | 2 | 11 | 8 | 5 | 3 | 11 | 7 | 5 | 5 | 10 | 20 | 17 | 16 | 7 | 5 | 5 | Pilra |
| Pilrb1 | 0 | 2 | 0 | 0 | 0 | 0 | 0 | 0 | 1 | 2 | 3 | 2 | 1 | 0 | 1 | 0 | 0 | 4 | 2 | 3 | 0 | 5 | 5 | 0 | 0 | 1 | 1 | 0 | Pilrb1 |
| Pilrb2 | 0 | 3 | 0 | 0 | 5 | 0 | 1 | 0 | 0 | 0 | 2 | 0 | 0 | 0 | 0 | 0 | 1 | 2 | 0 | 0 | 0 | 2 | 8 | 0 | 2 | 2 | 1 | 2 | Pilrb2 |
| Pim1 | 2546 | 2997 | 3894 | 3825 | 2818 | 2938 | 2951 | 1434 | 1752 | 1796 | 1786 | 2520 | 2684 | 2311 | 2574 | 2775 | 3541 | 3397 | 3187 | 2707 | 2948 | 1425 | 1599 | 2312 | 1832 | 3026 | 2528 | 2785 | Pim1 |
| Pim2 | 1455 | 1986 | 1850 | 2129 | 1807 | 1802 | 1798 | 893 | 751 | 963 | 814 | 943 | 920 | 965 | 1494 | 1618 | 1968 | 2069 | 1824 | 1489 | 1849 | 1218 | 781 | 963 | 1125 | 849 | 999 | 966 | Pim2 |
| Pim3 | 1288 | 1696 | 1501 | 1830 | 1541 | 1393 | 1334 | 1334 | 1446 | 1771 | 1410 | 1622 | 1261 | 1244 | 1201 | 1393 | 1752 | 1608 | 1570 | 1366 | 1437 | 1390 | 1496 | 1861 | 1587 | 1807 | 1261 | 1271 | Pim3 |
| Pimreg | 4418 | 5229 | 5160 | 3719 | 2810 | 3359 | 3519 | 3573 | 3756 | 3129 | 2945 | 5198 | 4220 | 5140 | 3766 | 3970 | 4400 | 3112 | 3808 | 3954 | 2988 | 2515 | 3216 | 4190 | 3415 | 6643 | 5905 | 5413 | Pimreg |
| Pin1 | 4306 | 5722 | 4978 | 5224 | 4148 | 4400 | 4292 | 2684 | 2768 | 2950 | 2774 | 3168 | 2894 | 3047 | 4308 | 4811 | 4877 | 4569 | 4918 | 4886 | 4586 | 2423 | 2827 | 3191 | 2963 | 3468 | 3283 | 3283 | Pin1 |
| Pin1rt1 | 3 | 3 | 2 | 5 | 7 | 1 | 6 | 2 | 1 | 3 | 3 | 1 | 0 | 0 | 0 | 0 | 0 | 3 | 3 | 5 | 2 | 2 | 0 | 2 | 0 | 0 | 1 | 0 | Pin1rt1 |
| Pin4 | 1547 | 1498 | 2361 | 2271 | 1521 | 1333 | 2052 | 1501 | 1030 | 1441 | 1342 | 1493 | 1538 | 2246 | 1807 | 1230 | 2002 | 1775 | 1672 | 1977 | 2056 | 1245 | 1098 | 1829 | 1470 | 1995 | 1782 | 2300 | Pin4 |
| Pink1 | 2804 | 3407 | 3557 | 3768 | 4050 | 3664 | 3939 | 4161 | 4301 | 4597 | 4158 | 3795 | 4001 | 4452 | 3210 | 2952 | 3541 | 3374 | 4011 | 3979 | 4658 | 4017 | 4419 | 4347 | 4617 | 3617 | 3946 | 4192 | Pink1 |
| Pinlyp | 5 | 15 | 2 | 1 | 9 | 1 | 1 | 1 | 0 | 6 | 3 | 2 | 3 | 1 | 1 | 3 | 5 | 8 | 1 | 2 | 5 | 3 | 1 | 2 | 0 | 1 | 6 | 8 | Pinlyp |
| Pinx1 | 946 | 1030 | 1146 | 984 | 835 | 916 | 945 | 1060 | 933 | 1035 | 820 | 1088 | 1016 | 1120 | 949 | 899 | 1151 | 905 | 947 | 1028 | 899 | 829 | 926 | 998 | 844 | 1388 | 1126 | 1262 | Pinx1 |
| Pip4k2a | 3325 | 3231 | 3585 | 4166 | 4077 | 3621 | 4232 | 1958 | 1935 | 2124 | 2126 | 1688 | 1935 | 1898 | 3541 | 3430 | 3541 | 4258 | 4017 | 3579 | 4156 | 2109 | 1999 | 1908 | 1935 | 1935 | 1833 | 1939 | Pip4k2a |
| Pip4k2b | 13021 | 15285 | 17168 | 18894 | 19784 | 16829 | 18206 | 10139 | 10932 | 12624 | 11914 | 11353 | 10829 | 11251 | 13229 | 13540 | 18338 | 19455 | 19425 | 15582 | 19570 | 13049 | 10375 | 12400 | 13328 | 11633 | 10212 | 10722 | Pip4k2b |
| Pip4k2c | 3247 | 3403 | 2964 | 3100 | 2920 | 2923 | 2721 | 1899 | 2395 | 2148 | 2311 | 2330 | 2216 | 1995 | 3038 | 3569 | 2872 | 3496 | 2710 | 2583 | 2788 | 2045 | 2378 | 2223 | 2113 | 2266 | 1952 | 2144 | Pip4k2c |
| Pip4p1 | 5353 | 5351 | 5129 | 5622 | 4847 | 4873 | 5568 | 3082 | 2845 | 3139 | 2976 | 3078 | 2845 | 3327 | 5265 | 4588 | 5256 | 5527 | 4933 | 4482 | 5611 | 3262 | 2787 | 3263 | 3210 | 3227 | 2939 | 3404 | Pip4p1 |
| Pip4p2 | 4300 | 3054 | 3928 | 3302 | 3508 | 3672 | 3774 | 6064 | 5490 | 4465 | 5391 | 6055 | 5942 | 6132 | 3833 | 3961 | 3788 | 3316 | 3512 | 3478 | 3488 | 5508 | 5590 | 5676 | 5117 | 7356 | 5857 | 6494 | Pip4p2 |
| Pip5k1a | 4750 | 4385 | 4581 | 4317 | 3877 | 4122 | 4242 | 3595 | 3460 | 3448 | 3515 | 4003 | 3700 | 3861 | 4189 | 4222 | 4398 | 4113 | 4194 | 4041 | 4218 | 3334 | 3594 | 3630 | 3499 | 4307 | 4033 | 4099 | Pip5k1a |
| Pip5k1b | 269 | 371 | 407 | 343 | 305 | 272 | 277 | 198 | 226 | 188 | 195 | 269 | 304 | 254 | 318 | 280 | 423 | 274 | 346 | 290 | 288 | 229 | 261 | 261 | 222 | 227 | 292 | 316 | Pip5k1b |
| Pip5k1c | 8804 | 11728 | 11477 | 14461 | 14594 | 13103 | 12277 | 8479 | 10544 | 12093 | 11431 | 11959 | 10102 | 11295 | 8414 | 10417 | 13027 | 14784 | 14679 | 10796 | 13044 | 12881 | 9607 | 12465 | 13700 | 11760 | 10040 | 10581 | Pip5k1c |
| Pip5kl1 | 294 | 353 | 283 | 356 | 394 | 369 | 336 | 137 | 162 | 178 | 133 | 142 | 122 | 130 | 227 | 329 | 362 | 328 | 436 | 386 | 298 | 187 | 153 | 156 | 175 | 129 | 123 | 145 | Pip5kl1 |
| Pipox | 354 | 394 | 462 | 317 | 315 | 379 | 391 | 663 | 623 | 638 | 501 | 1017 | 658 | 1139 | 376 | 313 | 477 | 315 | 391 | 402 | 340 | 708 | 482 | 931 | 839 | 1078 | 1041 | 1061 | Pipox |
| Pir | 172 | 158 | 205 | 112 | 100 | 112 | 92 | 139 | 168 | 133 | 158 | 228 | 248 | 183 | 174 | 173 | 124 | 85 | 97 | 117 | 93 | 135 | 208 | 178 | 104 | 233 | 200 | 228 | Pir |
| Pira12 | 0 | 0 | 0 | 0 | 0 | 0 | 0 | 0 | 2 | 0 | 0 | 2 | 0 | 0 | 0 | 1 | 0 | 0 | 0 | 0 | 0 | 0 | 2 | 0 | 0 | 1 | 0 | 0 | Pira12 |
| Pira13 | 0 | 0 | 0 | 1 | 0 | 0 | 0 | 1 | 0 | 0 | 0 | 0 | 0 | 0 | 0 | 0 | 0 | 0 | 0 | 0 | 3 | 0 | 1 | 0 | 0 | 0 | 0 | 0 | Pira13 |
| Pira2 | 1 | 0 | 0 | 0 | 0 | 0 | 1 | 0 | 0 | 0 | 0 | 0 | 0 | 0 | 0 | 0 | 0 | 0 | 0 | 0 | 0 | 0 | 3 | 2 | 0 | 2 | 0 | 0 | Pira2 |
| Pirb | 9 | 4 | 2 | 1 | 9 | 5 | 3 | 9 | 7 | 30 | 13 | 13 | 15 | 14 | 5 | 7 | 7 | 9 | 3 | 3 | 6 | 5 | 30 | 16 | 6 | 13 | 7 | 9 | Pirb |
| Pirt | 298 | 333 | 41 | 70 | 875 | 87 | 13 | 1329 | 1537 | 955 | 1456 | 73 | 1144 | 55 | 554 | 636 | 197 | 412 | 56 | 79 | 49 | 2214 | 928 | 69 | 932 | 79 | 85 | 71 | Pirt |
| Pisd | 1012 | 1126 | 1186 | 1265 | 1134 | 1000 | 1162 | 1010 | 918 | 1044 | 986 | 1000 | 956 | 1055 | 1130 | 979 | 1079 | 1131 | 1047 | 1084 | 1121 | 1155 | 948 | 1111 | 1016 | 1036 | 990 | 1110 | Pisd |
| Pithd1 | 1911 | 2110 | 2173 | 2152 | 2148 | 2042 | 1988 | 1406 | 1332 | 1446 | 1466 | 1578 | 1498 | 1569 | 1923 | 2049 | 2155 | 2092 | 2105 | 1738 | 1989 | 1604 | 1380 | 1579 | 1660 | 2049 | 1511 | 1698 | Pithd1 |
| Pitpna | 13068 | 12575 | 12196 | 14104 | 14341 | 13253 | 14613 | 9029 | 8843 | 9553 | 9139 | 9372 | 8842 | 9264 | 12999 | 12517 | 13348 | 13877 | 13847 | 12132 | 14022 | 10612 | 9200 | 9779 | 10174 | 9412 | 8646 | 9352 | Pitpna |
| Pitpnb | 10705 | 9849 | 11336 | 11515 | 10228 | 10723 | 10888 | 7275 | 7373 | 8113 | 7766 | 9290 | 8371 | 8969 | 10142 | 10236 | 10973 | 10690 | 10622 | 9440 | 10646 | 8027 | 7308 | 8561 | 8586 | 10198 | 9109 | 9743 | Pitpnb |
| Pitpnc1 | 4666 | 4189 | 4608 | 5337 | 6116 | 5805 | 6125 | 6536 | 6373 | 7618 | 7434 | 7392 | 6864 | 7535 | 4771 | 4447 | 4802 | 5821 | 5796 | 4978 | 6265 | 7797 | 6043 | 7846 | 7723 | 6542 | 7112 | 7424 | Pitpnc1 |
| Pitpnm1 | 0 | 0 | 0 | 0 | 0 | 0 | 0 | 0 | 0 | 0 | 0 | 0 | 0 | 0 | 0 | 0 | 0 | 0 | 0 | 0 | 0 | 0 | 1 | 0 | 0 | 1 | 0 | 0 | Pitpnm1 |
| Pitpnm2 | 1526 | 2127 | 2375 | 2324 | 3341 | 2714 | 2229 | 2982 | 4217 | 4970 | 4612 | 4785 | 4601 | 3862 | 1763 | 2161 | 2541 | 2637 | 3218 | 2193 | 2416 | 5173 | 3689 | 5112 | 5200 | 3991 | 3885 | 3838 | Pitpnm2 |
| Pitpnm3 | 100 | 149 | 180 | 163 | 512 | 316 | 287 | 460 | 523 | 598 | 506 | 384 | 407 | 344 | 145 | 137 | 172 | 310 | 322 | 333 | 377 | 722 | 443 | 389 | 667 | 144 | 400 | 267 | Pitpnm3 |
| Pitrm1 | 11953 | 11616 | 9281 | 8539 | 7329 | 8492 | 8043 | 7678 | 7982 | 6575 | 7313 | 7809 | 8222 | 7899 | 11251 | 11490 | 8181 | 8902 | 7720 | 8603 | 7878 | 6313 | 7958 | 6945 | 6786 | 8752 | 8275 | 8511 | Pitrm1 |
| Pitx1 | 206 | 58 | 19 | 88 | 92 | 32 | 18 | 225 | 367 | 286 | 785 | 45 | 104 | 65 | 54 | 58 | 46 | 67 | 43 | 31 | 18 | 214 | 317 | 412 | 44 | 561 | 85 | 61 | Pitx1 |
| Pitx2 | 930 | 767 | 1638 | 974 | 672 | 772 | 870 | 855 | 655 | 803 | 847 | 1267 | 725 | 1355 | 670 | 470 | 1081 | 769 | 963 | 1078 | 843 | 1103 | 911 | 1109 | 826 | 998 | 1590 | 1775 | Pitx2 |
| Pitx3 | 80 | 93 | 71 | 64 | 56 | 56 | 39 | 22 | 55 | 30 | 185 | 41 | 19 | 29 | 51 | 90 | 63 | 66 | 48 | 51 | 40 | 58 | 83 | 45 | 53 | 14 | 43 | 26 | Pitx3 |
| Piwil1 | 7 | 16 | 20 | 16 | 8 | 8 | 7 | 12 | 4 | 12 | 13 | 20 | 28 | 15 | 15 | 16 | 14 | 4 | 12 | 25 | 11 | 22 | 8 | 16 | 14 | 7 | 21 | 20 | Piwil1 |
| Piwil2 | 35 | 60 | 48 | 66 | 66 | 55 | 78 | 26 | 38 | 44 | 25 | 34 | 27 | 19 | 50 | 64 | 58 | 106 | 47 | 49 | 55 | 22 | 32 | 25 | 18 | 27 | 38 | 15 | Piwil2 |
| Piwil4 | 6 | 10 | 9 | 3 | 5 | 10 | 6 | 12 | 24 | 27 | 14 | 14 | 18 | 15 | 15 | 10 | 6 | 3 | 0 | 1 | 4 | 18 | 21 | 14 | 3 | 5 | 20 | 2 | Piwil4 |
| Pja1 | 13313 | 15905 | 13722 | 13757 | 14245 | 14274 | 15114 | 13468 | 13908 | 13040 | 12125 | 13170 | 11498 | 13418 | 14224 | 13169 | 14516 | 13627 | 15019 | 16347 | 14779 | 12775 | 13193 | 14050 | 14849 | 13734 | 13047 | 12845 | Pja1 |
| Pja2 | 26541 | 23983 | 20284 | 23936 | 26782 | 25973 | 26718 | 26714 | 22387 | 19457 | 20443 | 17672 | 18218 | 18251 | 29608 | 24252 | 21652 | 25534 | 23987 | 30904 | 26511 | 24463 | 22407 | 17547 | 20872 | 14474 | 18058 | 17349 | Pja2 |
| Pjvk | 24 | 16 | 36 | 18 | 24 | 8 | 12 | 24 | 27 | 38 | 30 | 48 | 36 | 30 | 8 | 10 | 24 | 7 | 21 | 16 | 18 | 33 | 38 | 28 | 41 | 72 | 40 | 64 | Pjvk |
| Pkd1 | 8410 | 10448 | 6307 | 10942 | 11706 | 11479 | 9702 | 7288 | 9651 | 9610 | 10995 | 9393 | 9922 | 8202 | 7416 | 9682 | 8167 | 11459 | 11215 | 8752 | 10397 | 9959 | 8886 | 8025 | 10714 | 5189 | 9697 | 7713 | Pkd1 |
| Pkd1l1 | 8 | 21 | 14 | 16 | 6 | 3 | 6 | 6 | 19 | 12 | 8 | 4 | 15 | 6 | 11 | 11 | 1 | 7 | 5 | 13 | 7 | 3 | 16 | 10 | 5 | 8 | 3 | 11 | Pkd1l1 |
| Pkd1l2 | 1 | 3 | 0 | 2 | 2 | 0 | 2 | 0 | 1 | 3 | 7 | 1 | 3 | 1 | 0 | 7 | 7 | 4 | 1 | 0 | 4 | 2 | 3 | 3 | 7 | 3 | 0 | 0 | Pkd1l2 |
| Pkd1l3 | 37 | 27 | 37 | 22 | 8 | 14 | 4 | 26 | 13 | 12 | 36 | 35 | 33 | 21 | 34 | 39 | 16 | 33 | 7 | 20 | 18 | 27 | 17 | 25 | 19 | 17 | 30 | 11 | Pkd1l3 |
| Pkd2 | 3399 | 2553 | 2960 | 2659 | 3147 | 3655 | 2627 | 7129 | 7836 | 8326 | 10616 | 7578 | 8123 | 7646 | 3051 | 3870 | 2765 | 3280 | 2672 | 2486 | 2533 | 8057 | 8343 | 7759 | 7306 | 7069 | 7372 | 6953 | Pkd2 |
| Pkd2l1 | 17 | 15 | 6 | 29 | 17 | 5 | 19 | 39 | 34 | 39 | 23 | 23 | 27 | 49 | 24 | 16 | 14 | 17 | 9 | 18 | 19 | 42 | 17 | 22 | 45 | 44 | 43 | 26 | Pkd2l1 |
| Pkd2l2 | 58 | 49 | 35 | 71 | 56 | 31 | 55 | 53 | 32 | 32 | 38 | 18 | 39 | 24 | 70 | 70 | 66 | 50 | 45 | 30 | 47 | 38 | 43 | 23 | 29 | 22 | 19 | 34 | Pkd2l2 |
| Pkdcc | 3049 | 3666 | 2703 | 2844 | 2284 | 2548 | 1849 | 2755 | 4219 | 3593 | 5147 | 3257 | 3650 | 2915 | 2771 | 4532 | 2746 | 3081 | 2102 | 1717 | 1684 | 3135 | 3790 | 3812 | 3238 | 5215 | 2989 | 2667 | Pkdcc |

| | | | | | | | | | | | | | | | | | | | | | | | | | | | | | |
|---|---|---|---|---|---|---|---|---|---|---|---|---|---|---|---|---|---|---|---|---|---|---|---|---|---|---|---|---|---|
| Pkdrej | 6 | 8 | 6 | 9 | 11 | 6 | 6 | 29 | 46 | 60 | 45 | 38 | 24 | 36 | 8 | 5 | 9 | 21 | 6 | 8 | 7 | 22 | 41 | 29 | 36 | 35 | 24 | 38 | Pkdrej |
| Pkhd1 | 7 | 4 | 0 | 0 | 4 | 3 | 0 | 2 | 11 | 2 | 32 | 4 | 16 | 5 | 8 | 2 | 7 | 1 | 1 | 2 | 0 | 7 | 41 | 0 | 0 | 1 | 0 | 4 | Pkhd1 |
| Pkhd1l1 | 295 | 375 | 197 | 79 | 44 | 75 | 56 | 123 | 101 | 56 | 42 | 162 | 611 | 380 | 290 | 325 | 88 | 47 | 53 | 103 | 34 | 35 | 92 | 37 | 39 | 186 | 360 | 720 | Pkhd1l1 |
| Pkia | 14956 | 12455 | 11509 | 17737 | 24649 | 22346 | 25107 | 23651 | 18929 | 18417 | 16852 | 14715 | 13794 | 16030 | 19726 | 13284 | 15058 | 19486 | 21685 | 24775 | 24153 | 22987 | 19506 | 15368 | 19765 | 12359 | 14234 | 15053 | Pkia |
| Pkib | 59 | 48 | 68 | 42 | 54 | 45 | 36 | 74 | 54 | 49 | 80 | 131 | 138 | 90 | 63 | 51 | 63 | 47 | 66 | 71 | 64 | 47 | 96 | 110 | 86 | 64 | 151 | 140 | Pkib |
| Pkig | 2147 | 2667 | 2663 | 2305 | 2210 | 2117 | 2508 | 1835 | 1943 | 1995 | 1952 | 2284 | 1893 | 2386 | 2090 | 2108 | 2553 | 2181 | 2473 | 2422 | 2513 | 1648 | 1855 | 2388 | 2125 | 2666 | 2067 | 2416 | Pkig |
| Pklr | 25 | 41 | 53 | 11 | 10 | 7 | 9 | 23 | 36 | 18 | 19 | 43 | 46 | 38 | 46 | 84 | 32 | 22 | 9 | 22 | 5 | 12 | 26 | 15 | 11 | 53 | 38 | 212 | Pklr |
| Pkm | 82060 | 95258 | 90877 | 87704 | 69900 | 75565 | 72846 | 66474 | 72188 | 75938 | 67459 | 86519 | 83722 | 96318 | 75689 | 73456 | 94264 | 74549 | 88402 | 77763 | 81103 | 54049 | 67379 | 87533 | 76714 | 100515 | 96241 | 100822 | Pkm |
| Pkmyt1 | 1732 | 1683 | 2015 | 1792 | 1141 | 1426 | 1519 | 1018 | 1313 | 1132 | 1237 | 1787 | 1436 | 1686 | 1266 | 1502 | 1860 | 1606 | 1571 | 1174 | 1322 | 988 | 1098 | 1481 | 1267 | 2187 | 1622 | 1661 | Pkmyt1 |
| Pkn1 | 6973 | 8356 | 8499 | 10426 | 10737 | 8798 | 9080 | 6105 | 7219 | 9236 | 8115 | 8730 | 7643 | 7827 | 6492 | 7668 | 9634 | 10823 | 11247 | 8012 | 10151 | 8971 | 6935 | 9926 | 9474 | 8363 | 7897 | 7917 | Pkn1 |
| Pkn2 | 12482 | 9714 | 10453 | 9934 | 9484 | 10624 | 11418 | 10345 | 9166 | 8396 | 8685 | 9933 | 9509 | 10225 | 11991 | 11062 | 9799 | 9928 | 9934 | 11343 | 10450 | 8238 | 8935 | 8478 | 8672 | 9488 | 10373 | 10329 | Pkn2 |
| Pkn3 | 511 | 616 | 862 | 535 | 458 | 428 | 372 | 492 | 541 | 602 | 617 | 855 | 877 | 724 | 409 | 548 | 745 | 483 | 567 | 450 | 399 | 449 | 530 | 696 | 570 | 918 | 832 | 763 | Pkn3 |
| Pknox1 | 2285 | 2538 | 2277 | 2704 | 2923 | 2609 | 2810 | 2567 | 2704 | 2805 | 2894 | 2533 | 2479 | 2579 | 2362 | 1979 | 2284 | 2835 | 2812 | 2425 | 2745 | 2545 | 2910 | 2612 | 2838 | 2545 | 2475 | 2459 | Pknox1 |
| Pknox2 | 3864 | 4560 | 4602 | 4413 | 4461 | 4142 | 3747 | 2847 | 2970 | 3543 | 3650 | 3658 | 3400 | 3484 | 4240 | 4164 | 4271 | 3979 | 4348 | 3836 | 3719 | 3808 | 3241 | 3391 | 3351 | 3896 | 3813 | 3774 | Pknox2 |
| Pkp1 | 113 | 67 | 38 | 9 | 5 | 51 | 1 | 342 | 684 | 301 | 231 | 65 | 153 | 84 | 57 | 45 | 9 | 61 | 9 | 2 | 0 | 10 | 801 | 186 | 37 | 22 | 65 | 96 | Pkp1 |
| Pkp2 | 1791 | 1595 | 1494 | 1112 | 663 | 760 | 807 | 325 | 243 | 186 | 181 | 169 | 240 | 193 | 1668 | 1627 | 1152 | 888 | 726 | 979 | 715 | 190 | 291 | 271 | 117 | 342 | 221 | 267 | Pkp2 |
| Pkp3 | 217 | 138 | 132 | 31 | 12 | 91 | 1 | 274 | 525 | 159 | 219 | 114 | 191 | 78 | 119 | 192 | 42 | 84 | 16 | 4 | 2 | 17 | 528 | 173 | 33 | 119 | 79 | 127 | Pkp3 |
| Pkp4 | 10048 | 9830 | 9787 | 11363 | 11338 | 10775 | 11270 | 5707 | 6075 | 6310 | 6168 | 6442 | 5896 | 5983 | 9480 | 9361 | 10153 | 11433 | 12047 | 10183 | 11180 | 5750 | 6091 | 6367 | 6161 | 6801 | 6144 | 5661 | Pkp4 |
| Pla1a | 9 | 11 | 12 | 12 | 21 | 8 | 15 | 22 | 28 | 16 | 13 | 14 | 20 | 17 | 16 | 9 | 15 | 22 | 9 | 15 | 15 | 18 | 23 | 7 | 27 | 15 | 24 | 19 | Pla1a |
| Pla2g10 | 0 | 1 | 1 | 0 | 2 | 0 | 6 | 0 | 7 | 1 | 11 | 9 | 2 | 3 | 1 | 2 | 1 | 11 | 1 | 0 | 0 | 2 | 10 | 7 | 3 | 2 | 7 | 2 | Pla2g10 |
| Pla2g12a | 978 | 914 | 793 | 964 | 848 | 794 | 1030 | 686 | 752 | 868 | 738 | 775 | 762 | 933 | 899 | 838 | 847 | 1017 | 898 | 947 | 843 | 747 | 741 | 805 | 819 | 950 | 757 | 876 | Pla2g12a |
| Pla2g12b | 0 | 0 | 1 | 0 | 0 | 0 | 0 | 0 | 2 | 3 | 2 | 1 | 3 | 0 | 0 | 0 | 0 | 0 | 0 | 0 | 0 | 2 | 0 | 0 | 0 | 2 | 2 | 0 | Pla2g12b |
| Pla2g15 | 1189 | 1514 | 1578 | 1648 | 1534 | 1337 | 1295 | 1009 | 1242 | 1228 | 1313 | 1266 | 1227 | 1154 | 1273 | 1447 | 1559 | 1784 | 1677 | 1330 | 1308 | 1233 | 1221 | 1312 | 1329 | 1502 | 1145 | 1188 | Pla2g15 |
| Pla2g1b | 0 | 6 | 1 | 0 | 0 | 4 | 2 | 0 | 1 | 0 | 0 | 0 | 1 | 1 | 3 | 5 | 3 | 1 | 2 | 1 | 3 | 2 | 1 | 0 | 0 | 0 | 1 | 0 | Pla2g1b |
| Pla2g2c | 18 | 19 | 8 | 15 | 10 | 12 | 13 | 29 | 16 | 10 | 3 | 12 | 35 | 27 | 24 | 12 | 9 | 20 | 20 | 14 | 9 | 18 | 21 | 12 | 12 | 19 | 23 | 21 | Pla2g2c |
| Pla2g2d | 2 | 6 | 3 | 6 | 11 | 9 | 3 | 13 | 19 | 27 | 21 | 10 | 25 | 10 | 7 | 4 | 20 | 0 | 6 | 8 | 2 | 10 | 17 | 9 | 8 | 2 | 12 | 12 | Pla2g2d |
| Pla2g2e | 0 | 1 | 0 | 0 | 0 | 0 | 0 | 0 | 0 | 1 | 0 | 0 | 0 | 0 | 3 | 2 | 0 | 0 | 2 | 0 | 0 | 0 | 1 | 0 | 1 | 1 | 2 | 0 | Pla2g2e |
| Pla2g2f | 15 | 16 | 38 | 1 | 1 | 9 | 6 | 13 | 23 | 13 | 10 | 18 | 8 | 11 | 7 | 11 | 13 | 15 | 4 | 22 | 2 | 5 | 47 | 9 | 6 | 4 | 10 | 14 | Pla2g2f |
| Pla2g3 | 131 | 163 | 106 | 109 | 164 | 141 | 153 | 112 | 135 | 131 | 125 | 118 | 141 | 95 | 129 | 108 | 79 | 130 | 125 | 142 | 153 | 117 | 106 | 154 | 175 | 182 | 117 | 134 | Pla2g3 |
| Pla2g4a | 720 | 430 | 281 | 180 | 173 | 273 | 115 | 553 | 659 | 407 | 618 | 384 | 449 | 353 | 539 | 695 | 222 | 255 | 103 | 142 | 107 | 353 | 828 | 415 | 186 | 411 | 315 | 328 | Pla2g4a |
| Pla2g4b | 338 | 333 | 414 | 489 | 519 | 502 | 498 | 477 | 467 | 696 | 527 | 654 | 648 | 651 | 261 | 314 | 457 | 464 | 566 | 402 | 463 | 692 | 460 | 713 | 678 | 175 | 668 | 573 | Pla2g4b |
| Pla2g4c | 653 | 602 | 724 | 776 | 806 | 790 | 939 | 458 | 369 | 530 | 459 | 595 | 500 | 603 | 599 | 777 | 721 | 795 | 839 | 760 | 879 | 428 | 326 | 500 | 520 | 550 | 594 | 586 | Pla2g4c |
| Pla2g4d | 8 | 2 | 0 | 1 | 1 | 1 | 1 | 1 | 3 | 2 | 3 | 5 | 3 | 4 | 4 | 4 | 3 | 4 | 1 | 0 | 2 | 3 | 5 | 6 | 2 | 6 | 0 | 0 | Pla2g4d |
| Pla2g4e | 2 | 7 | 8 | 19 | 77 | 61 | 34 | 72 | 111 | 80 | 99 | 39 | 43 | 40 | 16 | 9 | 5 | 46 | 31 | 48 | 38 | 114 | 80 | 37 | 80 | 24 | 33 | 18 | Pla2g4e |
| Pla2g4f | 12 | 29 | 9 | 22 | 27 | 23 | 29 | 10 | 7 | 3 | 1 | 5 | 7 | 10 | 28 | 40 | 36 | 15 | 25 | 14 | 19 | 3 | 19 | 6 | 3 | 16 | 2 | 7 | Pla2g4f |
| Pla2g5 | 9 | 5 | 6 | 4 | 10 | 3 | 11 | 16 | 21 | 13 | 23 | 20 | 24 | 14 | 5 | 3 | 13 | 8 | 12 | 14 | 13 | 2 | 18 | 10 | 6 | 15 | 12 | 19 | Pla2g5 |
| Pla2g6 | 1727 | 2659 | 2675 | 2903 | 2669 | 2377 | 2271 | 2003 | 2310 | 2656 | 2353 | 2672 | 2496 | 2409 | 1928 | 2016 | 2824 | 2731 | 3089 | 2245 | 2479 | 2859 | 2174 | 2817 | 2671 | 2320 | 2635 | 2477 | Pla2g6 |
| Pla2g7 | 296 | 338 | 287 | 239 | 853 | 374 | 254 | 725 | 830 | 653 | 831 | 222 | 448 | 251 | 488 | 372 | 265 | 384 | 294 | 339 | 242 | 974 | 1226 | 286 | 409 | 204 | 241 | 236 | Pla2g7 |
| Pla2r1 | 728 | 376 | 309 | 320 | 204 | 311 | 129 | 680 | 460 | 540 | 406 | 172 | 285 | 191 | 413 | 595 | 192 | 229 | 122 | 185 | 139 | 144 | 623 | 547 | 132 | 546 | 198 | 140 | Pla2r1 |
| Plaa | 10302 | 8235 | 9012 | 8582 | 8980 | 9136 | 9850 | 9987 | 8370 | 7605 | 7779 | 9715 | 9112 | 9528 | 10963 | 9512 | 8807 | 8853 | 8110 | 9880 | 9281 | 8936 | 8385 | 8442 | 8609 | 9580 | 9783 | 9772 | Plaa |
| Plaat1 | 76 | 77 | 114 | 90 | 151 | 97 | 129 | 142 | 118 | 97 | 104 | 87 | 90 | 73 | 102 | 83 | 87 | 95 | 111 | 114 | 94 | 149 | 134 | 106 | 129 | 104 | 108 | 65 | Plaat1 |
| Plaat3 | 442 | 381 | 308 | 273 | 271 | 357 | 325 | 316 | 379 | 291 | 318 | 265 | 316 | 280 | 364 | 433 | 307 | 282 | 306 | 398 | 291 | 264 | 417 | 337 | 324 | 444 | 296 | 244 | Plaat3 |
| Plaat5 | 2 | 1 | 0 | 4 | 2 | 1 | 4 | 0 | 0 | 0 | 0 | 0 | 0 | 0 | 0 | 2 | 2 | 1 | 1 | 3 | 3 | 0 | 5 | 2 | 0 | 3 | 0 | 0 | Plaat5 |
| Plac1 | 35 | 24 | 41 | 18 | 19 | 27 | 6 | 35 | 53 | 33 | 80 | 25 | 27 | 17 | 25 | 26 | 19 | 21 | 10 | 16 | 11 | 20 | 52 | 20 | 31 | 35 | 28 | 30 | Plac1 |
| Plac8 | 58 | 49 | 32 | 21 | 21 | 27 | 17 | 61 | 56 | 27 | 70 | 26 | 41 | 10 | 69 | 88 | 13 | 35 | 12 | 31 | 11 | 33 | 85 | 45 | 21 | 41 | 30 | 31 | Plac8 |
| Plac8l1 | 0 | 2 | 0 | 0 | 3 | 1 | 2 | 0 | 0 | 0 | 0 | 3 | 5 | 3 | 3 | 2 | 0 | 0 | 0 | 0 | 1 | 0 | 0 | 2 | 4 | 0 | 2 | 1 | Plac8l1 |
| Plac9a | 1 | 0 | 0 | 0 | 0 | 1 | 0 | 2 | 1 | 1 | 4 | 0 | 1 | 1 | 3 | 1 | 0 | 0 | 0 | 0 | 1 | 2 | 2 | 2 | 0 | 4 | 0 | 1 | Plac9a |
| Plag1 | 3284 | 2498 | 1880 | 2414 | 2257 | 2960 | 2708 | 1743 | 1734 | 1428 | 1806 | 1587 | 1786 | 1592 | 3078 | 2833 | 2048 | 2368 | 2113 | 2762 | 2565 | 1298 | 1608 | 1241 | 1291 | 851 | 1703 | 1561 | Plag1 |
| Plagl1 | 27633 | 24402 | 25665 | 23255 | 19581 | 24130 | 23711 | 17696 | 19550 | 16956 | 32752 | 22225 | 19635 | 21417 | 25423 | 22881 | 24987 | 23916 | 21646 | 22749 | 23109 | 18616 | 25873 | 17958 | 17505 | 21453 | 22984 | 22275 | Plagl1 |
| Plagl2 | 6413 | 6769 | 7463 | 6218 | 4840 | 5370 | 5569 | 4542 | 4358 | 4513 | 4286 | 6693 | 5822 | 6479 | 5885 | 6650 | 6778 | 5853 | 5657 | 4831 | 5474 | 4523 | 4037 | 5656 | 4788 | 6449 | 6326 | 7195 | Plagl2 |
| Plat | 5425 | 5855 | 5715 | 4837 | 5150 | 4724 | 3900 | 3285 | 3360 | 3185 | 3470 | 2994 | 3530 | 3034 | 5272 | 5839 | 5184 | 4830 | 4647 | 4926 | 4245 | 3293 | 4115 | 3166 | 2922 | 3687 | 3552 | 3037 | Plat |
| Platr25 | 2662 | 1976 | 2859 | 2507 | 2861 | 2682 | 3152 | 2904 | 2635 | 2539 | 2748 | 2682 | 2577 | 2709 | 2769 | 2599 | 2763 | 2601 | 2840 | 2678 | 2691 | 2755 | 2671 | 2533 | 2439 | 2826 | 2746 | 2693 | Platr25 |
| Plau | 371 | 338 | 363 | 193 | 203 | 238 | 161 | 261 | 367 | 256 | 469 | 202 | 306 | 239 | 345 | 434 | 324 | 322 | 226 | 255 | 172 | 282 | 494 | 330 | 232 | 295 | 245 | 229 | Plau |
| Plaur | 50 | 36 | 38 | 17 | 31 | 38 | 23 | 29 | 49 | 27 | 46 | 38 | 58 | 48 | 40 | 62 | 49 | 25 | 46 | 62 | 18 | 32 | 61 | 34 | 42 | 51 | 38 | 28 | Plaur |
| Plb1 | 9 | 17 | 6 | 2 | 9 | 8 | 0 | 10 | 38 | 52 | 13 | 18 | 25 | 17 | 22 | 23 | 0 | 13 | 3 | 5 | 3 | 13 | 32 | 16 | 4 | 8 | 19 | 11 | Plb1 |
| Plbd1 | 42 | 57 | 39 | 4 | 23 | 39 | 6 | 74 | 105 | 105 | 77 | 52 | 51 | 62 | 49 | 59 | 15 | 48 | 12 | 16 | 5 | 45 | 182 | 54 | 30 | 31 | 50 | 42 | Plbd1 |
| Plbd2 | 2922 | 4056 | 3542 | 3752 | 3288 | 3097 | 2851 | 1517 | 2169 | 2112 | 2307 | 2170 | 2229 | 1846 | 2941 | 3755 | 3541 | 3888 | 3250 | 2596 | 3023 | 2162 | 2300 | 2113 | 1978 | 2329 | 1986 | 1924 | Plbd2 |
| Plcb1 | 1429 | 1316 | 1035 | 2085 | 3602 | 2870 | 3174 | 2342 | 2482 | 2677 | 2763 | 1849 | 1762 | 1888 | 1797 | 1338 | 1328 | 2362 | 2364 | 2729 | 2966 | 2759 | 2479 | 2004 | 2955 | 1263 | 1533 | 1569 | Plcb1 |
| Plcb2 | 42 | 67 | 63 | 60 | 97 | 81 | 50 | 66 | 91 | 54 | 99 | 75 | 75 | 67 | 46 | 65 | 54 | 87 | 75 | 60 | 64 | 74 | 133 | 76 | 74 | 64 | 93 | 84 | Plcb2 |
| Plcb3 | 7481 | 7889 | 8213 | 7010 | 5397 | 5522 | 4550 | 2778 | 3702 | 3212 | 3749 | 3057 | 3453 | 2943 | 6285 | 9312 | 7167 | 6900 | 5459 | 4148 | 4602 | 2907 | 3566 | 3143 | 2631 | 3579 | 3097 | 2892 | Plcb3 |
| Plcb4 | 2145 | 1927 | 1866 | 2288 | 2817 | 2672 | 2484 | 3962 | 3533 | 3495 | 3373 | 2946 | 3417 | 2935 | 2647 | 2070 | 1951 | 2425 | 2223 | 2752 | 2492 | 4328 | 3633 | 2728 | 3576 | 2196 | 2907 | 2898 | Plcb4 |
| Plcd1 | 1854 | 2236 | 2102 | 1711 | 1546 | 1673 | 1176 | 1088 | 1296 | 1280 | 1518 | 1257 | 1633 | 1210 | 1821 | 2671 | 1763 | 1798 | 1355 | 1286 | 1190 | 1327 | 1549 | 1179 | 1082 | 1136 | 1474 | 1411 | Plcd1 |
| Plcd3 | 724 | 984 | 943 | 973 | 727 | 647 | 523 | 186 | 316 | 242 | 289 | 161 | 237 | 160 | 609 | 995 | 944 | 880 | 779 | 557 | 546 | 194 | 353 | 252 | 168 | 170 | 142 | 177 | Plcd3 |
| Plcd4 | 44 | 29 | 46 | 47 | 105 | 90 | 67 | 138 | 194 | 215 | 173 | 78 | 98 | 63 | 74 | 50 | 51 | 80 | 72 | 70 | 98 | 204 | 177 | 102 | 167 | 58 | 83 | 39 | Plcd4 |
| Plce1 | 2505 | 2544 | 1605 | 2037 | 2366 | 2537 | 2502 | 3607 | 3897 | 3592 | 4115 | 3330 | 3074 | 3186 | 2649 | 2422 | 2055 | 2221 | 2250 | 2424 | 2570 | 3098 | 3672 | 3128 | 3431 | 2355 | 3320 | 2775 | Plce1 |
| Plcg1 | 15684 | 16189 | 15013 | 18623 | 19248 | 18610 | 18867 | 16745 | 18154 | 20270 | 19426 | 20356 | 20137 | 20984 | 14599 | 15597 | 16584 | 18180 | 20074 | 15395 | 20070 | 20787 | 16613 | 20282 | 22106 | 18110 | 20409 | 20354 | Plcg1 |
| Plcg2 | 480 | 609 | 681 | 392 | 382 | 385 | 329 | 192 | 263 | 227 | 239 | 263 | 249 | 233 | 436 | 637 | 481 | 428 | 353 | 288 | 275 | 204 | 303 | 226 | 204 | 262 | 239 | 222 | Plcg2 |
| Plch1 | 6432 | 7043 | 5199 | 5667 | 5704 | 6316 | 5745 | 4759 | 3159 | 3229 | 2654 | 4643 | 4756 | 4299 | 7378 | 6004 | 5601 | 4778 | 5926 | 7282 | 5882 | 3461 | 3098 | 3253 | 3492 | 2866 | 5245 | 4866 | Plch1 |
| Plch2 | 479 | 687 | 625 | 920 | 1220 | 1160 | 1169 | 1059 | 1295 | 1514 | 1099 | 880 | 770 | 783 | 538 | 474 | 851 | 1066 | 1400 | 1012 | 1249 | 1389 | 1120 | 1367 | 1377 | 803 | 673 | 717 | Plch2 |
| Plcl1 | 1618 | 1481 | 1337 | 2037 | 2653 | 2536 | 2796 | 1916 | 1735 | 1788 | 1624 | 1768 | 1461 | 1699 | 1878 | 1547 | 1905 | 2243 | 2441 | 2480 | 2653 | 2142 | 1720 | 1715 | 2095 | 1357 | 1476 | 1532 | Plcl1 |
| Plcl2 | 1288 | 1198 | 1681 | 1397 | 1735 | 1741 | 1721 | 2919 | 2834 | 2787 | 2631 | 2677 | 2315 | 2676 | 1445 | 1047 | 1483 | 1289 | 1828 | 1708 | 1826 | 2971 | 2567 | 2497 | 3078 | 2677 | 2668 | 2650 | Plcl2 |
| Plcxd1 | 143 | 168 | 124 | 141 | 131 | 151 | 93 | 40 | 54 | 90 | 57 | 27 | 36 | 22 | 131 | 167 | 123 | 157 | 161 | 122 | 78 | 62 | 47 | 56 | 30 | 31 | 23 | 21 | Plcxd1 |
| Plcxd2 | 4327 | 4755 | 2976 | 4950 | 5949 | 5641 | 6236 | 3296 | 2782 | 2439 | 2172 | 2552 | 1977 | 2664 | 5243 | 3974 | 4143 | 5109 | 5415 | 6073 | 5977 | 2102 | 2230 | 2333 | 3089 | 1848 | 2273 | 2036 | Plcxd2 |
| Plcxd3 | 286 | 248 | 124 | 228 | 349 | 430 | 381 | 630 | 577 | 420 | 347 | 395 | 407 | 384 | 372 | 249 | 205 | 221 | 322 | 485 | 345 | 413 | 465 | 237 | 525 | 45 | 533 | 361 | Plcxd3 |
| Plcz1 | 2 | 0 | 0 | 1 | 8 | 5 | 8 | 4 | 8 | 5 | 5 | 4 | 5 | 2 | 1 | 1 | 0 | 3 | 6 | 1 | 2 | 3 | 6 | 4 | 5 | 5 | 0 | 4 | Plcz1 |
| Pld1 | 965 | 953 | 757 | 722 | 700 | 709 | 608 | 727 | 834 | 829 | 865 | 924 | 944 | 852 | 974 | 938 | 880 | 732 | 578 | 720 | 629 | 760 | 1046 | 579 | 664 | 707 | 822 | 710 | Pld1 |
| Pld2 | 647 | 683 | 817 | 665 | 1051 | 842 | 636 | 694 | 859 | 866 | 986 | 603 | 736 | 611 | 647 | 740 | 620 | 826 | 800 | 607 | 652 | 886 | 1083 | 750 | 810 | 591 | 718 | 593 | Pld2 |
| Pld3 | 4274 | 6005 | 4646 | 5380 | 5446 | 5150 | 4253 | 3121 | 3891 | 3707 | 3699 | 3106 | 3298 | 2770 | 4548 | 5312 | 5093 | 5438 | 5395 | 5015 | 4771 | 3636 | 4242 | 3422 | 3655 | 2974 | 3023 | 2734 | Pld3 |
| Pld4 | 7 | 38 | 14 | 3 | 23 | 4 | 2 | 26 | 10 | 6 | 8 | 7 | 12 | 1 | 7 | 2 | 3 | 18 | 47 | 41 | 4 | 23 | 10 | 31 | 4 | 29 | 6 | 15 | Pld4 |

| | | | | | | | | | | | | | | | | | | | | | | | | | | | | | |
|---|---|---|---|---|---|---|---|---|---|---|---|---|---|---|---|---|---|---|---|---|---|---|---|---|---|---|---|---|---|
| Pld5 | 35 | 41 | 45 | 52 | 155 | 68 | 44 | 192 | 214 | 291 | 285 | 191 | 299 | 175 | 78 | 50 | 38 | 73 | 72 | 70 | 73 | 373 | 287 | 166 | 228 | 131 | 143 | 184 | Pld5 |
| Pld6 | 57 | 27 | 47 | 39 | 41 | 49 | 31 | 50 | 59 | 47 | 43 | 51 | 48 | 34 | 58 | 50 | 44 | 51 | 45 | 23 | 44 | 42 | 56 | 53 | 71 | 59 | 51 | 61 | Pld6 |
| Plec | 6910 | 9604 | 10598 | 9591 | 8483 | 8437 | 5893 | 4501 | 5828 | 6303 | 7333 | 5598 | 6873 | 4731 | 6711 | 9444 | 10492 | 9682 | 7995 | 6352 | 6596 | 6259 | 6399 | 5762 | 5419 | 3291 | 5832 | 5375 | Plec |
| Plek | 796 | 504 | 652 | 412 | 452 | 480 | 407 | 1016 | 925 | 728 | 878 | 862 | 1174 | 852 | 708 | 880 | 447 | 454 | 346 | 564 | 407 | 999 | 1096 | 840 | 661 | 1084 | 952 | 779 | Plek |
| Plek2 | 105 | 103 | 75 | 64 | 38 | 63 | 61 | 78 | 97 | 82 | 52 | 88 | 332 | 142 | 80 | 99 | 66 | 46 | 61 | 78 | 48 | 53 | 103 | 69 | 56 | 135 | 131 | 218 | Plek2 |
| Plekha1 | 8412 | 8326 | 9451 | 9851 | 12078 | 10377 | 10451 | 11323 | 9269 | 10185 | 10019 | 8804 | 9647 | 8929 | 10486 | 9801 | 10039 | 10181 | 10402 | 11159 | 10681 | 12287 | 9514 | 8959 | 11172 | 6467 | 8881 | 8302 | Plekha1 |
| Plekha2 | 644 | 461 | 578 | 288 | 466 | 364 | 226 | 631 | 611 | 649 | 793 | 366 | 831 | 424 | 694 | 800 | 445 | 335 | 239 | 268 | 248 | 787 | 733 | 357 | 383 | 439 | 372 | 369 | Plekha2 |
| Plekha3 | 2400 | 2129 | 2503 | 2427 | 2467 | 2489 | 2605 | 1789 | 1735 | 1635 | 1795 | 1955 | 1772 | 2015 | 2391 | 2263 | 2328 | 2533 | 2433 | 2187 | 2522 | 1885 | 1651 | 1865 | 1789 | 2063 | 1887 | 1900 | Plekha3 |
| Plekha4 | 89 | 107 | 91 | 123 | 173 | 80 | 65 | 169 | 339 | 373 | 618 | 104 | 249 | 113 | 90 | 170 | 117 | 95 | 67 | 68 | 67 | 453 | 433 | 169 | 251 | 166 | 95 | 109 | Plekha4 |
| Plekha5 | 3262 | 3172 | 3750 | 3563 | 4584 | 4080 | 3871 | 6826 | 7003 | 7445 | 7397 | 7342 | 7485 | 7239 | 3303 | 3187 | 3586 | 3372 | 4187 | 3652 | 3949 | 7798 | 6321 | 7142 | 7545 | 6130 | 6972 | 7335 | Plekha5 |
| Plekha6 | 4279 | 5362 | 6218 | 6928 | 8782 | 7369 | 6814 | 8349 | 7523 | 10545 | 9078 | 11941 | 10637 | 11008 | 5006 | 4896 | 7130 | 6906 | 8265 | 6200 | 7443 | 12626 | 7306 | 10229 | 12023 | 6729 | 11766 | 11192 | Plekha6 |
| Plekha7 | 3683 | 4606 | 4621 | 4577 | 3842 | 4173 | 4008 | 1502 | 1721 | 1920 | 1522 | 2562 | 2254 | 2394 | 3580 | 3833 | 4898 | 4066 | 4437 | 3786 | 3995 | 1360 | 1726 | 2172 | 1918 | 2289 | 2488 | 2527 | Plekha7 |
| Plekha8 | 7934 | 6707 | 6436 | 6860 | 6102 | 6894 | 6888 | 5040 | 4907 | 4566 | 4625 | 5017 | 5158 | 5362 | 7195 | 7636 | 6446 | 6726 | 6151 | 6220 | 6871 | 4600 | 4343 | 4690 | 5117 | 4915 | 5398 | 5473 | Plekha8 |
| Plekhb1 | 193 | 249 | 97 | 137 | 234 | 145 | 101 | 267 | 381 | 293 | 380 | 205 | 268 | 148 | 305 | 229 | 129 | 185 | 171 | 147 | 114 | 369 | 369 | 195 | 210 | 221 | 202 | 163 | Plekhb1 |
| Plekhb2 | 4299 | 4978 | 4466 | 4701 | 5062 | 4540 | 4800 | 2927 | 2764 | 2851 | 2554 | 2953 | 2513 | 2918 | 4362 | 4171 | 4831 | 4524 | 5224 | 5184 | 4854 | 2949 | 2720 | 2813 | 2986 | 2770 | 2919 | 3010 | Plekhb2 |
| Plekhd1 | 293 | 333 | 310 | 333 | 533 | 312 | 201 | 350 | 290 | 304 | 359 | 238 | 456 | 261 | 336 | 414 | 367 | 387 | 341 | 357 | 275 | 530 | 261 | 209 | 385 | 57 | 292 | 299 | Plekhd1 |
| Plekhf1 | 591 | 725 | 644 | 649 | 431 | 467 | 380 | 93 | 161 | 129 | 135 | 121 | 113 | 76 | 484 | 668 | 597 | 540 | 509 | 544 | 413 | 87 | 148 | 105 | 110 | 125 | 117 | 85 | Plekhf1 |
| Plekhf2 | 2007 | 1680 | 1902 | 1848 | 1547 | 1596 | 1780 | 1660 | 1587 | 1522 | 1545 | 1538 | 1448 | 1546 | 1958 | 1850 | 1695 | 1661 | 1547 | 1548 | 1628 | 1275 | 1621 | 1644 | 1429 | 1790 | 1447 | 1658 | Plekhf2 |
| Plekhg1 | 2148 | 1658 | 1949 | 1376 | 1500 | 1689 | 1198 | 3075 | 3327 | 3194 | 3565 | 3240 | 4183 | 3472 | 1811 | 1920 | 1566 | 1407 | 1379 | 1510 | 1319 | 3130 | 3956 | 3027 | 2727 | 2778 | 3841 | 3674 | Plekhg1 |
| Plekhg2 | 4385 | 5840 | 7110 | 5989 | 5920 | 5489 | 4883 | 5192 | 6483 | 6751 | 7347 | 8593 | 8224 | 7526 | 4067 | 5503 | 7095 | 6119 | 6469 | 4540 | 5019 | 6154 | 6237 | 8088 | 6574 | 8978 | 7951 | 7852 | Plekhg2 |
| Plekhg3 | 1015 | 1053 | 1120 | 831 | 870 | 922 | 489 | 684 | 1092 | 975 | 1230 | 689 | 912 | 524 | 841 | 1182 | 991 | 973 | 778 | 582 | 573 | 794 | 1474 | 898 | 605 | 624 | 557 | 565 | Plekhg3 |
| Plekhg4 | 1115 | 1598 | 1393 | 2070 | 2376 | 2087 | 2044 | 382 | 413 | 581 | 462 | 492 | 371 | 476 | 1221 | 1425 | 1921 | 2447 | 2358 | 1896 | 2147 | 590 | 397 | 449 | 524 | 343 | 446 | 432 | Plekhg4 |
| Plekhg5 | 1445 | 1572 | 1762 | 1632 | 1739 | 1540 | 1362 | 1121 | 1468 | 1563 | 1696 | 1429 | 1399 | 1319 | 1120 | 1603 | 1737 | 1657 | 1932 | 1223 | 1425 | 1464 | 1465 | 1578 | 1326 | 1702 | 1219 | 1173 | Plekhg5 |
| Plekhg6 | 115 | 85 | 84 | 59 | 39 | 62 | 16 | 38 | 40 | 36 | 23 | 15 | 29 | 20 | 70 | 116 | 75 | 55 | 22 | 32 | 13 | 12 | 59 | 34 | 10 | 19 | 20 | 14 | Plekhg6 |
| Plekhh1 | 1747 | 1836 | 2218 | 1874 | 1983 | 1760 | 1607 | 1511 | 1489 | 1537 | 1673 | 2159 | 2190 | 1999 | 1725 | 1732 | 2027 | 1721 | 1946 | 1564 | 1694 | 1527 | 1325 | 1622 | 1706 | 1558 | 2260 | 2261 | Plekhh1 |
| Plekhh2 | 2748 | 2249 | 1913 | 1826 | 1787 | 2074 | 1819 | 1209 | 1519 | 1250 | 2181 | 956 | 1327 | 1020 | 2418 | 2848 | 1870 | 1723 | 1623 | 1944 | 1891 | 1076 | 1799 | 1068 | 988 | 972 | 1187 | 1044 | Plekhh2 |
| Plekhh3 | 1187 | 1707 | 1418 | 1886 | 1614 | 1514 | 1388 | 458 | 562 | 620 | 586 | 678 | 562 | 513 | 1195 | 1359 | 1859 | 1921 | 1747 | 1315 | 1490 | 695 | 649 | 626 | 628 | 661 | 575 | 540 | Plekhh3 |
| Plekhj1 | 1844 | 2217 | 2514 | 2369 | 2095 | 2050 | 2213 | 1887 | 2032 | 2560 | 2280 | 2569 | 2371 | 2849 | 1834 | 1717 | 2381 | 2041 | 2343 | 1906 | 2246 | 2012 | 2126 | 2826 | 2370 | 3026 | 2753 | 2864 | Plekhj1 |
| Plekhm1 | 3152 | 4141 | 3828 | 4177 | 3671 | 3501 | 3394 | 1727 | 2130 | 2351 | 2237 | 2488 | 2193 | 2260 | 2875 | 3881 | 4241 | 4357 | 3879 | 2766 | 3503 | 2722 | 1961 | 2377 | 2635 | 2151 | 2162 | 2258 | Plekhm1 |
| Plekhm2 | 1415 | 1708 | 2011 | 2049 | 2243 | 1949 | 1672 | 2179 | 2752 | 3018 | 3027 | 2813 | 2758 | 2478 | 1315 | 1749 | 1997 | 1955 | 2202 | 1528 | 1793 | 2968 | 2693 | 3045 | 3052 | 2857 | 2525 | 2553 | Plekhm2 |
| Plekhm3 | 1337 | 1533 | 927 | 1376 | 1585 | 1367 | 1363 | 1380 | 1551 | 1514 | 1390 | 1358 | 1606 | 1341 | 1422 | 1520 | 1229 | 1349 | 1322 | 1313 | 1467 | 1569 | 1299 | 1176 | 1546 | 744 | 1386 | 1246 | Plekhm3 |
| Plekhn1 | 5591 | 7061 | 5556 | 6874 | 7093 | 6902 | 6472 | 4880 | 5710 | 5952 | 5782 | 5687 | 5304 | 5071 | 5731 | 5863 | 6102 | 7069 | 7203 | 6653 | 6558 | 5701 | 5009 | 5493 | 5775 | 4371 | 5348 | 4774 | Plekhn1 |
| Plekho1 | 2074 | 2749 | 2857 | 2632 | 2966 | 2989 | 2546 | 4332 | 4880 | 4614 | 4388 | 5472 | 4613 | 5354 | 2124 | 2213 | 2986 | 2291 | 3439 | 2716 | 2813 | 4181 | 4574 | 5246 | 5184 | 6149 | 5108 | 5196 | Plekho1 |
| Plekho2 | 2600 | 3389 | 2805 | 2805 | 1997 | 2245 | 1804 | 865 | 1018 | 921 | 1236 | 1064 | 1089 | 979 | 2575 | 3485 | 2719 | 2861 | 2103 | 1632 | 1886 | 854 | 1202 | 945 | 833 | 1200 | 1117 | 1094 | Plekho2 |
| Plekhs1 | 15 | 4 | 5 | 10 | 21 | 3 | 18 | 28 | 36 | 45 | 27 | 11 | 28 | 23 | 12 | 11 | 14 | 7 | 2 | 6 | 18 | 37 | 37 | 37 | 14 | 34 | 23 | 11 | Plekhs1 |
| Plet1 | 7 | 15 | 14 | 7 | 12 | 8 | 9 | 12 | 25 | 30 | 29 | 4 | 10 | 6 | 18 | 16 | 2 | 1 | 14 | 5 | 5 | 17 | 13 | 11 | 25 | 9 | 30 | 11 | Plet1 |
| Plg | 6 | 1 | 3 | 2 | 13 | 6 | 3 | 6 | 12 | 0 | 1 | 1 | 8 | 5 | 3 | 6 | 12 | 8 | 4 | 3 | 6 | 5 | 4 | 7 | 12 | 13 | 0 | 4 | Plg |
| Plgrkt | 2718 | 2556 | 2328 | 2629 | 2375 | 2562 | 2839 | 2677 | 2314 | 2269 | 2094 | 2054 | 2064 | 2279 | 2813 | 2624 | 2396 | 2542 | 2512 | 2925 | 2693 | 2339 | 2184 | 2165 | 2437 | 2218 | 2123 | 2192 | Plgrkt |
| Plin1 | 3 | 3 | 4 | 6 | 9 | 0 | 10 | 5 | 7 | 3 | 2 | 4 | 3 | 4 | 8 | 5 | 0 | 12 | 2 | 6 | 8 | 5 | 4 | 2 | 2 | 7 | 1 | 4 | Plin1 |
| Plin2 | 1713 | 1748 | 1764 | 1126 | 1460 | 1374 | 1092 | 2949 | 3200 | 2521 | 2958 | 2824 | 3182 | 2733 | 1759 | 1899 | 1680 | 1126 | 1430 | 1501 | 1070 | 2595 | 3666 | 2673 | 2270 | 3125 | 3483 | 3114 | Plin2 |
| Plin3 | 4188 | 4261 | 4138 | 2968 | 2249 | 2693 | 2133 | 2295 | 2739 | 2045 | 2645 | 2165 | 2683 | 2101 | 3791 | 4298 | 3588 | 2826 | 2532 | 2488 | 2164 | 1728 | 3304 | 2025 | 1556 | 2786 | 2230 | 2270 | Plin3 |
| Plin4 | 7 | 6 | 14 | 11 | 11 | 10 | 9 | 6 | 15 | 3 | 9 | 5 | 11 | 2 | 7 | 9 | 5 | 14 | 3 | 12 | 10 | 10 | 10 | 8 | 4 | 4 | 7 | 0 | Plin4 |
| Plin5 | 6 | 17 | 7 | 23 | 10 | 15 | 13 | 5 | 7 | 8 | 8 | 3 | 2 | 2 | 13 | 14 | 23 | 15 | 6 | 12 | 4 | 12 | 14 | 9 | 13 | 3 | 9 | 1 | Plin5 |
| Plk1 | 2561 | 2934 | 3784 | 2712 | 2077 | 2259 | 2172 | 2712 | 3401 | 3410 | 3240 | 5755 | 4572 | 5392 | 2207 | 2630 | 3532 | 2129 | 2867 | 1939 | 2207 | 2705 | 2871 | 4819 | 3583 | 7379 | 5055 | 5620 | Plk1 |
| Plk2 | 6921 | 6021 | 4939 | 5978 | 5439 | 5940 | 5041 | 9856 | 9199 | 8720 | 8676 | 7306 | 8453 | 7085 | 7054 | 6891 | 4779 | 6309 | 5041 | 5069 | 5170 | 9608 | 10174 | 8505 | 8180 | 8792 | 6966 | 7575 | Plk2 |
| Plk3 | 899 | 1356 | 1454 | 1622 | 1448 | 1260 | 1127 | 768 | 804 | 950 | 821 | 1503 | 1364 | 1292 | 1076 | 1220 | 1568 | 1391 | 1463 | 1199 | 1196 | 956 | 744 | 1046 | 966 | 1325 | 1436 | 1423 | Plk3 |
| Plk4 | 603 | 116 | 131 | 170 | 70 | 158 | 108 | 670 | 145 | 106 | 145 | 442 | 176 | 295 | 394 | 202 | 76 | 103 | 122 | 190 | 93 | 162 | 233 | 118 | 167 | 248 | 238 | 529 | Plk4 |
| Plk5 | 135 | 103 | 76 | 120 | 79 | 69 | 114 | 34 | 29 | 40 | 20 | 17 | 20 | 20 | 90 | 121 | 103 | 102 | 131 | 86 | 73 | 40 | 35 | 34 | 23 | 20 | 17 | 20 | Plk5 |
| Pllp | 120 | 119 | 99 | 131 | 114 | 74 | 67 | 17 | 21 | 17 | 38 | 26 | 14 | 13 | 105 | 108 | 119 | 146 | 100 | 94 | 96 | 12 | 18 | 35 | 12 | 47 | 22 | 7 | Pllp |
| Pln | 0 | 1 | 0 | 6 | 2 | 4 | 3 | 4 | 6 | 3 | 0 | 0 | 0 | 0 | 1 | 0 | 0 | 3 | 2 | 0 | 2 | 2 | 2 | 2 | 0 | 0 | 1 | 1 | Pln |
| Plod1 | 3042 | 3683 | 3077 | 2796 | 2645 | 2777 | 1971 | 2077 | 3052 | 2742 | 3742 | 2071 | 2806 | 2055 | 2925 | 3978 | 3016 | 3126 | 2464 | 1908 | 2126 | 2669 | 3329 | 2446 | 2225 | 2555 | 2163 | 2057 | Plod1 |
| Plod2 | 7345 | 5110 | 5540 | 4678 | 3989 | 4537 | 3876 | 5448 | 6246 | 5383 | 8840 | 3838 | 4921 | 4061 | 7006 | 8081 | 4651 | 4864 | 3677 | 4380 | 3724 | 4969 | 6043 | 5006 | 3598 | 5646 | 4306 | 4484 | Plod2 |
| Plod3 | 2573 | 3297 | 3301 | 3245 | 3200 | 3154 | 2557 | 3309 | 4071 | 4332 | 4441 | 4249 | 3977 | 3965 | 2376 | 3166 | 3522 | 3231 | 3438 | 2447 | 2718 | 4380 | 4047 | 4476 | 4458 | 4338 | 4202 | 3817 | Plod3 |
| Plp1 | 1082 | 995 | 948 | 899 | 1267 | 906 | 1040 | 1811 | 1632 | 1810 | 1864 | 1480 | 1726 | 1447 | 1118 | 1327 | 963 | 1126 | 846 | 930 | 925 | 2182 | 1659 | 1520 | 1639 | 1420 | 1379 | 1354 | Plp1 |
| Plp2 | 888 | 832 | 781 | 554 | 581 | 724 | 622 | 1264 | 1619 | 1262 | 1577 | 1269 | 1479 | 1425 | 761 | 938 | 606 | 675 | 554 | 606 | 531 | 993 | 1651 | 1335 | 1180 | 1606 | 1311 | 1438 | Plp2 |
| Plpbp | 4916 | 5259 | 4091 | 4372 | 3874 | 4313 | 4145 | 3810 | 3550 | 3332 | 3434 | 3352 | 3511 | 3427 | 4603 | 4741 | 3859 | 4296 | 4150 | 4628 | 4379 | 3309 | 3384 | 3184 | 3268 | 3019 | 3507 | 3300 | Plpbp |
| Plpp1 | 813 | 767 | 571 | 615 | 801 | 682 | 751 | 935 | 1065 | 901 | 1089 | 697 | 844 | 733 | 815 | 827 | 676 | 753 | 729 | 802 | 699 | 1059 | 1212 | 776 | 806 | 727 | 728 | 782 | Plpp1 |
| Plpp2 | 732 | 651 | 667 | 487 | 497 | 502 | 340 | 402 | 468 | 411 | 400 | 350 | 420 | 351 | 587 | 683 | 517 | 439 | 455 | 477 | 378 | 389 | 534 | 411 | 311 | 484 | 424 | 407 | Plpp2 |
| Plpp3 | 3814 | 3562 | 3980 | 4367 | 4601 | 4584 | 4701 | 6740 | 7995 | 7510 | 8972 | 7667 | 8079 | 7870 | 4059 | 3960 | 4300 | 4519 | 4747 | 3874 | 4236 | 6988 | 8282 | 8586 | 7631 | 9060 | 7114 | 7242 | Plpp3 |
| Plpp4 | 55 | 29 | 58 | 67 | 79 | 54 | 39 | 110 | 129 | 157 | 112 | 109 | 153 | 118 | 30 | 53 | 58 | 51 | 45 | 76 | 56 | 177 | 107 | 141 | 112 | 74 | 150 | 167 | Plpp4 |
| Plpp5 | 768 | 727 | 676 | 686 | 671 | 728 | 640 | 1491 | 1943 | 1627 | 2110 | 1429 | 1733 | 1570 | 778 | 782 | 667 | 754 | 690 | 628 | 631 | 1795 | 1986 | 1570 | 1498 | 1814 | 1549 | 1534 | Plpp5 |
| Plpp6 | 1059 | 868 | 995 | 1070 | 1201 | 1105 | 1138 | 911 | 913 | 1029 | 928 | 805 | 895 | 893 | 1229 | 989 | 1028 | 1085 | 1019 | 971 | 1208 | 1103 | 943 | 911 | 948 | 892 | 809 | 803 | Plpp6 |
| Plpp7 | 126 | 174 | 151 | 177 | 307 | 185 | 166 | 327 | 400 | 388 | 448 | 329 | 262 | 251 | 162 | 164 | 184 | 175 | 222 | 225 | 205 | 423 | 392 | 298 | 342 | 299 | 300 | 260 | Plpp7 |
| Plppr1 | 872 | 880 | 949 | 1351 | 2044 | 1456 | 1751 | 1980 | 1660 | 1975 | 1693 | 1395 | 1745 | 1452 | 1155 | 884 | 1166 | 1562 | 1838 | 1692 | 1931 | 2398 | 1716 | 1642 | 2225 | 1310 | 1320 | 1411 | Plppr1 |
| Plppr2 | 3290 | 4647 | 4490 | 6565 | 7237 | 5912 | 5992 | 3138 | 3670 | 5157 | 3942 | 4018 | 3383 | 3657 | 3501 | 3828 | 6142 | 6781 | 7230 | 5089 | 6222 | 5242 | 3377 | 4386 | 4941 | 3622 | 3144 | 3299 | Plppr2 |
| Plppr3 | 8224 | 11562 | 10512 | 12137 | 13564 | 12430 | 10979 | 14747 | 16154 | 18603 | 15364 | 17403 | 15908 | 15970 | 8724 | 9249 | 13031 | 11581 | 14864 | 11983 | 11841 | 18927 | 14917 | 18281 | 20201 | 16595 | 16835 | 14873 | Plppr3 |
| Plppr4 | 211 | 232 | 144 | 486 | 1041 | 694 | 834 | 539 | 707 | 636 | 562 | 378 | 409 | 300 | 284 | 264 | 283 | 620 | 824 | 650 | 945 | 852 | 794 | 483 | 839 | 261 | 172 | 192 | Plppr4 |
| Plppr5 | 76 | 42 | 67 | 193 | 312 | 286 | 298 | 258 | 252 | 375 | 247 | 118 | 109 | 113 | 79 | 66 | 111 | 197 | 271 | 224 | 308 | 416 | 244 | 249 | 349 | 89 | 121 | 87 | Plppr5 |
| Plrg1 | 7345 | 7749 | 6797 | 6329 | 5246 | 5490 | 5815 | 5897 | 5406 | 4672 | 4239 | 5033 | 4907 | 5235 | 7630 | 7060 | 6525 | 5876 | 6058 | 6920 | 5817 | 4796 | 5278 | 4892 | 4935 | 5874 | 5591 | 5593 | Plrg1 |
| Pls1 | 1529 | 1110 | 1029 | 1053 | 893 | 862 | 898 | 620 | 481 | 448 | 414 | 345 | 433 | 351 | 1379 | 1355 | 961 | 916 | 778 | 791 | 791 | 314 | 464 | 477 | 288 | 539 | 392 | 388 | Pls1 |
| Pls3 | 11419 | 9120 | 9411 | 7237 | 6875 | 7548 | 8562 | 6657 | 6462 | 4950 | 6086 | 5628 | 6503 | 5823 | 11050 | 9431 | 7829 | 7517 | 7119 | 8535 | 7304 | 4782 | 6987 | 5185 | 4322 | 6446 | 5891 | 6407 | Pls3 |
| Plscr1 | 767 | 757 | 761 | 492 | 448 | 528 | 542 | 613 | 614 | 471 | 585 | 571 | 655 | 623 | 790 | 734 | 689 | 494 | 570 | 619 | 511 | 510 | 630 | 614 | 438 | 1004 | 714 | 735 | Plscr1 |
| Plscr2 | 30 | 31 | 16 | 13 | 10 | 17 | 19 | 37 | 49 | 23 | 87 | 37 | 56 | 33 | 22 | 25 | 15 | 11 | 7 | 12 | 15 | 53 | 63 | 49 | 27 | 72 | 63 | 64 | Plscr2 |
| Plscr3 | 2832 | 2983 | 3097 | 2328 | 2425 | 2377 | 2072 | 2727 | 3526 | 3167 | 3515 | 3182 | 3460 | 3093 | 2349 | 3188 | 2803 | 2511 | 2447 | 1911 | 2071 | 2831 | 3654 | 3359 | 2891 | 3803 | 3066 | 3198 | Plscr3 |
| Plscr4 | 113 | 66 | 128 | 97 | 112 | 113 | 60 | 165 | 233 | 188 | 402 | 161 | 179 | 170 | 92 | 113 | 102 | 69 | 69 | 91 | 72 | 174 | 247 | 228 | 120 | 201 | 158 | 172 | Plscr4 |
| Plscr5 | 3 | 1 | 0 | 7 | 15 | 13 | 8 | 7 | 10 | 3 | 6 | 2 | 2 | 9 | 8 | 5 | 8 | 4 | 5 | 8 | 13 | 5 | 4 | 2 | 7 | 5 | 3 | 6 | Plscr5 |
| Pltp | 2036 | 2757 | 3238 | 2262 | 2766 | 2124 | 1893 | 1809 | 2308 | 2334 | 2333 | 2723 | 3149 | 2302 | 2164 | 3065 | 2643 | 2436 | 2286 | 2392 | 2199 | 2299 | 2896 | 2232 | 2283 | 2866 | 3036 | 2522 | Pltp |
| Plvap | 879 | 1031 | 1065 | 762 | 578 | 581 | 455 | 612 | 840 | 841 | 1070 | 954 | 1054 | 850 | 918 | 1282 | 889 | 722 | 503 | 419 | 460 | 824 | 968 | 903 | 700 | 1319 | 906 | 1004 | Plvap |
| Plxdc1 | 50 | 53 | 34 | 60 | 52 | 40 | 38 | 31 | 46 | 30 | 36 | 42 | 8 | 12 | 44 | 84 | 36 | 100 | 55 | 41 | 36 | 37 | 51 | 27 | 55 | 32 | 14 | 33 | Plxdc1 |

| | | | | | | | | | | | | | | | | | | | | | | | | | | | | | |
|---|---|---|---|---|---|---|---|---|---|---|---|---|---|---|---|---|---|---|---|---|---|---|---|---|---|---|---|---|---|
| Plxdc2 | 1975 | 1681 | 1727 | 1232 | 1505 | 1510 | 1149 | 1950 | 2071 | 1837 | 2475 | 1551 | 2251 | 1403 | 1916 | 2069 | 1223 | 1457 | 1036 | 1271 | 1155 | 2035 | 2601 | 1424 | 1527 | 1445 | 1859 | 1767 | Plxdc2 |
| Plxna1 | 17168 | 22874 | 22999 | 28583 | 29355 | 26358 | 25228 | 16291 | 18947 | 23034 | 19508 | 24691 | 21975 | 21435 | 17689 | 22120 | 28763 | 29235 | 30400 | 22132 | 26476 | 26406 | 17547 | 23387 | 24818 | 14431 | 20549 | 21836 | Plxna1 |
| Plxna2 | 10663 | 11391 | 6576 | 13873 | 21042 | 19847 | 22087 | 17167 | 21754 | 23216 | 20406 | 19314 | 13110 | 20398 | 11819 | 10504 | 11143 | 15888 | 18955 | 15949 | 22685 | 19281 | 15360 | 19172 | 27832 | 11733 | 14559 | 13830 | Plxna2 |
| Plxna3 | 15759 | 20645 | 15086 | 22782 | 24209 | 21900 | 22910 | 14004 | 16270 | 18176 | 15013 | 16040 | 14870 | 14738 | 16736 | 17979 | 20296 | 25161 | 24068 | 19484 | 23077 | 20112 | 13740 | 16047 | 19193 | 9135 | 14405 | 13316 | Plxna3 |
| Plxna4 | 3377 | 3677 | 2136 | 5072 | 9624 | 7511 | 8079 | 8281 | 10567 | 11223 | 10199 | 7067 | 6763 | 7370 | 3932 | 3896 | 3177 | 6323 | 6770 | 5547 | 8191 | 10537 | 7538 | 7719 | 13664 | 3634 | 5721 | 5158 | Plxna4 |
| Plxnb1 | 8755 | 11148 | 13978 | 13223 | 14472 | 13554 | 11558 | 17319 | 18871 | 23440 | 18817 | 29170 | 23618 | 24362 | 9058 | 9111 | 14443 | 11909 | 15756 | 11279 | 12571 | 23009 | 16971 | 25908 | 25303 | 18749 | 25913 | 24525 | Plxnb1 |
| Plxnb2 | 24082 | 28273 | 27165 | 26744 | 24234 | 24540 | 21349 | 17723 | 22360 | 23395 | 22701 | 26003 | 24010 | 21910 | 21663 | 27951 | 26963 | 26311 | 26690 | 20292 | 22847 | 21697 | 23060 | 24512 | 22445 | 21891 | 22954 | 23193 | Plxnb2 |
| Plxnb3 | 18 | 14 | 9 | 9 | 41 | 22 | 7 | 40 | 80 | 66 | 151 | 30 | 75 | 21 | 13 | 24 | 2 | 22 | 5 | 7 | 13 | 95 | 143 | 37 | 39 | 25 | 20 | 17 | Plxnb3 |
| Plxnc1 | 2836 | 2725 | 2587 | 3438 | 5141 | 4461 | 5336 | 5268 | 5397 | 5597 | 4746 | 4926 | 3446 | 5247 | 2860 | 2398 | 3283 | 3857 | 4795 | 4389 | 5235 | 7088 | 5084 | 5498 | 6890 | 3729 | 4296 | 4358 | Plxnc1 |
| Plxnd1 | 5274 | 6994 | 6929 | 7189 | 7383 | 6793 | 6004 | 4258 | 6646 | 6665 | 7367 | 5960 | 6098 | 5131 | 4638 | 7302 | 6662 | 7980 | 7178 | 5402 | 6600 | 5897 | 6117 | 6955 | 6653 | 6474 | 5283 | 4536 | Plxnd1 |
| Pm20d1 | 74 | 71 | 121 | 29 | 50 | 32 | 28 | 16 | 37 | 18 | 18 | 52 | 41 | 35 | 65 | 96 | 60 | 37 | 22 | 29 | 20 | 22 | 38 | 25 | 18 | 35 | 48 | 42 | Pm20d1 |
| Pm20d2 | 110 | 74 | 51 | 53 | 109 | 120 | 99 | 156 | 177 | 154 | 148 | 133 | 159 | 107 | 65 | 58 | 34 | 47 | 61 | 74 | 86 | 135 | 197 | 135 | 173 | 72 | 128 | 120 | Pm20d2 |
| Pmaip1 | 1718 | 1193 | 1684 | 884 | 488 | 837 | 548 | 298 | 279 | 258 | 253 | 357 | 278 | 315 | 1325 | 1632 | 1055 | 1019 | 535 | 654 | 505 | 180 | 274 | 287 | 184 | 381 | 351 | 488 | Pmaip1 |
| Pmch | 51 | 37 | 15 | 115 | 77 | 129 | 192 | 136 | 96 | 60 | 66 | 101 | 12 | 109 | 44 | 31 | 57 | 90 | 117 | 210 | 162 | 137 | 105 | 96 | 114 | 125 | 49 | 66 | Pmch |
| Pmel | 194 | 87 | 55 | 17 | 29 | 92 | 11 | 125 | 309 | 124 | 215 | 93 | 124 | 36 | 123 | 164 | 30 | 70 | 30 | 13 | 22 | 69 | 230 | 111 | 64 | 94 | 63 | 50 | Pmel |
| Pmepa1 | 1250 | 1602 | 2008 | 2222 | 2284 | 2145 | 1857 | 1823 | 2050 | 2631 | 3046 | 2347 | 2211 | 2324 | 1363 | 1757 | 2199 | 2572 | 2192 | 1395 | 2095 | 2717 | 2227 | 2865 | 2884 | 2289 | 2112 | 2309 | Pmepa1 |
| Pmf1 | 1429 | 1620 | 1723 | 1326 | 1006 | 1213 | 1250 | 1652 | 1903 | 1726 | 1877 | 2566 | 2312 | 2743 | 1195 | 1371 | 1392 | 1078 | 1283 | 1233 | 1122 | 1370 | 1650 | 2297 | 2112 | 3265 | 2779 | 2993 | Pmf1 |
| Pmfbp1 | 240 | 385 | 367 | 467 | 645 | 538 | 304 | 606 | 452 | 572 | 476 | 407 | 756 | 364 | 434 | 477 | 498 | 413 | 429 | 544 | 375 | 707 | 411 | 305 | 541 | 122 | 466 | 396 | Pmfbp1 |
| Pmis2 | 0 | 0 | 0 | 0 | 0 | 0 | 1 | 0 | 0 | 0 | 2 | 0 | 0 | 0 | 0 | 0 | 0 | 0 | 0 | 0 | 0 | 0 | 0 | 0 | 0 | 0 | 0 | 0 | Pmis2 |
| Pml | 1644 | 1857 | 2103 | 1606 | 1410 | 1455 | 1301 | 1191 | 1587 | 1512 | 1765 | 1834 | 1768 | 1487 | 1485 | 2053 | 1875 | 1735 | 1820 | 1326 | 1361 | 1293 | 1654 | 1778 | 1401 | 2071 | 1527 | 1593 | Pml |
| Pmm1 | 2369 | 3098 | 2416 | 2589 | 2431 | 2459 | 2290 | 1288 | 1743 | 1419 | 1382 | 1257 | 1247 | 1284 | 2416 | 2799 | 2533 | 2678 | 2582 | 2642 | 2405 | 1297 | 1618 | 1511 | 1573 | 1512 | 1361 | 1247 | Pmm1 |
| Pmm2 | 1671 | 1615 | 1428 | 1295 | 1527 | 1707 | 1573 | 3094 | 3129 | 3107 | 3088 | 3312 | 3241 | 3338 | 1497 | 1319 | 1350 | 1343 | 1668 | 1660 | 1566 | 2959 | 2990 | 2981 | 3207 | 3069 | 3317 | 3309 | Pmm2 |
| Pmp22 | 1023 | 1079 | 1100 | 553 | 781 | 915 | 419 | 1784 | 2551 | 1707 | 2992 | 1289 | 2207 | 1225 | 1262 | 1314 | 753 | 917 | 554 | 674 | 420 | 2445 | 3105 | 1489 | 1691 | 1269 | 1583 | 1424 | Pmp22 |
| Pmpca | 9323 | 9377 | 9363 | 8611 | 7907 | 8079 | 8651 | 8774 | 8217 | 9027 | 8406 | 9648 | 8921 | 9520 | 8664 | 8963 | 9028 | 8636 | 8604 | 8019 | 8533 | 9546 | 8389 | 8945 | 8836 | 10072 | 9648 | 9875 | Pmpca |
| Pmpcb | 4771 | 4407 | 3973 | 3799 | 3542 | 3664 | 4216 | 5959 | 5011 | 4533 | 4308 | 5454 | 4568 | 5494 | 4638 | 3755 | 4118 | 3762 | 4005 | 4315 | 4123 | 4714 | 5060 | 5105 | 4814 | 5613 | 5176 | 5905 | Pmpcb |
| Pms1 | 1678 | 1158 | 1335 | 1107 | 1149 | 1211 | 1207 | 1236 | 1220 | 981 | 1081 | 1212 | 1198 | 1261 | 1742 | 1406 | 1285 | 1150 | 1166 | 1386 | 1197 | 993 | 1190 | 1222 | 1107 | 1522 | 1369 | 1297 | Pms1 |
| Pms2 | 3283 | 2989 | 2406 | 2567 | 2303 | 2580 | 2626 | 2209 | 1932 | 1906 | 2008 | 2146 | 1867 | 2136 | 2959 | 2718 | 2432 | 2415 | 2314 | 2564 | 2707 | 1902 | 1963 | 1844 | 2035 | 1843 | 2135 | 2209 | Pms2 |
| Pmvk | 2789 | 3694 | 3572 | 3518 | 3149 | 2781 | 2976 | 2249 | 2151 | 2352 | 2256 | 2641 | 2635 | 2479 | 3137 | 2963 | 3946 | 3416 | 3447 | 3502 | 2929 | 2421 | 2175 | 2814 | 2745 | 2825 | 2599 | 2648 | Pmvk |
| Pnck | 1476 | 1851 | 1484 | 2306 | 3203 | 2356 | 2600 | 2077 | 2328 | 2514 | 2159 | 1627 | 1453 | 1646 | 1586 | 1432 | 2126 | 2582 | 2756 | 2755 | 2744 | 2749 | 2289 | 2265 | 2667 | 1530 | 1464 | 1324 | Pnck |
| Pnisr | 16559 | 13924 | 16514 | 17705 | 20058 | 19898 | 20975 | 37530 | 31375 | 32903 | 34540 | 34976 | 33530 | 39960 | 17102 | 14741 | 17329 | 17019 | 19458 | 20930 | 21234 | 34086 | 29914 | 33748 | 34994 | 28040 | 35697 | 35251 | Pnisr |
| Pnkd | 4142 | 5245 | 4877 | 5537 | 5228 | 4461 | 4457 | 2425 | 3148 | 3014 | 3413 | 3272 | 3139 | 3040 | 4064 | 4719 | 5395 | 5560 | 4956 | 3914 | 4475 | 3078 | 2979 | 3179 | 3267 | 3617 | 2788 | 2815 | Pnkd |
| Pnkp | 1803 | 2286 | 2301 | 2184 | 1964 | 1959 | 1970 | 1307 | 1534 | 1603 | 1439 | 1730 | 1677 | 1657 | 1783 | 1960 | 2174 | 2125 | 2157 | 1893 | 1958 | 1440 | 1409 | 1664 | 1740 | 1782 | 1790 | 1785 | Pnkp |
| Pnldc1 | 167 | 191 | 185 | 202 | 242 | 272 | 240 | 258 | 304 | 325 | 241 | 292 | 304 | 309 | 164 | 190 | 217 | 253 | 229 | 192 | 186 | 276 | 201 | 302 | 280 | 279 | 286 | 288 | Pnldc1 |
| Pnlip | 0 | 0 | 0 | 3 | 1 | 3 | 2 | 5 | 5 | 1 | 10 | 1 | 2 | 1 | 0 | 3 | 0 | 4 | 0 | 2 | 4 | 5 | 11 | 2 | 0 | 1 | 1 | 1 | Pnlip |
| Pnliprp1 | 1 | 3 | 0 | 5 | 0 | 0 | 3 | 7 | 2 | 0 | 3 | 1 | 2 | 1 | 4 | 3 | 0 | 4 | 2 | 10 | 2 | 12 | 14 | 5 | 4 | 2 | 3 | 9 | Pnliprp1 |
| Pnliprp2 | 1 | 4 | 3 | 7 | 1 | 0 | 7 | 4 | 6 | 10 | 3 | 3 | 5 | 2 | 1 | 1 | 1 | 0 | 1 | 1 | 10 | 2 | 7 | 5 | 2 | 0 | 0 | 2 | Pnliprp2 |
| Pnma1 | 103 | 110 | 124 | 100 | 117 | 90 | 133 | 183 | 161 | 154 | 173 | 182 | 152 | 149 | 87 | 96 | 121 | 107 | 132 | 122 | 110 | 179 | 151 | 189 | 241 | 211 | 141 | 176 | Pnma1 |
| Pnma2 | 3662 | 4301 | 3733 | 4046 | 5021 | 4700 | 4512 | 2142 | 1848 | 1906 | 1668 | 1509 | 1455 | 1563 | 4258 | 3682 | 4422 | 4375 | 4537 | 4993 | 4824 | 2420 | 1733 | 1526 | 2221 | 1258 | 1399 | 1341 | Pnma2 |
| Pnma3 | 381 | 409 | 430 | 509 | 920 | 620 | 639 | 712 | 713 | 931 | 749 | 855 | 659 | 797 | 413 | 353 | 473 | 636 | 631 | 565 | 641 | 1046 | 682 | 818 | 1030 | 693 | 723 | 659 | Pnma3 |
| Pnma5 | 1 | 1 | 2 | 3 | 2 | 4 | 3 | 4 | 2 | 5 | 2 | 2 | 1 | 5 | 5 | 0 | 0 | 2 | 12 | 2 | 3 | 3 | 3 | 1 | 3 | 3 | 3 | 2 | Pnma5 |
| Pnmal1 | 2531 | 3092 | 2521 | 3804 | 4414 | 3992 | 4603 | 4437 | 3756 | 3993 | 3007 | 4011 | 2786 | 4093 | 2979 | 2217 | 3228 | 3714 | 4358 | 4440 | 4627 | 4752 | 3832 | 3927 | 4709 | 3384 | 3529 | 3849 | Pnmal1 |
| Pnmal2 | 6766 | 8940 | 10049 | 16386 | 19388 | 16237 | 16260 | 8674 | 10119 | 14445 | 12708 | 11084 | 8145 | 10710 | 7776 | 7328 | 14368 | 17574 | 18235 | 13610 | 17234 | 15493 | 10172 | 13548 | 16349 | 9779 | 8458 | 8923 | Pnmal2 |
| Pnmt | 5 | 4 | 0 | 0 | 16 | 1 | 2 | 10 | 25 | 14 | 49 | 0 | 12 | 3 | 3 | 2 | 2 | 4 | 4 | 5 | 0 | 17 | 22 | 0 | 4 | 0 | 1 | 0 | Pnmt |
| Pnn | 16766 | 15018 | 22670 | 20401 | 18128 | 17915 | 19945 | 20955 | 18276 | 22999 | 21804 | 24977 | 22654 | 29197 | 16763 | 15144 | 20702 | 18358 | 18529 | 18956 | 19164 | 20543 | 17582 | 25713 | 23149 | 27686 | 25330 | 26601 | Pnn |
| Pno1 | 3816 | 3320 | 3505 | 2827 | 2234 | 2530 | 2809 | 2164 | 1911 | 1618 | 1792 | 2152 | 2062 | 2489 | 3816 | 3248 | 3036 | 2360 | 2365 | 3067 | 2667 | 1724 | 1965 | 1967 | 1807 | 2597 | 2305 | 2656 | Pno1 |
| Pnoc | 138 | 193 | 189 | 247 | 417 | 345 | 337 | 710 | 597 | 820 | 689 | 983 | 932 | 1021 | 153 | 140 | 250 | 232 | 447 | 385 | 375 | 1068 | 547 | 926 | 1031 | 451 | 1022 | 994 | Pnoc |
| Pnp | 847 | 786 | 805 | 596 | 457 | 457 | 628 | 574 | 564 | 431 | 449 | 573 | 516 | 738 | 781 | 694 | 644 | 474 | 472 | 709 | 553 | 419 | 576 | 506 | 420 | 689 | 652 | 794 | Pnp |
| Pnp2 | 22 | 4 | 35 | 9 | 26 | 15 | 13 | 9 | 9 | 12 | 3 | 13 | 5 | 7 | 20 | 14 | 3 | 19 | 14 | 12 | 16 | 15 | 12 | 8 | 10 | 10 | 8 | 11 | Pnp2 |
| Pnpla1 | 6 | 7 | 0 | 14 | 1 | 4 | 3 | 10 | 7 | 7 | 1 | 2 | 3 | 0 | 7 | 12 | 10 | 7 | 4 | 2 | 1 | 0 | 11 | 1 | 0 | 3 | 1 | 4 | Pnpla1 |
| Pnpla2 | 1750 | 2035 | 1790 | 1821 | 1411 | 1418 | 1362 | 521 | 668 | 633 | 606 | 836 | 826 | 665 | 1444 | 1964 | 1710 | 1733 | 1490 | 1384 | 1422 | 613 | 627 | 699 | 650 | 854 | 673 | 663 | Pnpla2 |
| Pnpla3 | 377 | 515 | 376 | 519 | 456 | 408 | 448 | 134 | 97 | 111 | 142 | 106 | 129 | 111 | 389 | 448 | 417 | 521 | 415 | 457 | 453 | 160 | 162 | 122 | 152 | 106 | 109 | 109 | Pnpla3 |
| Pnpla5 | 0 | 0 | 0 | 0 | 0 | 0 | 1 | 0 | 0 | 0 | 0 | 0 | 0 | 2 | 0 | 0 | 0 | 0 | 0 | 0 | 0 | 0 | 0 | 1 | 3 | 0 | 0 | 1 | Pnpla5 |
| Pnpla6 | 2425 | 2966 | 2762 | 2911 | 3288 | 2899 | 2623 | 3251 | 3755 | 4016 | 3902 | 3512 | 3728 | 3192 | 2451 | 2406 | 2903 | 2812 | 3167 | 2769 | 2688 | 4102 | 3630 | 3444 | 3842 | 2758 | 3529 | 3042 | Pnpla6 |
| Pnpla7 | 474 | 536 | 390 | 486 | 434 | 433 | 373 | 443 | 615 | 620 | 670 | 504 | 648 | 458 | 454 | 485 | 357 | 487 | 397 | 393 | 434 | 511 | 603 | 541 | 551 | 578 | 513 | 545 | Pnpla7 |
| Pnpla8 | 7555 | 6682 | 6465 | 7917 | 8029 | 8101 | 9326 | 10148 | 8622 | 7779 | 7183 | 7722 | 7386 | 7760 | 9094 | 7297 | 7153 | 7772 | 7935 | 10196 | 8745 | 9110 | 8673 | 7487 | 8796 | 6739 | 8135 | 7787 | Pnpla8 |
| Pnpo | 1412 | 1717 | 1440 | 1438 | 1317 | 1281 | 1279 | 830 | 1153 | 788 | 824 | 793 | 1050 | 719 | 1505 | 1686 | 1348 | 1470 | 1461 | 1410 | 1239 | 829 | 1241 | 817 | 815 | 821 | 816 | 1073 | Pnpo |
| Pnpt1 | 5173 | 3829 | 4117 | 4129 | 3821 | 4182 | 4321 | 4367 | 3573 | 3104 | 3535 | 3506 | 3828 | 3691 | 5400 | 4446 | 3968 | 4103 | 3925 | 4750 | 4277 | 3684 | 3696 | 3434 | 3386 | 3903 | 3791 | 3756 | Pnpt1 |
| Pnrc1 | 2603 | 2229 | 2816 | 2933 | 2590 | 2626 | 2654 | 3339 | 3213 | 3306 | 3244 | 3350 | 3275 | 3882 | 2569 | 2270 | 2905 | 2670 | 2925 | 2269 | 2961 | 3599 | 3381 | 3865 | 3548 | 3760 | 3104 | 3663 | Pnrc1 |
| Pnrc2 | 7831 | 6396 | 7893 | 6451 | 5757 | 5993 | 7516 | 10291 | 8245 | 7420 | 7316 | 9212 | 8497 | 10215 | 8351 | 7038 | 7112 | 6077 | 6192 | 7406 | 6845 | 7854 | 8466 | 8652 | 7892 | 10990 | 10089 | 10890 | Pnrc2 |
| Poc1a | 1363 | 1634 | 1619 | 1162 | 721 | 815 | 986 | 552 | 680 | 568 | 576 | 744 | 692 | 718 | 1170 | 1400 | 1211 | 938 | 1051 | 853 | 833 | 433 | 636 | 732 | 610 | 1047 | 819 | 837 | Poc1a |
| Poc1b | 3849 | 3554 | 3914 | 3458 | 2903 | 3320 | 3314 | 2256 | 2253 | 2425 | 2321 | 3029 | 2844 | 3108 | 3373 | 3403 | 3544 | 3123 | 3434 | 2961 | 3514 | 2278 | 2277 | 2809 | 2468 | 3385 | 2884 | 3396 | Poc1b |
| Poc5 | 4862 | 4812 | 4009 | 4579 | 4466 | 4526 | 5041 | 3615 | 3339 | 3492 | 2922 | 3440 | 3013 | 3740 | 4787 | 4447 | 4108 | 4793 | 4306 | 4688 | 4938 | 3312 | 3178 | 3278 | 3455 | 3271 | 3525 | 3376 | Poc5 |
| Podn | 114 | 90 | 101 | 76 | 74 | 97 | 68 | 112 | 273 | 238 | 499 | 103 | 257 | 121 | 75 | 107 | 76 | 131 | 73 | 51 | 76 | 222 | 320 | 214 | 201 | 112 | 107 | 128 | Podn |
| Podnl1 | 5 | 5 | 7 | 9 | 4 | 3 | 1 | 20 | 17 | 35 | 53 | 24 | 23 | 20 | 8 | 1 | 1 | 3 | 4 | 2 | 0 | 28 | 27 | 20 | 21 | 13 | 13 | 5 | Podnl1 |
| Podxl | 7344 | 8017 | 8824 | 8219 | 8347 | 7911 | 8307 | 5576 | 5315 | 5886 | 5136 | 7171 | 6177 | 6814 | 7565 | 6826 | 8712 | 8359 | 9048 | 8244 | 8435 | 5843 | 5873 | 6377 | 6777 | 6772 | 6485 | 7050 | Podxl |
| Podxl2 | 5822 | 7975 | 7230 | 12249 | 13776 | 11687 | 10891 | 10408 | 11795 | 14301 | 12087 | 11603 | 10643 | 10918 | 6571 | 6371 | 10287 | 12141 | 13877 | 10522 | 12517 | 14265 | 11595 | 13132 | 15973 | 9705 | 9984 | 9577 | Podxl2 |
| Pof1b | 42 | 3 | 1 | 3 | 2 | 2 | 1 | 42 | 71 | 47 | 17 | 16 | 19 | 4 | 11 | 5 | 0 | 6 | 3 | 1 | 1 | 0 | 78 | 26 | 7 | 9 | 7 | 9 | Pof1b |
| Pofut1 | 3752 | 3375 | 3840 | 3569 | 3175 | 3673 | 3080 | 4444 | 4971 | 5117 | 5740 | 5768 | 5797 | 5387 | 3386 | 3819 | 3523 | 3560 | 3341 | 2852 | 3170 | 4774 | 4814 | 5459 | 5279 | 5227 | 5588 | 5915 | Pofut1 |
| Pofut2 | 3067 | 3989 | 4136 | 3420 | 2802 | 2874 | 2300 | 2619 | 3569 | 3497 | 4317 | 3719 | 3860 | 2938 | 2872 | 4146 | 3777 | 3583 | 2892 | 2249 | 2432 | 3395 | 4133 | 3735 | 3273 | 3998 | 3267 | 3157 | Pofut2 |
| Pogk | 13059 | 12594 | 9979 | 12971 | 14191 | 14953 | 14711 | 17986 | 16349 | 17431 | 15429 | 18195 | 16954 | 18823 | 12861 | 11397 | 11995 | 12740 | 14237 | 13918 | 15646 | 17630 | 15541 | 16006 | 18778 | 13573 | 18599 | 17977 | Pogk |
| Poglut1 | 5998 | 5230 | 4569 | 4605 | 4625 | 4881 | 5100 | 6219 | 5596 | 5266 | 4917 | 5343 | 5471 | 5048 | 5979 | 5270 | 4806 | 4521 | 4708 | 5394 | 4910 | 5178 | 5488 | 5033 | 4887 | 5285 | 5531 | 5186 | Poglut1 |
| Poglut2 | 3175 | 3181 | 2980 | 2566 | 2565 | 2541 | 2749 | 2175 | 2155 | 1925 | 2239 | 2250 | 2351 | 2088 | 3103 | 3113 | 2832 | 2896 | 2526 | 2938 | 2518 | 1865 | 2308 | 1973 | 1783 | 2491 | 2338 | 2230 | Poglut2 |
| Poglut3 | 456 | 350 | 550 | 359 | 302 | 430 | 212 | 1023 | 1299 | 1195 | 1658 | 931 | 1271 | 1008 | 493 | 618 | 368 | 429 | 200 | 272 | 237 | 1136 | 1625 | 1113 | 1008 | 1139 | 1052 | 1025 | Poglut3 |
| Pogz | 4415 | 4347 | 5453 | 5336 | 5913 | 5338 | 5574 | 10444 | 10710 | 11576 | 10591 | 12694 | 11019 | 12132 | 4092 | 4027 | 5524 | 4748 | 6566 | 5009 | 5498 | 12297 | 9281 | 12158 | 12188 | 10585 | 11545 | 11704 | Pogz |
| Pola1 | 7252 | 5419 | 5527 | 4780 | 3907 | 4511 | 6116 | 5587 | 4851 | 3758 | 3996 | 5461 | 4884 | 5853 | 6861 | 5582 | 5475 | 4661 | 4795 | 6050 | 5223 | 3415 | 4495 | 4681 | 4309 | 5239 | 5875 | 5993 | Pola1 |
| Pola2 | 6487 | 6819 | 6033 | 5614 | 3912 | 4704 | 5155 | 2909 | 3262 | 3042 | 2829 | 4057 | 3343 | 3680 | 5445 | 6189 | 5823 | 5296 | 4911 | 5021 | 4674 | 2333 | 2873 | 3335 | 2837 | 4358 | 4123 | 3949 | Pola2 |
| Polb | 4558 | 4384 | 3291 | 4452 | 4790 | 5115 | 5539 | 6234 | 5469 | 4984 | 5069 | 4630 | 4462 | 5166 | 4846 | 4505 | 3968 | 4464 | 4388 | 5851 | 5680 | 4906 | 5410 | 4151 | 6093 | 3586 | 5076 | 4414 | Polb |
| Pold1 | 4382 | 5629 | 6088 | 5314 | 3764 | 3965 | 3924 | 2894 | 4060 | 3870 | 3707 | 5449 | 4618 | 4359 | 3849 | 4785 | 5751 | 4782 | 4828 | 3784 | 3750 | 2841 | 3468 | 4698 | 3722 | 6515 | 5153 | 5020 | Pold1 |
| Pold2 | 2538 | 3302 | 2683 | 2655 | 1907 | 2200 | 2132 | 1667 | 2164 | 2118 | 1994 | 2578 | 2116 | 2048 | 2413 | 2611 | 2799 | 2335 | 2710 | 2176 | 2103 | 1516 | 2018 | 2207 | 1925 | 2919 | 2338 | 2205 | Pold2 |

| | | | | | | | | | | | | | | | | | | | | | | | | | | | | | |
|---|---|---|---|---|---|---|---|---|---|---|---|---|---|---|---|---|---|---|---|---|---|---|---|---|---|---|---|---|---|
| Pold3 | 6367 | 6070 | 5974 | 5057 | 5120 | 5419 | 5715 | 6018 | 5674 | 5611 | 5129 | 6972 | 5746 | 6927 | 5734 | 5563 | 5722 | 4927 | 5559 | 5644 | 5580 | 4968 | 5146 | 6279 | 5542 | 7760 | 7045 | 6965 | Pold3 |
| Pold4 | 5 | 1 | 1 | 2 | 1 | 2 | 1 | 0 | 2 | 0 | 3 | 0 | 3 | 0 | 3 | 2 | 1 | 3 | 2 | 1 | 2 | 0 | 1 | 0 | 1 | 1 | 5 | 1 | Pold4 |
| Poldip2 | 4861 | 5403 | 4775 | 4984 | 5184 | 5418 | 5427 | 5365 | 6139 | 6175 | 6251 | 6191 | 5880 | 6259 | 4659 | 4521 | 5215 | 4824 | 5479 | 5067 | 5400 | 5604 | 5855 | 6024 | 6317 | 6652 | 6143 | 6278 | Poldip2 |
| Poldip3 | 7472 | 8542 | 8408 | 8021 | 7696 | 7570 | 7616 | 9106 | 9704 | 9416 | 9279 | 10666 | 9612 | 10149 | 7142 | 7172 | 8565 | 7900 | 8691 | 7730 | 7704 | 9459 | 9569 | 10328 | 9886 | 11149 | 10550 | 11055 | Poldip3 |
| Pole | 8734 | 7665 | 5339 | 5586 | 4521 | 5969 | 6129 | 5327 | 5896 | 4508 | 4917 | 7477 | 6091 | 6866 | 7100 | 6812 | 5275 | 4944 | 5294 | 5975 | 5713 | 3695 | 4946 | 5083 | 4957 | 5576 | 7906 | 7056 | Pole |
| Pole2 | 1256 | 941 | 1192 | 789 | 633 | 715 | 805 | 985 | 887 | 789 | 776 | 1082 | 913 | 1115 | 1187 | 932 | 933 | 767 | 733 | 996 | 785 | 625 | 797 | 959 | 776 | 1285 | 1128 | 1221 | Pole2 |
| Pole3 | 8722 | 8815 | 9296 | 8044 | 6446 | 7188 | 7719 | 5412 | 4791 | 4756 | 4691 | 6127 | 5726 | 6392 | 8774 | 7656 | 8056 | 7208 | 7642 | 8587 | 7324 | 4177 | 4654 | 5460 | 5398 | 7159 | 6645 | 6761 | Pole3 |
| Pole4 | 1690 | 1422 | 1865 | 1562 | 1807 | 1566 | 1878 | 2969 | 2766 | 2483 | 2497 | 2631 | 2510 | 2900 | 1793 | 1439 | 1676 | 1714 | 1709 | 1740 | 1780 | 2765 | 2764 | 2735 | 2542 | 3127 | 2786 | 2913 | Pole4 |
| Polg | 8420 | 9717 | 8828 | 10245 | 8919 | 9314 | 9523 | 4218 | 4629 | 4821 | 4652 | 5460 | 5071 | 5166 | 8190 | 8737 | 9250 | 10255 | 9973 | 8008 | 9615 | 4674 | 4354 | 5285 | 5169 | 5242 | 4979 | 5122 | Polg |
| Polg2 | 890 | 853 | 776 | 939 | 692 | 726 | 765 | 552 | 583 | 680 | 601 | 596 | 685 | 627 | 787 | 794 | 790 | 919 | 739 | 742 | 703 | 633 | 599 | 584 | 600 | 673 | 589 | 605 | Polg2 |
| Polh | 2722 | 2642 | 2446 | 2299 | 1897 | 2156 | 2170 | 1964 | 1892 | 1830 | 1840 | 1942 | 2028 | 2109 | 2422 | 2739 | 2289 | 2534 | 1936 | 1915 | 2163 | 1546 | 1969 | 2041 | 1794 | 2319 | 2023 | 2095 | Polh |
| Poli | 890 | 960 | 781 | 946 | 803 | 905 | 1005 | 978 | 999 | 1128 | 949 | 1022 | 1001 | 1018 | 893 | 878 | 980 | 1022 | 931 | 948 | 932 | 971 | 983 | 923 | 986 | 1113 | 1049 | 943 | Poli |
| Polk | 2561 | 2056 | 1745 | 2184 | 1965 | 2165 | 2036 | 1326 | 1270 | 1066 | 1217 | 1033 | 1169 | 1077 | 2371 | 2412 | 1914 | 2238 | 1744 | 2225 | 2079 | 946 | 1234 | 1055 | 865 | 1021 | 1091 | 1104 | Polk |
| Poll | 873 | 1084 | 1125 | 807 | 767 | 703 | 745 | 813 | 1002 | 983 | 933 | 1122 | 1123 | 1127 | 818 | 970 | 977 | 726 | 882 | 768 | 760 | 812 | 958 | 1075 | 952 | 1408 | 1135 | 1129 | Poll |
| Polm | 915 | 1089 | 1136 | 978 | 985 | 999 | 957 | 701 | 897 | 802 | 855 | 1060 | 1044 | 879 | 839 | 1033 | 1131 | 859 | 1027 | 869 | 830 | 862 | 910 | 1017 | 798 | 1050 | 1011 | 980 | Polm |
| Poln | 326 | 306 | 173 | 237 | 153 | 231 | 254 | 88 | 110 | 49 | 71 | 94 | 101 | 102 | 289 | 323 | 184 | 215 | 156 | 227 | 208 | 84 | 79 | 68 | 73 | 91 | 107 | 68 | Poln |
| Polq | 3572 | 3078 | 2050 | 1991 | 1669 | 2342 | 2453 | 2586 | 2347 | 1872 | 1721 | 2532 | 2273 | 2763 | 3219 | 3184 | 2015 | 1746 | 1811 | 2852 | 2347 | 1255 | 1964 | 1672 | 2051 | 1774 | 2934 | 2504 | Polq |
| Polr1a | 6387 | 6664 | 5216 | 5572 | 4935 | 5735 | 5009 | 5299 | 6044 | 5495 | 5752 | 5911 | 6501 | 5797 | 5847 | 6796 | 5172 | 5438 | 5065 | 5142 | 5408 | 4769 | 5612 | 5378 | 5355 | 5325 | 6353 | 6442 | Polr1a |
| Polr1b | 3516 | 3488 | 3492 | 3665 | 3324 | 3428 | 3441 | 2773 | 3184 | 3130 | 3112 | 3339 | 3309 | 3248 | 3224 | 3306 | 3414 | 3562 | 3525 | 3008 | 3511 | 2901 | 2740 | 3174 | 3336 | 3601 | 3285 | 3405 | Polr1b |
| Polr1c | 3703 | 4412 | 4447 | 3984 | 3080 | 3172 | 2994 | 3957 | 3911 | 3997 | 3731 | 4325 | 3898 | 4063 | 3440 | 3709 | 4109 | 3627 | 4030 | 3749 | 3423 | 3188 | 4022 | 4092 | 3858 | 5108 | 4685 | 4345 | Polr1c |
| Polr1d | 4421 | 5819 | 4685 | 4740 | 4088 | 4177 | 4345 | 3046 | 3078 | 3046 | 3062 | 3293 | 3175 | 3211 | 4450 | 5012 | 4962 | 4734 | 4249 | 4941 | 4729 | 2381 | 3496 | 3595 | 3297 | 3622 | 3492 | 3685 | Polr1d |
| Polr1e | 3479 | 3907 | 2814 | 3250 | 2796 | 2966 | 3099 | 1487 | 1331 | 1293 | 1314 | 1493 | 1465 | 1441 | 3376 | 3382 | 3024 | 3156 | 2893 | 3198 | 3170 | 1138 | 1303 | 1361 | 1293 | 1654 | 1582 | 1663 | Polr1e |
| Polr1f | 4501 | 3197 | 3905 | 3471 | 2937 | 3281 | 3191 | 3295 | 2646 | 2369 | 2361 | 2702 | 2802 | 2617 | 4395 | 4224 | 3296 | 3241 | 2903 | 3640 | 3083 | 2622 | 2671 | 2613 | 2424 | 3011 | 2994 | 2941 | Polr1f |
| Polr1h | 1511 | 1792 | 1785 | 1545 | 1293 | 1313 | 1620 | 1432 | 1581 | 1381 | 1527 | 1549 | 1520 | 1831 | 1510 | 1441 | 1677 | 1448 | 1467 | 1694 | 1507 | 1203 | 1509 | 1691 | 1589 | 1853 | 1694 | 1989 | Polr1h |
| Polr1has | 280 | 224 | 354 | 231 | 233 | 224 | 268 | 271 | 251 | 302 | 246 | 369 | 319 | 378 | 319 | 220 | 379 | 237 | 272 | 286 | 213 | 212 | 252 | 359 | 262 | 394 | 302 | 396 | Polr1has |
| Polr2a | 16473 | 17162 | 19020 | 20139 | 17424 | 16795 | 14997 | 17070 | 20341 | 23592 | 21795 | 30981 | 25311 | 25472 | 15241 | 16916 | 19674 | 19425 | 19132 | 12228 | 16392 | 24132 | 18505 | 26652 | 23737 | 23354 | 24283 | 24163 | Polr2a |
| Polr2b | 25754 | 25093 | 20433 | 19766 | 16877 | 18634 | 21284 | 14889 | 12414 | 10323 | 9515 | 13309 | 11551 | 13137 | 27265 | 23226 | 19736 | 19550 | 18516 | 23353 | 20210 | 11351 | 11976 | 10846 | 11280 | 12876 | 13279 | 13481 | Polr2b |
| Polr2c | 3402 | 3883 | 3328 | 3231 | 3082 | 2996 | 3087 | 2635 | 3019 | 2763 | 2610 | 3092 | 2912 | 2994 | 3069 | 3283 | 3377 | 3022 | 3332 | 3248 | 3234 | 2572 | 2865 | 3120 | 2971 | 3523 | 3105 | 3114 | Polr2c |
| Polr2d | 3834 | 3814 | 3779 | 3468 | 2981 | 3253 | 3703 | 3027 | 2660 | 2473 | 2671 | 3407 | 2975 | 3460 | 3956 | 3524 | 3633 | 3309 | 3256 | 3789 | 3299 | 2630 | 2742 | 3068 | 2850 | 3910 | 3403 | 3690 | Polr2d |
| Polr2e | 4803 | 6499 | 5468 | 5024 | 3426 | 4064 | 3645 | 2815 | 3102 | 2970 | 2842 | 3353 | 3116 | 3160 | 4287 | 5271 | 4875 | 4223 | 4114 | 4663 | 3847 | 2206 | 3177 | 3189 | 2846 | 4152 | 3435 | 3316 | Polr2e |
| Polr2f | 2450 | 2972 | 2816 | 2913 | 2293 | 2075 | 2506 | 1793 | 1935 | 1853 | 2001 | 2134 | 2017 | 2366 | 2296 | 2266 | 2844 | 2631 | 2513 | 2545 | 2468 | 1475 | 1783 | 2398 | 2060 | 2669 | 2207 | 2370 | Polr2f |
| Polr2g | 4212 | 4161 | 4009 | 3934 | 3236 | 3366 | 3700 | 3518 | 3091 | 2978 | 2813 | 3284 | 3176 | 3323 | 4183 | 4012 | 4093 | 3682 | 3317 | 4171 | 3497 | 2973 | 3216 | 3331 | 2986 | 3841 | 3435 | 3405 | Polr2g |
| Polr2h | 2208 | 2456 | 2304 | 2295 | 1871 | 1874 | 2274 | 1801 | 1706 | 1562 | 1550 | 1699 | 1626 | 1756 | 2325 | 2230 | 2356 | 2161 | 2180 | 2762 | 2229 | 1282 | 1660 | 1699 | 1717 | 2079 | 1868 | 1854 | Polr2h |
| Polr2i | 746 | 988 | 919 | 929 | 778 | 760 | 889 | 1004 | 957 | 995 | 1080 | 1084 | 1111 | 1257 | 682 | 677 | 895 | 842 | 1001 | 1054 | 847 | 749 | 991 | 1192 | 1144 | 1453 | 1254 | 1158 | Polr2i |
| Polr2j | 3098 | 3497 | 3291 | 3455 | 2556 | 2836 | 3190 | 2231 | 2160 | 2222 | 2257 | 2361 | 2515 | 2916 | 3100 | 2869 | 3216 | 3013 | 2858 | 3444 | 3008 | 1785 | 2091 | 2621 | 2456 | 2991 | 2740 | 2807 | Polr2j |
| Polr2k | 1914 | 1754 | 2039 | 2172 | 1729 | 1491 | 2529 | 1931 | 1424 | 1833 | 1735 | 1847 | 1827 | 2987 | 2012 | 1504 | 2097 | 1631 | 1663 | 2177 | 2557 | 1501 | 1458 | 2303 | 1901 | 1957 | 2062 | 2881 | Polr2k |
| Polr2l | 2941 | 4100 | 3837 | 3827 | 2938 | 2328 | 3347 | 1622 | 1811 | 2080 | 2228 | 2232 | 2261 | 2389 | 2863 | 2928 | 3883 | 3425 | 2795 | 3142 | 3104 | 1469 | 1723 | 2323 | 2161 | 2453 | 2119 | 2524 | Polr2l |
| Polr2m | 13276 | 12481 | 15742 | 12382 | 12391 | 12020 | 12218 | 16861 | 17287 | 18868 | 18849 | 22227 | 19831 | 22723 | 12380 | 12069 | 14709 | 12022 | 12957 | 10603 | 12734 | 17044 | 17926 | 21355 | 18740 | 26399 | 22636 | 23644 | Polr2m |
| Polr3a | 3209 | 3408 | 1794 | 2004 | 2146 | 2793 | 2271 | 3213 | 3292 | 2537 | 2624 | 2603 | 2453 | 2769 | 3047 | 3175 | 1934 | 1955 | 1967 | 2929 | 2767 | 2169 | 2992 | 2307 | 2660 | 2297 | 3082 | 2624 | Polr3a |
| Polr3b | 2194 | 2251 | 1896 | 1888 | 2023 | 2046 | 1944 | 3467 | 3425 | 3037 | 3132 | 3194 | 3520 | 3543 | 2165 | 2036 | 1851 | 1812 | 2034 | 2104 | 2038 | 3232 | 3313 | 2926 | 3084 | 2844 | 3540 | 3572 | Polr3b |
| Polr3c | 3026 | 3190 | 2966 | 3072 | 2611 | 2637 | 2948 | 2545 | 2497 | 2814 | 2474 | 3144 | 2941 | 3245 | 2913 | 2713 | 2943 | 2791 | 2978 | 2816 | 2976 | 2465 | 2509 | 3015 | 2885 | 3583 | 3330 | 3493 | Polr3c |
| Polr3d | 1351 | 1475 | 1870 | 1758 | 1714 | 1598 | 1560 | 1829 | 2120 | 2123 | 2300 | 2147 | 2008 | 2491 | 1345 | 1386 | 1727 | 1701 | 1672 | 1354 | 1641 | 1902 | 2111 | 2351 | 2185 | 2749 | 2052 | 2528 | Polr3d |
| Polr3e | 3224 | 3335 | 3400 | 3323 | 3214 | 3617 | 3303 | 2932 | 3119 | 3015 | 3289 | 3378 | 3188 | 3393 | 2956 | 3167 | 3081 | 3303 | 3464 | 3102 | 3337 | 2772 | 2834 | 3140 | 3310 | 3335 | 3298 | 3192 | Polr3e |
| Polr3f | 5696 | 5114 | 4472 | 4696 | 4335 | 4765 | 5420 | 4941 | 4166 | 3800 | 3939 | 4643 | 4476 | 4883 | 5618 | 5202 | 4573 | 4588 | 4531 | 5076 | 5032 | 4116 | 4086 | 4172 | 4148 | 4248 | 4911 | 4641 | Polr3f |
| Polr3g | 3107 | 2432 | 2657 | 1736 | 1276 | 1497 | 1476 | 970 | 931 | 838 | 884 | 873 | 1077 | 910 | 2846 | 2999 | 2084 | 1888 | 1274 | 1757 | 1343 | 648 | 1061 | 735 | 546 | 1116 | 947 | 1043 | Polr3g |
| Polr3gl | 2708 | 3219 | 4053 | 4612 | 3844 | 3346 | 3766 | 1673 | 1926 | 2324 | 2189 | 2185 | 1975 | 2138 | 2859 | 3045 | 4569 | 4254 | 4190 | 3214 | 3738 | 2169 | 1934 | 2575 | 2347 | 2493 | 2205 | 2009 | Polr3gl |
| Polr3h | 2482 | 2925 | 2562 | 2675 | 2409 | 2627 | 2592 | 1523 | 1623 | 1514 | 1323 | 1602 | 1548 | 1509 | 2472 | 2538 | 2680 | 2742 | 2879 | 2756 | 2641 | 1295 | 1517 | 1618 | 1718 | 1763 | 1719 | 1524 | Polr3h |
| Polr3k | 8583 | 7186 | 8242 | 7534 | 8238 | 7802 | 9004 | 9955 | 7919 | 7955 | 7047 | 8300 | 7815 | 9173 | 8712 | 7506 | 7703 | 7646 | 7709 | 9504 | 8680 | 8326 | 7712 | 8478 | 9152 | 8564 | 8882 | 9105 | Polr3k |
| Polrmt | 1879 | 2382 | 2297 | 2722 | 2493 | 2359 | 1932 | 1704 | 2074 | 2284 | 1861 | 2184 | 1987 | 1889 | 1722 | 2136 | 2411 | 2445 | 2639 | 1956 | 2215 | 1947 | 1915 | 2402 | 2241 | 2505 | 2057 | 1983 | Polrmt |
| Pom121 | 7260 | 8759 | 11934 | 10043 | 8442 | 8226 | 7953 | 5991 | 7113 | 8311 | 8052 | 11375 | 8639 | 9736 | 6311 | 7751 | 11570 | 9592 | 9848 | 6205 | 8409 | 8674 | 6177 | 10283 | 9454 | 12357 | 8624 | 9941 | Pom121 |
| Pom121l2 | 0 | 0 | 9 | 1 | 2 | 8 | 4 | 1 | 1 | 6 | 3 | 0 | 6 | 2 | 8 | 8 | 10 | 1 | 3 | 3 | 4 | 0 | 3 | 2 | 10 | 3 | 5 | 5 | Pom121l2 |
| Pomc | 153 | 210 | 102 | 199 | 181 | 68 | 117 | 217 | 264 | 210 | 212 | 140 | 117 | 117 | 67 | 104 | 111 | 147 | 69 | 162 | 140 | 244 | 265 | 251 | 239 | 259 | 207 | 155 | Pomc |
| Pomgnt1 | 2170 | 2305 | 2439 | 2439 | 2747 | 2575 | 2447 | 3390 | 4094 | 4193 | 4115 | 4329 | 3848 | 3874 | 2230 | 2320 | 2465 | 2590 | 2576 | 2140 | 2550 | 4037 | 4273 | 3989 | 4195 | 4213 | 3998 | 3853 | Pomgnt1 |
| Pomgnt2 | 915 | 1415 | 1322 | 1372 | 1362 | 1326 | 1108 | 689 | 882 | 937 | 898 | 762 | 770 | 747 | 974 | 1319 | 1451 | 1524 | 1402 | 1049 | 1155 | 874 | 918 | 841 | 1084 | 780 | 698 | 700 | Pomgnt2 |
| Pomk | 1361 | 1308 | 1410 | 1420 | 1483 | 1396 | 1471 | 1938 | 1963 | 2041 | 2045 | 2078 | 1905 | 1940 | 1390 | 1192 | 1482 | 1399 | 1545 | 1363 | 1448 | 2067 | 1889 | 2032 | 2083 | 2053 | 2023 | 2109 | Pomk |
| Pomp | 7465 | 8374 | 8244 | 7917 | 6688 | 6706 | 7902 | 7603 | 6892 | 6659 | 6392 | 7589 | 7263 | 8101 | 7622 | 7163 | 8056 | 7065 | 7311 | 8521 | 7528 | 5805 | 6862 | 7573 | 7046 | 8484 | 7976 | 8591 | Pomp |
| Pomt1 | 1670 | 2121 | 1843 | 2103 | 2175 | 2308 | 2004 | 2158 | 2702 | 2880 | 2594 | 2549 | 2747 | 2804 | 1812 | 1878 | 2190 | 2190 | 2198 | 2143 | 2100 | 2760 | 2571 | 2712 | 2791 | 2596 | 2596 | 2635 | Pomt1 |
| Pomt2 | 1576 | 1711 | 1566 | 1537 | 1583 | 1568 | 1619 | 2373 | 2475 | 2206 | 2436 | 2634 | 2437 | 2449 | 1636 | 1472 | 1526 | 1655 | 1636 | 1714 | 1680 | 2448 | 2405 | 2296 | 2473 | 2310 | 2540 | 2580 | Pomt2 |
| Pon1 | 10 | 7 | 8 | 21 | 7 | 7 | 6 | 21 | 13 | 19 | 15 | 3 | 10 | 2 | 4 | 15 | 7 | 9 | 3 | 14 | 2 | 2 | 32 | 11 | 8 | 15 | 6 | 12 | Pon1 |
| Pon2 | 1000 | 939 | 1065 | 940 | 1064 | 963 | 962 | 1514 | 1668 | 1440 | 1504 | 2060 | 1944 | 1837 | 1040 | 1021 | 1021 | 891 | 905 | 952 | 879 | 1611 | 1658 | 1691 | 1662 | 2268 | 1796 | 2026 | Pon2 |
| Pon3 | 326 | 280 | 214 | 261 | 260 | 265 | 219 | 304 | 340 | 280 | 323 | 265 | 329 | 223 | 318 | 359 | 162 | 238 | 192 | 268 | 234 | 311 | 381 | 219 | 208 | 238 | 328 | 253 | Pon3 |
| Pop1 | 2698 | 2752 | 2738 | 2381 | 2010 | 2410 | 2404 | 2080 | 2116 | 1875 | 1838 | 2244 | 2129 | 2207 | 2413 | 2461 | 2483 | 2221 | 2392 | 2381 | 2255 | 1708 | 1999 | 2092 | 2117 | 2489 | 2288 | 2135 | Pop1 |
| Pop4 | 5832 | 6292 | 5870 | 5646 | 5047 | 5460 | 6170 | 4975 | 4622 | 4182 | 4139 | 5176 | 4982 | 5521 | 6343 | 5786 | 5770 | 5211 | 5640 | 7019 | 5530 | 4062 | 4405 | 4611 | 4786 | 5936 | 5268 | 5498 | Pop4 |
| Pop5 | 1548 | 1830 | 1460 | 1397 | 1090 | 1194 | 1371 | 887 | 725 | 765 | 784 | 800 | 821 | 829 | 1596 | 1275 | 1438 | 1319 | 1243 | 1380 | 1381 | 892 | 769 | 798 | 825 | 902 | 753 | 1042 | Pop5 |
| Pop7 | 336 | 436 | 412 | 434 | 435 | 484 | 430 | 774 | 885 | 945 | 954 | 1019 | 879 | 950 | 348 | 344 | 438 | 408 | 464 | 524 | 416 | 744 | 913 | 1068 | 1008 | 993 | 948 | 923 | Pop7 |
| Popdc2 | 12 | 22 | 35 | 38 | 24 | 28 | 25 | 23 | 37 | 24 | 93 | 13 | 44 | 16 | 15 | 48 | 42 | 36 | 39 | 18 | 23 | 25 | 67 | 44 | 13 | 26 | 20 | 26 | Popdc2 |
| Popdc3 | 8 | 24 | 8 | 19 | 33 | 11 | 15 | 17 | 40 | 28 | 42 | 25 | 27 | 12 | 24 | 18 | 15 | 33 | 24 | 15 | 13 | 40 | 45 | 17 | 16 | 6 | 12 | 13 | Popdc3 |
| Por | 8193 | 11118 | 11499 | 12380 | 9738 | 9588 | 8803 | 4269 | 4957 | 5310 | 5093 | 6551 | 5438 | 5990 | 8513 | 10961 | 11617 | 11886 | 10905 | 8099 | 9495 | 5591 | 4943 | 6260 | 5758 | 7413 | 5951 | 5994 | Por |
| Porcn | 1317 | 1717 | 1318 | 1736 | 2383 | 2022 | 2149 | 1644 | 1779 | 2004 | 1660 | 1767 | 1653 | 1854 | 1418 | 1432 | 1772 | 1893 | 2187 | 1995 | 2033 | 1990 | 1500 | 1812 | 2111 | 1400 | 1533 | 1518 | Porcn |
| Postn | 6982 | 3894 | 3082 | 3220 | 2869 | 3713 | 1311 | 10302 | 17526 | 9594 | 24174 | 4175 | 5669 | 3886 | 5556 | 7006 | 2745 | 4830 | 1309 | 1548 | 1257 | 10676 | 22612 | 7483 | 5716 | 4656 | 4699 | 3202 | Postn |
| Pot1a | 3778 | 3460 | 3079 | 3515 | 3250 | 3375 | 3676 | 3329 | 3190 | 2662 | 2733 | 3127 | 3175 | 3196 | 4034 | 3160 | 3096 | 3513 | 3307 | 4002 | 3436 | 2405 | 2989 | 2764 | 2800 | 3087 | 3166 | 3170 | Pot1a |
| Pot1b | 468 | 153 | 506 | 141 | 371 | 139 | 161 | 407 | 552 | 300 | 268 | 357 | 265 | 258 | 348 | 326 | 535 | 128 | 320 | 174 | 226 | 470 | 203 | 911 | 213 | 447 | 127 | 159 | Pot1b |
| Potegl | 28 | 13 | 16 | 47 | 50 | 37 | 46 | 11 | 2 | 6 | 8 | 2 | 12 | 15 | 12 | 23 | 29 | 23 | 41 | 31 | 36 | 25 | 3 | 7 | 13 | 5 | 8 | 11 | Potegl |
| Pou1f1 | 0 | 0 | 0 | 0 | 4 | 1 | 0 | 0 | 2 | 0 | 0 | 0 | 1 | 0 | 1 | 0 | 5 | 4 | 1 | 0 | 0 | 2 | 3 | 0 | 0 | 0 | 0 | 4 | Pou1f1 |
| Pou2af1 | 6 | 0 | 2 | 12 | 9 | 0 | 3 | 7 | 6 | 20 | 8 | 13 | 15 | 9 | 7 | 5 | 1 | 4 | 5 | 4 | 6 | 5 | 15 | 10 | 15 | 16 | 1 | 6 | Pou2af1 |
| Pou2f1 | 7268 | 5381 | 4856 | 5576 | 6640 | 7447 | 6805 | 8772 | 7834 | 7444 | 8172 | 8337 | 7791 | 9078 | 6730 | 6633 | 4883 | 5147 | 6036 | 6807 | 7427 | 6676 | 6671 | 6491 | 9127 | 5475 | 9927 | 7867 | Pou2f1 |
| Pou2f2 | 2205 | 2634 | 4909 | 4835 | 4935 | 4646 | 4060 | 3830 | 3070 | 5997 | 4403 | 6387 | 5103 | 6220 | 2635 | 2203 | 5229 | 4545 | 5305 | 4379 | 4302 | 6729 | 3181 | 6838 | 7037 | 5742 | 6410 | 6214 | Pou2f2 |
| Pou2f3 | 9 | 8 | 5 | 7 | 3 | 3 | 6 | 6 | 27 | 10 | 16 | 0 | 0 | 2 | 3 | 2 | 5 | 7 | 7 | 4 | 4 | 7 | 47 | 5 | 9 | 0 | 0 | 2 | Pou2f3 |
| Pou3f1 | 1324 | 1826 | 2128 | 2631 | 1808 | 1820 | 1812 | 1166 | 1012 | 1484 | 1172 | 1883 | 1348 | 1792 | 1491 | 1372 | 2593 | 2418 | 2092 | 1554 | 2008 | 1512 | 974 | 1816 | 1739 | 1612 | 1597 | 1532 | Pou3f1 |

| | | | | | | | | | | | | | | | | | | | | | | | | | | | | | |
|---|---|---|---|---|---|---|---|---|---|---|---|---|---|---|---|---|---|---|---|---|---|---|---|---|---|---|---|---|---|
| Pou3f2 | 6634 | 6458 | 12746 | 11851 | 10221 | 10731 | 10488 | 9387 | 6077 | 9203 | 6788 | 13944 | 11482 | 14683 | 8097 | 5720 | 12199 | 10281 | 11398 | 8595 | 11264 | 12074 | 5772 | 11121 | 11914 | 10470 | 15421 | 16581 | Pou3f2 |
| Pou3f3 | 12033 | 13504 | 14664 | 17595 | 14198 | 15067 | 16748 | 15069 | 13992 | 15481 | 11715 | 20411 | 15729 | 21616 | 12728 | 8682 | 17415 | 15353 | 16921 | 14567 | 16999 | 14170 | 10035 | 19371 | 18858 | 19102 | 18052 | 19355 | Pou3f3 |
| Pou3f4 | 3014 | 4265 | 5721 | 4334 | 3951 | 4191 | 3865 | 3444 | 2522 | 3100 | 2746 | 5307 | 4976 | 5418 | 3782 | 2918 | 5281 | 3651 | 4708 | 4208 | 4249 | 3823 | 2629 | 4196 | 4342 | 3271 | 6735 | 6380 | Pou3f4 |
| Pou4f1 | 2009 | 2008 | 3673 | 3501 | 4996 | 3367 | 2541 | 3777 | 2042 | 4735 | 4153 | 4881 | 5871 | 5272 | 2855 | 2355 | 3752 | 3505 | 4126 | 3210 | 3959 | 6082 | 3399 | 4585 | 5100 | 542 | 5302 | 5779 | Pou4f1 |
| Pou4f2 | 165 | 173 | 215 | 444 | 760 | 445 | 269 | 416 | 287 | 539 | 462 | 380 | 467 | 295 | 266 | 217 | 394 | 436 | 580 | 447 | 557 | 715 | 467 | 469 | 569 | 7 | 323 | 263 | Pou4f2 |
| Pou4f3 | 80 | 96 | 27 | 104 | 171 | 92 | 67 | 189 | 152 | 173 | 129 | 90 | 177 | 61 | 145 | 148 | 81 | 184 | 174 | 96 | 166 | 284 | 107 | 132 | 269 | 5 | 70 | 89 | Pou4f3 |
| Pou5f1 | 3 | 1 | 11 | 0 | 2 | 0 | 0 | 2 | 1 | 2 | 3 | 0 | 1 | 1 | 1 | 1 | 0 | 0 | 3 | 0 | 2 | 0 | 4 | 3 | 0 | 6 | 0 | 1 | Pou5f1 |
| Pou5f2 | 10 | 19 | 75 | 37 | 16 | 11 | 30 | 13 | 18 | 16 | 9 | 18 | 18 | 16 | 13 | 15 | 24 | 13 | 50 | 4 | 30 | 37 | 6 | 24 | 26 | 76 | 24 | 23 | Pou5f2 |
| Pou6f1 | 3084 | 3741 | 3808 | 5039 | 6928 | 6084 | 6007 | 6539 | 7210 | 9342 | 7418 | 9052 | 6815 | 8590 | 3302 | 2477 | 5459 | 5382 | 7354 | 4805 | 6374 | 9787 | 5878 | 9010 | 10524 | 6550 | 7587 | 7472 | Pou6f1 |
| Pou6f2 | 223 | 213 | 414 | 498 | 646 | 608 | 523 | 717 | 538 | 836 | 627 | 840 | 759 | 827 | 357 | 332 | 495 | 478 | 720 | 562 | 662 | 1054 | 454 | 843 | 889 | 485 | 846 | 758 | Pou6f2 |
| Pp2d1 | 1 | 0 | 0 | 0 | 1 | 1 | 1 | 1 | 1 | 0 | 4 | 0 | 2 | 1 | 1 | 0 | 0 | 2 | 2 | 1 | 4 | 0 | 1 | 1 | 4 | 1 | 1 | 1 | Pp2d1 |
| Ppa1 | 7729 | 7162 | 8085 | 6933 | 5893 | 6163 | 7510 | 6064 | 5629 | 5833 | 5214 | 6763 | 5697 | 7207 | 7330 | 6622 | 7473 | 6017 | 7221 | 7372 | 6998 | 5098 | 5456 | 6508 | 5928 | 8440 | 7043 | 7825 | Ppa1 |
| Ppa2 | 1369 | 1126 | 1346 | 1300 | 1423 | 1352 | 1568 | 1872 | 1804 | 1731 | 1765 | 1960 | 1714 | 2138 | 1383 | 1144 | 1350 | 1187 | 1485 | 1428 | 1387 | 1766 | 1885 | 1810 | 1801 | 2034 | 1972 | 2091 | Ppa2 |
| Ppan | 1869 | 2424 | 2216 | 2352 | 1633 | 1705 | 1631 | 1679 | 2062 | 2076 | 2262 | 2091 | 2058 | 2122 | 1767 | 2009 | 1977 | 1947 | 1840 | 1726 | 1716 | 1803 | 2192 | 2121 | 2002 | 2604 | 2205 | 2162 | Ppan |
| Ppara | 102 | 128 | 81 | 90 | 140 | 130 | 112 | 62 | 65 | 44 | 62 | 36 | 60 | 26 | 99 | 110 | 131 | 139 | 143 | 104 | 92 | 50 | 61 | 53 | 42 | 10 | 57 | 50 | Ppara |
| Ppard | 2336 | 3100 | 3547 | 3230 | 2547 | 2526 | 2274 | 951 | 1210 | 1729 | 1649 | 1395 | 1415 | 1239 | 2056 | 3009 | 3526 | 3130 | 2742 | 1955 | 2313 | 1327 | 1316 | 1478 | 1288 | 1593 | 1292 | 1190 | Ppard |
| Pparg | 32 | 29 | 26 | 61 | 47 | 51 | 56 | 24 | 20 | 13 | 47 | 1 | 10 | 6 | 21 | 30 | 25 | 68 | 51 | 32 | 58 | 17 | 47 | 21 | 11 | 3 | 5 | 6 | Pparg |
| Ppargc1a | 1483 | 1226 | 1027 | 1602 | 1762 | 1604 | 1669 | 1028 | 1102 | 943 | 980 | 830 | 746 | 675 | 1536 | 1274 | 1225 | 1683 | 1569 | 1673 | 1683 | 1193 | 896 | 765 | 981 | 458 | 772 | 618 | Ppargc1a |
| Ppargc1b | 213 | 283 | 194 | 274 | 286 | 269 | 225 | 68 | 154 | 108 | 220 | 74 | 105 | 71 | 208 | 269 | 233 | 314 | 276 | 174 | 207 | 82 | 159 | 94 | 88 | 54 | 103 | 74 | Ppargc1b |
| Ppat | 6298 | 5447 | 5544 | 4835 | 3915 | 4401 | 4939 | 3051 | 2592 | 2515 | 2568 | 3153 | 3002 | 3367 | 6186 | 5823 | 4985 | 5073 | 4325 | 5280 | 4618 | 2176 | 2715 | 2845 | 2473 | 4057 | 3961 | 3841 | Ppat |
| Ppbp | 271 | 195 | 340 | 99 | 54 | 72 | 68 | 162 | 149 | 135 | 155 | 226 | 323 | 153 | 194 | 321 | 130 | 119 | 50 | 146 | 79 | 155 | 146 | 150 | 62 | 324 | 272 | 157 | Ppbp |
| Ppcdc | 608 | 609 | 548 | 504 | 569 | 563 | 627 | 1071 | 1054 | 988 | 959 | 1068 | 1100 | 1017 | 574 | 566 | 508 | 533 | 620 | 639 | 575 | 907 | 1007 | 1020 | 1050 | 1162 | 963 | 1042 | Ppcdc |
| Ppcs | 476 | 485 | 461 | 422 | 383 | 433 | 491 | 513 | 547 | 442 | 424 | 466 | 448 | 396 | 563 | 470 | 439 | 437 | 365 | 504 | 377 | 458 | 526 | 507 | 386 | 583 | 497 | 500 | Ppcs |
| Ppdpf | 3016 | 3588 | 3405 | 3047 | 3391 | 3108 | 3101 | 3930 | 4254 | 4497 | 4132 | 5118 | 4374 | 4708 | 2830 | 2756 | 3571 | 3036 | 3644 | 3453 | 2951 | 3937 | 4122 | 5009 | 4765 | 5987 | 5063 | 4892 | Ppdpf |
| Ppef1 | 33 | 44 | 34 | 30 | 122 | 34 | 31 | 129 | 130 | 79 | 188 | 29 | 64 | 18 | 80 | 53 | 23 | 85 | 26 | 40 | 37 | 201 | 126 | 25 | 78 | 31 | 33 | 31 | Ppef1 |
| Ppef2 | 40 | 37 | 37 | 54 | 66 | 30 | 44 | 39 | 35 | 63 | 21 | 13 | 23 | 14 | 29 | 57 | 22 | 45 | 32 | 46 | 33 | 30 | 40 | 15 | 31 | 17 | 29 | 21 | Ppef2 |
| Ppfia1 | 8569 | 9208 | 7957 | 8305 | 7856 | 8238 | 8151 | 6577 | 6795 | 6982 | 6850 | 7849 | 7355 | 7586 | 7837 | 8364 | 8055 | 8490 | 8112 | 7108 | 8570 | 6375 | 6387 | 6940 | 7441 | 6918 | 7633 | 7414 | Ppfia1 |
| Ppfia2 | 2347 | 2770 | 1822 | 3867 | 5843 | 4728 | 4876 | 5448 | 4623 | 4465 | 3773 | 3119 | 2765 | 3243 | 3563 | 2412 | 2947 | 4151 | 4733 | 5561 | 5137 | 5812 | 4407 | 3434 | 5045 | 1990 | 2805 | 2613 | Ppfia2 |
| Ppfia3 | 2705 | 3545 | 4551 | 4595 | 5220 | 4369 | 4217 | 2450 | 2756 | 3586 | 2857 | 2867 | 2472 | 3041 | 2496 | 2742 | 4990 | 4682 | 5658 | 3754 | 4708 | 3816 | 2532 | 3361 | 3905 | 2671 | 2607 | 2766 | Ppfia3 |
| Ppfia4 | 1485 | 1822 | 1563 | 2066 | 2891 | 2405 | 1941 | 1180 | 1592 | 1729 | 2544 | 1069 | 1308 | 1013 | 1563 | 2093 | 1874 | 2464 | 2225 | 1891 | 2174 | 1770 | 1825 | 1061 | 1553 | 794 | 1078 | 921 | Ppfia4 |
| Ppfibp1 | 9352 | 7747 | 7306 | 5834 | 5106 | 6081 | 5950 | 4443 | 4973 | 3925 | 5533 | 3936 | 4749 | 3949 | 8387 | 9535 | 6195 | 6325 | 5137 | 6066 | 5115 | 3790 | 5465 | 3796 | 3345 | 4104 | 4048 | 4005 | Ppfibp1 |
| Ppfibp2 | 656 | 476 | 439 | 268 | 268 | 319 | 232 | 713 | 739 | 663 | 851 | 505 | 955 | 484 | 527 | 695 | 297 | 317 | 203 | 258 | 209 | 515 | 1018 | 539 | 390 | 727 | 523 | 553 | Ppfibp2 |
| Pphln1 | 8068 | 7272 | 8735 | 7443 | 6551 | 7258 | 7349 | 5584 | 5056 | 5228 | 4809 | 6483 | 6022 | 6296 | 8319 | 7739 | 7898 | 6963 | 6841 | 7023 | 6822 | 4774 | 4959 | 5565 | 5380 | 6589 | 6412 | 6763 | Pphln1 |
| Ppia | 49538 | 55900 | 53621 | 40667 | 38508 | 41217 | 43221 | 57074 | 58536 | 46846 | 51865 | 55542 | 61533 | 60909 | 50401 | 51976 | 45700 | 40132 | 43559 | 54690 | 41499 | 44037 | 59393 | 53363 | 52868 | 70092 | 65545 | 68443 | Ppia |
| Ppib | 12453 | 14017 | 13099 | 11680 | 10468 | 11284 | 11306 | 11913 | 13238 | 11805 | 13174 | 11994 | 12700 | 12094 | 12802 | 13545 | 12698 | 11580 | 11510 | 11906 | 11203 | 11212 | 14433 | 12669 | 11371 | 13272 | 12757 | 13238 | Ppib |
| Ppic | 3903 | 3289 | 3914 | 2497 | 1908 | 2543 | 1701 | 3483 | 4158 | 3349 | 4466 | 3268 | 4105 | 3390 | 3699 | 4442 | 2849 | 2773 | 1663 | 1954 | 1490 | 3337 | 5069 | 3618 | 2845 | 4164 | 3390 | 3623 | Ppic |
| Ppid | 10382 | 8046 | 9746 | 8659 | 8207 | 8616 | 10438 | 9912 | 8996 | 8516 | 8762 | 9142 | 8927 | 10401 | 10606 | 9544 | 8423 | 8340 | 8146 | 10053 | 9089 | 8612 | 9147 | 9367 | 8444 | 10218 | 9086 | 10099 | Ppid |
| Ppie | 2832 | 2929 | 2974 | 2460 | 2072 | 2266 | 2447 | 2363 | 2476 | 2193 | 2200 | 2735 | 2465 | 2736 | 2694 | 2546 | 2706 | 2225 | 2419 | 2704 | 2284 | 1927 | 2385 | 2478 | 2264 | 3263 | 2791 | 3050 | Ppie |
| Ppif | 2420 | 2748 | 2824 | 2922 | 2384 | 2466 | 2550 | 1546 | 1845 | 1846 | 1778 | 2066 | 1989 | 2076 | 2231 | 2643 | 2701 | 2872 | 2625 | 2219 | 2505 | 1536 | 1650 | 2015 | 1880 | 2374 | 2088 | 1976 | Ppif |
| Ppig | 13955 | 10555 | 14487 | 13338 | 11935 | 13053 | 12970 | 12737 | 11038 | 11654 | 11475 | 12181 | 12152 | 13238 | 14151 | 12110 | 13381 | 12123 | 12274 | 13100 | 12570 | 11307 | 10861 | 12527 | 11536 | 12942 | 13160 | 13258 | Ppig |
| Ppih | 1917 | 1738 | 1602 | 1274 | 1106 | 1435 | 1500 | 1964 | 1749 | 1741 | 1691 | 2166 | 1897 | 2533 | 1768 | 1684 | 1482 | 1088 | 1402 | 1770 | 1274 | 1592 | 1841 | 1839 | 1589 | 2334 | 2479 | 2563 | Ppih |
| Ppil1 | 3464 | 4139 | 3989 | 3646 | 2906 | 3400 | 3328 | 2663 | 2912 | 2931 | 2758 | 3367 | 3158 | 3418 | 3364 | 3252 | 3843 | 2979 | 3636 | 3511 | 3434 | 2398 | 2813 | 3365 | 2949 | 4179 | 3548 | 3898 | Ppil1 |
| Ppil2 | 8469 | 8937 | 7648 | 8195 | 7886 | 8118 | 8292 | 6988 | 7234 | 6509 | 6520 | 7072 | 6974 | 6961 | 8148 | 8230 | 8343 | 8311 | 8377 | 8635 | 8176 | 6854 | 7067 | 6856 | 7143 | 7149 | 7382 | 7007 | Ppil2 |
| Ppil3 | 1175 | 1015 | 1221 | 1051 | 1049 | 1135 | 1117 | 2081 | 1771 | 1692 | 1758 | 1768 | 1833 | 2054 | 1240 | 1103 | 1021 | 963 | 1098 | 1255 | 1126 | 1643 | 1790 | 1920 | 1700 | 2004 | 2091 | 2231 | Ppil3 |
| Ppil4 | 5479 | 4677 | 5417 | 4990 | 4844 | 5004 | 5434 | 6145 | 5066 | 5141 | 5042 | 5533 | 5138 | 5783 | 5624 | 5489 | 5015 | 4858 | 4797 | 5641 | 5023 | 5688 | 5119 | 5548 | 5323 | 5738 | 5678 | 5784 | Ppil4 |
| Ppil6 | 323 | 318 | 239 | 351 | 326 | 418 | 419 | 98 | 80 | 116 | 102 | 79 | 87 | 98 | 324 | 347 | 295 | 377 | 336 | 372 | 367 | 104 | 97 | 82 | 114 | 41 | 62 | 72 | Ppil6 |
| Ppip5k1 | 2045 | 2453 | 2026 | 2354 | 2508 | 2409 | 2395 | 2148 | 2179 | 2347 | 2240 | 1902 | 2032 | 1916 | 2048 | 2305 | 2306 | 2586 | 2400 | 2368 | 2645 | 2433 | 2104 | 2054 | 2330 | 1350 | 1990 | 1826 | Ppip5k1 |
| Ppip5k2 | 13532 | 10435 | 9248 | 10666 | 12363 | 13532 | 14443 | 14669 | 12543 | 11085 | 11170 | 11056 | 11281 | 12591 | 14292 | 11333 | 9952 | 11180 | 11060 | 14362 | 13515 | 12526 | 11661 | 10056 | 12179 | 8830 | 11404 | 11243 | Ppip5k2 |
| Ppl | 602 | 495 | 410 | 90 | 56 | 348 | 17 | 613 | 1000 | 567 | 461 | 365 | 471 | 300 | 479 | 590 | 103 | 262 | 48 | 22 | 14 | 33 | 1151 | 497 | 91 | 144 | 329 | 361 | Ppl |
| Ppm1a | 11778 | 10198 | 10229 | 12779 | 13404 | 12532 | 13722 | 13178 | 11398 | 11241 | 11268 | 11506 | 11100 | 11722 | 12283 | 10349 | 11332 | 12266 | 12567 | 13173 | 13622 | 12539 | 11207 | 11089 | 11959 | 10354 | 11351 | 11387 | Ppm1a |
| Ppm1b | 11863 | 10370 | 13407 | 12875 | 12039 | 12119 | 13101 | 8501 | 7717 | 8512 | 8327 | 9414 | 8487 | 9628 | 11466 | 10652 | 13347 | 12556 | 12894 | 11256 | 12507 | 9120 | 7350 | 9493 | 9583 | 10175 | 9190 | 9318 | Ppm1b |
| Ppm1d | 3413 | 2931 | 3130 | 3594 | 3184 | 3053 | 3532 | 2740 | 2299 | 2173 | 2171 | 2646 | 2420 | 2768 | 3302 | 3036 | 3246 | 3162 | 3514 | 3228 | 3465 | 2592 | 2353 | 2522 | 2513 | 3081 | 2850 | 2797 | Ppm1d |
| Ppm1e | 8977 | 8942 | 8422 | 12871 | 17255 | 15708 | 17232 | 9148 | 8206 | 9641 | 8155 | 8301 | 7421 | 8552 | 10537 | 8243 | 10828 | 13386 | 15192 | 15145 | 17416 | 10641 | 7471 | 8409 | 10789 | 4943 | 7618 | 7409 | Ppm1e |
| Ppm1f | 7147 | 7630 | 6775 | 6222 | 5680 | 6523 | 5662 | 5300 | 5293 | 4742 | 5393 | 5116 | 5494 | 4838 | 6898 | 6907 | 6216 | 6234 | 5992 | 6221 | 6285 | 5270 | 5541 | 5113 | 5268 | 4785 | 5108 | 5047 | Ppm1f |
| Ppm1g | 17165 | 18686 | 22076 | 20615 | 18036 | 17537 | 19132 | 9501 | 9639 | 11584 | 10706 | 13055 | 11560 | 13077 | 16009 | 16543 | 21411 | 19717 | 20202 | 15719 | 18838 | 10916 | 9271 | 12738 | 12400 | 14707 | 12492 | 13708 | Ppm1g |
| Ppm1h | 1295 | 1453 | 1569 | 1869 | 2624 | 1848 | 1826 | 1529 | 1299 | 1718 | 1829 | 1206 | 1535 | 1410 | 1582 | 1442 | 1925 | 2003 | 1905 | 1631 | 1996 | 2274 | 1426 | 1353 | 1678 | 943 | 1180 | 1244 | Ppm1h |
| Ppm1j | 531 | 575 | 647 | 606 | 396 | 368 | 337 | 55 | 68 | 81 | 95 | 55 | 66 | 56 | 414 | 719 | 620 | 603 | 422 | 325 | 396 | 52 | 76 | 77 | 64 | 108 | 49 | 31 | Ppm1j |
| Ppm1k | 2669 | 2432 | 3342 | 3992 | 3214 | 3137 | 3563 | 1343 | 1112 | 1329 | 1329 | 1196 | 1218 | 1332 | 2913 | 2616 | 3500 | 3810 | 3218 | 3003 | 3464 | 1472 | 1091 | 1402 | 1506 | 1467 | 1252 | 1315 | Ppm1k |
| Ppm1l | 7061 | 6239 | 5619 | 8916 | 10607 | 9961 | 10817 | 7433 | 6905 | 7947 | 7911 | 7941 | 7677 | 8304 | 7625 | 6260 | 7253 | 9468 | 9344 | 8909 | 11316 | 8651 | 6335 | 7386 | 9293 | 5366 | 8086 | 7694 | Ppm1l |
| Ppm1m | 1955 | 2163 | 2854 | 1918 | 1490 | 1633 | 1290 | 1305 | 1576 | 1609 | 1959 | 1714 | 1808 | 1531 | 1726 | 2379 | 2345 | 1859 | 1554 | 1531 | 1280 | 1479 | 1696 | 1617 | 1357 | 2417 | 1964 | 1678 | Ppm1m |
| Ppm1n | 13 | 3 | 13 | 11 | 5 | 1 | 3 | 11 | 3 | 10 | 7 | 10 | 7 | 5 | 3 | 4 | 3 | 12 | 7 | 2 | 9 | 22 | 8 | 8 | 6 | 15 | 3 | 11 | Ppm1n |
| Ppme1 | 8254 | 10032 | 8452 | 9244 | 9179 | 8794 | 9547 | 6981 | 6506 | 6777 | 5864 | 6787 | 6087 | 6469 | 8629 | 8006 | 8996 | 8544 | 9727 | 9616 | 9877 | 6784 | 6410 | 6726 | 7411 | 6528 | 7010 | 6683 | Ppme1 |
| Ppox | 5093 | 5659 | 5401 | 5517 | 5126 | 5036 | 5350 | 3301 | 3590 | 3652 | 3887 | 4137 | 3997 | 4027 | 4741 | 5296 | 5333 | 5241 | 5403 | 4881 | 5105 | 3417 | 3475 | 3831 | 3690 | 4493 | 4091 | 4141 | Ppox |
| Ppp1ca | 792 | 463 | 487 | 462 | 290 | 732 | 305 | 753 | 403 | 348 | 359 | 444 | 501 | 337 | 733 | 499 | 440 | 383 | 488 | 464 | 402 | 543 | 416 | 416 | 387 | 512 | 463 | 614 | Ppp1ca |
| Ppp1cb | 19858 | 14026 | 19928 | 18569 | 19111 | 19424 | 21358 | 31536 | 25623 | 24654 | 24158 | 26369 | 25424 | 27275 | 20886 | 16967 | 19328 | 17999 | 20192 | 21372 | 20000 | 27681 | 26346 | 27026 | 26750 | 28682 | 27044 | 28398 | Ppp1cb |
| Ppp1cc | 18142 | 17593 | 18810 | 17972 | 15557 | 16730 | 17310 | 15684 | 14060 | 14422 | 13262 | 17807 | 15403 | 16199 | 17693 | 16715 | 17763 | 16886 | 18502 | 17395 | 17115 | 14126 | 13383 | 15881 | 15571 | 19791 | 17295 | 17868 | Ppp1cc |
| Ppp1r10 | 5986 | 6213 | 7764 | 7200 | 6883 | 6075 | 6479 | 4732 | 5288 | 6483 | 5826 | 6920 | 6261 | 6991 | 5764 | 6415 | 7625 | 6892 | 7487 | 5645 | 6577 | 6132 | 5115 | 7498 | 6207 | 7382 | 6421 | 6819 | Ppp1r10 |
| Ppp1r11 | 3449 | 3608 | 3756 | 4118 | 3679 | 3509 | 4108 | 3684 | 3352 | 3665 | 3046 | 3472 | 3311 | 4141 | 3791 | 3005 | 3778 | 3748 | 4084 | 4113 | 4191 | 2994 | 3339 | 3718 | 3638 | 3871 | 3876 | 4156 | Ppp1r11 |
| Ppp1r12a | 8164 | 6438 | 7260 | 7927 | 8189 | 8356 | 8546 | 10630 | 9725 | 9866 | 10346 | 10125 | 10518 | 10827 | 8081 | 7020 | 7386 | 7542 | 8017 | 8173 | 8263 | 10497 | 9482 | 10100 | 10068 | 8998 | 10681 | 9873 | Ppp1r12a |
| Ppp1r12b | 3241 | 3465 | 2718 | 4923 | 5314 | 4747 | 4937 | 3560 | 3875 | 4558 | 4140 | 4164 | 3429 | 3781 | 3276 | 3428 | 3898 | 4946 | 4926 | 3950 | 5330 | 4400 | 3683 | 3722 | 4820 | 2337 | 3585 | 3464 | Ppp1r12b |
| Ppp1r12c | 7073 | 9028 | 8219 | 10097 | 8708 | 8172 | 8202 | 4700 | 4902 | 5277 | 5435 | 5829 | 5333 | 5447 | 6935 | 7910 | 8748 | 9647 | 8696 | 7463 | 8040 | 5282 | 4668 | 5572 | 5671 | 5839 | 5245 | 5334 | Ppp1r12c |
| Ppp1r13b | 2291 | 2677 | 2226 | 2970 | 3461 | 3078 | 2675 | 1998 | 2721 | 2629 | 2675 | 1867 | 1947 | 1644 | 2260 | 2876 | 2593 | 3356 | 3294 | 2363 | 2996 | 3068 | 2716 | 2274 | 2614 | 1633 | 1521 | 1526 | Ppp1r13b |
| Ppp1r13l | 1317 | 1792 | 1969 | 2074 | 1400 | 1346 | 1124 | 363 | 573 | 526 | 509 | 529 | 479 | 431 | 1159 | 1559 | 1969 | 1834 | 1612 | 1111 | 1341 | 329 | 618 | 589 | 410 | 624 | 377 | 446 | Ppp1r13l |
| Ppp1r14a | 228 | 225 | 249 | 191 | 190 | 190 | 185 | 210 | 198 | 249 | 233 | 255 | 359 | 281 | 231 | 193 | 245 | 254 | 249 | 195 | 197 | 124 | 138 | 307 | 223 | 297 | 263 | 297 | Ppp1r14a |
| Ppp1r14b | 5778 | 6432 | 6586 | 6547 | 6082 | 5532 | 7400 | 7035 | 7065 | 7351 | 7298 | 8162 | 6743 | 10251 | 5122 | 4876 | 6934 | 5464 | 6673 | 6270 | 7330 | 6458 | 6680 | 8863 | 8393 | 9838 | 8340 | 9589 | Ppp1r14b |
| Ppp1r14bl | 0 | 2 | 0 | 4 | 4 | 3 | 8 | 7 | 7 | 5 | 5 | 8 | 3 | 6 | 0 | 1 | 3 | 2 | 1 | 2 | 2 | 13 | 9 | 13 | 15 | 12 | 1 | 6 | Ppp1r14bl |
| Ppp1r14c | 326 | 289 | 330 | 508 | 1202 | 757 | 858 | 1175 | 1251 | 1244 | 1144 | 543 | 636 | 624 | 494 | 371 | 443 | 825 | 914 | 698 | 920 | 1415 | 1223 | 796 | 1261 | 321 | 402 | 402 | Ppp1r14c |
| Ppp1r14d | 2 | 2 | 0 | 0 | 0 | 0 | 0 | 2 | 3 | 0 | 4 | 0 | 1 | 3 | 3 | 5 | 0 | 1 | 0 | 0 | 0 | 0 | 3 | 5 | 0 | 7 | 0 | 0 | Ppp1r14d |
| Ppp1r15a | 1481 | 1938 | 1691 | 1889 | 1561 | 1706 | 1782 | 624 | 944 | 919 | 955 | 791 | 981 | 763 | 1564 | 1714 | 1749 | 2023 | 1777 | 1484 | 1773 | 662 | 1018 | 884 | 747 | 851 | 782 | 830 | Ppp1r15a |

| | | | | | | | | | | | | | | | | | | | | | | | | | | | | | |
|---|---|---|---|---|---|---|---|---|---|---|---|---|---|---|---|---|---|---|---|---|---|---|---|---|---|---|---|---|---|
| Ppp1r15b | 7243 | 7418 | 9060 | 9095 | 7309 | 7305 | 7394 | 6346 | 5927 | 6382 | 6590 | 7974 | 7498 | 7576 | 7106 | 7271 | 8473 | 8439 | 7716 | 6325 | 7537 | 7237 | 6064 | 7147 | 7077 | 8050 | 7941 | 8380 | Ppp1r15b |
| Ppp1r16a | 3556 | 4982 | 4382 | 4893 | 4363 | 4210 | 3800 | 2325 | 2860 | 2866 | 2749 | 3171 | 2947 | 2868 | 3626 | 3937 | 4971 | 4418 | 4925 | 3812 | 4002 | 3091 | 2913 | 3245 | 3079 | 2962 | 2915 | 2865 | Ppp1r16a |
| Ppp1r16b | 1160 | 1453 | 1277 | 1674 | 1980 | 1781 | 1793 | 629 | 806 | 813 | 797 | 545 | 578 | 606 | 1207 | 1247 | 1511 | 1574 | 1964 | 1639 | 1923 | 872 | 786 | 664 | 1093 | 416 | 643 | 541 | Ppp1r16b |
| Ppp1r17 | 632 | 681 | 655 | 423 | 660 | 404 | 289 | 778 | 629 | 647 | 662 | 672 | 1438 | 666 | 1106 | 1121 | 844 | 419 | 335 | 483 | 302 | 909 | 445 | 337 | 568 | 464 | 841 | 796 | Ppp1r17 |
| Ppp1r18 | 3722 | 4438 | 6004 | 4882 | 5326 | 4839 | 4171 | 4693 | 4825 | 5418 | 5632 | 5938 | 5763 | 5799 | 3758 | 4114 | 5648 | 4587 | 5408 | 4077 | 4413 | 5153 | 4732 | 5973 | 5947 | 6066 | 5600 | 5885 | Ppp1r18 |
| Ppp1r1a | 1705 | 2285 | 2112 | 2430 | 2509 | 2278 | 2223 | 1368 | 1317 | 1431 | 1228 | 1454 | 1484 | 1428 | 1832 | 1965 | 2274 | 2315 | 2588 | 2708 | 2298 | 1332 | 1373 | 1540 | 1552 | 1234 | 1493 | 1387 | Ppp1r1a |
| Ppp1r1b | 219 | 314 | 288 | 459 | 641 | 543 | 643 | 127 | 223 | 243 | 151 | 75 | 18 | 88 | 219 | 253 | 341 | 625 | 771 | 520 | 753 | 62 | 98 | 198 | 314 | 74 | 51 | 62 | Ppp1r1b |
| Ppp1r1c | 293 | 291 | 91 | 47 | 736 | 54 | 28 | 855 | 765 | 515 | 778 | 54 | 976 | 55 | 575 | 636 | 169 | 341 | 52 | 101 | 49 | 1238 | 499 | 47 | 541 | 4 | 56 | 72 | Ppp1r1c |
| Ppp1r2 | 8466 | 6604 | 8770 | 8321 | 9219 | 8168 | 8763 | 8145 | 7299 | 7297 | 8016 | 7607 | 8549 | 8364 | 9306 | 8601 | 8218 | 9027 | 7822 | 7483 | 8319 | 9608 | 7071 | 6966 | 8458 | 8609 | 7740 | 8546 | Ppp1r2 |
| Ppp1r21 | 3748 | 4213 | 3755 | 4636 | 5258 | 4953 | 5282 | 3526 | 3729 | 3993 | 3403 | 3633 | 3359 | 3613 | 4221 | 3907 | 4193 | 4796 | 5187 | 4741 | 5507 | 3897 | 3679 | 3630 | 4171 | 3332 | 3398 | 3532 | Ppp1r21 |
| Ppp1r26 | 1531 | 1824 | 1953 | 2028 | 2499 | 2053 | 1944 | 2256 | 2616 | 3136 | 2702 | 2646 | 2503 | 2578 | 1561 | 1615 | 2311 | 2187 | 2411 | 1889 | 1990 | 2762 | 2292 | 2808 | 2916 | 2407 | 2467 | 2429 | Ppp1r26 |
| Ppp1r27 | 0 | 0 | 0 | 0 | 0 | 0 | 0 | 0 | 0 | 0 | 6 | 0 | 0 | 0 | 0 | 1 | 0 | 0 | 2 | 0 | 2 | 0 | 11 | 0 | 0 | 0 | 0 | 0 | Ppp1r27 |
| Ppp1r32 | 76 | 102 | 83 | 89 | 111 | 117 | 80 | 66 | 84 | 83 | 61 | 111 | 118 | 73 | 98 | 98 | 139 | 90 | 126 | 96 | 105 | 62 | 87 | 68 | 86 | 84 | 73 | 61 | Ppp1r32 |
| Ppp1r35 | 1497 | 1925 | 1982 | 1951 | 1546 | 1490 | 1386 | 915 | 889 | 1054 | 860 | 1123 | 972 | 1031 | 1452 | 1313 | 2031 | 1745 | 1852 | 1642 | 1664 | 749 | 857 | 1129 | 958 | 1332 | 1164 | 1098 | Ppp1r35 |
| Ppp1r36 | 70 | 46 | 27 | 21 | 53 | 66 | 32 | 64 | 86 | 95 | 78 | 49 | 59 | 19 | 57 | 96 | 52 | 30 | 37 | 31 | 25 | 69 | 112 | 42 | 33 | 46 | 29 | 39 | Ppp1r36 |
| Ppp1r37 | 4602 | 6427 | 6745 | 7628 | 7146 | 6046 | 5768 | 3975 | 4634 | 5330 | 5128 | 4999 | 4461 | 4408 | 4733 | 5308 | 7109 | 7702 | 7602 | 5549 | 6424 | 5075 | 4859 | 5410 | 5410 | 5398 | 4284 | 4349 | Ppp1r37 |
| Ppp1r3a | 1 | 8 | 0 | 6 | 2 | 1 | 10 | 9 | 10 | 3 | 63 | 1 | 12 | 1 | 0 | 5 | 8 | 3 | 0 | 1 | 3 | 2 | 55 | 1 | 8 | 4 | 2 | 4 | Ppp1r3a |
| Ppp1r3b | 1599 | 1396 | 1395 | 1260 | 959 | 1144 | 672 | 1329 | 2136 | 1856 | 1978 | 989 | 1487 | 847 | 1236 | 1858 | 1270 | 1406 | 676 | 670 | 754 | 1773 | 2034 | 1611 | 1138 | 1458 | 1053 | 808 | Ppp1r3b |
| Ppp1r3c | 750 | 783 | 619 | 800 | 910 | 814 | 791 | 697 | 801 | 724 | 807 | 295 | 517 | 286 | 1044 | 815 | 687 | 884 | 715 | 821 | 752 | 695 | 1048 | 486 | 476 | 389 | 343 | 256 | Ppp1r3c |
| Ppp1r3d | 451 | 517 | 412 | 531 | 630 | 450 | 476 | 241 | 232 | 231 | 338 | 102 | 179 | 121 | 491 | 483 | 446 | 551 | 458 | 480 | 550 | 247 | 298 | 178 | 182 | 164 | 84 | 94 | Ppp1r3d |
| Ppp1r3e | 216 | 217 | 249 | 353 | 422 | 391 | 367 | 325 | 368 | 408 | 413 | 357 | 346 | 361 | 239 | 220 | 308 | 380 | 363 | 332 | 377 | 488 | 361 | 443 | 466 | 460 | 264 | 352 | Ppp1r3e |
| Ppp1r3f | 129 | 157 | 151 | 160 | 236 | 202 | 127 | 208 | 199 | 263 | 230 | 236 | 197 | 182 | 128 | 136 | 156 | 174 | 225 | 153 | 161 | 236 | 167 | 238 | 242 | 167 | 280 | 140 | Ppp1r3f |
| Ppp1r3g | 74 | 86 | 95 | 64 | 76 | 70 | 70 | 45 | 47 | 40 | 59 | 52 | 39 | 59 | 98 | 83 | 108 | 63 | 68 | 53 | 71 | 13 | 32 | 45 | 52 | 63 | 34 | 40 | Ppp1r3g |
| Ppp1r42 | 50 | 61 | 122 | 63 | 55 | 48 | 59 | 22 | 21 | 41 | 37 | 30 | 46 | 35 | 45 | 78 | 36 | 66 | 51 | 52 | 62 | 35 | 54 | 29 | 34 | 26 | 19 | 36 | Ppp1r42 |
| Ppp1r7 | 8712 | 8021 | 7014 | 7882 | 7313 | 7886 | 8638 | 5823 | 5719 | 6094 | 5717 | 7287 | 6525 | 7687 | 8359 | 7215 | 7496 | 7526 | 7157 | 7944 | 8019 | 5451 | 5407 | 5811 | 6339 | 6863 | 7677 | 7350 | Ppp1r7 |
| Ppp1r8 | 5170 | 5470 | 6070 | 5334 | 4942 | 5128 | 5062 | 5930 | 5526 | 5405 | 5107 | 6579 | 5629 | 6403 | 5239 | 4831 | 5859 | 4945 | 5525 | 5854 | 4897 | 5053 | 5273 | 6034 | 5769 | 7706 | 6919 | 7059 | Ppp1r8 |
| Ppp1r9a | 5869 | 5142 | 3504 | 5599 | 7245 | 7775 | 7190 | 12132 | 11963 | 11307 | 12801 | 10936 | 10314 | 11509 | 6080 | 5768 | 4494 | 5442 | 5843 | 6721 | 7504 | 11358 | 10957 | 8644 | 11781 | 6153 | 11302 | 8977 | Ppp1r9a |
| Ppp1r9b | 11227 | 14382 | 14942 | 18077 | 17232 | 15041 | 14208 | 10331 | 11037 | 13387 | 12642 | 12748 | 10473 | 11636 | 11054 | 11409 | 16560 | 17141 | 18654 | 13730 | 16696 | 12978 | 10958 | 14096 | 14925 | 13623 | 11375 | 12252 | Ppp1r9b |
| Ppp2ca | 25007 | 22786 | 22935 | 24496 | 25406 | 24918 | 27430 | 33208 | 28035 | 28297 | 25623 | 29393 | 27677 | 30251 | 26710 | 20350 | 23901 | 23942 | 26703 | 25948 | 28236 | 31001 | 28991 | 28077 | 29680 | 30778 | 30781 | 31623 | Ppp2ca |
| Ppp2cb | 5991 | 5763 | 6236 | 6072 | 5993 | 5519 | 6187 | 6566 | 6311 | 6101 | 5799 | 7076 | 6451 | 6921 | 5917 | 5539 | 6163 | 5721 | 6279 | 5972 | 6243 | 6081 | 6312 | 6457 | 6492 | 7208 | 6880 | 7173 | Ppp2cb |
| Ppp2r1a | 20251 | 24109 | 21385 | 23361 | 26189 | 24983 | 24061 | 23294 | 25911 | 25558 | 23600 | 28143 | 25753 | 26728 | 20343 | 19612 | 23059 | 22653 | 26974 | 22346 | 25070 | 27268 | 24227 | 25842 | 27436 | 28709 | 26260 | 27843 | Ppp2r1a |
| Ppp2r1b | 10219 | 10040 | 10474 | 9068 | 8235 | 8479 | 9485 | 6023 | 5772 | 5331 | 5451 | 6928 | 6455 | 6929 | 9777 | 9047 | 9603 | 8920 | 9082 | 9310 | 8704 | 4964 | 5348 | 6128 | 6053 | 7691 | 7325 | 7233 | Ppp2r1b |
| Ppp2r2a | 9127 | 7861 | 10317 | 10388 | 9694 | 8868 | 9666 | 7891 | 7380 | 7951 | 7674 | 9009 | 8181 | 8850 | 8987 | 8330 | 10210 | 10247 | 10006 | 7694 | 10337 | 8980 | 7234 | 8528 | 8396 | 10057 | 8566 | 9280 | Ppp2r2a |
| Ppp2r2b | 1986 | 2263 | 1670 | 3537 | 5475 | 4681 | 5662 | 4907 | 5607 | 5308 | 4351 | 3432 | 3006 | 3836 | 2643 | 1784 | 2418 | 4296 | 5001 | 5462 | 5517 | 4921 | 4458 | 4863 | 6411 | 3594 | 2489 | 2794 | Ppp2r2b |
| Ppp2r2c | 509 | 675 | 448 | 1286 | 3046 | 1888 | 2207 | 1993 | 2431 | 2502 | 2530 | 1189 | 951 | 1228 | 761 | 598 | 910 | 1818 | 2097 | 1865 | 2406 | 3008 | 2374 | 1883 | 2925 | 865 | 807 | 800 | Ppp2r2c |
| Ppp2r2d | 5132 | 4641 | 5018 | 5395 | 5434 | 5455 | 5795 | 5567 | 4940 | 5399 | 5063 | 5615 | 5240 | 5933 | 5160 | 4160 | 5109 | 5103 | 5568 | 4983 | 6047 | 5840 | 5000 | 5147 | 5686 | 5879 | 5616 | 6008 | Ppp2r2d |
| Ppp2r3a | 6584 | 6082 | 4773 | 6492 | 8715 | 8581 | 8311 | 9623 | 8370 | 8067 | 7575 | 6887 | 7377 | 7078 | 7764 | 5860 | 5964 | 7030 | 7644 | 8899 | 8616 | 8727 | 8348 | 6612 | 7879 | 4620 | 6809 | 6610 | Ppp2r3a |
| Ppp2r3c | 2309 | 1872 | 2161 | 2089 | 2747 | 2275 | 2731 | 4542 | 3551 | 3571 | 3303 | 3960 | 3881 | 4630 | 2641 | 2008 | 2224 | 2194 | 2420 | 2749 | 2702 | 3648 | 3398 | 3754 | 4049 | 4092 | 4267 | 4138 | Ppp2r3c |
| Ppp2r3d | 29 | 74 | 27 | 28 | 53 | 17 | 45 | 61 | 54 | 43 | 81 | 90 | 67 | 62 | 128 | 36 | 36 | 26 | 34 | 24 | 29 | 107 | 53 | 51 | 87 | 69 | 85 | 63 | Ppp2r3d |
| Ppp2r5a | 4410 | 3973 | 4305 | 3955 | 3398 | 3549 | 3727 | 2304 | 2456 | 2219 | 2541 | 2393 | 2369 | 2432 | 3990 | 3992 | 3783 | 3919 | 3626 | 3685 | 3514 | 1967 | 2650 | 2395 | 2083 | 2823 | 2366 | 2379 | Ppp2r5a |
| Ppp2r5b | 0 | 0 | 0 | 0 | 0 | 0 | 0 | 0 | 0 | 0 | 0 | 0 | 0 | 0 | 1 | 0 | 0 | 0 | 0 | 0 | 1 | 0 | 0 | 0 | 0 | 0 | 0 | 0 | Ppp2r5b |
| Ppp2r5c | 10671 | 10718 | 10510 | 10604 | 10324 | 9665 | 10690 | 10119 | 9667 | 9584 | 9276 | 9919 | 9552 | 9973 | 10933 | 10763 | 10798 | 10378 | 10162 | 10561 | 10409 | 9605 | 9785 | 9024 | 9338 | 10071 | 9879 | 10379 | Ppp2r5c |
| Ppp2r5d | 13998 | 16169 | 14029 | 13945 | 12024 | 12410 | 12201 | 6757 | 7420 | 7525 | 6823 | 8815 | 7917 | 7776 | 13226 | 13544 | 14336 | 13366 | 13986 | 11824 | 13083 | 6839 | 7304 | 7732 | 7154 | 9641 | 8367 | 8410 | Ppp2r5d |
| Ppp2r5e | 9566 | 7679 | 9342 | 9583 | 9793 | 9758 | 10683 | 9304 | 8201 | 9006 | 8554 | 10788 | 9187 | 10653 | 9430 | 8368 | 9427 | 9671 | 10256 | 8570 | 10523 | 10273 | 7717 | 9559 | 10178 | 9484 | 9718 | 10690 | Ppp2r5e |
| Ppp3ca | 10656 | 8317 | 10557 | 13211 | 14722 | 14052 | 15940 | 11803 | 10682 | 11878 | 11613 | 10101 | 9840 | 10952 | 10984 | 9037 | 11846 | 14047 | 14611 | 12283 | 16099 | 13926 | 10980 | 11684 | 13067 | 10044 | 9359 | 10611 | Ppp3ca |
| Ppp3cb | 11648 | 8695 | 10627 | 12190 | 15101 | 14265 | 16542 | 17315 | 15001 | 15672 | 15674 | 15946 | 14610 | 16973 | 12018 | 9780 | 11363 | 13091 | 14214 | 13143 | 16125 | 18842 | 15611 | 15806 | 17917 | 15264 | 15682 | 16667 | Ppp3cb |
| Ppp3cc | 490 | 403 | 438 | 458 | 491 | 531 | 443 | 663 | 710 | 616 | 553 | 533 | 497 | 543 | 521 | 498 | 480 | 445 | 352 | 456 | 398 | 710 | 700 | 476 | 613 | 478 | 516 | 441 | Ppp3cc |
| Ppp3r1 | 10156 | 7406 | 9628 | 11381 | 12393 | 11435 | 13892 | 9182 | 7622 | 8294 | 8411 | 7626 | 6946 | 8578 | 11072 | 8778 | 10276 | 11990 | 11608 | 11062 | 13633 | 10581 | 8071 | 8264 | 9228 | 7704 | 7377 | 8427 | Ppp3r1 |
| Ppp3r2 | 9 | 4 | 1 | 4 | 4 | 6 | 6 | 37 | 21 | 5 | 8 | 3 | 9 | 7 | 7 | 8 | 2 | 0 | 6 | 2 | 2 | 0 | 22 | 7 | 12 | 6 | 7 | 4 | Ppp3r2 |
| Ppp4c | 6508 | 7294 | 7686 | 6197 | 5364 | 5565 | 5890 | 6428 | 6749 | 6486 | 6173 | 7791 | 7292 | 7272 | 6220 | 6428 | 7094 | 5710 | 6324 | 5847 | 5797 | 5897 | 6771 | 7293 | 6599 | 8930 | 7907 | 7925 | Ppp4c |
| Ppp4r1 | 3722 | 3748 | 3793 | 3532 | 4343 | 3832 | 3680 | 6421 | 6298 | 6573 | 6357 | 7697 | 7077 | 7315 | 3728 | 3460 | 4048 | 3616 | 4005 | 3681 | 3731 | 7199 | 6130 | 6802 | 6998 | 7905 | 7151 | 7703 | Ppp4r1 |
| Ppp4r2 | 7179 | 5508 | 7982 | 7737 | 8040 | 8092 | 8812 | 11512 | 9542 | 10397 | 9732 | 11456 | 10638 | 11309 | 7762 | 7090 | 7726 | 7538 | 8068 | 8481 | 7613 | 11239 | 9769 | 10477 | 11126 | 12061 | 11515 | 12223 | Ppp4r2 |
| Ppp4r3a | 9115 | 7211 | 10791 | 9742 | 9723 | 9555 | 10712 | 12995 | 10785 | 11267 | 11993 | 14012 | 13201 | 15371 | 8871 | 8196 | 9933 | 9318 | 9955 | 9062 | 10137 | 12118 | 10195 | 13008 | 12754 | 13808 | 13955 | 15073 | Ppp4r3a |
| Ppp4r3b | 10017 | 7478 | 7644 | 7335 | 7841 | 8374 | 10059 | 12879 | 10797 | 8932 | 8870 | 11272 | 10661 | 11681 | 10357 | 8507 | 7993 | 7593 | 8267 | 10584 | 8961 | 9613 | 10585 | 9604 | 9562 | 10850 | 11943 | 11968 | Ppp4r3b |
| Ppp4r4 | 1916 | 1585 | 1752 | 1694 | 1643 | 1683 | 1847 | 816 | 690 | 756 | 598 | 796 | 759 | 771 | 1742 | 1568 | 1522 | 1492 | 1584 | 1678 | 1556 | 608 | 662 | 628 | 588 | 723 | 772 | 808 | Ppp4r4 |
| Ppp5c | 3643 | 4043 | 4458 | 4368 | 3890 | 3961 | 3802 | 3446 | 4164 | 4300 | 4318 | 4873 | 4602 | 4290 | 3500 | 3746 | 4446 | 4139 | 4618 | 3751 | 3832 | 4059 | 4317 | 4765 | 4464 | 5409 | 4450 | 4462 | Ppp5c |
| Ppp6c | 6473 | 5442 | 6158 | 5571 | 5600 | 5667 | 6549 | 6693 | 5553 | 4805 | 4480 | 5441 | 4935 | 5594 | 7457 | 6143 | 5749 | 5556 | 5711 | 6720 | 5966 | 5887 | 5642 | 5024 | 5088 | 5600 | 5653 | 5830 | Ppp6c |
| Ppp6r1 | 9145 | 11625 | 10285 | 10877 | 10103 | 9714 | 9894 | 7952 | 9173 | 9563 | 8806 | 10138 | 8929 | 9434 | 8983 | 9497 | 11214 | 10412 | 10922 | 8994 | 9804 | 8610 | 8535 | 9832 | 9572 | 9466 | 8930 | 9670 | Ppp6r1 |
| Ppp6r2 | 7391 | 7868 | 6764 | 8953 | 9973 | 9200 | 9680 | 7983 | 8282 | 8879 | 7888 | 8467 | 7945 | 8417 | 7552 | 6857 | 8194 | 9005 | 10017 | 8591 | 9858 | 9207 | 7394 | 8421 | 9899 | 6506 | 8191 | 8090 | Ppp6r2 |
| Ppp6r3 | 19912 | 19034 | 15729 | 17198 | 16184 | 17548 | 18074 | 17379 | 15362 | 14628 | 13490 | 16931 | 15159 | 15956 | 20094 | 17907 | 16388 | 17142 | 16454 | 18340 | 17694 | 14474 | 14748 | 14676 | 15158 | 14462 | 16870 | 15760 | Ppp6r3 |
| Pprc1 | 2646 | 3203 | 3380 | 3436 | 2940 | 2798 | 2596 | 3635 | 4411 | 4491 | 4854 | 5359 | 4966 | 5063 | 2362 | 3223 | 3321 | 3395 | 3150 | 2160 | 2808 | 4523 | 4073 | 4846 | 4542 | 5649 | 4962 | 5148 | Pprc1 |
| Ppt1 | 5015 | 4822 | 4209 | 4105 | 4112 | 4556 | 4672 | 5340 | 4792 | 4706 | 4761 | 4919 | 4917 | 5156 | 5176 | 4534 | 4144 | 4216 | 4018 | 4522 | 4456 | 5247 | 4723 | 4430 | 4756 | 4692 | 4780 | 5075 | Ppt1 |
| Ppt2 | 581 | 797 | 894 | 830 | 734 | 744 | 837 | 476 | 483 | 681 | 624 | 692 | 604 | 666 | 618 | 678 | 969 | 830 | 906 | 662 | 810 | 471 | 500 | 639 | 620 | 610 | 482 | 665 | Ppt2 |
| Pptc7 | 2805 | 2786 | 3141 | 3200 | 2572 | 2654 | 2833 | 1931 | 1827 | 1800 | 1831 | 1995 | 2057 | 2054 | 2719 | 3002 | 3089 | 3044 | 2749 | 2441 | 2888 | 1922 | 1766 | 1864 | 1988 | 2284 | 2195 | 2302 | Pptc7 |
| Ppwd1 | 2616 | 2021 | 2740 | 2045 | 1677 | 1900 | 2200 | 2434 | 2095 | 1984 | 2003 | 2457 | 2256 | 2474 | 2566 | 2499 | 2281 | 2037 | 1905 | 2270 | 1946 | 1778 | 2025 | 2304 | 1822 | 2808 | 2453 | 2436 | Ppwd1 |
| Ppy | 0 | 0 | 0 | 0 | 0 | 0 | 0 | 0 | 0 | 0 | 3 | 0 | 1 | 1 | 0 | 0 | 0 | 0 | 0 | 0 | 0 | 0 | 0 | 0 | 0 | 0 | 0 | 1 | Ppy |
| Pqbp1 | 3716 | 4373 | 4345 | 4140 | 4325 | 4203 | 3985 | 4041 | 4199 | 4277 | 4105 | 4403 | 4168 | 4428 | 3566 | 3325 | 4456 | 3985 | 4717 | 4212 | 4270 | 4149 | 4011 | 4838 | 4772 | 4917 | 4503 | 4332 | Pqbp1 |
| Pqlc3 | 208 | 143 | 168 | 91 | 92 | 112 | 65 | 186 | 214 | 155 | 201 | 94 | 144 | 118 | 174 | 236 | 109 | 118 | 51 | 96 | 47 | 92 | 269 | 151 | 77 | 117 | 123 | 144 | Pqlc3 |
| Pradc1 | 946 | 974 | 893 | 946 | 794 | 885 | 898 | 452 | 450 | 388 | 473 | 571 | 524 | 623 | 832 | 815 | 993 | 974 | 913 | 706 | 945 | 428 | 426 | 562 | 477 | 692 | 556 | 681 | Pradc1 |
| Praf2 | 1081 | 1222 | 830 | 1039 | 993 | 977 | 1319 | 1155 | 920 | 1092 | 917 | 876 | 940 | 1179 | 1064 | 743 | 977 | 852 | 921 | 1090 | 1144 | 1026 | 880 | 1133 | 978 | 954 | 947 | 1145 | Praf2 |
| Prag1 | 1614 | 2200 | 2428 | 3148 | 3657 | 2957 | 2629 | 2165 | 2761 | 3118 | 2990 | 3009 | 2868 | 2634 | 1683 | 2054 | 3052 | 3358 | 3706 | 2356 | 3052 | 3160 | 2742 | 3263 | 3318 | 2750 | 2351 | 2359 | Prag1 |
| Pram1 | 8 | 16 | 12 | 11 | 16 | 21 | 18 | 5 | 8 | 0 | 11 | 2 | 5 | 3 | 9 | 19 | 14 | 19 | 16 | 13 | 14 | 7 | 2 | 7 | 8 | 0 | 0 | 4 | Pram1 |
| Pramel12 | 0 | 0 | 0 | 0 | 0 | 0 | 0 | 0 | 1 | 0 | 0 | 0 | 0 | 0 | 0 | 0 | 0 | 0 | 0 | 0 | 0 | 0 | 0 | 0 | 0 | 0 | 0 | 0 | Pramel12 |
| Pramel13 | 7 | 3 | 7 | 0 | 0 | 11 | 0 | 15 | 22 | 6 | 9 | 9 | 5 | 4 | 7 | 3 | 7 | 11 | 0 | 0 | 0 | 7 | 21 | 7 | 7 | 1 | 3 | 5 | Pramel13 |
| Pramel18 | 0 | 3 | 0 | 0 | 0 | 0 | 0 | 0 | 0 | 0 | 0 | 0 | 0 | 0 | 0 | 1 | 0 | 0 | 0 | 0 | 0 | 0 | 0 | 0 | 0 | 0 | 0 | 0 | Pramel18 |
| Pramel19 | 0 | 0 | 0 | 0 | 0 | 0 | 0 | 0 | 0 | 0 | 1 | 0 | 0 | 0 | 0 | 0 | 0 | 0 | 0 | 0 | 0 | 0 | 0 | 0 | 0 | 0 | 0 | 0 | Pramel19 |
| Pramel3 | 0 | 0 | 0 | 0 | 0 | 0 | 0 | 0 | 0 | 0 | 2 | 0 | 0 | 0 | 0 | 2 | 0 | 0 | 0 | 0 | 0 | 0 | 0 | 0 | 0 | 0 | 0 | 0 | Pramel3 |
| Pramel5 | 0 | 0 | 0 | 0 | 0 | 0 | 0 | 0 | 0 | 0 | 0 | 0 | 2 | 0 | 0 | 0 | 0 | 2 | 0 | 0 | 0 | 0 | 1 | 1 | 0 | 0 | 0 | 0 | Pramel5 |
| Pramel6 | 0 | 0 | 0 | 0 | 0 | 0 | 0 | 0 | 0 | 1 | 0 | 0 | 0 | 0 | 0 | 0 | 0 | 0 | 0 | 0 | 0 | 0 | 0 | 0 | 0 | 0 | 0 | 0 | Pramel6 |

| | | | | | | | | | | | | | | | | | | | | | | | | | | | | | |
|---|---|---|---|---|---|---|---|---|---|---|---|---|---|---|---|---|---|---|---|---|---|---|---|---|---|---|---|---|---|
| Pramel7 | 0 | 0 | 0 | 0 | 0 | 0 | 0 | 0 | 0 | 0 | 0 | 0 | 0 | 0 | 0 | 0 | 0 | 0 | 0 | 0 | 0 | 0 | 0 | 2 | 0 | 0 | 0 | 0 | Pramel7 |
| Pramex1 | 0 | 0 | 0 | 0 | 0 | 0 | 0 | 2 | 0 | 1 | 7 | 1 | 1 | 0 | 0 | 0 | 0 | 0 | 2 | 0 | 0 | 0 | 5 | 0 | 0 | 0 | 0 | 0 | Pramex1 |
| Prap1 | 0 | 1 | 0 | 2 | 0 | 0 | 1 | 0 | 65 | 0 | 5 | 1 | 6 | 1 | 5 | 1 | 5 | 2 | 1 | 0 | 0 | 3 | 16 | 8 | 2 | 0 | 1 | 2 | Prap1 |
| Prc1 | 13162 | 12615 | 13546 | 10446 | 8138 | 9799 | 10865 | 11577 | 9727 | 9168 | 8323 | 14418 | 11530 | 14947 | 12891 | 11227 | 12199 | 9667 | 10411 | 11950 | 10233 | 7585 | 8755 | 11928 | 10550 | 15564 | 15102 | 15224 | Prc1 |
| Prcc | 3949 | 4685 | 6157 | 5751 | 5081 | 4345 | 4598 | 4041 | 4582 | 5130 | 4984 | 5575 | 4808 | 5320 | 3771 | 4111 | 5968 | 5399 | 5499 | 4307 | 4698 | 4852 | 4470 | 5892 | 5777 | 6645 | 4833 | 5466 | Prcc |
| Prcd | 10 | 15 | 13 | 8 | 25 | 17 | 6 | 6 | 13 | 14 | 33 | 10 | 17 | 12 | 12 | 3 | 12 | 10 | 12 | 22 | 26 | 12 | 17 | 8 | 23 | 6 | 6 | 11 | Prcd |
| Prcp | 2269 | 2064 | 2430 | 1624 | 1438 | 1566 | 1333 | 1646 | 1782 | 1438 | 1498 | 1412 | 1674 | 1507 | 2180 | 2484 | 1904 | 1733 | 1319 | 1600 | 1319 | 1477 | 1955 | 1486 | 1417 | 1686 | 1642 | 1544 | Prcp |
| Prdm1 | 414 | 179 | 266 | 191 | 189 | 191 | 166 | 274 | 345 | 324 | 239 | 153 | 179 | 147 | 211 | 247 | 232 | 167 | 196 | 176 | 145 | 140 | 338 | 250 | 149 | 297 | 157 | 145 | Prdm1 |
| Prdm10 | 867 | 864 | 1011 | 1251 | 1316 | 1276 | 1424 | 3819 | 3656 | 4293 | 4005 | 4710 | 4199 | 4809 | 926 | 946 | 1078 | 1229 | 1396 | 1156 | 1311 | 4752 | 3503 | 4207 | 4759 | 3733 | 4580 | 4527 | Prdm10 |
| Prdm11 | 329 | 385 | 243 | 264 | 542 | 513 | 313 | 772 | 1308 | 967 | 1322 | 880 | 907 | 755 | 243 | 385 | 286 | 337 | 400 | 411 | 371 | 810 | 1201 | 777 | 941 | 364 | 963 | 691 | Prdm11 |
| Prdm12 | 382 | 387 | 539 | 348 | 844 | 279 | 308 | 838 | 796 | 889 | 978 | 765 | 1047 | 783 | 402 | 523 | 627 | 470 | 429 | 199 | 397 | 1444 | 421 | 772 | 1033 | 603 | 783 | 942 | Prdm12 |
| Prdm13 | 76 | 263 | 22 | 328 | 216 | 262 | 80 | 230 | 231 | 217 | 190 | 85 | 490 | 96 | 251 | 338 | 334 | 163 | 85 | 198 | 63 | 262 | 293 | 126 | 177 | 156 | 87 | 64 | Prdm13 |
| Prdm14 | 3 | 10 | 1 | 3 | 3 | 0 | 2 | 4 | 4 | 1 | 1 | 1 | 1 | 1 | 0 | 12 | 2 | 7 | 4 | 3 | 4 | 7 | 2 | 2 | 4 | 0 | 0 | 4 | Prdm14 |
| Prdm15 | 1103 | 1250 | 1085 | 1052 | 1225 | 992 | 1158 | 3248 | 3111 | 2974 | 2766 | 3146 | 2825 | 3163 | 1025 | 968 | 1159 | 1162 | 1226 | 1167 | 1139 | 2984 | 2810 | 2971 | 3246 | 2473 | 2944 | 2922 | Prdm15 |
| Prdm16 | 1401 | 1748 | 1454 | 1905 | 2038 | 1934 | 1988 | 2646 | 3506 | 4128 | 3369 | 4286 | 3107 | 3910 | 1360 | 1397 | 1947 | 1910 | 2081 | 1553 | 1812 | 2664 | 2265 | 4178 | 3689 | 3922 | 3027 | 2940 | Prdm16 |
| Prdm2 | 8646 | 9842 | 10072 | 11953 | 11647 | 10378 | 10577 | 8466 | 8666 | 10126 | 9234 | 10094 | 9323 | 9982 | 8637 | 9119 | 11050 | 11312 | 11709 | 8747 | 11125 | 9829 | 8247 | 9792 | 10609 | 7937 | 9361 | 9373 | Prdm2 |
| Prdm4 | 6775 | 6504 | 6241 | 6029 | 5845 | 5816 | 5741 | 6064 | 5535 | 5436 | 4874 | 6015 | 5642 | 5926 | 6625 | 6122 | 6052 | 5905 | 5745 | 5551 | 6055 | 6168 | 5213 | 5204 | 5642 | 5723 | 5755 | 6036 | Prdm4 |
| Prdm5 | 429 | 422 | 595 | 347 | 338 | 438 | 248 | 830 | 907 | 720 | 970 | 785 | 978 | 796 | 480 | 542 | 411 | 311 | 262 | 387 | 302 | 805 | 1056 | 735 | 676 | 886 | 816 | 805 | Prdm5 |
| Prdm6 | 185 | 255 | 265 | 199 | 176 | 266 | 87 | 332 | 482 | 447 | 535 | 367 | 559 | 358 | 299 | 358 | 265 | 302 | 123 | 129 | 75 | 543 | 610 | 374 | 446 | 258 | 470 | 299 | Prdm6 |
| Prdm8 | 404 | 556 | 533 | 1273 | 1586 | 1285 | 1619 | 1340 | 1282 | 1819 | 1415 | 1614 | 991 | 1451 | 483 | 359 | 1004 | 1407 | 1782 | 1012 | 1608 | 1661 | 1055 | 1913 | 2347 | 1261 | 943 | 1076 | Prdm8 |
| Prdm9 | 1106 | 1219 | 1411 | 1760 | 1428 | 1395 | 1488 | 476 | 422 | 543 | 575 | 516 | 572 | 614 | 1146 | 1042 | 1504 | 1392 | 1375 | 1137 | 1500 | 551 | 379 | 599 | 540 | 443 | 601 | 572 | Prdm9 |
| Prdx1 | 23689 | 19422 | 18849 | 16015 | 12432 | 14386 | 16843 | 18387 | 16773 | 13226 | 14182 | 16755 | 17185 | 18500 | 23427 | 19865 | 16630 | 14279 | 14806 | 18752 | 15451 | 12818 | 17672 | 15545 | 13519 | 21709 | 18948 | 21500 | Prdx1 |
| Prdx2 | 40180 | 51079 | 44094 | 46622 | 42391 | 46849 | 46095 | 42636 | 43197 | 40926 | 37084 | 47141 | 45933 | 47109 | 42705 | 41033 | 46516 | 43565 | 50954 | 56660 | 45917 | 36095 | 40749 | 45989 | 46856 | 53149 | 50591 | 50618 | Prdx2 |
| Prdx3 | 4005 | 3782 | 3856 | 3088 | 2711 | 2940 | 3226 | 3926 | 3700 | 2817 | 3278 | 3405 | 3580 | 3545 | 4242 | 3956 | 3631 | 3059 | 3075 | 3695 | 2976 | 2969 | 3709 | 3121 | 3067 | 3770 | 3891 | 4036 | Prdx3 |
| Prdx4 | 5885 | 5070 | 5108 | 4075 | 3458 | 4141 | 4298 | 4456 | 4369 | 3422 | 4124 | 4425 | 4651 | 4611 | 5823 | 5354 | 4429 | 3791 | 3886 | 4700 | 4081 | 3461 | 4466 | 3829 | 3360 | 5267 | 4832 | 5622 | Prdx4 |
| Prdx5 | 3625 | 4292 | 4831 | 4461 | 4144 | 4004 | 3698 | 3549 | 3994 | 3942 | 3912 | 3816 | 3932 | 4035 | 3882 | 3913 | 4425 | 4391 | 4274 | 4282 | 3939 | 3233 | 4229 | 4242 | 4147 | 4296 | 4045 | 4241 | Prdx5 |
| Prdx6 | 9261 | 8393 | 7282 | 6392 | 5733 | 6462 | 7299 | 7211 | 6940 | 6099 | 6091 | 7232 | 7498 | 7799 | 8172 | 8252 | 7253 | 6247 | 6578 | 6906 | 6514 | 5147 | 7146 | 6676 | 6033 | 8917 | 7538 | 8679 | Prdx6 |
| Preb | 8628 | 9161 | 9350 | 9909 | 8961 | 9144 | 8912 | 6492 | 6736 | 7705 | 7047 | 7655 | 7040 | 7106 | 8268 | 8350 | 9667 | 9533 | 9334 | 8033 | 8880 | 7222 | 6348 | 7638 | 7592 | 7639 | 7313 | 7323 | Preb |
| Prelid1 | 9378 | 11501 | 10407 | 9537 | 8000 | 8734 | 7980 | 7612 | 8728 | 8275 | 8297 | 9358 | 9341 | 8076 | 8983 | 9984 | 9965 | 8482 | 9541 | 8907 | 7911 | 7359 | 9165 | 8534 | 8310 | 10784 | 10038 | 8968 | Prelid1 |
| Prelid2 | 493 | 327 | 379 | 222 | 155 | 297 | 206 | 267 | 264 | 229 | 282 | 250 | 282 | 329 | 510 | 504 | 229 | 216 | 220 | 263 | 227 | 214 | 331 | 171 | 138 | 284 | 288 | 343 | Prelid2 |
| Prelid3a | 843 | 948 | 1230 | 1120 | 1133 | 1172 | 1167 | 865 | 901 | 972 | 900 | 1101 | 988 | 1084 | 807 | 888 | 1102 | 1129 | 1272 | 1110 | 1079 | 894 | 784 | 1026 | 1146 | 941 | 1058 | 1057 | Prelid3a |
| Prelid3b | 3234 | 3533 | 3366 | 3272 | 3050 | 3030 | 3717 | 3226 | 3157 | 2847 | 2811 | 3289 | 3071 | 3830 | 3642 | 3226 | 3412 | 3056 | 3298 | 3770 | 3361 | 2770 | 3216 | 3331 | 3223 | 3730 | 3410 | 3580 | Prelid3b |
| Prelp | 175 | 255 | 173 | 246 | 352 | 299 | 150 | 600 | 778 | 656 | 898 | 434 | 709 | 374 | 245 | 235 | 189 | 266 | 345 | 210 | 214 | 1029 | 988 | 404 | 606 | 339 | 381 | 325 | Prelp |
| Prep | 7063 | 7090 | 7514 | 6429 | 6750 | 6757 | 6316 | 6620 | 6611 | 6724 | 7011 | 7405 | 7622 | 7435 | 6980 | 7404 | 7360 | 6643 | 6874 | 5971 | 6957 | 6818 | 6630 | 7080 | 7330 | 7430 | 7289 | 8014 | Prep |
| Prepl | 10997 | 11062 | 8909 | 10582 | 12368 | 11416 | 11977 | 10308 | 9568 | 9412 | 9228 | 8034 | 8523 | 8096 | 11350 | 10300 | 9318 | 10761 | 11404 | 12292 | 12292 | 10529 | 9572 | 7733 | 9571 | 5984 | 8280 | 7751 | Prepl |
| Prex1 | 6673 | 8489 | 9227 | 9270 | 9661 | 8242 | 7697 | 4587 | 5839 | 6028 | 5600 | 5537 | 5894 | 4734 | 6624 | 9514 | 9485 | 10072 | 9996 | 6711 | 8243 | 5583 | 5178 | 6262 | 6405 | 5040 | 4573 | 4485 | Prex1 |
| Prex2 | 1461 | 1036 | 826 | 706 | 839 | 1106 | 866 | 2000 | 2293 | 1322 | 1967 | 1694 | 2337 | 1822 | 1327 | 1390 | 708 | 711 | 729 | 1055 | 905 | 1357 | 2265 | 1172 | 1306 | 1165 | 2065 | 1653 | Prex2 |
| Prf1 | 0 | 0 | 0 | 0 | 0 | 0 | 0 | 0 | 0 | 0 | 1 | 0 | 0 | 0 | 0 | 1 | 0 | 0 | 0 | 0 | 0 | 0 | 1 | 0 | 0 | 0 | 0 | 0 | Prf1 |
| Prg2 | 16 | 12 | 28 | 13 | 2 | 4 | 3 | 2 | 0 | 6 | 40 | 29 | 9 | 21 | 5 | 18 | 10 | 8 | 7 | 7 | 6 | 2 | 0 | 10 | 0 | 24 | 1 | 2 | Prg2 |
| Prg3 | 2 | 0 | 3 | 0 | 0 | 0 | 1 | 0 | 1 | 0 | 3 | 0 | 1 | 1 | 3 | 1 | 0 | 0 | 2 | 0 | 0 | 0 | 0 | 0 | 0 | 1 | 0 | 0 | Prg3 |
| Prg4 | 8 | 4 | 16 | 2 | 5 | 2 | 2 | 13 | 25 | 14 | 190 | 0 | 5 | 7 | 1 | 5 | 2 | 0 | 5 | 5 | 2 | 7 | 27 | 11 | 1 | 5 | 6 | 4 | Prg4 |
| Prickle1 | 1910 | 1917 | 1733 | 1720 | 1803 | 1623 | 1208 | 1628 | 2128 | 1986 | 2620 | 1378 | 1888 | 1103 | 1792 | 2319 | 1601 | 1760 | 1317 | 1175 | 1281 | 1750 | 2208 | 1809 | 1406 | 1904 | 1538 | 1370 | Prickle1 |
| Prickle2 | 2700 | 2839 | 2783 | 3336 | 4377 | 3852 | 3313 | 3066 | 2911 | 3341 | 3351 | 2317 | 3316 | 2372 | 3162 | 3211 | 3146 | 3478 | 3538 | 3416 | 3858 | 4119 | 3180 | 2376 | 3347 | 1238 | 2821 | 2295 | Prickle2 |
| Prickle3 | 378 | 323 | 444 | 292 | 281 | 410 | 227 | 297 | 422 | 420 | 531 | 328 | 454 | 362 | 266 | 404 | 344 | 365 | 277 | 263 | 233 | 381 | 436 | 412 | 307 | 364 | 289 | 318 | Prickle3 |
| Prickle4 | 22 | 35 | 42 | 40 | 34 | 21 | 40 | 7 | 7 | 12 | 5 | 2 | 1 | 2 | 32 | 31 | 58 | 34 | 30 | 27 | 29 | 7 | 6 | 16 | 6 | 5 | 1 | 8 | Prickle4 |
| Prim1 | 6529 | 5010 | 5996 | 4829 | 3818 | 4794 | 5756 | 5294 | 4706 | 3944 | 4288 | 5561 | 4518 | 5773 | 6086 | 5176 | 5007 | 4303 | 5036 | 5771 | 4796 | 3542 | 4069 | 5108 | 4822 | 7061 | 6070 | 6438 | Prim1 |
| Prim2 | 4645 | 4545 | 4579 | 3931 | 2887 | 3041 | 3995 | 2611 | 2458 | 2139 | 2015 | 2810 | 2397 | 2823 | 4494 | 4051 | 4350 | 3626 | 3805 | 4242 | 3667 | 1751 | 2208 | 2469 | 2296 | 3283 | 2886 | 3017 | Prim2 |
| Prima1 | 81 | 96 | 207 | 112 | 115 | 126 | 21 | 84 | 95 | 67 | 106 | 116 | 170 | 101 | 80 | 101 | 137 | 116 | 128 | 60 | 97 | 107 | 111 | 74 | 114 | 37 | 207 | 167 | Prima1 |
| Primpol | 1356 | 1046 | 1066 | 1207 | 1021 | 1108 | 1199 | 1330 | 1548 | 1360 | 1438 | 1392 | 1349 | 1409 | 1357 | 1181 | 1107 | 1246 | 1105 | 1143 | 1171 | 1242 | 1522 | 1222 | 1246 | 1360 | 1362 | 1370 | Primpol |
| Prkaa1 | 3095 | 2334 | 2738 | 2684 | 2963 | 2831 | 3085 | 3666 | 3269 | 3113 | 3531 | 3267 | 3315 | 3245 | 3216 | 2816 | 2727 | 2642 | 2385 | 2864 | 2797 | 3621 | 3311 | 3097 | 3132 | 3082 | 3165 | 3313 | Prkaa1 |
| Prkaa2 | 1866 | 1503 | 1581 | 2292 | 3274 | 3069 | 3153 | 2538 | 2226 | 2602 | 2381 | 2002 | 2105 | 2106 | 2203 | 1758 | 1925 | 2432 | 2713 | 2637 | 3118 | 3175 | 2232 | 1887 | 2870 | 1001 | 1822 | 1775 | Prkaa2 |
| Prkab1 | 3011 | 3232 | 2659 | 2772 | 2337 | 2489 | 2439 | 2066 | 2400 | 2482 | 2247 | 2605 | 2474 | 2496 | 2701 | 2780 | 2633 | 2605 | 2368 | 2216 | 2342 | 2191 | 2353 | 2281 | 2309 | 2586 | 2352 | 2468 | Prkab1 |
| Prkab2 | 4436 | 4279 | 3899 | 4166 | 4010 | 4182 | 4179 | 2599 | 2319 | 2471 | 2684 | 2342 | 2510 | 2380 | 4601 | 4653 | 3897 | 4173 | 3575 | 3650 | 4046 | 2538 | 2447 | 2169 | 2485 | 2080 | 2395 | 2453 | Prkab2 |
| Prkaca | 12472 | 13491 | 14861 | 15358 | 13061 | 12146 | 13134 | 6130 | 6952 | 7076 | 8501 | 7957 | 7561 | 8448 | 11720 | 12442 | 15000 | 14574 | 13873 | 10438 | 13936 | 8224 | 7324 | 8346 | 8123 | 9018 | 7304 | 8491 | Prkaca |
| Prkacb | 13564 | 11776 | 11891 | 14058 | 15660 | 15443 | 16701 | 13854 | 13595 | 13768 | 12801 | 11989 | 11455 | 12929 | 13489 | 12523 | 12924 | 14857 | 15232 | 14290 | 17285 | 15313 | 13449 | 12314 | 14608 | 10915 | 11782 | 11794 | Prkacb |
| Prkag1 | 2767 | 2886 | 2940 | 2880 | 2542 | 2619 | 2816 | 3401 | 3509 | 3606 | 3154 | 3693 | 3409 | 3633 | 2777 | 2734 | 2859 | 2637 | 2782 | 2644 | 2689 | 3456 | 3551 | 3630 | 3578 | 4172 | 3668 | 3720 | Prkag1 |
| Prkag2 | 3539 | 3339 | 3750 | 3862 | 4877 | 4382 | 4844 | 4257 | 3592 | 4087 | 3635 | 4128 | 3662 | 4184 | 3603 | 3248 | 3994 | 4122 | 4750 | 4335 | 4921 | 4418 | 3627 | 3956 | 4584 | 3604 | 3905 | 4232 | Prkag2 |
| Prkag3 | 6 | 3 | 0 | 0 | 0 | 3 | 0 | 7 | 20 | 1 | 168 | 1 | 6 | 2 | 1 | 4 | 1 | 4 | 2 | 2 | 0 | 12 | 78 | 1 | 0 | 0 | 3 | 4 | Prkag3 |
| Prkar1a | 29121 | 26468 | 29823 | 31872 | 33738 | 31049 | 34030 | 35662 | 32098 | 32724 | 33519 | 32418 | 30640 | 33530 | 31018 | 26613 | 31382 | 33039 | 34542 | 32805 | 35226 | 37298 | 33179 | 34134 | 35690 | 32451 | 31749 | 33422 | Prkar1a |
| Prkar1b | 1243 | 1572 | 1188 | 2422 | 4547 | 3027 | 3285 | 4164 | 4643 | 5030 | 4105 | 2754 | 2982 | 2844 | 1704 | 1455 | 1869 | 3056 | 3421 | 3198 | 3816 | 6149 | 4454 | 3630 | 4800 | 2207 | 2458 | 2398 | Prkar1b |
| Prkar2a | 13064 | 12586 | 14320 | 14433 | 15002 | 14941 | 16063 | 10794 | 9934 | 11246 | 10614 | 11977 | 11183 | 11630 | 13375 | 11854 | 14994 | 14373 | 14744 | 14046 | 15236 | 11864 | 9432 | 10977 | 12034 | 10623 | 10906 | 11885 | Prkar2a |
| Prkar2b | 7465 | 6253 | 7669 | 9029 | 10945 | 10037 | 10746 | 10168 | 8221 | 9165 | 8100 | 7996 | 7463 | 8332 | 8290 | 7055 | 8157 | 9613 | 9222 | 9823 | 10325 | 11317 | 8356 | 8433 | 10219 | 7025 | 7406 | 7933 | Prkar2b |
| Prkca | 2433 | 2128 | 1641 | 3104 | 4534 | 3633 | 4368 | 2650 | 2774 | 2961 | 3104 | 1913 | 1797 | 1995 | 2471 | 2130 | 2373 | 3849 | 3832 | 3226 | 4653 | 3532 | 3109 | 2444 | 2973 | 1285 | 1823 | 1741 | Prkca |
| Prkcb | 1645 | 1744 | 1109 | 1983 | 2837 | 2694 | 2615 | 1962 | 1899 | 2163 | 1872 | 1555 | 1701 | 1560 | 1883 | 1604 | 1644 | 2135 | 2359 | 2478 | 2866 | 2181 | 1735 | 1363 | 2229 | 650 | 1527 | 1272 | Prkcb |
| Prkcd | 6040 | 6366 | 6562 | 6234 | 6217 | 5683 | 5683 | 2401 | 2834 | 2616 | 2899 | 2261 | 2734 | 2088 | 5712 | 6729 | 6488 | 6358 | 6152 | 5252 | 5591 | 2779 | 2718 | 2414 | 2322 | 2353 | 2024 | 2103 | Prkcd |
| Prkce | 2006 | 1828 | 1927 | 2978 | 5237 | 3489 | 4147 | 2924 | 3510 | 4002 | 3846 | 2029 | 2296 | 2163 | 2077 | 2238 | 2216 | 4252 | 3995 | 3257 | 4579 | 4695 | 3548 | 3004 | 4249 | 1726 | 1754 | 1830 | Prkce |
| Prkch | 944 | 750 | 728 | 570 | 718 | 748 | 506 | 785 | 888 | 833 | 949 | 752 | 824 | 673 | 795 | 962 | 663 | 764 | 597 | 549 | 528 | 872 | 979 | 757 | 654 | 817 | 674 | 627 | Prkch |
| Prkci | 12500 | 10242 | 10551 | 11219 | 9544 | 10213 | 11069 | 6224 | 5866 | 5513 | 5774 | 5843 | 5881 | 5853 | 11435 | 11173 | 10539 | 11205 | 9322 | 9746 | 10539 | 5447 | 5683 | 5725 | 5481 | 5864 | 5617 | 5656 | Prkci |
| Prkcq | 142 | 111 | 113 | 113 | 208 | 160 | 165 | 380 | 365 | 418 | 362 | 188 | 209 | 151 | 156 | 159 | 136 | 156 | 141 | 154 | 172 | 344 | 495 | 295 | 238 | 296 | 186 | 166 | Prkcq |
| Prkcsh | 6090 | 8230 | 8509 | 7816 | 6525 | 6583 | 6219 | 5376 | 6680 | 6795 | 7123 | 7075 | 7081 | 7460 | 5777 | 7809 | 7767 | 7270 | 7463 | 5929 | 6731 | 6527 | 6580 | 7385 | 6840 | 8385 | 6717 | 7477 | Prkcsh |
| Prkcz | 1556 | 1976 | 1740 | 2447 | 3347 | 2814 | 2493 | 2588 | 2783 | 3041 | 2570 | 2543 | 2211 | 2315 | 1898 | 1773 | 2071 | 2324 | 2887 | 2673 | 2869 | 3636 | 2585 | 2564 | 3543 | 1982 | 2112 | 2210 | Prkcz |
| Prkd1 | 1652 | 1420 | 1264 | 1793 | 1686 | 1533 | 2514 | 1130 | 1216 | 1297 | 1201 | 1463 | 1078 | 1529 | 1427 | 1196 | 1621 | 1834 | 2091 | 1615 | 2136 | 1026 | 1110 | 1488 | 1255 | 1681 | 1151 | 1222 | Prkd1 |
| Prkd2 | 1774 | 2208 | 2434 | 2197 | 1946 | 1934 | 1758 | 855 | 1173 | 1293 | 1369 | 1303 | 1242 | 1189 | 1642 | 2105 | 2388 | 2125 | 2265 | 1664 | 1902 | 1011 | 1285 | 1222 | 1162 | 1560 | 1198 | 1117 | Prkd2 |
| Prkd3 | 13499 | 11068 | 9422 | 9671 | 8876 | 10582 | 11158 | 12549 | 11028 | 9318 | 9937 | 12042 | 11474 | 12470 | 12530 | 11704 | 9368 | 9509 | 9426 | 11597 | 10467 | 8574 | 10661 | 9960 | 9841 | 10519 | 12452 | 12158 | Prkd3 |
| Prkdc | 5475 | 4779 | 2597 | 2838 | 2744 | 3725 | 3393 | 6052 | 5248 | 3374 | 3852 | 3947 | 4151 | 4157 | 4975 | 4867 | 2619 | 2616 | 2479 | 4107 | 3254 | 2785 | 4638 | 2820 | 3260 | 2715 | 4528 | 3690 | Prkdc |
| Prkg1 | 2969 | 2283 | 1901 | 1360 | 1309 | 1451 | 589 | 2269 | 2296 | 1900 | 2812 | 1733 | 3182 | 1686 | 3203 | 4322 | 1599 | 1520 | 781 | 778 | 681 | 2236 | 2641 | 1660 | 1716 | 1482 | 2055 | 2013 | Prkg1 |
| Prkg2 | 299 | 240 | 325 | 419 | 380 | 319 | 394 | 696 | 522 | 668 | 824 | 647 | 482 | 673 | 276 | 291 | 352 | 341 | 339 | 280 | 300 | 678 | 493 | 758 | 577 | 871 | 840 | 834 | Prkg2 |
| Prkn | 119 | 108 | 93 | 217 | 218 | 184 | 226 | 145 | 150 | 229 | 198 | 154 | 178 | 170 | 103 | 80 | 189 | 197 | 221 | 150 | 208 | 262 | 145 | 186 | 221 | 133 | 120 | 117 | Prkn |
| Prkra | 4424 | 4171 | 3575 | 3925 | 4021 | 3980 | 4375 | 4516 | 4004 | 3819 | 3473 | 4008 | 3834 | 4045 | 4781 | 4041 | 3912 | 3944 | 3882 | 4975 | 4115 | 4034 | 3924 | 3558 | 3936 | 4090 | 4011 | 4085 | Prkra |

| | | | | | | | | | | | | | | | | | | | | | | | | | | | | | |
|---|---|---|---|---|---|---|---|---|---|---|---|---|---|---|---|---|---|---|---|---|---|---|---|---|---|---|---|---|---|
| Prkrip1 | 2022 | 2374 | 2378 | 2465 | 2203 | 2217 | 2237 | 1901 | 1729 | 1981 | 1585 | 1993 | 1850 | 2009 | 2188 | 2099 | 2559 | 2402 | 2261 | 2376 | 2172 | 1744 | 1685 | 2058 | 1937 | 1834 | 2144 | 2148 | Prkrip1 |
| Prkx | 4516 | 3906 | 4941 | 6050 | 6019 | 5816 | 5830 | 3600 | 3342 | 3957 | 3674 | 3656 | 3623 | 3460 | 4615 | 4565 | 5561 | 6252 | 5834 | 4487 | 6018 | 4692 | 3078 | 3733 | 4417 | 3425 | 3185 | 3456 | Prkx |
| Prl | 0 | 0 | 0 | 0 | 0 | 0 | 0 | 1 | 0 | 0 | 0 | 0 | 0 | 0 | 0 | 0 | 0 | 0 | 0 | 0 | 0 | 0 | 1 | 0 | 0 | 0 | 0 | 0 | Prl |
| Prl2a1 | 0 | 0 | 0 | 0 | 0 | 0 | 0 | 0 | 9 | 0 | 0 | 0 | 0 | 0 | 0 | 0 | 0 | 0 | 0 | 0 | 0 | 0 | 0 | 0 | 0 | 0 | 0 | 0 | Prl2a1 |
| Prl2b1 | 0 | 0 | 0 | 0 | 0 | 0 | 0 | 0 | 13 | 0 | 0 | 0 | 0 | 0 | 0 | 0 | 0 | 0 | 0 | 0 | 0 | 0 | 6 | 0 | 0 | 0 | 0 | 0 | Prl2b1 |
| Prl2c2 | 0 | 0 | 0 | 0 | 0 | 0 | 2 | 0 | 3 | 0 | 0 | 0 | 0 | 0 | 0 | 0 | 0 | 1 | 0 | 0 | 0 | 0 | 0 | 0 | 0 | 0 | 0 | 0 | Prl2c2 |
| Prl2c3 | 0 | 0 | 0 | 0 | 0 | 0 | 0 | 0 | 9 | 0 | 0 | 0 | 0 | 0 | 0 | 0 | 0 | 0 | 0 | 0 | 0 | 0 | 5 | 0 | 0 | 0 | 0 | 0 | Prl2c3 |
| Prl2c5 | 0 | 0 | 0 | 0 | 0 | 0 | 0 | 0 | 5 | 0 | 0 | 0 | 0 | 0 | 0 | 0 | 0 | 0 | 0 | 0 | 0 | 0 | 0 | 0 | 0 | 0 | 0 | 0 | Prl2c5 |
| Prl3a1 | 0 | 0 | 0 | 0 | 0 | 0 | 0 | 0 | 2 | 0 | 0 | 0 | 0 | 0 | 0 | 0 | 0 | 0 | 0 | 0 | 0 | 0 | 1 | 0 | 0 | 0 | 0 | 0 | Prl3a1 |
| Prl3b1 | 0 | 1 | 0 | 0 | 0 | 0 | 0 | 1 | 35 | 0 | 0 | 4 | 3 | 0 | 0 | 2 | 0 | 0 | 0 | 0 | 0 | 0 | 12 | 0 | 1 | 3 | 2 | 1 | Prl3b1 |
| Prl3c1 | 0 | 0 | 0 | 0 | 0 | 0 | 0 | 0 | 1 | 0 | 0 | 0 | 0 | 0 | 0 | 0 | 0 | 0 | 0 | 0 | 0 | 0 | 0 | 0 | 0 | 0 | 0 | 0 | Prl3c1 |
| Prl3d1 | 0 | 0 | 0 | 0 | 0 | 0 | 0 | 0 | 0 | 0 | 2 | 0 | 0 | 2 | 1 | 1 | 0 | 0 | 0 | 0 | 0 | 0 | 0 | 0 | 0 | 0 | 0 | 0 | Prl3d1 |
| Prl3d2 | 0 | 0 | 0 | 0 | 1 | 0 | 0 | 0 | 0 | 2 | 0 | 0 | 0 | 0 | 1 | 0 | 3 | 0 | 0 | 0 | 0 | 0 | 0 | 0 | 0 | 1 | 0 | 0 | Prl3d2 |
| Prl3d3 | 0 | 0 | 0 | 0 | 0 | 0 | 0 | 0 | 0 | 0 | 0 | 0 | 1 | 0 | 0 | 0 | 0 | 0 | 0 | 0 | 0 | 0 | 0 | 0 | 0 | 0 | 0 | 0 | Prl3d3 |
| Prl5a1 | 0 | 0 | 0 | 0 | 3 | 0 | 0 | 0 | 0 | 0 | 0 | 0 | 0 | 0 | 0 | 0 | 0 | 0 | 0 | 0 | 0 | 0 | 0 | 0 | 0 | 0 | 0 | 0 | Prl5a1 |
| Prl6a1 | 1 | 0 | 0 | 0 | 2 | 0 | 0 | 0 | 1 | 0 | 0 | 0 | 0 | 0 | 0 | 0 | 0 | 0 | 0 | 0 | 0 | 0 | 0 | 0 | 0 | 0 | 0 | 0 | Prl6a1 |
| Prl7a1 | 0 | 0 | 0 | 0 | 0 | 0 | 0 | 0 | 0 | 0 | 0 | 0 | 0 | 0 | 0 | 0 | 0 | 0 | 0 | 0 | 0 | 0 | 2 | 0 | 0 | 0 | 0 | 0 | Prl7a1 |
| Prl7a2 | 0 | 0 | 0 | 0 | 0 | 0 | 0 | 0 | 13 | 0 | 0 | 0 | 0 | 0 | 0 | 0 | 0 | 1 | 0 | 0 | 0 | 0 | 5 | 0 | 0 | 0 | 0 | 0 | Prl7a2 |
| Prl7d1 | 0 | 0 | 0 | 0 | 0 | 0 | 0 | 0 | 2 | 0 | 0 | 0 | 0 | 0 | 0 | 0 | 0 | 0 | 0 | 0 | 0 | 0 | 1 | 0 | 0 | 0 | 0 | 0 | Prl7d1 |
| Prl8a1 | 0 | 0 | 0 | 0 | 0 | 0 | 0 | 0 | 1 | 0 | 0 | 0 | 0 | 0 | 0 | 0 | 0 | 0 | 0 | 0 | 0 | 0 | 0 | 0 | 0 | 0 | 0 | 0 | Prl8a1 |
| Prl8a2 | 0 | 0 | 0 | 2 | 0 | 0 | 12 | 2 | 8 | 2 | 0 | 0 | 0 | 0 | 0 | 0 | 0 | 2 | 0 | 2 | 0 | 0 | 3 | 0 | 0 | 0 | 0 | 0 | Prl8a2 |
| Prl8a6 | 0 | 0 | 0 | 0 | 0 | 0 | 0 | 0 | 1 | 0 | 0 | 0 | 0 | 0 | 0 | 0 | 0 | 0 | 0 | 0 | 0 | 0 | 0 | 0 | 0 | 0 | 0 | 0 | Prl8a6 |
| Prl8a8 | 0 | 0 | 0 | 0 | 0 | 0 | 0 | 1 | 6 | 0 | 0 | 1 | 0 | 0 | 0 | 0 | 0 | 0 | 0 | 0 | 0 | 0 | 6 | 0 | 0 | 0 | 0 | 0 | Prl8a8 |
| Prl8a9 | 0 | 0 | 0 | 0 | 0 | 0 | 0 | 0 | 30 | 0 | 0 | 0 | 0 | 0 | 0 | 0 | 0 | 0 | 0 | 0 | 0 | 0 | 6 | 0 | 0 | 0 | 0 | 0 | Prl8a9 |
| Prlh | 0 | 0 | 0 | 0 | 0 | 0 | 0 | 0 | 0 | 0 | 0 | 0 | 0 | 0 | 0 | 2 | 0 | 2 | 0 | 0 | 0 | 0 | 0 | 0 | 0 | 0 | 0 | 0 | Prlh |
| Prlhr | 29 | 34 | 30 | 51 | 56 | 38 | 42 | 4 | 8 | 9 | 15 | 4 | 3 | 5 | 15 | 29 | 49 | 43 | 57 | 37 | 63 | 7 | 3 | 9 | 1 | 1 | 5 | 4 | Prlhr |
| Prlr | 51 | 30 | 20 | 35 | 45 | 35 | 47 | 107 | 117 | 95 | 80 | 29 | 35 | 47 | 50 | 44 | 28 | 14 | 18 | 32 | 38 | 77 | 115 | 65 | 53 | 34 | 19 | 34 | Prlr |
| Prm1 | 0 | 0 | 0 | 0 | 0 | 0 | 0 | 0 | 0 | 0 | 0 | 0 | 0 | 0 | 0 | 1 | 0 | 0 | 0 | 0 | 0 | 0 | 1 | 0 | 0 | 0 | 0 | 0 | Prm1 |
| Prmt1 | 16296 | 19935 | 18604 | 17494 | 16154 | 16750 | 15811 | 19848 | 20725 | 20028 | 18108 | 23736 | 21177 | 21923 | 16150 | 16049 | 17980 | 16005 | 19530 | 19103 | 17051 | 17396 | 19527 | 22289 | 21710 | 27901 | 25679 | 24534 | Prmt1 |
| Prmt2 | 3224 | 4582 | 3994 | 5334 | 5821 | 5048 | 4930 | 5970 | 5726 | 7026 | 5804 | 5570 | 4859 | 5367 | 3690 | 3642 | 4949 | 5096 | 5861 | 5425 | 5168 | 6487 | 5768 | 6327 | 7071 | 5066 | 5108 | 5015 | Prmt2 |
| Prmt3 | 1378 | 1455 | 1453 | 1245 | 1176 | 1184 | 1309 | 2298 | 2042 | 1934 | 1956 | 2217 | 2175 | 2226 | 1382 | 1259 | 1350 | 1121 | 1112 | 1315 | 1151 | 1910 | 2048 | 1995 | 1918 | 2075 | 2366 | 2392 | Prmt3 |
| Prmt5 | 10813 | 12442 | 10408 | 10748 | 9186 | 9951 | 10086 | 8304 | 8286 | 7708 | 7037 | 8837 | 8410 | 9439 | 10582 | 10043 | 10337 | 9874 | 11321 | 10221 | 11112 | 7442 | 7972 | 8409 | 8259 | 10163 | 9556 | 10300 | Prmt5 |
| Prmt6 | 1879 | 1551 | 1754 | 1810 | 1701 | 1711 | 1758 | 2210 | 1964 | 1968 | 1930 | 2116 | 2165 | 2054 | 1825 | 1569 | 1622 | 1661 | 1763 | 1821 | 1589 | 2082 | 1973 | 1862 | 2062 | 2172 | 1969 | 2013 | Prmt6 |
| Prmt7 | 3524 | 4180 | 3521 | 3751 | 3451 | 3542 | 3226 | 4020 | 3896 | 3672 | 3621 | 4137 | 4325 | 4007 | 3571 | 3462 | 3593 | 3311 | 3833 | 3684 | 3461 | 3517 | 4105 | 3854 | 4129 | 4627 | 4485 | 4441 | Prmt7 |
| Prmt8 | 837 | 1223 | 1451 | 1552 | 2109 | 1614 | 1398 | 1939 | 1634 | 2355 | 1798 | 2372 | 2938 | 1986 | 1273 | 1124 | 1803 | 1391 | 1944 | 1723 | 1643 | 2585 | 1808 | 1999 | 2493 | 1812 | 2430 | 2262 | Prmt8 |
| Prmt9 | 3075 | 2986 | 2570 | 2786 | 2635 | 2795 | 2777 | 2881 | 2413 | 2047 | 2396 | 2073 | 2082 | 2243 | 3376 | 3205 | 2576 | 2674 | 2501 | 3009 | 2692 | 2309 | 2327 | 2097 | 2209 | 2247 | 2205 | 2148 | Prmt9 |
| Prn | 0 | 5 | 0 | 6 | 0 | 1 | 7 | 0 | 1 | 3 | 1 | 0 | 2 | 0 | 1 | 0 | 0 | 1 | 0 | 1 | 0 | 0 | 1 | 0 | 0 | 0 | 0 | 6 | Prn |
| Prnd | 0 | 1 | 0 | 2 | 1 | 5 | 3 | 0 | 3 | 0 | 0 | 0 | 0 | 0 | 0 | 0 | 0 | 0 | 0 | 0 | 1 | 2 | 1 | 0 | 5 | 0 | 0 | 0 | Prnd |
| Prnp | 5807 | 5566 | 4778 | 6227 | 6332 | 6039 | 6972 | 4515 | 4226 | 4307 | 3977 | 3707 | 3154 | 3693 | 6348 | 5247 | 5679 | 6269 | 6347 | 6595 | 6892 | 4963 | 4412 | 4037 | 4471 | 3770 | 3581 | 3402 | Prnp |
| Prob1 | 1013 | 1315 | 1554 | 1609 | 1374 | 1209 | 1178 | 552 | 547 | 662 | 928 | 677 | 609 | 633 | 1096 | 1206 | 1696 | 1502 | 1404 | 1074 | 1214 | 807 | 641 | 646 | 657 | 649 | 646 | 602 | Prob1 |
| Proc | 1 | 6 | 5 | 0 | 2 | 4 | 0 | 1 | 7 | 12 | 8 | 0 | 6 | 3 | 1 | 3 | 5 | 4 | 0 | 0 | 0 | 3 | 5 | 2 | 5 | 5 | 7 | 7 | Proc |
| Proca1 | 113 | 131 | 129 | 123 | 101 | 108 | 68 | 73 | 91 | 138 | 124 | 96 | 91 | 68 | 90 | 146 | 127 | 108 | 79 | 90 | 72 | 107 | 145 | 83 | 74 | 83 | 86 | 87 | Proca1 |
| Procr | 85 | 92 | 136 | 95 | 108 | 102 | 72 | 104 | 166 | 171 | 209 | 108 | 180 | 117 | 105 | 121 | 110 | 128 | 83 | 103 | 74 | 162 | 239 | 130 | 151 | 154 | 151 | 114 | Procr |
| Prodh2 | 0 | 2 | 0 | 0 | 7 | 2 | 2 | 0 | 5 | 0 | 0 | 0 | 0 | 0 | 1 | 5 | 2 | 0 | 1 | 1 | 1 | 0 | 0 | 0 | 0 | 0 | 0 | 0 | Prodh2 |
| Prok1 | 102 | 104 | 115 | 175 | 150 | 123 | 137 | 48 | 46 | 54 | 47 | 71 | 67 | 71 | 94 | 93 | 95 | 141 | 118 | 84 | 154 | 63 | 28 | 37 | 84 | 31 | 48 | 57 | Prok1 |
| Prok2 | 8 | 15 | 11 | 5 | 14 | 13 | 15 | 10 | 13 | 17 | 15 | 16 | 9 | 18 | 7 | 5 | 25 | 20 | 21 | 19 | 16 | 20 | 12 | 20 | 26 | 20 | 7 | 20 | Prok2 |
| Prokr1 | 1219 | 1400 | 1091 | 1059 | 1024 | 1088 | 1139 | 572 | 561 | 500 | 870 | 686 | 687 | 824 | 1294 | 1012 | 1161 | 1088 | 1203 | 1298 | 1143 | 383 | 733 | 537 | 615 | 650 | 702 | 748 | Prokr1 |
| Prokr2 | 601 | 492 | 454 | 549 | 744 | 756 | 834 | 545 | 425 | 623 | 621 | 404 | 254 | 427 | 600 | 472 | 516 | 647 | 726 | 736 | 849 | 707 | 497 | 554 | 772 | 625 | 443 | 452 | Prokr2 |
| Prom1 | 7451 | 7106 | 7482 | 4768 | 4608 | 6102 | 5122 | 8254 | 6411 | 6207 | 5308 | 8987 | 9082 | 8248 | 8461 | 7006 | 5825 | 4799 | 5375 | 6836 | 5284 | 5915 | 5949 | 6409 | 6114 | 9035 | 10191 | 9545 | Prom1 |
| Prom2 | 151 | 84 | 58 | 21 | 17 | 61 | 0 | 149 | 220 | 182 | 141 | 101 | 134 | 85 | 95 | 117 | 13 | 68 | 9 | 2 | 0 | 53 | 251 | 141 | 15 | 89 | 71 | 103 | Prom2 |
| Prop1 | 5 | 3 | 0 | 0 | 13 | 0 | 0 | 0 | 23 | 0 | 17 | 0 | 0 | 0 | 7 | 1 | 0 | 3 | 0 | 0 | 0 | 48 | 16 | 0 | 0 | 0 | 0 | 0 | Prop1 |
| Prorp | 859 | 735 | 818 | 605 | 711 | 628 | 772 | 1180 | 1126 | 984 | 1019 | 1187 | 1187 | 1260 | 807 | 697 | 630 | 632 | 650 | 753 | 702 | 1096 | 1164 | 1204 | 1010 | 1300 | 1320 | 1273 | Prorp |
| Prorsd1 | 355 | 258 | 403 | 261 | 286 | 231 | 304 | 438 | 441 | 444 | 503 | 471 | 500 | 561 | 390 | 291 | 328 | 251 | 207 | 235 | 229 | 366 | 531 | 469 | 351 | 591 | 526 | 590 | Prorsd1 |
| Pros1 | 1631 | 1154 | 1512 | 901 | 1012 | 1226 | 816 | 1768 | 1845 | 1382 | 1964 | 1527 | 1734 | 1518 | 1437 | 1659 | 1163 | 1055 | 819 | 975 | 771 | 1420 | 2128 | 1371 | 1180 | 1624 | 1629 | 1772 | Pros1 |
| Proser1 | 4364 | 4698 | 4944 | 5920 | 5231 | 5136 | 5321 | 3440 | 3845 | 4479 | 4143 | 5558 | 4509 | 4965 | 4072 | 4815 | 5917 | 5993 | 5465 | 4099 | 5402 | 4756 | 3407 | 4804 | 4941 | 5241 | 4656 | 4817 | Proser1 |
| Proser2 | 352 | 326 | 289 | 330 | 363 | 301 | 242 | 182 | 238 | 161 | 217 | 115 | 163 | 119 | 316 | 389 | 363 | 347 | 330 | 296 | 274 | 165 | 259 | 166 | 158 | 134 | 128 | 103 | Proser2 |
| Proser3 | 578 | 652 | 803 | 797 | 848 | 723 | 742 | 1231 | 1422 | 1606 | 1370 | 1804 | 1372 | 1712 | 447 | 562 | 913 | 750 | 926 | 633 | 768 | 1417 | 1202 | 1902 | 1624 | 1904 | 1577 | 1557 | Proser3 |
| Prox1 | 3325 | 3335 | 4083 | 4919 | 5192 | 5188 | 5613 | 3228 | 2805 | 3499 | 3510 | 4429 | 2866 | 4560 | 3767 | 3009 | 4919 | 4906 | 5143 | 3987 | 5612 | 4725 | 3161 | 3818 | 4298 | 3240 | 4070 | 4366 | Prox1 |
| Prox2 | 112 | 121 | 116 | 75 | 155 | 79 | 56 | 115 | 134 | 119 | 205 | 117 | 167 | 131 | 167 | 207 | 110 | 112 | 62 | 55 | 77 | 174 | 149 | 92 | 144 | 130 | 100 | 103 | Prox2 |
| Proz | 5 | 18 | 18 | 12 | 16 | 11 | 11 | 4 | 8 | 13 | 6 | 3 | 7 | 6 | 12 | 23 | 20 | 11 | 10 | 13 | 17 | 5 | 3 | 2 | 15 | 8 | 5 | 1 | Proz |
| Prpf18 | 6828 | 6694 | 6918 | 7344 | 6610 | 6423 | 7830 | 6283 | 5478 | 5514 | 5088 | 6099 | 5755 | 6413 | 7032 | 6398 | 6760 | 7056 | 6696 | 6956 | 7125 | 5842 | 5330 | 5955 | 5885 | 6090 | 5779 | 6006 | Prpf18 |
| Prpf19 | 21022 | 23013 | 19974 | 19108 | 17175 | 18238 | 17999 | 17396 | 16845 | 16212 | 15438 | 18508 | 17312 | 18633 | 20057 | 18913 | 19470 | 17353 | 19840 | 19572 | 18967 | 15373 | 17144 | 17680 | 17121 | 21033 | 19501 | 19988 | Prpf19 |
| Prpf3 | 5733 | 5808 | 5800 | 5515 | 4424 | 4689 | 4902 | 4455 | 3905 | 3706 | 3767 | 4553 | 4271 | 4518 | 5421 | 5267 | 5591 | 5042 | 4901 | 5343 | 4843 | 3910 | 3682 | 4453 | 4161 | 5142 | 4580 | 4700 | Prpf3 |
| Prpf31 | 7850 | 8526 | 8702 | 6820 | 6011 | 6324 | 6643 | 6851 | 6367 | 6629 | 5831 | 8125 | 7520 | 8567 | 7750 | 7273 | 7671 | 6170 | 6798 | 7219 | 6676 | 5746 | 6214 | 7269 | 6830 | 9480 | 8427 | 9176 | Prpf31 |
| Prpf38a | 5750 | 5228 | 5950 | 4674 | 4042 | 4463 | 4884 | 5718 | 4792 | 4450 | 4277 | 5086 | 4668 | 5137 | 5867 | 5220 | 5019 | 4556 | 4623 | 5535 | 4568 | 4286 | 4626 | 4759 | 4864 | 5538 | 5348 | 5700 | Prpf38a |
| Prpf38b | 12507 | 10563 | 14365 | 13584 | 12302 | 12762 | 13211 | 12887 | 11223 | 12484 | 12652 | 13702 | 12981 | 14874 | 12388 | 11465 | 13245 | 12882 | 12327 | 12107 | 12975 | 12562 | 11037 | 13603 | 12656 | 13619 | 13594 | 13794 | Prpf38b |
| Prpf39 | 11808 | 9056 | 11464 | 13338 | 11549 | 12567 | 14003 | 10220 | 8554 | 9529 | 10577 | 10198 | 9829 | 10528 | 12196 | 11291 | 11506 | 12923 | 11621 | 12611 | 12952 | 10098 | 8253 | 9983 | 9656 | 10380 | 9843 | 10537 | Prpf39 |
| Prpf4 | 5619 | 5148 | 5040 | 4671 | 4021 | 4119 | 4719 | 4051 | 3643 | 3381 | 3575 | 4328 | 4065 | 4389 | 4975 | 4976 | 4733 | 4479 | 4186 | 4455 | 4152 | 3594 | 3603 | 3758 | 3848 | 4648 | 4630 | 4794 | Prpf4 |
| Prpf40a | 18200 | 13106 | 15704 | 14591 | 14659 | 16225 | 17895 | 20539 | 17214 | 16397 | 16670 | 20234 | 19525 | 21817 | 17529 | 15105 | 14412 | 14311 | 14869 | 16783 | 16039 | 16446 | 16600 | 17784 | 17558 | 18952 | 21387 | 21447 | Prpf40a |
| Prpf40b | 5001 | 5922 | 6178 | 7155 | 7202 | 6718 | 6681 | 5350 | 6196 | 7485 | 6757 | 6887 | 6410 | 6548 | 4436 | 5329 | 7029 | 7042 | 7757 | 5997 | 6751 | 6605 | 5710 | 7028 | 7357 | 6352 | 6328 | 6093 | Prpf40b |
| Prpf4b | 23914 | 20500 | 19428 | 21495 | 21090 | 21998 | 23865 | 28298 | 22836 | 21488 | 23054 | 23998 | 22576 | 25533 | 24195 | 21732 | 20277 | 21389 | 20996 | 25559 | 23239 | 23088 | 22533 | 22496 | 22688 | 19860 | 23696 | 23917 | Prpf4b |
| Prpf6 | 11385 | 12705 | 11534 | 10351 | 9050 | 9664 | 9821 | 8061 | 7937 | 7271 | 7470 | 8110 | 7630 | 8349 | 11430 | 11390 | 10786 | 10599 | 10078 | 10803 | 10114 | 7441 | 7551 | 7846 | 7984 | 8555 | 8928 | 8348 | Prpf6 |
| Prpf8 | 40910 | 41452 | 30341 | 34036 | 32339 | 35902 | 34170 | 35077 | 40275 | 38323 | 34025 | 44127 | 40419 | 41728 | 37631 | 37608 | 33581 | 33066 | 36311 | 31409 | 37282 | 35790 | 35460 | 36470 | 38451 | 30650 | 42528 | 42464 | Prpf8 |
| Prph | 762 | 1146 | 214 | 578 | 4124 | 465 | 324 | 2571 | 3388 | 2667 | 4007 | 178 | 2315 | 167 | 1383 | 1611 | 803 | 1908 | 577 | 485 | 452 | 4226 | 2716 | 399 | 2345 | 217 | 218 | 188 | Prph |
| Prph2 | 1 | 0 | 0 | 0 | 3 | 3 | 0 | 0 | 0 | 7 | 0 | 0 | 0 | 0 | 1 | 0 | 0 | 0 | 2 | 0 | 4 | 0 | 3 | 0 | 4 | 3 | 0 | 0 | Prph2 |
| Prps1 | 5208 | 4417 | 4660 | 4103 | 4156 | 4105 | 4779 | 3971 | 3226 | 2899 | 2905 | 3465 | 3301 | 3777 | 5432 | 4139 | 4417 | 3986 | 4479 | 4864 | 4400 | 3288 | 3179 | 3206 | 3057 | 3859 | 3748 | 3852 | Prps1 |
| Prps1l3 | 2960 | 2404 | 2704 | 2170 | 2064 | 2097 | 2223 | 3390 | 2766 | 2238 | 2227 | 2581 | 2548 | 2781 | 2821 | 2482 | 2257 | 1988 | 2200 | 2541 | 2176 | 2232 | 2554 | 2625 | 2118 | 2691 | 2821 | 2860 | Prps1l3 |
| Prps2 | 2700 | 2158 | 2516 | 1895 | 2110 | 2087 | 2205 | 2220 | 2064 | 1848 | 1693 | 2401 | 2217 | 2231 | 2531 | 2496 | 2208 | 1877 | 1898 | 2123 | 1957 | 1953 | 2508 | 1965 | 1593 | 2612 | 2334 | 2331 | Prps2 |
| Prpsap1 | 5380 | 6265 | 5818 | 5725 | 5554 | 5309 | 5710 | 4289 | 4762 | 5303 | 4833 | 5326 | 5075 | 5379 | 4925 | 5948 | 6141 | 5710 | 5949 | 5014 | 5694 | 4654 | 4646 | 5288 | 5299 | 5497 | 5188 | 5536 | Prpsap1 |

| | | | | | | | | | | | | | | | | | | | | | | | | | | | | | |
|---|---|---|---|---|---|---|---|---|---|---|---|---|---|---|---|---|---|---|---|---|---|---|---|---|---|---|---|---|---|
| Prpsap2 | 4803 | 4807 | 4767 | 4540 | 4383 | 4490 | 5379 | 4323 | 3932 | 3713 | 3347 | 4594 | 3840 | 4772 | 5230 | 4332 | 5129 | 4138 | 4842 | 5116 | 4980 | 3725 | 3981 | 4183 | 3870 | 4643 | 4516 | 4682 | Prpsap2 |
| Prr11 | 3898 | 3527 | 4776 | 3553 | 2750 | 2963 | 3389 | 2873 | 2693 | 2853 | 2724 | 4116 | 3549 | 4268 | 3508 | 3348 | 4259 | 3287 | 3403 | 2673 | 3283 | 2607 | 2480 | 3584 | 3085 | 4736 | 3904 | 4171 | Prr11 |
| Prr12 | 4056 | 5472 | 7417 | 7628 | 7918 | 6016 | 5421 | 5561 | 7104 | 9403 | 8539 | 10012 | 8005 | 8754 | 3758 | 4335 | 7990 | 7356 | 8552 | 4526 | 6540 | 9870 | 6245 | 10739 | 10190 | 11333 | 7537 | 7971 | Prr12 |
| Prr13 | 1566 | 1897 | 1751 | 1757 | 1808 | 1449 | 1725 | 1834 | 1906 | 1779 | 1643 | 1887 | 2304 | 2229 | 1791 | 1780 | 1920 | 1834 | 1758 | 1774 | 1676 | 1878 | 1862 | 2072 | 2079 | 2040 | 1900 | 2243 | Prr13 |
| Prr14 | 3393 | 4063 | 4279 | 3976 | 3677 | 3482 | 3672 | 3385 | 3371 | 4015 | 3241 | 4341 | 3677 | 3992 | 3227 | 3249 | 4235 | 3744 | 4212 | 3462 | 3539 | 3649 | 3016 | 4194 | 3950 | 4312 | 4004 | 4003 | Prr14 |
| Prr14l | 5861 | 5472 | 3413 | 5082 | 5323 | 5673 | 5523 | 5351 | 4646 | 4648 | 4751 | 4743 | 4755 | 4670 | 5677 | 5494 | 4036 | 5116 | 4580 | 5458 | 5661 | 4654 | 4449 | 3824 | 4956 | 2868 | 4864 | 4402 | Prr14l |
| Prr15 | 689 | 639 | 570 | 825 | 578 | 570 | 856 | 253 | 228 | 361 | 214 | 338 | 381 | 468 | 542 | 433 | 657 | 604 | 717 | 723 | 705 | 124 | 245 | 446 | 287 | 673 | 318 | 602 | Prr15 |
| Prr15l | 29 | 6 | 7 | 16 | 8 | 1 | 0 | 37 | 33 | 38 | 20 | 5 | 9 | 3 | 16 | 19 | 0 | 3 | 0 | 0 | 0 | 12 | 39 | 41 | 0 | 30 | 3 | 8 | Prr15l |
| Prr16 | 269 | 131 | 164 | 146 | 148 | 150 | 105 | 385 | 476 | 338 | 484 | 149 | 265 | 178 | 182 | 207 | 139 | 200 | 89 | 135 | 135 | 252 | 434 | 306 | 199 | 186 | 136 | 179 | Prr16 |
| Prr18 | 90 | 134 | 149 | 164 | 435 | 252 | 278 | 772 | 855 | 1018 | 731 | 679 | 514 | 698 | 119 | 97 | 192 | 205 | 336 | 238 | 323 | 1125 | 808 | 873 | 1107 | 504 | 483 | 555 | Prr18 |
| Prr19 | 55 | 83 | 82 | 78 | 66 | 78 | 35 | 42 | 52 | 52 | 50 | 53 | 44 | 44 | 50 | 65 | 80 | 89 | 79 | 71 | 65 | 40 | 40 | 52 | 51 | 47 | 62 | 39 | Prr19 |
| Prr22 | 54 | 63 | 85 | 50 | 60 | 60 | 51 | 62 | 59 | 79 | 60 | 64 | 79 | 96 | 24 | 53 | 53 | 63 | 83 | 62 | 49 | 58 | 54 | 63 | 93 | 84 | 73 | 76 | Prr22 |
| Prr23a1 | 6 | 12 | 16 | 8 | 4 | 2 | 2 | 0 | 1 | 3 | 1 | 0 | 3 | 2 | 11 | 12 | 7 | 0 | 5 | 3 | 3 | 5 | 0 | 1 | 1 | 1 | 1 | 2 | Prr23a1 |
| Prr23a2 | 0 | 0 | 1 | 0 | 1 | 6 | 0 | 0 | 1 | 0 | 0 | 0 | 0 | 0 | 0 | 1 | 0 | 0 | 0 | 0 | 0 | 0 | 0 | 0 | 0 | 0 | 0 | 0 | Prr23a2 |
| Prr23a3 | 7 | 7 | 6 | 1 | 5 | 7 | 3 | 1 | 4 | 1 | 3 | 0 | 1 | 0 | 11 | 5 | 13 | 6 | 2 | 3 | 1 | 0 | 0 | 2 | 0 | 2 | 1 | 1 | Prr23a3 |
| Prr29 | 0 | 5 | 1 | 2 | 2 | 0 | 0 | 2 | 1 | 3 | 4 | 1 | 5 | 0 | 7 | 0 | 1 | 4 | 0 | 2 | 1 | 0 | 8 | 2 | 1 | 3 | 6 | 4 | Prr29 |
| Prr3 | 2825 | 3473 | 3161 | 3595 | 3045 | 3131 | 3063 | 2256 | 2688 | 2732 | 2675 | 2852 | 2384 | 2535 | 2822 | 3192 | 3369 | 3598 | 3034 | 2893 | 3001 | 2973 | 2445 | 2695 | 2707 | 2864 | 2508 | 2514 | Prr3 |
| Prr30 | 0 | 0 | 0 | 0 | 0 | 0 | 0 | 1 | 1 | 0 | 0 | 0 | 0 | 0 | 0 | 0 | 0 | 0 | 0 | 0 | 0 | 2 | 1 | 0 | 0 | 0 | 0 | 0 | Prr30 |
| Prr32 | 35 | 41 | 28 | 31 | 70 | 79 | 19 | 114 | 138 | 146 | 623 | 81 | 126 | 26 | 54 | 55 | 5 | 23 | 37 | 39 | 38 | 172 | 401 | 47 | 69 | 15 | 19 | 5 | Prr32 |
| Prr33 | 5 | 10 | 17 | 13 | 11 | 15 | 11 | 32 | 36 | 21 | 19 | 20 | 28 | 24 | 25 | 14 | 10 | 10 | 6 | 14 | 20 | 30 | 32 | 21 | 24 | 22 | 29 | 9 | Prr33 |
| Prr36 | 2320 | 2993 | 4151 | 4182 | 5429 | 4114 | 4100 | 5580 | 5946 | 8844 | 6499 | 9664 | 6793 | 8316 | 2280 | 2150 | 4795 | 4064 | 5876 | 3412 | 4345 | 10347 | 5377 | 9435 | 9853 | 9277 | 7174 | 7849 | Prr36 |
| Prr5 | 977 | 1374 | 1446 | 1181 | 1069 | 1016 | 755 | 355 | 548 | 523 | 703 | 411 | 552 | 383 | 885 | 1286 | 1276 | 1310 | 1070 | 830 | 868 | 556 | 653 | 473 | 471 | 543 | 493 | 436 | Prr5 |
| Prr5l | 536 | 454 | 555 | 281 | 334 | 387 | 289 | 866 | 852 | 763 | 832 | 1089 | 962 | 923 | 426 | 495 | 442 | 349 | 347 | 283 | 317 | 652 | 882 | 916 | 681 | 1240 | 1032 | 1178 | Prr5l |
| Prr7 | 271 | 345 | 344 | 417 | 395 | 344 | 316 | 469 | 558 | 634 | 613 | 726 | 523 | 554 | 264 | 238 | 366 | 414 | 418 | 309 | 366 | 668 | 484 | 721 | 709 | 752 | 636 | 541 | Prr7 |
| Prr9 | 0 | 0 | 0 | 0 | 0 | 0 | 0 | 0 | 1 | 0 | 0 | 0 | 0 | 1 | 0 | 1 | 0 | 0 | 0 | 0 | 0 | 0 | 0 | 0 | 0 | 0 | 0 | 0 | Prr9 |
| Prrc1 | 5313 | 5021 | 6125 | 6232 | 5677 | 5318 | 5046 | 3643 | 4467 | 4555 | 5489 | 4629 | 5012 | 4348 | 5073 | 5778 | 5873 | 6241 | 5335 | 4254 | 5089 | 4732 | 4608 | 4873 | 4604 | 5007 | 4431 | 4688 | Prrc1 |
| Prrc2a | 31238 | 43086 | 55368 | 54852 | 48195 | 43609 | 36105 | 28992 | 37592 | 48779 | 45090 | 56416 | 44430 | 47345 | 28288 | 36012 | 55963 | 52721 | 52786 | 32394 | 42290 | 45654 | 34354 | 55318 | 50660 | 60098 | 44337 | 44930 | Prrc2a |
| Prrc2b | 24314 | 27238 | 33334 | 41818 | 46590 | 40005 | 36930 | 32425 | 39113 | 52602 | 47223 | 53333 | 44100 | 46361 | 22826 | 27506 | 39591 | 42619 | 45550 | 29712 | 40712 | 56335 | 35474 | 51591 | 56367 | 32532 | 41946 | 43441 | Prrc2b |
| Prrc2c | 36440 | 33188 | 23552 | 31373 | 32540 | 37385 | 32962 | 33088 | 35104 | 34342 | 36935 | 36114 | 37426 | 36820 | 34888 | 36409 | 25540 | 30957 | 29346 | 32134 | 35466 | 27795 | 32212 | 27660 | 35826 | 23233 | 38420 | 31991 | Prrc2c |
| Prrg1 | 945 | 538 | 743 | 668 | 726 | 724 | 855 | 1071 | 1124 | 921 | 1176 | 876 | 1016 | 962 | 874 | 783 | 745 | 698 | 680 | 837 | 753 | 1008 | 1209 | 927 | 882 | 1050 | 932 | 964 | Prrg1 |
| Prrg2 | 291 | 331 | 256 | 266 | 251 | 273 | 227 | 554 | 625 | 594 | 663 | 603 | 679 | 672 | 280 | 303 | 319 | 298 | 221 | 238 | 234 | 523 | 603 | 661 | 535 | 707 | 651 | 662 | Prrg2 |
| Prrg3 | 2402 | 1991 | 2147 | 2176 | 2568 | 2775 | 2221 | 3228 | 3257 | 3279 | 3607 | 2864 | 3107 | 2758 | 2197 | 2252 | 2519 | 2270 | 2327 | 2290 | 2231 | 3768 | 3478 | 2878 | 3088 | 2227 | 2895 | 2737 | Prrg3 |
| Prrg4 | 556 | 364 | 268 | 231 | 195 | 221 | 166 | 137 | 153 | 98 | 116 | 85 | 133 | 76 | 411 | 506 | 273 | 229 | 181 | 202 | 220 | 70 | 186 | 95 | 66 | 61 | 64 | 81 | Prrg4 |
| Prrt1 | 945 | 1333 | 1650 | 2345 | 2553 | 2124 | 2385 | 1033 | 975 | 1742 | 1109 | 1449 | 976 | 1519 | 1052 | 929 | 2179 | 2539 | 2784 | 1914 | 2665 | 1920 | 992 | 1699 | 1649 | 1107 | 1216 | 1371 | Prrt1 |
| Prrt2 | 68 | 75 | 98 | 69 | 51 | 72 | 50 | 56 | 72 | 85 | 100 | 70 | 94 | 85 | 69 | 89 | 66 | 101 | 55 | 48 | 59 | 67 | 99 | 112 | 95 | 75 | 67 | 97 | Prrt2 |
| Prrt3 | 293 | 233 | 341 | 335 | 421 | 350 | 315 | 165 | 224 | 291 | 235 | 176 | 215 | 158 | 223 | 303 | 318 | 362 | 383 | 332 | 320 | 277 | 190 | 244 | 340 | 184 | 139 | 179 | Prrt3 |
| Prrt4 | 196 | 272 | 316 | 247 | 430 | 313 | 294 | 161 | 251 | 309 | 339 | 191 | 208 | 173 | 207 | 210 | 299 | 288 | 377 | 296 | 269 | 406 | 308 | 275 | 306 | 154 | 181 | 223 | Prrt4 |
| Prrx1 | 8126 | 3753 | 4259 | 2574 | 1501 | 4058 | 1821 | 11087 | 16341 | 11759 | 16510 | 7888 | 14376 | 8311 | 5893 | 6192 | 2980 | 2921 | 1831 | 1914 | 1578 | 10111 | 17410 | 10297 | 7553 | 10837 | 8341 | 8799 | Prrx1 |
| Prrx2 | 769 | 754 | 592 | 351 | 153 | 400 | 114 | 689 | 897 | 783 | 710 | 510 | 872 | 476 | 614 | 971 | 424 | 375 | 182 | 168 | 105 | 475 | 823 | 717 | 465 | 974 | 685 | 609 | Prrx2 |
| Prrxl1 | 209 | 251 | 228 | 163 | 786 | 170 | 59 | 607 | 615 | 657 | 876 | 201 | 1051 | 216 | 407 | 442 | 322 | 326 | 129 | 139 | 96 | 1064 | 539 | 144 | 466 | 40 | 261 | 370 | Prrxl1 |
| Prss1 | 0 | 0 | 0 | 0 | 0 | 0 | 0 | 0 | 1 | 0 | 0 | 0 | 0 | 0 | 0 | 1 | 0 | 0 | 0 | 0 | 0 | 0 | 0 | 0 | 0 | 0 | 0 | 0 | Prss1 |
| Prss12 | 716 | 607 | 610 | 556 | 726 | 456 | 258 | 824 | 1007 | 750 | 1097 | 414 | 785 | 434 | 653 | 850 | 652 | 639 | 441 | 308 | 297 | 1034 | 995 | 618 | 628 | 351 | 482 | 434 | Prss12 |
| Prss16 | 220 | 236 | 168 | 219 | 59 | 97 | 112 | 4 | 7 | 6 | 8 | 8 | 6 | 5 | 222 | 308 | 177 | 205 | 102 | 160 | 141 | 3 | 11 | 13 | 8 | 5 | 7 | 9 | Prss16 |
| Prss21 | 0 | 0 | 0 | 0 | 0 | 0 | 0 | 1 | 0 | 0 | 0 | 0 | 0 | 0 | 0 | 0 | 0 | 0 | 0 | 0 | 0 | 0 | 0 | 0 | 0 | 0 | 0 | 0 | Prss21 |
| Prss22 | 8 | 0 | 0 | 1 | 0 | 1 | 0 | 22 | 34 | 3 | 20 | 4 | 5 | 3 | 4 | 9 | 0 | 3 | 0 | 0 | 0 | 13 | 23 | 23 | 1 | 10 | 3 | 5 | Prss22 |
| Prss23 | 1358 | 940 | 1569 | 765 | 759 | 765 | 660 | 1586 | 1782 | 1221 | 1540 | 1263 | 1233 | 1148 | 1108 | 1402 | 991 | 968 | 656 | 828 | 592 | 1171 | 1985 | 1544 | 800 | 1858 | 1316 | 1275 | Prss23 |
| Prss27 | 0 | 0 | 0 | 0 | 0 | 0 | 0 | 1 | 1 | 0 | 0 | 0 | 0 | 2 | 0 | 0 | 0 | 0 | 0 | 0 | 0 | 0 | 1 | 0 | 4 | 0 | 1 | 0 | Prss27 |
| Prss28 | 0 | 0 | 0 | 0 | 0 | 0 | 0 | 0 | 1 | 0 | 0 | 0 | 0 | 0 | 0 | 0 | 0 | 0 | 0 | 0 | 0 | 0 | 1 | 0 | 0 | 0 | 0 | 0 | Prss28 |
| Prss3 | 6 | 6 | 4 | 0 | 4 | 0 | 2 | 2 | 2 | 0 | 0 | 0 | 31 | 1 | 9 | 12 | 8 | 14 | 0 | 0 | 0 | 22 | 3 | 2 | 5 | 5 | 6 | 4 | Prss3 |
| Prss30 | 2 | 0 | 0 | 0 | 0 | 0 | 0 | 0 | 1 | 0 | 0 | 0 | 0 | 0 | 0 | 2 | 0 | 0 | 0 | 0 | 0 | 3 | 1 | 0 | 4 | 0 | 0 | 0 | Prss30 |
| Prss32 | 1 | 0 | 1 | 4 | 3 | 0 | 1 | 5 | 2 | 0 | 0 | 5 | 1 | 1 | 1 | 0 | 0 | 0 | 0 | 0 | 0 | 0 | 1 | 1 | 1 | 0 | 1 | 1 | Prss32 |
| Prss33 | 5 | 5 | 2 | 1 | 10 | 0 | 3 | 22 | 21 | 34 | 14 | 8 | 18 | 14 | 3 | 5 | 6 | 1 | 3 | 6 | 2 | 10 | 17 | 15 | 3 | 23 | 15 | 11 | Prss33 |
| Prss34 | 1 | 0 | 0 | 0 | 0 | 0 | 0 | 0 | 0 | 1 | 5 | 3 | 0 | 0 | 0 | 0 | 0 | 0 | 0 | 0 | 0 | 0 | 5 | 2 | 1 | 1 | 0 | 1 | Prss34 |
| Prss35 | 942 | 726 | 565 | 444 | 464 | 713 | 270 | 641 | 849 | 904 | 1931 | 349 | 562 | 411 | 812 | 1005 | 503 | 863 | 315 | 221 | 262 | 869 | 2352 | 509 | 640 | 365 | 416 | 441 | Prss35 |
| Prss36 | 231 | 239 | 236 | 241 | 239 | 214 | 216 | 375 | 447 | 377 | 451 | 382 | 428 | 394 | 220 | 254 | 290 | 244 | 221 | 233 | 206 | 378 | 441 | 380 | 382 | 420 | 396 | 345 | Prss36 |
| Prss37 | 1 | 0 | 0 | 0 | 0 | 0 | 0 | 0 | 1 | 0 | 0 | 1 | 0 | 1 | 0 | 0 | 0 | 0 | 0 | 0 | 0 | 2 | 2 | 0 | 0 | 0 | 0 | 2 | Prss37 |
| Prss38 | 28 | 31 | 22 | 43 | 49 | 44 | 59 | 11 | 10 | 8 | 13 | 15 | 20 | 14 | 20 | 30 | 25 | 72 | 23 | 30 | 51 | 17 | 9 | 20 | 24 | 19 | 16 | 13 | Prss38 |
| Prss39 | 0 | 0 | 0 | 4 | 0 | 2 | 0 | 1 | 0 | 1 | 0 | 0 | 0 | 0 | 0 | 1 | 0 | 0 | 0 | 0 | 0 | 0 | 0 | 0 | 0 | 5 | 0 | 0 | Prss39 |
| Prss40 | 0 | 0 | 4 | 0 | 2 | 0 | 0 | 0 | 0 | 0 | 0 | 0 | 0 | 0 | 3 | 0 | 0 | 0 | 0 | 1 | 0 | 0 | 0 | 0 | 0 | 0 | 0 | 0 | Prss40 |
| Prss41 | 44 | 42 | 38 | 25 | 36 | 30 | 34 | 74 | 73 | 74 | 65 | 37 | 95 | 65 | 47 | 41 | 36 | 35 | 37 | 24 | 21 | 28 | 54 | 70 | 38 | 73 | 51 | 53 | Prss41 |
| Prss42 | 0 | 0 | 0 | 0 | 0 | 3 | 0 | 0 | 0 | 0 | 1 | 0 | 2 | 0 | 1 | 1 | 1 | 2 | 0 | 0 | 0 | 0 | 0 | 1 | 3 | 0 | 0 | 0 | Prss42 |
| Prss43 | 0 | 0 | 0 | 0 | 2 | 0 | 0 | 0 | 0 | 2 | 0 | 0 | 0 | 2 | 0 | 0 | 0 | 2 | 0 | 0 | 0 | 0 | 1 | 0 | 0 | 0 | 0 | 2 | Prss43 |
| Prss44 | 0 | 0 | 2 | 0 | 1 | 8 | 0 | 0 | 0 | 0 | 1 | 0 | 0 | 0 | 1 | 0 | 0 | 0 | 0 | 0 | 0 | 0 | 1 | 1 | 0 | 0 | 0 | 0 | Prss44 |
| Prss45 | 0 | 3 | 2 | 2 | 2 | 1 | 7 | 2 | 1 | 3 | 1 | 1 | 6 | 2 | 0 | 2 | 2 | 1 | 3 | 6 | 1 | 3 | 1 | 0 | 7 | 1 | 2 | 5 | Prss45 |
| Prss46 | 5 | 0 | 4 | 0 | 0 | 0 | 0 | 1 | 1 | 0 | 0 | 0 | 1 | 0 | 0 | 2 | 0 | 0 | 0 | 1 | 0 | 2 | 0 | 1 | 0 | 2 | 2 | 0 | Prss46 |
| Prss47 | 6 | 11 | 0 | 8 | 7 | 6 | 9 | 1 | 2 | 0 | 1 | 0 | 0 | 2 | 8 | 7 | 0 | 7 | 2 | 16 | 8 | 0 | 1 | 2 | 0 | 3 | 1 | 0 | Prss47 |
| Prss50 | 6 | 3 | 12 | 11 | 4 | 2 | 3 | 1 | 3 | 2 | 4 | 0 | 2 | 2 | 5 | 7 | 2 | 0 | 5 | 4 | 1 | 0 | 3 | 3 | 0 | 8 | 2 | 2 | Prss50 |
| Prss53 | 16 | 26 | 26 | 38 | 44 | 24 | 29 | 10 | 12 | 13 | 23 | 5 | 12 | 18 | 20 | 20 | 30 | 17 | 34 | 13 | 27 | 23 | 14 | 20 | 18 | 12 | 9 | 11 | Prss53 |
| Prss54 | 0 | 0 | 0 | 2 | 0 | 0 | 0 | 0 | 0 | 1 | 0 | 0 | 0 | 0 | 0 | 0 | 0 | 0 | 0 | 0 | 0 | 0 | 1 | 0 | 0 | 0 | 0 | 0 | Prss54 |
| Prss55 | 5 | 3 | 0 | 2 | 2 | 0 | 1 | 0 | 0 | 5 | 1 | 0 | 0 | 0 | 1 | 1 | 0 | 3 | 2 | 9 | 2 | 0 | 1 | 0 | 1 | 0 | 0 | 1 | Prss55 |
| Prss56 | 15 | 26 | 7 | 7 | 54 | 5 | 8 | 74 | 91 | 46 | 94 | 5 | 70 | 2 | 28 | 24 | 22 | 6 | 11 | 4 | 14 | 70 | 91 | 8 | 32 | 8 | 8 | 6 | Prss56 |
| Prss57 | 1 | 3 | 0 | 2 | 2 | 2 | 3 | 5 | 5 | 3 | 6 | 9 | 6 | 2 | 3 | 0 | 0 | 4 | 1 | 1 | 2 | 7 | 3 | 4 | 11 | 6 | 7 | 2 | Prss57 |
| Prss8 | 181 | 107 | 41 | 51 | 25 | 26 | 3 | 136 | 218 | 197 | 199 | 20 | 94 | 19 | 124 | 157 | 58 | 39 | 2 | 16 | 3 | 38 | 214 | 162 | 32 | 229 | 22 | 51 | Prss8 |
| Prtg | 862 | 763 | 1154 | 555 | 507 | 522 | 460 | 535 | 503 | 420 | 516 | 825 | 920 | 824 | 787 | 951 | 833 | 493 | 515 | 506 | 456 | 468 | 546 | 495 | 487 | 758 | 1118 | 1060 | Prtg |
| Prtn3 | 235 | 284 | 310 | 226 | 242 | 186 | 232 | 248 | 255 | 367 | 273 | 330 | 263 | 301 | 240 | 242 | 234 | 314 | 256 | 214 | 212 | 306 | 284 | 330 | 279 | 350 | 308 | 284 | Prtn3 |
| Prune1 | 2498 | 2611 | 2493 | 2446 | 3470 | 3034 | 3117 | 3982 | 3916 | 3914 | 3776 | 3600 | 3525 | 3513 | 2786 | 2365 | 2705 | 2807 | 3326 | 3102 | 3069 | 4690 | 3925 | 3758 | 4228 | 3418 | 3608 | 3446 | Prune1 |
| Prune2 | 1318 | 1283 | 443 | 609 | 1498 | 816 | 700 | 1812 | 1878 | 1037 | 1480 | 476 | 1171 | 534 | 1776 | 1828 | 635 | 831 | 545 | 815 | 664 | 1492 | 1358 | 431 | 1041 | 447 | 538 | 465 | Prune2 |
| Prx | 551 | 692 | 618 | 563 | 499 | 589 | 400 | 492 | 731 | 650 | 792 | 753 | 861 | 701 | 493 | 625 | 600 | 492 | 492 | 445 | 424 | 536 | 654 | 722 | 616 | 810 | 723 | 681 | Prx |
| Prxl2a | 2579 | 2591 | 2770 | 2827 | 2909 | 2838 | 2723 | 2673 | 2768 | 2947 | 2499 | 2490 | 2733 | 2504 | 2616 | 2505 | 2610 | 2656 | 2784 | 2657 | 2706 | 2488 | 2791 | 2833 | 2574 | 2840 | 2573 | 2600 | Prxl2a |

| | | | | | | | | | | | | | | | | | | | | | | | | | | | | | |
|---|---|---|---|---|---|---|---|---|---|---|---|---|---|---|---|---|---|---|---|---|---|---|---|---|---|---|---|---|---|
| Prxl2b | 1159 | 1461 | 1286 | 1261 | 1061 | 1068 | 915 | 657 | 690 | 712 | 690 | 630 | 695 | 575 | 1253 | 1293 | 1301 | 1128 | 1137 | 1253 | 965 | 541 | 691 | 821 | 622 | 691 | 817 | 706 | Prxl2b |
| Prxl2c | 1739 | 1241 | 1592 | 1417 | 1177 | 1246 | 1290 | 778 | 944 | 866 | 1147 | 771 | 897 | 816 | 1540 | 1646 | 1468 | 1402 | 1103 | 1194 | 1183 | 742 | 1144 | 851 | 764 | 935 | 790 | 888 | Prxl2c |
| Psap | 20964 | 25738 | 27159 | 26804 | 26575 | 25136 | 23652 | 22577 | 26858 | 28578 | 28428 | 26700 | 26080 | 26951 | 21854 | 24383 | 27527 | 27876 | 27115 | 23706 | 24690 | 27370 | 27829 | 28069 | 29159 | 28228 | 25451 | 26750 | Psap |
| Psapl1 | 0 | 0 | 0 | 0 | 0 | 0 | 0 | 5 | 19 | 12 | 2 | 0 | 2 | 0 | 0 | 1 | 0 | 0 | 0 | 0 | 0 | 0 | 10 | 2 | 0 | 1 | 0 | 7 | Psapl1 |
| Psat1 | 19893 | 19759 | 17333 | 22512 | 18384 | 21367 | 20449 | 14957 | 13224 | 14401 | 12691 | 16109 | 14664 | 16211 | 19690 | 16369 | 19705 | 19382 | 22576 | 24629 | 21089 | 11637 | 12784 | 15319 | 16253 | 18533 | 16823 | 16256 | Psat1 |
| Psca | 0 | 3 | 6 | 4 | 1 | 2 | 1 | 2 | 3 | 2 | 2 | 0 | 0 | 3 | 0 | 0 | 0 | 0 | 0 | 1 | 0 | 0 | 2 | 0 | 5 | 2 | 3 | 2 | Psca |
| Psd | 2323 | 3494 | 3163 | 6319 | 8784 | 6528 | 7007 | 4330 | 4932 | 6555 | 5253 | 5125 | 3631 | 4737 | 2764 | 2484 | 5538 | 7255 | 8178 | 6255 | 7773 | 8489 | 4514 | 6227 | 7922 | 3912 | 4138 | 3891 | Psd |
| Psd2 | 2226 | 2915 | 3235 | 4522 | 6289 | 4834 | 4976 | 4537 | 4448 | 6113 | 4565 | 5025 | 4190 | 4790 | 2683 | 2320 | 4201 | 4998 | 5972 | 4660 | 5237 | 7447 | 4346 | 5444 | 6955 | 3828 | 4202 | 4307 | Psd2 |
| Psd3 | 7151 | 5485 | 3857 | 4810 | 6046 | 7240 | 6755 | 6974 | 6714 | 5238 | 5895 | 5037 | 5362 | 5422 | 6980 | 6177 | 4049 | 5237 | 5100 | 6778 | 6722 | 4949 | 5883 | 3992 | 5465 | 2902 | 5704 | 4908 | Psd3 |
| Psd4 | 55 | 59 | 52 | 43 | 43 | 39 | 26 | 74 | 163 | 124 | 139 | 92 | 105 | 74 | 65 | 83 | 31 | 37 | 21 | 52 | 31 | 74 | 174 | 75 | 71 | 55 | 112 | 89 | Psd4 |
| Psen1 | 5787 | 6083 | 5216 | 5528 | 5665 | 5828 | 5686 | 5503 | 5536 | 5874 | 5616 | 5760 | 5625 | 5608 | 5703 | 5476 | 5116 | 5444 | 5767 | 5379 | 6214 | 5820 | 5309 | 5298 | 5950 | 4336 | 5794 | 5861 | Psen1 |
| Psen2 | 812 | 1010 | 957 | 1009 | 862 | 949 | 870 | 407 | 448 | 453 | 643 | 452 | 455 | 446 | 837 | 1061 | 948 | 912 | 840 | 832 | 884 | 500 | 530 | 498 | 463 | 468 | 427 | 433 | Psen2 |
| Psenen | 1345 | 1817 | 1836 | 1783 | 1816 | 1690 | 1930 | 1677 | 1929 | 1799 | 1928 | 1988 | 2206 | 2094 | 1391 | 1538 | 1845 | 1679 | 1916 | 2139 | 1737 | 1511 | 1964 | 2085 | 2058 | 2524 | 2427 | 2359 | Psenen |
| Psg16 | 13 | 7 | 29 | 12 | 36 | 17 | 22 | 35 | 54 | 67 | 65 | 13 | 27 | 40 | 8 | 9 | 13 | 25 | 13 | 14 | 35 | 75 | 49 | 27 | 65 | 25 | 17 | 12 | Psg16 |
| Psg17 | 32 | 48 | 19 | 42 | 26 | 40 | 26 | 7 | 26 | 8 | 15 | 27 | 24 | 11 | 44 | 44 | 21 | 42 | 26 | 28 | 31 | 23 | 10 | 23 | 13 | 23 | 14 | 11 | Psg17 |
| Psg18 | 0 | 0 | 0 | 0 | 0 | 0 | 0 | 0 | 14 | 0 | 0 | 0 | 0 | 0 | 0 | 0 | 0 | 0 | 0 | 0 | 0 | 0 | 5 | 0 | 0 | 0 | 0 | 0 | Psg18 |
| Psg19 | 12 | 16 | 31 | 28 | 20 | 14 | 7 | 0 | 2 | 0 | 1 | 0 | 7 | 0 | 16 | 10 | 16 | 35 | 12 | 14 | 16 | 0 | 3 | 3 | 0 | 3 | 0 | 4 | Psg19 |
| Psg21 | 0 | 0 | 0 | 0 | 0 | 0 | 0 | 0 | 15 | 0 | 0 | 0 | 0 | 0 | 0 | 0 | 0 | 0 | 0 | 0 | 0 | 0 | 12 | 0 | 0 | 0 | 0 | 0 | Psg21 |
| Psg23 | 0 | 0 | 0 | 0 | 0 | 0 | 1 | 0 | 16 | 0 | 0 | 0 | 0 | 0 | 0 | 0 | 0 | 0 | 0 | 0 | 0 | 0 | 1 | 0 | 0 | 0 | 0 | 0 | Psg23 |
| Psg25 | 0 | 0 | 0 | 0 | 0 | 0 | 0 | 0 | 7 | 0 | 0 | 0 | 0 | 0 | 0 | 0 | 0 | 0 | 0 | 0 | 0 | 0 | 1 | 1 | 0 | 0 | 0 | 0 | Psg25 |
| Psg26 | 0 | 0 | 0 | 0 | 0 | 0 | 0 | 0 | 2 | 0 | 0 | 0 | 0 | 0 | 0 | 0 | 0 | 0 | 0 | 0 | 0 | 0 | 0 | 0 | 0 | 0 | 0 | 0 | Psg26 |
| Psg27 | 0 | 0 | 0 | 0 | 0 | 0 | 0 | 0 | 16 | 0 | 0 | 0 | 0 | 0 | 0 | 0 | 0 | 0 | 0 | 0 | 0 | 0 | 5 | 0 | 0 | 0 | 0 | 0 | Psg27 |
| Psg28 | 0 | 0 | 0 | 0 | 0 | 0 | 0 | 0 | 4 | 0 | 1 | 0 | 0 | 0 | 0 | 0 | 0 | 2 | 0 | 0 | 0 | 0 | 3 | 0 | 0 | 0 | 0 | 0 | Psg28 |
| Psg29 | 0 | 0 | 0 | 0 | 2 | 13 | 4 | 0 | 11 | 6 | 6 | 2 | 2 | 0 | 0 | 0 | 0 | 0 | 4 | 0 | 4 | 3 | 1 | 10 | 0 | 0 | 0 | 1 | Psg29 |
| Psip1 | 47106 | 35685 | 48668 | 46532 | 41419 | 43548 | 47631 | 46868 | 35969 | 42755 | 35445 | 49542 | 45053 | 53124 | 46802 | 38224 | 46648 | 41531 | 43616 | 47115 | 44598 | 39966 | 34886 | 47887 | 44267 | 57260 | 50850 | 52027 | Psip1 |
| Pskh1 | 2208 | 2020 | 1968 | 1546 | 1227 | 1576 | 1297 | 1921 | 2618 | 2128 | 2580 | 2241 | 2563 | 1943 | 1838 | 2158 | 1570 | 1503 | 1300 | 1431 | 1148 | 1915 | 2636 | 2228 | 1708 | 2725 | 2181 | 2273 | Pskh1 |
| Psma1 | 13008 | 11679 | 12296 | 10496 | 9703 | 10194 | 12288 | 12432 | 10766 | 9682 | 9106 | 11379 | 10798 | 12458 | 13444 | 12161 | 11317 | 10314 | 10622 | 13520 | 11168 | 9560 | 10751 | 10834 | 10536 | 13500 | 12184 | 13079 | Psma1 |
| Psma2 | 13220 | 12675 | 12493 | 11456 | 10468 | 10556 | 12859 | 11105 | 9327 | 8578 | 8386 | 9246 | 9048 | 10822 | 14373 | 11982 | 12043 | 11183 | 11137 | 13618 | 12377 | 8988 | 9041 | 9637 | 8913 | 10773 | 10280 | 11450 | Psma2 |
| Psma3 | 10555 | 9257 | 9716 | 9446 | 8845 | 8834 | 11061 | 10415 | 8896 | 8376 | 8147 | 8643 | 8612 | 10914 | 11265 | 9004 | 9649 | 8635 | 9084 | 11290 | 10770 | 8477 | 8926 | 9381 | 8541 | 9794 | 9667 | 10801 | Psma3 |
| Psma4 | 11918 | 10137 | 9850 | 9337 | 8886 | 8864 | 10706 | 11771 | 9937 | 9024 | 9180 | 9576 | 9607 | 11098 | 12977 | 10720 | 9868 | 9256 | 9273 | 12095 | 10111 | 8961 | 10280 | 9729 | 9434 | 11367 | 10462 | 11427 | Psma4 |
| Psma5 | 10364 | 9605 | 10464 | 8620 | 7450 | 7973 | 8996 | 7886 | 7253 | 6498 | 6788 | 7220 | 6999 | 7953 | 10519 | 10023 | 9493 | 8603 | 8648 | 9940 | 8326 | 6241 | 7354 | 7571 | 7091 | 9252 | 8365 | 8600 | Psma5 |
| Psma6 | 12442 | 10826 | 11891 | 10191 | 9371 | 9536 | 11373 | 11339 | 9753 | 9397 | 9177 | 11044 | 10703 | 12267 | 12088 | 10742 | 10820 | 9367 | 10431 | 11511 | 10665 | 9513 | 10202 | 10841 | 10325 | 12778 | 11695 | 13218 | Psma6 |
| Psma7 | 12333 | 14707 | 14516 | 13525 | 11817 | 12174 | 13634 | 10831 | 11278 | 10426 | 9906 | 12000 | 11657 | 12499 | 12606 | 13398 | 14020 | 12591 | 13419 | 14873 | 13276 | 8886 | 11451 | 12137 | 11452 | 13978 | 12829 | 13630 | Psma7 |
| Psma8 | 35 | 34 | 39 | 9 | 20 | 10 | 17 | 39 | 47 | 27 | 47 | 21 | 40 | 24 | 26 | 39 | 19 | 21 | 14 | 18 | 16 | 65 | 38 | 25 | 23 | 17 | 44 | 21 | Psma8 |
| Psmb1 | 9788 | 10303 | 9980 | 9102 | 8415 | 8395 | 10216 | 8912 | 8882 | 8101 | 7825 | 8586 | 8698 | 9631 | 10011 | 9213 | 9589 | 8850 | 9277 | 10631 | 9598 | 7157 | 8963 | 8828 | 8445 | 10437 | 9597 | 10130 | Psmb1 |
| Psmb10 | 291 | 304 | 315 | 280 | 336 | 354 | 293 | 488 | 495 | 533 | 532 | 430 | 481 | 459 | 286 | 321 | 299 | 330 | 308 | 414 | 315 | 399 | 557 | 477 | 502 | 499 | 482 | 438 | Psmb10 |
| Psmb11 | 5 | 6 | 4 | 4 | 7 | 4 | 1 | 2 | 4 | 3 | 8 | 7 | 7 | 10 | 3 | 4 | 2 | 2 | 1 | 2 | 0 | 15 | 9 | 15 | 2 | 7 | 15 | 7 | Psmb11 |
| Psmb2 | 7131 | 8014 | 6938 | 6182 | 5402 | 5430 | 6831 | 5854 | 5671 | 4399 | 4770 | 5423 | 5641 | 6198 | 7446 | 7333 | 6608 | 5935 | 5668 | 7058 | 5918 | 4736 | 5847 | 5357 | 5375 | 6654 | 6266 | 6417 | Psmb2 |
| Psmb3 | 6120 | 8389 | 7075 | 7307 | 5668 | 6219 | 6162 | 4979 | 5922 | 5104 | 4928 | 5589 | 5480 | 5849 | 6142 | 7159 | 6969 | 6353 | 6807 | 7712 | 6374 | 3825 | 5923 | 5521 | 5366 | 6746 | 6426 | 6297 | Psmb3 |
| Psmb4 | 12560 | 13389 | 12588 | 12785 | 10738 | 11172 | 10841 | 10544 | 9495 | 9590 | 9374 | 10372 | 10181 | 9811 | 12642 | 11347 | 12604 | 11396 | 12506 | 12973 | 10822 | 8729 | 9776 | 10529 | 10262 | 11911 | 10892 | 10596 | Psmb4 |
| Psmb5 | 1993 | 2712 | 2460 | 2292 | 2243 | 2224 | 2263 | 2163 | 2263 | 2623 | 2272 | 2821 | 2569 | 2561 | 2020 | 2078 | 2348 | 2028 | 2465 | 2892 | 2148 | 1893 | 2264 | 2732 | 2488 | 3049 | 2903 | 2800 | Psmb5 |
| Psmb6 | 8957 | 10055 | 8578 | 7900 | 7345 | 7516 | 7778 | 9417 | 9471 | 8444 | 8220 | 9953 | 9628 | 10230 | 8479 | 8423 | 8691 | 7031 | 8456 | 8912 | 7520 | 7722 | 9470 | 9877 | 9109 | 11870 | 11035 | 11083 | Psmb6 |
| Psmb7 | 14627 | 13744 | 14064 | 12206 | 11345 | 10887 | 13437 | 14600 | 11899 | 11169 | 10674 | 13449 | 12598 | 14616 | 15227 | 12548 | 13505 | 11201 | 12109 | 14398 | 12681 | 11418 | 11593 | 13403 | 12304 | 15418 | 14556 | 15875 | Psmb7 |
| Psmb8 | 55 | 70 | 67 | 60 | 75 | 28 | 37 | 40 | 52 | 41 | 64 | 36 | 34 | 49 | 57 | 64 | 57 | 81 | 60 | 50 | 66 | 33 | 45 | 54 | 41 | 49 | 38 | 37 | Psmb8 |
| Psmb9 | 40 | 38 | 46 | 35 | 23 | 38 | 30 | 15 | 28 | 10 | 34 | 14 | 15 | 7 | 41 | 48 | 29 | 30 | 34 | 20 | 22 | 22 | 23 | 22 | 23 | 27 | 27 | 15 | Psmb9 |
| Psmc1 | 17055 | 16508 | 19123 | 15704 | 13808 | 13892 | 15665 | 13503 | 12218 | 12559 | 11721 | 14133 | 13925 | 15933 | 17840 | 15724 | 17208 | 14295 | 15197 | 17473 | 14019 | 10998 | 11978 | 13451 | 12898 | 17322 | 15743 | 16372 | Psmc1 |
| Psmc2 | 10584 | 11014 | 11118 | 9683 | 8492 | 8629 | 9618 | 9434 | 8209 | 7871 | 7788 | 9826 | 9188 | 10143 | 10609 | 9548 | 9970 | 9034 | 8888 | 10554 | 8931 | 7865 | 8143 | 9054 | 8619 | 10957 | 10507 | 10871 | Psmc2 |
| Psmc3 | 11608 | 13522 | 13352 | 10970 | 9717 | 10235 | 10014 | 10214 | 11390 | 10132 | 10332 | 11407 | 11031 | 11857 | 11832 | 11811 | 11976 | 10138 | 11321 | 12487 | 10494 | 8228 | 11339 | 11406 | 10519 | 14348 | 13041 | 12823 | Psmc3 |
| Psmc3ip | 1096 | 1168 | 1424 | 1081 | 878 | 1012 | 1307 | 982 | 969 | 972 | 857 | 1294 | 1215 | 1540 | 969 | 945 | 1323 | 900 | 1220 | 1400 | 1103 | 628 | 901 | 1356 | 1128 | 1673 | 1477 | 1453 | Psmc3ip |
| Psmc4 | 10430 | 12315 | 11792 | 10151 | 8623 | 9154 | 9518 | 9381 | 9255 | 8982 | 8070 | 10523 | 9400 | 10543 | 10257 | 9962 | 10761 | 9056 | 10495 | 10669 | 9568 | 7549 | 8969 | 10139 | 9350 | 12761 | 11125 | 11732 | Psmc4 |
| Psmc5 | 7571 | 8537 | 9019 | 7206 | 6749 | 6915 | 7396 | 8436 | 8118 | 8101 | 7703 | 9108 | 8873 | 10412 | 7911 | 7588 | 8018 | 6541 | 7452 | 7761 | 7441 | 7357 | 8407 | 9173 | 7805 | 11365 | 10516 | 11165 | Psmc5 |
| Psmc6 | 15777 | 11902 | 14229 | 12914 | 11775 | 13201 | 15355 | 16998 | 13846 | 11709 | 12203 | 15132 | 14048 | 16699 | 16945 | 13999 | 13079 | 12382 | 13116 | 16535 | 14261 | 12694 | 13688 | 13998 | 13484 | 16403 | 16363 | 16956 | Psmc6 |
| Psmd1 | 16085 | 15180 | 13755 | 12298 | 11618 | 12705 | 12697 | 12825 | 11514 | 10409 | 10303 | 12589 | 12132 | 12709 | 16026 | 13710 | 13301 | 11677 | 12179 | 13090 | 12110 | 11109 | 11683 | 11061 | 10872 | 13232 | 13042 | 13311 | Psmd1 |
| Psmd10 | 236 | 92 | 350 | 178 | 122 | 418 | 73 | 96 | 110 | 70 | 100 | 77 | 203 | 113 | 170 | 93 | 132 | 89 | 119 | 150 | 172 | 85 | 225 | 83 | 326 | 84 | 94 | 89 | Psmd10 |
| Psmd11 | 7952 | 7806 | 9639 | 8552 | 7344 | 7358 | 8275 | 7076 | 6908 | 7209 | 7000 | 8327 | 7529 | 8976 | 7787 | 7223 | 9028 | 8083 | 7817 | 6823 | 7957 | 7279 | 6730 | 8214 | 7781 | 10324 | 8284 | 9683 | Psmd11 |
| Psmd12 | 10208 | 9259 | 10601 | 9093 | 7979 | 8047 | 9571 | 7820 | 6728 | 6888 | 6475 | 8062 | 7659 | 8717 | 10354 | 9108 | 9700 | 8173 | 8856 | 10137 | 8492 | 6764 | 6783 | 7532 | 7092 | 8954 | 8605 | 8790 | Psmd12 |
| Psmd13 | 11339 | 12152 | 11442 | 10352 | 9155 | 9852 | 9629 | 9314 | 9290 | 8553 | 7997 | 10065 | 9414 | 9400 | 11087 | 11365 | 10885 | 9840 | 10563 | 11034 | 9767 | 7949 | 9001 | 9506 | 9204 | 11174 | 10766 | 10029 | Psmd13 |
| Psmd14 | 9303 | 7795 | 8716 | 7800 | 8219 | 8356 | 9688 | 8750 | 7364 | 7210 | 6808 | 8699 | 7888 | 9005 | 10146 | 8060 | 9057 | 8050 | 8877 | 9720 | 9089 | 7200 | 7287 | 7984 | 8269 | 9918 | 8990 | 9425 | Psmd14 |
| Psmd2 | 36131 | 39510 | 34552 | 32817 | 27204 | 28977 | 29290 | 27218 | 25728 | 23802 | 22373 | 27954 | 25937 | 26860 | 35554 | 33960 | 32850 | 29930 | 30901 | 31389 | 30071 | 22939 | 25072 | 25820 | 25181 | 30586 | 29900 | 28859 | Psmd2 |
| Psmd3 | 8200 | 10369 | 10933 | 10202 | 8354 | 7893 | 7751 | 5838 | 7310 | 7756 | 7633 | 9474 | 7570 | 8030 | 7346 | 9256 | 10404 | 9628 | 9373 | 7079 | 8372 | 7123 | 7200 | 8725 | 7730 | 10857 | 7920 | 8113 | Psmd3 |
| Psmd4 | 9770 | 12418 | 10637 | 10380 | 8555 | 9103 | 9138 | 7336 | 8018 | 7295 | 7361 | 8244 | 8274 | 8675 | 10015 | 10025 | 10520 | 10230 | 9803 | 9952 | 9422 | 6460 | 7831 | 8313 | 8239 | 10033 | 8856 | 9125 | Psmd4 |
| Psmd5 | 7063 | 7030 | 5765 | 6018 | 5050 | 5254 | 5909 | 5693 | 4856 | 4065 | 3989 | 4778 | 4415 | 4802 | 7598 | 6688 | 5509 | 5667 | 5332 | 6026 | 5379 | 4984 | 5006 | 4426 | 4493 | 5314 | 4693 | 5203 | Psmd5 |
| Psmd6 | 10317 | 9720 | 9846 | 8788 | 7913 | 7977 | 10030 | 9685 | 8277 | 7393 | 7177 | 8475 | 8400 | 9291 | 10764 | 9260 | 9014 | 7978 | 8525 | 10656 | 8866 | 7651 | 8389 | 8478 | 8038 | 9821 | 8959 | 9842 | Psmd6 |
| Psmd7 | 10568 | 11157 | 11347 | 9397 | 8157 | 8369 | 8939 | 8658 | 8464 | 8216 | 8209 | 9237 | 8853 | 9929 | 10407 | 9886 | 10605 | 9128 | 8432 | 9422 | 8520 | 7710 | 8297 | 9070 | 8465 | 10201 | 9375 | 9841 | Psmd7 |
| Psmd8 | 12012 | 13143 | 9567 | 9866 | 8552 | 8845 | 9334 | 9227 | 8625 | 7889 | 7175 | 8304 | 8083 | 8458 | 11757 | 11102 | 10166 | 9096 | 9763 | 10609 | 9443 | 8029 | 9048 | 8156 | 7701 | 9336 | 8606 | 8874 | Psmd8 |
| Psmd9 | 2810 | 3281 | 2758 | 3166 | 2384 | 2482 | 2731 | 1893 | 1990 | 2047 | 2128 | 2007 | 1870 | 2253 | 2794 | 2945 | 2867 | 2930 | 2584 | 2774 | 2842 | 1780 | 2056 | 2153 | 2106 | 2212 | 2136 | 2180 | Psmd9 |
| Psme1 | 2782 | 2907 | 2528 | 2971 | 3162 | 3183 | 3184 | 3334 | 3269 | 3604 | 3551 | 3367 | 3072 | 3562 | 2869 | 2633 | 2936 | 2921 | 3278 | 2979 | 3604 | 3205 | 3498 | 3684 | 3535 | 3715 | 3262 | 3711 | Psme1 |
| Psme2 | 3201 | 3152 | 3049 | 3005 | 2932 | 2834 | 3161 | 2757 | 2768 | 2862 | 2928 | 2941 | 3035 | 3282 | 2955 | 3055 | 3078 | 2954 | 3176 | 3536 | 3013 | 2510 | 2966 | 2932 | 2861 | 3211 | 3258 | 3431 | Psme2 |
| Psme2b | 1 | 0 | 0 | 0 | 0 | 0 | 0 | 0 | 0 | 1 | 0 | 0 | 0 | 0 | 0 | 0 | 0 | 0 | 2 | 0 | 0 | 0 | 0 | 0 | 0 | 1 | 1 | 2 | Psme2b |
| Psme3 | 11480 | 11057 | 11771 | 11074 | 9685 | 10055 | 11097 | 9537 | 9036 | 9544 | 9054 | 11304 | 10227 | 11547 | 10677 | 10146 | 11717 | 10649 | 11105 | 9275 | 10875 | 9799 | 8930 | 10844 | 10196 | 12421 | 10705 | 12420 | Psme3 |
| Psme3ip1 | 5249 | 5416 | 5929 | 5252 | 5001 | 4990 | 5673 | 4824 | 4295 | 4639 | 4127 | 5285 | 4687 | 5448 | 5349 | 4520 | 5778 | 5026 | 5664 | 5511 | 5243 | 4512 | 4089 | 4917 | 5089 | 5914 | 5361 | 5693 | Psme3ip1 |
| Psme4 | 33641 | 28557 | 25589 | 26903 | 25472 | 29864 | 31980 | 23000 | 18597 | 17759 | 17277 | 22901 | 22817 | 22594 | 33011 | 29735 | 26511 | 26316 | 26672 | 28453 | 30461 | 19522 | 18283 | 17918 | 19717 | 18125 | 24468 | 23922 | Psme4 |
| Psmf1 | 3162 | 3331 | 3151 | 3070 | 3134 | 3056 | 3147 | 2848 | 2927 | 2935 | 2846 | 2862 | 2947 | 3046 | 3198 | 3096 | 3157 | 2762 | 3030 | 3137 | 2984 | 2749 | 2931 | 2843 | 3078 | 3072 | 3013 | 3029 | Psmf1 |
| Psmg1 | 2210 | 2304 | 2196 | 1944 | 1780 | 1756 | 2031 | 1757 | 1560 | 1611 | 1597 | 1661 | 1779 | 1881 | 2027 | 2109 | 2243 | 1970 | 1944 | 2099 | 1852 | 1509 | 1614 | 1800 | 1618 | 1958 | 1962 | 2114 | Psmg1 |
| Psmg2 | 1979 | 2047 | 2162 | 2068 | 1807 | 1776 | 2265 | 1809 | 1927 | 1697 | 1789 | 1947 | 1968 | 2253 | 2163 | 2039 | 2119 | 1838 | 1950 | 2334 | 1988 | 1559 | 1994 | 1902 | 1817 | 2301 | 2099 | 2479 | Psmg2 |
| Psmg3 | 227 | 318 | 400 | 307 | 322 | 231 | 343 | 344 | 337 | 399 | 307 | 356 | 338 | 434 | 266 | 233 | 385 | 257 | 318 | 268 | 347 | 331 | 296 | 390 | 398 | 402 | 372 | 427 | Psmg3 |
| Psmg4 | 660 | 739 | 737 | 689 | 713 | 622 | 815 | 815 | 718 | 742 | 756 | 861 | 805 | 1037 | 596 | 484 | 761 | 643 | 652 | 738 | 741 | 571 | 694 | 941 | 859 | 1198 | 995 | 1111 | Psmg4 |
| Psors1c2 | 84 | 143 | 211 | 159 | 159 | 76 | 152 | 37 | 20 | 45 | 37 | 40 | 44 | 51 | 102 | 140 | 185 | 161 | 169 | 72 | 118 | 22 | 28 | 42 | 31 | 41 | 34 | 50 | Psors1c2 |
| Pspc1 | 4379 | 3841 | 5257 | 4149 | 5326 | 5165 | 5271 | 6627 | 6237 | 6587 | 6312 | 8080 | 7275 | 8792 | 4610 | 3777 | 4584 | 4434 | 4868 | 5502 | 4911 | 6772 | 5805 | 7142 | 7150 | 8328 | 7817 | 8053 | Pspc1 |

| | | | | | | | | | | | | | | | | | | | | | | | | | | | | | |
|---|---|---|---|---|---|---|---|---|---|---|---|---|---|---|---|---|---|---|---|---|---|---|---|---|---|---|---|---|---|
| Psph | 926 | 1000 | 865 | 1025 | 870 | 859 | 1068 | 1088 | 1033 | 975 | 861 | 1164 | 1151 | 1273 | 869 | 787 | 909 | 906 | 933 | 1004 | 1047 | 882 | 944 | 1244 | 1178 | 1303 | 1278 | 1239 | Psph |
| Pspn | 3 | 9 | 8 | 15 | 13 | 5 | 7 | 2 | 2 | 0 | 4 | 2 | 0 | 3 | 5 | 6 | 16 | 13 | 14 | 10 | 4 | 3 | 1 | 2 | 2 | 4 | 0 | 9 | Pspn |
| Psrc1 | 3635 | 4047 | 4785 | 3537 | 2743 | 3166 | 3320 | 2579 | 2818 | 2698 | 2447 | 4127 | 3495 | 4106 | 3328 | 3520 | 4232 | 3151 | 3670 | 3300 | 3166 | 1856 | 2416 | 3605 | 2947 | 5104 | 3947 | 4031 | Psrc1 |
| Pstk | 1127 | 954 | 1013 | 930 | 1050 | 1042 | 1031 | 1198 | 1281 | 1176 | 1299 | 1196 | 1252 | 1292 | 1149 | 938 | 931 | 980 | 1050 | 975 | 1014 | 1128 | 1292 | 1221 | 992 | 1238 | 1297 | 1272 | Pstk |
| Pstpip1 | 78 | 87 | 97 | 143 | 70 | 116 | 104 | 65 | 62 | 75 | 110 | 64 | 74 | 65 | 69 | 95 | 100 | 145 | 88 | 92 | 96 | 55 | 87 | 47 | 52 | 86 | 64 | 64 | Pstpip1 |
| Pstpip2 | 23 | 10 | 66 | 13 | 34 | 22 | 35 | 29 | 34 | 35 | 70 | 23 | 18 | 19 | 46 | 61 | 124 | 75 | 36 | 66 | 40 | 63 | 54 | 41 | 20 | 4 | 15 | 34 | Pstpip2 |
| Ptafr | 21 | 27 | 11 | 13 | 34 | 22 | 25 | 54 | 72 | 57 | 131 | 73 | 54 | 56 | 40 | 53 | 34 | 29 | 24 | 62 | 29 | 87 | 120 | 64 | 66 | 73 | 58 | 101 | Ptafr |
| Ptar1 | 8792 | 7479 | 7172 | 7813 | 7791 | 8959 | 9169 | 7740 | 6354 | 6079 | 6404 | 7753 | 7656 | 8166 | 8943 | 8205 | 7537 | 7416 | 7608 | 9459 | 8859 | 5730 | 6422 | 6007 | 6543 | 5736 | 8492 | 8270 | Ptar1 |
| Ptbp1 | 23236 | 24676 | 21303 | 17638 | 13425 | 16011 | 13866 | 17573 | 22795 | 19523 | 20350 | 22504 | 23516 | 20696 | 18676 | 24512 | 18789 | 16404 | 15820 | 16148 | 13921 | 14088 | 21920 | 21083 | 17583 | 29255 | 22907 | 22953 | Ptbp1 |
| Ptbp2 | 20390 | 16243 | 20599 | 19334 | 20868 | 22907 | 23673 | 35473 | 27825 | 26774 | 25056 | 36297 | 31369 | 38114 | 23375 | 17890 | 21049 | 18832 | 21688 | 26306 | 21932 | 32108 | 26177 | 31517 | 34098 | 36778 | 37048 | 38554 | Ptbp2 |
| Ptbp3 | 14090 | 9346 | 9601 | 8749 | 7972 | 9115 | 8581 | 8954 | 8283 | 6225 | 7670 | 6635 | 8263 | 6563 | 13444 | 13237 | 8835 | 8741 | 7388 | 10227 | 8012 | 7183 | 8774 | 6061 | 5937 | 6608 | 6789 | 7187 | Ptbp3 |
| Ptcd1 | 1577 | 2263 | 1677 | 1921 | 1780 | 1719 | 1703 | 1661 | 1975 | 1782 | 1710 | 1748 | 1738 | 1606 | 1713 | 2017 | 1649 | 2040 | 1818 | 1841 | 1724 | 1915 | 1897 | 1696 | 1798 | 1534 | 1743 | 1712 | Ptcd1 |
| Ptcd2 | 1899 | 2146 | 1832 | 1745 | 1657 | 1588 | 1973 | 2168 | 1906 | 1810 | 1663 | 1866 | 1684 | 2006 | 2037 | 1975 | 1785 | 1869 | 1760 | 2102 | 1809 | 1917 | 2008 | 1785 | 1783 | 2030 | 1966 | 2059 | Ptcd2 |
| Ptcd3 | 9943 | 8480 | 7111 | 7143 | 6528 | 7302 | 7907 | 8258 | 6910 | 5633 | 5864 | 7117 | 6914 | 6972 | 10056 | 9256 | 6810 | 6824 | 6643 | 8445 | 7381 | 6136 | 6984 | 5932 | 6047 | 6857 | 7524 | 7122 | Ptcd3 |
| Ptch1 | 7074 | 6579 | 7260 | 7348 | 8085 | 6898 | 7111 | 8716 | 9175 | 8251 | 8706 | 9850 | 7097 | 9008 | 6870 | 6653 | 7562 | 7411 | 6713 | 6080 | 6981 | 9728 | 7970 | 7975 | 7765 | 8240 | 9941 | 9406 | Ptch1 |
| Ptch2 | 1072 | 907 | 1216 | 922 | 1021 | 886 | 860 | 662 | 957 | 830 | 655 | 942 | 569 | 791 | 779 | 896 | 1252 | 1000 | 1019 | 700 | 852 | 799 | 801 | 798 | 726 | 988 | 896 | 781 | Ptch2 |
| Ptchd1 | 172 | 120 | 82 | 158 | 259 | 247 | 165 | 457 | 419 | 361 | 413 | 164 | 243 | 147 | 171 | 142 | 140 | 143 | 191 | 238 | 160 | 530 | 386 | 180 | 432 | 71 | 196 | 170 | Ptchd1 |
| Ptchd3 | 1 | 1 | 0 | 1 | 0 | 1 | 0 | 0 | 2 | 0 | 1 | 2 | 5 | 2 | 0 | 3 | 2 | 0 | 0 | 1 | 0 | 2 | 5 | 0 | 0 | 4 | 1 | 1 | Ptchd3 |
| Ptchd4 | 96 | 137 | 47 | 49 | 69 | 101 | 45 | 164 | 147 | 96 | 105 | 70 | 159 | 100 | 112 | 104 | 57 | 50 | 48 | 123 | 56 | 102 | 230 | 67 | 89 | 71 | 134 | 81 | Ptchd4 |
| Ptcra | 1 | 10 | 8 | 2 | 3 | 0 | 3 | 0 | 0 | 0 | 4 | 2 | 2 | 1 | 0 | 2 | 9 | 4 | 0 | 4 | 2 | 2 | 2 | 2 | 0 | 8 | 5 | 0 | Ptcra |
| Ptdss1 | 6567 | 6511 | 5238 | 5523 | 5454 | 5322 | 5491 | 5162 | 4820 | 4231 | 4159 | 3913 | 4153 | 3924 | 6931 | 5921 | 5203 | 5676 | 5543 | 5757 | 5762 | 4739 | 4708 | 4071 | 4187 | 3955 | 3943 | 4035 | Ptdss1 |
| Ptdss2 | 3418 | 3994 | 3484 | 3344 | 3228 | 3197 | 3169 | 3147 | 3367 | 3173 | 3221 | 3163 | 3188 | 3026 | 3398 | 3664 | 3551 | 3394 | 3292 | 3153 | 3099 | 3055 | 3414 | 3352 | 3293 | 3480 | 3270 | 3254 | Ptdss2 |
| Pten | 14704 | 10610 | 13154 | 16181 | 16450 | 16272 | 17654 | 12313 | 10574 | 11054 | 10851 | 10473 | 10377 | 11284 | 14991 | 12710 | 13580 | 16450 | 14825 | 14875 | 17276 | 12395 | 10572 | 10830 | 12161 | 9390 | 10261 | 11065 | Pten |
| Pter | 494 | 383 | 509 | 300 | 319 | 253 | 307 | 302 | 376 | 336 | 417 | 304 | 371 | 301 | 487 | 541 | 334 | 286 | 268 | 349 | 244 | 326 | 482 | 288 | 251 | 343 | 375 | 344 | Pter |
| Ptf1a | 8 | 70 | 1 | 23 | 25 | 22 | 15 | 32 | 35 | 26 | 18 | 1 | 58 | 6 | 51 | 62 | 43 | 12 | 13 | 37 | 6 | 35 | 24 | 5 | 19 | 10 | 6 | 1 | Ptf1a |
| Ptgdr | 15 | 14 | 8 | 13 | 29 | 28 | 19 | 21 | 38 | 39 | 35 | 13 | 19 | 20 | 8 | 13 | 7 | 24 | 16 | 19 | 13 | 32 | 56 | 25 | 39 | 12 | 5 | 9 | Ptgdr |
| Ptgdr2 | 10 | 2 | 3 | 1 | 2 | 0 | 2 | 12 | 7 | 16 | 6 | 12 | 12 | 5 | 0 | 6 | 6 | 3 | 5 | 3 | 2 | 18 | 13 | 11 | 5 | 4 | 9 | 5 | Ptgdr2 |
| Ptgds | 9 | 18 | 11 | 70 | 380 | 164 | 115 | 227 | 525 | 293 | 452 | 43 | 45 | 61 | 25 | 19 | 30 | 257 | 121 | 175 | 194 | 485 | 794 | 234 | 394 | 67 | 44 | 17 | Ptgds |
| Ptger1 | 202 | 322 | 329 | 429 | 296 | 253 | 247 | 84 | 82 | 187 | 85 | 80 | 61 | 100 | 214 | 307 | 467 | 480 | 347 | 267 | 290 | 69 | 74 | 162 | 100 | 116 | 79 | 59 | Ptger1 |
| Ptger2 | 48 | 22 | 25 | 33 | 51 | 56 | 20 | 24 | 42 | 34 | 28 | 9 | 15 | 17 | 29 | 34 | 35 | 33 | 19 | 27 | 19 | 43 | 43 | 25 | 28 | 22 | 15 | 12 | Ptger2 |
| Ptger3 | 239 | 190 | 143 | 161 | 200 | 156 | 120 | 247 | 214 | 173 | 314 | 113 | 132 | 115 | 200 | 312 | 96 | 235 | 121 | 132 | 116 | 202 | 285 | 174 | 138 | 137 | 139 | 98 | Ptger3 |
| Ptger4 | 270 | 241 | 203 | 196 | 137 | 106 | 74 | 70 | 110 | 119 | 147 | 41 | 68 | 51 | 258 | 409 | 148 | 240 | 104 | 95 | 99 | 77 | 91 | 76 | 48 | 89 | 52 | 59 | Ptger4 |
| Ptges | 1588 | 1710 | 1313 | 975 | 716 | 936 | 468 | 387 | 468 | 445 | 539 | 318 | 490 | 302 | 1283 | 2235 | 903 | 1186 | 558 | 596 | 474 | 334 | 571 | 416 | 281 | 515 | 334 | 352 | Ptges |
| Ptges2 | 1987 | 2364 | 1781 | 2134 | 1912 | 1940 | 1839 | 1252 | 1346 | 1222 | 1076 | 1127 | 1145 | 1009 | 1900 | 1879 | 1910 | 2040 | 2117 | 1927 | 2070 | 1265 | 1306 | 1176 | 1214 | 1151 | 1124 | 1013 | Ptges2 |
| Ptges3 | 15443 | 11050 | 17013 | 14321 | 12413 | 12513 | 15490 | 14734 | 12803 | 12758 | 12865 | 15082 | 15110 | 17110 | 15535 | 13879 | 14746 | 13115 | 13461 | 13718 | 14325 | 13658 | 13356 | 14964 | 13544 | 19192 | 15969 | 17864 | Ptges3 |
| Ptges3l | 47 | 41 | 48 | 27 | 43 | 44 | 45 | 50 | 56 | 64 | 133 | 61 | 68 | 67 | 50 | 68 | 50 | 54 | 30 | 26 | 38 | 38 | 84 | 60 | 53 | 70 | 43 | 49 | Ptges3l |
| Ptgfr | 53 | 29 | 46 | 27 | 56 | 37 | 16 | 215 | 242 | 296 | 283 | 156 | 150 | 138 | 37 | 38 | 22 | 40 | 13 | 25 | 21 | 175 | 287 | 267 | 146 | 128 | 107 | 107 | Ptgfr |
| Ptgfrn | 8215 | 7445 | 7996 | 7861 | 7906 | 7666 | 7774 | 6792 | 8834 | 8830 | 9684 | 9275 | 9212 | 8614 | 6624 | 7977 | 7759 | 8501 | 8656 | 7208 | 7887 | 6432 | 8961 | 9682 | 8394 | 9756 | 7775 | 8586 | Ptgfrn |
| Ptgir | 52 | 63 | 30 | 60 | 66 | 30 | 32 | 70 | 92 | 126 | 128 | 63 | 111 | 64 | 69 | 104 | 80 | 61 | 59 | 51 | 33 | 114 | 72 | 46 | 50 | 51 | 70 | 68 | Ptgir |
| Ptgis | 514 | 642 | 693 | 493 | 365 | 414 | 268 | 487 | 542 | 451 | 856 | 499 | 691 | 491 | 517 | 771 | 523 | 375 | 351 | 364 | 310 | 485 | 825 | 467 | 423 | 790 | 625 | 564 | Ptgis |
| Ptgr1 | 1317 | 1206 | 1524 | 644 | 512 | 769 | 439 | 942 | 969 | 788 | 902 | 1068 | 1162 | 1054 | 1319 | 1559 | 951 | 888 | 542 | 728 | 529 | 597 | 1109 | 881 | 617 | 1335 | 1177 | 1195 | Ptgr1 |
| Ptgr2 | 1517 | 1259 | 1603 | 1365 | 1403 | 1379 | 1414 | 1291 | 1307 | 1160 | 1139 | 1407 | 1401 | 1312 | 1580 | 1403 | 1499 | 1325 | 1331 | 1570 | 1348 | 1198 | 1265 | 1260 | 1285 | 1216 | 1334 | 1346 | Ptgr2 |
| Ptgs1 | 386 | 383 | 362 | 270 | 277 | 320 | 201 | 296 | 433 | 407 | 591 | 280 | 403 | 258 | 331 | 406 | 220 | 343 | 224 | 270 | 238 | 389 | 500 | 348 | 293 | 391 | 372 | 280 | Ptgs1 |
| Ptgs2 | 31 | 31 | 23 | 15 | 7 | 14 | 16 | 27 | 28 | 24 | 45 | 21 | 19 | 21 | 24 | 62 | 17 | 19 | 14 | 9 | 5 | 18 | 48 | 29 | 11 | 29 | 8 | 15 | Ptgs2 |
| Pth | 0 | 0 | 0 | 0 | 0 | 0 | 0 | 0 | 0 | 0 | 57 | 0 | 0 | 0 | 0 | 0 | 0 | 0 | 0 | 0 | 0 | 0 | 57 | 0 | 0 | 0 | 0 | 0 | Pth |
| Pth1r | 1456 | 1380 | 1437 | 832 | 797 | 1070 | 526 | 1470 | 2193 | 1643 | 2177 | 1428 | 1913 | 1138 | 1149 | 1547 | 1108 | 1043 | 840 | 652 | 566 | 1186 | 2186 | 1645 | 1032 | 1611 | 1563 | 1474 | Pth1r |
| Pth2 | 3 | 1 | 0 | 9 | 2 | 3 | 6 | 2 | 2 | 1 | 0 | 1 | 1 | 7 | 0 | 2 | 5 | 0 | 5 | 1 | 3 | 2 | 1 | 0 | 5 | 0 | 6 | 5 | Pth2 |
| Pth2r | 59 | 70 | 72 | 91 | 73 | 65 | 82 | 53 | 37 | 41 | 75 | 47 | 62 | 48 | 53 | 73 | 81 | 119 | 75 | 64 | 80 | 48 | 56 | 55 | 67 | 77 | 33 | 30 | Pth2r |
| Pthlh | 53 | 64 | 54 | 57 | 61 | 52 | 49 | 136 | 142 | 123 | 133 | 97 | 119 | 48 | 53 | 76 | 41 | 52 | 44 | 44 | 44 | 110 | 179 | 42 | 66 | 65 | 62 | 55 | Pthlh |
| Ptk2 | 10743 | 10034 | 9093 | 8881 | 9180 | 9665 | 9667 | 11000 | 10125 | 10075 | 9499 | 11444 | 10629 | 11632 | 10307 | 9749 | 9056 | 8696 | 9186 | 9434 | 9315 | 9895 | 9800 | 10237 | 10727 | 10558 | 11518 | 11352 | Ptk2 |
| Ptk2b | 208 | 226 | 150 | 157 | 141 | 113 | 89 | 89 | 113 | 93 | 109 | 47 | 109 | 61 | 170 | 185 | 123 | 187 | 125 | 84 | 122 | 72 | 132 | 77 | 109 | 61 | 76 | 63 | Ptk2b |
| Ptk6 | 5 | 18 | 6 | 22 | 29 | 14 | 13 | 4 | 1 | 12 | 2 | 2 | 3 | 2 | 17 | 10 | 9 | 14 | 16 | 17 | 12 | 15 | 4 | 0 | 5 | 0 | 5 | 1 | Ptk6 |
| Ptk7 | 11833 | 13419 | 14210 | 11637 | 9637 | 10492 | 8021 | 4860 | 9749 | 8576 | 10246 | 8754 | 9800 | 7619 | 10038 | 14713 | 12056 | 12154 | 10194 | 7850 | 8658 | 6582 | 10066 | 9421 | 7389 | 10132 | 7925 | 7919 | Ptk7 |
| Ptma | 96932 | 91802 | 113033 | 101647 | 86657 | 83895 | 109859 | 96231 | 87599 | 103567 | 92503 | 107394 | 103639 | 138117 | 92746 | 79295 | 106283 | 84746 | 94939 | 95719 | 103301 | 76575 | 81293 | 120156 | 103874 | 132325 | 110671 | 134009 | Ptma |
| Ptms | 13650 | 17195 | 18948 | 25699 | 26651 | 23036 | 20403 | 21070 | 23881 | 35206 | 26891 | 29756 | 24072 | 31609 | 13756 | 13687 | 23187 | 23955 | 28791 | 19858 | 25684 | 31030 | 24398 | 35858 | 34685 | 32876 | 26427 | 28734 | Ptms |
| Ptn | 14496 | 11609 | 12820 | 15916 | 19471 | 18215 | 16697 | 25460 | 25683 | 25887 | 32905 | 18811 | 19758 | 20522 | 16481 | 13972 | 13752 | 17938 | 17182 | 21276 | 17264 | 25218 | 36125 | 25977 | 25444 | 23183 | 18003 | 17553 | Ptn |
| Ptov1 | 12227 | 15971 | 15849 | 18991 | 16997 | 16556 | 13939 | 15042 | 16975 | 19756 | 17813 | 21349 | 17672 | 18478 | 12058 | 12952 | 18219 | 16666 | 19147 | 15140 | 15947 | 17400 | 16027 | 21283 | 20100 | 23153 | 18779 | 18052 | Ptov1 |
| Ptp4a1 | 68 | 60 | 79 | 115 | 50 | 62 | 55 | 104 | 129 | 84 | 86 | 109 | 79 | 79 | 63 | 44 | 61 | 75 | 86 | 74 | 64 | 110 | 99 | 128 | 104 | 122 | 96 | 125 | Ptp4a1 |
| Ptp4a1 | 0 | 1 | 0 | 5 | 0 | 0 | 1 | 0 | 0 | 0 | 3 | 4 | 1 | 1 | 0 | 2 | 6 | 0 | 1 | 1 | 0 | 2 | 2 | 6 | 3 | 1 | 1 | 1 | Ptp4a1 |
| Ptp4a2 | 32361 | 25365 | 29546 | 27815 | 25120 | 27782 | 32001 | 25745 | 21812 | 21148 | 22101 | 25528 | 24128 | 28369 | 31459 | 26929 | 28481 | 27097 | 27530 | 28756 | 30851 | 21626 | 22629 | 24585 | 23550 | 28195 | 26246 | 29834 | Ptp4a2 |
| Ptp4a3 | 971 | 1053 | 1191 | 1198 | 1320 | 1214 | 1036 | 1451 | 1893 | 1952 | 2205 | 1729 | 1918 | 1651 | 890 | 1091 | 1236 | 1153 | 1308 | 1094 | 1055 | 1721 | 1953 | 1897 | 1740 | 1952 | 1524 | 1881 | Ptp4a3 |
| Ptpa | 7668 | 8174 | 8484 | 8450 | 8437 | 8184 | 7420 | 9082 | 10191 | 10176 | 10321 | 11460 | 11119 | 10771 | 6926 | 7639 | 8364 | 7774 | 9120 | 7631 | 7860 | 9605 | 10281 | 11089 | 11033 | 12580 | 11303 | 11136 | Ptpa |
| Ptpdc1 | 2673 | 2425 | 2266 | 2600 | 2829 | 2877 | 2808 | 2944 | 3152 | 3344 | 3383 | 2854 | 2790 | 2893 | 2525 | 2442 | 2508 | 2584 | 2681 | 2483 | 2703 | 3050 | 3178 | 3108 | 2960 | 2821 | 2783 | 2662 | Ptpdc1 |
| Ptpn1 | 7354 | 9445 | 10111 | 10315 | 8915 | 8697 | 8980 | 4860 | 4828 | 5807 | 5416 | 6675 | 6127 | 6497 | 7189 | 8087 | 10639 | 9850 | 9976 | 7462 | 9183 | 6161 | 4623 | 6171 | 6774 | 6295 | 6279 | 6868 | Ptpn1 |
| Ptpn11 | 20075 | 20843 | 18777 | 19171 | 17807 | 18393 | 17966 | 13764 | 14380 | 13915 | 13542 | 15732 | 15119 | 15088 | 19235 | 18922 | 18773 | 18628 | 18332 | 18636 | 17455 | 13025 | 14006 | 14683 | 15125 | 14141 | 15983 | 15094 | Ptpn11 |
| Ptpn12 | 4862 | 4199 | 5044 | 5047 | 5536 | 5170 | 5679 | 7387 | 6505 | 6422 | 6493 | 7236 | 6969 | 7629 | 5008 | 4143 | 4954 | 4826 | 5292 | 5377 | 5602 | 7337 | 6472 | 6770 | 7269 | 7047 | 7074 | 7800 | Ptpn12 |
| Ptpn13 | 14207 | 15462 | 11352 | 11928 | 10091 | 11070 | 10247 | 5599 | 5907 | 5635 | 5152 | 5433 | 6320 | 5112 | 13193 | 15276 | 11488 | 11234 | 10404 | 10381 | 10544 | 4878 | 6733 | 4545 | 4497 | 3492 | 5768 | 5301 | Ptpn13 |
| Ptpn14 | 3576 | 3403 | 2704 | 2524 | 2234 | 2731 | 1434 | 3264 | 4569 | 3992 | 5172 | 2939 | 4273 | 2555 | 2981 | 4341 | 2588 | 2881 | 1740 | 1596 | 1613 | 3131 | 4814 | 3239 | 2820 | 2117 | 3218 | 2654 | Ptpn14 |
| Ptpn18 | 105 | 141 | 138 | 166 | 90 | 98 | 108 | 70 | 98 | 101 | 90 | 84 | 94 | 67 | 82 | 115 | 112 | 130 | 98 | 98 | 102 | 75 | 115 | 92 | 69 | 83 | 77 | 66 | Ptpn18 |
| Ptpn2 | 1895 | 1424 | 1799 | 1454 | 1629 | 1546 | 1846 | 3198 | 2818 | 2690 | 2744 | 2888 | 2812 | 3177 | 1850 | 1471 | 1637 | 1503 | 1597 | 1766 | 1712 | 2734 | 2966 | 2884 | 2930 | 3039 | 2898 | 3063 | Ptpn2 |
| Ptpn20 | 8 | 6 | 9 | 2 | 9 | 4 | 7 | 22 | 16 | 8 | 17 | 11 | 20 | 5 | 11 | 13 | 1 | 6 | 9 | 4 | 1 | 18 | 13 | 14 | 7 | 1 | 9 | 6 | Ptpn20 |
| Ptpn21 | 1183 | 1286 | 1906 | 1717 | 1513 | 1617 | 1543 | 1102 | 1214 | 1385 | 1543 | 1970 | 1463 | 1957 | 1031 | 1281 | 1785 | 1619 | 1870 | 1230 | 1576 | 1420 | 1233 | 1684 | 1611 | 1770 | 1683 | 1946 | Ptpn21 |
| Ptpn22 | 65 | 73 | 66 | 118 | 71 | 63 | 76 | 32 | 27 | 49 | 34 | 17 | 24 | 26 | 61 | 63 | 88 | 88 | 49 | 52 | 64 | 37 | 32 | 30 | 21 | 36 | 7 | 20 | Ptpn22 |
| Ptpn23 | 5742 | 7480 | 7975 | 8623 | 7917 | 7396 | 6603 | 4797 | 6297 | 7156 | 6762 | 7543 | 6432 | 6046 | 5436 | 6815 | 8228 | 8352 | 8548 | 5873 | 7167 | 7583 | 5933 | 7498 | 7304 | 7162 | 6373 | 6000 | Ptpn23 |
| Ptpn3 | 2155 | 2046 | 1881 | 2120 | 2621 | 1828 | 1863 | 1316 | 1338 | 1136 | 1353 | 394 | 1094 | 399 | 2256 | 2477 | 1990 | 2143 | 1880 | 1566 | 1886 | 1584 | 1028 | 642 | 835 | 444 | 400 | 399 | Ptpn3 |
| Ptpn4 | 11682 | 8661 | 6566 | 10205 | 11090 | 13317 | 13635 | 9959 | 8631 | 7002 | 7017 | 7981 | 7900 | 8246 | 12857 | 10471 | 7969 | 10482 | 10713 | 14274 | 14899 | 7389 | 8014 | 6010 | 8365 | 4952 | 8634 | 7734 | Ptpn4 |
| Ptpn5 | 1606 | 2196 | 1914 | 3514 | 5768 | 4135 | 3920 | 5047 | 5183 | 6787 | 5407 | 4885 | 4816 | 4641 | 2142 | 1812 | 2814 | 3505 | 4403 | 3625 | 4260 | 7660 | 4566 | 4732 | 6192 | 2764 | 3894 | 3734 | Ptpn5 |
| Ptpn6 | 231 | 219 | 246 | 152 | 186 | 186 | 120 | 254 | 329 | 271 | 374 | 243 | 246 | 180 | 235 | 282 | 141 | 162 | 143 | 144 | 140 | 219 | 441 | 276 | 192 | 365 | 252 | 220 | Ptpn6 |
| Ptpn7 | 45 | 54 | 80 | 13 | 38 | 51 | 20 | 73 | 72 | 55 | 70 | 72 | 94 | 33 | 57 | 60 | 43 | 26 | 34 | 30 | 24 | 47 | 100 | 52 | 61 | 80 | 107 | 52 | Ptpn7 |

| | | | | | | | | | | | | | | | | | | | | | | | | | | | | | |
|---|---|---|---|---|---|---|---|---|---|---|---|---|---|---|---|---|---|---|---|---|---|---|---|---|---|---|---|---|---|
| Ptpn9 | 3321 | 2485 | 2831 | 2666 | 3546 | 3206 | 3396 | 5628 | 5569 | 5416 | 5542 | 5663 | 5464 | 5860 | 2732 | 2630 | 2794 | 2790 | 3425 | 3102 | 3322 | 6057 | 5613 | 5447 | 5503 | 5874 | 5075 | 5488 | Ptpn9 |
| Ptpra | 9957 | 10549 | 8819 | 9848 | 9704 | 9509 | 10335 | 9384 | 9372 | 9495 | 8387 | 8612 | 8122 | 8833 | 10337 | 9536 | 9456 | 9843 | 9830 | 10023 | 10629 | 9115 | 9697 | 8762 | 9137 | 9034 | 8116 | 8737 | Ptpra |
| Ptprb | 728 | 655 | 723 | 724 | 999 | 1104 | 788 | 1000 | 1298 | 1471 | 1884 | 1427 | 1523 | 1496 | 793 | 779 | 571 | 787 | 807 | 777 | 1026 | 1569 | 1607 | 1224 | 1679 | 1078 | 1601 | 1364 | Ptprb |
| Ptprc | 224 | 191 | 175 | 156 | 210 | 149 | 141 | 222 | 256 | 197 | 331 | 207 | 226 | 193 | 218 | 292 | 134 | 182 | 132 | 256 | 177 | 254 | 366 | 192 | 150 | 223 | 186 | 197 | Ptprc |
| Ptprcap | 3 | 3 | 4 | 2 | 2 | 1 | 2 | 0 | 1 | 0 | 0 | 0 | 0 | 2 | 5 | 4 | 1 | 1 | 2 | 3 | 0 | 0 | 1 | 0 | 4 | 1 | 0 | 0 | Ptprcap |
| Ptprd | 12904 | 10572 | 7470 | 11873 | 14939 | 16348 | 14553 | 17894 | 19275 | 18024 | 21559 | 12892 | 14580 | 14303 | 13854 | 12324 | 9259 | 13424 | 11186 | 13719 | 14630 | 17587 | 20404 | 13862 | 17640 | 8562 | 13382 | 11882 | Ptprd |
| Ptpre | 276 | 257 | 198 | 422 | 625 | 523 | 552 | 343 | 476 | 502 | 478 | 234 | 332 | 275 | 314 | 343 | 257 | 517 | 519 | 456 | 559 | 568 | 472 | 324 | 512 | 194 | 214 | 239 | Ptpre |
| Ptprf | 19362 | 25001 | 27941 | 25404 | 24923 | 23126 | 19853 | 18807 | 23265 | 24657 | 21572 | 27141 | 24485 | 22580 | 18250 | 22538 | 28327 | 25457 | 26397 | 18770 | 21519 | 22640 | 21275 | 25376 | 22221 | 21783 | 23050 | 23883 | Ptprf |
| Ptprh | 9 | 19 | 23 | 49 | 79 | 40 | 27 | 43 | 49 | 77 | 70 | 38 | 45 | 21 | 17 | 27 | 23 | 44 | 50 | 56 | 42 | 47 | 47 | 16 | 67 | 21 | 40 | 24 | Ptprh |
| Ptprj | 1042 | 973 | 969 | 1038 | 1400 | 1032 | 1012 | 1164 | 1332 | 1431 | 1370 | 925 | 1078 | 940 | 1021 | 1129 | 956 | 1181 | 1183 | 938 | 1064 | 1470 | 1259 | 1052 | 1215 | 818 | 924 | 878 | Ptprj |
| Ptprk | 5287 | 4520 | 4008 | 4265 | 4511 | 4432 | 4257 | 4395 | 4799 | 4405 | 4531 | 4000 | 4350 | 3800 | 4729 | 4879 | 3949 | 4339 | 4228 | 4597 | 4500 | 4385 | 4770 | 4086 | 3948 | 4007 | 4085 | 4029 | Ptprk |
| Ptprm | 1311 | 1351 | 1156 | 1467 | 1699 | 1429 | 1165 | 1318 | 1712 | 1806 | 2172 | 1334 | 1520 | 1290 | 1283 | 1487 | 1235 | 1468 | 1284 | 1025 | 1397 | 1823 | 1862 | 1428 | 1657 | 1302 | 1238 | 1168 | Ptprm |
| Ptprn | 1696 | 2388 | 2815 | 3627 | 4403 | 3144 | 3162 | 1315 | 1395 | 1938 | 1779 | 1592 | 1464 | 1472 | 1900 | 2101 | 3845 | 3532 | 3998 | 3035 | 3315 | 2418 | 1405 | 1622 | 1832 | 1315 | 1617 | 1531 | Ptprn |
| Ptprn2 | 1737 | 2418 | 2165 | 3046 | 3362 | 3075 | 2568 | 2402 | 2380 | 2746 | 2257 | 3274 | 2437 | 2772 | 2018 | 1862 | 2868 | 2915 | 3268 | 2752 | 2732 | 3415 | 2090 | 2630 | 3317 | 2213 | 3022 | 2766 | Ptprn2 |
| Ptpro | 1505 | 1650 | 1336 | 3029 | 4471 | 4080 | 4298 | 5506 | 5318 | 5587 | 4868 | 4581 | 3948 | 4845 | 2073 | 1470 | 2067 | 3386 | 3849 | 3645 | 4694 | 7596 | 4185 | 5034 | 7277 | 2911 | 4155 | 4123 | Ptpro |
| Ptprq | 53 | 71 | 61 | 42 | 65 | 72 | 51 | 127 | 118 | 103 | 101 | 98 | 96 | 93 | 57 | 54 | 56 | 73 | 41 | 56 | 50 | 165 | 153 | 91 | 139 | 60 | 88 | 100 | Ptprq |
| Ptprr | 313 | 456 | 265 | 438 | 798 | 443 | 419 | 837 | 794 | 596 | 862 | 278 | 594 | 259 | 492 | 495 | 328 | 569 | 463 | 549 | 458 | 1123 | 625 | 305 | 648 | 154 | 236 | 290 | Ptprr |
| Ptprs | 31298 | 44273 | 45613 | 55460 | 55275 | 50412 | 48019 | 37002 | 45394 | 56836 | 48556 | 63721 | 48008 | 52134 | 31324 | 35772 | 54814 | 56286 | 61389 | 42437 | 49919 | 55032 | 41066 | 62535 | 64555 | 51157 | 51582 | 48635 | Ptprs |
| Ptprt | 1252 | 1177 | 526 | 2461 | 5071 | 3841 | 4132 | 3285 | 3340 | 3934 | 3678 | 1546 | 1951 | 1729 | 1743 | 1189 | 1207 | 3171 | 3441 | 3525 | 4829 | 4580 | 3422 | 1964 | 3971 | 623 | 1441 | 1248 | Ptprt |
| Ptpru | 5392 | 6613 | 6598 | 7277 | 6101 | 6505 | 5920 | 2694 | 4287 | 4388 | 3790 | 5197 | 3737 | 4290 | 4548 | 5722 | 7291 | 7212 | 7466 | 5570 | 6064 | 3355 | 3262 | 5037 | 4452 | 5237 | 4220 | 3826 | Ptpru |
| Ptprz1 | 25185 | 22570 | 14695 | 20426 | 25427 | 26975 | 31894 | 30140 | 26661 | 20410 | 17300 | 28449 | 13659 | 30089 | 27941 | 14530 | 20664 | 22459 | 25076 | 28599 | 30837 | 26065 | 20225 | 21217 | 24132 | 21934 | 26710 | 24426 | Ptprz1 |
| Ptrh1 | 354 | 435 | 416 | 493 | 268 | 333 | 344 | 158 | 198 | 172 | 199 | 163 | 220 | 163 | 357 | 369 | 403 | 412 | 346 | 345 | 268 | 145 | 178 | 195 | 188 | 197 | 206 | 153 | Ptrh1 |
| Ptrh2 | 2932 | 2528 | 2721 | 2393 | 2423 | 2339 | 2684 | 3505 | 3207 | 2791 | 2782 | 3552 | 3137 | 3865 | 2896 | 2403 | 2757 | 2308 | 2561 | 3046 | 2633 | 2924 | 3043 | 3483 | 3299 | 4198 | 3755 | 4016 | Ptrh2 |
| Ptrhd1 | 148 | 187 | 175 | 161 | 180 | 192 | 178 | 530 | 503 | 543 | 523 | 522 | 531 | 656 | 166 | 150 | 148 | 105 | 188 | 191 | 182 | 413 | 517 | 570 | 592 | 685 | 662 | 629 | Ptrhd1 |
| Pts | 2205 | 1894 | 2366 | 2054 | 2176 | 2260 | 2181 | 3048 | 2678 | 2872 | 2695 | 3170 | 3031 | 3297 | 2411 | 1811 | 2078 | 2057 | 2163 | 2305 | 2179 | 3166 | 2795 | 2997 | 2968 | 2783 | 2981 | 3189 | Pts |
| Pttg1 | 1513 | 1697 | 1782 | 1584 | 1289 | 1281 | 1415 | 764 | 803 | 688 | 744 | 748 | 945 | 889 | 1688 | 1675 | 1613 | 1579 | 1254 | 1393 | 1292 | 632 | 898 | 712 | 593 | 888 | 876 | 869 | Pttg1 |
| Pttg1ip | 6109 | 6480 | 5257 | 5440 | 4865 | 5043 | 4536 | 4205 | 4612 | 3886 | 4573 | 3806 | 4316 | 3632 | 6003 | 6587 | 5027 | 5332 | 4524 | 5316 | 4666 | 4114 | 5063 | 3884 | 4001 | 4487 | 3841 | 3890 | Pttg1ip |
| Pttg1ip2 | 8 | 5 | 1 | 6 | 3 | 7 | 10 | 0 | 0 | 0 | 0 | 5 | 0 | 2 | 5 | 6 | 1 | 8 | 5 | 10 | 6 | 0 | 1 | 2 | 1 | 0 | 2 | 2 | Pttg1ip2 |
| Ptx3 | 714 | 687 | 1236 | 503 | 548 | 778 | 537 | 2194 | 2169 | 2116 | 2276 | 2745 | 2544 | 2811 | 918 | 775 | 1027 | 568 | 684 | 687 | 564 | 1724 | 2362 | 3053 | 2037 | 3793 | 2783 | 3089 | Ptx3 |
| Ptx4 | 6 | 4 | 3 | 3 | 2 | 3 | 0 | 2 | 3 | 0 | 1 | 3 | 1 | 2 | 7 | 2 | 0 | 14 | 7 | 1 | 3 | 5 | 11 | 0 | 2 | 3 | 3 | 0 | Ptx4 |
| Pudp | 0 | 0 | 0 | 0 | 0 | 0 | 0 | 1 | 0 | 0 | 0 | 0 | 0 | 0 | 0 | 0 | 0 | 0 | 0 | 0 | 0 | 0 | 1 | 0 | 0 | 0 | 1 | 0 | Pudp |
| Puf60 | 6452 | 7884 | 7794 | 6252 | 5951 | 5961 | 6138 | 9898 | 10698 | 10330 | 9405 | 12202 | 10761 | 12204 | 6497 | 6326 | 7307 | 5709 | 6629 | 6267 | 6201 | 9606 | 10363 | 11679 | 10543 | 12967 | 12377 | 12715 | Puf60 |
| Pum1 | 6043 | 5713 | 8255 | 7906 | 7963 | 7429 | 8171 | 7761 | 8143 | 9942 | 9196 | 11448 | 9754 | 11799 | 6046 | 6052 | 8269 | 8282 | 8401 | 5981 | 8110 | 10661 | 7886 | 10806 | 10796 | 11692 | 9829 | 11568 | Pum1 |
| Pum2 | 22884 | 17691 | 17583 | 18709 | 21671 | 22517 | 24778 | 34054 | 28071 | 26586 | 25146 | 32104 | 29058 | 33662 | 23389 | 18552 | 19051 | 19226 | 21296 | 23685 | 23628 | 29955 | 26638 | 27710 | 29760 | 27853 | 32097 | 32442 | Pum2 |
| Pum3 | 7283 | 6797 | 7274 | 6081 | 5696 | 5829 | 6427 | 6417 | 5799 | 5515 | 5562 | 6114 | 6015 | 6659 | 7138 | 6650 | 6456 | 6023 | 5798 | 6889 | 5979 | 5101 | 5721 | 5884 | 5824 | 6944 | 6568 | 6643 | Pum3 |
| Pura | 2743 | 2116 | 2635 | 2656 | 2485 | 3259 | 3030 | 2409 | 2197 | 1718 | 1932 | 2032 | 2051 | 2043 | 2778 | 2643 | 2427 | 2591 | 2592 | 3064 | 3267 | 1778 | 2249 | 1942 | 2053 | 2170 | 2199 | 1984 | Pura |
| Purb | 16035 | 10817 | 12464 | 11992 | 11243 | 13321 | 13058 | 11138 | 10542 | 10156 | 13274 | 11104 | 11891 | 11484 | 13969 | 14214 | 11586 | 12420 | 10358 | 10823 | 12682 | 10510 | 11479 | 9617 | 10308 | 9722 | 12035 | 11744 | Purb |
| Purg | 2569 | 2807 | 3863 | 4095 | 3547 | 3226 | 3627 | 2579 | 2258 | 2882 | 2896 | 2590 | 2447 | 2862 | 2817 | 2815 | 4124 | 3979 | 3897 | 2947 | 3666 | 3200 | 2408 | 3058 | 3046 | 2953 | 2338 | 2803 | Purg |
| Pus1 | 1185 | 1576 | 1671 | 1495 | 1080 | 1171 | 1055 | 1044 | 1107 | 1286 | 1137 | 1263 | 1212 | 1253 | 1052 | 1394 | 1406 | 1271 | 1170 | 1101 | 1029 | 976 | 1178 | 1335 | 1190 | 1623 | 1373 | 1291 | Pus1 |
| Pus10 | 5547 | 4431 | 4189 | 4011 | 3447 | 4289 | 4722 | 3767 | 3225 | 2843 | 3242 | 3287 | 3869 | 3754 | 5575 | 5132 | 4403 | 3848 | 3754 | 4639 | 4205 | 2921 | 3383 | 3058 | 3096 | 3508 | 3647 | 4021 | Pus10 |
| Pus3 | 994 | 773 | 848 | 771 | 907 | 911 | 997 | 2111 | 1951 | 1497 | 1666 | 1963 | 1932 | 1994 | 1007 | 826 | 819 | 701 | 922 | 1125 | 854 | 1623 | 1847 | 1821 | 1652 | 2064 | 1948 | 2085 | Pus3 |
| Pus7 | 4840 | 4491 | 4429 | 4605 | 3741 | 4368 | 4257 | 2348 | 2399 | 2322 | 2615 | 2540 | 2605 | 2362 | 4551 | 4417 | 4152 | 4590 | 3949 | 4175 | 4202 | 2197 | 2367 | 2273 | 2352 | 2782 | 2456 | 2607 | Pus7 |
| Pus7l | 670 | 651 | 757 | 646 | 495 | 562 | 645 | 596 | 611 | 633 | 640 | 660 | 692 | 581 | 670 | 682 | 591 | 675 | 540 | 742 | 555 | 486 | 663 | 595 | 520 | 809 | 646 | 662 | Pus7l |
| Pusl1 | 1673 | 1976 | 1594 | 1844 | 1733 | 1851 | 1742 | 916 | 1060 | 1117 | 1222 | 1206 | 1207 | 1051 | 1493 | 1834 | 1688 | 1984 | 1830 | 1769 | 1734 | 1054 | 1129 | 1118 | 1040 | 1094 | 1095 | 1106 | Pusl1 |
| Pvalb | 3 | 20 | 0 | 13 | 92 | 26 | 3 | 122 | 101 | 63 | 55 | 1 | 66 | 3 | 20 | 35 | 15 | 15 | 21 | 32 | 7 | 99 | 102 | 2 | 98 | 0 | 1 | 1 | Pvalb |
| Pvr | 248 | 208 | 226 | 242 | 422 | 293 | 279 | 636 | 677 | 739 | 842 | 426 | 538 | 462 | 272 | 278 | 217 | 243 | 281 | 273 | 267 | 685 | 679 | 562 | 567 | 539 | 406 | 414 | Pvr |
| Pvrig | 6 | 3 | 8 | 1 | 1 | 0 | 3 | 10 | 0 | 11 | 5 | 3 | 0 | 6 | 1 | 4 | 0 | 3 | 4 | 7 | 5 | 7 | 1 | 2 | 5 | 0 | 3 | 8 | Pvrig |
| Pwp1 | 5523 | 5689 | 4615 | 4540 | 3953 | 4403 | 4775 | 4728 | 4373 | 3747 | 3663 | 4331 | 3997 | 4326 | 5759 | 5277 | 4509 | 4461 | 4253 | 5359 | 4564 | 3522 | 4198 | 3781 | 3885 | 4986 | 4344 | 4608 | Pwp1 |
| Pwp2 | 2128 | 2608 | 2591 | 2637 | 2170 | 2284 | 2050 | 2514 | 2650 | 2617 | 2634 | 3127 | 2866 | 2879 | 2087 | 2214 | 2332 | 2423 | 2275 | 2190 | 2137 | 2308 | 2424 | 2917 | 2910 | 3221 | 3109 | 2884 | Pwp2 |
| Pwwp2a | 3056 | 2849 | 4109 | 3728 | 3614 | 3528 | 3518 | 4150 | 3643 | 4471 | 4112 | 4714 | 4762 | 4736 | 3042 | 2967 | 3884 | 3483 | 3759 | 3254 | 3282 | 4707 | 3679 | 4651 | 4552 | 4953 | 4692 | 5057 | Pwwp2a |
| Pwwp2b | 42 | 22 | 14 | 6 | 3 | 12 | 18 | 2 | 6 | 5 | 1 | 8 | 1 | 0 | 8 | 11 | 14 | 47 | 14 | 5 | 4 | 3 | 4 | 6 | 6 | 4 | 2 | 4 | Pwwp2b |
| Pwwp3a | 7435 | 8304 | 10899 | 8326 | 6898 | 7316 | 6322 | 6069 | 5862 | 7128 | 6853 | 9336 | 8998 | 8983 | 6695 | 7519 | 9619 | 7530 | 8137 | 6534 | 7240 | 6475 | 6469 | 8755 | 7445 | 7170 | 10665 | 10448 | Pwwp3a |
| Pwwp3b | 6757 | 4753 | 5167 | 4566 | 3963 | 4496 | 4704 | 1866 | 2026 | 1580 | 2440 | 1791 | 2121 | 1788 | 6150 | 6259 | 4487 | 4568 | 3956 | 4668 | 4236 | 1594 | 2400 | 1602 | 1434 | 2017 | 1855 | 1932 | Pwwp3b |
| Pwwp4a | 2 | 0 | 0 | 2 | 2 | 5 | 1 | 2 | 2 | 2 | 3 | 5 | 2 | 2 | 5 | 2 | 1 | 2 | 0 | 7 | 1 | 2 | 17 | 3 | 9 | 2 | 7 | 11 | Pwwp4a |
| Pwwp4b | 2 | 3 | 12 | 8 | 2 | 3 | 6 | 4 | 5 | 11 | 3 | 10 | 3 | 5 | 7 | 2 | 9 | 2 | 10 | 6 | 7 | 10 | 8 | 16 | 9 | 0 | 12 | 4 | Pwwp4b |
| Pwwp4c | 2 | 2 | 0 | 1 | 2 | 12 | 6 | 6 | 2 | 11 | 3 | 10 | 3 | 5 | 0 | 6 | 5 | 4 | 9 | 10 | 7 | 12 | 5 | 6 | 7 | 7 | 9 | 6 | Pwwp4c |
| Pwwp4d | 0 | 2 | 3 | 1 | 2 | 1 | 3 | 0 | 4 | 2 | 2 | 0 | 3 | 1 | 1 | 2 | 0 | 4 | 4 | 2 | 3 | 5 | 3 | 1 | 1 | 0 | 0 | 0 | Pwwp4d |
| Pxdc1 | 355 | 341 | 406 | 266 | 303 | 323 | 230 | 416 | 383 | 377 | 503 | 290 | 308 | 283 | 297 | 376 | 346 | 326 | 265 | 230 | 221 | 371 | 447 | 314 | 328 | 370 | 354 | 356 | Pxdc1 |
| Pxdn | 15625 | 15308 | 13617 | 11808 | 12758 | 13691 | 9426 | 17043 | 20618 | 20376 | 24776 | 17688 | 20258 | 16090 | 13572 | 18608 | 12493 | 13644 | 11480 | 9766 | 9760 | 19567 | 20420 | 17638 | 17499 | 14563 | 17307 | 16247 | Pxdn |
| Pxk | 3806 | 4133 | 3479 | 4308 | 4500 | 4207 | 4796 | 2688 | 2780 | 2853 | 2853 | 2243 | 2236 | 2321 | 3956 | 4099 | 3843 | 4521 | 4398 | 4264 | 4837 | 3093 | 2738 | 2402 | 2988 | 2076 | 1904 | 2079 | Pxk |
| Pxmp2 | 467 | 496 | 453 | 420 | 391 | 376 | 393 | 173 | 154 | 205 | 211 | 201 | 210 | 229 | 455 | 506 | 391 | 404 | 423 | 413 | 390 | 142 | 169 | 175 | 144 | 219 | 207 | 191 | Pxmp2 |
| Pxmp4 | 1198 | 1550 | 1355 | 1339 | 1071 | 1168 | 969 | 749 | 883 | 807 | 976 | 751 | 934 | 682 | 1114 | 1340 | 1344 | 1248 | 1043 | 1068 | 956 | 734 | 947 | 823 | 719 | 866 | 807 | 760 | Pxmp4 |
| Pxn | 7932 | 9987 | 9354 | 10630 | 9037 | 8448 | 8212 | 3417 | 4172 | 4261 | 5477 | 3738 | 4095 | 3250 | 7569 | 9809 | 9521 | 10986 | 8890 | 7490 | 8841 | 4089 | 4994 | 4374 | 3690 | 4226 | 3546 | 3697 | Pxn |
| Pxt1 | 0 | 0 | 0 | 4 | 5 | 3 | 1 | 1 | 1 | 0 | 2 | 3 | 1 | 1 | 0 | 2 | 0 | 0 | 0 | 0 | 1 | 0 | 0 | 0 | 0 | 3 | 0 | 2 | Pxt1 |
| Pxylp1 | 3865 | 4467 | 4485 | 4546 | 4428 | 4502 | 4314 | 3203 | 2810 | 3260 | 2556 | 3408 | 3504 | 3108 | 4305 | 4076 | 4755 | 4189 | 4641 | 4534 | 4375 | 3481 | 2462 | 3006 | 3309 | 2927 | 3380 | 3249 | Pxylp1 |
| Pycard | 160 | 98 | 111 | 69 | 62 | 91 | 50 | 162 | 217 | 167 | 180 | 164 | 133 | 117 | 127 | 127 | 100 | 100 | 49 | 93 | 62 | 92 | 267 | 127 | 111 | 115 | 132 | 157 | Pycard |
| Pycr1 | 1408 | 1597 | 1485 | 1433 | 1118 | 1396 | 1101 | 960 | 1205 | 1081 | 1342 | 1218 | 1359 | 1244 | 1402 | 1505 | 1460 | 1438 | 1329 | 1236 | 1115 | 1033 | 1375 | 1215 | 1260 | 1394 | 1398 | 1431 | Pycr1 |
| Pycr2 | 2787 | 3258 | 3197 | 3051 | 2526 | 2503 | 2342 | 2022 | 2183 | 2048 | 2191 | 2465 | 2311 | 2222 | 2334 | 2863 | 3074 | 2748 | 2876 | 2621 | 2322 | 1853 | 2119 | 2580 | 2183 | 3006 | 2539 | 2487 | Pycr2 |
| Pycrl | 2158 | 2665 | 2131 | 2179 | 1866 | 2055 | 1811 | 2170 | 2707 | 2162 | 2160 | 2306 | 2327 | 2067 | 2122 | 2320 | 2291 | 2062 | 2397 | 2251 | 1996 | 1920 | 2599 | 2347 | 2188 | 2683 | 2572 | 2292 | Pycrl |
| Pygb | 18382 | 20506 | 16800 | 19492 | 17701 | 19525 | 18280 | 10275 | 12761 | 10525 | 11362 | 9737 | 10772 | 9466 | 17856 | 18448 | 17043 | 19914 | 19793 | 18316 | 18890 | 9606 | 13152 | 10707 | 10791 | 10149 | 9870 | 9602 | Pygb |
| Pygl | 1151 | 953 | 934 | 873 | 778 | 788 | 653 | 667 | 843 | 804 | 784 | 546 | 737 | 554 | 1004 | 1326 | 816 | 833 | 712 | 830 | 689 | 563 | 988 | 681 | 463 | 646 | 545 | 625 | Pygl |
| Pygo1 | 2624 | 1981 | 2187 | 2524 | 4428 | 3763 | 3551 | 8567 | 7793 | 8724 | 8196 | 7300 | 7446 | 7718 | 2883 | 2383 | 2768 | 2976 | 3536 | 3606 | 3652 | 8301 | 6912 | 7692 | 8582 | 5755 | 7797 | 7403 | Pygo1 |
| Pygo2 | 5154 | 6157 | 6391 | 6220 | 5105 | 4951 | 4787 | 4194 | 4278 | 4585 | 4950 | 4731 | 4736 | 4545 | 4911 | 6018 | 6017 | 6255 | 5442 | 4139 | 5074 | 4909 | 4398 | 4969 | 4493 | 5647 | 4445 | 4707 | Pygo2 |
| Pym1 | 890 | 1100 | 1289 | 1005 | 1114 | 1084 | 1044 | 1678 | 1978 | 1993 | 1956 | 2210 | 2177 | 1974 | 994 | 1035 | 1242 | 907 | 1131 | 1252 | 984 | 1529 | 1980 | 2026 | 1925 | 2731 | 2335 | 2182 | Pym1 |
| Pyroxd1 | 1203 | 1215 | 1097 | 1073 | 1050 | 1136 | 1196 | 2026 | 1854 | 1676 | 1622 | 2052 | 2036 | 2090 | 1344 | 1178 | 1117 | 1100 | 1096 | 1358 | 1141 | 1801 | 1827 | 1681 | 1912 | 1733 | 2069 | 1909 | Pyroxd1 |
| Pyroxd2 | 171 | 180 | 110 | 160 | 196 | 208 | 173 | 173 | 236 | 236 | 304 | 216 | 240 | 189 | 111 | 213 | 132 | 174 | 180 | 183 | 139 | 194 | 285 | 210 | 230 | 170 | 215 | 176 | Pyroxd2 |
| Pyurf | 604 | 608 | 568 | 607 | 567 | 564 | 626 | 681 | 517 | 574 | 526 | 482 | 625 | 602 | 636 | 491 | 571 | 517 | 514 | 598 | 590 | 500 | 537 | 512 | 423 | 486 | 529 | 633 | Pyurf |
| Pyy | 3 | 13 | 16 | 0 | 5 | 0 | 0 | 5 | 15 | 2 | 10 | 1 | 17 | 4 | 5 | 2 | 0 | 0 | 3 | 0 | 0 | 12 | 5 | 1 | 0 | 0 | 0 | 0 | Pyy |
| Pzp | 0 | 0 | 0 | 0 | 0 | 0 | 0 | 0 | 5 | 4 | 0 | 0 | 1 | 0 | 0 | 0 | 0 | 0 | 1 | 0 | 0 | 0 | 2 | 2 | 2 | 0 | 0 | 0 | Pzp |

| | | | | | | | | | | | | | | | | | | | | | | | | | | | | | |
|---|---|---|---|---|---|---|---|---|---|---|---|---|---|---|---|---|---|---|---|---|---|---|---|---|---|---|---|---|---|
| Qars | 12975 | 14581 | 11572 | 10753 | 8200 | 9790 | 9708 | 8288 | 8539 | 6677 | 7009 | 8027 | 8002 | 8023 | 12192 | 12534 | 11277 | 10280 | 9870 | 10550 | 9820 | 6510 | 8548 | 7616 | 7159 | 9270 | 8829 | 8992 | Qars |
| Qdpr | 6095 | 6176 | 5632 | 5785 | 4351 | 4412 | 4482 | 3563 | 3414 | 3408 | 3441 | 3281 | 3411 | 3240 | 5773 | 5814 | 5463 | 5263 | 4864 | 4921 | 4755 | 3003 | 3420 | 3828 | 2958 | 4448 | 3403 | 3541 | Qdpr |
| Qk | 22968 | 16983 | 23637 | 19824 | 20862 | 22054 | 24424 | 35222 | 31159 | 29725 | 31062 | 40815 | 36118 | 42498 | 23169 | 19170 | 22527 | 18946 | 21773 | 20770 | 22591 | 33640 | 29134 | 33694 | 33690 | 39124 | 40983 | 43107 | Qk |
| Qpct | 502 | 413 | 477 | 333 | 398 | 362 | 315 | 385 | 367 | 393 | 333 | 297 | 251 | 295 | 584 | 576 | 418 | 362 | 334 | 413 | 328 | 398 | 344 | 315 | 288 | 325 | 373 | 310 | Qpct |
| Qpctl | 2095 | 2548 | 2207 | 2277 | 1953 | 2103 | 2228 | 1172 | 1273 | 1289 | 1370 | 1431 | 1415 | 1469 | 2022 | 2305 | 2369 | 2340 | 2119 | 1954 | 2078 | 1460 | 1276 | 1391 | 1295 | 1402 | 1392 | 1409 | Qpctl |
| Qprt | 102 | 98 | 163 | 58 | 48 | 37 | 36 | 39 | 62 | 36 | 96 | 103 | 76 | 121 | 71 | 107 | 94 | 35 | 64 | 56 | 28 | 42 | 89 | 52 | 59 | 131 | 162 | 157 | Qprt |
| Qrfp | 2 | 1 | 0 | 1 | 2 | 1 | 6 | 2 | 4 | 5 | 7 | 14 | 3 | 4 | 5 | 1 | 7 | 1 | 2 | 8 | 3 | 0 | 5 | 9 | 2 | 1 | 5 | 6 | Qrfp |
| Qrfpr | 20 | 16 | 2 | 20 | 34 | 41 | 59 | 35 | 55 | 29 | 33 | 42 | 6 | 45 | 24 | 11 | 41 | 24 | 29 | 42 | 31 | 28 | 34 | 40 | 61 | 64 | 26 | 32 | Qrfpr |
| Qrfprl | 150 | 100 | 169 | 117 | 115 | 151 | 134 | 167 | 88 | 147 | 104 | 285 | 172 | 280 | 182 | 112 | 155 | 78 | 125 | 168 | 101 | 155 | 99 | 170 | 116 | 354 | 286 | 323 | Qrfprl |
| Qrich1 | 6308 | 6234 | 6719 | 6592 | 6433 | 5976 | 6669 | 7736 | 7232 | 7500 | 7076 | 8706 | 7741 | 8814 | 6348 | 5506 | 6895 | 6015 | 6960 | 5909 | 6476 | 8378 | 7349 | 8237 | 8080 | 9315 | 8447 | 8734 | Qrich1 |
| Qrich2 | 8 | 5 | 1 | 16 | 3 | 15 | 6 | 4 | 5 | 8 | 8 | 4 | 3 | 7 | 9 | 9 | 7 | 8 | 7 | 5 | 5 | 7 | 8 | 1 | 10 | 6 | 8 | 2 | Qrich2 |
| Qrsl1 | 603 | 577 | 532 | 524 | 558 | 570 | 587 | 1399 | 1330 | 1165 | 1182 | 1330 | 1304 | 1388 | 618 | 535 | 497 | 493 | 582 | 656 | 618 | 1118 | 1195 | 1109 | 1131 | 1188 | 1406 | 1453 | Qrsl1 |
| Qser1 | 9120 | 7197 | 7298 | 8565 | 9983 | 10974 | 11031 | 12760 | 11712 | 11761 | 11442 | 13274 | 12555 | 13762 | 8799 | 8081 | 7964 | 8282 | 9564 | 9016 | 10852 | 10611 | 10803 | 11213 | 12864 | 10819 | 13754 | 13027 | Qser1 |
| Qsox1 | 3982 | 5252 | 4895 | 4777 | 4679 | 4921 | 4033 | 2259 | 2786 | 2757 | 2854 | 2686 | 3333 | 2275 | 4135 | 5065 | 4869 | 4982 | 4668 | 4024 | 4260 | 2491 | 2964 | 2639 | 2655 | 2674 | 2574 | 2370 | Qsox1 |
| Qsox2 | 2458 | 2837 | 2772 | 2647 | 2135 | 2206 | 2294 | 1467 | 1664 | 1709 | 1723 | 2017 | 1909 | 1801 | 2352 | 2755 | 2675 | 2457 | 2251 | 2268 | 2157 | 1654 | 1681 | 1796 | 1707 | 1977 | 2040 | 1894 | Qsox2 |
| Qtrt1 | 1491 | 2097 | 1925 | 2179 | 1915 | 1671 | 1681 | 1425 | 1612 | 1606 | 1486 | 1681 | 1694 | 1654 | 1535 | 1497 | 2052 | 1931 | 2000 | 1836 | 1737 | 1514 | 1457 | 1761 | 1689 | 1805 | 1821 | 1690 | Qtrt1 |
| Qtrt2 | 3327 | 3165 | 2857 | 3230 | 2848 | 2601 | 3089 | 2947 | 2728 | 2624 | 2867 | 3090 | 2818 | 3100 | 3011 | 3178 | 3030 | 3266 | 2888 | 3050 | 2900 | 2674 | 2596 | 2879 | 2868 | 3408 | 3047 | 3096 | Qtrt2 |
| R3hcc1 | 2893 | 3176 | 3721 | 3327 | 3145 | 3074 | 2863 | 2412 | 2970 | 3082 | 3100 | 3259 | 3485 | 3401 | 2847 | 2974 | 3423 | 2960 | 3258 | 2970 | 3007 | 2473 | 2843 | 3330 | 2997 | 3946 | 3648 | 3787 | R3hcc1 |
| R3hcc1l | 2243 | 2073 | 2283 | 1832 | 1587 | 1512 | 1740 | 2179 | 1920 | 1637 | 1768 | 1951 | 1839 | 1986 | 2025 | 2104 | 2138 | 1754 | 1706 | 2013 | 1757 | 1805 | 1831 | 1975 | 1775 | 2080 | 1962 | 1963 | R3hcc1l |
| R3hdm1 | 13567 | 12463 | 11745 | 13365 | 14221 | 14466 | 15922 | 12064 | 11718 | 12038 | 11192 | 11941 | 11101 | 11980 | 13481 | 13241 | 12793 | 13428 | 13863 | 13546 | 15596 | 12325 | 10580 | 10715 | 12523 | 9910 | 11353 | 11153 | R3hdm1 |
| R3hdm2 | 4180 | 4336 | 4969 | 5784 | 6582 | 5802 | 5393 | 7272 | 8273 | 9776 | 9308 | 9640 | 8024 | 8876 | 4262 | 4113 | 5700 | 5996 | 6236 | 4480 | 5858 | 11067 | 7649 | 9847 | 10136 | 9509 | 7968 | 8200 | R3hdm2 |
| R3hdm4 | 6724 | 7882 | 9373 | 9469 | 8822 | 8027 | 8129 | 6449 | 7507 | 9177 | 8032 | 8543 | 7944 | 8588 | 6215 | 7046 | 9511 | 9093 | 9392 | 7064 | 8533 | 8174 | 6879 | 9491 | 8938 | 9051 | 7977 | 8515 | R3hdm4 |
| R3hdml | 1 | 3 | 3 | 4 | 0 | 0 | 0 | 0 | 0 | 0 | 3 | 0 | 1 | 3 | 1 | 2 | 0 | 1 | 0 | 0 | 2 | 2 | 1 | 2 | 1 | 8 | 0 | 1 | R3hdml |
| Rab10 | 18390 | 15825 | 17622 | 20363 | 21804 | 20746 | 23436 | 17862 | 15901 | 17095 | 16447 | 16308 | 15955 | 17476 | 19837 | 16889 | 18935 | 21523 | 20799 | 19444 | 23272 | 19584 | 16368 | 16726 | 18210 | 16138 | 15896 | 17368 | Rab10 |
| Rab11a | 8771 | 6976 | 7927 | 8369 | 9002 | 8887 | 10547 | 9371 | 7257 | 6916 | 6488 | 6697 | 6595 | 7762 | 10076 | 7562 | 8058 | 8291 | 8398 | 9911 | 10336 | 8204 | 7575 | 6777 | 7504 | 5953 | 6814 | 7503 | Rab11a |
| Rab11b | 3989 | 4973 | 4470 | 4907 | 5744 | 5169 | 5174 | 8910 | 10250 | 10624 | 9664 | 11155 | 9844 | 10632 | 4250 | 4196 | 4808 | 4819 | 5618 | 5283 | 5672 | 10322 | 9741 | 10797 | 10984 | 11344 | 10113 | 11015 | Rab11b |
| Rab11fip1 | 2455 | 2788 | 2598 | 2414 | 1246 | 1547 | 1348 | 418 | 416 | 436 | 427 | 478 | 510 | 458 | 2126 | 3361 | 2154 | 2053 | 1401 | 1363 | 1270 | 369 | 499 | 458 | 278 | 558 | 480 | 504 | Rab11fip1 |
| Rab11fip2 | 2573 | 1744 | 2061 | 2161 | 2832 | 2768 | 3010 | 5600 | 4936 | 4980 | 4802 | 4978 | 5229 | 4996 | 2507 | 1817 | 2117 | 2268 | 2553 | 2606 | 2722 | 5411 | 5106 | 4827 | 5245 | 3793 | 4817 | 4898 | Rab11fip2 |
| Rab11fip3 | 4683 | 5732 | 6365 | 7171 | 7155 | 6328 | 6138 | 5208 | 5975 | 7272 | 6628 | 7452 | 6224 | 7133 | 4381 | 5153 | 6711 | 7508 | 7260 | 5095 | 6529 | 7381 | 5304 | 7555 | 7780 | 6927 | 6306 | 6895 | Rab11fip3 |
| Rab11fip4 | 7387 | 8622 | 10471 | 10697 | 11622 | 10411 | 10098 | 8024 | 8881 | 10267 | 9201 | 9602 | 8850 | 9009 | 8019 | 8105 | 10629 | 10290 | 11433 | 9557 | 10283 | 10719 | 7522 | 9858 | 11866 | 7446 | 8969 | 9043 | Rab11fip4 |
| Rab11fip5 | 3697 | 4424 | 4679 | 4765 | 5428 | 5006 | 4449 | 1893 | 2328 | 2481 | 2623 | 2183 | 2194 | 1915 | 3724 | 4065 | 4949 | 5532 | 5382 | 3793 | 4853 | 2562 | 2515 | 2406 | 2449 | 1826 | 1841 | 1860 | Rab11fip5 |
| Rab12 | 5974 | 5469 | 6320 | 7041 | 6590 | 6419 | 7670 | 4829 | 4416 | 5270 | 4830 | 4936 | 4221 | 5267 | 6116 | 5430 | 6333 | 7000 | 6852 | 5631 | 7616 | 5549 | 4705 | 5374 | 5129 | 4958 | 4309 | 5052 | Rab12 |
| Rab13 | 1232 | 1323 | 1760 | 1253 | 1233 | 1242 | 1148 | 1152 | 1384 | 1400 | 1515 | 1387 | 1617 | 1469 | 1208 | 1435 | 1401 | 1248 | 1216 | 1429 | 1250 | 1233 | 1646 | 1495 | 1317 | 1742 | 1550 | 1486 | Rab13 |
| Rab14 | 15770 | 13006 | 15673 | 16823 | 17916 | 17197 | 19225 | 19825 | 16738 | 17498 | 17717 | 18380 | 17431 | 19015 | 16387 | 14088 | 15947 | 17607 | 17076 | 15789 | 18445 | 20546 | 17103 | 17723 | 18712 | 18425 | 17466 | 18724 | Rab14 |
| Rab15 | 1321 | 1159 | 1199 | 931 | 1550 | 1264 | 931 | 2237 | 2522 | 2438 | 2558 | 1441 | 2076 | 1381 | 1328 | 1369 | 1319 | 1136 | 1169 | 929 | 1014 | 2632 | 2598 | 1952 | 2109 | 1811 | 1392 | 1498 | Rab15 |
| Rab17 | 3 | 1 | 3 | 4 | 2 | 0 | 1 | 9 | 29 | 10 | 13 | 1 | 7 | 6 | 0 | 3 | 2 | 0 | 5 | 0 | 2 | 12 | 19 | 17 | 14 | 6 | 0 | 1 | Rab17 |
| Rab18 | 12491 | 9309 | 11636 | 12000 | 11490 | 10892 | 13118 | 10730 | 8946 | 9251 | 9621 | 9096 | 9029 | 9948 | 12853 | 10394 | 11638 | 11617 | 10676 | 11640 | 12137 | 9600 | 9086 | 9650 | 9788 | 9499 | 9024 | 9963 | Rab18 |
| Rab19 | 6 | 5 | 0 | 0 | 7 | 1 | 6 | 24 | 17 | 25 | 17 | 4 | 18 | 5 | 4 | 5 | 1 | 3 | 1 | 1 | 2 | 17 | 24 | 6 | 8 | 24 | 7 | 2 | Rab19 |
| Rab1a | 16959 | 14263 | 16302 | 15829 | 15144 | 14400 | 17245 | 17088 | 14496 | 14989 | 14603 | 15458 | 14700 | 16319 | 17766 | 15252 | 16443 | 15512 | 14581 | 15694 | 16367 | 17037 | 15386 | 16473 | 15899 | 17154 | 15466 | 16487 | Rab1a |
| Rab1b | 5307 | 6601 | 5478 | 5417 | 5144 | 5363 | 4822 | 7584 | 8740 | 8160 | 7804 | 9052 | 8219 | 8466 | 5126 | 5447 | 5748 | 4801 | 5718 | 5239 | 4989 | 7167 | 8718 | 8244 | 7952 | 9772 | 8667 | 8588 | Rab1b |
| Rab20 | 242 | 276 | 171 | 233 | 128 | 102 | 126 | 34 | 33 | 33 | 27 | 23 | 29 | 17 | 216 | 291 | 204 | 206 | 139 | 134 | 115 | 25 | 45 | 27 | 24 | 22 | 29 | 15 | Rab20 |
| Rab21 | 5026 | 4368 | 6540 | 6961 | 6335 | 5925 | 6328 | 4058 | 3617 | 4156 | 4375 | 4268 | 4180 | 4398 | 5169 | 5109 | 6315 | 6812 | 6039 | 4854 | 6592 | 5138 | 3867 | 4421 | 4852 | 4201 | 3942 | 4762 | Rab21 |
| Rab22a | 5526 | 5165 | 5046 | 5565 | 5257 | 5564 | 5563 | 5461 | 5395 | 5681 | 6431 | 5582 | 5619 | 6164 | 5083 | 5382 | 5335 | 5684 | 5249 | 4722 | 5499 | 5880 | 5831 | 5477 | 5991 | 5201 | 5781 | 6020 | Rab22a |
| Rab23 | 2192 | 2183 | 2528 | 2724 | 2891 | 2599 | 2825 | 2590 | 2653 | 2948 | 2747 | 3000 | 2860 | 2979 | 2078 | 2230 | 2861 | 2708 | 2805 | 2609 | 2794 | 2819 | 2654 | 2861 | 3034 | 3229 | 2803 | 2972 | Rab23 |
| Rab24 | 3392 | 3659 | 4252 | 3939 | 3379 | 3292 | 3396 | 3035 | 3015 | 3194 | 3296 | 3611 | 3375 | 3382 | 3175 | 3086 | 4278 | 3735 | 3711 | 3486 | 3304 | 3039 | 2952 | 3867 | 3356 | 4053 | 3183 | 3420 | Rab24 |
| Rab25 | 219 | 134 | 80 | 131 | 47 | 38 | 41 | 117 | 124 | 132 | 63 | 30 | 33 | 34 | 125 | 146 | 63 | 63 | 25 | 39 | 34 | 35 | 175 | 132 | 31 | 110 | 10 | 27 | Rab25 |
| Rab26 | 241 | 293 | 234 | 220 | 232 | 173 | 221 | 21 | 33 | 38 | 22 | 39 | 24 | 19 | 204 | 254 | 170 | 245 | 234 | 211 | 203 | 23 | 34 | 45 | 37 | 51 | 31 | 27 | Rab26 |
| Rab27a | 648 | 470 | 570 | 392 | 304 | 352 | 247 | 414 | 454 | 375 | 509 | 265 | 383 | 234 | 506 | 675 | 392 | 342 | 254 | 250 | 259 | 314 | 509 | 364 | 214 | 418 | 257 | 307 | Rab27a |
| Rab27b | 439 | 163 | 255 | 201 | 339 | 225 | 207 | 600 | 552 | 506 | 463 | 274 | 372 | 295 | 286 | 326 | 161 | 239 | 196 | 218 | 189 | 444 | 564 | 367 | 263 | 327 | 313 | 271 | Rab27b |
| Rab28 | 5457 | 4214 | 4732 | 4851 | 4061 | 4480 | 4979 | 4224 | 3639 | 3435 | 3501 | 4058 | 3625 | 4266 | 5405 | 4259 | 4868 | 4376 | 4485 | 4705 | 4773 | 4052 | 3660 | 3811 | 3875 | 4212 | 4295 | 4577 | Rab28 |
| Rab29 | 294 | 297 | 366 | 250 | 242 | 269 | 219 | 299 | 403 | 359 | 410 | 343 | 348 | 332 | 289 | 313 | 245 | 207 | 270 | 246 | 229 | 264 | 397 | 344 | 285 | 383 | 353 | 352 | Rab29 |
| Rab2a | 14643 | 13134 | 13960 | 14672 | 15415 | 14531 | 17462 | 17408 | 14645 | 14325 | 14111 | 14062 | 13066 | 14877 | 16847 | 13331 | 14523 | 14821 | 15371 | 17222 | 16609 | 16646 | 15321 | 14474 | 15337 | 14112 | 13613 | 14421 | Rab2a |
| Rab2b | 2429 | 2530 | 2476 | 2464 | 2677 | 2607 | 2867 | 3450 | 3101 | 3229 | 2903 | 3117 | 3212 | 3299 | 2767 | 2280 | 2477 | 2562 | 2627 | 2639 | 2711 | 4047 | 3102 | 3129 | 3394 | 2872 | 3197 | 3287 | Rab2b |
| Rab30 | 4689 | 3740 | 3455 | 4849 | 5609 | 5915 | 6080 | 4763 | 4470 | 3801 | 4329 | 3883 | 3715 | 4161 | 5251 | 4133 | 4034 | 4961 | 4839 | 5924 | 5888 | 4289 | 4347 | 3486 | 4409 | 3041 | 3803 | 3915 | Rab30 |
| Rab31 | 5424 | 5045 | 5070 | 4615 | 5412 | 5535 | 5197 | 6462 | 5900 | 5576 | 5552 | 5790 | 5651 | 6041 | 5687 | 4944 | 4837 | 4768 | 5527 | 5645 | 5356 | 6351 | 6263 | 5580 | 6064 | 6239 | 6334 | 6055 | Rab31 |
| Rab32 | 428 | 347 | 452 | 224 | 248 | 302 | 231 | 458 | 521 | 394 | 494 | 345 | 567 | 333 | 421 | 396 | 324 | 306 | 276 | 271 | 210 | 406 | 610 | 388 | 270 | 471 | 437 | 431 | Rab32 |
| Rab33a | 338 | 509 | 410 | 478 | 732 | 614 | 581 | 1038 | 940 | 875 | 810 | 880 | 736 | 825 | 473 | 294 | 496 | 519 | 659 | 713 | 654 | 1011 | 800 | 878 | 1138 | 849 | 733 | 819 | Rab33a |
| Rab33b | 1743 | 1373 | 1378 | 1871 | 1492 | 1592 | 1701 | 1562 | 1537 | 1434 | 1625 | 1338 | 1285 | 1456 | 1623 | 1738 | 1513 | 1791 | 1506 | 1654 | 1643 | 1623 | 1589 | 1469 | 1406 | 1459 | 1355 | 1354 | Rab33b |
| Rab34 | 1178 | 1033 | 1167 | 925 | 1073 | 1129 | 929 | 2456 | 2871 | 2379 | 2994 | 2301 | 2518 | 2239 | 1027 | 1233 | 1063 | 1002 | 1073 | 948 | 933 | 2461 | 2926 | 2588 | 2342 | 2892 | 2518 | 2328 | Rab34 |
| Rab35 | 4452 | 5465 | 5662 | 5960 | 5889 | 5450 | 5620 | 4501 | 4849 | 5143 | 5175 | 5738 | 4595 | 5423 | 4681 | 4778 | 6225 | 6026 | 6385 | 4828 | 5835 | 5207 | 4785 | 5472 | 5555 | 6070 | 4774 | 5364 | Rab35 |
| Rab36 | 1403 | 1553 | 1426 | 1762 | 1901 | 1707 | 1843 | 1110 | 1291 | 1643 | 1349 | 1669 | 1543 | 1589 | 1369 | 1588 | 1640 | 1818 | 1763 | 1557 | 1727 | 1602 | 1279 | 1516 | 1373 | 1412 | 1445 | 1402 | Rab36 |
| Rab37 | 325 | 342 | 325 | 244 | 200 | 171 | 212 | 116 | 102 | 100 | 104 | 127 | 142 | 92 | 332 | 400 | 271 | 256 | 155 | 206 | 201 | 62 | 105 | 106 | 98 | 127 | 95 | 141 | Rab37 |
| Rab38 | 215 | 144 | 152 | 59 | 56 | 79 | 34 | 142 | 223 | 132 | 132 | 77 | 115 | 69 | 131 | 148 | 73 | 96 | 46 | 61 | 19 | 92 | 274 | 126 | 57 | 135 | 106 | 109 | Rab38 |
| Rab39 | 995 | 876 | 1090 | 984 | 937 | 925 | 1111 | 326 | 215 | 240 | 251 | 294 | 291 | 308 | 1134 | 871 | 1125 | 917 | 920 | 1020 | 1023 | 282 | 265 | 295 | 268 | 258 | 407 | 356 | Rab39 |
| Rab39b | 2405 | 2185 | 2272 | 2630 | 2886 | 2655 | 3261 | 3507 | 2597 | 2863 | 2448 | 2941 | 2667 | 2941 | 2694 | 2118 | 2647 | 2585 | 3094 | 3234 | 3181 | 3225 | 2482 | 2566 | 3047 | 2439 | 3008 | 2921 | Rab39b |
| Rab3a | 9 | 7 | 3 | 9 | 7 | 1 | 3 | 4 | 2 | 5 | 113 | 2 | 1 | 2 | 4 | 6 | 2 | 3 | 7 | 5 | 6 | 5 | 3 | 1 | 9 | 3 | 0 | 5 | Rab3a |
| Rab3b | 735 | 1036 | 781 | 885 | 1213 | 837 | 963 | 874 | 910 | 857 | 732 | 677 | 716 | 684 | 968 | 966 | 842 | 924 | 941 | 1129 | 969 | 1059 | 926 | 806 | 872 | 756 | 697 | 735 | Rab3b |
| Rab3c | 9943 | 11647 | 8434 | 17994 | 25474 | 21809 | 21467 | 22917 | 16666 | 20478 | 16876 | 13151 | 16014 | 13892 | 16125 | 10666 | 13626 | 19873 | 20603 | 23753 | 23861 | 25502 | 16796 | 13746 | 21498 | 5308 | 13554 | 12346 | Rab3c |
| Rab3d | 3373 | 3533 | 3523 | 3549 | 3436 | 3341 | 3124 | 1159 | 1503 | 1407 | 1350 | 1200 | 1205 | 1081 | 3270 | 3591 | 3555 | 3551 | 3469 | 3191 | 3061 | 1285 | 1555 | 1262 | 1512 | 1410 | 1151 | 1145 | Rab3d |
| Rab3gap1 | 2377 | 2396 | 2294 | 2305 | 2406 | 2418 | 2289 | 3615 | 3521 | 3615 | 3572 | 3394 | 3329 | 3227 | 2352 | 2318 | 2429 | 2397 | 2423 | 2591 | 2409 | 3594 | 3746 | 3244 | 3442 | 2957 | 3448 | 3221 | Rab3gap1 |
| Rab3gap2 | 6855 | 6986 | 5058 | 6668 | 7019 | 7049 | 7030 | 5359 | 5512 | 5548 | 5350 | 5192 | 5202 | 5574 | 6861 | 6729 | 5760 | 6966 | 6129 | 6731 | 6943 | 5394 | 5185 | 4895 | 5763 | 3895 | 5081 | 5023 | Rab3gap2 |
| Rab3il1 | 2796 | 3304 | 2540 | 2262 | 2044 | 2281 | 1561 | 1555 | 2043 | 1653 | 1961 | 1604 | 2031 | 1518 | 2728 | 3772 | 1975 | 2549 | 1831 | 2003 | 1692 | 1744 | 2233 | 1617 | 1476 | 1844 | 1873 | 1705 | Rab3il1 |
| Rab3ip | 3868 | 4446 | 4570 | 5064 | 4787 | 4546 | 4874 | 2070 | 1963 | 2306 | 2081 | 2448 | 2194 | 2437 | 4175 | 4217 | 4784 | 4671 | 4990 | 3969 | 4810 | 2231 | 2011 | 2397 | 2163 | 2615 | 2221 | 2528 | Rab3ip |
| Rab40b | 624 | 660 | 635 | 701 | 812 | 747 | 808 | 230 | 311 | 280 | 449 | 230 | 204 | 192 | 724 | 818 | 790 | 850 | 747 | 667 | 711 | 309 | 397 | 295 | 244 | 186 | 167 | 204 | Rab40b |
| Rab40c | 3011 | 4324 | 3647 | 4678 | 3896 | 3670 | 3620 | 1983 | 2407 | 2573 | 2352 | 2610 | 2315 | 2248 | 3008 | 3867 | 4187 | 4456 | 4117 | 3136 | 3808 | 2580 | 2403 | 2437 | 2590 | 2682 | 2316 | 2324 | Rab40c |
| Rab42 | 27 | 37 | 55 | 44 | 31 | 35 | 17 | 9 | 16 | 15 | 21 | 15 | 9 | 12 | 42 | 38 | 36 | 31 | 34 | 39 | 29 | 17 | 23 | 13 | 9 | 16 | 26 | 6 | Rab42 |
| Rab43 | 83 | 105 | 117 | 92 | 83 | 79 | 59 | 65 | 64 | 56 | 63 | 58 | 65 | 68 | 88 | 74 | 80 | 73 | 88 | 100 | 93 | 60 | 56 | 47 | 63 | 40 | 62 | 51 | Rab43 |
| Rab44 | 8 | 5 | 7 | 0 | 3 | 3 | 0 | 13 | 14 | 19 | 9 | 13 | 20 | 4 | 7 | 19 | 2 | 8 | 5 | 4 | 3 | 3 | 25 | 19 | 30 | 17 | 22 | 12 | Rab44 |
| Rab4a | 1314 | 1522 | 1448 | 1490 | 1613 | 1398 | 1554 | 868 | 1055 | 1075 | 952 | 1058 | 1010 | 1041 | 1218 | 1441 | 1522 | 1478 | 1636 | 1375 | 1656 | 1071 | 1008 | 991 | 1117 | 1141 | 1069 | 1081 | Rab4a |

| | | | | | | | | | | | | | | | | | | | | | | | | | | | | | |
|---|---|---|---|---|---|---|---|---|---|---|---|---|---|---|---|---|---|---|---|---|---|---|---|---|---|---|---|---|---|
| Rab4b | 66 | 74 | 110 | 106 | 102 | 115 | 95 | 100 | 109 | 141 | 142 | 101 | 91 | 100 | 71 | 72 | 112 | 111 | 131 | 78 | 127 | 137 | 81 | 180 | 114 | 91 | 87 | 88 | Rab4b |
| Rab5a | 6852 | 5515 | 6912 | 6870 | 6932 | 7370 | 7801 | 6393 | 5586 | 6107 | 5774 | 6009 | 5883 | 6239 | 7253 | 6209 | 6685 | 6863 | 7190 | 7314 | 7205 | 6861 | 5828 | 5853 | 6133 | 6472 | 5993 | 6262 | Rab5a |
| Rab5b | 9259 | 9200 | 8897 | 8994 | 9485 | 8948 | 9814 | 9673 | 9526 | 8884 | 9075 | 9284 | 9110 | 9239 | 9315 | 8505 | 9188 | 9618 | 9421 | 9384 | 9298 | 10169 | 9607 | 9449 | 9939 | 9594 | 9231 | 9489 | Rab5b |
| Rab5c | 8017 | 10131 | 8857 | 9137 | 8977 | 9539 | 8373 | 7928 | 9143 | 8684 | 8450 | 8642 | 8869 | 8168 | 8331 | 8521 | 9441 | 9274 | 9798 | 9946 | 8935 | 7474 | 9230 | 8871 | 8868 | 9728 | 9019 | 8816 | Rab5c |
| Rab5if | 6090 | 6288 | 5215 | 5985 | 4437 | 4557 | 5618 | 3626 | 3250 | 3243 | 3050 | 3398 | 3282 | 3440 | 6100 | 5664 | 5465 | 5198 | 5103 | 5560 | 5358 | 3080 | 3393 | 3460 | 3214 | 3797 | 3492 | 3737 | Rab5if |
| Rab6a | 18748 | 14156 | 20153 | 21681 | 24094 | 21585 | 24259 | 18055 | 14713 | 17384 | 16927 | 15808 | 15974 | 16467 | 20368 | 16318 | 20234 | 21681 | 22097 | 19669 | 23545 | 20925 | 15165 | 16745 | 18242 | 16218 | 15567 | 17362 | Rab6a |
| Rab6b | 7757 | 9820 | 11480 | 14764 | 19312 | 15427 | 15840 | 12267 | 12648 | 16022 | 13685 | 13133 | 14816 | 12977 | 9781 | 9308 | 14263 | 15310 | 17596 | 13353 | 17204 | 19111 | 11582 | 14349 | 17498 | 9984 | 11859 | 12416 | Rab6b |
| Rab7 | 14783 | 15190 | 14867 | 14839 | 14964 | 13967 | 14733 | 14461 | 13294 | 13740 | 12956 | 13975 | 13191 | 14210 | 15664 | 13749 | 15035 | 14696 | 15172 | 15151 | 14862 | 15665 | 13806 | 13820 | 14429 | 14084 | 13696 | 13887 | Rab7 |
| Rab7b | 44 | 47 | 28 | 17 | 41 | 41 | 36 | 65 | 93 | 87 | 96 | 47 | 95 | 44 | 21 | 44 | 20 | 30 | 22 | 59 | 39 | 53 | 141 | 78 | 70 | 67 | 52 | 47 | Rab7b |
| Rab8a | 7098 | 7819 | 8027 | 7284 | 5818 | 6326 | 6765 | 5200 | 5146 | 4985 | 4978 | 5333 | 5499 | 5885 | 7142 | 6896 | 7460 | 6523 | 6728 | 7058 | 6670 | 4365 | 5414 | 5718 | 5301 | 6620 | 6172 | 6100 | Rab8a |
| Rab8b | 8772 | 7476 | 8352 | 8751 | 8191 | 8964 | 9877 | 6867 | 6384 | 5987 | 5803 | 7100 | 6143 | 7074 | 8905 | 7439 | 8536 | 8700 | 8266 | 8727 | 8670 | 6315 | 5887 | 6527 | 6872 | 6931 | 6639 | 6795 | Rab8b |
| Rab9 | 2963 | 2769 | 2510 | 2688 | 2242 | 2429 | 2900 | 2304 | 1893 | 1966 | 1813 | 2023 | 2155 | 2364 | 3173 | 2798 | 2599 | 2597 | 2365 | 2898 | 2658 | 2015 | 2265 | 1977 | 1933 | 2317 | 1973 | 2309 | Rab9 |
| Rab9b | 423 | 382 | 577 | 695 | 1006 | 893 | 850 | 1167 | 979 | 1233 | 1033 | 969 | 882 | 889 | 604 | 436 | 679 | 738 | 757 | 803 | 859 | 1669 | 988 | 1072 | 1252 | 644 | 803 | 838 | Rab9b |
| Rabac1 | 2098 | 3252 | 2613 | 2786 | 2495 | 2294 | 2099 | 1973 | 2169 | 2108 | 2195 | 1717 | 1808 | 1556 | 2458 | 2453 | 2745 | 2567 | 2753 | 2975 | 2247 | 1928 | 2418 | 1813 | 1971 | 1664 | 1779 | 1546 | Rabac1 |
| Rabep1 | 12936 | 11579 | 11579 | 13837 | 15414 | 13965 | 16509 | 12463 | 10144 | 10325 | 9508 | 9404 | 8910 | 10137 | 14600 | 11692 | 12736 | 14990 | 14269 | 15452 | 15639 | 12106 | 9848 | 9905 | 11631 | 8102 | 8793 | 9371 | Rabep1 |
| Rabep2 | 679 | 809 | 852 | 903 | 716 | 774 | 620 | 987 | 1091 | 1034 | 1176 | 1007 | 1090 | 1034 | 637 | 802 | 852 | 671 | 752 | 693 | 633 | 949 | 1131 | 1128 | 1066 | 1233 | 1058 | 967 | Rabep2 |
| Rabepk | 1527 | 1687 | 1520 | 1301 | 1207 | 1380 | 1258 | 834 | 952 | 830 | 843 | 754 | 858 | 873 | 1543 | 1571 | 1475 | 1396 | 1319 | 1463 | 1252 | 747 | 999 | 755 | 769 | 942 | 848 | 892 | Rabepk |
| Rabgap1 | 13701 | 12679 | 11050 | 13795 | 13389 | 13732 | 14701 | 12281 | 10556 | 10848 | 9762 | 10548 | 9778 | 11134 | 13967 | 11573 | 12497 | 13629 | 12987 | 14546 | 14754 | 11326 | 10523 | 10572 | 11173 | 9482 | 10584 | 10239 | Rabgap1 |
| Rabgap1l | 3459 | 2533 | 2367 | 3431 | 4912 | 4679 | 4811 | 6072 | 5582 | 5579 | 5129 | 4271 | 4179 | 4332 | 3779 | 3229 | 2768 | 4061 | 4109 | 4751 | 4833 | 6067 | 5620 | 4614 | 6087 | 2665 | 3848 | 3809 | Rabgap1l |
| Rabgef1 | 2103 | 2319 | 2582 | 2768 | 2366 | 2312 | 2598 | 1868 | 1652 | 1881 | 1812 | 1818 | 1944 | 1908 | 2334 | 2319 | 2745 | 2702 | 2603 | 2298 | 2528 | 2214 | 1663 | 1962 | 2091 | 1823 | 1747 | 1982 | Rabgef1 |
| Rabggta | 690 | 868 | 810 | 845 | 1068 | 828 | 888 | 1319 | 1494 | 1571 | 1447 | 1481 | 1462 | 1418 | 713 | 731 | 937 | 859 | 982 | 908 | 874 | 1614 | 1522 | 1506 | 1696 | 1584 | 1332 | 1422 | Rabggta |
| Rabggtb | 4005 | 4026 | 4015 | 4862 | 3553 | 3390 | 3713 | 4416 | 3756 | 3131 | 3380 | 3578 | 3509 | 3808 | 4292 | 3979 | 4018 | 4349 | 3581 | 4730 | 3468 | 3405 | 3808 | 3666 | 3388 | 4269 | 3642 | 3788 | Rabggtb |
| Rabif | 1488 | 1532 | 1514 | 1479 | 1579 | 1522 | 1467 | 2862 | 2711 | 2568 | 2470 | 2419 | 2473 | 2445 | 1644 | 1467 | 1534 | 1351 | 1536 | 1756 | 1341 | 2948 | 2752 | 2370 | 2681 | 2551 | 2430 | 2540 | Rabif |
| Rabl2 | 3456 | 3747 | 3612 | 3580 | 2971 | 2919 | 3210 | 2257 | 2056 | 2415 | 2137 | 2494 | 2452 | 2597 | 3319 | 3381 | 3577 | 3390 | 3116 | 2860 | 3069 | 2692 | 2241 | 2338 | 2244 | 2504 | 2471 | 2645 | Rabl2 |
| Rabl3 | 805 | 518 | 856 | 535 | 543 | 660 | 609 | 1625 | 1575 | 1229 | 1320 | 1801 | 1720 | 1924 | 741 | 581 | 673 | 471 | 534 | 720 | 558 | 1494 | 1415 | 1484 | 1324 | 1875 | 2185 | 2090 | Rabl3 |
| Rabl6 | 5298 | 5665 | 5820 | 5714 | 5578 | 5535 | 5302 | 5821 | 6469 | 7181 | 6828 | 7274 | 6865 | 6680 | 4970 | 5258 | 5691 | 5186 | 5926 | 5080 | 5420 | 6226 | 6388 | 7156 | 6978 | 7253 | 6930 | 6780 | Rabl6 |
| Rac1 | 17267 | 15847 | 19663 | 18684 | 17963 | 16884 | 19956 | 16261 | 16523 | 18193 | 17815 | 19616 | 17856 | 19300 | 17256 | 16851 | 20614 | 18575 | 18627 | 15445 | 18795 | 18927 | 17359 | 19090 | 17818 | 21400 | 16854 | 19490 | Rac1 |
| Rac2 | 127 | 189 | 165 | 137 | 113 | 153 | 111 | 149 | 236 | 199 | 207 | 176 | 191 | 153 | 131 | 246 | 145 | 157 | 94 | 155 | 118 | 182 | 335 | 177 | 170 | 219 | 185 | 144 | Rac2 |
| Rac3 | 1640 | 2285 | 1972 | 2846 | 3175 | 2597 | 2144 | 2185 | 2516 | 3005 | 2528 | 1828 | 2006 | 1607 | 1721 | 2093 | 2471 | 2968 | 2849 | 2477 | 2508 | 2749 | 2829 | 2175 | 2777 | 1676 | 1750 | 1489 | Rac3 |
| Racgap1 | 15052 | 16517 | 18046 | 16372 | 13598 | 14178 | 15825 | 10306 | 9943 | 11716 | 10024 | 16458 | 14239 | 16168 | 14210 | 14559 | 17919 | 15065 | 15438 | 12856 | 14957 | 9107 | 9114 | 13551 | 12744 | 18061 | 16214 | 16955 | Racgap1 |
| Rack1 | 52176 | 60908 | 47306 | 43422 | 37548 | 40035 | 42021 | 44506 | 50716 | 39239 | 39464 | 42832 | 42325 | 41929 | 50095 | 54193 | 44400 | 41882 | 44926 | 49364 | 41853 | 33632 | 51606 | 42628 | 39157 | 50129 | 45671 | 47895 | Rack1 |
| Rad1 | 1420 | 1372 | 1459 | 1335 | 1160 | 1236 | 1474 | 1280 | 1244 | 1125 | 1111 | 1447 | 1331 | 1345 | 1375 | 1302 | 1462 | 1171 | 1276 | 1247 | 1216 | 1110 | 1167 | 1346 | 1228 | 1567 | 1417 | 1415 | Rad1 |
| Rad17 | 2701 | 2293 | 2216 | 2227 | 2055 | 2008 | 2305 | 2538 | 2227 | 2155 | 2176 | 2160 | 2157 | 2396 | 2447 | 2748 | 2145 | 2140 | 2018 | 2456 | 2170 | 2179 | 2183 | 2218 | 2384 | 2375 | 2428 | 2449 | Rad17 |
| Rad18 | 2045 | 1727 | 1699 | 1414 | 1136 | 1378 | 1593 | 1737 | 1715 | 1448 | 1578 | 2068 | 1992 | 2096 | 2028 | 1680 | 1571 | 1194 | 1202 | 1617 | 1480 | 1155 | 1612 | 1584 | 1511 | 2177 | 2349 | 2122 | Rad18 |
| Rad21 | 22729 | 18745 | 22474 | 19856 | 20174 | 21246 | 23189 | 27909 | 25200 | 24743 | 23748 | 31348 | 27228 | 32843 | 21559 | 19452 | 20825 | 19086 | 21178 | 21386 | 22330 | 24593 | 24461 | 28452 | 26746 | 32912 | 30690 | 32120 | Rad21 |
| Rad21l | 39 | 42 | 39 | 53 | 79 | 46 | 45 | 10 | 18 | 9 | 12 | 8 | 7 | 14 | 45 | 40 | 44 | 39 | 24 | 55 | 60 | 13 | 19 | 7 | 17 | 7 | 6 | 7 | Rad21l |
| Rad23a | 2107 | 2658 | 2403 | 2256 | 2201 | 2088 | 2251 | 2194 | 2717 | 2873 | 2824 | 3379 | 2892 | 3130 | 1925 | 2147 | 2511 | 2284 | 2396 | 2252 | 2167 | 2236 | 2653 | 3128 | 2764 | 3771 | 3066 | 3290 | Rad23a |
| Rad23b | 27905 | 24604 | 29075 | 25837 | 24245 | 23718 | 28519 | 27307 | 23483 | 24367 | 23316 | 26881 | 25846 | 32843 | 28060 | 21656 | 27423 | 24126 | 26318 | 26197 | 29232 | 23616 | 23344 | 26986 | 25972 | 30612 | 27520 | 33460 | Rad23b |
| Rad50 | 3530 | 2839 | 2918 | 2683 | 2673 | 2851 | 3064 | 2899 | 2882 | 2544 | 2671 | 3075 | 2973 | 2963 | 3290 | 3049 | 2691 | 2545 | 2509 | 3102 | 2807 | 2141 | 2970 | 2700 | 2532 | 2879 | 3359 | 2842 | Rad50 |
| Rad51 | 4474 | 3466 | 4235 | 2765 | 2052 | 2509 | 3084 | 3324 | 2960 | 2862 | 2856 | 4469 | 3859 | 4396 | 3705 | 3339 | 3284 | 2371 | 2651 | 3080 | 2503 | 1962 | 2660 | 3646 | 2797 | 5346 | 4875 | 5040 | Rad51 |
| Rad51ap1 | 3220 | 2776 | 2887 | 2234 | 1723 | 2090 | 2709 | 2391 | 2397 | 1933 | 1938 | 2838 | 2402 | 2889 | 3017 | 2745 | 2506 | 2025 | 2162 | 2720 | 2216 | 1484 | 1990 | 2235 | 2175 | 3125 | 2886 | 3013 | Rad51ap1 |
| Rad51ap2 | 63 | 28 | 37 | 32 | 38 | 52 | 49 | 33 | 44 | 32 | 29 | 23 | 42 | 27 | 57 | 48 | 45 | 19 | 38 | 40 | 29 | 32 | 52 | 26 | 13 | 41 | 31 | 33 | Rad51ap2 |
| Rad51b | 412 | 416 | 316 | 289 | 235 | 293 | 312 | 354 | 310 | 260 | 243 | 352 | 313 | 349 | 359 | 359 | 345 | 273 | 281 | 355 | 273 | 201 | 290 | 335 | 270 | 378 | 407 | 374 | Rad51b |
| Rad51c | 781 | 734 | 1032 | 810 | 624 | 688 | 773 | 877 | 906 | 921 | 900 | 1082 | 1036 | 1194 | 728 | 699 | 897 | 711 | 754 | 778 | 717 | 754 | 864 | 1121 | 978 | 1469 | 1217 | 1138 | Rad51c |
| Rad51d | 858 | 997 | 876 | 918 | 927 | 1000 | 949 | 851 | 998 | 963 | 889 | 1016 | 1109 | 1084 | 920 | 856 | 972 | 931 | 1069 | 907 | 1013 | 911 | 959 | 1048 | 1026 | 1002 | 1081 | 1105 | Rad51d |
| Rad52 | 1293 | 1495 | 1544 | 1505 | 1361 | 1343 | 1483 | 1762 | 1464 | 1558 | 1440 | 1471 | 1565 | 1592 | 1431 | 1356 | 1546 | 1533 | 1505 | 1468 | 1568 | 1678 | 1416 | 1642 | 1623 | 1671 | 1564 | 1552 | Rad52 |
| Rad54b | 2010 | 1583 | 1680 | 1549 | 1106 | 1364 | 1711 | 1994 | 1803 | 1477 | 1689 | 2280 | 1968 | 2190 | 1655 | 1609 | 1435 | 1330 | 1336 | 1431 | 1443 | 1287 | 1553 | 1955 | 1652 | 2222 | 2289 | 2306 | Rad54b |
| Rad54l | 6771 | 5976 | 4552 | 3892 | 2886 | 4200 | 4482 | 3085 | 2968 | 2480 | 2390 | 4128 | 3501 | 4194 | 5496 | 5393 | 4274 | 3658 | 3871 | 4674 | 4272 | 1898 | 2509 | 2982 | 2955 | 4085 | 4620 | 4212 | Rad54l |
| Rad54l2 | 11152 | 10893 | 10248 | 12901 | 12815 | 12968 | 12754 | 8933 | 8755 | 10290 | 9724 | 11778 | 11162 | 11880 | 11087 | 10692 | 11972 | 11761 | 12742 | 10487 | 12791 | 9606 | 7810 | 10154 | 11489 | 8640 | 11705 | 11714 | Rad54l2 |
| Rad9a | 182 | 625 | 568 | 142 | 43 | 518 | 460 | 328 | 87 | 37 | 151 | 67 | 387 | 133 | 237 | 187 | 110 | 187 | 572 | 85 | 158 | 404 | 142 | 374 | 242 | 696 | 66 | 398 | Rad9a |
| Rad9b | 776 | 739 | 641 | 850 | 846 | 943 | 885 | 1119 | 1082 | 1139 | 1151 | 1026 | 899 | 1021 | 750 | 713 | 724 | 778 | 848 | 949 | 961 | 1071 | 1062 | 1043 | 1143 | 803 | 962 | 905 | Rad9b |
| Radil | 604 | 802 | 861 | 884 | 917 | 866 | 644 | 451 | 626 | 653 | 667 | 443 | 566 | 451 | 570 | 783 | 768 | 919 | 796 | 578 | 632 | 799 | 637 | 500 | 613 | 359 | 553 | 371 | Radil |
| Radx | 325 | 253 | 251 | 218 | 198 | 219 | 227 | 234 | 195 | 164 | 130 | 276 | 266 | 315 | 353 | 237 | 298 | 230 | 256 | 276 | 223 | 162 | 162 | 199 | 204 | 310 | 264 | 309 | Radx |
| Rae1 | 5007 | 5376 | 5111 | 5248 | 4534 | 4440 | 4797 | 4119 | 4029 | 4131 | 4080 | 4782 | 4077 | 4371 | 4838 | 4989 | 5034 | 4957 | 4817 | 4322 | 4689 | 3878 | 3952 | 4271 | 4385 | 5140 | 4373 | 4436 | Rae1 |
| Raet1d | 293 | 204 | 350 | 105 | 143 | 142 | 124 | 263 | 276 | 216 | 282 | 323 | 417 | 274 | 253 | 266 | 198 | 131 | 168 | 205 | 130 | 226 | 279 | 252 | 222 | 389 | 316 | 386 | Raet1d |
| Raet1e | 360 | 334 | 468 | 217 | 179 | 213 | 201 | 166 | 126 | 107 | 142 | 199 | 260 | 205 | 442 | 375 | 283 | 216 | 198 | 260 | 173 | 94 | 193 | 162 | 118 | 273 | 278 | 290 | Raet1e |
| Raf1 | 8905 | 8639 | 7804 | 7859 | 10071 | 9610 | 9503 | 15624 | 14545 | 13638 | 13383 | 15360 | 14051 | 14329 | 8615 | 7786 | 8689 | 8115 | 10195 | 9719 | 9528 | 15326 | 13560 | 13512 | 15312 | 14016 | 14492 | 14210 | Raf1 |
| Rag1 | 81 | 66 | 71 | 54 | 46 | 57 | 49 | 68 | 138 | 95 | 125 | 60 | 125 | 61 | 96 | 106 | 44 | 45 | 48 | 77 | 44 | 112 | 137 | 60 | 48 | 78 | 67 | 56 | Rag1 |
| Rag2 | 1 | 0 | 0 | 5 | 2 | 5 | 1 | 1 | 2 | 0 | 2 | 0 | 9 | 2 | 0 | 3 | 3 | 0 | 0 | 2 | 4 | 0 | 3 | 2 | 0 | 5 | 0 | 0 | Rag2 |
| Rai1 | 4593 | 5634 | 5873 | 6121 | 7908 | 6714 | 5863 | 7135 | 9564 | 9866 | 10299 | 10655 | 10112 | 9317 | 4406 | 5021 | 6610 | 6313 | 7441 | 5676 | 6120 | 10091 | 8380 | 10011 | 11160 | 7333 | 9362 | 8624 | Rai1 |
| Rai14 | 10468 | 9081 | 9104 | 6753 | 6262 | 7791 | 6351 | 11442 | 10795 | 10445 | 9808 | 12345 | 13218 | 11903 | 10275 | 10368 | 8418 | 6903 | 6903 | 6910 | 6401 | 7718 | 10617 | 11381 | 9297 | 10696 | 13646 | 12292 | Rai14 |
| Rai2 | 284 | 319 | 413 | 488 | 655 | 539 | 536 | 464 | 511 | 610 | 745 | 530 | 422 | 570 | 347 | 282 | 448 | 489 | 700 | 434 | 500 | 772 | 571 | 681 | 679 | 482 | 310 | 508 | Rai2 |
| Rala | 11780 | 9944 | 12010 | 12423 | 12518 | 11889 | 14711 | 12240 | 10214 | 10901 | 10205 | 10846 | 10084 | 12847 | 12194 | 9738 | 12187 | 12416 | 12197 | 12760 | 14561 | 11797 | 10044 | 11889 | 12434 | 11251 | 10777 | 12256 | Rala |
| Ralb | 4171 | 4254 | 4335 | 3759 | 3485 | 3692 | 3545 | 2429 | 2854 | 2638 | 3092 | 2492 | 2928 | 2578 | 4129 | 4801 | 4073 | 3762 | 3411 | 3855 | 3440 | 2267 | 3169 | 2520 | 2234 | 3075 | 2530 | 2674 | Ralb |
| Ralbp1 | 5490 | 5839 | 6354 | 5793 | 5492 | 5576 | 5188 | 4070 | 4785 | 4983 | 4835 | 4731 | 4779 | 4585 | 5186 | 6087 | 5992 | 5848 | 5621 | 4959 | 5259 | 4102 | 4641 | 4936 | 4674 | 5032 | 4611 | 4789 | Ralbp1 |
| Ralgapa1 | 9238 | 7741 | 4972 | 6242 | 7288 | 8269 | 8210 | 12419 | 10841 | 7978 | 8682 | 7908 | 8074 | 8314 | 9708 | 8146 | 5715 | 6430 | 6189 | 9374 | 8644 | 8793 | 10877 | 6510 | 8341 | 5462 | 8777 | 7700 | Ralgapa1 |
| Ralgapa2 | 5042 | 4536 | 3498 | 5171 | 5803 | 5887 | 6516 | 3880 | 3891 | 3654 | 3690 | 3705 | 3733 | 3920 | 4974 | 4414 | 3963 | 5045 | 5351 | 5493 | 6430 | 3781 | 3284 | 3452 | 4562 | 2435 | 3818 | 3302 | Ralgapa2 |
| Ralgapb | 12630 | 11928 | 8350 | 11479 | 12349 | 13204 | 13927 | 12416 | 11515 | 10359 | 10414 | 11140 | 10415 | 11621 | 13044 | 11232 | 9997 | 11976 | 11645 | 13258 | 14032 | 11035 | 10659 | 9774 | 11866 | 7659 | 11185 | 10568 | Ralgapb |
| Ralgds | 5980 | 7794 | 6235 | 7386 | 8306 | 7294 | 6505 | 6917 | 8187 | 8575 | 7581 | 7578 | 7403 | 6676 | 6399 | 7103 | 7156 | 7516 | 7940 | 6849 | 6940 | 8829 | 7847 | 7634 | 8554 | 6179 | 6826 | 6699 | Ralgds |
| Ralgps1 | 3297 | 4032 | 2873 | 5499 | 7475 | 6377 | 7226 | 6785 | 7204 | 8192 | 6747 | 6818 | 5920 | 6987 | 3902 | 3370 | 4441 | 5998 | 6616 | 6032 | 7381 | 9799 | 6141 | 6914 | 9451 | 4465 | 5912 | 5839 | Ralgps1 |
| Ralgps2 | 10127 | 8694 | 9087 | 9793 | 10969 | 11132 | 11454 | 11621 | 10398 | 11039 | 10604 | 11741 | 11432 | 11957 | 10779 | 9769 | 9386 | 10172 | 10545 | 10675 | 11637 | 11916 | 10460 | 10871 | 12407 | 8668 | 11380 | 10750 | Ralgps2 |
| Raly | 18079 | 21383 | 21191 | 20394 | 15191 | 15901 | 16380 | 11948 | 12727 | 13108 | 12433 | 15370 | 13438 | 15354 | 17439 | 17784 | 20300 | 18124 | 18118 | 17038 | 17066 | 9977 | 12963 | 14529 | 13160 | 18612 | 15788 | 16169 | Raly |
| Ralyl | 901 | 992 | 1035 | 1770 | 2320 | 1845 | 2201 | 1696 | 1484 | 1597 | 1470 | 1253 | 1145 | 1365 | 1333 | 933 | 1359 | 1684 | 1986 | 2405 | 2213 | 2064 | 1438 | 1535 | 1836 | 933 | 1288 | 1145 | Ralyl |
| Ramac | 4573 | 3548 | 4659 | 4394 | 4367 | 4274 | 5095 | 4636 | 3824 | 3734 | 3565 | 4132 | 3743 | 4366 | 4941 | 4090 | 4267 | 4076 | 4461 | 5268 | 4635 | 3882 | 4013 | 4292 | 3972 | 4841 | 4141 | 4600 | Ramac |
| Ramp1 | 18 | 33 | 20 | 18 | 20 | 26 | 17 | 27 | 39 | 23 | 78 | 11 | 20 | 9 | 21 | 17 | 14 | 29 | 11 | 30 | 22 | 50 | 76 | 17 | 18 | 9 | 21 | 11 | Ramp1 |
| Ramp2 | 379 | 401 | 428 | 319 | 458 | 463 | 363 | 847 | 932 | 1083 | 1067 | 929 | 954 | 977 | 397 | 392 | 425 | 447 | 394 | 429 | 414 | 939 | 1174 | 1028 | 911 | 1122 | 897 | 865 | Ramp2 |
| Ramp3 | 243 | 333 | 379 | 216 | 327 | 203 | 160 | 126 | 190 | 216 | 340 | 127 | 144 | 99 | 265 | 371 | 278 | 257 | 241 | 200 | 200 | 170 | 368 | 121 | 132 | 139 | 146 | 100 | Ramp3 |
| Ran | 26713 | 29039 | 29846 | 24300 | 20068 | 21531 | 24848 | 26956 | 25872 | 25691 | 23183 | 33981 | 29385 | 34542 | 25904 | 25051 | 27584 | 21721 | 24009 | 26838 | 22965 | 20493 | 24453 | 30357 | 26961 | 40192 | 37299 | 38525 | Ran |
| Ranbp1 | 0 | 0 | 1 | 0 | 0 | 0 | 0 | 0 | 0 | 0 | 0 | 0 | 0 | 2 | 1 | 0 | 0 | 1 | 0 | 0 | 0 | 0 | 0 | 0 | 0 | 0 | 0 | 0 | Ranbp1 |

| | | | | | | | | | | | | | | | | | | | | | | | | | | | | | |
|---|---|---|---|---|---|---|---|---|---|---|---|---|---|---|---|---|---|---|---|---|---|---|---|---|---|---|---|---|---|
| Ranbp10 | 5395 | 5825 | 5172 | 5897 | 5212 | 5191 | 5138 | 4693 | 4896 | 5337 | 4893 | 5964 | 6296 | 5945 | 5101 | 5572 | 5648 | 5677 | 5802 | 4911 | 5254 | 5792 | 4853 | 5574 | 5857 | 5410 | 5585 | 6162 | Ranbp10 |
| Ranbp17 | 833 | 802 | 712 | 586 | 785 | 699 | 703 | 1319 | 1043 | 994 | 1009 | 1287 | 1274 | 1221 | 964 | 887 | 654 | 540 | 588 | 846 | 638 | 1091 | 1023 | 868 | 1057 | 975 | 1364 | 1213 | Ranbp17 |
| Ranbp2 | 35317 | 27706 | 18124 | 20183 | 19633 | 25625 | 25245 | 29688 | 24797 | 18083 | 20130 | 21962 | 22679 | 22292 | 35336 | 30448 | 18247 | 19185 | 19216 | 30468 | 24157 | 16533 | 23836 | 15526 | 19015 | 14637 | 25635 | 21801 | Ranbp2 |
| Ranbp3 | 5475 | 6513 | 6289 | 6409 | 5746 | 5805 | 5464 | 5816 | 6692 | 6906 | 6440 | 7995 | 7089 | 7252 | 4809 | 5501 | 6034 | 5668 | 6191 | 5654 | 5597 | 6027 | 6299 | 7508 | 7065 | 8472 | 7627 | 7024 | Ranbp3 |
| Ranbp3l | 80 | 92 | 92 | 103 | 263 | 261 | 185 | 192 | 328 | 186 | 332 | 213 | 220 | 149 | 120 | 64 | 140 | 149 | 146 | 274 | 239 | 244 | 506 | 166 | 266 | 106 | 205 | 157 | Ranbp3l |
| Ranbp6 | 5888 | 5650 | 5639 | 5479 | 5280 | 5265 | 5758 | 6879 | 5201 | 4744 | 4480 | 4817 | 4647 | 4734 | 6646 | 6108 | 5140 | 5542 | 5164 | 6535 | 5387 | 6109 | 5168 | 4524 | 5096 | 4294 | 5418 | 4836 | Ranbp6 |
| Ranbp9 | 4768 | 3681 | 5284 | 5210 | 6016 | 5150 | 5849 | 5208 | 4795 | 5681 | 5427 | 5794 | 4944 | 5961 | 4716 | 4433 | 5402 | 5648 | 6046 | 4626 | 5975 | 6605 | 4966 | 5806 | 5808 | 5741 | 5152 | 5612 | Ranbp9 |
| Rangap1 | 18628 | 22646 | 25420 | 24570 | 20535 | 20259 | 20433 | 11909 | 13583 | 14678 | 12989 | 17864 | 15710 | 16677 | 17900 | 20416 | 25297 | 23242 | 24070 | 18815 | 20924 | 12733 | 12463 | 16526 | 15179 | 20034 | 17422 | 17969 | Rangap1 |
| Rangrf | 1051 | 1490 | 1579 | 1340 | 953 | 1058 | 1326 | 573 | 592 | 641 | 653 | 741 | 727 | 879 | 1088 | 1165 | 1471 | 1154 | 1160 | 1404 | 1125 | 443 | 656 | 755 | 702 | 996 | 867 | 871 | Rangrf |
| Rap1a | 5734 | 4215 | 5986 | 5366 | 4726 | 4947 | 5423 | 5004 | 4508 | 3808 | 4462 | 4511 | 4533 | 5250 | 6133 | 5517 | 5100 | 5061 | 4924 | 5539 | 5175 | 4482 | 4731 | 4514 | 4453 | 5105 | 4759 | 5295 | Rap1a |
| Rap1b | 13140 | 9446 | 12571 | 11806 | 11383 | 11620 | 11651 | 11466 | 9972 | 9359 | 10691 | 10497 | 10703 | 10529 | 13647 | 12252 | 11487 | 11432 | 11074 | 11153 | 11516 | 11353 | 11189 | 9965 | 10016 | 11655 | 10569 | 11208 | Rap1b |
| Rap1gap | 1110 | 1602 | 1490 | 1966 | 2851 | 1808 | 1589 | 1377 | 1762 | 1815 | 1786 | 988 | 1432 | 956 | 1344 | 1808 | 2011 | 2249 | 2196 | 1667 | 1748 | 2356 | 1588 | 1369 | 1677 | 891 | 949 | 853 | Rap1gap |
| Rap1gap2 | 1024 | 1174 | 1092 | 1469 | 2176 | 1563 | 1550 | 1989 | 2167 | 2464 | 2167 | 1438 | 1791 | 1546 | 1162 | 1136 | 1301 | 1521 | 1622 | 1572 | 1547 | 2468 | 2079 | 2053 | 2261 | 1271 | 1234 | 1392 | Rap1gap2 |
| Rap1gds1 | 6560 | 6457 | 6972 | 7054 | 7791 | 7185 | 7306 | 5929 | 5765 | 6069 | 5846 | 6243 | 6592 | 6298 | 6835 | 6523 | 7092 | 7164 | 7321 | 5947 | 7273 | 7312 | 5752 | 5920 | 6276 | 6256 | 6125 | 6687 | Rap1gds1 |
| Rap2a | 4803 | 5571 | 5920 | 6481 | 5998 | 5847 | 6107 | 3865 | 3899 | 4358 | 4013 | 4633 | 4122 | 4465 | 5023 | 5399 | 6572 | 6719 | 6458 | 5175 | 6253 | 4649 | 4127 | 4229 | 4979 | 4533 | 4379 | 4740 | Rap2a |
| Rap2b | 3667 | 3282 | 4178 | 4222 | 3404 | 3389 | 3737 | 2893 | 2861 | 2981 | 2988 | 3237 | 2952 | 3393 | 3571 | 3168 | 4171 | 3826 | 3375 | 3573 | 3593 | 2734 | 2986 | 3198 | 3157 | 4000 | 3151 | 3429 | Rap2b |
| Rap2c | 7998 | 5434 | 6865 | 5963 | 6175 | 6823 | 6873 | 8772 | 7662 | 7748 | 8344 | 8569 | 8308 | 8433 | 7730 | 7058 | 6642 | 6198 | 6311 | 6700 | 6294 | 8674 | 7938 | 7949 | 7925 | 8756 | 8168 | 8733 | Rap2c |
| Rapgef1 | 7210 | 8041 | 8460 | 9026 | 9680 | 9158 | 8263 | 9850 | 11476 | 13092 | 11918 | 14323 | 11492 | 12728 | 6562 | 7086 | 9184 | 9264 | 10553 | 7006 | 9330 | 13075 | 10409 | 13888 | 13697 | 12636 | 11935 | 12871 | Rapgef1 |
| Rapgef2 | 4124 | 4137 | 3530 | 4706 | 5009 | 5124 | 4719 | 4161 | 4397 | 5032 | 4858 | 4388 | 4454 | 4249 | 4098 | 4348 | 3901 | 4867 | 4858 | 4415 | 4905 | 4654 | 4328 | 4408 | 4865 | 3072 | 3903 | 3841 | Rapgef2 |
| Rapgef3 | 600 | 809 | 966 | 697 | 819 | 874 | 645 | 464 | 615 | 710 | 907 | 679 | 719 | 566 | 592 | 810 | 926 | 796 | 823 | 689 | 689 | 698 | 723 | 671 | 744 | 675 | 651 | 583 | Rapgef3 |
| Rapgef4 | 301 | 299 | 380 | 456 | 745 | 512 | 662 | 766 | 765 | 792 | 703 | 457 | 524 | 527 | 357 | 325 | 311 | 522 | 567 | 617 | 700 | 1089 | 754 | 585 | 866 | 373 | 417 | 477 | Rapgef4 |
| Rapgef5 | 3234 | 3349 | 2470 | 3385 | 6235 | 5109 | 4636 | 8194 | 6770 | 7176 | 5935 | 5243 | 6150 | 5375 | 4212 | 3115 | 3327 | 3919 | 4752 | 5362 | 4916 | 8569 | 7302 | 5269 | 7545 | 3482 | 5577 | 5015 | Rapgef5 |
| Rapgef6 | 7706 | 6489 | 4138 | 5119 | 5240 | 5685 | 5976 | 6222 | 5914 | 4459 | 5230 | 4445 | 5058 | 4669 | 7356 | 7016 | 4558 | 5190 | 4593 | 6225 | 5651 | 4736 | 5499 | 3725 | 4928 | 2893 | 4729 | 4272 | Rapgef6 |
| Rapgefl1 | 1937 | 2088 | 2261 | 2439 | 2810 | 2475 | 2272 | 922 | 1200 | 1256 | 1316 | 861 | 998 | 793 | 1667 | 2176 | 2635 | 2682 | 2730 | 1973 | 2486 | 1464 | 1212 | 1073 | 1245 | 707 | 801 | 866 | Rapgefl1 |
| Raph1 | 5790 | 4717 | 5171 | 6727 | 8120 | 8204 | 7968 | 7342 | 7591 | 8412 | 9048 | 7761 | 7510 | 8176 | 5781 | 5801 | 5911 | 7202 | 7377 | 6121 | 7881 | 10475 | 7022 | 7174 | 9461 | 5881 | 7184 | 7257 | Raph1 |
| Rara | 1425 | 2230 | 2107 | 2310 | 1794 | 1484 | 1188 | 1082 | 1585 | 1687 | 1904 | 1196 | 1533 | 965 | 1505 | 2223 | 2099 | 2238 | 1566 | 1263 | 1315 | 1298 | 1731 | 1378 | 1062 | 1626 | 1004 | 1053 | Rara |
| Rarb | 1120 | 794 | 737 | 593 | 658 | 416 | 421 | 1362 | 1551 | 1524 | 1769 | 1248 | 1407 | 1057 | 922 | 1133 | 499 | 576 | 396 | 424 | 399 | 1155 | 1796 | 1700 | 935 | 1936 | 1146 | 1180 | Rarb |
| Rarg | 1817 | 1716 | 1500 | 1064 | 740 | 1414 | 579 | 2246 | 3498 | 2884 | 3600 | 1620 | 2808 | 1672 | 1356 | 2164 | 1175 | 1345 | 796 | 590 | 569 | 2239 | 3351 | 2842 | 1699 | 3023 | 1930 | 1688 | Rarg |
| Rarres1 | 20 | 19 | 24 | 24 | 34 | 28 | 20 | 37 | 34 | 15 | 32 | 13 | 18 | 14 | 37 | 19 | 24 | 35 | 19 | 20 | 19 | 20 | 42 | 20 | 8 | 11 | 22 | 15 | Rarres1 |
| Rarres2 | 33 | 27 | 24 | 39 | 49 | 55 | 40 | 60 | 97 | 77 | 164 | 73 | 74 | 73 | 47 | 41 | 29 | 80 | 52 | 48 | 39 | 79 | 185 | 78 | 104 | 80 | 50 | 46 | Rarres2 |
| Rars | 11682 | 10754 | 11321 | 9018 | 7523 | 8186 | 8880 | 8132 | 7145 | 6527 | 6140 | 7577 | 7293 | 7777 | 12274 | 11135 | 9792 | 8719 | 7564 | 10323 | 8231 | 6266 | 7058 | 6778 | 6417 | 8498 | 7997 | 8434 | Rars |
| Rars2 | 2832 | 2439 | 2854 | 2509 | 2223 | 2448 | 2809 | 2914 | 2605 | 2348 | 2586 | 2779 | 2660 | 2913 | 2860 | 2696 | 2720 | 2349 | 2422 | 2725 | 2469 | 2633 | 2699 | 2613 | 2508 | 2990 | 3001 | 3049 | Rars2 |
| Rasa1 | 10409 | 8011 | 8974 | 10390 | 10915 | 12171 | 13398 | 12853 | 10910 | 9833 | 10049 | 11463 | 10605 | 12391 | 10849 | 8791 | 9506 | 10343 | 11042 | 11481 | 13034 | 11732 | 10659 | 10109 | 11325 | 10205 | 11250 | 11532 | Rasa1 |
| Rasa2 | 1703 | 1206 | 1418 | 1400 | 1814 | 1860 | 1842 | 3181 | 3016 | 2702 | 3085 | 2700 | 2975 | 2997 | 1681 | 1615 | 1355 | 1282 | 1674 | 1764 | 1735 | 2919 | 2933 | 2517 | 2886 | 2423 | 2779 | 2617 | Rasa2 |
| Rasa3 | 5306 | 5743 | 5291 | 5235 | 5478 | 5528 | 4898 | 3085 | 3506 | 3136 | 3494 | 3039 | 3291 | 2925 | 5425 | 5687 | 5276 | 5335 | 5372 | 4983 | 4946 | 3566 | 3609 | 2918 | 3204 | 3173 | 3192 | 2809 | Rasa3 |
| Rasa4 | 1971 | 2624 | 2397 | 2610 | 2725 | 2424 | 2123 | 932 | 1046 | 998 | 1360 | 846 | 954 | 807 | 2041 | 2324 | 2533 | 2788 | 2591 | 2350 | 2138 | 1415 | 1189 | 930 | 961 | 765 | 968 | 803 | Rasa4 |
| Rasal1 | 14 | 8 | 7 | 17 | 14 | 14 | 31 | 21 | 27 | 6 | 11 | 7 | 0 | 12 | 12 | 9 | 14 | 43 | 34 | 39 | 46 | 30 | 48 | 12 | 24 | 8 | 3 | 7 | Rasal1 |
| Rasal2 | 9131 | 9064 | 6461 | 9335 | 9519 | 10019 | 10410 | 10651 | 9011 | 8867 | 8073 | 8388 | 8610 | 9184 | 9742 | 7843 | 7413 | 8680 | 8845 | 10184 | 10626 | 8662 | 8378 | 6824 | 9987 | 4589 | 9411 | 8402 | Rasal2 |
| Rasal3 | 23 | 22 | 33 | 16 | 23 | 35 | 20 | 34 | 40 | 41 | 63 | 47 | 40 | 28 | 34 | 41 | 25 | 20 | 16 | 18 | 20 | 50 | 54 | 25 | 28 | 25 | 46 | 40 | Rasal3 |
| Rasd1 | 63 | 86 | 86 | 75 | 74 | 63 | 54 | 27 | 31 | 36 | 27 | 37 | 39 | 36 | 63 | 83 | 80 | 39 | 68 | 55 | 34 | 32 | 47 | 44 | 20 | 63 | 43 | 23 | Rasd1 |
| Rasd2 | 204 | 275 | 339 | 462 | 647 | 407 | 383 | 72 | 78 | 74 | 121 | 15 | 56 | 24 | 241 | 319 | 458 | 406 | 526 | 275 | 489 | 115 | 103 | 52 | 102 | 19 | 5 | 24 | Rasd2 |
| Rasef | 150 | 42 | 6 | 47 | 11 | 17 | 4 | 245 | 219 | 166 | 241 | 38 | 84 | 48 | 95 | 73 | 20 | 13 | 5 | 10 | 10 | 67 | 245 | 187 | 41 | 254 | 41 | 34 | Rasef |
| Rasgef1a | 87 | 121 | 123 | 247 | 335 | 249 | 292 | 115 | 206 | 203 | 164 | 96 | 91 | 112 | 141 | 123 | 154 | 259 | 259 | 185 | 306 | 226 | 144 | 159 | 213 | 110 | 101 | 68 | Rasgef1a |
| Rasgef1b | 1631 | 1805 | 1479 | 2802 | 3057 | 2717 | 3163 | 3091 | 2973 | 3315 | 2539 | 2821 | 1898 | 3226 | 1880 | 1328 | 1874 | 2951 | 3010 | 2764 | 3503 | 2904 | 2755 | 3390 | 4191 | 2252 | 2404 | 2541 | Rasgef1b |
| Rasgef1c | 53 | 75 | 18 | 91 | 154 | 137 | 108 | 193 | 253 | 207 | 197 | 151 | 142 | 126 | 65 | 77 | 41 | 111 | 127 | 103 | 115 | 274 | 223 | 137 | 256 | 93 | 71 | 78 | Rasgef1c |
| Rasgrf1 | 340 | 447 | 431 | 551 | 928 | 670 | 810 | 183 | 227 | 217 | 217 | 103 | 96 | 101 | 419 | 400 | 447 | 771 | 829 | 708 | 773 | 236 | 182 | 143 | 274 | 47 | 41 | 71 | Rasgrf1 |
| Rasgrf2 | 60 | 60 | 44 | 163 | 384 | 229 | 231 | 244 | 299 | 320 | 279 | 126 | 147 | 133 | 83 | 69 | 101 | 242 | 241 | 201 | 264 | 483 | 270 | 172 | 372 | 53 | 86 | 90 | Rasgrf2 |
| Rasgrp1 | 761 | 706 | 421 | 747 | 828 | 838 | 1043 | 812 | 779 | 559 | 439 | 975 | 612 | 957 | 692 | 591 | 712 | 646 | 817 | 920 | 835 | 560 | 384 | 699 | 630 | 1055 | 781 | 885 | Rasgrp1 |
| Rasgrp2 | 0 | 1 | 0 | 0 | 0 | 0 | 0 | 0 | 0 | 0 | 0 | 0 | 0 | 0 | 1 | 0 | 0 | 0 | 0 | 0 | 0 | 0 | 0 | 2 | 0 | 0 | 0 | 0 | Rasgrp2 |
| Rasgrp3 | 714 | 621 | 699 | 483 | 676 | 677 | 568 | 1017 | 991 | 1022 | 1369 | 957 | 1185 | 932 | 700 | 669 | 633 | 676 | 616 | 755 | 583 | 1140 | 1100 | 929 | 968 | 1007 | 1149 | 1037 | Rasgrp3 |
| Rasgrp4 | 159 | 169 | 210 | 167 | 162 | 197 | 131 | 123 | 152 | 202 | 184 | 188 | 220 | 188 | 117 | 196 | 188 | 172 | 163 | 115 | 148 | 174 | 179 | 203 | 151 | 251 | 196 | 198 | Rasgrp4 |
| Rasip1 | 287 | 302 | 424 | 370 | 467 | 403 | 329 | 628 | 915 | 917 | 985 | 771 | 775 | 692 | 293 | 404 | 419 | 417 | 429 | 323 | 358 | 947 | 948 | 885 | 798 | 885 | 703 | 673 | Rasip1 |
| Rasl10a | 24 | 22 | 25 | 21 | 15 | 19 | 8 | 5 | 7 | 1 | 22 | 14 | 9 | 6 | 15 | 41 | 19 | 10 | 16 | 17 | 19 | 7 | 20 | 3 | 0 | 18 | 3 | 5 | Rasl10a |
| Rasl10b | 2660 | 3504 | 3329 | 4599 | 5098 | 4270 | 4730 | 2116 | 2530 | 2898 | 2531 | 2335 | 1835 | 2273 | 2690 | 2881 | 4376 | 4887 | 5287 | 3654 | 5253 | 3031 | 2238 | 2801 | 3136 | 2222 | 1963 | 2083 | Rasl10b |
| Rasl11a | 178 | 179 | 260 | 204 | 240 | 193 | 225 | 256 | 213 | 267 | 250 | 308 | 315 | 333 | 196 | 176 | 239 | 156 | 235 | 234 | 201 | 231 | 170 | 303 | 263 | 401 | 294 | 411 | Rasl11a |
| Rasl11b | 1856 | 1891 | 1586 | 1654 | 1171 | 1267 | 1178 | 1528 | 1715 | 1538 | 1687 | 1113 | 1356 | 943 | 1569 | 2016 | 1455 | 1593 | 1172 | 1266 | 1106 | 1171 | 1858 | 1460 | 908 | 1656 | 1068 | 1149 | Rasl11b |
| Rasl12 | 366 | 388 | 363 | 321 | 382 | 352 | 344 | 361 | 489 | 491 | 457 | 445 | 477 | 447 | 378 | 478 | 322 | 374 | 422 | 322 | 340 | 424 | 444 | 383 | 455 | 462 | 440 | 439 | Rasl12 |
| Rasl2-9 | 12 | 1 | 4 | 5 | 6 | 0 | 4 | 5 | 4 | 4 | 6 | 4 | 8 | 6 | 8 | 2 | 7 | 8 | 5 | 4 | 7 | 10 | 10 | 8 | 0 | 4 | 7 | 4 | Rasl2-9 |
| Rassf1 | 2307 | 2640 | 2368 | 2413 | 1707 | 1843 | 1948 | 1931 | 1909 | 1709 | 1997 | 1885 | 2019 | 1975 | 2115 | 2332 | 2319 | 2229 | 1951 | 2098 | 1899 | 1659 | 2060 | 1949 | 1814 | 2165 | 2101 | 2090 | Rassf1 |
| Rassf10 | 287 | 292 | 226 | 233 | 272 | 233 | 193 | 203 | 202 | 232 | 250 | 93 | 156 | 116 | 269 | 273 | 190 | 281 | 189 | 183 | 222 | 194 | 246 | 203 | 187 | 146 | 109 | 93 | Rassf10 |
| Rassf2 | 6676 | 8307 | 8569 | 9200 | 9710 | 9509 | 8613 | 4951 | 4809 | 5712 | 5319 | 6008 | 5760 | 5918 | 7093 | 6928 | 9407 | 9073 | 9552 | 8357 | 8469 | 6044 | 5245 | 5920 | 6344 | 5396 | 6315 | 5675 | Rassf2 |
| Rassf3 | 2759 | 2384 | 2840 | 2148 | 1618 | 1622 | 1414 | 1181 | 1491 | 1445 | 1850 | 1022 | 1555 | 1115 | 2232 | 3464 | 1943 | 2225 | 1514 | 1424 | 1539 | 1121 | 1709 | 1347 | 1021 | 1532 | 1170 | 1107 | Rassf3 |
| Rassf4 | 9846 | 12519 | 7313 | 7825 | 7891 | 8776 | 7952 | 4322 | 3614 | 3082 | 3014 | 3646 | 5279 | 3178 | 11525 | 11988 | 8398 | 7913 | 7496 | 9918 | 7899 | 3200 | 4278 | 2611 | 3013 | 2087 | 3972 | 3739 | Rassf4 |
| Rassf5 | 210 | 197 | 200 | 228 | 245 | 193 | 159 | 134 | 141 | 153 | 216 | 50 | 136 | 99 | 253 | 247 | 220 | 261 | 147 | 151 | 180 | 201 | 193 | 92 | 142 | 71 | 93 | 87 | Rassf5 |
| Rassf6 | 44 | 53 | 26 | 43 | 47 | 37 | 16 | 26 | 49 | 26 | 49 | 13 | 40 | 21 | 54 | 43 | 17 | 29 | 12 | 24 | 26 | 30 | 43 | 47 | 30 | 42 | 28 | 44 | Rassf6 |
| Rassf7 | 1546 | 2060 | 1740 | 1709 | 1354 | 1526 | 1276 | 670 | 708 | 675 | 664 | 845 | 907 | 864 | 1485 | 1793 | 1814 | 1592 | 1559 | 1503 | 1448 | 632 | 743 | 821 | 634 | 1005 | 954 | 924 | Rassf7 |
| Rassf8 | 3362 | 3156 | 3573 | 3286 | 3275 | 3410 | 2851 | 1892 | 2419 | 2117 | 2770 | 2030 | 2402 | 1967 | 3257 | 3996 | 3183 | 3817 | 2869 | 2724 | 3015 | 2074 | 2760 | 2192 | 2110 | 1975 | 2058 | 2115 | Rassf8 |
| Rassf9 | 188 | 70 | 91 | 84 | 119 | 83 | 96 | 339 | 316 | 263 | 262 | 181 | 211 | 209 | 136 | 134 | 74 | 85 | 110 | 94 | 101 | 256 | 331 | 257 | 251 | 237 | 149 | 171 | Rassf9 |
| Raver2 | 3776 | 3757 | 3465 | 2879 | 2555 | 2659 | 2411 | 1884 | 2111 | 2041 | 1941 | 1892 | 2335 | 1782 | 3297 | 3753 | 3096 | 2971 | 2461 | 2590 | 2528 | 1584 | 1976 | 1593 | 1484 | 2087 | 1965 | 2012 | Raver2 |
| Rax | 120 | 147 | 38 | 73 | 65 | 42 | 32 | 77 | 64 | 36 | 45 | 100 | 41 | 95 | 86 | 86 | 65 | 99 | 48 | 82 | 70 | 104 | 98 | 101 | 108 | 160 | 92 | 44 | Rax |
| Rb1 | 10212 | 8932 | 7423 | 9633 | 10016 | 11291 | 12790 | 7921 | 6789 | 6318 | 6523 | 7335 | 6467 | 7306 | 11101 | 9008 | 8642 | 9261 | 9436 | 11372 | 12514 | 7252 | 6658 | 5897 | 7386 | 4870 | 6973 | 6589 | Rb1 |
| Rb1cc1 | 9173 | 7483 | 6741 | 8635 | 9235 | 9542 | 10264 | 12046 | 9644 | 8685 | 8627 | 8626 | 8740 | 9070 | 9959 | 8315 | 7497 | 8546 | 8812 | 10751 | 10044 | 10041 | 9728 | 8409 | 9945 | 6529 | 8871 | 8456 | Rb1cc1 |
| Rbak | 2643 | 2379 | 2097 | 2440 | 2208 | 2608 | 2543 | 2325 | 2222 | 2070 | 2240 | 2339 | 2169 | 2474 | 2507 | 2326 | 2126 | 2475 | 2153 | 2260 | 2473 | 2353 | 2152 | 2302 | 2305 | 2302 | 2209 | 2226 | Rbak |
| Rbbp4 | 35519 | 30347 | 34249 | 30692 | 26535 | 28574 | 31704 | 31731 | 27900 | 26708 | 25859 | 32729 | 29955 | 34047 | 33766 | 28245 | 32423 | 28751 | 31729 | 29559 | 31503 | 27215 | 26797 | 31212 | 30124 | 39035 | 34737 | 37667 | Rbbp4 |
| Rbbp5 | 6564 | 6785 | 5604 | 5984 | 5226 | 5717 | 5941 | 5953 | 4948 | 4708 | 4722 | 5207 | 4898 | 5321 | 6600 | 6196 | 5483 | 5689 | 5671 | 6218 | 6295 | 4938 | 4920 | 5006 | 5145 | 5145 | 5444 | 5548 | Rbbp5 |
| Rbbp6 | 13498 | 12337 | 12198 | 14832 | 13196 | 14250 | 13991 | 16080 | 14171 | 15578 | 15304 | 15989 | 15626 | 16773 | 13209 | 12623 | 13273 | 13668 | 13766 | 13300 | 14020 | 15141 | 13457 | 14993 | 15691 | 13794 | 15832 | 15482 | Rbbp6 |
| Rbbp7 | 15894 | 13859 | 16582 | 12597 | 11211 | 11867 | 12968 | 13578 | 13766 | 12370 | 13783 | 13840 | 13861 | 14816 | 15202 | 14394 | 13606 | 12768 | 11902 | 12715 | 11758 | 11834 | 14617 | 13547 | 11943 | 17491 | 14585 | 15548 | Rbbp7 |
| Rbbp8 | 3793 | 2649 | 3184 | 2407 | 2136 | 2623 | 2781 | 4172 | 3707 | 3062 | 2828 | 4123 | 3521 | 4295 | 3752 | 2752 | 2863 | 2100 | 2403 | 3128 | 2560 | 2806 | 3541 | 3313 | 3143 | 4727 | 4302 | 4226 | Rbbp8 |
| Rbbp8nl | 24 | 3 | 11 | 1 | 0 | 4 | 0 | 20 | 47 | 30 | 25 | 1 | 18 | 5 | 3 | 7 | 0 | 0 | 0 | 0 | 0 | 3 | 50 | 38 | 2 | 43 | 9 | 11 | Rbbp8nl |

| | | | | | | | | | | | | | | | | | | | | | | | | | | | | | |
|---|---|---|---|---|---|---|---|---|---|---|---|---|---|---|---|---|---|---|---|---|---|---|---|---|---|---|---|---|---|
| Rbbp9 | 3180 | 2514 | 2857 | 2399 | 2756 | 2629 | 2932 | 4796 | 4211 | 3894 | 4174 | 4304 | 4161 | 4492 | 3095 | 2569 | 2666 | 2446 | 2795 | 3212 | 2694 | 4324 | 4561 | 4170 | 4043 | 4760 | 4424 | 4676 | Rbbp9 |
| Rbck1 | 4342 | 5549 | 4604 | 5097 | 5018 | 4896 | 4571 | 4575 | 5425 | 5755 | 5264 | 5736 | 5261 | 5283 | 4189 | 4674 | 5307 | 5016 | 5483 | 4495 | 4919 | 5260 | 5066 | 5394 | 5575 | 5292 | 5328 | 5384 | Rbck1 |
| Rbfa | 1653 | 2313 | 1861 | 1768 | 1571 | 1534 | 1418 | 828 | 1040 | 1002 | 882 | 943 | 895 | 904 | 1582 | 1808 | 1800 | 1654 | 1671 | 1543 | 1448 | 909 | 1067 | 882 | 948 | 1122 | 948 | 915 | Rbfa |
| Rbfox1 | 1134 | 1442 | 1297 | 3252 | 5684 | 3837 | 3642 | 3596 | 3333 | 5018 | 4580 | 2427 | 3375 | 2554 | 2168 | 1562 | 2132 | 3713 | 3761 | 3766 | 4182 | 5484 | 3670 | 3153 | 4620 | 1173 | 1911 | 2040 | Rbfox1 |
| Rbfox2 | 20226 | 19547 | 21659 | 28965 | 33815 | 28216 | 27702 | 32110 | 32908 | 37783 | 37647 | 31433 | 30957 | 31124 | 21298 | 21121 | 25919 | 31486 | 30934 | 24983 | 29794 | 42847 | 31595 | 36407 | 42455 | 25442 | 28563 | 28944 | Rbfox2 |
| Rbfox3 | 789 | 1177 | 1219 | 2010 | 3296 | 2100 | 2129 | 1916 | 2202 | 3374 | 2673 | 2322 | 1994 | 2225 | 1069 | 906 | 1804 | 2478 | 2604 | 1846 | 2473 | 3213 | 2186 | 2761 | 3421 | 2125 | 1693 | 1774 | Rbfox3 |
| Rbis | 5173 | 4016 | 4761 | 4735 | 4061 | 3802 | 6198 | 4731 | 3526 | 3724 | 4060 | 3938 | 4257 | 5478 | 5384 | 4045 | 4517 | 4393 | 4275 | 5160 | 5672 | 3643 | 3587 | 4320 | 4112 | 4515 | 4424 | 5444 | Rbis |
| Rbks | 87 | 79 | 68 | 53 | 76 | 45 | 55 | 34 | 40 | 34 | 48 | 58 | 35 | 60 | 45 | 64 | 56 | 56 | 49 | 70 | 55 | 48 | 38 | 40 | 41 | 62 | 38 | 40 | Rbks |
| Rbl1 | 4050 | 3058 | 3864 | 3305 | 2963 | 3442 | 3962 | 4645 | 4213 | 4212 | 3719 | 6115 | 4880 | 6201 | 3704 | 3285 | 3566 | 2962 | 3418 | 3664 | 3323 | 3521 | 3722 | 4688 | 4265 | 5744 | 6099 | 5987 | Rbl1 |
| Rbl2 | 3624 | 3505 | 3066 | 3470 | 3774 | 4018 | 3888 | 2507 | 2580 | 2534 | 2697 | 2343 | 2326 | 2435 | 3506 | 3335 | 3055 | 3766 | 3685 | 3483 | 3869 | 2699 | 2515 | 2379 | 2543 | 2197 | 2292 | 2134 | Rbl2 |
| Rbm10 | 5014 | 6141 | 6250 | 5728 | 5889 | 5582 | 5058 | 7365 | 8374 | 8729 | 8163 | 9520 | 8532 | 9348 | 4805 | 5045 | 6189 | 5223 | 6350 | 5296 | 5328 | 7282 | 7875 | 9559 | 9147 | 9970 | 9883 | 9544 | Rbm10 |
| Rbm11 | 649 | 581 | 628 | 606 | 599 | 645 | 670 | 145 | 136 | 179 | 152 | 92 | 118 | 123 | 775 | 690 | 698 | 610 | 580 | 662 | 642 | 167 | 210 | 154 | 99 | 138 | 105 | 103 | Rbm11 |
| Rbm12 | 3276 | 2033 | 2871 | 2429 | 2486 | 2494 | 2514 | 4493 | 4158 | 3951 | 4023 | 5085 | 4588 | 4966 | 2970 | 2395 | 2611 | 2113 | 2318 | 2667 | 2253 | 4121 | 3952 | 4133 | 4503 | 5534 | 4875 | 4863 | Rbm12 |
| Rbm12b1 | 873 | 802 | 1094 | 971 | 1116 | 990 | 1071 | 1417 | 1230 | 992 | 1114 | 1153 | 1002 | 1242 | 894 | 874 | 1048 | 1053 | 1043 | 1228 | 1192 | 1298 | 1218 | 1057 | 1257 | 1043 | 1109 | 970 | Rbm12b1 |
| Rbm12b2 | 2588 | 2325 | 2977 | 3122 | 2802 | 2840 | 3047 | 3330 | 2477 | 2254 | 2472 | 2358 | 2307 | 2670 | 2745 | 2752 | 3177 | 3043 | 2907 | 3261 | 2944 | 2849 | 2572 | 2583 | 2607 | 2552 | 2496 | 2478 | Rbm12b2 |
| Rbm14 | 3047 | 3505 | 3053 | 3126 | 2923 | 2602 | 2711 | 2815 | 3113 | 3029 | 3095 | 3854 | 3191 | 3861 | 2689 | 2852 | 3329 | 2866 | 3206 | 2655 | 2907 | 2837 | 2816 | 3572 | 3217 | 3346 | 3352 | 3777 | Rbm14 |
| Rbm15 | 2067 | 1894 | 1670 | 1678 | 1348 | 1579 | 1719 | 2191 | 1704 | 1266 | 1382 | 1652 | 1537 | 1662 | 1924 | 2058 | 1628 | 1498 | 1449 | 2007 | 1752 | 1372 | 1712 | 1549 | 1589 | 2100 | 1927 | 1832 | Rbm15 |
| Rbm15b | 7892 | 8485 | 9432 | 9725 | 8816 | 8456 | 8739 | 9645 | 9412 | 9948 | 9215 | 11415 | 9757 | 10841 | 7559 | 7207 | 9851 | 9359 | 10085 | 7418 | 9199 | 10659 | 8579 | 11628 | 11298 | 12375 | 10901 | 11660 | Rbm15b |
| Rbm17 | 6692 | 6581 | 7982 | 6881 | 5702 | 5734 | 6395 | 6560 | 6063 | 7179 | 6856 | 7945 | 6922 | 8379 | 6620 | 6022 | 7434 | 6246 | 6067 | 6244 | 6139 | 6261 | 6275 | 7444 | 6852 | 9062 | 7711 | 8416 | Rbm17 |
| Rbm18 | 5996 | 5725 | 5704 | 6296 | 6061 | 6137 | 6886 | 8309 | 6652 | 6558 | 6423 | 7076 | 6117 | 6956 | 6626 | 5855 | 5986 | 5927 | 6316 | 7071 | 6693 | 7635 | 6699 | 6483 | 7233 | 6473 | 6777 | 6972 | Rbm18 |
| Rbm19 | 1464 | 1805 | 1766 | 1416 | 1258 | 1265 | 1226 | 2011 | 2438 | 2372 | 2304 | 2777 | 2790 | 2640 | 1295 | 1477 | 1510 | 1243 | 1524 | 1344 | 1200 | 2025 | 2233 | 2504 | 2534 | 3139 | 2866 | 2867 | Rbm19 |
| Rbm20 | 400 | 389 | 377 | 461 | 521 | 472 | 558 | 615 | 748 | 784 | 777 | 1025 | 906 | 975 | 359 | 401 | 542 | 514 | 557 | 543 | 519 | 505 | 885 | 1064 | 943 | 1061 | 834 | 945 | Rbm20 |
| Rbm22 | 3763 | 3746 | 4605 | 4577 | 4282 | 4053 | 4463 | 4656 | 4849 | 5537 | 5166 | 6701 | 5746 | 6663 | 3520 | 3780 | 4618 | 4296 | 4546 | 3528 | 4341 | 5559 | 4807 | 6147 | 5838 | 7276 | 5861 | 6605 | Rbm22 |
| Rbm24 | 551 | 396 | 756 | 635 | 718 | 606 | 716 | 688 | 608 | 697 | 1352 | 607 | 518 | 634 | 545 | 449 | 773 | 582 | 780 | 652 | 741 | 613 | 875 | 842 | 517 | 859 | 536 | 668 | Rbm24 |
| Rbm25 | 18754 | 14375 | 20180 | 20362 | 20261 | 22401 | 23177 | 27026 | 23865 | 29092 | 27972 | 31225 | 30132 | 34175 | 18948 | 16772 | 19766 | 19243 | 21002 | 19496 | 21905 | 27626 | 23172 | 29072 | 29091 | 28252 | 31164 | 31567 | Rbm25 |
| Rbm26 | 10869 | 8025 | 9585 | 9725 | 9619 | 10651 | 11962 | 11147 | 10280 | 10897 | 11693 | 11947 | 11721 | 12701 | 10370 | 9428 | 9791 | 9803 | 9382 | 9288 | 11227 | 11127 | 9759 | 11012 | 10859 | 10790 | 11998 | 12291 | Rbm26 |
| Rbm27 | 6801 | 5700 | 7197 | 8009 | 6995 | 7617 | 7849 | 6469 | 5492 | 6092 | 6064 | 6711 | 6255 | 7103 | 6870 | 6129 | 7149 | 7770 | 7183 | 6825 | 7542 | 6169 | 5238 | 6254 | 6790 | 6443 | 6678 | 6907 | Rbm27 |
| Rbm28 | 13144 | 13676 | 12726 | 12913 | 11400 | 11306 | 13021 | 9482 | 8778 | 8841 | 8904 | 9650 | 9029 | 10137 | 12955 | 13042 | 12725 | 12844 | 12273 | 11976 | 12541 | 8941 | 8497 | 9311 | 9221 | 8626 | 9616 | 9474 | Rbm28 |
| Rbm3 | 16576 | 14564 | 17013 | 15328 | 13979 | 16532 | 14702 | 21187 | 18926 | 16668 | 19801 | 23159 | 23578 | 26041 | 15672 | 14654 | 14932 | 13649 | 15385 | 19381 | 13761 | 15719 | 16877 | 20905 | 23140 | 28925 | 26503 | 27481 | Rbm3 |
| Rbm33 | 8450 | 8185 | 7396 | 9390 | 10902 | 10128 | 10579 | 13384 | 12756 | 14254 | 13448 | 15550 | 13283 | 14958 | 8410 | 7606 | 9113 | 9684 | 10798 | 9445 | 10793 | 14357 | 11413 | 14006 | 15433 | 9845 | 14214 | 13830 | Rbm33 |
| Rbm34 | 4037 | 3255 | 3810 | 3618 | 3299 | 3615 | 3927 | 2508 | 2225 | 2529 | 2570 | 3079 | 2851 | 3123 | 4054 | 3501 | 3502 | 3629 | 3117 | 3600 | 3540 | 2694 | 2336 | 2749 | 2566 | 3183 | 3029 | 3056 | Rbm34 |
| Rbm38 | 2418 | 3815 | 3467 | 2863 | 2266 | 2236 | 2161 | 1145 | 1388 | 1208 | 1143 | 1988 | 1946 | 1759 | 2314 | 3142 | 3310 | 2904 | 2906 | 2285 | 2091 | 1156 | 1205 | 1765 | 1404 | 2387 | 1796 | 2233 | Rbm38 |
| Rbm39 | 24220 | 18786 | 23040 | 20966 | 21966 | 23091 | 24939 | 49034 | 41837 | 39737 | 40950 | 44789 | 42135 | 45645 | 25440 | 21073 | 22556 | 20819 | 22946 | 26853 | 23324 | 40952 | 41730 | 42002 | 39849 | 44684 | 43985 | 44791 | Rbm39 |
| Rbm4 | 864 | 599 | 460 | 612 | 616 | 701 | 684 | 819 | 702 | 568 | 618 | 620 | 548 | 726 | 781 | 722 | 520 | 523 | 523 | 722 | 712 | 695 | 654 | 516 | 643 | 499 | 771 | 568 | Rbm4 |
| Rbm41 | 1629 | 1170 | 1179 | 1467 | 1582 | 1790 | 1783 | 2446 | 1910 | 1745 | 1840 | 1875 | 1818 | 2011 | 1820 | 1442 | 1356 | 1349 | 1492 | 1876 | 1769 | 1858 | 1860 | 1660 | 1948 | 1217 | 1835 | 1871 | Rbm41 |
| Rbm42 | 9 | 10 | 22 | 11 | 10 | 12 | 12 | 15 | 18 | 14 | 14 | 14 | 2 | 19 | 8 | 11 | 16 | 12 | 19 | 13 | 11 | 17 | 13 | 12 | 17 | 12 | 19 | 11 | Rbm42 |
| Rbm43 | 397 | 263 | 383 | 257 | 317 | 261 | 266 | 463 | 484 | 428 | 473 | 452 | 452 | 437 | 406 | 387 | 346 | 266 | 249 | 284 | 249 | 406 | 566 | 397 | 393 | 590 | 454 | 436 | Rbm43 |
| Rbm44 | 58 | 35 | 65 | 25 | 23 | 30 | 21 | 42 | 50 | 52 | 68 | 66 | 103 | 47 | 46 | 50 | 48 | 31 | 14 | 32 | 21 | 92 | 66 | 33 | 22 | 34 | 44 | 27 | Rbm44 |
| Rbm45 | 4119 | 4237 | 3995 | 4192 | 3863 | 3931 | 4422 | 4241 | 3868 | 3778 | 3646 | 4438 | 4073 | 4453 | 4130 | 4023 | 4174 | 3854 | 4062 | 4441 | 4060 | 3888 | 3854 | 4141 | 3967 | 4408 | 4595 | 4562 | Rbm45 |
| Rbm46 | 503 | 318 | 573 | 405 | 380 | 409 | 323 | 282 | 266 | 267 | 369 | 210 | 342 | 231 | 583 | 600 | 501 | 432 | 313 | 333 | 256 | 289 | 344 | 208 | 161 | 309 | 255 | 214 | Rbm46 |
| Rbm47 | 115 | 74 | 131 | 120 | 122 | 96 | 80 | 234 | 308 | 290 | 262 | 260 | 391 | 221 | 91 | 163 | 122 | 87 | 97 | 112 | 80 | 296 | 461 | 373 | 208 | 329 | 229 | 214 | Rbm47 |
| Rbm48 | 760 | 631 | 749 | 693 | 675 | 701 | 832 | 1044 | 815 | 833 | 835 | 908 | 688 | 956 | 799 | 695 | 727 | 774 | 653 | 724 | 746 | 931 | 778 | 917 | 884 | 1038 | 832 | 946 | Rbm48 |
| Rbm4b | 3211 | 3094 | 3349 | 3422 | 4449 | 4330 | 4281 | 9254 | 7458 | 7516 | 6701 | 8409 | 7117 | 8507 | 3694 | 2780 | 3845 | 3497 | 4583 | 4568 | 4480 | 8545 | 7351 | 7683 | 8456 | 7288 | 8126 | 8148 | Rbm4b |
| Rbm5 | 18175 | 15757 | 15293 | 16871 | 18819 | 18445 | 19088 | 33091 | 28716 | 29803 | 30280 | 28030 | 27041 | 29495 | 17922 | 15386 | 15288 | 16789 | 18503 | 19167 | 18628 | 31707 | 27513 | 28517 | 30692 | 22875 | 27935 | 26527 | Rbm5 |
| Rbm6 | 13433 | 12920 | 14570 | 13902 | 13214 | 13722 | 15008 | 15369 | 13261 | 15030 | 14029 | 15727 | 14787 | 16779 | 13863 | 12509 | 14139 | 13522 | 14210 | 14050 | 14220 | 15214 | 12885 | 15812 | 16032 | 15218 | 16154 | 16422 | Rbm6 |
| Rbm7 | 4210 | 3314 | 3833 | 3439 | 3174 | 3628 | 3940 | 5383 | 4214 | 3613 | 3687 | 4255 | 4211 | 4493 | 4278 | 3722 | 3387 | 3148 | 3237 | 4210 | 3504 | 4012 | 4465 | 3970 | 3822 | 4923 | 4485 | 4576 | Rbm7 |
| Rbm8a | 8360 | 7970 | 10916 | 9328 | 7744 | 7457 | 8861 | 6684 | 5839 | 7471 | 6704 | 8262 | 7572 | 9025 | 8323 | 7379 | 9814 | 8543 | 8270 | 8354 | 8529 | 6169 | 6038 | 8415 | 7930 | 10257 | 8765 | 9159 | Rbm8a |
| Rbm8a2 | 57 | 36 | 49 | 31 | 29 | 31 | 32 | 46 | 22 | 25 | 31 | 45 | 40 | 43 | 46 | 34 | 35 | 26 | 34 | 29 | 27 | 52 | 29 | 38 | 34 | 33 | 35 | 43 | Rbm8a2 |
| Rbms1 | 4831 | 3366 | 4210 | 2994 | 3897 | 3591 | 2320 | 6581 | 7296 | 6681 | 8962 | 4561 | 7614 | 4636 | 4574 | 5419 | 3328 | 3681 | 2369 | 2611 | 2390 | 6956 | 8209 | 5608 | 4953 | 5689 | 4910 | 4987 | Rbms1 |
| Rbms2 | 5274 | 5453 | 4907 | 4437 | 3340 | 3842 | 3624 | 2845 | 3296 | 2888 | 3467 | 2840 | 3393 | 3297 | 4824 | 5270 | 4604 | 4226 | 3645 | 3419 | 3924 | 2349 | 3598 | 2939 | 2479 | 3103 | 3141 | 3466 | Rbms2 |
| Rbms3 | 3511 | 2194 | 2258 | 2352 | 4196 | 2678 | 1830 | 4335 | 5177 | 4527 | 6315 | 2035 | 4093 | 1892 | 3319 | 4216 | 2275 | 3388 | 1992 | 1938 | 2085 | 5930 | 5379 | 2920 | 3370 | 2587 | 2045 | 1944 | Rbms3 |
| Rbmx | 17685 | 16030 | 18014 | 17901 | 19536 | 20021 | 20730 | 28427 | 24504 | 25780 | 23846 | 30068 | 26864 | 33138 | 18264 | 15687 | 18289 | 16801 | 19667 | 21207 | 20421 | 26978 | 22680 | 27262 | 29390 | 29529 | 29916 | 31454 | Rbmx |
| Rbmx2 | 1190 | 1115 | 1503 | 1155 | 1010 | 945 | 1336 | 1116 | 1014 | 962 | 999 | 1246 | 1091 | 1439 | 1222 | 1086 | 1191 | 1144 | 1112 | 1170 | 1227 | 1036 | 909 | 1213 | 1216 | 1508 | 1227 | 1457 | Rbmx2 |
| Rbmxl1 | 6726 | 7057 | 7733 | 6532 | 6412 | 6364 | 7111 | 7800 | 7237 | 6793 | 6555 | 8585 | 7886 | 9125 | 6887 | 6886 | 7802 | 6508 | 7029 | 7820 | 6447 | 5957 | 6925 | 7794 | 7146 | 9957 | 8918 | 9276 | Rbmxl1 |
| Rbmxl2 | 10 | 1 | 2 | 7 | 3 | 4 | 2 | 0 | 0 | 1 | 0 | 0 | 1 | 1 | 0 | 4 | 6 | 7 | 5 | 2 | 0 | 0 | 1 | 0 | 0 | 7 | 0 | 0 | Rbmxl2 |
| Rbmy | 0 | 0 | 0 | 0 | 0 | 0 | 0 | 0 | 0 | 1 | 0 | 0 | 0 | 0 | 0 | 0 | 0 | 0 | 0 | 0 | 0 | 0 | 0 | 0 | 0 | 0 | 0 | 0 | Rbmy |
| Rbp1 | 3160 | 2827 | 2142 | 3106 | 3051 | 2746 | 3467 | 3086 | 3785 | 3475 | 4318 | 3041 | 2137 | 3654 | 2793 | 2211 | 3086 | 2979 | 3170 | 2452 | 3417 | 2379 | 3503 | 4131 | 3375 | 4201 | 2503 | 2604 | Rbp1 |
| Rbp2 | 2 | 0 | 4 | 0 | 2 | 2 | 0 | 13 | 12 | 2 | 18 | 10 | 3 | 6 | 4 | 5 | 3 | 2 | 0 | 1 | 4 | 3 | 12 | 2 | 12 | 4 | 5 | 8 | Rbp2 |
| Rbp3 | 0 | 1 | 2 | 0 | 0 | 0 | 1 | 1 | 2 | 0 | 2 | 1 | 3 | 2 | 0 | 4 | 0 | 3 | 0 | 0 | 1 | 7 | 3 | 6 | 3 | 0 | 3 | 2 | Rbp3 |
| Rbp4 | 9 | 11 | 30 | 20 | 28 | 7 | 17 | 21 | 40 | 20 | 63 | 13 | 22 | 14 | 16 | 22 | 14 | 18 | 13 | 18 | 18 | 23 | 69 | 15 | 23 | 12 | 23 | 23 | Rbp4 |
| Rbp7 | 2 | 3 | 5 | 8 | 0 | 4 | 3 | 4 | 4 | 13 | 4 | 5 | 0 | 6 | 1 | 2 | 8 | 4 | 5 | 0 | 4 | 7 | 14 | 3 | 8 | 12 | 2 | 6 | Rbp7 |
| Rbpj | 10221 | 8386 | 9823 | 9834 | 10638 | 10348 | 11586 | 12676 | 11554 | 11972 | 11989 | 13896 | 12528 | 14001 | 9928 | 8893 | 9437 | 9819 | 10639 | 9526 | 10633 | 13089 | 10628 | 12401 | 13170 | 12999 | 13124 | 13567 | Rbpj |
| Rbpjl | 15 | 10 | 3 | 5 | 6 | 28 | 0 | 9 | 39 | 31 | 40 | 3 | 9 | 10 | 4 | 23 | 14 | 14 | 5 | 1 | 0 | 40 | 20 | 14 | 23 | 12 | 10 | 2 | Rbpjl |
| Rbpms | 1369 | 1107 | 1440 | 783 | 614 | 742 | 449 | 590 | 848 | 746 | 947 | 705 | 843 | 626 | 1208 | 1585 | 851 | 800 | 532 | 518 | 452 | 719 | 944 | 798 | 511 | 1029 | 632 | 644 | Rbpms |
| Rbpms2 | 912 | 891 | 1129 | 696 | 529 | 557 | 459 | 355 | 405 | 368 | 501 | 464 | 591 | 434 | 791 | 1030 | 859 | 567 | 509 | 394 | 411 | 401 | 457 | 418 | 296 | 570 | 525 | 516 | Rbpms2 |
| Rbsn | 4164 | 4074 | 3754 | 3787 | 3963 | 4111 | 3957 | 5667 | 5483 | 5326 | 5425 | 5471 | 5040 | 5161 | 3973 | 3765 | 3855 | 3616 | 4185 | 4058 | 3928 | 5457 | 5311 | 4966 | 5743 | 4780 | 5287 | 4990 | Rbsn |
| Rbx1 | 5796 | 5211 | 5822 | 5991 | 6103 | 5966 | 6792 | 9628 | 8300 | 8942 | 8182 | 9366 | 8465 | 10184 | 5943 | 5099 | 6245 | 5404 | 6420 | 7166 | 6370 | 8971 | 8213 | 9541 | 9595 | 9850 | 9596 | 10231 | Rbx1 |
| Rc3h1 | 7221 | 5918 | 5779 | 6466 | 6217 | 6966 | 7168 | 6704 | 5686 | 5817 | 6029 | 5805 | 5934 | 6388 | 6931 | 6757 | 6087 | 6852 | 6139 | 6292 | 7313 | 5835 | 5557 | 5356 | 6259 | 5125 | 6444 | 6462 | Rc3h1 |
| Rc3h2 | 9629 | 7881 | 9228 | 10758 | 11065 | 11911 | 12581 | 13948 | 11031 | 11888 | 11367 | 12829 | 12094 | 13691 | 10218 | 8521 | 10247 | 10612 | 10983 | 11796 | 12483 | 13526 | 10563 | 11800 | 13643 | 12311 | 13183 | 13381 | Rc3h2 |
| Rcan1 | 4051 | 4814 | 4832 | 5126 | 4969 | 4679 | 4533 | 2690 | 3152 | 3139 | 2846 | 2690 | 2532 | 2684 | 4180 | 4367 | 4889 | 5198 | 5146 | 4355 | 4818 | 3200 | 3214 | 3028 | 2993 | 2970 | 2686 | 2677 | Rcan1 |
| Rcan2 | 1606 | 1765 | 1867 | 2973 | 4985 | 3560 | 3323 | 4247 | 3886 | 4735 | 3793 | 3301 | 3675 | 3490 | 2736 | 1854 | 2373 | 3138 | 3261 | 3097 | 3713 | 6284 | 4167 | 3237 | 4245 | 2777 | 2724 | 3131 | Rcan2 |
| Rcan3 | 2997 | 2912 | 3087 | 3537 | 4123 | 3564 | 3633 | 2800 | 2901 | 3174 | 3183 | 2547 | 2431 | 2470 | 3021 | 3039 | 3574 | 3997 | 3809 | 3219 | 3506 | 3853 | 2871 | 2965 | 3360 | 2284 | 2277 | 2311 | Rcan3 |
| Rcbtb1 | 9449 | 8336 | 9388 | 8634 | 9345 | 9674 | 9628 | 10636 | 9724 | 9611 | 9267 | 12173 | 11071 | 12153 | 9228 | 8803 | 8898 | 8170 | 8955 | 9107 | 9323 | 9670 | 9081 | 10654 | 10600 | 11933 | 12248 | 12072 | Rcbtb1 |
| Rcbtb2 | 3365 | 2629 | 2929 | 2182 | 2299 | 2628 | 2282 | 9750 | 8419 | 7640 | 7996 | 9682 | 9456 | 9555 | 3340 | 3025 | 2669 | 1927 | 2336 | 2791 | 2257 | 7838 | 8405 | 8256 | 7932 | 10449 | 10127 | 10525 | Rcbtb2 |
| Rcc1 | 3864 | 4369 | 4759 | 3671 | 2688 | 3342 | 3222 | 3235 | 3397 | 3398 | 3243 | 5009 | 4397 | 4589 | 3406 | 3664 | 4082 | 3354 | 3455 | 3399 | 3055 | 2796 | 3123 | 4349 | 3754 | 6021 | 5317 | 5104 | Rcc1 |
| Rcc1l | 1057 | 1225 | 1268 | 997 | 883 | 768 | 895 | 1102 | 1187 | 1108 | 1284 | 1382 | 1332 | 1225 | 948 | 1153 | 1116 | 928 | 958 | 917 | 777 | 998 | 1166 | 1273 | 1268 | 1666 | 1447 | 1288 | Rcc1l |
| Rcc2 | 12364 | 14017 | 19470 | 15472 | 13870 | 13767 | 12649 | 14906 | 17693 | 19225 | 18301 | 25725 | 21711 | 23606 | 10406 | 12113 | 18593 | 14330 | 17317 | 11485 | 13870 | 15309 | 15151 | 24505 | 20558 | 30715 | 23480 | 24701 | Rcc2 |
| Rccd1 | 1839 | 1995 | 1812 | 1895 | 1865 | 1880 | 1873 | 1635 | 1736 | 1933 | 1625 | 1939 | 1717 | 1973 | 1853 | 1688 | 1871 | 1965 | 1976 | 1887 | 1901 | 1430 | 1616 | 1814 | 2056 | 1730 | 2070 | 1921 | Rccd1 |
| Rce1 | 1166 | 1450 | 1407 | 1525 | 1172 | 1199 | 1214 | 1276 | 1266 | 1318 | 1339 | 1435 | 1291 | 1464 | 1120 | 1176 | 1301 | 1294 | 1334 | 1228 | 1205 | 1302 | 1238 | 1462 | 1288 | 1719 | 1454 | 1523 | Rce1 |

| | | | | | | | | | | | | | | | | | | | | | | | | | | | | | |
|---|---|---|---|---|---|---|---|---|---|---|---|---|---|---|---|---|---|---|---|---|---|---|---|---|---|---|---|---|---|
| Rchy1 | 2716 | 2365 | 2591 | 2894 | 2757 | 2857 | 3027 | 3845 | 3419 | 3439 | 3492 | 3644 | 3359 | 3711 | 2851 | 2414 | 2621 | 2758 | 2918 | 2636 | 2807 | 3679 | 3529 | 3452 | 3669 | 3805 | 3525 | 3730 | Rchy1 |
| Rcl1 | 3110 | 3461 | 2649 | 2656 | 2144 | 2363 | 2392 | 1263 | 1336 | 1077 | 1221 | 1194 | 1369 | 1244 | 2773 | 3255 | 2602 | 2586 | 2439 | 2602 | 2423 | 894 | 1342 | 1199 | 1089 | 1706 | 1332 | 1302 | Rcl1 |
| Rcn1 | 12153 | 9810 | 11913 | 8094 | 6785 | 8622 | 7182 | 10574 | 11755 | 10397 | 13096 | 9925 | 11578 | 11099 | 11813 | 11603 | 9574 | 8797 | 6870 | 7495 | 6805 | 9356 | 13217 | 10732 | 9134 | 12997 | 10662 | 12140 | Rcn1 |
| Rcn2 | 8920 | 7917 | 9849 | 7901 | 7265 | 7858 | 8329 | 12125 | 10254 | 9984 | 10326 | 12852 | 11439 | 13921 | 9103 | 7962 | 9084 | 7312 | 7819 | 8103 | 7828 | 10960 | 10415 | 11660 | 10992 | 13513 | 12905 | 14222 | Rcn2 |
| Rcn3 | 4921 | 6226 | 6518 | 4636 | 2996 | 4107 | 2291 | 3940 | 6138 | 5265 | 7262 | 4070 | 5043 | 3873 | 4635 | 6331 | 5444 | 4909 | 3243 | 2935 | 2478 | 4328 | 7306 | 5351 | 4347 | 5340 | 4507 | 4033 | Rcn3 |
| Rcor1 | 5978 | 5516 | 6579 | 6993 | 6766 | 6599 | 6804 | 4811 | 5008 | 5526 | 5711 | 5552 | 5381 | 5501 | 5450 | 5927 | 6655 | 6785 | 6454 | 5020 | 6348 | 5152 | 4941 | 6127 | 5115 | 5671 | 4926 | 5342 | Rcor1 |
| Rcor2 | 6437 | 7705 | 10040 | 8624 | 8593 | 7824 | 7407 | 8101 | 8344 | 10648 | 8290 | 14659 | 10567 | 12134 | 6115 | 5892 | 10718 | 7896 | 10106 | 7261 | 7686 | 8791 | 6890 | 12995 | 11996 | 15579 | 12963 | 12912 | Rcor2 |
| Rcor3 | 573 | 473 | 344 | 459 | 579 | 632 | 564 | 1047 | 1036 | 785 | 922 | 855 | 887 | 827 | 607 | 514 | 392 | 387 | 520 | 656 | 526 | 804 | 924 | 628 | 832 | 601 | 938 | 741 | Rcor3 |
| Rcsd1 | 686 | 638 | 878 | 596 | 646 | 678 | 485 | 937 | 1051 | 1094 | 1540 | 1007 | 1143 | 899 | 682 | 829 | 658 | 727 | 544 | 547 | 534 | 1001 | 1554 | 1112 | 923 | 1133 | 952 | 871 | Rcsd1 |
| Rcvrn | 3 | 2 | 0 | 0 | 9 | 1 | 10 | 1 | 3 | 10 | 2 | 0 | 0 | 3 | 3 | 3 | 2 | 1 | 2 | 1 | 9 | 5 | 1 | 2 | 12 | 0 | 3 | 0 | Rcvrn |
| Rd3 | 0 | 0 | 0 | 0 | 11 | 0 | 0 | 4 | 2 | 48 | 9 | 1 | 3 | 1 | 5 | 2 | 0 | 1 | 2 | 0 | 1 | 3 | 4 | 5 | 1 | 5 | 2 | 0 | Rd3 |
| Rd3l | 2 | 0 | 0 | 0 | 0 | 0 | 0 | 0 | 0 | 1 | 2 | 0 | 0 | 0 | 0 | 0 | 0 | 0 | 0 | 0 | 0 | 0 | 1 | 0 | 0 | 0 | 0 | 0 | Rd3l |
| Rdh1 | 6 | 10 | 21 | 14 | 24 | 21 | 18 | 7 | 7 | 19 | 16 | 11 | 5 | 21 | 5 | 11 | 22 | 14 | 21 | 3 | 23 | 20 | 10 | 20 | 14 | 8 | 10 | 20 | Rdh1 |
| Rdh10 | 2042 | 1560 | 1825 | 1862 | 1988 | 1746 | 1886 | 1790 | 1949 | 1764 | 2264 | 1612 | 1878 | 1629 | 1861 | 1842 | 1725 | 2094 | 1769 | 1883 | 1857 | 2044 | 2179 | 1785 | 1763 | 1802 | 1576 | 1548 | Rdh10 |
| Rdh11 | 4910 | 4755 | 4390 | 3797 | 3980 | 4143 | 4403 | 3413 | 3143 | 2715 | 2712 | 2998 | 3158 | 3028 | 4809 | 4675 | 3890 | 4209 | 4222 | 4747 | 4350 | 2811 | 3193 | 2929 | 2804 | 3115 | 3077 | 3417 | Rdh11 |
| Rdh12 | 181 | 128 | 124 | 105 | 108 | 86 | | 72 | 105 | 85 | 85 | 99 | 81 | 114 | 166 | 157 | 172 | 139 | 131 | 131 | 113 | 77 | 103 | 92 | 68 | 88 | 70 | 70 | Rdh12 |
| Rdh13 | 3377 | 3711 | 3250 | 3166 | 2934 | 2797 | 3169 | 2650 | 2385 | 2368 | 2171 | 2337 | 2391 | 2402 | 3218 | 3157 | 3233 | 2870 | 3218 | 3143 | 3109 | 2379 | 2379 | 2257 | 2574 | 2527 | 2566 | 2441 | Rdh13 |
| Rdh14 | 595 | 419 | 458 | 435 | 574 | 554 | 646 | 1084 | 1171 | 862 | 1029 | 921 | 899 | 984 | 564 | 513 | 518 | 489 | 556 | 527 | 559 | 1105 | 1190 | 1060 | 918 | 1033 | 850 | 1011 | Rdh14 |
| Rdh16 | 2 | 3 | 4 | 8 | 6 | 1 | 4 | 1 | 2 | 1 | 0 | 0 | 1 | 0 | 5 | 2 | 6 | 2 | 2 | 9 | 3 | 0 | 1 | 0 | 1 | 3 | 0 | 0 | Rdh16 |
| Rdh16f2 | 1 | 2 | 1 | 1 | 1 | 0 | 2 | 0 | 0 | 1 | 0 | 0 | 0 | 0 | 0 | 5 | 0 | 2 | 0 | 2 | 3 | 0 | 1 | 0 | 0 | 0 | 0 | 0 | Rdh16f2 |
| Rdh19 | 0 | 1 | 0 | 0 | 0 | 0 | 0 | 0 | 0 | 0 | 1 | 0 | 0 | 0 | 0 | 1 | 0 | 0 | 0 | 0 | 1 | 0 | 0 | 0 | 0 | 0 | 0 | 0 | Rdh19 |
| Rdh5 | 204 | 269 | 218 | 215 | 214 | 208 | 189 | 206 | 241 | 271 | 237 | 272 | 242 | 287 | 203 | 242 | 270 | 263 | 292 | 195 | 174 | 286 | 271 | 241 | 262 | 180 | 294 | 258 | Rdh5 |
| Rdh7 | 0 | 0 | 0 | 0 | 0 | 0 | 0 | 0 | 1 | 0 | 0 | 0 | 0 | 0 | 0 | 0 | 0 | 0 | 0 | 0 | 0 | 0 | 0 | 1 | 0 | 0 | 0 | 0 | Rdh7 |
| Rdh9 | 0 | 2 | 0 | 7 | 1 | 2 | 2 | 0 | 2 | 0 | 2 | 2 | 1 | 3 | 5 | 4 | 0 | 3 | 0 | 0 | 7 | 2 | 2 | 0 | 4 | 0 | 2 | 0 | Rdh9 |
| Rdm1 | 850 | 989 | 863 | 877 | 738 | 895 | 965 | 1082 | 1109 | 936 | 964 | 1307 | 1055 | 1331 | 832 | 769 | 936 | 727 | 1023 | 983 | 911 | 812 | 1062 | 1209 | 1162 | 1429 | 1327 | 1283 | Rdm1 |
| Rdx | 26889 | 21771 | 25457 | 19570 | 19703 | 21806 | 21579 | 27667 | 24199 | 22079 | 22946 | 27147 | 26037 | 27457 | 27174 | 23929 | 22955 | 19513 | 20457 | 24715 | 20900 | 22546 | 24477 | 24201 | 23835 | 26832 | 29650 | 28078 | Rdx |
| Rec114 | 32 | 52 | 51 | 54 | 34 | 22 | 49 | 18 | 24 | 35 | 27 | 22 | 44 | 42 | 34 | 28 | 53 | 37 | 51 | 28 | 28 | 10 | 19 | 15 | 20 | 35 | 37 | 66 | Rec114 |
| Rec8 | 700 | 936 | 854 | 1018 | 1076 | 966 | 780 | 793 | 642 | 965 | 903 | 685 | 864 | 639 | 948 | 1083 | 979 | 835 | 873 | 863 | 1034 | 1105 | 784 | 666 | 821 | 805 | 650 | 749 | Rec8 |
| Reck | 4357 | 3959 | 3552 | 3146 | 2743 | 3848 | 2674 | 2768 | 3651 | 3091 | 4275 | 2980 | 3598 | 2922 | 4101 | 5184 | 3263 | 3759 | 2499 | 2587 | 2582 | 3023 | 4254 | 2951 | 2819 | 3010 | 3099 | 2822 | Reck |
| Recql | 5091 | 5217 | 4652 | 4729 | 4084 | 4088 | 4909 | 3709 | 3332 | 3144 | 3099 | 3774 | 3358 | 3961 | 4899 | 5042 | 4424 | 4604 | 4229 | 4673 | 4738 | 3110 | 3287 | 3406 | 3547 | 3622 | 4069 | 3961 | Recql |
| Recql4 | 1653 | 2113 | 1972 | 1708 | 1033 | 1188 | 1258 | 854 | 1063 | 1041 | 944 | 1447 | 1192 | 1302 | 1328 | 1698 | 1593 | 1460 | 1444 | 1275 | 1250 | 668 | 867 | 1280 | 1018 | 1975 | 1432 | 1402 | Recql4 |
| Recql5 | 2985 | 3566 | 3472 | 3675 | 3072 | 3217 | 3309 | 2170 | 2567 | 2750 | 2419 | 3101 | 2821 | 2944 | 2810 | 3044 | 3459 | 3279 | 3518 | 3057 | 3254 | 2580 | 2264 | 2997 | 2906 | 2824 | 3118 | 2947 | Recql5 |
| Reep1 | 6653 | 7219 | 6956 | 9348 | 14250 | 11152 | 11094 | 14247 | 13026 | 14732 | 12832 | 10843 | 10950 | 11030 | 8751 | 7011 | 8467 | 10740 | 11486 | 11251 | 11970 | 19006 | 12290 | 12026 | 16244 | 8150 | 9836 | 10275 | Reep1 |
| Reep2 | 1354 | 1856 | 1809 | 2578 | 3332 | 2534 | 2617 | 2033 | 2336 | 2917 | 2494 | 2573 | 2099 | 2497 | 1592 | 1456 | 2371 | 2716 | 3133 | 2360 | 2921 | 3240 | 2057 | 2596 | 3281 | 2099 | 2036 | 2071 | Reep2 |
| Reep3 | 6158 | 5583 | 6501 | 6145 | 6052 | 6326 | 6338 | 7675 | 6960 | 7589 | 7173 | 8244 | 7984 | 8508 | 6497 | 6386 | 6154 | 6048 | 5976 | 5445 | 6074 | 7461 | 7071 | 7633 | 7383 | 7798 | 7669 | 8444 | Reep3 |
| Reep4 | 1953 | 2477 | 2360 | 1916 | 1407 | 1438 | 1501 | 945 | 1254 | 934 | 1059 | 1372 | 1347 | 1177 | 1693 | 2441 | 2041 | 1994 | 1562 | 1592 | 1388 | 979 | 1218 | 1218 | 938 | 1716 | 1317 | 1304 | Reep4 |
| Reep5 | 7961 | 7784 | 8522 | 8095 | 12389 | 10201 | 9886 | 9785 | 9738 | 10523 | 10809 | 7261 | 8658 | 7296 | 8818 | 8576 | 8031 | 9987 | 9298 | 9975 | 9792 | 11946 | 10336 | 8610 | 10112 | 6698 | 7594 | 7135 | Reep5 |
| Reep6 | 1065 | 1598 | 1286 | 1386 | 796 | 881 | 622 | 153 | 199 | 197 | 158 | 150 | 152 | 121 | 1001 | 1436 | 1290 | 1306 | 938 | 815 | 725 | 117 | 184 | 212 | 120 | 192 | 167 | 171 | Reep6 |
| Reg1 | 0 | 0 | 0 | 0 | 0 | 0 | 0 | 0 | 1 | 0 | 0 | 0 | 0 | 0 | 0 | 0 | 0 | 0 | 0 | 0 | 0 | 0 | 0 | 0 | 1 | 0 | 0 | 0 | Reg1 |
| Reg3a | 0 | 0 | 0 | 0 | 0 | 1 | 0 | 0 | 1 | 0 | 0 | 0 | 0 | 0 | 0 | 0 | 0 | 0 | 0 | 0 | 0 | 0 | 0 | 0 | 0 | 0 | 0 | 0 | Reg3a |
| Reg3b | 0 | 0 | 0 | 0 | 2 | 0 | 1 | 0 | 1 | 1 | 0 | 1 | 1 | 0 | 0 | 0 | 0 | 0 | 4 | 0 | 1 | 2 | 1 | 0 | 3 | 1 | 2 | 0 | Reg3b |
| Reg3d | 0 | 0 | 0 | 0 | 0 | 0 | 0 | 0 | 0 | 0 | 0 | 0 | 0 | 0 | 0 | 0 | 0 | 0 | 0 | 0 | 0 | 0 | 0 | 0 | 1 | 0 | 0 | 0 | Reg3d |
| Reg3g | 0 | 0 | 0 | 0 | 0 | 0 | 0 | 26 | 14 | 43 | 6 | 0 | 0 | 0 | 0 | 0 | 0 | 0 | 0 | 0 | 0 | 0 | 25 | 21 | 0 | 3 | 0 | 0 | Reg3g |
| Reg4 | 2 | 0 | 0 | 2 | 0 | 0 | 0 | 1 | 0 | 0 | 0 | 0 | 0 | 4 | 0 | 0 | 0 | 1 | 0 | 0 | 0 | 2 | 0 | 0 | 0 | 0 | 0 | 0 | Reg4 |
| Rel | 807 | 745 | 568 | 771 | 614 | 756 | 700 | 338 | 297 | 300 | 402 | 216 | 316 | 239 | 742 | 818 | 636 | 811 | 591 | 598 | 652 | 282 | 375 | 279 | 274 | 197 | 234 | 247 | Rel |
| Rela | 1307 | 1635 | 1497 | 1621 | 1742 | 1608 | 1332 | 1885 | 2392 | 2426 | 2359 | 2443 | 2288 | 2110 | 1191 | 1556 | 1572 | 1688 | 1687 | 1253 | 1493 | 2234 | 2455 | 2506 | 2399 | 3046 | 2048 | 2331 | Rela |
| Relb | 204 | 260 | 265 | 213 | 177 | 199 | 150 | 181 | 217 | 197 | 252 | 157 | 222 | 165 | 182 | 258 | 234 | 239 | 136 | 164 | 171 | 234 | 270 | 186 | 180 | 175 | 158 | 195 | Relb |
| Relch | 8395 | 7292 | 5941 | 8175 | 7580 | 8343 | 8719 | 6729 | 5829 | 5018 | 5034 | 5441 | 5229 | 5177 | 9297 | 7742 | 6455 | 7948 | 7406 | 8716 | 8659 | 6413 | 5708 | 4698 | 5610 | 3869 | 5026 | 4932 | Relch |
| Rell1 | 3785 | 3194 | 3891 | 3364 | 3009 | 2986 | 3371 | 1379 | 1400 | 1378 | 1372 | 1491 | 1420 | 1561 | 3537 | 3605 | 3342 | 3422 | 3172 | 3049 | 3232 | 1161 | 1416 | 1577 | 1206 | 1886 | 1425 | 1495 | Rell1 |
| Rell2 | 843 | 1227 | 1205 | 1654 | 2365 | 1576 | 1724 | 752 | 1008 | 1233 | 1082 | 701 | 729 | 641 | 864 | 896 | 1612 | 1802 | 1852 | 1538 | 1801 | 1245 | 911 | 954 | 1171 | 591 | 560 | 517 | Rell2 |
| Reln | 8389 | 9115 | 3504 | 12472 | 16397 | 18221 | 19980 | 11731 | 13930 | 12273 | 11441 | 11649 | 7496 | 12072 | 8631 | 6305 | 6968 | 13761 | 16797 | 18515 | 21988 | 10373 | 11388 | 10026 | 16832 | 5575 | 10665 | 9257 | Reln |
| Relt | 424 | 512 | 573 | 594 | 592 | 515 | 504 | 205 | 198 | 218 | 242 | 205 | 181 | 213 | 442 | 517 | 557 | 579 | 567 | 505 | 510 | 204 | 211 | 209 | 227 | 226 | 198 | 148 | Relt |
| Rem1 | 50 | 47 | 31 | 36 | 37 | 31 | 13 | 12 | 20 | 16 | 36 | 10 | 14 | 13 | 63 | 76 | 43 | 41 | 34 | 30 | 15 | 20 | 30 | 13 | 18 | 8 | 9 | 6 | Rem1 |
| Rem2 | 752 | 1001 | 1071 | 915 | 1072 | 936 | 941 | 926 | 855 | 1028 | 817 | 1237 | 1176 | 1129 | 885 | 790 | 1161 | 869 | 1222 | 994 | 918 | 1155 | 876 | 1077 | 1035 | 965 | 1197 | 1098 | Rem2 |
| Ren1 | 0 | 1 | 0 | 0 | 44 | 3 | 1 | 24 | 57 | 49 | 103 | 0 | 20 | 0 | 18 | 2 | 3 | 2 | 0 | 1 | 0 | 55 | 87 | 0 | 6 | 0 | 0 | 0 | Ren1 |
| Renbp | 499 | 544 | 628 | 479 | 408 | 498 | 331 | 413 | 464 | 452 | 622 | 493 | 565 | 437 | 473 | 625 | 435 | 533 | 364 | 407 | 316 | 505 | 661 | 441 | 384 | 494 | 522 | 476 | Renbp |
| Rep15 | 113 | 85 | 121 | 124 | 97 | 79 | 131 | 60 | 56 | 63 | 42 | 48 | 32 | 32 | 83 | 104 | 100 | 85 | 116 | 78 | 58 | 72 | 80 | 44 | 37 | 71 | 42 | 62 | Rep15 |
| Repin1 | 1368 | 1642 | 1873 | 1886 | 1757 | 1591 | 1607 | 1491 | 1556 | 1752 | 1630 | 2072 | 1675 | 2123 | 1303 | 1347 | 2084 | 1520 | 1892 | 1478 | 1641 | 1646 | 1531 | 1968 | 1779 | 2239 | 2038 | 2085 | Repin1 |
| Reps1 | 5059 | 4786 | 4298 | 4617 | 5021 | 4817 | 4874 | 4683 | 4608 | 4786 | 4400 | 5437 | 4953 | 5135 | 4878 | 4489 | 4548 | 4541 | 5195 | 4317 | 5010 | 5091 | 4425 | 4737 | 4773 | 5249 | 4939 | 5253 | Reps1 |
| Reps2 | 1646 | 1090 | 901 | 1498 | 2004 | 2086 | 1969 | 1710 | 1799 | 1560 | 1746 | 1187 | 1134 | 1241 | 1444 | 1425 | 1241 | 2060 | 1654 | 1647 | 1979 | 1900 | 1936 | 1506 | 1706 | 790 | 1019 | 916 | Reps2 |
| Rer1 | 5421 | 5354 | 5299 | 4851 | 4482 | 4594 | 5051 | 5286 | 5120 | 4351 | 4914 | 4897 | 4820 | 4809 | 5732 | 5279 | 5014 | 4824 | 4722 | 5592 | 4869 | 4583 | 5135 | 4887 | 4640 | 5687 | 4709 | 5266 | Rer1 |
| Rere | 10972 | 12754 | 16559 | 19293 | 19350 | 16317 | 14928 | 9900 | 10787 | 15911 | 13918 | 16280 | 13397 | 13692 | 10668 | 12009 | 18354 | 19038 | 19575 | 12745 | 17077 | 16706 | 10100 | 15717 | 16588 | 12294 | 12856 | 13410 | Rere |
| Rerg | 309 | 210 | 142 | 280 | 232 | 272 | 208 | 460 | 410 | 335 | 340 | 174 | 151 | 160 | 240 | 262 | 191 | 330 | 229 | 218 | 224 | 162 | 440 | 409 | 174 | 366 | 132 | 134 | Rerg |
| Rergl | 0 | 0 | 0 | 0 | 0 | 0 | 0 | 0 | 0 | 0 | 0 | 0 | 0 | 0 | 0 | 0 | 0 | 0 | 0 | 0 | 0 | 0 | 1 | 0 | 0 | 0 | 0 | 0 | Rergl |
| Resf1 | 3586 | 2630 | 3489 | 2916 | 3190 | 3399 | 3076 | 5142 | 4339 | 4500 | 4658 | 4752 | 4860 | 4816 | 3376 | 3076 | 3130 | 2851 | 2999 | 3230 | 3091 | 4385 | 4402 | 4414 | 4483 | 4245 | 4696 | 4677 | Resf1 |
| Resp18 | 400 | 529 | 471 | 580 | 932 | 653 | 805 | 718 | 638 | 714 | 595 | 547 | 525 | 593 | 679 | 474 | 712 | 665 | 782 | 904 | 788 | 906 | 674 | 638 | 665 | 434 | 619 | 600 | Resp18 |
| Rest | 3240 | 2645 | 3213 | 3533 | 2786 | 3250 | 3023 | 2745 | 2944 | 2854 | 3386 | 2744 | 3308 | 2704 | 3091 | 3308 | 3177 | 3498 | 2935 | 2669 | 3048 | 2740 | 3153 | 2963 | 2363 | 3892 | 2880 | 2985 | Rest |
| Ret | 582 | 667 | 694 | 756 | 1871 | 769 | 571 | 833 | 1148 | 1033 | 1483 | 377 | 724 | 359 | 736 | 738 | 822 | 1054 | 775 | 719 | 690 | 1596 | 1106 | 401 | 940 | 226 | 411 | 381 | Ret |
| Retn | 6 | 16 | 8 | 18 | 17 | 17 | 22 | 9 | 3 | 9 | 6 | 15 | 9 | 16 | 13 | 13 | 10 | 31 | 16 | 16 | 21 | 7 | 6 | 11 | 1 | 5 | 9 | 6 | Retn |
| Retnla | 0 | 0 | 0 | 0 | 1 | 0 | 0 | 0 | 0 | 0 | 0 | 0 | 0 | 0 | 0 | 0 | 0 | 0 | 0 | 0 | 0 | 0 | 0 | 0 | 0 | 0 | 0 | 0 | Retnla |
| Retnlg | 0 | 0 | 0 | 2 | 0 | 0 | 2 | 0 | 2 | 8 | 5 | 3 | 5 | 2 | 0 | 0 | 0 | 7 | 5 | 0 | 1 | 0 | 8 | 3 | 0 | 3 | 5 | 1 | Retnlg |
| Retreg1 | 955 | 802 | 814 | 789 | 1088 | 903 | 884 | 885 | 828 | 917 | 760 | 453 | 644 | 557 | 1033 | 988 | 773 | 946 | 846 | 972 | 925 | 1061 | 913 | 737 | 732 | 735 | 571 | 599 | Retreg1 |
| Retreg2 | 4284 | 5158 | 5198 | 6659 | 6920 | 6088 | 5898 | 5012 | 5147 | 5585 | 5524 | 5807 | 5037 | 5199 | 4411 | 4410 | 6080 | 6285 | 7238 | 5526 | 6367 | 7031 | 5272 | 5528 | 6102 | 5592 | 4858 | 5375 | Retreg2 |
| Retreg3 | 8007 | 8928 | 7756 | 8618 | 7706 | 7726 | 7800 | 6504 | 6033 | 6120 | 5839 | 6598 | 5958 | 6006 | 8195 | 8079 | 7911 | 8944 | 7742 | 7689 | 7932 | 6879 | 6268 | 6431 | 6844 | 6496 | 5950 | 6259 | Retreg3 |
| Retsat | 294 | 270 | 270 | 214 | 242 | 287 | 178 | 315 | 475 | 324 | 415 | 278 | 373 | 310 | 252 | 342 | 254 | 266 | 196 | 269 | 237 | 312 | 467 | 282 | 329 | 433 | 380 | 330 | Retsat |
| Rev1 | 5554 | 4764 | 4451 | 5057 | 4719 | 4952 | 5561 | 5130 | 4490 | 4402 | 4002 | 4782 | 4293 | 4850 | 5275 | 4761 | 4610 | 4736 | 4848 | 5109 | 5106 | 4692 | 4356 | 4200 | 4602 | 4789 | 4382 | 4799 | Rev1 |
| Rev3l | 15384 | 11974 | 5914 | 10097 | 13862 | 15618 | 15090 | 22983 | 20187 | 14657 | 15873 | 12009 | 12944 | 12806 | 15955 | 13838 | 7165 | 10871 | 10670 | 18750 | 15691 | 15430 | 19497 | 10198 | 15410 | 7307 | 13372 | 10734 | Rev3l |
| Rex1bd | 1057 | 1273 | 1132 | 1230 | 1154 | 1108 | 967 | 1398 | 1389 | 1718 | 1423 | 1418 | 1395 | 1301 | 964 | 1009 | 1177 | 1109 | 1245 | 1232 | 1025 | 1297 | 1368 | 1728 | 1394 | 1785 | 1443 | 1571 | Rex1bd |
| Rex2 | 0 | 1 | 0 | 0 | 0 | 0 | 0 | 1 | 0 | 0 | 0 | 0 | 1 | 0 | 0 | 0 | 0 | 0 | 0 | 0 | 1 | 0 | 0 | 0 | 0 | 0 | 1 | 0 | Rex2 |

| | | | | | | | | | | | | | | | | | | | | | | | | | | | | | |
|---|---|---|---|---|---|---|---|---|---|---|---|---|---|---|---|---|---|---|---|---|---|---|---|---|---|---|---|---|---|
| Rexo1 | 3420 | 4668 | 5392 | 5078 | 5005 | 4479 | 3769 | 3816 | 4601 | 5585 | 4931 | 6473 | 4947 | 4964 | 3343 | 3911 | 5430 | 4692 | 5311 | 3562 | 4005 | 5417 | 4372 | 5938 | 5509 | 6378 | 5145 | 4841 | Rexo1 |
| Rexo2 | 2359 | 2006 | 2393 | 2359 | 2472 | 2115 | 2518 | 2160 | 2085 | 2164 | 2292 | 2093 | 2141 | 2297 | 2285 | 2089 | 2284 | 2334 | 2476 | 2340 | 2458 | 2087 | 2055 | 2285 | 2261 | 2295 | 2216 | 2427 | Rexo2 |
| Rexo4 | 2978 | 3139 | 2845 | 2943 | 2873 | 2870 | 2696 | 3551 | 3487 | 3492 | 3479 | 3433 | 3486 | 3408 | 2763 | 3054 | 2866 | 3093 | 2869 | 2964 | 2723 | 3182 | 3651 | 3508 | 3404 | 3473 | 3402 | 3344 | Rexo4 |
| Rexo5 | 2040 | 1559 | 1323 | 1218 | 1217 | 1420 | 1340 | 1417 | 1266 | 1095 | 1307 | 1315 | 1305 | 1437 | 1812 | 1775 | 1442 | 1223 | 1150 | 1394 | 1377 | 978 | 1328 | 998 | 991 | 1114 | 1382 | 1258 | Rexo5 |
| Rfc1 | 8868 | 8020 | 7606 | 6358 | 5189 | 6201 | 6139 | 7514 | 7654 | 6129 | 7085 | 7753 | 7475 | 7995 | 8227 | 8157 | 6487 | 6123 | 5715 | 6988 | 5597 | 5217 | 7249 | 6956 | 6023 | 8474 | 8458 | 8167 | Rfc1 |
| Rfc2 | 5585 | 6314 | 6237 | 5372 | 4907 | 5053 | 5226 | 4971 | 5441 | 5328 | 5081 | 6263 | 5691 | 5955 | 4977 | 5534 | 6166 | 5131 | 5834 | 5491 | 5217 | 4401 | 5155 | 5869 | 5599 | 7465 | 6527 | 6407 | Rfc2 |
| Rfc3 | 3162 | 3176 | 3179 | 2551 | 2528 | 2542 | 2963 | 3489 | 3444 | 2813 | 2635 | 3909 | 3225 | 3856 | 3134 | 2631 | 2788 | 2347 | 2894 | 3131 | 2628 | 2764 | 3018 | 3376 | 3032 | 4837 | 4194 | 4037 | Rfc3 |
| Rfc4 | 4149 | 3549 | 3873 | 3087 | 2167 | 2765 | 3205 | 3342 | 2745 | 2372 | 2130 | 3532 | 2975 | 3681 | 4026 | 3457 | 3175 | 2513 | 2856 | 3394 | 2877 | 1977 | 2564 | 2907 | 2642 | 4511 | 3858 | 4119 | Rfc4 |
| Rfc5 | 2344 | 2217 | 2309 | 1846 | 1716 | 1900 | 2149 | 2464 | 2726 | 2623 | 2397 | 3549 | 2862 | 3297 | 1995 | 1901 | 2257 | 1826 | 2109 | 1921 | 1910 | 2020 | 2495 | 3000 | 2549 | 4213 | 3595 | 3820 | Rfc5 |
| Rfesd | 1065 | 760 | 688 | 806 | 704 | 791 | 935 | 933 | 1044 | 802 | 844 | 967 | 976 | 990 | 1054 | 912 | 740 | 703 | 777 | 973 | 887 | 784 | 1068 | 851 | 865 | 870 | 1149 | 1164 | Rfesd |
| Rffl | 2893 | 2843 | 2413 | 2685 | 2095 | 1974 | 2683 | 1241 | 1268 | 1211 | 1110 | 1387 | 1036 | 1579 | 2653 | 2463 | 2415 | 2451 | 2358 | 2253 | 2467 | 1019 | 1046 | 1423 | 1129 | 1767 | 1221 | 1301 | Rffl |
| Rfk | 6171 | 5364 | 5931 | 5880 | 5026 | 4967 | 5951 | 3727 | 3338 | 3432 | 3516 | 3707 | 4042 | 4101 | 6291 | 5997 | 5750 | 5172 | 4945 | 5071 | 5218 | 3212 | 3555 | 3754 | 3534 | 4831 | 3698 | 4405 | Rfk |
| Rflna | 347 | 503 | 396 | 381 | 321 | 303 | 146 | 375 | 821 | 731 | 1049 | 302 | 490 | 301 | 434 | 632 | 348 | 418 | 224 | 151 | 197 | 633 | 706 | 477 | 367 | 513 | 314 | 313 | Rflna |
| Rflnb | 2773 | 2379 | 3599 | 2374 | 2032 | 2046 | 1728 | 1585 | 2065 | 1874 | 2648 | 1542 | 1825 | 1327 | 2509 | 3523 | 2697 | 2348 | 1968 | 1561 | 1524 | 1338 | 2207 | 1948 | 1119 | 1875 | 1338 | 1604 | Rflnb |
| Rfng | 1956 | 2626 | 2636 | 3168 | 3416 | 2656 | 2532 | 2350 | 2845 | 3303 | 3040 | 3125 | 2694 | 2845 | 1791 | 2377 | 3065 | 3208 | 3241 | 2365 | 2759 | 3561 | 2728 | 3338 | 3217 | 3018 | 2513 | 2476 | Rfng |
| Rfpl4 | 0 | 0 | 0 | 0 | 0 | 0 | 0 | 0 | 0 | 0 | 0 | 0 | 0 | 0 | 0 | 0 | 0 | 0 | 0 | 0 | 0 | 0 | 0 | 0 | 1 | 0 | 0 | 0 | Rfpl4 |
| Rfpl4b | 0 | 2 | 0 | 0 | 2 | 4 | 0 | 0 | 1 | 0 | 3 | 0 | 0 | 2 | 3 | 0 | 0 | 0 | 0 | 0 | 0 | 0 | 0 | 0 | 0 | 1 | 1 | 1 | Rfpl4b |
| Rft1 | 644 | 529 | 620 | 481 | 515 | 592 | 483 | 948 | 1129 | 989 | 1100 | 1100 | 1226 | 1006 | 591 | 550 | 552 | 513 | 524 | 542 | 497 | 916 | 1102 | 1064 | 923 | 1257 | 1113 | 1042 | Rft1 |
| Rftn1 | 459 | 568 | 561 | 559 | 781 | 640 | 757 | 537 | 831 | 824 | 790 | 747 | 421 | 689 | 456 | 495 | 699 | 677 | 713 | 601 | 635 | 777 | 768 | 785 | 868 | 810 | 549 | 529 | Rftn1 |
| Rftn2 | 1310 | 1085 | 1012 | 984 | 1002 | 1188 | 977 | 1606 | 1763 | 1713 | 1698 | 1265 | 1349 | 1298 | 1139 | 1192 | 1088 | 1020 | 1128 | 1102 | 1076 | 1432 | 1762 | 1703 | 1345 | 1612 | 1203 | 1220 | Rftn2 |
| Rfwd3 | 10154 | 9538 | 10212 | 8464 | 7614 | 8157 | 9163 | 8365 | 7853 | 7521 | 7124 | 9670 | 9017 | 10195 | 9869 | 9139 | 9251 | 8501 | 8341 | 8364 | 8647 | 6889 | 7167 | 8058 | 7956 | 11269 | 10104 | 10671 | Rfwd3 |
| Rfx1 | 1355 | 1694 | 1795 | 1981 | 2160 | 1778 | 1724 | 2077 | 2492 | 2932 | 2632 | 3049 | 2624 | 2621 | 1233 | 1431 | 2021 | 1902 | 2348 | 1716 | 1852 | 3182 | 2370 | 2957 | 2814 | 2688 | 2586 | 2533 | Rfx1 |
| Rfx2 | 1423 | 1994 | 2046 | 1788 | 1376 | 1395 | 1306 | 515 | 590 | 588 | 522 | 766 | 770 | 652 | 1358 | 1816 | 1924 | 1710 | 1588 | 1297 | 1380 | 610 | 671 | 643 | 636 | 876 | 658 | 679 | Rfx2 |
| Rfx3 | 4481 | 3721 | 3999 | 5071 | 4863 | 5323 | 5451 | 7618 | 6507 | 6597 | 5890 | 7449 | 6683 | 7756 | 4555 | 3709 | 4633 | 4545 | 5062 | 4933 | 5480 | 7209 | 6121 | 6623 | 7678 | 5409 | 7315 | 7316 | Rfx3 |
| Rfx4 | 1493 | 1837 | 2979 | 1653 | 1794 | 1822 | 1195 | 2881 | 2082 | 2577 | 2006 | 3957 | 4840 | 3716 | 2115 | 1924 | 2620 | 1443 | 1746 | 1521 | 1397 | 3065 | 2223 | 2594 | 2607 | 2792 | 4774 | 4685 | Rfx4 |
| Rfx5 | 2263 | 2150 | 2192 | 2355 | 2650 | 2540 | 2607 | 2455 | 2564 | 2451 | 2461 | 2217 | 2252 | 2265 | 2113 | 1970 | 2376 | 2423 | 2707 | 2343 | 2665 | 2496 | 2515 | 2633 | 2280 | 2090 | 2009 | 2185 | Rfx5 |
| Rfx6 | 0 | 1 | 0 | 0 | 0 | 0 | 0 | 1 | 0 | 0 | 6 | 0 | 0 | 0 | 0 | 0 | 0 | 0 | 0 | 0 | 0 | 0 | 1 | 0 | 0 | 0 | 0 | 0 | Rfx6 |
| Rfx7 | 10933 | 9093 | 8712 | 10655 | 11299 | 11899 | 12474 | 12612 | 10561 | 10723 | 10073 | 12530 | 11504 | 12353 | 10671 | 9401 | 9580 | 10441 | 11223 | 11278 | 12364 | 12101 | 9482 | 10003 | 12101 | 8828 | 11472 | 11744 | Rfx7 |
| Rfx8 | 13 | 14 | 13 | 2 | 2 | 3 | 3 | 22 | 86 | 5 | 100 | 20 | 39 | 13 | 20 | 7 | 17 | 14 | 3 | 10 | 4 | 42 | 54 | 8 | 10 | 27 | 31 | 28 | Rfx8 |
| Rfxank | 1407 | 1821 | 2011 | 1872 | 1709 | 1660 | 1561 | 1197 | 1329 | 1465 | 1455 | 1614 | 1514 | 1337 | 1211 | 1537 | 1951 | 1810 | 1907 | 1458 | 1702 | 1325 | 1311 | 1579 | 1436 | 1749 | 1342 | 1423 | Rfxank |
| Rfxap | 2083 | 2334 | 2339 | 2305 | 2215 | 1904 | 2263 | 1529 | 1370 | 1538 | 1524 | 1560 | 1418 | 1677 | 2090 | 2196 | 2326 | 2560 | 2159 | 2055 | 2239 | 1663 | 1458 | 1617 | 1657 | 1761 | 1367 | 1576 | Rfxap |
| Rgcc | 401 | 541 | 313 | 416 | 528 | 365 | 513 | 642 | 639 | 568 | 779 | 609 | 442 | 678 | 487 | 428 | 462 | 526 | 394 | 419 | 424 | 500 | 464 | 708 | 662 | 976 | 452 | 485 | Rgcc |
| Rgl1 | 3796 | 3832 | 3762 | 4212 | 3953 | 3625 | 3724 | 2235 | 2607 | 2630 | 2865 | 2384 | 2741 | 2282 | 3848 | 4044 | 3881 | 4324 | 3832 | 3249 | 3812 | 2744 | 2979 | 2533 | 2471 | 2995 | 2413 | 2608 | Rgl1 |
| Rgl2 | 4674 | 5904 | 6748 | 6781 | 6802 | 5840 | 5844 | 4573 | 5193 | 5949 | 5478 | 6244 | 5714 | 5910 | 4609 | 5207 | 7325 | 6302 | 7167 | 5362 | 6037 | 5538 | 4832 | 6388 | 5917 | 6373 | 5956 | 5846 | Rgl2 |
| Rgl3 | 630 | 860 | 1097 | 833 | 663 | 662 | 598 | 111 | 103 | 137 | 180 | 202 | 201 | 140 | 679 | 856 | 1071 | 804 | 692 | 554 | 614 | 179 | 154 | 181 | 171 | 221 | 173 | 198 | Rgl3 |
| Rgma | 3302 | 4842 | 6283 | 6026 | 6590 | 5348 | 4422 | 5159 | 5773 | 7560 | 6706 | 8357 | 7279 | 6741 | 3533 | 3751 | 6660 | 5657 | 6791 | 4033 | 5071 | 7869 | 5521 | 7453 | 7520 | 6475 | 6952 | 6647 | Rgma |
| Rgmb | 6276 | 7123 | 9501 | 12995 | 14142 | 11826 | 12034 | 11020 | 9323 | 12769 | 11654 | 12218 | 14049 | 11626 | 7675 | 7186 | 11770 | 13592 | 12654 | 9620 | 12499 | 17771 | 9635 | 11399 | 13240 | 11661 | 11698 | 12740 | Rgmb |
| Rgn | 18 | 9 | 14 | 1 | 2 | 4 | 9 | 11 | 19 | 7 | 5 | 11 | 11 | 23 | 21 | 7 | 9 | 4 | 11 | 7 | 8 | 5 | 15 | 16 | 11 | 18 | 15 | 14 | Rgn |
| Rgp1 | 6377 | 7451 | 8281 | 7394 | 6533 | 6525 | 6490 | 5003 | 5863 | 6636 | 6673 | 7479 | 6939 | 7078 | 6113 | 6988 | 7975 | 7201 | 6830 | 5329 | 6210 | 6335 | 5916 | 7044 | 6771 | 7056 | 6998 | 7334 | Rgp1 |
| Rgr | 0 | 0 | 0 | 0 | 0 | 0 | 0 | 2 | 2 | 7 | 9 | 0 | 0 | 1 | 0 | 0 | 0 | 0 | 1 | 0 | 0 | 2 | 1 | 4 | 6 | 0 | 0 | 1 | Rgr |
| Rgs1 | 0 | 0 | 0 | 3 | 2 | 0 | 0 | 2 | 2 | 0 | 0 | 2 | 5 | 0 | 0 | 2 | 0 | 0 | 1 | 1 | 0 | 2 | 4 | 5 | 3 | 2 | 1 | 4 | Rgs1 |
| Rgs10 | 654 | 693 | 627 | 683 | 835 | 534 | 824 | 475 | 434 | 415 | 491 | 257 | 294 | 271 | 630 | 621 | 679 | 691 | 666 | 725 | 802 | 643 | 504 | 356 | 370 | 286 | 315 | 295 | Rgs10 |
| Rgs11 | 422 | 260 | 213 | 347 | 444 | 348 | 163 | 335 | 400 | 507 | 476 | 88 | 193 | 112 | 314 | 404 | 260 | 379 | 284 | 198 | 216 | 297 | 422 | 454 | 169 | 321 | 77 | 81 | Rgs11 |
| Rgs12 | 5041 | 5522 | 6492 | 5903 | 6091 | 6130 | 5541 | 6964 | 8066 | 8616 | 7837 | 10948 | 9482 | 9773 | 4917 | 4961 | 6327 | 5702 | 6563 | 5010 | 5601 | 8442 | 7572 | 9476 | 9978 | 9412 | 10669 | 10116 | Rgs12 |
| Rgs13 | 52 | 46 | 89 | 60 | 51 | 85 | 111 | 198 | 56 | 72 | 65 | 138 | 104 | 212 | 99 | 38 | 67 | 69 | 82 | 118 | 100 | 142 | 121 | 138 | 124 | 188 | 246 | 263 | Rgs13 |
| Rgs14 | 25 | 22 | 16 | 8 | 7 | 38 | 11 | 16 | 30 | 38 | 50 | 10 | 17 | 14 | 11 | 19 | 5 | 13 | 9 | 6 | 17 | 25 | 26 | 13 | 19 | 24 | 9 | 18 | Rgs14 |
| Rgs16 | 2260 | 3316 | 3838 | 3916 | 3060 | 2968 | 3469 | 1757 | 1591 | 1824 | 1984 | 2760 | 1807 | 3057 | 2552 | 2301 | 3816 | 3725 | 3381 | 3169 | 3525 | 1873 | 1952 | 2589 | 2210 | 3178 | 3011 | 3398 | Rgs16 |
| Rgs17 | 1896 | 1469 | 1388 | 2014 | 2741 | 2624 | 2708 | 2105 | 1734 | 1545 | 1583 | 1351 | 1154 | 1401 | 2186 | 1596 | 1866 | 2218 | 2175 | 2638 | 2687 | 2232 | 1614 | 1244 | 1868 | 704 | 1133 | 1117 | Rgs17 |
| Rgs18 | 59 | 18 | 31 | 17 | 21 | 29 | 18 | 71 | 49 | 38 | 20 | 59 | 62 | 47 | 30 | 40 | 12 | 20 | 12 | 28 | 14 | 40 | 76 | 40 | 33 | 72 | 42 | 32 | Rgs18 |
| Rgs19 | 1137 | 1006 | 1177 | 1017 | 1065 | 1073 | 1073 | 1342 | 1599 | 1677 | 1618 | 1460 | 1588 | 1398 | 931 | 1037 | 1125 | 973 | 1189 | 858 | 1051 | 1385 | 1516 | 1522 | 1500 | 1774 | 1514 | 1612 | Rgs19 |
| Rgs2 | 4279 | 4363 | 4747 | 4911 | 4787 | 4478 | 5242 | 6349 | 4895 | 5309 | 4583 | 4832 | 4394 | 4888 | 5246 | 4129 | 4631 | 5037 | 5040 | 6290 | 5042 | 6156 | 5387 | 4966 | 5360 | 4821 | 5294 | 4859 | Rgs2 |
| Rgs20 | 491 | 456 | 530 | 717 | 945 | 896 | 1002 | 700 | 756 | 889 | 674 | 875 | 879 | 979 | 622 | 364 | 760 | 714 | 1018 | 844 | 1027 | 647 | 634 | 930 | 1027 | 746 | 723 | 758 | Rgs20 |
| Rgs21 | 0 | 1 | 0 | 0 | 0 | 0 | 0 | 0 | 0 | 0 | 5 | 1 | 1 | 2 | 1 | 0 | 0 | 0 | 0 | 0 | 0 | 0 | 0 | 0 | 0 | 0 | 0 | 0 | Rgs21 |
| Rgs22 | 59 | 70 | 53 | 37 | 61 | 106 | 36 | 169 | 119 | 106 | 92 | 135 | 145 | 69 | 70 | 94 | 49 | 66 | 51 | 66 | 70 | 94 | 163 | 73 | 93 | 30 | 81 | 108 | Rgs22 |
| Rgs3 | 1700 | 1951 | 2183 | 2173 | 2474 | 2284 | 1837 | 3019 | 3603 | 3938 | 3548 | 3925 | 3784 | 3531 | 1829 | 1892 | 2256 | 2140 | 2647 | 1973 | 2087 | 3910 | 3437 | 3670 | 3941 | 3786 | 3725 | 3424 | Rgs3 |
| Rgs4 | 1624 | 1703 | 1112 | 1621 | 2515 | 1746 | 1939 | 2280 | 1997 | 1832 | 2183 | 1552 | 1623 | 1576 | 2192 | 1860 | 1544 | 1915 | 1603 | 2215 | 1893 | 2679 | 2397 | 1369 | 2060 | 1479 | 1608 | 1615 | Rgs4 |
| Rgs5 | 2208 | 1887 | 1818 | 3114 | 4597 | 3943 | 4558 | 4539 | 4109 | 4295 | 4320 | 3182 | 2933 | 3590 | 2815 | 2097 | 2370 | 3904 | 3948 | 4573 | 4792 | 5583 | 4730 | 4376 | 5273 | 3116 | 2701 | 2846 | Rgs5 |
| Rgs6 | 398 | 276 | 244 | 617 | 957 | 964 | 645 | 817 | 737 | 930 | 629 | 534 | 513 | 527 | 360 | 312 | 364 | 910 | 902 | 769 | 973 | 966 | 779 | 818 | 1089 | 264 | 481 | 441 | Rgs6 |
| Rgs7 | 390 | 371 | 305 | 499 | 799 | 607 | 600 | 1014 | 936 | 794 | 728 | 501 | 534 | 567 | 497 | 409 | 381 | 506 | 591 | 630 | 593 | 1160 | 1024 | 575 | 813 | 428 | 565 | 449 | Rgs7 |
| Rgs7bp | 1240 | 991 | 929 | 1174 | 2381 | 1679 | 1769 | 1436 | 1847 | 1479 | 1859 | 718 | 1432 | 769 | 1473 | 1431 | 1151 | 1951 | 1446 | 1358 | 1705 | 2537 | 1517 | 821 | 1694 | 592 | 551 | 707 | Rgs7bp |
| Rgs8 | 3840 | 4299 | 4675 | 5022 | 6290 | 5710 | 5761 | 4515 | 4081 | 5507 | 4377 | 5841 | 3880 | 5675 | 4629 | 3090 | 5548 | 5260 | 6234 | 5853 | 6176 | 6615 | 4496 | 5148 | 6121 | 4269 | 6363 | 6107 | Rgs8 |
| Rgs9 | 948 | 954 | 284 | 781 | 1046 | 1206 | 797 | 898 | 901 | 780 | 887 | 558 | 695 | 556 | 980 | 1042 | 488 | 798 | 875 | 1443 | 1136 | 727 | 821 | 434 | 848 | 138 | 722 | 479 | Rgs9 |
| Rgs9bp | 1053 | 971 | 1192 | 819 | 695 | 670 | 644 | 98 | 115 | 102 | 132 | 101 | 88 | 87 | 898 | 1118 | 920 | 872 | 611 | 616 | 593 | 104 | 126 | 110 | 100 | 105 | 145 | 120 | Rgs9bp |
| Rgsl1 | 2 | 9 | 7 | 13 | 11 | 26 | 15 | 22 | 26 | 30 | 15 | 20 | 10 | 20 | 9 | 7 | 1 | 28 | 14 | 13 | 20 | 30 | 29 | 34 | 11 | 2 | 7 | 18 | Rgsl1 |
| Rhag | 102 | 117 | 177 | 20 | 10 | 16 | 16 | 27 | 31 | 58 | 31 | 86 | 270 | 211 | 104 | 152 | 29 | 15 | 16 | 35 | 13 | 30 | 30 | 25 | 6 | 112 | 142 | 490 | Rhag |
| Rhbdd1 | 1007 | 876 | 886 | 851 | 641 | 646 | 778 | 835 | 737 | 705 | 695 | 795 | 837 | 973 | 905 | 814 | 782 | 687 | 729 | 704 | 733 | 740 | 770 | 783 | 662 | 911 | 904 | 967 | Rhbdd1 |
| Rhbdd2 | 3731 | 3843 | 3470 | 3858 | 4097 | 3892 | 3976 | 3124 | 3439 | 3644 | 3348 | 3947 | 3506 | 3678 | 3620 | 3608 | 3990 | 3860 | 4027 | 3441 | 4028 | 3883 | 3322 | 3555 | 3875 | 3485 | 3636 | 3618 | Rhbdd2 |
| Rhbdd3 | 702 | 789 | 895 | 828 | 853 | 824 | 724 | 967 | 1280 | 1279 | 1450 | 1392 | 1352 | 1228 | 671 | 780 | 904 | 855 | 974 | 885 | 745 | 1270 | 1329 | 1445 | 1421 | 1564 | 1406 | 1273 | Rhbdd3 |
| Rhbdf1 | 1489 | 1916 | 2006 | 1735 | 1765 | 1554 | 1245 | 1331 | 1986 | 1921 | 2300 | 1374 | 1833 | 1200 | 1408 | 1974 | 1772 | 1871 | 1672 | 1203 | 1251 | 1714 | 2201 | 1481 | 1376 | 1628 | 1439 | 1326 | Rhbdf1 |
| Rhbdf2 | 947 | 1218 | 1109 | 914 | 727 | 577 | 552 | 175 | 260 | 196 | 275 | 161 | 336 | 153 | 845 | 1449 | 1000 | 845 | 632 | 597 | 531 | 187 | 269 | 221 | 187 | 278 | 160 | 177 | Rhbdf2 |
| Rhbdl1 | 272 | 292 | 401 | 414 | 280 | 207 | 206 | 90 | 73 | 144 | 120 | 136 | 138 | 102 | 204 | 287 | 358 | 357 | 368 | 210 | 215 | 104 | 82 | 142 | 134 | 124 | 100 | 94 | Rhbdl1 |
| Rhbdl2 | 21 | 16 | 25 | 16 | 31 | 24 | 29 | 62 | 46 | 64 | 65 | 53 | 27 | 38 | 12 | 13 | 20 | 33 | 23 | 28 | 29 | 30 | 60 | 48 | 23 | 57 | 28 | 42 | Rhbdl2 |
| Rhbdl3 | 2136 | 2553 | 3013 | 3984 | 4203 | 4186 | 4085 | 3058 | 2970 | 3557 | 2862 | 4059 | 3694 | 3932 | 2422 | 1887 | 3978 | 4212 | 4877 | 3694 | 4147 | 3252 | 2744 | 4692 | 4505 | 3267 | 3672 | 3833 | Rhbdl3 |
| Rhbg | 0 | 0 | 2 | 1 | 0 | 0 | 0 | 1 | 3 | 0 | 0 | 0 | 1 | 2 | 4 | 0 | 0 | 0 | 2 | 0 | 1 | 3 | 3 | 1 | 0 | 1 | 0 | 0 | Rhbg |
| Rhcg | 112 | 130 | 153 | 216 | 138 | 159 | 162 | 121 | 153 | 170 | 134 | 269 | 278 | 250 | 117 | 111 | 160 | 153 | 157 | 190 | 160 | 109 | 177 | 207 | 185 | 305 | 212 | 273 | Rhcg |
| Rhd | 111 | 188 | 132 | 47 | 43 | 49 | 38 | 53 | 80 | 45 | 37 | 139 | 288 | 184 | 117 | 192 | 76 | 35 | 19 | 58 | 32 | 38 | 85 | 38 | 33 | 114 | 160 | 433 | Rhd |
| Rheb | 6775 | 5085 | 5901 | 6179 | 5945 | 5603 | 6590 | 6272 | 5086 | 5020 | 5141 | 5285 | 5312 | 6130 | 6977 | 5680 | 5930 | 6046 | 5782 | 6578 | 6321 | 5195 | 5387 | 5366 | 5387 | 5843 | 5474 | 6169 | Rheb |
| Rhebl1 | 1285 | 1429 | 1519 | 1864 | 1636 | 1315 | 1673 | 1029 | 940 | 1038 | 931 | 942 | 948 | 978 | 1344 | 1332 | 1457 | 1586 | 1660 | 1540 | 1687 | 1110 | 906 | 991 | 1189 | 959 | 916 | 941 | Rhebl1 |
| Rhno1 | 5076 | 5404 | 5580 | 4885 | 4877 | 4853 | 5053 | 4970 | 4625 | 5050 | 4442 | 6222 | 5421 | 5718 | 4650 | 4575 | 5634 | 4601 | 5352 | 5218 | 4823 | 4470 | 4270 | 5614 | 5144 | 6458 | 6098 | 6083 | Rhno1 |

| | | | | | | | | | | | | | | | | | | | | | | | | | | | | | |
|---|---|---|---|---|---|---|---|---|---|---|---|---|---|---|---|---|---|---|---|---|---|---|---|---|---|---|---|---|---|
| Rho | 75 | 120 | 102 | 121 | 160 | 102 | 102 | 88 | 109 | 106 | 109 | 83 | 87 | 65 | 87 | 119 | 97 | 143 | 109 | 124 | 134 | 137 | 83 | 92 | 65 | 72 | 80 | 59 | Rho |
| Rhoa | 21940 | 18006 | 20972 | 17942 | 15995 | 16323 | 18759 | 20235 | 18107 | 16985 | 18484 | 18680 | 18621 | 21212 | 21742 | 19086 | 18934 | 18357 | 17206 | 16306 | 18932 | 19231 | 19434 | 18970 | 17638 | 22991 | 19703 | 23188 | Rhoa |
| Rhob | 4430 | 5334 | 6444 | 6983 | 6356 | 5689 | 5328 | 5769 | 6171 | 7467 | 7704 | 6438 | 6692 | 6552 | 4592 | 5297 | 6835 | 6985 | 6336 | 4682 | 5449 | 7379 | 7001 | 7442 | 8076 | 7404 | 6214 | 6793 | Rhob |
| Rhobtb1 | 587 | 574 | 877 | 605 | 753 | 713 | 607 | 1612 | 2017 | 2221 | 2510 | 2159 | 2236 | 2181 | 537 | 674 | 780 | 656 | 707 | 511 | 575 | 1943 | 2215 | 2102 | 1930 | 2333 | 2150 | 2284 | Rhobtb1 |
| Rhobtb2 | 4067 | 4947 | 5927 | 6534 | 6118 | 5329 | 5425 | 4064 | 3646 | 4829 | 4090 | 5099 | 3969 | 5187 | 3986 | 4418 | 6437 | 6415 | 6493 | 4669 | 5613 | 5396 | 3625 | 5194 | 5562 | 4911 | 4683 | 5114 | Rhobtb2 |
| Rhobtb3 | 10071 | 7005 | 8813 | 6590 | 6529 | 7424 | 7000 | 11302 | 9896 | 8975 | 10091 | 11811 | 10525 | 12182 | 9196 | 8032 | 7803 | 6396 | 6914 | 7582 | 6624 | 9160 | 9218 | 10117 | 9881 | 12081 | 12626 | 12617 | Rhobtb3 |
| Rhoc | 4221 | 4756 | 4453 | 3133 | 2459 | 3076 | 2316 | 2637 | 3652 | 3020 | 3981 | 2555 | 3176 | 2639 | 3988 | 4981 | 3651 | 3288 | 2773 | 2928 | 2277 | 2366 | 4325 | 2952 | 2412 | 3704 | 3013 | 2916 | Rhoc |
| Rhod | 661 | 688 | 563 | 381 | 309 | 380 | 208 | 403 | 438 | 331 | 466 | 263 | 323 | 253 | 535 | 832 | 461 | 407 | 291 | 296 | 212 | 252 | 515 | 355 | 203 | 420 | 311 | 323 | Rhod |
| Rhof | 667 | 758 | 636 | 648 | 697 | 658 | 552 | 209 | 274 | 323 | 265 | 219 | 205 | 204 | 640 | 749 | 649 | 675 | 642 | 663 | 602 | 306 | 263 | 273 | 278 | 183 | 201 | 166 | Rhof |
| Rhog | 935 | 1144 | 993 | 749 | 758 | 740 | 700 | 685 | 869 | 799 | 905 | 827 | 834 | 750 | 886 | 1308 | 933 | 953 | 814 | 694 | 704 | 605 | 1017 | 717 | 662 | 946 | 815 | 839 | Rhog |
| Rhoh | 21 | 19 | 11 | 39 | 22 | 31 | 27 | 26 | 40 | 5 | 53 | 20 | 18 | 19 | 22 | 21 | 21 | 25 | 21 | 22 | 30 | 22 | 56 | 20 | 18 | 22 | 27 | 7 | Rhoh |
| Rhoj | 649 | 701 | 772 | 530 | 609 | 632 | 514 | 1420 | 1760 | 1459 | 2710 | 1357 | 1783 | 1185 | 728 | 822 | 581 | 577 | 580 | 596 | 510 | 1684 | 2210 | 1438 | 1304 | 1515 | 1313 | 1232 | Rhoj |
| Rhoq | 7040 | 4987 | 6629 | 4862 | 4380 | 4590 | 4450 | 4554 | 4541 | 4423 | 6262 | 3897 | 4746 | 4168 | 6012 | 6340 | 5449 | 4789 | 4296 | 3871 | 4291 | 4739 | 5606 | 4340 | 3635 | 4795 | 4307 | 4691 | Rhoq |
| Rhot1 | 7732 | 7052 | 6791 | 6965 | 6551 | 7171 | 7626 | 6981 | 5938 | 5827 | 5659 | 5820 | 5495 | 6033 | 7936 | 7179 | 6431 | 7203 | 6785 | 7618 | 7371 | 6333 | 6018 | 5816 | 6097 | 5976 | 5886 | 6090 | Rhot1 |
| Rhot2 | 1811 | 2023 | 2297 | 2335 | 1677 | 1646 | 1935 | 1115 | 1096 | 1504 | 1282 | 1578 | 1236 | 1700 | 1486 | 1755 | 2321 | 2137 | 1810 | 1494 | 1949 | 1293 | 1125 | 1682 | 1360 | 1657 | 1453 | 1613 | Rhot2 |
| Rhou | 1983 | 1565 | 2461 | 1885 | 2898 | 2339 | 2215 | 3087 | 3004 | 3492 | 3442 | 3086 | 3402 | 3174 | 1998 | 2000 | 2270 | 2073 | 2385 | 1737 | 2193 | 4047 | 3018 | 3362 | 3668 | 3221 | 3101 | 3443 | Rhou |
| Rhov | 142 | 174 | 107 | 178 | 328 | 246 | 225 | 217 | 302 | 323 | 276 | 166 | 125 | 160 | 121 | 151 | 162 | 183 | 286 | 231 | 243 | 249 | 313 | 244 | 237 | 119 | 124 | 131 | Rhov |
| Rhox1 | 0 | 0 | 0 | 0 | 0 | 0 | 0 | 0 | 0 | 0 | 0 | 0 | 0 | 0 | 0 | 0 | 2 | 0 | 0 | 0 | 0 | 0 | 0 | 0 | 0 | 0 | 0 | 0 | Rhox1 |
| Rhox12 | 0 | 0 | 0 | 3 | 0 | 5 | 0 | 0 | 1 | 0 | 0 | 0 | 0 | 0 | 0 | 2 | 0 | 0 | 1 | 0 | 0 | 0 | 2 | 0 | 0 | 0 | 0 | 0 | Rhox12 |
| Rhox2a | 1 | 5 | 1 | 0 | 2 | 0 | 0 | 0 | 1 | 10 | 1 | 3 | 1 | 6 | 1 | 10 | 0 | 0 | 0 | 0 | 0 | 0 | 1 | 0 | 0 | 6 | 2 | 7 | Rhox2a |
| Rhox2b | 2 | 8 | 7 | 3 | 0 | 4 | 1 | 4 | 4 | 0 | 0 | 3 | 3 | 2 | 4 | 4 | 2 | 0 | 0 | 0 | 0 | 2 | 2 | 1 | 0 | 6 | 5 | 5 | Rhox2b |
| Rhox2c | 16 | 35 | 5 | 6 | 4 | 11 | 0 | 20 | 18 | 8 | 4 | 11 | 64 | 16 | 16 | 30 | 15 | 12 | 2 | 0 | 0 | 30 | 10 | 2 | 21 | 15 | 45 | 37 | Rhox2c |
| Rhox2d | 2 | 4 | 2 | 0 | 0 | 0 | 0 | 1 | 2 | 0 | 2 | 5 | 10 | 4 | 3 | 4 | 3 | 1 | 0 | 0 | 0 | 5 | 1 | 0 | 0 | 11 | 5 | 5 | Rhox2d |
| Rhox2e | 0 | 1 | 0 | 0 | 0 | 0 | 0 | 0 | 1 | 1 | 0 | 0 | 0 | 0 | 0 | 0 | 0 | 0 | 0 | 0 | 0 | 0 | 0 | 2 | 0 | 0 | 0 | 0 | Rhox2e |
| Rhox2f | 1 | 2 | 3 | 0 | 0 | 0 | 0 | 0 | 2 | 0 | 1 | 1 | 15 | 5 | 3 | 2 | 0 | 1 | 0 | 0 | 0 | 5 | 1 | 2 | 0 | 8 | 5 | 8 | Rhox2f |
| Rhox2g | 0 | 0 | 0 | 0 | 0 | 0 | 0 | 0 | 0 | 0 | 1 | 1 | 0 | 1 | 0 | 0 | 0 | 0 | 2 | 0 | 0 | 0 | 0 | 0 | 1 | 0 | 0 | 0 | Rhox2g |
| Rhox2h | 0 | 0 | 0 | 0 | 0 | 0 | 0 | 0 | 1 | 0 | 0 | 0 | 0 | 0 | 0 | 0 | 0 | 0 | 0 | 0 | 0 | 0 | 0 | 0 | 0 | 0 | 0 | 4 | Rhox2h |
| Rhox3c | 0 | 0 | 0 | 0 | 0 | 0 | 0 | 0 | 0 | 0 | 0 | 0 | 1 | 0 | 0 | 0 | 0 | 0 | 0 | 0 | 0 | 0 | 0 | 0 | 0 | 0 | 0 | 0 | Rhox3c |
| Rhox3f | 0 | 0 | 0 | 0 | 0 | 0 | 0 | 0 | 0 | 0 | 3 | 0 | 0 | 0 | 0 | 0 | 0 | 0 | 0 | 0 | 0 | 0 | 0 | 0 | 0 | 0 | 0 | 0 | Rhox3f |
| Rhox3g | 0 | 0 | 0 | 0 | 0 | 0 | 0 | 2 | 0 | 0 | 0 | 0 | 0 | 0 | 0 | 0 | 0 | 0 | 0 | 0 | 0 | 0 | 0 | 0 | 0 | 0 | 1 | 0 | Rhox3g |
| Rhox4a | 0 | 0 | 0 | 0 | 0 | 1 | 0 | 0 | 0 | 0 | 0 | 0 | 3 | 0 | 0 | 0 | 0 | 0 | 0 | 0 | 0 | 0 | 1 | 0 | 0 | 0 | 0 | 0 | Rhox4a |
| Rhox4a2 | 1 | 1 | 0 | 0 | 0 | 0 | 0 | 1 | 2 | 0 | 0 | 0 | 1 | 0 | 1 | 4 | 0 | 0 | 0 | 0 | 0 | 0 | 1 | 0 | 0 | 0 | 0 | 1 | Rhox4a2 |
| Rhox4b | 7 | 4 | 0 | 0 | 0 | 4 | 0 | 1 | 2 | 3 | 2 | 3 | 8 | 1 | 12 | 2 | 0 | 2 | 0 | 0 | 0 | 3 | 3 | 1 | 6 | 1 | 2 | 8 | Rhox4b |
| Rhox4c | 0 | 0 | 0 | 0 | 0 | 0 | 0 | 0 | 0 | 0 | 0 | 0 | 0 | 0 | 0 | 0 | 0 | 0 | 0 | 0 | 0 | 0 | 0 | 0 | 0 | 2 | 0 | 0 | Rhox4c |
| Rhox4d | 1 | 1 | 1 | 0 | 0 | 0 | 0 | 1 | 3 | 0 | 0 | 1 | 3 | 0 | 7 | 2 | 0 | 0 | 0 | 0 | 0 | 0 | 0 | 0 | 0 | 0 | 0 | 1 | Rhox4d |
| Rhox4e | 0 | 2 | 0 | 0 | 0 | 0 | 0 | 0 | 2 | 0 | 0 | 0 | 1 | 0 | 1 | 0 | 0 | 0 | 0 | 0 | 0 | 2 | 1 | 0 | 0 | 0 | 0 | 1 | Rhox4e |
| Rhox4f | 0 | 0 | 2 | 0 | 0 | 0 | 0 | 0 | 0 | 4 | 0 | 0 | 0 | 0 | 0 | 1 | 0 | 0 | 0 | 0 | 0 | 2 | 1 | 0 | 0 | 0 | 0 | 0 | Rhox4f |
| Rhox4g | 0 | 0 | 0 | 0 | 0 | 0 | 0 | 0 | 0 | 0 | 0 | 0 | 1 | 1 | 0 | 0 | 0 | 0 | 0 | 0 | 0 | 2 | 0 | 1 | 0 | 0 | 0 | 0 | Rhox4g |
| Rhpn1 | 2323 | 3709 | 3460 | 3427 | 2506 | 2440 | 2341 | 927 | 989 | 1032 | 893 | 1173 | 1097 | 1069 | 2380 | 3048 | 3493 | 3121 | 2813 | 2648 | 2264 | 1115 | 1018 | 1147 | 987 | 1286 | 1159 | 1098 | Rhpn1 |
| Rhpn2 | 1764 | 1811 | 1775 | 1657 | 1487 | 1499 | 1467 | 266 | 283 | 291 | 292 | 163 | 238 | 182 | 1528 | 1850 | 1699 | 1559 | 1475 | 1330 | 1422 | 147 | 344 | 257 | 165 | 219 | 179 | 171 | Rhpn2 |
| Ribc1 | 208 | 230 | 255 | 209 | 153 | 150 | 234 | 131 | 112 | 138 | 119 | 144 | 138 | 205 | 237 | 167 | 255 | 136 | 198 | 192 | 160 | 104 | 107 | 191 | 106 | 234 | 148 | 171 | Ribc1 |
| Ribc2 | 149 | 185 | 125 | 131 | 92 | 106 | 142 | 24 | 29 | 41 | 54 | 59 | 78 | 39 | 182 | 196 | 143 | 157 | 100 | 143 | 155 | 27 | 63 | 44 | 36 | 29 | 41 | 56 | Ribc2 |
| Ric1 | 11911 | 9572 | 7452 | 9540 | 9737 | 10790 | 10296 | 7867 | 7514 | 5997 | 7409 | 5767 | 6659 | 5716 | 10498 | 11062 | 8029 | 10258 | 9139 | 9539 | 10805 | 6458 | 7307 | 5604 | 6255 | 4562 | 5705 | 5720 | Ric1 |
| Ric3 | 4242 | 3844 | 3523 | 4079 | 3832 | 4206 | 4697 | 3761 | 3054 | 3407 | 2673 | 3875 | 3266 | 3978 | 4547 | 3396 | 3978 | 3810 | 4218 | 4278 | 4757 | 4012 | 2967 | 3275 | 3558 | 2935 | 3866 | 3686 | Ric3 |
| Ric8a | 4270 | 5333 | 3840 | 4665 | 4563 | 4581 | 4200 | 3938 | 4431 | 3870 | 3811 | 4121 | 4061 | 3773 | 4466 | 4625 | 4202 | 4292 | 4606 | 4387 | 4434 | 4278 | 4310 | 4001 | 4016 | 3709 | 3987 | 3719 | Ric8a |
| Ric8b | 4427 | 4586 | 4535 | 4449 | 4218 | 4122 | 4395 | 4673 | 4112 | 4175 | 4170 | 4041 | 3678 | 3954 | 4477 | 4443 | 4242 | 4215 | 4460 | 4541 | 4536 | 4639 | 4135 | 3798 | 4259 | 3722 | 3932 | 3896 | Ric8b |
| Rictor | 9393 | 8098 | 5174 | 7057 | 7606 | 8806 | 8789 | 10119 | 8549 | 7098 | 7755 | 7783 | 8140 | 8286 | 9506 | 8558 | 5460 | 7032 | 6798 | 9134 | 8690 | 7217 | 8404 | 6057 | 7928 | 4755 | 8535 | 7425 | Rictor |
| Rida | 972 | 764 | 1162 | 933 | 701 | 706 | 815 | 846 | 742 | 777 | 803 | 955 | 1001 | 979 | 931 | 953 | 973 | 737 | 740 | 1027 | 761 | 675 | 812 | 1008 | 771 | 1239 | 1054 | 1093 | Rida |
| Rif1 | 14634 | 11258 | 8550 | 10569 | 10356 | 12618 | 12511 | 14035 | 11543 | 9340 | 9508 | 11652 | 11510 | 12347 | 14227 | 12987 | 9177 | 10329 | 10601 | 14475 | 12543 | 8173 | 10795 | 8934 | 9817 | 8225 | 13316 | 11248 | Rif1 |
| Riiad1 | 413 | 455 | 501 | 633 | 533 | 506 | 706 | 304 | 214 | 258 | 234 | 282 | 319 | 339 | 469 | 400 | 560 | 579 | 476 | 482 | 618 | 262 | 194 | 289 | 224 | 286 | 306 | 293 | Riiad1 |
| Rilp | 42 | 49 | 40 | 84 | 82 | 76 | 60 | 76 | 66 | 60 | 66 | 74 | 67 | 56 | 22 | 65 | 64 | 76 | 55 | 43 | 64 | 97 | 80 | 60 | 59 | 50 | 63 | 64 | Rilp |
| Rilpl1 | 2174 | 2678 | 2809 | 2487 | 2112 | 1950 | 1907 | 890 | 1067 | 1017 | 1324 | 1060 | 1058 | 1080 | 2099 | 2519 | 2719 | 2176 | 2349 | 1985 | 1943 | 899 | 1100 | 1155 | 1018 | 1392 | 1029 | 1108 | Rilpl1 |
| Rilpl2 | 1696 | 1939 | 1647 | 1736 | 1311 | 1278 | 1427 | 673 | 790 | 608 | 693 | 627 | 621 | 587 | 1549 | 1766 | 1662 | 1611 | 1501 | 1485 | 1360 | 570 | 818 | 662 | 593 | 810 | 642 | 718 | Rilpl2 |
| Rimbp2 | 426 | 720 | 775 | 1046 | 1681 | 1159 | 1169 | 1313 | 1294 | 1605 | 1457 | 1109 | 1327 | 1069 | 729 | 678 | 1145 | 1130 | 1307 | 1087 | 1254 | 2329 | 1129 | 1222 | 1689 | 547 | 1032 | 1081 | Rimbp2 |
| Rimbp3 | 540 | 602 | 384 | 595 | 723 | 722 | 769 | 216 | 279 | 260 | 238 | 351 | 247 | 284 | 496 | 454 | 555 | 621 | 746 | 627 | 655 | 281 | 194 | 288 | 335 | 162 | 274 | 237 | Rimbp3 |
| Rimkla | 1072 | 1346 | 1000 | 1898 | 2700 | 2016 | 2259 | 1918 | 1853 | 1951 | 1564 | 1503 | 1277 | 1323 | 1416 | 1217 | 1365 | 2144 | 2201 | 2109 | 2355 | 2346 | 1485 | 1646 | 1988 | 1001 | 1170 | 1235 | Rimkla |
| Rimklb | 15077 | 11968 | 12375 | 17441 | 20140 | 19995 | 21397 | 19650 | 16311 | 18818 | 17711 | 19437 | 17834 | 21171 | 15870 | 12100 | 14847 | 17777 | 18781 | 19188 | 21373 | 19615 | 15151 | 17947 | 21285 | 14474 | 19133 | 18429 | Rimklb |
| Rims1 | 1 | 0 | 0 | 0 | 0 | 0 | 0 | 1 | 0 | 0 | 0 | 0 | 0 | 0 | 0 | 0 | 0 | 0 | 0 | 0 | 0 | 0 | 1 | 0 | 0 | 0 | 0 | 0 | Rims1 |
| Rims2 | 2070 | 1970 | 2323 | 2714 | 3844 | 3506 | 3854 | 3994 | 3381 | 4261 | 3700 | 4341 | 3797 | 4522 | 2583 | 1929 | 2659 | 2575 | 3393 | 3078 | 3799 | 5680 | 3322 | 3791 | 5122 | 3038 | 4535 | 4146 | Rims2 |
| Rims3 | 3988 | 5236 | 4790 | 5840 | 7565 | 5564 | 5670 | 2992 | 3255 | 3241 | 3015 | 2409 | 2581 | 2290 | 4706 | 4780 | 5654 | 5998 | 6229 | 5384 | 5704 | 4525 | 2634 | 2259 | 3737 | 1638 | 2327 | 2360 | Rims3 |
| Rims4 | 1043 | 1344 | 1849 | 2558 | 3444 | 2338 | 2473 | 1126 | 1176 | 1591 | 1289 | 1181 | 1058 | 1066 | 1213 | 1153 | 2235 | 2539 | 3091 | 2081 | 2787 | 2105 | 1116 | 1320 | 1715 | 1013 | 937 | 1175 | Rims4 |
| Rin1 | 406 | 587 | 594 | 622 | 472 | 475 | 428 | 205 | 266 | 304 | 244 | 305 | 303 | 281 | 393 | 558 | 504 | 558 | 491 | 397 | 458 | 304 | 239 | 370 | 293 | 272 | 309 | 268 | Rin1 |
| Rin2 | 1788 | 1613 | 1883 | 1544 | 1676 | 1655 | 1389 | 2659 | 2749 | 2767 | 3025 | 2227 | 2707 | 2129 | 1650 | 1930 | 1555 | 1657 | 1419 | 1294 | 1311 | 2799 | 2999 | 2795 | 2152 | 2374 | 2138 | 2142 | Rin2 |
| Rin3 | 176 | 280 | 312 | 230 | 283 | 251 | 133 | 509 | 743 | 726 | 824 | 638 | 792 | 488 | 208 | 347 | 290 | 232 | 231 | 134 | 145 | 792 | 906 | 643 | 582 | 815 | 710 | 534 | Rin3 |
| Ring1 | 1217 | 1607 | 2127 | 1815 | 1499 | 1433 | 1353 | 1953 | 2295 | 3062 | 2666 | 3869 | 3252 | 3905 | 1236 | 1154 | 2169 | 1600 | 1693 | 1398 | 1399 | 2425 | 2333 | 3304 | 2994 | 4094 | 4004 | 4073 | Ring1 |
| Rinl | 102 | 185 | 148 | 150 | 115 | 103 | 85 | 85 | 125 | 130 | 128 | 134 | 125 | 122 | 88 | 150 | 172 | 155 | 86 | 93 | 92 | 80 | 137 | 95 | 91 | 105 | 128 | 112 | Rinl |
| Rint1 | 2568 | 2665 | 2439 | 2620 | 2058 | 2338 | 2259 | 2347 | 2352 | 2172 | 2494 | 2266 | 2495 | 2281 | 2534 | 2734 | 2472 | 2564 | 2230 | 2252 | 2347 | 2186 | 2309 | 2306 | 2167 | 2188 | 2271 | 2357 | Rint1 |
| Riok1 | 4128 | 3342 | 3367 | 3285 | 2990 | 3200 | 3562 | 4075 | 3492 | 3383 | 3510 | 3570 | 3532 | 3824 | 4296 | 3545 | 3450 | 3219 | 2989 | 3700 | 3363 | 3345 | 3553 | 3363 | 3263 | 3422 | 3627 | 3719 | Riok1 |
| Riok2 | 6828 | 5671 | 5877 | 5735 | 4493 | 5546 | 6284 | 4285 | 4015 | 3544 | 3527 | 4216 | 4280 | 4552 | 6728 | 5997 | 5373 | 5312 | 5045 | 5964 | 5677 | 3599 | 3904 | 3812 | 3583 | 4478 | 4573 | 4485 | Riok2 |
| Riok3 | 11472 | 10204 | 9862 | 10351 | 9854 | 10465 | 11265 | 11164 | 9558 | 9338 | 9011 | 9634 | 9096 | 9809 | 12217 | 10474 | 10010 | 10186 | 9743 | 10724 | 10825 | 10808 | 9491 | 8958 | 9826 | 8709 | 9233 | 9940 | Riok3 |
| Riox1 | 1147 | 1394 | 1293 | 1297 | 1109 | 1080 | 969 | 902 | 974 | 831 | 879 | 933 | 951 | 1004 | 1027 | 1189 | 1314 | 1271 | 1298 | 1092 | 989 | 822 | 898 | 880 | 906 | 1186 | 917 | 902 | Riox1 |
| Riox2 | 1580 | 1548 | 1708 | 1439 | 1226 | 1329 | 1400 | 1335 | 1435 | 1286 | 1413 | 1506 | 1571 | 1557 | 1721 | 1627 | 1396 | 1428 | 1213 | 1457 | 1303 | 1041 | 1506 | 1479 | 1115 | 1648 | 1576 | 1600 | Riox2 |
| Ripk1 | 719 | 769 | 828 | 623 | 685 | 728 | 685 | 1526 | 1685 | 1427 | 1562 | 1355 | 1456 | 1365 | 767 | 706 | 716 | 689 | 756 | 734 | 676 | 1539 | 1893 | 1338 | 1242 | 1262 | 1327 | 1379 | Ripk1 |
| Ripk2 | 1792 | 1848 | 1637 | 1660 | 1603 | 1722 | 1700 | 661 | 631 | 565 | 547 | 576 | 646 | 634 | 1905 | 1814 | 1600 | 1721 | 1521 | 1542 | 1619 | 635 | 638 | 555 | 445 | 595 | 682 | 564 | Ripk2 |
| Ripk3 | 1501 | 986 | 905 | 640 | 362 | 513 | 285 | 335 | 558 | 334 | 623 | 209 | 409 | 184 | 906 | 1431 | 592 | 627 | 337 | 341 | 277 | 309 | 574 | 366 | 210 | 428 | 250 | 222 | Ripk3 |
| Ripk4 | 494 | 347 | 334 | 296 | 175 | 188 | 188 | 327 | 405 | 308 | 250 | 92 | 153 | 121 | 262 | 340 | 203 | 227 | 235 | 159 | 138 | 75 | 476 | 339 | 38 | 246 | 101 | 102 | Ripk4 |
| Ripor1 | 2806 | 3721 | 4159 | 4298 | 4507 | 3711 | 3080 | 2118 | 2567 | 3249 | 3049 | 3199 | 2743 | 2439 | 2707 | 3625 | 4333 | 4573 | 4409 | 2913 | 3663 | 3385 | 2691 | 3179 | 3276 | 3033 | 2553 | 2276 | Ripor1 |
| Ripor2 | 1884 | 2043 | 1643 | 2686 | 3678 | 3240 | 3251 | 2469 | 2698 | 3020 | 2644 | 2628 | 2349 | 2649 | 2119 | 1793 | 2506 | 3185 | 3381 | 2981 | 3555 | 3669 | 2491 | 2669 | 3356 | 1892 | 2528 | 2244 | Ripor2 |
| Ripor3 | 120 | 98 | 183 | 69 | 47 | 80 | 20 | 238 | 340 | 370 | 291 | 215 | 308 | 166 | 66 | 162 | 92 | 66 | 55 | 33 | 34 | 147 | 345 | 369 | 156 | 288 | 264 | 173 | Ripor3 |

| | | | | | | | | | | | | | | | | | | | | | | | | | | | | | |
|---|---|---|---|---|---|---|---|---|---|---|---|---|---|---|---|---|---|---|---|---|---|---|---|---|---|---|---|---|---|
| Ripply1 | 15 | 10 | 12 | 6 | 6 | 6 | 4 | 17 | 12 | 8 | 58 | 38 | 37 | 26 | 7 | 7 | 7 | 2 | 4 | 12 | 4 | 17 | 49 | 19 | 17 | 8 | 57 | 28 | Ripply1 |
| Ripply2 | 7 | 10 | 14 | 17 | 21 | 11 | 16 | 22 | 33 | 31 | 22 | 31 | 8 | 24 | 11 | 8 | 12 | 10 | 16 | 19 | 13 | 38 | 31 | 23 | 35 | 8 | 24 | 15 | Ripply2 |
| Ripply3 | 220 | 141 | 247 | 142 | 85 | 121 | 96 | 98 | 102 | 92 | 66 | 98 | 154 | 135 | 150 | 153 | 183 | 147 | 147 | 139 | 99 | 104 | 118 | 92 | 89 | 182 | 157 | 135 | Ripply3 |
| Rit1 | 2679 | 3207 | 2706 | 3142 | 2876 | 2839 | 3066 | 1984 | 1912 | 1613 | 1785 | 1682 | 1841 | 1758 | 2835 | 2671 | 2954 | 3106 | 3292 | 3076 | 3260 | 1975 | 2055 | 1754 | 1737 | 1868 | 1991 | 1844 | Rit1 |
| Rit2 | 209 | 246 | 283 | 445 | 1046 | 702 | 915 | 670 | 593 | 644 | 656 | 352 | 330 | 397 | 347 | 230 | 352 | 684 | 764 | 785 | 935 | 1024 | 715 | 545 | 790 | 263 | 363 | 310 | Rit2 |
| Rita1 | 1551 | 2338 | 1588 | 2159 | 1679 | 1589 | 1627 | 739 | 749 | 773 | 666 | 686 | 689 | 551 | 1559 | 2003 | 1773 | 1995 | 1817 | 1817 | 1781 | 703 | 750 | 747 | 738 | 695 | 652 | 552 | Rita1 |
| Rlbp1 | 257 | 291 | 110 | 454 | 759 | 720 | 948 | 1120 | 1406 | 1132 | 954 | 849 | 573 | 996 | 294 | 103 | 298 | 603 | 790 | 858 | 861 | 820 | 960 | 1219 | 1387 | 843 | 593 | 553 | Rlbp1 |
| Rlf | 8975 | 7753 | 6018 | 7194 | 6864 | 7751 | 7568 | 9123 | 7516 | 7249 | 6649 | 6692 | 6688 | 6871 | 9108 | 7957 | 6585 | 6710 | 6685 | 8424 | 7025 | 6928 | 7746 | 6034 | 6432 | 5120 | 6527 | 6319 | Rlf |
| Rlim | 8712 | 6488 | 7438 | 7497 | 7161 | 7645 | 7393 | 10135 | 8345 | 7719 | 8649 | 8263 | 8245 | 8315 | 8708 | 7802 | 7134 | 7258 | 6923 | 8261 | 7409 | 8604 | 8688 | 7817 | 8137 | 7531 | 8529 | 8476 | Rlim |
| Rln1 | 29 | 30 | 17 | 45 | 37 | 30 | 46 | 29 | 25 | 33 | 24 | 21 | 16 | 14 | 42 | 28 | 15 | 42 | 39 | 30 | 40 | 13 | 30 | 20 | 27 | 11 | 28 | 17 | Rln1 |
| Rln3 | 1 | 1 | 4 | 0 | 0 | 1 | 0 | 0 | 0 | 0 | 1 | 0 | 0 | 0 | 1 | 0 | 1 | 3 | 2 | 0 | 0 | 3 | 0 | 0 | 1 | 0 | 0 | 0 | Rln3 |
| Rmc1 | 3787 | 4219 | 2996 | 4114 | 3887 | 3957 | 4140 | 1915 | 2168 | 1744 | 1873 | 1703 | 2058 | 1641 | 3831 | 3562 | 3343 | 4044 | 4033 | 4373 | 4306 | 2095 | 2063 | 1745 | 1771 | 1498 | 1636 | 1712 | Rmc1 |
| Rmdn1 | 677 | 460 | 459 | 555 | 622 | 675 | 733 | 924 | 812 | 732 | 799 | 795 | 747 | 778 | 593 | 550 | 451 | 593 | 655 | 704 | 646 | 744 | 832 | 717 | 817 | 586 | 724 | 726 | Rmdn1 |
| Rmdn2 | 67 | 50 | 49 | 62 | 60 | 81 | 53 | 418 | 419 | 295 | 423 | 336 | 414 | 368 | 57 | 51 | 44 | 53 | 38 | 60 | 54 | 406 | 493 | 357 | 267 | 398 | 386 | 355 | Rmdn2 |
| Rmdn3 | 2880 | 3131 | 3477 | 3258 | 2754 | 2741 | 2951 | 2219 | 2099 | 2296 | 2263 | 2381 | 2402 | 2415 | 2842 | 2923 | 3170 | 3045 | 3200 | 2901 | 2949 | 2423 | 2080 | 2308 | 2387 | 2557 | 2269 | 2499 | Rmdn3 |
| Rmi1 | 1617 | 1316 | 1589 | 1423 | 1430 | 1451 | 1685 | 2172 | 1993 | 1898 | 2091 | 2305 | 2154 | 2173 | 1458 | 1435 | 1541 | 1472 | 1520 | 1565 | 1482 | 2156 | 1986 | 2185 | 2045 | 2251 | 2553 | 2369 | Rmi1 |
| Rmi2 | 423 | 402 | 609 | 513 | 295 | 374 | 455 | 457 | 439 | 489 | 559 | 792 | 589 | 705 | 365 | 435 | 618 | 401 | 460 | 319 | 430 | 349 | 396 | 579 | 507 | 947 | 744 | 767 | Rmi2 |
| Rmnd1 | 2261 | 2162 | 2204 | 2314 | 2201 | 2171 | 2456 | 2694 | 2558 | 2696 | 2604 | 2425 | 2325 | 2521 | 2379 | 2110 | 2278 | 2245 | 2276 | 2366 | 2315 | 2635 | 2782 | 2623 | 2564 | 2258 | 2523 | 2370 | Rmnd1 |
| Rmnd5a | 13883 | 11354 | 12289 | 12675 | 11307 | 11910 | 12671 | 8447 | 7462 | 7649 | 8162 | 8040 | 8350 | 8287 | 12982 | 13441 | 11499 | 12904 | 11228 | 10874 | 12293 | 8091 | 7939 | 7888 | 7757 | 7294 | 8534 | 8445 | Rmnd5a |
| Rmnd5b | 2495 | 3064 | 2341 | 2694 | 2412 | 2444 | 2272 | 1822 | 1912 | 1903 | 1873 | 2016 | 1907 | 1656 | 2430 | 2608 | 2415 | 2650 | 2566 | 2379 | 2394 | 1998 | 2010 | 1815 | 2131 | 1887 | 1667 | 1671 | Rmnd5b |
| Rnase1 | 5 | 0 | 9 | 0 | 0 | 4 | 0 | 2 | 2 | 0 | 7 | 0 | 0 | 2 | 0 | 0 | 1 | 2 | 0 | 4 | 0 | 2 | 3 | 2 | 0 | 0 | 2 | 2 | Rnase1 |
| Rnase10 | 1 | 0 | 0 | 0 | 0 | 0 | 0 | 1 | 5 | 0 | 6 | 2 | 0 | 0 | 0 | 0 | 0 | 0 | 0 | 0 | 0 | 0 | 4 | 0 | 0 | 1 | 1 | 0 | Rnase10 |
| Rnase11 | 0 | 0 | 0 | 0 | 0 | 0 | 0 | 0 | 0 | 0 | 0 | 0 | 0 | 0 | 1 | 0 | 0 | 0 | 1 | 0 | 0 | 0 | 0 | 0 | 0 | 0 | 0 | 0 | Rnase11 |
| Rnase12 | 1 | 0 | 0 | 0 | 5 | 0 | 1 | 1 | 2 | 1 | 3 | 1 | 0 | 2 | 0 | 2 | 0 | 2 | 0 | 0 | 2 | 2 | 3 | 3 | 1 | 0 | 2 | 5 | Rnase12 |
| Rnase4 | 1025 | 841 | 783 | 553 | 579 | 697 | 451 | 596 | 721 | 554 | 839 | 559 | 718 | 504 | 1029 | 1104 | 630 | 704 | 544 | 664 | 398 | 707 | 1107 | 550 | 526 | 570 | 495 | 515 | Rnase4 |
| Rnase6 | 8 | 7 | 4 | 3 | 11 | 9 | 6 | 7 | 12 | 13 | 13 | 5 | 12 | 7 | 7 | 2 | 5 | 0 | 5 | 18 | 4 | 8 | 12 | 13 | 5 | 6 | 7 | 6 | Rnase6 |
| Rnaseh1 | 1195 | 1355 | 1396 | 1517 | 1150 | 1335 | 1205 | 1211 | 1258 | 1438 | 1378 | 1501 | 1441 | 1367 | 1179 | 1214 | 1292 | 1366 | 1252 | 1261 | 1193 | 1135 | 1171 | 1439 | 1469 | 1552 | 1483 | 1445 | Rnaseh1 |
| Rnaseh2a | 3478 | 3831 | 3867 | 3693 | 2790 | 3129 | 3163 | 1629 | 1725 | 1653 | 1541 | 2269 | 1948 | 2188 | 3378 | 3391 | 3517 | 3373 | 3521 | 3570 | 3208 | 1268 | 1630 | 2029 | 1867 | 2795 | 2464 | 2262 | Rnaseh2a |
| Rnaseh2b | 1306 | 1193 | 1171 | 985 | 930 | 1004 | 1398 | 2507 | 2176 | 1802 | 1750 | 2536 | 1949 | 2917 | 1412 | 984 | 1060 | 934 | 1057 | 1323 | 1093 | 1648 | 1849 | 2194 | 2206 | 2948 | 2430 | 2667 | Rnaseh2b |
| Rnaseh2c | 2268 | 2985 | 3022 | 2835 | 1928 | 2205 | 2177 | 1546 | 1453 | 1779 | 1566 | 2156 | 1852 | 2273 | 2234 | 2165 | 2951 | 2245 | 2615 | 2563 | 2212 | 1088 | 1398 | 2078 | 1859 | 2395 | 2170 | 2307 | Rnaseh2c |
| Rnasek | 13 | 41 | 36 | 39 | 29 | 24 | 38 | 11 | 16 | 22 | 19 | 10 | 22 | 18 | 20 | 26 | 28 | 20 | 34 | 32 | 28 | 17 | 21 | 16 | 22 | 15 | 23 | 14 | Rnasek |
| Rnasel | 2228 | 2386 | 2028 | 2843 | 3689 | 3692 | 3253 | 8327 | 6873 | 7839 | 7280 | 8585 | 8568 | 9168 | 3059 | 2206 | 2720 | 2681 | 3257 | 3423 | 3463 | 9299 | 6910 | 7669 | 8874 | 5790 | 8989 | 8278 | Rnasel |
| Rnaset2a | 61 | 68 | 48 | 71 | 43 | 32 | 74 | 31 | 21 | 21 | 19 | 29 | 22 | 30 | 74 | 79 | 29 | 61 | 55 | 72 | 41 | 28 | 28 | 17 | 21 | 51 | 19 | 39 | Rnaset2a |
| Rnaset2b | 121 | 107 | 113 | 90 | 115 | 118 | 106 | 66 | 49 | 60 | 51 | 56 | 59 | 54 | 138 | 129 | 137 | 90 | 95 | 119 | 120 | 69 | 58 | 42 | 52 | 63 | 59 | 68 | Rnaset2b |
| Rnd1 | 1025 | 2031 | 1660 | 1913 | 1473 | 1587 | 1370 | 460 | 408 | 502 | 368 | 555 | 603 | 657 | 1238 | 1338 | 1778 | 1795 | 1510 | 1644 | 1605 | 488 | 469 | 554 | 543 | 510 | 601 | 673 | Rnd1 |
| Rnd2 | 5292 | 7479 | 7717 | 10215 | 9516 | 9514 | 9181 | 6167 | 5824 | 9028 | 6014 | 8413 | 8530 | 7826 | 6910 | 5580 | 10758 | 8779 | 10840 | 8884 | 9288 | 5008 | 4629 | 9422 | 9176 | 8211 | 8351 | 8098 | Rnd2 |
| Rnd3 | 12172 | 12130 | 14833 | 14689 | 14415 | 14426 | 13124 | 24389 | 16401 | 17993 | 15997 | 21062 | 23441 | 21865 | 15389 | 12506 | 16111 | 13511 | 14249 | 15228 | 13113 | 22877 | 16764 | 18658 | 20697 | 18274 | 24499 | 22536 | Rnd3 |
| Rnf10 | 21516 | 24770 | 27457 | 27544 | 24662 | 22635 | 23703 | 16957 | 18118 | 21084 | 18881 | 21645 | 20026 | 20616 | 20816 | 21246 | 28036 | 26599 | 27419 | 20427 | 25399 | 19905 | 17395 | 21785 | 21167 | 23314 | 19360 | 21430 | Rnf10 |
| Rnf103 | 2754 | 2118 | 2332 | 2717 | 2972 | 2826 | 2986 | 3389 | 2804 | 2840 | 2842 | 2742 | 2825 | 2777 | 2702 | 2364 | 2473 | 2834 | 2725 | 2880 | 2921 | 3309 | 2875 | 3105 | 3357 | 2915 | 2840 | 2928 | Rnf103 |
| Rnf11 | 6976 | 5244 | 6740 | 7681 | 8809 | 8256 | 9412 | 8819 | 7227 | 7224 | 7836 | 7494 | 7177 | 8298 | 7359 | 5807 | 7063 | 8313 | 8610 | 7600 | 9705 | 9663 | 7991 | 7767 | 8608 | 7372 | 6897 | 7806 | Rnf11 |
| Rnf111 | 4280 | 3778 | 5392 | 5590 | 6033 | 5236 | 5643 | 5655 | 4974 | 6105 | 5822 | 6783 | 6102 | 6971 | 4178 | 4292 | 5456 | 5503 | 5837 | 4560 | 5629 | 7377 | 5025 | 6638 | 6993 | 6263 | 5977 | 6950 | Rnf111 |
| Rnf112 | 630 | 761 | 602 | 1536 | 2060 | 1587 | 1772 | 1702 | 1751 | 2263 | 1691 | 1577 | 1038 | 1675 | 721 | 586 | 1161 | 1576 | 1923 | 1565 | 2041 | 2485 | 1555 | 2281 | 2454 | 1616 | 1223 | 1205 | Rnf112 |
| Rnf113a1 | 351 | 455 | 449 | 367 | 378 | 395 | 389 | 333 | 339 | 317 | 324 | 371 | 413 | 429 | 386 | 326 | 403 | 353 | 413 | 449 | 386 | 306 | 343 | 368 | 311 | 344 | 381 | 396 | Rnf113a1 |
| Rnf113a2 | 1175 | 1279 | 1338 | 1268 | 1082 | 1133 | 1195 | 866 | 873 | 866 | 880 | 892 | 1010 | 997 | 1216 | 1093 | 1227 | 1059 | 1288 | 1329 | 1135 | 710 | 791 | 1075 | 846 | 1102 | 1010 | 1005 | Rnf113a2 |
| Rnf114 | 6371 | 7855 | 6757 | 6906 | 6157 | 6402 | 5773 | 7113 | 6570 | 5987 | 5783 | 6977 | 6508 | 6526 | 6562 | 6310 | 6954 | 6444 | 6699 | 7039 | 5956 | 6453 | 6435 | 6819 | 6971 | 7571 | 7224 | 7008 | Rnf114 |
| Rnf115 | 6690 | 5605 | 6275 | 6071 | 5618 | 6173 | 7019 | 4631 | 3851 | 4390 | 3784 | 4549 | 4449 | 5132 | 6634 | 5796 | 6282 | 5558 | 6209 | 6325 | 6610 | 4475 | 3916 | 4355 | 4275 | 5077 | 4779 | 5345 | Rnf115 |
| Rnf121 | 3500 | 3438 | 3091 | 2936 | 3041 | 3111 | 2833 | 3366 | 3079 | 2785 | 2862 | 3030 | 2916 | 2715 | 3432 | 3283 | 2945 | 2849 | 3006 | 3234 | 2940 | 3029 | 2974 | 2718 | 3184 | 2913 | 3025 | 2856 | Rnf121 |
| Rnf122 | 2900 | 3280 | 2932 | 2697 | 3100 | 3052 | 2833 | 3639 | 3389 | 3492 | 3152 | 4044 | 3877 | 3778 | 2854 | 2684 | 3014 | 2664 | 3206 | 2952 | 2853 | 3395 | 3447 | 3628 | 3639 | 3937 | 4299 | 4319 | Rnf122 |
| Rnf123 | 3279 | 3978 | 3826 | 4256 | 4529 | 4194 | 3900 | 3591 | 4248 | 4494 | 4133 | 4502 | 4283 | 3901 | 3236 | 3731 | 4140 | 4199 | 4642 | 3534 | 4106 | 4430 | 4001 | 4327 | 4903 | 3483 | 4215 | 4022 | Rnf123 |
| Rnf125 | 50 | 44 | 50 | 36 | 66 | 41 | 31 | 57 | 74 | 65 | 91 | 39 | 94 | 44 | 62 | 76 | 35 | 45 | 31 | 44 | 34 | 57 | 75 | 41 | 41 | 49 | 44 | 44 | Rnf125 |
| Rnf126 | 2087 | 2648 | 2523 | 2721 | 2563 | 2081 | 2035 | 1782 | 1979 | 2296 | 2069 | 2587 | 1918 | 2085 | 1887 | 2246 | 2919 | 2691 | 2696 | 1916 | 2139 | 2286 | 1889 | 2450 | 2005 | 2903 | 2059 | 1944 | Rnf126 |
| Rnf128 | 450 | 213 | 244 | 219 | 206 | 245 | 242 | 217 | 199 | 188 | 211 | 97 | 134 | 98 | 326 | 348 | 176 | 235 | 193 | 189 | 215 | 130 | 241 | 184 | 102 | 173 | 63 | 127 | Rnf128 |
| Rnf13 | 4496 | 3946 | 3661 | 3657 | 3768 | 3903 | 4600 | 3612 | 3374 | 2745 | 2847 | 2874 | 2739 | 3002 | 5077 | 4149 | 3620 | 3959 | 3754 | 4723 | 4275 | 3178 | 3524 | 2794 | 2853 | 2978 | 2772 | 2997 | Rnf13 |
| Rnf130 | 10810 | 11415 | 8464 | 13638 | 17652 | 16121 | 17169 | 12283 | 11762 | 12804 | 11209 | 11116 | 9149 | 11206 | 11644 | 9467 | 11413 | 14696 | 15870 | 15223 | 17314 | 13859 | 10740 | 10712 | 14627 | 7549 | 9834 | 10005 | Rnf130 |
| Rnf133 | 3 | 0 | 0 | 2 | 2 | 0 | 0 | 0 | 2 | 0 | 0 | 0 | 0 | 2 | 1 | 1 | 0 | 0 | 0 | 0 | 0 | 0 | 0 | 0 | 0 | 3 | 0 | 2 | Rnf133 |
| Rnf135 | 118 | 128 | 145 | 103 | 117 | 108 | 68 | 187 | 272 | 229 | 240 | 177 | 279 | 179 | 142 | 151 | 117 | 128 | 84 | 100 | 77 | 236 | 307 | 197 | 190 | 255 | 200 | 182 | Rnf135 |
| Rnf138 | 4858 | 3435 | 5001 | 5627 | 5347 | 5614 | 5995 | 5211 | 4203 | 4698 | 4980 | 5476 | 5270 | 5682 | 4963 | 4400 | 5016 | 5132 | 5156 | 5069 | 5476 | 5922 | 4209 | 5302 | 5433 | 5053 | 5069 | 5882 | Rnf138 |
| Rnf138rt1 | 50 | 83 | 70 | 86 | 77 | 64 | 98 | 26 | 21 | 29 | 23 | 26 | 6 | 12 | 70 | 52 | 90 | 74 | 94 | 98 | 87 | 42 | 28 | 26 | 24 | 18 | 34 | 26 | Rnf138rt1 |
| Rnf139 | 3310 | 2724 | 2281 | 2427 | 2500 | 2882 | 2922 | 3667 | 3267 | 2432 | 2699 | 2888 | 2861 | 2944 | 3468 | 2896 | 2277 | 2311 | 2630 | 3257 | 3025 | 2879 | 3265 | 2576 | 2766 | 3184 | 3176 | 3021 | Rnf139 |
| Rnf14 | 7825 | 7589 | 5940 | 8556 | 10299 | 9211 | 10428 | 11467 | 9986 | 9363 | 9181 | 8449 | 8218 | 9158 | 9187 | 7372 | 7536 | 9061 | 9261 | 9962 | 10558 | 11894 | 10105 | 8599 | 10447 | 7736 | 8444 | 8857 | Rnf14 |
| Rnf141 | 3522 | 2846 | 2621 | 2709 | 3174 | 3153 | 3152 | 3662 | 3284 | 3020 | 2878 | 3143 | 3254 | 3126 | 3295 | 2999 | 2699 | 2835 | 2926 | 3588 | 3038 | 3088 | 3522 | 2917 | 3298 | 3137 | 2952 | 3113 | Rnf141 |
| Rnf144a | 4896 | 3852 | 5035 | 4485 | 5147 | 4920 | 4937 | 6172 | 6836 | 7111 | 7212 | 7863 | 7412 | 8019 | 4433 | 4588 | 4854 | 4589 | 5129 | 4091 | 4848 | 6572 | 6101 | 7428 | 7168 | 7953 | 7154 | 8015 | Rnf144a |
| Rnf144b | 542 | 444 | 465 | 307 | 349 | 374 | 287 | 299 | 438 | 330 | 580 | 282 | 355 | 345 | 435 | 483 | 392 | 397 | 302 | 310 | 298 | 324 | 436 | 299 | 275 | 302 | 338 | 329 | Rnf144b |
| Rnf145 | 11635 | 11170 | 11448 | 11540 | 11849 | 11949 | 11458 | 12471 | 11794 | 12430 | 11755 | 13984 | 14230 | 13786 | 11190 | 10132 | 11632 | 10994 | 11360 | 11336 | 11558 | 12606 | 11319 | 11987 | 12920 | 12957 | 14601 | 14563 | Rnf145 |
| Rnf146 | 5865 | 5224 | 5489 | 6277 | 5663 | 5591 | 6336 | 7060 | 5634 | 5483 | 5437 | 5887 | 5447 | 6165 | 6199 | 5228 | 5844 | 5903 | 5744 | 6310 | 6298 | 7033 | 5801 | 5901 | 6655 | 6083 | 6233 | 6518 | Rnf146 |
| Rnf148 | 2 | 0 | 0 | 1 | 1 | 0 | 0 | 1 | 0 | 0 | 0 | 0 | 0 | 2 | 4 | 0 | 0 | 0 | 0 | 2 | 0 | 0 | 0 | 0 | 0 | 0 | 0 | 0 | Rnf148 |
| Rnf149 | 1919 | 1629 | 1857 | 1578 | 1591 | 1546 | 1705 | 1542 | 1517 | 1377 | 1445 | 1437 | 1515 | 1460 | 1946 | 1672 | 1688 | 1581 | 1509 | 1604 | 1675 | 1352 | 1541 | 1442 | 1227 | 1452 | 1477 | 1434 | Rnf149 |
| Rnf150 | 1997 | 1575 | 1682 | 1760 | 2330 | 1788 | 1726 | 3854 | 3409 | 3440 | 3900 | 3023 | 3845 | 3283 | 1937 | 2167 | 1936 | 1744 | 1790 | 1731 | 1734 | 4264 | 3269 | 2969 | 3507 | 2135 | 3413 | 3122 | Rnf150 |
| Rnf151 | 25 | 22 | 10 | 8 | 10 | 4 | 7 | 13 | 8 | 3 | 7 | 21 | 19 | 9 | 17 | 10 | 3 | 28 | 15 | 6 | 9 | 13 | 9 | 16 | 17 | 16 | 14 | 8 | Rnf151 |
| Rnf152 | 1405 | 1146 | 1681 | 2131 | 4102 | 3230 | 3146 | 4352 | 3844 | 4162 | 4126 | 2960 | 3531 | 3076 | 1754 | 1363 | 1897 | 2409 | 3218 | 2887 | 3339 | 5919 | 3635 | 2665 | 4963 | 1407 | 2718 | 2763 | Rnf152 |
| Rnf157 | 5064 | 6360 | 6139 | 8613 | 10499 | 8530 | 7815 | 5047 | 5978 | 8248 | 7120 | 5759 | 5454 | 5334 | 5469 | 6277 | 7719 | 9444 | 9716 | 6950 | 8928 | 8087 | 5649 | 7086 | 8033 | 5242 | 4723 | 5408 | Rnf157 |
| Rnf165 | 4309 | 5083 | 8325 | 9622 | 9903 | 8230 | 8657 | 7746 | 7416 | 10792 | 8718 | 12991 | 9232 | 12353 | 4694 | 4382 | 9574 | 8906 | 10536 | 6631 | 9412 | 13409 | 6581 | 12600 | 13201 | 10096 | 9912 | 11598 | Rnf165 |
| Rnf166 | 1929 | 2515 | 2068 | 2327 | 1898 | 2042 | 2005 | 1296 | 1651 | 1587 | 1544 | 1575 | 1569 | 1484 | 1878 | 2258 | 2235 | 2242 | 2056 | 2017 | 1929 | 1450 | 1622 | 1593 | 1640 | 1711 | 1492 | 1482 | Rnf166 |
| Rnf167 | 3223 | 3648 | 3181 | 3346 | 3300 | 3168 | 3428 | 3661 | 3845 | 3576 | 3523 | 3819 | 3492 | 3779 | 3208 | 2948 | 3695 | 3421 | 3518 | 3379 | 3473 | 3785 | 3861 | 3864 | 3940 | 3914 | 3690 | 3893 | Rnf167 |
| Rnf168 | 4434 | 3874 | 4508 | 4162 | 3989 | 4225 | 4476 | 5252 | 4705 | 4761 | 4103 | 5360 | 4838 | 5657 | 4615 | 3885 | 4414 | 3704 | 4521 | 4173 | 4325 | 4259 | 4557 | 5090 | 4806 | 5924 | 5442 | 5772 | Rnf168 |
| Rnf169 | 3044 | 2773 | 3360 | 3183 | 3338 | 3370 | 3588 | 3559 | 3445 | 3990 | 3913 | 4456 | 4234 | 4515 | 2955 | 2955 | 3527 | 3193 | 3364 | 2914 | 3411 | 3761 | 3353 | 4039 | 4103 | 3785 | 4291 | 4643 | Rnf169 |
| Rnf17 | 256 | 132 | 113 | 127 | 116 | 181 | 92 | 225 | 238 | 162 | 235 | 209 | 212 | 134 | 160 | 238 | 115 | 80 | 48 | 116 | 71 | 349 | 232 | 93 | 160 | 157 | 174 | 154 | Rnf17 |
| Rnf170 | 1887 | 1189 | 1439 | 1495 | 1813 | 1765 | 1884 | 2215 | 1898 | 2002 | 2097 | 1934 | 2089 | 1975 | 1840 | 1531 | 1471 | 1542 | 1606 | 1920 | 1647 | 2129 | 2016 | 1843 | 1889 | 1937 | 1995 | 1829 | Rnf170 |
| Rnf180 | 430 | 371 | 414 | 466 | 748 | 636 | 769 | 1286 | 1283 | 1283 | 1000 | 1194 | 950 | 1206 | 530 | 442 | 578 | 498 | 630 | 683 | 743 | 1404 | 1037 | 1166 | 1343 | 1100 | 1011 | 1015 | Rnf180 |

| | | | | | | | | | | | | | | | | | | | | | | | | | | | | | |
|---|---|---|---|---|---|---|---|---|---|---|---|---|---|---|---|---|---|---|---|---|---|---|---|---|---|---|---|---|---|
| Rnf181 | 263 | 622 | 171 | 195 | 153 | 239 | 394 | 158 | 107 | 104 | 103 | 91 | 237 | 81 | 415 | 206 | 184 | 227 | 196 | 350 | 170 | 755 | 90 | 107 | 128 | 109 | 108 | 112 | Rnf181 |
| Rnf182 | 1926 | 1851 | 2111 | 2788 | 3487 | 2714 | 3103 | 1958 | 1920 | 2407 | 1937 | 1601 | 1538 | 1702 | 2116 | 1819 | 2699 | 2853 | 3188 | 2867 | 3275 | 2162 | 1978 | 2278 | 2238 | 2151 | 1445 | 1563 | Rnf182 |
| Rnf183 | 7 | 10 | 20 | 28 | 7 | 19 | 28 | 1 | 2 | 8 | 13 | 1 | 5 | 4 | 16 | 9 | 30 | 26 | 33 | 17 | 23 | 8 | 6 | 1 | 11 | 7 | 0 | 8 | Rnf183 |
| Rnf185 | 6133 | 6449 | 5750 | 5410 | 4912 | 4795 | 5209 | 3599 | 3215 | 3367 | 3142 | 3704 | 3339 | 3505 | 6298 | 5668 | 5730 | 5269 | 5401 | 5541 | 5232 | 3559 | 3198 | 3380 | 3486 | 3812 | 3785 | 3786 | Rnf185 |
| Rnf186 | 0 | 0 | 0 | 0 | 0 | 0 | 0 | 0 | 1 | 0 | 0 | 0 | 0 | 0 | 0 | 0 | 0 | 0 | 0 | 0 | 0 | 0 | 0 | 0 | 0 | 0 | 0 | 0 | Rnf186 |
| Rnf187 | 16368 | 20823 | 20077 | 22187 | 21692 | 19315 | 18859 | 13516 | 14991 | 16629 | 15072 | 16588 | 14550 | 14835 | 16429 | 18264 | 22489 | 21828 | 22285 | 18902 | 19842 | 16066 | 14904 | 17051 | 16866 | 17706 | 14476 | 15302 | Rnf187 |
| Rnf19a | 5702 | 5119 | 4787 | 5421 | 5710 | 5629 | 5921 | 6067 | 5184 | 4612 | 4617 | 4554 | 4736 | 4764 | 5988 | 5233 | 4816 | 5385 | 5391 | 5915 | 5834 | 5384 | 5207 | 4411 | 5019 | 4060 | 4903 | 4631 | Rnf19a |
| Rnf19b | 2623 | 2892 | 2463 | 3663 | 3700 | 3303 | 3398 | 2332 | 2244 | 2640 | 2281 | 2252 | 1970 | 2228 | 2595 | 2795 | 2946 | 3583 | 3732 | 3228 | 3653 | 2891 | 2460 | 2465 | 2939 | 2132 | 1901 | 2260 | Rnf19b |
| Rnf2 | 11470 | 8618 | 10148 | 9814 | 10429 | 10624 | 11534 | 13991 | 10874 | 9653 | 9949 | 11271 | 10947 | 12013 | 12129 | 10520 | 9677 | 10311 | 10493 | 11937 | 10765 | 11824 | 11250 | 10034 | 10429 | 12131 | 12112 | 12277 | Rnf2 |
| Rnf20 | 17786 | 16639 | 15314 | 15921 | 14838 | 15447 | 17497 | 15748 | 13277 | 12822 | 12856 | 13806 | 13477 | 15539 | 18315 | 15923 | 15316 | 16105 | 14284 | 16691 | 16615 | 13095 | 12605 | 13538 | 13574 | 12779 | 14091 | 14116 | Rnf20 |
| Rnf207 | 111 | 118 | 133 | 120 | 204 | 126 | 116 | 160 | 166 | 213 | 148 | 179 | 198 | 150 | 128 | 131 | 130 | 124 | 121 | 126 | 111 | 234 | 128 | 145 | 172 | 104 | 167 | 180 | Rnf207 |
| Rnf208 | 745 | 1014 | 1227 | 1289 | 1496 | 1278 | 1082 | 680 | 849 | 1122 | 895 | 838 | 655 | 617 | 740 | 745 | 1314 | 1197 | 1574 | 1073 | 1218 | 1186 | 809 | 929 | 1108 | 740 | 632 | 600 | Rnf208 |
| Rnf212 | 0 | 5 | 9 | 0 | 2 | 0 | 0 | 0 | 0 | 1 | 1 | 3 | 11 | 11 | 1 | 6 | 0 | 0 | 2 | 1 | 2 | 2 | 1 | 2 | 0 | 3 | 6 | 24 | Rnf212 |
| Rnf212b | 0 | 0 | 0 | 0 | 0 | 0 | 0 | 0 | 2 | 0 | 0 | 2 | 0 | 4 | 0 | 0 | 0 | 0 | 0 | 0 | 1 | 0 | 1 | 0 | 2 | 0 | 1 | 1 | Rnf212b |
| Rnf213 | 1751 | 1564 | 710 | 868 | 926 | 1332 | 856 | 1011 | 1147 | 822 | 1390 | 742 | 832 | 765 | 1456 | 1920 | 673 | 841 | 778 | 1972 | 1281 | 770 | 1315 | 720 | 719 | 615 | 1219 | 825 | Rnf213 |
| Rnf214 | 5344 | 5216 | 3704 | 5333 | 5929 | 6362 | 5840 | 6324 | 6323 | 7245 | 6612 | 6614 | 6489 | 6538 | 5619 | 5276 | 4497 | 5118 | 5620 | 5475 | 6595 | 6391 | 5848 | 5215 | 7421 | 3968 | 6744 | 6054 | Rnf214 |
| Rnf215 | 1458 | 1646 | 1481 | 1815 | 2050 | 1869 | 1590 | 2357 | 2604 | 2836 | 2578 | 2629 | 2409 | 2427 | 1429 | 1407 | 1668 | 1687 | 1922 | 1745 | 1752 | 2720 | 2563 | 2645 | 2894 | 2326 | 2435 | 2280 | Rnf215 |
| Rnf216 | 4470 | 4717 | 5395 | 5399 | 5483 | 4952 | 5418 | 3940 | 4435 | 4652 | 4569 | 5296 | 4791 | 5151 | 4261 | 4753 | 5562 | 5547 | 6194 | 4365 | 5365 | 4737 | 4516 | 4968 | 4837 | 5678 | 4700 | 5158 | Rnf216 |
| Rnf217 | 1226 | 928 | 984 | 1164 | 1730 | 1691 | 1734 | 2243 | 2312 | 1799 | 2488 | 1782 | 1858 | 1947 | 1275 | 1203 | 949 | 1129 | 1473 | 1735 | 1691 | 2077 | 2346 | 1586 | 1962 | 1469 | 1722 | 1799 | Rnf217 |
| Rnf220 | 4875 | 5860 | 6483 | 6418 | 6789 | 6425 | 6259 | 5671 | 6058 | 7554 | 6492 | 7818 | 6676 | 7207 | 5211 | 5206 | 6700 | 6554 | 7379 | 5864 | 6349 | 7319 | 6093 | 7786 | 7879 | 8305 | 6940 | 7175 | Rnf220 |
| Rnf222 | 3 | 4 | 15 | 8 | 4 | 8 | 6 | 9 | 12 | 11 | 11 | 9 | 11 | 4 | 5 | 4 | 13 | 11 | 11 | 3 | 11 | 12 | 10 | 2 | 4 | 10 | 6 | 6 | Rnf222 |
| Rnf223 | 276 | 491 | 440 | 462 | 405 | 331 | 412 | 96 | 165 | 142 | 144 | 159 | 150 | 159 | 305 | 393 | 593 | 432 | 465 | 367 | 374 | 165 | 117 | 169 | 170 | 189 | 173 | 140 | Rnf223 |
| Rnf224 | 0 | 0 | 0 | 0 | 1 | 0 | 0 | 0 | 0 | 0 | 0 | 0 | 0 | 0 | 0 | 1 | 0 | 0 | 0 | 0 | 0 | 2 | 0 | 0 | 3 | 0 | 0 | 0 | Rnf224 |
| Rnf225 | 218 | 286 | 240 | 303 | 229 | 214 | 248 | 131 | 127 | 125 | 134 | 189 | 135 | 162 | 195 | 259 | 270 | 269 | 205 | 203 | 220 | 124 | 126 | 150 | 143 | 126 | 162 | 186 | Rnf225 |
| Rnf227 | 3794 | 3506 | 3543 | 5728 | 6671 | 5896 | 7213 | 4857 | 3670 | 4085 | 3743 | 3079 | 2484 | 3319 | 4858 | 3377 | 4832 | 6217 | 6263 | 6982 | 7230 | 5125 | 3796 | 3651 | 4740 | 2708 | 2905 | 2890 | Rnf227 |
| Rnf24 | 6141 | 5244 | 4728 | 5303 | 6390 | 6261 | 6566 | 6293 | 5580 | 6084 | 5419 | 5691 | 5164 | 6019 | 5792 | 5154 | 5190 | 5708 | 6096 | 5494 | 6574 | 7210 | 5196 | 5431 | 6824 | 4488 | 5444 | 5749 | Rnf24 |
| Rnf25 | 2496 | 2804 | 2653 | 2614 | 2420 | 2517 | 2287 | 2684 | 2725 | 2892 | 2798 | 2755 | 2572 | 2528 | 2483 | 2625 | 2685 | 2545 | 2423 | 2500 | 2224 | 2939 | 2657 | 2719 | 2762 | 3022 | 2882 | 2676 | Rnf25 |
| Rnf26 | 711 | 714 | 785 | 638 | 536 | 579 | 788 | 525 | 536 | 519 | 748 | 670 | 722 | 1012 | 650 | 628 | 702 | 748 | 614 | 547 | 651 | 540 | 537 | 812 | 711 | 984 | 739 | 994 | Rnf26 |
| Rnf31 | 2208 | 2826 | 2086 | 2981 | 2644 | 2571 | 2361 | 1602 | 1939 | 2133 | 2118 | 2192 | 1965 | 2072 | 2051 | 2653 | 2549 | 2938 | 2590 | 2086 | 2661 | 2222 | 1935 | 2031 | 2228 | 1635 | 2055 | 1912 | Rnf31 |
| Rnf32 | 702 | 708 | 606 | 653 | 668 | 602 | 797 | 666 | 715 | 642 | 538 | 531 | 472 | 583 | 793 | 713 | 667 | 768 | 704 | 859 | 802 | 719 | 654 | 480 | 701 | 466 | 492 | 533 | Rnf32 |
| Rnf34 | 5901 | 6205 | 5479 | 5809 | 4626 | 4809 | 5256 | 4350 | 3700 | 3440 | 3089 | 3594 | 3612 | 3797 | 6405 | 5879 | 5289 | 5644 | 5003 | 5759 | 5365 | 3506 | 3576 | 3507 | 3657 | 4058 | 3850 | 4041 | Rnf34 |
| Rnf38 | 14378 | 11945 | 14272 | 16846 | 18202 | 18146 | 18423 | 18208 | 15947 | 18242 | 16938 | 20770 | 18978 | 20530 | 14654 | 13458 | 16545 | 17357 | 17147 | 15516 | 18457 | 21534 | 14533 | 18396 | 21057 | 17263 | 19414 | 19534 | Rnf38 |
| Rnf39 | 16 | 6 | 15 | 4 | 10 | 8 | 12 | 6 | 6 | 1 | 9 | 0 | 3 | 2 | 9 | 19 | 3 | 13 | 7 | 9 | 6 | 3 | 13 | 2 | 4 | 2 | 2 | 0 | Rnf39 |
| Rnf4 | 13976 | 13319 | 11755 | 10826 | 10533 | 10915 | 12178 | 12044 | 10654 | 8921 | 9447 | 10725 | 10440 | 11196 | 13776 | 12579 | 11456 | 10726 | 11287 | 12751 | 11689 | 9919 | 10738 | 9568 | 9596 | 11975 | 11211 | 11910 | Rnf4 |
| Rnf40 | 6895 | 8202 | 8561 | 9372 | 7647 | 7633 | 7785 | 5835 | 6756 | 7336 | 7252 | 8012 | 7223 | 8174 | 6163 | 7199 | 9041 | 8655 | 8567 | 6476 | 8234 | 6684 | 6314 | 8103 | 7914 | 8163 | 7710 | 8262 | Rnf40 |
| Rnf41 | 4601 | 5725 | 4186 | 5562 | 5632 | 5398 | 5237 | 4621 | 4647 | 4635 | 4173 | 4375 | 4075 | 4427 | 4632 | 4455 | 4919 | 5304 | 5710 | 5480 | 5641 | 4436 | 4577 | 4476 | 5007 | 4262 | 4328 | 4227 | Rnf41 |
| Rnf43 | 446 | 277 | 353 | 288 | 227 | 285 | 261 | 275 | 407 | 341 | 300 | 184 | 296 | 225 | 285 | 474 | 269 | 397 | 255 | 225 | 244 | 154 | 499 | 377 | 139 | 310 | 184 | 210 | Rnf43 |
| Rnf44 | 6290 | 8137 | 10694 | 9358 | 8695 | 7478 | 7614 | 7507 | 8319 | 10760 | 9431 | 12841 | 10061 | 12125 | 6057 | 7058 | 10948 | 8920 | 9432 | 6322 | 7603 | 10606 | 7804 | 12243 | 11416 | 13974 | 10758 | 12132 | Rnf44 |
| Rnf5 | 4254 | 5823 | 6029 | 5497 | 5067 | 4338 | 5330 | 3980 | 3798 | 4423 | 3665 | 4574 | 4045 | 4801 | 4683 | 4329 | 6508 | 5251 | 5669 | 5525 | 4873 | 3930 | 3550 | 5058 | 4770 | 5366 | 4811 | 4677 | Rnf5 |
| Rnf6 | 2150 | 2141 | 2139 | 2012 | 1742 | 1743 | 1989 | 1752 | 1646 | 1753 | 1909 | 1720 | 1606 | 1816 | 2089 | 2169 | 1986 | 2174 | 1695 | 2080 | 1992 | 1581 | 1918 | 1976 | 1772 | 1949 | 1726 | 1877 | Rnf6 |
| Rnf7 | 6100 | 5551 | 5977 | 5039 | 4974 | 4811 | 5510 | 6603 | 5876 | 5664 | 5677 | 6380 | 6301 | 6951 | 6132 | 5806 | 5460 | 4724 | 4999 | 5655 | 4932 | 5691 | 6325 | 6194 | 5536 | 7275 | 7032 | 7358 | Rnf7 |
| Rnf8 | 769 | 742 | 890 | 803 | 781 | 824 | 991 | 750 | 645 | 770 | 719 | 775 | 799 | 902 | 655 | 664 | 809 | 847 | 884 | 816 | 894 | 735 | 594 | 852 | 865 | 807 | 760 | 945 | Rnf8 |
| Rnft1 | 2675 | 1792 | 2260 | 1816 | 1822 | 1753 | 1744 | 1657 | 1593 | 1444 | 1817 | 1421 | 1630 | 1535 | 2411 | 2238 | 1949 | 2016 | 1699 | 2136 | 1640 | 1638 | 1836 | 1482 | 1429 | 1666 | 1461 | 1557 | Rnft1 |
| Rnft2 | 4180 | 5173 | 4315 | 5769 | 6200 | 5887 | 5830 | 4257 | 4361 | 4506 | 4366 | 4286 | 4366 | 4302 | 4406 | 4268 | 5233 | 5862 | 6248 | 5301 | 6249 | 5446 | 4124 | 4417 | 5312 | 3561 | 4336 | 4098 | Rnft2 |
| Rngtt | 7191 | 6424 | 6275 | 6304 | 5704 | 6192 | 7185 | 4407 | 3490 | 3448 | 3054 | 3770 | 3392 | 3772 | 7721 | 6738 | 6283 | 6103 | 5877 | 7060 | 6579 | 3603 | 3490 | 3352 | 3383 | 3460 | 3711 | 3790 | Rngtt |
| Rnh1 | 1752 | 1751 | 1425 | 1138 | 1132 | 1348 | 995 | 2549 | 3148 | 2431 | 2630 | 2508 | 2510 | 2305 | 1590 | 1664 | 1465 | 1209 | 1276 | 1224 | 1121 | 2291 | 3451 | 2522 | 2280 | 3126 | 2636 | 2580 | Rnh1 |
| Rnmt | 10416 | 9540 | 8659 | 9439 | 9326 | 9596 | 11203 | 8310 | 6817 | 6490 | 5904 | 7274 | 6346 | 7562 | 11234 | 9766 | 9128 | 9584 | 9168 | 11159 | 10621 | 7070 | 6490 | 6784 | 7496 | 5989 | 7275 | 7074 | Rnmt |
| Rnpc3 | 2050 | 1715 | 1632 | 2014 | 2408 | 2311 | 2465 | 4169 | 3344 | 3167 | 2928 | 3271 | 3064 | 3232 | 2421 | 1912 | 1898 | 1968 | 2336 | 2915 | 2518 | 3569 | 3203 | 2932 | 3262 | 2722 | 3251 | 2928 | Rnpc3 |
| Rnpep | 2889 | 2900 | 3271 | 2862 | 3208 | 3117 | 3177 | 3468 | 4060 | 4072 | 4068 | 4672 | 4007 | 3951 | 2636 | 2946 | 3201 | 2872 | 3546 | 2721 | 3279 | 4022 | 4145 | 4345 | 4226 | 4718 | 4250 | 4323 | Rnpep |
| Rnpepl1 | 1820 | 2154 | 2485 | 2316 | 2869 | 2490 | 2199 | 2329 | 2857 | 3277 | 3396 | 3070 | 2976 | 2723 | 1716 | 2064 | 2508 | 2437 | 2747 | 1964 | 2122 | 3110 | 2897 | 3340 | 3129 | 3282 | 2623 | 2775 | Rnpepl1 |
| Rnps1 | 13765 | 15749 | 16577 | 16469 | 14048 | 14250 | 15021 | 11582 | 12334 | 12909 | 12673 | 15534 | 14173 | 15473 | 13018 | 14112 | 16293 | 15224 | 15290 | 13918 | 14579 | 11971 | 11548 | 14470 | 14786 | 17045 | 15264 | 15666 | Rnps1 |
| Ro60 | 34231 | 26215 | 20601 | 25717 | 23339 | 29129 | 29323 | 18822 | 14796 | 11889 | 12099 | 15143 | 14528 | 16029 | 35699 | 31197 | 22627 | 24504 | 23338 | 31995 | 29317 | 12320 | 13690 | 11759 | 14047 | 10717 | 16625 | 15278 | Ro60 |
| Robo1 | 16055 | 16116 | 16441 | 18746 | 19431 | 19747 | 16718 | 16385 | 15858 | 19247 | 18002 | 18390 | 19681 | 17479 | 17003 | 18565 | 18140 | 19307 | 18965 | 14738 | 17956 | 20197 | 14795 | 18469 | 19195 | 12201 | 17270 | 17631 | Robo1 |
| Robo2 | 9393 | 8228 | 5678 | 10875 | 16239 | 15149 | 14201 | 17001 | 17663 | 17448 | 16334 | 12491 | 12452 | 13069 | 10639 | 9336 | 8047 | 12767 | 12387 | 13590 | 15630 | 17139 | 16889 | 14322 | 17661 | 7705 | 11111 | 10466 | Robo2 |
| Robo3 | 2990 | 6817 | 5833 | 6477 | 5781 | 5755 | 3357 | 7698 | 6604 | 8398 | 6746 | 7487 | 20515 | 6402 | 5784 | 7308 | 7962 | 4617 | 4476 | 4745 | 3007 | 9103 | 6776 | 3602 | 6180 | 4446 | 8479 | 8117 | Robo3 |
| Robo4 | 424 | 409 | 432 | 425 | 592 | 557 | 345 | 691 | 951 | 993 | 1158 | 980 | 1122 | 824 | 398 | 420 | 452 | 476 | 508 | 358 | 358 | 879 | 1101 | 994 | 862 | 1002 | 927 | 807 | Robo4 |
| Rock1 | 9721 | 6959 | 7960 | 8790 | 8270 | 9532 | 9902 | 9237 | 7978 | 7517 | 7983 | 8130 | 8418 | 8213 | 9728 | 9129 | 7995 | 8579 | 8402 | 9370 | 9405 | 7668 | 7851 | 7470 | 7752 | 6614 | 8349 | 7946 | Rock1 |
| Rock2 | 13175 | 10580 | 9291 | 11366 | 11562 | 12803 | 13095 | 11830 | 10345 | 9567 | 10086 | 9988 | 10190 | 10664 | 13684 | 12310 | 10331 | 11485 | 10872 | 13015 | 12851 | 10037 | 10299 | 8795 | 9972 | 6965 | 10168 | 9653 | Rock2 |
| Rogdi | 2332 | 3109 | 2768 | 3152 | 3089 | 2819 | 2960 | 2047 | 1980 | 2390 | 2110 | 2146 | 2161 | 2225 | 2583 | 2557 | 3242 | 3029 | 3162 | 2610 | 2907 | 2453 | 2017 | 2338 | 2381 | 2183 | 2079 | 2110 | Rogdi |
| Rom1 | 69 | 86 | 96 | 87 | 79 | 72 | 55 | 88 | 149 | 137 | 137 | 129 | 175 | 117 | 78 | 104 | 94 | 69 | 86 | 51 | 67 | 120 | 156 | 143 | 119 | 128 | 138 | 110 | Rom1 |
| Romo1 | 1132 | 1640 | 1664 | 1661 | 1347 | 1173 | 1428 | 1327 | 1216 | 1706 | 1608 | 1445 | 1364 | 1654 | 1137 | 953 | 1752 | 1318 | 1368 | 1658 | 1517 | 1044 | 1201 | 1886 | 1551 | 1774 | 1690 | 1923 | Romo1 |
| Ropn1 | 1 | 0 | 0 | 0 | 1 | 0 | 0 | 0 | 0 | 2 | 3 | 0 | 0 | 1 | 0 | 2 | 0 | 0 | 0 | 0 | 0 | 0 | 1 | 2 | 0 | 0 | 0 | 2 | Ropn1 |
| Ropn1l | 38 | 51 | 72 | 49 | 36 | 34 | 28 | 38 | 38 | 35 | 21 | 37 | 36 | 36 | 51 | 55 | 48 | 15 | 26 | 27 | 40 | 18 | 33 | 28 | 14 | 58 | 70 | 50 | Ropn1l |
| Ror1 | 1925 | 1201 | 1122 | 701 | 436 | 752 | 304 | 1475 | 2024 | 1341 | 2153 | 1341 | 1788 | 1242 | 1518 | 2246 | 870 | 752 | 360 | 379 | 327 | 1243 | 2023 | 1552 | 864 | 1639 | 1317 | 1295 | Ror1 |
| Ror2 | 1278 | 1404 | 1533 | 1081 | 952 | 1051 | 774 | 1039 | 1557 | 1577 | 1856 | 1919 | 2058 | 1451 | 948 | 1685 | 1173 | 1069 | 935 | 867 | 670 | 1068 | 1527 | 1781 | 1207 | 2626 | 1786 | 1742 | Ror2 |
| Rora | 1693 | 1187 | 984 | 1612 | 2034 | 2054 | 1886 | 1673 | 1442 | 1581 | 1585 | 1077 | 1080 | 1047 | 1718 | 1621 | 1021 | 1694 | 1505 | 2026 | 2023 | 1993 | 1908 | 1243 | 1897 | 714 | 929 | 900 | Rora |
| Rorb | 676 | 636 | 1197 | 1266 | 1388 | 949 | 1226 | 592 | 507 | 644 | 616 | 707 | 456 | 740 | 786 | 474 | 1382 | 1256 | 1339 | 907 | 1271 | 1041 | 585 | 897 | 848 | 920 | 661 | 668 | Rorb |
| Rorc | 400 | 439 | 243 | 476 | 407 | 464 | 588 | 200 | 220 | 189 | 178 | 146 | 158 | 213 | 335 | 329 | 333 | 533 | 581 | 639 | 540 | 119 | 247 | 298 | 154 | 251 | 145 | 158 | Rorc |
| Ros1 | 0 | 0 | 0 | 0 | 0 | 0 | 0 | 0 | 0 | 0 | 3 | 0 | 0 | 0 | 1 | 2 | 0 | 0 | 0 | 1 | 0 | 0 | 1 | 0 | 0 | 0 | 0 | 0 | Ros1 |
| Rp1 | 16 | 13 | 9 | 12 | 17 | 18 | 11 | 34 | 19 | 27 | 28 | 25 | 64 | 15 | 24 | 30 | 22 | 13 | 9 | 20 | 13 | 37 | 25 | 11 | 31 | 24 | 34 | 17 | Rp1 |
| Rp1l1 | 0 | 3 | 0 | 4 | 5 | 1 | 2 | 0 | 0 | 1 | 0 | 0 | 0 | 1 | 1 | 6 | 0 | 12 | 5 | 2 | 5 | 0 | 0 | 0 | 3 | 0 | 1 | 2 | Rp1l1 |
| Rp2 | 4418 | 2892 | 3657 | 3409 | 3346 | 3727 | 4238 | 2555 | 2261 | 1993 | 2059 | 2348 | 2384 | 2593 | 4535 | 3862 | 3342 | 3263 | 3373 | 3880 | 3841 | 2134 | 2534 | 2279 | 2134 | 2449 | 2606 | 2561 | Rp2 |
| Rp9 | 1923 | 1887 | 2238 | 2007 | 2105 | 1853 | 1876 | 1594 | 1720 | 1870 | 1757 | 1804 | 1877 | 1771 | 1644 | 1701 | 1983 | 1867 | 2140 | 1930 | 1899 | 1617 | 1797 | 2044 | 1944 | 2012 | 1725 | 1866 | Rp9 |
| Rpa1 | 17836 | 17415 | 16854 | 16780 | 12637 | 13591 | 16435 | 7747 | 7823 | 7291 | 7442 | 9613 | 8484 | 9694 | 16152 | 15787 | 15753 | 15594 | 15003 | 14622 | 15492 | 6269 | 7448 | 8582 | 7548 | 11478 | 9312 | 10360 | Rpa1 |
| Rpa2 | 5621 | 6231 | 5694 | 5159 | 3726 | 4221 | 4769 | 3090 | 3179 | 2615 | 2883 | 3914 | 3531 | 3925 | 5073 | 5051 | 5096 | 4572 | 4884 | 5105 | 4589 | 2115 | 3023 | 3510 | 3050 | 5183 | 4367 | 4408 | Rpa2 |
| Rpa3 | 3049 | 2569 | 3369 | 3156 | 2520 | 2677 | 3281 | 2218 | 1955 | 2052 | 1951 | 2552 | 2287 | 2804 | 2909 | 2969 | 3187 | 2938 | 2791 | 3544 | 2991 | 1567 | 1990 | 2439 | 2169 | 3170 | 2550 | 3069 | Rpa3 |
| Rpain | 1764 | 2160 | 1874 | 2062 | 1632 | 1576 | 1833 | 1423 | 1244 | 1272 | 1347 | 1416 | 1253 | 1480 | 1931 | 1900 | 1950 | 1963 | 1824 | 1930 | 1787 | 1193 | 1317 | 1392 | 1515 | 1388 | 1496 | 1703 | Rpain |
| Rpap1 | 1760 | 1993 | 2561 | 2039 | 1951 | 1989 | 1847 | 2847 | 3635 | 3809 | 3530 | 4855 | 4254 | 4339 | 1824 | 1870 | 2423 | 1886 | 2187 | 1717 | 1711 | 3474 | 3258 | 4325 | 3817 | 4486 | 4313 | 4577 | Rpap1 |
| Rpap2 | 1262 | 1247 | 1183 | 962 | 1013 | 1056 | 1191 | 1606 | 1448 | 1314 | 1370 | 1327 | 1407 | 1413 | 1387 | 1039 | 1201 | 957 | 1108 | 1295 | 1074 | 1358 | 1579 | 1392 | 1390 | 1524 | 1424 | 1435 | Rpap2 |

| | | | | | | | | | | | | | | | | | | | | | | | | | | | | | |
|---|---|---|---|---|---|---|---|---|---|---|---|---|---|---|---|---|---|---|---|---|---|---|---|---|---|---|---|---|---|
| Rpap3 | 3032 | 2747 | 3541 | 2990 | 2969 | 3047 | 3391 | 2431 | 2573 | 2599 | 2347 | 2964 | 2879 | 3279 | 2918 | 2886 | 3038 | 2771 | 2694 | 3081 | 3028 | 2403 | 2577 | 2770 | 2760 | 3101 | 3072 | 3321 | Rpap3 |
| Rpe | 4315 | 3653 | 4130 | 3437 | 3220 | 3225 | 3766 | 3638 | 3109 | 2679 | 2685 | 3266 | 2836 | 3235 | 4820 | 4059 | 3907 | 3358 | 3383 | 4238 | 3506 | 2789 | 3177 | 3030 | 2650 | 3427 | 3287 | 3457 | Rpe |
| Rpe65 | 1 | 1 | 0 | 0 | 0 | 0 | 0 | 0 | 0 | 2 | 0 | 4 | 0 | 0 | 0 | 0 | 0 | 0 | 0 | 0 | 0 | 0 | 3 | 0 | 0 | 0 | 0 | 0 | Rpe65 |
| Rpf1 | 1916 | 1811 | 2035 | 1745 | 1705 | 1724 | 1991 | 2964 | 2391 | 2513 | 2119 | 3032 | 2581 | 3147 | 2107 | 1636 | 2021 | 1636 | 1944 | 2005 | 1818 | 2378 | 2291 | 2800 | 2665 | 3436 | 3168 | 3069 | Rpf1 |
| Rpf2 | 2288 | 1921 | 2375 | 2322 | 2098 | 2019 | 2529 | 2772 | 2338 | 2140 | 2114 | 2409 | 2359 | 2900 | 2571 | 2185 | 2106 | 1975 | 2207 | 2749 | 2373 | 2379 | 2256 | 2473 | 2528 | 3002 | 2828 | 3072 | Rpf2 |
| Rpgr | 4194 | 3649 | 4016 | 3810 | 4047 | 4285 | 4507 | 5737 | 4341 | 4222 | 4095 | 4896 | 4841 | 5314 | 4619 | 3959 | 4015 | 3559 | 3926 | 5074 | 4286 | 4809 | 4471 | 4155 | 5012 | 4389 | 5112 | 4847 | Rpgr |
| Rpgrip1 | 66 | 60 | 45 | 65 | 100 | 92 | 85 | 154 | 144 | 167 | 130 | 106 | 108 | 90 | 74 | 68 | 90 | 87 | 110 | 73 | 73 | 112 | 121 | 122 | 137 | 88 | 113 | 104 | Rpgrip1 |
| Rpgrip1l | 5864 | 5109 | 4090 | 4093 | 3353 | 4010 | 4123 | 3596 | 3250 | 2659 | 2709 | 2944 | 2944 | 2809 | 5889 | 5304 | 4335 | 3832 | 3557 | 4551 | 4160 | 2642 | 3210 | 2570 | 2591 | 2063 | 2975 | 2813 | Rpgrip1l |
| Rph3a | 283 | 442 | 488 | 1067 | 1861 | 1148 | 1179 | 1161 | 1463 | 1795 | 1472 | 1019 | 949 | 1098 | 504 | 396 | 851 | 1157 | 1442 | 986 | 1344 | 2374 | 1135 | 1409 | 2037 | 1048 | 681 | 879 | Rph3a |
| Rph3al | 174 | 244 | 195 | 152 | 176 | 167 | 161 | 151 | 171 | 222 | 161 | 118 | 159 | 114 | 183 | 224 | 220 | 142 | 215 | 167 | 151 | 139 | 211 | 152 | 152 | 181 | 95 | 125 | Rph3al |
| Rpia | 2112 | 2168 | 2746 | 2793 | 2329 | 2275 | 2367 | 1788 | 1826 | 1914 | 1914 | 2149 | 1944 | 2190 | 2181 | 2121 | 2600 | 2651 | 2388 | 2261 | 2330 | 1963 | 1759 | 2090 | 2146 | 2842 | 2122 | 2213 | Rpia |
| Rpl10 | 23626 | 27775 | 25466 | 23865 | 21350 | 25511 | 23902 | 22517 | 27480 | 21501 | 25401 | 24083 | 26045 | 24621 | 24266 | 28241 | 23364 | 24223 | 25323 | 30561 | 23164 | 18229 | 28293 | 23430 | 23713 | 29486 | 27798 | 26320 | Rpl10 |
| Rpl10a | 21527 | 25952 | 21855 | 19916 | 15843 | 17096 | 17734 | 19075 | 22279 | 19407 | 20305 | 19891 | 20373 | 22009 | 19694 | 21387 | 19943 | 18127 | 18773 | 21111 | 18603 | 14641 | 22895 | 20958 | 18343 | 25623 | 21875 | 23292 | Rpl10a |
| Rpl10-ps3 | 130 | 82 | 68 | 70 | 62 | 90 | 106 | 138 | 73 | 61 | 75 | 91 | 83 | 83 | 125 | 87 | 56 | 87 | 83 | 115 | 99 | 112 | 72 | 86 | 100 | 82 | 100 | 97 | Rpl10-ps3 |
| Rpl11 | 21713 | 24396 | 24113 | 21838 | 17970 | 17949 | 22302 | 20508 | 21656 | 23245 | 22429 | 22637 | 22622 | 27746 | 21378 | 20718 | 22855 | 19805 | 19625 | 25044 | 21588 | 15649 | 22269 | 25522 | 21652 | 27143 | 24970 | 28656 | Rpl11 |
| Rpl12 | 21708 | 25543 | 24578 | 20787 | 16868 | 18189 | 17879 | 18523 | 21971 | 19618 | 22154 | 20751 | 22403 | 21401 | 19885 | 22338 | 21850 | 19558 | 19546 | 23891 | 17464 | 13341 | 22440 | 22452 | 19332 | 27272 | 23800 | 24141 | Rpl12 |
| Rpl13 | 29314 | 34138 | 28362 | 27167 | 23636 | 24388 | 24896 | 27645 | 30920 | 27916 | 29919 | 27620 | 29430 | 30352 | 27667 | 28100 | 28267 | 25543 | 26881 | 29618 | 24439 | 21031 | 31065 | 31016 | 26700 | 34427 | 30811 | 32554 | Rpl13 |
| Rpl13a | 1341 | 1355 | 1292 | 1301 | 1021 | 1028 | 1549 | 1630 | 1187 | 1714 | 1379 | 1556 | 1260 | 2225 | 1246 | 833 | 1245 | 1096 | 1170 | 1174 | 1699 | 1242 | 1284 | 1919 | 1424 | 1506 | 1521 | 2127 | Rpl13a |
| Rpl14 | 24112 | 27153 | 29041 | 23284 | 19302 | 20245 | 22911 | 21550 | 23529 | 24299 | 25224 | 24484 | 25382 | 27940 | 23114 | 25221 | 25190 | 21206 | 20440 | 25854 | 21036 | 16177 | 24419 | 25691 | 23119 | 32308 | 27632 | 30725 | Rpl14 |
| Rpl15 | 29389 | 30803 | 29155 | 29338 | 26867 | 28763 | 29513 | 28955 | 30735 | 26066 | 27772 | 27002 | 28233 | 28237 | 29590 | 29718 | 28292 | 28630 | 29862 | 33413 | 29387 | 22583 | 32668 | 27356 | 27852 | 33390 | 30341 | 30617 | Rpl15 |
| Rpl17 | 7474 | 14715 | 17778 | 13422 | 13864 | 11032 | 16553 | 9558 | 14559 | 15419 | 13564 | 15393 | 11637 | 17492 | 8831 | 14039 | 12381 | 12319 | 16096 | 16994 | 15128 | 3603 | 17713 | 12828 | 9405 | 16459 | 18091 | 16720 | Rpl17 |
| Rpl18 | 22300 | 28998 | 25134 | 25414 | 19467 | 21080 | 20533 | 18890 | 21791 | 21519 | 22323 | 21721 | 22160 | 23263 | 21415 | 21855 | 24886 | 22148 | 24758 | 26720 | 22739 | 14352 | 23127 | 24114 | 21264 | 27477 | 25356 | 24932 | Rpl18 |
| Rpl18a | 16278 | 21255 | 18175 | 18642 | 15633 | 18064 | 15008 | 15819 | 19298 | 16223 | 17823 | 17107 | 19781 | 16179 | 16561 | 19013 | 17700 | 16549 | 19671 | 23549 | 16087 | 12216 | 19926 | 20149 | 17497 | 21456 | 21835 | 18838 | Rpl18a |
| Rpl19 | 6534 | 8016 | 5581 | 6636 | 5267 | 5704 | 7366 | 5992 | 6096 | 7172 | 6452 | 6047 | 6273 | 10128 | 6297 | 5913 | 6551 | 5434 | 5422 | 6948 | 8150 | 4878 | 6313 | 7678 | 5886 | 6864 | 6722 | 9433 | Rpl19 |
| Rpl21 | 6730 | 8260 | 8293 | 7422 | 7521 | 6348 | 7266 | 7477 | 6625 | 7774 | 8566 | 6720 | 6397 | 10651 | 6600 | 5245 | 5891 | 5683 | 5338 | 6392 | 9013 | 5096 | 7680 | 9370 | 7181 | 8282 | 6120 | 10219 | Rpl21 |
| Rpl22 | 23822 | 22110 | 23922 | 22720 | 19067 | 19583 | 22338 | 20421 | 20338 | 19174 | 20378 | 19345 | 20610 | 22288 | 23004 | 21461 | 22051 | 20393 | 19830 | 24092 | 21194 | 15884 | 20895 | 22043 | 19502 | 24086 | 21367 | 23799 | Rpl22 |
| Rpl22l1 | 8397 | 5713 | 8414 | 8736 | 6774 | 7444 | 9888 | 8361 | 7150 | 8560 | 8636 | 8571 | 9365 | 9582 | 8043 | 6694 | 7715 | 6927 | 7449 | 9357 | 8438 | 5890 | 7534 | 9580 | 7854 | 10266 | 9577 | 11097 | Rpl22l1 |
| Rpl23 | 24322 | 23500 | 28466 | 25320 | 20211 | 18565 | 27306 | 23132 | 20689 | 24451 | 25406 | 22803 | 23853 | 34187 | 24005 | 20422 | 26445 | 22740 | 20578 | 25216 | 25855 | 17741 | 21557 | 27307 | 23775 | 29189 | 24474 | 32739 | Rpl23 |
| Rpl23a | 20882 | 28179 | 24492 | 24511 | 19726 | 19133 | 18439 | 20458 | 23282 | 24712 | 23057 | 22134 | 21349 | 23839 | 21096 | 20814 | 23526 | 20657 | 22158 | 23769 | 21475 | 15401 | 25023 | 24905 | 22021 | 26461 | 23095 | 23629 | Rpl23a |
| Rpl24 | 60 | 56 | 60 | 95 | 61 | 22 | 56 | 88 | 71 | 82 | 66 | 72 | 64 | 78 | 54 | 69 | 56 | 85 | 51 | 74 | 66 | 70 | 73 | 60 | 60 | 90 | 63 | 89 | Rpl24 |
| Rpl26 | 31881 | 31682 | 28706 | 29163 | 25570 | 28086 | 29433 | 29264 | 30814 | 28070 | 29292 | 28176 | 31547 | 29793 | 32762 | 30799 | 27787 | 26798 | 29914 | 37319 | 28213 | 23983 | 31924 | 28945 | 27057 | 34859 | 32913 | 32149 | Rpl26 |
| Rpl27 | 16957 | 18527 | 20750 | 20742 | 17019 | 15841 | 20128 | 16452 | 17623 | 17695 | 20712 | 19318 | 21361 | 23918 | 15654 | 15351 | 19966 | 18153 | 19830 | 21134 | 18943 | 12477 | 17374 | 21596 | 20650 | 25222 | 22353 | 24954 | Rpl27 |
| Rpl27a | 22912 | 25461 | 22601 | 22275 | 18023 | 19560 | 22009 | 21498 | 21156 | 22158 | 23516 | 20680 | 22857 | 27438 | 22598 | 19501 | 21983 | 20329 | 21362 | 25001 | 22322 | 15924 | 22605 | 23772 | 20731 | 26251 | 24346 | 27108 | Rpl27a |
| Rpl28 | 18403 | 25409 | 24832 | 23510 | 18072 | 17668 | 18728 | 16472 | 20294 | 22944 | 23737 | 21027 | 22363 | 24911 | 16361 | 17550 | 24278 | 21053 | 20851 | 20246 | 21077 | 14375 | 20463 | 25918 | 21272 | 28532 | 24539 | 26119 | Rpl28 |
| Rpl29 | 9840 | 13003 | 9812 | 10712 | 8784 | 8585 | 9400 | 9191 | 16136 | 11123 | 10626 | 10465 | 11387 | 11302 | 15177 | 19186 | 10209 | 9044 | 13651 | 12068 | 8888 | 10164 | 14417 | 10968 | 11004 | 12512 | 11217 | 11325 | Rpl29 |
| Rpl3 | 61554 | 67617 | 58550 | 52412 | 45236 | 49139 | 52346 | 52567 | 56729 | 45472 | 50845 | 51692 | 51877 | 54073 | 60488 | 62082 | 53340 | 50815 | 49984 | 57322 | 51859 | 40200 | 58016 | 51569 | 46946 | 61327 | 56428 | 60392 | Rpl3 |
| Rpl30 | 18120 | 14737 | 19567 | 17480 | 14840 | 4935 | 20031 | 17061 | 16855 | 18101 | 16194 | 9670 | 16299 | 18337 | 17571 | 14163 | 12645 | 14635 | 14145 | 15405 | 17493 | 7484 | 14185 | 14477 | 18395 | 12986 | 18441 | 27442 | Rpl30 |
| Rpl31 | 27299 | 26631 | 24165 | 24093 | 19150 | 17677 | 28861 | 26203 | 23521 | 22394 | 23301 | 20631 | 22311 | 31482 | 27051 | 20510 | 23995 | 21200 | 20979 | 24976 | 28047 | 19460 | 23761 | 26041 | 22479 | 25567 | 23928 | 33276 | Rpl31 |
| Rpl32 | 26550 | 28982 | 25274 | 24908 | 21018 | 22347 | 26759 | 24483 | 24478 | 24641 | 25946 | 23836 | 26863 | 31173 | 25897 | 22786 | 25095 | 22850 | 24199 | 28096 | 25732 | 18615 | 25009 | 27729 | 24398 | 29459 | 27778 | 32257 | Rpl32 |
| Rpl34 | 17684 | 20636 | 20366 | 22406 | 18924 | 18770 | 23608 | 17591 | 19770 | 20373 | 23921 | 19655 | 21647 | 23570 | 16656 | 15767 | 21205 | 19893 | 20775 | 23635 | 21731 | 12582 | 19575 | 23234 | 22258 | 23497 | 21454 | 25047 | Rpl34 |
| Rpl35 | 17085 | 23678 | 22352 | 24297 | 19136 | 18922 | 22819 | 16455 | 20106 | 21881 | 22620 | 20588 | 22712 | 25679 | 17609 | 18408 | 22425 | 20966 | 22093 | 26240 | 24036 | 12053 | 21347 | 24315 | 22733 | 25236 | 23850 | 26915 | Rpl35 |
| Rpl35a | 15541 | 15075 | 16767 | 16769 | 12870 | 10949 | 18508 | 15376 | 13038 | 15595 | 17111 | 14124 | 14848 | 22435 | 14919 | 10626 | 16901 | 14732 | 14468 | 15569 | 17205 | 10758 | 13337 | 17679 | 15900 | 18288 | 16114 | 22483 | Rpl35a |
| Rpl36 | 9103 | 11882 | 10014 | 11721 | 8876 | 7940 | 10910 | 8495 | 9357 | 11104 | 12351 | 9616 | 10951 | 12334 | 8156 | 6774 | 10776 | 9215 | 10173 | 10314 | 11138 | 6288 | 9346 | 12300 | 11332 | 11991 | 10851 | 12936 | Rpl36 |
| Rpl36a | 16932 | 14559 | 15760 | 16500 | 12715 | 11718 | 19144 | 16442 | 14450 | 15272 | 19011 | 14393 | 16667 | 24244 | 16250 | 11461 | 16026 | 13380 | 13558 | 16409 | 18351 | 11466 | 14214 | 20038 | 16438 | 18092 | 16381 | 24194 | Rpl36a |
| Rpl36al | 32 | 30 | 32 | 19 | 28 | 22 | 18 | 13 | 21 | 34 | 17 | 29 | 12 | 34 | 25 | 21 | 31 | 32 | 29 | 27 | 21 | 20 | 17 | 37 | 24 | 29 | 15 | 21 | Rpl36al |
| Rpl36a-ps1 | 189 | 92 | 106 | 105 | 83 | 68 | 150 | 155 | 85 | 135 | 150 | 110 | 158 | 149 | 137 | 86 | 110 | 98 | 101 | 128 | 141 | 115 | 98 | 134 | 125 | 137 | 126 | 163 | Rpl36a-ps1 |
| Rpl37 | 9541 | 10977 | 11444 | 12356 | 9784 | 6960 | 14781 | 9779 | 9362 | 13068 | 13149 | 10358 | 10998 | 19587 | 8987 | 6892 | 12249 | 9593 | 9564 | 11109 | 14420 | 7038 | 9532 | 15118 | 12348 | 12711 | 11345 | 18523 | Rpl37 |
| Rpl37a | 15076 | 19442 | 18864 | 20882 | 15369 | 12106 | 24701 | 14803 | 14480 | 20878 | 20397 | 16977 | 16856 | 29273 | 14562 | 11403 | 20478 | 16370 | 15569 | 17601 | 23999 | 10363 | 15288 | 23080 | 19015 | 19916 | 17674 | 28659 | Rpl37a |
| Rpl38 | 6253 | 7254 | 7095 | 9092 | 6122 | 4918 | 12002 | 6546 | 5520 | 9135 | 9096 | 6533 | 7318 | 18484 | 5782 | 3681 | 8479 | 6678 | 6053 | 7685 | 12908 | 4386 | 5523 | 10828 | 8481 | 7858 | 7110 | 15136 | Rpl38 |
| Rpl39 | 15847 | 14552 | 13950 | 17641 | 13560 | 11456 | 24687 | 16799 | 12571 | 18105 | 19337 | 14201 | 15324 | 30346 | 14733 | 8188 | 15549 | 13107 | 12678 | 14755 | 24623 | 11221 | 12363 | 21391 | 17125 | 16345 | 15016 | 28796 | Rpl39 |
| Rpl39l | 15 | 10 | 6 | 12 | 7 | 13 | 1 | 24 | 46 | 22 | 45 | 16 | 25 | 33 | 24 | 17 | 9 | 40 | 15 | 14 | 11 | 38 | 10 | 19 | 36 | 11 | 21 | 26 | Rpl39l |
| Rpl3l | 0 | 1 | 0 | 2 | 0 | 0 | 3 | 1 | 1 | 1 | 8 | 0 | 2 | 3 | 5 | 2 | 0 | 1 | 0 | 1 | 1 | 0 | 2 | 1 | 3 | 0 | 1 | 1 | Rpl3l |
| Rpl4 | 75873 | 77399 | 71767 | 62320 | 54154 | 56385 | 63490 | 69060 | 69216 | 61050 | 65808 | 66102 | 66141 | 70566 | 74282 | 73445 | 66174 | 61533 | 60114 | 73411 | 60660 | 53013 | 71317 | 67037 | 60555 | 79872 | 70541 | 73640 | Rpl4 |
| Rpl41 | 29991 | 40694 | 35303 | 37308 | 28142 | 25599 | 44283 | 29286 | 32508 | 37254 | 39243 | 32288 | 34962 | 51775 | 28581 | 24491 | 36398 | 30480 | 31038 | 34121 | 43164 | 19793 | 32556 | 40500 | 35662 | 39819 | 34767 | 52707 | Rpl41 |
| Rpl5 | 58302 | 54662 | 55453 | 48392 | 41708 | 42583 | 55569 | 56304 | 54372 | 46520 | 47854 | 49835 | 48088 | 62209 | 59480 | 55621 | 51183 | 45997 | 44307 | 53971 | 52175 | 41834 | 55893 | 52386 | 46274 | 60510 | 53093 | 66773 | Rpl5 |
| Rpl6 | 40775 | 43058 | 45858 | 43157 | 35551 | 37334 | 38091 | 37637 | 41318 | 45064 | 42774 | 41285 | 43323 | 42616 | 41233 | 42818 | 42229 | 38149 | 39766 | 45832 | 35776 | 30156 | 42331 | 41711 | 39410 | 50956 | 52350 | 45369 | Rpl6 |
| Rpl7 | 74495 | 64812 | 60335 | 54595 | 47013 | 50418 | 61753 | 70650 | 63010 | 51355 | 52857 | 57213 | 58064 | 65313 | 76490 | 66212 | 56730 | 52297 | 52718 | 69947 | 56145 | 54266 | 66040 | 57046 | 52849 | 70594 | 60921 | 69466 | Rpl7 |
| Rpl7a | 14595 | 28546 | 14970 | 14466 | 12653 | 14047 | 12662 | 13080 | 16336 | 19641 | 13204 | 19895 | 15738 | 16526 | 21629 | 17656 | 13848 | 19513 | 16471 | 28826 | 12217 | 11535 | 13858 | 15556 | 14528 | 19904 | 15846 | 15273 | Rpl7a |
| Rpl7l1 | 6840 | 6841 | 8046 | 6315 | 5452 | 5758 | 6439 | 7272 | 6995 | 7039 | 7018 | 7812 | 7727 | 8665 | 6485 | 6557 | 7047 | 6017 | 6489 | 6342 | 6064 | 5721 | 7434 | 8255 | 7447 | 10167 | 9207 | 9696 | Rpl7l1 |
| Rpl8 | 41655 | 54472 | 45742 | 43736 | 35390 | 38959 | 37392 | 32681 | 39401 | 36087 | 39018 | 39989 | 41294 | 38757 | 38121 | 42684 | 44938 | 39635 | 43917 | 42195 | 38170 | 27591 | 39328 | 41351 | 37365 | 50259 | 44666 | 42972 | Rpl8 |
| Rpl9 | 29215 | 29263 | 28987 | 27230 | 23204 | 22648 | 30042 | 27158 | 26089 | 24304 | 27792 | 26052 | 27073 | 32011 | 28208 | 25080 | 27452 | 24710 | 25090 | 28102 | 27655 | 20865 | 26373 | 27246 | 25563 | 31873 | 29529 | 34902 | Rpl9 |
| Rpl9-ps6 | 3757 | 3422 | 4279 | 3578 | 2442 | 2282 | 3351 | 3463 | 2841 | 3145 | 3517 | 2833 | 3144 | 4477 | 3690 | 2807 | 3650 | 3134 | 2761 | 3106 | 3367 | 2562 | 3045 | 3352 | 3211 | 3781 | 3263 | 4406 | Rpl9-ps6 |
| Rplp0 | 64974 | 80161 | 64016 | 62374 | 53425 | 56661 | 57349 | 57984 | 70798 | 58287 | 62549 | 58978 | 61595 | 62979 | 61021 | 67040 | 60913 | 58659 | 62532 | 66802 | 60357 | 45434 | 73256 | 63444 | 59238 | 72382 | 64539 | 67872 | Rplp0 |
| Rplp1 | 29617 | 37709 | 34293 | 31950 | 25908 | 27597 | 27975 | 26633 | 32219 | 29643 | 34147 | 28854 | 32804 | 30552 | 27426 | 27550 | 32296 | 29059 | 31212 | 33799 | 28103 | 19402 | 31559 | 34388 | 30252 | 37299 | 34789 | 34992 | Rplp1 |
| Rplp2 | 14760 | 19876 | 22885 | 21743 | 16671 | 14032 | 17584 | 11866 | 13691 | 18560 | 18517 | 18174 | 18901 | 19822 | 12456 | 12072 | 22739 | 17299 | 18837 | 13924 | 18477 | 11256 | 13779 | 22024 | 18183 | 22768 | 19765 | 23471 | Rplp2 |
| Rpn1 | 15175 | 16582 | 13455 | 13382 | 11576 | 11711 | 11989 | 11347 | 12493 | 10765 | 11989 | 9621 | 10549 | 9761 | 15151 | 15637 | 13319 | 13059 | 12752 | 12528 | 12289 | 10420 | 12474 | 10483 | 9832 | 10819 | 10213 | 10562 | Rpn1 |
| Rpn2 | 15941 | 18317 | 14971 | 13752 | 12273 | 13199 | 13132 | 13583 | 14258 | 12752 | 14026 | 13833 | 13882 | 13745 | 15709 | 16375 | 14157 | 13904 | 13379 | 12753 | 13120 | 13354 | 14334 | 13180 | 12379 | 15280 | 13930 | 14348 | Rpn2 |
| Rpp14 | 2399 | 2307 | 2303 | 2450 | 2211 | 2398 | 2486 | 2994 | 2677 | 2745 | 2547 | 2870 | 2942 | 3130 | 2616 | 2328 | 2363 | 2426 | 2375 | 2818 | 2328 | 2552 | 2673 | 2882 | 2665 | 3280 | 2880 | 3058 | Rpp14 |
| Rpp21 | 352 | 444 | 387 | 454 | 417 | 422 | 381 | 583 | 670 | 713 | 640 | 737 | 731 | 617 | 357 | 307 | 366 | 432 | 429 | 361 | 349 | 647 | 628 | 681 | 588 | 727 | 721 | 659 | Rpp21 |
| Rpp25 | 173 | 244 | 205 | 255 | 290 | 218 | 303 | 249 | 194 | 284 | 142 | 323 | 201 | 284 | 206 | 143 | 235 | 302 | 330 | 237 | 332 | 311 | 190 | 324 | 252 | 111 | 286 | 273 | Rpp25 |
| Rpp25l | 1096 | 1716 | 1364 | 1293 | 1149 | 1214 | 1212 | 1222 | 1341 | 1262 | 1198 | 1396 | 1427 | 1355 | 1062 | 1193 | 1389 | 1180 | 1308 | 1484 | 1224 | 911 | 1333 | 1480 | 1249 | 1802 | 1470 | 1488 | Rpp25l |
| Rpp30 | 1458 | 1285 | 1441 | 1298 | 1286 | 1343 | 1665 | 1818 | 1721 | 1356 | 1349 | 1833 | 1729 | 2166 | 1490 | 1306 | 1363 | 1173 | 1384 | 1906 | 1428 | 1465 | 1467 | 1661 | 1520 | 1995 | 2077 | 2188 | Rpp30 |
| Rpp38 | 249 | 251 | 307 | 246 | 254 | 207 | 231 | 476 | 474 | 333 | 409 | 433 | 373 | 461 | 289 | 223 | 282 | 207 | 251 | 310 | 230 | 328 | 509 | 422 | 372 | 460 | 425 | 513 | Rpp38 |
| Rpp40 | 1493 | 1389 | 1194 | 989 | 931 | 848 | 1215 | 569 | 588 | 534 | 519 | 538 | 553 | 636 | 1416 | 1428 | 1130 | 949 | 957 | 1118 | 1126 | 508 | 622 | 485 | 486 | 732 | 650 | 679 | Rpp40 |
| Rprd1a | 10560 | 9453 | 9112 | 9156 | 8634 | 9153 | 10140 | 7239 | 6141 | 5297 | 5452 | 6889 | 6292 | 6903 | 10429 | 9823 | 8961 | 9295 | 9187 | 9810 | 9640 | 6122 | 5769 | 6250 | 6236 | 6306 | 6959 | 6737 | Rprd1a |
| Rprd1b | 4174 | 4266 | 5149 | 4340 | 4169 | 3967 | 4264 | 3664 | 3647 | 4022 | 3645 | 5238 | 4443 | 4936 | 3920 | 4269 | 4662 | 4319 | 4455 | 3517 | 4094 | 4136 | 3408 | 4522 | 4129 | 5594 | 4608 | 5368 | Rprd1b |
| Rprd2 | 2308 | 2017 | 2893 | 2849 | 3589 | 2766 | 3039 | 5478 | 5256 | 5874 | 5648 | 6089 | 5769 | 6183 | 2259 | 1944 | 2859 | 2591 | 3334 | 2671 | 3161 | 6353 | 5029 | 6087 | 6189 | 5260 | 5604 | 6000 | Rprd2 |
| Rprm | 1459 | 2071 | 1848 | 2174 | 3235 | 2295 | 2659 | 3199 | 3624 | 3969 | 3167 | 2923 | 3171 | 3081 | 1807 | 1610 | 2268 | 2485 | 2805 | 2947 | 2648 | 2951 | 3075 | 3893 | 4138 | 3409 | 2450 | 2712 | Rprm |
| Rprml | 13 | 7 | 7 | 4 | 18 | 11 | 7 | 5 | 9 | 5 | 3 | 1 | 1 | 2 | 3 | 16 | 6 | 22 | 8 | 19 | 20 | 5 | 6 | 13 | 9 | 2 | 0 | 0 | Rprml |

| | | | | | | | | | | | | | | | | | | | | | | | | | | | | | |
|---|---|---|---|---|---|---|---|---|---|---|---|---|---|---|---|---|---|---|---|---|---|---|---|---|---|---|---|---|---|
| Rps10 | 478 | 669 | 574 | 692 | 412 | 477 | 448 | 519 | 492 | 448 | 469 | 527 | 449 | 466 | 423 | 629 | 627 | 605 | 569 | 482 | 450 | 451 | 484 | 502 | 456 | 624 | 568 | 530 | Rps10 |
| Rps11 | 38514 | 35922 | 35273 | 32333 | 26160 | 28359 | 32918 | 34899 | 30268 | 29312 | 30992 | 31997 | 34429 | 41616 | 35311 | 28561 | 32969 | 29101 | 31789 | 31325 | 33213 | 29698 | 31725 | 35678 | 32213 | 41414 | 39252 | 43155 | Rps11 |
| Rps12 | 14731 | 17624 | 17125 | 14745 | 15227 | 11001 | 16479 | 13496 | 16586 | 13776 | 17628 | 12706 | 11904 | 17113 | 17198 | 14895 | 19316 | 13337 | 13989 | 13876 | 14370 | 8540 | 17662 | 20251 | 16724 | 19692 | 17572 | 16714 | Rps12 |
| Rps13 | 17985 | 23177 | 20216 | 19025 | 17566 | 17184 | 21596 | 17307 | 18634 | 18272 | 20347 | 20546 | 22651 | 24141 | 16664 | 17803 | 19607 | 17398 | 19417 | 20626 | 20142 | 14437 | 18797 | 22197 | 19914 | 25496 | 23692 | 26985 | Rps13 |
| Rps14 | 45065 | 45863 | 58688 | 57600 | 44368 | 48747 | 43914 | 43840 | 46669 | 49042 | 51190 | 49524 | 52031 | 50497 | 42617 | 45011 | 51282 | 50741 | 57578 | 57117 | 48088 | 30901 | 49624 | 53453 | 50115 | 62819 | 57632 | 56048 | Rps14 |
| Rps15 | 25745 | 34127 | 30387 | 30535 | 23401 | 25549 | 25533 | 22304 | 26679 | 27731 | 28055 | 27381 | 29374 | 29890 | 24204 | 24857 | 30553 | 27533 | 29379 | 31713 | 27726 | 17985 | 27023 | 29786 | 27745 | 35524 | 32681 | 31506 | Rps15 |
| Rps15a | 37171 | 33778 | 31452 | 33993 | 30202 | 29140 | 42390 | 36777 | 31045 | 32034 | 36034 | 31023 | 33318 | 42094 | 36646 | 27392 | 32321 | 33598 | 32289 | 37709 | 41830 | 26859 | 31051 | 34455 | 34799 | 37032 | 34443 | 45258 | Rps15a |
| Rps16 | 20805 | 28149 | 24581 | 23323 | 17784 | 18194 | 20911 | 17456 | 19249 | 19488 | 20022 | 18750 | 19995 | 24126 | 19093 | 20508 | 24118 | 19788 | 20123 | 24238 | 21805 | 12472 | 19944 | 22248 | 18616 | 24978 | 22585 | 26107 | Rps16 |
| Rps17 | 28108 | 31881 | 31078 | 28321 | 23247 | 23357 | 29356 | 25415 | 26412 | 26102 | 28367 | 26561 | 29995 | 34496 | 27187 | 24585 | 29191 | 25222 | 27048 | 31753 | 28946 | 19519 | 26168 | 30000 | 26840 | 34478 | 32450 | 36844 | Rps17 |
| Rps18 | 29784 | 42057 | 35133 | 33188 | 27754 | 30148 | 30900 | 26848 | 32164 | 28947 | 32648 | 31805 | 34329 | 34285 | 28963 | 32008 | 33559 | 30422 | 34233 | 37987 | 30146 | 19554 | 33207 | 33508 | 30128 | 38714 | 37379 | 37244 | Rps18 |
| Rps18-ps5 | 899 | 1513 | 658 | 740 | 542 | 699 | 619 | 788 | 644 | 688 | 700 | 737 | 785 | 751 | 790 | 688 | 995 | 632 | 933 | 1009 | 628 | 480 | 673 | 811 | 687 | 784 | 831 | 802 | Rps18-ps5 |
| Rps18-ps6 | 100 | 207 | 70 | 62 | 41 | 32 | 69 | 90 | 58 | 84 | 73 | 66 | 48 | 80 | 102 | 58 | 108 | 53 | 79 | 60 | 71 | 52 | 53 | 87 | 70 | 61 | 66 | 77 | Rps18-ps6 |
| Rps19 | 20336 | 26184 | 25156 | 22626 | 19046 | 19138 | 23062 | 17778 | 21161 | 20046 | 22758 | 22573 | 23498 | 23504 | 19316 | 21016 | 23142 | 20854 | 22337 | 25310 | 21318 | 13207 | 21407 | 24657 | 21356 | 27541 | 25421 | 26608 | Rps19 |
| Rps19bp1 | 1385 | 1631 | 1457 | 1653 | 1330 | 1273 | 1458 | 1128 | 1077 | 1220 | 1158 | 1133 | 1134 | 1327 | 1305 | 1343 | 1542 | 1413 | 1457 | 1618 | 1465 | 1023 | 1088 | 1393 | 1259 | 1258 | 1237 | 1239 | Rps19bp1 |
| Rps2 | 49397 | 60245 | 63551 | 52179 | 41025 | 42484 | 40400 | 33273 | 40611 | 37467 | 40314 | 46879 | 46962 | 40272 | 45254 | 54247 | 53164 | 46992 | 48985 | 48514 | 40307 | 29245 | 41547 | 43968 | 38987 | 57900 | 51309 | 48184 | Rps2 |
| Rps20 | 17656 | 19955 | 16179 | 15885 | 13114 | 12882 | 18521 | 16745 | 16548 | 17701 | 17182 | 16877 | 17958 | 22156 | 16663 | 15127 | 16312 | 14069 | 14394 | 17480 | 17757 | 12123 | 17037 | 18767 | 15484 | 19653 | 17903 | 23715 | Rps20 |
| Rps21 | 9781 | 11976 | 11640 | 13664 | 9757 | 7395 | 17442 | 9972 | 9052 | 14524 | 14067 | 10691 | 11513 | 22977 | 8645 | 6622 | 13196 | 9960 | 9854 | 11430 | 16421 | 6410 | 9148 | 16284 | 12660 | 13086 | 11329 | 21065 | Rps21 |
| Rps23 | 29899 | 31343 | 32308 | 30477 | 25089 | 22646 | 32964 | 28256 | 27779 | 31110 | 31064 | 30561 | 30874 | 42070 | 28393 | 25211 | 31076 | 26646 | 27562 | 33017 | 31840 | 21246 | 27594 | 34496 | 29182 | 37688 | 33128 | 43110 | Rps23 |
| Rps23rg1 | 63 | 89 | 99 | 61 | 25 | 40 | 41 | 49 | 47 | 55 | 35 | 70 | 84 | 82 | 70 | 48 | 68 | 30 | 26 | 46 | 35 | 33 | 57 | 69 | 50 | 124 | 105 | 138 | Rps23rg1 |
| Rps24 | 59158 | 55543 | 57866 | 54853 | 47463 | 50716 | 58069 | 54986 | 53359 | 50105 | 56926 | 53916 | 58857 | 61920 | 62122 | 54089 | 53850 | 51000 | 53220 | 67233 | 54357 | 44264 | 54298 | 55958 | 51851 | 64862 | 60022 | 65124 | Rps24 |
| Rps25 | 26366 | 26479 | 24209 | 23765 | 20215 | 20462 | 28052 | 25959 | 23550 | 21945 | 26876 | 22995 | 24891 | 29005 | 25581 | 20747 | 23720 | 21754 | 23615 | 27948 | 25707 | 19136 | 23746 | 27682 | 23981 | 28930 | 27380 | 32465 | Rps25 |
| Rps26 | 23458 | 27655 | 29689 | 29520 | 23078 | 22518 | 27028 | 21989 | 24401 | 27011 | 32182 | 25998 | 29423 | 31645 | 23194 | 21592 | 27954 | 26803 | 26239 | 29459 | 25674 | 17146 | 23934 | 30777 | 28657 | 33784 | 29440 | 32246 | Rps26 |
| Rps27 | 22780 | 22193 | 16670 | 19317 | 16393 | 12313 | 31021 | 23798 | 15769 | 21303 | 23067 | 16988 | 18604 | 39980 | 20367 | 7375 | 19087 | 16218 | 16364 | 18035 | 30775 | 12935 | 16905 | 25656 | 20996 | 19059 | 18095 | 36191 | Rps27 |
| Rps27a | 27609 | 25287 | 26609 | 27126 | 21525 | 15258 | 30350 | 25203 | 26954 | 28126 | 27647 | 28759 | 25362 | 34098 | 22105 | 24680 | 26052 | 16088 | 23148 | 30010 | 27995 | 18698 | 28420 | 24997 | 17568 | 31891 | 12401 | 35318 | Rps27a |
| Rps27l | 5039 | 4274 | 5413 | 4843 | 3306 | 3258 | 5559 | 3938 | 3269 | 3530 | 3578 | 3791 | 3924 | 5459 | 4803 | 3544 | 4780 | 3856 | 3906 | 4509 | 5160 | 2526 | 3239 | 4572 | 3547 | 4632 | 4069 | 6041 | Rps27l |
| Rps27rt | 7199 | 6164 | 5694 | 8620 | 3887 | 3776 | 9386 | 6891 | 6663 | 5440 | 6438 | 4361 | 4373 | 8790 | 7483 | 7124 | 6951 | 4221 | 4173 | 4271 | 7117 | 7782 | 4907 | 6966 | 5218 | 4969 | 5269 | 10186 | Rps27rt |
| Rps28 | 8611 | 11630 | 10991 | 11860 | 8527 | 6303 | 15003 | 8613 | 7743 | 12651 | 11294 | 9242 | 9260 | 22748 | 7949 | 5379 | 12100 | 8220 | 8364 | 9335 | 15985 | 6462 | 7833 | 15624 | 10663 | 11419 | 10565 | 20181 | Rps28 |
| Rps29 | 8154 | 11548 | 9610 | 12473 | 8885 | 6648 | 16290 | 9417 | 8740 | 14087 | 12647 | 11035 | 10739 | 27270 | 7618 | 5237 | 12227 | 8755 | 7833 | 10660 | 17088 | 6015 | 8770 | 18275 | 11989 | 12492 | 11488 | 23425 | Rps29 |
| Rps3 | 54612 | 56571 | 59402 | 53866 | 44705 | 46218 | 45590 | 49906 | 50355 | 47622 | 50776 | 51622 | 51058 | 51168 | 52830 | 54114 | 54647 | 49280 | 51176 | 58155 | 46586 | 35906 | 53942 | 54039 | 47310 | 63154 | 56128 | 56185 | Rps3 |
| Rps3a1 | 51938 | 47811 | 50512 | 42128 | 35664 | 37474 | 45587 | 48147 | 46854 | 42240 | 44765 | 44309 | 45650 | 50298 | 52878 | 50478 | 44086 | 40292 | 39894 | 51429 | 41493 | 36741 | 50375 | 45653 | 40990 | 55372 | 48758 | 55796 | Rps3a1 |
| Rps4x | 74790 | 65361 | 63122 | 54340 | 46383 | 52020 | 58967 | 71938 | 66993 | 52310 | 56660 | 59562 | 60087 | 64894 | 75707 | 67574 | 55332 | 53764 | 53296 | 67100 | 55075 | 54719 | 66579 | 59954 | 54814 | 74960 | 64494 | 73076 | Rps4x |
| Rps5 | 30847 | 39311 | 29445 | 28952 | 24826 | 26121 | 26703 | 27523 | 32646 | 28665 | 28691 | 28278 | 28791 | 27130 | 29727 | 32774 | 28828 | 26510 | 29562 | 33127 | 27694 | 21363 | 34070 | 29170 | 26901 | 33240 | 31502 | 31138 | Rps5 |
| Rps6 | 46822 | 46336 | 44693 | 38326 | 33552 | 35590 | 39409 | 44711 | 41547 | 35791 | 41566 | 40456 | 42200 | 44446 | 46252 | 42238 | 40387 | 35365 | 38143 | 42774 | 37533 | 34968 | 43610 | 40693 | 39280 | 48881 | 45198 | 49293 | Rps6 |
| Rps6ka1 | 3959 | 4574 | 5063 | 4656 | 3908 | 3672 | 3429 | 1156 | 1505 | 1496 | 1537 | 1643 | 1669 | 1288 | 3516 | 5302 | 4671 | 4614 | 4383 | 2993 | 3689 | 1430 | 1665 | 1653 | 1462 | 1940 | 1479 | 1584 | Rps6ka1 |
| Rps6ka2 | 6111 | 7070 | 6543 | 7341 | 7340 | 6842 | 6739 | 2938 | 3687 | 3537 | 3840 | 4564 | 4148 | 3949 | 6099 | 7345 | 7227 | 7507 | 7294 | 5240 | 6821 | 4214 | 3226 | 3524 | 4088 | 3356 | 3970 | 4047 | Rps6ka2 |
| Rps6ka3 | 15160 | 11365 | 11172 | 10989 | 12692 | 13594 | 13800 | 11162 | 10305 | 9189 | 10668 | 10252 | 11499 | 10580 | 14876 | 13921 | 10974 | 11700 | 11252 | 12817 | 13337 | 10066 | 9963 | 9050 | 9842 | 8159 | 10884 | 10681 | Rps6ka3 |
| Rps6ka5 | 2533 | 2219 | 1972 | 3023 | 3240 | 3320 | 3807 | 2159 | 2181 | 1911 | 1761 | 1729 | 1543 | 1864 | 2685 | 2245 | 2217 | 3168 | 3015 | 3180 | 3749 | 2177 | 1952 | 1705 | 2035 | 1379 | 1547 | 1613 | Rps6ka5 |
| Rps6ka6 | 2 | 2 | 1 | 0 | 0 | 4 | 2 | 2 | 1 | 1 | 0 | 2 | 7 | 2 | 1 | 1 | 0 | 1 | 2 | 3 | 1 | 3 | 2 | 0 | 1 | 0 | 9 | 1 | Rps6ka6 |
| Rps6kb1 | 8530 | 6155 | 6907 | 7282 | 6532 | 7343 | 7987 | 8606 | 7782 | 7588 | 8124 | 8200 | 8053 | 9008 | 8132 | 7691 | 6624 | 7162 | 6543 | 7313 | 7209 | 8006 | 7425 | 7691 | 8041 | 7993 | 8230 | 8737 | Rps6kb1 |
| Rps6kb2 | 370 | 543 | 112 | 1036 | 646 | 135 | 405 | 442 | 681 | 667 | 503 | 328 | 877 | 934 | 204 | 296 | 269 | 321 | 1005 | 386 | 139 | 413 | 214 | 581 | 466 | 332 | 593 | 240 | Rps6kb2 |
| Rps6kc1 | 2871 | 3252 | 2968 | 3384 | 3475 | 3146 | 3143 | 3132 | 3227 | 3216 | 3351 | 3115 | 3009 | 3371 | 2981 | 2897 | 3124 | 3235 | 3402 | 3249 | 3356 | 3240 | 3118 | 3454 | 3479 | 3022 | 3337 | 3269 | Rps6kc1 |
| Rps6kl1 | 490 | 741 | 559 | 922 | 1305 | 901 | 920 | 573 | 734 | 796 | 818 | 514 | 522 | 473 | 644 | 681 | 866 | 1115 | 1127 | 931 | 982 | 962 | 766 | 589 | 848 | 471 | 476 | 387 | Rps6kl1 |
| Rps7 | 27639 | 28175 | 25979 | 27623 | 23800 | 24726 | 29188 | 28325 | 30105 | 26158 | 27250 | 25328 | 27877 | 29668 | 27662 | 27024 | 27817 | 23830 | 27343 | 33871 | 29238 | 21345 | 31719 | 28417 | 26153 | 30799 | 28145 | 32287 | Rps7 |
| Rps8 | 35234 | 42732 | 41361 | 39470 | 33606 | 36545 | 37386 | 33904 | 38418 | 36630 | 41189 | 37366 | 41489 | 47495 | 34393 | 36965 | 38418 | 36945 | 39016 | 43517 | 38433 | 25367 | 39977 | 43499 | 38888 | 47765 | 43262 | 48265 | Rps8 |
| Rps9 | 30620 | 39652 | 40652 | 36545 | 30757 | 30177 | 33907 | 27385 | 32754 | 35535 | 36904 | 38203 | 38270 | 40489 | 28317 | 33711 | 40248 | 34782 | 34436 | 35225 | 31841 | 23562 | 34725 | 41479 | 34871 | 45851 | 40778 | 42007 | Rps9 |
| Rpsa | 76582 | 92306 | 72742 | 66009 | 57309 | 63826 | 67251 | 67303 | 77825 | 60145 | 64614 | 70350 | 71754 | 73116 | 75798 | 80563 | 68082 | 63661 | 68170 | 79679 | 66243 | 50532 | 78109 | 68045 | 63378 | 84661 | 78025 | 80402 | Rpsa |
| Rptor | 3336 | 4072 | 4438 | 4435 | 4245 | 4241 | 3998 | 3886 | 4537 | 5117 | 5008 | 5793 | 4798 | 5031 | 3303 | 3780 | 4667 | 4253 | 4894 | 3252 | 4205 | 4749 | 4370 | 5201 | 5361 | 4773 | 4988 | 5350 | Rptor |
| Rpusd1 | 777 | 984 | 729 | 769 | 717 | 811 | 602 | 696 | 871 | 778 | 844 | 853 | 770 | 773 | 666 | 804 | 910 | 790 | 774 | 818 | 714 | 703 | 874 | 824 | 821 | 974 | 824 | 852 | Rpusd1 |
| Rpusd2 | 905 | 977 | 891 | 983 | 810 | 814 | 892 | 648 | 644 | 614 | 626 | 856 | 909 | 863 | 816 | 857 | 1021 | 828 | 949 | 771 | 912 | 648 | 568 | 792 | 708 | 686 | 925 | 911 | Rpusd2 |
| Rpusd3 | 403 | 395 | 464 | 357 | 376 | 404 | 339 | 387 | 472 | 424 | 419 | 571 | 569 | 479 | 339 | 343 | 374 | 305 | 467 | 402 | 360 | 384 | 436 | 534 | 551 | 675 | 566 | 627 | Rpusd3 |
| Rpusd4 | 937 | 1160 | 1087 | 913 | 887 | 759 | 897 | 1077 | 1094 | 1013 | 964 | 1199 | 1059 | 1060 | 936 | 935 | 998 | 726 | 898 | 858 | 860 | 954 | 1083 | 998 | 999 | 1225 | 1178 | 1184 | Rpusd4 |
| Rrad | 156 | 140 | 214 | 128 | 112 | 120 | 117 | 68 | 90 | 86 | 240 | 70 | 56 | 55 | 179 | 157 | 174 | 149 | 135 | 99 | 133 | 82 | 152 | 92 | 90 | 70 | 60 | 61 | Rrad |
| Rraga | 7471 | 7373 | 7235 | 6822 | 6402 | 6495 | 7483 | 7392 | 6567 | 6351 | 6072 | 6748 | 6645 | 7008 | 7949 | 6731 | 7130 | 6721 | 6717 | 7325 | 7070 | 7088 | 6829 | 7025 | 6508 | 7476 | 6979 | 7577 | Rraga |
| Rragb | 3319 | 2991 | 3086 | 3167 | 3562 | 3587 | 4089 | 2419 | 2154 | 2138 | 1713 | 2350 | 2056 | 2457 | 3679 | 2631 | 3191 | 3129 | 3603 | 4208 | 3871 | 2311 | 2048 | 1962 | 2150 | 2699 | 2427 | 2216 | Rragb |
| Rragc | 4924 | 4809 | 5069 | 4849 | 4311 | 4851 | 4640 | 4056 | 3817 | 4221 | 4322 | 4492 | 4136 | 4488 | 4716 | 5062 | 5074 | 4825 | 4584 | 3946 | 4654 | 4609 | 4123 | 3992 | 4247 | 4610 | 4186 | 4470 | Rragc |
| Rragd | 6569 | 5784 | 6867 | 6150 | 6983 | 7209 | 7021 | 6095 | 4697 | 5086 | 5367 | 6165 | 6827 | 6100 | 7114 | 5849 | 6489 | 5817 | 6884 | 7242 | 6744 | 6149 | 5381 | 5445 | 5692 | 5658 | 6650 | 6725 | Rragd |
| Rras | 851 | 887 | 822 | 622 | 517 | 568 | 470 | 346 | 489 | 405 | 505 | 373 | 403 | 382 | 662 | 906 | 648 | 664 | 506 | 435 | 489 | 426 | 498 | 369 | 331 | 473 | 417 | 464 | Rras |
| Rras2 | 4579 | 3534 | 3861 | 3678 | 3335 | 3613 | 4031 | 2589 | 2104 | 2008 | 2219 | 2006 | 1850 | 2142 | 4378 | 3879 | 3717 | 3685 | 3668 | 4045 | 4004 | 2242 | 2238 | 2003 | 2196 | 2579 | 2013 | 2257 | Rras2 |
| Rrbp1 | 14759 | 15729 | 21070 | 18081 | 14592 | 15844 | 12838 | 9436 | 12687 | 13410 | 19628 | 12135 | 14787 | 11830 | 13266 | 17837 | 19153 | 17650 | 14681 | 11253 | 13464 | 12343 | 14987 | 14896 | 11920 | 14001 | 12944 | 11871 | Rrbp1 |
| Rreb1 | 1548 | 1520 | 1900 | 1578 | 1730 | 1593 | 1368 | 1220 | 1880 | 1741 | 2519 | 1626 | 1728 | 1335 | 1449 | 1769 | 1658 | 1791 | 1655 | 1347 | 1540 | 1616 | 2205 | 1657 | 1371 | 1103 | 1383 | 1359 | Rreb1 |
| Rrh | 75 | 51 | 25 | 34 | 56 | 78 | 72 | 44 | 45 | 29 | 39 | 59 | 35 | 36 | 75 | 65 | 24 | 56 | 48 | 77 | 79 | 32 | 36 | 42 | 44 | 35 | 35 | 42 | Rrh |
| Rrm1 | 24810 | 21704 | 20388 | 15915 | 13922 | 16939 | 17581 | 21351 | 19146 | 15249 | 14558 | 23089 | 20016 | 23346 | 23379 | 19359 | 18256 | 14854 | 17183 | 19706 | 17045 | 13648 | 16949 | 18950 | 16717 | 26769 | 25542 | 25421 | Rrm1 |
| Rrm2 | 10593 | 8886 | 12462 | 7933 | 6652 | 8115 | 9443 | 12606 | 10995 | 10312 | 9649 | 21015 | 15680 | 23085 | 9228 | 7561 | 10684 | 6747 | 8997 | 8684 | 8189 | 9351 | 8652 | 16044 | 13257 | 25565 | 21632 | 23571 | Rrm2 |
| Rrm2b | 1931 | 1715 | 2038 | 2100 | 2023 | 2068 | 2252 | 1480 | 1292 | 1218 | 1326 | 1361 | 1295 | 1419 | 2083 | 1813 | 2071 | 2031 | 2064 | 2096 | 2053 | 1469 | 1302 | 1276 | 1380 | 1298 | 1368 | 1340 | Rrm2b |
| Rrn3 | 9028 | 9070 | 9187 | 9215 | 9550 | 9287 | 9980 | 10927 | 9049 | 9746 | 9108 | 10823 | 11116 | 11509 | 9539 | 8165 | 9288 | 8930 | 9834 | 9610 | 9788 | 10206 | 9013 | 10013 | 10966 | 10367 | 11581 | 12003 | Rrn3 |
| Rrnad1 | 2666 | 3184 | 3112 | 3261 | 2890 | 2611 | 3046 | 2925 | 3031 | 3259 | 3443 | 3536 | 3150 | 3691 | 2524 | 2687 | 3301 | 3024 | 2954 | 2606 | 2935 | 3389 | 2926 | 3621 | 3439 | 3235 | 3345 | 3296 | Rrnad1 |
| Rrp1 | 15779 | 18852 | 19514 | 18470 | 16694 | 16603 | 17362 | 12351 | 13074 | 13670 | 12950 | 14394 | 13702 | 14713 | 15423 | 15639 | 19568 | 17668 | 18642 | 17391 | 17325 | 12541 | 12571 | 15115 | 14783 | 16185 | 14487 | 14916 | Rrp1 |
| Rrp12 | 4651 | 5882 | 4151 | 4871 | 3585 | 4020 | 4034 | 2313 | 2653 | 2724 | 2679 | 2852 | 2877 | 2958 | 4199 | 4748 | 4356 | 4557 | 3904 | 3614 | 4077 | 2174 | 2397 | 2693 | 2720 | 3022 | 2918 | 3166 | Rrp12 |
| Rrp15 | 1855 | 1793 | 1986 | 1636 | 1255 | 1386 | 1502 | 1588 | 1725 | 1435 | 1631 | 1901 | 1703 | 2073 | 1788 | 1823 | 1663 | 1555 | 1333 | 1740 | 1356 | 1372 | 1649 | 1784 | 1747 | 2477 | 1916 | 1976 | Rrp15 |
| Rrp1b | 4349 | 4386 | 3951 | 4233 | 3382 | 3708 | 3551 | 2821 | 3302 | 3239 | 3521 | 3892 | 3633 | 3608 | 3782 | 4541 | 4018 | 4228 | 3711 | 3514 | 3763 | 2824 | 2950 | 3506 | 3356 | 3658 | 3901 | 3948 | Rrp1b |
| Rrp36 | 1528 | 1871 | 1806 | 1792 | 1332 | 1672 | 1596 | 1178 | 1473 | 1433 | 1520 | 1610 | 1490 | 1643 | 1398 | 1688 | 1826 | 1729 | 1615 | 1463 | 1531 | 1105 | 1387 | 1526 | 1388 | 1791 | 1569 | 1683 | Rrp36 |
| Rrp7a | 4696 | 6011 | 6399 | 5597 | 5117 | 5466 | 4740 | 3734 | 4537 | 4598 | 4562 | 5033 | 4685 | 4391 | 4681 | 5135 | 5901 | 5382 | 5894 | 4899 | 5238 | 4350 | 4724 | 4734 | 4606 | 5419 | 4926 | 4859 | Rrp7a |
| Rrp8 | 2008 | 1995 | 1942 | 1775 | 1516 | 1491 | 1505 | 2288 | 2077 | 1934 | 1930 | 2028 | 2003 | 1985 | 1941 | 1861 | 1725 | 1536 | 1449 | 1598 | 1432 | 1863 | 2145 | 1980 | 1859 | 2390 | 2166 | 2254 | Rrp8 |
| Rrp9 | 2992 | 4166 | 3359 | 3603 | 2474 | 2603 | 2643 | 1239 | 1548 | 1503 | 1466 | 1737 | 1508 | 1760 | 2788 | 3318 | 3573 | 3365 | 2986 | 2538 | 2891 | 1312 | 1489 | 1869 | 1544 | 2191 | 1836 | 1724 | Rrp9 |
| Rrs1 | 2775 | 2779 | 3918 | 3177 | 2548 | 2543 | 2573 | 2000 | 2029 | 2374 | 2280 | 2965 | 2650 | 2946 | 2474 | 2843 | 3480 | 2917 | 2759 | 2488 | 2434 | 1910 | 2035 | 2715 | 2458 | 3625 | 2783 | 2941 | Rrs1 |
| Rs1 | 3 | 1 | 0 | 0 | 9 | 4 | 4 | 7 | 5 | 3 | 7 | 3 | 11 | 0 | 0 | 3 | 0 | 1 | 1 | 0 | 1 | 5 | 6 | 0 | 2 | 3 | 7 | 0 | Rs1 |
| Rsad1 | 662 | 590 | 750 | 614 | 714 | 599 | 591 | 614 | 696 | 782 | 785 | 791 | 751 | 778 | 601 | 640 | 746 | 599 | 683 | 663 | 651 | 770 | 704 | 796 | 708 | 787 | 672 | 827 | Rsad1 |
| Rsad2 | 47 | 50 | 31 | 26 | 29 | 27 | 18 | 56 | 55 | 54 | 92 | 52 | 73 | 74 | 58 | 74 | 41 | 32 | 26 | 60 | 38 | 52 | 89 | 47 | 23 | 68 | 62 | 80 | Rsad2 |
| Rsbn1 | 9491 | 7520 | 7293 | 9118 | 9074 | 9852 | 10296 | 9561 | 7135 | 7089 | 6708 | 7251 | 6643 | 7385 | 10010 | 8312 | 8127 | 9234 | 9206 | 10269 | 10316 | 8316 | 6750 | 6567 | 7960 | 5817 | 7345 | 7513 | Rsbn1 |

| | | | | | | | | | | | | | | | | | | | | | | | | | | | | | |
|---|---|---|---|---|---|---|---|---|---|---|---|---|---|---|---|---|---|---|---|---|---|---|---|---|---|---|---|---|---|
| Rsbn1l | 3925 | 2844 | 3775 | 3980 | 4078 | 4036 | 4298 | 3729 | 2978 | 3429 | 3344 | 3737 | 3570 | 3698 | 3915 | 3234 | 3672 | 3927 | 3961 | 4093 | 4307 | 3456 | 2979 | 3349 | 3619 | 3297 | 3790 | 3827 | Rsbn1l |
| Rsf1 | 7251 | 6037 | 5821 | 6610 | 7571 | 8689 | 8472 | 11986 | 10064 | 9410 | 9615 | 9941 | 9639 | 10784 | 7615 | 6592 | 6167 | 6444 | 7376 | 9022 | 8476 | 9078 | 9616 | 8652 | 10517 | 8099 | 10161 | 9486 | Rsf1 |
| Rskr | 159 | 162 | 151 | 141 | 130 | 130 | 153 | 71 | 58 | 59 | 67 | 63 | 75 | 72 | 125 | 103 | 177 | 168 | 178 | 161 | 155 | 65 | 63 | 56 | 69 | 51 | 79 | 65 | Rskr |
| Rsl1 | 713 | 539 | 627 | 512 | 685 | 577 | 678 | 767 | 681 | 558 | 653 | 544 | 510 | 548 | 691 | 536 | 598 | 572 | 604 | 669 | 581 | 648 | 747 | 547 | 599 | 639 | 560 | 568 | Rsl1 |
| Rsl1d1 | 11219 | 9386 | 11043 | 8659 | 8242 | 9268 | 9947 | 11088 | 9579 | 9098 | 8808 | 11067 | 10584 | 10869 | 11418 | 10125 | 9707 | 8440 | 8935 | 11906 | 8888 | 8809 | 9586 | 9520 | 9610 | 12656 | 11441 | 11515 | Rsl1d1 |
| Rsl24d1 | 4099 | 3795 | 4313 | 3641 | 2734 | 2837 | 3399 | 2958 | 2683 | 2724 | 2636 | 3177 | 2914 | 3156 | 3870 | 3653 | 4018 | 3416 | 3156 | 3403 | 3088 | 2550 | 2733 | 3103 | 2879 | 3840 | 3029 | 3519 | Rsl24d1 |
| Rslcan18 | 32 | 17 | 24 | 18 | 19 | 21 | 25 | 27 | 25 | 34 | 22 | 30 | 31 | 19 | 21 | 31 | 23 | 11 | 30 | 16 | 21 | 12 | 21 | 13 | 26 | 20 | 48 | 23 | Rslcan18 |
| Rsph1 | 128 | 229 | 162 | 240 | 229 | 232 | 196 | 133 | 128 | 131 | 139 | 153 | 192 | 118 | 240 | 219 | 242 | 196 | 171 | 224 | 181 | 135 | 224 | 112 | 111 | 122 | 121 | 131 | Rsph1 |
| Rsph10b | 657 | 738 | 458 | 921 | 791 | 736 | 816 | 364 | 367 | 441 | 418 | 294 | 382 | 254 | 781 | 682 | 680 | 1007 | 774 | 732 | 853 | 461 | 414 | 266 | 336 | 186 | 230 | 193 | Rsph10b |
| Rsph14 | 20 | 28 | 11 | 21 | 14 | 25 | 6 | 15 | 21 | 12 | 10 | 13 | 12 | 12 | 17 | 30 | 8 | 13 | 12 | 25 | 14 | 18 | 23 | 11 | 12 | 12 | 1 | 7 | Rsph14 |
| Rsph3a | 1181 | 1302 | 1318 | 1297 | 1105 | 1077 | 1341 | 1174 | 988 | 1056 | 999 | 1099 | 1132 | 1345 | 1350 | 1079 | 1304 | 1179 | 1274 | 1168 | 1248 | 1168 | 1102 | 1147 | 1190 | 1135 | 1097 | 1137 | Rsph3a |
| Rsph3b | 813 | 829 | 885 | 819 | 691 | 757 | 767 | 670 | 629 | 647 | 603 | 651 | 567 | 643 | 803 | 693 | 812 | 728 | 859 | 727 | 780 | 725 | 725 | 647 | 609 | 707 | 703 | 694 | Rsph3b |
| Rsph4a | 262 | 304 | 211 | 227 | 319 | 282 | 315 | 151 | 182 | 156 | 123 | 220 | 237 | 144 | 303 | 327 | 236 | 326 | 291 | 337 | 323 | 237 | 382 | 151 | 110 | 103 | 91 | 116 | Rsph4a |
| Rsph6a | 2 | 13 | 12 | 16 | 4 | 15 | 7 | 2 | 1 | 0 | 0 | 1 | 1 | 2 | 8 | 9 | 6 | 13 | 4 | 2 | 8 | 2 | 0 | 1 | 2 | 3 | 3 | 1 | Rsph6a |
| Rsph9 | 618 | 770 | 494 | 614 | 711 | 619 | 640 | 479 | 554 | 416 | 455 | 413 | 481 | 392 | 696 | 672 | 570 | 651 | 704 | 819 | 671 | 471 | 581 | 443 | 439 | 371 | 467 | 419 | Rsph9 |
| Rspo1 | 820 | 1016 | 1320 | 765 | 402 | 554 | 469 | 634 | 545 | 702 | 548 | 1263 | 1254 | 1093 | 793 | 976 | 877 | 646 | 478 | 482 | 424 | 433 | 846 | 959 | 622 | 1748 | 1332 | 1659 | Rspo1 |
| Rspo2 | 1247 | 749 | 1436 | 788 | 657 | 637 | 906 | 1707 | 966 | 1128 | 1293 | 1882 | 1670 | 1949 | 895 | 1141 | 934 | 629 | 841 | 943 | 769 | 957 | 1675 | 1580 | 1177 | 3165 | 1978 | 2541 | Rspo2 |
| Rspo3 | 1562 | 970 | 1720 | 965 | 1078 | 1190 | 1368 | 2141 | 1636 | 1659 | 1568 | 2732 | 1989 | 2800 | 1571 | 1179 | 1307 | 866 | 1269 | 1695 | 1267 | 1900 | 2585 | 2146 | 2131 | 3603 | 2659 | 3420 | Rspo3 |
| Rspo4 | 21 | 29 | 11 | 18 | 21 | 26 | 11 | 12 | 16 | 21 | 16 | 2 | 17 | 4 | 26 | 36 | 14 | 8 | 16 | 12 | 11 | 10 | 25 | 10 | 5 | 3 | 1 | 7 | Rspo4 |
| Rspry1 | 6373 | 5511 | 5479 | 5484 | 5004 | 5425 | 5747 | 5464 | 4519 | 4249 | 4299 | 4552 | 4493 | 4808 | 6551 | 5558 | 5533 | 5382 | 5223 | 6190 | 5631 | 4720 | 4625 | 4350 | 4423 | 4828 | 4997 | 4723 | Rspry1 |
| Rsrc1 | 4870 | 4380 | 5164 | 3794 | 3826 | 4107 | 3967 | 5688 | 4490 | 4500 | 4300 | 5131 | 5050 | 5581 | 5004 | 4060 | 4244 | 3812 | 3743 | 4447 | 4073 | 4719 | 4815 | 4931 | 4752 | 5317 | 5875 | 5685 | Rsrc1 |
| Rsrc2 | 8433 | 7077 | 9963 | 9215 | 9291 | 8960 | 9544 | 16552 | 13303 | 14491 | 14559 | 14928 | 14705 | 16322 | 9655 | 7904 | 9635 | 8715 | 8956 | 10947 | 9298 | 14681 | 13570 | 15044 | 14923 | 15512 | 15931 | 15918 | Rsrc2 |
| Rsrp1 | 30910 | 24593 | 28735 | 35133 | 33490 | 32184 | 35119 | 30382 | 27811 | 32414 | 33267 | 31460 | 30670 | 32896 | 31275 | 26066 | 31081 | 33826 | 31392 | 30299 | 34520 | 36054 | 28657 | 32539 | 33920 | 31079 | 29530 | 30505 | Rsrp1 |
| Rsu1 | 3187 | 2856 | 2682 | 2043 | 1874 | 1988 | 1849 | 2380 | 2585 | 1929 | 2501 | 1808 | 2515 | 1993 | 3196 | 3041 | 2179 | 2178 | 1787 | 2238 | 1793 | 2194 | 2957 | 1942 | 1587 | 2359 | 2064 | 2117 | Rsu1 |
| Rtbdn | 108 | 136 | 169 | 215 | 287 | 246 | 276 | 109 | 108 | 169 | 107 | 126 | 115 | 82 | 160 | 107 | 203 | 272 | 270 | 230 | 284 | 189 | 80 | 117 | 140 | 83 | 126 | 90 | Rtbdn |
| Rtca | 4693 | 4944 | 4548 | 4485 | 3933 | 4242 | 4832 | 3676 | 3227 | 3277 | 3012 | 3686 | 3405 | 3931 | 4975 | 4364 | 4339 | 4199 | 4191 | 4604 | 4379 | 3347 | 3103 | 3416 | 3233 | 4042 | 3858 | 4132 | Rtca |
| Rtcb | 15755 | 16825 | 15986 | 14546 | 13044 | 14286 | 13266 | 12511 | 12525 | 11318 | 11674 | 12488 | 12191 | 11990 | 16105 | 15747 | 14732 | 13885 | 14114 | 15612 | 13174 | 10410 | 12508 | 12028 | 11714 | 14383 | 13675 | 12803 | Rtcb |
| Rtel1 | 3963 | 4282 | 3569 | 3420 | 3085 | 3527 | 3283 | 4593 | 4837 | 4376 | 4394 | 5447 | 5074 | 5079 | 3898 | 3777 | 3423 | 3055 | 3243 | 3523 | 3299 | 4380 | 4330 | 4537 | 4640 | 4820 | 5573 | 5385 | Rtel1 |
| Rtf1 | 14492 | 13303 | 15446 | 15653 | 14135 | 14642 | 15346 | 10386 | 9799 | 11219 | 10133 | 11957 | 10679 | 12005 | 13901 | 13537 | 14676 | 15113 | 14504 | 14327 | 14844 | 11175 | 9277 | 11551 | 11692 | 12387 | 11670 | 11765 | Rtf1 |
| Rtf2 | 5657 | 7329 | 5910 | 5934 | 4925 | 5251 | 5508 | 4239 | 4727 | 4414 | 4158 | 4901 | 4511 | 4706 | 5726 | 5808 | 5891 | 5748 | 5806 | 5843 | 5371 | 4162 | 4500 | 4689 | 4819 | 5271 | 5097 | 5088 | Rtf2 |
| Rtkn | 1164 | 1472 | 1650 | 1032 | 891 | 1020 | 664 | 1630 | 2371 | 2157 | 2696 | 2323 | 2768 | 1967 | 1087 | 1408 | 1417 | 935 | 1001 | 812 | 744 | 2084 | 2233 | 2134 | 1652 | 2358 | 2568 | 2238 | Rtkn |
| Rtkn2 | 1418 | 1446 | 1712 | 1349 | 954 | 1201 | 1338 | 1006 | 1010 | 1097 | 871 | 1840 | 1284 | 1905 | 1324 | 1209 | 1511 | 1155 | 1265 | 1191 | 1265 | 919 | 877 | 1346 | 1336 | 2010 | 1878 | 1811 | Rtkn2 |
| Rtl10 | 172 | 224 | 277 | 194 | 199 | 174 | 144 | 159 | 211 | 230 | 229 | 241 | 206 | 187 | 206 | 261 | 202 | 176 | 161 | 175 | 192 | 266 | 185 | 218 | 207 | 241 | 198 | 223 | Rtl10 |
| Rtl3 | 389 | 339 | 225 | 269 | 151 | 246 | 64 | 920 | 1444 | 1239 | 3322 | 539 | 638 | 576 | 385 | 460 | 309 | 377 | 72 | 69 | 48 | 1285 | 2402 | 1191 | 810 | 773 | 803 | 433 | Rtl3 |
| Rtl4 | 24 | 16 | 24 | 11 | 45 | 32 | 28 | 64 | 72 | 60 | 108 | 38 | 40 | 42 | 22 | 31 | 23 | 41 | 34 | 23 | 35 | 94 | 74 | 34 | 37 | 21 | 41 | 20 | Rtl4 |
| Rtl5 | 5019 | 5999 | 5093 | 6891 | 7263 | 7040 | 8136 | 6439 | 6054 | 6998 | 5762 | 8048 | 5532 | 8548 | 5090 | 4404 | 6960 | 6628 | 7874 | 7454 | 7977 | 7790 | 5352 | 7715 | 8650 | 8534 | 7631 | 7695 | Rtl5 |
| Rtl6 | 2806 | 3234 | 3623 | 4167 | 4643 | 4291 | 4005 | 4406 | 5572 | 6536 | 5649 | 6387 | 5525 | 5793 | 2901 | 2999 | 4215 | 4031 | 5075 | 3313 | 4195 | 6662 | 4989 | 6589 | 6817 | 6086 | 5617 | 5524 | Rtl6 |
| Rtl8a | 0 | 1 | 0 | 0 | 0 | 0 | 2 | 2 | 0 | 0 | 1 | 0 | 0 | 0 | 0 | 2 | 1 | 0 | 0 | 0 | 0 | 0 | 3 | 0 | 1 | 0 | 0 | 0 | Rtl8a |
| Rtl8b | 6 | 1 | 0 | 2 | 2 | 3 | 1 | 4 | 2 | 2 | 4 | 1 | 1 | 0 | 0 | 1 | 1 | 3 | 1 | 0 | 4 | 0 | 6 | 2 | 3 | 1 | 5 | 0 | Rtl8b |
| Rtl9 | 175 | 166 | 83 | 185 | 120 | 141 | 166 | 117 | 193 | 156 | 209 | 102 | 126 | 106 | 173 | 146 | 98 | 150 | 168 | 141 | 150 | 140 | 173 | 109 | 119 | 88 | 128 | 70 | Rtl9 |
| Rtn1 | 8543 | 10000 | 9009 | 17663 | 28079 | 20039 | 23373 | 22674 | 22352 | 30624 | 24011 | 21403 | 20882 | 23274 | 12262 | 8816 | 14776 | 19876 | 22040 | 19653 | 24908 | 36270 | 21892 | 25396 | 31032 | 16977 | 16747 | 19939 | Rtn1 |
| Rtn2 | 2481 | 3669 | 3699 | 4255 | 4320 | 3698 | 3831 | 2961 | 3188 | 3912 | 3040 | 3801 | 2959 | 3362 | 2633 | 2626 | 4146 | 3996 | 4826 | 3322 | 4191 | 4288 | 3010 | 3679 | 4312 | 3504 | 3447 | 3211 | Rtn2 |
| Rtn3 | 38738 | 38852 | 39904 | 44850 | 48201 | 43468 | 51561 | 47887 | 37388 | 41535 | 37439 | 44778 | 43095 | 48103 | 44978 | 32909 | 42789 | 44636 | 46014 | 48825 | 48826 | 51521 | 35819 | 43285 | 47361 | 41539 | 47109 | 47680 | Rtn3 |
| Rtn4 | 19954 | 16028 | 22528 | 22437 | 27747 | 23974 | 25999 | 39796 | 33765 | 38265 | 36906 | 38282 | 34607 | 40669 | 21592 | 17728 | 22265 | 22742 | 24476 | 21145 | 26118 | 45761 | 33456 | 38227 | 42186 | 36083 | 36660 | 37855 | Rtn4 |
| Rtn4ip1 | 929 | 919 | 1033 | 920 | 917 | 869 | 934 | 985 | 1103 | 956 | 976 | 1168 | 1038 | 1064 | 1000 | 917 | 944 | 916 | 959 | 894 | 926 | 946 | 977 | 996 | 1127 | 1148 | 1109 | 1081 | Rtn4ip1 |
| Rtn4rl1 | 496 | 523 | 568 | 572 | 867 | 626 | 529 | 479 | 549 | 597 | 578 | 412 | 519 | 433 | 430 | 601 | 654 | 672 | 754 | 504 | 641 | 777 | 603 | 550 | 689 | 172 | 397 | 450 | Rtn4rl1 |
| Rtn4rl2 | 454 | 691 | 646 | 778 | 715 | 615 | 477 | 299 | 337 | 339 | 369 | 289 | 270 | 243 | 442 | 529 | 765 | 694 | 839 | 596 | 561 | 326 | 287 | 413 | 414 | 334 | 222 | 198 | Rtn4rl2 |
| Rtp1 | 36 | 16 | 8 | 25 | 8 | 12 | 8 | 143 | 75 | 182 | 77 | 28 | 50 | 29 | 22 | 22 | 10 | 1 | 6 | 5 | 9 | 3 | 138 | 212 | 15 | 155 | 2 | 23 | Rtp1 |
| Rtp2 | 24 | 12 | 0 | 31 | 2 | 0 | 2 | 57 | 60 | 113 | 59 | 21 | 31 | 16 | 11 | 16 | 3 | 1 | 0 | 0 | 4 | 2 | 62 | 156 | 3 | 70 | 2 | 2 | Rtp2 |
| Rtp3 | 10 | 13 | 5 | 2 | 4 | 10 | 1 | 1 | 7 | 3 | 10 | 7 | 0 | 3 | 12 | 5 | 1 | 6 | 7 | 2 | 4 | 5 | 8 | 7 | 2 | 0 | 7 | 12 | Rtp3 |
| Rtp4 | 43 | 27 | 32 | 46 | 58 | 36 | 69 | 107 | 83 | 143 | 91 | 92 | 96 | 81 | 46 | 52 | 44 | 35 | 58 | 98 | 88 | 127 | 104 | 128 | 81 | 146 | 169 | 126 | Rtp4 |
| Rtraf | 14657 | 13694 | 11660 | 11824 | 10460 | 11798 | 13867 | 11125 | 9299 | 8528 | 8889 | 9865 | 9901 | 10777 | 15231 | 12337 | 12214 | 11429 | 11865 | 14281 | 13355 | 8519 | 9443 | 9892 | 8877 | 11238 | 10400 | 11618 | Rtraf |
| Rttn | 2817 | 2600 | 1420 | 1776 | 1717 | 2047 | 2075 | 2698 | 2483 | 2011 | 2002 | 2323 | 2351 | 2383 | 2683 | 2452 | 1610 | 1612 | 1784 | 2502 | 1944 | 1763 | 2302 | 1897 | 1831 | 1836 | 2557 | 2204 | Rttn |
| Rubcn | 5678 | 6766 | 5853 | 7387 | 7435 | 6863 | 7020 | 3968 | 4413 | 4740 | 4685 | 4569 | 4059 | 4096 | 5822 | 6087 | 6888 | 7579 | 7564 | 6198 | 7229 | 5192 | 4236 | 4414 | 4945 | 3630 | 4071 | 3875 | Rubcn |
| Rubcnl | 72 | 27 | 34 | 26 | 41 | 74 | 34 | 42 | 43 | 46 | 19 | 30 | 39 | 19 | 44 | 51 | 50 | 40 | 49 | 36 | 34 | 28 | 54 | 34 | 27 | 41 | 41 | 32 | Rubcnl |
| Rufy1 | 1570 | 1576 | 1649 | 1442 | 1718 | 1505 | 1658 | 1997 | 2029 | 2234 | 2004 | 2318 | 2005 | 2376 | 1548 | 1378 | 1836 | 1511 | 1822 | 1718 | 1657 | 2030 | 1885 | 2217 | 2188 | 2181 | 2299 | 2377 | Rufy1 |
| Rufy2 | 4706 | 4104 | 3569 | 4581 | 5224 | 4935 | 5316 | 5197 | 4579 | 4482 | 4446 | 4190 | 4058 | 4515 | 4891 | 4141 | 4122 | 4602 | 4709 | 5114 | 5348 | 5185 | 4435 | 4267 | 4926 | 3284 | 3801 | 4132 | Rufy2 |
| Rufy3 | 16678 | 18196 | 17222 | 23990 | 30603 | 25257 | 30249 | 28694 | 22270 | 26490 | 22008 | 24909 | 23159 | 27212 | 21343 | 15825 | 22013 | 25003 | 27300 | 27314 | 30181 | 32527 | 21885 | 24755 | 29315 | 18688 | 24116 | 25661 | Rufy3 |
| Rufy4 | 55 | 102 | 102 | 89 | 70 | 77 | 59 | 117 | 74 | 86 | 78 | 92 | 92 | 96 | 108 | 104 | 80 | 100 | 79 | 78 | 67 | 89 | 74 | 91 | 87 | 53 | 109 | 62 | Rufy4 |
| Rundc1 | 2778 | 3587 | 3184 | 3508 | 2918 | 3146 | 2923 | 1926 | 1960 | 2105 | 2004 | 2142 | 2058 | 2121 | 2715 | 3111 | 3260 | 3316 | 3298 | 2797 | 3091 | 2244 | 1856 | 2066 | 2216 | 2214 | 2080 | 2123 | Rundc1 |
| Rundc3a | 5027 | 8247 | 6864 | 10238 | 10089 | 8674 | 9013 | 6459 | 6560 | 8211 | 6140 | 7273 | 6244 | 7292 | 6438 | 5306 | 9518 | 9552 | 10873 | 8659 | 10164 | 8061 | 6060 | 7808 | 8694 | 6390 | 6686 | 6444 | Rundc3a |
| Rundc3b | 2143 | 1779 | 2099 | 2529 | 3079 | 2626 | 3098 | 3333 | 2794 | 2980 | 2711 | 2612 | 2532 | 2726 | 2330 | 1738 | 2318 | 2441 | 2759 | 2809 | 2918 | 3928 | 2664 | 2772 | 3348 | 2143 | 2323 | 2497 | Rundc3b |
| Runx1 | 355 | 198 | 180 | 196 | 226 | 126 | 42 | 642 | 802 | 875 | 926 | 283 | 440 | 254 | 265 | 417 | 146 | 272 | 64 | 56 | 69 | 700 | 731 | 709 | 393 | 707 | 313 | 279 | Runx1 |
| Runx1t1 | 3024 | 2745 | 4156 | 6028 | 8651 | 6666 | 6614 | 7929 | 7196 | 10971 | 10295 | 8063 | 8471 | 8518 | 3943 | 3147 | 5771 | 6183 | 7275 | 5466 | 7178 | 12699 | 7950 | 9408 | 11606 | 5505 | 7626 | 8053 | Runx1t1 |
| Runx2 | 624 | 345 | 359 | 322 | 143 | 436 | 91 | 1199 | 1303 | 1590 | 1661 | 1017 | 1335 | 925 | 402 | 602 | 374 | 485 | 132 | 52 | 64 | 1340 | 1218 | 1664 | 899 | 1471 | 872 | 964 | Runx2 |
| Runx3 | 187 | 77 | 49 | 21 | 48 | 30 | 7 | 114 | 165 | 129 | 180 | 26 | 110 | 20 | 69 | 139 | 35 | 41 | 9 | 17 | 10 | 92 | 133 | 82 | 46 | 42 | 27 | 20 | Runx3 |
| Rusc1 | 1922 | 2305 | 2055 | 2868 | 3278 | 2602 | 3125 | 1730 | 1805 | 2300 | 2037 | 1742 | 1691 | 2021 | 1866 | 1598 | 2748 | 2821 | 2883 | 2314 | 3382 | 2943 | 1569 | 2116 | 2463 | 1424 | 1610 | 1952 | Rusc1 |
| Rusc2 | 1993 | 2671 | 3108 | 3964 | 6553 | 4733 | 3854 | 5991 | 7302 | 9932 | 8465 | 8033 | 6893 | 6634 | 2164 | 2270 | 3911 | 4431 | 6096 | 3894 | 4442 | 11364 | 6836 | 8992 | 11090 | 6889 | 6175 | 6062 | Rusc2 |
| Rusf1 | 1933 | 2182 | 2409 | 2165 | 1935 | 1809 | 1713 | 1366 | 1643 | 1822 | 1815 | 1699 | 1621 | 1694 | 1730 | 2153 | 2158 | 2093 | 1892 | 1660 | 1853 | 1706 | 1726 | 1774 | 1537 | 1765 | 1553 | 1644 | Rusf1 |
| Ruvbl1 | 4236 | 4507 | 4792 | 3760 | 3256 | 3272 | 3417 | 4078 | 4010 | 3824 | 3849 | 4499 | 4351 | 4889 | 3787 | 4160 | 4431 | 3470 | 3665 | 3827 | 3473 | 3314 | 4082 | 4624 | 4365 | 5863 | 5025 | 5196 | Ruvbl1 |
| Ruvbl2 | 5687 | 7301 | 6737 | 5821 | 4680 | 5084 | 4814 | 4352 | 4954 | 5444 | 4417 | 6437 | 5368 | 5774 | 5159 | 5651 | 6530 | 5286 | 5839 | 5417 | 5146 | 4199 | 4904 | 5681 | 5288 | 7227 | 6009 | 6272 | Ruvbl2 |
| Rwdd1 | 3825 | 3214 | 3218 | 3064 | 2871 | 3031 | 3409 | 3499 | 3143 | 2952 | 3008 | 3203 | 3244 | 3289 | 3847 | 3424 | 3070 | 2718 | 2913 | 3431 | 3164 | 2603 | 3326 | 3098 | 3039 | 3687 | 3287 | 3415 | Rwdd1 |
| Rwdd2a | 722 | 807 | 803 | 725 | 673 | 660 | 771 | 118 | 132 | 111 | 102 | 59 | 103 | 94 | 882 | 775 | 746 | 743 | 696 | 844 | 819 | 110 | 133 | 127 | 114 | 123 | 89 | 103 | Rwdd2a |
| Rwdd2b | 234 | 232 | 205 | 189 | 245 | 187 | 197 | 815 | 716 | 676 | 709 | 735 | 751 | 733 | 198 | 180 | 209 | 162 | 219 | 220 | 188 | 693 | 717 | 686 | 656 | 797 | 751 | 779 | Rwdd2b |
| Rwdd3 | 443 | 320 | 382 | 382 | 489 | 456 | 591 | 640 | 733 | 554 | 573 | 608 | 445 | 724 | 434 | 321 | 402 | 361 | 552 | 546 | 503 | 448 | 551 | 727 | 761 | 719 | 429 | 508 | Rwdd3 |
| Rwdd4a | 7647 | 6713 | 6928 | 7541 | 5510 | 5959 | 6540 | 3687 | 3247 | 3144 | 3164 | 3822 | 3567 | 3815 | 7603 | 6822 | 7274 | 6824 | 5884 | 6645 | 6235 | 3314 | 3167 | 3662 | 3248 | 4223 | 3711 | 3952 | Rwdd4a |
| Rxfp1 | 81 | 8 | 1 | 11 | 26 | 21 | 26 | 164 | 120 | 40 | 72 | 25 | 19 | 16 | 13 | 12 | 6 | 11 | 15 | 26 | 18 | 33 | 230 | 20 | 20 | 8 | 5 | 13 | Rxfp1 |
| Rxfp2 | 36 | 26 | 27 | 46 | 41 | 48 | 35 | 37 | 32 | 38 | 38 | 24 | 26 | 22 | 46 | 34 | 42 | 26 | 75 | 75 | 68 | 38 | 63 | 26 | 45 | 29 | 38 | 21 | Rxfp2 |
| Rxfp3 | 97 | 66 | 137 | 64 | 110 | 91 | 41 | 221 | 274 | 235 | 289 | 250 | 364 | 219 | 88 | 104 | 63 | 102 | 56 | 27 | 43 | 226 | 341 | 208 | 205 | 235 | 232 | 260 | Rxfp3 |

| | | | | | | | | | | | | | | | | | | | | | | | | | | | | | |
|---|---|---|---|---|---|---|---|---|---|---|---|---|---|---|---|---|---|---|---|---|---|---|---|---|---|---|---|---|---|
| Rxfp4 | 0 | 0 | 0 | 0 | 1 | 0 | 0 | 0 | 0 | 0 | 0 | 0 | 0 | 0 | 0 | 0 | 0 | 0 | 0 | 0 | 0 | 0 | 0 | 0 | 0 | 0 | 0 | 0 | Rxfp4 |
| Rxra | 3746 | 4170 | 5477 | 4042 | 3834 | 3801 | 2816 | 3378 | 4790 | 4347 | 5025 | 4738 | 5065 | 3874 | 3398 | 4352 | 4706 | 3772 | 3779 | 2751 | 2933 | 3863 | 4836 | 4657 | 3385 | 5359 | 4430 | 4459 | Rxra |
| Rxrb | 3286 | 4208 | 4149 | 3999 | 3698 | 3658 | 3403 | 4297 | 4993 | 4862 | 4887 | 5047 | 4764 | 4719 | 3038 | 3736 | 4397 | 3828 | 4297 | 3664 | 3588 | 4558 | 4787 | 5235 | 5293 | 5478 | 5139 | 5008 | Rxrb |
| Rxrg | 120 | 148 | 121 | 147 | 229 | 165 | 172 | 233 | 393 | 234 | 495 | 119 | 147 | 112 | 115 | 136 | 119 | 198 | 221 | 172 | 203 | 331 | 401 | 172 | 251 | 206 | 119 | 114 | Rxrg |
| Rxylt1 | 2832 | 2240 | 2322 | 2410 | 2360 | 2257 | 2668 | 1917 | 1692 | 1478 | 1625 | 1522 | 1643 | 1621 | 2864 | 2289 | 2332 | 2103 | 2268 | 2618 | 2545 | 1753 | 1729 | 1529 | 1420 | 1692 | 1495 | 1676 | Rxylt1 |
| Rybp | 3488 | 2964 | 3355 | 4367 | 4031 | 4041 | 4114 | 4009 | 3680 | 4028 | 4328 | 3481 | 3597 | 3727 | 3196 | 3689 | 3203 | 4315 | 3919 | 3430 | 4139 | 4027 | 3700 | 4155 | 4401 | 3849 | 3503 | 3673 | Rybp |
| Ryk | 8999 | 7778 | 8157 | 6969 | 6708 | 7109 | 7414 | 6687 | 6665 | 5837 | 6280 | 6674 | 6583 | 6416 | 8536 | 7911 | 7722 | 7228 | 7356 | 7256 | 6983 | 5584 | 6754 | 6369 | 5949 | 7742 | 6823 | 7064 | Ryk |
| Ryr1 | 1241 | 1297 | 1019 | 1501 | 1496 | 1297 | 1282 | 971 | 1452 | 1149 | 3742 | 892 | 863 | 907 | 990 | 1668 | 1199 | 1444 | 1416 | 1056 | 1275 | 1260 | 2260 | 950 | 1169 | 516 | 788 | 663 | Ryr1 |
| Ryr2 | 600 | 602 | 257 | 809 | 1570 | 1413 | 1307 | 1621 | 1567 | 1433 | 1390 | 647 | 762 | 759 | 638 | 471 | 506 | 1018 | 1051 | 1380 | 1508 | 1639 | 1245 | 766 | 1367 | 397 | 659 | 494 | Ryr2 |
| Ryr3 | 886 | 989 | 499 | 724 | 1316 | 1232 | 1212 | 1589 | 1590 | 1096 | 1425 | 1048 | 1117 | 1056 | 1265 | 884 | 588 | 969 | 867 | 1349 | 1427 | 1400 | 1452 | 711 | 1460 | 497 | 1307 | 951 | Ryr3 |
| S100a1 | 144 | 116 | 183 | 121 | 129 | 117 | 103 | 132 | 163 | 170 | 176 | 104 | 134 | 136 | 113 | 117 | 97 | 150 | 160 | 104 | 135 | 102 | 189 | 121 | 134 | 103 | 106 | 133 | S100a1 |
| S100a10 | 3384 | 3044 | 3142 | 1846 | 1972 | 2304 | 1203 | 3388 | 4361 | 3148 | 4498 | 2568 | 4098 | 2948 | 3291 | 3632 | 2290 | 2500 | 1414 | 1512 | 1398 | 2690 | 5128 | 3331 | 2605 | 3130 | 3303 | 3636 | S100a10 |
| S100a11 | 3710 | 3658 | 3185 | 2012 | 1177 | 2271 | 1091 | 3125 | 4191 | 2924 | 4476 | 2661 | 4107 | 3163 | 3253 | 3545 | 2041 | 2316 | 1285 | 1295 | 1172 | 2254 | 4872 | 3482 | 2487 | 3254 | 3201 | 3687 | S100a11 |
| S100a13 | 181 | 147 | 164 | 144 | 128 | 112 | 115 | 210 | 191 | 175 | 217 | 105 | 208 | 182 | 158 | 160 | 129 | 160 | 96 | 116 | 116 | 142 | 208 | 179 | 149 | 172 | 153 | 173 | S100a13 |
| S100a14 | 33 | 3 | 1 | 4 | 7 | 14 | 3 | 88 | 178 | 131 | 117 | 12 | 24 | 10 | 15 | 5 | 0 | 21 | 3 | 0 | 0 | 5 | 257 | 87 | 16 | 8 | 6 | 27 | S100a14 |
| S100a16 | 439 | 457 | 428 | 333 | 313 | 362 | 299 | 723 | 961 | 740 | 879 | 744 | 921 | 741 | 397 | 449 | 426 | 277 | 278 | 397 | 335 | 680 | 1040 | 838 | 791 | 780 | 858 | 794 | S100a16 |
| S100a2 | 0 | 2 | 1 | 0 | 0 | 0 | 0 | 4 | 1 | 7 | 10 | 2 | 0 | 0 | 0 | 0 | 0 | 0 | 0 | 0 | 0 | 3 | 1 | 0 | 1 | 3 | 0 | 0 | S100a2 |
| S100a3 | 0 | 0 | 0 | 0 | 0 | 0 | 0 | 0 | 0 | 5 | 5 | 0 | 0 | 0 | 0 | 2 | 0 | 0 | 0 | 0 | 0 | 3 | 2 | 0 | 0 | 0 | 0 | 0 | S100a3 |
| S100a4 | 0 | 0 | 0 | 0 | 0 | 0 | 0 | 4 | 0 | 0 | 0 | 0 | 0 | 0 | 0 | 0 | 0 | 0 | 0 | 0 | 0 | 0 | 0 | 5 | 3 | 0 | 0 | 0 | S100a4 |
| S100a5 | 2 | 3 | 0 | 2 | 0 | 0 | 0 | 7 | 7 | 1 | 7 | 0 | 2 | 0 | 4 | 0 | 0 | 0 | 0 | 0 | 0 | 0 | 1 | 10 | 2 | 5 | 0 | 0 | S100a5 |
| S100a6 | 437 | 408 | 494 | 331 | 329 | 390 | 397 | 440 | 512 | 587 | 594 | 468 | 567 | 425 | 392 | 485 | 328 | 371 | 306 | 359 | 367 | 331 | 570 | 499 | 447 | 493 | 406 | 548 | S100a6 |
| S100a7a | 2 | 6 | 7 | 4 | 6 | 8 | 6 | 2 | 3 | 2 | 14 | 0 | 14 | 1 | 5 | 12 | 14 | 9 | 10 | 1 | 3 | 3 | 14 | 11 | 6 | 1 | 5 | 2 | S100a7a |
| S100a8 | 62 | 23 | 55 | 64 | 34 | 43 | 89 | 46 | 96 | 85 | 239 | 73 | 178 | 62 | 36 | 66 | 64 | 85 | 55 | 24 | 39 | 75 | 142 | 131 | 39 | 93 | 146 | 176 | S100a8 |
| S100a9 | 40 | 22 | 39 | 52 | 31 | 58 | 23 | 24 | 73 | 47 | 170 | 72 | 147 | 62 | 45 | 91 | 74 | 58 | 34 | 18 | 38 | 60 | 104 | 132 | 52 | 114 | 96 | 104 | S100a9 |
| S100b | 18 | 22 | 3 | 18 | 35 | 24 | 1 | 88 | 209 | 202 | 306 | 45 | 83 | 46 | 32 | 21 | 12 | 36 | 11 | 6 | 2 | 221 | 171 | 148 | 133 | 74 | 37 | 38 | S100b |
| S100g | 1 | 1 | 0 | 1 | 3 | 0 | 0 | 2 | 18 | 0 | 4 | 3 | 2 | 0 | 4 | 0 | 0 | 0 | 0 | 5 | 1 | 3 | 8 | 1 | 1 | 5 | 0 | 1 | S100g |
| S100pbp | 6369 | 5224 | 6821 | 6233 | 6027 | 5855 | 6420 | 5521 | 4921 | 5159 | 5501 | 5851 | 5456 | 6270 | 5830 | 5627 | 6461 | 6318 | 5965 | 5407 | 6363 | 5571 | 5015 | 5738 | 5389 | 6085 | 5739 | 6284 | S100pbp |
| S100z | 35 | 12 | 0 | 35 | 0 | 0 | 0 | 54 | 31 | 58 | 34 | 3 | 27 | 21 | 21 | 24 | 6 | 1 | 0 | 0 | 1 | 0 | 28 | 80 | 2 | 72 | 1 | 6 | S100z |
| S1pr1 | 1801 | 1576 | 1687 | 1601 | 1744 | 1809 | 1770 | 2071 | 2005 | 2089 | 1892 | 2467 | 2022 | 2425 | 1685 | 1421 | 1690 | 1678 | 1807 | 1774 | 1725 | 2227 | 2367 | 2305 | 2175 | 2599 | 2213 | 2346 | S1pr1 |
| S1pr2 | 2320 | 2474 | 2298 | 1835 | 1774 | 1788 | 1431 | 1556 | 2136 | 1691 | 2205 | 1530 | 1955 | 1390 | 1875 | 2541 | 2019 | 2056 | 1777 | 1626 | 1473 | 1372 | 2263 | 1629 | 1231 | 1905 | 1639 | 1677 | S1pr2 |
| S1pr3 | 5027 | 4466 | 4002 | 2136 | 1285 | 2680 | 1026 | 3642 | 5019 | 3719 | 4813 | 3528 | 5054 | 2866 | 3889 | 5739 | 2503 | 2687 | 1334 | 1441 | 1069 | 2692 | 5546 | 3645 | 2544 | 4120 | 3989 | 3578 | S1pr3 |
| S1pr4 | 8 | 11 | 8 | 20 | 14 | 32 | 15 | 7 | 14 | 43 | 24 | 27 | 18 | 10 | 21 | 18 | 17 | 14 | 14 | 17 | 9 | 13 | 18 | 25 | 15 | 18 | 12 | 17 | S1pr4 |
| S1pr5 | 20 | 21 | 11 | 35 | 86 | 43 | 34 | 57 | 73 | 87 | 74 | 35 | 69 | 39 | 37 | 23 | 37 | 23 | 50 | 26 | 28 | 119 | 78 | 36 | 91 | 28 | 27 | 18 | S1pr5 |
| Saa1 | 1 | 1 | 0 | 3 | 0 | 0 | 0 | 1 | 5 | 0 | 9 | 1 | 0 | 2 | 3 | 1 | 3 | 0 | 12 | 0 | 0 | 5 | 1 | 0 | 4 | 2 | 6 | 1 | Saa1 |
| Saa2 | 0 | 0 | 0 | 2 | 2 | 0 | 0 | 0 | 1 | 1 | 4 | 0 | 1 | 0 | 3 | 5 | 0 | 0 | 1 | 0 | 0 | 2 | 0 | 2 | 0 | 2 | 0 | 2 | Saa2 |
| Saa3 | 2 | 1 | 0 | 1 | 0 | 0 | 2 | 2 | 2 | 3 | 2 | 0 | 0 | 2 | 0 | 3 | 0 | 0 | 2 | 2 | 0 | 2 | 3 | 2 | 2 | 2 | 0 | 6 | Saa3 |
| Saa4 | 0 | 1 | 1 | 0 | 0 | 0 | 0 | 0 | 1 | 0 | 0 | 0 | 0 | 0 | 0 | 0 | 0 | 0 | 0 | 0 | 0 | 0 | 0 | 0 | 0 | 0 | 0 | 1 | Saa4 |
| Saal1 | 3140 | 3099 | 2696 | 2573 | 2281 | 2540 | 2538 | 2920 | 2881 | 2579 | 2394 | 2948 | 2719 | 2971 | 2964 | 3015 | 2626 | 2485 | 2538 | 2921 | 2508 | 2291 | 2678 | 2660 | 2416 | 3065 | 2974 | 3177 | Saal1 |
| Sac3d1 | 767 | 957 | 798 | 815 | 825 | 735 | 736 | 753 | 831 | 797 | 732 | 882 | 856 | 901 | 688 | 761 | 948 | 796 | 947 | 764 | 794 | 680 | 729 | 869 | 805 | 1023 | 805 | 979 | Sac3d1 |
| Sacm1l | 10791 | 8360 | 9663 | 9166 | 8516 | 8694 | 9336 | 7095 | 5709 | 5161 | 5998 | 5932 | 6050 | 5586 | 11006 | 10193 | 9158 | 9163 | 8473 | 10007 | 9008 | 6423 | 6103 | 5566 | 5776 | 5534 | 5760 | 5918 | Sacm1l |
| Sacs | 11974 | 10055 | 5367 | 7896 | 9493 | 12235 | 11705 | 12759 | 11363 | 6931 | 7525 | 7000 | 7290 | 7584 | 12617 | 10555 | 6362 | 7913 | 8313 | 14453 | 12790 | 7310 | 10066 | 5552 | 8323 | 4181 | 8693 | 6506 | Sacs |
| Sae1 | 15693 | 14530 | 16799 | 14107 | 12872 | 12907 | 14074 | 11726 | 11412 | 10811 | 11021 | 13787 | 12776 | 14096 | 14288 | 13714 | 15587 | 13466 | 14583 | 12662 | 13547 | 10119 | 10908 | 13177 | 11420 | 17046 | 12934 | 14621 | Sae1 |
| Safb | 8308 | 9710 | 13435 | 12885 | 12423 | 11279 | 10823 | 12664 | 13374 | 17371 | 16384 | 18979 | 16498 | 18826 | 8201 | 8842 | 13309 | 12006 | 13490 | 10199 | 11396 | 15730 | 12384 | 19465 | 18870 | 19761 | 17278 | 18513 | Safb |
| Safb2 | 6431 | 8290 | 9552 | 10710 | 9705 | 8658 | 8472 | 7889 | 8260 | 10855 | 10783 | 11538 | 10607 | 11293 | 6043 | 8118 | 9545 | 9976 | 9848 | 7610 | 8960 | 9680 | 7631 | 11418 | 12044 | 10418 | 10879 | 10804 | Safb2 |
| Sag | 257 | 308 | 256 | 399 | 272 | 328 | 296 | 49 | 59 | 89 | 65 | 90 | 51 | 99 | 252 | 291 | 307 | 382 | 274 | 209 | 311 | 75 | 60 | 67 | 64 | 80 | 50 | 38 | Sag |
| Sall1 | 1942 | 2292 | 3521 | 2986 | 3491 | 2956 | 3386 | 3737 | 4317 | 5359 | 3967 | 7908 | 5517 | 7033 | 2143 | 1717 | 3653 | 2785 | 4309 | 2869 | 3110 | 3689 | 2646 | 6626 | 6111 | 7301 | 5399 | 6116 | Sall1 |
| Sall2 | 5682 | 6066 | 6782 | 5996 | 7155 | 6651 | 6392 | 9640 | 9776 | 10640 | 9367 | 13227 | 11311 | 13063 | 5337 | 4967 | 6920 | 6040 | 7966 | 6087 | 6629 | 11820 | 9068 | 11816 | 11122 | 13609 | 12805 | 14031 | Sall2 |
| Sall3 | 907 | 1256 | 1883 | 1540 | 1919 | 1198 | 1342 | 1730 | 1474 | 2261 | 2244 | 2665 | 1977 | 2493 | 1146 | 757 | 1938 | 1412 | 1578 | 1005 | 1345 | 2428 | 1652 | 2269 | 2192 | 2809 | 2307 | 2585 | Sall3 |
| Sall4 | 747 | 751 | 1658 | 841 | 555 | 486 | 454 | 181 | 176 | 202 | 189 | 555 | 511 | 507 | 618 | 808 | 1159 | 583 | 638 | 495 | 518 | 194 | 151 | 302 | 254 | 432 | 597 | 711 | Sall4 |
| Samd1 | 1829 | 2036 | 2634 | 2193 | 2362 | 2017 | 2132 | 2450 | 2922 | 3444 | 3227 | 3250 | 2814 | 3239 | 1545 | 1798 | 2421 | 2265 | 2461 | 1732 | 2233 | 2623 | 2937 | 3681 | 3050 | 3964 | 2738 | 3375 | Samd1 |
| Samd10 | 1065 | 1418 | 1086 | 1656 | 2256 | 1991 | 1973 | 3891 | 3600 | 3885 | 2946 | 3312 | 3185 | 3628 | 1454 | 997 | 1477 | 1690 | 1917 | 2083 | 2073 | 4565 | 3339 | 3430 | 4064 | 2900 | 3000 | 3325 | Samd10 |
| Samd11 | 52 | 113 | 115 | 135 | 163 | 139 | 96 | 182 | 166 | 197 | 205 | 243 | 301 | 252 | 86 | 79 | 127 | 135 | 125 | 118 | 118 | 276 | 180 | 208 | 236 | 92 | 298 | 318 | Samd11 |
| Samd12 | 543 | 496 | 290 | 551 | 1122 | 810 | 917 | 1374 | 1248 | 1085 | 1046 | 603 | 644 | 666 | 692 | 517 | 374 | 715 | 765 | 863 | 1042 | 1320 | 1263 | 588 | 1077 | 432 | 673 | 568 | Samd12 |
| Samd13 | 13 | 12 | 12 | 4 | 11 | 13 | 11 | 4 | 7 | 7 | 7 | 8 | 8 | 7 | 8 | 14 | 21 | 13 | 24 | 28 | 4 | 7 | 5 | 5 | 10 | 8 | 5 | 15 | Samd13 |
| Samd14 | 5549 | 6835 | 7032 | 8585 | 11164 | 9044 | 8537 | 11246 | 11815 | 14131 | 12383 | 13632 | 11783 | 12579 | 5734 | 5395 | 8707 | 9651 | 11481 | 8161 | 9590 | 15214 | 10686 | 14972 | 15526 | 13715 | 11821 | 11568 | Samd14 |
| Samd15 | 20 | 41 | 21 | 39 | 38 | 44 | 44 | 78 | 59 | 63 | 80 | 60 | 81 | 70 | 42 | 29 | 35 | 41 | 36 | 68 | 45 | 82 | 56 | 86 | 101 | 60 | 45 | 80 | Samd15 |
| Samd3 | 221 | 146 | 179 | 181 | 229 | 226 | 327 | 604 | 592 | 368 | 297 | 767 | 102 | 863 | 215 | 118 | 210 | 191 | 268 | 329 | 256 | 468 | 323 | 538 | 597 | 1193 | 539 | 846 | Samd3 |
| Samd4 | 1927 | 1708 | 2540 | 1987 | 1987 | 1961 | 1925 | 1750 | 2076 | 2131 | 2983 | 2116 | 2219 | 2313 | 1755 | 1896 | 2296 | 1918 | 2013 | 1578 | 1963 | 1917 | 2339 | 2366 | 2136 | 2082 | 2069 | 2294 | Samd4 |
| Samd4b | 7356 | 9125 | 9855 | 9315 | 7437 | 7381 | 7099 | 4567 | 5388 | 5665 | 5587 | 6539 | 5664 | 6189 | 6757 | 8861 | 9383 | 9039 | 8303 | 6039 | 7477 | 5606 | 5039 | 6449 | 5730 | 7428 | 5162 | 6069 | Samd4b |
| Samd5 | 1841 | 1113 | 2005 | 1284 | 1598 | 1605 | 1497 | 3187 | 2890 | 2758 | 2666 | 2973 | 2943 | 3181 | 1622 | 1607 | 1472 | 1384 | 1467 | 1476 | 1576 | 2861 | 2867 | 2880 | 2824 | 2509 | 3412 | 3189 | Samd5 |
| Samd7 | 0 | 0 | 0 | 0 | 1 | 0 | 0 | 1 | 1 | 0 | 0 | 0 | 0 | 0 | 0 | 0 | 0 | 2 | 0 | 0 | 1 | 2 | 1 | 0 | 0 | 0 | 0 | 0 | Samd7 |
| Samd8 | 7002 | 6151 | 6754 | 9413 | 8482 | 8607 | 9576 | 4851 | 4308 | 4504 | 4527 | 4563 | 4544 | 5007 | 7233 | 6574 | 7273 | 9303 | 8259 | 8357 | 9369 | 5043 | 4183 | 4576 | 5650 | 4025 | 4803 | 4803 | Samd8 |
| Samd9l | 170 | 108 | 145 | 111 | 156 | 209 | 105 | 318 | 331 | 281 | 377 | 214 | 227 | 238 | 190 | 176 | 129 | 169 | 106 | 238 | 149 | 424 | 565 | 257 | 312 | 174 | 242 | 271 | Samd9l |
| Samhd1 | 6531 | 6011 | 7495 | 6228 | 5463 | 5421 | 5703 | 3319 | 3650 | 3930 | 3809 | 4494 | 4340 | 4456 | 5931 | 6360 | 6903 | 6259 | 5768 | 5253 | 5617 | 3293 | 3528 | 4141 | 3475 | 5031 | 4328 | 4711 | Samhd1 |
| Samm50 | 3978 | 3878 | 4507 | 3783 | 3548 | 3266 | 3737 | 5233 | 5057 | 5659 | 5310 | 5830 | 5283 | 6400 | 3774 | 3409 | 4200 | 3717 | 3926 | 3435 | 3871 | 5142 | 5089 | 6115 | 6020 | 6400 | 5663 | 6705 | Samm50 |
| Samsn1 | 106 | 83 | 127 | 86 | 72 | 62 | 92 | 82 | 103 | 90 | 86 | 68 | 102 | 73 | 120 | 119 | 95 | 83 | 44 | 109 | 65 | 80 | 128 | 97 | 46 | 102 | 81 | 89 | Samsn1 |
| Sanbr | 7175 | 5888 | 6202 | 7796 | 9976 | 9309 | 10467 | 13517 | 10778 | 10205 | 9149 | 10267 | 8768 | 10138 | 8985 | 6256 | 6922 | 8226 | 8936 | 10741 | 9632 | 12653 | 10336 | 9764 | 11763 | 8362 | 9769 | 9272 | Sanbr |
| Sap130 | 7855 | 8606 | 8178 | 8682 | 7683 | 7936 | 7695 | 4945 | 5804 | 6137 | 5960 | 6307 | 6270 | 6084 | 7456 | 7995 | 8165 | 8382 | 8507 | 6576 | 7735 | 5897 | 5404 | 6067 | 6063 | 6397 | 6152 | 6041 | Sap130 |
| Sap18 | 5935 | 6195 | 7189 | 6385 | 5495 | 5579 | 6322 | 4534 | 4647 | 4798 | 5019 | 5355 | 5296 | 5732 | 5803 | 6007 | 6544 | 6061 | 5995 | 6103 | 6025 | 3999 | 4849 | 5456 | 5020 | 6523 | 5554 | 5708 | Sap18 |
| Sap18b | 1043 | 1002 | 1366 | 1032 | 838 | 790 | 862 | 739 | 714 | 833 | 813 | 885 | 858 | 870 | 973 | 906 | 1132 | 969 | 915 | 931 | 911 | 633 | 789 | 864 | 813 | 1111 | 894 | 973 | Sap18b |
| Sap30 | 2804 | 2886 | 3086 | 3280 | 2523 | 2441 | 3099 | 2749 | 2084 | 2440 | 1875 | 3017 | 2337 | 3528 | 2925 | 2294 | 3154 | 2726 | 2924 | 3271 | 2861 | 2047 | 1834 | 2960 | 2701 | 3493 | 2893 | 3535 | Sap30 |
| Sap30bp | 5425 | 6176 | 5581 | 5634 | 4808 | 5059 | 5061 | 5475 | 5276 | 5234 | 4691 | 5641 | 5246 | 5695 | 5669 | 5082 | 5496 | 4937 | 5565 | 5361 | 5207 | 4976 | 4939 | 5462 | 5583 | 5859 | 5824 | 5752 | Sap30bp |
| Sap30l | 1470 | 1795 | 1398 | 1596 | 1563 | 1356 | 1532 | 1005 | 1043 | 1053 | 1013 | 1079 | 966 | 1162 | 1481 | 1419 | 1545 | 1611 | 1508 | 1534 | 1677 | 921 | 929 | 1214 | 1099 | 1018 | 1084 | 1093 | Sap30l |
| Sapcd1 | 160 | 216 | 267 | 228 | 174 | 165 | 253 | 120 | 139 | 136 | 159 | 229 | 122 | 218 | 177 | 196 | 306 | 255 | 242 | 198 | 212 | 77 | 148 | 181 | 187 | 246 | 192 | 215 | Sapcd1 |
| Sapcd2 | 1749 | 1983 | 1983 | 1690 | 1221 | 1546 | 1410 | 1034 | 1243 | 1158 | 970 | 1815 | 1428 | 1578 | 1439 | 1510 | 1898 | 1413 | 1624 | 1468 | 1353 | 790 | 981 | 1466 | 1213 | 2189 | 1822 | 1682 | Sapcd2 |
| Sar1a | 18007 | 16303 | 17238 | 16200 | 13107 | 14646 | 14375 | 13953 | 12795 | 12743 | 13476 | 13042 | 12343 | 12464 | 17934 | 17392 | 15932 | 15839 | 14152 | 16272 | 13939 | 12049 | 13764 | 12883 | 12554 | 14194 | 13063 | 12977 | Sar1a |
| Sar1b | 3071 | 2845 | 3151 | 2871 | 2946 | 2838 | 3433 | 3318 | 2892 | 2746 | 2805 | 3057 | 2947 | 3239 | 3422 | 3083 | 3173 | 2743 | 2897 | 3628 | 3198 | 2617 | 3262 | 2977 | 3038 | 3032 | 2944 | 3348 | Sar1b |
| Saraf | 10860 | 12405 | 9815 | 11598 | 10150 | 9364 | 9789 | 7287 | 6899 | 6309 | 6558 | 5697 | 5797 | 5368 | 11621 | 11989 | 10561 | 11756 | 10137 | 11078 | 10175 | 6806 | 7489 | 6617 | 6434 | 6324 | 5944 | 5453 | Saraf |
| Sardh | 321 | 457 | 355 | 290 | 377 | 381 | 185 | 493 | 807 | 656 | 920 | 453 | 775 | 425 | 405 | 534 | 279 | 324 | 246 | 283 | 245 | 795 | 941 | 526 | 569 | 618 | 580 | 444 | Sardh |

| | | | | | | | | | | | | | | | | | | | | | | | | | | | | | |
|---|---|---|---|---|---|---|---|---|---|---|---|---|---|---|---|---|---|---|---|---|---|---|---|---|---|---|---|---|---|
| Sarm1 | 2015 | 2855 | 2521 | 3493 | 4606 | 3471 | 3850 | 2466 | 2738 | 3374 | 2592 | 3051 | 2595 | 3003 | 2441 | 2156 | 3639 | 3672 | 4547 | 3393 | 4060 | 4157 | 2407 | 3231 | 3979 | 2219 | 2537 | 2830 | Sarm1 |
| Sarnp | 3245 | 2833 | 4213 | 3891 | 2872 | 2936 | 2663 | 2958 | 3066 | 3318 | 1556 | 2952 | 3480 | 3722 | 3343 | 3105 | 1826 | 2810 | 3451 | 3452 | 3558 | 2637 | 3133 | 4117 | 3325 | 4567 | 4106 | 3421 | Sarnp |
| Sars | 13654 | 16164 | 15005 | 15710 | 14470 | 14663 | 14081 | 11554 | 11399 | 11911 | 10862 | 12620 | 12177 | 13123 | 14186 | 12993 | 15137 | 13764 | 16346 | 15648 | 15731 | 11317 | 11596 | 12421 | 13248 | 13828 | 13529 | 13624 | Sars |
| Sars2 | 403 | 548 | 598 | 429 | 434 | 446 | 327 | 850 | 1103 | 1078 | 1085 | 1361 | 1311 | 1277 | 385 | 478 | 531 | 403 | 497 | 460 | 392 | 969 | 1002 | 1198 | 1013 | 1603 | 1261 | 1265 | Sars2 |
| Sart1 | 6055 | 7335 | 10419 | 9216 | 7340 | 6976 | 7354 | 5003 | 5904 | 7509 | 6904 | 8592 | 7042 | 8291 | 5752 | 6548 | 10102 | 8176 | 8458 | 6127 | 7438 | 6395 | 5559 | 8539 | 7583 | 9468 | 7453 | 8326 | Sart1 |
| Sart3 | 6392 | 7413 | 8668 | 8471 | 7321 | 7044 | 6778 | 6173 | 6757 | 7827 | 7130 | 9385 | 7837 | 8185 | 5827 | 6695 | 8568 | 7844 | 8344 | 6160 | 7097 | 6998 | 6273 | 8983 | 8311 | 9695 | 8459 | 8624 | Sart3 |
| Sash1 | 3970 | 4000 | 4721 | 4753 | 4021 | 4255 | 3750 | 3127 | 3408 | 3653 | 4434 | 3638 | 3976 | 3406 | 3686 | 4674 | 4430 | 4590 | 4105 | 3162 | 3956 | 3584 | 4011 | 3543 | 3476 | 2682 | 3403 | 3482 | Sash1 |
| Sash3 | 66 | 54 | 64 | 29 | 37 | 37 | 20 | 83 | 101 | 63 | 88 | 80 | 75 | 50 | 38 | 51 | 35 | 28 | 25 | 40 | 34 | 77 | 170 | 61 | 75 | 96 | 74 | 69 | Sash3 |
| Sass6 | 2978 | 2494 | 2710 | 2622 | 2756 | 2843 | 3076 | 2880 | 2553 | 2536 | 2482 | 3330 | 3244 | 3450 | 3032 | 2611 | 2738 | 2487 | 2734 | 3024 | 3023 | 2398 | 2258 | 2803 | 2677 | 3227 | 3477 | 3588 | Sass6 |
| Sat1 | 2639 | 2483 | 2808 | 2408 | 2401 | 2408 | 2286 | 2240 | 1965 | 1918 | 2059 | 1789 | 1814 | 1887 | 3022 | 2850 | 2296 | 2340 | 2374 | 2951 | 2246 | 1736 | 2283 | 1914 | 1684 | 2199 | 1840 | 2000 | Sat1 |
| Sat2 | 842 | 872 | 823 | 764 | 641 | 868 | 771 | 554 | 547 | 582 | 559 | 565 | 599 | 569 | 997 | 835 | 743 | 748 | 714 | 824 | 780 | 541 | 555 | 540 | 581 | 561 | 623 | 592 | Sat2 |
| Satb1 | 4848 | 3949 | 5517 | 5491 | 6275 | 5759 | 6473 | 5084 | 4777 | 5735 | 5284 | 6642 | 5664 | 7017 | 4847 | 4231 | 6204 | 5468 | 5999 | 4890 | 6134 | 6417 | 4493 | 6508 | 6420 | 5846 | 5705 | 6497 | Satb1 |
| Satb2 | 660 | 580 | 807 | 1029 | 956 | 882 | 1040 | 972 | 1065 | 1103 | 966 | 716 | 677 | 784 | 518 | 567 | 773 | 976 | 1029 | 667 | 1032 | 745 | 749 | 1183 | 1345 | 679 | 669 | 725 | Satb2 |
| Sav1 | 5723 | 4687 | 5947 | 5652 | 4930 | 5082 | 5331 | 3990 | 3687 | 4130 | 4626 | 4567 | 4350 | 4699 | 5180 | 4904 | 5686 | 5504 | 4934 | 4261 | 5482 | 4131 | 3961 | 4485 | 4228 | 5224 | 4124 | 5110 | Sav1 |
| Saxo1 | 1 | 3 | 0 | 0 | 0 | 0 | 0 | 0 | 2 | 0 | 6 | 8 | 1 | 0 | 0 | 1 | 0 | 0 | 0 | 2 | 0 | 0 | 4 | 0 | 0 | 0 | 2 | 0 | Saxo1 |
| Saxo2 | 404 | 304 | 321 | 326 | 332 | 353 | 361 | 335 | 283 | 301 | 206 | 251 | 265 | 191 | 398 | 452 | 279 | 305 | 252 | 325 | 368 | 314 | 354 | 173 | 230 | 217 | 167 | 211 | Saxo2 |
| Saysd1 | 221 | 283 | 238 | 297 | 320 | 307 | 322 | 473 | 475 | 437 | 490 | 420 | 418 | 441 | 222 | 207 | 279 | 264 | 335 | 312 | 298 | 481 | 417 | 489 | 473 | 487 | 406 | 434 | Saysd1 |
| Sbds | 2699 | 3196 | 3237 | 3008 | 2703 | 2701 | 2737 | 2347 | 2405 | 2340 | 2227 | 2593 | 2376 | 2653 | 2794 | 2787 | 3115 | 2775 | 2858 | 3088 | 2573 | 2112 | 2272 | 2415 | 2344 | 2871 | 2561 | 2861 | Sbds |
| Sbf1 | 11246 | 15073 | 15030 | 18086 | 16400 | 15395 | 14267 | 7651 | 9279 | 10155 | 9860 | 11387 | 9403 | 9517 | 10952 | 12996 | 16766 | 17601 | 18223 | 12162 | 15427 | 10444 | 8424 | 10242 | 11189 | 9070 | 9735 | 9113 | Sbf1 |
| Sbf2 | 11851 | 12126 | 7423 | 9700 | 11739 | 12496 | 12409 | 15428 | 13960 | 12838 | 11456 | 13134 | 11668 | 12739 | 11892 | 10140 | 9090 | 10462 | 10925 | 13262 | 12735 | 12521 | 12747 | 11074 | 13901 | 7669 | 12740 | 11521 | Sbf2 |
| Sbk1 | 24410 | 29400 | 30769 | 39757 | 46327 | 38837 | 36998 | 36395 | 39391 | 50041 | 39994 | 49822 | 38865 | 44446 | 25669 | 23883 | 38683 | 39741 | 46791 | 35847 | 40632 | 52119 | 34429 | 50617 | 55381 | 45913 | 38652 | 41425 | Sbk1 |
| Sbk2 | 9 | 2 | 0 | 3 | 2 | 2 | 3 | 9 | 7 | 8 | 11 | 1 | 6 | 2 | 4 | 3 | 3 | 2 | 5 | 5 | 4 | 2 | 7 | 1 | 6 | 0 | 1 | 0 | Sbk2 |
| Sbk3 | 80 | 117 | 84 | 220 | 242 | 236 | 256 | 53 | 49 | 93 | 70 | 54 | 35 | 65 | 83 | 73 | 180 | 265 | 250 | 180 | 216 | 97 | 60 | 59 | 74 | 39 | 24 | 47 | Sbk3 |
| Sbno1 | 17901 | 16857 | 13270 | 16414 | 15037 | 16297 | 17429 | 17520 | 15087 | 14546 | 15341 | 15865 | 15517 | 16639 | 17762 | 16454 | 14655 | 16403 | 14825 | 17237 | 17503 | 15695 | 14688 | 13992 | 16392 | 11800 | 16823 | 15875 | Sbno1 |
| Sbno2 | 933 | 911 | 1123 | 925 | 1183 | 962 | 808 | 1608 | 2148 | 2200 | 1964 | 2404 | 2206 | 1865 | 749 | 918 | 1099 | 913 | 1165 | 812 | 809 | 2246 | 1980 | 2194 | 2269 | 2094 | 1897 | 1969 | Sbno2 |
| Sbp | 0 | 0 | 0 | 0 | 0 | 2 | 0 | 0 | 0 | 0 | 0 | 0 | 0 | 0 | 0 | 0 | 0 | 0 | 2 | 0 | 0 | 0 | 0 | 0 | 0 | 0 | 0 | 1 | Sbp |
| Sbpl | 0 | 0 | 0 | 0 | 0 | 0 | 0 | 0 | 0 | 0 | 3 | 0 | 0 | 0 | 0 | 0 | 0 | 0 | 0 | 0 | 0 | 0 | 0 | 0 | 0 | 0 | 0 | 0 | Sbpl |
| Sbsn | 600 | 667 | 972 | 1017 | 1019 | 864 | 948 | 600 | 637 | 758 | 825 | 740 | 759 | 937 | 611 | 553 | 1132 | 1011 | 1057 | 758 | 961 | 822 | 601 | 914 | 949 | 710 | 757 | 845 | Sbsn |
| Sbspon | 37 | 43 | 6 | 27 | 14 | 68 | 8 | 73 | 148 | 76 | 105 | 34 | 64 | 40 | 47 | 33 | 32 | 37 | 9 | 3 | 12 | 55 | 126 | 57 | 37 | 29 | 46 | 43 | Sbspon |
| Sc5d | 9356 | 9042 | 9355 | 9327 | 8512 | 9217 | 9836 | 5763 | 5019 | 5150 | 5023 | 5564 | 5853 | 6499 | 9703 | 9658 | 8776 | 9421 | 8506 | 9511 | 9297 | 5574 | 4925 | 5167 | 5704 | 5204 | 6316 | 6258 | Sc5d |
| Scaf1 | 7794 | 9628 | 12141 | 11897 | 11817 | 10119 | 8532 | 8521 | 10363 | 12992 | 11896 | 13700 | 11543 | 11249 | 7059 | 8469 | 12419 | 11253 | 12908 | 8143 | 9719 | 12355 | 9862 | 13512 | 12847 | 14454 | 11755 | 10773 | Scaf1 |
| Scaf11 | 12480 | 10464 | 10707 | 10873 | 11427 | 11895 | 11887 | 12058 | 11707 | 11661 | 12355 | 13768 | 13086 | 13689 | 12144 | 11249 | 11012 | 10633 | 10852 | 10895 | 11652 | 10243 | 11507 | 11782 | 11275 | 11854 | 13909 | 13494 | Scaf11 |
| Scaf4 | 5496 | 6272 | 6913 | 7443 | 6958 | 6567 | 6773 | 4267 | 4341 | 5052 | 4813 | 6236 | 5051 | 5934 | 5359 | 5979 | 7380 | 7231 | 7310 | 5722 | 7057 | 5073 | 3897 | 5519 | 5463 | 5477 | 5497 | 5503 | Scaf4 |
| Scaf8 | 4855 | 4106 | 4658 | 4665 | 5174 | 4835 | 5411 | 6979 | 6714 | 6754 | 6314 | 7169 | 6473 | 7421 | 4644 | 4277 | 4655 | 4540 | 5097 | 4194 | 5242 | 7033 | 6214 | 6758 | 6728 | 6251 | 6396 | 7392 | Scaf8 |
| Scai | 8742 | 5827 | 4500 | 7268 | 7824 | 9682 | 9484 | 12658 | 10760 | 8080 | 10359 | 8566 | 8997 | 9290 | 8672 | 7906 | 5442 | 7211 | 7596 | 9488 | 9801 | 9207 | 10409 | 7702 | 9768 | 5907 | 9081 | 8109 | Scai |
| Scamp1 | 8418 | 7021 | 8337 | 9875 | 11581 | 10294 | 11061 | 8683 | 7442 | 8090 | 7455 | 6657 | 6867 | 6710 | 9079 | 8011 | 8902 | 10329 | 10459 | 10684 | 10596 | 10225 | 7547 | 7383 | 8743 | 5761 | 5999 | 6057 | Scamp1 |
| Scamp2 | 2237 | 2526 | 2321 | 2131 | 2055 | 2244 | 2077 | 2805 | 2951 | 2916 | 3058 | 3137 | 3221 | 3115 | 2184 | 2334 | 2286 | 2100 | 2138 | 2074 | 2074 | 3018 | 3125 | 2850 | 3033 | 3086 | 3248 | 3237 | Scamp2 |
| Scamp3 | 2308 | 2852 | 2854 | 2778 | 2774 | 2652 | 2459 | 1984 | 2631 | 2644 | 2723 | 2807 | 2789 | 2401 | 2279 | 2424 | 2958 | 2849 | 2858 | 2596 | 2479 | 2633 | 2557 | 2806 | 2757 | 2858 | 2676 | 2409 | Scamp3 |
| Scamp4 | 24 | 12 | 16 | 32 | 17 | 14 | 17 | 17 | 19 | 16 | 13 | 22 | 20 | 27 | 17 | 17 | 35 | 22 | 27 | 24 | 31 | 25 | 20 | 26 | 26 | 26 | 13 | 32 | Scamp4 |
| Scamp5 | 4798 | 5537 | 5245 | 6881 | 8587 | 8057 | 7582 | 6185 | 6292 | 6777 | 5848 | 6417 | 5610 | 5880 | 5248 | 4628 | 6224 | 7379 | 8171 | 7018 | 7797 | 8380 | 5722 | 6357 | 7845 | 4692 | 5858 | 5592 | Scamp5 |
| Scand1 | 989 | 1219 | 1066 | 1228 | 1249 | 1207 | 835 | 1421 | 1340 | 1562 | 1473 | 1496 | 1523 | 1140 | 886 | 846 | 1307 | 1095 | 1474 | 1187 | 903 | 1496 | 1367 | 1695 | 1574 | 1727 | 1519 | 1201 | Scand1 |
| Scap | 4382 | 5624 | 5338 | 6093 | 5447 | 5036 | 4892 | 3535 | 4394 | 4588 | 4481 | 4596 | 4090 | 3965 | 4122 | 4998 | 5530 | 6124 | 5726 | 4575 | 5068 | 4436 | 4297 | 4689 | 4636 | 4886 | 4105 | 4044 | Scap |
| Scaper | 2616 | 2425 | 1839 | 2724 | 3708 | 3754 | 3771 | 6135 | 5337 | 5176 | 4899 | 4119 | 4273 | 4528 | 3385 | 2547 | 2186 | 3108 | 3223 | 4048 | 3812 | 6311 | 5465 | 4456 | 5733 | 3179 | 4195 | 3925 | Scaper |
| Scara3 | 647 | 441 | 476 | 405 | 418 | 555 | 337 | 1282 | 1870 | 1432 | 1937 | 1127 | 1229 | 1018 | 537 | 604 | 433 | 532 | 337 | 306 | 279 | 1145 | 2124 | 1381 | 1030 | 1702 | 1166 | 1057 | Scara3 |
| Scara5 | 286 | 304 | 449 | 269 | 115 | 277 | 79 | 404 | 557 | 578 | 1180 | 470 | 545 | 493 | 310 | 369 | 282 | 296 | 67 | 126 | 94 | 439 | 1450 | 506 | 370 | 537 | 562 | 453 | Scara5 |
| Scarb1 | 4309 | 5147 | 5109 | 4988 | 3776 | 4166 | 3765 | 1375 | 1790 | 1762 | 1940 | 1836 | 1850 | 1678 | 3907 | 5116 | 4880 | 5218 | 4382 | 3248 | 3949 | 1836 | 1927 | 1929 | 1758 | 2277 | 1858 | 1849 | Scarb1 |
| Scarb2 | 9069 | 8096 | 8378 | 10126 | 12183 | 11268 | 11990 | 13953 | 11966 | 12831 | 12246 | 12217 | 11005 | 12997 | 9932 | 7763 | 9093 | 10106 | 11217 | 11828 | 12020 | 13851 | 11343 | 12530 | 14245 | 11039 | 11824 | 12182 | Scarb2 |
| Scarf1 | 218 | 339 | 262 | 276 | 287 | 230 | 204 | 275 | 329 | 353 | 368 | 404 | 400 | 313 | 214 | 274 | 227 | 295 | 273 | 257 | 244 | 361 | 452 | 364 | 381 | 414 | 417 | 299 | Scarf1 |
| Scarf2 | 1293 | 1359 | 1332 | 770 | 475 | 842 | 296 | 1656 | 2464 | 2179 | 3369 | 1673 | 2674 | 1291 | 985 | 1607 | 1068 | 873 | 460 | 313 | 321 | 1960 | 2626 | 2278 | 1486 | 2420 | 1898 | 1487 | Scarf2 |
| Scart1 | 0 | 0 | 0 | 1 | 0 | 0 | 0 | 0 | 0 | 3 | 5 | 1 | 0 | 0 | 0 | 0 | 0 | 1 | 0 | 0 | 0 | 0 | 2 | 1 | 0 | 1 | 0 | 0 | Scart1 |
| Scart2 | 0 | 0 | 0 | 0 | 0 | 0 | 0 | 0 | 0 | 0 | 0 | 0 | 0 | 0 | 0 | 0 | 0 | 0 | 0 | 0 | 0 | 0 | 0 | 2 | 0 | 0 | 0 | 0 | Scart2 |
| Sccpdh | 3591 | 3288 | 3351 | 3189 | 4170 | 3723 | 3741 | 4621 | 3849 | 3518 | 3687 | 3275 | 3556 | 3549 | 4387 | 3468 | 3440 | 3230 | 3611 | 4119 | 3583 | 4316 | 3976 | 3359 | 3617 | 3413 | 3452 | 3760 | Sccpdh |
| Scd1 | 6249 | 5794 | 6441 | 4203 | 3397 | 4057 | 3666 | 1856 | 1851 | 1822 | 1705 | 2757 | 3113 | 3003 | 5374 | 6489 | 5147 | 3921 | 3218 | 3281 | 3331 | 1723 | 1737 | 1917 | 1755 | 3530 | 3420 | 3895 | Scd1 |
| Scd2 | 57142 | 64526 | 69661 | 63650 | 57293 | 56688 | 60373 | 56313 | 55831 | 62031 | 58344 | 75735 | 71321 | 76063 | 57614 | 66423 | 69339 | 63585 | 58201 | 50248 | 56474 | 61705 | 52668 | 65681 | 66231 | 79369 | 77170 | 83018 | Scd2 |
| Scd3 | 54 | 101 | 74 | 88 | 124 | 122 | 120 | 73 | 72 | 86 | 111 | 51 | 86 | 66 | 79 | 67 | 79 | 122 | 103 | 93 | 131 | 115 | 91 | 65 | 77 | 41 | 48 | 49 | Scd3 |
| Scd4 | 110 | 95 | 50 | 67 | 67 | 87 | 87 | 24 | 33 | 39 | 32 | 21 | 19 | 20 | 84 | 86 | 52 | 78 | 82 | 61 | 70 | 35 | 32 | 43 | 16 | 15 | 21 | 14 | Scd4 |
| Scel | 57 | 48 | 40 | 19 | 28 | 35 | 31 | 70 | 87 | 69 | 67 | 40 | 57 | 61 | 74 | 53 | 48 | 63 | 27 | 37 | 19 | 23 | 98 | 47 | 31 | 40 | 40 | 38 | Scel |
| Scfd1 | 2049 | 1555 | 1722 | 1554 | 1611 | 1742 | 1610 | 4347 | 4139 | 3361 | 3643 | 3704 | 3804 | 3760 | 2321 | 1826 | 1619 | 1358 | 1509 | 2031 | 1646 | 3698 | 4229 | 3424 | 3305 | 3818 | 3863 | 3711 | Scfd1 |
| Scfd2 | 1080 | 1127 | 862 | 1118 | 1174 | 961 | 1186 | 1053 | 1105 | 1030 | 1017 | 1040 | 966 | 1028 | 1162 | 1174 | 975 | 1161 | 1081 | 1046 | 1193 | 1064 | 1092 | 972 | 1101 | 980 | 1076 | 998 | Scfd2 |
| Scg2 | 1322 | 1481 | 1167 | 1835 | 3401 | 2367 | 2418 | 2505 | 2433 | 2269 | 1919 | 1128 | 1136 | 1155 | 2073 | 1231 | 1572 | 2610 | 2271 | 2846 | 2588 | 3050 | 2717 | 1532 | 2148 | 1063 | 930 | 866 | Scg2 |
| Scg3 | 5044 | 5881 | 5493 | 7803 | 9679 | 8746 | 8470 | 7800 | 6390 | 8233 | 6428 | 7753 | 6693 | 7649 | 7032 | 4896 | 7131 | 7984 | 8547 | 8034 | 8960 | 10328 | 6108 | 7299 | 9055 | 6728 | 7180 | 7164 | Scg3 |
| Scg5 | 1514 | 1905 | 1493 | 2525 | 3023 | 2725 | 3035 | 2970 | 2426 | 2508 | 2082 | 2560 | 2100 | 2798 | 2168 | 1456 | 2121 | 2488 | 2950 | 3199 | 3354 | 3033 | 2385 | 2393 | 3218 | 2011 | 2409 | 2521 | Scg5 |
| Scgb1a1 | 0 | 0 | 0 | 0 | 0 | 0 | 0 | 0 | 0 | 0 | 0 | 0 | 0 | 0 | 0 | 0 | 0 | 1 | 0 | 0 | 0 | 2 | 0 | 0 | 0 | 0 | 0 | 0 | Scgb1a1 |
| Scgb2b2 | 0 | 0 | 0 | 0 | 0 | 0 | 0 | 0 | 0 | 0 | 0 | 0 | 0 | 0 | 0 | 0 | 0 | 0 | 0 | 0 | 2 | 0 | 0 | 0 | 0 | 0 | 0 | 0 | Scgb2b2 |
| Scgb3a1 | 1 | 0 | 0 | 0 | 0 | 0 | 0 | 0 | 0 | 0 | 2 | 1 | 0 | 0 | 1 | 0 | 0 | 0 | 0 | 0 | 0 | 2 | 0 | 0 | 4 | 0 | 0 | 2 | Scgb3a1 |
| Scgb3a2 | 0 | 2 | 0 | 1 | 0 | 4 | 1 | 1 | 4 | 0 | 6 | 0 | 0 | 0 | 0 | 0 | 0 | 0 | 0 | 0 | 3 | 2 | 1 | 1 | 0 | 0 | 1 | 0 | Scgb3a2 |
| Scgn | 78 | 68 | 60 | 49 | 67 | 88 | 106 | 49 | 59 | 59 | 33 | 46 | 16 | 72 | 91 | 37 | 68 | 111 | 90 | 118 | 88 | 62 | 58 | 42 | 46 | 60 | 52 | 40 | Scgn |
| Schip1 | 73 | 82 | 75 | 67 | 63 | 53 | 67 | 71 | 91 | 57 | 35 | 58 | 61 | 67 | 76 | 72 | 96 | 80 | 48 | 65 | 86 | 47 | 76 | 60 | 81 | 45 | 62 | 61 | Schip1 |
| Scimp | 1 | 2 | 4 | 0 | 0 | 0 | 1 | 2 | 2 | 0 | 3 | 0 | 2 | 5 | 0 | 0 | 0 | 0 | 1 | 1 | 3 | 0 | 1 | 4 | 2 | 2 | 6 | 5 | Scimp |
| Scin | 17 | 18 | 14 | 15 | 10 | 44 | 2 | 67 | 222 | 202 | 465 | 9 | 14 | 10 | 15 | 16 | 2 | 39 | 28 | 9 | 10 | 182 | 179 | 110 | 99 | 20 | 3 | 9 | Scin |
| Sclt1 | 3913 | 3309 | 3228 | 3272 | 3127 | 3463 | 3536 | 3082 | 2815 | 2372 | 2384 | 2735 | 2764 | 2725 | 4059 | 3720 | 3151 | 3274 | 3151 | 3946 | 3447 | 2202 | 2602 | 2357 | 2474 | 2633 | 2869 | 2746 | Sclt1 |
| Scly | 1543 | 1939 | 1911 | 1859 | 1649 | 1653 | 1507 | 630 | 693 | 783 | 764 | 843 | 810 | 817 | 1436 | 1766 | 2117 | 1750 | 1822 | 1506 | 1526 | 647 | 730 | 784 | 741 | 942 | 845 | 851 | Scly |
| Scmh1 | 2408 | 2731 | 3102 | 3551 | 4164 | 3898 | 3806 | 3090 | 3381 | 4165 | 3841 | 3854 | 3856 | 3688 | 2536 | 2683 | 3598 | 3528 | 4264 | 3085 | 3734 | 4380 | 3286 | 3662 | 4267 | 3341 | 3332 | 3383 | Scmh1 |
| Scml2 | 540 | 386 | 484 | 443 | 543 | 591 | 510 | 950 | 1001 | 798 | 1216 | 883 | 978 | 943 | 516 | 532 | 422 | 442 | 536 | 483 | 503 | 896 | 1018 | 879 | 878 | 978 | 812 | 900 | Scml2 |
| Scml4 | 444 | 522 | 367 | 837 | 1015 | 933 | 830 | 782 | 672 | 791 | 678 | 603 | 480 | 568 | 522 | 461 | 600 | 861 | 882 | 838 | 1083 | 976 | 665 | 697 | 1071 | 358 | 442 | 412 | Scml4 |
| Scn10a | 133 | 210 | 2 | 8 | 554 | 19 | 18 | 581 | 810 | 330 | 742 | 17 | 479 | 10 | 298 | 343 | 80 | 273 | 17 | 10 | 14 | 993 | 386 | 10 | 483 | 12 | 1 | 6 | Scn10a |
| Scn11a | 96 | 81 | 64 | 56 | 56 | 36 | 80 | 100 | 85 | 71 | 82 | 202 | 84 | 180 | 67 | 40 | 59 | 109 | 90 | 72 | 61 | 57 | 54 | 105 | 81 | 207 | 106 | 132 | Scn11a |
| Scn1a | 848 | 654 | 621 | 878 | 1744 | 1370 | 1342 | 1397 | 1207 | 1191 | 1230 | 783 | 773 | 909 | 943 | 700 | 825 | 933 | 1197 | 1421 | 1336 | 1307 | 1080 | 811 | 1207 | 508 | 775 | 678 | Scn1a |
| Scn1b | 190 | 166 | 157 | 149 | 245 | 215 | 140 | 109 | 145 | 194 | 212 | 85 | 108 | 59 | 136 | 206 | 222 | 234 | 168 | 111 | 164 | 165 | 201 | 134 | 160 | 69 | 67 | 45 | Scn1b |

| | | | | | | | | | | | | | | | | | | | | | | | | | | | | | |
|---|---|---|---|---|---|---|---|---|---|---|---|---|---|---|---|---|---|---|---|---|---|---|---|---|---|---|---|---|---|
| Scn2a | 1404 | 1475 | 1006 | 1919 | 3634 | 3112 | 3046 | 4030 | 3302 | 3466 | 3079 | 1975 | 2311 | 2358 | 2259 | 1388 | 1590 | 2243 | 2553 | 3294 | 3195 | 3853 | 3166 | 2228 | 3452 | 1007 | 2119 | 1928 | Scn2a |
| Scn2b | 458 | 518 | 933 | 718 | 939 | 620 | 695 | 843 | 852 | 1122 | 1213 | 760 | 982 | 885 | 455 | 563 | 927 | 768 | 802 | 656 | 816 | 1183 | 948 | 996 | 1092 | 804 | 868 | 908 | Scn2b |
| Scn3a | 4567 | 4424 | 2743 | 5136 | 7983 | 7811 | 7483 | 15404 | 11873 | 10711 | 10379 | 10554 | 10552 | 11034 | 6360 | 4102 | 3950 | 5379 | 6502 | 8740 | 7918 | 13540 | 10897 | 8641 | 13051 | 4981 | 11089 | 9770 | Scn3a |
| Scn3b | 5121 | 5823 | 3896 | 8271 | 13920 | 11058 | 11931 | 9166 | 8029 | 8413 | 7757 | 5009 | 5214 | 5247 | 7697 | 4647 | 6181 | 9698 | 10880 | 12072 | 12399 | 11643 | 7695 | 5678 | 8575 | 3168 | 4272 | 4377 | Scn3b |
| Scn4a | 32 | 34 | 36 | 37 | 12 | 3 | 20 | 16 | 18 | 7 | 115 | 10 | 15 | 15 | 17 | 30 | 37 | 56 | 16 | 6 | 22 | 20 | 75 | 7 | 19 | 13 | 20 | 15 | Scn4a |
| Scn4b | 83 | 73 | 64 | 46 | 52 | 42 | 42 | 51 | 38 | 63 | 50 | 80 | 78 | 76 | 57 | 72 | 50 | 33 | 48 | 55 | 58 | 37 | 66 | 83 | 58 | 92 | 83 | 75 | Scn4b |
| Scn5a | 1488 | 1541 | 1425 | 1911 | 1440 | 1264 | 1713 | 509 | 663 | 625 | 585 | 911 | 797 | 827 | 1098 | 1233 | 1737 | 1585 | 1721 | 1414 | 1348 | 485 | 360 | 789 | 607 | 932 | 608 | 769 | Scn5a |
| Scn7a | 165 | 157 | 55 | 30 | 555 | 55 | 18 | 607 | 676 | 381 | 721 | 60 | 344 | 35 | 413 | 364 | 86 | 237 | 53 | 63 | 37 | 830 | 484 | 30 | 340 | 7 | 71 | 34 | Scn7a |
| Scn8a | 3715 | 4263 | 3457 | 5256 | 6631 | 6182 | 5880 | 4352 | 4388 | 5601 | 4606 | 5217 | 4614 | 5174 | 4025 | 3937 | 4692 | 5410 | 6007 | 4927 | 6408 | 5464 | 3671 | 4767 | 6068 | 3098 | 5340 | 4675 | Scn8a |
| Scn9a | 1610 | 1378 | 450 | 836 | 2993 | 1220 | 983 | 4885 | 4624 | 2957 | 3930 | 944 | 3256 | 989 | 2533 | 2867 | 830 | 1459 | 890 | 1273 | 1090 | 4737 | 2955 | 1107 | 2729 | 656 | 1110 | 794 | Scn9a |
| Scnm1 | 1908 | 1973 | 2369 | 2165 | 1813 | 2014 | 1929 | 2676 | 2447 | 2979 | 2520 | 2877 | 2583 | 3078 | 1960 | 1709 | 2453 | 1929 | 2276 | 2393 | 2036 | 2336 | 2513 | 3000 | 2779 | 3129 | 2988 | 3067 | Scnm1 |
| Scnn1a | 60 | 42 | 39 | 49 | 31 | 38 | 19 | 71 | 105 | 83 | 86 | 51 | 66 | 49 | 47 | 51 | 58 | 37 | 33 | 26 | 37 | 97 | 83 | 80 | 54 | 80 | 65 | 62 | Scnn1a |
| Scnn1b | 14 | 15 | 2 | 2 | 3 | 4 | 2 | 5 | 7 | 5 | 5 | 0 | 0 | 10 | 1 | 9 | 7 | 1 | 6 | 1 | 1 | 12 | 11 | 6 | 8 | 13 | 0 | 2 | Scnn1b |
| Scnn1g | 28 | 14 | 3 | 0 | 2 | 0 | 0 | 9 | 26 | 2 | 2 | 2 | 7 | 4 | 12 | 24 | 1 | 6 | 0 | 0 | 3 | 2 | 18 | 6 | 2 | 4 | 2 | 2 | Scnn1g |
| Sco1 | 1563 | 1515 | 1596 | 1507 | 1235 | 1495 | 1484 | 1254 | 1155 | 1075 | 1269 | 1313 | 1195 | 1315 | 1622 | 1561 | 1435 | 1456 | 1396 | 1413 | 1354 | 1190 | 1184 | 1178 | 1118 | 1418 | 1455 | 1466 | Sco1 |
| Sco2 | 13 | 10 | 12 | 24 | 11 | 3 | 7 | 34 | 23 | 44 | 35 | 26 | 24 | 29 | 9 | 2 | 14 | 8 | 9 | 14 | 12 | 38 | 28 | 42 | 18 | 29 | 30 | 47 | Sco2 |
| Scoc | 6987 | 5897 | 7087 | 7118 | 6870 | 6881 | 8375 | 7731 | 6512 | 5818 | 6195 | 6414 | 6910 | 7550 | 8004 | 7185 | 6614 | 7273 | 6936 | 7941 | 7381 | 7123 | 6138 | 6286 | 6557 | 6656 | 6994 | 7190 | Scoc |
| Scp2 | 6216 | 5532 | 6710 | 5615 | 5193 | 5175 | 6478 | 6483 | 5529 | 4724 | 4986 | 5779 | 5593 | 6278 | 7259 | 5910 | 6182 | 5407 | 5644 | 7007 | 5778 | 5243 | 5721 | 5503 | 5304 | 6609 | 6352 | 6687 | Scp2 |
| Scp2d1 | 0 | 0 | 0 | 0 | 0 | 0 | 0 | 0 | 0 | 0 | 0 | 0 | 0 | 0 | 0 | 0 | 0 | 0 | 4 | 0 | 0 | 0 | 0 | 0 | 0 | 0 | 0 | 0 | Scp2d1 |
| Scpep1 | 4148 | 3924 | 4459 | 3244 | 2826 | 2953 | 2775 | 2378 | 2518 | 2198 | 2573 | 2566 | 2782 | 2533 | 3787 | 4166 | 4016 | 3342 | 3068 | 3138 | 2670 | 2186 | 2725 | 2530 | 2183 | 3457 | 3091 | 2686 | Scpep1 |
| Scrg1 | 13 | 5 | 7 | 13 | 11 | 12 | 15 | 33 | 39 | 35 | 63 | 9 | 14 | 7 | 18 | 14 | 6 | 12 | 9 | 25 | 32 | 63 | 34 | 35 | 29 | 32 | 15 | 1 | Scrg1 |
| Scrib | 9325 | 13381 | 15721 | 15095 | 13192 | 12263 | 10934 | 7838 | 9322 | 10616 | 10057 | 13579 | 10917 | 11934 | 9274 | 11894 | 15574 | 14224 | 14767 | 10203 | 11995 | 10532 | 8398 | 12410 | 11434 | 13097 | 11884 | 12269 | Scrib |
| Scrn1 | 13399 | 14692 | 11195 | 15361 | 17117 | 16624 | 18633 | 13319 | 13659 | 13573 | 10510 | 15083 | 11742 | 15616 | 14569 | 11210 | 14465 | 15449 | 18466 | 16645 | 19050 | 13067 | 10120 | 13571 | 16088 | 13303 | 13466 | 13795 | Scrn1 |
| Scrn2 | 599 | 660 | 717 | 496 | 442 | 489 | 403 | 744 | 818 | 785 | 786 | 1060 | 1016 | 885 | 513 | 595 | 644 | 490 | 605 | 536 | 390 | 570 | 838 | 865 | 829 | 1183 | 1191 | 990 | Scrn2 |
| Scrn3 | 443 | 371 | 421 | 425 | 387 | 438 | 553 | 950 | 841 | 760 | 836 | 873 | 845 | 930 | 504 | 396 | 411 | 430 | 391 | 487 | 433 | 765 | 950 | 840 | 670 | 1001 | 869 | 890 | Scrn3 |
| Scrt1 | 1197 | 1933 | 2424 | 3113 | 4060 | 2913 | 2516 | 3722 | 3642 | 6315 | 4887 | 6204 | 5801 | 5666 | 1539 | 1284 | 3296 | 2950 | 4058 | 2346 | 3083 | 7113 | 3379 | 6129 | 7023 | 4990 | 5054 | 5237 | Scrt1 |
| Scrt2 | 915 | 1438 | 1667 | 2045 | 3584 | 2230 | 2000 | 4280 | 4384 | 6483 | 5078 | 6156 | 6338 | 6075 | 1625 | 1250 | 2516 | 2020 | 2853 | 1735 | 2246 | 6691 | 3726 | 6338 | 6896 | 5135 | 4637 | 5479 | Scrt2 |
| Sct | 12 | 19 | 17 | 18 | 16 | 11 | 8 | 6 | 15 | 8 | 20 | 11 | 18 | 9 | 12 | 19 | 8 | 10 | 9 | 9 | 8 | 10 | 20 | 15 | 3 | 17 | 1 | 9 | Sct |
| Sctr | 46 | 75 | 58 | 33 | 87 | 65 | 63 | 61 | 65 | 57 | 90 | 97 | 125 | 81 | 53 | 67 | 37 | 53 | 89 | 65 | 62 | 90 | 97 | 74 | 88 | 70 | 92 | 70 | Sctr |
| Scube1 | 5832 | 6855 | 6951 | 6329 | 6115 | 5356 | 4452 | 2947 | 3583 | 4339 | 4864 | 4312 | 4998 | 3683 | 4909 | 7870 | 6313 | 5744 | 5797 | 4480 | 4875 | 3982 | 3100 | 4181 | 3650 | 3714 | 4266 | 3763 | Scube1 |
| Scube2 | 151 | 238 | 238 | 129 | 135 | 225 | 134 | 231 | 238 | 156 | 368 | 226 | 263 | 200 | 216 | 241 | 210 | 207 | 218 | 185 | 154 | 212 | 279 | 212 | 218 | 148 | 244 | 280 | Scube2 |
| Scube3 | 4395 | 5461 | 7414 | 4093 | 3738 | 4464 | 2847 | 5083 | 5924 | 7184 | 6786 | 7688 | 8043 | 7259 | 4378 | 5772 | 5704 | 4278 | 3785 | 3209 | 3080 | 7600 | 5447 | 7709 | 6546 | 6989 | 7587 | 8088 | Scube3 |
| Scx | 59 | 59 | 34 | 45 | 45 | 28 | 27 | 154 | 235 | 211 | 434 | 79 | 175 | 55 | 63 | 65 | 63 | 36 | 39 | 15 | 22 | 177 | 287 | 122 | 107 | 221 | 88 | 76 | Scx |
| Scyl1 | 3915 | 5054 | 4471 | 5016 | 4553 | 4373 | 4142 | 3214 | 4228 | 4325 | 4086 | 4117 | 3876 | 3618 | 3866 | 4781 | 4889 | 4676 | 4720 | 4323 | 4307 | 3820 | 4323 | 4092 | 4250 | 4457 | 3694 | 3801 | Scyl1 |
| Scyl2 | 9945 | 9388 | 8325 | 9059 | 8825 | 9043 | 9624 | 8156 | 6850 | 6301 | 6100 | 6799 | 6634 | 6779 | 10485 | 9493 | 8393 | 9084 | 8762 | 9904 | 9397 | 7072 | 6697 | 6037 | 6727 | 6043 | 6838 | 6699 | Scyl2 |
| Scyl3 | 3028 | 2538 | 2340 | 2533 | 2437 | 2684 | 2635 | 2593 | 2427 | 2305 | 2260 | 2499 | 2463 | 2511 | 2879 | 2805 | 2351 | 2292 | 2403 | 2402 | 2637 | 2593 | 2398 | 2291 | 2386 | 2242 | 2474 | 2365 | Scyl3 |
| Sdad1 | 6802 | 6522 | 6329 | 5600 | 5276 | 5884 | 5322 | 5648 | 5466 | 4914 | 5430 | 5622 | 5704 | 5847 | 6493 | 6269 | 5791 | 5426 | 5390 | 5944 | 5318 | 4725 | 5011 | 5100 | 5471 | 5408 | 6213 | 6092 | Sdad1 |
| Sdc1 | 6472 | 6360 | 6392 | 4390 | 3549 | 4595 | 2954 | 4004 | 5138 | 3888 | 4517 | 4157 | 4659 | 3777 | 5196 | 6867 | 4620 | 4429 | 3772 | 3761 | 3231 | 2884 | 5187 | 4106 | 3146 | 5411 | 4742 | 4353 | Sdc1 |
| Sdc2 | 3603 | 2753 | 3282 | 1555 | 1717 | 2134 | 1206 | 3519 | 4185 | 2994 | 3946 | 2879 | 4092 | 3190 | 3535 | 4213 | 2213 | 2177 | 1282 | 1440 | 1221 | 3270 | 4610 | 3034 | 2606 | 3367 | 3550 | 3514 | Sdc2 |
| Sdc3 | 6251 | 7510 | 7237 | 9783 | 13875 | 11760 | 9821 | 12379 | 15565 | 17418 | 16998 | 12950 | 11595 | 11638 | 6769 | 7387 | 8441 | 11392 | 12432 | 9728 | 11424 | 19590 | 14912 | 15475 | 18917 | 12490 | 11240 | 10423 | Sdc3 |
| Sdc4 | 1562 | 1107 | 1545 | 1199 | 815 | 982 | 760 | 799 | 1144 | 1056 | 1265 | 648 | 780 | 599 | 1298 | 1469 | 1239 | 1093 | 787 | 800 | 734 | 693 | 1193 | 944 | 603 | 1051 | 754 | 705 | Sdc4 |
| Sdcbp | 23028 | 20064 | 18290 | 18836 | 16134 | 17350 | 19330 | 15242 | 12883 | 10536 | 10777 | 12086 | 12809 | 12509 | 25939 | 22332 | 19262 | 18358 | 17256 | 21638 | 18206 | 12482 | 14215 | 10793 | 10758 | 13283 | 12478 | 13602 | Sdcbp |
| Sdcbp2 | 42 | 45 | 26 | 61 | 38 | 29 | 54 | 40 | 47 | 52 | 42 | 58 | 32 | 43 | 46 | 66 | 38 | 56 | 58 | 38 | 51 | 32 | 40 | 30 | 21 | 34 | 29 | 23 | Sdcbp2 |
| Sdccag8 | 1653 | 1831 | 1783 | 1857 | 1687 | 1565 | 1767 | 1211 | 1091 | 1253 | 1075 | 1083 | 1191 | 1160 | 1775 | 1586 | 1866 | 1746 | 1753 | 1768 | 1774 | 1041 | 1033 | 1113 | 1180 | 995 | 1104 | 1105 | Sdccag8 |
| Sde2 | 3748 | 3834 | 3805 | 3711 | 3088 | 3153 | 3016 | 2895 | 2766 | 2864 | 3065 | 3041 | 2854 | 2968 | 3422 | 3487 | 3413 | 3630 | 3274 | 3372 | 3144 | 2744 | 2823 | 2957 | 3101 | 3265 | 3098 | 2995 | Sde2 |
| Sdf2 | 2272 | 2590 | 2569 | 2382 | 2074 | 1834 | 1896 | 3174 | 3026 | 2963 | 2753 | 2712 | 2749 | 2736 | 2425 | 2437 | 2284 | 2271 | 2222 | 2478 | 1995 | 2991 | 3330 | 2951 | 2893 | 2943 | 2924 | 2789 | Sdf2 |
| Sdf2l1 | 1589 | 1918 | 2351 | 2017 | 1289 | 1309 | 1326 | 880 | 751 | 935 | 867 | 905 | 806 | 807 | 1659 | 1695 | 2255 | 1617 | 1659 | 1308 | 1365 | 668 | 805 | 1154 | 785 | 1143 | 895 | 991 | Sdf2l1 |
| Sdf4 | 6427 | 6071 | 6034 | 5582 | 5465 | 5606 | 5493 | 7978 | 7687 | 7450 | 7620 | 7910 | 7578 | 8062 | 6692 | 6069 | 5781 | 5399 | 5699 | 5555 | 5512 | 8079 | 8106 | 7906 | 7557 | 8734 | 8112 | 8472 | Sdf4 |
| Sdha | 15428 | 16381 | 16064 | 14080 | 13506 | 13720 | 14086 | 14959 | 14956 | 14302 | 13890 | 16232 | 15316 | 16313 | 15033 | 14447 | 15410 | 13537 | 14662 | 14513 | 14329 | 13800 | 14630 | 15496 | 14662 | 17713 | 16707 | 16624 | Sdha |
| Sdhaf1 | 1213 | 1398 | 1270 | 1258 | 1035 | 941 | 1133 | 827 | 833 | 778 | 771 | 913 | 848 | 1114 | 1216 | 1119 | 1240 | 922 | 1084 | 1038 | 1171 | 735 | 788 | 963 | 883 | 1186 | 954 | 1102 | Sdhaf1 |
| Sdhaf2 | 2868 | 2693 | 2747 | 2552 | 2361 | 2556 | 2680 | 3093 | 2718 | 2673 | 2553 | 2521 | 2751 | 2841 | 2987 | 2839 | 2562 | 2319 | 2427 | 2727 | 2467 | 2918 | 2874 | 2773 | 2513 | 2763 | 2751 | 2980 | Sdhaf2 |
| Sdhaf3 | 1223 | 891 | 967 | 968 | 996 | 1024 | 1113 | 1047 | 883 | 781 | 897 | 997 | 937 | 1070 | 1303 | 986 | 1008 | 935 | 994 | 1143 | 1015 | 847 | 893 | 905 | 894 | 1042 | 926 | 1050 | Sdhaf3 |
| Sdhaf4 | 953 | 852 | 879 | 982 | 1017 | 891 | 1341 | 1139 | 1107 | 1012 | 1076 | 958 | 954 | 1278 | 1072 | 771 | 979 | 950 | 1044 | 1131 | 1312 | 1021 | 1056 | 1177 | 1063 | 1132 | 968 | 1164 | Sdhaf4 |
| Sdhb | 6412 | 7719 | 6245 | 6304 | 5789 | 6377 | 6421 | 6408 | 6532 | 6096 | 5467 | 6500 | 6017 | 6334 | 6799 | 6437 | 6163 | 6270 | 7034 | 7320 | 6785 | 5090 | 6572 | 6331 | 6278 | 7225 | 6801 | 6810 | Sdhb |
| Sdhc | 7476 | 9075 | 8728 | 7789 | 7471 | 7669 | 7762 | 7279 | 7742 | 7549 | 7438 | 8635 | 8214 | 8141 | 7482 | 7785 | 8447 | 7412 | 7994 | 7602 | 7436 | 6958 | 8336 | 8297 | 7910 | 9721 | 9036 | 9176 | Sdhc |
| Sdhd | 6515 | 6682 | 6199 | 5201 | 4717 | 4768 | 4976 | 5816 | 5461 | 4516 | 4975 | 5118 | 5237 | 5044 | 6641 | 6335 | 5873 | 5130 | 5128 | 6293 | 4775 | 4508 | 5641 | 5006 | 4748 | 6115 | 5396 | 5529 | Sdhd |
| Sdk1 | 1331 | 1747 | 1433 | 1625 | 1946 | 1605 | 1037 | 1586 | 2019 | 2045 | 2836 | 1539 | 2011 | 1325 | 1386 | 2094 | 1483 | 2056 | 1390 | 1202 | 1293 | 2358 | 2298 | 1902 | 1825 | 1286 | 1342 | 1233 | Sdk1 |
| Sdk2 | 1239 | 1566 | 2136 | 1697 | 2240 | 2009 | 1164 | 1570 | 1973 | 2637 | 3346 | 2338 | 2882 | 1895 | 1250 | 1793 | 2085 | 1935 | 1878 | 1229 | 1493 | 2918 | 2071 | 2412 | 2434 | 1027 | 2444 | 2139 | Sdk2 |
| Sdr16c5 | 0 | 0 | 0 | 0 | 0 | 0 | 0 | 0 | 2 | 0 | 0 | 0 | 0 | 0 | 0 | 0 | 0 | 0 | 0 | 0 | 0 | 0 | 2 | 0 | 0 | 0 | 0 | 0 | Sdr16c5 |
| Sdr16c6 | 0 | 0 | 0 | 0 | 0 | 0 | 0 | 5 | 16 | 7 | 2 | 2 | 0 | 0 | 0 | 0 | 0 | 0 | 0 | 0 | 0 | 0 | 6 | 0 | 0 | 0 | 0 | 0 | Sdr16c6 |
| Sdr39u1 | 871 | 837 | 653 | 862 | 1216 | 1112 | 1066 | 1263 | 1626 | 1567 | 1585 | 1146 | 1221 | 1189 | 816 | 685 | 749 | 962 | 1087 | 1114 | 1067 | 1459 | 1761 | 1376 | 1601 | 1375 | 1113 | 1167 | Sdr39u1 |
| Sdr42e1 | 161 | 130 | 121 | 141 | 108 | 67 | 89 | 92 | 105 | 80 | 106 | 89 | 67 | 63 | 131 | 144 | 137 | 133 | 84 | 84 | 86 | 74 | 131 | 94 | 55 | 92 | 48 | 78 | Sdr42e1 |
| Sdr9c7 | 0 | 3 | 0 | 0 | 0 | 0 | 0 | 0 | 1 | 0 | 0 | 1 | 0 | 0 | 1 | 0 | 0 | 0 | 0 | 0 | 0 | 0 | 2 | 0 | 0 | 0 | 0 | 0 | Sdr9c7 |
| Sds | 0 | 0 | 0 | 4 | 0 | 3 | 0 | 1 | 0 | 5 | 0 | 1 | 0 | 0 | 0 | 0 | 1 | 0 | 1 | 0 | 0 | 0 | 1 | 1 | 0 | 3 | 1 | 0 | Sds |
| Sdsl | 107 | 177 | 191 | 192 | 229 | 156 | 164 | 187 | 166 | 227 | 217 | 281 | 242 | 241 | 142 | 143 | 170 | 182 | 183 | 198 | 174 | 222 | 201 | 255 | 229 | 216 | 268 | 254 | Sdsl |
| Sebox | 2 | 9 | 17 | 5 | 14 | 5 | 10 | 10 | 5 | 3 | 5 | 5 | 3 | 10 | 1 | 0 | 8 | 8 | 10 | 4 | 4 | 7 | 5 | 6 | 10 | 7 | 12 | 8 | Sebox |
| Sec1 | 31 | 25 | 36 | 16 | 30 | 25 | 16 | 62 | 126 | 124 | 128 | 50 | 96 | 42 | 21 | 44 | 14 | 23 | 11 | 22 | 13 | 38 | 119 | 99 | 41 | 89 | 48 | 51 | Sec1 |
| Sec11a | 6391 | 6050 | 6753 | 5663 | 4671 | 4961 | 5041 | 4925 | 4633 | 4386 | 4909 | 4667 | 4943 | 4550 | 6675 | 6312 | 5782 | 5471 | 5047 | 6146 | 4758 | 4299 | 4710 | 4673 | 4548 | 5391 | 4824 | 4940 | Sec11a |
| Sec11c | 2931 | 2763 | 2291 | 2680 | 2520 | 2384 | 2958 | 1606 | 1428 | 1412 | 1574 | 1160 | 1109 | 1384 | 2850 | 2684 | 2432 | 2632 | 2467 | 2715 | 2768 | 1457 | 1494 | 1449 | 1398 | 1201 | 1115 | 1233 | Sec11c |
| Sec13 | 6159 | 7013 | 5986 | 5319 | 4260 | 4698 | 4869 | 5926 | 6423 | 5201 | 5594 | 5908 | 5666 | 5953 | 6003 | 6231 | 5642 | 4758 | 4850 | 5628 | 4849 | 4769 | 6629 | 5754 | 5574 | 6657 | 6186 | 6524 | Sec13 |
| Sec14l1 | 12696 | 16602 | 16954 | 18887 | 16494 | 15380 | 15758 | 7124 | 8447 | 9331 | 9182 | 9565 | 8850 | 8978 | 12899 | 15471 | 18512 | 19008 | 17681 | 13612 | 16303 | 9611 | 8334 | 10163 | 10373 | 9249 | 8526 | 8850 | Sec14l1 |
| Sec14l2 | 390 | 418 | 484 | 500 | 571 | 457 | 398 | 365 | 391 | 416 | 347 | 268 | 329 | 316 | 404 | 483 | 468 | 419 | 493 | 476 | 416 | 299 | 373 | 385 | 317 | 447 | 328 | 318 | Sec14l2 |
| Sec14l3 | 3 | 5 | 3 | 3 | 7 | 5 | 4 | 22 | 12 | 16 | 8 | 8 | 0 | 16 | 1 | 2 | 2 | 4 | 2 | 5 | 4 | 13 | 23 | 20 | 8 | 20 | 2 | 4 | Sec14l3 |
| Sec14l4 | 0 | 3 | 0 | 1 | 11 | 0 | 0 | 5 | 9 | 16 | 11 | 0 | 6 | 6 | 0 | 1 | 0 | 0 | 0 | 0 | 0 | 2 | 19 | 12 | 1 | 17 | 0 | 0 | Sec14l4 |
| Sec14l5 | 945 | 1394 | 1299 | 1509 | 1478 | 1370 | 1107 | 220 | 182 | 319 | 256 | 277 | 400 | 324 | 1133 | 1283 | 1477 | 1428 | 1438 | 1315 | 1314 | 286 | 232 | 249 | 259 | 74 | 351 | 286 | Sec14l5 |
| Sec16a | 14110 | 15136 | 11810 | 15832 | 14236 | 15031 | 14996 | 9360 | 10915 | 11593 | 11802 | 11855 | 11590 | 11300 | 13589 | 14967 | 13662 | 16325 | 14657 | 11666 | 15691 | 11277 | 10207 | 10414 | 11819 | 7671 | 11381 | 11057 | Sec16a |
| Sec16b | 173 | 155 | 191 | 183 | 146 | 162 | 153 | 155 | 263 | 240 | 445 | 164 | 211 | 137 | 210 | 213 | 159 | 186 | 169 | 161 | 126 | 175 | 297 | 174 | 152 | 109 | 120 | 114 | Sec16b |
| Sec22a | 2513 | 2230 | 1891 | 1942 | 2003 | 2212 | 2332 | 2331 | 2078 | 1816 | 1734 | 1859 | 1714 | 1827 | 2769 | 2180 | 2017 | 2035 | 2033 | 2386 | 2155 | 1843 | 2194 | 1688 | 1717 | 1977 | 1882 | 2057 | Sec22a |
| Sec22b | 7199 | 6625 | 6310 | 6298 | 5357 | 5893 | 5944 | 6050 | 5416 | 4766 | 5550 | 5171 | 5151 | 5072 | 7423 | 6718 | 6277 | 5973 | 5350 | 6066 | 5220 | 5623 | 5510 | 5002 | 4835 | 5869 | 5027 | 5343 | Sec22b |

| | | | | | | | | | | | | | | | | | | | | | | | | | | | | | |
|---|---|---|---|---|---|---|---|---|---|---|---|---|---|---|---|---|---|---|---|---|---|---|---|---|---|---|---|---|---|
| Sec22c | 4336 | 4710 | 5220 | 6595 | 6199 | 5948 | 6257 | 3621 | 3795 | 4432 | 4022 | 4760 | 4234 | 4644 | 4311 | 4463 | 5880 | 6498 | 6225 | 4958 | 6289 | 5083 | 3357 | 4747 | 4912 | 4274 | 4428 | 4799 | Sec22c |
| Sec23a | 15174 | 13646 | 12687 | 13380 | 13404 | 14263 | 14357 | 13081 | 11912 | 10011 | 11395 | 10432 | 10470 | 10587 | 16264 | 14018 | 13059 | 13760 | 13108 | 15921 | 14299 | 12373 | 12095 | 10239 | 11168 | 9366 | 10466 | 10372 | Sec23a |
| Sec23b | 9529 | 10271 | 8557 | 7919 | 6788 | 7203 | 7588 | 6433 | 5622 | 4878 | 4887 | 5973 | 5725 | 5995 | 9650 | 8416 | 8270 | 7259 | 7646 | 8375 | 8030 | 4705 | 5726 | 5434 | 5066 | 6518 | 6566 | 6710 | Sec23b |
| Sec23ip | 8502 | 9016 | 8602 | 8490 | 7780 | 7753 | 8492 | 6685 | 6620 | 6769 | 6666 | 7580 | 7106 | 8095 | 8558 | 8491 | 8613 | 8386 | 7528 | 7805 | 8179 | 7046 | 6308 | 6683 | 6897 | 7558 | 7673 | 7805 | Sec23ip |
| Sec24a | 3509 | 3052 | 3899 | 4163 | 4118 | 4213 | 4842 | 2761 | 2763 | 2893 | 3461 | 3254 | 3151 | 3228 | 3493 | 3529 | 3888 | 4242 | 3933 | 3313 | 4543 | 3419 | 2733 | 2862 | 3107 | 2676 | 2876 | 3071 | Sec24a |
| Sec24b | 5037 | 5151 | 5802 | 5832 | 6656 | 5941 | 6265 | 5923 | 6292 | 7263 | 6785 | 7274 | 6632 | 7119 | 4894 | 5155 | 5837 | 5841 | 6399 | 4764 | 6262 | 7417 | 6104 | 6910 | 6996 | 6592 | 6177 | 7115 | Sec24b |
| Sec24c | 15379 | 15633 | 17486 | 16740 | 15185 | 14632 | 15024 | 12169 | 12561 | 12937 | 12894 | 13599 | 13116 | 13531 | 14747 | 15283 | 17398 | 16314 | 16184 | 13167 | 15272 | 14292 | 11829 | 13411 | 13001 | 14232 | 12753 | 13205 | Sec24c |
| Sec24d | 3718 | 3141 | 2879 | 2425 | 2265 | 2515 | 2050 | 1899 | 3014 | 2111 | 3619 | 1664 | 2315 | 1542 | 3137 | 3847 | 2577 | 2839 | 2123 | 1953 | 2043 | 2044 | 3632 | 1982 | 1801 | 1969 | 1868 | 1634 | Sec24d |
| Sec31a | 13055 | 13054 | 12057 | 11555 | 10292 | 10902 | 10800 | 10086 | 12021 | 11812 | 12821 | 11291 | 11247 | 10666 | 11710 | 12878 | 11652 | 12025 | 10977 | 9662 | 10617 | 11956 | 12485 | 11140 | 11001 | 10849 | 10528 | 10948 | Sec31a |
| Sec31b | 130 | 155 | 157 | 162 | 129 | 58 | 83 | 42 | 65 | 35 | 78 | 61 | 66 | 45 | 105 | 186 | 154 | 108 | 132 | 132 | 95 | 50 | 49 | 50 | 42 | 44 | 86 | 58 | Sec31b |
| Sec61a1 | 14245 | 16565 | 14085 | 13364 | 12130 | 12784 | 11833 | 10408 | 12428 | 11026 | 12349 | 11429 | 12038 | 10380 | 13843 | 15704 | 13282 | 13617 | 12970 | 12645 | 11754 | 10901 | 12442 | 11265 | 11376 | 12405 | 11898 | 10964 | Sec61a1 |
| Sec61a2 | 7982 | 6398 | 5846 | 5926 | 6658 | 7860 | 7096 | 6831 | 6555 | 6094 | 6910 | 6151 | 6454 | 6049 | 7512 | 7372 | 5676 | 6236 | 6265 | 7750 | 7132 | 6152 | 6356 | 5695 | 6131 | 5509 | 6148 | 5821 | Sec61a2 |
| Sec61b | 2887 | 3534 | 3295 | 3038 | 2447 | 2341 | 2813 | 2405 | 2459 | 2418 | 2727 | 2487 | 2755 | 3034 | 2832 | 2794 | 3330 | 2860 | 2603 | 2979 | 2707 | 1801 | 2504 | 2938 | 2503 | 3175 | 2743 | 3252 | Sec61b |
| Sec61g | 4176 | 4000 | 3770 | 4120 | 3268 | 2848 | 5374 | 4092 | 3269 | 4123 | 4362 | 3587 | 3768 | 5916 | 3967 | 2759 | 4194 | 3207 | 3317 | 3814 | 5028 | 3208 | 3486 | 4534 | 3906 | 4204 | 3810 | 6044 | Sec61g |
| Sec62 | 8397 | 6996 | 8657 | 7819 | 7829 | 7859 | 8487 | 11615 | 9648 | 9610 | 10044 | 9677 | 9668 | 9807 | 9175 | 8308 | 7789 | 7704 | 7678 | 9200 | 8026 | 10870 | 10508 | 9671 | 9613 | 10368 | 9874 | 10007 | Sec62 |
| Sec63 | 14762 | 11231 | 13845 | 14516 | 13215 | 14321 | 15525 | 13283 | 11284 | 12339 | 12540 | 14207 | 12703 | 15128 | 14277 | 13030 | 13969 | 13777 | 13344 | 12378 | 14430 | 13272 | 11086 | 13473 | 13101 | 13761 | 13763 | 14938 | Sec63 |
| Secisbp2 | 781 | 935 | 982 | 961 | 1109 | 993 | 977 | 1970 | 2115 | 2172 | 2147 | 2497 | 2384 | 2258 | 901 | 769 | 950 | 851 | 1160 | 1031 | 869 | 2107 | 2137 | 2344 | 2304 | 2432 | 2501 | 2409 | Secisbp2 |
| Secisbp2l | 10010 | 8935 | 8898 | 10920 | 11126 | 11303 | 12240 | 9116 | 8680 | 9055 | 9712 | 9479 | 9615 | 9961 | 10626 | 9583 | 9826 | 11157 | 10649 | 11107 | 11614 | 9804 | 8376 | 8936 | 10351 | 7153 | 9635 | 9327 | Secisbp2l |
| Sectm1a | 1 | 0 | 0 | 0 | 1 | 2 | 0 | 0 | 5 | 0 | 2 | 4 | 0 | 0 | 0 | 0 | 0 | 0 | 0 | 0 | 0 | 0 | 9 | 0 | 0 | 1 | 2 | 2 | Sectm1a |
| Sectm1b | 0 | 0 | 0 | 0 | 0 | 0 | 0 | 1 | 2 | 0 | 1 | 0 | 0 | 0 | 0 | 0 | 0 | 0 | 0 | 0 | 0 | 2 | 5 | 0 | 0 | 0 | 0 | 0 | Sectm1b |
| Seh1l | 3775 | 2641 | 3038 | 2618 | 2828 | 2952 | 3222 | 7646 | 7011 | 6235 | 6641 | 7281 | 7193 | 7794 | 3456 | 2868 | 2960 | 2735 | 2929 | 3187 | 2784 | 6005 | 6829 | 6848 | 6347 | 8347 | 7645 | 7734 | Seh1l |
| Sel1l | 14671 | 14463 | 11551 | 14198 | 13113 | 13686 | 14250 | 10220 | 10385 | 11198 | 11601 | 10547 | 10553 | 10666 | 14004 | 14932 | 12217 | 15153 | 13167 | 11876 | 14885 | 11575 | 10049 | 10508 | 11524 | 8052 | 10197 | 10361 | Sel1l |
| Sel1l2 | 0 | 2 | 5 | 0 | 0 | 0 | 1 | 0 | 0 | 1 | 3 | 1 | 1 | 0 | 0 | 2 | 0 | 2 | 0 | 0 | 2 | 0 | 1 | 0 | 0 | 2 | 0 | 1 | Sel1l2 |
| Sel1l3 | 975 | 1020 | 880 | 1133 | 1680 | 1458 | 1785 | 1762 | 2105 | 1944 | 1842 | 2175 | 1413 | 2208 | 1093 | 806 | 1191 | 1385 | 1707 | 1441 | 1630 | 1913 | 1733 | 2075 | 2394 | 2126 | 1514 | 1762 | Sel1l3 |
| Sele | 0 | 0 | 0 | 0 | 0 | 3 | 0 | 1 | 0 | 0 | 7 | 0 | 0 | 0 | 1 | 2 | 1 | 1 | 0 | 0 | 1 | 0 | 4 | 2 | 0 | 0 | 0 | 0 | Sele |
| Selenbp1 | 268 | 313 | 240 | 322 | 353 | 237 | 240 | 396 | 560 | 462 | 548 | 275 | 314 | 219 | 247 | 350 | 251 | 318 | 300 | 270 | 292 | 364 | 632 | 380 | 358 | 288 | 238 | 274 | Selenbp1 |
| Selenbp2 | 2 | 0 | 0 | 0 | 1 | 1 | 1 | 5 | 3 | 2 | 5 | 0 | 1 | 0 | 1 | 4 | 0 | 0 | 0 | 1 | 0 | 0 | 5 | 5 | 2 | 0 | 0 | 0 | Selenbp2 |
| Selenof | 16237 | 14299 | 14539 | 15097 | 12773 | 14260 | 15535 | 14493 | 13984 | 11930 | 12192 | 11905 | 11953 | 13050 | 17664 | 15888 | 14764 | 13874 | 14251 | 18052 | 15689 | 11150 | 14956 | 12463 | 11712 | 13432 | 12604 | 13992 | Selenof |
| Selenoh | 5341 | 6338 | 6252 | 5870 | 4610 | 5182 | 5599 | 4493 | 4641 | 4637 | 4533 | 6348 | 5787 | 6924 | 5070 | 4996 | 6189 | 5039 | 5675 | 6778 | 5168 | 3205 | 4154 | 6104 | 5335 | 8055 | 7089 | 6754 | Selenoh |
| Selenoi | 8035 | 7603 | 6133 | 7995 | 8080 | 8393 | 7577 | 7697 | 6481 | 6033 | 6155 | 6206 | 6009 | 6067 | 8327 | 7718 | 7010 | 7998 | 7534 | 9123 | 7576 | 6706 | 6260 | 5600 | 7128 | 5180 | 6359 | 5891 | Selenoi |
| Selenok | 5995 | 5918 | 6159 | 5753 | 5470 | 5006 | 6604 | 6699 | 5353 | 5466 | 5160 | 5015 | 5235 | 6424 | 6558 | 5122 | 5817 | 5477 | 5834 | 6368 | 6520 | 5857 | 5353 | 5726 | 5693 | 5687 | 5510 | 6342 | Selenok |
| Selenom | 1171 | 1456 | 1484 | 1407 | 981 | 1121 | 746 | 729 | 1089 | 921 | 1339 | 765 | 816 | 707 | 1261 | 1555 | 1327 | 1173 | 1050 | 959 | 827 | 722 | 1208 | 1028 | 666 | 1108 | 798 | 705 | Selenom |
| Selenon | 3160 | 4016 | 3982 | 3101 | 3193 | 3479 | 2686 | 4835 | 6298 | 6612 | 6825 | 7292 | 6688 | 6363 | 3210 | 4220 | 3845 | 3536 | 3497 | 2486 | 2747 | 6704 | 6354 | 7019 | 6255 | 7259 | 6251 | 6810 | Selenon |
| Selenoo | 1914 | 2088 | 1882 | 1904 | 1703 | 1947 | 1986 | 1881 | 1885 | 1930 | 1715 | 2044 | 1759 | 1845 | 1928 | 1767 | 1732 | 1837 | 2041 | 1966 | 1889 | 1765 | 1784 | 1940 | 1981 | 1950 | 1973 | 1850 | Selenoo |
| Selenop | 4984 | 4665 | 4944 | 4344 | 5458 | 5106 | 4399 | 5796 | 6208 | 5313 | 7379 | 4805 | 5706 | 4600 | 5806 | 6232 | 3975 | 5522 | 3921 | 5867 | 4690 | 6515 | 9552 | 4943 | 4928 | 5127 | 4998 | 4250 | Selenop |
| Selenos | 2291 | 2488 | 2212 | 2374 | 2068 | 2196 | 2304 | 1960 | 2125 | 1940 | 2253 | 1756 | 1927 | 1884 | 2443 | 2407 | 2187 | 2474 | 1894 | 2309 | 2081 | 1928 | 2238 | 2006 | 1843 | 2068 | 1740 | 1978 | Selenos |
| Selenot | 9357 | 7431 | 10652 | 9739 | 10690 | 9852 | 10966 | 12270 | 9619 | 9387 | 10357 | 10943 | 11186 | 11968 | 10259 | 8420 | 10009 | 9021 | 9933 | 10343 | 10372 | 12156 | 10372 | 10743 | 11491 | 12034 | 11685 | 12256 | Selenot |
| Selenov | 50 | 76 | 79 | 66 | 79 | 91 | 55 | 53 | 40 | 27 | 52 | 72 | 101 | 62 | 57 | 72 | 66 | 85 | 64 | 64 | 65 | 47 | 45 | 43 | 49 | 20 | 60 | 94 | Selenov |
| Selenow | 12222 | 15967 | 15507 | 16822 | 15855 | 14255 | 17511 | 12688 | 13648 | 14774 | 14345 | 14459 | 13888 | 18825 | 11772 | 11506 | 16474 | 15835 | 16476 | 14395 | 18283 | 11814 | 13215 | 17115 | 15831 | 16337 | 14367 | 18176 | Selenow |
| Sell | 2 | 0 | 2 | 3 | 1 | 5 | 0 | 4 | 2 | 2 | 9 | 3 | 5 | 2 | 3 | 2 | 7 | 7 | 2 | 0 | 2 | 0 | 8 | 8 | 3 | 0 | 1 | 2 | Sell |
| Selp | 0 | 6 | 0 | 1 | 2 | 0 | 1 | 1 | 2 | 1 | 8 | 0 | 0 | 0 | 0 | 2 | 0 | 3 | 0 | 1 | 0 | 2 | 1 | 3 | 2 | 2 | 2 | 1 | Selp |
| Selplg | 33 | 54 | 65 | 44 | 37 | 42 | 22 | 22 | 63 | 53 | 59 | 35 | 40 | 28 | 34 | 51 | 58 | 58 | 41 | 41 | 29 | 50 | 57 | 43 | 36 | 34 | 35 | 34 | Selplg |
| Sem1 | 5638 | 6154 | 6743 | 6227 | 4914 | 3923 | 7469 | 5410 | 4759 | 6260 | 5596 | 5954 | 5408 | 8812 | 5354 | 3817 | 6934 | 5072 | 5134 | 5861 | 7073 | 3778 | 4653 | 7750 | 5864 | 7338 | 6327 | 8812 | Sem1 |
| Sema3a | 3037 | 1948 | 1341 | 1410 | 1565 | 1978 | 1431 | 4735 | 5090 | 4221 | 4868 | 3028 | 3672 | 2796 | 2666 | 2970 | 1284 | 1577 | 1471 | 1633 | 1346 | 2912 | 5176 | 3395 | 2844 | 3587 | 2468 | 2514 | Sema3a |
| Sema3b | 181 | 212 | 173 | 153 | 239 | 186 | 129 | 206 | 311 | 263 | 435 | 231 | 289 | 182 | 212 | 234 | 149 | 180 | 120 | 157 | 128 | 309 | 407 | 232 | 212 | 239 | 225 | 177 | Sema3b |
| Sema3c | 2490 | 2109 | 2232 | 2062 | 1992 | 3371 | 2110 | 3980 | 4529 | 4157 | 6092 | 3968 | 3601 | 4292 | 3066 | 3066 | 1856 | 3061 | 1731 | 2041 | 1896 | 4306 | 6490 | 4166 | 4632 | 2678 | 3871 | 3967 | Sema3c |
| Sema3d | 1096 | 586 | 414 | 386 | 305 | 780 | 249 | 1871 | 2261 | 1584 | 2054 | 674 | 963 | 653 | 919 | 1044 | 407 | 611 | 275 | 304 | 251 | 1111 | 2487 | 1144 | 837 | 581 | 525 | 582 | Sema3d |
| Sema3e | 576 | 595 | 629 | 479 | 797 | 588 | 523 | 644 | 873 | 901 | 839 | 417 | 603 | 367 | 737 | 770 | 553 | 599 | 603 | 587 | 548 | 735 | 823 | 422 | 498 | 275 | 424 | 342 | Sema3e |
| Sema3f | 4509 | 5300 | 5592 | 5379 | 5033 | 4701 | 3623 | 3944 | 4525 | 5028 | 4929 | 4305 | 4648 | 3710 | 3778 | 6014 | 5380 | 5630 | 5051 | 3534 | 4018 | 5043 | 4709 | 5082 | 4612 | 3973 | 3977 | 3865 | Sema3f |
| Sema3g | 368 | 468 | 530 | 430 | 451 | 462 | 398 | 484 | 623 | 795 | 785 | 765 | 820 | 812 | 344 | 427 | 542 | 492 | 475 | 435 | 432 | 714 | 590 | 822 | 723 | 921 | 853 | 881 | Sema3g |
| Sema4a | 784 | 926 | 1089 | 1204 | 2014 | 1545 | 1506 | 1752 | 1795 | 2333 | 1828 | 2170 | 2036 | 2079 | 848 | 850 | 1374 | 1369 | 1796 | 1412 | 1503 | 2852 | 1581 | 2364 | 2558 | 1516 | 1923 | 1920 | Sema4a |
| Sema4b | 1242 | 1494 | 1781 | 1831 | 1928 | 1821 | 1728 | 1463 | 1749 | 2004 | 1797 | 2068 | 1919 | 1955 | 1175 | 1441 | 1750 | 1622 | 1945 | 1556 | 1648 | 1885 | 1668 | 2093 | 2148 | 2277 | 1814 | 1924 | Sema4b |
| Sema4c | 2753 | 2938 | 3748 | 3644 | 4174 | 3697 | 2998 | 5273 | 5833 | 6531 | 6170 | 7517 | 6538 | 6161 | 2397 | 2736 | 3596 | 3451 | 4414 | 2928 | 3105 | 7127 | 5299 | 7258 | 7377 | 7372 | 6968 | 6112 | Sema4c |
| Sema4d | 1628 | 1665 | 2047 | 2438 | 2630 | 2139 | 2312 | 957 | 1237 | 1289 | 1272 | 1092 | 924 | 1038 | 1402 | 1842 | 2255 | 2631 | 2356 | 1844 | 2288 | 1188 | 964 | 1322 | 1475 | 1173 | 851 | 892 | Sema4d |
| Sema4f | 961 | 1011 | 893 | 1302 | 2133 | 1358 | 1241 | 1202 | 1440 | 1688 | 1375 | 1102 | 1133 | 960 | 843 | 1035 | 1166 | 1462 | 1634 | 1162 | 1364 | 1811 | 1212 | 1262 | 1670 | 882 | 739 | 865 | Sema4f |
| Sema4g | 5938 | 6836 | 7234 | 8416 | 10696 | 8698 | 8341 | 5210 | 6192 | 7890 | 6823 | 6340 | 6357 | 6137 | 5730 | 7242 | 7967 | 8785 | 9725 | 6802 | 8655 | 8311 | 5515 | 6940 | 8034 | 5267 | 5441 | 5616 | Sema4g |
| Sema5a | 10337 | 6724 | 4104 | 7283 | 6950 | 9173 | 9635 | 9340 | 11187 | 9721 | 11512 | 10214 | 7944 | 10373 | 7686 | 7711 | 5432 | 7766 | 7367 | 6333 | 8415 | 6565 | 9248 | 9193 | 9401 | 8115 | 8635 | 7538 | Sema5a |
| Sema5b | 2884 | 4120 | 4596 | 4881 | 4990 | 4857 | 5078 | 4167 | 4544 | 5363 | 4304 | 8369 | 5916 | 7252 | 3302 | 2948 | 5180 | 4727 | 5851 | 4166 | 4803 | 5347 | 3733 | 6576 | 6279 | 7530 | 7785 | 7280 | Sema5b |
| Sema6a | 3873 | 3907 | 5139 | 5478 | 5897 | 5278 | 4680 | 5446 | 5801 | 6957 | 6494 | 6719 | 6774 | 6005 | 3936 | 4257 | 5370 | 5270 | 5978 | 4385 | 4885 | 8552 | 5858 | 6675 | 7107 | 6231 | 5958 | 6338 | Sema6a |
| Sema6b | 1812 | 2907 | 2768 | 3486 | 3819 | 2890 | 2637 | 1451 | 1726 | 2032 | 1802 | 1359 | 1285 | 1152 | 1924 | 2259 | 3212 | 3652 | 3801 | 2832 | 2998 | 2217 | 1731 | 1744 | 2146 | 1097 | 1177 | 1067 | Sema6b |
| Sema6c | 5105 | 6231 | 7106 | 8378 | 9277 | 8102 | 7014 | 6056 | 5998 | 8305 | 7208 | 7834 | 7000 | 6988 | 5052 | 5559 | 8357 | 8390 | 9491 | 6406 | 7637 | 9553 | 5641 | 7969 | 8802 | 6645 | 6749 | 6565 | Sema6c |
| Sema6d | 3393 | 3050 | 2763 | 3835 | 7309 | 5415 | 5548 | 7960 | 9001 | 8109 | 8093 | 6319 | 6151 | 6438 | 4040 | 3495 | 3363 | 4576 | 5333 | 4965 | 5689 | 8074 | 7075 | 6617 | 9301 | 3982 | 5255 | 4512 | Sema6d |
| Sema7a | 907 | 1312 | 1407 | 1797 | 2193 | 1813 | 1856 | 1420 | 1657 | 2003 | 1738 | 1701 | 1735 | 1445 | 1175 | 1126 | 1714 | 1700 | 2240 | 1599 | 1961 | 1982 | 1527 | 1942 | 2219 | 1686 | 1282 | 1438 | Sema7a |
| Senp1 | 5794 | 5552 | 5154 | 4622 | 4549 | 4900 | 4766 | 5361 | 5057 | 4654 | 4519 | 5916 | 5234 | 5514 | 5272 | 5139 | 5062 | 4463 | 4750 | 4513 | 4790 | 4953 | 4739 | 4868 | 4921 | 5291 | 5788 | 5853 | Senp1 |
| Senp2 | 10751 | 9929 | 10786 | 10739 | 9579 | 9905 | 11402 | 6436 | 6077 | 6649 | 6701 | 7379 | 6887 | 7512 | 9846 | 10017 | 10422 | 10271 | 10169 | 9022 | 10743 | 6348 | 5798 | 6938 | 6858 | 7076 | 6918 | 7618 | Senp2 |
| Senp3 | 9846 | 10210 | 9520 | 9727 | 8620 | 8867 | 9543 | 7690 | 7385 | 7552 | 7227 | 7331 | 7039 | 7282 | 9711 | 9330 | 9248 | 9709 | 9263 | 8649 | 9168 | 7441 | 7144 | 7738 | 7327 | 8065 | 7241 | 7541 | Senp3 |
| Senp5 | 8050 | 7286 | 6799 | 7658 | 7670 | 8069 | 8238 | 7353 | 6461 | 6636 | 6915 | 7332 | 7648 | 8061 | 7904 | 7433 | 6976 | 7491 | 7467 | 7447 | 8123 | 7553 | 6355 | 6824 | 7856 | 6123 | 7614 | 7787 | Senp5 |
| Senp6 | 5881 | 4772 | 6005 | 5140 | 6851 | 6331 | 6821 | 15863 | 13263 | 13640 | 13414 | 14329 | 14142 | 15087 | 6290 | 5379 | 5716 | 5311 | 6533 | 6718 | 6344 | 14603 | 13161 | 13723 | 13862 | 14395 | 14478 | 15075 | Senp6 |
| Senp7 | 6651 | 5645 | 5323 | 6361 | 6192 | 6744 | 6844 | 9006 | 7203 | 6409 | 6716 | 6547 | 6538 | 7022 | 6881 | 6175 | 5465 | 5688 | 6122 | 7472 | 6942 | 7524 | 7269 | 6342 | 7285 | 5212 | 7130 | 6603 | Senp7 |
| Senp8 | 1907 | 1670 | 2403 | 1775 | 1555 | 1552 | 1706 | 1763 | 1412 | 1455 | 1343 | 1727 | 1589 | 1693 | 2296 | 1875 | 2234 | 1697 | 1684 | 2007 | 1661 | 1617 | 1436 | 1642 | 1419 | 1740 | 1780 | 1879 | Senp8 |
| Sephs1 | 7008 | 5722 | 6579 | 5547 | 5585 | 5998 | 5727 | 13211 | 11511 | 11729 | 10503 | 14431 | 12819 | 14604 | 6618 | 5740 | 6093 | 5083 | 6152 | 6130 | 5622 | 11217 | 11005 | 12541 | 12108 | 15846 | 14612 | 14834 | Sephs1 |
| Sephs2 | 3908 | 3581 | 4639 | 4487 | 3810 | 3752 | 4198 | 2099 | 2103 | 2512 | 2571 | 2726 | 2669 | 2813 | 3541 | 3575 | 4217 | 4079 | 3873 | 3422 | 4128 | 2034 | 2121 | 2951 | 2566 | 3454 | 2667 | 3201 | Sephs2 |
| Sepsecs | 3395 | 2839 | 2429 | 2878 | 2481 | 2901 | 2856 | 2216 | 2227 | 1871 | 2160 | 2285 | 2082 | 2226 | 3243 | 2983 | 2497 | 2869 | 2471 | 2909 | 2792 | 2032 | 2119 | 2040 | 1898 | 1912 | 2309 | 2160 | Sepsecs |
| Septin1 | 571 | 626 | 877 | 649 | 549 | 621 | 602 | 753 | 761 | 864 | 739 | 1150 | 1074 | 1180 | 553 | 482 | 723 | 521 | 742 | 505 | 620 | 767 | 667 | 1179 | 1004 | 1374 | 1192 | 1286 | Septin1 |
| Septin10 | 2899 | 2025 | 2335 | 1995 | 2044 | 2128 | 2504 | 2401 | 2466 | 2159 | 2214 | 2336 | 2449 | 2541 | 2712 | 2478 | 2023 | 2210 | 2107 | 2529 | 2263 | 1825 | 2519 | 2250 | 2109 | 2634 | 2302 | 2474 | Septin10 |
| Septin11 | 38444 | 34998 | 37408 | 33999 | 31305 | 33002 | 33829 | 31528 | 31127 | 32130 | 31732 | 37496 | 36844 | 37595 | 36415 | 36190 | 36177 | 33594 | 32862 | 31096 | 32425 | 30014 | 30165 | 34107 | 31632 | 37281 | 37116 | 36827 | Septin11 |
| Septin12 | 0 | 1 | 0 | 0 | 1 | 0 | 0 | 0 | 0 | 0 | 0 | 1 | 0 | 0 | 0 | 0 | 0 | 1 | 1 | 0 | 0 | 0 | 2 | 2 | 0 | 1 | 7 | 1 | Septin12 |
| Septin14 | 0 | 1 | 0 | 0 | 0 | 3 | 0 | 0 | 0 | 0 | 0 | 0 | 1 | 0 | 1 | 0 | 0 | 0 | 0 | 0 | 0 | 0 | 0 | 0 | 0 | 0 | 0 | 0 | Septin14 |
| Septin2 | 442 | 282 | 357 | 432 | 443 | 432 | 428 | 402 | 391 | 472 | 524 | 428 | 425 | 499 | 352 | 345 | 333 | 459 | 362 | 406 | 492 | 383 | 365 | 446 | 380 | 322 | 414 | 417 | Septin2 |

| | | | | | | | | | | | | | | | | | | | | | | | | | | | | | |
|---|---|---|---|---|---|---|---|---|---|---|---|---|---|---|---|---|---|---|---|---|---|---|---|---|---|---|---|---|---|
| Septin3 | 25517 | 26521 | 26225 | 35127 | 41581 | 37824 | 39378 | 36494 | 32991 | 41539 | 31912 | 40502 | 33670 | 40404 | 27920 | 22145 | 31616 | 35550 | 40999 | 34066 | 41259 | 47861 | 29119 | 39302 | 48015 | 31104 | 36193 | 37858 | Septin3 |
| Septin4 | 967 | 1113 | 1276 | 1072 | 1112 | 924 | 925 | 364 | 470 | 440 | 791 | 472 | 473 | 459 | 1117 | 1102 | 1279 | 1038 | 1187 | 1094 | 1034 | 550 | 638 | 485 | 444 | 378 | 487 | 475 | Septin4 |
| Septin5 | 295 | 416 | 308 | 462 | 490 | 328 | 396 | 380 | 363 | 719 | 415 | 363 | 315 | 431 | 342 | 219 | 444 | 409 | 410 | 294 | 508 | 515 | 290 | 515 | 459 | 298 | 310 | 361 | Septin5 |
| Septin6 | 8809 | 8604 | 7800 | 7289 | 8341 | 7682 | 8047 | 8618 | 8017 | 7383 | 7811 | 7022 | 7489 | 7472 | 9171 | 8492 | 7499 | 7363 | 7773 | 8190 | 7679 | 8433 | 7920 | 6705 | 7366 | 7012 | 7469 | 7244 | Septin6 |
| Septin7 | 20431 | 14301 | 18508 | 16316 | 15795 | 17020 | 18605 | 27091 | 23236 | 19982 | 21724 | 21272 | 21358 | 23824 | 22337 | 18405 | 16581 | 15926 | 15966 | 20521 | 17077 | 22311 | 24377 | 22723 | 21282 | 23862 | 22813 | 24305 | Septin7 |
| Septin8 | 6704 | 6256 | 7435 | 6155 | 6473 | 6757 | 6186 | 4246 | 6158 | 5728 | 6872 | 5407 | 5384 | 5160 | 5905 | 6969 | 6693 | 7174 | 6859 | 4767 | 6008 | 4971 | 6049 | 5979 | 5454 | 5571 | 4775 | 5165 | Septin8 |
| Septin9 | 12190 | 12413 | 14012 | 10942 | 10143 | 10550 | 9533 | 11280 | 14014 | 13968 | 13875 | 16076 | 14893 | 15439 | 10580 | 12507 | 11549 | 10220 | 10929 | 9391 | 9319 | 11122 | 14025 | 15315 | 12766 | 19645 | 14876 | 15723 | Septin9 |
| Serac1 | 3338 | 3839 | 2228 | 3206 | 3022 | 3213 | 3371 | 2058 | 1992 | 1993 | 1825 | 1803 | 1786 | 1848 | 3380 | 3479 | 2598 | 3043 | 2831 | 3347 | 3547 | 1873 | 1839 | 1547 | 2069 | 1183 | 1762 | 1557 | Serac1 |
| Serbp1 | 40067 | 35947 | 42994 | 37239 | 33400 | 34592 | 39634 | 42923 | 38037 | 36125 | 36607 | 42888 | 40364 | 48826 | 41577 | 37401 | 38893 | 35064 | 35409 | 40755 | 37897 | 33411 | 37799 | 42576 | 39131 | 48662 | 45240 | 50633 | Serbp1 |
| Serf1 | 2173 | 2043 | 2053 | 2846 | 2802 | 2201 | 3604 | 2836 | 2191 | 3127 | 2709 | 2339 | 2352 | 3948 | 2294 | 1663 | 2467 | 2525 | 2413 | 2855 | 3560 | 2740 | 2372 | 3268 | 3194 | 2628 | 2388 | 3264 | Serf1 |
| Serf2 | 12265 | 14424 | 12946 | 12359 | 10470 | 11033 | 11533 | 10142 | 11087 | 9880 | 11796 | 10756 | 11345 | 10964 | 12254 | 12889 | 12645 | 12032 | 11331 | 12706 | 10784 | 8898 | 11695 | 10566 | 10350 | 12206 | 11554 | 11405 | Serf2 |
| Sergef | 675 | 967 | 818 | 914 | 1215 | 1006 | 979 | 782 | 786 | 779 | 743 | 817 | 657 | 617 | 736 | 778 | 1032 | 1041 | 1124 | 1019 | 1010 | 794 | 742 | 745 | 860 | 803 | 750 | 635 | Sergef |
| Serhl | 649 | 694 | 732 | 521 | 380 | 519 | 391 | 563 | 689 | 599 | 760 | 615 | 677 | 600 | 599 | 732 | 670 | 500 | 444 | 429 | 398 | 498 | 698 | 637 | 460 | 831 | 710 | 644 | Serhl |
| Serinc1 | 22691 | 19913 | 18389 | 21587 | 24436 | 22869 | 25463 | 31269 | 25338 | 22051 | 21996 | 21891 | 22003 | 22636 | 27216 | 22100 | 20042 | 22404 | 22508 | 28278 | 24883 | 29721 | 25955 | 21084 | 24554 | 19442 | 21976 | 22297 | Serinc1 |
| Serinc2 | 2052 | 1871 | 1538 | 1478 | 1714 | 1668 | 1621 | 3011 | 3839 | 3540 | 3603 | 3508 | 3155 | 3295 | 1622 | 1972 | 1566 | 1493 | 1844 | 1732 | 1558 | 2998 | 3519 | 3683 | 3568 | 4286 | 3310 | 3398 | Serinc2 |
| Serinc3 | 11727 | 11180 | 11282 | 9711 | 10440 | 10120 | 10987 | 12677 | 10467 | 9914 | 10019 | 10581 | 11011 | 11630 | 13388 | 11712 | 10434 | 10555 | 9559 | 12449 | 10735 | 12519 | 11843 | 10315 | 10229 | 10702 | 10693 | 12450 | Serinc3 |
| Serinc4 | 33 | 31 | 38 | 33 | 47 | 35 | 61 | 197 | 190 | 205 | 188 | 142 | 212 | 233 | 32 | 34 | 19 | 25 | 51 | 59 | 70 | 217 | 181 | 196 | 215 | 167 | 194 | 152 | Serinc4 |
| Serinc5 | 6552 | 6470 | 5417 | 6277 | 8216 | 7273 | 7666 | 3778 | 3865 | 3554 | 4424 | 3040 | 3463 | 3130 | 6733 | 6474 | 6043 | 7371 | 7226 | 7957 | 7591 | 3716 | 4376 | 3194 | 4023 | 2238 | 2873 | 2854 | Serinc5 |
| Serp1 | 18316 | 13243 | 15750 | 13708 | 10894 | 13557 | 14196 | 12220 | 11206 | 9364 | 11392 | 11059 | 11644 | 11948 | 17900 | 16242 | 13610 | 13309 | 12151 | 12805 | 13728 | 10009 | 12444 | 10436 | 9899 | 13165 | 11864 | 13176 | Serp1 |
| Serp2 | 468 | 618 | 505 | 580 | 752 | 543 | 667 | 637 | 595 | 702 | 543 | 531 | 472 | 602 | 579 | 474 | 533 | 538 | 624 | 690 | 639 | 703 | 609 | 641 | 613 | 453 | 556 | 481 | Serp2 |
| Serpina10 | 1 | 0 | 0 | 0 | 0 | 0 | 0 | 1 | 2 | 0 | 0 | 0 | 0 | 0 | 0 | 2 | 0 | 0 | 0 | 0 | 0 | 0 | 4 | 1 | 0 | 0 | 0 | 0 | Serpina10 |
| Serpina12 | 0 | 0 | 0 | 0 | 0 | 0 | 0 | 0 | 0 | 0 | 0 | 0 | 0 | 0 | 0 | 0 | 0 | 0 | 0 | 0 | 0 | 0 | 1 | 0 | 6 | 0 | 0 | 0 | Serpina12 |
| Serpina1a | 0 | 1 | 0 | 0 | 0 | 0 | 0 | 0 | 2 | 0 | 0 | 0 | 0 | 2 | 0 | 0 | 0 | 2 | 0 | 0 | 0 | 0 | 0 | 2 | 0 | 0 | 0 | 0 | Serpina1a |
| Serpina1b | 0 | 0 | 0 | 0 | 1 | 0 | 1 | 0 | 12 | 0 | 2 | 1 | 0 | 0 | 0 | 6 | 0 | 2 | 0 | 0 | 0 | 0 | 2 | 0 | 0 | 0 | 1 | 0 | Serpina1b |
| Serpina1c | 0 | 0 | 0 | 0 | 2 | 0 | 0 | 1 | 2 | 2 | 0 | 0 | 0 | 0 | 0 | 0 | 0 | 0 | 0 | 0 | 0 | 0 | 0 | 0 | 0 | 0 | 0 | 0 | Serpina1c |
| Serpina1d | 0 | 1 | 0 | 0 | 0 | 0 | 0 | 0 | 1 | 0 | 0 | 0 | 0 | 0 | 0 | 0 | 0 | 0 | 0 | 0 | 0 | 0 | 0 | 0 | 0 | 0 | 0 | 0 | Serpina1d |
| Serpina1e | 0 | 0 | 0 | 0 | 0 | 0 | 0 | 0 | 3 | 0 | 0 | 0 | 0 | 0 | 0 | 0 | 0 | 0 | 0 | 0 | 0 | 0 | 4 | 1 | 0 | 0 | 0 | 0 | Serpina1e |
| Serpina3b | 0 | 0 | 0 | 0 | 0 | 0 | 0 | 0 | 0 | 0 | 0 | 0 | 0 | 0 | 0 | 0 | 1 | 0 | 2 | 0 | 0 | 0 | 0 | 0 | 0 | 0 | 0 | 0 | Serpina3b |
| Serpina3f | 2 | 1 | 0 | 3 | 0 | 2 | 3 | 12 | 7 | 7 | 11 | 20 | 25 | 15 | 4 | 7 | 6 | 1 | 8 | 1 | 3 | 17 | 6 | 13 | 9 | 18 | 22 | 12 | Serpina3f |
| Serpina3g | 1 | 2 | 0 | 1 | 15 | 0 | 0 | 11 | 11 | 9 | 7 | 5 | 9 | 0 | 7 | 10 | 6 | 3 | 0 | 1 | 1 | 15 | 8 | 1 | 3 | 2 | 6 | 4 | Serpina3g |
| Serpina3i | 3 | 3 | 2 | 0 | 2 | 0 | 0 | 2 | 9 | 4 | 8 | 0 | 7 | 11 | 0 | 2 | 3 | 2 | 0 | 0 | 1 | 2 | 4 | 1 | 5 | 1 | 3 | 5 | Serpina3i |
| Serpina3m | 0 | 0 | 0 | 0 | 0 | 0 | 0 | 0 | 0 | 0 | 0 | 0 | 0 | 0 | 0 | 0 | 0 | 0 | 0 | 0 | 0 | 0 | 0 | 0 | 0 | 0 | 0 | 1 | Serpina3m |
| Serpina3n | 1 | 6 | 0 | 0 | 7 | 0 | 0 | 11 | 17 | 12 | 63 | 1 | 0 | 0 | 1 | 2 | 0 | 6 | 0 | 0 | 0 | 15 | 16 | 2 | 20 | 0 | 0 | 1 | Serpina3n |
| Serpina6 | 0 | 0 | 0 | 0 | 0 | 0 | 0 | 0 | 6 | 0 | 0 | 0 | 0 | 1 | 0 | 0 | 0 | 0 | 0 | 0 | 0 | 2 | 1 | 0 | 0 | 0 | 0 | 0 | Serpina6 |
| Serpina7 | 13 | 9 | 0 | 0 | 3 | 3 | 0 | 4 | 2 | 4 | 0 | 2 | 2 | 1 | 1 | 8 | 2 | 2 | 3 | 1 | 0 | 2 | 5 | 7 | 5 | 5 | 8 | 1 | Serpina7 |
| Serpina9 | 0 | 1 | 0 | 0 | 0 | 0 | 1 | 0 | 2 | 2 | 0 | 1 | 1 | 1 | 0 | 1 | 5 | 0 | 0 | 0 | 0 | 0 | 1 | 0 | 0 | 0 | 0 | 0 | Serpina9 |
| Serpinb11 | 2 | 0 | 0 | 0 | 0 | 0 | 0 | 23 | 41 | 38 | 2 | 0 | 0 | 0 | 1 | 0 | 0 | 0 | 0 | 0 | 0 | 0 | 46 | 18 | 0 | 0 | 0 | 0 | Serpinb11 |
| Serpinb12 | 0 | 0 | 0 | 0 | 0 | 0 | 0 | 0 | 4 | 3 | 5 | 0 | 0 | 0 | 1 | 0 | 0 | 0 | 0 | 0 | 0 | 2 | 3 | 2 | 0 | 3 | 1 | 0 | Serpinb12 |
| Serpinb1a | 67 | 48 | 69 | 55 | 45 | 89 | 25 | 62 | 127 | 76 | 142 | 56 | 58 | 47 | 57 | 44 | 43 | 91 | 64 | 35 | 30 | 112 | 183 | 85 | 77 | 36 | 77 | 57 | Serpinb1a |
| Serpinb1b | 47 | 44 | 31 | 33 | 92 | 69 | 65 | 125 | 113 | 137 | 84 | 85 | 134 | 51 | 67 | 79 | 21 | 43 | 72 | 61 | 63 | 127 | 250 | 51 | 93 | 34 | 31 | 49 | Serpinb1b |
| Serpinb1c | 2 | 0 | 0 | 0 | 0 | 0 | 1 | 0 | 1 | 0 | 3 | 0 | 0 | 1 | 0 | 1 | 0 | 0 | 0 | 0 | 4 | 0 | 0 | 0 | 0 | 1 | 0 | 0 | Serpinb1c |
| Serpinb2 | 7 | 8 | 10 | 6 | 2 | 12 | 22 | 33 | 39 | 27 | 27 | 52 | 31 | 53 | 13 | 10 | 1 | 11 | 13 | 22 | 11 | 18 | 44 | 60 | 45 | 41 | 48 | 57 | Serpinb2 |
| Serpinb3a | 2 | 0 | 0 | 0 | 0 | 0 | 0 | 28 | 40 | 44 | 0 | 0 | 0 | 0 | 0 | 2 | 0 | 0 | 0 | 0 | 0 | 0 | 27 | 0 | 11 | 0 | 0 | 0 | Serpinb3a |
| Serpinb3b | 0 | 0 | 0 | 0 | 0 | 0 | 0 | 0 | 0 | 0 | 0 | 0 | 0 | 0 | 0 | 0 | 0 | 0 | 0 | 0 | 0 | 0 | 1 | 0 | 0 | 0 | 0 | 1 | Serpinb3b |
| Serpinb3c | 0 | 0 | 0 | 0 | 0 | 0 | 0 | 0 | 2 | 5 | 0 | 0 | 0 | 0 | 0 | 0 | 0 | 0 | 0 | 0 | 0 | 0 | 3 | 1 | 0 | 0 | 0 | 0 | Serpinb3c |
| Serpinb3d | 0 | 0 | 0 | 0 | 1 | 0 | 0 | 0 | 1 | 2 | 0 | 0 | 0 | 0 | 0 | 0 | 0 | 0 | 0 | 0 | 0 | 3 | 0 | 0 | 0 | 0 | 0 | 0 | Serpinb3d |
| Serpinb5 | 10 | 0 | 3 | 4 | 1 | 5 | 2 | 23 | 71 | 27 | 22 | 2 | 6 | 3 | 3 | 9 | 1 | 3 | 0 | 1 | 0 | 3 | 87 | 6 | 0 | 8 | 2 | 1 | Serpinb5 |
| Serpinb6a | 2513 | 2002 | 2016 | 1711 | 1330 | 1853 | 1477 | 2328 | 2782 | 1941 | 2655 | 2137 | 2614 | 2345 | 2471 | 2491 | 1691 | 1756 | 1327 | 1618 | 1339 | 2151 | 3251 | 2095 | 1884 | 2357 | 2349 | 2341 | Serpinb6a |
| Serpinb6b | 46 | 48 | 46 | 30 | 47 | 34 | 45 | 74 | 82 | 65 | 62 | 59 | 115 | 50 | 40 | 43 | 22 | 20 | 26 | 54 | 30 | 79 | 98 | 48 | 52 | 46 | 41 | 44 | Serpinb6b |
| Serpinb6c | 30 | 34 | 15 | 16 | 41 | 36 | 44 | 13 | 11 | 19 | 11 | 15 | 6 | 11 | 38 | 32 | 23 | 26 | 24 | 29 | 40 | 20 | 8 | 11 | 17 | 16 | 5 | 2 | Serpinb6c |
| Serpinb6e | 10 | 10 | 7 | 11 | 16 | 21 | 13 | 2 | 5 | 0 | 0 | 8 | 2 | 0 | 13 | 6 | 6 | 28 | 11 | 5 | 22 | 3 | 1 | 2 | 3 | 0 | 1 | 4 | Serpinb6e |
| Serpinb7 | 1 | 0 | 0 | 0 | 0 | 0 | 0 | 44 | 34 | 49 | 28 | 2 | 1 | 3 | 0 | 0 | 0 | 0 | 2 | 0 | 0 | 0 | 46 | 27 | 0 | 2 | 0 | 2 | Serpinb7 |
| Serpinb8 | 91 | 86 | 50 | 29 | 110 | 44 | 8 | 125 | 135 | 109 | 149 | 17 | 138 | 24 | 105 | 187 | 36 | 59 | 32 | 24 | 26 | 89 | 140 | 38 | 58 | 32 | 17 | 36 | Serpinb8 |
| Serpinb9 | 65 | 31 | 55 | 19 | 49 | 46 | 50 | 167 | 222 | 161 | 202 | 123 | 108 | 110 | 46 | 61 | 20 | 40 | 29 | 53 | 36 | 224 | 247 | 105 | 140 | 123 | 100 | 108 | Serpinb9 |
| Serpinb9b | 2 | 2 | 0 | 6 | 1 | 0 | 3 | 10 | 16 | 10 | 13 | 9 | 11 | 9 | 3 | 0 | 8 | 0 | 1 | 5 | 4 | 10 | 10 | 14 | 20 | 8 | 14 | 7 | Serpinb9b |
| Serpinb9c | 0 | 0 | 0 | 0 | 0 | 0 | 0 | 0 | 0 | 0 | 0 | 0 | 0 | 0 | 0 | 0 | 0 | 0 | 0 | 0 | 0 | 0 | 1 | 0 | 0 | 0 | 0 | 0 | Serpinb9c |
| Serpinb9d | 0 | 0 | 0 | 0 | 0 | 0 | 0 | 0 | 2 | 0 | 0 | 0 | 0 | 0 | 0 | 0 | 0 | 0 | 0 | 0 | 0 | 0 | 1 | 0 | 0 | 0 | 0 | 0 | Serpinb9d |
| Serpinb9e | 1 | 0 | 0 | 0 | 0 | 0 | 0 | 0 | 4 | 0 | 0 | 0 | 1 | 0 | 0 | 0 | 0 | 0 | 0 | 0 | 0 | 0 | 0 | 0 | 0 | 0 | 0 | 0 | Serpinb9e |
| Serpinb9f | 0 | 0 | 0 | 0 | 0 | 0 | 0 | 0 | 1 | 0 | 0 | 0 | 0 | 0 | 0 | 0 | 0 | 0 | 0 | 0 | 0 | 0 | 0 | 0 | 0 | 0 | 0 | 0 | Serpinb9f |
| Serpinb9g | 0 | 0 | 0 | 0 | 0 | 0 | 0 | 0 | 4 | 0 | 0 | 0 | 0 | 0 | 0 | 0 | 0 | 0 | 0 | 0 | 0 | 0 | 3 | 0 | 0 | 0 | 0 | 0 | Serpinb9g |
| Serpinc1 | 47 | 41 | 58 | 87 | 52 | 50 | 60 | 33 | 38 | 24 | 49 | 34 | 68 | 36 | 46 | 58 | 66 | 85 | 86 | 61 | 71 | 27 | 41 | 47 | 42 | 15 | 44 | 37 | Serpinc1 |
| Serpind1 | 33 | 29 | 27 | 36 | 56 | 27 | 56 | 29 | 30 | 27 | 21 | 21 | 18 | 9 | 40 | 20 | 49 | 70 | 34 | 37 | 42 | 43 | 50 | 25 | 28 | 8 | 9 | 14 | Serpind1 |
| Serpine1 | 84 | 98 | 101 | 126 | 111 | 82 | 88 | 85 | 150 | 148 | 151 | 66 | 98 | 101 | 100 | 97 | 81 | 87 | 100 | 78 | 97 | 132 | 153 | 103 | 93 | 111 | 86 | 101 | Serpine1 |
| Serpine2 | 5445 | 4222 | 3476 | 2942 | 4043 | 3276 | 2056 | 5742 | 5633 | 4884 | 6102 | 2868 | 4554 | 2537 | 4760 | 5483 | 3194 | 3713 | 2625 | 2726 | 2496 | 5193 | 6688 | 3635 | 3196 | 3439 | 3052 | 2890 | Serpine2 |
| Serpine3 | 1 | 2 | 0 | 2 | 5 | 5 | 9 | 2 | 4 | 0 | 0 | 0 | 0 | 2 | 7 | 2 | 1 | 12 | 3 | 3 | 4 | 2 | 3 | 1 | 0 | 1 | 0 | 0 | Serpine3 |
| Serpinf1 | 2234 | 1998 | 1620 | 1180 | 874 | 1462 | 518 | 1723 | 2462 | 1983 | 2594 | 1298 | 1983 | 1275 | 1836 | 2653 | 1133 | 1749 | 653 | 759 | 593 | 1798 | 2877 | 1913 | 1629 | 1930 | 1374 | 1442 | Serpinf1 |
| Serpinf2 | 10 | 25 | 39 | 27 | 13 | 21 | 21 | 4 | 16 | 16 | 6 | 10 | 7 | 3 | 9 | 43 | 38 | 40 | 21 | 18 | 35 | 22 | 13 | 11 | 14 | 4 | 5 | 11 | Serpinf2 |
| Serping1 | 683 | 851 | 794 | 575 | 618 | 765 | 439 | 1233 | 1361 | 1308 | 1393 | 1287 | 1537 | 1284 | 928 | 862 | 692 | 753 | 533 | 641 | 494 | 1313 | 1760 | 1277 | 1111 | 1377 | 1486 | 1328 | Serping1 |
| Serpinh1 | 32290 | 36934 | 34799 | 24283 | 17357 | 25482 | 14517 | 28228 | 42339 | 33436 | 49013 | 30036 | 38474 | 28549 | 29728 | 42031 | 28456 | 28274 | 17907 | 18235 | 14968 | 27262 | 50138 | 34500 | 27722 | 38601 | 32969 | 31424 | Serpinh1 |
| Serpini1 | 6036 | 6162 | 5668 | 8239 | 11397 | 9028 | 11092 | 8642 | 6814 | 6694 | 5794 | 4524 | 5185 | 5187 | 9621 | 7057 | 7628 | 9741 | 9248 | 11986 | 10996 | 8423 | 7095 | 5806 | 7522 | 3594 | 4581 | 4729 | Serpini1 |
| Serpini2 | 0 | 0 | 0 | 0 | 0 | 2 | 0 | 1 | 2 | 0 | 0 | 0 | 0 | 0 | 0 | 0 | 0 | 0 | 0 | 0 | 0 | 0 | 0 | 0 | 2 | 0 | 0 | 0 | Serpini2 |
| Sertad1 | 287 | 412 | 395 | 309 | 254 | 256 | 207 | 261 | 390 | 257 | 363 | 234 | 348 | 243 | 302 | 316 | 289 | 285 | 262 | 326 | 194 | 246 | 363 | 299 | 172 | 303 | 384 | 233 | Sertad1 |
| Sertad2 | 4357 | 4615 | 4986 | 5398 | 4791 | 4851 | 5344 | 2615 | 2907 | 3235 | 3159 | 3510 | 3526 | 3487 | 4499 | 4791 | 5128 | 5469 | 5058 | 4166 | 5201 | 2829 | 2838 | 3487 | 3067 | 3527 | 3403 | 3763 | Sertad2 |
| Sertad3 | 1159 | 1380 | 1307 | 1256 | 944 | 1075 | 960 | 470 | 564 | 473 | 576 | 521 | 635 | 557 | 1064 | 1204 | 1272 | 1159 | 1086 | 1071 | 984 | 426 | 611 | 546 | 479 | 678 | 654 | 615 | Sertad3 |
| Sertad4 | 2461 | 1845 | 2237 | 2324 | 2595 | 2565 | 2191 | 4107 | 4292 | 4967 | 5727 | 3877 | 4183 | 3893 | 2116 | 2262 | 2216 | 2485 | 2432 | 1868 | 2339 | 5127 | 4743 | 4929 | 4574 | 4147 | 3620 | 3557 | Sertad4 |
| Sertm1 | 1032 | 994 | 1077 | 1349 | 1931 | 1776 | 1861 | 725 | 667 | 839 | 589 | 594 | 712 | 669 | 1234 | 911 | 1262 | 1602 | 1811 | 1913 | 1908 | 799 | 871 | 738 | 891 | 737 | 568 | 633 | Sertm1 |
| Sesn1 | 4847 | 4608 | 3860 | 4566 | 4785 | 4744 | 5082 | 3516 | 3711 | 3304 | 3355 | 2971 | 2910 | 3097 | 4956 | 4757 | 4415 | 5128 | 4682 | 5128 | 5138 | 3334 | 3706 | 3265 | 3332 | 2960 | 2477 | 2840 | Sesn1 |
| Sesn2 | 2614 | 3817 | 3867 | 3852 | 3036 | 2906 | 2675 | 576 | 702 | 721 | 718 | 773 | 756 | 830 | 2544 | 3263 | 3751 | 3623 | 3441 | 2452 | 2929 | 703 | 755 | 739 | 582 | 915 | 786 | 879 | Sesn2 |
| Sesn3 | 23006 | 21320 | 18255 | 22827 | 21944 | 24529 | 24246 | 13676 | 13493 | 13441 | 14862 | 13318 | 15431 | 13612 | 22808 | 23907 | 20025 | 23091 | 20459 | 20258 | 24335 | 13734 | 13705 | 12008 | 13002 | 9628 | 14190 | 12861 | Sesn3 |

| | | | | | | | | | | | | | | | | | | | | | | | | | | | | | |
|---|---|---|---|---|---|---|---|---|---|---|---|---|---|---|---|---|---|---|---|---|---|---|---|---|---|---|---|---|---|
| Sestd1 | 9840 | 8189 | 7415 | 10131 | 13704 | 12875 | 12850 | 17434 | 15227 | 15919 | 15199 | 14389 | 13454 | 14391 | 10646 | 8627 | 9021 | 11047 | 12016 | 12416 | 12919 | 18290 | 13279 | 13557 | 17890 | 10404 | 14451 | 13046 | Sestd1 |
| Set | 36369 | 29575 | 44484 | 33040 | 30309 | 30702 | 37793 | 46132 | 39035 | 43859 | 41029 | 55940 | 51030 | 63406 | 34305 | 29897 | 38374 | 30261 | 32805 | 33780 | 34230 | 39746 | 38200 | 52567 | 47755 | 67812 | 57711 | 66669 | Set |
| Setbp1 | 3704 | 3346 | 3149 | 4401 | 5161 | 5268 | 5460 | 11852 | 12060 | 14436 | 13014 | 17389 | 14111 | 17098 | 3703 | 3204 | 4020 | 4420 | 5331 | 4354 | 5435 | 14302 | 10395 | 14243 | 16939 | 10326 | 16269 | 15721 | Setbp1 |
| Setd1a | 1819 | 2154 | 2627 | 2350 | 2302 | 2212 | 2239 | 1785 | 2230 | 2557 | 2698 | 2986 | 2635 | 2804 | 1652 | 1912 | 2624 | 2431 | 2514 | 2012 | 2239 | 2361 | 2057 | 3097 | 2551 | 2973 | 2774 | 2619 | Setd1a |
| Setd1b | 1724 | 2150 | 3272 | 2918 | 3664 | 2668 | 2448 | 3947 | 4458 | 6273 | 5853 | 7022 | 5082 | 5814 | 1590 | 1738 | 3591 | 2794 | 3609 | 1978 | 2893 | 6545 | 3996 | 7017 | 6708 | 6234 | 5217 | 5744 | Setd1b |
| Setd2 | 15031 | 12883 | 9339 | 13109 | 14496 | 16185 | 15901 | 17982 | 16776 | 15362 | 15991 | 16634 | 16314 | 17479 | 14869 | 13671 | 11249 | 12679 | 13450 | 14743 | 16098 | 15303 | 16106 | 12964 | 16413 | 9985 | 16712 | 15151 | Setd2 |
| Setd3 | 12076 | 11990 | 11999 | 12256 | 11730 | 12078 | 12971 | 8394 | 7765 | 8428 | 8598 | 8409 | 8475 | 9350 | 12096 | 11467 | 12012 | 12108 | 12126 | 12444 | 12783 | 8051 | 7882 | 8404 | 8932 | 8317 | 8450 | 8897 | Setd3 |
| Setd4 | 371 | 416 | 508 | 543 | 591 | 441 | 505 | 806 | 859 | 975 | 916 | 917 | 957 | 806 | 400 | 393 | 505 | 519 | 548 | 427 | 493 | 864 | 896 | 1054 | 896 | 852 | 858 | 888 | Setd4 |
| Setd5 | 19775 | 17089 | 18002 | 19948 | 20478 | 20751 | 19636 | 19274 | 19051 | 20740 | 21724 | 22503 | 21524 | 20921 | 18770 | 18397 | 19782 | 19648 | 19913 | 17204 | 20154 | 20386 | 18478 | 19775 | 20421 | 17547 | 21792 | 20689 | Setd5 |
| Setd6 | 997 | 927 | 1186 | 1055 | 991 | 916 | 1124 | 1067 | 920 | 989 | 950 | 1138 | 1038 | 1237 | 949 | 793 | 1029 | 1086 | 1047 | 1065 | 1047 | 917 | 784 | 1181 | 1189 | 1371 | 1104 | 1218 | Setd6 |
| Setd7 | 3060 | 2630 | 3363 | 3192 | 3733 | 3684 | 3651 | 4553 | 4853 | 4944 | 5594 | 5214 | 5163 | 5268 | 2917 | 2753 | 3522 | 3275 | 3876 | 3247 | 3533 | 4841 | 4576 | 5323 | 5284 | 4802 | 5218 | 5188 | Setd7 |
| Setdb1 | 12762 | 12524 | 12217 | 12128 | 11713 | 11942 | 13198 | 12190 | 11993 | 11821 | 11271 | 12917 | 12027 | 13029 | 12595 | 11830 | 12318 | 12034 | 12564 | 11432 | 12690 | 12248 | 11001 | 12151 | 11933 | 12637 | 12265 | 13021 | Setdb1 |
| Setdb2 | 1940 | 1446 | 1208 | 1315 | 1183 | 1401 | 1422 | 1010 | 861 | 824 | 903 | 910 | 846 | 990 | 1911 | 1599 | 1074 | 1199 | 1160 | 1414 | 1275 | 732 | 916 | 806 | 770 | 916 | 902 | 850 | Setdb2 |
| Setmar | 183 | 267 | 273 | 209 | 218 | 161 | 209 | 460 | 400 | 450 | 372 | 403 | 502 | 431 | 248 | 242 | 254 | 199 | 234 | 276 | 200 | 428 | 463 | 422 | 340 | 506 | 518 | 412 | Setmar |
| Setx | 19484 | 16819 | 11627 | 14977 | 16312 | 19644 | 19608 | 18437 | 15568 | 13467 | 14153 | 14606 | 14832 | 15742 | 19711 | 16897 | 12356 | 14714 | 14816 | 19897 | 19612 | 12121 | 14593 | 11281 | 15901 | 9862 | 16418 | 14012 | Setx |
| Sez6 | 3700 | 6000 | 4451 | 12177 | 14484 | 12154 | 13689 | 5991 | 8161 | 10765 | 7765 | 8309 | 5139 | 7289 | 4415 | 3678 | 9425 | 13774 | 15264 | 9745 | 14692 | 10866 | 6835 | 10914 | 13557 | 6550 | 5007 | 5234 | Sez6 |
| Sez6l | 1978 | 2664 | 2962 | 5954 | 9236 | 7172 | 7816 | 4455 | 5444 | 8213 | 6044 | 6289 | 4657 | 6179 | 2567 | 2361 | 4888 | 7087 | 8486 | 6395 | 8449 | 9987 | 5255 | 7755 | 10077 | 3914 | 4444 | 4742 | Sez6l |
| Sez6l2 | 1512 | 2262 | 2183 | 3897 | 5654 | 4095 | 3870 | 3055 | 3620 | 4817 | 3658 | 3709 | 3122 | 3415 | 1850 | 1846 | 3560 | 4471 | 5062 | 3387 | 4649 | 5601 | 3155 | 4139 | 5093 | 2828 | 2907 | 3082 | Sez6l2 |
| Sf1 | 0 | 3 | 0 | 0 | 0 | 0 | 1 | 1 | 0 | 1 | 2 | 0 | 0 | 1 | 0 | 0 | 0 | 0 | 0 | 0 | 0 | 0 | 1 | 0 | 1 | 0 | 1 | 0 | Sf1 |
| Sf3a1 | 8053 | 8120 | 10865 | 9908 | 8665 | 7872 | 7630 | 9964 | 10540 | 11735 | 11207 | 15287 | 12105 | 14094 | 7251 | 8213 | 10699 | 9425 | 9913 | 6205 | 8418 | 13369 | 9655 | 14212 | 13414 | 16678 | 12187 | 14551 | Sf3a1 |
| Sf3a2 | 2323 | 3144 | 4388 | 3628 | 3306 | 2778 | 2509 | 2762 | 3438 | 4016 | 4082 | 4835 | 3868 | 4558 | 1975 | 2520 | 4219 | 3334 | 3862 | 2258 | 2831 | 3867 | 3161 | 5176 | 4245 | 6277 | 3998 | 4434 | Sf3a2 |
| Sf3a3 | 7530 | 7284 | 8651 | 6149 | 5301 | 5743 | 6566 | 7465 | 6815 | 6672 | 6667 | 8258 | 7508 | 9470 | 7030 | 6724 | 7066 | 5281 | 6169 | 6688 | 6024 | 5863 | 6627 | 7992 | 7178 | 9973 | 9230 | 9538 | Sf3a3 |
| Sf3b1 | 49876 | 42618 | 36702 | 38893 | 41889 | 43975 | 48137 | 62008 | 53446 | 44168 | 46339 | 51640 | 48036 | 52021 | 52339 | 44176 | 38199 | 39748 | 40816 | 50475 | 44564 | 49501 | 50910 | 46193 | 47712 | 43670 | 50465 | 49039 | Sf3b1 |
| Sf3b2 | 24264 | 27459 | 29081 | 26549 | 23940 | 23601 | 25652 | 23709 | 25938 | 27933 | 24581 | 32525 | 28161 | 32692 | 23165 | 24179 | 29071 | 24883 | 26633 | 22329 | 25816 | 24645 | 24388 | 29426 | 28222 | 34309 | 30303 | 33831 | Sf3b2 |
| Sf3b3 | 27358 | 27750 | 22485 | 19963 | 18119 | 20338 | 20240 | 25442 | 22612 | 19624 | 19445 | 25237 | 22999 | 23044 | 25288 | 23576 | 21606 | 19459 | 20374 | 21581 | 20653 | 20296 | 21904 | 21363 | 21270 | 25423 | 26097 | 24930 | Sf3b3 |
| Sf3b4 | 4083 | 4983 | 5598 | 4953 | 4498 | 4087 | 4123 | 3303 | 3938 | 3922 | 3984 | 4642 | 4052 | 4743 | 3835 | 4519 | 5514 | 4644 | 4894 | 3605 | 4278 | 3900 | 3751 | 4658 | 4218 | 5981 | 4242 | 4558 | Sf3b4 |
| Sf3b5 | 2382 | 3162 | 2514 | 2401 | 1675 | 1828 | 1872 | 1500 | 1557 | 1418 | 1552 | 1590 | 1602 | 1490 | 2292 | 2190 | 2393 | 2041 | 2067 | 2295 | 1854 | 1158 | 1582 | 1763 | 1494 | 2026 | 1741 | 1645 | Sf3b5 |
| Sf3b6 | 7125 | 6889 | 6938 | 7148 | 6089 | 6536 | 8152 | 5951 | 5160 | 4845 | 4458 | 5402 | 5251 | 6444 | 7800 | 6851 | 6661 | 6392 | 6756 | 8553 | 7554 | 4370 | 4974 | 5289 | 5060 | 5920 | 5742 | 6419 | Sf3b6 |
| Sfi1 | 1088 | 1242 | 1291 | 1220 | 1173 | 1028 | 1130 | 1738 | 1945 | 1796 | 1827 | 1454 | 2038 | 1611 | 1048 | 1168 | 1102 | 1276 | 1156 | 935 | 1137 | 1683 | 1685 | 1630 | 1839 | 1614 | 1991 | 1720 | Sfi1 |
| Sfmbt1 | 3564 | 2963 | 3362 | 3942 | 4309 | 4268 | 4635 | 4197 | 3976 | 3936 | 3806 | 4544 | 3677 | 4710 | 3527 | 2951 | 3685 | 3688 | 4274 | 4116 | 4395 | 4075 | 3634 | 4250 | 4760 | 4427 | 4491 | 4488 | Sfmbt1 |
| Sfmbt2 | 901 | 696 | 668 | 799 | 988 | 921 | 822 | 485 | 567 | 493 | 552 | 380 | 411 | 364 | 824 | 769 | 847 | 785 | 771 | 899 | 821 | 535 | 608 | 331 | 448 | 298 | 408 | 325 | Sfmbt2 |
| Sfn | 667 | 561 | 438 | 265 | 323 | 412 | 212 | 688 | 1208 | 651 | 634 | 459 | 608 | 396 | 448 | 513 | 319 | 428 | 316 | 208 | 224 | 177 | 1247 | 651 | 208 | 343 | 389 | 504 | Sfn |
| Sfpq | 39362 | 31848 | 40389 | 36670 | 35727 | 36170 | 40208 | 59501 | 53093 | 55310 | 55148 | 62301 | 55801 | 71514 | 35102 | 33272 | 38348 | 32981 | 38559 | 35005 | 41834 | 51210 | 47945 | 57266 | 60310 | 58196 | 62046 | 64938 | Sfpq |
| Sfr1 | 11242 | 10324 | 12540 | 10361 | 9464 | 9684 | 10891 | 11707 | 10458 | 10413 | 10723 | 11907 | 11575 | 13378 | 11747 | 10051 | 11807 | 10002 | 10203 | 12041 | 10200 | 9648 | 10764 | 12217 | 11059 | 14460 | 12658 | 13848 | Sfr1 |
| Sfrp1 | 24480 | 23844 | 22281 | 22007 | 16248 | 18292 | 22148 | 12482 | 13228 | 12836 | 12877 | 17267 | 16097 | 18138 | 22362 | 22260 | 20965 | 19860 | 19002 | 17938 | 18940 | 8637 | 10008 | 15586 | 13052 | 22643 | 15150 | 17757 | Sfrp1 |
| Sfrp2 | 3842 | 3286 | 4192 | 2933 | 2655 | 3563 | 2481 | 5144 | 9321 | 6024 | 10520 | 3227 | 3593 | 3374 | 3467 | 3836 | 3741 | 3673 | 2865 | 2160 | 2364 | 6829 | 11691 | 4826 | 4239 | 4529 | 3127 | 3489 | Sfrp2 |
| Sfrp4 | 358 | 314 | 377 | 345 | 390 | 284 | 437 | 194 | 194 | 190 | 183 | 171 | 156 | 178 | 343 | 264 | 351 | 320 | 325 | 364 | 382 | 180 | 177 | 168 | 187 | 186 | 230 | 151 | Sfrp4 |
| Sfrp5 | 21 | 32 | 3 | 18 | 116 | 3 | 2 | 106 | 166 | 130 | 222 | 11 | 79 | 9 | 12 | 37 | 22 | 64 | 2 | 1 | 7 | 206 | 155 | 16 | 91 | 19 | 5 | 9 | Sfrp5 |
| Sfswap | 8711 | 8612 | 7862 | 8988 | 9803 | 9838 | 9131 | 9729 | 9798 | 10837 | 10180 | 10407 | 9611 | 10159 | 8500 | 8175 | 8324 | 8921 | 9604 | 8859 | 9578 | 9984 | 9007 | 9842 | 10460 | 8956 | 10072 | 9258 | Sfswap |
| Sft2d1 | 2646 | 2587 | 2504 | 2886 | 2990 | 2875 | 3136 | 2633 | 2386 | 2267 | 2461 | 2456 | 2445 | 2596 | 2596 | 2721 | 2787 | 3101 | 2793 | 2719 | 3044 | 2841 | 2232 | 2295 | 2633 | 2043 | 2219 | 2300 | Sft2d1 |
| Sft2d2 | 5338 | 4436 | 3714 | 3631 | 2891 | 3457 | 3129 | 2232 | 2564 | 2153 | 2751 | 2335 | 2698 | 2245 | 4149 | 5162 | 3186 | 3882 | 3158 | 2932 | 3035 | 2229 | 2834 | 2350 | 2056 | 2517 | 2392 | 2404 | Sft2d2 |
| Sft2d3 | 621 | 447 | 565 | 463 | 483 | 462 | 518 | 691 | 656 | 670 | 631 | 704 | 722 | 715 | 564 | 466 | 518 | 449 | 514 | 551 | 520 | 685 | 704 | 605 | 629 | 642 | 830 | 806 | Sft2d3 |
| Sfta2 | 2 | 2 | 0 | 0 | 0 | 0 | 1 | 1 | 1 | 3 | 1 | 7 | 1 | 4 | 0 | 0 | 0 | 0 | 0 | 0 | 0 | 0 | 6 | 2 | 1 | 2 | 0 | 0 | Sfta2 |
| Sftpa1 | 0 | 0 | 0 | 0 | 0 | 0 | 0 | 0 | 0 | 0 | 0 | 0 | 0 | 0 | 0 | 0 | 0 | 1 | 0 | 0 | 1 | 2 | 1 | 0 | 0 | 2 | 0 | 0 | Sftpa1 |
| Sftpb | 0 | 0 | 0 | 0 | 0 | 0 | 0 | 0 | 0 | 0 | 0 | 0 | 0 | 0 | 0 | 0 | 0 | 0 | 0 | 2 | 0 | 0 | 0 | 0 | 0 | 0 | 0 | 0 | Sftpb |
| Sftpc | 0 | 0 | 0 | 0 | 0 | 0 | 0 | 0 | 1 | 3 | 0 | 1 | 0 | 0 | 0 | 1 | 0 | 0 | 1 | 0 | 0 | 0 | 0 | 0 | 0 | 4 | 0 | 0 | Sftpc |
| Sftpd | 0 | 0 | 0 | 0 | 0 | 0 | 0 | 0 | 0 | 3 | 0 | 0 | 0 | 0 | 0 | 0 | 0 | 0 | 0 | 0 | 0 | 0 | 0 | 0 | 2 | 0 | 0 | 0 | Sftpd |
| Sfxn1 | 5112 | 4759 | 5522 | 4820 | 5181 | 4997 | 5340 | 5790 | 5712 | 5862 | 5157 | 6132 | 6653 | 6288 | 5375 | 4868 | 5136 | 4745 | 5079 | 5621 | 4895 | 5706 | 5216 | 5607 | 6000 | 6710 | 6840 | 6497 | Sfxn1 |
| Sfxn2 | 4226 | 4447 | 4015 | 4903 | 3608 | 4023 | 4075 | 1800 | 2167 | 2185 | 2623 | 2170 | 2198 | 2144 | 3814 | 4541 | 3789 | 4957 | 3674 | 3442 | 3908 | 2166 | 2100 | 2276 | 2254 | 2209 | 2379 | 2390 | Sfxn2 |
| Sfxn3 | 2686 | 2948 | 2660 | 3478 | 4749 | 4070 | 4253 | 3582 | 3961 | 4039 | 3616 | 3330 | 3322 | 3353 | 3189 | 2811 | 3307 | 3877 | 4210 | 3684 | 4282 | 4814 | 3624 | 3681 | 4366 | 3223 | 2965 | 3109 | Sfxn3 |
| Sfxn4 | 702 | 632 | 605 | 578 | 542 | 679 | 594 | 838 | 779 | 651 | 771 | 683 | 875 | 746 | 765 | 759 | 677 | 541 | 563 | 656 | 580 | 708 | 751 | 641 | 681 | 670 | 752 | 782 | Sfxn4 |
| Sfxn5 | 971 | 1065 | 1136 | 1402 | 1397 | 1369 | 1621 | 1197 | 1412 | 1382 | 1332 | 1534 | 1380 | 1580 | 885 | 886 | 1300 | 1451 | 1525 | 1209 | 1557 | 1151 | 1310 | 1679 | 1590 | 1614 | 1348 | 1609 | Sfxn5 |
| Sgca | 2 | 8 | 1 | 12 | 2 | 7 | 4 | 11 | 24 | 4 | 114 | 3 | 6 | 1 | 7 | 5 | 14 | 8 | 6 | 1 | 3 | 20 | 68 | 7 | 0 | 2 | 1 | 1 | Sgca |
| Sgcb | 3377 | 2825 | 3083 | 2703 | 2495 | 2774 | 2598 | 2425 | 2249 | 2082 | 2691 | 2092 | 2210 | 2049 | 3323 | 3418 | 2792 | 2840 | 2512 | 2597 | 2619 | 2565 | 2616 | 1992 | 2185 | 2104 | 2023 | 2131 | Sgcb |
| Sgcd | 261 | 177 | 36 | 104 | 147 | 199 | 92 | 304 | 404 | 246 | 416 | 187 | 279 | 172 | 236 | 277 | 54 | 134 | 79 | 114 | 108 | 165 | 523 | 142 | 257 | 66 | 250 | 129 | Sgcd |
| Sgce | 1299 | 1261 | 1520 | 1186 | 1426 | 1401 | 1266 | 2514 | 2444 | 2406 | 2541 | 2602 | 2404 | 2566 | 1385 | 1275 | 1626 | 1207 | 1292 | 1408 | 1249 | 2988 | 2626 | 2482 | 2115 | 2841 | 2629 | 2659 | Sgce |
| Sgcg | 17 | 2 | 4 | 0 | 0 | 3 | 1 | 23 | 63 | 21 | 316 | 7 | 18 | 3 | 8 | 9 | 2 | 1 | 0 | 4 | 4 | 17 | 192 | 29 | 9 | 12 | 1 | 1 | Sgcg |
| Sgcz | 55 | 40 | 18 | 48 | 66 | 78 | 59 | 100 | 73 | 59 | 72 | 24 | 42 | 34 | 55 | 50 | 35 | 53 | 65 | 89 | 54 | 95 | 76 | 42 | 39 | 17 | 56 | 38 | Sgcz |
| Sgf29 | 553 | 711 | 663 | 582 | 560 | 548 | 555 | 1060 | 1160 | 1083 | 1018 | 1096 | 1073 | 1213 | 579 | 526 | 619 | 463 | 622 | 693 | 582 | 929 | 1127 | 1202 | 1134 | 1294 | 1189 | 1297 | Sgf29 |
| Sgip1 | 2004 | 1994 | 1738 | 3068 | 4880 | 4098 | 4167 | 3588 | 3289 | 4121 | 3460 | 3263 | 3248 | 3327 | 2548 | 2156 | 2322 | 3483 | 3783 | 3768 | 4353 | 4729 | 3108 | 3116 | 4428 | 2185 | 2767 | 2919 | Sgip1 |
| Sgk1 | 1684 | 1543 | 1426 | 1373 | 1380 | 1360 | 1373 | 1755 | 1505 | 1362 | 1473 | 1277 | 1742 | 1270 | 1630 | 1924 | 1279 | 1579 | 1206 | 1347 | 1277 | 1755 | 1728 | 1346 | 1412 | 1287 | 1403 | 1311 | Sgk1 |
| Sgk2 | 0 | 8 | 2 | 8 | 3 | 8 | 2 | 0 | 3 | 3 | 2 | 1 | 3 | 0 | 3 | 5 | 6 | 6 | 2 | 1 | 2 | 5 | 0 | 0 | 3 | 1 | 0 | 5 | Sgk2 |
| Sgk3 | 682 | 311 | 418 | 220 | 489 | 442 | 434 | 956 | 1095 | 709 | 929 | 784 | 764 | 713 | 526 | 544 | 301 | 381 | 362 | 528 | 344 | 804 | 1169 | 742 | 584 | 717 | 675 | 765 | Sgk3 |
| Sgms1 | 1427 | 1113 | 1202 | 1129 | 1080 | 1172 | 1019 | 929 | 1055 | 908 | 1137 | 840 | 1061 | 794 | 1316 | 1403 | 954 | 1115 | 1013 | 990 | 961 | 932 | 1298 | 963 | 824 | 976 | 707 | 866 | Sgms1 |
| Sgms2 | 0 | 0 | 1 | 0 | 0 | 0 | 0 | 0 | 0 | 0 | 0 | 0 | 0 | 0 | 0 | 2 | 0 | 0 | 0 | 0 | 0 | 0 | 2 | 0 | 0 | 0 | 0 | 0 | Sgms2 |
| Sgo1 | 2870 | 2261 | 2404 | 1869 | 1537 | 1712 | 2215 | 3435 | 2750 | 2569 | 2260 | 3958 | 2992 | 4023 | 2454 | 2062 | 2179 | 1650 | 1845 | 1980 | 1855 | 2130 | 2551 | 3151 | 2592 | 4419 | 3980 | 4044 | Sgo1 |
| Sgo2a | 5335 | 3989 | 4900 | 3641 | 2738 | 3660 | 3500 | 4196 | 3737 | 3386 | 3045 | 4897 | 4265 | 4680 | 4661 | 4555 | 4297 | 3367 | 3605 | 3965 | 3220 | 2821 | 3348 | 3935 | 3622 | 5085 | 5205 | 4832 | Sgo2a |
| Sgo2b | 2 | 1 | 0 | 1 | 0 | 0 | 0 | 1 | 2 | 0 | 3 | 0 | 3 | 1 | 1 | 0 | 0 | 0 | 0 | 0 | 0 | 3 | 8 | 4 | 5 | 8 | 1 | 1 | Sgo2b |
| Sgpl1 | 15943 | 15017 | 12516 | 12619 | 11965 | 13031 | 12722 | 10490 | 9178 | 8756 | 8958 | 9416 | 9593 | 9197 | 15360 | 14221 | 12439 | 12940 | 12405 | 12322 | 12496 | 9899 | 9509 | 8891 | 9318 | 8976 | 9167 | 9463 | Sgpl1 |
| Sgpp1 | 2045 | 1400 | 1695 | 1969 | 1874 | 2130 | 2085 | 1368 | 1397 | 1233 | 1378 | 1139 | 1017 | 1179 | 2062 | 1807 | 1673 | 2209 | 1810 | 2047 | 1976 | 1235 | 1397 | 1171 | 1129 | 1329 | 1140 | 1087 | Sgpp1 |
| Sgpp2 | 837 | 545 | 491 | 428 | 443 | 266 | 248 | 269 | 224 | 311 | 239 | 133 | 210 | 113 | 591 | 784 | 422 | 380 | 293 | 357 | 272 | 125 | 256 | 283 | 96 | 327 | 121 | 148 | Sgpp2 |
| Sgsh | 625 | 677 | 902 | 565 | 590 | 524 | 506 | 413 | 531 | 559 | 552 | 754 | 657 | 534 | 533 | 667 | 715 | 560 | 482 | 424 | 434 | 513 | 608 | 556 | 517 | 689 | 697 | 605 | Sgsh |
| Sgsm1 | 2593 | 3554 | 2724 | 3877 | 3952 | 3534 | 3569 | 1832 | 1865 | 2320 | 1886 | 2151 | 1777 | 2142 | 2724 | 3037 | 3453 | 3839 | 3890 | 3319 | 3758 | 2770 | 1699 | 2216 | 2560 | 1762 | 1951 | 2070 | Sgsm1 |
| Sgsm2 | 3651 | 4805 | 4970 | 6041 | 6172 | 5598 | 5180 | 3154 | 3606 | 4149 | 3644 | 4254 | 3556 | 3637 | 3575 | 4546 | 5597 | 5864 | 6148 | 4877 | 5392 | 4801 | 3249 | 3997 | 4593 | 3019 | 3596 | 3599 | Sgsm2 |
| Sgsm3 | 2573 | 3669 | 3375 | 3274 | 3380 | 3039 | 3165 | 1628 | 2016 | 1989 | 1865 | 1788 | 1736 | 1762 | 2789 | 3240 | 3197 | 3212 | 3558 | 3069 | 3161 | 2064 | 1909 | 1821 | 2094 | 1865 | 1706 | 1734 | Sgsm3 |
| Sgta | 8639 | 11365 | 9572 | 10652 | 9198 | 9281 | 8900 | 6754 | 8460 | 8516 | 7714 | 8657 | 8166 | 8383 | 8377 | 9379 | 10966 | 9896 | 10484 | 9021 | 9374 | 7498 | 8014 | 9042 | 8634 | 9673 | 8434 | 8421 | Sgta |
| Sgtb | 6642 | 7561 | 5871 | 7553 | 7352 | 7396 | 7315 | 3380 | 2598 | 2739 | 2469 | 2624 | 2626 | 2730 | 7941 | 7032 | 6765 | 7545 | 7220 | 7807 | 7813 | 3205 | 2738 | 2594 | 3171 | 2147 | 2552 | 2693 | Sgtb |
| Sh2b1 | 5637 | 6929 | 8149 | 7455 | 6810 | 6409 | 6440 | 5356 | 6334 | 6931 | 7035 | 7331 | 6550 | 7195 | 5103 | 6541 | 7765 | 7526 | 7499 | 5680 | 6514 | 6933 | 5785 | 7403 | 7325 | 7422 | 6491 | 6890 | Sh2b1 |
| Sh2b2 | 1433 | 1684 | 1752 | 2068 | 2257 | 1886 | 1692 | 1248 | 1313 | 1527 | 1456 | 1549 | 1239 | 1277 | 1407 | 1573 | 2036 | 2059 | 2482 | 1578 | 1908 | 1851 | 1291 | 1646 | 1730 | 1487 | 1318 | 1262 | Sh2b2 |

| | | | | | | | | | | | | | | | | | | | | | | | | | | | | | |
|---|---|---|---|---|---|---|---|---|---|---|---|---|---|---|---|---|---|---|---|---|---|---|---|---|---|---|---|---|---|
| Sh2b3 | 3151 | 3498 | 3360 | 2975 | 2472 | 2431 | 2261 | 977 | 1255 | 1235 | 1603 | 1246 | 1418 | 1174 | 2939 | 3572 | 3243 | 2969 | 2323 | 2117 | 2356 | 1360 | 1381 | 1373 | 1163 | 1573 | 1367 | 1277 | Sh2b3 |
| Sh2d1a | 2 | 0 | 3 | 11 | 2 | 4 | 8 | 5 | 5 | 2 | 3 | 4 | 2 | 2 | 3 | 1 | 2 | 6 | 3 | 1 | 3 | 3 | 4 | 10 | 3 | 7 | 9 | 2 | Sh2d1a |
| Sh2d1b1 | 0 | 4 | 0 | 1 | 0 | 3 | 0 | 5 | 2 | 0 | 9 | 1 | 5 | 2 | 3 | 4 | 3 | 2 | 2 | 5 | 0 | 0 | 7 | 2 | 7 | 7 | 5 | 1 | Sh2d1b1 |
| Sh2d1b2 | 0 | 0 | 0 | 0 | 0 | 0 | 0 | 0 | 0 | 0 | 0 | 0 | 0 | 0 | 0 | 0 | 0 | 0 | 0 | 0 | 0 | 0 | 1 | 0 | 1 | 0 | 0 | 0 | Sh2d1b2 |
| Sh2d2a | 5 | 0 | 1 | 5 | 1 | 0 | 0 | 0 | 1 | 0 | 0 | 0 | 1 | 2 | 3 | 5 | 2 | 1 | 0 | 1 | 0 | 0 | 2 | 1 | 0 | 2 | 3 | 0 | Sh2d2a |
| Sh2d3c | 1373 | 2027 | 1726 | 1871 | 1685 | 1419 | 1264 | 1045 | 1261 | 1464 | 1439 | 1502 | 1479 | 1196 | 1350 | 1759 | 1913 | 1631 | 1645 | 1250 | 1375 | 1340 | 1308 | 1452 | 1349 | 1805 | 1413 | 1314 | Sh2d3c |
| Sh2d4a | 17 | 25 | 8 | 1 | 17 | 2 | 6 | 72 | 81 | 53 | 56 | 18 | 58 | 22 | 18 | 25 | 5 | 26 | 15 | 9 | 4 | 37 | 71 | 17 | 22 | 4 | 35 | 36 | Sh2d4a |
| Sh2d4b | 1177 | 1246 | 1161 | 2076 | 1975 | 1795 | 2331 | 504 | 456 | 649 | 583 | 539 | 468 | 583 | 1178 | 1173 | 1688 | 2368 | 2031 | 1458 | 2217 | 720 | 392 | 613 | 713 | 397 | 409 | 415 | Sh2d4b |
| Sh2d5 | 989 | 1302 | 1184 | 1372 | 1763 | 1472 | 1386 | 542 | 591 | 693 | 643 | 470 | 607 | 492 | 976 | 1280 | 1423 | 1620 | 1644 | 1429 | 1446 | 864 | 579 | 554 | 670 | 347 | 437 | 356 | Sh2d5 |
| Sh2d6 | 46 | 78 | 68 | 81 | 93 | 101 | 60 | 94 | 66 | 67 | 76 | 101 | 101 | 122 | 78 | 61 | 70 | 63 | 73 | 61 | 80 | 95 | 65 | 78 | 84 | 76 | 93 | 90 | Sh2d6 |
| Sh2d7 | 2 | 1 | 2 | 1 | 15 | 11 | 8 | 4 | 20 | 5 | 4 | 8 | 5 | 13 | 3 | 2 | 5 | 13 | 9 | 5 | 12 | 2 | 3 | 7 | 11 | 11 | 3 | 5 | Sh2d7 |
| Sh3bgr | 175 | 137 | 239 | 193 | 170 | 152 | 177 | 73 | 65 | 80 | 282 | 66 | 79 | 66 | 182 | 169 | 161 | 173 | 179 | 152 | 193 | 90 | 187 | 73 | 63 | 70 | 60 | 68 | Sh3bgr |
| Sh3bgrl | 25262 | 17157 | 24805 | 22289 | 22309 | 23836 | 27178 | 28234 | 23238 | 20196 | 20783 | 26183 | 24188 | 27737 | 27677 | 22181 | 22477 | 21709 | 22551 | 27786 | 24769 | 24105 | 23295 | 23808 | 24234 | 29372 | 27021 | 28147 | Sh3bgrl |
| Sh3bgrl2 | 3547 | 3308 | 2939 | 2786 | 2918 | 2812 | 3093 | 1429 | 1412 | 1289 | 1134 | 1156 | 1107 | 1092 | 3654 | 3324 | 2875 | 3266 | 2929 | 3372 | 3235 | 1135 | 1460 | 1167 | 1309 | 1186 | 1040 | 1061 | Sh3bgrl2 |
| Sh3bgrl3 | 2393 | 3104 | 2447 | 2161 | 2102 | 2134 | 1872 | 1923 | 2372 | 2367 | 2401 | 2155 | 2489 | 2151 | 2293 | 2655 | 2365 | 2128 | 2099 | 2192 | 2107 | 2024 | 2537 | 2199 | 2166 | 2574 | 2166 | 2303 | Sh3bgrl3 |
| Sh3bp1 | 354 | 346 | 407 | 296 | 169 | 247 | 173 | 160 | 205 | 210 | 287 | 126 | 164 | 128 | 208 | 421 | 290 | 283 | 192 | 167 | 151 | 137 | 242 | 185 | 82 | 259 | 114 | 146 | Sh3bp1 |
| Sh3bp2 | 845 | 869 | 1209 | 1208 | 1713 | 1380 | 1281 | 2078 | 2464 | 2669 | 2201 | 2588 | 2209 | 2245 | 771 | 863 | 1169 | 1213 | 1633 | 1216 | 1314 | 2767 | 2481 | 2633 | 2962 | 2585 | 2294 | 2293 | Sh3bp2 |
| Sh3bp4 | 4523 | 4964 | 4196 | 4259 | 4105 | 4140 | 3718 | 2282 | 3002 | 2506 | 2698 | 2754 | 2784 | 2447 | 4236 | 4889 | 4580 | 4185 | 4390 | 3632 | 3706 | 2237 | 3128 | 2364 | 2508 | 2923 | 2769 | 2623 | Sh3bp4 |
| Sh3bp5 | 2407 | 2208 | 2669 | 3023 | 4997 | 3881 | 3774 | 6149 | 6363 | 7592 | 7241 | 5702 | 5818 | 6208 | 2767 | 2657 | 2814 | 3728 | 3820 | 3142 | 4058 | 8734 | 6524 | 6959 | 8508 | 5401 | 5069 | 5478 | Sh3bp5 |
| Sh3bp5l | 3862 | 5038 | 4400 | 4787 | 4304 | 4112 | 3999 | 3284 | 3425 | 3713 | 3529 | 3995 | 3705 | 3723 | 3703 | 3953 | 4485 | 4612 | 4446 | 4269 | 4236 | 3436 | 3289 | 3892 | 4124 | 4021 | 3995 | 3881 | Sh3bp5l |
| Sh3d19 | 1911 | 1625 | 1757 | 1329 | 1633 | 1627 | 1038 | 2093 | 2352 | 2471 | 2788 | 1900 | 2299 | 1909 | 1673 | 2293 | 1387 | 1699 | 1216 | 1075 | 1045 | 2378 | 2807 | 2097 | 2037 | 2128 | 1869 | 1790 | Sh3d19 |
| Sh3d21 | 212 | 188 | 118 | 167 | 65 | 111 | 58 | 173 | 265 | 171 | 342 | 148 | 308 | 137 | 160 | 211 | 108 | 158 | 86 | 69 | 66 | 162 | 271 | 204 | 143 | 210 | 159 | 139 | Sh3d21 |
| Sh3gl1 | 4452 | 5732 | 5374 | 4879 | 4217 | 4335 | 3867 | 3322 | 4226 | 3943 | 4227 | 4289 | 4038 | 3841 | 4146 | 5196 | 5060 | 4796 | 4545 | 3891 | 3816 | 3549 | 4361 | 4193 | 3870 | 4974 | 3812 | 3986 | Sh3gl1 |
| Sh3gl2 | 1034 | 1091 | 806 | 1439 | 2645 | 1825 | 2018 | 2287 | 2340 | 2315 | 1961 | 1466 | 1496 | 1560 | 1396 | 1026 | 1342 | 1808 | 1948 | 2125 | 2197 | 2525 | 2256 | 1839 | 2410 | 1227 | 1335 | 1318 | Sh3gl2 |
| Sh3gl3 | 1381 | 1131 | 990 | 1211 | 1501 | 1378 | 1560 | 1414 | 1207 | 1124 | 1160 | 925 | 944 | 1027 | 1447 | 1379 | 1301 | 1324 | 1326 | 1524 | 1541 | 1343 | 1170 | 1078 | 1198 | 925 | 797 | 882 | Sh3gl3 |
| Sh3glb1 | 13093 | 11410 | 12355 | 15932 | 18722 | 17131 | 18710 | 15478 | 14294 | 16316 | 16663 | 14672 | 14666 | 15081 | 14126 | 12025 | 14442 | 17164 | 16818 | 14815 | 19006 | 18870 | 15305 | 14309 | 16865 | 12292 | 13716 | 14658 | Sh3glb1 |
| Sh3glb2 | 4231 | 6020 | 5184 | 6745 | 6985 | 6250 | 6050 | 4123 | 4969 | 5159 | 4832 | 4644 | 4622 | 4515 | 4456 | 5337 | 6519 | 7141 | 7335 | 6299 | 5881 | 5150 | 4699 | 4905 | 5643 | 4629 | 4468 | 3925 | Sh3glb2 |
| Sh3kbp1 | 2426 | 2233 | 2429 | 2585 | 3427 | 2662 | 2842 | 3396 | 3170 | 3591 | 3300 | 2910 | 3002 | 2823 | 2633 | 2414 | 2687 | 2630 | 2840 | 3050 | 2796 | 3703 | 3510 | 3151 | 3386 | 2658 | 2683 | 2747 | Sh3kbp1 |
| Sh3pxd2a | 7623 | 7924 | 8451 | 9138 | 8571 | 9081 | 7287 | 8197 | 11385 | 12222 | 17008 | 11511 | 11578 | 10097 | 6674 | 10008 | 8413 | 10124 | 8507 | 6111 | 7513 | 10808 | 12951 | 12936 | 11177 | 8199 | 9646 | 9378 | Sh3pxd2a |
| Sh3pxd2b | 4710 | 5357 | 6702 | 5958 | 7333 | 6500 | 5392 | 7910 | 9264 | 10819 | 11479 | 11519 | 10929 | 10522 | 4627 | 5629 | 6983 | 6714 | 7174 | 4716 | 5870 | 11939 | 9499 | 11122 | 11163 | 8890 | 10696 | 11065 | Sh3pxd2b |
| Sh3rf1 | 3291 | 3455 | 3797 | 5103 | 6125 | 5408 | 5556 | 4174 | 4414 | 5437 | 4300 | 5395 | 4101 | 5151 | 3339 | 3026 | 4653 | 5418 | 5947 | 4815 | 5596 | 5746 | 3955 | 5489 | 6030 | 5153 | 4120 | 4923 | Sh3rf1 |
| Sh3rf2 | 45 | 28 | 15 | 7 | 9 | 41 | 4 | 73 | 143 | 86 | 114 | 30 | 70 | 44 | 32 | 37 | 12 | 14 | 0 | 8 | 7 | 23 | 142 | 38 | 42 | 44 | 37 | 36 | Sh3rf2 |
| Sh3rf3 | 497 | 572 | 289 | 883 | 1267 | 955 | 1207 | 979 | 1226 | 1427 | 1222 | 1147 | 494 | 1060 | 514 | 431 | 680 | 1059 | 1216 | 864 | 1271 | 1414 | 888 | 1477 | 1677 | 918 | 608 | 606 | Sh3rf3 |
| Sh3tc1 | 216 | 240 | 256 | 199 | 236 | 221 | 148 | 212 | 304 | 337 | 362 | 339 | 373 | 270 | 190 | 283 | 220 | 250 | 246 | 179 | 190 | 314 | 381 | 310 | 242 | 324 | 352 | 255 | Sh3tc1 |
| Sh3tc2 | 113 | 109 | 86 | 50 | 83 | 97 | 50 | 222 | 290 | 263 | 265 | 176 | 344 | 223 | 100 | 159 | 82 | 51 | 56 | 61 | 50 | 212 | 306 | 190 | 199 | 203 | 215 | 271 | Sh3tc2 |
| Sh3yl1 | 1257 | 1292 | 1543 | 1713 | 1938 | 1572 | 1797 | 1040 | 1004 | 1277 | 1023 | 979 | 1083 | 1119 | 1275 | 1257 | 1664 | 1573 | 1869 | 1663 | 1820 | 1245 | 1022 | 1074 | 1232 | 992 | 883 | 1102 | Sh3yl1 |
| Shank1 | 3693 | 5061 | 5452 | 9236 | 12842 | 9150 | 9524 | 7013 | 8700 | 12388 | 9735 | 8246 | 7091 | 7619 | 3885 | 3991 | 7967 | 10385 | 11727 | 7752 | 10652 | 14775 | 7130 | 11003 | 12670 | 5488 | 6394 | 6421 | Shank1 |
| Shank2 | 676 | 707 | 941 | 1386 | 2099 | 1466 | 1558 | 2394 | 2778 | 3807 | 2830 | 2860 | 2134 | 2701 | 674 | 673 | 1218 | 1473 | 1838 | 1323 | 1725 | 4423 | 2454 | 3210 | 4175 | 1924 | 2244 | 2312 | Shank2 |
| Shank3 | 1768 | 2421 | 3154 | 2932 | 3379 | 2521 | 2359 | 1921 | 2491 | 3022 | 3066 | 2482 | 2383 | 2118 | 1614 | 2192 | 3203 | 3006 | 3173 | 2007 | 2450 | 3472 | 2431 | 2943 | 3172 | 2744 | 2296 | 2009 | Shank3 |
| Sharpin | 1880 | 2323 | 2526 | 2823 | 2491 | 2216 | 2414 | 1801 | 1865 | 2127 | 2018 | 2196 | 1966 | 2084 | 1876 | 1964 | 2619 | 2597 | 2627 | 2294 | 2578 | 2146 | 1781 | 2247 | 2282 | 2099 | 2050 | 1958 | Sharpin |
| Shb | 1510 | 2031 | 1728 | 2820 | 2548 | 2159 | 2405 | 1051 | 1328 | 1569 | 1256 | 1452 | 687 | 1537 | 1542 | 1455 | 2421 | 2649 | 2544 | 1927 | 2716 | 1399 | 1041 | 1776 | 1788 | 1791 | 938 | 1138 | Shb |
| Shbg | 3 | 1 | 3 | 6 | 0 | 2 | 0 | 0 | 0 | 0 | 0 | 0 | 1 | 3 | 0 | 2 | 0 | 6 | 3 | 0 | 0 | 2 | 0 | 0 | 0 | 0 | 0 | 0 | Shbg |
| Shc1 | 5980 | 5596 | 6008 | 4419 | 3533 | 3638 | 2653 | 3604 | 4631 | 4279 | 5851 | 3447 | 4547 | 3299 | 5003 | 7208 | 4845 | 4735 | 3098 | 2917 | 2767 | 3887 | 5310 | 4102 | 3404 | 4742 | 3799 | 3646 | Shc1 |
| Shc2 | 1715 | 1709 | 2324 | 2050 | 2714 | 2092 | 1771 | 2843 | 2974 | 3150 | 3219 | 2620 | 3357 | 2285 | 1809 | 2152 | 2283 | 2155 | 2155 | 1789 | 1860 | 3758 | 2731 | 2582 | 3089 | 2419 | 2791 | 2667 | Shc2 |
| Shc3 | 606 | 551 | 214 | 563 | 552 | 658 | 518 | 640 | 798 | 647 | 523 | 683 | 584 | 713 | 454 | 483 | 338 | 536 | 496 | 554 | 689 | 476 | 627 | 382 | 786 | 288 | 790 | 623 | Shc3 |
| Shc4 | 235 | 228 | 230 | 236 | 373 | 212 | 216 | 424 | 430 | 480 | 460 | 283 | 480 | 364 | 285 | 253 | 287 | 233 | 279 | 217 | 224 | 617 | 389 | 340 | 448 | 327 | 274 | 278 | Shc4 |
| Shcbp1 | 4035 | 2612 | 4015 | 2396 | 1999 | 2720 | 2990 | 3197 | 2946 | 2403 | 2346 | 3894 | 3521 | 4173 | 3520 | 3280 | 3082 | 2310 | 2549 | 3404 | 2540 | 2090 | 2573 | 3290 | 2744 | 5168 | 4136 | 4139 | Shcbp1 |
| Shcbp1l | 60 | 74 | 51 | 40 | 84 | 45 | 56 | 39 | 37 | 41 | 44 | 25 | 24 | 28 | 62 | 66 | 44 | 63 | 33 | 58 | 45 | 22 | 33 | 29 | 43 | 20 | 17 | 19 | Shcbp1l |
| Shd | 1692 | 2507 | 2495 | 2207 | 2934 | 2707 | 2024 | 3335 | 3129 | 3516 | 3056 | 3764 | 3999 | 3381 | 2099 | 1858 | 2608 | 2224 | 2878 | 2756 | 2372 | 3855 | 2947 | 3679 | 3725 | 3230 | 3865 | 3494 | Shd |
| She | 318 | 278 | 376 | 265 | 392 | 434 | 258 | 563 | 571 | 669 | 854 | 587 | 716 | 640 | 323 | 386 | 351 | 318 | 347 | 301 | 299 | 685 | 765 | 629 | 580 | 609 | 572 | 687 | She |
| Shf | 2186 | 3137 | 2813 | 3285 | 3617 | 3251 | 2657 | 1688 | 1908 | 2122 | 1917 | 1908 | 2037 | 1685 | 2432 | 2488 | 3310 | 3354 | 3755 | 3023 | 3155 | 2047 | 1770 | 1810 | 2103 | 1748 | 1697 | 1580 | Shf |
| Shfl | 277 | 269 | 284 | 265 | 201 | 228 | 145 | 95 | 119 | 177 | 181 | 101 | 103 | 97 | 161 | 274 | 304 | 337 | 204 | 195 | 226 | 160 | 163 | 164 | 88 | 124 | 86 | 119 | Shfl |
| Shh | 811 | 904 | 1810 | 847 | 641 | 645 | 660 | 816 | 618 | 642 | 538 | 1080 | 820 | 1177 | 940 | 893 | 1286 | 635 | 620 | 681 | 675 | 934 | 475 | 761 | 579 | 1498 | 1397 | 1310 | Shh |
| Shisa2 | 3827 | 3655 | 2898 | 3753 | 2814 | 3158 | 3993 | 2859 | 3367 | 3079 | 3577 | 4015 | 2825 | 4166 | 3361 | 3407 | 3657 | 3635 | 3724 | 3053 | 3208 | 2085 | 2324 | 4048 | 3235 | 5816 | 3332 | 3350 | Shisa2 |
| Shisa3 | 499 | 466 | 762 | 492 | 606 | 599 | 603 | 1629 | 1491 | 1103 | 1503 | 1344 | 1724 | 1485 | 501 | 635 | 513 | 551 | 784 | 947 | 763 | 2543 | 1360 | 1352 | 1376 | 229 | 1987 | 1996 | Shisa3 |
| Shisa4 | 1368 | 1476 | 1764 | 1547 | 1709 | 1641 | 1621 | 1556 | 1599 | 2164 | 1823 | 2197 | 1864 | 2333 | 1375 | 1361 | 1849 | 1610 | 1803 | 1487 | 1739 | 1908 | 1748 | 2243 | 2251 | 2435 | 2300 | 2336 | Shisa4 |
| Shisa5 | 1227 | 1261 | 1441 | 1269 | 1144 | 1243 | 906 | 1476 | 1877 | 1816 | 2147 | 1517 | 1657 | 1302 | 1211 | 1483 | 1195 | 1212 | 1029 | 1031 | 877 | 1750 | 2155 | 1781 | 1502 | 1755 | 1407 | 1332 | Shisa5 |
| Shisa6 | 213 | 215 | 147 | 425 | 711 | 582 | 636 | 865 | 835 | 1051 | 882 | 922 | 764 | 914 | 320 | 254 | 318 | 514 | 597 | 501 | 672 | 1036 | 728 | 931 | 1155 | 585 | 695 | 756 | Shisa6 |
| Shisa7 | 3565 | 4647 | 3658 | 5566 | 8296 | 6136 | 6537 | 5950 | 5712 | 7258 | 5827 | 5139 | 4574 | 5244 | 4110 | 3546 | 4984 | 5963 | 7112 | 5673 | 6613 | 9184 | 4993 | 5494 | 7369 | 3409 | 4278 | 4333 | Shisa7 |
| Shisa8 | 46 | 60 | 56 | 40 | 73 | 42 | 54 | 43 | 53 | 53 | 26 | 72 | 70 | 52 | 40 | 36 | 46 | 37 | 58 | 52 | 47 | 42 | 74 | 56 | 62 | 72 | 59 | 53 | Shisa8 |
| Shisa9 | 104 | 188 | 105 | 321 | 692 | 411 | 457 | 581 | 639 | 670 | 656 | 324 | 272 | 344 | 227 | 151 | 198 | 437 | 468 | 407 | 533 | 1008 | 734 | 503 | 748 | 216 | 234 | 266 | Shisa9 |
| Shisal1 | 912 | 932 | 1203 | 1418 | 2501 | 1896 | 1831 | 2877 | 2946 | 3558 | 2985 | 2788 | 2832 | 2436 | 1149 | 921 | 1482 | 1541 | 2134 | 1571 | 1845 | 4094 | 2645 | 3040 | 3817 | 1953 | 2658 | 2445 | Shisal1 |
| Shisal2a | 25 | 28 | 30 | 42 | 36 | 22 | 27 | 43 | 59 | 72 | 93 | 42 | 45 | 45 | 32 | 49 | 49 | 19 | 19 | 28 | 25 | 52 | 54 | 76 | 48 | 56 | 81 | 42 | Shisal2a |
| Shisal2b | 3 | 4 | 6 | 3 | 11 | 1 | 6 | 10 | 6 | 16 | 12 | 7 | 23 | 21 | 8 | 2 | 10 | 1 | 12 | 15 | 4 | 8 | 15 | 9 | 5 | 3 | 8 | 15 | Shisal2b |
| Shkbp1 | 2949 | 3448 | 3406 | 3506 | 3372 | 3105 | 3157 | 2285 | 2451 | 2859 | 2382 | 2911 | 2461 | 2702 | 2777 | 3022 | 3402 | 3319 | 3849 | 2769 | 3223 | 2864 | 2282 | 2995 | 2927 | 2938 | 2912 | 2752 | Shkbp1 |
| Shld1 | 196 | 193 | 142 | 180 | 224 | 232 | 234 | 688 | 601 | 563 | 527 | 601 | 576 | 558 | 210 | 164 | 222 | 235 | 215 | 274 | 219 | 625 | 612 | 585 | 562 | 696 | 557 | 552 | Shld1 |
| Shld2 | 438 | 351 | 301 | 408 | 330 | 409 | 421 | 699 | 629 | 577 | 614 | 658 | 718 | 674 | 473 | 389 | 334 | 379 | 352 | 406 | 399 | 485 | 667 | 538 | 559 | 676 | 733 | 655 | Shld2 |
| Shld3 | 332 | 200 | 210 | 267 | 236 | 275 | 325 | 267 | 185 | 206 | 152 | 232 | 239 | 253 | 289 | 233 | 204 | 191 | 210 | 232 | 246 | 241 | 254 | 212 | 236 | 268 | 221 | 298 | Shld3 |
| Shmt1 | 3298 | 2939 | 2711 | 2191 | 1600 | 1952 | 2217 | 1765 | 1973 | 1831 | 1609 | 2498 | 2368 | 2495 | 2608 | 2660 | 2356 | 2013 | 2051 | 2295 | 2034 | 1078 | 1698 | 2057 | 1723 | 3154 | 2715 | 2801 | Shmt1 |
| Shmt2 | 5108 | 6499 | 6518 | 6802 | 5199 | 5552 | 5529 | 3219 | 4313 | 4344 | 4092 | 5186 | 4513 | 5263 | 4739 | 5472 | 6743 | 6117 | 6706 | 5352 | 5786 | 3212 | 4073 | 5119 | 4583 | 6425 | 4981 | 5635 | Shmt2 |
| Shoc1 | 0 | 0 | 0 | 0 | 0 | 0 | 0 | 0 | 0 | 1 | 0 | 0 | 0 | 4 | 0 | 0 | 0 | 0 | 3 | 0 | 0 | 0 | 0 | 2 | 0 | 2 | 0 | 0 | Shoc1 |
| Shoc2 | 11720 | 9869 | 10215 | 10421 | 10690 | 10812 | 12103 | 10425 | 8305 | 7544 | 7515 | 8264 | 8359 | 9085 | 12535 | 10181 | 9632 | 10573 | 10365 | 12440 | 11396 | 9225 | 8719 | 8086 | 8152 | 8687 | 8677 | 9464 | Shoc2 |
| Shox2 | 2031 | 1395 | 3489 | 2282 | 3090 | 2639 | 2352 | 4305 | 2693 | 4770 | 3739 | 5689 | 5976 | 6596 | 2380 | 1305 | 2788 | 2258 | 3248 | 1893 | 3078 | 6351 | 3164 | 5188 | 4690 | 3784 | 7009 | 8060 | Shox2 |
| Shpk | 396 | 517 | 401 | 434 | 376 | 335 | 360 | 149 | 166 | 149 | 199 | 174 | 147 | 126 | 430 | 456 | 430 | 469 | 264 | 357 | 306 | 159 | 190 | 150 | 170 | 149 | 128 | 127 | Shpk |
| Shprh | 12211 | 10367 | 7264 | 9814 | 9981 | 11080 | 11850 | 10209 | 8883 | 7083 | 7493 | 8525 | 7686 | 8533 | 12699 | 10827 | 8205 | 9628 | 9595 | 12741 | 11919 | 7718 | 8418 | 6892 | 8168 | 5677 | 8644 | 7751 | Shprh |
| Shq1 | 1634 | 1808 | 1619 | 1530 | 1347 | 1411 | 1460 | 813 | 720 | 754 | 741 | 845 | 780 | 747 | 1565 | 1611 | 1646 | 1464 | 1536 | 1619 | 1309 | 610 | 764 | 762 | 813 | 828 | 915 | 895 | Shq1 |
| Shroom1 | 138 | 129 | 126 | 110 | 116 | 128 | 76 | 158 | 299 | 302 | 332 | 131 | 215 | 116 | 115 | 143 | 131 | 127 | 112 | 90 | 99 | 229 | 260 | 245 | 231 | 297 | 152 | 127 | Shroom1 |
| Shroom2 | 3135 | 2666 | 2455 | 3236 | 4590 | 3918 | 3698 | 5200 | 5193 | 4768 | 4708 | 3455 | 3370 | 3496 | 3432 | 2977 | 2810 | 3383 | 3535 | 3539 | 3796 | 5902 | 4833 | 3690 | 4674 | 2612 | 3074 | 3157 | Shroom2 |
| Shroom3 | 3246 | 4344 | 5213 | 4729 | 3455 | 3368 | 3057 | 1642 | 1875 | 2197 | 2030 | 2974 | 2357 | 2412 | 2960 | 4307 | 5172 | 3894 | 4164 | 2947 | 3236 | 1770 | 1904 | 2787 | 1949 | 3634 | 2509 | 2661 | Shroom3 |

| | | | | | | | | | | | | | | | | | | | | | | | | | | | | | |
|---|---|---|---|---|---|---|---|---|---|---|---|---|---|---|---|---|---|---|---|---|---|---|---|---|---|---|---|---|---|
| Shroom4 | 2456 | 1887 | 1674 | 1655 | 1433 | 1912 | 1330 | 2225 | 2426 | 2311 | 2553 | 2135 | 2827 | 2055 | 1904 | 2303 | 1499 | 1647 | 1399 | 1255 | 1364 | 1793 | 2346 | 1985 | 1870 | 1729 | 2396 | 2181 | Shroom4 |
| Shtn1 | 1983 | 1915 | 912 | 3140 | 4943 | 3285 | 4883 | 4076 | 4384 | 4463 | 3680 | 2250 | 1482 | 2397 | 2289 | 1548 | 2589 | 4043 | 4012 | 3506 | 4720 | 5185 | 3579 | 3888 | 5012 | 2986 | 1548 | 1608 | Shtn1 |
| Siae | 1608 | 1832 | 1410 | 1511 | 1363 | 1537 | 1552 | 1299 | 1247 | 1191 | 1162 | 1057 | 1052 | 1053 | 1991 | 1691 | 1358 | 1588 | 1361 | 1618 | 1491 | 1201 | 1350 | 888 | 939 | 969 | 1065 | 1013 | Siae |
| Siah1a | 1675 | 1409 | 1993 | 1962 | 2140 | 1851 | 2138 | 3226 | 2699 | 3198 | 2846 | 3331 | 2986 | 3397 | 1767 | 1477 | 1865 | 1763 | 2193 | 1855 | 1974 | 3507 | 2895 | 3244 | 3432 | 3476 | 3323 | 3404 | Siah1a |
| Siah1b | 942 | 907 | 1056 | 686 | 504 | 536 | 745 | 953 | 866 | 866 | 837 | 1227 | 1090 | 1253 | 831 | 769 | 858 | 527 | 657 | 784 | 632 | 750 | 822 | 1058 | 909 | 1476 | 1238 | 1266 | Siah1b |
| Siah2 | 935 | 1094 | 913 | 1107 | 981 | 923 | 974 | 540 | 539 | 637 | 569 | 512 | 538 | 529 | 961 | 977 | 1046 | 1124 | 1100 | 815 | 1014 | 590 | 550 | 545 | 572 | 712 | 488 | 522 | Siah2 |
| Siah3 | 1031 | 1159 | 859 | 1461 | 1856 | 1831 | 1901 | 1633 | 1769 | 1914 | 1538 | 2149 | 1589 | 2121 | 1229 | 827 | 1291 | 1547 | 1955 | 1931 | 1906 | 1671 | 1480 | 2232 | 2390 | 1968 | 1578 | 1805 | Siah3 |
| Sidt1 | 40 | 41 | 34 | 61 | 155 | 105 | 91 | 147 | 201 | 219 | 208 | 84 | 99 | 98 | 58 | 57 | 60 | 94 | 106 | 157 | 140 | 227 | 324 | 100 | 151 | 75 | 51 | 55 | Sidt1 |
| Sidt2 | 5785 | 6681 | 6586 | 6902 | 6551 | 6400 | 6053 | 4158 | 5321 | 5163 | 5822 | 4780 | 5172 | 4251 | 5483 | 6418 | 6291 | 7287 | 6710 | 5216 | 5999 | 5668 | 5422 | 5082 | 4900 | 4503 | 4443 | 4330 | Sidt2 |
| Sigirr | 46 | 48 | 46 | 39 | 39 | 28 | 42 | 74 | 89 | 98 | 105 | 70 | 94 | 90 | 37 | 51 | 52 | 51 | 61 | 39 | 42 | 79 | 129 | 123 | 74 | 125 | 85 | 70 | Sigirr |
| Siglec1 | 27 | 52 | 40 | 81 | 65 | 42 | 49 | 23 | 58 | 56 | 116 | 78 | 61 | 63 | 42 | 61 | 38 | 96 | 30 | 62 | 84 | 77 | 113 | 96 | 107 | 31 | 65 | 58 | Siglec1 |
| Siglece | 13 | 10 | 2 | 7 | 7 | 9 | 8 | 17 | 26 | 9 | 22 | 13 | 19 | 16 | 17 | 9 | 5 | 19 | 2 | 13 | 13 | 15 | 30 | 27 | 6 | 23 | 23 | 14 | Siglece |
| Siglecf | 6 | 3 | 9 | 2 | 1 | 1 | 1 | 9 | 8 | 6 | 13 | 7 | 19 | 6 | 4 | 7 | 1 | 7 | 0 | 5 | 0 | 12 | 7 | 11 | 6 | 8 | 14 | 12 | Siglecf |
| Siglecg | 1 | 1 | 2 | 0 | 2 | 1 | 0 | 1 | 7 | 1 | 16 | 1 | 9 | 10 | 0 | 2 | 3 | 1 | 1 | 5 | 4 | 3 | 6 | 7 | 0 | 4 | 6 | 0 | Siglecg |
| Siglech | 50 | 68 | 50 | 38 | 64 | 64 | 61 | 87 | 85 | 67 | 86 | 60 | 93 | 62 | 59 | 45 | 28 | 88 | 75 | 93 | 60 | 74 | 80 | 85 | 58 | 82 | 84 | 77 | Siglech |
| Sigmar1 | 3051 | 3806 | 3639 | 3243 | 2531 | 2861 | 2437 | 1716 | 2075 | 1966 | 2143 | 2038 | 2175 | 1943 | 2736 | 3502 | 3271 | 2941 | 2632 | 2444 | 2399 | 1813 | 2155 | 2136 | 1728 | 2666 | 2148 | 2308 | Sigmar1 |
| Sik1 | 2225 | 2977 | 3121 | 3657 | 3257 | 3025 | 2607 | 1878 | 2251 | 2611 | 2877 | 2501 | 2282 | 2270 | 2230 | 2854 | 3497 | 3807 | 3277 | 2547 | 2829 | 2451 | 2267 | 2526 | 2643 | 2545 | 2116 | 2140 | Sik1 |
| Sik2 | 3047 | 3128 | 2261 | 3227 | 3342 | 3294 | 3344 | 1839 | 2230 | 1863 | 1910 | 1861 | 1761 | 1836 | 2852 | 3141 | 2680 | 3122 | 2999 | 2769 | 3382 | 1803 | 1946 | 1766 | 2044 | 948 | 1654 | 1717 | Sik2 |
| Sik3 | 4008 | 4647 | 4983 | 5546 | 5610 | 5199 | 4952 | 3153 | 3834 | 4489 | 4488 | 4553 | 4155 | 4254 | 3699 | 4588 | 5481 | 5355 | 5813 | 4161 | 5220 | 4647 | 3823 | 4434 | 4486 | 4054 | 4140 | 4315 | Sik3 |
| Sike1 | 9812 | 9316 | 8864 | 9861 | 9745 | 10066 | 10445 | 8626 | 7633 | 8269 | 7685 | 9121 | 8930 | 9394 | 9948 | 9002 | 9471 | 9255 | 9369 | 9396 | 10467 | 9337 | 7642 | 7951 | 8899 | 7165 | 9581 | 8837 | Sike1 |
| Sil1 | 592 | 657 | 592 | 526 | 620 | 574 | 550 | 761 | 824 | 762 | 792 | 875 | 917 | 806 | 580 | 581 | 546 | 585 | 614 | 588 | 588 | 772 | 882 | 829 | 798 | 902 | 891 | 849 | Sil1 |
| Sim1 | 1225 | 824 | 1159 | 1040 | 804 | 1175 | 1509 | 1366 | 1220 | 941 | 933 | 1715 | 629 | 1566 | 1092 | 537 | 1133 | 989 | 967 | 1404 | 1127 | 1322 | 1157 | 1396 | 1327 | 1387 | 1883 | 1772 | Sim1 |
| Sim2 | 160 | 141 | 351 | 85 | 90 | 66 | 135 | 245 | 406 | 186 | 620 | 307 | 67 | 283 | 80 | 188 | 236 | 107 | 120 | 133 | 92 | 346 | 339 | 220 | 137 | 397 | 421 | 313 | Sim2 |
| Simc1 | 2933 | 2845 | 2182 | 2143 | 2875 | 3025 | 2789 | 3388 | 3172 | 2784 | 2722 | 2725 | 2608 | 2612 | 2956 | 2517 | 2356 | 2486 | 2707 | 3046 | 2958 | 3253 | 2862 | 2659 | 2899 | 2390 | 2521 | 2359 | Simc1 |
| Sin3a | 17935 | 18382 | 15795 | 16437 | 14162 | 15435 | 16355 | 9774 | 9014 | 9062 | 8358 | 10766 | 9816 | 10851 | 16870 | 16333 | 15302 | 15418 | 15007 | 15984 | 16562 | 9214 | 8617 | 9446 | 9401 | 10570 | 11023 | 10710 | Sin3a |
| Sin3b | 6339 | 8033 | 6976 | 7557 | 7189 | 6877 | 6281 | 6662 | 7097 | 6823 | 7012 | 6982 | 6690 | 6780 | 5880 | 6779 | 7249 | 7613 | 7366 | 7104 | 6606 | 7167 | 6894 | 7209 | 7809 | 6680 | 6839 | 6694 | Sin3b |
| Sinhcaf | 6521 | 6049 | 7592 | 5850 | 5558 | 5894 | 5729 | 8120 | 6367 | 5699 | 5566 | 9517 | 8536 | 9787 | 6727 | 5830 | 6920 | 5351 | 6081 | 6963 | 5647 | 6137 | 5854 | 7547 | 7165 | 10617 | 10379 | 10237 | Sinhcaf |
| Sipa1 | 2708 | 3302 | 3649 | 3169 | 2450 | 2391 | 2125 | 1588 | 1949 | 2245 | 2877 | 2310 | 2469 | 1901 | 2211 | 3338 | 3320 | 3007 | 2689 | 1981 | 2047 | 2343 | 2421 | 2408 | 2141 | 2715 | 2255 | 1830 | Sipa1 |
| Sipa1l1 | 6384 | 6713 | 6323 | 6058 | 6229 | 6025 | 6697 | 5428 | 6337 | 6215 | 5919 | 7352 | 6734 | 7055 | 5672 | 6300 | 6451 | 6343 | 6770 | 5822 | 6157 | 5374 | 5829 | 6452 | 6107 | 5748 | 7044 | 7085 | Sipa1l1 |
| Sipa1l2 | 9231 | 11487 | 10907 | 14030 | 12931 | 12160 | 11863 | 6278 | 6680 | 8174 | 6958 | 8233 | 6923 | 7960 | 9415 | 10618 | 12429 | 13861 | 13683 | 10262 | 13030 | 7554 | 6539 | 8067 | 8631 | 5630 | 7160 | 7554 | Sipa1l2 |
| Sipa1l3 | 3516 | 4755 | 5395 | 5813 | 5657 | 4436 | 4024 | 2010 | 2632 | 3228 | 2862 | 3082 | 2485 | 2353 | 3231 | 4245 | 5867 | 5753 | 5452 | 3628 | 4618 | 3081 | 2598 | 3310 | 3065 | 2684 | 2183 | 2204 | Sipa1l3 |
| Sirpa | 1504 | 1568 | 1921 | 1929 | 2557 | 2325 | 2399 | 2464 | 3174 | 3303 | 3507 | 3058 | 2954 | 2748 | 1535 | 1673 | 1950 | 1946 | 2596 | 2106 | 2323 | 3444 | 3124 | 3392 | 3263 | 3248 | 2666 | 2582 | Sirpa |
| Sirpb1b | 0 | 0 | 0 | 0 | 0 | 0 | 0 | 0 | 0 | 0 | 2 | 0 | 0 | 0 | 0 | 1 | 0 | 0 | 0 | 0 | 0 | 2 | 0 | 0 | 0 | 0 | 0 | 0 | Sirpb1b |
| Sirpb1c | 0 | 0 | 0 | 0 | 0 | 0 | 0 | 0 | 0 | 0 | 0 | 0 | 0 | 0 | 0 | 0 | 0 | 0 | 0 | 0 | 0 | 0 | 0 | 0 | 0 | 6 | 0 | 0 | Sirpb1c |
| Sirt1 | 2873 | 2198 | 2962 | 2564 | 2525 | 2869 | 2887 | 4412 | 3704 | 3462 | 3693 | 4367 | 4187 | 4221 | 2927 | 2700 | 2641 | 2732 | 2608 | 2674 | 2806 | 4304 | 3624 | 3645 | 3872 | 4107 | 4495 | 4196 | Sirt1 |
| Sirt2 | 5936 | 7862 | 6613 | 7190 | 6814 | 7277 | 6643 | 4593 | 5059 | 5227 | 5134 | 4993 | 4788 | 4980 | 6306 | 6655 | 6942 | 6888 | 7355 | 6980 | 7234 | 5041 | 4994 | 5212 | 5283 | 4613 | 4822 | 4755 | Sirt2 |
| Sirt3 | 1249 | 1284 | 1127 | 1194 | 1244 | 1359 | 1158 | 1373 | 1320 | 1264 | 1264 | 1266 | 1438 | 1297 | 1212 | 1156 | 1218 | 1140 | 1281 | 1289 | 1181 | 1190 | 1357 | 1174 | 1342 | 1135 | 1463 | 1246 | Sirt3 |
| Sirt4 | 839 | 1122 | 1487 | 1507 | 1334 | 1345 | 1347 | 839 | 910 | 1204 | 1229 | 1044 | 1112 | 1111 | 931 | 1057 | 1610 | 1595 | 1542 | 1158 | 1366 | 1091 | 975 | 1334 | 1169 | 1187 | 1091 | 1072 | Sirt4 |
| Sirt5 | 511 | 416 | 363 | 407 | 549 | 567 | 489 | 906 | 1021 | 930 | 939 | 919 | 903 | 930 | 463 | 505 | 459 | 427 | 510 | 561 | 514 | 759 | 872 | 901 | 822 | 879 | 904 | 889 | Sirt5 |
| Sirt6 | 1359 | 1591 | 2078 | 1905 | 2002 | 1684 | 1673 | 2049 | 2385 | 2765 | 2789 | 3125 | 2874 | 2948 | 1280 | 1357 | 2089 | 1747 | 2253 | 1543 | 1646 | 2677 | 2244 | 3331 | 3060 | 3520 | 2960 | 2887 | Sirt6 |
| Sirt7 | 2996 | 3777 | 3420 | 4141 | 3242 | 3192 | 3431 | 1944 | 2140 | 2326 | 2021 | 2387 | 2241 | 2224 | 3029 | 3421 | 3658 | 3760 | 3478 | 3001 | 3204 | 2288 | 2005 | 2355 | 2543 | 2193 | 2272 | 2301 | Sirt7 |
| Sis | 1 | 0 | 0 | 0 | 2 | 3 | 0 | 5 | 2 | 1 | 0 | 0 | 0 | 0 | 1 | 1 | 0 | 0 | 0 | 0 | 2 | 0 | 0 | 0 | 0 | 1 | 1 | 2 | Sis |
| Sit1 | 0 | 2 | 0 | 0 | 0 | 0 | 0 | 0 | 0 | 0 | 1 | 0 | 0 | 0 | 1 | 1 | 0 | 1 | 0 | 0 | 0 | 0 | 0 | 0 | 0 | 0 | 0 | 0 | Sit1 |
| Siva1 | 3089 | 3860 | 3251 | 2835 | 2046 | 2408 | 2475 | 1684 | 1903 | 1628 | 1668 | 2130 | 1997 | 1984 | 2745 | 3291 | 2971 | 2752 | 2586 | 3148 | 2308 | 1158 | 1774 | 1959 | 1610 | 2657 | 2273 | 2249 | Siva1 |
| Six1 | 1937 | 1147 | 1043 | 922 | 552 | 456 | 54 | 2127 | 2160 | 2145 | 3188 | 1018 | 2176 | 1117 | 1561 | 2172 | 822 | 800 | 104 | 73 | 63 | 2251 | 2476 | 2032 | 1154 | 2634 | 1328 | 1232 | Six1 |
| Six2 | 2992 | 2443 | 1858 | 1428 | 352 | 673 | 156 | 2032 | 2364 | 2408 | 2435 | 1757 | 2663 | 1675 | 1907 | 3702 | 1417 | 1100 | 129 | 94 | 116 | 1726 | 1865 | 2947 | 1181 | 5216 | 1857 | 2114 | Six2 |
| Six3 | 1761 | 1597 | 2198 | 2412 | 1829 | 1583 | 2265 | 1578 | 1429 | 2023 | 1316 | 2560 | 933 | 2684 | 1229 | 849 | 2484 | 2133 | 2381 | 1480 | 2428 | 2191 | 967 | 2764 | 2281 | 3152 | 2133 | 2467 | Six3 |
| Six4 | 1707 | 1230 | 1337 | 1215 | 1019 | 1050 | 883 | 1034 | 959 | 925 | 1216 | 573 | 1014 | 669 | 1389 | 1840 | 1320 | 1140 | 844 | 770 | 826 | 834 | 1060 | 908 | 494 | 1089 | 636 | 673 | Six4 |
| Six5 | 1034 | 1236 | 1432 | 1065 | 791 | 936 | 826 | 1638 | 2150 | 1873 | 2011 | 1894 | 2033 | 1787 | 902 | 1061 | 1373 | 1029 | 1012 | 801 | 810 | 1602 | 2050 | 2005 | 1461 | 2639 | 1994 | 1953 | Six5 |
| Six6 | 288 | 167 | 5 | 169 | 184 | 9 | 103 | 203 | 358 | 111 | 189 | 11 | 8 | 5 | 117 | 127 | 10 | 182 | 32 | 85 | 130 | 495 | 192 | 288 | 189 | 191 | 16 | 6 | Six6 |
| Ska1 | 896 | 704 | 882 | 621 | 512 | 646 | 781 | 1018 | 950 | 892 | 815 | 1289 | 1063 | 1392 | 777 | 682 | 783 | 523 | 746 | 748 | 695 | 648 | 825 | 1195 | 987 | 1645 | 1324 | 1391 | Ska1 |
| Ska2 | 5592 | 4438 | 5654 | 5030 | 4734 | 5069 | 5705 | 8392 | 6522 | 6559 | 6566 | 8896 | 7580 | 9747 | 5652 | 4787 | 5437 | 4912 | 5375 | 5702 | 5290 | 6879 | 6209 | 8612 | 7799 | 10704 | 8986 | 9778 | Ska2 |
| Ska3 | 2601 | 2147 | 2448 | 1969 | 1500 | 1709 | 2075 | 1542 | 1447 | 1383 | 1409 | 2164 | 1995 | 2315 | 2124 | 2266 | 2176 | 1769 | 1824 | 1891 | 1857 | 1185 | 1504 | 1865 | 1657 | 2787 | 2113 | 2339 | Ska3 |
| Skap1 | 113 | 126 | 112 | 135 | 184 | 159 | 201 | 79 | 75 | 95 | 125 | 89 | 104 | 105 | 131 | 121 | 153 | 133 | 147 | 155 | 154 | 99 | 105 | 87 | 107 | 102 | 98 | 140 | Skap1 |
| Skap2 | 677 | 755 | 673 | 578 | 1379 | 667 | 507 | 622 | 826 | 708 | 1137 | 494 | 653 | 478 | 1084 | 936 | 618 | 691 | 496 | 620 | 516 | 581 | 1219 | 498 | 412 | 399 | 453 | 534 | Skap2 |
| Ski | 7996 | 9272 | 9722 | 10093 | 8529 | 8366 | 8119 | 5961 | 7763 | 8034 | 8677 | 8150 | 7978 | 7795 | 7534 | 9382 | 9756 | 10532 | 8865 | 6695 | 8333 | 8221 | 7789 | 8724 | 7918 | 9005 | 7023 | 7603 | Ski |
| Skida1 | 4092 | 3556 | 4544 | 4085 | 3579 | 4145 | 4393 | 4929 | 4194 | 5105 | 4349 | 6339 | 5750 | 6985 | 4138 | 3343 | 4407 | 3668 | 4422 | 3820 | 4245 | 4801 | 4132 | 5687 | 5307 | 6762 | 6737 | 7339 | Skida1 |
| Skil | 8701 | 8429 | 7333 | 8600 | 9056 | 8679 | 9261 | 9591 | 7523 | 7189 | 7751 | 8412 | 8443 | 8691 | 9713 | 8907 | 8362 | 8890 | 8359 | 9641 | 9338 | 7753 | 7603 | 7321 | 8017 | 5977 | 9258 | 8717 | Skil |
| Skint3 | 0 | 0 | 0 | 0 | 0 | 0 | 0 | 0 | 2 | 0 | 1 | 0 | 0 | 0 | 1 | 2 | 0 | 0 | 0 | 0 | 0 | 3 | 0 | 1 | 0 | 0 | 0 | 0 | Skint3 |
| Skint8 | 0 | 1 | 0 | 2 | 0 | 0 | 0 | 0 | 0 | 0 | 0 | 0 | 0 | 0 | 1 | 1 | 0 | 0 | 0 | 0 | 0 | 0 | 2 | 0 | 6 | 0 | 0 | 0 | Skint8 |
| Skiv2l | 4891 | 5995 | 5738 | 5816 | 5692 | 5548 | 5117 | 5378 | 6298 | 6666 | 6343 | 7318 | 6391 | 6604 | 4640 | 5277 | 6178 | 5604 | 6141 | 4993 | 5336 | 6274 | 5754 | 6971 | 6756 | 6824 | 6702 | 6762 | Skiv2l |
| Skor1 | 47 | 147 | 39 | 205 | 651 | 272 | 64 | 532 | 575 | 903 | 907 | 99 | 837 | 66 | 165 | 179 | 263 | 202 | 148 | 177 | 69 | 775 | 838 | 99 | 421 | 90 | 46 | 64 | Skor1 |
| Skor2 | 29 | 143 | 10 | 354 | 524 | 289 | 37 | 287 | 349 | 484 | 436 | 34 | 408 | 29 | 146 | 188 | 219 | 305 | 143 | 210 | 29 | 645 | 399 | 22 | 417 | 14 | 21 | 12 | Skor2 |
| Skp1 | 33428 | 26977 | 32898 | 31479 | 28986 | 28900 | 36904 | 32282 | 26169 | 26800 | 25882 | 29335 | 27533 | 32082 | 35256 | 29914 | 31219 | 30897 | 30440 | 35986 | 33255 | 26532 | 26761 | 29836 | 28235 | 33698 | 29849 | 33367 | Skp1 |
| Skp2 | 0 | 0 | 0 | 0 | 1 | 1 | 1 | 0 | 1 | 0 | 0 | 1 | 0 | 1 | 1 | 0 | 0 | 3 | 2 | 0 | 0 | 0 | 0 | 1 | 0 | 0 | 0 | 0 | Skp2 |
| Sla | 50 | 30 | 60 | 79 | 183 | 141 | 148 | 259 | 355 | 319 | 235 | 137 | 179 | 154 | 40 | 65 | 44 | 118 | 140 | 101 | 201 | 321 | 368 | 281 | 463 | 186 | 173 | 155 | Sla |
| Sla2 | 6 | 20 | 2 | 17 | 10 | 19 | 1 | 5 | 9 | 6 | 8 | 1 | 15 | 7 | 4 | 16 | 8 | 14 | 16 | 4 | 14 | 8 | 10 | 8 | 0 | 22 | 8 | 7 | Sla2 |
| Slain1 | 2023 | 2426 | 2838 | 3466 | 3849 | 3120 | 3957 | 2251 | 2057 | 2470 | 2143 | 3036 | 1802 | 3115 | 2217 | 1875 | 3323 | 3436 | 4024 | 2867 | 4054 | 3056 | 1992 | 3027 | 3285 | 2755 | 2249 | 2693 | Slain1 |
| Slain2 | 4985 | 4158 | 4878 | 4785 | 5571 | 5344 | 5589 | 5908 | 5142 | 5429 | 5260 | 5727 | 5432 | 5793 | 4812 | 4102 | 4804 | 5039 | 5562 | 5122 | 5661 | 6154 | 5334 | 5495 | 5931 | 5858 | 5656 | 5815 | Slain2 |
| Slamf1 | 7 | 5 | 6 | 6 | 10 | 15 | 13 | 16 | 9 | 10 | 11 | 9 | 16 | 11 | 11 | 7 | 8 | 7 | 4 | 10 | 8 | 12 | 14 | 20 | 5 | 24 | 8 | 18 | Slamf1 |
| Slamf6 | 1 | 0 | 1 | 0 | 2 | 3 | 0 | 7 | 1 | 0 | 7 | 4 | 5 | 1 | 1 | 2 | 0 | 0 | 0 | 5 | 3 | 0 | 3 | 2 | 0 | 3 | 9 | 1 | Slamf6 |
| Slamf7 | 0 | 0 | 0 | 0 | 0 | 2 | 0 | 2 | 4 | 0 | 3 | 1 | 0 | 0 | 0 | 0 | 0 | 0 | 0 | 0 | 0 | 0 | 8 | 6 | 3 | 4 | 0 | 1 | Slamf7 |
| Slamf8 | 3 | 5 | 3 | 3 | 2 | 2 | 0 | 0 | 2 | 3 | 1 | 0 | 1 | 1 | 0 | 2 | 2 | 1 | 1 | 2 | 1 | 0 | 1 | 0 | 0 | 0 | 2 | 0 | Slamf8 |
| Slamf9 | 9 | 6 | 11 | 1 | 10 | 4 | 4 | 7 | 12 | 12 | 5 | 3 | 10 | 7 | 13 | 9 | 10 | 0 | 6 | 6 | 0 | 7 | 13 | 7 | 3 | 10 | 6 | 11 | Slamf9 |
| Slbp | 9374 | 8102 | 9365 | 8388 | 6271 | 7183 | 7970 | 6333 | 5600 | 4823 | 5099 | 6853 | 6297 | 6709 | 8968 | 8495 | 8471 | 7734 | 7444 | 9104 | 6921 | 4744 | 5594 | 6032 | 5374 | 8762 | 7377 | 7544 | Slbp |
| Slc10a1 | 0 | 0 | 0 | 0 | 0 | 0 | 0 | 2 | 0 | 0 | 0 | 0 | 1 | 2 | 0 | 0 | 0 | 2 | 0 | 0 | 0 | 0 | 0 | 0 | 0 | 0 | 0 | 0 | Slc10a1 |
| Slc10a2 | 0 | 0 | 0 | 0 | 0 | 0 | 0 | 0 | 0 | 0 | 2 | 0 | 0 | 0 | 0 | 0 | 0 | 0 | 0 | 0 | 0 | 0 | 0 | 0 | 0 | 0 | 0 | 0 | Slc10a2 |
| Slc10a3 | 308 | 285 | 208 | 187 | 187 | 235 | 204 | 267 | 272 | 233 | 229 | 241 | 257 | 225 | 320 | 298 | 233 | 264 | 190 | 226 | 173 | 251 | 337 | 226 | 165 | 250 | 257 | 247 | Slc10a3 |
| Slc10a4 | 208 | 243 | 241 | 346 | 548 | 347 | 355 | 488 | 587 | 660 | 508 | 611 | 534 | 606 | 232 | 220 | 404 | 366 | 428 | 336 | 387 | 805 | 396 | 568 | 627 | 537 | 504 | 500 | Slc10a4 |

| | | | | | | | | | | | | | | | | | | | | | | | | | | | | | |
|---|---|---|---|---|---|---|---|---|---|---|---|---|---|---|---|---|---|---|---|---|---|---|---|---|---|---|---|---|---|
| Slc10a4-ps | 74 | 71 | 129 | 92 | 104 | 70 | 92 | 106 | 72 | 66 | 102 | 99 | 90 | 74 | 87 | 86 | 97 | 76 | 79 | 101 | 80 | 135 | 72 | 84 | 68 | 95 | 127 | 91 | Slc10a4-ps |
| Slc10a5 | 1 | 9 | 0 | 7 | 4 | 10 | 4 | 10 | 2 | 3 | 1 | 0 | 0 | 3 | 8 | 6 | 6 | 8 | 1 | 4 | 3 | 0 | 6 | 5 | 0 | 0 | 3 | 0 | Slc10a5 |
| Slc10a6 | 6 | 3 | 0 | 1 | 7 | 5 | 3 | 7 | 9 | 14 | 20 | 3 | 8 | 2 | 16 | 9 | 2 | 12 | 5 | 6 | 3 | 12 | 21 | 1 | 3 | 7 | 6 | 1 | Slc10a6 |
| Slc10a7 | 2145 | 1896 | 1939 | 1792 | 1685 | 1674 | 1507 | 1688 | 1740 | 1510 | 1840 | 1377 | 1564 | 1497 | 2077 | 2051 | 1802 | 1843 | 1585 | 1734 | 1483 | 1542 | 1726 | 1638 | 1494 | 1491 | 1597 | 1475 | Slc10a7 |
| Slc11a1 | 63 | 97 | 46 | 66 | 92 | 66 | 66 | 81 | 110 | 74 | 111 | 85 | 64 | 69 | 66 | 90 | 93 | 91 | 44 | 101 | 63 | 100 | 151 | 89 | 96 | 88 | 113 | 82 | Slc11a1 |
| Slc11a2 | 4805 | 6025 | 4557 | 4944 | 5044 | 5071 | 4671 | 2950 | 3063 | 2901 | 2959 | 3104 | 3269 | 2938 | 5021 | 5462 | 4717 | 4845 | 4763 | 4958 | 4975 | 3014 | 2708 | 2799 | 3224 | 2812 | 3205 | 2890 | Slc11a2 |
| Slc12a1 | 5 | 4 | 1 | 1 | 18 | 7 | 6 | 6 | 16 | 16 | 12 | 3 | 8 | 1 | 5 | 3 | 1 | 4 | 4 | 4 | 10 | 7 | 9 | 3 | 11 | 1 | 8 | 2 | Slc12a1 |
| Slc12a2 | 11557 | 8524 | 8296 | 8955 | 8500 | 9684 | 8948 | 9297 | 9245 | 7680 | 8783 | 8525 | 9069 | 8520 | 10925 | 11009 | 7946 | 9575 | 7733 | 8448 | 8896 | 8199 | 8862 | 7817 | 7435 | 7558 | 8498 | 7937 | Slc12a2 |
| Slc12a3 | 115 | 193 | 75 | 208 | 174 | 161 | 152 | 31 | 28 | 34 | 36 | 26 | 36 | 20 | 187 | 206 | 170 | 177 | 140 | 117 | 160 | 57 | 20 | 16 | 24 | 17 | 20 | 20 | Slc12a3 |
| Slc12a4 | 1661 | 1542 | 1415 | 1135 | 1013 | 1144 | 763 | 1376 | 2097 | 1865 | 2486 | 1372 | 1579 | 1041 | 1185 | 1981 | 1217 | 1330 | 990 | 643 | 761 | 1432 | 2235 | 1615 | 1159 | 1467 | 1226 | 1182 | Slc12a4 |
| Slc12a5 | 1393 | 2075 | 2259 | 3925 | 5960 | 4277 | 4242 | 2998 | 3040 | 4516 | 3572 | 2628 | 2584 | 2683 | 1731 | 1688 | 3406 | 4725 | 5260 | 3777 | 4837 | 5650 | 2703 | 3604 | 4794 | 1594 | 2277 | 2099 | Slc12a5 |
| Slc12a6 | 3593 | 3207 | 3174 | 3717 | 4115 | 3943 | 4580 | 3559 | 3261 | 3520 | 3592 | 3338 | 3590 | 3750 | 3659 | 3343 | 3415 | 3970 | 3634 | 3807 | 4552 | 4231 | 3219 | 3326 | 3803 | 3127 | 3141 | 3475 | Slc12a6 |
| Slc12a7 | 7636 | 8082 | 7037 | 6646 | 6513 | 6487 | 5761 | 4284 | 4879 | 4690 | 4309 | 6402 | 6179 | 5756 | 6672 | 8273 | 6769 | 6091 | 6694 | 5665 | 5909 | 4050 | 4104 | 4814 | 4623 | 5720 | 6316 | 5658 | Slc12a7 |
| Slc12a8 | 66 | 52 | 53 | 43 | 43 | 37 | 58 | 57 | 68 | 61 | 48 | 54 | 71 | 63 | 34 | 47 | 57 | 76 | 48 | 49 | 44 | 37 | 65 | 58 | 39 | 89 | 89 | 69 | Slc12a8 |
| Slc12a9 | 2555 | 3770 | 3677 | 4320 | 3744 | 3428 | 3023 | 1492 | 1859 | 1949 | 1968 | 1956 | 2056 | 1735 | 2623 | 3422 | 4035 | 4043 | 3963 | 2989 | 3128 | 2151 | 1689 | 2025 | 2143 | 1785 | 2007 | 1626 | Slc12a9 |
| Slc13a1 | 0 | 0 | 0 | 0 | 2 | 0 | 0 | 2 | 3 | 0 | 1 | 1 | 0 | 0 | 1 | 0 | 0 | 0 | 1 | 0 | 2 | 2 | 1 | 0 | 0 | 0 | 0 | 0 | Slc13a1 |
| Slc13a2 | 1 | 0 | 0 | 0 | 0 | 0 | 0 | 0 | 1 | 0 | 0 | 0 | 1 | 0 | 1 | 0 | 0 | 3 | 0 | 1 | 0 | 5 | 1 | 1 | 0 | 1 | 1 | 4 | Slc13a2 |
| Slc13a3 | 59 | 97 | 60 | 90 | 159 | 100 | 106 | 118 | 131 | 144 | 133 | 112 | 67 | 121 | 59 | 65 | 107 | 120 | 129 | 110 | 108 | 132 | 159 | 133 | 143 | 119 | 121 | 119 | Slc13a3 |
| Slc13a4 | 20 | 71 | 51 | 54 | 204 | 112 | 69 | 103 | 171 | 106 | 226 | 64 | 78 | 20 | 58 | 65 | 51 | 123 | 72 | 119 | 120 | 184 | 290 | 83 | 144 | 27 | 21 | 37 | Slc13a4 |
| Slc13a5 | 135 | 156 | 168 | 108 | 151 | 204 | 152 | 56 | 80 | 89 | 105 | 62 | 75 | 29 | 142 | 164 | 149 | 131 | 155 | 182 | 135 | 105 | 69 | 62 | 96 | 24 | 48 | 39 | Slc13a5 |
| Slc14a1 | 12 | 7 | 8 | 3 | 11 | 8 | 4 | 4 | 10 | 1 | 18 | 5 | 18 | 13 | 21 | 13 | 9 | 8 | 7 | 14 | 13 | 3 | 78 | 12 | 6 | 19 | 20 | 11 | Slc14a1 |
| Slc14a2 | 66 | 85 | 44 | 83 | 90 | 78 | 95 | 123 | 161 | 135 | 110 | 164 | 129 | 161 | 55 | 63 | 49 | 97 | 91 | 106 | 99 | 132 | 179 | 186 | 187 | 180 | 121 | 123 | Slc14a2 |
| Slc15a1 | 3 | 5 | 4 | 4 | 2 | 2 | 1 | 5 | 4 | 1 | 5 | 5 | 5 | 2 | 5 | 2 | 0 | 0 | 4 | 4 | 1 | 10 | 5 | 0 | 5 | 8 | 1 | 4 | Slc15a1 |
| Slc15a2 | 1037 | 1679 | 1092 | 713 | 1359 | 2569 | 2954 | 2794 | 784 | 1798 | 2091 | 807 | 1812 | 1044 | 849 | 1467 | 1873 | 438 | 388 | 1622 | 1459 | 1103 | 1139 | 1800 | 1154 | 1214 | 178 | 695 | Slc15a2 |
| Slc15a3 | 50 | 34 | 41 | 47 | 30 | 37 | 32 | 64 | 78 | 93 | 105 | 67 | 77 | 77 | 46 | 58 | 29 | 36 | 40 | 37 | 38 | 48 | 124 | 70 | 63 | 68 | 58 | 47 | Slc15a3 |
| Slc15a4 | 1444 | 2050 | 1783 | 1801 | 1654 | 1557 | 1519 | 1470 | 1291 | 1193 | 1239 | 1322 | 1313 | 1192 | 1410 | 1769 | 1833 | 1753 | 1771 | 1718 | 1660 | 1337 | 1233 | 1277 | 1434 | 1199 | 1325 | 1267 | Slc15a4 |
| Slc15a5 | 1 | 1 | 0 | 0 | 0 | 0 | 0 | 6 | 2 | 6 | 0 | 0 | 1 | 0 | 4 | 2 | 0 | 0 | 1 | 0 | 0 | 2 | 3 | 1 | 0 | 4 | 0 | 2 | Slc15a5 |
| Slc16a1 | 4866 | 4367 | 4627 | 3552 | 2408 | 3040 | 2911 | 2209 | 2656 | 1949 | 2165 | 2558 | 2808 | 2456 | 4457 | 4812 | 3576 | 3452 | 2652 | 3266 | 2601 | 1686 | 2558 | 2451 | 1924 | 3297 | 2721 | 2784 | Slc16a1 |
| Slc16a10 | 1208 | 1063 | 1146 | 965 | 1249 | 1006 | 893 | 686 | 1026 | 892 | 795 | 953 | 1284 | 581 | 1200 | 1524 | 941 | 1077 | 922 | 882 | 1041 | 917 | 1407 | 510 | 529 | 609 | 535 | 726 | Slc16a10 |
| Slc16a11 | 29 | 16 | 16 | 16 | 25 | 15 | 12 | 18 | 33 | 21 | 22 | 16 | 8 | 7 | 24 | 33 | 19 | 24 | 17 | 18 | 14 | 35 | 43 | 38 | 18 | 18 | 10 | 5 | Slc16a11 |
| Slc16a12 | 83 | 95 | 105 | 81 | 160 | 135 | 105 | 291 | 280 | 270 | 290 | 343 | 356 | 286 | 136 | 150 | 78 | 118 | 107 | 116 | 113 | 304 | 419 | 274 | 248 | 255 | 274 | 246 | Slc16a12 |
| Slc16a13 | 807 | 983 | 1044 | 878 | 919 | 918 | 837 | 927 | 1011 | 889 | 907 | 1139 | 1212 | 1117 | 938 | 822 | 963 | 828 | 934 | 898 | 766 | 994 | 971 | 1072 | 1004 | 1143 | 1191 | 1186 | Slc16a13 |
| Slc16a14 | 313 | 301 | 343 | 364 | 512 | 566 | 539 | 944 | 898 | 987 | 791 | 1093 | 1016 | 1155 | 364 | 227 | 298 | 340 | 522 | 457 | 466 | 1044 | 721 | 895 | 1107 | 512 | 1204 | 1057 | Slc16a14 |
| Slc16a2 | 3267 | 3155 | 3455 | 3590 | 4981 | 4177 | 4302 | 2906 | 3564 | 3576 | 3841 | 3211 | 3130 | 2949 | 3220 | 3457 | 3603 | 4006 | 4550 | 3761 | 4543 | 4032 | 4442 | 3201 | 3757 | 2898 | 2682 | 2995 | Slc16a2 |
| Slc16a3 | 2638 | 2853 | 2888 | 1847 | 1289 | 1735 | 1095 | 729 | 1065 | 961 | 1261 | 806 | 1041 | 873 | 2298 | 2896 | 2575 | 1864 | 1472 | 1464 | 1140 | 662 | 1211 | 1016 | 776 | 1251 | 997 | 1015 | Slc16a3 |
| Slc16a4 | 62 | 56 | 65 | 95 | 82 | 109 | 82 | 151 | 246 | 164 | 382 | 126 | 172 | 106 | 99 | 85 | 58 | 117 | 65 | 97 | 64 | 224 | 382 | 104 | 150 | 110 | 114 | 94 | Slc16a4 |
| Slc16a5 | 1 | 1 | 0 | 3 | 0 | 0 | 2 | 5 | 2 | 14 | 6 | 0 | 1 | 0 | 0 | 0 | 0 | 0 | 0 | 0 | 0 | 3 | 1 | 1 | 0 | 19 | 0 | 0 | Slc16a5 |
| Slc16a6 | 907 | 867 | 1078 | 1011 | 854 | 901 | 813 | 689 | 740 | 823 | 878 | 947 | 1022 | 909 | 935 | 872 | 956 | 965 | 733 | 873 | 702 | 657 | 885 | 852 | 844 | 913 | 900 | 907 | Slc16a6 |
| Slc16a7 | 1043 | 921 | 888 | 1013 | 1150 | 1214 | 1116 | 1274 | 1242 | 1032 | 1311 | 723 | 1060 | 815 | 1224 | 1113 | 793 | 1034 | 964 | 1308 | 1090 | 1237 | 1310 | 716 | 980 | 565 | 779 | 781 | Slc16a7 |
| Slc16a8 | 0 | 4 | 3 | 7 | 5 | 6 | 0 | 6 | 7 | 1 | 6 | 1 | 3 | 2 | 1 | 2 | 3 | 0 | 3 | 3 | 3 | 8 | 15 | 3 | 1 | 0 | 0 | 0 | Slc16a8 |
| Slc16a9 | 131 | 133 | 199 | 155 | 189 | 168 | 115 | 285 | 308 | 209 | 412 | 143 | 276 | 212 | 185 | 168 | 160 | 185 | 93 | 106 | 118 | 463 | 440 | 267 | 343 | 145 | 200 | 189 | Slc16a9 |
| Slc17a1 | 0 | 0 | 26 | 8 | 1 | 2 | 4 | 4 | 2 | 0 | 3 | 1 | 6 | 6 | 1 | 0 | 8 | 6 | 5 | 6 | 1 | 5 | 4 | 11 | 8 | 0 | 0 | 17 | Slc17a1 |
| Slc17a2 | 3 | 1 | 3 | 6 | 0 | 1 | 6 | 0 | 2 | 1 | 0 | 0 | 2 | 0 | 8 | 5 | 0 | 2 | 0 | 2 | 2 | 0 | 1 | 0 | 2 | 0 | 0 | 0 | Slc17a2 |
| Slc17a3 | 1 | 0 | 0 | 0 | 0 | 0 | 0 | 0 | 0 | 0 | 0 | 0 | 0 | 2 | 0 | 3 | 0 | 0 | 0 | 0 | 2 | 0 | 2 | 0 | 1 | 2 | 0 | 0 | Slc17a3 |
| Slc17a4 | 0 | 0 | 0 | 0 | 1 | 0 | 0 | 0 | 0 | 0 | 2 | 0 | 0 | 0 | 0 | 1 | 0 | 0 | 0 | 0 | 0 | 0 | 0 | 0 | 0 | 0 | 0 | 2 | Slc17a4 |
| Slc17a5 | 1527 | 1458 | 1318 | 1628 | 1652 | 1749 | 1571 | 1202 | 1243 | 1278 | 1313 | 1170 | 1193 | 1130 | 1495 | 1505 | 1277 | 1618 | 1559 | 1595 | 1582 | 1203 | 1339 | 1218 | 1309 | 1097 | 1169 | 1065 | Slc17a5 |
| Slc17a6 | 2249 | 1989 | 2727 | 2757 | 5107 | 4310 | 4605 | 9835 | 6814 | 8282 | 7106 | 8660 | 7571 | 9015 | 3175 | 1674 | 3271 | 2838 | 4886 | 4532 | 4292 | 10418 | 7723 | 7951 | 9448 | 6210 | 8725 | 9322 | Slc17a6 |
| Slc17a7 | 656 | 791 | 614 | 1063 | 1226 | 1129 | 1428 | 595 | 829 | 669 | 500 | 686 | 371 | 587 | 609 | 614 | 824 | 1412 | 1565 | 1235 | 1534 | 446 | 546 | 1054 | 1108 | 729 | 440 | 464 | Slc17a7 |
| Slc17a8 | 181 | 244 | 363 | 297 | 430 | 381 | 283 | 684 | 406 | 482 | 433 | 388 | 972 | 391 | 390 | 313 | 351 | 303 | 276 | 308 | 266 | 719 | 423 | 265 | 531 | 292 | 493 | 444 | Slc17a8 |
| Slc17a9 | 290 | 389 | 319 | 562 | 295 | 328 | 409 | 51 | 45 | 45 | 76 | 66 | 76 | 83 | 268 | 371 | 431 | 544 | 408 | 309 | 410 | 45 | 45 | 96 | 38 | 44 | 70 | 68 | Slc17a9 |
| Slc18a1 | 22 | 10 | 15 | 11 | 22 | 18 | 13 | 32 | 25 | 20 | 23 | 33 | 28 | 9 | 5 | 16 | 24 | 9 | 21 | 35 | 13 | 10 | 49 | 18 | 23 | 18 | 21 | 19 | Slc18a1 |
| Slc18a2 | 2180 | 2257 | 2100 | 2751 | 2960 | 3091 | 4255 | 2640 | 2190 | 2492 | 2007 | 2757 | 1312 | 2761 | 2686 | 2055 | 2947 | 2895 | 3467 | 2784 | 3702 | 3070 | 1643 | 2924 | 3478 | 2514 | 2194 | 1843 | Slc18a2 |
| Slc18a3 | 60 | 55 | 59 | 83 | 176 | 58 | 47 | 54 | 96 | 98 | 102 | 20 | 104 | 22 | 103 | 78 | 92 | 70 | 80 | 44 | 55 | 95 | 87 | 30 | 34 | 11 | 15 | 12 | Slc18a3 |
| Slc18b1 | 1359 | 1056 | 1482 | 1090 | 1336 | 1371 | 993 | 2142 | 1889 | 1965 | 1847 | 1626 | 1754 | 1642 | 1300 | 1363 | 1162 | 1175 | 1345 | 1058 | 1008 | 2032 | 1904 | 1918 | 1783 | 1898 | 1542 | 1641 | Slc18b1 |
| Slc19a1 | 1069 | 1210 | 1174 | 1162 | 1239 | 1289 | 1074 | 1082 | 1410 | 1326 | 1314 | 1449 | 1351 | 1330 | 1034 | 1196 | 1151 | 1170 | 1262 | 1110 | 1156 | 1290 | 1399 | 1229 | 1420 | 1281 | 1375 | 1271 | Slc19a1 |
| Slc19a2 | 2301 | 2314 | 2084 | 2210 | 1685 | 1867 | 1960 | 939 | 915 | 856 | 964 | 1112 | 933 | 872 | 2176 | 2347 | 2130 | 2309 | 1883 | 1787 | 1885 | 917 | 961 | 847 | 877 | 1078 | 1038 | 941 | Slc19a2 |
| Slc19a3 | 37 | 29 | 9 | 15 | 20 | 43 | 23 | 45 | 62 | 34 | 54 | 41 | 32 | 32 | 28 | 28 | 22 | 31 | 27 | 48 | 21 | 52 | 43 | 38 | 47 | 27 | 30 | 28 | Slc19a3 |
| Slc1a1 | 743 | 782 | 615 | 995 | 1567 | 1229 | 1425 | 929 | 900 | 909 | 826 | 465 | 452 | 497 | 765 | 745 | 716 | 1263 | 1357 | 1317 | 1576 | 1041 | 827 | 708 | 1093 | 300 | 295 | 282 | Slc1a1 |
| Slc1a2 | 5358 | 5510 | 6571 | 9039 | 12712 | 10892 | 11645 | 12490 | 9314 | 12839 | 11176 | 11961 | 9408 | 12355 | 7378 | 4639 | 8516 | 9496 | 11010 | 10399 | 12141 | 15530 | 11157 | 11146 | 13846 | 7772 | 13235 | 12651 | Slc1a2 |
| Slc1a3 | 5772 | 5867 | 4754 | 6561 | 10610 | 8381 | 8425 | 10253 | 11131 | 9845 | 9597 | 7023 | 8165 | 7126 | 7536 | 6030 | 5767 | 7943 | 8037 | 8371 | 8824 | 10601 | 11204 | 7069 | 8968 | 6021 | 6122 | 6005 | Slc1a3 |
| Slc1a4 | 4992 | 5510 | 5258 | 5998 | 6424 | 6191 | 6098 | 3899 | 4011 | 4448 | 4267 | 3985 | 4417 | 4401 | 5058 | 5002 | 5394 | 6259 | 6385 | 5904 | 6201 | 4639 | 4232 | 4372 | 5131 | 4375 | 3808 | 4405 | Slc1a4 |
| Slc1a5 | 5659 | 6690 | 7110 | 5881 | 4233 | 4613 | 4903 | 1462 | 1518 | 1628 | 1688 | 2166 | 2159 | 2390 | 5172 | 6083 | 6258 | 5800 | 5160 | 4746 | 4614 | 1260 | 1520 | 1991 | 1601 | 2550 | 2546 | 2554 | Slc1a5 |
| Slc1a6 | 374 | 370 | 357 | 271 | 417 | 447 | 297 | 198 | 244 | 277 | 208 | 163 | 175 | 122 | 315 | 317 | 331 | 406 | 365 | 302 | 290 | 261 | 282 | 184 | 180 | 154 | 151 | 150 | Slc1a6 |
| Slc1a7 | 1 | 6 | 0 | 0 | 2 | 2 | 0 | 6 | 3 | 5 | 1 | 0 | 1 | 2 | 0 | 3 | 0 | 0 | 2 | 4 | 0 | 2 | 5 | 7 | 8 | 2 | 5 | 1 | Slc1a7 |
| Slc20a1 | 5260 | 4842 | 4634 | 4303 | 5133 | 5209 | 5372 | 10009 | 8884 | 7991 | 7672 | 9598 | 9624 | 9496 | 5486 | 4294 | 4614 | 4141 | 5140 | 5766 | 5097 | 8976 | 8177 | 8155 | 9265 | 8516 | 9717 | 9881 | Slc20a1 |
| Slc20a2 | 1390 | 1296 | 1270 | 1324 | 1204 | 1280 | 1036 | 1922 | 2342 | 1968 | 2358 | 2408 | 2375 | 2134 | 1299 | 1396 | 1137 | 1184 | 1128 | 1148 | 1101 | 1788 | 2504 | 2138 | 1971 | 2687 | 2349 | 2194 | Slc20a2 |
| Slc22a1 | 7 | 16 | 15 | 12 | 24 | 9 | 13 | 6 | 1 | 5 | 12 | 0 | 9 | 0 | 7 | 6 | 8 | 7 | 12 | 9 | 12 | 8 | 6 | 0 | 8 | 1 | 2 | 1 | Slc22a1 |
| Slc22a13 | 8 | 5 | 2 | 15 | 11 | 8 | 13 | 1 | 1 | 3 | 2 | 2 | 0 | 0 | 7 | 9 | 6 | 20 | 16 | 6 | 8 | 2 | 1 | 1 | 2 | 0 | 2 | 0 | Slc22a13 |
| Slc22a14 | 13 | 13 | 8 | 14 | 18 | 11 | 16 | 5 | 5 | 3 | 5 | 3 | 3 | 3 | 11 | 11 | 31 | 22 | 11 | 8 | 13 | 0 | 6 | 2 | 7 | 0 | 2 | 4 | Slc22a14 |
| Slc22a15 | 639 | 631 | 634 | 707 | 899 | 796 | 708 | 1446 | 1371 | 1514 | 1480 | 1580 | 1826 | 1545 | 723 | 746 | 759 | 687 | 789 | 739 | 760 | 1537 | 1468 | 1283 | 1495 | 1152 | 1463 | 1494 | Slc22a15 |
| Slc22a16 | 0 | 0 | 0 | 0 | 0 | 0 | 1 | 0 | 0 | 0 | 0 | 0 | 0 | 0 | 0 | 0 | 0 | 0 | 0 | 0 | 0 | 0 | 0 | 0 | 0 | 0 | 0 | 0 | Slc22a16 |
| Slc22a17 | 7251 | 8961 | 8639 | 10615 | 11783 | 9631 | 10429 | 8930 | 9876 | 10686 | 10562 | 9736 | 9243 | 9126 | 7007 | 7595 | 9590 | 11221 | 11847 | 9837 | 11093 | 11575 | 10095 | 11161 | 11346 | 9596 | 8867 | 8577 | Slc22a17 |
| Slc22a18 | 42 | 26 | 19 | 26 | 39 | 28 | 10 | 53 | 56 | 27 | 57 | 25 | 35 | 15 | 18 | 40 | 13 | 18 | 15 | 8 | 14 | 17 | 60 | 49 | 25 | 24 | 20 | 18 | Slc22a18 |
| Slc22a19 | 3 | 1 | 0 | 0 | 7 | 5 | 1 | 10 | 8 | 9 | 0 | 7 | 8 | 5 | 1 | 5 | 0 | 2 | 2 | 0 | 1 | 5 | 11 | 2 | 1 | 0 | 2 | 2 | Slc22a19 |
| Slc22a2 | 1 | 3 | 0 | 3 | 11 | 4 | 8 | 23 | 30 | 16 | 35 | 5 | 9 | 5 | 5 | 0 | 1 | 4 | 12 | 20 | 21 | 47 | 50 | 21 | 26 | 6 | 8 | 5 | Slc22a2 |
| Slc22a20 | 0 | 0 | 0 | 0 | 0 | 0 | 0 | 0 | 0 | 0 | 0 | 0 | 0 | 0 | 0 | 1 | 0 | 0 | 0 | 0 | 0 | 0 | 0 | 0 | 0 | 0 | 0 | 0 | Slc22a20 |
| Slc22a21 | 736 | 594 | 628 | 546 | 615 | 677 | 611 | 245 | 310 | 278 | 319 | 324 | 327 | 239 | 723 | 741 | 650 | 574 | 641 | 507 | 520 | 326 | 359 | 185 | 173 | 284 | 246 | 235 | Slc22a21 |
| Slc22a22 | 2 | 3 | 0 | 7 | 1 | 0 | 0 | 4 | 4 | 0 | 1 | 2 | 6 | 5 | 3 | 1 | 0 | 2 | 4 | 1 | 6 | 2 | 3 | 0 | 0 | 1 | 0 | 0 | Slc22a22 |
| Slc22a23 | 5143 | 5928 | 6214 | 6852 | 7745 | 7438 | 7096 | 5417 | 5286 | 6257 | 5297 | 7310 | 5883 | 7066 | 5649 | 4631 | 6696 | 6661 | 7670 | 6678 | 7517 | 6654 | 5458 | 6521 | 6618 | 6264 | 6829 | 7225 | Slc22a23 |

| | | | | | | | | | | | | | | | | | | | | | | | | | | | | | |
|---|---|---|---|---|---|---|---|---|---|---|---|---|---|---|---|---|---|---|---|---|---|---|---|---|---|---|---|---|---|
| Slc22a27 | 0 | 0 | 0 | 0 | 0 | 0 | 0 | 0 | 0 | 0 | 0 | 0 | 0 | 0 | 1 | 0 | 0 | 0 | 0 | 0 | 0 | 0 | 0 | 1 | 0 | 0 | 0 | 0 | Slc22a27 |
| Slc22a28 | 0 | 0 | 0 | 0 | 0 | 2 | 0 | 1 | 0 | 0 | 0 | 0 | 1 | 0 | 0 | 0 | 0 | 0 | 0 | 0 | 0 | 0 | 0 | 2 | 0 | 1 | 3 | 1 | Slc22a28 |
| Slc22a29 | 6 | 3 | 4 | 1 | 0 | 2 | 2 | 0 | 5 | 1 | 6 | 2 | 9 | 2 | 3 | 3 | 1 | 8 | 4 | 2 | 4 | 8 | 8 | 2 | 5 | 4 | 0 | 1 | Slc22a29 |
| Slc22a3 | 14 | 24 | 27 | 13 | 24 | 24 | 9 | 82 | 93 | 83 | 93 | 46 | 101 | 30 | 25 | 30 | 36 | 31 | 24 | 40 | 31 | 65 | 117 | 46 | 54 | 24 | 50 | 14 | Slc22a3 |
| Slc22a30 | 0 | 1 | 5 | 0 | 0 | 0 | 1 | 0 | 1 | 0 | 3 | 3 | 3 | 2 | 1 | 3 | 0 | 0 | 1 | 1 | 0 | 2 | 2 | 3 | 2 | 5 | 1 | 2 | Slc22a30 |
| Slc22a4 | 72 | 69 | 62 | 53 | 65 | 46 | 38 | 84 | 90 | 78 | 74 | 79 | 178 | 136 | 74 | 63 | 52 | 51 | 25 | 49 | 35 | 62 | 107 | 58 | 59 | 67 | 85 | 123 | Slc22a4 |
| Slc22a5 | 1390 | 1485 | 1336 | 1242 | 1342 | 1468 | 1226 | 977 | 1058 | 914 | 1064 | 985 | 1034 | 822 | 1294 | 1629 | 1232 | 1350 | 1265 | 1253 | 1202 | 1056 | 1045 | 891 | 887 | 907 | 897 | 873 | Slc22a5 |
| Slc22a6 | 134 | 138 | 78 | 240 | 415 | 331 | 247 | 501 | 624 | 561 | 540 | 354 | 382 | 326 | 111 | 114 | 149 | 275 | 307 | 400 | 352 | 770 | 809 | 479 | 623 | 231 | 432 | 243 | Slc22a6 |
| Slc22a7 | 17 | 23 | 14 | 36 | 15 | 31 | 26 | 5 | 5 | 9 | 0 | 7 | 5 | 12 | 33 | 8 | 27 | 23 | 33 | 19 | 43 | 12 | 7 | 5 | 9 | 7 | 14 | 7 | Slc22a7 |
| Slc22a8 | 18 | 37 | 56 | 66 | 142 | 94 | 112 | 143 | 161 | 203 | 202 | 138 | 150 | 121 | 36 | 24 | 43 | 80 | 123 | 130 | 138 | 322 | 221 | 192 | 217 | 84 | 128 | 83 | Slc22a8 |
| Slc23a1 | 1 | 1 | 3 | 7 | 0 | 2 | 3 | 1 | 4 | 1 | 2 | 0 | 1 | 1 | 3 | 5 | 1 | 1 | 0 | 4 | 0 | 5 | 1 | 1 | 0 | 0 | 0 | 0 | Slc23a1 |
| Slc23a2 | 4439 | 4250 | 3359 | 4184 | 5446 | 4967 | 4819 | 4404 | 4762 | 4059 | 4280 | 3686 | 3811 | 3695 | 4722 | 4169 | 3847 | 4664 | 4718 | 5106 | 4953 | 4903 | 4766 | 3675 | 4677 | 2775 | 3514 | 3266 | Slc23a2 |
| Slc23a3 | 15 | 18 | 5 | 36 | 8 | 19 | 9 | 11 | 16 | 6 | 11 | 9 | 12 | 10 | 12 | 23 | 21 | 32 | 14 | 20 | 13 | 5 | 17 | 7 | 8 | 8 | 9 | 14 | Slc23a3 |
| Slc23a4 | 144 | 138 | 84 | 163 | 177 | 174 | 121 | 96 | 90 | 113 | 117 | 97 | 99 | 121 | 108 | 166 | 104 | 166 | 158 | 161 | 162 | 82 | 99 | 86 | 103 | 85 | 100 | 87 | Slc23a4 |
| Slc24a1 | 2 | 6 | 7 | 12 | 11 | 16 | 6 | 22 | 16 | 25 | 21 | 11 | 23 | 12 | 5 | 6 | 10 | 4 | 12 | 5 | 10 | 22 | 27 | 20 | 15 | 18 | 3 | 4 | Slc24a1 |
| Slc24a2 | 245 | 181 | 146 | 293 | 458 | 465 | 371 | 556 | 466 | 442 | 458 | 319 | 335 | 357 | 261 | 231 | 168 | 303 | 327 | 483 | 421 | 373 | 478 | 373 | 467 | 212 | 385 | 309 | Slc24a2 |
| Slc24a3 | 292 | 365 | 338 | 617 | 1127 | 772 | 797 | 920 | 923 | 1121 | 1365 | 741 | 759 | 846 | 371 | 363 | 450 | 760 | 796 | 709 | 861 | 1516 | 1159 | 960 | 1197 | 640 | 805 | 750 | Slc24a3 |
| Slc24a4 | 24 | 19 | 20 | 41 | 98 | 74 | 51 | 117 | 118 | 126 | 127 | 74 | 105 | 55 | 22 | 36 | 21 | 48 | 65 | 89 | 59 | 110 | 167 | 93 | 127 | 32 | 45 | 47 | Slc24a4 |
| Slc24a5 | 37 | 31 | 61 | 38 | 34 | 25 | 25 | 54 | 80 | 47 | 62 | 58 | 74 | 53 | 24 | 30 | 30 | 33 | 29 | 33 | 11 | 72 | 90 | 103 | 59 | 58 | 79 | 30 | Slc24a5 |
| Slc25a1 | 4685 | 6043 | 5935 | 5771 | 5419 | 5392 | 4748 | 2740 | 3115 | 3559 | 2932 | 3454 | 3144 | 3004 | 4714 | 5396 | 6217 | 5437 | 5903 | 4683 | 4980 | 3487 | 3279 | 3225 | 3262 | 3640 | 3190 | 3221 | Slc25a1 |
| Slc25a10 | 3550 | 4622 | 3704 | 3919 | 3018 | 3317 | 2947 | 1198 | 1565 | 1363 | 1486 | 1652 | 1610 | 1398 | 3185 | 3880 | 3559 | 3431 | 3369 | 3193 | 2865 | 1158 | 1456 | 1474 | 1399 | 2067 | 1596 | 1467 | Slc25a10 |
| Slc25a11 | 4131 | 4676 | 3616 | 3831 | 3705 | 3789 | 3933 | 4288 | 4526 | 4120 | 3972 | 4235 | 4065 | 4349 | 4200 | 4016 | 4012 | 3710 | 3861 | 3829 | 4028 | 4104 | 4446 | 4046 | 3996 | 4754 | 4194 | 4339 | Slc25a11 |
| Slc25a12 | 5597 | 5982 | 4945 | 5740 | 6356 | 5844 | 6785 | 4509 | 4562 | 4542 | 3965 | 4615 | 3530 | 4088 | 5757 | 5234 | 5840 | 5862 | 6163 | 6178 | 6292 | 4884 | 4535 | 4019 | 4579 | 4524 | 4128 | 3980 | Slc25a12 |
| Slc25a13 | 1912 | 1816 | 1958 | 1348 | 1050 | 1247 | 1349 | 1342 | 1257 | 1149 | 1053 | 1655 | 1320 | 1671 | 1797 | 1645 | 1691 | 1168 | 1242 | 1268 | 1157 | 1008 | 1159 | 1301 | 1045 | 2198 | 1725 | 1942 | Slc25a13 |
| Slc25a14 | 1628 | 1610 | 1640 | 1866 | 1754 | 1844 | 2150 | 1616 | 1275 | 1261 | 1348 | 1446 | 1210 | 1481 | 1977 | 1460 | 1672 | 1910 | 1912 | 2169 | 1989 | 1594 | 1269 | 1437 | 1458 | 1361 | 1336 | 1441 | Slc25a14 |
| Slc25a15 | 2377 | 2187 | 2642 | 1916 | 1734 | 1909 | 1902 | 1304 | 1347 | 1178 | 1262 | 1250 | 1322 | 1205 | 2177 | 2143 | 2045 | 1950 | 1896 | 1854 | 1747 | 1044 | 1228 | 1154 | 1191 | 1458 | 1276 | 1301 | Slc25a15 |
| Slc25a16 | 2397 | 1745 | 1955 | 1982 | 1798 | 2048 | 2391 | 1268 | 1186 | 1171 | 1181 | 1389 | 1284 | 1417 | 2144 | 1825 | 1717 | 2029 | 1772 | 1946 | 2018 | 1392 | 1193 | 1335 | 1315 | 1380 | 1287 | 1292 | Slc25a16 |
| Slc25a17 | 5976 | 5367 | 5441 | 5487 | 5431 | 5193 | 5944 | 6281 | 5498 | 5292 | 5142 | 5254 | 5378 | 5678 | 6281 | 5468 | 5597 | 5344 | 5446 | 6414 | 5563 | 5521 | 5680 | 5387 | 5353 | 5727 | 5559 | 5543 | Slc25a17 |
| Slc25a18 | 37 | 70 | 45 | 104 | 146 | 95 | 99 | 111 | 150 | 123 | 125 | 85 | 144 | 95 | 47 | 54 | 96 | 83 | 142 | 81 | 132 | 127 | 157 | 71 | 161 | 56 | 112 | 125 | Slc25a18 |
| Slc25a19 | 1516 | 2090 | 2034 | 2246 | 2151 | 1864 | 1929 | 1194 | 1364 | 1490 | 1343 | 1439 | 1289 | 1326 | 1520 | 1754 | 2118 | 2278 | 2387 | 1805 | 2139 | 1340 | 1250 | 1586 | 1668 | 1307 | 1313 | 1391 | Slc25a19 |
| Slc25a2 | 29 | 30 | 57 | 49 | 37 | 23 | 51 | 18 | 27 | 37 | 12 | 41 | 37 | 35 | 30 | 20 | 72 | 36 | 27 | 40 | 46 | 28 | 23 | 31 | 40 | 44 | 19 | 51 | Slc25a2 |
| Slc25a20 | 3223 | 3347 | 3278 | 3101 | 3103 | 3090 | 3162 | 2310 | 2216 | 1887 | 2123 | 1967 | 2088 | 2379 | 3356 | 2859 | 3094 | 3160 | 3403 | 3481 | 3433 | 1977 | 2226 | 2292 | 2016 | 2367 | 2135 | 2461 | Slc25a20 |
| Slc25a21 | 23 | 16 | 27 | 2 | 3 | 8 | 7 | 20 | 19 | 16 | 9 | 21 | 34 | 21 | 34 | 43 | 29 | 3 | 9 | 12 | 6 | 17 | 19 | 28 | 10 | 28 | 23 | 57 | Slc25a21 |
| Slc25a22 | 3324 | 4499 | 4805 | 5777 | 5587 | 5049 | 4871 | 2662 | 3252 | 3901 | 3524 | 3953 | 3553 | 3720 | 3400 | 4167 | 5737 | 5720 | 5626 | 4509 | 4917 | 3912 | 3016 | 4033 | 4216 | 4040 | 3188 | 3336 | Slc25a22 |
| Slc25a23 | 10139 | 12839 | 11721 | 13032 | 14108 | 12951 | 12149 | 6251 | 6946 | 8224 | 6432 | 8072 | 6582 | 6629 | 9653 | 10172 | 13626 | 13755 | 14486 | 11995 | 12961 | 8502 | 6337 | 7542 | 8175 | 7573 | 6746 | 6408 | Slc25a23 |
| Slc25a24 | 2556 | 1748 | 2487 | 1195 | 992 | 1516 | 905 | 2967 | 3133 | 2816 | 3998 | 3208 | 3916 | 3236 | 2440 | 2866 | 1610 | 1341 | 864 | 940 | 815 | 2851 | 3715 | 2762 | 2282 | 3769 | 3436 | 3680 | Slc25a24 |
| Slc25a25 | 3012 | 4346 | 3630 | 5204 | 4550 | 4421 | 4472 | 2740 | 2954 | 3070 | 2839 | 2885 | 2641 | 3041 | 3320 | 3515 | 4512 | 4803 | 4647 | 4140 | 4652 | 2973 | 2655 | 3012 | 3400 | 2844 | 2989 | 3008 | Slc25a25 |
| Slc25a26 | 149 | 164 | 155 | 148 | 120 | 113 | 112 | 419 | 460 | 469 | 655 | 604 | 602 | 592 | 112 | 145 | 115 | 130 | 116 | 133 | 127 | 463 | 489 | 538 | 518 | 541 | 638 | 593 | Slc25a26 |
| Slc25a27 | 6636 | 6071 | 5178 | 7864 | 8822 | 8978 | 9507 | 6929 | 6257 | 7251 | 6634 | 7379 | 6652 | 7572 | 6339 | 5784 | 6562 | 7772 | 8510 | 7981 | 9370 | 7591 | 5748 | 6875 | 8200 | 4866 | 6675 | 6605 | Slc25a27 |
| Slc25a28 | 2765 | 3286 | 2987 | 3631 | 2762 | 2997 | 3084 | 1689 | 1849 | 1817 | 1922 | 2297 | 1936 | 2076 | 2502 | 2818 | 3022 | 3193 | 3004 | 2724 | 3027 | 1750 | 1735 | 2328 | 2007 | 2467 | 1961 | 2086 | Slc25a28 |
| Slc25a29 | 791 | 1059 | 938 | 1112 | 1031 | 1098 | 1022 | 331 | 461 | 511 | 461 | 487 | 395 | 450 | 737 | 850 | 1073 | 1205 | 1174 | 764 | 1016 | 480 | 375 | 567 | 535 | 586 | 452 | 394 | Slc25a29 |
| Slc25a3 | 26272 | 25235 | 26495 | 24690 | 22163 | 20336 | 24029 | 20262 | 19514 | 19447 | 20914 | 22685 | 21913 | 23088 | 24747 | 24203 | 25394 | 23311 | 23270 | 20377 | 23257 | 21709 | 19514 | 21966 | 20379 | 25629 | 21665 | 24230 | Slc25a3 |
| Slc25a30 | 3466 | 3448 | 3382 | 3966 | 3776 | 3605 | 4247 | 1988 | 1823 | 1966 | 1610 | 1915 | 2000 | 2120 | 3510 | 3396 | 3635 | 3774 | 3662 | 3953 | 4023 | 1928 | 1675 | 1902 | 1934 | 2086 | 2011 | 2079 | Slc25a30 |
| Slc25a31 | 1 | 2 | 0 | 2 | 2 | 13 | 0 | 1 | 2 | 3 | 3 | 0 | 3 | 0 | 1 | 1 | 1 | 2 | 1 | 13 | 0 | 2 | 1 | 0 | 1 | 0 | 0 | 0 | Slc25a31 |
| Slc25a32 | 55 | 21 | 36 | 128 | 25 | 58 | 37 | 54 | 206 | 56 | 21 | 24 | 69 | 87 | 103 | 43 | 63 | 41 | 25 | 41 | 20 | 58 | 67 | 62 | 31 | 18 | 34 | 24 | Slc25a32 |
| Slc25a33 | 794 | 941 | 881 | 971 | 902 | 805 | 915 | 531 | 554 | 508 | 623 | 503 | 528 | 568 | 944 | 957 | 880 | 1034 | 836 | 905 | 919 | 570 | 579 | 494 | 535 | 494 | 508 | 555 | Slc25a33 |
| Slc25a34 | 13 | 27 | 24 | 46 | 31 | 27 | 26 | 32 | 13 | 25 | 43 | 11 | 35 | 39 | 20 | 35 | 42 | 35 | 23 | 27 | 26 | 35 | 24 | 28 | 18 | 16 | 28 | 26 | Slc25a34 |
| Slc25a35 | 540 | 561 | 441 | 651 | 462 | 421 | 458 | 216 | 220 | 278 | 225 | 131 | 178 | 177 | 504 | 635 | 520 | 583 | 432 | 426 | 434 | 152 | 242 | 299 | 143 | 206 | 126 | 125 | Slc25a35 |
| Slc25a36 | 13162 | 9203 | 10466 | 11592 | 11596 | 12464 | 12875 | 11828 | 11145 | 9764 | 11717 | 9217 | 9986 | 9323 | 13367 | 12183 | 9945 | 11340 | 10459 | 12134 | 11959 | 10579 | 11258 | 8853 | 9515 | 7748 | 9648 | 8832 | Slc25a36 |
| Slc25a37 | 5477 | 5017 | 5372 | 3539 | 3392 | 3421 | 3596 | 2282 | 2500 | 2498 | 2685 | 2886 | 4740 | 4344 | 4759 | 5598 | 4348 | 3822 | 3195 | 3133 | 3405 | 2015 | 2544 | 2366 | 1887 | 3306 | 3496 | 7233 | Slc25a37 |
| Slc25a38 | 818 | 1040 | 1062 | 922 | 945 | 1001 | 857 | 1106 | 1313 | 1294 | 1204 | 1266 | 1347 | 1274 | 851 | 967 | 1042 | 867 | 1043 | 1049 | 956 | 1305 | 1289 | 1334 | 1311 | 1428 | 1307 | 1481 | Slc25a38 |
| Slc25a39 | 5369 | 7176 | 6281 | 5838 | 4972 | 4825 | 5026 | 2666 | 3561 | 3275 | 3096 | 3699 | 3513 | 3362 | 5045 | 6428 | 6253 | 5554 | 5666 | 4819 | 5119 | 3004 | 3684 | 3504 | 3122 | 4251 | 3440 | 3712 | Slc25a39 |
| Slc25a4 | 22835 | 26073 | 24466 | 24229 | 24439 | 22157 | 25679 | 22366 | 22496 | 21898 | 22433 | 20775 | 22744 | 23968 | 24929 | 24229 | 24450 | 24230 | 23149 | 27932 | 24424 | 20404 | 23290 | 23352 | 22881 | 22622 | 21725 | 23695 | Slc25a4 |
| Slc25a40 | 2503 | 1623 | 1718 | 1955 | 1807 | 2309 | 2296 | 2121 | 2059 | 1694 | 1959 | 1714 | 1676 | 1821 | 2490 | 2143 | 1807 | 1940 | 1927 | 2462 | 2106 | 1820 | 1856 | 1695 | 1609 | 1626 | 1786 | 1693 | Slc25a40 |
| Slc25a41 | 9 | 27 | 26 | 20 | 20 | 16 | 22 | 7 | 2 | 0 | 5 | 0 | 3 | 2 | 13 | 19 | 21 | 41 | 25 | 14 | 21 | 8 | 1 | 2 | 4 | 6 | 3 | 5 | Slc25a41 |
| Slc25a42 | 1326 | 1735 | 1704 | 2154 | 1667 | 1591 | 1446 | 584 | 796 | 928 | 1053 | 760 | 713 | 691 | 1163 | 1918 | 1640 | 2293 | 1635 | 1386 | 1512 | 1066 | 848 | 840 | 933 | 694 | 590 | 633 | Slc25a42 |
| Slc25a43 | 16 | 11 | 13 | 8 | 7 | 15 | 9 | 16 | 30 | 19 | 19 | 17 | 28 | 32 | 5 | 10 | 8 | 9 | 10 | 5 | 6 | 22 | 32 | 29 | 15 | 16 | 17 | 27 | Slc25a43 |
| Slc25a44 | 4482 | 5301 | 4289 | 5024 | 4526 | 4722 | 4785 | 2884 | 3155 | 3205 | 2984 | 3138 | 2860 | 2953 | 4361 | 4360 | 4509 | 4986 | 4945 | 4594 | 5115 | 3340 | 2974 | 3092 | 3565 | 2852 | 2841 | 3108 | Slc25a44 |
| Slc25a45 | 55 | 62 | 102 | 61 | 76 | 65 | 60 | 120 | 195 | 191 | 249 | 133 | 197 | 113 | 74 | 79 | 46 | 47 | 53 | 59 | 65 | 152 | 284 | 185 | 159 | 216 | 187 | 142 | Slc25a45 |
| Slc25a46 | 5787 | 4642 | 5609 | 6051 | 5787 | 5609 | 6652 | 6069 | 4884 | 5049 | 4890 | 5702 | 5319 | 5679 | 6186 | 5051 | 5702 | 5624 | 5863 | 5680 | 6101 | 6731 | 5014 | 5575 | 5823 | 6028 | 5453 | 5895 | Slc25a46 |
| Slc25a47 | 654 | 706 | 581 | 560 | 499 | 646 | 553 | 525 | 634 | 613 | 558 | 751 | 612 | 657 | 551 | 576 | 679 | 609 | 634 | 573 | 571 | 551 | 597 | 687 | 723 | 672 | 695 | 710 | Slc25a47 |
| Slc25a48 | 57 | 26 | 30 | 27 | 4 | 35 | 3 | 27 | 40 | 14 | 20 | 5 | 12 | 9 | 38 | 26 | 17 | 34 | 6 | 3 | 3 | 7 | 36 | 22 | 11 | 10 | 12 | 7 | Slc25a48 |
| Slc25a5 | 19158 | 23453 | 20661 | 17728 | 16691 | 18307 | 19005 | 19848 | 18999 | 17415 | 16544 | 23157 | 21996 | 24846 | 19622 | 17935 | 21118 | 16677 | 19260 | 22299 | 18805 | 15580 | 18149 | 21663 | 20286 | 26079 | 25306 | 25386 | Slc25a5 |
| Slc25a51 | 7913 | 8129 | 8236 | 8712 | 7909 | 7504 | 8396 | 7566 | 7327 | 7796 | 7533 | 7888 | 7317 | 8318 | 7477 | 7269 | 8583 | 8273 | 8214 | 7270 | 8335 | 8168 | 6888 | 8259 | 8348 | 7938 | 7599 | 8350 | Slc25a51 |
| Slc25a53 | 1143 | 1102 | 1093 | 1188 | 1064 | 1175 | 1233 | 1760 | 1547 | 1288 | 1325 | 1619 | 1572 | 1703 | 1307 | 988 | 1077 | 1029 | 975 | 1263 | 1203 | 1494 | 1421 | 1410 | 1476 | 1484 | 1689 | 1623 | Slc25a53 |
| Slc25a54 | 0 | 0 | 0 | 0 | 0 | 0 | 0 | 0 | 0 | 0 | 0 | 0 | 0 | 0 | 0 | 0 | 0 | 0 | 0 | 0 | 2 | 0 | 0 | 0 | 0 | 0 | 0 | 0 | Slc25a54 |
| Slc26a1 | 18 | 16 | 17 | 17 | 25 | 32 | 13 | 173 | 214 | 174 | 177 | 142 | 184 | 125 | 12 | 19 | 19 | 11 | 27 | 24 | 29 | 216 | 178 | 119 | 173 | 186 | 153 | 104 | Slc26a1 |
| Slc26a10 | 183 | 228 | 341 | 273 | 232 | 210 | 183 | 93 | 93 | 137 | 144 | 128 | 142 | 111 | 144 | 233 | 242 | 254 | 212 | 132 | 178 | 94 | 97 | 141 | 125 | 172 | 128 | 133 | Slc26a10 |
| Slc26a11 | 1023 | 1167 | 1076 | 1070 | 941 | 1056 | 944 | 448 | 576 | 478 | 689 | 633 | 625 | 565 | 870 | 1160 | 1072 | 1104 | 1027 | 957 | 835 | 560 | 551 | 566 | 572 | 705 | 604 | 576 | Slc26a11 |
| Slc26a2 | 3876 | 3035 | 2619 | 2707 | 2065 | 2873 | 2316 | 1806 | 2209 | 2354 | 2679 | 1831 | 2083 | 1622 | 3075 | 3857 | 2391 | 2968 | 2026 | 2184 | 2506 | 1836 | 1968 | 1997 | 1759 | 1428 | 1786 | 1715 | Slc26a2 |
| Slc26a3 | 0 | 1 | 3 | 1 | 4 | 2 | 7 | 2 | 3 | 4 | 2 | 1 | 2 | 1 | 1 | 0 | 2 | 1 | 1 | 0 | 1 | 3 | 2 | 1 | 0 | 2 | 1 | 1 | Slc26a3 |
| Slc26a4 | 2 | 2 | 0 | 0 | 3 | 4 | 1 | 4 | 14 | 3 | 21 | 0 | 11 | 0 | 5 | 2 | 2 | 3 | 6 | 0 | 4 | 18 | 10 | 12 | 11 | 7 | 8 | 0 | Slc26a4 |
| Slc26a5 | 23 | 17 | 7 | 22 | 24 | 19 | 16 | 17 | 15 | 10 | 16 | 9 | 22 | 28 | 18 | 16 | 0 | 34 | 22 | 25 | 46 | 15 | 12 | 13 | 20 | 8 | 17 | 23 | Slc26a5 |
| Slc26a6 | 5 | 0 | 4 | 3 | 2 | 2 | 1 | 2 | 1 | 3 | 0 | 0 | 0 | 0 | 1 | 1 | 2 | 1 | 2 | 1 | 4 | 0 | 1 | 3 | 2 | 0 | 1 | 2 | Slc26a6 |
| Slc26a7 | 280 | 129 | 102 | 209 | 254 | 284 | 363 | 628 | 592 | 456 | 555 | 100 | 104 | 94 | 174 | 171 | 125 | 276 | 212 | 304 | 332 | 289 | 924 | 378 | 206 | 166 | 34 | 76 | Slc26a7 |
| Slc26a8 | 89 | 100 | 104 | 131 | 160 | 147 | 165 | 93 | 128 | 159 | 105 | 133 | 120 | 116 | 102 | 104 | 103 | 122 | 113 | 124 | 139 | 157 | 122 | 132 | 154 | 98 | 152 | 113 | Slc26a8 |
| Slc26a9 | 7 | 9 | 7 | 11 | 5 | 21 | 6 | 4 | 18 | 8 | 22 | 12 | 11 | 3 | 1 | 9 | 1 | 6 | 2 | 10 | 3 | 10 | 11 | 13 | 5 | 1 | 0 | 1 | Slc26a9 |
| Slc27a1 | 3990 | 5450 | 5346 | 7037 | 7213 | 6352 | 6891 | 3552 | 4349 | 4717 | 4473 | 4177 | 4002 | 3447 | 4146 | 4645 | 6751 | 7679 | 7708 | 6183 | 6381 | 4652 | 4203 | 4957 | 4713 | 3802 | 3554 | 3163 | Slc27a1 |
| Slc27a2 | 314 | 204 | 158 | 342 | 319 | 212 | 189 | 559 | 451 | 626 | 465 | 255 | 299 | 317 | 233 | 241 | 161 | 160 | 266 | 235 | 189 | 480 | 519 | 704 | 323 | 562 | 331 | 288 | Slc27a2 |

| | | | | | | | | | | | | | | | | | | | | | | | | | | | | | |
|---|---|---|---|---|---|---|---|---|---|---|---|---|---|---|---|---|---|---|---|---|---|---|---|---|---|---|---|---|---|
| Slc27a3 | 618 | 796 | 801 | 841 | 838 | 764 | 599 | 1354 | 1801 | 1810 | 1939 | 1405 | 1666 | 1208 | 686 | 872 | 827 | 854 | 863 | 576 | 594 | 1760 | 1726 | 1534 | 1731 | 1523 | 1389 | 1203 | Slc27a3 |
| Slc27a4 | 3099 | 3972 | 4004 | 4273 | 4289 | 3954 | 3452 | 3313 | 3861 | 4093 | 3723 | 3738 | 3825 | 3398 | 3343 | 3727 | 4159 | 4160 | 4523 | 3548 | 4047 | 4553 | 3809 | 4015 | 4078 | 3418 | 3519 | 3438 | Slc27a4 |
| Slc27a5 | 20 | 17 | 18 | 39 | 49 | 53 | 32 | 26 | 35 | 32 | 34 | 22 | 16 | 29 | 24 | 26 | 72 | 48 | 49 | 17 | 32 | 23 | 17 | 37 | 23 | 39 | 17 | 14 | Slc27a5 |
| Slc27a6 | 247 | 199 | 264 | 139 | 99 | 155 | 60 | 175 | 255 | 260 | 330 | 197 | 274 | 206 | 204 | 301 | 153 | 142 | 81 | 76 | 63 | 192 | 278 | 202 | 233 | 260 | 187 | 215 | Slc27a6 |
| Slc28a1 | 1 | 5 | 0 | 10 | 2 | 4 | 6 | 0 | 1 | 2 | 5 | 0 | 12 | 2 | 1 | 9 | 6 | 6 | 5 | 2 | 7 | 0 | 13 | 1 | 9 | 0 | 1 | 12 | Slc28a1 |
| Slc28a2 | 7 | 4 | 4 | 0 | 1 | 5 | 1 | 4 | 5 | 2 | 49 | 0 | 10 | 1 | 3 | 2 | 1 | 1 | 5 | 1 | 4 | 0 | 21 | 7 | 3 | 6 | 8 | 2 | Slc28a2 |
| Slc28a2b | 0 | 0 | 0 | 0 | 0 | 0 | 0 | 0 | 0 | 0 | 1 | 0 | 0 | 0 | 0 | 0 | 0 | 1 | 0 | 1 | 1 | 0 | 0 | 0 | 0 | 0 | 0 | 0 | Slc28a2b |
| Slc28a3 | 22 | 10 | 3 | 5 | 11 | 9 | 3 | 15 | 16 | 14 | 16 | 0 | 5 | 5 | 12 | 13 | 10 | 4 | 2 | 6 | 2 | 7 | 17 | 15 | 1 | 4 | 1 | 1 | Slc28a3 |
| Slc29a1 | 3002 | 3338 | 3233 | 2667 | 2554 | 2700 | 2626 | 1243 | 1714 | 1422 | 1750 | 1560 | 1471 | 1428 | 2661 | 3271 | 2986 | 2886 | 2870 | 2837 | 2522 | 1333 | 1779 | 1679 | 1471 | 1855 | 1618 | 1671 | Slc29a1 |
| Slc29a2 | 518 | 694 | 552 | 512 | 718 | 580 | 580 | 673 | 752 | 711 | 700 | 752 | 676 | 696 | 508 | 638 | 672 | 536 | 663 | 611 | 562 | 755 | 662 | 728 | 833 | 742 | 758 | 685 | Slc29a2 |
| Slc29a3 | 1434 | 1570 | 1459 | 1289 | 1603 | 1511 | 1569 | 1796 | 2217 | 1968 | 2025 | 2153 | 2046 | 1957 | 1618 | 1413 | 1574 | 1400 | 1570 | 1302 | 1487 | 2094 | 2016 | 1785 | 1862 | 1826 | 2087 | 2077 | Slc29a3 |
| Slc29a4 | 3069 | 4266 | 3972 | 5056 | 5471 | 4619 | 4235 | 2419 | 2880 | 3320 | 2651 | 3184 | 2625 | 2681 | 3252 | 3407 | 4789 | 5160 | 5911 | 3984 | 4830 | 3711 | 2483 | 3076 | 3564 | 2660 | 2463 | 2421 | Slc29a4 |
| Slc2a1 | 6373 | 7979 | 7936 | 6561 | 5763 | 6176 | 5463 | 5356 | 6517 | 7072 | 5976 | 7335 | 7185 | 6818 | 6091 | 6926 | 6839 | 6440 | 6544 | 5832 | 5883 | 5180 | 6543 | 7409 | 6259 | 7780 | 7160 | 7524 | Slc2a1 |
| Slc2a10 | 309 | 282 | 291 | 225 | 194 | 280 | 178 | 502 | 557 | 430 | 528 | 431 | 474 | 420 | 315 | 315 | 272 | 263 | 185 | 170 | 147 | 500 | 530 | 493 | 365 | 520 | 401 | 389 | Slc2a10 |
| Slc2a12 | 379 | 363 | 303 | 277 | 340 | 329 | 189 | 482 | 553 | 447 | 453 | 556 | 586 | 352 | 367 | 461 | 234 | 285 | 300 | 313 | 280 | 471 | 794 | 395 | 383 | 350 | 439 | 394 | Slc2a12 |
| Slc2a13 | 2074 | 1793 | 1388 | 2020 | 2287 | 2168 | 2331 | 1323 | 1402 | 1275 | 1394 | 1020 | 1145 | 1031 | 2234 | 2069 | 1570 | 2334 | 1988 | 2358 | 2260 | 1479 | 1500 | 979 | 1220 | 772 | 1015 | 960 | Slc2a13 |
| Slc2a2 | 21 | 16 | 10 | 19 | 29 | 49 | 38 | 5 | 2 | 2 | 6 | 1 | 2 | 1 | 13 | 13 | 15 | 40 | 21 | 25 | 55 | 7 | 8 | 0 | 1 | 8 | 2 | 0 | Slc2a2 |
| Slc2a3 | 8889 | 9566 | 9784 | 7238 | 6973 | 8296 | 6569 | 3326 | 4247 | 2798 | 3198 | 3292 | 4024 | 3124 | 8890 | 8517 | 7130 | 7933 | 6566 | 7590 | 6371 | 2075 | 4448 | 3134 | 2221 | 1987 | 3339 | 3682 | Slc2a3 |
| Slc2a4 | 1637 | 1790 | 1614 | 1526 | 1168 | 1458 | 1543 | 630 | 902 | 727 | 709 | 840 | 819 | 826 | 1416 | 1600 | 1545 | 1467 | 1340 | 1380 | 1341 | 498 | 731 | 736 | 576 | 1179 | 811 | 846 | Slc2a4 |
| Slc2a5 | 138 | 233 | 187 | 132 | 187 | 131 | 163 | 138 | 124 | 109 | 156 | 106 | 162 | 123 | 164 | 210 | 260 | 191 | 227 | 154 | 211 | 164 | 146 | 119 | 101 | 215 | 159 | 176 | Slc2a5 |
| Slc2a6 | 235 | 351 | 246 | 230 | 345 | 270 | 277 | 95 | 164 | 146 | 123 | 105 | 144 | 114 | 294 | 346 | 273 | 265 | 243 | 316 | 293 | 179 | 137 | 115 | 127 | 106 | 129 | 108 | Slc2a6 |
| Slc2a7 | 0 | 0 | 1 | 0 | 0 | 3 | 0 | 0 | 0 | 0 | 0 | 0 | 0 | 0 | 0 | 5 | 3 | 3 | 0 | 0 | 0 | 0 | 1 | 0 | 0 | 0 | 0 | 0 | Slc2a7 |
| Slc2a8 | 1553 | 2046 | 1769 | 1991 | 1735 | 1795 | 1605 | 597 | 764 | 776 | 720 | 762 | 739 | 577 | 1381 | 1881 | 1856 | 2107 | 1732 | 1556 | 1579 | 652 | 734 | 690 | 687 | 757 | 767 | 627 | Slc2a8 |
| Slc2a9 | 53 | 51 | 40 | 29 | 42 | 39 | 39 | 57 | 98 | 43 | 93 | 74 | 76 | 40 | 57 | 61 | 38 | 31 | 49 | 40 | 37 | 47 | 79 | 50 | 59 | 86 | 53 | 53 | Slc2a9 |
| Slc30a1 | 77 | 34 | 48 | 72 | 71 | 62 | 70 | 72 | 51 | 46 | 43 | 53 | 60 | 36 | 76 | 64 | 74 | 70 | 64 | 79 | 50 | 52 | 65 | 61 | 54 | 57 | 40 | 50 | Slc30a1 |
| Slc30a10 | 918 | 739 | 820 | 1463 | 2736 | 2490 | 3242 | 2142 | 2588 | 2953 | 2171 | 2326 | 1828 | 2397 | 920 | 780 | 969 | 1750 | 2734 | 2386 | 3125 | 1743 | 2214 | 2917 | 3589 | 1730 | 1558 | 2013 | Slc30a10 |
| Slc30a2 | 17 | 16 | 13 | 12 | 11 | 4 | 7 | 1 | 4 | 1 | 6 | 1 | 2 | 2 | 17 | 14 | 6 | 12 | 3 | 6 | 9 | 5 | 14 | 2 | 0 | 7 | 2 | 6 | Slc30a2 |
| Slc30a3 | 346 | 695 | 686 | 705 | 862 | 752 | 432 | 557 | 469 | 791 | 541 | 569 | 950 | 392 | 525 | 536 | 765 | 691 | 786 | 605 | 510 | 847 | 373 | 365 | 523 | 131 | 602 | 471 | Slc30a3 |
| Slc30a5 | 6719 | 5951 | 5801 | 5784 | 4940 | 5723 | 5760 | 4419 | 4186 | 3749 | 4201 | 4135 | 3828 | 3972 | 6794 | 6284 | 5373 | 5869 | 5035 | 5628 | 5399 | 4096 | 4159 | 4236 | 4053 | 4574 | 3883 | 4306 | Slc30a5 |
| Slc30a6 | 1136 | 958 | 1066 | 1013 | 896 | 959 | 927 | 1160 | 1196 | 1085 | 1098 | 1137 | 1232 | 1230 | 1039 | 994 | 997 | 957 | 921 | 1010 | 1023 | 1093 | 1267 | 1173 | 1117 | 1340 | 1141 | 1132 | Slc30a6 |
| Slc30a7 | 6345 | 5974 | 4095 | 5029 | 4970 | 5442 | 5547 | 4686 | 4331 | 3921 | 4199 | 3876 | 3997 | 3965 | 6257 | 6212 | 4392 | 5620 | 4615 | 5564 | 5669 | 3967 | 4438 | 3367 | 3878 | 3088 | 4085 | 3900 | Slc30a7 |
| Slc30a8 | 6 | 3 | 8 | 0 | 5 | 1 | 8 | 2 | 1 | 6 | 2 | 0 | 2 | 0 | 1 | 3 | 0 | 7 | 5 | 3 | 2 | 12 | 1 | 3 | 2 | 2 | 0 | 1 | Slc30a8 |
| Slc30a9 | 12854 | 12156 | 11353 | 12354 | 11737 | 11942 | 13009 | 10014 | 8540 | 8688 | 8054 | 8788 | 8197 | 8685 | 13829 | 12206 | 12290 | 12532 | 11628 | 12820 | 12366 | 9580 | 8442 | 8237 | 9090 | 7900 | 8686 | 8529 | Slc30a9 |
| Slc31a1 | 11477 | 10595 | 9615 | 9253 | 8494 | 8995 | 10013 | 5961 | 5217 | 4865 | 4987 | 5428 | 5384 | 5354 | 11814 | 10668 | 9396 | 9480 | 8611 | 9946 | 9355 | 4988 | 5825 | 4872 | 4866 | 5697 | 5527 | 5880 | Slc31a1 |
| Slc31a2 | 976 | 1050 | 1092 | 1040 | 1166 | 1142 | 954 | 564 | 597 | 584 | 766 | 518 | 668 | 549 | 936 | 1116 | 1065 | 1165 | 1089 | 1055 | 991 | 764 | 734 | 603 | 607 | 575 | 557 | 508 | Slc31a2 |
| Slc32a1 | 514 | 685 | 649 | 1451 | 1926 | 1382 | 1396 | 1310 | 1439 | 2118 | 1637 | 1749 | 1349 | 1792 | 508 | 352 | 1433 | 1524 | 1908 | 1069 | 1749 | 2197 | 1111 | 2024 | 2229 | 1591 | 1416 | 1498 | Slc32a1 |
| Slc33a1 | 3295 | 2811 | 2607 | 2869 | 2615 | 2430 | 2928 | 2014 | 1808 | 1734 | 1789 | 1632 | 1690 | 1666 | 3345 | 3018 | 2605 | 2969 | 2681 | 2867 | 2649 | 1988 | 1923 | 1798 | 1889 | 1783 | 1683 | 1688 | Slc33a1 |
| Slc34a1 | 0 | 3 | 0 | 0 | 3 | 0 | 0 | 1 | 2 | 0 | 1 | 0 | 7 | 1 | 4 | 0 | 1 | 1 | 2 | 1 | 3 | 0 | 1 | 0 | 0 | 1 | 1 | 1 | Slc34a1 |
| Slc34a2 | 1 | 0 | 0 | 0 | 0 | 0 | 0 | 7 | 13 | 8 | 3 | 0 | 3 | 0 | 1 | 1 | 2 | 0 | 2 | 1 | 1 | 0 | 8 | 3 | 0 | 2 | 0 | 0 | Slc34a2 |
| Slc34a3 | 1 | 1 | 0 | 0 | 0 | 3 | 0 | 0 | 0 | 0 | 0 | 0 | 0 | 0 | 0 | 0 | 0 | 0 | 0 | 0 | 0 | 0 | 0 | 0 | 0 | 0 | 1 | 0 | Slc34a3 |
| Slc35a1 | 1413 | 1234 | 1766 | 1332 | 1585 | 1656 | 1592 | 2864 | 2429 | 2649 | 2607 | 3098 | 3005 | 3047 | 1598 | 1457 | 1654 | 1360 | 1697 | 1571 | 1576 | 2887 | 2590 | 2812 | 2571 | 3023 | 3060 | 3021 | Slc35a1 |
| Slc35a2 | 2170 | 2553 | 2318 | 2405 | 2015 | 1856 | 1980 | 1304 | 1495 | 1544 | 1765 | 1642 | 1556 | 1442 | 1921 | 2231 | 2431 | 2308 | 2067 | 1990 | 1948 | 1634 | 1539 | 1640 | 1622 | 1616 | 1539 | 1556 | Slc35a2 |
| Slc35a3 | 2189 | 1482 | 2028 | 1872 | 1932 | 1749 | 1884 | 2076 | 1892 | 1663 | 2079 | 1807 | 2165 | 2023 | 2217 | 2091 | 1895 | 1716 | 1739 | 1806 | 1747 | 1880 | 1985 | 1789 | 1671 | 1813 | 1843 | 1982 | Slc35a3 |
| Slc35a4 | 4083 | 4323 | 4192 | 3387 | 3071 | 3234 | 3099 | 4136 | 4944 | 4480 | 4784 | 5210 | 5328 | 4847 | 3579 | 4056 | 3633 | 3327 | 3358 | 3181 | 2979 | 4044 | 4603 | 4569 | 4603 | 6032 | 5189 | 5352 | Slc35a4 |
| Slc35a5 | 1884 | 1534 | 1665 | 1655 | 2076 | 2054 | 2168 | 3021 | 2438 | 2166 | 2210 | 2083 | 2167 | 2198 | 2126 | 1583 | 1698 | 1709 | 2090 | 2408 | 2071 | 2717 | 2396 | 2107 | 2424 | 1778 | 2073 | 2105 | Slc35a5 |
| Slc35b1 | 4347 | 4369 | 4039 | 3587 | 3449 | 3478 | 4034 | 4991 | 4302 | 3890 | 4100 | 4243 | 4072 | 4426 | 4404 | 3793 | 3892 | 3524 | 3907 | 4172 | 3908 | 4380 | 4422 | 4265 | 4414 | 4518 | 4398 | 4711 | Slc35b1 |
| Slc35b2 | 3930 | 4736 | 4134 | 4335 | 3515 | 4196 | 3552 | 3154 | 3441 | 3082 | 3068 | 3344 | 3539 | 3042 | 3957 | 4205 | 4073 | 4059 | 4060 | 4097 | 3604 | 2583 | 3601 | 3171 | 3170 | 3723 | 3737 | 3205 | Slc35b2 |
| Slc35b3 | 2081 | 1602 | 1522 | 1704 | 1716 | 1817 | 1840 | 1779 | 1734 | 1700 | 1691 | 1781 | 1669 | 1687 | 2070 | 1833 | 1649 | 1650 | 1608 | 1795 | 1633 | 1825 | 1743 | 1604 | 1573 | 1625 | 1643 | 1565 | Slc35b3 |
| Slc35b4 | 4684 | 4531 | 3939 | 3843 | 3810 | 3918 | 3956 | 4042 | 4102 | 3866 | 4049 | 4107 | 3799 | 3716 | 4272 | 4498 | 3876 | 4292 | 3359 | 3886 | 3959 | 4390 | 3960 | 3833 | 3771 | 3630 | 4095 | 4095 | Slc35b4 |
| Slc35c1 | 1059 | 1468 | 1447 | 1386 | 1172 | 1236 | 1163 | 581 | 723 | 779 | 757 | 741 | 818 | 776 | 1129 | 1429 | 1520 | 1467 | 1296 | 1086 | 1166 | 769 | 797 | 760 | 784 | 865 | 679 | 834 | Slc35c1 |
| Slc35c2 | 2900 | 4260 | 3517 | 3359 | 2874 | 2864 | 3117 | 2006 | 2338 | 2302 | 2216 | 2436 | 2321 | 2264 | 2763 | 3086 | 3461 | 2999 | 3319 | 3035 | 3031 | 2122 | 2284 | 2325 | 2480 | 2644 | 2524 | 2339 | Slc35c2 |
| Slc35d1 | 1257 | 991 | 1468 | 1269 | 1429 | 1240 | 1253 | 1480 | 1399 | 1867 | 1980 | 1333 | 1484 | 1297 | 990 | 1090 | 1290 | 1278 | 1378 | 1132 | 1231 | 1673 | 1460 | 2029 | 1337 | 2147 | 1428 | 1314 | Slc35d1 |
| Slc35d2 | 138 | 106 | 154 | 87 | 109 | 90 | 68 | 136 | 189 | 157 | 211 | 175 | 249 | 140 | 196 | 161 | 116 | 92 | 50 | 86 | 81 | 155 | 223 | 122 | 144 | 136 | 193 | 136 | Slc35d2 |
| Slc35d3 | 74 | 79 | 137 | 104 | 124 | 75 | 107 | 73 | 59 | 68 | 86 | 52 | 83 | 64 | 79 | 90 | 124 | 97 | 119 | 98 | 128 | 102 | 80 | 74 | 86 | 73 | 41 | 49 | Slc35d3 |
| Slc35e1 | 6493 | 6702 | 5920 | 6593 | 5253 | 6167 | 5579 | 3496 | 3969 | 4044 | 4474 | 4498 | 4031 | 4387 | 5738 | 6368 | 5917 | 6694 | 5920 | 5057 | 6104 | 4092 | 3950 | 4236 | 4094 | 4235 | 4479 | 4399 | Slc35e1 |
| Slc35e2 | 5666 | 5672 | 6119 | 6794 | 5626 | 5608 | 5827 | 4167 | 3627 | 4293 | 4155 | 4544 | 4265 | 4439 | 5846 | 5644 | 6209 | 6707 | 5743 | 5342 | 5726 | 4968 | 3563 | 4583 | 4589 | 4203 | 4388 | 4628 | Slc35e2 |
| Slc35e3 | 5208 | 4689 | 5752 | 5724 | 5555 | 5598 | 5802 | 3897 | 3896 | 4524 | 4404 | 4225 | 4119 | 4425 | 5082 | 4986 | 5623 | 5873 | 5688 | 4479 | 5764 | 4440 | 3772 | 4312 | 4647 | 4355 | 3948 | 4386 | Slc35e3 |
| Slc35e4 | 1105 | 1489 | 970 | 1266 | 1249 | 1324 | 1150 | 425 | 455 | 412 | 370 | 408 | 405 | 371 | 1051 | 1138 | 1221 | 1409 | 1400 | 1231 | 1293 | 458 | 498 | 407 | 476 | 428 | 393 | 318 | Slc35e4 |
| Slc35f1 | 5350 | 4877 | 5473 | 7501 | 8781 | 8377 | 9534 | 8609 | 7051 | 8256 | 6689 | 8716 | 7191 | 9670 | 6577 | 3921 | 6750 | 7613 | 8770 | 8547 | 9614 | 9451 | 7066 | 8755 | 10266 | 7481 | 8219 | 8681 | Slc35f1 |
| Slc35f2 | 337 | 242 | 300 | 276 | 448 | 370 | 341 | 366 | 386 | 410 | 401 | 287 | 247 | 292 | 268 | 288 | 279 | 288 | 372 | 314 | 355 | 311 | 346 | 376 | 386 | 393 | 258 | 295 | Slc35f2 |
| Slc35f3 | 69 | 50 | 64 | 66 | 160 | 74 | 80 | 57 | 96 | 98 | 119 | 50 | 83 | 54 | 49 | 67 | 59 | 109 | 94 | 76 | 89 | 110 | 110 | 56 | 111 | 44 | 42 | 58 | Slc35f3 |
| Slc35f4 | 24 | 16 | 12 | 31 | 81 | 41 | 49 | 122 | 143 | 128 | 90 | 48 | 56 | 40 | 25 | 19 | 13 | 30 | 37 | 39 | 51 | 107 | 109 | 51 | 114 | 13 | 44 | 32 | Slc35f4 |
| Slc35f5 | 2851 | 2680 | 2653 | 2262 | 2371 | 2419 | 2366 | 2055 | 2163 | 1897 | 2225 | 1680 | 1957 | 1774 | 2891 | 2826 | 2403 | 2485 | 2293 | 2514 | 2188 | 2184 | 2344 | 1717 | 1791 | 1787 | 1919 | 1805 | Slc35f5 |
| Slc35f6 | 2237 | 2925 | 1851 | 2174 | 2073 | 2168 | 2135 | 1219 | 1326 | 1315 | 1306 | 1151 | 1195 | 1200 | 2247 | 2554 | 2315 | 2195 | 1940 | 2049 | 2179 | 1240 | 1325 | 1146 | 1240 | 1170 | 1270 | 1221 | Slc35f6 |
| Slc35g1 | 829 | 801 | 1005 | 787 | 664 | 802 | 822 | 630 | 461 | 450 | 511 | 702 | 614 | 766 | 825 | 744 | 688 | 705 | 679 | 738 | 775 | 428 | 484 | 589 | 555 | 672 | 838 | 843 | Slc35g1 |
| Slc35g2 | 554 | 575 | 593 | 573 | 695 | 532 | 568 | 343 | 265 | 214 | 216 | 213 | 232 | 211 | 657 | 541 | 538 | 574 | 562 | 804 | 626 | 299 | 287 | 244 | 192 | 250 | 256 | 223 | Slc35g2 |
| Slc35g3 | 15 | 26 | 18 | 12 | 47 | 35 | 17 | 51 | 44 | 61 | 29 | 45 | 67 | 22 | 42 | 51 | 20 | 17 | 36 | 32 | 13 | 52 | 85 | 18 | 41 | 41 | 16 | 15 | Slc35g3 |
| Slc36a1 | 1982 | 2278 | 2511 | 2763 | 3161 | 2592 | 2487 | 1524 | 1869 | 2312 | 2183 | 2017 | 1919 | 1711 | 1919 | 2413 | 2923 | 3218 | 2893 | 1911 | 2647 | 2425 | 1872 | 2190 | 2228 | 1743 | 1820 | 1657 | Slc36a1 |
| Slc36a2 | 6 | 22 | 9 | 7 | 27 | 16 | 7 | 23 | 29 | 24 | 36 | 3 | 18 | 4 | 15 | 26 | 13 | 24 | 7 | 8 | 10 | 45 | 21 | 8 | 16 | 6 | 10 | 2 | Slc36a2 |
| Slc36a3 | 0 | 0 | 0 | 0 | 0 | 0 | 0 | 2 | 0 | 0 | 0 | 0 | 0 | 0 | 0 | 0 | 0 | 0 | 0 | 0 | 0 | 0 | 0 | 0 | 0 | 0 | 0 | 0 | Slc36a3 |
| Slc36a4 | 4902 | 4183 | 3583 | 4659 | 5593 | 5441 | 5924 | 5934 | 5446 | 5110 | 5157 | 5056 | 4625 | 5094 | 5267 | 4183 | 4092 | 4707 | 5106 | 5434 | 5758 | 5432 | 4899 | 4712 | 5530 | 4367 | 4693 | 4384 | Slc36a4 |
| Slc37a1 | 315 | 431 | 485 | 571 | 753 | 502 | 567 | 449 | 579 | 612 | 513 | 582 | 566 | 589 | 390 | 456 | 644 | 529 | 640 | 478 | 536 | 792 | 460 | 657 | 659 | 595 | 565 | 462 | Slc37a1 |
| Slc37a2 | 428 | 484 | 550 | 426 | 448 | 465 | 408 | 253 | 266 | 331 | 294 | 272 | 319 | 309 | 448 | 526 | 399 | 450 | 390 | 436 | 409 | 312 | 390 | 294 | 333 | 284 | 307 | 246 | Slc37a2 |
| Slc37a3 | 6575 | 6000 | 6433 | 6162 | 6890 | 6518 | 6402 | 5473 | 5337 | 5442 | 5487 | 5147 | 5346 | 5098 | 6889 | 5974 | 6282 | 6061 | 6631 | 7066 | 6502 | 5444 | 5427 | 5203 | 5540 | 5218 | 5274 | 5061 | Slc37a3 |
| Slc37a4 | 1607 | 1884 | 1570 | 1759 | 1719 | 1585 | 1801 | 1230 | 1176 | 1256 | 1211 | 1149 | 1183 | 1003 | 1763 | 1543 | 1750 | 1739 | 1702 | 1769 | 1806 | 1445 | 1408 | 1088 | 1297 | 1045 | 1110 | 1072 | Slc37a4 |
| Slc38a1 | 11954 | 10461 | 13250 | 13244 | 17095 | 15465 | 14938 | 21336 | 19744 | 24231 | 22210 | 25882 | 25306 | 26621 | 11587 | 10956 | 14201 | 13278 | 16583 | 13612 | 15491 | 24227 | 17953 | 23894 | 27781 | 20826 | 25861 | 25898 | Slc38a1 |
| Slc38a10 | 11618 | 14484 | 14770 | 15428 | 13571 | 13233 | 11864 | 6814 | 8402 | 8983 | 8855 | 8850 | 9027 | 7832 | 11315 | 12894 | 15309 | 14740 | 14816 | 11150 | 12860 | 8607 | 8822 | 8611 | 9226 | 8979 | 8629 | 8412 | Slc38a10 |
| Slc38a11 | 58 | 67 | 63 | 59 | 74 | 62 | 56 | 50 | 41 | 40 | 47 | 66 | 60 | 70 | 55 | 43 | 41 | 54 | 40 | 88 | 74 | 90 | 63 | 48 | 36 | 50 | 110 | 85 | Slc38a11 |
| Slc38a2 | 18676 | 16485 | 16035 | 16517 | 19154 | 18282 | 19130 | 30682 | 26105 | 24677 | 25338 | 27443 | 25715 | 29116 | 17868 | 16718 | 16351 | 16718 | 19543 | 19723 | 19169 | 29494 | 24536 | 26297 | 29065 | 25537 | 28262 | 29145 | Slc38a2 |

| | | | | | | | | | | | | | | | | | | | | | | | | | | | | | |
|---|---|---|---|---|---|---|---|---|---|---|---|---|---|---|---|---|---|---|---|---|---|---|---|---|---|---|---|---|---|
| Slc38a3 | 1772 | 2363 | 2357 | 2160 | 2488 | 2096 | 2010 | 1259 | 1554 | 1672 | 1458 | 1438 | 1555 | 1343 | 1843 | 2176 | 2216 | 2590 | 2299 | 2333 | 2275 | 1711 | 1814 | 1571 | 1731 | 1242 | 1327 | 1197 | Slc38a3 |
| Slc38a4 | 4750 | 3195 | 3529 | 1633 | 996 | 1804 | 771 | 1844 | 2547 | 1904 | 3818 | 1486 | 2783 | 1517 | 4200 | 5108 | 2005 | 2037 | 923 | 1172 | 788 | 1866 | 3787 | 1789 | 1341 | 1891 | 1941 | 1851 | Slc38a4 |
| Slc38a5 | 113 | 209 | 133 | 269 | 533 | 477 | 475 | 455 | 452 | 497 | 450 | 355 | 289 | 296 | 170 | 131 | 163 | 329 | 468 | 437 | 502 | 592 | 469 | 431 | 574 | 203 | 242 | 301 | Slc38a5 |
| Slc38a6 | 1754 | 1508 | 1626 | 1656 | 1393 | 1631 | 1275 | 1606 | 1600 | 1891 | 2328 | 1534 | 1774 | 1663 | 1513 | 2042 | 1586 | 1563 | 1122 | 1245 | 1259 | 1865 | 1779 | 1557 | 1664 | 1455 | 1682 | 1413 | Slc38a6 |
| Slc38a7 | 2568 | 3325 | 2947 | 3453 | 2806 | 2786 | 3142 | 1773 | 2109 | 2306 | 1924 | 2068 | 2038 | 1928 | 2403 | 2919 | 3160 | 3593 | 3412 | 3120 | 3258 | 2366 | 2021 | 2062 | 2464 | 2176 | 2192 | 2182 | Slc38a7 |
| Slc38a8 | 13 | 8 | 3 | 3 | 4 | 5 | 0 | 5 | 7 | 2 | 13 | 39 | 2 | 10 | 4 | 2 | 6 | 3 | 11 | 6 | 5 | 15 | 8 | 2 | 4 | 7 | 7 | 7 | Slc38a8 |
| Slc38a9 | 5037 | 4849 | 4226 | 5263 | 5021 | 5341 | 5330 | 3163 | 2789 | 2825 | 2759 | 2925 | 2931 | 3047 | 5083 | 4818 | 4847 | 5304 | 4821 | 5184 | 5570 | 2944 | 2717 | 2750 | 3037 | 2201 | 3067 | 2991 | Slc38a9 |
| Slc39a1 | 5191 | 5286 | 5081 | 4529 | 3741 | 4009 | 4140 | 4096 | 4128 | 3711 | 4092 | 3930 | 4500 | 4173 | 4750 | 4986 | 4475 | 4092 | 4090 | 4012 | 4034 | 3725 | 4442 | 3970 | 3756 | 4909 | 4541 | 4867 | Slc39a1 |
| Slc39a10 | 9748 | 8299 | 8010 | 7524 | 8818 | 8596 | 9180 | 16153 | 13509 | 11986 | 11307 | 12898 | 13051 | 12902 | 10158 | 8568 | 7935 | 7814 | 8827 | 10588 | 8873 | 13485 | 14012 | 11946 | 12512 | 11474 | 13763 | 13377 | Slc39a10 |
| Slc39a11 | 1100 | 1274 | 979 | 1224 | 1275 | 1431 | 1225 | 1249 | 1301 | 1359 | 1238 | 1228 | 1328 | 1266 | 1205 | 1166 | 1210 | 1420 | 1330 | 1225 | 1242 | 1228 | 1381 | 1338 | 1386 | 1287 | 1141 | 1262 | Slc39a11 |
| Slc39a12 | 47 | 67 | 58 | 29 | 102 | 65 | 42 | 165 | 150 | 146 | 88 | 125 | 186 | 118 | 88 | 65 | 66 | 51 | 51 | 77 | 65 | 165 | 219 | 84 | 104 | 76 | 144 | 132 | Slc39a12 |
| Slc39a13 | 1421 | 1744 | 1643 | 1503 | 1528 | 1506 | 1255 | 1280 | 1649 | 1570 | 1906 | 1397 | 1498 | 1278 | 1407 | 1643 | 1705 | 1595 | 1481 | 1393 | 1269 | 1537 | 1744 | 1487 | 1500 | 1317 | 1384 | 1240 | Slc39a13 |
| Slc39a14 | 3095 | 2845 | 2671 | 2836 | 2608 | 2818 | 2773 | 1668 | 2155 | 2063 | 2377 | 1856 | 2194 | 1855 | 2469 | 3071 | 2892 | 2726 | 2614 | 2639 | 2783 | 1960 | 1901 | 1849 | 2085 | 1557 | 1866 | 1860 | Slc39a14 |
| Slc39a2 | 727 | 962 | 801 | 1363 | 1330 | 1173 | 1293 | 691 | 692 | 768 | 729 | 609 | 551 | 698 | 744 | 738 | 1073 | 1460 | 1305 | 1018 | 1339 | 889 | 621 | 854 | 837 | 535 | 562 | 657 | Slc39a2 |
| Slc39a3 | 1343 | 1887 | 1702 | 2294 | 2648 | 2145 | 2091 | 2717 | 3213 | 3743 | 3148 | 3085 | 2948 | 2962 | 1476 | 1488 | 2104 | 2395 | 2708 | 2064 | 2233 | 3626 | 2959 | 3639 | 3823 | 3417 | 3221 | 2885 | Slc39a3 |
| Slc39a4 | 12 | 17 | 4 | 22 | 13 | 12 | 16 | 13 | 21 | 21 | 17 | 5 | 19 | 11 | 16 | 29 | 5 | 17 | 13 | 18 | 17 | 23 | 18 | 7 | 22 | 9 | 12 | 0 | Slc39a4 |
| Slc39a5 | 10 | 18 | 14 | 5 | 16 | 10 | 10 | 5 | 8 | 7 | 9 | 3 | 7 | 1 | 16 | 10 | 8 | 21 | 5 | 2 | 4 | 5 | 8 | 0 | 4 | 5 | 2 | 5 | Slc39a5 |
| Slc39a6 | 10864 | 10023 | 10321 | 11923 | 13863 | 13007 | 12952 | 11223 | 10590 | 11423 | 11353 | 11317 | 10713 | 11034 | 11381 | 9668 | 11780 | 12378 | 13054 | 13317 | 13297 | 12074 | 10333 | 10939 | 11785 | 11591 | 10689 | 10872 | Slc39a6 |
| Slc39a7 | 120 | 168 | 127 | 162 | 106 | 56 | 141 | 73 | 78 | 131 | 113 | 108 | 76 | 188 | 94 | 50 | 188 | 92 | 66 | 76 | 189 | 65 | 81 | 200 | 104 | 113 | 95 | 169 | Slc39a7 |
| Slc39a8 | 1225 | 1134 | 1480 | 988 | 1022 | 996 | 1040 | 1351 | 1161 | 1203 | 1128 | 1513 | 1799 | 1658 | 1374 | 1320 | 1052 | 966 | 999 | 1085 | 1037 | 1196 | 1278 | 1304 | 1187 | 1503 | 1655 | 2039 | Slc39a8 |
| Slc39a9 | 6993 | 6791 | 5456 | 5783 | 5361 | 5797 | 5910 | 5417 | 5018 | 4612 | 4746 | 5097 | 5029 | 4966 | 6637 | 6138 | 5266 | 5773 | 5471 | 5963 | 6266 | 5036 | 5136 | 4619 | 4844 | 4170 | 5106 | 5277 | Slc39a9 |
| Slc3a1 | 554 | 640 | 549 | 1089 | 779 | 721 | 982 | 142 | 119 | 154 | 102 | 166 | 230 | 166 | 505 | 648 | 744 | 1039 | 893 | 557 | 950 | 192 | 115 | 144 | 183 | 122 | 192 | 189 | Slc3a1 |
| Slc3a2 | 3986 | 4426 | 4617 | 4511 | 4602 | 4326 | 4183 | 4090 | 4376 | 4838 | 4407 | 5095 | 4750 | 5447 | 3935 | 3953 | 4263 | 4582 | 4744 | 3949 | 4626 | 4585 | 4585 | 5090 | 4723 | 5578 | 4967 | 5394 | Slc3a2 |
| Slc40a1 | 1807 | 1635 | 1832 | 1476 | 1321 | 1511 | 1358 | 1607 | 1596 | 1329 | 1725 | 1599 | 1746 | 1718 | 1847 | 2108 | 1341 | 1732 | 1015 | 1415 | 1503 | 1858 | 2148 | 1470 | 1676 | 1612 | 1547 | 1564 | Slc40a1 |
| Slc41a1 | 2633 | 3295 | 3400 | 3295 | 3199 | 2960 | 3076 | 2242 | 2758 | 2472 | 2866 | 2580 | 2548 | 2516 | 2604 | 2991 | 3204 | 3318 | 3435 | 2948 | 3182 | 2697 | 3141 | 2880 | 2775 | 2601 | 2600 | 2467 | Slc41a1 |
| Slc41a2 | 114 | 79 | 82 | 108 | 237 | 154 | 168 | 326 | 391 | 434 | 471 | 227 | 271 | 248 | 131 | 108 | 110 | 167 | 131 | 131 | 146 | 456 | 379 | 324 | 392 | 163 | 231 | 210 | Slc41a2 |
| Slc41a3 | 280 | 324 | 269 | 286 | 271 | 277 | 193 | 115 | 174 | 177 | 188 | 184 | 237 | 158 | 290 | 341 | 315 | 250 | 262 | 219 | 218 | 172 | 212 | 154 | 117 | 155 | 202 | 183 | Slc41a3 |
| Slc43a1 | 991 | 790 | 934 | 592 | 545 | 644 | 572 | 697 | 791 | 796 | 872 | 966 | 976 | 954 | 825 | 900 | 801 | 560 | 627 | 582 | 521 | 503 | 750 | 917 | 628 | 1317 | 1022 | 1042 | Slc43a1 |
| Slc43a2 | 1036 | 1188 | 1411 | 1675 | 2695 | 1955 | 1804 | 2558 | 2806 | 3740 | 3465 | 3108 | 3255 | 2880 | 1229 | 1195 | 1708 | 1991 | 2268 | 1709 | 2022 | 4204 | 2597 | 3399 | 3993 | 2114 | 2804 | 2625 | Slc43a2 |
| Slc43a3 | 1618 | 1252 | 1196 | 742 | 713 | 773 | 574 | 1049 | 1174 | 1053 | 1420 | 1103 | 1422 | 941 | 1358 | 1763 | 866 | 913 | 572 | 611 | 506 | 996 | 1367 | 1058 | 722 | 1343 | 1080 | 1140 | Slc43a3 |
| Slc44a1 | 5233 | 3591 | 4076 | 3760 | 3386 | 3976 | 3676 | 3820 | 3903 | 3444 | 4101 | 2746 | 3109 | 2922 | 4742 | 4836 | 3526 | 4039 | 3398 | 3467 | 3564 | 3389 | 4419 | 3277 | 3022 | 3391 | 2836 | 2922 | Slc44a1 |
| Slc44a2 | 11147 | 11083 | 9775 | 10599 | 9456 | 9898 | 9354 | 5213 | 6160 | 6090 | 6571 | 5332 | 5761 | 5266 | 10052 | 10856 | 10182 | 10513 | 9690 | 8707 | 9285 | 5681 | 6088 | 5828 | 5210 | 5793 | 5165 | 5142 | Slc44a2 |
| Slc44a3 | 75 | 31 | 18 | 51 | 12 | 21 | 10 | 57 | 55 | 52 | 64 | 26 | 20 | 11 | 36 | 40 | 15 | 13 | 28 | 22 | 24 | 32 | 71 | 48 | 9 | 48 | 14 | 8 | Slc44a3 |
| Slc44a4 | 8 | 9 | 6 | 11 | 3 | 3 | 2 | 4 | 7 | 9 | 11 | 5 | 7 | 4 | 11 | 10 | 6 | 4 | 2 | 1 | 2 | 8 | 11 | 10 | 7 | 7 | 0 | 1 | Slc44a4 |
| Slc44a5 | 857 | 942 | 567 | 1100 | 1559 | 1485 | 1385 | 2209 | 1898 | 1654 | 1498 | 1488 | 1698 | 1478 | 1083 | 1231 | 934 | 1147 | 1338 | 1894 | 1632 | 1978 | 1885 | 1427 | 2232 | 765 | 1425 | 1260 | Slc44a5 |
| Slc45a1 | 442 | 678 | 628 | 957 | 1314 | 1029 | 914 | 784 | 906 | 1061 | 880 | 857 | 831 | 793 | 618 | 575 | 899 | 990 | 1146 | 932 | 975 | 1186 | 789 | 939 | 1180 | 723 | 685 | 656 | Slc45a1 |
| Slc45a2 | 17 | 15 | 13 | 2 | 2 | 13 | 1 | 13 | 40 | 21 | 21 | 18 | 20 | 10 | 9 | 8 | 6 | 4 | 2 | 1 | 1 | 5 | 34 | 15 | 5 | 5 | 8 | 6 | Slc45a2 |
| Slc45a3 | 919 | 1271 | 1438 | 1004 | 742 | 772 | 725 | 423 | 326 | 416 | 371 | 631 | 667 | 653 | 913 | 945 | 1160 | 861 | 891 | 808 | 871 | 490 | 456 | 445 | 458 | 378 | 872 | 917 | Slc45a3 |
| Slc45a4 | 3698 | 4285 | 3875 | 4822 | 4592 | 4264 | 4155 | 1713 | 2085 | 2228 | 2299 | 2342 | 2223 | 2238 | 3562 | 4189 | 4428 | 4791 | 4361 | 3388 | 4253 | 2374 | 2038 | 2376 | 2300 | 2165 | 2077 | 2268 | Slc45a4 |
| Slc46a1 | 299 | 342 | 313 | 296 | 335 | 270 | 256 | 396 | 431 | 357 | 398 | 384 | 483 | 403 | 261 | 252 | 360 | 278 | 346 | 342 | 302 | 431 | 452 | 421 | 405 | 499 | 463 | 476 | Slc46a1 |
| Slc46a2 | 0 | 2 | 0 | 0 | 0 | 7 | 0 | 1 | 2 | 3 | 0 | 0 | 0 | 0 | 0 | 0 | 0 | 1 | 2 | 1 | 0 | 3 | 1 | 1 | 1 | 0 | 0 | 0 | Slc46a2 |
| Slc46a3 | 675 | 645 | 668 | 551 | 596 | 556 | 568 | 171 | 152 | 178 | 152 | 129 | 195 | 180 | 638 | 646 | 688 | 518 | 541 | 567 | 516 | 169 | 233 | 184 | 130 | 152 | 171 | 191 | Slc46a3 |
| Slc47a1 | 8 | 10 | 0 | 1 | 6 | 5 | 1 | 10 | 19 | 29 | 14 | 1 | 8 | 14 | 8 | 9 | 1 | 4 | 1 | 5 | 8 | 5 | 9 | 11 | 6 | 0 | 0 | 7 | Slc47a1 |
| Slc47a2 | 5 | 3 | 4 | 0 | 6 | 1 | 0 | 1 | 6 | 3 | 4 | 0 | 0 | 1 | 3 | 6 | 5 | 0 | 1 | 2 | 4 | 3 | 3 | 0 | 0 | 1 | 6 | 0 | Slc47a2 |
| Slc48a1 | 2098 | 2472 | 2247 | 2065 | 2169 | 2211 | 2220 | 1903 | 2208 | 1871 | 1926 | 1988 | 2004 | 1932 | 2127 | 2291 | 2220 | 2222 | 2268 | 2419 | 2079 | 2000 | 2250 | 1941 | 2087 | 2024 | 2049 | 2028 | Slc48a1 |
| Slc49a4 | 2390 | 2505 | 2231 | 2663 | 3547 | 3408 | 3074 | 3425 | 3200 | 3309 | 3061 | 2676 | 2725 | 2544 | 2868 | 2311 | 2519 | 2902 | 3293 | 3232 | 3379 | 3471 | 3087 | 2928 | 3508 | 2590 | 2410 | 2470 | Slc49a4 |
| Slc4a1 | 2245 | 3981 | 3623 | 1282 | 451 | 1022 | 933 | 1231 | 2048 | 1404 | 1083 | 2628 | 8894 | 4415 | 2487 | 3340 | 2017 | 876 | 908 | 1480 | 661 | 688 | 1935 | 1151 | 1031 | 3223 | 4212 | 9863 | Slc4a1 |
| Slc4a10 | 580 | 640 | 323 | 747 | 1618 | 1392 | 1187 | 1542 | 1619 | 1430 | 1215 | 903 | 1110 | 610 | 707 | 792 | 501 | 961 | 1069 | 1227 | 1327 | 1972 | 2040 | 820 | 1326 | 355 | 474 | 437 | Slc4a10 |
| Slc4a11 | 356 | 437 | 416 | 561 | 369 | 381 | 391 | 106 | 130 | 167 | 122 | 162 | 128 | 126 | 276 | 333 | 394 | 432 | 452 | 390 | 386 | 130 | 157 | 156 | 150 | 171 | 117 | 108 | Slc4a11 |
| Slc4a1ap | 2765 | 2647 | 2949 | 2534 | 2428 | 2426 | 2475 | 3374 | 2873 | 3004 | 2803 | 3074 | 2969 | 3261 | 2781 | 2417 | 2724 | 2447 | 2442 | 2575 | 2566 | 2841 | 2825 | 3008 | 3304 | 3210 | 3235 | 3210 | Slc4a1ap |
| Slc4a2 | 5410 | 7011 | 6935 | 6765 | 5968 | 5516 | 5282 | 3208 | 4154 | 3752 | 4030 | 4240 | 4199 | 3285 | 5342 | 6577 | 7355 | 6544 | 6515 | 5157 | 5389 | 3494 | 4570 | 4035 | 3287 | 4013 | 3727 | 3597 | Slc4a2 |
| Slc4a3 | 1281 | 1405 | 1529 | 1582 | 2181 | 1857 | 1458 | 3445 | 4477 | 4809 | 4969 | 4275 | 4314 | 3838 | 1201 | 1523 | 1640 | 1807 | 1927 | 1530 | 1652 | 4919 | 4308 | 4497 | 4664 | 3835 | 3782 | 3589 | Slc4a3 |
| Slc4a4 | 902 | 722 | 867 | 668 | 1048 | 950 | 743 | 999 | 1185 | 1026 | 1322 | 991 | 1277 | 1036 | 898 | 920 | 794 | 872 | 768 | 713 | 865 | 1275 | 1378 | 856 | 1132 | 802 | 1102 | 968 | Slc4a4 |
| Slc4a5 | 255 | 404 | 85 | 114 | 555 | 301 | 117 | 459 | 728 | 658 | 550 | 673 | 1116 | 127 | 262 | 607 | 141 | 144 | 118 | 181 | 172 | 800 | 1298 | 159 | 291 | 110 | 129 | 165 | Slc4a5 |
| Slc4a7 | 9983 | 6700 | 6144 | 8315 | 8308 | 10005 | 10075 | 11771 | 9699 | 9255 | 9377 | 11549 | 11007 | 11841 | 9558 | 7310 | 7052 | 7699 | 8150 | 9519 | 9416 | 10562 | 8801 | 9346 | 11078 | 7608 | 12358 | 11193 | Slc4a7 |
| Slc4a8 | 5364 | 5365 | 2760 | 5801 | 8354 | 8240 | 8247 | 4879 | 5319 | 5674 | 4845 | 4315 | 4405 | 4568 | 5774 | 5087 | 4164 | 6644 | 7093 | 7502 | 9013 | 5658 | 4216 | 3908 | 6265 | 2403 | 4565 | 3599 | Slc4a8 |
| Slc4a9 | 10 | 13 | 10 | 0 | 5 | 9 | 3 | 16 | 18 | 4 | 21 | 4 | 23 | 7 | 11 | 10 | 6 | 13 | 9 | 4 | 5 | 18 | 19 | 11 | 25 | 2 | 16 | 8 | Slc4a9 |
| Slc50a1 | 923 | 1027 | 1040 | 992 | 970 | 881 | 857 | 896 | 1096 | 1095 | 1135 | 819 | 1077 | 859 | 930 | 1076 | 1041 | 1024 | 1035 | 966 | 874 | 902 | 1155 | 992 | 930 | 1016 | 939 | 884 | Slc50a1 |
| Slc51a | 1 | 0 | 0 | 0 | 1 | 0 | 0 | 0 | 1 | 0 | 6 | 8 | 0 | 1 | 1 | 0 | 0 | 2 | 0 | 3 | 0 | 0 | 1 | 2 | 3 | 4 | 1 | 0 | Slc51a |
| Slc51b | 0 | 0 | 0 | 1 | 0 | 0 | 0 | 0 | 0 | 0 | 0 | 0 | 0 | 0 | 0 | 0 | 0 | 0 | 0 | 0 | 0 | 0 | 0 | 0 | 0 | 0 | 0 | 0 | Slc51b |
| Slc52a2 | 1609 | 2184 | 1951 | 1946 | 1843 | 1946 | 1731 | 1064 | 1333 | 1187 | 1182 | 1448 | 1347 | 1239 | 1671 | 1878 | 1934 | 1948 | 1982 | 1798 | 1672 | 1210 | 1274 | 1212 | 1160 | 1354 | 1300 | 1257 | Slc52a2 |
| Slc52a3 | 93 | 148 | 104 | 99 | 76 | 95 | 61 | 33 | 65 | 36 | 53 | 34 | 77 | 50 | 111 | 128 | 53 | 72 | 84 | 56 | 71 | 50 | 80 | 56 | 46 | 58 | 55 | 53 | Slc52a3 |
| Slc5a1 | 2 | 5 | 0 | 0 | 7 | 0 | 1 | 7 | 9 | 10 | 5 | 0 | 1 | 2 | 1 | 5 | 0 | 4 | 0 | 0 | 0 | 12 | 8 | 1 | 1 | 0 | 0 | 0 | Slc5a1 |
| Slc5a10 | 20 | 16 | 10 | 22 | 41 | 36 | 20 | 0 | 2 | 0 | 0 | 0 | 3 | 4 | 15 | 30 | 14 | 26 | 21 | 24 | 19 | 5 | 0 | 3 | 5 | 9 | 0 | 1 | Slc5a10 |
| Slc5a11 | 8 | 4 | 10 | 3 | 3 | 2 | 4 | 9 | 7 | 2 | 12 | 7 | 1 | 10 | 4 | 4 | 5 | 4 | 12 | 4 | 8 | 10 | 9 | 8 | 5 | 2 | 7 | 7 | Slc5a11 |
| Slc5a12 | 2 | 1 | 0 | 0 | 0 | 0 | 0 | 0 | 0 | 0 | 0 | 0 | 7 | 0 | 1 | 0 | 0 | 0 | 0 | 0 | 0 | 3 | 4 | 0 | 0 | 0 | 2 | 0 | Slc5a12 |
| Slc5a2 | 1 | 1 | 0 | 0 | 0 | 4 | 1 | 0 | 2 | 1 | 3 | 4 | 1 | 11 | 1 | 2 | 0 | 2 | 1 | 1 | 0 | 0 | 1 | 1 | 1 | 1 | 0 | 6 | Slc5a2 |
| Slc5a3 | 3109 | 2782 | 3854 | 3774 | 4601 | 3878 | 3918 | 3814 | 3600 | 3782 | 3956 | 3352 | 3424 | 2909 | 3451 | 3593 | 4293 | 4468 | 4216 | 3748 | 3823 | 5583 | 4060 | 3577 | 3815 | 3039 | 2974 | 3095 | Slc5a3 |
| Slc5a4a | 2 | 1 | 2 | 3 | 5 | 6 | 3 | 5 | 5 | 6 | 6 | 4 | 0 | 2 | 3 | 3 | 3 | 2 | 2 | 6 | 1 | 5 | 3 | 8 | 9 | 0 | 1 | 5 | Slc5a4a |
| Slc5a4b | 0 | 1 | 0 | 0 | 0 | 4 | 0 | 1 | 4 | 4 | 0 | 0 | 0 | 0 | 1 | 0 | 0 | 0 | 1 | 0 | 0 | 0 | 1 | 3 | 2 | 4 | 2 | 0 | Slc5a4b |
| Slc5a5 | 127 | 217 | 189 | 235 | 200 | 152 | 116 | 49 | 53 | 51 | 72 | 71 | 69 | 63 | 98 | 178 | 182 | 209 | 161 | 135 | 115 | 87 | 54 | 83 | 54 | 69 | 86 | 59 | Slc5a5 |
| Slc5a6 | 1080 | 1145 | 1184 | 1227 | 1264 | 1174 | 1063 | 1202 | 1298 | 1358 | 1348 | 1157 | 1264 | 1143 | 1120 | 1142 | 1123 | 1163 | 1222 | 1190 | 1059 | 1579 | 1459 | 1151 | 1396 | 1123 | 1110 | 1015 | Slc5a6 |
| Slc5a7 | 168 | 96 | 138 | 132 | 421 | 241 | 160 | 553 | 489 | 602 | 597 | 276 | 585 | 245 | 240 | 168 | 180 | 166 | 142 | 207 | 147 | 633 | 495 | 246 | 344 | 192 | 361 | 285 | Slc5a7 |
| Slc5a8 | 12 | 3 | 0 | 4 | 8 | 2 | 3 | 20 | 9 | 8 | 5 | 3 | 15 | 3 | 7 | 9 | 0 | 10 | 2 | 7 | 4 | 2 | 17 | 5 | 6 | 11 | 0 | 6 | Slc5a8 |
| Slc5a9 | 6 | 2 | 0 | 2 | 1 | 2 | 2 | 12 | 22 | 3 | 13 | 1 | 1 | 0 | 0 | 2 | 1 | 0 | 1 | 0 | 0 | 8 | 12 | 7 | 0 | 8 | 3 | 1 | Slc5a9 |
| Slc66a1 | 1368 | 1653 | 1501 | 1790 | 1576 | 1533 | 1382 | 669 | 704 | 772 | 851 | 723 | 814 | 714 | 1259 | 1469 | 1559 | 1859 | 1561 | 1406 | 1438 | 812 | 650 | 780 | 793 | 644 | 711 | 689 | Slc66a1 |
| Slc66a2 | 1506 | 1967 | 1436 | 1742 | 1947 | 1768 | 1483 | 1053 | 1200 | 1095 | 1114 | 875 | 1000 | 767 | 1606 | 1929 | 1505 | 1853 | 1814 | 1714 | 1599 | 1185 | 1231 | 930 | 1240 | 923 | 822 | 761 | Slc66a2 |
| Slc6a1 | 3696 | 4872 | 3755 | 7724 | 9679 | 8415 | 9623 | 5805 | 6234 | 7824 | 6283 | 7814 | 5045 | 7756 | 4527 | 3180 | 6684 | 8514 | 9098 | 7303 | 9324 | 9954 | 5086 | 8044 | 9475 | 5405 | 5926 | 5955 | Slc6a1 |
| Slc6a11 | 72 | 99 | 79 | 157 | 718 | 248 | 265 | 429 | 620 | 659 | 669 | 283 | 301 | 251 | 132 | 78 | 110 | 320 | 294 | 211 | 345 | 618 | 845 | 299 | 525 | 172 | 205 | 199 | Slc6a11 |

| | | | | | | | | | | | | | | | | | | | | | | | | | | | | | |
|---|---|---|---|---|---|---|---|---|---|---|---|---|---|---|---|---|---|---|---|---|---|---|---|---|---|---|---|---|---|
| Slc6a12 | 5 | 3 | 10 | 7 | 10 | 15 | 22 | 15 | 17 | 32 | 38 | 20 | 18 | 17 | 8 | 5 | 6 | 25 | 23 | 17 | 19 | 55 | 52 | 21 | 25 | 24 | 31 | 14 | Slc6a12 |
| Slc6a13 | 239 | 237 | 268 | 474 | 992 | 604 | 524 | 524 | 737 | 688 | 927 | 395 | 485 | 403 | 278 | 238 | 324 | 557 | 637 | 882 | 769 | 789 | 1278 | 634 | 718 | 448 | 371 | 309 | Slc6a13 |
| Slc6a14 | 5 | 1 | 0 | 0 | 0 | 2 | 2 | 23 | 21 | 15 | 32 | 14 | 22 | 18 | 7 | 3 | 0 | 6 | 3 | 0 | 0 | 3 | 33 | 30 | 2 | 3 | 21 | 27 | Slc6a14 |
| Slc6a15 | 2907 | 2674 | 2649 | 2942 | 4744 | 4310 | 4140 | 3908 | 3872 | 3624 | 3025 | 3434 | 3633 | 2870 | 3597 | 3137 | 2786 | 3307 | 4034 | 4878 | 4091 | 4435 | 5741 | 2757 | 3724 | 2466 | 2751 | 2668 | Slc6a15 |
| Slc6a16 | 24 | 24 | 12 | 38 | 53 | 30 | 49 | 16 | 17 | 10 | 21 | 8 | 16 | 6 | 15 | 26 | 29 | 37 | 37 | 21 | 34 | 22 | 36 | 13 | 13 | 8 | 7 | 6 | Slc6a16 |
| Slc6a17 | 2021 | 1976 | 1890 | 3308 | 4365 | 3744 | 3805 | 3142 | 3807 | 4146 | 3473 | 2775 | 2160 | 2713 | 1873 | 2168 | 2334 | 4225 | 4010 | 3555 | 4398 | 4385 | 3466 | 4086 | 4834 | 2769 | 1990 | 2133 | Slc6a17 |
| Slc6a19 | 8 | 4 | 1 | 3 | 8 | 6 | 7 | 7 | 18 | 9 | 16 | 11 | 16 | 6 | 11 | 3 | 0 | 2 | 12 | 2 | 3 | 25 | 12 | 13 | 20 | 3 | 12 | 9 | Slc6a19 |
| Slc6a2 | 10 | 71 | 37 | 197 | 188 | 232 | 19 | 199 | 176 | 244 | 303 | 34 | 303 | 9 | 99 | 101 | 206 | 200 | 76 | 133 | 69 | 291 | 317 | 35 | 195 | 39 | 15 | 17 | Slc6a2 |
| Slc6a20a | 38 | 41 | 69 | 59 | 124 | 96 | 77 | 143 | 197 | 216 | 210 | 185 | 220 | 193 | 47 | 54 | 74 | 76 | 103 | 156 | 116 | 249 | 300 | 171 | 215 | 230 | 273 | 155 | Slc6a20a |
| Slc6a20b | 1 | 4 | 12 | 7 | 7 | 6 | 8 | 6 | 1 | 5 | 5 | 5 | 3 | 9 | 3 | 6 | 10 | 1 | 10 | 6 | 5 | 3 | 5 | 3 | 6 | 2 | 15 | 11 | Slc6a20b |
| Slc6a21 | 0 | 0 | 0 | 0 | 0 | 0 | 0 | 0 | 0 | 0 | 1 | 0 | 0 | 0 | 0 | 0 | 0 | 1 | 0 | 0 | 0 | 0 | 0 | 0 | 0 | 0 | 0 | 0 | Slc6a21 |
| Slc6a3 | 21 | 42 | 53 | 36 | 52 | 51 | 60 | 57 | 77 | 27 | 45 | 37 | 28 | 24 | 58 | 54 | 30 | 56 | 78 | 61 | 55 | 45 | 79 | 56 | 58 | 19 | 22 | 21 | Slc6a3 |
| Slc6a4 | 76 | 168 | 104 | 191 | 242 | 233 | 86 | 304 | 276 | 215 | 154 | 109 | 346 | 72 | 216 | 227 | 178 | 213 | 143 | 228 | 137 | 326 | 312 | 64 | 181 | 72 | 136 | 64 | Slc6a4 |
| Slc6a5 | 10 | 69 | 19 | 77 | 857 | 195 | 13 | 928 | 927 | 1131 | 1461 | 12 | 1065 | 18 | 247 | 146 | 74 | 111 | 26 | 54 | 13 | 1233 | 1002 | 15 | 179 | 3 | 22 | 23 | Slc6a5 |
| Slc6a6 | 6380 | 6671 | 7127 | 7305 | 8413 | 8023 | 7241 | 7245 | 8054 | 9941 | 9091 | 10354 | 9889 | 9379 | 5931 | 7270 | 7093 | 7732 | 7986 | 6327 | 7222 | 11025 | 8074 | 10168 | 11392 | 8118 | 9609 | 8937 | Slc6a6 |
| Slc6a7 | 58 | 108 | 94 | 207 | 386 | 323 | 331 | 154 | 146 | 225 | 213 | 109 | 100 | 86 | 83 | 100 | 165 | 284 | 405 | 365 | 407 | 309 | 149 | 168 | 383 | 66 | 86 | 65 | Slc6a7 |
| Slc6a8 | 6172 | 6334 | 7182 | 6977 | 7097 | 6708 | 6766 | 3559 | 3899 | 4039 | 3892 | 4314 | 3904 | 3880 | 5777 | 5923 | 7357 | 7178 | 7649 | 5850 | 6817 | 4288 | 3535 | 4298 | 4225 | 4494 | 3921 | 4017 | Slc6a8 |
| Slc6a9 | 603 | 745 | 929 | 964 | 1144 | 899 | 778 | 753 | 1030 | 1068 | 1215 | 897 | 1181 | 847 | 603 | 753 | 913 | 855 | 1082 | 834 | 844 | 1004 | 1169 | 938 | 863 | 919 | 845 | 1137 | Slc6a9 |
| Slc7a1 | 4479 | 4879 | 5053 | 6403 | 7209 | 6488 | 6422 | 3626 | 4214 | 5006 | 4780 | 4609 | 4086 | 4928 | 4540 | 4590 | 5510 | 6776 | 7018 | 5448 | 7122 | 5295 | 4260 | 5056 | 5923 | 3765 | 4011 | 4593 | Slc7a1 |
| Slc7a10 | 75 | 120 | 91 | 109 | 209 | 195 | 117 | 138 | 221 | 223 | 183 | 92 | 125 | 94 | 61 | 127 | 137 | 168 | 156 | 144 | 141 | 224 | 333 | 122 | 149 | 82 | 48 | 95 | Slc7a10 |
| Slc7a11 | 419 | 223 | 245 | 237 | 444 | 359 | 310 | 1178 | 887 | 945 | 1440 | 981 | 761 | 1074 | 342 | 217 | 229 | 226 | 348 | 462 | 380 | 692 | 1792 | 857 | 966 | 808 | 883 | 966 | Slc7a11 |
| Slc7a13 | 0 | 3 | 0 | 7 | 0 | 0 | 0 | 1 | 4 | 6 | 3 | 1 | 3 | 0 | 1 | 2 | 0 | 3 | 0 | 0 | 0 | 0 | 1 | 1 | 0 | 0 | 0 | 6 | Slc7a13 |
| Slc7a14 | 434 | 560 | 331 | 851 | 1527 | 1055 | 924 | 584 | 668 | 703 | 692 | 437 | 539 | 438 | 611 | 540 | 578 | 923 | 934 | 954 | 1156 | 871 | 683 | 426 | 706 | 183 | 304 | 366 | Slc7a14 |
| Slc7a15 | 0 | 0 | 0 | 0 | 0 | 0 | 0 | 0 | 0 | 0 | 0 | 0 | 0 | 0 | 0 | 0 | 0 | 0 | 0 | 0 | 0 | 0 | 0 | 0 | 0 | 0 | 0 | 4 | Slc7a15 |
| Slc7a2 | 3119 | 2066 | 1460 | 1630 | 1657 | 2200 | 1856 | 1357 | 1679 | 1412 | 1976 | 1121 | 1268 | 1231 | 2205 | 2738 | 1575 | 2224 | 1615 | 1769 | 1845 | 1145 | 1656 | 1360 | 1199 | 971 | 1219 | 1130 | Slc7a2 |
| Slc7a3 | 423 | 424 | 504 | 551 | 431 | 399 | 466 | 143 | 153 | 163 | 182 | 179 | 150 | 195 | 386 | 454 | 424 | 594 | 551 | 439 | 455 | 154 | 165 | 203 | 141 | 181 | 144 | 193 | Slc7a3 |
| Slc7a4 | 240 | 339 | 276 | 412 | 629 | 469 | 501 | 333 | 498 | 540 | 430 | 294 | 286 | 246 | 272 | 291 | 336 | 465 | 517 | 442 | 512 | 568 | 493 | 335 | 526 | 269 | 261 | 223 | Slc7a4 |
| Slc7a5 | 5508 | 6691 | 8738 | 7943 | 7082 | 6369 | 6260 | 3085 | 3692 | 4627 | 4266 | 5482 | 4569 | 5164 | 5453 | 6160 | 7475 | 8178 | 7606 | 5220 | 6962 | 4767 | 3744 | 5181 | 5284 | 6307 | 4471 | 4952 | Slc7a5 |
| Slc7a6 | 6936 | 6449 | 6756 | 6575 | 7234 | 7040 | 6978 | 7275 | 6105 | 6619 | 6176 | 6992 | 7225 | 7180 | 6750 | 5944 | 6568 | 6402 | 7391 | 7040 | 6723 | 7010 | 5726 | 6692 | 7272 | 7047 | 7232 | 7255 | Slc7a6 |
| Slc7a6os | 3485 | 3226 | 3447 | 3794 | 3183 | 3246 | 3457 | 3229 | 2881 | 3220 | 3138 | 3593 | 3300 | 3653 | 3227 | 3093 | 3566 | 3563 | 3225 | 3164 | 3381 | 3118 | 3024 | 3371 | 3488 | 4063 | 3416 | 3786 | Slc7a6os |
| Slc7a7 | 144 | 223 | 266 | 263 | 247 | 187 | 217 | 226 | 339 | 281 | 378 | 380 | 337 | 358 | 236 | 179 | 277 | 274 | 256 | 212 | 230 | 391 | 339 | 330 | 406 | 406 | 387 | 443 | Slc7a7 |
| Slc7a8 | 667 | 858 | 763 | 1013 | 1220 | 1048 | 1059 | 558 | 748 | 807 | 760 | 709 | 760 | 635 | 822 | 903 | 874 | 1084 | 1130 | 979 | 1249 | 957 | 937 | 735 | 857 | 566 | 618 | 511 | Slc7a8 |
| Slc7a9 | 3 | 2 | 0 | 5 | 4 | 2 | 1 | 4 | 11 | 8 | 10 | 5 | 7 | 2 | 1 | 6 | 12 | 2 | 2 | 6 | 5 | 10 | 6 | 0 | 7 | 20 | 6 | 7 | Slc7a9 |
| Slc8a1 | 1951 | 1719 | 1174 | 2124 | 4369 | 3541 | 3082 | 8214 | 7364 | 6426 | 6573 | 3946 | 4863 | 3893 | 2657 | 2345 | 1635 | 2584 | 2707 | 3852 | 3562 | 7544 | 6937 | 4143 | 6537 | 2283 | 3768 | 3385 | Slc8a1 |
| Slc8a2 | 3885 | 4956 | 4410 | 6179 | 6262 | 5195 | 5111 | 2159 | 2408 | 3057 | 2653 | 2624 | 2563 | 2605 | 3993 | 3762 | 5933 | 5853 | 6131 | 4361 | 5884 | 2923 | 2211 | 2691 | 2849 | 2425 | 2308 | 2194 | Slc8a2 |
| Slc8a3 | 431 | 422 | 490 | 584 | 883 | 713 | 700 | 725 | 784 | 1085 | 974 | 663 | 658 | 525 | 517 | 463 | 590 | 722 | 679 | 642 | 721 | 1198 | 775 | 758 | 1177 | 405 | 545 | 511 | Slc8a3 |
| Slc8b1 | 1467 | 1937 | 1730 | 1654 | 904 | 1207 | 929 | 269 | 291 | 304 | 359 | 377 | 454 | 288 | 1249 | 2139 | 1499 | 1597 | 1117 | 1088 | 1013 | 302 | 417 | 360 | 319 | 387 | 385 | 320 | Slc8b1 |
| Slc9a1 | 2307 | 3119 | 3138 | 2814 | 2719 | 2551 | 2152 | 1810 | 2460 | 2754 | 2690 | 2848 | 2561 | 2584 | 2196 | 3111 | 2898 | 3055 | 2735 | 1899 | 2180 | 2775 | 2456 | 2736 | 2608 | 2864 | 2461 | 2447 | Slc9a1 |
| Slc9a2 | 27 | 24 | 8 | 26 | 21 | 43 | 10 | 11 | 56 | 38 | 34 | 12 | 11 | 10 | 28 | 22 | 32 | 34 | 20 | 10 | 15 | 45 | 32 | 35 | 41 | 16 | 9 | 24 | Slc9a2 |
| Slc9a3 | 0 | 0 | 0 | 0 | 2 | 0 | 1 | 1 | 2 | 2 | 6 | 0 | 0 | 1 | 0 | 0 | 0 | 0 | 0 | 1 | 0 | 2 | 4 | 0 | 0 | 0 | 1 | 0 | Slc9a3 |
| Slc9a3r1 | 4533 | 5977 | 5089 | 4755 | 3467 | 3396 | 3366 | 1476 | 1698 | 1623 | 1381 | 1281 | 1411 | 1183 | 4478 | 5453 | 4869 | 4338 | 3794 | 3905 | 3465 | 1332 | 1916 | 1447 | 1254 | 1742 | 1278 | 1341 | Slc9a3r1 |
| Slc9a3r2 | 1514 | 1624 | 1612 | 1129 | 995 | 1082 | 664 | 990 | 1414 | 1354 | 1450 | 1257 | 1419 | 982 | 1303 | 1934 | 1224 | 1250 | 933 | 846 | 754 | 1238 | 1570 | 1301 | 1083 | 1473 | 1285 | 1076 | Slc9a3r2 |
| Slc9a4 | 0 | 0 | 0 | 0 | 4 | 0 | 2 | 4 | 5 | 2 | 11 | 0 | 0 | 2 | 0 | 0 | 0 | 0 | 1 | 0 | 0 | 0 | 1 | 0 | 7 | 0 | 2 | 0 | Slc9a4 |
| Slc9a5 | 3488 | 4120 | 3977 | 4604 | 4696 | 4877 | 4362 | 1970 | 2480 | 2594 | 2492 | 2864 | 2734 | 2480 | 3239 | 3925 | 4314 | 4510 | 5094 | 3848 | 4383 | 2613 | 2097 | 2808 | 2814 | 2585 | 2808 | 2402 | Slc9a5 |
| Slc9a6 | 7981 | 7417 | 6875 | 8726 | 10247 | 9801 | 10296 | 8545 | 6916 | 6840 | 6541 | 5849 | 5856 | 6224 | 9260 | 7178 | 7494 | 9579 | 9333 | 10345 | 10373 | 9037 | 6955 | 6056 | 7757 | 4383 | 5692 | 5950 | Slc9a6 |
| Slc9a7 | 300 | 280 | 72 | 247 | 360 | 435 | 400 | 365 | 299 | 292 | 260 | 168 | 206 | 163 | 290 | 274 | 107 | 246 | 313 | 458 | 400 | 206 | 315 | 135 | 248 | 38 | 145 | 120 | Slc9a7 |
| Slc9a8 | 782 | 850 | 806 | 871 | 1060 | 927 | 879 | 2279 | 2739 | 2778 | 2650 | 3093 | 2950 | 2852 | 736 | 796 | 935 | 784 | 983 | 881 | 802 | 2640 | 2480 | 2813 | 2716 | 2717 | 3067 | 2852 | Slc9a8 |
| Slc9a9 | 566 | 600 | 395 | 434 | 653 | 446 | 425 | 718 | 675 | 649 | 644 | 398 | 572 | 369 | 658 | 657 | 408 | 519 | 437 | 465 | 453 | 802 | 638 | 407 | 555 | 420 | 425 | 382 | Slc9a9 |
| Slc9b1 | 16 | 8 | 9 | 5 | 5 | 8 | 10 | 5 | 7 | 9 | 10 | 13 | 9 | 1 | 7 | 19 | 6 | 7 | 7 | 6 | 7 | 2 | 5 | 2 | 5 | 2 | 2 | 11 | Slc9b1 |
| Slc9b2 | 220 | 192 | 191 | 97 | 72 | 94 | 74 | 133 | 119 | 150 | 90 | 143 | 147 | 90 | 133 | 172 | 158 | 84 | 72 | 106 | 55 | 115 | 140 | 131 | 83 | 183 | 172 | 151 | Slc9b2 |
| Slc9c1 | 0 | 0 | 0 | 0 | 0 | 0 | 0 | 0 | 0 | 0 | 0 | 0 | 0 | 0 | 0 | 0 | 0 | 3 | 0 | 0 | 0 | 0 | 0 | 0 | 0 | 0 | 0 | 0 | Slc9c1 |
| Slco1a4 | 5 | 5 | 3 | 18 | 87 | 52 | 57 | 132 | 151 | 133 | 107 | 35 | 59 | 33 | 12 | 4 | 1 | 34 | 71 | 56 | 111 | 182 | 267 | 102 | 160 | 11 | 22 | 25 | Slco1a4 |
| Slco1a5 | 17 | 10 | 10 | 4 | 3 | 9 | 11 | 17 | 46 | 36 | 30 | 27 | 74 | 20 | 15 | 18 | 12 | 4 | 5 | 4 | 7 | 37 | 42 | 25 | 32 | 29 | 31 | 30 | Slco1a5 |
| Slco1a6 | 5 | 6 | 0 | 0 | 2 | 2 | 0 | 5 | 9 | 0 | 3 | 10 | 12 | 0 | 7 | 9 | 0 | 0 | 2 | 0 | 1 | 12 | 11 | 1 | 0 | 0 | 0 | 1 | Slco1a6 |
| Slco1b2 | 6 | 3 | 3 | 10 | 5 | 9 | 2 | 4 | 9 | 4 | 10 | 10 | 6 | 1 | 4 | 16 | 3 | 0 | 0 | 4 | 4 | 15 | 3 | 1 | 2 | 8 | 6 | 5 | Slco1b2 |
| Slco1c1 | 213 | 238 | 152 | 147 | 509 | 317 | 295 | 609 | 653 | 834 | 563 | 559 | 650 | 294 | 243 | 290 | 112 | 259 | 305 | 292 | 393 | 966 | 981 | 448 | 657 | 237 | 231 | 291 | Slco1c1 |
| Slco2a1 | 145 | 112 | 100 | 131 | 172 | 115 | 111 | 311 | 409 | 342 | 460 | 163 | 276 | 194 | 123 | 134 | 103 | 145 | 121 | 56 | 158 | 274 | 439 | 388 | 271 | 296 | 205 | 160 | Slco2a1 |
| Slco2b1 | 204 | 280 | 273 | 306 | 408 | 368 | 296 | 394 | 564 | 565 | 635 | 496 | 621 | 539 | 247 | 307 | 301 | 371 | 333 | 277 | 312 | 638 | 691 | 508 | 635 | 537 | 533 | 428 | Slco2b1 |
| Slco3a1 | 795 | 852 | 765 | 917 | 1266 | 1200 | 1159 | 1695 | 1897 | 1893 | 2009 | 1525 | 1533 | 1575 | 840 | 986 | 823 | 1148 | 1254 | 1162 | 1331 | 1783 | 1721 | 1736 | 2095 | 1369 | 1592 | 1411 | Slco3a1 |
| Slco4a1 | 531 | 858 | 780 | 599 | 557 | 498 | 469 | 199 | 172 | 148 | 176 | 198 | 268 | 194 | 667 | 738 | 771 | 494 | 493 | 528 | 407 | 157 | 126 | 152 | 148 | 236 | 253 | 221 | Slco4a1 |
| Slco4c1 | 213 | 73 | 42 | 111 | 119 | 63 | 60 | 182 | 116 | 165 | 128 | 10 | 46 | 33 | 102 | 93 | 29 | 76 | 71 | 66 | 49 | 58 | 165 | 215 | 29 | 160 | 15 | 21 | Slco4c1 |
| Slco5a1 | 3022 | 2683 | 2400 | 3086 | 4535 | 4243 | 3813 | 9011 | 7480 | 7754 | 6695 | 7728 | 7335 | 7668 | 3609 | 2589 | 3028 | 2933 | 3718 | 4308 | 4012 | 8570 | 6607 | 6260 | 8743 | 4575 | 7438 | 6961 | Slco5a1 |
| Slco6c1 | 2 | 3 | 0 | 2 | 2 | 2 | 9 | 0 | 0 | 2 | 0 | 1 | 1 | 3 | 1 | 1 | 2 | 1 | 0 | 2 | 1 | 2 | 2 | 1 | 2 | 3 | 0 | 0 | Slco6c1 |
| Slf1 | 4950 | 3869 | 4466 | 4409 | 4200 | 4611 | 5502 | 4738 | 4051 | 3664 | 3590 | 4415 | 3603 | 4507 | 5246 | 4386 | 4314 | 3910 | 4328 | 5534 | 4755 | 3992 | 4212 | 4277 | 4200 | 4562 | 4625 | 4584 | Slf1 |
| Slf2 | 9014 | 8448 | 7614 | 8785 | 9206 | 9243 | 9597 | 9303 | 8239 | 8076 | 9608 | 8562 | 8197 | 9061 | 8578 | 8474 | 8428 | 9286 | 8521 | 8860 | 9560 | 9439 | 8262 | 8028 | 9230 | 6539 | 8268 | 8484 | Slf2 |
| Slfn14 | 13 | 15 | 13 | 2 | 0 | 0 | 1 | 18 | 10 | 4 | 1 | 26 | 103 | 32 | 8 | 9 | 5 | 0 | 2 | 4 | 3 | 0 | 16 | 5 | 7 | 41 | 50 | 96 | Slfn14 |
| Slfn2 | 3 | 1 | 3 | 4 | 2 | 2 | 1 | 6 | 17 | 8 | 12 | 9 | 11 | 7 | 1 | 1 | 0 | 1 | 2 | 3 | 0 | 13 | 15 | 5 | 21 | 6 | 8 | 0 | Slfn2 |
| Slfn3 | 12 | 11 | 2 | 9 | 1 | 16 | 2 | 15 | 17 | 16 | 19 | 12 | 22 | 9 | 13 | 12 | 5 | 7 | 5 | 8 | 3 | 5 | 26 | 7 | 6 | 8 | 3 | 9 | Slfn3 |
| Slfn4 | 0 | 1 | 6 | 1 | 0 | 3 | 0 | 10 | 7 | 9 | 13 | 1 | 1 | 0 | 0 | 4 | 0 | 4 | 0 | 0 | 0 | 13 | 5 | 3 | 11 | 1 | 0 | 0 | Slfn4 |
| Slfn5 | 200 | 124 | 146 | 178 | 167 | 166 | 143 | 118 | 105 | 126 | 194 | 100 | 163 | 115 | 146 | 192 | 125 | 132 | 132 | 190 | 148 | 160 | 181 | 160 | 116 | 120 | 167 | 131 | Slfn5 |
| Slfn8 | 0 | 0 | 2 | 0 | 0 | 0 | 0 | 6 | 2 | 0 | 0 | 0 | 0 | 1 | 1 | 0 | 0 | 0 | 1 | 0 | 0 | 0 | 5 | 8 | 0 | 0 | 0 | 1 | Slfn8 |
| Slfn9 | 2089 | 1524 | 1781 | 1235 | 1262 | 1733 | 1435 | 2332 | 2405 | 2093 | 2088 | 2320 | 2447 | 2113 | 1853 | 1979 | 1460 | 1139 | 1422 | 1652 | 1310 | 1419 | 2168 | 2061 | 1891 | 2348 | 2266 | 2176 | Slfn9 |
| Slfnl1 | 1 | 6 | 3 | 11 | 11 | 13 | 1 | 13 | 11 | 5 | 1 | 15 | 10 | 5 | 7 | 3 | 16 | 4 | 8 | 9 | 12 | 3 | 7 | 5 | 5 | 8 | 5 | 1 | Slfnl1 |
| Slirp | 1877 | 1830 | 2037 | 2138 | 1894 | 1604 | 2652 | 2649 | 1949 | 2466 | 2708 | 2482 | 2671 | 3388 | 1973 | 1401 | 2143 | 1718 | 1842 | 2268 | 2278 | 1890 | 2043 | 3163 | 2679 | 2852 | 2664 | 3591 | Slirp |
| Slit1 | 2455 | 4538 | 3498 | 6063 | 8743 | 6268 | 6134 | 4538 | 5254 | 6399 | 5242 | 5378 | 5278 | 4425 | 4342 | 3972 | 5368 | 6725 | 7527 | 5742 | 6747 | 7346 | 4563 | 5696 | 6540 | 2504 | 4227 | 3583 | Slit1 |
| Slit2 | 12221 | 11808 | 10574 | 11698 | 11591 | 12597 | 12538 | 9095 | 8610 | 8062 | 8922 | 8840 | 11263 | 8526 | 13511 | 13302 | 11641 | 11405 | 11106 | 12192 | 12135 | 9194 | 9401 | 7193 | 8412 | 5377 | 10005 | 8901 | Slit2 |
| Slit3 | 1780 | 2173 | 1652 | 883 | 903 | 1292 | 458 | 1215 | 2181 | 1747 | 2192 | 1324 | 2047 | 963 | 1691 | 2654 | 1258 | 1326 | 593 | 468 | 531 | 1856 | 2139 | 1317 | 1341 | 805 | 1461 | 1139 | Slit3 |
| Slitrk1 | 925 | 1021 | 1284 | 1673 | 2140 | 1877 | 1818 | 2053 | 2140 | 2639 | 2418 | 2059 | 2087 | 2096 | 1159 | 878 | 1522 | 1515 | 2025 | 1592 | 1938 | 2876 | 2027 | 2217 | 2982 | 1790 | 1689 | 1982 | Slitrk1 |
| Slitrk2 | 1558 | 1371 | 1295 | 1662 | 2120 | 1929 | 2430 | 2147 | 1943 | 1920 | 2042 | 1619 | 1479 | 1687 | 1652 | 1295 | 1669 | 1690 | 1937 | 2218 | 2306 | 2214 | 2098 | 1778 | 2313 | 1497 | 1385 | 1685 | Slitrk2 |

| | | | | | | | | | | | | | | | | | | | | | | | | | | | | | |
|---|---|---|---|---|---|---|---|---|---|---|---|---|---|---|---|---|---|---|---|---|---|---|---|---|---|---|---|---|---|
| Slitrk3 | 787 | 953 | 859 | 1242 | 2082 | 1658 | 1675 | 1722 | 1567 | 1814 | 1670 | 1071 | 1102 | 1074 | 1014 | 785 | 1355 | 1347 | 1820 | 1567 | 1814 | 2411 | 1423 | 1248 | 1747 | 717 | 903 | 952 | Slitrk3 |
| Slitrk4 | 1453 | 1018 | 883 | 1336 | 1656 | 1688 | 1753 | 1122 | 1009 | 835 | 820 | 770 | 695 | 794 | 1457 | 1228 | 1114 | 1250 | 1487 | 1716 | 1755 | 1123 | 887 | 732 | 933 | 587 | 769 | 801 | Slitrk4 |
| Slitrk5 | 1431 | 1435 | 1564 | 1456 | 2087 | 1839 | 1966 | 2832 | 2482 | 2858 | 2368 | 2523 | 2531 | 2803 | 1569 | 1319 | 1854 | 1587 | 1948 | 1800 | 1866 | 3298 | 2233 | 2549 | 3074 | 2295 | 2786 | 2776 | Slitrk5 |
| Slitrk6 | 760 | 352 | 216 | 256 | 367 | 311 | 185 | 1677 | 1456 | 894 | 945 | 419 | 803 | 247 | 475 | 668 | 272 | 296 | 256 | 400 | 196 | 740 | 1437 | 588 | 372 | 576 | 296 | 298 | Slitrk6 |
| Slk | 14798 | 12024 | 10694 | 11914 | 11475 | 12409 | 11996 | 8934 | 8485 | 7348 | 8382 | 8228 | 8028 | 8413 | 14311 | 13395 | 10790 | 11813 | 10880 | 12861 | 11851 | 7082 | 8413 | 7570 | 7824 | 6478 | 8176 | 7889 | Slk |
| Slmap | 9135 | 6942 | 7920 | 8007 | 8106 | 8511 | 8634 | 6000 | 5537 | 5725 | 5915 | 5965 | 5987 | 6123 | 8375 | 8060 | 7995 | 8010 | 7443 | 7772 | 8147 | 5919 | 5678 | 5632 | 5939 | 5581 | 6211 | 5916 | Slmap |
| Sln | 18 | 1 | 0 | 7 | 1 | 5 | 4 | 31 | 82 | 11 | 454 | 1 | 16 | 2 | 4 | 13 | 0 | 10 | 12 | 5 | 9 | 48 | 345 | 12 | 18 | 13 | 0 | 2 | Sln |
| Slpi | 8 | 10 | 6 | 24 | 15 | 44 | 1 | 26 | 89 | 139 | 225 | 31 | 39 | 17 | 13 | 16 | 3 | 55 | 12 | 0 | 0 | 202 | 62 | 76 | 117 | 24 | 22 | 24 | Slpi |
| Sltm | 8002 | 6928 | 8977 | 8449 | 8991 | 8536 | 9295 | 13015 | 10706 | 12158 | 11354 | 12965 | 12247 | 13773 | 8310 | 7303 | 8712 | 7985 | 8893 | 9024 | 8821 | 12524 | 10403 | 12357 | 13006 | 12352 | 12925 | 13556 | Sltm |
| Slu7 | 5457 | 5702 | 5359 | 5118 | 4671 | 4596 | 5039 | 5768 | 5037 | 5030 | 4672 | 5154 | 4712 | 5563 | 5803 | 5501 | 5215 | 4905 | 4828 | 5311 | 5011 | 5178 | 5009 | 5259 | 5164 | 5214 | 5096 | 5556 | Slu7 |
| Slurp1 | 1 | 0 | 0 | 0 | 0 | 0 | 0 | 2 | 2 | 1 | 3 | 0 | 0 | 0 | 0 | 0 | 0 | 0 | 0 | 0 | 0 | 0 | 4 | 1 | 0 | 0 | 0 | 0 | Slurp1 |
| Slurp2 | 0 | 1 | 0 | 0 | 0 | 0 | 0 | 0 | 0 | 0 | 0 | 0 | 0 | 0 | 0 | 1 | 0 | 0 | 0 | 0 | 2 | 0 | 0 | 0 | 0 | 0 | 0 | 0 | Slurp2 |
| Slx1b | 1426 | 1553 | 1761 | 2139 | 2077 | 1924 | 2166 | 1336 | 1293 | 1512 | 1409 | 1565 | 1414 | 1637 | 1573 | 1341 | 2181 | 2010 | 2254 | 2045 | 2194 | 1477 | 1205 | 1622 | 1583 | 1709 | 1587 | 1593 | Slx1b |
| Slx4 | 3400 | 3994 | 3492 | 4388 | 4277 | 4093 | 3761 | 3331 | 3971 | 4144 | 4114 | 4147 | 3992 | 4055 | 3248 | 3894 | 3772 | 4360 | 4020 | 3463 | 3850 | 4124 | 3673 | 3962 | 4283 | 2795 | 3926 | 3711 | Slx4 |
| Slx4ip | 1876 | 1742 | 2019 | 1758 | 2030 | 1753 | 1871 | 3574 | 2955 | 2952 | 2884 | 2804 | 2851 | 3298 | 2032 | 1750 | 1885 | 1586 | 1858 | 1950 | 1874 | 3322 | 2930 | 2897 | 2900 | 2417 | 3140 | 3067 | Slx4ip |
| Smad1 | 2773 | 2541 | 2968 | 3497 | 3809 | 3425 | 3302 | 4267 | 3974 | 4445 | 4331 | 4511 | 4228 | 4313 | 2763 | 2565 | 3312 | 3218 | 3782 | 2903 | 3408 | 4906 | 4060 | 4339 | 4646 | 4491 | 3934 | 4186 | Smad1 |
| Smad2 | 8329 | 7463 | 6730 | 7634 | 8629 | 8718 | 8460 | 11323 | 9787 | 8955 | 9006 | 9482 | 9394 | 10033 | 9133 | 7231 | 7258 | 7561 | 8207 | 8884 | 8706 | 10101 | 9227 | 8307 | 10208 | 7825 | 10387 | 9376 | Smad2 |
| Smad3 | 4547 | 4201 | 4673 | 3636 | 3827 | 4231 | 2872 | 5796 | 5890 | 5824 | 6558 | 7513 | 7622 | 6640 | 4205 | 4683 | 4155 | 3854 | 3691 | 3193 | 3039 | 5150 | 6242 | 6290 | 5223 | 7920 | 7709 | 7738 | Smad3 |
| Smad4 | 8981 | 8013 | 7453 | 8146 | 8579 | 9098 | 8805 | 8764 | 8725 | 8362 | 8313 | 10051 | 9305 | 9326 | 8280 | 7953 | 7787 | 8067 | 9143 | 8216 | 9096 | 7419 | 8598 | 8884 | 9083 | 8120 | 9892 | 9725 | Smad4 |
| Smad5 | 14819 | 11849 | 11756 | 12413 | 13358 | 13860 | 13974 | 20316 | 19029 | 17021 | 18276 | 19881 | 19426 | 19055 | 13705 | 12639 | 12169 | 12769 | 13403 | 12750 | 13633 | 18092 | 17301 | 18393 | 19833 | 16808 | 19901 | 19300 | Smad5 |
| Smad6 | 657 | 650 | 622 | 531 | 371 | 445 | 270 | 455 | 512 | 423 | 538 | 404 | 519 | 360 | 514 | 650 | 537 | 561 | 345 | 438 | 302 | 356 | 605 | 522 | 332 | 687 | 465 | 446 | Smad6 |
| Smad7 | 746 | 684 | 951 | 845 | 757 | 656 | 561 | 852 | 820 | 887 | 857 | 837 | 906 | 765 | 592 | 734 | 847 | 763 | 736 | 561 | 638 | 897 | 935 | 1004 | 758 | 1055 | 683 | 730 | Smad7 |
| Smad9 | 499 | 384 | 378 | 344 | 407 | 448 | 296 | 376 | 450 | 405 | 518 | 163 | 293 | 161 | 394 | 526 | 415 | 442 | 362 | 346 | 348 | 376 | 487 | 261 | 284 | 241 | 179 | 170 | Smad9 |
| Smagp | 170 | 186 | 194 | 134 | 133 | 106 | 107 | 93 | 100 | 121 | 122 | 92 | 119 | 78 | 161 | 225 | 143 | 176 | 104 | 115 | 107 | 80 | 161 | 106 | 80 | 100 | 96 | 76 | Smagp |
| Smap1 | 6910 | 6731 | 7918 | 7683 | 7878 | 7513 | 7911 | 6295 | 6309 | 6484 | 6014 | 6480 | 6430 | 6601 | 7346 | 6946 | 7313 | 7990 | 8002 | 6916 | 8034 | 7006 | 6261 | 6457 | 6604 | 6804 | 6111 | 6802 | Smap1 |
| Smap2 | 5538 | 5932 | 6161 | 8042 | 8799 | 7747 | 8427 | 4633 | 4884 | 5820 | 4924 | 4820 | 4280 | 4974 | 5968 | 5563 | 7266 | 8335 | 8074 | 6446 | 8685 | 6061 | 4678 | 5506 | 5479 | 4568 | 4166 | 4575 | Smap2 |
| Smarca1 | 6034 | 4267 | 4920 | 4425 | 4874 | 5539 | 4967 | 7805 | 7458 | 6026 | 6563 | 5609 | 6293 | 5813 | 6244 | 5667 | 4538 | 4518 | 4701 | 5601 | 4816 | 7386 | 7512 | 5460 | 5904 | 4988 | 5621 | 5543 | Smarca1 |
| Smarca2 | 10532 | 9489 | 10925 | 13399 | 13909 | 13135 | 14079 | 9238 | 10995 | 11468 | 12255 | 7926 | 7928 | 8306 | 10320 | 10324 | 11454 | 13937 | 13801 | 11877 | 13874 | 11159 | 11830 | 10891 | 10780 | 8134 | 6962 | 7162 | Smarca2 |
| Smarca4 | 16070 | 19606 | 24498 | 25232 | 25939 | 23883 | 22753 | 25753 | 29952 | 39151 | 33474 | 45869 | 36067 | 40939 | 16613 | 17988 | 26416 | 24662 | 28554 | 18940 | 24895 | 37224 | 27911 | 42331 | 41949 | 41512 | 35404 | 40075 | Smarca4 |
| Smarca5 | 19750 | 14982 | 20917 | 18120 | 15884 | 17499 | 19501 | 20855 | 15830 | 16915 | 16123 | 22993 | 20729 | 24297 | 19568 | 16733 | 18756 | 16423 | 17259 | 17886 | 18093 | 16967 | 15865 | 19938 | 18329 | 24478 | 23575 | 24696 | Smarca5 |
| Smarcad1 | 7344 | 5843 | 7313 | 6817 | 7715 | 7395 | 8356 | 15875 | 12319 | 12046 | 11276 | 15069 | 13743 | 15495 | 7705 | 6500 | 7249 | 6376 | 7652 | 8239 | 7700 | 14232 | 12063 | 12841 | 13335 | 13226 | 15077 | 15424 | Smarcad1 |
| Smarcal1 | 2536 | 3136 | 2262 | 2422 | 2647 | 2614 | 2576 | 4190 | 4439 | 3734 | 3874 | 4137 | 3895 | 4178 | 2774 | 2490 | 2371 | 2360 | 2697 | 2916 | 2782 | 3741 | 4198 | 3804 | 3879 | 3914 | 4335 | 4004 | Smarcal1 |
| Smarcb1 | 4508 | 4984 | 5972 | 5600 | 5800 | 5413 | 5099 | 7021 | 8251 | 9085 | 8245 | 10110 | 8917 | 9799 | 4538 | 4789 | 6417 | 5547 | 6530 | 4684 | 5440 | 9082 | 7733 | 10014 | 9691 | 11490 | 9002 | 9927 | Smarcb1 |
| Smarcc1 | 18919 | 16605 | 20419 | 17617 | 17220 | 17715 | 17820 | 23146 | 22652 | 24524 | 22427 | 29501 | 25952 | 27815 | 17962 | 17485 | 19543 | 17388 | 18453 | 15320 | 17325 | 24249 | 20942 | 26921 | 24462 | 28680 | 26427 | 28603 | Smarcc1 |
| Smarcc2 | 17144 | 17877 | 19869 | 22497 | 25825 | 23797 | 22847 | 30003 | 30986 | 37498 | 31425 | 37019 | 30379 | 35573 | 17677 | 16488 | 22059 | 22877 | 26338 | 20973 | 24298 | 38160 | 28508 | 39119 | 39969 | 34623 | 31777 | 33272 | Smarcc2 |
| Smarcd1 | 4915 | 6004 | 8104 | 7776 | 8229 | 6932 | 7152 | 9213 | 10869 | 13338 | 11340 | 16553 | 12740 | 14512 | 4999 | 5103 | 8508 | 7201 | 9080 | 6010 | 7519 | 14484 | 9708 | 14942 | 15078 | 17137 | 13109 | 14656 | Smarcd1 |
| Smarcd2 | 2686 | 2762 | 2668 | 2545 | 2307 | 2200 | 2059 | 2264 | 2871 | 2939 | 3166 | 2065 | 2428 | 2096 | 2498 | 2913 | 2602 | 2561 | 2237 | 1912 | 2073 | 1882 | 3374 | 2992 | 1888 | 3225 | 2107 | 2085 | Smarcd2 |
| Smarcd3 | 3353 | 3874 | 4074 | 4958 | 5462 | 4993 | 5259 | 4207 | 4788 | 5820 | 5277 | 6070 | 4111 | 6054 | 3112 | 2865 | 4614 | 4762 | 6108 | 4707 | 5506 | 5761 | 4891 | 6681 | 6360 | 6557 | 5586 | 5777 | Smarcd3 |
| Smarce1 | 18345 | 16453 | 19894 | 17811 | 17874 | 18237 | 20202 | 29441 | 24973 | 27099 | 24956 | 29713 | 27024 | 34287 | 18846 | 14943 | 19028 | 17139 | 19251 | 19818 | 20148 | 26759 | 24118 | 29996 | 29855 | 33762 | 30976 | 34505 | Smarce1 |
| Smbd1 | 0 | 3 | 0 | 0 | 0 | 5 | 0 | 0 | 2 | 6 | 0 | 0 | 0 | 2 | 0 | 0 | 0 | 0 | 0 | 0 | 0 | 0 | 2 | 1 | 0 | 0 | 0 | 1 | Smbd1 |
| Smc1a | 18010 | 16123 | 16674 | 13909 | 13360 | 14768 | 15812 | 23526 | 21021 | 19078 | 18086 | 22778 | 19764 | 24566 | 18155 | 15123 | 15323 | 13008 | 14121 | 18015 | 14931 | 18449 | 19318 | 21038 | 19575 | 23914 | 23937 | 23010 | Smc1a |
| Smc1b | 31 | 22 | 34 | 30 | 26 | 54 | 28 | 51 | 38 | 30 | 51 | 50 | 50 | 35 | 36 | 71 | 39 | 8 | 24 | 38 | 28 | 50 | 50 | 29 | 32 | 57 | 55 | 47 | Smc1b |
| Smc2 | 21679 | 16431 | 15349 | 12901 | 10856 | 13830 | 14972 | 16254 | 13798 | 10628 | 11005 | 17010 | 14410 | 17214 | 20653 | 18143 | 14443 | 12215 | 12494 | 17468 | 13325 | 9586 | 12505 | 12702 | 11778 | 15076 | 18169 | 16962 | Smc2 |
| Smc3 | 26180 | 22562 | 26115 | 24412 | 20728 | 22756 | 26232 | 21162 | 18262 | 18158 | 11065 | 21214 | 19165 | 22872 | 27576 | 24898 | 24809 | 23473 | 21655 | 24889 | 23788 | 18095 | 17133 | 20042 | 18992 | 22434 | 21847 | 21614 | Smc3 |
| Smc4 | 25806 | 18467 | 24783 | 19220 | 15234 | 18993 | 20798 | 21901 | 18821 | 17745 | 15966 | 26714 | 22745 | 25982 | 25996 | 21865 | 21554 | 17926 | 19253 | 23537 | 18790 | 14776 | 16973 | 21572 | 19006 | 29894 | 28081 | 27023 | Smc4 |
| Smc5 | 16318 | 12660 | 13034 | 14104 | 13363 | 14959 | 16652 | 12810 | 10489 | 8932 | 10549 | 11772 | 11313 | 12344 | 16793 | 14612 | 13070 | 13431 | 13493 | 15797 | 15359 | 10144 | 10012 | 10105 | 10284 | 9443 | 12699 | 12294 | Smc5 |
| Smc6 | 16948 | 12748 | 14883 | 14060 | 12666 | 14585 | 15996 | 13335 | 11396 | 10705 | 11134 | 12984 | 11718 | 13900 | 16172 | 14270 | 13412 | 13244 | 13552 | 15898 | 14599 | 10672 | 11222 | 11722 | 11020 | 12396 | 13192 | 13092 | Smc6 |
| Smchd1 | 24401 | 19257 | 14945 | 16827 | 15832 | 19429 | 19646 | 15706 | 13677 | 12201 | 12554 | 13988 | 13380 | 14176 | 23373 | 20133 | 16275 | 16443 | 16518 | 20490 | 19145 | 10948 | 12657 | 11963 | 12491 | 11465 | 14942 | 13454 | Smchd1 |
| Smco1 | 5 | 0 | 0 | 4 | 5 | 4 | 1 | 6 | 1 | 0 | 3 | 1 | 3 | 3 | 9 | 1 | 2 | 3 | 2 | 4 | 2 | 5 | 1 | 2 | 1 | 3 | 2 | 0 | Smco1 |
| Smco2 | 7 | 10 | 6 | 10 | 7 | 6 | 13 | 0 | 0 | 0 | 0 | 0 | 3 | 0 | 7 | 2 | 13 | 8 | 5 | 4 | 6 | 0 | 0 | 0 | 0 | 0 | 0 | 1 | Smco2 |
| Smco3 | 215 | 153 | 180 | 141 | 92 | 153 | 67 | 129 | 122 | 111 | 105 | 84 | 156 | 90 | 222 | 267 | 104 | 177 | 36 | 78 | 41 | 97 | 157 | 97 | 66 | 122 | 74 | 89 | Smco3 |
| Smco4 | 218 | 192 | 307 | 270 | 233 | 229 | 239 | 319 | 400 | 494 | 586 | 487 | 540 | 650 | 175 | 191 | 277 | 229 | 245 | 165 | 224 | 277 | 428 | 623 | 433 | 656 | 485 | 547 | Smco4 |
| Smcr8 | 5253 | 5386 | 3217 | 3734 | 3830 | 4441 | 4093 | 4608 | 4277 | 3831 | 3827 | 4387 | 4107 | 4164 | 4904 | 4643 | 3386 | 3701 | 3854 | 4593 | 4487 | 3770 | 4178 | 3411 | 4203 | 2658 | 4823 | 4302 | Smcr8 |
| Smdt1 | 2531 | 2920 | 3345 | 3333 | 3073 | 3046 | 2935 | 2634 | 2858 | 3445 | 3506 | 3681 | 3525 | 3811 | 2488 | 2361 | 3398 | 3090 | 3122 | 2980 | 2997 | 2892 | 2926 | 3833 | 3705 | 4220 | 3847 | 3915 | Smdt1 |
| Smg1 | 36657 | 34511 | 17932 | 23686 | 23231 | 33280 | 30328 | 26280 | 23389 | 15469 | 18666 | 16937 | 17669 | 17991 | 35273 | 34714 | 19476 | 24569 | 21599 | 34629 | 33583 | 16066 | 21514 | 14032 | 18149 | 11476 | 20231 | 15836 | Smg1 |
| Smg5 | 7763 | 8980 | 7598 | 7790 | 7685 | 8392 | 7773 | 9890 | 10053 | 9939 | 9061 | 12299 | 11220 | 11153 | 7348 | 7460 | 8384 | 7180 | 8626 | 7959 | 7696 | 9874 | 9555 | 9993 | 10840 | 11265 | 12485 | 11895 | Smg5 |
| Smg6 | 6191 | 6532 | 6453 | 7176 | 6509 | 6474 | 6973 | 5499 | 5568 | 5950 | 5696 | 6660 | 6283 | 6652 | 6028 | 6220 | 6934 | 6585 | 7125 | 5752 | 7063 | 6545 | 5047 | 6030 | 6539 | 5341 | 6480 | 6273 | Smg6 |
| Smg7 | 9570 | 9640 | 7796 | 9194 | 9934 | 9946 | 9890 | 11703 | 10931 | 10679 | 10679 | 11801 | 11310 | 12183 | 9728 | 8753 | 8970 | 8912 | 9650 | 9778 | 9642 | 11867 | 10101 | 10676 | 11806 | 9377 | 11929 | 11152 | Smg7 |
| Smg8 | 1817 | 1992 | 1685 | 1685 | 1564 | 1587 | 1601 | 2230 | 2095 | 1960 | 1781 | 2031 | 1915 | 2008 | 1828 | 1826 | 1639 | 1499 | 1540 | 1593 | 1482 | 2025 | 2110 | 1919 | 1976 | 2149 | 2081 | 2086 | Smg8 |
| Smg9 | 504 | 566 | 662 | 615 | 801 | 741 | 625 | 1203 | 1502 | 1654 | 1492 | 1661 | 1549 | 1543 | 489 | 517 | 689 | 665 | 778 | 611 | 633 | 1681 | 1232 | 1513 | 1684 | 1599 | 1484 | 1440 | Smg9 |
| Smgc | 0 | 0 | 0 | 0 | 0 | 0 | 0 | 1 | 0 | 3 | 4 | 0 | 0 | 0 | 7 | 1 | 0 | 0 | 0 | 0 | 0 | 3 | 1 | 0 | 1 | 0 | 0 | 0 | Smgc |
| Smim1 | 287 | 257 | 298 | 234 | 247 | 179 | 121 | 173 | 275 | 224 | 295 | 187 | 227 | 155 | 222 | 336 | 225 | 220 | 122 | 146 | 161 | 199 | 368 | 244 | 149 | 217 | 170 | 185 | Smim1 |
| Smim10l1 | 1783 | 1359 | 2349 | 2145 | 2859 | 2511 | 2644 | 6872 | 5777 | 7287 | 7094 | 6596 | 6848 | 7725 | 2159 | 1698 | 2299 | 2323 | 2463 | 2496 | 2617 | 7267 | 6308 | 7062 | 7198 | 7154 | 6258 | 7588 | Smim10l1 |
| Smim10l2a | 373 | 363 | 307 | 318 | 646 | 451 | 364 | 532 | 728 | 730 | 651 | 495 | 424 | 534 | 369 | 291 | 357 | 332 | 486 | 395 | 394 | 708 | 681 | 509 | 694 | 532 | 395 | 430 | Smim10l2a |
| Smim11 | 902 | 1025 | 1375 | 1031 | 977 | 868 | 1057 | 911 | 867 | 1095 | 1202 | 1073 | 1065 | 1450 | 789 | 755 | 1312 | 916 | 951 | 1044 | 1030 | 759 | 862 | 1484 | 1175 | 1519 | 1332 | 1417 | Smim11 |
| Smim12 | 1160 | 1500 | 1279 | 1328 | 927 | 966 | 1048 | 633 | 704 | 726 | 736 | 845 | 786 | 810 | 984 | 1189 | 1357 | 1075 | 1092 | 1103 | 1040 | 628 | 762 | 833 | 700 | 975 | 754 | 831 | Smim12 |
| Smim13 | 4062 | 3177 | 3503 | 4012 | 5083 | 4661 | 5203 | 4321 | 3906 | 4014 | 4236 | 3809 | 3652 | 3662 | 4030 | 3530 | 4027 | 4769 | 4955 | 4285 | 5006 | 4622 | 3922 | 3955 | 4338 | 3581 | 3543 | 3570 | Smim13 |
| Smim14 | 3753 | 2972 | 2476 | 3909 | 4434 | 4125 | 4450 | 3924 | 3557 | 3832 | 3432 | 3181 | 3120 | 3190 | 4025 | 3166 | 3279 | 3908 | 3938 | 4297 | 4359 | 4328 | 3640 | 3365 | 3990 | 2591 | 2972 | 3116 | Smim14 |
| Smim15 | 6234 | 5197 | 6357 | 5343 | 4435 | 4496 | 5492 | 4628 | 4018 | 3509 | 3827 | 4259 | 4251 | 4578 | 6956 | 6090 | 5867 | 5235 | 4697 | 6171 | 5277 | 3679 | 4206 | 4240 | 3712 | 4985 | 4546 | 5114 | Smim15 |
| Smim17 | 196 | 216 | 251 | 367 | 488 | 381 | 434 | 285 | 285 | 326 | 295 | 144 | 225 | 212 | 249 | 196 | 264 | 421 | 458 | 406 | 422 | 338 | 265 | 227 | 319 | 152 | 195 | 195 | Smim17 |
| Smim18 | 54 | 82 | 38 | 99 | 226 | 155 | 189 | 557 | 454 | 449 | 415 | 352 | 457 | 450 | 113 | 72 | 118 | 121 | 177 | 203 | 159 | 576 | 462 | 471 | 590 | 282 | 399 | 386 | Smim18 |
| Smim19 | 774 | 723 | 948 | 712 | 707 | 708 | 790 | 788 | 794 | 803 | 904 | 933 | 855 | 951 | 708 | 666 | 743 | 649 | 715 | 658 | 735 | 839 | 796 | 822 | 791 | 978 | 816 | 962 | Smim19 |
| Smim20 | 882 | 999 | 798 | 711 | 785 | 772 | 902 | 1089 | 1011 | 930 | 945 | 1020 | 1074 | 994 | 843 | 822 | 696 | 753 | 845 | 866 | 802 | 867 | 966 | 874 | 917 | 957 | 1181 | 1005 | Smim20 |
| Smim22 | 9 | 1 | 0 | 2 | 13 | 7 | 6 | 13 | 21 | 29 | 15 | 14 | 18 | 4 | 5 | 17 | 14 | 2 | 5 | 2 | 3 | 5 | 26 | 19 | 1 | 5 | 3 | 2 | Smim22 |
| Smim24 | 121 | 181 | 119 | 174 | 142 | 153 | 121 | 105 | 129 | 90 | 131 | 111 | 102 | 136 | 131 | 156 | 152 | 176 | 139 | 111 | 133 | 58 | 131 | 140 | 70 | 181 | 107 | 153 | Smim24 |
| Smim26 | 281 | 276 | 378 | 393 | 412 | 299 | 474 | 390 | 334 | 440 | 430 | 383 | 405 | 573 | 264 | 190 | 387 | 317 | 316 | 483 | 474 | 312 | 403 | 476 | 442 | 445 | 422 | 515 | Smim26 |
| Smim27 | 97 | 84 | 72 | 103 | 62 | 66 | 98 | 127 | 97 | 142 | 113 | 156 | 103 | 247 | 79 | 41 | 75 | 64 | 85 | 61 | 103 | 63 | 105 | 149 | 125 | 150 | 163 | 218 | Smim27 |
| Smim3 | 370 | 282 | 368 | 200 | 239 | 221 | 171 | 374 | 430 | 309 | 346 | 387 | 501 | 366 | 338 | 347 | 287 | 196 | 203 | 235 | 174 | 222 | 452 | 371 | 236 | 566 | 399 | 438 | Smim3 |
| Smim31 | 0 | 0 | 0 | 0 | 0 | 0 | 0 | 0 | 1 | 0 | 2 | 0 | 0 | 1 | 0 | 1 | 0 | 0 | 0 | 1 | 0 | 0 | 1 | 0 | 0 | 0 | 0 | 1 | Smim31 |

| | | | | | | | | | | | | | | | | | | | | | | | | | | | | | |
|---|---|---|---|---|---|---|---|---|---|---|---|---|---|---|---|---|---|---|---|---|---|---|---|---|---|---|---|---|---|
| Smim38 | 60 | 75 | 97 | 76 | 72 | 74 | 41 | 39 | 49 | 52 | 55 | 56 | 75 | 29 | 41 | 95 | 105 | 95 | 57 | 35 | 60 | 52 | 28 | 51 | 43 | 47 | 45 | 43 | Smim38 |
| Smim4 | 582 | 653 | 601 | 680 | 651 | 548 | 807 | 394 | 335 | 412 | 307 | 382 | 408 | 557 | 485 | 488 | 664 | 625 | 651 | 799 | 830 | 247 | 313 | 494 | 390 | 410 | 407 | 484 | Smim4 |
| Smim40 | 20 | 14 | 7 | 20 | 15 | 21 | 18 | 1 | 3 | 6 | 6 | 3 | 9 | 5 | 8 | 15 | 21 | 17 | 22 | 20 | 23 | 8 | 5 | 5 | 6 | 3 | 0 | 6 | Smim40 |
| Smim40-ps | 8 | 3 | 2 | 1 | 2 | 4 | 3 | 2 | 1 | 2 | 0 | 0 | 0 | 0 | 1 | 3 | 0 | 0 | 5 | 3 | 4 | 0 | 0 | 1 | 1 | 0 | 5 | 1 | Smim40-ps |
| Smim41 | 15 | 42 | 17 | 33 | 19 | 30 | 28 | 10 | 8 | 14 | 13 | 5 | 10 | 11 | 26 | 15 | 35 | 28 | 17 | 13 | 27 | 25 | 12 | 3 | 12 | 0 | 2 | 11 | Smim41 |
| Smim43 | 69 | 60 | 83 | 46 | 63 | 50 | 91 | 83 | 63 | 46 | 69 | 102 | 46 | 109 | 79 | 37 | 90 | 48 | 73 | 70 | 52 | 99 | 60 | 55 | 63 | 117 | 156 | 145 | Smim43 |
| Smim5 | 9 | 11 | 5 | 8 | 13 | 12 | 6 | 20 | 36 | 30 | 69 | 26 | 35 | 13 | 8 | 19 | 10 | 4 | 4 | 12 | 5 | 27 | 74 | 11 | 8 | 22 | 14 | 12 | Smim5 |
| Smim6 | 1 | 3 | 0 | 0 | 2 | 1 | 1 | 2 | 1 | 0 | 6 | 0 | 1 | 0 | 4 | 3 | 0 | 0 | 0 | 1 | 4 | 2 | 5 | 2 | 1 | 2 | 2 | 4 | Smim6 |
| Smim7 | 6361 | 5993 | 5609 | 5753 | 5545 | 5936 | 6251 | 7990 | 7166 | 6507 | 6649 | 7073 | 6628 | 6993 | 6727 | 5175 | 5836 | 5498 | 5892 | 6484 | 6067 | 7220 | 7345 | 6622 | 6918 | 7625 | 7254 | 7428 | Smim7 |
| Smim8 | 621 | 466 | 580 | 642 | 637 | 670 | 809 | 918 | 828 | 939 | 760 | 941 | 953 | 1060 | 764 | 491 | 679 | 555 | 605 | 804 | 723 | 866 | 889 | 929 | 936 | 1032 | 1047 | 993 | Smim8 |
| Smim9 | 3 | 2 | 13 | 8 | 0 | 3 | 3 | 7 | 6 | 14 | 3 | 4 | 9 | 10 | 8 | 5 | 0 | 2 | 5 | 6 | 3 | 5 | 4 | 1 | 5 | 3 | 12 | 4 | Smim9 |
| Smlr1 | 2 | 1 | 2 | 0 | 4 | 1 | 2 | 0 | 2 | 1 | 1 | 3 | 3 | 0 | 1 | 4 | 0 | 0 | 12 | 3 | 6 | 0 | 4 | 0 | 5 | 1 | 1 | 1 | Smlr1 |
| Smn1 | 2332 | 2580 | 3004 | 2508 | 1826 | 1849 | 2191 | 1871 | 1776 | 2139 | 1891 | 2494 | 2287 | 2554 | 2190 | 2315 | 2625 | 2030 | 2118 | 1936 | 2193 | 2114 | 1733 | 2222 | 2117 | 2747 | 2377 | 2707 | Smn1 |
| Smndc1 | 4327 | 3217 | 4873 | 4365 | 3830 | 3960 | 4470 | 4495 | 3760 | 4476 | 4070 | 5364 | 4601 | 5510 | 4133 | 3414 | 4328 | 4022 | 4278 | 3578 | 4346 | 4720 | 3775 | 4815 | 4622 | 5956 | 4993 | 5842 | Smndc1 |
| Smo | 6180 | 6789 | 7069 | 4809 | 4228 | 5195 | 4282 | 7603 | 8691 | 7847 | 8487 | 10098 | 10258 | 9338 | 5571 | 6252 | 6258 | 4664 | 5205 | 4239 | 4120 | 7264 | 8967 | 9143 | 7426 | 12395 | 10669 | 10456 | Smo |
| Smoc1 | 3324 | 3556 | 4607 | 3949 | 3841 | 3486 | 2196 | 2321 | 2595 | 2741 | 2709 | 2607 | 3725 | 2475 | 3282 | 4438 | 4792 | 3642 | 3377 | 2304 | 2891 | 3439 | 2753 | 2516 | 2582 | 2043 | 2920 | 2767 | Smoc1 |
| Smoc2 | 1940 | 1419 | 978 | 826 | 209 | 1027 | 75 | 1575 | 3494 | 2128 | 4493 | 1084 | 2256 | 961 | 1626 | 1977 | 869 | 1268 | 198 | 118 | 65 | 2022 | 4040 | 1871 | 1259 | 1117 | 1275 | 1027 | Smoc2 |
| Smox | 1775 | 1875 | 1947 | 2036 | 2269 | 2043 | 1840 | 1939 | 2344 | 2559 | 2412 | 2573 | 2363 | 2208 | 1534 | 1905 | 2070 | 2030 | 2367 | 1726 | 1918 | 2249 | 2214 | 2589 | 2633 | 2888 | 2432 | 2059 | Smox |
| Smpd1 | 2763 | 3284 | 2531 | 2966 | 3363 | 3504 | 2883 | 2651 | 2932 | 3131 | 2651 | 2723 | 2603 | 2365 | 2771 | 2995 | 2855 | 3142 | 3249 | 3257 | 3061 | 2625 | 3070 | 2618 | 2729 | 2586 | 2642 | 2497 | Smpd1 |
| Smpd2 | 2460 | 2867 | 2535 | 2407 | 2014 | 2101 | 2130 | 1540 | 1654 | 1607 | 1381 | 1690 | 1504 | 1619 | 2333 | 2313 | 2192 | 2366 | 2268 | 2331 | 2067 | 1422 | 1658 | 1616 | 1410 | 2115 | 1705 | 1716 | Smpd2 |
| Smpd3 | 3310 | 4781 | 4881 | 6175 | 7257 | 5718 | 6166 | 3894 | 4361 | 5501 | 4342 | 4540 | 3567 | 4413 | 4017 | 3740 | 6391 | 6504 | 7130 | 5383 | 6697 | 6059 | 3763 | 5082 | 6201 | 3561 | 3583 | 3942 | Smpd3 |
| Smpd4 | 5736 | 5857 | 5008 | 4596 | 4279 | 4411 | 4856 | 5528 | 5890 | 5533 | 5283 | 6311 | 5999 | 6247 | 4952 | 5373 | 4813 | 4217 | 4788 | 4942 | 4602 | 5297 | 5333 | 6030 | 5893 | 7358 | 6328 | 6534 | Smpd4 |
| Smpd5 | 45 | 35 | 28 | 30 | 37 | 44 | 21 | 46 | 100 | 71 | 65 | 65 | 62 | 48 | 32 | 61 | 34 | 21 | 30 | 33 | 20 | 70 | 80 | 61 | 68 | 89 | 51 | 40 | Smpd5 |
| Smpdl3a | 573 | 610 | 483 | 468 | 702 | 526 | 513 | 949 | 929 | 726 | 718 | 634 | 849 | 575 | 789 | 674 | 538 | 530 | 526 | 621 | 552 | 846 | 954 | 598 | 684 | 634 | 662 | 634 | Smpdl3a |
| Smpdl3b | 48 | 33 | 53 | 21 | 15 | 55 | 32 | 129 | 130 | 113 | 126 | 113 | 116 | 132 | 32 | 42 | 27 | 46 | 38 | 51 | 25 | 80 | 151 | 148 | 144 | 190 | 122 | 153 | Smpdl3b |
| Smpx | 7 | 0 | 0 | 0 | 2 | 3 | 0 | 20 | 28 | 1 | 261 | 0 | 0 | 3 | 5 | 3 | 0 | 3 | 0 | 5 | 0 | 23 | 229 | 2 | 5 | 0 | 0 | 2 | Smpx |
| Smr2l | 0 | 0 | 0 | 0 | 0 | 0 | 0 | 0 | 0 | 3 | 0 | 0 | 0 | 0 | 0 | 0 | 0 | 0 | 0 | 0 | 0 | 0 | 0 | 0 | 0 | 0 | 0 | 0 | Smr2l |
| Sms | 7309 | 6031 | 7495 | 7357 | 8668 | 8021 | 9284 | 9607 | 8179 | 8949 | 7905 | 9204 | 8763 | 9958 | 7752 | 6809 | 7616 | 7542 | 8228 | 9052 | 8671 | 10176 | 8352 | 9034 | 9706 | 9211 | 8951 | 9956 | Sms |
| Smtn | 2644 | 3697 | 3537 | 3146 | 2389 | 2552 | 1881 | 1469 | 2025 | 2116 | 2790 | 2552 | 2738 | 2150 | 2574 | 3780 | 3327 | 3067 | 2582 | 1830 | 2041 | 1917 | 2377 | 2353 | 2071 | 2619 | 2398 | 2239 | Smtn |
| Smtnl1 | 1 | 4 | 2 | 3 | 3 | 0 | 2 | 5 | 2 | 2 | 1 | 0 | 1 | 3 | 0 | 0 | 0 | 0 | 4 | 0 | 1 | 3 | 6 | 3 | 0 | 1 | 0 | 5 | Smtnl1 |
| Smtnl2 | 852 | 888 | 1014 | 771 | 507 | 589 | 455 | 249 | 426 | 386 | 612 | 179 | 271 | 193 | 702 | 1112 | 870 | 778 | 503 | 386 | 400 | 322 | 590 | 325 | 167 | 349 | 156 | 172 | Smtnl2 |
| Smu1 | 11480 | 10351 | 9849 | 8413 | 7264 | 7992 | 8861 | 9624 | 8050 | 6775 | 6806 | 8662 | 7791 | 8441 | 11151 | 10064 | 9378 | 8207 | 8429 | 10332 | 8415 | 7085 | 7991 | 7533 | 7473 | 9310 | 8914 | 8945 | Smu1 |
| Smug1 | 1607 | 1587 | 1615 | 1585 | 2225 | 1732 | 1747 | 2483 | 2455 | 2696 | 3413 | 2599 | 2381 | 2618 | 1770 | 1368 | 1860 | 1562 | 1839 | 1757 | 1770 | 2578 | 2856 | 2519 | 2693 | 2528 | 2509 | 2749 | Smug1 |
| Smurf1 | 3899 | 4285 | 5137 | 5706 | 6943 | 6134 | 5688 | 4632 | 5404 | 6176 | 6053 | 6090 | 5608 | 5519 | 3745 | 3825 | 5565 | 5939 | 7065 | 4829 | 6071 | 6909 | 5290 | 5954 | 6928 | 4933 | 5531 | 5533 | Smurf1 |
| Smurf2 | 8483 | 7508 | 6887 | 7300 | 7185 | 7454 | 7345 | 9813 | 8566 | 7731 | 7816 | 8039 | 8094 | 8527 | 8296 | 7794 | 6581 | 7436 | 7102 | 8179 | 7832 | 8071 | 8564 | 7480 | 8324 | 6979 | 8311 | 8420 | Smurf2 |
| Smyd1 | 181 | 119 | 93 | 252 | 153 | 87 | 169 | 143 | 340 | 92 | 1946 | 50 | 161 | 57 | 181 | 138 | 146 | 164 | 139 | 105 | 175 | 241 | 1145 | 79 | 75 | 44 | 51 | 50 | Smyd1 |
| Smyd2 | 1899 | 1841 | 1651 | 1592 | 1762 | 1610 | 1422 | 2070 | 2484 | 2096 | 2323 | 1877 | 1988 | 1749 | 1601 | 1822 | 1702 | 1652 | 1605 | 1493 | 1473 | 1968 | 2419 | 2064 | 2050 | 2226 | 1835 | 1918 | Smyd2 |
| Smyd3 | 1732 | 1534 | 1542 | 1713 | 1868 | 1453 | 1542 | 1136 | 1130 | 1074 | 1069 | 818 | 979 | 915 | 1506 | 1520 | 1461 | 1809 | 1588 | 1395 | 1516 | 1131 | 1126 | 1052 | 1148 | 962 | 908 | 900 | Smyd3 |
| Smyd4 | 839 | 917 | 691 | 715 | 734 | 906 | 849 | 1055 | 999 | 962 | 979 | 1069 | 862 | 1042 | 756 | 850 | 721 | 809 | 760 | 817 | 830 | 849 | 966 | 994 | 951 | 1089 | 1099 | 1066 | Smyd4 |
| Smyd5 | 1961 | 2068 | 2097 | 1819 | 1861 | 1942 | 1677 | 2452 | 2698 | 2612 | 2795 | 2870 | 2999 | 2689 | 1905 | 1959 | 1833 | 1723 | 1923 | 1722 | 1783 | 2789 | 2861 | 2638 | 2790 | 3007 | 2788 | 3026 | Smyd5 |
| Snai1 | 723 | 958 | 701 | 596 | 275 | 471 | 154 | 622 | 779 | 738 | 724 | 416 | 559 | 341 | 542 | 1329 | 560 | 476 | 224 | 167 | 135 | 518 | 907 | 755 | 334 | 777 | 459 | 412 | Snai1 |
| Snai2 | 1225 | 745 | 928 | 454 | 206 | 453 | 151 | 992 | 1384 | 945 | 1275 | 737 | 1081 | 722 | 865 | 1206 | 495 | 540 | 178 | 207 | 137 | 700 | 1450 | 888 | 501 | 941 | 726 | 744 | Snai2 |
| Snai3 | 25 | 10 | 0 | 3 | 6 | 1 | 1 | 4 | 9 | 9 | 17 | 0 | 5 | 0 | 5 | 7 | 3 | 0 | 2 | 8 | 0 | 7 | 25 | 6 | 0 | 8 | 0 | 1 | Snai3 |
| Snap23 | 7328 | 5719 | 7537 | 5749 | 4160 | 5040 | 4944 | 3990 | 3681 | 3628 | 4408 | 4009 | 4671 | 4343 | 6451 | 6748 | 5872 | 5426 | 4597 | 4774 | 4653 | 3663 | 4088 | 4010 | 3549 | 4998 | 4302 | 4697 | Snap23 |
| Snap25 | 5568 | 5403 | 5521 | 8183 | 10611 | 8521 | 10047 | 5283 | 4748 | 5301 | 4691 | 4116 | 4238 | 4457 | 7391 | 5966 | 7136 | 8648 | 8704 | 8372 | 9896 | 7235 | 4739 | 4527 | 5952 | 3215 | 3771 | 3839 | Snap25 |
| Snap29 | 5887 | 5826 | 5070 | 5812 | 5477 | 5627 | 6106 | 4749 | 4116 | 4105 | 3974 | 3729 | 3780 | 3852 | 6265 | 5430 | 5515 | 5965 | 5577 | 5508 | 6113 | 4809 | 4094 | 3957 | 4260 | 3670 | 3699 | 4012 | Snap29 |
| Snap47 | 5842 | 6758 | 4613 | 5951 | 8468 | 7123 | 7787 | 9297 | 8829 | 8163 | 7639 | 6213 | 6408 | 6502 | 6589 | 5912 | 5903 | 7039 | 7904 | 8485 | 8194 | 9245 | 8958 | 7297 | 8159 | 6293 | 6198 | 6086 | Snap47 |
| Snap91 | 4702 | 5256 | 4602 | 8416 | 12579 | 9595 | 10561 | 7984 | 7185 | 8070 | 7353 | 5857 | 5736 | 5588 | 6129 | 4939 | 6397 | 9865 | 10015 | 9650 | 11296 | 10574 | 6717 | 6620 | 9125 | 3843 | 4775 | 4927 | Snap91 |
| Snapc1 | 4894 | 4462 | 4160 | 5486 | 5219 | 5352 | 5766 | 3623 | 3242 | 3098 | 2971 | 3247 | 2887 | 3385 | 5451 | 4739 | 4715 | 5646 | 4913 | 5588 | 5815 | 3811 | 3114 | 3130 | 3420 | 2635 | 3155 | 3090 | Snapc1 |
| Snapc2 | 1807 | 2108 | 2196 | 2040 | 1775 | 1790 | 1998 | 1508 | 1727 | 1676 | 1590 | 1857 | 1762 | 1828 | 1664 | 1899 | 2065 | 1935 | 2223 | 1713 | 1943 | 1771 | 1693 | 1763 | 1807 | 2008 | 1812 | 1907 | Snapc2 |
| Snapc3 | 4637 | 3372 | 2835 | 3378 | 3933 | 4783 | 4644 | 6693 | 5620 | 4489 | 4519 | 5113 | 4834 | 5161 | 4510 | 3748 | 3136 | 3335 | 3984 | 5143 | 4858 | 4908 | 5357 | 4096 | 5112 | 3513 | 5382 | 4588 | Snapc3 |
| Snapc4 | 4497 | 5286 | 4245 | 5306 | 4961 | 5195 | 4840 | 3103 | 3441 | 3466 | 3589 | 3700 | 3497 | 3394 | 4183 | 5230 | 4558 | 5366 | 4915 | 4180 | 5002 | 3589 | 3071 | 3537 | 3750 | 2876 | 3453 | 3375 | Snapc4 |
| Snapc5 | 967 | 856 | 1251 | 1175 | 911 | 706 | 1089 | 1423 | 1033 | 1351 | 1060 | 1412 | 1371 | 2421 | 1089 | 651 | 1132 | 793 | 853 | 921 | 1185 | 1242 | 1080 | 1740 | 1345 | 1692 | 1515 | 2296 | Snapc5 |
| Snapin | 2008 | 1869 | 1949 | 1648 | 1755 | 1760 | 1836 | 2669 | 2487 | 2260 | 2329 | 2475 | 2427 | 2692 | 2235 | 1804 | 1881 | 1532 | 1648 | 1831 | 1749 | 2448 | 2678 | 2446 | 2303 | 2638 | 2497 | 2584 | Snapin |
| Snca | 1037 | 1151 | 1019 | 1446 | 2225 | 1461 | 2150 | 1711 | 1910 | 1659 | 1300 | 912 | 1433 | 1077 | 1396 | 1234 | 1267 | 1655 | 1875 | 2169 | 1951 | 1567 | 1847 | 1307 | 1634 | 988 | 956 | 1311 | Snca |
| Sncaip | 1712 | 1546 | 1358 | 1607 | 2119 | 1919 | 1872 | 3764 | 4218 | 4169 | 3795 | 3285 | 2758 | 2983 | 1705 | 1426 | 1572 | 1736 | 1894 | 1807 | 1755 | 3840 | 4007 | 3589 | 3749 | 2938 | 2602 | 2445 | Sncaip |
| Sncb | 272 | 397 | 459 | 816 | 1145 | 819 | 951 | 238 | 429 | 585 | 443 | 245 | 128 | 250 | 347 | 281 | 712 | 859 | 977 | 678 | 994 | 483 | 408 | 412 | 561 | 217 | 179 | 160 | Sncb |
| Sncg | 855 | 1551 | 769 | 658 | 2365 | 599 | 293 | 2135 | 2478 | 1976 | 2590 | 547 | 2577 | 725 | 1764 | 2076 | 1006 | 1212 | 481 | 719 | 475 | 2775 | 2014 | 491 | 1392 | 141 | 875 | 833 | Sncg |
| Snd1 | 11796 | 13042 | 13227 | 12536 | 10397 | 10270 | 10257 | 7477 | 9531 | 9338 | 9481 | 9860 | 9400 | 9002 | 10653 | 13013 | 13190 | 12205 | 12158 | 8731 | 10860 | 8413 | 9587 | 9609 | 8808 | 11288 | 8952 | 9522 | Snd1 |
| Sned1 | 3944 | 3853 | 3386 | 4468 | 3850 | 4294 | 2073 | 3677 | 5398 | 5656 | 8478 | 4832 | 6606 | 4355 | 3214 | 5809 | 3497 | 4568 | 2094 | 2773 | 2231 | 6386 | 6734 | 5033 | 4599 | 3615 | 4943 | 3580 | Sned1 |
| Snf8 | 2940 | 3204 | 3737 | 3586 | 3273 | 3113 | 3347 | 2518 | 2577 | 3024 | 2955 | 3251 | 3268 | 3525 | 2933 | 3103 | 3599 | 3528 | 3399 | 3124 | 3439 | 2857 | 2647 | 3233 | 3119 | 3373 | 3254 | 3375 | Snf8 |
| Snhg11 | 9959 | 11955 | 12437 | 22619 | 49438 | 30036 | 28981 | 31265 | 28501 | 41573 | 40472 | 17105 | 29452 | 19819 | 19094 | 14808 | 18070 | 29211 | 26734 | 25014 | 30795 | 54864 | 23648 | 21527 | 34972 | 7834 | 16418 | 17610 | Snhg11 |
| Snhg4 | 7 | 16 | 7 | 4 | 13 | 7 | 7 | 7 | 7 | 10 | 10 | 13 | 10 | 6 | 17 | 16 | 17 | 25 | 25 | 14 | 15 | 15 | 6 | 10 | 5 | 16 | 7 | 9 | Snhg4 |
| Snip1 | 2430 | 2710 | 2245 | 2177 | 1805 | 2173 | 1964 | 2056 | 1940 | 1679 | 1699 | 1947 | 1864 | 1831 | 2337 | 2458 | 2111 | 2291 | 1979 | 2089 | 1912 | 1803 | 1933 | 1954 | 1734 | 2350 | 2000 | 1934 | Snip1 |
| Snn | 7848 | 7601 | 7440 | 7470 | 9549 | 9074 | 9650 | 16342 | 15659 | 15417 | 14431 | 16747 | 15707 | 18114 | 8590 | 6455 | 8011 | 7560 | 8821 | 9074 | 9182 | 16088 | 14352 | 16307 | 18066 | 16616 | 16163 | 16709 | Snn |
| Snorc | 1 | 1 | 0 | 0 | 2 | 3 | 1 | 4 | 11 | 10 | 70 | 0 | 0 | 4 | 0 | 0 | 0 | 7 | 0 | 0 | 2 | 3 | 6 | 2 | 3 | 3 | 0 | 0 | Snorc |
| Snph | 653 | 940 | 972 | 1325 | 2157 | 1494 | 1449 | 943 | 1131 | 1426 | 1240 | 790 | 782 | 732 | 844 | 856 | 1204 | 1670 | 1709 | 1421 | 1541 | 1534 | 988 | 925 | 1341 | 488 | 691 | 621 | Snph |
| Snrk | 3932 | 4752 | 4148 | 5080 | 5365 | 5089 | 4738 | 3825 | 4341 | 4526 | 4507 | 4087 | 3851 | 3970 | 3978 | 4405 | 4634 | 5361 | 4938 | 4685 | 4996 | 4405 | 4225 | 4172 | 4719 | 3699 | 3907 | 3808 | Snrk |
| Snrnp200 | 27319 | 28295 | 18166 | 19229 | 17586 | 20919 | 18726 | 27505 | 27029 | 22936 | 21487 | 27004 | 24808 | 25830 | 24885 | 24064 | 19186 | 17896 | 19838 | 21153 | 20468 | 20468 | 24708 | 22048 | 23677 | 18969 | 28775 | 26925 | Snrnp200 |
| Snrnp25 | 1299 | 1684 | 1578 | 1408 | 1019 | 1201 | 1131 | 883 | 987 | 761 | 841 | 974 | 1052 | 1010 | 1315 | 1553 | 1341 | 1182 | 1333 | 1460 | 1116 | 556 | 924 | 938 | 876 | 1322 | 1183 | 1152 | Snrnp25 |
| Snrnp27 | 2515 | 2483 | 2594 | 2465 | 1893 | 2162 | 2522 | 2948 | 2568 | 2667 | 2689 | 2691 | 2425 | 3165 | 2699 | 2305 | 2400 | 2121 | 2064 | 2502 | 2487 | 2373 | 2591 | 2999 | 2844 | 3128 | 2797 | 3261 | Snrnp27 |
| Snrnp35 | 442 | 470 | 719 | 602 | 575 | 529 | 552 | 1022 | 1032 | 1192 | 1090 | 1347 | 1332 | 1581 | 400 | 420 | 743 | 547 | 691 | 682 | 589 | 869 | 992 | 1472 | 1333 | 1488 | 1607 | 1609 | Snrnp35 |
| Snrnp40 | 532 | 166 | 128 | 126 | 108 | 123 | 95 | 204 | 129 | 98 | 112 | 114 | 156 | 114 | 241 | 210 | 118 | 112 | 101 | 179 | 93 | 162 | 104 | 118 | 126 | 158 | 150 | 163 | Snrnp40 |
| Snrnp48 | 8337 | 7235 | 7494 | 7842 | 6622 | 6729 | 7526 | 6890 | 5553 | 5562 | 6143 | 6449 | 6093 | 6628 | 8116 | 7749 | 7124 | 7328 | 6588 | 7486 | 7249 | 5790 | 5642 | 5925 | 6104 | 5884 | 6305 | 6482 | Snrnp48 |
| Snrnp70 | 24641 | 28481 | 33603 | 35469 | 31747 | 31243 | 29109 | 29222 | 30686 | 39223 | 34799 | 41811 | 36585 | 41121 | 23782 | 22936 | 36539 | 32914 | 35004 | 29191 | 30577 | 32891 | 28997 | 44285 | 41703 | 43035 | 40680 | 40745 | Snrnp70 |
| Snrpa | 5819 | 7351 | 6799 | 5800 | 5173 | 5272 | 5787 | 6270 | 6720 | 6691 | 6407 | 7841 | 7251 | 7708 | 5439 | 5811 | 6584 | 5223 | 6216 | 5789 | 5574 | 5528 | 6439 | 7829 | 6830 | 9141 | 7835 | 8079 | Snrpa |
| Snrpa1 | 3879 | 3966 | 3932 | 3565 | 2786 | 2905 | 3610 | 2679 | 2487 | 2738 | 2260 | 2842 | 2814 | 3277 | 3843 | 3439 | 3846 | 3177 | 2929 | 3549 | 3326 | 2114 | 2400 | 2843 | 2371 | 3330 | 2910 | 3612 | Snrpa1 |
| Snrpb | 9161 | 12025 | 11644 | 9956 | 8049 | 8384 | 8736 | 7709 | 8347 | 8078 | 8244 | 10507 | 9664 | 10124 | 8458 | 9607 | 10552 | 8595 | 9551 | 8970 | 8199 | 6308 | 8377 | 9655 | 8757 | 12347 | 11072 | 11112 | Snrpb |
| Snrpb2 | 9037 | 7150 | 8870 | 7644 | 5997 | 6616 | 7705 | 7571 | 6100 | 6023 | 5631 | 7079 | 7046 | 8374 | 9403 | 8052 | 7919 | 6660 | 7028 | 8594 | 7144 | 5800 | 6162 | 6971 | 6203 | 8541 | 7931 | 8880 | Snrpb2 |

| | | | | | | | | | | | | | | | | | | | | | | | | | | | | | |
|---|---|---|---|---|---|---|---|---|---|---|---|---|---|---|---|---|---|---|---|---|---|---|---|---|---|---|---|---|---|
| Snrpc | 4127 | 5263 | 5452 | 4917 | 3791 | 4224 | 4465 | 3505 | 3930 | 3645 | 3861 | 4443 | 4347 | 5667 | 4001 | 4412 | 5107 | 4123 | 4690 | 4307 | 4511 | 3285 | 3916 | 4873 | 4205 | 5841 | 4708 | 5110 | Snrpc |
| Snrpd1 | 7999 | 7039 | 9513 | 8288 | 7018 | 8016 | 9065 | 9017 | 8190 | 8547 | 8692 | 9797 | 9799 | 12683 | 8301 | 6775 | 8671 | 7013 | 8184 | 9785 | 8316 | 7265 | 7787 | 10366 | 9889 | 13076 | 11353 | 12957 | Snrpd1 |
| Snrpd2 | 8143 | 9686 | 9188 | 9333 | 7381 | 8120 | 8371 | 6328 | 6645 | 6639 | 7178 | 7172 | 8402 | 8747 | 7721 | 7299 | 9114 | 8107 | 9097 | 9917 | 8796 | 4782 | 6616 | 8228 | 7554 | 9055 | 9397 | 9375 | Snrpd2 |
| Snrpd3 | 12157 | 11688 | 14688 | 13147 | 10441 | 10843 | 12436 | 9701 | 8592 | 8586 | 9138 | 10489 | 10204 | 12043 | 12511 | 10244 | 13486 | 11820 | 11988 | 13095 | 11963 | 8134 | 8341 | 11083 | 10426 | 13765 | 12119 | 12721 | Snrpd3 |
| Snrpe | 6262 | 5319 | 6739 | 6137 | 5342 | 5199 | 8417 | 6836 | 5249 | 6576 | 6414 | 7239 | 7431 | 10043 | 6254 | 4911 | 6399 | 5187 | 5863 | 7113 | 7454 | 4662 | 5503 | 8087 | 6969 | 9111 | 8165 | 10792 | Snrpe |
| Snrpf | 3073 | 3327 | 4277 | 3626 | 2482 | 2337 | 3548 | 2876 | 2378 | 3054 | 2985 | 3338 | 3178 | 4959 | 2880 | 2146 | 3881 | 2999 | 2831 | 3118 | 3317 | 2201 | 2153 | 4264 | 3334 | 4227 | 3568 | 4603 | Snrpf |
| Snrpg | 6147 | 6208 | 6987 | 6721 | 4675 | 3986 | 8451 | 5401 | 4294 | 5714 | 5533 | 5787 | 5530 | 9452 | 5848 | 3831 | 6694 | 5093 | 5181 | 6491 | 7043 | 3688 | 3998 | 7302 | 5717 | 7321 | 6401 | 9600 | Snrpg |
| Snrpn | 2705 | 2917 | 2672 | 3598 | 4023 | 2755 | 4613 | 3894 | 2873 | 3822 | 2867 | 3295 | 2339 | 4203 | 3104 | 1689 | 3472 | 3350 | 3548 | 3078 | 5243 | 4428 | 2738 | 3536 | 3987 | 2544 | 2607 | 3540 | Snrpn |
| Snta1 | 1009 | 1275 | 1198 | 1137 | 1003 | 874 | 931 | 366 | 431 | 388 | 446 | 333 | 414 | 329 | 881 | 1174 | 1166 | 1088 | 990 | 936 | 949 | 433 | 526 | 427 | 418 | 433 | 375 | 407 | Snta1 |
| Sntb1 | 268 | 163 | 183 | 160 | 182 | 168 | 66 | 401 | 456 | 385 | 790 | 292 | 481 | 253 | 174 | 273 | 130 | 169 | 104 | 111 | 104 | 386 | 680 | 284 | 227 | 368 | 266 | 304 | Sntb1 |
| Sntb2 | 3680 | 3050 | 2539 | 3098 | 2677 | 3206 | 2552 | 2174 | 2403 | 2162 | 3106 | 2085 | 2237 | 2328 | 3150 | 3472 | 2481 | 2947 | 2346 | 2480 | 2736 | 2281 | 2602 | 1890 | 2107 | 1656 | 2229 | 2108 | Sntb2 |
| Sntg1 | 235 | 242 | 173 | 326 | 439 | 404 | 501 | 434 | 421 | 335 | 304 | 275 | 194 | 318 | 327 | 145 | 256 | 285 | 408 | 476 | 497 | 475 | 284 | 265 | 437 | 235 | 259 | 298 | Sntg1 |
| Sntg2 | 35 | 38 | 27 | 57 | 64 | 70 | 64 | 78 | 54 | 49 | 52 | 50 | 44 | 33 | 42 | 37 | 60 | 59 | 64 | 70 | 59 | 58 | 65 | 40 | 41 | 21 | 55 | 40 | Sntg2 |
| Sntn | 0 | 0 | 0 | 0 | 0 | 0 | 0 | 0 | 0 | 0 | 0 | 0 | 0 | 0 | 0 | 0 | 0 | 0 | 0 | 0 | 1 | 0 | 0 | 0 | 0 | 0 | 0 | 1 | Sntn |
| Snu13 | 9823 | 9694 | 8176 | 8060 | 6908 | 7660 | 8888 | 7320 | 7479 | 5712 | 6022 | 6460 | 6370 | 7241 | 9542 | 9271 | 8066 | 7823 | 7904 | 9859 | 8590 | 6010 | 7253 | 6347 | 6172 | 7950 | 7232 | 7735 | Snu13 |
| Snupn | 1403 | 1534 | 1608 | 1572 | 1291 | 1345 | 1483 | 1058 | 987 | 966 | 1038 | 1100 | 1002 | 1103 | 1419 | 1445 | 1523 | 1426 | 1431 | 1507 | 1496 | 967 | 959 | 1057 | 1129 | 1081 | 1026 | 1143 | Snupn |
| Snw1 | 7413 | 6764 | 7830 | 6492 | 6098 | 6539 | 7097 | 11685 | 9713 | 9798 | 9071 | 10896 | 10054 | 11804 | 7763 | 6525 | 6868 | 5790 | 6422 | 7492 | 6513 | 9075 | 9565 | 10581 | 9645 | 12526 | 11273 | 12129 | Snw1 |
| Snx1 | 8008 | 8593 | 8045 | 6933 | 5967 | 6439 | 6617 | 6789 | 6386 | 5799 | 5980 | 6204 | 6062 | 6234 | 8178 | 7671 | 7426 | 6691 | 6996 | 7610 | 6678 | 5559 | 6669 | 6098 | 5869 | 6820 | 6740 | 6527 | Snx1 |
| Snx10 | 2413 | 2136 | 2925 | 2565 | 3256 | 2956 | 3155 | 2739 | 2115 | 2366 | 2203 | 2301 | 2312 | 2674 | 2894 | 2479 | 2896 | 2678 | 2985 | 3120 | 3101 | 2642 | 2179 | 2430 | 2689 | 2393 | 2730 | 2811 | Snx10 |
| Snx11 | 4207 | 5033 | 4695 | 5795 | 5343 | 5028 | 4568 | 2958 | 2985 | 3795 | 3682 | 3625 | 3327 | 3351 | 4038 | 4212 | 5375 | 5317 | 5195 | 4443 | 4729 | 3412 | 3009 | 3482 | 3762 | 3340 | 3479 | 3279 | Snx11 |
| Snx12 | 3982 | 3861 | 4481 | 3917 | 3969 | 4166 | 4222 | 4644 | 4200 | 4504 | 4477 | 5447 | 5094 | 5362 | 3993 | 3351 | 4244 | 3634 | 4392 | 4089 | 4195 | 4602 | 4264 | 5114 | 5375 | 5723 | 5953 | 5892 | Snx12 |
| Snx13 | 3516 | 2803 | 2454 | 2871 | 3521 | 3431 | 3711 | 4555 | 4112 | 3761 | 3797 | 3590 | 3483 | 3696 | 3820 | 3008 | 2675 | 3060 | 2907 | 3684 | 3249 | 4152 | 4215 | 3404 | 3620 | 2812 | 3499 | 3523 | Snx13 |
| Snx14 | 4340 | 3891 | 3841 | 4186 | 4153 | 4144 | 4803 | 2545 | 2338 | 2364 | 2534 | 2393 | 2415 | 2364 | 4366 | 4092 | 3851 | 4391 | 3970 | 4200 | 4343 | 2638 | 2432 | 2159 | 2245 | 2117 | 2352 | 2239 | Snx14 |
| Snx15 | 559 | 730 | 769 | 656 | 893 | 824 | 730 | 1042 | 1255 | 1545 | 1330 | 1149 | 1148 | 1197 | 568 | 634 | 858 | 772 | 874 | 676 | 768 | 1206 | 1260 | 1258 | 1359 | 1159 | 1139 | 1127 | Snx15 |
| Snx16 | 1813 | 1706 | 1775 | 2080 | 2493 | 2210 | 2620 | 2417 | 2054 | 1799 | 1945 | 1403 | 1359 | 1560 | 2134 | 1851 | 2015 | 2517 | 2266 | 2657 | 2575 | 2433 | 2152 | 1684 | 2032 | 1355 | 1385 | 1428 | Snx16 |
| Snx17 | 5980 | 6989 | 6549 | 5862 | 5347 | 5549 | 5540 | 5801 | 6096 | 5478 | 5621 | 6826 | 5909 | 6076 | 5847 | 5670 | 6408 | 5431 | 6287 | 5531 | 5673 | 5681 | 6183 | 6196 | 6098 | 7166 | 6958 | 6750 | Snx17 |
| Snx18 | 4557 | 4576 | 4572 | 4360 | 3748 | 3683 | 3465 | 3059 | 3395 | 3249 | 3750 | 2765 | 3295 | 2746 | 4412 | 5314 | 4397 | 4653 | 3504 | 3180 | 3640 | 3153 | 3546 | 3099 | 2839 | 3491 | 2912 | 3052 | Snx18 |
| Snx19 | 4146 | 3984 | 3838 | 3648 | 3892 | 3733 | 3536 | 4586 | 4835 | 4898 | 4782 | 5399 | 5125 | 4928 | 4162 | 3998 | 3833 | 3895 | 3711 | 3423 | 3686 | 5081 | 4746 | 4725 | 4982 | 4687 | 5367 | 5298 | Snx19 |
| Snx2 | 10567 | 8601 | 9454 | 8105 | 6856 | 7780 | 7844 | 7364 | 7032 | 5967 | 6720 | 6403 | 7021 | 7250 | 10439 | 9899 | 8187 | 8502 | 6697 | 8377 | 7274 | 6834 | 7855 | 6633 | 6095 | 7383 | 7008 | 6977 | Snx2 |
| Snx20 | 44 | 54 | 27 | 42 | 36 | 37 | 29 | 43 | 36 | 34 | 50 | 23 | 57 | 20 | 37 | 39 | 39 | 34 | 37 | 50 | 33 | 47 | 72 | 56 | 24 | 31 | 28 | 42 | Snx20 |
| Snx21 | 1007 | 1143 | 1014 | 957 | 1015 | 893 | 873 | 552 | 693 | 642 | 700 | 505 | 516 | 517 | 901 | 1045 | 941 | 978 | 909 | 949 | 902 | 645 | 776 | 590 | 573 | 575 | 533 | 479 | Snx21 |
| Snx22 | 696 | 828 | 774 | 715 | 668 | 641 | 610 | 201 | 264 | 282 | 298 | 186 | 229 | 138 | 698 | 873 | 700 | 634 | 711 | 669 | 566 | 216 | 257 | 243 | 232 | 244 | 178 | 147 | Snx22 |
| Snx24 | 945 | 767 | 1081 | 587 | 551 | 619 | 470 | 1082 | 1018 | 961 | 986 | 1133 | 1279 | 1139 | 945 | 963 | 788 | 599 | 512 | 529 | 432 | 956 | 1061 | 1008 | 962 | 1319 | 1223 | 1215 | Snx24 |
| Snx25 | 4000 | 3672 | 4032 | 3675 | 3542 | 3550 | 3898 | 1961 | 1746 | 1851 | 2005 | 1686 | 1715 | 1589 | 3960 | 3727 | 3590 | 3895 | 3677 | 3922 | 3873 | 2005 | 1874 | 1694 | 1747 | 1692 | 1603 | 1620 | Snx25 |
| Snx27 | 11340 | 10751 | 10275 | 11669 | 13237 | 12590 | 13685 | 10903 | 9718 | 10566 | 10160 | 10892 | 9652 | 10818 | 11298 | 9963 | 11296 | 12569 | 12651 | 11730 | 13846 | 11718 | 9578 | 10598 | 11753 | 8425 | 10382 | 10913 | Snx27 |
| Snx29 | 1271 | 1495 | 1015 | 1495 | 1586 | 1513 | 1418 | 1229 | 1278 | 1270 | 1237 | 1175 | 1192 | 1215 | 1237 | 1293 | 1431 | 1510 | 1634 | 1463 | 1564 | 1312 | 1235 | 1028 | 1528 | 528 | 1292 | 1061 | Snx29 |
| Snx3 | 5523 | 5722 | 6069 | 5607 | 5132 | 5042 | 6360 | 4281 | 4610 | 4803 | 5019 | 4700 | 4571 | 5006 | 5304 | 5102 | 6154 | 5498 | 5631 | 5220 | 5958 | 4761 | 4970 | 5331 | 4545 | 5779 | 4522 | 5328 | Snx3 |
| Snx30 | 6990 | 7074 | 6574 | 8187 | 8691 | 8967 | 8142 | 3959 | 3643 | 4056 | 4099 | 3808 | 4065 | 3942 | 7212 | 7259 | 7609 | 8348 | 8136 | 7638 | 8594 | 4381 | 3811 | 3493 | 4328 | 2755 | 3747 | 3768 | Snx30 |
| Snx31 | 98 | 75 | 123 | 41 | 28 | 67 | 22 | 57 | 80 | 49 | 36 | 26 | 45 | 26 | 128 | 129 | 57 | 77 | 27 | 30 | 12 | 17 | 86 | 42 | 11 | 9 | 26 | 38 | Snx31 |
| Snx32 | 2077 | 3080 | 2719 | 3408 | 3925 | 3403 | 3628 | 2450 | 2359 | 2792 | 2331 | 2157 | 1990 | 2367 | 2600 | 2451 | 3412 | 3880 | 3822 | 3554 | 3795 | 3145 | 2249 | 2522 | 3005 | 1961 | 1885 | 2008 | Snx32 |
| Snx33 | 686 | 652 | 814 | 660 | 796 | 684 | 531 | 815 | 985 | 866 | 1213 | 973 | 1144 | 845 | 614 | 713 | 729 | 643 | 734 | 537 | 567 | 725 | 1132 | 1099 | 866 | 1258 | 1077 | 991 | Snx33 |
| Snx4 | 6598 | 5399 | 5557 | 5068 | 5114 | 5480 | 5816 | 6177 | 5603 | 5056 | 5056 | 5245 | 5420 | 5237 | 6956 | 5731 | 5628 | 5382 | 4987 | 6409 | 5519 | 5695 | 5829 | 5037 | 4890 | 5537 | 5318 | 5423 | Snx4 |
| Snx5 | 11605 | 8727 | 10598 | 8840 | 7211 | 7800 | 8525 | 11267 | 10852 | 9366 | 10690 | 11041 | 11252 | 11225 | 11236 | 10260 | 8773 | 8098 | 7793 | 8117 | 7633 | 9752 | 11698 | 10243 | 9193 | 13339 | 11196 | 11605 | Snx5 |
| Snx6 | 10716 | 9482 | 10401 | 8281 | 8613 | 8239 | 10029 | 10796 | 9816 | 8731 | 8742 | 9102 | 9095 | 10429 | 11314 | 9953 | 9352 | 8782 | 8299 | 10052 | 9062 | 9067 | 10078 | 9047 | 8878 | 10605 | 10091 | 10633 | Snx6 |
| Snx7 | 450 | 459 | 589 | 413 | 483 | 408 | 383 | 517 | 592 | 445 | 643 | 495 | 523 | 445 | 567 | 589 | 428 | 495 | 353 | 392 | 328 | 523 | 706 | 473 | 447 | 543 | 502 | 489 | Snx7 |
| Snx8 | 2569 | 3657 | 3268 | 3646 | 3152 | 3018 | 2762 | 1481 | 1767 | 1708 | 2023 | 1883 | 1862 | 1788 | 2772 | 3154 | 3246 | 3430 | 3314 | 2719 | 2926 | 1810 | 1911 | 1770 | 1887 | 1980 | 1973 | 1832 | Snx8 |
| Snx9 | 856 | 803 | 944 | 597 | 656 | 696 | 422 | 1621 | 2157 | 1909 | 2420 | 1685 | 2003 | 1827 | 774 | 1044 | 644 | 679 | 521 | 501 | 456 | 1753 | 2349 | 1884 | 1603 | 2316 | 1879 | 2039 | Snx9 |
| Soat1 | 6912 | 6150 | 8230 | 6958 | 5798 | 5991 | 6211 | 2732 | 2799 | 2897 | 3469 | 3268 | 3332 | 3211 | 6319 | 6701 | 7373 | 7180 | 5891 | 5404 | 6437 | 3029 | 2926 | 3157 | 2956 | 3673 | 3612 | 3582 | Soat1 |
| Soat2 | 1 | 1 | 0 | 0 | 6 | 0 | 4 | 1 | 5 | 1 | 4 | 11 | 8 | 7 | 1 | 3 | 7 | 1 | 0 | 1 | 2 | 8 | 8 | 6 | 0 | 10 | 5 | 7 | Soat2 |
| Sobp | 2441 | 2427 | 3366 | 4435 | 5627 | 4336 | 4168 | 3869 | 3617 | 5970 | 4732 | 4481 | 3527 | 4188 | 2595 | 2538 | 4225 | 4645 | 5641 | 3594 | 4976 | 7067 | 3492 | 5621 | 6283 | 4265 | 3588 | 3755 | Sobp |
| Socs1 | 100 | 81 | 134 | 82 | 90 | 91 | 44 | 79 | 106 | 98 | 134 | 75 | 113 | 62 | 66 | 89 | 82 | 59 | 69 | 47 | 46 | 97 | 161 | 104 | 77 | 105 | 56 | 46 | Socs1 |
| Socs2 | 2697 | 2222 | 2320 | 2956 | 3504 | 3318 | 3727 | 3506 | 3002 | 3533 | 3089 | 2471 | 2683 | 2661 | 2943 | 2317 | 2527 | 2793 | 3320 | 3642 | 3774 | 3282 | 3387 | 3370 | 3268 | 3086 | 2192 | 2682 | Socs2 |
| Socs3 | 638 | 752 | 922 | 627 | 820 | 755 | 641 | 629 | 737 | 647 | 679 | 701 | 641 | 551 | 550 | 662 | 753 | 682 | 846 | 633 | 604 | 660 | 656 | 735 | 633 | 825 | 562 | 669 | Socs3 |
| Socs4 | 5050 | 4002 | 4654 | 4973 | 4622 | 4652 | 5650 | 5843 | 4653 | 4349 | 4525 | 5236 | 5142 | 5947 | 5018 | 4550 | 4923 | 4654 | 4658 | 5186 | 5171 | 5061 | 4546 | 5160 | 5120 | 5427 | 5726 | 5624 | Socs4 |
| Socs5 | 2749 | 2842 | 2792 | 2923 | 3391 | 3170 | 3168 | 2760 | 2703 | 2595 | 2754 | 2300 | 2182 | 2270 | 2751 | 2872 | 2796 | 3054 | 3306 | 2974 | 3069 | 2577 | 2843 | 2406 | 2413 | 2128 | 2264 | 2131 | Socs5 |
| Socs6 | 5583 | 4308 | 5210 | 4785 | 4498 | 4719 | 5192 | 5467 | 4598 | 4286 | 4790 | 5645 | 5852 | 6179 | 5421 | 4847 | 4905 | 4316 | 4448 | 5010 | 4726 | 4441 | 4591 | 4721 | 4661 | 5813 | 6280 | 6501 | Socs6 |
| Socs7 | 9498 | 11334 | 10000 | 14609 | 14188 | 12999 | 13917 | 8398 | 7726 | 9096 | 8161 | 8737 | 7954 | 8905 | 9941 | 9057 | 12568 | 14786 | 14691 | 11676 | 15202 | 10325 | 7102 | 8943 | 10503 | 6312 | 8220 | 8772 | Socs7 |
| Sod1 | 10100 | 10995 | 10410 | 10010 | 8251 | 8140 | 9881 | 7421 | 7294 | 7453 | 7848 | 7841 | 8021 | 9376 | 9908 | 9027 | 10615 | 9043 | 8782 | 9867 | 9642 | 6126 | 7651 | 8517 | 7831 | 9647 | 8723 | 9894 | Sod1 |
| Sod2 | 9696 | 9553 | 9384 | 10462 | 9441 | 9209 | 10018 | 4628 | 4135 | 4482 | 4132 | 4192 | 4119 | 4289 | 9661 | 8854 | 9405 | 10136 | 9006 | 9632 | 9681 | 4565 | 3842 | 4113 | 4472 | 3995 | 3963 | 4050 | Sod2 |
| Sod3 | 13 | 9 | 14 | 13 | 27 | 9 | 13 | 17 | 40 | 36 | 59 | 34 | 31 | 22 | 25 | 19 | 15 | 21 | 12 | 15 | 25 | 42 | 78 | 33 | 20 | 24 | 14 | 7 | Sod3 |
| Soga1 | 5730 | 7020 | 1930 | 7988 | 10760 | 9389 | 7920 | 8818 | 10225 | 11545 | 11097 | 11553 | 10381 | 10124 | 5354 | 6094 | 5209 | 8616 | 10096 | 7559 | 9473 | 11184 | 8855 | 7803 | 14059 | 1823 | 10398 | 8686 | Soga1 |
| Soga3 | 2163 | 2933 | 2591 | 3612 | 4877 | 4137 | 4031 | 6163 | 6286 | 7667 | 6488 | 6680 | 5675 | 7274 | 2462 | 2129 | 3447 | 3706 | 4552 | 3983 | 4594 | 7056 | 5357 | 7183 | 8841 | 4292 | 6430 | 6369 | Soga3 |
| Sohlh1 | 0 | 0 | 0 | 1 | 0 | 0 | 0 | 0 | 1 | 0 | 0 | 0 | 0 | 0 | 0 | 0 | 0 | 0 | 0 | 0 | 0 | 0 | 1 | 0 | 0 | 0 | 0 | 0 | Sohlh1 |
| Sohlh2 | 1 | 2 | 0 | 0 | 6 | 1 | 1 | 5 | 0 | 0 | 4 | 1 | 0 | 1 | 1 | 5 | 1 | 1 | 0 | 3 | 1 | 0 | 1 | 0 | 4 | 0 | 0 | 1 | Sohlh2 |
| Son | 63997 | 58684 | 35392 | 46546 | 51676 | 58495 | 55578 | 85022 | 74462 | 62078 | 66966 | 65645 | 67140 | 69298 | 64353 | 53537 | 40185 | 47249 | 48538 | 62702 | 59528 | 58967 | 69178 | 52290 | 67857 | 42585 | 73338 | 63726 | Son |
| Sorbs1 | 3735 | 4591 | 3302 | 4154 | 5614 | 5337 | 5323 | 3866 | 4736 | 4774 | 5018 | 3750 | 3927 | 3832 | 4172 | 4554 | 3533 | 4864 | 4777 | 4797 | 5652 | 4943 | 4163 | 3667 | 5162 | 1896 | 3847 | 3245 | Sorbs1 |
| Sorbs2 | 5269 | 5021 | 5061 | 6164 | 7810 | 7818 | 8382 | 12908 | 13078 | 12799 | 12206 | 13428 | 11118 | 13603 | 6105 | 4658 | 5830 | 6417 | 7579 | 8286 | 8262 | 13363 | 10457 | 12674 | 15037 | 10254 | 11327 | 11527 | Sorbs2 |
| Sorbs3 | 1278 | 1254 | 1518 | 855 | 1004 | 1025 | 758 | 1286 | 1840 | 1584 | 2000 | 1552 | 1788 | 1344 | 1085 | 1542 | 1141 | 1016 | 973 | 729 | 685 | 1450 | 2039 | 1714 | 1242 | 2073 | 1579 | 1474 | Sorbs3 |
| Sorcs1 | 1407 | 1555 | 729 | 1787 | 2190 | 2016 | 2171 | 2257 | 2243 | 2173 | 2090 | 1602 | 1669 | 1758 | 1753 | 1439 | 1469 | 1892 | 1753 | 1870 | 2132 | 2832 | 1986 | 1669 | 2281 | 1267 | 1414 | 1284 | Sorcs1 |
| Sorcs2 | 2082 | 2472 | 2364 | 2401 | 2631 | 2780 | 2198 | 2910 | 3466 | 3670 | 4096 | 3432 | 3047 | 2828 | 2294 | 2164 | 2371 | 2574 | 2703 | 2379 | 2305 | 4259 | 4213 | 3259 | 3676 | 2764 | 3191 | 3141 | Sorcs2 |
| Sorcs3 | 648 | 780 | 745 | 1158 | 1490 | 1281 | 1354 | 1170 | 1149 | 1358 | 1218 | 1119 | 1089 | 1161 | 820 | 758 | 1000 | 1268 | 1286 | 1393 | 1380 | 1744 | 1050 | 1088 | 1501 | 791 | 952 | 1003 | Sorcs3 |
| Sord | 3640 | 3881 | 3348 | 3197 | 2822 | 3032 | 2879 | 1723 | 1917 | 1588 | 1460 | 1757 | 1788 | 1846 | 3389 | 3287 | 3452 | 3021 | 3229 | 3007 | 2968 | 1704 | 1990 | 1611 | 1498 | 2031 | 1964 | 2028 | Sord |
| Sorl1 | 3838 | 3580 | 2100 | 3746 | 6073 | 5403 | 5370 | 4658 | 5786 | 5868 | 5458 | 4649 | 4589 | 4533 | 3775 | 3717 | 2802 | 4629 | 4798 | 4173 | 5511 | 4102 | 4752 | 4492 | 5131 | 3179 | 3853 | 3420 | Sorl1 |
| Sort1 | 20690 | 22136 | 19008 | 22548 | 21865 | 21836 | 23126 | 12044 | 12259 | 13608 | 12579 | 14682 | 13417 | 14533 | 21402 | 22013 | 21402 | 22782 | 22070 | 19559 | 23504 | 14492 | 11833 | 12888 | 14766 | 9654 | 14554 | 14431 | Sort1 |
| Sos1 | 9366 | 7788 | 7904 | 11054 | 11588 | 11547 | 12231 | 8074 | 7081 | 8297 | 8272 | 8423 | 8247 | 8675 | 9179 | 9127 | 9151 | 11061 | 10647 | 9654 | 12275 | 9513 | 6733 | 7285 | 8886 | 6137 | 8404 | 8217 | Sos1 |
| Sos2 | 3058 | 2655 | 2981 | 2950 | 4229 | 3688 | 3508 | 4264 | 3948 | 3962 | 4310 | 3926 | 3935 | 4035 | 3272 | 2916 | 3211 | 3314 | 3617 | 3391 | 3719 | 4421 | 3897 | 3612 | 4348 | 3087 | 3956 | 3955 | Sos2 |
| Sost | 20 | 20 | 13 | 12 | 16 | 8 | 12 | 40 | 48 | 44 | 21 | 59 | 87 | 35 | 30 | 24 | 29 | 11 | 25 | 14 | 12 | 40 | 49 | 26 | 30 | 52 | 44 | 61 | Sost |
| Sostdc1 | 473 | 307 | 314 | 189 | 344 | 334 | 270 | 1021 | 1473 | 998 | 660 | 297 | 548 | 281 | 414 | 451 | 175 | 233 | 265 | 251 | 314 | 495 | 1659 | 690 | 282 | 365 | 252 | 249 | Sostdc1 |
| Sowaha | 156 | 122 | 137 | 208 | 226 | 176 | 227 | 109 | 122 | 154 | 112 | 119 | 62 | 89 | 153 | 114 | 187 | 163 | 206 | 127 | 199 | 52 | 71 | 202 | 100 | 210 | 55 | 56 | Sowaha |

| | | | | | | | | | | | | | | | | | | | | | | | | | | | | | |
|---|---|---|---|---|---|---|---|---|---|---|---|---|---|---|---|---|---|---|---|---|---|---|---|---|---|---|---|---|---|
| Sowahb | 88 | 92 | 10 | 45 | 25 | 22 | 21 | 38 | 36 | 32 | 28 | 5 | 31 | 11 | 57 | 83 | 24 | 55 | 23 | 10 | 23 | 3 | 32 | 34 | 1 | 43 | 9 | 9 | Sowahb |
| Sowahc | 3186 | 3162 | 3005 | 3653 | 3016 | 3206 | 3416 | 1958 | 1931 | 1872 | 1848 | 1948 | 2031 | 2049 | 3277 | 3293 | 3176 | 3711 | 3090 | 3164 | 3407 | 2037 | 2051 | 1860 | 1926 | 1981 | 2027 | 2098 | Sowahc |
| Sowahd | 1 | 2 | 3 | 6 | 2 | 0 | 2 | 0 | 0 | 1 | 0 | 0 | 0 | 0 | 4 | 0 | 1 | 2 | 0 | 7 | 3 | 0 | 1 | 1 | 0 | 0 | 0 | 0 | Sowahd |
| Sox1 | 2957 | 2973 | 4750 | 4808 | 4204 | 3569 | 5015 | 3662 | 3267 | 4311 | 3144 | 6859 | 3382 | 7107 | 3000 | 1812 | 5517 | 4163 | 5284 | 3092 | 5265 | 4831 | 2172 | 6310 | 5431 | 8049 | 5391 | 6378 | Sox1 |
| Sox10 | 305 | 351 | 119 | 128 | 513 | 58 | 21 | 540 | 801 | 680 | 1078 | 135 | 667 | 105 | 384 | 614 | 256 | 199 | 53 | 22 | 39 | 792 | 645 | 394 | 340 | 390 | 129 | 106 | Sox10 |
| Sox11 | 35722 | 30233 | 60570 | 66277 | 61057 | 54869 | 65269 | 61505 | 52012 | 72934 | 54355 | 81508 | 62723 | 84813 | 38469 | 32062 | 67554 | 63345 | 65876 | 47252 | 63768 | 84288 | 44575 | 91089 | 90399 | 112676 | 72235 | 84591 | Sox11 |
| Sox12 | 6256 | 7663 | 8174 | 7787 | 8632 | 8204 | 7717 | 10050 | 11002 | 13432 | 11054 | 14010 | 11908 | 12434 | 6117 | 6463 | 9071 | 7657 | 9521 | 7348 | 8190 | 12389 | 10002 | 13460 | 13099 | 14797 | 11950 | 12248 | Sox12 |
| Sox13 | 1694 | 2249 | 3465 | 2252 | 2093 | 1776 | 1330 | 1737 | 1730 | 2140 | 1861 | 2948 | 2969 | 2597 | 1534 | 2223 | 2928 | 1800 | 2183 | 1556 | 1600 | 2146 | 1684 | 2484 | 2112 | 2613 | 3039 | 3183 | Sox13 |
| Sox14 | 280 | 289 | 433 | 682 | 808 | 679 | 494 | 766 | 470 | 1033 | 655 | 819 | 982 | 686 | 332 | 248 | 585 | 588 | 886 | 699 | 748 | 1288 | 595 | 954 | 1155 | 205 | 734 | 662 | Sox14 |
| Sox15 | 9 | 10 | 0 | 3 | 8 | 10 | 7 | 4 | 3 | 11 | 6 | 11 | 3 | 3 | 1 | 9 | 9 | 8 | 9 | 10 | 8 | 3 | 4 | 5 | 10 | 11 | 14 | 6 | Sox15 |
| Sox17 | 195 | 189 | 345 | 289 | 345 | 285 | 264 | 325 | 362 | 396 | 455 | 434 | 505 | 458 | 212 | 232 | 273 | 294 | 309 | 235 | 307 | 394 | 403 | 462 | 568 | 537 | 493 | 398 | Sox17 |
| Sox18 | 393 | 339 | 346 | 232 | 267 | 232 | 179 | 493 | 696 | 504 | 696 | 452 | 627 | 375 | 284 | 372 | 290 | 249 | 283 | 173 | 149 | 510 | 832 | 558 | 445 | 624 | 459 | 395 | Sox18 |
| Sox2 | 6620 | 5943 | 11009 | 9976 | 8716 | 7968 | 10810 | 8802 | 6671 | 9978 | 7611 | 14846 | 9050 | 16602 | 6618 | 4123 | 11470 | 8694 | 10382 | 7588 | 10513 | 9825 | 6313 | 14267 | 11741 | 18945 | 12487 | 15497 | Sox2 |
| Sox21 | 2558 | 2407 | 4261 | 3488 | 3309 | 2987 | 3341 | 3217 | 2590 | 3229 | 2300 | 5130 | 3751 | 4791 | 2401 | 1878 | 4086 | 3133 | 3985 | 2887 | 3513 | 3283 | 2025 | 4569 | 3780 | 4430 | 4213 | 4736 | Sox21 |
| Sox3 | 1939 | 2847 | 3738 | 3640 | 2189 | 2205 | 2167 | 1269 | 1271 | 1767 | 1395 | 3448 | 2113 | 2920 | 1548 | 1550 | 3897 | 2999 | 3115 | 2131 | 2417 | 1384 | 1011 | 2839 | 2245 | 3946 | 2621 | 2804 | Sox3 |
| Sox30 | 68 | 75 | 76 | 80 | 82 | 65 | 66 | 37 | 43 | 74 | 36 | 70 | 54 | 46 | 71 | 79 | 71 | 63 | 79 | 53 | 68 | 48 | 45 | 38 | 44 | 59 | 46 | 40 | Sox30 |
| Sox4 | 31847 | 30130 | 38130 | 49572 | 49361 | 45613 | 56331 | 55371 | 49326 | 57233 | 51480 | 60183 | 45012 | 67249 | 34862 | 26810 | 42559 | 49489 | 51670 | 44408 | 58400 | 64280 | 47012 | 66573 | 69997 | 59624 | 54522 | 63146 | Sox4 |
| Sox5 | 4735 | 4270 | 3423 | 4996 | 5368 | 5921 | 6822 | 5650 | 6575 | 6476 | 5638 | 5555 | 3695 | 5984 | 4983 | 3626 | 4422 | 4855 | 5719 | 5712 | 6557 | 5690 | 4700 | 6511 | 8461 | 4972 | 4991 | 4651 | Sox5 |
| Sox6 | 4593 | 4323 | 3486 | 4302 | 4554 | 6506 | 6642 | 5217 | 6366 | 6217 | 5964 | 7811 | 5082 | 7772 | 4549 | 3326 | 4238 | 4343 | 5315 | 5723 | 6125 | 5673 | 5114 | 6462 | 7484 | 5784 | 7229 | 6524 | Sox6 |
| Sox7 | 270 | 286 | 420 | 293 | 353 | 276 | 285 | 482 | 470 | 583 | 662 | 485 | 627 | 501 | 291 | 326 | 293 | 328 | 305 | 234 | 291 | 643 | 544 | 580 | 531 | 651 | 492 | 476 | Sox7 |
| Sox8 | 761 | 926 | 650 | 1147 | 1010 | 905 | 923 | 793 | 850 | 937 | 967 | 754 | 413 | 698 | 737 | 879 | 983 | 1219 | 1079 | 747 | 1047 | 872 | 741 | 920 | 913 | 746 | 493 | 438 | Sox8 |
| Sox9 | 3066 | 3409 | 5615 | 5002 | 3573 | 3818 | 3605 | 4628 | 4949 | 6986 | 6782 | 7363 | 6775 | 6811 | 3276 | 3766 | 5227 | 4429 | 4235 | 2654 | 3484 | 6460 | 4285 | 7895 | 6177 | 10109 | 6186 | 7252 | Sox9 |
| Sp1 | 9047 | 7184 | 7351 | 7391 | 7261 | 7755 | 8077 | 10481 | 10352 | 8365 | 9522 | 9562 | 9871 | 9720 | 8408 | 7663 | 7609 | 7287 | 7381 | 7886 | 7956 | 8478 | 9828 | 8648 | 8996 | 8572 | 10599 | 10003 | Sp1 |
| Sp100 | 35 | 22 | 11 | 22 | 21 | 17 | 11 | 27 | 40 | 49 | 92 | 38 | 29 | 33 | 26 | 39 | 13 | 25 | 10 | 36 | 24 | 65 | 49 | 42 | 27 | 61 | 71 | 57 | Sp100 |
| Sp110 | 29 | 26 | 32 | 27 | 35 | 32 | 27 | 51 | 32 | 44 | 73 | 48 | 62 | 34 | 54 | 44 | 22 | 34 | 17 | 44 | 36 | 58 | 67 | 30 | 61 | 67 | 52 | 51 | Sp110 |
| Sp140 | 46 | 46 | 27 | 46 | 21 | 36 | 20 | 42 | 53 | 15 | 57 | 56 | 36 | 45 | 44 | 79 | 22 | 28 | 28 | 31 | 31 | 43 | 104 | 38 | 36 | 55 | 28 | 38 | Sp140 |
| Sp2 | 787 | 985 | 1475 | 1081 | 977 | 838 | 808 | 1028 | 1299 | 1551 | 1374 | 1921 | 1469 | 1726 | 749 | 847 | 1424 | 987 | 1232 | 638 | 884 | 1390 | 1071 | 1748 | 1561 | 2082 | 1548 | 1634 | Sp2 |
| Sp3 | 8987 | 6608 | 7404 | 6939 | 7502 | 7799 | 8838 | 14084 | 11905 | 9712 | 10081 | 11904 | 10875 | 12829 | 8918 | 7258 | 7290 | 6937 | 7811 | 8599 | 8254 | 11282 | 11174 | 10918 | 11069 | 12759 | 12386 | 12716 | Sp3 |
| Sp4 | 5819 | 4997 | 3830 | 5363 | 5332 | 6046 | 6574 | 7339 | 5760 | 4953 | 4557 | 5681 | 4813 | 5731 | 5992 | 4687 | 4757 | 5378 | 5440 | 6602 | 6316 | 5855 | 5240 | 4712 | 5924 | 3983 | 5898 | 5268 | Sp4 |
| Sp5 | 497 | 521 | 903 | 573 | 443 | 566 | 363 | 289 | 278 | 282 | 228 | 450 | 348 | 379 | 426 | 368 | 552 | 482 | 689 | 495 | 495 | 292 | 328 | 423 | 362 | 419 | 407 | 485 | Sp5 |
| Sp6 | 152 | 110 | 133 | 93 | 78 | 98 | 68 | 100 | 98 | 86 | 62 | 67 | 86 | 53 | 98 | 125 | 86 | 84 | 61 | 67 | 60 | 37 | 136 | 65 | 41 | 62 | 49 | 82 | Sp6 |
| Sp7 | 130 | 79 | 58 | 107 | 70 | 97 | 29 | 307 | 276 | 425 | 347 | 88 | 129 | 92 | 84 | 125 | 53 | 138 | 42 | 29 | 34 | 378 | 316 | 392 | 360 | 65 | 71 | 63 | Sp7 |
| Sp8 | 1275 | 794 | 1360 | 1480 | 990 | 1075 | 1639 | 1542 | 1235 | 1325 | 987 | 1937 | 1378 | 2279 | 1051 | 858 | 1447 | 917 | 1413 | 906 | 1412 | 976 | 773 | 2062 | 1362 | 3055 | 1554 | 2129 | Sp8 |
| Sp9 | 911 | 921 | 918 | 1738 | 1422 | 1453 | 2016 | 1143 | 982 | 1474 | 1024 | 1870 | 327 | 1920 | 828 | 306 | 1583 | 1786 | 1956 | 1090 | 2036 | 1785 | 702 | 2145 | 1849 | 2227 | 1239 | 1316 | Sp9 |
| Spa17 | 335 | 436 | 413 | 475 | 362 | 372 | 447 | 437 | 328 | 382 | 378 | 487 | 538 | 624 | 404 | 353 | 436 | 355 | 399 | 459 | 435 | 322 | 323 | 470 | 424 | 534 | 459 | 529 | Spa17 |
| Spaar | 3 | 4 | 0 | 0 | 2 | 5 | 0 | 11 | 6 | 8 | 22 | 0 | 2 | 3 | 0 | 9 | 6 | 10 | 2 | 0 | 4 | 5 | 15 | 6 | 0 | 5 | 6 | 7 | Spaar |
| Spaca1 | 85 | 62 | 72 | 46 | 62 | 60 | 48 | 43 | 66 | 56 | 114 | 62 | 99 | 51 | 51 | 73 | 37 | 31 | 46 | 38 | 27 | 35 | 82 | 44 | 52 | 38 | 73 | 47 | Spaca1 |
| Spaca3 | 0 | 0 | 0 | 0 | 0 | 1 | 0 | 0 | 1 | 1 | 0 | 0 | 0 | 0 | 3 | 0 | 0 | 0 | 1 | 1 | 0 | 0 | 0 | 0 | 3 | 0 | 5 | 4 | Spaca3 |
| Spaca5 | 3 | 0 | 1 | 0 | 1 | 0 | 3 | 0 | 0 | 4 | 0 | 1 | 0 | 1 | 0 | 2 | 1 | 0 | 0 | 0 | 3 | 0 | 0 | 6 | 1 | 0 | 0 | 0 | Spaca5 |
| Spaca6 | 2475 | 1960 | 2318 | 2158 | 2583 | 2761 | 2368 | 7618 | 7044 | 8517 | 9444 | 9155 | 10270 | 10086 | 2203 | 2084 | 2015 | 1924 | 2258 | 2268 | 2330 | 8256 | 6434 | 7956 | 8533 | 5313 | 9783 | 9032 | Spaca6 |
| Spaca7 | 0 | 2 | 2 | 1 | 1 | 0 | 0 | 0 | 1 | 2 | 3 | 0 | 2 | 0 | 1 | 1 | 0 | 1 | 0 | 0 | 1 | 0 | 1 | 1 | 0 | 4 | 0 | 1 | Spaca7 |
| Spaca9 | 21 | 40 | 11 | 32 | 32 | 25 | 22 | 37 | 36 | 21 | 24 | 45 | 39 | 19 | 26 | 48 | 21 | 18 | 26 | 21 | 16 | 27 | 32 | 15 | 32 | 18 | 27 | 14 | Spaca9 |
| Spag1 | 1389 | 1565 | 1232 | 1570 | 1436 | 1498 | 1581 | 1430 | 1177 | 1324 | 1160 | 1143 | 1209 | 1191 | 1402 | 1410 | 1426 | 1508 | 1541 | 1528 | 1609 | 1457 | 1181 | 1261 | 1358 | 1119 | 1252 | 1104 | Spag1 |
| Spag11a | 0 | 0 | 0 | 0 | 0 | 0 | 0 | 0 | 0 | 0 | 0 | 0 | 0 | 0 | 0 | 0 | 0 | 0 | 0 | 1 | 0 | 0 | 0 | 0 | 0 | 0 | 0 | 0 | Spag11a |
| Spag11b | 0 | 0 | 0 | 0 | 0 | 3 | 0 | 0 | 1 | 0 | 0 | 0 | 3 | 0 | 0 | 0 | 0 | 0 | 0 | 0 | 1 | 0 | 0 | 0 | 0 | 0 | 0 | 0 | Spag11b |
| Spag16 | 226 | 190 | 263 | 149 | 175 | 189 | 173 | 177 | 164 | 124 | 128 | 238 | 296 | 212 | 223 | 235 | 165 | 118 | 146 | 182 | 171 | 179 | 174 | 122 | 184 | 127 | 200 | 179 | Spag16 |
| Spag17 | 80 | 68 | 27 | 42 | 92 | 75 | 40 | 85 | 147 | 90 | 80 | 80 | 125 | 54 | 80 | 123 | 24 | 36 | 45 | 48 | 44 | 90 | 211 | 60 | 41 | 26 | 56 | 63 | Spag17 |
| Spag4 | 573 | 650 | 797 | 733 | 828 | 666 | 645 | 375 | 410 | 423 | 439 | 533 | 468 | 446 | 510 | 537 | 808 | 701 | 884 | 704 | 717 | 485 | 365 | 471 | 486 | 586 | 501 | 492 | Spag4 |
| Spag5 | 9085 | 8859 | 8350 | 7529 | 5195 | 6167 | 7445 | 6826 | 6227 | 5482 | 5169 | 8922 | 7073 | 8825 | 8090 | 7393 | 7276 | 6345 | 6713 | 7193 | 6637 | 4466 | 5262 | 7074 | 6366 | 9249 | 8851 | 8641 | Spag5 |
| Spag6 | 18 | 25 | 34 | 13 | 7 | 24 | 8 | 23 | 6 | 19 | 17 | 16 | 18 | 18 | 28 | 13 | 16 | 10 | 8 | 13 | 11 | 3 | 6 | 6 | 19 | 8 | 15 | 21 | Spag6 |
| Spag6l | 444 | 503 | 324 | 490 | 668 | 663 | 685 | 364 | 314 | 414 | 293 | 257 | 329 | 231 | 579 | 530 | 448 | 574 | 639 | 777 | 652 | 449 | 482 | 226 | 265 | 154 | 309 | 206 | Spag6l |
| Spag7 | 1444 | 1630 | 2031 | 1679 | 1518 | 1541 | 1715 | 3245 | 3236 | 3501 | 3077 | 4053 | 3636 | 4335 | 1476 | 1428 | 1845 | 1499 | 1663 | 1665 | 1487 | 3253 | 3421 | 4200 | 3560 | 4907 | 4517 | 4448 | Spag7 |
| Spag8 | 159 | 269 | 293 | 237 | 261 | 227 | 239 | 173 | 215 | 211 | 182 | 256 | 286 | 165 | 179 | 238 | 289 | 245 | 285 | 259 | 264 | 175 | 256 | 206 | 180 | 216 | 206 | 210 | Spag8 |
| Spag9 | 33116 | 29615 | 21357 | 28633 | 32878 | 34296 | 34330 | 50060 | 39603 | 36839 | 35897 | 40339 | 36894 | 42878 | 37238 | 30056 | 25577 | 31147 | 28922 | 35318 | 35633 | 40653 | 38230 | 34444 | 41793 | 25109 | 42736 | 39647 | Spag9 |
| Sparc | 23801 | 26594 | 25622 | 20204 | 16760 | 24180 | 12776 | 23875 | 32163 | 27863 | 39591 | 25791 | 31615 | 24998 | 25354 | 30904 | 22069 | 26114 | 14719 | 14026 | 13451 | 28194 | 42208 | 28001 | 26822 | 27017 | 26819 | 26210 | Sparc |
| Sparcl1 | 3873 | 3868 | 4084 | 4367 | 7137 | 5001 | 4120 | 7544 | 8809 | 7941 | 9262 | 5280 | 7478 | 5395 | 4804 | 4827 | 4045 | 5426 | 4663 | 6088 | 4838 | 9782 | 10774 | 6289 | 7705 | 6291 | 5530 | 4824 | Sparcl1 |
| Spast | 15879 | 12458 | 13914 | 17737 | 19140 | 19043 | 21083 | 18417 | 15587 | 17690 | 15990 | 17557 | 15723 | 18529 | 16862 | 13451 | 15593 | 17652 | 18569 | 17953 | 20670 | 20449 | 14940 | 16923 | 19578 | 15611 | 16494 | 17978 | Spast |
| Spata1 | 803 | 545 | 695 | 739 | 815 | 920 | 892 | 631 | 622 | 518 | 612 | 559 | 586 | 565 | 856 | 795 | 549 | 677 | 661 | 891 | 862 | 485 | 638 | 465 | 499 | 393 | 523 | 476 | Spata1 |
| Spata13 | 9808 | 11454 | 11106 | 12679 | 11001 | 11451 | 12597 | 5821 | 6078 | 6575 | 5481 | 10718 | 8594 | 9870 | 10026 | 9216 | 12210 | 12325 | 12387 | 10383 | 12347 | 5701 | 4692 | 7695 | 7638 | 8277 | 9516 | 9884 | Spata13 |
| Spata16 | 0 | 0 | 0 | 0 | 1 | 2 | 0 | 0 | 0 | 0 | 0 | 2 | 0 | 0 | 3 | 0 | 0 | 0 | 0 | 0 | 1 | 0 | 1 | 1 | 0 | 1 | 0 | 4 | Spata16 |
| Spata17 | 43 | 59 | 23 | 19 | 48 | 49 | 64 | 64 | 77 | 48 | 44 | 73 | 88 | 55 | 58 | 80 | 31 | 37 | 41 | 54 | 30 | 70 | 83 | 46 | 45 | 49 | 65 | 53 | Spata17 |
| Spata18 | 33 | 35 | 12 | 22 | 47 | 19 | 20 | 56 | 77 | 40 | 60 | 47 | 75 | 9 | 33 | 62 | 23 | 36 | 52 | 50 | 28 | 67 | 110 | 22 | 34 | 7 | 34 | 19 | Spata18 |
| Spata19 | 12 | 5 | 6 | 21 | 28 | 16 | 22 | 2 | 1 | 0 | 0 | 4 | 2 | 1 | 13 | 7 | 20 | 37 | 26 | 14 | 28 | 3 | 0 | 3 | 1 | 2 | 0 | 1 | Spata19 |
| Spata2 | 2771 | 3600 | 4287 | 4357 | 3699 | 3522 | 3545 | 1734 | 2075 | 2221 | 2169 | 2482 | 2155 | 2216 | 2726 | 3264 | 4059 | 4253 | 4214 | 2896 | 3684 | 2436 | 1920 | 2329 | 2606 | 2450 | 2142 | 2248 | Spata2 |
| Spata20 | 8 | 14 | 9 | 7 | 12 | 12 | 8 | 5 | 13 | 21 | 8 | 1 | 11 | 14 | 8 | 14 | 13 | 17 | 14 | 13 | 15 | 7 | 21 | 9 | 9 | 5 | 14 | 8 | Spata20 |
| Spata21 | 0 | 1 | 0 | 0 | 0 | 0 | 0 | 4 | 2 | 0 | 0 | 0 | 0 | 0 | 0 | 1 | 0 | 0 | 0 | 0 | 0 | 5 | 2 | 3 | 3 | 0 | 0 | 1 | Spata21 |
| Spata22 | 0 | 0 | 0 | 0 | 0 | 0 | 0 | 0 | 0 | 0 | 0 | 0 | 0 | 0 | 0 | 0 | 0 | 0 | 0 | 0 | 0 | 2 | 0 | 0 | 0 | 0 | 0 | 0 | Spata22 |
| Spata24 | 219 | 236 | 123 | 229 | 232 | 245 | 244 | 112 | 144 | 164 | 162 | 181 | 196 | 151 | 227 | 231 | 177 | 263 | 235 | 186 | 226 | 192 | 159 | 141 | 165 | 102 | 146 | 158 | Spata24 |
| Spata25 | 21 | 23 | 23 | 14 | 20 | 17 | 23 | 11 | 14 | 14 | 11 | 8 | 28 | 17 | 15 | 26 | 22 | 39 | 35 | 33 | 20 | 10 | 14 | 16 | 34 | 20 | 28 | 19 | Spata25 |
| Spata2l | 258 | 326 | 286 | 314 | 279 | 233 | 242 | 100 | 115 | 129 | 92 | 77 | 83 | 69 | 228 | 371 | 283 | 384 | 254 | 237 | 260 | 142 | 135 | 105 | 96 | 83 | 74 | 81 | Spata2l |
| Spata3 | 1 | 2 | 0 | 0 | 0 | 0 | 0 | 0 | 1 | 0 | 0 | 0 | 0 | 2 | 0 | 0 | 2 | 0 | 2 | 0 | 1 | 0 | 3 | 0 | 3 | 0 | 0 | 1 | Spata3 |
| Spata31 | 0 | 0 | 0 | 0 | 0 | 0 | 0 | 0 | 2 | 0 | 0 | 2 | 0 | 1 | 0 | 0 | 0 | 0 | 0 | 0 | 0 | 0 | 0 | 0 | 0 | 0 | 0 | 0 | Spata31 |
| Spata31d1a | 10 | 8 | 18 | 8 | 3 | 24 | 18 | 0 | 3 | 0 | 0 | 0 | 2 | 0 | 5 | 16 | 8 | 12 | 13 | 8 | 10 | 0 | 6 | 2 | 5 | 0 | 0 | 1 | Spata31d1a |
| Spata31d1b | 0 | 5 | 2 | 12 | 2 | 1 | 3 | 2 | 2 | 5 | 5 | 4 | 3 | 6 | 4 | 2 | 1 | 0 | 2 | 2 | 3 | 0 | 3 | 2 | 1 | 5 | 3 | 2 | Spata31d1b |
| Spata31d1c | 0 | 0 | 0 | 0 | 2 | 0 | 0 | 1 | 0 | 2 | 3 | 0 | 1 | 0 | 1 | 0 | 0 | 0 | 0 | 0 | 0 | 0 | 0 | 0 | 0 | 0 | 2 | 2 | Spata31d1c |
| Spata31d1d | 7 | 10 | 9 | 7 | 9 | 15 | 13 | 2 | 2 | 3 | 3 | 3 | 7 | 1 | 9 | 12 | 10 | 7 | 12 | 16 | 7 | 0 | 3 | 8 | 0 | 3 | 0 | 1 | Spata31d1d |
| Spata32 | 7 | 3 | 0 | 8 | 10 | 1 | 4 | 9 | 13 | 10 | 7 | 3 | 3 | 1 | 4 | 4 | 8 | 11 | 4 | 13 | 4 | 15 | 7 | 4 | 12 | 0 | 5 | 5 | Spata32 |
| Spata33 | 683 | 685 | 506 | 756 | 694 | 693 | 792 | 485 | 460 | 351 | 419 | 429 | 471 | 487 | 818 | 689 | 693 | 851 | 715 | 714 | 738 | 520 | 454 | 413 | 410 | 563 | 468 | 552 | Spata33 |
| Spata4 | 9 | 4 | 3 | 0 | 2 | 0 | 3 | 7 | 6 | 8 | 11 | 0 | 7 | 2 | 3 | 9 | 3 | 0 | 0 | 0 | 8 | 8 | 5 | 6 | 0 | 4 | 3 | 4 | Spata4 |

| | | | | | | | | | | | | | | | | | | | | | | | | | | | | | |
|---|---|---|---|---|---|---|---|---|---|---|---|---|---|---|---|---|---|---|---|---|---|---|---|---|---|---|---|---|---|
| Spata45 | 0 | 2 | 1 | 5 | 0 | 0 | 0 | 0 | 1 | 4 | 0 | 0 | 0 | 1 | 3 | 0 | 0 | 1 | 1 | 1 | 0 | 0 | 0 | 1 | 0 | 0 | 0 | 0 | Spata45 |
| Spata46 | 0 | 2 | 0 | 0 | 0 | 0 | 0 | 0 | 0 | 0 | 0 | 0 | 0 | 0 | 0 | 0 | 0 | 0 | 0 | 0 | 0 | 0 | 0 | 0 | 0 | 0 | 0 | 0 | Spata46 |
| Spata48 | 13 | 7 | 14 | 15 | 21 | 28 | 11 | 60 | 41 | 33 | 60 | 52 | 61 | 64 | 9 | 12 | 24 | 12 | 10 | 22 | 15 | 52 | 54 | 44 | 68 | 55 | 64 | 39 | Spata48 |
| Spata5 | 3964 | 3524 | 3278 | 3424 | 2853 | 3168 | 3472 | 2673 | 2107 | 1835 | 1927 | 2279 | 2087 | 2319 | 3985 | 3378 | 2982 | 3378 | 2780 | 3424 | 3363 | 2172 | 2170 | 2151 | 2084 | 2523 | 2368 | 2480 | Spata5 |
| Spata6 | 3660 | 3127 | 3369 | 2548 | 2165 | 2332 | 2297 | 1292 | 1271 | 1070 | 1275 | 1357 | 1589 | 1288 | 3684 | 3562 | 2870 | 2287 | 2085 | 2518 | 2244 | 1263 | 1399 | 1067 | 988 | 1466 | 1427 | 1491 | Spata6 |
| Spata6l | 38 | 19 | 17 | 17 | 27 | 32 | 28 | 64 | 55 | 60 | 60 | 42 | 46 | 29 | 34 | 34 | 21 | 24 | 40 | 31 | 29 | 37 | 66 | 25 | 42 | 33 | 40 | 23 | Spata6l |
| Spata7 | 1858 | 1869 | 1906 | 1890 | 1880 | 1833 | 2036 | 1198 | 1134 | 1189 | 1011 | 1300 | 1101 | 1200 | 2058 | 1878 | 1896 | 1844 | 1801 | 2098 | 1979 | 1183 | 1003 | 1112 | 1076 | 1331 | 1238 | 1270 | Spata7 |
| Spata9 | 8 | 5 | 12 | 9 | 19 | 7 | 16 | 17 | 15 | 10 | 25 | 21 | 10 | 10 | 16 | 8 | 21 | 10 | 8 | 16 | 19 | 27 | 15 | 9 | 20 | 11 | 10 | 7 | Spata9 |
| Spatc1 | 0 | 0 | 6 | 0 | 0 | 0 | 0 | 0 | 0 | 0 | 0 | 0 | 0 | 0 | 0 | 0 | 0 | 0 | 0 | 0 | 1 | 0 | 0 | 0 | 0 | 0 | 0 | 0 | Spatc1 |
| Spatc1l | 9 | 31 | 21 | 10 | 20 | 13 | 12 | 4 | 9 | 10 | 12 | 4 | 1 | 5 | 12 | 30 | 12 | 13 | 16 | 18 | 9 | 13 | 6 | 18 | 6 | 4 | 8 | 9 | Spatc1l |
| Spats1 | 16 | 22 | 18 | 22 | 11 | 16 | 35 | 34 | 29 | 51 | 40 | 22 | 23 | 40 | 26 | 22 | 17 | 21 | 24 | 24 | 33 | 22 | 42 | 23 | 23 | 31 | 33 | 27 | Spats1 |
| Spats2 | 7630 | 7314 | 8681 | 8096 | 8084 | 7678 | 8272 | 5190 | 4803 | 5494 | 5096 | 5930 | 5812 | 6189 | 7551 | 7331 | 8317 | 7957 | 8066 | 6811 | 8017 | 5516 | 4747 | 6008 | 5681 | 6749 | 5487 | 6449 | Spats2 |
| Spats2l | 1103 | 1062 | 938 | 822 | 1026 | 1228 | 755 | 2766 | 3899 | 3150 | 3868 | 2611 | 2929 | 2398 | 1303 | 1240 | 853 | 1082 | 868 | 914 | 750 | 3058 | 4339 | 2843 | 2599 | 2431 | 2467 | 2442 | Spats2l |
| Spc24 | 1747 | 1682 | 2259 | 1602 | 1046 | 1388 | 1426 | 1849 | 1973 | 2261 | 2104 | 3005 | 2558 | 3095 | 1448 | 1528 | 1905 | 1293 | 1606 | 1445 | 1287 | 1078 | 1690 | 2868 | 2193 | 4265 | 2972 | 3265 | Spc24 |
| Spc25 | 4702 | 3534 | 4850 | 3560 | 2749 | 3202 | 4209 | 4739 | 3985 | 3604 | 3416 | 5680 | 4295 | 6091 | 4867 | 4138 | 4400 | 3280 | 3349 | 4175 | 3507 | 2891 | 3394 | 4820 | 4081 | 6625 | 6064 | 6467 | Spc25 |
| Spcs1 | 4429 | 4107 | 4024 | 3879 | 3360 | 3520 | 4266 | 4318 | 3863 | 3488 | 3780 | 3312 | 3545 | 3674 | 4622 | 4216 | 3854 | 3703 | 3907 | 4606 | 4110 | 3581 | 4292 | 3825 | 3402 | 4081 | 3572 | 3990 | Spcs1 |
| Spcs2 | 9236 | 7986 | 9155 | 7201 | 6763 | 6738 | 7464 | 9442 | 7982 | 7717 | 7541 | 8241 | 8205 | 8925 | 9563 | 8281 | 8027 | 7112 | 7323 | 8203 | 7380 | 8246 | 8567 | 8025 | 7829 | 9615 | 8823 | 9563 | Spcs2 |
| Spcs3 | 8922 | 7138 | 7910 | 7630 | 6810 | 7383 | 8223 | 7717 | 6672 | 5970 | 6634 | 6434 | 6402 | 6702 | 8831 | 7868 | 7398 | 7381 | 7069 | 8527 | 7501 | 6331 | 6919 | 6396 | 6181 | 7216 | 6845 | 7180 | Spcs3 |
| Spdef | 87 | 124 | 73 | 134 | 133 | 121 | 129 | 18 | 30 | 43 | 22 | 31 | 45 | 29 | 105 | 119 | 89 | 156 | 110 | 89 | 108 | 32 | 13 | 58 | 26 | 23 | 19 | 21 | Spdef |
| Spdl1 | 1614 | 1485 | 1973 | 1243 | 993 | 1107 | 1295 | 1778 | 1592 | 1424 | 1349 | 2126 | 1766 | 2160 | 1605 | 1589 | 1547 | 1012 | 1232 | 1384 | 1067 | 1106 | 1414 | 1770 | 1516 | 2459 | 2447 | 2308 | Spdl1 |
| Spdya | 106 | 55 | 126 | 125 | 91 | 108 | 104 | 79 | 68 | 57 | 73 | 100 | 70 | 100 | 94 | 94 | 97 | 96 | 103 | 107 | 72 | 69 | 75 | 90 | 88 | 125 | 139 | 100 | Spdya |
| Spdye4a | 7 | 10 | 3 | 6 | 3 | 0 | 7 | 1 | 4 | 0 | 3 | 1 | 7 | 1 | 5 | 9 | 8 | 6 | 5 | 4 | 9 | 0 | 1 | 0 | 0 | 0 | 5 | 1 | Spdye4a |
| Spdye4b | 0 | 2 | 0 | 0 | 1 | 0 | 0 | 5 | 0 | 0 | 0 | 0 | 0 | 0 | 0 | 0 | 1 | 0 | 0 | 0 | 0 | 0 | 1 | 0 | 0 | 0 | 0 | 0 | Spdye4b |
| Spdye4c | 0 | 0 | 0 | 0 | 0 | 0 | 0 | 0 | 0 | 0 | 0 | 0 | 0 | 0 | 0 | 1 | 0 | 0 | 0 | 0 | 0 | 0 | 0 | 0 | 0 | 0 | 0 | 0 | Spdye4c |
| Specc1 | 7097 | 6940 | 6417 | 7336 | 7018 | 7010 | 6712 | 3882 | 4144 | 3968 | 3919 | 5511 | 4783 | 5120 | 6763 | 6598 | 7217 | 6959 | 7423 | 6092 | 6544 | 4309 | 3814 | 4417 | 4474 | 4429 | 5002 | 4932 | Specc1 |
| Specc1l | 5353 | 6214 | 5078 | 5999 | 6705 | 6422 | 6348 | 11052 | 11534 | 11138 | 10641 | 11039 | 10624 | 11197 | 6086 | 5429 | 5729 | 6325 | 6537 | 6152 | 6646 | 10388 | 10864 | 10560 | 11361 | 8683 | 11243 | 10524 | Specc1l |
| Speer2 | 0 | 0 | 0 | 0 | 0 | 0 | 0 | 1 | 4 | 2 | 0 | 2 | 1 | 0 | 0 | 1 | 0 | 0 | 0 | 1 | 1 | 0 | 0 | 1 | 0 | 0 | 0 | 1 | Speer2 |
| Speer4a | 0 | 0 | 0 | 0 | 0 | 0 | 0 | 2 | 2 | 1 | 0 | 4 | 1 | 2 | 0 | 0 | 0 | 0 | 0 | 0 | 1 | 2 | 1 | 0 | 0 | 0 | 0 | 0 | Speer4a |
| Speer4b | 0 | 0 | 0 | 0 | 0 | 0 | 0 | 0 | 0 | 0 | 0 | 0 | 0 | 1 | 0 | 0 | 0 | 0 | 0 | 0 | 0 | 0 | 0 | 4 | 0 | 0 | 0 | 0 | Speer4b |
| Speer4c | 0 | 0 | 0 | 0 | 0 | 0 | 0 | 1 | 0 | 0 | 0 | 0 | 0 | 0 | 0 | 0 | 0 | 0 | 0 | 0 | 0 | 0 | 0 | 0 | 0 | 0 | 0 | 0 | Speer4c |
| Speer4e | 0 | 0 | 0 | 0 | 0 | 0 | 0 | 0 | 0 | 0 | 0 | 0 | 2 | 0 | 0 | 0 | 0 | 0 | 0 | 0 | 0 | 0 | 0 | 0 | 0 | 0 | 0 | 0 | Speer4e |
| Speer4f1 | 0 | 0 | 0 | 1 | 0 | 0 | 0 | 0 | 0 | 0 | 0 | 0 | 0 | 0 | 0 | 1 | 0 | 0 | 0 | 0 | 0 | 0 | 0 | 1 | 0 | 0 | 0 | 0 | Speer4f1 |
| Speer4f2 | 0 | 0 | 0 | 0 | 0 | 0 | 0 | 0 | 1 | 0 | 0 | 0 | 0 | 0 | 0 | 0 | 0 | 0 | 0 | 0 | 0 | 0 | 0 | 0 | 0 | 0 | 0 | 0 | Speer4f2 |
| Spef1 | 2648 | 3473 | 4275 | 4786 | 4474 | 3666 | 4884 | 2120 | 2317 | 3143 | 2515 | 3492 | 2808 | 3405 | 2825 | 3037 | 5064 | 4802 | 5150 | 3601 | 4827 | 2969 | 2339 | 3652 | 3427 | 3504 | 2873 | 3131 | Spef1 |
| Spef1l | 25 | 41 | 32 | 47 | 27 | 36 | 32 | 24 | 28 | 12 | 30 | 40 | 37 | 23 | 32 | 37 | 39 | 30 | 23 | 21 | 27 | 15 | 37 | 16 | 51 | 41 | 37 | 38 | Spef1l |
| Spef2 | 193 | 162 | 101 | 140 | 240 | 180 | 164 | 293 | 232 | 236 | 208 | 226 | 271 | 143 | 199 | 319 | 145 | 178 | 122 | 257 | 185 | 252 | 398 | 132 | 190 | 144 | 153 | 142 | Spef2 |
| Speg | 2706 | 3536 | 4124 | 3110 | 3238 | 3022 | 2216 | 1772 | 2160 | 2469 | 2755 | 2095 | 2429 | 1658 | 2306 | 3571 | 3688 | 3408 | 3183 | 2245 | 2373 | 2876 | 2148 | 2357 | 2295 | 1645 | 2113 | 1749 | Speg |
| Spem1 | 111 | 212 | 153 | 268 | 242 | 221 | 256 | 59 | 35 | 106 | 38 | 76 | 54 | 63 | 116 | 139 | 239 | 306 | 265 | 175 | 257 | 90 | 28 | 73 | 66 | 60 | 59 | 82 | Spem1 |
| Spem2 | 0 | 0 | 8 | 0 | 1 | 0 | 0 | 0 | 0 | 0 | 2 | 0 | 0 | 0 | 3 | 2 | 5 | 0 | 0 | 0 | 1 | 0 | 0 | 0 | 0 | 0 | 0 | 0 | Spem2 |
| Spen | 8433 | 11109 | 10959 | 14825 | 15186 | 13288 | 11063 | 8275 | 9248 | 12480 | 11537 | 14143 | 10910 | 11978 | 7655 | 10142 | 12860 | 15331 | 14977 | 9738 | 13019 | 13593 | 8136 | 12519 | 14850 | 5790 | 11364 | 10892 | Spen |
| Spertl | 0 | 2 | 0 | 1 | 0 | 3 | 0 | 0 | 1 | 0 | 0 | 1 | 0 | 0 | 0 | 0 | 6 | 0 | 1 | 0 | 2 | 3 | 3 | 1 | 6 | 0 | 3 | 0 | Spertl |
| Spesp1 | 2 | 1 | 0 | 0 | 3 | 0 | 3 | 1 | 0 | 0 | 0 | 3 | 0 | 1 | 4 | 0 | 1 | 0 | 0 | 0 | 1 | 0 | 0 | 0 | 3 | 1 | 1 | 2 | Spesp1 |
| Spg11 | 3751 | 3724 | 2261 | 2779 | 3088 | 3565 | 3286 | 5556 | 5567 | 4610 | 4465 | 5427 | 5201 | 4992 | 3849 | 3224 | 2598 | 2660 | 2975 | 3458 | 3359 | 4338 | 5317 | 4314 | 5024 | 3280 | 5854 | 4779 | Spg11 |
| Spg20 | 4496 | 3868 | 4050 | 2890 | 2650 | 2998 | 2853 | 4435 | 5347 | 4316 | 5372 | 4472 | 4906 | 4686 | 4007 | 4323 | 3450 | 3240 | 2764 | 2869 | 2817 | 3639 | 5388 | 4628 | 3742 | 5431 | 4462 | 5179 | Spg20 |
| Spg21 | 4676 | 4495 | 3508 | 4094 | 3438 | 3577 | 4164 | 4380 | 4036 | 3642 | 3835 | 3547 | 3309 | 3818 | 4767 | 4380 | 4238 | 3528 | 3788 | 4207 | 4035 | 4010 | 4354 | 3434 | 3584 | 3771 | 3606 | 3881 | Spg21 |
| Spg7 | 2185 | 2742 | 2849 | 2993 | 2857 | 2788 | 2539 | 3633 | 3840 | 4310 | 4028 | 4643 | 4309 | 4566 | 2198 | 2614 | 2907 | 2829 | 3027 | 2506 | 2726 | 4045 | 3611 | 4480 | 4498 | 4426 | 4695 | 4411 | Spg7 |
| Sphk1 | 197 | 236 | 219 | 133 | 142 | 193 | 66 | 147 | 171 | 157 | 165 | 135 | 272 | 122 | 181 | 283 | 148 | 157 | 97 | 95 | 81 | 189 | 187 | 144 | 163 | 162 | 213 | 185 | Sphk1 |
| Sphk2 | 1791 | 2731 | 2434 | 2812 | 1941 | 1963 | 1719 | 1547 | 1865 | 1846 | 1949 | 2130 | 2011 | 1851 | 1672 | 2220 | 2545 | 2453 | 2237 | 1751 | 1879 | 1850 | 1855 | 2146 | 2010 | 2394 | 1954 | 1828 | Sphk2 |
| Sphkap | 484 | 483 | 208 | 509 | 1261 | 902 | 1082 | 1432 | 1596 | 1189 | 1084 | 762 | 583 | 825 | 647 | 401 | 411 | 647 | 906 | 1070 | 1045 | 1205 | 1351 | 928 | 1474 | 538 | 643 | 501 | Sphkap |
| Spi1 | 36 | 79 | 52 | 57 | 53 | 49 | 26 | 96 | 170 | 114 | 143 | 153 | 111 | 90 | 40 | 78 | 53 | 47 | 38 | 38 | 55 | 129 | 189 | 112 | 96 | 140 | 126 | 87 | Spi1 |
| Spib | 10 | 17 | 27 | 24 | 16 | 5 | 7 | 7 | 16 | 3 | 9 | 11 | 8 | 11 | 13 | 26 | 31 | 57 | 26 | 16 | 13 | 5 | 21 | 11 | 10 | 12 | 22 | 7 | Spib |
| Spic | 0 | 0 | 8 | 2 | 0 | 0 | 0 | 2 | 8 | 1 | 1 | 2 | 7 | 0 | 3 | 5 | 3 | 8 | 1 | 0 | 0 | 3 | 5 | 7 | 4 | 0 | 3 | 2 | Spic |
| Spice1 | 2633 | 2607 | 2880 | 2494 | 2201 | 2088 | 2534 | 2385 | 2274 | 2154 | 2091 | 2438 | 2380 | 2394 | 2676 | 2435 | 2902 | 2510 | 2325 | 2447 | 2457 | 2112 | 2323 | 2321 | 1930 | 2488 | 2597 | 2586 | Spice1 |
| Spidr | 3307 | 2752 | 2728 | 2254 | 1684 | 2128 | 2364 | 2602 | 2291 | 2004 | 2103 | 2517 | 2391 | 2532 | 3011 | 2904 | 2460 | 2049 | 2014 | 2273 | 1983 | 1619 | 2387 | 2182 | 1828 | 2532 | 2613 | 2586 | Spidr |
| Spin1 | 10129 | 7908 | 10009 | 10533 | 13016 | 11880 | 13006 | 20069 | 18732 | 19635 | 19129 | 20012 | 18844 | 20503 | 9723 | 8728 | 10564 | 10666 | 13000 | 10986 | 12522 | 23761 | 18305 | 20537 | 22139 | 20053 | 19282 | 20063 | Spin1 |
| Spin2c | 652 | 461 | 464 | 379 | 518 | 503 | 523 | 672 | 653 | 519 | 489 | 525 | 506 | 546 | 576 | 531 | 406 | 428 | 460 | 573 | 494 | 496 | 572 | 589 | 493 | 661 | 495 | 540 | Spin2c |
| Spin4 | 885 | 557 | 642 | 550 | 547 | 633 | 650 | 707 | 680 | 462 | 651 | 663 | 643 | 701 | 872 | 795 | 520 | 500 | 547 | 669 | 610 | 556 | 720 | 592 | 504 | 667 | 725 | 841 | Spin4 |
| Spindoc | 3086 | 3887 | 4394 | 4432 | 3957 | 4164 | 3724 | 2869 | 3158 | 3749 | 3386 | 3781 | 3708 | 3512 | 3009 | 3252 | 4639 | 4205 | 4878 | 3626 | 3858 | 3345 | 3087 | 3765 | 3855 | 4398 | 3691 | 3709 | Spindoc |
| Spink1 | 0 | 1 | 0 | 0 | 0 | 0 | 0 | 4 | 7 | 0 | 1 | 0 | 0 | 0 | 0 | 0 | 0 | 0 | 0 | 0 | 0 | 0 | 3 | 0 | 1 | 0 | 0 | 0 | Spink1 |
| Spink10 | 1152 | 1071 | 649 | 1119 | 1171 | 1229 | 1279 | 520 | 537 | 517 | 507 | 402 | 494 | 412 | 1320 | 1335 | 947 | 1176 | 916 | 1175 | 1266 | 600 | 528 | 350 | 470 | 302 | 443 | 388 | Spink10 |
| Spink12 | 1 | 0 | 0 | 0 | 0 | 0 | 0 | 0 | 0 | 0 | 0 | 0 | 0 | 0 | 0 | 0 | 0 | 0 | 0 | 0 | 0 | 0 | 1 | 0 | 0 | 0 | 0 | 0 | Spink12 |
| Spink13 | 5 | 3 | 0 | 13 | 7 | 3 | 12 | 1 | 9 | 0 | 3 | 4 | 0 | 11 | 1 | 3 | 1 | 19 | 12 | 8 | 13 | 0 | 1 | 17 | 13 | 1 | 0 | 0 | Spink13 |
| Spink2 | 9 | 9 | 10 | 10 | 8 | 9 | 8 | 2 | 15 | 14 | 23 | 9 | 19 | 5 | 16 | 12 | 14 | 4 | 8 | 4 | 7 | 8 | 10 | 12 | 8 | 7 | 13 | 8 | Spink2 |
| Spink4 | 0 | 0 | 0 | 0 | 0 | 0 | 0 | 0 | 0 | 0 | 0 | 0 | 0 | 0 | 3 | 1 | 0 | 0 | 0 | 0 | 0 | 0 | 0 | 0 | 0 | 0 | 0 | 0 | Spink4 |
| Spink5 | 1 | 0 | 0 | 0 | 0 | 0 | 0 | 2 | 3 | 5 | 21 | 0 | 0 | 0 | 1 | 0 | 0 | 0 | 0 | 0 | 0 | 2 | 11 | 2 | 0 | 3 | 0 | 0 | Spink5 |
| Spink7 | 0 | 0 | 0 | 0 | 2 | 0 | 1 | 0 | 0 | 0 | 0 | 0 | 0 | 0 | 0 | 1 | 0 | 0 | 1 | 0 | 0 | 0 | 0 | 0 | 0 | 0 | 0 | 0 | Spink7 |
| Spink8 | 1 | 0 | 0 | 0 | 0 | 10 | 1 | 0 | 1 | 0 | 1 | 0 | 3 | 1 | 4 | 1 | 0 | 1 | 0 | 0 | 0 | 0 | 5 | 0 | 2 | 0 | 0 | 2 | Spink8 |
| Spint1 | 290 | 179 | 127 | 70 | 78 | 85 | 21 | 230 | 281 | 227 | 193 | 74 | 146 | 88 | 160 | 234 | 83 | 96 | 36 | 32 | 28 | 70 | 314 | 158 | 23 | 199 | 107 | 103 | Spint1 |
| Spint2 | 1351 | 1057 | 937 | 1016 | 1006 | 959 | 854 | 1144 | 1503 | 1536 | 1394 | 979 | 1134 | 914 | 955 | 1139 | 791 | 975 | 1012 | 867 | 883 | 856 | 1671 | 1488 | 830 | 1760 | 774 | 868 | Spint2 |
| Spint4 | 0 | 0 | 0 | 0 | 0 | 0 | 0 | 0 | 0 | 0 | 0 | 0 | 0 | 1 | 0 | 0 | 0 | 0 | 0 | 0 | 0 | 0 | 1 | 0 | 0 | 0 | 0 | 0 | Spint4 |
| Spire1 | 11108 | 11491 | 11358 | 13187 | 14306 | 13865 | 14884 | 8135 | 7458 | 8847 | 7660 | 8994 | 9055 | 9418 | 12057 | 11150 | 12447 | 13391 | 14062 | 12748 | 14682 | 8794 | 6845 | 8581 | 9566 | 7467 | 8612 | 9149 | Spire1 |
| Spire2 | 1058 | 1536 | 1537 | 1942 | 2056 | 1685 | 1573 | 1153 | 1218 | 1467 | 1268 | 1349 | 1164 | 1171 | 1193 | 1129 | 1884 | 1766 | 2204 | 1623 | 1744 | 1649 | 1207 | 1326 | 1635 | 1272 | 1252 | 1206 | Spire2 |
| Spn | 16 | 25 | 15 | 28 | 10 | 5 | 17 | 18 | 27 | 16 | 27 | 27 | 23 | 13 | 34 | 25 | 28 | 20 | 7 | 14 | 12 | 12 | 26 | 25 | 13 | 11 | 16 | 28 | Spn |
| Spns1 | 3170 | 4059 | 3849 | 3742 | 3798 | 3525 | 3127 | 3046 | 3699 | 3781 | 3670 | 3733 | 3868 | 3245 | 3025 | 3483 | 4087 | 3653 | 4107 | 3396 | 3257 | 3663 | 3455 | 3865 | 3759 | 3873 | 3753 | 3302 | Spns1 |
| Spns2 | 1215 | 1678 | 1470 | 1916 | 2333 | 1983 | 1735 | 1669 | 1932 | 2335 | 1783 | 1742 | 1822 | 1524 | 1357 | 1499 | 1897 | 1960 | 2007 | 1821 | 1796 | 2446 | 1655 | 1785 | 2144 | 1551 | 1594 | 1553 | Spns2 |
| Spns3 | 5 | 3 | 4 | 2 | 3 | 2 | 2 | 6 | 7 | 1 | 8 | 10 | 9 | 2 | 5 | 8 | 3 | 3 | 8 | 4 | 2 | 8 | 7 | 9 | 5 | 12 | 8 | 5 | Spns3 |
| Spo11 | 51 | 26 | 22 | 32 | 48 | 78 | 41 | 126 | 192 | 95 | 117 | 89 | 96 | 62 | 67 | 48 | 28 | 48 | 49 | 56 | 42 | 58 | 187 | 105 | 69 | 94 | 57 | 88 | Spo11 |
| Spocd1 | 20 | 22 | 27 | 40 | 16 | 19 | 19 | 4 | 9 | 4 | 12 | 14 | 14 | 9 | 9 | 9 | 17 | 32 | 23 | 12 | 29 | 17 | 6 | 7 | 6 | 3 | 6 | 1 | Spocd1 |
| Spock1 | 1761 | 1734 | 2252 | 3248 | 5980 | 3673 | 3106 | 3326 | 3362 | 5216 | 4886 | 2952 | 3982 | 2943 | 2272 | 2079 | 3101 | 4074 | 3878 | 2911 | 4211 | 6916 | 3736 | 3731 | 5055 | 1869 | 2553 | 2966 | Spock1 |

| | | | | | | | | | | | | | | | | | | | | | | | | | | | | | |
|---|---|---|---|---|---|---|---|---|---|---|---|---|---|---|---|---|---|---|---|---|---|---|---|---|---|---|---|---|---|
| Spock2 | 2902 | 3528 | 4144 | 6162 | 10494 | 7325 | 6729 | 8005 | 9110 | 11742 | 10708 | 7609 | 8589 | 6714 | 3700 | 3619 | 5336 | 7483 | 8339 | 6419 | 7973 | 14567 | 8569 | 8759 | 12482 | 5469 | 6458 | 6427 | Spock2 |
| Spock3 | 1275 | 994 | 912 | 1360 | 2167 | 1888 | 1333 | 2654 | 2274 | 1846 | 2385 | 1316 | 1970 | 1447 | 1925 | 1547 | 1050 | 1684 | 1372 | 1731 | 1481 | 3068 | 2760 | 1251 | 2061 | 931 | 1249 | 1310 | Spock3 |
| Spon1 | 3160 | 5179 | 4499 | 3832 | 5974 | 4292 | 2718 | 6843 | 5919 | 5926 | 5924 | 3666 | 9009 | 3076 | 7474 | 7064 | 4642 | 4165 | 3223 | 3716 | 3006 | 7147 | 5391 | 2915 | 4273 | 2435 | 4780 | 3542 | Spon1 |
| Spon2 | 48 | 104 | 66 | 54 | 74 | 100 | 49 | 131 | 199 | 125 | 216 | 135 | 139 | 122 | 61 | 76 | 52 | 80 | 58 | 42 | 63 | 139 | 315 | 155 | 138 | 135 | 158 | 131 | Spon2 |
| Spop | 9596 | 9411 | 13811 | 11814 | 11131 | 10020 | 11448 | 8636 | 7978 | 9744 | 8699 | 10085 | 9539 | 10338 | 9792 | 9800 | 13236 | 11348 | 11130 | 10517 | 10701 | 9678 | 8403 | 10156 | 10051 | 10975 | 10258 | 10452 | Spop |
| Spopfm3 | 0 | 0 | 0 | 0 | 0 | 0 | 0 | 0 | 0 | 0 | 0 | 0 | 0 | 0 | 0 | 0 | 0 | 0 | 0 | 0 | 0 | 0 | 0 | 0 | 1 | 0 | 0 | 0 | Spopfm3 |
| Spopl | 1970 | 1484 | 1905 | 2182 | 2312 | 2425 | 2591 | 2866 | 2548 | 2618 | 3072 | 3067 | 3234 | 3185 | 1923 | 1987 | 1947 | 2046 | 2159 | 2208 | 2284 | 2976 | 2695 | 2711 | 2928 | 2382 | 2823 | 3163 | Spopl |
| Spout1 | 3245 | 4059 | 4683 | 4769 | 3465 | 3576 | 3351 | 2097 | 2736 | 3181 | 2928 | 2959 | 2994 | 2822 | 3086 | 3738 | 4475 | 3998 | 4168 | 3392 | 3529 | 2321 | 2745 | 3163 | 2830 | 3842 | 3018 | 3026 | Spout1 |
| Spp1 | 331 | 317 | 212 | 207 | 155 | 219 | 102 | 122 | 288 | 153 | 255 | 113 | 314 | 146 | 446 | 438 | 169 | 219 | 142 | 106 | 117 | 328 | 403 | 137 | 177 | 79 | 152 | 150 | Spp1 |
| Spp2 | 1 | 2 | 0 | 0 | 6 | 3 | 1 | 2 | 14 | 2 | 0 | 3 | 1 | 4 | 0 | 3 | 0 | 2 | 3 | 3 | 4 | 3 | 9 | 2 | 8 | 1 | 2 | 2 | Spp2 |
| Sppl2a | 5206 | 3845 | 4051 | 4146 | 4165 | 4589 | 4712 | 7598 | 6809 | 5917 | 6380 | 6364 | 6398 | 6849 | 5036 | 4432 | 3940 | 4377 | 4347 | 5150 | 4455 | 6403 | 6965 | 6378 | 6592 | 5987 | 6746 | 6724 | Sppl2a |
| Sppl2b | 3092 | 4377 | 3831 | 4030 | 3619 | 3422 | 3067 | 2152 | 2779 | 2667 | 2809 | 2613 | 2552 | 2132 | 2995 | 3812 | 3834 | 3661 | 3746 | 3273 | 3006 | 2642 | 2718 | 2638 | 2724 | 2773 | 2307 | 2279 | Sppl2b |
| Sppl2c | 0 | 0 | 0 | 0 | 0 | 0 | 0 | 0 | 1 | 0 | 0 | 0 | 2 | 0 | 0 | 0 | 1 | 0 | 0 | 0 | 2 | 2 | 0 | 0 | 0 | 0 | 0 | 0 | Sppl2c |
| Sppl3 | 4295 | 4366 | 5159 | 5017 | 5093 | 4568 | 4757 | 5013 | 5197 | 5362 | 5555 | 5887 | 5350 | 5443 | 3767 | 4396 | 5029 | 5016 | 5026 | 3985 | 4664 | 5989 | 5095 | 5727 | 5760 | 6312 | 5285 | 5633 | Sppl3 |
| Spr | 890 | 997 | 1299 | 822 | 818 | 953 | 811 | 1094 | 1326 | 1099 | 1155 | 1338 | 1436 | 1196 | 815 | 917 | 958 | 842 | 958 | 1004 | 783 | 866 | 1327 | 1289 | 1247 | 1866 | 1505 | 1443 | Spr |
| Spred1 | 8691 | 5608 | 7367 | 7576 | 7670 | 7747 | 7965 | 9408 | 8763 | 8117 | 9001 | 8687 | 8962 | 8718 | 7713 | 7466 | 7435 | 7878 | 7279 | 6764 | 7543 | 8863 | 8375 | 8336 | 8589 | 8602 | 8488 | 8799 | Spred1 |
| Spred2 | 2030 | 1688 | 2579 | 2402 | 2676 | 2211 | 2345 | 2047 | 2580 | 2710 | 2861 | 2837 | 2388 | 2903 | 1792 | 1855 | 2343 | 2156 | 2530 | 1741 | 2314 | 2689 | 2256 | 2857 | 2577 | 3429 | 2293 | 2730 | Spred2 |
| Spred3 | 3142 | 2820 | 3005 | 3717 | 3881 | 3457 | 3504 | 3446 | 3294 | 3952 | 3619 | 3698 | 3531 | 3858 | 2686 | 2718 | 3570 | 3423 | 3599 | 3073 | 3646 | 4455 | 2998 | 3805 | 4237 | 3577 | 3612 | 3368 | Spred3 |
| Spring1 | 5580 | 6043 | 5276 | 6961 | 6485 | 6121 | 5955 | 3219 | 3289 | 3507 | 3425 | 3675 | 3333 | 3297 | 5629 | 5527 | 5926 | 6722 | 6652 | 6305 | 6462 | 3325 | 3042 | 3577 | 3876 | 3103 | 3245 | 3241 | Spring1 |
| Sprn | 90 | 116 | 76 | 96 | 322 | 196 | 168 | 547 | 611 | 591 | 488 | 310 | 341 | 310 | 133 | 87 | 104 | 144 | 156 | 167 | 190 | 744 | 575 | 424 | 487 | 263 | 308 | 214 | Sprn |
| Sprr1a | 2 | 1 | 0 | 0 | 0 | 0 | 0 | 27 | 34 | 34 | 3 | 0 | 0 | 0 | 0 | 0 | 0 | 0 | 1 | 0 | 0 | 0 | 29 | 9 | 0 | 0 | 0 | 0 | Sprr1a |
| Sprr1b | 0 | 0 | 0 | 0 | 0 | 0 | 0 | 2 | 2 | 0 | 0 | 0 | 0 | 0 | 0 | 0 | 0 | 0 | 0 | 0 | 0 | 0 | 5 | 0 | 0 | 0 | 0 | 0 | Sprr1b |
| Sprr2a1 | 0 | 0 | 0 | 0 | 0 | 0 | 0 | 0 | 0 | 0 | 1 | 0 | 0 | 0 | 0 | 0 | 0 | 0 | 0 | 0 | 0 | 0 | 1 | 2 | 0 | 0 | 0 | 0 | Sprr2a1 |
| Sprr2a3 | 0 | 0 | 0 | 0 | 0 | 0 | 0 | 0 | 0 | 0 | 0 | 0 | 0 | 0 | 0 | 0 | 0 | 0 | 0 | 0 | 0 | 0 | 3 | 0 | 0 | 0 | 0 | 0 | Sprr2a3 |
| Sprr2b | 0 | 1 | 5 | 0 | 0 | 0 | 0 | 0 | 2 | 0 | 0 | 0 | 10 | 0 | 0 | 2 | 0 | 0 | 0 | 0 | 0 | 0 | 1 | 0 | 0 | 0 | 0 | 0 | Sprr2b |
| Sprr2f | 0 | 0 | 0 | 0 | 0 | 0 | 0 | 0 | 1 | 0 | 0 | 0 | 0 | 0 | 0 | 0 | 0 | 0 | 0 | 0 | 0 | 0 | 0 | 0 | 0 | 0 | 0 | 0 | Sprr2f |
| Sprr2h | 0 | 0 | 0 | 0 | 0 | 0 | 0 | 0 | 0 | 0 | 0 | 0 | 0 | 0 | 0 | 0 | 0 | 0 | 0 | 0 | 0 | 0 | 1 | 0 | 0 | 0 | 0 | 0 | Sprr2h |
| Sprr2k | 0 | 0 | 0 | 0 | 0 | 0 | 0 | 0 | 0 | 0 | 0 | 0 | 0 | 0 | 0 | 0 | 0 | 0 | 0 | 0 | 0 | 0 | 1 | 0 | 0 | 0 | 0 | 0 | Sprr2k |
| Sprr3 | 0 | 0 | 0 | 0 | 0 | 0 | 0 | 0 | 0 | 5 | 0 | 0 | 0 | 0 | 0 | 0 | 0 | 0 | 0 | 0 | 0 | 0 | 1 | 0 | 0 | 0 | 0 | 0 | Sprr3 |
| Sprr4 | 0 | 0 | 0 | 0 | 0 | 0 | 0 | 0 | 1 | 0 | 0 | 0 | 0 | 0 | 0 | 0 | 0 | 0 | 0 | 0 | 0 | 0 | 0 | 0 | 0 | 0 | 0 | 0 | Sprr4 |
| Sprtn | 3926 | 4173 | 4475 | 4618 | 3593 | 3911 | 3764 | 2811 | 2812 | 2933 | 3017 | 3272 | 3138 | 3239 | 3700 | 4012 | 4305 | 4479 | 3884 | 3294 | 3978 | 3150 | 2762 | 3057 | 3366 | 3214 | 3391 | 3379 | Sprtn |
| Spry1 | 1069 | 948 | 1358 | 1047 | 992 | 860 | 907 | 1154 | 1191 | 1163 | 1390 | 1754 | 1574 | 1600 | 820 | 905 | 1172 | 906 | 1054 | 821 | 872 | 1099 | 1223 | 1457 | 1115 | 2109 | 1427 | 1683 | Spry1 |
| Spry2 | 1053 | 1020 | 1436 | 1267 | 1368 | 1130 | 1006 | 1239 | 1316 | 1335 | 2048 | 1219 | 1415 | 1211 | 972 | 1043 | 1334 | 1147 | 1240 | 926 | 1020 | 1521 | 1505 | 1319 | 1225 | 1427 | 1130 | 1169 | Spry2 |
| Spry4 | 649 | 446 | 459 | 461 | 517 | 387 | 353 | 462 | 538 | 611 | 916 | 340 | 439 | 359 | 392 | 571 | 452 | 436 | 390 | 291 | 363 | 588 | 646 | 480 | 409 | 541 | 322 | 329 | Spry4 |
| Spryd3 | 1220 | 1411 | 1632 | 1969 | 2351 | 2066 | 1667 | 1544 | 1930 | 2228 | 2228 | 1647 | 1826 | 1469 | 1258 | 1492 | 1938 | 2088 | 2180 | 1870 | 1834 | 2181 | 2136 | 1869 | 2276 | 1732 | 1626 | 1518 | Spryd3 |
| Spryd4 | 1035 | 1114 | 929 | 836 | 785 | 758 | 700 | 747 | 819 | 731 | 737 | 792 | 836 | 726 | 945 | 1064 | 844 | 807 | 705 | 845 | 709 | 637 | 849 | 686 | 734 | 945 | 888 | 794 | Spryd4 |
| Spryd7 | 1991 | 1553 | 1632 | 1983 | 2274 | 1876 | 2562 | 1749 | 1469 | 1395 | 1391 | 1260 | 1105 | 1377 | 2392 | 1726 | 1885 | 2210 | 2097 | 2682 | 2389 | 1778 | 1515 | 1362 | 1548 | 1288 | 1316 | 1269 | Spryd7 |
| Spsb1 | 837 | 1002 | 727 | 660 | 692 | 634 | 560 | 470 | 548 | 506 | 738 | 464 | 717 | 441 | 791 | 1114 | 882 | 737 | 677 | 636 | 547 | 496 | 614 | 501 | 414 | 658 | 579 | 555 | Spsb1 |
| Spsb2 | 397 | 588 | 603 | 658 | 525 | 468 | 387 | 136 | 220 | 168 | 247 | 227 | 222 | 205 | 485 | 505 | 633 | 587 | 516 | 528 | 443 | 184 | 260 | 226 | 205 | 229 | 217 | 212 | Spsb2 |
| Spsb3 | 2241 | 2931 | 2498 | 2986 | 2491 | 2685 | 2318 | 1748 | 2141 | 2023 | 1801 | 1990 | 1919 | 1862 | 2197 | 2565 | 2497 | 2818 | 2872 | 2420 | 2455 | 1937 | 2049 | 1970 | 2091 | 2260 | 2023 | 1778 | Spsb3 |
| Spsb4 | 1401 | 1707 | 2413 | 1529 | 1413 | 1533 | 1252 | 1867 | 1811 | 2054 | 1787 | 3293 | 3046 | 3017 | 1557 | 1438 | 2052 | 1161 | 1643 | 1410 | 1298 | 2214 | 1566 | 2272 | 2380 | 3330 | 3541 | 3501 | Spsb4 |
| Spta1 | 323 | 416 | 391 | 124 | 55 | 127 | 133 | 208 | 238 | 150 | 142 | 371 | 1082 | 613 | 406 | 539 | 198 | 100 | 115 | 192 | 83 | 110 | 270 | 105 | 128 | 310 | 528 | 1239 | Spta1 |
| Sptan1 | 72472 | 86660 | 44349 | 75196 | 87123 | 86731 | 87900 | 71836 | 72120 | 66091 | 64168 | 55362 | 55318 | 56395 | 80665 | 73364 | 57515 | 81081 | 80734 | 87628 | 98257 | 63634 | 67966 | 52487 | 73828 | 30693 | 55035 | 49623 | Sptan1 |
| Sptb | 1015 | 1504 | 1561 | 1138 | 2280 | 1298 | 1150 | 553 | 1090 | 1264 | 1254 | 811 | 1894 | 1159 | 1108 | 1657 | 1359 | 1395 | 1541 | 1133 | 1373 | 1414 | 1419 | 721 | 1097 | 584 | 1031 | 2346 | Sptb |
| Sptbn1 | 33299 | 33777 | 25791 | 31168 | 40168 | 38419 | 33339 | 39782 | 42243 | 41167 | 46752 | 34387 | 38425 | 34125 | 34502 | 35367 | 28240 | 35315 | 32083 | 33181 | 36560 | 42509 | 43019 | 31663 | 42418 | 20450 | 35895 | 30807 | Sptbn1 |
| Sptbn2 | 5201 | 6931 | 7963 | 10566 | 14942 | 12408 | 10151 | 13431 | 15783 | 20463 | 16068 | 19728 | 16887 | 17495 | 6444 | 5923 | 10608 | 10439 | 14024 | 9578 | 12236 | 20986 | 13198 | 18065 | 23413 | 10499 | 15416 | 16822 | Sptbn2 |
| Sptbn4 | 920 | 1376 | 1420 | 2601 | 3637 | 2496 | 2495 | 1000 | 1309 | 1892 | 1555 | 828 | 771 | 744 | 1151 | 1168 | 2057 | 3168 | 3049 | 2001 | 2712 | 2002 | 1052 | 1267 | 1768 | 440 | 551 | 545 | Sptbn4 |
| Sptbn5 | 99 | 87 | 106 | 111 | 122 | 96 | 91 | 54 | 74 | 56 | 81 | 70 | 116 | 76 | 87 | 125 | 79 | 94 | 83 | 83 | 91 | 55 | 74 | 74 | 45 | 75 | 99 | 76 | Sptbn5 |
| Sptlc1 | 4513 | 3886 | 3769 | 3569 | 2673 | 3045 | 2958 | 3147 | 2785 | 2895 | 2770 | 2847 | 2935 | 2846 | 4065 | 4159 | 3167 | 3280 | 2745 | 3674 | 2913 | 2548 | 2972 | 2743 | 2743 | 2884 | 2923 | 2841 | Sptlc1 |
| Sptlc2 | 12716 | 11496 | 9293 | 9903 | 9895 | 10961 | 10760 | 6259 | 5558 | 5104 | 5281 | 5194 | 5474 | 5393 | 12765 | 12289 | 9243 | 10618 | 8765 | 11115 | 10435 | 6124 | 5443 | 4500 | 5438 | 4163 | 5022 | 4939 | Sptlc2 |
| Sptlc3 | 22 | 3 | 3 | 0 | 1 | 2 | 1 | 44 | 47 | 25 | 12 | 11 | 5 | 9 | 18 | 9 | 1 | 0 | 4 | 1 | 0 | 7 | 48 | 13 | 8 | 24 | 6 | 5 | Sptlc3 |
| Sptssa | 3430 | 2760 | 3589 | 2962 | 2921 | 2776 | 3467 | 3612 | 3010 | 2800 | 2742 | 2937 | 2892 | 3629 | 3562 | 3001 | 3381 | 2846 | 2886 | 3562 | 3313 | 2854 | 3043 | 3076 | 2783 | 3522 | 3191 | 3736 | Sptssa |
| Sptssb | 22 | 16 | 13 | 11 | 18 | 13 | 18 | 9 | 30 | 10 | 34 | 12 | 18 | 23 | 17 | 23 | 6 | 12 | 11 | 23 | 24 | 27 | 30 | 16 | 30 | 14 | 21 | 15 | Sptssb |
| Spty2d1 | 3372 | 3067 | 2953 | 3302 | 2316 | 2641 | 2735 | 2114 | 2252 | 2102 | 2402 | 1770 | 2239 | 2016 | 3033 | 3516 | 2708 | 3075 | 2302 | 2401 | 2633 | 2005 | 2334 | 2223 | 1992 | 1909 | 2054 | 2000 | Spty2d1 |
| Spx | 302 | 355 | 257 | 460 | 422 | 490 | 460 | 267 | 250 | 237 | 232 | 248 | 256 | 170 | 313 | 375 | 363 | 503 | 384 | 451 | 448 | 277 | 191 | 195 | 297 | 138 | 195 | 165 | Spx |
| Sqle | 15791 | 17807 | 15685 | 15614 | 13969 | 14431 | 15184 | 10365 | 9631 | 9880 | 8454 | 10799 | 11321 | 11835 | 17069 | 16566 | 14887 | 15989 | 13584 | 14842 | 14550 | 10525 | 9137 | 9942 | 10559 | 11845 | 11861 | 12728 | Sqle |
| Sqor | 149 | 131 | 148 | 108 | 83 | 142 | 121 | 154 | 163 | 161 | 147 | 128 | 133 | 152 | 165 | 184 | 109 | 156 | 92 | 150 | 139 | 139 | 173 | 163 | 107 | 120 | 179 | 128 | Sqor |
| Sqstm1 | 10889 | 13390 | 11536 | 13045 | 12188 | 12070 | 12370 | 7855 | 8988 | 8490 | 8247 | 6985 | 7497 | 7103 | 11366 | 11254 | 12380 | 12777 | 13327 | 12831 | 12883 | 8530 | 9788 | 7580 | 8076 | 7128 | 6839 | 7135 | Sqstm1 |
| Sra1 | 2059 | 2236 | 2230 | 1949 | 2127 | 1934 | 2158 | 2589 | 2550 | 2546 | 2599 | 2619 | 2465 | 2754 | 2109 | 1998 | 2231 | 2003 | 2007 | 1988 | 2007 | 2413 | 2823 | 2666 | 2381 | 2869 | 2795 | 2821 | Sra1 |
| Srarp | 1 | 3 | 0 | 7 | 0 | 0 | 3 | 5 | 7 | 7 | 3 | 5 | 5 | 2 | 0 | 1 | 0 | 4 | 0 | 6 | 0 | 7 | 6 | 5 | 2 | 8 | 2 | 9 | Srarp |
| Srbd1 | 4083 | 3909 | 4127 | 3514 | 3215 | 3211 | 3738 | 3555 | 3272 | 3336 | 3307 | 3735 | 3486 | 4143 | 3922 | 3516 | 3970 | 3217 | 3208 | 3465 | 3617 | 3222 | 3185 | 3769 | 3702 | 4170 | 4187 | 4157 | Srbd1 |
| Src | 2265 | 2729 | 3259 | 3322 | 4565 | 3747 | 3312 | 6622 | 8107 | 9045 | 8273 | 9522 | 7771 | 8406 | 2301 | 2531 | 3819 | 3407 | 4333 | 3190 | 3440 | 9005 | 7324 | 9800 | 9961 | 9938 | 8142 | 8102 | Src |
| Srcap | 153 | 153 | 109 | 154 | 228 | 203 | 169 | 541 | 562 | 686 | 580 | 710 | 643 | 579 | 160 | 112 | 147 | 141 | 196 | 183 | 214 | 475 | 438 | 624 | 711 | 203 | 715 | 598 | Srcap |
| Srcin1 | 3016 | 4118 | 4983 | 6856 | 8335 | 6836 | 6434 | 5904 | 6681 | 9425 | 7750 | 8895 | 6665 | 7893 | 2962 | 2901 | 6495 | 7126 | 8828 | 5661 | 7335 | 10634 | 5718 | 10325 | 11954 | 6904 | 7158 | 7048 | Srcin1 |
| Srd5a1 | 2861 | 3213 | 2845 | 2887 | 2348 | 2279 | 2784 | 856 | 909 | 921 | 715 | 887 | 868 | 956 | 2839 | 2646 | 2853 | 2549 | 2705 | 2369 | 2750 | 682 | 808 | 874 | 810 | 916 | 867 | 940 | Srd5a1 |
| Srd5a2 | 10 | 10 | 21 | 12 | 17 | 11 | 21 | 4 | 6 | 7 | 23 | 8 | 9 | 18 | 15 | 6 | 7 | 8 | 20 | 3 | 10 | 5 | 8 | 7 | 9 | 16 | 2 | 2 | Srd5a2 |
| Srd5a3 | 363 | 278 | 315 | 245 | 320 | 283 | 293 | 606 | 711 | 612 | 727 | 647 | 722 | 607 | 331 | 338 | 312 | 308 | 301 | 319 | 281 | 663 | 695 | 684 | 627 | 749 | 669 | 686 | Srd5a3 |
| Srebf1 | 4489 | 6576 | 5559 | 5905 | 4111 | 4147 | 4009 | 2913 | 3796 | 3457 | 3328 | 4305 | 3843 | 3557 | 4229 | 5420 | 5499 | 5186 | 4602 | 3789 | 3766 | 3237 | 3443 | 3875 | 3546 | 5045 | 4058 | 3737 | Srebf1 |
| Srebf2 | 14439 | 19625 | 20779 | 19741 | 17660 | 16313 | 15726 | 11872 | 14024 | 15775 | 15066 | 19071 | 16636 | 18000 | 14580 | 17403 | 20999 | 18737 | 19420 | 13685 | 16577 | 15465 | 13045 | 16851 | 17075 | 17970 | 17632 | 18688 | Srebf2 |
| Srek1 | 4658 | 3472 | 4690 | 4804 | 6017 | 5808 | 5629 | 13769 | 12069 | 14241 | 14584 | 15215 | 14562 | 15430 | 4398 | 3875 | 4883 | 4770 | 5681 | 5009 | 5403 | 14121 | 11884 | 14327 | 14560 | 13768 | 14814 | 14324 | Srek1 |
| Srek1ip1 | 2481 | 1848 | 2768 | 2147 | 2065 | 2063 | 2309 | 3002 | 2701 | 2362 | 2553 | 2868 | 3009 | 2878 | 2600 | 2189 | 2252 | 1951 | 2348 | 2671 | 2223 | 2511 | 2703 | 2796 | 2482 | 3024 | 3058 | 3186 | Srek1ip1 |
| Srf | 2374 | 3162 | 2880 | 2811 | 2385 | 2316 | 1793 | 1558 | 2023 | 1956 | 2124 | 2046 | 2095 | 1700 | 1975 | 2849 | 2713 | 2776 | 2408 | 1963 | 1958 | 1830 | 2034 | 1966 | 2020 | 2244 | 2011 | 1708 | Srf |
| Srfbp1 | 1998 | 1823 | 1859 | 1698 | 1410 | 1661 | 1685 | 1782 | 1798 | 1562 | 1497 | 1727 | 1685 | 1700 | 2062 | 1888 | 1699 | 1690 | 1416 | 2003 | 1535 | 1358 | 1632 | 1565 | 1508 | 1800 | 1801 | 1712 | Srfbp1 |
| Srgap1 | 2842 | 2973 | 3537 | 5025 | 6808 | 5496 | 4836 | 6495 | 6953 | 11345 | 9322 | 8866 | 8604 | 8152 | 2985 | 3170 | 4442 | 5563 | 5801 | 3916 | 5647 | 13418 | 6489 | 9224 | 10658 | 5675 | 7240 | 7522 | Srgap1 |
| Srgap2 | 11766 | 11192 | 10397 | 12948 | 13791 | 13387 | 14583 | 8361 | 8824 | 9292 | 9043 | 9079 | 8566 | 8825 | 11718 | 11516 | 11305 | 13029 | 13654 | 11846 | 14754 | 9461 | 8753 | 8727 | 10144 | 6345 | 8571 | 8014 | Srgap2 |
| Srgap3 | 22002 | 26307 | 25041 | 30264 | 32989 | 32189 | 29266 | 26837 | 23079 | 29152 | 24430 | 34160 | 33417 | 32252 | 25732 | 23147 | 29599 | 29196 | 31690 | 28446 | 32966 | 33690 | 22811 | 27593 | 33745 | 18187 | 35570 | 33791 | Srgap3 |
| Srgn | 146 | 115 | 188 | 110 | 151 | 145 | 132 | 115 | 160 | 144 | 156 | 108 | 169 | 145 | 113 | 136 | 117 | 150 | 101 | 155 | 128 | 137 | 201 | 140 | 131 | 125 | 149 | 140 | Srgn |
| Sri | 3118 | 2535 | 3220 | 2537 | 2745 | 2807 | 2684 | 3276 | 3304 | 3176 | 3442 | 2885 | 3386 | 3445 | 3125 | 2986 | 2523 | 2638 | 2331 | 2574 | 2417 | 3075 | 3552 | 3157 | 2772 | 3499 | 3149 | 3429 | Sri |
| Srl | 4090 | 3749 | 2778 | 3640 | 3616 | 3537 | 3933 | 3178 | 3377 | 2732 | 5228 | 2662 | 2642 | 2668 | 3769 | 3298 | 3245 | 3265 | 3596 | 3661 | 3818 | 3113 | 4513 | 2494 | 2625 | 2029 | 2701 | 2466 | Srl |

| | | | | | | | | | | | | | | | | | | | | | | | | | | | | | |
|---|---|---|---|---|---|---|---|---|---|---|---|---|---|---|---|---|---|---|---|---|---|---|---|---|---|---|---|---|---|
| Srm | 2158 | 2883 | 2607 | 2441 | 2114 | 2046 | 2034 | 2102 | 2718 | 2368 | 2752 | 2531 | 2261 | 2370 | 2152 | 2481 | 2525 | 2315 | 2243 | 1980 | 2195 | 2326 | 2653 | 2650 | 2556 | 3246 | 2556 | 2480 | Srm |
| Srms | 10 | 20 | 16 | 26 | 20 | 21 | 10 | 4 | 11 | 10 | 13 | 7 | 20 | 14 | 8 | 12 | 10 | 24 | 18 | 5 | 13 | 17 | 14 | 4 | 11 | 4 | 13 | 8 | Srms |
| Srp14 | 4941 | 5717 | 5407 | 5953 | 5133 | 4679 | 5747 | 4805 | 4402 | 4394 | 4127 | 4595 | 4427 | 4965 | 5563 | 5041 | 5381 | 5239 | 5275 | 6199 | 5706 | 3898 | 4469 | 4555 | 4628 | 4795 | 4527 | 4747 | Srp14 |
| Srp19 | 3199 | 2582 | 2988 | 2869 | 2386 | 2256 | 2929 | 3096 | 2673 | 2672 | 2824 | 2916 | 2971 | 3493 | 3033 | 2678 | 2948 | 2507 | 2568 | 2720 | 2702 | 2779 | 2879 | 3221 | 2665 | 3251 | 2874 | 3430 | Srp19 |
| Srp54a | 315 | 392 | 456 | 379 | 245 | 181 | 375 | 366 | 281 | 347 | 138 | 360 | 314 | 328 | 286 | 198 | 492 | 271 | 119 | 246 | 214 | 251 | 293 | 458 | 366 | 364 | 295 | 453 | Srp54a |
| Srp54b | 510 | 315 | 401 | 219 | 295 | 343 | 310 | 530 | 312 | 217 | 220 | 294 | 243 | 441 | 493 | 308 | 548 | 503 | 330 | 340 | 283 | 463 | 239 | 236 | 330 | 317 | 336 | 349 | Srp54b |
| Srp54c | 1791 | 1253 | 1362 | 1307 | 1277 | 1288 | 1458 | 1631 | 1486 | 1313 | 1585 | 1434 | 1210 | 1518 | 1957 | 1581 | 1516 | 1557 | 1323 | 1655 | 1146 | 1765 | 1540 | 1347 | 1391 | 1426 | 1386 | 1272 | Srp54c |
| Srp68 | 6053 | 7051 | 6492 | 5723 | 5101 | 5555 | 5203 | 5954 | 6632 | 6463 | 6626 | 6857 | 6753 | 6613 | 5854 | 6256 | 6050 | 5569 | 5679 | 5293 | 5097 | 5661 | 6669 | 6950 | 6458 | 7972 | 7304 | 6844 | Srp68 |
| Srp72 | 14893 | 13807 | 12744 | 12137 | 11838 | 12209 | 13116 | 13411 | 12724 | 12325 | 12093 | 11574 | 12187 | 12092 | 14711 | 12934 | 11973 | 12485 | 12080 | 12682 | 13117 | 12148 | 12803 | 12031 | 12206 | 12473 | 12342 | 12723 | Srp72 |
| Srp9 | 4778 | 4299 | 4493 | 4357 | 3612 | 3628 | 4468 | 4279 | 3851 | 3312 | 3491 | 3671 | 3655 | 4176 | 5056 | 4502 | 4472 | 3916 | 3960 | 4587 | 4233 | 3437 | 3988 | 3818 | 3640 | 4316 | 3834 | 4297 | Srp9 |
| Srpk1 | 11704 | 11931 | 12044 | 12355 | 12080 | 11357 | 12699 | 12073 | 10971 | 11227 | 10439 | 13039 | 12107 | 13552 | 11512 | 11080 | 11991 | 11955 | 12931 | 12313 | 12518 | 11416 | 10125 | 12055 | 12088 | 12538 | 13054 | 13200 | Srpk1 |
| Srpk2 | 13779 | 11489 | 12986 | 14062 | 16643 | 15242 | 17926 | 13432 | 11762 | 12627 | 12387 | 12339 | 11578 | 13461 | 14388 | 12242 | 14484 | 15172 | 15647 | 15184 | 17531 | 14018 | 11914 | 12490 | 13651 | 10939 | 12311 | 12741 | Srpk2 |
| Srpk3 | 13 | 7 | 8 | 0 | 20 | 13 | 2 | 40 | 78 | 13 | 491 | 3 | 24 | 4 | 16 | 18 | 0 | 7 | 0 | 2 | 3 | 75 | 314 | 12 | 15 | 12 | 2 | 8 | Srpk3 |
| Srpr | 6211 | 6163 | 6118 | 5206 | 4268 | 4722 | 4456 | 5757 | 6081 | 5651 | 6754 | 5763 | 6212 | 5519 | 5751 | 6173 | 5226 | 4870 | 4486 | 4413 | 4427 | 5763 | 6581 | 5534 | 5325 | 6317 | 6342 | 6092 | Srpr |
| Srprb | 4809 | 5850 | 5321 | 5065 | 4665 | 4877 | 4539 | 3635 | 4097 | 3881 | 4170 | 3936 | 4082 | 3800 | 4701 | 5254 | 5381 | 5335 | 5199 | 4599 | 4816 | 3908 | 4168 | 4069 | 3964 | 4637 | 4283 | 4290 | Srprb |
| Srpx | 343 | 356 | 305 | 334 | 166 | 404 | 166 | 568 | 925 | 673 | 878 | 387 | 617 | 440 | 386 | 389 | 327 | 441 | 158 | 158 | 161 | 603 | 1010 | 608 | 540 | 455 | 430 | 407 | Srpx |
| Srpx2 | 98 | 75 | 126 | 43 | 62 | 80 | 53 | 85 | 96 | 96 | 172 | 93 | 127 | 101 | 108 | 125 | 70 | 77 | 39 | 75 | 60 | 62 | 177 | 77 | 115 | 79 | 84 | 100 | Srpx2 |
| Srr | 5622 | 5503 | 3930 | 5315 | 5479 | 5835 | 5801 | 4252 | 4246 | 3788 | 3838 | 3644 | 3303 | 3343 | 6010 | 4854 | 4862 | 5470 | 5280 | 6053 | 5869 | 4488 | 4194 | 3268 | 3952 | 2767 | 3446 | 3129 | Srr |
| Srrd | 899 | 1070 | 1073 | 1003 | 849 | 791 | 875 | 629 | 729 | 711 | 767 | 1021 | 787 | 941 | 779 | 1014 | 1075 | 902 | 982 | 706 | 972 | 745 | 763 | 953 | 769 | 990 | 798 | 860 | Srrd |
| Srrm1 | 18521 | 16260 | 20772 | 20383 | 19895 | 19916 | 18704 | 17702 | 18771 | 21284 | 22460 | 23750 | 21643 | 22713 | 17023 | 17522 | 20683 | 19775 | 20455 | 16459 | 19085 | 21631 | 17516 | 22107 | 21597 | 22987 | 21291 | 21600 | Srrm1 |
| Srrm2 | 65227 | 70315 | 56736 | 80832 | 86051 | 83648 | 83230 | 78704 | 78293 | 92129 | 83070 | 99914 | 90679 | 92769 | 63241 | 65236 | 73268 | 82812 | 86547 | 70884 | 86071 | 98012 | 70910 | 85485 | 95617 | 48495 | 92625 | 86783 | Srrm2 |
| Srrm3 | 1350 | 2162 | 2554 | 3530 | 4066 | 3017 | 2919 | 2633 | 2383 | 3883 | 3155 | 3479 | 3483 | 3407 | 1851 | 1744 | 3234 | 3341 | 3794 | 2681 | 3363 | 4700 | 2350 | 3384 | 4081 | 1967 | 3238 | 3186 | Srrm3 |
| Srrm4 | 3674 | 4688 | 5953 | 7963 | 10582 | 8584 | 9096 | 6530 | 6565 | 8756 | 6946 | 10272 | 7595 | 9648 | 4751 | 4055 | 7701 | 8460 | 9729 | 6843 | 9601 | 10639 | 5520 | 9655 | 10631 | 6811 | 7704 | 8959 | Srrm4 |
| Srrt | 11505 | 13352 | 17002 | 15137 | 13476 | 13220 | 13961 | 12698 | 14346 | 16948 | 15753 | 19625 | 16771 | 19591 | 10512 | 11921 | 16143 | 13620 | 15252 | 11892 | 13766 | 14781 | 12943 | 19614 | 17378 | 21545 | 17920 | 19673 | Srrt |
| Srsf1 | 1925 | 1089 | 1487 | 1238 | 1064 | 1379 | 1045 | 2447 | 1711 | 1356 | 1465 | 1957 | 1761 | 1838 | 1869 | 1315 | 1267 | 1108 | 1446 | 1424 | 1067 | 2151 | 1393 | 1669 | 1781 | 1902 | 1943 | 1756 | Srsf1 |
| Srsf10 | 27413 | 18939 | 28818 | 26487 | 24164 | 24544 | 28547 | 30884 | 23690 | 24117 | 25015 | 28758 | 27481 | 32107 | 27170 | 23241 | 25902 | 25632 | 25955 | 28505 | 26002 | 25011 | 22557 | 28398 | 27793 | 31323 | 30351 | 31468 | Srsf10 |
| Srsf11 | 34538 | 28055 | 28214 | 30316 | 28769 | 31390 | 33275 | 31382 | 27928 | 28756 | 30925 | 30499 | 29877 | 31481 | 33676 | 29321 | 28246 | 29719 | 29725 | 31162 | 31251 | 30393 | 27082 | 30687 | 29954 | 27871 | 30204 | 30674 | Srsf11 |
| Srsf12 | 839 | 757 | 1099 | 1080 | 1514 | 1268 | 1263 | 1225 | 1243 | 1502 | 1371 | 882 | 993 | 991 | 982 | 943 | 1139 | 1246 | 1223 | 1170 | 1230 | 1656 | 1272 | 1114 | 1481 | 697 | 793 | 915 | Srsf12 |
| Srsf2 | 30428 | 25701 | 30739 | 29288 | 25873 | 25344 | 31687 | 28355 | 25857 | 27759 | 27463 | 34206 | 30758 | 36961 | 29376 | 25709 | 29501 | 27436 | 27198 | 25698 | 29408 | 28289 | 24840 | 32683 | 29663 | 40248 | 32722 | 38352 | Srsf2 |
| Srsf3 | 45548 | 37719 | 45972 | 41806 | 35304 | 36951 | 41642 | 57268 | 49767 | 47657 | 46001 | 56870 | 53518 | 58426 | 46649 | 42563 | 42490 | 38874 | 39937 | 47334 | 37694 | 45308 | 46768 | 53749 | 49534 | 65827 | 59420 | 59267 | Srsf3 |
| Srsf4 | 5343 | 6093 | 7047 | 6766 | 6544 | 6037 | 6192 | 6705 | 7476 | 8732 | 7755 | 10325 | 7952 | 10067 | 4927 | 5074 | 7262 | 6315 | 7356 | 5620 | 6509 | 7551 | 6620 | 9785 | 8780 | 11652 | 9103 | 9463 | Srsf4 |
| Srsf5 | 29103 | 26042 | 37566 | 33283 | 29720 | 28439 | 31161 | 39438 | 33395 | 36569 | 39208 | 41592 | 40580 | 44915 | 30241 | 26664 | 33378 | 30681 | 31179 | 33801 | 29662 | 36330 | 33479 | 42904 | 39944 | 47391 | 43776 | 43518 | Srsf5 |
| Srsf6 | 22945 | 17431 | 25726 | 22148 | 22189 | 23322 | 23752 | 45143 | 39218 | 42692 | 43014 | 50703 | 47035 | 51955 | 21402 | 17900 | 23779 | 20251 | 24908 | 21581 | 23290 | 46486 | 37971 | 47625 | 46373 | 53633 | 51702 | 53128 | Srsf6 |
| Srsf7 | 21659 | 18135 | 22186 | 20706 | 19041 | 19325 | 22481 | 34397 | 28702 | 27977 | 27022 | 32136 | 29120 | 33029 | 21593 | 18914 | 21598 | 19553 | 21655 | 24315 | 21177 | 26964 | 27544 | 31814 | 29179 | 36612 | 32669 | 34458 | Srsf7 |
| Srsf9 | 4077 | 4460 | 6213 | 5392 | 4725 | 4166 | 4527 | 2851 | 3040 | 3631 | 3793 | 4371 | 3915 | 4505 | 3676 | 4072 | 5988 | 5010 | 4861 | 4150 | 4718 | 2839 | 3058 | 4511 | 3870 | 5195 | 4348 | 4542 | Srsf9 |
| Srxn1 | 1698 | 1957 | 2097 | 1578 | 1600 | 1438 | 1500 | 904 | 901 | 805 | 888 | 936 | 995 | 914 | 1973 | 1725 | 1961 | 1503 | 1607 | 1408 | 1415 | 973 | 862 | 838 | 879 | 1147 | 1035 | 1067 | Srxn1 |
| Ss18 | 9535 | 9103 | 9535 | 8845 | 7837 | 8135 | 8233 | 7519 | 7266 | 6653 | 7479 | 7061 | 7668 | 7274 | 9700 | 10033 | 8809 | 8632 | 7920 | 8766 | 7853 | 6609 | 7431 | 7147 | 6735 | 8047 | 7071 | 7568 | Ss18 |
| Ss18l1 | 4012 | 4296 | 3807 | 6191 | 6931 | 6381 | 6220 | 5746 | 5299 | 6147 | 5095 | 5484 | 4603 | 5240 | 4478 | 3881 | 5040 | 6840 | 6457 | 6006 | 6598 | 8081 | 4948 | 6030 | 7271 | 4432 | 4860 | 4736 | Ss18l1 |
| Ss18l2 | 1555 | 1526 | 1415 | 1771 | 1398 | 1347 | 1848 | 1109 | 886 | 1064 | 1052 | 901 | 950 | 1199 | 1751 | 1319 | 1605 | 1482 | 1340 | 1658 | 1740 | 1021 | 926 | 1088 | 1018 | 935 | 931 | 1047 | Ss18l2 |
| Ssb | 30993 | 24390 | 33784 | 28749 | 24971 | 26298 | 30961 | 26843 | 21864 | 22490 | 21448 | 26999 | 24900 | 29527 | 34309 | 28243 | 30525 | 26290 | 27115 | 33468 | 28323 | 23039 | 22162 | 26118 | 23726 | 31380 | 28143 | 30835 | Ssb |
| Ssbp1 | 3952 | 3531 | 3957 | 3766 | 3198 | 3600 | 4130 | 4754 | 3812 | 3926 | 3805 | 4992 | 4609 | 5064 | 4146 | 3444 | 3902 | 3360 | 3646 | 4369 | 3594 | 3534 | 3861 | 4762 | 4172 | 5320 | 5244 | 5389 | Ssbp1 |
| Ssbp2 | 9931 | 9868 | 7384 | 10037 | 10106 | 10336 | 11732 | 7460 | 7287 | 6612 | 6147 | 6283 | 5725 | 6643 | 10439 | 8506 | 9156 | 9881 | 10695 | 11962 | 11299 | 6647 | 6853 | 6472 | 6475 | 6068 | 6209 | 6508 | Ssbp2 |
| Ssbp3 | 18692 | 21199 | 26215 | 26876 | 25444 | 22898 | 22952 | 18371 | 19343 | 25360 | 22450 | 25628 | 24057 | 26066 | 18977 | 20108 | 28439 | 25974 | 26684 | 19636 | 24897 | 24179 | 18848 | 26720 | 25843 | 27345 | 24145 | 25481 | Ssbp3 |
| Ssbp4 | 2909 | 4095 | 3031 | 3838 | 3505 | 3236 | 2721 | 2201 | 2437 | 2416 | 2231 | 2380 | 2391 | 2023 | 2879 | 3348 | 3644 | 3531 | 3697 | 3164 | 2769 | 2669 | 2354 | 2317 | 2292 | 2611 | 2177 | 1883 | Ssbp4 |
| Ssc4d | 113 | 106 | 80 | 64 | 53 | 78 | 27 | 123 | 190 | 172 | 167 | 70 | 103 | 94 | 113 | 171 | 48 | 64 | 21 | 33 | 33 | 135 | 164 | 150 | 82 | 199 | 83 | 97 | Ssc4d |
| Ssc5d | 386 | 491 | 549 | 478 | 478 | 476 | 334 | 408 | 596 | 605 | 1105 | 639 | 559 | 469 | 353 | 522 | 562 | 408 | 496 | 357 | 274 | 752 | 925 | 601 | 459 | 644 | 580 | 509 | Ssc5d |
| Ssh1 | 1715 | 1867 | 2119 | 2192 | 2305 | 1985 | 1998 | 2114 | 2548 | 2898 | 2822 | 3179 | 2987 | 3132 | 1536 | 1801 | 2241 | 2075 | 2245 | 1670 | 2022 | 3212 | 2408 | 3057 | 2898 | 3231 | 2811 | 3077 | Ssh1 |
| Ssh2 | 4938 | 4378 | 3788 | 5899 | 6798 | 7026 | 6614 | 6970 | 6555 | 7592 | 6867 | 7139 | 6058 | 7128 | 5079 | 4658 | 4784 | 5924 | 6456 | 5813 | 7118 | 7822 | 6090 | 6372 | 8612 | 5266 | 6512 | 6459 | Ssh2 |
| Ssh3 | 1848 | 2150 | 2462 | 2772 | 3076 | 2649 | 2361 | 1185 | 1493 | 1806 | 1609 | 1224 | 1371 | 1186 | 1780 | 2393 | 2968 | 3044 | 2967 | 2188 | 2446 | 1965 | 1441 | 1449 | 1561 | 904 | 1074 | 1038 | Ssh3 |
| Ssmem1 | 9 | 16 | 8 | 21 | 14 | 9 | 13 | 17 | 12 | 4 | 6 | 12 | 26 | 9 | 13 | 9 | 10 | 13 | 12 | 10 | 10 | 8 | 12 | 14 | 5 | 9 | 23 | 21 | Ssmem1 |
| Ssna1 | 2866 | 3758 | 3236 | 3095 | 2612 | 2805 | 3073 | 2721 | 3009 | 2566 | 2841 | 3151 | 3063 | 3011 | 2823 | 2889 | 3161 | 2873 | 3170 | 3668 | 2842 | 2247 | 2948 | 3143 | 2858 | 3492 | 3274 | 3313 | Ssna1 |
| Sspn | 466 | 416 | 491 | 363 | 367 | 512 | 239 | 806 | 1016 | 918 | 1384 | 690 | 958 | 681 | 534 | 537 | 336 | 483 | 235 | 237 | 252 | 946 | 1285 | 910 | 729 | 750 | 820 | 726 | Sspn |
| Sspo | 87 | 67 | 90 | 66 | 91 | 65 | 61 | 106 | 120 | 130 | 182 | 114 | 197 | 76 | 78 | 118 | 103 | 73 | 81 | 41 | 80 | 155 | 159 | 128 | 99 | 95 | 138 | 83 | Sspo |
| Ssr1 | 18424 | 15275 | 16803 | 15291 | 13269 | 14882 | 15037 | 14770 | 14566 | 14317 | 16218 | 15902 | 16071 | 16591 | 17537 | 17519 | 15434 | 15520 | 13377 | 14161 | 14672 | 14237 | 15578 | 14961 | 15126 | 17234 | 16966 | 17573 | Ssr1 |
| Ssr2 | 9404 | 10024 | 8049 | 7297 | 6005 | 6976 | 6288 | 6068 | 6576 | 5327 | 6911 | 5483 | 6044 | 5608 | 8650 | 9606 | 7850 | 7335 | 6900 | 7268 | 6278 | 5320 | 6879 | 5690 | 5364 | 7090 | 6313 | 5959 | Ssr2 |
| Ssr3 | 32590 | 25639 | 31724 | 25700 | 24095 | 25743 | 27597 | 26247 | 23167 | 20806 | 24336 | 23345 | 25697 | 25129 | 33695 | 29923 | 28566 | 27177 | 24491 | 29315 | 25540 | 22231 | 24728 | 23424 | 22456 | 25883 | 26164 | 26407 | Ssr3 |
| Ssr4 | 2336 | 2709 | 2367 | 2292 | 2230 | 2154 | 2325 | 2334 | 2718 | 2563 | 2757 | 2412 | 2545 | 2521 | 2495 | 2422 | 2465 | 2236 | 2201 | 2647 | 2346 | 1980 | 2721 | 2447 | 2477 | 2667 | 2411 | 2695 | Ssr4 |
| Ssrp1 | 22238 | 20756 | 21751 | 17488 | 16195 | 17717 | 18177 | 25495 | 23896 | 23254 | 22079 | 27090 | 24824 | 29190 | 20663 | 18640 | 19374 | 16286 | 17987 | 18247 | 17493 | 21587 | 22645 | 25871 | 23436 | 32010 | 28476 | 30155 | Ssrp1 |
| Sst | 257 | 333 | 397 | 559 | 1145 | 579 | 707 | 883 | 1155 | 1506 | 1154 | 916 | 600 | 906 | 245 | 232 | 503 | 644 | 783 | 761 | 807 | 1258 | 1277 | 1244 | 1454 | 635 | 765 | 776 | Sst |
| Sstr1 | 58 | 78 | 42 | 93 | 223 | 120 | 148 | 153 | 228 | 181 | 196 | 136 | 78 | 102 | 87 | 70 | 105 | 140 | 195 | 226 | 121 | 224 | 193 | 159 | 213 | 123 | 127 | 135 | Sstr1 |
| Sstr2 | 323 | 560 | 322 | 583 | 865 | 716 | 713 | 1109 | 1269 | 1327 | 871 | 942 | 1042 | 935 | 493 | 502 | 589 | 602 | 787 | 736 | 782 | 849 | 1063 | 1361 | 1346 | 792 | 760 | 681 | Sstr2 |
| Sstr2 | 114 | 137 | 255 | 123 | 110 | 133 | 65 | 132 | 55 | 76 | 60 | 123 | 281 | 103 | 169 | 202 | 159 | 129 | 76 | 79 | 68 | 120 | 41 | 73 | 116 | 29 | 170 | 209 | Sstr2 |
| Sstr3 | 3 | 3 | 0 | 10 | 36 | 15 | 23 | 96 | 183 | 136 | 100 | 117 | 74 | 130 | 11 | 7 | 12 | 10 | 23 | 19 | 27 | 90 | 112 | 199 | 225 | 95 | 48 | 55 | Sstr3 |
| Sstr4 | 28 | 42 | 27 | 17 | 25 | 45 | 17 | 59 | 82 | 60 | 97 | 52 | 69 | 38 | 24 | 56 | 42 | 35 | 30 | 25 | 38 | 112 | 93 | 48 | 74 | 24 | 66 | 36 | Sstr4 |
| Sstr5 | 16 | 17 | 10 | 9 | 19 | 15 | 10 | 7 | 17 | 8 | 11 | 1 | 2 | 3 | 9 | 19 | 16 | 15 | 11 | 14 | 9 | 8 | 6 | 5 | 5 | 6 | 3 | 2 | Sstr5 |
| Ssu2 | 5 | 1 | 1 | 0 | 0 | 0 | 0 | 24 | 27 | 11 | 36 | 10 | 20 | 13 | 15 | 7 | 2 | 2 | 5 | 4 | 1 | 18 | 21 | 14 | 10 | 28 | 7 | 7 | Ssu2 |
| Ssu72 | 3720 | 3990 | 3722 | 3662 | 3251 | 3248 | 3236 | 3568 | 3283 | 3152 | 3158 | 3343 | 3445 | 3275 | 3777 | 3734 | 3635 | 3404 | 3416 | 3879 | 3314 | 3003 | 3535 | 3269 | 3243 | 3800 | 3665 | 3682 | Ssu72 |
| Ssx2ip | 6223 | 5367 | 6245 | 6287 | 5078 | 5290 | 5927 | 3107 | 2635 | 3118 | 2834 | 3423 | 2985 | 3647 | 5838 | 4972 | 5909 | 5711 | 5259 | 4949 | 5871 | 2573 | 2671 | 3559 | 2772 | 4339 | 3096 | 3592 | Ssx2ip |
| Ssxa1 | 0 | 1 | 0 | 0 | 2 | 0 | 1 | 0 | 0 | 0 | 0 | 0 | 0 | 0 | 0 | 2 | 0 | 0 | 0 | 0 | 0 | 0 | 0 | 0 | 0 | 0 | 1 | 0 | Ssxa1 |
| St13 | 25915 | 21709 | 27018 | 23550 | 20638 | 21735 | 23807 | 23491 | 19828 | 21262 | 20505 | 21688 | 20860 | 24418 | 25452 | 22068 | 24510 | 22198 | 21903 | 24223 | 22552 | 20753 | 20249 | 22994 | 21582 | 25699 | 22789 | 24872 | St13 |
| St14 | 603 | 330 | 203 | 223 | 137 | 142 | 55 | 346 | 448 | 345 | 304 | 73 | 158 | 88 | 306 | 423 | 138 | 209 | 108 | 77 | 76 | 105 | 516 | 389 | 68 | 365 | 83 | 101 | St14 |
| St18 | 2315 | 2384 | 2225 | 2259 | 2467 | 2840 | 2654 | 6116 | 3758 | 3830 | 3255 | 6017 | 5124 | 6046 | 3108 | 1949 | 2611 | 1949 | 2458 | 3002 | 2629 | 5167 | 3453 | 4278 | 4912 | 4309 | 6630 | 6336 | St18 |
| St3gal1 | 2246 | 2306 | 2521 | 2255 | 2673 | 2342 | 2207 | 2263 | 2934 | 3033 | 3833 | 2255 | 2756 | 2248 | 2116 | 2492 | 2379 | 2477 | 2286 | 1913 | 2261 | 3280 | 3088 | 2634 | 2916 | 2539 | 2019 | 2218 | St3gal1 |
| St3gal2 | 3190 | 3670 | 3770 | 3677 | 4213 | 3843 | 3292 | 3403 | 4060 | 4485 | 4757 | 4204 | 4273 | 4127 | 2991 | 3547 | 3765 | 3986 | 3899 | 3289 | 3749 | 4289 | 4095 | 4346 | 4514 | 4306 | 3950 | 4187 | St3gal2 |
| St3gal3 | 1758 | 2155 | 2291 | 2288 | 2302 | 1989 | 1890 | 1503 | 1655 | 1749 | 1847 | 1716 | 1532 | 1786 | 1785 | 1944 | 2199 | 2376 | 2265 | 1761 | 2202 | 1711 | 1728 | 1838 | 1741 | 1936 | 1557 | 1691 | St3gal3 |
| St3gal4 | 1068 | 758 | 919 | 708 | 790 | 783 | 738 | 1145 | 1265 | 1226 | 1498 | 998 | 1014 | 882 | 810 | 889 | 713 | 791 | 734 | 583 | 686 | 1044 | 1474 | 1209 | 1027 | 1359 | 930 | 971 | St3gal4 |
| St3gal5 | 2475 | 3008 | 3028 | 3055 | 3669 | 3173 | 3429 | 2152 | 2160 | 2132 | 2334 | 2191 | 2691 | 2383 | 2798 | 2758 | 3099 | 3286 | 3448 | 3058 | 3202 | 2348 | 2205 | 2185 | 2299 | 2267 | 2260 | 2589 | St3gal5 |
| St3gal6 | 390 | 282 | 441 | 269 | 386 | 357 | 265 | 501 | 505 | 500 | 591 | 528 | 629 | 478 | 385 | 376 | 304 | 270 | 284 | 340 | 325 | 535 | 640 | 508 | 429 | 560 | 528 | 623 | St3gal6 |
| St6gal1 | 2635 | 2503 | 2752 | 2232 | 3005 | 2795 | 2618 | 5177 | 4825 | 5607 | 4955 | 5415 | 5057 | 5182 | 2541 | 2494 | 2534 | 2338 | 2883 | 2574 | 2611 | 5492 | 4961 | 5449 | 5545 | 4873 | 5073 | 5374 | St6gal1 |

| | | | | | | | | | | | | | | | | | | | | | | | | | | | | | |
|---|---|---|---|---|---|---|---|---|---|---|---|---|---|---|---|---|---|---|---|---|---|---|---|---|---|---|---|---|---|
| St6gal2 | 1 | 0 | 0 | 0 | 0 | 0 | 0 | 0 | 0 | 0 | 0 | 0 | 0 | 0 | 0 | 0 | 0 | 0 | 2 | 0 | 0 | 0 | 0 | 0 | 0 | 0 | 0 | 0 | St6gal2 |
| St6galnac1 | 6 | 10 | 0 | 10 | 20 | 6 | 10 | 5 | 5 | 0 | 0 | 0 | 0 | 0 | 18 | 16 | 6 | 13 | 9 | 8 | 10 | 0 | 0 | 2 | 0 | 0 | 0 | 0 | St6galnac1 |
| St6galnac2 | 57 | 54 | 60 | 57 | 61 | 36 | 20 | 66 | 110 | 122 | 154 | 40 | 92 | 68 | 70 | 55 | 48 | 62 | 55 | 15 | 41 | 53 | 104 | 96 | 38 | 85 | 57 | 40 | St6galnac2 |
| St6galnac2 | 3 | 3 | 10 | 7 | 3 | 6 | 3 | 7 | 12 | 8 | 11 | 9 | 6 | 10 | 8 | 5 | 1 | 4 | 0 | 3 | 6 | 13 | 14 | 6 | 6 | 6 | 7 | 2 | St6galnac2 |
| St6galnac3 | 2382 | 1682 | 1772 | 1524 | 1403 | 1702 | 1387 | 2281 | 2141 | 1903 | 2122 | 1732 | 1957 | 1674 | 1966 | 2346 | 1442 | 1620 | 1270 | 1690 | 1477 | 1875 | 2233 | 1695 | 1756 | 1938 | 1808 | 1832 | St6galnac3 |
| St6galnac4 | 2278 | 2458 | 2259 | 2277 | 2197 | 2192 | 2039 | 828 | 1135 | 1132 | 1305 | 949 | 1060 | 818 | 1979 | 2658 | 2284 | 2594 | 2307 | 1732 | 2067 | 1019 | 1141 | 1050 | 1015 | 1046 | 962 | 803 | St6galnac4 |
| St6galnac5 | 998 | 1095 | 1148 | 1731 | 2831 | 2202 | 1843 | 1716 | 1594 | 2265 | 1743 | 1222 | 1718 | 1057 | 1416 | 1231 | 1674 | 2110 | 2261 | 1864 | 2147 | 2652 | 1548 | 1495 | 2140 | 572 | 995 | 1013 | St6galnac5 |
| St6galnac6 | 2443 | 2703 | 2711 | 2580 | 2417 | 2322 | 2337 | 981 | 1124 | 1237 | 1372 | 964 | 1063 | 816 | 2132 | 2577 | 2661 | 2760 | 2462 | 2072 | 2171 | 981 | 1269 | 1115 | 1017 | 1169 | 890 | 871 | St6galnac6 |
| St7 | 1504 | 1572 | 1264 | 1404 | 1922 | 1763 | 1992 | 3162 | 2643 | 2630 | 2425 | 2462 | 2198 | 2451 | 1812 | 1377 | 1420 | 1541 | 1848 | 1878 | 2016 | 3070 | 2769 | 2353 | 2856 | 2018 | 2245 | 2506 | St7 |
| St7l | 1434 | 1408 | 1405 | 1433 | 1379 | 1329 | 1344 | 2420 | 1971 | 1951 | 1880 | 1855 | 1676 | 1878 | 1636 | 1513 | 1374 | 1374 | 1418 | 1671 | 1408 | 2196 | 2080 | 1891 | 1981 | 1833 | 1834 | 1768 | St7l |
| St8sia1 | 2298 | 2326 | 2167 | 3049 | 4476 | 4061 | 3816 | 5763 | 4854 | 5500 | 4671 | 5142 | 5278 | 4831 | 2993 | 2379 | 2897 | 3265 | 4027 | 3809 | 4262 | 6913 | 4388 | 4524 | 6377 | 2578 | 4917 | 4582 | St8sia1 |
| St8sia2 | 4555 | 4317 | 4516 | 6482 | 10860 | 8685 | 9219 | 11429 | 12274 | 15216 | 13083 | 13298 | 11267 | 13123 | 5169 | 4051 | 5852 | 7449 | 9330 | 6916 | 9456 | 19938 | 11180 | 13308 | 16615 | 9861 | 9993 | 11432 | St8sia2 |
| St8sia3 | 444 | 383 | 472 | 527 | 1113 | 807 | 997 | 3435 | 2745 | 3306 | 2780 | 2552 | 2282 | 2673 | 570 | 349 | 637 | 569 | 890 | 910 | 924 | 4351 | 2750 | 2723 | 3433 | 1814 | 2524 | 2470 | St8sia3 |
| St8sia4 | 5569 | 4954 | 4785 | 6050 | 7408 | 6923 | 7751 | 7707 | 5643 | 5288 | 4736 | 4606 | 4467 | 5024 | 7374 | 5531 | 5319 | 6119 | 6800 | 8491 | 7432 | 7018 | 6242 | 4659 | 6415 | 3516 | 4522 | 4643 | St8sia4 |
| St8sia5 | 99 | 175 | 125 | 223 | 299 | 325 | 301 | 99 | 96 | 96 | 79 | 91 | 44 | 103 | 148 | 121 | 169 | 254 | 341 | 268 | 298 | 115 | 73 | 87 | 159 | 58 | 84 | 70 | St8sia5 |
| St8sia6 | 269 | 109 | 42 | 123 | 80 | 106 | 78 | 244 | 180 | 267 | 228 | 55 | 132 | 92 | 144 | 156 | 92 | 73 | 58 | 105 | 84 | 84 | 231 | 233 | 72 | 349 | 62 | 71 | St8sia6 |
| Stab1 | 3140 | 4769 | 4626 | 3589 | 3166 | 3747 | 2360 | 2612 | 4649 | 4368 | 6052 | 5621 | 6200 | 4415 | 3261 | 4835 | 3896 | 4022 | 2992 | 2803 | 2910 | 4993 | 6778 | 4967 | 4563 | 3621 | 5450 | 4455 | Stab1 |
| Stab2 | 16 | 9 | 32 | 9 | 3 | 19 | 3 | 10 | 27 | 21 | 31 | 24 | 24 | 16 | 25 | 30 | 10 | 10 | 4 | 14 | 9 | 23 | 31 | 20 | 13 | 25 | 33 | 12 | Stab2 |
| Stac | 88 | 90 | 43 | 75 | 140 | 105 | 101 | 156 | 185 | 164 | 281 | 136 | 95 | 199 | 80 | 51 | 61 | 102 | 108 | 96 | 104 | 206 | 199 | 173 | 140 | 199 | 143 | 97 | Stac |
| Stac2 | 1239 | 1609 | 1752 | 2096 | 2500 | 1951 | 1931 | 239 | 395 | 462 | 455 | 338 | 201 | 366 | 1422 | 1488 | 2108 | 2282 | 2308 | 1514 | 1986 | 543 | 356 | 322 | 397 | 427 | 244 | 248 | Stac2 |
| Stac3 | 248 | 290 | 363 | 505 | 496 | 440 | 450 | 390 | 445 | 638 | 975 | 506 | 526 | 535 | 278 | 304 | 443 | 515 | 547 | 370 | 433 | 779 | 636 | 544 | 652 | 460 | 474 | 471 | Stac3 |
| Stag1 | 5744 | 4244 | 4391 | 4181 | 4574 | 5414 | 5409 | 13423 | 11532 | 10573 | 10619 | 12191 | 11394 | 12535 | 5498 | 4754 | 4318 | 4130 | 4609 | 5286 | 5108 | 10853 | 10989 | 10837 | 11476 | 10087 | 12491 | 12232 | Stag1 |
| Stag2 | 12911 | 9208 | 9046 | 8945 | 9943 | 11377 | 11871 | 19378 | 16704 | 13094 | 13937 | 16762 | 15956 | 17854 | 12976 | 10502 | 9374 | 8649 | 10123 | 13097 | 11819 | 12711 | 15833 | 14372 | 14387 | 14089 | 17451 | 17280 | Stag2 |
| Stag3 | 1356 | 1110 | 1229 | 1186 | 1363 | 1393 | 1427 | 995 | 765 | 742 | 984 | 658 | 726 | 636 | 1251 | 1161 | 1116 | 1111 | 964 | 1000 | 1116 | 1176 | 798 | 535 | 610 | 487 | 549 | 578 | Stag3 |
| Stam | 6457 | 6287 | 5892 | 6808 | 7459 | 7428 | 7810 | 7754 | 6877 | 6143 | 6211 | 6277 | 5720 | 6609 | 7087 | 6684 | 6179 | 6736 | 7182 | 7516 | 7712 | 6945 | 6729 | 6119 | 6654 | 5830 | 6092 | 6424 | Stam |
| Stam2 | 5819 | 6006 | 5453 | 5371 | 5178 | 5588 | 5852 | 5482 | 4675 | 3984 | 4212 | 4583 | 4196 | 4446 | 6467 | 5516 | 5422 | 5116 | 5320 | 6150 | 5729 | 4620 | 4838 | 4242 | 4567 | 3990 | 4610 | 4577 | Stam2 |
| Stambp | 1732 | 1698 | 1623 | 1795 | 1920 | 1792 | 1954 | 2187 | 2098 | 2020 | 2071 | 1811 | 1735 | 2041 | 1836 | 1612 | 1686 | 1802 | 1752 | 1937 | 2000 | 2393 | 2083 | 1999 | 2217 | 1795 | 1879 | 1882 | Stambp |
| Stambpl1 | 475 | 413 | 493 | 473 | 439 | 411 | 353 | 354 | 297 | 352 | 341 | 202 | 279 | 192 | 402 | 476 | 386 | 392 | 391 | 332 | 420 | 307 | 348 | 240 | 228 | 170 | 177 | 153 | Stambpl1 |
| Stap1 | 7 | 8 | 10 | 3 | 5 | 10 | 6 | 9 | 5 | 5 | 6 | 7 | 1 | 2 | 3 | 8 | 10 | 4 | 8 | 10 | 5 | 15 | 15 | 5 | 2 | 7 | 0 | 2 | Stap1 |
| Stap2 | 145 | 200 | 149 | 103 | 146 | 126 | 124 | 197 | 231 | 161 | 250 | 187 | 236 | 178 | 167 | 219 | 139 | 134 | 149 | 159 | 115 | 184 | 232 | 201 | 151 | 209 | 167 | 202 | Stap2 |
| Star | 526 | 595 | 405 | 475 | 359 | 507 | 451 | 162 | 142 | 128 | 154 | 232 | 198 | 223 | 537 | 488 | 394 | 534 | 415 | 508 | 519 | 162 | 153 | 189 | 178 | 205 | 203 | 166 | Star |
| Stard10 | 904 | 980 | 830 | 830 | 872 | 826 | 607 | 569 | 639 | 612 | 559 | 407 | 446 | 326 | 878 | 921 | 929 | 790 | 920 | 808 | 686 | 465 | 698 | 505 | 379 | 596 | 431 | 371 | Stard10 |
| Stard13 | 1022 | 1197 | 1007 | 982 | 965 | 791 | 705 | 628 | 870 | 885 | 1370 | 583 | 888 | 595 | 1025 | 1463 | 909 | 1024 | 746 | 646 | 668 | 894 | 1014 | 629 | 797 | 646 | 593 | 590 | Stard13 |
| Stard3 | 1777 | 2691 | 2445 | 2395 | 2009 | 2034 | 1905 | 1873 | 2070 | 2032 | 1881 | 2143 | 2083 | 2056 | 1861 | 2230 | 2395 | 2325 | 2322 | 2170 | 1946 | 1987 | 1963 | 2175 | 2173 | 2416 | 2223 | 2071 | Stard3 |
| Stard3nl | 3701 | 3911 | 3205 | 3769 | 3791 | 3677 | 4095 | 3798 | 3542 | 3286 | 3230 | 3126 | 2841 | 3094 | 3965 | 3731 | 3473 | 3681 | 3609 | 4275 | 4144 | 3541 | 3669 | 3206 | 3379 | 3196 | 3249 | 3278 | Stard3nl |
| Stard4 | 8145 | 7873 | 6619 | 7265 | 5648 | 6454 | 7119 | 4549 | 4135 | 3732 | 4163 | 5065 | 5144 | 5617 | 7855 | 7482 | 6521 | 7174 | 5959 | 7162 | 6758 | 3679 | 3900 | 4286 | 4194 | 5002 | 5645 | 5737 | Stard4 |
| Stard5 | 2408 | 2193 | 1957 | 1959 | 1703 | 1979 | 1593 | 813 | 799 | 708 | 830 | 663 | 689 | 637 | 2147 | 2349 | 2055 | 1708 | 1915 | 1635 | 1719 | 592 | 880 | 733 | 558 | 855 | 782 | 676 | Stard5 |
| Stard6 | 42 | 36 | 12 | 53 | 29 | 16 | 45 | 74 | 73 | 85 | 71 | 58 | 87 | 92 | 42 | 36 | 38 | 44 | 30 | 41 | 36 | 60 | 88 | 65 | 77 | 63 | 105 | 98 | Stard6 |
| Stard7 | 12157 | 11066 | 10331 | 11114 | 11204 | 11498 | 12840 | 10718 | 9403 | 8680 | 8490 | 10129 | 8541 | 9860 | 12179 | 10575 | 11293 | 11569 | 11713 | 12309 | 12253 | 10098 | 9117 | 9218 | 9945 | 10752 | 9910 | 10223 | Stard7 |
| Stard8 | 2227 | 2959 | 2954 | 1600 | 1164 | 1772 | 879 | 928 | 1297 | 1314 | 1707 | 1476 | 2218 | 1193 | 2469 | 3568 | 1820 | 1692 | 1140 | 1093 | 936 | 1472 | 1848 | 1070 | 1208 | 1123 | 1594 | 1409 | Stard8 |
| Stard9 | 5608 | 5694 | 2211 | 3354 | 4268 | 5416 | 3951 | 3773 | 4376 | 2861 | 4469 | 2405 | 3200 | 2455 | 5350 | 5693 | 2526 | 3396 | 3321 | 5398 | 4865 | 2391 | 4412 | 1993 | 3119 | 1513 | 3271 | 2058 | Stard9 |
| Stat1 | 979 | 879 | 947 | 972 | 779 | 777 | 888 | 408 | 419 | 524 | 497 | 426 | 480 | 554 | 998 | 959 | 955 | 969 | 775 | 867 | 893 | 476 | 471 | 431 | 417 | 597 | 458 | 520 | Stat1 |
| Stat2 | 3854 | 4336 | 4709 | 4629 | 4398 | 4194 | 4032 | 2136 | 2291 | 2676 | 2523 | 2452 | 2684 | 2395 | 3753 | 4650 | 4519 | 5117 | 4238 | 3955 | 4209 | 2630 | 2318 | 2452 | 2504 | 2140 | 2465 | 2370 | Stat2 |
| Stat3 | 7393 | 8409 | 8008 | 9274 | 9782 | 8500 | 8769 | 2696 | 3289 | 3488 | 3993 | 2754 | 3180 | 2662 | 7665 | 8382 | 8727 | 9994 | 9076 | 7126 | 8514 | 3800 | 3571 | 3269 | 3364 | 2711 | 2636 | 2762 | Stat3 |
| Stat4 | 138 | 111 | 113 | 61 | 95 | 107 | 121 | 81 | 85 | 39 | 48 | 130 | 115 | 175 | 156 | 100 | 94 | 65 | 78 | 126 | 84 | 27 | 36 | 98 | 72 | 204 | 139 | 148 | Stat4 |
| Stat5a | 168 | 219 | 204 | 124 | 135 | 196 | 111 | 112 | 152 | 164 | 222 | 122 | 159 | 121 | 158 | 223 | 130 | 135 | 113 | 98 | 106 | 132 | 246 | 149 | 119 | 162 | 145 | 119 | Stat5a |
| Stat5b | 2087 | 2302 | 2632 | 2277 | 2337 | 2355 | 2062 | 1585 | 1843 | 1891 | 2103 | 1783 | 2013 | 1816 | 1899 | 2197 | 2493 | 2195 | 2427 | 1979 | 2052 | 1860 | 1896 | 1778 | 1843 | 1874 | 1962 | 2061 | Stat5b |
| Stat6 | 677 | 615 | 833 | 463 | 468 | 521 | 360 | 812 | 1131 | 1009 | 1182 | 820 | 1161 | 758 | 596 | 815 | 594 | 516 | 413 | 435 | 390 | 906 | 1504 | 890 | 711 | 968 | 926 | 878 | Stat6 |
| Stau1 | 8586 | 9173 | 9958 | 10069 | 9940 | 9618 | 9712 | 7982 | 8437 | 9624 | 8974 | 9619 | 8650 | 9731 | 8352 | 8779 | 9834 | 9823 | 10024 | 8353 | 10305 | 9152 | 8534 | 9728 | 9397 | 9180 | 8788 | 9835 | Stau1 |
| Stau2 | 3857 | 3247 | 3504 | 3829 | 5012 | 4504 | 4768 | 6425 | 5944 | 6041 | 5431 | 5179 | 5029 | 5285 | 4048 | 3284 | 3508 | 4006 | 4547 | 4687 | 4778 | 7294 | 5829 | 5784 | 6185 | 5085 | 5043 | 5376 | Stau2 |
| Stbd1 | 1145 | 965 | 1029 | 772 | 819 | 798 | 598 | 399 | 480 | 410 | 539 | 330 | 463 | 319 | 935 | 1045 | 883 | 820 | 718 | 767 | 648 | 391 | 625 | 368 | 271 | 361 | 385 | 325 | Stbd1 |
| Stc1 | 669 | 485 | 612 | 652 | 690 | 634 | 685 | 490 | 574 | 491 | 635 | 364 | 338 | 379 | 658 | 540 | 640 | 647 | 573 | 509 | 612 | 541 | 652 | 414 | 444 | 401 | 311 | 317 | Stc1 |
| Stc2 | 580 | 622 | 694 | 954 | 704 | 741 | 819 | 219 | 220 | 210 | 374 | 279 | 227 | 278 | 674 | 526 | 683 | 791 | 793 | 667 | 836 | 201 | 289 | 346 | 219 | 338 | 242 | 310 | Stc2 |
| Steap1 | 276 | 238 | 273 | 240 | 150 | 119 | 151 | 62 | 76 | 65 | 63 | 60 | 79 | 74 | 269 | 228 | 222 | 194 | 149 | 207 | 185 | 80 | 88 | 81 | 59 | 93 | 69 | 69 | Steap1 |
| Steap2 | 765 | 608 | 799 | 508 | 662 | 740 | 616 | 761 | 854 | 803 | 779 | 712 | 752 | 697 | 712 | 702 | 674 | 591 | 605 | 663 | 575 | 714 | 808 | 690 | 635 | 718 | 743 | 727 | Steap2 |
| Steap3 | 255 | 357 | 352 | 226 | 205 | 268 | 161 | 144 | 243 | 148 | 246 | 210 | 297 | 210 | 334 | 420 | 227 | 298 | 228 | 176 | 200 | 184 | 371 | 221 | 186 | 239 | 201 | 298 | Steap3 |
| Steap4 | 14 | 6 | 31 | 11 | 7 | 7 | 10 | 50 | 71 | 70 | 92 | 16 | 49 | 37 | 15 | 14 | 8 | 13 | 5 | 9 | 6 | 18 | 120 | 78 | 31 | 41 | 38 | 39 | Steap4 |
| Steep1 | 4953 | 3962 | 4457 | 3787 | 3657 | 3790 | 4431 | 4362 | 4185 | 3426 | 3924 | 4258 | 4275 | 4188 | 4944 | 4410 | 4191 | 3955 | 3705 | 4175 | 3909 | 3673 | 4071 | 3685 | 3620 | 4312 | 4056 | 4285 | Steep1 |
| Stfa1 | 13 | 17 | 17 | 27 | 7 | 33 | 12 | 17 | 35 | 38 | 114 | 34 | 61 | 45 | 12 | 50 | 24 | 26 | 15 | 13 | 20 | 40 | 70 | 57 | 22 | 61 | 53 | 65 | Stfa1 |
| Stfa2 | 0 | 1 | 2 | 6 | 2 | 0 | 8 | 7 | 11 | 19 | 21 | 15 | 39 | 5 | 1 | 8 | 0 | 7 | 2 | 2 | 4 | 12 | 8 | 20 | 17 | 10 | 10 | 12 | Stfa2 |
| Stfa2l1 | 0 | 1 | 5 | 4 | 2 | 3 | 8 | 4 | 16 | 0 | 24 | 4 | 20 | 10 | 4 | 5 | 10 | 11 | 2 | 3 | 2 | 7 | 12 | 24 | 7 | 8 | 19 | 8 | Stfa2l1 |
| Stfa3 | 0 | 3 | 2 | 13 | 5 | 10 | 4 | 11 | 11 | 27 | 25 | 15 | 37 | 14 | 5 | 10 | 14 | 6 | 4 | 9 | 8 | 2 | 29 | 40 | 3 | 21 | 26 | 26 | Stfa3 |
| Stil | 6696 | 5722 | 4625 | 4355 | 3214 | 4189 | 4596 | 3560 | 3247 | 2742 | 2754 | 4296 | 3886 | 4307 | 5764 | 5510 | 4505 | 3912 | 3887 | 4199 | 4058 | 2316 | 2926 | 3385 | 2805 | 4414 | 4457 | 4145 | Stil |
| Stim1 | 4211 | 4195 | 4310 | 4457 | 4017 | 3838 | 3578 | 1578 | 1961 | 1736 | 2285 | 1702 | 1893 | 1619 | 3545 | 4407 | 4370 | 4517 | 4131 | 3144 | 3800 | 1932 | 2115 | 1749 | 1596 | 1609 | 1673 | 1651 | Stim1 |
| Stim2 | 3045 | 3022 | 3200 | 3767 | 4148 | 3586 | 4299 | 2461 | 2410 | 2691 | 2684 | 2485 | 2460 | 2567 | 3034 | 2965 | 3415 | 3826 | 3819 | 3467 | 4215 | 2806 | 2498 | 2554 | 2566 | 2633 | 2201 | 2503 | Stim2 |
| Stimate | 2132 | 2240 | 2365 | 2467 | 2428 | 2356 | 2209 | 1326 | 1554 | 1548 | 1753 | 1489 | 1670 | 1507 | 1961 | 2403 | 2144 | 2426 | 2360 | 2079 | 2196 | 1434 | 1531 | 1577 | 1642 | 1501 | 1579 | 1517 | Stimate |
| Sting1 | 111 | 112 | 64 | 70 | 83 | 58 | 46 | 140 | 266 | 177 | 243 | 126 | 180 | 133 | 91 | 136 | 61 | 83 | 65 | 67 | 61 | 144 | 316 | 156 | 119 | 183 | 145 | 140 | Sting1 |
| Stip1 | 25967 | 27666 | 29789 | 26459 | 21681 | 21455 | 23176 | 18036 | 16024 | 18460 | 16317 | 20012 | 17359 | 20668 | 24991 | 22151 | 28543 | 23624 | 25606 | 21724 | 24317 | 16065 | 15817 | 20625 | 18106 | 23289 | 20650 | 21965 | Stip1 |
| Stk10 | 884 | 1098 | 1250 | 1241 | 1160 | 980 | 1041 | 420 | 547 | 685 | 709 | 592 | 655 | 626 | 903 | 1167 | 1144 | 1231 | 1161 | 817 | 1102 | 590 | 634 | 731 | 562 | 590 | 585 | 565 | Stk10 |
| Stk11 | 6151 | 7980 | 9759 | 9134 | 8221 | 7846 | 7350 | 4756 | 5808 | 6724 | 6590 | 7587 | 6641 | 7277 | 5945 | 6367 | 9872 | 8474 | 9740 | 6960 | 7957 | 5285 | 5517 | 7938 | 7102 | 8993 | 7239 | 7271 | Stk11 |
| Stk11ip | 3785 | 4729 | 4053 | 4406 | 3251 | 3357 | 3295 | 3072 | 3441 | 3450 | 3380 | 3850 | 3851 | 3553 | 3603 | 4240 | 4224 | 3900 | 3698 | 3283 | 3229 | 3278 | 3271 | 3570 | 3466 | 3668 | 3724 | 3725 | Stk11ip |
| Stk16 | 4517 | 4902 | 4313 | 4416 | 4119 | 3953 | 4722 | 4545 | 4507 | 4237 | 4531 | 4817 | 4652 | 4806 | 4391 | 4237 | 4481 | 4277 | 4453 | 4307 | 4418 | 4441 | 4355 | 4521 | 4687 | 5093 | 4803 | 4934 | Stk16 |
| Stk17b | 2252 | 1856 | 1945 | 1808 | 1335 | 1592 | 1781 | 1040 | 914 | 784 | 779 | 919 | 1047 | 955 | 2371 | 2219 | 1742 | 1779 | 1533 | 2052 | 1605 | 790 | 1031 | 848 | 833 | 1110 | 1002 | 1086 | Stk17b |
| Stk19 | 1578 | 1937 | 1820 | 2264 | 2225 | 2087 | 1949 | 1519 | 1519 | 1834 | 1895 | 1828 | 1852 | 1922 | 1535 | 1589 | 2275 | 2205 | 2177 | 1983 | 1984 | 1770 | 1549 | 1993 | 1940 | 1792 | 1837 | 1804 | Stk19 |
| Stk24 | 4411 | 3657 | 3820 | 3748 | 3494 | 3684 | 3772 | 3883 | 3930 | 3979 | 4079 | 4441 | 4067 | 4762 | 3808 | 3757 | 3924 | 3738 | 3825 | 3236 | 3737 | 3928 | 3829 | 4272 | 4081 | 4500 | 4238 | 4868 | Stk24 |
| Stk25 | 14177 | 15944 | 14065 | 15116 | 15565 | 15185 | 14921 | 12334 | 12306 | 12080 | 11998 | 12217 | 11581 | 11566 | 14827 | 14231 | 14777 | 15221 | 15974 | 15524 | 15039 | 13463 | 12178 | 12499 | 12903 | 12226 | 11893 | 11665 | Stk25 |
| Stk26 | 2098 | 929 | 1008 | 666 | 475 | 820 | 506 | 1271 | 2233 | 1308 | 3347 | 615 | 1060 | 651 | 1623 | 2072 | 696 | 817 | 457 | 601 | 452 | 1399 | 2299 | 1215 | 809 | 1217 | 722 | 808 | Stk26 |
| Stk3 | 3276 | 2234 | 2692 | 2397 | 2086 | 2323 | 2407 | 2387 | 2283 | 1943 | 2255 | 2082 | 2377 | 2143 | 3066 | 2830 | 2199 | 2324 | 2178 | 2415 | 2179 | 1871 | 2519 | 2141 | 1929 | 2571 | 2250 | 2544 | Stk3 |
| Stk31 | 18 | 19 | 18 | 35 | 21 | 16 | 50 | 9 | 9 | 5 | 10 | 8 | 10 | 6 | 34 | 12 | 20 | 21 | 15 | 5 | 29 | 5 | 12 | 5 | 14 | 2 | 10 | 4 | Stk31 |

| | | | | | | | | | | | | | | | | | | | | | | | | | | | | | |
|---|---|---|---|---|---|---|---|---|---|---|---|---|---|---|---|---|---|---|---|---|---|---|---|---|---|---|---|---|---|
| Stk32a | 255 | 170 | 208 | 252 | 229 | 177 | 184 | 324 | 275 | 381 | 439 | 215 | 172 | 170 | 218 | 208 | 187 | 182 | 187 | 211 | 190 | 267 | 344 | 374 | 202 | 284 | 198 | 199 | Stk32a |
| Stk32b | 1240 | 1145 | 952 | 1745 | 2676 | 2322 | 2213 | 2387 | 2859 | 3043 | 2612 | 1950 | 1844 | 1876 | 1262 | 1144 | 1428 | 2196 | 2327 | 2067 | 2596 | 3537 | 2473 | 2437 | 3034 | 1380 | 1518 | 1615 | Stk32b |
| Stk32c | 28 | 173 | 39 | 65 | 451 | 262 | 27 | 171 | 30 | 348 | 61 | 26 | 48 | 144 | 26 | 84 | 35 | 213 | 108 | 107 | 31 | 38 | 28 | 63 | 366 | 16 | 86 | 14 | Stk32c |
| Stk33 | 584 | 614 | 840 | 490 | 483 | 568 | 506 | 699 | 517 | 616 | 484 | 937 | 779 | 949 | 666 | 523 | 689 | 457 | 600 | 555 | 535 | 682 | 522 | 705 | 653 | 682 | 1181 | 1044 | Stk33 |
| Stk35 | 2791 | 2597 | 3218 | 3083 | 2995 | 2758 | 3006 | 2876 | 3179 | 3117 | 3354 | 4058 | 3393 | 3781 | 2378 | 2494 | 3246 | 2860 | 3102 | 2381 | 3063 | 3285 | 2977 | 3742 | 3544 | 4037 | 3625 | 3961 | Stk35 |
| Stk36 | 2749 | 3472 | 3020 | 3228 | 3335 | 2926 | 3358 | 2650 | 2675 | 2847 | 2539 | 3009 | 2791 | 2797 | 2732 | 3079 | 3490 | 3214 | 3458 | 3025 | 3267 | 2976 | 2646 | 2780 | 3124 | 2551 | 2932 | 2887 | Stk36 |
| Stk38 | 15326 | 14195 | 15018 | 15215 | 14604 | 15020 | 16108 | 11054 | 10257 | 11390 | 10947 | 13002 | 11922 | 13028 | 15054 | 14020 | 15223 | 14681 | 14855 | 13849 | 15309 | 11774 | 10023 | 11718 | 12425 | 12163 | 12731 | 12955 | Stk38 |
| Stk38l | 3992 | 3664 | 3762 | 3710 | 2928 | 3234 | 3260 | 2065 | 1788 | 1583 | 1776 | 1777 | 1941 | 1716 | 4187 | 3788 | 3351 | 3516 | 2946 | 3578 | 3097 | 1654 | 1861 | 1567 | 1687 | 1824 | 1755 | 1841 | Stk38l |
| Stk39 | 4842 | 4614 | 5673 | 4467 | 4909 | 4767 | 4662 | 5539 | 5038 | 5077 | 4927 | 6146 | 6643 | 6263 | 5182 | 4721 | 5032 | 4381 | 4597 | 4388 | 4331 | 5750 | 5143 | 5033 | 5136 | 6255 | 6199 | 6487 | Stk39 |
| Stk4 | 11929 | 11474 | 9352 | 10066 | 8693 | 9553 | 10156 | 7238 | 6306 | 6165 | 5775 | 7290 | 6480 | 7118 | 11639 | 10168 | 9778 | 9859 | 9277 | 10417 | 10196 | 6385 | 6008 | 6612 | 7017 | 6353 | 7241 | 7390 | Stk4 |
| Stk40 | 6316 | 8155 | 8053 | 7347 | 5929 | 5949 | 5515 | 3251 | 3700 | 3858 | 4002 | 4288 | 4179 | 4081 | 6005 | 7615 | 7743 | 6848 | 6361 | 5422 | 5936 | 3673 | 3638 | 4199 | 3805 | 4468 | 3685 | 4307 | Stk40 |
| Stkld1 | 3 | 5 | 1 | 7 | 8 | 10 | 11 | 13 | 12 | 15 | 23 | 29 | 14 | 11 | 9 | 8 | 2 | 11 | 5 | 9 | 11 | 17 | 19 | 16 | 16 | 17 | 28 | 8 | Stkld1 |
| Stmn1 | 51365 | 54771 | 55053 | 66577 | 70268 | 62750 | 78360 | 74802 | 66954 | 78128 | 64103 | 76815 | 64753 | 80736 | 58587 | 45196 | 62066 | 66966 | 73300 | 70596 | 80837 | 80021 | 62174 | 81896 | 84579 | 75138 | 69301 | 78782 | Stmn1 |
| Stmn2 | 18204 | 21475 | 17093 | 24723 | 36387 | 29012 | 32407 | 45878 | 36359 | 37142 | 32402 | 28979 | 33962 | 31902 | 26570 | 19195 | 22393 | 28051 | 30276 | 35052 | 32470 | 47912 | 35481 | 32141 | 40744 | 23180 | 29141 | 30524 | Stmn2 |
| Stmn3 | 5376 | 8037 | 5971 | 9613 | 12378 | 9565 | 11644 | 12234 | 11830 | 11967 | 9507 | 9407 | 8788 | 10182 | 7923 | 5691 | 8633 | 9613 | 11752 | 12394 | 12949 | 12325 | 11585 | 10161 | 12945 | 8424 | 9549 | 9597 | Stmn3 |
| Stmn4 | 5424 | 7266 | 6675 | 9594 | 11146 | 8749 | 10920 | 8243 | 7674 | 9064 | 7328 | 6495 | 5338 | 7914 | 7713 | 5321 | 9013 | 9516 | 10776 | 10612 | 11608 | 10169 | 7519 | 8219 | 9335 | 6508 | 6723 | 7155 | Stmn4 |
| Stmnd1 | 0 | 3 | 8 | 4 | 0 | 0 | 0 | 0 | 9 | 7 | 17 | 1 | 5 | 0 | 4 | 3 | 1 | 0 | 2 | 1 | 0 | 3 | 5 | 0 | 7 | 0 | 3 | 2 | Stmnd1 |
| Stn1 | 1510 | 1480 | 1529 | 1069 | 1204 | 1345 | 1150 | 2045 | 2118 | 1986 | 2100 | 2207 | 2301 | 2056 | 1298 | 1438 | 1314 | 1197 | 1248 | 1247 | 1147 | 1761 | 2034 | 1874 | 1847 | 2520 | 2393 | 2200 | Stn1 |
| Stom | 3606 | 3314 | 3111 | 2779 | 2553 | 2623 | 2452 | 1072 | 1198 | 963 | 1215 | 813 | 1012 | 802 | 3413 | 3593 | 3011 | 2747 | 2177 | 2458 | 2348 | 1053 | 1552 | 993 | 897 | 1100 | 894 | 989 | Stom |
| Stoml1 | 2591 | 3174 | 2429 | 3618 | 3512 | 3339 | 3125 | 2623 | 2927 | 2638 | 2493 | 2624 | 2459 | 2371 | 2860 | 2759 | 3006 | 3356 | 3642 | 3438 | 3158 | 3146 | 2849 | 2709 | 3065 | 2360 | 2486 | 2281 | Stoml1 |
| Stoml2 | 4738 | 5399 | 5118 | 4341 | 3607 | 4095 | 4099 | 4612 | 4769 | 4210 | 4343 | 5099 | 4884 | 5191 | 4555 | 4529 | 4527 | 3779 | 4387 | 5074 | 3954 | 3684 | 4709 | 4878 | 4518 | 6200 | 5833 | 5749 | Stoml2 |
| Stoml3 | 24 | 20 | 24 | 41 | 15 | 26 | 44 | 9 | 3 | 5 | 8 | 8 | 3 | 6 | 17 | 29 | 31 | 50 | 32 | 17 | 39 | 3 | 6 | 2 | 1 | 6 | 8 | 1 | Stoml3 |
| Ston1 | 1316 | 1373 | 1383 | 1079 | 1163 | 1280 | 1093 | 923 | 1364 | 1012 | 1403 | 1086 | 1280 | 928 | 1234 | 1515 | 1365 | 1274 | 1256 | 979 | 1066 | 886 | 1399 | 1126 | 902 | 931 | 1010 | 943 | Ston1 |
| Ston2 | 4331 | 4399 | 4435 | 6119 | 5375 | 4855 | 6301 | 2500 | 2823 | 2823 | 2712 | 3810 | 3679 | 3830 | 4178 | 4339 | 5212 | 5885 | 5488 | 4809 | 5854 | 2102 | 2666 | 3287 | 3096 | 2926 | 3517 | 3819 | Ston2 |
| Stox1 | 294 | 276 | 252 | 247 | 192 | 212 | 235 | 211 | 180 | 251 | 175 | 204 | 193 | 213 | 274 | 313 | 221 | 223 | 232 | 180 | 212 | 172 | 211 | 239 | 244 | 207 | 155 | 179 | Stox1 |
| Stox2 | 15606 | 14194 | 15091 | 19242 | 21315 | 20907 | 21131 | 19723 | 18488 | 22387 | 19216 | 25046 | 21368 | 25990 | 15329 | 13738 | 17616 | 17865 | 21378 | 17901 | 21337 | 22363 | 16204 | 20212 | 25970 | 13749 | 24205 | 24143 | Stox2 |
| Stpg1 | 8 | 17 | 25 | 15 | 20 | 24 | 15 | 39 | 26 | 38 | 35 | 63 | 82 | 43 | 22 | 27 | 30 | 26 | 16 | 19 | 21 | 38 | 47 | 28 | 45 | 40 | 55 | 61 | Stpg1 |
| Stpg2 | 5 | 4 | 2 | 4 | 10 | 8 | 8 | 5 | 7 | 1 | 1 | 8 | 6 | 3 | 4 | 3 | 6 | 0 | 4 | 3 | 4 | 5 | 8 | 12 | 3 | 5 | 3 | 7 | Stpg2 |
| Stpg3 | 1 | 2 | 0 | 1 | 2 | 1 | 0 | 0 | 1 | 1 | 4 | 0 | 0 | 0 | 0 | 1 | 0 | 1 | 2 | 1 | 6 | 0 | 0 | 2 | 3 | 4 | 2 | 0 | Stpg3 |
| Stpg4 | 0 | 3 | 0 | 7 | 1 | 15 | 9 | 1 | 13 | 5 | 6 | 2 | 2 | 1 | 3 | 2 | 9 | 7 | 5 | 10 | 9 | 8 | 5 | 0 | 4 | 7 | 8 | 2 | Stpg4 |
| Stra6 | 713 | 590 | 642 | 469 | 894 | 488 | 523 | 1084 | 1266 | 1255 | 1646 | 948 | 1318 | 932 | 564 | 769 | 564 | 668 | 571 | 511 | 549 | 1447 | 1450 | 1334 | 994 | 1929 | 810 | 954 | Stra6 |
| Stra6l | 178 | 154 | 122 | 212 | 268 | 335 | 272 | 267 | 253 | 355 | 309 | 275 | 180 | 287 | 171 | 114 | 163 | 216 | 253 | 302 | 358 | 289 | 210 | 261 | 409 | 152 | 227 | 173 | Stra6l |
| Stra8 | 0 | 0 | 0 | 0 | 0 | 0 | 0 | 0 | 0 | 0 | 0 | 0 | 0 | 0 | 0 | 2 | 0 | 0 | 0 | 0 | 0 | 0 | 0 | 0 | 0 | 2 | 0 | 0 | Stra8 |
| Strada | 2388 | 2642 | 2353 | 2284 | 2144 | 2590 | 2129 | 2386 | 2584 | 2359 | 2511 | 2599 | 2583 | 2705 | 2441 | 2533 | 2195 | 2165 | 2240 | 2602 | 2333 | 2159 | 2530 | 2434 | 2580 | 2728 | 2907 | 2632 | Strada |
| Stradb | 2844 | 2744 | 2460 | 3111 | 3704 | 3411 | 3850 | 3384 | 3198 | 2915 | 2927 | 2615 | 2633 | 2775 | 3041 | 2602 | 2968 | 3288 | 3420 | 3812 | 4031 | 3457 | 3176 | 2857 | 3059 | 2244 | 2575 | 2676 | Stradb |
| Strap | 22885 | 18489 | 21866 | 18552 | 17487 | 17819 | 20046 | 17653 | 14576 | 14367 | 13381 | 17582 | 16023 | 17403 | 22971 | 19767 | 19288 | 17539 | 18227 | 20449 | 18628 | 14800 | 14863 | 15561 | 15261 | 19665 | 18204 | 18823 | Strap |
| Strbp | 15252 | 12552 | 15693 | 17058 | 18547 | 17611 | 19703 | 16838 | 13392 | 16146 | 14403 | 16728 | 15492 | 17958 | 15587 | 12920 | 16465 | 17146 | 18651 | 16344 | 19104 | 19231 | 13345 | 17074 | 18685 | 16157 | 16066 | 17751 | Strbp |
| Strc | 2 | 3 | 0 | 8 | 8 | 6 | 2 | 5 | 6 | 0 | 5 | 3 | 8 | 5 | 0 | 2 | 10 | 12 | 11 | 3 | 4 | 7 | 5 | 11 | 1 | 8 | 9 | 4 | Strc |
| Strip1 | 9388 | 10549 | 8445 | 10216 | 10738 | 10355 | 10860 | 11093 | 11277 | 11155 | 9848 | 11714 | 10237 | 11324 | 9393 | 8828 | 9655 | 10294 | 11036 | 10441 | 11021 | 12333 | 10550 | 11438 | 12246 | 10377 | 11068 | 11160 | Strip1 |
| Strip2 | 1187 | 1074 | 879 | 1066 | 1235 | 1088 | 1158 | 275 | 313 | 282 | 251 | 188 | 278 | 173 | 1187 | 1323 | 1035 | 1158 | 944 | 1136 | 1111 | 309 | 351 | 181 | 270 | 147 | 213 | 184 | Strip2 |
| Strit1 | 0 | 0 | 0 | 0 | 0 | 0 | 0 | 0 | 0 | 0 | 0 | 0 | 0 | 0 | 1 | 0 | 0 | 0 | 0 | 0 | 0 | 0 | 0 | 0 | 0 | 0 | 0 | 0 | Strit1 |
| Strn | 5494 | 4620 | 4550 | 5444 | 5605 | 5821 | 5612 | 4714 | 4473 | 4050 | 4580 | 4964 | 4797 | 4802 | 5474 | 5400 | 4933 | 5416 | 5156 | 5073 | 5745 | 4610 | 4409 | 3940 | 4792 | 3782 | 5054 | 4379 | Strn |
| Strn3 | 6158 | 4358 | 5312 | 5192 | 6251 | 6358 | 6519 | 9738 | 8721 | 7947 | 8557 | 9634 | 8946 | 9640 | 6626 | 5288 | 5817 | 5576 | 6273 | 6287 | 6229 | 9354 | 8708 | 8272 | 8750 | 8775 | 9860 | 9630 | Strn3 |
| Strn4 | 6717 | 8352 | 7688 | 8785 | 9970 | 9108 | 9733 | 10345 | 11555 | 12541 | 11413 | 13203 | 11378 | 12232 | 6711 | 6818 | 9049 | 9134 | 10715 | 8682 | 9688 | 12624 | 11110 | 12879 | 13359 | 12135 | 12221 | 11714 | Strn4 |
| Stt3a | 12423 | 10568 | 10335 | 8274 | 7050 | 7696 | 7383 | 9649 | 8708 | 8077 | 9610 | 8220 | 8811 | 7989 | 11501 | 11612 | 8764 | 8905 | 7314 | 8225 | 7200 | 8900 | 9563 | 8193 | 7289 | 9298 | 8407 | 8594 | Stt3a |
| Stt3b | 14935 | 14068 | 14991 | 14601 | 13812 | 13602 | 14152 | 14067 | 13188 | 12541 | 13605 | 12784 | 13113 | 12962 | 15140 | 15217 | 13906 | 13970 | 13764 | 13342 | 14580 | 13966 | 13502 | 12258 | 12750 | 12947 | 12831 | 13604 | Stt3b |
| Stub1 | 5022 | 6374 | 5680 | 5793 | 5139 | 5185 | 5243 | 3423 | 4418 | 4549 | 4570 | 4718 | 4581 | 4531 | 4812 | 5274 | 6135 | 5302 | 5598 | 4884 | 5371 | 3873 | 4326 | 4790 | 4521 | 5360 | 4596 | 4778 | Stub1 |
| Stum | 662 | 749 | 570 | 737 | 1209 | 775 | 761 | 307 | 417 | 468 | 683 | 173 | 251 | 148 | 758 | 889 | 673 | 850 | 871 | 754 | 862 | 551 | 498 | 275 | 366 | 215 | 145 | 155 | Stum |
| Stx11 | 602 | 485 | 589 | 299 | 322 | 410 | 238 | 159 | 167 | 113 | 173 | 148 | 214 | 116 | 553 | 581 | 458 | 415 | 303 | 251 | 253 | 155 | 202 | 109 | 94 | 153 | 172 | 155 | Stx11 |
| Stx12 | 11397 | 9798 | 9522 | 11161 | 12119 | 11545 | 13784 | 12772 | 10421 | 10422 | 9434 | 10284 | 9395 | 10398 | 12381 | 10249 | 10673 | 11565 | 12036 | 14358 | 12999 | 12198 | 10613 | 10432 | 11392 | 10111 | 9917 | 10250 | Stx12 |
| Stx16 | 5376 | 5199 | 6253 | 6566 | 6664 | 6246 | 6886 | 7306 | 7492 | 8556 | 8907 | 8587 | 7730 | 8533 | 4981 | 5062 | 6638 | 6867 | 7052 | 5754 | 6806 | 8283 | 7121 | 9605 | 9200 | 8410 | 8378 | 7862 | Stx16 |
| Stx17 | 3952 | 3244 | 3072 | 2951 | 3043 | 3286 | 3414 | 3452 | 3293 | 2768 | 3278 | 2807 | 3071 | 3241 | 3835 | 3649 | 2866 | 3022 | 2763 | 3277 | 3118 | 3008 | 3335 | 2759 | 2674 | 3034 | 3026 | 2896 | Stx17 |
| Stx18 | 1077 | 1057 | 1128 | 785 | 731 | 831 | 759 | 1464 | 1439 | 1387 | 1351 | 1506 | 1567 | 1412 | 1058 | 1016 | 907 | 789 | 792 | 855 | 721 | 1240 | 1546 | 1473 | 1249 | 1607 | 1644 | 1518 | Stx18 |
| Stx19 | 1 | 1 | 0 | 0 | 0 | 0 | 0 | 4 | 20 | 5 | 8 | 2 | 6 | 1 | 0 | 2 | 0 | 0 | 0 | 0 | 0 | 3 | 22 | 9 | 0 | 3 | 0 | 1 | Stx19 |
| Stx1a | 2317 | 3355 | 2919 | 3735 | 4165 | 3466 | 3256 | 2277 | 2742 | 3086 | 2816 | 1941 | 2159 | 1836 | 2411 | 3004 | 3161 | 4384 | 3982 | 3445 | 3544 | 3180 | 2688 | 2655 | 3019 | 2217 | 2056 | 1733 | Stx1a |
| Stx1b | 3177 | 3309 | 3928 | 5669 | 7798 | 5934 | 5908 | 4938 | 5559 | 7071 | 6029 | 5766 | 5160 | 5370 | 3268 | 3140 | 4971 | 6266 | 7159 | 4853 | 6499 | 8702 | 4846 | 6586 | 7914 | 4482 | 4561 | 4780 | Stx1b |
| Stx2 | 3633 | 4082 | 3501 | 3568 | 3472 | 3572 | 3206 | 2351 | 2556 | 2583 | 2627 | 2525 | 2631 | 2429 | 3657 | 4255 | 3612 | 3548 | 3233 | 3417 | 3212 | 2525 | 2553 | 2290 | 2268 | 2677 | 2589 | 2470 | Stx2 |
| Stx3 | 2054 | 1574 | 1904 | 1541 | 1978 | 1951 | 1818 | 1556 | 1223 | 1343 | 1154 | 1210 | 1236 | 1446 | 1899 | 1512 | 1486 | 1610 | 1852 | 2071 | 1889 | 1460 | 1265 | 1131 | 1449 | 798 | 1422 | 1415 | Stx3 |
| Stx4a | 1911 | 2087 | 2372 | 1758 | 1580 | 1807 | 1611 | 1279 | 1710 | 1797 | 2020 | 1564 | 1721 | 1621 | 1738 | 1944 | 2009 | 1819 | 1662 | 1582 | 1556 | 1360 | 2022 | 1809 | 1499 | 2110 | 1649 | 1656 | Stx4a |
| Stx5a | 7183 | 8428 | 6597 | 7603 | 6815 | 7062 | 7119 | 6719 | 6692 | 6801 | 7214 | 6232 | 6224 | 6242 | 6964 | 7473 | 6938 | 7701 | 7105 | 7329 | 7396 | 6330 | 6727 | 6387 | 6376 | 6063 | 6093 | 6052 | Stx5a |
| Stx6 | 8597 | 9573 | 7294 | 8873 | 8629 | 9263 | 9303 | 5675 | 5941 | 5256 | 5938 | 6417 | 5979 | 6470 | 8528 | 8501 | 8545 | 9045 | 8936 | 8536 | 9832 | 6034 | 5747 | 6207 | 6491 | 5982 | 6150 | 6120 | Stx6 |
| Stx7 | 9961 | 9106 | 9211 | 11267 | 13426 | 11748 | 13045 | 9474 | 8426 | 8850 | 8062 | 7343 | 6922 | 7170 | 11462 | 9830 | 10299 | 12224 | 12445 | 12376 | 12333 | 10940 | 8904 | 8007 | 9469 | 6886 | 6816 | 6886 | Stx7 |
| Stx8 | 1416 | 1490 | 1350 | 1203 | 951 | 1211 | 1220 | 1390 | 1256 | 1300 | 1117 | 1164 | 1220 | 1344 | 1468 | 1210 | 1238 | 1173 | 1253 | 1315 | 1203 | 1255 | 1271 | 1172 | 1172 | 1339 | 1189 | 1491 | Stx8 |
| Stxbp1 | 7170 | 8627 | 9502 | 13654 | 16836 | 13417 | 13491 | 7974 | 8919 | 12605 | 10201 | 9384 | 7986 | 8878 | 7871 | 8318 | 12228 | 14896 | 15706 | 11155 | 15319 | 13948 | 8370 | 10721 | 13481 | 7689 | 7596 | 8082 | Stxbp1 |
| Stxbp2 | 1626 | 1685 | 1636 | 1563 | 1724 | 1595 | 1753 | 1544 | 1809 | 1771 | 1460 | 1812 | 1393 | 1658 | 1375 | 1474 | 1674 | 1645 | 1693 | 1623 | 1652 | 1616 | 1705 | 1975 | 1734 | 2091 | 1564 | 1616 | Stxbp2 |
| Stxbp3 | 1735 | 1396 | 1612 | 1554 | 1322 | 1352 | 1350 | 1338 | 1313 | 1123 | 1356 | 1253 | 1370 | 1352 | 1621 | 1879 | 1444 | 1382 | 1230 | 1576 | 1263 | 1213 | 1521 | 1177 | 1057 | 1392 | 1318 | 1284 | Stxbp3 |
| Stxbp4 | 3385 | 3055 | 2654 | 2715 | 2694 | 2769 | 2977 | 4665 | 4271 | 3997 | 3882 | 4785 | 4841 | 4685 | 3312 | 3056 | 2804 | 2689 | 2703 | 2840 | 2834 | 4121 | 4392 | 4221 | 4141 | 3602 | 4782 | 4850 | Stxbp4 |
| Stxbp5 | 4850 | 4306 | 5024 | 5665 | 6807 | 6482 | 6899 | 3750 | 3718 | 4185 | 4110 | 3816 | 3916 | 3998 | 4912 | 4329 | 5405 | 6007 | 6138 | 5719 | 6718 | 4416 | 3591 | 3630 | 4116 | 2788 | 3930 | 3732 | Stxbp5 |
| Stxbp5l | 982 | 1008 | 351 | 664 | 844 | 1253 | 1028 | 592 | 405 | 344 | 352 | 342 | 303 | 367 | 1183 | 742 | 448 | 609 | 687 | 1361 | 1179 | 334 | 500 | 159 | 383 | 55 | 420 | 249 | Stxbp5l |
| Stxbp6 | 798 | 636 | 954 | 732 | 1078 | 895 | 1029 | 1419 | 1296 | 1256 | 1136 | 1427 | 1245 | 1392 | 827 | 739 | 885 | 716 | 899 | 941 | 943 | 1305 | 1289 | 1395 | 1206 | 1418 | 1610 | 1466 | Stxbp6 |
| Styk1 | 107 | 74 | 143 | 61 | 130 | 100 | 120 | 116 | 138 | 137 | 173 | 128 | 120 | 115 | 79 | 91 | 48 | 86 | 91 | 89 | 122 | 150 | 120 | 101 | 149 | 159 | 144 | 140 | Styk1 |
| Styx | 3037 | 2051 | 2558 | 3170 | 3291 | 3338 | 3655 | 2306 | 2182 | 2155 | 2406 | 2198 | 2414 | 2146 | 2859 | 3146 | 2634 | 3342 | 2877 | 3053 | 3491 | 2410 | 2163 | 1961 | 2345 | 1969 | 2151 | 2156 | Styx |
| Styxl1 | 16 | 24 | 13 | 22 | 21 | 11 | 17 | 9 | 21 | 14 | 11 | 11 | 16 | 14 | 33 | 19 | 17 | 22 | 12 | 22 | 16 | 8 | 13 | 8 | 8 | 11 | 24 | 18 | Styxl1 |
| Sub1 | 19104 | 16468 | 19104 | 20247 | 19307 | 18358 | 21617 | 17462 | 13945 | 14596 | 15052 | 14617 | 14673 | 16417 | 22030 | 17914 | 19109 | 19089 | 18713 | 24674 | 20360 | 15206 | 14680 | 15204 | 15516 | 16383 | 15937 | 16881 | Sub1 |
| Sucla2 | 6016 | 4721 | 5387 | 5055 | 4970 | 4886 | 5981 | 6535 | 5359 | 5256 | 5282 | 5621 | 5594 | 6350 | 6138 | 5146 | 4793 | 5064 | 4908 | 6085 | 5272 | 5691 | 5745 | 5487 | 5312 | 6509 | 5682 | 6329 | Sucla2 |
| Suclg1 | 6668 | 6833 | 6280 | 5728 | 5303 | 5418 | 5972 | 4301 | 4254 | 3818 | 3873 | 4252 | 4259 | 4382 | 6501 | 6520 | 6057 | 5853 | 5597 | 6625 | 5610 | 3676 | 4332 | 4015 | 3938 | 4804 | 4436 | 4721 | Suclg1 |
| Suclg2 | 2294 | 1989 | 2161 | 1479 | 1394 | 1528 | 1352 | 2387 | 2396 | 2198 | 2429 | 2499 | 2401 | 2673 | 2113 | 2242 | 1567 | 1411 | 1294 | 1575 | 1417 | 1653 | 2643 | 2490 | 1995 | 3351 | 2572 | 2725 | Suclg2 |
| Sucnr1 | 0 | 1 | 1 | 0 | 0 | 0 | 1 | 1 | 2 | 1 | 6 | 0 | 2 | 0 | 4 | 2 | 1 | 0 | 2 | 0 | 2 | 0 | 3 | 1 | 6 | 0 | 6 | 1 | Sucnr1 |
| Suco | 10568 | 7743 | 8224 | 8614 | 8523 | 9291 | 9333 | 8426 | 7558 | 6910 | 7234 | 7791 | 7745 | 8026 | 10309 | 9284 | 8192 | 8587 | 8055 | 9392 | 8810 | 7411 | 7226 | 7111 | 7084 | 7293 | 7620 | 7886 | Suco |

| | | | | | | | | | | | | | | | | | | | | | | | | | | | | | |
|---|---|---|---|---|---|---|---|---|---|---|---|---|---|---|---|---|---|---|---|---|---|---|---|---|---|---|---|---|---|
| Suds3 | 12389 | 13499 | 13483 | 14418 | 12401 | 12273 | 13020 | 7860 | 7725 | 8567 | 7622 | 8432 | 7896 | 9038 | 11859 | 11335 | 13935 | 13537 | 14294 | 12385 | 13932 | 7626 | 7382 | 8646 | 8694 | 8690 | 8621 | 9053 | Suds3 |
| Sufu | 4531 | 5340 | 5657 | 5723 | 4971 | 4905 | 4562 | 3274 | 3904 | 4081 | 4029 | 4766 | 4257 | 4567 | 4032 | 5134 | 5490 | 5604 | 5297 | 3925 | 5072 | 3745 | 3816 | 4560 | 4193 | 5047 | 4227 | 4580 | Sufu |
| Sugct | 70 | 41 | 95 | 77 | 52 | 75 | 58 | 61 | 76 | 93 | 69 | 77 | 62 | 66 | 76 | 79 | 71 | 57 | 55 | 56 | 59 | 90 | 90 | 56 | 60 | 66 | 49 | 72 | Sugct |
| Sugp1 | 3873 | 4112 | 4004 | 4678 | 4269 | 4349 | 4081 | 3530 | 4163 | 4169 | 4255 | 4593 | 4341 | 4591 | 4030 | 3717 | 4406 | 4394 | 4495 | 3638 | 4129 | 4181 | 3878 | 4335 | 4719 | 4419 | 4190 | 4314 | Sugp1 |
| Sugp2 | 8450 | 9993 | 8809 | 11578 | 11454 | 10797 | 11688 | 8999 | 8912 | 10319 | 9553 | 10713 | 9486 | 10375 | 8665 | 8508 | 10262 | 11274 | 11956 | 9976 | 11709 | 11214 | 8016 | 10377 | 11875 | 8250 | 9735 | 9975 | Sugp2 |
| Sugt1 | 10589 | 9005 | 8697 | 8340 | 8049 | 8646 | 9637 | 8767 | 7429 | 6824 | 6461 | 7910 | 7204 | 8008 | 10941 | 9693 | 8981 | 8419 | 8352 | 9977 | 9332 | 7406 | 7671 | 7258 | 7120 | 7993 | 8229 | 8254 | Sugt1 |
| Sulf1 | 10823 | 10718 | 10188 | 6420 | 7643 | 8130 | 5111 | 15299 | 14954 | 11487 | 13444 | 9906 | 17361 | 8989 | 14213 | 15457 | 8641 | 7410 | 5761 | 7660 | 5749 | 11887 | 15800 | 8368 | 8464 | 9106 | 11923 | 10383 | Sulf1 |
| Sulf2 | 11066 | 12938 | 13660 | 9971 | 9810 | 10167 | 5655 | 12146 | 15177 | 15278 | 19655 | 14220 | 20181 | 12312 | 11424 | 16957 | 11444 | 11197 | 7614 | 5757 | 6142 | 16883 | 16717 | 13872 | 12803 | 16619 | 14645 | 14512 | Sulf2 |
| Sult1a1 | 31 | 30 | 45 | 46 | 60 | 35 | 57 | 35 | 80 | 69 | 123 | 27 | 23 | 23 | 32 | 30 | 42 | 62 | 62 | 50 | 75 | 84 | 135 | 56 | 62 | 11 | 17 | 15 | Sult1a1 |
| Sult1b1 | 12 | 3 | 0 | 3 | 0 | 0 | 0 | 6 | 7 | 14 | 2 | 0 | 0 | 1 | 5 | 2 | 2 | 0 | 1 | 0 | 0 | 0 | 16 | 11 | 0 | 26 | 0 | 0 | Sult1b1 |
| Sult1c1 | 3 | 0 | 0 | 2 | 0 | 0 | 0 | 5 | 12 | 21 | 7 | 0 | 0 | 0 | 3 | 1 | 0 | 0 | 0 | 0 | 1 | 0 | 20 | 16 | 1 | 5 | 0 | 0 | Sult1c1 |
| Sult1d1 | 9 | 1 | 0 | 4 | 0 | 0 | 0 | 16 | 23 | 21 | 14 | 0 | 2 | 1 | 4 | 1 | 0 | 0 | 0 | 0 | 0 | 0 | 40 | 54 | 1 | 35 | 0 | 0 | Sult1d1 |
| Sult1e1 | 143 | 41 | 0 | 139 | 0 | 1 | 1 | 276 | 124 | 177 | 137 | 0 | 61 | 29 | 70 | 82 | 8 | 0 | 0 | 1 | 0 | 2 | 215 | 440 | 0 | 489 | 2 | 0 | Sult1e1 |
| Sult2a7 | 0 | 0 | 0 | 0 | 0 | 1 | 0 | 0 | 0 | 0 | 0 | 0 | 0 | 0 | 1 | 1 | 0 | 3 | 0 | 0 | 0 | 0 | 0 | 0 | 0 | 0 | 0 | 0 | Sult2a7 |
| Sult2a8 | 0 | 0 | 0 | 0 | 0 | 0 | 1 | 0 | 0 | 1 | 0 | 0 | 0 | 0 | 0 | 0 | 0 | 0 | 0 | 3 | 0 | 0 | 0 | 0 | 0 | 0 | 0 | 0 | Sult2a8 |
| Sult2b1 | 75 | 84 | 78 | 80 | 67 | 61 | 75 | 88 | 82 | 117 | 68 | 72 | 59 | 53 | 69 | 57 | 58 | 56 | 56 | 62 | 55 | 89 | 81 | 84 | 76 | 74 | 37 | 53 | Sult2b1 |
| Sult3a1 | 0 | 0 | 0 | 0 | 0 | 0 | 0 | 1 | 0 | 0 | 0 | 0 | 1 | 0 | 0 | 0 | 0 | 0 | 0 | 0 | 0 | 0 | 0 | 0 | 0 | 0 | 0 | 0 | Sult3a1 |
| Sult4a1 | 1825 | 2366 | 2297 | 3101 | 3915 | 2893 | 2997 | 1981 | 2480 | 2889 | 2334 | 1657 | 1505 | 1709 | 2020 | 2126 | 2656 | 3220 | 3597 | 2900 | 3253 | 2916 | 2331 | 2454 | 2826 | 1756 | 1462 | 1269 | Sult4a1 |
| Sult5a1 | 163 | 154 | 84 | 63 | 44 | 89 | 35 | 181 | 315 | 198 | 326 | 101 | 195 | 95 | 129 | 180 | 87 | 134 | 30 | 17 | 13 | 160 | 368 | 136 | 101 | 136 | 170 | 158 | Sult5a1 |
| Sult6b1 | 5 | 7 | 2 | 1 | 11 | 4 | 8 | 7 | 15 | 5 | 12 | 0 | 0 | 3 | 11 | 9 | 2 | 19 | 7 | 4 | 4 | 2 | 14 | 14 | 8 | 10 | 1 | 6 | Sult6b1 |
| Sult6b2 | 23 | 22 | 19 | 33 | 31 | 36 | 38 | 11 | 11 | 8 | 14 | 7 | 10 | 6 | 13 | 21 | 24 | 39 | 39 | 22 | 30 | 20 | 16 | 10 | 13 | 8 | 16 | 9 | Sult6b2 |
| Sumf1 | 916 | 942 | 1241 | 866 | 776 | 865 | 830 | 1142 | 1219 | 1117 | 1251 | 1267 | 1338 | 1315 | 1017 | 1075 | 956 | 887 | 844 | 816 | 739 | 1153 | 1379 | 1260 | 1095 | 1661 | 1350 | 1256 | Sumf1 |
| Sumf2 | 1438 | 1633 | 1458 | 1299 | 1192 | 1322 | 1095 | 1570 | 1870 | 1520 | 1855 | 1565 | 1758 | 1541 | 1483 | 1574 | 1360 | 1315 | 1137 | 1260 | 1137 | 1375 | 1999 | 1609 | 1529 | 1742 | 1720 | 1640 | Sumf2 |
| Sumo1 | 12689 | 9963 | 13049 | 11239 | 9973 | 10653 | 13748 | 11655 | 9800 | 9169 | 9076 | 11354 | 10651 | 12469 | 13710 | 11883 | 11704 | 10517 | 11475 | 14598 | 12261 | 9272 | 9956 | 10625 | 10333 | 13684 | 12121 | 13659 | Sumo1 |
| Sumo2 | 11900 | 11410 | 5823 | 11432 | 10222 | 10495 | 7077 | 13824 | 10518 | 12540 | 11997 | 14984 | 10669 | 17055 | 8834 | 11810 | 12430 | 9530 | 12064 | 15120 | 12238 | 8813 | 11361 | 13594 | 14374 | 10443 | 16636 | 18278 | Sumo2 |
| Sumo3 | 6108 | 4998 | 5882 | 5292 | 4718 | 5289 | 5306 | 8013 | 7182 | 6967 | 6850 | 7827 | 7419 | 8102 | 5919 | 4727 | 5146 | 4602 | 5403 | 5460 | 5547 | 6727 | 7228 | 7644 | 7102 | 9407 | 8320 | 8967 | Sumo3 |
| Sun1 | 10338 | 11625 | 8456 | 9592 | 8922 | 10257 | 9189 | 5887 | 6061 | 5352 | 5822 | 6019 | 5854 | 5651 | 10255 | 10191 | 8867 | 9866 | 9158 | 10058 | 9663 | 5544 | 5763 | 5578 | 6142 | 5111 | 6255 | 5553 | Sun1 |
| Sun2 | 5231 | 6386 | 7359 | 7984 | 7721 | 7101 | 7415 | 4593 | 4993 | 6295 | 5749 | 5963 | 5581 | 5978 | 5124 | 5549 | 7631 | 7949 | 8613 | 5902 | 7507 | 5720 | 4768 | 6474 | 6958 | 5934 | 5262 | 5917 | Sun2 |
| Sun3 | 1 | 2 | 3 | 8 | 2 | 3 | 0 | 9 | 2 | 1 | 4 | 3 | 1 | 5 | 8 | 3 | 5 | 12 | 2 | 3 | 4 | 3 | 7 | 2 | 2 | 3 | 1 | 6 | Sun3 |
| Sun5 | 0 | 0 | 0 | 0 | 0 | 0 | 0 | 0 | 1 | 0 | 0 | 0 | 0 | 0 | 0 | 0 | 0 | 0 | 0 | 0 | 0 | 0 | 0 | 0 | 0 | 0 | 0 | 0 | Sun5 |
| Suox | 2073 | 2357 | 2081 | 1896 | 1601 | 1739 | 1496 | 957 | 950 | 848 | 907 | 778 | 852 | 776 | 2016 | 2302 | 1804 | 1776 | 1625 | 1690 | 1611 | 817 | 1069 | 809 | 770 | 929 | 897 | 877 | Suox |
| Supt16 | 25452 | 22702 | 24690 | 21375 | 18780 | 21755 | 22787 | 23243 | 20372 | 18819 | 18145 | 25867 | 22885 | 25833 | 25358 | 22535 | 23114 | 19992 | 20620 | 24429 | 21249 | 17296 | 19045 | 21199 | 20783 | 25589 | 26935 | 25488 | Supt16 |
| Supt20 | 10304 | 9487 | 9359 | 9294 | 9270 | 9194 | 10073 | 9589 | 9199 | 8790 | 9635 | 9537 | 9200 | 9892 | 10147 | 9703 | 9108 | 9089 | 9308 | 10188 | 9909 | 9155 | 8842 | 9211 | 8679 | 9073 | 9094 | 8795 | Supt20 |
| Supt3 | 728 | 685 | 723 | 582 | 512 | 473 | 510 | 832 | 908 | 849 | 907 | 849 | 963 | 1018 | 593 | 699 | 628 | 512 | 559 | 452 | 600 | 904 | 908 | 764 | 853 | 1056 | 919 | 1025 | Supt3 |
| Supt4a | 2451 | 2806 | 2713 | 2704 | 2401 | 2188 | 2544 | 2282 | 2437 | 2140 | 2500 | 2526 | 2592 | 2611 | 2673 | 2607 | 2560 | 2335 | 2453 | 2880 | 2507 | 2022 | 2383 | 2469 | 2308 | 2793 | 2672 | 2766 | Supt4a |
| Supt5 | 17710 | 20352 | 20115 | 20794 | 18185 | 18151 | 17404 | 11930 | 13282 | 13634 | 12868 | 14685 | 13151 | 13599 | 16742 | 18405 | 19751 | 19430 | 20011 | 16811 | 17889 | 12995 | 12479 | 14566 | 14031 | 14820 | 13959 | 13991 | Supt5 |
| Supt6 | 19474 | 20609 | 17055 | 19115 | 16878 | 17654 | 18105 | 12974 | 13867 | 14803 | 14274 | 14668 | 14274 | 14953 | 18670 | 19613 | 17649 | 18652 | 16858 | 15680 | 18164 | 14509 | 13093 | 14060 | 15381 | 11222 | 14916 | 14089 | Supt6 |
| Supt7l | 1247 | 1307 | 1408 | 1218 | 1355 | 1266 | 1309 | 2038 | 2078 | 2097 | 2046 | 2629 | 2335 | 2424 | 1313 | 1205 | 1409 | 1136 | 1384 | 1317 | 1261 | 2054 | 2084 | 2435 | 2219 | 2697 | 2480 | 2622 | Supt7l |
| Supv3l1 | 1955 | 1886 | 1905 | 1557 | 1489 | 1775 | 1789 | 2127 | 2122 | 1979 | 1935 | 2401 | 2240 | 2392 | 1789 | 1651 | 1743 | 1533 | 1683 | 1693 | 1764 | 2057 | 2030 | 2048 | 2204 | 2320 | 2219 | 2397 | Supv3l1 |
| Surf1 | 1301 | 1511 | 1437 | 1308 | 1307 | 1341 | 1402 | 1419 | 1523 | 1459 | 1486 | 1509 | 1529 | 1510 | 1366 | 1301 | 1304 | 1356 | 1389 | 1521 | 1377 | 1457 | 1542 | 1532 | 1512 | 1642 | 1557 | 1477 | Surf1 |
| Surf2 | 2194 | 2824 | 2602 | 2852 | 2408 | 2498 | 2256 | 2566 | 2190 | 2452 | 2210 | 2518 | 2439 | 2544 | 2487 | 2414 | 2593 | 2632 | 2524 | 2775 | 2376 | 2453 | 2084 | 2506 | 2483 | 2637 | 2625 | 2356 | Surf2 |
| Surf4 | 12537 | 12487 | 12537 | 9765 | 8715 | 9611 | 8410 | 10538 | 11848 | 10203 | 12487 | 10622 | 11616 | 9981 | 11339 | 12699 | 11276 | 9598 | 9470 | 8730 | 8557 | 10860 | 13041 | 10783 | 9865 | 12747 | 11391 | 11190 | Surf4 |
| Surf6 | 2374 | 2870 | 2692 | 2819 | 2549 | 2594 | 2277 | 2503 | 2791 | 2829 | 2805 | 2815 | 2833 | 2937 | 2236 | 2502 | 2904 | 2737 | 2583 | 2548 | 2425 | 2506 | 2603 | 2907 | 2920 | 3106 | 3217 | 2962 | Surf6 |
| Susd1 | 239 | 187 | 207 | 151 | 141 | 174 | 103 | 249 | 293 | 256 | 285 | 222 | 280 | 220 | 227 | 225 | 108 | 177 | 119 | 143 | 110 | 229 | 322 | 239 | 209 | 284 | 224 | 195 | Susd1 |
| Susd2 | 258 | 447 | 335 | 404 | 342 | 305 | 304 | 216 | 229 | 273 | 321 | 260 | 243 | 235 | 311 | 396 | 387 | 491 | 364 | 270 | 349 | 296 | 311 | 252 | 249 | 300 | 238 | 256 | Susd2 |
| Susd3 | 116 | 96 | 68 | 90 | 62 | 66 | 30 | 20 | 20 | 18 | 23 | 26 | 22 | 15 | 78 | 143 | 61 | 76 | 48 | 51 | 44 | 12 | 27 | 18 | 34 | 37 | 16 | 18 | Susd3 |
| Susd4 | 1700 | 1759 | 2130 | 2443 | 2964 | 2185 | 2302 | 2052 | 2178 | 2532 | 2073 | 2033 | 2108 | 1881 | 1873 | 1861 | 2377 | 2363 | 2713 | 2227 | 2490 | 3083 | 2084 | 2147 | 2496 | 1734 | 1888 | 1812 | Susd4 |
| Susd5 | 59 | 59 | 17 | 82 | 44 | 187 | 7 | 396 | 817 | 1098 | 1891 | 71 | 141 | 67 | 58 | 76 | 30 | 114 | 83 | 6 | 6 | 819 | 548 | 794 | 388 | 197 | 72 | 56 | Susd5 |
| Susd6 | 3658 | 3548 | 4027 | 4090 | 3671 | 3483 | 3818 | 2584 | 2604 | 2713 | 2936 | 2954 | 2869 | 2844 | 3563 | 3712 | 3986 | 4002 | 3811 | 3043 | 3542 | 3038 | 2774 | 2739 | 2994 | 2791 | 2895 | 3113 | Susd6 |
| Suv39h1 | 5371 | 5268 | 5157 | 4388 | 4222 | 4575 | 4721 | 5157 | 5229 | 4918 | 4833 | 6446 | 5708 | 6171 | 4941 | 4637 | 4785 | 4208 | 4729 | 4613 | 4499 | 4659 | 4801 | 5441 | 5119 | 7380 | 6370 | 6547 | Suv39h1 |
| Suv39h2 | 3042 | 2472 | 2916 | 2597 | 2467 | 2691 | 3261 | 2789 | 2307 | 2161 | 2028 | 3642 | 3020 | 3522 | 2996 | 2504 | 2799 | 2385 | 2634 | 2818 | 2740 | 1791 | 1963 | 2569 | 2545 | 3828 | 3562 | 3716 | Suv39h2 |
| Suz12 | 14293 | 10979 | 14965 | 15825 | 14398 | 14787 | 18518 | 11203 | 9716 | 11186 | 10005 | 14442 | 12217 | 14945 | 13694 | 11439 | 15054 | 15403 | 15751 | 14357 | 17032 | 10657 | 9506 | 13874 | 12395 | 15257 | 13705 | 14878 | Suz12 |
| Sv2a | 5171 | 5788 | 4626 | 7708 | 10900 | 8848 | 8422 | 7914 | 7797 | 8942 | 7009 | 6049 | 5863 | 5798 | 5782 | 5189 | 6437 | 8863 | 9772 | 8708 | 9422 | 10305 | 6955 | 7228 | 9683 | 4270 | 5438 | 4956 | Sv2a |
| Sv2b | 339 | 312 | 251 | 493 | 900 | 731 | 983 | 833 | 999 | 840 | 956 | 641 | 467 | 681 | 410 | 310 | 319 | 560 | 775 | 806 | 912 | 871 | 1170 | 768 | 960 | 523 | 569 | 528 | Sv2b |
| Sv2c | 367 | 367 | 465 | 520 | 1612 | 760 | 592 | 945 | 1130 | 1460 | 1651 | 298 | 908 | 286 | 580 | 419 | 430 | 714 | 723 | 518 | 658 | 1751 | 1090 | 409 | 616 | 208 | 285 | 271 | Sv2c |
| Sval1 | 0 | 0 | 0 | 0 | 0 | 0 | 0 | 1 | 0 | 0 | 0 | 0 | 0 | 0 | 0 | 0 | 0 | 0 | 0 | 0 | 0 | 0 | 1 | 0 | 0 | 0 | 0 | 0 | Sval1 |
| Svbp | 2239 | 2794 | 2670 | 2449 | 2636 | 2356 | 2976 | 2303 | 2539 | 2806 | 2579 | 2348 | 2229 | 2550 | 2395 | 2227 | 2858 | 2467 | 2939 | 2935 | 2777 | 2141 | 2594 | 2541 | 2430 | 2330 | 2352 | 2535 | Svbp |
| Svep1 | 2019 | 2592 | 1415 | 1237 | 488 | 1671 | 241 | 1988 | 3131 | 2725 | 3694 | 2265 | 3401 | 2171 | 2250 | 2987 | 1156 | 1922 | 394 | 445 | 300 | 1699 | 3567 | 2236 | 2178 | 1369 | 2771 | 1931 | Svep1 |
| Svil | 5740 | 5900 | 5193 | 4264 | 4598 | 5166 | 4050 | 4802 | 6788 | 5110 | 8418 | 5103 | 6208 | 5200 | 5372 | 6694 | 4543 | 5049 | 4026 | 4085 | 4163 | 4502 | 8630 | 5241 | 4521 | 3950 | 5201 | 4688 | Svil |
| Svip | 3249 | 2934 | 3024 | 3421 | 3626 | 3683 | 4408 | 2049 | 1798 | 1672 | 1521 | 1909 | 1826 | 1941 | 4061 | 3174 | 3170 | 3372 | 3489 | 4480 | 4158 | 1813 | 1827 | 1821 | 1897 | 1755 | 1735 | 1952 | Svip |
| Svop | 1209 | 1670 | 1475 | 2489 | 3784 | 2954 | 2807 | 2864 | 3157 | 4101 | 3109 | 3501 | 2929 | 3164 | 1531 | 1208 | 2622 | 2765 | 3609 | 2662 | 3229 | 4390 | 2786 | 3680 | 4489 | 3004 | 2974 | 2995 | Svop |
| Svopl | 107 | 101 | 52 | 157 | 90 | 48 | 99 | 156 | 158 | 189 | 109 | 62 | 120 | 31 | 102 | 131 | 64 | 117 | 77 | 76 | 79 | 55 | 190 | 199 | 36 | 136 | 37 | 28 | Svopl |
| Svs3a | 0 | 0 | 0 | 0 | 0 | 0 | 1 | 0 | 0 | 0 | 0 | 0 | 0 | 0 | 0 | 0 | 0 | 0 | 0 | 0 | 0 | 0 | 0 | 0 | 0 | 0 | 0 | 0 | Svs3a |
| Svs4 | 0 | 0 | 0 | 0 | 0 | 0 | 0 | 0 | 0 | 0 | 0 | 0 | 0 | 0 | 0 | 0 | 0 | 0 | 0 | 0 | 0 | 0 | 1 | 0 | 0 | 0 | 0 | 0 | Svs4 |
| Swap70 | 3417 | 3070 | 3636 | 3093 | 2621 | 3068 | 3091 | 1918 | 2422 | 2273 | 2687 | 2187 | 2457 | 2396 | 3086 | 3490 | 3193 | 3109 | 2904 | 2624 | 2806 | 1927 | 2366 | 2401 | 2017 | 2619 | 2353 | 2385 | Swap70 |
| Swi5 | 13411 | 14698 | 15323 | 13953 | 12335 | 13355 | 13264 | 11269 | 11598 | 10532 | 11010 | 11790 | 12238 | 13243 | 13640 | 13661 | 14241 | 12952 | 13612 | 15893 | 13351 | 9247 | 11817 | 11762 | 11523 | 13941 | 12783 | 13582 | Swi5 |
| Swsap1 | 669 | 713 | 686 | 681 | 625 | 589 | 711 | 1187 | 1056 | 1061 | 913 | 1166 | 1074 | 1330 | 690 | 543 | 655 | 530 | 711 | 663 | 665 | 1073 | 1070 | 1243 | 1141 | 1386 | 1284 | 1413 | Swsap1 |
| Swt1 | 3708 | 3392 | 3365 | 3216 | 2552 | 3184 | 3179 | 2317 | 1895 | 1681 | 1663 | 2010 | 1868 | 1754 | 3981 | 3840 | 3069 | 3006 | 2876 | 3565 | 3011 | 1703 | 1916 | 1737 | 1714 | 1771 | 1950 | 2079 | Swt1 |
| Syap1 | 6304 | 4802 | 5177 | 4716 | 4224 | 4414 | 4994 | 3646 | 3305 | 2840 | 3296 | 2899 | 3163 | 2949 | 6533 | 6168 | 4626 | 4802 | 3821 | 4951 | 4530 | 3452 | 3911 | 3170 | 2881 | 3346 | 3115 | 3171 | Syap1 |
| Sybu | 1089 | 1345 | 1206 | 1673 | 2443 | 2280 | 2291 | 2887 | 2338 | 2532 | 1968 | 2540 | 1974 | 2594 | 1454 | 942 | 1503 | 1866 | 2477 | 2124 | 2504 | 3678 | 2179 | 2598 | 3201 | 1960 | 2538 | 2448 | Sybu |
| Syce1 | 14 | 11 | 12 | 16 | 11 | 19 | 9 | 5 | 7 | 15 | 30 | 12 | 16 | 3 | 9 | 9 | 20 | 13 | 6 | 10 | 4 | 8 | 6 | 7 | 7 | 3 | 5 | 4 | Syce1 |
| Syce1l | 33 | 59 | 23 | 80 | 55 | 44 | 83 | 21 | 19 | 10 | 12 | 16 | 17 | 12 | 45 | 38 | 56 | 98 | 75 | 54 | 76 | 25 | 15 | 34 | 15 | 14 | 24 | 14 | Syce1l |
| Syce2 | 1683 | 1559 | 1249 | 1481 | 1263 | 1338 | 1767 | 1021 | 961 | 949 | 790 | 932 | 851 | 1077 | 1345 | 1291 | 1253 | 1338 | 1528 | 1792 | 1604 | 632 | 861 | 1159 | 989 | 1274 | 924 | 1097 | Syce2 |
| Syce3 | 0 | 0 | 0 | 0 | 0 | 0 | 0 | 0 | 0 | 0 | 0 | 0 | 1 | 1 | 0 | 0 | 0 | 2 | 0 | 0 | 0 | 0 | 0 | 0 | 0 | 0 | 0 | 0 | Syce3 |
| Sycn | 0 | 0 | 0 | 0 | 0 | 0 | 0 | 0 | 0 | 0 | 0 | 0 | 0 | 0 | 0 | 0 | 0 | 0 | 0 | 0 | 0 | 0 | 2 | 0 | 0 | 0 | 0 | 0 | Sycn |
| Sycp1 | 0 | 0 | 0 | 0 | 2 | 0 | 0 | 0 | 2 | 3 | 0 | 0 | 0 | 1 | 0 | 0 | 0 | 0 | 1 | 0 | 1 | 0 | 1 | 0 | 0 | 0 | 0 | 0 | Sycp1 |
| Sycp2 | 59 | 41 | 34 | 65 | 92 | 86 | 77 | 81 | 76 | 76 | 89 | 26 | 54 | 28 | 76 | 62 | 45 | 79 | 72 | 79 | 100 | 84 | 82 | 48 | 84 | 40 | 41 | 19 | Sycp2 |
| Sycp2l | 7 | 4 | 13 | 4 | 3 | 4 | 4 | 9 | 5 | 10 | 16 | 8 | 1 | 9 | 9 | 10 | 1 | 8 | 9 | 4 | 4 | 12 | 6 | 7 | 5 | 19 | 13 | 14 | Sycp2l |
| Sycp3 | 22 | 28 | 40 | 22 | 37 | 29 | 28 | 21 | 22 | 19 | 34 | 18 | 31 | 9 | 22 | 19 | 32 | 24 | 12 | 18 | 27 | 28 | 22 | 12 | 24 | 8 | 17 | 13 | Sycp3 |

| | | | | | | | | | | | | | | | | | | | | | | | | | | | | | |
|---|---|---|---|---|---|---|---|---|---|---|---|---|---|---|---|---|---|---|---|---|---|---|---|---|---|---|---|---|---|
| Syde1 | 1638 | 1690 | 1765 | 1112 | 964 | 1365 | 977 | 1730 | 2198 | 1852 | 2269 | 1927 | 2239 | 1742 | 1444 | 1643 | 1352 | 1116 | 1079 | 1003 | 905 | 1512 | 2139 | 1904 | 1517 | 2423 | 2019 | 1798 | Syde1 |
| Syde2 | 1283 | 1105 | 1096 | 1166 | 1145 | 1160 | 1048 | 670 | 778 | 636 | 744 | 683 | 684 | 666 | 1092 | 1293 | 1117 | 1148 | 1113 | 929 | 977 | 587 | 680 | 664 | 545 | 624 | 589 | 625 | Syde2 |
| Syf2 | 2957 | 3240 | 3168 | 2853 | 2598 | 2513 | 3007 | 2988 | 2814 | 2621 | 2649 | 2657 | 2530 | 3476 | 3247 | 2897 | 3038 | 2914 | 2692 | 3294 | 3155 | 2575 | 2798 | 3023 | 2900 | 3333 | 2790 | 3252 | Syf2 |
| Syk | 539 | 644 | 584 | 497 | 466 | 535 | 313 | 519 | 714 | 629 | 716 | 622 | 848 | 498 | 547 | 843 | 619 | 467 | 452 | 476 | 318 | 623 | 827 | 538 | 652 | 659 | 600 | 567 | Syk |
| Sympk | 12923 | 15938 | 18021 | 16407 | 14369 | 13828 | 13058 | 9550 | 11281 | 13230 | 11873 | 15010 | 12303 | 12825 | 11755 | 14590 | 17206 | 16002 | 16747 | 11037 | 14226 | 12109 | 10978 | 13866 | 12965 | 15624 | 12280 | 13091 | Sympk |
| Syn1 | 2121 | 2685 | 3299 | 4753 | 6119 | 4269 | 4400 | 2489 | 2763 | 3905 | 3145 | 3040 | 2079 | 2508 | 2461 | 2188 | 4298 | 5229 | 5644 | 3301 | 5210 | 5569 | 2452 | 3224 | 3928 | 2503 | 1833 | 2191 | Syn1 |
| Syn2 | 1186 | 1495 | 1074 | 2319 | 3881 | 2673 | 2739 | 1461 | 1773 | 1806 | 1672 | 937 | 904 | 901 | 1483 | 1399 | 2045 | 2795 | 2905 | 2485 | 3174 | 2336 | 1457 | 1289 | 2056 | 652 | 687 | 686 | Syn2 |
| Syn3 | 346 | 504 | 166 | 625 | 1027 | 841 | 781 | 747 | 926 | 827 | 776 | 695 | 650 | 619 | 458 | 422 | 284 | 603 | 667 | 729 | 840 | 984 | 855 | 532 | 947 | 83 | 712 | 431 | Syn3 |
| Syna | 2 | 0 | 0 | 0 | 2 | 1 | 0 | 1 | 1 | 0 | 1 | 1 | 0 | 0 | 5 | 1 | 0 | 0 | 2 | 2 | 0 | 0 | 3 | 2 | 0 | 6 | 0 | 0 | Syna |
| Sync | 104 | 144 | 130 | 111 | 121 | 114 | 86 | 81 | 104 | 67 | 348 | 59 | 117 | 48 | 88 | 134 | 94 | 138 | 82 | 118 | 83 | 112 | 236 | 51 | 49 | 65 | 38 | 49 | Sync |
| Syncrip | 20556 | 13761 | 17193 | 16515 | 16436 | 18550 | 20749 | 29775 | 25092 | 22901 | 23699 | 27094 | 25203 | 29715 | 19852 | 15949 | 15733 | 15738 | 17478 | 19215 | 19349 | 24371 | 24203 | 25279 | 26743 | 28406 | 28416 | 29374 | Syncrip |
| Syndig1 | 279 | 327 | 315 | 253 | 319 | 330 | 244 | 355 | 410 | 445 | 479 | 386 | 425 | 336 | 309 | 340 | 321 | 296 | 287 | 242 | 216 | 394 | 494 | 382 | 355 | 333 | 365 | 439 | Syndig1 |
| Syndig1l | 25 | 35 | 73 | 71 | 230 | 102 | 96 | 169 | 156 | 208 | 160 | 83 | 125 | 85 | 62 | 51 | 66 | 98 | 120 | 106 | 114 | 267 | 189 | 144 | 215 | 59 | 62 | 66 | Syndig1l |
| Syne1 | 2262 | 1854 | 825 | 1161 | 2157 | 1996 | 1409 | 3488 | 4372 | 2728 | 3439 | 1679 | 2452 | 1614 | 1913 | 2523 | 907 | 1451 | 1279 | 2150 | 1732 | 2542 | 3981 | 1494 | 2460 | 830 | 2091 | 1463 | Syne1 |
| Syne2 | 19904 | 19963 | 9509 | 12832 | 14850 | 18151 | 17585 | 20395 | 22645 | 15029 | 14808 | 18595 | 15631 | 19129 | 18592 | 18801 | 11550 | 13372 | 15256 | 22296 | 19443 | 10709 | 17642 | 14278 | 18134 | 11706 | 21104 | 15829 | Syne2 |
| Syne3 | 602 | 636 | 316 | 594 | 428 | 496 | 379 | 184 | 203 | 179 | 405 | 88 | 104 | 84 | 513 | 652 | 379 | 533 | 411 | 372 | 381 | 117 | 311 | 167 | 116 | 64 | 71 | 68 | Syne3 |
| Syne4 | 1149 | 1280 | 1071 | 1295 | 1049 | 859 | 926 | 644 | 798 | 847 | 1082 | 530 | 706 | 568 | 959 | 1393 | 1136 | 1379 | 938 | 841 | 885 | 683 | 796 | 738 | 541 | 869 | 526 | 478 | Syne4 |
| Syngap1 | 7656 | 9519 | 11648 | 13814 | 14138 | 11558 | 12695 | 8234 | 9676 | 12659 | 11378 | 12004 | 10680 | 11673 | 7035 | 7992 | 13924 | 13569 | 15084 | 10255 | 13522 | 13525 | 8834 | 13782 | 14395 | 10985 | 10443 | 10964 | Syngap1 |
| Syngr1 | 2076 | 2631 | 2528 | 4035 | 5420 | 4122 | 4161 | 1441 | 1802 | 2308 | 1711 | 1287 | 1112 | 1226 | 2238 | 2174 | 3490 | 4768 | 5088 | 3869 | 4670 | 2293 | 1674 | 1863 | 2449 | 1014 | 1082 | 1056 | Syngr1 |
| Syngr2 | 1405 | 1695 | 1376 | 1083 | 835 | 1000 | 875 | 459 | 570 | 458 | 554 | 481 | 589 | 483 | 1225 | 1554 | 1128 | 936 | 960 | 1018 | 897 | 429 | 724 | 449 | 444 | 549 | 474 | 564 | Syngr2 |
| Syngr3 | 925 | 1183 | 1264 | 1411 | 1669 | 1311 | 1436 | 1185 | 1267 | 1562 | 1362 | 1023 | 780 | 894 | 914 | 955 | 1348 | 1293 | 1633 | 1330 | 1575 | 2017 | 1339 | 1379 | 1626 | 908 | 837 | 866 | Syngr3 |
| Syngr4 | 16 | 15 | 9 | 14 | 16 | 7 | 16 | 12 | 10 | 9 | 8 | 21 | 22 | 17 | 16 | 15 | 16 | 10 | 8 | 8 | 11 | 13 | 22 | 10 | 12 | 11 | 14 | 19 | Syngr4 |
| Synj1 | 9004 | 8811 | 7875 | 12118 | 14037 | 13604 | 14148 | 8054 | 7700 | 8733 | 7616 | 7481 | 7235 | 7685 | 10108 | 8463 | 10349 | 12810 | 12295 | 12036 | 14332 | 9787 | 6942 | 7124 | 9804 | 4359 | 7140 | 7136 | Synj1 |
| Synj2 | 320 | 326 | 300 | 180 | 191 | 200 | 111 | 388 | 426 | 329 | 638 | 294 | 545 | 329 | 314 | 471 | 203 | 175 | 126 | 101 | 107 | 366 | 562 | 361 | 293 | 365 | 373 | 290 | Synj2 |
| Synj2bp | 5046 | 3820 | 4583 | 4431 | 3926 | 4033 | 4464 | 3714 | 2856 | 2787 | 3106 | 3107 | 2924 | 3303 | 4845 | 4452 | 4493 | 4307 | 4011 | 4323 | 4521 | 3145 | 2991 | 2831 | 3217 | 2680 | 3176 | 3513 | Synj2bp |
| Synm | 1016 | 1144 | 795 | 1321 | 2386 | 1253 | 1288 | 1238 | 1396 | 1193 | 1622 | 558 | 1428 | 509 | 1473 | 1596 | 1067 | 1837 | 1180 | 1156 | 1298 | 1900 | 1074 | 596 | 1407 | 366 | 479 | 436 | Synm |
| Synpo | 2775 | 3026 | 2929 | 1851 | 1027 | 1762 | 622 | 2986 | 4192 | 4497 | 5905 | 2799 | 5325 | 2513 | 2494 | 4421 | 2188 | 1979 | 934 | 663 | 677 | 3706 | 3861 | 4001 | 2335 | 4884 | 3373 | 3297 | Synpo |
| Synpo2 | 2768 | 2002 | 2096 | 1216 | 936 | 1153 | 768 | 883 | 1144 | 860 | 1671 | 948 | 1507 | 897 | 2302 | 3288 | 1388 | 1445 | 733 | 855 | 717 | 740 | 1474 | 891 | 620 | 1034 | 1340 | 951 | Synpo2 |
| Synpo2l | 45 | 16 | 6 | 18 | 21 | 13 | 7 | 95 | 247 | 67 | 1720 | 45 | 81 | 36 | 20 | 40 | 25 | 21 | 9 | 19 | 12 | 221 | 847 | 39 | 53 | 39 | 43 | 15 | Synpo2l |
| Synpr | 484 | 394 | 420 | 563 | 977 | 800 | 822 | 1194 | 1018 | 921 | 866 | 659 | 857 | 788 | 841 | 581 | 458 | 700 | 695 | 941 | 825 | 1295 | 1236 | 598 | 868 | 706 | 762 | 750 | Synpr |
| Synrg | 5072 | 5193 | 4795 | 6539 | 7051 | 6589 | 6885 | 7669 | 7272 | 8450 | 7649 | 8895 | 7665 | 8463 | 5089 | 4594 | 5950 | 6373 | 7024 | 5674 | 7209 | 8961 | 6866 | 8051 | 9543 | 5964 | 8287 | 8018 | Synrg |
| Syp | 4986 | 6304 | 6636 | 9621 | 11072 | 8741 | 9164 | 6920 | 6590 | 8424 | 7068 | 5977 | 5404 | 6328 | 6078 | 4656 | 8492 | 9741 | 10328 | 8533 | 10413 | 8651 | 6066 | 7156 | 9270 | 5326 | 5345 | 5647 | Syp |
| Sypl | 6003 | 4418 | 4856 | 4706 | 4062 | 4467 | 4811 | 4354 | 4057 | 3274 | 4078 | 3748 | 4241 | 3786 | 5501 | 5751 | 4107 | 4576 | 3981 | 4692 | 4177 | 3735 | 4431 | 3622 | 3748 | 4189 | 3769 | 4085 | Sypl |
| Sypl2 | 280 | 257 | 200 | 242 | 357 | 264 | 354 | 167 | 236 | 254 | 336 | 330 | 204 | 375 | 200 | 199 | 341 | 337 | 427 | 314 | 424 | 185 | 247 | 363 | 374 | 234 | 274 | 268 | Sypl2 |
| Sys1 | 978 | 999 | 919 | 861 | 901 | 833 | 952 | 1125 | 1112 | 1180 | 1109 | 1058 | 1072 | 1308 | 988 | 826 | 902 | 800 | 881 | 911 | 1028 | 1003 | 1039 | 1057 | 1177 | 987 | 1069 | 1256 | Sys1 |
| Syt1 | 4827 | 5393 | 3937 | 7693 | 10858 | 8599 | 10387 | 8707 | 8017 | 8236 | 7191 | 6306 | 5162 | 6383 | 6344 | 4187 | 6318 | 8444 | 9173 | 9572 | 10408 | 10926 | 7419 | 7111 | 9748 | 4492 | 5783 | 5459 | Syt1 |
| Syt10 | 18 | 35 | 7 | 22 | 48 | 25 | 31 | 24 | 22 | 27 | 34 | 30 | 10 | 6 | 30 | 29 | 16 | 31 | 25 | 43 | 32 | 22 | 27 | 21 | 13 | 5 | 21 | 8 | Syt10 |
| Syt11 | 13275 | 15558 | 16154 | 16484 | 17766 | 16415 | 17824 | 20019 | 19816 | 21250 | 17834 | 25456 | 20593 | 24515 | 15202 | 11992 | 16607 | 15964 | 18798 | 17348 | 17956 | 21383 | 17962 | 22323 | 24421 | 23308 | 24889 | 24744 | Syt11 |
| Syt12 | 108 | 160 | 158 | 141 | 165 | 111 | 101 | 98 | 136 | 204 | 182 | 152 | 174 | 166 | 128 | 186 | 126 | 144 | 139 | 140 | 122 | 147 | 168 | 188 | 166 | 255 | 158 | 163 | Syt12 |
| Syt13 | 2122 | 2743 | 2831 | 4087 | 5178 | 4486 | 4986 | 2845 | 2128 | 4350 | 3683 | 3180 | 2699 | 2896 | 2782 | 2858 | 3607 | 4667 | 5824 | 3771 | 5967 | 6119 | 3194 | 4339 | 5331 | 2255 | 2780 | 2682 | Syt13 |
| Syt14 | 5905 | 5402 | 3716 | 5154 | 5849 | 6142 | 6129 | 4929 | 4124 | 3604 | 3362 | 4119 | 3814 | 4338 | 5916 | 5448 | 4568 | 5252 | 5347 | 6041 | 6114 | 4274 | 3646 | 3299 | 4160 | 2765 | 4373 | 3836 | Syt14 |
| Syt15 | 17 | 15 | 10 | 24 | 29 | 31 | 16 | 40 | 53 | 62 | 75 | 55 | 62 | 63 | 15 | 5 | 21 | 21 | 28 | 16 | 20 | 72 | 69 | 67 | 45 | 85 | 56 | 56 | Syt15 |
| Syt16 | 856 | 783 | 521 | 1204 | 2287 | 1781 | 2064 | 2626 | 2532 | 2563 | 2061 | 1582 | 1380 | 1632 | 1027 | 717 | 778 | 1654 | 1592 | 1914 | 2063 | 3180 | 2279 | 1915 | 2846 | 1235 | 1409 | 1196 | Syt16 |
| Syt17 | 288 | 272 | 396 | 289 | 403 | 262 | 270 | 242 | 277 | 261 | 293 | 205 | 218 | 263 | 264 | 246 | 322 | 277 | 316 | 303 | 258 | 322 | 269 | 223 | 253 | 212 | 225 | 203 | Syt17 |
| Syt2 | 375 | 423 | 606 | 370 | 961 | 455 | 316 | 488 | 510 | 566 | 776 | 463 | 684 | 426 | 517 | 583 | 538 | 534 | 463 | 363 | 390 | 737 | 419 | 345 | 530 | 299 | 479 | 522 | Syt2 |
| Syt3 | 261 | 394 | 471 | 499 | 882 | 631 | 563 | 390 | 569 | 655 | 569 | 375 | 394 | 344 | 302 | 317 | 416 | 486 | 803 | 583 | 650 | 682 | 463 | 532 | 628 | 323 | 315 | 314 | Syt3 |
| Syt4 | 4845 | 5255 | 4718 | 8576 | 14753 | 11362 | 12367 | 13606 | 11471 | 11094 | 10182 | 6903 | 7912 | 7202 | 8433 | 5297 | 6843 | 9927 | 11386 | 14105 | 12641 | 14365 | 11631 | 8661 | 12177 | 3913 | 6974 | 6560 | Syt4 |
| Syt5 | 547 | 709 | 506 | 878 | 1248 | 1131 | 995 | 1567 | 1884 | 1894 | 1566 | 1474 | 1216 | 1468 | 559 | 447 | 691 | 866 | 1076 | 985 | 1104 | 2237 | 1614 | 1708 | 2020 | 1151 | 1231 | 1262 | Syt5 |
| Syt6 | 568 | 556 | 487 | 777 | 1173 | 1077 | 1116 | 1054 | 969 | 1369 | 1037 | 1083 | 1032 | 1111 | 566 | 475 | 729 | 868 | 1168 | 764 | 1160 | 1547 | 830 | 1146 | 1551 | 635 | 1008 | 1021 | Syt6 |
| Syt7 | 3713 | 4736 | 6539 | 7361 | 9061 | 7356 | 7108 | 4483 | 4392 | 6068 | 5258 | 5686 | 5332 | 5451 | 4352 | 4210 | 7181 | 7647 | 8836 | 6684 | 7783 | 7641 | 4418 | 6377 | 7276 | 4455 | 5149 | 5470 | Syt7 |
| Syt8 | 7 | 3 | 3 | 0 | 0 | 9 | 0 | 21 | 37 | 12 | 14 | 13 | 12 | 20 | 1 | 5 | 0 | 0 | 2 | 0 | 3 | 7 | 21 | 20 | 2 | 8 | 9 | 12 | Syt8 |
| Syt9 | 822 | 847 | 617 | 718 | 1151 | 867 | 896 | 945 | 981 | 846 | 848 | 816 | 870 | 789 | 854 | 817 | 932 | 846 | 904 | 856 | 851 | 1151 | 892 | 752 | 935 | 737 | 736 | 694 | Syt9 |
| Sytl1 | 95 | 225 | 175 | 196 | 149 | 155 | 92 | 117 | 158 | 71 | 135 | 102 | 141 | 100 | 96 | 179 | 212 | 243 | 162 | 132 | 143 | 84 | 176 | 145 | 93 | 89 | 132 | 109 | Sytl1 |
| Sytl2 | 472 | 442 | 436 | 284 | 466 | 398 | 320 | 363 | 675 | 430 | 1289 | 243 | 369 | 211 | 556 | 614 | 342 | 341 | 332 | 462 | 317 | 561 | 896 | 263 | 329 | 190 | 261 | 221 | Sytl2 |
| Sytl3 | 13 | 6 | 3 | 3 | 20 | 4 | 6 | 13 | 21 | 21 | 18 | 22 | 15 | 13 | 13 | 11 | 5 | 2 | 4 | 4 | 4 | 10 | 28 | 19 | 10 | 16 | 2 | 6 | Sytl3 |
| Sytl4 | 383 | 299 | 377 | 264 | 178 | 244 | 151 | 302 | 294 | 278 | 433 | 206 | 316 | 218 | 323 | 413 | 243 | 313 | 179 | 242 | 183 | 229 | 627 | 256 | 181 | 199 | 253 | 202 | Sytl4 |
| Sytl5 | 91 | 81 | 8 | 40 | 32 | 107 | 83 | 134 | 152 | 73 | 84 | 105 | 53 | 143 | 99 | 51 | 28 | 45 | 46 | 105 | 83 | 55 | 69 | 60 | 73 | 50 | 113 | 71 | Sytl5 |
| Syvn1 | 3056 | 4226 | 4668 | 4523 | 4012 | 3350 | 3303 | 2560 | 2965 | 3528 | 3590 | 3458 | 3322 | 2981 | 2840 | 3815 | 4394 | 4500 | 4224 | 2865 | 3413 | 3768 | 2964 | 3684 | 3345 | 3694 | 2991 | 3076 | Syvn1 |
| Szrd1 | 5459 | 6411 | 8349 | 7195 | 5718 | 5869 | 5141 | 5477 | 6681 | 7103 | 7552 | 7729 | 7979 | 7377 | 5206 | 6066 | 7821 | 6729 | 6597 | 5193 | 5548 | 6183 | 6968 | 8349 | 7248 | 9744 | 8100 | 8152 | Szrd1 |
| Szt2 | 5244 | 6326 | 4513 | 7120 | 7478 | 6723 | 6386 | 6125 | 7369 | 8143 | 7510 | 8894 | 7737 | 7727 | 4751 | 5812 | 6058 | 7058 | 7493 | 5353 | 6744 | 7919 | 6279 | 7311 | 8486 | 4072 | 7975 | 7001 | Szt2 |
| T | 106 | 113 | 138 | 144 | 92 | 70 | 83 | 45 | 50 | 35 | 64 | 60 | 64 | 71 | 79 | 152 | 114 | 164 | 97 | 72 | 81 | 53 | 39 | 67 | 48 | 67 | 64 | 77 | T |
| T2 | 197 | 195 | 181 | 220 | 113 | 165 | 152 | 45 | 33 | 82 | 58 | 58 | 61 | 39 | 160 | 280 | 222 | 223 | 164 | 127 | 133 | 57 | 47 | 46 | 56 | 44 | 29 | 57 | T2 |
| Taar1 | 0 | 0 | 0 | 0 | 0 | 0 | 0 | 0 | 0 | 0 | 0 | 0 | 0 | 1 | 0 | 0 | 0 | 0 | 0 | 0 | 0 | 0 | 0 | 0 | 0 | 0 | 0 | 1 | Taar1 |
| Taar2 | 0 | 1 | 0 | 0 | 0 | 0 | 0 | 2 | 0 | 4 | 0 | 0 | 0 | 0 | 0 | 0 | 0 | 0 | 0 | 0 | 0 | 0 | 1 | 0 | 0 | 3 | 0 | 0 | Taar2 |
| Taar3 | 0 | 0 | 0 | 0 | 0 | 1 | 0 | 0 | 0 | 0 | 3 | 0 | 0 | 1 | 0 | 0 | 0 | 0 | 0 | 0 | 0 | 0 | 1 | 0 | 0 | 0 | 0 | 0 | Taar3 |
| Taar4 | 0 | 0 | 0 | 0 | 0 | 0 | 0 | 0 | 0 | 0 | 0 | 0 | 0 | 0 | 0 | 0 | 0 | 0 | 0 | 0 | 0 | 0 | 2 | 0 | 0 | 0 | 0 | 0 | Taar4 |
| Taar5 | 1 | 1 | 0 | 2 | 1 | 10 | 0 | 4 | 21 | 5 | 9 | 3 | 6 | 0 | 0 | 1 | 2 | 3 | 4 | 2 | 1 | 12 | 4 | 2 | 2 | 1 | 0 | 1 | Taar5 |
| Taar6 | 0 | 0 | 0 | 0 | 0 | 0 | 0 | 0 | 1 | 0 | 0 | 0 | 0 | 0 | 0 | 0 | 0 | 0 | 0 | 0 | 0 | 0 | 1 | 0 | 0 | 0 | 0 | 0 | Taar6 |
| Taar7a | 0 | 0 | 0 | 0 | 0 | 0 | 0 | 0 | 0 | 0 | 0 | 0 | 0 | 0 | 0 | 0 | 0 | 0 | 0 | 0 | 0 | 0 | 2 | 5 | 0 | 0 | 0 | 0 | Taar7a |
| Taar7b | 0 | 0 | 0 | 0 | 0 | 0 | 0 | 0 | 0 | 0 | 0 | 0 | 0 | 0 | 0 | 0 | 0 | 0 | 0 | 0 | 0 | 0 | 1 | 0 | 0 | 0 | 0 | 0 | Taar7b |
| Taar7d | 0 | 0 | 0 | 0 | 0 | 0 | 0 | 9 | 2 | 0 | 0 | 0 | 0 | 0 | 0 | 0 | 0 | 0 | 0 | 0 | 0 | 0 | 0 | 2 | 0 | 7 | 0 | 0 | Taar7d |
| Taar7f | 0 | 0 | 0 | 0 | 0 | 0 | 0 | 1 | 0 | 3 | 1 | 0 | 0 | 0 | 0 | 0 | 0 | 0 | 0 | 0 | 0 | 0 | 0 | 0 | 0 | 0 | 0 | 0 | Taar7f |
| Taar9 | 0 | 0 | 0 | 0 | 0 | 0 | 0 | 0 | 0 | 0 | 0 | 1 | 0 | 1 | 0 | 0 | 0 | 0 | 0 | 0 | 0 | 0 | 0 | 4 | 3 | 0 | 0 | 0 | Taar9 |
| Tab1 | 1020 | 1131 | 1191 | 1062 | 1169 | 1149 | 1170 | 2621 | 3278 | 3022 | 2817 | 3352 | 3031 | 2819 | 1014 | 1001 | 1022 | 1060 | 1273 | 1133 | 1094 | 2747 | 3062 | 3094 | 2871 | 3550 | 3362 | 3062 | Tab1 |
| Tab2 | 8759 | 7343 | 7935 | 7432 | 7756 | 7715 | 7505 | 10568 | 10353 | 10566 | 10992 | 11021 | 10401 | 10676 | 8284 | 7731 | 7803 | 7710 | 7701 | 6766 | 7277 | 11139 | 10289 | 10185 | 10376 | 11110 | 10574 | 10682 | Tab2 |
| Tab3 | 3953 | 3047 | 2866 | 3872 | 4104 | 4390 | 4666 | 3954 | 3510 | 3233 | 3223 | 3226 | 3375 | 3539 | 4118 | 3406 | 3086 | 3766 | 3705 | 4055 | 4177 | 3848 | 3494 | 2993 | 3614 | 2596 | 3076 | 3233 | Tab3 |
| Tac1 | 456 | 409 | 412 | 549 | 691 | 608 | 647 | 431 | 455 | 473 | 562 | 301 | 285 | 381 | 500 | 396 | 546 | 494 | 630 | 669 | 758 | 456 | 504 | 446 | 469 | 288 | 287 | 314 | Tac1 |
| Tac2 | 5 | 9 | 0 | 4 | 1 | 2 | 0 | 1 | 2 | 0 | 12 | 1 | 2 | 3 | 3 | 6 | 6 | 3 | 2 | 2 | 4 | 2 | 9 | 3 | 13 | 0 | 0 | 4 | Tac2 |
| Tac4 | 2 | 1 | 0 | 0 | 1 | 0 | 0 | 0 | 0 | 1 | 0 | 0 | 1 | 0 | 0 | 1 | 0 | 0 | 0 | 0 | 0 | 0 | 0 | 2 | 0 | 0 | 0 | 0 | Tac4 |

| | | | | | | | | | | | | | | | | | | | | | | | | | | | | | |
|---|---|---|---|---|---|---|---|---|---|---|---|---|---|---|---|---|---|---|---|---|---|---|---|---|---|---|---|---|---|
| Tacc1 | 4778 | 5429 | 6027 | 5994 | 6196 | 5860 | 6118 | 4610 | 4467 | 5101 | 4997 | 6117 | 5584 | 6363 | 4507 | 4689 | 6127 | 5939 | 6063 | 5134 | 6419 | 5138 | 4222 | 5395 | 5858 | 4712 | 6362 | 6357 | Tacc1 |
| Tacc2 | 4368 | 4480 | 5860 | 6252 | 7184 | 6344 | 5546 | 7017 | 7989 | 9493 | 8536 | 9437 | 8658 | 8733 | 4387 | 4359 | 6044 | 5909 | 7554 | 5089 | 6186 | 9605 | 7080 | 9059 | 9595 | 7685 | 8326 | 8699 | Tacc2 |
| Tacc3 | 11136 | 10718 | 11553 | 8553 | 5655 | 7263 | 8557 | 6611 | 6000 | 5543 | 5414 | 9181 | 7968 | 9553 | 9699 | 9656 | 9852 | 7408 | 7847 | 7426 | 7321 | 4664 | 5380 | 7491 | 6208 | 11609 | 9284 | 10090 | Tacc3 |
| Taco1 | 965 | 1093 | 1134 | 831 | 860 | 854 | 748 | 920 | 946 | 829 | 863 | 1024 | 1066 | 1030 | 963 | 945 | 1090 | 821 | 965 | 1023 | 775 | 750 | 916 | 972 | 906 | 1129 | 1063 | 1009 | Taco1 |
| Tacr1 | 123 | 211 | 143 | 222 | 498 | 327 | 259 | 412 | 390 | 469 | 442 | 288 | 406 | 327 | 224 | 157 | 218 | 261 | 283 | 197 | 314 | 560 | 350 | 270 | 355 | 122 | 321 | 287 | Tacr1 |
| Tacr2 | 0 | 0 | 3 | 0 | 0 | 0 | 0 | 2 | 0 | 0 | 1 | 0 | 1 | 0 | 0 | 3 | 2 | 1 | 0 | 0 | 0 | 0 | 1 | 0 | 0 | 0 | 0 | 0 | Tacr2 |
| Tacr3 | 158 | 151 | 144 | 149 | 311 | 183 | 226 | 250 | 204 | 254 | 203 | 147 | 192 | 110 | 185 | 145 | 158 | 227 | 263 | 279 | 190 | 246 | 189 | 139 | 162 | 97 | 92 | 154 | Tacr3 |
| Tacstd2 | 148 | 126 | 103 | 14 | 5 | 81 | 6 | 220 | 405 | 183 | 172 | 125 | 194 | 145 | 148 | 121 | 17 | 37 | 18 | 10 | 3 | 20 | 450 | 174 | 37 | 68 | 132 | 195 | Tacstd2 |
| Tada1 | 2299 | 2103 | 2149 | 2302 | 2774 | 2512 | 2773 | 4153 | 3947 | 3586 | 3350 | 3924 | 3494 | 3724 | 2363 | 2122 | 2329 | 2354 | 2915 | 2997 | 2854 | 3948 | 3897 | 3822 | 4142 | 3654 | 3794 | 3757 | Tada1 |
| Tada2a | 3041 | 2634 | 3368 | 3356 | 2941 | 2817 | 3693 | 2585 | 2497 | 2916 | 2886 | 2892 | 3104 | 3409 | 2889 | 2707 | 3072 | 3356 | 2915 | 2908 | 3098 | 2660 | 2471 | 3201 | 2897 | 3467 | 2833 | 3337 | Tada2a |
| Tada2b | 2222 | 2262 | 2735 | 3027 | 3091 | 2850 | 3063 | 1992 | 1767 | 2286 | 2108 | 2533 | 2240 | 2636 | 2136 | 2179 | 2770 | 3167 | 2918 | 2207 | 3113 | 2628 | 1945 | 2350 | 2431 | 2499 | 2271 | 2713 | Tada2b |
| Tada3 | 955 | 1330 | 1662 | 1480 | 1552 | 1261 | 1269 | 2004 | 2619 | 3029 | 2831 | 3178 | 2715 | 2826 | 882 | 1048 | 1501 | 1314 | 1651 | 1146 | 1306 | 2578 | 2524 | 3253 | 2963 | 3521 | 2504 | 2878 | Tada3 |
| Taf1 | 15610 | 13643 | 10556 | 12726 | 12265 | 14085 | 15179 | 11681 | 10224 | 9710 | 9724 | 11373 | 9928 | 12137 | 14927 | 13238 | 12077 | 12400 | 12239 | 14496 | 14629 | 9371 | 9864 | 9310 | 10593 | 8196 | 11237 | 10723 | Taf1 |
| Taf10 | 5166 | 5927 | 4936 | 5807 | 4839 | 4813 | 6182 | 3226 | 2872 | 2997 | 2927 | 3337 | 3053 | 3959 | 5169 | 4218 | 5451 | 5369 | 5337 | 4858 | 5997 | 2924 | 2826 | 3348 | 3304 | 3712 | 3477 | 3821 | Taf10 |
| Taf11 | 3690 | 3970 | 4413 | 3685 | 3224 | 3163 | 3962 | 3675 | 3544 | 3894 | 3545 | 4419 | 4224 | 4711 | 3640 | 3653 | 3892 | 3582 | 3387 | 3783 | 3485 | 3529 | 3404 | 4270 | 4165 | 4544 | 4584 | 4883 | Taf11 |
| Taf12 | 2924 | 2934 | 3203 | 2790 | 2387 | 2568 | 2746 | 1873 | 1753 | 1867 | 1597 | 2181 | 1975 | 2297 | 2740 | 2595 | 3079 | 2494 | 2527 | 2703 | 2599 | 1731 | 1808 | 2038 | 1935 | 2418 | 2135 | 2339 | Taf12 |
| Taf13 | 6454 | 6510 | 8553 | 7627 | 5551 | 5938 | 5955 | 4209 | 3206 | 3261 | 3308 | 4122 | 4288 | 4377 | 8231 | 7284 | 7650 | 6691 | 5866 | 7447 | 5813 | 3571 | 3082 | 4073 | 3918 | 4364 | 4647 | 4609 | Taf13 |
| Taf15 | 11993 | 12029 | 20764 | 18713 | 14267 | 13860 | 14657 | 12312 | 12706 | 17775 | 16513 | 20783 | 18250 | 21054 | 11257 | 11657 | 19600 | 14915 | 15952 | 13271 | 13659 | 13525 | 12312 | 21467 | 18283 | 26238 | 21264 | 19451 | Taf15 |
| Taf1a | 1835 | 1982 | 1467 | 1977 | 1753 | 1868 | 1847 | 1701 | 1667 | 1586 | 1821 | 1780 | 1814 | 1934 | 1940 | 2038 | 1452 | 1743 | 1612 | 1776 | 1809 | 1678 | 1658 | 1711 | 1801 | 1520 | 1771 | 1701 | Taf1a |
| Taf1b | 1944 | 1694 | 1723 | 1784 | 1629 | 1797 | 1893 | 2457 | 2169 | 2110 | 1989 | 1967 | 1923 | 2193 | 1884 | 1477 | 1635 | 1521 | 1621 | 2060 | 1905 | 2134 | 2073 | 1964 | 2212 | 2038 | 2131 | 2083 | Taf1b |
| Taf1c | 1100 | 1589 | 1556 | 1586 | 1389 | 1211 | 1184 | 1138 | 1380 | 1386 | 1197 | 1505 | 1300 | 1428 | 982 | 1431 | 1539 | 1315 | 1581 | 1154 | 1162 | 1268 | 1137 | 1422 | 1414 | 1663 | 1325 | 1307 | Taf1c |
| Taf1d | 5696 | 4694 | 5434 | 6227 | 4595 | 4712 | 5195 | 5472 | 4636 | 4521 | 5086 | 5226 | 5161 | 6157 | 5654 | 5308 | 5572 | 5461 | 4611 | 5279 | 5190 | 4520 | 4682 | 5305 | 4921 | 5481 | 5535 | 6083 | Taf1d |
| Taf2 | 10049 | 8972 | 8245 | 8415 | 7834 | 8782 | 9036 | 8436 | 7594 | 7238 | 7086 | 8832 | 8122 | 8673 | 9738 | 8874 | 7890 | 8433 | 7667 | 8205 | 8527 | 7439 | 7221 | 7354 | 7593 | 7370 | 8764 | 8624 | Taf2 |
| Taf3 | 2618 | 2825 | 3055 | 3414 | 2678 | 2835 | 2759 | 1627 | 1781 | 1984 | 1743 | 2259 | 2017 | 2255 | 2681 | 2755 | 3094 | 2990 | 2915 | 2545 | 2872 | 1878 | 1532 | 2021 | 2029 | 2294 | 2170 | 2277 | Taf3 |
| Taf4 | 3524 | 3889 | 4466 | 5113 | 4599 | 4310 | 4812 | 2762 | 2463 | 3431 | 3175 | 3797 | 3062 | 3737 | 3233 | 3678 | 4921 | 4908 | 4599 | 3714 | 4759 | 3367 | 2309 | 3445 | 3565 | 3796 | 3367 | 3533 | Taf4 |
| Taf4b | 873 | 720 | 706 | 549 | 456 | 487 | 510 | 252 | 246 | 212 | 261 | 216 | 340 | 217 | 878 | 910 | 539 | 479 | 404 | 517 | 383 | 307 | 320 | 163 | 195 | 195 | 223 | 233 | Taf4b |
| Taf5 | 2709 | 2299 | 2396 | 2472 | 2225 | 2375 | 2743 | 2268 | 1762 | 1784 | 1759 | 2070 | 1838 | 2286 | 2527 | 2248 | 2449 | 2409 | 2427 | 2408 | 2551 | 2134 | 1735 | 1849 | 2204 | 2234 | 2030 | 2214 | Taf5 |
| Taf5l | 1390 | 1280 | 1113 | 1092 | 991 | 1168 | 976 | 2089 | 2052 | 1838 | 2066 | 2215 | 2118 | 2159 | 1043 | 1146 | 1110 | 1093 | 1107 | 1055 | 1017 | 1724 | 2133 | 2134 | 1862 | 2591 | 2358 | 2222 | Taf5l |
| Taf6 | 6264 | 7573 | 7090 | 6303 | 5118 | 5433 | 5741 | 4584 | 4856 | 4606 | 4321 | 5576 | 4857 | 5286 | 6001 | 6390 | 6672 | 5659 | 6071 | 5873 | 5655 | 4406 | 4552 | 5321 | 4831 | 5843 | 5406 | 5587 | Taf6 |
| Taf6l | 1990 | 2616 | 2329 | 2481 | 1936 | 1989 | 1782 | 1180 | 1341 | 1218 | 1314 | 1460 | 1382 | 1296 | 1925 | 2102 | 2355 | 2360 | 2233 | 1829 | 1821 | 1215 | 1311 | 1331 | 1261 | 1599 | 1443 | 1356 | Taf6l |
| Taf7 | 3652 | 3238 | 3549 | 3332 | 2723 | 2852 | 3375 | 3040 | 2637 | 2360 | 2489 | 3128 | 2578 | 2982 | 3957 | 3537 | 3321 | 3204 | 2840 | 3566 | 2878 | 2369 | 2557 | 2606 | 2492 | 3102 | 2919 | 2893 | Taf7 |
| Taf7l | 14 | 15 | 16 | 15 | 18 | 4 | 4 | 4 | 6 | 9 | 11 | 11 | 2 | 4 | 15 | 18 | 14 | 21 | 5 | 2 | 6 | 10 | 11 | 1 | 4 | 5 | 10 | 1 | Taf7l |
| Taf7l2 | 0 | 0 | 0 | 0 | 0 | 0 | 0 | 0 | 0 | 0 | 0 | 0 | 0 | 0 | 0 | 0 | 1 | 1 | 0 | 3 | 0 | 0 | 2 | 0 | 0 | 0 | 0 | 2 | Taf7l2 |
| Taf8 | 2052 | 2437 | 2032 | 1985 | 1687 | 1742 | 1619 | 2086 | 2347 | 2223 | 2159 | 2533 | 2435 | 2477 | 1742 | 2008 | 2006 | 1892 | 1842 | 1701 | 1639 | 2117 | 2163 | 2517 | 2084 | 2732 | 2506 | 2453 | Taf8 |
| Taf9 | 28 | 10 | 42 | 49 | 17 | 19 | 7 | 10 | 11 | 16 | 13 | 11 | 10 | 29 | 21 | 18 | 28 | 18 | 23 | 38 | 19 | 20 | 12 | 22 | 21 | 16 | 19 | 13 | Taf9 |
| Taf9b | 194 | 160 | 187 | 144 | 241 | 163 | 202 | 926 | 858 | 786 | 843 | 879 | 780 | 861 | 228 | 176 | 162 | 152 | 203 | 214 | 139 | 966 | 858 | 791 | 819 | 1021 | 873 | 985 | Taf9b |
| Tafa1 | 59 | 40 | 77 | 154 | 323 | 185 | 270 | 204 | 248 | 212 | 255 | 61 | 85 | 95 | 83 | 48 | 93 | 210 | 196 | 268 | 337 | 281 | 295 | 129 | 218 | 46 | 83 | 46 | Tafa1 |
| Tafa2 | 732 | 760 | 568 | 1096 | 2104 | 1415 | 1512 | 1991 | 1998 | 1985 | 2004 | 1130 | 1370 | 1112 | 1207 | 889 | 1087 | 1473 | 1370 | 1493 | 1639 | 3006 | 2061 | 1410 | 2335 | 930 | 974 | 943 | Tafa2 |
| Tafa3 | 24 | 18 | 14 | 13 | 11 | 5 | 7 | 16 | 30 | 17 | 35 | 5 | 18 | 16 | 28 | 17 | 17 | 15 | 10 | 8 | 4 | 20 | 27 | 21 | 20 | 13 | 7 | 9 | Tafa3 |
| Tafa4 | 369 | 575 | 604 | 487 | 348 | 368 | 232 | 208 | 124 | 144 | 121 | 167 | 239 | 184 | 563 | 596 | 555 | 393 | 426 | 466 | 369 | 182 | 194 | 156 | 171 | 62 | 248 | 221 | Tafa4 |
| Tafa5 | 828 | 1035 | 990 | 1438 | 1924 | 1514 | 1381 | 1706 | 1717 | 2394 | 1719 | 1654 | 1429 | 1525 | 974 | 715 | 1467 | 1376 | 1601 | 1315 | 1532 | 2065 | 1629 | 1815 | 2064 | 1595 | 1304 | 1386 | Tafa5 |
| Tagap | 35 | 29 | 17 | 26 | 21 | 16 | 13 | 53 | 39 | 42 | 40 | 37 | 40 | 28 | 28 | 37 | 31 | 20 | 18 | 20 | 13 | 37 | 52 | 40 | 19 | 35 | 40 | 25 | Tagap |
| Tagap1 | 2158 | 1900 | 1931 | 2006 | 2077 | 2108 | 2095 | 2595 | 2538 | 2375 | 2596 | 2878 | 2637 | 2498 | 2101 | 1729 | 1855 | 1986 | 2220 | 2204 | 1960 | 2508 | 2454 | 2659 | 2574 | 3073 | 2841 | 2611 | Tagap1 |
| Tagln | 1535 | 2219 | 1735 | 744 | 311 | 1008 | 248 | 891 | 1336 | 822 | 1592 | 1851 | 2522 | 1904 | 2097 | 2308 | 1540 | 1102 | 302 | 304 | 247 | 617 | 1783 | 1278 | 870 | 1361 | 2662 | 2294 | Tagln |
| Tagln2 | 2944 | 3234 | 3207 | 2682 | 2610 | 2491 | 2433 | 1898 | 2688 | 2246 | 2942 | 1805 | 2222 | 1727 | 2480 | 3633 | 2636 | 2785 | 2575 | 2721 | 2503 | 1820 | 2855 | 2192 | 2185 | 2372 | 1839 | 1842 | Tagln2 |
| Tagln3 | 3300 | 4866 | 3904 | 5221 | 6941 | 6050 | 5972 | 10097 | 9152 | 10053 | 8638 | 10465 | 10075 | 11215 | 4909 | 3674 | 4997 | 5014 | 6524 | 6102 | 6637 | 9434 | 8685 | 10193 | 11471 | 8784 | 10982 | 11163 | Tagln3 |
| Tal1 | 843 | 967 | 1943 | 1447 | 1694 | 1580 | 1111 | 2922 | 1790 | 2776 | 2215 | 3347 | 4846 | 3428 | 974 | 805 | 1539 | 1265 | 1819 | 1261 | 1499 | 3763 | 1848 | 2891 | 3320 | 890 | 4195 | 4481 | Tal1 |
| Tal2 | 1683 | 1429 | 4001 | 1652 | 1727 | 2020 | 1278 | 4742 | 1795 | 3186 | 2275 | 6680 | 6907 | 7526 | 1762 | 1012 | 2723 | 1269 | 2410 | 1854 | 1988 | 4380 | 2667 | 4680 | 4293 | 1078 | 9559 | 9779 | Tal2 |
| Taldo1 | 8368 | 12057 | 10252 | 10174 | 8885 | 8971 | 9724 | 6582 | 7678 | 7681 | 6890 | 8288 | 8207 | 8377 | 8525 | 9552 | 10505 | 9562 | 10050 | 10779 | 9541 | 5663 | 7567 | 7826 | 7745 | 9229 | 8494 | 8775 | Taldo1 |
| Tamalin | 1039 | 1079 | 1314 | 1201 | 954 | 936 | 809 | 573 | 610 | 638 | 715 | 669 | 728 | 595 | 939 | 1065 | 1254 | 1010 | 1149 | 733 | 872 | 453 | 655 | 779 | 445 | 1069 | 703 | 687 | Tamalin |
| Tamm41 | 2099 | 2234 | 1784 | 1898 | 1456 | 1545 | 1850 | 1434 | 1495 | 1351 | 1279 | 1641 | 1447 | 1625 | 1829 | 1834 | 1747 | 1669 | 1774 | 1796 | 1801 | 1272 | 1327 | 1472 | 1421 | 1751 | 1541 | 1621 | Tamm41 |
| Tanc1 | 4538 | 4160 | 3491 | 3541 | 3326 | 3772 | 3562 | 2397 | 3147 | 2906 | 3187 | 2876 | 3001 | 2739 | 3516 | 4494 | 3467 | 3737 | 3402 | 2908 | 3153 | 1970 | 3112 | 2963 | 2303 | 2695 | 2753 | 2724 | Tanc1 |
| Tanc2 | 5435 | 5101 | 3305 | 5908 | 9578 | 8924 | 8682 | 8328 | 8945 | 9120 | 9830 | 7206 | 7173 | 7187 | 6057 | 6099 | 4388 | 7097 | 7228 | 7552 | 9207 | 9633 | 8510 | 6789 | 10402 | 4051 | 6878 | 5834 | Tanc2 |
| Tango6 | 294 | 318 | 316 | 228 | 241 | 263 | 221 | 669 | 829 | 730 | 769 | 1004 | 941 | 876 | 287 | 329 | 302 | 239 | 266 | 250 | 232 | 635 | 790 | 769 | 728 | 825 | 931 | 1013 | Tango6 |
| Tank | 3756 | 3247 | 3225 | 3647 | 3381 | 3596 | 4250 | 2291 | 2146 | 1748 | 2002 | 2229 | 2160 | 2221 | 4151 | 3537 | 3311 | 3714 | 3509 | 4221 | 3750 | 1988 | 2012 | 1895 | 1957 | 1951 | 2345 | 2232 | Tank |
| Taok1 | 11026 | 8343 | 7794 | 8986 | 10714 | 12389 | 12307 | 18014 | 15154 | 13268 | 13889 | 15428 | 14603 | 16005 | 11513 | 8942 | 8480 | 8836 | 10098 | 12807 | 12346 | 14028 | 14329 | 12433 | 15529 | 11303 | 16845 | 14939 | Taok1 |
| Taok2 | 13251 | 16964 | 17051 | 18715 | 16578 | 16318 | 15374 | 12350 | 13681 | 15753 | 14148 | 16632 | 14914 | 15449 | 12589 | 14390 | 18374 | 18077 | 18191 | 14295 | 16265 | 15276 | 12923 | 16480 | 16507 | 13355 | 15926 | 15564 | Taok2 |
| Taok3 | 3813 | 4724 | 5327 | 5786 | 5208 | 4474 | 4963 | 2409 | 2293 | 3148 | 2726 | 2359 | 2100 | 2549 | 3922 | 4199 | 5313 | 5801 | 5507 | 4365 | 5266 | 3309 | 2234 | 2809 | 3268 | 2084 | 2205 | 2334 | Taok3 |
| Tap1 | 654 | 550 | 517 | 451 | 390 | 452 | 377 | 139 | 195 | 153 | 184 | 111 | 129 | 136 | 480 | 618 | 482 | 467 | 429 | 387 | 368 | 120 | 237 | 127 | 111 | 197 | 110 | 146 | Tap1 |
| Tap2 | 262 | 246 | 367 | 286 | 335 | 230 | 189 | 31 | 67 | 64 | 88 | 39 | 61 | 48 | 224 | 301 | 351 | 296 | 322 | 230 | 258 | 65 | 90 | 42 | 59 | 65 | 48 | 43 | Tap2 |
| Tapbp | 1308 | 1477 | 1761 | 1537 | 1536 | 1499 | 1367 | 1122 | 1367 | 1694 | 1561 | 1281 | 1377 | 1291 | 1411 | 1534 | 1619 | 1702 | 1460 | 1363 | 1396 | 1524 | 1543 | 1513 | 1363 | 1238 | 1247 | 1398 | Tapbp |
| Tapbpl | 52 | 64 | 57 | 39 | 105 | 90 | 63 | 126 | 210 | 227 | 162 | 124 | 141 | 133 | 78 | 62 | 44 | 69 | 86 | 67 | 61 | 180 | 219 | 152 | 154 | 117 | 120 | 128 | Tapbpl |
| Tapt1 | 3463 | 3105 | 2804 | 2886 | 2946 | 3225 | 3182 | 4231 | 3759 | 3629 | 3691 | 3523 | 3637 | 3364 | 3384 | 3287 | 2724 | 3062 | 3033 | 3325 | 3118 | 3781 | 3921 | 3265 | 3500 | 3463 | 3543 | 3469 | Tapt1 |
| Tarbp1 | 1187 | 1060 | 843 | 1087 | 1112 | 1110 | 1048 | 1244 | 1256 | 1261 | 1255 | 1132 | 1169 | 1095 | 1007 | 1219 | 998 | 1067 | 1003 | 1048 | 1062 | 1029 | 1234 | 1111 | 1145 | 1029 | 1073 | 1136 | Tarbp1 |
| Tarbp2 | 1163 | 1491 | 1517 | 1195 | 1303 | 1279 | 1077 | 2671 | 3130 | 2841 | 2720 | 3131 | 3108 | 3041 | 1263 | 1347 | 1421 | 1082 | 1314 | 1275 | 1091 | 2697 | 2922 | 3065 | 3139 | 3742 | 3435 | 3211 | Tarbp2 |
| Tardbp | 60422 | 49325 | 56590 | 50964 | 45213 | 49476 | 54828 | 59542 | 49453 | 46627 | 48128 | 59124 | 55697 | 61724 | 59038 | 52666 | 52422 | 46662 | 49233 | 58583 | 50315 | 45616 | 47656 | 54611 | 52890 | 59752 | 62865 | 61561 | Tardbp |
| Tarm1 | 2 | 0 | 0 | 5 | 0 | 0 | 2 | 0 | 0 | 1 | 2 | 2 | 1 | 2 | 0 | 4 | 1 | 3 | 2 | 2 | 5 | 0 | 5 | 0 | 0 | 2 | 0 | 2 | Tarm1 |
| Tars | 9421 | 10727 | 9844 | 8578 | 6461 | 7078 | 7526 | 6066 | 6137 | 5433 | 5198 | 6440 | 6344 | 7101 | 9618 | 10140 | 8712 | 7978 | 7134 | 8404 | 7004 | 4652 | 5948 | 6017 | 5491 | 6942 | 6830 | 7174 | Tars |
| Tars2 | 1525 | 1656 | 1585 | 1468 | 1852 | 1485 | 1523 | 2886 | 3222 | 3067 | 3075 | 3408 | 3172 | 3280 | 1389 | 1421 | 1661 | 1576 | 1850 | 1565 | 1532 | 2839 | 2891 | 3073 | 3445 | 3086 | 3633 | 3259 | Tars2 |
| Tarsl2 | 1008 | 860 | 874 | 895 | 980 | 872 | 1005 | 1486 | 1358 | 1389 | 1379 | 1379 | 1201 | 1426 | 1076 | 897 | 890 | 869 | 931 | 1022 | 1000 | 1277 | 1413 | 1220 | 1316 | 1388 | 1334 | 1430 | Tarsl2 |
| Tas1r1 | 6 | 4 | 4 | 0 | 5 | 2 | 1 | 7 | 21 | 16 | 33 | 11 | 12 | 9 | 3 | 2 | 1 | 1 | 5 | 7 | 4 | 15 | 30 | 8 | 12 | 8 | 17 | 7 | Tas1r1 |
| Tas1r2 | 9 | 14 | 0 | 23 | 13 | 12 | 15 | 2 | 0 | 0 | 3 | 1 | 2 | 2 | 11 | 18 | 8 | 19 | 12 | 5 | 13 | 0 | 5 | 1 | 0 | 0 | 0 | 6 | Tas1r2 |
| Tas1r3 | 128 | 174 | 132 | 206 | 188 | 147 | 177 | 57 | 78 | 85 | 77 | 101 | 101 | 95 | 108 | 141 | 183 | 198 | 185 | 115 | 163 | 79 | 72 | 118 | 117 | 41 | 89 | 85 | Tas1r3 |
| Tas2r108 | 16 | 14 | 6 | 23 | 32 | 34 | 34 | 11 | 14 | 13 | 2 | 11 | 7 | 12 | 8 | 14 | 21 | 18 | 12 | 21 | 18 | 10 | 8 | 9 | 7 | 8 | 17 | 5 | Tas2r108 |
| Tas2r119 | 0 | 0 | 0 | 0 | 0 | 0 | 0 | 0 | 0 | 0 | 0 | 0 | 0 | 0 | 0 | 0 | 0 | 0 | 0 | 0 | 0 | 0 | 2 | 0 | 0 | 0 | 0 | 0 | Tas2r119 |
| Tas2r126 | 0 | 1 | 0 | 0 | 2 | 0 | 1 | 0 | 0 | 0 | 0 | 0 | 0 | 0 | 0 | 1 | 0 | 2 | 2 | 0 | 1 | 0 | 0 | 1 | 0 | 0 | 0 | 0 | Tas2r126 |
| Tas2r134 | 0 | 2 | 0 | 0 | 0 | 0 | 2 | 0 | 0 | 0 | 0 | 0 | 0 | 0 | 1 | 0 | 0 | 0 | 0 | 0 | 0 | 0 | 0 | 0 | 0 | 0 | 0 | 0 | Tas2r134 |
| Tas2r135 | 14 | 7 | 16 | 16 | 22 | 6 | 9 | 1 | 0 | 1 | 7 | 0 | 5 | 3 | 7 | 3 | 6 | 12 | 13 | 14 | 10 | 2 | 3 | 5 | 4 | 2 | 3 | 1 | Tas2r135 |
| Tas2r137 | 17 | 25 | 8 | 20 | 16 | 24 | 20 | 11 | 3 | 10 | 4 | 8 | 14 | 12 | 17 | 23 | 2 | 15 | 16 | 33 | 28 | 5 | 13 | 1 | 9 | 2 | 14 | 11 | Tas2r137 |

| | | | | | | | | | | | | | | | | | | | | | | | | | | | | | |
|---|---|---|---|---|---|---|---|---|---|---|---|---|---|---|---|---|---|---|---|---|---|---|---|---|---|---|---|---|---|
| Tas2r143 | 3 | 3 | 1 | 3 | 7 | 6 | 0 | 0 | 1 | 2 | 1 | 3 | 1 | 2 | 7 | 5 | 0 | 0 | 9 | 2 | 4 | 0 | 2 | 1 | 2 | 3 | 0 | 1 | Tas2r143 |
| Tasl | 5 | 6 | 8 | 6 | 3 | 1 | 3 | 1 | 9 | 8 | 8 | 4 | 14 | 9 | 5 | 3 | 1 | 3 | 5 | 4 | 4 | 10 | 16 | 8 | 7 | 4 | 3 | 6 | Tasl |
| Tasor | 4609 | 3302 | 4669 | 5286 | 5831 | 5964 | 6140 | 8209 | 7047 | 7606 | 8262 | 7580 | 7646 | 7679 | 4759 | 4272 | 4902 | 5290 | 5592 | 5021 | 5695 | 9157 | 7075 | 7549 | 8022 | 5805 | 7649 | 7344 | Tasor |
| Tasor2 | 18453 | 15308 | 10311 | 14130 | 14590 | 17614 | 18338 | 15599 | 13560 | 11513 | 12249 | 12527 | 11911 | 12010 | 18697 | 16787 | 12293 | 14978 | 14018 | 18026 | 16925 | 11817 | 12873 | 10145 | 12104 | 7787 | 13335 | 11689 | Tasor2 |
| Tasp1 | 979 | 807 | 794 | 782 | 786 | 694 | 886 | 863 | 768 | 819 | 761 | 755 | 838 | 880 | 895 | 794 | 831 | 828 | 758 | 784 | 826 | 678 | 812 | 766 | 722 | 928 | 755 | 871 | Tasp1 |
| Tat | 0 | 0 | 0 | 7 | 2 | 0 | 0 | 0 | 2 | 0 | 0 | 0 | 0 | 2 | 0 | 0 | 1 | 2 | 0 | 2 | 0 | 0 | 1 | 1 | 0 | 0 | 0 | 2 | Tat |
| Tatdn1 | 1779 | 1534 | 1644 | 1792 | 1674 | 1654 | 2131 | 1672 | 1497 | 1395 | 1523 | 1628 | 1623 | 1697 | 1937 | 1742 | 1741 | 1690 | 1571 | 1906 | 1805 | 1724 | 1532 | 1602 | 1535 | 1627 | 1617 | 1768 | Tatdn1 |
| Tatdn2 | 5236 | 5508 | 3928 | 4677 | 3814 | 4359 | 4217 | 3158 | 2909 | 2670 | 2537 | 2721 | 2578 | 2721 | 5246 | 4625 | 4132 | 4240 | 4328 | 4195 | 4631 | 2811 | 2685 | 2569 | 2678 | 2656 | 2811 | 2940 | Tatdn2 |
| Tatdn3 | 444 | 413 | 356 | 395 | 472 | 510 | 352 | 1037 | 952 | 845 | 1085 | 858 | 961 | 879 | 414 | 402 | 340 | 419 | 394 | 481 | 458 | 979 | 970 | 803 | 1029 | 687 | 880 | 853 | Tatdn3 |
| Tax1bp1 | 16290 | 14470 | 14844 | 13886 | 13703 | 13911 | 15294 | 14477 | 12660 | 11380 | 11255 | 12643 | 12327 | 12956 | 17735 | 15208 | 14832 | 13936 | 13868 | 16746 | 14425 | 12616 | 12908 | 12060 | 12091 | 12727 | 12938 | 13144 | Tax1bp1 |
| Tax1bp3 | 1459 | 1799 | 1770 | 1606 | 1671 | 1787 | 1585 | 2049 | 2267 | 2139 | 2413 | 2284 | 2519 | 2303 | 1454 | 1734 | 1603 | 1629 | 1778 | 1797 | 1591 | 1868 | 2450 | 2355 | 2254 | 2677 | 2508 | 2417 | Tax1bp3 |
| Taz | 3892 | 3915 | 3564 | 3828 | 3557 | 3767 | 3813 | 4154 | 3806 | 3925 | 3958 | 4005 | 3952 | 4043 | 3982 | 3599 | 3783 | 3789 | 3776 | 3774 | 3797 | 4218 | 3959 | 3733 | 4080 | 3522 | 4091 | 3881 | Taz |
| Tbata | 0 | 1 | 0 | 3 | 9 | 2 | 11 | 6 | 16 | 17 | 16 | 2 | 14 | 13 | 1 | 2 | 1 | 10 | 3 | 8 | 11 | 13 | 10 | 14 | 23 | 9 | 10 | 13 | Tbata |
| Tbc1d1 | 6000 | 5789 | 5272 | 5617 | 4928 | 5043 | 4969 | 3156 | 3361 | 3564 | 3681 | 3919 | 3746 | 3755 | 5611 | 6090 | 5366 | 5604 | 5163 | 4580 | 4805 | 3096 | 3245 | 3609 | 3326 | 3998 | 3595 | 3529 | Tbc1d1 |
| Tbc1d10a | 1105 | 1478 | 1355 | 1409 | 1259 | 1167 | 1029 | 819 | 1094 | 1022 | 1039 | 1014 | 965 | 793 | 1129 | 1347 | 1325 | 1381 | 1374 | 1154 | 1045 | 830 | 1150 | 1008 | 1063 | 1019 | 919 | 840 | Tbc1d10a |
| Tbc1d10b | 5530 | 7024 | 7276 | 8748 | 7617 | 7046 | 6968 | 4095 | 4547 | 5381 | 5094 | 5517 | 4787 | 5476 | 5414 | 6135 | 7838 | 8871 | 8489 | 6235 | 7676 | 5287 | 4339 | 5658 | 5475 | 6025 | 4815 | 5305 | Tbc1d10b |
| Tbc1d10c | 0 | 0 | 0 | 0 | 0 | 1 | 0 | 0 | 1 | 0 | 2 | 2 | 2 | 0 | 0 | 0 | 0 | 0 | 0 | 0 | 0 | 0 | 0 | 0 | 2 | 1 | 1 | 1 | Tbc1d10c |
| Tbc1d12 | 2795 | 2215 | 2564 | 2589 | 3354 | 3122 | 3188 | 2444 | 2176 | 1898 | 1999 | 1837 | 1998 | 1934 | 3187 | 2534 | 2568 | 2861 | 2904 | 3351 | 3135 | 2298 | 2146 | 1751 | 2080 | 1672 | 1771 | 1849 | Tbc1d12 |
| Tbc1d13 | 5900 | 6406 | 5754 | 5968 | 5771 | 5836 | 5863 | 5978 | 6105 | 5936 | 5936 | 6160 | 6036 | 5912 | 5658 | 5217 | 6023 | 5982 | 6420 | 5845 | 5975 | 6589 | 5972 | 6347 | 6132 | 6182 | 6337 | 6051 | Tbc1d13 |
| Tbc1d14 | 8263 | 7713 | 6555 | 8019 | 9102 | 8833 | 9221 | 9056 | 8983 | 8382 | 8105 | 7924 | 7343 | 8202 | 8010 | 6995 | 6719 | 8225 | 8779 | 9256 | 9350 | 8824 | 8385 | 8072 | 9127 | 7349 | 7426 | 7595 | Tbc1d14 |
| Tbc1d15 | 6356 | 4852 | 4726 | 4826 | 3972 | 4717 | 4683 | 4502 | 4036 | 3492 | 4074 | 4001 | 3977 | 4183 | 6215 | 5755 | 4330 | 4940 | 3956 | 5034 | 4450 | 3547 | 4245 | 3684 | 3680 | 4027 | 3979 | 4262 | Tbc1d15 |
| Tbc1d16 | 20612 | 26145 | 30707 | 34002 | 34218 | 30195 | 30412 | 16601 | 18782 | 23600 | 21914 | 23020 | 21175 | 21999 | 20836 | 23294 | 32705 | 34807 | 35714 | 24892 | 33553 | 23277 | 17744 | 24020 | 26902 | 17953 | 20944 | 21981 | Tbc1d16 |
| Tbc1d17 | 2643 | 3530 | 3313 | 4039 | 3729 | 3406 | 3393 | 2201 | 2885 | 2863 | 3084 | 3003 | 2897 | 2700 | 2557 | 3171 | 3592 | 4039 | 4163 | 2749 | 3538 | 3115 | 2847 | 3223 | 3229 | 3016 | 2759 | 2664 | Tbc1d17 |
| Tbc1d19 | 3275 | 3340 | 2685 | 2960 | 3104 | 3275 | 3461 | 4272 | 3569 | 3106 | 3232 | 3249 | 3188 | 3427 | 3705 | 3162 | 2916 | 2801 | 2909 | 3541 | 3416 | 3676 | 3527 | 2945 | 3172 | 2794 | 3402 | 3375 | Tbc1d19 |
| Tbc1d2 | 381 | 496 | 364 | 389 | 380 | 340 | 316 | 133 | 147 | 151 | 150 | 93 | 128 | 99 | 364 | 530 | 385 | 363 | 430 | 300 | 314 | 105 | 186 | 128 | 96 | 103 | 110 | 93 | Tbc1d2 |
| Tbc1d20 | 5944 | 6701 | 5504 | 6187 | 5061 | 5572 | 5633 | 4317 | 4218 | 4065 | 4265 | 3964 | 3961 | 4012 | 5657 | 6245 | 5452 | 5951 | 5220 | 5095 | 5585 | 4286 | 4750 | 4082 | 4128 | 4380 | 4013 | 4321 | Tbc1d20 |
| Tbc1d21 | 0 | 0 | 0 | 0 | 0 | 0 | 0 | 0 | 0 | 0 | 0 | 0 | 0 | 0 | 0 | 1 | 0 | 0 | 0 | 0 | 0 | 0 | 1 | 0 | 0 | 0 | 0 | 0 | Tbc1d21 |
| Tbc1d22a | 806 | 938 | 795 | 858 | 793 | 791 | 628 | 1138 | 1288 | 1051 | 1116 | 1162 | 1165 | 1129 | 839 | 917 | 794 | 866 | 829 | 803 | 707 | 1096 | 1232 | 1015 | 1236 | 1175 | 1168 | 1137 | Tbc1d22a |
| Tbc1d22b | 2827 | 3468 | 3829 | 3841 | 3854 | 3362 | 3160 | 2174 | 2589 | 3099 | 2706 | 3182 | 2617 | 2952 | 2672 | 2972 | 3865 | 3787 | 3899 | 2768 | 3578 | 3034 | 2392 | 3101 | 3171 | 3090 | 2658 | 2886 | Tbc1d22b |
| Tbc1d23 | 6218 | 4992 | 4524 | 4742 | 5036 | 5337 | 5623 | 5577 | 4957 | 4371 | 4784 | 4380 | 4403 | 4738 | 6070 | 5540 | 4266 | 5040 | 4651 | 5605 | 5582 | 4588 | 5071 | 4088 | 4651 | 3879 | 4381 | 4281 | Tbc1d23 |
| Tbc1d24 | 10830 | 11440 | 9024 | 14525 | 14424 | 14087 | 14441 | 6629 | 6409 | 7201 | 6767 | 5838 | 5857 | 5761 | 11646 | 11116 | 11449 | 15436 | 14110 | 12592 | 14949 | 8577 | 6379 | 6154 | 7567 | 3642 | 5449 | 5482 | Tbc1d24 |
| Tbc1d25 | 909 | 1004 | 876 | 1064 | 1050 | 961 | 1027 | 912 | 986 | 966 | 868 | 890 | 822 | 825 | 799 | 950 | 986 | 957 | 1054 | 934 | 1044 | 962 | 903 | 856 | 916 | 824 | 702 | 718 | Tbc1d25 |
| Tbc1d2b | 6647 | 7014 | 6306 | 5612 | 4604 | 5774 | 4443 | 2926 | 3518 | 3306 | 3856 | 3205 | 3810 | 3014 | 6098 | 7374 | 5295 | 6020 | 4563 | 4054 | 4376 | 2695 | 3860 | 3233 | 2797 | 2791 | 2932 | 3082 | Tbc1d2b |
| Tbc1d30 | 1328 | 1410 | 1423 | 1957 | 3021 | 2094 | 2501 | 2484 | 2493 | 3082 | 2417 | 1934 | 1836 | 1987 | 1522 | 1312 | 2130 | 2088 | 2490 | 2013 | 2461 | 3425 | 2353 | 2283 | 3030 | 1797 | 1763 | 1641 | Tbc1d30 |
| Tbc1d31 | 6623 | 6284 | 4745 | 5429 | 4147 | 5009 | 5518 | 2550 | 2280 | 1977 | 2208 | 2343 | 2205 | 2503 | 6142 | 5944 | 4675 | 5129 | 4447 | 5593 | 5403 | 1795 | 2174 | 2132 | 2197 | 2252 | 2645 | 2347 | Tbc1d31 |
| Tbc1d32 | 1204 | 940 | 726 | 880 | 884 | 984 | 971 | 1561 | 1561 | 1369 | 1379 | 1357 | 1384 | 1434 | 1142 | 1142 | 812 | 813 | 838 | 950 | 978 | 1414 | 1517 | 1252 | 1295 | 1228 | 1508 | 1308 | Tbc1d32 |
| Tbc1d4 | 1247 | 1087 | 1001 | 1139 | 1546 | 1383 | 1326 | 1617 | 1714 | 1846 | 2113 | 1574 | 1681 | 1421 | 1291 | 1372 | 982 | 1302 | 1336 | 1169 | 1323 | 1703 | 1977 | 1767 | 1779 | 1435 | 1432 | 1335 | Tbc1d4 |
| Tbc1d5 | 1294 | 1221 | 1097 | 1077 | 1193 | 1281 | 1095 | 1758 | 1917 | 1817 | 1944 | 2006 | 1956 | 1807 | 1244 | 1295 | 1099 | 1110 | 1068 | 1054 | 1155 | 1724 | 1827 | 1682 | 1699 | 1741 | 2059 | 2031 | Tbc1d5 |
| Tbc1d7 | 773 | 715 | 652 | 599 | 697 | 703 | 724 | 1569 | 1455 | 1365 | 1297 | 1436 | 1431 | 1450 | 750 | 679 | 659 | 679 | 690 | 871 | 816 | 1482 | 1480 | 1414 | 1590 | 1552 | 1484 | 1654 | Tbc1d7 |
| Tbc1d8 | 1306 | 1544 | 1367 | 1751 | 2214 | 1766 | 1842 | 849 | 1093 | 998 | 1280 | 933 | 845 | 791 | 1395 | 1560 | 1459 | 1999 | 1784 | 1579 | 1977 | 1205 | 1134 | 873 | 1106 | 869 | 830 | 833 | Tbc1d8 |
| Tbc1d8b | 1878 | 1338 | 1508 | 1282 | 1235 | 1454 | 1431 | 1263 | 1387 | 1062 | 1342 | 1075 | 1086 | 1099 | 1692 | 1518 | 1385 | 1393 | 1381 | 1549 | 1353 | 981 | 1441 | 1064 | 960 | 954 | 1097 | 992 | Tbc1d8b |
| Tbc1d9 | 4057 | 4317 | 5100 | 5818 | 9247 | 6567 | 7005 | 7624 | 7510 | 9449 | 8133 | 9382 | 8984 | 8818 | 4985 | 4417 | 5935 | 6540 | 7635 | 6314 | 7355 | 10840 | 7809 | 8381 | 10316 | 6738 | 8228 | 8956 | Tbc1d9 |
| Tbc1d9b | 2623 | 3233 | 3036 | 2854 | 3193 | 3078 | 2553 | 4659 | 5700 | 5729 | 5394 | 5345 | 5393 | 4667 | 2856 | 3293 | 2826 | 2952 | 3120 | 2651 | 2773 | 5521 | 5891 | 5101 | 5223 | 4980 | 4966 | 4616 | Tbc1d9b |
| Tbca | 6834 | 5844 | 6564 | 7009 | 6003 | 5523 | 7774 | 6708 | 5474 | 6715 | 6317 | 5968 | 5460 | 7810 | 7090 | 5050 | 6916 | 6216 | 6216 | 7418 | 7813 | 5419 | 5728 | 6965 | 6427 | 7369 | 6008 | 7725 | Tbca |
| Tbcb | 4555 | 6357 | 6037 | 6494 | 5999 | 5984 | 5555 | 4205 | 4699 | 5304 | 4837 | 5247 | 5044 | 4983 | 4717 | 5013 | 6186 | 5889 | 6670 | 6600 | 5759 | 4348 | 4691 | 5449 | 5403 | 5689 | 5162 | 5198 | Tbcb |
| Tbcc | 1237 | 1487 | 1362 | 1440 | 1070 | 1115 | 1155 | 1050 | 1023 | 1143 | 1090 | 1194 | 1127 | 1152 | 1253 | 1209 | 1440 | 1338 | 1308 | 1213 | 1113 | 974 | 1004 | 1192 | 1102 | 1342 | 1260 | 1203 | Tbcc |
| Tbccd1 | 2203 | 2428 | 2010 | 1701 | 1672 | 1776 | 1799 | 1762 | 1771 | 1577 | 1479 | 1674 | 1651 | 1680 | 2201 | 2084 | 1803 | 1931 | 1683 | 2057 | 1776 | 1462 | 1726 | 1576 | 1479 | 1738 | 1713 | 1834 | Tbccd1 |
| Tbcd | 8371 | 8715 | 6707 | 7497 | 7456 | 7910 | 6955 | 5908 | 6030 | 5073 | 5112 | 5986 | 6199 | 5631 | 8277 | 7759 | 6947 | 7582 | 7943 | 7365 | 7341 | 5559 | 5591 | 5249 | 6125 | 5647 | 6147 | 5732 | Tbcd |
| Tbce | 2703 | 2277 | 2350 | 2595 | 3053 | 3049 | 3162 | 6868 | 5738 | 6099 | 5967 | 6137 | 6132 | 6596 | 2996 | 2290 | 2476 | 2675 | 2950 | 3227 | 3181 | 6741 | 5521 | 5696 | 6588 | 5417 | 6629 | 6127 | Tbce |
| Tbcel | 3412 | 3730 | 3857 | 4075 | 3733 | 3500 | 3652 | 2966 | 2968 | 3339 | 3265 | 3597 | 3728 | 3464 | 3509 | 3815 | 4079 | 4001 | 3640 | 3509 | 3618 | 3415 | 3064 | 3378 | 3525 | 3250 | 3430 | 3513 | Tbcel |
| Tbck | 3135 | 2917 | 2804 | 2708 | 2601 | 2805 | 2890 | 3551 | 3573 | 3465 | 3539 | 3252 | 3153 | 3200 | 3173 | 3023 | 2826 | 2826 | 2449 | 2694 | 2787 | 3691 | 3408 | 3326 | 3571 | 2844 | 3335 | 3304 | Tbck |
| Tbk1 | 3579 | 3753 | 3283 | 3869 | 3403 | 3507 | 3755 | 3665 | 3511 | 3736 | 3417 | 3831 | 3587 | 4035 | 3649 | 3724 | 3473 | 3868 | 3324 | 3800 | 3680 | 3252 | 3563 | 3457 | 3763 | 3380 | 3686 | 3790 | Tbk1 |
| Tbkbp1 | 1047 | 1414 | 1340 | 1597 | 2212 | 1722 | 1682 | 2756 | 3129 | 3914 | 3973 | 3065 | 2944 | 2571 | 1046 | 1185 | 1679 | 1841 | 2200 | 1616 | 1673 | 3723 | 3120 | 3467 | 3693 | 2983 | 2551 | 2536 | Tbkbp1 |
| Tbl1x | 10183 | 8342 | 8851 | 7602 | 8371 | 9073 | 8859 | 14090 | 13502 | 13439 | 14042 | 15775 | 14579 | 15330 | 9356 | 8669 | 8168 | 7650 | 8405 | 7921 | 8463 | 13272 | 13263 | 13633 | 13793 | 13984 | 15265 | 15333 | Tbl1x |
| Tbl1xr1 | 3463 | 2410 | 4039 | 4047 | 3821 | 3288 | 3553 | 4832 | 4522 | 5116 | 5180 | 5110 | 5193 | 5036 | 3260 | 3148 | 3935 | 3811 | 3394 | 2963 | 3405 | 5559 | 4554 | 5337 | 5293 | 6501 | 5012 | 5261 | Tbl1xr1 |
| Tbl2 | 7630 | 7930 | 6128 | 7293 | 5711 | 6430 | 5873 | 3029 | 3222 | 2927 | 3632 | 3262 | 2936 | 3196 | 7399 | 7557 | 6358 | 7156 | 5872 | 5854 | 6285 | 3048 | 3119 | 3228 | 3090 | 3314 | 3056 | 3302 | Tbl2 |
| Tbl3 | 5357 | 6661 | 5161 | 5157 | 4282 | 4683 | 4295 | 3202 | 3694 | 3405 | 3010 | 4260 | 3984 | 3498 | 4883 | 5143 | 5076 | 4921 | 5219 | 4936 | 4307 | 2787 | 3271 | 3697 | 3620 | 4794 | 4472 | 4025 | Tbl3 |
| Tbp | 5693 | 5801 | 5087 | 5476 | 5033 | 5149 | 5546 | 4945 | 3957 | 4164 | 3581 | 4600 | 4157 | 4558 | 5799 | 5132 | 5179 | 5236 | 5081 | 6291 | 5630 | 4091 | 3894 | 4011 | 4485 | 4477 | 4568 | 4352 | Tbp |
| Tbpl1 | 6029 | 4676 | 5762 | 5413 | 5841 | 5639 | 6083 | 6851 | 5479 | 5741 | 5480 | 6697 | 6543 | 7188 | 5930 | 4837 | 5541 | 5105 | 5728 | 6083 | 5845 | 7072 | 5481 | 6340 | 6832 | 6703 | 7181 | 7333 | Tbpl1 |
| Tbr1 | 1283 | 1036 | 1104 | 1655 | 2533 | 2212 | 3629 | 4892 | 6011 | 5190 | 3697 | 6619 | 2193 | 7471 | 1199 | 673 | 1687 | 1690 | 3204 | 2401 | 3005 | 3425 | 3393 | 7495 | 8303 | 7801 | 3777 | 5250 | Tbr1 |
| Tbrg1 | 6213 | 6388 | 5786 | 5550 | 4448 | 4934 | 5175 | 4379 | 5248 | 4385 | 5266 | 4839 | 5422 | 5244 | 6207 | 6344 | 5274 | 5148 | 4822 | 4763 | 4868 | 5125 | 5589 | 4732 | 4593 | 5336 | 5258 | 5597 | Tbrg1 |
| Tbrg4 | 2714 | 2949 | 3139 | 2806 | 2616 | 2633 | 2539 | 2640 | 2992 | 3009 | 3063 | 3809 | 3565 | 3532 | 2400 | 2482 | 2797 | 2463 | 2848 | 2379 | 2586 | 3006 | 2818 | 3286 | 3273 | 3998 | 3675 | 3809 | Tbrg4 |
| Tbx1 | 140 | 130 | 119 | 112 | 71 | 72 | 80 | 154 | 344 | 148 | 462 | 80 | 163 | 69 | 129 | 224 | 85 | 89 | 78 | 56 | 40 | 169 | 363 | 155 | 97 | 149 | 89 | 71 | Tbx1 |
| Tbx10 | 0 | 1 | 0 | 0 | 0 | 3 | 0 | 4 | 0 | 12 | 2 | 0 | 18 | 1 | 17 | 7 | 0 | 6 | 0 | 0 | 0 | 10 | 3 | 0 | 0 | 0 | 3 | 1 | Tbx10 |
| Tbx15 | 1798 | 1490 | 1403 | 825 | 257 | 1369 | 180 | 1231 | 2759 | 1514 | 2896 | 1498 | 2520 | 1360 | 1891 | 2178 | 1058 | 1268 | 352 | 217 | 166 | 1216 | 2989 | 1791 | 1128 | 1607 | 1476 | 1495 | Tbx15 |
| Tbx18 | 1234 | 824 | 1608 | 777 | 831 | 1020 | 863 | 1086 | 1594 | 1362 | 1447 | 1667 | 1963 | 1606 | 1158 | 1396 | 1101 | 868 | 747 | 923 | 819 | 1374 | 1647 | 1347 | 1316 | 1698 | 1751 | 1657 | Tbx18 |
| Tbx19 | 5 | 4 | 0 | 2 | 12 | 0 | 0 | 7 | 7 | 2 | 16 | 0 | 2 | 2 | 3 | 1 | 0 | 10 | 1 | 4 | 6 | 18 | 16 | 2 | 6 | 1 | 5 | 0 | Tbx19 |
| Tbx2 | 1029 | 304 | 245 | 281 | 359 | 174 | 125 | 1564 | 1354 | 1462 | 1182 | 377 | 719 | 276 | 384 | 565 | 195 | 228 | 237 | 134 | 192 | 834 | 1305 | 1075 | 462 | 1423 | 263 | 285 | Tbx2 |
| Tbx20 | 31 | 58 | 31 | 39 | 191 | 54 | 17 | 288 | 264 | 220 | 183 | 23 | 456 | 15 | 276 | 132 | 41 | 37 | 4 | 18 | 26 | 321 | 144 | 17 | 31 | 20 | 13 | 11 | Tbx20 |
| Tbx21 | 46 | 55 | 106 | 70 | 37 | 27 | 39 | 7 | 12 | 8 | 7 | 11 | 5 | 19 | 41 | 45 | 115 | 28 | 40 | 42 | 34 | 15 | 13 | 7 | 22 | 8 | 7 | 17 | Tbx21 |
| Tbx22 | 331 | 22 | 29 | 37 | 5 | 12 | 3 | 642 | 560 | 280 | 409 | 85 | 88 | 63 | 33 | 138 | 12 | 4 | 4 | 1 | 3 | 43 | 383 | 368 | 43 | 667 | 28 | 45 | Tbx22 |
| Tbx3 | 880 | 239 | 131 | 239 | 442 | 102 | 108 | 1192 | 1065 | 856 | 988 | 152 | 523 | 132 | 290 | 424 | 178 | 294 | 91 | 112 | 110 | 847 | 1075 | 573 | 419 | 704 | 88 | 128 | Tbx3 |
| Tbx4 | 47 | 24 | 18 | 16 | 9 | 24 | 10 | 76 | 89 | 73 | 117 | 35 | 50 | 36 | 26 | 47 | 30 | 15 | 8 | 21 | 13 | 35 | 76 | 57 | 27 | 60 | 20 | 23 | Tbx4 |
| Tbx5 | 9 | 0 | 0 | 0 | 0 | 0 | 1 | 11 | 15 | 16 | 73 | 15 | 5 | 18 | 0 | 2 | 3 | 7 | 0 | 0 | 0 | 5 | 134 | 11 | 3 | 16 | 15 | 15 | Tbx5 |
| Tbx6 | 107 | 193 | 177 | 214 | 174 | 156 | 163 | 51 | 107 | 108 | 87 | 116 | 110 | 105 | 112 | 181 | 202 | 235 | 178 | 154 | 172 | 89 | 74 | 154 | 98 | 76 | 65 | 85 | Tbx6 |
| Tbxa2r | 43 | 60 | 54 | 78 | 76 | 95 | 70 | 68 | 103 | 128 | 119 | 97 | 90 | 78 | 62 | 51 | 68 | 79 | 79 | 62 | 82 | 90 | 114 | 117 | 155 | 109 | 105 | 81 | Tbxa2r |
| Tbxas1 | 76 | 58 | 52 | 56 | 55 | 74 | 54 | 112 | 120 | 135 | 134 | 141 | 187 | 101 | 79 | 83 | 52 | 74 | 58 | 69 | 42 | 135 | 173 | 103 | 127 | 125 | 157 | 134 | Tbxas1 |
| Tc2n | 102 | 57 | 74 | 98 | 72 | 63 | 53 | 44 | 51 | 38 | 68 | 10 | 24 | 37 | 105 | 93 | 85 | 77 | 54 | 56 | 63 | 48 | 69 | 56 | 27 | 35 | 10 | 17 | Tc2n |
| Tcaf1 | 22423 | 22060 | 20857 | 23826 | 29794 | 28391 | 28546 | 38008 | 31410 | 33851 | 30475 | 34168 | 31967 | 34676 | 24965 | 20526 | 23947 | 24796 | 27976 | 28482 | 28603 | 38621 | 30783 | 33073 | 37012 | 27099 | 34987 | 33019 | Tcaf1 |

| | | | | | | | | | | | | | | | | | | | | | | | | | | | | | |
|---|---|---|---|---|---|---|---|---|---|---|---|---|---|---|---|---|---|---|---|---|---|---|---|---|---|---|---|---|---|
| Tcaf2 | 2070 | 2049 | 2034 | 1381 | 1238 | 1381 | 1196 | 666 | 658 | 563 | 864 | 838 | 1009 | 709 | 1994 | 2216 | 1548 | 1301 | 1230 | 1234 | 1113 | 551 | 743 | 533 | 533 | 768 | 897 | 741 | Tcaf2 |
| Tcaf3 | 0 | 0 | 0 | 0 | 0 | 0 | 0 | 0 | 1 | 0 | 0 | 0 | 0 | 0 | 3 | 0 | 0 | 0 | 0 | 1 | 0 | 0 | 0 | 3 | 0 | 0 | 1 | 0 | Tcaf3 |
| Tcaim | 601 | 580 | 659 | 642 | 667 | 666 | 746 | 793 | 754 | 774 | 787 | 744 | 799 | 753 | 597 | 584 | 703 | 648 | 576 | 628 | 700 | 735 | 805 | 816 | 702 | 801 | 794 | 756 | Tcaim |
| Tcam1 | 29 | 23 | 34 | 34 | 16 | 22 | 26 | 40 | 41 | 30 | 41 | 41 | 39 | 20 | 32 | 32 | 27 | 30 | 21 | 23 | 21 | 22 | 23 | 30 | 20 | 38 | 20 | 18 | Tcam1 |
| Tcap | 17 | 35 | 36 | 38 | 28 | 21 | 36 | 18 | 18 | 32 | 78 | 14 | 26 | 23 | 22 | 22 | 46 | 41 | 20 | 20 | 27 | 33 | 67 | 15 | 13 | 24 | 16 | 12 | Tcap |
| Tcea1 | 8310 | 6125 | 7721 | 7018 | 6686 | 7429 | 8320 | 8485 | 7214 | 7023 | 7025 | 7771 | 7473 | 8010 | 8636 | 7784 | 6934 | 7049 | 7109 | 8113 | 7691 | 7005 | 7767 | 7241 | 7289 | 8540 | 7778 | 8119 | Tcea1 |
| Tcea2 | 759 | 733 | 739 | 792 | 1091 | 921 | 999 | 1226 | 1284 | 1215 | 1206 | 1279 | 1156 | 1219 | 762 | 703 | 765 | 813 | 1116 | 955 | 932 | 1450 | 1243 | 1340 | 1440 | 1205 | 1121 | 1099 | Tcea2 |
| Tcea3 | 239 | 268 | 285 | 203 | 152 | 176 | 91 | 83 | 157 | 120 | 208 | 108 | 163 | 100 | 290 | 361 | 248 | 212 | 126 | 126 | 88 | 145 | 231 | 120 | 117 | 114 | 98 | 115 | Tcea3 |
| Tceal1 | 693 | 574 | 649 | 516 | 818 | 711 | 876 | 1503 | 1228 | 1163 | 1179 | 1239 | 1196 | 1296 | 748 | 570 | 692 | 602 | 868 | 983 | 894 | 1312 | 1290 | 1232 | 1195 | 1239 | 1247 | 1365 | Tceal1 |
| Tceal3 | 1419 | 1876 | 2212 | 2097 | 2215 | 1938 | 2219 | 1269 | 1386 | 1647 | 1729 | 1351 | 1430 | 1652 | 1662 | 1494 | 2285 | 2071 | 2282 | 2035 | 2110 | 1519 | 1431 | 1597 | 1794 | 1373 | 1293 | 1335 | Tceal3 |
| Tceal5 | 1113 | 1321 | 1513 | 1862 | 1699 | 1538 | 1867 | 613 | 658 | 1079 | 984 | 647 | 652 | 805 | 1222 | 1186 | 1563 | 1655 | 1592 | 1573 | 1818 | 697 | 795 | 868 | 918 | 694 | 558 | 689 | Tceal5 |
| Tceal6 | 886 | 1053 | 1096 | 1346 | 1412 | 1238 | 1544 | 498 | 579 | 811 | 808 | 414 | 445 | 554 | 886 | 738 | 1358 | 1368 | 1288 | 1271 | 1581 | 585 | 649 | 668 | 720 | 366 | 413 | 415 | Tceal6 |
| Tceal7 | 27 | 10 | 23 | 9 | 35 | 23 | 23 | 101 | 122 | 47 | 1268 | 51 | 36 | 50 | 33 | 29 | 28 | 22 | 24 | 25 | 24 | 104 | 733 | 43 | 45 | 30 | 19 | 42 | Tceal7 |
| Tceal8 | 4403 | 3472 | 4170 | 3491 | 3713 | 3680 | 4139 | 6589 | 5835 | 5158 | 6019 | 5474 | 6090 | 6346 | 4894 | 3874 | 3992 | 3323 | 3960 | 5049 | 3770 | 5419 | 6226 | 5709 | 5100 | 6606 | 6272 | 6572 | Tceal8 |
| Tceal9 | 14497 | 11317 | 14138 | 11832 | 10604 | 11409 | 13914 | 14566 | 13185 | 11344 | 13029 | 13029 | 13159 | 15833 | 14944 | 12428 | 13201 | 11244 | 11561 | 15019 | 12684 | 11165 | 13925 | 14461 | 12069 | 16282 | 13476 | 15974 | Tceal9 |
| Tceanc | 871 | 615 | 826 | 674 | 622 | 660 | 715 | 938 | 817 | 829 | 910 | 1111 | 976 | 1018 | 874 | 727 | 862 | 627 | 616 | 694 | 651 | 856 | 858 | 924 | 918 | 1187 | 1044 | 1067 | Tceanc |
| Tceanc2 | 1765 | 1831 | 1728 | 2073 | 2125 | 1818 | 2239 | 2417 | 2402 | 2488 | 2540 | 2168 | 2189 | 2330 | 1679 | 1712 | 1852 | 2035 | 2144 | 1950 | 2195 | 2237 | 2178 | 2397 | 2533 | 2062 | 2214 | 2281 | Tceanc2 |
| Tcerg1 | 13027 | 10146 | 13370 | 12776 | 13543 | 13858 | 15051 | 13982 | 13273 | 14521 | 14011 | 17040 | 15957 | 18023 | 12575 | 10970 | 12918 | 12367 | 13869 | 12157 | 14174 | 14616 | 12096 | 15547 | 15527 | 17105 | 16819 | 17912 | Tcerg1 |
| Tcerg1l | 178 | 244 | 336 | 369 | 721 | 352 | 374 | 453 | 470 | 689 | 825 | 381 | 475 | 386 | 287 | 301 | 381 | 390 | 447 | 361 | 446 | 872 | 559 | 507 | 620 | 260 | 285 | 385 | Tcerg1l |
| Tcf12 | 28575 | 26325 | 24878 | 20277 | 19811 | 22318 | 19263 | 30283 | 26097 | 23840 | 23890 | 27953 | 30620 | 27393 | 29700 | 27452 | 22907 | 19201 | 19387 | 22922 | 19477 | 23788 | 26240 | 23556 | 22389 | 25059 | 31403 | 30495 | Tcf12 |
| Tcf15 | 209 | 214 | 189 | 229 | 114 | 126 | 142 | 37 | 44 | 67 | 70 | 48 | 90 | 48 | 179 | 220 | 209 | 149 | 141 | 104 | 121 | 53 | 54 | 54 | 66 | 82 | 45 | 46 | Tcf15 |
| Tcf19 | 5182 | 6030 | 4943 | 4629 | 3229 | 3839 | 4377 | 1713 | 1991 | 1466 | 1678 | 2284 | 1810 | 2255 | 4725 | 5060 | 4792 | 4566 | 3939 | 4370 | 3970 | 1145 | 1769 | 1926 | 1761 | 2671 | 2264 | 2273 | Tcf19 |
| Tcf20 | 13321 | 12781 | 10414 | 13185 | 12724 | 13630 | 14142 | 10723 | 10645 | 11197 | 10454 | 10812 | 9965 | 11364 | 12443 | 11597 | 11906 | 13193 | 12669 | 12753 | 14178 | 10301 | 9655 | 10979 | 12286 | 7818 | 10537 | 10567 | Tcf20 |
| Tcf21 | 8 | 7 | 3 | 6 | 11 | 0 | 0 | 5 | 7 | 7 | 44 | 2 | 9 | 2 | 1 | 9 | 0 | 7 | 0 | 2 | 2 | 17 | 17 | 1 | 4 | 8 | 0 | 0 | Tcf21 |
| Tcf23 | 0 | 4 | 0 | 4 | 2 | 1 | 0 | 0 | 0 | 2 | 5 | 1 | 1 | 0 | 0 | 0 | 0 | 4 | 2 | 0 | 3 | 2 | 0 | 0 | 0 | 0 | 0 | 0 | Tcf23 |
| Tcf24 | 120 | 234 | 148 | 186 | 236 | 225 | 105 | 221 | 142 | 144 | 430 | 18 | 269 | 21 | 303 | 360 | 181 | 205 | 104 | 83 | 103 | 227 | 171 | 25 | 120 | 39 | 22 | 15 | Tcf24 |
| Tcf25 | 14391 | 16795 | 14365 | 15778 | 18216 | 16427 | 17273 | 16972 | 18280 | 17497 | 17162 | 16944 | 16827 | 16842 | 15008 | 14239 | 15811 | 16143 | 17585 | 16626 | 17451 | 18489 | 17968 | 17441 | 18613 | 16229 | 16719 | 16770 | Tcf25 |
| Tcf3 | 7934 | 9400 | 10400 | 8454 | 7668 | 7441 | 6436 | 9377 | 11533 | 11559 | 11093 | 15140 | 12919 | 12888 | 6782 | 8073 | 9895 | 7535 | 8757 | 6782 | 6693 | 9301 | 10459 | 13776 | 11335 | 18115 | 13672 | 13283 | Tcf3 |
| Tcf4 | 17553 | 12847 | 14955 | 19851 | 21122 | 22565 | 24913 | 28953 | 31087 | 30492 | 29468 | 28932 | 28210 | 30037 | 16905 | 15400 | 16886 | 19070 | 21990 | 22118 | 23455 | 23688 | 29627 | 32766 | 34447 | 27984 | 26005 | 26366 | Tcf4 |
| Tcf7 | 2102 | 2128 | 3259 | 2144 | 1773 | 1788 | 1630 | 651 | 601 | 710 | 727 | 841 | 773 | 692 | 1787 | 2408 | 2367 | 2053 | 1974 | 1570 | 1763 | 708 | 727 | 748 | 701 | 819 | 737 | 806 | Tcf7 |
| Tcf7l1 | 2482 | 2932 | 3220 | 2770 | 1906 | 2087 | 1635 | 1914 | 2634 | 2477 | 2767 | 2842 | 2460 | 2374 | 2076 | 2927 | 3011 | 2617 | 2360 | 1591 | 1738 | 1875 | 2448 | 3236 | 1938 | 4056 | 2331 | 2315 | Tcf7l1 |
| Tcf7l2 | 3405 | 3312 | 8814 | 6638 | 7130 | 6601 | 9598 | 8802 | 4296 | 12508 | 9460 | 16989 | 8873 | 17954 | 3700 | 2446 | 7225 | 7009 | 9593 | 5689 | 9472 | 18098 | 6551 | 18460 | 17305 | 14981 | 16426 | 19504 | Tcf7l2 |
| Tcfl5 | 431 | 429 | 447 | 351 | 365 | 334 | 349 | 85 | 115 | 85 | 121 | 68 | 88 | 49 | 373 | 469 | 323 | 385 | 289 | 365 | 300 | 104 | 102 | 67 | 87 | 60 | 63 | 62 | Tcfl5 |
| Tchh | 399 | 328 | 545 | 443 | 402 | 495 | 450 | 81 | 122 | 125 | 157 | 114 | 178 | 132 | 386 | 353 | 518 | 385 | 422 | 422 | 448 | 109 | 141 | 146 | 131 | 92 | 145 | 171 | Tchh |
| Tchp | 2952 | 3871 | 3885 | 4385 | 3361 | 3443 | 3481 | 1697 | 1832 | 2315 | 2113 | 2467 | 2346 | 2683 | 2946 | 3530 | 4062 | 4127 | 3519 | 2971 | 3544 | 2050 | 1672 | 2512 | 2284 | 2631 | 2694 | 2595 | Tchp |
| Tcim | 1111 | 821 | 819 | 699 | 671 | 617 | 684 | 749 | 852 | 762 | 891 | 501 | 643 | 531 | 804 | 1101 | 687 | 643 | 506 | 638 | 628 | 565 | 995 | 769 | 560 | 886 | 435 | 441 | Tcim |
| Tcirg1 | 2531 | 2707 | 2321 | 1790 | 1666 | 1728 | 1300 | 1130 | 1498 | 1358 | 1768 | 1152 | 1410 | 979 | 1928 | 3232 | 2079 | 2250 | 1528 | 1455 | 1348 | 1258 | 1630 | 1316 | 1088 | 1349 | 1074 | 1021 | Tcirg1 |
| Tcl1 | 0 | 0 | 0 | 0 | 0 | 0 | 0 | 0 | 0 | 0 | 0 | 0 | 1 | 0 | 0 | 0 | 0 | 0 | 0 | 0 | 0 | 0 | 0 | 0 | 0 | 0 | 0 | 0 | Tcl1 |
| Tcl1b1 | 0 | 2 | 0 | 14 | 0 | 1 | 4 | 0 | 0 | 0 | 0 | 4 | 1 | 0 | 1 | 3 | 0 | 2 | 1 | 4 | 4 | 2 | 0 | 0 | 1 | 0 | 5 | 0 | Tcl1b1 |
| Tcl1b2 | 10 | 9 | 1 | 2 | 3 | 1 | 7 | 1 | 2 | 0 | 0 | 1 | 2 | 0 | 4 | 5 | 5 | 7 | 0 | 6 | 8 | 0 | 1 | 1 | 1 | 4 | 2 | 0 | Tcl1b2 |
| Tcl1b3 | 0 | 0 | 0 | 0 | 0 | 0 | 0 | 0 | 1 | 0 | 0 | 0 | 0 | 0 | 0 | 0 | 0 | 0 | 0 | 0 | 0 | 0 | 0 | 0 | 0 | 0 | 0 | 0 | Tcl1b3 |
| Tcl1b5 | 0 | 0 | 0 | 1 | 0 | 0 | 0 | 0 | 0 | 0 | 0 | 0 | 0 | 0 | 0 | 0 | 0 | 0 | 0 | 0 | 0 | 0 | 0 | 0 | 0 | 0 | 0 | 0 | Tcl1b5 |
| Tcn2 | 2204 | 2663 | 2211 | 1862 | 1777 | 1941 | 1551 | 1342 | 1789 | 1461 | 1654 | 1437 | 1906 | 1408 | 2167 | 2774 | 1895 | 1951 | 1764 | 1842 | 1522 | 1489 | 2047 | 1325 | 1208 | 1526 | 1590 | 1542 | Tcn2 |
| Tcof1 | 4142 | 5325 | 7039 | 5644 | 4403 | 4466 | 3932 | 3315 | 4649 | 5358 | 4999 | 6347 | 5096 | 5590 | 3861 | 5101 | 6522 | 5519 | 5374 | 3699 | 4104 | 4299 | 4434 | 5859 | 4781 | 7451 | 5252 | 5447 | Tcof1 |
| Tcp1 | 37272 | 34007 | 33226 | 28508 | 26716 | 28261 | 32551 | 38085 | 32180 | 28350 | 27948 | 34132 | 32241 | 34327 | 39146 | 33370 | 31283 | 27927 | 29828 | 34672 | 30789 | 30338 | 31844 | 30841 | 29552 | 39281 | 35705 | 37676 | Tcp1 |
| Tcp10a | 30 | 33 | 22 | 40 | 32 | 54 | 61 | 0 | 0 | 5 | 1 | 3 | 1 | 1 | 34 | 30 | 27 | 55 | 40 | 27 | 47 | 2 | 5 | 1 | 8 | 0 | 2 | 0 | Tcp10a |
| Tcp10b | 53 | 47 | 60 | 90 | 51 | 47 | 66 | 5 | 6 | 10 | 19 | 3 | 14 | 1 | 57 | 41 | 61 | 62 | 38 | 30 | 59 | 22 | 10 | 4 | 8 | 8 | 7 | 2 | Tcp10b |
| Tcp10c | 2 | 2 | 0 | 2 | 2 | 0 | 2 | 5 | 7 | 3 | 0 | 0 | 0 | 0 | 4 | 2 | 1 | 4 | 1 | 2 | 1 | 0 | 1 | 0 | 5 | 0 | 0 | 1 | Tcp10c |
| Tcp11 | 315 | 400 | 476 | 458 | 424 | 374 | 336 | 93 | 85 | 125 | 111 | 121 | 126 | 114 | 306 | 423 | 438 | 478 | 450 | 331 | 384 | 95 | 117 | 127 | 109 | 112 | 95 | 107 | Tcp11 |
| Tcp11l1 | 1651 | 1676 | 846 | 1014 | 1327 | 1784 | 1306 | 2245 | 2217 | 1769 | 1966 | 1565 | 1616 | 1722 | 1577 | 1561 | 1041 | 1136 | 1082 | 1615 | 1595 | 1424 | 1906 | 1369 | 1875 | 1070 | 1809 | 1575 | Tcp11l1 |
| Tcp11l2 | 3156 | 3632 | 2636 | 3425 | 3131 | 2914 | 3390 | 1171 | 1151 | 1066 | 1187 | 783 | 1275 | 907 | 3239 | 3119 | 3023 | 3451 | 3003 | 3079 | 3394 | 1260 | 1257 | 986 | 1030 | 750 | 918 | 964 | Tcp11l2 |
| Tcp11x2 | 0 | 0 | 0 | 0 | 0 | 2 | 0 | 0 | 0 | 1 | 0 | 0 | 0 | 0 | 0 | 0 | 0 | 0 | 0 | 2 | 0 | 0 | 0 | 0 | 0 | 0 | 0 | 0 | Tcp11x2 |
| Tcta | 940 | 993 | 833 | 817 | 920 | 920 | 879 | 1465 | 1528 | 1496 | 1430 | 1309 | 1376 | 1454 | 1117 | 974 | 778 | 790 | 813 | 934 | 799 | 1596 | 1567 | 1226 | 1371 | 1372 | 1440 | 1385 | Tcta |
| Tcte1 | 13 | 29 | 19 | 20 | 55 | 39 | 26 | 31 | 35 | 21 | 29 | 50 | 54 | 18 | 20 | 27 | 31 | 11 | 35 | 30 | 28 | 18 | 49 | 35 | 61 | 41 | 22 | 33 | Tcte1 |
| Tcte2 | 200 | 219 | 203 | 258 | 306 | 313 | 326 | 353 | 386 | 427 | 397 | 427 | 384 | 464 | 237 | 201 | 262 | 245 | 249 | 311 | 308 | 419 | 346 | 364 | 490 | 299 | 388 | 351 | Tcte2 |
| Tctn1 | 3926 | 4947 | 5225 | 5640 | 4781 | 4821 | 4717 | 2648 | 2848 | 3393 | 3298 | 3670 | 3648 | 3695 | 3823 | 4608 | 5393 | 5267 | 5308 | 4419 | 4916 | 3447 | 2778 | 3535 | 3490 | 3212 | 3733 | 3792 | Tctn1 |
| Tctn3 | 3058 | 3372 | 1959 | 2819 | 2216 | 2696 | 2661 | 1440 | 1441 | 1374 | 1317 | 1424 | 1330 | 1337 | 2871 | 3024 | 2227 | 2797 | 2198 | 2582 | 2591 | 1338 | 1432 | 1206 | 1344 | 1124 | 1418 | 1316 | Tctn3 |
| Tdg | 6802 | 7125 | 8102 | 7050 | 6783 | 7496 | 6729 | 9484 | 9447 | 9824 | 9416 | 13207 | 12443 | 12369 | 6795 | 6849 | 7631 | 6733 | 7424 | 6921 | 6351 | 9386 | 8745 | 10909 | 11489 | 13923 | 13237 | 12844 | Tdg |
| Tdgf1 | 2 | 3 | 4 | 0 | 7 | 3 | 0 | 4 | 2 | 4 | 8 | 1 | 7 | 2 | 3 | 1 | 2 | 0 | 0 | 3 | 2 | 5 | 3 | 1 | 1 | 0 | 0 | 6 | Tdgf1 |
| Tdh | 0 | 2 | 0 | 0 | 2 | 3 | 1 | 4 | 4 | 8 | 10 | 4 | 11 | 3 | 0 | 1 | 0 | 0 | 4 | 0 | 5 | 5 | 11 | 1 | 4 | 6 | 1 | 2 | Tdh |
| Tdo2 | 2 | 0 | 0 | 0 | 2 | 3 | 2 | 4 | 0 | 5 | 0 | 1 | 1 | 2 | 0 | 0 | 2 | 0 | 4 | 4 | 5 | 2 | 5 | 2 | 0 | 0 | 0 | 0 | Tdo2 |
| Tdp1 | 3988 | 4147 | 3429 | 3640 | 3329 | 3747 | 3787 | 3121 | 3085 | 2832 | 2740 | 3306 | 3102 | 3404 | 3703 | 3403 | 3500 | 3626 | 3751 | 3727 | 3848 | 2757 | 2794 | 3092 | 3028 | 3719 | 3521 | 3272 | Tdp1 |
| Tdp2 | 5743 | 4152 | 5519 | 5198 | 4228 | 4508 | 5145 | 4500 | 3919 | 3956 | 3882 | 4901 | 4369 | 5122 | 5470 | 5116 | 4957 | 4779 | 4354 | 4924 | 4748 | 4049 | 3809 | 4426 | 4077 | 5447 | 4756 | 5204 | Tdp2 |
| Tdrd1 | 212 | 230 | 130 | 123 | 147 | 222 | 102 | 112 | 234 | 204 | 428 | 135 | 211 | 139 | 179 | 252 | 114 | 160 | 62 | 107 | 80 | 144 | 365 | 97 | 146 | 65 | 106 | 78 | Tdrd1 |
| Tdrd12 | 28 | 27 | 25 | 42 | 38 | 25 | 29 | 20 | 38 | 24 | 18 | 20 | 25 | 16 | 36 | 27 | 12 | 33 | 44 | 27 | 44 | 27 | 25 | 16 | 27 | 10 | 15 | 14 | Tdrd12 |
| Tdrd3 | 2902 | 2272 | 2447 | 2427 | 2903 | 2712 | 2853 | 4842 | 4870 | 4548 | 4602 | 4422 | 4332 | 4622 | 2838 | 2499 | 2275 | 2488 | 2509 | 2873 | 2715 | 4597 | 4905 | 4496 | 4444 | 4813 | 4415 | 4614 | Tdrd3 |
| Tdrd5 | 35 | 30 | 27 | 23 | 46 | 36 | 37 | 76 | 57 | 63 | 49 | 41 | 41 | 36 | 47 | 61 | 38 | 50 | 25 | 53 | 42 | 57 | 65 | 42 | 68 | 42 | 30 | 40 | Tdrd5 |
| Tdrd6 | 473 | 456 | 171 | 591 | 448 | 603 | 640 | 73 | 88 | 101 | 71 | 71 | 99 | 98 | 447 | 440 | 328 | 654 | 421 | 498 | 738 | 90 | 71 | 49 | 105 | 25 | 46 | 57 | Tdrd6 |
| Tdrd7 | 2838 | 3098 | 2704 | 2432 | 2704 | 2690 | 2547 | 1423 | 1528 | 1347 | 1590 | 1325 | 1437 | 1249 | 3065 | 2720 | 2726 | 2436 | 2533 | 2789 | 2417 | 1648 | 1637 | 1257 | 1436 | 1217 | 1292 | 1208 | Tdrd7 |
| Tdrd9 | 93 | 98 | 120 | 77 | 89 | 71 | 70 | 114 | 111 | 110 | 121 | 129 | 155 | 134 | 66 | 98 | 110 | 106 | 70 | 129 | 83 | 112 | 98 | 118 | 107 | 210 | 137 | 148 | Tdrd9 |
| Tdrkh | 6342 | 6963 | 6750 | 7958 | 8337 | 7941 | 9770 | 6792 | 5061 | 5440 | 4615 | 5839 | 5054 | 6283 | 7961 | 5655 | 7743 | 7827 | 8394 | 8652 | 8932 | 6376 | 4983 | 5485 | 6035 | 4936 | 5836 | 5978 | Tdrkh |
| Tdrp | 867 | 991 | 1032 | 1016 | 845 | 911 | 832 | 318 | 381 | 321 | 291 | 483 | 473 | 443 | 877 | 1028 | 933 | 900 | 851 | 867 | 866 | 379 | 504 | 401 | 433 | 440 | 449 | 530 | Tdrp |
| Tead1 | 13340 | 11550 | 12234 | 13703 | 12413 | 13238 | 13788 | 10437 | 10743 | 12322 | 12620 | 14020 | 13761 | 13812 | 12232 | 13618 | 12459 | 13376 | 11963 | 9765 | 13426 | 10477 | 11119 | 11671 | 10938 | 10983 | 13330 | 13241 | Tead1 |
| Tead2 | 6867 | 7368 | 8501 | 5785 | 5115 | 5798 | 5558 | 11580 | 12356 | 12449 | 12095 | 17531 | 15079 | 17471 | 5834 | 6163 | 8278 | 5034 | 6632 | 5436 | 5643 | 9641 | 11802 | 15875 | 12239 | 20373 | 17426 | 18691 | Tead2 |
| Tead3 | 3878 | 4722 | 4756 | 4396 | 3758 | 3756 | 3739 | 1939 | 2506 | 2293 | 2716 | 2367 | 2740 | 2152 | 3312 | 4170 | 4726 | 4122 | 4303 | 3146 | 3403 | 1681 | 2531 | 2633 | 1695 | 3087 | 2348 | 2127 | Tead3 |
| Tead4 | 203 | 217 | 225 | 144 | 102 | 161 | 74 | 177 | 248 | 181 | 484 | 225 | 227 | 170 | 228 | 253 | 156 | 142 | 108 | 72 | 77 | 174 | 346 | 208 | 121 | 182 | 205 | 196 | Tead4 |
| Tec | 794 | 910 | 824 | 835 | 785 | 721 | 673 | 374 | 342 | 405 | 518 | 362 | 497 | 351 | 926 | 1060 | 625 | 949 | 563 | 853 | 724 | 376 | 640 | 394 | 346 | 311 | 390 | 345 | Tec |
| Tecpr1 | 5158 | 6927 | 7222 | 8963 | 8211 | 7682 | 6907 | 3654 | 4845 | 5540 | 5341 | 5214 | 4620 | 4474 | 5058 | 6652 | 7983 | 9145 | 8811 | 6212 | 7368 | 5653 | 4548 | 5351 | 6009 | 4480 | 4477 | 4218 | Tecpr1 |
| Tecpr2 | 3340 | 3833 | 3186 | 4414 | 4294 | 3786 | 3739 | 2622 | 2950 | 3343 | 3208 | 3100 | 3068 | 2871 | 3178 | 3857 | 3868 | 4472 | 4216 | 3437 | 3855 | 3302 | 2771 | 3277 | 3333 | 2110 | 2866 | 2822 | Tecpr2 |

| | | | | | | | | | | | | | | | | | | | | | | | | | | | | | |
|---|---|---|---|---|---|---|---|---|---|---|---|---|---|---|---|---|---|---|---|---|---|---|---|---|---|---|---|---|---|
| Tecr | 13025 | 18307 | 15543 | 16112 | 15234 | 13535 | 15292 | 12863 | 13621 | 13725 | 11988 | 13059 | 13041 | 13917 | 13872 | 14109 | 16048 | 15311 | 16506 | 15234 | 16100 | 13164 | 13470 | 13255 | 13212 | 14246 | 13132 | 14310 | Tecr |
| Tecrl | 5 | 2 | 0 | 4 | 4 | 8 | 3 | 13 | 41 | 16 | 129 | 16 | 17 | 11 | 8 | 5 | 0 | 2 | 2 | 3 | 0 | 47 | 97 | 11 | 11 | 18 | 23 | 23 | Tecrl |
| Tecta | 91 | 124 | 62 | 149 | 99 | 100 | 73 | 125 | 198 | 283 | 423 | 99 | 206 | 56 | 105 | 160 | 70 | 119 | 69 | 90 | 86 | 222 | 189 | 87 | 101 | 82 | 113 | 74 | Tecta |
| Tectb | 5 | 18 | 6 | 11 | 2 | 6 | 7 | 13 | 14 | 55 | 94 | 18 | 14 | 9 | 13 | 4 | 12 | 12 | 2 | 15 | 5 | 48 | 16 | 19 | 20 | 12 | 8 | 20 | Tectb |
| Tedc1 | 3742 | 4504 | 4135 | 3940 | 2960 | 3014 | 3152 | 1500 | 1767 | 1617 | 1765 | 2414 | 2265 | 2107 | 3228 | 4034 | 3712 | 3593 | 3325 | 2810 | 3047 | 1377 | 1596 | 2175 | 1678 | 3073 | 2401 | 2500 | Tedc1 |
| Tedc2 | 2119 | 2560 | 2314 | 2166 | 1624 | 1742 | 1831 | 1235 | 1367 | 1349 | 1390 | 1603 | 1505 | 1653 | 1982 | 2263 | 2314 | 2093 | 1852 | 1991 | 1834 | 1176 | 1286 | 1555 | 1370 | 1882 | 1693 | 1823 | Tedc2 |
| Teddm1b | 0 | 0 | 0 | 0 | 0 | 0 | 0 | 0 | 1 | 0 | 0 | 0 | 1 | 1 | 0 | 0 | 0 | 3 | 0 | 0 | 0 | 0 | 2 | 0 | 2 | 0 | 0 | 0 | Teddm1b |
| Teddm2 | 36 | 22 | 20 | 27 | 38 | 29 | 42 | 48 | 66 | 94 | 50 | 56 | 49 | 63 | 29 | 23 | 48 | 35 | 48 | 41 | 47 | 57 | 78 | 77 | 78 | 60 | 41 | 45 | Teddm2 |
| Teddm3 | 0 | 0 | 0 | 0 | 0 | 0 | 0 | 0 | 0 | 0 | 0 | 0 | 0 | 0 | 0 | 0 | 0 | 0 | 0 | 0 | 0 | 0 | 1 | 0 | 0 | 0 | 0 | 0 | Teddm3 |
| Tef | 6023 | 5810 | 4563 | 6341 | 5221 | 6099 | 5850 | 3998 | 4395 | 4217 | 4605 | 4167 | 4011 | 4148 | 5641 | 5689 | 5278 | 6214 | 5612 | 5264 | 5784 | 4406 | 4526 | 4108 | 4536 | 3522 | 4043 | 3993 | Tef |
| Tefm | 486 | 434 | 639 | 476 | 461 | 377 | 497 | 742 | 819 | 612 | 688 | 836 | 765 | 749 | 522 | 470 | 502 | 437 | 444 | 577 | 467 | 638 | 805 | 748 | 689 | 911 | 754 | 932 | Tefm |
| Tek | 1129 | 1064 | 1195 | 1036 | 1190 | 1166 | 1184 | 1594 | 1641 | 1719 | 1975 | 1611 | 1930 | 1761 | 1222 | 1283 | 962 | 1143 | 959 | 1271 | 1097 | 1800 | 2276 | 1678 | 1731 | 1710 | 1683 | 1622 | Tek |
| Tekt1 | 437 | 458 | 268 | 382 | 384 | 361 | 449 | 341 | 306 | 267 | 275 | 320 | 306 | 274 | 439 | 460 | 438 | 388 | 407 | 448 | 421 | 274 | 328 | 260 | 269 | 292 | 223 | 205 | Tekt1 |
| Tekt2 | 298 | 353 | 287 | 428 | 366 | 365 | 346 | 394 | 551 | 559 | 591 | 437 | 485 | 386 | 276 | 365 | 364 | 484 | 330 | 270 | 340 | 398 | 539 | 589 | 444 | 565 | 363 | 418 | Tekt2 |
| Tekt3 | 2 | 0 | 6 | 3 | 0 | 1 | 0 | 0 | 2 | 0 | 0 | 0 | 1 | 0 | 0 | 9 | 1 | 0 | 0 | 0 | 1 | 2 | 0 | 0 | 0 | 0 | 0 | 0 | Tekt3 |
| Tekt4 | 8 | 22 | 15 | 22 | 29 | 28 | 18 | 37 | 40 | 22 | 33 | 56 | 49 | 21 | 18 | 31 | 17 | 11 | 16 | 28 | 22 | 42 | 52 | 21 | 26 | 25 | 23 | 20 | Tekt4 |
| Tekt5 | 7 | 16 | 13 | 6 | 24 | 13 | 20 | 31 | 41 | 63 | 38 | 9 | 17 | 15 | 17 | 9 | 19 | 23 | 19 | 21 | 18 | 67 | 48 | 26 | 33 | 24 | 10 | 14 | Tekt5 |
| Telo2 | 2310 | 2746 | 2488 | 2743 | 2787 | 2739 | 2584 | 2809 | 3234 | 3347 | 3304 | 3522 | 3185 | 3179 | 2041 | 2459 | 2787 | 2894 | 2899 | 2436 | 2634 | 3228 | 2995 | 3402 | 3521 | 3243 | 3550 | 3275 | Telo2 |
| Ten1 | 475 | 614 | 551 | 479 | 605 | 567 | 592 | 1005 | 1116 | 1084 | 1088 | 1132 | 1144 | 1108 | 553 | 477 | 525 | 591 | 634 | 605 | 588 | 973 | 1149 | 1184 | 1212 | 1191 | 1120 | 1175 | Ten1 |
| Tenm1 | 1412 | 1228 | 581 | 1682 | 2557 | 2751 | 2794 | 3116 | 2740 | 2249 | 2068 | 1798 | 1280 | 1521 | 1578 | 1002 | 1088 | 1797 | 1960 | 3276 | 3013 | 2438 | 2163 | 1336 | 2516 | 594 | 1386 | 1210 | Tenm1 |
| Tenm2 | 2647 | 3514 | 3768 | 7121 | 9259 | 7780 | 7164 | 4601 | 5080 | 7712 | 5600 | 5311 | 5914 | 5006 | 3737 | 3527 | 5080 | 7980 | 8391 | 6156 | 8693 | 8338 | 5299 | 5927 | 8688 | 2361 | 4883 | 4663 | Tenm2 |
| Tenm3 | 15282 | 17467 | 11451 | 13880 | 14857 | 17426 | 12855 | 14309 | 15489 | 17167 | 19147 | 17217 | 20248 | 16274 | 16192 | 19260 | 12173 | 15610 | 13560 | 12294 | 14589 | 14472 | 16333 | 13643 | 16067 | 8548 | 19496 | 16738 | Tenm3 |
| Tenm4 | 12064 | 11961 | 10591 | 14949 | 15652 | 15570 | 16025 | 10147 | 12344 | 14049 | 13142 | 15363 | 14914 | 13640 | 10578 | 14545 | 12222 | 15940 | 16988 | 13670 | 16938 | 11645 | 10861 | 12814 | 15939 | 10387 | 13364 | 13766 | Tenm4 |
| Tent2 | 5356 | 3835 | 4671 | 3935 | 4033 | 4826 | 4914 | 5159 | 4820 | 4193 | 4597 | 5182 | 5042 | 5095 | 5325 | 4467 | 4040 | 3925 | 4215 | 4507 | 4631 | 4933 | 5082 | 4579 | 4389 | 5401 | 5254 | 5312 | Tent2 |
| Tent4a | 3182 | 3763 | 4067 | 4376 | 4193 | 3967 | 3803 | 2933 | 3206 | 3715 | 3418 | 3807 | 3442 | 3601 | 3082 | 3592 | 4153 | 4417 | 4443 | 3462 | 4019 | 3688 | 2974 | 3558 | 3833 | 3836 | 3270 | 3715 | Tent4a |
| Tent4b | 8778 | 8073 | 9300 | 11103 | 9986 | 10133 | 11001 | 5307 | 4433 | 5115 | 5180 | 5397 | 5180 | 5364 | 9634 | 8792 | 10013 | 11101 | 10369 | 9605 | 11113 | 5706 | 4457 | 5202 | 5472 | 5213 | 5417 | 5458 | Tent4b |
| Tent5a | 1509 | 1422 | 1577 | 1435 | 1317 | 1395 | 1172 | 1489 | 1611 | 1598 | 2480 | 1506 | 1878 | 1483 | 1585 | 1653 | 1487 | 1578 | 1084 | 1151 | 1182 | 2134 | 2232 | 1540 | 1595 | 1184 | 1945 | 1742 | Tent5a |
| Tent5b | 286 | 374 | 357 | 301 | 155 | 215 | 121 | 24 | 38 | 41 | 35 | 25 | 31 | 27 | 268 | 434 | 262 | 229 | 175 | 131 | 118 | 32 | 67 | 32 | 19 | 41 | 31 | 27 | Tent5b |
| Tent5c | 2224 | 2515 | 2134 | 1842 | 1136 | 1330 | 1330 | 879 | 1190 | 936 | 783 | 1245 | 3472 | 1444 | 2483 | 2337 | 1863 | 1615 | 1455 | 1724 | 1318 | 702 | 1348 | 873 | 856 | 1143 | 1589 | 2410 | Tent5c |
| Tent5d | 37 | 9 | 3 | 2 | 7 | 3 | 4 | 12 | 11 | 9 | 1 | 9 | 8 | 6 | 13 | 16 | 5 | 11 | 0 | 13 | 2 | 3 | 10 | 11 | 1 | 5 | 3 | 5 | Tent5d |
| Tep1 | 596 | 693 | 637 | 620 | 565 | 629 | 390 | 902 | 1093 | 1101 | 1182 | 1206 | 1322 | 1103 | 585 | 819 | 537 | 519 | 535 | 436 | 497 | 983 | 1191 | 913 | 921 | 729 | 1185 | 1060 | Tep1 |
| Tepp | 30 | 48 | 39 | 39 | 51 | 42 | 44 | 15 | 21 | 38 | 24 | 10 | 23 | 14 | 49 | 39 | 32 | 24 | 41 | 29 | 35 | 32 | 39 | 22 | 13 | 14 | 9 | 18 | Tepp |
| Tepsin | 1617 | 2419 | 2204 | 2193 | 1991 | 1873 | 1939 | 1165 | 1439 | 1406 | 1481 | 1574 | 1378 | 1503 | 1478 | 1998 | 2342 | 2114 | 2205 | 1736 | 1762 | 1335 | 1364 | 1581 | 1498 | 1736 | 1539 | 1351 | Tepsin |
| Terb1 | 76 | 67 | 79 | 104 | 133 | 114 | 96 | 67 | 55 | 41 | 57 | 21 | 69 | 27 | 107 | 107 | 101 | 109 | 111 | 110 | 103 | 42 | 52 | 38 | 58 | 31 | 29 | 44 | Terb1 |
| Terb2 | 0 | 0 | 0 | 0 | 0 | 0 | 0 | 0 | 0 | 0 | 0 | 0 | 0 | 0 | 0 | 0 | 0 | 0 | 0 | 0 | 0 | 0 | 0 | 2 | 0 | 0 | 0 | 1 | Terb2 |
| Terf1 | 1709 | 1370 | 1851 | 1591 | 1256 | 1332 | 1429 | 1892 | 1589 | 1731 | 1658 | 2056 | 1827 | 2207 | 1673 | 1453 | 1700 | 1330 | 1503 | 1531 | 1376 | 1445 | 1573 | 2019 | 1586 | 2508 | 2049 | 2134 | Terf1 |
| Terf2 | 3999 | 4242 | 5297 | 5181 | 4978 | 4513 | 5161 | 4368 | 4581 | 5243 | 4716 | 5279 | 4698 | 5565 | 4155 | 4218 | 5159 | 5229 | 5381 | 4444 | 5244 | 5307 | 4390 | 5426 | 5323 | 5560 | 4706 | 5212 | Terf2 |
| Terf2ip | 4736 | 5135 | 5320 | 6244 | 5546 | 5229 | 5850 | 3330 | 3185 | 3596 | 3237 | 3697 | 3598 | 3758 | 4845 | 4937 | 5650 | 6029 | 5670 | 5175 | 6042 | 4017 | 3124 | 3880 | 3848 | 3825 | 3335 | 3760 | Terf2ip |
| Tert | 976 | 1052 | 800 | 906 | 1024 | 892 | 836 | 1662 | 1740 | 1690 | 1753 | 1606 | 1673 | 1553 | 742 | 1016 | 841 | 863 | 934 | 824 | 745 | 1733 | 1608 | 1685 | 1644 | 1128 | 1692 | 1444 | Tert |
| Tes | 5913 | 4374 | 4045 | 3266 | 3121 | 3023 | 2407 | 3449 | 3503 | 2840 | 3174 | 2028 | 3114 | 2134 | 5069 | 6124 | 3365 | 3252 | 2493 | 2991 | 2498 | 2672 | 3728 | 2524 | 2162 | 2608 | 2120 | 2302 | Tes |
| Tes3-ps | 180 | 82 | 95 | 155 | 155 | 144 | 152 | 188 | 93 | 109 | 90 | 99 | 117 | 104 | 195 | 125 | 148 | 179 | 142 | 159 | 135 | 155 | 106 | 95 | 97 | 120 | 116 | 129 | Tes3-ps |
| Tesc | 203 | 263 | 175 | 293 | 281 | 202 | 259 | 103 | 69 | 180 | 77 | 50 | 66 | 31 | 183 | 199 | 249 | 262 | 288 | 225 | 277 | 40 | 150 | 154 | 48 | 166 | 19 | 32 | Tesc |
| Tescl | 1 | 2 | 0 | 0 | 2 | 2 | 0 | 1 | 1 | 0 | 0 | 0 | 0 | 1 | 4 | 1 | 7 | 1 | 0 | 0 | 4 | 2 | 3 | 2 | 1 | 3 | 0 | 0 | Tescl |
| Tesk1 | 2625 | 3302 | 2865 | 3438 | 3697 | 3084 | 2996 | 3284 | 3810 | 4260 | 3730 | 3913 | 3212 | 3384 | 2288 | 2763 | 3574 | 3358 | 3890 | 2809 | 3202 | 4293 | 3545 | 4244 | 4280 | 4135 | 3263 | 3292 | Tesk1 |
| Tesk2 | 1538 | 1373 | 1419 | 1386 | 976 | 1083 | 1217 | 455 | 420 | 449 | 451 | 513 | 442 | 498 | 1415 | 1376 | 1358 | 1302 | 1066 | 1110 | 1172 | 428 | 462 | 534 | 379 | 546 | 488 | 520 | Tesk2 |
| Tesl1 | 0 | 1 | 0 | 0 | 0 | 0 | 2 | 0 | 0 | 0 | 2 | 0 | 1 | 1 | 1 | 1 | 0 | 0 | 0 | 0 | 0 | 0 | 2 | 2 | 4 | 0 | 3 | 0 | Tesl1 |
| Tesmin | 16 | 25 | 30 | 39 | 22 | 26 | 13 | 27 | 29 | 38 | 36 | 16 | 26 | 32 | 20 | 37 | 22 | 44 | 42 | 21 | 27 | 35 | 30 | 19 | 23 | 23 | 28 | 26 | Tesmin |
| Tespa1 | 13 | 6 | 2 | 3 | 2 | 3 | 0 | 5 | 8 | 8 | 14 | 3 | 17 | 4 | 7 | 7 | 8 | 2 | 2 | 5 | 4 | 2 | 16 | 4 | 2 | 9 | 8 | 4 | Tespa1 |
| Tet1 | 4525 | 4164 | 3375 | 3561 | 3517 | 4697 | 4123 | 4956 | 4666 | 3947 | 4017 | 3908 | 3901 | 4202 | 4560 | 4464 | 3311 | 3382 | 3529 | 4546 | 4333 | 3801 | 4295 | 3341 | 4348 | 3201 | 4172 | 3622 | Tet1 |
| Tet2 | 5855 | 4776 | 3862 | 4942 | 5926 | 6579 | 5832 | 8267 | 7212 | 6842 | 7504 | 7848 | 7606 | 7960 | 5687 | 4625 | 4244 | 4840 | 5438 | 6056 | 6197 | 7060 | 6724 | 6595 | 7873 | 4799 | 8020 | 7168 | Tet2 |
| Tet3 | 6785 | 6985 | 9174 | 10701 | 11343 | 9910 | 9865 | 9677 | 10877 | 13495 | 12526 | 15461 | 13634 | 14353 | 6498 | 7015 | 10250 | 10869 | 11306 | 7944 | 9963 | 13809 | 9989 | 14475 | 15694 | 13549 | 13739 | 14733 | Tet3 |
| Tex10 | 7462 | 6935 | 6068 | 5710 | 5042 | 5393 | 6191 | 5275 | 4668 | 3569 | 3718 | 4435 | 4094 | 4403 | 7647 | 6918 | 5619 | 5768 | 5341 | 6540 | 5731 | 4246 | 4458 | 3921 | 3782 | 4450 | 4552 | 4357 | Tex10 |
| Tex101 | 0 | 1 | 1 | 0 | 0 | 0 | 0 | 0 | 0 | 0 | 0 | 0 | 0 | 0 | 1 | 0 | 0 | 0 | 0 | 0 | 1 | 0 | 1 | 0 | 0 | 0 | 0 | 0 | Tex101 |
| Tex11 | 5 | 10 | 1 | 1 | 9 | 3 | 6 | 4 | 7 | 10 | 4 | 3 | 11 | 15 | 8 | 12 | 8 | 10 | 0 | 4 | 19 | 7 | 8 | 3 | 4 | 4 | 8 | 4 | Tex11 |
| Tex12 | 0 | 0 | 0 | 0 | 0 | 0 | 0 | 1 | 0 | 0 | 9 | 2 | 0 | 1 | 0 | 0 | 1 | 0 | 0 | 3 | 0 | 0 | 0 | 0 | 0 | 0 | 2 | 0 | Tex12 |
| Tex13a | 0 | 0 | 0 | 0 | 0 | 0 | 0 | 0 | 0 | 0 | 0 | 0 | 0 | 0 | 0 | 0 | 0 | 0 | 0 | 0 | 1 | 0 | 0 | 0 | 0 | 0 | 0 | 0 | Tex13a |
| Tex13c1 | 0 | 0 | 0 | 0 | 0 | 0 | 0 | 0 | 0 | 0 | 0 | 1 | 0 | 0 | 0 | 0 | 0 | 0 | 0 | 0 | 0 | 0 | 0 | 0 | 2 | 0 | 0 | 0 | Tex13c1 |
| Tex14 | 32 | 24 | 34 | 23 | 28 | 21 | 13 | 38 | 30 | 28 | 63 | 21 | 42 | 21 | 29 | 26 | 21 | 36 | 23 | 30 | 25 | 38 | 56 | 16 | 15 | 29 | 35 | 13 | Tex14 |
| Tex15 | 5637 | 3007 | 1907 | 1853 | 1506 | 2820 | 1829 | 2717 | 2197 | 1853 | 2088 | 976 | 1639 | 1135 | 4880 | 4738 | 1643 | 1883 | 1630 | 3275 | 2158 | 760 | 3227 | 1618 | 873 | 1308 | 1367 | 997 | Tex15 |
| Tex16 | 159 | 130 | 53 | 55 | 29 | 117 | 66 | 78 | 78 | 14 | 51 | 74 | 26 | 51 | 177 | 86 | 25 | 19 | 23 | 152 | 106 | 10 | 83 | 15 | 47 | 32 | 95 | 42 | Tex16 |
| Tex19.1 | 0 | 1 | 1 | 0 | 0 | 2 | 1 | 1 | 1 | 0 | 2 | 0 | 0 | 0 | 1 | 5 | 0 | 0 | 0 | 0 | 0 | 2 | 0 | 0 | 0 | 0 | 0 | 0 | Tex19.1 |
| Tex19.2 | 0 | 0 | 0 | 0 | 0 | 0 | 0 | 1 | 0 | 2 | 0 | 0 | 0 | 0 | 0 | 0 | 0 | 1 | 0 | 0 | 0 | 0 | 1 | 0 | 0 | 0 | 0 | 1 | Tex19.2 |
| Tex2 | 2970 | 3805 | 3628 | 4639 | 4935 | 3837 | 4480 | 2140 | 2267 | 2504 | 2459 | 1953 | 2116 | 2028 | 3392 | 3438 | 4143 | 4852 | 4400 | 3679 | 4782 | 3014 | 2224 | 2169 | 2656 | 1612 | 1777 | 1904 | Tex2 |
| Tex21 | 5 | 1 | 4 | 4 | 0 | 0 | 0 | 1 | 1 | 3 | 5 | 0 | 5 | 4 | 7 | 5 | 3 | 3 | 2 | 4 | 1 | 0 | 3 | 3 | 0 | 3 | 0 | 0 | Tex21 |
| Tex22 | 2 | 6 | 4 | 14 | 11 | 5 | 7 | 0 | 6 | 0 | 2 | 2 | 1 | 4 | 1 | 4 | 17 | 11 | 7 | 13 | 5 | 7 | 2 | 6 | 0 | 1 | 12 | 6 | Tex22 |
| Tex24 | 0 | 0 | 1 | 4 | 0 | 0 | 0 | 0 | 0 | 0 | 0 | 0 | 0 | 0 | 0 | 2 | 0 | 2 | 0 | 1 | 0 | 0 | 2 | 0 | 0 | 0 | 0 | 0 | Tex24 |
| Tex26 | 168 | 114 | 124 | 121 | 121 | 113 | 152 | 95 | 103 | 110 | 76 | 152 | 120 | 117 | 149 | 103 | 94 | 77 | 115 | 130 | 132 | 69 | 83 | 109 | 126 | 162 | 143 | 85 | Tex26 |
| Tex261 | 5443 | 5722 | 4809 | 4641 | 4138 | 4563 | 4340 | 3705 | 4127 | 3596 | 3713 | 4006 | 3874 | 3706 | 4967 | 5165 | 4780 | 4500 | 4533 | 4262 | 4444 | 3651 | 4141 | 3586 | 3585 | 4088 | 3798 | 3833 | Tex261 |
| Tex264 | 1361 | 1810 | 1289 | 1607 | 1789 | 1781 | 1591 | 1097 | 1472 | 1296 | 1523 | 915 | 1196 | 990 | 1469 | 1734 | 1544 | 1796 | 1715 | 1618 | 1622 | 1193 | 1603 | 1188 | 1167 | 1084 | 994 | 910 | Tex264 |
| Tex28 | 1 | 0 | 0 | 0 | 0 | 0 | 0 | 0 | 2 | 0 | 0 | 0 | 0 | 0 | 0 | 1 | 1 | 0 | 0 | 0 | 0 | 0 | 0 | 0 | 0 | 0 | 0 | 0 | Tex28 |
| Tex29 | 21 | 28 | 22 | 13 | 11 | 31 | 25 | 21 | 28 | 13 | 28 | 24 | 8 | 26 | 5 | 20 | 14 | 25 | 33 | 39 | 27 | 18 | 22 | 19 | 33 | 6 | 29 | 8 | Tex29 |
| Tex30 | 1804 | 1315 | 1722 | 1699 | 1301 | 1403 | 1805 | 1023 | 671 | 761 | 612 | 995 | 868 | 1208 | 1817 | 1542 | 1544 | 1559 | 1509 | 1712 | 1670 | 688 | 647 | 912 | 742 | 1146 | 955 | 1213 | Tex30 |
| Tex33 | 0 | 0 | 11 | 0 | 1 | 2 | 1 | 0 | 0 | 1 | 0 | 0 | 0 | 0 | 4 | 0 | 0 | 0 | 0 | 0 | 0 | 0 | 0 | 0 | 0 | 3 | 0 | 0 | Tex33 |
| Tex35 | 80 | 82 | 26 | 105 | 80 | 135 | 106 | 16 | 23 | 16 | 18 | 35 | 40 | 32 | 82 | 49 | 68 | 78 | 90 | 104 | 112 | 13 | 14 | 25 | 34 | 18 | 60 | 21 | Tex35 |
| Tex36 | 1 | 2 | 0 | 0 | 2 | 3 | 0 | 1 | 2 | 5 | 5 | 0 | 0 | 5 | 0 | 0 | 0 | 3 | 2 | 12 | 3 | 0 | 3 | 1 | 5 | 0 | 0 | 1 | Tex36 |
| Tex37 | 0 | 0 | 0 | 0 | 0 | 0 | 0 | 0 | 0 | 0 | 0 | 0 | 0 | 0 | 0 | 0 | 0 | 0 | 0 | 0 | 0 | 0 | 1 | 0 | 0 | 0 | 0 | 0 | Tex37 |
| Tex38 | 13 | 10 | 6 | 3 | 19 | 10 | 8 | 9 | 15 | 41 | 30 | 23 | 24 | 23 | 15 | 12 | 13 | 13 | 12 | 3 | 4 | 22 | 19 | 15 | 23 | 33 | 17 | 18 | Tex38 |
| Tex43 | 7 | 10 | 7 | 3 | 6 | 9 | 3 | 2 | 7 | 5 | 6 | 7 | 0 | 1 | 4 | 9 | 3 | 7 | 6 | 12 | 6 | 5 | 1 | 2 | 0 | 2 | 15 | 2 | Tex43 |
| Tex44 | 1 | 0 | 0 | 13 | 2 | 2 | 1 | 0 | 0 | 3 | 0 | 0 | 3 | 5 | 1 | 0 | 0 | 0 | 5 | 0 | 2 | 8 | 1 | 0 | 0 | 0 | 0 | 0 | Tex44 |

| | | | | | | | | | | | | | | | | | | | | | | | | | | | | | |
|---|---|---|---|---|---|---|---|---|---|---|---|---|---|---|---|---|---|---|---|---|---|---|---|---|---|---|---|---|---|
| Tex45 | 42 | 59 | 28 | 73 | 44 | 60 | 34 | 9 | 5 | 14 | 10 | 8 | 7 | 6 | 46 | 44 | 59 | 61 | 55 | 41 | 46 | 12 | 6 | 8 | 5 | 11 | 3 | 12 | Tex45 |
| Tex46 | 0 | 0 | 0 | 0 | 0 | 0 | 0 | 0 | 0 | 0 | 1 | 0 | 0 | 1 | 0 | 0 | 0 | 0 | 2 | 0 | 0 | 0 | 1 | 0 | 0 | 0 | 0 | 0 | Tex46 |
| Tex48 | 0 | 0 | 0 | 0 | 0 | 0 | 0 | 1 | 0 | 0 | 0 | 0 | 0 | 4 | 0 | 0 | 0 | 0 | 0 | 0 | 1 | 0 | 0 | 0 | 0 | 0 | 0 | 0 | Tex48 |
| Tex49 | 6 | 7 | 0 | 1 | 6 | 0 | 1 | 5 | 0 | 3 | 2 | 4 | 2 | 1 | 1 | 2 | 0 | 0 | 0 | 5 | 3 | 0 | 2 | 3 | 2 | 6 | 0 | 4 | Tex49 |
| Tex50 | 3 | 11 | 2 | 8 | 6 | 4 | 13 | 1 | 2 | 3 | 0 | 0 | 1 | 0 | 5 | 5 | 16 | 11 | 11 | 0 | 11 | 2 | 1 | 0 | 0 | 0 | 2 | 1 | Tex50 |
| Tex52 | 130 | 156 | 108 | 129 | 80 | 90 | 135 | 116 | 111 | 109 | 78 | 104 | 91 | 100 | 96 | 129 | 132 | 110 | 136 | 103 | 115 | 79 | 68 | 160 | 99 | 114 | 122 | 90 | Tex52 |
| Tex55 | 0 | 0 | 0 | 0 | 0 | 0 | 0 | 0 | 0 | 0 | 0 | 0 | 0 | 0 | 0 | 0 | 0 | 0 | 0 | 0 | 0 | 0 | 1 | 0 | 0 | 0 | 0 | 0 | Tex55 |
| Tex9 | 7448 | 6081 | 6146 | 6314 | 5863 | 6635 | 7004 | 4098 | 3699 | 3993 | 3485 | 4964 | 4765 | 5373 | 7481 | 6143 | 6205 | 5832 | 6022 | 7203 | 6786 | 3618 | 3452 | 4022 | 4091 | 4400 | 5194 | 5001 | Tex9 |
| Tfam | 4832 | 4251 | 4589 | 4496 | 4015 | 4062 | 4578 | 4588 | 4177 | 4053 | 4000 | 4759 | 4465 | 4776 | 4963 | 4575 | 4524 | 4182 | 4274 | 4708 | 4137 | 3957 | 4073 | 4474 | 4355 | 5560 | 4944 | 5122 | Tfam |
| Tfap2a | 529 | 496 | 642 | 845 | 1240 | 945 | 305 | 1287 | 1358 | 1837 | 1385 | 745 | 1892 | 761 | 637 | 734 | 816 | 840 | 662 | 506 | 400 | 1910 | 1440 | 1031 | 1297 | 671 | 648 | 858 | Tfap2a |
| Tfap2b | 2353 | 2185 | 2821 | 4362 | 6442 | 5448 | 3831 | 9685 | 7237 | 13113 | 9693 | 9827 | 16491 | 11018 | 2843 | 2923 | 4341 | 4092 | 6072 | 4194 | 5228 | 15014 | 8166 | 13196 | 12972 | 4657 | 9298 | 11364 | Tfap2b |
| Tfap2c | 1209 | 1142 | 1251 | 1296 | 1064 | 1170 | 1580 | 1097 | 1270 | 1237 | 795 | 1731 | 925 | 1705 | 811 | 756 | 1257 | 1214 | 1570 | 1434 | 1404 | 451 | 916 | 1668 | 1622 | 2142 | 1119 | 1459 | Tfap2c |
| Tfap2d | 145 | 155 | 368 | 180 | 357 | 316 | 260 | 909 | 487 | 919 | 544 | 1085 | 1000 | 1107 | 178 | 101 | 305 | 222 | 493 | 382 | 434 | 1118 | 674 | 1142 | 1091 | 237 | 1275 | 1291 | Tfap2d |
| Tfap2e | 60 | 67 | 207 | 167 | 177 | 122 | 152 | 106 | 106 | 157 | 110 | 323 | 277 | 221 | 88 | 59 | 189 | 140 | 199 | 139 | 140 | 137 | 110 | 279 | 189 | 116 | 198 | 272 | Tfap2e |
| Tfap4 | 514 | 470 | 800 | 607 | 628 | 524 | 620 | 744 | 911 | 1024 | 955 | 1207 | 956 | 1231 | 423 | 500 | 721 | 534 | 637 | 481 | 535 | 566 | 852 | 1213 | 1044 | 1530 | 940 | 1240 | Tfap4 |
| Tfb1m | 827 | 808 | 794 | 679 | 519 | 529 | 659 | 894 | 818 | 745 | 782 | 809 | 809 | 850 | 881 | 777 | 713 | 615 | 582 | 630 | 622 | 668 | 827 | 733 | 701 | 902 | 891 | 930 | Tfb1m |
| Tfb2m | 2135 | 1836 | 1711 | 1874 | 1706 | 1748 | 1925 | 1906 | 1702 | 1532 | 1593 | 1693 | 1616 | 1609 | 2194 | 2125 | 1662 | 1935 | 1760 | 2067 | 1913 | 1589 | 1753 | 1521 | 1622 | 1755 | 1723 | 1640 | Tfb2m |
| Tfcp2 | 3886 | 4050 | 4287 | 4894 | 4574 | 4320 | 4727 | 3406 | 3216 | 3870 | 3680 | 3696 | 3745 | 3859 | 3909 | 3971 | 4749 | 4529 | 4275 | 4083 | 4589 | 3811 | 3122 | 3676 | 3672 | 3477 | 3941 | 3919 | Tfcp2 |
| Tfcp2l1 | 811 | 733 | 559 | 618 | 531 | 539 | 532 | 114 | 153 | 102 | 155 | 100 | 102 | 67 | 723 | 797 | 547 | 600 | 473 | 530 | 533 | 102 | 159 | 86 | 114 | 52 | 84 | 62 | Tfcp2l1 |
| Tfdp1 | 11197 | 11576 | 12381 | 11195 | 10286 | 10544 | 12421 | 9608 | 9860 | 9263 | 9448 | 11898 | 10468 | 12610 | 10941 | 10609 | 12393 | 10939 | 11274 | 11148 | 11540 | 8482 | 9453 | 11324 | 10497 | 13964 | 12163 | 13118 | Tfdp1 |
| Tfdp2 | 11114 | 9772 | 9268 | 8816 | 8746 | 9595 | 9934 | 8172 | 6336 | 6225 | 5720 | 9533 | 9349 | 9488 | 10572 | 9851 | 9010 | 8389 | 9199 | 9669 | 9904 | 6644 | 6357 | 6853 | 7295 | 6124 | 11579 | 10551 | Tfdp2 |
| Tfe3 | 4601 | 5228 | 5139 | 5627 | 4971 | 4826 | 4863 | 2258 | 2275 | 2718 | 2553 | 2986 | 2819 | 2997 | 4356 | 4555 | 5284 | 5213 | 5183 | 3869 | 4970 | 2682 | 2375 | 2796 | 2639 | 2864 | 2829 | 3012 | Tfe3 |
| Tfeb | 621 | 837 | 785 | 746 | 547 | 663 | 476 | 123 | 200 | 149 | 199 | 108 | 163 | 94 | 541 | 787 | 871 | 723 | 628 | 455 | 507 | 127 | 190 | 152 | 154 | 185 | 132 | 104 | Tfeb |
| Tfec | 23 | 14 | 36 | 8 | 16 | 23 | 11 | 48 | 54 | 42 | 69 | 62 | 66 | 60 | 25 | 23 | 20 | 13 | 12 | 16 | 15 | 45 | 90 | 39 | 54 | 72 | 60 | 61 | Tfec |
| Tff2 | 0 | 0 | 0 | 0 | 0 | 0 | 0 | 0 | 0 | 3 | 0 | 0 | 0 | 1 | 0 | 1 | 0 | 0 | 0 | 0 | 0 | 0 | 0 | 0 | 0 | 0 | 0 | 0 | Tff2 |
| Tff3 | 3 | 4 | 5 | 9 | 12 | 5 | 2 | 9 | 0 | 14 | 8 | 13 | 7 | 10 | 3 | 0 | 0 | 4 | 9 | 1 | 3 | 2 | 11 | 2 | 1 | 19 | 9 | 13 | Tff3 |
| Tfg | 6867 | 7042 | 7396 | 6428 | 6677 | 6377 | 7012 | 7590 | 6637 | 6668 | 6621 | 7875 | 7334 | 8193 | 7681 | 6390 | 6886 | 6325 | 6773 | 7084 | 6660 | 7132 | 6829 | 7302 | 7175 | 8444 | 8007 | 8572 | Tfg |
| Tfip11 | 4197 | 5288 | 5231 | 4991 | 4044 | 4198 | 3952 | 2743 | 3783 | 3920 | 3683 | 4452 | 4115 | 3989 | 3779 | 4825 | 5131 | 4593 | 4381 | 3604 | 3936 | 3638 | 3397 | 4149 | 4245 | 4851 | 4188 | 4206 | Tfip11 |
| Tfpi | 1751 | 1327 | 1764 | 1250 | 1279 | 1596 | 1124 | 1407 | 1807 | 1485 | 2348 | 1312 | 1712 | 1421 | 1871 | 2088 | 1311 | 1630 | 992 | 1475 | 1182 | 1753 | 2539 | 1448 | 1454 | 1527 | 1648 | 1524 | Tfpi |
| Tfpi2 | 35 | 14 | 27 | 17 | 12 | 31 | 10 | 39 | 36 | 22 | 54 | 36 | 19 | 33 | 40 | 19 | 13 | 20 | 9 | 9 | 13 | 28 | 55 | 20 | 18 | 46 | 23 | 39 | Tfpi2 |
| Tfpt | 1082 | 1158 | 1201 | 1181 | 1208 | 1194 | 1095 | 912 | 928 | 1142 | 1170 | 1152 | 1173 | 1327 | 994 | 935 | 1239 | 1069 | 1153 | 1169 | 1168 | 961 | 1024 | 1273 | 1156 | 1295 | 1120 | 1270 | Tfpt |
| Tfr2 | 401 | 650 | 464 | 987 | 989 | 875 | 726 | 178 | 220 | 267 | 181 | 252 | 271 | 244 | 468 | 529 | 863 | 1073 | 994 | 754 | 861 | 302 | 175 | 207 | 286 | 112 | 205 | 190 | Tfr2 |
| Tfrc | 18089 | 13757 | 12261 | 12806 | 13044 | 16524 | 16568 | 14691 | 11491 | 8986 | 9964 | 12527 | 11989 | 13529 | 19043 | 15535 | 11148 | 12834 | 13150 | 20750 | 15579 | 12089 | 11437 | 9105 | 13340 | 10665 | 13619 | 13011 | Tfrc |
| Tg | 21 | 15 | 13 | 18 | 18 | 22 | 10 | 26 | 47 | 31 | 52 | 24 | 56 | 30 | 37 | 47 | 19 | 32 | 15 | 12 | 13 | 47 | 59 | 16 | 33 | 8 | 69 | 32 | Tg |
| Tgds | 2287 | 2252 | 2207 | 2059 | 1610 | 1932 | 1863 | 1895 | 1886 | 1494 | 1921 | 1782 | 1723 | 1645 | 2232 | 2411 | 2051 | 1976 | 1748 | 2073 | 1769 | 1564 | 1855 | 1690 | 1581 | 1896 | 1807 | 1726 | Tgds |
| Tgfa | 589 | 428 | 493 | 583 | 1114 | 806 | 838 | 539 | 743 | 991 | 925 | 461 | 371 | 487 | 514 | 502 | 657 | 738 | 786 | 678 | 770 | 847 | 628 | 695 | 783 | 518 | 342 | 395 | Tgfa |
| Tgfb1 | 340 | 445 | 464 | 385 | 343 | 371 | 241 | 348 | 462 | 461 | 511 | 405 | 526 | 353 | 342 | 440 | 396 | 452 | 340 | 229 | 294 | 436 | 514 | 436 | 385 | 561 | 370 | 310 | Tgfb1 |
| Tgfb1i1 | 570 | 817 | 764 | 372 | 405 | 597 | 274 | 1083 | 1502 | 1407 | 1870 | 1643 | 2124 | 1496 | 604 | 904 | 508 | 529 | 367 | 297 | 296 | 1365 | 1935 | 1481 | 1332 | 1774 | 1811 | 1697 | Tgfb1i1 |
| Tgfb2 | 1558 | 1194 | 1541 | 1370 | 1199 | 1382 | 1065 | 2898 | 3165 | 3660 | 5032 | 2295 | 4017 | 2376 | 1547 | 1760 | 1256 | 1335 | 1002 | 942 | 1079 | 2837 | 3834 | 3470 | 2025 | 3909 | 2377 | 2433 | Tgfb2 |
| Tgfb3 | 947 | 824 | 940 | 773 | 595 | 819 | 447 | 1111 | 1809 | 1691 | 2841 | 1037 | 1354 | 1007 | 845 | 1144 | 876 | 884 | 563 | 346 | 420 | 1611 | 2264 | 1443 | 1193 | 1338 | 1159 | 927 | Tgfb3 |
| Tgfbi | 3876 | 3736 | 3263 | 1955 | 1140 | 1662 | 650 | 2354 | 3539 | 3099 | 6445 | 2368 | 4338 | 2239 | 3349 | 6011 | 2114 | 2113 | 719 | 557 | 571 | 3278 | 4125 | 3027 | 2046 | 5048 | 2651 | 2549 | Tgfbi |
| Tgfbr1 | 24646 | 18749 | 20308 | 19951 | 17692 | 19618 | 20227 | 19974 | 15045 | 14563 | 14737 | 19788 | 19415 | 19674 | 25293 | 22171 | 20065 | 18899 | 17809 | 20050 | 19420 | 16688 | 14922 | 16175 | 15702 | 18139 | 21783 | 20939 | Tgfbr1 |
| Tgfbr2 | 1374 | 1195 | 1307 | 976 | 970 | 1304 | 732 | 2177 | 3301 | 2792 | 4062 | 2263 | 2724 | 1976 | 1294 | 1487 | 1101 | 1267 | 908 | 842 | 756 | 2811 | 4237 | 2555 | 2429 | 2389 | 2404 | 1968 | Tgfbr2 |
| Tgfbr3 | 1460 | 1188 | 1493 | 737 | 832 | 920 | 503 | 953 | 1609 | 1084 | 1761 | 1008 | 1372 | 858 | 1234 | 1748 | 973 | 1040 | 550 | 533 | 608 | 1089 | 1906 | 999 | 812 | 830 | 973 | 1002 | Tgfbr3 |
| Tgfbr3l | 235 | 228 | 317 | 355 | 421 | 380 | 387 | 413 | 432 | 534 | 491 | 506 | 438 | 554 | 273 | 219 | 319 | 392 | 341 | 304 | 323 | 528 | 397 | 575 | 524 | 549 | 533 | 492 | Tgfbr3l |
| Tgfbrap1 | 3329 | 3778 | 3080 | 4196 | 4427 | 4267 | 4124 | 3705 | 3920 | 4608 | 4475 | 4550 | 4230 | 4445 | 3672 | 3547 | 4020 | 4422 | 4383 | 3917 | 4338 | 4550 | 3836 | 4359 | 4672 | 3487 | 4091 | 4186 | Tgfbrap1 |
| Tgif1 | 2360 | 2116 | 2798 | 1847 | 1540 | 1517 | 1574 | 1951 | 1788 | 1756 | 1923 | 1947 | 2682 | 2068 | 2193 | 2493 | 2428 | 1687 | 1630 | 1534 | 1589 | 1442 | 2186 | 2082 | 1391 | 2644 | 2198 | 2451 | Tgif1 |
| Tgif2 | 3931 | 4563 | 4896 | 4352 | 3385 | 3644 | 3695 | 4364 | 4748 | 4430 | 4418 | 6026 | 5148 | 5426 | 3603 | 4253 | 4580 | 4101 | 4195 | 3290 | 3751 | 3728 | 4335 | 5352 | 4816 | 7177 | 5675 | 5770 | Tgif2 |
| Tgm1 | 58 | 57 | 25 | 27 | 34 | 58 | 28 | 85 | 130 | 36 | 63 | 47 | 67 | 42 | 49 | 83 | 35 | 54 | 26 | 30 | 35 | 32 | 135 | 38 | 23 | 28 | 51 | 74 | Tgm1 |
| Tgm2 | 963 | 1234 | 1256 | 1021 | 1168 | 1205 | 673 | 1158 | 1729 | 1598 | 1834 | 1224 | 1419 | 1017 | 1069 | 1509 | 1019 | 1382 | 801 | 1218 | 935 | 1552 | 2306 | 1726 | 1313 | 1424 | 1389 | 1143 | Tgm2 |
| Tgm3 | 29 | 19 | 23 | 25 | 22 | 44 | 56 | 26 | 39 | 17 | 14 | 10 | 23 | 7 | 25 | 30 | 10 | 25 | 18 | 29 | 47 | 20 | 33 | 12 | 21 | 13 | 33 | 12 | Tgm3 |
| Tgm4 | 22 | 20 | 19 | 32 | 24 | 23 | 17 | 33 | 37 | 25 | 44 | 22 | 49 | 34 | 30 | 14 | 23 | 32 | 22 | 22 | 14 | 43 | 30 | 34 | 46 | 24 | 40 | 27 | Tgm4 |
| Tgm5 | 1 | 6 | 3 | 0 | 7 | 0 | 0 | 1 | 4 | 1 | 11 | 2 | 5 | 0 | 1 | 3 | 0 | 10 | 5 | 4 | 1 | 2 | 3 | 3 | 3 | 0 | 0 | 5 | Tgm5 |
| Tgm6 | 1 | 0 | 1 | 0 | 1 | 2 | 4 | 6 | 2 | 2 | 8 | 0 | 3 | 5 | 0 | 5 | 0 | 0 | 6 | 3 | 4 | 5 | 5 | 11 | 15 | 1 | 5 | 4 | Tgm6 |
| Tgm7 | 6 | 22 | 8 | 3 | 9 | 8 | 20 | 6 | 9 | 8 | 1 | 10 | 7 | 4 | 7 | 23 | 16 | 20 | 8 | 3 | 29 | 18 | 1 | 4 | 6 | 7 | 2 | 9 | Tgm7 |
| Tgoln1 | 10216 | 10757 | 8370 | 9348 | 7234 | 8250 | 7784 | 6537 | 5301 | 4696 | 4638 | 4964 | 4864 | 4858 | 11192 | 11066 | 8596 | 9708 | 6886 | 9402 | 8166 | 5815 | 5559 | 4595 | 4879 | 4372 | 5474 | 4944 | Tgoln1 |
| Tgs1 | 10807 | 9084 | 8748 | 8547 | 8210 | 8137 | 9077 | 9973 | 7805 | 6711 | 7112 | 8481 | 8227 | 8552 | 11457 | 10142 | 8824 | 8381 | 7945 | 10148 | 8217 | 7959 | 7689 | 7518 | 7276 | 8103 | 8758 | 8475 | Tgs1 |
| Tgtp1 | 0 | 0 | 0 | 0 | 0 | 0 | 0 | 0 | 1 | 0 | 1 | 2 | 0 | 0 | 4 | 0 | 0 | 0 | 0 | 1 | 0 | 5 | 0 | 3 | 2 | 0 | 0 | 0 | Tgtp1 |
| Th | 302 | 476 | 295 | 365 | 569 | 325 | 361 | 543 | 723 | 536 | 603 | 522 | 434 | 460 | 347 | 318 | 518 | 415 | 370 | 393 | 335 | 912 | 577 | 437 | 638 | 474 | 540 | 530 | Th |
| Tha1 | 241 | 326 | 425 | 299 | 280 | 249 | 253 | 101 | 158 | 172 | 195 | 210 | 175 | 162 | 212 | 262 | 403 | 311 | 303 | 274 | 205 | 189 | 149 | 176 | 199 | 246 | 215 | 191 | Tha1 |
| Thada | 2668 | 2498 | 1419 | 1778 | 1820 | 1981 | 1994 | 2503 | 2638 | 2120 | 2352 | 2346 | 2613 | 2404 | 2356 | 2288 | 1612 | 1732 | 1797 | 1964 | 2013 | 1811 | 2435 | 1736 | 2402 | 1399 | 2573 | 2273 | Thada |
| Thap1 | 512 | 485 | 645 | 498 | 717 | 490 | 738 | 885 | 812 | 803 | 782 | 933 | 869 | 965 | 491 | 474 | 687 | 620 | 699 | 596 | 715 | 998 | 794 | 1046 | 979 | 985 | 831 | 961 | Thap1 |
| Thap11 | 3465 | 3927 | 2950 | 3068 | 2617 | 2551 | 2851 | 2480 | 2259 | 1983 | 1985 | 2337 | 2081 | 2150 | 3347 | 3231 | 3095 | 2950 | 2892 | 2919 | 2694 | 2207 | 2194 | 2303 | 2026 | 2723 | 2306 | 2257 | Thap11 |
| Thap12 | 15216 | 12428 | 13215 | 11379 | 9991 | 11634 | 11313 | 11434 | 10313 | 9884 | 10152 | 12132 | 11767 | 11838 | 14541 | 13788 | 12385 | 11055 | 10199 | 11504 | 10628 | 9561 | 9912 | 10124 | 10135 | 12470 | 12156 | 12164 | Thap12 |
| Thap2 | 2620 | 2512 | 2973 | 3048 | 3158 | 3091 | 3199 | 3802 | 3251 | 3861 | 3498 | 3914 | 3775 | 4148 | 2789 | 2695 | 3109 | 3070 | 3381 | 3000 | 3497 | 3888 | 3180 | 3762 | 3943 | 3759 | 3856 | 4225 | Thap2 |
| Thap3 | 645 | 892 | 1034 | 1018 | 922 | 806 | 744 | 795 | 1029 | 1120 | 1270 | 1184 | 1108 | 1097 | 687 | 805 | 1122 | 851 | 992 | 782 | 750 | 929 | 1052 | 1396 | 1108 | 1403 | 1271 | 1174 | Thap3 |
| Thap4 | 2124 | 2154 | 2078 | 2204 | 1899 | 1922 | 1859 | 1419 | 1657 | 1537 | 1492 | 1526 | 1516 | 1359 | 1874 | 2039 | 2087 | 1958 | 2087 | 1776 | 1930 | 1521 | 1596 | 1576 | 1419 | 1798 | 1379 | 1479 | Thap4 |
| Thap7 | 519 | 613 | 586 | 573 | 593 | 699 | 532 | 834 | 982 | 955 | 987 | 1050 | 974 | 916 | 471 | 534 | 700 | 580 | 677 | 563 | 621 | 944 | 904 | 1160 | 869 | 1140 | 960 | 901 | Thap7 |
| Thbd | 1102 | 999 | 1559 | 992 | 981 | 1025 | 411 | 1238 | 1552 | 1220 | 1967 | 1485 | 2074 | 1578 | 1261 | 1281 | 1409 | 1326 | 454 | 431 | 482 | 2057 | 2031 | 1377 | 1378 | 1418 | 2187 | 1440 | Thbd |
| Thbs1 | 7413 | 8199 | 7366 | 3705 | 1970 | 4725 | 1589 | 5878 | 10500 | 8726 | 12211 | 7925 | 14073 | 7487 | 7291 | 10390 | 5531 | 5011 | 2198 | 1682 | 1516 | 9984 | 10792 | 7352 | 7032 | 7852 | 11567 | 9424 | Thbs1 |
| Thbs2 | 1944 | 2352 | 1666 | 1754 | 672 | 2478 | 200 | 2430 | 3539 | 3683 | 5052 | 2447 | 3368 | 2562 | 2232 | 3037 | 1299 | 2824 | 365 | 286 | 226 | 5534 | 5448 | 3843 | 4593 | 1384 | 2514 | 2466 | Thbs2 |
| Thbs3 | 3400 | 4098 | 4060 | 4206 | 4443 | 4465 | 3884 | 4603 | 4748 | 5321 | 5230 | 5893 | 5780 | 5526 | 3281 | 4092 | 4399 | 4540 | 4535 | 4027 | 4035 | 6181 | 4846 | 5616 | 6092 | 4462 | 6057 | 5565 | Thbs3 |
| Thbs4 | 322 | 360 | 408 | 286 | 269 | 270 | 230 | 269 | 470 | 323 | 2324 | 374 | 460 | 301 | 345 | 458 | 284 | 375 | 248 | 286 | 241 | 543 | 872 | 274 | 260 | 397 | 447 | 358 | Thbs4 |
| Theg | 0 | 1 | 0 | 0 | 1 | 0 | 0 | 1 | 0 | 0 | 0 | 0 | 0 | 0 | 0 | 0 | 0 | 0 | 0 | 0 | 0 | 0 | 0 | 0 | 0 | 0 | 0 | 0 | Theg |
| Thegl | 82 | 117 | 56 | 100 | 90 | 105 | 117 | 9 | 7 | 10 | 4 | 15 | 17 | 9 | 116 | 90 | 101 | 117 | 77 | 122 | 85 | 10 | 14 | 13 | 2 | 8 | 13 | 2 | Thegl |
| Them4 | 1129 | 1009 | 753 | 1059 | 1540 | 1419 | 1529 | 1292 | 1083 | 1042 | 976 | 783 | 792 | 833 | 1370 | 939 | 1005 | 1284 | 1329 | 1668 | 1520 | 1337 | 1130 | 868 | 993 | 645 | 645 | 712 | Them4 |
| Them5 | 3 | 6 | 7 | 0 | 7 | 0 | 2 | 11 | 15 | 22 | 10 | 15 | 12 | 9 | 4 | 5 | 0 | 3 | 5 | 3 | 4 | 7 | 27 | 11 | 13 | 24 | 7 | 9 | Them5 |
| Them6 | 314 | 416 | 389 | 394 | 385 | 280 | 302 | 84 | 134 | 180 | 108 | 110 | 135 | 144 | 276 | 329 | 396 | 349 | 377 | 306 | 323 | 122 | 118 | 130 | 155 | 152 | 127 | 122 | Them6 |
| Them7 | 1 | 6 | 8 | 4 | 0 | 0 | 1 | 0 | 5 | 8 | 3 | 2 | 5 | 3 | 5 | 3 | 1 | 0 | 0 | 0 | 0 | 5 | 3 | 2 | 2 | 2 | 0 | 0 | Them7 |

| | | | | | | | | | | | | | | | | | | | | | | | | | | | | | |
|---|---|---|---|---|---|---|---|---|---|---|---|---|---|---|---|---|---|---|---|---|---|---|---|---|---|---|---|---|---|
| Themis | 1 | 0 | 0 | 0 | 0 | 0 | 1 | 0 | 2 | 0 | 2 | 2 | 5 | 0 | 0 | 0 | 0 | 0 | 0 | 0 | 0 | 0 | 4 | 0 | 0 | 0 | 0 | 2 | Themis |
| Themis2 | 119 | 97 | 165 | 90 | 78 | 98 | 66 | 53 | 86 | 86 | 111 | 157 | 108 | 68 | 84 | 128 | 97 | 119 | 62 | 78 | 47 | 125 | 161 | 69 | 103 | 126 | 94 | 77 | Themis2 |
| Themis3 | 0 | 3 | 0 | 1 | 2 | 1 | 6 | 2 | 1 | 3 | 2 | 2 | 0 | 0 | 0 | 3 | 0 | 0 | 0 | 3 | 5 | 2 | 5 | 4 | 2 | 2 | 0 | 2 | Themis3 |
| Thg1l | 781 | 710 | 949 | 630 | 539 | 550 | 555 | 1314 | 1261 | 1207 | 1161 | 1377 | 1302 | 1545 | 820 | 758 | 683 | 519 | 610 | 731 | 491 | 1220 | 1346 | 1307 | 1191 | 1625 | 1546 | 1704 | Thg1l |
| Thnsl1 | 2963 | 2835 | 2492 | 2560 | 2338 | 2481 | 2693 | 2348 | 2034 | 1537 | 1647 | 1657 | 1547 | 1879 | 3559 | 2643 | 2491 | 2419 | 2293 | 3142 | 2602 | 1855 | 1852 | 1705 | 1748 | 1774 | 1963 | 1734 | Thnsl1 |
| Thnsl2 | 118 | 167 | 143 | 114 | 109 | 157 | 88 | 164 | 161 | 141 | 150 | 202 | 211 | 164 | 169 | 164 | 104 | 88 | 122 | 157 | 101 | 165 | 225 | 142 | 152 | 173 | 172 | 165 | Thnsl2 |
| Thoc1 | 6754 | 5379 | 6506 | 5478 | 5488 | 5800 | 6397 | 9570 | 7778 | 6913 | 7073 | 8190 | 8056 | 8477 | 6923 | 6291 | 5877 | 5560 | 5758 | 7330 | 5934 | 7855 | 7839 | 7618 | 7453 | 8415 | 8496 | 8556 | Thoc1 |
| Thoc2 | 17844 | 12710 | 11272 | 13841 | 13724 | 16141 | 16941 | 21126 | 17876 | 13710 | 14762 | 16191 | 16418 | 17122 | 18642 | 15932 | 11930 | 13222 | 13400 | 18096 | 16399 | 14674 | 17049 | 13378 | 15173 | 11368 | 17806 | 15378 | Thoc2 |
| Thoc2l | 7742 | 6290 | 5447 | 7235 | 6227 | 7431 | 7767 | 7261 | 5884 | 5191 | 5656 | 6164 | 6095 | 6348 | 7580 | 7608 | 5699 | 6655 | 6064 | 7393 | 7531 | 5147 | 5745 | 4797 | 5402 | 4587 | 6742 | 6036 | Thoc2l |
| Thoc3 | 4674 | 4818 | 5032 | 4732 | 3963 | 4179 | 3696 | 3223 | 3364 | 3596 | 3624 | 4528 | 4153 | 3985 | 4383 | 4552 | 4953 | 4464 | 4316 | 3992 | 3885 | 3387 | 3526 | 3938 | 3611 | 5164 | 4659 | 4367 | Thoc3 |
| Thoc5 | 3719 | 4146 | 3745 | 3690 | 3657 | 4083 | 3691 | 4637 | 4520 | 4746 | 3968 | 4820 | 4320 | 4732 | 3907 | 3218 | 3615 | 3606 | 4265 | 4253 | 4071 | 4348 | 4201 | 4740 | 4841 | 4925 | 4690 | 4904 | Thoc5 |
| Thoc6 | 1089 | 1431 | 1527 | 1253 | 1181 | 1145 | 1105 | 1631 | 1773 | 1714 | 1779 | 1831 | 1996 | 1801 | 1031 | 1152 | 1458 | 1130 | 1280 | 1228 | 1007 | 1322 | 1667 | 1846 | 1768 | 2252 | 2122 | 1957 | Thoc6 |
| Thoc7 | 4328 | 3491 | 4763 | 3680 | 4258 | 4200 | 4689 | 6464 | 5351 | 5847 | 5636 | 6874 | 6569 | 7223 | 4404 | 3754 | 4176 | 3536 | 4244 | 4919 | 4371 | 5743 | 5560 | 6814 | 6099 | 7406 | 6916 | 7844 | Thoc7 |
| Thop1 | 5405 | 6480 | 5410 | 5039 | 3973 | 4299 | 4130 | 2712 | 3469 | 3289 | 3135 | 3379 | 3300 | 2960 | 4763 | 5525 | 5297 | 4833 | 4772 | 4226 | 4396 | 2806 | 3449 | 3286 | 3065 | 3951 | 3242 | 3326 | Thop1 |
| Thpo | 112 | 124 | 144 | 135 | 144 | 148 | 125 | 81 | 77 | 93 | 105 | 61 | 75 | 62 | 91 | 118 | 143 | 118 | 138 | 130 | 105 | 110 | 78 | 80 | 75 | 57 | 72 | 51 | Thpo |
| Thra | 7699 | 10252 | 8296 | 14284 | 17349 | 15076 | 14911 | 13915 | 16476 | 20365 | 16692 | 14055 | 12316 | 14392 | 8773 | 8032 | 12167 | 14905 | 17410 | 13730 | 16757 | 19893 | 16450 | 18417 | 21190 | 12652 | 11538 | 11900 | Thra |
| Thrap3 | 28114 | 26618 | 29293 | 27848 | 24144 | 25902 | 27840 | 23320 | 24072 | 24672 | 25306 | 28487 | 27568 | 30831 | 26352 | 24779 | 27340 | 26143 | 26587 | 24914 | 27294 | 23181 | 23037 | 27286 | 25722 | 30907 | 28286 | 30584 | Thrap3 |
| Thrb | 181 | 167 | 135 | 133 | 126 | 127 | 121 | 239 | 283 | 266 | 202 | 192 | 185 | 180 | 170 | 180 | 149 | 150 | 98 | 128 | 125 | 130 | 224 | 274 | 139 | 114 | 167 | 139 | Thrb |
| Thrsp | 5 | 4 | 1 | 17 | 21 | 13 | 20 | 11 | 20 | 19 | 7 | 1 | 7 | 7 | 4 | 3 | 5 | 7 | 21 | 24 | 22 | 15 | 17 | 26 | 17 | 4 | 10 | 2 | Thrsp |
| Thsd1 | 887 | 992 | 1204 | 947 | 1183 | 1079 | 930 | 695 | 793 | 956 | 979 | 950 | 1081 | 893 | 944 | 998 | 1023 | 1049 | 1037 | 961 | 995 | 921 | 886 | 874 | 884 | 1063 | 941 | 845 | Thsd1 |
| Thsd4 | 1589 | 1729 | 1381 | 1650 | 1695 | 1974 | 1529 | 2464 | 3046 | 2636 | 3843 | 2465 | 2387 | 2411 | 1813 | 1369 | 1759 | 2061 | 1682 | 1413 | 1594 | 2904 | 2976 | 2387 | 2715 | 1579 | 2668 | 1998 | Thsd4 |
| Thsd7a | 2668 | 3099 | 3250 | 4635 | 6581 | 6766 | 4560 | 7018 | 5749 | 7693 | 7325 | 6139 | 10842 | 6314 | 4768 | 4520 | 4131 | 4725 | 4493 | 4836 | 5445 | 9927 | 6160 | 4683 | 8119 | 2283 | 7244 | 6478 | Thsd7a |
| Thsd7b | 1356 | 1394 | 551 | 928 | 1737 | 997 | 874 | 1888 | 1784 | 1553 | 1842 | 715 | 1638 | 814 | 1946 | 2275 | 844 | 1368 | 833 | 905 | 903 | 2151 | 1404 | 1075 | 1344 | 769 | 729 | 738 | Thsd7b |
| Thtpa | 676 | 742 | 701 | 774 | 722 | 696 | 713 | 819 | 906 | 1041 | 852 | 959 | 908 | 1003 | 618 | 499 | 764 | 590 | 781 | 712 | 824 | 867 | 851 | 1069 | 979 | 983 | 962 | 990 | Thtpa |
| Thumpd1 | 6676 | 6131 | 7006 | 6631 | 6199 | 6095 | 6289 | 7798 | 6325 | 6990 | 6847 | 7258 | 7195 | 7190 | 6771 | 5637 | 6839 | 5996 | 6177 | 6644 | 6085 | 7038 | 6286 | 7329 | 7076 | 7864 | 7262 | 7408 | Thumpd1 |
| Thumpd2 | 510 | 449 | 434 | 470 | 562 | 510 | 537 | 626 | 709 | 540 | 672 | 740 | 672 | 685 | 484 | 434 | 414 | 489 | 553 | 591 | 515 | 635 | 707 | 681 | 592 | 723 | 737 | 730 | Thumpd2 |
| Thumpd3 | 3712 | 3242 | 3501 | 3138 | 3017 | 3204 | 3698 | 4302 | 3448 | 3205 | 3191 | 3778 | 3441 | 3872 | 4127 | 3433 | 3330 | 3031 | 3099 | 3817 | 3238 | 3372 | 3493 | 3388 | 3672 | 3943 | 3873 | 3984 | Thumpd3 |
| Thy1 | 371 | 237 | 222 | 216 | 274 | 285 | 242 | 61 | 201 | 129 | 251 | 12 | 24 | 24 | 229 | 257 | 217 | 276 | 304 | 271 | 298 | 63 | 248 | 81 | 101 | 17 | 15 | 7 | Thy1 |
| Thyn1 | 2870 | 2654 | 2915 | 2499 | 2302 | 2513 | 2759 | 3127 | 2893 | 2706 | 2652 | 2831 | 3032 | 3364 | 2951 | 2602 | 2614 | 2322 | 2548 | 2917 | 2557 | 2410 | 2676 | 2984 | 2883 | 3340 | 2987 | 3419 | Thyn1 |
| Tia1 | 16461 | 11817 | 10779 | 13498 | 16858 | 18721 | 18799 | 39591 | 33339 | 29861 | 32682 | 32254 | 30257 | 33774 | 16944 | 13785 | 11941 | 13072 | 15389 | 19512 | 18786 | 31904 | 31894 | 28781 | 33079 | 22993 | 32247 | 28814 | Tia1 |
| Tial1 | 14198 | 10632 | 14121 | 16025 | 16565 | 17096 | 19480 | 15732 | 14075 | 16374 | 16357 | 17144 | 15815 | 18154 | 14364 | 12435 | 15378 | 16595 | 16453 | 15166 | 18247 | 18063 | 13785 | 16592 | 16885 | 15959 | 16612 | 16897 | Tial1 |
| Tiam1 | 4774 | 4426 | 5227 | 5354 | 5294 | 4842 | 4740 | 3729 | 4387 | 4956 | 5180 | 4796 | 4829 | 4505 | 3906 | 5203 | 5207 | 5207 | 5240 | 3969 | 4768 | 4896 | 4250 | 4962 | 5034 | 3897 | 4043 | 4194 | Tiam1 |
| Tiam2 | 1855 | 1410 | 1028 | 1580 | 2618 | 2292 | 2677 | 2376 | 3640 | 3300 | 2800 | 1414 | 1024 | 1455 | 1602 | 2002 | 1031 | 2179 | 2491 | 2039 | 2867 | 2032 | 2792 | 3099 | 4181 | 1116 | 851 | 773 | Tiam2 |
| Ticam1 | 767 | 805 | 736 | 675 | 466 | 564 | 475 | 320 | 428 | 324 | 430 | 379 | 327 | 285 | 649 | 787 | 676 | 631 | 524 | 526 | 425 | 287 | 480 | 364 | 310 | 419 | 354 | 336 | Ticam1 |
| Ticam2 | 84 | 56 | 53 | 78 | 58 | 64 | 54 | 76 | 55 | 78 | 61 | 61 | 82 | 65 | 82 | 115 | 64 | 54 | 46 | 62 | 63 | 53 | 76 | 64 | 41 | 54 | 88 | 84 | Ticam2 |
| Ticrr | 5628 | 5436 | 4919 | 5189 | 3620 | 4273 | 4960 | 2396 | 2625 | 2498 | 2460 | 4353 | 3823 | 4046 | 4714 | 5094 | 5231 | 4967 | 4436 | 3501 | 4426 | 2308 | 2144 | 3272 | 2850 | 3721 | 3814 | 4226 | Ticrr |
| Tie1 | 1819 | 2244 | 2665 | 2065 | 2055 | 2013 | 1592 | 1601 | 2236 | 2467 | 2788 | 2427 | 3009 | 2188 | 1617 | 2267 | 2126 | 2155 | 1968 | 1450 | 1678 | 2201 | 2452 | 2421 | 2321 | 3036 | 2636 | 2286 | Tie1 |
| Tifa | 359 | 295 | 431 | 333 | 377 | 372 | 485 | 303 | 278 | 221 | 248 | 301 | 247 | 282 | 456 | 263 | 346 | 305 | 441 | 486 | 435 | 241 | 277 | 257 | 286 | 279 | 277 | 310 | Tifa |
| Tifab | 48 | 43 | 48 | 53 | 43 | 48 | 39 | 100 | 129 | 93 | 147 | 92 | 133 | 95 | 36 | 55 | 45 | 30 | 29 | 50 | 29 | 105 | 168 | 99 | 92 | 139 | 91 | 104 | Tifab |
| Tigar | 2482 | 3303 | 3434 | 3864 | 3254 | 3022 | 3346 | 1561 | 1536 | 1674 | 1631 | 1757 | 1769 | 1982 | 2674 | 2716 | 3525 | 3650 | 3192 | 3137 | 3516 | 1843 | 1429 | 1911 | 2064 | 1728 | 1744 | 1784 | Tigar |
| Tigd2 | 1398 | 1030 | 1325 | 1068 | 1152 | 1231 | 1323 | 1076 | 934 | 931 | 968 | 1020 | 1002 | 995 | 1395 | 1164 | 1262 | 1100 | 1065 | 1260 | 1233 | 1023 | 990 | 962 | 877 | 1205 | 1012 | 1054 | Tigd2 |
| Tigd3 | 505 | 579 | 740 | 697 | 693 | 712 | 695 | 719 | 732 | 702 | 613 | 799 | 686 | 824 | 454 | 524 | 671 | 657 | 791 | 680 | 667 | 710 | 616 | 821 | 896 | 867 | 775 | 865 | Tigd3 |
| Tigd4 | 13 | 11 | 5 | 16 | 7 | 2 | 13 | 10 | 18 | 3 | 92 | 4 | 14 | 2 | 15 | 11 | 8 | 13 | 5 | 7 | 9 | 13 | 67 | 9 | 4 | 10 | 3 | 5 | Tigd4 |
| Tigd5 | 844 | 1167 | 833 | 1255 | 1067 | 992 | 948 | 581 | 676 | 642 | 681 | 808 | 662 | 665 | 807 | 942 | 1048 | 1185 | 1164 | 920 | 1113 | 851 | 629 | 762 | 781 | 663 | 698 | 614 | Tigd5 |
| Tigit | 0 | 0 | 3 | 3 | 2 | 0 | 0 | 1 | 0 | 0 | 1 | 1 | 1 | 1 | 0 | 0 | 1 | 0 | 0 | 0 | 0 | 7 | 0 | 1 | 3 | 3 | 7 | 1 | Tigit |
| Timd2 | 2 | 0 | 7 | 1 | 3 | 3 | 2 | 0 | 7 | 0 | 0 | 0 | 0 | 1 | 3 | 1 | 0 | 0 | 0 | 0 | 1 | 3 | 3 | 0 | 2 | 8 | 0 | 1 | Timd2 |
| Timd4 | 17 | 13 | 6 | 11 | 14 | 19 | 7 | 17 | 8 | 9 | 26 | 20 | 11 | 13 | 20 | 12 | 7 | 17 | 5 | 20 | 9 | 15 | 23 | 21 | 14 | 14 | 14 | 18 | Timd4 |
| Timeless | 3121 | 3223 | 3536 | 3115 | 2911 | 3007 | 2994 | 3519 | 4402 | 4272 | 4113 | 6044 | 5200 | 5753 | 2794 | 2820 | 3354 | 2998 | 3141 | 2786 | 2814 | 3207 | 3520 | 5132 | 4059 | 6944 | 5972 | 5707 | Timeless |
| Timm10 | 854 | 1213 | 1068 | 1106 | 1010 | 1037 | 1130 | 746 | 870 | 726 | 840 | 854 | 857 | 941 | 948 | 1055 | 1053 | 1104 | 1013 | 1316 | 1127 | 675 | 885 | 945 | 947 | 1019 | 951 | 939 | Timm10 |
| Timm10b | 652 | 556 | 589 | 730 | 644 | 649 | 732 | 608 | 471 | 564 | 591 | 542 | 507 | 667 | 728 | 615 | 622 | 594 | 586 | 595 | 768 | 632 | 547 | 644 | 570 | 550 | 502 | 595 | Timm10b |
| Timm13 | 1683 | 2067 | 2157 | 1896 | 1707 | 1568 | 1726 | 2132 | 2267 | 2576 | 2374 | 2620 | 2499 | 2732 | 1668 | 1548 | 2077 | 1662 | 2007 | 1848 | 1713 | 1878 | 2329 | 2622 | 2459 | 2833 | 2732 | 2669 | Timm13 |
| Timm17a | 2694 | 2889 | 2536 | 2672 | 2667 | 2747 | 2770 | 2983 | 2765 | 2902 | 2595 | 2876 | 2793 | 3100 | 2714 | 2746 | 2577 | 2507 | 2603 | 3061 | 2789 | 2475 | 2812 | 2731 | 2911 | 2775 | 3267 | 3014 | Timm17a |
| Timm17b | 2451 | 3272 | 3094 | 2797 | 2250 | 2536 | 2425 | 2224 | 2417 | 2419 | 2355 | 2620 | 2227 | 2450 | 2600 | 2579 | 3011 | 2532 | 2853 | 2702 | 2460 | 2015 | 2441 | 2567 | 2471 | 3076 | 2796 | 2500 | Timm17b |
| Timm21 | 1871 | 1414 | 1769 | 1432 | 1626 | 1502 | 1840 | 3108 | 2495 | 2533 | 2463 | 2872 | 2679 | 3183 | 1858 | 1422 | 1577 | 1247 | 1665 | 1968 | 1562 | 2420 | 2643 | 2838 | 2553 | 2920 | 2904 | 3077 | Timm21 |
| Timm22 | 3422 | 3601 | 3053 | 2972 | 2480 | 2688 | 2771 | 2500 | 2464 | 2382 | 2233 | 2609 | 2507 | 2464 | 3498 | 3259 | 3026 | 2917 | 2719 | 3067 | 2889 | 2186 | 2731 | 2352 | 2157 | 2886 | 2680 | 2797 | Timm22 |
| Timm23 | 2910 | 2969 | 2639 | 1732 | 1670 | 1860 | 1858 | 1793 | 1827 | 1482 | 2117 | 1929 | 2257 | 1950 | 2144 | 3325 | 2378 | 2059 | 1725 | 2834 | 1933 | 1370 | 1860 | 2509 | 1782 | 2470 | 2180 | 2403 | Timm23 |
| Timm29 | 7321 | 6991 | 7197 | 7199 | 7113 | 6933 | 7176 | 6891 | 5894 | 5813 | 5363 | 6121 | 5558 | 5942 | 7402 | 6092 | 7148 | 6863 | 7782 | 6977 | 7349 | 7234 | 5590 | 6007 | 6688 | 6387 | 6001 | 6270 | Timm29 |
| Timm44 | 2669 | 3037 | 3059 | 3112 | 2924 | 2920 | 3013 | 3368 | 3826 | 3808 | 3800 | 4001 | 3767 | 3936 | 2475 | 2725 | 3091 | 3055 | 3035 | 2777 | 2724 | 3382 | 3643 | 4015 | 3701 | 4245 | 4083 | 3978 | Timm44 |
| Timm50 | 3855 | 4992 | 4016 | 3612 | 3430 | 3493 | 3626 | 3317 | 3728 | 3374 | 3158 | 4165 | 3835 | 3976 | 3731 | 4063 | 4041 | 3297 | 4111 | 3855 | 3637 | 2872 | 3473 | 3735 | 3606 | 4605 | 4251 | 4105 | Timm50 |
| Timm8a1 | 1385 | 1077 | 1268 | 1084 | 1028 | 1098 | 1339 | 959 | 863 | 767 | 750 | 947 | 869 | 841 | 1404 | 1260 | 1187 | 1144 | 1110 | 1410 | 1199 | 697 | 878 | 748 | 758 | 1167 | 1076 | 1035 | Timm8a1 |
| Timm8a2 | 7 | 14 | 7 | 13 | 3 | 16 | 16 | 2 | 8 | 8 | 3 | 3 | 6 | 3 | 7 | 15 | 21 | 22 | 11 | 12 | 18 | 3 | 5 | 6 | 1 | 1 | 1 | 14 | Timm8a2 |
| Timm8b | 2618 | 3119 | 2818 | 3119 | 2801 | 2667 | 3400 | 3656 | 3303 | 3534 | 3313 | 3446 | 3376 | 4417 | 3032 | 2415 | 2924 | 2617 | 2967 | 3856 | 3487 | 2822 | 3300 | 3886 | 3556 | 4019 | 3971 | 4548 | Timm8b |
| Timm9 | 2345 | 2302 | 1975 | 2301 | 2146 | 2467 | 2605 | 1723 | 1900 | 1672 | 1838 | 1675 | 1903 | 1872 | 2308 | 2331 | 2011 | 2146 | 2094 | 2528 | 2492 | 1514 | 1878 | 1714 | 1829 | 1791 | 1766 | 1936 | Timm9 |
| Timmdc1 | 2299 | 2580 | 3123 | 2718 | 2325 | 2322 | 2656 | 1882 | 1816 | 2227 | 1919 | 2104 | 2019 | 2357 | 2443 | 2338 | 2861 | 2492 | 2511 | 2556 | 2503 | 1755 | 1928 | 2347 | 2207 | 2383 | 2237 | 2328 | Timmdc1 |
| Timp1 | 146 | 181 | 167 | 113 | 88 | 111 | 57 | 125 | 226 | 158 | 297 | 167 | 244 | 165 | 108 | 185 | 94 | 147 | 76 | 70 | 55 | 207 | 295 | 157 | 175 | 213 | 251 | 208 | Timp1 |
| Timp2 | 4682 | 4174 | 4365 | 3848 | 4289 | 4952 | 3491 | 6530 | 8143 | 7458 | 9416 | 6011 | 6691 | 6212 | 4259 | 5290 | 4011 | 5074 | 3710 | 2903 | 3677 | 8211 | 9270 | 7806 | 6908 | 7140 | 5689 | 6241 | Timp2 |
| Timp3 | 7503 | 6285 | 5930 | 5198 | 5514 | 5376 | 4377 | 5008 | 6835 | 5400 | 8690 | 3792 | 5830 | 3682 | 7919 | 8607 | 5477 | 6722 | 3861 | 4477 | 4298 | 6689 | 8141 | 4413 | 4544 | 4213 | 4065 | 3571 | Timp3 |
| Timp4 | 6 | 4 | 0 | 10 | 79 | 50 | 8 | 138 | 142 | 85 | 96 | 21 | 54 | 3 | 13 | 12 | 3 | 21 | 29 | 33 | 27 | 109 | 163 | 27 | 68 | 4 | 12 | 7 | Timp4 |
| Tinag | 12 | 5 | 4 | 0 | 0 | 0 | 0 | 10 | 11 | 10 | 6 | 5 | 15 | 12 | 18 | 9 | 1 | 0 | 0 | 0 | 0 | 3 | 14 | 2 | 4 | 24 | 15 | 7 | Tinag |
| Tinagl1 | 249 | 409 | 428 | 181 | 192 | 195 | 85 | 169 | 322 | 254 | 324 | 264 | 365 | 188 | 276 | 539 | 241 | 209 | 122 | 94 | 109 | 226 | 431 | 172 | 180 | 202 | 198 | 281 | Tinagl1 |
| Tincr | 376 | 617 | 604 | 754 | 623 | 595 | 572 | 203 | 187 | 295 | 245 | 273 | 264 | 243 | 297 | 538 | 727 | 841 | 718 | 523 | 548 | 343 | 213 | 312 | 278 | 429 | 271 | 262 | Tincr |
| Tinf2 | 4718 | 4879 | 4100 | 3774 | 2806 | 3238 | 3732 | 2524 | 2510 | 2417 | 2215 | 3266 | 2751 | 3076 | 4217 | 3972 | 3765 | 3334 | 3333 | 3674 | 3337 | 2074 | 2207 | 2545 | 2396 | 3602 | 3397 | 3304 | Tinf2 |
| Tiparp | 5139 | 4576 | 4387 | 4494 | 3890 | 4034 | 4364 | 3066 | 2408 | 2436 | 2460 | 2460 | 2631 | 2407 | 5501 | 5228 | 4441 | 4510 | 4044 | 4607 | 4119 | 2699 | 2601 | 2426 | 2652 | 2389 | 2707 | 2417 | Tiparp |
| Tipin | 5424 | 4067 | 5159 | 3893 | 3053 | 3677 | 4469 | 4534 | 4031 | 3560 | 3548 | 5321 | 4635 | 6218 | 5297 | 4354 | 4429 | 3510 | 3619 | 4585 | 3904 | 3182 | 3700 | 4436 | 3891 | 7010 | 5681 | 6617 | Tipin |
| Tiprl | 5621 | 5203 | 5675 | 6120 | 5610 | 5518 | 6093 | 4886 | 4369 | 4298 | 4074 | 4489 | 4112 | 4624 | 5942 | 5424 | 5886 | 5820 | 5697 | 6306 | 5538 | 4492 | 4435 | 4577 | 4544 | 5102 | 4398 | 4681 | Tiprl |
| Tirap | 1172 | 1097 | 1335 | 862 | 829 | 934 | 754 | 733 | 791 | 732 | 852 | 907 | 1101 | 1108 | 1142 | 1128 | 1212 | 803 | 976 | 899 | 859 | 734 | 805 | 817 | 705 | 1095 | 1144 | 1107 | Tirap |
| Tjap1 | 2859 | 4235 | 5114 | 4381 | 3585 | 3153 | 3197 | 2535 | 2843 | 3253 | 3524 | 4188 | 3839 | 3832 | 2714 | 3838 | 4913 | 4118 | 4032 | 2690 | 3154 | 2852 | 2863 | 4020 | 3221 | 4957 | 3588 | 4002 | Tjap1 |
| Tjp1 | 16515 | 14854 | 15585 | 15156 | 14575 | 14870 | 15657 | 12565 | 12947 | 12422 | 13359 | 13643 | 14469 | 13967 | 15893 | 16035 | 14579 | 15097 | 14054 | 14222 | 15294 | 12009 | 13778 | 12122 | 12294 | 10758 | 13927 | 13413 | Tjp1 |
| Tjp2 | 4916 | 5148 | 6697 | 5387 | 4853 | 4887 | 4430 | 2619 | 3015 | 3393 | 3005 | 4136 | 3666 | 3829 | 4367 | 5101 | 6220 | 5381 | 5142 | 4107 | 4523 | 2732 | 2970 | 3920 | 3244 | 4838 | 4019 | 4198 | Tjp2 |

| | | | | | | | | | | | | | | | | | | | | | | | | | | | | | |
|---|---|---|---|---|---|---|---|---|---|---|---|---|---|---|---|---|---|---|---|---|---|---|---|---|---|---|---|---|---|
| Tjp3 | 397 | 417 | 471 | 475 | 356 | 287 | 337 | 299 | 362 | 455 | 291 | 354 | 263 | 321 | 326 | 296 | 466 | 375 | 441 | 345 | 345 | 251 | 358 | 564 | 317 | 546 | 298 | 314 | Tjp3 |
| Tk1 | 1390 | 1399 | 1422 | 1142 | 961 | 1207 | 1161 | 2241 | 2843 | 2318 | 2447 | 3780 | 3103 | 3382 | 1160 | 1079 | 1285 | 991 | 1287 | 1350 | 1063 | 1465 | 2420 | 3202 | 2633 | 4048 | 3690 | 3408 | Tk1 |
| Tk2 | 2649 | 2930 | 2827 | 3213 | 2860 | 2882 | 3062 | 1754 | 1949 | 2080 | 1867 | 2112 | 1977 | 2291 | 2720 | 2821 | 3057 | 2978 | 2950 | 2573 | 3194 | 2008 | 1838 | 2002 | 2325 | 2130 | 2022 | 2277 | Tk2 |
| Tkfc | 1538 | 1838 | 1840 | 1670 | 1724 | 1680 | 1665 | 1787 | 2181 | 2049 | 2351 | 2640 | 2232 | 2698 | 1458 | 1543 | 1877 | 1728 | 2001 | 1647 | 1791 | 2152 | 1952 | 2498 | 2322 | 2950 | 2454 | 2608 | Tkfc |
| Tkt | 12204 | 15426 | 12625 | 12582 | 11608 | 12540 | 11500 | 9594 | 12397 | 11702 | 10345 | 12640 | 11644 | 11481 | 10843 | 12799 | 12974 | 11793 | 14138 | 12332 | 12289 | 8995 | 11941 | 12636 | 11883 | 15224 | 12077 | 12708 | Tkt |
| Tktl1 | 1 | 0 | 1 | 2 | 0 | 0 | 1 | 2 | 0 | 3 | 0 | 1 | 0 | 0 | 0 | 1 | 0 | 0 | 0 | 0 | 2 | 0 | 0 | 0 | 0 | 0 | 3 | 0 | Tktl1 |
| Tktl2 | 1 | 6 | 8 | 8 | 7 | 31 | 12 | 1 | 4 | 5 | 3 | 0 | 3 | 1 | 5 | 1 | 5 | 4 | 6 | 2 | 4 | 3 | 3 | 2 | 2 | 0 | 1 | 0 | Tktl2 |
| Tlcd1 | 1017 | 983 | 963 | 1080 | 780 | 844 | 872 | 427 | 430 | 434 | 487 | 479 | 378 | 565 | 919 | 931 | 1140 | 968 | 837 | 781 | 924 | 416 | 515 | 429 | 361 | 550 | 438 | 418 | Tlcd1 |
| Tlcd2 | 180 | 124 | 180 | 92 | 61 | 115 | 61 | 203 | 294 | 232 | 264 | 210 | 235 | 152 | 104 | 172 | 97 | 103 | 70 | 54 | 61 | 159 | 323 | 213 | 131 | 290 | 188 | 163 | Tlcd2 |
| Tlcd3a | 2261 | 2243 | 2229 | 1914 | 1962 | 1979 | 1901 | 1542 | 1504 | 1612 | 1523 | 1549 | 1614 | 1569 | 2178 | 2284 | 2103 | 1952 | 2060 | 1950 | 1966 | 1734 | 1504 | 1430 | 1669 | 1688 | 1608 | 1717 | Tlcd3a |
| Tlcd3b | 2187 | 3446 | 3502 | 5534 | 7113 | 5403 | 5634 | 4395 | 4869 | 6397 | 5259 | 5143 | 4583 | 5317 | 2652 | 2401 | 4985 | 5690 | 7097 | 4673 | 6347 | 7595 | 4281 | 6136 | 6988 | 4539 | 4465 | 4719 | Tlcd3b |
| Tlcd4 | 738 | 500 | 511 | 623 | 696 | 728 | 808 | 655 | 590 | 489 | 561 | 556 | 284 | 524 | 804 | 501 | 627 | 638 | 809 | 923 | 832 | 466 | 482 | 499 | 616 | 416 | 413 | 519 | Tlcd4 |
| Tlcd5 | 3096 | 2794 | 3301 | 3136 | 2908 | 3032 | 3245 | 2032 | 1902 | 2012 | 1963 | 2268 | 2189 | 2518 | 3154 | 2805 | 3060 | 3042 | 2862 | 2847 | 3136 | 2386 | 1869 | 2080 | 2256 | 2147 | 2187 | 2244 | Tlcd5 |
| Tldc2 | 7 | 10 | 17 | 5 | 4 | 0 | 6 | 7 | 3 | 9 | 3 | 5 | 8 | 12 | 7 | 4 | 8 | 14 | 19 | 5 | 5 | 3 | 2 | 4 | 10 | 8 | 3 | 4 | Tldc2 |
| Tle1 | 4854 | 5275 | 5413 | 5770 | 5900 | 5222 | 5273 | 7132 | 6757 | 7069 | 6463 | 7557 | 6361 | 7244 | 5118 | 4925 | 5769 | 5668 | 6010 | 5376 | 5253 | 7424 | 6590 | 7170 | 7489 | 8031 | 6563 | 7153 | Tle1 |
| Tle2 | 1835 | 1942 | 2083 | 2411 | 2388 | 2145 | 1743 | 1789 | 2581 | 3014 | 3375 | 2002 | 2301 | 1570 | 1538 | 2100 | 2344 | 2663 | 2233 | 1671 | 2087 | 2861 | 2363 | 2761 | 2450 | 1975 | 1897 | 1587 | Tle2 |
| Tle3 | 5914 | 7882 | 10194 | 9742 | 7706 | 6542 | 6450 | 7461 | 8361 | 9967 | 8984 | 11773 | 10210 | 11106 | 5636 | 7026 | 10004 | 8776 | 8259 | 5516 | 7363 | 8863 | 7274 | 12223 | 10165 | 11065 | 11168 | 11330 | Tle3 |
| Tle4 | 6729 | 6536 | 6906 | 7850 | 8102 | 7016 | 8113 | 7493 | 7016 | 7008 | 6616 | 8569 | 7165 | 8307 | 6752 | 6631 | 7463 | 7674 | 7850 | 7077 | 7822 | 8858 | 6064 | 7546 | 8045 | 7611 | 7527 | 8441 | Tle4 |
| Tle5 | 9926 | 12228 | 11979 | 11825 | 11512 | 10789 | 9686 | 10864 | 12349 | 14323 | 13246 | 13786 | 12513 | 13091 | 9432 | 9189 | 12266 | 11548 | 12320 | 10366 | 10135 | 12078 | 12124 | 14978 | 13732 | 15430 | 13269 | 12914 | Tle5 |
| Tle6 | 178 | 202 | 178 | 192 | 151 | 171 | 93 | 134 | 190 | 147 | 218 | 117 | 161 | 88 | 166 | 236 | 212 | 172 | 150 | 132 | 107 | 160 | 200 | 159 | 155 | 162 | 137 | 127 | Tle6 |
| Tle7 | 0 | 0 | 3 | 0 | 1 | 0 | 0 | 0 | 0 | 0 | 0 | 0 | 1 | 0 | 0 | 2 | 1 | 0 | 0 | 1 | 2 | 0 | 0 | 0 | 0 | 0 | 0 | 0 | Tle7 |
| Tlk1 | 6226 | 4722 | 6049 | 5861 | 7081 | 6574 | 7315 | 8816 | 8062 | 8342 | 8397 | 10207 | 9554 | 10099 | 6179 | 5156 | 6302 | 5815 | 6894 | 6341 | 6902 | 9225 | 7627 | 8673 | 8881 | 10025 | 9862 | 10118 | Tlk1 |
| Tlk2 | 5171 | 4706 | 5343 | 4766 | 5146 | 4974 | 5291 | 8074 | 6913 | 6465 | 6535 | 7300 | 7017 | 7828 | 5596 | 4394 | 5217 | 4525 | 5336 | 5281 | 5138 | 7195 | 6528 | 6981 | 6981 | 7687 | 7649 | 7988 | Tlk2 |
| Tll1 | 248 | 174 | 209 | 203 | 281 | 278 | 255 | 468 | 442 | 489 | 632 | 307 | 327 | 307 | 202 | 212 | 233 | 210 | 233 | 244 | 229 | 543 | 637 | 356 | 446 | 258 | 321 | 253 | Tll1 |
| Tll2 | 193 | 298 | 243 | 281 | 347 | 271 | 292 | 200 | 225 | 205 | 270 | 238 | 212 | 196 | 255 | 262 | 258 | 398 | 334 | 242 | 378 | 307 | 263 | 250 | 311 | 133 | 255 | 171 | Tll2 |
| Tln1 | 11674 | 12610 | 9963 | 8924 | 7198 | 8692 | 5530 | 8679 | 12216 | 11054 | 12839 | 10396 | 12875 | 9449 | 10028 | 15193 | 8869 | 9453 | 7143 | 5825 | 6325 | 9431 | 13221 | 10811 | 9265 | 8143 | 11073 | 9808 | Tln1 |
| Tln2 | 6508 | 6532 | 4517 | 4825 | 5221 | 5145 | 4072 | 4557 | 6039 | 5130 | 6854 | 4535 | 5988 | 4146 | 5879 | 7547 | 4177 | 5478 | 4163 | 3937 | 4514 | 4924 | 6376 | 3839 | 4765 | 3212 | 5348 | 4311 | Tln2 |
| Tlnrd1 | 2222 | 2167 | 2118 | 1847 | 1969 | 1813 | 1455 | 1936 | 2063 | 1997 | 2307 | 2066 | 2303 | 1861 | 1792 | 2230 | 2110 | 1784 | 1969 | 1558 | 1661 | 2097 | 2054 | 2074 | 1824 | 2433 | 1981 | 1880 | Tlnrd1 |
| Tlr1 | 17 | 16 | 15 | 3 | 21 | 13 | 20 | 56 | 57 | 67 | 85 | 73 | 40 | 48 | 17 | 16 | 13 | 15 | 12 | 22 | 16 | 75 | 81 | 62 | 57 | 69 | 64 | 47 | Tlr1 |
| Tlr11 | 0 | 0 | 0 | 0 | 0 | 0 | 0 | 0 | 0 | 2 | 1 | 0 | 0 | 0 | 0 | 0 | 0 | 0 | 0 | 0 | 0 | 0 | 2 | 3 | 0 | 0 | 0 | 0 | Tlr11 |
| Tlr12 | 44 | 65 | 83 | 71 | 100 | 71 | 99 | 99 | 91 | 176 | 124 | 151 | 110 | 136 | 51 | 54 | 109 | 90 | 129 | 70 | 109 | 152 | 108 | 172 | 159 | 169 | 89 | 97 | Tlr12 |
| Tlr13 | 25 | 34 | 22 | 16 | 24 | 25 | 21 | 64 | 99 | 49 | 87 | 60 | 65 | 52 | 32 | 32 | 9 | 42 | 30 | 31 | 24 | 58 | 102 | 68 | 48 | 59 | 43 | 59 | Tlr13 |
| Tlr2 | 316 | 218 | 262 | 164 | 160 | 208 | 139 | 192 | 199 | 189 | 234 | 74 | 147 | 80 | 231 | 292 | 166 | 200 | 159 | 141 | 137 | 132 | 230 | 113 | 124 | 125 | 126 | 81 | Tlr2 |
| Tlr3 | 149 | 100 | 119 | 80 | 149 | 122 | 129 | 212 | 208 | 225 | 200 | 229 | 188 | 219 | 104 | 101 | 109 | 106 | 118 | 131 | 121 | 289 | 217 | 224 | 185 | 211 | 241 | 179 | Tlr3 |
| Tlr4 | 165 | 103 | 89 | 81 | 126 | 115 | 82 | 156 | 158 | 145 | 189 | 131 | 161 | 134 | 141 | 199 | 85 | 87 | 84 | 143 | 107 | 165 | 225 | 104 | 151 | 113 | 143 | 103 | Tlr4 |
| Tlr5 | 205 | 212 | 121 | 129 | 118 | 166 | 101 | 83 | 148 | 117 | 148 | 98 | 166 | 87 | 239 | 247 | 105 | 171 | 85 | 95 | 86 | 134 | 199 | 80 | 88 | 57 | 110 | 103 | Tlr5 |
| Tlr6 | 47 | 42 | 73 | 41 | 20 | 33 | 11 | 38 | 91 | 60 | 104 | 76 | 109 | 86 | 44 | 80 | 37 | 34 | 31 | 30 | 11 | 62 | 114 | 80 | 78 | 71 | 81 | 69 | Tlr6 |
| Tlr7 | 30 | 25 | 39 | 17 | 30 | 35 | 35 | 116 | 104 | 117 | 166 | 108 | 94 | 103 | 49 | 52 | 44 | 29 | 24 | 50 | 31 | 127 | 175 | 115 | 83 | 91 | 130 | 115 | Tlr7 |
| Tlr8 | 9 | 9 | 9 | 6 | 7 | 17 | 7 | 31 | 41 | 29 | 45 | 18 | 22 | 21 | 22 | 15 | 9 | 14 | 12 | 25 | 10 | 45 | 59 | 17 | 20 | 30 | 17 | 21 | Tlr8 |
| Tlr9 | 6 | 8 | 5 | 4 | 11 | 16 | 2 | 15 | 26 | 19 | 43 | 22 | 19 | 26 | 11 | 11 | 8 | 14 | 3 | 9 | 12 | 23 | 34 | 20 | 28 | 12 | 21 | 19 | Tlr9 |
| Tlx1 | 23 | 96 | 15 | 23 | 100 | 65 | 1 | 137 | 133 | 137 | 138 | 17 | 356 | 18 | 165 | 147 | 44 | 77 | 6 | 4 | 4 | 177 | 118 | 9 | 50 | 4 | 17 | 11 | Tlx1 |
| Tlx2 | 37 | 60 | 6 | 1 | 74 | 4 | 0 | 110 | 89 | 56 | 88 | 0 | 110 | 2 | 59 | 83 | 19 | 33 | 0 | 1 | 1 | 105 | 66 | 2 | 37 | 4 | 0 | 0 | Tlx2 |
| Tlx3 | 52 | 117 | 89 | 75 | 249 | 73 | 34 | 134 | 132 | 205 | 297 | 26 | 244 | 26 | 107 | 147 | 138 | 92 | 68 | 65 | 27 | 184 | 189 | 23 | 98 | 1 | 27 | 19 | Tlx3 |
| Tm2d1 | 950 | 660 | 963 | 948 | 1161 | 1038 | 1161 | 1773 | 1431 | 1413 | 1459 | 1478 | 1505 | 1757 | 998 | 854 | 931 | 921 | 1086 | 1218 | 1170 | 1644 | 1592 | 1584 | 1555 | 1552 | 1599 | 1546 | Tm2d1 |
| Tm2d2 | 3623 | 4251 | 2976 | 3442 | 3066 | 3181 | 3264 | 2595 | 2377 | 1979 | 2083 | 2178 | 2359 | 2299 | 3574 | 3556 | 3191 | 3601 | 3324 | 3754 | 3363 | 2187 | 2330 | 2250 | 2210 | 2400 | 2264 | 2275 | Tm2d2 |
| Tm2d3 | 1019 | 1373 | 1130 | 1186 | 1078 | 1072 | 1117 | 1205 | 1279 | 1111 | 1128 | 1234 | 1085 | 1083 | 1106 | 1234 | 1138 | 1106 | 1133 | 1284 | 1020 | 1155 | 1303 | 1172 | 1239 | 1265 | 1220 | 1095 | Tm2d3 |
| Tm4sf1 | 300 | 215 | 302 | 194 | 300 | 321 | 187 | 592 | 723 | 568 | 760 | 487 | 807 | 378 | 355 | 323 | 286 | 305 | 241 | 301 | 228 | 643 | 1042 | 554 | 599 | 502 | 773 | 528 | Tm4sf1 |
| Tm4sf19 | 7 | 8 | 8 | 12 | 12 | 18 | 15 | 11 | 14 | 6 | 4 | 8 | 9 | 7 | 13 | 15 | 5 | 14 | 12 | 14 | 11 | 10 | 3 | 2 | 9 | 7 | 8 | 5 | Tm4sf19 |
| Tm4sf20 | 0 | 0 | 0 | 0 | 1 | 0 | 0 | 0 | 0 | 0 | 0 | 0 | 0 | 0 | 0 | 0 | 0 | 0 | 0 | 0 | 0 | 0 | 0 | 0 | 0 | 0 | 0 | 0 | Tm4sf20 |
| Tm4sf4 | 0 | 1 | 0 | 0 | 3 | 0 | 1 | 1 | 4 | 3 | 5 | 2 | 0 | 0 | 1 | 1 | 7 | 0 | 4 | 0 | 3 | 2 | 3 | 5 | 1 | 0 | 0 | 1 | Tm4sf4 |
| Tm4sf5 | 7 | 5 | 0 | 2 | 10 | 10 | 6 | 2 | 7 | 2 | 3 | 5 | 15 | 3 | 4 | 3 | 6 | 4 | 4 | 5 | 4 | 13 | 8 | 5 | 6 | 8 | 16 | 5 | Tm4sf5 |
| Tm6sf1 | 797 | 732 | 765 | 562 | 667 | 700 | 689 | 672 | 664 | 664 | 1159 | 691 | 824 | 782 | 885 | 759 | 645 | 566 | 652 | 704 | 678 | 747 | 1021 | 667 | 592 | 692 | 783 | 679 | Tm6sf1 |
| Tm6sf2 | 250 | 475 | 456 | 686 | 538 | 420 | 370 | 278 | 260 | 308 | 239 | 234 | 251 | 229 | 277 | 415 | 594 | 675 | 581 | 372 | 432 | 383 | 236 | 271 | 301 | 194 | 241 | 252 | Tm6sf2 |
| Tm7sf2 | 2110 | 2488 | 2620 | 2730 | 2547 | 2437 | 2348 | 2025 | 1952 | 2285 | 2264 | 2515 | 2454 | 2440 | 1932 | 2110 | 2556 | 2516 | 2698 | 2264 | 2239 | 2692 | 1902 | 2394 | 2664 | 2322 | 2650 | 2444 | Tm7sf2 |
| Tm7sf3 | 1880 | 1612 | 1720 | 1738 | 2020 | 1856 | 1892 | 2055 | 2280 | 2171 | 2481 | 1907 | 2023 | 1964 | 1832 | 1698 | 1683 | 1843 | 1789 | 1649 | 1717 | 1982 | 2293 | 1940 | 1932 | 2381 | 2016 | 1933 | Tm7sf3 |
| Tm9sf1 | 2631 | 3107 | 2921 | 2771 | 2360 | 2414 | 2112 | 2191 | 2683 | 2685 | 2814 | 2835 | 2643 | 2520 | 2447 | 2882 | 3110 | 2714 | 2722 | 2188 | 2162 | 2296 | 2742 | 2685 | 2772 | 2826 | 2781 | 2643 | Tm9sf1 |
| Tm9sf2 | 12422 | 9555 | 10870 | 8968 | 8757 | 9386 | 9566 | 10271 | 9225 | 8736 | 9597 | 9292 | 9626 | 9269 | 12398 | 10828 | 9692 | 9765 | 8715 | 9020 | 9047 | 10014 | 10325 | 8938 | 8460 | 9940 | 9519 | 10009 | Tm9sf2 |
| Tm9sf3 | 19912 | 14943 | 19527 | 17733 | 17680 | 18797 | 19217 | 22129 | 19907 | 18884 | 21191 | 20708 | 20832 | 20418 | 20274 | 18866 | 18178 | 18412 | 16737 | 17591 | 18148 | 21301 | 21187 | 19128 | 20395 | 20362 | 20499 | 20906 | Tm9sf3 |
| Tm9sf4 | 4154 | 4561 | 4164 | 4172 | 4111 | 4078 | 3891 | 5379 | 5960 | 5502 | 5706 | 5782 | 5410 | 5219 | 4146 | 4260 | 4260 | 4201 | 4144 | 3788 | 3971 | 6076 | 5913 | 5261 | 5641 | 5287 | 5566 | 5542 | Tm9sf4 |
| Tm9sf5 | 0 | 0 | 4 | 0 | 5 | 0 | 0 | 0 | 1 | 0 | 0 | 2 | 0 | 1 | 0 | 0 | 2 | 0 | 1 | 2 | 1 | 0 | 1 | 0 | 0 | 2 | 0 | 0 | Tm9sf5 |
| Tma16 | 2487 | 2162 | 2117 | 2374 | 2192 | 2207 | 2520 | 1883 | 1794 | 1764 | 1747 | 2118 | 2104 | 2190 | 2409 | 2089 | 2275 | 2205 | 2093 | 2382 | 2347 | 1619 | 1802 | 1995 | 1975 | 2185 | 2130 | 2302 | Tma16 |
| Tma7 | 6877 | 7044 | 8352 | 8132 | 6559 | 6284 | 8610 | 6174 | 5657 | 7262 | 6911 | 6889 | 6873 | 9276 | 7067 | 6673 | 8183 | 7072 | 7052 | 8110 | 8221 | 5257 | 6051 | 7285 | 6855 | 8226 | 7424 | 8823 | Tma7 |
| Tmbim1 | 353 | 353 | 329 | 269 | 240 | 338 | 155 | 592 | 866 | 612 | 1028 | 484 | 696 | 524 | 327 | 362 | 276 | 313 | 223 | 214 | 186 | 655 | 1071 | 632 | 542 | 586 | 607 | 577 | Tmbim1 |
| Tmbim4 | 2062 | 2200 | 2150 | 1983 | 1438 | 1562 | 1580 | 1419 | 1276 | 1211 | 1293 | 1259 | 1328 | 1299 | 2229 | 2090 | 1839 | 1762 | 1648 | 1850 | 1601 | 1484 | 1291 | 1332 | 1319 | 1332 | 1183 | 1425 | Tmbim4 |
| Tmbim6 | 15357 | 18321 | 19930 | 19598 | 16585 | 16622 | 14800 | 11264 | 13419 | 14300 | 13762 | 14139 | 14801 | 12823 | 15708 | 17828 | 19670 | 18588 | 17924 | 15976 | 14521 | 12257 | 13989 | 14882 | 13845 | 16143 | 14664 | 13874 | Tmbim6 |
| Tmbim7 | 1 | 0 | 0 | 0 | 0 | 1 | 0 | 0 | 0 | 0 | 0 | 0 | 0 | 0 | 0 | 0 | 0 | 0 | 2 | 0 | 0 | 0 | 0 | 2 | 0 | 0 | 0 | 0 | Tmbim7 |
| Tmc1 | 43 | 39 | 21 | 14 | 19 | 25 | 11 | 37 | 42 | 16 | 11 | 21 | 44 | 23 | 37 | 48 | 39 | 14 | 5 | 18 | 12 | 22 | 37 | 11 | 17 | 15 | 24 | 27 | Tmc1 |
| Tmc2 | 1 | 3 | 6 | 7 | 0 | 4 | 3 | 0 | 0 | 3 | 5 | 18 | 6 | 4 | 1 | 2 | 2 | 4 | 2 | 1 | 0 | 0 | 5 | 2 | 5 | 0 | 5 | 12 | Tmc2 |
| Tmc3 | 241 | 245 | 161 | 120 | 116 | 145 | 78 | 57 | 51 | 76 | 177 | 63 | 136 | 48 | 177 | 257 | 177 | 143 | 143 | 147 | 102 | 38 | 121 | 47 | 30 | 2 | 115 | 69 | Tmc3 |
| Tmc4 | 1578 | 1917 | 2300 | 2776 | 2526 | 2042 | 2242 | 584 | 640 | 932 | 865 | 607 | 711 | 656 | 1461 | 2020 | 2423 | 3021 | 2336 | 1731 | 2099 | 909 | 633 | 842 | 629 | 590 | 648 | 558 | Tmc4 |
| Tmc5 | 27 | 42 | 36 | 31 | 47 | 41 | 36 | 44 | 37 | 47 | 45 | 64 | 52 | 72 | 50 | 28 | 30 | 32 | 25 | 49 | 43 | 47 | 69 | 72 | 53 | 63 | 76 | 78 | Tmc5 |
| Tmc6 | 994 | 941 | 1049 | 723 | 509 | 598 | 430 | 355 | 473 | 493 | 638 | 373 | 575 | 265 | 733 | 1134 | 712 | 761 | 561 | 442 | 359 | 409 | 529 | 404 | 277 | 442 | 368 | 415 | Tmc6 |
| Tmc7 | 2227 | 2068 | 1910 | 1939 | 1767 | 1968 | 2045 | 1679 | 1442 | 1446 | 1307 | 1946 | 1666 | 1816 | 2040 | 1909 | 1913 | 1790 | 1947 | 2037 | 1898 | 1506 | 1389 | 1624 | 1711 | 1660 | 1747 | 1774 | Tmc7 |
| Tmc8 | 73 | 125 | 197 | 89 | 68 | 109 | 66 | 64 | 77 | 104 | 105 | 93 | 120 | 90 | 70 | 132 | 145 | 100 | 79 | 68 | 77 | 102 | 114 | 78 | 112 | 105 | 98 | 107 | Tmc8 |
| Tmcc1 | 5548 | 6260 | 6107 | 6933 | 7398 | 6657 | 7454 | 6771 | 5580 | 6319 | 5762 | 6548 | 5644 | 6239 | 6033 | 5464 | 6859 | 6762 | 7261 | 6565 | 7336 | 7792 | 5395 | 5924 | 6615 | 5713 | 5962 | 6616 | Tmcc1 |
| Tmcc2 | 2543 | 3863 | 3834 | 4646 | 5142 | 4253 | 3846 | 3632 | 4197 | 4906 | 4126 | 4718 | 6102 | 4476 | 3195 | 3214 | 4772 | 4513 | 5002 | 3773 | 4362 | 4807 | 4062 | 4795 | 5154 | 4556 | 4470 | 5040 | Tmcc2 |
| Tmcc3 | 1527 | 1538 | 1347 | 1580 | 1523 | 1543 | 1438 | 1032 | 1114 | 956 | 1392 | 736 | 827 | 813 | 1341 | 1466 | 1501 | 1724 | 1501 | 1374 | 1524 | 1026 | 1295 | 1071 | 924 | 849 | 705 | 757 | Tmcc3 |
| Tmco1 | 3002 | 3081 | 3172 | 2916 | 2548 | 2475 | 3264 | 3248 | 2958 | 2730 | 2740 | 2805 | 2804 | 3251 | 3417 | 3118 | 3097 | 2832 | 2736 | 3236 | 3128 | 2689 | 3112 | 2933 | 2751 | 3010 | 3045 | 3373 | Tmco1 |
| Tmco2 | 6 | 0 | 2 | 0 | 2 | 0 | 2 | 0 | 1 | 6 | 1 | 0 | 0 | 2 | 4 | 0 | 0 | 1 | 0 | 1 | 1 | 0 | 1 | 4 | 0 | 2 | 0 | 1 | Tmco2 |

| | | | | | | | | | | | | | | | | | | | | | | | | | | | | | |
|---|---|---|---|---|---|---|---|---|---|---|---|---|---|---|---|---|---|---|---|---|---|---|---|---|---|---|---|---|---|
| Tmco3 | 2586 | 2962 | 2424 | 2662 | 2980 | 3004 | 2751 | 2732 | 2792 | 2592 | 2814 | 2357 | 2719 | 2551 | 2651 | 2962 | 2468 | 2732 | 2890 | 2758 | 2888 | 2592 | 2896 | 2592 | 2575 | 2355 | 2613 | 2591 | Tmco3 |
| Tmco4 | 46 | 45 | 59 | 53 | 55 | 43 | 18 | 93 | 127 | 132 | 125 | 108 | 129 | 96 | 47 | 82 | 52 | 36 | 48 | 30 | 33 | 95 | 182 | 112 | 81 | 154 | 115 | 96 | Tmco4 |
| Tmco5 | 0 | 0 | 0 | 0 | 1 | 0 | 0 | 1 | 0 | 0 | 0 | 0 | 0 | 0 | 0 | 0 | 0 | 0 | 0 | 0 | 0 | 0 | 0 | 0 | 0 | 0 | 0 | 1 | Tmco5 |
| Tmco5b | 0 | 0 | 0 | 0 | 1 | 0 | 0 | 0 | 0 | 0 | 0 | 2 | 0 | 0 | 0 | 2 | 0 | 1 | 3 | 0 | 4 | 0 | 1 | 1 | 5 | 2 | 0 | 1 | Tmco5b |
| Tmco6 | 837 | 980 | 1140 | 1056 | 810 | 859 | 778 | 1072 | 1070 | 1138 | 1047 | 1313 | 1128 | 1195 | 785 | 823 | 1053 | 875 | 960 | 832 | 789 | 1041 | 1052 | 1200 | 1202 | 1236 | 1303 | 1280 | Tmco6 |
| Tmdd1 | 0 | 3 | 2 | 0 | 1 | 0 | 1 | 0 | 1 | 0 | 3 | 0 | 5 | 0 | 4 | 1 | 5 | 4 | 1 | 0 | 0 | 0 | 1 | 2 | 0 | 0 | 1 | 0 | Tmdd1 |
| Tmed1 | 487 | 536 | 627 | 366 | 375 | 430 | 389 | 1257 | 1482 | 1242 | 1351 | 1539 | 1492 | 1379 | 516 | 558 | 479 | 372 | 443 | 511 | 352 | 1163 | 1501 | 1301 | 1154 | 1661 | 1490 | 1432 | Tmed1 |
| Tmed10 | 12140 | 10509 | 12531 | 11485 | 11407 | 11487 | 12222 | 11883 | 11852 | 11929 | 12122 | 11918 | 11969 | 12366 | 12493 | 11566 | 12397 | 11731 | 11337 | 11709 | 11720 | 12457 | 12258 | 11978 | 11570 | 13247 | 12121 | 12819 | Tmed10 |
| Tmed2 | 11506 | 9509 | 10606 | 9186 | 8107 | 8663 | 9021 | 7795 | 7187 | 6642 | 7732 | 6959 | 7499 | 6942 | 11190 | 10581 | 9068 | 9157 | 8641 | 9088 | 8262 | 6985 | 8148 | 6840 | 6936 | 8451 | 7592 | 7346 | Tmed2 |
| Tmed3 | 828 | 873 | 730 | 696 | 722 | 749 | 639 | 1364 | 1753 | 1388 | 1720 | 1288 | 1319 | 1221 | 916 | 811 | 712 | 864 | 629 | 696 | 690 | 1422 | 1889 | 1468 | 1228 | 1370 | 1235 | 1212 | Tmed3 |
| Tmed4 | 5088 | 5637 | 5223 | 5122 | 4912 | 4665 | 5002 | 5041 | 4682 | 4642 | 4391 | 5044 | 4801 | 5434 | 5124 | 5122 | 5201 | 5036 | 5108 | 5133 | 5034 | 4809 | 4809 | 5203 | 5065 | 5629 | 5363 | 5474 | Tmed4 |
| Tmed5 | 5755 | 3803 | 5651 | 4898 | 3835 | 4919 | 4975 | 3047 | 2712 | 2151 | 2880 | 2833 | 3138 | 3421 | 5836 | 5495 | 4836 | 4307 | 4161 | 4813 | 4748 | 2665 | 3008 | 2683 | 2661 | 3411 | 3166 | 3440 | Tmed5 |
| Tmed6 | 0 | 0 | 1 | 3 | 1 | 0 | 4 | 1 | 1 | 0 | 0 | 0 | 0 | 0 | 0 | 0 | 0 | 1 | 0 | 1 | 2 | 2 | 0 | 2 | 0 | 0 | 2 | 0 | Tmed6 |
| Tmed7 | 15546 | 11619 | 13032 | 12586 | 11005 | 12525 | 13144 | 12770 | 11226 | 9728 | 10517 | 10465 | 10461 | 10792 | 15647 | 13676 | 12232 | 12279 | 11603 | 12866 | 12962 | 11262 | 11678 | 10253 | 10365 | 11099 | 10871 | 11444 | Tmed7 |
| Tmed8 | 5493 | 5204 | 4177 | 4821 | 5570 | 5578 | 5385 | 3609 | 3227 | 3113 | 3038 | 2936 | 3269 | 3035 | 5612 | 5534 | 4704 | 5185 | 5005 | 5431 | 5684 | 3494 | 3018 | 2521 | 3228 | 2115 | 3212 | 2940 | Tmed8 |
| Tmed9 | 7046 | 7936 | 8754 | 7939 | 6075 | 6568 | 6297 | 6655 | 6746 | 7503 | 7866 | 7008 | 7473 | 7272 | 6587 | 6848 | 8234 | 6884 | 6958 | 7133 | 6230 | 5720 | 7191 | 8590 | 7026 | 8714 | 7951 | 7980 | Tmed9 |
| Tmeff1 | 11885 | 10432 | 13206 | 14399 | 17412 | 15651 | 16084 | 21943 | 15678 | 18956 | 16566 | 18058 | 18850 | 19346 | 14867 | 11275 | 14286 | 14997 | 15746 | 16491 | 15740 | 21973 | 15952 | 18431 | 20646 | 15523 | 18792 | 19365 | Tmeff1 |
| Tmeff2 | 1505 | 1460 | 1065 | 2156 | 3492 | 2346 | 2362 | 1603 | 1705 | 1608 | 2020 | 812 | 1236 | 819 | 2049 | 1609 | 1803 | 2556 | 2472 | 2457 | 2508 | 2425 | 1865 | 987 | 1551 | 711 | 746 | 767 | Tmeff2 |
| Tmem100 | 701 | 447 | 514 | 419 | 434 | 467 | 364 | 414 | 588 | 460 | 644 | 276 | 408 | 346 | 543 | 576 | 435 | 413 | 369 | 449 | 384 | 393 | 727 | 410 | 386 | 472 | 346 | 298 | Tmem100 |
| Tmem101 | 1305 | 1455 | 1542 | 1348 | 1311 | 1299 | 1188 | 1294 | 1467 | 1565 | 1565 | 1686 | 1562 | 1471 | 1170 | 1181 | 1647 | 1295 | 1523 | 1435 | 1256 | 1362 | 1450 | 1844 | 1497 | 1977 | 1873 | 1716 | Tmem101 |
| Tmem102 | 104 | 75 | 39 | 61 | 69 | 28 | 35 | 35 | 35 | 35 | 27 | 12 | 20 | 7 | 90 | 86 | 35 | 63 | 27 | 28 | 42 | 8 | 46 | 24 | 8 | 30 | 7 | 4 | Tmem102 |
| Tmem104 | 929 | 1278 | 1161 | 1214 | 1013 | 970 | 935 | 586 | 732 | 820 | 818 | 665 | 710 | 582 | 886 | 1248 | 1144 | 1197 | 987 | 820 | 970 | 739 | 711 | 795 | 759 | 776 | 630 | 637 | Tmem104 |
| Tmem106a | 98 | 107 | 83 | 98 | 125 | 113 | 84 | 109 | 160 | 141 | 170 | 126 | 180 | 132 | 142 | 150 | 124 | 108 | 104 | 113 | 105 | 147 | 230 | 144 | 150 | 151 | 151 | 167 | Tmem106a |
| Tmem106b | 6234 | 5182 | 4453 | 5219 | 5579 | 5699 | 5541 | 8272 | 7459 | 6084 | 6970 | 6130 | 6377 | 5957 | 6924 | 5941 | 4668 | 5334 | 4749 | 6261 | 5689 | 7304 | 8150 | 5641 | 6442 | 5033 | 6091 | 6073 | Tmem106b |
| Tmem106c | 1441 | 1822 | 2008 | 1830 | 1648 | 1667 | 1681 | 1653 | 1648 | 1708 | 1526 | 1715 | 1601 | 1604 | 1636 | 1655 | 1905 | 1716 | 1750 | 1646 | 1642 | 1750 | 1777 | 1763 | 1805 | 1946 | 1806 | 1758 | Tmem106c |
| Tmem107 | 790 | 1063 | 1020 | 913 | 695 | 726 | 968 | 481 | 550 | 532 | 534 | 534 | 629 | 598 | 749 | 811 | 1044 | 905 | 887 | 855 | 880 | 465 | 576 | 565 | 517 | 802 | 558 | 631 | Tmem107 |
| Tmem108 | 1162 | 1221 | 1374 | 1836 | 1733 | 1379 | 1549 | 740 | 935 | 1086 | 910 | 1057 | 846 | 1107 | 1135 | 1276 | 1531 | 1729 | 1718 | 1128 | 1520 | 1108 | 927 | 1140 | 1207 | 1261 | 868 | 822 | Tmem108 |
| Tmem109 | 1488 | 1702 | 1694 | 1354 | 1340 | 1322 | 1148 | 1780 | 2267 | 1965 | 2369 | 1744 | 1949 | 1674 | 1302 | 1677 | 1277 | 1403 | 1245 | 1214 | 1079 | 1776 | 2421 | 2017 | 1743 | 2114 | 1806 | 1836 | Tmem109 |
| Tmem11 | 1503 | 1681 | 1813 | 1673 | 1467 | 1417 | 1473 | 1421 | 1522 | 1558 | 1635 | 1754 | 1640 | 1839 | 1435 | 1426 | 1681 | 1333 | 1673 | 1400 | 1532 | 1330 | 1586 | 1830 | 1638 | 2203 | 1858 | 2015 | Tmem11 |
| Tmem114 | 5 | 7 | 4 | 4 | 4 | 1 | 3 | 0 | 4 | 4 | 1 | 5 | 0 | 2 | 11 | 5 | 2 | 2 | 4 | 4 | 2 | 7 | 3 | 2 | 3 | 5 | 1 | 1 | Tmem114 |
| Tmem115 | 2467 | 3537 | 2662 | 2705 | 2177 | 2492 | 2057 | 1568 | 1780 | 1689 | 1611 | 1569 | 1583 | 1525 | 2469 | 2929 | 2750 | 2633 | 2441 | 2367 | 2418 | 1606 | 1763 | 1568 | 1612 | 1758 | 1641 | 1626 | Tmem115 |
| Tmem116 | 35 | 42 | 37 | 34 | 52 | 45 | 27 | 46 | 41 | 45 | 58 | 35 | 43 | 34 | 25 | 36 | 25 | 33 | 39 | 36 | 21 | 43 | 40 | 41 | 41 | 36 | 55 | 43 | Tmem116 |
| Tmem117 | 51 | 48 | 42 | 28 | 90 | 57 | 51 | 159 | 149 | 137 | 164 | 67 | 107 | 76 | 87 | 79 | 71 | 63 | 60 | 70 | 70 | 137 | 164 | 95 | 124 | 118 | 77 | 77 | Tmem117 |
| Tmem119 | 924 | 835 | 630 | 475 | 250 | 492 | 107 | 1053 | 1505 | 1073 | 1768 | 899 | 1412 | 843 | 662 | 1262 | 516 | 627 | 202 | 156 | 109 | 1029 | 1843 | 1160 | 898 | 1443 | 1082 | 961 | Tmem119 |
| Tmem120a | 2684 | 3569 | 2651 | 2829 | 2532 | 2729 | 2675 | 1303 | 1346 | 1303 | 1258 | 1178 | 1142 | 1212 | 2571 | 2811 | 2726 | 2905 | 3101 | 2997 | 2953 | 1258 | 1439 | 1202 | 1345 | 1245 | 1267 | 1244 | Tmem120a |
| Tmem120b | 937 | 1272 | 1059 | 1389 | 1141 | 1101 | 1103 | 382 | 517 | 494 | 577 | 497 | 439 | 441 | 1023 | 1201 | 1203 | 1347 | 1170 | 1073 | 1249 | 570 | 540 | 503 | 466 | 385 | 381 | 408 | Tmem120b |
| Tmem121 | 1343 | 1774 | 1295 | 1380 | 1239 | 1388 | 1003 | 520 | 609 | 521 | 463 | 682 | 616 | 462 | 1352 | 1379 | 1493 | 1265 | 1479 | 1329 | 1126 | 566 | 595 | 550 | 622 | 668 | 733 | 522 | Tmem121 |
| Tmem121b | 1339 | 1545 | 1653 | 2128 | 2657 | 2170 | 2257 | 1292 | 1431 | 1949 | 1455 | 1634 | 1540 | 1554 | 1416 | 1247 | 2108 | 2256 | 2439 | 2109 | 2407 | 1978 | 1236 | 1573 | 1930 | 1395 | 1330 | 1405 | Tmem121b |
| Tmem123 | 13024 | 10233 | 12665 | 8263 | 6322 | 7858 | 7825 | 6594 | 5369 | 4484 | 5274 | 6865 | 7030 | 7152 | 13217 | 12370 | 9897 | 7962 | 6700 | 8140 | 7074 | 4517 | 5978 | 5586 | 4514 | 7479 | 8022 | 8158 | Tmem123 |
| Tmem125 | 31 | 14 | 9 | 4 | 6 | 9 | 3 | 16 | 30 | 10 | 30 | 4 | 25 | 13 | 9 | 17 | 6 | 9 | 5 | 2 | 4 | 10 | 38 | 17 | 2 | 26 | 10 | 8 | Tmem125 |
| Tmem126a | 1339 | 1130 | 1337 | 1127 | 1291 | 1217 | 1486 | 1711 | 1393 | 1646 | 1486 | 1572 | 1758 | 1923 | 1313 | 1210 | 1385 | 1179 | 1076 | 1373 | 1361 | 1626 | 1407 | 1780 | 1479 | 1931 | 1947 | 1832 | Tmem126a |
| Tmem126b | 2418 | 2267 | 2683 | 2201 | 2182 | 2211 | 2315 | 2800 | 2392 | 2427 | 2533 | 2565 | 2561 | 2618 | 2542 | 2560 | 2337 | 2202 | 2200 | 2787 | 2380 | 2575 | 2665 | 2525 | 2347 | 2701 | 2626 | 2854 | Tmem126b |
| Tmem127 | 6717 | 6669 | 6838 | 7565 | 7533 | 7154 | 7197 | 6510 | 6328 | 6353 | 6576 | 6402 | 6148 | 6521 | 6451 | 5998 | 7248 | 7440 | 7362 | 6649 | 7567 | 7610 | 6394 | 6392 | 7131 | 6129 | 6255 | 6408 | Tmem127 |
| Tmem128 | 2072 | 2094 | 2224 | 1914 | 1838 | 1863 | 2183 | 2972 | 2436 | 2344 | 2189 | 2676 | 2618 | 2893 | 2446 | 2059 | 2107 | 1791 | 2005 | 2507 | 1868 | 2538 | 2527 | 2579 | 2658 | 2821 | 2867 | 2985 | Tmem128 |
| Tmem129 | 2605 | 3017 | 2440 | 2375 | 1696 | 2102 | 1980 | 1530 | 1575 | 1309 | 1395 | 1442 | 1556 | 1440 | 2521 | 2578 | 2406 | 2168 | 2217 | 2351 | 2099 | 1399 | 1586 | 1408 | 1355 | 1763 | 1533 | 1480 | Tmem129 |
| Tmem130 | 2663 | 3708 | 2540 | 6313 | 9366 | 7590 | 7990 | 6599 | 6602 | 8364 | 6257 | 4979 | 3935 | 4880 | 3647 | 2461 | 4833 | 7304 | 8468 | 7843 | 9104 | 10181 | 5898 | 6561 | 8520 | 3831 | 4134 | 4264 | Tmem130 |
| Tmem131 | 11683 | 10490 | 10527 | 12023 | 12546 | 12282 | 12506 | 8576 | 9923 | 10721 | 10729 | 11668 | 10290 | 10377 | 10440 | 12080 | 11783 | 11833 | 12101 | 10115 | 12410 | 10317 | 9040 | 9889 | 10609 | 8306 | 9912 | 9670 | Tmem131 |
| Tmem131l | 10156 | 9744 | 6914 | 9705 | 10942 | 11220 | 12196 | 12461 | 13357 | 13230 | 11629 | 14897 | 11174 | 15070 | 9581 | 8774 | 9036 | 10059 | 11288 | 10085 | 11845 | 12860 | 10310 | 13855 | 15211 | 12792 | 12992 | 12540 | Tmem131l |
| Tmem132a | 8393 | 10223 | 12696 | 13563 | 16257 | 12900 | 11085 | 12549 | 14534 | 18149 | 16343 | 16370 | 14713 | 14310 | 8400 | 9771 | 13133 | 13745 | 15790 | 11461 | 12364 | 18120 | 14348 | 18701 | 18651 | 16695 | 14695 | 14005 | Tmem132a |
| Tmem132b | 655 | 593 | 469 | 1054 | 1784 | 1336 | 1554 | 1581 | 1879 | 2074 | 1647 | 1164 | 1090 | 1240 | 721 | 657 | 684 | 1166 | 1287 | 1218 | 1441 | 2204 | 1441 | 1338 | 2397 | 657 | 853 | 926 | Tmem132b |
| Tmem132c | 7286 | 6548 | 7727 | 4813 | 3864 | 5398 | 3879 | 3519 | 4413 | 3685 | 4010 | 4289 | 5861 | 4027 | 5648 | 7857 | 5349 | 4941 | 4269 | 4150 | 3873 | 2161 | 5110 | 4067 | 2917 | 4323 | 4574 | 4756 | Tmem132c |
| Tmem132d | 198 | 174 | 174 | 237 | 424 | 321 | 263 | 318 | 383 | 426 | 373 | 275 | 223 | 170 | 185 | 237 | 200 | 259 | 238 | 249 | 284 | 426 | 390 | 267 | 337 | 176 | 193 | 195 | Tmem132d |
| Tmem132e | 803 | 979 | 1088 | 1191 | 1731 | 1263 | 1065 | 470 | 550 | 673 | 771 | 415 | 521 | 301 | 819 | 904 | 1265 | 1154 | 1662 | 1000 | 1207 | 760 | 594 | 404 | 564 | 327 | 342 | 337 | Tmem132e |
| Tmem134 | 228 | 136 | 79 | 81 | 64 | 63 | 77 | 94 | 56 | 82 | 89 | 66 | 211 | 49 | 80 | 90 | 73 | 58 | 60 | 65 | 63 | 122 | 83 | 92 | 54 | 140 | 84 | 80 | Tmem134 |
| Tmem135 | 3006 | 2750 | 2532 | 2659 | 2963 | 2820 | 3053 | 2229 | 2051 | 1839 | 2043 | 1943 | 1915 | 1906 | 2976 | 2738 | 2466 | 2984 | 2639 | 2814 | 2909 | 2384 | 2138 | 1866 | 2034 | 1812 | 1825 | 1772 | Tmem135 |
| Tmem138 | 1194 | 1226 | 1223 | 966 | 1061 | 945 | 1207 | 1431 | 1353 | 1154 | 1193 | 1504 | 1535 | 1536 | 1249 | 1074 | 1150 | 1004 | 1077 | 1064 | 1053 | 1444 | 1251 | 1396 | 1163 | 1550 | 1626 | 1667 | Tmem138 |
| Tmem139 | 46 | 36 | 18 | 23 | 16 | 6 | 9 | 18 | 28 | 30 | 27 | 11 | 22 | 11 | 32 | 44 | 17 | 32 | 14 | 8 | 10 | 17 | 30 | 18 | 9 | 28 | 9 | 20 | Tmem139 |
| Tmem140 | 28 | 23 | 17 | 33 | 16 | 15 | 16 | 28 | 57 | 46 | 65 | 50 | 52 | 37 | 25 | 35 | 14 | 26 | 16 | 21 | 20 | 40 | 66 | 34 | 28 | 40 | 29 | 39 | Tmem140 |
| Tmem141 | 243 | 276 | 244 | 206 | 320 | 268 | 242 | 441 | 474 | 409 | 412 | 321 | 366 | 325 | 295 | 263 | 253 | 243 | 311 | 304 | 249 | 409 | 453 | 382 | 375 | 336 | 299 | 393 | Tmem141 |
| Tmem143 | 307 | 368 | 431 | 385 | 450 | 411 | 356 | 481 | 664 | 657 | 592 | 600 | 586 | 536 | 323 | 351 | 565 | 387 | 457 | 359 | 410 | 592 | 640 | 588 | 644 | 513 | 550 | 586 | Tmem143 |
| Tmem144 | 157 | 169 | 217 | 205 | 214 | 159 | 236 | 245 | 208 | 243 | 188 | 324 | 289 | 358 | 165 | 129 | 168 | 155 | 231 | 196 | 197 | 134 | 254 | 305 | 216 | 337 | 261 | 311 | Tmem144 |
| Tmem145 | 1425 | 2074 | 2191 | 3732 | 4340 | 3097 | 3131 | 2036 | 1871 | 2929 | 2184 | 2273 | 2108 | 2017 | 1814 | 1629 | 3399 | 3715 | 3986 | 2856 | 3554 | 3803 | 1766 | 2516 | 2998 | 1585 | 1988 | 1893 | Tmem145 |
| Tmem147 | 2373 | 2877 | 2401 | 2431 | 2458 | 2164 | 2550 | 2759 | 2652 | 2664 | 2422 | 2290 | 2449 | 2499 | 2313 | 2266 | 2497 | 2334 | 2561 | 2459 | 2660 | 2476 | 2832 | 2593 | 2476 | 2592 | 2504 | 2711 | Tmem147 |
| Tmem14a | 90 | 87 | 142 | 94 | 139 | 131 | 154 | 466 | 395 | 407 | 430 | 432 | 432 | 477 | 142 | 104 | 188 | 120 | 155 | 193 | 171 | 429 | 425 | 386 | 382 | 425 | 481 | 511 | Tmem14a |
| Tmem14c | 3946 | 4400 | 4265 | 3986 | 3594 | 3641 | 3448 | 2838 | 3258 | 2829 | 3180 | 3262 | 3458 | 3034 | 3680 | 3998 | 3986 | 3776 | 3852 | 3968 | 3339 | 2481 | 3199 | 3210 | 2739 | 3734 | 3717 | 3319 | Tmem14c |
| Tmem150a | 28 | 165 | 20 | 13 | 19 | 149 | 70 | 62 | 37 | 22 | 18 | 25 | 12 | 11 | 53 | 20 | 22 | 18 | 58 | 121 | 11 | 28 | 20 | 68 | 89 | 11 | 8 | 8 | Tmem150a |
| Tmem150b | 1 | 1 | 0 | 6 | 3 | 2 | 6 | 0 | 2 | 3 | 10 | 9 | 2 | 18 | 4 | 3 | 1 | 0 | 5 | 5 | 6 | 5 | 4 | 4 | 1 | 5 | 6 | 0 | Tmem150b |
| Tmem150c | 408 | 454 | 565 | 672 | 1106 | 765 | 860 | 1073 | 1218 | 1537 | 1168 | 988 | 929 | 1020 | 522 | 383 | 571 | 738 | 871 | 846 | 1012 | 1470 | 1209 | 1221 | 1617 | 675 | 860 | 873 | Tmem150c |
| Tmem151a | 200 | 284 | 230 | 264 | 437 | 294 | 264 | 122 | 175 | 234 | 160 | 106 | 81 | 100 | 211 | 258 | 204 | 357 | 307 | 276 | 301 | 241 | 200 | 122 | 181 | 106 | 87 | 95 | Tmem151a |
| Tmem151b | 1834 | 2204 | 2817 | 2954 | 3782 | 3128 | 3075 | 3715 | 4096 | 4658 | 4071 | 4539 | 3704 | 3963 | 1855 | 1738 | 3213 | 3268 | 4189 | 2929 | 3325 | 4986 | 3704 | 5035 | 5401 | 4599 | 3828 | 3832 | Tmem151b |
| Tmem154 | 62 | 45 | 46 | 23 | 46 | 47 | 41 | 90 | 115 | 84 | 66 | 46 | 42 | 20 | 50 | 60 | 36 | 35 | 37 | 48 | 29 | 45 | 104 | 69 | 35 | 34 | 33 | 32 | Tmem154 |
| Tmem156 | 7 | 3 | 5 | 0 | 4 | 8 | 2 | 12 | 10 | 7 | 14 | 7 | 5 | 4 | 12 | 5 | 9 | 9 | 16 | 7 | 9 | 7 | 15 | 2 | 5 | 6 | 12 | 13 | Tmem156 |
| Tmem158 | 548 | 612 | 574 | 729 | 959 | 778 | 902 | 596 | 566 | 734 | 670 | 451 | 369 | 467 | 550 | 501 | 740 | 778 | 920 | 776 | 927 | 722 | 660 | 667 | 752 | 508 | 351 | 441 | Tmem158 |
| Tmem159 | 1814 | 1950 | 2107 | 1741 | 1384 | 1324 | 1372 | 607 | 649 | 582 | 666 | 490 | 777 | 642 | 1789 | 1977 | 1869 | 1667 | 1308 | 1361 | 1394 | 551 | 770 | 590 | 404 | 619 | 729 | 739 | Tmem159 |
| Tmem160 | 747 | 869 | 851 | 769 | 749 | 654 | 569 | 438 | 414 | 431 | 428 | 456 | 494 | 394 | 703 | 637 | 852 | 790 | 781 | 713 | 565 | 371 | 429 | 464 | 469 | 499 | 503 | 400 | Tmem160 |
| Tmem161a | 1571 | 1863 | 2181 | 1714 | 1635 | 1586 | 1543 | 1909 | 2077 | 2049 | 2064 | 2117 | 2024 | 1980 | 1502 | 1639 | 2041 | 1681 | 1839 | 1633 | 1502 | 1796 | 2019 | 2222 | 2042 | 2481 | 2133 | 2103 | Tmem161a |
| Tmem161b | 3325 | 2628 | 3336 | 3591 | 3296 | 3537 | 4196 | 2586 | 2108 | 2327 | 2083 | 2708 | 2426 | 2659 | 3810 | 2961 | 3475 | 3551 | 3434 | 3990 | 3568 | 2814 | 2101 | 2487 | 2437 | 2832 | 2438 | 2843 | Tmem161b |
| Tmem163 | 765 | 928 | 950 | 1103 | 1577 | 1202 | 1138 | 1701 | 1391 | 1810 | 1512 | 1972 | 2042 | 1968 | 978 | 759 | 1136 | 1132 | 1543 | 1336 | 1250 | 2189 | 1470 | 1860 | 2009 | 1627 | 2229 | 2301 | Tmem163 |
| Tmem164 | 7996 | 7085 | 7740 | 6543 | 6316 | 6442 | 5501 | 6432 | 6670 | 6966 | 8034 | 7489 | 8818 | 6738 | 7191 | 8279 | 7173 | 6951 | 6410 | 5223 | 5755 | 6383 | 6890 | 7960 | 6568 | 9073 | 7620 | 7488 | Tmem164 |

| | | | | | | | | | | | | | | | | | | | | | | | | | | | | | |
|---|---|---|---|---|---|---|---|---|---|---|---|---|---|---|---|---|---|---|---|---|---|---|---|---|---|---|---|---|---|
| Tmem165 | 7728 | 5793 | 6224 | 5882 | 5353 | 5951 | 5893 | 6099 | 5572 | 4966 | 5697 | 5517 | 5838 | 6168 | 7402 | 7136 | 5617 | 6077 | 5241 | 5911 | 5683 | 5274 | 6082 | 5462 | 4796 | 6575 | 5932 | 6502 | Tmem165 |
| Tmem167 | 14587 | 10551 | 9153 | 9181 | 9079 | 11163 | 12046 | 13945 | 11942 | 9492 | 10497 | 10974 | 11077 | 11830 | 14493 | 12278 | 9087 | 9180 | 9164 | 14023 | 11382 | 9981 | 12033 | 9674 | 10224 | 10647 | 11941 | 11769 | Tmem167 |
| Tmem167b | 7495 | 6801 | 6681 | 6739 | 5977 | 6599 | 7016 | 6787 | 5515 | 5460 | 5377 | 5876 | 5313 | 6122 | 7907 | 6741 | 6906 | 6897 | 6385 | 6919 | 6902 | 6355 | 5941 | 5784 | 6051 | 6098 | 6009 | 6048 | Tmem167b |
| Tmem168 | 2435 | 1729 | 1959 | 1782 | 1924 | 1841 | 2002 | 3566 | 3399 | 2617 | 3013 | 2981 | 3154 | 2991 | 2397 | 2009 | 1692 | 1794 | 1809 | 2111 | 1771 | 2769 | 3521 | 2836 | 2724 | 3128 | 3163 | 3203 | Tmem168 |
| Tmem169 | 2110 | 2128 | 2238 | 2658 | 3202 | 2907 | 3273 | 2341 | 1933 | 2310 | 2174 | 2177 | 2036 | 2322 | 2776 | 2008 | 2744 | 2746 | 2741 | 2855 | 3069 | 2884 | 1966 | 2194 | 2280 | 1807 | 1958 | 2194 | Tmem169 |
| Tmem17 | 105 | 107 | 151 | 97 | 185 | 121 | 123 | 530 | 477 | 374 | 399 | 451 | 431 | 431 | 169 | 98 | 104 | 100 | 168 | 179 | 154 | 436 | 504 | 394 | 458 | 473 | 463 | 458 | Tmem17 |
| Tmem170 | 1678 | 1689 | 1952 | 1833 | 1305 | 1373 | 1698 | 1072 | 943 | 936 | 1007 | 1063 | 1175 | 1100 | 1513 | 1734 | 1823 | 1621 | 1526 | 1303 | 1500 | 954 | 970 | 1042 | 937 | 1239 | 1061 | 1219 | Tmem170 |
| Tmem170b | 5447 | 3854 | 4914 | 6475 | 7131 | 7006 | 7771 | 5174 | 4268 | 5286 | 4736 | 4560 | 4433 | 4952 | 5583 | 4555 | 5552 | 6864 | 6749 | 6512 | 7579 | 6269 | 4119 | 4886 | 6056 | 3548 | 4645 | 4584 | Tmem170b |
| Tmem171 | 0 | 0 | 0 | 0 | 0 | 0 | 2 | 0 | 2 | 0 | 0 | 0 | 1 | 2 | 1 | 2 | 2 | 0 | 0 | 2 | 0 | 3 | 4 | 2 | 1 | 0 | 0 | 0 | Tmem171 |
| Tmem174 | 0 | 0 | 0 | 0 | 0 | 0 | 2 | 0 | 0 | 0 | 0 | 0 | 0 | 0 | 1 | 0 | 0 | 4 | 0 | 0 | 0 | 0 | 0 | 0 | 0 | 0 | 0 | 0 | Tmem174 |
| Tmem175 | 858 | 877 | 825 | 821 | 870 | 935 | 976 | 1566 | 1795 | 1402 | 1548 | 1458 | 1431 | 1284 | 944 | 864 | 875 | 819 | 876 | 886 | 855 | 1606 | 1722 | 1374 | 1536 | 1317 | 1489 | 1225 | Tmem175 |
| Tmem176a | 1240 | 1095 | 1103 | 991 | 928 | 977 | 855 | 1167 | 1384 | 1093 | 1383 | 972 | 1281 | 930 | 1126 | 1368 | 931 | 1093 | 765 | 890 | 776 | 1213 | 1442 | 1164 | 910 | 1398 | 1110 | 1048 | Tmem176a |
| Tmem176b | 2178 | 2300 | 1831 | 1951 | 2351 | 2321 | 2240 | 3173 | 3648 | 3038 | 3565 | 2552 | 3142 | 2462 | 2167 | 2389 | 2079 | 2246 | 2275 | 2441 | 2035 | 3177 | 3864 | 3210 | 2824 | 3182 | 2709 | 2617 | Tmem176b |
| Tmem177 | 1216 | 1488 | 1225 | 1282 | 1346 | 1425 | 1427 | 1279 | 1275 | 1184 | 1062 | 1215 | 1200 | 1089 | 1353 | 1134 | 1315 | 1206 | 1508 | 1665 | 1536 | 939 | 1192 | 1149 | 1228 | 1332 | 1148 | 1216 | Tmem177 |
| Tmem178 | 457 | 359 | 404 | 641 | 1072 | 774 | 916 | 1261 | 1210 | 1341 | 1079 | 1165 | 961 | 1249 | 487 | 326 | 549 | 613 | 1002 | 730 | 936 | 1303 | 1059 | 1432 | 1403 | 1027 | 980 | 1003 | Tmem178 |
| Tmem178b | 3939 | 4590 | 3685 | 7308 | 9727 | 8793 | 9281 | 7238 | 6779 | 8051 | 7128 | 6147 | 6467 | 6723 | 5283 | 4125 | 5409 | 7772 | 8011 | 7775 | 9819 | 9660 | 6036 | 5838 | 9264 | 3426 | 6104 | 5814 | Tmem178b |
| Tmem179 | 419 | 542 | 357 | 779 | 1201 | 850 | 882 | 771 | 938 | 1136 | 946 | 727 | 628 | 653 | 566 | 463 | 776 | 792 | 1068 | 886 | 918 | 1255 | 867 | 941 | 1283 | 614 | 550 | 527 | Tmem179 |
| Tmem179b | 591 | 592 | 519 | 475 | 392 | 439 | 574 | 576 | 494 | 511 | 577 | 466 | 590 | 496 | 526 | 510 | 594 | 524 | 479 | 486 | 501 | 521 | 608 | 580 | 515 | 689 | 559 | 579 | Tmem179b |
| Tmem18 | 3403 | 3001 | 3195 | 2981 | 2751 | 2890 | 2777 | 2699 | 2539 | 2383 | 2437 | 2364 | 2575 | 2382 | 3516 | 3382 | 3012 | 2821 | 2764 | 3172 | 2673 | 2199 | 2584 | 2327 | 2257 | 2571 | 2778 | 2585 | Tmem18 |
| Tmem181a | 4344 | 4016 | 4354 | 4513 | 4716 | 4662 | 4611 | 3171 | 3297 | 3128 | 3429 | 3114 | 3275 | 3270 | 4634 | 4428 | 4077 | 4848 | 4257 | 4508 | 4259 | 3574 | 3004 | 2837 | 3275 | 3003 | 3120 | 3151 | Tmem181a |
| Tmem182 | 2 | 1 | 0 | 2 | 1 | 4 | 4 | 13 | 16 | 15 | 122 | 1 | 12 | 7 | 3 | 5 | 7 | 0 | 5 | 0 | 2 | 27 | 117 | 8 | 12 | 7 | 2 | 6 | Tmem182 |
| Tmem183a | 12314 | 11811 | 10508 | 11787 | 11189 | 11072 | 12592 | 10603 | 8786 | 9331 | 8016 | 9750 | 8848 | 9202 | 12897 | 10999 | 11239 | 11283 | 11992 | 12990 | 12284 | 9523 | 9132 | 9156 | 9534 | 10143 | 9941 | 9443 | Tmem183a |
| Tmem184a | 47 | 38 | 38 | 53 | 36 | 16 | 27 | 45 | 48 | 78 | 83 | 16 | 41 | 21 | 29 | 45 | 28 | 33 | 35 | 32 | 24 | 30 | 91 | 96 | 10 | 84 | 6 | 25 | Tmem184a |
| Tmem184b | 4746 | 6100 | 6115 | 5496 | 4506 | 4580 | 4178 | 2659 | 3330 | 3330 | 3903 | 3526 | 3457 | 3269 | 4233 | 5987 | 5935 | 5285 | 4747 | 3668 | 4337 | 3096 | 3278 | 3387 | 3247 | 3916 | 3022 | 3185 | Tmem184b |
| Tmem184c | 7158 | 6745 | 5983 | 6192 | 6507 | 6274 | 6835 | 4708 | 4124 | 4220 | 4021 | 3999 | 3917 | 4204 | 7102 | 6656 | 6465 | 6730 | 6519 | 6413 | 6885 | 4654 | 3840 | 4006 | 4135 | 4043 | 3969 | 3884 | Tmem184c |
| Tmem185a | 3061 | 2959 | 2860 | 2791 | 2906 | 2715 | 2680 | 2611 | 2422 | 2072 | 2098 | 2533 | 2361 | 2217 | 3033 | 2618 | 2913 | 2732 | 2851 | 3175 | 2735 | 2557 | 2372 | 2374 | 2385 | 2619 | 2350 | 2436 | Tmem185a |
| Tmem185b | 632 | 631 | 804 | 562 | 621 | 668 | 594 | 1211 | 1195 | 1194 | 1135 | 1194 | 1312 | 1152 | 673 | 639 | 598 | 558 | 655 | 690 | 603 | 1173 | 1206 | 1124 | 1118 | 1292 | 1364 | 1224 | Tmem185b |
| Tmem186 | 1857 | 1823 | 2133 | 1678 | 1856 | 1689 | 1792 | 2660 | 2555 | 2599 | 2453 | 2839 | 2573 | 2776 | 1863 | 1652 | 1906 | 1590 | 1896 | 1962 | 1826 | 2361 | 2362 | 2796 | 2627 | 2924 | 2931 | 2905 | Tmem186 |
| Tmem189 | 2331 | 2653 | 2542 | 2598 | 2267 | 2249 | 2039 | 912 | 1061 | 1110 | 1102 | 1004 | 1083 | 1070 | 2300 | 2587 | 2520 | 2459 | 2222 | 2004 | 2229 | 954 | 997 | 989 | 1021 | 1170 | 998 | 1027 | Tmem189 |
| Tmem19 | 3224 | 3249 | 3213 | 3547 | 3120 | 3240 | 3503 | 2157 | 2181 | 2321 | 2181 | 2295 | 2311 | 2368 | 3221 | 3419 | 3457 | 3418 | 3162 | 2919 | 3355 | 2341 | 2273 | 2344 | 2435 | 2426 | 2284 | 2528 | Tmem19 |
| Tmem190 | 0 | 1 | 0 | 0 | 0 | 0 | 0 | 2 | 1 | 1 | 0 | 0 | 0 | 4 | 0 | 0 | 0 | 0 | 1 | 0 | 0 | 0 | 0 | 0 | 0 | 1 | 1 | 0 | Tmem190 |
| Tmem191c | 450 | 621 | 683 | 862 | 883 | 735 | 701 | 358 | 489 | 727 | 572 | 605 | 548 | 585 | 422 | 555 | 788 | 843 | 846 | 691 | 752 | 650 | 375 | 631 | 641 | 588 | 569 | 553 | Tmem191c |
| Tmem192 | 776 | 745 | 498 | 652 | 476 | 581 | 559 | 381 | 413 | 322 | 443 | 357 | 379 | 338 | 683 | 660 | 559 | 631 | 575 | 591 | 571 | 348 | 438 | 356 | 372 | 339 | 352 | 329 | Tmem192 |
| Tmem196 | 131 | 100 | 156 | 162 | 396 | 305 | 346 | 304 | 306 | 349 | 299 | 285 | 271 | 303 | 175 | 91 | 207 | 216 | 295 | 269 | 331 | 433 | 318 | 256 | 329 | 234 | 307 | 325 | Tmem196 |
| Tmem198 | 1028 | 1685 | 1378 | 2055 | 1907 | 1681 | 1491 | 805 | 879 | 998 | 728 | 936 | 761 | 796 | 1188 | 1239 | 1898 | 2054 | 2182 | 1522 | 1767 | 1223 | 818 | 1010 | 1101 | 799 | 882 | 726 | Tmem198 |
| Tmem198b | 2344 | 2496 | 2562 | 2105 | 2060 | 1859 | 2084 | 1685 | 1765 | 1745 | 1898 | 1981 | 2132 | 1973 | 2196 | 2155 | 2467 | 2108 | 2144 | 1937 | 1970 | 1741 | 1880 | 1879 | 1728 | 2268 | 2034 | 2104 | Tmem198b |
| Tmem199 | 1954 | 1970 | 2145 | 2110 | 1969 | 1777 | 2066 | 2348 | 1877 | 1907 | 1774 | 2334 | 2073 | 2439 | 2097 | 1688 | 2150 | 1887 | 2113 | 2393 | 1994 | 2102 | 1985 | 2177 | 2116 | 2337 | 2346 | 2540 | Tmem199 |
| Tmem200a | 720 | 633 | 956 | 771 | 1148 | 988 | 1005 | 2043 | 1905 | 2306 | 2077 | 2055 | 1701 | 1950 | 808 | 544 | 983 | 795 | 1022 | 947 | 976 | 2648 | 1995 | 2187 | 2196 | 1854 | 1864 | 1984 | Tmem200a |
| Tmem200b | 85 | 68 | 85 | 60 | 32 | 50 | 21 | 147 | 213 | 165 | 239 | 94 | 126 | 65 | 75 | 87 | 61 | 80 | 33 | 16 | 18 | 134 | 223 | 188 | 95 | 123 | 83 | 84 | Tmem200b |
| Tmem200c | 281 | 290 | 366 | 349 | 422 | 406 | 348 | 226 | 227 | 299 | 264 | 321 | 243 | 256 | 301 | 252 | 436 | 275 | 401 | 337 | 341 | 204 | 269 | 331 | 256 | 323 | 286 | 258 | Tmem200c |
| Tmem201 | 4409 | 6040 | 6482 | 7248 | 6352 | 5777 | 5425 | 3477 | 4260 | 4958 | 5294 | 5325 | 4826 | 4758 | 4005 | 5354 | 6919 | 7628 | 6223 | 4753 | 5759 | 5238 | 4047 | 5277 | 5323 | 5329 | 4733 | 4837 | Tmem201 |
| Tmem202 | 1 | 2 | 1 | 3 | 4 | 2 | 2 | 1 | 3 | 6 | 8 | 2 | 2 | 3 | 3 | 2 | 0 | 0 | 5 | 4 | 5 | 2 | 10 | 2 | 11 | 3 | 7 | 0 | Tmem202 |
| Tmem203 | 562 | 613 | 590 | 518 | 506 | 486 | 540 | 405 | 517 | 405 | 518 | 543 | 533 | 513 | 585 | 485 | 560 | 610 | 520 | 533 | 452 | 453 | 560 | 505 | 493 | 532 | 550 | 457 | Tmem203 |
| Tmem204 | 271 | 316 | 354 | 300 | 283 | 303 | 258 | 271 | 320 | 289 | 353 | 263 | 346 | 296 | 290 | 288 | 299 | 246 | 304 | 350 | 236 | 287 | 444 | 270 | 302 | 381 | 320 | 273 | Tmem204 |
| Tmem205 | 707 | 785 | 688 | 815 | 652 | 706 | 689 | 622 | 623 | 563 | 637 | 563 | 545 | 585 | 644 | 713 | 677 | 763 | 683 | 840 | 755 | 523 | 693 | 574 | 591 | 557 | 664 | 563 | Tmem205 |
| Tmem207 | 3 | 5 | 0 | 7 | 2 | 2 | 6 | 9 | 9 | 4 | 1 | 15 | 9 | 10 | 0 | 5 | 10 | 3 | 0 | 1 | 4 | 2 | 3 | 7 | 11 | 8 | 3 | 9 | Tmem207 |
| Tmem208 | 1168 | 1481 | 1495 | 1470 | 1453 | 1480 | 1606 | 1363 | 1563 | 1213 | 1422 | 1320 | 1477 | 1327 | 1360 | 1473 | 1425 | 1544 | 1632 | 1796 | 1397 | 1273 | 1528 | 1396 | 1477 | 1495 | 1428 | 1428 | Tmem208 |
| Tmem209 | 3127 | 2710 | 2458 | 2506 | 2647 | 2584 | 3176 | 4822 | 4317 | 4004 | 3528 | 5132 | 4267 | 4988 | 3281 | 2511 | 2666 | 2517 | 2687 | 3030 | 2927 | 4204 | 4187 | 4140 | 4276 | 5000 | 4937 | 4829 | Tmem209 |
| Tmem210 | 1 | 0 | 3 | 0 | 1 | 0 | 4 | 0 | 2 | 0 | 1 | 1 | 0 | 0 | 0 | 1 | 6 | 0 | 0 | 2 | 2 | 0 | 0 | 1 | 4 | 0 | 0 | 1 | Tmem210 |
| Tmem211 | 20 | 10 | 6 | 7 | 7 | 5 | 6 | 37 | 25 | 43 | 30 | 7 | 26 | 12 | 18 | 6 | 19 | 4 | 8 | 5 | 3 | 10 | 28 | 68 | 8 | 45 | 23 | 7 | Tmem211 |
| Tmem212 | 0 | 0 | 0 | 0 | 0 | 0 | 0 | 0 | 0 | 0 | 1 | 0 | 0 | 0 | 0 | 0 | 0 | 0 | 0 | 0 | 0 | 0 | 0 | 0 | 0 | 0 | 0 | 0 | Tmem212 |
| Tmem213 | 1 | 0 | 0 | 0 | 0 | 3 | 0 | 5 | 2 | 11 | 2 | 0 | 0 | 0 | 1 | 0 | 0 | 1 | 3 | 0 | 1 | 0 | 11 | 14 | 0 | 8 | 0 | 0 | Tmem213 |
| Tmem214 | 5072 | 6048 | 4692 | 6162 | 6205 | 5892 | 6089 | 4655 | 5152 | 5014 | 5350 | 4683 | 5007 | 4428 | 4923 | 5485 | 5330 | 6410 | 6195 | 5229 | 6352 | 5646 | 5190 | 4741 | 5107 | 4333 | 4637 | 4646 | Tmem214 |
| Tmem215 | 10 | 10 | 12 | 7 | 11 | 26 | 13 | 11 | 26 | 23 | 23 | 9 | 8 | 17 | 9 | 9 | 6 | 7 | 6 | 16 | 10 | 15 | 28 | 20 | 11 | 9 | 2 | 12 | Tmem215 |
| Tmem216 | 1325 | 1335 | 1352 | 1142 | 933 | 952 | 1044 | 1050 | 1023 | 879 | 916 | 954 | 991 | 1074 | 1353 | 1340 | 1191 | 1029 | 1005 | 1345 | 1038 | 1001 | 1040 | 1022 | 941 | 1154 | 1056 | 1226 | Tmem216 |
| Tmem217 | 1 | 2 | 12 | 2 | 0 | 4 | 2 | 2 | 0 | 2 | 0 | 1 | 3 | 0 | 0 | 2 | 2 | 1 | 0 | 1 | 5 | 0 | 3 | 0 | 0 | 1 | 0 | 2 | Tmem217 |
| Tmem218 | 1440 | 1322 | 1701 | 1340 | 1028 | 1164 | 1187 | 890 | 799 | 776 | 727 | 962 | 934 | 893 | 1485 | 1363 | 1578 | 1182 | 1186 | 1296 | 1095 | 724 | 786 | 835 | 667 | 964 | 990 | 914 | Tmem218 |
| Tmem219 | 659 | 765 | 635 | 670 | 705 | 634 | 639 | 666 | 717 | 651 | 774 | 823 | 846 | 852 | 653 | 544 | 701 | 616 | 651 | 742 | 668 | 675 | 748 | 683 | 792 | 776 | 966 | 857 | Tmem219 |
| Tmem220 | 103 | 111 | 124 | 88 | 77 | 75 | 57 | 84 | 71 | 123 | 140 | 114 | 145 | 120 | 132 | 114 | 129 | 69 | 66 | 84 | 85 | 69 | 116 | 125 | 68 | 105 | 138 | 107 | Tmem220 |
| Tmem221 | 48 | 33 | 15 | 18 | 26 | 29 | 11 | 10 | 21 | 6 | 19 | 2 | 2 | 0 | 40 | 32 | 27 | 37 | 5 | 26 | 14 | 5 | 14 | 9 | 4 | 18 | 6 | 5 | Tmem221 |
| Tmem222 | 2354 | 2726 | 2503 | 2323 | 2338 | 2371 | 2439 | 2690 | 2744 | 2281 | 2364 | 2509 | 2358 | 2414 | 2476 | 2406 | 2391 | 2221 | 2555 | 2613 | 2424 | 2692 | 2708 | 2562 | 2466 | 2815 | 2500 | 2617 | Tmem222 |
| Tmem223 | 1233 | 1649 | 1531 | 1589 | 1526 | 1596 | 1512 | 2501 | 2884 | 2645 | 2671 | 3325 | 3401 | 3165 | 1341 | 1324 | 1619 | 1384 | 1521 | 1605 | 1281 | 1806 | 2682 | 2700 | 2570 | 2746 | 3406 | 2782 | Tmem223 |
| Tmem229a | 180 | 145 | 221 | 188 | 295 | 256 | 251 | 471 | 465 | 425 | 434 | 379 | 369 | 423 | 214 | 157 | 246 | 147 | 323 | 293 | 270 | 521 | 419 | 337 | 427 | 480 | 461 | 399 | Tmem229a |
| Tmem229b | 1903 | 2147 | 1789 | 3225 | 5176 | 4159 | 3969 | 4113 | 4568 | 5359 | 4630 | 3614 | 3325 | 3548 | 2267 | 1937 | 2640 | 3818 | 4631 | 3550 | 4432 | 6141 | 4003 | 4684 | 5825 | 2530 | 2997 | 3048 | Tmem229b |
| Tmem230 | 1373 | 1158 | 1050 | 1082 | 1231 | 1187 | 1226 | 2097 | 1912 | 1828 | 2059 | 1991 | 2135 | 2109 | 1451 | 1235 | 1153 | 1099 | 1136 | 1278 | 1143 | 1992 | 2153 | 1828 | 1816 | 1937 | 2173 | 2162 | Tmem230 |
| Tmem231 | 2118 | 2112 | 2124 | 1975 | 1559 | 1679 | 1533 | 1211 | 1331 | 1382 | 1305 | 1340 | 1377 | 1325 | 2000 | 2176 | 1968 | 1723 | 1662 | 1664 | 1530 | 1338 | 1443 | 1381 | 1284 | 1577 | 1409 | 1419 | Tmem231 |
| Tmem232 | 12 | 18 | 8 | 13 | 29 | 22 | 13 | 7 | 25 | 27 | 3 | 14 | 19 | 9 | 11 | 30 | 12 | 10 | 7 | 29 | 23 | 37 | 30 | 17 | 21 | 8 | 9 | 14 | Tmem232 |
| Tmem233 | 0 | 0 | 3 | 0 | 0 | 0 | 0 | 1 | 0 | 0 | 3 | 2 | 3 | 0 | 0 | 0 | 0 | 0 | 2 | 0 | 0 | 0 | 3 | 0 | 2 | 0 | 0 | 0 | Tmem233 |
| Tmem234 | 5419 | 5565 | 6391 | 5403 | 4817 | 5278 | 4736 | 5283 | 5291 | 5846 | 5751 | 6572 | 6145 | 6531 | 4990 | 5476 | 5589 | 5105 | 5533 | 5281 | 4769 | 5422 | 5693 | 6090 | 6041 | 7064 | 6951 | 6713 | Tmem234 |
| Tmem235 | 30 | 40 | 6 | 37 | 16 | 26 | 16 | 5 | 5 | 7 | 6 | 3 | 2 | 9 | 15 | 28 | 28 | 50 | 25 | 27 | 23 | 8 | 8 | 10 | 6 | 12 | 6 | 9 | Tmem235 |
| Tmem236 | 39 | 37 | 48 | 48 | 49 | 69 | 94 | 5 | 15 | 11 | 16 | 20 | 5 | 16 | 30 | 25 | 41 | 62 | 62 | 31 | 63 | 20 | 6 | 12 | 10 | 12 | 9 | 6 | Tmem236 |
| Tmem237 | 1919 | 1812 | 1886 | 1594 | 1613 | 1811 | 1868 | 2381 | 2183 | 1870 | 1930 | 2220 | 2054 | 2458 | 2086 | 1745 | 1839 | 1624 | 1889 | 2082 | 1877 | 2074 | 2108 | 2151 | 2076 | 2335 | 2266 | 2417 | Tmem237 |
| Tmem238 | 27 | 22 | 17 | 22 | 24 | 22 | 19 | 51 | 81 | 76 | 81 | 56 | 52 | 49 | 21 | 17 | 19 | 18 | 25 | 22 | 21 | 43 | 77 | 60 | 44 | 104 | 43 | 46 | Tmem238 |
| Tmem238l | 0 | 3 | 0 | 0 | 2 | 0 | 1 | 1 | 6 | 7 | 4 | 0 | 0 | 0 | 3 | 5 | 0 | 2 | 0 | 0 | 0 | 5 | 1 | 0 | 3 | 3 | 0 | 1 | Tmem238l |
| Tmem240 | 208 | 338 | 277 | 505 | 763 | 566 | 571 | 593 | 576 | 795 | 644 | 484 | 459 | 481 | 286 | 250 | 433 | 494 | 768 | 600 | 656 | 909 | 577 | 611 | 819 | 399 | 454 | 413 | Tmem240 |
| Tmem241 | 917 | 737 | 670 | 724 | 826 | 775 | 792 | 635 | 596 | 508 | 581 | 398 | 496 | 441 | 865 | 805 | 687 | 742 | 762 | 837 | 769 | 551 | 631 | 394 | 481 | 465 | 497 | 519 | Tmem241 |
| Tmem242 | 2658 | 3029 | 3137 | 2750 | 2422 | 2552 | 2411 | 3168 | 3311 | 3258 | 3489 | 3556 | 3464 | 3671 | 2661 | 2994 | 3080 | 2538 | 2638 | 3027 | 2472 | 2460 | 3642 | 3555 | 3319 | 4449 | 3998 | 3939 | Tmem242 |
| Tmem243 | 806 | 657 | 853 | 789 | 695 | 778 | 951 | 372 | 315 | 375 | 277 | 442 | 420 | 536 | 880 | 688 | 812 | 739 | 762 | 850 | 791 | 334 | 314 | 369 | 310 | 475 | 475 | 463 | Tmem243 |
| Tmem245 | 11127 | 9506 | 6067 | 7679 | 7249 | 10471 | 9750 | 7566 | 7359 | 6253 | 7201 | 6403 | 6855 | 7075 | 9944 | 10674 | 6762 | 7714 | 7442 | 9164 | 10423 | 5656 | 6936 | 5108 | 6481 | 4386 | 7542 | 6159 | Tmem245 |

| | | | | | | | | | | | | | | | | | | | | | | | | | | | | | |
|---|---|---|---|---|---|---|---|---|---|---|---|---|---|---|---|---|---|---|---|---|---|---|---|---|---|---|---|---|---|
| Tmem248 | 3878 | 3485 | 4294 | 3697 | 3523 | 3308 | 3247 | 2989 | 3244 | 3203 | 3591 | 3753 | 3770 | 3600 | 3551 | 3993 | 3844 | 3393 | 3448 | 3071 | 3344 | 3715 | 3243 | 3507 | 3473 | 4429 | 3689 | 3909 | Tmem248 |
| Tmem249 | 21 | 32 | 37 | 39 | 25 | 23 | 29 | 11 | 21 | 1 | 6 | 21 | 7 | 11 | 30 | 16 | 34 | 28 | 43 | 27 | 20 | 13 | 9 | 9 | 15 | 8 | 14 | 11 | Tmem249 |
| Tmem25 | 490 | 575 | 521 | 631 | 746 | 565 | 567 | 325 | 451 | 462 | 542 | 462 | 477 | 357 | 502 | 469 | 643 | 536 | 694 | 588 | 592 | 553 | 409 | 360 | 471 | 372 | 384 | 354 | Tmem25 |
| Tmem250-ps | 934 | 1001 | 1181 | 882 | 905 | 895 | 919 | 1445 | 1873 | 1712 | 1972 | 2016 | 1827 | 1741 | 884 | 900 | 959 | 988 | 1072 | 852 | 928 | 1641 | 1942 | 2004 | 1708 | 2373 | 1864 | 1974 | Tmem250-ps |
| Tmem251 | 1335 | 1259 | 1112 | 1150 | 937 | 1061 | 1215 | 1137 | 943 | 818 | 880 | 1084 | 905 | 1104 | 1358 | 1234 | 1165 | 1137 | 969 | 1340 | 1011 | 961 | 924 | 977 | 905 | 1197 | 1106 | 1129 | Tmem251 |
| Tmem252 | 17 | 16 | 15 | 22 | 24 | 32 | 28 | 55 | 60 | 75 | 110 | 59 | 60 | 61 | 33 | 10 | 17 | 22 | 35 | 28 | 36 | 62 | 73 | 88 | 86 | 76 | 73 | 61 | Tmem252 |
| Tmem253 | 50 | 34 | 48 | 58 | 47 | 72 | 55 | 51 | 63 | 64 | 67 | 67 | 45 | 82 | 38 | 32 | 64 | 63 | 96 | 48 | 57 | 47 | 47 | 58 | 49 | 59 | 67 | 74 | Tmem253 |
| Tmem254a | 0 | 0 | 0 | 0 | 0 | 0 | 1 | 0 | 0 | 0 | 0 | 0 | 0 | 0 | 0 | 0 | 0 | 0 | 0 | 0 | 0 | 2 | 0 | 0 | 0 | 0 | 0 | 0 | Tmem254a |
| Tmem255a | 1746 | 1392 | 1174 | 1409 | 2011 | 1664 | 1697 | 1796 | 1863 | 1397 | 2004 | 907 | 1111 | 941 | 1614 | 1537 | 1363 | 1508 | 1418 | 1977 | 1531 | 1813 | 2253 | 1176 | 1291 | 1300 | 955 | 1049 | Tmem255a |
| Tmem255b | 3 | 2 | 2 | 9 | 2 | 4 | 2 | 10 | 16 | 3 | 18 | 0 | 6 | 4 | 7 | 5 | 1 | 0 | 5 | 10 | 0 | 8 | 9 | 2 | 7 | 8 | 1 | 5 | Tmem255b |
| Tmem256 | 2685 | 3255 | 3089 | 3179 | 2670 | 2524 | 3585 | 2082 | 1926 | 2149 | 2479 | 2186 | 2361 | 2860 | 2652 | 2103 | 3155 | 2993 | 2967 | 3181 | 3412 | 1519 | 1988 | 2586 | 2265 | 2648 | 2287 | 2712 | Tmem256 |
| Tmem258 | 2438 | 2308 | 2657 | 2588 | 2074 | 1794 | 2702 | 2789 | 2394 | 2619 | 2686 | 2256 | 2459 | 3882 | 2463 | 1502 | 2557 | 2089 | 2035 | 2309 | 2723 | 2090 | 2370 | 2974 | 2795 | 2901 | 2566 | 3537 | Tmem258 |
| Tmem259 | 4907 | 6430 | 6465 | 6409 | 5859 | 5156 | 5120 | 3932 | 4559 | 4994 | 4745 | 4975 | 4574 | 4535 | 4553 | 5243 | 6792 | 6146 | 6512 | 5043 | 5442 | 4714 | 4404 | 5227 | 4866 | 5267 | 4618 | 4309 | Tmem259 |
| Tmem26 | 3049 | 1620 | 1308 | 945 | 822 | 1448 | 534 | 3645 | 3044 | 2133 | 2537 | 1299 | 2161 | 1180 | 2028 | 2314 | 1119 | 1317 | 683 | 743 | 573 | 1415 | 3475 | 1560 | 1050 | 1747 | 1332 | 1441 | Tmem26 |
| Tmem260 | 4622 | 4050 | 3846 | 3609 | 3170 | 3343 | 3274 | 3794 | 3673 | 3545 | 3714 | 3384 | 3751 | 3335 | 4392 | 3977 | 3668 | 3617 | 3145 | 3551 | 3292 | 3527 | 3731 | 3325 | 3164 | 2993 | 3557 | 3401 | Tmem260 |
| Tmem262 | 2 | 4 | 2 | 2 | 2 | 5 | 4 | 6 | 5 | 8 | 5 | 12 | 22 | 13 | 5 | 2 | 0 | 2 | 2 | 2 | 4 | 18 | 9 | 8 | 15 | 19 | 27 | 8 | Tmem262 |
| Tmem263 | 7897 | 5581 | 7756 | 7670 | 6441 | 7311 | 7906 | 4756 | 4433 | 4096 | 5473 | 4427 | 4831 | 4835 | 7441 | 7504 | 7008 | 7697 | 6356 | 6357 | 7430 | 4896 | 4724 | 4782 | 4615 | 5009 | 4338 | 4866 | Tmem263 |
| Tmem266 | 295 | 448 | 452 | 517 | 585 | 439 | 426 | 232 | 234 | 289 | 225 | 224 | 249 | 195 | 360 | 414 | 424 | 437 | 514 | 428 | 508 | 282 | 277 | 244 | 359 | 163 | 194 | 209 | Tmem266 |
| Tmem267 | 619 | 470 | 928 | 799 | 882 | 1024 | 980 | 528 | 470 | 714 | 599 | 716 | 610 | 683 | 764 | 528 | 936 | 571 | 657 | 687 | 1218 | 907 | 733 | 793 | 739 | 628 | 860 | 656 | Tmem267 |
| Tmem268 | 516 | 454 | 656 | 495 | 430 | 457 | 418 | 470 | 583 | 569 | 500 | 533 | 497 | 530 | 407 | 461 | 575 | 453 | 500 | 394 | 410 | 399 | 511 | 548 | 493 | 680 | 450 | 570 | Tmem268 |
| Tmem269 | 157 | 245 | 180 | 334 | 240 | 249 | 249 | 56 | 68 | 51 | 36 | 59 | 90 | 72 | 195 | 209 | 233 | 337 | 259 | 232 | 242 | 52 | 40 | 56 | 71 | 50 | 72 | 44 | Tmem269 |
| Tmem270 | 0 | 0 | 0 | 1 | 0 | 0 | 0 | 0 | 0 | 1 | 2 | 0 | 0 | 0 | 0 | 0 | 0 | 0 | 0 | 0 | 2 | 0 | 0 | 0 | 0 | 0 | 0 | 0 | Tmem270 |
| Tmem273 | 9 | 6 | 11 | 17 | 9 | 8 | 4 | 12 | 24 | 19 | 21 | 16 | 18 | 6 | 13 | 12 | 8 | 15 | 10 | 10 | 14 | 18 | 29 | 26 | 10 | 23 | 21 | 8 | Tmem273 |
| Tmem274 | 0 | 0 | 0 | 0 | 0 | 0 | 0 | 2 | 0 | 0 | 0 | 0 | 0 | 0 | 0 | 0 | 0 | 0 | 0 | 0 | 1 | 0 | 0 | 0 | 0 | 0 | 0 | 0 | Tmem274 |
| Tmem28 | 938 | 1228 | 1291 | 1562 | 1717 | 1567 | 1472 | 608 | 644 | 715 | 641 | 772 | 662 | 661 | 910 | 886 | 1467 | 1530 | 1831 | 1250 | 1656 | 735 | 593 | 770 | 835 | 773 | 750 | 707 | Tmem28 |
| Tmem29 | 1283 | 1153 | 851 | 1095 | 1191 | 1401 | 1402 | 2807 | 2176 | 1840 | 1940 | 1984 | 1955 | 2104 | 1502 | 1188 | 995 | 1117 | 1232 | 1629 | 1384 | 2134 | 2313 | 1833 | 2199 | 1733 | 2100 | 1918 | Tmem29 |
| Tmem30a | 16125 | 12407 | 14095 | 15550 | 14574 | 15780 | 17639 | 14476 | 11618 | 10088 | 10757 | 11533 | 11523 | 12554 | 18753 | 16331 | 13844 | 15367 | 14371 | 17062 | 16830 | 13785 | 12301 | 11000 | 11784 | 10542 | 11577 | 12349 | Tmem30a |
| Tmem30b | 960 | 520 | 384 | 318 | 129 | 195 | 29 | 390 | 386 | 438 | 390 | 157 | 268 | 138 | 616 | 879 | 312 | 306 | 30 | 21 | 24 | 317 | 405 | 396 | 145 | 469 | 166 | 126 | Tmem30b |
| Tmem30c | 61 | 56 | 82 | 95 | 111 | 89 | 94 | 62 | 43 | 32 | 63 | 50 | 56 | 77 | 54 | 72 | 51 | 87 | 97 | 52 | 130 | 79 | 49 | 65 | 55 | 66 | 64 | 52 | Tmem30c |
| Tmem33 | 11749 | 9430 | 10253 | 10564 | 11972 | 12150 | 13019 | 14161 | 11006 | 10251 | 10200 | 11116 | 11027 | 11563 | 13181 | 10647 | 11049 | 10382 | 11330 | 13613 | 12042 | 12183 | 11248 | 10335 | 11537 | 9678 | 11577 | 11487 | Tmem33 |
| Tmem35a | 3283 | 3377 | 3043 | 3865 | 5182 | 4498 | 5310 | 5140 | 4191 | 4209 | 3859 | 4036 | 2904 | 4078 | 3969 | 2597 | 3971 | 4147 | 5160 | 5537 | 5169 | 5691 | 4337 | 4279 | 5011 | 3695 | 3841 | 3507 | Tmem35a |
| Tmem35b | 523 | 515 | 508 | 516 | 591 | 582 | 577 | 493 | 489 | 572 | 566 | 571 | 565 | 480 | 467 | 475 | 535 | 578 | 581 | 544 | 550 | 585 | 491 | 472 | 489 | 536 | 503 | 511 | Tmem35b |
| Tmem37 | 223 | 327 | 245 | 170 | 123 | 158 | 110 | 99 | 140 | 102 | 125 | 85 | 137 | 139 | 220 | 338 | 210 | 155 | 125 | 154 | 110 | 90 | 209 | 104 | 88 | 165 | 122 | 106 | Tmem37 |
| Tmem38a | 1722 | 1902 | 1771 | 2186 | 2140 | 2105 | 1887 | 1066 | 1200 | 1166 | 1679 | 1049 | 1059 | 971 | 1619 | 1749 | 2067 | 2165 | 2367 | 1738 | 2078 | 1165 | 1514 | 1225 | 1091 | 1078 | 1046 | 1055 | Tmem38a |
| Tmem38b | 1533 | 1561 | 1901 | 1605 | 1283 | 1333 | 1501 | 833 | 695 | 699 | 791 | 897 | 805 | 942 | 1510 | 1373 | 1717 | 1556 | 1449 | 1439 | 1478 | 750 | 730 | 861 | 785 | 820 | 1078 | 1044 | Tmem38b |
| Tmem39a | 2468 | 2344 | 2201 | 1888 | 1676 | 1878 | 1824 | 2828 | 3134 | 2842 | 3551 | 2558 | 2971 | 2639 | 2111 | 2280 | 2126 | 2104 | 1884 | 2005 | 1680 | 2916 | 3214 | 2786 | 2461 | 2780 | 2787 | 2716 | Tmem39a |
| Tmem39b | 655 | 923 | 859 | 846 | 741 | 752 | 697 | 1386 | 1543 | 1549 | 1456 | 1804 | 1574 | 1821 | 655 | 665 | 822 | 810 | 859 | 744 | 799 | 1549 | 1423 | 1678 | 1710 | 1984 | 1865 | 1896 | Tmem39b |
| Tmem40 | 25 | 28 | 30 | 5 | 17 | 6 | 18 | 32 | 54 | 28 | 52 | 64 | 70 | 35 | 24 | 48 | 24 | 21 | 12 | 23 | 18 | 45 | 56 | 41 | 51 | 91 | 53 | 42 | Tmem40 |
| Tmem41a | 1444 | 1609 | 1653 | 1600 | 1559 | 1477 | 1481 | 763 | 789 | 681 | 856 | 710 | 838 | 660 | 1494 | 1444 | 1519 | 1379 | 1612 | 1555 | 1389 | 705 | 752 | 670 | 707 | 720 | 759 | 727 | Tmem41a |
| Tmem41b | 5396 | 4797 | 4234 | 4643 | 4024 | 4518 | 4866 | 3938 | 3403 | 2783 | 3134 | 3207 | 3244 | 3283 | 5773 | 5531 | 4204 | 4549 | 3832 | 5101 | 4627 | 3432 | 3451 | 2869 | 3254 | 3206 | 3407 | 3204 | Tmem41b |
| Tmem42 | 37 | 60 | 40 | 50 | 70 | 51 | 75 | 189 | 205 | 201 | 214 | 188 | 144 | 203 | 49 | 55 | 41 | 26 | 45 | 58 | 58 | 202 | 195 | 187 | 190 | 198 | 181 | 249 | Tmem42 |
| Tmem43 | 5283 | 6452 | 5310 | 5209 | 4637 | 5422 | 4898 | 3896 | 4397 | 3949 | 3966 | 4766 | 4332 | 4242 | 5210 | 5436 | 5202 | 5270 | 4960 | 5093 | 4937 | 3910 | 4344 | 4074 | 4396 | 4851 | 4955 | 4658 | Tmem43 |
| Tmem44 | 2212 | 2352 | 2486 | 3528 | 5490 | 4395 | 4402 | 7010 | 7636 | 9466 | 8481 | 8145 | 7564 | 7990 | 2661 | 2225 | 3362 | 3915 | 5160 | 3668 | 4922 | 9909 | 6803 | 8509 | 10226 | 6058 | 7626 | 7243 | Tmem44 |
| Tmem45a | 340 | 241 | 268 | 200 | 163 | 292 | 134 | 460 | 620 | 437 | 719 | 257 | 415 | 265 | 257 | 392 | 232 | 318 | 139 | 168 | 113 | 433 | 748 | 324 | 324 | 373 | 281 | 316 | Tmem45a |
| Tmem45a2 | 0 | 0 | 0 | 0 | 0 | 1 | 0 | 1 | 3 | 10 | 2 | 0 | 2 | 0 | 0 | 2 | 0 | 2 | 0 | 0 | 0 | 3 | 0 | 3 | 1 | 4 | 8 | 1 | Tmem45a2 |
| Tmem45b | 1 | 0 | 0 | 1 | 1 | 0 | 0 | 6 | 3 | 0 | 5 | 2 | 1 | 2 | 1 | 1 | 0 | 0 | 4 | 0 | 1 | 0 | 9 | 2 | 0 | 4 | 0 | 1 | Tmem45b |
| Tmem47 | 6680 | 4847 | 6993 | 5579 | 5416 | 5793 | 6093 | 6282 | 4700 | 4267 | 4651 | 6298 | 5425 | 6322 | 6881 | 6118 | 6131 | 5497 | 5557 | 6222 | 5603 | 6696 | 5807 | 5394 | 5211 | 7139 | 6875 | 6984 | Tmem47 |
| Tmem50a | 4086 | 4741 | 5063 | 4952 | 4538 | 4454 | 4626 | 3213 | 3449 | 3399 | 3544 | 3414 | 3539 | 3534 | 4195 | 4611 | 5010 | 4782 | 4815 | 5041 | 4782 | 3283 | 3840 | 3809 | 3598 | 3720 | 3626 | 3564 | Tmem50a |
| Tmem50b | 6030 | 6736 | 6404 | 6957 | 6725 | 6238 | 7110 | 2485 | 2329 | 2458 | 2195 | 2046 | 2225 | 2039 | 6552 | 6295 | 6519 | 7001 | 6819 | 7305 | 7058 | 2478 | 2293 | 2292 | 2400 | 2016 | 2323 | 2156 | Tmem50b |
| Tmem51 | 8 | 3 | 9 | 1 | 4 | 0 | 9 | 5 | 7 | 2 | 5 | 3 | 3 | 2 | 1 | 2 | 8 | 2 | 2 | 0 | 1 | 8 | 5 | 2 | 1 | 8 | 2 | 2 | Tmem51 |
| Tmem52 | 2 | 4 | 6 | 4 | 9 | 15 | 1 | 2 | 1 | 3 | 4 | 3 | 8 | 2 | 4 | 9 | 17 | 7 | 6 | 9 | 3 | 2 | 6 | 2 | 3 | 1 | 1 | 1 | Tmem52 |
| Tmem52b | 0 | 2 | 0 | 2 | 5 | 5 | 1 | 1 | 2 | 0 | 1 | 5 | 1 | 2 | 1 | 1 | 0 | 0 | 2 | 5 | 0 | 3 | 1 | 0 | 3 | 5 | 0 | 0 | Tmem52b |
| Tmem53 | 182 | 234 | 199 | 224 | 146 | 114 | 168 | 120 | 136 | 161 | 150 | 155 | 154 | 94 | 200 | 183 | 238 | 168 | 172 | 177 | 119 | 147 | 181 | 172 | 101 | 188 | 181 | 140 | Tmem53 |
| Tmem54 | 92 | 43 | 30 | 31 | 16 | 30 | 4 | 46 | 68 | 60 | 35 | 26 | 31 | 19 | 46 | 58 | 20 | 12 | 4 | 5 | 5 | 3 | 87 | 63 | 13 | 89 | 14 | 9 | Tmem54 |
| Tmem59 | 8358 | 7824 | 7852 | 8101 | 8222 | 7808 | 8561 | 8709 | 7959 | 7936 | 7692 | 7605 | 7998 | 7629 | 8468 | 7734 | 7771 | 7907 | 8119 | 7692 | 8430 | 8306 | 8721 | 7629 | 7731 | 7979 | 7518 | 7976 | Tmem59 |
| Tmem59l | 747 | 668 | 625 | 757 | 1123 | 783 | 831 | 808 | 994 | 1002 | 930 | 657 | 716 | 721 | 604 | 640 | 730 | 878 | 967 | 852 | 852 | 1038 | 1070 | 843 | 1139 | 602 | 599 | 610 | Tmem59l |
| Tmem60 | 1706 | 1676 | 1611 | 1615 | 1325 | 1485 | 1868 | 1305 | 1069 | 1009 | 960 | 1227 | 1109 | 1177 | 1837 | 1779 | 1643 | 1574 | 1440 | 1927 | 1787 | 1039 | 1089 | 1061 | 1027 | 1326 | 1106 | 1309 | Tmem60 |
| Tmem61 | 0 | 0 | 0 | 0 | 0 | 2 | 0 | 0 | 1 | 0 | 0 | 0 | 0 | 0 | 1 | 0 | 0 | 0 | 0 | 1 | 0 | 7 | 0 | 0 | 0 | 0 | 0 | 0 | Tmem61 |
| Tmem62 | 280 | 226 | 241 | 224 | 388 | 302 | 249 | 834 | 656 | 756 | 657 | 551 | 553 | 581 | 324 | 253 | 308 | 281 | 344 | 305 | 355 | 961 | 664 | 580 | 732 | 510 | 599 | 553 | Tmem62 |
| Tmem63a | 923 | 903 | 974 | 711 | 462 | 585 | 418 | 386 | 607 | 573 | 743 | 384 | 634 | 343 | 757 | 1150 | 779 | 745 | 537 | 471 | 411 | 491 | 677 | 447 | 374 | 484 | 404 | 437 | Tmem63a |
| Tmem63b | 5812 | 7201 | 8671 | 9728 | 10605 | 9053 | 9245 | 6903 | 7376 | 9772 | 8185 | 9813 | 8587 | 9207 | 5921 | 6510 | 9429 | 9525 | 11097 | 8145 | 9340 | 10494 | 7124 | 9800 | 10616 | 8386 | 8707 | 8926 | Tmem63b |
| Tmem63c | 227 | 296 | 273 | 423 | 853 | 559 | 569 | 570 | 581 | 613 | 599 | 399 | 370 | 366 | 328 | 301 | 340 | 609 | 732 | 634 | 672 | 949 | 508 | 462 | 768 | 197 | 315 | 268 | Tmem63c |
| Tmem64 | 5186 | 3997 | 4389 | 3584 | 2875 | 3394 | 2906 | 3065 | 3057 | 2401 | 3311 | 2437 | 3061 | 2539 | 4970 | 5431 | 3458 | 4035 | 2529 | 2765 | 2838 | 2988 | 3629 | 2348 | 2260 | 2576 | 2650 | 2726 | Tmem64 |
| Tmem65 | 6181 | 4452 | 6015 | 6178 | 7351 | 6586 | 7199 | 5961 | 5704 | 5358 | 5743 | 5260 | 5629 | 5582 | 6562 | 5814 | 6050 | 6722 | 6430 | 5992 | 6664 | 7143 | 5677 | 5494 | 6053 | 5447 | 5239 | 5505 | Tmem65 |
| Tmem67 | 2335 | 2277 | 2026 | 2325 | 2345 | 2422 | 2646 | 2585 | 2175 | 2012 | 2106 | 2228 | 2096 | 2350 | 2537 | 2286 | 2203 | 2369 | 2364 | 3072 | 2561 | 2346 | 2181 | 2084 | 2072 | 2067 | 2195 | 2135 | Tmem67 |
| Tmem68 | 1588 | 1145 | 1542 | 1546 | 1622 | 1583 | 1890 | 3429 | 2770 | 2941 | 2607 | 3007 | 2620 | 3158 | 1743 | 1348 | 1455 | 1531 | 1760 | 1818 | 1698 | 3160 | 2823 | 2813 | 3044 | 2838 | 2997 | 3128 | Tmem68 |
| Tmem69 | 1238 | 1180 | 1556 | 1206 | 1182 | 1168 | 1265 | 1690 | 1385 | 1512 | 1481 | 1663 | 1638 | 1868 | 1295 | 1137 | 1417 | 1193 | 1266 | 1246 | 1273 | 1534 | 1304 | 1684 | 1437 | 1697 | 1617 | 2016 | Tmem69 |
| Tmem70 | 2486 | 2027 | 2382 | 2330 | 2267 | 2277 | 2682 | 2470 | 2257 | 2100 | 2278 | 2411 | 2332 | 2533 | 2502 | 2306 | 2407 | 2376 | 2353 | 2464 | 2339 | 2374 | 2511 | 2394 | 2395 | 2684 | 2344 | 2485 | Tmem70 |
| Tmem71 | 17 | 8 | 26 | 17 | 11 | 12 | 4 | 18 | 18 | 11 | 23 | 14 | 15 | 5 | 11 | 17 | 7 | 10 | 12 | 22 | 22 | 22 | 19 | 11 | 22 | 8 | 21 | 17 | Tmem71 |
| Tmem72 | 7 | 5 | 0 | 1 | 107 | 37 | 12 | 104 | 207 | 143 | 136 | 41 | 36 | 3 | 9 | 16 | 2 | 15 | 42 | 19 | 48 | 264 | 445 | 7 | 73 | 1 | 3 | 1 | Tmem72 |
| Tmem74 | 794 | 751 | 684 | 948 | 1216 | 1024 | 1149 | 858 | 777 | 708 | 626 | 533 | 477 | 590 | 993 | 727 | 870 | 1024 | 1015 | 1175 | 1274 | 734 | 654 | 565 | 948 | 396 | 511 | 515 | Tmem74 |
| Tmem74b | 141 | 143 | 115 | 149 | 405 | 338 | 294 | 510 | 643 | 637 | 518 | 483 | 418 | 506 | 156 | 121 | 148 | 163 | 355 | 308 | 252 | 638 | 557 | 448 | 771 | 297 | 415 | 345 | Tmem74b |
| Tmem74bos | 82 | 132 | 95 | 215 | 136 | 171 | 177 | 27 | 35 | 23 | 26 | 36 | 26 | 49 | 100 | 110 | 136 | 196 | 147 | 132 | 230 | 40 | 37 | 46 | 39 | 36 | 29 | 27 | Tmem74bos |
| Tmem79 | 179 | 241 | 260 | 189 | 156 | 204 | 177 | 127 | 211 | 148 | 187 | 176 | 187 | 178 | 166 | 206 | 217 | 232 | 185 | 160 | 177 | 79 | 206 | 200 | 93 | 169 | 163 | 173 | Tmem79 |
| Tmem80 | 721 | 813 | 881 | 874 | 1028 | 1079 | 959 | 2037 | 1904 | 2254 | 2170 | 2452 | 2430 | 2515 | 816 | 759 | 875 | 796 | 1025 | 999 | 950 | 2271 | 1848 | 2530 | 2432 | 2366 | 2604 | 2487 | Tmem80 |
| Tmem81 | 264 | 229 | 273 | 297 | 232 | 231 | 201 | 175 | 173 | 189 | 232 | 227 | 181 | 172 | 261 | 232 | 212 | 234 | 248 | 196 | 221 | 206 | 214 | 189 | 212 | 271 | 214 | 184 | Tmem81 |
| Tmem82 | 40 | 29 | 38 | 37 | 38 | 38 | 16 | 12 | 22 | 31 | 19 | 20 | 16 | 21 | 21 | 50 | 49 | 25 | 26 | 25 | 42 | 20 | 27 | 23 | 12 | 24 | 19 | 19 | Tmem82 |
| Tmem86a | 873 | 996 | 832 | 990 | 818 | 850 | 825 | 700 | 826 | 723 | 844 | 575 | 671 | 657 | 873 | 918 | 825 | 997 | 803 | 852 | 847 | 752 | 923 | 717 | 693 | 699 | 621 | 565 | Tmem86a |
| Tmem86b | 1023 | 1327 | 1525 | 1625 | 1704 | 1373 | 1516 | 982 | 950 | 1160 | 967 | 1314 | 1149 | 1193 | 1011 | 1097 | 1648 | 1568 | 1742 | 1425 | 1450 | 1245 | 788 | 1219 | 1448 | 1212 | 1334 | 1216 | Tmem86b |

| | | | | | | | | | | | | | | | | | | | | | | | | | | | | | |
|---|---|---|---|---|---|---|---|---|---|---|---|---|---|---|---|---|---|---|---|---|---|---|---|---|---|---|---|---|---|
| Tmem87a | 3504 | 2688 | 3247 | 3152 | 3112 | 3184 | 3505 | 3968 | 3691 | 3619 | 3700 | 3747 | 3790 | 3821 | 3516 | 3115 | 3226 | 3201 | 3264 | 3461 | 3048 | 3801 | 3693 | 3507 | 3722 | 3745 | 3691 | 3763 | Tmem87a |
| Tmem87b | 4715 | 3945 | 3981 | 4038 | 3976 | 4388 | 4256 | 4860 | 3962 | 4034 | 4359 | 4138 | 4276 | 4258 | 4739 | 4609 | 3928 | 3782 | 3940 | 4440 | 4295 | 4426 | 4288 | 3694 | 4110 | 3400 | 4269 | 3972 | Tmem87b |
| Tmem88 | 248 | 348 | 328 | 164 | 177 | 214 | 155 | 162 | 187 | 181 | 217 | 215 | 225 | 243 | 211 | 336 | 264 | 186 | 182 | 169 | 163 | 180 | 244 | 202 | 160 | 284 | 215 | 266 | Tmem88 |
| Tmem88b | 20 | 22 | 20 | 25 | 52 | 52 | 28 | 87 | 80 | 104 | 154 | 58 | 74 | 57 | 28 | 18 | 20 | 28 | 34 | 17 | 27 | 107 | 115 | 77 | 81 | 55 | 48 | 47 | Tmem88b |
| Tmem89 | 0 | 0 | 2 | 0 | 0 | 1 | 0 | 0 | 0 | 0 | 0 | 0 | 0 | 0 | 0 | 1 | 0 | 0 | 0 | 0 | 0 | 0 | 0 | 1 | 0 | 0 | 0 | 0 | Tmem89 |
| Tmem8b | 4061 | 4545 | 4801 | 6135 | 6948 | 5845 | 5870 | 3983 | 4146 | 5166 | 4427 | 4868 | 4433 | 4497 | 4237 | 4192 | 6005 | 6272 | 7049 | 5071 | 6266 | 5710 | 3703 | 4877 | 5414 | 4335 | 4284 | 4371 | Tmem8b |
| Tmem9 | 4426 | 5029 | 4987 | 4175 | 3939 | 4046 | 3758 | 4084 | 4700 | 4081 | 4546 | 4188 | 4552 | 4278 | 4212 | 4611 | 4544 | 3940 | 4338 | 4285 | 3830 | 3811 | 5121 | 4237 | 4049 | 4886 | 4597 | 4446 | Tmem9 |
| Tmem91 | 46 | 45 | 40 | 68 | 133 | 117 | 46 | 62 | 61 | 56 | 79 | 35 | 17 | 40 | 28 | 76 | 44 | 64 | 94 | 85 | 68 | 62 | 51 | 60 | 57 | 39 | 20 | 39 | Tmem91 |
| Tmem94 | 2127 | 2752 | 3355 | 2962 | 3190 | 2714 | 2366 | 3675 | 4790 | 5350 | 4984 | 6097 | 5058 | 5204 | 2097 | 2602 | 3405 | 2796 | 3128 | 2215 | 2579 | 5259 | 4279 | 5708 | 5268 | 5916 | 4779 | 5385 | Tmem94 |
| Tmem95 | 74 | 65 | 80 | 57 | 46 | 34 | 31 | 16 | 25 | 29 | 46 | 28 | 20 | 22 | 62 | 67 | 46 | 75 | 29 | 45 | 49 | 30 | 27 | 34 | 23 | 19 | 19 | 18 | Tmem95 |
| Tmem97 | 1503 | 1782 | 1566 | 1309 | 911 | 1154 | 1060 | 1304 | 1331 | 1306 | 1311 | 1341 | 1429 | 1498 | 1423 | 1509 | 1348 | 1110 | 1124 | 1393 | 1077 | 1029 | 1504 | 1258 | 1229 | 1694 | 1622 | 1639 | Tmem97 |
| Tmem98 | 1005 | 967 | 1263 | 732 | 713 | 721 | 736 | 1751 | 1957 | 1697 | 1674 | 2213 | 2146 | 2080 | 960 | 1101 | 1060 | 673 | 814 | 657 | 668 | 1469 | 1723 | 2047 | 1662 | 2951 | 2246 | 2410 | Tmem98 |
| Tmem9b | 1937 | 1937 | 1692 | 1850 | 2121 | 1863 | 2024 | 2207 | 1920 | 1809 | 1787 | 1483 | 1655 | 1638 | 2259 | 1784 | 1854 | 1847 | 1776 | 2215 | 2074 | 2095 | 2076 | 1620 | 1772 | 1554 | 1569 | 1647 | Tmem9b |
| Tmf1 | 5463 | 4444 | 4953 | 5429 | 4947 | 5253 | 5559 | 4752 | 3996 | 4181 | 4220 | 4305 | 4397 | 4708 | 5304 | 4684 | 5076 | 5133 | 4930 | 5279 | 5524 | 4477 | 4122 | 4220 | 4628 | 3864 | 4291 | 4609 | Tmf1 |
| Tmie | 44 | 52 | 68 | 39 | 53 | 23 | 42 | 51 | 64 | 80 | 66 | 26 | 56 | 16 | 58 | 53 | 32 | 37 | 40 | 44 | 47 | 58 | 67 | 38 | 42 | 34 | 13 | 20 | Tmie |
| Tmigd1 | 1 | 0 | 0 | 0 | 1 | 1 | 1 | 0 | 0 | 2 | 5 | 0 | 0 | 0 | 1 | 0 | 2 | 0 | 0 | 0 | 0 | 0 | 2 | 1 | 2 | 0 | 0 | 0 | Tmigd1 |
| Tmigd3 | 0 | 2 | 0 | 0 | 0 | 0 | 0 | 0 | 0 | 0 | 0 | 0 | 5 | 0 | 0 | 2 | 0 | 0 | 0 | 1 | 0 | 0 | 0 | 0 | 0 | 0 | 0 | 0 | Tmigd3 |
| Tmod1 | 216 | 249 | 182 | 236 | 187 | 170 | 179 | 92 | 152 | 93 | 460 | 84 | 200 | 109 | 191 | 293 | 166 | 220 | 151 | 154 | 147 | 139 | 392 | 74 | 60 | 101 | 98 | 171 | Tmod1 |
| Tmod2 | 14715 | 13731 | 9733 | 15124 | 20742 | 20695 | 21425 | 20914 | 17745 | 16837 | 16592 | 15209 | 13397 | 16550 | 16391 | 12950 | 11957 | 16126 | 17309 | 22034 | 21510 | 19524 | 16071 | 13657 | 21117 | 9312 | 15254 | 13897 | Tmod2 |
| Tmod3 | 16533 | 13824 | 13750 | 13863 | 12887 | 13528 | 14967 | 9432 | 8496 | 7137 | 7927 | 8443 | 8450 | 8816 | 16719 | 15432 | 13603 | 14270 | 12961 | 15033 | 14220 | 7518 | 8744 | 8037 | 7604 | 8899 | 8606 | 9143 | Tmod3 |
| Tmod4 | 107 | 112 | 140 | 120 | 103 | 108 | 127 | 82 | 82 | 96 | 114 | 89 | 87 | 97 | 78 | 98 | 141 | 130 | 146 | 79 | 109 | 104 | 95 | 98 | 100 | 113 | 94 | 95 | Tmod4 |
| Tmpo | 30496 | 22560 | 29175 | 23944 | 24044 | 25989 | 29033 | 40819 | 35446 | 33119 | 33697 | 45970 | 39531 | 47264 | 28397 | 24157 | 27194 | 23022 | 27274 | 27480 | 26815 | 32042 | 33048 | 39648 | 36707 | 49314 | 47236 | 49100 | Tmpo |
| Tmppe | 479 | 643 | 702 | 667 | 670 | 631 | 502 | 343 | 418 | 425 | 496 | 461 | 476 | 388 | 418 | 588 | 658 | 657 | 628 | 468 | 576 | 444 | 423 | 417 | 450 | 238 | 429 | 412 | Tmppe |
| Tmprss11a | 6 | 0 | 3 | 0 | 0 | 0 | 0 | 9 | 25 | 4 | 10 | 2 | 1 | 0 | 0 | 7 | 0 | 2 | 0 | 0 | 0 | 3 | 28 | 1 | 9 | 0 | 0 | 0 | Tmprss11a |
| Tmprss11b | 7 | 0 | 0 | 0 | 2 | 0 | 0 | 24 | 51 | 22 | 66 | 3 | 0 | 2 | 4 | 2 | 0 | 0 | 0 | 0 | 0 | 2 | 61 | 15 | 6 | 2 | 0 | 0 | Tmprss11b |
| Tmprss11d | 6 | 2 | 3 | 2 | 0 | 2 | 4 | 5 | 8 | 0 | 1 | 1 | 0 | 3 | 5 | 1 | 0 | 6 | 2 | 4 | 4 | 2 | 5 | 4 | 0 | 1 | 2 | 2 | Tmprss11d |
| Tmprss11e | 0 | 0 | 0 | 0 | 0 | 0 | 0 | 1 | 6 | 1 | 2 | 0 | 7 | 0 | 3 | 0 | 2 | 1 | 2 | 0 | 0 | 2 | 4 | 2 | 0 | 3 | 0 | 0 | Tmprss11e |
| Tmprss11f | 7 | 1 | 0 | 0 | 0 | 0 | 0 | 1 | 0 | 0 | 5 | 2 | 1 | 2 | 1 | 0 | 0 | 3 | 0 | 0 | 0 | 0 | 1 | 0 | 0 | 3 | 0 | 0 | Tmprss11f |
| Tmprss11g | 0 | 0 | 0 | 0 | 0 | 0 | 0 | 0 | 2 | 0 | 2 | 0 | 0 | 0 | 0 | 0 | 0 | 0 | 0 | 1 | 0 | 3 | 1 | 0 | 0 | 0 | 0 | 0 | Tmprss11g |
| Tmprss12 | 0 | 0 | 0 | 0 | 0 | 0 | 1 | 1 | 0 | 0 | 0 | 0 | 0 | 0 | 1 | 0 | 3 | 0 | 0 | 0 | 0 | 2 | 3 | 0 | 1 | 0 | 0 | 1 | Tmprss12 |
| Tmprss13 | 18 | 10 | 8 | 1 | 0 | 1 | 0 | 40 | 81 | 37 | 16 | 16 | 24 | 6 | 12 | 9 | 5 | 3 | 7 | 0 | 2 | 2 | 83 | 35 | 2 | 13 | 10 | 7 | Tmprss13 |
| Tmprss15 | 1 | 3 | 0 | 1 | 5 | 2 | 0 | 6 | 2 | 0 | 0 | 0 | 1 | 4 | 5 | 1 | 2 | 2 | 1 | 1 | 0 | 0 | 1 | 3 | 1 | 4 | 8 | 0 | Tmprss15 |
| Tmprss2 | 58 | 56 | 47 | 21 | 11 | 38 | 0 | 116 | 138 | 99 | 112 | 48 | 69 | 42 | 90 | 68 | 30 | 41 | 8 | 4 | 4 | 12 | 142 | 118 | 12 | 44 | 38 | 42 | Tmprss2 |
| Tmprss3 | 2 | 0 | 0 | 0 | 0 | 0 | 1 | 5 | 7 | 3 | 2 | 0 | 3 | 2 | 1 | 2 | 0 | 0 | 0 | 0 | 1 | 5 | 5 | 2 | 0 | 0 | 1 | 1 | Tmprss3 |
| Tmprss4 | 12 | 3 | 0 | 8 | 2 | 3 | 0 | 40 | 26 | 37 | 44 | 1 | 11 | 2 | 7 | 3 | 1 | 2 | 5 | 0 | 1 | 2 | 52 | 77 | 0 | 33 | 0 | 1 | Tmprss4 |
| Tmprss5 | 84 | 80 | 73 | 69 | 81 | 88 | 67 | 115 | 129 | 131 | 177 | 163 | 176 | 161 | 76 | 72 | 105 | 87 | 97 | 84 | 54 | 149 | 135 | 140 | 181 | 154 | 164 | 140 | Tmprss5 |
| Tmprss6 | 44 | 16 | 0 | 40 | 7 | 4 | 1 | 89 | 63 | 153 | 92 | 5 | 45 | 13 | 26 | 14 | 7 | 1 | 2 | 0 | 3 | 7 | 107 | 161 | 16 | 158 | 3 | 7 | Tmprss6 |
| Tmprss7 | 127 | 139 | 106 | 140 | 179 | 167 | 166 | 101 | 105 | 95 | 89 | 166 | 92 | 126 | 131 | 127 | 167 | 213 | 172 | 154 | 176 | 100 | 87 | 113 | 124 | 80 | 148 | 120 | Tmprss7 |
| Tmprss9 | 55 | 76 | 58 | 138 | 103 | 74 | 70 | 13 | 15 | 23 | 19 | 16 | 26 | 16 | 51 | 69 | 114 | 180 | 81 | 44 | 75 | 47 | 23 | 23 | 28 | 10 | 12 | 27 | Tmprss9 |
| Tmsb10 | 37527 | 50555 | 51037 | 58091 | 56819 | 48096 | 41604 | 46137 | 50696 | 66703 | 59182 | 51879 | 54191 | 52016 | 40788 | 36633 | 55784 | 56126 | 56676 | 49444 | 50860 | 52472 | 53369 | 63992 | 65101 | 59167 | 53843 | 50717 | Tmsb10 |
| Tmsb15a | 0 | 0 | 0 | 0 | 0 | 0 | 0 | 2 | 0 | 1 | 6 | 1 | 5 | 1 | 0 | 0 | 1 | 0 | 1 | 0 | 2 | 2 | 4 | 6 | 3 | 0 | 0 | 5 | Tmsb15a |
| Tmsb15b1 | 30 | 29 | 15 | 12 | 20 | 12 | 47 | 22 | 20 | 10 | 18 | 2 | 14 | 27 | 36 | 16 | 9 | 9 | 16 | 25 | 22 | 18 | 13 | 18 | 16 | 19 | 13 | 15 | Tmsb15b1 |
| Tmsb15b2 | 9 | 8 | 13 | 26 | 7 | 11 | 9 | 6 | 7 | 8 | 4 | 4 | 5 | 3 | 8 | 3 | 12 | 7 | 12 | 18 | 16 | 5 | 10 | 20 | 9 | 3 | 8 | 0 | Tmsb15b2 |
| Tmsb15l | 9 | 10 | 2 | 21 | 15 | 12 | 20 | 17 | 4 | 3 | 1 | 7 | 9 | 9 | 12 | 10 | 14 | 14 | 10 | 23 | 18 | 5 | 5 | 15 | 5 | 2 | 2 | 11 | Tmsb15l |
| Tmsb4x | 59926 | 63017 | 62632 | 66837 | 69494 | 69471 | 83356 | 80156 | 69613 | 71153 | 66584 | 76983 | 78831 | 89593 | 73984 | 61707 | 66329 | 62714 | 71729 | 95876 | 78287 | 65993 | 72247 | 78131 | 79505 | 83481 | 87088 | 90956 | Tmsb4x |
| Tmtc1 | 1351 | 1256 | 961 | 1213 | 1906 | 1533 | 1182 | 1570 | 2015 | 1788 | 2249 | 1339 | 1786 | 1192 | 1439 | 1650 | 1010 | 1569 | 1027 | 1246 | 1296 | 2430 | 2151 | 1268 | 1733 | 757 | 1333 | 1078 | Tmtc1 |
| Tmtc2 | 3244 | 2745 | 2766 | 2984 | 2918 | 2726 | 2940 | 3761 | 3648 | 3505 | 3705 | 2727 | 3393 | 2766 | 3389 | 3066 | 3072 | 2793 | 2582 | 2376 | 2691 | 4027 | 3224 | 3120 | 3067 | 2953 | 2722 | 2636 | Tmtc2 |
| Tmtc3 | 6737 | 4801 | 4538 | 4770 | 4139 | 5118 | 5127 | 4338 | 4021 | 3357 | 4064 | 4057 | 4238 | 4355 | 6045 | 6214 | 4183 | 4952 | 4036 | 4820 | 4766 | 3723 | 4133 | 3428 | 3419 | 3653 | 4154 | 4144 | Tmtc3 |
| Tmtc4 | 3767 | 3380 | 3384 | 3471 | 3850 | 3571 | 3798 | 4209 | 3876 | 3841 | 3661 | 3641 | 3731 | 3662 | 4180 | 3453 | 3610 | 3646 | 3590 | 4012 | 3746 | 4226 | 3952 | 3344 | 3576 | 3392 | 3837 | 3706 | Tmtc4 |
| Tmub1 | 773 | 1025 | 896 | 1005 | 904 | 798 | 829 | 662 | 728 | 758 | 743 | 735 | 687 | 698 | 732 | 776 | 968 | 956 | 994 | 880 | 979 | 719 | 738 | 626 | 724 | 860 | 672 | 680 | Tmub1 |
| Tmub2 | 3111 | 3893 | 3538 | 3483 | 3697 | 3576 | 3429 | 2860 | 3418 | 3461 | 3362 | 3479 | 3497 | 3266 | 3212 | 3381 | 3789 | 3168 | 3965 | 3605 | 3504 | 3390 | 3486 | 3559 | 3600 | 3411 | 3744 | 3393 | Tmub2 |
| Tmx1 | 11185 | 8822 | 11617 | 10985 | 11488 | 11034 | 12525 | 10793 | 8466 | 7756 | 8577 | 9160 | 8912 | 9805 | 12792 | 10761 | 10932 | 11308 | 10856 | 13448 | 11722 | 9752 | 9041 | 9262 | 9237 | 10003 | 9610 | 10119 | Tmx1 |
| Tmx2 | 6073 | 6280 | 5642 | 5952 | 5564 | 5503 | 6496 | 5733 | 4874 | 4648 | 4309 | 4770 | 4676 | 5272 | 6724 | 6111 | 5621 | 5862 | 5509 | 6237 | 6156 | 5803 | 4882 | 4605 | 4883 | 5101 | 4825 | 5461 | Tmx2 |
| Tmx3 | 8232 | 5360 | 6848 | 7084 | 7409 | 8208 | 8154 | 9495 | 8155 | 6995 | 8234 | 8551 | 8520 | 9186 | 8223 | 7343 | 6292 | 6958 | 6743 | 8155 | 7796 | 8245 | 8251 | 7526 | 7746 | 7241 | 8500 | 8513 | Tmx3 |
| Tmx4 | 13320 | 11882 | 11963 | 15660 | 16167 | 15503 | 17411 | 16352 | 12672 | 13342 | 12081 | 12762 | 11467 | 13679 | 15187 | 11301 | 13410 | 15517 | 15196 | 17231 | 16513 | 17313 | 12524 | 13053 | 15728 | 10461 | 12614 | 12946 | Tmx4 |
| Tnc | 4196 | 5218 | 3001 | 3131 | 3778 | 6122 | 1154 | 6491 | 10745 | 9258 | 14509 | 5201 | 7470 | 5017 | 4135 | 5746 | 3012 | 6377 | 2233 | 1109 | 1923 | 12205 | 14592 | 5936 | 9009 | 4438 | 5781 | 5257 | Tnc |
| Tnf | 2 | 6 | 3 | 1 | 0 | 0 | 0 | 4 | 3 | 3 | 8 | 5 | 6 | 2 | 0 | 2 | 9 | 3 | 1 | 4 | 1 | 2 | 4 | 0 | 5 | 3 | 2 | 7 | Tnf |
| Tnfaip1 | 9254 | 9006 | 8303 | 7630 | 7283 | 7751 | 7953 | 7238 | 7603 | 7240 | 7036 | 8175 | 7472 | 8056 | 8475 | 8256 | 7967 | 7696 | 7770 | 7309 | 7799 | 7152 | 7765 | 7422 | 6953 | 8982 | 8172 | 8454 | Tnfaip1 |
| Tnfaip2 | 758 | 613 | 659 | 434 | 214 | 322 | 188 | 288 | 367 | 342 | 299 | 172 | 177 | 136 | 549 | 881 | 417 | 379 | 244 | 229 | 221 | 120 | 425 | 350 | 105 | 385 | 137 | 178 | Tnfaip2 |
| Tnfaip3 | 505 | 440 | 390 | 449 | 479 | 525 | 483 | 318 | 336 | 338 | 426 | 237 | 286 | 278 | 427 | 435 | 444 | 497 | 440 | 428 | 480 | 314 | 426 | 315 | 294 | 218 | 192 | 211 | Tnfaip3 |
| Tnfaip6 | 1031 | 711 | 890 | 500 | 433 | 576 | 535 | 812 | 696 | 545 | 534 | 701 | 696 | 663 | 982 | 857 | 670 | 506 | 517 | 471 | 504 | 364 | 728 | 704 | 423 | 1020 | 629 | 729 | Tnfaip6 |
| Tnfaip8 | 3511 | 3148 | 3483 | 2385 | 1795 | 2161 | 2247 | 2120 | 2263 | 1839 | 2380 | 1795 | 2443 | 1795 | 3494 | 3826 | 2756 | 2415 | 1999 | 2413 | 1880 | 1581 | 2561 | 1941 | 1487 | 2687 | 2194 | 2255 | Tnfaip8 |
| Tnfaip8l1 | 751 | 890 | 539 | 746 | 628 | 573 | 644 | 418 | 504 | 445 | 472 | 461 | 438 | 435 | 827 | 807 | 649 | 837 | 687 | 718 | 705 | 371 | 517 | 431 | 441 | 429 | 437 | 409 | Tnfaip8l1 |
| Tnfaip8l2 | 13 | 29 | 17 | 31 | 26 | 19 | 18 | 26 | 49 | 35 | 59 | 56 | 32 | 22 | 21 | 51 | 8 | 15 | 31 | 38 | 29 | 30 | 91 | 51 | 17 | 45 | 45 | 53 | Tnfaip8l2 |
| Tnfaip8l3 | 22 | 20 | 37 | 39 | 87 | 35 | 57 | 71 | 47 | 37 | 58 | 27 | 36 | 17 | 41 | 31 | 31 | 43 | 41 | 46 | 46 | 42 | 45 | 22 | 57 | 27 | 30 | 11 | Tnfaip8l3 |
| Tnfrsf10b | 1380 | 1265 | 1252 | 971 | 664 | 794 | 670 | 405 | 497 | 459 | 664 | 333 | 514 | 370 | 1126 | 1558 | 1075 | 1034 | 733 | 681 | 645 | 376 | 553 | 440 | 344 | 554 | 414 | 375 | Tnfrsf10b |
| Tnfrsf11a | 70 | 63 | 74 | 70 | 71 | 115 | 70 | 98 | 100 | 104 | 146 | 100 | 101 | 70 | 76 | 79 | 56 | 107 | 72 | 72 | 65 | 112 | 161 | 116 | 83 | 114 | 79 | 90 | Tnfrsf11a |
| Tnfrsf11b | 113 | 83 | 145 | 94 | 120 | 128 | 88 | 184 | 176 | 140 | 183 | 118 | 171 | 105 | 100 | 92 | 83 | 112 | 106 | 135 | 108 | 169 | 162 | 102 | 105 | 112 | 138 | 145 | Tnfrsf11b |
| Tnfrsf12a | 490 | 644 | 522 | 683 | 389 | 418 | 418 | 100 | 98 | 111 | 190 | 97 | 78 | 92 | 365 | 544 | 527 | 638 | 425 | 449 | 395 | 57 | 131 | 140 | 71 | 107 | 70 | 78 | Tnfrsf12a |
| Tnfrsf13b | 24 | 29 | 54 | 20 | 34 | 35 | 37 | 46 | 66 | 46 | 63 | 54 | 64 | 73 | 50 | 30 | 41 | 17 | 43 | 33 | 22 | 58 | 93 | 64 | 43 | 76 | 67 | 68 | Tnfrsf13b |
| Tnfrsf13c | 66 | 87 | 106 | 66 | 52 | 67 | 65 | 18 | 4 | 16 | 15 | 29 | 6 | 21 | 82 | 65 | 108 | 95 | 76 | 65 | 53 | 15 | 6 | 16 | 27 | 24 | 21 | 23 | Tnfrsf13c |
| Tnfrsf14 | 32 | 48 | 45 | 45 | 24 | 29 | 16 | 56 | 68 | 48 | 62 | 110 | 176 | 119 | 30 | 57 | 28 | 19 | 18 | 22 | 12 | 63 | 69 | 69 | 64 | 72 | 122 | 202 | Tnfrsf14 |
| Tnfrsf17 | 2 | 2 | 1 | 6 | 5 | 7 | 3 | 0 | 3 | 1 | 6 | 2 | 12 | 2 | 5 | 7 | 1 | 2 | 5 | 6 | 2 | 2 | 4 | 2 | 7 | 0 | 0 | 4 | Tnfrsf17 |
| Tnfrsf18 | 2 | 2 | 0 | 0 | 2 | 0 | 2 | 6 | 5 | 6 | 5 | 1 | 1 | 1 | 0 | 0 | 0 | 0 | 1 | 4 | 0 | 10 | 10 | 6 | 9 | 1 | 3 | 0 | Tnfrsf18 |
| Tnfrsf19 | 11404 | 10404 | 10979 | 8918 | 7499 | 10169 | 9343 | 8889 | 9035 | 7794 | 7195 | 11401 | 11928 | 11279 | 10741 | 9805 | 8847 | 9286 | 9277 | 11214 | 8476 | 5491 | 10054 | 9302 | 8410 | 11039 | 11848 | 12397 | Tnfrsf19 |
| Tnfrsf1a | 2141 | 2241 | 2454 | 1890 | 1734 | 1878 | 1503 | 1121 | 1620 | 1417 | 1807 | 1205 | 1556 | 1213 | 1940 | 2700 | 2126 | 1944 | 1781 | 1299 | 1525 | 1235 | 1766 | 1488 | 1014 | 1862 | 1156 | 1275 | Tnfrsf1a |
| Tnfrsf1b | 245 | 337 | 353 | 278 | 265 | 265 | 211 | 184 | 277 | 269 | 301 | 262 | 294 | 275 | 236 | 327 | 199 | 318 | 230 | 188 | 226 | 286 | 364 | 203 | 251 | 294 | 241 | 190 | Tnfrsf1b |
| Tnfrsf21 | 2136 | 2140 | 2433 | 2569 | 3176 | 2494 | 2573 | 2535 | 2598 | 3249 | 2857 | 2566 | 2233 | 2455 | 2080 | 2152 | 2502 | 2719 | 2726 | 2382 | 2544 | 3227 | 2632 | 2870 | 3035 | 2923 | 2200 | 2357 | Tnfrsf21 |
| Tnfrsf22 | 1614 | 1465 | 1329 | 1115 | 965 | 1151 | 982 | 602 | 591 | 636 | 674 | 630 | 738 | 599 | 1273 | 1611 | 1159 | 1041 | 926 | 1001 | 919 | 576 | 629 | 582 | 532 | 781 | 683 | 661 | Tnfrsf22 |

| | | | | | | | | | | | | | | | | | | | | | | | | | | | | | |
|---|---|---|---|---|---|---|---|---|---|---|---|---|---|---|---|---|---|---|---|---|---|---|---|---|---|---|---|---|---|
| Tnfrsf23 | 181 | 181 | 162 | 223 | 270 | 225 | 215 | 89 | 105 | 81 | 131 | 56 | 70 | 70 | 227 | 229 | 168 | 271 | 200 | 232 | 239 | 115 | 131 | 74 | 87 | 51 | 65 | 50 | Tnfrsf23 |
| Tnfrsf25 | 283 | 383 | 536 | 527 | 466 | 382 | 321 | 195 | 176 | 286 | 331 | 174 | 223 | 179 | 231 | 470 | 552 | 600 | 570 | 395 | 386 | 269 | 214 | 279 | 282 | 236 | 198 | 202 | Tnfrsf25 |
| Tnfrsf26 | 125 | 114 | 136 | 125 | 111 | 120 | 126 | 18 | 23 | 25 | 34 | 52 | 53 | 40 | 135 | 144 | 151 | 166 | 118 | 115 | 101 | 55 | 41 | 48 | 23 | 41 | 43 | 58 | Tnfrsf26 |
| Tnfrsf4 | 76 | 91 | 82 | 109 | 99 | 78 | 111 | 74 | 96 | 120 | 121 | 146 | 107 | 143 | 78 | 76 | 88 | 78 | 103 | 70 | 65 | 100 | 72 | 99 | 125 | 126 | 131 | 113 | Tnfrsf4 |
| Tnfrsf8 | 0 | 3 | 9 | 3 | 6 | 6 | 2 | 11 | 17 | 14 | 16 | 11 | 11 | 14 | 4 | 4 | 10 | 11 | 12 | 4 | 4 | 22 | 16 | 10 | 7 | 23 | 7 | 7 | Tnfrsf8 |
| Tnfrsf9 | 9 | 17 | 19 | 24 | 16 | 15 | 17 | 5 | 24 | 19 | 25 | 5 | 15 | 17 | 18 | 21 | 9 | 30 | 23 | 9 | 7 | 10 | 21 | 32 | 18 | 22 | 10 | 17 | Tnfrsf9 |
| Tnfsf10 | 5 | 3 | 7 | 2 | 8 | 5 | 1 | 4 | 7 | 25 | 28 | 10 | 7 | 11 | 3 | 5 | 5 | 6 | 9 | 4 | 7 | 15 | 13 | 10 | 13 | 5 | 1 | 7 | Tnfsf10 |
| Tnfsf11 | 23 | 14 | 29 | 9 | 10 | 17 | 16 | 17 | 28 | 12 | 31 | 15 | 3 | 16 | 33 | 18 | 17 | 20 | 9 | 15 | 9 | 13 | 19 | 37 | 12 | 34 | 27 | 7 | Tnfsf11 |
| Tnfsf12 | 18 | 49 | 44 | 36 | 43 | 30 | 22 | 66 | 85 | 78 | 120 | 52 | 66 | 54 | 40 | 45 | 34 | 39 | 12 | 16 | 27 | 65 | 89 | 83 | 65 | 57 | 56 | 55 | Tnfsf12 |
| Tnfsf13 | 7 | 9 | 6 | 2 | 10 | 0 | 9 | 9 | 17 | 7 | 19 | 4 | 5 | 6 | 5 | 8 | 1 | 8 | 3 | 2 | 8 | 10 | 16 | 11 | 4 | 9 | 8 | 8 | Tnfsf13 |
| Tnfsf13b | 70 | 95 | 55 | 107 | 65 | 101 | 89 | 38 | 33 | 38 | 34 | 35 | 39 | 44 | 79 | 68 | 71 | 96 | 69 | 84 | 66 | 62 | 31 | 19 | 26 | 34 | 43 | 51 | Tnfsf13b |
| Tnfsf14 | 9 | 4 | 4 | 3 | 2 | 2 | 2 | 7 | 10 | 14 | 7 | 10 | 12 | 4 | 5 | 9 | 19 | 7 | 2 | 2 | 7 | 7 | 8 | 5 | 6 | 20 | 14 | 14 | Tnfsf14 |
| Tnfsf15 | 10 | 10 | 3 | 6 | 0 | 0 | 1 | 2 | 7 | 8 | 9 | 9 | 20 | 6 | 1 | 12 | 7 | 4 | 0 | 1 | 2 | 5 | 17 | 5 | 4 | 3 | 8 | 8 | Tnfsf15 |
| Tnfsf18 | 0 | 0 | 0 | 1 | 2 | 0 | 4 | 2 | 10 | 1 | 14 | 1 | 0 | 4 | 0 | 2 | 0 | 4 | 2 | 0 | 1 | 15 | 11 | 9 | 3 | 6 | 5 | 2 | Tnfsf18 |
| Tnfsf4 | 0 | 0 | 1 | 0 | 3 | 0 | 1 | 0 | 2 | 4 | 1 | 0 | 6 | 0 | 3 | 0 | 3 | 0 | 2 | 0 | 1 | 2 | 2 | 0 | 2 | 2 | 0 | 4 | Tnfsf4 |
| Tnfsf8 | 0 | 1 | 0 | 0 | 3 | 3 | 4 | 4 | 2 | 7 | 2 | 0 | 2 | 1 | 0 | 0 | 0 | 3 | 7 | 3 | 4 | 2 | 5 | 7 | 3 | 6 | 1 | 0 | Tnfsf8 |
| Tnfsf9 | 275 | 278 | 322 | 259 | 320 | 273 | 304 | 164 | 138 | 165 | 164 | 215 | 181 | 234 | 240 | 222 | 345 | 253 | 351 | 301 | 341 | 145 | 144 | 174 | 179 | 247 | 199 | 159 | Tnfsf9 |
| Tnfsfm13 | 1 | 2 | 1 | 4 | 0 | 2 | 1 | 4 | 6 | 10 | 2 | 2 | 1 | 9 | 0 | 5 | 0 | 1 | 1 | 2 | 1 | 2 | 3 | 2 | 2 | 1 | 1 | 7 | Tnfsfm13 |
| Tnik | 5174 | 5122 | 4089 | 6543 | 10065 | 9013 | 9342 | 11053 | 10683 | 11903 | 10398 | 8657 | 9801 | 9548 | 6398 | 5811 | 5353 | 7339 | 7877 | 8606 | 9211 | 13094 | 9599 | 9392 | 12557 | 4527 | 8692 | 7630 | Tnik |
| Tnip1 | 633 | 771 | 879 | 793 | 752 | 595 | 571 | 872 | 1172 | 1076 | 1265 | 959 | 1122 | 875 | 683 | 856 | 877 | 863 | 767 | 594 | 572 | 1111 | 1218 | 1111 | 958 | 1108 | 945 | 946 | Tnip1 |
| Tnip2 | 807 | 849 | 873 | 793 | 781 | 794 | 674 | 368 | 504 | 478 | 609 | 396 | 432 | 410 | 749 | 830 | 816 | 815 | 703 | 734 | 701 | 421 | 543 | 475 | 411 | 461 | 476 | 444 | Tnip2 |
| Tnip3 | 2 | 1 | 0 | 1 | 0 | 0 | 0 | 0 | 2 | 1 | 0 | 9 | 1 | 1 | 1 | 0 | 0 | 0 | 0 | 0 | 0 | 0 | 1 | 2 | 6 | 3 | 3 | 2 | Tnip3 |
| Tnk1 | 100 | 46 | 16 | 21 | 11 | 9 | 7 | 74 | 119 | 93 | 92 | 24 | 51 | 15 | 34 | 45 | 15 | 23 | 7 | 7 | 3 | 20 | 113 | 88 | 1 | 114 | 21 | 27 | Tnk1 |
| Tnk2 | 6312 | 8502 | 8937 | 10521 | 9905 | 9106 | 8773 | 4770 | 5900 | 7033 | 6560 | 7005 | 6004 | 6054 | 5978 | 6920 | 9936 | 10789 | 10922 | 7770 | 9089 | 6293 | 5339 | 7258 | 7153 | 7760 | 6513 | 6037 | Tnk2 |
| Tnks | 20891 | 18846 | 13237 | 18222 | 21368 | 21923 | 22550 | 22313 | 20821 | 18978 | 19156 | 17826 | 18548 | 18790 | 20969 | 19355 | 16112 | 18959 | 19214 | 20787 | 23285 | 21031 | 18724 | 15604 | 21003 | 11282 | 18121 | 17255 | Tnks |
| Tnks1bp1 | 7251 | 9417 | 10152 | 10306 | 9576 | 9159 | 8328 | 5145 | 6615 | 6850 | 7379 | 6675 | 6901 | 6641 | 6767 | 8530 | 10445 | 10309 | 10304 | 7672 | 8779 | 6251 | 6325 | 6821 | 6802 | 6207 | 6548 | 6579 | Tnks1bp1 |
| Tnks2 | 39233 | 34741 | 33178 | 35005 | 33900 | 36043 | 35624 | 37742 | 30565 | 28528 | 29913 | 31207 | 30953 | 30334 | 40081 | 36443 | 34068 | 35050 | 34249 | 38202 | 35827 | 32392 | 31713 | 28381 | 30506 | 26033 | 32891 | 30882 | Tnks2 |
| Tnmd | 722 | 650 | 402 | 466 | 36 | 723 | 8 | 613 | 679 | 520 | 662 | 760 | 890 | 918 | 1030 | 648 | 552 | 716 | 35 | 44 | 12 | 628 | 1153 | 710 | 552 | 397 | 880 | 777 | Tnmd |
| Tnn | 115 | 15 | 6 | 187 | 15 | 362 | 2 | 198 | 395 | 461 | 402 | 111 | 102 | 171 | 21 | 19 | 21 | 384 | 41 | 7 | 2 | 531 | 572 | 417 | 607 | 38 | 80 | 52 | Tnn |
| Tnnc1 | 136 | 39 | 51 | 52 | 94 | 53 | 53 | 155 | 442 | 131 | 2682 | 52 | 132 | 48 | 54 | 111 | 68 | 57 | 49 | 38 | 52 | 307 | 1591 | 98 | 68 | 46 | 66 | 37 | Tnnc1 |
| Tnnc2 | 60 | 32 | 14 | 32 | 47 | 19 | 41 | 66 | 126 | 14 | 1399 | 9 | 34 | 4 | 26 | 52 | 41 | 37 | 26 | 26 | 30 | 77 | 783 | 25 | 27 | 12 | 8 | 11 | Tnnc2 |
| Tnni1 | 100 | 17 | 12 | 13 | 33 | 13 | 13 | 122 | 339 | 38 | 2557 | 22 | 75 | 19 | 51 | 95 | 25 | 17 | 5 | 10 | 10 | 202 | 1576 | 51 | 41 | 17 | 17 | 9 | Tnni1 |
| Tnni2 | 10 | 2 | 17 | 5 | 13 | 6 | 8 | 26 | 62 | 16 | 529 | 8 | 9 | 11 | 16 | 14 | 0 | 1 | 4 | 10 | 2 | 18 | 316 | 33 | 9 | 14 | 3 | 20 | Tnni2 |
| Tnni3 | 746 | 1069 | 1350 | 1068 | 547 | 672 | 703 | 138 | 133 | 169 | 136 | 174 | 204 | 234 | 719 | 974 | 1088 | 802 | 615 | 571 | 570 | 150 | 164 | 172 | 151 | 251 | 164 | 233 | Tnni3 |
| Tnni3k | 2 | 2 | 2 | 0 | 2 | 0 | 7 | 1 | 7 | 3 | 3 | 4 | 7 | 2 | 3 | 2 | 0 | 2 | 1 | 14 | 2 | 12 | 10 | 2 | 2 | 3 | 5 | 4 | Tnni3k |
| Tnnt1 | 262 | 187 | 215 | 154 | 128 | 118 | 120 | 333 | 637 | 227 | 3719 | 230 | 380 | 184 | 191 | 310 | 190 | 130 | 114 | 138 | 103 | 448 | 2057 | 247 | 174 | 211 | 200 | 228 | Tnnt1 |
| Tnnt2 | 103 | 28 | 8 | 20 | 32 | 14 | 12 | 126 | 341 | 64 | 2655 | 16 | 107 | 17 | 44 | 84 | 23 | 9 | 16 | 5 | 10 | 196 | 1440 | 56 | 37 | 15 | 27 | 17 | Tnnt2 |
| Tnnt3 | 39 | 34 | 32 | 8 | 38 | 21 | 7 | 83 | 169 | 76 | 1659 | 54 | 75 | 32 | 36 | 43 | 24 | 33 | 19 | 10 | 11 | 125 | 981 | 47 | 53 | 24 | 45 | 42 | Tnnt3 |
| Tnpo1 | 33322 | 27479 | 22368 | 24920 | 24414 | 28122 | 26832 | 38844 | 30838 | 26341 | 28199 | 32505 | 32447 | 35276 | 33367 | 28893 | 22791 | 23112 | 23610 | 30616 | 26930 | 28600 | 30327 | 25884 | 30040 | 25345 | 36839 | 35087 | Tnpo1 |
| Tnpo2 | 10912 | 12731 | 14576 | 16811 | 16520 | 14345 | 16036 | 10911 | 11478 | 13577 | 13641 | 14037 | 13385 | 15224 | 10669 | 10554 | 16890 | 16693 | 17861 | 13110 | 16698 | 13607 | 10930 | 15119 | 15647 | 13718 | 13739 | 15165 | Tnpo2 |
| Tnpo3 | 15052 | 14000 | 12869 | 11374 | 11240 | 12058 | 12987 | 15245 | 13008 | 11685 | 10943 | 13688 | 13027 | 13919 | 15217 | 12763 | 12448 | 11119 | 11871 | 13705 | 12028 | 12641 | 12653 | 12498 | 12825 | 14154 | 14633 | 14356 | Tnpo3 |
| Tnr | 340 | 400 | 155 | 762 | 1385 | 1265 | 1210 | 1054 | 1350 | 1259 | 935 | 686 | 417 | 622 | 438 | 364 | 381 | 877 | 1087 | 1207 | 1375 | 1407 | 1023 | 789 | 1576 | 312 | 506 | 440 | Tnr |
| Tnrc18 | 6409 | 7978 | 9697 | 10453 | 11618 | 9900 | 8390 | 14867 | 19722 | 24224 | 22163 | 25930 | 21045 | 21635 | 5952 | 7137 | 10857 | 10318 | 12124 | 8114 | 9192 | 22688 | 17366 | 27489 | 25463 | 23733 | 20382 | 19490 | Tnrc18 |
| Tnrc6a | 12048 | 11902 | 9426 | 11728 | 13375 | 13435 | 13123 | 17002 | 15788 | 16764 | 17135 | 17302 | 17476 | 17875 | 12228 | 11445 | 11324 | 11375 | 12157 | 12733 | 13739 | 15151 | 14687 | 14153 | 17358 | 11193 | 18439 | 16400 | Tnrc6a |
| Tnrc6b | 7048 | 6499 | 5795 | 7452 | 8283 | 8817 | 7842 | 10418 | 10112 | 9732 | 11086 | 10220 | 10713 | 10465 | 7109 | 6801 | 6486 | 7353 | 7014 | 7925 | 8868 | 9727 | 9281 | 8806 | 11391 | 7278 | 11916 | 9634 | Tnrc6b |
| Tnrc6c | 10293 | 11540 | 11477 | 15976 | 16233 | 15410 | 14477 | 11351 | 12763 | 14819 | 13868 | 14478 | 13277 | 13171 | 9844 | 10963 | 13874 | 15597 | 16100 | 12989 | 15152 | 14840 | 10905 | 14442 | 16730 | 9120 | 13476 | 12104 | Tnrc6c |
| Tns1 | 3319 | 3432 | 3484 | 3015 | 3184 | 3479 | 2152 | 3832 | 5256 | 5164 | 6779 | 4784 | 5784 | 4147 | 2960 | 4159 | 3211 | 3861 | 2844 | 2067 | 2309 | 4871 | 5956 | 5110 | 4362 | 3892 | 4431 | 4440 | Tns1 |
| Tns2 | 612 | 642 | 756 | 741 | 668 | 728 | 450 | 1454 | 2471 | 2754 | 3574 | 1634 | 2082 | 1376 | 525 | 744 | 706 | 731 | 648 | 470 | 466 | 2366 | 2585 | 2441 | 1733 | 2192 | 1653 | 1404 | Tns2 |
| Tns3 | 3617 | 3181 | 3136 | 2414 | 2309 | 2760 | 1787 | 5859 | 7992 | 7236 | 8855 | 5553 | 7841 | 5224 | 3061 | 4184 | 2734 | 2742 | 2144 | 1706 | 1855 | 5748 | 7921 | 6876 | 5474 | 6255 | 6393 | 5525 | Tns3 |
| Tns4 | 77 | 94 | 100 | 10 | 7 | 21 | 11 | 44 | 49 | 29 | 29 | 40 | 51 | 32 | 96 | 115 | 24 | 39 | 5 | 4 | 2 | 12 | 88 | 39 | 19 | 24 | 44 | 57 | Tns4 |
| Tnxb | 24 | 29 | 33 | 66 | 129 | 72 | 40 | 59 | 60 | 80 | 186 | 30 | 62 | 51 | 20 | 37 | 30 | 64 | 52 | 33 | 60 | 95 | 179 | 58 | 97 | 53 | 35 | 32 | Tnxb |
| Tob1 | 2366 | 2231 | 2846 | 3401 | 2603 | 2536 | 3102 | 2093 | 2027 | 2195 | 2206 | 2463 | 2278 | 2936 | 2352 | 2317 | 2712 | 3034 | 2668 | 2222 | 3067 | 2568 | 2226 | 2723 | 2570 | 2598 | 2301 | 2894 | Tob1 |
| Tob2 | 1864 | 2442 | 2967 | 3326 | 2417 | 2332 | 2126 | 1324 | 1542 | 1787 | 1873 | 2026 | 1727 | 1751 | 1742 | 2164 | 3133 | 3110 | 2526 | 1918 | 2237 | 1623 | 1671 | 1997 | 1694 | 2235 | 1775 | 1686 | Tob2 |
| Toe1 | 811 | 966 | 1038 | 928 | 840 | 824 | 797 | 1581 | 1631 | 1325 | 1423 | 1754 | 1515 | 1574 | 833 | 836 | 860 | 800 | 933 | 996 | 781 | 1273 | 1687 | 1623 | 1581 | 1887 | 1635 | 1658 | Toe1 |
| Togaram1 | 6530 | 5905 | 4687 | 5564 | 4858 | 5550 | 5774 | 4742 | 4208 | 3982 | 4229 | 3933 | 4333 | 4163 | 6138 | 6330 | 4765 | 5673 | 4643 | 5420 | 5454 | 3965 | 4098 | 3509 | 3863 | 3125 | 4078 | 4024 | Togaram1 |
| Togaram2 | 17 | 6 | 6 | 13 | 30 | 24 | 16 | 16 | 30 | 45 | 43 | 21 | 42 | 12 | 17 | 9 | 9 | 21 | 19 | 29 | 17 | 25 | 40 | 28 | 27 | 24 | 10 | 7 | Togaram2 |
| Tollip | 6572 | 6759 | 6936 | 7660 | 7386 | 7475 | 7335 | 5994 | 6284 | 6263 | 6023 | 7070 | 6459 | 6993 | 6724 | 6457 | 7116 | 7274 | 7476 | 6340 | 7201 | 6918 | 6012 | 6764 | 7267 | 7114 | 6610 | 7010 | Tollip |
| Tom1 | 648 | 777 | 646 | 682 | 595 | 635 | 613 | 397 | 524 | 369 | 530 | 441 | 505 | 333 | 632 | 890 | 607 | 670 | 606 | 582 | 619 | 471 | 562 | 456 | 372 | 487 | 533 | 401 | Tom1 |
| Tom1l1 | 2253 | 1786 | 1707 | 1482 | 1603 | 1627 | 1421 | 1638 | 1589 | 1343 | 1461 | 1326 | 1496 | 1213 | 2180 | 2176 | 1479 | 1785 | 1387 | 1843 | 1530 | 1181 | 1822 | 1428 | 1072 | 1572 | 1284 | 1445 | Tom1l1 |
| Tom1l2 | 8483 | 10507 | 7989 | 11005 | 11414 | 10435 | 10203 | 4721 | 5498 | 6419 | 6395 | 5130 | 5155 | 4646 | 9026 | 10087 | 9812 | 11946 | 11087 | 8979 | 10408 | 6433 | 5315 | 5222 | 6346 | 4093 | 4664 | 4460 | Tom1l2 |
| Tomm20 | 11911 | 10830 | 10020 | 11889 | 13604 | 13087 | 13245 | 10650 | 9826 | 9350 | 9221 | 9290 | 9103 | 9292 | 12311 | 10355 | 10839 | 12316 | 12805 | 12982 | 13775 | 11010 | 9467 | 8800 | 10359 | 9145 | 9921 | 9147 | Tomm20 |
| Tomm20l | 0 | 0 | 0 | 0 | 0 | 0 | 0 | 0 | 0 | 3 | 0 | 0 | 0 | 1 | 0 | 0 | 0 | 0 | 0 | 0 | 0 | 0 | 0 | 0 | 0 | 0 | 0 | 0 | Tomm20l |
| Tomm22 | 6612 | 6150 | 6555 | 5888 | 5099 | 5141 | 6296 | 6099 | 5141 | 4928 | 5074 | 5643 | 5650 | 6610 | 6531 | 5530 | 6050 | 5335 | 5382 | 6090 | 5959 | 5132 | 5159 | 6037 | 5629 | 6783 | 6251 | 6967 | Tomm22 |
| Tomm34 | 3479 | 3916 | 3472 | 3466 | 2810 | 3002 | 2995 | 2292 | 2494 | 2304 | 2379 | 2371 | 2300 | 2351 | 3231 | 3537 | 3369 | 3301 | 3277 | 3140 | 3011 | 2236 | 2355 | 2571 | 2204 | 2918 | 2231 | 2296 | Tomm34 |
| Tomm40 | 3993 | 4769 | 4967 | 4674 | 4071 | 3847 | 3572 | 2494 | 3218 | 3200 | 3044 | 3829 | 3111 | 3382 | 3624 | 4174 | 4962 | 4402 | 4521 | 3658 | 3577 | 2757 | 2947 | 3435 | 3172 | 4174 | 3402 | 3260 | Tomm40 |
| Tomm40l | 1742 | 1756 | 1563 | 1604 | 1803 | 1899 | 1730 | 2072 | 2374 | 2310 | 2092 | 2397 | 2149 | 2355 | 1770 | 1695 | 1827 | 1642 | 1949 | 1783 | 1827 | 2085 | 2248 | 2383 | 2394 | 2690 | 2400 | 2333 | Tomm40l |
| Tomm5 | 2344 | 2395 | 2266 | 2092 | 1633 | 1571 | 2175 | 1651 | 1723 | 1608 | 1641 | 1562 | 1676 | 2071 | 2236 | 2068 | 2196 | 1766 | 1779 | 2211 | 1961 | 1210 | 1774 | 1650 | 1590 | 2028 | 1902 | 2029 | Tomm5 |
| Tomm6 | 2 | 1 | 6 | 0 | 0 | 2 | 1 | 0 | 1 | 0 | 0 | 1 | 1 | 0 | 1 | 2 | 0 | 7 | 1 | 1 | 4 | 3 | 0 | 0 | 0 | 0 | 0 | 0 | Tomm6 |
| Tomm7 | 2487 | 2634 | 3461 | 3194 | 2392 | 1740 | 3558 | 2720 | 2351 | 3647 | 2832 | 3049 | 2673 | 5403 | 2308 | 1535 | 3384 | 2317 | 2339 | 2786 | 3660 | 1841 | 2175 | 4220 | 3054 | 3422 | 3076 | 5187 | Tomm7 |
| Tomm70a | 19286 | 15637 | 16427 | 16093 | 15715 | 16443 | 18051 | 16714 | 13719 | 13316 | 12464 | 14613 | 13306 | 14694 | 20408 | 17618 | 15959 | 15650 | 15082 | 17883 | 16383 | 14629 | 13499 | 13068 | 13799 | 14468 | 14377 | 14827 | Tomm70a |
| Tomt | 6 | 5 | 5 | 5 | 2 | 2 | 6 | 5 | 7 | 3 | 9 | 15 | 3 | 12 | 5 | 5 | 5 | 0 | 9 | 1 | 8 | 13 | 4 | 18 | 2 | 9 | 5 | 1 | Tomt |
| Tonsl | 1535 | 1839 | 1958 | 1752 | 1468 | 1595 | 1509 | 1271 | 1713 | 1796 | 1647 | 2079 | 1914 | 1818 | 1349 | 1823 | 1859 | 1744 | 1739 | 1262 | 1415 | 1469 | 1430 | 1959 | 1611 | 2225 | 1986 | 1895 | Tonsl |
| Top1 | 25929 | 21669 | 28556 | 26924 | 25954 | 26339 | 30841 | 33321 | 26645 | 31331 | 28093 | 34407 | 29899 | 37325 | 27459 | 22039 | 27634 | 26187 | 27837 | 27885 | 28784 | 29352 | 25572 | 35876 | 34313 | 37264 | 33941 | 36997 | Top1 |
| Top1mt | 504 | 680 | 737 | 754 | 629 | 741 | 606 | 518 | 676 | 555 | 680 | 479 | 539 | 548 | 625 | 680 | 673 | 632 | 649 | 760 | 574 | 523 | 627 | 552 | 603 | 538 | 514 | 444 | Top1mt |
| Top2a | 63149 | 48445 | 45475 | 34894 | 26722 | 34827 | 38471 | 55879 | 47098 | 38040 | 35015 | 60045 | 49227 | 61354 | 54974 | 49456 | 40972 | 31577 | 34601 | 44803 | 35371 | 30694 | 39849 | 47121 | 43002 | 57102 | 65921 | 62949 | Top2a |
| Top2b | 47205 | 35364 | 37919 | 44189 | 48071 | 51533 | 58108 | 63093 | 50692 | 52369 | 47824 | 60148 | 51781 | 63151 | 48185 | 38096 | 42960 | 44076 | 47968 | 48106 | 54517 | 59277 | 47739 | 54141 | 60813 | 48802 | 58676 | 58256 | Top2b |
| Top3a | 3245 | 3862 | 2940 | 3037 | 2381 | 2677 | 2752 | 1677 | 1807 | 1683 | 1650 | 1852 | 1833 | 1732 | 2931 | 3221 | 2971 | 2896 | 2822 | 2799 | 2878 | 1484 | 1718 | 1740 | 1700 | 1797 | 2073 | 1825 | Top3a |
| Top3b | 3058 | 3072 | 2595 | 2723 | 3004 | 3012 | 2839 | 4279 | 4627 | 4094 | 4256 | 4413 | 4530 | 4148 | 2931 | 2918 | 2881 | 2709 | 2947 | 2996 | 2996 | 4346 | 4612 | 4332 | 4320 | 4156 | 4379 | 4307 | Top3b |
| Topaz1 | 0 | 0 | 0 | 0 | 0 | 2 | 0 | 1 | 5 | 0 | 0 | 0 | 0 | 2 | 0 | 0 | 6 | 0 | 0 | 0 | 0 | 0 | 0 | 2 | 1 | 0 | 0 | 0 | Topaz1 |

| | | | | | | | | | | | | | | | | | | | | | | | | | | | | | |
|---|---|---|---|---|---|---|---|---|---|---|---|---|---|---|---|---|---|---|---|---|---|---|---|---|---|---|---|---|---|
| Topbp1 | 19162 | 16951 | 14266 | 14670 | 12366 | 14238 | 16794 | 13514 | 13047 | 11221 | 11191 | 16483 | 13464 | 16303 | 17567 | 14657 | 14559 | 13921 | 14314 | 16239 | 14875 | 9399 | 11404 | 13018 | 12157 | 15176 | 16621 | 16376 | Topbp1 |
| Topors | 4046 | 3394 | 3970 | 3872 | 3808 | 3809 | 3806 | 5824 | 4848 | 4477 | 4917 | 5586 | 5201 | 5560 | 4179 | 3757 | 3639 | 3561 | 3619 | 3909 | 3417 | 4943 | 5009 | 4964 | 4572 | 5744 | 5564 | 5230 | Topors |
| Toporsl | 1 | 0 | 4 | 0 | 0 | 1 | 0 | 0 | 1 | 0 | 1 | 0 | 0 | 0 | 1 | 2 | 0 | 0 | 2 | 0 | 0 | 0 | 1 | 0 | 0 | 0 | 0 | 0 | Toporsl |
| Tor1a | 2201 | 2704 | 2036 | 2262 | 2077 | 2254 | 2120 | 1845 | 1842 | 1783 | 1760 | 2007 | 1737 | 1901 | 2119 | 2334 | 2287 | 2355 | 2293 | 2422 | 2216 | 1541 | 1996 | 1858 | 1931 | 2104 | 2092 | 2076 | Tor1a |
| Tor1aip1 | 1578 | 1305 | 1659 | 1611 | 1486 | 1553 | 1707 | 3027 | 3211 | 3574 | 3461 | 3895 | 3731 | 3844 | 1428 | 1367 | 1810 | 1589 | 1704 | 1525 | 1717 | 3158 | 3247 | 3751 | 3332 | 4270 | 3706 | 3980 | Tor1aip1 |
| Tor1aip2 | 14922 | 13394 | 12677 | 11750 | 9831 | 10629 | 11145 | 10916 | 9950 | 8757 | 9665 | 10041 | 10292 | 10420 | 14817 | 14122 | 11507 | 11488 | 10269 | 11736 | 10799 | 9068 | 10261 | 9641 | 8504 | 10788 | 10442 | 10759 | Tor1aip2 |
| Tor1b | 4201 | 4441 | 3905 | 4050 | 3499 | 3865 | 3927 | 3628 | 3722 | 3556 | 3953 | 3845 | 3654 | 4091 | 3973 | 4248 | 3909 | 4087 | 3855 | 4027 | 3799 | 3415 | 3671 | 3973 | 3831 | 3949 | 3949 | 3887 | Tor1b |
| Tor2a | 1107 | 1363 | 1414 | 1172 | 1178 | 1176 | 1136 | 1730 | 1975 | 1768 | 1754 | 1741 | 1914 | 1719 | 1233 | 1127 | 1446 | 1087 | 1330 | 1362 | 1180 | 1699 | 1941 | 1872 | 1637 | 1976 | 1870 | 1915 | Tor2a |
| Tor3a | 626 | 710 | 729 | 579 | 623 | 610 | 532 | 272 | 332 | 315 | 274 | 327 | 363 | 283 | 651 | 805 | 674 | 581 | 554 | 668 | 550 | 221 | 397 | 308 | 248 | 341 | 352 | 288 | Tor3a |
| Tor4a | 182 | 173 | 185 | 144 | 128 | 179 | 108 | 178 | 197 | 150 | 227 | 118 | 176 | 132 | 124 | 237 | 127 | 171 | 125 | 134 | 99 | 159 | 298 | 159 | 115 | 176 | 157 | 115 | Tor4a |
| Tox | 916 | 870 | 1201 | 1180 | 1322 | 1293 | 1153 | 2453 | 2407 | 2970 | 2601 | 3486 | 3312 | 3555 | 1124 | 897 | 1177 | 1001 | 1307 | 1040 | 1300 | 3517 | 2444 | 3201 | 3310 | 3060 | 3079 | 3550 | Tox |
| Tox2 | 2532 | 3080 | 3742 | 3528 | 3121 | 2940 | 3688 | 1356 | 1301 | 1611 | 1262 | 1996 | 1522 | 2224 | 2632 | 2421 | 4025 | 3157 | 3567 | 2682 | 3808 | 2461 | 1188 | 1861 | 2078 | 2094 | 2038 | 2200 | Tox2 |
| Tox3 | 2668 | 2692 | 4326 | 3898 | 4191 | 3567 | 3890 | 5063 | 4013 | 5044 | 4223 | 6939 | 5585 | 7015 | 3008 | 2186 | 4442 | 3407 | 4781 | 3285 | 4016 | 6119 | 3874 | 6284 | 6210 | 6185 | 6373 | 6942 | Tox3 |
| Tox4 | 11818 | 12082 | 10605 | 11383 | 9790 | 10079 | 10977 | 8308 | 7829 | 7515 | 7441 | 8014 | 7307 | 8243 | 11661 | 10460 | 10931 | 11146 | 10785 | 10499 | 11228 | 7688 | 7607 | 7897 | 8090 | 8496 | 7818 | 8240 | Tox4 |
| Tpbg | 2228 | 2307 | 2931 | 2218 | 2072 | 2063 | 2233 | 2052 | 1830 | 1878 | 1866 | 2408 | 2532 | 2456 | 2296 | 2192 | 2373 | 2151 | 2057 | 2139 | 2068 | 1768 | 2483 | 2269 | 1737 | 2554 | 2662 | 2752 | Tpbg |
| Tpbgl | 104 | 94 | 132 | 73 | 164 | 90 | 96 | 89 | 99 | 85 | 108 | 56 | 85 | 36 | 120 | 113 | 111 | 121 | 108 | 54 | 97 | 147 | 111 | 69 | 84 | 26 | 43 | 44 | Tpbgl |
| Tpbpa | 0 | 0 | 0 | 0 | 0 | 0 | 1 | 0 | 85 | 0 | 0 | 0 | 0 | 0 | 0 | 0 | 0 | 0 | 0 | 0 | 0 | 0 | 28 | 0 | 0 | 0 | 0 | 5 | Tpbpa |
| Tpbpb | 0 | 1 | 0 | 0 | 0 | 0 | 0 | 0 | 30 | 0 | 0 | 0 | 0 | 0 | 0 | 0 | 0 | 0 | 0 | 0 | 0 | 0 | 9 | 0 | 0 | 0 | 0 | 0 | Tpbpb |
| Tpcn1 | 4331 | 5264 | 6293 | 6267 | 5239 | 5460 | 4803 | 2218 | 3044 | 3793 | 4093 | 3389 | 3474 | 2980 | 4102 | 5508 | 5941 | 6696 | 5601 | 4084 | 5131 | 3275 | 3410 | 3605 | 3419 | 3368 | 2796 | 2855 | Tpcn1 |
| Tpcn2 | 1068 | 1091 | 1252 | 1005 | 903 | 955 | 725 | 827 | 1135 | 1171 | 1264 | 1149 | 1271 | 954 | 943 | 1160 | 1009 | 913 | 808 | 811 | 645 | 954 | 1166 | 1014 | 867 | 1021 | 1123 | 939 | Tpcn2 |
| Tpd52 | 2699 | 2062 | 2044 | 2224 | 2465 | 1993 | 2016 | 2193 | 2005 | 2080 | 2107 | 1421 | 1918 | 1400 | 2925 | 2776 | 1989 | 2335 | 1880 | 2158 | 1991 | 1875 | 2310 | 1907 | 1543 | 2491 | 1362 | 1402 | Tpd52 |
| Tpd52l1 | 776 | 799 | 866 | 445 | 526 | 502 | 415 | 553 | 666 | 566 | 665 | 438 | 712 | 451 | 740 | 975 | 473 | 599 | 351 | 523 | 445 | 545 | 718 | 449 | 480 | 524 | 497 | 501 | Tpd52l1 |
| Tpd52l2 | 6905 | 6604 | 6761 | 5766 | 5335 | 5698 | 5896 | 6118 | 5912 | 5315 | 5519 | 5724 | 5646 | 5973 | 6803 | 6372 | 6139 | 5617 | 5845 | 6450 | 5603 | 5167 | 6058 | 5749 | 5243 | 6663 | 5949 | 6389 | Tpd52l2 |
| Tpgs1 | 328 | 406 | 447 | 337 | 432 | 402 | 318 | 895 | 1007 | 993 | 873 | 910 | 948 | 773 | 323 | 276 | 465 | 313 | 466 | 417 | 309 | 804 | 945 | 891 | 916 | 1033 | 1052 | 799 | Tpgs1 |
| Tpgs2 | 6124 | 6254 | 4917 | 5971 | 6207 | 5782 | 7105 | 5321 | 5021 | 4696 | 4215 | 5003 | 4138 | 4975 | 6782 | 5787 | 5762 | 5834 | 6331 | 6755 | 6889 | 5389 | 4592 | 4762 | 4980 | 4706 | 4752 | 4638 | Tpgs2 |
| Tph1 | 43 | 26 | 47 | 16 | 18 | 28 | 17 | 31 | 33 | 52 | 17 | 24 | 26 | 49 | 40 | 39 | 29 | 36 | 5 | 20 | 18 | 25 | 22 | 25 | 33 | 47 | 37 | 24 | Tph1 |
| Tph2 | 69 | 142 | 66 | 128 | 217 | 129 | 75 | 208 | 147 | 113 | 112 | 76 | 211 | 43 | 189 | 213 | 161 | 151 | 121 | 169 | 115 | 170 | 204 | 51 | 175 | 16 | 70 | 25 | Tph2 |
| Tpi1 | 25853 | 25908 | 23676 | 21174 | 16872 | 20180 | 18182 | 18389 | 22155 | 18930 | 19659 | 19300 | 19878 | 19050 | 23952 | 23701 | 22164 | 19882 | 21698 | 22930 | 18964 | 15025 | 23528 | 20287 | 17866 | 25616 | 21065 | 22198 | Tpi1 |
| Tpk1 | 276 | 212 | 199 | 143 | 168 | 237 | 145 | 465 | 469 | 370 | 454 | 374 | 475 | 411 | 319 | 283 | 148 | 134 | 129 | 242 | 131 | 406 | 545 | 384 | 331 | 394 | 399 | 401 | Tpk1 |
| Tpm1 | 21206 | 18396 | 18227 | 12105 | 8768 | 13323 | 7768 | 16411 | 19373 | 17155 | 22947 | 16611 | 21363 | 17173 | 20151 | 23755 | 13748 | 14716 | 7557 | 7492 | 7775 | 13951 | 23361 | 18059 | 12862 | 19422 | 18784 | 18488 | Tpm1 |
| Tpm2 | 2621 | 2602 | 2874 | 1641 | 932 | 1654 | 705 | 1780 | 2663 | 1844 | 7571 | 2085 | 2935 | 2011 | 2455 | 3327 | 2260 | 1872 | 873 | 780 | 806 | 1671 | 5075 | 2270 | 1424 | 2394 | 2402 | 2265 | Tpm2 |
| Tpm3 | 23670 | 24221 | 28309 | 25275 | 22754 | 23800 | 26060 | 16882 | 18786 | 18873 | 20190 | 22523 | 22837 | 24074 | 23489 | 23701 | 26578 | 24406 | 24723 | 23118 | 25024 | 16962 | 18938 | 22401 | 21453 | 25921 | 22609 | 25103 | Tpm3 |
| Tpm3-rs7 | 2871 | 2363 | 3159 | 2293 | 1962 | 1881 | 2477 | 2042 | 1761 | 1819 | 1806 | 2030 | 1896 | 2621 | 2751 | 1991 | 2599 | 2158 | 2117 | 1968 | 2683 | 1831 | 1717 | 2093 | 1931 | 2467 | 2076 | 2680 | Tpm3-rs7 |
| Tpm4 | 24218 | 18572 | 20381 | 13122 | 12439 | 15401 | 14039 | 25976 | 24219 | 20750 | 23584 | 25999 | 27632 | 27026 | 22322 | 21295 | 16136 | 13554 | 12880 | 15055 | 12766 | 19629 | 26404 | 23719 | 20093 | 31005 | 28544 | 29146 | Tpm4 |
| Tpmt | 2955 | 2792 | 2793 | 2556 | 2366 | 2533 | 2723 | 1715 | 1334 | 1119 | 1205 | 1546 | 1615 | 1766 | 3131 | 2708 | 2692 | 2544 | 2354 | 2741 | 2590 | 1455 | 1269 | 1252 | 1261 | 1568 | 1819 | 1810 | Tpmt |
| Tpo | 10 | 0 | 0 | 13 | 11 | 14 | 9 | 12 | 2 | 8 | 11 | 2 | 5 | 6 | 3 | 6 | 2 | 1 | 5 | 4 | 4 | 7 | 8 | 7 | 7 | 3 | 3 | 0 | Tpo |
| Tpp1 | 4525 | 5196 | 4221 | 3893 | 3845 | 3715 | 4019 | 3717 | 3692 | 3385 | 3491 | 3736 | 3796 | 3608 | 4857 | 4766 | 4259 | 4536 | 3715 | 4037 | 3688 | 3969 | 3994 | 3419 | 3314 | 3789 | 3725 | 3726 | Tpp1 |
| Tpp2 | 16153 | 13411 | 12047 | 11839 | 11377 | 12174 | 13609 | 10417 | 8912 | 7690 | 8242 | 8727 | 8732 | 9089 | 16163 | 14492 | 11189 | 12042 | 10872 | 13448 | 12410 | 8462 | 8895 | 7969 | 7888 | 8131 | 9173 | 9036 | Tpp2 |
| Tppp | 1921 | 2146 | 1737 | 1951 | 2041 | 2006 | 2167 | 1291 | 1052 | 1079 | 976 | 762 | 856 | 867 | 2011 | 1801 | 1896 | 1973 | 2035 | 2344 | 2058 | 1308 | 979 | 857 | 1000 | 752 | 787 | 760 | Tppp |
| Tppp2 | 0 | 0 | 0 | 0 | 0 | 0 | 0 | 0 | 0 | 0 | 0 | 0 | 1 | 0 | 1 | 0 | 2 | 6 | 0 | 0 | 0 | 0 | 0 | 0 | 0 | 0 | 0 | 0 | Tppp2 |
| Tppp3 | 1263 | 2389 | 2091 | 1848 | 2147 | 1656 | 1253 | 1545 | 1729 | 1614 | 1981 | 1205 | 2072 | 1096 | 1834 | 2245 | 2109 | 1729 | 1718 | 1853 | 1366 | 1689 | 1946 | 1022 | 1367 | 1117 | 1482 | 1372 | Tppp3 |
| Tpr | 33933 | 28042 | 25081 | 28599 | 27330 | 31881 | 30574 | 25382 | 23731 | 23056 | 22938 | 25684 | 25181 | 25366 | 33366 | 31338 | 26795 | 27870 | 26682 | 29771 | 29799 | 21681 | 23397 | 21334 | 22819 | 18839 | 26990 | 24011 | Tpr |
| Tpra1 | 1716 | 2319 | 2234 | 2238 | 2130 | 1924 | 1914 | 1396 | 1646 | 1542 | 1705 | 1478 | 1539 | 1361 | 1689 | 2169 | 2384 | 2168 | 2326 | 1910 | 1944 | 1532 | 1588 | 1655 | 1421 | 1544 | 1478 | 1373 | Tpra1 |
| Tprg | 3 | 1 | 0 | 0 | 3 | 0 | 3 | 6 | 2 | 0 | 1 | 2 | 0 | 1 | 0 | 0 | 0 | 4 | 0 | 0 | 0 | 3 | 4 | 0 | 0 | 2 | 0 | 0 | Tprg |
| Tprgl | 5199 | 5869 | 5122 | 4939 | 4879 | 4789 | 4788 | 4554 | 4420 | 4546 | 4232 | 4245 | 4406 | 4284 | 5156 | 5189 | 5224 | 4966 | 4956 | 4769 | 4975 | 4428 | 4888 | 4403 | 4326 | 4315 | 4099 | 4427 | Tprgl |
| Tprkb | 1684 | 1396 | 1687 | 1674 | 2014 | 1838 | 1947 | 2242 | 2122 | 2028 | 2135 | 2094 | 2167 | 2414 | 1694 | 1445 | 1780 | 1665 | 1838 | 1861 | 1763 | 2216 | 2259 | 2176 | 2303 | 2332 | 2196 | 2410 | Tprkb |
| Tprn | 729 | 1025 | 848 | 1134 | 958 | 925 | 856 | 460 | 488 | 590 | 577 | 475 | 484 | 440 | 815 | 871 | 1012 | 1147 | 959 | 715 | 898 | 637 | 551 | 564 | 546 | 517 | 436 | 475 | Tprn |
| Tpsab1 | 0 | 0 | 0 | 0 | 0 | 0 | 0 | 0 | 0 | 0 | 0 | 0 | 0 | 1 | 0 | 0 | 0 | 0 | 0 | 0 | 0 | 0 | 0 | 1 | 0 | 0 | 0 | 0 | Tpsab1 |
| Tpsb2 | 0 | 0 | 0 | 0 | 0 | 1 | 0 | 0 | 7 | 10 | 11 | 0 | 0 | 0 | 0 | 0 | 0 | 0 | 0 | 0 | 0 | 10 | 18 | 2 | 2 | 0 | 2 | 0 | Tpsb2 |
| Tpsg1 | 0 | 0 | 0 | 0 | 0 | 0 | 0 | 0 | 6 | 1 | 12 | 0 | 0 | 0 | 0 | 0 | 0 | 2 | 0 | 0 | 0 | 2 | 6 | 1 | 0 | 0 | 0 | 0 | Tpsg1 |
| Tpst1 | 1738 | 1769 | 1747 | 1524 | 1713 | 1886 | 1571 | 2052 | 2411 | 2085 | 2259 | 2128 | 2225 | 2146 | 1693 | 1809 | 1775 | 1593 | 1684 | 1655 | 1577 | 1863 | 2508 | 2134 | 1998 | 2479 | 2265 | 2305 | Tpst1 |
| Tpst2 | 1433 | 2035 | 1713 | 1952 | 1537 | 1378 | 1322 | 634 | 813 | 730 | 711 | 819 | 765 | 701 | 1387 | 1778 | 1795 | 1938 | 1461 | 1403 | 1372 | 705 | 822 | 831 | 742 | 892 | 794 | 779 | Tpst2 |
| Tpt1 | 93068 | 90624 | 88726 | 90075 | 79583 | 79377 | 100321 | 92713 | 85198 | 89018 | 89226 | 82755 | 86948 | 104236 | 96985 | 81988 | 89227 | 81290 | 89870 | 95367 | 98909 | 75185 | 94057 | 93434 | 90489 | 103045 | 92969 | 108034 | Tpt1 |
| Tpte | 0 | 1 | 0 | 1 | 0 | 0 | 2 | 0 | 0 | 1 | 0 | 0 | 0 | 0 | 4 | 0 | 3 | 1 | 0 | 1 | 0 | 2 | 3 | 3 | 0 | 0 | 3 | 0 | Tpte |
| Tpx2 | 15554 | 13559 | 14760 | 10832 | 7888 | 9274 | 11184 | 13501 | 11801 | 10969 | 9985 | 16582 | 13464 | 17183 | 13647 | 12146 | 12516 | 9610 | 10320 | 11106 | 9436 | 8798 | 10409 | 14149 | 12020 | 19730 | 17646 | 17162 | Tpx2 |
| Tra2a | 28410 | 26907 | 31051 | 31681 | 27042 | 28204 | 31848 | 30440 | 24054 | 25149 | 26924 | 26091 | 26439 | 30323 | 27933 | 27485 | 30145 | 30793 | 28708 | 32380 | 31728 | 23190 | 22719 | 26522 | 28040 | 28140 | 28624 | 28321 | Tra2a |
| Tra2b | 19796 | 16959 | 21758 | 19897 | 17416 | 18056 | 20995 | 24651 | 20256 | 20071 | 20669 | 24643 | 24299 | 26039 | 20611 | 18718 | 19904 | 18671 | 19716 | 22634 | 18797 | 20187 | 20042 | 24228 | 23015 | 30658 | 26858 | 29309 | Tra2b |
| Trabd | 3601 | 4649 | 4334 | 4179 | 3390 | 3716 | 3200 | 2320 | 2656 | 2591 | 2744 | 3318 | 2884 | 2725 | 3290 | 4131 | 4439 | 3889 | 3875 | 3401 | 3319 | 2485 | 2686 | 3033 | 2667 | 3477 | 3006 | 2948 | Trabd |
| Trabd2b | 460 | 534 | 733 | 400 | 331 | 440 | 197 | 699 | 873 | 811 | 1115 | 825 | 1204 | 822 | 549 | 671 | 502 | 519 | 287 | 338 | 229 | 1048 | 997 | 784 | 855 | 682 | 952 | 957 | Trabd2b |
| Tradd | 637 | 727 | 719 | 628 | 574 | 532 | 460 | 228 | 299 | 317 | 313 | 257 | 239 | 237 | 554 | 634 | 683 | 757 | 534 | 543 | 473 | 287 | 373 | 300 | 274 | 315 | 265 | 275 | Tradd |
| Traf1 | 37 | 54 | 50 | 44 | 33 | 49 | 15 | 79 | 102 | 85 | 138 | 72 | 109 | 49 | 58 | 79 | 42 | 55 | 26 | 12 | 23 | 149 | 151 | 69 | 77 | 33 | 77 | 62 | Traf1 |
| Traf2 | 1678 | 2264 | 1993 | 2137 | 1906 | 2042 | 2064 | 1743 | 1833 | 1745 | 1635 | 2039 | 1808 | 1950 | 1750 | 1836 | 2160 | 2239 | 2208 | 2040 | 1975 | 1417 | 1871 | 1969 | 1953 | 2181 | 2170 | 1989 | Traf2 |
| Traf3 | 7388 | 8162 | 7457 | 8543 | 10059 | 9195 | 8943 | 5326 | 5732 | 6367 | 5605 | 6865 | 6175 | 6509 | 7457 | 7896 | 8936 | 9167 | 9520 | 8463 | 9112 | 6721 | 5229 | 6240 | 6583 | 4571 | 6259 | 6284 | Traf3 |
| Traf3ip1 | 2317 | 2636 | 3082 | 3129 | 2716 | 2620 | 2730 | 1652 | 1692 | 2078 | 2091 | 2238 | 2142 | 2423 | 2116 | 2543 | 3132 | 2824 | 2718 | 2379 | 2504 | 2050 | 1586 | 2394 | 2350 | 2425 | 2231 | 2338 | Traf3ip1 |
| Traf3ip2 | 99 | 65 | 63 | 29 | 48 | 58 | 30 | 116 | 188 | 194 | 162 | 87 | 108 | 53 | 57 | 77 | 43 | 57 | 26 | 30 | 25 | 77 | 192 | 137 | 54 | 172 | 56 | 61 | Traf3ip2 |
| Traf3ip3 | 17 | 10 | 27 | 9 | 18 | 8 | 3 | 81 | 107 | 69 | 260 | 63 | 74 | 70 | 18 | 28 | 12 | 22 | 14 | 36 | 15 | 84 | 217 | 59 | 59 | 101 | 64 | 61 | Traf3ip3 |
| Traf4 | 4092 | 5586 | 4717 | 4893 | 5148 | 4682 | 4259 | 6447 | 6945 | 7032 | 6462 | 7688 | 7514 | 7177 | 4050 | 4549 | 5159 | 4789 | 5511 | 4769 | 4553 | 6933 | 6771 | 7071 | 7512 | 8066 | 7903 | 7242 | Traf4 |
| Traf5 | 20 | 7 | 11 | 3 | 3 | 4 | 34 | 9 | 7 | 10 | 6 | 1 | 7 | 7 | 11 | 14 | 37 | 8 | 27 | 7 | 4 | 27 | 5 | 5 | 2 | 7 | 3 | 32 | Traf5 |
| Traf6 | 7198 | 6482 | 5019 | 5283 | 4556 | 5260 | 5338 | 4280 | 3741 | 3428 | 3241 | 4078 | 3839 | 4158 | 6957 | 5840 | 5176 | 4961 | 4924 | 5740 | 5385 | 3200 | 3514 | 3330 | 3503 | 3411 | 4327 | 4189 | Traf6 |
| Traf7 | 10378 | 12639 | 11330 | 11489 | 9369 | 10381 | 9775 | 6332 | 7521 | 7787 | 7220 | 8634 | 7765 | 7771 | 9705 | 11052 | 11170 | 11225 | 11145 | 9373 | 9847 | 7150 | 7174 | 8246 | 7787 | 8996 | 8094 | 8070 | Traf7 |
| Trafd1 | 6667 | 8304 | 7884 | 8869 | 7949 | 8129 | 7698 | 5823 | 5944 | 6666 | 5964 | 6550 | 6049 | 6324 | 6783 | 7417 | 8492 | 8886 | 8436 | 7820 | 7947 | 6739 | 5781 | 6720 | 7107 | 6167 | 6207 | 6221 | Trafd1 |
| Traip | 2575 | 2737 | 2241 | 1950 | 1470 | 1740 | 1830 | 1551 | 1523 | 1387 | 1329 | 1773 | 1627 | 1845 | 2207 | 2359 | 1975 | 1821 | 1781 | 1963 | 1762 | 1128 | 1343 | 1669 | 1489 | 2292 | 1979 | 1915 | Traip |
| Trak1 | 6382 | 8062 | 9540 | 10894 | 11030 | 9700 | 9955 | 5376 | 6548 | 8338 | 7325 | 8243 | 6507 | 7250 | 6417 | 6996 | 10434 | 11194 | 11512 | 8071 | 10900 | 7897 | 6143 | 8165 | 9260 | 7335 | 6703 | 7040 | Trak1 |
| Trak2 | 6388 | 6727 | 5736 | 7288 | 7723 | 7254 | 7618 | 5950 | 6015 | 5996 | 5933 | 5536 | 5703 | 5743 | 6682 | 6877 | 6468 | 7783 | 7126 | 7350 | 7772 | 6527 | 5983 | 5499 | 6866 | 4207 | 5223 | 5497 | Trak2 |
| Tram1 | 4831 | 3487 | 4365 | 3389 | 3446 | 3686 | 3688 | 3964 | 3894 | 3467 | 4425 | 3839 | 4222 | 3789 | 4589 | 4379 | 4005 | 3716 | 3451 | 3498 | 3344 | 4007 | 4404 | 3863 | 3300 | 4643 | 3855 | 4123 | Tram1 |
| Tram1l1 | 3189 | 3325 | 3120 | 3774 | 3863 | 3638 | 3730 | 3605 | 2714 | 2882 | 2501 | 2750 | 2687 | 2995 | 3807 | 2805 | 3539 | 3730 | 3940 | 4084 | 3932 | 3554 | 2864 | 2911 | 3085 | 2573 | 2891 | 2887 | Tram1l1 |
| Tram2 | 3272 | 3516 | 4096 | 3283 | 2471 | 2868 | 2335 | 1432 | 1728 | 1794 | 2206 | 1702 | 2123 | 1523 | 3054 | 4115 | 3442 | 3507 | 2574 | 2127 | 2393 | 1734 | 1849 | 1714 | 1460 | 1774 | 1727 | 1701 | Tram2 |
| Trank1 | 1962 | 1922 | 828 | 2770 | 4858 | 4264 | 4691 | 3149 | 3335 | 3190 | 2861 | 1928 | 1014 | 2068 | 1996 | 1516 | 1767 | 3390 | 4018 | 4889 | 5511 | 3948 | 3415 | 2333 | 4396 | 1051 | 1610 | 1307 | Trank1 |
| Trap1 | 5486 | 5771 | 4921 | 4541 | 3804 | 4449 | 4392 | 4588 | 5033 | 4587 | 4756 | 4914 | 4961 | 5006 | 4836 | 5374 | 4657 | 4340 | 4354 | 4467 | 4239 | 3837 | 5022 | 4616 | 4392 | 5373 | 4886 | 5173 | Trap1 |

| | | | | | | | | | | | | | | | | | | | | | | | | | | | | | |
|---|---|---|---|---|---|---|---|---|---|---|---|---|---|---|---|---|---|---|---|---|---|---|---|---|---|---|---|---|---|
| Trap1a | 2 | 1 | 0 | 0 | 1 | 0 | 1 | 1 | 5 | 4 | 0 | 2 | 1 | 0 | 0 | 7 | 10 | 0 | 4 | 3 | 1 | 0 | 3 | 1 | 9 | 3 | 2 | 0 | Trap1a |
| Trappc1 | 6950 | 6669 | 6082 | 6223 | 5105 | 5166 | 6615 | 5168 | 4294 | 3892 | 3949 | 4290 | 4588 | 5077 | 7636 | 5949 | 6164 | 5930 | 5777 | 7151 | 6270 | 4478 | 4181 | 4463 | 4139 | 5048 | 5229 | 5211 | Trappc1 |
| Trappc10 | 6580 | 7524 | 5301 | 7483 | 7445 | 7653 | 7487 | 5688 | 5857 | 6040 | 5885 | 6133 | 5574 | 5566 | 6393 | 6200 | 6431 | 7578 | 7381 | 6751 | 7994 | 6253 | 5364 | 5700 | 6795 | 4628 | 5871 | 5676 | Trappc10 |
| Trappc11 | 10135 | 11204 | 8182 | 9335 | 8206 | 8478 | 9250 | 6878 | 6635 | 5327 | 5856 | 6093 | 6136 | 6179 | 10437 | 10101 | 8713 | 9183 | 8170 | 9806 | 9187 | 6142 | 6417 | 5687 | 5941 | 5778 | 6202 | 6165 | Trappc11 |
| Trappc12 | 2742 | 3117 | 2784 | 2742 | 2675 | 2627 | 2742 | 3351 | 3896 | 3469 | 3385 | 3775 | 3432 | 3438 | 2680 | 2739 | 2799 | 2671 | 2895 | 2757 | 2789 | 3135 | 3829 | 3253 | 3590 | 3611 | 3550 | 3366 | Trappc12 |
| Trappc13 | 4411 | 4074 | 4646 | 4702 | 4160 | 3927 | 4819 | 4197 | 3364 | 3441 | 3385 | 3775 | 3460 | 4029 | 4807 | 4449 | 4475 | 4524 | 4237 | 4735 | 4383 | 3417 | 3592 | 3838 | 3665 | 4019 | 3793 | 4062 | Trappc13 |
| Trappc14 | 3448 | 5037 | 4371 | 5435 | 5447 | 4362 | 5045 | 3306 | 3533 | 3993 | 3569 | 3656 | 3345 | 3757 | 3653 | 3637 | 5304 | 5463 | 5480 | 4287 | 5046 | 3940 | 3178 | 3936 | 4236 | 4020 | 3521 | 3501 | Trappc14 |
| Trappc2 | 1886 | 1562 | 1759 | 2082 | 2113 | 1988 | 2472 | 2175 | 1729 | 1503 | 1592 | 1662 | 1744 | 1828 | 2314 | 1852 | 1842 | 2152 | 2014 | 2615 | 2264 | 2132 | 1710 | 1837 | 2041 | 1596 | 1672 | 1912 | Trappc2 |
| Trappc2l | 1353 | 1450 | 1235 | 1432 | 1546 | 1338 | 1573 | 1671 | 1565 | 1481 | 1515 | 1412 | 1496 | 1669 | 1630 | 1210 | 1457 | 1413 | 1451 | 1654 | 1596 | 1564 | 1647 | 1600 | 1523 | 1455 | 1489 | 1708 | Trappc2l |
| Trappc3 | 4755 | 5186 | 5374 | 5433 | 5218 | 4858 | 5291 | 4466 | 4437 | 4747 | 4523 | 4398 | 4361 | 4517 | 5018 | 4910 | 5500 | 5302 | 5509 | 5942 | 5437 | 4286 | 4697 | 4829 | 5102 | 4899 | 4567 | 4812 | Trappc3 |
| Trappc3l | 1 | 1 | 0 | 0 | 4 | 0 | 0 | 5 | 1 | 3 | 4 | 3 | 0 | 0 | 1 | 1 | 0 | 0 | 0 | 0 | 1 | 3 | 3 | 3 | 7 | 0 | 9 | 1 | Trappc3l |
| Trappc4 | 3222 | 3867 | 3291 | 3717 | 3547 | 3398 | 3553 | 4427 | 3996 | 3990 | 3618 | 4316 | 4141 | 4452 | 3578 | 3104 | 3612 | 3394 | 3613 | 4138 | 3697 | 4172 | 3955 | 4098 | 4128 | 4168 | 4443 | 4600 | Trappc4 |
| Trappc5 | 2962 | 3393 | 3135 | 3640 | 3641 | 3465 | 3277 | 3151 | 3055 | 3600 | 3244 | 3477 | 3525 | 3397 | 3137 | 2709 | 3525 | 3252 | 3630 | 3758 | 3393 | 2996 | 3043 | 3678 | 3443 | 3767 | 3588 | 3452 | Trappc5 |
| Trappc6a | 568 | 613 | 557 | 572 | 501 | 500 | 402 | 431 | 490 | 497 | 457 | 535 | 518 | 434 | 475 | 594 | 570 | 541 | 523 | 556 | 413 | 361 | 498 | 550 | 438 | 638 | 554 | 483 | Trappc6a |
| Trappc6b | 2233 | 1923 | 2691 | 2645 | 2740 | 2570 | 2813 | 2888 | 2611 | 3343 | 2811 | 2999 | 2768 | 3451 | 2342 | 1826 | 2671 | 2431 | 2583 | 2614 | 2716 | 3218 | 2643 | 3317 | 3151 | 3210 | 2961 | 3256 | Trappc6b |
| Trappc8 | 6568 | 5833 | 5260 | 5594 | 5852 | 5964 | 6090 | 7072 | 6305 | 5829 | 6123 | 6104 | 5871 | 6189 | 6613 | 5738 | 5282 | 5743 | 5577 | 5918 | 6134 | 6625 | 6188 | 5531 | 6436 | 4996 | 6011 | 5860 | Trappc8 |
| Trappc9 | 2809 | 3539 | 2826 | 3463 | 3507 | 3334 | 2991 | 2402 | 3047 | 3357 | 2912 | 3239 | 3047 | 2982 | 2558 | 2984 | 3164 | 3425 | 3638 | 2732 | 3238 | 3272 | 2951 | 3090 | 3217 | 3282 | 2676 | 2959 | Trappc9 |
| Trarg1 | 17 | 32 | 26 | 30 | 56 | 31 | 10 | 40 | 56 | 20 | 37 | 2 | 28 | 2 | 21 | 29 | 32 | 23 | 16 | 4 | 6 | 45 | 36 | 5 | 31 | 5 | 2 | 0 | Trarg1 |
| Trat1 | 50 | 78 | 29 | 86 | 31 | 62 | 72 | 9 | 8 | 10 | 15 | 8 | 15 | 4 | 91 | 77 | 65 | 129 | 48 | 36 | 94 | 18 | 12 | 5 | 1 | 2 | 14 | 9 | Trat1 |
| Trdmt1 | 917 | 703 | 607 | 728 | 772 | 800 | 866 | 915 | 822 | 706 | 846 | 847 | 797 | 910 | 989 | 733 | 670 | 634 | 620 | 753 | 869 | 809 | 874 | 853 | 771 | 754 | 853 | 858 | Trdmt1 |
| Trdn | 5 | 2 | 1 | 1 | 1 | 0 | 1 | 10 | 38 | 10 | 210 | 2 | 3 | 7 | 7 | 4 | 1 | 2 | 6 | 3 | 3 | 18 | 156 | 2 | 5 | 2 | 3 | 2 | Trdn |
| Treh | 0 | 0 | 0 | 0 | 0 | 2 | 0 | 0 | 0 | 0 | 0 | 0 | 0 | 0 | 0 | 0 | 0 | 0 | 0 | 0 | 0 | 0 | 0 | 0 | 0 | 0 | 0 | 0 | Treh |
| Trem1 | 0 | 0 | 0 | 0 | 0 | 0 | 0 | 0 | 0 | 0 | 0 | 0 | 0 | 0 | 0 | 0 | 0 | 1 | 0 | 0 | 0 | 0 | 0 | 0 | 0 | 0 | 1 | 0 | Trem1 |
| Trem2 | 2 | 2 | 3 | 12 | 0 | 2 | 2 | 6 | 6 | 0 | 6 | 3 | 2 | 4 | 4 | 11 | 1 | 6 | 2 | 7 | 13 | 28 | 3 | 2 | 2 | 2 | 30 | 4 | Trem2 |
| Trem3 | 0 | 0 | 0 | 0 | 0 | 0 | 0 | 0 | 0 | 0 | 0 | 0 | 0 | 0 | 0 | 2 | 0 | 0 | 0 | 0 | 0 | 0 | 0 | 0 | 0 | 0 | 0 | 0 | Trem3 |
| Treml1 | 2 | 5 | 1 | 0 | 0 | 0 | 2 | 1 | 5 | 0 | 3 | 14 | 6 | 0 | 1 | 23 | 0 | 1 | 1 | 2 | 1 | 7 | 2 | 20 | 0 | 6 | 5 | 0 | Treml1 |
| Treml2 | 1 | 3 | 8 | 0 | 1 | 1 | 0 | 2 | 2 | 1 | 6 | 9 | 2 | 2 | 0 | 10 | 1 | 0 | 1 | 1 | 1 | 2 | 0 | 0 | 1 | 17 | 2 | 8 | Treml2 |
| Trerf1 | 0 | 0 | 0 | 0 | 0 | 0 | 0 | 0 | 0 | 1 | 0 | 0 | 0 | 0 | 0 | 0 | 0 | 0 | 0 | 0 | 0 | 0 | 0 | 0 | 0 | 0 | 0 | 0 | Trerf1 |
| Trex2 | 2 | 0 | 0 | 1 | 0 | 0 | 0 | 1 | 1 | 0 | 0 | 11 | 2 | 0 | 0 | 4 | 0 | 0 | 0 | 0 | 1 | 0 | 3 | 1 | 0 | 0 | 0 | 0 | Trex2 |
| Trf | 689 | 914 | 737 | 665 | 905 | 916 | 684 | 844 | 1108 | 915 | 1065 | 894 | 961 | 775 | 766 | 1046 | 664 | 812 | 755 | 884 | 740 | 1086 | 1251 | 797 | 993 | 874 | 881 | 746 | Trf |
| Trh | 55 | 57 | 56 | 45 | 95 | 43 | 70 | 92 | 104 | 89 | 63 | 89 | 35 | 113 | 74 | 34 | 68 | 78 | 84 | 67 | 87 | 129 | 119 | 80 | 117 | 109 | 119 | 102 | Trh |
| Trhde | 325 | 314 | 170 | 317 | 650 | 528 | 413 | 813 | 694 | 606 | 626 | 267 | 474 | 335 | 429 | 431 | 302 | 435 | 449 | 412 | 460 | 909 | 598 | 313 | 610 | 263 | 300 | 293 | Trhde |
| Trhr | 8 | 19 | 20 | 29 | 46 | 16 | 29 | 45 | 68 | 49 | 77 | 28 | 53 | 15 | 28 | 17 | 21 | 35 | 48 | 38 | 46 | 55 | 78 | 28 | 54 | 8 | 35 | 26 | Trhr |
| Trhr2 | 3 | 0 | 0 | 0 | 3 | 5 | 0 | 7 | 15 | 15 | 53 | 4 | 6 | 5 | 0 | 5 | 1 | 1 | 3 | 0 | 1 | 25 | 9 | 14 | 7 | 8 | 3 | 9 | Trhr2 |
| Triap1 | 1688 | 2038 | 2052 | 1867 | 1397 | 1418 | 1721 | 1116 | 1081 | 1076 | 1054 | 1234 | 1157 | 1398 | 1615 | 1701 | 1839 | 1734 | 1444 | 1744 | 1465 | 959 | 1237 | 1308 | 1262 | 1543 | 1339 | 1372 | Triap1 |
| Trib1 | 1057 | 1002 | 1539 | 1452 | 1576 | 1341 | 1278 | 1215 | 1233 | 1501 | 1533 | 1597 | 1588 | 1527 | 916 | 1004 | 1438 | 1350 | 1606 | 1174 | 1333 | 1477 | 1271 | 1589 | 1385 | 1882 | 1548 | 1604 | Trib1 |
| Trib2 | 4541 | 4161 | 5298 | 6166 | 8461 | 6836 | 7591 | 10390 | 10615 | 11742 | 11638 | 9765 | 9837 | 10658 | 4950 | 4264 | 6036 | 6458 | 8013 | 6673 | 7519 | 13754 | 9890 | 11154 | 12921 | 9756 | 8915 | 9700 | Trib2 |
| Trib3 | 93 | 125 | 88 | 107 | 81 | 114 | 82 | 81 | 108 | 97 | 94 | 122 | 338 | 160 | 59 | 121 | 98 | 83 | 108 | 115 | 75 | 40 | 111 | 96 | 63 | 134 | 134 | 206 | Trib3 |
| Tril | 2422 | 2128 | 2860 | 1155 | 1136 | 1685 | 877 | 2652 | 3483 | 3093 | 4142 | 3575 | 5040 | 3091 | 2033 | 3062 | 1593 | 1413 | 1240 | 889 | 869 | 3055 | 4253 | 2898 | 2193 | 4647 | 3958 | 4326 | Tril |
| Trim10 | 235 | 289 | 289 | 131 | 86 | 140 | 108 | 147 | 215 | 142 | 113 | 321 | 1013 | 427 | 227 | 278 | 183 | 95 | 129 | 206 | 128 | 97 | 223 | 101 | 137 | 458 | 523 | 898 | Trim10 |
| Trim11 | 2478 | 3318 | 2854 | 3062 | 2455 | 2789 | 2462 | 1661 | 1912 | 2006 | 2077 | 1976 | 2098 | 1964 | 2147 | 3027 | 2855 | 3134 | 2670 | 2364 | 2388 | 1890 | 1832 | 2108 | 2044 | 2206 | 2101 | 1915 | Trim11 |
| Trim12a | 276 | 192 | 222 | 166 | 135 | 266 | 107 | 310 | 407 | 377 | 418 | 263 | 396 | 266 | 244 | 307 | 124 | 250 | 109 | 188 | 113 | 334 | 491 | 213 | 233 | 252 | 285 | 256 | Trim12a |
| Trim12c | 756 | 481 | 559 | 505 | 468 | 521 | 430 | 794 | 769 | 662 | 803 | 560 | 786 | 620 | 649 | 818 | 501 | 529 | 457 | 571 | 439 | 663 | 1042 | 633 | 659 | 678 | 607 | 660 | Trim12c |
| Trim13 | 2148 | 1830 | 1480 | 1814 | 1999 | 1922 | 2275 | 2243 | 1924 | 1643 | 1473 | 1567 | 1486 | 1621 | 2463 | 1709 | 1749 | 1812 | 2014 | 2538 | 2232 | 2052 | 1892 | 1600 | 1768 | 1532 | 1421 | 1620 | Trim13 |
| Trim14 | 38 | 28 | 23 | 20 | 39 | 24 | 18 | 15 | 21 | 28 | 46 | 7 | 15 | 24 | 26 | 49 | 24 | 24 | 16 | 24 | 14 | 15 | 38 | 34 | 23 | 30 | 34 | 17 | Trim14 |
| Trim15 | 5 | 10 | 1 | 8 | 1 | 1 | 1 | 6 | 8 | 0 | 5 | 10 | 9 | 6 | 0 | 7 | 0 | 2 | 2 | 0 | 3 | 5 | 10 | 2 | 2 | 4 | 3 | 13 | Trim15 |
| Trim16 | 616 | 410 | 431 | 331 | 484 | 437 | 332 | 794 | 804 | 754 | 820 | 587 | 685 | 501 | 584 | 537 | 382 | 436 | 412 | 337 | 343 | 612 | 893 | 680 | 522 | 571 | 581 | 503 | Trim16 |
| Trim17 | 98 | 129 | 73 | 294 | 200 | 165 | 301 | 84 | 108 | 125 | 107 | 80 | 67 | 85 | 141 | 92 | 184 | 278 | 158 | 206 | 257 | 105 | 116 | 123 | 105 | 54 | 58 | 53 | Trim17 |
| Trim2 | 14056 | 14146 | 13517 | 15879 | 21012 | 20239 | 20195 | 21623 | 18299 | 19892 | 16443 | 19461 | 19771 | 20096 | 17205 | 14207 | 15667 | 16733 | 19124 | 19607 | 21081 | 21286 | 16454 | 17678 | 22821 | 11263 | 21169 | 19263 | Trim2 |
| Trim21 | 236 | 187 | 274 | 167 | 178 | 187 | 102 | 267 | 302 | 267 | 340 | 263 | 367 | 286 | 237 | 249 | 174 | 166 | 147 | 207 | 153 | 244 | 390 | 293 | 219 | 349 | 293 | 293 | Trim21 |
| Trim23 | 4947 | 3814 | 4409 | 4593 | 4644 | 4694 | 5453 | 5440 | 4519 | 4069 | 4434 | 4387 | 4223 | 4648 | 5366 | 4610 | 4242 | 4665 | 4415 | 5456 | 4973 | 4928 | 4752 | 4328 | 4632 | 4408 | 4489 | 4514 | Trim23 |
| Trim24 | 6278 | 5614 | 6216 | 5786 | 6541 | 6242 | 6754 | 10209 | 8528 | 9265 | 8528 | 10658 | 9936 | 11003 | 6505 | 5245 | 6324 | 5358 | 6357 | 6179 | 6501 | 9598 | 8469 | 9030 | 9183 | 8917 | 11622 | 11271 | Trim24 |
| Trim25 | 1009 | 797 | 1010 | 810 | 1006 | 787 | 746 | 650 | 732 | 902 | 1084 | 796 | 855 | 743 | 857 | 1059 | 797 | 894 | 773 | 652 | 759 | 787 | 1003 | 852 | 864 | 776 | 898 | 713 | Trim25 |
| Trim26 | 3685 | 4112 | 4524 | 4750 | 4571 | 4271 | 4057 | 3036 | 3292 | 3669 | 3743 | 3704 | 3937 | 3776 | 3708 | 3803 | 4376 | 4419 | 4330 | 3446 | 4155 | 3526 | 3239 | 3581 | 3567 | 3641 | 3344 | 3748 | Trim26 |
| Trim27 | 6181 | 6387 | 7363 | 7141 | 5951 | 5824 | 6016 | 4532 | 4425 | 4694 | 4613 | 5478 | 4905 | 5429 | 5718 | 5963 | 7108 | 6837 | 6440 | 5069 | 6226 | 4542 | 4442 | 4951 | 4547 | 6417 | 5059 | 5609 | Trim27 |
| Trim28 | 26746 | 30100 | 34571 | 31593 | 27536 | 26390 | 26284 | 27919 | 31347 | 31941 | 30125 | 40843 | 33898 | 35999 | 24437 | 26211 | 33941 | 29212 | 33556 | 23281 | 28427 | 31163 | 29380 | 37239 | 33524 | 47579 | 35652 | 39237 | Trim28 |
| Trim29 | 349 | 176 | 96 | 19 | 13 | 147 | 7 | 294 | 756 | 303 | 395 | 116 | 191 | 142 | 150 | 223 | 44 | 153 | 41 | 7 | 0 | 8 | 861 | 275 | 42 | 63 | 135 | 183 | Trim29 |
| Trim3 | 4369 | 5328 | 5096 | 6185 | 6724 | 6132 | 5482 | 3602 | 4065 | 4884 | 4656 | 4214 | 3974 | 3743 | 4607 | 5234 | 5487 | 6475 | 6442 | 5331 | 5715 | 4705 | 4017 | 4230 | 4795 | 3715 | 3912 | 3604 | Trim3 |
| Trim30a | 118 | 76 | 104 | 49 | 94 | 55 | 65 | 125 | 108 | 150 | 201 | 126 | 117 | 152 | 115 | 116 | 75 | 86 | 76 | 180 | 90 | 157 | 190 | 144 | 145 | 175 | 166 | 169 | Trim30a |
| Trim30b | 3 | 5 | 0 | 0 | 3 | 3 | 2 | 2 | 7 | 5 | 5 | 3 | 10 | 3 | 5 | 5 | 15 | 1 | 0 | 15 | 3 | 5 | 7 | 8 | 18 | 2 | 7 | 2 | Trim30b |
| Trim30c | 3 | 1 | 0 | 0 | 0 | 0 | 1 | 10 | 7 | 10 | 13 | 4 | 9 | 17 | 3 | 4 | 3 | 1 | 0 | 1 | 2 | 7 | 15 | 6 | 7 | 4 | 15 | 5 | Trim30c |
| Trim30d | 25 | 11 | 9 | 19 | 20 | 31 | 23 | 39 | 34 | 21 | 51 | 50 | 40 | 40 | 22 | 34 | 17 | 32 | 18 | 39 | 28 | 37 | 47 | 35 | 23 | 30 | 49 | 34 | Trim30d |
| Trim31 | 0 | 0 | 0 | 0 | 0 | 0 | 0 | 1 | 1 | 0 | 0 | 2 | 1 | 2 | 0 | 0 | 5 | 1 | 0 | 2 | 0 | 2 | 1 | 1 | 0 | 0 | 0 | 1 | Trim31 |
| Trim32 | 7130 | 8147 | 7192 | 8578 | 9603 | 8288 | 9453 | 8252 | 8460 | 7818 | 7057 | 6820 | 6066 | 6683 | 7843 | 7186 | 7982 | 9095 | 9684 | 9011 | 9444 | 9175 | 8091 | 7336 | 8320 | 6482 | 6219 | 6445 | Trim32 |
| Trim33 | 10381 | 8098 | 8216 | 9391 | 10573 | 11043 | 11416 | 14719 | 12142 | 11652 | 11284 | 13122 | 12557 | 13623 | 10864 | 9124 | 8481 | 9369 | 10124 | 10730 | 10877 | 13541 | 11723 | 11210 | 12976 | 10593 | 12965 | 13161 | Trim33 |
| Trim34a | 47 | 44 | 87 | 45 | 52 | 47 | 32 | 114 | 126 | 146 | 159 | 135 | 162 | 115 | 57 | 68 | 67 | 77 | 33 | 66 | 41 | 179 | 168 | 152 | 105 | 228 | 178 | 141 | Trim34a |
| Trim34b | 0 | 0 | 0 | 0 | 2 | 2 | 4 | 1 | 0 | 6 | 1 | 3 | 1 | 2 | 0 | 0 | 0 | 0 | 0 | 0 | 0 | 0 | 3 | 2 | 1 | 0 | 0 | 0 | Trim34b |
| Trim35 | 19547 | 20688 | 19719 | 22130 | 23365 | 22356 | 22589 | 21252 | 19913 | 19971 | 19482 | 22058 | 19949 | 20662 | 20441 | 17562 | 21461 | 22511 | 24794 | 22236 | 23660 | 24231 | 19358 | 21285 | 23178 | 20866 | 20959 | 20902 | Trim35 |
| Trim36 | 2947 | 2842 | 3248 | 4308 | 5133 | 4268 | 4240 | 4659 | 3633 | 4345 | 4060 | 4245 | 4081 | 4436 | 3593 | 2953 | 3892 | 4230 | 4503 | 4059 | 4396 | 5454 | 3537 | 3980 | 4621 | 3254 | 3997 | 4174 | Trim36 |
| Trim37 | 19778 | 18955 | 17696 | 19532 | 20380 | 20070 | 22227 | 16127 | 13497 | 14040 | 11627 | 15160 | 13282 | 15556 | 20803 | 16369 | 19375 | 19623 | 20774 | 21578 | 22141 | 15827 | 12732 | 14029 | 15642 | 13407 | 15507 | 15035 | Trim37 |
| Trim38 | 14 | 8 | 12 | 13 | 9 | 3 | 9 | 5 | 1 | 0 | 2 | 1 | 2 | 3 | 9 | 9 | 9 | 0 | 12 | 3 | 7 | 0 | 4 | 1 | 1 | 5 | 1 | 0 | Trim38 |
| Trim39 | 2682 | 2747 | 3154 | 3199 | 3508 | 3288 | 3417 | 5432 | 4927 | 6167 | 6172 | 5885 | 5578 | 6218 | 2780 | 2614 | 3306 | 3113 | 3443 | 3076 | 3372 | 6198 | 4798 | 5650 | 6325 | 5151 | 5489 | 6057 | Trim39 |
| Trim40 | 0 | 0 | 0 | 0 | 0 | 2 | 0 | 4 | 2 | 0 | 0 | 0 | 0 | 0 | 0 | 2 | 0 | 0 | 0 | 0 | 2 | 0 | 1 | 0 | 0 | 0 | 0 | 0 | Trim40 |
| Trim41 | 3564 | 4334 | 5080 | 4790 | 4223 | 4195 | 4008 | 4264 | 4813 | 5663 | 5671 | 5920 | 5632 | 5666 | 3506 | 3664 | 5033 | 4450 | 4771 | 4195 | 4119 | 4943 | 4633 | 6316 | 5757 | 6454 | 6046 | 5967 | Trim41 |
| Trim42 | 1 | 1 | 0 | 0 | 0 | 4 | 2 | 0 | 0 | 1 | 0 | 0 | 0 | 0 | 0 | 0 | 3 | 4 | 0 | 1 | 1 | 0 | 0 | 0 | 0 | 0 | 0 | 0 | Trim42 |
| Trim43a | 53 | 79 | 39 | 124 | 78 | 94 | 106 | 26 | 14 | 20 | 17 | 18 | 15 | 11 | 47 | 68 | 58 | 94 | 104 | 108 | 88 | 23 | 19 | 9 | 23 | 4 | 17 | 7 | Trim43a |
| Trim43b | 1 | 1 | 0 | 0 | 1 | 0 | 0 | 0 | 0 | 0 | 0 | 2 | 1 | 0 | 0 | 0 | 0 | 3 | 0 | 0 | 0 | 0 | 0 | 0 | 0 | 0 | 0 | 0 | Trim43b |
| Trim43c | 0 | 0 | 0 | 0 | 0 | 0 | 0 | 2 | 0 | 5 | 0 | 0 | 0 | 0 | 0 | 0 | 0 | 0 | 0 | 0 | 1 | 0 | 0 | 0 | 0 | 0 | 0 | 0 | Trim43c |
| Trim44 | 19352 | 17554 | 17724 | 21059 | 23521 | 22868 | 23926 | 24715 | 22393 | 24684 | 24194 | 24686 | 23395 | 25156 | 20305 | 17541 | 20124 | 21712 | 22379 | 20163 | 24749 | 26893 | 22234 | 23442 | 26575 | 19566 | 23484 | 24100 | Trim44 |

| | | | | | | | | | | | | | | | | | | | | | | | | | | | | | |
|---|---|---|---|---|---|---|---|---|---|---|---|---|---|---|---|---|---|---|---|---|---|---|---|---|---|---|---|---|---|
| Trim45 | 1205 | 1410 | 1225 | 1456 | 1355 | 1369 | 1219 | 1406 | 1476 | 1584 | 1429 | 1490 | 1627 | 1395 | 1171 | 1200 | 1451 | 1369 | 1401 | 1181 | 1294 | 1497 | 1471 | 1472 | 1510 | 1535 | 1562 | 1418 | Trim45 |
| Trim46 | 1615 | 2216 | 2274 | 3291 | 4120 | 3270 | 3086 | 3197 | 3903 | 4725 | 3908 | 3794 | 3113 | 3192 | 1646 | 1796 | 2927 | 3287 | 4032 | 2768 | 3272 | 5320 | 3446 | 4622 | 5397 | 3089 | 3201 | 2948 | Trim46 |
| Trim47 | 119 | 140 | 162 | 141 | 164 | 147 | 120 | 118 | 198 | 201 | 308 | 99 | 154 | 146 | 109 | 174 | 114 | 195 | 150 | 96 | 128 | 162 | 225 | 162 | 141 | 131 | 113 | 119 | Trim47 |
| Trim5 | 32 | 16 | 36 | 8 | 20 | 26 | 7 | 26 | 26 | 35 | 43 | 20 | 39 | 23 | 38 | 34 | 16 | 25 | 12 | 31 | 9 | 32 | 71 | 15 | 33 | 17 | 29 | 38 | Trim5 |
| Trim50 | 0 | 2 | 0 | 0 | 0 | 0 | 0 | 1 | 0 | 2 | 1 | 0 | 0 | 5 | 0 | 0 | 3 | 1 | 0 | 0 | 0 | 0 | 1 | 1 | 0 | 2 | 0 | 0 | Trim50 |
| Trim52 | 0 | 0 | 0 | 0 | 0 | 1 | 0 | 0 | 0 | 0 | 0 | 0 | 0 | 0 | 0 | 0 | 0 | 0 | 0 | 0 | 0 | 0 | 0 | 0 | 0 | 0 | 0 | 0 | Trim52 |
| Trim54 | 2 | 1 | 1 | 0 | 0 | 0 | 0 | 2 | 18 | 3 | 115 | 0 | 2 | 0 | 1 | 2 | 0 | 0 | 0 | 0 | 0 | 15 | 71 | 1 | 4 | 0 | 0 | 0 | Trim54 |
| Trim55 | 36 | 4 | 0 | 0 | 6 | 4 | 1 | 37 | 124 | 6 | 890 | 0 | 18 | 0 | 9 | 25 | 0 | 4 | 0 | 4 | 0 | 60 | 524 | 10 | 11 | 0 | 1 | 0 | Trim55 |
| Trim56 | 1440 | 1260 | 1341 | 1342 | 1479 | 1383 | 1269 | 1570 | 1757 | 1858 | 2088 | 2105 | 2418 | 2395 | 1199 | 1411 | 1198 | 1263 | 1386 | 1188 | 1269 | 1342 | 1836 | 1965 | 1816 | 1577 | 2238 | 2182 | Trim56 |
| Trim58 | 2 | 5 | 5 | 0 | 5 | 4 | 3 | 12 | 21 | 8 | 18 | 30 | 103 | 28 | 7 | 4 | 5 | 4 | 2 | 6 | 8 | 10 | 10 | 11 | 3 | 17 | 35 | 72 | Trim58 |
| Trim59 | 7973 | 5315 | 8216 | 6588 | 5967 | 6503 | 7381 | 9149 | 7303 | 7650 | 7104 | 11144 | 9821 | 10917 | 7001 | 6459 | 7406 | 6576 | 7303 | 6569 | 7062 | 7454 | 7034 | 9316 | 8386 | 13280 | 11665 | 12394 | Trim59 |
| Trim6 | 323 | 276 | 389 | 119 | 145 | 171 | 122 | 61 | 82 | 62 | 113 | 113 | 147 | 94 | 322 | 363 | 228 | 141 | 111 | 129 | 117 | 97 | 102 | 113 | 37 | 169 | 169 | 123 | Trim6 |
| Trim60 | 16 | 10 | 7 | 13 | 7 | 1 | 15 | 9 | 3 | 4 | 0 | 1 | 0 | 1 | 16 | 9 | 9 | 20 | 5 | 4 | 13 | 2 | 1 | 0 | 0 | 0 | 0 | 2 | Trim60 |
| Trim61 | 27 | 15 | 26 | 16 | 20 | 23 | 29 | 7 | 3 | 5 | 7 | 2 | 1 | 6 | 28 | 10 | 24 | 23 | 25 | 12 | 16 | 2 | 8 | 5 | 1 | 17 | 2 | 11 | Trim61 |
| Trim62 | 3854 | 4717 | 4063 | 5041 | 5607 | 4884 | 4314 | 1905 | 2096 | 2552 | 2311 | 2265 | 2422 | 1861 | 3819 | 4888 | 4887 | 5477 | 5054 | 3774 | 4689 | 2797 | 1956 | 2207 | 2277 | 2288 | 1921 | 1860 | Trim62 |
| Trim63 | 16 | 6 | 8 | 6 | 11 | 11 | 2 | 32 | 29 | 36 | 194 | 26 | 31 | 11 | 7 | 12 | 2 | 9 | 3 | 3 | 3 | 47 | 124 | 22 | 14 | 25 | 8 | 23 | Trim63 |
| Trim65 | 114 | 131 | 219 | 132 | 155 | 137 | 130 | 220 | 270 | 359 | 345 | 258 | 268 | 319 | 82 | 104 | 174 | 178 | 150 | 100 | 139 | 339 | 323 | 351 | 316 | 360 | 253 | 272 | Trim65 |
| Trim66 | 397 | 349 | 184 | 681 | 553 | 504 | 465 | 658 | 489 | 887 | 635 | 173 | 307 | 235 | 373 | 367 | 346 | 450 | 484 | 506 | 536 | 197 | 658 | 1041 | 166 | 586 | 184 | 155 | Trim66 |
| Trim67 | 2568 | 3620 | 3544 | 6548 | 9959 | 7079 | 7755 | 5674 | 5745 | 7923 | 6361 | 4869 | 4765 | 4756 | 3463 | 2816 | 5711 | 7553 | 8455 | 5881 | 8671 | 10196 | 5148 | 5741 | 8626 | 3139 | 3932 | 4130 | Trim67 |
| Trim68 | 904 | 739 | 703 | 591 | 623 | 599 | 558 | 935 | 1006 | 944 | 1086 | 863 | 1134 | 788 | 802 | 842 | 608 | 616 | 495 | 584 | 505 | 884 | 983 | 829 | 730 | 895 | 915 | 782 | Trim68 |
| Trim69 | 16 | 18 | 23 | 20 | 7 | 28 | 23 | 10 | 7 | 7 | 13 | 1 | 8 | 2 | 12 | 33 | 38 | 23 | 9 | 20 | 17 | 5 | 3 | 7 | 7 | 9 | 1 | 6 | Trim69 |
| Trim7 | 120 | 86 | 65 | 79 | 140 | 160 | 132 | 35 | 61 | 53 | 95 | 59 | 79 | 49 | 116 | 86 | 95 | 81 | 141 | 98 | 117 | 47 | 68 | 48 | 94 | 40 | 62 | 39 | Trim7 |
| Trim71 | 442 | 481 | 871 | 546 | 331 | 267 | 203 | 206 | 146 | 149 | 165 | 260 | 354 | 217 | 426 | 564 | 616 | 299 | 277 | 220 | 181 | 157 | 107 | 155 | 131 | 343 | 310 | 352 | Trim71 |
| Trim72 | 102 | 179 | 298 | 459 | 322 | 283 | 284 | 40 | 91 | 93 | 718 | 99 | 98 | 93 | 128 | 150 | 380 | 479 | 395 | 157 | 279 | 134 | 366 | 78 | 100 | 77 | 98 | 69 | Trim72 |
| Trim75 | 13 | 3 | 15 | 2 | 2 | 1 | 3 | 5 | 1 | 0 | 4 | 0 | 0 | 1 | 8 | 4 | 2 | 4 | 2 | 8 | 3 | 0 | 0 | 0 | 3 | 2 | 0 | 0 | Trim75 |
| Trim8 | 1465 | 1900 | 2687 | 2372 | 2703 | 2081 | 1969 | 3101 | 3255 | 4097 | 3868 | 4432 | 3776 | 4224 | 1515 | 1583 | 2771 | 2168 | 2781 | 1850 | 2141 | 4243 | 3355 | 4555 | 4771 | 4968 | 4154 | 4187 | Trim8 |
| Trim80 | 3 | 3 | 1 | 2 | 7 | 1 | 2 | 1 | 2 | 2 | 0 | 2 | 7 | 7 | 4 | 1 | 1 | 1 | 1 | 4 | 8 | 10 | 2 | 7 | 2 | 14 | 6 | 15 | Trim80 |
| Trim9 | 3350 | 3867 | 4294 | 5258 | 7334 | 6262 | 6174 | 3998 | 3647 | 4347 | 3787 | 3589 | 3563 | 3604 | 4240 | 3590 | 4573 | 5843 | 6511 | 6397 | 7378 | 5342 | 3902 | 3601 | 4893 | 2004 | 3516 | 3432 | Trim9 |
| Triml1 | 0 | 0 | 0 | 0 | 0 | 0 | 0 | 0 | 1 | 0 | 2 | 0 | 0 | 0 | 0 | 0 | 0 | 0 | 0 | 0 | 0 | 0 | 0 | 0 | 0 | 0 | 0 | 0 | Triml1 |
| Triml2 | 0 | 0 | 0 | 0 | 0 | 0 | 0 | 0 | 2 | 0 | 0 | 0 | 0 | 0 | 0 | 0 | 1 | 0 | 2 | 0 | 0 | 3 | 0 | 0 | 0 | 0 | 0 | 1 | Triml2 |
| Trio | 10439 | 11431 | 8256 | 13232 | 16994 | 14960 | 15414 | 14169 | 16993 | 18780 | 19840 | 16931 | 15748 | 15826 | 10086 | 11174 | 10706 | 14127 | 15112 | 12908 | 16241 | 16798 | 15905 | 14931 | 19741 | 10174 | 15659 | 14406 | Trio |
| Triobp | 5192 | 6918 | 7181 | 7860 | 7180 | 6488 | 5653 | 3262 | 4524 | 5145 | 4491 | 5125 | 4534 | 4346 | 4833 | 6071 | 7496 | 7690 | 7974 | 5573 | 6075 | 5183 | 4043 | 5295 | 5303 | 4527 | 4458 | 4112 | Triobp |
| Trip10 | 2587 | 3239 | 3782 | 3529 | 3015 | 3101 | 2750 | 1054 | 1163 | 1118 | 1300 | 1364 | 1312 | 1277 | 2403 | 2823 | 3651 | 3354 | 3175 | 2726 | 2895 | 1009 | 1206 | 1353 | 1110 | 1499 | 1312 | 1406 | Trip10 |
| Trip11 | 4021 | 3073 | 2907 | 3687 | 4237 | 4490 | 4416 | 5107 | 5072 | 5219 | 6119 | 5171 | 5296 | 5514 | 3821 | 3596 | 3140 | 3695 | 3599 | 3913 | 4285 | 5005 | 5259 | 4467 | 5533 | 3678 | 5613 | 4761 | Trip11 |
| Trip12 | 36440 | 31399 | 26488 | 30108 | 31767 | 35600 | 35121 | 39718 | 35461 | 33782 | 34632 | 39059 | 38081 | 37875 | 36224 | 32387 | 27438 | 28595 | 30939 | 33974 | 34067 | 35087 | 34582 | 30714 | 34974 | 27245 | 41576 | 36914 | Trip12 |
| Trip13 | 1827 | 1562 | 2030 | 1273 | 1088 | 1111 | 1381 | 1638 | 1720 | 1440 | 1645 | 2583 | 2195 | 2651 | 1602 | 1759 | 1657 | 1228 | 1223 | 1176 | 1180 | 1374 | 1618 | 2015 | 1677 | 3121 | 2382 | 2861 | Trip13 |
| Trip4 | 4692 | 4042 | 3932 | 4456 | 4597 | 4584 | 5022 | 4671 | 4324 | 4100 | 4520 | 4256 | 4447 | 4360 | 5250 | 4395 | 4204 | 4639 | 4392 | 4730 | 4706 | 4508 | 4675 | 4033 | 4087 | 4230 | 4353 | 4347 | Trip4 |
| Trip6 | 2786 | 3083 | 3321 | 3063 | 2473 | 2692 | 2297 | 2500 | 3111 | 2805 | 3107 | 2917 | 2834 | 2570 | 2367 | 2931 | 3498 | 3047 | 2872 | 2641 | 2271 | 2216 | 3207 | 3058 | 2527 | 3752 | 2914 | 2756 | Trip6 |
| Triqk | 802 | 590 | 638 | 855 | 794 | 833 | 914 | 843 | 783 | 737 | 657 | 583 | 611 | 611 | 944 | 721 | 797 | 798 | 774 | 935 | 911 | 857 | 901 | 588 | 689 | 544 | 648 | 649 | Triqk |
| Trir | 4380 | 4997 | 4816 | 4837 | 4414 | 3757 | 5139 | 4244 | 3749 | 4667 | 4224 | 4598 | 4190 | 5668 | 4278 | 3477 | 5106 | 4512 | 4736 | 4221 | 4924 | 3706 | 3660 | 5161 | 4691 | 5500 | 4795 | 5366 | Trir |
| Trit1 | 1718 | 1812 | 1552 | 1985 | 1709 | 1802 | 2016 | 2114 | 1847 | 1966 | 1777 | 2084 | 1944 | 2105 | 1817 | 1572 | 1663 | 1782 | 1824 | 1749 | 1876 | 1997 | 1934 | 1879 | 2092 | 1964 | 2141 | 2083 | Trit1 |
| Trmo | 140 | 117 | 188 | 131 | 161 | 124 | 153 | 361 | 335 | 354 | 284 | 339 | 328 | 365 | 136 | 93 | 153 | 147 | 118 | 168 | 147 | 302 | 394 | 355 | 375 | 501 | 366 | 395 | Trmo |
| Trmt1 | 4668 | 5240 | 5095 | 6341 | 5830 | 5544 | 5342 | 3486 | 3658 | 3958 | 3979 | 4023 | 3743 | 3877 | 4656 | 4914 | 5778 | 5685 | 6238 | 4932 | 5603 | 3780 | 3646 | 3809 | 3761 | 4070 | 3944 | 3946 | Trmt1 |
| Trmt10a | 878 | 667 | 781 | 620 | 560 | 620 | 629 | 1305 | 1217 | 1136 | 1124 | 1540 | 1377 | 1574 | 803 | 700 | 709 | 583 | 607 | 700 | 567 | 1211 | 1265 | 1348 | 1120 | 1684 | 1558 | 1634 | Trmt10a |
| Trmt10b | 1301 | 1215 | 1141 | 1324 | 1097 | 1051 | 1198 | 1132 | 988 | 893 | 1015 | 972 | 1007 | 901 | 1291 | 1266 | 1291 | 1177 | 1004 | 1257 | 1256 | 1081 | 965 | 934 | 1095 | 990 | 896 | 922 | Trmt10b |
| Trmt10c | 2008 | 1795 | 1823 | 1682 | 1279 | 1548 | 1750 | 2312 | 1879 | 1529 | 1575 | 1689 | 1665 | 1744 | 2004 | 1965 | 1575 | 1591 | 1362 | 1859 | 1591 | 1601 | 1797 | 1696 | 1728 | 1904 | 1845 | 1818 | Trmt10c |
| Trmt11 | 1141 | 789 | 872 | 901 | 790 | 927 | 962 | 1055 | 941 | 787 | 841 | 1094 | 979 | 1037 | 1110 | 1048 | 823 | 843 | 864 | 1037 | 826 | 907 | 857 | 988 | 928 | 1272 | 1061 | 1168 | Trmt11 |
| Trmt112 | 1498 | 1903 | 2157 | 1867 | 1620 | 1576 | 1578 | 1534 | 2215 | 2136 | 2331 | 2353 | 2581 | 2574 | 1352 | 1479 | 1909 | 1657 | 1827 | 1723 | 1560 | 1576 | 1952 | 2573 | 2217 | 3001 | 2613 | 2744 | Trmt112 |
| Trmt12 | 1400 | 1437 | 1197 | 1306 | 1252 | 1214 | 1287 | 905 | 953 | 848 | 809 | 1026 | 1027 | 992 | 1472 | 1379 | 1167 | 1251 | 1159 | 1402 | 1239 | 934 | 908 | 838 | 889 | 918 | 1141 | 1003 | Trmt12 |
| Trmt13 | 1897 | 1570 | 1413 | 1695 | 1624 | 1862 | 1866 | 1937 | 1804 | 1762 | 1916 | 1921 | 1810 | 2009 | 1983 | 1826 | 1423 | 1648 | 1497 | 1882 | 1754 | 1793 | 1843 | 1687 | 1847 | 1570 | 1929 | 1810 | Trmt13 |
| Trmt1l | 2451 | 2725 | 3132 | 3102 | 3237 | 2695 | 2925 | 2889 | 2663 | 3025 | 2815 | 3258 | 2995 | 3103 | 2606 | 2839 | 3471 | 3121 | 3183 | 2717 | 3063 | 3051 | 2802 | 3162 | 3141 | 3409 | 2962 | 3167 | Trmt1l |
| Trmt2a | 0 | 0 | 0 | 0 | 1 | 0 | 0 | 0 | 0 | 0 | 0 | 0 | 0 | 0 | 0 | 0 | 0 | 0 | 0 | 0 | 0 | 2 | 0 | 0 | 0 | 0 | 0 | 0 | Trmt2a |
| Trmt2b | 1437 | 1198 | 1283 | 1147 | 1028 | 1164 | 1240 | 1078 | 1097 | 886 | 964 | 866 | 898 | 854 | 1389 | 1222 | 1088 | 1270 | 1226 | 1191 | 1087 | 866 | 1094 | 865 | 767 | 1115 | 868 | 902 | Trmt2b |
| Trmt44 | 901 | 1157 | 641 | 846 | 848 | 876 | 789 | 950 | 875 | 816 | 845 | 882 | 911 | 862 | 843 | 970 | 744 | 877 | 922 | 837 | 940 | 846 | 840 | 920 | 961 | 715 | 989 | 909 | Trmt44 |
| Trmt5 | 1190 | 1056 | 1041 | 838 | 923 | 1032 | 1029 | 1417 | 1339 | 1136 | 1148 | 1262 | 1221 | 1170 | 1278 | 1143 | 1108 | 917 | 990 | 1168 | 986 | 1064 | 1280 | 1129 | 1001 | 1326 | 1416 | 1269 | Trmt5 |
| Trmt6 | 5015 | 4090 | 3973 | 3742 | 3443 | 3905 | 4185 | 3621 | 3286 | 3376 | 3267 | 3763 | 3750 | 3773 | 4804 | 4368 | 3574 | 3449 | 3292 | 3937 | 3720 | 3024 | 3245 | 3393 | 3385 | 4108 | 4051 | 3913 | Trmt6 |
| Trmt61a | 783 | 1017 | 1012 | 887 | 736 | 811 | 605 | 747 | 924 | 827 | 994 | 1234 | 1179 | 956 | 723 | 728 | 975 | 923 | 902 | 758 | 683 | 682 | 800 | 1172 | 1043 | 1431 | 1303 | 1137 | Trmt61a |
| Trmt9b | 584 | 504 | 600 | 524 | 677 | 657 | 638 | 338 | 303 | 345 | 316 | 255 | 346 | 328 | 579 | 485 | 519 | 524 | 634 | 699 | 584 | 333 | 348 | 304 | 298 | 298 | 310 | 282 | Trmt9b |
| Trmu | 1192 | 1289 | 1484 | 1362 | 1393 | 1338 | 1343 | 1513 | 1578 | 1583 | 1682 | 1736 | 1664 | 1612 | 1207 | 1126 | 1522 | 1337 | 1505 | 1375 | 1366 | 1589 | 1496 | 1865 | 1707 | 1872 | 1749 | 1810 | Trmu |
| Trnau1ap | 2036 | 2363 | 1929 | 2186 | 1960 | 1933 | 2106 | 2154 | 2250 | 1919 | 1769 | 2025 | 1961 | 1904 | 2039 | 2224 | 2113 | 1986 | 2300 | 2421 | 2043 | 1795 | 2130 | 2055 | 1809 | 2308 | 2098 | 2066 | Trnau1ap |
| Trnp1 | 382 | 384 | 292 | 378 | 473 | 342 | 316 | 287 | 308 | 330 | 331 | 147 | 235 | 134 | 369 | 398 | 341 | 376 | 312 | 309 | 323 | 336 | 337 | 257 | 235 | 197 | 93 | 117 | Trnp1 |
| Trnt1 | 6109 | 4932 | 5217 | 5397 | 4659 | 4899 | 5899 | 5261 | 4494 | 4010 | 4136 | 4678 | 4540 | 4852 | 6464 | 5577 | 4923 | 5283 | 4960 | 6189 | 5303 | 4246 | 4629 | 4270 | 4184 | 4965 | 4818 | 5177 | Trnt1 |
| Tro | 14072 | 16422 | 9411 | 13597 | 15701 | 16115 | 13748 | 17524 | 16752 | 14800 | 15012 | 13459 | 13020 | 12615 | 14577 | 13481 | 12028 | 13766 | 14635 | 17439 | 14963 | 16207 | 16017 | 12553 | 16609 | 7895 | 13969 | 10866 | Tro |
| Troap | 1427 | 1765 | 2206 | 1656 | 1050 | 1190 | 1257 | 1018 | 1232 | 1261 | 1193 | 1943 | 1649 | 1711 | 1137 | 1565 | 1918 | 1389 | 1578 | 1148 | 1194 | 904 | 1082 | 1714 | 1345 | 2341 | 1847 | 1816 | Troap |
| Trp53 | 6343 | 6679 | 6680 | 5901 | 5367 | 5617 | 5650 | 4437 | 5849 | 5386 | 5229 | 6510 | 6269 | 6909 | 5397 | 6104 | 6354 | 5790 | 6078 | 5071 | 5601 | 4812 | 5467 | 6499 | 5348 | 8083 | 6058 | 6998 | Trp53 |
| Trp53bp1 | 27501 | 29020 | 20128 | 28542 | 32434 | 33036 | 32913 | 37063 | 32118 | 31670 | 29428 | 32851 | 31541 | 33459 | 30203 | 24489 | 24656 | 27300 | 30209 | 33285 | 33943 | 33906 | 29093 | 29154 | 35861 | 21645 | 34092 | 31018 | Trp53bp1 |
| Trp53bp2 | 6389 | 6841 | 6918 | 7878 | 7157 | 6702 | 7052 | 4566 | 4788 | 5285 | 5028 | 6281 | 5457 | 5802 | 6173 | 6773 | 7528 | 7386 | 7548 | 6080 | 7157 | 5076 | 4603 | 5670 | 5609 | 5800 | 5241 | 5599 | Trp53bp2 |
| Trp53i11 | 6798 | 9589 | 12365 | 10519 | 11751 | 10202 | 9430 | 8413 | 9251 | 12137 | 9339 | 15168 | 13710 | 12927 | 7635 | 9282 | 13220 | 9902 | 12409 | 8358 | 9421 | 13179 | 9255 | 13429 | 13433 | 14290 | 13300 | 13945 | Trp53i11 |
| Trp53i13 | 1276 | 1577 | 1520 | 1559 | 1211 | 1265 | 1171 | 542 | 822 | 596 | 871 | 805 | 790 | 631 | 1242 | 1423 | 1477 | 1687 | 1274 | 1269 | 1043 | 617 | 770 | 805 | 636 | 1140 | 741 | 708 | Trp53i13 |
| Trp53inp1 | 12907 | 10230 | 9785 | 10940 | 10692 | 10984 | 12783 | 10952 | 8525 | 7150 | 6970 | 7913 | 7413 | 7902 | 14223 | 12272 | 10768 | 11727 | 10756 | 13862 | 11745 | 8599 | 8695 | 7385 | 8658 | 7143 | 7524 | 7449 | Trp53inp1 |
| Trp53inp2 | 7315 | 8672 | 10554 | 10729 | 12165 | 10006 | 10471 | 13996 | 13235 | 16381 | 14785 | 16694 | 16406 | 17222 | 9119 | 9078 | 11724 | 11113 | 11967 | 10991 | 11381 | 19766 | 13621 | 16372 | 18696 | 14646 | 15301 | 17565 | Trp53inp2 |
| Trp53rka | 1924 | 2059 | 2478 | 2278 | 1914 | 2008 | 1931 | 1562 | 1541 | 1703 | 1754 | 1947 | 1926 | 1822 | 1929 | 2139 | 2225 | 2148 | 2127 | 1861 | 1827 | 1549 | 1584 | 1931 | 1844 | 2349 | 2040 | 2035 | Trp53rka |
| Trp53rkb | 758 | 819 | 826 | 932 | 749 | 825 | 798 | 552 | 524 | 570 | 599 | 683 | 609 | 626 | 767 | 727 | 804 | 843 | 874 | 718 | 840 | 618 | 511 | 578 | 656 | 672 | 633 | 652 | Trp53rkb |
| Trp53tg5 | 0 | 0 | 0 | 0 | 0 | 0 | 0 | 1 | 0 | 0 | 0 | 0 | 0 | 0 | 1 | 0 | 0 | 0 | 0 | 0 | 0 | 0 | 1 | 0 | 0 | 0 | 0 | 0 | Trp53tg5 |
| Trp63 | 976 | 284 | 277 | 18 | 33 | 230 | 4 | 931 | 1495 | 591 | 710 | 257 | 370 | 252 | 363 | 622 | 64 | 253 | 32 | 21 | 4 | 47 | 1590 | 503 | 92 | 223 | 237 | 273 | Trp63 |
| Trp73 | 469 | 516 | 643 | 730 | 637 | 673 | 818 | 752 | 934 | 876 | 592 | 1422 | 802 | 1243 | 414 | 510 | 689 | 584 | 877 | 627 | 752 | 568 | 1011 | 1151 | 1097 | 1458 | 1005 | 1151 | Trp73 |
| Trpa1 | 102 | 78 | 69 | 64 | 48 | 44 | 70 | 54 | 73 | 45 | 44 | 97 | 57 | 71 | 92 | 70 | 51 | 43 | 49 | 100 | 73 | 42 | 45 | 52 | 40 | 90 | 86 | 70 | Trpa1 |
| Trpc1 | 1646 | 1412 | 1415 | 1648 | 1730 | 1680 | 1673 | 1073 | 981 | 1021 | 883 | 859 | 882 | 818 | 1576 | 1481 | 1525 | 1593 | 1637 | 1556 | 1783 | 1059 | 940 | 874 | 982 | 747 | 815 | 812 | Trpc1 |
| Trpc3 | 375 | 380 | 500 | 401 | 638 | 404 | 437 | 611 | 629 | 654 | 806 | 476 | 589 | 474 | 389 | 424 | 368 | 374 | 494 | 424 | 445 | 794 | 687 | 556 | 632 | 555 | 553 | 545 | Trpc3 |

| | | | | | | | | | | | | | | | | | | | | | | | | | | | | | |
|---|---|---|---|---|---|---|---|---|---|---|---|---|---|---|---|---|---|---|---|---|---|---|---|---|---|---|---|---|---|
| Trpc4 | 314 | 273 | 325 | 300 | 358 | 386 | 355 | 401 | 485 | 380 | 338 | 219 | 206 | 266 | 286 | 277 | 318 | 270 | 398 | 400 | 377 | 470 | 300 | 324 | 433 | 175 | 263 | 203 | Trpc4 |
| Trpc4ap | 11259 | 12295 | 11230 | 11938 | 11757 | 11644 | 11599 | 10810 | 11402 | 10899 | 10707 | 11339 | 11058 | 11309 | 10789 | 11459 | 11503 | 11739 | 12022 | 10791 | 12170 | 11608 | 11563 | 10783 | 11806 | 10874 | 10712 | 10993 | Trpc4ap |
| Trpc5 | 65 | 65 | 24 | 76 | 158 | 131 | 121 | 165 | 218 | 131 | 218 | 68 | 144 | 103 | 99 | 64 | 45 | 80 | 82 | 157 | 142 | 177 | 215 | 67 | 147 | 5 | 74 | 89 | Trpc5 |
| Trpc5os | 24 | 8 | 27 | 24 | 19 | 31 | 48 | 1 | 4 | 5 | 1 | 1 | 5 | 4 | 13 | 21 | 17 | 18 | 19 | 19 | 21 | 0 | 3 | 5 | 2 | 14 | 3 | 4 | Trpc5os |
| Trpc6 | 63 | 38 | 65 | 10 | 57 | 57 | 30 | 85 | 57 | 43 | 66 | 73 | 95 | 76 | 41 | 65 | 51 | 21 | 39 | 70 | 24 | 45 | 78 | 87 | 63 | 123 | 108 | 89 | Trpc6 |
| Trpc7 | 39 | 49 | 13 | 80 | 145 | 97 | 136 | 120 | 129 | 151 | 116 | 78 | 58 | 84 | 54 | 23 | 70 | 109 | 149 | 130 | 109 | 197 | 104 | 116 | 194 | 75 | 63 | 65 | Trpc7 |
| Trpm1 | 103 | 16 | 16 | 5 | 20 | 17 | 9 | 46 | 75 | 62 | 68 | 63 | 45 | 57 | 24 | 24 | 9 | 23 | 21 | 19 | 9 | 50 | 76 | 61 | 47 | 74 | 44 | 49 | Trpm1 |
| Trpm2 | 129 | 148 | 82 | 188 | 398 | 269 | 275 | 177 | 264 | 280 | 239 | 163 | 243 | 188 | 173 | 188 | 192 | 325 | 294 | 235 | 290 | 251 | 223 | 203 | 346 | 79 | 156 | 116 | Trpm2 |
| Trpm3 | 1517 | 1474 | 939 | 1479 | 2931 | 2631 | 2192 | 4036 | 4397 | 3933 | 3904 | 3804 | 4196 | 2740 | 1940 | 2087 | 1048 | 1801 | 1941 | 3028 | 2717 | 4314 | 7035 | 2736 | 3549 | 2011 | 2664 | 2718 | Trpm3 |
| Trpm4 | 2036 | 2428 | 2595 | 2977 | 2369 | 2298 | 2271 | 929 | 1373 | 1191 | 1593 | 968 | 1344 | 930 | 1998 | 2433 | 2525 | 2972 | 2491 | 1911 | 2228 | 1123 | 1435 | 1167 | 1075 | 1034 | 1046 | 974 | Trpm4 |
| Trpm5 | 54 | 51 | 30 | 79 | 106 | 112 | 53 | 289 | 597 | 416 | 1152 | 177 | 236 | 131 | 63 | 69 | 61 | 108 | 65 | 86 | 68 | 508 | 1045 | 344 | 325 | 157 | 142 | 112 | Trpm5 |
| Trpm6 | 135 | 108 | 105 | 79 | 57 | 89 | 85 | 89 | 93 | 94 | 97 | 108 | 115 | 92 | 115 | 116 | 81 | 53 | 54 | 93 | 63 | 82 | 111 | 65 | 87 | 56 | 87 | 94 | Trpm6 |
| Trpm7 | 15169 | 10263 | 8777 | 10083 | 10488 | 12866 | 12553 | 13133 | 12353 | 10295 | 11772 | 11457 | 11397 | 11797 | 14249 | 12833 | 9434 | 10230 | 10357 | 12270 | 12173 | 10910 | 11863 | 10119 | 10749 | 8984 | 12270 | 10993 | Trpm7 |
| Trpm8 | 0 | 1 | 1 | 1 | 2 | 6 | 10 | 4 | 9 | 8 | 7 | 5 | 10 | 4 | 9 | 2 | 8 | 0 | 7 | 10 | 4 | 7 | 6 | 5 | 9 | 3 | 7 | 5 | Trpm8 |
| Trps1 | 3678 | 2179 | 2518 | 2282 | 2192 | 2431 | 2372 | 6582 | 6286 | 4313 | 4556 | 3301 | 3512 | 3389 | 2763 | 2473 | 2231 | 2024 | 2293 | 2658 | 2237 | 3212 | 6455 | 3729 | 2977 | 3652 | 3741 | 3712 | Trps1 |
| Trpt1 | 453 | 537 | 534 | 547 | 502 | 522 | 540 | 399 | 419 | 440 | 454 | 436 | 449 | 421 | 471 | 458 | 526 | 548 | 570 | 506 | 526 | 391 | 428 | 490 | 471 | 513 | 427 | 425 | Trpt1 |
| Trpv1 | 9 | 10 | 0 | 24 | 71 | 6 | 15 | 33 | 86 | 53 | 147 | 4 | 19 | 0 | 18 | 9 | 9 | 46 | 7 | 7 | 5 | 122 | 44 | 2 | 61 | 1 | 2 | 4 | Trpv1 |
| Trpv2 | 288 | 315 | 306 | 261 | 479 | 386 | 311 | 689 | 718 | 693 | 670 | 503 | 708 | 502 | 319 | 323 | 255 | 297 | 391 | 345 | 317 | 891 | 733 | 648 | 727 | 488 | 621 | 465 | Trpv2 |
| Trpv3 | 8 | 6 | 5 | 0 | 11 | 2 | 6 | 18 | 35 | 32 | 42 | 17 | 28 | 9 | 9 | 9 | 1 | 1 | 5 | 2 | 2 | 30 | 46 | 29 | 24 | 9 | 23 | 9 | Trpv3 |
| Trpv4 | 216 | 156 | 166 | 170 | 142 | 168 | 94 | 121 | 257 | 315 | 421 | 61 | 81 | 47 | 154 | 206 | 156 | 164 | 140 | 151 | 144 | 234 | 230 | 245 | 109 | 134 | 43 | 62 | Trpv4 |
| Trpv5 | 0 | 3 | 0 | 3 | 2 | 0 | 4 | 7 | 5 | 0 | 3 | 0 | 6 | 2 | 0 | 0 | 0 | 3 | 4 | 0 | 2 | 3 | 6 | 1 | 3 | 7 | 2 | 2 | Trpv5 |
| Trpv6 | 3 | 10 | 2 | 8 | 2 | 2 | 0 | 21 | 21 | 8 | 35 | 4 | 18 | 2 | 9 | 1 | 0 | 4 | 2 | 0 | 1 | 7 | 16 | 6 | 6 | 14 | 2 | 0 | Trpv6 |
| Trrap | 12824 | 14537 | 10823 | 14549 | 16257 | 16796 | 14937 | 16618 | 19997 | 22132 | 21136 | 24539 | 23157 | 22264 | 11954 | 14164 | 12758 | 14922 | 16377 | 13427 | 16292 | 18939 | 18272 | 17539 | 23384 | 13580 | 23835 | 20595 | Trrap |
| Trub1 | 2220 | 1988 | 1918 | 2064 | 1955 | 2108 | 2101 | 1993 | 2086 | 2192 | 2348 | 2202 | 2294 | 2201 | 2027 | 2088 | 2118 | 2092 | 1859 | 2010 | 2061 | 2134 | 2137 | 2260 | 2036 | 2003 | 2020 | 2169 | Trub1 |
| Trub2 | 2718 | 2566 | 1852 | 2157 | 2488 | 2531 | 2347 | 3053 | 3282 | 2915 | 2944 | 3799 | 3759 | 3336 | 2589 | 2375 | 2016 | 2182 | 2472 | 2641 | 2422 | 2650 | 3006 | 2897 | 3499 | 3205 | 3934 | 3640 | Trub2 |
| Try5 | 1 | 2 | 0 | 0 | 3 | 12 | 1 | 4 | 2 | 0 | 0 | 11 | 10 | 0 | 1 | 2 | 0 | 3 | 1 | 1 | 6 | 0 | 2 | 3 | 5 | 7 | 0 | 4 | Try5 |
| Tsacc | 55 | 46 | 63 | 43 | 44 | 37 | 45 | 99 | 77 | 89 | 83 | 92 | 75 | 82 | 50 | 29 | 42 | 36 | 51 | 55 | 61 | 85 | 77 | 96 | 71 | 63 | 78 | 85 | Tsacc |
| Tsc1 | 8945 | 9792 | 7719 | 9120 | 10301 | 10005 | 10017 | 9180 | 8725 | 8965 | 8595 | 8845 | 8444 | 9041 | 9126 | 8528 | 8553 | 9458 | 10111 | 9562 | 10705 | 9463 | 8054 | 8329 | 10127 | 5740 | 8937 | 8395 | Tsc1 |
| Tsc2 | 11532 | 14024 | 12648 | 14799 | 13484 | 12985 | 13037 | 8200 | 10180 | 11236 | 10645 | 12079 | 10720 | 11050 | 10722 | 13785 | 14002 | 15033 | 14643 | 10413 | 13887 | 11448 | 9482 | 11205 | 11615 | 9285 | 10268 | 10619 | Tsc2 |
| Tsc22d1 | 22231 | 22984 | 24814 | 24968 | 27956 | 23570 | 25781 | 35457 | 34199 | 36981 | 36160 | 36229 | 35249 | 36050 | 23488 | 22792 | 25515 | 25431 | 26707 | 25150 | 26026 | 41271 | 34575 | 37194 | 37466 | 36759 | 34070 | 37193 | Tsc22d1 |
| Tsc22d2 | 6518 | 5910 | 7378 | 9876 | 7972 | 8624 | 9313 | 5163 | 4412 | 4747 | 5147 | 5238 | 5559 | 5619 | 7152 | 7048 | 7883 | 10045 | 7925 | 6943 | 8956 | 6497 | 4622 | 5389 | 5750 | 5025 | 5138 | 5372 | Tsc22d2 |
| Tsc22d3 | 1794 | 2141 | 2540 | 1891 | 2073 | 1768 | 1532 | 3038 | 2722 | 3055 | 2873 | 3010 | 3161 | 2912 | 2022 | 2221 | 2265 | 1775 | 1889 | 2091 | 1687 | 2812 | 2848 | 2975 | 2824 | 3089 | 3233 | 3273 | Tsc22d3 |
| Tsc22d4 | 1982 | 2550 | 2807 | 2678 | 2518 | 2426 | 1880 | 1668 | 2059 | 2103 | 2548 | 2044 | 2002 | 1976 | 1865 | 2414 | 2529 | 2809 | 2557 | 1772 | 2255 | 1942 | 2277 | 2569 | 2023 | 2651 | 1816 | 1765 | Tsc22d4 |
| Tsen15 | 1353 | 1117 | 1021 | 1050 | 825 | 934 | 988 | 1290 | 1345 | 1120 | 1329 | 1303 | 1169 | 1378 | 1254 | 1309 | 1029 | 995 | 919 | 942 | 1028 | 1110 | 1364 | 1355 | 1069 | 1519 | 1203 | 1351 | Tsen15 |
| Tsen2 | 809 | 941 | 779 | 854 | 915 | 972 | 980 | 1301 | 1441 | 1444 | 1335 | 1304 | 1234 | 1400 | 793 | 794 | 810 | 913 | 983 | 897 | 1029 | 1198 | 1398 | 1454 | 1353 | 1313 | 1384 | 1390 | Tsen2 |
| Tsen34 | 6284 | 7159 | 6699 | 6470 | 6151 | 5829 | 5302 | 4598 | 4961 | 4757 | 4415 | 4638 | 4351 | 4280 | 5813 | 6196 | 6953 | 6478 | 6917 | 5882 | 5722 | 4132 | 5170 | 5046 | 4757 | 5583 | 4912 | 4423 | Tsen34 |
| Tsen54 | 1789 | 2352 | 2063 | 2200 | 1701 | 1724 | 1694 | 1098 | 1098 | 1016 | 1038 | 1086 | 1181 | 1077 | 1817 | 2043 | 2136 | 1944 | 1876 | 1831 | 1796 | 1116 | 1117 | 1191 | 1135 | 1321 | 1227 | 1102 | Tsen54 |
| Tsfm | 1788 | 1830 | 2139 | 1865 | 1516 | 1608 | 1672 | 1070 | 1120 | 1143 | 1169 | 1426 | 1484 | 1597 | 1725 | 1667 | 2023 | 1782 | 1723 | 1540 | 1505 | 1191 | 1082 | 1498 | 1276 | 1716 | 1497 | 1683 | Tsfm |
| Tsg101 | 7987 | 7118 | 6765 | 7339 | 7823 | 7557 | 8914 | 10898 | 8753 | 9144 | 8229 | 9571 | 8537 | 10586 | 8780 | 6646 | 7345 | 7668 | 7983 | 8254 | 8658 | 9910 | 8804 | 9599 | 9896 | 10011 | 9836 | 10370 | Tsg101 |
| Tsga10 | 1772 | 1750 | 1879 | 2108 | 2024 | 2091 | 2145 | 3718 | 2798 | 2867 | 2696 | 2791 | 2840 | 2972 | 2243 | 1914 | 1902 | 1913 | 1981 | 2586 | 2028 | 3319 | 2972 | 2627 | 3240 | 2186 | 2967 | 2860 | Tsga10 |
| Tsga10ip | 3 | 3 | 2 | 2 | 6 | 2 | 0 | 6 | 4 | 2 | 2 | 0 | 16 | 1 | 3 | 5 | 1 | 1 | 0 | 0 | 0 | 3 | 9 | 7 | 9 | 3 | 1 | 2 | Tsga10ip |
| Tsga13 | 27 | 31 | 18 | 50 | 27 | 30 | 47 | 22 | 17 | 38 | 28 | 14 | 31 | 20 | 32 | 19 | 41 | 42 | 47 | 23 | 41 | 53 | 23 | 18 | 46 | 39 | 17 | 19 | Tsga13 |
| Tshb | 0 | 0 | 0 | 4 | 7 | 3 | 1 | 2 | 9 | 0 | 2 | 2 | 7 | 4 | 0 | 2 | 0 | 0 | 0 | 3 | 0 | 18 | 9 | 0 | 0 | 0 | 0 | 0 | Tshb |
| Tshr | 158 | 161 | 74 | 222 | 244 | 194 | 304 | 76 | 121 | 80 | 92 | 89 | 48 | 97 | 140 | 67 | 122 | 193 | 245 | 247 | 235 | 43 | 39 | 112 | 102 | 80 | 57 | 65 | Tshr |
| Tshz1 | 3599 | 3296 | 3985 | 4971 | 6073 | 4563 | 4693 | 3657 | 4107 | 5879 | 6484 | 4625 | 4124 | 4359 | 3440 | 3401 | 4757 | 4655 | 5195 | 4084 | 4836 | 4478 | 5155 | 5080 | 4376 | 5540 | 3750 | 4195 | Tshz1 |
| Tshz2 | 1491 | 1663 | 1434 | 2342 | 4336 | 2594 | 2282 | 2862 | 3631 | 4419 | 5432 | 2211 | 2456 | 2323 | 1921 | 1610 | 1995 | 2535 | 2536 | 2295 | 2433 | 4009 | 4499 | 3001 | 3217 | 2224 | 1988 | 1893 | Tshz2 |
| Tshz3 | 1433 | 1461 | 1860 | 2174 | 2738 | 2127 | 2044 | 1281 | 1529 | 1942 | 2041 | 1157 | 1613 | 1269 | 1667 | 1466 | 1926 | 2051 | 2346 | 1890 | 2137 | 1790 | 1619 | 1262 | 1372 | 1160 | 1029 | 1191 | Tshz3 |
| Tsks | 2 | 8 | 5 | 12 | 0 | 13 | 8 | 6 | 2 | 0 | 7 | 3 | 2 | 2 | 1 | 8 | 9 | 1 | 15 | 13 | 4 | 3 | 6 | 7 | 4 | 3 | 10 | 1 | Tsks |
| Tsku | 1270 | 1496 | 1293 | 1123 | 1035 | 1066 | 851 | 990 | 1380 | 1235 | 1484 | 1055 | 1118 | 1066 | 1192 | 1463 | 1183 | 1109 | 966 | 1072 | 867 | 1074 | 1626 | 1273 | 1123 | 1353 | 1031 | 1108 | Tsku |
| Tslp | 50 | 22 | 31 | 28 | 27 | 23 | 25 | 15 | 21 | 21 | 56 | 10 | 14 | 6 | 30 | 37 | 24 | 33 | 9 | 27 | 23 | 37 | 36 | 22 | 29 | 16 | 16 | 14 | Tslp |
| Tsn | 16252 | 15512 | 17534 | 15745 | 15820 | 15405 | 17693 | 17009 | 14504 | 14747 | 13960 | 16854 | 16125 | 17706 | 16866 | 14748 | 16979 | 15334 | 15999 | 18399 | 16492 | 15014 | 14022 | 16421 | 16798 | 18109 | 17872 | 17673 | Tsn |
| Tsnax | 8772 | 8415 | 8861 | 8319 | 8617 | 8046 | 7871 | 9900 | 9141 | 8990 | 9219 | 9208 | 8569 | 8987 | 8668 | 7926 | 8590 | 8270 | 8457 | 8784 | 7973 | 9703 | 9159 | 9358 | 9674 | 8781 | 9588 | 8942 | Tsnax |
| Tsnaxip1 | 42 | 41 | 23 | 42 | 34 | 37 | 49 | 49 | 31 | 32 | 46 | 91 | 94 | 33 | 50 | 72 | 27 | 35 | 34 | 63 | 60 | 60 | 94 | 59 | 63 | 34 | 41 | 21 | Tsnaxip1 |
| Tspan1 | 14 | 2 | 0 | 6 | 0 | 4 | 0 | 6 | 7 | 23 | 8 | 2 | 6 | 2 | 3 | 5 | 0 | 1 | 7 | 3 | 0 | 3 | 16 | 17 | 5 | 7 | 0 | 1 | Tspan1 |
| Tspan10 | 3 | 0 | 2 | 0 | 0 | 2 | 1 | 5 | 5 | 4 | 2 | 2 | 12 | 3 | 0 | 2 | 0 | 1 | 0 | 0 | 0 | 3 | 11 | 2 | 3 | 0 | 2 | 2 | Tspan10 |
| Tspan11 | 715 | 878 | 769 | 654 | 505 | 735 | 473 | 1376 | 1388 | 1549 | 1310 | 1565 | 1975 | 1354 | 806 | 962 | 802 | 546 | 744 | 656 | 556 | 1320 | 1439 | 1452 | 1401 | 1520 | 1549 | 1599 | Tspan11 |
| Tspan12 | 1726 | 1478 | 1835 | 1850 | 2052 | 1713 | 2068 | 2259 | 1893 | 1795 | 1793 | 1934 | 1937 | 1986 | 1783 | 1951 | 1814 | 1771 | 1867 | 2309 | 1889 | 1788 | 1977 | 1909 | 1807 | 2344 | 1792 | 2086 | Tspan12 |
| Tspan13 | 10634 | 10267 | 10402 | 12450 | 13857 | 11937 | 15099 | 12002 | 9489 | 10510 | 9037 | 9570 | 9044 | 10395 | 12274 | 9729 | 12154 | 12687 | 13503 | 14020 | 13949 | 12567 | 10037 | 9895 | 11353 | 9423 | 9749 | 10579 | Tspan13 |
| Tspan14 | 5382 | 5914 | 5683 | 4929 | 4722 | 4855 | 4488 | 5490 | 5621 | 5392 | 4907 | 5371 | 5261 | 5510 | 5300 | 5300 | 5299 | 4743 | 4842 | 4812 | 4192 | 4993 | 5548 | 5629 | 5226 | 6011 | 5487 | 5432 | Tspan14 |
| Tspan15 | 1857 | 1830 | 1817 | 1825 | 1888 | 1925 | 1764 | 707 | 818 | 717 | 708 | 673 | 802 | 740 | 1826 | 1890 | 1785 | 1891 | 1884 | 1749 | 1885 | 673 | 802 | 717 | 706 | 974 | 736 | 757 | Tspan15 |
| Tspan17 | 1087 | 1566 | 1880 | 1851 | 1979 | 1582 | 1446 | 459 | 609 | 628 | 609 | 584 | 730 | 533 | 1182 | 1386 | 1965 | 1924 | 1854 | 1522 | 1411 | 769 | 567 | 502 | 603 | 452 | 580 | 549 | Tspan17 |
| Tspan18 | 2662 | 3512 | 4351 | 3637 | 3892 | 3687 | 3277 | 3907 | 4097 | 5189 | 5341 | 6085 | 6133 | 5605 | 3297 | 3588 | 4297 | 3716 | 4032 | 2853 | 3184 | 5947 | 4979 | 5544 | 5160 | 6810 | 5649 | 6106 | Tspan18 |
| Tspan2 | 2285 | 1695 | 2374 | 1675 | 1894 | 1923 | 1760 | 1319 | 1061 | 1064 | 1129 | 1061 | 1247 | 1226 | 2258 | 2110 | 2042 | 1695 | 1655 | 1826 | 1799 | 1191 | 1038 | 1039 | 1030 | 1051 | 1216 | 1196 | Tspan2 |
| Tspan3 | 14237 | 14694 | 14939 | 13595 | 13516 | 13131 | 15559 | 15218 | 14170 | 13250 | 13678 | 14844 | 14937 | 16564 | 14960 | 14130 | 14686 | 13606 | 14012 | 14172 | 15134 | 14313 | 15055 | 14789 | 13805 | 15983 | 15218 | 17857 | Tspan3 |
| Tspan31 | 5571 | 6157 | 5767 | 5129 | 4734 | 5064 | 5040 | 5264 | 5187 | 4704 | 4966 | 5421 | 5433 | 5194 | 5588 | 5396 | 5689 | 4802 | 5421 | 5233 | 5098 | 4513 | 5223 | 5405 | 4913 | 6264 | 5817 | 5719 | Tspan31 |
| Tspan32 | 55 | 46 | 42 | 22 | 12 | 14 | 19 | 59 | 51 | 27 | 33 | 54 | 94 | 70 | 44 | 53 | 34 | 21 | 9 | 13 | 17 | 37 | 43 | 33 | 30 | 101 | 131 | 125 | Tspan32 |
| Tspan33 | 1210 | 1423 | 1740 | 1578 | 1478 | 1373 | 1599 | 313 | 330 | 420 | 378 | 394 | 561 | 488 | 1228 | 1234 | 1672 | 1454 | 1583 | 1301 | 1403 | 371 | 407 | 389 | 388 | 478 | 547 | 749 | Tspan33 |
| Tspan4 | 1203 | 1541 | 1311 | 1083 | 1136 | 1035 | 859 | 730 | 923 | 807 | 1129 | 715 | 940 | 585 | 1241 | 1460 | 1167 | 1231 | 1001 | 1004 | 912 | 824 | 1060 | 818 | 726 | 871 | 850 | 724 | Tspan4 |
| Tspan5 | 3031 | 2936 | 3269 | 3911 | 5028 | 4228 | 4386 | 3915 | 3841 | 4422 | 4033 | 3706 | 3628 | 3712 | 3374 | 3146 | 4041 | 4355 | 4580 | 4127 | 4460 | 4590 | 4010 | 4264 | 4781 | 3788 | 3455 | 3762 | Tspan5 |
| Tspan6 | 8459 | 6332 | 7085 | 6421 | 7069 | 7325 | 7611 | 10476 | 8998 | 6912 | 6730 | 7722 | 8077 | 8055 | 8975 | 7739 | 6377 | 6753 | 7125 | 9164 | 7085 | 8955 | 9295 | 7701 | 7786 | 8922 | 8552 | 8314 | Tspan6 |
| Tspan7 | 5060 | 5028 | 4955 | 6402 | 7755 | 6301 | 7505 | 5048 | 5650 | 6059 | 5332 | 5189 | 4324 | 5160 | 4989 | 4316 | 5909 | 6802 | 7712 | 6538 | 7638 | 6975 | 5515 | 5802 | 6582 | 5671 | 4522 | 4808 | Tspan7 |
| Tspan8 | 25 | 14 | 20 | 10 | 10 | 10 | 7 | 48 | 77 | 53 | 75 | 20 | 43 | 23 | 18 | 27 | 6 | 36 | 3 | 1 | 2 | 30 | 117 | 40 | 32 | 14 | 30 | 33 | Tspan8 |
| Tspan9 | 4918 | 5236 | 5305 | 4134 | 3714 | 4038 | 2834 | 3052 | 4254 | 3404 | 5183 | 2875 | 3856 | 2517 | 4596 | 6498 | 4275 | 4388 | 3574 | 2807 | 3064 | 3265 | 5129 | 3170 | 2748 | 3242 | 2973 | 2749 | Tspan9 |
| Tspear | 85 | 126 | 130 | 90 | 67 | 69 | 73 | 48 | 71 | 56 | 46 | 52 | 75 | 68 | 103 | 117 | 82 | 65 | 86 | 91 | 73 | 32 | 69 | 56 | 51 | 79 | 72 | 90 | Tspear |
| Tspo | 230 | 280 | 239 | 159 | 191 | 174 | 122 | 153 | 215 | 172 | 156 | 172 | 261 | 163 | 211 | 301 | 245 | 193 | 178 | 232 | 142 | 109 | 238 | 219 | 156 | 233 | 260 | 224 | Tspo |
| Tspo2 | 0 | 6 | 0 | 1 | 9 | 1 | 3 | 0 | 2 | 1 | 13 | 1 | 6 | 7 | 20 | 1 | 0 | 3 | 0 | 2 | 0 | 3 | 17 | 2 | 7 | 1 | 0 | 0 | Tspo2 |
| Tspoap1 | 4895 | 6934 | 6230 | 12128 | 15015 | 12868 | 12279 | 6909 | 7154 | 9365 | 7515 | 9291 | 7994 | 9006 | 5843 | 5428 | 9845 | 12739 | 14158 | 11532 | 13232 | 11782 | 5965 | 9134 | 11650 | 4998 | 8108 | 7819 | Tspoap1 |
| Tspyl1 | 11764 | 11611 | 12490 | 14059 | 13626 | 12429 | 14056 | 8376 | 7710 | 8907 | 8613 | 7455 | 7749 | 8459 | 12601 | 11898 | 12837 | 14079 | 13377 | 12131 | 14283 | 9757 | 8041 | 8417 | 9458 | 7985 | 7510 | 8520 | Tspyl1 |
| Tspyl2 | 4538 | 4800 | 5279 | 6124 | 5767 | 5464 | 5739 | 4896 | 4514 | 5398 | 4978 | 5513 | 4982 | 5547 | 4531 | 4252 | 5641 | 5778 | 5764 | 4826 | 5803 | 5798 | 4221 | 5515 | 5641 | 5336 | 5011 | 5433 | Tspyl2 |

| | | | | | | | | | | | | | | | | | | | | | | | | | | | | | |
|---|---|---|---|---|---|---|---|---|---|---|---|---|---|---|---|---|---|---|---|---|---|---|---|---|---|---|---|---|---|
| Tspyl3 | 597 | 460 | 474 | 630 | 907 | 906 | 716 | 1196 | 1402 | 1449 | 1353 | 924 | 942 | 964 | 510 | 491 | 498 | 632 | 786 | 673 | 736 | 1624 | 1449 | 1170 | 1370 | 1034 | 865 | 888 | Tspyl3 |
| Tspyl4 | 13024 | 15088 | 11068 | 16111 | 18411 | 15917 | 17914 | 14044 | 12122 | 12303 | 10812 | 9779 | 9559 | 10120 | 15725 | 12544 | 14147 | 16956 | 16971 | 19024 | 18449 | 15264 | 12189 | 10458 | 13133 | 7967 | 9473 | 9664 | Tspyl4 |
| Tspyl5 | 1523 | 1500 | 1501 | 1921 | 2392 | 2172 | 2227 | 1613 | 1506 | 1608 | 1453 | 1249 | 1170 | 1253 | 1888 | 1399 | 1615 | 1873 | 2205 | 2100 | 2282 | 1955 | 1621 | 1393 | 1720 | 1133 | 1198 | 1188 | Tspyl5 |
| Tsr1 | 5406 | 4971 | 4906 | 4484 | 4183 | 4173 | 4748 | 5155 | 4552 | 4161 | 4535 | 5133 | 4766 | 5321 | 5414 | 4759 | 4317 | 4220 | 4256 | 4827 | 4524 | 3880 | 4568 | 4712 | 4349 | 5323 | 5163 | 5285 | Tsr1 |
| Tsr2 | 2759 | 2566 | 2557 | 2566 | 2234 | 2513 | 2638 | 2696 | 2455 | 2464 | 2361 | 2476 | 2499 | 2795 | 2718 | 2282 | 2428 | 2619 | 2386 | 2726 | 2455 | 2369 | 2359 | 2527 | 2507 | 2770 | 2629 | 2796 | Tsr2 |
| Tsr3 | 1631 | 1987 | 2020 | 1855 | 1634 | 1624 | 1701 | 1674 | 1679 | 1717 | 1767 | 1909 | 2031 | 2043 | 1611 | 1682 | 1938 | 1783 | 1769 | 1851 | 1692 | 1594 | 1587 | 2004 | 1830 | 2381 | 2188 | 1952 | Tsr3 |
| Tssc4 | 2046 | 2632 | 2436 | 2394 | 2110 | 2196 | 1893 | 2140 | 2512 | 2455 | 2170 | 2682 | 2392 | 2624 | 1917 | 2164 | 2425 | 2099 | 2512 | 2178 | 2078 | 1965 | 2302 | 2672 | 2475 | 3228 | 2723 | 2655 | Tssc4 |
| Tssk1 | 5 | 13 | 6 | 4 | 9 | 13 | 9 | 10 | 12 | 14 | 8 | 10 | 9 | 13 | 11 | 8 | 15 | 18 | 9 | 7 | 13 | 8 | 13 | 9 | 10 | 5 | 15 | 8 | Tssk1 |
| Tssk2 | 0 | 2 | 1 | 0 | 1 | 0 | 0 | 2 | 0 | 0 | 0 | 0 | 0 | 0 | 4 | 2 | 0 | 0 | 4 | 0 | 0 | 0 | 1 | 0 | 0 | 3 | 1 | 0 | Tssk2 |
| Tssk3 | 2 | 1 | 2 | 12 | 11 | 4 | 3 | 0 | 2 | 1 | 2 | 4 | 1 | 0 | 3 | 2 | 2 | 2 | 2 | 4 | 4 | 2 | 3 | 1 | 5 | 0 | 1 | 0 | Tssk3 |
| Tssk4 | 232 | 183 | 191 | 157 | 186 | 190 | 158 | 127 | 148 | 141 | 200 | 128 | 124 | 144 | 171 | 206 | 197 | 157 | 136 | 165 | 159 | 84 | 131 | 135 | 111 | 136 | 117 | 139 | Tssk4 |
| Tssk5 | 1 | 0 | 0 | 0 | 0 | 0 | 0 | 0 | 0 | 0 | 0 | 0 | 0 | 1 | 0 | 0 | 0 | 0 | 0 | 0 | 0 | 0 | 1 | 0 | 0 | 0 | 0 | 0 | Tssk5 |
| Tssk6 | 12 | 15 | 15 | 18 | 11 | 8 | 3 | 13 | 7 | 7 | 9 | 13 | 10 | 12 | 8 | 12 | 25 | 8 | 8 | 6 | 13 | 5 | 11 | 7 | 15 | 11 | 23 | 8 | Tssk6 |
| Tst | 469 | 600 | 464 | 499 | 473 | 448 | 420 | 299 | 411 | 471 | 321 | 298 | 306 | 287 | 409 | 602 | 510 | 484 | 553 | 525 | 461 | 236 | 491 | 365 | 352 | 388 | 275 | 280 | Tst |
| Tstd1 | 145 | 180 | 272 | 212 | 209 | 165 | 235 | 173 | 160 | 228 | 189 | 215 | 179 | 249 | 141 | 132 | 314 | 198 | 279 | 189 | 216 | 180 | 168 | 279 | 187 | 270 | 229 | 295 | Tstd1 |
| Tstd2 | 4380 | 4730 | 4605 | 4283 | 3856 | 4074 | 4060 | 4523 | 4397 | 4116 | 4424 | 4804 | 4758 | 4987 | 4207 | 4377 | 4335 | 4012 | 4118 | 4151 | 4106 | 4131 | 4380 | 4428 | 4499 | 4984 | 5003 | 5165 | Tstd2 |
| Tstd3 | 824 | 690 | 1004 | 861 | 1043 | 903 | 963 | 993 | 802 | 923 | 838 | 1147 | 1084 | 1324 | 1092 | 826 | 985 | 813 | 953 | 1030 | 974 | 993 | 893 | 914 | 986 | 1048 | 1158 | 1311 | Tstd3 |
| Tsx | 58 | 55 | 39 | 124 | 59 | 70 | 83 | 35 | 22 | 24 | 43 | 38 | 42 | 44 | 55 | 58 | 82 | 123 | 78 | 88 | 105 | 38 | 20 | 27 | 60 | 37 | 13 | 38 | Tsx |
| Ttbk1 | 4297 | 6488 | 7362 | 9628 | 9517 | 7647 | 7238 | 4001 | 4374 | 5998 | 4818 | 5816 | 4152 | 5471 | 4444 | 4363 | 9153 | 9292 | 9563 | 6515 | 8453 | 6518 | 3563 | 6130 | 7232 | 4810 | 4936 | 4742 | Ttbk1 |
| Ttbk2 | 12768 | 10436 | 9471 | 13116 | 14933 | 15819 | 17154 | 9572 | 7700 | 8301 | 7983 | 7856 | 7731 | 8386 | 13351 | 11316 | 11167 | 14048 | 13695 | 14781 | 17377 | 9302 | 7476 | 6972 | 9716 | 5712 | 7842 | 7713 | Ttbk2 |
| Ttc1 | 4072 | 4129 | 4213 | 3992 | 3557 | 3753 | 3881 | 3284 | 3021 | 2827 | 2781 | 3017 | 3048 | 3104 | 4453 | 4171 | 4055 | 3873 | 3582 | 4198 | 3792 | 2704 | 3019 | 3087 | 3077 | 3196 | 3126 | 3318 | Ttc1 |
| Ttc12 | 664 | 734 | 642 | 554 | 525 | 568 | 453 | 391 | 340 | 325 | 332 | 374 | 459 | 322 | 761 | 746 | 599 | 538 | 503 | 519 | 529 | 391 | 447 | 239 | 309 | 355 | 408 | 426 | Ttc12 |
| Ttc13 | 5569 | 5916 | 5277 | 5585 | 5740 | 5851 | 5903 | 4528 | 4538 | 4678 | 4447 | 4727 | 4701 | 4557 | 5545 | 5673 | 5102 | 5798 | 5559 | 5667 | 6114 | 4777 | 4208 | 4359 | 5210 | 4365 | 4750 | 4727 | Ttc13 |
| Ttc14 | 16679 | 11863 | 9743 | 12709 | 12930 | 16090 | 16288 | 17363 | 14672 | 12080 | 13851 | 12917 | 13292 | 14081 | 17453 | 14871 | 10596 | 12544 | 11652 | 17915 | 16234 | 12552 | 14197 | 11016 | 12690 | 8937 | 13415 | 12504 | Ttc14 |
| Ttc16 | 142 | 172 | 141 | 266 | 187 | 141 | 156 | 77 | 90 | 61 | 76 | 111 | 111 | 85 | 129 | 175 | 158 | 240 | 233 | 140 | 189 | 114 | 95 | 73 | 73 | 57 | 93 | 78 | Ttc16 |
| Ttc17 | 2148 | 2110 | 1865 | 2188 | 2549 | 2331 | 2566 | 2945 | 3249 | 3090 | 2994 | 2777 | 2791 | 2911 | 2041 | 2077 | 2046 | 2328 | 2395 | 2127 | 2467 | 3290 | 3020 | 2845 | 3135 | 2105 | 2626 | 2676 | Ttc17 |
| Ttc19 | 3106 | 2611 | 2004 | 2296 | 2636 | 2776 | 2934 | 2507 | 2389 | 1907 | 2169 | 1574 | 1837 | 1959 | 3025 | 2843 | 2277 | 2359 | 2456 | 3193 | 3054 | 2356 | 2346 | 1664 | 1962 | 1542 | 1683 | 1661 | Ttc19 |
| Ttc21a | 185 | 233 | 102 | 200 | 255 | 172 | 149 | 223 | 297 | 306 | 236 | 253 | 314 | 182 | 186 | 274 | 141 | 168 | 178 | 223 | 174 | 246 | 410 | 232 | 195 | 147 | 210 | 191 | Ttc21a |
| Ttc21b | 5677 | 5663 | 4638 | 4472 | 4284 | 4437 | 4802 | 2920 | 2688 | 2357 | 2495 | 2325 | 2516 | 2696 | 5835 | 5416 | 4828 | 4635 | 4217 | 4981 | 4698 | 2301 | 2682 | 2264 | 2382 | 2325 | 2673 | 2582 | Ttc21b |
| Ttc22 | 0 | 10 | 10 | 7 | 20 | 17 | 6 | 33 | 66 | 81 | 62 | 23 | 75 | 43 | 7 | 6 | 20 | 14 | 24 | 10 | 11 | 69 | 61 | 53 | 60 | 42 | 56 | 37 | Ttc22 |
| Ttc23 | 2441 | 2246 | 2261 | 1735 | 1487 | 1691 | 1721 | 1904 | 2065 | 1701 | 2018 | 2079 | 2244 | 2087 | 2247 | 2323 | 1949 | 1822 | 1709 | 2060 | 1712 | 1293 | 2166 | 1915 | 1629 | 2133 | 2337 | 2438 | Ttc23 |
| Ttc23l | 6 | 4 | 0 | 5 | 2 | 1 | 1 | 4 | 2 | 5 | 2 | 1 | 7 | 7 | 3 | 6 | 1 | 2 | 2 | 7 | 3 | 3 | 3 | 4 | 6 | 0 | 5 | 0 | Ttc23l |
| Ttc24 | 25 | 3 | 1 | 7 | 7 | 0 | 0 | 38 | 32 | 49 | 28 | 0 | 6 | 2 | 3 | 6 | 1 | 6 | 0 | 0 | 0 | 10 | 39 | 78 | 0 | 76 | 0 | 0 | Ttc24 |
| Ttc26 | 3127 | 2611 | 2872 | 2496 | 2372 | 2545 | 2699 | 2049 | 1827 | 1588 | 1595 | 1829 | 1664 | 2035 | 3120 | 2896 | 2661 | 2322 | 2500 | 2934 | 2633 | 1649 | 1759 | 1764 | 1645 | 1811 | 1922 | 1975 | Ttc26 |
| Ttc27 | 1929 | 1715 | 1587 | 1325 | 1263 | 1302 | 1339 | 1976 | 1853 | 1612 | 1482 | 1673 | 1670 | 1749 | 1808 | 1732 | 1411 | 1339 | 1276 | 1413 | 1230 | 1472 | 1816 | 1668 | 1616 | 1785 | 1872 | 1969 | Ttc27 |
| Ttc28 | 6786 | 8813 | 8617 | 13457 | 16598 | 14011 | 12334 | 8929 | 12236 | 14964 | 13101 | 13533 | 11463 | 11882 | 7249 | 8231 | 11391 | 15141 | 15875 | 11184 | 14503 | 14634 | 10830 | 15299 | 16306 | 8165 | 10323 | 10281 | Ttc28 |
| Ttc29 | 18 | 32 | 22 | 12 | 29 | 27 | 19 | 35 | 32 | 22 | 19 | 31 | 49 | 16 | 20 | 46 | 52 | 36 | 51 | 39 | 21 | 32 | 54 | 23 | 31 | 18 | 31 | 30 | Ttc29 |
| Ttc3 | 54689 | 54795 | 36535 | 50511 | 63290 | 62868 | 66001 | 95979 | 82100 | 72759 | 67133 | 68460 | 65314 | 71582 | 64570 | 50659 | 45833 | 53820 | 57859 | 78466 | 65817 | 78408 | 80452 | 62846 | 80503 | 43541 | 69841 | 64743 | Ttc3 |
| Ttc30a1 | 1360 | 1114 | 1176 | 1092 | 1046 | 1136 | 1199 | 1196 | 1135 | 910 | 1071 | 1094 | 1102 | 998 | 1313 | 1234 | 1088 | 990 | 1120 | 1275 | 1086 | 1044 | 1138 | 1035 | 927 | 1191 | 1109 | 1176 | Ttc30a1 |
| Ttc30a2 | 118 | 116 | 111 | 110 | 112 | 129 | 134 | 64 | 71 | 106 | 51 | 55 | 79 | 63 | 140 | 114 | 144 | 194 | 124 | 135 | 150 | 80 | 67 | 74 | 65 | 57 | 71 | 63 | Ttc30a2 |
| Ttc30b | 851 | 796 | 760 | 699 | 909 | 801 | 826 | 1034 | 1035 | 855 | 844 | 791 | 920 | 782 | 1036 | 787 | 839 | 725 | 934 | 966 | 878 | 999 | 1196 | 780 | 699 | 794 | 734 | 775 | Ttc30b |
| Ttc32 | 119 | 106 | 191 | 110 | 126 | 120 | 112 | 519 | 391 | 435 | 457 | 472 | 457 | 600 | 137 | 146 | 129 | 89 | 127 | 126 | 135 | 341 | 364 | 409 | 404 | 482 | 433 | 558 | Ttc32 |
| Ttc33 | 1945 | 1632 | 1944 | 1953 | 1891 | 1746 | 1985 | 2700 | 2255 | 2123 | 2189 | 2111 | 2119 | 2366 | 2080 | 1932 | 1873 | 1976 | 1926 | 2473 | 1986 | 2288 | 2259 | 2180 | 2237 | 2356 | 2285 | 2279 | Ttc33 |
| Ttc34 | 12 | 10 | 1 | 11 | 9 | 9 | 2 | 11 | 40 | 36 | 21 | 29 | 36 | 22 | 12 | 7 | 1 | 0 | 2 | 4 | 4 | 25 | 45 | 21 | 21 | 22 | 14 | 17 | Ttc34 |
| Ttc36 | 9 | 12 | 22 | 12 | 18 | 21 | 20 | 4 | 12 | 16 | 26 | 9 | 10 | 16 | 8 | 11 | 19 | 14 | 22 | 22 | 23 | 15 | 6 | 20 | 15 | 10 | 14 | 9 | Ttc36 |
| Ttc37 | 6772 | 5859 | 4365 | 5329 | 5119 | 5777 | 5945 | 4716 | 4374 | 3535 | 3991 | 3545 | 3485 | 3764 | 6878 | 6201 | 4852 | 5290 | 4787 | 6530 | 5806 | 3335 | 4203 | 3192 | 3559 | 2632 | 3796 | 3500 | Ttc37 |
| Ttc38 | 594 | 511 | 556 | 481 | 453 | 446 | 380 | 546 | 615 | 608 | 686 | 484 | 591 | 442 | 553 | 604 | 452 | 518 | 401 | 456 | 399 | 429 | 760 | 605 | 546 | 580 | 465 | 436 | Ttc38 |
| Ttc39a | 111 | 76 | 72 | 33 | 49 | 56 | 50 | 99 | 115 | 93 | 152 | 106 | 119 | 100 | 86 | 83 | 58 | 45 | 51 | 46 | 38 | 65 | 102 | 68 | 74 | 136 | 117 | 126 | Ttc39a |
| Ttc39b | 2403 | 1752 | 1177 | 1951 | 2325 | 2593 | 2691 | 1846 | 1840 | 1786 | 1480 | 1218 | 1194 | 1208 | 2182 | 2009 | 1447 | 1841 | 2092 | 2787 | 2581 | 1140 | 1489 | 1674 | 1861 | 1074 | 992 | 910 | Ttc39b |
| Ttc39c | 798 | 670 | 743 | 710 | 1042 | 915 | 863 | 1207 | 1161 | 1154 | 1071 | 1031 | 1007 | 1043 | 810 | 759 | 717 | 774 | 851 | 790 | 937 | 1318 | 1093 | 1129 | 1278 | 1045 | 1031 | 1049 | Ttc39c |
| Ttc39d | 21 | 22 | 23 | 25 | 12 | 21 | 23 | 15 | 21 | 14 | 17 | 26 | 18 | 28 | 17 | 24 | 16 | 19 | 8 | 22 | 23 | 10 | 18 | 28 | 15 | 33 | 17 | 21 | Ttc39d |
| Ttc4 | 5052 | 5192 | 4893 | 4784 | 5053 | 4855 | 5393 | 6692 | 6385 | 5860 | 5556 | 5872 | 5584 | 5951 | 5203 | 4800 | 5295 | 4949 | 5037 | 5866 | 5115 | 6134 | 6040 | 5929 | 5890 | 6328 | 5848 | 6151 | Ttc4 |
| Ttc41 | 235 | 206 | 208 | 194 | 224 | 189 | 207 | 427 | 398 | 464 | 370 | 500 | 505 | 562 | 222 | 231 | 220 | 162 | 238 | 225 | 216 | 455 | 344 | 417 | 437 | 362 | 543 | 474 | Ttc41 |
| Ttc5 | 6667 | 6341 | 6781 | 6927 | 7216 | 7074 | 7005 | 6950 | 6813 | 6599 | 6241 | 7167 | 6417 | 7115 | 6550 | 6113 | 6911 | 6893 | 6950 | 7238 | 7195 | 6527 | 6503 | 6815 | 6864 | 6526 | 7131 | 7106 | Ttc5 |
| Ttc6 | 176 | 117 | 226 | 146 | 152 | 154 | 123 | 322 | 222 | 199 | 197 | 236 | 217 | 241 | 187 | 162 | 165 | 139 | 143 | 213 | 138 | 201 | 177 | 230 | 169 | 170 | 361 | 293 | Ttc6 |
| Ttc7 | 1081 | 1430 | 1263 | 1262 | 1336 | 1110 | 1120 | 375 | 554 | 546 | 502 | 331 | 361 | 348 | 988 | 1278 | 1293 | 1313 | 1320 | 996 | 1213 | 486 | 555 | 492 | 445 | 358 | 291 | 330 | Ttc7 |
| Ttc7b | 1245 | 1221 | 1346 | 1079 | 1468 | 1315 | 1131 | 1761 | 2143 | 2043 | 1927 | 2190 | 1921 | 2069 | 1129 | 1197 | 1197 | 1087 | 1266 | 1091 | 1171 | 2077 | 2104 | 1882 | 2005 | 2131 | 2092 | 1983 | Ttc7b |
| Ttc8 | 1954 | 1879 | 2035 | 1953 | 1681 | 1794 | 1966 | 2142 | 1983 | 1944 | 1870 | 2309 | 2075 | 2432 | 2122 | 1743 | 1909 | 1686 | 1696 | 2005 | 1768 | 2037 | 1964 | 1948 | 1842 | 2231 | 2256 | 2445 | Ttc8 |
| Ttc9 | 923 | 942 | 1122 | 1185 | 1912 | 1432 | 1732 | 1690 | 1679 | 1805 | 1657 | 1679 | 1170 | 1651 | 953 | 661 | 1107 | 1220 | 1780 | 1295 | 1746 | 2057 | 1542 | 1821 | 2318 | 1421 | 1326 | 1374 | Ttc9 |
| Ttc9b | 375 | 561 | 422 | 907 | 1172 | 869 | 991 | 631 | 770 | 961 | 760 | 784 | 530 | 847 | 417 | 276 | 895 | 930 | 1248 | 750 | 1134 | 991 | 635 | 968 | 1097 | 800 | 535 | 574 | Ttc9b |
| Ttc9c | 8163 | 7510 | 7263 | 8046 | 6942 | 7395 | 8141 | 7585 | 6330 | 6536 | 6087 | 7495 | 6952 | 8035 | 8174 | 7255 | 7864 | 7925 | 7737 | 7815 | 8085 | 6044 | 6400 | 6990 | 7026 | 7985 | 7671 | 8191 | Ttc9c |
| Ttf1 | 2486 | 2529 | 2622 | 2485 | 2045 | 2331 | 2395 | 2445 | 2235 | 2331 | 2396 | 2747 | 2514 | 2733 | 2534 | 2393 | 2545 | 2121 | 2304 | 2317 | 2323 | 2334 | 2252 | 2304 | 2513 | 2325 | 2623 | 2579 | Ttf1 |
| Ttf2 | 5485 | 5428 | 4973 | 4624 | 3562 | 4078 | 4167 | 2976 | 2925 | 2781 | 2585 | 4285 | 3563 | 3918 | 5000 | 4833 | 4486 | 4265 | 4076 | 4272 | 4063 | 2264 | 2695 | 3050 | 3008 | 4113 | 4292 | 4075 | Ttf2 |
| Tti1 | 2442 | 2509 | 2340 | 2026 | 1852 | 1963 | 2044 | 3369 | 3143 | 2820 | 2841 | 3392 | 3150 | 3282 | 2338 | 2164 | 2294 | 1941 | 2253 | 2208 | 2117 | 2687 | 3014 | 3118 | 2986 | 3563 | 3740 | 3522 | Tti1 |
| Tti2 | 2415 | 2467 | 2441 | 2538 | 2357 | 2347 | 2562 | 2144 | 1997 | 1898 | 1861 | 2361 | 1992 | 2271 | 2436 | 2098 | 2579 | 2509 | 2400 | 2461 | 2414 | 2100 | 2019 | 2218 | 2026 | 2283 | 2122 | 2176 | Tti2 |
| Ttk | 5361 | 4548 | 4898 | 3590 | 2680 | 3352 | 4266 | 3532 | 3147 | 2687 | 2578 | 4339 | 3461 | 4379 | 5021 | 4761 | 4262 | 3213 | 3553 | 4401 | 3590 | 2197 | 2680 | 3447 | 3003 | 4805 | 4458 | 4423 | Ttk |
| Ttl | 3888 | 3877 | 3512 | 4047 | 4540 | 4396 | 4394 | 3518 | 4077 | 4068 | 4494 | 3820 | 3680 | 3623 | 3695 | 3815 | 3708 | 4162 | 4221 | 4277 | 4354 | 3644 | 4066 | 3671 | 4472 | 3699 | 3376 | 3463 | Ttl |
| Ttll1 | 2543 | 3088 | 2782 | 2907 | 3216 | 2847 | 2972 | 3047 | 2767 | 3183 | 2590 | 2849 | 2693 | 2835 | 2718 | 2600 | 2927 | 2906 | 3179 | 3121 | 3203 | 3578 | 2959 | 3076 | 3365 | 2742 | 2936 | 2862 | Ttll1 |
| Ttll10 | 90 | 37 | 11 | 76 | 11 | 10 | 0 | 64 | 82 | 67 | 62 | 4 | 33 | 11 | 46 | 46 | 3 | 0 | 3 | 1 | 0 | 18 | 78 | 74 | 6 | 52 | 5 | 12 | Ttll10 |
| Ttll11 | 332 | 383 | 365 | 507 | 518 | 408 | 398 | 294 | 275 | 249 | 294 | 313 | 215 | 225 | 386 | 384 | 430 | 437 | 406 | 356 | 414 | 319 | 261 | 242 | 305 | 229 | 249 | 210 | Ttll11 |
| Ttll12 | 5324 | 6574 | 6163 | 5826 | 5281 | 5167 | 4850 | 3436 | 4243 | 4130 | 3836 | 4502 | 4092 | 3729 | 5255 | 5654 | 6309 | 5348 | 6231 | 5050 | 5020 | 3795 | 3871 | 4050 | 3938 | 4857 | 3926 | 4196 | Ttll12 |
| Ttll13 | 77 | 86 | 43 | 66 | 92 | 111 | 97 | 66 | 72 | 77 | 85 | 60 | 101 | 84 | 84 | 61 | 85 | 75 | 92 | 65 | 83 | 65 | 79 | 88 | 94 | 52 | 85 | 51 | Ttll13 |
| Ttll2 | 0 | 0 | 0 | 0 | 0 | 0 | 0 | 0 | 2 | 0 | 0 | 0 | 0 | 0 | 0 | 0 | 0 | 0 | 0 | 0 | 0 | 0 | 0 | 0 | 1 | 0 | 0 | 1 | Ttll2 |
| Ttll3 | 1119 | 1437 | 1108 | 1785 | 1457 | 1365 | 1361 | 793 | 844 | 1053 | 1118 | 992 | 981 | 884 | 1033 | 1476 | 1320 | 1858 | 1397 | 1256 | 1396 | 1252 | 933 | 917 | 1017 | 821 | 894 | 780 | Ttll3 |
| Ttll4 | 11779 | 10639 | 8718 | 9217 | 9665 | 11061 | 11292 | 10487 | 9576 | 9487 | 8966 | 12691 | 11650 | 12668 | 10764 | 10191 | 9747 | 9253 | 9968 | 10427 | 11193 | 8667 | 8514 | 9685 | 10520 | 9093 | 13096 | 12481 | Ttll4 |
| Ttll5 | 2347 | 2406 | 2166 | 2247 | 2207 | 1977 | 2221 | 1939 | 1923 | 2061 | 1888 | 1992 | 1982 | 1765 | 2304 | 2287 | 2265 | 2423 | 2274 | 2199 | 2305 | 1856 | 1916 | 1794 | 2026 | 1707 | 1880 | 1782 | Ttll5 |
| Ttll6 | 38 | 38 | 8 | 32 | 62 | 27 | 39 | 73 | 70 | 117 | 83 | 45 | 64 | 57 | 30 | 34 | 31 | 43 | 44 | 56 | 55 | 63 | 125 | 111 | 61 | 71 | 37 | 21 | Ttll6 |
| Ttll7 | 5268 | 5262 | 3557 | 5173 | 6377 | 5185 | 6015 | 4499 | 3769 | 3025 | 3118 | 2110 | 3079 | 2333 | 7017 | 6244 | 4495 | 5986 | 4868 | 6243 | 5713 | 4236 | 2917 | 2148 | 3007 | 1579 | 2210 | 2123 | Ttll7 |
| Ttll8 | 9 | 5 | 5 | 4 | 10 | 10 | 15 | 15 | 34 | 19 | 17 | 18 | 7 | 11 | 5 | 11 | 0 | 6 | 19 | 14 | 16 | 15 | 32 | 16 | 33 | 19 | 23 | 15 | Ttll8 |

| | | | | | | | | | | | | | | | | | | | | | | | | | | | | | |
|---|---|---|---|---|---|---|---|---|---|---|---|---|---|---|---|---|---|---|---|---|---|---|---|---|---|---|---|---|---|
| Ttll9 | 78 | 129 | 66 | 137 | 125 | 109 | 103 | 90 | 119 | 107 | 131 | 101 | 100 | 104 | 92 | 87 | 83 | 103 | 113 | 122 | 114 | 189 | 151 | 84 | 142 | 65 | 130 | 81 | Ttll9 |
| Ttn | 619 | 86 | 7 | 15 | 74 | 70 | 45 | 773 | 2439 | 202 | 10163 | 83 | 297 | 28 | 255 | 651 | 20 | 70 | 23 | 56 | 37 | 944 | 8189 | 143 | 184 | 21 | 58 | 36 | Ttn |
| Ttpa | 221 | 84 | 175 | 77 | 160 | 103 | 104 | 277 | 246 | 306 | 246 | 143 | 257 | 155 | 177 | 174 | 109 | 84 | 85 | 89 | 76 | 137 | 294 | 217 | 109 | 270 | 160 | 203 | Ttpa |
| Ttpal | 4881 | 4786 | 4271 | 4372 | 4555 | 4864 | 4785 | 6289 | 6075 | 5786 | 5835 | 5973 | 5856 | 5920 | 5077 | 4487 | 4473 | 4544 | 4651 | 4459 | 4542 | 6542 | 6236 | 5574 | 6009 | 5515 | 5930 | 6039 | Ttpal |
| Ttr | 5076 | 10001 | 2011 | 5446 | 20728 | 11607 | 10573 | 11347 | 17663 | 16184 | 12706 | 14307 | 18608 | 5307 | 7034 | 10824 | 4190 | 7670 | 10749 | 12192 | 12129 | 17938 | 33175 | 8540 | 11136 | 5402 | 4597 | 4703 | Ttr |
| Ttyh1 | 6030 | 7423 | 8102 | 9487 | 11671 | 9907 | 10723 | 11717 | 11158 | 13308 | 10817 | 15275 | 14247 | 15399 | 7304 | 5440 | 9265 | 9263 | 10834 | 8629 | 9857 | 12330 | 9848 | 13573 | 14763 | 13857 | 12029 | 14150 | Ttyh1 |
| Ttyh2 | 2479 | 2553 | 2504 | 2458 | 2822 | 2538 | 2397 | 1265 | 1530 | 1714 | 2089 | 1231 | 1545 | 1310 | 2302 | 2766 | 2313 | 2640 | 2569 | 2244 | 2540 | 1601 | 1658 | 1575 | 1745 | 1214 | 1110 | 1123 | Ttyh2 |
| Ttyh3 | 18108 | 24303 | 26342 | 28892 | 28752 | 25391 | 21703 | 19069 | 25131 | 29941 | 26359 | 32092 | 26215 | 25039 | 16868 | 21059 | 29030 | 29453 | 31170 | 20886 | 23526 | 27681 | 22938 | 33151 | 31960 | 34678 | 24279 | 25352 | Ttyh3 |
| Tub | 8051 | 8919 | 7842 | 12838 | 16952 | 14110 | 15744 | 11775 | 11632 | 13025 | 11234 | 11565 | 9169 | 11400 | 9308 | 7418 | 12019 | 13699 | 15543 | 12040 | 15311 | 17400 | 9576 | 11473 | 14734 | 8190 | 9151 | 9425 | Tub |
| Tuba1a | 138636 | 175043 | 113694 | 165612 | 180260 | 171545 | 169767 | 205367 | 171297 | 160708 | 133298 | 152004 | 143758 | 158019 | 178902 | 136361 | 147625 | 158223 | 184215 | 198775 | 192157 | 196277 | 170693 | 158132 | 182387 | 135362 | 158916 | 165171 | Tuba1a |
| Tuba1b | 31668 | 37204 | 33352 | 26604 | 22812 | 26520 | 24082 | 31372 | 32862 | 27265 | 29516 | 39111 | 37054 | 37874 | 29218 | 27694 | 30080 | 24555 | 28499 | 27544 | 25344 | 21383 | 32059 | 36471 | 32406 | 48168 | 43742 | 42515 | Tuba1b |
| Tuba1c | 7650 | 8279 | 6271 | 5587 | 4286 | 5071 | 5178 | 5381 | 5338 | 4391 | 3872 | 5538 | 5146 | 4854 | 6883 | 7223 | 6084 | 5036 | 5882 | 6082 | 4941 | 3902 | 5229 | 5257 | 4528 | 7382 | 5882 | 6025 | Tuba1c |
| Tuba3a | 0 | 2 | 0 | 1 | 0 | 1 | 1 | 2 | 2 | 2 | 5 | 4 | 0 | 3 | 0 | 3 | 0 | 0 | 0 | 0 | 3 | 3 | 4 | 1 | 0 | 2 | 0 | 1 | Tuba3a |
| Tuba3b | 0 | 0 | 0 | 0 | 0 | 0 | 0 | 0 | 0 | 0 | 0 | 0 | 0 | 0 | 0 | 2 | 0 | 0 | 0 | 0 | 0 | 0 | 0 | 0 | 0 | 0 | 0 | 0 | Tuba3b |
| Tuba4a | 1495 | 1627 | 1042 | 1245 | 1123 | 1161 | 1195 | 415 | 590 | 414 | 631 | 330 | 457 | 294 | 1416 | 1584 | 1079 | 1294 | 1305 | 1467 | 1201 | 276 | 645 | 538 | 422 | 563 | 329 | 317 | Tuba4a |
| Tuba8 | 17 | 10 | 7 | 27 | 25 | 22 | 15 | 11 | 10 | 3 | 8 | 7 | 22 | 7 | 15 | 17 | 17 | 22 | 7 | 21 | 15 | 7 | 16 | 7 | 6 | 11 | 8 | 15 | Tuba8 |
| Tubal3 | 3 | 4 | 0 | 3 | 2 | 2 | 2 | 0 | 3 | 5 | 0 | 2 | 1 | 1 | 4 | 4 | 9 | 0 | 4 | 1 | 7 | 0 | 1 | 5 | 6 | 0 | 2 | 1 | Tubal3 |
| Tubb1 | 120 | 72 | 158 | 47 | 38 | 57 | 30 | 100 | 102 | 80 | 79 | 112 | 228 | 78 | 70 | 115 | 63 | 33 | 33 | 79 | 45 | 99 | 91 | 73 | 68 | 228 | 200 | 103 | Tubb1 |
| Tubb2a | 12345 | 20460 | 15270 | 21009 | 24414 | 21438 | 20122 | 18208 | 19421 | 19581 | 16525 | 12601 | 14855 | 13952 | 16833 | 14866 | 19432 | 20145 | 21618 | 25870 | 23299 | 20102 | 19465 | 15224 | 19508 | 9758 | 13038 | 11860 | Tubb2a |
| Tubb2b | 96963 | 141312 | 122420 | 166635 | 169604 | 161414 | 152713 | 138745 | 131886 | 154973 | 121155 | 140905 | 126090 | 147967 | 124213 | 96680 | 151941 | 157887 | 183103 | 177742 | 185750 | 139738 | 133108 | 161613 | 176659 | 130647 | 151091 | 146591 | Tubb2b |
| Tubb3 | 33400 | 53165 | 38774 | 53867 | 63964 | 58011 | 51447 | 64489 | 64189 | 69177 | 51843 | 67436 | 62732 | 62487 | 44213 | 36572 | 52026 | 52735 | 65307 | 59358 | 58243 | 73915 | 61624 | 66575 | 76982 | 59286 | 66494 | 62246 | Tubb3 |
| Tubb4a | 5828 | 7756 | 6337 | 8462 | 11188 | 9505 | 8275 | 8636 | 8802 | 10310 | 8404 | 9866 | 9586 | 8788 | 7090 | 5889 | 8041 | 8823 | 10557 | 9355 | 9397 | 12614 | 7693 | 9180 | 10604 | 7330 | 8894 | 8936 | Tubb4a |
| Tubb4b | 19962 | 24310 | 21072 | 18175 | 14735 | 16248 | 15370 | 13198 | 14937 | 12877 | 12922 | 17798 | 17608 | 16154 | 18742 | 19077 | 19513 | 16096 | 17861 | 17356 | 15638 | 11428 | 14498 | 15334 | 14015 | 21276 | 18899 | 18443 | Tubb4b |
| Tubb5 | 123531 | 149183 | 116698 | 123750 | 130217 | 130057 | 128633 | 145026 | 155956 | 152264 | 129141 | 171101 | 156548 | 165143 | 126630 | 114617 | 128310 | 121595 | 144232 | 134527 | 136548 | 150124 | 142408 | 160201 | 164825 | 183756 | 168137 | 174109 | Tubb5 |
| Tubb6 | 2352 | 2664 | 2700 | 1502 | 1049 | 1588 | 981 | 1172 | 1576 | 1066 | 2673 | 1441 | 1765 | 1316 | 2275 | 2751 | 2268 | 1498 | 1163 | 1368 | 1019 | 1120 | 2554 | 1215 | 1064 | 1428 | 1940 | 1536 | Tubb6 |
| Tubd1 | 569 | 537 | 483 | 541 | 431 | 471 | 478 | 603 | 577 | 594 | 565 | 648 | 576 | 598 | 601 | 534 | 567 | 503 | 518 | 586 | 516 | 565 | 638 | 646 | 573 | 669 | 638 | 610 | Tubd1 |
| Tube1 | 2010 | 1614 | 1418 | 1385 | 1288 | 1539 | 1642 | 1515 | 1402 | 1180 | 1304 | 1382 | 1591 | 1452 | 1862 | 1860 | 1347 | 1325 | 1332 | 1819 | 1597 | 1096 | 1276 | 1243 | 1341 | 1438 | 1712 | 1578 | Tube1 |
| Tubg1 | 3091 | 4163 | 4267 | 3951 | 3355 | 3433 | 3270 | 2006 | 2565 | 2842 | 2564 | 3646 | 2739 | 3159 | 2931 | 3566 | 4336 | 3624 | 3915 | 3074 | 3165 | 2578 | 2676 | 3200 | 3032 | 4048 | 3172 | 3297 | Tubg1 |
| Tubg2 | 203 | 233 | 203 | 346 | 506 | 418 | 339 | 431 | 496 | 559 | 501 | 381 | 353 | 335 | 227 | 206 | 265 | 336 | 411 | 349 | 453 | 675 | 503 | 462 | 584 | 384 | 277 | 280 | Tubg2 |
| Tubgcp2 | 6236 | 6572 | 4722 | 5698 | 5866 | 6267 | 6184 | 5542 | 6155 | 5405 | 5463 | 6415 | 5831 | 5818 | 6045 | 6048 | 5049 | 5428 | 5687 | 6278 | 5979 | 4996 | 5536 | 5357 | 6096 | 6425 | 6270 | 5814 | Tubgcp2 |
| Tubgcp3 | 4735 | 5309 | 4111 | 3952 | 4160 | 4257 | 4571 | 5794 | 5682 | 4777 | 4580 | 5734 | 5095 | 5664 | 4639 | 4203 | 4288 | 4058 | 4535 | 5126 | 4538 | 4893 | 5307 | 5115 | 5316 | 5530 | 5710 | 5745 | Tubgcp3 |
| Tubgcp4 | 8043 | 7650 | 6205 | 6337 | 6275 | 6710 | 7016 | 6829 | 5825 | 5128 | 5253 | 5907 | 5821 | 5977 | 7934 | 7503 | 6401 | 6292 | 6304 | 7569 | 7138 | 5212 | 5797 | 5441 | 5472 | 5077 | 6229 | 5973 | Tubgcp4 |
| Tubgcp5 | 3596 | 3007 | 2651 | 2892 | 3328 | 3568 | 3869 | 4975 | 4494 | 3677 | 3867 | 4124 | 3999 | 4353 | 3864 | 3000 | 2898 | 2791 | 3140 | 3932 | 3514 | 4007 | 4361 | 3662 | 3826 | 3604 | 4279 | 4199 | Tubgcp5 |
| Tubgcp6 | 8390 | 10222 | 7612 | 10317 | 8656 | 8914 | 8827 | 4517 | 5006 | 5559 | 5093 | 6036 | 5562 | 5613 | 7875 | 9138 | 8989 | 10433 | 9488 | 7924 | 9546 | 4986 | 4782 | 5478 | 5819 | 4120 | 5820 | 5811 | Tubgcp6 |
| Tufm | 7238 | 9001 | 8744 | 7511 | 6559 | 6588 | 6461 | 4730 | 5438 | 5405 | 5308 | 5618 | 5587 | 5643 | 6560 | 7514 | 8530 | 6923 | 7377 | 6608 | 6627 | 4527 | 5091 | 5728 | 5380 | 6917 | 5901 | 6065 | Tufm |
| Tuft1 | 1380 | 1408 | 1434 | 1303 | 1189 | 1146 | 1063 | 272 | 384 | 306 | 243 | 253 | 383 | 255 | 1321 | 1441 | 1287 | 1310 | 1116 | 1143 | 1216 | 266 | 410 | 267 | 200 | 334 | 270 | 389 | Tuft1 |
| Tug1 | 13437 | 11035 | 13992 | 12579 | 13104 | 12974 | 15073 | 16569 | 14997 | 15786 | 14679 | 15741 | 15111 | 16577 | 13510 | 12150 | 14069 | 12866 | 13358 | 11704 | 14033 | 18182 | 14961 | 15976 | 15830 | 17939 | 15873 | 16665 | Tug1 |
| Tulp1 | 166 | 190 | 201 | 246 | 150 | 228 | 154 | 144 | 149 | 193 | 167 | 175 | 202 | 154 | 140 | 187 | 190 | 160 | 203 | 123 | 151 | 109 | 180 | 203 | 109 | 226 | 217 | 211 | Tulp1 |
| Tulp2 | 10 | 9 | 0 | 3 | 4 | 3 | 6 | 10 | 25 | 19 | 30 | 2 | 8 | 2 | 5 | 12 | 1 | 2 | 6 | 1 | 4 | 7 | 19 | 10 | 7 | 8 | 5 | 6 | Tulp2 |
| Tulp3 | 4896 | 4980 | 5564 | 5344 | 3914 | 4508 | 3901 | 2713 | 3277 | 3194 | 3543 | 3263 | 3337 | 2903 | 4330 | 5505 | 5246 | 5191 | 4287 | 3418 | 4051 | 3160 | 3251 | 3351 | 2961 | 3597 | 2933 | 3176 | Tulp3 |
| Tulp4 | 13451 | 12705 | 10747 | 17252 | 21585 | 21453 | 21246 | 20584 | 20185 | 22136 | 21843 | 20845 | 21144 | 21749 | 14467 | 13664 | 13424 | 17765 | 18873 | 17579 | 21565 | 22082 | 18429 | 17947 | 27247 | 13805 | 21088 | 19066 | Tulp4 |
| Tunar | 205 | 257 | 188 | 455 | 923 | 599 | 507 | 572 | 553 | 794 | 567 | 368 | 431 | 365 | 253 | 262 | 392 | 555 | 598 | 592 | 608 | 944 | 553 | 525 | 791 | 269 | 265 | 287 | Tunar |
| Tusc1 | 118 | 105 | 118 | 150 | 192 | 130 | 162 | 341 | 356 | 390 | 394 | 336 | 314 | 400 | 141 | 93 | 143 | 167 | 152 | 128 | 171 | 386 | 350 | 346 | 353 | 331 | 330 | 405 | Tusc1 |
| Tusc2 | 3959 | 4758 | 3512 | 3706 | 3128 | 3321 | 3742 | 2252 | 2209 | 2269 | 2097 | 2367 | 2243 | 2599 | 4131 | 3705 | 3872 | 3575 | 3338 | 3597 | 3595 | 2358 | 2190 | 2229 | 2354 | 2506 | 2321 | 2676 | Tusc2 |
| Tut1 | 2476 | 2824 | 3020 | 2981 | 2612 | 2593 | 2556 | 1976 | 2280 | 2298 | 2416 | 2728 | 2573 | 2632 | 2404 | 2602 | 3001 | 2604 | 2791 | 2533 | 2610 | 2154 | 2086 | 2602 | 2553 | 3190 | 2611 | 2645 | Tut1 |
| Tut4 | 13493 | 10367 | 9679 | 10584 | 10716 | 12512 | 12139 | 15259 | 14452 | 13325 | 14631 | 16025 | 15415 | 16237 | 12171 | 11702 | 10199 | 10366 | 10869 | 11405 | 12029 | 13954 | 13713 | 12507 | 13985 | 11984 | 16646 | 15659 | Tut4 |
| Tut7 | 10665 | 10215 | 7869 | 9375 | 8888 | 9655 | 10166 | 9000 | 8394 | 7838 | 8048 | 8074 | 7849 | 8763 | 10399 | 10087 | 8204 | 9394 | 8556 | 9346 | 9902 | 7683 | 8154 | 7455 | 8423 | 6091 | 8430 | 8190 | Tut7 |
| Tvp23a | 3399 | 3710 | 3274 | 3773 | 4196 | 4222 | 3879 | 3716 | 3862 | 4067 | 3466 | 3665 | 3560 | 3432 | 3236 | 3123 | 3695 | 3861 | 4295 | 4006 | 3978 | 4040 | 3306 | 3818 | 4383 | 3589 | 3674 | 3578 | Tvp23a |
| Tvp23b | 1899 | 1726 | 2141 | 1924 | 1761 | 1700 | 1598 | 2238 | 2172 | 1828 | 2121 | 1887 | 1887 | 1734 | 2151 | 1965 | 2029 | 1804 | 1860 | 2059 | 1554 | 2030 | 2273 | 1928 | 1977 | 1985 | 1947 | 1883 | Tvp23b |
| Twf1 | 8773 | 8307 | 9165 | 8937 | 7927 | 8356 | 9316 | 7555 | 6517 | 6933 | 6737 | 8137 | 7555 | 8455 | 9274 | 8358 | 9156 | 8307 | 8138 | 8611 | 8554 | 6998 | 6694 | 7669 | 7597 | 9113 | 8315 | 8565 | Twf1 |
| Twf2 | 2252 | 3191 | 2429 | 2316 | 1991 | 2210 | 1890 | 1470 | 1707 | 1368 | 1583 | 1473 | 1473 | 1376 | 2251 | 2647 | 2372 | 2485 | 2177 | 2438 | 2019 | 1300 | 1749 | 1441 | 1338 | 1637 | 1483 | 1410 | Twf2 |
| Twist1 | 3400 | 2214 | 2227 | 1042 | 331 | 1230 | 194 | 2667 | 3060 | 2289 | 2853 | 1948 | 3399 | 2028 | 2294 | 3400 | 1253 | 1164 | 394 | 279 | 196 | 1420 | 3172 | 2514 | 1228 | 2652 | 2051 | 2255 | Twist1 |
| Twist2 | 1027 | 719 | 490 | 224 | 52 | 468 | 31 | 821 | 1587 | 803 | 1077 | 601 | 1039 | 578 | 729 | 1005 | 337 | 524 | 107 | 71 | 29 | 239 | 1438 | 901 | 334 | 574 | 593 | 608 | Twist2 |
| Twnk | 3086 | 3492 | 3559 | 3671 | 3017 | 2934 | 3203 | 2395 | 2408 | 2722 | 2571 | 2890 | 2652 | 2854 | 2988 | 2981 | 3361 | 3423 | 3270 | 2898 | 3279 | 2323 | 2259 | 2875 | 3027 | 3143 | 2898 | 2911 | Twnk |
| Twsg1 | 7820 | 5585 | 6213 | 5477 | 5492 | 6082 | 5561 | 9780 | 9855 | 9085 | 9924 | 9443 | 9774 | 9170 | 7007 | 7031 | 5390 | 5590 | 5354 | 5852 | 5383 | 8397 | 10453 | 9248 | 8355 | 10242 | 9546 | 9640 | Twsg1 |
| Txk | 5 | 10 | 2 | 7 | 2 | 1 | 4 | 9 | 6 | 3 | 5 | 7 | 8 | 6 | 0 | 6 | 1 | 14 | 3 | 12 | 13 | 3 | 8 | 9 | 7 | 4 | 0 | 4 | Txk |
| Txlna | 10822 | 12337 | 11188 | 11261 | 9959 | 10378 | 10327 | 11657 | 12450 | 12159 | 12582 | 13484 | 12820 | 13568 | 10081 | 10661 | 11681 | 10942 | 11489 | 10361 | 10477 | 11490 | 12249 | 14234 | 13062 | 14089 | 13775 | 14002 | Txlna |
| Txlnb | 333 | 466 | 557 | 365 | 313 | 298 | 229 | 541 | 430 | 424 | 738 | 597 | 905 | 689 | 442 | 434 | 516 | 291 | 254 | 304 | 279 | 587 | 733 | 474 | 393 | 275 | 852 | 769 | Txlnb |
| Txlng | 4579 | 3762 | 4361 | 4487 | 4070 | 4255 | 4891 | 2695 | 2202 | 2321 | 2298 | 2635 | 2604 | 2688 | 4759 | 4036 | 4342 | 4473 | 4219 | 4415 | 4542 | 2648 | 2067 | 2556 | 2541 | 2495 | 2631 | 2680 | Txlng |
| Txn1 | 8511 | 8751 | 9328 | 8229 | 6915 | 7111 | 9324 | 6616 | 6550 | 6377 | 7236 | 8115 | 8275 | 9838 | 7973 | 7569 | 9013 | 7120 | 7404 | 8661 | 8267 | 5370 | 6587 | 8466 | 7109 | 10396 | 8990 | 11122 | Txn1 |
| Txn2 | 4429 | 5590 | 4764 | 4218 | 3735 | 3773 | 3826 | 4586 | 5155 | 4598 | 4579 | 5018 | 5018 | 5091 | 4036 | 4643 | 4574 | 4009 | 4055 | 4065 | 3902 | 4411 | 5177 | 4886 | 4739 | 5408 | 5281 | 5266 | Txn2 |
| Txndc11 | 3119 | 2953 | 2712 | 2639 | 2597 | 2711 | 2778 | 2374 | 2529 | 2470 | 2883 | 2319 | 2628 | 2247 | 3001 | 3010 | 2494 | 2774 | 2572 | 2507 | 2733 | 2582 | 2385 | 2474 | 2409 | 2272 | 2436 | 2434 | Txndc11 |
| Txndc12 | 4471 | 4605 | 4501 | 4231 | 3532 | 3929 | 4050 | 3750 | 3552 | 3334 | 3367 | 3481 | 3399 | 3770 | 4342 | 4360 | 4234 | 3820 | 4032 | 4381 | 4001 | 3116 | 3748 | 3485 | 3147 | 4105 | 3707 | 3766 | Txndc12 |
| Txndc15 | 1998 | 1902 | 2033 | 2010 | 2591 | 2285 | 2520 | 3279 | 3276 | 3504 | 3293 | 3276 | 3317 | 3388 | 2097 | 1924 | 2174 | 2063 | 2586 | 2443 | 2433 | 3283 | 3412 | 3331 | 3336 | 3115 | 3062 | 3469 | Txndc15 |
| Txndc16 | 10153 | 9628 | 7590 | 9573 | 9831 | 9838 | 10699 | 5327 | 4868 | 4494 | 4270 | 3961 | 3717 | 4157 | 11107 | 9034 | 8734 | 9946 | 9241 | 10544 | 10823 | 5006 | 4664 | 4008 | 4380 | 3356 | 3829 | 3678 | Txndc16 |
| Txndc17 | 5046 | 4956 | 5226 | 4421 | 3782 | 3550 | 4512 | 3462 | 3173 | 3047 | 3193 | 3192 | 3554 | 3827 | 4946 | 4397 | 4675 | 4382 | 3754 | 4522 | 4210 | 2787 | 3522 | 3479 | 3071 | 3971 | 3514 | 3927 | Txndc17 |
| Txndc2 | 15 | 17 | 6 | 10 | 24 | 15 | 21 | 13 | 15 | 12 | 14 | 16 | 27 | 13 | 7 | 16 | 23 | 14 | 12 | 20 | 8 | 13 | 12 | 14 | 9 | 6 | 12 | 7 | Txndc2 |
| Txndc5 | 10377 | 10475 | 9464 | 7871 | 6645 | 8462 | 6427 | 7391 | 9277 | 8048 | 9992 | 7161 | 8247 | 6621 | 9721 | 10724 | 8349 | 8386 | 7111 | 6834 | 6236 | 7526 | 10436 | 7227 | 6844 | 8798 | 7674 | 7188 | Txndc5 |
| Txndc8 | 2 | 0 | 0 | 0 | 0 | 0 | 0 | 0 | 0 | 0 | 0 | 0 | 0 | 0 | 0 | 0 | 0 | 0 | 0 | 0 | 0 | 0 | 0 | 0 | 0 | 0 | 0 | 0 | Txndc8 |
| Txndc9 | 4527 | 4333 | 4385 | 4003 | 3692 | 3720 | 4362 | 7464 | 6282 | 6006 | 5732 | 6775 | 6575 | 7389 | 4953 | 4328 | 4239 | 3732 | 3971 | 4657 | 4167 | 6293 | 6386 | 6481 | 6036 | 7067 | 7021 | 7887 | Txndc9 |
| Txnip | 8878 | 8196 | 11271 | 9457 | 8374 | 7402 | 7492 | 7040 | 9451 | 7227 | 8959 | 6198 | 6778 | 5206 | 8640 | 8420 | 9620 | 10586 | 8293 | 7796 | 7425 | 6717 | 10481 | 7302 | 7037 | 6974 | 5826 | 5007 | Txnip |
| Txnl1 | 11814 | 9635 | 10853 | 9876 | 8948 | 9557 | 10885 | 9308 | 8383 | 8121 | 7902 | 9131 | 8545 | 9436 | 11967 | 10440 | 9999 | 9694 | 9239 | 10715 | 9977 | 8430 | 8476 | 8725 | 8114 | 10423 | 9504 | 10314 | Txnl1 |
| Txnl4a | 3015 | 3676 | 2644 | 3358 | 2793 | 3049 | 3629 | 2137 | 2036 | 2161 | 1811 | 2116 | 1969 | 2220 | 3224 | 2746 | 3116 | 3318 | 3478 | 3207 | 3781 | 1831 | 1959 | 2304 | 2373 | 2143 | 2030 | 2389 | Txnl4a |
| Txnl4b | 309 | 312 | 295 | 228 | 277 | 275 | 306 | 736 | 674 | 671 | 646 | 664 | 713 | 767 | 342 | 282 | 320 | 217 | 314 | 298 | 317 | 698 | 805 | 703 | 630 | 835 | 794 | 700 | Txnl4b |
| Txnrd1 | 15423 | 14044 | 18275 | 16006 | 14857 | 14816 | 16091 | 11970 | 11744 | 13478 | 12549 | 17330 | 14548 | 16497 | 14089 | 14664 | 17072 | 15606 | 16615 | 13079 | 15645 | 13399 | 10996 | 15926 | 15632 | 18278 | 15982 | 17322 | Txnrd1 |
| Txnrd2 | 33 | 105 | 129 | 110 | 115 | 185 | 177 | 418 | 168 | 212 | 198 | 290 | 43 | 33 | 142 | 265 | 20 | 294 | 113 | 203 | 21 | 50 | 261 | 324 | 240 | 59 | 316 | 128 | Txnrd2 |
| Txnrd3 | 1431 | 1341 | 1809 | 1396 | 1292 | 1379 | 1294 | 981 | 907 | 888 | 845 | 1071 | 1071 | 1020 | 1428 | 1296 | 1557 | 1334 | 1233 | 1139 | 1286 | 1125 | 924 | 887 | 932 | 1053 | 1096 | 1044 | Txnrd3 |
| Tyk2 | 1216 | 1649 | 1665 | 1696 | 1642 | 1526 | 1360 | 1566 | 1945 | 2032 | 1955 | 1864 | 1872 | 1708 | 1361 | 1618 | 1738 | 1825 | 1679 | 1312 | 1367 | 1965 | 1930 | 2036 | 1950 | 1765 | 1879 | 1784 | Tyk2 |
| Tymp | 1 | 1 | 1 | 1 | 6 | 6 | 4 | 6 | 8 | 6 | 5 | 12 | 3 | 10 | 3 | 1 | 7 | 2 | 6 | 1 | 7 | 3 | 11 | 6 | 7 | 6 | 3 | 4 | Tymp |

| | | | | | | | | | | | | | | | | | | | | | | | | | | | | | |
|---|---|---|---|---|---|---|---|---|---|---|---|---|---|---|---|---|---|---|---|---|---|---|---|---|---|---|---|---|---|
| Tyms | 4376 | 3829 | 3975 | 2913 | 2304 | 2888 | 3419 | 4093 | 4044 | 3336 | 3309 | 4501 | 3781 | 4742 | 3835 | 3711 | 3657 | 2666 | 3023 | 3554 | 3080 | 2513 | 3579 | 4232 | 3372 | 5679 | 4694 | 5180 | Tyms |
| Tyr | 30 | 7 | 2 | 0 | 0 | 12 | 3 | 35 | 63 | 44 | 48 | 33 | 31 | 11 | 16 | 5 | 0 | 25 | 5 | 6 | 0 | 20 | 55 | 33 | 14 | 11 | 10 | 6 | Tyr |
| Tyro3 | 4080 | 4522 | 4748 | 3900 | 4151 | 4061 | 3635 | 3670 | 4164 | 3706 | 3645 | 4299 | 4139 | 3840 | 3877 | 4122 | 4672 | 3765 | 4701 | 3692 | 3774 | 3354 | 3972 | 4301 | 3578 | 4555 | 4177 | 4118 | Tyro3 |
| Tyrobp | 172 | 251 | 220 | 195 | 182 | 156 | 142 | 206 | 288 | 249 | 288 | 216 | 214 | 189 | 146 | 222 | 182 | 227 | 151 | 206 | 152 | 217 | 389 | 226 | 255 | 211 | 258 | 190 | Tyrobp |
| Tyrp1 | 13 | 4 | 7 | 1 | 16 | 12 | 16 | 15 | 40 | 17 | 24 | 7 | 5 | 13 | 3 | 3 | 9 | 17 | 6 | 7 | 13 | 23 | 22 | 13 | 23 | 23 | 9 | 17 | Tyrp1 |
| Tysnd1 | 2031 | 2496 | 2131 | 2128 | 1697 | 1688 | 1534 | 1180 | 1322 | 1264 | 1284 | 1342 | 1326 | 1257 | 1795 | 2133 | 2195 | 2129 | 1938 | 1592 | 1614 | 1315 | 1328 | 1405 | 1434 | 1576 | 1263 | 1285 | Tysnd1 |
| Tyw1 | 2480 | 2935 | 2618 | 2489 | 2264 | 2169 | 2199 | 1926 | 1919 | 1835 | 1828 | 2052 | 2012 | 1907 | 2306 | 2518 | 2431 | 2382 | 2427 | 2256 | 2242 | 1634 | 1987 | 1858 | 1801 | 2074 | 2201 | 2028 | Tyw1 |
| Tyw3 | 693 | 575 | 636 | 604 | 661 | 678 | 663 | 1160 | 1070 | 1259 | 1082 | 1218 | 1077 | 1230 | 545 | 507 | 510 | 511 | 781 | 620 | 593 | 986 | 1083 | 1199 | 1060 | 1240 | 1189 | 1239 | Tyw3 |
| Tyw5 | 433 | 327 | 509 | 389 | 428 | 435 | 515 | 976 | 922 | 792 | 908 | 973 | 896 | 976 | 516 | 390 | 454 | 316 | 503 | 496 | 408 | 952 | 986 | 882 | 808 | 943 | 951 | 983 | Tyw5 |
| U2af1 | 7556 | 8300 | 8293 | 7358 | 6473 | 6843 | 7254 | 8328 | 8174 | 7763 | 8078 | 9632 | 8963 | 9932 | 7370 | 7097 | 7934 | 7074 | 7356 | 8086 | 7227 | 6863 | 7890 | 8810 | 8533 | 10425 | 10187 | 10385 | U2af1 |
| U2af1l4 | 1 | 4 | 0 | 5 | 0 | 0 | 1 | 0 | 1 | 10 | 2 | 9 | 0 | 4 | 0 | 0 | 0 | 4 | 3 | 3 | 4 | 3 | 3 | 9 | 0 | 2 | 0 | 4 | U2af1l4 |
| U2af2 | 13551 | 16363 | 17571 | 16374 | 14755 | 14670 | 14580 | 17077 | 19439 | 21375 | 19360 | 24560 | 21825 | 24609 | 12678 | 12238 | 17198 | 14317 | 16937 | 14044 | 15408 | 17398 | 17583 | 23920 | 21994 | 28736 | 23538 | 25590 | U2af2 |
| U2surp | 20013 | 15460 | 16284 | 15622 | 14883 | 16427 | 18026 | 19643 | 17178 | 15522 | 15840 | 17558 | 16960 | 18446 | 20356 | 16395 | 16131 | 15084 | 15173 | 19290 | 16687 | 15567 | 16209 | 16069 | 16370 | 15557 | 18836 | 17575 | U2surp |
| Uaca | 6412 | 7005 | 9161 | 6449 | 5345 | 5711 | 4976 | 4007 | 4700 | 5310 | 5625 | 5712 | 6238 | 5617 | 6505 | 7745 | 7384 | 6058 | 5331 | 5081 | 4962 | 4482 | 5471 | 5374 | 5053 | 5995 | 6452 | 5779 | Uaca |
| Uap1 | 5105 | 4844 | 4421 | 4309 | 3895 | 3922 | 4562 | 3197 | 3005 | 2830 | 2861 | 2797 | 2772 | 2855 | 4971 | 4974 | 4463 | 4475 | 3888 | 4631 | 4195 | 3056 | 3001 | 2883 | 2739 | 2862 | 2767 | 2859 | Uap1 |
| Uap1l1 | 354 | 395 | 344 | 339 | 275 | 321 | 193 | 181 | 252 | 169 | 229 | 139 | 194 | 115 | 331 | 429 | 358 | 343 | 282 | 317 | 203 | 152 | 353 | 217 | 180 | 195 | 167 | 132 | Uap1l1 |
| Uba1 | 51401 | 59553 | 48415 | 55265 | 54933 | 57685 | 52404 | 52595 | 55132 | 57181 | 50656 | 58689 | 55281 | 54818 | 51444 | 46128 | 52223 | 52755 | 61886 | 52137 | 58157 | 57685 | 51572 | 55209 | 60767 | 55839 | 59080 | 57512 | Uba1 |
| Uba1y | 6 | 10 | 4 | 3 | 22 | 10 | 13 | 10 | 16 | 5 | 9 | 15 | 20 | 9 | 0 | 0 | 0 | 0 | 0 | 0 | 0 | 0 | 0 | 0 | 0 | 0 | 0 | 0 | Uba1y |
| Uba2 | 17704 | 16136 | 14923 | 14539 | 13120 | 13514 | 15922 | 15370 | 13428 | 12595 | 11741 | 14933 | 13150 | 15490 | 17682 | 14920 | 14237 | 13753 | 13794 | 15439 | 14908 | 12770 | 13685 | 13612 | 13529 | 16249 | 14518 | 16465 | Uba2 |
| Uba3 | 7732 | 5645 | 6721 | 6640 | 6322 | 6380 | 7293 | 11235 | 9167 | 8899 | 8714 | 9647 | 9256 | 10299 | 7945 | 6833 | 6407 | 6399 | 6547 | 7893 | 6814 | 9361 | 9593 | 9594 | 9393 | 10379 | 9624 | 10154 | Uba3 |
| Uba5 | 5915 | 5752 | 5224 | 5607 | 5188 | 5325 | 5804 | 6705 | 6126 | 5627 | 6295 | 5839 | 5772 | 5894 | 5916 | 6380 | 5234 | 5548 | 4902 | 5596 | 5432 | 6027 | 6295 | 5867 | 6042 | 5148 | 5827 | 5781 | Uba5 |
| Uba52 | 225 | 260 | 303 | 322 | 192 | 170 | 207 | 170 | 157 | 240 | 211 | 276 | 208 | 226 | 142 | 164 | 341 | 252 | 255 | 193 | 202 | 139 | 156 | 340 | 221 | 310 | 244 | 329 | Uba52 |
| Uba6 | 12927 | 10318 | 8616 | 9272 | 7681 | 9412 | 9647 | 7592 | 6446 | 5067 | 5513 | 6070 | 6120 | 6323 | 12939 | 12291 | 8719 | 8912 | 7741 | 10477 | 9012 | 5765 | 6449 | 5059 | 5619 | 4615 | 6564 | 6355 | Uba6 |
| Uba7 | 120 | 154 | 131 | 46 | 52 | 58 | 64 | 68 | 105 | 161 | 164 | 149 | 193 | 138 | 148 | 187 | 102 | 50 | 45 | 88 | 59 | 99 | 168 | 85 | 93 | 137 | 157 | 284 | Uba7 |
| Ubac1 | 3460 | 3710 | 2771 | 3058 | 3105 | 3262 | 3260 | 2943 | 3175 | 2668 | 2832 | 2721 | 2724 | 2687 | 3492 | 3383 | 3155 | 3051 | 3141 | 3183 | 3159 | 2891 | 2951 | 2744 | 2840 | 2902 | 2421 | 2903 | Ubac1 |
| Ubac2 | 2756 | 2804 | 2768 | 2506 | 2273 | 2246 | 2214 | 1944 | 2089 | 1881 | 1965 | 2362 | 2245 | 2142 | 2552 | 2593 | 2773 | 2272 | 2508 | 2465 | 2297 | 1851 | 2227 | 2225 | 2095 | 2690 | 2291 | 2375 | Ubac2 |
| Ubald1 | 1446 | 1885 | 1275 | 1853 | 1840 | 1832 | 1652 | 1485 | 1845 | 1968 | 1890 | 1641 | 1657 | 1602 | 1383 | 1555 | 1618 | 1930 | 1860 | 1677 | 1782 | 1968 | 1790 | 1831 | 1729 | 1599 | 1438 | 1450 | Ubald1 |
| Ubald2 | 1416 | 1963 | 1665 | 1795 | 1799 | 1668 | 1360 | 2286 | 2478 | 2673 | 2269 | 2627 | 2485 | 2204 | 1435 | 1287 | 1875 | 1878 | 2202 | 1948 | 1574 | 2174 | 2234 | 2866 | 2835 | 2970 | 2816 | 2302 | Ubald2 |
| Ubap1 | 4207 | 4644 | 4923 | 4247 | 4019 | 4012 | 4155 | 3858 | 3504 | 3920 | 3277 | 4058 | 3884 | 4142 | 4183 | 4018 | 4727 | 3903 | 4575 | 4103 | 4116 | 3705 | 3370 | 3801 | 3828 | 4656 | 4326 | 4489 | Ubap1 |
| Ubap1l | 174 | 215 | 167 | 395 | 493 | 326 | 306 | 204 | 197 | 190 | 200 | 112 | 201 | 117 | 199 | 250 | 239 | 383 | 425 | 311 | 336 | 317 | 173 | 153 | 211 | 83 | 156 | 97 | Ubap1l |
| Ubap2 | 8989 | 11143 | 12902 | 12375 | 10667 | 9976 | 9724 | 7376 | 9007 | 10681 | 9823 | 11203 | 9475 | 10275 | 8801 | 10511 | 12462 | 12175 | 11543 | 8184 | 10702 | 9640 | 8121 | 11342 | 10233 | 11877 | 9358 | 10098 | Ubap2 |
| Ubap2l | 17334 | 17229 | 16967 | 17249 | 16499 | 17095 | 16841 | 16825 | 17915 | 18225 | 18298 | 20240 | 19469 | 19559 | 16481 | 16400 | 17021 | 16421 | 17482 | 15330 | 17314 | 18229 | 17506 | 19030 | 18724 | 20209 | 19805 | 20093 | Ubap2l |
| Ubash3a | 6 | 2 | 4 | 0 | 5 | 0 | 1 | 6 | 5 | 12 | 7 | 1 | 10 | 5 | 1 | 12 | 5 | 0 | 0 | 3 | 0 | 7 | 8 | 6 | 0 | 8 | 1 | 11 | Ubash3a |
| Ubash3b | 2392 | 2203 | 2121 | 2315 | 3521 | 2580 | 2768 | 2353 | 2328 | 2210 | 2369 | 1564 | 1998 | 1444 | 2714 | 2314 | 2343 | 2872 | 2627 | 2348 | 2819 | 3103 | 1904 | 1580 | 2343 | 1569 | 1565 | 1513 | Ubash3b |
| Ubb | 47843 | 59456 | 48976 | 51292 | 53560 | 52318 | 55297 | 54461 | 60158 | 51798 | 55294 | 57933 | 57200 | 61911 | 50225 | 49763 | 53438 | 51352 | 58811 | 56242 | 56772 | 54094 | 60582 | 61167 | 60859 | 66682 | 64896 | 60163 | Ubb |
| Ubc | 6888 | 7431 | 9086 | 8708 | 8264 | 7724 | 8085 | 8006 | 9756 | 11616 | 13140 | 13058 | 11853 | 12704 | 6948 | 8090 | 9489 | 9554 | 10305 | 5930 | 8243 | 11655 | 10139 | 13008 | 12588 | 14685 | 12794 | 11843 | Ubc |
| Ubd | 0 | 0 | 0 | 0 | 0 | 0 | 0 | 0 | 0 | 0 | 0 | 0 | 0 | 0 | 0 | 0 | 0 | 0 | 0 | 0 | 0 | 0 | 0 | 2 | 0 | 0 | 0 | 1 | Ubd |
| Ube2a | 3290 | 2985 | 3155 | 3316 | 3032 | 3011 | 3511 | 2442 | 2164 | 2179 | 2120 | 2338 | 2171 | 2551 | 3522 | 3089 | 3182 | 3122 | 2890 | 3128 | 3329 | 2371 | 2311 | 2285 | 2204 | 2461 | 2083 | 2500 | Ube2a |
| Ube2b | 13279 | 10974 | 10970 | 13104 | 12002 | 12019 | 14707 | 11870 | 9716 | 9442 | 9997 | 9650 | 9719 | 10615 | 14372 | 12522 | 11768 | 12616 | 12357 | 14760 | 13783 | 10457 | 9935 | 10457 | 10155 | 9906 | 9642 | 10377 | Ube2b |
| Ube2c | 3378 | 3961 | 4586 | 3369 | 2295 | 2631 | 3006 | 3698 | 4180 | 4121 | 3990 | 6599 | 5332 | 7026 | 3032 | 3132 | 4079 | 2496 | 3323 | 2944 | 2898 | 2864 | 3717 | 5909 | 4760 | 8839 | 6701 | 7245 | Ube2c |
| Ube2cbp | 90 | 109 | 113 | 75 | 122 | 65 | 106 | 302 | 311 | 278 | 289 | 311 | 310 | 322 | 82 | 73 | 107 | 84 | 92 | 84 | 94 | 289 | 286 | 306 | 247 | 359 | 349 | 348 | Ube2cbp |
| Ube2d1 | 5174 | 5034 | 6490 | 6891 | 7648 | 6847 | 7420 | 6591 | 5975 | 7592 | 6613 | 7382 | 6518 | 7756 | 5540 | 5165 | 6901 | 6650 | 7248 | 6192 | 7409 | 8417 | 5781 | 7548 | 8111 | 7288 | 6695 | 7614 | Ube2d1 |
| Ube2d2a | 9283 | 6807 | 9806 | 10071 | 10469 | 10617 | 11670 | 10679 | 8511 | 10396 | 10008 | 11486 | 10675 | 11995 | 9546 | 8001 | 9682 | 10069 | 10474 | 9825 | 10735 | 11571 | 8686 | 10963 | 11377 | 12181 | 10982 | 12060 | Ube2d2a |
| Ube2d3 | 24724 | 18646 | 22191 | 21650 | 22191 | 22550 | 25647 | 30157 | 25359 | 23370 | 25057 | 27410 | 26468 | 28938 | 25776 | 21029 | 22080 | 21734 | 21669 | 23664 | 24229 | 27902 | 26083 | 26474 | 25727 | 29784 | 27761 | 30720 | Ube2d3 |
| Ube2e1 | 6651 | 5251 | 7006 | 6159 | 6510 | 6232 | 7387 | 8888 | 7043 | 7497 | 7020 | 7679 | 8010 | 9114 | 6831 | 5803 | 6627 | 5887 | 6606 | 7136 | 6772 | 7723 | 7323 | 8180 | 7998 | 8888 | 8513 | 9137 | Ube2e1 |
| Ube2e2 | 1316 | 1145 | 1259 | 1498 | 1712 | 1536 | 1606 | 1883 | 1731 | 1895 | 1825 | 1551 | 1564 | 1680 | 1431 | 1199 | 1336 | 1537 | 1477 | 1465 | 1567 | 1871 | 1957 | 1745 | 1888 | 1537 | 1571 | 1550 | Ube2e2 |
| Ube2e3 | 7384 | 6315 | 7390 | 7102 | 8759 | 8408 | 8789 | 13854 | 11670 | 12775 | 11277 | 13380 | 12644 | 14294 | 8466 | 6317 | 7463 | 7066 | 8259 | 8383 | 8835 | 13595 | 11509 | 12303 | 13058 | 13362 | 12936 | 14584 | Ube2e3 |
| Ube2f | 4463 | 4458 | 3658 | 3690 | 3101 | 3456 | 4089 | 3134 | 2618 | 2412 | 2333 | 2465 | 2361 | 2667 | 4429 | 3987 | 3669 | 3648 | 3186 | 4097 | 3800 | 2540 | 2582 | 2396 | 2476 | 2701 | 2615 | 2908 | Ube2f |
| Ube2g1 | 8637 | 6267 | 8349 | 7944 | 8593 | 8277 | 9185 | 10616 | 8462 | 8561 | 8203 | 10034 | 9593 | 10249 | 8399 | 7213 | 7873 | 7747 | 8678 | 7948 | 9005 | 10291 | 8877 | 9046 | 9766 | 10347 | 9387 | 10942 | Ube2g1 |
| Ube2g2 | 3053 | 3587 | 3586 | 3666 | 3118 | 3185 | 2856 | 2568 | 2578 | 2539 | 2538 | 2840 | 2667 | 2603 | 3040 | 3137 | 3540 | 3503 | 3470 | 3155 | 2947 | 2363 | 2662 | 3042 | 2978 | 3333 | 2848 | 2785 | Ube2g2 |
| Ube2h | 11465 | 10870 | 13458 | 14803 | 14305 | 12807 | 13742 | 9122 | 8253 | 9990 | 9887 | 10000 | 9317 | 10095 | 11322 | 11136 | 13534 | 14662 | 13913 | 11571 | 13477 | 10724 | 8576 | 10689 | 10963 | 9537 | 8966 | 9909 | Ube2h |
| Ube2i | 17800 | 16232 | 16524 | 16022 | 15913 | 15892 | 16894 | 18237 | 16918 | 17197 | 17530 | 21038 | 19870 | 21560 | 16578 | 15479 | 15880 | 15846 | 16623 | 15466 | 16557 | 17137 | 15995 | 19244 | 19511 | 20249 | 21256 | 22126 | Ube2i |
| Ube2j1 | 4571 | 4355 | 4972 | 5042 | 6089 | 5434 | 5979 | 6233 | 5052 | 5580 | 5441 | 5498 | 5167 | 5648 | 5368 | 4283 | 4979 | 5321 | 5675 | 5799 | 5702 | 6036 | 5268 | 5560 | 5852 | 5394 | 5455 | 5798 | Ube2j1 |
| Ube2j2 | 3171 | 3281 | 3796 | 3569 | 3158 | 3091 | 3317 | 3207 | 3121 | 3654 | 3347 | 4022 | 3498 | 4212 | 3115 | 2958 | 3741 | 3299 | 3461 | 2899 | 3347 | 3501 | 2970 | 3775 | 3414 | 4456 | 3856 | 4087 | Ube2j2 |
| Ube2k | 12102 | 10291 | 13730 | 13189 | 13060 | 13134 | 14142 | 12451 | 10514 | 11966 | 11266 | 11922 | 11345 | 12525 | 12803 | 10565 | 12741 | 13622 | 13471 | 14349 | 13716 | 12120 | 10570 | 12646 | 12965 | 13063 | 12221 | 12873 | Ube2k |
| Ube2l3 | 12832 | 13711 | 14340 | 12630 | 11474 | 12005 | 13116 | 11883 | 12201 | 12414 | 11182 | 13940 | 13477 | 14703 | 13239 | 12560 | 13355 | 11656 | 12599 | 12976 | 12660 | 11949 | 12122 | 12871 | 12900 | 16379 | 14014 | 15078 | Ube2l3 |
| Ube2l6 | 1143 | 1287 | 1265 | 881 | 719 | 936 | 868 | 819 | 975 | 749 | 681 | 701 | 1444 | 889 | 1267 | 1247 | 1087 | 919 | 889 | 1133 | 884 | 483 | 1082 | 704 | 693 | 874 | 951 | 1443 | Ube2l6 |
| Ube2m | 5051 | 5814 | 6287 | 6201 | 5803 | 5255 | 5480 | 5157 | 5558 | 6175 | 5934 | 6095 | 5762 | 6695 | 4965 | 4761 | 6186 | 5844 | 6356 | 5010 | 6192 | 5904 | 5622 | 6718 | 6235 | 7215 | 5712 | 6608 | Ube2m |
| Ube2n | 14042 | 12734 | 15816 | 15165 | 15162 | 14900 | 17074 | 14818 | 12473 | 13295 | 12192 | 14836 | 13590 | 15888 | 14977 | 12592 | 15463 | 14697 | 15641 | 15347 | 15733 | 14531 | 12244 | 14142 | 14542 | 16324 | 14920 | 15796 | Ube2n |
| Ube2o | 7018 | 9416 | 10379 | 11639 | 10968 | 9611 | 8916 | 5241 | 6707 | 7449 | 6730 | 7319 | 7419 | 6773 | 7159 | 8911 | 11126 | 11205 | 11099 | 8099 | 9550 | 7456 | 6113 | 7208 | 7950 | 7003 | 6195 | 6880 | Ube2o |
| Ube2q1 | 7379 | 7811 | 7791 | 8006 | 8068 | 7721 | 8141 | 8760 | 7701 | 8633 | 7792 | 9115 | 8097 | 9611 | 7308 | 6618 | 8304 | 7954 | 8179 | 7236 | 8379 | 8985 | 7610 | 8917 | 8853 | 9316 | 8939 | 9743 | Ube2q1 |
| Ube2q2 | 3711 | 2862 | 3830 | 3208 | 3607 | 3382 | 3571 | 4887 | 4219 | 4408 | 4511 | 4809 | 4779 | 4970 | 3840 | 3593 | 3634 | 3350 | 3324 | 3454 | 3308 | 4705 | 3959 | 4638 | 4727 | 5124 | 4585 | 5280 | Ube2q2 |
| Ube2ql1 | 1070 | 1603 | 1307 | 2090 | 4206 | 2619 | 2506 | 3002 | 3158 | 3883 | 3528 | 2537 | 2728 | 2397 | 1636 | 1372 | 1842 | 2660 | 3053 | 2437 | 2962 | 4523 | 2880 | 2989 | 3853 | 1726 | 2022 | 2122 | Ube2ql1 |
| Ube2r2 | 7116 | 6430 | 9073 | 7866 | 7718 | 7013 | 7582 | 11251 | 10274 | 14189 | 12256 | 14491 | 12786 | 15033 | 6912 | 6495 | 8749 | 7158 | 8451 | 5788 | 7997 | 13217 | 10672 | 15109 | 13017 | 16981 | 13278 | 15535 | Ube2r2 |
| Ube2s | 9036 | 11675 | 10118 | 10260 | 8056 | 8684 | 8268 | 6875 | 7749 | 6794 | 6792 | 9141 | 7878 | 8453 | 9216 | 10350 | 9729 | 9236 | 9857 | 9565 | 7970 | 5904 | 7385 | 8164 | 7747 | 11433 | 9404 | 8605 | Ube2s |
| Ube2t | 2000 | 2147 | 2238 | 2166 | 1471 | 1706 | 1942 | 1045 | 966 | 991 | 898 | 1355 | 1243 | 1421 | 2003 | 2047 | 2311 | 2056 | 1919 | 2073 | 1807 | 780 | 927 | 1162 | 1067 | 1473 | 1192 | 1354 | Ube2t |
| Ube2u | 150 | 155 | 128 | 184 | 97 | 110 | 120 | 42 | 35 | 36 | 49 | 65 | 65 | 42 | 150 | 180 | 125 | 136 | 133 | 119 | 88 | 37 | 39 | 33 | 47 | 60 | 55 | 58 | Ube2u |
| Ube2v1 | 57 | 29 | 28 | 56 | 37 | 68 | 49 | 51 | 50 | 52 | 29 | 62 | 81 | 62 | 49 | 35 | 50 | 43 | 41 | 40 | 51 | 63 | 43 | 65 | 29 | 29 | 87 | 45 | Ube2v1 |
| Ube2v2 | 2660 | 2673 | 2394 | 2533 | 3014 | 3444 | 2814 | 2908 | 3599 | 2549 | 2176 | 2446 | 2876 | 2881 | 3051 | 2705 | 2840 | 2730 | 2763 | 4424 | 2815 | 2956 | 2648 | 2674 | 3131 | 2795 | 2431 | 2500 | Ube2v2 |
| Ube2w | 4182 | 3221 | 3964 | 4119 | 4618 | 4546 | 4903 | 4358 | 3741 | 3319 | 3636 | 3288 | 3285 | 3797 | 4704 | 3888 | 3958 | 4430 | 4190 | 4948 | 4699 | 4079 | 3744 | 3510 | 3569 | 3529 | 3436 | 3745 | Ube2w |
| Ube2z | 6754 | 6543 | 7116 | 7063 | 7155 | 6643 | 6575 | 6146 | 6580 | 6577 | 7186 | 6234 | 6061 | 6089 | 6345 | 6823 | 7170 | 7587 | 7138 | 5585 | 6948 | 7080 | 6763 | 6675 | 6658 | 7197 | 5610 | 6330 | Ube2z |
| Ube3a | 15955 | 12053 | 11518 | 14011 | 15100 | 16202 | 16919 | 23908 | 19131 | 14867 | 16081 | 16259 | 15939 | 16926 | 18303 | 14911 | 13506 | 13881 | 14703 | 18190 | 17069 | 17891 | 19100 | 14424 | 17366 | 13105 | 17512 | 16743 | Ube3a |
| Ube3b | 6424 | 7743 | 7054 | 7131 | 7283 | 6804 | 6885 | 6119 | 7990 | 8523 | 7977 | 9318 | 8780 | 8228 | 6221 | 7584 | 7858 | 7165 | 7675 | 5609 | 6762 | 8011 | 7504 | 8536 | 8656 | 8671 | 8049 | 8744 | Ube3b |
| Ube3c | 8784 | 8344 | 6743 | 7171 | 7301 | 7623 | 8356 | 8965 | 7927 | 6867 | 6830 | 7894 | 6894 | 7776 | 8974 | 7518 | 6804 | 7159 | 7575 | 8143 | 8410 | 8006 | 7754 | 7032 | 7767 | 6824 | 7779 | 7626 | Ube3c |
| Ube4a | 15418 | 15368 | 10988 | 12565 | 11858 | 12540 | 13634 | 12560 | 11137 | 9548 | 9736 | 9750 | 9967 | 9441 | 16249 | 14233 | 11811 | 13244 | 11836 | 13694 | 13047 | 11032 | 10754 | 9302 | 10333 | 7908 | 10365 | 9610 | Ube4a |
| Ube4b | 11749 | 12117 | 13438 | 13701 | 13958 | 12912 | 13357 | 11187 | 11587 | 13600 | 12657 | 14955 | 12948 | 13952 | 11380 | 11802 | 14066 | 13033 | 13824 | 11157 | 13772 | 14051 | 11126 | 13237 | 13820 | 12840 | 12953 | 14242 | Ube4b |
| Ubfd1 | 6640 | 6001 | 6971 | 6691 | 6266 | 6228 | 5754 | 9145 | 8518 | 8365 | 9095 | 7978 | 8003 | 7469 | 6519 | 6186 | 6900 | 6529 | 6555 | 5778 | 5991 | 9670 | 8984 | 8670 | 9224 | 9082 | 8421 | 8224 | Ubfd1 |
| Ubiad1 | 759 | 804 | 843 | 656 | 710 | 719 | 675 | 1055 | 1154 | 1195 | 1035 | 1398 | 1158 | 1036 | 761 | 762 | 881 | 661 | 871 | 703 | 667 | 1131 | 1054 | 1235 | 1147 | 1326 | 1195 | 1241 | Ubiad1 |
| Ubl3 | 3756 | 3859 | 4329 | 4778 | 4982 | 4439 | 5141 | 2867 | 3074 | 3389 | 3146 | 3452 | 3133 | 3508 | 3649 | 3696 | 4591 | 4503 | 4785 | 4227 | 5038 | 3402 | 3153 | 3544 | 3322 | 3808 | 2862 | 3463 | Ubl3 |

| | | | | | | | | | | | | | | | | | | | | | | | | | | | | | |
|---|---|---|---|---|---|---|---|---|---|---|---|---|---|---|---|---|---|---|---|---|---|---|---|---|---|---|---|---|---|
| Ubl4a | 2696 | 2345 | 2346 | 2301 | 2407 | 2543 | 2663 | 3350 | 3170 | 3082 | 2924 | 3701 | 3308 | 3626 | 2520 | 2203 | 2334 | 2170 | 2618 | 2364 | 2539 | 3342 | 3263 | 3291 | 3174 | 3914 | 3626 | 3882 | Ubl4a |
| Ubl4b | 17 | 10 | 19 | 7 | 7 | 1 | 2 | 23 | 13 | 14 | 14 | 18 | 17 | 15 | 9 | 9 | 17 | 2 | 0 | 5 | 4 | 8 | 17 | 34 | 17 | 41 | 17 | 18 | Ubl4b |
| Ubl5 | 5560 | 6430 | 6065 | 6433 | 5998 | 5267 | 7771 | 5503 | 4578 | 5644 | 5677 | 5441 | 5905 | 7070 | 5731 | 4274 | 6544 | 6022 | 6313 | 6126 | 7530 | 5285 | 4592 | 6251 | 6058 | 6280 | 5847 | 7275 | Ubl5 |
| Ubl7 | 2397 | 2802 | 2808 | 3255 | 2923 | 2778 | 2961 | 2434 | 2810 | 3101 | 2775 | 3050 | 2745 | 3177 | 2471 | 2518 | 2845 | 2989 | 3160 | 2500 | 3073 | 3103 | 2707 | 3201 | 3015 | 3422 | 2779 | 3170 | Ubl7 |
| Ublcp1 | 4683 | 4132 | 4414 | 4005 | 3454 | 3512 | 4114 | 4187 | 3545 | 3163 | 3398 | 3761 | 3842 | 4199 | 4853 | 4315 | 4008 | 3656 | 3698 | 4053 | 3927 | 3347 | 3493 | 3446 | 3399 | 3904 | 4113 | 4167 | Ublcp1 |
| Ubn1 | 10649 | 11195 | 9482 | 11334 | 9398 | 10466 | 9715 | 9048 | 8525 | 9073 | 9257 | 9674 | 9409 | 9561 | 10163 | 11089 | 10210 | 10631 | 9489 | 8829 | 9952 | 8911 | 8495 | 9151 | 9527 | 8064 | 9903 | 9676 | Ubn1 |
| Ubn2 | 13229 | 10974 | 9688 | 11633 | 12334 | 15070 | 14534 | 17075 | 14368 | 12868 | 13654 | 13675 | 14232 | 15026 | 12612 | 12009 | 10804 | 11378 | 11875 | 13848 | 15538 | 13488 | 13045 | 11942 | 14753 | 10000 | 15336 | 13655 | Ubn2 |
| Ubox5 | 1553 | 1585 | 1227 | 1062 | 1037 | 1167 | 1259 | 1095 | 1331 | 994 | 1024 | 1240 | 1228 | 1276 | 1400 | 1345 | 1269 | 1047 | 1153 | 1276 | 1257 | 1043 | 1156 | 1219 | 1129 | 1464 | 1249 | 1375 | Ubox5 |
| Ubp1 | 15430 | 13984 | 15351 | 16505 | 16579 | 16741 | 17857 | 12225 | 11829 | 13931 | 12835 | 15122 | 13275 | 15035 | 14927 | 13886 | 15950 | 16372 | 16971 | 14688 | 17734 | 13739 | 11094 | 14149 | 14005 | 13905 | 13928 | 14571 | Ubp1 |
| Ubqln1 | 14479 | 14423 | 18669 | 18033 | 18056 | 16398 | 18372 | 12323 | 12297 | 14813 | 13495 | 15610 | 13768 | 15094 | 14479 | 13440 | 19060 | 18306 | 19286 | 14291 | 18303 | 14713 | 12007 | 15810 | 15486 | 16399 | 14256 | 15692 | Ubqln1 |
| Ubqln2 | 10746 | 10547 | 16011 | 14663 | 14974 | 13054 | 13692 | 12708 | 11683 | 16141 | 14060 | 18957 | 13615 | 18288 | 10744 | 10326 | 15552 | 14764 | 15520 | 10355 | 14550 | 19403 | 11687 | 17510 | 17854 | 20176 | 15080 | 17827 | Ubqln2 |
| Ubqln3 | 0 | 0 | 0 | 0 | 0 | 0 | 0 | 1 | 2 | 11 | 1 | 0 | 0 | 0 | 0 | 0 | 0 | 0 | 0 | 0 | 0 | 0 | 6 | 3 | 0 | 0 | 0 | 0 | Ubqln3 |
| Ubqln4 | 8140 | 9020 | 10240 | 9447 | 9428 | 8489 | 9093 | 7525 | 8307 | 9197 | 8263 | 10268 | 8966 | 10288 | 7777 | 8129 | 10819 | 9034 | 10394 | 7867 | 9359 | 8988 | 7655 | 10030 | 9596 | 11318 | 9726 | 10554 | Ubqln4 |
| Ubqln5 | 0 | 0 | 0 | 0 | 0 | 0 | 0 | 2 | 0 | 5 | 0 | 0 | 0 | 0 | 0 | 0 | 0 | 0 | 0 | 0 | 0 | 0 | 1 | 0 | 0 | 1 | 0 | 0 | Ubqln5 |
| Ubqlnl | 0 | 0 | 0 | 0 | 0 | 0 | 0 | 7 | 5 | 24 | 2 | 0 | 0 | 0 | 0 | 0 | 0 | 0 | 0 | 0 | 0 | 0 | 10 | 10 | 0 | 0 | 0 | 0 | Ubqlnl |
| Ubr1 | 5027 | 4075 | 3469 | 4417 | 5427 | 5460 | 5475 | 7956 | 7256 | 6427 | 6884 | 6862 | 6634 | 7151 | 5313 | 4251 | 3768 | 4261 | 4481 | 5619 | 5353 | 6843 | 6970 | 5704 | 6988 | 4886 | 7424 | 6563 | Ubr1 |
| Ubr2 | 10813 | 10803 | 9191 | 11664 | 12108 | 11691 | 11990 | 7574 | 7987 | 8482 | 8830 | 8285 | 8240 | 8526 | 9972 | 10554 | 10652 | 11523 | 11982 | 10166 | 12648 | 8751 | 7655 | 8228 | 8846 | 5100 | 7968 | 7944 | Ubr2 |
| Ubr3 | 16702 | 13669 | 11047 | 13993 | 16176 | 17334 | 17349 | 15880 | 14100 | 12609 | 13377 | 13876 | 14350 | 15281 | 17180 | 13981 | 12147 | 14087 | 13938 | 16789 | 17799 | 14001 | 13459 | 11534 | 14138 | 9602 | 14578 | 13981 | Ubr3 |
| Ubr4 | 19927 | 24062 | 14314 | 21971 | 25265 | 26367 | 23474 | 16751 | 20239 | 21377 | 21406 | 18661 | 20579 | 18714 | 18834 | 23601 | 16337 | 22920 | 22509 | 20130 | 27140 | 17161 | 18533 | 14792 | 22928 | 11000 | 20828 | 16231 | Ubr4 |
| Ubr5 | 27811 | 24750 | 17592 | 22691 | 26323 | 29193 | 29129 | 34166 | 31646 | 29637 | 29412 | 34207 | 32493 | 35766 | 27167 | 24707 | 19890 | 22922 | 24333 | 26712 | 29487 | 28754 | 29341 | 26269 | 32920 | 21361 | 36185 | 32325 | Ubr5 |
| Ubr7 | 13064 | 12788 | 10700 | 10411 | 8073 | 9592 | 10261 | 7184 | 6337 | 5514 | 5204 | 6745 | 5964 | 6886 | 12951 | 11456 | 10418 | 9738 | 9806 | 10870 | 10061 | 5718 | 6099 | 6201 | 5924 | 7904 | 7380 | 7195 | Ubr7 |
| Ubtd1 | 919 | 1285 | 977 | 1340 | 983 | 992 | 768 | 432 | 564 | 522 | 654 | 399 | 488 | 405 | 945 | 1099 | 1109 | 1121 | 1019 | 941 | 886 | 436 | 589 | 485 | 412 | 573 | 440 | 414 | Ubtd1 |
| Ubtd2 | 2790 | 2871 | 3308 | 2791 | 2887 | 2951 | 2778 | 2530 | 2538 | 2342 | 2638 | 2928 | 2877 | 2897 | 2821 | 2631 | 2856 | 2744 | 3068 | 2592 | 2741 | 2643 | 2607 | 2781 | 2872 | 3025 | 2886 | 2968 | Ubtd2 |
| Ubtf | 5601 | 5924 | 7960 | 6631 | 6762 | 6533 | 6331 | 11258 | 12301 | 13886 | 12987 | 14106 | 12420 | 14192 | 5145 | 5270 | 7260 | 6099 | 7305 | 6141 | 6603 | 12064 | 12264 | 15274 | 14318 | 15987 | 13154 | 14587 | Ubtf |
| Ubtfl1 | 1 | 0 | 2 | 0 | 0 | 2 | 0 | 1 | 0 | 0 | 0 | 0 | 0 | 0 | 0 | 0 | 0 | 0 | 0 | 0 | 2 | 0 | 0 | 0 | 0 | 0 | 2 | 0 | Ubtfl1 |
| Ubxn1 | 6786 | 7836 | 8137 | 7787 | 6365 | 6600 | 6857 | 5361 | 5885 | 6186 | 5907 | 6627 | 6151 | 6983 | 6502 | 6551 | 8233 | 7060 | 7139 | 7149 | 6847 | 5148 | 5758 | 7282 | 6406 | 8129 | 6593 | 7466 | Ubxn1 |
| Ubxn10 | 223 | 318 | 257 | 196 | 253 | 284 | 285 | 206 | 214 | 240 | 173 | 267 | 286 | 182 | 286 | 317 | 256 | 244 | 295 | 300 | 266 | 234 | 307 | 145 | 204 | 150 | 187 | 166 | Ubxn10 |
| Ubxn11 | 571 | 644 | 592 | 634 | 718 | 659 | 514 | 503 | 590 | 621 | 603 | 508 | 581 | 512 | 488 | 667 | 584 | 599 | 632 | 545 | 648 | 708 | 675 | 489 | 534 | 444 | 529 | 478 | Ubxn11 |
| Ubxn2a | 2731 | 2097 | 2290 | 2709 | 3691 | 3403 | 3930 | 2617 | 2308 | 2368 | 2420 | 2306 | 2208 | 2520 | 2937 | 2290 | 2407 | 3119 | 3321 | 3617 | 3699 | 2164 | 2385 | 2545 | 2639 | 2019 | 2170 | 2174 | Ubxn2a |
| Ubxn2b | 5928 | 5704 | 4968 | 6561 | 6213 | 6208 | 7382 | 4609 | 3751 | 3613 | 3463 | 4199 | 3818 | 4450 | 6494 | 5432 | 5480 | 6522 | 6162 | 6801 | 7158 | 4244 | 3660 | 3739 | 4055 | 3319 | 3946 | 4208 | Ubxn2b |
| Ubxn4 | 8920 | 8053 | 9333 | 9062 | 8811 | 8871 | 9456 | 9050 | 7806 | 8635 | 8538 | 8781 | 8899 | 9188 | 9525 | 8255 | 9254 | 8652 | 8864 | 8971 | 8610 | 8938 | 8381 | 8762 | 8910 | 8533 | 8794 | 9314 | Ubxn4 |
| Ubxn6 | 1912 | 2405 | 1968 | 2302 | 2790 | 2376 | 2443 | 3485 | 3685 | 3669 | 3440 | 3512 | 3334 | 3357 | 2028 | 1886 | 2365 | 2586 | 2884 | 2445 | 2585 | 3659 | 3750 | 3617 | 3869 | 3250 | 3516 | 3377 | Ubxn6 |
| Ubxn7 | 11900 | 9211 | 10447 | 11203 | 11936 | 12272 | 12351 | 16603 | 13196 | 12301 | 12505 | 13248 | 12826 | 14001 | 11586 | 10361 | 10244 | 10813 | 11413 | 12145 | 12387 | 14028 | 12705 | 12706 | 14505 | 12375 | 13702 | 14055 | Ubxn7 |
| Ubxn8 | 100 | 84 | 110 | 93 | 162 | 119 | 118 | 758 | 799 | 826 | 955 | 849 | 951 | 974 | 156 | 96 | 124 | 106 | 143 | 75 | 117 | 737 | 823 | 840 | 834 | 859 | 949 | 967 | Ubxn8 |
| Uchl1 | 12442 | 17404 | 17212 | 21829 | 25215 | 19861 | 22696 | 13572 | 15681 | 18213 | 15843 | 15660 | 15071 | 16782 | 14483 | 13952 | 20952 | 22210 | 24639 | 20833 | 22863 | 18563 | 14683 | 17594 | 21138 | 14723 | 14027 | 15507 | Uchl1 |
| Uchl3 | 5134 | 4600 | 4704 | 5130 | 4793 | 4521 | 6088 | 3684 | 2944 | 3032 | 2561 | 2747 | 2785 | 3168 | 5656 | 4513 | 4870 | 4676 | 5000 | 5957 | 5653 | 3329 | 3119 | 3058 | 3281 | 2990 | 2891 | 3070 | Uchl3 |
| Uchl4 | 133 | 149 | 153 | 135 | 116 | 134 | 125 | 88 | 60 | 74 | 74 | 56 | 61 | 93 | 148 | 150 | 115 | 101 | 158 | 187 | 130 | 95 | 68 | 58 | 71 | 70 | 77 | 61 | Uchl4 |
| Uchl5 | 4290 | 3387 | 4862 | 4464 | 3391 | 3557 | 4427 | 2637 | 2092 | 2428 | 2261 | 2861 | 2698 | 3100 | 4342 | 4009 | 4556 | 4096 | 3763 | 4228 | 3796 | 2321 | 2067 | 2956 | 2516 | 3540 | 3155 | 3140 | Uchl5 |
| Uck1 | 2979 | 3689 | 3085 | 3182 | 2773 | 2668 | 2696 | 2329 | 2530 | 2391 | 2344 | 2488 | 2279 | 2399 | 2736 | 3040 | 3147 | 2921 | 3081 | 2789 | 2756 | 2293 | 2433 | 2537 | 2672 | 3115 | 2496 | 2519 | Uck1 |
| Uck2 | 10754 | 10870 | 12371 | 10166 | 9773 | 9845 | 8823 | 9152 | 8811 | 9723 | 8567 | 11907 | 10760 | 11358 | 10300 | 10604 | 11443 | 9523 | 10555 | 9283 | 9153 | 8778 | 8300 | 10242 | 9931 | 13000 | 11802 | 11838 | Uck2 |
| Uckl1 | 1028 | 1297 | 1215 | 1379 | 1286 | 1262 | 1250 | 1733 | 1772 | 1908 | 1772 | 2069 | 1749 | 1948 | 1063 | 1055 | 1360 | 1345 | 1364 | 1220 | 1222 | 1831 | 1762 | 1995 | 1719 | 1815 | 1908 | 1857 | Uckl1 |
| Ucma | 3 | 14 | 6 | 7 | 3 | 17 | 6 | 10 | 24 | 22 | 22 | 7 | 15 | 11 | 17 | 9 | 7 | 9 | 8 | 3 | 11 | 15 | 30 | 25 | 25 | 8 | 15 | 11 | Ucma |
| Ucn | 0 | 1 | 2 | 0 | 4 | 5 | 0 | 0 | 0 | 0 | 0 | 0 | 0 | 0 | 1 | 4 | 0 | 2 | 0 | 0 | 0 | 0 | 0 | 0 | 3 | 0 | 0 | 0 | Ucn |
| Ucn2 | 23 | 38 | 39 | 38 | 32 | 45 | 34 | 16 | 30 | 27 | 11 | 16 | 22 | 19 | 22 | 51 | 32 | 37 | 41 | 7 | 28 | 12 | 27 | 31 | 6 | 17 | 29 | 25 | Ucn2 |
| Ucn3 | 0 | 0 | 0 | 0 | 1 | 0 | 0 | 0 | 0 | 0 | 0 | 0 | 0 | 0 | 0 | 0 | 0 | 0 | 0 | 2 | 0 | 0 | 1 | 0 | 0 | 0 | 0 | 0 | Ucn3 |
| Ucp1 | 29 | 22 | 14 | 42 | 55 | 32 | 34 | 26 | 24 | 13 | 26 | 15 | 19 | 21 | 24 | 27 | 42 | 51 | 45 | 26 | 49 | 35 | 20 | 11 | 15 | 23 | 10 | 9 | Ucp1 |
| Ucp2 | 3984 | 4488 | 4447 | 3478 | 2544 | 2829 | 2375 | 1833 | 2313 | 2578 | 2553 | 2052 | 2700 | 1995 | 3558 | 5238 | 3979 | 2960 | 2614 | 2270 | 2288 | 1788 | 3108 | 2392 | 1558 | 2977 | 1905 | 2428 | Ucp2 |
| Ucp3 | 5 | 3 | 7 | 7 | 5 | 2 | 10 | 2 | 1 | 3 | 5 | 1 | 5 | 2 | 7 | 12 | 8 | 2 | 10 | 0 | 5 | 2 | 3 | 2 | 0 | 6 | 2 | 0 | Ucp3 |
| Uevld | 2938 | 2256 | 2539 | 2148 | 1964 | 2419 | 2453 | 2329 | 2177 | 1935 | 2008 | 2322 | 2260 | 2689 | 2552 | 2332 | 2343 | 2195 | 2115 | 2443 | 2245 | 1684 | 2056 | 2178 | 2046 | 2236 | 2594 | 2506 | Uevld |
| Ufc1 | 1911 | 2105 | 2134 | 1911 | 2027 | 1878 | 2095 | 3515 | 3587 | 3828 | 3819 | 4028 | 3955 | 4549 | 1965 | 1820 | 2125 | 1862 | 2075 | 2120 | 2061 | 3215 | 3694 | 3976 | 3979 | 4645 | 4388 | 4755 | Ufc1 |
| Ufd1 | 6134 | 5913 | 5483 | 5245 | 4955 | 5269 | 6037 | 5407 | 4916 | 4572 | 4359 | 4952 | 4484 | 5387 | 6343 | 5541 | 5041 | 5346 | 5245 | 5987 | 5715 | 4465 | 4826 | 4728 | 4868 | 4737 | 4929 | 5277 | Ufd1 |
| Ufl1 | 3902 | 3145 | 3180 | 3551 | 3323 | 3394 | 3741 | 4252 | 3676 | 3535 | 3687 | 3699 | 3726 | 3990 | 3965 | 3488 | 3466 | 3472 | 3306 | 3718 | 3671 | 3659 | 3749 | 3555 | 3614 | 3690 | 3668 | 3992 | Ufl1 |
| Ufm1 | 7990 | 7092 | 7771 | 7343 | 6710 | 7034 | 7799 | 6565 | 5483 | 5537 | 5797 | 6225 | 5874 | 6653 | 8446 | 7353 | 7421 | 6781 | 6584 | 7814 | 7676 | 5979 | 5681 | 5792 | 6151 | 6300 | 6689 | 6805 | Ufm1 |
| Ufsp1 | 262 | 386 | 359 | 373 | 333 | 269 | 301 | 171 | 208 | 196 | 256 | 166 | 174 | 236 | 330 | 340 | 368 | 351 | 323 | 344 | 272 | 184 | 188 | 174 | 240 | 197 | 198 | 205 | Ufsp1 |
| Ufsp2 | 3586 | 2953 | 3256 | 2806 | 2581 | 2867 | 3089 | 2687 | 2576 | 2296 | 2611 | 2586 | 2491 | 2514 | 3415 | 3203 | 2846 | 2672 | 2617 | 2453 | 2749 | 2241 | 2677 | 2382 | 2487 | 2589 | 2563 | 2619 | Ufsp2 |
| Ugcg | 6140 | 4547 | 6140 | 6158 | 6314 | 6106 | 6368 | 5773 | 4915 | 4886 | 5161 | 5296 | 5187 | 5462 | 6310 | 5544 | 5876 | 6410 | 5819 | 6060 | 6053 | 6094 | 4660 | 5249 | 5663 | 5592 | 5290 | 5376 | Ugcg |
| Ugdh | 7019 | 5765 | 5816 | 5579 | 5129 | 5768 | 5931 | 5554 | 5657 | 5372 | 5641 | 4840 | 4872 | 5197 | 6657 | 6109 | 5875 | 5410 | 4941 | 5846 | 5493 | 5365 | 5418 | 5409 | 4781 | 6033 | 4799 | 4828 | Ugdh |
| Uggt1 | 14078 | 13790 | 11683 | 12541 | 11902 | 13051 | 12722 | 11103 | 10452 | 10818 | 11057 | 11281 | 11331 | 11719 | 13502 | 13312 | 11862 | 12482 | 12044 | 10907 | 13255 | 11560 | 10488 | 9817 | 11485 | 8608 | 11943 | 11335 | Uggt1 |
| Uggt2 | 4008 | 3386 | 2272 | 3168 | 4007 | 4517 | 3983 | 5113 | 4527 | 4086 | 4867 | 4015 | 4733 | 4445 | 4286 | 3795 | 2724 | 3347 | 3300 | 4696 | 4263 | 4045 | 4606 | 3206 | 4379 | 1941 | 4749 | 3687 | Uggt2 |
| Ugp2 | 8179 | 6030 | 7580 | 6626 | 6699 | 6454 | 7577 | 6035 | 5670 | 4912 | 6148 | 5059 | 5289 | 5289 | 8335 | 7361 | 6889 | 6753 | 6574 | 7094 | 6795 | 5609 | 6491 | 4954 | 4808 | 5796 | 5299 | 5675 | Ugp2 |
| Ugt1a1 | 0 | 0 | 0 | 0 | 0 | 0 | 0 | 0 | 0 | 1 | 0 | 0 | 1 | 6 | 0 | 0 | 3 | 0 | 0 | 0 | 0 | 0 | 0 | 0 | 0 | 0 | 1 | 0 | Ugt1a1 |
| Ugt1a10 | 0 | 0 | 0 | 0 | 0 | 0 | 0 | 0 | 0 | 0 | 0 | 1 | 0 | 0 | 0 | 0 | 0 | 0 | 0 | 0 | 0 | 0 | 0 | 0 | 0 | 0 | 0 | 0 | Ugt1a10 |
| Ugt1a2 | 0 | 0 | 0 | 0 | 0 | 0 | 0 | 0 | 0 | 0 | 0 | 1 | 0 | 0 | 0 | 0 | 0 | 0 | 0 | 0 | 0 | 0 | 1 | 0 | 0 | 0 | 0 | 0 | Ugt1a2 |
| Ugt1a5 | 0 | 0 | 0 | 0 | 0 | 0 | 0 | 0 | 0 | 7 | 0 | 0 | 0 | 0 | 0 | 0 | 0 | 0 | 0 | 0 | 0 | 0 | 0 | 0 | 0 | 0 | 0 | 1 | Ugt1a5 |
| Ugt1a6a | 0 | 0 | 0 | 3 | 9 | 0 | 1 | 11 | 12 | 3 | 9 | 5 | 3 | 3 | 1 | 0 | 2 | 0 | 0 | 1 | 3 | 7 | 12 | 5 | 3 | 4 | 6 | 5 | Ugt1a6a |
| Ugt1a6b | 0 | 3 | 0 | 0 | 12 | 0 | 6 | 4 | 7 | 5 | 3 | 0 | 1 | 4 | 4 | 4 | 0 | 3 | 0 | 4 | 5 | 8 | 11 | 1 | 1 | 0 | 0 | 0 | Ugt1a6b |
| Ugt1a7c | 21 | 19 | 25 | 13 | 11 | 36 | 26 | 22 | 36 | 27 | 46 | 24 | 48 | 27 | 20 | 19 | 10 | 29 | 14 | 35 | 13 | 27 | 52 | 38 | 28 | 22 | 38 | 25 | Ugt1a7c |
| Ugt1a9 | 0 | 0 | 0 | 0 | 0 | 0 | 0 | 0 | 0 | 0 | 3 | 0 | 0 | 0 | 0 | 0 | 0 | 0 | 0 | 0 | 0 | 0 | 0 | 0 | 0 | 0 | 0 | 0 | Ugt1a9 |
| Ugt2a2 | 0 | 0 | 0 | 0 | 0 | 0 | 0 | 4 | 2 | 9 | 2 | 0 | 0 | 0 | 0 | 0 | 0 | 0 | 0 | 0 | 0 | 0 | 0 | 0 | 0 | 6 | 0 | 0 | Ugt2a2 |
| Ugt2a3 | 1 | 2 | 0 | 0 | 0 | 0 | 0 | 1 | 0 | 0 | 0 | 0 | 0 | 0 | 0 | 0 | 0 | 0 | 0 | 0 | 0 | 0 | 0 | 0 | 0 | 2 | 0 | 0 | Ugt2a3 |
| Ugt2b34 | 0 | 0 | 0 | 0 | 0 | 0 | 0 | 1 | 1 | 0 | 0 | 0 | 0 | 0 | 0 | 0 | 0 | 0 | 0 | 0 | 0 | 0 | 2 | 0 | 0 | 0 | 0 | 0 | Ugt2b34 |
| Ugt2b35 | 0 | 0 | 0 | 0 | 0 | 2 | 0 | 2 | 4 | 0 | 0 | 5 | 0 | 0 | 0 | 0 | 0 | 0 | 0 | 0 | 0 | 0 | 1 | 3 | 0 | 0 | 0 | 0 | Ugt2b35 |
| Ugt2b36 | 0 | 0 | 0 | 0 | 0 | 0 | 0 | 0 | 0 | 0 | 0 | 0 | 0 | 0 | 0 | 0 | 0 | 0 | 0 | 0 | 0 | 0 | 3 | 0 | 0 | 0 | 0 | 1 | Ugt2b36 |
| Ugt2b37 | 0 | 0 | 0 | 0 | 1 | 0 | 0 | 0 | 0 | 0 | 0 | 0 | 0 | 0 | 0 | 0 | 0 | 0 | 0 | 0 | 0 | 0 | 0 | 0 | 0 | 0 | 0 | 0 | Ugt2b37 |
| Ugt2b38 | 0 | 0 | 0 | 0 | 0 | 0 | 0 | 0 | 0 | 0 | 0 | 0 | 0 | 0 | 0 | 0 | 0 | 0 | 0 | 0 | 0 | 0 | 0 | 0 | 0 | 0 | 0 | 1 | Ugt2b38 |
| Ugt2b5 | 1 | 0 | 0 | 0 | 0 | 0 | 0 | 0 | 2 | 0 | 6 | 0 | 6 | 0 | 0 | 2 | 0 | 0 | 0 | 0 | 2 | 5 | 1 | 2 | 1 | 0 | 0 | 0 | Ugt2b5 |
| Ugt3a1 | 1 | 1 | 0 | 0 | 0 | 0 | 0 | 0 | 0 | 0 | 0 | 0 | 0 | 0 | 0 | 1 | 0 | 0 | 3 | 2 | 0 | 0 | 0 | 0 | 0 | 0 | 0 | 0 | Ugt3a1 |
| Ugt3a2 | 0 | 0 | 0 | 0 | 1 | 0 | 0 | 0 | 0 | 0 | 0 | 0 | 0 | 0 | 0 | 0 | 0 | 0 | 0 | 0 | 2 | 0 | 1 | 0 | 0 | 0 | 0 | 0 | Ugt3a2 |
| Ugt8a | 1084 | 1063 | 1038 | 1181 | 1231 | 1047 | 1357 | 509 | 414 | 467 | 410 | 458 | 372 | 546 | 1486 | 956 | 1146 | 1160 | 1069 | 1306 | 1203 | 486 | 372 | 389 | 453 | 536 | 536 | 482 | Ugt8a |

| | | | | | | | | | | | | | | | | | | | | | | | | | | | | | |
|---|---|---|---|---|---|---|---|---|---|---|---|---|---|---|---|---|---|---|---|---|---|---|---|---|---|---|---|---|---|
| Uhmk1 | 9661 | 8229 | 7184 | 8240 | 8516 | 8717 | 8651 | 10667 | 8993 | 8672 | 9108 | 9091 | 10098 | 9396 | 9807 | 8794 | 7645 | 7791 | 7648 | 8531 | 8420 | 9117 | 8904 | 7871 | 8882 | 6643 | 10060 | 9439 | Uhmk1 |
| Uhrf1 | 13615 | 15247 | 18020 | 13114 | 9611 | 10935 | 11145 | 8375 | 10444 | 9016 | 8808 | 14767 | 11995 | 13868 | 12028 | 13794 | 15586 | 11872 | 12970 | 11408 | 10581 | 6495 | 8570 | 11832 | 9714 | 18192 | 15092 | 15469 | Uhrf1 |
| Uhrf1bp1 | 6429 | 7403 | 6056 | 8154 | 7662 | 7122 | 6968 | 3642 | 4128 | 4451 | 4154 | 4399 | 4433 | 4126 | 6125 | 7363 | 6684 | 7920 | 7542 | 6214 | 7242 | 4204 | 3777 | 3872 | 4165 | 3177 | 4386 | 4199 | Uhrf1bp1 |
| Uhrf1bp1l | 4599 | 4118 | 3716 | 4224 | 5105 | 4991 | 5439 | 6609 | 5737 | 6507 | 6192 | 6910 | 6345 | 6584 | 4474 | 4252 | 4081 | 4524 | 4696 | 4502 | 5172 | 6470 | 5565 | 5624 | 6678 | 4828 | 6656 | 6376 | Uhrf1bp1l |
| Uhrf2 | 9780 | 9140 | 9595 | 10198 | 9447 | 9395 | 10229 | 12320 | 10254 | 9815 | 9733 | 10636 | 9815 | 10982 | 9948 | 9461 | 9800 | 9645 | 10066 | 10178 | 10218 | 10193 | 10400 | 10474 | 10498 | 10769 | 10815 | 10853 | Uhrf2 |
| Uimc1 | 2337 | 1938 | 2461 | 2214 | 2130 | 2251 | 2387 | 2884 | 2609 | 2504 | 2530 | 2746 | 2544 | 3015 | 2335 | 2036 | 2153 | 2138 | 2251 | 2345 | 2342 | 2572 | 2512 | 2780 | 2528 | 3072 | 2695 | 2828 | Uimc1 |
| Ulbp1 | 23 | 9 | 6 | 7 | 12 | 12 | 13 | 12 | 2 | 6 | 6 | 0 | 6 | 4 | 42 | 12 | 9 | 8 | 7 | 9 | 13 | 12 | 8 | 7 | 10 | 4 | 12 | 30 | Ulbp1 |
| Ulk1 | 6989 | 9062 | 8236 | 11377 | 10716 | 9674 | 9243 | 6487 | 7518 | 8772 | 7993 | 9910 | 8053 | 8196 | 6610 | 8541 | 10213 | 11994 | 11144 | 7665 | 10094 | 9728 | 6673 | 9470 | 10099 | 8452 | 8075 | 8270 | Ulk1 |
| Ulk2 | 7087 | 6365 | 6890 | 7230 | 9056 | 8229 | 8267 | 13061 | 11504 | 11339 | 10929 | 11058 | 10998 | 11070 | 8057 | 6711 | 7260 | 7912 | 7864 | 8195 | 8437 | 13886 | 11343 | 10544 | 12935 | 9329 | 10802 | 11127 | Ulk2 |
| Ulk3 | 1225 | 1170 | 1499 | 1660 | 1790 | 1610 | 1793 | 1308 | 1498 | 1832 | 1733 | 1743 | 1505 | 1628 | 1116 | 1092 | 1765 | 1742 | 1815 | 1380 | 1583 | 2037 | 1436 | 1843 | 1897 | 1783 | 1570 | 1556 | Ulk3 |
| Ulk4 | 833 | 967 | 796 | 772 | 676 | 725 | 638 | 597 | 665 | 609 | 533 | 654 | 795 | 599 | 818 | 824 | 775 | 691 | 794 | 853 | 753 | 501 | 650 | 668 | 615 | 640 | 774 | 685 | Ulk4 |
| Umad1 | 1311 | 1120 | 1208 | 1494 | 1536 | 1385 | 1630 | 1154 | 924 | 912 | 1028 | 842 | 849 | 917 | 1543 | 1326 | 1254 | 1459 | 1304 | 1503 | 1587 | 1295 | 1015 | 948 | 1012 | 838 | 817 | 945 | Umad1 |
| Umod | 0 | 0 | 0 | 0 | 0 | 0 | 0 | 0 | 0 | 0 | 0 | 0 | 0 | 1 | 0 | 0 | 0 | 0 | 0 | 0 | 0 | 0 | 0 | 0 | 0 | 0 | 0 | 0 | Umod |
| Umodl1 | 13 | 5 | 3 | 3 | 3 | 1 | 6 | 16 | 12 | 35 | 11 | 1 | 15 | 3 | 5 | 12 | 2 | 8 | 1 | 9 | 5 | 5 | 14 | 17 | 6 | 33 | 3 | 5 | Umodl1 |
| Umps | 5424 | 5314 | 4833 | 3903 | 3066 | 3718 | 3869 | 3975 | 3678 | 3240 | 3255 | 3918 | 3809 | 4124 | 4900 | 4831 | 4446 | 3781 | 3815 | 4405 | 3666 | 2707 | 3607 | 3715 | 3338 | 5075 | 4487 | 4679 | Umps |
| Unc119 | 2222 | 2574 | 2315 | 2576 | 2401 | 2414 | 2204 | 2076 | 2349 | 2388 | 2156 | 2607 | 2268 | 2368 | 2011 | 2245 | 2676 | 2351 | 2752 | 2452 | 2401 | 2122 | 2308 | 2344 | 2361 | 2946 | 2432 | 2397 | Unc119 |
| Unc119b | 8173 | 9606 | 9589 | 8703 | 6769 | 7290 | 7365 | 4731 | 4837 | 4798 | 4509 | 6169 | 5753 | 5679 | 7544 | 8588 | 9070 | 8194 | 8623 | 7510 | 7121 | 4037 | 4666 | 5682 | 5154 | 6525 | 6223 | 5967 | Unc119b |
| Unc13a | 8213 | 11934 | 10564 | 19433 | 20228 | 16206 | 18625 | 5388 | 5717 | 8461 | 6617 | 6501 | 4862 | 6097 | 9066 | 9386 | 15518 | 20314 | 20097 | 13873 | 19605 | 9705 | 5229 | 7351 | 9548 | 3258 | 5196 | 5131 | Unc13a |
| Unc13b | 10070 | 10750 | 8630 | 10834 | 11315 | 11884 | 11895 | 9640 | 8864 | 8853 | 7441 | 9556 | 8802 | 9655 | 10934 | 9559 | 9555 | 10797 | 10949 | 11394 | 12066 | 9570 | 8770 | 8066 | 9568 | 6904 | 9902 | 9181 | Unc13b |
| Unc13c | 102 | 125 | 101 | 197 | 449 | 396 | 298 | 460 | 468 | 327 | 495 | 215 | 227 | 218 | 196 | 131 | 90 | 227 | 253 | 398 | 312 | 466 | 513 | 229 | 433 | 154 | 193 | 167 | Unc13c |
| Unc13d | 53 | 97 | 97 | 78 | 85 | 47 | 51 | 60 | 105 | 67 | 92 | 88 | 104 | 84 | 45 | 92 | 100 | 99 | 113 | 85 | 60 | 114 | 118 | 116 | 111 | 93 | 120 | 89 | Unc13d |
| Unc45a | 4839 | 5662 | 5552 | 6429 | 5810 | 5524 | 5336 | 3381 | 3931 | 4669 | 4267 | 4380 | 4427 | 4050 | 4519 | 5143 | 5925 | 6212 | 6121 | 4636 | 5554 | 4279 | 3748 | 4624 | 4382 | 4363 | 4112 | 4232 | Unc45a |
| Unc45b | 48 | 33 | 55 | 29 | 57 | 66 | 36 | 67 | 197 | 82 | 612 | 43 | 76 | 31 | 53 | 65 | 35 | 43 | 52 | 52 | 46 | 152 | 432 | 83 | 36 | 35 | 24 | 49 | Unc45b |
| Unc50 | 2934 | 2737 | 2532 | 2530 | 2353 | 2443 | 2501 | 2945 | 2583 | 2343 | 2543 | 2590 | 2536 | 2471 | 3070 | 2595 | 2749 | 2398 | 2480 | 2812 | 2403 | 2516 | 2763 | 2651 | 2559 | 2811 | 2753 | 2537 | Unc50 |
| Unc5a | 820 | 1240 | 1251 | 1772 | 2099 | 1581 | 1626 | 662 | 901 | 996 | 877 | 722 | 493 | 605 | 901 | 903 | 1890 | 1919 | 2021 | 1536 | 1713 | 1166 | 861 | 922 | 1159 | 758 | 526 | 497 | Unc5a |
| Unc5b | 4346 | 4650 | 6199 | 5272 | 4167 | 4095 | 3941 | 2120 | 2470 | 2749 | 2522 | 3463 | 2950 | 3104 | 3621 | 4106 | 5474 | 4828 | 4766 | 3501 | 4012 | 2620 | 2334 | 3083 | 2598 | 3416 | 3540 | 3862 | Unc5b |
| Unc5c | 3787 | 3614 | 2841 | 2498 | 2928 | 3590 | 2046 | 7739 | 7668 | 6909 | 8050 | 4541 | 7367 | 4674 | 4487 | 4871 | 2359 | 2484 | 1993 | 3144 | 2156 | 8163 | 8478 | 3921 | 5946 | 3030 | 5746 | 4650 | Unc5c |
| Unc5cl | 1 | 1 | 0 | 0 | 0 | 0 | 0 | 0 | 5 | 0 | 0 | 0 | 0 | 0 | 1 | 1 | 0 | 0 | 0 | 0 | 0 | 0 | 0 | 0 | 0 | 0 | 0 | 0 | Unc5cl |
| Unc5d | 1069 | 813 | 591 | 1825 | 2689 | 2324 | 2461 | 2053 | 2034 | 2064 | 1810 | 1144 | 1359 | 1346 | 1094 | 903 | 1166 | 1979 | 2339 | 2228 | 2813 | 2284 | 1721 | 1649 | 2615 | 699 | 1047 | 910 | Unc5d |
| Unc79 | 1043 | 1173 | 777 | 1381 | 2246 | 2038 | 2235 | 3546 | 3522 | 4040 | 3460 | 3515 | 2987 | 3428 | 1275 | 909 | 1059 | 1501 | 1866 | 1670 | 2243 | 4338 | 2785 | 3193 | 4581 | 1516 | 3309 | 2908 | Unc79 |
| Unc80 | 321 | 428 | 271 | 524 | 977 | 874 | 781 | 837 | 933 | 887 | 955 | 538 | 526 | 535 | 473 | 408 | 397 | 592 | 588 | 807 | 904 | 1029 | 822 | 556 | 927 | 331 | 543 | 430 | Unc80 |
| Unc93a | 1 | 0 | 0 | 1 | 1 | 0 | 2 | 0 | 1 | 0 | 0 | 2 | 0 | 4 | 0 | 1 | 0 | 0 | 0 | 1 | 0 | 0 | 3 | 0 | 0 | 0 | 0 | 1 | Unc93a |
| Unc93a2 | 0 | 0 | 0 | 0 | 0 | 2 | 0 | 0 | 0 | 0 | 0 | 0 | 0 | 0 | 0 | 0 | 0 | 0 | 0 | 0 | 0 | 0 | 0 | 0 | 0 | 0 | 0 | 0 | Unc93a2 |
| Unc93b1 | 324 | 396 | 360 | 286 | 331 | 294 | 217 | 286 | 424 | 346 | 426 | 351 | 380 | 293 | 320 | 467 | 302 | 361 | 237 | 281 | 244 | 336 | 548 | 378 | 317 | 363 | 338 | 277 | Unc93b1 |
| Uncx | 621 | 1024 | 1350 | 1273 | 919 | 960 | 448 | 1072 | 834 | 1279 | 987 | 1347 | 1709 | 1051 | 758 | 772 | 1604 | 874 | 1035 | 989 | 572 | 1445 | 808 | 1244 | 1257 | 1269 | 1361 | 1360 | Uncx |
| Ung | 1615 | 1900 | 2132 | 2231 | 1523 | 1517 | 1749 | 883 | 1188 | 1038 | 1093 | 1931 | 1379 | 1606 | 1469 | 1621 | 1900 | 1856 | 1792 | 1626 | 1560 | 732 | 977 | 1517 | 1184 | 2590 | 1720 | 1889 | Ung |
| Unk | 2370 | 2860 | 3307 | 3301 | 3313 | 3028 | 2927 | 3037 | 3792 | 4390 | 4197 | 5142 | 4394 | 4550 | 2211 | 2514 | 3526 | 3126 | 3744 | 2598 | 3006 | 4062 | 3596 | 4842 | 4603 | 4763 | 4528 | 4446 | Unk |
| Unkl | 1295 | 1466 | 1646 | 1645 | 1649 | 1547 | 1292 | 1023 | 1267 | 1392 | 1316 | 1265 | 1401 | 1133 | 1139 | 1406 | 1523 | 1639 | 1532 | 1231 | 1358 | 1355 | 1221 | 1264 | 1444 | 1130 | 1204 | 1116 | Unkl |
| Uox | 0 | 0 | 0 | 1 | 0 | 1 | 1 | 0 | 3 | 0 | 0 | 0 | 0 | 0 | 0 | 0 | 0 | 0 | 0 | 0 | 0 | 0 | 1 | 2 | 0 | 0 | 0 | 0 | Uox |
| Upb1 | 5 | 6 | 8 | 7 | 2 | 3 | 1 | 4 | 16 | 11 | 9 | 4 | 16 | 2 | 7 | 16 | 0 | 15 | 9 | 3 | 5 | 8 | 16 | 12 | 7 | 6 | 5 | 7 | Upb1 |
| Upf1 | 5905 | 7074 | 8981 | 8910 | 7576 | 6710 | 6326 | 5023 | 6126 | 7099 | 7100 | 8564 | 6856 | 7232 | 5076 | 6690 | 8869 | 8298 | 8067 | 5506 | 6958 | 7072 | 5644 | 8173 | 7303 | 9088 | 6842 | 7160 | Upf1 |
| Upf2 | 9250 | 8203 | 8852 | 8946 | 8183 | 8376 | 9614 | 7355 | 6529 | 6047 | 6256 | 6738 | 6434 | 7439 | 9245 | 8770 | 8820 | 8442 | 8249 | 9436 | 8539 | 6151 | 6204 | 6547 | 6022 | 6141 | 6743 | 6565 | Upf2 |
| Upf3a | 1024 | 900 | 1056 | 1002 | 1186 | 1008 | 1069 | 2906 | 2874 | 3653 | 3262 | 3598 | 3476 | 3853 | 959 | 855 | 1133 | 986 | 1110 | 1029 | 1114 | 3021 | 2932 | 3662 | 3420 | 4035 | 3574 | 3805 | Upf3a |
| Upf3b | 6123 | 4793 | 6410 | 6924 | 6536 | 6517 | 7677 | 5848 | 4959 | 6002 | 5179 | 6273 | 5818 | 7193 | 6115 | 5285 | 6584 | 6729 | 6726 | 6715 | 7098 | 5959 | 4550 | 6423 | 6284 | 6250 | 6254 | 6761 | Upf3b |
| Upk1a | 161 | 226 | 212 | 241 | 217 | 154 | 193 | 77 | 73 | 45 | 38 | 87 | 84 | 76 | 157 | 178 | 234 | 202 | 184 | 178 | 187 | 77 | 52 | 78 | 87 | 78 | 57 | 102 | Upk1a |
| Upk1b | 116 | 168 | 119 | 14 | 6 | 162 | 1 | 159 | 207 | 82 | 229 | 174 | 231 | 203 | 219 | 174 | 39 | 113 | 30 | 13 | 0 | 7 | 293 | 129 | 32 | 29 | 150 | 260 | Upk1b |
| Upk2 | 20 | 7 | 3 | 1 | 2 | 4 | 0 | 13 | 26 | 19 | 18 | 10 | 3 | 1 | 11 | 7 | 6 | 0 | 1 | 1 | 1 | 2 | 30 | 16 | 0 | 5 | 0 | 5 | Upk2 |
| Upk3a | 3 | 9 | 0 | 12 | 6 | 11 | 4 | 2 | 9 | 10 | 8 | 5 | 3 | 4 | 9 | 5 | 3 | 2 | 9 | 3 | 12 | 0 | 19 | 4 | 5 | 7 | 6 | 1 | Upk3a |
| Upk3b | 7 | 4 | 10 | 7 | 1 | 11 | 6 | 5 | 8 | 6 | 6 | 2 | 9 | 12 | 3 | 12 | 1 | 6 | 3 | 3 | 0 | 2 | 2 | 3 | 5 | 1 | 3 | 6 | Upk3b |
| Upk3bl | 35 | 39 | 15 | 4 | 1 | 36 | 3 | 43 | 97 | 20 | 44 | 54 | 48 | 19 | 32 | 34 | 5 | 30 | 0 | 5 | 2 | 13 | 82 | 45 | 5 | 39 | 15 | 40 | Upk3bl |
| Upp1 | 150 | 191 | 147 | 126 | 79 | 114 | 83 | 96 | 91 | 96 | 88 | 81 | 95 | 62 | 170 | 157 | 123 | 97 | 98 | 114 | 97 | 90 | 83 | 88 | 54 | 105 | 113 | 113 | Upp1 |
| Upp2 | 220 | 233 | 222 | 228 | 245 | 322 | 289 | 303 | 297 | 304 | 291 | 225 | 230 | 271 | 252 | 223 | 225 | 230 | 241 | 242 | 269 | 296 | 294 | 284 | 264 | 182 | 228 | 185 | Upp2 |
| Uprt | 621 | 594 | 339 | 612 | 767 | 683 | 678 | 637 | 627 | 561 | 668 | 504 | 494 | 561 | 595 | 642 | 481 | 636 | 664 | 668 | 724 | 496 | 583 | 449 | 631 | 189 | 526 | 466 | Uprt |
| Uqcc1 | 1945 | 1854 | 1924 | 1653 | 1596 | 1706 | 1855 | 2787 | 2718 | 2498 | 2523 | 2728 | 2739 | 2995 | 1946 | 1752 | 1706 | 1597 | 1887 | 1849 | 1779 | 2508 | 2663 | 2741 | 2775 | 3067 | 2874 | 3030 | Uqcc1 |
| Uqcc2 | 1626 | 2070 | 1832 | 1918 | 1651 | 1643 | 1926 | 2254 | 2294 | 2473 | 2403 | 2271 | 2362 | 3025 | 1625 | 1558 | 1987 | 1696 | 1954 | 1910 | 2188 | 1883 | 2291 | 2802 | 2314 | 2732 | 2574 | 3140 | Uqcc2 |
| Uqcc3 | 641 | 746 | 727 | 633 | 646 | 643 | 613 | 834 | 906 | 906 | 911 | 916 | 957 | 885 | 632 | 668 | 785 | 592 | 618 | 730 | 578 | 672 | 858 | 1051 | 858 | 1048 | 996 | 932 | Uqcc3 |
| Uqcr10 | 2381 | 3221 | 3036 | 2703 | 2180 | 1989 | 3037 | 2147 | 2339 | 2727 | 2718 | 2519 | 2361 | 3332 | 2194 | 2023 | 3193 | 2457 | 2354 | 2901 | 2938 | 1623 | 2448 | 2987 | 2576 | 2971 | 2702 | 3547 | Uqcr10 |
| Uqcr11 | 1946 | 2337 | 2297 | 2878 | 2195 | 1826 | 2419 | 1711 | 1819 | 2037 | 2094 | 1987 | 2117 | 2240 | 1863 | 1580 | 2495 | 2168 | 2277 | 2568 | 2297 | 1367 | 1746 | 2452 | 2248 | 2222 | 2172 | 2358 | Uqcr11 |
| Uqcrb | 3314 | 2515 | 3214 | 3117 | 2817 | 2542 | 4611 | 4372 | 3587 | 3849 | 4174 | 3836 | 3848 | 6058 | 3807 | 2313 | 2927 | 2659 | 2858 | 3377 | 4041 | 3878 | 3722 | 4562 | 4020 | 4854 | 4267 | 5901 | Uqcrb |
| Uqcrc1 | 589 | 284 | 290 | 299 | 206 | 254 | 196 | 343 | 433 | 267 | 237 | 286 | 348 | 215 | 406 | 290 | 271 | 215 | 261 | 195 | 194 | 403 | 234 | 284 | 248 | 293 | 246 | 439 | Uqcrc1 |
| Uqcrc2 | 14144 | 12402 | 11419 | 10951 | 10555 | 10842 | 11903 | 15099 | 13059 | 11451 | 10660 | 11970 | 11500 | 12497 | 14206 | 11745 | 11202 | 10285 | 11395 | 13288 | 12110 | 11800 | 12851 | 11856 | 11580 | 13655 | 12566 | 13621 | Uqcrc2 |
| Uqcrfs1 | 6715 | 6693 | 6073 | 5906 | 5591 | 5493 | 6064 | 6476 | 5720 | 5310 | 5099 | 5843 | 5621 | 6221 | 6759 | 5969 | 5983 | 5327 | 5817 | 6053 | 5902 | 5748 | 5840 | 5940 | 5705 | 6529 | 6100 | 6399 | Uqcrfs1 |
| Uqcrh | 11042 | 10669 | 10072 | 10385 | 8649 | 8355 | 11180 | 8789 | 7643 | 7366 | 7809 | 7213 | 7181 | 8872 | 11641 | 8754 | 10184 | 9131 | 9622 | 11258 | 11017 | 6838 | 8128 | 8300 | 7662 | 8297 | 8158 | 9813 | Uqcrh |
| Uqcrq | 3189 | 4081 | 3629 | 3630 | 3066 | 2872 | 3672 | 3030 | 2874 | 2923 | 3155 | 3006 | 3013 | 3647 | 3179 | 2771 | 3775 | 3014 | 3442 | 3647 | 3666 | 2435 | 2792 | 3456 | 3107 | 3680 | 3245 | 3875 | Uqcrq |
| Urah | 2 | 2 | 0 | 0 | 0 | 0 | 0 | 2 | 12 | 10 | 10 | 2 | 0 | 2 | 1 | 5 | 0 | 2 | 0 | 1 | 0 | 2 | 6 | 2 | 0 | 0 | 3 | 4 | Urah |
| Urb1 | 3854 | 4917 | 3064 | 4115 | 3067 | 3463 | 3462 | 2281 | 2737 | 2594 | 2513 | 2938 | 2597 | 2555 | 3587 | 4370 | 3326 | 4081 | 3437 | 3492 | 3637 | 2351 | 2404 | 2321 | 2577 | 1676 | 2697 | 2496 | Urb1 |
| Urb2 | 4368 | 4748 | 3194 | 3552 | 2767 | 3264 | 3311 | 2523 | 2508 | 2332 | 2363 | 2779 | 2460 | 2488 | 3827 | 4351 | 3282 | 3411 | 2768 | 3339 | 3286 | 2226 | 2467 | 2270 | 2285 | 2343 | 2939 | 2655 | Urb2 |
| Urgcp | 1355 | 1701 | 1719 | 1864 | 1778 | 1634 | 1639 | 1765 | 1985 | 2157 | 2037 | 1925 | 1863 | 1900 | 1447 | 1533 | 1753 | 1784 | 1788 | 1397 | 1818 | 2094 | 1934 | 2090 | 2159 | 2079 | 1818 | 1956 | Urgcp |
| Uri1 | 6590 | 5326 | 6747 | 6333 | 5602 | 5787 | 6836 | 4068 | 3826 | 3867 | 3868 | 4442 | 4211 | 4559 | 6630 | 5829 | 6387 | 5752 | 5703 | 5857 | 6285 | 4126 | 3822 | 4120 | 4041 | 4967 | 4640 | 4912 | Uri1 |
| Urm1 | 2064 | 2326 | 2372 | 2160 | 1878 | 1888 | 1750 | 2129 | 2162 | 2107 | 2062 | 2202 | 2338 | 2226 | 1990 | 2231 | 2337 | 1881 | 2090 | 2178 | 1822 | 1890 | 2271 | 2224 | 2188 | 2274 | 2260 | 2403 | Urm1 |
| Uroc1 | 0 | 0 | 0 | 0 | 2 | 0 | 0 | 0 | 3 | 1 | 2 | 0 | 0 | 3 | 4 | 0 | 1 | 0 | 0 | 0 | 0 | 0 | 1 | 0 | 0 | 0 | 0 | 1 | Uroc1 |
| Urod | 4637 | 5462 | 4652 | 4184 | 3368 | 3395 | 3629 | 2414 | 2531 | 2256 | 2195 | 2323 | 2910 | 2539 | 4246 | 4868 | 4283 | 3859 | 3938 | 4190 | 3636 | 1923 | 2597 | 2554 | 2212 | 2895 | 2717 | 3075 | Urod |
| Uros | 1723 | 1824 | 1458 | 1731 | 1694 | 1538 | 1871 | 1931 | 1725 | 1622 | 1579 | 1490 | 1456 | 1606 | 1718 | 1626 | 1648 | 1745 | 1686 | 1791 | 1798 | 1750 | 1706 | 1679 | 1636 | 1534 | 1353 | 1710 | Uros |
| Usb1 | 1210 | 1551 | 1467 | 1534 | 1387 | 1314 | 1403 | 1331 | 1632 | 1321 | 1420 | 1697 | 1389 | 1493 | 1280 | 1371 | 1355 | 1598 | 1481 | 1539 | 1358 | 1404 | 1612 | 1510 | 1449 | 1802 | 1433 | 1595 | Usb1 |
| Use1 | 1969 | 2395 | 2097 | 2394 | 2406 | 2421 | 2138 | 2680 | 3302 | 3291 | 3271 | 2925 | 2734 | 2969 | 2061 | 2119 | 2204 | 2229 | 2468 | 2621 | 2305 | 2456 | 3477 | 3220 | 3012 | 3161 | 3175 | 2994 | Use1 |
| Usf1 | 3402 | 4208 | 5135 | 5147 | 5538 | 5032 | 4527 | 5808 | 6096 | 7500 | 6601 | 8302 | 7012 | 7261 | 3364 | 3569 | 5308 | 4407 | 5816 | 5127 | 4463 | 6592 | 5934 | 7915 | 8260 | 8483 | 7868 | 6963 | Usf1 |
| Usf2 | 1143 | 1322 | 1440 | 1574 | 1775 | 1483 | 1573 | 2243 | 2398 | 2880 | 2590 | 2771 | 2400 | 2684 | 1079 | 1090 | 1598 | 1407 | 1815 | 1369 | 1545 | 2553 | 2158 | 3080 | 2803 | 2974 | 2489 | 2573 | Usf2 |
| Usf3 | 6414 | 5841 | 3540 | 4413 | 5131 | 6479 | 5957 | 6357 | 5823 | 4382 | 4871 | 4951 | 5190 | 4863 | 6084 | 5689 | 3739 | 4390 | 4606 | 6438 | 6306 | 4268 | 5223 | 3759 | 4978 | 3294 | 5635 | 4351 | Usf3 |
| Ush1c | 98 | 138 | 109 | 83 | 92 | 63 | 95 | 117 | 151 | 150 | 170 | 114 | 127 | 101 | 107 | 76 | 118 | 83 | 104 | 97 | 95 | 174 | 100 | 91 | 104 | 138 | 148 | 167 | Ush1c |

| | | | | | | | | | | | | | | | | | | | | | | | | | | | | | |
|---|---|---|---|---|---|---|---|---|---|---|---|---|---|---|---|---|---|---|---|---|---|---|---|---|---|---|---|---|---|
| Ush1g | 18 | 35 | 30 | 26 | 31 | 16 | 11 | 0 | 2 | 7 | 2 | 5 | 7 | 4 | 25 | 42 | 39 | 15 | 29 | 13 | 20 | 5 | 3 | 0 | 2 | 1 | 1 | 2 | Ush1g |
| Ush2a | 32 | 35 | 22 | 15 | 29 | 21 | 18 | 45 | 47 | 32 | 28 | 14 | 44 | 19 | 28 | 40 | 19 | 21 | 13 | 36 | 27 | 38 | 40 | 7 | 31 | 7 | 20 | 13 | Ush2a |
| Ushbp1 | 156 | 185 | 193 | 163 | 194 | 182 | 163 | 405 | 521 | 559 | 744 | 547 | 685 | 495 | 144 | 176 | 190 | 199 | 192 | 190 | 157 | 576 | 647 | 556 | 600 | 594 | 500 | 482 | Ushbp1 |
| Uso1 | 11003 | 10100 | 10593 | 10084 | 9979 | 10559 | 10861 | 8954 | 8319 | 8393 | 8846 | 8806 | 9126 | 8801 | 11333 | 10316 | 9926 | 10529 | 10246 | 9713 | 10301 | 9326 | 8739 | 8366 | 8573 | 8130 | 9223 | 8878 | Uso1 |
| Usp1 | 16958 | 12436 | 15416 | 12163 | 10177 | 12152 | 14096 | 12389 | 10853 | 9320 | 9424 | 14781 | 12445 | 15822 | 15841 | 13277 | 13123 | 11568 | 11759 | 13773 | 12242 | 8246 | 9382 | 12191 | 10543 | 17437 | 15316 | 16338 | Usp1 |
| Usp10 | 5567 | 6520 | 6987 | 7011 | 6104 | 5799 | 6451 | 5211 | 6316 | 6370 | 6120 | 8113 | 7243 | 7465 | 5297 | 6301 | 7064 | 6631 | 6887 | 5458 | 6542 | 6346 | 5997 | 7210 | 6701 | 8416 | 6866 | 7833 | Usp10 |
| Usp11 | 13034 | 14462 | 14417 | 16296 | 14278 | 14378 | 14881 | 10335 | 11294 | 12250 | 11035 | 12492 | 11428 | 11780 | 13457 | 13119 | 15325 | 15600 | 15582 | 12362 | 15153 | 13540 | 10771 | 12079 | 12041 | 11111 | 11352 | 11430 | Usp11 |
| Usp12 | 5038 | 5069 | 5438 | 5564 | 5744 | 5471 | 5140 | 4540 | 4472 | 4772 | 4726 | 4576 | 4817 | 4541 | 5186 | 5136 | 5577 | 5496 | 5211 | 5494 | 5354 | 4458 | 4238 | 4268 | 4654 | 4273 | 4357 | 4555 | Usp12 |
| Usp13 | 784 | 733 | 896 | 913 | 1434 | 1132 | 977 | 1465 | 1443 | 1604 | 1593 | 1361 | 1382 | 1405 | 938 | 870 | 918 | 924 | 1175 | 982 | 1071 | 1591 | 1276 | 1200 | 1536 | 1131 | 1348 | 1244 | Usp13 |
| Usp14 | 12514 | 10420 | 11292 | 11251 | 11520 | 11235 | 13366 | 10243 | 8843 | 7698 | 8302 | 8954 | 8974 | 9451 | 13813 | 11499 | 11089 | 11361 | 11083 | 13169 | 12240 | 9504 | 8638 | 8060 | 8504 | 9360 | 9793 | 9606 | Usp14 |
| Usp15 | 11318 | 9649 | 8391 | 9284 | 8639 | 8897 | 9802 | 9300 | 8174 | 7084 | 7116 | 6913 | 6854 | 7097 | 11207 | 10390 | 8025 | 8942 | 8193 | 10427 | 9556 | 7653 | 7836 | 6321 | 7249 | 6019 | 6901 | 6577 | Usp15 |
| Usp16 | 6216 | 5091 | 5045 | 5092 | 4660 | 5059 | 5244 | 5561 | 4463 | 4403 | 4737 | 4548 | 4369 | 4845 | 6345 | 5829 | 4664 | 5033 | 4431 | 5435 | 5061 | 4898 | 4669 | 4439 | 4577 | 4524 | 4580 | 4519 | Usp16 |
| Usp17lb | 0 | 0 | 0 | 0 | 2 | 0 | 0 | 1 | 1 | 0 | 0 | 0 | 0 | 0 | 0 | 0 | 0 | 0 | 0 | 0 | 0 | 0 | 0 | 0 | 0 | 0 | 0 | 0 | Usp17lb |
| Usp17lc | 0 | 1 | 0 | 0 | 0 | 0 | 0 | 0 | 0 | 0 | 0 | 0 | 0 | 0 | 0 | 0 | 0 | 0 | 0 | 0 | 0 | 0 | 0 | 0 | 0 | 0 | 0 | 0 | Usp17lc |
| Usp17ld | 0 | 0 | 0 | 0 | 0 | 0 | 0 | 0 | 0 | 0 | 0 | 0 | 0 | 0 | 0 | 0 | 0 | 0 | 0 | 0 | 0 | 0 | 0 | 0 | 0 | 1 | 0 | 0 | Usp17ld |
| Usp17le | 0 | 0 | 0 | 0 | 0 | 0 | 0 | 0 | 0 | 0 | 0 | 0 | 0 | 0 | 0 | 0 | 0 | 0 | 0 | 0 | 0 | 0 | 1 | 0 | 1 | 0 | 0 | 0 | Usp17le |
| Usp18 | 45 | 55 | 54 | 52 | 55 | 46 | 47 | 23 | 27 | 25 | 37 | 30 | 33 | 35 | 67 | 63 | 76 | 45 | 38 | 86 | 80 | 33 | 40 | 48 | 33 | 60 | 41 | 38 | Usp18 |
| Usp19 | 9881 | 11760 | 12468 | 12623 | 12077 | 11188 | 10977 | 10595 | 12466 | 13663 | 12350 | 14893 | 13140 | 13183 | 9286 | 10576 | 13354 | 12555 | 13465 | 10248 | 11309 | 13042 | 11857 | 14982 | 14022 | 14029 | 13391 | 13347 | Usp19 |
| Usp2 | 1009 | 1168 | 1345 | 1205 | 831 | 844 | 896 | 260 | 337 | 299 | 393 | 371 | 357 | 354 | 994 | 1150 | 1187 | 1030 | 997 | 773 | 861 | 286 | 393 | 347 | 284 | 456 | 357 | 390 | Usp2 |
| Usp20 | 5742 | 7872 | 6675 | 8248 | 7031 | 6845 | 6464 | 2751 | 3139 | 3239 | 3034 | 3207 | 2994 | 2842 | 5591 | 6615 | 7553 | 8214 | 7638 | 6106 | 6806 | 3614 | 3054 | 3166 | 3468 | 2937 | 2751 | 2740 | Usp20 |
| Usp21 | 2510 | 2910 | 3250 | 3336 | 3623 | 3134 | 3303 | 4183 | 4642 | 5078 | 4765 | 5261 | 4797 | 5089 | 2462 | 2517 | 3458 | 3323 | 3956 | 2688 | 3455 | 5085 | 4410 | 5610 | 5098 | 6114 | 4921 | 5150 | Usp21 |
| Usp22 | 21191 | 21572 | 19460 | 22729 | 26543 | 24516 | 24868 | 31051 | 32238 | 32039 | 30135 | 32197 | 30199 | 31257 | 21473 | 19644 | 21811 | 23896 | 25535 | 22681 | 25111 | 37460 | 29679 | 30951 | 36061 | 28146 | 30932 | 31507 | Usp22 |
| Usp24 | 6827 | 5824 | 4560 | 4985 | 6752 | 7416 | 6876 | 11195 | 11117 | 9428 | 10256 | 10671 | 10622 | 10779 | 6721 | 6287 | 4622 | 5340 | 6227 | 7165 | 7284 | 8597 | 10372 | 8081 | 10651 | 6753 | 11390 | 9843 | Usp24 |
| Usp25 | 8198 | 6953 | 7151 | 7356 | 6486 | 6952 | 7230 | 3107 | 2961 | 2778 | 2877 | 3341 | 2998 | 3208 | 7590 | 7653 | 7238 | 6902 | 6704 | 6239 | 7089 | 2453 | 2851 | 2982 | 2609 | 3361 | 3356 | 3437 | Usp25 |
| Usp26 | 5 | 2 | 0 | 0 | 0 | 1 | 1 | 1 | 1 | 0 | 1 | 0 | 1 | 1 | 12 | 6 | 5 | 0 | 1 | 2 | 0 | 0 | 3 | 3 | 6 | 3 | 8 | 0 | Usp26 |
| Usp27x | 1144 | 1100 | 1163 | 1446 | 1419 | 1234 | 1419 | 1167 | 1116 | 1182 | 1168 | 1037 | 1016 | 1112 | 1077 | 1006 | 1228 | 1323 | 1400 | 1195 | 1387 | 1445 | 997 | 1119 | 1299 | 1061 | 1024 | 1108 | Usp27x |
| Usp28 | 10207 | 9896 | 8889 | 9594 | 8663 | 9538 | 9677 | 4513 | 4209 | 3992 | 4462 | 4738 | 4662 | 4752 | 9901 | 10273 | 9013 | 9708 | 8839 | 8512 | 9481 | 4582 | 4025 | 4113 | 4367 | 4401 | 4953 | 4828 | Usp28 |
| Usp29 | 4456 | 4821 | 3417 | 6487 | 7248 | 7043 | 7752 | 5948 | 5167 | 5467 | 5296 | 4249 | 3901 | 4786 | 5330 | 3935 | 4810 | 6660 | 6618 | 7053 | 7873 | 7055 | 4807 | 4998 | 6147 | 3295 | 3920 | 3863 | Usp29 |
| Usp3 | 7478 | 6486 | 6101 | 6719 | 7231 | 7581 | 8209 | 9703 | 8408 | 8501 | 7258 | 9488 | 8324 | 9823 | 7547 | 5882 | 6829 | 6666 | 8020 | 7518 | 8448 | 8819 | 8000 | 8845 | 9420 | 9214 | 9284 | 9890 | Usp3 |
| Usp30 | 2610 | 2923 | 2831 | 3099 | 2873 | 2837 | 2670 | 2017 | 2235 | 2394 | 2296 | 2412 | 2184 | 2286 | 2420 | 2574 | 2825 | 2899 | 2999 | 2549 | 2848 | 2313 | 2174 | 2463 | 2463 | 2423 | 2395 | 2362 | Usp30 |
| Usp31 | 6061 | 5757 | 4969 | 6583 | 6560 | 6225 | 6498 | 4439 | 4120 | 4038 | 4104 | 4176 | 3926 | 4179 | 5916 | 5877 | 5320 | 6398 | 5912 | 5671 | 6638 | 4587 | 3600 | 3545 | 4682 | 3125 | 4278 | 3986 | Usp31 |
| Usp32 | 10843 | 9607 | 6930 | 10571 | 11885 | 12329 | 13504 | 9384 | 7618 | 7707 | 7174 | 7332 | 7245 | 7538 | 11743 | 9544 | 8793 | 10736 | 10682 | 11830 | 13470 | 8722 | 7535 | 6537 | 8429 | 4538 | 7224 | 7066 | Usp32 |
| Usp33 | 15034 | 13136 | 11874 | 13176 | 14192 | 14306 | 16024 | 11416 | 9460 | 9231 | 9074 | 8843 | 8529 | 9308 | 16751 | 14240 | 13004 | 14093 | 13711 | 16083 | 15214 | 10955 | 9569 | 8817 | 9940 | 7871 | 8906 | 8947 | Usp33 |
| Usp34 | 21339 | 16669 | 10292 | 12908 | 16007 | 19898 | 18877 | 30602 | 27107 | 20266 | 21066 | 20822 | 21836 | 22161 | 21149 | 18700 | 11400 | 12935 | 13944 | 21120 | 19661 | 19308 | 25009 | 15872 | 21573 | 13524 | 23585 | 19719 | Usp34 |
| Usp35 | 1321 | 1477 | 1584 | 1761 | 1964 | 1825 | 1697 | 801 | 1063 | 1132 | 1101 | 1001 | 1065 | 938 | 1346 | 1551 | 1703 | 1898 | 1861 | 1438 | 1691 | 1168 | 977 | 1103 | 1286 | 756 | 938 | 949 | Usp35 |
| Usp36 | 3456 | 3994 | 4044 | 4577 | 3959 | 3824 | 3245 | 2646 | 3187 | 3353 | 3526 | 3650 | 3317 | 3220 | 2988 | 3809 | 4052 | 4468 | 4025 | 3215 | 3539 | 3469 | 2939 | 3588 | 3650 | 3491 | 3382 | 3405 | Usp36 |
| Usp37 | 8416 | 7490 | 6777 | 7483 | 6144 | 6902 | 7841 | 6721 | 5852 | 5364 | 5184 | 6358 | 5668 | 6150 | 8304 | 7807 | 6991 | 7657 | 6245 | 7707 | 7244 | 5350 | 5461 | 5646 | 5651 | 5766 | 6574 | 6566 | Usp37 |
| Usp38 | 7698 | 7607 | 6382 | 6612 | 5393 | 6561 | 6823 | 4220 | 3927 | 3497 | 3409 | 4397 | 4010 | 4293 | 7453 | 7313 | 6388 | 6431 | 5587 | 6158 | 6342 | 3850 | 3659 | 3660 | 3834 | 4181 | 4135 | 4271 | Usp38 |
| Usp39 | 370 | 195 | 163 | 192 | 175 | 199 | 184 | 222 | 297 | 106 | 118 | 130 | 120 | 123 | 368 | 217 | 174 | 503 | 179 | 334 | 153 | 167 | 235 | 143 | 266 | 156 | 146 | 127 | Usp39 |
| Usp4 | 13652 | 14329 | 11753 | 12803 | 12458 | 13592 | 13118 | 13280 | 12591 | 12572 | 11459 | 12801 | 12433 | 13365 | 13634 | 12486 | 11947 | 12915 | 13266 | 13080 | 13729 | 12880 | 11951 | 12072 | 13438 | 12524 | 13710 | 13416 | Usp4 |
| Usp40 | 2719 | 2123 | 1854 | 1788 | 2086 | 2217 | 2169 | 5163 | 5371 | 4445 | 4855 | 5004 | 4985 | 4827 | 2391 | 2092 | 1859 | 1675 | 2006 | 2341 | 2026 | 4318 | 5308 | 4191 | 4401 | 3949 | 5216 | 4480 | Usp40 |
| Usp42 | 3751 | 4160 | 4135 | 4467 | 4480 | 4338 | 4347 | 4731 | 4957 | 5528 | 5428 | 6024 | 5700 | 5636 | 3737 | 4058 | 4497 | 4470 | 4732 | 3882 | 4630 | 5725 | 4704 | 5594 | 6027 | 5368 | 5703 | 5835 | Usp42 |
| Usp43 | 212 | 181 | 176 | 181 | 123 | 119 | 92 | 117 | 137 | 140 | 101 | 141 | 132 | 144 | 144 | 227 | 172 | 150 | 126 | 76 | 88 | 132 | 156 | 156 | 84 | 153 | 116 | 140 | Usp43 |
| Usp44 | 1547 | 1748 | 1787 | 1146 | 737 | 787 | 884 | 237 | 163 | 143 | 156 | 387 | 357 | 423 | 1452 | 1375 | 1648 | 978 | 810 | 1066 | 903 | 160 | 124 | 257 | 179 | 360 | 673 | 540 | Usp44 |
| Usp45 | 5404 | 4344 | 4564 | 4741 | 4583 | 4901 | 5116 | 3563 | 3228 | 2868 | 3206 | 3569 | 3706 | 3686 | 4961 | 4807 | 4745 | 4486 | 4432 | 4802 | 4941 | 3163 | 3183 | 3163 | 3314 | 3013 | 3611 | 3724 | Usp45 |
| Usp46 | 4960 | 3806 | 3988 | 3498 | 4111 | 4093 | 3784 | 5716 | 5497 | 4836 | 5005 | 4889 | 5059 | 4855 | 4688 | 4715 | 3584 | 3854 | 3887 | 3797 | 3860 | 5227 | 5519 | 4895 | 4935 | 5266 | 4997 | 5102 | Usp46 |
| Usp47 | 24119 | 20793 | 17333 | 18712 | 20909 | 21711 | 23258 | 28050 | 24175 | 21328 | 20946 | 23834 | 22299 | 23911 | 24788 | 20531 | 18665 | 19570 | 19999 | 24112 | 21966 | 22760 | 23282 | 20732 | 23187 | 19491 | 23530 | 22653 | Usp47 |
| Usp48 | 13913 | 12888 | 11820 | 13450 | 13449 | 13959 | 14757 | 12664 | 11781 | 12591 | 13597 | 12702 | 12073 | 13385 | 12965 | 12587 | 12633 | 13246 | 13348 | 12456 | 14944 | 12549 | 11516 | 11737 | 13754 | 10034 | 12937 | 12447 | Usp48 |
| Usp49 | 5098 | 4881 | 5751 | 6153 | 6010 | 5822 | 6835 | 4073 | 3737 | 5129 | 4219 | 5515 | 4349 | 5964 | 4950 | 4733 | 6284 | 6256 | 6198 | 4795 | 6752 | 5867 | 3442 | 5303 | 5361 | 5584 | 5004 | 5760 | Usp49 |
| Usp5 | 10819 | 12090 | 13721 | 12851 | 12466 | 12425 | 11618 | 11446 | 13399 | 14207 | 13595 | 15589 | 14606 | 14831 | 10706 | 10640 | 13558 | 12184 | 13975 | 11502 | 11940 | 12975 | 12848 | 14988 | 15165 | 16193 | 15268 | 15499 | Usp5 |
| Usp50 | 5 | 2 | 2 | 2 | 2 | 11 | 9 | 6 | 2 | 0 | 1 | 3 | 0 | 5 | 7 | 3 | 1 | 7 | 7 | 3 | 8 | 2 | 3 | 1 | 2 | 6 | 1 | 0 | Usp50 |
| Usp51 | 549 | 485 | 563 | 403 | 356 | 445 | 426 | 304 | 300 | 299 | 289 | 518 | 331 | 490 | 444 | 421 | 569 | 319 | 415 | 443 | 416 | 353 | 197 | 310 | 360 | 490 | 420 | 475 | Usp51 |
| Usp53 | 2091 | 1658 | 1294 | 1735 | 1773 | 2029 | 2073 | 1029 | 1014 | 877 | 1075 | 882 | 1021 | 834 | 2019 | 1941 | 1457 | 2010 | 1685 | 2167 | 1895 | 904 | 1045 | 917 | 850 | 717 | 894 | 920 | Usp53 |
| Usp54 | 5057 | 5116 | 4405 | 4356 | 4570 | 4839 | 4603 | 4184 | 3959 | 3660 | 3522 | 4316 | 4148 | 4205 | 4919 | 4404 | 4127 | 4311 | 4297 | 4663 | 4790 | 4348 | 3793 | 3746 | 4174 | 3364 | 4528 | 4169 | Usp54 |
| Usp6nl | 2147 | 1666 | 2177 | 2075 | 2257 | 2084 | 1905 | 3893 | 3961 | 4204 | 3964 | 4655 | 4280 | 4637 | 1842 | 2007 | 1983 | 1862 | 2211 | 1995 | 2136 | 4044 | 3845 | 3956 | 4205 | 4231 | 4749 | 4667 | Usp6nl |
| Usp7 | 19553 | 16817 | 17745 | 19263 | 20356 | 20529 | 21978 | 21238 | 18700 | 19285 | 19263 | 21954 | 19599 | 22542 | 18815 | 17063 | 18574 | 19431 | 19647 | 18468 | 21374 | 20673 | 17479 | 19418 | 20953 | 18999 | 20872 | 21370 | Usp7 |
| Usp8 | 7944 | 7920 | 7097 | 7054 | 6169 | 6402 | 6881 | 6328 | 6000 | 5515 | 5911 | 5675 | 5517 | 6014 | 7981 | 7647 | 6643 | 6784 | 6020 | 6968 | 6507 | 5476 | 6148 | 5395 | 5445 | 5307 | 6015 | 5919 | Usp8 |
| Usp9x | 31883 | 27559 | 12349 | 19849 | 21898 | 29349 | 27755 | 36097 | 31525 | 22404 | 23473 | 24447 | 24107 | 24258 | 32592 | 27402 | 15986 | 19897 | 20065 | 31824 | 28110 | 20413 | 29584 | 17006 | 24940 | 11020 | 27524 | 21342 | Usp9x |
| Usp9y | 1 | 0 | 2 | 2 | 1 | 0 | 0 | 1 | 0 | 0 | 1 | 1 | 0 | 0 | 0 | 0 | 0 | 0 | 0 | 0 | 0 | 0 | 0 | 0 | 0 | 0 | 0 | 0 | Usp9y |
| Uspl1 | 7764 | 7481 | 6281 | 6676 | 6352 | 6765 | 7044 | 7371 | 6391 | 5743 | 5736 | 6355 | 6072 | 6639 | 8165 | 6899 | 6458 | 6326 | 6424 | 7529 | 7178 | 5643 | 5994 | 5786 | 5905 | 5935 | 7133 | 6916 | Uspl1 |
| Ust | 428 | 355 | 390 | 378 | 510 | 437 | 344 | 543 | 623 | 560 | 794 | 422 | 452 | 438 | 380 | 431 | 336 | 495 | 365 | 349 | 428 | 617 | 773 | 549 | 557 | 483 | 417 | 417 | Ust |
| Utf1 | 18 | 21 | 9 | 13 | 5 | 10 | 1 | 4 | 3 | 0 | 2 | 2 | 0 | 0 | 11 | 31 | 9 | 12 | 2 | 14 | 1 | 0 | 3 | 0 | 0 | 1 | 1 | 0 | Utf1 |
| Utp11 | 4154 | 3987 | 5058 | 4035 | 3051 | 3381 | 3768 | 2729 | 2487 | 2932 | 2958 | 2771 | 2912 | 3152 | 4146 | 4171 | 4298 | 3925 | 3415 | 3682 | 3429 | 2608 | 2813 | 3303 | 2680 | 3480 | 3038 | 3226 | Utp11 |
| Utp14a | 5226 | 4887 | 4720 | 4652 | 3968 | 4238 | 5171 | 4172 | 3896 | 3743 | 3659 | 3912 | 3904 | 4328 | 5527 | 4917 | 4599 | 4812 | 4353 | 5061 | 4645 | 3609 | 3788 | 3975 | 3866 | 4421 | 4120 | 4313 | Utp14a |
| Utp14b | 2776 | 2251 | 2835 | 2389 | 2004 | 2012 | 2093 | 1987 | 1827 | 1747 | 1891 | 2006 | 2144 | 2419 | 2391 | 2597 | 2408 | 2045 | 1809 | 1904 | 1810 | 1659 | 1655 | 2001 | 1925 | 1889 | 2303 | 2363 | Utp14b |
| Utp15 | 5079 | 4861 | 4526 | 4534 | 3862 | 4335 | 4491 | 4078 | 3598 | 3344 | 3603 | 3914 | 3715 | 4065 | 5069 | 4784 | 4386 | 4474 | 3968 | 4767 | 4345 | 3265 | 3398 | 3497 | 3673 | 3926 | 4006 | 4022 | Utp15 |
| Utp18 | 4564 | 4093 | 4615 | 4719 | 4171 | 4371 | 5296 | 3622 | 3027 | 2913 | 2967 | 3655 | 3277 | 3820 | 4919 | 4284 | 4519 | 4578 | 4579 | 5164 | 5053 | 3111 | 2949 | 3433 | 3355 | 4025 | 3575 | 3840 | Utp18 |
| Utp20 | 3791 | 3390 | 2235 | 2110 | 2319 | 2620 | 2548 | 6152 | 5978 | 4535 | 5333 | 5798 | 5687 | 5757 | 3257 | 3324 | 2152 | 2155 | 2198 | 2829 | 2609 | 3877 | 5623 | 4503 | 4863 | 4324 | 6837 | 5715 | Utp20 |
| Utp23 | 1194 | 1287 | 1314 | 1327 | 1222 | 1160 | 1352 | 1318 | 1150 | 1226 | 1173 | 1218 | 1225 | 1417 | 1276 | 1077 | 1299 | 1259 | 1213 | 1256 | 1430 | 1395 | 1208 | 1217 | 1339 | 1328 | 1163 | 1513 | Utp23 |
| Utp25 | 2440 | 2592 | 2337 | 2611 | 2466 | 2396 | 2480 | 2982 | 2795 | 2728 | 2854 | 2865 | 2851 | 3193 | 2437 | 2297 | 2291 | 2270 | 2387 | 2468 | 2624 | 2797 | 2638 | 2766 | 3267 | 2741 | 3076 | 3074 | Utp25 |
| Utp3 | 4094 | 4211 | 4699 | 4141 | 3080 | 3378 | 3713 | 3673 | 3732 | 3623 | 3556 | 4284 | 4118 | 4623 | 4003 | 3782 | 4113 | 3422 | 3423 | 4126 | 3479 | 3011 | 3671 | 4287 | 3657 | 5313 | 4548 | 4620 | Utp3 |
| Utp4 | 5989 | 6466 | 5917 | 5321 | 4255 | 4514 | 4685 | 2950 | 2890 | 2513 | 2735 | 3000 | 2905 | 2941 | 5606 | 5868 | 4979 | 5152 | 4443 | 5231 | 4734 | 2510 | 2932 | 2737 | 2607 | 3614 | 3232 | 3121 | Utp4 |
| Utp6 | 10010 | 9505 | 10275 | 9197 | 8367 | 9098 | 10114 | 8534 | 7434 | 7504 | 7490 | 8851 | 8067 | 9402 | 10263 | 9012 | 9131 | 8918 | 9016 | 10210 | 9298 | 6935 | 7659 | 8595 | 8197 | 9674 | 9157 | 9522 | Utp6 |
| Utrn | 6892 | 5856 | 4251 | 3537 | 3753 | 4875 | 3085 | 5997 | 7279 | 5461 | 7789 | 5200 | 6903 | 5296 | 6156 | 8170 | 3536 | 3662 | 2844 | 3641 | 3392 | 4610 | 7753 | 4125 | 4867 | 3980 | 6047 | 5063 | Utrn |
| Uts2b | 6 | 6 | 0 | 4 | 30 | 7 | 1 | 34 | 30 | 52 | 56 | 0 | 33 | 4 | 17 | 24 | 2 | 8 | 0 | 2 | 1 | 38 | 30 | 4 | 8 | 0 | 1 | 5 | Uts2b |
| Uts2r | 1 | 3 | 7 | 5 | 10 | 10 | 2 | 5 | 4 | 3 | 30 | 5 | 7 | 1 | 9 | 9 | 6 | 1 | 4 | 2 | 4 | 7 | 17 | 2 | 4 | 1 | 0 | 1 | Uts2r |
| Uty | 2646 | 2022 | 1856 | 2174 | 2392 | 2587 | 2640 | 3236 | 3507 | 2806 | 2566 | 3214 | 3375 | 3463 | 1 | 0 | 0 | 0 | 0 | 0 | 0 | 0 | 14 | 1 | 0 | 0 | 2 | 0 | Uty |
| Uvrag | 3098 | 3019 | 2704 | 3098 | 3121 | 2909 | 2889 | 1614 | 1673 | 1641 | 1703 | 1643 | 1631 | 1534 | 3024 | 3083 | 2833 | 3095 | 2855 | 2657 | 2858 | 1813 | 1728 | 1647 | 1640 | 1682 | 1440 | 1472 | Uvrag |
| Uvssa | 3541 | 3184 | 2451 | 2910 | 2603 | 3555 | 2659 | 3085 | 2854 | 2781 | 3093 | 2782 | 2948 | 2702 | 3377 | 3449 | 2748 | 3064 | 2512 | 3007 | 3019 | 2503 | 2797 | 2491 | 2965 | 1991 | 2943 | 2489 | Uvssa |

| | | | | | | | | | | | | | | | | | | | | | | | | | | | | | |
|---|---|---|---|---|---|---|---|---|---|---|---|---|---|---|---|---|---|---|---|---|---|---|---|---|---|---|---|---|---|
| Uxs1 | 2307 | 1972 | 2084 | 1923 | 1847 | 1921 | 1979 | 1912 | 1801 | 1813 | 2044 | 1705 | 1859 | 1977 | 2151 | 2225 | 2022 | 2029 | 1545 | 1919 | 1828 | 1808 | 2001 | 1929 | 1891 | 2037 | 1675 | 1914 | Uxs1 |
| Uxt | 452 | 507 | 414 | 406 | 344 | 406 | 469 | 673 | 594 | 506 | 534 | 577 | 665 | 765 | 443 | 393 | 414 | 307 | 424 | 496 | 454 | 434 | 541 | 640 | 551 | 719 | 816 | 828 | Uxt |
| V1ra8 | 0 | 0 | 0 | 0 | 0 | 0 | 0 | 0 | 1 | 0 | 0 | 0 | 0 | 0 | 0 | 0 | 0 | 0 | 0 | 0 | 0 | 0 | 0 | 0 | 0 | 0 | 0 | 0 | V1ra8 |
| Vac14 | 2694 | 3490 | 3269 | 3298 | 3210 | 3138 | 3229 | 2434 | 2679 | 2838 | 2622 | 3281 | 2990 | 3279 | 2735 | 3019 | 3621 | 3329 | 3678 | 2785 | 3306 | 2856 | 2642 | 2792 | 3222 | 3063 | 3109 | 3122 | Vac14 |
| Vamp1 | 2169 | 2345 | 2136 | 2467 | 2317 | 2170 | 2225 | 1037 | 980 | 1131 | 1218 | 1085 | 1266 | 1109 | 2123 | 2079 | 2296 | 2345 | 2121 | 2120 | 2107 | 1140 | 1029 | 1012 | 1075 | 952 | 1194 | 1146 | Vamp1 |
| Vamp2 | 7988 | 8689 | 9943 | 14350 | 15144 | 12280 | 13502 | 8223 | 8170 | 11474 | 11200 | 9465 | 8096 | 10078 | 7900 | 8329 | 12681 | 15407 | 13913 | 9761 | 15160 | 13416 | 8225 | 10645 | 12152 | 9207 | 7398 | 8848 | Vamp2 |
| Vamp3 | 8623 | 7515 | 8413 | 6733 | 4814 | 5845 | 6296 | 5442 | 5309 | 4663 | 5159 | 5569 | 5443 | 5607 | 8297 | 8412 | 6825 | 6155 | 5615 | 6278 | 5907 | 4562 | 5834 | 5233 | 4513 | 6417 | 5948 | 5876 | Vamp3 |
| Vamp4 | 4675 | 3578 | 3887 | 3830 | 4056 | 4158 | 4568 | 3489 | 3152 | 2849 | 3233 | 2988 | 3138 | 3066 | 4916 | 4291 | 3899 | 4000 | 4063 | 4416 | 4381 | 3367 | 3382 | 3133 | 3222 | 2937 | 3004 | 3165 | Vamp4 |
| Vamp5 | 3 | 1 | 1 | 5 | 1 | 3 | 1 | 27 | 3 | 8 | 3 | 3 | 7 | 4 | 7 | 6 | 8 | 1 | 6 | 5 | 6 | 8 | 25 | 7 | 3 | 3 | 8 | 5 | Vamp5 |
| Vamp8 | 25 | 14 | 12 | 9 | 8 | 18 | 2 | 37 | 29 | 23 | 24 | 21 | 24 | 18 | 29 | 51 | 3 | 8 | 9 | 14 | 6 | 17 | 32 | 31 | 10 | 180 | 16 | 23 | Vamp8 |
| Vamp9 | 2 | 6 | 0 | 5 | 8 | 8 | 6 | 10 | 8 | 19 | 11 | 5 | 11 | 11 | 3 | 5 | 2 | 4 | 10 | 6 | 4 | 5 | 11 | 4 | 5 | 7 | 2 | 13 | Vamp9 |
| Vangl1 | 944 | 872 | 993 | 823 | 837 | 771 | 660 | 696 | 943 | 953 | 1122 | 659 | 1040 | 695 | 916 | 1078 | 782 | 777 | 680 | 668 | 694 | 1051 | 1205 | 764 | 784 | 779 | 808 | 751 | Vangl1 |
| Vangl2 | 8463 | 9876 | 13964 | 13084 | 13234 | 11398 | 11746 | 10549 | 10062 | 13140 | 10314 | 15371 | 13904 | 14793 | 8925 | 8826 | 14325 | 11996 | 14385 | 11228 | 11982 | 12952 | 9499 | 14452 | 14670 | 14204 | 14292 | 15021 | Vangl2 |
| Vapa | 6547 | 5768 | 6882 | 7430 | 7004 | 6344 | 7917 | 6942 | 6466 | 7276 | 6634 | 7090 | 6787 | 8705 | 6531 | 6486 | 7057 | 6967 | 7337 | 6980 | 7402 | 7200 | 6319 | 7254 | 7276 | 7902 | 6524 | 7937 | Vapa |
| Vapb | 7428 | 8257 | 9180 | 9448 | 9551 | 8760 | 8882 | 7591 | 7785 | 9460 | 8828 | 9112 | 8349 | 9017 | 7642 | 7537 | 9392 | 9141 | 9044 | 7614 | 9020 | 9028 | 8095 | 8966 | 9506 | 7887 | 8958 | 8791 | Vapb |
| Vars | 7147 | 8823 | 11419 | 9454 | 8089 | 7859 | 6900 | 6165 | 7911 | 8796 | 8678 | 12332 | 9426 | 10014 | 6341 | 8082 | 10515 | 9136 | 10065 | 6369 | 7721 | 7693 | 7497 | 11137 | 9088 | 14146 | 9929 | 10531 | Vars |
| Vars2 | 1980 | 2518 | 2801 | 2943 | 2815 | 2405 | 2352 | 1926 | 2408 | 2469 | 2351 | 2711 | 2479 | 2425 | 1944 | 2223 | 2941 | 2725 | 2804 | 2199 | 2338 | 2264 | 2148 | 2788 | 2526 | 2734 | 2392 | 2371 | Vars2 |
| Vash1 | 2848 | 4054 | 4253 | 4505 | 5205 | 4734 | 3979 | 3736 | 4449 | 5305 | 4713 | 5224 | 4393 | 4390 | 2933 | 3360 | 4546 | 4604 | 5647 | 4177 | 4356 | 4904 | 4311 | 5802 | 6074 | 4728 | 4650 | 4555 | Vash1 |
| Vash2 | 3203 | 2828 | 3246 | 3104 | 4116 | 3728 | 3737 | 5400 | 5419 | 5777 | 6189 | 5038 | 5358 | 5228 | 3072 | 2845 | 3501 | 3266 | 3725 | 3242 | 3578 | 6380 | 5671 | 5257 | 6227 | 5286 | 4951 | 5239 | Vash2 |
| Vasn | 731 | 1057 | 761 | 644 | 421 | 697 | 356 | 641 | 1100 | 927 | 1240 | 686 | 1141 | 618 | 744 | 1185 | 625 | 801 | 398 | 383 | 359 | 884 | 1143 | 889 | 667 | 948 | 853 | 698 | Vasn |
| Vasp | 4864 | 5544 | 6522 | 5890 | 4956 | 4713 | 4640 | 2838 | 3164 | 3274 | 3993 | 3326 | 3366 | 3329 | 4458 | 4965 | 5886 | 5474 | 5470 | 3960 | 4776 | 2984 | 3510 | 3568 | 3314 | 4312 | 3377 | 3582 | Vasp |
| Vat1 | 12557 | 16088 | 16404 | 16453 | 14304 | 13859 | 13080 | 9039 | 10135 | 10671 | 10282 | 9937 | 10331 | 10535 | 12924 | 12756 | 17040 | 15250 | 16760 | 12630 | 14817 | 10611 | 9990 | 11212 | 12127 | 9708 | 9630 | 10531 | Vat1 |
| Vat1l | 6060 | 6550 | 6172 | 6419 | 8674 | 6763 | 7701 | 4769 | 4806 | 5052 | 4303 | 4457 | 4439 | 3864 | 6440 | 5529 | 6670 | 6839 | 7531 | 7520 | 8351 | 6401 | 6378 | 3871 | 4544 | 3321 | 4035 | 4093 | Vat1l |
| Vav1 | 106 | 108 | 159 | 65 | 92 | 106 | 61 | 116 | 138 | 145 | 179 | 152 | 129 | 101 | 80 | 114 | 80 | 113 | 64 | 94 | 72 | 117 | 190 | 131 | 102 | 192 | 141 | 123 | Vav1 |
| Vav2 | 5159 | 6411 | 7354 | 8144 | 8790 | 7842 | 7323 | 4560 | 4419 | 5788 | 5549 | 6595 | 6087 | 5968 | 5287 | 5461 | 8020 | 7987 | 9192 | 6600 | 7870 | 6231 | 4620 | 5819 | 6325 | 4991 | 6448 | 6165 | Vav2 |
| Vav3 | 1583 | 1375 | 1741 | 1634 | 1904 | 1968 | 1705 | 3126 | 1898 | 2443 | 2362 | 2578 | 2999 | 2627 | 1953 | 1478 | 1627 | 1554 | 1865 | 1760 | 1752 | 3332 | 2574 | 2495 | 2663 | 1678 | 2994 | 3084 | Vav3 |
| Vax1 | 157 | 153 | 27 | 221 | 227 | 163 | 337 | 216 | 325 | 339 | 289 | 509 | 28 | 417 | 158 | 69 | 213 | 200 | 279 | 158 | 230 | 393 | 172 | 642 | 409 | 798 | 260 | 249 | Vax1 |
| Vax2 | 35 | 54 | 83 | 45 | 34 | 43 | 35 | 1 | 2 | 1 | 1 | 4 | 1 | 3 | 41 | 48 | 102 | 52 | 54 | 43 | 45 | 5 | 4 | 7 | 2 | 2 | 3 | 5 | Vax2 |
| Vbp1 | 7303 | 5620 | 7200 | 7499 | 7664 | 7676 | 9308 | 9968 | 7674 | 7885 | 7108 | 8805 | 8351 | 9766 | 8884 | 6421 | 7345 | 7567 | 7618 | 9037 | 8619 | 9454 | 7992 | 8545 | 8794 | 9236 | 8808 | 9824 | Vbp1 |
| Vcam1 | 4137 | 3997 | 3573 | 3037 | 3309 | 4260 | 3007 | 6075 | 5779 | 5310 | 6072 | 5182 | 7563 | 5417 | 4832 | 4810 | 3581 | 3596 | 3498 | 4040 | 3419 | 5429 | 6745 | 5396 | 4927 | 5317 | 5828 | 6217 | Vcam1 |
| Vcan | 36727 | 26510 | 13052 | 16882 | 18638 | 29708 | 20072 | 66100 | 71753 | 41622 | 56623 | 39890 | 44673 | 42279 | 31302 | 34057 | 13903 | 18944 | 16404 | 26011 | 21001 | 35735 | 65920 | 34186 | 40658 | 31892 | 42961 | 33729 | Vcan |
| Vcl | 17353 | 15867 | 13568 | 10174 | 7894 | 10143 | 7916 | 4971 | 6143 | 4832 | 6999 | 5123 | 6717 | 4878 | 15609 | 19745 | 10277 | 11239 | 7611 | 7800 | 8073 | 4944 | 7265 | 4471 | 4124 | 4722 | 6200 | 5292 | Vcl |
| Vcp | 50033 | 45671 | 46355 | 42256 | 41154 | 43414 | 41915 | 47096 | 40706 | 42264 | 40262 | 47534 | 46899 | 49656 | 47178 | 42670 | 42900 | 43493 | 46076 | 39593 | 43980 | 42745 | 42529 | 45002 | 44644 | 55136 | 50498 | 53235 | Vcp |
| Vcpip1 | 9498 | 8178 | 7827 | 10361 | 9372 | 9847 | 10351 | 7823 | 6490 | 6752 | 6939 | 7419 | 7265 | 7271 | 10130 | 9598 | 8611 | 10534 | 8785 | 9427 | 10273 | 7615 | 6518 | 6278 | 7579 | 5841 | 7442 | 6991 | Vcpip1 |
| Vcpkmt | 917 | 808 | 937 | 948 | 884 | 943 | 976 | 1059 | 1000 | 901 | 1097 | 961 | 961 | 1043 | 939 | 831 | 845 | 953 | 871 | 921 | 975 | 944 | 917 | 963 | 1143 | 1024 | 997 | 1044 | Vcpkmt |
| Vdac1 | 21648 | 22020 | 22243 | 21155 | 20073 | 19528 | 21325 | 18075 | 17752 | 16848 | 17239 | 17766 | 17517 | 19100 | 21859 | 20295 | 21345 | 20860 | 21231 | 20955 | 21489 | 18314 | 18188 | 18166 | 18414 | 19973 | 18193 | 19536 | Vdac1 |
| Vdac2 | 14076 | 13504 | 12975 | 12277 | 11292 | 11719 | 12797 | 10552 | 9222 | 8977 | 9262 | 10270 | 9844 | 10459 | 14377 | 13043 | 12889 | 12193 | 12323 | 12929 | 12439 | 9586 | 9692 | 9568 | 9662 | 10939 | 10831 | 11049 | Vdac2 |
| Vdac3 | 11003 | 9227 | 10165 | 9505 | 9426 | 9468 | 11073 | 11251 | 9870 | 9373 | 9864 | 10490 | 9815 | 10894 | 11388 | 9749 | 10185 | 9760 | 9940 | 10820 | 10236 | 9966 | 9736 | 10087 | 9958 | 11765 | 11203 | 12167 | Vdac3 |
| Vdr | 510 | 386 | 342 | 241 | 165 | 187 | 174 | 51 | 63 | 54 | 106 | 18 | 18 | 11 | 414 | 600 | 289 | 318 | 150 | 180 | 159 | 48 | 55 | 67 | 30 | 58 | 27 | 27 | Vdr |
| Vegfa | 6114 | 5467 | 7506 | 6981 | 7036 | 6887 | 5886 | 5576 | 6283 | 7579 | 7290 | 6211 | 6118 | 6965 | 5185 | 5798 | 7740 | 7222 | 7308 | 5720 | 6334 | 6271 | 6536 | 7645 | 6965 | 7082 | 6301 | 6641 | Vegfa |
| Vegfb | 1261 | 1419 | 1610 | 1469 | 1234 | 1198 | 1210 | 1067 | 1274 | 1509 | 1492 | 1502 | 1331 | 1345 | 1033 | 1217 | 1670 | 1465 | 1548 | 1133 | 1382 | 1250 | 1261 | 1587 | 1283 | 1650 | 1285 | 1299 | Vegfb |
| Vegfc | 331 | 232 | 342 | 315 | 291 | 245 | 292 | 381 | 407 | 423 | 512 | 521 | 416 | 445 | 302 | 308 | 265 | 218 | 341 | 259 | 247 | 398 | 507 | 439 | 371 | 633 | 452 | 549 | Vegfc |
| Vegfd | 92 | 68 | 97 | 55 | 65 | 89 | 37 | 115 | 185 | 150 | 283 | 113 | 179 | 87 | 133 | 102 | 60 | 90 | 48 | 45 | 37 | 152 | 253 | 122 | 116 | 119 | 122 | 96 | Vegfd |
| Veph1 | 1122 | 1183 | 1412 | 658 | 452 | 1197 | 406 | 1515 | 1357 | 1246 | 1243 | 1815 | 2056 | 1912 | 1398 | 1474 | 954 | 930 | 588 | 552 | 491 | 1300 | 1691 | 1407 | 1392 | 1178 | 2090 | 2040 | Veph1 |
| Vezf1 | 430 | 242 | 359 | 390 | 304 | 349 | 341 | 481 | 323 | 320 | 360 | 408 | 418 | 847 | 421 | 319 | 324 | 458 | 383 | 411 | 315 | 521 | 324 | 394 | 425 | 409 | 450 | 406 | Vezf1 |
| Vezt | 9797 | 9900 | 7950 | 9199 | 10316 | 10430 | 11318 | 11057 | 9852 | 8916 | 7990 | 8421 | 8378 | 9020 | 11068 | 9337 | 9029 | 9018 | 9825 | 12755 | 11346 | 10210 | 9282 | 8120 | 9890 | 6747 | 8536 | 8179 | Vezt |
| Vgf | 144 | 190 | 163 | 294 | 412 | 277 | 260 | 106 | 157 | 217 | 172 | 98 | 112 | 65 | 156 | 181 | 214 | 353 | 301 | 220 | 282 | 252 | 136 | 131 | 253 | 57 | 49 | 55 | Vgf |
| Vgll1 | 5 | 11 | 0 | 0 | 0 | 16 | 1 | 13 | 17 | 12 | 9 | 16 | 37 | 16 | 8 | 9 | 0 | 8 | 0 | 0 | 0 | 0 | 28 | 11 | 2 | 8 | 22 | 20 | Vgll1 |
| Vgll2 | 46 | 21 | 13 | 44 | 42 | 30 | 25 | 42 | 77 | 43 | 407 | 33 | 2 | 48 | 16 | 9 | 51 | 21 | 34 | 9 | 19 | 82 | 259 | 49 | 33 | 56 | 24 | 27 | Vgll2 |
| Vgll3 | 1474 | 752 | 630 | 513 | 577 | 889 | 148 | 1870 | 2784 | 1600 | 2283 | 1070 | 1748 | 1042 | 1247 | 1084 | 484 | 892 | 233 | 218 | 131 | 864 | 3218 | 1569 | 849 | 718 | 1033 | 910 | Vgll3 |
| Vgll4 | 3817 | 4118 | 4234 | 3948 | 4025 | 4052 | 3306 | 3936 | 4521 | 4037 | 4509 | 4507 | 4917 | 3955 | 3402 | 3995 | 4193 | 3940 | 4169 | 4012 | 3421 | 3654 | 4449 | 4741 | 4155 | 5101 | 4644 | 4198 | Vgll4 |
| Vhl | 4683 | 4300 | 4341 | 4158 | 4322 | 4509 | 4624 | 4949 | 4302 | 3866 | 3657 | 3892 | 3638 | 3949 | 5170 | 4231 | 4492 | 4390 | 4529 | 4697 | 4583 | 4298 | 4355 | 4091 | 4225 | 4274 | 3907 | 4183 | Vhl |
| Vil1 | 7 | 22 | 10 | 27 | 7 | 22 | 23 | 4 | 7 | 11 | 7 | 7 | 1 | 3 | 4 | 11 | 23 | 41 | 24 | 24 | 30 | 12 | 7 | 10 | 12 | 6 | 5 | 13 | Vil1 |
| Vill | 912 | 956 | 880 | 634 | 443 | 610 | 330 | 692 | 1085 | 825 | 1177 | 626 | 917 | 569 | 818 | 1290 | 715 | 675 | 417 | 477 | 358 | 714 | 906 | 825 | 545 | 993 | 654 | 488 | Vill |
| Vim | 70708 | 84564 | 78624 | 56696 | 47087 | 54970 | 50860 | 63306 | 63508 | 56563 | 55442 | 74548 | 75206 | 73197 | 75814 | 72525 | 71086 | 54688 | 55058 | 59752 | 49536 | 54209 | 66699 | 64334 | 57209 | 81719 | 81148 | 81051 | Vim |
| Vinac1 | 108 | 98 | 54 | 126 | 81 | 117 | 68 | 81 | 83 | 54 | 93 | 86 | 71 | 62 | 138 | 112 | 79 | 80 | 80 | 112 | 90 | 87 | 71 | 60 | 62 | 44 | 69 | 52 | Vinac1 |
| Vip | 17 | 35 | 8 | 26 | 25 | 22 | 19 | 33 | 21 | 17 | 31 | 11 | 39 | 10 | 34 | 45 | 17 | 19 | 10 | 12 | 9 | 42 | 15 | 25 | 22 | 0 | 21 | 34 | Vip |
| Vipas39 | 5140 | 4987 | 4188 | 4264 | 4125 | 4657 | 4741 | 6001 | 5500 | 5075 | 4766 | 5497 | 5129 | 5387 | 5259 | 4351 | 4065 | 3803 | 4240 | 4916 | 4691 | 5160 | 5114 | 4964 | 5492 | 5637 | 5318 | 5539 | Vipas39 |
| Vipr1 | 167 | 168 | 196 | 246 | 295 | 222 | 221 | 71 | 85 | 113 | 71 | 86 | 103 | 59 | 141 | 200 | 192 | 278 | 277 | 237 | 251 | 77 | 71 | 90 | 101 | 60 | 74 | 56 | Vipr1 |
| Vipr2 | 226 | 223 | 272 | 176 | 160 | 219 | 189 | 331 | 330 | 330 | 312 | 502 | 445 | 482 | 170 | 192 | 241 | 133 | 192 | 160 | 191 | 316 | 259 | 304 | 334 | 426 | 493 | 498 | Vipr2 |
| Virma | 13897 | 13212 | 10022 | 9875 | 8987 | 10190 | 10807 | 11218 | 10631 | 8791 | 9147 | 10581 | 9950 | 10688 | 13270 | 12116 | 9579 | 9793 | 9724 | 10690 | 10688 | 8711 | 10024 | 8942 | 9195 | 8407 | 11147 | 10250 | Virma |
| Vit | 802 | 1158 | 970 | 940 | 1008 | 1102 | 1063 | 1318 | 1263 | 1108 | 987 | 1362 | 1410 | 1517 | 1125 | 828 | 1169 | 996 | 1108 | 1143 | 1224 | 1283 | 919 | 1205 | 1281 | 1070 | 1477 | 1463 | Vit |
| Vkorc1 | 22 | 43 | 21 | 32 | 29 | 21 | 36 | 24 | 26 | 38 | 11 | 18 | 23 | 31 | 24 | 28 | 53 | 31 | 26 | 17 | 30 | 42 | 23 | 26 | 26 | 17 | 42 | 49 | Vkorc1 |
| Vkorc1l1 | 10077 | 8691 | 9517 | 9654 | 9077 | 9761 | 10347 | 6535 | 5749 | 5978 | 5987 | 6539 | 6561 | 6560 | 10297 | 9044 | 9117 | 9812 | 9282 | 9102 | 10007 | 7396 | 5746 | 5890 | 6821 | 6411 | 6743 | 7084 | Vkorc1l1 |
| Vldlr | 4317 | 3156 | 2694 | 3251 | 4789 | 4834 | 4491 | 7041 | 5611 | 5655 | 5278 | 4576 | 4009 | 4903 | 4048 | 3414 | 3065 | 3310 | 4208 | 4731 | 4492 | 6156 | 5558 | 4660 | 6158 | 2794 | 4683 | 4383 | Vldlr |
| Vma21 | 6390 | 4680 | 5884 | 5640 | 5938 | 6383 | 6505 | 6978 | 5553 | 5075 | 5501 | 5941 | 5901 | 6156 | 7166 | 5671 | 5744 | 5849 | 5907 | 6995 | 6143 | 6634 | 5890 | 5302 | 6133 | 5583 | 6062 | 6448 | Vma21 |
| Vmac | 2196 | 2520 | 2084 | 2369 | 2010 | 2127 | 2237 | 938 | 1089 | 946 | 1057 | 1067 | 970 | 1048 | 2122 | 2385 | 2055 | 2139 | 2186 | 2147 | 2049 | 971 | 1232 | 992 | 929 | 1125 | 1011 | 979 | Vmac |
| Vmn1r1 | 0 | 0 | 0 | 0 | 0 | 0 | 0 | 0 | 0 | 0 | 0 | 0 | 0 | 0 | 1 | 0 | 0 | 1 | 0 | 5 | 0 | 0 | 0 | 0 | 0 | 0 | 0 | 0 | Vmn1r1 |
| Vmn1r10 | 0 | 0 | 0 | 0 | 0 | 1 | 0 | 0 | 0 | 0 | 0 | 0 | 0 | 0 | 0 | 0 | 0 | 3 | 0 | 0 | 0 | 0 | 0 | 0 | 0 | 0 | 0 | 0 | Vmn1r10 |
| Vmn1r14 | 0 | 0 | 0 | 0 | 0 | 0 | 0 | 0 | 0 | 0 | 0 | 0 | 0 | 0 | 0 | 0 | 0 | 0 | 0 | 1 | 0 | 0 | 0 | 0 | 0 | 0 | 0 | 0 | Vmn1r14 |
| Vmn1r175 | 0 | 0 | 0 | 0 | 0 | 0 | 0 | 1 | 0 | 0 | 0 | 0 | 0 | 0 | 0 | 0 | 0 | 0 | 0 | 0 | 0 | 0 | 0 | 0 | 0 | 0 | 0 | 0 | Vmn1r175 |
| Vmn1r178 | 0 | 0 | 0 | 0 | 0 | 0 | 0 | 2 | 0 | 0 | 0 | 0 | 0 | 0 | 0 | 0 | 0 | 0 | 0 | 0 | 0 | 0 | 0 | 0 | 0 | 0 | 0 | 0 | Vmn1r178 |
| Vmn1r180 | 0 | 0 | 0 | 0 | 0 | 0 | 0 | 0 | 2 | 3 | 0 | 0 | 0 | 0 | 0 | 0 | 0 | 0 | 0 | 0 | 0 | 0 | 3 | 0 | 0 | 1 | 0 | 0 | Vmn1r180 |
| Vmn1r181 | 0 | 0 | 0 | 0 | 0 | 0 | 0 | 0 | 0 | 0 | 0 | 0 | 0 | 0 | 0 | 0 | 0 | 0 | 0 | 0 | 0 | 0 | 1 | 0 | 0 | 0 | 0 | 0 | Vmn1r181 |
| Vmn1r183 | 0 | 0 | 4 | 0 | 0 | 0 | 0 | 0 | 0 | 0 | 0 | 0 | 0 | 0 | 0 | 0 | 0 | 0 | 0 | 0 | 0 | 0 | 0 | 0 | 0 | 0 | 0 | 0 | Vmn1r183 |
| Vmn1r184 | 0 | 0 | 0 | 0 | 0 | 0 | 0 | 1 | 0 | 0 | 0 | 0 | 0 | 0 | 0 | 0 | 0 | 0 | 0 | 0 | 0 | 0 | 1 | 0 | 0 | 0 | 0 | 0 | Vmn1r184 |
| Vmn1r188 | 0 | 0 | 0 | 0 | 0 | 5 | 1 | 0 | 0 | 3 | 0 | 2 | 0 | 0 | 0 | 0 | 1 | 0 | 0 | 0 | 1 | 2 | 0 | 1 | 0 | 4 | 0 | 0 | Vmn1r188 |
| Vmn1r193 | 1 | 0 | 0 | 0 | 0 | 0 | 0 | 0 | 1 | 0 | 0 | 0 | 0 | 0 | 0 | 0 | 0 | 0 | 0 | 0 | 0 | 0 | 0 | 0 | 0 | 0 | 0 | 0 | Vmn1r193 |
| Vmn1r199 | 0 | 0 | 0 | 0 | 0 | 0 | 0 | 0 | 1 | 2 | 0 | 0 | 0 | 0 | 0 | 0 | 0 | 0 | 0 | 0 | 0 | 0 | 0 | 0 | 0 | 0 | 0 | 0 | Vmn1r199 |

| | | | | | | | | | | | | | | | | | | | | | | | | | | | | | |
|---|---|---|---|---|---|---|---|---|---|---|---|---|---|---|---|---|---|---|---|---|---|---|---|---|---|---|---|---|---|
| Vmn1r200 | 0 | 0 | 0 | 2 | 0 | 0 | 0 | 0 | 1 | 3 | 0 | 0 | 0 | 0 | 1 | 0 | 1 | 0 | 2 | 1 | 0 | 2 | 3 | 2 | 0 | 0 | 3 | 1 | Vmn1r200 |
| Vmn1r206 | 0 | 0 | 0 | 0 | 0 | 0 | 0 | 1 | 1 | 0 | 0 | 0 | 0 | 3 | 0 | 0 | 0 | 0 | 0 | 0 | 0 | 0 | 0 | 0 | 0 | 0 | 0 | 0 | Vmn1r206 |
| Vmn1r207 | 0 | 0 | 0 | 0 | 0 | 0 | 0 | 0 | 0 | 0 | 1 | 0 | 0 | 0 | 0 | 0 | 0 | 0 | 0 | 0 | 0 | 0 | 0 | 0 | 0 | 0 | 0 | 0 | Vmn1r207 |
| Vmn1r210 | 0 | 0 | 0 | 0 | 0 | 0 | 0 | 0 | 0 | 0 | 0 | 0 | 0 | 0 | 1 | 0 | 0 | 0 | 0 | 0 | 0 | 0 | 0 | 0 | 0 | 0 | 0 | 0 | Vmn1r210 |
| Vmn1r211 | 0 | 0 | 0 | 0 | 0 | 0 | 0 | 1 | 0 | 0 | 0 | 0 | 0 | 0 | 0 | 0 | 0 | 0 | 0 | 0 | 0 | 0 | 0 | 0 | 0 | 0 | 0 | 0 | Vmn1r211 |
| Vmn1r218 | 0 | 0 | 0 | 0 | 0 | 0 | 0 | 0 | 1 | 0 | 0 | 0 | 0 | 0 | 0 | 0 | 0 | 0 | 0 | 0 | 0 | 0 | 0 | 0 | 0 | 0 | 0 | 0 | Vmn1r218 |
| Vmn1r219 | 0 | 0 | 0 | 0 | 0 | 0 | 0 | 0 | 0 | 1 | 0 | 0 | 0 | 0 | 0 | 0 | 0 | 0 | 0 | 0 | 0 | 0 | 0 | 0 | 0 | 0 | 0 | 0 | Vmn1r219 |
| Vmn1r220 | 1 | 2 | 7 | 2 | 0 | 0 | 3 | 0 | 0 | 0 | 0 | 0 | 0 | 0 | 5 | 0 | 0 | 0 | 4 | 4 | 0 | 0 | 0 | 0 | 0 | 0 | 0 | 0 | Vmn1r220 |
| Vmn1r222 | 1 | 1 | 2 | 1 | 0 | 2 | 0 | 0 | 1 | 0 | 0 | 1 | 0 | 2 | 0 | 0 | 0 | 0 | 0 | 0 | 0 | 2 | 0 | 3 | 0 | 0 | 0 | 0 | Vmn1r222 |
| Vmn1r223 | 0 | 0 | 0 | 0 | 0 | 0 | 0 | 0 | 0 | 0 | 0 | 0 | 0 | 0 | 0 | 0 | 2 | 0 | 0 | 0 | 0 | 0 | 0 | 0 | 0 | 0 | 0 | 0 | Vmn1r223 |
| Vmn1r224 | 0 | 0 | 0 | 0 | 0 | 0 | 0 | 0 | 0 | 0 | 0 | 0 | 0 | 0 | 0 | 0 | 0 | 0 | 0 | 0 | 0 | 0 | 1 | 0 | 0 | 0 | 0 | 0 | Vmn1r224 |
| Vmn1r227 | 0 | 0 | 0 | 0 | 0 | 0 | 0 | 1 | 0 | 0 | 0 | 0 | 0 | 0 | 0 | 0 | 0 | 0 | 0 | 0 | 0 | 0 | 0 | 0 | 0 | 0 | 0 | 0 | Vmn1r227 |
| Vmn1r228 | 5 | 0 | 0 | 0 | 0 | 0 | 0 | 0 | 2 | 0 | 0 | 0 | 0 | 0 | 0 | 0 | 0 | 3 | 0 | 0 | 0 | 0 | 1 | 2 | 0 | 0 | 0 | 0 | Vmn1r228 |
| Vmn1r232 | 0 | 0 | 0 | 0 | 3 | 0 | 0 | 1 | 1 | 0 | 0 | 0 | 0 | 1 | 0 | 0 | 1 | 0 | 0 | 0 | 0 | 2 | 2 | 0 | 0 | 0 | 0 | 0 | Vmn1r232 |
| Vmn1r233 | 0 | 1 | 0 | 0 | 2 | 0 | 0 | 1 | 0 | 0 | 2 | 0 | 0 | 0 | 0 | 0 | 0 | 1 | 0 | 0 | 0 | 0 | 0 | 0 | 0 | 0 | 0 | 0 | Vmn1r233 |
| Vmn1r234 | 0 | 1 | 0 | 0 | 0 | 0 | 0 | 0 | 0 | 0 | 0 | 0 | 0 | 0 | 0 | 0 | 0 | 0 | 0 | 0 | 0 | 0 | 0 | 0 | 0 | 0 | 0 | 0 | Vmn1r234 |
| Vmn1r30 | 0 | 0 | 0 | 0 | 0 | 0 | 0 | 0 | 2 | 0 | 0 | 0 | 0 | 0 | 1 | 0 | 0 | 0 | 2 | 0 | 0 | 0 | 1 | 0 | 0 | 0 | 0 | 0 | Vmn1r30 |
| Vmn1r31 | 0 | 0 | 0 | 0 | 0 | 0 | 0 | 0 | 0 | 0 | 0 | 0 | 0 | 0 | 0 | 0 | 0 | 0 | 0 | 1 | 0 | 0 | 0 | 0 | 0 | 0 | 0 | 0 | Vmn1r31 |
| Vmn1r32 | 0 | 0 | 0 | 0 | 0 | 0 | 0 | 0 | 0 | 0 | 1 | 0 | 0 | 0 | 0 | 1 | 0 | 0 | 0 | 0 | 0 | 0 | 1 | 1 | 1 | 0 | 0 | 0 | Vmn1r32 |
| Vmn1r34 | 0 | 0 | 0 | 0 | 0 | 0 | 0 | 2 | 0 | 0 | 0 | 0 | 0 | 0 | 0 | 1 | 0 | 0 | 0 | 0 | 0 | 0 | 0 | 0 | 0 | 0 | 2 | 0 | Vmn1r34 |
| Vmn1r35 | 0 | 0 | 0 | 0 | 0 | 0 | 0 | 0 | 0 | 0 | 0 | 0 | 0 | 0 | 0 | 0 | 0 | 0 | 0 | 0 | 0 | 2 | 0 | 2 | 0 | 0 | 0 | 0 | Vmn1r35 |
| Vmn1r36 | 0 | 0 | 0 | 0 | 0 | 1 | 0 | 0 | 0 | 0 | 0 | 0 | 0 | 0 | 0 | 0 | 1 | 0 | 0 | 0 | 0 | 0 | 0 | 0 | 0 | 0 | 0 | 0 | Vmn1r36 |
| Vmn1r37 | 0 | 0 | 0 | 0 | 0 | 0 | 0 | 0 | 0 | 0 | 0 | 0 | 0 | 0 | 0 | 0 | 0 | 0 | 0 | 0 | 0 | 0 | 0 | 2 | 0 | 0 | 0 | 0 | Vmn1r37 |
| Vmn1r38 | 0 | 0 | 0 | 0 | 2 | 0 | 0 | 0 | 0 | 0 | 1 | 0 | 0 | 0 | 0 | 0 | 0 | 0 | 0 | 0 | 0 | 0 | 0 | 0 | 0 | 0 | 0 | 0 | Vmn1r38 |
| Vmn1r39 | 0 | 1 | 0 | 0 | 2 | 4 | 0 | 0 | 0 | 0 | 0 | 0 | 0 | 0 | 0 | 0 | 0 | 0 | 0 | 0 | 0 | 0 | 0 | 0 | 0 | 0 | 0 | 0 | Vmn1r39 |
| Vmn1r4 | 10 | 12 | 16 | 24 | 30 | 10 | 26 | 40 | 44 | 33 | 22 | 35 | 42 | 50 | 28 | 9 | 29 | 19 | 24 | 43 | 21 | 28 | 41 | 34 | 36 | 38 | 45 | 36 | Vmn1r4 |
| Vmn1r40 | 0 | 2 | 3 | 0 | 0 | 0 | 1 | 0 | 0 | 3 | 4 | 0 | 0 | 0 | 0 | 1 | 6 | 3 | 2 | 0 | 7 | 2 | 1 | 0 | 4 | 0 | 0 | 2 | Vmn1r40 |
| Vmn1r41 | 0 | 0 | 0 | 0 | 0 | 0 | 1 | 0 | 0 | 0 | 2 | 2 | 0 | 0 | 0 | 0 | 0 | 4 | 1 | 0 | 2 | 0 | 1 | 1 | 3 | 0 | 0 | 0 | Vmn1r41 |
| Vmn1r43 | 0 | 0 | 0 | 0 | 0 | 0 | 0 | 0 | 1 | 0 | 0 | 0 | 0 | 0 | 0 | 0 | 0 | 0 | 0 | 0 | 0 | 0 | 1 | 0 | 0 | 0 | 0 | 0 | Vmn1r43 |
| Vmn1r45 | 0 | 0 | 0 | 0 | 0 | 0 | 0 | 0 | 0 | 0 | 0 | 0 | 0 | 0 | 0 | 2 | 0 | 0 | 0 | 0 | 0 | 0 | 0 | 0 | 0 | 0 | 0 | 0 | Vmn1r45 |
| Vmn1r49 | 0 | 0 | 0 | 0 | 0 | 0 | 0 | 0 | 1 | 0 | 0 | 0 | 0 | 0 | 0 | 0 | 0 | 0 | 0 | 0 | 0 | 0 | 0 | 0 | 0 | 0 | 0 | 0 | Vmn1r49 |
| Vmn1r5 | 1 | 0 | 0 | 0 | 0 | 0 | 0 | 0 | 0 | 0 | 0 | 2 | 2 | 0 | 3 | 0 | 0 | 1 | 1 | 0 | 0 | 0 | 0 | 0 | 0 | 0 | 0 | 0 | Vmn1r5 |
| Vmn1r52 | 0 | 0 | 0 | 0 | 0 | 0 | 0 | 0 | 0 | 0 | 2 | 0 | 0 | 0 | 0 | 0 | 0 | 0 | 0 | 0 | 0 | 0 | 0 | 0 | 0 | 0 | 0 | 0 | Vmn1r52 |
| Vmn1r53 | 1 | 0 | 0 | 0 | 0 | 0 | 0 | 0 | 0 | 0 | 0 | 1 | 0 | 0 | 0 | 0 | 1 | 0 | 0 | 0 | 0 | 2 | 1 | 0 | 0 | 0 | 1 | 0 | Vmn1r53 |
| Vmn1r54 | 0 | 0 | 0 | 0 | 0 | 0 | 0 | 0 | 1 | 0 | 0 | 0 | 0 | 0 | 0 | 0 | 0 | 0 | 0 | 0 | 0 | 0 | 1 | 0 | 0 | 0 | 0 | 0 | Vmn1r54 |
| Vmn1r56 | 0 | 0 | 0 | 0 | 0 | 0 | 0 | 0 | 0 | 0 | 0 | 0 | 0 | 0 | 0 | 0 | 0 | 0 | 0 | 0 | 0 | 0 | 0 | 0 | 0 | 2 | 0 | 0 | Vmn1r56 |
| Vmn1r57 | 2 | 1 | 0 | 2 | 2 | 4 | 1 | 0 | 0 | 0 | 0 | 0 | 0 | 0 | 5 | 2 | 0 | 0 | 1 | 1 | 3 | 2 | 1 | 0 | 0 | 0 | 0 | 0 | Vmn1r57 |
| Vmn1r58 | 0 | 0 | 0 | 0 | 0 | 0 | 0 | 0 | 0 | 0 | 0 | 0 | 0 | 0 | 0 | 0 | 0 | 0 | 0 | 0 | 0 | 0 | 1 | 0 | 0 | 0 | 0 | 0 | Vmn1r58 |
| Vmn1r6 | 1 | 2 | 1 | 0 | 0 | 0 | 0 | 0 | 0 | 0 | 0 | 0 | 3 | 0 | 1 | 0 | 0 | 2 | 0 | 4 | 2 | 0 | 0 | 0 | 0 | 1 | 3 | 0 | Vmn1r6 |
| Vmn1r65 | 0 | 0 | 0 | 0 | 0 | 0 | 3 | 0 | 0 | 0 | 0 | 0 | 0 | 1 | 0 | 0 | 0 | 0 | 0 | 0 | 0 | 0 | 0 | 0 | 0 | 0 | 0 | 0 | Vmn1r65 |
| Vmn1r71 | 0 | 0 | 0 | 0 | 0 | 0 | 0 | 0 | 2 | 0 | 0 | 1 | 0 | 0 | 0 | 0 | 0 | 0 | 0 | 0 | 0 | 0 | 0 | 0 | 0 | 0 | 0 | 0 | Vmn1r71 |
| Vmn1r72 | 1 | 0 | 0 | 0 | 0 | 0 | 0 | 0 | 0 | 0 | 0 | 0 | 0 | 0 | 0 | 0 | 0 | 0 | 0 | 0 | 0 | 0 | 0 | 0 | 0 | 0 | 0 | 0 | Vmn1r72 |
| Vmn1r75 | 0 | 0 | 0 | 0 | 0 | 0 | 0 | 0 | 1 | 0 | 0 | 0 | 0 | 0 | 0 | 0 | 0 | 0 | 0 | 0 | 0 | 0 | 0 | 0 | 0 | 0 | 0 | 0 | Vmn1r75 |
| Vmn1r82 | 0 | 0 | 0 | 0 | 0 | 0 | 0 | 0 | 0 | 1 | 0 | 0 | 0 | 0 | 0 | 0 | 0 | 0 | 0 | 0 | 0 | 0 | 0 | 0 | 0 | 0 | 0 | 0 | Vmn1r82 |
| Vmn1r84 | 1 | 0 | 0 | 0 | 0 | 0 | 0 | 0 | 0 | 2 | 1 | 0 | 0 | 0 | 0 | 0 | 0 | 0 | 0 | 0 | 0 | 0 | 1 | 1 | 0 | 0 | 0 | 0 | Vmn1r84 |
| Vmn1r85 | 0 | 0 | 0 | 0 | 2 | 5 | 6 | 4 | 2 | 12 | 9 | 3 | 6 | 0 | 3 | 0 | 0 | 7 | 1 | 6 | 7 | 13 | 5 | 0 | 1 | 2 | 5 | 0 | Vmn1r85 |
| Vmn1r86 | 0 | 0 | 0 | 0 | 0 | 1 | 1 | 0 | 0 | 3 | 0 | 0 | 1 | 0 | 3 | 0 | 0 | 0 | 0 | 0 | 3 | 0 | 0 | 2 | 1 | 0 | 0 | 2 | Vmn1r86 |
| Vmn1r87 | 0 | 0 | 0 | 0 | 0 | 0 | 0 | 0 | 0 | 0 | 0 | 0 | 0 | 0 | 0 | 0 | 0 | 0 | 0 | 0 | 0 | 0 | 0 | 4 | 0 | 0 | 0 | 0 | Vmn1r87 |
| Vmn1r88 | 0 | 0 | 0 | 0 | 0 | 0 | 0 | 2 | 0 | 0 | 0 | 0 | 0 | 0 | 1 | 0 | 0 | 0 | 0 | 0 | 0 | 0 | 0 | 0 | 0 | 0 | 0 | 0 | Vmn1r88 |
| Vmn1r9 | 0 | 1 | 0 | 0 | 0 | 0 | 1 | 0 | 0 | 0 | 0 | 0 | 0 | 0 | 0 | 0 | 0 | 1 | 0 | 1 | 0 | 3 | 0 | 1 | 2 | 0 | 0 | 0 | Vmn1r9 |
| Vmn1r90 | 30 | 34 | 11 | 24 | 41 | 84 | 56 | 28 | 39 | 51 | 80 | 56 | 59 | 54 | 75 | 64 | 60 | 54 | 34 | 51 | 55 | 99 | 70 | 58 | 84 | 28 | 53 | 61 | Vmn1r90 |
| Vmn2r1 | 142 | 110 | 123 | 216 | 218 | 219 | 188 | 57 | 63 | 62 | 74 | 70 | 91 | 78 | 179 | 129 | 114 | 143 | 164 | 151 | 175 | 125 | 59 | 57 | 82 | 56 | 67 | 59 | Vmn2r1 |
| Vmn2r100 | 0 | 0 | 0 | 0 | 0 | 0 | 0 | 0 | 0 | 0 | 0 | 1 | 0 | 0 | 0 | 0 | 0 | 0 | 0 | 0 | 0 | 0 | 0 | 0 | 0 | 0 | 0 | 0 | Vmn2r100 |
| Vmn2r110 | 0 | 0 | 0 | 0 | 0 | 0 | 0 | 0 | 0 | 0 | 0 | 0 | 0 | 0 | 0 | 0 | 1 | 0 | 0 | 0 | 0 | 0 | 1 | 0 | 0 | 0 | 0 | 0 | Vmn2r110 |
| Vmn2r111 | 0 | 1 | 0 | 0 | 0 | 0 | 0 | 0 | 0 | 0 | 0 | 0 | 0 | 0 | 1 | 0 | 0 | 0 | 0 | 0 | 0 | 0 | 0 | 0 | 0 | 0 | 0 | 0 | Vmn2r111 |
| Vmn2r118 | 0 | 0 | 1 | 0 | 1 | 0 | 0 | 0 | 1 | 0 | 3 | 0 | 0 | 0 | 0 | 0 | 0 | 0 | 1 | 0 | 0 | 0 | 0 | 0 | 0 | 0 | 0 | 0 | Vmn2r118 |
| Vmn2r129 | 0 | 0 | 0 | 0 | 0 | 0 | 0 | 0 | 0 | 0 | 0 | 0 | 0 | 0 | 0 | 0 | 0 | 0 | 0 | 0 | 0 | 0 | 1 | 0 | 2 | 0 | 0 | 0 | Vmn2r129 |
| Vmn2r130 | 0 | 0 | 0 | 0 | 0 | 0 | 0 | 0 | 0 | 0 | 0 | 0 | 0 | 0 | 0 | 1 | 0 | 0 | 0 | 0 | 0 | 0 | 0 | 0 | 0 | 0 | 0 | 0 | Vmn2r130 |
| Vmn2r15 | 1 | 0 | 0 | 0 | 0 | 0 | 2 | 1 | 0 | 2 | 0 | 0 | 0 | 2 | 0 | 0 | 0 | 0 | 0 | 0 | 0 | 0 | 0 | 0 | 0 | 0 | 1 | 2 | Vmn2r15 |
| Vmn2r18 | 40 | 34 | 19 | 16 | 16 | 25 | 12 | 5 | 7 | 3 | 1 | 4 | 1 | 9 | 26 | 16 | 10 | 17 | 9 | 19 | 17 | 2 | 5 | 10 | 8 | 14 | 17 | 6 | Vmn2r18 |
| Vmn2r2 | 0 | 0 | 4 | 3 | 0 | 3 | 2 | 2 | 2 | 5 | 3 | 3 | 2 | 4 | 1 | 2 | 2 | 0 | 0 | 1 | 1 | 3 | 3 | 0 | 1 | 0 | 1 | 1 | Vmn2r2 |
| Vmn2r20 | 0 | 0 | 0 | 0 | 0 | 0 | 0 | 0 | 0 | 4 | 1 | 0 | 0 | 0 | 0 | 0 | 0 | 0 | 0 | 0 | 0 | 0 | 1 | 0 | 0 | 2 | 0 | 0 | Vmn2r20 |
| Vmn2r21 | 0 | 0 | 0 | 0 | 0 | 0 | 0 | 0 | 1 | 0 | 0 | 0 | 0 | 0 | 0 | 0 | 0 | 0 | 0 | 0 | 0 | 0 | 0 | 0 | 0 | 0 | 0 | 0 | Vmn2r21 |
| Vmn2r22 | 0 | 0 | 0 | 0 | 0 | 0 | 0 | 0 | 0 | 3 | 0 | 0 | 0 | 0 | 0 | 0 | 0 | 0 | 0 | 0 | 0 | 0 | 0 | 0 | 0 | 0 | 0 | 0 | Vmn2r22 |
| Vmn2r23 | 0 | 0 | 0 | 0 | 0 | 0 | 0 | 2 | 0 | 0 | 0 | 0 | 0 | 0 | 0 | 0 | 0 | 0 | 0 | 0 | 0 | 0 | 0 | 0 | 0 | 0 | 0 | 0 | Vmn2r23 |
| Vmn2r24 | 0 | 0 | 0 | 0 | 0 | 0 | 0 | 0 | 0 | 0 | 0 | 0 | 0 | 0 | 0 | 0 | 0 | 0 | 0 | 0 | 0 | 2 | 0 | 0 | 0 | 0 | 0 | 0 | Vmn2r24 |
| Vmn2r25 | 0 | 0 | 0 | 0 | 0 | 0 | 0 | 0 | 0 | 0 | 0 | 0 | 0 | 0 | 0 | 0 | 0 | 0 | 0 | 0 | 0 | 0 | 1 | 0 | 0 | 0 | 0 | 0 | Vmn2r25 |
| Vmn2r26 | 0 | 0 | 0 | 0 | 0 | 0 | 0 | 0 | 1 | 2 | 0 | 0 | 1 | 0 | 1 | 0 | 0 | 0 | 0 | 0 | 0 | 0 | 0 | 2 | 0 | 0 | 2 | 5 | Vmn2r26 |
| Vmn2r27 | 0 | 0 | 0 | 0 | 0 | 0 | 0 | 2 | 0 | 0 | 5 | 0 | 0 | 0 | 0 | 0 | 0 | 0 | 0 | 0 | 0 | 0 | 0 | 0 | 0 | 0 | 0 | 0 | Vmn2r27 |
| Vmn2r29 | 7 | 7 | 3 | 2 | 10 | 5 | 11 | 34 | 30 | 37 | 27 | 36 | 16 | 10 | 5 | 10 | 5 | 10 | 15 | 13 | 4 | 33 | 50 | 22 | 13 | 9 | 24 | 8 | Vmn2r29 |
| Vmn2r3 | 0 | 0 | 0 | 0 | 0 | 0 | 0 | 2 | 0 | 0 | 0 | 0 | 0 | 0 | 0 | 0 | 0 | 0 | 1 | 0 | 0 | 0 | 0 | 2 | 2 | 0 | 1 | 1 | Vmn2r3 |
| Vmn2r30 | 1 | 1 | 0 | 0 | 0 | 3 | 0 | 0 | 0 | 0 | 0 | 1 | 0 | 1 | 3 | 2 | 3 | 2 | 0 | 5 | 1 | 0 | 0 | 0 | 0 | 0 | 0 | 0 | Vmn2r30 |
| Vmn2r31 | 0 | 0 | 0 | 0 | 0 | 0 | 0 | 0 | 0 | 0 | 0 | 0 | 0 | 0 | 0 | 0 | 0 | 0 | 0 | 0 | 0 | 2 | 0 | 0 | 0 | 0 | 0 | 0 | Vmn2r31 |
| Vmn2r4 | 0 | 0 | 0 | 0 | 2 | 0 | 0 | 0 | 0 | 0 | 0 | 0 | 0 | 0 | 0 | 0 | 0 | 0 | 0 | 0 | 0 | 0 | 0 | 0 | 0 | 0 | 0 | 0 | Vmn2r4 |
| Vmn2r5 | 1 | 0 | 0 | 0 | 0 | 0 | 0 | 0 | 0 | 0 | 0 | 0 | 0 | 0 | 0 | 0 | 0 | 0 | 0 | 0 | 0 | 0 | 4 | 5 | 0 | 1 | 0 | 0 | Vmn2r5 |
| Vmn2r52 | 0 | 0 | 0 | 2 | 0 | 0 | 0 | 0 | 0 | 0 | 0 | 0 | 0 | 0 | 0 | 0 | 0 | 0 | 0 | 0 | 0 | 0 | 0 | 0 | 0 | 0 | 0 | 0 | Vmn2r52 |
| Vmn2r53 | 0 | 0 | 0 | 0 | 1 | 0 | 1 | 0 | 2 | 1 | 1 | 0 | 1 | 0 | 0 | 3 | 1 | 4 | 0 | 0 | 3 | 0 | 0 | 3 | 9 | 0 | 5 | 0 | Vmn2r53 |
| Vmn2r54 | 0 | 0 | 0 | 0 | 0 | 0 | 2 | 0 | 0 | 0 | 2 | 1 | 0 | 1 | 0 | 0 | 1 | 0 | 0 | 1 | 2 | 0 | 1 | 0 | 0 | 0 | 0 | 0 | Vmn2r54 |
| Vmn2r55 | 0 | 0 | 0 | 0 | 0 | 9 | 1 | 0 | 0 | 0 | 0 | 0 | 0 | 0 | 0 | 4 | 0 | 0 | 2 | 0 | 2 | 0 | 0 | 0 | 0 | 0 | 0 | 0 | Vmn2r55 |

| | | | | | | | | | | | | | | | | | | | | | | | | | | | | | |
|---|---|---|---|---|---|---|---|---|---|---|---|---|---|---|---|---|---|---|---|---|---|---|---|---|---|---|---|---|---|
| Vmn2r56 | 1 | 2 | 0 | 4 | 0 | 1 | 2 | 0 | 0 | 0 | 1 | 1 | 1 | 0 | 5 | 1 | 1 | 2 | 1 | 0 | 4 | 0 | 1 | 2 | 0 | 0 | 0 | 2 | Vmn2r56 |
| Vmn2r57 | 63 | 38 | 26 | 76 | 50 | 45 | 57 | 45 | 48 | 60 | 81 | 28 | 48 | 52 | 51 | 48 | 71 | 53 | 52 | 42 | 59 | 50 | 42 | 53 | 36 | 60 | 48 | 45 | Vmn2r57 |
| Vmn2r6 | 0 | 0 | 0 | 0 | 0 | 0 | 0 | 1 | 0 | 0 | 0 | 0 | 0 | 0 | 0 | 0 | 0 | 0 | 0 | 0 | 0 | 0 | 3 | 0 | 0 | 0 | 0 | 0 | Vmn2r6 |
| Vmn2r7 | 1 | 0 | 0 | 0 | 0 | 0 | 1 | 4 | 0 | 0 | 0 | 0 | 0 | 0 | 0 | 1 | 0 | 2 | 1 | 0 | 0 | 0 | 0 | 0 | 0 | 0 | 0 | 0 | Vmn2r7 |
| Vmn2r73 | 0 | 0 | 0 | 0 | 0 | 0 | 0 | 0 | 0 | 0 | 0 | 0 | 0 | 0 | 0 | 0 | 0 | 0 | 0 | 0 | 0 | 0 | 0 | 2 | 0 | 0 | 0 | 0 | Vmn2r73 |
| Vmn2r79 | 0 | 0 | 0 | 0 | 0 | 0 | 0 | 0 | 0 | 1 | 0 | 0 | 0 | 0 | 0 | 0 | 0 | 0 | 0 | 0 | 0 | 0 | 0 | 0 | 0 | 0 | 0 | 0 | Vmn2r79 |
| Vmn2r81 | 1 | 0 | 0 | 0 | 0 | 0 | 0 | 0 | 0 | 0 | 0 | 0 | 0 | 0 | 0 | 0 | 0 | 0 | 0 | 0 | 0 | 0 | 0 | 0 | 0 | 0 | 0 | 0 | Vmn2r81 |
| Vmn2r82 | 0 | 0 | 0 | 4 | 0 | 0 | 1 | 1 | 0 | 0 | 0 | 0 | 1 | 0 | 3 | 0 | 0 | 0 | 0 | 0 | 0 | 0 | 0 | 0 | 2 | 0 | 0 | 0 | Vmn2r82 |
| Vmn2r83 | 0 | 0 | 0 | 0 | 0 | 4 | 0 | 0 | 0 | 0 | 0 | 0 | 0 | 0 | 0 | 0 | 2 | 0 | 0 | 0 | 0 | 0 | 0 | 0 | 0 | 0 | 0 | 0 | Vmn2r83 |
| Vmn2r84 | 6 | 5 | 5 | 7 | 21 | 14 | 31 | 5 | 1 | 8 | 2 | 2 | 5 | 4 | 21 | 12 | 17 | 43 | 19 | 6 | 36 | 7 | 6 | 6 | 3 | 4 | 6 | 2 | Vmn2r84 |
| Vmn2r85 | 2 | 10 | 5 | 13 | 10 | 13 | 22 | 15 | 7 | 12 | 5 | 2 | 10 | 2 | 9 | 9 | 13 | 14 | 9 | 19 | 20 | 7 | 7 | 5 | 8 | 4 | 9 | 4 | Vmn2r85 |
| Vmn2r86 | 15 | 6 | 5 | 22 | 19 | 9 | 10 | 15 | 8 | 8 | 14 | 3 | 11 | 5 | 7 | 11 | 5 | 7 | 19 | 4 | 13 | 7 | 4 | 7 | 3 | 5 | 7 | 7 | Vmn2r86 |
| Vmn2r87 | 55 | 43 | 59 | 62 | 111 | 106 | 104 | 77 | 73 | 52 | 72 | 72 | 82 | 57 | 67 | 47 | 63 | 114 | 103 | 62 | 93 | 62 | 50 | 55 | 71 | 37 | 65 | 76 | Vmn2r87 |
| Vmn2r90 | 0 | 0 | 0 | 0 | 1 | 0 | 0 | 0 | 0 | 3 | 0 | 5 | 0 | 0 | 0 | 0 | 0 | 0 | 0 | 0 | 4 | 0 | 1 | 2 | 0 | 0 | 0 | 1 | Vmn2r90 |
| Vmn2r96 | 0 | 0 | 0 | 0 | 0 | 0 | 0 | 0 | 0 | 0 | 0 | 0 | 0 | 0 | 0 | 0 | 0 | 0 | 0 | 0 | 0 | 0 | 0 | 0 | 0 | 1 | 0 | 0 | Vmn2r96 |
| Vmo1 | 0 | 1 | 3 | 2 | 0 | 1 | 0 | 6 | 2 | 5 | 8 | 1 | 7 | 1 | 4 | 8 | 5 | 4 | 3 | 0 | 1 | 7 | 11 | 2 | 0 | 4 | 3 | 6 | Vmo1 |
| Vmp1 | 4597 | 4164 | 4189 | 3765 | 3580 | 3408 | 3957 | 4030 | 3378 | 3122 | 2896 | 3069 | 3159 | 3409 | 5098 | 4575 | 3823 | 3909 | 3438 | 4371 | 3704 | 3429 | 3774 | 3134 | 3012 | 3457 | 3218 | 3408 | Vmp1 |
| Vnn1 | 50 | 68 | 104 | 19 | 16 | 27 | 12 | 34 | 72 | 38 | 73 | 50 | 98 | 64 | 80 | 95 | 23 | 46 | 14 | 5 | 8 | 45 | 61 | 47 | 33 | 51 | 83 | 69 | Vnn1 |
| Vnn3 | 1 | 0 | 0 | 2 | 2 | 5 | 3 | 0 | 2 | 8 | 2 | 0 | 0 | 0 | 3 | 3 | 0 | 0 | 1 | 6 | 0 | 5 | 2 | 3 | 5 | 0 | 1 | 1 | Vnn3 |
| Vopp1 | 8395 | 8066 | 8573 | 10047 | 13011 | 10782 | 11534 | 12563 | 10696 | 12014 | 10645 | 10660 | 10309 | 11477 | 9918 | 7142 | 9769 | 10551 | 11482 | 11307 | 11617 | 13521 | 10223 | 11039 | 12736 | 10520 | 10144 | 10981 | Vopp1 |
| Vpreb1 | 1 | 2 | 4 | 1 | 1 | 9 | 6 | 33 | 26 | 30 | 46 | 17 | 27 | 20 | 3 | 1 | 3 | 2 | 2 | 1 | 2 | 23 | 26 | 20 | 23 | 28 | 23 | 21 | Vpreb1 |
| Vpreb2 | 0 | 0 | 0 | 0 | 0 | 1 | 0 | 1 | 0 | 0 | 0 | 2 | 2 | 0 | 0 | 0 | 0 | 0 | 0 | 0 | 1 | 0 | 1 | 0 | 0 | 0 | 0 | 0 | Vpreb2 |
| Vpreb3 | 2 | 5 | 0 | 6 | 3 | 4 | 8 | 4 | 1 | 0 | 3 | 0 | 0 | 1 | 0 | 8 | 2 | 4 | 2 | 1 | 3 | 0 | 3 | 2 | 0 | 5 | 0 | 0 | Vpreb3 |
| Vps11 | 4139 | 5148 | 3743 | 4245 | 4064 | 4093 | 4152 | 3863 | 3766 | 3481 | 3309 | 3875 | 3633 | 3452 | 4134 | 4076 | 4273 | 4218 | 4327 | 4415 | 4380 | 3623 | 3897 | 3538 | 3876 | 3407 | 3817 | 3647 | Vps11 |
| Vps13a | 10105 | 7952 | 4657 | 5790 | 5291 | 7351 | 7063 | 5333 | 4563 | 3011 | 3285 | 3523 | 3932 | 3764 | 10077 | 9387 | 4969 | 5663 | 5086 | 8925 | 6999 | 2700 | 4517 | 2836 | 3433 | 2115 | 4114 | 3478 | Vps13a |
| Vps13b | 5632 | 4278 | 1776 | 2224 | 2877 | 3754 | 3004 | 6427 | 6257 | 3817 | 4594 | 3321 | 4227 | 3475 | 5230 | 4997 | 1858 | 2392 | 2291 | 4095 | 3475 | 3225 | 6218 | 2752 | 3763 | 2187 | 4068 | 3188 | Vps13b |
| Vps13c | 7690 | 8111 | 3497 | 5600 | 6622 | 8145 | 7451 | 6896 | 6576 | 4241 | 4993 | 4405 | 4735 | 4750 | 8294 | 7615 | 4446 | 5827 | 5517 | 9314 | 8313 | 4224 | 5734 | 3257 | 5118 | 2182 | 5452 | 3757 | Vps13c |
| Vps13d | 9081 | 8719 | 3754 | 5388 | 6102 | 8107 | 6492 | 7421 | 7414 | 5202 | 6479 | 4570 | 4985 | 4797 | 8591 | 8261 | 3912 | 5273 | 5029 | 8176 | 7784 | 4742 | 6850 | 3783 | 6229 | 2818 | 5861 | 4184 | Vps13d |
| Vps16 | 5375 | 5955 | 5597 | 5863 | 5317 | 5517 | 5782 | 5506 | 5644 | 5601 | 5188 | 6197 | 5642 | 6087 | 5222 | 4950 | 5777 | 5328 | 5732 | 5730 | 5567 | 4859 | 5410 | 5829 | 5714 | 6734 | 6240 | 6224 | Vps16 |
| Vps18 | 1944 | 2625 | 2868 | 2828 | 2806 | 2422 | 2126 | 1842 | 2318 | 2336 | 2404 | 2652 | 2248 | 2326 | 1843 | 2356 | 2807 | 2715 | 2936 | 1978 | 2425 | 2232 | 2174 | 2504 | 2424 | 2709 | 2328 | 2340 | Vps18 |
| Vps25 | 36 | 21 | 41 | 23 | 28 | 46 | 34 | 29 | 40 | 68 | 25 | 50 | 32 | 27 | 9 | 48 | 46 | 40 | 15 | 21 | 29 | 48 | 29 | 31 | 57 | 62 | 41 | 44 | Vps25 |
| Vps26a | 8728 | 7446 | 8502 | 7379 | 6567 | 7186 | 8617 | 5150 | 4345 | 4414 | 4199 | 5344 | 4835 | 5312 | 8931 | 8235 | 7478 | 7520 | 7215 | 8180 | 7648 | 4238 | 4500 | 4987 | 4439 | 5648 | 5417 | 5680 | Vps26a |
| Vps26b | 9296 | 8826 | 8821 | 12048 | 11980 | 11095 | 11808 | 8435 | 7753 | 9085 | 8364 | 8649 | 7837 | 8593 | 8950 | 8161 | 10594 | 11988 | 12520 | 9639 | 13271 | 9651 | 7991 | 9042 | 10052 | 7711 | 7717 | 8532 | Vps26b |
| Vps26c | 5039 | 5856 | 4764 | 4412 | 4093 | 4044 | 4101 | 3813 | 3565 | 3139 | 3161 | 2859 | 3032 | 3078 | 5219 | 5355 | 4464 | 4314 | 4321 | 4963 | 4224 | 3521 | 3705 | 3009 | 3265 | 3072 | 3131 | 3078 | Vps26c |
| Vps28 | 2933 | 3341 | 2639 | 3054 | 3290 | 2877 | 3339 | 3748 | 3264 | 3538 | 2874 | 2999 | 3111 | 3393 | 3363 | 2515 | 3354 | 3110 | 3150 | 3442 | 3220 | 3554 | 3240 | 3389 | 3422 | 3066 | 3276 | 2969 | Vps28 |
| Vps29 | 7367 | 6612 | 7058 | 6921 | 6667 | 7100 | 7654 | 5735 | 5145 | 4988 | 4969 | 4739 | 5011 | 5158 | 7439 | 6855 | 6918 | 6965 | 6846 | 7656 | 7653 | 5511 | 5303 | 4998 | 5412 | 4917 | 5136 | 5293 | Vps29 |
| Vps33a | 4111 | 4525 | 3721 | 4456 | 5375 | 4794 | 4912 | 5475 | 5586 | 5491 | 5411 | 5592 | 5610 | 5633 | 4491 | 4076 | 4263 | 4540 | 4950 | 4652 | 5217 | 6037 | 5222 | 5197 | 6366 | 4393 | 5671 | 5579 | Vps33a |
| Vps33b | 2491 | 2747 | 2740 | 3094 | 2819 | 2828 | 2733 | 2027 | 1978 | 2219 | 2039 | 2259 | 2116 | 2157 | 2498 | 2475 | 2895 | 2987 | 3108 | 2433 | 2742 | 2692 | 1906 | 2241 | 2249 | 2126 | 2331 | 2146 | Vps33b |
| Vps35 | 25202 | 21009 | 20176 | 19096 | 17743 | 18721 | 22274 | 20548 | 16637 | 14100 | 14021 | 14872 | 14983 | 16225 | 27077 | 22797 | 19091 | 19173 | 18434 | 24412 | 20673 | 15938 | 16977 | 14016 | 14663 | 15186 | 15543 | 16211 | Vps35 |
| Vps35l | 4133 | 4635 | 3708 | 3715 | 3904 | 3971 | 4007 | 6332 | 6369 | 5887 | 5187 | 5943 | 5608 | 5596 | 4321 | 3985 | 3509 | 3812 | 3920 | 4276 | 4144 | 6066 | 6274 | 5585 | 6204 | 5517 | 5986 | 5420 | Vps35l |
| Vps36 | 4277 | 3809 | 4298 | 4103 | 4458 | 4036 | 4626 | 6161 | 5487 | 5628 | 4987 | 5928 | 5329 | 6402 | 4261 | 3669 | 4406 | 4008 | 4283 | 4507 | 4371 | 5177 | 5176 | 5569 | 5662 | 6458 | 6105 | 6189 | Vps36 |
| Vps37a | 8887 | 7083 | 5864 | 7580 | 7697 | 8651 | 8887 | 6974 | 5862 | 5254 | 5679 | 5337 | 5410 | 5848 | 8500 | 8122 | 6654 | 7644 | 7161 | 8897 | 8796 | 6009 | 5874 | 4911 | 5645 | 4154 | 5545 | 5253 | Vps37a |
| Vps37b | 2704 | 3587 | 3924 | 4367 | 3823 | 3290 | 3619 | 2507 | 2796 | 3313 | 2922 | 3506 | 2851 | 3178 | 2823 | 3133 | 4245 | 4271 | 4024 | 3196 | 3869 | 3263 | 2779 | 3489 | 3761 | 3513 | 2831 | 3131 | Vps37b |
| Vps37c | 1004 | 1286 | 1586 | 1476 | 1568 | 1286 | 1406 | 1255 | 1818 | 1968 | 1768 | 1934 | 1715 | 1723 | 1106 | 1216 | 1749 | 1579 | 1755 | 1138 | 1440 | 1826 | 1731 | 1868 | 1960 | 2229 | 1729 | 1708 | Vps37c |
| Vps37d | 845 | 1111 | 1192 | 1204 | 1564 | 1318 | 1262 | 1503 | 1543 | 1808 | 1590 | 1629 | 1442 | 1621 | 829 | 805 | 1353 | 1196 | 1638 | 1266 | 1273 | 1641 | 1564 | 1752 | 1850 | 1556 | 1550 | 1457 | Vps37d |
| Vps39 | 6343 | 6941 | 5361 | 6624 | 6599 | 6944 | 6550 | 7350 | 7264 | 6274 | 6098 | 6523 | 6457 | 6389 | 6367 | 5939 | 5638 | 6287 | 6453 | 7040 | 6859 | 6781 | 6899 | 6320 | 7145 | 5426 | 6754 | 6262 | Vps39 |
| Vps41 | 7553 | 7503 | 6109 | 6595 | 7499 | 7356 | 7729 | 10944 | 9172 | 7921 | 7729 | 8073 | 8125 | 8283 | 9112 | 7546 | 6478 | 6851 | 6899 | 8417 | 7654 | 9789 | 9412 | 7661 | 8912 | 7592 | 8648 | 8204 | Vps41 |
| Vps45 | 2647 | 2655 | 2267 | 2559 | 2380 | 2383 | 2633 | 2982 | 2562 | 2393 | 2283 | 2241 | 2053 | 2382 | 2759 | 2281 | 2406 | 2424 | 2484 | 2777 | 2675 | 2518 | 2640 | 2344 | 2504 | 2178 | 2156 | 2493 | Vps45 |
| Vps4a | 5860 | 6110 | 5982 | 6134 | 5954 | 5417 | 6353 | 6075 | 5564 | 5667 | 5337 | 5389 | 4927 | 5964 | 6070 | 5374 | 5838 | 6015 | 6196 | 5990 | 6321 | 6139 | 5385 | 5855 | 6294 | 5771 | 5341 | 5921 | Vps4a |
| Vps4b | 7922 | 6645 | 6602 | 5797 | 5831 | 6113 | 6849 | 6559 | 5852 | 4995 | 5267 | 5957 | 5790 | 6295 | 8099 | 7132 | 6191 | 5951 | 5814 | 7191 | 6547 | 5197 | 5966 | 5465 | 5262 | 6097 | 5935 | 6179 | Vps4b |
| Vps50 | 5878 | 5267 | 5357 | 5418 | 5778 | 6002 | 6455 | 6110 | 4987 | 4429 | 4456 | 4934 | 4639 | 5305 | 6852 | 5428 | 5446 | 5202 | 5675 | 6820 | 6079 | 5078 | 4849 | 4552 | 5220 | 4975 | 5069 | 5213 | Vps50 |
| Vps51 | 568 | 703 | 527 | 572 | 718 | 587 | 542 | 1760 | 2057 | 1974 | 1856 | 2045 | 1804 | 1867 | 559 | 603 | 553 | 549 | 670 | 555 | 598 | 2050 | 1999 | 1968 | 1991 | 1980 | 2048 | 1876 | Vps51 |
| Vps52 | 2233 | 2488 | 2843 | 2639 | 2928 | 2639 | 2579 | 3627 | 4192 | 4377 | 4063 | 4785 | 4314 | 4382 | 2318 | 2241 | 2715 | 2658 | 3123 | 2399 | 2690 | 4318 | 4075 | 4660 | 4432 | 4730 | 4144 | 4462 | Vps52 |
| Vps53 | 3420 | 3953 | 3992 | 4009 | 3976 | 3830 | 4453 | 3127 | 3101 | 3775 | 3147 | 3318 | 3232 | 3767 | 3492 | 3404 | 4065 | 3986 | 4158 | 3514 | 4349 | 3444 | 3074 | 3783 | 3761 | 3392 | 3419 | 3589 | Vps53 |
| Vps54 | 4037 | 3045 | 4118 | 3692 | 4134 | 4044 | 4105 | 5658 | 4951 | 5165 | 5514 | 5870 | 5966 | 6237 | 4061 | 3606 | 3768 | 3639 | 3869 | 3602 | 3883 | 5763 | 5050 | 5258 | 5399 | 5826 | 5607 | 6190 | Vps54 |
| Vps72 | 4300 | 4751 | 5370 | 5069 | 4417 | 4539 | 4752 | 5533 | 5742 | 6423 | 5548 | 7428 | 6005 | 7579 | 4316 | 3941 | 5257 | 4818 | 5422 | 4469 | 4958 | 6089 | 5586 | 7653 | 6987 | 8455 | 7202 | 7841 | Vps72 |
| Vps8 | 3893 | 4594 | 3555 | 3872 | 4498 | 4687 | 4235 | 4323 | 4420 | 3788 | 3694 | 4027 | 3766 | 3951 | 4588 | 3886 | 3859 | 4196 | 4524 | 4512 | 4703 | 4331 | 4168 | 3621 | 4353 | 2926 | 4081 | 3849 | Vps8 |
| Vps9d1 | 1397 | 1759 | 1829 | 1920 | 2302 | 2002 | 1711 | 2551 | 3177 | 3571 | 3332 | 3318 | 3100 | 3031 | 1403 | 1734 | 2099 | 2023 | 2362 | 1675 | 1906 | 3776 | 2836 | 3358 | 3608 | 3194 | 2933 | 3059 | Vps9d1 |
| Vrk1 | 6172 | 5510 | 5479 | 5461 | 4648 | 5191 | 6109 | 4169 | 4416 | 4111 | 3836 | 5269 | 4626 | 5597 | 5598 | 5062 | 5158 | 4846 | 5156 | 5602 | 5425 | 3342 | 3829 | 4840 | 5045 | 5813 | 5619 | 5641 | Vrk1 |
| Vrk2 | 427 | 267 | 436 | 229 | 227 | 245 | 225 | 537 | 503 | 407 | 442 | 619 | 586 | 585 | 359 | 406 | 312 | 201 | 217 | 273 | 222 | 391 | 521 | 555 | 388 | 726 | 786 | 776 | Vrk2 |
| Vrk3 | 1662 | 2056 | 1779 | 1563 | 1443 | 1455 | 1425 | 1473 | 1603 | 1478 | 1442 | 1842 | 1746 | 1709 | 1571 | 1496 | 1709 | 1308 | 1744 | 1489 | 1344 | 1420 | 1702 | 1647 | 1550 | 1991 | 1892 | 1755 | Vrk3 |
| Vrtn | 0 | 1 | 1 | 0 | 0 | 0 | 0 | 2 | 0 | 2 | 0 | 1 | 1 | 0 | 0 | 0 | 0 | 0 | 0 | 0 | 0 | 0 | 0 | 0 | 0 | 0 | 0 | 0 | Vrtn |
| Vsig1 | 0 | 0 | 0 | 0 | 0 | 0 | 0 | 0 | 0 | 0 | 0 | 0 | 0 | 0 | 0 | 0 | 0 | 0 | 0 | 0 | 0 | 0 | 1 | 0 | 0 | 0 | 0 | 0 | Vsig1 |
| Vsig10 | 501 | 438 | 699 | 432 | 408 | 461 | 341 | 529 | 564 | 574 | 719 | 465 | 616 | 473 | 442 | 495 | 430 | 401 | 416 | 337 | 298 | 396 | 662 | 679 | 435 | 798 | 636 | 559 | Vsig10 |
| Vsig10l | 397 | 441 | 495 | 453 | 588 | 545 | 534 | 276 | 370 | 399 | 339 | 333 | 294 | 235 | 365 | 439 | 494 | 474 | 598 | 409 | 496 | 349 | 336 | 289 | 293 | 275 | 278 | 225 | Vsig10l |
| Vsig2 | 80 | 100 | 109 | 130 | 183 | 111 | 134 | 117 | 151 | 271 | 271 | 237 | 268 | 177 | 87 | 109 | 88 | 146 | 156 | 82 | 124 | 237 | 160 | 186 | 251 | 141 | 205 | 184 | Vsig2 |
| Vsig4 | 1 | 1 | 0 | 3 | 5 | 0 | 0 | 0 | 3 | 0 | 9 | 0 | 2 | 7 | 0 | 0 | 0 | 1 | 0 | 3 | 1 | 8 | 5 | 0 | 4 | 0 | 1 | 5 | Vsig4 |
| Vsig8 | 6 | 3 | 3 | 12 | 10 | 1 | 6 | 9 | 7 | 1 | 4 | 0 | 2 | 5 | 7 | 4 | 1 | 6 | 7 | 5 | 1 | 5 | 9 | 0 | 6 | 2 | 0 | 1 | Vsig8 |
| Vsir | 429 | 389 | 429 | 411 | 372 | 350 | 249 | 247 | 356 | 380 | 519 | 328 | 348 | 261 | 332 | 503 | 337 | 326 | 295 | 312 | 222 | 354 | 460 | 357 | 294 | 347 | 354 | 311 | Vsir |
| Vsnl1 | 459 | 413 | 450 | 699 | 1235 | 870 | 925 | 485 | 536 | 672 | 549 | 307 | 306 | 341 | 549 | 417 | 510 | 810 | 774 | 860 | 1009 | 824 | 520 | 395 | 616 | 208 | 302 | 272 | Vsnl1 |
| Vstm2a | 521 | 448 | 427 | 932 | 1453 | 1270 | 1244 | 1823 | 1823 | 1639 | 1514 | 1032 | 815 | 1093 | 682 | 421 | 776 | 1044 | 1271 | 1386 | 1309 | 2206 | 1805 | 1388 | 2033 | 1026 | 805 | 894 | Vstm2a |
| Vstm2b | 614 | 785 | 931 | 822 | 1087 | 972 | 794 | 724 | 974 | 1174 | 1175 | 870 | 870 | 765 | 584 | 811 | 873 | 742 | 953 | 705 | 720 | 1021 | 856 | 919 | 904 | 763 | 668 | 714 | Vstm2b |
| Vstm2l | 403 | 600 | 400 | 838 | 1438 | 917 | 788 | 1132 | 1008 | 1442 | 1069 | 799 | 854 | 705 | 596 | 498 | 731 | 1011 | 1272 | 1040 | 1034 | 1592 | 1052 | 1017 | 1161 | 413 | 609 | 555 | Vstm2l |
| Vstm4 | 148 | 129 | 125 | 115 | 174 | 146 | 112 | 355 | 490 | 355 | 696 | 295 | 349 | 261 | 137 | 178 | 129 | 141 | 136 | 113 | 116 | 389 | 768 | 307 | 316 | 299 | 325 | 280 | Vstm4 |
| Vstm5 | 43 | 60 | 38 | 55 | 192 | 83 | 77 | 399 | 446 | 449 | 420 | 226 | 302 | 230 | 91 | 70 | 35 | 89 | 79 | 98 | 80 | 511 | 377 | 293 | 374 | 175 | 207 | 244 | Vstm5 |
| Vsx1 | 17 | 10 | 27 | 15 | 14 | 7 | 10 | 28 | 24 | 41 | 35 | 22 | 102 | 26 | 22 | 29 | 20 | 21 | 5 | 2 | 16 | 13 | 21 | 16 | 5 | 11 | 51 | 18 | Vsx1 |
| Vsx2 | 45 | 97 | 82 | 127 | 234 | 130 | 22 | 171 | 136 | 268 | 208 | 97 | 420 | 50 | 138 | 136 | 197 | 125 | 85 | 72 | 68 | 259 | 90 | 109 | 147 | 15 | 102 | 47 | Vsx2 |
| Vta1 | 3783 | 3841 | 4066 | 3769 | 3616 | 3281 | 3898 | 3999 | 3329 | 3270 | 2820 | 3896 | 3361 | 3978 | 4243 | 3573 | 3954 | 3444 | 3550 | 4093 | 3847 | 3526 | 3282 | 3344 | 3379 | 3805 | 4005 | 4181 | Vta1 |
| Vtcn1 | 6 | 5 | 2 | 0 | 0 | 4 | 0 | 21 | 22 | 4 | 22 | 14 | 11 | 24 | 4 | 8 | 0 | 8 | 0 | 0 | 0 | 12 | 21 | 6 | 1 | 1 | 8 | 24 | Vtcn1 |

| | | | | | | | | | | | | | | | | | | | | | | | | | | | | | |
|---|---|---|---|---|---|---|---|---|---|---|---|---|---|---|---|---|---|---|---|---|---|---|---|---|---|---|---|---|---|
| Vti1a | 1747 | 1757 | 1795 | 1591 | 1834 | 1742 | 1817 | 2501 | 2277 | 2489 | 2159 | 2208 | 2344 | 2402 | 1705 | 1476 | 1691 | 1640 | 1811 | 1918 | 1808 | 2403 | 2289 | 2352 | 2410 | 2312 | 2529 | 2622 | Vti1a |
| Vti1b | 3419 | 3415 | 3290 | 3346 | 3239 | 3042 | 3622 | 4140 | 3715 | 3831 | 3538 | 3164 | 3280 | 3725 | 3749 | 3207 | 3465 | 3374 | 3251 | 3783 | 3474 | 3813 | 3710 | 3814 | 3771 | 3595 | 3232 | 3507 | Vti1b |
| Vtn | 1971 | 3078 | 3528 | 2897 | 3584 | 3061 | 2642 | 2677 | 2656 | 3024 | 2862 | 2937 | 4280 | 2805 | 3133 | 3306 | 2980 | 3078 | 3097 | 4438 | 3042 | 2896 | 3480 | 2688 | 3051 | 3222 | 3526 | 2987 | Vtn |
| Vwa1 | 774 | 898 | 919 | 923 | 1051 | 1042 | 803 | 916 | 1106 | 1211 | 1175 | 1089 | 1069 | 897 | 704 | 814 | 921 | 890 | 1117 | 1015 | 906 | 1143 | 1190 | 1191 | 1108 | 1089 | 1052 | 946 | Vwa1 |
| Vwa2 | 218 | 164 | 82 | 88 | 101 | 119 | 54 | 193 | 361 | 223 | 279 | 126 | 214 | 120 | 127 | 199 | 85 | 160 | 58 | 40 | 50 | 99 | 317 | 178 | 105 | 259 | 106 | 148 | Vwa2 |
| Vwa3a | 166 | 178 | 85 | 144 | 226 | 259 | 169 | 205 | 273 | 233 | 146 | 230 | 274 | 151 | 164 | 259 | 107 | 189 | 187 | 238 | 201 | 206 | 394 | 150 | 176 | 82 | 139 | 122 | Vwa3a |
| Vwa3b | 76 | 54 | 39 | 89 | 50 | 61 | 76 | 51 | 71 | 34 | 45 | 26 | 58 | 29 | 58 | 74 | 29 | 57 | 48 | 71 | 82 | 60 | 95 | 57 | 30 | 31 | 22 | 19 | Vwa3b |
| Vwa5a | 2414 | 1767 | 1439 | 1127 | 1341 | 1416 | 1059 | 1783 | 1972 | 1426 | 1958 | 1288 | 1723 | 1245 | 2106 | 2298 | 1227 | 1436 | 1084 | 1454 | 997 | 1542 | 2647 | 1275 | 1213 | 1253 | 1416 | 1477 | Vwa5a |
| Vwa5b1 | 87 | 140 | 54 | 104 | 253 | 186 | 131 | 171 | 272 | 243 | 238 | 247 | 312 | 149 | 99 | 201 | 71 | 152 | 168 | 197 | 172 | 366 | 357 | 148 | 306 | 107 | 156 | 136 | Vwa5b1 |
| Vwa5b2 | 532 | 595 | 601 | 893 | 1597 | 956 | 742 | 876 | 952 | 1162 | 1103 | 660 | 1080 | 581 | 671 | 723 | 852 | 966 | 1052 | 924 | 842 | 1482 | 818 | 765 | 1001 | 358 | 667 | 606 | Vwa5b2 |
| Vwa7 | 145 | 209 | 227 | 313 | 234 | 190 | 244 | 62 | 59 | 134 | 92 | 109 | 102 | 65 | 131 | 225 | 326 | 366 | 280 | 195 | 213 | 105 | 91 | 111 | 102 | 58 | 87 | 114 | Vwa7 |
| Vwa8 | 1711 | 1612 | 1379 | 1242 | 1263 | 1405 | 1120 | 2022 | 2292 | 1966 | 2115 | 1903 | 2123 | 1918 | 1514 | 1798 | 1181 | 1150 | 1146 | 1209 | 1222 | 1818 | 2217 | 1835 | 1925 | 1606 | 1961 | 1892 | Vwa8 |
| Vwc2 | 119 | 128 | 119 | 90 | 223 | 194 | 208 | 315 | 274 | 319 | 363 | 203 | 162 | 219 | 138 | 95 | 139 | 144 | 166 | 211 | 213 | 401 | 343 | 248 | 372 | 161 | 165 | 173 | Vwc2 |
| Vwc2l | 91 | 99 | 59 | 120 | 342 | 241 | 198 | 519 | 448 | 440 | 359 | 163 | 265 | 154 | 170 | 145 | 115 | 153 | 243 | 252 | 258 | 525 | 397 | 199 | 430 | 85 | 205 | 151 | Vwc2l |
| Vwce | 76 | 50 | 47 | 43 | 43 | 71 | 56 | 85 | 129 | 113 | 162 | 117 | 115 | 69 | 26 | 71 | 60 | 44 | 65 | 24 | 51 | 94 | 146 | 96 | 120 | 90 | 65 | 49 | Vwce |
| Vwde | 24 | 10 | 2 | 13 | 19 | 8 | 11 | 23 | 26 | 9 | 19 | 0 | 7 | 7 | 8 | 14 | 3 | 10 | 10 | 12 | 16 | 12 | 27 | 3 | 9 | 19 | 3 | 7 | Vwde |
| Vwf | 1498 | 1973 | 2638 | 2226 | 1810 | 2074 | 1373 | 895 | 1737 | 2131 | 2306 | 2840 | 2993 | 2826 | 1805 | 2160 | 2532 | 2479 | 1669 | 1358 | 1566 | 3019 | 2333 | 2546 | 2858 | 2354 | 3276 | 3093 | Vwf |
| Vxn | 308 | 384 | 319 | 434 | 794 | 644 | 472 | 2288 | 1670 | 2023 | 1564 | 1701 | 2903 | 1536 | 675 | 500 | 513 | 528 | 538 | 646 | 607 | 2513 | 1620 | 1387 | 1971 | 691 | 1715 | 1435 | Vxn |
| Wac | 9866 | 8827 | 9264 | 9351 | 9917 | 9546 | 10782 | 14760 | 11994 | 12330 | 11616 | 13221 | 12452 | 13889 | 10433 | 8173 | 9396 | 9097 | 10006 | 10122 | 10114 | 14105 | 11909 | 12594 | 13180 | 13845 | 13389 | 14030 | Wac |
| Wapl | 14910 | 11573 | 12940 | 12869 | 11607 | 13169 | 13508 | 12454 | 10434 | 10000 | 10458 | 12400 | 12039 | 12934 | 14660 | 13210 | 12193 | 11994 | 11976 | 13224 | 12605 | 9783 | 10321 | 10802 | 10390 | 11024 | 13123 | 13082 | Wapl |
| Wars | 6436 | 8243 | 8096 | 7518 | 6505 | 6384 | 6315 | 4570 | 5201 | 5587 | 5002 | 6982 | 5954 | 6396 | 6057 | 6591 | 7742 | 6728 | 7118 | 5535 | 6826 | 5105 | 5034 | 5803 | 5928 | 6980 | 6172 | 6713 | Wars |
| Wars2 | 487 | 421 | 384 | 366 | 335 | 418 | 282 | 1133 | 1208 | 991 | 1158 | 1094 | 1083 | 1115 | 448 | 469 | 356 | 362 | 302 | 332 | 325 | 993 | 1192 | 1064 | 903 | 883 | 1128 | 1117 | Wars2 |
| Was | 43 | 46 | 43 | 37 | 34 | 29 | 36 | 68 | 73 | 79 | 83 | 41 | 90 | 51 | 34 | 69 | 53 | 63 | 34 | 30 | 28 | 60 | 100 | 77 | 64 | 80 | 73 | 62 | Was |
| Wasf1 | 4460 | 4717 | 6896 | 7806 | 7889 | 6488 | 7214 | 4551 | 4184 | 5981 | 5179 | 6391 | 4942 | 6283 | 4675 | 4586 | 7363 | 7439 | 7822 | 5142 | 8102 | 7392 | 4103 | 6195 | 6523 | 5648 | 4861 | 6258 | Wasf1 |
| Wasf2 | 4547 | 4276 | 5428 | 4295 | 3580 | 3623 | 3609 | 4196 | 5497 | 5260 | 6255 | 4941 | 5439 | 4759 | 3790 | 4952 | 4605 | 4164 | 3819 | 2982 | 3548 | 4273 | 5579 | 5605 | 4516 | 6441 | 4393 | 4739 | Wasf2 |
| Wasf3 | 5971 | 6362 | 8546 | 8678 | 8430 | 7740 | 7479 | 4217 | 4176 | 4668 | 4727 | 5096 | 4947 | 4847 | 6204 | 6835 | 8485 | 8253 | 8306 | 6744 | 7559 | 5249 | 3818 | 4570 | 5079 | 4279 | 4764 | 4904 | Wasf3 |
| Washc1 | 2712 | 3594 | 2891 | 3664 | 3409 | 3001 | 3022 | 2214 | 2673 | 2622 | 2633 | 2682 | 2414 | 2500 | 2695 | 3114 | 3447 | 3684 | 3490 | 3011 | 3174 | 2675 | 2503 | 2819 | 2640 | 2794 | 2542 | 2546 | Washc1 |
| Washc2 | 9056 | 9048 | 7958 | 8138 | 8420 | 8502 | 9219 | 8587 | 7899 | 7898 | 8135 | 8081 | 8411 | 8715 | 9476 | 9108 | 8398 | 8318 | 8210 | 8474 | 8825 | 8485 | 7883 | 7589 | 8011 | 6574 | 8237 | 8246 | Washc2 |
| Washc3 | 391 | 365 | 346 | 300 | 306 | 317 | 332 | 1036 | 962 | 843 | 992 | 899 | 844 | 1020 | 426 | 382 | 397 | 340 | 340 | 341 | 342 | 884 | 1040 | 936 | 906 | 1029 | 877 | 980 | Washc3 |
| Washc4 | 7093 | 5575 | 4683 | 6021 | 5539 | 6532 | 6279 | 7090 | 6058 | 5212 | 6296 | 5603 | 5887 | 5860 | 7183 | 6855 | 5251 | 5562 | 5036 | 6145 | 6148 | 5549 | 6227 | 4817 | 5406 | 4131 | 5625 | 5207 | Washc4 |
| Washc5 | 8449 | 8557 | 6854 | 6459 | 6452 | 6764 | 6993 | 7220 | 7164 | 6522 | 6409 | 7382 | 6792 | 7465 | 8144 | 8224 | 6831 | 6697 | 6095 | 7174 | 6870 | 6122 | 6753 | 6151 | 6629 | 6850 | 7471 | 7351 | Washc5 |
| Wasl | 6413 | 5205 | 7309 | 7262 | 6910 | 7057 | 7350 | 5715 | 5015 | 5575 | 5812 | 6114 | 5435 | 6382 | 6231 | 6368 | 6946 | 7451 | 6807 | 5428 | 7237 | 6709 | 5145 | 5859 | 6066 | 6541 | 5774 | 6113 | Wasl |
| Wbp1 | 7121 | 9352 | 7733 | 8876 | 7330 | 7469 | 7695 | 7220 | 7317 | 7530 | 6421 | 8218 | 7428 | 7800 | 7044 | 7540 | 8763 | 7657 | 8500 | 7916 | 7986 | 6562 | 7048 | 7816 | 7519 | 9034 | 8724 | 7817 | Wbp1 |
| Wbp11 | 6339 | 6517 | 9339 | 7761 | 6411 | 6087 | 6329 | 6174 | 6709 | 8119 | 7446 | 9624 | 7905 | 9111 | 5869 | 6340 | 8247 | 6825 | 7129 | 5309 | 6528 | 7319 | 6405 | 8773 | 7723 | 11507 | 8012 | 9596 | Wbp11 |
| Wbp1l | 5944 | 7126 | 7986 | 7611 | 6723 | 6548 | 5719 | 3634 | 4696 | 5081 | 5200 | 5718 | 5214 | 4749 | 5658 | 7009 | 7559 | 7469 | 7141 | 5235 | 6019 | 5116 | 4772 | 5523 | 5379 | 5830 | 5127 | 5184 | Wbp1l |
| Wbp2 | 1361 | 1611 | 1815 | 1943 | 2712 | 2034 | 2344 | 3555 | 4027 | 5209 | 4486 | 5277 | 4381 | 5235 | 1523 | 1544 | 2117 | 2358 | 2683 | 1767 | 2277 | 5259 | 4062 | 5520 | 5506 | 5329 | 4013 | 5064 | Wbp2 |
| Wbp2nl | 7 | 10 | 5 | 22 | 5 | 7 | 8 | 2 | 4 | 3 | 4 | 14 | 9 | 6 | 17 | 14 | 10 | 2 | 5 | 7 | 10 | 8 | 3 | 5 | 2 | 9 | 12 | 5 | Wbp2nl |
| Wbp4 | 5313 | 5014 | 5830 | 5602 | 5574 | 5414 | 5869 | 4846 | 4169 | 4626 | 4286 | 4680 | 4500 | 4849 | 5608 | 4945 | 5611 | 5465 | 5624 | 5669 | 5718 | 4547 | 4196 | 4645 | 4494 | 5007 | 4690 | 4864 | Wbp4 |
| Wdcp | 2640 | 3245 | 2703 | 3160 | 2563 | 2500 | 2710 | 1237 | 1130 | 1210 | 1253 | 1253 | 1200 | 1286 | 2521 | 2850 | 3311 | 3099 | 2722 | 2479 | 2851 | 1056 | 1111 | 1267 | 1282 | 1343 | 1290 | 1351 | Wdcp |
| Wdfy1 | 2215 | 1901 | 1902 | 2621 | 2822 | 2752 | 2807 | 2402 | 2151 | 2432 | 2505 | 2292 | 2248 | 2330 | 2291 | 1805 | 2396 | 2511 | 2513 | 2435 | 2796 | 3090 | 2243 | 2265 | 2669 | 2173 | 2206 | 2256 | Wdfy1 |
| Wdfy2 | 8877 | 8115 | 6721 | 7229 | 6703 | 7354 | 7253 | 4110 | 3749 | 3698 | 3586 | 3993 | 3730 | 3775 | 8401 | 8006 | 6901 | 7394 | 6685 | 6880 | 7324 | 3746 | 3848 | 3676 | 3905 | 3477 | 4058 | 3891 | Wdfy2 |
| Wdfy3 | 6717 | 7196 | 5106 | 7641 | 11254 | 10189 | 9270 | 14377 | 15777 | 18299 | 17937 | 16789 | 17503 | 17239 | 6852 | 7608 | 5975 | 8403 | 8955 | 8014 | 10460 | 15224 | 14544 | 13688 | 20401 | 9852 | 17331 | 15276 | Wdfy3 |
| Wdfy4 | 122 | 144 | 127 | 107 | 66 | 137 | 79 | 120 | 187 | 139 | 211 | 89 | 141 | 103 | 136 | 150 | 74 | 122 | 90 | 107 | 88 | 104 | 234 | 119 | 183 | 90 | 141 | 110 | Wdfy4 |
| Wdhd1 | 9090 | 7464 | 7862 | 6608 | 6065 | 6927 | 7082 | 5931 | 5141 | 4724 | 4695 | 6806 | 6090 | 6684 | 7950 | 7956 | 7194 | 6291 | 6446 | 7538 | 6608 | 4117 | 4599 | 5501 | 5113 | 6837 | 6960 | 7011 | Wdhd1 |
| Wdpcp | 1093 | 1056 | 965 | 875 | 676 | 797 | 870 | 960 | 857 | 673 | 769 | 919 | 839 | 834 | 1094 | 985 | 840 | 735 | 847 | 964 | 830 | 795 | 863 | 764 | 741 | 843 | 922 | 949 | Wdpcp |
| Wdr1 | 12335 | 12346 | 10944 | 10238 | 10194 | 10327 | 9735 | 9971 | 10752 | 9494 | 10223 | 9267 | 10356 | 9257 | 11802 | 12063 | 10347 | 10705 | 10318 | 10225 | 9954 | 9499 | 11021 | 9591 | 9375 | 10413 | 10133 | 9647 | Wdr1 |
| Wdr11 | 3839 | 3317 | 3113 | 2964 | 3257 | 3553 | 3176 | 6356 | 6295 | 5638 | 6108 | 5993 | 5757 | 5865 | 3687 | 3446 | 2933 | 2945 | 3038 | 3320 | 3097 | 5633 | 6300 | 5738 | 5778 | 5484 | 6140 | 5879 | Wdr11 |
| Wdr12 | 6434 | 6255 | 5349 | 5987 | 5285 | 5780 | 6488 | 3753 | 3098 | 2913 | 3006 | 3032 | 3320 | 3586 | 6599 | 6126 | 5583 | 5932 | 5640 | 6780 | 6488 | 2822 | 3086 | 3044 | 3306 | 3357 | 3362 | 3527 | Wdr12 |
| Wdr13 | 4229 | 4573 | 4088 | 5947 | 7984 | 7013 | 6968 | 10265 | 9723 | 11422 | 10277 | 10893 | 9677 | 10343 | 4646 | 3945 | 5365 | 6501 | 7112 | 6407 | 7243 | 12810 | 9098 | 10301 | 11762 | 8007 | 9518 | 9086 | Wdr13 |
| Wdr17 | 2080 | 2003 | 1580 | 2447 | 2545 | 2359 | 3014 | 1089 | 938 | 946 | 833 | 880 | 616 | 955 | 2302 | 1728 | 2100 | 2358 | 2304 | 2886 | 2850 | 1108 | 922 | 771 | 1085 | 657 | 740 | 840 | Wdr17 |
| Wdr18 | 4738 | 6096 | 5282 | 4919 | 4228 | 4508 | 4059 | 3709 | 3963 | 3581 | 3810 | 4382 | 4100 | 3706 | 4594 | 5108 | 4884 | 4541 | 4739 | 4540 | 4189 | 3656 | 3828 | 4093 | 4132 | 4738 | 4296 | 4182 | Wdr18 |
| Wdr19 | 5479 | 5441 | 4011 | 4403 | 4269 | 4589 | 4484 | 3087 | 2950 | 2579 | 2419 | 2605 | 2744 | 2429 | 5333 | 5359 | 4223 | 4364 | 3991 | 4795 | 4374 | 2645 | 2864 | 2277 | 2360 | 1835 | 2660 | 2527 | Wdr19 |
| Wdr20 | 2488 | 2572 | 2348 | 2613 | 2333 | 2366 | 2305 | 2396 | 2191 | 2200 | 2263 | 2315 | 2342 | 2412 | 2463 | 2315 | 2099 | 2181 | 2384 | 2490 | 2319 | 2182 | 2294 | 2118 | 2348 | 2232 | 2289 | 2331 | Wdr20 |
| Wdr20rt | 0 | 0 | 0 | 3 | 0 | 0 | 0 | 0 | 0 | 0 | 0 | 0 | 0 | 0 | 0 | 1 | 0 | 2 | 0 | 0 | 0 | 0 | 1 | 0 | 0 | 0 | 0 | 0 | Wdr20rt |
| Wdr24 | 1643 | 2160 | 1775 | 1778 | 1627 | 1547 | 1476 | 1791 | 2031 | 1804 | 1825 | 2022 | 1727 | 1783 | 1510 | 1681 | 1794 | 1812 | 1802 | 1786 | 1642 | 1753 | 2058 | 1988 | 2022 | 2207 | 1895 | 1883 | Wdr24 |
| Wdr25 | 269 | 336 | 377 | 289 | 322 | 284 | 267 | 430 | 470 | 512 | 442 | 526 | 531 | 425 | 286 | 326 | 391 | 280 | 384 | 235 | 299 | 381 | 439 | 462 | 425 | 563 | 513 | 479 | Wdr25 |
| Wdr26 | 25343 | 21537 | 26110 | 29995 | 28406 | 29105 | 31004 | 23154 | 20208 | 23723 | 22705 | 25192 | 24149 | 26103 | 25022 | 22887 | 27532 | 28745 | 28144 | 23747 | 30264 | 24383 | 20014 | 23953 | 24679 | 22060 | 24452 | 25023 | Wdr26 |
| Wdr27 | 91 | 86 | 83 | 97 | 124 | 121 | 122 | 76 | 69 | 91 | 56 | 72 | 101 | 96 | 91 | 92 | 96 | 86 | 81 | 108 | 100 | 80 | 93 | 80 | 92 | 51 | 94 | 95 | Wdr27 |
| Wdr3 | 5920 | 5828 | 5463 | 4714 | 4238 | 4404 | 4891 | 5178 | 4761 | 3922 | 4101 | 4483 | 4397 | 4604 | 6013 | 5756 | 4711 | 4850 | 4254 | 5203 | 4372 | 4176 | 4621 | 4453 | 4278 | 4778 | 4740 | 4631 | Wdr3 |
| Wdr31 | 661 | 551 | 307 | 396 | 475 | 564 | 507 | 567 | 659 | 565 | 614 | 657 | 594 | 596 | 588 | 569 | 331 | 407 | 414 | 607 | 552 | 438 | 650 | 507 | 707 | 564 | 726 | 536 | Wdr31 |
| Wdr33 | 7286 | 7481 | 7051 | 7443 | 7236 | 7344 | 7383 | 7751 | 7559 | 7591 | 7358 | 8256 | 7908 | 8310 | 6895 | 7199 | 7568 | 7030 | 7813 | 6959 | 7465 | 7267 | 7309 | 7827 | 8028 | 8692 | 8453 | 8569 | Wdr33 |
| Wdr35 | 6810 | 6658 | 5983 | 5821 | 4997 | 5501 | 6341 | 4163 | 3897 | 3643 | 3672 | 4090 | 3886 | 4278 | 6864 | 6488 | 5434 | 6214 | 5273 | 6624 | 6036 | 3491 | 3950 | 3635 | 3699 | 3664 | 4274 | 4218 | Wdr35 |
| Wdr36 | 4872 | 4209 | 4098 | 3380 | 3103 | 3511 | 3698 | 4820 | 4317 | 3804 | 4008 | 4360 | 4289 | 4670 | 4598 | 4226 | 3917 | 3471 | 3356 | 3709 | 3472 | 3979 | 4423 | 4089 | 3886 | 4779 | 4833 | 4802 | Wdr36 |
| Wdr37 | 10416 | 9460 | 9715 | 10871 | 10947 | 11072 | 11478 | 8418 | 7060 | 7368 | 6873 | 7841 | 7638 | 7839 | 11112 | 9194 | 10509 | 10506 | 10459 | 11668 | 11127 | 8624 | 6899 | 7040 | 8039 | 7028 | 8022 | 7586 | Wdr37 |
| Wdr38 | 72 | 82 | 53 | 78 | 65 | 65 | 88 | 48 | 59 | 57 | 75 | 45 | 74 | 149 | 73 | 72 | 68 | 80 | 50 | 85 | 93 | 43 | 64 | 67 | 50 | 52 | 76 | 63 | Wdr38 |
| Wdr4 | 2043 | 2610 | 2464 | 2619 | 2655 | 2730 | 2289 | 3129 | 3429 | 3534 | 3366 | 3900 | 4034 | 3568 | 2023 | 2247 | 2678 | 2545 | 2809 | 2460 | 2406 | 3604 | 3225 | 3537 | 3838 | 3677 | 3826 | 3502 | Wdr4 |
| Wdr41 | 707 | 740 | 720 | 817 | 958 | 931 | 869 | 2256 | 2235 | 2280 | 2299 | 2053 | 2119 | 2271 | 795 | 752 | 743 | 789 | 860 | 857 | 863 | 2246 | 2271 | 2154 | 2189 | 2289 | 2138 | 2150 | Wdr41 |
| Wdr43 | 6326 | 5405 | 5768 | 5415 | 4795 | 4949 | 5399 | 6175 | 5609 | 5130 | 5419 | 6085 | 5769 | 6247 | 6224 | 5631 | 5055 | 5132 | 4534 | 5713 | 4903 | 4998 | 5460 | 5431 | 5494 | 6787 | 6238 | 6280 | Wdr43 |
| Wdr44 | 1584 | 1376 | 1461 | 2106 | 2232 | 2184 | 2294 | 1334 | 1074 | 1345 | 1411 | 1186 | 1057 | 1262 | 1832 | 1519 | 1860 | 2091 | 2049 | 1933 | 2317 | 1531 | 1201 | 1200 | 1358 | 1015 | 1011 | 1069 | Wdr44 |
| Wdr45 | 396 | 381 | 289 | 437 | 341 | 300 | 352 | 391 | 334 | 370 | 361 | 316 | 319 | 476 | 324 | 285 | 283 | 349 | 307 | 257 | 376 | 344 | 331 | 412 | 334 | 267 | 327 | 358 | Wdr45 |
| Wdr45b | 5278 | 5420 | 4441 | 3915 | 3950 | 4254 | 4322 | 5068 | 4684 | 3819 | 3918 | 4199 | 3991 | 4328 | 5169 | 5035 | 4463 | 4099 | 4306 | 4937 | 4078 | 4233 | 4902 | 4224 | 4073 | 4620 | 4296 | 4538 | Wdr45b |
| Wdr46 | 3410 | 4098 | 3849 | 3878 | 2999 | 3049 | 3214 | 1672 | 1941 | 2019 | 1890 | 2433 | 2232 | 2368 | 2910 | 3539 | 4107 | 3575 | 3458 | 2867 | 3107 | 1430 | 1859 | 2393 | 2172 | 3044 | 2391 | 2567 | Wdr46 |
| Wdr47 | 10380 | 11100 | 9021 | 12098 | 14010 | 13296 | 13680 | 10875 | 8897 | 8850 | 7271 | 8177 | 7524 | 8253 | 12330 | 9671 | 10576 | 12692 | 13150 | 14426 | 13763 | 10602 | 8467 | 8047 | 10134 | 5687 | 7746 | 7395 | Wdr47 |
| Wdr48 | 8195 | 8552 | 6946 | 7649 | 7202 | 7350 | 8370 | 10088 | 8906 | 8857 | 8476 | 9281 | 8424 | 9745 | 8300 | 7477 | 7467 | 7104 | 7681 | 7991 | 8645 | 8789 | 8816 | 8938 | 9223 | 8272 | 8969 | 9119 | Wdr48 |
| Wdr49 | 3 | 3 | 1 | 1 | 10 | 2 | 4 | 6 | 9 | 3 | 2 | 3 | 2 | 6 | 5 | 1 | 0 | 1 | 2 | 1 | 3 | 7 | 8 | 4 | 2 | 4 | 2 | 2 | Wdr49 |
| Wdr5 | 9055 | 9150 | 9152 | 8520 | 6659 | 7303 | 7400 | 5203 | 5179 | 5055 | 4995 | 6155 | 5432 | 5963 | 7919 | 8130 | 8489 | 7826 | 7607 | 7302 | 7337 | 4809 | 5118 | 5794 | 5437 | 7007 | 6195 | 6446 | Wdr5 |
| Wdr53 | 555 | 534 | 580 | 490 | 513 | 518 | 438 | 608 | 634 | 633 | 544 | 696 | 644 | 712 | 605 | 548 | 530 | 467 | 494 | 549 | 486 | 583 | 646 | 837 | 672 | 727 | 773 | 768 | Wdr53 |
| Wdr54 | 974 | 1247 | 1003 | 1226 | 1070 | 998 | 1115 | 524 | 527 | 622 | 481 | 636 | 602 | 569 | 1014 | 1007 | 1181 | 1093 | 1180 | 1164 | 1068 | 515 | 531 | 641 | 607 | 711 | 608 | 600 | Wdr54 |
| Wdr55 | 1788 | 2230 | 1999 | 1892 | 1559 | 1667 | 1694 | 1492 | 1508 | 1395 | 1355 | 1401 | 1421 | 1627 | 1709 | 1840 | 1807 | 1645 | 1587 | 1870 | 1708 | 1180 | 1463 | 1453 | 1334 | 1812 | 1639 | 1718 | Wdr55 |

| | | | | | | | | | | | | | | | | | | | | | | | | | | | | | |
|---|---|---|---|---|---|---|---|---|---|---|---|---|---|---|---|---|---|---|---|---|---|---|---|---|---|---|---|---|---|
| Wdr59 | 2467 | 2562 | 2203 | 2339 | 2588 | 2499 | 2417 | 2395 | 2530 | 2405 | 2425 | 2159 | 2352 | 2072 | 2304 | 2210 | 2389 | 2561 | 2489 | 2368 | 2439 | 2567 | 2370 | 2251 | 2449 | 2106 | 2258 | 2174 | Wdr59 |
| Wdr5b | 869 | 706 | 749 | 723 | 830 | 758 | 872 | 1138 | 1046 | 991 | 908 | 1138 | 997 | 1135 | 831 | 626 | 771 | 770 | 861 | 912 | 769 | 989 | 990 | 1008 | 1031 | 1185 | 1124 | 1046 | Wdr5b |
| Wdr6 | 19295 | 23894 | 24900 | 28027 | 30703 | 28749 | 24370 | 24099 | 28599 | 30742 | 28440 | 37518 | 32488 | 30843 | 19372 | 21107 | 28126 | 28030 | 31910 | 25002 | 26879 | 34228 | 27575 | 33698 | 35350 | 37493 | 33755 | 31495 | Wdr6 |
| Wdr61 | 4786 | 4636 | 4909 | 4122 | 3746 | 3750 | 3989 | 3583 | 3353 | 2943 | 3038 | 3674 | 3433 | 3615 | 4770 | 4911 | 4584 | 3913 | 3796 | 4631 | 3719 | 2941 | 3416 | 3475 | 3180 | 4164 | 3885 | 3832 | Wdr61 |
| Wdr62 | 2813 | 3512 | 3408 | 3333 | 2334 | 2503 | 2851 | 819 | 1131 | 1055 | 1122 | 1981 | 1475 | 1622 | 2451 | 3175 | 3388 | 3250 | 3020 | 2238 | 2672 | 994 | 990 | 1385 | 1290 | 1834 | 1572 | 1618 | Wdr62 |
| Wdr64 | 0 | 0 | 0 | 3 | 0 | 0 | 0 | 2 | 2 | 1 | 0 | 2 | 2 | 3 | 0 | 1 | 1 | 0 | 0 | 1 | 3 | 3 | 5 | 2 | 2 | 0 | 0 | 6 | Wdr64 |
| Wdr66 | 336 | 331 | 187 | 376 | 356 | 293 | 322 | 249 | 290 | 332 | 226 | 211 | 347 | 192 | 311 | 419 | 298 | 377 | 344 | 309 | 370 | 247 | 305 | 335 | 196 | 313 | 174 | 150 | Wdr66 |
| Wdr7 | 6033 | 5977 | 4783 | 7085 | 8627 | 7996 | 7965 | 6405 | 6387 | 7014 | 5958 | 5912 | 5574 | 6066 | 7046 | 6012 | 6249 | 7554 | 7511 | 7099 | 8315 | 7817 | 6055 | 5761 | 7454 | 3682 | 5781 | 5569 | Wdr7 |
| Wdr70 | 4082 | 4311 | 4210 | 3739 | 2929 | 3196 | 3117 | 2882 | 2725 | 2605 | 2640 | 3144 | 2872 | 2917 | 4068 | 3794 | 3923 | 3406 | 3316 | 3476 | 3126 | 2680 | 2720 | 2924 | 2828 | 3448 | 3149 | 3152 | Wdr70 |
| Wdr72 | 32 | 54 | 57 | 43 | 59 | 18 | 42 | 55 | 66 | 76 | 30 | 35 | 54 | 26 | 86 | 68 | 24 | 21 | 35 | 49 | 26 | 69 | 82 | 20 | 17 | 34 | 42 | 28 | Wdr72 |
| Wdr73 | 2135 | 2337 | 2218 | 2203 | 2047 | 2261 | 2061 | 2656 | 2632 | 2690 | 2397 | 3053 | 2946 | 2750 | 2081 | 2160 | 2427 | 2224 | 2240 | 2251 | 2069 | 2401 | 2472 | 2806 | 2873 | 3144 | 3001 | 3023 | Wdr73 |
| Wdr74 | 2135 | 2417 | 2033 | 2125 | 1892 | 1959 | 2044 | 1910 | 2063 | 2052 | 2003 | 2164 | 2007 | 2126 | 2168 | 2102 | 2160 | 2105 | 2164 | 2042 | 2074 | 1699 | 1998 | 2116 | 2066 | 2409 | 2238 | 2181 | Wdr74 |
| Wdr75 | 6719 | 6167 | 5343 | 5863 | 4856 | 5164 | 5721 | 5678 | 5327 | 4513 | 4491 | 5238 | 5330 | 5692 | 6821 | 6037 | 5302 | 5304 | 5045 | 6400 | 5474 | 4562 | 4873 | 4788 | 4868 | 5819 | 5613 | 5802 | Wdr75 |
| Wdr76 | 3177 | 2938 | 3857 | 2847 | 2293 | 2597 | 3124 | 2757 | 2241 | 2276 | 2130 | 3783 | 3020 | 3904 | 3101 | 2717 | 2997 | 2589 | 2909 | 3107 | 2845 | 1918 | 2036 | 2826 | 2634 | 4101 | 3876 | 4150 | Wdr76 |
| Wdr77 | 5885 | 4743 | 5323 | 5227 | 4822 | 5222 | 5984 | 4913 | 3987 | 4125 | 4178 | 4882 | 4623 | 5152 | 5987 | 4918 | 4715 | 5028 | 5121 | 4909 | 5690 | 4791 | 4032 | 4418 | 4670 | 5104 | 4702 | 5295 | Wdr77 |
| Wdr81 | 2127 | 2905 | 2949 | 3378 | 3066 | 2749 | 2461 | 1613 | 2046 | 2359 | 2298 | 2232 | 2036 | 1916 | 1991 | 2558 | 3206 | 3354 | 3126 | 2053 | 2800 | 2267 | 2130 | 2402 | 2410 | 1698 | 1916 | 2003 | Wdr81 |
| Wdr82 | 17762 | 16394 | 15316 | 16729 | 16283 | 16659 | 17325 | 22319 | 19144 | 18316 | 17194 | 19670 | 16689 | 19656 | 17884 | 14438 | 16223 | 16463 | 17115 | 17718 | 17288 | 21250 | 18183 | 18968 | 20625 | 19732 | 19205 | 19409 | Wdr82 |
| Wdr83 | 1834 | 2116 | 2194 | 2119 | 1919 | 2025 | 1915 | 2056 | 2174 | 2175 | 2159 | 2424 | 2327 | 2325 | 1768 | 1836 | 2060 | 1915 | 2295 | 2103 | 1817 | 2079 | 2011 | 2622 | 2348 | 2479 | 2651 | 2436 | Wdr83 |
| Wdr83os | 1799 | 2144 | 1964 | 1912 | 1799 | 1865 | 2119 | 1762 | 1907 | 1724 | 1830 | 2101 | 2057 | 2172 | 1721 | 1752 | 1820 | 2147 | 2022 | 1940 | 2017 | 1756 | 1840 | 1881 | 1905 | 2502 | 2231 | 2290 | Wdr83os |
| Wdr86 | 1930 | 1557 | 1208 | 923 | 752 | 853 | 434 | 1533 | 1826 | 1316 | 1358 | 1010 | 1386 | 752 | 1292 | 2163 | 926 | 922 | 521 | 599 | 416 | 790 | 1827 | 1347 | 648 | 1621 | 1092 | 819 | Wdr86 |
| Wdr88 | 0 | 3 | 0 | 0 | 0 | 0 | 0 | 2 | 2 | 0 | 3 | 0 | 0 | 2 | 1 | 2 | 0 | 0 | 1 | 4 | 1 | 0 | 1 | 0 | 0 | 0 | 0 | 1 | Wdr88 |
| Wdr89 | 247 | 238 | 294 | 213 | 172 | 255 | 228 | 276 | 304 | 287 | 348 | 331 | 348 | 329 | 239 | 245 | 247 | 191 | 222 | 166 | 211 | 247 | 295 | 351 | 314 | 463 | 310 | 352 | Wdr89 |
| Wdr90 | 4367 | 5401 | 5263 | 5821 | 4564 | 4762 | 4500 | 2854 | 3623 | 3826 | 3496 | 5409 | 4434 | 4118 | 3982 | 5052 | 5786 | 5359 | 5542 | 4295 | 4261 | 3367 | 3423 | 4433 | 3962 | 4102 | 4602 | 4430 | Wdr90 |
| Wdr91 | 1967 | 2611 | 2267 | 2126 | 2017 | 1880 | 1838 | 1310 | 1543 | 1729 | 1584 | 1691 | 1721 | 1530 | 1908 | 2272 | 2158 | 2259 | 2131 | 1794 | 1849 | 1651 | 1453 | 1561 | 1619 | 1676 | 1539 | 1594 | Wdr91 |
| Wdr92 | 1977 | 1881 | 2053 | 1661 | 1339 | 1546 | 1514 | 1661 | 1580 | 1351 | 1365 | 1435 | 1408 | 1498 | 1878 | 2053 | 1730 | 1614 | 1474 | 1821 | 1413 | 1290 | 1568 | 1507 | 1427 | 1568 | 1635 | 1585 | Wdr92 |
| Wdr93 | 24 | 38 | 10 | 28 | 52 | 33 | 27 | 34 | 50 | 16 | 30 | 25 | 58 | 26 | 24 | 47 | 23 | 35 | 31 | 49 | 34 | 38 | 67 | 37 | 35 | 29 | 45 | 46 | Wdr93 |
| Wdr95 | 25 | 35 | 11 | 35 | 52 | 60 | 38 | 21 | 33 | 55 | 46 | 51 | 40 | 38 | 20 | 14 | 36 | 36 | 35 | 40 | 38 | 43 | 39 | 32 | 42 | 30 | 27 | 46 | Wdr95 |
| Wdsub1 | 1207 | 1084 | 1246 | 917 | 912 | 840 | 894 | 852 | 895 | 805 | 942 | 822 | 873 | 691 | 1056 | 1119 | 880 | 930 | 893 | 1018 | 877 | 655 | 1058 | 829 | 626 | 904 | 809 | 833 | Wdsub1 |
| Wdtc1 | 5182 | 6869 | 7090 | 7191 | 7319 | 6304 | 6231 | 3695 | 4333 | 4893 | 4208 | 4655 | 4549 | 4312 | 5095 | 6156 | 7740 | 7156 | 7758 | 5674 | 6795 | 4503 | 3979 | 4859 | 4746 | 4814 | 4303 | 4149 | Wdtc1 |
| Wee1 | 3796 | 3334 | 4428 | 3639 | 2857 | 2873 | 3452 | 2965 | 2949 | 2637 | 2807 | 4233 | 3382 | 3985 | 3281 | 3267 | 3654 | 3210 | 3500 | 2860 | 3191 | 2460 | 2712 | 3427 | 2819 | 4859 | 3811 | 4303 | Wee1 |
| Wee2 | 10 | 13 | 10 | 22 | 27 | 6 | 20 | 11 | 21 | 25 | 6 | 12 | 17 | 12 | 16 | 13 | 12 | 31 | 23 | 17 | 22 | 12 | 25 | 11 | 28 | 14 | 30 | 7 | Wee2 |
| Wfdc1 | 22 | 35 | 41 | 39 | 24 | 37 | 28 | 34 | 36 | 35 | 37 | 39 | 36 | 30 | 37 | 37 | 31 | 30 | 34 | 41 | 27 | 37 | 61 | 39 | 50 | 24 | 52 | 38 | Wfdc1 |
| Wfdc10 | 1 | 0 | 0 | 0 | 0 | 0 | 1 | 0 | 0 | 5 | 5 | 0 | 0 | 0 | 0 | 0 | 0 | 0 | 0 | 0 | 0 | 2 | 2 | 2 | 0 | 0 | 0 | 2 | Wfdc10 |
| Wfdc11 | 0 | 0 | 0 | 0 | 0 | 0 | 0 | 0 | 1 | 0 | 0 | 0 | 0 | 0 | 0 | 0 | 0 | 0 | 0 | 0 | 0 | 0 | 0 | 0 | 0 | 0 | 0 | 0 | Wfdc11 |
| Wfdc12 | 0 | 0 | 0 | 0 | 0 | 0 | 0 | 0 | 1 | 0 | 8 | 0 | 1 | 0 | 0 | 1 | 3 | 0 | 2 | 0 | 0 | 2 | 3 | 3 | 2 | 0 | 0 | 1 | Wfdc12 |
| Wfdc13 | 3 | 2 | 1 | 4 | 0 | 0 | 0 | 0 | 0 | 0 | 1 | 0 | 3 | 0 | 0 | 3 | 0 | 0 | 0 | 0 | 0 | 0 | 0 | 0 | 6 | 0 | 0 | 0 | Wfdc13 |
| Wfdc15a | 1 | 0 | 0 | 0 | 0 | 0 | 0 | 0 | 0 | 0 | 0 | 0 | 0 | 0 | 0 | 0 | 0 | 0 | 0 | 0 | 2 | 0 | 0 | 0 | 0 | 0 | 0 | 0 | Wfdc15a |
| Wfdc17 | 0 | 2 | 1 | 3 | 0 | 2 | 0 | 1 | 4 | 1 | 11 | 0 | 0 | 1 | 0 | 1 | 0 | 0 | 1 | 0 | 4 | 0 | 3 | 0 | 0 | 3 | 2 | 5 | Wfdc17 |
| Wfdc18 | 0 | 0 | 0 | 0 | 0 | 0 | 0 | 0 | 0 | 0 | 0 | 0 | 0 | 0 | 0 | 0 | 0 | 0 | 0 | 3 | 0 | 0 | 3 | 0 | 0 | 0 | 0 | 0 | Wfdc18 |
| Wfdc2 | 401 | 392 | 256 | 255 | 122 | 193 | 204 | 278 | 376 | 312 | 336 | 297 | 406 | 337 | 307 | 301 | 264 | 174 | 208 | 229 | 155 | 185 | 421 | 409 | 227 | 596 | 385 | 385 | Wfdc2 |
| Wfdc21 | 0 | 0 | 0 | 0 | 0 | 3 | 0 | 0 | 2 | 3 | 0 | 1 | 3 | 1 | 0 | 0 | 6 | 0 | 5 | 0 | 0 | 0 | 3 | 2 | 0 | 0 | 1 | 1 | Wfdc21 |
| Wfdc3 | 60 | 87 | 55 | 68 | 59 | 38 | 37 | 32 | 24 | 38 | 52 | 31 | 41 | 37 | 80 | 64 | 98 | 67 | 63 | 74 | 43 | 47 | 57 | 47 | 32 | 64 | 37 | 44 | Wfdc3 |
| Wfdc5 | 0 | 0 | 3 | 0 | 0 | 0 | 0 | 0 | 0 | 0 | 0 | 0 | 0 | 0 | 0 | 0 | 1 | 0 | 0 | 0 | 0 | 0 | 0 | 0 | 0 | 0 | 0 | 0 | Wfdc5 |
| Wfdc6a | 0 | 0 | 0 | 0 | 1 | 0 | 0 | 1 | 0 | 0 | 0 | 1 | 0 | 0 | 0 | 0 | 0 | 0 | 0 | 0 | 0 | 2 | 0 | 0 | 3 | 0 | 0 | 0 | Wfdc6a |
| Wfdc6b | 0 | 0 | 0 | 3 | 0 | 0 | 0 | 0 | 0 | 0 | 0 | 0 | 0 | 0 | 0 | 2 | 0 | 0 | 0 | 0 | 0 | 0 | 0 | 0 | 2 | 0 | 0 | 0 | Wfdc6b |
| Wfdc8 | 0 | 0 | 0 | 0 | 0 | 0 | 1 | 1 | 1 | 0 | 1 | 3 | 0 | 6 | 0 | 0 | 0 | 0 | 0 | 0 | 1 | 0 | 1 | 0 | 0 | 1 | 1 | 1 | Wfdc8 |
| Wfikkn1 | 95 | 136 | 123 | 156 | 89 | 126 | 121 | 40 | 34 | 46 | 33 | 46 | 53 | 39 | 104 | 107 | 126 | 140 | 109 | 113 | 116 | 38 | 39 | 51 | 42 | 73 | 46 | 66 | Wfikkn1 |
| Wfikkn2 | 336 | 420 | 355 | 399 | 508 | 623 | 350 | 699 | 1043 | 932 | 1113 | 874 | 1008 | 573 | 306 | 490 | 338 | 426 | 387 | 413 | 481 | 924 | 1498 | 867 | 654 | 848 | 723 | 611 | Wfikkn2 |
| Wfs1 | 1617 | 2003 | 1973 | 1982 | 1829 | 1654 | 1672 | 586 | 791 | 834 | 880 | 793 | 733 | 656 | 1545 | 1735 | 2135 | 2127 | 1922 | 1601 | 1756 | 690 | 873 | 817 | 678 | 1016 | 836 | 796 | Wfs1 |
| Whamm | 946 | 1088 | 1091 | 1245 | 1119 | 1073 | 1121 | 675 | 714 | 803 | 714 | 690 | 712 | 707 | 940 | 960 | 1144 | 1185 | 1163 | 925 | 1086 | 692 | 671 | 735 | 845 | 780 | 588 | 665 | Whamm |
| Whrn | 3255 | 4637 | 4506 | 4982 | 5164 | 4917 | 4074 | 1403 | 1685 | 2028 | 1886 | 2387 | 1868 | 1800 | 3066 | 3802 | 5144 | 5133 | 5321 | 4087 | 4720 | 1975 | 1612 | 1950 | 2046 | 1767 | 2357 | 1920 | Whrn |
| Wif1 | 1083 | 724 | 783 | 608 | 531 | 689 | 565 | 673 | 934 | 713 | 752 | 562 | 778 | 541 | 750 | 966 | 639 | 684 | 545 | 703 | 568 | 531 | 1131 | 597 | 453 | 662 | 550 | 621 | Wif1 |
| Wipf1 | 1527 | 1155 | 1772 | 1150 | 1100 | 1072 | 924 | 2017 | 2431 | 2252 | 2587 | 2003 | 2425 | 1907 | 1451 | 1692 | 1473 | 1198 | 981 | 880 | 883 | 1840 | 2512 | 2230 | 1588 | 2531 | 1964 | 2193 | Wipf1 |
| Wipf2 | 1926 | 2160 | 3390 | 3357 | 2949 | 2555 | 2670 | 2586 | 2576 | 3479 | 3151 | 3634 | 3246 | 3456 | 1967 | 2182 | 3417 | 3328 | 3220 | 2127 | 2910 | 3920 | 2473 | 3956 | 3532 | 3878 | 3173 | 3723 | Wipf2 |
| Wipf3 | 610 | 599 | 706 | 768 | 1253 | 908 | 997 | 667 | 936 | 1180 | 1043 | 853 | 721 | 804 | 605 | 604 | 681 | 1049 | 1146 | 779 | 978 | 1342 | 874 | 1039 | 1186 | 883 | 642 | 614 | Wipf3 |
| Wipi1 | 5726 | 6023 | 6276 | 6199 | 4760 | 5131 | 4779 | 1564 | 1926 | 1864 | 2415 | 1602 | 2003 | 1602 | 4927 | 6271 | 5940 | 6105 | 4955 | 3990 | 4970 | 1556 | 2345 | 1683 | 1467 | 1574 | 1515 | 1533 | Wipi1 |
| Wipi2 | 12524 | 13453 | 12230 | 12506 | 11005 | 11804 | 11485 | 6541 | 7021 | 7021 | 6818 | 7627 | 7652 | 7402 | 12010 | 12657 | 12388 | 12838 | 11623 | 10842 | 11498 | 6711 | 6829 | 7340 | 7513 | 7369 | 7681 | 7613 | Wipi2 |
| Wiz | 4711 | 6045 | 7192 | 7134 | 7252 | 6424 | 5554 | 6437 | 8173 | 9509 | 8490 | 11011 | 8707 | 9405 | 4453 | 5432 | 7422 | 6656 | 7621 | 5192 | 6168 | 9528 | 7192 | 10288 | 9821 | 11621 | 8839 | 9025 | Wiz |
| Wls | 14521 | 11657 | 14262 | 8072 | 7988 | 9522 | 7148 | 11428 | 10256 | 8478 | 9617 | 10688 | 13668 | 10043 | 14838 | 14983 | 9513 | 8623 | 7189 | 9243 | 7413 | 9413 | 13622 | 8389 | 7210 | 8819 | 12376 | 12265 | Wls |
| Wnk1 | 35142 | 31507 | 26213 | 30543 | 30287 | 34148 | 33297 | 28496 | 27713 | 25408 | 29403 | 29069 | 30509 | 30145 | 33792 | 34656 | 27128 | 29458 | 28208 | 30023 | 34044 | 26348 | 27370 | 23249 | 27404 | 20767 | 31665 | 27763 | Wnk1 |
| Wnk2 | 4392 | 6147 | 5946 | 8731 | 10775 | 8386 | 7420 | 6531 | 7757 | 10341 | 8542 | 9614 | 7461 | 7613 | 4709 | 5288 | 7431 | 9368 | 10412 | 6952 | 8787 | 11436 | 6519 | 9407 | 11789 | 6735 | 7073 | 6673 | Wnk2 |
| Wnk3 | 6247 | 5552 | 3303 | 4874 | 5851 | 6999 | 6724 | 9652 | 7506 | 5840 | 5471 | 6087 | 5881 | 6476 | 7266 | 5634 | 4092 | 4736 | 5365 | 7735 | 6904 | 6726 | 6805 | 4719 | 6966 | 3515 | 6684 | 5561 | Wnk3 |
| Wnk4 | 851 | 961 | 1178 | 976 | 785 | 818 | 530 | 409 | 566 | 780 | 929 | 603 | 735 | 541 | 708 | 1135 | 1134 | 1010 | 757 | 561 | 656 | 784 | 711 | 628 | 536 | 646 | 578 | 589 | Wnk4 |
| Wnt1 | 36 | 66 | 101 | 23 | 67 | 32 | 20 | 76 | 57 | 98 | 72 | 151 | 226 | 89 | 41 | 68 | 70 | 20 | 25 | 55 | 45 | 94 | 137 | 85 | 64 | 11 | 162 | 170 | Wnt1 |
| Wnt10a | 37 | 60 | 52 | 21 | 29 | 52 | 10 | 50 | 88 | 31 | 72 | 72 | 84 | 83 | 44 | 37 | 70 | 46 | 26 | 22 | 28 | 27 | 109 | 89 | 45 | 43 | 89 | 95 | Wnt10a |
| Wnt10b | 135 | 63 | 59 | 22 | 6 | 84 | 8 | 256 | 379 | 140 | 169 | 129 | 201 | 129 | 92 | 103 | 45 | 45 | 33 | 13 | 9 | 25 | 425 | 230 | 58 | 54 | 155 | 169 | Wnt10b |
| Wnt11 | 376 | 365 | 297 | 215 | 141 | 289 | 98 | 264 | 615 | 276 | 688 | 202 | 425 | 208 | 378 | 470 | 265 | 349 | 161 | 101 | 80 | 246 | 777 | 324 | 268 | 153 | 178 | 178 | Wnt11 |
| Wnt16 | 279 | 304 | 239 | 218 | 101 | 96 | 102 | 81 | 97 | 100 | 107 | 37 | 39 | 33 | 169 | 371 | 219 | 241 | 94 | 103 | 107 | 28 | 124 | 96 | 16 | 78 | 27 | 44 | Wnt16 |
| Wnt2 | 37 | 38 | 32 | 15 | 7 | 57 | 7 | 62 | 132 | 58 | 143 | 47 | 76 | 62 | 34 | 44 | 27 | 48 | 9 | 9 | 6 | 42 | 181 | 86 | 32 | 21 | 34 | 68 | Wnt2 |
| Wnt2b | 549 | 580 | 600 | 436 | 354 | 520 | 285 | 554 | 498 | 563 | 637 | 724 | 795 | 615 | 564 | 646 | 439 | 462 | 436 | 371 | 400 | 306 | 942 | 686 | 506 | 394 | 694 | 713 | Wnt2b |
| Wnt3 | 469 | 620 | 836 | 556 | 545 | 559 | 383 | 761 | 573 | 615 | 543 | 699 | 790 | 760 | 520 | 520 | 674 | 495 | 618 | 559 | 497 | 662 | 797 | 695 | 602 | 579 | 955 | 1013 | Wnt3 |
| Wnt3a | 95 | 107 | 163 | 60 | 49 | 85 | 47 | 220 | 279 | 292 | 210 | 391 | 489 | 326 | 105 | 107 | 88 | 76 | 78 | 83 | 56 | 165 | 399 | 289 | 255 | 424 | 409 | 437 | Wnt3a |
| Wnt4 | 486 | 402 | 538 | 236 | 384 | 382 | 197 | 686 | 996 | 719 | 766 | 350 | 535 | 398 | 477 | 556 | 290 | 424 | 298 | 278 | 227 | 234 | 1234 | 596 | 288 | 392 | 370 | 427 | Wnt4 |
| Wnt5a | 2984 | 1542 | 2382 | 1531 | 2081 | 1844 | 1465 | 8276 | 6516 | 6792 | 5186 | 3332 | 5408 | 2901 | 2033 | 2049 | 1853 | 1578 | 1594 | 1375 | 1429 | 3407 | 7805 | 5655 | 2824 | 4562 | 3251 | 3171 | Wnt5a |
| Wnt5b | 600 | 513 | 619 | 424 | 380 | 418 | 365 | 506 | 494 | 517 | 619 | 659 | 626 | 536 | 422 | 532 | 549 | 453 | 443 | 402 | 313 | 326 | 489 | 553 | 422 | 718 | 653 | 570 | Wnt5b |
| Wnt6 | 303 | 254 | 269 | 85 | 113 | 221 | 67 | 446 | 719 | 420 | 505 | 364 | 433 | 312 | 240 | 243 | 131 | 182 | 101 | 91 | 61 | 134 | 873 | 448 | 194 | 217 | 329 | 332 | Wnt6 |
| Wnt7a | 887 | 1068 | 1176 | 1339 | 1549 | 1523 | 1307 | 1148 | 1307 | 1510 | 1092 | 1802 | 1129 | 1659 | 862 | 662 | 1489 | 1174 | 1865 | 1300 | 1476 | 1034 | 1078 | 1676 | 1679 | 1597 | 1417 | 1477 | Wnt7a |
| Wnt7b | 2219 | 2856 | 3175 | 2694 | 3195 | 3460 | 3214 | 4537 | 5319 | 4668 | 3573 | 6604 | 4280 | 5998 | 2495 | 1859 | 3359 | 2761 | 3783 | 3265 | 3199 | 4438 | 5227 | 5642 | 5516 | 7652 | 5963 | 6467 | Wnt7b |
| Wnt8a | 2 | 3 | 0 | 3 | 2 | 0 | 1 | 1 | 2 | 0 | 5 | 0 | 7 | 0 | 1 | 7 | 1 | 1 | 3 | 8 | 0 | 3 | 0 | 2 | 2 | 0 | 1 | 0 | Wnt8a |

| | | | | | | | | | | | | | | | | | | | | | | | | | | | | | |
|---|---|---|---|---|---|---|---|---|---|---|---|---|---|---|---|---|---|---|---|---|---|---|---|---|---|---|---|---|---|
| Wnt8b | 1167 | 1056 | 2327 | 754 | 568 | 911 | 872 | 948 | 830 | 934 | 681 | 2700 | 1963 | 2489 | 1059 | 878 | 1313 | 616 | 885 | 1136 | 716 | 662 | 1048 | 1494 | 1159 | 3254 | 2868 | 3704 | Wnt8b |
| Wnt9a | 444 | 530 | 774 | 446 | 347 | 380 | 249 | 396 | 452 | 485 | 619 | 690 | 677 | 540 | 421 | 543 | 574 | 426 | 351 | 352 | 243 | 455 | 649 | 579 | 449 | 609 | 604 | 694 | Wnt9a |
| Wnt9b | 354 | 192 | 97 | 17 | 30 | 86 | 16 | 256 | 258 | 106 | 87 | 149 | 181 | 134 | 120 | 228 | 56 | 109 | 23 | 23 | 19 | 53 | 338 | 174 | 70 | 116 | 112 | 170 | Wnt9b |
| Wrap53 | 2466 | 3275 | 2622 | 2724 | 2207 | 2400 | 2434 | 1342 | 1427 | 1314 | 1237 | 1616 | 1351 | 1550 | 2502 | 2674 | 2582 | 2532 | 2518 | 2405 | 2328 | 1248 | 1339 | 1415 | 1374 | 1731 | 1565 | 1565 | Wrap53 |
| Wrap73 | 1922 | 2553 | 2173 | 2150 | 1958 | 2034 | 2000 | 1408 | 1596 | 1594 | 1413 | 1663 | 1433 | 1438 | 1662 | 1981 | 2102 | 2106 | 2376 | 2041 | 2078 | 1415 | 1620 | 1638 | 1742 | 1963 | 1715 | 1637 | Wrap73 |
| Wrn | 2339 | 1885 | 1925 | 1717 | 1670 | 1911 | 1791 | 2141 | 2040 | 1701 | 2172 | 2067 | 2197 | 2105 | 2271 | 2322 | 1913 | 1680 | 1627 | 1763 | 1644 | 1669 | 2073 | 1825 | 1792 | 1607 | 2295 | 2023 | Wrn |
| Wrnip1 | 3718 | 3676 | 3799 | 3999 | 4269 | 3707 | 3869 | 3149 | 3173 | 3138 | 3239 | 3125 | 2742 | 2950 | 3742 | 3522 | 4066 | 3987 | 4404 | 4092 | 4046 | 3374 | 3221 | 3444 | 3346 | 3298 | 2760 | 2963 | Wrnip1 |
| Wsb1 | 28617 | 19098 | 22776 | 24432 | 29374 | 28134 | 29262 | 39892 | 34406 | 33404 | 36951 | 36177 | 37394 | 38886 | 30064 | 23855 | 23229 | 25757 | 23820 | 28437 | 27551 | 41540 | 34308 | 33424 | 36618 | 30822 | 36146 | 34539 | Wsb1 |
| Wsb2 | 6000 | 6938 | 6047 | 7281 | 8839 | 7415 | 7853 | 5606 | 5721 | 6505 | 5893 | 5718 | 5226 | 5305 | 6132 | 5841 | 7320 | 7947 | 8299 | 7058 | 8453 | 7364 | 5738 | 5652 | 6662 | 5141 | 4879 | 5260 | Wsb2 |
| Wscd1 | 2909 | 4247 | 3948 | 3822 | 4313 | 4011 | 3825 | 2928 | 2801 | 3407 | 2438 | 4574 | 3783 | 3886 | 3171 | 2931 | 4177 | 3747 | 4669 | 3972 | 3990 | 3217 | 3020 | 3530 | 3624 | 3336 | 4600 | 4594 | Wscd1 |
| Wscd2 | 488 | 716 | 844 | 775 | 789 | 824 | 619 | 734 | 872 | 1031 | 991 | 1121 | 1224 | 1018 | 570 | 664 | 811 | 733 | 806 | 660 | 662 | 1034 | 883 | 1072 | 985 | 927 | 1119 | 1162 | Wscd2 |
| Wt1 | 2 | 4 | 3 | 5 | 11 | 7 | 9 | 12 | 16 | 23 | 69 | 7 | 35 | 6 | 4 | 13 | 0 | 4 | 1 | 2 | 4 | 10 | 41 | 4 | 0 | 0 | 7 | 11 | Wt1 |
| Wtap | 7762 | 6525 | 7391 | 7463 | 7586 | 7353 | 8468 | 10490 | 9208 | 8624 | 8753 | 9489 | 9507 | 10239 | 7863 | 6520 | 7158 | 6871 | 7690 | 8224 | 7981 | 9436 | 8903 | 9119 | 9577 | 10192 | 9910 | 10342 | Wtap |
| Wtip | 972 | 1040 | 1021 | 953 | 1064 | 893 | 827 | 1108 | 1352 | 1334 | 1419 | 1147 | 1226 | 1040 | 840 | 966 | 1022 | 983 | 1036 | 876 | 886 | 1073 | 1347 | 1302 | 1098 | 1299 | 1128 | 1108 | Wtip |
| Wwc1 | 3913 | 4765 | 5795 | 5869 | 6177 | 5377 | 5292 | 3554 | 4223 | 4970 | 3777 | 4678 | 3931 | 4283 | 3953 | 4461 | 5895 | 5781 | 6340 | 4925 | 5675 | 3987 | 4202 | 5043 | 5166 | 4370 | 4124 | 4155 | Wwc1 |
| Wwc2 | 5777 | 5973 | 5670 | 5358 | 5285 | 5288 | 4932 | 3283 | 3755 | 3624 | 3953 | 3637 | 4289 | 3393 | 5488 | 6613 | 5464 | 5620 | 4930 | 4527 | 4883 | 3263 | 3873 | 3348 | 3063 | 3325 | 3635 | 3598 | Wwc2 |
| Wwox | 963 | 879 | 1011 | 1017 | 966 | 927 | 904 | 626 | 643 | 701 | 679 | 782 | 793 | 843 | 901 | 945 | 1014 | 843 | 968 | 872 | 995 | 739 | 652 | 772 | 710 | 895 | 715 | 826 | Wwox |
| Wwp1 | 6685 | 4707 | 3926 | 4253 | 4709 | 5430 | 5214 | 5548 | 4856 | 3716 | 4474 | 4351 | 4688 | 4431 | 6564 | 5861 | 3809 | 4260 | 4230 | 5589 | 4941 | 4182 | 5166 | 3479 | 3919 | 3251 | 4642 | 4349 | Wwp1 |
| Wwp2 | 4362 | 4926 | 4070 | 4720 | 3814 | 4738 | 3446 | 7184 | 12424 | 12409 | 15326 | 6542 | 7404 | 5910 | 4094 | 4454 | 4071 | 4086 | 4408 | 3795 | 3952 | 11114 | 9027 | 10105 | 7809 | 9842 | 6653 | 6080 | Wwp2 |
| Wwtr1 | 7680 | 6180 | 6703 | 5680 | 5330 | 5875 | 5496 | 4595 | 4965 | 4630 | 5518 | 4649 | 4801 | 4817 | 7392 | 7309 | 6036 | 6034 | 5389 | 5014 | 5543 | 4069 | 5712 | 4595 | 4381 | 4908 | 4761 | 4828 | Wwtr1 |
| Xab2 | 4654 | 6056 | 5135 | 5049 | 4032 | 4568 | 4223 | 3378 | 4095 | 3907 | 3373 | 4347 | 3941 | 3920 | 4396 | 4972 | 4904 | 4622 | 4934 | 4276 | 4377 | 3302 | 3845 | 4163 | 3887 | 4749 | 4435 | 4027 | Xab2 |
| Xaf1 | 14 | 13 | 24 | 15 | 30 | 22 | 20 | 29 | 31 | 96 | 94 | 50 | 49 | 66 | 20 | 15 | 10 | 8 | 13 | 56 | 38 | 89 | 87 | 79 | 59 | 67 | 70 | 71 | Xaf1 |
| Xbp1 | 4693 | 4661 | 5299 | 4302 | 4230 | 4208 | 4312 | 5305 | 5041 | 4692 | 5493 | 3879 | 4225 | 3774 | 4447 | 4620 | 4715 | 4376 | 4826 | 4254 | 4026 | 5212 | 5491 | 4966 | 4758 | 4545 | 3842 | 4400 | Xbp1 |
| Xcl1 | 0 | 0 | 0 | 0 | 0 | 0 | 0 | 0 | 0 | 0 | 0 | 0 | 0 | 0 | 0 | 1 | 0 | 0 | 0 | 0 | 0 | 0 | 0 | 0 | 0 | 0 | 0 | 0 | Xcl1 |
| Xdh | 13 | 6 | 4 | 20 | 17 | 17 | 15 | 2 | 22 | 13 | 60 | 2 | 18 | 7 | 11 | 5 | 12 | 23 | 21 | 4 | 6 | 8 | 33 | 5 | 14 | 8 | 12 | 8 | Xdh |
| Xiap | 5880 | 3869 | 4917 | 5260 | 6325 | 6471 | 6549 | 11535 | 10495 | 10264 | 11315 | 11615 | 11522 | 12311 | 5772 | 4886 | 5193 | 5271 | 5945 | 5700 | 6128 | 11075 | 10331 | 10502 | 10687 | 9421 | 12007 | 11354 | Xiap |
| Xirp1 | 0 | 0 | 0 | 2 | 0 | 0 | 0 | 9 | 23 | 8 | 193 | 1 | 8 | 0 | 1 | 4 | 0 | 0 | 5 | 0 | 1 | 18 | 129 | 2 | 9 | 4 | 0 | 4 | Xirp1 |
| Xirp2 | 0 | 1 | 0 | 6 | 1 | 0 | 0 | 5 | 2 | 0 | 0 | 0 | 1 | 0 | 7 | 2 | 0 | 0 | 0 | 0 | 1 | 3 | 5 | 0 | 1 | 0 | 0 | 0 | Xirp2 |
| Xk | 810 | 696 | 648 | 721 | 946 | 839 | 1041 | 818 | 714 | 770 | 546 | 816 | 1192 | 954 | 785 | 664 | 666 | 900 | 893 | 1079 | 958 | 772 | 645 | 623 | 640 | 754 | 779 | 1151 | Xk |
| Xkr4 | 967 | 931 | 107 | 542 | 947 | 1435 | 1121 | 2440 | 2104 | 1153 | 1261 | 791 | 1066 | 950 | 1296 | 934 | 205 | 474 | 508 | 1498 | 1514 | 692 | 1715 | 363 | 1458 | 160 | 1144 | 655 | Xkr4 |
| Xkr5 | 164 | 153 | 166 | 100 | 85 | 95 | 73 | 264 | 391 | 312 | 354 | 500 | 448 | 403 | 111 | 161 | 127 | 70 | 83 | 110 | 84 | 229 | 297 | 315 | 240 | 579 | 481 | 406 | Xkr5 |
| Xkr6 | 1916 | 1666 | 2067 | 1991 | 1887 | 1877 | 2001 | 1657 | 1360 | 1793 | 1501 | 1893 | 1809 | 1820 | 1791 | 1604 | 1961 | 1822 | 1905 | 1664 | 1844 | 1910 | 1272 | 1705 | 1799 | 1777 | 1764 | 1900 | Xkr6 |
| Xkr7 | 637 | 879 | 1087 | 1490 | 1820 | 1332 | 1323 | 977 | 1121 | 1480 | 1198 | 1067 | 1091 | 943 | 791 | 692 | 1294 | 1442 | 1652 | 1195 | 1567 | 1343 | 926 | 1297 | 1668 | 425 | 977 | 903 | Xkr7 |
| Xkr8 | 837 | 969 | 755 | 759 | 697 | 755 | 627 | 536 | 539 | 485 | 578 | 472 | 482 | 506 | 845 | 924 | 768 | 757 | 647 | 791 | 693 | 481 | 562 | 507 | 510 | 521 | 560 | 522 | Xkr8 |
| Xkr9 | 6 | 0 | 0 | 0 | 0 | 4 | 1 | 1 | 6 | 4 | 5 | 0 | 2 | 0 | 0 | 2 | 1 | 2 | 2 | 1 | 0 | 3 | 4 | 2 | 0 | 1 | 0 | 2 | Xkr9 |
| Xkrx | 133 | 89 | 67 | 150 | 187 | 166 | 174 | 28 | 21 | 40 | 25 | 31 | 31 | 12 | 111 | 96 | 139 | 157 | 158 | 129 | 151 | 37 | 31 | 29 | 18 | 42 | 22 | 14 | Xkrx |
| Xlr | 52 | 53 | 98 | 62 | 45 | 25 | 29 | 38 | 63 | 32 | 66 | 54 | 79 | 48 | 73 | 65 | 53 | 81 | 70 | 69 | 32 | 67 | 107 | 55 | 66 | 43 | 55 | 55 | Xlr |
| Xlr3a | 227 | 274 | 344 | 230 | 125 | 160 | 274 | 73 | 130 | 207 | 227 | 160 | 120 | 213 | 248 | 111 | 155 | 129 | 71 | 201 | 194 | 391 | 275 | 168 | 226 | 212 | 123 | 220 | Xlr3a |
| Xlr3b | 929 | 1313 | 1835 | 1298 | 1258 | 846 | 980 | 79 | 662 | 552 | 766 | 566 | 677 | 841 | 1199 | 231 | 712 | 1004 | 380 | 1151 | 945 | 1491 | 1030 | 1325 | 1524 | 885 | 675 | 957 | Xlr3b |
| Xlr3c | 6 | 10 | 15 | 8 | 7 | 4 | 4 | 9 | 9 | 10 | 16 | 15 | 7 | 7 | 3 | 2 | 5 | 3 | 2 | 12 | 16 | 28 | 8 | 7 | 7 | 10 | 2 | 5 | Xlr3c |
| Xlr4a | 58 | 73 | 213 | 59 | 19 | 8 | 34 | 9 | 34 | 46 | 142 | 41 | 15 | 151 | 198 | 13 | 29 | 62 | 12 | 104 | 86 | 130 | 71 | 58 | 92 | 81 | 5 | 195 | Xlr4a |
| Xlr4b | 48 | 67 | 76 | 70 | 43 | 47 | 36 | 2 | 36 | 44 | 46 | 35 | 52 | 63 | 46 | 9 | 24 | 50 | 15 | 58 | 19 | 182 | 32 | 61 | 135 | 110 | 21 | 76 | Xlr4b |
| Xlr4c | 0 | 5 | 11 | 5 | 1 | 3 | 4 | 0 | 0 | 3 | 4 | 1 | 1 | 7 | 8 | 2 | 0 | 4 | 0 | 7 | 1 | 13 | 3 | 2 | 3 | 0 | 1 | 14 | Xlr4c |
| Xlr5a | 1 | 1 | 1 | 0 | 16 | 0 | 0 | 0 | 1 | 0 | 0 | 0 | 0 | 0 | 7 | 0 | 0 | 0 | 0 | 2 | 1 | 2 | 3 | 0 | 0 | 1 | 0 | 0 | Xlr5a |
| Xlr5b | 1 | 0 | 0 | 2 | 0 | 0 | 0 | 0 | 2 | 0 | 3 | 0 | 1 | 0 | 0 | 0 | 0 | 0 | 0 | 0 | 0 | 2 | 1 | 0 | 4 | 0 | 0 | 1 | Xlr5b |
| Xlr5c | 0 | 0 | 2 | 0 | 0 | 0 | 0 | 0 | 1 | 0 | 0 | 0 | 0 | 0 | 0 | 0 | 0 | 0 | 0 | 0 | 0 | 0 | 0 | 0 | 0 | 0 | 0 | 1 | Xlr5c |
| Xndc1 | 593 | 489 | 455 | 593 | 593 | 608 | 708 | 1073 | 990 | 1233 | 1128 | 1234 | 1054 | 1503 | 484 | 384 | 496 | 614 | 623 | 557 | 701 | 996 | 991 | 1197 | 1202 | 1045 | 1148 | 1317 | Xndc1 |
| Xntrpc | 36 | 60 | 50 | 80 | 52 | 25 | 64 | 65 | 56 | 74 | 63 | 85 | 56 | 60 | 50 | 35 | 56 | 54 | 56 | 20 | 36 | 74 | 40 | 61 | 90 | 52 | 71 | 90 | Xntrpc |
| Xpa | 1151 | 974 | 883 | 1093 | 1068 | 943 | 1238 | 1121 | 899 | 990 | 862 | 1007 | 907 | 1126 | 1150 | 916 | 954 | 1037 | 967 | 1229 | 1052 | 976 | 874 | 1092 | 1032 | 898 | 984 | 991 | Xpa |
| Xpc | 3801 | 4238 | 3458 | 3928 | 3499 | 3606 | 3612 | 2540 | 2627 | 2464 | 2539 | 2777 | 2872 | 2839 | 3915 | 3842 | 3777 | 3902 | 3770 | 3858 | 3660 | 2358 | 2500 | 2648 | 2976 | 2741 | 2989 | 2794 | Xpc |
| Xpnpep1 | 5906 | 6576 | 5178 | 4896 | 4618 | 5161 | 4226 | 4279 | 5118 | 4435 | 4722 | 4899 | 4769 | 4300 | 5562 | 6108 | 5098 | 4814 | 5001 | 5281 | 4505 | 3893 | 5192 | 4484 | 4300 | 5159 | 5015 | 4694 | Xpnpep1 |
| Xpnpep2 | 194 | 211 | 288 | 126 | 70 | 194 | 65 | 182 | 269 | 181 | 317 | 253 | 397 | 231 | 215 | 255 | 122 | 171 | 63 | 91 | 84 | 239 | 409 | 172 | 164 | 326 | 346 | 307 | Xpnpep2 |
| Xpnpep3 | 2532 | 2484 | 2254 | 2302 | 1996 | 2048 | 2251 | 1627 | 1599 | 1624 | 1751 | 1888 | 1675 | 1623 | 2352 | 2562 | 2274 | 2360 | 1911 | 2123 | 2191 | 1536 | 1635 | 1624 | 1629 | 1640 | 1622 | 1698 | Xpnpep3 |
| Xpo1 | 33300 | 23993 | 29551 | 28229 | 29336 | 30415 | 34925 | 37800 | 33310 | 31091 | 32240 | 36554 | 34665 | 38628 | 34756 | 29052 | 29628 | 28883 | 29267 | 33292 | 31741 | 32410 | 31769 | 33313 | 31883 | 36153 | 35620 | 37976 | Xpo1 |
| Xpo4 | 6198 | 4648 | 3997 | 3976 | 3828 | 4431 | 4260 | 6169 | 5609 | 4570 | 5367 | 5638 | 5582 | 5758 | 5976 | 5627 | 3821 | 3867 | 3711 | 4857 | 4043 | 4568 | 5629 | 4578 | 4832 | 4497 | 6086 | 5586 | Xpo4 |
| Xpo5 | 16891 | 17925 | 13529 | 13704 | 12561 | 14079 | 14228 | 12298 | 11693 | 10527 | 10063 | 12667 | 11775 | 12405 | 16312 | 14574 | 14308 | 13009 | 13808 | 14204 | 14037 | 10662 | 10804 | 10951 | 11303 | 11129 | 13418 | 13184 | Xpo5 |
| Xpo6 | 9904 | 10534 | 10150 | 11511 | 11322 | 10648 | 11534 | 7455 | 8410 | 9186 | 8467 | 10212 | 9160 | 9393 | 9397 | 10382 | 11179 | 11329 | 11643 | 9285 | 11223 | 9399 | 7900 | 8818 | 9357 | 9111 | 8887 | 9172 | Xpo6 |
| Xpo7 | 11886 | 10531 | 9698 | 9393 | 9750 | 9987 | 10695 | 11704 | 11040 | 10415 | 9812 | 11190 | 11438 | 11862 | 11091 | 9513 | 9978 | 9149 | 10121 | 10223 | 10943 | 10684 | 10392 | 10369 | 11263 | 10576 | 11423 | 12617 | Xpo7 |
| Xpot | 10169 | 9347 | 9153 | 8966 | 9037 | 9150 | 9555 | 8056 | 7786 | 7392 | 8005 | 8355 | 8277 | 8646 | 9637 | 9522 | 8980 | 9282 | 9082 | 8595 | 9269 | 7454 | 7961 | 7893 | 7920 | 7976 | 8335 | 8662 | Xpot |
| Xpr1 | 26280 | 25391 | 20955 | 35692 | 39361 | 39332 | 45645 | 23935 | 22209 | 24556 | 20887 | 22409 | 18277 | 23381 | 28846 | 23554 | 28072 | 36805 | 37667 | 37558 | 45904 | 29244 | 20135 | 21695 | 31261 | 12388 | 20934 | 20109 | Xpr1 |
| Xrcc1 | 4633 | 5759 | 5715 | 5453 | 4534 | 4901 | 4494 | 3706 | 4473 | 4783 | 4484 | 5374 | 4949 | 4990 | 4142 | 4653 | 5674 | 5026 | 5481 | 4502 | 4453 | 3649 | 4095 | 5359 | 4434 | 6428 | 5131 | 5035 | Xrcc1 |
| Xrcc2 | 1463 | 1164 | 1197 | 1060 | 875 | 1011 | 1134 | 1380 | 1260 | 1195 | 982 | 1488 | 1263 | 1483 | 1171 | 1137 | 997 | 898 | 1142 | 1276 | 985 | 802 | 1059 | 1325 | 1273 | 1550 | 1407 | 1550 | Xrcc2 |
| Xrcc3 | 1726 | 2055 | 1927 | 1891 | 1312 | 1552 | 1574 | 540 | 646 | 592 | 547 | 979 | 747 | 819 | 1390 | 1861 | 2070 | 1763 | 1626 | 1440 | 1454 | 438 | 570 | 756 | 638 | 1250 | 974 | 938 | Xrcc3 |
| Xrcc4 | 313 | 290 | 234 | 241 | 323 | 367 | 359 | 812 | 692 | 548 | 585 | 654 | 652 | 652 | 336 | 291 | 284 | 269 | 348 | 408 | 342 | 688 | 736 | 590 | 578 | 795 | 610 | 706 | Xrcc4 |
| Xrcc5 | 2868 | 2344 | 2269 | 1565 | 1671 | 1787 | 1637 | 3559 | 3291 | 2608 | 2948 | 3213 | 3271 | 3269 | 2716 | 2558 | 1993 | 1533 | 1545 | 2047 | 1600 | 2473 | 3290 | 2816 | 2593 | 3874 | 3584 | 3544 | Xrcc5 |
| Xrcc6 | 5054 | 4450 | 4275 | 4262 | 3527 | 4021 | 4329 | 3204 | 3208 | 2964 | 3103 | 3089 | 3037 | 3323 | 4772 | 4420 | 3928 | 4241 | 3776 | 4207 | 4237 | 2558 | 3147 | 3356 | 3111 | 3759 | 3039 | 3354 | Xrcc6 |
| Xrn1 | 9255 | 7387 | 5113 | 6596 | 7463 | 9137 | 8567 | 9978 | 8604 | 6738 | 7944 | 8128 | 8057 | 8402 | 9051 | 7932 | 5848 | 6490 | 6980 | 9155 | 8887 | 7232 | 8165 | 6094 | 7947 | 5004 | 9159 | 7779 | Xrn1 |
| Xrn2 | 14282 | 12248 | 12463 | 11493 | 11149 | 11859 | 12970 | 14040 | 11898 | 10717 | 10280 | 12811 | 11736 | 13222 | 14561 | 11538 | 11160 | 11038 | 11433 | 13519 | 12070 | 11622 | 11617 | 11086 | 11595 | 13282 | 13124 | 12860 | Xrn2 |
| Xrra1 | 45 | 54 | 43 | 27 | 47 | 38 | 44 | 24 | 35 | 43 | 26 | 28 | 29 | 28 | 59 | 41 | 52 | 36 | 26 | 49 | 26 | 33 | 34 | 34 | 19 | 22 | 34 | 28 | Xrra1 |
| Xxylt1 | 2509 | 2440 | 2614 | 2021 | 2058 | 1981 | 1847 | 1589 | 1838 | 1765 | 2024 | 1753 | 1903 | 1789 | 2222 | 2593 | 2233 | 2221 | 1854 | 1563 | 1851 | 1719 | 1871 | 1627 | 1515 | 1895 | 1729 | 1844 | Xxylt1 |
| Xylb | 1087 | 1032 | 1070 | 1112 | 1024 | 1016 | 1078 | 503 | 592 | 699 | 623 | 885 | 649 | 805 | 827 | 966 | 1128 | 1088 | 1194 | 835 | 1035 | 587 | 524 | 741 | 600 | 952 | 752 | 813 | Xylb |
| Xylt1 | 1625 | 1662 | 1558 | 2078 | 2003 | 1911 | 2305 | 1723 | 2210 | 2407 | 2507 | 2701 | 2470 | 2692 | 1551 | 1565 | 1722 | 2133 | 2058 | 1916 | 1997 | 2486 | 1872 | 2201 | 2476 | 2032 | 2288 | 2611 | Xylt1 |
| Xylt2 | 2097 | 2575 | 2271 | 2349 | 2296 | 2142 | 1833 | 1511 | 2069 | 1990 | 2284 | 1890 | 1914 | 1775 | 1911 | 2368 | 2348 | 2354 | 2194 | 1839 | 1849 | 1778 | 2076 | 1964 | 1842 | 1985 | 2023 | 1753 | Xylt2 |
| Yae1d1 | 4756 | 4314 | 3293 | 4247 | 4291 | 4779 | 5237 | 4560 | 3903 | 3417 | 3291 | 3881 | 3686 | 4158 | 5014 | 4478 | 3809 | 4197 | 4053 | 5561 | 5178 | 3661 | 3787 | 3429 | 3862 | 2985 | 4179 | 3887 | Yae1d1 |
| Yaf2 | 2678 | 2224 | 2796 | 2531 | 2734 | 2620 | 3041 | 2809 | 2288 | 2177 | 2220 | 2634 | 2489 | 2830 | 2869 | 2395 | 2697 | 2691 | 2948 | 2913 | 3083 | 2847 | 2394 | 2417 | 2415 | 2712 | 2665 | 2838 | Yaf2 |
| Yap1 | 8522 | 6340 | 9570 | 6481 | 5510 | 6342 | 5456 | 10455 | 10884 | 10811 | 12255 | 12231 | 12775 | 12790 | 6949 | 7587 | 7168 | 5949 | 6204 | 4809 | 5336 | 8589 | 11240 | 12039 | 9193 | 15497 | 11827 | 14074 | Yap1 |
| Yars | 8935 | 9762 | 8867 | 9325 | 8683 | 9349 | 9069 | 6342 | 6356 | 6661 | 6181 | 7472 | 7059 | 7596 | 8720 | 8661 | 9111 | 9168 | 9577 | 8590 | 9809 | 6610 | 6419 | 6880 | 7454 | 7608 | 7876 | 8052 | Yars |
| Yars2 | 909 | 773 | 1024 | 787 | 922 | 882 | 922 | 1388 | 1205 | 1146 | 1119 | 1307 | 1209 | 1475 | 903 | 873 | 927 | 778 | 839 | 1002 | 925 | 1160 | 1218 | 1140 | 1218 | 1549 | 1366 | 1453 | Yars2 |
| Ybey | 377 | 365 | 524 | 398 | 409 | 385 | 411 | 537 | 471 | 590 | 552 | 659 | 575 | 629 | 404 | 341 | 499 | 429 | 351 | 422 | 416 | 453 | 516 | 575 | 558 | 669 | 568 | 671 | Ybey |

| | | | | | | | | | | | | | | | | | | | | | | | | | | | | | |
|---|---|---|---|---|---|---|---|---|---|---|---|---|---|---|---|---|---|---|---|---|---|---|---|---|---|---|---|---|---|
| Ybx1 | 55063 | 57861 | 69908 | 65267 | 59441 | 55876 | 59251 | 46989 | 47723 | 56719 | 49667 | 58105 | 51551 | 61241 | 53655 | 52167 | 65446 | 60203 | 63832 | 56024 | 62544 | 45475 | 48011 | 58605 | 57492 | 67795 | 57701 | 61248 | Ybx1 |
| Ybx2 | 5008 | 5483 | 5555 | 6377 | 4979 | 4770 | 5259 | 1320 | 1285 | 1503 | 1435 | 1479 | 1330 | 1293 | 4278 | 5135 | 5800 | 6107 | 5801 | 4305 | 4604 | 1546 | 1322 | 1604 | 1353 | 1818 | 1343 | 1280 | Ybx2 |
| Ybx3 | 5828 | 6296 | 7287 | 6283 | 5646 | 5470 | 4973 | 4670 | 5811 | 5824 | 6588 | 6154 | 5925 | 6095 | 5309 | 6084 | 6531 | 6064 | 6104 | 4809 | 5419 | 4677 | 5954 | 6355 | 5437 | 7408 | 5998 | 6214 | Ybx3 |
| Ydjc | 631 | 774 | 628 | 651 | 830 | 652 | 619 | 401 | 569 | 542 | 551 | 286 | 352 | 301 | 636 | 621 | 669 | 676 | 696 | 727 | 654 | 565 | 467 | 389 | 469 | 268 | 302 | 273 | Ydjc |
| Yeats2 | 8120 | 7525 | 6417 | 7306 | 7183 | 7982 | 7168 | 10000 | 10921 | 10244 | 10583 | 10759 | 10656 | 10564 | 7279 | 7872 | 6511 | 7123 | 7183 | 6852 | 7329 | 9687 | 10156 | 9825 | 10379 | 8209 | 10922 | 10059 | Yeats2 |
| Yeats4 | 2994 | 3030 | 3420 | 3125 | 3062 | 2810 | 3336 | 5851 | 4899 | 5774 | 4897 | 6554 | 5364 | 7114 | 3030 | 2295 | 3262 | 2750 | 3441 | 3216 | 3464 | 5360 | 4717 | 6333 | 5950 | 7281 | 6420 | 7162 | Yeats4 |
| Yes1 | 5999 | 4583 | 5795 | 6011 | 5693 | 5867 | 6606 | 4059 | 3442 | 3444 | 3864 | 3898 | 3909 | 4305 | 6384 | 5955 | 5539 | 5829 | 5327 | 6134 | 6124 | 3623 | 3610 | 3610 | 3624 | 3880 | 3811 | 4179 | Yes1 |
| Yif1a | 1630 | 2019 | 1524 | 1392 | 1136 | 1239 | 1222 | 1072 | 1309 | 1108 | 1087 | 1107 | 1212 | 1081 | 1609 | 1629 | 1493 | 1264 | 1128 | 1398 | 1248 | 964 | 1341 | 1088 | 944 | 1185 | 1230 | 1157 | Yif1a |
| Yif1b | 1336 | 1772 | 1522 | 1402 | 1292 | 1308 | 1026 | 882 | 1128 | 1020 | 1170 | 1158 | 1154 | 875 | 1278 | 1493 | 1545 | 1225 | 1493 | 1172 | 1180 | 866 | 1238 | 1095 | 1009 | 1296 | 1178 | 994 | Yif1b |
| Yipf1 | 1658 | 1867 | 1329 | 1510 | 1532 | 1434 | 1465 | 1680 | 1708 | 1335 | 1513 | 1283 | 1357 | 1252 | 1763 | 1783 | 1373 | 1442 | 1615 | 1803 | 1521 | 1397 | 1768 | 1240 | 1466 | 1256 | 1361 | 1282 | Yipf1 |
| Yipf2 | 376 | 380 | 405 | 314 | 419 | 378 | 292 | 849 | 864 | 853 | 871 | 862 | 872 | 865 | 331 | 341 | 350 | 344 | 284 | 313 | 312 | 755 | 914 | 781 | 837 | 903 | 859 | 862 | Yipf2 |
| Yipf3 | 2659 | 3568 | 3161 | 3457 | 2707 | 2879 | 2690 | 2155 | 2573 | 2663 | 2629 | 2586 | 2486 | 2423 | 2734 | 2925 | 3326 | 3254 | 3352 | 2660 | 2864 | 2481 | 2586 | 2720 | 2561 | 2699 | 2622 | 2398 | Yipf3 |
| Yipf4 | 2397 | 1836 | 2236 | 2391 | 1975 | 2182 | 2408 | 2748 | 2120 | 2175 | 2223 | 2235 | 2278 | 2336 | 2540 | 2221 | 2125 | 2189 | 2110 | 2416 | 2255 | 2563 | 2345 | 2488 | 2509 | 2390 | 2418 | 2438 | Yipf4 |
| Yipf5 | 4564 | 3652 | 4062 | 3859 | 3646 | 3677 | 3930 | 4827 | 4439 | 4085 | 4386 | 4152 | 4123 | 3977 | 4609 | 4206 | 3866 | 3957 | 3683 | 3966 | 3881 | 4659 | 4779 | 3973 | 4049 | 4406 | 4291 | 4174 | Yipf5 |
| Yipf6 | 3938 | 3048 | 3489 | 3888 | 4001 | 3922 | 4225 | 4651 | 3662 | 3908 | 3991 | 4313 | 4266 | 4516 | 4032 | 3382 | 3564 | 3713 | 3775 | 4036 | 3969 | 4515 | 3695 | 3934 | 3997 | 3850 | 4302 | 4145 | Yipf6 |
| Yipf7 | 2 | 3 | 4 | 0 | 5 | 7 | 3 | 1 | 2 | 0 | 15 | 7 | 0 | 0 | 7 | 9 | 6 | 4 | 4 | 3 | 2 | 0 | 5 | 2 | 4 | 2 | 1 | 4 | Yipf7 |
| Yjefn3 | 137 | 219 | 264 | 301 | 242 | 191 | 184 | 128 | 123 | 175 | 153 | 163 | 192 | 126 | 191 | 172 | 289 | 303 | 309 | 208 | 212 | 244 | 148 | 234 | 193 | 173 | 207 | 158 | Yjefn3 |
| Yju2 | 517 | 600 | 665 | 724 | 632 | 630 | 654 | 960 | 1083 | 1077 | 962 | 1228 | 1178 | 1365 | 535 | 455 | 651 | 598 | 680 | 616 | 660 | 916 | 986 | 1270 | 1244 | 1407 | 1361 | 1416 | Yju2 |
| Ykt6 | 13609 | 14120 | 14265 | 14209 | 12337 | 12229 | 12245 | 10572 | 10640 | 10679 | 10328 | 10839 | 10211 | 10842 | 12988 | 12872 | 13917 | 13439 | 12950 | 12290 | 12200 | 10494 | 9958 | 10924 | 11255 | 11951 | 11238 | 11261 | Ykt6 |
| Ylpm1 | 10980 | 10033 | 12197 | 13198 | 13758 | 13617 | 13422 | 12841 | 13146 | 15281 | 14266 | 16385 | 14402 | 16518 | 10747 | 11156 | 12698 | 12734 | 13411 | 10497 | 13893 | 14674 | 11417 | 15141 | 15881 | 12324 | 14319 | 15276 | Ylpm1 |
| Yme1l1 | 18765 | 14309 | 16555 | 14691 | 14064 | 14771 | 16748 | 18104 | 15658 | 13646 | 14746 | 15896 | 15645 | 16400 | 19440 | 17614 | 15433 | 14520 | 14686 | 18222 | 15292 | 15244 | 16573 | 14849 | 14627 | 16399 | 16706 | 17289 | Yme1l1 |
| Yod1 | 1998 | 1436 | 1712 | 1930 | 1761 | 2045 | 2284 | 2313 | 2129 | 1936 | 1774 | 2234 | 2209 | 2442 | 2116 | 1792 | 1962 | 1697 | 1876 | 2120 | 2055 | 2017 | 2048 | 2112 | 1921 | 2122 | 2323 | 2537 | Yod1 |
| Ypel1 | 2149 | 2120 | 2018 | 1936 | 2492 | 2623 | 2737 | 5120 | 4696 | 4733 | 3980 | 4952 | 4312 | 5486 | 2235 | 1764 | 2019 | 1906 | 2583 | 2696 | 2806 | 4939 | 4553 | 4855 | 5474 | 5226 | 5281 | 5185 | Ypel1 |
| Ypel2 | 2587 | 2871 | 2564 | 3747 | 2675 | 3058 | 3379 | 1551 | 1461 | 1862 | 1828 | 1574 | 1546 | 1822 | 2740 | 2736 | 3118 | 3554 | 3063 | 2635 | 3287 | 1882 | 1353 | 2115 | 1790 | 1800 | 1655 | 1812 | Ypel2 |
| Ypel3 | 1882 | 2645 | 2194 | 2851 | 2803 | 2562 | 2633 | 1850 | 2179 | 2391 | 2206 | 1966 | 2137 | 2097 | 2060 | 2293 | 2889 | 2946 | 3052 | 2636 | 2684 | 2309 | 2153 | 2336 | 2437 | 2119 | 1909 | 1854 | Ypel3 |
| Ypel4 | 768 | 909 | 937 | 1222 | 1453 | 1117 | 1292 | 1280 | 1240 | 1796 | 1409 | 1724 | 1881 | 1920 | 778 | 703 | 1207 | 1205 | 1491 | 1116 | 1412 | 1803 | 1147 | 1872 | 1840 | 1346 | 1769 | 1869 | Ypel4 |
| Ypel5 | 3963 | 3515 | 4014 | 4283 | 4291 | 4320 | 4949 | 4629 | 3587 | 3518 | 3328 | 3649 | 3774 | 3972 | 4667 | 3839 | 4278 | 4291 | 4263 | 5019 | 4690 | 4299 | 3988 | 3652 | 4270 | 3599 | 3624 | 4082 | Ypel5 |
| Yrdc | 1931 | 1890 | 2321 | 2212 | 2011 | 1743 | 2150 | 1476 | 1336 | 1532 | 1329 | 1606 | 1390 | 1794 | 1782 | 1615 | 2135 | 2126 | 2225 | 1725 | 2251 | 1572 | 1295 | 1665 | 1593 | 1722 | 1495 | 1729 | Yrdc |
| Ythdc1 | 9370 | 7866 | 10547 | 11341 | 10258 | 10283 | 11901 | 11342 | 9722 | 11708 | 11111 | 11948 | 11324 | 13502 | 9233 | 8888 | 10585 | 10830 | 10470 | 9351 | 11022 | 11729 | 9417 | 12549 | 12043 | 11859 | 11747 | 12893 | Ythdc1 |
| Ythdc2 | 3945 | 2951 | 2757 | 3303 | 3818 | 4014 | 3936 | 4644 | 3666 | 3044 | 3587 | 3416 | 3559 | 3654 | 4085 | 3377 | 3167 | 3311 | 3513 | 4412 | 4083 | 3801 | 3696 | 3108 | 3621 | 2559 | 3665 | 3616 | Ythdc2 |
| Ythdf1 | 8194 | 8654 | 8629 | 8762 | 7376 | 7233 | 7623 | 6245 | 6113 | 5729 | 5909 | 6738 | 6386 | 6740 | 7796 | 8056 | 8324 | 8244 | 7580 | 7038 | 7706 | 6147 | 5979 | 6579 | 6373 | 7598 | 6735 | 7253 | Ythdf1 |
| Ythdf2 | 6998 | 6094 | 7270 | 6476 | 7407 | 6827 | 7717 | 9104 | 7666 | 8178 | 7677 | 9203 | 8744 | 9753 | 6887 | 6227 | 7342 | 6534 | 7564 | 7032 | 7234 | 8732 | 7402 | 8541 | 9022 | 10120 | 9415 | 10151 | Ythdf2 |
| Ythdf3 | 14324 | 10327 | 15128 | 15052 | 14492 | 15241 | 16474 | 13623 | 11804 | 12355 | 13782 | 15504 | 15080 | 16091 | 14057 | 13330 | 14397 | 14786 | 14621 | 12897 | 15152 | 14639 | 11804 | 14028 | 14235 | 14674 | 15591 | 15991 | Ythdf3 |
| Ywhab | 32987 | 31007 | 33221 | 33432 | 34845 | 32810 | 35113 | 29634 | 27774 | 29545 | 27969 | 29990 | 28207 | 28638 | 33534 | 30931 | 33237 | 33905 | 34379 | 31659 | 34937 | 32946 | 28622 | 28320 | 29734 | 31061 | 28755 | 30134 | Ywhab |
| Ywhae | 63789 | 55620 | 69545 | 61786 | 63509 | 58728 | 72167 | 82857 | 70361 | 78204 | 70434 | 85388 | 78118 | 94676 | 66851 | 53112 | 67183 | 60005 | 65449 | 63004 | 69316 | 84725 | 71741 | 87009 | 82586 | 97473 | 84158 | 97735 | Ywhae |
| Ywhag | 22429 | 27076 | 31264 | 35483 | 41042 | 33812 | 34719 | 31072 | 33169 | 39377 | 36530 | 35705 | 33539 | 37152 | 24857 | 24114 | 33940 | 36716 | 39121 | 31187 | 37040 | 43758 | 31321 | 39374 | 45011 | 33413 | 32060 | 36256 | Ywhag |
| Ywhah | 10200 | 9268 | 11088 | 10390 | 12980 | 10589 | 12664 | 16425 | 14627 | 15447 | 15052 | 15487 | 15766 | 17223 | 10833 | 9104 | 11218 | 10907 | 11933 | 11045 | 12780 | 17619 | 14879 | 15904 | 16143 | 17082 | 15177 | 18002 | Ywhah |
| Ywhaq | 32746 | 30307 | 28751 | 27648 | 28804 | 28823 | 33255 | 40646 | 34279 | 29670 | 29759 | 34349 | 33806 | 36771 | 36415 | 30968 | 29270 | 28064 | 30124 | 36992 | 31009 | 34920 | 34934 | 31711 | 33373 | 37703 | 35958 | 38911 | Ywhaq |
| Ywhaz | 35275 | 31029 | 38894 | 35881 | 37018 | 35479 | 40534 | 51933 | 46143 | 45470 | 44141 | 52527 | 49277 | 53836 | 36540 | 30746 | 37298 | 34614 | 37640 | 39217 | 38544 | 51523 | 46368 | 50314 | 52052 | 57231 | 52632 | 55890 | Ywhaz |
| Yy1 | 9853 | 8860 | 11045 | 10611 | 10073 | 9970 | 10813 | 10304 | 9277 | 9969 | 10007 | 11625 | 10411 | 11539 | 9455 | 9338 | 10851 | 10402 | 10546 | 8909 | 10887 | 10325 | 9222 | 10511 | 10669 | 11585 | 10771 | 11858 | Yy1 |
| Zadh2 | 1884 | 1563 | 2439 | 1760 | 2146 | 1790 | 1876 | 1961 | 2039 | 2206 | 3269 | 2063 | 2453 | 2164 | 1685 | 1802 | 2204 | 1806 | 1901 | 1661 | 1708 | 2201 | 2503 | 2303 | 2157 | 2572 | 2060 | 2265 | Zadh2 |
| Zan | 39 | 24 | 31 | 22 | 46 | 31 | 18 | 56 | 112 | 65 | 112 | 67 | 59 | 52 | 17 | 32 | 21 | 44 | 37 | 16 | 26 | 94 | 87 | 70 | 92 | 55 | 50 | 43 | Zan |
| Zap70 | 23 | 9 | 20 | 28 | 29 | 21 | 25 | 31 | 31 | 32 | 33 | 25 | 51 | 24 | 17 | 15 | 20 | 21 | 32 | 18 | 21 | 50 | 42 | 49 | 22 | 39 | 31 | 24 | Zap70 |
| Zar1 | 39 | 33 | 48 | 38 | 38 | 46 | 25 | 59 | 66 | 44 | 60 | 51 | 98 | 65 | 36 | 33 | 36 | 23 | 37 | 51 | 36 | 62 | 76 | 55 | 81 | 29 | 57 | 72 | Zar1 |
| Zar1l | 0 | 0 | 0 | 0 | 1 | 0 | 0 | 0 | 0 | 0 | 0 | 0 | 0 | 0 | 3 | 0 | 0 | 0 | 2 | 0 | 3 | 0 | 0 | 2 | 0 | 0 | 0 | 1 | Zar1l |
| Zbbx | 28 | 19 | 7 | 23 | 43 | 28 | 30 | 40 | 37 | 42 | 9 | 30 | 31 | 35 | 44 | 44 | 16 | 12 | 23 | 35 | 40 | 30 | 51 | 30 | 41 | 31 | 26 | 40 | Zbbx |
| Zbed3 | 1132 | 1184 | 1616 | 1379 | 1405 | 1297 | 1158 | 1737 | 1980 | 2068 | 2212 | 2312 | 2129 | 2344 | 988 | 1200 | 1534 | 1275 | 1478 | 1073 | 1170 | 2012 | 2043 | 2244 | 1921 | 3103 | 2035 | 2217 | Zbed3 |
| Zbed4 | 4531 | 4519 | 4129 | 4098 | 3849 | 4145 | 4573 | 4332 | 3681 | 3279 | 3092 | 4076 | 3958 | 4290 | 4683 | 4536 | 4145 | 4232 | 4050 | 4233 | 4172 | 3885 | 3423 | 3471 | 3991 | 3730 | 4432 | 4398 | Zbed4 |
| Zbed5 | 997 | 967 | 787 | 930 | 718 | 855 | 798 | 1175 | 1017 | 949 | 948 | 769 | 944 | 881 | 1021 | 1048 | 803 | 978 | 822 | 949 | 842 | 1111 | 1012 | 852 | 934 | 721 | 783 | 833 | Zbed5 |
| Zbed6 | 6105 | 4907 | 287 | 2117 | 2959 | 4930 | 3739 | 8987 | 8104 | 4304 | 6738 | 4329 | 5444 | 5041 | 5273 | 5649 | 773 | 2047 | 2085 | 5322 | 4839 | 2237 | 7763 | 1478 | 4855 | 191 | 6154 | 3315 | Zbed6 |
| Zbp1 | 6 | 8 | 8 | 12 | 7 | 25 | 10 | 10 | 10 | 4 | 14 | 12 | 8 | 13 | 17 | 10 | 2 | 8 | 5 | 25 | 5 | 8 | 12 | 9 | 14 | 10 | 23 | 11 | Zbp1 |
| Zbtb1 | 1609 | 1151 | 1516 | 1400 | 1609 | 1687 | 1561 | 3171 | 2960 | 2835 | 3297 | 2713 | 2958 | 3040 | 1581 | 1427 | 1505 | 1362 | 1624 | 1548 | 1494 | 2874 | 3044 | 2845 | 2865 | 2935 | 2924 | 3014 | Zbtb1 |
| Zbtb10 | 3120 | 2703 | 4135 | 4033 | 3706 | 3792 | 3724 | 3418 | 2578 | 3116 | 3338 | 3557 | 3772 | 3836 | 3142 | 2958 | 3942 | 3481 | 3504 | 3061 | 3552 | 3522 | 2552 | 3306 | 3672 | 3308 | 3758 | 4074 | Zbtb10 |
| Zbtb11 | 3306 | 2999 | 3252 | 3762 | 3267 | 3313 | 3727 | 2796 | 2419 | 2680 | 2824 | 2600 | 2704 | 2826 | 3312 | 3202 | 3186 | 3888 | 3223 | 2880 | 3419 | 3397 | 2419 | 2712 | 2774 | 2513 | 2637 | 2646 | Zbtb11 |
| Zbtb12 | 5140 | 6809 | 7495 | 7265 | 6856 | 6371 | 6823 | 6312 | 6693 | 7899 | 6527 | 8637 | 7198 | 8696 | 5387 | 4874 | 8230 | 6640 | 8089 | 6737 | 7337 | 6953 | 6159 | 9282 | 8366 | 9426 | 8836 | 8768 | Zbtb12 |
| Zbtb14 | 1945 | 1785 | 2284 | 1916 | 2431 | 2188 | 2273 | 2726 | 2760 | 3144 | 2819 | 3340 | 2947 | 3152 | 1999 | 1725 | 2253 | 2068 | 2573 | 2267 | 2304 | 2814 | 2619 | 3236 | 2976 | 3584 | 3195 | 3101 | Zbtb14 |
| Zbtb16 | 769 | 993 | 2176 | 1222 | 989 | 1114 | 755 | 410 | 411 | 645 | 572 | 786 | 829 | 662 | 916 | 958 | 1613 | 1071 | 1237 | 946 | 1019 | 643 | 410 | 609 | 661 | 406 | 968 | 864 | Zbtb16 |
| Zbtb17 | 1775 | 2380 | 2402 | 2457 | 2189 | 2023 | 1862 | 2827 | 3173 | 3326 | 3022 | 3842 | 3268 | 3316 | 1747 | 2034 | 2297 | 2151 | 2429 | 1942 | 1985 | 3113 | 3058 | 3595 | 3597 | 4021 | 3552 | 3427 | Zbtb17 |
| Zbtb18 | 10441 | 10271 | 11216 | 13350 | 14967 | 14409 | 17309 | 16063 | 15208 | 14695 | 12651 | 16606 | 12621 | 17015 | 10770 | 8670 | 12629 | 13305 | 16627 | 15678 | 17023 | 12940 | 13253 | 18275 | 20073 | 13912 | 14082 | 14725 | Zbtb18 |
| Zbtb2 | 737 | 982 | 940 | 927 | 1008 | 912 | 1037 | 1694 | 1672 | 1727 | 1563 | 1970 | 1693 | 1956 | 796 | 759 | 949 | 886 | 1093 | 907 | 1024 | 1666 | 1467 | 1828 | 1762 | 1788 | 1827 | 2060 | Zbtb2 |
| Zbtb20 | 3735 | 3400 | 2491 | 3475 | 4429 | 5175 | 4509 | 13085 | 11967 | 8462 | 8252 | 9486 | 6997 | 10413 | 3857 | 3425 | 3007 | 3378 | 4053 | 5146 | 5450 | 8626 | 9182 | 7876 | 11176 | 5697 | 10132 | 7698 | Zbtb20 |
| Zbtb21 | 1555 | 1589 | 1735 | 1708 | 1708 | 1609 | 1830 | 1958 | 2008 | 2072 | 2208 | 2149 | 2085 | 2030 | 1577 | 1511 | 1749 | 1779 | 1824 | 1731 | 1722 | 1888 | 1926 | 1966 | 2081 | 1968 | 2090 | 2053 | Zbtb21 |
| Zbtb22 | 1293 | 1603 | 1473 | 1512 | 1484 | 1445 | 1325 | 1763 | 2052 | 1972 | 1788 | 2420 | 2127 | 2291 | 1286 | 1182 | 1581 | 1314 | 1519 | 1338 | 1398 | 1988 | 1845 | 2192 | 2132 | 2586 | 2381 | 2328 | Zbtb22 |
| Zbtb24 | 2043 | 2300 | 2045 | 2244 | 1784 | 1784 | 1817 | 1689 | 1515 | 1637 | 1597 | 1604 | 1636 | 1605 | 1948 | 2094 | 1950 | 2060 | 1839 | 1725 | 1826 | 1629 | 1627 | 1804 | 1684 | 1669 | 1594 | 1647 | Zbtb24 |
| Zbtb25 | 1340 | 1175 | 1246 | 1290 | 1161 | 1222 | 1387 | 1102 | 1125 | 1133 | 1122 | 1194 | 1154 | 1240 | 1271 | 1099 | 1306 | 1240 | 1183 | 1185 | 1314 | 1255 | 1074 | 1254 | 1153 | 1101 | 1274 | 1250 | Zbtb25 |
| Zbtb26 | 2450 | 2033 | 1993 | 2055 | 2055 | 2249 | 2196 | 3307 | 2609 | 2289 | 2507 | 2542 | 2555 | 2716 | 2488 | 2212 | 2024 | 1950 | 1945 | 2367 | 2216 | 2438 | 2520 | 2336 | 2371 | 2122 | 2932 | 2739 | Zbtb26 |
| Zbtb3 | 167 | 223 | 255 | 167 | 221 | 220 | 182 | 437 | 433 | 363 | 376 | 321 | 400 | 440 | 198 | 177 | 198 | 193 | 234 | 187 | 200 | 368 | 452 | 445 | 454 | 342 | 371 | 367 | Zbtb3 |
| Zbtb32 | 18 | 41 | 46 | 35 | 20 | 13 | 19 | 11 | 14 | 24 | 12 | 18 | 18 | 16 | 30 | 23 | 30 | 45 | 34 | 28 | 29 | 8 | 17 | 22 | 17 | 8 | 23 | 18 | Zbtb32 |
| Zbtb33 | 7720 | 5843 | 6613 | 7020 | 7376 | 7850 | 8582 | 9683 | 7097 | 6642 | 6258 | 8147 | 7288 | 8639 | 8157 | 6449 | 6963 | 7125 | 7409 | 8432 | 8306 | 8181 | 6980 | 7179 | 7925 | 6669 | 8464 | 8485 | Zbtb33 |
| Zbtb34 | 3235 | 3111 | 4148 | 4628 | 4197 | 3718 | 3682 | 2242 | 2288 | 2654 | 2761 | 2717 | 2706 | 2598 | 3070 | 3477 | 4085 | 4552 | 3762 | 3040 | 3674 | 3068 | 2284 | 2790 | 2865 | 2471 | 2496 | 2540 | Zbtb34 |
| Zbtb37 | 5394 | 4422 | 3359 | 4745 | 5121 | 5869 | 5612 | 7649 | 6745 | 5934 | 6456 | 5813 | 5426 | 6250 | 5227 | 4917 | 4127 | 4909 | 4789 | 5794 | 5915 | 6163 | 6483 | 5279 | 6542 | 3945 | 6456 | 5614 | Zbtb37 |
| Zbtb38 | 692 | 636 | 714 | 750 | 1431 | 1159 | 1136 | 2324 | 2183 | 2396 | 2049 | 1813 | 1559 | 1735 | 827 | 570 | 776 | 814 | 1284 | 1215 | 1232 | 2446 | 2111 | 2036 | 2511 | 1254 | 1593 | 1533 | Zbtb38 |
| Zbtb39 | 2726 | 3081 | 3869 | 3780 | 3346 | 3151 | 2913 | 2591 | 2742 | 3274 | 2759 | 3987 | 3395 | 3681 | 2666 | 2845 | 3925 | 3368 | 3728 | 2554 | 3315 | 3090 | 2548 | 3476 | 3500 | 3413 | 3445 | 3968 | Zbtb39 |
| Zbtb4 | 1103 | 1190 | 1048 | 1099 | 1647 | 1564 | 1246 | 1707 | 1894 | 1916 | 2254 | 1281 | 1607 | 1197 | 1112 | 1163 | 1068 | 1300 | 1340 | 1267 | 1464 | 1860 | 2155 | 1668 | 1743 | 1098 | 1417 | 1243 | Zbtb4 |
| Zbtb40 | 662 | 792 | 844 | 788 | 689 | 607 | 727 | 1882 | 1974 | 2526 | 2179 | 2193 | 1983 | 2177 | 603 | 631 | 822 | 684 | 1046 | 678 | 703 | 2107 | 1839 | 2554 | 2564 | 1604 | 2100 | 2218 | Zbtb40 |
| Zbtb41 | 9042 | 7705 | 7450 | 10351 | 10037 | 11218 | 11933 | 10025 | 7455 | 6868 | 6804 | 8107 | 6812 | 8485 | 10501 | 8513 | 8780 | 10436 | 10301 | 12397 | 11432 | 8839 | 7633 | 7071 | 8450 | 6296 | 8772 | 8431 | Zbtb41 |
| Zbtb42 | 644 | 648 | 798 | 699 | 679 | 735 | 651 | 242 | 242 | 255 | 206 | 380 | 364 | 331 | 724 | 625 | 825 | 620 | 722 | 610 | 556 | 244 | 302 | 300 | 248 | 408 | 365 | 420 | Zbtb42 |
| Zbtb43 | 3816 | 4251 | 3541 | 4075 | 3730 | 3869 | 3723 | 3046 | 2909 | 2906 | 2767 | 3249 | 2959 | 3270 | 3767 | 4052 | 3648 | 4297 | 3762 | 3585 | 4014 | 3021 | 2647 | 2997 | 3195 | 2835 | 3083 | 3161 | Zbtb43 |
| Zbtb44 | 2592 | 2031 | 3383 | 3330 | 3077 | 2901 | 3101 | 2833 | 2412 | 2827 | 2939 | 3168 | 2993 | 3324 | 2432 | 2441 | 3453 | 3309 | 3243 | 2596 | 3051 | 3392 | 2425 | 3283 | 3350 | 3784 | 2996 | 3291 | Zbtb44 |

| | | | | | | | | | | | | | | | | | | | | | | | | | | | | | |
|---|---|---|---|---|---|---|---|---|---|---|---|---|---|---|---|---|---|---|---|---|---|---|---|---|---|---|---|---|---|
| Zbtb45 | 692 | 703 | 986 | 934 | 965 | 873 | 838 | 1220 | 1353 | 1697 | 1444 | 1803 | 1354 | 1704 | 588 | 590 | 993 | 829 | 1064 | 736 | 856 | 1711 | 1344 | 1898 | 1661 | 1940 | 1404 | 1608 | Zbtb45 |
| Zbtb46 | 3498 | 4038 | 3897 | 4002 | 4058 | 3788 | 3836 | 2014 | 1611 | 1808 | 1793 | 1874 | 1830 | 1744 | 3949 | 3697 | 4051 | 3881 | 3760 | 3986 | 3792 | 2511 | 1612 | 1655 | 2121 | 1434 | 2038 | 1805 | Zbtb46 |
| Zbtb48 | 1415 | 2136 | 1790 | 2256 | 1359 | 1573 | 1476 | 1065 | 1145 | 1190 | 1204 | 1333 | 1237 | 1178 | 1369 | 1794 | 1987 | 1819 | 1633 | 1608 | 1387 | 1206 | 1190 | 1279 | 1242 | 1358 | 1428 | 1191 | Zbtb48 |
| Zbtb49 | 1375 | 1626 | 1110 | 1451 | 1433 | 1435 | 1381 | 1249 | 1264 | 1255 | 1165 | 1302 | 1266 | 1237 | 1302 | 1530 | 1308 | 1566 | 1317 | 1437 | 1385 | 1151 | 1115 | 1119 | 1369 | 947 | 1314 | 1044 | Zbtb49 |
| Zbtb5 | 5660 | 6464 | 5228 | 5958 | 5971 | 5674 | 5580 | 5343 | 4970 | 4947 | 4381 | 5106 | 4812 | 4839 | 5559 | 5566 | 5628 | 5765 | 6131 | 6015 | 6104 | 4913 | 4667 | 4971 | 5538 | 4786 | 5016 | 5079 | Zbtb5 |
| Zbtb6 | 2925 | 2462 | 2828 | 3429 | 3373 | 3327 | 3741 | 4185 | 3485 | 3419 | 3522 | 3410 | 3277 | 3608 | 3241 | 2722 | 2970 | 3506 | 3183 | 3841 | 3655 | 3723 | 3592 | 3640 | 3770 | 3390 | 3469 | 3743 | Zbtb6 |
| Zbtb7a | 1233 | 1473 | 1784 | 2028 | 1831 | 1489 | 1528 | 1371 | 1551 | 1905 | 1920 | 1731 | 1690 | 1798 | 1192 | 1219 | 1990 | 1829 | 1897 | 1362 | 1490 | 1658 | 1626 | 2093 | 1808 | 1989 | 1658 | 1673 | Zbtb7a |
| Zbtb7b | 738 | 1041 | 1009 | 1016 | 720 | 696 | 640 | 270 | 282 | 368 | 306 | 186 | 208 | 172 | 618 | 857 | 946 | 899 | 895 | 677 | 690 | 134 | 394 | 357 | 126 | 401 | 202 | 186 | Zbtb7b |
| Zbtb7c | 271 | 342 | 327 | 478 | 542 | 417 | 387 | 464 | 512 | 574 | 574 | 470 | 301 | 436 | 281 | 304 | 385 | 473 | 469 | 440 | 423 | 772 | 566 | 664 | 601 | 471 | 313 | 368 | Zbtb7c |
| Zbtb8a | 376 | 388 | 432 | 362 | 536 | 440 | 421 | 797 | 825 | 722 | 804 | 792 | 803 | 735 | 436 | 377 | 467 | 362 | 521 | 450 | 456 | 947 | 903 | 766 | 777 | 696 | 798 | 749 | Zbtb8a |
| Zbtb8b | 712 | 897 | 539 | 977 | 1013 | 914 | 910 | 611 | 621 | 510 | 441 | 355 | 407 | 287 | 884 | 965 | 737 | 1005 | 850 | 1089 | 873 | 673 | 606 | 442 | 479 | 255 | 279 | 271 | Zbtb8b |
| Zbtb8os | 1517 | 1316 | 1558 | 1448 | 1222 | 1395 | 1397 | 1366 | 1234 | 1258 | 1347 | 1576 | 1557 | 1657 | 1444 | 1336 | 1326 | 1351 | 1315 | 1428 | 1336 | 1257 | 1349 | 1588 | 1389 | 1578 | 1743 | 1717 | Zbtb8os |
| Zbtb9 | 306 | 326 | 299 | 448 | 324 | 369 | 386 | 182 | 129 | 179 | 149 | 176 | 170 | 229 | 307 | 285 | 336 | 384 | 370 | 308 | 396 | 247 | 132 | 199 | 217 | 164 | 206 | 214 | Zbtb9 |
| Zc2hc1a | 7917 | 6519 | 6658 | 8670 | 9460 | 9269 | 9744 | 7863 | 6422 | 6143 | 5642 | 5627 | 5464 | 5809 | 9505 | 7527 | 7594 | 8876 | 8499 | 10509 | 9369 | 7553 | 6421 | 5779 | 6682 | 5325 | 5251 | 5428 | Zc2hc1a |
| Zc2hc1b | 8 | 10 | 4 | 4 | 2 | 3 | 1 | 1 | 2 | 0 | 9 | 2 | 0 | 1 | 3 | 3 | 7 | 8 | 0 | 1 | 5 | 0 | 1 | 1 | 1 | 4 | 1 | 7 | Zc2hc1b |
| Zc2hc1c | 224 | 285 | 308 | 270 | 268 | 276 | 247 | 122 | 100 | 113 | 124 | 166 | 168 | 116 | 256 | 301 | 299 | 249 | 309 | 269 | 275 | 95 | 152 | 123 | 116 | 138 | 137 | 147 | Zc2hc1c |
| Zc3h10 | 1472 | 1832 | 1784 | 1792 | 1577 | 1412 | 1562 | 1398 | 1397 | 1707 | 1345 | 1737 | 1580 | 1588 | 1246 | 1559 | 1708 | 1665 | 1808 | 1393 | 1570 | 1409 | 1284 | 1729 | 1552 | 1711 | 1625 | 1710 | Zc3h10 |
| Zc3h11a | 1040 | 720 | 545 | 835 | 775 | 1063 | 1049 | 1700 | 1433 | 934 | 1510 | 1176 | 1260 | 1483 | 1085 | 1011 | 542 | 759 | 775 | 930 | 1140 | 1170 | 1490 | 1025 | 1178 | 470 | 1273 | 1275 | Zc3h11a |
| Zc3h12a | 106 | 172 | 138 | 144 | 93 | 95 | 104 | 24 | 54 | 47 | 42 | 28 | 43 | 29 | 79 | 227 | 194 | 142 | 103 | 79 | 99 | 32 | 52 | 42 | 23 | 60 | 48 | 40 | Zc3h12a |
| Zc3h12b | 751 | 542 | 478 | 498 | 598 | 633 | 535 | 651 | 648 | 566 | 696 | 629 | 575 | 598 | 653 | 626 | 574 | 532 | 548 | 622 | 549 | 653 | 613 | 514 | 541 | 476 | 646 | 538 | Zc3h12b |
| Zc3h12c | 4012 | 3771 | 2726 | 3725 | 3688 | 4418 | 4421 | 4774 | 3652 | 3464 | 3372 | 4720 | 4384 | 5240 | 4385 | 3096 | 3233 | 3158 | 3778 | 4867 | 4567 | 3698 | 3359 | 3642 | 4495 | 2973 | 5605 | 4808 | Zc3h12c |
| Zc3h12d | 25 | 24 | 32 | 70 | 42 | 50 | 44 | 10 | 26 | 32 | 49 | 25 | 26 | 14 | 38 | 23 | 44 | 70 | 33 | 27 | 33 | 40 | 25 | 25 | 15 | 23 | 24 | 15 | Zc3h12d |
| Zc3h13 | 12908 | 12925 | 12597 | 15027 | 13287 | 14002 | 15072 | 9236 | 9409 | 10721 | 10001 | 11192 | 10486 | 12086 | 12330 | 12065 | 13451 | 14399 | 13410 | 13669 | 14437 | 9087 | 8211 | 10878 | 11138 | 9448 | 10667 | 11039 | Zc3h13 |
| Zc3h14 | 18468 | 15455 | 14415 | 13517 | 12487 | 13564 | 15414 | 13676 | 11920 | 10317 | 10360 | 12159 | 11426 | 12401 | 18969 | 15397 | 13426 | 13168 | 12886 | 15751 | 14132 | 11304 | 11729 | 10587 | 10369 | 12700 | 12260 | 12506 | Zc3h14 |
| Zc3h15 | 13560 | 12232 | 14618 | 13479 | 12553 | 12459 | 14729 | 12660 | 10306 | 10819 | 10383 | 11745 | 11218 | 13415 | 14884 | 12863 | 13383 | 12776 | 13012 | 13963 | 13286 | 11754 | 10384 | 11806 | 11414 | 13414 | 12085 | 12648 | Zc3h15 |
| Zc3h18 | 6392 | 7474 | 9205 | 8155 | 6717 | 6615 | 6452 | 5729 | 6673 | 8165 | 7362 | 9110 | 8031 | 9102 | 5884 | 6685 | 8321 | 7275 | 7424 | 5982 | 6768 | 6447 | 6084 | 8654 | 7804 | 9875 | 8077 | 9022 | Zc3h18 |
| Zc3h3 | 1318 | 2084 | 2362 | 2409 | 2018 | 1710 | 1619 | 1095 | 1295 | 1628 | 1573 | 1618 | 1322 | 1347 | 1213 | 1837 | 2175 | 2506 | 2138 | 1383 | 1787 | 1491 | 1359 | 1574 | 1587 | 1949 | 1318 | 1377 | Zc3h3 |
| Zc3h4 | 3149 | 3270 | 5198 | 3716 | 4222 | 3617 | 3566 | 7330 | 8993 | 11119 | 9362 | 12189 | 10046 | 11048 | 3026 | 3016 | 4868 | 3620 | 4885 | 3081 | 3613 | 9859 | 8000 | 11811 | 11198 | 10965 | 10068 | 10966 | Zc3h4 |
| Zc3h6 | 1047 | 688 | 767 | 1119 | 992 | 1035 | 1236 | 954 | 854 | 887 | 840 | 1128 | 983 | 1084 | 1094 | 746 | 863 | 889 | 881 | 1024 | 1113 | 978 | 891 | 1019 | 920 | 859 | 939 | 1009 | Zc3h6 |
| Zc3h7a | 11300 | 9228 | 9664 | 9809 | 10184 | 10979 | 11559 | 9355 | 8232 | 8238 | 8862 | 8348 | 8716 | 8572 | 11239 | 10449 | 9631 | 9993 | 10198 | 11688 | 11508 | 8793 | 8287 | 8005 | 8276 | 7100 | 8435 | 8153 | Zc3h7a |
| Zc3h7b | 8155 | 8815 | 11476 | 11358 | 13921 | 11582 | 10724 | 13515 | 16202 | 19517 | 18266 | 20407 | 16380 | 18397 | 8156 | 8158 | 12265 | 11717 | 13296 | 9504 | 12274 | 19153 | 15850 | 20956 | 20349 | 20247 | 17053 | 17813 | Zc3h7b |
| Zc3h8 | 716 | 587 | 845 | 932 | 943 | 853 | 969 | 855 | 751 | 801 | 792 | 810 | 770 | 781 | 816 | 646 | 898 | 878 | 886 | 985 | 852 | 800 | 732 | 865 | 976 | 957 | 679 | 765 | Zc3h8 |
| Zc3hav1 | 391 | 388 | 458 | 267 | 319 | 310 | 247 | 305 | 376 | 371 | 475 | 360 | 786 | 431 | 435 | 500 | 345 | 296 | 282 | 409 | 272 | 329 | 414 | 367 | 300 | 354 | 573 | 722 | Zc3hav1 |
| Zc3hav1l | 2512 | 1762 | 2660 | 2187 | 2718 | 2521 | 2378 | 5388 | 5463 | 5964 | 6040 | 6201 | 6002 | 6067 | 2325 | 2402 | 2520 | 2178 | 2540 | 2045 | 2199 | 6411 | 5350 | 6042 | 5905 | 6047 | 6019 | 5952 | Zc3hav1l |
| Zc3hc1 | 2853 | 3207 | 2564 | 2602 | 2362 | 2479 | 2452 | 2922 | 2899 | 2394 | 2357 | 2843 | 2843 | 2756 | 2934 | 2778 | 2760 | 2360 | 2513 | 2795 | 2582 | 2496 | 2951 | 2453 | 2439 | 3396 | 3002 | 2922 | Zc3hc1 |
| Zc4h2 | 2774 | 2351 | 2587 | 2668 | 3096 | 2985 | 3437 | 6017 | 4359 | 4149 | 3555 | 4442 | 3718 | 5143 | 3417 | 2415 | 2717 | 2820 | 3243 | 3875 | 3480 | 5407 | 4295 | 4683 | 4846 | 4470 | 4371 | 4702 | Zc4h2 |
| Zcchc10 | 490 | 419 | 640 | 394 | 391 | 387 | 486 | 797 | 644 | 661 | 764 | 910 | 797 | 984 | 446 | 444 | 505 | 398 | 419 | 440 | 405 | 719 | 702 | 793 | 711 | 1000 | 1049 | 970 | Zcchc10 |
| Zcchc12 | 7124 | 8303 | 7434 | 9561 | 10099 | 9478 | 10487 | 6630 | 5958 | 6821 | 5062 | 5251 | 4118 | 5547 | 8400 | 6847 | 9084 | 9710 | 10188 | 10549 | 11160 | 8776 | 5773 | 6436 | 7481 | 3922 | 4862 | 4812 | Zcchc12 |
| Zcchc13 | 0 | 0 | 0 | 0 | 1 | 1 | 3 | 2 | 2 | 0 | 6 | 4 | 5 | 0 | 1 | 0 | 1 | 10 | 3 | 2 | 4 | 3 | 0 | 2 | 5 | 3 | 3 | 1 | Zcchc13 |
| Zcchc14 | 10577 | 13221 | 13202 | 16231 | 15338 | 13967 | 14500 | 8896 | 10150 | 12662 | 11501 | 13222 | 11024 | 12534 | 10007 | 11869 | 15425 | 16862 | 15895 | 11196 | 15638 | 12323 | 9073 | 12871 | 13690 | 10854 | 11133 | 11662 | Zcchc14 |
| Zcchc17 | 4146 | 4582 | 4393 | 4158 | 3540 | 3755 | 3950 | 2939 | 2823 | 2978 | 2756 | 2837 | 2601 | 2919 | 4450 | 4328 | 4226 | 4127 | 3955 | 4305 | 3886 | 2535 | 2804 | 2987 | 2894 | 3238 | 2909 | 3071 | Zcchc17 |
| Zcchc18 | 4075 | 4316 | 4071 | 4624 | 6713 | 5621 | 6645 | 7193 | 6697 | 6812 | 5852 | 6159 | 5468 | 6502 | 4486 | 3593 | 4799 | 4918 | 6179 | 6543 | 6227 | 8219 | 6635 | 6793 | 7956 | 6067 | 5504 | 5953 | Zcchc18 |
| Zcchc2 | 3468 | 3550 | 3847 | 4337 | 3701 | 3507 | 3616 | 1704 | 1547 | 1762 | 1721 | 1851 | 1880 | 1671 | 3314 | 3689 | 3891 | 4134 | 3690 | 2864 | 3556 | 1943 | 1544 | 1624 | 1715 | 1702 | 1715 | 1753 | Zcchc2 |
| Zcchc24 | 6127 | 5311 | 5363 | 4518 | 3422 | 4518 | 3174 | 3441 | 4354 | 3739 | 5881 | 3596 | 4625 | 3810 | 5297 | 7244 | 4541 | 5179 | 3289 | 3220 | 3396 | 3938 | 5357 | 4263 | 3406 | 4297 | 3767 | 3653 | Zcchc24 |
| Zcchc3 | 2264 | 2623 | 3727 | 2821 | 2155 | 2426 | 2016 | 2340 | 2742 | 2884 | 3242 | 3628 | 3443 | 3375 | 2048 | 2664 | 3078 | 2514 | 2572 | 2074 | 2021 | 2461 | 2731 | 3409 | 2966 | 4918 | 3737 | 3719 | Zcchc3 |
| Zcchc4 | 1151 | 953 | 1012 | 1030 | 1039 | 1152 | 1079 | 1746 | 1788 | 1760 | 1773 | 2053 | 1917 | 2154 | 1009 | 953 | 1061 | 854 | 1044 | 1122 | 1060 | 1686 | 1711 | 1752 | 1670 | 1811 | 2029 | 2009 | Zcchc4 |
| Zcchc7 | 3645 | 3039 | 3092 | 3316 | 3524 | 3576 | 3881 | 4813 | 4261 | 3883 | 4045 | 3747 | 3833 | 3944 | 3571 | 3241 | 3114 | 3498 | 3409 | 3864 | 3704 | 4736 | 4152 | 3773 | 4129 | 3018 | 3688 | 3826 | Zcchc7 |
| Zcchc8 | 5267 | 5560 | 5979 | 6044 | 5505 | 5575 | 5921 | 5007 | 5145 | 5760 | 5552 | 6221 | 5477 | 6423 | 4851 | 5378 | 5889 | 5989 | 5891 | 5365 | 5755 | 5259 | 4942 | 6203 | 6207 | 6561 | 6080 | 6087 | Zcchc8 |
| Zcchc9 | 1504 | 1367 | 1438 | 1412 | 1410 | 1388 | 1433 | 2273 | 1939 | 2017 | 2018 | 2313 | 2298 | 2521 | 1391 | 1268 | 1381 | 1282 | 1499 | 1467 | 1401 | 2189 | 1926 | 2321 | 2078 | 2502 | 2382 | 2442 | Zcchc9 |
| Zcrb1 | 5297 | 4849 | 4716 | 5073 | 5190 | 5165 | 6302 | 5511 | 4788 | 5017 | 5116 | 5387 | 5068 | 6114 | 5258 | 4747 | 4979 | 4945 | 5146 | 5386 | 5770 | 5137 | 4688 | 5410 | 5434 | 4852 | 5624 | 5798 | Zcrb1 |
| Zcwpw1 | 1441 | 1085 | 1126 | 1118 | 965 | 1218 | 1020 | 1512 | 1217 | 1207 | 1670 | 1208 | 1272 | 1233 | 1178 | 1266 | 1109 | 1066 | 929 | 1044 | 927 | 1280 | 1385 | 1231 | 1053 | 1398 | 1204 | 1094 | Zcwpw1 |
| Zcwpw2 | 8 | 14 | 17 | 20 | 21 | 15 | 18 | 48 | 49 | 58 | 65 | 40 | 44 | 39 | 9 | 11 | 25 | 11 | 13 | 7 | 25 | 72 | 53 | 49 | 48 | 44 | 30 | 43 | Zcwpw2 |
| Zdbf2 | 15574 | 12653 | 11451 | 17667 | 16231 | 21083 | 22134 | 10727 | 9746 | 8437 | 10645 | 11355 | 9205 | 12846 | 15672 | 14395 | 13641 | 16699 | 16721 | 18368 | 22557 | 11102 | 9468 | 8537 | 11788 | 8749 | 13318 | 13008 | Zdbf2 |
| Zdhhc1 | 2693 | 3105 | 2967 | 3306 | 3134 | 3207 | 2754 | 2481 | 2795 | 3025 | 3205 | 2825 | 3149 | 2521 | 2598 | 2967 | 3020 | 3045 | 3080 | 2806 | 2851 | 3001 | 2821 | 2694 | 2843 | 2260 | 2696 | 2500 | Zdhhc1 |
| Zdhhc11 | 1 | 2 | 0 | 3 | 4 | 5 | 2 | 1 | 2 | 0 | 7 | 0 | 0 | 0 | 4 | 5 | 0 | 3 | 2 | 6 | 6 | 0 | 1 | 2 | 1 | 0 | 0 | 1 | Zdhhc11 |
| Zdhhc12 | 967 | 1343 | 1152 | 1043 | 829 | 934 | 932 | 438 | 573 | 412 | 557 | 520 | 558 | 526 | 1000 | 1134 | 1233 | 988 | 947 | 890 | 847 | 475 | 502 | 579 | 529 | 720 | 589 | 538 | Zdhhc12 |
| Zdhhc13 | 3567 | 3239 | 2816 | 3003 | 2900 | 3009 | 3380 | 2849 | 2357 | 2077 | 1993 | 2182 | 2234 | 2292 | 3580 | 3215 | 2837 | 2893 | 2745 | 3490 | 3021 | 2252 | 2358 | 2002 | 2186 | 2153 | 2181 | 2292 | Zdhhc13 |
| Zdhhc14 | 438 | 490 | 398 | 623 | 803 | 583 | 587 | 399 | 447 | 583 | 442 | 375 | 321 | 301 | 436 | 474 | 475 | 753 | 717 | 601 | 634 | 660 | 441 | 437 | 508 | 358 | 259 | 310 | Zdhhc14 |
| Zdhhc15 | 2814 | 2079 | 2278 | 1580 | 1695 | 2026 | 2078 | 2800 | 2701 | 2172 | 2525 | 2329 | 2457 | 2446 | 2912 | 2523 | 1914 | 1825 | 1564 | 2149 | 1759 | 2237 | 2804 | 2030 | 2031 | 2375 | 2708 | 2705 | Zdhhc15 |
| Zdhhc16 | 3505 | 3704 | 3072 | 3451 | 2876 | 3404 | 3378 | 2749 | 2921 | 2779 | 2690 | 2934 | 2869 | 2932 | 3253 | 3220 | 3390 | 3358 | 3283 | 3418 | 3377 | 2568 | 2790 | 2883 | 2986 | 3346 | 2807 | 2909 | Zdhhc16 |
| Zdhhc17 | 4689 | 4057 | 3963 | 4831 | 6041 | 5591 | 5875 | 9732 | 8211 | 7416 | 7905 | 7797 | 7534 | 8097 | 5536 | 4154 | 4330 | 4902 | 5497 | 6296 | 5653 | 9025 | 7888 | 7650 | 8855 | 6349 | 7919 | 7164 | Zdhhc17 |
| Zdhhc18 | 1964 | 2411 | 3209 | 3026 | 2941 | 2739 | 2400 | 1650 | 1968 | 2486 | 2110 | 2506 | 2101 | 2063 | 1884 | 2321 | 3292 | 3051 | 3144 | 2246 | 2599 | 2269 | 1770 | 2684 | 2448 | 2818 | 2142 | 2168 | Zdhhc18 |
| Zdhhc19 | 1 | 2 | 0 | 2 | 2 | 0 | 1 | 2 | 0 | 1 | 0 | 5 | 1 | 4 | 0 | 0 | 0 | 0 | 1 | 3 | 0 | 3 | 3 | 1 | 2 | 0 | 1 | 0 | Zdhhc19 |
| Zdhhc2 | 2881 | 2159 | 2162 | 2487 | 3424 | 2634 | 3653 | 3968 | 3310 | 3365 | 3055 | 3480 | 2444 | 3520 | 3170 | 2219 | 2666 | 2648 | 2962 | 3045 | 3174 | 4114 | 3363 | 3755 | 3393 | 4405 | 3291 | 3437 | Zdhhc2 |
| Zdhhc20 | 10565 | 7809 | 9849 | 9775 | 9278 | 10114 | 10087 | 7605 | 6749 | 7131 | 7342 | 7669 | 7641 | 8044 | 10539 | 10129 | 9279 | 9532 | 9138 | 9606 | 9959 | 7182 | 7270 | 6876 | 7259 | 6967 | 7333 | 7440 | Zdhhc20 |
| Zdhhc21 | 7276 | 5312 | 5196 | 6446 | 6556 | 7375 | 7194 | 6901 | 6386 | 5787 | 6210 | 6656 | 6470 | 6828 | 7204 | 6641 | 5847 | 6031 | 5973 | 7090 | 7371 | 5892 | 6184 | 5485 | 6711 | 4823 | 6746 | 6535 | Zdhhc21 |
| Zdhhc22 | 75 | 114 | 86 | 144 | 396 | 175 | 162 | 360 | 465 | 415 | 516 | 119 | 313 | 126 | 116 | 129 | 124 | 160 | 201 | 189 | 193 | 693 | 342 | 151 | 448 | 64 | 108 | 95 | Zdhhc22 |
| Zdhhc23 | 497 | 452 | 387 | 626 | 861 | 704 | 754 | 315 | 402 | 352 | 333 | 253 | 239 | 213 | 527 | 498 | 528 | 710 | 648 | 656 | 827 | 401 | 390 | 366 | 422 | 169 | 195 | 206 | Zdhhc23 |
| Zdhhc24 | 1600 | 2065 | 1795 | 2066 | 1789 | 1669 | 1675 | 1299 | 1142 | 1373 | 1319 | 1319 | 1430 | 1339 | 1618 | 1657 | 1992 | 1953 | 1667 | 1748 | 1778 | 1452 | 1232 | 1326 | 1528 | 1263 | 1284 | 1404 | Zdhhc24 |
| Zdhhc3 | 6244 | 6378 | 5598 | 5449 | 5677 | 6442 | 6010 | 7217 | 7200 | 7222 | 6838 | 7522 | 7541 | 7679 | 5947 | 5687 | 5708 | 5236 | 5732 | 5776 | 5928 | 6968 | 6909 | 6949 | 7220 | 7536 | 7877 | 7918 | Zdhhc3 |
| Zdhhc4 | 1346 | 1526 | 1308 | 1292 | 1353 | 1290 | 1260 | 1485 | 1743 | 1571 | 1624 | 1841 | 1736 | 1642 | 1172 | 1389 | 1191 | 1278 | 1238 | 1435 | 1325 | 1576 | 1652 | 1597 | 1663 | 1774 | 1734 | 1697 | Zdhhc4 |
| Zdhhc5 | 6921 | 6842 | 6026 | 6208 | 5369 | 5646 | 5741 | 4973 | 5221 | 5181 | 5213 | 5251 | 5116 | 4994 | 6194 | 6688 | 5839 | 6213 | 5870 | 5462 | 5756 | 5160 | 5267 | 4995 | 4874 | 5032 | 5166 | 5025 | Zdhhc5 |
| Zdhhc6 | 4670 | 4420 | 4164 | 3588 | 3193 | 3475 | 3832 | 3832 | 3472 | 2944 | 2993 | 3642 | 3615 | 3740 | 4704 | 3971 | 3844 | 3217 | 3324 | 4114 | 3632 | 2959 | 3417 | 3252 | 3090 | 3891 | 3905 | 4043 | Zdhhc6 |
| Zdhhc7 | 1844 | 2357 | 2282 | 2479 | 1852 | 1826 | 1626 | 1596 | 1671 | 1780 | 1766 | 1859 | 1902 | 1729 | 1756 | 2171 | 2439 | 2138 | 2024 | 1659 | 1751 | 1771 | 1790 | 1874 | 1912 | 2265 | 1894 | 1760 | Zdhhc7 |
| Zdhhc8 | 0 | 0 | 0 | 0 | 0 | 0 | 0 | 0 | 0 | 0 | 0 | 0 | 0 | 0 | 0 | 0 | 0 | 0 | 2 | 0 | 0 | 0 | 0 | 0 | 0 | 0 | 0 | 0 | Zdhhc8 |
| Zdhhc9 | 3931 | 4412 | 4241 | 4618 | 4972 | 4666 | 4937 | 3511 | 3545 | 4017 | 3631 | 3721 | 3475 | 3638 | 3977 | 4111 | 4774 | 5007 | 5013 | 4181 | 5218 | 4918 | 3480 | 3743 | 4055 | 3696 | 3289 | 3519 | Zdhhc9 |
| Zeb1 | 8772 | 9357 | 9308 | 9858 | 11072 | 10803 | 10587 | 11497 | 10270 | 11066 | 11057 | 12749 | 11679 | 12704 | 10097 | 8529 | 9046 | 9169 | 10600 | 11581 | 10834 | 11446 | 10963 | 10716 | 12003 | 10307 | 13606 | 12546 | Zeb1 |
| Zeb2 | 5383 | 4533 | 4730 | 4926 | 6211 | 6266 | 6609 | 13663 | 13693 | 11265 | 10488 | 11270 | 9433 | 11952 | 5916 | 4554 | 5077 | 4859 | 5968 | 6308 | 6297 | 10417 | 11306 | 11838 | 12839 | 9350 | 11183 | 10528 | Zeb2 |
| Zer1 | 4111 | 4904 | 4695 | 6032 | 6164 | 5579 | 5557 | 3953 | 4977 | 5715 | 5369 | 5344 | 4884 | 4832 | 4114 | 4576 | 5262 | 5928 | 6477 | 4958 | 5796 | 5763 | 4622 | 5359 | 5598 | 4041 | 4828 | 4696 | Zer1 |

| | | | | | | | | | | | | | | | | | | | | | | | | | | | | | |
|---|---|---|---|---|---|---|---|---|---|---|---|---|---|---|---|---|---|---|---|---|---|---|---|---|---|---|---|---|---|
| Zfand1 | 2954 | 2496 | 2678 | 2510 | 2411 | 2550 | 2854 | 2492 | 2340 | 2173 | 2314 | 2676 | 2592 | 2629 | 3057 | 2762 | 2544 | 2538 | 2360 | 2993 | 2590 | 2246 | 2335 | 2134 | 2179 | 2645 | 2754 | 2496 | Zfand1 |
| Zfand2a | 2067 | 2384 | 1924 | 2266 | 2247 | 2251 | 2147 | 1871 | 2074 | 2153 | 2049 | 2185 | 2091 | 1923 | 1902 | 2041 | 2169 | 2265 | 2247 | 2076 | 2040 | 2154 | 2035 | 2108 | 2294 | 2152 | 1998 | 2051 | Zfand2a |
| Zfand2b | 821 | 1148 | 875 | 966 | 830 | 825 | 822 | 742 | 874 | 794 | 789 | 818 | 720 | 764 | 882 | 904 | 892 | 756 | 940 | 914 | 784 | 705 | 859 | 806 | 675 | 981 | 718 | 833 | Zfand2b |
| Zfand3 | 6734 | 8776 | 9601 | 9217 | 8836 | 8137 | 7456 | 7188 | 7814 | 9000 | 8006 | 9607 | 9003 | 8701 | 6996 | 7238 | 10045 | 8644 | 9587 | 7823 | 7784 | 7656 | 7745 | 9748 | 9029 | 10915 | 9386 | 9003 | Zfand3 |
| Zfand4 | 1927 | 1390 | 1851 | 1612 | 1644 | 1686 | 1964 | 1104 | 917 | 941 | 809 | 1351 | 1185 | 1359 | 1895 | 1498 | 1702 | 1676 | 1760 | 1881 | 1755 | 954 | 929 | 1093 | 1058 | 1374 | 1323 | 1444 | Zfand4 |
| Zfand5 | 11565 | 8849 | 10536 | 12785 | 12260 | 12256 | 12930 | 16339 | 13528 | 13367 | 13564 | 15135 | 14596 | 15860 | 12002 | 10328 | 10684 | 11968 | 11620 | 11700 | 12786 | 17662 | 13796 | 14274 | 15777 | 14353 | 15012 | 15892 | Zfand5 |
| Zfand6 | 13898 | 10908 | 14274 | 14173 | 13322 | 12750 | 15088 | 10695 | 8922 | 8752 | 8976 | 9993 | 9530 | 9944 | 15217 | 13261 | 13928 | 14155 | 13485 | 14584 | 13446 | 9899 | 9427 | 9295 | 9063 | 9986 | 9503 | 10263 | Zfand6 |
| Zfat | 85 | 131 | 91 | 74 | 123 | 108 | 106 | 692 | 723 | 761 | 702 | 691 | 710 | 801 | 80 | 99 | 137 | 57 | 148 | 113 | 90 | 662 | 759 | 826 | 741 | 833 | 826 | 794 | Zfat |
| Zfc3h1 | 18334 | 15933 | 11905 | 17047 | 15501 | 18107 | 17541 | 12384 | 10397 | 9574 | 10356 | 10087 | 10223 | 9949 | 18046 | 17175 | 13667 | 15936 | 14923 | 17223 | 17815 | 10863 | 9898 | 8208 | 9887 | 6316 | 10425 | 9077 | Zfc3h1 |
| Zfhx2 | 5132 | 7080 | 8585 | 11074 | 12229 | 9805 | 9673 | 6618 | 6958 | 10782 | 8767 | 12386 | 9104 | 10747 | 5280 | 5556 | 11047 | 10817 | 12424 | 8044 | 10654 | 12773 | 6141 | 11860 | 11422 | 7562 | 9793 | 9781 | Zfhx2 |
| Zfhx3 | 3893 | 3731 | 6703 | 6287 | 6250 | 6351 | 4906 | 7028 | 7004 | 13044 | 10680 | 12402 | 13213 | 11247 | 3646 | 4347 | 6889 | 6155 | 6560 | 4281 | 5879 | 13785 | 7058 | 13216 | 13042 | 9894 | 11024 | 11896 | Zfhx3 |
| Zfhx4 | 6819 | 5538 | 5927 | 5415 | 6391 | 7563 | 5965 | 19838 | 18325 | 19017 | 19189 | 18309 | 24621 | 19053 | 7030 | 6893 | 5478 | 4787 | 5379 | 6284 | 6479 | 18544 | 16643 | 14684 | 17950 | 12544 | 22331 | 19190 | Zfhx4 |
| Zfp1 | 879 | 793 | 898 | 1030 | 1104 | 1081 | 1122 | 1523 | 1495 | 1492 | 1642 | 1497 | 1472 | 1538 | 822 | 758 | 875 | 975 | 1128 | 1000 | 1042 | 1601 | 1441 | 1478 | 1478 | 1618 | 1488 | 1442 | Zfp1 |
| Zfp101 | 5542 | 3835 | 4034 | 3525 | 3034 | 3421 | 3537 | 4704 | 4268 | 3699 | 3825 | 4279 | 4240 | 4399 | 5098 | 4864 | 3442 | 3131 | 2927 | 4127 | 3344 | 3604 | 3987 | 3515 | 3456 | 3808 | 4735 | 4348 | Zfp101 |
| Zfp105 | 1428 | 1218 | 1470 | 1248 | 1248 | 1222 | 1386 | 2201 | 1772 | 1895 | 1716 | 2171 | 1946 | 2332 | 1316 | 1210 | 1331 | 1043 | 1188 | 1321 | 1214 | 1928 | 1693 | 2095 | 1960 | 2401 | 2409 | 2479 | Zfp105 |
| Zfp106 | 27472 | 24955 | 17687 | 19603 | 18567 | 21238 | 20466 | 20783 | 19442 | 17322 | 19950 | 19806 | 19085 | 18933 | 25322 | 24411 | 18149 | 19848 | 18417 | 21382 | 20471 | 16407 | 19621 | 16459 | 18058 | 13908 | 20828 | 18358 | Zfp106 |
| Zfp108 | 363 | 357 | 246 | 251 | 389 | 403 | 345 | 524 | 459 | 423 | 413 | 301 | 341 | 393 | 363 | 310 | 255 | 310 | 295 | 386 | 331 | 386 | 498 | 307 | 427 | 240 | 374 | 280 | Zfp108 |
| Zfp109 | 379 | 358 | 339 | 337 | 385 | 390 | 382 | 484 | 516 | 507 | 469 | 454 | 481 | 454 | 377 | 379 | 392 | 330 | 459 | 332 | 346 | 418 | 467 | 440 | 442 | 308 | 413 | 446 | Zfp109 |
| Zfp11 | 829 | 856 | 977 | 844 | 954 | 956 | 948 | 1010 | 957 | 839 | 866 | 920 | 855 | 832 | 940 | 735 | 897 | 869 | 877 | 1071 | 866 | 949 | 836 | 938 | 855 | 911 | 952 | 946 | Zfp11 |
| Zfp110 | 2942 | 2632 | 2779 | 2277 | 2824 | 2840 | 2966 | 3676 | 3415 | 2955 | 2806 | 2852 | 2761 | 2853 | 3005 | 2509 | 2728 | 2355 | 3010 | 3319 | 2792 | 2976 | 3443 | 3075 | 2880 | 2913 | 2962 | 2909 | Zfp110 |
| Zfp111 | 1311 | 1055 | 1332 | 1026 | 1198 | 1253 | 1240 | 2083 | 1941 | 1657 | 1976 | 2059 | 2005 | 2035 | 1146 | 1113 | 1177 | 1138 | 1148 | 1253 | 1220 | 1785 | 2065 | 1834 | 1894 | 1880 | 2223 | 2209 | Zfp111 |
| Zfp112 | 501 | 492 | 314 | 424 | 551 | 605 | 525 | 485 | 606 | 489 | 481 | 419 | 425 | 424 | 524 | 467 | 459 | 432 | 444 | 647 | 600 | 476 | 533 | 406 | 493 | 351 | 442 | 371 | Zfp112 |
| Zfp113 | 2874 | 2621 | 2866 | 2803 | 2788 | 2649 | 3070 | 4686 | 3752 | 3639 | 3614 | 4439 | 3948 | 4838 | 2900 | 2507 | 2939 | 2699 | 3010 | 3027 | 2943 | 3778 | 3676 | 4196 | 4503 | 3952 | 4756 | 4710 | Zfp113 |
| Zfp114 | 210 | 179 | 84 | 134 | 195 | 187 | 203 | 391 | 423 | 355 | 377 | 354 | 356 | 385 | 196 | 150 | 161 | 156 | 252 | 263 | 237 | 281 | 386 | 348 | 407 | 255 | 447 | 339 | Zfp114 |
| Zfp119a | 617 | 371 | 274 | 355 | 488 | 571 | 596 | 865 | 783 | 559 | 615 | 720 | 767 | 700 | 562 | 488 | 378 | 359 | 483 | 683 | 592 | 620 | 710 | 553 | 692 | 451 | 776 | 698 | Zfp119a |
| Zfp119b | 287 | 262 | 315 | 273 | 290 | 284 | 324 | 466 | 438 | 333 | 360 | 384 | 401 | 453 | 322 | 227 | 304 | 237 | 339 | 320 | 323 | 505 | 441 | 400 | 414 | 439 | 404 | 433 | Zfp119b |
| Zfp12 | 3173 | 3549 | 3441 | 3451 | 3437 | 3514 | 3361 | 4399 | 3864 | 3936 | 3605 | 3898 | 3657 | 3988 | 3129 | 3068 | 3356 | 3220 | 3536 | 3402 | 3600 | 4034 | 4043 | 4026 | 4087 | 3513 | 3780 | 3855 | Zfp12 |
| Zfp120 | 1525 | 1195 | 1656 | 1527 | 1664 | 1803 | 1943 | 1975 | 1561 | 1564 | 1356 | 1606 | 1471 | 1638 | 1858 | 1521 | 1662 | 1555 | 1698 | 2174 | 1761 | 1669 | 1639 | 1570 | 1644 | 1549 | 1567 | 1709 | Zfp120 |
| Zfp128 | 1151 | 1170 | 967 | 1087 | 1240 | 1152 | 1268 | 1611 | 1299 | 1380 | 1088 | 1153 | 1048 | 1219 | 1242 | 938 | 1148 | 1029 | 1262 | 1342 | 1297 | 1432 | 1331 | 1206 | 1411 | 1129 | 1167 | 1214 | Zfp128 |
| Zfp13 | 1614 | 1953 | 1900 | 1951 | 1794 | 1791 | 1563 | 1535 | 1786 | 2012 | 1977 | 2134 | 2021 | 1971 | 1524 | 1775 | 1889 | 1741 | 1964 | 1707 | 1659 | 1708 | 1576 | 1996 | 2028 | 2064 | 2030 | 1949 | Zfp13 |
| Zfp131 | 6381 | 5189 | 5602 | 5670 | 5399 | 5836 | 6357 | 7757 | 6163 | 5687 | 5453 | 6649 | 6074 | 7101 | 6472 | 5544 | 5567 | 5678 | 5695 | 6618 | 6195 | 6066 | 5873 | 6250 | 6386 | 6830 | 6580 | 6579 | Zfp131 |
| Zfp14 | 852 | 870 | 839 | 915 | 1007 | 979 | 1033 | 1139 | 984 | 1101 | 835 | 969 | 764 | 955 | 974 | 790 | 874 | 879 | 1100 | 1004 | 1075 | 1173 | 979 | 968 | 960 | 817 | 829 | 847 | Zfp14 |
| Zfp141 | 2400 | 1859 | 2246 | 2383 | 2680 | 2686 | 2806 | 2383 | 2149 | 2017 | 2035 | 2081 | 2106 | 2315 | 2407 | 2081 | 2395 | 2439 | 2607 | 2729 | 2525 | 2209 | 2110 | 2202 | 2134 | 2203 | 2107 | 2115 | Zfp141 |
| Zfp142 | 4236 | 5125 | 3822 | 5298 | 4642 | 4907 | 4449 | 2666 | 3011 | 3301 | 3151 | 3451 | 3196 | 3075 | 3938 | 4566 | 4225 | 4997 | 4648 | 4039 | 5042 | 2991 | 2581 | 3049 | 3436 | 2446 | 3331 | 3081 | Zfp142 |
| Zfp143 | 2970 | 2805 | 2837 | 2435 | 2302 | 2315 | 2786 | 2982 | 2663 | 2774 | 2271 | 3213 | 2626 | 3012 | 2900 | 2433 | 2680 | 2453 | 2576 | 2960 | 2575 | 2558 | 2405 | 2772 | 2543 | 3166 | 2986 | 3125 | Zfp143 |
| Zfp146 | 6414 | 6173 | 6282 | 5455 | 6018 | 5953 | 6223 | 10462 | 9788 | 8773 | 8141 | 10125 | 9738 | 10156 | 6434 | 6119 | 6289 | 5733 | 6260 | 7309 | 5822 | 8808 | 9236 | 9211 | 9314 | 11561 | 10656 | 10581 | Zfp146 |
| Zfp148 | 9471 | 7593 | 6485 | 7764 | 9085 | 9903 | 10502 | 12082 | 10827 | 9333 | 9868 | 10118 | 9424 | 10494 | 9643 | 8034 | 7392 | 7911 | 8676 | 10025 | 9945 | 9961 | 10423 | 9243 | 10150 | 8223 | 10235 | 10066 | Zfp148 |
| Zfp157 | 5224 | 4380 | 4242 | 4992 | 5533 | 5444 | 5833 | 9126 | 7359 | 6974 | 6540 | 6847 | 6561 | 7171 | 5898 | 4670 | 4826 | 4842 | 5376 | 6112 | 5655 | 8560 | 7244 | 6834 | 7715 | 5954 | 6943 | 6806 | Zfp157 |
| Zfp160 | 4071 | 2835 | 3694 | 3529 | 3599 | 4025 | 4205 | 4847 | 4237 | 3956 | 4136 | 4299 | 4160 | 4550 | 4019 | 3597 | 3530 | 3415 | 3741 | 4061 | 3869 | 4453 | 3932 | 4016 | 4128 | 4149 | 4364 | 4467 | Zfp160 |
| Zfp169 | 1311 | 1621 | 535 | 1373 | 1283 | 1408 | 1236 | 1070 | 970 | 857 | 912 | 766 | 816 | 908 | 1311 | 1208 | 992 | 1356 | 1227 | 1399 | 1537 | 983 | 940 | 725 | 991 | 254 | 992 | 805 | Zfp169 |
| Zfp174 | 830 | 792 | 507 | 684 | 776 | 816 | 734 | 702 | 663 | 609 | 559 | 459 | 550 | 425 | 857 | 750 | 552 | 694 | 626 | 833 | 784 | 618 | 683 | 484 | 570 | 416 | 497 | 417 | Zfp174 |
| Zfp180 | 2593 | 2591 | 2789 | 2604 | 3161 | 2674 | 2835 | 4247 | 3667 | 3521 | 3279 | 3826 | 3708 | 3820 | 2603 | 2366 | 2607 | 2487 | 3024 | 3079 | 2830 | 4044 | 3730 | 3802 | 3832 | 3868 | 3949 | 3780 | Zfp180 |
| Zfp182 | 1275 | 1113 | 1229 | 1483 | 1727 | 1709 | 1753 | 2351 | 1862 | 2058 | 1901 | 1729 | 1958 | 1924 | 1501 | 1285 | 1443 | 1405 | 1577 | 1713 | 1623 | 2212 | 1776 | 1880 | 1867 | 1527 | 1891 | 1728 | Zfp182 |
| Zfp184 | 2461 | 2193 | 1982 | 2145 | 1853 | 2013 | 2101 | 2481 | 2167 | 1964 | 1675 | 2284 | 2033 | 2229 | 2517 | 2107 | 2114 | 2130 | 2039 | 2478 | 2003 | 2027 | 2008 | 2176 | 2117 | 2439 | 2238 | 2187 | Zfp184 |
| Zfp185 | 771 | 588 | 527 | 385 | 205 | 583 | 211 | 653 | 827 | 598 | 1039 | 506 | 778 | 480 | 824 | 873 | 319 | 513 | 243 | 204 | 189 | 580 | 1218 | 498 | 398 | 468 | 494 | 516 | Zfp185 |
| Zfp189 | 2025 | 1933 | 1832 | 1968 | 1958 | 1960 | 2086 | 2004 | 1554 | 1476 | 1351 | 1461 | 1499 | 1487 | 2210 | 1938 | 1936 | 1970 | 1862 | 2388 | 1984 | 1683 | 1551 | 1314 | 1525 | 1448 | 1531 | 1544 | Zfp189 |
| Zfp2 | 1945 | 1489 | 1644 | 1658 | 2386 | 2172 | 2456 | 4423 | 3508 | 3398 | 2884 | 3732 | 3317 | 3804 | 2334 | 1629 | 1825 | 1692 | 2130 | 2612 | 2277 | 3985 | 3302 | 3447 | 3792 | 3119 | 3539 | 3482 | Zfp2 |
| Zfp202 | 1204 | 1231 | 1109 | 1218 | 1175 | 1325 | 1380 | 1391 | 1357 | 1337 | 1386 | 1325 | 1283 | 1258 | 1226 | 1076 | 1065 | 1304 | 1362 | 1236 | 1270 | 1462 | 1284 | 1248 | 1228 | 1263 | 1211 | 1234 | Zfp202 |
| Zfp207 | 25494 | 21469 | 23343 | 24620 | 22698 | 24785 | 25807 | 34467 | 27698 | 26825 | 27641 | 30313 | 29165 | 31257 | 26170 | 22805 | 23432 | 23270 | 25024 | 28180 | 25425 | 28973 | 27171 | 30512 | 29923 | 32055 | 31479 | 32201 | Zfp207 |
| Zfp212 | 2139 | 2363 | 2384 | 2443 | 2044 | 2053 | 2209 | 1918 | 1956 | 2109 | 1937 | 2250 | 2141 | 2173 | 2086 | 2058 | 2533 | 2330 | 2380 | 2069 | 2074 | 1893 | 2039 | 2286 | 2066 | 2573 | 2282 | 2224 | Zfp212 |
| Zfp213 | 1243 | 1506 | 1351 | 1285 | 1068 | 1199 | 1059 | 1109 | 1210 | 1181 | 1004 | 1274 | 1090 | 1058 | 1171 | 1236 | 1409 | 1161 | 1284 | 1231 | 1105 | 954 | 1079 | 1193 | 984 | 1399 | 1168 | 1040 | Zfp213 |
| Zfp217 | 3750 | 3744 | 5053 | 4772 | 3976 | 3722 | 4004 | 2332 | 2476 | 2793 | 2832 | 2870 | 3005 | 3159 | 3381 | 3586 | 4552 | 4306 | 4132 | 3309 | 3980 | 2003 | 2550 | 3201 | 2398 | 3386 | 2712 | 3133 | Zfp217 |
| Zfp219 | 4084 | 5212 | 6190 | 5327 | 4997 | 4487 | 4358 | 7049 | 7152 | 7906 | 6799 | 10756 | 8327 | 9893 | 3757 | 3952 | 6236 | 4522 | 6090 | 4440 | 4573 | 7449 | 6270 | 10497 | 8813 | 12003 | 9776 | 9976 | Zfp219 |
| Zfp229 | 1433 | 1094 | 1423 | 1500 | 1399 | 1581 | 1579 | 2236 | 1889 | 2076 | 1759 | 2099 | 2085 | 2073 | 1485 | 1313 | 1262 | 1361 | 1387 | 1532 | 1359 | 1913 | 2041 | 1813 | 1982 | 1606 | 1984 | 2002 | Zfp229 |
| Zfp235 | 1220 | 1359 | 967 | 1275 | 1330 | 1333 | 1227 | 1596 | 1339 | 1291 | 1366 | 1262 | 1227 | 1152 | 1167 | 1208 | 1275 | 1385 | 1299 | 1416 | 1314 | 1638 | 1259 | 1286 | 1433 | 1123 | 1146 | 1070 | Zfp235 |
| Zfp236 | 2129 | 2165 | 1972 | 2434 | 3096 | 2686 | 2835 | 5014 | 5360 | 5590 | 5729 | 5753 | 5126 | 5488 | 2130 | 2157 | 2199 | 2524 | 3041 | 2566 | 2784 | 5285 | 5269 | 5069 | 6022 | 3779 | 5445 | 5061 | Zfp236 |
| Zfp239 | 1264 | 1217 | 1201 | 1180 | 1202 | 1171 | 1340 | 1049 | 1055 | 1074 | 928 | 986 | 1116 | 1162 | 1209 | 1172 | 1275 | 1214 | 1216 | 1297 | 1143 | 946 | 942 | 1041 | 989 | 1105 | 980 | 1092 | Zfp239 |
| Zfp24 | 6116 | 5621 | 6045 | 5802 | 6088 | 6158 | 6273 | 11611 | 8814 | 8216 | 7821 | 10553 | 9596 | 11190 | 7083 | 5715 | 6159 | 5667 | 5994 | 7199 | 6278 | 9588 | 8679 | 9018 | 9788 | 10120 | 11037 | 11179 | Zfp24 |
| Zfp248 | 965 | 926 | 1083 | 1071 | 1150 | 1061 | 1154 | 1566 | 1303 | 1527 | 1449 | 1367 | 1480 | 1479 | 953 | 995 | 1079 | 1019 | 1182 | 1107 | 1113 | 1694 | 1324 | 1399 | 1451 | 1516 | 1293 | 1406 | Zfp248 |
| Zfp251 | 6018 | 6509 | 5668 | 6631 | 6374 | 6523 | 6305 | 4412 | 4512 | 4963 | 4545 | 4625 | 4262 | 4320 | 6039 | 5940 | 6136 | 6683 | 6667 | 6152 | 6478 | 4786 | 4535 | 4639 | 5171 | 4707 | 4571 | 4507 | Zfp251 |
| Zfp26 | 6200 | 4199 | 4496 | 4832 | 4944 | 5294 | 5068 | 8225 | 7412 | 5752 | 6368 | 6777 | 6829 | 6773 | 6223 | 4932 | 4627 | 4665 | 4747 | 5664 | 5169 | 6654 | 6752 | 6198 | 6376 | 5598 | 7403 | 6597 | Zfp26 |
| Zfp260 | 6522 | 4935 | 5674 | 5046 | 5809 | 5839 | 6043 | 13082 | 11014 | 9456 | 9874 | 10107 | 9932 | 10084 | 6465 | 6059 | 5522 | 5277 | 5668 | 6800 | 5769 | 10858 | 11003 | 10055 | 10105 | 10565 | 10308 | 10900 | Zfp260 |
| Zfp263 | 3744 | 3496 | 3234 | 3440 | 3210 | 3281 | 3423 | 3866 | 3553 | 3436 | 3341 | 3624 | 3269 | 3688 | 3348 | 3258 | 3057 | 3308 | 3155 | 3292 | 3387 | 3648 | 3421 | 3405 | 3813 | 3413 | 3488 | 3684 | Zfp263 |
| Zfp266 | 19484 | 15177 | 14216 | 15878 | 17356 | 18435 | 19412 | 23637 | 19618 | 17701 | 18455 | 18641 | 18671 | 20372 | 18780 | 16045 | 15122 | 16901 | 16948 | 19577 | 19442 | 18808 | 18788 | 18228 | 19446 | 15887 | 20075 | 19949 | Zfp266 |
| Zfp267 | 4094 | 3806 | 3023 | 3795 | 3890 | 4013 | 3981 | 3777 | 3412 | 2940 | 2818 | 2776 | 2844 | 2810 | 4461 | 3916 | 3080 | 3705 | 3511 | 4126 | 3811 | 3367 | 3086 | 2530 | 2959 | 2068 | 2896 | 2746 | Zfp267 |
| Zfp268 | 84 | 50 | 120 | 32 | 52 | 71 | 49 | 76 | 70 | 81 | 105 | 92 | 126 | 55 | 102 | 118 | 65 | 64 | 48 | 41 | 50 | 77 | 135 | 71 | 32 | 105 | 112 | 102 | Zfp268 |
| Zfp27 | 1630 | 1654 | 1293 | 1467 | 1628 | 1742 | 1698 | 2158 | 2037 | 1885 | 1873 | 1944 | 1854 | 1859 | 1816 | 1528 | 1491 | 1598 | 1640 | 1689 | 1579 | 2070 | 1959 | 2073 | 2021 | 1692 | 2160 | 1773 | Zfp27 |
| Zfp273 | 574 | 320 | 487 | 436 | 596 | 564 | 649 | 1051 | 986 | 890 | 816 | 871 | 839 | 858 | 611 | 453 | 468 | 465 | 600 | 675 | 584 | 882 | 950 | 862 | 900 | 970 | 833 | 857 | Zfp273 |
| Zfp275 | 5729 | 5256 | 4391 | 4599 | 4561 | 5392 | 4966 | 5405 | 5544 | 4643 | 5631 | 4825 | 4812 | 4646 | 5437 | 5460 | 4421 | 4940 | 4864 | 5083 | 5062 | 4722 | 5444 | 4246 | 4719 | 4174 | 5019 | 5097 | Zfp275 |
| Zfp276 | 2714 | 3628 | 2707 | 3017 | 2898 | 2769 | 2558 | 2142 | 2353 | 2469 | 2314 | 2395 | 2328 | 2171 | 2487 | 3208 | 3037 | 3010 | 2975 | 2731 | 2610 | 2500 | 2267 | 2373 | 2591 | 2097 | 2337 | 2160 | Zfp276 |
| Zfp277 | 4849 | 4209 | 3755 | 3777 | 3301 | 3524 | 3934 | 2514 | 2282 | 1891 | 1889 | 1912 | 1852 | 1848 | 5061 | 4534 | 3595 | 3778 | 3599 | 4224 | 3818 | 2074 | 2378 | 1849 | 1881 | 2312 | 1966 | 2092 | Zfp277 |
| Zfp28 | 1907 | 1691 | 1383 | 1449 | 1613 | 1598 | 1736 | 2219 | 1991 | 1871 | 1878 | 2082 | 1970 | 2065 | 1855 | 1673 | 1501 | 1427 | 1459 | 1662 | 1770 | 2064 | 1820 | 1867 | 1941 | 1703 | 2141 | 1908 | Zfp28 |
| Zfp280b | 3041 | 3181 | 3578 | 2932 | 2403 | 2495 | 2701 | 2181 | 1996 | 2054 | 2096 | 2596 | 2455 | 2521 | 2760 | 3114 | 3238 | 2619 | 2614 | 2492 | 2422 | 2042 | 1968 | 2373 | 2128 | 2585 | 3059 | 2720 | Zfp280b |
| Zfp280c | 4422 | 2878 | 2996 | 2978 | 3741 | 4041 | 3940 | 5352 | 4708 | 4764 | 4834 | 5684 | 5374 | 5648 | 4129 | 3288 | 2920 | 2996 | 3240 | 3644 | 3598 | 4998 | 4403 | 4333 | 4634 | 4532 | 5775 | 5220 | Zfp280c |
| Zfp280d | 4357 | 3222 | 4506 | 4328 | 4947 | 4624 | 4971 | 9165 | 7785 | 7652 | 7277 | 8504 | 8138 | 9005 | 4723 | 3720 | 4436 | 3925 | 4789 | 4773 | 4580 | 9167 | 7512 | 7943 | 8057 | 8403 | 8653 | 8802 | Zfp280d |
| Zfp281 | 7621 | 6474 | 6555 | 6694 | 6298 | 7177 | 7168 | 7569 | 5931 | 5521 | 5507 | 6535 | 6311 | 6797 | 7499 | 6787 | 6793 | 6625 | 6355 | 7134 | 7138 | 6657 | 5505 | 5875 | 6771 | 5753 | 7217 | 6916 | Zfp281 |
| Zfp282 | 2485 | 2598 | 2758 | 3022 | 3498 | 3097 | 2771 | 3756 | 4172 | 4081 | 3976 | 4098 | 3818 | 3987 | 2389 | 2294 | 3080 | 2775 | 3527 | 2847 | 3099 | 4261 | 3956 | 4464 | 4571 | 4067 | 3756 | 4069 | Zfp282 |
| Zfp286 | 1780 | 1651 | 1455 | 1639 | 1941 | 1833 | 1976 | 2827 | 2574 | 2431 | 2368 | 2159 | 2203 | 2255 | 1857 | 1536 | 1586 | 1705 | 2104 | 2228 | 1999 | 2567 | 2434 | 2240 | 2563 | 2279 | 2334 | 2214 | Zfp286 |
| Zfp287 | 1581 | 1508 | 1298 | 1614 | 1720 | 1825 | 1741 | 1909 | 1879 | 1855 | 1829 | 1689 | 1571 | 1640 | 1659 | 1491 | 1336 | 1566 | 1548 | 1872 | 1762 | 1870 | 1758 | 1510 | 2005 | 1272 | 1711 | 1518 | Zfp287 |

| | | | | | | | | | | | | | | | | | | | | | | | | | | | | | |
|---|---|---|---|---|---|---|---|---|---|---|---|---|---|---|---|---|---|---|---|---|---|---|---|---|---|---|---|---|---|
| Zfp292 | 12980 | 10172 | 6583 | 9374 | 10512 | 12647 | 12320 | 23132 | 18743 | 15402 | 14683 | 16986 | 15842 | 17325 | 13842 | 10573 | 8357 | 8604 | 10319 | 14783 | 12122 | 15299 | 17189 | 13569 | 17005 | 10255 | 18928 | 15957 | Zfp292 |
| Zfp296 | 24 | 29 | 62 | 14 | 20 | 4 | 12 | 33 | 26 | 42 | 19 | 29 | 22 | 27 | 9 | 19 | 24 | 26 | 21 | 19 | 13 | 30 | 32 | 44 | 31 | 33 | 35 | 19 | Zfp296 |
| Zfp3 | 685 | 650 | 686 | 777 | 814 | 778 | 742 | 805 | 670 | 762 | 702 | 846 | 725 | 724 | 728 | 594 | 686 | 728 | 768 | 745 | 825 | 810 | 707 | 706 | 753 | 586 | 716 | 741 | Zfp3 |
| Zfp30 | 1741 | 1664 | 1706 | 1855 | 2047 | 1925 | 1994 | 1690 | 1550 | 1774 | 1629 | 1701 | 1500 | 1628 | 1691 | 1478 | 1781 | 1787 | 2209 | 1931 | 2060 | 1860 | 1562 | 1800 | 1746 | 1612 | 1592 | 1557 | Zfp30 |
| Zfp300 | 429 | 353 | 375 | 399 | 583 | 570 | 562 | 525 | 465 | 454 | 365 | 382 | 401 | 380 | 492 | 378 | 415 | 425 | 401 | 614 | 541 | 463 | 434 | 287 | 356 | 284 | 353 | 377 | Zfp300 |
| Zfp316 | 2781 | 3609 | 4214 | 4763 | 4566 | 4054 | 4021 | 3482 | 3989 | 4786 | 4222 | 4740 | 4106 | 4457 | 2952 | 3012 | 5076 | 4768 | 5071 | 3480 | 4295 | 4687 | 3625 | 5149 | 5374 | 4347 | 4338 | 4348 | Zfp316 |
| Zfp317 | 4592 | 4363 | 4058 | 4328 | 4675 | 4559 | 5026 | 6544 | 5675 | 5382 | 5074 | 5681 | 5372 | 5902 | 5003 | 4470 | 4403 | 4483 | 4493 | 5127 | 4970 | 5803 | 5541 | 5476 | 5810 | 5323 | 5839 | 5518 | Zfp317 |
| Zfp318 | 9172 | 7871 | 6845 | 8444 | 9717 | 9839 | 9659 | 9646 | 8596 | 9050 | 9385 | 8826 | 9088 | 9032 | 9171 | 8495 | 7903 | 8176 | 8859 | 8809 | 10034 | 9722 | 8231 | 7852 | 9805 | 5943 | 8685 | 8693 | Zfp318 |
| Zfp319 | 3697 | 4686 | 5243 | 6531 | 6313 | 5340 | 5123 | 4043 | 4285 | 5377 | 4719 | 5277 | 4458 | 4815 | 3927 | 4157 | 6311 | 6571 | 6987 | 4187 | 5952 | 6109 | 3961 | 5592 | 5763 | 5048 | 4430 | 4948 | Zfp319 |
| Zfp322a | 9185 | 7671 | 8502 | 8700 | 7927 | 8235 | 9397 | 8529 | 6943 | 6512 | 6793 | 7022 | 6922 | 7744 | 9090 | 9134 | 8281 | 8353 | 7849 | 9403 | 8592 | 7046 | 6800 | 6987 | 7298 | 6781 | 6995 | 7376 | Zfp322a |
| Zfp324 | 2348 | 2795 | 2290 | 2710 | 2534 | 2454 | 2496 | 2130 | 2144 | 1948 | 2157 | 1832 | 1814 | 1947 | 2474 | 2326 | 2606 | 2574 | 2608 | 2510 | 2598 | 2117 | 2058 | 1923 | 2040 | 1601 | 1765 | 1918 | Zfp324 |
| Zfp326 | 6608 | 4736 | 6721 | 6084 | 5939 | 6227 | 6876 | 8675 | 7449 | 7491 | 8094 | 8486 | 8653 | 9762 | 6367 | 5344 | 6107 | 5454 | 5995 | 6384 | 6060 | 7984 | 7041 | 8419 | 8531 | 8350 | 8929 | 9517 | Zfp326 |
| Zfp329 | 6330 | 4769 | 4969 | 5780 | 6431 | 7322 | 7201 | 8672 | 7035 | 6957 | 6602 | 6839 | 6878 | 7694 | 6521 | 5382 | 5205 | 5654 | 6072 | 6825 | 7479 | 7536 | 6644 | 6480 | 7809 | 4952 | 7554 | 7005 | Zfp329 |
| Zfp330 | 4523 | 4241 | 4328 | 4385 | 3724 | 3914 | 4040 | 5258 | 4484 | 4173 | 4005 | 4573 | 4276 | 4672 | 4797 | 4511 | 4461 | 4090 | 3967 | 4428 | 3824 | 4291 | 4411 | 4615 | 4186 | 4837 | 4682 | 4906 | Zfp330 |
| Zfp334 | 5218 | 5175 | 5309 | 5180 | 4928 | 5421 | 4998 | 6592 | 6054 | 6443 | 5676 | 7444 | 7058 | 7368 | 5102 | 4689 | 5482 | 4718 | 5723 | 4905 | 5101 | 6316 | 5652 | 6596 | 6780 | 6696 | 8344 | 7752 | Zfp334 |
| Zfp335 | 1538 | 1968 | 1659 | 2032 | 2181 | 1895 | 1799 | 2271 | 2886 | 3123 | 2776 | 3354 | 2784 | 2629 | 1449 | 1680 | 1820 | 2040 | 2258 | 1722 | 1804 | 3080 | 2669 | 3102 | 3100 | 2836 | 2739 | 2497 | Zfp335 |
| Zfp341 | 113 | 136 | 198 | 208 | 295 | 192 | 171 | 658 | 854 | 922 | 824 | 871 | 705 | 808 | 131 | 161 | 196 | 254 | 241 | 153 | 236 | 946 | 744 | 981 | 1010 | 1013 | 757 | 711 | Zfp341 |
| Zfp345 | 1 | 3 | 2 | 5 | 7 | 4 | 2 | 1 | 9 | 10 | 8 | 1 | 10 | 3 | 8 | 2 | 1 | 11 | 10 | 9 | 8 | 3 | 14 | 2 | 2 | 1 | 14 | 1 | Zfp345 |
| Zfp346 | 4555 | 5075 | 4317 | 4571 | 3925 | 4283 | 4329 | 2610 | 2810 | 2626 | 2969 | 2824 | 2941 | 2627 | 4603 | 4597 | 4187 | 4579 | 4216 | 4130 | 4174 | 2734 | 2876 | 2765 | 2693 | 2896 | 2775 | 2677 | Zfp346 |
| Zfp35 | 3098 | 2961 | 2667 | 2420 | 2606 | 2588 | 2778 | 4040 | 3078 | 2804 | 2447 | 3069 | 2891 | 2930 | 3282 | 2914 | 2656 | 2469 | 2465 | 3356 | 2752 | 3106 | 3150 | 2882 | 2859 | 3407 | 3039 | 3090 | Zfp35 |
| Zfp352 | 0 | 0 | 0 | 1 | 0 | 0 | 0 | 0 | 0 | 0 | 0 | 0 | 0 | 0 | 0 | 0 | 0 | 0 | 0 | 0 | 0 | 0 | 0 | 1 | 0 | 0 | 0 | 0 | Zfp352 |
| Zfp354a | 1113 | 966 | 1091 | 1192 | 1313 | 1248 | 1179 | 1446 | 1221 | 1110 | 1241 | 1113 | 1140 | 1105 | 1221 | 1060 | 1162 | 1201 | 1222 | 1270 | 1213 | 1409 | 1225 | 1222 | 1355 | 1068 | 1154 | 1194 | Zfp354a |
| Zfp354b | 234 | 243 | 225 | 231 | 331 | 349 | 278 | 300 | 243 | 216 | 222 | 203 | 155 | 187 | 297 | 251 | 216 | 300 | 281 | 329 | 311 | 304 | 237 | 219 | 233 | 185 | 180 | 160 | Zfp354b |
| Zfp354c | 10942 | 8854 | 8082 | 10174 | 11734 | 12006 | 12247 | 12469 | 10729 | 10429 | 11552 | 9454 | 9417 | 9924 | 11041 | 9436 | 9749 | 10953 | 11162 | 11538 | 12619 | 11528 | 10736 | 9992 | 11339 | 8194 | 9490 | 9702 | Zfp354c |
| Zfp358 | 2057 | 2522 | 2011 | 1881 | 2153 | 1925 | 1997 | 3480 | 3459 | 3327 | 2878 | 3224 | 3034 | 3264 | 2025 | 1843 | 2232 | 1625 | 2342 | 2106 | 2080 | 3405 | 3190 | 3496 | 3153 | 3498 | 3400 | 3208 | Zfp358 |
| Zfp36 | 355 | 477 | 389 | 387 | 328 | 349 | 292 | 209 | 363 | 401 | 368 | 336 | 356 | 279 | 327 | 495 | 335 | 482 | 345 | 244 | 297 | 307 | 388 | 382 | 275 | 401 | 244 | 259 | Zfp36 |
| Zfp362 | 2461 | 2801 | 4070 | 3282 | 3614 | 3047 | 2558 | 4388 | 5526 | 6771 | 6263 | 7797 | 6289 | 6670 | 2273 | 2540 | 3964 | 2835 | 3841 | 2299 | 2716 | 6370 | 5022 | 7654 | 6798 | 9201 | 6306 | 6547 | Zfp362 |
| Zfp365 | 573 | 447 | 508 | 429 | 586 | 497 | 504 | 375 | 391 | 437 | 417 | 530 | 517 | 474 | 493 | 526 | 599 | 568 | 517 | 395 | 523 | 418 | 432 | 452 | 464 | 487 | 473 | 494 | Zfp365 |
| Zfp366 | 227 | 234 | 374 | 240 | 401 | 323 | 275 | 419 | 476 | 515 | 525 | 508 | 557 | 500 | 229 | 241 | 219 | 305 | 403 | 352 | 355 | 592 | 515 | 538 | 502 | 471 | 476 | 483 | Zfp366 |
| Zfp367 | 3493 | 2503 | 3330 | 3071 | 2476 | 2721 | 3305 | 2199 | 2203 | 1958 | 1791 | 2917 | 2403 | 3069 | 3212 | 2798 | 3094 | 2937 | 2586 | 3070 | 2894 | 1693 | 1924 | 2595 | 2137 | 3202 | 2931 | 3030 | Zfp367 |
| Zfp369 | 2654 | 2153 | 2850 | 2491 | 2839 | 2872 | 2596 | 3690 | 3395 | 3042 | 3349 | 3327 | 3295 | 2750 | 2719 | 2365 | 2847 | 2600 | 3055 | 2875 | 2437 | 3457 | 3029 | 3086 | 3341 | 3332 | 3241 | 2651 | Zfp369 |
| Zfp36l1 | 9372 | 7988 | 14142 | 10566 | 7733 | 7221 | 7558 | 11176 | 11426 | 11384 | 13436 | 14144 | 13681 | 14422 | 8375 | 9584 | 11596 | 9198 | 8257 | 6742 | 7780 | 10664 | 12835 | 14296 | 10473 | 19436 | 13323 | 16154 | Zfp36l1 |
| Zfp36l2 | 2646 | 2480 | 3307 | 2554 | 2142 | 2184 | 1780 | 2964 | 3385 | 3001 | 3773 | 3230 | 3773 | 2946 | 2541 | 3195 | 2629 | 2593 | 1858 | 1629 | 1779 | 3228 | 3587 | 3381 | 2666 | 4092 | 3069 | 3350 | Zfp36l2 |
| Zfp36l3 | 3 | 1 | 2 | 1 | 0 | 1 | 1 | 0 | 0 | 0 | 2 | 0 | 2 | 0 | 0 | 0 | 1 | 1 | 4 | 1 | 0 | 0 | 1 | 2 | 0 | 0 | 1 | 0 | Zfp36l3 |
| Zfp37 | 2392 | 1935 | 2591 | 2200 | 2669 | 2460 | 2464 | 4063 | 3210 | 3262 | 3029 | 3460 | 3538 | 3413 | 2685 | 2118 | 2423 | 2183 | 2660 | 2907 | 2220 | 4092 | 3205 | 3368 | 3298 | 3359 | 3782 | 3548 | Zfp37 |
| Zfp382 | 801 | 878 | 322 | 787 | 817 | 830 | 891 | 723 | 712 | 714 | 583 | 613 | 623 | 712 | 816 | 791 | 509 | 763 | 782 | 787 | 886 | 608 | 627 | 479 | 652 | 139 | 676 | 500 | Zfp382 |
| Zfp383 | 2222 | 1842 | 2157 | 2232 | 2428 | 2214 | 2500 | 3920 | 3522 | 3535 | 3818 | 3668 | 3772 | 4053 | 2338 | 1902 | 2370 | 2163 | 2387 | 2428 | 2480 | 4114 | 3603 | 3939 | 3910 | 3774 | 4007 | 3870 | Zfp383 |
| Zfp384 | 4081 | 4372 | 3973 | 4329 | 4561 | 4586 | 4427 | 6036 | 6171 | 6562 | 6422 | 6574 | 5877 | 6178 | 4189 | 3776 | 4344 | 4255 | 4671 | 4118 | 4533 | 6911 | 5795 | 6427 | 6474 | 6119 | 5950 | 5838 | Zfp384 |
| Zfp385a | 3790 | 5212 | 6211 | 5295 | 4656 | 4179 | 3454 | 2105 | 2335 | 2687 | 2689 | 3001 | 2701 | 2591 | 3614 | 4388 | 5839 | 4884 | 5179 | 3159 | 4036 | 2887 | 2426 | 3001 | 2605 | 3232 | 2635 | 2838 | Zfp385a |
| Zfp385b | 140 | 128 | 129 | 194 | 231 | 215 | 199 | 199 | 274 | 303 | 419 | 186 | 158 | 176 | 148 | 111 | 146 | 179 | 227 | 154 | 186 | 376 | 304 | 305 | 242 | 186 | 179 | 209 | Zfp385b |
| Zfp385c | 52 | 47 | 23 | 15 | 26 | 25 | 17 | 45 | 58 | 32 | 48 | 24 | 18 | 20 | 32 | 35 | 21 | 17 | 23 | 14 | 12 | 40 | 32 | 29 | 27 | 36 | 16 | 6 | Zfp385c |
| Zfp386 | 4486 | 3334 | 3538 | 3600 | 3833 | 4379 | 4273 | 5199 | 4500 | 4410 | 4060 | 4773 | 4478 | 4375 | 4709 | 3924 | 3736 | 3554 | 3881 | 4966 | 4279 | 4505 | 4467 | 4204 | 4229 | 4610 | 4835 | 4421 | Zfp386 |
| Zfp39 | 1333 | 1252 | 1168 | 1332 | 996 | 1146 | 1315 | 1147 | 1160 | 1027 | 1010 | 1109 | 978 | 1024 | 1374 | 1140 | 1016 | 1140 | 990 | 1175 | 1086 | 1156 | 1040 | 1136 | 1025 | 1165 | 952 | 1013 | Zfp39 |
| Zfp395 | 3301 | 3059 | 3863 | 3449 | 2455 | 2675 | 2249 | 1663 | 2398 | 2289 | 2834 | 2073 | 2185 | 1960 | 2569 | 3667 | 3333 | 3316 | 2817 | 1962 | 2409 | 1990 | 2386 | 2354 | 1753 | 2657 | 1863 | 2086 | Zfp395 |
| Zfp397 | 12243 | 9827 | 8852 | 9865 | 10131 | 11909 | 12260 | 13842 | 11372 | 11344 | 11053 | 12906 | 12264 | 13915 | 12159 | 10641 | 8894 | 9673 | 9902 | 11647 | 12068 | 11306 | 11084 | 10753 | 12151 | 9488 | 12921 | 12462 | Zfp397 |
| Zfp398 | 5106 | 4488 | 3057 | 4432 | 4874 | 5219 | 5370 | 4621 | 4252 | 3995 | 4103 | 3503 | 3655 | 3686 | 5167 | 4414 | 3888 | 4385 | 4517 | 5317 | 5606 | 4461 | 3857 | 3486 | 4356 | 2421 | 3671 | 3566 | Zfp398 |
| Zfp40 | 2846 | 2230 | 3097 | 3103 | 3550 | 3493 | 3686 | 3377 | 2898 | 3231 | 2808 | 3052 | 2639 | 3076 | 2934 | 2522 | 3160 | 3297 | 3451 | 3566 | 3479 | 3603 | 2698 | 2872 | 3026 | 2814 | 2730 | 2937 | Zfp40 |
| Zfp407 | 1669 | 1601 | 1230 | 1374 | 1478 | 1688 | 1563 | 3606 | 3326 | 2710 | 2914 | 3170 | 3064 | 3299 | 1774 | 1593 | 1381 | 1216 | 1513 | 1653 | 1507 | 2896 | 3265 | 2693 | 3057 | 1740 | 3392 | 2975 | Zfp407 |
| Zfp408 | 2283 | 2484 | 2945 | 2831 | 2409 | 2533 | 2338 | 1997 | 2252 | 2812 | 2595 | 2979 | 2884 | 2913 | 1941 | 2228 | 2690 | 2527 | 2786 | 2097 | 2449 | 2319 | 2127 | 3130 | 2984 | 3173 | 3133 | 2927 | Zfp408 |
| Zfp41 | 14794 | 16415 | 13327 | 12798 | 12523 | 13503 | 13373 | 16668 | 13869 | 13922 | 11481 | 19326 | 16677 | 18936 | 15634 | 13274 | 14949 | 11948 | 13690 | 14295 | 13590 | 14222 | 13082 | 14538 | 15509 | 14151 | 20694 | 19574 | Zfp41 |
| Zfp410 | 7690 | 7790 | 6654 | 6705 | 5774 | 6224 | 7175 | 6708 | 5708 | 4768 | 4945 | 5856 | 5577 | 6359 | 7789 | 6923 | 6602 | 6239 | 6435 | 7147 | 6842 | 5411 | 5351 | 5395 | 5472 | 6102 | 6571 | 6715 | Zfp410 |
| Zfp414 | 442 | 523 | 572 | 507 | 599 | 453 | 529 | 895 | 1012 | 1173 | 1065 | 1219 | 984 | 1150 | 414 | 329 | 532 | 474 | 602 | 521 | 504 | 1091 | 951 | 1159 | 1070 | 1336 | 1196 | 1205 | Zfp414 |
| Zfp418 | 451 | 478 | 306 | 491 | 469 | 486 | 403 | 570 | 475 | 429 | 393 | 415 | 404 | 454 | 475 | 414 | 367 | 461 | 462 | 465 | 472 | 541 | 469 | 398 | 478 | 315 | 436 | 403 | Zfp418 |
| Zfp42 | 2 | 2 | 4 | 1 | 2 | 3 | 0 | 0 | 2 | 3 | 1 | 0 | 0 | 1 | 8 | 5 | 0 | 0 | 2 | 0 | 0 | 0 | 1 | 0 | 0 | 1 | 3 | 0 | Zfp42 |
| Zfp420 | 1353 | 1346 | 1389 | 1532 | 1730 | 1572 | 1857 | 2125 | 1678 | 1908 | 1896 | 1640 | 1564 | 1725 | 1472 | 1441 | 1698 | 1922 | 1687 | 1516 | 1826 | 2495 | 1711 | 1877 | 2055 | 1512 | 1582 | 1747 | Zfp420 |
| Zfp422 | 7381 | 6221 | 7944 | 5937 | 6223 | 6463 | 7196 | 13522 | 11682 | 10903 | 9949 | 14370 | 12517 | 15425 | 6974 | 6368 | 7086 | 5263 | 6756 | 7162 | 6340 | 11411 | 11057 | 12777 | 11917 | 16338 | 14970 | 15989 | Zfp422 |
| Zfp423 | 1392 | 1779 | 2444 | 2167 | 2412 | 2034 | 1703 | 2889 | 3273 | 4230 | 4002 | 4399 | 4628 | 3942 | 1460 | 1845 | 2401 | 2014 | 2492 | 1593 | 1863 | 4568 | 3223 | 4461 | 4240 | 4521 | 4238 | 4438 | Zfp423 |
| Zfp426 | 6046 | 5642 | 4790 | 5240 | 5181 | 5413 | 5915 | 5853 | 4563 | 4333 | 4403 | 4707 | 4525 | 5049 | 6231 | 5449 | 5018 | 4989 | 5311 | 6186 | 5907 | 5053 | 4702 | 4323 | 4726 | 4380 | 4891 | 4851 | Zfp426 |
| Zfp428 | 700 | 906 | 899 | 1078 | 1415 | 1263 | 1048 | 2323 | 2474 | 2954 | 2491 | 2711 | 2239 | 2623 | 736 | 593 | 1037 | 1077 | 1446 | 1206 | 1273 | 2894 | 2311 | 3040 | 3172 | 2666 | 2343 | 2352 | Zfp428 |
| Zfp429 | 400 | 289 | 392 | 236 | 333 | 349 | 311 | 480 | 390 | 338 | 386 | 348 | 354 | 308 | 356 | 357 | 379 | 331 | 319 | 375 | 289 | 426 | 502 | 317 | 331 | 421 | 338 | 388 | Zfp429 |
| Zfp433 | 157 | 136 | 137 | 99 | 119 | 169 | 149 | 288 | 306 | 236 | 251 | 268 | 382 | 305 | 189 | 142 | 140 | 121 | 132 | 201 | 125 | 322 | 279 | 257 | 197 | 219 | 339 | 268 | Zfp433 |
| Zfp438 | 164 | 180 | 221 | 160 | 182 | 172 | 184 | 353 | 353 | 363 | 281 | 391 | 359 | 349 | 174 | 143 | 176 | 147 | 232 | 178 | 142 | 349 | 366 | 378 | 352 | 305 | 330 | 379 | Zfp438 |
| Zfp442 | 690 | 596 | 283 | 478 | 557 | 641 | 698 | 641 | 571 | 465 | 517 | 469 | 504 | 467 | 785 | 732 | 422 | 402 | 488 | 755 | 604 | 358 | 561 | 326 | 449 | 124 | 503 | 428 | Zfp442 |
| Zfp444 | 3397 | 3980 | 4238 | 4922 | 4556 | 4006 | 4507 | 2672 | 2521 | 2964 | 2565 | 2813 | 2625 | 2713 | 3432 | 3413 | 4899 | 4867 | 5082 | 3912 | 4765 | 3141 | 2506 | 3222 | 3202 | 2958 | 2708 | 2910 | Zfp444 |
| Zfp445 | 20780 | 18063 | 15576 | 18204 | 21728 | 22110 | 22521 | 32615 | 27652 | 26717 | 27074 | 27238 | 26822 | 28512 | 21601 | 18485 | 17176 | 18329 | 19537 | 23325 | 21949 | 28047 | 25989 | 24552 | 28848 | 18845 | 28293 | 26040 | Zfp445 |
| Zfp446 | 1096 | 1316 | 1380 | 1273 | 1237 | 1210 | 1328 | 1726 | 1673 | 1887 | 1669 | 1979 | 1833 | 1967 | 1071 | 1084 | 1475 | 1310 | 1428 | 1280 | 1350 | 1978 | 1458 | 1855 | 1727 | 1651 | 2038 | 2006 | Zfp446 |
| Zfp449 | 1272 | 963 | 1133 | 1105 | 1065 | 1052 | 989 | 1188 | 1056 | 987 | 1081 | 915 | 1008 | 1001 | 1279 | 1247 | 1027 | 984 | 1005 | 1099 | 1012 | 1160 | 1189 | 948 | 997 | 1021 | 973 | 977 | Zfp449 |
| Zfp451 | 6499 | 5405 | 6725 | 7230 | 8909 | 7571 | 8853 | 10517 | 9190 | 9875 | 9407 | 9690 | 9161 | 10294 | 6593 | 5523 | 7405 | 7784 | 8514 | 7621 | 8283 | 11789 | 8631 | 9917 | 10903 | 9110 | 9614 | 10181 | Zfp451 |
| Zfp454 | 1163 | 1130 | 1039 | 1058 | 1308 | 1237 | 1127 | 1303 | 1144 | 1073 | 1076 | 1201 | 1244 | 1137 | 1178 | 951 | 1145 | 1092 | 1269 | 1377 | 1179 | 1235 | 1094 | 1218 | 1299 | 1225 | 1433 | 1308 | Zfp454 |
| Zfp455 | 89 | 60 | 86 | 79 | 147 | 136 | 153 | 324 | 289 | 246 | 212 | 176 | 227 | 191 | 104 | 76 | 116 | 101 | 156 | 139 | 118 | 274 | 296 | 210 | 236 | 230 | 181 | 203 | Zfp455 |
| Zfp456 | 129 | 101 | 112 | 101 | 110 | 152 | 103 | 217 | 173 | 171 | 188 | 118 | 137 | 125 | 142 | 161 | 104 | 98 | 116 | 143 | 85 | 139 | 192 | 102 | 123 | 123 | 139 | 155 | Zfp456 |
| Zfp457 | 85 | 66 | 45 | 76 | 116 | 90 | 69 | 38 | 49 | 69 | 57 | 17 | 56 | 48 | 58 | 48 | 48 | 102 | 74 | 96 | 72 | 90 | 52 | 59 | 44 | 43 | 41 | 33 | Zfp457 |
| Zfp458 | 766 | 656 | 441 | 612 | 762 | 697 | 826 | 822 | 682 | 603 | 543 | 516 | 488 | 554 | 885 | 675 | 596 | 633 | 686 | 887 | 755 | 730 | 695 | 512 | 555 | 355 | 566 | 478 | Zfp458 |
| Zfp459 | 171 | 119 | 172 | 161 | 167 | 162 | 174 | 112 | 123 | 87 | 105 | 103 | 98 | 107 | 137 | 160 | 151 | 140 | 164 | 181 | 192 | 69 | 101 | 126 | 95 | 169 | 103 | 109 | Zfp459 |
| Zfp46 | 4026 | 4689 | 4492 | 5039 | 4999 | 4754 | 4329 | 4280 | 4140 | 4387 | 4324 | 4107 | 4097 | 3746 | 4147 | 4025 | 4749 | 5238 | 5429 | 4622 | 4916 | 4756 | 4225 | 4339 | 4971 | 3419 | 3846 | 3789 | Zfp46 |
| Zfp462 | 6819 | 6214 | 8136 | 8227 | 10369 | 9159 | 9095 | 18608 | 19237 | 23935 | 21113 | 28630 | 26752 | 28810 | 6687 | 6593 | 8338 | 7405 | 10174 | 7624 | 9069 | 22491 | 16518 | 24898 | 28396 | 21127 | 28574 | 27681 | Zfp462 |
| Zfp467 | 618 | 625 | 663 | 748 | 994 | 851 | 772 | 2050 | 2267 | 2575 | 2547 | 2496 | 2462 | 2395 | 541 | 590 | 695 | 691 | 912 | 670 | 806 | 2257 | 2206 | 2693 | 2288 | 2642 | 2159 | 2324 | Zfp467 |
| Zfp469 | 181 | 320 | 406 | 294 | 241 | 340 | 145 | 271 | 722 | 928 | 1624 | 617 | 898 | 482 | 186 | 372 | 357 | 391 | 189 | 101 | 139 | 1094 | 1001 | 755 | 850 | 428 | 681 | 622 | Zfp469 |
| Zfp472 | 1354 | 927 | 1169 | 1078 | 918 | 903 | 1158 | 1038 | 977 | 792 | 882 | 928 | 971 | 1036 | 1299 | 1259 | 1080 | 1069 | 895 | 1167 | 917 | 1001 | 995 | 1004 | 967 | 1359 | 1066 | 1116 | Zfp472 |

| | | | | | | | | | | | | | | | | | | | | | | | | | | | | | |
|---|---|---|---|---|---|---|---|---|---|---|---|---|---|---|---|---|---|---|---|---|---|---|---|---|---|---|---|---|---|
| Zfp473 | 400 | 356 | 368 | 235 | 206 | 222 | 209 | 245 | 238 | 210 | 197 | 263 | 213 | 270 | 357 | 300 | 321 | 209 | 249 | 277 | 216 | 172 | 230 | 267 | 221 | 296 | 303 | 297 | Zfp473 |
| Zfp474 | 3 | 5 | 0 | 2 | 9 | 0 | 1 | 12 | 17 | 19 | 14 | 14 | 32 | 1 | 4 | 9 | 3 | 1 | 5 | 7 | 4 | 23 | 28 | 3 | 11 | 5 | 5 | 6 | Zfp474 |
| Zfp488 | 131 | 126 | 98 | 79 | 106 | 124 | 55 | 67 | 92 | 78 | 82 | 33 | 60 | 36 | 108 | 178 | 119 | 101 | 92 | 82 | 79 | 82 | 92 | 65 | 70 | 35 | 29 | 34 | Zfp488 |
| Zfp493 | 631 | 619 | 674 | 556 | 677 | 615 | 615 | 551 | 479 | 447 | 470 | 447 | 458 | 471 | 733 | 670 | 576 | 598 | 577 | 637 | 557 | 426 | 496 | 429 | 432 | 440 | 393 | 515 | Zfp493 |
| Zfp503 | 2802 | 2491 | 4248 | 2920 | 3027 | 2259 | 2321 | 3055 | 3114 | 3325 | 4595 | 2763 | 3166 | 2836 | 2739 | 2307 | 3306 | 2603 | 2809 | 1768 | 2505 | 3115 | 3674 | 3161 | 2539 | 3143 | 2626 | 2959 | Zfp503 |
| Zfp507 | 4749 | 4292 | 3577 | 3860 | 3941 | 4387 | 4199 | 5490 | 5583 | 5256 | 5462 | 5801 | 5551 | 5659 | 4425 | 4101 | 3918 | 3752 | 3909 | 4063 | 4332 | 4659 | 5277 | 5234 | 5239 | 4295 | 5724 | 5577 | Zfp507 |
| Zfp51 | 771 | 553 | 686 | 657 | 661 | 712 | 700 | 1186 | 928 | 899 | 652 | 872 | 874 | 864 | 803 | 731 | 620 | 609 | 701 | 874 | 680 | 976 | 1010 | 855 | 789 | 896 | 729 | 879 | Zfp51 |
| Zfp511 | 836 | 1037 | 1105 | 1062 | 1057 | 940 | 1029 | 1188 | 1213 | 1379 | 1317 | 1429 | 1262 | 1402 | 795 | 863 | 1169 | 941 | 1088 | 924 | 995 | 1348 | 1186 | 1521 | 1424 | 1534 | 1319 | 1304 | Zfp511 |
| Zfp512 | 6614 | 6701 | 7195 | 7891 | 8093 | 7759 | 8042 | 9424 | 8655 | 9294 | 8559 | 9484 | 8523 | 9594 | 7006 | 5948 | 7645 | 7630 | 7849 | 7838 | 8105 | 9979 | 8242 | 9386 | 9830 | 8975 | 9209 | 9193 | Zfp512 |
| Zfp512b | 7022 | 7415 | 7952 | 9502 | 10725 | 10162 | 9183 | 10247 | 11810 | 13703 | 13344 | 13009 | 11932 | 11636 | 6804 | 7013 | 9027 | 9756 | 10460 | 8315 | 9513 | 13832 | 11072 | 13348 | 14026 | 11065 | 11656 | 10987 | Zfp512b |
| Zfp513 | 1052 | 1195 | 1261 | 1332 | 1349 | 1236 | 1157 | 1546 | 1763 | 1973 | 1828 | 2139 | 1926 | 1872 | 965 | 893 | 1416 | 1296 | 1382 | 1159 | 1186 | 1883 | 1652 | 2000 | 1913 | 2291 | 1937 | 1875 | Zfp513 |
| Zfp516 | 8947 | 9528 | 9474 | 9727 | 8826 | 8498 | 8030 | 3979 | 4818 | 4884 | 5207 | 5589 | 5248 | 5100 | 7762 | 9111 | 10020 | 9376 | 9696 | 7507 | 8370 | 4069 | 4536 | 5425 | 4531 | 5218 | 4928 | 5040 | Zfp516 |
| Zfp518a | 2841 | 1868 | 2154 | 2047 | 2423 | 2443 | 2432 | 4667 | 4280 | 3773 | 3678 | 4322 | 4384 | 4130 | 2772 | 2229 | 2170 | 1817 | 2390 | 2819 | 2364 | 3561 | 4233 | 3650 | 3483 | 3294 | 4416 | 4081 | Zfp518a |
| Zfp518b | 5891 | 5284 | 5871 | 6216 | 7348 | 7201 | 7201 | 9649 | 9039 | 10025 | 9259 | 11031 | 10401 | 11490 | 5480 | 5254 | 5947 | 6113 | 7337 | 6236 | 7356 | 10059 | 8307 | 10160 | 11231 | 8223 | 11338 | 11337 | Zfp518b |
| Zfp52 | 660 | 464 | 665 | 527 | 583 | 710 | 601 | 347 | 368 | 309 | 360 | 265 | 348 | 285 | 616 | 674 | 421 | 585 | 577 | 616 | 517 | 289 | 349 | 336 | 317 | 300 | 285 | 241 | Zfp52 |
| Zfp521 | 3907 | 4102 | 4049 | 3813 | 4342 | 4422 | 3803 | 7155 | 7761 | 7805 | 7454 | 7500 | 8852 | 7341 | 4168 | 4314 | 3970 | 3861 | 3930 | 3784 | 3554 | 7992 | 7586 | 7363 | 7320 | 7391 | 7553 | 7861 | Zfp521 |
| Zfp523 | 2249 | 2827 | 2830 | 3192 | 3571 | 3104 | 2615 | 3324 | 3886 | 4447 | 3922 | 4449 | 4165 | 3739 | 2359 | 2425 | 3281 | 3166 | 3559 | 2647 | 2890 | 4286 | 3599 | 4243 | 4294 | 4669 | 4206 | 3607 | Zfp523 |
| Zfp524 | 16 | 44 | 34 | 40 | 34 | 34 | 36 | 59 | 72 | 77 | 95 | 85 | 105 | 65 | 32 | 24 | 42 | 47 | 44 | 43 | 51 | 50 | 78 | 83 | 76 | 91 | 65 | 66 | Zfp524 |
| Zfp526 | 959 | 1104 | 1339 | 1075 | 893 | 926 | 850 | 808 | 887 | 958 | 989 | 1087 | 1017 | 1001 | 778 | 929 | 1119 | 1099 | 991 | 726 | 878 | 934 | 937 | 1068 | 1052 | 908 | 955 | 915 | Zfp526 |
| Zfp53 | 1135 | 929 | 1030 | 940 | 907 | 972 | 1104 | 868 | 812 | 740 | 770 | 811 | 787 | 777 | 1098 | 895 | 890 | 945 | 962 | 1042 | 960 | 747 | 812 | 770 | 676 | 718 | 686 | 774 | Zfp53 |
| Zfp532 | 3715 | 3709 | 4206 | 4001 | 5026 | 4684 | 4075 | 10815 | 11814 | 13404 | 11515 | 13563 | 12529 | 12089 | 3779 | 3566 | 4609 | 3609 | 5306 | 4268 | 4226 | 12746 | 10389 | 12957 | 13743 | 11035 | 12595 | 12221 | Zfp532 |
| Zfp534 | 0 | 2 | 4 | 7 | 0 | 3 | 1 | 1 | 3 | 1 | 1 | 3 | 3 | 0 | 3 | 5 | 2 | 2 | 2 | 0 | 1 | 3 | 1 | 0 | 1 | 0 | 0 | 1 | Zfp534 |
| Zfp536 | 2150 | 2547 | 3828 | 4634 | 5122 | 4555 | 4361 | 2757 | 3019 | 4494 | 3520 | 4323 | 3875 | 4201 | 2648 | 2347 | 4253 | 4414 | 4807 | 3298 | 4324 | 4289 | 2775 | 3742 | 4581 | 3479 | 3934 | 4006 | Zfp536 |
| Zfp54 | 250 | 179 | 270 | 200 | 337 | 330 | 324 | 264 | 305 | 280 | 274 | 239 | 253 | 243 | 297 | 234 | 268 | 223 | 249 | 264 | 228 | 281 | 298 | 232 | 223 | 201 | 246 | 228 | Zfp54 |
| Zfp541 | 13 | 27 | 12 | 13 | 34 | 29 | 27 | 7 | 17 | 16 | 22 | 12 | 9 | 5 | 15 | 22 | 20 | 40 | 23 | 20 | 20 | 13 | 21 | 9 | 14 | 3 | 6 | 8 | Zfp541 |
| Zfp551 | 309 | 282 | 154 | 275 | 488 | 407 | 335 | 761 | 622 | 573 | 518 | 467 | 488 | 411 | 284 | 172 | 253 | 224 | 401 | 471 | 390 | 685 | 610 | 422 | 538 | 220 | 433 | 349 | Zfp551 |
| Zfp553 | 2708 | 3203 | 4499 | 4140 | 4091 | 3662 | 3422 | 3375 | 3603 | 4680 | 4484 | 5503 | 4501 | 5296 | 2499 | 2900 | 4515 | 3876 | 4527 | 2834 | 3766 | 4306 | 3482 | 5363 | 5191 | 6263 | 4871 | 5504 | Zfp553 |
| Zfp558 | 1480 | 1304 | 1117 | 1625 | 2311 | 2243 | 2272 | 3184 | 2851 | 2919 | 2519 | 2502 | 2088 | 2482 | 1577 | 1211 | 1403 | 1811 | 1993 | 2150 | 2031 | 3501 | 2698 | 2317 | 3312 | 1771 | 2377 | 2176 | Zfp558 |
| Zfp560 | 814 | 625 | 777 | 747 | 1190 | 1121 | 1168 | 1856 | 1795 | 1729 | 1785 | 1568 | 1694 | 1725 | 854 | 710 | 829 | 847 | 1029 | 1002 | 995 | 1977 | 1526 | 1679 | 1877 | 1426 | 1576 | 1582 | Zfp560 |
| Zfp563 | 1742 | 1475 | 1572 | 1546 | 2044 | 2178 | 2204 | 3352 | 2811 | 2568 | 2487 | 2294 | 2121 | 2401 | 1929 | 1463 | 1577 | 1615 | 1879 | 2399 | 2235 | 3133 | 2699 | 2437 | 2878 | 2012 | 2316 | 2243 | Zfp563 |
| Zfp566 | 593 | 526 | 597 | 573 | 476 | 477 | 581 | 706 | 564 | 624 | 648 | 764 | 616 | 733 | 546 | 551 | 579 | 500 | 519 | 553 | 570 | 750 | 550 | 719 | 763 | 720 | 686 | 756 | Zfp566 |
| Zfp568 | 5285 | 4534 | 3656 | 3258 | 2832 | 3176 | 2538 | 3013 | 3041 | 2768 | 4265 | 3026 | 3614 | 2891 | 4080 | 5072 | 3322 | 3027 | 2579 | 2701 | 2686 | 2665 | 3703 | 2573 | 2444 | 2426 | 3491 | 3172 | Zfp568 |
| Zfp57 | 6174 | 5372 | 5977 | 7300 | 11547 | 9292 | 9874 | 17039 | 15592 | 18715 | 16688 | 15603 | 13899 | 16715 | 6446 | 5351 | 7258 | 7806 | 9685 | 8966 | 9501 | 21231 | 14637 | 17808 | 20211 | 13968 | 13796 | 13944 | Zfp57 |
| Zfp574 | 1450 | 1938 | 2056 | 2262 | 1816 | 1596 | 1368 | 1456 | 1705 | 1880 | 1847 | 2236 | 1831 | 1868 | 1353 | 1798 | 2155 | 2078 | 1926 | 1326 | 1630 | 1910 | 1633 | 2063 | 1885 | 2264 | 1805 | 1841 | Zfp574 |
| Zfp575 | 161 | 171 | 194 | 246 | 321 | 263 | 202 | 239 | 274 | 284 | 316 | 212 | 200 | 143 | 161 | 164 | 233 | 298 | 252 | 175 | 231 | 296 | 258 | 258 | 326 | 221 | 143 | 172 | Zfp575 |
| Zfp579 | 2184 | 2920 | 3045 | 3473 | 3377 | 3056 | 2777 | 2539 | 2811 | 3512 | 3178 | 3471 | 3108 | 3351 | 2167 | 2254 | 3588 | 3463 | 3850 | 2485 | 2927 | 3952 | 2557 | 3852 | 3670 | 3726 | 3396 | 3239 | Zfp579 |
| Zfp58 | 641 | 542 | 586 | 546 | 554 | 486 | 574 | 587 | 466 | 510 | 534 | 513 | 492 | 526 | 622 | 541 | 546 | 528 | 591 | 555 | 571 | 460 | 481 | 508 | 522 | 457 | 556 | 511 | Zfp58 |
| Zfp580 | 1085 | 1071 | 1364 | 1458 | 1393 | 1094 | 1167 | 933 | 860 | 1167 | 1031 | 1074 | 897 | 1157 | 953 | 801 | 1384 | 1228 | 1377 | 1038 | 1306 | 1063 | 893 | 1180 | 1175 | 1142 | 987 | 1100 | Zfp580 |
| Zfp583 | 403 | 365 | 374 | 381 | 483 | 402 | 485 | 950 | 864 | 865 | 851 | 759 | 753 | 822 | 425 | 386 | 380 | 373 | 431 | 408 | 427 | 979 | 780 | 830 | 913 | 761 | 807 | 784 | Zfp583 |
| Zfp59 | 1768 | 1524 | 1650 | 1518 | 1769 | 1728 | 1751 | 1856 | 1676 | 1633 | 1537 | 1724 | 1762 | 1693 | 1854 | 1671 | 1570 | 1607 | 1569 | 1871 | 1528 | 1636 | 1733 | 1606 | 1717 | 1726 | 1764 | 1720 | Zfp59 |
| Zfp592 | 2186 | 2139 | 2391 | 2416 | 3112 | 2802 | 2522 | 3379 | 3706 | 4096 | 4273 | 4373 | 4080 | 4088 | 1985 | 2433 | 2487 | 2590 | 3008 | 2189 | 2508 | 4679 | 3667 | 4156 | 4223 | 3531 | 3811 | 4093 | Zfp592 |
| Zfp593 | 1107 | 1462 | 1356 | 1251 | 978 | 1064 | 1063 | 572 | 673 | 582 | 663 | 649 | 730 | 688 | 1088 | 1294 | 1451 | 1231 | 1113 | 1098 | 1021 | 515 | 650 | 738 | 731 | 965 | 764 | 724 | Zfp593 |
| Zfp595 | 457 | 322 | 282 | 295 | 358 | 359 | 399 | 707 | 680 | 563 | 589 | 550 | 616 | 528 | 423 | 401 | 330 | 291 | 344 | 448 | 395 | 583 | 684 | 494 | 502 | 501 | 542 | 487 | Zfp595 |
| Zfp597 | 2103 | 1953 | 1770 | 2253 | 2495 | 2522 | 2496 | 2920 | 2317 | 2315 | 2242 | 2282 | 2141 | 2322 | 2229 | 2083 | 2021 | 2304 | 2184 | 2637 | 2543 | 2740 | 2406 | 2087 | 2667 | 1694 | 2323 | 2243 | Zfp597 |
| Zfp598 | 4726 | 6215 | 6491 | 6527 | 5635 | 5102 | 5068 | 3700 | 4232 | 4917 | 4562 | 4980 | 4420 | 4333 | 4238 | 5544 | 6334 | 6050 | 6086 | 4676 | 4965 | 4430 | 3957 | 5301 | 4768 | 5571 | 4627 | 4551 | Zfp598 |
| Zfp599 | 1096 | 883 | 970 | 848 | 1091 | 1116 | 1215 | 1164 | 1000 | 998 | 852 | 931 | 920 | 974 | 1108 | 1009 | 956 | 930 | 1148 | 1314 | 1159 | 994 | 978 | 956 | 1014 | 917 | 989 | 968 | Zfp599 |
| Zfp60 | 9239 | 8109 | 8614 | 9351 | 10919 | 10937 | 11734 | 16617 | 12788 | 12822 | 11520 | 13495 | 11863 | 14155 | 10787 | 8502 | 9943 | 9599 | 10668 | 12124 | 11075 | 14999 | 12387 | 12351 | 14655 | 11088 | 13089 | 12907 | Zfp60 |
| Zfp600 | 0 | 1 | 6 | 0 | 2 | 0 | 0 | 1 | 1 | 0 | 0 | 0 | 0 | 0 | 1 | 0 | 0 | 0 | 0 | 0 | 1 | 5 | 0 | 0 | 0 | 0 | 0 | 0 | Zfp600 |
| Zfp605 | 1062 | 952 | 1164 | 1243 | 1474 | 1452 | 1494 | 2153 | 1846 | 1933 | 1622 | 1987 | 1794 | 2017 | 1389 | 951 | 1240 | 1306 | 1486 | 1472 | 1469 | 2120 | 1929 | 1934 | 2118 | 1612 | 1734 | 1906 | Zfp605 |
| Zfp606 | 5745 | 4758 | 5366 | 5253 | 6129 | 6616 | 6655 | 8800 | 7078 | 6596 | 5889 | 6787 | 6735 | 7172 | 6773 | 5192 | 5452 | 5066 | 6308 | 7563 | 6391 | 7693 | 6977 | 6411 | 6971 | 6258 | 7669 | 7122 | Zfp606 |
| Zfp607a | 621 | 498 | 301 | 399 | 375 | 481 | 436 | 557 | 440 | 489 | 434 | 488 | 459 | 556 | 510 | 447 | 284 | 328 | 350 | 454 | 409 | 317 | 484 | 330 | 384 | 414 | 579 | 424 | Zfp607a |
| Zfp607b | 1506 | 1372 | 1383 | 1751 | 1942 | 1981 | 2084 | 1669 | 1328 | 1558 | 1360 | 1427 | 1291 | 1671 | 1576 | 1305 | 1785 | 1960 | 2110 | 2148 | 2129 | 1666 | 1238 | 1548 | 1696 | 1284 | 1350 | 1397 | Zfp607b |
| Zfp608 | 4241 | 4046 | 4896 | 5418 | 5736 | 4970 | 5337 | 6608 | 6573 | 7790 | 7077 | 7661 | 6679 | 7505 | 4218 | 3808 | 5474 | 5203 | 5725 | 4845 | 5317 | 7611 | 6453 | 7946 | 7868 | 6322 | 7059 | 7093 | Zfp608 |
| Zfp609 | 8886 | 9240 | 11087 | 12729 | 13383 | 11966 | 11972 | 9506 | 10368 | 11827 | 11158 | 14164 | 11522 | 13446 | 8115 | 8335 | 11988 | 12509 | 14043 | 9159 | 12667 | 13364 | 9139 | 13058 | 13495 | 9905 | 11524 | 12838 | Zfp609 |
| Zfp61 | 1588 | 1471 | 1695 | 1597 | 1946 | 1788 | 1729 | 2290 | 2289 | 2117 | 2274 | 2501 | 2328 | 2289 | 1635 | 1428 | 1646 | 1425 | 1894 | 1740 | 1637 | 2289 | 2142 | 2250 | 2331 | 2578 | 2330 | 2287 | Zfp61 |
| Zfp612 | 4132 | 4357 | 3829 | 4624 | 4301 | 4043 | 4878 | 2491 | 2400 | 2483 | 2336 | 2335 | 2156 | 2378 | 4298 | 4052 | 4233 | 4607 | 4225 | 4419 | 4687 | 2483 | 2353 | 2317 | 2565 | 2042 | 2352 | 2118 | Zfp612 |
| Zfp616 | 0 | 0 | 0 | 0 | 0 | 0 | 0 | 0 | 0 | 2 | 0 | 0 | 0 | 0 | 0 | 0 | 0 | 0 | 0 | 0 | 0 | 0 | 0 | 0 | 0 | 1 | 0 | 0 | Zfp616 |
| Zfp617 | 3501 | 2591 | 3143 | 2879 | 3032 | 3387 | 3624 | 3650 | 3343 | 3115 | 2943 | 3505 | 3436 | 3644 | 3773 | 3144 | 3240 | 3011 | 3311 | 3394 | 3308 | 3150 | 3287 | 3282 | 3207 | 3570 | 3598 | 3669 | Zfp617 |
| Zfp618 | 2445 | 2484 | 2294 | 2194 | 2808 | 2780 | 2029 | 8607 | 8820 | 10512 | 10011 | 11083 | 11590 | 10306 | 2413 | 2664 | 2680 | 2289 | 2681 | 1979 | 2063 | 10609 | 8601 | 9624 | 10257 | 6775 | 11251 | 10563 | Zfp618 |
| Zfp619 | 601 | 413 | 292 | 523 | 502 | 563 | 608 | 1667 | 1438 | 1057 | 997 | 1205 | 1069 | 1175 | 673 | 456 | 396 | 372 | 472 | 664 | 521 | 1176 | 1313 | 1014 | 1084 | 682 | 1030 | 1087 | Zfp619 |
| Zfp62 | 7336 | 5573 | 6376 | 6373 | 6764 | 7063 | 7451 | 11414 | 9050 | 8501 | 8318 | 9572 | 9215 | 9981 | 7942 | 6485 | 6779 | 6489 | 6648 | 8060 | 7040 | 9105 | 8857 | 8905 | 9273 | 8915 | 10024 | 9660 | Zfp62 |
| Zfp622 | 2843 | 3180 | 3143 | 3546 | 2909 | 2771 | 2730 | 2508 | 2316 | 2203 | 2415 | 2454 | 2135 | 2448 | 2864 | 2910 | 3303 | 3357 | 3067 | 2688 | 2800 | 2603 | 2378 | 2469 | 2282 | 2623 | 2329 | 2458 | Zfp622 |
| Zfp623 | 1528 | 1640 | 1471 | 1505 | 1367 | 1415 | 1491 | 1978 | 1658 | 1479 | 1540 | 1631 | 1525 | 1722 | 1600 | 1427 | 1474 | 1391 | 1496 | 1489 | 1518 | 1808 | 1690 | 1684 | 1792 | 1867 | 1806 | 1758 | Zfp623 |
| Zfp626 | 2914 | 2272 | 2693 | 2741 | 2515 | 2740 | 3125 | 3015 | 2511 | 2485 | 2524 | 2773 | 2708 | 3063 | 3021 | 2374 | 2763 | 2491 | 2727 | 2748 | 2935 | 2585 | 2457 | 2647 | 2799 | 2466 | 3017 | 3105 | Zfp626 |
| Zfp628 | 448 | 536 | 743 | 655 | 673 | 488 | 470 | 1001 | 1132 | 1587 | 1338 | 1674 | 1201 | 1346 | 384 | 410 | 810 | 529 | 756 | 424 | 542 | 1412 | 1002 | 1662 | 1444 | 1833 | 1319 | 1295 | Zfp628 |
| Zfp629 | 10057 | 10896 | 12572 | 11506 | 11221 | 11022 | 10324 | 9056 | 8296 | 9457 | 8846 | 10977 | 10497 | 10766 | 10229 | 10094 | 12660 | 10938 | 12535 | 9622 | 11231 | 10574 | 7780 | 9963 | 10264 | 9224 | 10643 | 11578 | Zfp629 |
| Zfp637 | 1306 | 1226 | 1131 | 1183 | 1347 | 1246 | 1383 | 2806 | 2720 | 2574 | 2513 | 2621 | 2464 | 2595 | 1282 | 1071 | 1271 | 1154 | 1303 | 1380 | 1378 | 2618 | 2626 | 2893 | 2528 | 3037 | 2575 | 2528 | Zfp637 |
| Zfp638 | 13396 | 11212 | 9627 | 10872 | 12749 | 13586 | 13974 | 27026 | 21635 | 18064 | 17855 | 19564 | 18540 | 20565 | 14358 | 12237 | 10630 | 10702 | 12630 | 16195 | 13591 | 19373 | 21083 | 17048 | 19557 | 14196 | 19410 | 18613 | Zfp638 |
| Zfp639 | 4628 | 4301 | 3713 | 4249 | 4090 | 4273 | 4461 | 3975 | 3670 | 3159 | 3327 | 3287 | 3513 | 3462 | 4723 | 4444 | 3985 | 4018 | 3966 | 4739 | 4217 | 3529 | 3577 | 3120 | 3474 | 3345 | 3473 | 3605 | Zfp639 |
| Zfp64 | 1391 | 1576 | 1599 | 1537 | 1560 | 1422 | 1146 | 1178 | 1469 | 1347 | 1312 | 1389 | 1382 | 1222 | 1302 | 1397 | 1591 | 1287 | 1599 | 1388 | 1319 | 1088 | 1469 | 1438 | 1257 | 1446 | 1417 | 1276 | Zfp64 |
| Zfp641 | 333 | 295 | 439 | 401 | 807 | 564 | 520 | 862 | 753 | 993 | 738 | 656 | 637 | 718 | 390 | 300 | 379 | 408 | 575 | 526 | 494 | 1240 | 757 | 715 | 956 | 511 | 762 | 667 | Zfp641 |
| Zfp644 | 5613 | 4548 | 4685 | 4650 | 5167 | 5478 | 5551 | 10794 | 8877 | 8166 | 7995 | 8867 | 8742 | 9309 | 6050 | 4903 | 4715 | 4571 | 4846 | 6129 | 5498 | 8490 | 8655 | 7969 | 8387 | 7092 | 9316 | 9100 | Zfp644 |
| Zfp646 | 4414 | 5098 | 4656 | 4973 | 4981 | 4786 | 4736 | 5511 | 5869 | 6564 | 5997 | 7686 | 6668 | 7198 | 4400 | 4599 | 5005 | 4917 | 5449 | 4374 | 4851 | 6549 | 5335 | 6813 | 7149 | 5140 | 7496 | 7444 | Zfp646 |
| Zfp647 | 912 | 1047 | 1126 | 978 | 973 | 904 | 903 | 1281 | 1364 | 1317 | 1291 | 1530 | 1351 | 1386 | 808 | 848 | 956 | 823 | 1044 | 1008 | 936 | 1325 | 1333 | 1526 | 1476 | 1585 | 1427 | 1483 | Zfp647 |
| Zfp648 | 8 | 4 | 1 | 1 | 7 | 3 | 1 | 20 | 56 | 37 | 88 | 15 | 33 | 4 | 5 | 5 | 1 | 3 | 2 | 1 | 1 | 47 | 36 | 22 | 17 | 3 | 1 | 0 | Zfp648 |
| Zfp65 | 2231 | 1913 | 2523 | 2373 | 2491 | 2423 | 2753 | 2510 | 2258 | 2362 | 2157 | 2311 | 2257 | 2503 | 2411 | 2065 | 2525 | 2259 | 2377 | 2644 | 2548 | 2346 | 2310 | 2472 | 2458 | 2483 | 2437 | 2395 | Zfp65 |
| Zfp651 | 1510 | 1797 | 1893 | 2275 | 2497 | 2324 | 1990 | 2285 | 2895 | 3022 | 3215 | 2725 | 2558 | 2281 | 1400 | 1550 | 2006 | 2410 | 2514 | 1942 | 2213 | 2689 | 2862 | 2866 | 2942 | 2792 | 2461 | 2407 | Zfp651 |
| Zfp652 | 6579 | 5894 | 7101 | 8611 | 8941 | 8426 | 8785 | 8124 | 7825 | 9605 | 9570 | 9064 | 8769 | 9258 | 6326 | 6030 | 8046 | 8552 | 8916 | 6627 | 8731 | 9926 | 7524 | 9391 | 10308 | 8062 | 8300 | 8632 | Zfp652 |
| Zfp653 | 388 | 568 | 550 | 655 | 603 | 666 | 594 | 924 | 1032 | 1160 | 1044 | 1156 | 1084 | 1233 | 414 | 396 | 641 | 583 | 651 | 509 | 612 | 1093 | 952 | 1259 | 1261 | 1215 | 1181 | 1169 | Zfp653 |

| | | | | | | | | | | | | | | | | | | | | | | | | | | | | | |
|---|---|---|---|---|---|---|---|---|---|---|---|---|---|---|---|---|---|---|---|---|---|---|---|---|---|---|---|---|---|
| Zfp654 | 4445 | 3539 | 3476 | 4711 | 3944 | 4621 | 4632 | 3793 | 3263 | 2959 | 3191 | 3232 | 3111 | 3316 | 4577 | 4491 | 3729 | 4492 | 3790 | 4212 | 4590 | 3568 | 3181 | 3045 | 3419 | 2507 | 3452 | 3274 | Zfp654 |
| Zfp655 | 4684 | 4084 | 3626 | 3910 | 3464 | 3763 | 3796 | 4645 | 3594 | 3215 | 3127 | 3254 | 3315 | 3303 | 4937 | 4505 | 3394 | 4011 | 3311 | 4363 | 3747 | 3653 | 3728 | 3072 | 3479 | 3302 | 3526 | 3253 | Zfp655 |
| Zfp658 | 1160 | 1293 | 1313 | 1433 | 1438 | 1333 | 1405 | 1607 | 1319 | 1323 | 1310 | 1412 | 1174 | 1461 | 1175 | 1108 | 1467 | 1368 | 1453 | 1308 | 1414 | 1738 | 1308 | 1384 | 1587 | 1367 | 1391 | 1368 | Zfp658 |
| Zfp661 | 1679 | 1749 | 1469 | 1552 | 1556 | 1513 | 1690 | 1319 | 1128 | 1189 | 1129 | 1261 | 1106 | 1138 | 1689 | 1637 | 1614 | 1533 | 1566 | 1595 | 1459 | 1218 | 1141 | 1177 | 1267 | 1080 | 1232 | 1240 | Zfp661 |
| Zfp663 | 939 | 1198 | 1131 | 1252 | 1248 | 1135 | 1134 | 1027 | 921 | 974 | 887 | 991 | 985 | 1043 | 1230 | 962 | 1330 | 1193 | 1233 | 1224 | 1125 | 1245 | 772 | 1045 | 1133 | 902 | 963 | 951 | Zfp663 |
| Zfp664 | 24120 | 22965 | 24812 | 22483 | 21298 | 21511 | 22914 | 23250 | 18582 | 20690 | 18410 | 24892 | 23200 | 24969 | 24062 | 20770 | 24404 | 21532 | 22977 | 21693 | 22876 | 20732 | 17860 | 22670 | 22515 | 25117 | 25134 | 26304 | Zfp664 |
| Zfp667 | 3579 | 3410 | 3407 | 3706 | 3217 | 3444 | 3557 | 1992 | 1538 | 1514 | 1388 | 1461 | 1394 | 1518 | 3938 | 3441 | 3290 | 3515 | 3113 | 4127 | 3351 | 1721 | 1644 | 1497 | 1492 | 1477 | 1542 | 1585 | Zfp667 |
| Zfp668 | 1420 | 1596 | 1922 | 1924 | 2059 | 2035 | 1939 | 2138 | 2214 | 2678 | 2391 | 2771 | 2535 | 2722 | 1437 | 1407 | 2079 | 1885 | 2205 | 1508 | 1859 | 3061 | 1897 | 2872 | 3236 | 2570 | 2481 | 2813 | Zfp668 |
| Zfp672 | 2560 | 2633 | 1912 | 2229 | 2314 | 2262 | 2356 | 2337 | 2431 | 2456 | 2532 | 2259 | 2252 | 2348 | 2491 | 2446 | 2124 | 2217 | 2354 | 2416 | 2403 | 2400 | 2389 | 2367 | 2439 | 2182 | 2245 | 2262 | Zfp672 |
| Zfp677 | 599 | 422 | 423 | 547 | 632 | 639 | 578 | 542 | 449 | 475 | 402 | 436 | 527 | 449 | 460 | 488 | 589 | 508 | 404 | 542 | 464 | 493 | 443 | 397 | 425 | 309 | 294 | 406 | Zfp677 |
| Zfp68 | 5638 | 5078 | 5332 | 5491 | 6036 | 6172 | 6773 | 9914 | 8119 | 7201 | 6912 | 7176 | 6918 | 7322 | 6670 | 5735 | 5574 | 5807 | 6257 | 7152 | 6460 | 8450 | 8081 | 7375 | 8230 | 6918 | 7391 | 7454 | Zfp68 |
| Zfp683 | 0 | 1 | 5 | 0 | 1 | 0 | 0 | 1 | 2 | 0 | 0 | 0 | 1 | 0 | 3 | 2 | 0 | 1 | 0 | 4 | 2 | 0 | 1 | 0 | 0 | 0 | 0 | 2 | Zfp683 |
| Zfp687 | 6584 | 8339 | 7319 | 8205 | 7060 | 7421 | 7401 | 3792 | 3975 | 4138 | 4027 | 5233 | 4900 | 4915 | 6399 | 6771 | 7631 | 7650 | 7695 | 6716 | 7697 | 3860 | 3710 | 4511 | 4510 | 4943 | 5466 | 5277 | Zfp687 |
| Zfp688 | 684 | 765 | 874 | 737 | 650 | 695 | 647 | 466 | 530 | 609 | 545 | 681 | 595 | 633 | 632 | 623 | 782 | 735 | 722 | 662 | 656 | 570 | 553 | 632 | 572 | 777 | 647 | 679 | Zfp688 |
| Zfp689 | 977 | 910 | 1218 | 1129 | 1267 | 1132 | 1133 | 924 | 1060 | 1313 | 1138 | 1383 | 1235 | 1311 | 960 | 836 | 1202 | 1029 | 1237 | 1010 | 1239 | 1198 | 895 | 1369 | 1230 | 1517 | 1263 | 1359 | Zfp689 |
| Zfp69 | 737 | 850 | 689 | 817 | 664 | 764 | 656 | 431 | 441 | 440 | 472 | 432 | 493 | 483 | 712 | 827 | 722 | 723 | 680 | 600 | 742 | 414 | 445 | 395 | 404 | 380 | 516 | 447 | Zfp69 |
| Zfp691 | 745 | 1075 | 1241 | 944 | 766 | 899 | 863 | 650 | 588 | 780 | 648 | 888 | 853 | 918 | 814 | 928 | 1145 | 867 | 985 | 792 | 845 | 665 | 605 | 798 | 754 | 959 | 852 | 939 | Zfp691 |
| Zfp692 | 1756 | 2081 | 2053 | 2200 | 2126 | 1906 | 2126 | 1937 | 2072 | 2287 | 2201 | 2443 | 2092 | 2249 | 1725 | 1860 | 2279 | 2317 | 2320 | 1841 | 1992 | 2161 | 2052 | 2554 | 2260 | 2624 | 2274 | 2216 | Zfp692 |
| Zfp697 | 2008 | 1934 | 2464 | 2558 | 2802 | 2331 | 2531 | 2982 | 2430 | 3080 | 2851 | 2858 | 2616 | 3036 | 2032 | 2053 | 2445 | 2495 | 2618 | 2224 | 2684 | 3588 | 2404 | 2941 | 3587 | 2598 | 2671 | 2954 | Zfp697 |
| Zfp7 | 1729 | 1638 | 1546 | 1411 | 1286 | 1284 | 1336 | 1226 | 1148 | 1133 | 1054 | 1137 | 1143 | 1197 | 1539 | 1560 | 1409 | 1303 | 1358 | 1470 | 1276 | 1130 | 1122 | 1225 | 1065 | 1264 | 1214 | 1258 | Zfp7 |
| Zfp703 | 1847 | 2169 | 2150 | 2315 | 2458 | 1945 | 1733 | 1917 | 2573 | 2590 | 3229 | 2168 | 2231 | 1774 | 1775 | 2199 | 2101 | 2370 | 1992 | 1450 | 1861 | 2498 | 2933 | 2187 | 2154 | 2573 | 1463 | 1679 | Zfp703 |
| Zfp704 | 8495 | 7520 | 5396 | 7387 | 10105 | 11762 | 10838 | 9418 | 10828 | 9783 | 11570 | 8335 | 8348 | 9395 | 7851 | 8644 | 5845 | 8278 | 8617 | 9346 | 11936 | 8014 | 10045 | 7727 | 10900 | 6035 | 9031 | 7136 | Zfp704 |
| Zfp706 | 14937 | 12510 | 14712 | 15255 | 15320 | 14795 | 16376 | 13650 | 12082 | 13105 | 12256 | 14203 | 12976 | 14100 | 14736 | 12495 | 14921 | 15038 | 15609 | 14659 | 16578 | 14417 | 12255 | 13385 | 13567 | 15070 | 13396 | 15101 | Zfp706 |
| Zfp707 | 726 | 922 | 824 | 879 | 730 | 713 | 761 | 579 | 697 | 732 | 674 | 627 | 633 | 602 | 719 | 843 | 809 | 795 | 842 | 741 | 736 | 627 | 641 | 760 | 674 | 662 | 637 | 589 | Zfp707 |
| Zfp708 | 1 | 0 | 0 | 0 | 0 | 0 | 0 | 0 | 0 | 0 | 1 | 2 | 0 | 1 | 3 | 0 | 0 | 0 | 0 | 0 | 1 | 0 | 0 | 0 | 0 | 0 | 0 | 1 | Zfp708 |
| Zfp709 | 1301 | 921 | 1031 | 942 | 1307 | 1477 | 1398 | 1774 | 1595 | 1472 | 1374 | 1176 | 1381 | 1371 | 1316 | 1051 | 1094 | 930 | 1226 | 1436 | 1263 | 1544 | 1547 | 1218 | 1442 | 1135 | 1332 | 1266 | Zfp709 |
| Zfp710 | 1173 | 1573 | 2317 | 1699 | 1972 | 1788 | 1551 | 2155 | 2631 | 3156 | 2699 | 3811 | 3047 | 3467 | 1241 | 1232 | 2301 | 1645 | 2243 | 1362 | 1569 | 2802 | 2382 | 3801 | 3248 | 4343 | 3619 | 3640 | Zfp710 |
| Zfp711 | 2961 | 2046 | 3436 | 3536 | 4255 | 3918 | 4348 | 5385 | 4430 | 4693 | 4668 | 5145 | 4716 | 5149 | 3371 | 2516 | 3206 | 3444 | 3956 | 3917 | 4032 | 5581 | 4393 | 4681 | 5408 | 4351 | 5174 | 5046 | Zfp711 |
| Zfp712 | 903 | 776 | 810 | 929 | 890 | 974 | 1040 | 1112 | 932 | 781 | 733 | 728 | 700 | 746 | 1101 | 890 | 965 | 986 | 903 | 1135 | 1057 | 957 | 896 | 759 | 761 | 734 | 674 | 748 | Zfp712 |
| Zfp715 | 3706 | 3347 | 2980 | 3522 | 3683 | 3739 | 3798 | 3472 | 3027 | 2568 | 2679 | 2501 | 2583 | 2665 | 3878 | 3527 | 3019 | 3509 | 3319 | 3807 | 3691 | 3362 | 2954 | 2382 | 2727 | 2207 | 2308 | 2444 | Zfp715 |
| Zfp719 | 2028 | 1792 | 1733 | 1552 | 1824 | 1825 | 1818 | 2494 | 2277 | 1905 | 1886 | 1786 | 1929 | 1970 | 2076 | 1822 | 1686 | 1667 | 1710 | 2100 | 1902 | 2054 | 2172 | 1758 | 1910 | 1879 | 2045 | 1949 | Zfp719 |
| Zfp72 | 633 | 476 | 683 | 557 | 691 | 753 | 677 | 777 | 793 | 620 | 717 | 478 | 575 | 529 | 708 | 618 | 591 | 576 | 560 | 734 | 594 | 775 | 796 | 523 | 561 | 562 | 511 | 511 | Zfp72 |
| Zfp729a | 1802 | 1342 | 1130 | 1475 | 1807 | 1857 | 1938 | 2532 | 2344 | 2053 | 1834 | 1919 | 1718 | 1845 | 1945 | 1568 | 1410 | 1405 | 1680 | 2141 | 1864 | 1972 | 2297 | 1754 | 1949 | 1528 | 1887 | 1781 | Zfp729a |
| Zfp729b | 698 | 466 | 660 | 660 | 721 | 684 | 724 | 1025 | 958 | 831 | 761 | 741 | 745 | 761 | 715 | 641 | 643 | 693 | 690 | 722 | 714 | 869 | 983 | 804 | 730 | 855 | 726 | 832 | Zfp729b |
| Zfp735 | 0 | 0 | 0 | 0 | 0 | 0 | 0 | 0 | 0 | 0 | 0 | 0 | 0 | 0 | 0 | 0 | 0 | 0 | 0 | 0 | 0 | 0 | 0 | 0 | 0 | 0 | 0 | 2 | Zfp735 |
| Zfp738 | 3624 | 2610 | 2430 | 2807 | 3142 | 3474 | 3365 | 5059 | 4445 | 3611 | 3935 | 3459 | 3477 | 3285 | 3786 | 3414 | 2801 | 3009 | 2996 | 4047 | 3488 | 3833 | 4086 | 3206 | 3876 | 2771 | 3543 | 3002 | Zfp738 |
| Zfp74 | 825 | 610 | 910 | 814 | 1066 | 863 | 955 | 1850 | 1639 | 1944 | 1691 | 1962 | 1723 | 1746 | 770 | 754 | 876 | 778 | 1022 | 768 | 974 | 2095 | 1757 | 1848 | 1829 | 1861 | 1804 | 1888 | Zfp74 |
| Zfp740 | 14200 | 13984 | 14860 | 16814 | 16910 | 16552 | 17013 | 14488 | 14754 | 16748 | 15815 | 18264 | 16987 | 18064 | 13321 | 13218 | 16249 | 16263 | 17649 | 14063 | 17120 | 16243 | 14120 | 18016 | 18215 | 18000 | 17290 | 17131 | Zfp740 |
| Zfp746 | 1985 | 1971 | 2353 | 2256 | 2495 | 2400 | 2285 | 2628 | 2903 | 3121 | 2873 | 3231 | 2888 | 3045 | 2105 | 1894 | 2374 | 2433 | 2710 | 2041 | 2246 | 2944 | 2731 | 3153 | 2967 | 3430 | 2702 | 3163 | Zfp746 |
| Zfp747 | 706 | 960 | 899 | 859 | 815 | 790 | 683 | 482 | 595 | 559 | 510 | 575 | 614 | 524 | 673 | 787 | 811 | 815 | 783 | 817 | 721 | 496 | 568 | 604 | 534 | 656 | 561 | 571 | Zfp747 |
| Zfp748 | 2554 | 2127 | 1820 | 1953 | 2052 | 2325 | 2404 | 3339 | 2980 | 2210 | 2363 | 2395 | 2261 | 2323 | 2801 | 2414 | 1810 | 1871 | 2011 | 2825 | 2389 | 2573 | 2898 | 2020 | 2241 | 2241 | 2392 | 2301 | Zfp748 |
| Zfp750 | 50 | 10 | 9 | 2 | 1 | 12 | 0 | 128 | 194 | 115 | 65 | 41 | 74 | 18 | 26 | 30 | 8 | 14 | 1 | 0 | 2 | 8 | 230 | 80 | 6 | 42 | 31 | 31 | Zfp750 |
| Zfp758 | 1181 | 967 | 1351 | 1224 | 1101 | 1223 | 1129 | 1330 | 1024 | 1092 | 946 | 1059 | 1102 | 1049 | 1361 | 1340 | 1345 | 1085 | 1188 | 1354 | 1213 | 1081 | 1122 | 981 | 972 | 969 | 1178 | 1148 | Zfp758 |
| Zfp759 | 614 | 420 | 486 | 425 | 645 | 645 | 689 | 1083 | 957 | 787 | 782 | 627 | 672 | 676 | 676 | 525 | 545 | 533 | 614 | 736 | 630 | 872 | 966 | 816 | 804 | 695 | 652 | 630 | Zfp759 |
| Zfp760 | 4487 | 3476 | 4131 | 4333 | 4199 | 4556 | 4948 | 5584 | 4482 | 4478 | 4104 | 5272 | 5026 | 5619 | 4342 | 4001 | 3946 | 4013 | 4131 | 4834 | 4034 | 4655 | 4334 | 4649 | 4510 | 4790 | 4998 | 5320 | Zfp760 |
| Zfp763 | 3188 | 2547 | 2949 | 3111 | 3164 | 3155 | 3508 | 3745 | 2987 | 2698 | 2554 | 2657 | 2545 | 2775 | 3549 | 3021 | 3062 | 3095 | 3017 | 3994 | 3550 | 3171 | 2887 | 2781 | 2854 | 2742 | 2696 | 3089 | Zfp763 |
| Zfp764 | 385 | 387 | 455 | 379 | 422 | 443 | 399 | 509 | 535 | 562 | 470 | 496 | 570 | 512 | 413 | 375 | 457 | 339 | 479 | 482 | 359 | 396 | 509 | 526 | 504 | 622 | 485 | 554 | Zfp764 |
| Zfp768 | 1901 | 2325 | 2601 | 2495 | 2211 | 2148 | 1829 | 2023 | 2193 | 2669 | 2417 | 2748 | 2621 | 2593 | 1925 | 1956 | 2754 | 2092 | 2486 | 1941 | 2040 | 2249 | 2177 | 2780 | 2522 | 3047 | 2590 | 2772 | Zfp768 |
| Zfp770 | 2694 | 2149 | 2460 | 2595 | 2653 | 2749 | 2956 | 2467 | 2426 | 2290 | 2394 | 2229 | 2391 | 2280 | 2641 | 2287 | 2394 | 2666 | 2543 | 2814 | 2751 | 2483 | 2302 | 2265 | 2420 | 2348 | 2320 | 2328 | Zfp770 |
| Zfp771 | 1009 | 1234 | 937 | 1115 | 1013 | 835 | 885 | 871 | 902 | 949 | 907 | 910 | 826 | 749 | 823 | 888 | 999 | 1015 | 1181 | 901 | 931 | 867 | 918 | 986 | 897 | 1029 | 900 | 758 | Zfp771 |
| Zfp772 | 1887 | 1883 | 1680 | 1928 | 1462 | 1646 | 1581 | 1005 | 835 | 867 | 847 | 868 | 710 | 799 | 1949 | 1922 | 1596 | 1650 | 1554 | 1562 | 1567 | 1089 | 927 | 899 | 947 | 891 | 840 | 854 | Zfp772 |
| Zfp773 | 446 | 407 | 494 | 594 | 936 | 834 | 806 | 1744 | 1265 | 1454 | 1099 | 1360 | 1193 | 1303 | 476 | 349 | 721 | 574 | 881 | 888 | 824 | 1865 | 1241 | 1487 | 1677 | 1142 | 1313 | 1272 | Zfp773 |
| Zfp775 | 900 | 1149 | 1035 | 1161 | 1181 | 1027 | 972 | 876 | 765 | 938 | 823 | 984 | 779 | 759 | 969 | 1023 | 1195 | 1109 | 1257 | 1000 | 1020 | 817 | 763 | 945 | 1009 | 1069 | 833 | 720 | Zfp775 |
| Zfp777 | 1647 | 1651 | 1849 | 1828 | 1914 | 1912 | 1517 | 1761 | 2057 | 2186 | 2037 | 2429 | 2063 | 2139 | 1552 | 1583 | 1796 | 1727 | 1968 | 1580 | 1661 | 2085 | 1910 | 2289 | 2212 | 2549 | 2228 | 2231 | Zfp777 |
| Zfp78 | 504 | 504 | 511 | 598 | 543 | 553 | 562 | 337 | 272 | 348 | 289 | 211 | 226 | 225 | 542 | 502 | 480 | 596 | 495 | 576 | 475 | 277 | 330 | 240 | 298 | 218 | 227 | 269 | Zfp78 |
| Zfp780b | 2566 | 2294 | 2112 | 2489 | 2533 | 2439 | 2640 | 2661 | 2292 | 2094 | 2317 | 2154 | 2093 | 2286 | 2570 | 2546 | 2360 | 2621 | 2420 | 2675 | 2567 | 2508 | 2370 | 1977 | 2263 | 1677 | 2334 | 2169 | Zfp780b |
| Zfp781 | 165 | 182 | 148 | 125 | 149 | 200 | 152 | 287 | 246 | 199 | 156 | 176 | 183 | 212 | 136 | 163 | 146 | 154 | 148 | 224 | 144 | 251 | 240 | 187 | 194 | 188 | 156 | 183 | Zfp781 |
| Zfp784 | 118 | 97 | 112 | 119 | 146 | 135 | 122 | 343 | 375 | 462 | 392 | 393 | 379 | 359 | 116 | 88 | 126 | 125 | 150 | 100 | 132 | 426 | 434 | 375 | 440 | 358 | 334 | 354 | Zfp784 |
| Zfp786 | 674 | 733 | 634 | 848 | 982 | 826 | 932 | 964 | 948 | 941 | 768 | 836 | 847 | 918 | 814 | 643 | 835 | 853 | 934 | 944 | 967 | 1073 | 847 | 981 | 979 | 751 | 847 | 802 | Zfp786 |
| Zfp787 | 825 | 959 | 949 | 874 | 743 | 695 | 707 | 1188 | 1148 | 1394 | 1311 | 1328 | 1209 | 1297 | 752 | 652 | 944 | 657 | 856 | 859 | 779 | 1006 | 1130 | 1243 | 1168 | 1363 | 1374 | 1311 | Zfp787 |
| Zfp788 | 2731 | 2124 | 3174 | 3089 | 3763 | 3550 | 3796 | 5244 | 4542 | 4544 | 4207 | 4380 | 4206 | 4549 | 3182 | 2382 | 3161 | 3240 | 3591 | 3503 | 3499 | 5651 | 4148 | 4193 | 4464 | 4303 | 4303 | 4460 | Zfp788 |
| Zfp790 | 2693 | 2955 | 2336 | 2975 | 2695 | 2929 | 2778 | 3042 | 2656 | 2457 | 2335 | 2165 | 2151 | 2335 | 2917 | 2803 | 2549 | 2731 | 2427 | 3156 | 2907 | 2677 | 2532 | 2140 | 2515 | 1930 | 2158 | 2298 | Zfp790 |
| Zfp791 | 364 | 290 | 393 | 346 | 449 | 396 | 387 | 374 | 352 | 449 | 346 | 380 | 345 | 302 | 340 | 363 | 382 | 306 | 369 | 428 | 424 | 449 | 366 | 386 | 341 | 373 | 311 | 335 | Zfp791 |
| Zfp799 | 4886 | 4228 | 3388 | 4327 | 4086 | 4628 | 4505 | 5551 | 4665 | 4386 | 4324 | 4379 | 4350 | 4476 | 4986 | 4202 | 3775 | 4436 | 3998 | 4504 | 4658 | 4634 | 4620 | 4140 | 4610 | 3838 | 4606 | 4226 | Zfp799 |
| Zfp800 | 2240 | 1876 | 1675 | 2024 | 1825 | 2074 | 1950 | 1696 | 1397 | 1322 | 1399 | 1617 | 1598 | 1633 | 2229 | 1942 | 1670 | 1924 | 1675 | 2081 | 1910 | 1517 | 1465 | 1419 | 1413 | 1219 | 1762 | 1666 | Zfp800 |
| Zfp804a | 243 | 270 | 214 | 378 | 718 | 495 | 432 | 942 | 906 | 976 | 803 | 492 | 564 | 495 | 392 | 281 | 286 | 393 | 448 | 543 | 480 | 1343 | 711 | 522 | 955 | 259 | 483 | 462 | Zfp804a |
| Zfp804b | 48 | 46 | 14 | 28 | 88 | 94 | 63 | 87 | 111 | 34 | 62 | 67 | 73 | 62 | 69 | 55 | 15 | 34 | 41 | 117 | 98 | 70 | 115 | 25 | 66 | 12 | 59 | 42 | Zfp804b |
| Zfp808 | 591 | 333 | 655 | 415 | 451 | 560 | 474 | 663 | 695 | 662 | 649 | 744 | 811 | 618 | 629 | 550 | 491 | 409 | 456 | 447 | 400 | 598 | 618 | 639 | 570 | 797 | 800 | 731 | Zfp808 |
| Zfp809 | 622 | 536 | 472 | 520 | 604 | 659 | 587 | 1003 | 991 | 855 | 884 | 808 | 916 | 825 | 587 | 508 | 591 | 487 | 638 | 640 | 582 | 837 | 1032 | 904 | 831 | 766 | 916 | 777 | Zfp809 |
| Zfp81 | 6187 | 4840 | 4710 | 6695 | 7030 | 7202 | 7930 | 7852 | 6158 | 5932 | 5770 | 5853 | 5425 | 6246 | 6717 | 5605 | 5514 | 6771 | 6930 | 8019 | 7900 | 6901 | 6183 | 5858 | 6680 | 4457 | 5954 | 5819 | Zfp81 |
| Zfp810 | 2042 | 1592 | 1705 | 1557 | 1960 | 2021 | 2020 | 3886 | 3347 | 3601 | 3280 | 3612 | 3606 | 3647 | 2097 | 1807 | 1676 | 1483 | 1914 | 1866 | 1991 | 3852 | 3270 | 3433 | 3530 | 3507 | 3833 | 3570 | Zfp810 |
| Zfp811 | 5842 | 5249 | 7055 | 7114 | 8035 | 8182 | 8615 | 9078 | 6605 | 8658 | 7025 | 8893 | 8439 | 9919 | 6808 | 5083 | 7159 | 7108 | 8191 | 8002 | 8412 | 10358 | 6701 | 8296 | 9519 | 7247 | 9949 | 10434 | Zfp811 |
| Zfp819 | 114 | 133 | 89 | 88 | 34 | 40 | 73 | 34 | 22 | 35 | 18 | 31 | 17 | 26 | 102 | 98 | 86 | 53 | 76 | 83 | 62 | 30 | 35 | 39 | 23 | 41 | 52 | 59 | Zfp819 |
| Zfp82 | 111 | 93 | 92 | 162 | 154 | 128 | 127 | 137 | 111 | 127 | 166 | 126 | 102 | 134 | 128 | 90 | 121 | 106 | 117 | 132 | 106 | 137 | 128 | 141 | 158 | 115 | 128 | 122 | Zfp82 |
| Zfp820 | 1621 | 983 | 1442 | 1105 | 1010 | 1398 | 1342 | 949 | 944 | 652 | 730 | 958 | 995 | 970 | 1447 | 1278 | 1363 | 1072 | 1224 | 1352 | 1132 | 703 | 855 | 821 | 706 | 1102 | 1137 | 1030 | Zfp820 |
| Zfp821 | 3406 | 3576 | 3898 | 4161 | 5043 | 4654 | 4618 | 6781 | 6171 | 7122 | 6209 | 7064 | 6168 | 7157 | 3512 | 2972 | 4009 | 4289 | 5264 | 4571 | 4698 | 7344 | 5778 | 7396 | 8126 | 6705 | 6693 | 6855 | Zfp821 |
| Zfp825 | 732 | 630 | 922 | 684 | 680 | 708 | 754 | 1060 | 957 | 941 | 945 | 1178 | 1154 | 1050 | 767 | 750 | 868 | 753 | 757 | 728 | 622 | 1079 | 965 | 1058 | 901 | 1209 | 1099 | 1072 | Zfp825 |
| Zfp827 | 1829 | 1714 | 2262 | 2074 | 2852 | 2648 | 2338 | 4825 | 5037 | 6382 | 6355 | 6156 | 6178 | 5885 | 1841 | 1820 | 2391 | 2247 | 2863 | 1951 | 2475 | 6208 | 4834 | 5979 | 6211 | 4848 | 5718 | 5573 | Zfp827 |
| Zfp830 | 1782 | 1580 | 1871 | 1940 | 1462 | 1488 | 1614 | 1481 | 1345 | 1505 | 1339 | 1574 | 1405 | 1574 | 1737 | 1587 | 1782 | 1559 | 1629 | 1444 | 1609 | 1362 | 1323 | 1706 | 1392 | 1785 | 1500 | 1720 | Zfp830 |

| | | | | | | | | | | | | | | | | | | | | | | | | | | | | | |
|---|---|---|---|---|---|---|---|---|---|---|---|---|---|---|---|---|---|---|---|---|---|---|---|---|---|---|---|---|---|
| Zfp831 | 2 | 4 | 6 | 5 | 2 | 1 | 6 | 4 | 8 | 2 | 13 | 3 | 2 | 1 | 1 | 5 | 2 | 1 | 5 | 0 | 4 | 7 | 14 | 8 | 10 | 1 | 2 | 2 | Zfp831 |
| Zfp839 | 709 | 638 | 979 | 711 | 706 | 680 | 618 | 611 | 859 | 815 | 958 | 791 | 875 | 669 | 604 | 706 | 797 | 721 | 729 | 532 | 584 | 737 | 790 | 771 | 777 | 953 | 782 | 663 | Zfp839 |
| Zfp84 | 2343 | 2609 | 2065 | 2455 | 2603 | 2521 | 2505 | 3226 | 2597 | 2694 | 2582 | 2490 | 2245 | 2406 | 2503 | 2270 | 2440 | 2578 | 2388 | 2871 | 2666 | 2709 | 2761 | 2462 | 2863 | 2095 | 2428 | 2384 | Zfp84 |
| Zfp846 | 2963 | 2667 | 1892 | 2659 | 3634 | 3817 | 3657 | 6613 | 5766 | 5072 | 4727 | 4467 | 4395 | 4651 | 3237 | 2521 | 2211 | 2666 | 3171 | 3921 | 4122 | 5320 | 5294 | 3774 | 5548 | 2989 | 4522 | 4101 | Zfp846 |
| Zfp85 | 946 | 738 | 891 | 977 | 943 | 973 | 1034 | 917 | 748 | 683 | 627 | 701 | 628 | 634 | 1149 | 922 | 922 | 925 | 1008 | 1226 | 974 | 797 | 822 | 672 | 629 | 718 | 723 | 783 | Zfp85 |
| Zfp850 | 2842 | 2436 | 2311 | 2240 | 2274 | 2443 | 2541 | 2066 | 1712 | 1511 | 1442 | 1650 | 1716 | 1808 | 3038 | 2698 | 2268 | 2220 | 2095 | 3004 | 2297 | 1459 | 1543 | 1325 | 1519 | 1643 | 1669 | 1828 | Zfp850 |
| Zfp853 | 88 | 97 | 87 | 78 | 92 | 110 | 60 | 99 | 166 | 146 | 140 | 98 | 94 | 66 | 63 | 111 | 107 | 103 | 103 | 73 | 79 | 154 | 153 | 105 | 115 | 63 | 63 | 76 | Zfp853 |
| Zfp865 | 3009 | 3334 | 2785 | 3700 | 3289 | 3293 | 2928 | 2844 | 3268 | 3532 | 3610 | 4021 | 3540 | 3496 | 2664 | 2838 | 3187 | 3333 | 3386 | 2716 | 3218 | 3489 | 2973 | 3501 | 3583 | 3168 | 3815 | 3443 | Zfp865 |
| Zfp866 | 4170 | 4184 | 3591 | 4130 | 4428 | 4721 | 4217 | 8261 | 6888 | 6147 | 5989 | 6310 | 6112 | 6388 | 4348 | 3667 | 4001 | 3849 | 4491 | 4789 | 4455 | 6986 | 6594 | 5949 | 6713 | 4850 | 6656 | 6273 | Zfp866 |
| Zfp867 | 2632 | 2416 | 2200 | 2163 | 2128 | 2515 | 2516 | 1961 | 1907 | 1567 | 1497 | 1710 | 1444 | 1671 | 2980 | 2389 | 2271 | 2326 | 2225 | 2812 | 2454 | 1763 | 1917 | 1634 | 1637 | 1611 | 1527 | 1723 | Zfp867 |
| Zfp868 | 2017 | 1712 | 1830 | 1940 | 1901 | 2115 | 2025 | 3197 | 2609 | 2405 | 2144 | 2574 | 2334 | 2507 | 2106 | 1735 | 1852 | 1656 | 2134 | 2212 | 1957 | 2787 | 2642 | 2495 | 2569 | 2643 | 2721 | 2673 | Zfp868 |
| Zfp869 | 2915 | 2486 | 2588 | 2704 | 2788 | 2768 | 3058 | 4725 | 4271 | 3733 | 3759 | 4187 | 4051 | 4146 | 3101 | 2598 | 2781 | 2663 | 3037 | 2808 | 2815 | 4096 | 3955 | 3998 | 4083 | 4081 | 4453 | 4221 | Zfp869 |
| Zfp87 | 1921 | 1757 | 1678 | 1775 | 1725 | 1746 | 1982 | 2137 | 1782 | 1435 | 1422 | 1588 | 1479 | 1612 | 2008 | 1805 | 1792 | 1754 | 1883 | 2179 | 1811 | 1975 | 1725 | 1599 | 1709 | 1423 | 1644 | 1659 | Zfp87 |
| Zfp870 | 1969 | 1837 | 1580 | 1787 | 1910 | 1946 | 2114 | 1671 | 1443 | 1479 | 1292 | 1460 | 1435 | 1507 | 1904 | 1635 | 1635 | 1795 | 1977 | 2213 | 1985 | 1414 | 1448 | 1427 | 1450 | 1402 | 1632 | 1479 | Zfp870 |
| Zfp871 | 12641 | 10850 | 11343 | 12852 | 11875 | 12854 | 12421 | 14581 | 12621 | 11858 | 12504 | 13475 | 12813 | 13267 | 11910 | 12632 | 11359 | 13074 | 11939 | 12052 | 12343 | 13697 | 11945 | 11959 | 13609 | 11501 | 13837 | 12422 | Zfp871 |
| Zfp872 | 77 | 83 | 126 | 100 | 54 | 76 | 65 | 9 | 11 | 5 | 3 | 5 | 19 | 23 | 92 | 121 | 119 | 91 | 74 | 92 | 74 | 8 | 22 | 19 | 10 | 11 | 9 | 12 | Zfp872 |
| Zfp873 | 459 | 425 | 558 | 437 | 615 | 591 | 603 | 1605 | 1386 | 1184 | 1240 | 1300 | 1192 | 1330 | 525 | 400 | 606 | 487 | 571 | 701 | 583 | 1414 | 1377 | 1285 | 1326 | 1152 | 1274 | 1332 | Zfp873 |
| Zfp874a | 1158 | 790 | 900 | 843 | 785 | 766 | 949 | 1012 | 993 | 717 | 836 | 681 | 860 | 691 | 1114 | 1085 | 882 | 823 | 869 | 1135 | 804 | 827 | 1029 | 764 | 717 | 774 | 765 | 794 | Zfp874a |
| Zfp874b | 897 | 729 | 895 | 907 | 843 | 815 | 943 | 852 | 761 | 755 | 863 | 517 | 658 | 636 | 907 | 970 | 921 | 1045 | 909 | 1023 | 888 | 804 | 880 | 734 | 655 | 599 | 636 | 724 | Zfp874b |
| Zfp879 | 76 | 55 | 77 | 87 | 101 | 81 | 70 | 237 | 199 | 181 | 148 | 160 | 159 | 134 | 94 | 58 | 74 | 68 | 84 | 110 | 88 | 209 | 175 | 168 | 146 | 113 | 124 | 140 | Zfp879 |
| Zfp882 | 1405 | 1184 | 1056 | 1244 | 1467 | 1518 | 1506 | 2865 | 2209 | 2034 | 2088 | 2041 | 1955 | 2089 | 1486 | 1072 | 1265 | 1130 | 1693 | 1766 | 1522 | 2060 | 2040 | 1976 | 2153 | 1276 | 1985 | 1909 | Zfp882 |
| Zfp9 | 2064 | 1599 | 1771 | 1826 | 2214 | 2164 | 1852 | 2566 | 2667 | 2644 | 2815 | 2070 | 2422 | 2111 | 1941 | 2076 | 1992 | 2169 | 1790 | 1912 | 1699 | 3018 | 2853 | 2240 | 2498 | 2111 | 1988 | 1926 | Zfp9 |
| Zfp90 | 2187 | 1821 | 1941 | 1792 | 2509 | 2475 | 2370 | 4444 | 4139 | 4311 | 3791 | 4644 | 3911 | 4293 | 2189 | 1678 | 2227 | 1906 | 2586 | 2356 | 2313 | 4508 | 3978 | 4358 | 4765 | 4078 | 4051 | 4237 | Zfp90 |
| Zfp91 | 8797 | 6613 | 8223 | 8034 | 8542 | 8994 | 9867 | 12993 | 10467 | 10353 | 10573 | 11979 | 11620 | 13568 | 8864 | 7080 | 7938 | 7689 | 8310 | 9365 | 9103 | 11570 | 10108 | 11153 | 11392 | 12098 | 12181 | 13180 | Zfp91 |
| Zfp92 | 93 | 90 | 86 | 85 | 168 | 95 | 141 | 99 | 110 | 127 | 85 | 77 | 82 | 66 | 73 | 77 | 98 | 87 | 101 | 124 | 125 | 119 | 100 | 96 | 149 | 30 | 48 | 64 | Zfp92 |
| Zfp93 | 761 | 713 | 633 | 642 | 828 | 758 | 792 | 973 | 854 | 800 | 771 | 746 | 793 | 792 | 703 | 637 | 706 | 662 | 706 | 795 | 703 | 792 | 870 | 962 | 732 | 933 | 791 | 722 | Zfp93 |
| Zfp930 | 1677 | 1372 | 1780 | 1691 | 1589 | 1612 | 1677 | 1849 | 1738 | 1561 | 1189 | 1760 | 1605 | 1643 | 1849 | 1742 | 1703 | 1471 | 1578 | 1962 | 1733 | 1554 | 1730 | 1635 | 1472 | 1919 | 1829 | 1677 | Zfp930 |
| Zfp931 | 1111 | 704 | 945 | 755 | 841 | 964 | 798 | 985 | 876 | 669 | 800 | 716 | 756 | 705 | 1147 | 1057 | 742 | 793 | 761 | 1024 | 794 | 600 | 944 | 724 | 707 | 792 | 841 | 780 | Zfp931 |
| Zfp932 | 3971 | 2945 | 3549 | 3815 | 4389 | 4688 | 4780 | 3968 | 3438 | 3596 | 3549 | 3519 | 3424 | 3818 | 3939 | 3604 | 3773 | 3803 | 4404 | 4415 | 4602 | 4141 | 3404 | 3537 | 3647 | 3523 | 3764 | 3429 | Zfp932 |
| Zfp933 | 2269 | 1882 | 2316 | 2179 | 2105 | 2300 | 2364 | 2446 | 2149 | 2208 | 2171 | 2229 | 2188 | 2369 | 2425 | 2031 | 2400 | 2221 | 2261 | 2223 | 2160 | 2413 | 2149 | 2383 | 2491 | 2467 | 2203 | 2292 | Zfp933 |
| Zfp934 | 729 | 462 | 871 | 569 | 603 | 699 | 731 | 904 | 842 | 813 | 888 | 895 | 1038 | 1004 | 736 | 677 | 705 | 571 | 577 | 683 | 578 | 942 | 845 | 914 | 752 | 1203 | 963 | 1075 | Zfp934 |
| Zfp935 | 247 | 191 | 230 | 217 | 197 | 213 | 236 | 276 | 259 | 281 | 293 | 266 | 261 | 298 | 185 | 210 | 154 | 196 | 212 | 276 | 254 | 222 | 257 | 304 | 257 | 258 | 268 | 316 | Zfp935 |
| Zfp936 | 1 | 1 | 0 | 0 | 0 | 1 | 1 | 4 | 0 | 1 | 0 | 0 | 0 | 0 | 0 | 3 | 0 | 0 | 1 | 1 | 0 | 2 | 0 | 0 | 0 | 0 | 6 | 0 | Zfp936 |
| Zfp937 | 497 | 426 | 422 | 442 | 563 | 527 | 477 | 780 | 652 | 581 | 632 | 569 | 684 | 606 | 633 | 540 | 484 | 447 | 580 | 608 | 487 | 662 | 704 | 555 | 559 | 515 | 619 | 574 | Zfp937 |
| Zfp938 | 1646 | 1227 | 1579 | 1480 | 1597 | 1547 | 1840 | 1633 | 1430 | 1461 | 1316 | 1366 | 1372 | 1441 | 1863 | 1530 | 1524 | 1446 | 1563 | 1845 | 1673 | 1427 | 1476 | 1261 | 1555 | 1325 | 1438 | 1588 | Zfp938 |
| Zfp94 | 514 | 570 | 611 | 641 | 664 | 608 | 634 | 760 | 873 | 923 | 839 | 886 | 831 | 855 | 560 | 519 | 592 | 670 | 711 | 704 | 626 | 856 | 818 | 948 | 970 | 863 | 868 | 835 | Zfp94 |
| Zfp940 | 1694 | 1481 | 1360 | 1355 | 1539 | 1570 | 1882 | 1605 | 1460 | 1398 | 1198 | 1431 | 1159 | 1360 | 1664 | 1276 | 1527 | 1536 | 1582 | 1832 | 1770 | 1384 | 1330 | 1362 | 1385 | 1318 | 1392 | 1346 | Zfp940 |
| Zfp941 | 1324 | 1180 | 1068 | 1320 | 1429 | 1496 | 1293 | 1254 | 1083 | 1008 | 859 | 930 | 846 | 892 | 1501 | 1160 | 1182 | 1153 | 1267 | 1462 | 1307 | 1395 | 1046 | 823 | 989 | 653 | 869 | 777 | Zfp941 |
| Zfp942 | 3675 | 2500 | 3221 | 3100 | 2882 | 3428 | 3579 | 2754 | 2506 | 2242 | 2083 | 2417 | 2279 | 2340 | 3857 | 3481 | 3174 | 2933 | 2885 | 3530 | 3391 | 2401 | 2410 | 2271 | 2064 | 2409 | 2407 | 2366 | Zfp942 |
| Zfp943 | 2092 | 1417 | 1968 | 1937 | 1976 | 2271 | 2370 | 2386 | 2009 | 2131 | 1983 | 2240 | 2258 | 2206 | 2372 | 2079 | 2214 | 1798 | 2014 | 2346 | 2194 | 2254 | 2076 | 2276 | 2133 | 2282 | 2234 | 2307 | Zfp943 |
| Zfp944 | 2719 | 2204 | 2809 | 3136 | 2771 | 3062 | 3612 | 1789 | 1556 | 1650 | 1512 | 1821 | 1590 | 1855 | 3103 | 2801 | 3085 | 3068 | 2826 | 3354 | 3143 | 1801 | 1632 | 1696 | 1829 | 1748 | 1773 | 1855 | Zfp944 |
| Zfp945 | 4841 | 2971 | 3134 | 3595 | 4408 | 5077 | 4900 | 5929 | 5548 | 4209 | 4479 | 4735 | 4923 | 5089 | 5070 | 3932 | 3310 | 3345 | 3824 | 5734 | 4610 | 4495 | 5188 | 3903 | 4278 | 3509 | 5088 | 4571 | Zfp945 |
| Zfp946 | 4851 | 3340 | 5051 | 5112 | 5962 | 5982 | 6781 | 4920 | 3845 | 4302 | 3640 | 4360 | 4003 | 4559 | 5521 | 4293 | 5366 | 5226 | 5828 | 5989 | 5935 | 4609 | 3876 | 4266 | 4239 | 4172 | 4312 | 4333 | Zfp946 |
| Zfp947 | 332 | 161 | 280 | 227 | 284 | 254 | 305 | 258 | 257 | 205 | 236 | 234 | 281 | 202 | 235 | 267 | 265 | 216 | 238 | 278 | 248 | 197 | 266 | 218 | 160 | 245 | 277 | 214 | Zfp947 |
| Zfp948 | 3054 | 2594 | 2711 | 3329 | 2781 | 3090 | 3026 | 1981 | 1888 | 2064 | 1786 | 1870 | 1691 | 1796 | 2996 | 2886 | 2487 | 3097 | 2580 | 3137 | 2851 | 2077 | 1941 | 1670 | 1838 | 1761 | 1628 | 1604 | Zfp948 |
| Zfp949 | 2006 | 1686 | 2055 | 1990 | 2150 | 2110 | 2197 | 1832 | 1548 | 1729 | 1751 | 1396 | 1683 | 1379 | 2069 | 1823 | 1962 | 1962 | 2254 | 2026 | 2137 | 1907 | 1493 | 1558 | 1701 | 1381 | 1426 | 1321 | Zfp949 |
| Zfp950 | 3843 | 2673 | 2263 | 2386 | 2643 | 3279 | 2939 | 4787 | 4596 | 3516 | 4644 | 3209 | 3722 | 3675 | 3429 | 3778 | 2371 | 2609 | 2390 | 3336 | 2930 | 3551 | 4570 | 3246 | 3434 | 2672 | 3348 | 3088 | Zfp950 |
| Zfp951 | 240 | 184 | 330 | 166 | 313 | 270 | 215 | 436 | 389 | 318 | 430 | 306 | 547 | 400 | 297 | 312 | 212 | 195 | 208 | 294 | 236 | 500 | 517 | 322 | 354 | 341 | 447 | 362 | Zfp951 |
| Zfp952 | 2799 | 2614 | 2515 | 2639 | 2886 | 2899 | 3127 | 3094 | 2685 | 2475 | 2472 | 2581 | 2364 | 2669 | 2970 | 2641 | 2656 | 2852 | 2816 | 3096 | 2930 | 2846 | 2700 | 2523 | 2543 | 2377 | 2545 | 2617 | Zfp952 |
| Zfp953 | 517 | 352 | 395 | 370 | 367 | 376 | 332 | 409 | 367 | 402 | 386 | 348 | 404 | 344 | 504 | 424 | 401 | 319 | 327 | 500 | 366 | 378 | 434 | 350 | 337 | 344 | 393 | 370 | Zfp953 |
| Zfp954 | 1521 | 1646 | 1152 | 1450 | 1342 | 1504 | 1369 | 1349 | 1121 | 1030 | 904 | 928 | 838 | 862 | 1618 | 1471 | 1415 | 1437 | 1444 | 1549 | 1401 | 1088 | 1106 | 950 | 1093 | 950 | 883 | 939 | Zfp954 |
| Zfp955a | 3398 | 2945 | 2976 | 3053 | 2727 | 3151 | 3169 | 3352 | 2948 | 2543 | 2398 | 2487 | 2521 | 2519 | 3899 | 3290 | 2979 | 3179 | 2850 | 3842 | 2943 | 2811 | 3016 | 2427 | 2623 | 2607 | 2599 | 2573 | Zfp955a |
| Zfp955b | 4899 | 4426 | 3673 | 4686 | 4294 | 4541 | 4835 | 4246 | 3879 | 3476 | 3444 | 3295 | 3334 | 3257 | 5280 | 4607 | 3867 | 4645 | 4050 | 5488 | 4678 | 3838 | 3795 | 3070 | 3537 | 2690 | 3350 | 3396 | Zfp955b |
| Zfp956 | 1044 | 1046 | 995 | 949 | 1201 | 1155 | 1083 | 1044 | 1157 | 1170 | 1369 | 834 | 1046 | 832 | 1058 | 974 | 1109 | 1199 | 1202 | 1244 | 1109 | 1203 | 1155 | 1117 | 1169 | 1039 | 900 | 849 | Zfp956 |
| Zfp957 | 0 | 0 | 0 | 0 | 0 | 0 | 2 | 0 | 0 | 0 | 0 | 0 | 0 | 0 | 0 | 0 | 0 | 0 | 0 | 0 | 0 | 3 | 0 | 0 | 3 | 0 | 0 | 0 | Zfp957 |
| Zfp958 | 2202 | 1919 | 2074 | 1938 | 1712 | 2055 | 1975 | 1657 | 1438 | 1204 | 1291 | 1333 | 1359 | 1352 | 2387 | 2236 | 2162 | 2008 | 1862 | 2337 | 2032 | 1521 | 1543 | 1445 | 1395 | 1229 | 1347 | 1351 | Zfp958 |
| Zfp959 | 689 | 501 | 621 | 630 | 666 | 623 | 745 | 1106 | 859 | 822 | 823 | 1071 | 858 | 927 | 653 | 611 | 688 | 578 | 716 | 771 | 651 | 1018 | 865 | 917 | 917 | 1018 | 959 | 1082 | Zfp959 |
| Zfp960 | 307 | 203 | 269 | 182 | 197 | 223 | 208 | 200 | 205 | 131 | 149 | 182 | 193 | 211 | 307 | 262 | 213 | 196 | 195 | 299 | 197 | 175 | 221 | 174 | 127 | 201 | 189 | 201 | Zfp960 |
| Zfp961 | 2638 | 1969 | 2689 | 2070 | 2294 | 2157 | 2527 | 2223 | 1833 | 1885 | 1700 | 2281 | 1934 | 2316 | 2451 | 2249 | 2359 | 2246 | 2357 | 2539 | 2239 | 2015 | 1716 | 2251 | 2034 | 2504 | 2037 | 2190 | Zfp961 |
| Zfp963 | 277 | 237 | 333 | 352 | 338 | 341 | 437 | 372 | 307 | 445 | 411 | 442 | 428 | 530 | 287 | 224 | 329 | 343 | 343 | 313 | 379 | 568 | 379 | 500 | 433 | 514 | 420 | 522 | Zfp963 |
| Zfp964 | 952 | 840 | 682 | 813 | 790 | 861 | 850 | 695 | 684 | 553 | 587 | 570 | 492 | 521 | 1023 | 841 | 693 | 894 | 737 | 878 | 861 | 597 | 685 | 613 | 588 | 562 | 508 | 548 | Zfp964 |
| Zfp965 | 75 | 30 | 36 | 66 | 51 | 76 | 45 | 93 | 64 | 50 | 54 | 59 | 92 | 49 | 71 | 70 | 29 | 68 | 27 | 75 | 52 | 32 | 55 | 46 | 68 | 24 | 57 | 43 | Zfp965 |
| Zfp966 | 10 | 0 | 7 | 9 | 9 | 9 | 35 | 10 | 11 | 8 | 6 | 5 | 18 | 12 | 30 | 16 | 1 | 22 | 5 | 14 | 8 | 7 | 17 | 32 | 11 | 8 | 21 | 15 | Zfp966 |
| Zfp967 | 3 | 2 | 0 | 1 | 4 | 4 | 2 | 1 | 4 | 8 | 3 | 5 | 2 | 1 | 3 | 5 | 5 | 1 | 2 | 2 | 1 | 8 | 4 | 5 | 1 | 2 | 1 | 1 | Zfp967 |
| Zfp968 | 22 | 32 | 4 | 14 | 18 | 32 | 12 | 33 | 26 | 9 | 13 | 21 | 24 | 17 | 25 | 29 | 13 | 10 | 17 | 43 | 26 | 22 | 28 | 10 | 23 | 3 | 27 | 17 | Zfp968 |
| Zfp968-ps | 7 | 1 | 7 | 6 | 0 | 8 | 4 | 6 | 7 | 3 | 1 | 3 | 5 | 2 | 12 | 6 | 1 | 4 | 3 | 3 | 5 | 7 | 5 | 7 | 6 | 2 | 7 | 0 | Zfp968-ps |
| Zfp969 | 163 | 54 | 18 | 46 | 101 | 125 | 48 | 144 | 82 | 40 | 111 | 127 | 65 | 132 | 145 | 65 | 43 | 26 | 41 | 47 | 38 | 89 | 144 | 93 | 40 | 73 | 38 | 97 | Zfp969 |
| Zfp97 | 262 | 193 | 192 | 150 | 168 | 253 | 153 | 165 | 180 | 183 | 198 | 168 | 202 | 192 | 262 | 246 | 229 | 127 | 181 | 257 | 191 | 154 | 246 | 189 | 181 | 164 | 163 | 154 | Zfp97 |
| Zfp970 | 1265 | 919 | 778 | 849 | 912 | 1045 | 1009 | 937 | 1009 | 872 | 1032 | 903 | 1124 | 899 | 1404 | 1266 | 900 | 828 | 918 | 1140 | 936 | 1031 | 1048 | 885 | 944 | 788 | 952 | 869 | Zfp970 |
| Zfp971 | 1172 | 950 | 577 | 700 | 866 | 970 | 815 | 1141 | 1114 | 841 | 920 | 827 | 885 | 792 | 1280 | 1185 | 695 | 664 | 806 | 1083 | 778 | 785 | 1056 | 619 | 768 | 294 | 926 | 693 | Zfp971 |
| Zfp972 | 7 | 10 | 12 | 15 | 12 | 10 | 15 | 5 | 9 | 41 | 25 | 23 | 23 | 15 | 15 | 11 | 13 | 28 | 9 | 18 | 18 | 10 | 23 | 17 | 20 | 30 | 24 | 17 | Zfp972 |
| Zfp973 | 283 | 222 | 133 | 154 | 198 | 214 | 184 | 226 | 252 | 139 | 306 | 159 | 174 | 214 | 364 | 183 | 180 | 107 | 138 | 193 | 227 | 155 | 246 | 128 | 173 | 89 | 189 | 227 | Zfp973 |
| Zfp974 | 2224 | 2138 | 1702 | 2134 | 2036 | 2238 | 2181 | 2629 | 1954 | 1760 | 1719 | 1883 | 1742 | 1936 | 2352 | 2227 | 1880 | 2089 | 1869 | 2483 | 2164 | 2087 | 1950 | 1577 | 2027 | 1286 | 1915 | 1854 | Zfp974 |
| Zfp975 | 624 | 477 | 863 | 887 | 889 | 670 | 733 | 774 | 703 | 860 | 819 | 781 | 892 | 851 | 761 | 801 | 846 | 766 | 792 | 684 | 649 | 906 | 707 | 788 | 792 | 807 | 698 | 876 | Zfp975 |
| Zfp976 | 776 | 573 | 936 | 895 | 1012 | 943 | 1000 | 868 | 761 | 691 | 696 | 797 | 822 | 751 | 800 | 809 | 941 | 821 | 1019 | 860 | 963 | 815 | 744 | 713 | 690 | 657 | 760 | 657 | Zfp976 |
| Zfp977 | 112 | 102 | 102 | 126 | 143 | 147 | 149 | 129 | 110 | 125 | 129 | 90 | 115 | 83 | 120 | 132 | 133 | 183 | 173 | 161 | 101 | 147 | 137 | 112 | 113 | 108 | 102 | 100 | Zfp977 |
| Zfp978 | 3 | 0 | 0 | 0 | 1 | 0 | 1 | 0 | 1 | 0 | 3 | 0 | 1 | 1 | 0 | 2 | 0 | 2 | 2 | 0 | 0 | 2 | 2 | 1 | 0 | 0 | 0 | 1 | Zfp978 |
| Zfp979 | 99 | 58 | 163 | 67 | 136 | 112 | 73 | 285 | 253 | 232 | 294 | 215 | 319 | 242 | 141 | 140 | 83 | 111 | 95 | 112 | 79 | 314 | 325 | 203 | 233 | 304 | 308 | 272 | Zfp979 |
| Zfp980 | 3 | 0 | 7 | 1 | 2 | 0 | 2 | 2 | 6 | 2 | 3 | 1 | 3 | 1 | 0 | 4 | 1 | 1 | 0 | 4 | 0 | 0 | 5 | 1 | 4 | 2 | 1 | 2 | Zfp980 |

| | | | | | | | | | | | | | | | | | | | | | | | | | | | | | |
|---|---|---|---|---|---|---|---|---|---|---|---|---|---|---|---|---|---|---|---|---|---|---|---|---|---|---|---|---|---|
| Zfp981 | 23 | 24 | 37 | 61 | 68 | 26 | 49 | 13 | 16 | 19 | 21 | 17 | 11 | 19 | 58 | 53 | 28 | 37 | 29 | 50 | 38 | 48 | 21 | 18 | 17 | 20 | 17 | 17 | Zfp981 |
| Zfp982 | 42 | 46 | 72 | 52 | 47 | 41 | 29 | 180 | 131 | 139 | 113 | 104 | 203 | 135 | 51 | 72 | 48 | 37 | 48 | 62 | 39 | 130 | 175 | 89 | 107 | 128 | 177 | 142 | Zfp982 |
| Zfp983 | 1322 | 1358 | 1472 | 1754 | 1774 | 1751 | 1926 | 2023 | 1490 | 1617 | 1582 | 1498 | 1450 | 1540 | 1437 | 1316 | 1559 | 1641 | 1615 | 1824 | 1654 | 2003 | 1544 | 1539 | 1760 | 1268 | 1446 | 1496 | Zfp983 |
| Zfp984 | 881 | 578 | 1006 | 949 | 875 | 827 | 1021 | 775 | 768 | 715 | 651 | 820 | 792 | 752 | 1004 | 900 | 912 | 951 | 828 | 975 | 869 | 832 | 849 | 695 | 650 | 981 | 803 | 909 | Zfp984 |
| Zfp985 | 6 | 3 | 7 | 8 | 0 | 4 | 9 | 2 | 12 | 8 | 9 | 5 | 7 | 6 | 3 | 4 | 12 | 1 | 5 | 5 | 3 | 7 | 11 | 7 | 12 | 9 | 2 | 13 | Zfp985 |
| Zfp986 | 6 | 2 | 0 | 17 | 15 | 18 | 8 | 5 | 2 | 8 | 1 | 3 | 3 | 1 | 15 | 0 | 3 | 14 | 8 | 0 | 15 | 12 | 3 | 7 | 6 | 0 | 0 | 1 | Zfp986 |
| Zfp987 | 0 | 0 | 0 | 0 | 0 | 0 | 0 | 0 | 0 | 0 | 0 | 0 | 0 | 0 | 0 | 0 | 0 | 0 | 0 | 1 | 1 | 0 | 0 | 0 | 0 | 0 | 0 | 0 | Zfp987 |
| Zfp988 | 0 | 0 | 3 | 3 | 10 | 1 | 7 | 2 | 1 | 17 | 0 | 0 | 0 | 1 | 8 | 5 | 5 | 8 | 1 | 0 | 7 | 3 | 0 | 0 | 0 | 3 | 0 | 1 | Zfp988 |
| Zfp989 | 1 | 1 | 1 | 1 | 1 | 0 | 0 | 1 | 0 | 0 | 1 | 0 | 1 | 1 | 5 | 2 | 1 | 2 | 0 | 6 | 0 | 0 | 2 | 0 | 2 | 0 | 2 | 0 | Zfp989 |
| Zfp990 | 5 | 1 | 3 | 4 | 10 | 10 | 3 | 4 | 4 | 14 | 6 | 0 | 0 | 3 | 3 | 12 | 8 | 9 | 2 | 12 | 10 | 2 | 2 | 2 | 0 | 3 | 2 | 2 | Zfp990 |
| Zfp991 | 431 | 238 | 489 | 312 | 313 | 399 | 305 | 425 | 419 | 307 | 342 | 391 | 374 | 404 | 464 | 541 | 333 | 306 | 358 | 454 | 321 | 371 | 461 | 330 | 352 | 377 | 439 | 530 | Zfp991 |
| Zfp992 | 309 | 160 | 288 | 148 | 182 | 219 | 116 | 263 | 242 | 185 | 179 | 139 | 218 | 143 | 361 | 389 | 111 | 116 | 118 | 283 | 166 | 155 | 294 | 105 | 168 | 171 | 229 | 169 | Zfp992 |
| Zfp993 | 112 | 49 | 196 | 139 | 74 | 85 | 83 | 101 | 129 | 117 | 141 | 48 | 227 | 70 | 107 | 157 | 71 | 76 | 113 | 88 | 80 | 94 | 177 | 69 | 110 | 112 | 114 | 69 | Zfp993 |
| Zfp994 | 810 | 557 | 812 | 862 | 1006 | 854 | 894 | 1230 | 1088 | 1077 | 880 | 1072 | 1015 | 1001 | 845 | 765 | 892 | 765 | 863 | 1088 | 857 | 1089 | 1126 | 1045 | 1089 | 921 | 1097 | 1025 | Zfp994 |
| Zfp995 | 912 | 625 | 951 | 1125 | 1103 | 1144 | 1322 | 1011 | 924 | 1031 | 979 | 1103 | 987 | 996 | 1106 | 829 | 991 | 1089 | 1097 | 1146 | 1189 | 1023 | 845 | 954 | 980 | 1076 | 1080 | 997 | Zfp995 |
| Zfpl1 | 1024 | 1202 | 1324 | 1288 | 1010 | 1054 | 1069 | 939 | 1054 | 1272 | 1099 | 1077 | 1064 | 1191 | 949 | 1020 | 1264 | 1242 | 1133 | 1019 | 1134 | 899 | 1062 | 1180 | 1105 | 1268 | 1067 | 1151 | Zfpl1 |
| Zfpm1 | 1633 | 2222 | 2201 | 2163 | 1969 | 1748 | 1209 | 548 | 666 | 732 | 732 | 743 | 981 | 734 | 1629 | 1981 | 2299 | 2038 | 2145 | 1451 | 1514 | 824 | 676 | 710 | 675 | 542 | 793 | 763 | Zfpm1 |
| Zfpm2 | 1053 | 1169 | 1205 | 1480 | 2176 | 1881 | 1838 | 1862 | 1594 | 1960 | 1700 | 1733 | 1924 | 1914 | 1167 | 1026 | 1395 | 1815 | 1954 | 1753 | 2157 | 2438 | 1585 | 2049 | 2349 | 970 | 1988 | 1828 | Zfpm2 |
| Zfr | 15190 | 12971 | 13262 | 13819 | 15277 | 14640 | 15756 | 21426 | 17549 | 16296 | 16883 | 16940 | 15940 | 17060 | 16118 | 13113 | 13818 | 14198 | 14496 | 16192 | 15683 | 18997 | 17988 | 16496 | 17031 | 15800 | 17066 | 17451 | Zfr |
| Zfr2 | 2307 | 2930 | 2933 | 4029 | 3792 | 3351 | 3131 | 1428 | 1846 | 2105 | 2017 | 1627 | 1531 | 1464 | 2421 | 2700 | 3509 | 4084 | 3845 | 2919 | 3355 | 2398 | 1655 | 1894 | 1948 | 1180 | 1412 | 1321 | Zfr2 |
| Zfta | 6614 | 7348 | 9524 | 9225 | 11178 | 9304 | 9562 | 21058 | 20618 | 26096 | 24516 | 27303 | 23605 | 28222 | 6597 | 5686 | 10527 | 9370 | 11925 | 8787 | 9950 | 25308 | 18485 | 28845 | 29672 | 26268 | 25543 | 25299 | Zfta |
| Zfx | 2766 | 2027 | 2570 | 2378 | 2811 | 2830 | 2744 | 7347 | 6387 | 5284 | 6306 | 6622 | 6330 | 6854 | 2742 | 2309 | 2378 | 2111 | 2724 | 2975 | 2611 | 6274 | 6273 | 6076 | 6073 | 5338 | 6968 | 6928 | Zfx |
| Zfy1 | 0 | 1 | 2 | 0 | 3 | 6 | 2 | 0 | 2 | 0 | 4 | 2 | 7 | 11 | 0 | 0 | 0 | 0 | 0 | 0 | 0 | 0 | 0 | 0 | 0 | 0 | 0 | 0 | Zfy1 |
| Zfyve1 | 4346 | 4904 | 4526 | 4870 | 5257 | 4866 | 4805 | 4798 | 4630 | 4806 | 4688 | 5109 | 4820 | 4906 | 4251 | 4158 | 4770 | 4803 | 5520 | 4709 | 5009 | 5015 | 4566 | 5044 | 5374 | 4391 | 5187 | 4895 | Zfyve1 |
| Zfyve16 | 5018 | 4108 | 3368 | 4074 | 4057 | 4662 | 4354 | 5244 | 4805 | 3911 | 4573 | 3678 | 3892 | 4181 | 5246 | 4936 | 3644 | 4173 | 3624 | 4914 | 4576 | 4278 | 5042 | 3631 | 4227 | 3003 | 3942 | 3923 | Zfyve16 |
| Zfyve19 | 1799 | 2181 | 1991 | 2009 | 1798 | 1938 | 1837 | 1463 | 1773 | 1574 | 1623 | 1543 | 1591 | 1567 | 1683 | 1901 | 1916 | 1839 | 1958 | 1888 | 1719 | 1491 | 1814 | 1580 | 1556 | 1751 | 1683 | 1576 | Zfyve19 |
| Zfyve21 | 1810 | 2137 | 1905 | 1665 | 1061 | 1363 | 1254 | 983 | 1210 | 985 | 1179 | 1086 | 1068 | 895 | 1640 | 2033 | 1774 | 1598 | 1252 | 1353 | 1214 | 952 | 1144 | 1119 | 943 | 1307 | 1089 | 1016 | Zfyve21 |
| Zfyve26 | 2020 | 2271 | 1541 | 1901 | 2112 | 2130 | 2048 | 2536 | 2992 | 2735 | 2963 | 3249 | 3079 | 2974 | 1970 | 2065 | 1881 | 2072 | 2162 | 1905 | 2154 | 2720 | 2699 | 2519 | 2864 | 1978 | 3206 | 2917 | Zfyve26 |
| Zfyve27 | 6817 | 8499 | 6981 | 9752 | 9423 | 8527 | 8936 | 4780 | 4857 | 5240 | 5073 | 4846 | 4749 | 4768 | 7639 | 7191 | 8615 | 10118 | 9453 | 8558 | 9137 | 5858 | 4393 | 5017 | 5665 | 3784 | 4692 | 4586 | Zfyve27 |
| Zfyve28 | 1044 | 1361 | 1360 | 1435 | 1602 | 1313 | 1293 | 294 | 412 | 315 | 384 | 304 | 327 | 269 | 1113 | 1205 | 1301 | 1471 | 1372 | 1277 | 1318 | 411 | 342 | 340 | 429 | 177 | 259 | 243 | Zfyve28 |
| Zfyve9 | 2959 | 2236 | 2077 | 2182 | 2833 | 2776 | 2665 | 6305 | 5759 | 5286 | 5138 | 4954 | 5196 | 5224 | 2819 | 2406 | 2369 | 2226 | 2655 | 2806 | 2699 | 5431 | 5478 | 4796 | 5244 | 3985 | 5199 | 5161 | Zfyve9 |
| Zg16 | 1 | 0 | 0 | 0 | 0 | 0 | 1 | 0 | 0 | 0 | 0 | 0 | 2 | 2 | 3 | 0 | 0 | 0 | 2 | 0 | 0 | 0 | 0 | 0 | 0 | 1 | 1 | 0 | Zg16 |
| Zglp1 | 0 | 0 | 0 | 0 | 0 | 0 | 0 | 0 | 0 | 0 | 0 | 0 | 2 | 0 | 0 | 0 | 0 | 1 | 0 | 0 | 0 | 0 | 2 | 0 | 1 | 0 | 0 | 0 | Zglp1 |
| Zgpat | 3147 | 4024 | 3289 | 4452 | 4610 | 4005 | 3880 | 3679 | 3816 | 4028 | 4022 | 3919 | 3615 | 3726 | 3227 | 3338 | 3904 | 4614 | 4490 | 3865 | 4241 | 4274 | 3599 | 4106 | 4428 | 2747 | 3640 | 3584 | Zgpat |
| Zgrf1 | 5651 | 4073 | 3085 | 3367 | 2478 | 3600 | 3979 | 3008 | 3044 | 2412 | 2400 | 3720 | 3316 | 3909 | 5116 | 3971 | 3390 | 3101 | 2974 | 4180 | 3432 | 1744 | 2519 | 2622 | 2452 | 3047 | 4038 | 3574 | Zgrf1 |
| Zhx2 | 671 | 816 | 1110 | 992 | 1162 | 895 | 841 | 1371 | 1858 | 1990 | 1864 | 2209 | 1414 | 1829 | 790 | 833 | 1166 | 1062 | 1193 | 791 | 915 | 1755 | 1563 | 2095 | 2018 | 2350 | 1571 | 1608 | Zhx2 |
| Zhx3 | 3354 | 3579 | 3192 | 3935 | 4193 | 4109 | 4172 | 2656 | 2931 | 2910 | 3047 | 2791 | 2485 | 2780 | 3517 | 3406 | 3719 | 4058 | 4202 | 3860 | 4482 | 3141 | 2680 | 2559 | 3519 | 2249 | 2721 | 2529 | Zhx3 |
| Zic1 | 13906 | 14299 | 23177 | 19310 | 19617 | 18194 | 20021 | 17837 | 16294 | 19311 | 18557 | 22769 | 19367 | 26285 | 16764 | 12930 | 19546 | 16527 | 18763 | 17406 | 18888 | 23255 | 20995 | 21046 | 22915 | 23194 | 23178 | 25648 | Zic1 |
| Zic2 | 3812 | 4326 | 8208 | 5637 | 5479 | 5004 | 5474 | 5255 | 4803 | 5701 | 5426 | 7617 | 6843 | 8132 | 4876 | 4599 | 6217 | 5437 | 5622 | 4928 | 4915 | 7001 | 5902 | 6699 | 7112 | 8435 | 7475 | 7977 | Zic2 |
| Zic3 | 3876 | 4220 | 6559 | 5573 | 4709 | 5374 | 5001 | 4426 | 3376 | 4186 | 3709 | 5999 | 5459 | 6259 | 4953 | 3717 | 5657 | 4723 | 5359 | 4466 | 5006 | 5852 | 4084 | 4864 | 4887 | 4602 | 5958 | 6697 | Zic3 |
| Zic4 | 4387 | 4239 | 6653 | 4891 | 5920 | 5742 | 6387 | 6309 | 5669 | 6470 | 5975 | 8450 | 7099 | 9207 | 4874 | 3585 | 5390 | 4714 | 6274 | 5738 | 5996 | 7855 | 7180 | 6654 | 7517 | 7203 | 8808 | 8850 | Zic4 |
| Zic5 | 2615 | 2553 | 5515 | 3676 | 3386 | 3205 | 3691 | 2734 | 2125 | 2794 | 2494 | 3928 | 3395 | 4326 | 3289 | 2817 | 3923 | 3153 | 3493 | 3159 | 3578 | 3716 | 2858 | 3385 | 3485 | 4313 | 3850 | 4546 | Zic5 |
| Zik1 | 7128 | 6643 | 6246 | 7007 | 7634 | 8427 | 8558 | 11017 | 8646 | 7451 | 6517 | 9091 | 7542 | 8928 | 8325 | 6650 | 6924 | 7340 | 8124 | 10210 | 8110 | 8843 | 8074 | 7669 | 9102 | 8435 | 9082 | 9016 | Zik1 |
| Zim1 | 3510 | 3085 | 1594 | 2616 | 3239 | 4307 | 3015 | 2828 | 5288 | 3716 | 12532 | 1808 | 2617 | 1925 | 3479 | 3566 | 1965 | 3952 | 2365 | 2999 | 3053 | 5093 | 6661 | 2453 | 3316 | 1545 | 1739 | 1353 | Zim1 |
| Zkscan1 | 7605 | 6643 | 6232 | 6955 | 7118 | 7533 | 7649 | 10359 | 9105 | 8398 | 8098 | 8787 | 8417 | 9137 | 8058 | 6330 | 6672 | 6687 | 6943 | 8194 | 7702 | 9403 | 8501 | 8510 | 9638 | 7572 | 9302 | 9200 | Zkscan1 |
| Zkscan14 | 689 | 866 | 563 | 861 | 640 | 632 | 549 | 542 | 564 | 512 | 506 | 540 | 431 | 476 | 733 | 738 | 666 | 692 | 620 | 694 | 591 | 597 | 550 | 530 | 495 | 410 | 535 | 475 | Zkscan14 |
| Zkscan16 | 1285 | 1233 | 388 | 753 | 725 | 1131 | 855 | 602 | 503 | 412 | 607 | 407 | 535 | 408 | 1120 | 1211 | 538 | 733 | 634 | 1171 | 976 | 348 | 537 | 315 | 399 | 122 | 427 | 333 | Zkscan16 |
| Zkscan17 | 2595 | 2725 | 3302 | 2515 | 2531 | 2611 | 2563 | 2378 | 2778 | 2932 | 2777 | 4020 | 3576 | 3703 | 2383 | 2588 | 3142 | 2505 | 2916 | 2082 | 2517 | 2635 | 2606 | 3536 | 3170 | 4407 | 3570 | 3906 | Zkscan17 |
| Zkscan2 | 4812 | 4321 | 3646 | 5415 | 7017 | 6835 | 7062 | 6099 | 5716 | 6323 | 5912 | 6163 | 5342 | 5943 | 5015 | 4391 | 4616 | 5892 | 5866 | 5594 | 7184 | 7583 | 5120 | 5171 | 6786 | 3377 | 5688 | 5041 | Zkscan2 |
| Zkscan3 | 13610 | 13468 | 10507 | 11566 | 11221 | 12045 | 12481 | 12838 | 11456 | 11109 | 10742 | 11263 | 10646 | 11704 | 13337 | 12822 | 11792 | 11707 | 12029 | 13038 | 12689 | 10828 | 10954 | 10607 | 11054 | 9256 | 11459 | 11584 | Zkscan3 |
| Zkscan4 | 550 | 535 | 548 | 617 | 625 | 604 | 616 | 579 | 543 | 608 | 508 | 608 | 542 | 575 | 521 | 441 | 667 | 578 | 666 | 674 | 558 | 638 | 478 | 580 | 644 | 600 | 546 | 523 | Zkscan4 |
| Zkscan5 | 2321 | 2503 | 1984 | 2276 | 2108 | 2170 | 2118 | 2141 | 2024 | 2007 | 1960 | 2181 | 2088 | 2087 | 2298 | 2114 | 1917 | 1982 | 2075 | 2127 | 2019 | 1775 | 1911 | 2026 | 1865 | 2109 | 2196 | 2074 | Zkscan5 |
| Zkscan6 | 1226 | 1519 | 1234 | 1632 | 1314 | 1286 | 1421 | 1104 | 1177 | 1160 | 1086 | 991 | 1060 | 1001 | 1303 | 1221 | 1223 | 1473 | 1319 | 1495 | 1463 | 1071 | 1175 | 1065 | 1228 | 1079 | 1044 | 1035 | Zkscan6 |
| Zkscan7 | 2240 | 2204 | 2214 | 2270 | 2084 | 2328 | 2259 | 2174 | 2113 | 1962 | 2011 | 2131 | 2166 | 2152 | 2360 | 2185 | 2214 | 2233 | 2043 | 2150 | 2189 | 1983 | 2016 | 1897 | 2105 | 1712 | 2159 | 2192 | Zkscan7 |
| Zkscan8 | 5337 | 4076 | 2899 | 3701 | 3879 | 4749 | 4615 | 8459 | 7264 | 5591 | 5873 | 6306 | 6530 | 6477 | 5525 | 4407 | 3403 | 3746 | 3628 | 5137 | 4438 | 5230 | 7025 | 5014 | 5929 | 4183 | 7074 | 5897 | Zkscan8 |
| Zmat1 | 989 | 759 | 1147 | 1054 | 1050 | 1004 | 1026 | 1077 | 1016 | 1211 | 1284 | 1191 | 1217 | 1055 | 940 | 863 | 1102 | 1069 | 976 | 961 | 938 | 1235 | 1019 | 1087 | 1106 | 1042 | 1081 | 1028 | Zmat1 |
| Zmat2 | 10690 | 9841 | 11844 | 11037 | 9342 | 9386 | 10910 | 10039 | 8562 | 9579 | 8714 | 9843 | 9740 | 11135 | 11147 | 9303 | 11091 | 9651 | 10258 | 11095 | 10712 | 8448 | 8739 | 10399 | 9684 | 11530 | 10727 | 11583 | Zmat2 |
| Zmat3 | 4312 | 4455 | 7192 | 6514 | 5986 | 5545 | 5785 | 2782 | 2708 | 3637 | 3059 | 4336 | 3587 | 3811 | 4298 | 4679 | 7305 | 6477 | 6548 | 4172 | 5911 | 3923 | 2716 | 4019 | 3994 | 4860 | 3688 | 4264 | Zmat3 |
| Zmat4 | 661 | 517 | 292 | 686 | 1079 | 685 | 668 | 780 | 653 | 703 | 643 | 369 | 432 | 398 | 587 | 458 | 506 | 703 | 793 | 789 | 743 | 927 | 770 | 482 | 700 | 371 | 363 | 400 | Zmat4 |
| Zmat5 | 858 | 1177 | 1049 | 1081 | 938 | 940 | 974 | 933 | 906 | 751 | 854 | 972 | 1013 | 1056 | 951 | 874 | 1070 | 974 | 1000 | 1088 | 1029 | 810 | 868 | 1014 | 1026 | 1064 | 1078 | 1028 | Zmat5 |
| Zmiz1 | 10681 | 13530 | 16039 | 17872 | 19889 | 17161 | 14643 | 16189 | 19331 | 25980 | 22324 | 28533 | 22535 | 24125 | 10748 | 12305 | 18528 | 17865 | 19409 | 12732 | 16765 | 28201 | 17305 | 27039 | 27078 | 21045 | 21861 | 22820 | Zmiz1 |
| Zmiz2 | 9194 | 12811 | 13152 | 14807 | 15496 | 12677 | 12172 | 9639 | 11527 | 14625 | 11522 | 15631 | 11743 | 12782 | 8675 | 10426 | 14901 | 14641 | 16302 | 10695 | 13262 | 14947 | 10142 | 14988 | 16014 | 14695 | 12343 | 12418 | Zmiz2 |
| Zmpste24 | 4015 | 3193 | 3835 | 3227 | 3526 | 3640 | 3502 | 4058 | 3437 | 3167 | 3358 | 3454 | 3773 | 3656 | 4046 | 3393 | 3328 | 3333 | 3216 | 3977 | 3514 | 3514 | 3592 | 3326 | 3393 | 3613 | 3810 | 3890 | Zmpste24 |
| Zmym1 | 3838 | 3496 | 3193 | 3350 | 2644 | 3129 | 3475 | 2737 | 2440 | 2285 | 2119 | 2507 | 2502 | 2723 | 3645 | 3807 | 3187 | 3210 | 2950 | 3735 | 3281 | 2216 | 2362 | 2277 | 2347 | 2608 | 2625 | 2627 | Zmym1 |
| Zmym2 | 15416 | 12730 | 11590 | 12323 | 14110 | 14670 | 15954 | 25342 | 19966 | 17516 | 15873 | 19920 | 18102 | 19512 | 16746 | 13288 | 12792 | 12272 | 14379 | 17198 | 14712 | 19773 | 18768 | 17065 | 18983 | 15823 | 20149 | 19695 | Zmym2 |
| Zmym3 | 7148 | 7594 | 6260 | 8335 | 10391 | 9366 | 9601 | 10285 | 11500 | 11300 | 11347 | 10100 | 10164 | 9929 | 7270 | 6923 | 7738 | 9148 | 9914 | 8816 | 9803 | 12731 | 10930 | 10426 | 11950 | 8349 | 9440 | 9077 | Zmym3 |
| Zmym4 | 11174 | 9681 | 8827 | 9173 | 10368 | 11163 | 10972 | 17210 | 14703 | 14215 | 13706 | 16521 | 15627 | 17039 | 11281 | 9306 | 9510 | 9160 | 9982 | 11104 | 10656 | 14442 | 14013 | 13611 | 14870 | 12462 | 17402 | 16359 | Zmym4 |
| Zmym5 | 9048 | 7667 | 8984 | 8835 | 9704 | 10030 | 10391 | 8021 | 6835 | 7338 | 7433 | 7920 | 7735 | 8118 | 9030 | 7997 | 8741 | 9403 | 9338 | 9709 | 10077 | 8184 | 6616 | 7447 | 7796 | 7410 | 8040 | 8091 | Zmym5 |
| Zmym6 | 4460 | 4094 | 3431 | 3911 | 4590 | 4478 | 4492 | 5263 | 4668 | 4261 | 4573 | 4183 | 3979 | 4246 | 4834 | 4259 | 3998 | 4207 | 4226 | 4952 | 4679 | 4899 | 4708 | 4011 | 4443 | 3166 | 4099 | 3794 | Zmym6 |
| Zmynd10 | 242 | 445 | 310 | 468 | 371 | 335 | 297 | 166 | 199 | 225 | 133 | 215 | 200 | 142 | 237 | 375 | 421 | 480 | 437 | 321 | 374 | 254 | 237 | 204 | 237 | 185 | 186 | 172 | Zmynd10 |
| Zmynd11 | 8308 | 7381 | 8111 | 8882 | 11119 | 9604 | 10761 | 17434 | 16434 | 17938 | 16833 | 17000 | 15866 | 18211 | 8339 | 7358 | 8378 | 9190 | 10398 | 10159 | 10943 | 18102 | 15931 | 17338 | 18211 | 15576 | 16497 | 16870 | Zmynd11 |
| Zmynd12 | 14 | 24 | 10 | 23 | 28 | 33 | 17 | 16 | 19 | 7 | 14 | 17 | 27 | 16 | 18 | 16 | 15 | 19 | 28 | 23 | 23 | 3 | 25 | 11 | 6 | 10 | 24 | 15 | Zmynd12 |
| Zmynd15 | 60 | 74 | 91 | 114 | 115 | 85 | 112 | 68 | 80 | 101 | 142 | 90 | 100 | 93 | 53 | 88 | 110 | 113 | 108 | 69 | 86 | 117 | 89 | 119 | 100 | 96 | 59 | 66 | Zmynd15 |
| Zmynd19 | 6931 | 7480 | 6824 | 7188 | 6166 | 6923 | 7022 | 4549 | 4208 | 4212 | 4144 | 4884 | 4386 | 5090 | 6657 | 6096 | 6862 | 6963 | 7006 | 6987 | 7221 | 4029 | 3979 | 4862 | 4757 | 5518 | 4896 | 5219 | Zmynd19 |
| Zmynd8 | 4267 | 4609 | 5700 | 5078 | 6457 | 5385 | 5034 | 8275 | 9962 | 11974 | 10217 | 10825 | 9748 | 10023 | 4168 | 4421 | 6261 | 5197 | 6248 | 4880 | 5355 | 11132 | 8935 | 11609 | 11397 | 9348 | 9759 | 9563 | Zmynd8 |
| Znfx1 | 2217 | 2805 | 2399 | 3137 | 3396 | 2871 | 2582 | 1866 | 2146 | 2541 | 2679 | 2611 | 2372 | 2357 | 1977 | 2656 | 2949 | 3240 | 3070 | 2270 | 2852 | 2714 | 2242 | 2431 | 2764 | 1760 | 2374 | 2345 | Znfx1 |
| Znhit1 | 889 | 1011 | 1004 | 998 | 956 | 1014 | 929 | 1568 | 1830 | 1898 | 1763 | 1784 | 1656 | 1815 | 894 | 864 | 1014 | 1021 | 1070 | 1002 | 1012 | 1666 | 1832 | 1852 | 2030 | 2102 | 1735 | 1866 | Znhit1 |
| Znhit2 | 564 | 730 | 579 | 631 | 520 | 547 | 568 | 496 | 571 | 489 | 503 | 557 | 502 | 587 | 576 | 538 | 588 | 559 | 564 | 628 | 532 | 518 | 563 | 589 | 467 | 696 | 589 | 630 | Znhit2 |
| Znhit3 | 1708 | 1684 | 1685 | 1520 | 1316 | 1446 | 1552 | 1324 | 1196 | 1044 | 1039 | 1077 | 1058 | 1259 | 1762 | 1503 | 1684 | 1404 | 1499 | 1688 | 1480 | 1021 | 1104 | 1182 | 1145 | 1384 | 1325 | 1269 | Znhit3 |

| | | | | | | | | | | | | | | | | | | | | | | | | | | | | | |
|---|---|---|---|---|---|---|---|---|---|---|---|---|---|---|---|---|---|---|---|---|---|---|---|---|---|---|---|---|---|
| Znhit6 | 2606 | 2600 | 2407 | 2518 | 2113 | 2237 | 2328 | 2138 | 2121 | 2175 | 2209 | 2084 | 2068 | 2252 | 2637 | 2518 | 2461 | 2474 | 1905 | 2183 | 2192 | 2059 | 2047 | 2210 | 2150 | 2504 | 2165 | 2517 | Znhit6 |
| Znrd2 | 40 | 25 | 22 | 32 | 29 | 15 | 40 | 24 | 21 | 27 | 32 | 23 | 24 | 32 | 26 | 33 | 35 | 28 | 22 | 21 | 35 | 40 | 18 | 26 | 41 | 34 | 19 | 20 | Znrd2 |
| Znrf1 | 5345 | 6336 | 7184 | 9393 | 11003 | 8650 | 9614 | 7880 | 8464 | 10993 | 9766 | 9817 | 8410 | 10176 | 5583 | 5557 | 9202 | 9567 | 10796 | 7549 | 10197 | 11673 | 7969 | 11042 | 11445 | 9836 | 8380 | 9231 | Znrf1 |
| Znrf2 | 4840 | 4199 | 3581 | 6159 | 6995 | 7015 | 8532 | 4428 | 4205 | 3957 | 3679 | 3369 | 2956 | 3818 | 4912 | 4238 | 4560 | 7122 | 6714 | 6936 | 8563 | 3930 | 3875 | 4160 | 5322 | 2748 | 3049 | 3087 | Znrf2 |
| Znrf3 | 2520 | 2137 | 2426 | 2358 | 1956 | 2368 | 2244 | 1447 | 1437 | 1487 | 1585 | 1917 | 1819 | 1978 | 1948 | 2394 | 2097 | 2409 | 2084 | 1930 | 2205 | 1499 | 1618 | 1602 | 1647 | 1747 | 1971 | 2027 | Znrf3 |
| Znrf4 | 0 | 3 | 5 | 1 | 0 | 0 | 1 | 0 | 1 | 0 | 0 | 0 | 1 | 0 | 0 | 0 | 0 | 6 | 0 | 0 | 0 | 0 | 1 | 0 | 0 | 2 | 0 | 0 | Znrf4 |
| Zp1 | 0 | 2 | 4 | 0 | 4 | 0 | 0 | 4 | 2 | 1 | 1 | 0 | 1 | 1 | 0 | 3 | 2 | 0 | 4 | 0 | 4 | 2 | 3 | 7 | 0 | 1 | 0 | 0 | Zp1 |
| Zp2 | 3 | 1 | 2 | 4 | 1 | 2 | 2 | 1 | 0 | 1 | 18 | 1 | 2 | 0 | 1 | 7 | 0 | 0 | 3 | 3 | 2 | 2 | 8 | 0 | 0 | 2 | 0 | 0 | Zp2 |
| Zp3 | 0 | 1 | 0 | 2 | 2 | 2 | 0 | 4 | 2 | 5 | 0 | 0 | 2 | 2 | 1 | 0 | 0 | 0 | 2 | 1 | 3 | 2 | 7 | 3 | 11 | 0 | 6 | 2 | Zp3 |
| Zp3r | 0 | 0 | 0 | 0 | 0 | 0 | 0 | 0 | 0 | 0 | 3 | 0 | 0 | 1 | 0 | 1 | 1 | 0 | 0 | 0 | 0 | 0 | 0 | 0 | 0 | 0 | 0 | 0 | Zp3r |
| Zpbp | 23 | 15 | 17 | 48 | 17 | 19 | 30 | 89 | 73 | 93 | 83 | 59 | 76 | 42 | 25 | 11 | 17 | 25 | 35 | 48 | 29 | 107 | 71 | 73 | 74 | 41 | 88 | 80 | Zpbp |
| Zpbp2 | 17 | 16 | 23 | 23 | 12 | 13 | 9 | 7 | 10 | 10 | 15 | 7 | 8 | 9 | 20 | 16 | 25 | 18 | 9 | 4 | 12 | 8 | 14 | 7 | 4 | 9 | 7 | 9 | Zpbp2 |
| Zpld1 | 110 | 118 | 74 | 86 | 36 | 174 | 17 | 228 | 353 | 183 | 327 | 111 | 396 | 181 | 249 | 142 | 41 | 125 | 79 | 22 | 9 | 299 | 438 | 131 | 165 | 57 | 103 | 76 | Zpld1 |
| Zpld2 | 0 | 0 | 0 | 0 | 0 | 0 | 0 | 0 | 0 | 0 | 1 | 0 | 0 | 1 | 0 | 0 | 0 | 1 | 0 | 0 | 2 | 0 | 0 | 0 | 0 | 0 | 0 | 0 | Zpld2 |
| Zpr1 | 3780 | 4563 | 3502 | 3926 | 3269 | 3431 | 3446 | 3136 | 3115 | 2768 | 2737 | 3032 | 2714 | 2917 | 3866 | 3833 | 3532 | 3715 | 3289 | 3698 | 3525 | 2704 | 2873 | 3031 | 2919 | 3378 | 3220 | 3001 | Zpr1 |
| Zranb1 | 9260 | 7332 | 7448 | 9567 | 8789 | 9345 | 10331 | 8742 | 7063 | 6971 | 6436 | 6671 | 6489 | 7125 | 9659 | 7585 | 7675 | 8853 | 8953 | 9993 | 9986 | 8153 | 7018 | 6855 | 7770 | 6639 | 6929 | 6947 | Zranb1 |
| Zranb2 | 20213 | 13882 | 16723 | 16359 | 16703 | 18151 | 20051 | 23720 | 19244 | 17881 | 18454 | 21223 | 20927 | 22593 | 20214 | 17249 | 15973 | 16011 | 17005 | 20836 | 18412 | 19348 | 19260 | 19430 | 19356 | 20807 | 22111 | 21887 | Zranb2 |
| Zranb3 | 2555 | 2497 | 2270 | 1819 | 1461 | 1773 | 1881 | 2263 | 2101 | 1772 | 1829 | 2342 | 2235 | 2436 | 2465 | 2309 | 2074 | 1585 | 1792 | 2096 | 1878 | 1462 | 1847 | 1907 | 1780 | 2360 | 2589 | 2493 | Zranb3 |
| Zrsr1 | 1226 | 1318 | 1153 | 1627 | 1527 | 1220 | 1363 | 1417 | 1218 | 1339 | 1207 | 1006 | 1033 | 1128 | 1329 | 1230 | 1306 | 1482 | 1476 | 1354 | 1445 | 1439 | 1293 | 1265 | 1316 | 1182 | 1126 | 1065 | Zrsr1 |
| Zrsr2 | 2832 | 2341 | 2735 | 2951 | 2899 | 2898 | 3086 | 3942 | 3427 | 3717 | 3940 | 4122 | 3771 | 4153 | 2709 | 2386 | 2722 | 2890 | 2850 | 2869 | 2866 | 3696 | 3424 | 3931 | 3846 | 4121 | 4076 | 3993 | Zrsr2 |
| Zscan10 | 24 | 31 | 43 | 12 | 6 | 7 | 0 | 9 | 9 | 13 | 17 | 13 | 23 | 13 | 11 | 33 | 8 | 12 | 0 | 5 | 6 | 15 | 14 | 14 | 6 | 13 | 3 | 9 | Zscan10 |
| Zscan12 | 2079 | 2021 | 2041 | 1903 | 1930 | 1886 | 2069 | 2541 | 2218 | 1972 | 1862 | 2328 | 2284 | 2517 | 2048 | 1925 | 1919 | 1623 | 2014 | 2302 | 2009 | 1828 | 1908 | 2174 | 2227 | 2306 | 2502 | 2461 | Zscan12 |
| Zscan18 | 2067 | 2084 | 1737 | 2114 | 2567 | 2348 | 2106 | 2186 | 2038 | 2108 | 2177 | 1706 | 1768 | 1559 | 2174 | 2044 | 1893 | 2195 | 2291 | 2622 | 2097 | 2436 | 2008 | 1740 | 2138 | 1551 | 1655 | 1461 | Zscan18 |
| Zscan2 | 717 | 949 | 1000 | 1216 | 1308 | 1063 | 879 | 1331 | 1379 | 1686 | 1383 | 1438 | 1156 | 1230 | 771 | 906 | 1136 | 1166 | 1355 | 1054 | 1063 | 1365 | 1418 | 1589 | 1582 | 1523 | 1242 | 1259 | Zscan2 |
| Zscan20 | 1226 | 1541 | 1068 | 1166 | 1112 | 1278 | 1016 | 1154 | 1007 | 842 | 1070 | 997 | 949 | 897 | 1138 | 1357 | 993 | 1177 | 1068 | 1217 | 1370 | 921 | 1044 | 824 | 1185 | 770 | 1192 | 889 | Zscan20 |
| Zscan21 | 6579 | 7014 | 6476 | 6377 | 6748 | 6983 | 6795 | 8728 | 7421 | 6884 | 6368 | 8077 | 7304 | 8151 | 7286 | 6381 | 6942 | 6260 | 7162 | 7756 | 6875 | 7611 | 7222 | 7339 | 7830 | 8390 | 8573 | 8520 | Zscan21 |
| Zscan22 | 1658 | 1719 | 1452 | 1382 | 1320 | 1248 | 1323 | 1473 | 1280 | 1376 | 1188 | 1233 | 1183 | 1174 | 1560 | 1534 | 1398 | 1412 | 1390 | 1435 | 1409 | 1083 | 1204 | 1345 | 1087 | 1436 | 1175 | 1337 | Zscan22 |
| Zscan25 | 596 | 612 | 665 | 616 | 708 | 716 | 656 | 841 | 990 | 917 | 932 | 1160 | 1033 | 979 | 587 | 525 | 751 | 578 | 785 | 616 | 611 | 964 | 1013 | 1150 | 1000 | 1121 | 1073 | 1049 | Zscan25 |
| Zscan26 | 6854 | 6591 | 6116 | 6202 | 6329 | 6595 | 6849 | 7061 | 6665 | 6565 | 7659 | 6479 | 6767 | 7211 | 6454 | 6795 | 6529 | 6324 | 6242 | 6635 | 6627 | 7026 | 6814 | 6749 | 6805 | 5761 | 6310 | 6372 | Zscan26 |
| Zscan29 | 2234 | 2123 | 2654 | 2778 | 2403 | 2652 | 2739 | 2242 | 1900 | 2300 | 2139 | 2559 | 2315 | 2603 | 2115 | 2238 | 2595 | 2719 | 2666 | 2268 | 2668 | 2262 | 1919 | 2378 | 2525 | 2450 | 2533 | 2641 | Zscan29 |
| Zscan5b | 0 | 0 | 0 | 0 | 0 | 0 | 0 | 0 | 0 | 1 | 0 | 0 | 1 | 0 | 0 | 0 | 0 | 0 | 0 | 0 | 2 | 0 | 0 | 0 | 0 | 0 | 0 | 0 | Zscan5b |
| Zswim1 | 768 | 834 | 900 | 825 | 903 | 723 | 919 | 733 | 833 | 860 | 772 | 1030 | 984 | 993 | 700 | 823 | 1045 | 993 | 1018 | 741 | 890 | 942 | 731 | 907 | 888 | 1160 | 848 | 976 | Zswim1 |
| Zswim2 | 2 | 3 | 0 | 3 | 7 | 0 | 3 | 1 | 0 | 0 | 1 | 2 | 3 | 0 | 1 | 2 | 2 | 0 | 3 | 0 | 2 | 0 | 5 | 2 | 3 | 10 | 3 | 0 | Zswim2 |
| Zswim3 | 1105 | 1430 | 1118 | 1111 | 1026 | 1107 | 1071 | 702 | 727 | 750 | 641 | 582 | 657 | 719 | 1004 | 1105 | 1389 | 1140 | 1042 | 1068 | 1170 | 779 | 686 | 757 | 783 | 594 | 665 | 748 | Zswim3 |
| Zswim4 | 4992 | 6108 | 5569 | 5750 | 5662 | 5172 | 5330 | 5671 | 5981 | 6532 | 5389 | 7079 | 6438 | 6734 | 4700 | 4992 | 6070 | 5197 | 6211 | 5197 | 5386 | 6532 | 5456 | 6755 | 7034 | 6923 | 6696 | 6759 | Zswim4 |
| Zswim5 | 6051 | 7577 | 6270 | 7349 | 8354 | 7699 | 8172 | 6489 | 5830 | 6869 | 5272 | 8895 | 5708 | 8357 | 7034 | 5160 | 8624 | 7140 | 8943 | 7086 | 8663 | 8031 | 4794 | 7340 | 8357 | 7350 | 7209 | 7970 | Zswim5 |
| Zswim6 | 5138 | 5833 | 5798 | 7953 | 7459 | 7027 | 7137 | 5152 | 4750 | 5764 | 5186 | 5942 | 5388 | 5835 | 5545 | 5260 | 6792 | 7669 | 7091 | 5707 | 7353 | 7038 | 4656 | 5368 | 6891 | 4717 | 5278 | 5663 | Zswim6 |
| Zswim7 | 165 | 187 | 129 | 108 | 114 | 124 | 126 | 249 | 259 | 243 | 256 | 234 | 240 | 231 | 166 | 142 | 121 | 111 | 130 | 138 | 115 | 224 | 234 | 208 | 203 | 298 | 286 | 280 | Zswim7 |
| Zswim8 | 16319 | 21053 | 21408 | 24677 | 23159 | 21782 | 20002 | 12118 | 14599 | 17265 | 16219 | 17250 | 15605 | 15307 | 15883 | 19038 | 23434 | 24594 | 23704 | 17353 | 21225 | 17406 | 13707 | 16601 | 17816 | 13813 | 14898 | 14790 | Zswim8 |
| Zswim9 | 1609 | 1710 | 1907 | 1855 | 1811 | 1889 | 1801 | 2353 | 2465 | 2794 | 2556 | 2635 | 2487 | 2786 | 1631 | 1677 | 1746 | 1701 | 1824 | 2017 | 1823 | 2234 | 2331 | 2679 | 2562 | 2623 | 2834 | 2588 | Zswim9 |
| Zup1 | 1970 | 1533 | 2041 | 2125 | 2001 | 2232 | 2110 | 2648 | 2143 | 2074 | 2048 | 2221 | 2292 | 2336 | 2018 | 1826 | 1982 | 1931 | 1946 | 2299 | 2122 | 2105 | 2060 | 2179 | 2025 | 2050 | 2200 | 2112 | Zup1 |
| Zw10 | 6050 | 6175 | 5724 | 5181 | 4528 | 4661 | 5471 | 4347 | 3791 | 3894 | 3587 | 4546 | 4118 | 4553 | 6041 | 5421 | 5473 | 4870 | 5202 | 5582 | 5055 | 3511 | 3693 | 4041 | 3753 | 4717 | 4453 | 4675 | Zw10 |
| Zwilch | 4640 | 3838 | 4435 | 3369 | 2227 | 2852 | 3353 | 2811 | 2696 | 2551 | 2494 | 3620 | 3378 | 3746 | 3990 | 3857 | 3808 | 3258 | 2916 | 3204 | 2991 | 2162 | 2365 | 3095 | 2518 | 4154 | 3603 | 3642 | Zwilch |
| Zwint | 22863 | 23908 | 21866 | 21875 | 21917 | 21153 | 23931 | 23957 | 20036 | 18423 | 18228 | 20601 | 19828 | 21804 | 25134 | 20690 | 22503 | 20852 | 22853 | 26613 | 23110 | 20954 | 19480 | 19305 | 20976 | 21745 | 21063 | 22037 | Zwint |
| Zxdb | 654 | 352 | 711 | 668 | 704 | 740 | 730 | 953 | 868 | 690 | 785 | 743 | 781 | 859 | 778 | 555 | 607 | 671 | 670 | 660 | 741 | 1059 | 784 | 774 | 804 | 843 | 805 | 843 | Zxdb |
| Zxdc | 2361 | 2692 | 2146 | 2996 | 2688 | 2640 | 2706 | 2088 | 2019 | 2016 | 2022 | 2123 | 1814 | 2005 | 2268 | 2231 | 2427 | 3017 | 2877 | 2576 | 2830 | 2319 | 1916 | 1814 | 2202 | 1725 | 1930 | 1890 | Zxdc |
| Zyg11a | 31 | 29 | 3 | 18 | 23 | 33 | 17 | 5 | 5 | 5 | 8 | 13 | 6 | 3 | 34 | 53 | 20 | 17 | 3 | 20 | 21 | 7 | 13 | 3 | 11 | 0 | 7 | 5 | Zyg11a |
| Zyg11b | 20269 | 18849 | 11722 | 17978 | 22243 | 23874 | 24674 | 23448 | 19764 | 17808 | 17380 | 17254 | 16895 | 18233 | 21493 | 17700 | 14589 | 18647 | 20304 | 24454 | 24886 | 19515 | 19388 | 14617 | 20494 | 9986 | 19033 | 16880 | Zyg11b |
| Zyx | 3084 | 3938 | 4303 | 3148 | 3346 | 3054 | 2271 | 3174 | 4182 | 3953 | 5241 | 4765 | 4635 | 3910 | 2823 | 3852 | 3604 | 3203 | 3397 | 2102 | 2532 | 3618 | 4684 | 4372 | 3699 | 5890 | 4391 | 4188 | Zyx |
| Zzef1 | 9481 | 9771 | 4639 | 7075 | 8010 | 9236 | 8001 | 7173 | 7781 | 6001 | 6874 | 5641 | 5957 | 5624 | 9336 | 9456 | 5357 | 7170 | 6959 | 8928 | 8783 | 5548 | 6890 | 4615 | 6661 | 3168 | 6335 | 4826 | Zzef1 |
| Zzz3 | 5419 | 4416 | 4644 | 4485 | 4961 | 4995 | 4720 | 10748 | 9540 | 8687 | 8963 | 9951 | 9668 | 9678 | 5429 | 4867 | 4650 | 4277 | 4646 | 5012 | 4480 | 9635 | 9178 | 8804 | 9578 | 8771 | 10004 | 9644 | Zzz3 |
