## Supplemental Table S2 for "Rapid effects of valproic acid on the fetal brain transcriptome: Implications for brain development and autism"

| Gene | Counts | Male | | | Female | | |
|---|---|---|---|---|---|---|---|
| | | FC | LFC | $p_{FDR}$ | FC | LFC | $p_{FDR}$ |
| 1110004F10Rik | 10227 | -1.21 | -0.28 | 1.1E-03 | -1.29 | -0.37 | 1.6E-05 |
| 1110017D15Rik | 366 | 3.55 | 1.83 | 8.6E-25 | 3.61 | 1.85 | 2.2E-25 |
| 1110038F14Rik | 1540 | -1.54 | -0.62 | 4.0E-09 | -1.73 | -0.79 | 5.8E-14 |
| 1110059E24Rik | 3275 | 1.34 | 0.42 | 9.4E-05 | 1.38 | 0.47 | 1.3E-05 |
| 1110059G10Rik | 1092 | -1.68 | -0.75 | 5.0E-15 | -1.68 | -0.75 | 4.8E-15 |
| 1190007I07Rik | 446 | -1.50 | -0.59 | 5.3E-08 | -1.72 | -0.78 | 4.2E-13 |
| 1500009C09Rik | 1708 | 1.64 | 0.71 | 1.2E-03 | 1.69 | 0.76 | 5.1E-04 |
| 1500009L16Rik | 1141 | 1.74 | 0.80 | 1.6E-32 | 1.66 | 0.73 | 3.5E-27 |
| 1600014C10Rik | 1018 | -1.80 | -0.85 | 3.0E-28 | -1.79 | -0.84 | 1.6E-27 |
| 1700001O22Rik | 185 | 2.86 | 1.51 | 1.5E-18 | 2.53 | 1.34 | 8.7E-15 |
| 1700003E16Rik | 238 | 4.56 | 2.19 | 2.5E-33 | 4.03 | 2.01 | 1.5E-28 |
| 1700017B05Rik | 1189 | 1.56 | 0.64 | 1.1E-04 | 1.44 | 0.53 | 1.8E-03 |
| 1700019D03Rik | 360 | 3.51 | 1.81 | 4.4E-11 | 3.41 | 1.77 | 1.4E-10 |
| 1700029J07Rik | 352 | -1.87 | -0.90 | 1.1E-19 | -1.75 | -0.81 | 4.3E-16 |
| 1700030J22Rik | 533 | 1.90 | 0.93 | 4.2E-19 | 1.87 | 0.90 | 3.8E-18 |
| 1700030K09Rik | 823 | -5.92 | -2.57 | 2.8E-83 | -5.85 | -2.55 | 2.3E-82 |
| 1700037C18Rik | 116 | -2.51 | -1.33 | 3.2E-16 | -2.77 | -1.47 | 2.5E-19 |
| 1700037H04Rik | 3120 | 1.90 | 0.92 | 2.0E-12 | 1.71 | 0.77 | 5.5E-09 |
| 1700066M21Rik | 1664 | 1.35 | 0.44 | 1.7E-03 | 1.40 | 0.48 | 4.7E-04 |
| 1700088E04Rik | 521 | 2.56 | 1.36 | 1.8E-39 | 2.73 | 1.45 | 9.3E-45 |
| 1700093K21Rik | 119 | 4.46 | 2.16 | 2.7E-13 | 6.25 | 2.64 | 7.4E-19 |
| 1700094D03Rik | 356 | 1.85 | 0.89 | 1.8E-13 | 1.81 | 0.85 | 1.5E-12 |
| 1700102P08Rik | 274 | 1.88 | 0.91 | 6.0E-10 | 1.90 | 0.93 | 2.8E-10 |
| 1810010H24Rik | 193 | 1.79 | 0.84 | 9.5E-08 | 1.93 | 0.95 | 1.6E-09 |
| 1810024B03Rik | 123 | -2.57 | -1.36 | 9.0E-14 | -2.62 | -1.39 | 3.7E-14 |
| 1810055G02Rik | 2101 | 1.65 | 0.73 | 3.2E-09 | 1.60 | 0.68 | 2.9E-08 |
| 2210016L21Rik | 2072 | -1.29 | -0.37 | 4.0E-08 | -1.31 | -0.39 | 4.2E-09 |
| 2310011J03Rik | 1664 | 1.77 | 0.82 | 3.7E-11 | 1.59 | 0.67 | 8.7E-08 |
| 2310022B05Rik | 6485 | 1.28 | 0.36 | 4.5E-04 | 1.23 | 0.30 | 3.3E-03 |
| 2410002F23Rik | 6002 | -1.43 | -0.51 | 1.2E-35 | -1.47 | -0.55 | 7.9E-41 |
| 2410004B18Rik | 2022 | 1.51 | 0.59 | 2.0E-21 | 1.50 | 0.59 | 5.3E-21 |
| 2510002D24Rik | 507 | -1.43 | -0.52 | 8.8E-10 | -1.47 | -0.56 | 4.6E-11 |
| 2510039O18Rik | 2533 | 1.56 | 0.64 | 6.6E-08 | 1.42 | 0.51 | 2.6E-05 |
| 2610008E11Rik | 7868 | -1.45 | -0.54 | 3.6E-12 | -1.37 | -0.46 | 6.1E-09 |
| 2610021A01Rik | 2839 | 1.24 | 0.31 | 6.4E-05 | 1.32 | 0.40 | 2.1E-07 |
| 2610044O15Rik8 | 2098 | -1.26 | -0.33 | 4.1E-05 | -1.26 | -0.33 | 4.9E-05 |
| 2610301B20Rik | 3491 | 1.40 | 0.49 | 1.0E-09 | 1.43 | 0.52 | 6.0E-11 |
| 2610507B11Rik | 19385 | 1.31 | 0.39 | 5.8E-07 | 1.35 | 0.43 | 2.9E-08 |
| 2700049A03Rik | 3410 | 1.24 | 0.30 | 1.7E-04 | 1.34 | 0.42 | 1.2E-07 |
| 2700062C07Rik | 729 | -1.73 | -0.79 | 3.2E-14 | -1.89 | -0.92 | 6.2E-19 |
| 2700097O09Rik | 2207 | 1.32 | 0.40 | 9.0E-09 | 1.39 | 0.47 | 1.4E-11 |
| 2900026A02Rik | 3248 | 1.98 | 0.98 | 6.5E-17 | 2.00 | 1.00 | 1.9E-17 |
| 3110009E18Rik | 124 | -1.63 | -0.71 | 4.0E-04 | -2.03 | -1.02 | 1.9E-07 |
| 3110040N11Rik | 681 | -1.89 | -0.92 | 2.6E-28 | -1.90 | -0.93 | 1.1E-28 |

| | | | | | | | |
|---|---|---|---|---|---|---|---|
| 3110082I17Rik | 1166 | 1.58 | 0.66 | 1.4E-04 | 1.31 | 0.39 | 3.3E-02 |
| 3110082J24Rik | 140 | -2.58 | -1.37 | 2.6E-11 | -2.49 | -1.32 | 1.3E-10 |
| 3830406C13Rik | 812 | -1.29 | -0.37 | 5.4E-03 | -1.31 | -0.39 | 3.5E-03 |
| 4921507P07Rik | 215 | 1.39 | 0.48 | 1.6E-03 | 1.43 | 0.51 | 6.9E-04 |
| 4921524J17Rik | 1645 | 1.69 | 0.76 | 3.8E-18 | 1.68 | 0.75 | 9.6E-18 |
| 4921536K21Rik | 210 | 1.50 | 0.58 | 2.3E-04 | 1.29 | 0.36 | 2.8E-02 |
| 4930430F08Rik | 1323 | 1.56 | 0.65 | 1.0E-16 | 1.64 | 0.71 | 3.0E-20 |
| 4930522L14Rik | 384 | 1.67 | 0.74 | 5.2E-12 | 1.80 | 0.85 | 1.4E-15 |
| 4930563E22Rik | 428 | -1.54 | -0.62 | 6.7E-09 | -1.43 | -0.51 | 2.2E-06 |
| 4930579G24Rik | 703 | 1.77 | 0.83 | 2.4E-07 | 1.74 | 0.80 | 6.0E-07 |
| 4930590J08Rik | 403 | 1.53 | 0.61 | 4.4E-05 | 1.71 | 0.77 | 2.0E-07 |
| 4931406C07Rik | 2467 | -1.28 | -0.36 | 1.8E-10 | -1.22 | -0.29 | 4.1E-07 |
| 4933407L21Rik | 155 | -1.74 | -0.80 | 1.3E-03 | -1.72 | -0.78 | 1.8E-03 |
| 5031439G07Rik | 16940 | 1.81 | 0.86 | 6.9E-20 | 1.77 | 0.83 | 1.4E-18 |
| 5430403G16Rik | 254 | 1.40 | 0.48 | 3.3E-05 | 1.67 | 0.74 | 1.4E-10 |
| 5730409E04Rik | 732 | -2.89 | -1.53 | 5.0E-11 | -2.79 | -1.48 | 2.3E-10 |
| 5730455P16Rik | 3267 | -1.37 | -0.46 | 9.0E-12 | -1.42 | -0.50 | 4.2E-14 |
| 6030458C11Rik | 3419 | -1.30 | -0.38 | 2.4E-22 | -1.22 | -0.29 | 6.6E-13 |
| 6330403K07Rik | 13572 | -1.41 | -0.50 | 4.6E-08 | -1.41 | -0.50 | 5.1E-08 |
| 6330409D20Rik | 174 | 1.62 | 0.69 | 1.5E-06 | 1.44 | 0.52 | 3.4E-04 |
| 6430548M08Rik | 4719 | 2.41 | 1.27 | 3.2E-26 | 2.40 | 1.26 | 8.8E-26 |
| 6430550D23Rik | 223 | 2.21 | 1.14 | 4.0E-20 | 2.43 | 1.28 | 1.1E-24 |
| 6430571L13Rik | 352 | 1.33 | 0.41 | 1.1E-02 | 1.40 | 0.49 | 2.2E-03 |
| 9030025P20Rik | 127 | -1.74 | -0.80 | 7.4E-10 | -1.74 | -0.80 | 5.7E-10 |
| 9130019O22Rik | 628 | 1.51 | 0.59 | 7.5E-08 | 1.67 | 0.74 | 2.1E-11 |
| 9530068E07Rik | 7749 | -1.23 | -0.30 | 7.8E-04 | -1.31 | -0.39 | 7.4E-06 |
| 9530077C05Rik | 513 | 1.48 | 0.57 | 5.6E-08 | 1.29 | 0.37 | 5.8E-04 |
| 9930021J03Rik | 5543 | -1.31 | -0.39 | 3.7E-05 | -1.23 | -0.30 | 2.2E-03 |
| 9930104L06Rik | 2337 | 1.34 | 0.43 | 6.6E-11 | 1.28 | 0.35 | 1.0E-07 |
| A130010J15Rik | 930 | -1.20 | -0.27 | 1.2E-04 | -1.26 | -0.33 | 1.1E-06 |
| A2m | 717 | -1.38 | -0.46 | 1.5E-02 | -1.50 | -0.58 | 1.7E-03 |
| A3galt2 | 1683 | 6.97 | 2.80 | 3.7E-70 | 7.64 | 2.93 | 1.1E-76 |
| A430033K04Rik | 1960 | 1.32 | 0.41 | 1.1E-07 | 1.38 | 0.46 | 1.5E-09 |
| A630001G21Rik | 105 | -3.08 | -1.62 | 3.0E-17 | -3.32 | -1.73 | 9.0E-19 |
| A830018L16Rik | 169 | -2.39 | -1.25 | 4.0E-07 | -2.35 | -1.23 | 7.9E-07 |
| A930004D18Rik | 1042 | -1.59 | -0.67 | 5.1E-08 | -1.64 | -0.72 | 5.5E-09 |
| A930033H14Rik | 144 | -2.63 | -1.39 | 1.9E-21 | -2.72 | -1.44 | 1.1E-22 |
| AA986860 | 139 | 1.99 | 1.00 | 1.8E-08 | 2.84 | 1.50 | 1.5E-17 |
| Aaas | 3111 | 1.61 | 0.69 | 5.9E-06 | 1.40 | 0.49 | 1.8E-03 |
| Aacs | 3674 | 2.02 | 1.01 | 1.4E-32 | 1.86 | 0.90 | 1.4E-25 |
| Aagab | 6156 | 1.76 | 0.82 | 1.0E-50 | 1.75 | 0.81 | 1.8E-49 |
| Aak1 | 15896 | 1.46 | 0.54 | 1.2E-04 | 1.62 | 0.70 | 5.1E-07 |
| Aard | 215 | -1.56 | -0.64 | 6.0E-05 | -1.71 | -0.77 | 1.1E-06 |
| Aars2 | 1417 | -1.35 | -0.44 | 4.1E-04 | -1.49 | -0.57 | 2.6E-06 |
| Aasdh | 1572 | -1.65 | -0.72 | 3.7E-16 | -1.58 | -0.66 | 7.4E-14 |
| Aasdhppt | 2945 | 1.31 | 0.39 | 2.0E-07 | 1.35 | 0.44 | 5.7E-09 |
| Aass | 396 | 2.23 | 1.15 | 1.7E-04 | 2.28 | 1.19 | 1.1E-04 |

| | | | | | | | |
|---|---|---|---|---|---|---|---|
| Aatk | 1600 | -2.10 | -1.07 | 7.9E-04 | -2.11 | -1.07 | 8.0E-04 |
| Abat | 8490 | 1.58 | 0.66 | 2.4E-08 | 1.68 | 0.75 | 1.7E-10 |
| Abca3 | 6623 | 1.32 | 0.40 | 3.3E-03 | 1.34 | 0.43 | 1.7E-03 |
| Abca4 | 342 | -1.60 | -0.68 | 6.6E-03 | -1.47 | -0.56 | 3.0E-02 |
| Abca5 | 3173 | 2.02 | 1.02 | 3.1E-09 | 2.52 | 1.33 | 4.1E-15 |
| Abca7 | 4344 | 2.59 | 1.37 | 3.7E-24 | 2.58 | 1.37 | 7.2E-24 |
| Abca9 | 1695 | -1.86 | -0.90 | 6.9E-06 | -1.71 | -0.78 | 1.2E-04 |
| Abcb10 | 2247 | -1.24 | -0.30 | 1.3E-03 | -1.33 | -0.41 | 1.0E-05 |
| Abcb6 | 130 | 1.52 | 0.60 | 5.4E-03 | 1.36 | 0.44 | 4.8E-02 |
| Abcb7 | 4367 | -1.46 | -0.55 | 3.7E-08 | -1.41 | -0.50 | 6.6E-07 |
| Abcb8 | 1836 | -2.93 | -1.55 | 1.2E-109 | -3.17 | -1.67 | 1.9E-126 |
| Abcb9 | 338 | 1.78 | 0.83 | 2.6E-08 | 1.73 | 0.79 | 1.3E-07 |
| Abcc5 | 13864 | -1.55 | -0.63 | 1.8E-04 | -1.31 | -0.39 | 2.7E-02 |
| Abcc9 | 1497 | -1.57 | -0.65 | 4.0E-05 | -1.51 | -0.60 | 1.8E-04 |
| Abcd2 | 4781 | 1.82 | 0.86 | 5.0E-11 | 1.72 | 0.79 | 2.5E-09 |
| Abcd4 | 2037 | 1.37 | 0.45 | 6.0E-04 | 1.33 | 0.41 | 1.8E-03 |
| Abce1 | 14484 | -1.25 | -0.32 | 2.2E-03 | -1.20 | -0.26 | 1.3E-02 |
| Abcf1 | 11952 | 1.56 | 0.64 | 3.4E-11 | 1.44 | 0.53 | 7.0E-08 |
| Abcf3 | 5371 | 1.49 | 0.58 | 7.5E-22 | 1.46 | 0.54 | 2.8E-19 |
| Abcg5 | 148 | -2.12 | -1.08 | 2.2E-07 | -2.08 | -1.06 | 4.3E-07 |
| Abhd10 | 2112 | -1.33 | -0.41 | 2.8E-08 | -1.35 | -0.43 | 8.5E-09 |
| Abhd11 | 1511 | 1.31 | 0.39 | 2.4E-10 | 1.29 | 0.37 | 3.2E-09 |
| Abhd12 | 3115 | 1.31 | 0.39 | 3.8E-10 | 1.31 | 0.38 | 1.0E-09 |
| Abhd13 | 3905 | 1.25 | 0.33 | 3.9E-05 | 1.25 | 0.32 | 5.3E-05 |
| Abhd14a | 1728 | 1.46 | 0.54 | 7.3E-11 | 1.41 | 0.49 | 5.1E-09 |
| Abhd14b | 827 | 1.95 | 0.96 | 3.1E-07 | 2.14 | 1.10 | 5.2E-09 |
| Abhd15 | 321 | 6.55 | 2.71 | 7.9E-82 | 6.24 | 2.64 | 5.7E-77 |
| Abhd16a | 1764 | -2.00 | -1.00 | 1.9E-26 | -2.21 | -1.14 | 5.6E-34 |
| Abhd17a | 3178 | 1.55 | 0.63 | 6.7E-11 | 1.48 | 0.56 | 1.0E-08 |
| Abhd17b | 3631 | 1.25 | 0.32 | 1.1E-04 | 1.20 | 0.26 | 2.3E-03 |
| Abhd18 | 124 | 2.70 | 1.43 | 9.7E-04 | 2.44 | 1.29 | 3.3E-03 |
| Abhd2 | 8026 | 1.26 | 0.34 | 9.8E-04 | 1.29 | 0.37 | 3.0E-04 |
| Abhd3 | 263 | 3.84 | 1.94 | 2.2E-29 | 3.78 | 1.92 | 1.4E-28 |
| Abhd4 | 3131 | 1.34 | 0.42 | 3.3E-06 | 1.31 | 0.39 | 1.8E-05 |
| Abhd5 | 1217 | -1.31 | -0.39 | 1.0E-03 | -1.36 | -0.44 | 1.9E-04 |
| Abhd6 | 1224 | 1.86 | 0.90 | 2.5E-23 | 1.93 | 0.95 | 7.3E-26 |
| Abhd8 | 5131 | 1.90 | 0.93 | 3.5E-32 | 1.83 | 0.87 | 8.2E-29 |
| Abi3 | 103 | -1.42 | -0.51 | 1.5E-02 | -1.43 | -0.52 | 1.4E-02 |
| Abl2 | 6428 | -1.33 | -0.42 | 1.3E-03 | -1.25 | -0.33 | 1.4E-02 |
| Ablim1 | 4502 | -1.34 | -0.43 | 2.6E-03 | -1.25 | -0.33 | 2.4E-02 |
| Ablim2 | 250 | 2.11 | 1.08 | 9.5E-03 | 2.25 | 1.17 | 4.8E-03 |
| Abr | 12513 | -1.60 | -0.68 | 5.8E-06 | -1.66 | -0.73 | 1.3E-06 |
| Abracl | 3562 | -1.63 | -0.70 | 9.6E-06 | -1.72 | -0.78 | 7.0E-07 |
| Abraxas1 | 745 | -1.24 | -0.31 | 3.1E-03 | -1.26 | -0.33 | 1.6E-03 |
| Abt1 | 1337 | 1.36 | 0.45 | 8.3E-05 | 1.28 | 0.35 | 2.1E-03 |
| Abtb2 | 2483 | 2.20 | 1.14 | 1.8E-38 | 2.19 | 1.13 | 8.4E-38 |
| Acaa1b | 135 | 2.59 | 1.37 | 2.2E-14 | 2.91 | 1.54 | 2.0E-17 |

| | | | | | | | |
|---|---|---|---|---|---|---|---|
| Acaa2 | 2091 | 1.42 | 0.51 | 2.5E-04 | 1.41 | 0.50 | 3.3E-04 |
| Acacb | 599 | 3.69 | 1.88 | 2.6E-51 | 3.64 | 1.86 | 5.1E-50 |
| Acad12 | 162 | -1.35 | -0.43 | 4.5E-04 | -1.54 | -0.62 | 2.6E-07 |
| Acadm | 4036 | 1.80 | 0.85 | 7.8E-53 | 1.92 | 0.94 | 3.2E-64 |
| Acan | 1045 | -6.91 | -2.79 | 4.2E-03 | -5.36 | -2.42 | 1.4E-02 |
| Acap2 | 8163 | 1.68 | 0.75 | 6.4E-24 | 1.79 | 0.84 | 7.9E-30 |
| Acap3 | 6900 | 1.42 | 0.50 | 2.2E-04 | 1.35 | 0.43 | 1.8E-03 |
| Acbd4 | 451 | 3.12 | 1.64 | 5.8E-30 | 3.09 | 1.63 | 2.9E-29 |
| Accs | 1681 | -1.34 | -0.42 | 1.1E-05 | -1.31 | -0.39 | 6.0E-05 |
| Acd | 3697 | 1.91 | 0.93 | 2.1E-34 | 1.82 | 0.86 | 1.4E-29 |
| Acer3 | 1362 | 1.49 | 0.57 | 5.2E-08 | 1.56 | 0.64 | 9.1E-10 |
| Ache | 892 | 1.73 | 0.79 | 1.1E-03 | 1.82 | 0.87 | 3.1E-04 |
| Acin1 | 25484 | -1.41 | -0.49 | 4.6E-05 | -1.52 | -0.60 | 5.6E-07 |
| Ackr1 | 1403 | 2.20 | 1.14 | 8.0E-09 | 2.19 | 1.13 | 1.0E-08 |
| Ackr3 | 2346 | -1.31 | -0.39 | 9.9E-03 | -1.40 | -0.49 | 1.0E-03 |
| Acly | 42598 | 1.52 | 0.60 | 2.4E-17 | 1.49 | 0.58 | 5.5E-16 |
| Aco1 | 9220 | 1.81 | 0.86 | 7.4E-27 | 1.74 | 0.80 | 2.2E-23 |
| Acot1 | 1481 | -1.26 | -0.34 | 6.0E-05 | -1.24 | -0.31 | 2.1E-04 |
| Acot13 | 852 | 1.40 | 0.48 | 1.3E-05 | 1.43 | 0.51 | 3.4E-06 |
| Acot2 | 211 | -2.95 | -1.56 | 1.6E-26 | -2.72 | -1.45 | 6.1E-23 |
| Acot6 | 253 | -3.52 | -1.81 | 7.4E-35 | -3.63 | -1.86 | 1.9E-36 |
| Acot7 | 4558 | 1.22 | 0.28 | 2.8E-02 | 1.27 | 0.34 | 8.2E-03 |
| Acot8 | 990 | 1.57 | 0.65 | 7.1E-12 | 1.51 | 0.60 | 3.7E-10 |
| Acot9 | 1147 | -1.46 | -0.55 | 7.6E-05 | -1.37 | -0.46 | 1.3E-03 |
| Acox1 | 7899 | 1.33 | 0.41 | 5.2E-11 | 1.32 | 0.40 | 2.4E-10 |
| Acp2 | 4785 | -1.74 | -0.80 | 2.5E-32 | -1.73 | -0.79 | 2.1E-31 |
| Acp6 | 1651 | 2.05 | 1.04 | 3.9E-48 | 2.10 | 1.07 | 9.5E-51 |
| Acrbp | 689 | -1.47 | -0.55 | 9.7E-05 | -1.47 | -0.56 | 9.8E-05 |
| Acsbg1 | 622 | -1.39 | -0.47 | 8.5E-03 | -1.62 | -0.70 | 7.1E-05 |
| Acsf3 | 819 | -2.85 | -1.51 | 1.8E-57 | -3.08 | -1.62 | 7.1E-66 |
| Acsl1 | 2150 | 3.11 | 1.64 | 6.5E-40 | 3.28 | 1.72 | 1.3E-43 |
| Acsl3 | 9041 | 1.51 | 0.59 | 3.9E-11 | 1.47 | 0.55 | 8.6E-10 |
| Acsl4 | 4014 | 1.62 | 0.69 | 1.5E-12 | 1.66 | 0.73 | 6.7E-14 |
| Acsl5 | 1132 | -1.93 | -0.95 | 1.9E-10 | -1.92 | -0.94 | 2.3E-10 |
| Acss1 | 3013 | -1.30 | -0.38 | 1.2E-02 | -1.33 | -0.41 | 7.0E-03 |
| Acss2 | 2917 | 2.04 | 1.03 | 5.3E-76 | 2.05 | 1.03 | 2.3E-76 |
| Acta1 | 391 | -15.56 | -3.96 | 1.5E-04 | -9.63 | -3.27 | 2.1E-03 |
| Actb | 107135 | -1.36 | -0.44 | 2.4E-08 | -1.48 | -0.57 | 4.5E-13 |
| Actc1 | 906 | -21.26 | -4.41 | 7.1E-04 | -12.85 | -3.68 | 5.4E-03 |
| Actl6a | 6126 | -1.28 | -0.36 | 1.1E-02 | -1.41 | -0.50 | 3.6E-04 |
| Actn4 | 15736 | 1.43 | 0.52 | 3.2E-05 | 1.37 | 0.46 | 2.6E-04 |
| Actr10 | 9766 | 1.26 | 0.33 | 1.7E-06 | 1.28 | 0.36 | 1.5E-07 |
| Actr1b | 10314 | 1.34 | 0.42 | 2.0E-11 | 1.25 | 0.33 | 3.5E-07 |
| Actr3b | 1001 | 1.98 | 0.99 | 3.9E-11 | 1.89 | 0.92 | 8.7E-10 |
| Actr5 | 1604 | 1.53 | 0.61 | 2.4E-13 | 1.52 | 0.60 | 8.0E-13 |
| Actr6 | 2081 | 1.36 | 0.44 | 3.1E-08 | 1.41 | 0.49 | 4.1E-10 |
| Actr8 | 3093 | -1.56 | -0.64 | 3.6E-27 | -1.61 | -0.68 | 2.0E-30 |

| | | | | | | | |
|---|---|---|---|---|---|---|---|
| Acvr1c | 155 | 2.57 | 1.36 | 5.6E-09 | 2.50 | 1.32 | 1.6E-08 |
| Acvr2b | 2847 | -2.10 | -1.07 | 3.0E-05 | -1.80 | -0.85 | 1.1E-03 |
| Acy1 | 1770 | 1.70 | 0.77 | 2.3E-17 | 1.68 | 0.75 | 1.2E-16 |
| Acyp1 | 881 | 1.26 | 0.34 | 2.2E-03 | 1.19 | 0.25 | 2.9E-02 |
| Ada | 284 | 1.30 | 0.38 | 1.9E-02 | 1.35 | 0.44 | 7.7E-03 |
| Adal | 1824 | -1.54 | -0.62 | 3.2E-17 | -1.40 | -0.49 | 5.7E-11 |
| Adam11 | 2711 | -1.52 | -0.61 | 1.0E-03 | -1.37 | -0.46 | 1.5E-02 |
| Adam12 | 911 | -2.63 | -1.39 | 5.6E-07 | -2.60 | -1.38 | 7.7E-07 |
| Adam15 | 3573 | 3.00 | 1.58 | 9.2E-52 | 2.93 | 1.55 | 1.2E-49 |
| Adam19 | 3857 | -1.57 | -0.65 | 1.3E-07 | -1.46 | -0.55 | 1.3E-05 |
| Adam22 | 4153 | -1.60 | -0.68 | 1.7E-04 | -1.45 | -0.54 | 3.2E-03 |
| Adam33 | 231 | -2.35 | -1.23 | 1.8E-03 | -2.49 | -1.32 | 8.6E-04 |
| Adam8 | 1522 | 2.38 | 1.25 | 5.7E-17 | 2.21 | 1.14 | 2.4E-14 |
| Adam9 | 4828 | 1.66 | 0.73 | 1.5E-44 | 1.65 | 0.72 | 1.2E-43 |
| Adamts1 | 1953 | -1.77 | -0.82 | 4.0E-04 | -1.74 | -0.80 | 6.3E-04 |
| Adamts12 | 2424 | -1.77 | -0.83 | 2.4E-04 | -1.65 | -0.72 | 1.5E-03 |
| Adamts14 | 272 | -1.71 | -0.77 | 3.0E-02 | -1.75 | -0.81 | 2.4E-02 |
| Adamts15 | 1767 | -1.70 | -0.76 | 1.0E-02 | -1.76 | -0.82 | 5.8E-03 |
| Adamts18 | 484 | -3.16 | -1.66 | 2.9E-12 | -2.65 | -1.41 | 4.3E-09 |
| Adamts2 | 893 | -2.09 | -1.06 | 5.1E-03 | -2.11 | -1.07 | 4.8E-03 |
| Adamts20 | 2579 | 1.69 | 0.76 | 3.0E-07 | 1.91 | 0.94 | 1.9E-10 |
| Adamts3 | 803 | -1.39 | -0.48 | 1.4E-02 | -1.32 | -0.40 | 4.5E-02 |
| Adamts5 | 815 | -2.10 | -1.07 | 4.1E-04 | -2.04 | -1.03 | 7.0E-04 |
| Adamts8 | 138 | -1.90 | -0.93 | 1.4E-04 | -1.96 | -0.97 | 7.6E-05 |
| Adamtsl1 | 1146 | -1.93 | -0.95 | 3.5E-03 | -1.71 | -0.77 | 2.0E-02 |
| Adamtsl2 | 585 | -1.51 | -0.59 | 7.3E-06 | -1.63 | -0.71 | 6.6E-08 |
| Adamtsl3 | 186 | -3.07 | -1.62 | 1.8E-04 | -2.33 | -1.22 | 6.1E-03 |
| Adap2 | 138 | -1.48 | -0.57 | 1.0E-05 | -1.52 | -0.60 | 2.5E-06 |
| Adar | 8383 | 1.32 | 0.40 | 3.5E-04 | 1.33 | 0.41 | 1.9E-04 |
| Adarb1 | 2417 | 1.89 | 0.92 | 2.6E-18 | 1.90 | 0.92 | 2.4E-18 |
| Adck1 | 1097 | -1.58 | -0.66 | 8.4E-16 | -1.54 | -0.63 | 3.0E-14 |
| Adcy3 | 1057 | -1.69 | -0.76 | 1.4E-08 | -1.81 | -0.86 | 1.1E-10 |
| Adcy4 | 713 | -1.48 | -0.57 | 1.9E-02 | -1.47 | -0.56 | 2.3E-02 |
| Adcy7 | 424 | -1.57 | -0.65 | 5.6E-04 | -1.46 | -0.55 | 4.6E-03 |
| Adcy9 | 825 | 1.95 | 0.96 | 8.7E-06 | 2.04 | 1.03 | 1.8E-06 |
| Adcyap1 | 725 | -2.68 | -1.42 | 2.1E-15 | -2.78 | -1.48 | 1.6E-16 |
| Add2 | 16044 | -1.89 | -0.92 | 2.8E-06 | -1.87 | -0.90 | 4.8E-06 |
| Add3 | 8147 | 1.95 | 0.96 | 4.8E-14 | 1.96 | 0.97 | 3.6E-14 |
| Adgra1 | 539 | -1.85 | -0.89 | 4.2E-06 | -1.71 | -0.78 | 6.8E-05 |
| Adgra3 | 7782 | -1.32 | -0.41 | 1.3E-08 | -1.38 | -0.46 | 7.6E-11 |
| Adgrb1 | 4790 | -1.43 | -0.52 | 3.8E-02 | -1.52 | -0.60 | 1.5E-02 |
| Adgrb2 | 7542 | 1.56 | 0.64 | 2.6E-04 | 1.58 | 0.66 | 1.8E-04 |
| Adgre5 | 1171 | -1.53 | -0.61 | 4.6E-04 | -1.62 | -0.69 | 6.3E-05 |
| Adgrf5 | 2348 | -1.37 | -0.45 | 2.5E-05 | -1.27 | -0.34 | 1.7E-03 |
| Adgrg1 | 13224 | -2.36 | -1.24 | 6.4E-16 | -2.52 | -1.33 | 2.9E-18 |
| Adgrg2 | 336 | -2.54 | -1.34 | 4.5E-08 | -2.07 | -1.05 | 2.6E-05 |
| Adgrg6 | 1070 | -2.33 | -1.22 | 2.5E-04 | -2.01 | -1.00 | 3.1E-03 |

| | | | | | | | |
|---|---|---|---|---|---|---|---|
| Adgrl2 | 16496 | -1.34 | -0.42 | 2.9E-05 | -1.32 | -0.40 | 8.8E-05 |
| Adgrv1 | 9048 | 1.65 | 0.73 | 2.2E-03 | 1.93 | 0.95 | 5.2E-05 |
| Adi1 | 1022 | -2.04 | -1.03 | 1.2E-23 | -2.31 | -1.21 | 3.8E-32 |
| Adipor2 | 9283 | 1.50 | 0.59 | 8.5E-31 | 1.43 | 0.52 | 2.1E-24 |
| Adk | 3332 | 1.30 | 0.37 | 1.4E-02 | 1.30 | 0.38 | 1.4E-02 |
| Adnp | 233 | -1.63 | -0.71 | 4.0E-07 | -1.58 | -0.66 | 3.0E-06 |
| Adora2a | 296 | -1.25 | -0.32 | 4.9E-02 | -1.36 | -0.45 | 4.9E-03 |
| Adprs | 1967 | -1.31 | -0.39 | 2.8E-06 | -1.36 | -0.44 | 6.9E-08 |
| Adra2a | 747 | -1.66 | -0.73 | 2.3E-09 | -2.01 | -1.01 | 1.1E-16 |
| Adrb1 | 136 | 2.96 | 1.57 | 1.6E-19 | 2.57 | 1.36 | 4.2E-15 |
| Adss | 9487 | 1.32 | 0.40 | 2.7E-07 | 1.33 | 0.41 | 1.2E-07 |
| Aen | 4571 | 1.97 | 0.98 | 5.6E-47 | 1.90 | 0.93 | 3.4E-42 |
| Afap1l1 | 1070 | -1.37 | -0.45 | 2.9E-03 | -1.30 | -0.38 | 1.4E-02 |
| Aff1 | 2450 | 2.00 | 1.00 | 1.3E-17 | 2.09 | 1.06 | 1.3E-19 |
| Aff3 | 5158 | -1.41 | -0.49 | 2.2E-05 | -1.35 | -0.43 | 2.3E-04 |
| Afg1l | 270 | -1.72 | -0.78 | 1.5E-12 | -1.59 | -0.67 | 1.7E-09 |
| Afg3l1 | 4163 | -1.56 | -0.64 | 4.5E-12 | -1.75 | -0.80 | 1.9E-18 |
| Afg3l2 | 6049 | 1.76 | 0.82 | 6.4E-25 | 1.82 | 0.86 | 1.9E-27 |
| Afmid | 234 | 2.34 | 1.23 | 3.4E-15 | 2.19 | 1.13 | 5.6E-13 |
| Aftph | 6785 | 1.51 | 0.59 | 2.3E-10 | 1.64 | 0.71 | 1.3E-14 |
| Aga | 449 | -1.96 | -0.97 | 3.0E-23 | -1.92 | -0.94 | 5.7E-22 |
| Agbl5 | 4182 | -1.54 | -0.62 | 4.6E-13 | -1.60 | -0.68 | 2.2E-15 |
| Agfg1 | 11006 | 1.30 | 0.38 | 3.7E-05 | 1.39 | 0.48 | 1.5E-07 |
| Agfg2 | 2354 | 2.43 | 1.28 | 1.1E-24 | 2.10 | 1.07 | 1.9E-17 |
| Aggf1 | 6715 | 1.26 | 0.33 | 4.6E-06 | 1.17 | 0.23 | 1.9E-03 |
| Agk | 1373 | -1.77 | -0.83 | 2.0E-36 | -1.76 | -0.81 | 4.3E-35 |
| Agl | 6683 | 1.28 | 0.36 | 3.8E-05 | 1.39 | 0.48 | 2.6E-08 |
| Ago3 | 5603 | 1.60 | 0.68 | 3.5E-07 | 1.90 | 0.92 | 1.9E-12 |
| Agpat1 | 6364 | 1.40 | 0.48 | 1.4E-07 | 1.31 | 0.39 | 2.9E-05 |
| Agpat2 | 179 | -1.97 | -0.98 | 3.5E-05 | -2.09 | -1.06 | 6.7E-06 |
| Agpat3 | 4554 | -1.46 | -0.54 | 3.6E-05 | -1.49 | -0.58 | 1.0E-05 |
| Agpat5 | 5028 | 1.37 | 0.46 | 4.5E-11 | 1.41 | 0.50 | 6.3E-13 |
| Agtr2 | 720 | -6.44 | -2.69 | 2.7E-04 | -4.45 | -2.15 | 4.1E-03 |
| Agtrap | 1685 | 2.30 | 1.20 | 3.0E-18 | 2.08 | 1.06 | 2.0E-14 |
| Ahcyl2 | 4631 | 1.49 | 0.57 | 2.9E-28 | 1.51 | 0.59 | 4.6E-30 |
| Ahi1 | 10423 | 1.57 | 0.65 | 1.1E-04 | 1.60 | 0.68 | 5.6E-05 |
| Ahnak | 9132 | -3.07 | -1.62 | 1.2E-07 | -2.64 | -1.40 | 5.6E-06 |
| Ahrr | 201 | -3.20 | -1.68 | 4.7E-26 | -2.64 | -1.40 | 1.8E-18 |
| Ahsa2 | 8656 | -1.26 | -0.33 | 6.6E-04 | -1.22 | -0.29 | 3.4E-03 |
| AI413582 | 586 | 2.61 | 1.38 | 1.1E-50 | 2.78 | 1.48 | 4.0E-57 |
| AI429214 | 364 | 1.39 | 0.47 | 3.9E-07 | 1.44 | 0.52 | 2.1E-08 |
| AI661453 | 130 | -2.15 | -1.10 | 3.0E-03 | -2.03 | -1.02 | 6.7E-03 |
| AI837181 | 1415 | -1.70 | -0.77 | 1.5E-15 | -1.85 | -0.89 | 1.5E-20 |
| AI987944 | 964 | 1.70 | 0.76 | 8.1E-15 | 1.79 | 0.84 | 1.3E-17 |
| Aida | 7488 | 1.42 | 0.50 | 1.4E-14 | 1.46 | 0.54 | 1.3E-16 |
| Aifm1 | 5036 | 1.53 | 0.62 | 8.5E-11 | 1.46 | 0.55 | 1.0E-08 |
| Aifm2 | 725 | 2.47 | 1.30 | 1.0E-17 | 2.10 | 1.07 | 2.9E-12 |

| | | | | | | | |
|---|---|---|---|---|---|---|---|
| Aifm3 | 718 | -1.38 | -0.47 | 4.1E-03 | -1.60 | -0.68 | 2.1E-05 |
| Aim2 | 1185 | -1.50 | -0.59 | 1.6E-03 | -1.79 | -0.84 | 3.7E-06 |
| Aimp2 | 1731 | -1.23 | -0.30 | 9.2E-04 | -1.32 | -0.40 | 8.4E-06 |
| Ak1 | 1321 | -1.46 | -0.54 | 2.0E-04 | -1.41 | -0.50 | 7.1E-04 |
| Ak2 | 7116 | 1.41 | 0.49 | 2.2E-04 | 1.27 | 0.34 | 1.2E-02 |
| Ak3 | 4309 | -1.28 | -0.35 | 1.5E-04 | -1.34 | -0.42 | 4.4E-06 |
| Ak6 | 1427 | 1.50 | 0.58 | 1.4E-07 | 1.48 | 0.57 | 3.3E-07 |
| Akap1 | 5233 | 1.78 | 0.83 | 3.8E-28 | 1.68 | 0.75 | 3.4E-23 |
| Akap10 | 1782 | -1.54 | -0.62 | 3.6E-14 | -1.47 | -0.55 | 2.4E-11 |
| Akap13 | 5952 | 1.25 | 0.32 | 1.5E-02 | 1.46 | 0.55 | 1.8E-05 |
| Akap17b | 2484 | 1.38 | 0.46 | 3.0E-07 | 1.43 | 0.51 | 1.3E-08 |
| Akap5 | 1493 | 1.53 | 0.62 | 2.5E-07 | 1.53 | 0.61 | 3.1E-07 |
| Akap7 | 976 | -1.99 | -0.99 | 9.9E-22 | -2.16 | -1.11 | 5.8E-27 |
| Akap8 | 17422 | -1.26 | -0.34 | 3.2E-05 | -1.29 | -0.37 | 6.1E-06 |
| Akap8l | 8525 | 1.60 | 0.67 | 5.9E-12 | 1.61 | 0.69 | 2.6E-12 |
| Akap9 | 18776 | -2.30 | -1.20 | 5.3E-09 | -1.85 | -0.89 | 2.4E-05 |
| Akip1 | 342 | -1.26 | -0.34 | 1.6E-02 | -1.50 | -0.58 | 1.5E-05 |
| Akirin1 | 7540 | 1.59 | 0.67 | 2.3E-16 | 1.54 | 0.62 | 2.1E-14 |
| Akr1b10 | 542 | 1.79 | 0.84 | 9.7E-07 | 1.70 | 0.77 | 8.8E-06 |
| Akr1b3 | 3895 | 1.43 | 0.51 | 1.5E-03 | 1.34 | 0.43 | 1.0E-02 |
| Akr7a5 | 1487 | 1.51 | 0.60 | 1.6E-07 | 1.41 | 0.50 | 1.4E-05 |
| Akt1 | 15951 | -1.40 | -0.48 | 5.5E-09 | -1.45 | -0.54 | 4.7E-11 |
| Akt1s1 | 2273 | -2.05 | -1.03 | 7.0E-38 | -2.20 | -1.14 | 1.3E-45 |
| Akt2 | 4906 | -2.03 | -1.02 | 2.9E-50 | -2.08 | -1.06 | 5.6E-54 |
| Aktip | 5460 | 1.47 | 0.56 | 2.3E-17 | 1.47 | 0.55 | 3.8E-17 |
| Aldh16a1 | 1460 | -1.64 | -0.72 | 4.0E-08 | -1.77 | -0.82 | 2.7E-10 |
| Aldh18a1 | 9896 | 1.58 | 0.66 | 5.9E-12 | 1.45 | 0.54 | 2.7E-08 |
| Aldh1b1 | 643 | 2.12 | 1.08 | 1.4E-15 | 2.13 | 1.09 | 8.6E-16 |
| Aldh2 | 6283 | 1.35 | 0.43 | 1.2E-03 | 1.30 | 0.38 | 4.9E-03 |
| Aldh3a2 | 1949 | -1.37 | -0.46 | 1.4E-16 | -1.37 | -0.46 | 2.5E-16 |
| Aldh4a1 | 1721 | 2.23 | 1.16 | 6.8E-42 | 2.16 | 1.11 | 1.6E-38 |
| Aldh6a1 | 3320 | 1.56 | 0.64 | 6.0E-09 | 1.63 | 0.71 | 1.4E-10 |
| Aldh7a1 | 5826 | 1.28 | 0.36 | 5.4E-04 | 1.24 | 0.31 | 3.3E-03 |
| Aldoa | 19582 | 1.32 | 0.40 | 1.1E-03 | 1.50 | 0.59 | 1.5E-04 |
| Aldoa | 19582 | 1.32 | 0.40 | 1.1E-03 | 1.25 | 0.32 | 1.2E-02 |
| Aldoa | 251 | 1.64 | 0.71 | 3.3E-06 | 1.50 | 0.59 | 1.5E-04 |
| Aldoa | 251 | 1.64 | 0.71 | 3.3E-06 | 1.25 | 0.32 | 1.2E-02 |
| Alg13 | 3561 | 2.06 | 1.05 | 1.3E-25 | 1.97 | 0.98 | 2.2E-22 |
| Alg3 | 980 | -1.23 | -0.30 | 3.8E-04 | -1.33 | -0.41 | 5.3E-07 |
| Alg8 | 1473 | -1.74 | -0.80 | 4.9E-13 | -1.78 | -0.83 | 5.6E-14 |
| Alkbh1 | 2871 | -1.37 | -0.46 | 5.8E-20 | -1.40 | -0.49 | 1.7E-22 |
| Alkbh3 | 782 | -1.69 | -0.76 | 5.4E-14 | -1.52 | -0.60 | 3.5E-09 |
| Alkbh4 | 826 | 1.41 | 0.50 | 5.5E-08 | 1.23 | 0.29 | 1.9E-03 |
| Alkbh5 | 7600 | 1.28 | 0.35 | 2.0E-03 | 1.18 | 0.24 | 4.6E-02 |
| Alkbh6 | 1091 | 1.46 | 0.55 | 2.1E-09 | 1.37 | 0.45 | 1.1E-06 |
| Alpl | 811 | -1.66 | -0.73 | 1.2E-05 | -1.64 | -0.71 | 2.1E-05 |
| Als2 | 6429 | 1.63 | 0.70 | 3.2E-30 | 1.72 | 0.78 | 4.6E-37 |

| | | | | | | | |
|---|---|---|---|---|---|---|---|
| Als2cl | 363 | 1.41 | 0.50 | 1.0E-02 | 1.53 | 0.61 | 1.5E-03 |
| Alx4 | 2332 | -2.78 | -1.47 | 2.2E-03 | -3.42 | -1.77 | 1.9E-04 |
| Alyref | 6003 | -1.29 | -0.36 | 5.2E-04 | -1.39 | -0.47 | 5.0E-06 |
| Amacr | 350 | -2.04 | -1.03 | 1.3E-19 | -2.04 | -1.03 | 1.7E-19 |
| Amer3 | 1277 | -2.39 | -1.26 | 7.9E-05 | -2.40 | -1.26 | 7.3E-05 |
| Amfr | 7509 | -1.43 | -0.52 | 1.6E-12 | -1.50 | -0.58 | 8.6E-16 |
| Amigo1 | 3006 | 1.68 | 0.75 | 1.2E-08 | 1.76 | 0.82 | 4.0E-10 |
| Amigo3 | 338 | 2.13 | 1.09 | 4.8E-22 | 2.06 | 1.04 | 4.4E-20 |
| Ammecr1 | 2133 | -1.26 | -0.33 | 1.1E-03 | -1.24 | -0.31 | 2.6E-03 |
| Ammecr1l | 5365 | -1.22 | -0.29 | 7.3E-04 | -1.26 | -0.34 | 5.8E-05 |
| Amn1 | 2385 | -1.55 | -0.63 | 2.5E-09 | -1.42 | -0.50 | 2.9E-06 |
| Amotl1 | 8448 | -1.60 | -0.68 | 2.3E-08 | -1.48 | -0.56 | 4.7E-06 |
| Amotl2 | 6480 | -1.60 | -0.68 | 8.2E-11 | -1.72 | -0.78 | 4.1E-14 |
| Ampd3 | 174 | -3.19 | -1.67 | 4.5E-11 | -3.04 | -1.61 | 2.9E-10 |
| Anapc1 | 19417 | -1.36 | -0.45 | 1.8E-04 | -1.27 | -0.35 | 4.3E-03 |
| Anapc11 | 3450 | -1.20 | -0.26 | 6.8E-03 | -1.31 | -0.39 | 3.1E-05 |
| Anapc15 | 2482 | -1.85 | -0.88 | 1.6E-29 | -1.85 | -0.89 | 7.2E-30 |
| Anapc7 | 6014 | 1.27 | 0.34 | 9.3E-08 | 1.24 | 0.31 | 1.8E-06 |
| Angel1 | 1761 | 2.10 | 1.07 | 4.7E-41 | 2.11 | 1.08 | 9.2E-42 |
| Angel2 | 4953 | -2.17 | -1.12 | 1.7E-48 | -2.19 | -1.13 | 2.2E-49 |
| Angpt1 | 1607 | -1.57 | -0.65 | 5.2E-05 | -1.48 | -0.56 | 6.1E-04 |
| Angpt2 | 223 | -2.28 | -1.19 | 6.3E-10 | -2.33 | -1.22 | 1.6E-10 |
| Angptl1 | 297 | -7.05 | -2.82 | 8.4E-05 | -5.19 | -2.38 | 1.1E-03 |
| Angptl4 | 809 | -2.71 | -1.44 | 1.8E-11 | -2.77 | -1.47 | 6.0E-12 |
| Ank1 | 889 | -1.75 | -0.80 | 3.2E-04 | -1.81 | -0.86 | 1.2E-04 |
| Ankhd1 | 9952 | 1.34 | 0.42 | 2.7E-02 | 1.63 | 0.70 | 1.3E-04 |
| Ankib1 | 7029 | 1.34 | 0.42 | 1.1E-05 | 1.41 | 0.49 | 2.0E-07 |
| Ankle2 | 4642 | -1.42 | -0.50 | 3.2E-24 | -1.46 | -0.55 | 1.8E-28 |
| Ankrd12 | 8990 | 1.43 | 0.52 | 7.3E-03 | 1.69 | 0.76 | 6.4E-05 |
| Ankrd13a | 9540 | 2.21 | 1.15 | 3.1E-40 | 2.08 | 1.06 | 1.8E-34 |
| Ankrd13d | 1571 | 1.41 | 0.50 | 1.1E-04 | 1.40 | 0.49 | 1.6E-04 |
| Ankrd24 | 1925 | -1.27 | -0.34 | 1.3E-02 | -1.25 | -0.32 | 2.5E-02 |
| Ankrd27 | 3783 | 1.48 | 0.56 | 4.2E-19 | 1.49 | 0.58 | 4.6E-20 |
| Ankrd28 | 8082 | 1.30 | 0.38 | 1.1E-04 | 1.33 | 0.42 | 1.9E-05 |
| Ankrd33b | 129 | 1.81 | 0.86 | 1.2E-06 | 1.79 | 0.84 | 2.7E-06 |
| Ankrd34a | 518 | 2.58 | 1.37 | 2.6E-11 | 2.73 | 1.45 | 1.6E-12 |
| Ankrd34b | 450 | 5.40 | 2.43 | 7.8E-20 | 6.84 | 2.77 | 2.8E-25 |
| Ankrd35 | 295 | 4.05 | 2.02 | 5.4E-21 | 3.63 | 1.86 | 5.3E-18 |
| Ankrd39 | 785 | 1.77 | 0.83 | 6.4E-21 | 1.78 | 0.83 | 3.4E-21 |
| Ankrd42 | 1551 | 2.04 | 1.03 | 1.1E-33 | 2.25 | 1.17 | 2.4E-43 |
| Ankrd44 | 2666 | -1.88 | -0.91 | 1.8E-11 | -1.73 | -0.79 | 8.1E-09 |
| Ankrd54 | 2350 | 1.55 | 0.63 | 4.7E-11 | 1.42 | 0.51 | 1.2E-07 |
| Ankrd9 | 290 | 1.82 | 0.86 | 2.2E-09 | 1.65 | 0.72 | 8.4E-07 |
| Anks6 | 1065 | 1.54 | 0.62 | 2.2E-08 | 1.42 | 0.50 | 7.9E-06 |
| Ankzf1 | 2571 | 1.33 | 0.42 | 2.0E-12 | 1.37 | 0.45 | 2.9E-14 |
| Anln | 4143 | -1.43 | -0.52 | 7.3E-04 | -1.42 | -0.50 | 1.1E-03 |
| Ano1 | 1278 | -2.06 | -1.04 | 6.9E-10 | -2.29 | -1.20 | 9.7E-13 |

| | | | | | | | |
|---|---|---|---|---|---|---|---|
| Ano10 | 3264 | 1.41 | 0.50 | 2.7E-09 | 1.47 | 0.56 | 2.2E-11 |
| Ano4 | 413 | -1.56 | -0.64 | 8.1E-03 | -1.41 | -0.50 | 4.5E-02 |
| Ano8 | 6678 | -1.61 | -0.69 | 4.3E-05 | -1.73 | -0.79 | 2.4E-06 |
| Anp32b | 15767 | -1.75 | -0.81 | 2.0E-06 | -2.04 | -1.03 | 9.7E-10 |
| Antkmt | 1730 | 1.65 | 0.72 | 6.9E-05 | 1.43 | 0.52 | 5.3E-03 |
| Antxr1 | 3034 | -2.63 | -1.39 | 1.6E-11 | -2.62 | -1.39 | 1.8E-11 |
| Antxr2 | 1034 | -1.92 | -0.94 | 7.5E-09 | -1.96 | -0.97 | 2.0E-09 |
| Anxa1 | 480 | -4.00 | -2.00 | 4.4E-05 | -3.16 | -1.66 | 8.3E-04 |
| Aoc2 | 1338 | 1.90 | 0.93 | 2.2E-15 | 1.92 | 0.94 | 1.0E-15 |
| Ap1ar | 7400 | 1.31 | 0.39 | 1.1E-11 | 1.25 | 0.32 | 3.9E-08 |
| Ap1m1 | 7303 | -1.57 | -0.65 | 1.4E-24 | -1.66 | -0.73 | 8.4E-31 |
| Ap1s2 | 5451 | 1.35 | 0.43 | 1.5E-03 | 1.33 | 0.41 | 2.7E-03 |
| Ap1s3 | 359 | 3.17 | 1.66 | 4.5E-11 | 2.97 | 1.57 | 5.8E-10 |
| Ap2a1 | 9022 | 1.36 | 0.45 | 6.8E-04 | 1.29 | 0.37 | 6.2E-03 |
| Ap2a2 | 15202 | 1.38 | 0.46 | 3.0E-18 | 1.32 | 0.40 | 3.8E-14 |
| Ap3d1 | 14765 | 1.28 | 0.36 | 1.4E-08 | 1.24 | 0.31 | 1.2E-06 |
| Ap3m1 | 10524 | 1.32 | 0.40 | 1.2E-04 | 1.37 | 0.46 | 1.1E-05 |
| Ap3m2 | 4202 | 1.37 | 0.45 | 1.4E-05 | 1.45 | 0.54 | 1.7E-07 |
| Ap3s1 | 7782 | 1.36 | 0.45 | 2.4E-07 | 1.38 | 0.46 | 9.3E-08 |
| Ap4m1 | 4211 | -2.05 | -1.03 | 3.1E-83 | -2.15 | -1.10 | 5.6E-94 |
| Ap4s1 | 970 | -1.99 | -0.99 | 4.5E-21 | -2.05 | -1.03 | 8.7E-23 |
| Ap5s1 | 904 | 1.45 | 0.53 | 1.1E-11 | 1.40 | 0.49 | 5.6E-10 |
| Apaf1 | 8393 | 1.69 | 0.76 | 1.3E-37 | 1.77 | 0.82 | 4.7E-44 |
| Apba2 | 6951 | -1.57 | -0.65 | 2.4E-04 | -1.59 | -0.67 | 1.3E-04 |
| Apbb1ip | 142 | -1.73 | -0.79 | 1.1E-04 | -1.57 | -0.65 | 1.7E-03 |
| Apc2 | 25582 | -1.41 | -0.49 | 3.9E-02 | -1.48 | -0.57 | 1.7E-02 |
| Apeh | 4262 | 1.48 | 0.57 | 1.9E-09 | 1.46 | 0.55 | 7.3E-09 |
| Apex2 | 1280 | -1.55 | -0.63 | 5.1E-15 | -1.55 | -0.63 | 6.2E-15 |
| Aph1a | 4926 | 1.43 | 0.52 | 2.2E-08 | 1.31 | 0.39 | 2.6E-05 |
| Aph1b | 4461 | 1.42 | 0.50 | 2.7E-06 | 1.44 | 0.53 | 1.0E-06 |
| Aph1c | 811 | 5.81 | 2.54 | 8.3E-124 | 6.10 | 2.61 | 9.0E-130 |
| Api5 | 17264 | 1.38 | 0.47 | 5.7E-06 | 1.35 | 0.43 | 3.3E-05 |
| Aplf | 1775 | 2.52 | 1.33 | 1.2E-96 | 2.52 | 1.33 | 6.5E-96 |
| Aplnr | 1799 | -2.28 | -1.19 | 1.5E-24 | -2.46 | -1.30 | 7.7E-29 |
| Aplp2 | 22093 | 1.67 | 0.74 | 3.3E-41 | 1.73 | 0.79 | 2.3E-46 |
| Apmap | 2812 | 1.50 | 0.58 | 8.0E-20 | 1.44 | 0.53 | 2.1E-16 |
| Apobec3 | 247 | -1.83 | -0.88 | 1.5E-13 | -1.73 | -0.79 | 4.6E-11 |
| Apold1 | 534 | 1.52 | 0.60 | 3.0E-07 | 1.52 | 0.61 | 2.3E-07 |
| Apoo | 2242 | 1.27 | 0.34 | 5.5E-04 | 1.33 | 0.41 | 3.5E-05 |
| Appl2 | 4832 | 1.21 | 0.28 | 2.7E-07 | 1.27 | 0.35 | 6.3E-11 |
| Aptx | 2949 | -1.28 | -0.36 | 2.4E-05 | -1.33 | -0.42 | 9.1E-07 |
| Aqp1 | 1564 | -1.90 | -0.93 | 3.5E-02 | -2.05 | -1.04 | 1.8E-02 |
| Aqp11 | 570 | 2.21 | 1.15 | 2.4E-22 | 2.13 | 1.09 | 2.5E-20 |
| Araf | 8449 | -1.33 | -0.41 | 2.4E-08 | -1.30 | -0.38 | 2.5E-07 |
| Arap2 | 360 | -1.70 | -0.77 | 3.4E-04 | -1.44 | -0.52 | 1.8E-02 |
| Arap3 | 1599 | -1.61 | -0.69 | 2.3E-06 | -1.76 | -0.82 | 1.6E-08 |
| Arc | 170 | 1.64 | 0.72 | 1.7E-04 | 1.66 | 0.73 | 1.4E-04 |

| | | | | | | | |
|---|---|---|---|---|---|---|---|
| Arf2 | 10786 | 1.55 | 0.63 | 4.6E-19 | 1.53 | 0.62 | 4.2E-18 |
| Arf3 | 12109 | 1.27 | 0.35 | 5.6E-06 | 1.20 | 0.26 | 9.6E-04 |
| Arfgap2 | 5954 | -1.56 | -0.64 | 1.2E-19 | -1.67 | -0.74 | 6.8E-26 |
| Arfgap3 | 2660 | 1.43 | 0.52 | 1.8E-25 | 1.34 | 0.42 | 7.3E-17 |
| Arfgef2 | 3514 | 1.70 | 0.77 | 3.0E-19 | 1.78 | 0.83 | 4.9E-22 |
| Arfip2 | 6408 | 1.42 | 0.50 | 9.9E-25 | 1.41 | 0.49 | 7.9E-24 |
| Arfrp1 | 3761 | -1.27 | -0.34 | 1.5E-05 | -1.32 | -0.40 | 4.0E-07 |
| Arglu1 | 25853 | -1.21 | -0.28 | 2.3E-02 | -1.25 | -0.32 | 7.1E-03 |
| Arhgap18 | 2849 | 1.30 | 0.38 | 1.8E-06 | 1.33 | 0.41 | 2.4E-07 |
| Arhgap20 | 2673 | 1.66 | 0.73 | 3.4E-08 | 1.71 | 0.77 | 4.9E-09 |
| Arhgap21 | 15502 | -1.42 | -0.51 | 1.1E-04 | -1.31 | -0.39 | 3.3E-03 |
| Arhgap22 | 170 | 2.12 | 1.08 | 1.6E-04 | 2.69 | 1.43 | 5.8E-07 |
| Arhgap23 | 3685 | 1.44 | 0.53 | 2.1E-05 | 1.41 | 0.50 | 6.4E-05 |
| Arhgap24 | 398 | -1.80 | -0.85 | 4.3E-04 | -1.67 | -0.74 | 2.4E-03 |
| Arhgap26 | 817 | 1.81 | 0.85 | 2.4E-08 | 1.85 | 0.89 | 7.1E-09 |
| Arhgap27 | 432 | -1.52 | -0.61 | 1.9E-05 | -1.63 | -0.71 | 5.5E-07 |
| Arhgap30 | 138 | -2.88 | -1.53 | 1.6E-17 | -2.93 | -1.55 | 4.6E-18 |
| Arhgap31 | 2522 | -1.75 | -0.81 | 9.8E-08 | -1.75 | -0.81 | 9.7E-08 |
| Arhgap35 | 20557 | -1.31 | -0.39 | 3.6E-04 | -1.27 | -0.34 | 2.0E-03 |
| Arhgap39 | 4939 | 1.51 | 0.59 | 7.1E-11 | 1.51 | 0.60 | 5.2E-11 |
| Arhgap4 | 571 | 1.65 | 0.73 | 1.5E-12 | 1.81 | 0.85 | 7.0E-17 |
| Arhgap42 | 1837 | -3.86 | -1.95 | 6.0E-65 | -3.41 | -1.77 | 9.9E-54 |
| Arhgap6 | 177 | -1.78 | -0.83 | 3.5E-04 | -1.61 | -0.69 | 3.5E-03 |
| Arhgdig | 193 | -1.88 | -0.91 | 3.1E-02 | -2.05 | -1.03 | 1.4E-02 |
| Arhgef10l | 2812 | -1.57 | -0.65 | 2.7E-08 | -1.69 | -0.76 | 8.7E-11 |
| Arhgef11 | 7475 | 1.47 | 0.56 | 9.8E-08 | 1.48 | 0.56 | 8.2E-08 |
| Arhgef15 | 851 | -2.42 | -1.27 | 4.5E-24 | -2.73 | -1.45 | 1.3E-30 |
| Arhgef18 | 4572 | 1.46 | 0.54 | 2.6E-16 | 1.43 | 0.52 | 8.1E-15 |
| Arhgef2 | 9820 | -1.95 | -0.96 | 1.2E-15 | -2.08 | -1.06 | 9.6E-19 |
| Arhgef28 | 730 | -1.73 | -0.79 | 9.6E-06 | -1.50 | -0.58 | 1.3E-03 |
| Arhgef39 | 947 | -1.82 | -0.86 | 1.7E-06 | -2.04 | -1.03 | 9.7E-09 |
| Arhgef5 | 758 | -3.56 | -1.83 | 1.8E-14 | -3.47 | -1.79 | 6.8E-14 |
| Arhgef7 | 12176 | -1.30 | -0.38 | 7.5E-08 | -1.35 | -0.43 | 6.8E-10 |
| Arid1a | 20363 | -1.49 | -0.58 | 4.8E-04 | -1.58 | -0.66 | 5.9E-05 |
| Arid1b | 5659 | -2.13 | -1.09 | 1.2E-16 | -2.17 | -1.12 | 2.7E-17 |
| Arid2 | 11217 | -1.62 | -0.70 | 2.8E-15 | -1.59 | -0.67 | 4.7E-14 |
| Arid3b | 1991 | 1.71 | 0.78 | 9.3E-07 | 1.47 | 0.56 | 6.1E-04 |
| Arid4a | 3784 | 1.23 | 0.30 | 2.1E-06 | 1.28 | 0.35 | 2.5E-08 |
| Arid5a | 365 | 2.65 | 1.41 | 2.9E-14 | 2.94 | 1.56 | 3.9E-17 |
| Arid5b | 4505 | 1.59 | 0.67 | 8.1E-13 | 1.81 | 0.86 | 2.5E-20 |
| Arih2 | 5232 | 1.35 | 0.44 | 9.7E-15 | 1.31 | 0.39 | 4.1E-12 |
| Arl10 | 1376 | -1.26 | -0.33 | 9.8E-03 | -1.26 | -0.33 | 1.1E-02 |
| Arl13b | 3545 | 2.03 | 1.02 | 1.9E-43 | 2.15 | 1.11 | 1.4E-50 |
| Arl15 | 3996 | 1.26 | 0.34 | 1.0E-03 | 1.29 | 0.36 | 3.9E-04 |
| Arl16 | 1235 | 1.49 | 0.58 | 2.2E-15 | 1.44 | 0.53 | 3.6E-13 |
| Arl2 | 1908 | -1.61 | -0.68 | 1.2E-30 | -1.65 | -0.72 | 5.3E-34 |
| Arl3 | 5824 | 1.55 | 0.63 | 2.0E-12 | 1.43 | 0.52 | 1.1E-08 |

| | | | | | | | |
|---|---|---|---|---|---|---|---|
| Arl4a | 5987 | 1.26 | 0.33 | 9.3E-04 | 1.25 | 0.32 | 1.7E-03 |
| Arl5c | 218 | 4.28 | 2.10 | 3.2E-49 | 4.10 | 2.04 | 2.1E-45 |
| Arl6 | 2653 | 1.84 | 0.88 | 9.4E-21 | 1.90 | 0.93 | 1.1E-22 |
| Arl6ip1 | 26887 | 1.42 | 0.50 | 5.7E-09 | 1.39 | 0.47 | 5.9E-08 |
| Arl6ip5 | 4917 | 1.36 | 0.45 | 4.7E-11 | 1.42 | 0.51 | 7.1E-14 |
| Arl6ip6 | 5014 | 1.65 | 0.72 | 2.9E-08 | 1.48 | 0.57 | 1.6E-05 |
| Arl8b | 8363 | 1.30 | 0.37 | 7.1E-11 | 1.27 | 0.34 | 2.8E-09 |
| Armc1 | 6660 | 1.30 | 0.38 | 6.7E-07 | 1.28 | 0.36 | 2.1E-06 |
| Armc2 | 481 | 1.50 | 0.58 | 1.8E-03 | 1.59 | 0.67 | 2.8E-04 |
| Armc5 | 927 | -1.88 | -0.91 | 1.1E-13 | -1.91 | -0.93 | 3.2E-14 |
| Armc6 | 1666 | -1.47 | -0.56 | 8.8E-06 | -1.43 | -0.51 | 5.0E-05 |
| Armc7 | 682 | 1.34 | 0.42 | 2.8E-04 | 1.30 | 0.37 | 1.3E-03 |
| Armc9 | 3088 | 2.29 | 1.20 | 9.0E-84 | 2.26 | 1.18 | 3.9E-81 |
| Armcx1 | 4223 | -2.14 | -1.10 | 3.3E-52 | -2.12 | -1.08 | 1.4E-50 |
| Armcx3 | 4569 | 1.46 | 0.54 | 1.2E-17 | 1.47 | 0.55 | 3.9E-18 |
| Armh3 | 1724 | -1.56 | -0.64 | 1.9E-25 | -1.69 | -0.76 | 4.2E-35 |
| Armh4 | 3827 | 1.38 | 0.47 | 1.9E-02 | 1.54 | 0.62 | 1.3E-03 |
| Arnt2 | 18880 | -1.34 | -0.42 | 5.6E-03 | -1.41 | -0.50 | 1.1E-03 |
| Arntl | 1203 | 2.17 | 1.12 | 8.8E-45 | 2.07 | 1.05 | 1.6E-39 |
| Arntl2 | 511 | 1.71 | 0.78 | 3.1E-06 | 2.17 | 1.12 | 1.2E-11 |
| Arpc2 | 12830 | 1.25 | 0.32 | 8.6E-12 | 1.24 | 0.31 | 3.9E-11 |
| Arpc5l | 4951 | 1.37 | 0.46 | 8.4E-14 | 1.29 | 0.36 | 4.6E-09 |
| Arpin | 1002 | -1.62 | -0.70 | 6.5E-09 | -1.76 | -0.82 | 8.3E-12 |
| Arpp19 | 15783 | 1.31 | 0.39 | 3.8E-09 | 1.33 | 0.41 | 5.4E-10 |
| Arpp21 | 589 | -3.31 | -1.73 | 2.8E-06 | -2.75 | -1.46 | 9.2E-05 |
| Arrb1 | 7323 | -1.38 | -0.47 | 2.7E-03 | -1.33 | -0.42 | 8.5E-03 |
| Arrb2 | 3883 | 1.37 | 0.45 | 8.6E-06 | 1.29 | 0.37 | 3.4E-04 |
| Arrdc1 | 1143 | 2.55 | 1.35 | 1.7E-38 | 2.40 | 1.26 | 6.2E-34 |
| Arrdc3 | 6108 | -1.47 | -0.56 | 6.9E-06 | -1.37 | -0.45 | 3.6E-04 |
| Arsb | 1585 | -1.66 | -0.74 | 4.0E-12 | -1.68 | -0.75 | 1.3E-12 |
| Arsi | 441 | -2.46 | -1.30 | 2.2E-05 | -2.61 | -1.38 | 6.1E-06 |
| Arsk | 783 | -1.26 | -0.33 | 1.1E-03 | -1.18 | -0.24 | 2.2E-02 |
| As3mt | 858 | -1.86 | -0.90 | 1.1E-27 | -1.68 | -0.75 | 1.7E-19 |
| Asah1 | 7276 | 1.75 | 0.80 | 3.8E-18 | 1.76 | 0.82 | 9.3E-19 |
| Asah2 | 331 | -2.27 | -1.18 | 1.2E-06 | -2.09 | -1.06 | 1.6E-05 |
| Asap3 | 1872 | 1.98 | 0.98 | 5.5E-12 | 2.03 | 1.02 | 7.9E-13 |
| Asb1 | 3892 | 1.55 | 0.64 | 9.9E-16 | 1.61 | 0.69 | 3.7E-18 |
| Asb13 | 1451 | -1.37 | -0.45 | 1.9E-05 | -1.53 | -0.62 | 2.8E-09 |
| Asb3 | 1497 | -1.48 | -0.57 | 7.9E-10 | -1.42 | -0.50 | 6.2E-08 |
| Asb6 | 1892 | 1.27 | 0.35 | 1.4E-04 | 1.16 | 0.22 | 2.2E-02 |
| Asb7 | 2931 | 1.43 | 0.52 | 3.3E-10 | 1.38 | 0.47 | 2.1E-08 |
| Ascc1 | 1461 | -1.30 | -0.38 | 4.4E-12 | -1.33 | -0.41 | 1.1E-13 |
| Asf1b | 2507 | 2.77 | 1.47 | 2.3E-13 | 2.31 | 1.21 | 2.6E-09 |
| Ash1l | 9264 | -1.69 | -0.75 | 1.4E-05 | -1.40 | -0.48 | 6.9E-03 |
| Ash2l | 7989 | -1.37 | -0.46 | 7.0E-14 | -1.46 | -0.55 | 2.3E-19 |
| Asl | 1544 | 2.09 | 1.06 | 7.5E-42 | 1.93 | 0.95 | 1.2E-33 |
| Asns | 2610 | 1.56 | 0.65 | 3.7E-09 | 1.57 | 0.65 | 2.4E-09 |

| | | | | | | | |
|---|---|---|---|---|---|---|---|
| Asphd2 | 3058 | 1.61 | 0.68 | 4.7E-12 | 1.67 | 0.74 | 8.4E-14 |
| Aspn | 763 | -7.74 | -2.95 | 2.8E-05 | -7.45 | -2.90 | 4.1E-05 |
| Aspscr1 | 3308 | -1.36 | -0.45 | 2.3E-13 | -1.43 | -0.51 | 1.8E-17 |
| Asrgl1 | 3173 | 1.79 | 0.84 | 8.6E-24 | 1.77 | 0.82 | 8.8E-23 |
| Aste1 | 987 | 1.35 | 0.44 | 8.0E-06 | 1.48 | 0.57 | 4.6E-09 |
| Astn1 | 10065 | -1.39 | -0.48 | 6.5E-03 | -1.34 | -0.43 | 1.7E-02 |
| Asxl1 | 9564 | -1.39 | -0.48 | 7.4E-06 | -1.42 | -0.51 | 1.8E-06 |
| Asxl3 | 5208 | -1.75 | -0.81 | 1.5E-10 | -1.57 | -0.65 | 2.7E-07 |
| Atad2b | 7614 | -1.87 | -0.91 | 6.3E-25 | -1.78 | -0.83 | 5.1E-21 |
| Atad3a | 4669 | 1.36 | 0.45 | 1.1E-06 | 1.28 | 0.36 | 1.2E-04 |
| Atad5 | 6034 | 1.31 | 0.39 | 9.0E-03 | 1.39 | 0.48 | 1.0E-03 |
| Atf1 | 1643 | -2.42 | -1.28 | 6.1E-20 | -2.75 | -1.46 | 9.4E-26 |
| Atf3 | 165 | 2.93 | 1.55 | 1.4E-10 | 2.24 | 1.16 | 2.3E-06 |
| Atf4 | 7357 | -1.27 | -0.34 | 6.0E-06 | -1.37 | -0.45 | 1.2E-09 |
| Atf5 | 2343 | -1.41 | -0.50 | 1.6E-02 | -1.69 | -0.75 | 1.6E-04 |
| Atf6 | 4850 | 1.97 | 0.98 | 4.7E-32 | 2.09 | 1.06 | 1.8E-37 |
| Atf7ip | 13847 | -1.60 | -0.68 | 2.0E-13 | -1.56 | -0.64 | 4.3E-12 |
| Atf7ip2 | 124 | 1.89 | 0.91 | 1.2E-02 | 1.71 | 0.77 | 3.7E-02 |
| Atg10 | 362 | -1.59 | -0.67 | 1.1E-10 | -1.63 | -0.71 | 7.8E-12 |
| Atg101 | 1809 | 1.46 | 0.54 | 1.2E-07 | 1.38 | 0.47 | 5.8E-06 |
| Atg12 | 8267 | 1.53 | 0.61 | 4.6E-15 | 1.60 | 0.68 | 2.5E-18 |
| Atg13 | 4511 | 1.36 | 0.45 | 1.5E-17 | 1.35 | 0.44 | 8.9E-17 |
| Atg16l2 | 961 | 1.84 | 0.88 | 3.2E-27 | 1.70 | 0.76 | 7.5E-21 |
| Atg2b | 5427 | 1.30 | 0.38 | 4.4E-04 | 1.44 | 0.52 | 9.4E-07 |
| Atg4b | 7841 | 1.39 | 0.47 | 4.0E-21 | 1.42 | 0.51 | 2.4E-24 |
| Atg4d | 2225 | 1.45 | 0.54 | 6.9E-10 | 1.31 | 0.39 | 1.2E-05 |
| Atg5 | 2643 | 1.25 | 0.32 | 8.8E-04 | 1.32 | 0.41 | 2.2E-05 |
| Atl1 | 2686 | -2.63 | -1.39 | 2.5E-14 | -2.61 | -1.39 | 3.4E-14 |
| Atl3 | 4777 | -1.75 | -0.81 | 2.3E-12 | -1.60 | -0.68 | 6.1E-09 |
| Atmin | 4631 | 1.44 | 0.52 | 2.0E-23 | 1.44 | 0.52 | 2.9E-23 |
| Atn1 | 628 | -1.57 | -0.65 | 4.7E-02 | -1.78 | -0.83 | 9.9E-03 |
| Atoh8 | 272 | -1.46 | -0.54 | 9.1E-03 | -1.34 | -0.43 | 4.6E-02 |
| Atox1 | 1991 | 1.44 | 0.53 | 2.4E-08 | 1.41 | 0.50 | 1.5E-07 |
| Atp10a | 473 | -1.89 | -0.92 | 3.3E-16 | -1.89 | -0.92 | 4.2E-16 |
| Atp10b | 100 | -4.83 | -2.27 | 2.0E-04 | -3.02 | -1.59 | 1.2E-02 |
| Atp13a2 | 8534 | 1.45 | 0.54 | 7.3E-05 | 1.41 | 0.49 | 3.4E-04 |
| Atp13a3 | 9720 | 1.67 | 0.74 | 7.0E-22 | 1.72 | 0.78 | 1.0E-24 |
| Atp1a2 | 4737 | -1.68 | -0.75 | 6.9E-03 | -1.63 | -0.71 | 1.1E-02 |
| Atp1b2 | 3050 | 1.81 | 0.86 | 3.5E-08 | 1.69 | 0.75 | 1.6E-06 |
| Atp1b3 | 9061 | 1.32 | 0.40 | 8.7E-17 | 1.26 | 0.33 | 1.2E-11 |
| Atp23 | 769 | 2.30 | 1.20 | 4.7E-33 | 2.29 | 1.20 | 9.9E-33 |
| Atp2b4 | 8446 | 1.45 | 0.54 | 3.0E-05 | 1.60 | 0.68 | 1.2E-07 |
| Atp4a | 133 | 3.99 | 2.00 | 7.4E-52 | 3.56 | 1.83 | 6.5E-42 |
| Atp6ap2 | 6746 | 1.43 | 0.52 | 3.2E-13 | 1.50 | 0.59 | 1.2E-16 |
| Atp6v0a2 | 6136 | 1.42 | 0.51 | 2.4E-14 | 1.40 | 0.49 | 2.6E-13 |
| Atp6v0b | 6380 | 1.68 | 0.74 | 1.2E-20 | 1.70 | 0.77 | 5.7E-22 |
| Atp6v0d1 | 7619 | 1.29 | 0.37 | 2.4E-07 | 1.23 | 0.30 | 2.6E-05 |

| | | | | | | | |
|---|---|---|---|---|---|---|---|
| Atp6v0e | 2228 | 1.22 | 0.28 | 4.1E-04 | 1.28 | 0.36 | 5.5E-06 |
| Atp6v1a | 14716 | 1.74 | 0.80 | 1.7E-37 | 1.81 | 0.85 | 5.8E-43 |
| Atp6v1d | 6816 | 1.42 | 0.51 | 2.8E-16 | 1.46 | 0.55 | 4.4E-19 |
| Atp6v1e1 | 7661 | 1.21 | 0.28 | 2.6E-05 | 1.26 | 0.33 | 3.0E-07 |
| Atp6v1h | 8350 | 1.23 | 0.30 | 2.2E-03 | 1.30 | 0.38 | 6.6E-05 |
| Atp7b | 677 | 1.90 | 0.93 | 7.2E-24 | 1.81 | 0.86 | 2.5E-20 |
| Atp8a1 | 9431 | 1.84 | 0.88 | 1.1E-10 | 2.00 | 1.00 | 1.5E-13 |
| Atp8b2 | 6504 | 1.44 | 0.52 | 6.7E-12 | 1.42 | 0.51 | 3.6E-11 |
| Atp8b3 | 122 | 1.34 | 0.43 | 1.3E-02 | 1.29 | 0.37 | 3.5E-02 |
| Atpaf1 | 881 | -1.49 | -0.58 | 6.9E-13 | -1.56 | -0.64 | 8.5E-16 |
| Atpsckmt | 1169 | -1.80 | -0.85 | 9.0E-08 | -1.68 | -0.75 | 3.2E-06 |
| Atrx | 29927 | -1.80 | -0.85 | 2.5E-06 | -1.44 | -0.52 | 5.0E-03 |
| Atxn3 | 3712 | 1.22 | 0.29 | 3.1E-04 | 1.26 | 0.33 | 2.9E-05 |
| Atxn7 | 2642 | -1.81 | -0.85 | 1.8E-14 | -1.85 | -0.89 | 1.6E-15 |
| Atxn7l2 | 2048 | -2.73 | -1.45 | 6.1E-23 | -2.93 | -1.55 | 5.1E-26 |
| AU021092 | 225 | -1.57 | -0.65 | 2.8E-04 | -1.54 | -0.62 | 5.7E-04 |
| AU022252 | 8011 | 1.90 | 0.93 | 6.5E-31 | 1.92 | 0.94 | 2.0E-31 |
| AU041133 | 486 | -1.91 | -0.93 | 1.1E-07 | -1.58 | -0.66 | 2.5E-04 |
| Auh | 1414 | -1.25 | -0.32 | 1.5E-07 | -1.28 | -0.36 | 5.7E-09 |
| Aven | 440 | -3.43 | -1.78 | 2.0E-36 | -3.41 | -1.77 | 7.0E-36 |
| Avpi1 | 314 | 2.04 | 1.03 | 4.3E-08 | 1.78 | 0.83 | 1.2E-05 |
| AW209491 | 1182 | -1.26 | -0.33 | 6.3E-06 | -1.23 | -0.30 | 4.6E-05 |
| Azin1 | 23358 | 1.33 | 0.41 | 2.7E-08 | 1.34 | 0.43 | 7.6E-09 |
| Azin2 | 1208 | 2.86 | 1.52 | 6.9E-31 | 3.06 | 1.61 | 9.4E-35 |
| B230219D22Rik | 12291 | -1.27 | -0.35 | 9.5E-07 | -1.28 | -0.35 | 8.5E-07 |
| B3galnt2 | 4658 | 1.44 | 0.52 | 7.5E-10 | 1.38 | 0.46 | 6.5E-08 |
| B3galt2 | 876 | -1.75 | -0.81 | 3.1E-03 | -1.50 | -0.59 | 3.7E-02 |
| B3galt4 | 182 | -2.31 | -1.21 | 7.3E-14 | -2.33 | -1.22 | 4.2E-14 |
| B3galt6 | 813 | -1.37 | -0.46 | 1.6E-09 | -1.42 | -0.51 | 1.3E-11 |
| B3gat3 | 4425 | 1.25 | 0.32 | 8.4E-05 | 1.24 | 0.31 | 1.1E-04 |
| B3glct | 3008 | -1.29 | -0.37 | 5.0E-04 | -1.29 | -0.37 | 4.3E-04 |
| B3gnt4 | 109 | 1.43 | 0.51 | 4.0E-04 | 1.26 | 0.34 | 2.4E-02 |
| B3gnt5 | 6584 | -1.38 | -0.47 | 2.3E-06 | -1.34 | -0.42 | 3.1E-05 |
| B3gnt7 | 252 | 1.47 | 0.56 | 1.3E-02 | 1.47 | 0.56 | 1.3E-02 |
| B3gnt9 | 319 | -2.48 | -1.31 | 7.4E-09 | -2.60 | -1.38 | 1.2E-09 |
| B4galnt4 | 4625 | -1.58 | -0.66 | 1.9E-13 | -1.64 | -0.71 | 1.1E-15 |
| B4galt2 | 4167 | -1.53 | -0.62 | 5.9E-13 | -1.57 | -0.65 | 1.9E-14 |
| B4galt5 | 3724 | 1.57 | 0.65 | 8.0E-10 | 1.44 | 0.53 | 7.8E-07 |
| B4galt6 | 4780 | 1.33 | 0.41 | 8.6E-05 | 1.34 | 0.43 | 4.5E-05 |
| B9d1 | 779 | -1.49 | -0.57 | 9.2E-08 | -1.51 | -0.59 | 2.9E-08 |
| B9d2 | 594 | -1.66 | -0.73 | 4.5E-13 | -1.59 | -0.67 | 2.3E-11 |
| Bace1 | 6271 | 1.83 | 0.88 | 4.3E-22 | 1.90 | 0.92 | 2.0E-24 |
| Bag4 | 4451 | 1.43 | 0.52 | 3.5E-13 | 1.44 | 0.52 | 1.9E-13 |
| Bahd1 | 5131 | 2.45 | 1.29 | 2.3E-18 | 2.18 | 1.12 | 4.6E-14 |
| Baiap2 | 1884 | 2.79 | 1.48 | 6.0E-73 | 2.98 | 1.57 | 2.4E-82 |
| Baiap2l1 | 386 | 3.78 | 1.92 | 2.5E-09 | 3.63 | 1.86 | 8.1E-09 |
| Baiap2l2 | 100 | 2.75 | 1.46 | 2.4E-10 | 2.11 | 1.08 | 3.7E-06 |

| | | | | | | | |
|---|---|---|---|---|---|---|---|
| Baiap3 | 573 | 2.07 | 1.05 | 2.5E-08 | 2.04 | 1.03 | 5.1E-08 |
| Bak1 | 1703 | -1.74 | -0.80 | 8.9E-08 | -1.87 | -0.91 | 1.0E-09 |
| Banf1 | 4611 | -1.32 | -0.40 | 3.1E-03 | -1.50 | -0.58 | 9.9E-06 |
| Barhl1 | 964 | -1.66 | -0.73 | 2.1E-02 | -1.95 | -0.97 | 1.8E-03 |
| Basp1 | 35657 | -1.40 | -0.48 | 1.5E-03 | -1.46 | -0.55 | 3.2E-04 |
| Bax | 1829 | 1.37 | 0.45 | 1.8E-04 | 1.24 | 0.32 | 1.1E-02 |
| Baz2a | 13887 | 1.76 | 0.82 | 1.2E-12 | 1.77 | 0.82 | 8.3E-13 |
| Baz2b | 10640 | -1.60 | -0.68 | 9.1E-08 | -1.41 | -0.50 | 1.2E-04 |
| Bbc3 | 762 | 2.22 | 1.15 | 5.6E-19 | 2.14 | 1.10 | 1.9E-17 |
| Bbip1 | 3597 | -1.29 | -0.37 | 6.9E-06 | -1.25 | -0.32 | 1.4E-04 |
| Bbln | 846 | 1.71 | 0.77 | 2.6E-23 | 1.56 | 0.64 | 1.8E-16 |
| Bbof1 | 478 | 1.38 | 0.46 | 2.1E-02 | 1.61 | 0.68 | 4.8E-04 |
| Bbs1 | 2679 | -1.27 | -0.35 | 1.6E-05 | -1.31 | -0.39 | 9.6E-07 |
| Bbs10 | 554 | -1.25 | -0.32 | 1.5E-02 | -1.39 | -0.48 | 2.6E-04 |
| Bbs2 | 2236 | -1.32 | -0.41 | 3.9E-14 | -1.25 | -0.32 | 2.1E-09 |
| Bbs5 | 984 | 1.21 | 0.27 | 9.4E-05 | 1.28 | 0.35 | 3.7E-07 |
| Bbx | 4976 | 1.48 | 0.56 | 2.4E-09 | 1.74 | 0.80 | 9.1E-18 |
| BC024063 | 540 | -1.50 | -0.59 | 2.7E-06 | -1.21 | -0.27 | 4.1E-02 |
| BC024139 | 1571 | -1.85 | -0.89 | 3.0E-05 | -1.46 | -0.55 | 1.3E-02 |
| BC024978 | 706 | -2.49 | -1.32 | 1.0E-17 | -2.68 | -1.42 | 2.2E-20 |
| BC025920 | 665 | -1.66 | -0.73 | 3.2E-11 | -1.69 | -0.75 | 7.9E-12 |
| BC028528 | 213 | -1.84 | -0.88 | 1.5E-08 | -1.82 | -0.86 | 3.7E-08 |
| BC029722 | 645 | -1.33 | -0.41 | 3.5E-10 | -1.26 | -0.34 | 2.8E-07 |
| BC034090 | 663 | -3.26 | -1.71 | 1.6E-48 | -2.98 | -1.57 | 3.0E-41 |
| BC035947 | 114 | -2.57 | -1.36 | 1.3E-07 | -3.09 | -1.63 | 3.3E-10 |
| Bcap31 | 3970 | 1.34 | 0.42 | 3.5E-07 | 1.28 | 0.36 | 1.4E-05 |
| Bcar1 | 5039 | 1.29 | 0.36 | 2.5E-03 | 1.24 | 0.31 | 1.0E-02 |
| Bcas1 | 279 | -1.78 | -0.83 | 1.5E-03 | -1.52 | -0.60 | 2.4E-02 |
| Bcas2 | 6556 | -1.19 | -0.25 | 3.2E-03 | -1.27 | -0.35 | 2.4E-05 |
| Bcat1 | 5539 | -1.32 | -0.40 | 1.3E-04 | -1.37 | -0.46 | 1.0E-05 |
| Bcat2 | 1499 | -1.61 | -0.69 | 1.5E-10 | -1.77 | -0.83 | 8.1E-15 |
| Bcdin3d | 987 | -1.23 | -0.30 | 4.4E-03 | -1.40 | -0.49 | 1.6E-06 |
| Bckdha | 1962 | 1.26 | 0.34 | 1.7E-03 | 1.22 | 0.28 | 1.1E-02 |
| Bckdhb | 1363 | 1.32 | 0.40 | 8.1E-05 | 1.26 | 0.33 | 1.3E-03 |
| Bckdk | 2521 | -1.23 | -0.29 | 4.6E-03 | -1.34 | -0.43 | 2.9E-05 |
| Bcl10 | 2338 | 1.68 | 0.75 | 2.6E-19 | 1.59 | 0.67 | 1.4E-15 |
| Bcl11a | 10078 | -1.59 | -0.67 | 6.6E-05 | -1.70 | -0.76 | 5.2E-06 |
| Bcl11b | 7112 | -1.58 | -0.66 | 1.7E-02 | -1.67 | -0.74 | 8.1E-03 |
| Bcl2 | 2372 | -1.32 | -0.40 | 3.9E-03 | -1.30 | -0.37 | 7.5E-03 |
| Bcl2l1 | 5311 | -1.76 | -0.81 | 3.0E-17 | -1.91 | -0.93 | 3.2E-22 |
| Bcl2l11 | 2437 | 1.34 | 0.43 | 5.0E-06 | 1.35 | 0.43 | 5.0E-06 |
| Bcl2l12 | 1325 | -1.43 | -0.52 | 1.8E-03 | -1.60 | -0.68 | 3.6E-05 |
| Bcl7a | 8903 | -1.89 | -0.92 | 4.0E-08 | -1.95 | -0.97 | 6.2E-09 |
| Bcl7b | 879 | -2.11 | -1.08 | 4.2E-18 | -2.32 | -1.22 | 8.9E-23 |
| Bcl9 | 11054 | -1.44 | -0.52 | 1.1E-04 | -1.52 | -0.60 | 7.8E-06 |
| Bcl9l | 7001 | 1.33 | 0.41 | 5.5E-03 | 1.23 | 0.30 | 4.6E-02 |
| Bclaf1 | 28096 | -1.53 | -0.61 | 6.9E-13 | -1.44 | -0.53 | 7.4E-10 |

| | | | | | | | |
|---|---|---|---|---|---|---|---|
| Bcorl1 | 4918 | 1.89 | 0.92 | 1.9E-10 | 1.80 | 0.85 | 5.5E-09 |
| Bcr | 9406 | 1.36 | 0.44 | 4.2E-03 | 1.35 | 0.43 | 5.0E-03 |
| Bdh1 | 2911 | -1.55 | -0.63 | 3.9E-06 | -1.60 | -0.68 | 7.1E-07 |
| Bdh2 | 169 | -1.82 | -0.87 | 6.5E-04 | -1.66 | -0.73 | 4.7E-03 |
| Bdkrb2 | 181 | 1.57 | 0.65 | 3.4E-02 | 1.82 | 0.86 | 4.9E-03 |
| Bdnf | 423 | 2.57 | 1.36 | 9.7E-12 | 3.10 | 1.63 | 3.3E-16 |
| Bdp1 | 6087 | 1.23 | 0.30 | 5.0E-02 | 1.43 | 0.51 | 5.8E-04 |
| Bean1 | 143 | -3.65 | -1.87 | 3.4E-08 | -3.24 | -1.70 | 6.3E-07 |
| Bend3 | 1944 | -3.55 | -1.83 | 3.6E-26 | -4.02 | -2.01 | 2.3E-31 |
| Bend4 | 3105 | 1.68 | 0.75 | 2.7E-19 | 1.82 | 0.87 | 1.8E-25 |
| Bend5 | 1357 | -1.84 | -0.88 | 3.8E-18 | -1.83 | -0.87 | 8.0E-18 |
| Bex1 | 3571 | 1.43 | 0.52 | 7.1E-05 | 1.36 | 0.44 | 8.7E-04 |
| Bex4 | 3020 | 1.52 | 0.60 | 5.9E-07 | 1.40 | 0.48 | 7.7E-05 |
| Bgn | 6732 | -1.47 | -0.55 | 1.5E-02 | -1.54 | -0.62 | 6.2E-03 |
| Bhlhe22 | 1614 | -2.33 | -1.22 | 9.0E-03 | -2.04 | -1.03 | 3.0E-02 |
| Bhlhe40 | 487 | -1.44 | -0.53 | 6.3E-05 | -1.45 | -0.53 | 5.8E-05 |
| Bhlhe41 | 341 | 1.95 | 0.96 | 1.2E-07 | 1.93 | 0.95 | 1.6E-07 |
| Bid | 1333 | 2.75 | 1.46 | 5.9E-73 | 2.86 | 1.52 | 6.5E-78 |
| Bin1 | 5487 | -1.41 | -0.49 | 2.8E-05 | -1.34 | -0.42 | 3.7E-04 |
| Bin3 | 1758 | 1.54 | 0.62 | 6.1E-11 | 1.45 | 0.54 | 2.0E-08 |
| Birc3 | 260 | -3.35 | -1.74 | 7.0E-11 | -2.97 | -1.57 | 5.6E-09 |
| Bivm | 2916 | -1.62 | -0.70 | 4.6E-26 | -1.55 | -0.63 | 9.0E-22 |
| Bloc1s6 | 3335 | -1.33 | -0.41 | 9.3E-10 | -1.41 | -0.50 | 1.2E-13 |
| Blvra | 600 | -1.52 | -0.60 | 4.5E-12 | -1.41 | -0.50 | 1.6E-08 |
| Blvrb | 723 | 2.00 | 1.00 | 4.8E-07 | 1.85 | 0.89 | 1.0E-05 |
| Bmerb1 | 3258 | -1.45 | -0.54 | 2.0E-02 | -1.39 | -0.48 | 4.1E-02 |
| Bmf | 5588 | 2.71 | 1.44 | 2.6E-26 | 2.81 | 1.49 | 3.8E-28 |
| Bmi1 | 7346 | -2.08 | -1.05 | 4.9E-29 | -2.12 | -1.08 | 1.7E-30 |
| Bmp1 | 6263 | -1.80 | -0.85 | 1.8E-09 | -1.70 | -0.77 | 5.9E-08 |
| Bmp2 | 291 | -1.49 | -0.58 | 3.6E-02 | -1.68 | -0.74 | 6.3E-03 |
| Bmp2k | 2466 | 1.55 | 0.63 | 2.6E-25 | 1.55 | 0.64 | 1.2E-25 |
| Bmp3 | 418 | -2.22 | -1.15 | 3.4E-05 | -2.17 | -1.12 | 6.5E-05 |
| Bmper | 990 | -2.37 | -1.24 | 6.5E-10 | -2.22 | -1.15 | 1.2E-08 |
| Bmpr1b | 1162 | -1.94 | -0.96 | 3.6E-14 | -1.71 | -0.77 | 1.3E-09 |
| Bmt2 | 4931 | 1.66 | 0.73 | 2.2E-25 | 1.64 | 0.71 | 3.6E-24 |
| Bnc2 | 799 | -3.32 | -1.73 | 1.3E-06 | -2.34 | -1.22 | 8.3E-04 |
| Bnip1 | 1686 | -1.49 | -0.57 | 1.3E-17 | -1.52 | -0.61 | 1.9E-19 |
| Bnip3 | 4078 | 1.79 | 0.84 | 4.4E-15 | 1.98 | 0.98 | 2.4E-20 |
| Boc | 4150 | -1.69 | -0.76 | 1.4E-11 | -1.67 | -0.74 | 4.6E-11 |
| Bok | 962 | -1.98 | -0.99 | 2.2E-13 | -2.06 | -1.05 | 7.1E-15 |
| Bola1 | 708 | -1.35 | -0.43 | 3.3E-03 | -1.41 | -0.50 | 6.7E-04 |
| Bola2 | 2477 | 1.59 | 0.67 | 2.6E-05 | 1.48 | 0.56 | 5.2E-04 |
| Bop1 | 4752 | 1.96 | 0.97 | 1.3E-24 | 1.84 | 0.88 | 2.8E-20 |
| Bora | 3772 | 1.79 | 0.84 | 5.4E-08 | 1.66 | 0.73 | 2.9E-06 |
| Borcs5 | 959 | 1.32 | 0.40 | 1.3E-06 | 1.19 | 0.25 | 2.9E-03 |
| Borcs6 | 1007 | 2.28 | 1.19 | 1.5E-35 | 2.19 | 1.13 | 2.1E-32 |
| Borcs7 | 761 | -4.14 | -2.05 | 1.8E-95 | -4.37 | -2.13 | 1.7E-102 |

| | | | | | | | |
|---|---|---|---|---|---|---|---|
| Bphl | 856 | -1.27 | -0.34 | 1.5E-04 | -1.26 | -0.34 | 2.4E-04 |
| Bpnt1 | 1925 | 1.34 | 0.42 | 8.5E-11 | 1.31 | 0.39 | 2.7E-09 |
| Bpnt2 | 17949 | 1.31 | 0.39 | 2.7E-08 | 1.31 | 0.39 | 2.2E-08 |
| Bptf | 13737 | -1.93 | -0.95 | 4.8E-18 | -1.73 | -0.79 | 6.8E-13 |
| Brap | 4740 | 1.39 | 0.48 | 5.0E-16 | 1.33 | 0.41 | 7.2E-12 |
| Brat1 | 2214 | 1.34 | 0.42 | 1.1E-06 | 1.29 | 0.36 | 3.2E-05 |
| Brca1 | 5043 | 1.43 | 0.52 | 4.0E-03 | 1.48 | 0.56 | 1.7E-03 |
| Brd1 | 7073 | -1.39 | -0.48 | 1.9E-04 | -1.47 | -0.55 | 1.5E-05 |
| Brd2 | 19021 | -1.26 | -0.33 | 1.3E-04 | -1.35 | -0.43 | 6.2E-07 |
| Brd3 | 16321 | -1.22 | -0.29 | 3.1E-02 | -1.32 | -0.40 | 2.8E-03 |
| Brd8 | 12694 | -1.89 | -0.92 | 4.6E-39 | -1.87 | -0.91 | 8.6E-38 |
| Brd9 | 7341 | -1.79 | -0.84 | 4.6E-40 | -1.73 | -0.79 | 3.3E-35 |
| Brdt | 662 | 2.46 | 1.30 | 1.3E-38 | 2.56 | 1.36 | 1.1E-41 |
| Brf1 | 5730 | 1.40 | 0.49 | 1.4E-07 | 1.33 | 0.42 | 1.0E-05 |
| Bri3bp | 6156 | 1.63 | 0.70 | 5.2E-20 | 1.57 | 0.65 | 2.2E-17 |
| Brinp2 | 2925 | 2.82 | 1.49 | 2.7E-12 | 2.61 | 1.38 | 1.2E-10 |
| Brinp3 | 271 | -1.66 | -0.74 | 3.7E-02 | -1.74 | -0.80 | 2.4E-02 |
| Brpf1 | 5325 | 1.41 | 0.50 | 3.4E-07 | 1.31 | 0.39 | 9.2E-05 |
| Brpf3 | 2554 | -4.32 | -2.11 | 2.5E-97 | -4.82 | -2.27 | 7.1E-112 |
| Brsk1 | 6395 | -1.46 | -0.55 | 1.0E-02 | -1.45 | -0.54 | 1.3E-02 |
| Brsk2 | 7140 | -1.74 | -0.80 | 2.6E-04 | -1.86 | -0.89 | 4.1E-05 |
| Brwd1 | 13743 | -1.37 | -0.45 | 5.2E-06 | -1.29 | -0.37 | 2.3E-04 |
| Brwd3 | 6365 | 1.42 | 0.50 | 2.2E-03 | 1.66 | 0.73 | 4.9E-06 |
| Bsdc1 | 3880 | 1.42 | 0.50 | 2.2E-11 | 1.38 | 0.46 | 9.5E-10 |
| Btaf1 | 10163 | 1.41 | 0.49 | 1.9E-02 | 1.62 | 0.70 | 6.9E-04 |
| Btbd11 | 1822 | 1.72 | 0.79 | 3.2E-12 | 1.76 | 0.81 | 5.0E-13 |
| Btbd17 | 734 | -4.75 | -2.25 | 6.4E-24 | -5.87 | -2.55 | 1.6E-30 |
| Btbd19 | 161 | -2.27 | -1.18 | 1.6E-12 | -2.31 | -1.21 | 5.6E-13 |
| Btbd6 | 759 | -2.37 | -1.25 | 9.8E-26 | -2.28 | -1.19 | 1.8E-23 |
| Btd | 733 | -1.99 | -0.99 | 4.2E-20 | -2.02 | -1.02 | 5.7E-21 |
| Btf3l4 | 7993 | -1.56 | -0.64 | 7.6E-10 | -1.58 | -0.66 | 2.1E-10 |
| Btg3 | 1481 | 1.92 | 0.94 | 1.5E-22 | 1.75 | 0.81 | 5.6E-17 |
| Btn2a2 | 187 | 6.94 | 2.80 | 7.0E-27 | 7.96 | 2.99 | 2.5E-30 |
| Bves | 348 | 1.43 | 0.52 | 2.7E-03 | 1.86 | 0.89 | 1.1E-07 |
| Bzw2 | 17444 | 1.54 | 0.62 | 4.4E-20 | 1.56 | 0.64 | 2.0E-21 |
| C130074G19Rik | 948 | -1.97 | -0.98 | 3.0E-18 | -2.07 | -1.05 | 1.0E-20 |
| C1d | 4742 | -1.56 | -0.64 | 3.1E-11 | -1.52 | -0.60 | 4.2E-10 |
| C1galt1 | 3395 | 1.41 | 0.49 | 3.6E-04 | 1.57 | 0.65 | 1.5E-06 |
| C1galt1c1 | 859 | -1.41 | -0.49 | 5.0E-08 | -1.31 | -0.39 | 2.2E-05 |
| C1qa | 319 | -1.50 | -0.58 | 1.6E-03 | -1.52 | -0.60 | 1.2E-03 |
| C1qb | 574 | -1.29 | -0.37 | 2.9E-02 | -1.31 | -0.39 | 2.2E-02 |
| C1qc | 389 | -1.33 | -0.42 | 1.0E-02 | -1.40 | -0.49 | 2.4E-03 |
| C1ql1 | 439 | 1.80 | 0.85 | 8.0E-05 | 1.60 | 0.68 | 2.0E-03 |
| C1qtnf1 | 444 | -1.45 | -0.53 | 2.7E-04 | -1.43 | -0.52 | 4.5E-04 |
| C1qtnf2 | 139 | -1.99 | -0.99 | 1.2E-04 | -2.13 | -1.09 | 2.5E-05 |
| C1qtnf3 | 468 | -6.03 | -2.59 | 4.3E-03 | -5.12 | -2.35 | 1.0E-02 |
| C1qtnf6 | 999 | -2.16 | -1.11 | 2.3E-06 | -2.26 | -1.18 | 5.4E-07 |

| | | | | | | | |
|---|---|---|---|---|---|---|---|
| C2cd2 | 2170 | 1.43 | 0.52 | 5.8E-15 | 1.43 | 0.51 | 1.8E-14 |
| C2cd2l | 2917 | 2.23 | 1.16 | 9.0E-17 | 1.99 | 0.99 | 1.3E-12 |
| C2cd3 | 5598 | -1.95 | -0.97 | 2.3E-29 | -1.78 | -0.83 | 3.8E-22 |
| C2cd4c | 507 | 3.89 | 1.96 | 1.7E-25 | 3.76 | 1.91 | 3.0E-24 |
| C2cd5 | 6100 | -1.54 | -0.62 | 1.9E-20 | -1.50 | -0.59 | 3.3E-18 |
| C330018D20Rik | 1208 | -1.57 | -0.66 | 1.9E-18 | -1.57 | -0.65 | 2.5E-18 |
| C3ar1 | 150 | -3.43 | -1.78 | 1.9E-16 | -4.18 | -2.06 | 1.9E-21 |
| C77080 | 1982 | 1.87 | 0.90 | 2.1E-13 | 1.69 | 0.76 | 9.7E-10 |
| C9orf72 | 4763 | 1.74 | 0.80 | 6.8E-17 | 1.79 | 0.84 | 2.4E-18 |
| Cabin1 | 8714 | 1.32 | 0.40 | 1.3E-05 | 1.33 | 0.42 | 5.1E-06 |
| Cables1 | 400 | -1.40 | -0.49 | 4.5E-05 | -1.35 | -0.43 | 3.6E-04 |
| Cabp1 | 1881 | 2.68 | 1.42 | 1.2E-09 | 3.27 | 1.71 | 1.9E-13 |
| Cabp7 | 196 | 1.59 | 0.67 | 8.2E-03 | 1.59 | 0.67 | 8.2E-03 |
| Cacna1b | 6214 | 1.34 | 0.42 | 4.7E-02 | 1.42 | 0.50 | 1.7E-02 |
| Cacna1c | 3446 | -1.47 | -0.55 | 3.1E-03 | -1.38 | -0.47 | 1.5E-02 |
| Cacna2d1 | 9393 | -1.84 | -0.88 | 2.7E-08 | -1.59 | -0.67 | 3.0E-05 |
| Cacnb2 | 498 | -2.08 | -1.06 | 9.5E-13 | -1.80 | -0.85 | 1.2E-08 |
| Cacng4 | 9218 | -1.41 | -0.50 | 4.0E-04 | -1.39 | -0.48 | 6.8E-04 |
| Cacng5 | 350 | 2.70 | 1.44 | 7.1E-09 | 2.95 | 1.56 | 3.2E-10 |
| Cacul1 | 8840 | 1.25 | 0.33 | 6.2E-06 | 1.20 | 0.26 | 4.4E-04 |
| Cacybp | 11670 | 1.44 | 0.53 | 9.8E-10 | 1.45 | 0.54 | 4.1E-10 |
| Cadm1 | 12690 | 1.25 | 0.32 | 1.3E-02 | 1.28 | 0.36 | 5.3E-03 |
| Cadm3 | 12942 | 1.41 | 0.50 | 3.4E-02 | 1.46 | 0.55 | 2.0E-02 |
| Calcoco1 | 6000 | 1.69 | 0.75 | 5.4E-25 | 1.65 | 0.72 | 6.7E-23 |
| Calcrl | 1048 | -1.37 | -0.46 | 8.9E-03 | -1.40 | -0.49 | 5.1E-03 |
| Calm1 | 52234 | 1.44 | 0.52 | 1.0E-23 | 1.44 | 0.53 | 2.4E-24 |
| Calml4 | 151 | -1.90 | -0.93 | 5.4E-03 | -1.71 | -0.77 | 2.4E-02 |
| Camk1 | 1971 | -1.61 | -0.69 | 5.9E-24 | -1.60 | -0.67 | 5.7E-23 |
| Camk2a | 4142 | 1.94 | 0.95 | 1.6E-07 | 1.84 | 0.88 | 1.4E-06 |
| Camk2n2 | 1252 | 1.58 | 0.66 | 3.0E-03 | 1.57 | 0.65 | 4.0E-03 |
| Camkk1 | 2077 | 2.66 | 1.41 | 7.2E-41 | 2.53 | 1.34 | 6.9E-37 |
| Camkk2 | 5909 | 1.61 | 0.68 | 2.2E-08 | 1.59 | 0.67 | 4.2E-08 |
| Camkv | 1980 | 2.50 | 1.32 | 3.4E-28 | 2.30 | 1.20 | 1.1E-23 |
| Camsap1 | 13218 | -1.26 | -0.33 | 1.6E-02 | -1.27 | -0.35 | 9.7E-03 |
| Cand1 | 17691 | -1.20 | -0.26 | 8.7E-06 | -1.26 | -0.33 | 1.0E-08 |
| Capn1 | 489 | -1.80 | -0.85 | 2.4E-03 | -1.84 | -0.88 | 1.6E-03 |
| Capn2 | 7231 | -1.71 | -0.78 | 3.3E-17 | -1.73 | -0.79 | 1.1E-17 |
| Capn5 | 2088 | 1.56 | 0.64 | 1.6E-07 | 1.64 | 0.72 | 4.5E-09 |
| Capn7 | 8313 | 1.25 | 0.32 | 2.3E-03 | 1.31 | 0.39 | 1.3E-04 |
| Capns1 | 8743 | 1.34 | 0.42 | 3.7E-07 | 1.28 | 0.36 | 1.7E-05 |
| Caprin1 | 55348 | -1.58 | -0.66 | 1.3E-21 | -1.59 | -0.67 | 1.8E-22 |
| Caprin2 | 3184 | 1.25 | 0.32 | 1.4E-03 | 1.31 | 0.39 | 1.0E-04 |
| Car14 | 812 | -1.43 | -0.51 | 1.9E-02 | -1.65 | -0.72 | 6.9E-04 |
| Car15 | 130 | 1.72 | 0.78 | 1.2E-03 | 1.64 | 0.71 | 3.3E-03 |
| Car5b | 228 | 5.57 | 2.48 | 5.3E-32 | 5.97 | 2.58 | 3.6E-34 |
| Car8 | 332 | 2.53 | 1.34 | 2.6E-19 | 2.40 | 1.26 | 3.0E-17 |
| Card10 | 814 | 2.42 | 1.28 | 3.3E-14 | 2.34 | 1.23 | 3.4E-13 |

| | | | | | | | |
|---|---|---|---|---|---|---|---|
| Card19 | 435 | 2.20 | 1.13 | 1.2E-12 | 2.43 | 1.28 | 1.1E-15 |
| Card6 | 193 | -1.36 | -0.44 | 5.6E-03 | -1.27 | -0.34 | 3.5E-02 |
| Carf | 1365 | -1.32 | -0.40 | 8.0E-05 | -1.21 | -0.28 | 8.2E-03 |
| Carhsp1 | 20590 | 1.83 | 0.87 | 1.9E-25 | 1.65 | 0.72 | 9.2E-18 |
| Carm1 | 6783 | -1.40 | -0.48 | 1.9E-06 | -1.52 | -0.61 | 1.4E-09 |
| Carmil1 | 1707 | -1.35 | -0.43 | 4.6E-08 | -1.37 | -0.46 | 9.1E-09 |
| Cars | 3887 | 1.29 | 0.37 | 2.9E-06 | 1.19 | 0.25 | 2.4E-03 |
| Casc1 | 491 | 3.30 | 1.72 | 1.3E-43 | 3.90 | 1.96 | 1.1E-55 |
| Casd1 | 4406 | -2.83 | -1.50 | 1.1E-69 | -2.67 | -1.42 | 2.8E-62 |
| Cask | 10174 | -1.30 | -0.37 | 4.7E-03 | -1.24 | -0.31 | 2.3E-02 |
| Casp3 | 16566 | 1.37 | 0.45 | 1.3E-13 | 1.40 | 0.49 | 1.0E-15 |
| Casp6 | 1948 | 1.32 | 0.40 | 1.1E-03 | 1.21 | 0.28 | 2.8E-02 |
| Casp7 | 707 | -3.14 | -1.65 | 6.0E-76 | -3.16 | -1.66 | 1.8E-76 |
| Casp8 | 570 | -2.31 | -1.21 | 1.3E-09 | -2.43 | -1.28 | 1.0E-10 |
| Casp8ap2 | 8583 | 1.32 | 0.40 | 1.6E-03 | 1.51 | 0.59 | 1.6E-06 |
| Casp9 | 3031 | 1.53 | 0.62 | 4.9E-34 | 1.56 | 0.64 | 1.2E-36 |
| Casq2 | 132 | -14.04 | -3.81 | 5.9E-04 | -7.48 | -2.90 | 1.1E-02 |
| Castor1 | 129 | 1.99 | 0.99 | 1.3E-05 | 2.06 | 1.04 | 5.6E-06 |
| Castor2 | 10331 | 2.01 | 1.00 | 1.3E-11 | 1.90 | 0.92 | 5.4E-10 |
| Casz1 | 1306 | 2.43 | 1.28 | 2.5E-08 | 3.00 | 1.59 | 3.2E-12 |
| Catsper2 | 311 | -2.49 | -1.31 | 4.5E-21 | -2.32 | -1.21 | 4.0E-18 |
| Catsperg1 | 507 | 1.77 | 0.82 | 3.0E-05 | 1.97 | 0.98 | 5.1E-07 |
| Cav1 | 1285 | -2.23 | -1.16 | 1.1E-05 | -2.08 | -1.05 | 7.4E-05 |
| Cav2 | 468 | -1.73 | -0.79 | 3.3E-07 | -1.60 | -0.68 | 1.7E-05 |
| Cavin2 | 477 | -2.96 | -1.56 | 8.5E-07 | -2.44 | -1.29 | 6.2E-05 |
| Cavin3 | 596 | -2.31 | -1.21 | 6.8E-15 | -2.62 | -1.39 | 3.7E-19 |
| Cavin4 | 374 | -1.36 | -0.44 | 4.9E-03 | -1.41 | -0.50 | 1.6E-03 |
| Cbfa2t3 | 2680 | -2.22 | -1.15 | 6.2E-05 | -2.21 | -1.14 | 7.2E-05 |
| Cbfb | 7215 | -1.39 | -0.48 | 3.6E-07 | -1.37 | -0.46 | 1.2E-06 |
| Cbln2 | 2008 | -1.82 | -0.87 | 1.5E-05 | -1.71 | -0.78 | 1.2E-04 |
| Cbr4 | 986 | -1.63 | -0.70 | 4.3E-12 | -1.68 | -0.75 | 1.2E-13 |
| Cbs | 736 | 5.02 | 2.33 | 2.1E-38 | 5.08 | 2.35 | 7.7E-39 |
| Cbwd1 | 1852 | 1.62 | 0.70 | 1.5E-14 | 1.58 | 0.66 | 4.1E-13 |
| Cbx3 | 15310 | -1.55 | -0.63 | 4.3E-07 | -1.63 | -0.70 | 1.4E-08 |
| Cbx4 | 1849 | 1.40 | 0.48 | 1.1E-06 | 1.32 | 0.41 | 5.5E-05 |
| Cbx5 | 44773 | -1.40 | -0.48 | 2.2E-08 | -1.40 | -0.49 | 1.9E-08 |
| Cbx7 | 1830 | 4.93 | 2.30 | 1.3E-101 | 4.72 | 2.24 | 7.4E-96 |
| Cbx8 | 1833 | 1.49 | 0.58 | 5.5E-05 | 1.39 | 0.48 | 1.0E-03 |
| Cc2d1a | 2014 | -1.38 | -0.47 | 3.6E-03 | -1.44 | -0.52 | 1.1E-03 |
| Cc2d1b | 3193 | -1.42 | -0.51 | 5.1E-15 | -1.41 | -0.50 | 2.5E-14 |
| Cc2d2a | 1640 | -2.47 | -1.30 | 3.1E-60 | -2.35 | -1.23 | 8.5E-54 |
| Ccar1 | 17537 | -1.57 | -0.65 | 3.0E-13 | -1.60 | -0.68 | 2.9E-14 |
| Ccar2 | 12647 | -1.19 | -0.25 | 1.5E-02 | -1.30 | -0.38 | 1.3E-04 |
| Ccdc102a | 794 | -2.77 | -1.47 | 4.1E-17 | -3.02 | -1.59 | 6.1E-20 |
| Ccdc103 | 151 | 2.25 | 1.17 | 2.1E-18 | 1.98 | 0.98 | 2.2E-13 |
| Ccdc107 | 973 | 1.42 | 0.51 | 2.7E-25 | 1.46 | 0.55 | 5.4E-29 |
| Ccdc112 | 3027 | 1.72 | 0.78 | 4.7E-39 | 1.64 | 0.72 | 3.3E-33 |

| | | | | | | | |
|---|---|---|---|---|---|---|---|
| Ccdc117 | 3769 | 1.28 | 0.35 | 4.2E-07 | 1.23 | 0.30 | 2.3E-05 |
| Ccdc12 | 1430 | -2.35 | -1.23 | 5.2E-17 | -2.50 | -1.32 | 2.3E-19 |
| Ccdc125 | 149 | -2.06 | -1.04 | 6.0E-10 | -2.25 | -1.17 | 5.6E-12 |
| Ccdc127 | 6622 | 1.26 | 0.34 | 6.5E-12 | 1.32 | 0.40 | 6.4E-16 |
| Ccdc13 | 328 | 1.69 | 0.76 | 3.7E-08 | 1.61 | 0.69 | 6.3E-07 |
| Ccdc130 | 666 | -1.15 | -0.20 | 1.2E-02 | -1.25 | -0.32 | 5.5E-05 |
| Ccdc134 | 2442 | 1.86 | 0.90 | 6.7E-29 | 1.81 | 0.86 | 1.5E-26 |
| Ccdc14 | 1920 | 1.28 | 0.36 | 3.4E-03 | 1.35 | 0.43 | 3.9E-04 |
| Ccdc148 | 411 | 1.88 | 0.91 | 2.7E-08 | 1.86 | 0.89 | 5.4E-08 |
| Ccdc149 | 486 | -3.36 | -1.75 | 4.8E-22 | -3.59 | -1.84 | 3.1E-24 |
| Ccdc158 | 447 | 3.90 | 1.96 | 7.6E-48 | 3.97 | 1.99 | 1.3E-48 |
| Ccdc160 | 275 | 1.76 | 0.82 | 1.3E-10 | 1.81 | 0.86 | 1.5E-11 |
| Ccdc166 | 479 | -1.30 | -0.38 | 2.0E-04 | -1.31 | -0.39 | 1.7E-04 |
| Ccdc167 | 1711 | -1.80 | -0.85 | 1.4E-16 | -1.78 | -0.83 | 5.9E-16 |
| Ccdc171 | 970 | 1.68 | 0.75 | 8.3E-10 | 1.89 | 0.92 | 3.5E-14 |
| Ccdc173 | 710 | 1.34 | 0.42 | 2.8E-04 | 1.34 | 0.42 | 3.0E-04 |
| Ccdc174 | 2762 | -1.16 | -0.22 | 3.9E-03 | -1.25 | -0.32 | 1.5E-05 |
| Ccdc177 | 2538 | -1.77 | -0.82 | 2.7E-05 | -1.86 | -0.89 | 4.6E-06 |
| Ccdc18 | 1388 | 1.36 | 0.45 | 9.4E-03 | 1.62 | 0.70 | 2.9E-05 |
| Ccdc181 | 1227 | -1.27 | -0.35 | 6.5E-10 | -1.34 | -0.42 | 5.9E-14 |
| Ccdc184 | 1968 | 1.81 | 0.85 | 5.1E-05 | 1.77 | 0.82 | 9.9E-05 |
| Ccdc186 | 6306 | 1.34 | 0.43 | 2.7E-04 | 1.54 | 0.62 | 6.7E-08 |
| Ccdc187 | 525 | -2.65 | -1.41 | 3.0E-10 | -2.45 | -1.29 | 8.9E-09 |
| Ccdc189 | 542 | 1.67 | 0.74 | 1.4E-18 | 1.70 | 0.76 | 1.9E-19 |
| Ccdc190 | 141 | -2.71 | -1.44 | 1.3E-10 | -3.21 | -1.68 | 4.7E-14 |
| Ccdc191 | 852 | -2.36 | -1.24 | 4.3E-42 | -2.42 | -1.28 | 1.9E-44 |
| Ccdc25 | 3989 | 1.40 | 0.49 | 1.5E-07 | 1.24 | 0.31 | 1.3E-03 |
| Ccdc28a | 576 | 2.16 | 1.11 | 1.2E-23 | 2.26 | 1.18 | 3.8E-26 |
| Ccdc28b | 2212 | 1.31 | 0.39 | 5.8E-11 | 1.29 | 0.36 | 1.3E-09 |
| Ccdc3 | 720 | -1.78 | -0.83 | 2.7E-04 | -1.42 | -0.51 | 3.3E-02 |
| Ccdc30 | 1291 | 1.61 | 0.69 | 5.4E-13 | 1.66 | 0.73 | 1.5E-14 |
| Ccdc32 | 2118 | 1.96 | 0.97 | 7.1E-77 | 1.99 | 0.99 | 3.2E-79 |
| Ccdc39 | 1972 | 1.21 | 0.28 | 2.7E-02 | 1.41 | 0.49 | 4.6E-05 |
| Ccdc40 | 1640 | -1.32 | -0.40 | 1.1E-04 | -1.31 | -0.39 | 1.5E-04 |
| Ccdc50 | 15638 | -1.34 | -0.43 | 8.4E-07 | -1.22 | -0.28 | 1.6E-03 |
| Ccdc57 | 660 | -1.25 | -0.32 | 3.4E-05 | -1.28 | -0.36 | 4.4E-06 |
| Ccdc58 | 1615 | -1.63 | -0.70 | 1.6E-07 | -1.64 | -0.71 | 1.2E-07 |
| Ccdc59 | 2734 | -1.24 | -0.31 | 4.7E-07 | -1.27 | -0.34 | 2.0E-08 |
| Ccdc61 | 2023 | 2.16 | 1.11 | 1.4E-15 | 2.04 | 1.03 | 2.2E-13 |
| Ccdc62 | 466 | -1.28 | -0.36 | 5.4E-04 | -1.33 | -0.41 | 6.5E-05 |
| Ccdc65 | 140 | 1.47 | 0.56 | 8.7E-04 | 1.72 | 0.78 | 2.3E-06 |
| Ccdc71l | 1909 | -1.35 | -0.43 | 2.1E-04 | -1.34 | -0.42 | 3.2E-04 |
| Ccdc77 | 1897 | -1.25 | -0.33 | 8.9E-03 | -1.25 | -0.32 | 1.2E-02 |
| Ccdc8 | 3091 | -1.52 | -0.61 | 1.2E-05 | -1.66 | -0.73 | 1.0E-07 |
| Ccdc80 | 1279 | -1.83 | -0.87 | 3.5E-02 | -1.93 | -0.95 | 2.1E-02 |
| Ccdc81 | 105 | 1.72 | 0.78 | 3.2E-03 | 2.32 | 1.21 | 3.2E-06 |
| Ccdc82 | 4387 | 1.77 | 0.82 | 3.2E-24 | 1.88 | 0.91 | 1.3E-29 |

| | | | | | | | |
|---|---|---|---|---|---|---|---|
| Ccdc84 | 1702 | 1.76 | 0.82 | 1.2E-36 | 1.76 | 0.81 | 5.5E-36 |
| Ccdc85a | 698 | -2.16 | -1.11 | 1.1E-05 | -2.08 | -1.05 | 3.1E-05 |
| Ccdc85b | 1811 | 1.66 | 0.73 | 2.3E-18 | 1.56 | 0.64 | 2.7E-14 |
| Ccdc85c | 8705 | 1.90 | 0.93 | 4.9E-09 | 1.67 | 0.74 | 3.8E-06 |
| Ccdc89 | 319 | 2.27 | 1.19 | 3.1E-16 | 2.35 | 1.23 | 1.9E-17 |
| Ccdc9 | 3731 | -1.25 | -0.32 | 3.2E-02 | -1.40 | -0.48 | 8.4E-04 |
| Ccdc90b | 2338 | -1.40 | -0.48 | 2.8E-08 | -1.43 | -0.51 | 3.6E-09 |
| Ccdc91 | 1888 | -1.95 | -0.96 | 2.9E-49 | -1.82 | -0.87 | 4.0E-40 |
| Ccdc92 | 1216 | 1.71 | 0.77 | 4.3E-07 | 1.67 | 0.74 | 1.5E-06 |
| Ccdc93 | 5120 | 1.41 | 0.49 | 4.3E-07 | 1.57 | 0.65 | 1.2E-11 |
| Ccdc96 | 916 | 2.36 | 1.24 | 9.6E-26 | 2.25 | 1.17 | 5.4E-23 |
| Ccdc9b | 471 | -3.05 | -1.61 | 6.3E-20 | -3.50 | -1.81 | 7.4E-25 |
| Cchcr1 | 2513 | 1.51 | 0.59 | 2.9E-05 | 1.41 | 0.49 | 6.1E-04 |
| Ccm2 | 3078 | 1.42 | 0.51 | 6.5E-14 | 1.36 | 0.44 | 9.3E-11 |
| Ccm2l | 212 | -2.42 | -1.28 | 2.5E-18 | -2.59 | -1.38 | 3.3E-21 |
| Ccn3 | 158 | -1.97 | -0.98 | 9.4E-03 | -2.01 | -1.01 | 7.7E-03 |
| Ccn4 | 1193 | -2.74 | -1.46 | 2.5E-05 | -2.72 | -1.44 | 3.1E-05 |
| Ccnb1 | 10225 | -1.36 | -0.44 | 3.9E-02 | -1.49 | -0.58 | 6.1E-03 |
| Ccnb2 | 2837 | -1.51 | -0.60 | 3.1E-03 | -1.70 | -0.76 | 1.3E-04 |
| Ccnd1 | 17325 | -1.54 | -0.63 | 7.0E-03 | -1.93 | -0.95 | 2.8E-05 |
| Ccne1 | 1957 | 1.85 | 0.89 | 1.6E-08 | 1.67 | 0.74 | 3.2E-06 |
| Ccne2 | 2220 | -1.32 | -0.40 | 1.7E-02 | -1.46 | -0.55 | 9.0E-04 |
| Ccng1 | 5463 | 1.49 | 0.57 | 2.9E-08 | 1.58 | 0.66 | 1.4E-10 |
| Ccng2 | 9036 | -1.43 | -0.52 | 2.1E-09 | -1.44 | -0.53 | 1.2E-09 |
| Ccnh | 4084 | 1.61 | 0.69 | 1.3E-14 | 1.63 | 0.70 | 5.7E-15 |
| Ccnj | 3077 | -1.80 | -0.85 | 3.1E-08 | -2.09 | -1.06 | 2.8E-12 |
| Ccnjl | 1366 | 1.32 | 0.40 | 5.5E-03 | 1.26 | 0.33 | 2.7E-02 |
| Ccnl1 | 7139 | -1.48 | -0.57 | 3.4E-21 | -1.45 | -0.53 | 9.0E-19 |
| Ccnl2 | 14250 | -1.53 | -0.62 | 1.4E-22 | -1.56 | -0.64 | 2.4E-24 |
| Ccnyl1 | 4081 | 1.64 | 0.71 | 1.9E-20 | 1.52 | 0.61 | 4.0E-15 |
| Ccpg1 | 364 | 1.28 | 0.35 | 1.4E-02 | 1.30 | 0.38 | 8.6E-03 |
| Ccsap | 878 | 2.02 | 1.01 | 4.5E-17 | 1.95 | 0.96 | 1.9E-15 |
| Ccz1 | 4162 | -1.33 | -0.41 | 4.0E-08 | -1.34 | -0.43 | 8.4E-09 |
| Cd1d1 | 1511 | -2.80 | -1.49 | 9.0E-33 | -2.75 | -1.46 | 1.6E-31 |
| Cd200 | 4220 | -2.05 | -1.03 | 1.3E-04 | -2.05 | -1.04 | 1.2E-04 |
| Cd24a | 41102 | -1.71 | -0.77 | 2.9E-09 | -1.62 | -0.70 | 9.9E-08 |
| Cd276 | 3140 | -2.13 | -1.09 | 5.0E-36 | -2.18 | -1.13 | 2.4E-38 |
| Cd2bp2 | 8250 | -1.41 | -0.50 | 1.0E-08 | -1.55 | -0.63 | 3.5E-13 |
| Cd320 | 1616 | 1.42 | 0.50 | 1.8E-06 | 1.46 | 0.54 | 2.4E-07 |
| Cd33 | 184 | -2.17 | -1.12 | 2.4E-16 | -2.06 | -1.05 | 1.2E-14 |
| Cd36 | 285 | -2.06 | -1.04 | 1.3E-03 | -1.67 | -0.74 | 2.8E-02 |
| Cd38 | 232 | -1.84 | -0.88 | 8.7E-05 | -1.90 | -0.93 | 3.5E-05 |
| Cd3eap | 2035 | 1.75 | 0.81 | 1.2E-17 | 1.61 | 0.69 | 5.0E-13 |
| Cd40 | 239 | 1.34 | 0.42 | 2.4E-03 | 1.22 | 0.29 | 4.6E-02 |
| Cd44 | 597 | -1.79 | -0.84 | 9.1E-03 | -1.65 | -0.72 | 2.8E-02 |
| Cd46 | 481 | -1.70 | -0.77 | 9.4E-06 | -1.71 | -0.78 | 8.2E-06 |
| Cd55 | 250 | -2.57 | -1.36 | 4.2E-05 | -2.23 | -1.16 | 5.9E-04 |

| | | | | | | | |
|---|---|---|---|---|---|---|---|
| Cd68 | 471 | -1.25 | -0.33 | 6.6E-04 | -1.48 | -0.57 | 8.0E-10 |
| Cd83 | 264 | -1.94 | -0.96 | 1.4E-05 | -1.82 | -0.86 | 1.1E-04 |
| Cd93 | 3630 | -1.56 | -0.65 | 1.4E-14 | -1.59 | -0.67 | 1.2E-15 |
| Cdan1 | 3766 | 1.34 | 0.42 | 2.6E-09 | 1.31 | 0.39 | 7.8E-08 |
| Cdc14b | 2432 | 1.80 | 0.85 | 1.0E-13 | 1.81 | 0.86 | 6.2E-14 |
| Cdc20 | 4773 | -1.38 | -0.46 | 2.0E-02 | -1.63 | -0.71 | 2.3E-04 |
| Cdc23 | 10337 | -1.28 | -0.36 | 1.2E-09 | -1.34 | -0.42 | 1.2E-12 |
| Cdc26 | 5117 | -1.31 | -0.39 | 3.3E-05 | -1.35 | -0.43 | 3.8E-06 |
| Cdc37 | 12052 | 1.31 | 0.39 | 7.2E-04 | 1.22 | 0.29 | 1.4E-02 |
| Cdc40 | 4695 | 1.41 | 0.49 | 4.2E-10 | 1.45 | 0.54 | 9.1E-12 |
| Cdc42bpb | 14607 | 1.34 | 0.42 | 2.8E-04 | 1.31 | 0.39 | 9.7E-04 |
| Cdc42ep1 | 456 | -1.82 | -0.87 | 8.3E-07 | -1.93 | -0.95 | 6.5E-08 |
| Cdc42ep2 | 579 | 1.33 | 0.41 | 4.2E-02 | 1.42 | 0.51 | 1.2E-02 |
| Cdc42ep3 | 1376 | -1.42 | -0.51 | 4.9E-06 | -1.44 | -0.53 | 2.1E-06 |
| Cdca2 | 3963 | -1.48 | -0.56 | 7.6E-04 | -1.52 | -0.61 | 2.8E-04 |
| Cdca4 | 4171 | -1.25 | -0.32 | 2.6E-02 | -1.41 | -0.49 | 4.2E-04 |
| Cdh11 | 12820 | -2.40 | -1.26 | 2.1E-06 | -2.37 | -1.24 | 3.3E-06 |
| Cdh2 | 18934 | -1.42 | -0.51 | 1.1E-06 | -1.53 | -0.62 | 2.5E-09 |
| Cdh5 | 3922 | -1.44 | -0.53 | 9.4E-05 | -1.57 | -0.65 | 1.2E-06 |
| Cdin1 | 1528 | -1.24 | -0.31 | 4.3E-04 | -1.25 | -0.32 | 3.3E-04 |
| Cdip1 | 7843 | 1.37 | 0.45 | 1.3E-23 | 1.34 | 0.42 | 5.0E-20 |
| Cdk1 | 5461 | -1.27 | -0.34 | 4.9E-02 | -1.31 | -0.39 | 2.4E-02 |
| Cdk12 | 4941 | -1.37 | -0.45 | 3.4E-13 | -1.34 | -0.42 | 9.2E-12 |
| Cdk18 | 156 | 2.59 | 1.37 | 2.5E-04 | 2.52 | 1.33 | 3.9E-04 |
| Cdk2 | 6622 | 1.41 | 0.49 | 2.7E-04 | 1.34 | 0.42 | 2.0E-03 |
| Cdk20 | 833 | 1.51 | 0.59 | 3.0E-06 | 1.36 | 0.45 | 5.6E-04 |
| Cdk2ap1 | 6508 | -2.04 | -1.03 | 7.8E-18 | -2.29 | -1.19 | 1.9E-23 |
| Cdk4 | 28254 | -1.36 | -0.45 | 6.5E-04 | -1.52 | -0.60 | 3.1E-06 |
| Cdk5 | 4008 | 1.32 | 0.40 | 5.4E-11 | 1.28 | 0.36 | 4.3E-09 |
| Cdk5r1 | 11497 | -1.50 | -0.58 | 2.3E-02 | -1.52 | -0.60 | 1.9E-02 |
| Cdk5r2 | 2504 | 1.78 | 0.83 | 3.4E-05 | 1.72 | 0.78 | 1.1E-04 |
| Cdk5rap1 | 577 | -1.31 | -0.39 | 1.7E-02 | -1.56 | -0.64 | 4.9E-05 |
| Cdk5rap2 | 8016 | -1.37 | -0.45 | 6.2E-04 | -1.27 | -0.34 | 1.2E-02 |
| Cdk6 | 3329 | -1.68 | -0.75 | 4.5E-07 | -1.72 | -0.78 | 1.6E-07 |
| Cdkal1 | 1026 | -1.29 | -0.36 | 1.9E-06 | -1.34 | -0.42 | 2.1E-08 |
| Cdkl1 | 110 | -1.52 | -0.60 | 1.2E-02 | -1.46 | -0.55 | 2.4E-02 |
| Cdkl3 | 1310 | 1.30 | 0.38 | 4.0E-05 | 1.36 | 0.45 | 1.5E-06 |
| Cdkl5 | 3056 | 1.71 | 0.77 | 5.9E-08 | 1.95 | 0.96 | 9.5E-12 |
| Cdkn1a | 2143 | 2.19 | 1.13 | 3.0E-16 | 2.27 | 1.18 | 9.8E-18 |
| Cdkn1c | 22953 | -1.44 | -0.53 | 5.6E-03 | -1.43 | -0.51 | 7.3E-03 |
| Cdkn2d | 2103 | 1.74 | 0.80 | 1.5E-25 | 1.81 | 0.85 | 5.3E-29 |
| Cdkn3 | 834 | 1.63 | 0.70 | 7.2E-04 | 1.55 | 0.63 | 2.5E-03 |
| Cdo1 | 3567 | -1.51 | -0.59 | 5.1E-06 | -1.44 | -0.53 | 5.5E-05 |
| Cdon | 13736 | -1.37 | -0.45 | 4.4E-04 | -1.20 | -0.27 | 4.7E-02 |
| Cdpf1 | 498 | 1.42 | 0.51 | 4.7E-06 | 1.38 | 0.46 | 3.8E-05 |
| Cdr2 | 3434 | 1.61 | 0.69 | 2.1E-21 | 1.61 | 0.68 | 3.9E-21 |
| Cds2 | 13091 | 1.52 | 0.61 | 1.7E-07 | 1.58 | 0.66 | 1.1E-08 |

| | | | | | | | |
|---|---|---|---|---|---|---|---|
| Cdyl2 | 247 | -1.56 | -0.64 | 5.1E-04 | -1.35 | -0.44 | 2.3E-02 |
| Cebpa | 309 | 2.22 | 1.15 | 9.5E-05 | 2.31 | 1.21 | 3.9E-05 |
| Celf6 | 1504 | 1.51 | 0.60 | 1.1E-03 | 1.44 | 0.53 | 4.8E-03 |
| Cemip | 1137 | -1.97 | -0.98 | 1.5E-03 | -2.09 | -1.06 | 5.6E-04 |
| Cenpa | 5877 | -1.41 | -0.50 | 3.5E-03 | -1.51 | -0.59 | 4.5E-04 |
| Cenpj | 3591 | 1.54 | 0.62 | 1.2E-06 | 1.52 | 0.60 | 3.0E-06 |
| Cenpp | 917 | 1.59 | 0.67 | 1.3E-03 | 1.59 | 0.67 | 1.3E-03 |
| Cenps | 654 | -2.01 | -1.00 | 5.9E-10 | -2.27 | -1.18 | 2.4E-13 |
| Cenpu | 1602 | 1.76 | 0.82 | 3.9E-05 | 1.65 | 0.72 | 3.3E-04 |
| Cenpx | 1493 | 1.36 | 0.44 | 1.3E-06 | 1.25 | 0.32 | 4.6E-04 |
| Cep126 | 1964 | 1.72 | 0.78 | 4.1E-28 | 1.95 | 0.96 | 3.6E-42 |
| Cep135 | 2358 | -1.51 | -0.60 | 7.3E-09 | -1.38 | -0.46 | 9.7E-06 |
| Cep152 | 2567 | 1.45 | 0.54 | 1.2E-03 | 1.52 | 0.61 | 2.7E-04 |
| Cep162 | 4241 | 1.45 | 0.54 | 5.3E-07 | 1.60 | 0.68 | 1.4E-10 |
| Cep164 | 5199 | 1.84 | 0.88 | 8.2E-20 | 1.80 | 0.85 | 1.2E-18 |
| Cep170 | 22187 | -1.49 | -0.58 | 1.9E-04 | -1.31 | -0.39 | 1.6E-02 |
| Cep192 | 9075 | 1.50 | 0.59 | 3.7E-04 | 1.70 | 0.77 | 2.1E-06 |
| Cep20 | 3063 | -1.50 | -0.59 | 9.8E-16 | -1.54 | -0.62 | 1.6E-17 |
| Cep250 | 8284 | -1.40 | -0.48 | 2.0E-05 | -1.40 | -0.48 | 1.8E-05 |
| Cep41 | 1580 | -1.91 | -0.93 | 2.3E-44 | -1.94 | -0.96 | 1.8E-46 |
| Cep44 | 2453 | 1.28 | 0.35 | 1.7E-07 | 1.36 | 0.44 | 6.8E-11 |
| Cep57l1 | 853 | -1.99 | -1.00 | 9.0E-30 | -1.94 | -0.96 | 1.8E-27 |
| Cep63 | 2728 | -1.97 | -0.98 | 4.3E-57 | -1.95 | -0.97 | 1.5E-55 |
| Cep68 | 4986 | -1.29 | -0.37 | 5.3E-12 | -1.28 | -0.36 | 3.3E-11 |
| Cep72 | 1922 | -1.46 | -0.54 | 3.4E-06 | -1.41 | -0.50 | 2.3E-05 |
| Cep78 | 4427 | 1.38 | 0.47 | 3.5E-07 | 1.33 | 0.41 | 9.9E-06 |
| Cep83 | 5442 | 1.54 | 0.62 | 3.0E-21 | 1.56 | 0.65 | 8.8E-23 |
| Cep85 | 3229 | 1.79 | 0.84 | 3.1E-37 | 1.78 | 0.84 | 1.5E-36 |
| Cep97 | 3828 | -1.35 | -0.43 | 1.5E-04 | -1.26 | -0.34 | 3.7E-03 |
| Cept1 | 2373 | -1.33 | -0.41 | 1.1E-09 | -1.31 | -0.39 | 7.5E-09 |
| Cerk | 2810 | -1.27 | -0.34 | 4.6E-03 | -1.25 | -0.33 | 7.6E-03 |
| Cers2 | 3488 | -2.42 | -1.27 | 1.1E-71 | -2.49 | -1.32 | 1.5E-76 |
| Cers5 | 4273 | -1.42 | -0.51 | 1.6E-12 | -1.37 | -0.46 | 2.7E-10 |
| Cetn3 | 7685 | -1.39 | -0.48 | 2.7E-05 | -1.37 | -0.45 | 8.6E-05 |
| Cfap126 | 172 | -1.70 | -0.77 | 1.5E-03 | -1.59 | -0.67 | 6.8E-03 |
| Cfap20 | 4759 | 1.52 | 0.60 | 5.2E-27 | 1.40 | 0.48 | 9.6E-18 |
| Cfap206 | 296 | 2.51 | 1.33 | 3.1E-10 | 2.84 | 1.51 | 8.6E-13 |
| Cfap298 | 916 | -1.56 | -0.64 | 1.4E-12 | -1.66 | -0.73 | 6.3E-16 |
| Cfap300 | 1539 | 1.66 | 0.73 | 2.5E-23 | 1.70 | 0.77 | 1.3E-25 |
| Cfap36 | 4695 | 1.46 | 0.55 | 8.3E-15 | 1.47 | 0.56 | 2.7E-15 |
| Cfap410 | 1559 | 1.53 | 0.62 | 7.3E-19 | 1.46 | 0.55 | 6.2E-15 |
| Cfap45 | 282 | 2.89 | 1.53 | 2.4E-25 | 2.78 | 1.47 | 2.3E-23 |
| Cfap53 | 101 | 1.79 | 0.84 | 4.2E-05 | 1.78 | 0.83 | 5.4E-05 |
| Cfap54 | 468 | 1.55 | 0.63 | 6.9E-03 | 1.81 | 0.85 | 2.0E-04 |
| Cfap69 | 2479 | 2.92 | 1.55 | 1.3E-45 | 3.42 | 1.77 | 1.9E-59 |
| Cfap70 | 206 | 2.50 | 1.32 | 1.3E-13 | 2.45 | 1.29 | 4.4E-13 |
| Cfap74 | 568 | -1.40 | -0.48 | 1.7E-06 | -1.39 | -0.48 | 2.5E-06 |

| | | | | | | | |
|---|---|---|---|---|---|---|---|
| Cfap77 | 122 | 3.23 | 1.69 | 1.5E-26 | 3.01 | 1.59 | 3.7E-23 |
| Cfap97 | 2423 | -1.46 | -0.54 | 3.7E-16 | -1.37 | -0.45 | 1.5E-11 |
| Cfl2 | 11017 | -1.27 | -0.34 | 1.2E-03 | -1.31 | -0.39 | 2.2E-04 |
| Cflar | 1690 | -1.35 | -0.43 | 1.9E-06 | -1.26 | -0.34 | 2.8E-04 |
| Cfp | 1881 | 1.60 | 0.68 | 3.8E-09 | 1.55 | 0.64 | 4.4E-08 |
| Cgn | 2200 | 3.58 | 1.84 | 8.6E-68 | 3.79 | 1.92 | 1.4E-73 |
| Cgref1 | 306 | -2.15 | -1.10 | 4.0E-06 | -1.99 | -0.99 | 4.0E-05 |
| Cgrrf1 | 981 | 2.19 | 1.13 | 2.0E-27 | 2.01 | 1.00 | 8.4E-22 |
| Chac1 | 122 | 2.27 | 1.18 | 1.6E-07 | 2.17 | 1.12 | 6.5E-07 |
| Chac2 | 562 | 1.40 | 0.48 | 1.2E-05 | 1.39 | 0.48 | 1.7E-05 |
| Chadl | 173 | -3.87 | -1.95 | 1.1E-06 | -3.53 | -1.82 | 6.6E-06 |
| Chaf1b | 3562 | 2.01 | 1.01 | 9.2E-10 | 1.77 | 0.83 | 6.5E-07 |
| Champ1 | 6108 | -1.18 | -0.24 | 1.0E-06 | -1.25 | -0.33 | 2.1E-11 |
| Chchd1 | 2480 | -1.43 | -0.51 | 1.4E-07 | -1.48 | -0.56 | 8.2E-09 |
| Chchd10 | 361 | 3.40 | 1.77 | 2.6E-37 | 3.34 | 1.74 | 6.1E-36 |
| Chchd5 | 727 | -1.29 | -0.37 | 1.1E-04 | -1.31 | -0.39 | 5.1E-05 |
| Chchd7 | 3824 | 3.11 | 1.64 | 4.1E-86 | 3.15 | 1.65 | 6.6E-88 |
| Chd1l | 784 | -3.66 | -1.87 | 4.5E-33 | -4.14 | -2.05 | 2.2E-39 |
| Chd2 | 12282 | 1.21 | 0.28 | 2.5E-02 | 1.35 | 0.43 | 3.5E-04 |
| Chd3 | 39510 | -1.98 | -0.99 | 9.7E-21 | -2.00 | -1.00 | 3.8E-21 |
| Chd7 | 13012 | -1.35 | -0.44 | 1.4E-02 | -1.33 | -0.42 | 2.0E-02 |
| Chfr | 8151 | 1.31 | 0.39 | 1.1E-11 | 1.32 | 0.40 | 7.9E-12 |
| Chga | 3469 | 1.71 | 0.78 | 5.8E-05 | 1.76 | 0.82 | 2.3E-05 |
| Chka | 6804 | 1.63 | 0.71 | 1.0E-27 | 1.73 | 0.79 | 3.2E-34 |
| Chmp2a | 3720 | -1.31 | -0.39 | 1.4E-12 | -1.32 | -0.40 | 1.9E-13 |
| Chmp2b | 4359 | 1.74 | 0.80 | 2.0E-08 | 1.73 | 0.79 | 2.7E-08 |
| Chmp3 | 3350 | -2.78 | -1.48 | 1.4E-199 | -2.68 | -1.42 | 5.3E-185 |
| Chmp6 | 1978 | -1.30 | -0.38 | 4.5E-04 | -1.41 | -0.50 | 3.7E-06 |
| Chn2 | 1592 | -1.27 | -0.34 | 9.9E-03 | -1.35 | -0.43 | 1.1E-03 |
| Chodl | 420 | -2.04 | -1.03 | 3.2E-03 | -1.70 | -0.76 | 3.3E-02 |
| Chp1 | 9369 | 2.04 | 1.03 | 8.9E-62 | 2.00 | 1.00 | 9.8E-59 |
| Chpf | 2567 | 1.82 | 0.87 | 5.5E-13 | 1.71 | 0.77 | 1.6E-10 |
| Chrac1 | 769 | -2.63 | -1.39 | 1.4E-41 | -2.75 | -1.46 | 2.6E-45 |
| Chrm4 | 208 | 11.36 | 3.51 | 2.0E-48 | 8.27 | 3.05 | 5.5E-38 |
| Chrna3 | 653 | -3.04 | -1.60 | 1.9E-19 | -2.56 | -1.35 | 3.8E-14 |
| Chrna5 | 623 | 3.27 | 1.71 | 6.1E-13 | 4.22 | 2.08 | 1.7E-18 |
| Chrna7 | 408 | 1.28 | 0.36 | 9.1E-03 | 1.42 | 0.51 | 1.9E-04 |
| Chrnb2 | 4806 | -1.36 | -0.44 | 2.1E-02 | -1.31 | -0.39 | 4.6E-02 |
| Chrnb4 | 184 | 1.99 | 0.99 | 1.6E-07 | 2.11 | 1.07 | 1.2E-08 |
| Chst11 | 2891 | 1.29 | 0.37 | 4.1E-03 | 1.40 | 0.48 | 1.6E-04 |
| Chst12 | 580 | 1.57 | 0.65 | 4.0E-04 | 1.48 | 0.57 | 2.2E-03 |
| Chst14 | 715 | -1.49 | -0.58 | 2.8E-02 | -1.61 | -0.69 | 8.8E-03 |
| Chst15 | 2424 | -1.96 | -0.97 | 1.4E-22 | -2.03 | -1.02 | 6.0E-25 |
| Chst3 | 1295 | -1.36 | -0.44 | 6.1E-05 | -1.43 | -0.52 | 2.1E-06 |
| Chst7 | 181 | 2.41 | 1.27 | 1.9E-19 | 2.33 | 1.22 | 5.7E-18 |
| Chst8 | 639 | 1.32 | 0.40 | 3.0E-04 | 1.29 | 0.37 | 9.5E-04 |
| Chtf18 | 2217 | 1.61 | 0.69 | 8.3E-04 | 1.36 | 0.44 | 3.9E-02 |

| | | | | | | | |
|---|---|---|---|---|---|---|---|
| Chtop | 18206 | -1.44 | -0.53 | 8.7E-23 | -1.45 | -0.53 | 3.6E-23 |
| Churc1 | 154 | -1.31 | -0.39 | 2.0E-02 | -1.26 | -0.34 | 4.9E-02 |
| Ciao1 | 3298 | 1.32 | 0.40 | 1.4E-14 | 1.32 | 0.41 | 7.6E-15 |
| Ciao2b | 1229 | 1.32 | 0.40 | 7.2E-05 | 1.18 | 0.24 | 2.4E-02 |
| Cib1 | 1177 | 2.12 | 1.09 | 8.8E-21 | 1.88 | 0.91 | 6.9E-15 |
| Cibar1 | 3133 | -1.70 | -0.76 | 3.3E-18 | -1.66 | -0.73 | 1.3E-16 |
| Cilk1 | 5801 | -1.24 | -0.31 | 9.9E-03 | -1.30 | -0.38 | 1.6E-03 |
| Cilp2 | 132 | 1.94 | 0.96 | 4.0E-04 | 2.18 | 1.13 | 3.0E-05 |
| Cinp | 2804 | 1.55 | 0.63 | 2.2E-14 | 1.51 | 0.59 | 8.5E-13 |
| Cipc | 5271 | -1.29 | -0.37 | 3.6E-03 | -1.38 | -0.47 | 1.8E-04 |
| Cisd2 | 5288 | -1.41 | -0.50 | 1.0E-06 | -1.53 | -0.61 | 1.0E-09 |
| Cisd3 | 485 | 2.01 | 1.00 | 5.9E-28 | 1.84 | 0.88 | 1.0E-21 |
| Cish | 654 | 1.36 | 0.44 | 5.7E-04 | 1.28 | 0.36 | 5.4E-03 |
| Cited1 | 266 | 2.30 | 1.20 | 4.0E-22 | 2.14 | 1.10 | 8.1E-19 |
| Ckap2 | 4326 | -1.24 | -0.31 | 3.8E-02 | -1.29 | -0.36 | 1.3E-02 |
| Ckap2l | 4485 | -1.61 | -0.69 | 1.2E-05 | -1.77 | -0.82 | 1.2E-07 |
| Ckap4 | 7252 | -1.41 | -0.50 | 1.7E-04 | -1.52 | -0.60 | 5.0E-06 |
| Clba1 | 343 | 2.73 | 1.45 | 6.4E-61 | 2.53 | 1.34 | 4.6E-53 |
| Clcn1 | 1327 | 1.73 | 0.79 | 6.8E-08 | 1.74 | 0.80 | 4.8E-08 |
| Clcn2 | 4232 | 1.41 | 0.49 | 4.1E-08 | 1.38 | 0.46 | 2.8E-07 |
| Clcn3 | 10982 | 1.40 | 0.49 | 1.8E-07 | 1.46 | 0.54 | 4.1E-09 |
| Clcn4 | 10609 | -1.57 | -0.65 | 1.3E-06 | -1.48 | -0.56 | 3.5E-05 |
| Clcn6 | 8009 | 1.34 | 0.42 | 1.2E-06 | 1.31 | 0.39 | 6.2E-06 |
| Clcn7 | 4232 | 1.25 | 0.32 | 5.5E-07 | 1.25 | 0.32 | 3.9E-07 |
| Cldn15 | 125 | 3.24 | 1.70 | 2.4E-18 | 3.38 | 1.76 | 2.0E-19 |
| Cldn34c1 | 671 | -1.54 | -0.63 | 2.4E-06 | -1.55 | -0.64 | 1.6E-06 |
| Cldn5 | 1246 | -1.61 | -0.69 | 4.5E-07 | -1.76 | -0.82 | 1.6E-09 |
| Cldnd1 | 5844 | 1.33 | 0.41 | 2.2E-19 | 1.32 | 0.40 | 3.2E-18 |
| Clec14a | 759 | -3.29 | -1.72 | 3.6E-27 | -3.43 | -1.78 | 6.6E-29 |
| Clec16a | 5184 | 1.68 | 0.75 | 6.9E-18 | 1.69 | 0.76 | 1.8E-18 |
| Clec1a | 118 | -1.94 | -0.95 | 1.7E-05 | -1.72 | -0.78 | 5.0E-04 |
| Clec2d | 138 | -2.73 | -1.45 | 9.0E-10 | -2.87 | -1.52 | 1.4E-10 |
| Clec2l | 312 | -1.62 | -0.69 | 1.3E-03 | -1.62 | -0.70 | 1.3E-03 |
| Clic6 | 1509 | -1.80 | -0.85 | 3.4E-04 | -1.58 | -0.66 | 6.3E-03 |
| Clip1 | 8820 | -1.76 | -0.81 | 2.0E-17 | -1.70 | -0.76 | 2.4E-15 |
| Clk1 | 14069 | -1.55 | -0.63 | 2.0E-18 | -1.47 | -0.56 | 1.3E-14 |
| Clk4 | 7583 | -1.41 | -0.49 | 1.1E-08 | -1.34 | -0.42 | 1.1E-06 |
| Clmn | 1658 | -1.54 | -0.62 | 5.0E-04 | -1.39 | -0.47 | 9.7E-03 |
| Clmp | 6553 | -2.55 | -1.35 | 9.1E-41 | -2.47 | -1.30 | 6.7E-38 |
| Cln6 | 704 | -1.48 | -0.56 | 2.7E-05 | -1.62 | -0.70 | 1.8E-07 |
| Cln8 | 2506 | 1.77 | 0.82 | 2.3E-31 | 1.68 | 0.75 | 4.8E-26 |
| Clptm1 | 6600 | 1.28 | 0.35 | 3.2E-07 | 1.20 | 0.27 | 1.2E-04 |
| Clspn | 3913 | -1.40 | -0.49 | 8.2E-03 | -1.59 | -0.67 | 2.2E-04 |
| Cltb | 2795 | 1.51 | 0.60 | 1.9E-03 | 1.61 | 0.68 | 3.4E-04 |
| Cluap1 | 1198 | -1.30 | -0.37 | 2.9E-06 | -1.32 | -0.41 | 3.8E-07 |
| Cluh | 5623 | 1.87 | 0.90 | 1.6E-20 | 1.84 | 0.88 | 2.2E-19 |
| Clvs1 | 2618 | -1.44 | -0.53 | 2.3E-03 | -1.36 | -0.44 | 1.1E-02 |

| | | | | | | | |
|---|---|---|---|---|---|---|---|
| Clvs2 | 420 | -2.15 | -1.10 | 8.9E-08 | -2.02 | -1.01 | 1.0E-06 |
| Cmc2 | 1014 | 1.68 | 0.75 | 4.6E-10 | 1.69 | 0.76 | 3.6E-10 |
| Cmc4 | 460 | -1.27 | -0.34 | 2.4E-03 | -1.31 | -0.39 | 4.4E-04 |
| Cmklr1 | 181 | -1.51 | -0.59 | 1.3E-03 | -1.60 | -0.67 | 2.3E-04 |
| Cmpk1 | 6914 | -1.59 | -0.67 | 9.0E-17 | -1.64 | -0.71 | 1.5E-18 |
| Cmpk2 | 785 | 1.26 | 0.33 | 2.5E-02 | 1.26 | 0.33 | 2.4E-02 |
| Cmss1 | 735 | -1.21 | -0.27 | 2.5E-02 | -1.27 | -0.34 | 3.8E-03 |
| Cmtm3 | 1767 | -2.68 | -1.42 | 1.6E-13 | -2.72 | -1.45 | 6.9E-14 |
| Cmtm4 | 9730 | 1.69 | 0.76 | 9.8E-31 | 1.80 | 0.85 | 2.4E-38 |
| Cmtm6 | 2645 | 1.70 | 0.76 | 5.0E-21 | 1.64 | 0.71 | 2.3E-18 |
| Cmtm8 | 340 | 1.77 | 0.83 | 2.6E-07 | 2.06 | 1.04 | 7.7E-11 |
| Cmtr1 | 1355 | 1.43 | 0.51 | 2.0E-09 | 1.38 | 0.46 | 8.8E-08 |
| Cmtr2 | 951 | -1.61 | -0.68 | 1.0E-14 | -1.53 | -0.61 | 7.0E-12 |
| Cnbd2 | 267 | 1.27 | 0.34 | 1.5E-02 | 1.22 | 0.28 | 4.8E-02 |
| Cndp2 | 5492 | 1.77 | 0.82 | 5.6E-22 | 1.67 | 0.74 | 4.3E-18 |
| Cnih4 | 3610 | 1.38 | 0.47 | 4.7E-09 | 1.31 | 0.39 | 1.0E-06 |
| Cnksr3 | 908 | 1.58 | 0.66 | 5.5E-09 | 1.44 | 0.53 | 3.9E-06 |
| Cnmd | 909 | -4.16 | -2.06 | 3.9E-03 | -3.51 | -1.81 | 1.2E-02 |
| Cnnm2 | 1491 | 2.06 | 1.04 | 1.6E-18 | 2.25 | 1.17 | 6.5E-23 |
| Cnnm4 | 1746 | 2.91 | 1.54 | 2.7E-34 | 2.79 | 1.48 | 9.9E-32 |
| Cnot2 | 7847 | -1.52 | -0.61 | 7.8E-21 | -1.53 | -0.61 | 5.1E-21 |
| Cnot6 | 21529 | -1.36 | -0.44 | 2.0E-05 | -1.41 | -0.49 | 1.7E-06 |
| Cnot7 | 13952 | -1.26 | -0.33 | 6.4E-04 | -1.31 | -0.39 | 6.4E-05 |
| Cnot8 | 4283 | -1.24 | -0.31 | 1.1E-07 | -1.28 | -0.35 | 1.2E-09 |
| Cnpy2 | 3064 | -1.54 | -0.63 | 7.0E-12 | -1.53 | -0.62 | 1.3E-11 |
| Cnpy4 | 2653 | -1.45 | -0.54 | 1.1E-07 | -1.60 | -0.68 | 1.9E-11 |
| Cnr1 | 6596 | -2.09 | -1.07 | 6.8E-05 | -1.84 | -0.88 | 1.2E-03 |
| Cnrip1 | 1388 | -1.74 | -0.80 | 3.6E-04 | -1.68 | -0.75 | 8.8E-04 |
| Cntd1 | 112 | -1.36 | -0.44 | 7.8E-03 | -1.42 | -0.51 | 2.7E-03 |
| Cntln | 2662 | -1.76 | -0.82 | 4.9E-14 | -1.69 | -0.75 | 4.3E-12 |
| Cntn2 | 19589 | -1.55 | -0.63 | 9.7E-03 | -1.46 | -0.55 | 2.7E-02 |
| Cntnap2 | 11429 | 1.42 | 0.51 | 7.8E-04 | 1.26 | 0.33 | 3.6E-02 |
| Cntrl | 6499 | -1.34 | -0.42 | 1.6E-03 | -1.24 | -0.31 | 2.4E-02 |
| Coa6 | 515 | 1.22 | 0.29 | 5.0E-04 | 1.34 | 0.42 | 2.0E-07 |
| Coa7 | 1488 | -1.47 | -0.56 | 8.8E-14 | -1.55 | -0.63 | 2.1E-17 |
| Cobl | 961 | -1.46 | -0.55 | 8.8E-03 | -1.44 | -0.52 | 1.3E-02 |
| Coch | 684 | 1.53 | 0.61 | 1.6E-04 | 1.61 | 0.69 | 2.1E-05 |
| Cog1 | 2647 | -1.82 | -0.86 | 2.4E-47 | -1.85 | -0.89 | 4.9E-50 |
| Cog4 | 3889 | -1.22 | -0.29 | 1.5E-03 | -1.31 | -0.39 | 1.5E-05 |
| Cog6 | 4067 | -1.25 | -0.32 | 5.5E-06 | -1.20 | -0.27 | 1.5E-04 |
| Cog7 | 1381 | -1.24 | -0.31 | 2.4E-08 | -1.28 | -0.35 | 7.3E-11 |
| Coil | 2073 | -2.22 | -1.15 | 1.3E-47 | -2.36 | -1.24 | 7.5E-55 |
| Col12a1 | 13954 | -2.79 | -1.48 | 6.4E-03 | -2.53 | -1.34 | 1.5E-02 |
| Col14a1 | 1956 | -2.72 | -1.44 | 2.3E-02 | -2.62 | -1.39 | 3.0E-02 |
| Col15a1 | 1651 | -2.06 | -1.04 | 3.0E-05 | -2.04 | -1.03 | 3.9E-05 |
| Col16a1 | 3628 | -2.17 | -1.12 | 1.3E-02 | -2.44 | -1.29 | 3.8E-03 |
| Col17a1 | 289 | -4.53 | -2.18 | 1.8E-02 | -4.98 | -2.32 | 1.2E-02 |

| | | | | | | | |
|---|---|---|---|---|---|---|---|
| Col20a1 | 387 | 1.87 | 0.90 | 3.3E-04 | 2.41 | 1.27 | 3.2E-07 |
| Col27a1 | 2555 | -2.36 | -1.24 | 4.8E-05 | -2.07 | -1.05 | 6.6E-04 |
| Col2a1 | 22191 | -2.13 | -1.09 | 1.7E-03 | -2.00 | -1.00 | 4.6E-03 |
| Col3a1 | 56698 | -1.71 | -0.77 | 1.7E-02 | -1.80 | -0.85 | 9.1E-03 |
| Col5a2 | 17331 | -1.65 | -0.72 | 1.3E-02 | -1.75 | -0.81 | 5.2E-03 |
| Col6a2 | 6048 | -1.76 | -0.82 | 2.1E-02 | -1.75 | -0.81 | 2.3E-02 |
| Col6a3 | 8071 | -2.05 | -1.04 | 3.6E-03 | -1.85 | -0.89 | 1.4E-02 |
| Col6a6 | 212 | -4.15 | -2.05 | 3.8E-04 | -4.13 | -2.05 | 4.2E-04 |
| Col8a1 | 1772 | -3.32 | -1.73 | 1.7E-03 | -2.95 | -1.56 | 5.1E-03 |
| Col9a1 | 3311 | -3.79 | -1.92 | 6.7E-03 | -2.85 | -1.51 | 3.7E-02 |
| Col9a2 | 2538 | -3.06 | -1.62 | 2.7E-04 | -2.57 | -1.36 | 2.5E-03 |
| Col9a3 | 3814 | -3.25 | -1.70 | 5.2E-04 | -2.85 | -1.51 | 2.3E-03 |
| Commd3 | 3042 | -1.28 | -0.36 | 1.7E-05 | -1.22 | -0.29 | 5.1E-04 |
| Commd4 | 2514 | -1.29 | -0.37 | 3.6E-05 | -1.33 | -0.41 | 3.0E-06 |
| Commd7 | 2340 | -1.36 | -0.45 | 3.8E-13 | -1.42 | -0.51 | 9.3E-17 |
| Copa | 36394 | 1.28 | 0.35 | 2.5E-04 | 1.33 | 0.41 | 2.1E-05 |
| Coprs | 1510 | 1.44 | 0.52 | 1.1E-22 | 1.41 | 0.49 | 2.6E-20 |
| Cops3 | 7303 | -1.50 | -0.58 | 1.9E-10 | -1.51 | -0.60 | 6.2E-11 |
| Cops7a | 2650 | -1.42 | -0.51 | 1.1E-10 | -1.50 | -0.59 | 4.2E-14 |
| Coq10b | 1729 | 1.44 | 0.52 | 9.3E-13 | 1.44 | 0.53 | 6.0E-13 |
| Coq2 | 2309 | -1.51 | -0.59 | 1.4E-31 | -1.52 | -0.61 | 4.9E-33 |
| Coq3 | 770 | -1.20 | -0.26 | 3.2E-03 | -1.26 | -0.34 | 1.3E-04 |
| Coq7 | 1063 | -2.45 | -1.29 | 2.6E-22 | -2.89 | -1.53 | 8.6E-31 |
| Coq8a | 206 | 2.22 | 1.15 | 3.2E-06 | 2.46 | 1.30 | 1.2E-07 |
| Coro1c | 16337 | 1.54 | 0.62 | 1.6E-07 | 1.38 | 0.46 | 1.3E-04 |
| Coro2b | 4077 | -1.92 | -0.94 | 2.7E-08 | -1.96 | -0.97 | 8.4E-09 |
| Coro7 | 4182 | 2.14 | 1.10 | 1.1E-33 | 2.07 | 1.05 | 6.5E-31 |
| Cotl1 | 9672 | 1.60 | 0.67 | 1.7E-10 | 1.48 | 0.57 | 1.1E-07 |
| Cox10 | 978 | -1.58 | -0.66 | 3.2E-08 | -1.66 | -0.73 | 8.6E-10 |
| Cox15 | 1961 | -1.52 | -0.60 | 2.7E-18 | -1.63 | -0.71 | 7.6E-25 |
| Cox5a | 6087 | 1.34 | 0.42 | 4.6E-04 | 1.25 | 0.32 | 1.1E-02 |
| Cox6a1 | 5372 | 1.35 | 0.44 | 6.6E-06 | 1.23 | 0.30 | 2.5E-03 |
| Cox6b2 | 1118 | 2.93 | 1.55 | 3.4E-12 | 2.39 | 1.26 | 2.4E-08 |
| Cpa4 | 183 | -2.84 | -1.51 | 2.0E-11 | -3.27 | -1.71 | 2.1E-14 |
| Cpd | 10366 | 1.55 | 0.63 | 8.9E-07 | 1.69 | 0.76 | 2.9E-09 |
| Cpeb1 | 415 | 2.14 | 1.10 | 3.4E-08 | 2.44 | 1.29 | 8.7E-11 |
| Cpeb3 | 1400 | 1.63 | 0.71 | 4.0E-03 | 1.63 | 0.70 | 4.5E-03 |
| Cpeb4 | 8073 | 1.60 | 0.68 | 9.9E-05 | 1.56 | 0.64 | 2.5E-04 |
| Cplane2 | 256 | 1.54 | 0.62 | 1.2E-07 | 1.52 | 0.61 | 2.2E-07 |
| Cplx2 | 9065 | -1.54 | -0.62 | 3.9E-06 | -1.57 | -0.65 | 1.5E-06 |
| Cpne2 | 688 | -1.75 | -0.81 | 7.2E-12 | -1.99 | -0.99 | 3.5E-17 |
| Cpne3 | 5698 | 1.80 | 0.85 | 2.5E-13 | 1.98 | 0.98 | 1.4E-17 |
| Cpox | 1946 | 1.43 | 0.51 | 5.2E-06 | 1.31 | 0.39 | 6.6E-04 |
| Cpped1 | 1131 | -2.31 | -1.21 | 3.4E-47 | -2.30 | -1.20 | 2.1E-46 |
| Cpsf4 | 3745 | 1.32 | 0.40 | 8.0E-10 | 1.29 | 0.37 | 9.1E-09 |
| Cpsf7 | 11294 | -1.50 | -0.58 | 5.6E-09 | -1.61 | -0.69 | 5.6E-12 |
| Cpt1a | 5622 | 2.78 | 1.47 | 2.8E-81 | 2.79 | 1.48 | 1.2E-81 |

| | | | | | | | |
|---|---|---|---|---|---|---|---|
| Cpt1c | 3037 | -1.41 | -0.50 | 6.1E-05 | -1.46 | -0.55 | 8.5E-06 |
| Cpt2 | 1271 | 1.38 | 0.46 | 4.1E-04 | 1.29 | 0.37 | 5.1E-03 |
| Cptp | 1411 | 1.59 | 0.67 | 1.1E-23 | 1.42 | 0.50 | 7.4E-14 |
| Cpxm1 | 2037 | -1.64 | -0.72 | 8.2E-03 | -1.74 | -0.80 | 3.1E-03 |
| Cr1l | 1897 | -2.47 | -1.30 | 3.7E-66 | -2.48 | -1.31 | 9.2E-67 |
| Crabp2 | 5842 | 1.65 | 0.72 | 1.1E-02 | 1.54 | 0.62 | 3.1E-02 |
| Cracd | 9972 | -2.63 | -1.40 | 5.7E-15 | -2.59 | -1.37 | 1.6E-14 |
| Cracr2a | 123 | -1.45 | -0.53 | 3.5E-04 | -1.25 | -0.32 | 4.0E-02 |
| Cradd | 337 | -2.41 | -1.27 | 7.6E-29 | -2.67 | -1.42 | 1.1E-35 |
| Crcp | 2318 | -1.41 | -0.50 | 3.3E-10 | -1.45 | -0.54 | 8.6E-12 |
| Creb3 | 2520 | 1.42 | 0.51 | 4.5E-15 | 1.43 | 0.52 | 1.2E-15 |
| Crebbp | 5496 | -1.93 | -0.95 | 1.7E-10 | -2.06 | -1.04 | 2.0E-12 |
| Crebl2 | 736 | 1.50 | 0.58 | 4.5E-06 | 1.55 | 0.64 | 5.1E-07 |
| Creg1 | 1453 | 2.29 | 1.20 | 9.4E-16 | 2.33 | 1.22 | 3.0E-16 |
| Creg2 | 635 | 2.15 | 1.10 | 1.1E-05 | 2.50 | 1.32 | 1.2E-07 |
| Creld1 | 1982 | -1.33 | -0.42 | 1.1E-03 | -1.39 | -0.48 | 1.6E-04 |
| Creld2 | 2382 | 1.51 | 0.60 | 3.8E-08 | 1.42 | 0.51 | 3.4E-06 |
| Crem | 689 | -1.46 | -0.55 | 5.5E-11 | -1.33 | -0.41 | 1.2E-06 |
| Crip2 | 6183 | 1.73 | 0.79 | 4.8E-11 | 1.60 | 0.68 | 1.9E-08 |
| Crip3 | 184 | 1.55 | 0.63 | 2.9E-02 | 1.77 | 0.82 | 3.9E-03 |
| Crispld1 | 1486 | -2.10 | -1.07 | 2.3E-04 | -2.05 | -1.04 | 3.6E-04 |
| Crlf2 | 205 | -2.49 | -1.31 | 1.3E-10 | -2.23 | -1.16 | 1.7E-08 |
| Crlf3 | 2219 | 1.35 | 0.44 | 1.5E-06 | 1.28 | 0.36 | 1.1E-04 |
| Crnkl1 | 5053 | 1.36 | 0.44 | 5.4E-06 | 1.37 | 0.46 | 2.5E-06 |
| Crppa | 339 | -4.16 | -2.06 | 6.4E-39 | -4.42 | -2.14 | 6.6E-42 |
| Crtac1 | 730 | 1.35 | 0.44 | 3.8E-02 | 1.40 | 0.49 | 2.1E-02 |
| Crtap | 1364 | -1.77 | -0.82 | 2.0E-03 | -1.84 | -0.88 | 9.6E-04 |
| Crtc1 | 3923 | -1.86 | -0.89 | 2.8E-07 | -1.99 | -1.00 | 8.4E-09 |
| Crtc3 | 1065 | -2.32 | -1.21 | 2.3E-15 | -2.44 | -1.29 | 3.7E-17 |
| Cry1 | 3590 | 2.57 | 1.36 | 1.2E-98 | 2.41 | 1.27 | 5.4E-86 |
| Cry2 | 3881 | 2.01 | 1.00 | 2.8E-27 | 1.94 | 0.95 | 1.1E-24 |
| Cryab | 123 | -3.46 | -1.79 | 1.5E-04 | -3.23 | -1.69 | 3.9E-04 |
| Crybb3 | 192 | 6.07 | 2.60 | 1.2E-24 | 4.63 | 2.21 | 2.5E-18 |
| Cryzl2 | 408 | -1.24 | -0.31 | 2.0E-03 | -1.28 | -0.36 | 3.3E-04 |
| Cs | 22561 | 1.44 | 0.52 | 9.5E-15 | 1.37 | 0.45 | 3.1E-11 |
| Csad | 1597 | -2.54 | -1.34 | 5.5E-62 | -2.69 | -1.43 | 6.2E-70 |
| Csdc2 | 3302 | 1.69 | 0.76 | 6.3E-04 | 1.70 | 0.76 | 6.4E-04 |
| Csde1 | 31901 | -1.98 | -0.99 | 6.1E-40 | -2.01 | -1.00 | 3.0E-41 |
| Csf1 | 731 | -1.61 | -0.68 | 1.2E-07 | -1.80 | -0.85 | 4.1E-11 |
| Csf1r | 1501 | -1.39 | -0.47 | 8.8E-04 | -1.46 | -0.54 | 1.2E-04 |
| Csf2ra | 1229 | -1.87 | -0.90 | 4.1E-08 | -1.79 | -0.84 | 4.2E-07 |
| Csgalnact1 | 179 | -4.40 | -2.14 | 2.5E-08 | -3.40 | -1.76 | 5.4E-06 |
| Csnk1a1 | 29607 | -1.40 | -0.48 | 4.7E-15 | -1.39 | -0.47 | 1.5E-14 |
| Csnk1e | 14294 | -1.44 | -0.52 | 1.2E-06 | -1.54 | -0.62 | 6.5E-09 |
| Csnk1g3 | 6823 | 1.27 | 0.34 | 9.6E-07 | 1.28 | 0.36 | 2.8E-07 |
| Csnk2a1 | 16382 | -1.27 | -0.34 | 5.8E-08 | -1.26 | -0.34 | 8.2E-08 |
| Csnk2a2 | 7023 | 1.29 | 0.37 | 2.4E-06 | 1.19 | 0.25 | 1.6E-03 |

| | | | | | | | |
|---|---|---|---|---|---|---|---|
| Csnk2b | 5717 | 1.39 | 0.48 | 1.0E-11 | 1.32 | 0.40 | 1.4E-08 |
| Cspg4b | 833 | -2.63 | -1.39 | 4.2E-07 | -2.09 | -1.06 | 1.6E-04 |
| Csrnp1 | 316 | 2.98 | 1.57 | 1.1E-18 | 3.18 | 1.67 | 1.2E-20 |
| Csrp1 | 4290 | 1.40 | 0.49 | 1.2E-02 | 1.46 | 0.55 | 4.7E-03 |
| Cstf1 | 2832 | -1.19 | -0.25 | 1.7E-02 | -1.28 | -0.35 | 4.5E-04 |
| Cstf2 | 8569 | -1.27 | -0.34 | 1.5E-04 | -1.22 | -0.29 | 1.7E-03 |
| Cstf3 | 8664 | 1.54 | 0.62 | 1.4E-18 | 1.51 | 0.59 | 7.3E-17 |
| Ctbp1 | 16292 | -1.57 | -0.65 | 9.3E-10 | -1.72 | -0.78 | 1.4E-13 |
| Ctbp2 | 5708 | -1.37 | -0.45 | 4.3E-08 | -1.49 | -0.57 | 2.5E-12 |
| Ctc1 | 6242 | -1.46 | -0.55 | 3.8E-14 | -1.50 | -0.59 | 3.3E-16 |
| Ctcf | 9148 | -2.34 | -1.23 | 1.8E-38 | -2.56 | -1.36 | 6.6E-47 |
| Ctdnep1 | 3625 | -1.31 | -0.39 | 1.6E-04 | -1.45 | -0.54 | 1.1E-07 |
| Ctdp1 | 2854 | 1.45 | 0.54 | 4.2E-06 | 1.29 | 0.37 | 2.1E-03 |
| Ctdsp1 | 3876 | -1.31 | -0.39 | 1.1E-03 | -1.37 | -0.45 | 1.6E-04 |
| Ctdspl | 1742 | -1.35 | -0.43 | 2.2E-05 | -1.40 | -0.48 | 1.9E-06 |
| Ctf1 | 217 | -1.96 | -0.97 | 7.0E-10 | -1.82 | -0.86 | 6.6E-08 |
| Cth | 764 | 2.80 | 1.48 | 3.5E-15 | 2.58 | 1.37 | 4.4E-13 |
| Ctnnal1 | 3594 | 1.38 | 0.47 | 1.6E-05 | 1.36 | 0.45 | 3.8E-05 |
| Ctnnbip1 | 5198 | 1.49 | 0.58 | 7.0E-14 | 1.42 | 0.50 | 1.1E-10 |
| Ctnnbl1 | 5721 | 1.36 | 0.45 | 4.6E-05 | 1.29 | 0.37 | 8.7E-04 |
| Ctns | 2248 | 1.62 | 0.70 | 1.4E-24 | 1.57 | 0.65 | 2.5E-21 |
| Ctps | 6445 | -1.23 | -0.30 | 3.5E-04 | -1.28 | -0.35 | 2.2E-05 |
| Ctr9 | 13165 | 1.54 | 0.62 | 1.2E-27 | 1.54 | 0.62 | 1.4E-27 |
| Ctsf | 2119 | 1.51 | 0.59 | 2.2E-14 | 1.58 | 0.66 | 1.1E-17 |
| Ctsk | 939 | -1.80 | -0.85 | 1.6E-04 | -1.73 | -0.79 | 4.9E-04 |
| Ctsl | 7274 | 1.76 | 0.82 | 2.5E-17 | 1.84 | 0.88 | 5.9E-20 |
| Ctso | 456 | -1.88 | -0.91 | 2.0E-13 | -1.86 | -0.90 | 5.1E-13 |
| Ctu2 | 1465 | 1.42 | 0.51 | 1.7E-06 | 1.29 | 0.37 | 7.8E-04 |
| Ctxn1 | 4659 | -1.61 | -0.68 | 1.4E-07 | -1.67 | -0.74 | 1.1E-08 |
| Cul1 | 12269 | -1.66 | -0.74 | 2.9E-19 | -1.71 | -0.78 | 2.3E-21 |
| Cul4b | 6130 | -1.37 | -0.46 | 4.5E-08 | -1.34 | -0.43 | 4.0E-07 |
| Cul5 | 5670 | -1.59 | -0.67 | 5.2E-13 | -1.59 | -0.67 | 5.0E-13 |
| Cul7 | 10139 | -1.69 | -0.75 | 4.5E-14 | -1.80 | -0.85 | 1.2E-17 |
| Cuta | 2284 | 1.88 | 0.91 | 8.5E-17 | 1.78 | 0.83 | 4.3E-14 |
| Cutc | 309 | -1.96 | -0.97 | 4.9E-10 | -1.88 | -0.91 | 5.8E-09 |
| Cwc22 | 2985 | -1.61 | -0.69 | 3.1E-17 | -1.71 | -0.77 | 3.1E-21 |
| Cwc27 | 1565 | -1.52 | -0.60 | 1.8E-07 | -1.57 | -0.65 | 2.1E-08 |
| Cwf19l2 | 2645 | -1.26 | -0.34 | 1.1E-03 | -1.25 | -0.32 | 1.9E-03 |
| Cx3cr1 | 687 | -2.05 | -1.04 | 1.2E-17 | -2.28 | -1.19 | 1.3E-22 |
| Cxcl12 | 7565 | -1.87 | -0.90 | 4.5E-06 | -1.77 | -0.82 | 3.3E-05 |
| Cxcl14 | 1869 | -3.84 | -1.94 | 2.6E-04 | -3.31 | -1.73 | 1.3E-03 |
| Cxcl16 | 107 | -1.57 | -0.66 | 1.2E-02 | -1.53 | -0.61 | 2.2E-02 |
| Cyb561 | 1603 | 1.77 | 0.82 | 3.0E-09 | 1.88 | 0.91 | 5.1E-11 |
| Cyb561d1 | 1050 | -1.83 | -0.87 | 2.6E-13 | -2.07 | -1.05 | 8.3E-19 |
| Cyb5d2 | 286 | -2.36 | -1.24 | 5.6E-29 | -2.09 | -1.06 | 1.7E-21 |
| Cyb5r1 | 2107 | 2.19 | 1.13 | 1.2E-27 | 2.07 | 1.05 | 6.6E-24 |
| Cyb5r3 | 8245 | 1.32 | 0.40 | 5.2E-04 | 1.19 | 0.26 | 3.2E-02 |

| | | | | | | | |
|---|---|---|---|---|---|---|---|
| Cyb5rl | 715 | 2.41 | 1.27 | 4.5E-21 | 2.17 | 1.12 | 1.3E-16 |
| Cybc1 | 3699 | 1.28 | 0.36 | 5.3E-13 | 1.29 | 0.37 | 1.8E-13 |
| Cyfip1 | 6506 | -1.45 | -0.54 | 1.4E-06 | -1.45 | -0.54 | 1.6E-06 |
| Cyfip2 | 14832 | 1.38 | 0.46 | 3.9E-03 | 1.38 | 0.47 | 3.5E-03 |
| Cyld | 4587 | 1.98 | 0.98 | 2.2E-37 | 2.11 | 1.08 | 1.1E-44 |
| Cyp1b1 | 971 | -1.68 | -0.75 | 1.1E-04 | -1.57 | -0.65 | 8.5E-04 |
| Cyp20a1 | 1058 | -1.96 | -0.97 | 1.0E-23 | -1.92 | -0.94 | 2.7E-22 |
| Cyp2j9 | 105 | -1.97 | -0.97 | 8.6E-07 | -1.64 | -0.71 | 4.8E-04 |
| Cyp2u1 | 339 | 3.79 | 1.92 | 2.5E-30 | 3.25 | 1.70 | 5.2E-24 |
| Cyp39a1 | 2465 | 1.65 | 0.72 | 8.5E-09 | 1.61 | 0.68 | 4.8E-08 |
| Cyp46a1 | 2009 | 3.45 | 1.79 | 4.9E-32 | 3.13 | 1.65 | 2.8E-27 |
| Cyp4f16 | 1154 | -1.62 | -0.70 | 5.5E-23 | -1.51 | -0.60 | 5.1E-17 |
| Cyp4f17 | 1326 | 1.41 | 0.49 | 3.5E-07 | 1.41 | 0.50 | 2.5E-07 |
| Cyp4x1 | 281 | -2.15 | -1.10 | 3.5E-04 | -1.95 | -0.96 | 2.1E-03 |
| Cyp7b1 | 188 | -3.56 | -1.83 | 4.3E-19 | -2.86 | -1.52 | 1.7E-13 |
| Cyria | 5583 | 1.59 | 0.67 | 7.6E-03 | 1.78 | 0.83 | 7.5E-04 |
| Cys1 | 127 | -1.85 | -0.88 | 8.1E-06 | -1.55 | -0.64 | 1.9E-03 |
| Cystm1 | 331 | 1.68 | 0.74 | 2.3E-05 | 1.76 | 0.81 | 3.4E-06 |
| Cyth2 | 6637 | -2.00 | -1.00 | 6.1E-19 | -2.08 | -1.06 | 5.1E-21 |
| Cyth3 | 4357 | 1.54 | 0.62 | 2.3E-16 | 1.55 | 0.63 | 4.8E-17 |
| Cyth4 | 285 | -2.70 | -1.43 | 1.2E-21 | -2.85 | -1.51 | 1.3E-23 |
| Cyyr1 | 443 | -2.12 | -1.08 | 4.0E-15 | -2.11 | -1.08 | 6.5E-15 |
| Czib | 2721 | -2.23 | -1.15 | 9.0E-93 | -2.22 | -1.15 | 6.9E-92 |
| D030056L22Rik | 4096 | 1.41 | 0.50 | 2.4E-07 | 1.39 | 0.47 | 1.1E-06 |
| D11Wsu47e | 1846 | 1.38 | 0.46 | 4.6E-07 | 1.44 | 0.52 | 1.1E-08 |
| D130043K22Rik | 412 | 1.69 | 0.76 | 6.1E-04 | 1.98 | 0.99 | 6.0E-06 |
| D16Ertd472e | 2077 | 1.33 | 0.41 | 4.6E-07 | 1.30 | 0.37 | 4.5E-06 |
| D2hgdh | 868 | -2.34 | -1.23 | 1.2E-86 | -2.15 | -1.11 | 2.8E-70 |
| D3Ertd751e | 1750 | 1.32 | 0.40 | 1.2E-03 | 1.37 | 0.46 | 1.8E-04 |
| D430041D05Rik | 2958 | -1.62 | -0.70 | 9.4E-03 | -1.49 | -0.58 | 3.6E-02 |
| D5Ertd579e | 8205 | -1.31 | -0.39 | 1.3E-03 | -1.19 | -0.26 | 4.2E-02 |
| D630003M21Rik | 489 | -3.06 | -1.61 | 4.7E-14 | -2.85 | -1.51 | 2.0E-12 |
| D630045J12Rik | 6799 | -1.84 | -0.88 | 2.8E-08 | -1.91 | -0.93 | 3.4E-09 |
| D8Ertd738e | 3382 | 1.30 | 0.38 | 4.9E-04 | 1.24 | 0.31 | 5.9E-03 |
| Daam1 | 10455 | 1.41 | 0.50 | 2.0E-06 | 1.47 | 0.55 | 1.1E-07 |
| Daam2 | 2596 | -2.17 | -1.12 | 3.1E-08 | -1.88 | -0.91 | 8.9E-06 |
| Dab2 | 6440 | -1.69 | -0.76 | 1.0E-02 | -1.63 | -0.70 | 1.8E-02 |
| Dach1 | 7194 | -1.35 | -0.43 | 1.8E-02 | -1.45 | -0.54 | 2.5E-03 |
| Dach2 | 949 | 1.29 | 0.37 | 9.3E-03 | 1.34 | 0.42 | 2.4E-03 |
| Dact3 | 3383 | 2.51 | 1.33 | 2.8E-37 | 2.43 | 1.28 | 1.1E-34 |
| Dad1 | 4263 | 1.38 | 0.47 | 4.2E-11 | 1.36 | 0.44 | 6.1E-10 |
| Dag1 | 13119 | 1.54 | 0.62 | 1.1E-10 | 1.46 | 0.55 | 1.7E-08 |
| Dalrd3 | 2680 | 1.39 | 0.48 | 1.7E-08 | 1.36 | 0.44 | 2.1E-07 |
| Dand5 | 279 | -1.26 | -0.33 | 1.3E-02 | -1.31 | -0.39 | 3.1E-03 |
| Dap3 | 6410 | 1.46 | 0.54 | 1.9E-24 | 1.41 | 0.50 | 8.5E-21 |
| Dapk3 | 1594 | -1.34 | -0.42 | 5.2E-05 | -1.45 | -0.54 | 1.4E-07 |
| Dars2 | 2508 | 1.29 | 0.37 | 5.2E-04 | 1.27 | 0.35 | 1.2E-03 |

| | | | | | | | |
|---|---|---|---|---|---|---|---|
| Dbn1 | 29718 | -1.63 | -0.70 | 3.2E-08 | -1.72 | -0.78 | 5.9E-10 |
| Dbndd1 | 1386 | -1.32 | -0.40 | 1.5E-02 | -1.27 | -0.35 | 3.6E-02 |
| Dbndd2 | 793 | 1.56 | 0.64 | 1.2E-03 | 1.56 | 0.64 | 1.3E-03 |
| Dbp | 617 | 2.06 | 1.04 | 6.1E-15 | 2.07 | 1.05 | 4.9E-15 |
| Dbr1 | 1966 | 1.34 | 0.42 | 2.1E-04 | 1.25 | 0.32 | 7.1E-03 |
| Dbt | 2528 | 1.68 | 0.75 | 7.7E-27 | 1.66 | 0.73 | 6.3E-26 |
| Dcaf10 | 3951 | 1.37 | 0.45 | 2.0E-08 | 1.39 | 0.47 | 4.1E-09 |
| Dcaf11 | 6593 | 1.40 | 0.49 | 1.3E-15 | 1.39 | 0.48 | 3.5E-15 |
| Dcaf12 | 6873 | 1.29 | 0.37 | 3.4E-08 | 1.21 | 0.27 | 5.3E-05 |
| Dcaf12l1 | 807 | -1.67 | -0.74 | 4.7E-11 | -1.74 | -0.80 | 1.3E-12 |
| Dcaf15 | 3609 | -1.59 | -0.67 | 4.4E-11 | -1.70 | -0.76 | 3.5E-14 |
| Dcaf17 | 2623 | -1.51 | -0.59 | 3.7E-12 | -1.46 | -0.54 | 1.9E-10 |
| Dcaf6 | 3572 | 1.53 | 0.61 | 1.2E-11 | 1.54 | 0.62 | 7.1E-12 |
| Dcaf8 | 10297 | -1.66 | -0.73 | 1.4E-45 | -1.67 | -0.74 | 5.8E-47 |
| Dcbld1 | 1946 | 1.71 | 0.78 | 2.1E-30 | 1.60 | 0.68 | 1.4E-23 |
| Dcbld2 | 9176 | 1.74 | 0.80 | 2.6E-25 | 1.68 | 0.75 | 3.3E-22 |
| Dchs1 | 14620 | -2.27 | -1.18 | 1.2E-19 | -2.24 | -1.16 | 4.8E-19 |
| Dclre1a | 861 | -2.32 | -1.22 | 2.6E-36 | -2.13 | -1.09 | 2.0E-29 |
| Dclre1c | 2028 | 1.82 | 0.87 | 1.3E-18 | 1.97 | 0.98 | 2.5E-23 |
| Dcn | 3091 | -2.03 | -1.02 | 4.2E-04 | -1.80 | -0.85 | 4.0E-03 |
| Dcp1a | 3802 | -1.27 | -0.35 | 3.3E-08 | -1.28 | -0.36 | 8.4E-09 |
| Dcp1b | 2376 | -1.31 | -0.39 | 1.6E-03 | -1.36 | -0.45 | 2.6E-04 |
| Dcps | 2121 | -1.27 | -0.34 | 3.8E-03 | -1.34 | -0.42 | 3.2E-04 |
| Dct | 331 | -2.12 | -1.09 | 2.9E-04 | -1.62 | -0.69 | 2.6E-02 |
| Dctn3 | 3745 | 1.47 | 0.55 | 1.4E-11 | 1.37 | 0.46 | 3.5E-08 |
| Dctn4 | 8294 | -1.38 | -0.47 | 3.3E-11 | -1.38 | -0.46 | 7.9E-11 |
| Dctpp1 | 3129 | 1.58 | 0.66 | 8.0E-06 | 1.43 | 0.52 | 5.9E-04 |
| Dcun1d2 | 3015 | 1.30 | 0.37 | 1.2E-09 | 1.34 | 0.42 | 7.9E-12 |
| Dcun1d4 | 7950 | 1.63 | 0.70 | 5.2E-24 | 1.59 | 0.67 | 1.1E-21 |
| Dcxr | 432 | 4.29 | 2.10 | 7.8E-33 | 3.82 | 1.93 | 5.0E-28 |
| Ddb2 | 1301 | 1.33 | 0.41 | 4.3E-13 | 1.34 | 0.42 | 1.2E-13 |
| Ddc | 823 | -1.30 | -0.38 | 1.8E-02 | -1.35 | -0.43 | 6.1E-03 |
| Ddhd1 | 5364 | 1.50 | 0.59 | 2.7E-06 | 1.57 | 0.65 | 1.4E-07 |
| Ddi2 | 7416 | 1.26 | 0.34 | 6.9E-05 | 1.23 | 0.30 | 5.0E-04 |
| Ddit3 | 543 | 1.56 | 0.64 | 2.9E-09 | 1.46 | 0.54 | 7.1E-07 |
| Ddn | 163 | 2.13 | 1.09 | 1.0E-03 | 2.21 | 1.14 | 5.9E-04 |
| Ddr1 | 11079 | -1.86 | -0.90 | 5.0E-10 | -2.09 | -1.06 | 1.2E-13 |
| Ddr2 | 5445 | -2.47 | -1.31 | 2.6E-03 | -2.11 | -1.08 | 1.5E-02 |
| Ddrgk1 | 1924 | -1.73 | -0.79 | 2.3E-15 | -1.86 | -0.89 | 3.8E-19 |
| Ddx1 | 13757 | -1.24 | -0.31 | 2.2E-04 | -1.31 | -0.39 | 2.8E-06 |
| Ddx10 | 4240 | 1.69 | 0.76 | 6.1E-22 | 1.65 | 0.73 | 2.8E-20 |
| Ddx11 | 3633 | 1.54 | 0.62 | 1.4E-05 | 1.41 | 0.50 | 6.2E-04 |
| Ddx18 | 3951 | -1.18 | -0.24 | 4.2E-02 | -1.28 | -0.35 | 2.3E-03 |
| Ddx31 | 1299 | 1.29 | 0.37 | 5.9E-07 | 1.25 | 0.32 | 1.7E-05 |
| Ddx39b | 23977 | -1.46 | -0.55 | 3.8E-07 | -1.63 | -0.71 | 3.7E-11 |
| Ddx3x | 52188 | -1.58 | -0.66 | 5.5E-09 | -1.63 | -0.71 | 4.1E-10 |
| Ddx4 | 255 | 1.62 | 0.69 | 8.2E-11 | 1.92 | 0.94 | 3.5E-18 |

| | | | | | | | |
|---|---|---|---|---|---|---|---|
| Ddx43 | 183 | 4.49 | 2.17 | 3.7E-40 | 4.36 | 2.12 | 3.2E-38 |
| Ddx46 | 18570 | 1.33 | 0.41 | 9.8E-09 | 1.40 | 0.49 | 5.5E-12 |
| Ddx47 | 5207 | -1.26 | -0.33 | 1.3E-08 | -1.30 | -0.38 | 5.9E-11 |
| Ddx5 | 96299 | -1.55 | -0.63 | 3.2E-16 | -1.54 | -0.62 | 6.8E-16 |
| Ddx52 | 4427 | -1.39 | -0.47 | 6.5E-05 | -1.38 | -0.47 | 7.7E-05 |
| Ddx59 | 785 | -1.38 | -0.46 | 4.6E-07 | -1.42 | -0.50 | 4.5E-08 |
| Ddx6 | 31286 | -1.65 | -0.72 | 1.5E-13 | -1.65 | -0.72 | 1.2E-13 |
| Deaf1 | 3240 | -1.47 | -0.55 | 4.7E-07 | -1.57 | -0.65 | 2.5E-09 |
| Decr1 | 1576 | -1.29 | -0.37 | 5.7E-03 | -1.35 | -0.43 | 1.0E-03 |
| Decr2 | 1465 | -1.41 | -0.49 | 4.0E-18 | -1.48 | -0.56 | 4.3E-23 |
| Dedd2 | 2426 | 2.40 | 1.26 | 2.0E-34 | 2.27 | 1.18 | 2.8E-30 |
| Def6 | 165 | -2.25 | -1.17 | 3.0E-13 | -2.37 | -1.25 | 6.7E-15 |
| Def8 | 3147 | -1.66 | -0.73 | 3.4E-30 | -1.77 | -0.82 | 3.4E-38 |
| Dele1 | 3442 | -2.00 | -1.00 | 3.5E-38 | -2.12 | -1.08 | 1.8E-44 |
| Dennd1b | 2370 | -1.61 | -0.69 | 5.6E-09 | -1.54 | -0.62 | 1.7E-07 |
| Dennd2c | 724 | 1.57 | 0.65 | 2.2E-03 | 1.69 | 0.75 | 3.3E-04 |
| Dennd3 | 722 | 3.25 | 1.70 | 6.8E-49 | 3.32 | 1.73 | 4.3E-50 |
| Dennd4a | 3995 | 1.70 | 0.76 | 7.5E-10 | 1.95 | 0.96 | 4.8E-15 |
| Dennd4b | 1868 | -1.35 | -0.43 | 8.8E-04 | -1.42 | -0.51 | 8.1E-05 |
| Dennd4c | 3654 | 1.31 | 0.39 | 4.8E-03 | 1.54 | 0.62 | 3.5E-06 |
| Depdc1a | 2152 | -1.61 | -0.68 | 3.2E-05 | -1.58 | -0.66 | 5.9E-05 |
| Depdc5 | 6414 | 1.60 | 0.68 | 1.5E-25 | 1.66 | 0.73 | 5.1E-29 |
| Depdc7 | 341 | 3.69 | 1.88 | 5.7E-45 | 2.96 | 1.57 | 9.6E-32 |
| Deptor | 656 | 2.82 | 1.49 | 3.6E-08 | 3.42 | 1.77 | 4.7E-11 |
| Desi1 | 5067 | 1.87 | 0.90 | 2.9E-22 | 1.72 | 0.78 | 5.3E-17 |
| Desi2 | 156 | 2.14 | 1.10 | 1.5E-03 | 1.65 | 0.72 | 4.6E-02 |
| Det1 | 761 | -1.56 | -0.64 | 5.1E-13 | -1.63 | -0.70 | 1.6E-15 |
| Deup1 | 152 | -2.16 | -1.11 | 5.9E-05 | -1.90 | -0.93 | 9.2E-04 |
| Dexi | 1203 | 1.73 | 0.79 | 2.2E-28 | 1.73 | 0.79 | 7.7E-28 |
| Dffa | 2410 | 1.46 | 0.54 | 1.7E-07 | 1.39 | 0.47 | 7.1E-06 |
| Dffb | 721 | -1.22 | -0.28 | 4.0E-04 | -1.34 | -0.42 | 9.8E-08 |
| Dgat2 | 654 | 1.83 | 0.88 | 3.1E-15 | 1.75 | 0.81 | 3.9E-13 |
| Dgke | 4603 | 1.70 | 0.77 | 2.4E-11 | 1.93 | 0.95 | 9.4E-17 |
| Dgkh | 2714 | 1.35 | 0.43 | 2.3E-02 | 1.65 | 0.72 | 8.2E-05 |
| Dgkq | 4510 | 1.47 | 0.56 | 3.0E-10 | 1.43 | 0.52 | 6.1E-09 |
| Dgkz | 5474 | 1.29 | 0.36 | 1.2E-04 | 1.21 | 0.27 | 5.1E-03 |
| Dglucy | 138 | -2.10 | -1.07 | 7.0E-08 | -1.77 | -0.82 | 5.3E-05 |
| Dguok | 1403 | -1.42 | -0.50 | 5.4E-21 | -1.43 | -0.52 | 6.9E-22 |
| Dhcr24 | 11186 | 1.71 | 0.77 | 6.2E-07 | 1.53 | 0.62 | 9.2E-05 |
| Dhcr7 | 4286 | 1.56 | 0.64 | 6.0E-08 | 1.38 | 0.46 | 1.2E-04 |
| Dhdds | 4891 | 1.45 | 0.53 | 4.8E-12 | 1.35 | 0.43 | 3.1E-08 |
| Dhdh | 392 | -2.06 | -1.04 | 5.1E-10 | -2.38 | -1.25 | 6.0E-14 |
| Dhfr | 4614 | 1.56 | 0.64 | 3.7E-04 | 1.38 | 0.47 | 1.1E-02 |
| Dhrs1 | 3342 | 1.39 | 0.47 | 5.1E-10 | 1.37 | 0.45 | 3.9E-09 |
| Dhrs11 | 413 | 1.60 | 0.67 | 2.0E-04 | 1.57 | 0.65 | 3.7E-04 |
| Dhrs13 | 824 | -1.49 | -0.57 | 3.0E-06 | -1.56 | -0.64 | 1.3E-07 |
| Dhrs3 | 373 | -2.92 | -1.54 | 1.4E-07 | -2.74 | -1.46 | 8.1E-07 |

| | | | | | | | |
|---|---|---|---|---|---|---|---|
| Dhrs4 | 1450 | -1.80 | -0.85 | 4.4E-15 | -1.86 | -0.90 | 9.1E-17 |
| Dhrs7 | 292 | 1.40 | 0.49 | 1.1E-05 | 1.34 | 0.42 | 2.1E-04 |
| Dhtkd1 | 2346 | 2.25 | 1.17 | 1.1E-34 | 2.45 | 1.29 | 3.8E-42 |
| Dhx16 | 4807 | 1.45 | 0.54 | 1.6E-07 | 1.31 | 0.39 | 1.7E-04 |
| Dhx29 | 6663 | 1.59 | 0.67 | 1.0E-18 | 1.60 | 0.68 | 4.3E-19 |
| Dhx30 | 9045 | -1.55 | -0.63 | 3.6E-20 | -1.65 | -0.72 | 7.1E-26 |
| Dhx32 | 3663 | 1.46 | 0.54 | 2.5E-06 | 1.27 | 0.35 | 3.8E-03 |
| Dhx34 | 1346 | -1.48 | -0.57 | 3.7E-05 | -1.58 | -0.66 | 1.4E-06 |
| Dhx35 | 1732 | -1.39 | -0.48 | 2.0E-08 | -1.47 | -0.55 | 6.8E-11 |
| Dhx38 | 4641 | -1.31 | -0.39 | 4.5E-04 | -1.47 | -0.56 | 2.6E-07 |
| Dhx40 | 7658 | 1.49 | 0.58 | 7.0E-09 | 1.36 | 0.44 | 1.6E-05 |
| Dhx58 | 230 | 1.68 | 0.75 | 1.2E-06 | 1.69 | 0.76 | 1.1E-06 |
| Diaph1 | 5191 | 1.37 | 0.46 | 6.0E-07 | 1.29 | 0.36 | 9.0E-05 |
| Dido1 | 10630 | -1.25 | -0.32 | 1.7E-04 | -1.18 | -0.24 | 6.3E-03 |
| Dip2b | 6915 | -1.26 | -0.34 | 3.0E-03 | -1.28 | -0.36 | 1.7E-03 |
| Dip2c | 3203 | -2.00 | -1.00 | 2.1E-11 | -1.94 | -0.95 | 1.9E-10 |
| Dipk1a | 1834 | 1.25 | 0.32 | 1.1E-04 | 1.24 | 0.31 | 2.3E-04 |
| Dipk1b | 877 | -1.64 | -0.71 | 6.3E-10 | -1.72 | -0.79 | 7.5E-12 |
| Dipk1c | 399 | -2.10 | -1.07 | 6.2E-10 | -2.52 | -1.33 | 1.2E-14 |
| Diras1 | 1287 | -1.61 | -0.69 | 1.0E-02 | -1.50 | -0.58 | 3.2E-02 |
| Dis3l | 1606 | -4.79 | -2.26 | 9.0E-221 | -5.27 | -2.40 | 1.1E-246 |
| Dis3l2 | 2406 | -1.30 | -0.38 | 6.4E-12 | -1.31 | -0.39 | 1.3E-12 |
| Disc1 | 150 | -2.17 | -1.12 | 8.3E-13 | -1.71 | -0.78 | 1.1E-06 |
| Disp3 | 1853 | 1.66 | 0.73 | 4.6E-03 | 1.86 | 0.89 | 5.0E-04 |
| Dixdc1 | 6724 | 1.41 | 0.50 | 1.4E-06 | 1.50 | 0.59 | 1.0E-08 |
| Dlat | 4593 | 1.26 | 0.33 | 1.9E-07 | 1.20 | 0.26 | 8.0E-05 |
| Dlec1 | 384 | 2.92 | 1.55 | 7.0E-15 | 3.07 | 1.62 | 4.1E-16 |
| Dleu7 | 201 | -2.27 | -1.19 | 1.0E-04 | -2.42 | -1.27 | 2.8E-05 |
| Dlg5 | 6522 | -2.95 | -1.56 | 3.9E-61 | -2.96 | -1.57 | 1.3E-61 |
| Dll3 | 1323 | -1.96 | -0.97 | 2.5E-04 | -2.48 | -1.31 | 4.6E-07 |
| Dll4 | 467 | -1.23 | -0.29 | 9.3E-03 | -1.30 | -0.38 | 7.5E-04 |
| Dmac2 | 961 | -1.59 | -0.67 | 4.6E-32 | -1.66 | -0.74 | 2.7E-38 |
| Dmac2l | 751 | 1.27 | 0.34 | 3.6E-06 | 1.35 | 0.44 | 3.4E-09 |
| Dmc1 | 448 | 1.52 | 0.61 | 2.9E-06 | 1.53 | 0.61 | 2.6E-06 |
| Dmpk | 2879 | 1.32 | 0.40 | 4.3E-03 | 1.26 | 0.33 | 2.2E-02 |
| Dmrt3 | 463 | -1.61 | -0.69 | 3.2E-02 | -1.95 | -0.96 | 2.2E-03 |
| Dmrtb1 | 808 | -1.66 | -0.73 | 5.7E-04 | -1.86 | -0.89 | 2.0E-05 |
| Dna2 | 2001 | -1.73 | -0.79 | 1.3E-09 | -1.81 | -0.86 | 4.5E-11 |
| Dnaaf2 | 349 | 1.88 | 0.91 | 1.5E-07 | 1.71 | 0.77 | 1.1E-05 |
| Dnaaf3 | 789 | 5.09 | 2.35 | 1.9E-32 | 4.34 | 2.12 | 1.6E-26 |
| Dnaaf9 | 13174 | -1.27 | -0.35 | 2.2E-03 | -1.22 | -0.29 | 1.4E-02 |
| Dnah1 | 1064 | -1.18 | -0.24 | 4.2E-02 | -1.35 | -0.43 | 2.0E-04 |
| Dnah10 | 231 | 2.35 | 1.23 | 4.2E-06 | 2.49 | 1.32 | 8.6E-07 |
| Dnah17 | 157 | 5.09 | 2.35 | 4.6E-23 | 5.86 | 2.55 | 7.3E-27 |
| Dnah2 | 868 | 1.43 | 0.51 | 2.1E-02 | 1.71 | 0.77 | 3.7E-04 |
| Dnah7a | 128 | -1.32 | -0.40 | 8.3E-03 | -1.35 | -0.44 | 4.9E-03 |
| Dnai1 | 162 | 2.08 | 1.05 | 3.7E-12 | 2.28 | 1.19 | 5.6E-15 |

| | | | | | | | |
|---|---|---|---|---|---|---|---|
| Dnai4 | 1831 | 1.64 | 0.72 | 2.0E-11 | 1.76 | 0.81 | 2.2E-14 |
| Dnaja1 | 33803 | -1.39 | -0.47 | 9.3E-09 | -1.37 | -0.46 | 2.9E-08 |
| Dnajb11 | 5675 | 1.34 | 0.43 | 6.2E-10 | 1.23 | 0.30 | 1.6E-05 |
| Dnajb12 | 1385 | -2.19 | -1.13 | 3.3E-27 | -2.49 | -1.31 | 3.7E-36 |
| Dnajb14 | 3074 | -1.60 | -0.68 | 1.4E-09 | -1.46 | -0.55 | 1.6E-06 |
| Dnajb3 | 233 | -1.25 | -0.32 | 3.5E-02 | -1.37 | -0.46 | 2.7E-03 |
| Dnajb4 | 3434 | 1.49 | 0.58 | 7.3E-22 | 1.51 | 0.59 | 6.6E-23 |
| Dnajc1 | 4220 | 1.42 | 0.50 | 1.9E-06 | 1.46 | 0.55 | 1.9E-07 |
| Dnajc12 | 1458 | 4.40 | 2.14 | 5.0E-179 | 4.66 | 2.22 | 1.6E-192 |
| Dnajc15 | 2498 | 1.45 | 0.54 | 3.7E-09 | 1.48 | 0.57 | 3.9E-10 |
| Dnajc16 | 3084 | 1.34 | 0.43 | 1.1E-07 | 1.36 | 0.45 | 2.3E-08 |
| Dnajc17 | 720 | -1.80 | -0.85 | 5.3E-11 | -1.91 | -0.94 | 3.7E-13 |
| Dnajc19 | 1159 | -1.35 | -0.44 | 1.9E-05 | -1.46 | -0.55 | 7.3E-08 |
| Dnajc21 | 2760 | 1.29 | 0.36 | 1.1E-04 | 1.24 | 0.31 | 1.3E-03 |
| Dnajc27 | 1872 | -1.48 | -0.57 | 1.1E-20 | -1.51 | -0.59 | 5.3E-22 |
| Dnajc28 | 827 | 1.56 | 0.64 | 6.2E-14 | 1.64 | 0.71 | 3.5E-17 |
| Dnajc3 | 5172 | 1.39 | 0.48 | 1.1E-05 | 1.49 | 0.58 | 9.0E-08 |
| Dnajc7 | 16769 | -1.41 | -0.49 | 1.5E-16 | -1.44 | -0.52 | 1.4E-18 |
| Dnal1 | 4156 | 1.22 | 0.29 | 1.0E-03 | 1.33 | 0.41 | 3.3E-06 |
| Dnase2a | 1349 | 2.87 | 1.52 | 2.5E-30 | 2.61 | 1.39 | 2.5E-25 |
| Dnhd1 | 1087 | 2.57 | 1.36 | 1.4E-14 | 3.22 | 1.69 | 1.1E-21 |
| Dnlz | 2018 | -1.70 | -0.76 | 1.6E-19 | -1.84 | -0.88 | 1.0E-25 |
| Dnm2 | 8421 | 1.83 | 0.88 | 3.1E-27 | 1.74 | 0.80 | 1.2E-22 |
| Dnmt3a | 15809 | -2.22 | -1.15 | 4.4E-38 | -2.11 | -1.07 | 3.4E-33 |
| Dnmt3b | 1342 | 2.05 | 1.04 | 3.0E-09 | 1.94 | 0.96 | 5.0E-08 |
| Dnpep | 5429 | 1.56 | 0.64 | 5.8E-19 | 1.43 | 0.51 | 2.0E-12 |
| Doc2a | 401 | 1.74 | 0.80 | 3.8E-08 | 1.56 | 0.64 | 1.5E-05 |
| Dock1 | 4066 | -1.49 | -0.58 | 7.5E-04 | -1.38 | -0.47 | 7.5E-03 |
| Dock11 | 5389 | -1.52 | -0.60 | 6.0E-08 | -1.40 | -0.49 | 1.4E-05 |
| Dock5 | 1163 | -1.66 | -0.73 | 3.3E-04 | -1.59 | -0.67 | 1.1E-03 |
| Dock6 | 4532 | 1.37 | 0.46 | 8.8E-03 | 1.29 | 0.37 | 3.8E-02 |
| Dock8 | 232 | -2.26 | -1.18 | 2.3E-04 | -2.08 | -1.06 | 1.0E-03 |
| Dok1 | 427 | 1.81 | 0.86 | 2.5E-07 | 1.62 | 0.70 | 3.4E-05 |
| Dok2 | 207 | -2.13 | -1.09 | 1.0E-14 | -2.02 | -1.02 | 6.3E-13 |
| Dok3 | 943 | 1.93 | 0.95 | 3.9E-10 | 1.81 | 0.85 | 2.0E-08 |
| Dolk | 1049 | 1.29 | 0.37 | 1.9E-04 | 1.22 | 0.28 | 5.4E-03 |
| Donson | 2658 | 1.28 | 0.35 | 5.2E-07 | 1.24 | 0.31 | 9.9E-06 |
| Dop1b | 4122 | 1.41 | 0.50 | 4.7E-03 | 1.47 | 0.56 | 1.5E-03 |
| Dot1l | 8282 | -1.24 | -0.31 | 1.4E-02 | -1.25 | -0.32 | 1.4E-02 |
| Dpf3 | 2106 | 2.45 | 1.29 | 5.1E-06 | 2.85 | 1.51 | 7.5E-08 |
| Dph1 | 786 | 1.57 | 0.65 | 5.9E-09 | 1.55 | 0.63 | 1.9E-08 |
| Dph2 | 1551 | 1.56 | 0.64 | 8.5E-10 | 1.37 | 0.45 | 2.0E-05 |
| Dph3 | 2461 | -1.86 | -0.90 | 1.4E-37 | -1.96 | -0.97 | 8.2E-44 |
| Dph5 | 733 | -1.35 | -0.44 | 2.0E-07 | -1.39 | -0.48 | 1.3E-08 |
| Dph6 | 1587 | -1.49 | -0.57 | 1.2E-15 | -1.56 | -0.64 | 2.6E-19 |
| Dpm1 | 1536 | 1.52 | 0.60 | 2.2E-11 | 1.49 | 0.58 | 1.7E-10 |
| Dpp4 | 163 | -1.87 | -0.90 | 5.5E-03 | -1.69 | -0.75 | 2.3E-02 |

| | | | | | | | |
|---|---|---|---|---|---|---|---|
| Dpp7 | 1375 | 1.33 | 0.41 | 1.4E-10 | 1.38 | 0.46 | 7.7E-13 |
| Dpy19l3 | 4852 | 1.73 | 0.79 | 2.6E-32 | 1.81 | 0.85 | 1.8E-37 |
| Dpy19l4 | 5668 | 1.22 | 0.28 | 2.8E-03 | 1.33 | 0.41 | 1.1E-05 |
| Dpysl2 | 11973 | -1.40 | -0.48 | 2.7E-02 | -1.70 | -0.77 | 2.8E-04 |
| Dpysl3 | 81721 | -1.46 | -0.55 | 4.0E-03 | -1.42 | -0.51 | 8.2E-03 |
| Dqx1 | 269 | 2.72 | 1.44 | 1.9E-17 | 2.83 | 1.50 | 1.0E-18 |
| Dr1 | 6353 | 1.43 | 0.52 | 6.5E-09 | 1.31 | 0.39 | 1.7E-05 |
| Draxin | 7093 | -1.46 | -0.54 | 1.0E-02 | -1.48 | -0.56 | 8.0E-03 |
| Drd2 | 191 | 1.98 | 0.98 | 7.8E-06 | 1.92 | 0.94 | 2.1E-05 |
| Drp2 | 4079 | -1.67 | -0.74 | 1.2E-04 | -1.67 | -0.74 | 1.3E-04 |
| Dsc2 | 211 | -4.09 | -2.03 | 1.9E-03 | -4.13 | -2.05 | 1.9E-03 |
| Dscc1 | 692 | -1.34 | -0.42 | 1.9E-02 | -1.44 | -0.53 | 3.0E-03 |
| Dse | 2031 | -1.82 | -0.86 | 5.7E-04 | -1.85 | -0.89 | 4.1E-04 |
| Dsn1 | 2055 | -1.29 | -0.37 | 7.6E-03 | -1.35 | -0.43 | 1.8E-03 |
| Dtd1 | 889 | -1.44 | -0.53 | 7.4E-07 | -1.42 | -0.51 | 1.8E-06 |
| Dtd2 | 894 | 1.56 | 0.64 | 2.9E-09 | 1.48 | 0.56 | 2.8E-07 |
| Dtna | 2903 | 2.01 | 1.00 | 9.3E-15 | 1.91 | 0.93 | 7.3E-13 |
| Dtnbp1 | 2370 | 1.52 | 0.60 | 3.6E-21 | 1.49 | 0.58 | 1.6E-19 |
| Dtwd1 | 519 | -1.30 | -0.37 | 6.3E-05 | -1.24 | -0.31 | 1.3E-03 |
| Dtwd2 | 221 | -4.39 | -2.13 | 4.5E-49 | -5.13 | -2.36 | 1.5E-57 |
| Dtx3l | 382 | -2.88 | -1.53 | 1.7E-23 | -2.71 | -1.44 | 6.4E-21 |
| Dtx4 | 7804 | -1.27 | -0.34 | 4.4E-03 | -1.31 | -0.39 | 9.2E-04 |
| Dtymk | 5211 | -1.31 | -0.39 | 1.1E-05 | -1.41 | -0.49 | 2.1E-08 |
| Dus2 | 902 | -1.70 | -0.77 | 2.1E-16 | -1.82 | -0.86 | 2.3E-20 |
| Dus3l | 2652 | -1.91 | -0.94 | 1.2E-16 | -1.94 | -0.96 | 2.6E-17 |
| Dusp1 | 1299 | -1.64 | -0.71 | 5.0E-09 | -1.77 | -0.82 | 9.9E-12 |
| Dusp10 | 751 | 1.47 | 0.55 | 2.4E-06 | 1.39 | 0.48 | 5.5E-05 |
| Dusp14 | 524 | 1.97 | 0.98 | 6.0E-14 | 2.02 | 1.01 | 8.1E-15 |
| Dusp15 | 477 | 2.05 | 1.03 | 1.4E-05 | 2.24 | 1.16 | 9.4E-07 |
| Dusp22 | 1088 | -1.40 | -0.49 | 5.1E-09 | -1.29 | -0.37 | 1.5E-05 |
| Dusp23 | 361 | 2.06 | 1.04 | 8.3E-19 | 2.10 | 1.07 | 7.5E-20 |
| Dusp27 | 234 | -2.23 | -1.15 | 1.3E-04 | -2.54 | -1.35 | 6.9E-06 |
| Dusp28 | 458 | 1.52 | 0.60 | 2.1E-08 | 1.45 | 0.54 | 7.5E-07 |
| Dusp3 | 1149 | -1.29 | -0.37 | 1.1E-05 | -1.28 | -0.36 | 2.1E-05 |
| Dusp5 | 207 | 3.15 | 1.65 | 2.6E-17 | 2.63 | 1.39 | 1.3E-12 |
| Duxbl1 | 200 | -3.71 | -1.89 | 7.7E-19 | -4.26 | -2.09 | 2.1E-22 |
| Dvl1 | 5782 | 1.40 | 0.48 | 2.0E-06 | 1.32 | 0.40 | 9.8E-05 |
| Dvl2 | 4139 | 1.46 | 0.55 | 6.6E-06 | 1.35 | 0.43 | 4.4E-04 |
| Dym | 1916 | 1.28 | 0.36 | 1.1E-08 | 1.18 | 0.24 | 1.9E-04 |
| Dynll1 | 18125 | 1.46 | 0.55 | 6.3E-11 | 1.41 | 0.49 | 5.2E-09 |
| Dynll2 | 41071 | 1.50 | 0.59 | 1.2E-03 | 1.36 | 0.44 | 1.8E-02 |
| Dynlt1f | 2100 | -1.60 | -0.68 | 1.2E-17 | -1.62 | -0.69 | 3.5E-18 |
| Dyrk1a | 7472 | -1.44 | -0.52 | 2.6E-08 | -1.54 | -0.62 | 2.1E-11 |
| Dyrk1b | 1249 | -2.21 | -1.14 | 3.1E-12 | -2.12 | -1.09 | 3.7E-11 |
| Dyrk3 | 1300 | 1.38 | 0.47 | 1.7E-04 | 1.33 | 0.42 | 1.0E-03 |
| Dzank1 | 2463 | 1.87 | 0.90 | 5.0E-08 | 2.02 | 1.02 | 6.4E-10 |
| Dzip1l | 2016 | -1.26 | -0.33 | 5.5E-05 | -1.22 | -0.28 | 6.3E-04 |

| | | | | | | | |
|---|---|---|---|---|---|---|---|
| E130114P18Rik | 1187 | -2.86 | -1.52 | 1.5E-18 | -2.71 | -1.44 | 1.0E-16 |
| E130311K13Rik | 149 | -2.95 | -1.56 | 4.8E-23 | -2.42 | -1.28 | 9.7E-16 |
| E4f1 | 2294 | -1.24 | -0.31 | 1.3E-02 | -1.30 | -0.37 | 2.9E-03 |
| Eaf2 | 184 | 3.27 | 1.71 | 1.0E-22 | 2.80 | 1.48 | 2.2E-17 |
| Eapp | 2895 | -1.47 | -0.55 | 3.6E-12 | -1.48 | -0.57 | 1.1E-12 |
| Ears2 | 1104 | -1.37 | -0.46 | 1.6E-08 | -1.36 | -0.45 | 4.7E-08 |
| Ebf1 | 6893 | -1.54 | -0.62 | 6.9E-05 | -1.34 | -0.42 | 9.0E-03 |
| Ebf3 | 12814 | -1.68 | -0.75 | 2.6E-04 | -1.60 | -0.68 | 1.1E-03 |
| Ebpl | 917 | -1.78 | -0.83 | 2.6E-56 | -1.96 | -0.97 | 8.2E-75 |
| Ecd | 3036 | -2.25 | -1.17 | 3.7E-66 | -2.43 | -1.28 | 9.2E-79 |
| Ece1 | 4433 | 1.48 | 0.56 | 1.7E-07 | 1.41 | 0.50 | 4.5E-06 |
| Ech1 | 1513 | -1.73 | -0.79 | 1.0E-10 | -1.72 | -0.78 | 2.2E-10 |
| Echdc2 | 513 | -1.29 | -0.37 | 1.0E-02 | -1.34 | -0.42 | 3.9E-03 |
| Echs1 | 4538 | -1.19 | -0.25 | 3.1E-03 | -1.25 | -0.33 | 8.1E-05 |
| Eci1 | 764 | -1.64 | -0.72 | 5.1E-11 | -1.90 | -0.93 | 1.4E-17 |
| Ecpas | 9645 | -1.62 | -0.70 | 2.0E-10 | -1.50 | -0.58 | 1.6E-07 |
| Ecscr | 463 | -1.77 | -0.82 | 8.0E-15 | -1.77 | -0.82 | 8.7E-15 |
| Eda | 935 | 1.60 | 0.68 | 9.6E-10 | 1.54 | 0.63 | 2.4E-08 |
| Eda2r | 381 | 2.03 | 1.02 | 1.5E-18 | 2.07 | 1.05 | 1.4E-19 |
| Edaradd | 238 | -1.78 | -0.83 | 4.3E-05 | -1.87 | -0.90 | 1.1E-05 |
| Edem1 | 3750 | 1.35 | 0.43 | 6.5E-11 | 1.31 | 0.39 | 3.9E-09 |
| Edem3 | 10282 | 1.57 | 0.65 | 9.5E-11 | 1.72 | 0.78 | 3.4E-15 |
| Edn3 | 547 | -2.36 | -1.24 | 1.4E-10 | -2.12 | -1.08 | 2.8E-08 |
| Ednra | 1262 | -2.67 | -1.42 | 8.0E-05 | -2.60 | -1.38 | 1.4E-04 |
| Ednrb | 7012 | -2.12 | -1.09 | 3.8E-15 | -1.87 | -0.90 | 8.9E-11 |
| Edrf1 | 4278 | 1.29 | 0.37 | 1.1E-05 | 1.35 | 0.43 | 2.4E-07 |
| Eea1 | 5899 | 1.27 | 0.34 | 3.7E-03 | 1.46 | 0.55 | 2.1E-06 |
| Eef1akmt2 | 1990 | -1.47 | -0.55 | 1.8E-06 | -1.44 | -0.53 | 5.8E-06 |
| Eef1akmt4 | 244 | 1.80 | 0.85 | 1.7E-07 | 1.60 | 0.68 | 3.8E-05 |
| Eef1d | 11624 | 1.35 | 0.44 | 7.9E-05 | 1.25 | 0.33 | 3.9E-03 |
| Eef1e1 | 2925 | -1.34 | -0.42 | 7.1E-07 | -1.27 | -0.34 | 6.7E-05 |
| Eef2k | 3029 | -1.43 | -0.52 | 4.0E-10 | -1.50 | -0.58 | 1.4E-12 |
| Eepd1 | 753 | 1.91 | 0.94 | 1.6E-11 | 1.93 | 0.95 | 8.0E-12 |
| Efcab14 | 4204 | 1.31 | 0.39 | 3.9E-08 | 1.32 | 0.41 | 1.2E-08 |
| Efcab2 | 998 | -1.67 | -0.74 | 3.2E-18 | -1.53 | -0.61 | 1.4E-12 |
| Efcab5 | 152 | -1.85 | -0.89 | 6.5E-05 | -1.73 | -0.79 | 4.6E-04 |
| Efcab6 | 111 | 2.66 | 1.41 | 7.2E-09 | 2.66 | 1.41 | 6.8E-09 |
| Efhb | 184 | 1.75 | 0.81 | 1.8E-07 | 1.77 | 0.82 | 1.5E-07 |
| Efhd1 | 206 | 3.86 | 1.95 | 8.1E-44 | 3.21 | 1.68 | 2.5E-33 |
| Efl1 | 1990 | 1.38 | 0.46 | 2.0E-08 | 1.37 | 0.46 | 3.6E-08 |
| Efna3 | 1500 | 2.61 | 1.38 | 1.2E-19 | 2.86 | 1.51 | 3.1E-23 |
| Efna5 | 4440 | -1.71 | -0.78 | 9.9E-07 | -1.90 | -0.92 | 4.3E-09 |
| Efr3a | 5605 | 1.74 | 0.80 | 1.0E-15 | 1.91 | 0.94 | 5.7E-21 |
| Efs | 1843 | -3.12 | -1.64 | 6.1E-42 | -3.18 | -1.67 | 4.0E-43 |
| Egfl6 | 1461 | -2.63 | -1.40 | 6.5E-03 | -2.18 | -1.13 | 3.2E-02 |
| Egfl7 | 1234 | -1.34 | -0.42 | 4.1E-04 | -1.43 | -0.52 | 1.1E-05 |
| Egflam | 1500 | -1.66 | -0.73 | 2.2E-03 | -1.70 | -0.77 | 1.4E-03 |

| | | | | | | | |
|---|---|---|---|---|---|---|---|
| Egfr | 1567 | -1.89 | -0.92 | 1.1E-02 | -1.69 | -0.76 | 3.9E-02 |
| Egln2 | 3902 | -1.51 | -0.59 | 7.8E-10 | -1.59 | -0.67 | 2.3E-12 |
| Egln3 | 1020 | 2.18 | 1.13 | 4.0E-23 | 2.35 | 1.23 | 2.3E-27 |
| Egr1 | 400 | 1.74 | 0.80 | 2.8E-04 | 1.71 | 0.77 | 4.8E-04 |
| Ehd2 | 1794 | -1.51 | -0.59 | 1.2E-02 | -1.49 | -0.58 | 1.6E-02 |
| Ehd3 | 1653 | 1.41 | 0.50 | 1.8E-08 | 1.38 | 0.47 | 1.4E-07 |
| Ehd4 | 1799 | 1.95 | 0.96 | 1.5E-07 | 1.79 | 0.84 | 5.0E-06 |
| Ehhadh | 313 | 1.60 | 0.68 | 1.1E-03 | 1.58 | 0.66 | 1.6E-03 |
| Ehmt1 | 11001 | -1.32 | -0.40 | 3.2E-07 | -1.44 | -0.53 | 5.5E-12 |
| Ei24 | 10552 | 1.34 | 0.42 | 4.3E-12 | 1.30 | 0.38 | 5.6E-10 |
| Eif1 | 19564 | -1.35 | -0.44 | 1.9E-05 | -1.47 | -0.56 | 3.8E-08 |
| Eif1a | 6290 | 1.38 | 0.46 | 4.7E-05 | 1.38 | 0.47 | 3.8E-05 |
| Eif1b | 5344 | -1.24 | -0.32 | 2.6E-02 | -1.33 | -0.41 | 3.4E-03 |
| Eif2a | 5182 | -1.26 | -0.33 | 8.3E-04 | -1.19 | -0.25 | 1.4E-02 |
| Eif2ak1 | 6552 | 1.40 | 0.49 | 1.8E-14 | 1.37 | 0.45 | 2.3E-12 |
| Eif2ak3 | 3360 | 1.66 | 0.74 | 1.2E-36 | 1.64 | 0.72 | 9.9E-35 |
| Eif2ak4 | 3967 | 1.65 | 0.73 | 3.6E-28 | 1.76 | 0.82 | 1.6E-35 |
| Eif2b2 | 3102 | 1.34 | 0.42 | 6.5E-08 | 1.19 | 0.25 | 2.5E-03 |
| Eif2d | 2628 | -1.32 | -0.40 | 1.4E-10 | -1.36 | -0.45 | 9.0E-13 |
| Eif3b | 17521 | 1.30 | 0.38 | 5.4E-06 | 1.20 | 0.26 | 2.3E-03 |
| Eif4a1 | 54504 | -1.46 | -0.54 | 2.9E-08 | -1.54 | -0.62 | 1.4E-10 |
| Eif4a2 | 34220 | -1.54 | -0.63 | 2.4E-11 | -1.41 | -0.50 | 1.5E-07 |
| Eif4b | 28381 | -1.25 | -0.33 | 3.4E-13 | -1.27 | -0.34 | 2.5E-14 |
| Eif4ebp1 | 706 | 1.61 | 0.69 | 2.7E-03 | 1.45 | 0.54 | 2.1E-02 |
| Eif4ebp2 | 26725 | 1.59 | 0.67 | 4.9E-06 | 1.41 | 0.50 | 9.0E-04 |
| Eif4g2 | 151975 | -1.58 | -0.66 | 2.1E-18 | -1.59 | -0.67 | 1.8E-18 |
| Eif5 | 26189 | -1.47 | -0.56 | 6.0E-10 | -1.55 | -0.63 | 2.3E-12 |
| Eif5a2 | 3776 | 2.20 | 1.13 | 2.3E-26 | 2.41 | 1.27 | 1.2E-32 |
| Eif5b | 19023 | 1.30 | 0.38 | 4.0E-06 | 1.27 | 0.35 | 3.3E-05 |
| Eif6 | 5069 | 1.57 | 0.65 | 4.5E-11 | 1.43 | 0.52 | 1.9E-07 |
| Elac1 | 1298 | -1.29 | -0.36 | 1.5E-08 | -1.27 | -0.34 | 1.1E-07 |
| Elac2 | 3479 | 1.44 | 0.53 | 4.5E-12 | 1.32 | 0.40 | 2.5E-07 |
| Elavl4 | 13824 | -1.58 | -0.66 | 1.1E-05 | -1.45 | -0.54 | 4.3E-04 |
| Elf1 | 1314 | -1.54 | -0.63 | 5.5E-14 | -1.64 | -0.72 | 6.2E-18 |
| Elf4 | 973 | 1.37 | 0.46 | 1.9E-02 | 1.31 | 0.39 | 4.7E-02 |
| Elf5 | 101 | -10.13 | -3.34 | 1.0E-02 | -14.82 | -3.89 | 2.8E-03 |
| Elk1 | 4408 | 1.40 | 0.49 | 8.7E-06 | 1.47 | 0.56 | 3.7E-07 |
| Elk3 | 2200 | -1.68 | -0.75 | 6.9E-07 | -1.60 | -0.68 | 8.7E-06 |
| Ell2 | 1029 | 2.67 | 1.42 | 6.1E-20 | 2.67 | 1.42 | 5.9E-20 |
| Elmo2 | 11087 | 1.89 | 0.92 | 1.1E-50 | 1.89 | 0.92 | 8.8E-51 |
| Elmod3 | 1141 | 1.49 | 0.58 | 4.7E-17 | 1.56 | 0.64 | 1.3E-20 |
| Eln | 1922 | -3.09 | -1.63 | 5.3E-04 | -2.83 | -1.50 | 1.5E-03 |
| Eloa | 5844 | 1.39 | 0.48 | 1.4E-07 | 1.36 | 0.44 | 1.4E-06 |
| Elovl4 | 3897 | 1.60 | 0.68 | 3.2E-13 | 1.64 | 0.71 | 1.7E-14 |
| Elovl5 | 8373 | 2.20 | 1.14 | 1.2E-38 | 2.13 | 1.09 | 9.9E-36 |
| Elovl7 | 200 | 2.09 | 1.06 | 3.7E-07 | 2.02 | 1.01 | 1.6E-06 |
| Elp3 | 7801 | -1.30 | -0.38 | 2.7E-17 | -1.35 | -0.43 | 4.6E-22 |

| | | | | | | | |
|---|---|---|---|---|---|---|---|
| Elp5 | 5319 | 1.51 | 0.60 | 8.5E-18 | 1.50 | 0.58 | 6.3E-17 |
| Elp6 | 1035 | -3.00 | -1.59 | 1.0E-81 | -3.22 | -1.69 | 7.7E-92 |
| Emc10 | 3890 | -1.48 | -0.56 | 2.3E-12 | -1.58 | -0.66 | 1.0E-16 |
| Emc2 | 3514 | 1.29 | 0.36 | 6.1E-05 | 1.29 | 0.37 | 4.7E-05 |
| Emc8 | 5610 | 1.60 | 0.68 | 5.8E-17 | 1.50 | 0.58 | 9.8E-13 |
| Emc9 | 270 | 1.54 | 0.63 | 8.8E-05 | 1.55 | 0.63 | 8.0E-05 |
| Emcn | 748 | -2.03 | -1.02 | 2.7E-06 | -2.03 | -1.02 | 2.8E-06 |
| Eme1 | 1560 | -1.42 | -0.51 | 3.2E-02 | -1.63 | -0.71 | 2.2E-03 |
| Emilin1 | 4377 | -2.01 | -1.01 | 1.7E-02 | -2.20 | -1.13 | 7.0E-03 |
| Emilin3 | 724 | -1.78 | -0.83 | 1.6E-02 | -1.70 | -0.77 | 2.8E-02 |
| Eml2 | 2644 | 1.77 | 0.82 | 1.3E-09 | 1.64 | 0.71 | 1.8E-07 |
| Eml3 | 2022 | 1.27 | 0.34 | 1.2E-03 | 1.22 | 0.28 | 8.9E-03 |
| Eml4 | 15217 | 1.40 | 0.49 | 2.8E-17 | 1.48 | 0.57 | 2.3E-23 |
| Eml5 | 16668 | 1.95 | 0.97 | 7.4E-08 | 2.10 | 1.07 | 2.0E-09 |
| Emp1 | 4540 | -3.07 | -1.62 | 4.5E-08 | -2.63 | -1.39 | 3.0E-06 |
| Emp2 | 457 | -1.49 | -0.57 | 4.9E-02 | -1.62 | -0.70 | 1.6E-02 |
| Emp3 | 587 | -1.83 | -0.87 | 2.6E-04 | -1.74 | -0.80 | 8.3E-04 |
| Enah | 27791 | -1.27 | -0.35 | 5.4E-05 | -1.31 | -0.39 | 7.1E-06 |
| Endod1 | 886 | -1.92 | -0.94 | 4.3E-12 | -1.92 | -0.94 | 5.9E-12 |
| Endog | 173 | -1.42 | -0.51 | 2.5E-05 | -1.50 | -0.58 | 1.1E-06 |
| Endov | 3296 | -1.36 | -0.44 | 4.5E-06 | -1.43 | -0.52 | 7.6E-08 |
| Eng | 1751 | -1.31 | -0.39 | 1.5E-02 | -1.40 | -0.49 | 2.3E-03 |
| Engase | 898 | 2.13 | 1.09 | 2.2E-20 | 2.01 | 1.00 | 1.7E-17 |
| Enho | 1731 | -1.53 | -0.61 | 3.6E-06 | -1.39 | -0.48 | 3.8E-04 |
| Enkd1 | 1285 | 1.61 | 0.68 | 3.6E-05 | 1.42 | 0.50 | 3.1E-03 |
| Eno1 | 41339 | 1.30 | 0.38 | 1.9E-03 | 1.23 | 0.30 | 1.7E-02 |
| Eno1b | 1567 | 1.33 | 0.41 | 1.1E-03 | 1.30 | 0.38 | 2.9E-03 |
| Eno2 | 2586 | 2.76 | 1.46 | 2.9E-16 | 2.82 | 1.49 | 7.1E-17 |
| Enoph1 | 2270 | 1.73 | 0.79 | 8.1E-29 | 1.64 | 0.71 | 8.5E-24 |
| Enox2 | 1188 | -1.37 | -0.45 | 2.7E-05 | -1.27 | -0.35 | 1.5E-03 |
| Enpep | 313 | -2.79 | -1.48 | 1.9E-04 | -2.55 | -1.35 | 7.4E-04 |
| Enpp4 | 875 | 1.33 | 0.41 | 6.2E-09 | 1.42 | 0.50 | 1.0E-12 |
| Ensa | 9994 | -1.46 | -0.54 | 5.2E-26 | -1.49 | -0.57 | 9.7E-29 |
| Entpd1 | 1019 | -2.00 | -1.00 | 3.6E-33 | -1.92 | -0.94 | 3.2E-29 |
| Entpd5 | 2041 | 1.41 | 0.49 | 1.2E-08 | 1.44 | 0.52 | 1.1E-09 |
| Entpd6 | 2345 | 1.30 | 0.38 | 4.7E-06 | 1.32 | 0.40 | 1.4E-06 |
| Entpd7 | 1194 | 1.48 | 0.57 | 1.8E-06 | 1.32 | 0.40 | 9.3E-04 |
| Eola1 | 1192 | 1.38 | 0.46 | 2.3E-08 | 1.39 | 0.47 | 1.2E-08 |
| Eomes | 3938 | -1.81 | -0.85 | 1.6E-02 | -1.86 | -0.89 | 1.1E-02 |
| Ep300 | 10587 | -1.39 | -0.48 | 1.2E-05 | -1.31 | -0.39 | 4.0E-04 |
| Epb41l1 | 5748 | -2.00 | -1.00 | 4.9E-09 | -2.08 | -1.06 | 6.4E-10 |
| Epb41l2 | 9234 | 1.33 | 0.42 | 6.0E-04 | 1.40 | 0.49 | 4.6E-05 |
| Epb41l3 | 7883 | -1.36 | -0.45 | 1.8E-05 | -1.26 | -0.33 | 1.7E-03 |
| Epb41l5 | 5238 | 1.94 | 0.96 | 4.1E-16 | 1.84 | 0.88 | 9.9E-14 |
| Epc1 | 5451 | -1.97 | -0.98 | 1.6E-20 | -2.09 | -1.06 | 3.7E-24 |
| Epc2 | 5022 | -3.53 | -1.82 | 7.0E-57 | -3.45 | -1.79 | 8.2E-55 |
| Epdr1 | 1016 | -1.45 | -0.54 | 5.7E-09 | -1.55 | -0.63 | 4.6E-12 |

| | | | | | | | |
|---|---|---|---|---|---|---|---|
| Epha10 | 2594 | 2.50 | 1.32 | 1.6E-07 | 2.58 | 1.37 | 5.6E-08 |
| Epha2 | 858 | 2.11 | 1.07 | 4.7E-07 | 2.00 | 1.00 | 3.2E-06 |
| Epha8 | 1177 | 1.69 | 0.75 | 3.5E-03 | 1.46 | 0.54 | 4.1E-02 |
| Ephb2 | 25546 | 1.44 | 0.52 | 1.4E-03 | 1.29 | 0.37 | 2.8E-02 |
| Ephb6 | 1846 | -1.44 | -0.53 | 6.5E-04 | -1.56 | -0.64 | 2.8E-05 |
| Ephx1 | 339 | 2.87 | 1.52 | 4.9E-08 | 3.13 | 1.64 | 3.6E-09 |
| Epm2a | 334 | -3.79 | -1.92 | 1.3E-36 | -3.98 | -1.99 | 4.4E-39 |
| Epop | 1631 | 3.78 | 1.92 | 8.7E-28 | 3.08 | 1.62 | 3.6E-20 |
| Epor | 184 | 2.54 | 1.35 | 4.0E-12 | 2.64 | 1.40 | 6.6E-13 |
| Eprs | 22041 | 1.32 | 0.40 | 6.4E-07 | 1.40 | 0.48 | 1.3E-09 |
| Eps8 | 1184 | 1.27 | 0.35 | 3.0E-02 | 1.32 | 0.40 | 1.2E-02 |
| Epyc | 353 | -2.80 | -1.48 | 5.5E-03 | -2.61 | -1.39 | 1.0E-02 |
| Eral1 | 1356 | -1.39 | -0.48 | 1.6E-15 | -1.49 | -0.58 | 2.5E-22 |
| Erap1 | 425 | -1.99 | -0.99 | 4.2E-10 | -2.11 | -1.07 | 1.2E-11 |
| Erbin | 14418 | -2.02 | -1.02 | 2.6E-26 | -1.88 | -0.91 | 3.6E-21 |
| Erc1 | 11179 | 1.36 | 0.45 | 1.9E-03 | 1.50 | 0.58 | 4.0E-05 |
| Ercc2 | 5173 | 1.59 | 0.66 | 2.0E-11 | 1.47 | 0.55 | 3.4E-08 |
| Ercc3 | 2601 | -1.21 | -0.27 | 9.5E-05 | -1.27 | -0.35 | 3.8E-07 |
| Ercc6 | 3104 | -2.05 | -1.04 | 3.7E-45 | -1.92 | -0.94 | 3.0E-37 |
| Ercc6l2 | 2908 | -1.79 | -0.84 | 1.2E-15 | -1.69 | -0.76 | 7.3E-13 |
| Ercc8 | 1754 | 1.30 | 0.38 | 3.6E-05 | 1.31 | 0.39 | 2.8E-05 |
| Erg | 1097 | -2.37 | -1.25 | 4.0E-09 | -2.44 | -1.29 | 1.2E-09 |
| Ergic2 | 6972 | -1.39 | -0.48 | 2.8E-08 | -1.35 | -0.44 | 5.2E-07 |
| Eri1 | 5184 | 1.54 | 0.62 | 2.7E-21 | 1.57 | 0.65 | 2.7E-23 |
| Eri2 | 3184 | -1.51 | -0.59 | 1.1E-10 | -1.50 | -0.59 | 2.0E-10 |
| Erich1 | 884 | 1.67 | 0.74 | 2.2E-21 | 1.57 | 0.65 | 8.7E-17 |
| Erich2 | 208 | 2.30 | 1.20 | 8.8E-19 | 2.12 | 1.08 | 1.9E-15 |
| Erich3 | 262 | -1.67 | -0.74 | 2.2E-04 | -1.38 | -0.46 | 2.5E-02 |
| Erlec1 | 2417 | -1.50 | -0.58 | 4.2E-08 | -1.39 | -0.47 | 1.0E-05 |
| Erlin1 | 2755 | 1.27 | 0.35 | 4.8E-07 | 1.25 | 0.32 | 4.4E-06 |
| Ermap | 432 | -2.82 | -1.49 | 8.0E-03 | -3.09 | -1.63 | 3.9E-03 |
| Ermard | 630 | -2.02 | -1.01 | 6.6E-38 | -1.85 | -0.89 | 3.0E-29 |
| Ern1 | 3334 | 2.07 | 1.05 | 2.2E-28 | 2.16 | 1.11 | 6.6E-32 |
| Ero1a | 4175 | 1.46 | 0.54 | 3.8E-11 | 1.51 | 0.60 | 3.3E-13 |
| Ero1b | 1506 | 2.60 | 1.38 | 2.1E-44 | 2.60 | 1.38 | 2.3E-44 |
| Erp44 | 4425 | 1.39 | 0.48 | 2.3E-16 | 1.39 | 0.47 | 5.7E-16 |
| Esam | 977 | -1.91 | -0.94 | 8.4E-27 | -2.07 | -1.05 | 3.2E-33 |
| Esd | 8994 | 1.36 | 0.45 | 6.3E-06 | 1.31 | 0.39 | 8.5E-05 |
| Esf1 | 5083 | 1.29 | 0.36 | 8.6E-04 | 1.25 | 0.32 | 3.3E-03 |
| Esrp2 | 159 | -2.97 | -1.57 | 2.7E-02 | -3.92 | -1.97 | 5.0E-03 |
| Esrrg | 3114 | 1.54 | 0.62 | 8.5E-03 | 1.67 | 0.74 | 1.6E-03 |
| Esyt1 | 4184 | 1.68 | 0.75 | 3.1E-07 | 1.63 | 0.70 | 1.5E-06 |
| Esyt2 | 5362 | 1.23 | 0.30 | 2.3E-04 | 1.31 | 0.39 | 1.5E-06 |
| Esyt3 | 111 | 5.12 | 2.36 | 8.5E-21 | 6.66 | 2.74 | 3.8E-26 |
| Etf1 | 14961 | 1.27 | 0.35 | 2.4E-04 | 1.23 | 0.30 | 1.9E-03 |
| Etfa | 4000 | 1.28 | 0.36 | 1.6E-03 | 1.21 | 0.28 | 1.7E-02 |
| Etfbkmt | 315 | -1.43 | -0.52 | 2.2E-04 | -1.51 | -0.59 | 2.1E-05 |

| | | | | | | | |
|---|---|---|---|---|---|---|---|
| Etfdh | 4623 | 1.64 | 0.71 | 1.1E-27 | 1.66 | 0.73 | 3.6E-29 |
| Etfrf1 | 1202 | -1.35 | -0.44 | 2.0E-05 | -1.31 | -0.38 | 2.2E-04 |
| Ethe1 | 687 | 2.65 | 1.41 | 3.5E-40 | 2.51 | 1.33 | 6.9E-36 |
| Etl4 | 2994 | -1.90 | -0.92 | 8.4E-15 | -1.81 | -0.86 | 6.8E-13 |
| Etnk1 | 20437 | 1.43 | 0.52 | 5.8E-06 | 1.64 | 0.72 | 2.6E-10 |
| Etnk2 | 538 | 3.78 | 1.92 | 2.7E-42 | 3.47 | 1.79 | 5.1E-37 |
| Ets2 | 2942 | -1.45 | -0.54 | 8.6E-03 | -1.42 | -0.50 | 1.5E-02 |
| Etv4 | 452 | -1.41 | -0.49 | 2.5E-02 | -1.36 | -0.45 | 4.7E-02 |
| Eva1b | 400 | -1.75 | -0.80 | 2.5E-03 | -1.82 | -0.86 | 1.2E-03 |
| Evc2 | 1346 | -1.48 | -0.57 | 1.7E-05 | -1.63 | -0.71 | 7.2E-08 |
| Evi5l | 7872 | 1.50 | 0.58 | 5.0E-06 | 1.45 | 0.53 | 3.6E-05 |
| Ewsr1 | 37253 | -1.31 | -0.39 | 2.3E-11 | -1.36 | -0.45 | 1.4E-14 |
| Exd1 | 115 | 1.77 | 0.82 | 3.0E-06 | 1.83 | 0.87 | 1.0E-06 |
| Exoc3 | 8530 | -1.26 | -0.33 | 7.8E-06 | -1.23 | -0.30 | 4.7E-05 |
| Exoc3l2 | 133 | -1.70 | -0.77 | 3.9E-04 | -1.70 | -0.76 | 4.4E-04 |
| Exoc6 | 1031 | -1.67 | -0.74 | 5.4E-12 | -1.60 | -0.68 | 2.1E-10 |
| Exosc10 | 4863 | -2.34 | -1.23 | 1.6E-128 | -2.43 | -1.28 | 5.7E-139 |
| Exosc7 | 1863 | -1.31 | -0.39 | 5.0E-04 | -1.40 | -0.49 | 7.8E-06 |
| Exosc8 | 3595 | -1.28 | -0.36 | 8.1E-04 | -1.29 | -0.36 | 7.1E-04 |
| Exosc9 | 4861 | 1.27 | 0.35 | 3.1E-03 | 1.25 | 0.32 | 6.2E-03 |
| Ext1 | 5543 | 1.28 | 0.35 | 3.9E-04 | 1.29 | 0.37 | 2.6E-04 |
| Extl1 | 449 | 5.64 | 2.50 | 1.8E-16 | 5.82 | 2.54 | 5.8E-17 |
| Extl2 | 2463 | 1.65 | 0.72 | 1.1E-25 | 1.76 | 0.81 | 4.8E-32 |
| Eya1 | 1127 | -2.29 | -1.19 | 9.7E-07 | -2.10 | -1.07 | 1.4E-05 |
| Eya3 | 5720 | 1.27 | 0.34 | 1.4E-05 | 1.27 | 0.34 | 1.5E-05 |
| Eya4 | 1779 | -2.09 | -1.06 | 3.7E-05 | -1.89 | -0.92 | 4.2E-04 |
| Ezr | 9712 | 1.47 | 0.55 | 1.6E-07 | 1.39 | 0.48 | 7.4E-06 |
| F13a1 | 1708 | -1.39 | -0.47 | 1.5E-02 | -1.40 | -0.48 | 1.4E-02 |
| F2rl1 | 139 | 2.31 | 1.21 | 2.3E-02 | 2.17 | 1.12 | 3.9E-02 |
| F8 | 193 | -2.40 | -1.26 | 6.2E-07 | -2.06 | -1.04 | 5.3E-05 |
| Faap20 | 1527 | -1.54 | -0.62 | 3.1E-12 | -1.70 | -0.77 | 6.6E-18 |
| Fabp5 | 12171 | -1.45 | -0.54 | 1.2E-07 | -1.46 | -0.55 | 7.7E-08 |
| Fabp7 | 24237 | -1.40 | -0.48 | 5.0E-04 | -1.29 | -0.37 | 9.7E-03 |
| Fads2 | 13692 | -1.50 | -0.59 | 1.0E-07 | -1.76 | -0.81 | 9.2E-14 |
| Faf2 | 5627 | -1.25 | -0.33 | 2.0E-15 | -1.30 | -0.37 | 5.1E-20 |
| Fahd1 | 725 | -1.41 | -0.49 | 4.7E-07 | -1.41 | -0.49 | 5.4E-07 |
| Fahd2a | 715 | -1.25 | -0.33 | 7.8E-06 | -1.29 | -0.37 | 4.5E-07 |
| Faim | 2662 | 1.46 | 0.55 | 2.4E-15 | 1.42 | 0.51 | 2.3E-13 |
| Fam102a | 2258 | 1.52 | 0.60 | 4.3E-08 | 1.50 | 0.59 | 1.1E-07 |
| Fam102b | 6586 | 1.45 | 0.54 | 9.7E-06 | 1.53 | 0.61 | 4.5E-07 |
| Fam104a | 2897 | 1.50 | 0.58 | 4.5E-20 | 1.38 | 0.47 | 3.1E-13 |
| Fam107a | 219 | -3.39 | -1.76 | 9.2E-05 | -3.53 | -1.82 | 5.4E-05 |
| Fam107b | 6022 | 1.35 | 0.43 | 1.1E-11 | 1.32 | 0.40 | 2.1E-10 |
| Fam111a | 4379 | 1.55 | 0.63 | 3.7E-05 | 1.57 | 0.65 | 2.1E-05 |
| Fam114a1 | 1545 | -1.70 | -0.77 | 8.8E-03 | -1.75 | -0.81 | 5.9E-03 |
| Fam117a | 1339 | 2.73 | 1.45 | 1.9E-13 | 2.69 | 1.43 | 5.3E-13 |
| Fam117b | 9506 | 1.31 | 0.39 | 2.2E-02 | 1.29 | 0.36 | 3.3E-02 |

| | | | | | | | |
|---|---|---|---|---|---|---|---|
| Fam118a | 1972 | 1.58 | 0.66 | 5.0E-20 | 1.50 | 0.58 | 1.2E-15 |
| Fam118b | 3527 | 1.41 | 0.50 | 9.7E-09 | 1.39 | 0.48 | 4.0E-08 |
| Fam120b | 6222 | 1.22 | 0.29 | 2.6E-04 | 1.27 | 0.35 | 1.0E-05 |
| Fam122b | 1710 | 1.50 | 0.58 | 1.1E-09 | 1.54 | 0.62 | 7.4E-11 |
| Fam126a | 12417 | 1.68 | 0.75 | 6.5E-18 | 1.82 | 0.87 | 8.3E-24 |
| Fam131b | 3839 | 1.27 | 0.35 | 5.7E-03 | 1.22 | 0.29 | 2.5E-02 |
| Fam131c | 183 | 2.05 | 1.04 | 2.2E-04 | 1.65 | 0.72 | 1.2E-02 |
| Fam135a | 9395 | 1.28 | 0.36 | 1.9E-03 | 1.44 | 0.52 | 3.3E-06 |
| Fam13a | 1152 | 1.86 | 0.90 | 2.3E-20 | 1.91 | 0.93 | 1.0E-21 |
| Fam13c | 1917 | -1.61 | -0.69 | 2.4E-11 | -1.62 | -0.70 | 1.2E-11 |
| Fam149a | 829 | -1.62 | -0.70 | 7.9E-08 | -1.56 | -0.64 | 9.6E-07 |
| Fam149b | 3470 | -1.58 | -0.66 | 2.8E-35 | -1.60 | -0.68 | 1.5E-36 |
| Fam151b | 114 | -2.12 | -1.09 | 1.3E-09 | -2.20 | -1.14 | 1.9E-10 |
| Fam160a1 | 194 | 2.05 | 1.03 | 1.6E-02 | 2.65 | 1.41 | 8.8E-04 |
| Fam160a2 | 6107 | 1.38 | 0.46 | 1.3E-09 | 1.40 | 0.48 | 3.1E-10 |
| Fam161a | 329 | -1.46 | -0.55 | 1.7E-05 | -1.36 | -0.44 | 6.9E-04 |
| Fam161b | 1588 | 1.36 | 0.44 | 2.3E-07 | 1.36 | 0.44 | 2.0E-07 |
| Fam163a | 413 | -1.86 | -0.89 | 2.4E-04 | -1.93 | -0.95 | 9.2E-05 |
| Fam168a | 15927 | -1.28 | -0.35 | 1.5E-02 | -1.23 | -0.30 | 4.5E-02 |
| Fam168b | 28932 | -1.47 | -0.55 | 2.2E-09 | -1.54 | -0.62 | 1.6E-11 |
| Fam169a | 2275 | -1.31 | -0.39 | 7.2E-03 | -1.29 | -0.36 | 1.4E-02 |
| Fam171a1 | 4578 | -1.95 | -0.96 | 2.3E-29 | -1.99 | -0.99 | 3.4E-31 |
| Fam171a2 | 4922 | -1.31 | -0.39 | 3.7E-03 | -1.35 | -0.44 | 9.8E-04 |
| Fam174b | 1868 | -1.51 | -0.59 | 8.4E-05 | -1.42 | -0.51 | 9.2E-04 |
| Fam177a2 | 339 | 1.93 | 0.95 | 3.5E-08 | 1.68 | 0.75 | 1.6E-05 |
| Fam184a | 2615 | 1.67 | 0.74 | 4.7E-13 | 1.89 | 0.92 | 1.3E-19 |
| Fam185a | 723 | 1.60 | 0.68 | 1.3E-20 | 1.58 | 0.66 | 3.2E-19 |
| Fam189a2 | 598 | 1.65 | 0.72 | 1.2E-05 | 1.66 | 0.73 | 9.5E-06 |
| Fam193a | 7160 | -1.31 | -0.39 | 3.0E-04 | -1.37 | -0.45 | 3.1E-05 |
| Fam199x | 3512 | 1.66 | 0.73 | 1.5E-14 | 1.74 | 0.80 | 7.2E-17 |
| Fam204a | 2660 | -1.47 | -0.56 | 1.5E-08 | -1.38 | -0.46 | 3.5E-06 |
| Fam20a | 144 | -1.98 | -0.98 | 1.4E-07 | -2.11 | -1.08 | 6.1E-09 |
| Fam210b | 4597 | 2.98 | 1.58 | 1.3E-37 | 3.08 | 1.62 | 8.7E-40 |
| Fam214a | 3614 | -1.26 | -0.33 | 3.0E-03 | -1.19 | -0.25 | 3.2E-02 |
| Fam217b | 976 | 1.60 | 0.68 | 8.8E-04 | 1.76 | 0.82 | 5.6E-05 |
| Fam219a | 5230 | 1.87 | 0.91 | 5.6E-09 | 1.75 | 0.81 | 2.4E-07 |
| Fam219b | 1632 | -1.66 | -0.73 | 1.9E-21 | -1.73 | -0.79 | 6.3E-25 |
| Fam221a | 677 | 3.24 | 1.70 | 1.2E-122 | 3.43 | 1.78 | 9.2E-131 |
| Fam222a | 973 | 1.71 | 0.77 | 7.3E-05 | 1.41 | 0.50 | 1.3E-02 |
| Fam228a | 125 | -2.93 | -1.55 | 4.6E-18 | -2.28 | -1.19 | 2.7E-11 |
| Fam228b | 258 | -3.29 | -1.72 | 8.0E-17 | -2.76 | -1.46 | 1.8E-12 |
| Fam229b | 532 | -2.07 | -1.05 | 6.8E-20 | -2.05 | -1.04 | 1.6E-19 |
| Fam234a | 2112 | 1.45 | 0.54 | 1.3E-07 | 1.39 | 0.48 | 3.4E-06 |
| Fam241a | 671 | 2.62 | 1.39 | 2.3E-28 | 2.96 | 1.57 | 1.3E-35 |
| Fam241b | 954 | -1.84 | -0.88 | 1.7E-05 | -1.68 | -0.75 | 2.9E-04 |
| Fam32a | 7381 | -1.33 | -0.41 | 2.3E-08 | -1.27 | -0.35 | 2.2E-06 |
| Fam3a | 992 | -2.33 | -1.22 | 2.6E-81 | -2.37 | -1.24 | 3.5E-84 |

| | | | | | | | |
|---|---|---|---|---|---|---|---|
| Fam50a | 3391 | 1.52 | 0.60 | 1.3E-10 | 1.38 | 0.47 | 9.0E-07 |
| Fam53a | 2076 | 1.27 | 0.35 | 7.3E-04 | 1.18 | 0.24 | 2.4E-02 |
| Fam53c | 6809 | 1.59 | 0.67 | 2.6E-19 | 1.39 | 0.47 | 3.7E-10 |
| Fam76a | 2920 | 1.43 | 0.51 | 6.3E-23 | 1.31 | 0.38 | 2.8E-13 |
| Fam78a | 234 | 1.99 | 0.99 | 1.1E-03 | 2.17 | 1.12 | 2.3E-04 |
| Fam81a | 711 | 1.30 | 0.38 | 2.2E-03 | 1.31 | 0.39 | 2.0E-03 |
| Fam83g | 141 | 4.13 | 2.05 | 7.6E-17 | 3.90 | 1.96 | 1.9E-15 |
| Fam83h | 667 | 1.60 | 0.68 | 1.9E-08 | 1.59 | 0.67 | 2.8E-08 |
| Fam98b | 7756 | -1.32 | -0.41 | 2.1E-03 | -1.47 | -0.56 | 1.7E-05 |
| Fam98c | 976 | 2.59 | 1.37 | 4.5E-43 | 2.47 | 1.31 | 5.2E-39 |
| Fanca | 3134 | 1.64 | 0.71 | 7.8E-06 | 1.63 | 0.71 | 9.9E-06 |
| Fancc | 1041 | -1.99 | -0.99 | 2.1E-24 | -2.03 | -1.02 | 1.2E-25 |
| Fancd2 | 1668 | -1.85 | -0.89 | 3.7E-07 | -1.79 | -0.84 | 2.0E-06 |
| Fance | 1589 | 1.35 | 0.43 | 1.5E-07 | 1.27 | 0.34 | 4.0E-05 |
| Fancg | 1810 | 1.57 | 0.66 | 1.5E-26 | 1.43 | 0.51 | 1.4E-16 |
| Fanci | 1724 | -1.33 | -0.41 | 4.1E-02 | -1.40 | -0.48 | 1.4E-02 |
| Fancl | 966 | -1.54 | -0.62 | 1.6E-06 | -1.59 | -0.67 | 2.1E-07 |
| Fancm | 5418 | 2.10 | 1.07 | 7.9E-16 | 2.27 | 1.18 | 4.3E-19 |
| Fank1 | 216 | 1.71 | 0.77 | 4.6E-08 | 2.03 | 1.02 | 3.6E-13 |
| Fap | 313 | -3.11 | -1.64 | 2.7E-03 | -2.64 | -1.40 | 1.2E-02 |
| Far1 | 14928 | 1.24 | 0.31 | 1.8E-03 | 1.38 | 0.47 | 1.6E-06 |
| Far2 | 180 | -2.03 | -1.02 | 2.3E-02 | -2.09 | -1.07 | 1.8E-02 |
| Farp1 | 6684 | -1.72 | -0.78 | 5.0E-11 | -1.72 | -0.78 | 7.1E-11 |
| Farp2 | 1216 | 2.44 | 1.29 | 1.7E-10 | 2.99 | 1.58 | 3.7E-15 |
| Fars2 | 691 | -1.61 | -0.68 | 1.2E-15 | -1.65 | -0.73 | 1.4E-17 |
| Fastk | 3314 | -1.28 | -0.35 | 5.9E-07 | -1.34 | -0.43 | 1.3E-09 |
| Fat2 | 174 | -5.11 | -2.35 | 4.0E-04 | -4.15 | -2.05 | 2.3E-03 |
| Fat4 | 13221 | -1.48 | -0.56 | 1.0E-05 | -1.32 | -0.40 | 2.4E-03 |
| Faxc | 6099 | -2.28 | -1.19 | 4.8E-12 | -2.08 | -1.05 | 1.1E-09 |
| Fbln7 | 142 | -1.80 | -0.84 | 5.9E-04 | -2.16 | -1.11 | 4.6E-06 |
| Fbxl12 | 1954 | 1.29 | 0.37 | 8.4E-07 | 1.20 | 0.26 | 7.5E-04 |
| Fbxl15 | 588 | 1.28 | 0.36 | 3.5E-04 | 1.20 | 0.26 | 1.2E-02 |
| Fbxl17 | 2918 | -1.41 | -0.50 | 1.1E-05 | -1.37 | -0.45 | 7.3E-05 |
| Fbxl19 | 11669 | -1.45 | -0.53 | 6.8E-05 | -1.52 | -0.60 | 6.1E-06 |
| Fbxl21 | 373 | 3.18 | 1.67 | 2.4E-56 | 3.23 | 1.69 | 2.4E-57 |
| Fbxl22 | 405 | 1.29 | 0.37 | 1.0E-02 | 1.26 | 0.33 | 2.2E-02 |
| Fbxl3 | 4323 | -1.77 | -0.82 | 5.5E-43 | -1.70 | -0.77 | 2.2E-37 |
| Fbxl6 | 2515 | 1.36 | 0.45 | 8.2E-08 | 1.24 | 0.31 | 3.5E-04 |
| Fbxl7 | 1539 | -2.08 | -1.06 | 1.4E-17 | -2.13 | -1.09 | 1.3E-18 |
| Fbxo10 | 3410 | 1.60 | 0.68 | 2.3E-24 | 1.47 | 0.56 | 1.1E-16 |
| Fbxo11 | 13248 | -1.37 | -0.46 | 3.5E-04 | -1.41 | -0.49 | 1.1E-04 |
| Fbxo2 | 214 | 3.52 | 1.81 | 1.3E-13 | 3.87 | 1.95 | 2.1E-15 |
| Fbxo22 | 3710 | -1.47 | -0.55 | 3.5E-10 | -1.56 | -0.64 | 1.7E-13 |
| Fbxo25 | 3316 | 1.33 | 0.41 | 1.3E-05 | 1.33 | 0.41 | 1.1E-05 |
| Fbxo27 | 347 | 1.47 | 0.55 | 8.3E-05 | 1.39 | 0.48 | 8.8E-04 |
| Fbxo31 | 2805 | 2.07 | 1.05 | 7.4E-16 | 1.94 | 0.96 | 2.2E-13 |
| Fbxo33 | 1584 | 1.59 | 0.67 | 1.5E-17 | 1.51 | 0.59 | 6.6E-14 |

| | | | | | | | |
|---|---|---|---|---|---|---|---|
| Fbxo36 | 343 | 3.19 | 1.67 | 3.1E-30 | 2.87 | 1.52 | 4.4E-25 |
| Fbxo4 | 284 | -1.59 | -0.67 | 9.7E-12 | -1.47 | -0.56 | 1.6E-08 |
| Fbxo42 | 4700 | 1.43 | 0.52 | 2.0E-09 | 1.39 | 0.48 | 4.0E-08 |
| Fbxo46 | 1348 | 1.31 | 0.39 | 2.3E-04 | 1.28 | 0.35 | 9.7E-04 |
| Fbxo7 | 624 | -2.04 | -1.03 | 1.5E-11 | -2.18 | -1.12 | 1.6E-13 |
| Fbxo8 | 1641 | 1.32 | 0.40 | 3.2E-08 | 1.33 | 0.41 | 2.0E-08 |
| Fbxw17 | 1384 | -1.44 | -0.52 | 1.7E-14 | -1.43 | -0.52 | 3.6E-14 |
| Fbxw4 | 698 | -1.86 | -0.90 | 6.8E-22 | -2.07 | -1.05 | 1.3E-29 |
| Fbxw7 | 5430 | 1.62 | 0.70 | 7.8E-20 | 1.59 | 0.67 | 3.6E-18 |
| Fcgr3 | 267 | -1.59 | -0.66 | 7.3E-04 | -1.55 | -0.63 | 1.4E-03 |
| Fcho1 | 4276 | 1.86 | 0.89 | 1.5E-07 | 1.70 | 0.77 | 7.5E-06 |
| Fchsd1 | 2990 | 3.28 | 1.71 | 1.2E-54 | 3.24 | 1.70 | 1.4E-53 |
| Fchsd2 | 5823 | -1.31 | -0.39 | 3.3E-03 | -1.29 | -0.36 | 6.2E-03 |
| Fcrls | 592 | -1.55 | -0.63 | 9.4E-05 | -1.50 | -0.59 | 2.9E-04 |
| Fcsk | 1452 | 1.39 | 0.48 | 3.2E-07 | 1.26 | 0.33 | 5.5E-04 |
| Fdxacb1 | 471 | -2.83 | -1.50 | 1.9E-55 | -3.04 | -1.61 | 3.6E-63 |
| Fdxr | 1181 | 1.25 | 0.32 | 1.4E-02 | 1.26 | 0.33 | 1.1E-02 |
| Fem1b | 27041 | 1.31 | 0.39 | 2.7E-05 | 1.32 | 0.40 | 2.4E-05 |
| Fermt1 | 105 | -5.84 | -2.55 | 4.8E-04 | -5.28 | -2.40 | 1.1E-03 |
| Fes | 460 | -2.58 | -1.37 | 2.7E-23 | -2.77 | -1.47 | 1.4E-26 |
| Fez2 | 1807 | 1.40 | 0.48 | 1.6E-18 | 1.36 | 0.44 | 9.9E-16 |
| Fezf2 | 1417 | -1.73 | -0.79 | 5.8E-03 | -1.99 | -1.00 | 4.1E-04 |
| Fgd3 | 1058 | -1.33 | -0.41 | 1.0E-02 | -1.43 | -0.51 | 1.1E-03 |
| Fgd4 | 4039 | -2.39 | -1.26 | 3.5E-15 | -2.16 | -1.11 | 4.8E-12 |
| Fgd5 | 1667 | -1.42 | -0.50 | 2.1E-04 | -1.38 | -0.46 | 7.7E-04 |
| Fgd6 | 3623 | 2.21 | 1.14 | 3.2E-34 | 2.39 | 1.26 | 4.2E-41 |
| Fgf11 | 2991 | 2.83 | 1.50 | 2.1E-29 | 2.81 | 1.49 | 5.0E-29 |
| Fgf15 | 1934 | -1.55 | -0.64 | 5.4E-04 | -1.80 | -0.85 | 2.5E-06 |
| Fgf18 | 123 | -2.31 | -1.21 | 3.5E-05 | -2.11 | -1.08 | 2.5E-04 |
| Fgf2 | 295 | -2.94 | -1.56 | 3.9E-15 | -2.79 | -1.48 | 1.1E-13 |
| Fgf3 | 153 | -1.90 | -0.92 | 1.7E-02 | -2.55 | -1.35 | 3.6E-04 |
| Fgfbp3 | 1992 | -1.61 | -0.68 | 6.1E-04 | -1.74 | -0.80 | 5.3E-05 |
| Fgfr1op2 | 6435 | 1.26 | 0.33 | 5.9E-05 | 1.31 | 0.38 | 3.4E-06 |
| Fgfr3 | 7687 | -1.49 | -0.57 | 4.4E-04 | -1.69 | -0.76 | 2.3E-06 |
| Fgfrl1 | 1853 | 1.59 | 0.67 | 6.8E-05 | 1.47 | 0.55 | 1.1E-03 |
| Fhdc1 | 667 | 1.32 | 0.40 | 2.6E-03 | 1.46 | 0.55 | 3.0E-05 |
| Fhl2 | 217 | -1.51 | -0.59 | 2.1E-02 | -1.74 | -0.80 | 1.6E-03 |
| Fhod1 | 1722 | 2.00 | 1.00 | 2.7E-09 | 1.74 | 0.80 | 2.7E-06 |
| Fibin | 898 | -4.54 | -2.18 | 3.6E-04 | -3.82 | -1.93 | 1.8E-03 |
| Fig4 | 3204 | 2.15 | 1.10 | 4.9E-48 | 2.21 | 1.15 | 1.8E-51 |
| Fitm2 | 1371 | 1.22 | 0.28 | 1.5E-07 | 1.25 | 0.33 | 1.1E-09 |
| Fiz1 | 2808 | -2.09 | -1.06 | 9.1E-28 | -2.25 | -1.17 | 1.9E-33 |
| Fjx1 | 2433 | -1.32 | -0.40 | 1.6E-03 | -1.42 | -0.51 | 5.1E-05 |
| Fkbp11 | 289 | 1.65 | 0.73 | 7.9E-05 | 1.56 | 0.64 | 6.1E-04 |
| Fkbp14 | 3396 | 2.54 | 1.34 | 2.1E-33 | 2.71 | 1.44 | 4.8E-38 |
| Fkbp5 | 5988 | 1.62 | 0.69 | 2.1E-10 | 1.50 | 0.58 | 1.3E-07 |
| Fkbpl | 574 | 1.24 | 0.31 | 4.2E-03 | 1.28 | 0.36 | 9.7E-04 |

| | | | | | | | |
|---|---|---|---|---|---|---|---|
| Fkrp | 1757 | -2.59 | -1.38 | 4.4E-61 | -2.66 | -1.41 | 1.7E-64 |
| Fktn | 3633 | -1.32 | -0.40 | 2.5E-06 | -1.29 | -0.37 | 2.2E-05 |
| Flad1 | 1338 | -1.49 | -0.57 | 6.5E-14 | -1.53 | -0.62 | 6.9E-16 |
| Fli1 | 720 | -2.11 | -1.08 | 9.2E-08 | -2.34 | -1.22 | 1.0E-09 |
| Flot1 | 2825 | 1.57 | 0.65 | 2.2E-12 | 1.42 | 0.51 | 6.3E-08 |
| Flrt2 | 7693 | -1.73 | -0.79 | 2.6E-08 | -1.52 | -0.60 | 3.0E-05 |
| Flvcr1 | 2493 | 1.91 | 0.94 | 2.4E-29 | 1.76 | 0.82 | 1.7E-22 |
| Flvcr2 | 242 | 1.53 | 0.61 | 6.7E-05 | 1.37 | 0.45 | 3.9E-03 |
| Fmc1 | 463 | -1.39 | -0.48 | 4.4E-03 | -1.39 | -0.47 | 5.1E-03 |
| Fmn1 | 217 | -2.81 | -1.49 | 7.2E-04 | -2.01 | -1.01 | 2.7E-02 |
| Fmo1 | 1524 | -2.21 | -1.15 | 1.8E-05 | -2.37 | -1.24 | 3.1E-06 |
| Fn3k | 279 | 1.68 | 0.75 | 2.1E-14 | 1.71 | 0.78 | 2.9E-15 |
| Fn3krp | 1919 | 1.83 | 0.87 | 9.1E-37 | 1.73 | 0.79 | 1.4E-30 |
| Fnbp1l | 43753 | -1.68 | -0.75 | 6.0E-06 | -1.60 | -0.68 | 5.4E-05 |
| Fndc3b | 3394 | -1.38 | -0.46 | 2.6E-03 | -1.25 | -0.33 | 4.1E-02 |
| Fndc4 | 3380 | -1.30 | -0.38 | 7.1E-09 | -1.37 | -0.45 | 2.9E-12 |
| Fnip1 | 5694 | 1.57 | 0.65 | 6.2E-11 | 1.69 | 0.76 | 1.2E-14 |
| Fnip2 | 4024 | 1.38 | 0.47 | 3.1E-05 | 1.41 | 0.49 | 1.1E-05 |
| Fntb | 1654 | -1.75 | -0.81 | 2.8E-25 | -1.82 | -0.86 | 1.9E-28 |
| Focad | 2776 | 1.50 | 0.59 | 2.5E-11 | 1.55 | 0.63 | 7.3E-13 |
| Folr1 | 192 | -1.60 | -0.68 | 1.6E-02 | -1.70 | -0.76 | 6.8E-03 |
| Fos | 108 | 1.92 | 0.94 | 1.0E-03 | 2.49 | 1.32 | 3.2E-06 |
| Foxa1 | 549 | -1.47 | -0.55 | 2.2E-02 | -1.75 | -0.81 | 5.7E-04 |
| Foxc1 | 1824 | -1.79 | -0.84 | 6.9E-04 | -2.13 | -1.09 | 7.9E-06 |
| Foxf2 | 516 | -1.34 | -0.42 | 8.5E-03 | -1.35 | -0.43 | 7.6E-03 |
| Foxj1 | 1534 | 1.24 | 0.31 | 4.0E-02 | 1.30 | 0.38 | 1.1E-02 |
| Foxj2 | 1949 | 1.29 | 0.36 | 8.3E-03 | 1.26 | 0.34 | 1.5E-02 |
| Foxk1 | 6300 | 1.42 | 0.51 | 2.6E-04 | 1.33 | 0.41 | 3.6E-03 |
| Foxl2 | 300 | -1.93 | -0.95 | 6.5E-03 | -2.40 | -1.26 | 2.5E-04 |
| Foxm1 | 5223 | -1.42 | -0.50 | 4.9E-03 | -1.62 | -0.69 | 8.1E-05 |
| Foxo1 | 2595 | 1.84 | 0.88 | 2.8E-13 | 1.80 | 0.85 | 1.8E-12 |
| Foxo3 | 4755 | 1.66 | 0.74 | 1.9E-11 | 1.70 | 0.77 | 2.6E-12 |
| Foxo4 | 3776 | 1.37 | 0.45 | 1.8E-05 | 1.36 | 0.44 | 3.1E-05 |
| Foxo6 | 990 | 1.56 | 0.64 | 1.2E-02 | 1.50 | 0.59 | 2.3E-02 |
| Foxp2 | 6899 | -1.49 | -0.58 | 2.2E-08 | -1.58 | -0.66 | 1.6E-10 |
| Foxq1 | 238 | -1.28 | -0.35 | 2.2E-02 | -1.40 | -0.49 | 1.2E-03 |
| Fpgt | 1526 | -1.27 | -0.35 | 4.3E-04 | -1.17 | -0.23 | 2.5E-02 |
| Frat1 | 1231 | 1.85 | 0.89 | 2.6E-09 | 1.57 | 0.65 | 1.8E-05 |
| Frat2 | 1496 | 1.65 | 0.73 | 9.8E-08 | 1.49 | 0.57 | 3.2E-05 |
| Frg2f1 | 271 | -1.33 | -0.41 | 2.7E-02 | -1.35 | -0.44 | 2.0E-02 |
| Frk | 338 | -3.44 | -1.78 | 2.6E-06 | -3.42 | -1.77 | 3.2E-06 |
| Frmd3 | 746 | -2.44 | -1.29 | 1.1E-05 | -2.05 | -1.04 | 5.0E-04 |
| Frmd4a | 12232 | -2.17 | -1.12 | 1.5E-26 | -2.10 | -1.07 | 2.1E-24 |
| Frmd4b | 5483 | -1.30 | -0.38 | 3.1E-09 | -1.30 | -0.38 | 4.9E-09 |
| Frmd8 | 2840 | 1.94 | 0.96 | 7.5E-18 | 1.90 | 0.92 | 1.0E-16 |
| Frmpd1 | 237 | -1.38 | -0.47 | 1.4E-02 | -1.35 | -0.43 | 2.5E-02 |
| Frrs1 | 547 | -1.68 | -0.75 | 1.6E-07 | -1.75 | -0.81 | 1.4E-08 |

| | | | | | | | |
|---|---|---|---|---|---|---|---|
| Fryl | 10889 | 1.65 | 0.73 | 2.4E-09 | 1.74 | 0.80 | 4.0E-11 |
| Fsbp | 229 | -1.98 | -0.99 | 1.0E-08 | -2.01 | -1.00 | 6.7E-09 |
| Fsd1 | 2961 | 1.42 | 0.50 | 1.8E-04 | 1.36 | 0.44 | 1.1E-03 |
| Fst | 1006 | -1.41 | -0.49 | 2.3E-02 | -1.83 | -0.87 | 3.1E-05 |
| Fstl3 | 146 | 3.32 | 1.73 | 1.3E-09 | 2.98 | 1.58 | 4.5E-08 |
| Ftsj1 | 2682 | 1.47 | 0.56 | 4.7E-19 | 1.42 | 0.51 | 8.5E-16 |
| Ftsj3 | 8106 | 1.50 | 0.58 | 1.9E-08 | 1.42 | 0.50 | 1.7E-06 |
| Fubp1 | 40486 | -1.33 | -0.41 | 7.9E-08 | -1.27 | -0.34 | 7.3E-06 |
| Fuca1 | 2707 | 1.96 | 0.97 | 5.8E-98 | 1.80 | 0.85 | 7.0E-75 |
| Fundc1 | 2603 | -1.71 | -0.77 | 1.1E-19 | -1.63 | -0.70 | 2.4E-16 |
| Fundc2 | 3228 | -1.31 | -0.39 | 1.1E-06 | -1.31 | -0.39 | 1.6E-06 |
| Fus | 58452 | -1.32 | -0.40 | 6.0E-03 | -1.49 | -0.58 | 5.2E-05 |
| Fut10 | 3079 | 2.19 | 1.13 | 1.4E-16 | 2.16 | 1.11 | 3.9E-16 |
| Fut4 | 139 | -1.42 | -0.50 | 2.4E-02 | -1.51 | -0.59 | 8.1E-03 |
| Fuz | 1216 | -1.23 | -0.30 | 5.2E-03 | -1.32 | -0.40 | 1.4E-04 |
| Fv1 | 1634 | 2.05 | 1.04 | 1.8E-51 | 2.05 | 1.04 | 3.0E-51 |
| Fxn | 615 | -1.96 | -0.97 | 1.2E-17 | -2.04 | -1.03 | 1.4E-19 |
| Fyb | 173 | -2.30 | -1.20 | 1.2E-08 | -2.25 | -1.17 | 3.3E-08 |
| Fyco1 | 2757 | 1.46 | 0.55 | 6.1E-07 | 1.51 | 0.59 | 6.2E-08 |
| Fyttd1 | 12474 | 1.30 | 0.38 | 1.9E-06 | 1.35 | 0.43 | 5.6E-08 |
| Fzd5 | 608 | 2.35 | 1.24 | 1.0E-10 | 1.80 | 0.85 | 1.3E-05 |
| G0s2 | 110 | -2.25 | -1.17 | 1.6E-06 | -2.30 | -1.20 | 7.9E-07 |
| Gaa | 4614 | 1.31 | 0.39 | 1.1E-02 | 1.32 | 0.41 | 8.5E-03 |
| Gab3 | 120 | 2.53 | 1.34 | 5.1E-17 | 2.09 | 1.07 | 1.9E-11 |
| Gabarapl1 | 8427 | 1.45 | 0.53 | 2.4E-12 | 1.40 | 0.49 | 1.9E-10 |
| Gabarapl2 | 6142 | 1.32 | 0.40 | 7.4E-06 | 1.32 | 0.40 | 5.2E-06 |
| Gabbr2 | 570 | -2.47 | -1.30 | 7.9E-03 | -2.25 | -1.17 | 1.9E-02 |
| Gabpa | 6018 | -1.47 | -0.56 | 1.2E-15 | -1.48 | -0.57 | 5.0E-16 |
| Gabpb1 | 3868 | -1.53 | -0.62 | 1.9E-09 | -1.69 | -0.76 | 6.4E-14 |
| Gabpb2 | 5710 | -2.39 | -1.26 | 9.2E-30 | -2.20 | -1.14 | 1.6E-24 |
| Gabra1 | 591 | -1.75 | -0.81 | 2.6E-03 | -1.68 | -0.75 | 5.9E-03 |
| Gabra2 | 1504 | -1.66 | -0.73 | 1.8E-02 | -1.77 | -0.83 | 7.2E-03 |
| Gabra4 | 739 | 2.46 | 1.30 | 3.0E-08 | 2.63 | 1.39 | 2.6E-09 |
| Gabre | 736 | 2.44 | 1.29 | 1.4E-36 | 2.45 | 1.29 | 7.5E-37 |
| Gabrg1 | 541 | 3.41 | 1.77 | 5.3E-16 | 3.10 | 1.63 | 7.7E-14 |
| Gabrg2 | 3554 | -2.47 | -1.30 | 5.0E-07 | -2.25 | -1.17 | 7.1E-06 |
| Gabrq | 1039 | 3.52 | 1.81 | 1.1E-36 | 3.63 | 1.86 | 1.9E-38 |
| Gad1 | 3462 | -2.50 | -1.32 | 1.0E-04 | -2.54 | -1.35 | 7.7E-05 |
| Gad2 | 5532 | -3.18 | -1.67 | 2.9E-08 | -3.38 | -1.76 | 4.9E-09 |
| Gadd45gip1 | 1310 | -1.72 | -0.78 | 2.2E-16 | -1.74 | -0.80 | 4.7E-17 |
| Gal3st3 | 791 | -1.33 | -0.41 | 8.2E-03 | -1.47 | -0.56 | 2.5E-04 |
| Galc | 2296 | -1.30 | -0.38 | 1.1E-07 | -1.32 | -0.40 | 2.5E-08 |
| Gale | 1183 | 1.89 | 0.92 | 1.8E-20 | 1.73 | 0.79 | 1.6E-15 |
| Galns | 568 | 1.50 | 0.59 | 3.9E-04 | 1.44 | 0.53 | 1.7E-03 |
| Galnt1 | 7835 | -1.43 | -0.52 | 1.3E-08 | -1.43 | -0.52 | 1.1E-08 |
| Galnt10 | 4324 | 1.96 | 0.97 | 2.5E-16 | 1.91 | 0.93 | 4.3E-15 |
| Galnt12 | 352 | 2.15 | 1.10 | 3.3E-06 | 1.85 | 0.89 | 2.4E-04 |

| | | | | | | | |
|---|---|---|---|---|---|---|---|
| Galnt14 | 565 | -2.75 | -1.46 | 7.3E-21 | -2.53 | -1.34 | 1.2E-17 |
| Galnt5 | 124 | -3.70 | -1.89 | 2.6E-03 | -3.74 | -1.90 | 2.6E-03 |
| Galnt7 | 2010 | -1.47 | -0.56 | 1.3E-10 | -1.45 | -0.53 | 7.3E-10 |
| Gamt | 468 | 1.43 | 0.52 | 5.3E-03 | 1.42 | 0.50 | 7.4E-03 |
| Gan | 3930 | 1.39 | 0.47 | 5.0E-05 | 1.40 | 0.48 | 3.5E-05 |
| Garem1 | 2204 | 1.92 | 0.94 | 1.6E-12 | 2.20 | 1.14 | 1.2E-17 |
| Garnl3 | 2398 | 1.85 | 0.89 | 2.1E-10 | 2.05 | 1.04 | 7.2E-14 |
| Gart | 10721 | 1.96 | 0.97 | 9.5E-18 | 1.77 | 0.82 | 4.6E-13 |
| Gas1 | 5642 | -1.78 | -0.83 | 7.0E-06 | -1.92 | -0.94 | 3.1E-07 |
| Gas2 | 1719 | -1.55 | -0.63 | 2.7E-04 | -1.45 | -0.54 | 2.1E-03 |
| Gas2l1 | 2396 | 2.68 | 1.42 | 1.2E-27 | 2.51 | 1.33 | 3.1E-24 |
| Gas2l3 | 2510 | 1.30 | 0.38 | 2.3E-04 | 1.32 | 0.40 | 7.5E-05 |
| Gas6 | 1399 | 1.51 | 0.59 | 8.8E-05 | 1.41 | 0.50 | 1.2E-03 |
| Gas7 | 1662 | -1.64 | -0.71 | 6.9E-03 | -1.67 | -0.74 | 4.8E-03 |
| Gas8 | 3924 | 1.49 | 0.58 | 2.1E-10 | 1.40 | 0.49 | 1.1E-07 |
| Gask1b | 385 | -3.66 | -1.87 | 9.1E-10 | -3.38 | -1.76 | 1.0E-08 |
| Gata2 | 1069 | -1.56 | -0.64 | 5.1E-03 | -1.74 | -0.80 | 4.6E-04 |
| Gatad2a | 7326 | 1.45 | 0.54 | 2.5E-07 | 1.32 | 0.40 | 1.6E-04 |
| Gatb | 739 | -1.90 | -0.92 | 1.5E-20 | -1.78 | -0.83 | 5.3E-17 |
| Gatd1 | 1870 | -1.44 | -0.53 | 1.2E-13 | -1.59 | -0.67 | 2.1E-21 |
| Gba | 2522 | 1.47 | 0.55 | 4.2E-14 | 1.46 | 0.55 | 8.1E-14 |
| Gba2 | 6864 | 1.51 | 0.59 | 2.2E-13 | 1.45 | 0.53 | 5.0E-11 |
| Gbe1 | 851 | -1.62 | -0.70 | 4.7E-07 | -1.67 | -0.74 | 7.3E-08 |
| Gbp7 | 141 | -2.29 | -1.20 | 2.9E-12 | -2.25 | -1.17 | 9.1E-12 |
| Gbp9 | 205 | -2.37 | -1.24 | 2.7E-08 | -2.59 | -1.37 | 6.4E-10 |
| Gbx1 | 311 | -2.29 | -1.20 | 3.5E-03 | -2.39 | -1.26 | 2.2E-03 |
| Gca | 1493 | 3.99 | 2.00 | 2.3E-40 | 3.93 | 1.97 | 1.9E-39 |
| Gcat | 161 | 2.19 | 1.13 | 2.0E-07 | 2.25 | 1.17 | 8.8E-08 |
| Gcdh | 3383 | 2.08 | 1.06 | 7.6E-69 | 1.95 | 0.96 | 3.5E-57 |
| Gch1 | 460 | 5.58 | 2.48 | 7.6E-65 | 5.25 | 2.39 | 1.8E-60 |
| Gck | 180 | -1.50 | -0.59 | 1.3E-03 | -1.80 | -0.85 | 2.0E-06 |
| Gclc | 5236 | 1.89 | 0.92 | 7.7E-41 | 1.93 | 0.95 | 2.0E-43 |
| Gcsh | 4319 | 1.49 | 0.57 | 1.6E-06 | 1.41 | 0.50 | 4.2E-05 |
| Gdap2 | 2521 | 1.47 | 0.56 | 2.0E-10 | 1.48 | 0.56 | 1.5E-10 |
| Gde1 | 4093 | 1.41 | 0.49 | 6.3E-10 | 1.42 | 0.51 | 1.9E-10 |
| Gdf10 | 847 | -2.94 | -1.56 | 1.1E-04 | -2.43 | -1.28 | 1.7E-03 |
| Gdf11 | 5930 | 1.88 | 0.91 | 2.4E-15 | 1.64 | 0.72 | 7.1E-10 |
| Gdf5 | 103 | -3.07 | -1.62 | 8.0E-12 | -2.66 | -1.41 | 3.7E-09 |
| Gdpd1 | 4982 | -1.60 | -0.67 | 1.5E-05 | -1.58 | -0.66 | 2.7E-05 |
| Gdpd2 | 998 | -1.39 | -0.48 | 5.4E-03 | -1.43 | -0.52 | 2.6E-03 |
| Gdpd5 | 3727 | 1.47 | 0.55 | 2.2E-04 | 1.41 | 0.50 | 1.0E-03 |
| Gdpgp1 | 755 | 2.34 | 1.22 | 7.5E-40 | 2.27 | 1.18 | 5.4E-37 |
| Gem | 273 | 1.98 | 0.98 | 3.5E-16 | 1.90 | 0.92 | 2.2E-14 |
| Gemin5 | 6237 | 1.82 | 0.86 | 1.5E-23 | 1.83 | 0.87 | 6.5E-24 |
| Gemin6 | 770 | -1.37 | -0.46 | 1.4E-03 | -1.39 | -0.48 | 8.7E-04 |
| Get4 | 2170 | -1.25 | -0.32 | 2.9E-04 | -1.31 | -0.38 | 1.4E-05 |
| Gfer | 1341 | -1.40 | -0.49 | 7.1E-11 | -1.45 | -0.53 | 9.2E-13 |

| | | | | | | | |
|---|---|---|---|---|---|---|---|
| Gfod1 | 1416 | 1.47 | 0.56 | 9.5E-03 | 1.54 | 0.62 | 3.8E-03 |
| Gfpt2 | 1177 | 1.67 | 0.74 | 1.2E-07 | 1.78 | 0.83 | 2.5E-09 |
| Gfra1 | 3622 | -1.55 | -0.63 | 2.1E-04 | -1.28 | -0.36 | 4.4E-02 |
| Gfra4 | 183 | -2.28 | -1.19 | 5.0E-06 | -2.08 | -1.05 | 6.2E-05 |
| Gga1 | 3531 | -1.46 | -0.54 | 6.5E-05 | -1.56 | -0.64 | 1.8E-06 |
| Gga2 | 8276 | 1.33 | 0.41 | 3.9E-09 | 1.25 | 0.32 | 5.8E-06 |
| Ggact | 204 | -1.29 | -0.36 | 9.5E-03 | -1.27 | -0.34 | 1.7E-02 |
| Ggct | 639 | 1.83 | 0.87 | 4.2E-37 | 1.96 | 0.97 | 1.9E-45 |
| Ggcx | 132 | 2.65 | 1.41 | 6.9E-05 | -1.68 | -0.75 | 4.4E-02 |
| Ggn | 146 | 2.21 | 1.14 | 1.2E-18 | 2.04 | 1.03 | 1.9E-15 |
| Ggt5 | 149 | -1.64 | -0.71 | 1.5E-04 | -2.05 | -1.04 | 1.6E-08 |
| Ggta1 | 834 | -1.34 | -0.42 | 1.3E-02 | -1.35 | -0.43 | 1.0E-02 |
| Ghdc | 476 | 1.40 | 0.49 | 1.6E-02 | 1.34 | 0.42 | 4.2E-02 |
| Ghr | 1697 | -1.85 | -0.88 | 9.4E-04 | -1.60 | -0.68 | 1.4E-02 |
| Gid4 | 5221 | 1.66 | 0.73 | 3.4E-16 | 1.59 | 0.67 | 9.9E-14 |
| Gimap4 | 212 | -2.02 | -1.01 | 3.3E-11 | -2.00 | -1.00 | 5.6E-11 |
| Gimap6 | 412 | -1.64 | -0.71 | 1.5E-05 | -1.36 | -0.44 | 9.9E-03 |
| Gimap8 | 132 | -2.59 | -1.37 | 2.0E-19 | -2.54 | -1.35 | 1.6E-18 |
| Gin1 | 729 | -1.71 | -0.78 | 1.1E-17 | -1.60 | -0.67 | 1.2E-13 |
| Gins1 | 1687 | -1.49 | -0.58 | 9.9E-04 | -1.63 | -0.70 | 5.1E-05 |
| Gins2 | 2081 | 1.55 | 0.63 | 4.5E-04 | 1.39 | 0.48 | 9.5E-03 |
| Gins3 | 1696 | 1.68 | 0.75 | 2.4E-07 | 1.63 | 0.70 | 1.4E-06 |
| Gins4 | 2733 | 1.39 | 0.48 | 4.0E-04 | 1.29 | 0.37 | 8.4E-03 |
| Gipc1 | 4619 | 1.42 | 0.51 | 2.3E-06 | 1.28 | 0.36 | 9.8E-04 |
| Gipc2 | 110 | 1.63 | 0.70 | 3.4E-05 | 1.76 | 0.81 | 1.4E-06 |
| Gipr | 596 | 2.14 | 1.10 | 1.2E-07 | 2.07 | 1.05 | 3.9E-07 |
| Git1 | 8435 | 1.65 | 0.72 | 1.2E-09 | 1.55 | 0.63 | 1.5E-07 |
| Git2 | 4258 | 1.31 | 0.39 | 2.4E-10 | 1.36 | 0.44 | 8.0E-13 |
| Gja1 | 9093 | -2.66 | -1.41 | 7.9E-22 | -2.96 | -1.57 | 1.3E-26 |
| Gja4 | 397 | -1.39 | -0.47 | 3.6E-04 | -1.50 | -0.59 | 7.4E-06 |
| Gja5 | 313 | -3.74 | -1.90 | 5.9E-11 | -3.94 | -1.98 | 9.2E-12 |
| Gjb6 | 465 | -5.42 | -2.44 | 4.9E-20 | -5.69 | -2.51 | 5.1E-21 |
| Gjc1 | 5342 | -1.22 | -0.29 | 4.4E-02 | -1.33 | -0.41 | 2.9E-03 |
| Gjd2 | 871 | -1.98 | -0.98 | 2.2E-04 | -1.93 | -0.95 | 3.8E-04 |
| Gk | 1714 | 2.06 | 1.04 | 6.2E-32 | 2.06 | 1.05 | 4.3E-32 |
| Gk5 | 932 | 1.74 | 0.80 | 1.9E-06 | 2.11 | 1.08 | 9.5E-11 |
| Gkap1 | 2822 | 1.43 | 0.51 | 9.3E-09 | 1.36 | 0.44 | 9.4E-07 |
| Gla | 1237 | 2.24 | 1.17 | 1.5E-33 | 2.08 | 1.05 | 1.8E-27 |
| Glb1l | 1778 | -1.33 | -0.42 | 2.4E-07 | -1.32 | -0.40 | 7.8E-07 |
| Glb1l2 | 428 | 1.34 | 0.42 | 1.3E-03 | 1.27 | 0.34 | 1.1E-02 |
| Glce | 4012 | 1.43 | 0.51 | 1.4E-04 | 1.57 | 0.65 | 1.1E-06 |
| Gli2 | 2735 | -1.79 | -0.84 | 1.0E-07 | -1.89 | -0.92 | 5.9E-09 |
| Gli3 | 7586 | -1.36 | -0.44 | 6.1E-03 | -1.44 | -0.53 | 9.1E-04 |
| Glis3 | 662 | -1.49 | -0.57 | 6.5E-04 | -1.44 | -0.52 | 2.0E-03 |
| Glmp | 2725 | 1.48 | 0.57 | 1.1E-07 | 1.44 | 0.53 | 8.9E-07 |
| Glrx | 1131 | 1.88 | 0.91 | 4.3E-22 | 2.01 | 1.01 | 1.5E-26 |
| Glrx5 | 3014 | 1.77 | 0.82 | 1.4E-14 | 1.60 | 0.68 | 3.5E-10 |

| | | | | | | | |
|---|---|---|---|---|---|---|---|
| Gls2 | 319 | 4.06 | 2.02 | 2.3E-62 | 4.84 | 2.28 | 4.0E-75 |
| Glt1d1 | 409 | -2.54 | -1.34 | 8.5E-15 | -2.94 | -1.56 | 2.3E-19 |
| Glt8d1 | 4240 | -1.42 | -0.50 | 2.8E-21 | -1.44 | -0.53 | 2.5E-23 |
| Glt8d2 | 236 | -3.39 | -1.76 | 2.4E-06 | -2.98 | -1.57 | 2.9E-05 |
| Glyctk | 135 | -2.01 | -1.01 | 4.5E-06 | -2.18 | -1.12 | 3.5E-07 |
| Gm10037 | 225 | -1.41 | -0.49 | 8.4E-03 | -1.39 | -0.48 | 1.2E-02 |
| Gm10226 | 301 | 1.28 | 0.36 | 6.1E-03 | 1.30 | 0.38 | 3.8E-03 |
| Gm10406 | 198 | -3.41 | -1.77 | 1.8E-22 | -4.09 | -2.03 | 2.2E-28 |
| Gm1141 | 166 | 2.05 | 1.03 | 8.4E-11 | 1.97 | 0.98 | 6.1E-10 |
| Gm11627 | 194 | -2.44 | -1.29 | 3.0E-09 | -2.58 | -1.37 | 3.2E-10 |
| Gm12184 | 863 | -1.91 | -0.93 | 2.9E-11 | -2.11 | -1.08 | 1.2E-14 |
| Gm13199 | 201 | 3.05 | 1.61 | 1.1E-16 | 2.56 | 1.36 | 2.6E-12 |
| Gm13889 | 360 | -1.49 | -0.57 | 4.6E-02 | -1.64 | -0.71 | 1.2E-02 |
| Gm14403 | 283 | 1.42 | 0.50 | 5.0E-03 | 1.70 | 0.77 | 1.3E-05 |
| Gm14443 | 290 | 1.45 | 0.53 | 3.1E-05 | 1.52 | 0.60 | 2.4E-06 |
| Gm14698 | 269 | 1.28 | 0.35 | 3.7E-02 | 1.31 | 0.39 | 2.0E-02 |
| Gm15446 | 2257 | 1.25 | 0.33 | 4.2E-03 | 1.36 | 0.44 | 8.4E-05 |
| Gm17655 | 130 | 1.46 | 0.55 | 5.5E-04 | 1.44 | 0.53 | 8.3E-04 |
| Gm19426 | 193 | -2.31 | -1.21 | 3.1E-12 | -2.52 | -1.33 | 2.0E-14 |
| Gm20075 | 1385 | -1.74 | -0.80 | 2.5E-09 | -1.76 | -0.82 | 8.2E-10 |
| Gm2115 | 108 | -2.38 | -1.25 | 8.6E-08 | -2.52 | -1.33 | 1.0E-08 |
| Gm21992 | 810 | 2.55 | 1.35 | 5.5E-08 | 2.18 | 1.12 | 7.5E-06 |
| Gm2237 | 528 | -1.70 | -0.77 | 1.8E-10 | -1.73 | -0.79 | 6.3E-11 |
| Gm2897 | 350 | -1.55 | -0.63 | 5.9E-05 | -1.50 | -0.59 | 2.3E-04 |
| Gm29695 | 1260 | -3.27 | -1.71 | 1.4E-22 | -3.49 | -1.80 | 5.6E-25 |
| Gm2974 | 142 | -3.33 | -1.74 | 2.1E-08 | -3.31 | -1.73 | 2.2E-08 |
| Gm2a | 1586 | 1.89 | 0.92 | 4.4E-28 | 1.92 | 0.94 | 3.2E-29 |
| Gm3055 | 181 | -1.65 | -0.72 | 1.4E-06 | -1.52 | -0.60 | 7.9E-05 |
| Gm3252 | 108 | -3.00 | -1.59 | 4.3E-09 | -3.33 | -1.74 | 1.4E-10 |
| Gm32687 | 231 | -2.16 | -1.11 | 3.1E-15 | -2.18 | -1.13 | 2.3E-15 |
| Gm3468 | 248 | -2.62 | -1.39 | 6.3E-18 | -2.47 | -1.30 | 8.6E-16 |
| Gm35339 | 1143 | 1.34 | 0.42 | 1.7E-02 | 1.36 | 0.45 | 1.2E-02 |
| Gm3636 | 685 | -1.64 | -0.71 | 2.8E-12 | -1.71 | -0.78 | 2.1E-14 |
| Gm3667 | 153 | -3.80 | -1.93 | 2.8E-16 | -3.58 | -1.84 | 9.5E-15 |
| Gm3696 | 169 | -3.32 | -1.73 | 9.1E-26 | -3.10 | -1.63 | 7.4E-23 |
| Gm3739 | 365 | -1.70 | -0.77 | 3.1E-09 | -1.67 | -0.74 | 1.2E-08 |
| Gm3752 | 139 | -2.99 | -1.58 | 1.1E-08 | -3.93 | -1.97 | 6.6E-13 |
| Gm3854 | 398 | -2.85 | -1.51 | 9.7E-29 | -2.65 | -1.41 | 2.6E-25 |
| Gm42372 | 3027 | 2.41 | 1.27 | 4.9E-34 | 2.40 | 1.26 | 1.0E-33 |
| Gm42688 | 115 | 1.55 | 0.63 | 2.9E-04 | 1.63 | 0.71 | 5.1E-05 |
| Gm42715 | 288 | -2.72 | -1.44 | 3.8E-07 | -3.55 | -1.83 | 8.3E-11 |
| Gm45902 | 1531 | 1.21 | 0.27 | 3.1E-03 | 1.31 | 0.39 | 2.1E-05 |
| Gm4924 | 793 | -2.05 | -1.03 | 8.3E-16 | -2.03 | -1.02 | 1.6E-15 |
| Gm49322 | 172 | -1.49 | -0.58 | 2.5E-03 | -1.68 | -0.75 | 6.3E-05 |
| Gm49336 | 7744 | 1.34 | 0.42 | 4.1E-06 | 1.36 | 0.44 | 1.1E-06 |
| Gm49396 | 191 | -2.29 | -1.19 | 9.4E-14 | -2.51 | -1.33 | 1.4E-16 |
| Gm50387 | 243 | 1.54 | 0.62 | 2.1E-05 | 1.45 | 0.54 | 2.4E-04 |

| | | | | | | | |
|---|---|---|---|---|---|---|---|
| Gm5148 | 214 | 1.38 | 0.47 | 2.6E-02 | 1.49 | 0.58 | 5.2E-03 |
| Gm6169 | 103 | -1.57 | -0.65 | 1.6E-03 | -1.74 | -0.80 | 8.9E-05 |
| Gm7072 | 2178 | -1.27 | -0.35 | 1.5E-05 | -1.20 | -0.26 | 1.6E-03 |
| Gm7694 | 1043 | 1.48 | 0.57 | 2.2E-05 | 1.30 | 0.38 | 5.8E-03 |
| Gm8113 | 439 | -1.77 | -0.83 | 4.1E-08 | -1.39 | -0.48 | 2.4E-03 |
| Gm8251 | 473 | 4.56 | 2.19 | 3.6E-16 | 6.85 | 2.78 | 3.6E-25 |
| Gm8281 | 342 | -2.96 | -1.56 | 3.2E-23 | -2.69 | -1.43 | 1.4E-19 |
| Gm8773 | 153 | -1.91 | -0.93 | 8.4E-15 | -1.77 | -0.83 | 7.8E-12 |
| Gm960 | 252 | 3.27 | 1.71 | 2.1E-12 | 3.13 | 1.65 | 1.4E-11 |
| Gm973 | 140 | -2.27 | -1.18 | 5.4E-08 | -1.72 | -0.78 | 4.1E-04 |
| Gm9833 | 2800 | -1.48 | -0.57 | 2.4E-08 | -1.43 | -0.51 | 5.7E-07 |
| Gmeb1 | 3965 | -1.74 | -0.80 | 5.1E-33 | -1.80 | -0.85 | 5.8E-37 |
| Gmeb2 | 2345 | -2.62 | -1.39 | 1.7E-34 | -2.80 | -1.48 | 2.9E-39 |
| Gmfg | 119 | -1.36 | -0.45 | 2.2E-02 | -1.34 | -0.42 | 3.3E-02 |
| Gmnc | 454 | -5.89 | -2.56 | 6.1E-21 | -6.17 | -2.62 | 5.6E-22 |
| Gmppb | 1101 | 1.48 | 0.57 | 1.9E-08 | 1.51 | 0.60 | 3.3E-09 |
| Gmpr | 738 | 2.05 | 1.04 | 1.2E-19 | 2.01 | 1.00 | 2.0E-18 |
| Gmpr2 | 2079 | -1.55 | -0.63 | 1.2E-15 | -1.55 | -0.63 | 1.9E-15 |
| Gna11 | 6584 | 1.62 | 0.70 | 1.0E-16 | 1.47 | 0.56 | 3.9E-11 |
| Gna12 | 7711 | 1.25 | 0.32 | 8.8E-04 | 1.17 | 0.22 | 2.8E-02 |
| Gnai1 | 4191 | 1.42 | 0.50 | 4.4E-03 | 1.43 | 0.51 | 3.8E-03 |
| Gnai2 | 27733 | 1.59 | 0.67 | 1.2E-07 | 1.42 | 0.50 | 9.8E-05 |
| Gnai3 | 20244 | 1.24 | 0.31 | 1.6E-03 | 1.27 | 0.34 | 4.6E-04 |
| Gnaq | 15834 | -1.32 | -0.40 | 4.5E-03 | -1.38 | -0.46 | 1.1E-03 |
| Gnaz | 1957 | 1.67 | 0.74 | 1.4E-06 | 1.64 | 0.71 | 3.2E-06 |
| Gnb1l | 483 | 1.57 | 0.65 | 2.1E-15 | 1.60 | 0.68 | 1.3E-16 |
| Gnb2 | 12033 | -1.18 | -0.24 | 2.2E-02 | -1.26 | -0.33 | 1.2E-03 |
| Gnb4 | 3977 | 1.21 | 0.27 | 2.8E-02 | 1.32 | 0.40 | 1.1E-03 |
| Gnb5 | 2854 | 2.17 | 1.12 | 1.5E-102 | 2.09 | 1.07 | 2.3E-93 |
| Gne | 1958 | 1.21 | 0.27 | 4.3E-03 | 1.29 | 0.36 | 9.5E-05 |
| Gng11 | 698 | -1.75 | -0.80 | 1.1E-12 | -1.84 | -0.88 | 5.4E-15 |
| Gng12 | 3927 | 1.73 | 0.79 | 6.0E-11 | 1.76 | 0.82 | 1.5E-11 |
| Gng2 | 25537 | -1.38 | -0.47 | 5.3E-03 | -1.38 | -0.46 | 6.3E-03 |
| Gng4 | 880 | 1.56 | 0.64 | 4.9E-02 | 1.66 | 0.74 | 2.2E-02 |
| Gng7 | 1266 | 1.64 | 0.71 | 3.6E-10 | 1.64 | 0.71 | 3.2E-10 |
| Gnl1 | 5018 | 1.61 | 0.68 | 7.8E-08 | 1.45 | 0.54 | 3.1E-05 |
| Gnmt | 141 | 4.21 | 2.07 | 7.8E-35 | 4.20 | 2.07 | 3.1E-34 |
| Gnptab | 4426 | 1.25 | 0.32 | 3.1E-03 | 1.29 | 0.37 | 6.9E-04 |
| Gnrh1 | 223 | 2.28 | 1.19 | 1.3E-13 | 2.09 | 1.07 | 3.5E-11 |
| Gns | 7927 | 1.66 | 0.73 | 1.2E-17 | 1.62 | 0.70 | 4.2E-16 |
| Golga1 | 4085 | -1.53 | -0.62 | 4.5E-16 | -1.45 | -0.54 | 2.2E-12 |
| Golgb1 | 13900 | 1.52 | 0.60 | 2.2E-04 | 1.80 | 0.85 | 1.1E-07 |
| Golm2 | 8324 | 1.42 | 0.51 | 2.5E-08 | 1.35 | 0.43 | 3.1E-06 |
| Golph3l | 3575 | 1.31 | 0.39 | 1.2E-05 | 1.41 | 0.50 | 2.5E-08 |
| Gosr1 | 2574 | -1.57 | -0.65 | 7.1E-28 | -1.66 | -0.73 | 3.5E-34 |
| Got1 | 4602 | 2.17 | 1.12 | 1.3E-48 | 2.01 | 1.00 | 1.7E-39 |
| Gpaa1 | 3100 | 1.31 | 0.39 | 2.1E-05 | 1.28 | 0.36 | 1.0E-04 |

| | | | | | | | |
|---|---|---|---|---|---|---|---|
| Gpam | 5132 | 1.37 | 0.45 | 2.1E-14 | 1.40 | 0.48 | 3.9E-16 |
| Gpank1 | 1132 | 1.40 | 0.49 | 2.6E-06 | 1.29 | 0.37 | 5.6E-04 |
| Gpat4 | 12130 | 1.28 | 0.36 | 1.3E-15 | 1.22 | 0.28 | 4.7E-10 |
| Gpatch3 | 462 | -1.23 | -0.30 | 1.6E-02 | -1.39 | -0.48 | 6.7E-05 |
| Gpc1 | 11787 | -1.31 | -0.39 | 3.3E-03 | -1.47 | -0.56 | 1.5E-05 |
| Gpcpd1 | 5896 | -1.55 | -0.63 | 1.3E-09 | -1.47 | -0.55 | 1.5E-07 |
| Gpd1l | 3737 | 1.89 | 0.91 | 1.0E-31 | 1.88 | 0.91 | 1.2E-31 |
| Gper1 | 147 | -1.95 | -0.97 | 2.4E-04 | -2.02 | -1.02 | 1.1E-04 |
| Gpkow | 5534 | 1.29 | 0.37 | 2.7E-12 | 1.32 | 0.40 | 2.9E-14 |
| Gpn2 | 827 | 1.26 | 0.34 | 3.1E-03 | 1.18 | 0.24 | 4.2E-02 |
| Gpn3 | 1664 | 1.25 | 0.32 | 5.1E-04 | 1.18 | 0.24 | 8.9E-03 |
| Gpr135 | 446 | 1.62 | 0.69 | 7.2E-07 | 1.74 | 0.80 | 9.0E-09 |
| Gpr137c | 3083 | 1.35 | 0.43 | 1.6E-04 | 1.29 | 0.37 | 1.5E-03 |
| Gpr153 | 1775 | -1.27 | -0.34 | 3.6E-02 | -1.37 | -0.46 | 4.8E-03 |
| Gpr155 | 1773 | 2.54 | 1.35 | 2.4E-23 | 2.85 | 1.51 | 5.2E-29 |
| Gpr157 | 312 | 2.70 | 1.43 | 1.1E-13 | 2.56 | 1.35 | 2.7E-12 |
| Gpr161 | 7389 | -1.59 | -0.67 | 3.4E-05 | -1.61 | -0.69 | 1.9E-05 |
| Gpr162 | 3077 | -1.63 | -0.70 | 6.0E-03 | -1.68 | -0.75 | 3.5E-03 |
| Gpr165 | 242 | -1.59 | -0.67 | 1.2E-02 | -1.49 | -0.57 | 3.5E-02 |
| Gpr17 | 126 | -7.49 | -2.90 | 4.6E-21 | -6.79 | -2.76 | 4.2E-19 |
| Gpr180 | 1468 | 1.40 | 0.49 | 5.6E-09 | 1.38 | 0.46 | 3.8E-08 |
| Gpr21 | 504 | -4.20 | -2.07 | 6.5E-12 | -4.37 | -2.13 | 1.8E-12 |
| Gpr27 | 1524 | -1.63 | -0.71 | 1.1E-02 | -1.56 | -0.64 | 2.3E-02 |
| Gpr3 | 118 | 8.60 | 3.10 | 1.4E-40 | 12.12 | 3.60 | 3.2E-50 |
| Gpr35 | 178 | 3.29 | 1.72 | 1.0E-14 | 2.91 | 1.54 | 3.8E-12 |
| Gpr4 | 302 | 3.41 | 1.77 | 7.1E-31 | 3.56 | 1.83 | 8.0E-33 |
| Gpr45 | 762 | -1.90 | -0.93 | 3.8E-12 | -1.78 | -0.83 | 5.2E-10 |
| Gpr50 | 342 | 2.18 | 1.12 | 1.7E-05 | 1.56 | 0.64 | 1.9E-02 |
| Gpr88 | 221 | -2.28 | -1.19 | 3.7E-03 | -2.10 | -1.07 | 9.6E-03 |
| Gprc5a | 115 | 1.85 | 0.89 | 3.1E-03 | 1.75 | 0.81 | 7.8E-03 |
| Gprin2 | 3251 | 1.58 | 0.66 | 1.9E-03 | 1.67 | 0.74 | 4.2E-04 |
| Gpsm1 | 8087 | -1.40 | -0.49 | 1.5E-04 | -1.48 | -0.57 | 7.8E-06 |
| Gpt | 1344 | 1.71 | 0.78 | 1.5E-07 | 1.64 | 0.71 | 2.0E-06 |
| Gpt2 | 4155 | 1.87 | 0.91 | 3.4E-42 | 1.80 | 0.84 | 1.0E-36 |
| Gpx7 | 796 | -2.06 | -1.04 | 1.7E-05 | -2.19 | -1.13 | 2.8E-06 |
| Gramd1a | 13884 | -1.45 | -0.54 | 2.0E-05 | -1.46 | -0.54 | 1.8E-05 |
| Gramd1c | 308 | -1.75 | -0.81 | 2.0E-06 | -1.71 | -0.77 | 6.8E-06 |
| Gramd2 | 314 | 2.60 | 1.38 | 1.8E-05 | 2.76 | 1.47 | 4.8E-06 |
| Gramd3 | 1346 | 1.66 | 0.73 | 2.8E-05 | 1.69 | 0.76 | 1.4E-05 |
| Gramd4 | 4172 | -1.25 | -0.32 | 1.7E-02 | -1.29 | -0.37 | 5.3E-03 |
| Grap2 | 239 | -2.41 | -1.27 | 2.8E-03 | -2.39 | -1.25 | 3.3E-03 |
| Grb10 | 45862 | -1.51 | -0.59 | 5.4E-03 | -1.49 | -0.58 | 6.7E-03 |
| Grb14 | 635 | 1.47 | 0.55 | 2.9E-03 | 1.68 | 0.75 | 4.1E-05 |
| Grb2 | 9573 | 1.44 | 0.53 | 5.9E-08 | 1.37 | 0.46 | 3.4E-06 |
| Grcc10 | 360 | -1.82 | -0.86 | 1.6E-13 | -1.80 | -0.85 | 3.3E-13 |
| Grem2 | 471 | -2.21 | -1.14 | 6.5E-04 | -1.76 | -0.81 | 1.9E-02 |
| Grhl1 | 214 | 1.92 | 0.94 | 2.5E-03 | 1.97 | 0.98 | 1.7E-03 |

| | | | | | | | |
|---|---|---|---|---|---|---|---|
| Grhpr | 879 | 1.38 | 0.46 | 3.4E-03 | 1.30 | 0.38 | 1.7E-02 |
| Grid2ip | 169 | 2.70 | 1.43 | 6.3E-24 | 3.40 | 1.76 | 1.4E-34 |
| Grik4 | 520 | -1.44 | -0.53 | 1.2E-03 | -1.38 | -0.47 | 4.7E-03 |
| Grik5 | 7313 | -1.31 | -0.39 | 2.2E-02 | -1.40 | -0.49 | 3.6E-03 |
| Grin2a | 270 | -2.28 | -1.19 | 1.5E-08 | -2.27 | -1.18 | 2.0E-08 |
| Grk1 | 127 | 6.57 | 2.72 | 2.6E-52 | 7.74 | 2.95 | 6.3E-57 |
| Grk2 | 4821 | -1.39 | -0.48 | 4.8E-06 | -1.50 | -0.58 | 1.5E-08 |
| Grk5 | 2884 | 1.76 | 0.81 | 2.1E-52 | 1.79 | 0.84 | 9.1E-56 |
| Grk6 | 6916 | 1.34 | 0.42 | 3.6E-04 | 1.25 | 0.32 | 7.9E-03 |
| Grm2 | 321 | -1.39 | -0.47 | 2.8E-02 | -1.49 | -0.57 | 7.1E-03 |
| Grm3 | 394 | -1.84 | -0.88 | 2.7E-03 | -1.59 | -0.67 | 2.7E-02 |
| Grn | 4795 | 1.71 | 0.77 | 8.7E-09 | 1.61 | 0.69 | 3.9E-07 |
| Grpel1 | 3863 | 1.42 | 0.51 | 1.0E-05 | 1.39 | 0.48 | 3.6E-05 |
| Grtp1 | 1292 | 2.34 | 1.22 | 6.8E-57 | 2.28 | 1.19 | 8.5E-54 |
| Gsap | 411 | -2.09 | -1.07 | 1.1E-05 | -1.84 | -0.88 | 3.5E-04 |
| Gsdmd | 232 | -2.82 | -1.49 | 1.4E-23 | -2.57 | -1.36 | 1.3E-19 |
| Gsdme | 636 | -2.27 | -1.19 | 1.8E-24 | -2.23 | -1.16 | 2.1E-23 |
| Gse1 | 7420 | -1.26 | -0.34 | 4.5E-02 | -1.36 | -0.44 | 7.4E-03 |
| Gspt2 | 1761 | 1.39 | 0.47 | 2.3E-07 | 1.36 | 0.44 | 1.6E-06 |
| Gsr | 1705 | -1.55 | -0.63 | 5.0E-08 | -1.64 | -0.71 | 7.0E-10 |
| Gsta4 | 2086 | 2.41 | 1.27 | 1.7E-10 | 2.19 | 1.13 | 1.4E-08 |
| Gstk1 | 486 | 1.46 | 0.54 | 9.9E-05 | 1.61 | 0.69 | 5.3E-07 |
| Gsto1 | 1957 | 1.60 | 0.68 | 4.5E-21 | 1.57 | 0.65 | 4.3E-19 |
| Gstt1 | 316 | 1.80 | 0.84 | 5.0E-06 | 1.62 | 0.70 | 1.9E-04 |
| Gstt2 | 285 | 1.88 | 0.91 | 1.8E-08 | 1.62 | 0.70 | 2.1E-05 |
| Gstt3 | 130 | 2.88 | 1.53 | 1.2E-03 | 2.60 | 1.38 | 3.7E-03 |
| Gstz1 | 2191 | -1.43 | -0.52 | 3.4E-13 | -1.39 | -0.48 | 2.6E-11 |
| Gsx1 | 239 | -2.00 | -1.00 | 7.3E-04 | -2.44 | -1.29 | 1.0E-05 |
| Gtf2a1 | 4806 | -1.42 | -0.51 | 1.0E-03 | -1.64 | -0.71 | 2.7E-06 |
| Gtf2e2 | 5060 | 1.57 | 0.65 | 1.3E-17 | 1.58 | 0.66 | 5.0E-18 |
| Gtf2f2 | 2453 | 1.52 | 0.61 | 1.9E-09 | 1.54 | 0.63 | 5.5E-10 |
| Gtf2h1 | 4471 | 1.24 | 0.31 | 6.4E-04 | 1.25 | 0.32 | 4.0E-04 |
| Gtf2h2 | 2883 | -1.28 | -0.36 | 1.4E-05 | -1.30 | -0.38 | 4.2E-06 |
| Gtf2i | 23958 | -2.36 | -1.24 | 1.2E-34 | -2.54 | -1.35 | 8.0E-41 |
| Gtf2ird1 | 4286 | -1.54 | -0.62 | 5.8E-18 | -1.67 | -0.74 | 1.1E-24 |
| Gtf2ird2 | 1253 | -2.52 | -1.33 | 7.4E-30 | -2.63 | -1.40 | 1.1E-32 |
| Gtf3c3 | 3562 | -1.90 | -0.93 | 1.3E-31 | -1.85 | -0.89 | 4.4E-29 |
| Gtf3c4 | 12558 | 1.31 | 0.39 | 7.5E-07 | 1.37 | 0.46 | 5.2E-09 |
| Gtf3c5 | 3162 | -2.05 | -1.03 | 2.9E-32 | -2.17 | -1.12 | 2.4E-37 |
| Gtf3c6 | 3797 | -2.77 | -1.47 | 7.5E-40 | -2.96 | -1.57 | 2.8E-45 |
| Gtpbp1 | 6758 | 1.33 | 0.41 | 2.1E-09 | 1.22 | 0.29 | 3.2E-05 |
| Gtpbp10 | 2227 | -1.28 | -0.36 | 8.1E-04 | -1.30 | -0.38 | 3.6E-04 |
| Gtpbp3 | 2274 | 2.13 | 1.09 | 1.1E-20 | 1.92 | 0.94 | 1.2E-15 |
| Gtpbp8 | 2089 | -1.39 | -0.48 | 1.3E-07 | -1.36 | -0.45 | 7.8E-07 |
| Gucy1b1 | 2626 | 2.27 | 1.18 | 5.9E-13 | 2.44 | 1.28 | 4.3E-15 |
| Gucy1b2 | 147 | -2.85 | -1.51 | 1.0E-07 | -2.64 | -1.40 | 1.0E-06 |
| Guf1 | 7414 | -1.39 | -0.47 | 1.2E-09 | -1.27 | -0.34 | 1.5E-05 |

| | | | | | | | |
|---|---|---|---|---|---|---|---|
| Gulp1 | 2021 | -1.45 | -0.53 | 1.4E-02 | -1.46 | -0.55 | 1.2E-02 |
| Gusb | 1547 | -1.38 | -0.46 | 7.6E-04 | -1.43 | -0.52 | 1.5E-04 |
| Gxylt1 | 7302 | 1.48 | 0.57 | 2.0E-07 | 1.55 | 0.63 | 8.2E-09 |
| Gypc | 922 | -2.16 | -1.11 | 8.9E-08 | -2.21 | -1.14 | 3.7E-08 |
| Gzf1 | 4642 | 1.56 | 0.64 | 4.8E-14 | 1.56 | 0.64 | 8.2E-14 |
| Gzmm | 159 | 2.08 | 1.06 | 2.3E-13 | 1.99 | 1.00 | 5.4E-12 |
| H1f0 | 57016 | 3.20 | 1.68 | 1.7E-28 | 2.83 | 1.50 | 5.8E-23 |
| H1f10 | 4974 | 1.70 | 0.77 | 2.0E-05 | 1.45 | 0.53 | 3.9E-03 |
| H1f2 | 993 | 3.77 | 1.91 | 2.1E-31 | 4.02 | 2.01 | 2.3E-34 |
| H2-Ab1 | 245 | 17.04 | 4.09 | 1.2E-73 | 13.41 | 3.75 | 4.0E-63 |
| H2ac10 | 154 | -1.57 | -0.65 | 9.6E-03 | -1.67 | -0.74 | 3.2E-03 |
| H2aj | 2580 | 1.93 | 0.95 | 9.2E-18 | 1.75 | 0.81 | 3.1E-13 |
| H2bc6 | 131 | 2.08 | 1.06 | 5.7E-12 | 1.99 | 0.99 | 9.6E-11 |
| H2bu2 | 3332 | -1.35 | -0.43 | 1.4E-03 | -1.33 | -0.41 | 2.7E-03 |
| H2-D1 | 2189 | 3.05 | 1.61 | 5.7E-53 | 2.71 | 1.44 | 2.2E-42 |
| H2-DMa | 345 | 2.02 | 1.01 | 2.1E-30 | 1.92 | 0.94 | 6.9E-26 |
| H2-K1 | 1897 | 5.53 | 2.47 | 1.4E-29 | 4.36 | 2.12 | 3.1E-22 |
| H2-M5 | 875 | -1.67 | -0.74 | 1.4E-04 | -1.69 | -0.76 | 8.3E-05 |
| H2-Q4 | 756 | 3.41 | 1.77 | 3.7E-18 | 2.61 | 1.38 | 1.8E-11 |
| H2-Q6 | 102 | 12.35 | 3.63 | 1.1E-43 | 7.20 | 2.85 | 1.1E-30 |
| H2-Q7 | 283 | 3.27 | 1.71 | 3.3E-13 | 2.63 | 1.40 | 3.5E-09 |
| H2-T23 | 263 | -1.29 | -0.36 | 3.2E-02 | -1.31 | -0.39 | 2.1E-02 |
| H3f3b | 103547 | 1.44 | 0.53 | 3.5E-13 | 1.43 | 0.51 | 1.9E-12 |
| H60b | 205 | 2.00 | 1.00 | 1.5E-05 | 2.09 | 1.07 | 4.1E-06 |
| H6pd | 838 | -1.60 | -0.68 | 3.9E-03 | -1.68 | -0.75 | 1.4E-03 |
| Habp4 | 3625 | 2.54 | 1.34 | 4.3E-40 | 2.51 | 1.33 | 4.3E-39 |
| Hacd1 | 399 | -2.20 | -1.14 | 5.1E-16 | -2.02 | -1.01 | 6.5E-13 |
| Hacd2 | 1259 | 1.75 | 0.81 | 1.3E-31 | 1.52 | 0.60 | 3.5E-18 |
| Hadh | 4271 | 1.88 | 0.91 | 1.5E-08 | 1.80 | 0.84 | 1.9E-07 |
| Hadha | 10854 | 1.47 | 0.55 | 1.6E-09 | 1.45 | 0.54 | 5.1E-09 |
| Hadhb | 4314 | 1.64 | 0.72 | 2.3E-08 | 1.65 | 0.72 | 1.9E-08 |
| Hagh | 1445 | 2.73 | 1.45 | 1.7E-84 | 2.72 | 1.44 | 1.9E-83 |
| Hap1 | 12514 | 1.96 | 0.97 | 2.8E-08 | 1.72 | 0.78 | 1.1E-05 |
| Hapln1 | 1964 | -4.31 | -2.11 | 1.6E-05 | -3.45 | -1.79 | 3.0E-04 |
| Hapln4 | 275 | 27.66 | 4.79 | 3.9E-126 | 23.71 | 4.57 | 2.8E-119 |
| Has2 | 670 | -2.71 | -1.44 | 1.9E-07 | -2.01 | -1.01 | 3.6E-04 |
| Haspin | 987 | 1.46 | 0.54 | 1.2E-03 | 1.38 | 0.46 | 6.6E-03 |
| Hat1 | 8050 | 1.38 | 0.47 | 4.1E-03 | 1.32 | 0.40 | 1.6E-02 |
| Haus3 | 2447 | 1.30 | 0.38 | 5.2E-03 | 1.27 | 0.35 | 1.1E-02 |
| Haus6 | 6494 | 1.41 | 0.49 | 3.9E-04 | 1.34 | 0.42 | 3.1E-03 |
| Hax1 | 3290 | -1.22 | -0.29 | 5.4E-04 | -1.29 | -0.37 | 1.0E-05 |
| Hbegf | 484 | 1.45 | 0.53 | 3.9E-04 | 1.37 | 0.46 | 2.7E-03 |
| Hbp1 | 6532 | -1.41 | -0.50 | 1.1E-08 | -1.34 | -0.42 | 1.8E-06 |
| Hccs | 2971 | 1.30 | 0.38 | 1.0E-05 | 1.30 | 0.38 | 1.1E-05 |
| Hcfc1 | 17725 | -2.30 | -1.20 | 2.7E-29 | -2.29 | -1.20 | 4.5E-29 |
| Hcls1 | 187 | -1.39 | -0.48 | 7.2E-03 | -1.65 | -0.72 | 3.2E-05 |
| Hcn2 | 509 | 4.94 | 2.30 | 1.9E-20 | 4.59 | 2.20 | 1.0E-18 |

| | | | | | | | |
|---|---|---|---|---|---|---|---|
| Hcn3 | 2701 | 2.06 | 1.04 | 1.5E-07 | 1.98 | 0.99 | 6.8E-07 |
| Hdac1 | 12630 | 1.67 | 0.74 | 3.1E-11 | 1.53 | 0.61 | 5.4E-08 |
| Hdac10 | 1291 | -1.29 | -0.37 | 7.8E-07 | -1.28 | -0.35 | 2.2E-06 |
| Hdac11 | 4312 | 1.46 | 0.54 | 5.7E-08 | 1.49 | 0.57 | 9.5E-09 |
| Hdac3 | 9789 | 1.27 | 0.34 | 1.1E-08 | 1.19 | 0.25 | 3.6E-05 |
| Hdac5 | 8709 | 1.59 | 0.67 | 4.5E-09 | 1.52 | 0.61 | 1.4E-07 |
| Hdac6 | 10541 | 1.56 | 0.64 | 1.2E-15 | 1.43 | 0.52 | 1.7E-10 |
| Hdac7 | 4354 | -2.07 | -1.05 | 5.5E-18 | -2.11 | -1.07 | 8.5E-19 |
| Hdac9 | 1503 | -1.37 | -0.46 | 2.9E-11 | -1.28 | -0.36 | 2.2E-07 |
| Hddc3 | 865 | -1.48 | -0.57 | 3.4E-12 | -1.47 | -0.56 | 9.2E-12 |
| Hdgfl2 | 10422 | 1.68 | 0.74 | 2.0E-09 | 1.51 | 0.59 | 2.5E-06 |
| Hdgfl3 | 17974 | -1.70 | -0.77 | 8.9E-07 | -1.64 | -0.71 | 5.7E-06 |
| Hdhd3 | 360 | 1.90 | 0.93 | 5.1E-14 | 1.79 | 0.84 | 1.2E-11 |
| Hdhd5 | 1737 | 1.38 | 0.47 | 1.5E-17 | 1.37 | 0.45 | 2.7E-16 |
| Hdlbp | 25696 | -1.29 | -0.37 | 9.1E-03 | -1.25 | -0.32 | 2.5E-02 |
| Hdx | 1392 | -1.61 | -0.69 | 1.4E-07 | -1.37 | -0.45 | 7.0E-04 |
| Heatr1 | 8066 | 1.48 | 0.57 | 4.6E-05 | 1.59 | 0.67 | 1.2E-06 |
| Heatr5a | 4914 | 1.31 | 0.39 | 8.5E-04 | 1.38 | 0.46 | 5.2E-05 |
| Heatr5b | 5065 | 1.23 | 0.30 | 1.9E-02 | 1.33 | 0.41 | 1.2E-03 |
| Heatr6 | 3708 | -1.59 | -0.67 | 5.8E-16 | -1.62 | -0.69 | 6.4E-17 |
| Hebp1 | 408 | 1.58 | 0.66 | 1.8E-03 | 1.47 | 0.56 | 9.6E-03 |
| Heca | 4999 | 1.57 | 0.65 | 1.5E-10 | 1.55 | 0.63 | 7.3E-10 |
| Hectd1 | 13159 | -2.01 | -1.01 | 1.5E-13 | -1.74 | -0.80 | 6.0E-09 |
| Hectd3 | 5325 | 1.91 | 0.93 | 5.9E-29 | 1.88 | 0.91 | 2.4E-27 |
| Hecw1 | 4482 | -2.00 | -1.00 | 2.2E-03 | -1.70 | -0.76 | 2.3E-02 |
| Hecw2 | 1165 | -2.41 | -1.27 | 4.0E-22 | -2.08 | -1.05 | 1.7E-15 |
| Heg1 | 5703 | -1.59 | -0.67 | 2.3E-05 | -1.72 | -0.78 | 6.5E-07 |
| Helb | 1918 | 2.87 | 1.52 | 1.7E-93 | 2.79 | 1.48 | 2.9E-88 |
| Helq | 2316 | 1.33 | 0.41 | 1.2E-09 | 1.34 | 0.42 | 7.6E-10 |
| Helz | 6671 | 1.25 | 0.32 | 8.3E-04 | 1.29 | 0.36 | 1.3E-04 |
| Hemk1 | 1325 | -1.48 | -0.57 | 4.2E-08 | -1.41 | -0.50 | 1.9E-06 |
| Herc3 | 3540 | 1.81 | 0.85 | 6.3E-13 | 1.89 | 0.92 | 8.5E-15 |
| Herc6 | 176 | -1.28 | -0.36 | 2.0E-02 | -1.30 | -0.38 | 1.3E-02 |
| Hes1 | 1561 | -1.52 | -0.60 | 3.5E-06 | -1.70 | -0.77 | 2.6E-09 |
| Hes7 | 133 | 15.80 | 3.98 | 1.4E-59 | 12.83 | 3.68 | 2.3E-53 |
| Hexb | 1597 | 1.36 | 0.44 | 5.3E-05 | 1.40 | 0.49 | 7.9E-06 |
| Hexim2 | 362 | -1.51 | -0.59 | 6.0E-05 | -1.52 | -0.60 | 4.7E-05 |
| Heyl | 818 | -1.59 | -0.67 | 7.4E-05 | -1.61 | -0.69 | 4.4E-05 |
| Hfm1 | 1011 | -1.80 | -0.85 | 1.3E-11 | -1.50 | -0.58 | 4.4E-06 |
| Hgfac | 114 | 1.56 | 0.64 | 3.3E-04 | 1.85 | 0.89 | 4.6E-07 |
| Hhat | 198 | -4.83 | -2.27 | 1.3E-29 | -4.91 | -2.30 | 3.1E-30 |
| Hhipl1 | 540 | -1.95 | -0.96 | 1.2E-06 | -2.28 | -1.19 | 1.5E-09 |
| Hhipl2 | 159 | 3.25 | 1.70 | 6.5E-22 | 3.83 | 1.94 | 1.1E-27 |
| Hibadh | 2808 | 1.33 | 0.41 | 5.1E-10 | 1.33 | 0.41 | 7.2E-10 |
| Hibch | 2245 | 1.36 | 0.45 | 1.9E-14 | 1.45 | 0.53 | 4.5E-20 |
| Hic1 | 634 | -3.80 | -1.93 | 1.5E-08 | -3.26 | -1.70 | 6.6E-07 |
| Hid1 | 4424 | 2.44 | 1.29 | 4.4E-34 | 2.21 | 1.14 | 4.9E-27 |

| | | | | | | | |
|---|---|---|---|---|---|---|---|
| Hif1a | 12695 | -1.47 | -0.55 | 1.9E-15 | -1.47 | -0.55 | 2.7E-15 |
| Hif3a | 2892 | 1.31 | 0.39 | 1.4E-04 | 1.39 | 0.48 | 2.9E-06 |
| Higd1a | 3706 | 1.48 | 0.56 | 2.2E-10 | 1.43 | 0.52 | 7.1E-09 |
| Hinfp | 4876 | 1.51 | 0.59 | 4.6E-08 | 1.61 | 0.69 | 1.6E-10 |
| Hint3 | 411 | -1.62 | -0.70 | 7.2E-06 | -1.57 | -0.65 | 3.4E-05 |
| Hip1r | 3544 | 2.26 | 1.18 | 4.8E-18 | 2.31 | 1.21 | 6.6E-19 |
| Hipk1 | 11888 | 1.27 | 0.35 | 9.8E-06 | 1.30 | 0.38 | 1.4E-06 |
| Hipk3 | 3357 | -2.01 | -1.01 | 1.4E-18 | -1.88 | -0.91 | 2.9E-15 |
| Hirip3 | 5800 | -2.09 | -1.06 | 9.5E-10 | -2.44 | -1.29 | 8.1E-14 |
| Hivep1 | 7862 | 1.29 | 0.37 | 3.0E-03 | 1.35 | 0.43 | 4.7E-04 |
| Hjurp | 22332 | -1.52 | -0.61 | 1.0E-14 | -1.46 | -0.55 | 3.3E-12 |
| Hk1 | 8782 | 1.89 | 0.92 | 5.8E-18 | 1.85 | 0.89 | 5.1E-17 |
| Hk2 | 3714 | 2.13 | 1.09 | 1.5E-12 | 1.93 | 0.95 | 1.1E-09 |
| Hlf | 1314 | 1.69 | 0.76 | 1.5E-13 | 1.61 | 0.68 | 2.7E-11 |
| Hltf | 4546 | 1.28 | 0.36 | 8.3E-07 | 1.36 | 0.44 | 6.8E-10 |
| Hmbox1 | 6274 | -1.35 | -0.43 | 9.4E-08 | -1.29 | -0.36 | 9.1E-06 |
| Hmbs | 1550 | -1.94 | -0.95 | 1.4E-06 | -2.12 | -1.08 | 3.4E-08 |
| Hmces | 1681 | 1.31 | 0.39 | 5.1E-07 | 1.16 | 0.22 | 7.1E-03 |
| Hmg20a | 6866 | -1.82 | -0.87 | 3.0E-61 | -1.77 | -0.83 | 6.9E-56 |
| Hmg20b | 3967 | 1.66 | 0.73 | 2.1E-06 | 1.52 | 0.60 | 1.1E-04 |
| Hmgb1 | 26362 | -1.34 | -0.43 | 4.7E-03 | -1.44 | -0.52 | 4.7E-04 |
| Hmgb2 | 24394 | -1.43 | -0.52 | 1.3E-02 | -1.62 | -0.69 | 7.5E-04 |
| Hmgb3 | 18826 | -1.41 | -0.50 | 3.7E-04 | -1.60 | -0.68 | 9.1E-07 |
| Hmgcl | 2057 | 2.00 | 1.00 | 7.4E-38 | 1.85 | 0.89 | 6.3E-30 |
| Hmgcll1 | 1224 | -2.45 | -1.29 | 1.6E-23 | -2.48 | -1.31 | 4.2E-24 |
| Hmgcs2 | 417 | -2.10 | -1.07 | 1.6E-04 | -1.80 | -0.85 | 3.3E-03 |
| Hmgn2 | 31781 | -1.38 | -0.46 | 1.0E-03 | -1.56 | -0.64 | 4.3E-06 |
| Hmgn3 | 4436 | 1.56 | 0.64 | 2.0E-10 | 1.50 | 0.58 | 7.0E-09 |
| Hmgxb4 | 4217 | -1.80 | -0.84 | 3.7E-29 | -1.85 | -0.88 | 6.7E-32 |
| Hmox1 | 1128 | 1.40 | 0.49 | 5.2E-03 | 1.37 | 0.46 | 9.3E-03 |
| Hmx2 | 110 | -1.63 | -0.70 | 5.9E-04 | -1.53 | -0.62 | 3.1E-03 |
| Hnrnpa2b1 | 162044 | -1.26 | -0.33 | 5.7E-03 | -1.36 | -0.45 | 1.3E-04 |
| Hnrnpab | 47049 | -1.53 | -0.61 | 2.7E-05 | -1.75 | -0.81 | 1.5E-08 |
| Hnrnph1 | 93936 | -1.60 | -0.68 | 9.0E-14 | -1.61 | -0.68 | 5.2E-14 |
| Hnrnph3 | 14582 | -1.34 | -0.42 | 4.3E-03 | -1.53 | -0.62 | 1.8E-05 |
| Hnrnpk | 69304 | -1.30 | -0.38 | 9.0E-05 | -1.36 | -0.45 | 3.4E-06 |
| Hnrnpl | 25633 | -1.47 | -0.56 | 9.6E-09 | -1.61 | -0.68 | 1.2E-12 |
| Hnrnpll | 11084 | -1.45 | -0.54 | 5.3E-09 | -1.50 | -0.59 | 1.4E-10 |
| Hnrnpm | 33103 | -1.43 | -0.52 | 6.8E-05 | -1.62 | -0.69 | 5.7E-08 |
| Hnrnpr | 28816 | -1.23 | -0.29 | 1.6E-03 | -1.26 | -0.34 | 2.7E-04 |
| Hnrnpu | 3018 | -1.59 | -0.67 | 3.3E-04 | -1.59 | -0.67 | 3.4E-04 |
| Hnrnpul1 | 16418 | -1.25 | -0.32 | 3.5E-04 | -1.34 | -0.42 | 1.6E-06 |
| Homer1 | 5300 | 1.61 | 0.69 | 9.6E-20 | 1.63 | 0.71 | 7.6E-21 |
| Homer3 | 3874 | -1.27 | -0.35 | 1.0E-02 | -1.43 | -0.51 | 9.4E-05 |
| Homez | 1824 | 1.89 | 0.92 | 1.5E-51 | 1.70 | 0.77 | 5.8E-36 |
| Hook1 | 8347 | 1.27 | 0.34 | 6.6E-03 | 1.43 | 0.52 | 2.3E-05 |
| Hook2 | 1648 | 1.25 | 0.33 | 3.9E-03 | 1.19 | 0.26 | 2.7E-02 |

| | | | | | | | |
|---|---|---|---|---|---|---|---|
| Hp1bp3 | 20976 | -1.82 | -0.86 | 8.3E-50 | -1.88 | -0.91 | 3.2E-55 |
| Hpcal1 | 1994 | 1.74 | 0.80 | 9.1E-29 | 1.79 | 0.84 | 2.5E-31 |
| Hpgds | 611 | -1.44 | -0.52 | 1.8E-03 | -1.33 | -0.41 | 1.8E-02 |
| Hps1 | 1466 | -1.36 | -0.45 | 1.8E-05 | -1.44 | -0.53 | 3.7E-07 |
| Hps3 | 2160 | 1.39 | 0.48 | 5.0E-09 | 1.44 | 0.53 | 9.8E-11 |
| Hps4 | 2656 | 1.34 | 0.43 | 2.4E-06 | 1.34 | 0.43 | 2.5E-06 |
| Hps6 | 307 | 2.52 | 1.33 | 7.4E-15 | 2.41 | 1.27 | 1.7E-13 |
| Hrh1 | 125 | 12.12 | 3.60 | 1.8E-67 | 13.08 | 3.71 | 4.0E-67 |
| Hrh3 | 823 | 2.58 | 1.37 | 5.1E-15 | 2.34 | 1.23 | 2.6E-12 |
| Hs1bp3 | 1314 | 1.47 | 0.56 | 9.1E-07 | 1.41 | 0.49 | 1.9E-05 |
| Hs2st1 | 12086 | 1.56 | 0.64 | 2.4E-17 | 1.47 | 0.56 | 2.2E-13 |
| Hs3st3a1 | 299 | -1.68 | -0.75 | 1.0E-02 | -1.70 | -0.76 | 9.7E-03 |
| Hs6st1 | 7939 | 1.60 | 0.68 | 1.0E-07 | 1.48 | 0.56 | 1.4E-05 |
| Hs6st2 | 2902 | -1.33 | -0.41 | 3.6E-03 | -1.36 | -0.44 | 1.6E-03 |
| Hscb | 393 | 1.64 | 0.71 | 1.4E-08 | 1.48 | 0.57 | 7.9E-06 |
| Hsd17b11 | 777 | -1.30 | -0.37 | 8.4E-03 | -1.36 | -0.45 | 1.5E-03 |
| Hsd17b7 | 3103 | 1.90 | 0.93 | 6.8E-16 | 1.94 | 0.96 | 7.5E-17 |
| Hsdl1 | 7041 | 1.38 | 0.46 | 5.2E-05 | 1.32 | 0.41 | 4.4E-04 |
| Hsdl2 | 3424 | 2.16 | 1.11 | 6.0E-73 | 2.31 | 1.21 | 2.4E-86 |
| Hspa12b | 1268 | -2.05 | -1.04 | 2.3E-06 | -2.16 | -1.11 | 4.2E-07 |
| Hspa14 | 3991 | 1.65 | 0.72 | 2.6E-10 | 1.54 | 0.62 | 5.6E-08 |
| Hspa1a | 235 | 6.27 | 2.65 | 1.4E-31 | 5.59 | 2.48 | 6.7E-28 |
| Hspa1b | 412 | 5.93 | 2.57 | 3.1E-35 | 5.20 | 2.38 | 1.9E-30 |
| Hspa2 | 996 | 6.34 | 2.66 | 1.0E-96 | 6.09 | 2.61 | 1.8E-92 |
| Hspa4 | 35409 | 1.28 | 0.35 | 8.5E-07 | 1.24 | 0.31 | 1.6E-05 |
| Hspbap1 | 1511 | 1.91 | 0.93 | 6.4E-44 | 1.80 | 0.85 | 1.0E-36 |
| Hspbp1 | 1929 | -1.19 | -0.25 | 3.6E-03 | -1.27 | -0.35 | 3.4E-05 |
| Hsph1 | 17576 | 1.48 | 0.56 | 1.2E-13 | 1.41 | 0.50 | 6.5E-11 |
| Htr2a | 167 | -1.63 | -0.70 | 9.8E-04 | -1.59 | -0.67 | 1.8E-03 |
| Htra2 | 2586 | 1.64 | 0.72 | 8.4E-15 | 1.48 | 0.57 | 1.1E-09 |
| Hunk | 4047 | -1.60 | -0.68 | 3.0E-12 | -1.72 | -0.78 | 5.3E-16 |
| Hvcn1 | 308 | -3.11 | -1.64 | 7.2E-13 | -3.41 | -1.77 | 7.8E-15 |
| Hyal1 | 396 | 1.50 | 0.58 | 6.6E-05 | 1.28 | 0.36 | 1.7E-02 |
| Hykk | 1143 | 1.83 | 0.87 | 4.0E-14 | 1.82 | 0.87 | 4.9E-14 |
| Hyou1 | 13989 | 1.52 | 0.60 | 2.3E-10 | 1.45 | 0.53 | 2.4E-08 |
| Hypk | 437 | 1.72 | 0.78 | 1.4E-14 | 1.71 | 0.78 | 3.3E-14 |
| Iba57 | 547 | -2.57 | -1.36 | 2.7E-23 | -2.68 | -1.42 | 3.0E-25 |
| Ibsp | 120 | -8.11 | -3.02 | 2.3E-02 | -7.07 | -2.82 | 3.6E-02 |
| Ibtk | 6460 | -1.63 | -0.71 | 2.9E-33 | -1.60 | -0.67 | 3.8E-30 |
| Ica1 | 848 | -1.41 | -0.49 | 2.3E-04 | -1.30 | -0.38 | 5.0E-03 |
| Ica1l | 573 | -2.37 | -1.25 | 1.1E-03 | -2.19 | -1.13 | 3.3E-03 |
| Icam2 | 259 | -2.22 | -1.15 | 1.1E-10 | -2.23 | -1.16 | 7.9E-11 |
| Icam5 | 104 | 16.47 | 4.04 | 6.4E-39 | 16.73 | 4.06 | 3.0E-37 |
| Ice2 | 2118 | -1.69 | -0.76 | 1.2E-11 | -1.47 | -0.56 | 8.2E-07 |
| Icmt | 4406 | 1.33 | 0.41 | 1.3E-08 | 1.30 | 0.38 | 1.6E-07 |
| Icosl | 192 | 1.48 | 0.57 | 9.5E-03 | 1.45 | 0.54 | 1.4E-02 |
| Id2 | 3994 | -1.42 | -0.51 | 1.7E-03 | -1.33 | -0.41 | 1.2E-02 |

| | | | | | | | |
|---|---|---|---|---|---|---|---|
| Id3 | 3101 | -1.50 | -0.58 | 4.0E-02 | -1.76 | -0.82 | 3.3E-03 |
| Id4 | 21596 | -1.43 | -0.52 | 3.5E-02 | -1.49 | -0.58 | 1.8E-02 |
| Idh2 | 9459 | -1.21 | -0.27 | 2.6E-03 | -1.29 | -0.37 | 3.2E-05 |
| Idh3a | 7492 | 1.95 | 0.96 | 7.7E-63 | 1.87 | 0.90 | 4.7E-55 |
| Idh3g | 5378 | -1.25 | -0.32 | 1.0E-06 | -1.28 | -0.36 | 2.1E-08 |
| Idi1 | 5613 | 1.48 | 0.56 | 3.6E-03 | 1.34 | 0.42 | 3.4E-02 |
| Idnk | 2715 | 2.11 | 1.08 | 3.9E-38 | 2.02 | 1.02 | 3.9E-34 |
| Idua | 719 | -1.59 | -0.67 | 9.4E-12 | -1.74 | -0.80 | 3.1E-16 |
| Ier2 | 1311 | 1.69 | 0.76 | 8.4E-07 | 1.61 | 0.69 | 9.0E-06 |
| Ier3 | 242 | 2.27 | 1.18 | 2.6E-17 | 2.36 | 1.24 | 6.6E-19 |
| Ier3ip1 | 3017 | 1.24 | 0.31 | 6.8E-04 | 1.29 | 0.36 | 7.1E-05 |
| Ier5 | 3457 | 1.29 | 0.36 | 5.8E-03 | 1.21 | 0.28 | 4.0E-02 |
| Iffo1 | 2039 | 1.39 | 0.47 | 8.9E-05 | 1.29 | 0.37 | 3.2E-03 |
| Iffo2 | 1872 | 1.96 | 0.97 | 1.8E-17 | 1.89 | 0.92 | 1.1E-15 |
| Ifi27 | 240 | -2.18 | -1.12 | 8.8E-05 | -2.21 | -1.14 | 6.5E-05 |
| Ifit2 | 673 | -1.91 | -0.93 | 9.0E-14 | -1.89 | -0.91 | 2.5E-13 |
| Ifitm1 | 167 | -1.64 | -0.72 | 1.6E-02 | -1.54 | -0.63 | 3.8E-02 |
| Ifnar2 | 657 | -2.25 | -1.17 | 9.4E-56 | -2.45 | -1.30 | 4.4E-67 |
| Ifngr1 | 1370 | 1.49 | 0.58 | 2.2E-06 | 1.43 | 0.51 | 3.2E-05 |
| Ifngr2 | 2471 | 1.40 | 0.48 | 2.6E-08 | 1.39 | 0.47 | 5.9E-08 |
| Ifrd1 | 4234 | 1.27 | 0.35 | 2.9E-03 | 1.20 | 0.26 | 2.7E-02 |
| Ift122 | 2446 | -1.72 | -0.78 | 5.9E-67 | -1.69 | -0.76 | 2.2E-62 |
| Ift140 | 5845 | 1.69 | 0.76 | 1.8E-25 | 1.71 | 0.78 | 1.4E-26 |
| Ift20 | 3147 | 1.53 | 0.61 | 1.1E-28 | 1.49 | 0.57 | 3.6E-25 |
| Ift22 | 2643 | 1.41 | 0.50 | 3.1E-13 | 1.30 | 0.38 | 2.9E-08 |
| Ift27 | 2598 | 1.60 | 0.68 | 1.0E-09 | 1.41 | 0.50 | 1.0E-05 |
| Ift52 | 2607 | -1.36 | -0.44 | 2.2E-07 | -1.45 | -0.54 | 2.7E-10 |
| Ift57 | 2127 | 1.82 | 0.86 | 8.1E-56 | 1.81 | 0.85 | 1.3E-54 |
| Ift80 | 4843 | 1.51 | 0.59 | 5.6E-13 | 1.63 | 0.70 | 6.9E-18 |
| Igdcc4 | 4547 | -1.87 | -0.90 | 4.1E-16 | -1.81 | -0.86 | 1.7E-14 |
| Igf1 | 2781 | -3.30 | -1.72 | 8.9E-04 | -3.18 | -1.67 | 1.3E-03 |
| Igf2 | 1793 | -2.59 | -1.37 | 1.7E-03 | -2.82 | -1.50 | 6.0E-04 |
| Igf2bp2 | 7288 | -1.42 | -0.51 | 7.5E-04 | -1.59 | -0.67 | 6.5E-06 |
| Igf2r | 18313 | 1.42 | 0.51 | 2.4E-03 | 1.64 | 0.72 | 1.4E-05 |
| Igfbp3 | 1355 | -2.79 | -1.48 | 6.3E-05 | -2.34 | -1.23 | 1.1E-03 |
| Igfbp5 | 21794 | -2.57 | -1.36 | 5.4E-09 | -2.37 | -1.24 | 1.2E-07 |
| Ighmbp2 | 1000 | -1.70 | -0.76 | 1.7E-15 | -1.80 | -0.85 | 8.5E-19 |
| Igsf3 | 10994 | 1.44 | 0.52 | 1.8E-05 | 1.51 | 0.60 | 9.0E-07 |
| Igsf8 | 7797 | 1.67 | 0.74 | 5.1E-07 | 1.59 | 0.67 | 7.6E-06 |
| Igsf9b | 7558 | 1.92 | 0.94 | 1.2E-09 | 1.97 | 0.98 | 2.5E-10 |
| Iho1 | 199 | 1.40 | 0.49 | 1.0E-04 | 1.22 | 0.29 | 2.7E-02 |
| Ikbip | 2039 | 1.45 | 0.53 | 2.7E-04 | 1.58 | 0.66 | 4.7E-06 |
| Ikbkb | 7244 | 1.40 | 0.48 | 4.4E-14 | 1.45 | 0.54 | 5.9E-17 |
| Ikbkg | 3453 | -1.29 | -0.37 | 7.3E-05 | -1.22 | -0.29 | 2.3E-03 |
| Ikzf1 | 271 | -1.58 | -0.66 | 1.5E-05 | -1.59 | -0.67 | 1.2E-05 |
| Ikzf4 | 1884 | -1.58 | -0.66 | 3.7E-04 | -1.68 | -0.75 | 5.0E-05 |
| Il10rb | 191 | -1.99 | -0.99 | 1.2E-12 | -2.03 | -1.02 | 5.0E-13 |

| | | | | | | | |
|---|---|---|---|---|---|---|---|
| Il16 | 121 | -2.78 | -1.47 | 2.0E-07 | -2.40 | -1.27 | 1.1E-05 |
| Il17d | 192 | -1.77 | -0.82 | 8.3E-08 | -1.88 | -0.91 | 2.3E-09 |
| Il17ra | 1921 | 3.40 | 1.76 | 1.7E-57 | 3.83 | 1.94 | 7.9E-69 |
| Il17rd | 5562 | 1.54 | 0.63 | 2.7E-06 | 1.70 | 0.77 | 6.3E-09 |
| Il18 | 351 | -1.75 | -0.81 | 2.1E-06 | -1.89 | -0.92 | 7.2E-08 |
| Il1rap | 1176 | 1.50 | 0.59 | 6.1E-04 | 1.73 | 0.79 | 3.1E-06 |
| Il1rl2 | 235 | -2.05 | -1.04 | 6.7E-05 | -1.86 | -0.90 | 6.9E-04 |
| Il20rb | 249 | 1.78 | 0.83 | 1.1E-08 | 1.60 | 0.68 | 4.2E-06 |
| Ilf3 | 19523 | -1.64 | -0.71 | 2.8E-19 | -1.73 | -0.79 | 1.4E-23 |
| Ilrun | 7355 | 1.28 | 0.36 | 1.1E-04 | 1.20 | 0.26 | 6.4E-03 |
| Ilvbl | 2182 | 1.30 | 0.38 | 3.9E-06 | 1.31 | 0.39 | 1.7E-06 |
| Immp1l | 2141 | 1.34 | 0.42 | 1.5E-04 | 1.41 | 0.49 | 6.9E-06 |
| Imp3 | 1956 | 1.27 | 0.34 | 4.0E-05 | 1.23 | 0.30 | 4.0E-04 |
| Imp4 | 5720 | 1.59 | 0.67 | 1.8E-20 | 1.55 | 0.63 | 3.9E-18 |
| Impa2 | 1044 | 2.29 | 1.19 | 1.2E-17 | 2.24 | 1.16 | 1.0E-16 |
| Impdh1 | 3231 | 1.62 | 0.69 | 1.2E-09 | 1.48 | 0.57 | 8.2E-07 |
| Impdh2 | 8677 | 1.32 | 0.40 | 3.5E-04 | 1.21 | 0.27 | 1.8E-02 |
| Impg2 | 400 | 2.09 | 1.07 | 9.0E-12 | 2.40 | 1.26 | 6.0E-16 |
| Inafm1 | 1121 | 1.25 | 0.32 | 3.3E-04 | 1.20 | 0.26 | 3.3E-03 |
| Inafm2 | 2534 | 2.04 | 1.03 | 4.2E-20 | 1.86 | 0.90 | 1.6E-15 |
| Ing1 | 5645 | 1.28 | 0.36 | 2.5E-06 | 1.17 | 0.22 | 4.6E-03 |
| Ing2 | 1279 | 1.68 | 0.75 | 2.1E-27 | 1.55 | 0.63 | 1.1E-19 |
| Ing3 | 2264 | -1.26 | -0.34 | 1.9E-04 | -1.30 | -0.38 | 2.4E-05 |
| Ing4 | 221 | -2.15 | -1.10 | 6.4E-12 | -1.91 | -0.93 | 1.0E-08 |
| Ing5 | 1826 | -2.42 | -1.27 | 3.8E-47 | -2.41 | -1.27 | 1.2E-46 |
| Inip | 5102 | 1.43 | 0.51 | 1.2E-06 | 1.35 | 0.43 | 5.3E-05 |
| Inka2 | 980 | 1.91 | 0.93 | 5.7E-26 | 1.91 | 0.93 | 9.2E-26 |
| Ino80 | 2782 | -1.76 | -0.82 | 1.6E-36 | -1.74 | -0.80 | 1.7E-34 |
| Ino80b | 1537 | 1.57 | 0.65 | 1.7E-13 | 1.51 | 0.59 | 2.4E-11 |
| Ino80c | 1239 | -2.98 | -1.57 | 3.8E-84 | -2.99 | -1.58 | 1.3E-84 |
| Ino80d | 4072 | -1.79 | -0.84 | 3.2E-09 | -1.81 | -0.86 | 1.3E-09 |
| Inpp1 | 1129 | -2.01 | -1.00 | 6.8E-32 | -2.01 | -1.01 | 8.4E-32 |
| Inpp4a | 5834 | 1.63 | 0.71 | 1.2E-06 | 1.61 | 0.69 | 2.6E-06 |
| Inpp5a | 2354 | 1.54 | 0.63 | 3.1E-12 | 1.53 | 0.61 | 1.3E-11 |
| Inpp5b | 3385 | 1.73 | 0.79 | 1.4E-32 | 1.75 | 0.80 | 1.8E-33 |
| Inpp5d | 366 | -1.78 | -0.83 | 1.4E-09 | -1.82 | -0.86 | 3.2E-10 |
| Inpp5e | 2670 | -1.19 | -0.25 | 2.0E-05 | -1.26 | -0.34 | 5.9E-09 |
| Inpp5k | 3089 | 1.58 | 0.66 | 2.6E-16 | 1.60 | 0.68 | 3.3E-17 |
| Insig2 | 2536 | 1.42 | 0.51 | 1.1E-16 | 1.43 | 0.51 | 5.7E-17 |
| Insl5 | 114 | 8.84 | 3.14 | 2.4E-42 | 9.24 | 3.21 | 1.1E-44 |
| Insl6 | 170 | 5.11 | 2.35 | 8.7E-23 | 6.14 | 2.62 | 1.3E-27 |
| Insm1 | 4447 | -1.38 | -0.47 | 3.7E-02 | -1.53 | -0.61 | 5.6E-03 |
| Insyn1 | 1174 | -1.42 | -0.51 | 1.6E-02 | -1.44 | -0.53 | 1.2E-02 |
| Insyn2b | 577 | -1.58 | -0.66 | 5.6E-03 | -1.45 | -0.54 | 2.7E-02 |
| Ints10 | 5797 | 1.20 | 0.26 | 3.0E-04 | 1.25 | 0.33 | 5.4E-06 |
| Ints12 | 2087 | -1.47 | -0.56 | 5.9E-12 | -1.50 | -0.58 | 6.1E-13 |
| Ints14 | 2799 | -2.49 | -1.32 | 3.0E-42 | -2.59 | -1.37 | 1.2E-45 |

| | | | | | | | |
|---|---|---|---|---|---|---|---|
| Ints2 | 5845 | -1.49 | -0.58 | 3.3E-13 | -1.41 | -0.49 | 7.3E-10 |
| Ints3 | 7990 | -1.19 | -0.25 | 1.5E-03 | -1.28 | -0.35 | 5.0E-06 |
| Ints5 | 2837 | 1.33 | 0.41 | 3.6E-05 | 1.21 | 0.27 | 8.5E-03 |
| Ints6 | 3398 | -1.53 | -0.62 | 3.8E-20 | -1.47 | -0.55 | 2.7E-16 |
| Ints6l | 7039 | -1.46 | -0.55 | 1.8E-06 | -1.30 | -0.38 | 1.1E-03 |
| Ints7 | 5990 | -1.89 | -0.92 | 1.2E-40 | -1.95 | -0.96 | 3.7E-44 |
| Ints9 | 3292 | 1.47 | 0.56 | 7.9E-08 | 1.32 | 0.40 | 1.6E-04 |
| Invs | 1538 | -1.37 | -0.45 | 5.6E-13 | -1.27 | -0.35 | 5.6E-08 |
| Ip6k2 | 1224 | -1.63 | -0.70 | 3.5E-04 | -1.78 | -0.83 | 1.9E-05 |
| Ipcef1 | 153 | -3.60 | -1.85 | 1.3E-13 | -3.91 | -1.97 | 3.5E-15 |
| Ipmk | 4642 | 1.68 | 0.74 | 1.3E-27 | 1.58 | 0.66 | 5.4E-22 |
| Ipo13 | 7274 | 1.83 | 0.87 | 7.3E-34 | 1.77 | 0.82 | 2.3E-30 |
| Ipo5 | 30086 | 1.32 | 0.40 | 1.4E-03 | 1.25 | 0.32 | 1.2E-02 |
| Ipo7 | 26828 | 1.22 | 0.28 | 5.5E-04 | 1.27 | 0.35 | 2.0E-05 |
| Iqcc | 1401 | -1.26 | -0.34 | 2.2E-08 | -1.27 | -0.34 | 2.0E-08 |
| Iqcg | 764 | 2.20 | 1.14 | 1.7E-23 | 2.18 | 1.13 | 5.8E-23 |
| Iqgap3 | 4143 | 1.40 | 0.49 | 1.9E-03 | 1.39 | 0.47 | 2.5E-03 |
| Iqsec2 | 1627 | 1.47 | 0.56 | 8.2E-05 | 1.40 | 0.49 | 7.0E-04 |
| Irag1 | 160 | -3.10 | -1.63 | 1.2E-18 | -2.75 | -1.46 | 4.0E-15 |
| Irak1 | 3461 | -1.98 | -0.99 | 1.1E-53 | -2.18 | -1.13 | 2.3E-69 |
| Irak2 | 155 | -1.82 | -0.87 | 2.3E-05 | -2.11 | -1.07 | 1.3E-07 |
| Irak4 | 345 | 1.62 | 0.70 | 7.7E-07 | 1.62 | 0.69 | 1.1E-06 |
| Irf1 | 346 | 1.81 | 0.85 | 7.1E-07 | 1.56 | 0.64 | 2.7E-04 |
| Irf2bpl | 5211 | -1.30 | -0.38 | 2.7E-02 | -1.52 | -0.60 | 3.4E-04 |
| Irf3 | 3445 | 1.56 | 0.64 | 6.3E-15 | 1.55 | 0.63 | 2.2E-14 |
| Irf8 | 271 | 2.06 | 1.05 | 2.5E-09 | 1.87 | 0.90 | 3.8E-07 |
| Irf9 | 2001 | 1.39 | 0.48 | 1.1E-06 | 1.29 | 0.37 | 2.4E-04 |
| Irgm2 | 123 | -1.62 | -0.69 | 3.6E-03 | -1.57 | -0.65 | 7.0E-03 |
| Irs2 | 3664 | 1.30 | 0.38 | 6.3E-04 | 1.22 | 0.29 | 1.1E-02 |
| Irs4 | 831 | 2.25 | 1.17 | 2.2E-11 | 1.68 | 0.74 | 3.1E-05 |
| Isca1 | 5555 | 1.69 | 0.76 | 5.1E-39 | 1.72 | 0.78 | 3.5E-41 |
| Iscu | 1586 | 1.36 | 0.44 | 2.5E-06 | 1.32 | 0.40 | 2.7E-05 |
| Isl1 | 3389 | -1.54 | -0.62 | 1.4E-03 | -1.45 | -0.53 | 6.9E-03 |
| Islr | 2279 | -1.93 | -0.95 | 2.4E-04 | -1.85 | -0.89 | 6.5E-04 |
| Isoc2a | 1669 | 2.02 | 1.01 | 6.4E-21 | 1.82 | 0.87 | 1.5E-15 |
| Isy1 | 1105 | -1.79 | -0.84 | 2.6E-18 | -1.96 | -0.97 | 3.4E-24 |
| Isyna1 | 4556 | 1.55 | 0.63 | 3.6E-06 | 1.43 | 0.52 | 1.6E-04 |
| Itfg2 | 1573 | -2.14 | -1.10 | 5.3E-38 | -2.35 | -1.23 | 2.1E-47 |
| Itga10 | 613 | 1.41 | 0.50 | 4.3E-03 | 1.49 | 0.58 | 8.8E-04 |
| Itga2 | 669 | -2.23 | -1.16 | 2.5E-13 | -1.94 | -0.96 | 1.8E-09 |
| Itga3 | 1680 | 1.42 | 0.50 | 2.6E-03 | 1.53 | 0.62 | 1.7E-04 |
| Itga6 | 7402 | 1.47 | 0.55 | 2.0E-18 | 1.48 | 0.57 | 2.3E-19 |
| Itga7 | 472 | 1.79 | 0.84 | 2.7E-02 | 2.16 | 1.11 | 3.1E-03 |
| Itgav | 3580 | -1.29 | -0.37 | 4.2E-03 | -1.21 | -0.27 | 4.2E-02 |
| Itgb1 | 18966 | -1.39 | -0.48 | 2.2E-03 | -1.30 | -0.38 | 1.6E-02 |
| Itgb1bp2 | 259 | -1.83 | -0.87 | 2.2E-02 | -1.75 | -0.81 | 3.7E-02 |
| Itgb4 | 216 | -3.23 | -1.69 | 1.3E-03 | -2.90 | -1.54 | 3.9E-03 |

| | | | | | | | |
|---|---|---|---|---|---|---|---|
| Itgb8 | 6530 | -1.44 | -0.53 | 5.0E-03 | -1.33 | -0.41 | 3.2E-02 |
| Itgbl1 | 163 | -1.98 | -0.98 | 3.0E-03 | -2.11 | -1.07 | 1.1E-03 |
| Itpk1 | 2507 | 2.59 | 1.37 | 4.0E-31 | 2.55 | 1.35 | 5.0E-30 |
| Itpka | 175 | 17.28 | 4.11 | 1.1E-58 | 10.83 | 3.44 | 3.3E-44 |
| Itpkc | 1319 | 1.35 | 0.43 | 1.7E-06 | 1.29 | 0.37 | 4.9E-05 |
| Itpr1 | 4428 | 1.77 | 0.82 | 6.2E-07 | 2.01 | 1.01 | 6.7E-10 |
| Itpr3 | 1378 | 1.60 | 0.67 | 3.0E-03 | 1.68 | 0.75 | 9.9E-04 |
| Itprid2 | 2892 | 1.64 | 0.72 | 4.2E-10 | 1.75 | 0.81 | 1.3E-12 |
| Itprip | 1837 | 1.58 | 0.66 | 2.3E-06 | 1.45 | 0.53 | 1.6E-04 |
| Itpripl1 | 1070 | 1.32 | 0.40 | 1.3E-05 | 1.20 | 0.26 | 5.5E-03 |
| Itpripl2 | 1637 | -2.01 | -1.01 | 2.7E-08 | -1.97 | -0.98 | 6.7E-08 |
| Itsn2 | 2301 | -1.77 | -0.83 | 5.7E-11 | -1.52 | -0.61 | 2.2E-06 |
| Iws1 | 8227 | -1.56 | -0.64 | 3.9E-15 | -1.50 | -0.58 | 1.6E-12 |
| Jade1 | 2510 | -1.59 | -0.67 | 7.2E-12 | -1.61 | -0.69 | 1.6E-12 |
| Jag2 | 3207 | 1.48 | 0.56 | 2.4E-04 | 1.45 | 0.54 | 4.7E-04 |
| Jak1 | 8991 | 1.25 | 0.32 | 5.9E-03 | 1.31 | 0.39 | 7.7E-04 |
| Jazf1 | 715 | -1.96 | -0.97 | 5.4E-07 | -1.82 | -0.86 | 9.7E-06 |
| Jdp2 | 392 | -1.34 | -0.43 | 2.5E-02 | -1.36 | -0.45 | 2.0E-02 |
| Jmjd4 | 3149 | 1.34 | 0.42 | 4.6E-05 | 1.42 | 0.51 | 8.2E-07 |
| Jmjd8 | 2988 | -1.64 | -0.72 | 3.8E-31 | -1.74 | -0.80 | 4.3E-38 |
| Josd1 | 4103 | 1.53 | 0.62 | 1.6E-21 | 1.52 | 0.60 | 1.2E-20 |
| Jph2 | 121 | -3.23 | -1.69 | 1.4E-04 | -2.35 | -1.23 | 6.8E-03 |
| Jrkl | 2004 | 1.25 | 0.32 | 4.0E-04 | 1.30 | 0.38 | 2.2E-05 |
| Junb | 206 | 2.19 | 1.13 | 5.8E-10 | 2.20 | 1.14 | 5.4E-10 |
| Kank3 | 1409 | 1.41 | 0.50 | 1.5E-08 | 1.29 | 0.36 | 5.1E-05 |
| Kansl1 | 8932 | -1.43 | -0.51 | 1.0E-14 | -1.47 | -0.56 | 3.6E-17 |
| Kansl1l | 2089 | -1.80 | -0.85 | 1.7E-19 | -1.61 | -0.68 | 4.1E-13 |
| Kansl2 | 5771 | -1.35 | -0.43 | 1.8E-10 | -1.43 | -0.51 | 2.0E-14 |
| Kantr | 1925 | 1.61 | 0.69 | 6.1E-07 | 1.49 | 0.58 | 3.2E-05 |
| Kars | 12303 | 1.32 | 0.40 | 3.2E-05 | 1.25 | 0.32 | 1.1E-03 |
| Kat14 | 2292 | -1.55 | -0.64 | 3.2E-11 | -1.68 | -0.74 | 6.1E-15 |
| Kat2a | 3607 | -1.46 | -0.54 | 3.7E-06 | -1.53 | -0.61 | 1.8E-07 |
| Kat2b | 1923 | -1.71 | -0.78 | 1.0E-09 | -1.73 | -0.79 | 4.2E-10 |
| Kat5 | 3136 | -2.02 | -1.02 | 3.0E-52 | -2.04 | -1.03 | 2.5E-53 |
| Kat6b | 7883 | -2.68 | -1.42 | 1.9E-23 | -2.43 | -1.28 | 3.8E-19 |
| Kat7 | 5227 | -1.29 | -0.36 | 4.0E-08 | -1.38 | -0.46 | 2.5E-12 |
| Katnal1 | 2730 | -1.97 | -0.98 | 2.7E-24 | -2.04 | -1.03 | 9.5E-27 |
| Katnb1 | 3482 | 1.81 | 0.85 | 6.1E-23 | 1.69 | 0.75 | 4.7E-18 |
| Katnip | 3767 | 1.32 | 0.40 | 1.3E-06 | 1.35 | 0.43 | 1.6E-07 |
| Kbtbd11 | 3918 | -1.56 | -0.64 | 8.4E-07 | -1.45 | -0.54 | 4.3E-05 |
| Kbtbd2 | 4187 | -1.24 | -0.31 | 1.2E-05 | -1.31 | -0.39 | 2.0E-08 |
| Kbtbd8 | 1457 | 1.41 | 0.50 | 1.5E-05 | 1.45 | 0.54 | 2.5E-06 |
| Kcmf1 | 4316 | -1.26 | -0.33 | 3.4E-07 | -1.28 | -0.36 | 2.8E-08 |
| Kcna1 | 358 | 1.97 | 0.97 | 1.1E-03 | 2.34 | 1.22 | 3.3E-05 |
| Kcna5 | 191 | -1.64 | -0.71 | 3.9E-05 | -1.93 | -0.95 | 4.1E-08 |
| Kcna6 | 2547 | 1.69 | 0.76 | 4.5E-06 | 1.79 | 0.84 | 4.2E-07 |
| Kcnab1 | 333 | -1.75 | -0.81 | 1.5E-03 | -1.62 | -0.69 | 7.4E-03 |

| | | | | | | | |
|---|---|---|---|---|---|---|---|
| Kcnab2 | 2430 | 2.11 | 1.08 | 6.4E-17 | 1.96 | 0.97 | 7.1E-14 |
| Kcnab3 | 1102 | 4.03 | 2.01 | 7.6E-61 | 3.63 | 1.86 | 3.9E-52 |
| Kcnb1 | 1903 | 1.44 | 0.52 | 4.3E-03 | 1.49 | 0.58 | 1.6E-03 |
| Kcnb2 | 823 | -2.71 | -1.44 | 3.4E-07 | -2.29 | -1.20 | 2.8E-05 |
| Kcnc3 | 5043 | 5.14 | 2.36 | 3.4E-42 | 4.74 | 2.24 | 3.4E-38 |
| Kcnd1 | 719 | 1.59 | 0.67 | 3.5E-09 | 1.60 | 0.67 | 2.7E-09 |
| Kcne1l | 432 | -1.50 | -0.58 | 9.8E-04 | -1.38 | -0.46 | 1.1E-02 |
| Kcne2 | 425 | -2.54 | -1.35 | 3.1E-03 | -2.49 | -1.32 | 4.0E-03 |
| Kcne4 | 181 | -2.35 | -1.23 | 1.5E-03 | -2.33 | -1.22 | 1.8E-03 |
| Kcng1 | 529 | 2.01 | 1.01 | 3.1E-04 | 2.08 | 1.05 | 1.6E-04 |
| Kcnh1 | 270 | -2.05 | -1.03 | 2.8E-03 | -1.85 | -0.89 | 1.2E-02 |
| Kcnh3 | 315 | 2.13 | 1.09 | 2.8E-03 | 1.95 | 0.97 | 8.8E-03 |
| Kcnh8 | 391 | -2.58 | -1.37 | 3.1E-15 | -2.40 | -1.26 | 5.3E-13 |
| Kcnip2 | 325 | 1.65 | 0.72 | 1.9E-03 | 1.63 | 0.71 | 2.4E-03 |
| Kcnj10 | 1277 | 2.72 | 1.44 | 9.7E-28 | 2.66 | 1.41 | 1.8E-26 |
| Kcnj5 | 154 | -2.10 | -1.07 | 1.1E-05 | -2.11 | -1.08 | 9.2E-06 |
| Kcnj8 | 268 | -4.32 | -2.11 | 5.2E-17 | -4.71 | -2.24 | 6.2E-19 |
| Kcnk10 | 2769 | -1.47 | -0.55 | 1.1E-03 | -1.68 | -0.75 | 7.7E-06 |
| Kcnk2 | 1372 | -2.84 | -1.51 | 8.1E-19 | -3.12 | -1.64 | 4.1E-22 |
| Kcnk5 | 925 | 2.04 | 1.03 | 3.1E-06 | 1.72 | 0.78 | 5.2E-04 |
| Kcnk6 | 220 | -1.65 | -0.72 | 6.6E-03 | -1.48 | -0.56 | 4.0E-02 |
| Kcnmb4 | 843 | -1.68 | -0.75 | 6.3E-18 | -1.71 | -0.77 | 5.2E-19 |
| Kcnn3 | 2098 | 2.69 | 1.43 | 5.6E-09 | 3.26 | 1.71 | 2.2E-12 |
| Kcnq5 | 265 | -2.13 | -1.09 | 9.7E-07 | -1.70 | -0.77 | 8.3E-04 |
| Kcnrg | 233 | 1.70 | 0.77 | 7.0E-05 | 1.76 | 0.82 | 2.2E-05 |
| Kctd12 | 13048 | -1.60 | -0.67 | 3.3E-03 | -1.51 | -0.60 | 1.0E-02 |
| Kctd12b | 1142 | -1.53 | -0.61 | 8.6E-06 | -1.42 | -0.51 | 2.8E-04 |
| Kctd14 | 139 | -2.40 | -1.26 | 1.2E-15 | -2.37 | -1.25 | 4.1E-15 |
| Kctd15 | 3599 | -2.13 | -1.09 | 7.3E-16 | -2.26 | -1.17 | 4.0E-18 |
| Kctd18 | 2322 | -1.26 | -0.33 | 7.3E-05 | -1.15 | -0.21 | 1.6E-02 |
| Kctd21 | 700 | 1.37 | 0.45 | 1.8E-08 | 1.25 | 0.32 | 1.1E-04 |
| Kctd9 | 2584 | 1.40 | 0.48 | 4.6E-06 | 1.38 | 0.47 | 8.9E-06 |
| Kdelr1 | 5091 | -1.31 | -0.39 | 1.5E-11 | -1.36 | -0.44 | 1.2E-14 |
| Kdm1a | 15192 | -1.26 | -0.33 | 6.0E-05 | -1.39 | -0.47 | 5.8E-09 |
| Kdm1b | 5673 | 2.26 | 1.18 | 3.4E-130 | 2.28 | 1.19 | 4.1E-133 |
| Kdm4a | 11601 | -1.61 | -0.69 | 2.1E-13 | -1.70 | -0.77 | 2.2E-16 |
| Kdm5c | 12138 | -1.40 | -0.48 | 1.1E-14 | -1.53 | -0.61 | 3.7E-23 |
| Kdm6b | 8176 | -1.30 | -0.38 | 4.8E-02 | -1.51 | -0.59 | 1.4E-03 |
| Kdm8 | 397 | -3.65 | -1.87 | 2.5E-33 | -3.86 | -1.95 | 4.9E-36 |
| Kdr | 5366 | -1.35 | -0.43 | 9.8E-08 | -1.37 | -0.46 | 1.6E-08 |
| Kdsr | 2243 | 1.48 | 0.57 | 7.9E-15 | 1.50 | 0.59 | 6.2E-16 |
| Keap1 | 2944 | -1.44 | -0.52 | 5.1E-14 | -1.55 | -0.63 | 5.1E-20 |
| Khdc4 | 9694 | -1.26 | -0.33 | 3.3E-11 | -1.27 | -0.34 | 1.3E-11 |
| Khk | 296 | 1.37 | 0.46 | 5.6E-03 | 1.30 | 0.38 | 2.6E-02 |
| Khnyn | 4786 | 1.32 | 0.40 | 2.1E-04 | 1.31 | 0.39 | 4.0E-04 |
| Khsrp | 12134 | -1.35 | -0.43 | 1.1E-02 | -1.53 | -0.61 | 1.9E-04 |
| Kif13a | 3851 | -1.99 | -0.99 | 6.3E-38 | -2.02 | -1.01 | 1.6E-39 |

| | | | | | | | |
|---|---|---|---|---|---|---|---|
| Kif13b | 3411 | 1.29 | 0.36 | 4.0E-03 | 1.48 | 0.56 | 4.3E-06 |
| Kif14 | 2071 | -1.46 | -0.54 | 4.8E-04 | -1.48 | -0.56 | 2.9E-04 |
| Kif17 | 387 | 4.17 | 2.06 | 2.5E-27 | 4.83 | 2.27 | 1.1E-32 |
| Kif1c | 6763 | 1.43 | 0.51 | 1.3E-04 | 1.33 | 0.41 | 2.6E-03 |
| Kif20a | 6033 | -1.41 | -0.49 | 1.3E-02 | -1.53 | -0.61 | 1.8E-03 |
| Kif24 | 2027 | 1.69 | 0.76 | 9.8E-05 | 1.76 | 0.81 | 2.9E-05 |
| Kif26b | 5534 | -2.50 | -1.32 | 1.3E-11 | -2.26 | -1.18 | 2.1E-09 |
| Kif27 | 755 | 2.09 | 1.07 | 3.6E-29 | 2.25 | 1.17 | 9.1E-35 |
| Kif3b | 7784 | 1.40 | 0.48 | 3.7E-08 | 1.44 | 0.52 | 2.1E-09 |
| Kif4 | 5927 | -1.35 | -0.43 | 2.8E-02 | -1.42 | -0.51 | 8.5E-03 |
| Kif6 | 249 | 1.66 | 0.74 | 7.6E-08 | 1.84 | 0.88 | 1.2E-10 |
| Kif7 | 2665 | 2.22 | 1.15 | 4.7E-21 | 2.09 | 1.07 | 3.0E-18 |
| Kifap3 | 11972 | -1.30 | -0.38 | 1.6E-03 | -1.24 | -0.31 | 1.0E-02 |
| Kifc2 | 2883 | 2.76 | 1.46 | 5.5E-16 | 2.75 | 1.46 | 6.8E-16 |
| Kifc3 | 1352 | 1.55 | 0.64 | 2.0E-06 | 1.59 | 0.67 | 5.4E-07 |
| Kifc5b | 1917 | 1.56 | 0.65 | 3.3E-04 | 1.37 | 0.46 | 1.4E-02 |
| Kiss1r | 156 | 1.93 | 0.95 | 9.5E-07 | 2.00 | 1.00 | 2.7E-07 |
| Klc4 | 3631 | 1.34 | 0.43 | 1.2E-05 | 1.27 | 0.35 | 3.8E-04 |
| Klf11 | 1422 | -1.62 | -0.70 | 6.2E-36 | -1.67 | -0.74 | 2.2E-40 |
| Klf15 | 380 | 2.00 | 1.00 | 1.4E-08 | 2.02 | 1.02 | 7.9E-09 |
| Klf16 | 1425 | 2.00 | 1.00 | 1.5E-16 | 1.78 | 0.83 | 8.3E-12 |
| Klf3 | 3631 | -1.51 | -0.60 | 2.9E-15 | -1.46 | -0.55 | 7.9E-13 |
| Klf4 | 882 | -1.56 | -0.64 | 1.3E-02 | -1.59 | -0.67 | 9.9E-03 |
| Klf7 | 6968 | -1.26 | -0.34 | 2.5E-02 | -1.27 | -0.34 | 2.3E-02 |
| Klf9 | 778 | 4.16 | 2.06 | 4.4E-45 | 3.97 | 1.99 | 4.3E-42 |
| Klhdc1 | 283 | -1.47 | -0.56 | 1.1E-02 | -1.45 | -0.54 | 1.5E-02 |
| Klhdc10 | 18793 | 1.28 | 0.35 | 1.6E-03 | 1.24 | 0.31 | 5.7E-03 |
| Klhdc2 | 13166 | -1.32 | -0.40 | 6.1E-07 | -1.33 | -0.41 | 2.5E-07 |
| Klhdc3 | 8405 | 1.30 | 0.38 | 1.1E-06 | 1.22 | 0.28 | 3.8E-04 |
| Klhdc7a | 156 | -4.69 | -2.23 | 6.7E-20 | -4.23 | -2.08 | 1.3E-17 |
| Klhdc8b | 3415 | 1.27 | 0.35 | 6.3E-05 | 1.17 | 0.22 | 1.4E-02 |
| Klhl13 | 8680 | -1.37 | -0.46 | 1.8E-06 | -1.39 | -0.48 | 6.0E-07 |
| Klhl15 | 1585 | 1.56 | 0.65 | 5.0E-10 | 1.46 | 0.54 | 2.3E-07 |
| Klhl21 | 994 | 2.11 | 1.08 | 5.1E-09 | 1.99 | 0.99 | 8.9E-08 |
| Klhl23 | 5853 | -1.73 | -0.79 | 1.4E-09 | -1.79 | -0.84 | 1.1E-10 |
| Klhl24 | 11392 | -1.71 | -0.78 | 2.2E-10 | -1.53 | -0.62 | 6.6E-07 |
| Klhl25 | 3590 | 2.04 | 1.03 | 1.6E-47 | 1.97 | 0.98 | 4.3E-43 |
| Klhl26 | 1693 | 1.63 | 0.71 | 1.4E-18 | 1.58 | 0.66 | 2.2E-16 |
| Klhl28 | 5847 | 1.48 | 0.57 | 2.7E-06 | 1.70 | 0.77 | 1.4E-10 |
| Klhl3 | 898 | -2.18 | -1.12 | 1.0E-08 | -2.16 | -1.11 | 1.6E-08 |
| Klhl32 | 1209 | 1.94 | 0.95 | 2.1E-11 | 2.09 | 1.06 | 6.6E-14 |
| Klhl35 | 324 | -6.75 | -2.75 | 1.8E-27 | -7.46 | -2.90 | 2.9E-30 |
| Klhl36 | 744 | 2.71 | 1.44 | 2.0E-47 | 2.56 | 1.35 | 2.8E-42 |
| Klhl4 | 635 | -1.77 | -0.82 | 3.8E-04 | -1.61 | -0.69 | 3.4E-03 |
| Klhl42 | 1888 | -1.30 | -0.38 | 7.5E-08 | -1.30 | -0.38 | 9.6E-08 |
| Klhl5 | 5569 | 1.26 | 0.33 | 1.0E-04 | 1.24 | 0.31 | 2.9E-04 |
| Klhl6 | 383 | -1.37 | -0.45 | 1.0E-03 | -1.46 | -0.54 | 7.0E-05 |

| | | | | | | | |
|---|---|---|---|---|---|---|---|
| Klhl8 | 1637 | 1.25 | 0.33 | 1.7E-02 | 1.30 | 0.38 | 4.9E-03 |
| Kmt2e | 18204 | -1.33 | -0.41 | 3.4E-04 | -1.36 | -0.45 | 8.9E-05 |
| Kmt5b | 6543 | -6.45 | -2.69 | 2.2E-62 | -6.59 | -2.72 | 9.4E-64 |
| Knop1 | 15778 | 1.57 | 0.65 | 7.1E-13 | 1.52 | 0.61 | 2.9E-11 |
| Kpna3 | 14148 | 1.27 | 0.34 | 5.0E-05 | 1.28 | 0.35 | 2.4E-05 |
| Kpnb1 | 44199 | -1.47 | -0.56 | 8.8E-07 | -1.62 | -0.69 | 6.2E-10 |
| Kptn | 941 | -2.00 | -1.00 | 4.4E-45 | -2.16 | -1.11 | 2.4E-55 |
| Kras | 16185 | 1.38 | 0.46 | 2.1E-07 | 1.32 | 0.40 | 6.2E-06 |
| Kremen1 | 4278 | 2.41 | 1.27 | 1.0E-34 | 2.46 | 1.30 | 3.4E-36 |
| Krt1 | 134 | -3.54 | -1.82 | 5.3E-05 | -3.14 | -1.65 | 2.9E-04 |
| Krt10 | 289 | -2.75 | -1.46 | 9.3E-13 | -2.61 | -1.39 | 1.5E-11 |
| Krt14 | 629 | -8.89 | -3.15 | 2.2E-03 | -9.84 | -3.30 | 1.4E-03 |
| Krt15 | 501 | -5.82 | -2.54 | 9.5E-03 | -4.33 | -2.11 | 3.4E-02 |
| Krt17 | 127 | -10.18 | -3.35 | 1.3E-04 | -7.14 | -2.84 | 1.4E-03 |
| Krtcap2 | 3095 | 1.35 | 0.43 | 5.8E-05 | 1.25 | 0.32 | 3.3E-03 |
| Kxd1 | 1417 | -1.30 | -0.38 | 1.0E-06 | -1.40 | -0.48 | 3.1E-10 |
| Kyat3 | 996 | 2.74 | 1.45 | 4.8E-26 | 2.99 | 1.58 | 2.1E-30 |
| L2hgdh | 2708 | 2.54 | 1.35 | 3.4E-100 | 2.56 | 1.35 | 4.1E-101 |
| L3hypdh | 561 | 1.45 | 0.53 | 1.0E-05 | 1.56 | 0.64 | 1.0E-07 |
| L3mbtl1 | 236 | 2.23 | 1.15 | 1.1E-05 | 2.42 | 1.28 | 1.1E-06 |
| Lactb | 1520 | 2.50 | 1.32 | 1.6E-60 | 2.52 | 1.34 | 9.3E-62 |
| Lactb2 | 1288 | 1.66 | 0.73 | 6.8E-08 | 1.83 | 0.87 | 8.5E-11 |
| Lage3 | 1477 | -1.23 | -0.30 | 1.1E-02 | -1.39 | -0.48 | 2.4E-05 |
| Lair1 | 178 | -1.92 | -0.94 | 9.0E-08 | -1.75 | -0.81 | 5.7E-06 |
| Lama3 | 271 | -3.18 | -1.67 | 8.9E-06 | -2.09 | -1.07 | 6.3E-03 |
| Lama4 | 4671 | -1.49 | -0.57 | 1.6E-03 | -1.38 | -0.47 | 1.2E-02 |
| Lama5 | 3349 | 2.39 | 1.26 | 2.7E-09 | 2.73 | 1.45 | 6.1E-12 |
| Lamb1 | 8515 | -1.45 | -0.54 | 4.2E-03 | -1.39 | -0.48 | 1.3E-02 |
| Lamc1 | 11716 | 1.25 | 0.32 | 2.2E-02 | 1.33 | 0.41 | 2.9E-03 |
| Lamp2 | 10284 | 1.26 | 0.33 | 2.0E-02 | 1.35 | 0.43 | 2.2E-03 |
| Lamp5 | 587 | -6.05 | -2.60 | 9.0E-04 | -4.36 | -2.13 | 7.7E-03 |
| Lamtor1 | 2306 | -2.19 | -1.13 | 1.7E-49 | -2.29 | -1.20 | 6.3E-55 |
| Lamtor2 | 4312 | 1.52 | 0.60 | 3.5E-05 | 1.40 | 0.49 | 1.0E-03 |
| Lamtor3 | 3349 | 1.34 | 0.42 | 2.2E-13 | 1.31 | 0.39 | 1.3E-11 |
| Lancl1 | 4659 | 1.22 | 0.29 | 5.6E-04 | 1.25 | 0.33 | 8.5E-05 |
| Lancl2 | 4527 | -2.54 | -1.34 | 4.9E-49 | -2.70 | -1.43 | 1.2E-55 |
| Laptm4b | 3095 | 1.53 | 0.62 | 4.7E-11 | 1.59 | 0.67 | 7.5E-13 |
| Larp1b | 901 | 2.87 | 1.52 | 2.6E-47 | 2.71 | 1.44 | 3.1E-42 |
| Larp4b | 11106 | 1.38 | 0.46 | 7.8E-18 | 1.39 | 0.48 | 5.8E-19 |
| Larp6 | 215 | -2.27 | -1.18 | 1.4E-07 | -2.29 | -1.20 | 9.7E-08 |
| Larp7 | 4216 | -2.29 | -1.20 | 1.2E-14 | -2.53 | -1.34 | 3.8E-18 |
| Las1l | 6968 | 1.40 | 0.49 | 4.1E-24 | 1.37 | 0.46 | 1.2E-21 |
| Lat2 | 610 | 3.03 | 1.60 | 5.1E-19 | 2.64 | 1.40 | 6.3E-15 |
| Lax1 | 162 | 4.73 | 2.24 | 1.2E-23 | 6.18 | 2.63 | 2.9E-31 |
| Layn | 181 | 3.02 | 1.60 | 2.8E-16 | 2.68 | 1.42 | 4.9E-13 |
| Lbh | 9353 | -1.31 | -0.39 | 6.9E-06 | -1.39 | -0.47 | 3.5E-08 |
| Lbhd1 | 915 | 1.37 | 0.46 | 6.5E-06 | 1.29 | 0.36 | 4.1E-04 |

| | | | | | | | |
|---|---|---|---|---|---|---|---|
| Lbp | 122 | -1.62 | -0.70 | 2.9E-02 | -1.85 | -0.88 | 5.2E-03 |
| Lck | 258 | -1.33 | -0.41 | 8.4E-04 | -1.38 | -0.47 | 1.1E-04 |
| Lclat1 | 2797 | 1.53 | 0.61 | 5.2E-25 | 1.55 | 0.63 | 3.2E-26 |
| Lcmt1 | 3502 | 1.71 | 0.77 | 6.8E-36 | 1.76 | 0.82 | 2.8E-40 |
| Lcp2 | 129 | -2.55 | -1.35 | 6.3E-13 | -2.19 | -1.13 | 1.5E-09 |
| Lct | 129 | 2.20 | 1.14 | 4.8E-11 | 2.62 | 1.39 | 2.5E-15 |
| Ldah | 2921 | -1.40 | -0.48 | 8.0E-13 | -1.39 | -0.47 | 2.9E-12 |
| Ldb1 | 15935 | -1.30 | -0.38 | 6.7E-05 | -1.40 | -0.49 | 3.0E-07 |
| Ldb3 | 118 | -4.27 | -2.09 | 1.0E-03 | -3.63 | -1.86 | 4.0E-03 |
| Ldlr | 2768 | -1.36 | -0.44 | 3.2E-02 | -1.57 | -0.65 | 1.1E-03 |
| Ldlrap1 | 1021 | 1.80 | 0.85 | 8.7E-04 | 1.67 | 0.74 | 3.9E-03 |
| Lekr1 | 546 | 1.29 | 0.37 | 2.3E-06 | 1.25 | 0.32 | 5.2E-05 |
| Lemd1 | 426 | -2.08 | -1.06 | 7.0E-08 | -1.93 | -0.95 | 1.6E-06 |
| Leng1 | 1294 | 1.84 | 0.88 | 8.2E-14 | 1.72 | 0.79 | 3.0E-11 |
| Leng8 | 22520 | -1.25 | -0.32 | 1.4E-02 | -1.34 | -0.42 | 1.2E-03 |
| Letm1 | 3858 | -1.46 | -0.55 | 1.3E-12 | -1.53 | -0.62 | 9.3E-16 |
| Lfng | 1615 | -1.64 | -0.72 | 8.8E-08 | -1.69 | -0.76 | 1.3E-08 |
| Lgals7 | 208 | -1.83 | -0.87 | 2.7E-03 | -1.52 | -0.61 | 4.4E-02 |
| Lgals8 | 1760 | 1.66 | 0.73 | 1.4E-20 | 1.72 | 0.78 | 2.3E-23 |
| Lgals9 | 363 | -1.66 | -0.73 | 1.5E-03 | -1.65 | -0.72 | 2.0E-03 |
| Lgi1 | 361 | -3.37 | -1.75 | 1.6E-06 | -2.98 | -1.57 | 1.9E-05 |
| Lgi4 | 135 | -2.19 | -1.13 | 1.5E-03 | -1.81 | -0.85 | 2.0E-02 |
| Lgmn | 3063 | 1.31 | 0.38 | 7.3E-04 | 1.32 | 0.40 | 4.2E-04 |
| Lgr4 | 5083 | 1.38 | 0.47 | 7.1E-04 | 1.46 | 0.55 | 6.4E-05 |
| Lgr6 | 150 | -2.42 | -1.28 | 1.0E-02 | -3.09 | -1.63 | 9.2E-04 |
| Lhfpl2 | 3650 | -1.69 | -0.75 | 1.5E-04 | -1.71 | -0.77 | 1.0E-04 |
| Lhx8 | 610 | -2.58 | -1.37 | 7.4E-03 | -2.81 | -1.49 | 3.5E-03 |
| Lhx9 | 7621 | -1.85 | -0.89 | 1.5E-03 | -2.00 | -1.00 | 3.3E-04 |
| Lias | 2575 | -1.22 | -0.29 | 4.3E-05 | -1.27 | -0.35 | 9.5E-07 |
| Lig3 | 7660 | -1.35 | -0.43 | 8.4E-10 | -1.36 | -0.44 | 3.6E-10 |
| Lig4 | 2648 | 1.29 | 0.36 | 1.5E-07 | 1.26 | 0.33 | 1.9E-06 |
| Limch1 | 6122 | -1.62 | -0.69 | 1.6E-10 | -1.43 | -0.52 | 2.1E-06 |
| Limd1 | 5555 | 1.28 | 0.36 | 1.7E-04 | 1.24 | 0.32 | 1.0E-03 |
| Limd2 | 4598 | -1.98 | -0.99 | 9.5E-19 | -2.17 | -1.12 | 9.7E-24 |
| Limk1 | 7085 | 1.66 | 0.73 | 1.4E-10 | 1.58 | 0.66 | 8.8E-09 |
| Limk2 | 10289 | 1.28 | 0.36 | 1.3E-05 | 1.31 | 0.39 | 2.5E-06 |
| Lims1 | 9873 | 1.40 | 0.49 | 1.2E-04 | 1.45 | 0.54 | 2.2E-05 |
| Lin37 | 976 | 1.46 | 0.54 | 1.4E-10 | 1.32 | 0.40 | 4.0E-06 |
| Lin52 | 1684 | 1.30 | 0.38 | 1.4E-06 | 1.25 | 0.32 | 5.2E-05 |
| Lin7b | 298 | 6.34 | 2.67 | 4.5E-69 | 6.45 | 2.69 | 1.5E-69 |
| Lin9 | 1581 | -1.81 | -0.86 | 5.3E-15 | -1.81 | -0.85 | 7.9E-15 |
| Lingo1 | 1683 | -1.58 | -0.66 | 2.3E-02 | -1.52 | -0.61 | 4.0E-02 |
| Lingo3 | 150 | 2.55 | 1.35 | 4.5E-15 | 2.34 | 1.23 | 1.9E-12 |
| Lins1 | 1541 | -1.54 | -0.62 | 1.6E-35 | -1.51 | -0.59 | 5.3E-32 |
| Lipa | 3263 | 1.69 | 0.76 | 1.3E-08 | 1.64 | 0.71 | 1.2E-07 |
| Lipe | 975 | 1.97 | 0.98 | 4.4E-20 | 1.90 | 0.92 | 6.8E-18 |
| Lipo3 | 689 | -2.83 | -1.50 | 6.6E-45 | -3.13 | -1.64 | 2.0E-53 |

| | | | | | | | |
|---|---|---|---|---|---|---|---|
| Lipt2 | 411 | -1.44 | -0.53 | 2.1E-05 | -1.42 | -0.51 | 4.4E-05 |
| Litaf | 2654 | 1.46 | 0.54 | 6.8E-07 | 1.50 | 0.59 | 7.7E-08 |
| Lix1l | 12076 | 1.36 | 0.44 | 6.4E-07 | 1.28 | 0.36 | 6.7E-05 |
| Llgl2 | 368 | 2.79 | 1.48 | 9.8E-07 | 2.83 | 1.50 | 7.5E-07 |
| Llph | 3168 | 1.35 | 0.43 | 4.9E-03 | 1.24 | 0.32 | 4.5E-02 |
| Lmbr1l | 4606 | 1.54 | 0.62 | 4.1E-11 | 1.47 | 0.55 | 5.6E-09 |
| Lmbrd2 | 164 | 1.49 | 0.58 | 9.5E-05 | 1.41 | 0.50 | 9.5E-04 |
| Lmf1 | 769 | -1.75 | -0.81 | 8.0E-14 | -1.77 | -0.83 | 2.3E-14 |
| Lmo1 | 1246 | -1.39 | -0.47 | 3.2E-02 | -1.54 | -0.62 | 3.9E-03 |
| Lmo2 | 1194 | -1.86 | -0.90 | 4.5E-14 | -2.14 | -1.10 | 1.5E-20 |
| Lmtk2 | 4256 | 1.29 | 0.37 | 7.0E-03 | 1.26 | 0.33 | 1.8E-02 |
| Lmx1a | 1011 | -1.57 | -0.65 | 1.5E-03 | -1.81 | -0.86 | 1.9E-05 |
| Lnpep | 6564 | 1.21 | 0.28 | 7.6E-04 | 1.25 | 0.32 | 8.7E-05 |
| Lnpk | 5745 | 1.28 | 0.35 | 3.6E-03 | 1.46 | 0.55 | 3.3E-06 |
| Lnx2 | 1049 | 1.51 | 0.59 | 3.1E-10 | 1.48 | 0.56 | 2.5E-09 |
| Lonp1 | 8249 | 1.49 | 0.58 | 1.4E-08 | 1.35 | 0.43 | 3.0E-05 |
| Lonp2 | 7629 | 1.57 | 0.65 | 1.5E-30 | 1.63 | 0.71 | 2.4E-36 |
| Lonrf3 | 714 | 3.43 | 1.78 | 1.6E-15 | 4.34 | 2.12 | 1.8E-21 |
| Lox | 5374 | -2.76 | -1.46 | 1.5E-05 | -2.58 | -1.37 | 6.0E-05 |
| Lpar1 | 2608 | -1.97 | -0.98 | 5.8E-06 | -1.99 | -0.99 | 4.4E-06 |
| Lpar2 | 1612 | 1.27 | 0.34 | 2.5E-07 | 1.22 | 0.29 | 2.0E-05 |
| Lpar6 | 1124 | -1.83 | -0.87 | 2.3E-10 | -1.86 | -0.90 | 6.0E-11 |
| Lpcat2 | 491 | 1.34 | 0.42 | 2.6E-02 | 1.56 | 0.64 | 6.1E-04 |
| Lpcat3 | 1550 | 1.31 | 0.39 | 4.7E-05 | 1.31 | 0.38 | 5.1E-05 |
| Lpgat1 | 9502 | -1.52 | -0.60 | 7.8E-11 | -1.43 | -0.51 | 3.3E-08 |
| Lpin1 | 2745 | 1.49 | 0.58 | 1.1E-08 | 1.53 | 0.61 | 1.3E-09 |
| Lrfn1 | 2114 | 1.32 | 0.40 | 2.5E-03 | 1.24 | 0.31 | 2.5E-02 |
| Lrfn3 | 1428 | 1.47 | 0.56 | 1.0E-05 | 1.35 | 0.43 | 8.8E-04 |
| Lrfn4 | 2999 | -2.18 | -1.13 | 9.9E-20 | -2.31 | -1.21 | 1.2E-22 |
| Lrguk | 610 | 1.82 | 0.86 | 5.3E-14 | 2.25 | 1.17 | 2.3E-24 |
| Lrif1 | 2501 | 1.26 | 0.34 | 3.3E-06 | 1.29 | 0.36 | 4.3E-07 |
| Lrp10 | 3105 | 1.91 | 0.93 | 9.3E-21 | 1.71 | 0.77 | 1.4E-14 |
| Lrp2 | 2856 | 1.43 | 0.52 | 3.2E-02 | 1.64 | 0.71 | 2.8E-03 |
| Lrp3 | 5457 | -1.37 | -0.45 | 1.1E-04 | -1.50 | -0.58 | 4.7E-07 |
| Lrp4 | 4657 | -1.50 | -0.59 | 6.6E-10 | -1.55 | -0.63 | 3.0E-11 |
| Lrp6 | 10654 | -1.26 | -0.33 | 1.0E-04 | -1.14 | -0.19 | 3.4E-02 |
| Lrpap1 | 6185 | 1.44 | 0.53 | 9.5E-23 | 1.43 | 0.52 | 6.0E-22 |
| Lrpprc | 8740 | 1.63 | 0.71 | 1.8E-21 | 1.61 | 0.69 | 2.4E-20 |
| Lrr1 | 597 | 1.70 | 0.76 | 6.7E-05 | 1.49 | 0.58 | 3.2E-03 |
| Lrrc1 | 2446 | 1.64 | 0.71 | 7.0E-12 | 1.59 | 0.67 | 9.9E-11 |
| Lrrc10b | 118 | 2.31 | 1.21 | 2.9E-13 | 1.73 | 0.79 | 2.9E-06 |
| Lrrc17 | 1709 | -2.33 | -1.22 | 4.0E-03 | -2.22 | -1.15 | 7.0E-03 |
| Lrrc20 | 2243 | 1.31 | 0.39 | 2.3E-06 | 1.20 | 0.26 | 2.1E-03 |
| Lrrc23 | 263 | -1.33 | -0.41 | 1.3E-02 | -1.38 | -0.46 | 5.7E-03 |
| Lrrc28 | 1998 | 1.88 | 0.91 | 1.7E-44 | 1.90 | 0.92 | 7.3E-46 |
| Lrrc3 | 1388 | 2.38 | 1.25 | 1.2E-29 | 2.60 | 1.38 | 1.4E-35 |
| Lrrc4 | 888 | -3.27 | -1.71 | 1.1E-12 | -3.64 | -1.86 | 8.2E-15 |

| | | | | | | | |
|---|---|---|---|---|---|---|---|
| Lrrc40 | 5790 | 1.31 | 0.38 | 7.8E-04 | 1.31 | 0.39 | 6.7E-04 |
| Lrrc41 | 3730 | -1.41 | -0.50 | 3.8E-14 | -1.40 | -0.49 | 1.0E-13 |
| Lrrc42 | 4775 | 1.83 | 0.87 | 9.5E-30 | 1.68 | 0.75 | 2.1E-22 |
| Lrrc45 | 6089 | 1.34 | 0.42 | 3.7E-05 | 1.29 | 0.37 | 3.9E-04 |
| Lrrc49 | 4403 | -1.92 | -0.94 | 7.1E-19 | -1.84 | -0.88 | 1.9E-16 |
| Lrrc4b | 4217 | -2.74 | -1.46 | 6.3E-12 | -2.98 | -1.58 | 8.4E-14 |
| Lrrc51 | 184 | 1.29 | 0.37 | 1.8E-02 | 1.31 | 0.39 | 1.1E-02 |
| Lrrc58 | 7447 | -1.24 | -0.31 | 2.1E-02 | -1.36 | -0.44 | 6.2E-04 |
| Lrrc59 | 14255 | 1.31 | 0.39 | 1.3E-07 | 1.21 | 0.27 | 2.8E-04 |
| Lrrc61 | 1302 | -2.43 | -1.28 | 1.5E-40 | -2.85 | -1.51 | 6.2E-56 |
| Lrrc73 | 581 | 2.51 | 1.33 | 5.2E-17 | 2.35 | 1.24 | 7.8E-15 |
| Lrrc75a | 1592 | 2.01 | 1.01 | 6.6E-43 | 1.96 | 0.97 | 8.3E-40 |
| Lrrc75b | 1622 | -3.53 | -1.82 | 2.0E-11 | -3.80 | -1.93 | 1.1E-12 |
| Lrrc8b | 3769 | -1.51 | -0.60 | 9.8E-09 | -1.45 | -0.54 | 3.2E-07 |
| Lrrc8d | 2753 | -1.33 | -0.41 | 2.6E-04 | -1.43 | -0.52 | 2.4E-06 |
| Lrrc9 | 597 | 1.48 | 0.57 | 1.9E-02 | 1.97 | 0.98 | 2.8E-05 |
| Lrrfip1 | 2511 | 1.27 | 0.35 | 4.8E-04 | 1.37 | 0.45 | 4.8E-06 |
| Lrrk1 | 709 | -2.99 | -1.58 | 2.8E-13 | -2.85 | -1.51 | 3.1E-12 |
| Lrrk2 | 647 | -2.40 | -1.26 | 1.1E-08 | -2.20 | -1.14 | 2.9E-07 |
| Lrrn4cl | 131 | 1.84 | 0.88 | 2.9E-07 | 1.91 | 0.94 | 5.2E-08 |
| Lrrtm1 | 551 | -2.92 | -1.55 | 3.0E-22 | -2.75 | -1.46 | 6.4E-20 |
| Lrrtm4 | 422 | -1.66 | -0.73 | 1.5E-03 | -1.53 | -0.61 | 8.7E-03 |
| Lrtm2 | 305 | 1.88 | 0.91 | 3.4E-02 | 2.02 | 1.01 | 1.9E-02 |
| Lrwd1 | 2507 | -1.32 | -0.40 | 1.9E-05 | -1.47 | -0.56 | 9.1E-10 |
| Lsm1 | 1777 | 1.33 | 0.42 | 2.8E-08 | 1.28 | 0.35 | 3.1E-06 |
| Lsm10 | 531 | -1.47 | -0.56 | 9.1E-14 | -1.58 | -0.66 | 4.8E-19 |
| Lsm11 | 4238 | 1.29 | 0.37 | 2.7E-02 | 1.31 | 0.38 | 2.1E-02 |
| Lsm12 | 3551 | -1.18 | -0.24 | 1.6E-03 | -1.31 | -0.38 | 3.1E-07 |
| Lsm6 | 4424 | -1.48 | -0.56 | 2.8E-06 | -1.63 | -0.70 | 3.3E-09 |
| Lsp1 | 523 | -2.44 | -1.29 | 3.1E-06 | -2.54 | -1.35 | 1.0E-06 |
| Lss | 7158 | 1.25 | 0.32 | 1.7E-02 | 1.23 | 0.30 | 2.7E-02 |
| Lta4h | 6868 | 1.56 | 0.64 | 2.7E-16 | 1.53 | 0.61 | 5.1E-15 |
| Ltbp1 | 2596 | -1.92 | -0.94 | 1.9E-03 | -1.64 | -0.71 | 2.2E-02 |
| Ltbp2 | 136 | -3.60 | -1.85 | 9.4E-03 | -3.17 | -1.66 | 2.1E-02 |
| Ltbp3 | 3021 | -1.66 | -0.74 | 1.4E-04 | -1.57 | -0.65 | 7.9E-04 |
| Ltbp4 | 3390 | 1.74 | 0.80 | 1.8E-06 | 1.61 | 0.69 | 5.1E-05 |
| Ltbr | 659 | -2.63 | -1.40 | 5.9E-24 | -2.73 | -1.45 | 1.4E-25 |
| Luc7l | 15156 | -1.34 | -0.42 | 6.6E-08 | -1.29 | -0.37 | 2.7E-06 |
| Luc7l2 | 24711 | -1.62 | -0.70 | 1.7E-08 | -1.65 | -0.72 | 5.4E-09 |
| Lum | 7596 | -1.94 | -0.95 | 1.1E-03 | -1.92 | -0.94 | 1.3E-03 |
| Lurap1 | 660 | 3.74 | 1.90 | 2.6E-89 | 3.80 | 1.92 | 3.3E-90 |
| Lxn | 1279 | -1.84 | -0.88 | 2.5E-10 | -1.68 | -0.75 | 9.4E-08 |
| Ly6e | 2255 | -1.27 | -0.34 | 1.1E-07 | -1.32 | -0.40 | 4.3E-10 |
| Ly75 | 515 | 1.83 | 0.87 | 1.0E-05 | 2.05 | 1.03 | 1.6E-07 |
| Ly86 | 164 | -1.52 | -0.61 | 8.7E-04 | -1.44 | -0.53 | 4.5E-03 |
| Lyl1 | 193 | -1.77 | -0.82 | 5.9E-06 | -1.72 | -0.78 | 2.0E-05 |
| Lyn | 618 | -1.26 | -0.34 | 1.8E-02 | -1.34 | -0.42 | 2.7E-03 |

| | | | | | | | |
|---|---|---|---|---|---|---|---|
| Lynx1 | 536 | 4.01 | 2.00 | 3.1E-30 | 3.76 | 1.91 | 1.6E-27 |
| Lypd6 | 1468 | 1.77 | 0.82 | 6.8E-17 | 1.69 | 0.76 | 1.3E-14 |
| Lypd6b | 164 | -2.04 | -1.03 | 1.9E-04 | -2.09 | -1.06 | 1.3E-04 |
| Lyplal1 | 543 | -1.66 | -0.73 | 1.0E-11 | -1.61 | -0.69 | 2.0E-10 |
| Lyrm2 | 1993 | -1.69 | -0.76 | 1.9E-14 | -1.72 | -0.79 | 2.2E-15 |
| Lyrm9 | 2462 | 1.86 | 0.89 | 1.6E-13 | 1.88 | 0.91 | 5.3E-14 |
| Lysmd2 | 268 | -2.63 | -1.40 | 6.8E-15 | -2.71 | -1.44 | 1.3E-15 |
| Lysmd3 | 3487 | 1.43 | 0.51 | 2.3E-05 | 1.55 | 0.63 | 1.6E-07 |
| Lyst | 3193 | 1.41 | 0.50 | 2.9E-02 | 1.80 | 0.85 | 1.3E-04 |
| Lyve1 | 462 | -1.85 | -0.89 | 3.0E-07 | -1.93 | -0.95 | 3.9E-08 |
| Lyz2 | 356 | -1.45 | -0.54 | 2.1E-02 | -1.60 | -0.68 | 3.1E-03 |
| Lzic | 2336 | -1.20 | -0.26 | 2.4E-07 | -1.27 | -0.34 | 5.6E-12 |
| Lztr1 | 7298 | -1.32 | -0.40 | 3.1E-15 | -1.33 | -0.41 | 6.8E-16 |
| Lzts2 | 3586 | -1.20 | -0.27 | 3.2E-02 | -1.33 | -0.42 | 5.7E-04 |
| Lzts3 | 2386 | 2.00 | 1.00 | 6.1E-16 | 1.84 | 0.88 | 1.9E-12 |
| M6pr | 6568 | 1.48 | 0.57 | 6.5E-07 | 1.42 | 0.51 | 9.3E-06 |
| Mab21l1 | 3550 | -1.65 | -0.72 | 5.9E-04 | -1.51 | -0.59 | 5.3E-03 |
| Mab21l3 | 237 | 1.90 | 0.92 | 9.2E-07 | 1.91 | 0.94 | 7.1E-07 |
| Macir | 3330 | -1.23 | -0.30 | 2.5E-02 | -1.30 | -0.38 | 3.4E-03 |
| Maco1 | 10206 | -1.27 | -0.34 | 1.5E-04 | -1.33 | -0.41 | 3.7E-06 |
| Macrod1 | 174 | 1.57 | 0.65 | 9.6E-05 | 1.48 | 0.56 | 8.5E-04 |
| Macroh2a2 | 11646 | -2.12 | -1.08 | 1.9E-34 | -2.39 | -1.26 | 3.1E-46 |
| Mad2l2 | 2438 | 1.28 | 0.35 | 3.3E-07 | 1.24 | 0.31 | 7.7E-06 |
| Madd | 10647 | 1.63 | 0.71 | 2.1E-05 | 1.63 | 0.70 | 2.6E-05 |
| Maea | 8647 | 1.32 | 0.40 | 3.1E-07 | 1.23 | 0.30 | 1.6E-04 |
| Maf | 3212 | -1.63 | -0.70 | 3.8E-06 | -1.64 | -0.71 | 2.8E-06 |
| Mafa | 182 | 1.95 | 0.97 | 5.3E-03 | 2.98 | 1.57 | 3.4E-06 |
| Mafb | 999 | -1.57 | -0.65 | 8.5E-05 | -1.71 | -0.78 | 2.2E-06 |
| Maff | 259 | 1.61 | 0.69 | 6.0E-07 | 1.55 | 0.63 | 5.4E-06 |
| Maged2 | 13167 | -1.29 | -0.36 | 2.5E-03 | -1.26 | -0.33 | 6.8E-03 |
| Magi1 | 9802 | -1.96 | -0.97 | 2.7E-20 | -2.01 | -1.01 | 1.4E-21 |
| Magi3 | 4987 | 1.43 | 0.51 | 3.7E-10 | 1.53 | 0.61 | 7.5E-14 |
| Magohb | 733 | -1.46 | -0.55 | 1.5E-05 | -1.42 | -0.51 | 6.7E-05 |
| Mak | 566 | 5.75 | 2.52 | 4.7E-48 | 6.92 | 2.79 | 1.1E-57 |
| Malt1 | 1476 | 2.13 | 1.09 | 8.0E-14 | 2.48 | 1.31 | 2.3E-19 |
| Mamdc2 | 168 | -1.92 | -0.94 | 4.9E-07 | -2.11 | -1.08 | 8.2E-09 |
| Mamdc4 | 859 | -1.55 | -0.63 | 5.9E-07 | -1.56 | -0.64 | 3.6E-07 |
| Maml1 | 3053 | -1.49 | -0.57 | 1.3E-08 | -1.57 | -0.65 | 9.3E-11 |
| Maml2 | 1738 | -1.27 | -0.34 | 5.0E-08 | -1.29 | -0.37 | 5.1E-09 |
| Man1c1 | 1997 | 2.20 | 1.14 | 1.2E-21 | 2.02 | 1.01 | 2.5E-17 |
| Man2b1 | 3263 | 1.69 | 0.76 | 1.0E-16 | 1.58 | 0.66 | 8.6E-13 |
| Man2b2 | 852 | -1.35 | -0.43 | 8.6E-03 | -1.43 | -0.52 | 1.4E-03 |
| Man2c1 | 3909 | -1.36 | -0.45 | 1.8E-11 | -1.41 | -0.50 | 7.0E-14 |
| Manba | 721 | 1.92 | 0.94 | 1.1E-13 | 2.10 | 1.07 | 2.8E-17 |
| Manbal | 1636 | -1.48 | -0.57 | 3.9E-13 | -1.59 | -0.67 | 4.4E-18 |
| Manf | 6389 | 1.53 | 0.61 | 3.9E-12 | 1.46 | 0.55 | 5.6E-10 |
| Maoa | 3840 | 1.36 | 0.44 | 9.8E-05 | 1.39 | 0.48 | 2.0E-05 |

| | | | | | | | |
|---|---|---|---|---|---|---|---|
| Map10 | 976 | 1.39 | 0.47 | 1.4E-15 | 1.36 | 0.45 | 8.4E-14 |
| Map1a | 10875 | 1.62 | 0.69 | 2.0E-04 | 1.61 | 0.69 | 2.3E-04 |
| Map1lc3b | 12691 | 1.56 | 0.64 | 4.5E-30 | 1.58 | 0.66 | 2.3E-31 |
| Map1s | 2513 | -1.42 | -0.50 | 4.4E-05 | -1.54 | -0.62 | 4.0E-07 |
| Map2 | 68999 | -1.59 | -0.67 | 2.1E-04 | -1.41 | -0.49 | 7.8E-03 |
| Map2k1 | 4241 | 1.70 | 0.77 | 2.3E-18 | 1.67 | 0.74 | 3.6E-17 |
| Map2k2 | 5166 | -1.19 | -0.25 | 9.3E-03 | -1.31 | -0.39 | 3.1E-05 |
| Map2k3 | 2130 | 1.60 | 0.68 | 1.6E-05 | 1.53 | 0.62 | 8.9E-05 |
| Map2k5 | 1956 | -1.25 | -0.32 | 1.2E-06 | -1.35 | -0.43 | 4.4E-11 |
| Map2k6 | 3115 | 1.62 | 0.70 | 3.2E-24 | 1.59 | 0.67 | 1.6E-22 |
| Map2k7 | 2563 | -2.07 | -1.05 | 6.2E-18 | -2.22 | -1.15 | 3.6E-21 |
| Map3k10 | 4529 | 1.54 | 0.63 | 1.2E-06 | 1.48 | 0.56 | 1.5E-05 |
| Map3k13 | 3556 | 1.29 | 0.37 | 4.1E-02 | 1.34 | 0.42 | 2.0E-02 |
| Map3k14 | 709 | 1.82 | 0.86 | 5.6E-10 | 1.64 | 0.72 | 3.7E-07 |
| Map3k15 | 270 | 2.08 | 1.06 | 1.2E-12 | 1.98 | 0.98 | 3.9E-11 |
| Map3k20 | 1614 | 1.27 | 0.35 | 1.6E-04 | 1.29 | 0.36 | 9.5E-05 |
| Map3k3 | 3929 | 1.29 | 0.37 | 1.7E-08 | 1.23 | 0.30 | 5.4E-06 |
| Map3k5 | 380 | -1.77 | -0.83 | 6.1E-04 | -1.75 | -0.81 | 9.3E-04 |
| Map3k7 | 13156 | 1.34 | 0.42 | 5.0E-10 | 1.35 | 0.43 | 1.4E-10 |
| Map4k1 | 311 | -1.27 | -0.35 | 6.1E-03 | -1.37 | -0.45 | 3.2E-04 |
| Map4k4 | 38454 | -1.65 | -0.72 | 1.7E-19 | -1.67 | -0.74 | 3.9E-20 |
| Map7d2 | 2946 | 1.74 | 0.80 | 9.0E-09 | 1.71 | 0.77 | 2.4E-08 |
| Map7d3 | 405 | 2.19 | 1.13 | 1.1E-08 | 2.28 | 1.19 | 2.2E-09 |
| Mapk1 | 20963 | 1.33 | 0.41 | 3.2E-15 | 1.29 | 0.37 | 7.0E-13 |
| Mapk11 | 3675 | -1.26 | -0.33 | 2.9E-02 | -1.41 | -0.49 | 9.9E-04 |
| Mapk12 | 1570 | 1.81 | 0.85 | 2.5E-07 | 1.58 | 0.66 | 9.8E-05 |
| Mapk14 | 7022 | 1.55 | 0.64 | 4.6E-40 | 1.52 | 0.60 | 9.9E-36 |
| Mapk15 | 391 | 3.12 | 1.64 | 3.3E-28 | 2.79 | 1.48 | 4.8E-23 |
| Mapk1ip1l | 10734 | -1.23 | -0.30 | 6.1E-03 | -1.32 | -0.40 | 1.9E-04 |
| Mapk6 | 16863 | -1.38 | -0.46 | 5.1E-09 | -1.44 | -0.52 | 3.1E-11 |
| Mapk8ip1 | 12031 | 1.45 | 0.54 | 1.9E-07 | 1.37 | 0.46 | 1.0E-05 |
| Mapk9 | 10625 | 1.65 | 0.72 | 1.4E-14 | 1.80 | 0.85 | 1.0E-19 |
| Mapkapk2 | 8710 | 1.49 | 0.58 | 2.1E-07 | 1.33 | 0.41 | 3.7E-04 |
| Mapkapk3 | 645 | 3.87 | 1.95 | 1.5E-23 | 3.78 | 1.92 | 1.1E-22 |
| Mapre1 | 23742 | -1.32 | -0.40 | 7.0E-08 | -1.33 | -0.41 | 2.3E-08 |
| Mapre2 | 14965 | 1.35 | 0.43 | 1.6E-05 | 1.37 | 0.46 | 4.6E-06 |
| Mapre3 | 4964 | 1.36 | 0.44 | 1.3E-02 | 1.31 | 0.39 | 3.0E-02 |
| Marchf2 | 2921 | 2.06 | 1.04 | 5.8E-39 | 2.15 | 1.10 | 3.3E-43 |
| Marchf3 | 280 | 2.09 | 1.06 | 2.0E-09 | 2.03 | 1.02 | 9.6E-09 |
| Marchf7 | 11402 | -1.29 | -0.36 | 9.0E-04 | -1.21 | -0.28 | 1.4E-02 |
| Marcks | 64056 | -2.10 | -1.07 | 3.1E-25 | -2.26 | -1.18 | 2.7E-30 |
| Marcksl1 | 49573 | -1.44 | -0.53 | 3.9E-11 | -1.55 | -0.63 | 2.4E-15 |
| Mark1 | 5757 | -1.39 | -0.48 | 3.0E-03 | -1.42 | -0.50 | 1.7E-03 |
| Mark4 | 5821 | 1.52 | 0.60 | 5.5E-05 | 1.40 | 0.49 | 1.2E-03 |
| Mars1 | 6851 | 1.25 | 0.32 | 8.5E-05 | 1.18 | 0.24 | 3.8E-03 |
| Mars2 | 3923 | 2.16 | 1.11 | 2.3E-27 | 2.29 | 1.20 | 1.5E-31 |
| Marveld1 | 1808 | -1.59 | -0.66 | 1.3E-03 | -1.56 | -0.64 | 1.9E-03 |

| | | | | | | | |
|---|---|---|---|---|---|---|---|
| Marveld2 | 138 | -2.05 | -1.04 | 9.7E-04 | -2.58 | -1.37 | 1.1E-05 |
| Marveld3 | 145 | 3.70 | 1.89 | 1.2E-22 | 3.18 | 1.67 | 5.6E-18 |
| Maskbp3 | 102 | 1.44 | 0.53 | 2.1E-02 | 1.73 | 0.79 | 4.1E-04 |
| Masp1 | 3860 | -2.88 | -1.52 | 2.5E-18 | -2.81 | -1.49 | 1.5E-17 |
| Mast3 | 433 | 2.89 | 1.53 | 1.1E-05 | 2.18 | 1.13 | 1.5E-03 |
| Mast4 | 5669 | 1.35 | 0.43 | 6.9E-03 | 1.51 | 0.59 | 1.7E-04 |
| Matk | 460 | 2.03 | 1.02 | 3.9E-06 | 2.13 | 1.09 | 8.7E-07 |
| Matn1 | 333 | -15.73 | -3.98 | 2.2E-03 | -11.44 | -3.52 | 7.4E-03 |
| Matn2 | 712 | -2.14 | -1.09 | 3.0E-03 | -2.04 | -1.03 | 5.8E-03 |
| Matn4 | 1519 | -6.04 | -2.59 | 6.0E-03 | -5.27 | -2.40 | 1.2E-02 |
| Matr3 | 23931 | -1.27 | -0.34 | 3.7E-03 | -1.21 | -0.28 | 2.0E-02 |
| Mavs | 1242 | 1.63 | 0.70 | 1.7E-05 | 1.58 | 0.66 | 5.4E-05 |
| Max | 5377 | 1.73 | 0.79 | 2.7E-37 | 1.58 | 0.66 | 7.3E-26 |
| Maz | 16904 | -1.71 | -0.77 | 4.9E-09 | -1.90 | -0.92 | 2.2E-12 |
| Mb21d2 | 2045 | -1.25 | -0.32 | 2.0E-02 | -1.23 | -0.30 | 3.2E-02 |
| Mbd2 | 2535 | 1.31 | 0.39 | 3.7E-07 | 1.19 | 0.25 | 1.8E-03 |
| Mbd3 | 6414 | -2.01 | -1.01 | 1.9E-13 | -2.27 | -1.18 | 3.7E-18 |
| Mbd4 | 2576 | 1.30 | 0.37 | 1.3E-05 | 1.33 | 0.41 | 1.8E-06 |
| Mblac2 | 1590 | 2.37 | 1.24 | 5.2E-14 | 2.25 | 1.17 | 1.4E-12 |
| Mbnl2 | 6663 | 1.25 | 0.32 | 4.7E-02 | 1.36 | 0.44 | 5.7E-03 |
| Mboat2 | 6334 | 1.25 | 0.33 | 1.3E-05 | 1.28 | 0.36 | 1.3E-06 |
| Mboat7 | 5265 | 1.93 | 0.95 | 3.0E-26 | 1.91 | 0.94 | 2.5E-25 |
| Mbp | 2022 | 1.26 | 0.33 | 3.0E-04 | 1.19 | 0.26 | 6.0E-03 |
| Mbtps1 | 9913 | -1.40 | -0.48 | 9.0E-08 | -1.39 | -0.47 | 1.9E-07 |
| Mc4r | 165 | 1.45 | 0.54 | 6.3E-03 | 1.60 | 0.67 | 6.3E-04 |
| Mcam | 4295 | 1.79 | 0.84 | 4.6E-17 | 1.65 | 0.72 | 5.1E-13 |
| Mcat | 1582 | -1.28 | -0.36 | 9.0E-06 | -1.38 | -0.46 | 5.5E-09 |
| Mcee | 606 | 1.35 | 0.43 | 1.7E-06 | 1.38 | 0.46 | 2.9E-07 |
| Mcf2l | 7583 | -2.10 | -1.07 | 2.1E-07 | -2.16 | -1.11 | 7.2E-08 |
| Mcfd2 | 3212 | 1.29 | 0.37 | 9.2E-05 | 1.28 | 0.35 | 2.2E-04 |
| Mcidas | 133 | -4.10 | -2.04 | 1.0E-13 | -3.80 | -1.93 | 1.4E-12 |
| Mcl1 | 13988 | -1.20 | -0.26 | 6.8E-05 | -1.26 | -0.34 | 2.8E-07 |
| Mcm3ap | 5291 | -3.54 | -1.83 | 1.2E-114 | -3.59 | -1.85 | 5.1E-117 |
| Mcm9 | 1058 | -1.31 | -0.39 | 4.3E-06 | -1.29 | -0.37 | 2.3E-05 |
| Mcoln1 | 4522 | 1.35 | 0.43 | 1.2E-05 | 1.29 | 0.37 | 1.8E-04 |
| Mcrip2 | 211 | 3.23 | 1.69 | 2.1E-26 | 2.97 | 1.57 | 9.2E-23 |
| Mcrs1 | 4070 | -1.89 | -0.92 | 8.1E-32 | -2.11 | -1.08 | 2.3E-43 |
| Mcts1 | 3739 | -1.31 | -0.39 | 6.6E-05 | -1.24 | -0.32 | 1.4E-03 |
| Mcub | 136 | 2.23 | 1.15 | 4.6E-22 | 1.96 | 0.97 | 5.5E-16 |
| Mdfic | 1291 | -2.33 | -1.22 | 7.0E-17 | -2.50 | -1.32 | 1.3E-19 |
| Mdh1 | 19514 | 1.53 | 0.62 | 7.9E-14 | 1.63 | 0.70 | 9.6E-18 |
| Mdk | 9601 | -1.57 | -0.65 | 4.2E-03 | -1.77 | -0.82 | 2.3E-04 |
| Mdm1 | 7281 | 2.48 | 1.31 | 4.4E-50 | 2.41 | 1.27 | 5.8E-47 |
| Mdm2 | 5593 | -1.29 | -0.37 | 4.5E-04 | -1.41 | -0.49 | 1.9E-06 |
| Mdm4 | 17786 | -1.36 | -0.44 | 1.7E-05 | -1.19 | -0.25 | 1.9E-02 |
| Mdn1 | 11812 | 1.59 | 0.67 | 1.6E-03 | 1.93 | 0.95 | 5.1E-06 |
| Mdp1 | 2372 | -1.28 | -0.36 | 7.1E-07 | -1.28 | -0.36 | 5.5E-07 |

| | | | | | | | |
|---|---|---|---|---|---|---|---|
| Me1 | 1309 | -1.46 | -0.55 | 8.6E-05 | -1.39 | -0.47 | 9.3E-04 |
| Meaf6 | 4406 | -2.01 | -1.01 | 5.3E-12 | -2.16 | -1.11 | 1.8E-14 |
| Mecom | 1546 | -1.69 | -0.76 | 5.4E-03 | -1.53 | -0.61 | 2.8E-02 |
| Mecp2 | 11258 | -1.59 | -0.67 | 2.3E-05 | -1.54 | -0.63 | 7.6E-05 |
| Mecr | 1644 | 1.95 | 0.96 | 2.9E-23 | 1.86 | 0.90 | 2.0E-20 |
| Med10 | 3468 | 1.47 | 0.56 | 3.9E-16 | 1.38 | 0.47 | 1.1E-11 |
| Med11 | 678 | -1.95 | -0.97 | 7.8E-22 | -1.89 | -0.92 | 9.4E-20 |
| Med12 | 7067 | -1.57 | -0.65 | 7.3E-11 | -1.45 | -0.53 | 1.6E-07 |
| Med14 | 9015 | 1.22 | 0.29 | 1.0E-02 | 1.30 | 0.38 | 4.8E-04 |
| Med16 | 2471 | -2.03 | -1.02 | 6.5E-41 | -2.18 | -1.12 | 2.3E-49 |
| Med22 | 6460 | 1.33 | 0.41 | 1.0E-09 | 1.26 | 0.33 | 1.2E-06 |
| Med24 | 6283 | -1.62 | -0.69 | 1.1E-26 | -1.65 | -0.72 | 5.1E-29 |
| Med6 | 2093 | -1.57 | -0.65 | 1.2E-09 | -1.62 | -0.70 | 6.8E-11 |
| Med9 | 1312 | -1.19 | -0.25 | 1.6E-02 | -1.26 | -0.33 | 1.2E-03 |
| Mef2a | 6251 | -1.34 | -0.42 | 3.2E-04 | -1.28 | -0.35 | 3.3E-03 |
| Mef2d | 6108 | 1.54 | 0.62 | 1.3E-05 | 1.47 | 0.56 | 1.1E-04 |
| Megf10 | 3621 | -1.96 | -0.97 | 6.1E-22 | -1.80 | -0.84 | 9.4E-17 |
| Megf9 | 12905 | 1.41 | 0.49 | 4.8E-07 | 1.43 | 0.52 | 1.2E-07 |
| Meioc | 118 | 2.85 | 1.51 | 1.8E-03 | 4.19 | 2.07 | 1.6E-05 |
| Meis2 | 15231 | -1.49 | -0.57 | 9.6E-04 | -1.39 | -0.47 | 7.0E-03 |
| Memo1 | 4132 | -1.22 | -0.29 | 1.6E-03 | -1.28 | -0.36 | 6.5E-05 |
| Meox2 | 259 | -3.50 | -1.81 | 7.2E-03 | -2.77 | -1.47 | 3.3E-02 |
| Mepce | 2915 | -1.45 | -0.54 | 1.1E-06 | -1.54 | -0.63 | 8.9E-09 |
| Mest | 36402 | -1.38 | -0.47 | 3.6E-02 | -1.45 | -0.53 | 1.7E-02 |
| Metap1 | 5356 | 1.39 | 0.47 | 4.4E-15 | 1.44 | 0.53 | 8.6E-19 |
| Metap1d | 2494 | -1.43 | -0.51 | 8.3E-09 | -1.23 | -0.30 | 1.4E-03 |
| Metrn | 4092 | 2.39 | 1.26 | 5.0E-10 | 2.07 | 1.05 | 2.6E-07 |
| Mettl14 | 3357 | -1.18 | -0.24 | 1.8E-02 | -1.26 | -0.34 | 9.1E-04 |
| Mettl15 | 259 | -1.58 | -0.66 | 1.3E-07 | -1.61 | -0.69 | 5.8E-08 |
| Mettl16 | 4340 | -1.46 | -0.55 | 6.6E-33 | -1.44 | -0.53 | 1.9E-30 |
| Mettl17 | 2775 | 1.34 | 0.42 | 2.0E-10 | 1.30 | 0.38 | 9.5E-09 |
| Mettl21a | 888 | -1.51 | -0.59 | 2.9E-13 | -1.53 | -0.62 | 2.8E-14 |
| Mettl22 | 504 | -2.07 | -1.05 | 5.4E-16 | -2.19 | -1.13 | 2.3E-18 |
| Mettl25 | 505 | -2.81 | -1.49 | 3.5E-46 | -2.85 | -1.51 | 4.1E-47 |
| Mettl27 | 770 | 2.27 | 1.19 | 1.1E-17 | 2.28 | 1.19 | 1.1E-17 |
| Mettl4 | 3171 | -2.09 | -1.06 | 1.3E-46 | -1.90 | -0.93 | 6.6E-36 |
| Mettl6 | 4156 | 1.30 | 0.37 | 5.8E-06 | 1.27 | 0.35 | 3.0E-05 |
| Mettl8 | 1816 | 1.49 | 0.57 | 2.5E-14 | 1.62 | 0.70 | 8.7E-21 |
| Mettl9 | 8475 | -1.38 | -0.47 | 1.2E-06 | -1.46 | -0.55 | 1.1E-08 |
| Mex3a | 23844 | -1.76 | -0.81 | 1.1E-07 | -2.00 | -1.00 | 4.7E-11 |
| Mex3b | 7788 | -2.08 | -1.06 | 1.4E-23 | -2.26 | -1.18 | 5.5E-29 |
| Mfap1a | 3409 | 1.28 | 0.35 | 2.4E-09 | 1.29 | 0.36 | 6.8E-10 |
| Mfap1b | 1919 | 1.26 | 0.33 | 1.2E-05 | 1.26 | 0.33 | 8.8E-06 |
| Mfap2 | 5735 | -1.57 | -0.65 | 3.9E-05 | -1.58 | -0.66 | 2.9E-05 |
| Mfap3 | 9210 | 1.28 | 0.36 | 1.3E-05 | 1.36 | 0.44 | 7.5E-08 |
| Mfap3l | 1004 | 2.22 | 1.15 | 3.2E-18 | 2.68 | 1.42 | 4.1E-27 |
| Mfap4 | 4350 | -2.10 | -1.07 | 2.9E-04 | -1.97 | -0.98 | 1.0E-03 |

| | | | | | | | |
|---|---|---|---|---|---|---|---|
| Mfge8 | 3006 | 1.48 | 0.57 | 2.8E-05 | 1.35 | 0.44 | 1.7E-03 |
| Mfhas1 | 3184 | 1.27 | 0.35 | 4.4E-03 | 1.22 | 0.29 | 1.9E-02 |
| Mfng | 3161 | -2.22 | -1.15 | 4.6E-08 | -2.64 | -1.40 | 1.9E-11 |
| Mfsd1 | 5510 | 1.64 | 0.71 | 2.0E-14 | 1.60 | 0.68 | 3.1E-13 |
| Mfsd10 | 1380 | -1.93 | -0.95 | 2.7E-21 | -2.18 | -1.12 | 3.5E-29 |
| Mfsd11 | 2408 | 1.45 | 0.53 | 5.2E-21 | 1.51 | 0.59 | 1.9E-25 |
| Mfsd12 | 2585 | 4.22 | 2.08 | 2.9E-55 | 3.84 | 1.94 | 1.8E-48 |
| Mfsd13a | 472 | 1.91 | 0.94 | 3.5E-11 | 1.82 | 0.86 | 1.4E-09 |
| Mfsd14b | 4325 | 1.30 | 0.38 | 4.5E-06 | 1.34 | 0.42 | 3.9E-07 |
| Mfsd2b | 505 | 1.50 | 0.58 | 1.6E-04 | 1.52 | 0.60 | 9.2E-05 |
| Mfsd3 | 268 | 1.43 | 0.52 | 6.7E-06 | 1.38 | 0.46 | 6.7E-05 |
| Mfsd4b4 | 1123 | -1.39 | -0.48 | 9.5E-13 | -1.26 | -0.33 | 1.2E-06 |
| Mfsd6 | 1074 | -2.97 | -1.57 | 7.5E-21 | -3.04 | -1.60 | 1.2E-21 |
| Mfsd9 | 638 | 3.03 | 1.60 | 4.9E-41 | 3.06 | 1.61 | 2.2E-41 |
| Mgarp | 268 | 1.96 | 0.97 | 7.6E-03 | 1.83 | 0.87 | 1.8E-02 |
| Mgat1 | 3069 | -1.35 | -0.44 | 1.4E-07 | -1.44 | -0.52 | 2.4E-10 |
| Mgat3 | 4905 | -1.61 | -0.69 | 9.8E-09 | -1.67 | -0.74 | 5.6E-10 |
| Mgat4a | 1876 | 1.31 | 0.39 | 2.7E-02 | 1.42 | 0.51 | 3.4E-03 |
| Mgat5b | 3614 | -1.79 | -0.84 | 7.9E-04 | -1.85 | -0.89 | 3.9E-04 |
| Mgme1 | 683 | -1.97 | -0.98 | 4.3E-19 | -1.98 | -0.98 | 2.3E-19 |
| Mgp | 202 | -5.29 | -2.40 | 6.4E-06 | -3.90 | -1.96 | 2.8E-04 |
| Mgrn1 | 8129 | 1.26 | 0.33 | 1.7E-06 | 1.22 | 0.29 | 3.4E-05 |
| Mgst3 | 1564 | 1.59 | 0.67 | 3.8E-11 | 1.68 | 0.75 | 7.9E-14 |
| Mib2 | 3792 | 1.39 | 0.48 | 1.5E-04 | 1.31 | 0.39 | 1.9E-03 |
| Mical1 | 2455 | -1.84 | -0.88 | 7.2E-19 | -2.06 | -1.05 | 3.2E-26 |
| Mical2 | 885 | 2.42 | 1.27 | 6.4E-15 | 2.31 | 1.21 | 1.5E-13 |
| Mical3 | 6409 | 1.48 | 0.57 | 3.2E-04 | 1.44 | 0.53 | 8.1E-04 |
| Micall1 | 4284 | -1.33 | -0.41 | 1.8E-04 | -1.31 | -0.39 | 3.8E-04 |
| Micall2 | 1258 | 2.10 | 1.07 | 2.5E-20 | 2.01 | 1.00 | 5.7E-18 |
| Micu1 | 5433 | 1.87 | 0.90 | 4.9E-71 | 1.86 | 0.89 | 2.2E-69 |
| Micu2 | 3704 | -2.11 | -1.08 | 2.6E-46 | -2.13 | -1.09 | 2.3E-47 |
| Mideas | 3080 | 1.29 | 0.37 | 2.1E-04 | 1.28 | 0.36 | 4.1E-04 |
| Midn | 15309 | -1.91 | -0.93 | 3.1E-09 | -2.18 | -1.12 | 5.8E-13 |
| Mier3 | 3964 | -1.28 | -0.36 | 1.4E-06 | -1.22 | -0.28 | 2.1E-04 |
| Miga2 | 2929 | 1.72 | 0.78 | 1.2E-36 | 1.70 | 0.77 | 1.2E-35 |
| Miip | 3843 | 1.34 | 0.42 | 1.9E-05 | 1.20 | 0.26 | 1.1E-02 |
| Minar1 | 411 | 1.86 | 0.89 | 5.2E-03 | 2.71 | 1.44 | 4.2E-06 |
| Minar2 | 3572 | -1.49 | -0.58 | 9.5E-04 | -1.42 | -0.51 | 4.1E-03 |
| Mindy1 | 2377 | -1.64 | -0.72 | 1.4E-44 | -1.69 | -0.76 | 9.4E-50 |
| Mindy2 | 5630 | 1.45 | 0.54 | 6.3E-05 | 1.56 | 0.64 | 1.5E-06 |
| Mindy3 | 3678 | 1.36 | 0.45 | 5.0E-09 | 1.39 | 0.47 | 4.8E-10 |
| Mindy4 | 560 | -5.12 | -2.36 | 1.1E-97 | -4.78 | -2.26 | 3.3E-90 |
| Mink1 | 6672 | 1.33 | 0.41 | 6.4E-04 | 1.25 | 0.32 | 9.3E-03 |
| Mios | 2877 | -1.30 | -0.38 | 2.3E-06 | -1.19 | -0.25 | 2.1E-03 |
| Mis18a | 1946 | 1.57 | 0.65 | 5.2E-05 | 1.46 | 0.54 | 8.4E-04 |
| Misp | 245 | 2.97 | 1.57 | 1.2E-07 | 3.14 | 1.65 | 2.5E-08 |
| Mitf | 516 | 2.23 | 1.15 | 3.3E-14 | 2.43 | 1.28 | 3.5E-17 |

| | | | | | | | |
|---|---|---|---|---|---|---|---|
| Mki67 | 37376 | -1.42 | -0.50 | 4.5E-03 | -1.33 | -0.41 | 2.3E-02 |
| Mknk1 | 3209 | 2.16 | 1.11 | 2.2E-70 | 2.23 | 1.16 | 2.9E-76 |
| Mknk2 | 8080 | 1.76 | 0.82 | 1.5E-10 | 1.64 | 0.71 | 3.2E-08 |
| Mks1 | 1023 | -2.03 | -1.02 | 1.4E-25 | -2.08 | -1.06 | 3.7E-27 |
| Mlc1 | 600 | -2.97 | -1.57 | 1.8E-14 | -3.17 | -1.67 | 3.5E-16 |
| Mlec | 23501 | 1.44 | 0.53 | 1.3E-06 | 1.46 | 0.54 | 6.8E-07 |
| Mlf1 | 943 | 1.63 | 0.71 | 3.3E-06 | 1.79 | 0.84 | 2.9E-08 |
| Mlf2 | 14065 | 1.26 | 0.34 | 1.1E-04 | 1.21 | 0.28 | 1.9E-03 |
| Mlh1 | 2263 | -1.47 | -0.56 | 8.7E-09 | -1.57 | -0.65 | 2.1E-11 |
| Mlh3 | 1742 | -1.77 | -0.82 | 6.2E-28 | -1.57 | -0.65 | 5.7E-18 |
| Mllt10 | 5619 | -1.47 | -0.56 | 2.0E-18 | -1.50 | -0.59 | 1.8E-20 |
| Mllt11 | 16031 | -1.56 | -0.65 | 2.5E-03 | -1.49 | -0.57 | 8.0E-03 |
| Mllt3 | 7786 | -1.52 | -0.61 | 4.1E-11 | -1.47 | -0.56 | 1.3E-09 |
| Mlst8 | 2354 | -2.67 | -1.42 | 8.3E-74 | -2.96 | -1.57 | 6.3E-90 |
| Mlxip | 9877 | 2.09 | 1.06 | 2.1E-28 | 2.00 | 1.00 | 2.7E-25 |
| Mlxipl | 515 | 4.55 | 2.19 | 5.5E-31 | 3.73 | 1.90 | 1.1E-23 |
| Mmab | 1482 | -1.85 | -0.88 | 6.7E-26 | -1.90 | -0.93 | 2.9E-28 |
| Mmachc | 1470 | -1.62 | -0.70 | 1.1E-09 | -1.74 | -0.80 | 2.4E-12 |
| Mmd | 6246 | 1.48 | 0.57 | 2.4E-06 | 1.48 | 0.57 | 2.6E-06 |
| Mmgt2 | 1037 | -1.33 | -0.41 | 2.3E-05 | -1.45 | -0.53 | 3.2E-08 |
| Mmp14 | 5686 | -2.88 | -1.52 | 6.7E-19 | -2.89 | -1.53 | 5.1E-19 |
| Mmp17 | 2242 | 2.88 | 1.52 | 1.5E-29 | 2.57 | 1.36 | 7.3E-24 |
| Mmp23 | 259 | 1.97 | 0.98 | 1.6E-06 | 2.07 | 1.05 | 2.2E-07 |
| Mmp24 | 2663 | -1.59 | -0.67 | 2.4E-02 | -1.54 | -0.62 | 3.8E-02 |
| Mmp25 | 191 | -1.45 | -0.54 | 5.4E-03 | -1.39 | -0.47 | 1.6E-02 |
| Mmrn2 | 804 | -1.33 | -0.41 | 2.3E-05 | -1.34 | -0.43 | 1.2E-05 |
| Mms19 | 6431 | 1.64 | 0.71 | 8.3E-20 | 1.70 | 0.77 | 1.1E-22 |
| Mms22l | 3477 | 1.34 | 0.42 | 6.8E-03 | 1.33 | 0.41 | 8.9E-03 |
| Mob2 | 1818 | 1.39 | 0.47 | 1.9E-14 | 1.40 | 0.49 | 4.7E-15 |
| Mob3a | 3676 | 1.55 | 0.63 | 2.5E-09 | 1.48 | 0.57 | 1.3E-07 |
| Mob3c | 423 | -1.48 | -0.56 | 4.6E-09 | -1.71 | -0.77 | 6.8E-16 |
| Mocs2 | 4439 | 1.58 | 0.66 | 2.6E-26 | 1.62 | 0.69 | 7.4E-29 |
| Mocs3 | 562 | 1.25 | 0.32 | 6.2E-06 | 1.29 | 0.36 | 3.9E-07 |
| Mogat2 | 213 | 2.29 | 1.20 | 1.3E-05 | 1.78 | 0.83 | 3.1E-03 |
| Mok | 770 | 1.68 | 0.75 | 6.5E-21 | 1.80 | 0.85 | 2.6E-26 |
| Mon1a | 907 | -1.30 | -0.38 | 3.3E-04 | -1.39 | -0.48 | 3.8E-06 |
| Mon1b | 2895 | 1.36 | 0.44 | 9.1E-05 | 1.28 | 0.36 | 1.6E-03 |
| Mon2 | 9326 | 1.28 | 0.36 | 3.3E-05 | 1.35 | 0.44 | 2.5E-07 |
| Morc3 | 3727 | -1.29 | -0.36 | 9.7E-13 | -1.24 | -0.31 | 1.1E-09 |
| Morc4 | 1402 | 1.49 | 0.58 | 2.7E-05 | 1.50 | 0.59 | 2.0E-05 |
| Morf4l1 | 20022 | -1.42 | -0.51 | 1.4E-15 | -1.48 | -0.56 | 5.8E-19 |
| Morn1 | 629 | 2.37 | 1.25 | 3.8E-47 | 2.23 | 1.16 | 1.3E-40 |
| Mospd2 | 2650 | 1.27 | 0.34 | 4.1E-04 | 1.35 | 0.43 | 5.8E-06 |
| Mov10 | 1177 | -1.70 | -0.77 | 5.7E-05 | -1.84 | -0.88 | 3.5E-06 |
| Mpc1 | 1384 | 1.32 | 0.40 | 3.5E-04 | 1.40 | 0.49 | 1.2E-05 |
| Mpc2 | 2092 | 1.24 | 0.31 | 1.5E-03 | 1.30 | 0.38 | 9.2E-05 |

| | | | | | | | |
|---|---|---|---|---|---|---|---|
| Mpdu1 | 2495 | -2.10 | -1.07 | 2.0E-36 | -2.28 | -1.19 | 1.1E-44 |
| Mpeg1 | 670 | -1.43 | -0.52 | 2.5E-03 | -1.32 | -0.40 | 2.0E-02 |
| Mphosph8 | 7814 | 1.32 | 0.40 | 2.6E-07 | 1.30 | 0.38 | 1.4E-06 |
| Mphosph9 | 3551 | -1.49 | -0.57 | 3.2E-06 | -1.41 | -0.49 | 6.6E-05 |
| Mpi | 3535 | 1.41 | 0.49 | 4.2E-14 | 1.35 | 0.43 | 3.6E-11 |
| Mpp2 | 4567 | 1.88 | 0.91 | 4.0E-12 | 1.75 | 0.81 | 7.9E-10 |
| Mpp3 | 3135 | 1.28 | 0.36 | 8.7E-03 | 1.29 | 0.36 | 7.6E-03 |
| Mpp4 | 216 | 2.66 | 1.41 | 2.6E-10 | 3.07 | 1.62 | 4.0E-13 |
| Mpp7 | 2944 | 1.78 | 0.83 | 6.8E-19 | 1.74 | 0.80 | 1.3E-17 |
| Mppe1 | 364 | -2.72 | -1.45 | 5.3E-15 | -2.59 | -1.37 | 1.3E-13 |
| Mpped2 | 4714 | -2.58 | -1.37 | 8.2E-12 | -2.59 | -1.37 | 6.9E-12 |
| Mpst | 1986 | 1.50 | 0.58 | 9.2E-11 | 1.35 | 0.43 | 2.3E-06 |
| Mpv17l | 761 | 2.04 | 1.03 | 1.2E-10 | 2.17 | 1.12 | 2.0E-12 |
| Mpzl1 | 9286 | 1.61 | 0.69 | 7.5E-40 | 1.60 | 0.68 | 5.8E-39 |
| Mpzl3 | 134 | -4.67 | -2.22 | 5.0E-42 | -4.85 | -2.28 | 3.0E-43 |
| Mrc1 | 2082 | -1.32 | -0.40 | 1.5E-02 | -1.34 | -0.42 | 1.1E-02 |
| Mrc2 | 4882 | -1.66 | -0.73 | 1.4E-04 | -1.81 | -0.86 | 6.4E-06 |
| Mreg | 417 | 2.03 | 1.02 | 3.4E-04 | 3.03 | 1.60 | 9.6E-09 |
| Mrfap1 | 18037 | -1.27 | -0.34 | 9.8E-04 | -1.38 | -0.46 | 7.0E-06 |
| Mrgpre | 425 | -1.50 | -0.59 | 5.4E-06 | -1.55 | -0.63 | 9.1E-07 |
| Mrm1 | 852 | 1.88 | 0.91 | 1.9E-20 | 1.90 | 0.93 | 3.6E-21 |
| Mrm3 | 468 | -1.20 | -0.27 | 4.8E-03 | -1.35 | -0.43 | 2.3E-06 |
| Mro | 349 | 1.59 | 0.67 | 3.8E-07 | 1.58 | 0.66 | 5.4E-07 |
| Mroh1 | 5037 | 2.01 | 1.00 | 4.9E-20 | 2.01 | 1.01 | 3.3E-20 |
| Mrpl11 | 2893 | -1.35 | -0.43 | 1.9E-06 | -1.40 | -0.48 | 8.0E-08 |
| Mrpl14 | 1093 | 1.30 | 0.37 | 1.9E-05 | 1.23 | 0.30 | 6.8E-04 |
| Mrpl17 | 2490 | 1.31 | 0.40 | 5.8E-10 | 1.23 | 0.30 | 3.0E-06 |
| Mrpl18 | 2767 | -1.19 | -0.25 | 2.6E-02 | -1.28 | -0.35 | 1.6E-03 |
| Mrpl21 | 1573 | -1.42 | -0.50 | 2.3E-06 | -1.52 | -0.60 | 1.4E-08 |
| Mrpl23 | 1564 | -1.43 | -0.52 | 3.5E-07 | -1.53 | -0.62 | 1.0E-09 |
| Mrpl24 | 6353 | 1.30 | 0.38 | 8.3E-10 | 1.27 | 0.34 | 2.9E-08 |
| Mrpl27 | 1124 | -2.18 | -1.13 | 6.8E-36 | -2.22 | -1.15 | 1.6E-37 |
| Mrpl28 | 3257 | 1.41 | 0.50 | 1.8E-06 | 1.28 | 0.35 | 1.0E-03 |
| Mrpl3 | 3732 | -1.49 | -0.57 | 9.9E-22 | -1.58 | -0.66 | 2.0E-28 |
| Mrpl34 | 1952 | 1.37 | 0.45 | 1.3E-05 | 1.20 | 0.26 | 1.5E-02 |
| Mrpl35 | 1437 | -1.51 | -0.59 | 6.3E-11 | -1.48 | -0.56 | 5.6E-10 |
| Mrpl37 | 3517 | -1.30 | -0.37 | 6.0E-08 | -1.36 | -0.44 | 1.5E-10 |
| Mrpl40 | 1394 | -1.38 | -0.46 | 3.1E-05 | -1.44 | -0.53 | 1.8E-06 |
| Mrpl41 | 2210 | 1.36 | 0.45 | 2.9E-07 | 1.39 | 0.47 | 5.2E-08 |
| Mrpl43 | 2286 | 1.53 | 0.61 | 5.6E-13 | 1.40 | 0.49 | 1.4E-08 |
| Mrpl44 | 1438 | -1.47 | -0.55 | 4.6E-09 | -1.52 | -0.60 | 1.8E-10 |
| Mrpl45 | 1841 | -1.90 | -0.92 | 2.6E-39 | -2.03 | -1.02 | 1.7E-47 |
| Mrpl46 | 1168 | -1.26 | -0.33 | 3.0E-03 | -1.30 | -0.38 | 6.7E-04 |
| Mrpl48 | 1790 | -1.90 | -0.93 | 1.5E-54 | -1.85 | -0.89 | 5.3E-50 |
| Mrpl49 | 3480 | 1.53 | 0.61 | 1.7E-09 | 1.40 | 0.49 | 2.3E-06 |
| Mrpl54 | 740 | -2.53 | -1.34 | 1.6E-19 | -2.62 | -1.39 | 8.1E-21 |

| | | | | | | | |
|---|---|---|---|---|---|---|---|
| Mrpl55 | 1554 | 1.36 | 0.44 | 4.5E-06 | 1.29 | 0.36 | 2.0E-04 |
| Mrpl58 | 2394 | -1.50 | -0.58 | 9.8E-08 | -1.69 | -0.76 | 2.2E-12 |
| Mrps12 | 1130 | 1.32 | 0.40 | 4.0E-04 | 1.23 | 0.30 | 9.2E-03 |
| Mrps15 | 1902 | -1.48 | -0.56 | 7.7E-18 | -1.47 | -0.56 | 1.9E-17 |
| Mrps16 | 1164 | -1.30 | -0.38 | 2.7E-04 | -1.37 | -0.45 | 1.1E-05 |
| Mrps18a | 1426 | -1.53 | -0.62 | 1.7E-24 | -1.60 | -0.68 | 3.2E-29 |
| Mrps22 | 2285 | 1.26 | 0.34 | 2.7E-04 | 1.30 | 0.38 | 3.2E-05 |
| Mrps26 | 1920 | 1.27 | 0.35 | 1.5E-07 | 1.17 | 0.22 | 1.3E-03 |
| Mrps28 | 457 | -1.59 | -0.67 | 9.3E-06 | -1.60 | -0.68 | 6.0E-06 |
| Mrps30 | 1973 | -1.32 | -0.40 | 6.5E-07 | -1.37 | -0.46 | 1.6E-08 |
| Mrps35 | 1439 | -1.73 | -0.79 | 1.9E-26 | -1.90 | -0.93 | 1.7E-35 |
| Mrps5 | 4667 | -1.31 | -0.39 | 1.8E-12 | -1.35 | -0.43 | 1.3E-14 |
| Mrps6 | 1363 | 1.55 | 0.63 | 1.2E-07 | 1.48 | 0.56 | 3.1E-06 |
| Mrrf | 1458 | -1.64 | -0.72 | 6.2E-40 | -1.67 | -0.74 | 3.2E-42 |
| Mrs2 | 8554 | 1.23 | 0.30 | 4.0E-04 | 1.32 | 0.40 | 2.3E-06 |
| Mrtfa | 3292 | 1.38 | 0.46 | 1.4E-04 | 1.23 | 0.30 | 1.7E-02 |
| Mrtfb | 3074 | -1.45 | -0.54 | 6.7E-10 | -1.39 | -0.48 | 5.4E-08 |
| Mrto4 | 2130 | -1.22 | -0.29 | 3.1E-03 | -1.36 | -0.45 | 2.4E-06 |
| Ms4a6b | 163 | -1.99 | -1.00 | 4.1E-06 | -2.21 | -1.15 | 1.0E-07 |
| Ms4a6c | 182 | -2.07 | -1.05 | 1.1E-07 | -2.11 | -1.08 | 4.2E-08 |
| Msantd1 | 237 | 1.47 | 0.56 | 1.9E-05 | 1.45 | 0.53 | 5.5E-05 |
| Msantd3 | 2656 | -1.62 | -0.70 | 8.5E-10 | -1.79 | -0.84 | 1.1E-13 |
| Msh2 | 10003 | 1.36 | 0.45 | 1.9E-06 | 1.29 | 0.36 | 1.3E-04 |
| Msh3 | 3059 | -1.36 | -0.45 | 7.2E-08 | -1.37 | -0.46 | 3.6E-08 |
| Msh5 | 739 | 2.41 | 1.27 | 1.7E-16 | 2.29 | 1.19 | 9.1E-15 |
| Msh6 | 10340 | 1.26 | 0.33 | 9.7E-03 | 1.20 | 0.26 | 4.7E-02 |
| Msi1 | 10774 | -2.33 | -1.22 | 5.8E-07 | -2.84 | -1.51 | 5.1E-10 |
| Msl1 | 15044 | -1.86 | -0.89 | 2.0E-14 | -2.02 | -1.01 | 3.0E-18 |
| Msl3l2 | 386 | -1.94 | -0.96 | 1.6E-31 | -2.03 | -1.02 | 4.4E-35 |
| Msmo1 | 7657 | 1.46 | 0.55 | 1.1E-03 | 1.41 | 0.49 | 3.7E-03 |
| Msrb1 | 1392 | 2.02 | 1.02 | 4.7E-27 | 1.94 | 0.95 | 5.6E-24 |
| Msrb2 | 231 | -3.77 | -1.91 | 1.4E-29 | -3.82 | -1.93 | 5.9E-30 |
| Mt1 | 609 | 1.63 | 0.71 | 4.2E-05 | 1.61 | 0.69 | 6.5E-05 |
| Mta1 | 7459 | -1.24 | -0.31 | 4.9E-02 | -1.40 | -0.49 | 1.4E-03 |
| Mta3 | 1599 | -1.65 | -0.72 | 6.7E-18 | -1.70 | -0.77 | 5.2E-20 |
| Mtap | 5054 | 1.58 | 0.66 | 1.5E-05 | 1.43 | 0.52 | 9.2E-04 |
| Mtarc2 | 2559 | 1.44 | 0.52 | 1.7E-08 | 1.52 | 0.60 | 7.6E-11 |
| Mtch2 | 3929 | 1.29 | 0.37 | 9.8E-08 | 1.29 | 0.37 | 1.2E-07 |
| Mterf3 | 2886 | -1.39 | -0.48 | 3.2E-07 | -1.34 | -0.42 | 7.0E-06 |
| Mtfmt | 1768 | 1.52 | 0.61 | 1.2E-38 | 1.53 | 0.61 | 5.8E-39 |
| Mtfp1 | 351 | 2.31 | 1.21 | 6.1E-23 | 2.27 | 1.18 | 4.8E-22 |
| Mtfr1 | 4828 | 1.43 | 0.52 | 5.5E-09 | 1.39 | 0.47 | 1.6E-07 |
| Mtfr1l | 6647 | 1.58 | 0.66 | 5.0E-36 | 1.57 | 0.65 | 4.0E-35 |
| Mtfr2 | 863 | 1.84 | 0.88 | 3.5E-07 | 1.67 | 0.74 | 2.0E-05 |
| Mtg1 | 878 | -1.96 | -0.97 | 1.4E-30 | -1.93 | -0.95 | 2.1E-29 |
| Mtg2 | 1525 | -1.22 | -0.28 | 1.3E-03 | -1.27 | -0.35 | 7.0E-05 |

| | | | | | | | |
|---|---|---|---|---|---|---|---|
| Mthfd1 | 4316 | 1.28 | 0.36 | 4.3E-03 | 1.19 | 0.26 | 4.7E-02 |
| Mthfd1l | 881 | -1.83 | -0.87 | 5.5E-05 | -1.98 | -0.98 | 5.0E-06 |
| Mthfd2 | 2200 | 1.64 | 0.71 | 3.6E-07 | 1.53 | 0.61 | 1.6E-05 |
| Mthfd2l | 2867 | -1.72 | -0.78 | 7.9E-13 | -1.66 | -0.73 | 2.6E-11 |
| Mthfsl | 295 | -2.34 | -1.22 | 7.3E-19 | -2.06 | -1.04 | 6.8E-14 |
| Mtif2 | 4422 | -1.27 | -0.34 | 1.2E-05 | -1.26 | -0.33 | 2.3E-05 |
| Mtm1 | 1136 | 2.88 | 1.53 | 2.2E-42 | 3.11 | 1.64 | 2.6E-48 |
| Mtmr1 | 2726 | -1.58 | -0.66 | 2.8E-06 | -1.45 | -0.53 | 1.8E-04 |
| Mtmr10 | 2554 | 1.55 | 0.63 | 1.6E-16 | 1.61 | 0.68 | 3.2E-19 |
| Mtmr11 | 1131 | 1.25 | 0.32 | 8.0E-03 | 1.31 | 0.39 | 1.1E-03 |
| Mtmr12 | 2972 | 1.49 | 0.58 | 4.8E-18 | 1.49 | 0.58 | 7.4E-18 |
| Mtmr14 | 3162 | 1.56 | 0.64 | 1.4E-14 | 1.45 | 0.53 | 2.7E-10 |
| Mtmr2 | 4932 | -1.58 | -0.66 | 1.5E-16 | -1.57 | -0.65 | 2.3E-16 |
| Mtmr6 | 8264 | 1.35 | 0.43 | 2.7E-08 | 1.41 | 0.50 | 1.6E-10 |
| Mtmr9 | 5920 | -1.26 | -0.33 | 7.4E-04 | -1.22 | -0.28 | 4.4E-03 |
| Mtor | 10534 | 1.48 | 0.57 | 7.4E-11 | 1.56 | 0.65 | 1.1E-13 |
| Mtpap | 2047 | -1.22 | -0.29 | 1.2E-03 | -1.25 | -0.32 | 2.9E-04 |
| Mtrf1 | 404 | -1.89 | -0.92 | 1.6E-30 | -1.70 | -0.77 | 9.6E-22 |
| Mtrf1l | 1065 | 1.46 | 0.54 | 2.6E-17 | 1.50 | 0.58 | 1.4E-19 |
| Mtrr | 1171 | -1.54 | -0.62 | 7.2E-26 | -1.67 | -0.74 | 8.2E-36 |
| Mtss2 | 5444 | -1.40 | -0.48 | 5.0E-03 | -1.48 | -0.56 | 1.0E-03 |
| Mturn | 13174 | 1.56 | 0.64 | 7.3E-05 | 1.65 | 0.72 | 7.8E-06 |
| Mtus1 | 4353 | -2.09 | -1.06 | 1.1E-42 | -2.22 | -1.15 | 6.9E-50 |
| Mtx1 | 1889 | -1.38 | -0.46 | 2.9E-06 | -1.52 | -0.60 | 7.6E-10 |
| Mtx2 | 3879 | -1.28 | -0.35 | 4.3E-05 | -1.23 | -0.30 | 6.2E-04 |
| Mtx3 | 3934 | -2.82 | -1.49 | 3.3E-51 | -2.51 | -1.33 | 8.9E-41 |
| Mul1 | 2768 | 1.51 | 0.60 | 1.8E-12 | 1.39 | 0.47 | 2.9E-08 |
| Mvb12a | 1557 | 1.66 | 0.73 | 1.4E-29 | 1.57 | 0.65 | 1.0E-23 |
| Mvb12b | 9118 | -1.92 | -0.94 | 2.8E-08 | -2.02 | -1.02 | 1.9E-09 |
| Mvd | 2606 | 1.87 | 0.90 | 8.7E-17 | 1.77 | 0.83 | 3.0E-14 |
| Mvk | 2925 | 1.82 | 0.87 | 5.6E-16 | 1.68 | 0.75 | 3.7E-12 |
| Mvp | 706 | 1.65 | 0.72 | 1.9E-12 | 1.48 | 0.56 | 5.3E-08 |
| Mxd1 | 3518 | 1.83 | 0.87 | 4.4E-15 | 1.77 | 0.82 | 1.2E-13 |
| Mxi1 | 5443 | 1.44 | 0.52 | 1.2E-09 | 1.40 | 0.48 | 3.0E-08 |
| Mxra7 | 2547 | 1.90 | 0.93 | 1.9E-18 | 1.88 | 0.91 | 9.9E-18 |
| Mybl1 | 2585 | 1.55 | 0.63 | 1.9E-06 | 1.70 | 0.77 | 5.0E-09 |
| Mybph | 185 | -33.70 | -5.07 | 4.2E-05 | -15.23 | -3.93 | 1.9E-03 |
| Myc | 840 | -2.08 | -1.05 | 1.1E-07 | -2.04 | -1.03 | 2.4E-07 |
| Mycbp | 1104 | 1.69 | 0.76 | 1.3E-08 | 1.72 | 0.78 | 4.8E-09 |
| Mycbpap | 316 | 1.51 | 0.60 | 2.1E-04 | 1.49 | 0.58 | 3.7E-04 |
| Mycn | 3287 | -1.40 | -0.48 | 2.1E-03 | -1.51 | -0.59 | 1.4E-04 |
| Myct1 | 154 | -2.06 | -1.05 | 6.6E-08 | -2.28 | -1.19 | 9.7E-10 |
| Myd88 | 307 | -1.39 | -0.48 | 1.5E-04 | -1.45 | -0.54 | 1.7E-05 |
| Myef2 | 23440 | -1.45 | -0.53 | 5.0E-09 | -1.41 | -0.49 | 6.5E-08 |
| Myh3 | 1854 | -21.90 | -4.45 | 8.0E-05 | -12.37 | -3.63 | 1.6E-03 |
| Myh7 | 338 | -7.07 | -2.82 | 1.4E-04 | -4.58 | -2.20 | 3.7E-03 |
| Myh8 | 737 | -9.69 | -3.28 | 9.6E-04 | -8.45 | -3.08 | 2.1E-03 |

| | | | | | | | |
|---|---|---|---|---|---|---|---|
| Myl12b | 341 | 1.54 | 0.63 | 3.8E-12 | 1.46 | 0.54 | 2.3E-09 |
| Myl4 | 243 | -15.83 | -3.98 | 1.5E-05 | -7.79 | -2.96 | 1.6E-03 |
| Myl6b | 933 | 1.44 | 0.52 | 1.5E-02 | 1.43 | 0.52 | 1.6E-02 |
| Mylk | 1785 | -1.20 | -0.27 | 4.4E-02 | -1.25 | -0.32 | 1.5E-02 |
| Mylk4 | 118 | -10.43 | -3.38 | 1.3E-04 | -6.26 | -2.65 | 3.4E-03 |
| Myo1a | 639 | 5.90 | 2.56 | 5.0E-60 | 5.49 | 2.46 | 1.8E-55 |
| Myo1b | 11754 | -2.12 | -1.08 | 1.4E-42 | -2.08 | -1.06 | 7.9E-41 |
| Myo1e | 3165 | 3.35 | 1.74 | 1.4E-64 | 3.28 | 1.71 | 2.9E-62 |
| Myo1f | 179 | -1.59 | -0.67 | 3.7E-04 | -1.56 | -0.64 | 8.2E-04 |
| Myo5b | 385 | -1.89 | -0.92 | 2.7E-04 | -1.92 | -0.94 | 1.9E-04 |
| Myo6 | 5438 | 1.90 | 0.92 | 3.4E-28 | 2.05 | 1.04 | 3.5E-35 |
| Myo7a | 2242 | -1.76 | -0.81 | 1.5E-12 | -1.75 | -0.81 | 2.0E-12 |
| Myo9b | 12403 | 1.43 | 0.51 | 3.5E-08 | 1.52 | 0.61 | 5.1E-11 |
| Myog | 193 | -29.97 | -4.91 | 1.3E-03 | -16.74 | -4.07 | 8.7E-03 |
| Myom1 | 412 | -2.44 | -1.29 | 3.6E-03 | -1.90 | -0.93 | 4.3E-02 |
| Myorg | 2371 | 3.42 | 1.77 | 1.3E-67 | 3.30 | 1.72 | 9.2E-64 |
| Mypop | 765 | 1.92 | 0.94 | 2.4E-17 | 1.76 | 0.82 | 4.0E-13 |
| Myrip | 448 | 1.53 | 0.61 | 1.2E-02 | 1.71 | 0.77 | 1.4E-03 |
| Myt1 | 8901 | -1.76 | -0.81 | 1.9E-07 | -1.81 | -0.85 | 4.2E-08 |
| Mzf1 | 456 | -4.50 | -2.17 | 9.2E-55 | -4.22 | -2.08 | 1.3E-50 |
| N4bp2 | 9983 | -1.56 | -0.64 | 4.6E-13 | -1.46 | -0.54 | 1.4E-09 |
| N4bp2l1 | 706 | 2.22 | 1.15 | 4.7E-13 | 2.33 | 1.22 | 1.3E-14 |
| N4bp2l2 | 6276 | -1.29 | -0.37 | 3.1E-06 | -1.23 | -0.30 | 1.7E-04 |
| N6amt1 | 946 | -4.03 | -2.01 | 1.1E-128 | -4.09 | -2.03 | 1.3E-130 |
| Naa16 | 1973 | -1.31 | -0.39 | 6.6E-07 | -1.29 | -0.37 | 2.7E-06 |
| Naa50 | 13698 | 1.45 | 0.53 | 3.4E-07 | 1.34 | 0.42 | 6.6E-05 |
| Naalad2 | 290 | -1.85 | -0.89 | 1.1E-05 | -1.85 | -0.89 | 1.1E-05 |
| Nabp1 | 1079 | 2.44 | 1.29 | 2.6E-51 | 2.61 | 1.38 | 6.6E-59 |
| Nacad | 4515 | -2.17 | -1.12 | 6.8E-07 | -2.32 | -1.21 | 7.2E-08 |
| Naga | 1940 | 1.85 | 0.89 | 6.5E-18 | 1.80 | 0.85 | 2.3E-16 |
| Nagk | 1716 | -1.44 | -0.52 | 1.9E-09 | -1.51 | -0.59 | 9.1E-12 |
| Naglu | 690 | 1.42 | 0.51 | 4.3E-05 | 1.42 | 0.50 | 6.4E-05 |
| Naif1 | 341 | -2.56 | -1.35 | 2.8E-34 | -2.98 | -1.58 | 2.4E-45 |
| Nanos2 | 219 | 3.81 | 1.93 | 2.2E-10 | 3.19 | 1.67 | 4.3E-08 |
| Nans | 4872 | 1.76 | 0.82 | 7.4E-31 | 1.65 | 0.73 | 1.4E-24 |
| Nap1l5 | 3791 | 1.90 | 0.93 | 4.5E-09 | 2.04 | 1.03 | 5.8E-11 |
| Napa | 6805 | 1.33 | 0.41 | 2.0E-10 | 1.24 | 0.31 | 1.5E-06 |
| Napepld | 1868 | 2.18 | 1.12 | 3.1E-08 | 1.73 | 0.79 | 1.3E-04 |
| Nars2 | 1491 | 1.89 | 0.92 | 1.3E-20 | 1.94 | 0.96 | 3.3E-22 |
| Nasp | 36412 | -1.30 | -0.38 | 1.7E-02 | -1.39 | -0.48 | 2.7E-03 |
| Nat10 | 4931 | 1.30 | 0.38 | 1.8E-07 | 1.24 | 0.31 | 2.6E-05 |
| Nat14 | 696 | -1.32 | -0.40 | 4.5E-03 | -1.51 | -0.59 | 1.4E-05 |
| Nat8f4 | 594 | -1.43 | -0.52 | 2.7E-07 | -1.48 | -0.57 | 1.8E-08 |
| Natd1 | 5000 | 1.26 | 0.33 | 3.8E-07 | 1.20 | 0.26 | 8.7E-05 |
| Naxd | 2877 | -1.31 | -0.39 | 2.3E-08 | -1.35 | -0.43 | 6.1E-10 |
| Nbeal1 | 6207 | 1.52 | 0.60 | 3.3E-03 | 1.89 | 0.92 | 3.7E-06 |
| Nbeal2 | 1295 | 2.27 | 1.19 | 3.1E-20 | 2.11 | 1.08 | 5.9E-17 |

| | | | | | | | |
|---|---|---|---|---|---|---|---|
| Nbl1 | 1625 | -1.81 | -0.85 | 1.3E-03 | -1.72 | -0.79 | 3.3E-03 |
| Nbn | 1557 | -3.74 | -1.90 | 7.6E-65 | -3.81 | -1.93 | 1.8E-66 |
| Ncald | 3248 | 1.87 | 0.90 | 3.2E-09 | 1.74 | 0.80 | 1.7E-07 |
| Ncapd3 | 5583 | 1.43 | 0.52 | 7.3E-05 | 1.52 | 0.61 | 2.9E-06 |
| Ncbp1 | 12054 | 1.31 | 0.39 | 6.3E-06 | 1.30 | 0.37 | 1.4E-05 |
| Ncdn | 4818 | 1.34 | 0.42 | 3.0E-03 | 1.28 | 0.35 | 1.4E-02 |
| Nceh1 | 1038 | 3.19 | 1.67 | 8.8E-90 | 3.22 | 1.69 | 1.8E-90 |
| Ncf1 | 181 | -2.08 | -1.06 | 9.7E-11 | -2.22 | -1.15 | 2.0E-12 |
| Nck2 | 2584 | -1.21 | -0.27 | 1.5E-04 | -1.34 | -0.43 | 1.8E-09 |
| Ncl | 55506 | -1.28 | -0.35 | 3.9E-03 | -1.38 | -0.46 | 1.4E-04 |
| Ncln | 4700 | 1.44 | 0.52 | 2.4E-05 | 1.32 | 0.40 | 1.7E-03 |
| Ncoa2 | 6533 | -1.45 | -0.53 | 3.8E-09 | -1.36 | -0.44 | 1.4E-06 |
| Ncoa3 | 5325 | 1.35 | 0.43 | 1.9E-06 | 1.38 | 0.46 | 3.7E-07 |
| Ncoa5 | 9033 | -1.33 | -0.41 | 6.2E-05 | -1.44 | -0.53 | 1.4E-07 |
| Ncoa6 | 9034 | -1.38 | -0.46 | 7.4E-09 | -1.33 | -0.41 | 3.7E-07 |
| Ncoa7 | 1481 | -2.09 | -1.06 | 7.8E-22 | -2.05 | -1.03 | 9.9E-21 |
| Ncs1 | 6683 | -2.01 | -1.01 | 3.8E-11 | -2.09 | -1.07 | 2.5E-12 |
| Ncstn | 6281 | -1.57 | -0.66 | 1.5E-30 | -1.62 | -0.69 | 3.8E-34 |
| Ndel1 | 3903 | 1.24 | 0.31 | 2.1E-07 | 1.26 | 0.34 | 1.6E-08 |
| Ndfip1 | 16911 | 1.25 | 0.32 | 2.4E-03 | 1.25 | 0.32 | 2.2E-03 |
| Ndor1 | 767 | 1.37 | 0.46 | 6.7E-06 | 1.37 | 0.45 | 8.7E-06 |
| Ndp | 239 | -2.13 | -1.09 | 5.0E-10 | -2.13 | -1.09 | 5.4E-10 |
| Ndrg3 | 6322 | 1.73 | 0.79 | 7.9E-29 | 1.72 | 0.78 | 3.8E-28 |
| Ndrg4 | 7781 | 1.44 | 0.53 | 1.1E-03 | 1.48 | 0.57 | 4.5E-04 |
| Ndst1 | 10114 | -1.30 | -0.38 | 6.7E-06 | -1.25 | -0.32 | 1.3E-04 |
| Ndst2 | 2010 | -1.33 | -0.41 | 1.0E-07 | -1.38 | -0.47 | 9.5E-10 |
| Ndst3 | 2340 | -1.64 | -0.72 | 9.5E-05 | -1.47 | -0.55 | 3.0E-03 |
| Ndufa2 | 2265 | -1.29 | -0.37 | 1.1E-02 | -1.36 | -0.44 | 2.0E-03 |
| Ndufa7 | 3458 | -1.25 | -0.32 | 5.0E-05 | -1.28 | -0.36 | 5.6E-06 |
| Ndufa9 | 5537 | -1.26 | -0.33 | 6.7E-05 | -1.30 | -0.37 | 4.5E-06 |
| Ndufaf1 | 1120 | -1.74 | -0.79 | 9.5E-19 | -1.88 | -0.91 | 5.0E-24 |
| Ndufaf2 | 1728 | 1.48 | 0.56 | 3.8E-06 | 1.44 | 0.53 | 1.7E-05 |
| Ndufaf4 | 3275 | 1.33 | 0.41 | 2.8E-09 | 1.38 | 0.47 | 1.3E-11 |
| Ndufaf6 | 453 | 1.36 | 0.44 | 6.5E-07 | 1.16 | 0.22 | 2.1E-02 |
| Ndufaf7 | 1775 | -1.58 | -0.66 | 2.1E-33 | -1.52 | -0.60 | 4.2E-28 |
| Ndufaf8 | 326 | -1.99 | -0.99 | 1.2E-08 | -2.01 | -1.01 | 7.1E-09 |
| Ndufb3 | 2690 | -1.27 | -0.34 | 1.3E-02 | -1.25 | -0.32 | 2.1E-02 |
| Ndufb6 | 2727 | 1.30 | 0.38 | 1.3E-03 | 1.29 | 0.37 | 2.0E-03 |
| Ndufs7 | 1435 | -1.26 | -0.33 | 4.7E-04 | -1.37 | -0.46 | 8.4E-07 |
| Ndufs8 | 2218 | -1.46 | -0.55 | 4.6E-08 | -1.53 | -0.62 | 7.3E-10 |
| Ndufv3 | 3865 | 1.55 | 0.63 | 1.4E-19 | 1.52 | 0.60 | 7.6E-18 |
| Neb | 476 | -8.89 | -3.15 | 8.6E-04 | -6.33 | -2.66 | 5.6E-03 |
| Necab3 | 636 | 1.30 | 0.38 | 1.9E-02 | 1.35 | 0.43 | 6.4E-03 |
| Necap2 | 2857 | 1.37 | 0.46 | 6.9E-07 | 1.24 | 0.31 | 1.2E-03 |
| Nectin2 | 3683 | 1.84 | 0.88 | 1.7E-15 | 1.83 | 0.87 | 3.9E-15 |
| Nedd4 | 141961 | -1.35 | -0.43 | 3.0E-06 | -1.32 | -0.40 | 1.4E-05 |
| Nedd4l | 8509 | -1.28 | -0.36 | 4.7E-02 | -1.32 | -0.41 | 2.4E-02 |

| | | | | | | | |
|---|---|---|---|---|---|---|---|
| Nedd9 | 3178 | 1.41 | 0.49 | 8.3E-12 | 1.38 | 0.46 | 1.7E-10 |
| Neil1 | 491 | -2.31 | -1.21 | 2.0E-27 | -2.33 | -1.22 | 4.2E-28 |
| Neil3 | 1825 | -1.61 | -0.68 | 5.5E-05 | -1.66 | -0.73 | 1.7E-05 |
| Nek1 | 5685 | 1.59 | 0.67 | 6.8E-17 | 1.77 | 0.82 | 8.1E-25 |
| Nek3 | 465 | -1.67 | -0.74 | 1.2E-08 | -1.84 | -0.88 | 7.2E-12 |
| Nek4 | 2231 | -1.41 | -0.49 | 5.2E-11 | -1.51 | -0.59 | 1.9E-15 |
| Nek9 | 9825 | -1.36 | -0.44 | 1.7E-14 | -1.33 | -0.41 | 5.1E-13 |
| Nelfa | 2468 | -1.37 | -0.46 | 3.0E-07 | -1.52 | -0.61 | 4.5E-12 |
| Nelfb | 5282 | -1.26 | -0.34 | 1.8E-05 | -1.39 | -0.47 | 1.1E-09 |
| Nelfe | 3000 | 1.53 | 0.61 | 1.1E-06 | 1.37 | 0.45 | 4.6E-04 |
| Nemf | 8657 | 1.24 | 0.31 | 2.1E-07 | 1.30 | 0.37 | 1.9E-10 |
| Nemp1 | 3795 | 2.39 | 1.26 | 1.5E-27 | 2.38 | 1.25 | 1.9E-27 |
| Nenf | 818 | -1.63 | -0.71 | 1.4E-22 | -1.63 | -0.70 | 3.2E-22 |
| Net1 | 3061 | -1.46 | -0.55 | 1.7E-02 | -1.48 | -0.57 | 1.4E-02 |
| Neto2 | 6922 | -1.66 | -0.73 | 1.1E-03 | -1.56 | -0.65 | 4.5E-03 |
| Neu1 | 1849 | 2.16 | 1.11 | 2.8E-37 | 2.09 | 1.06 | 3.8E-34 |
| Neu3 | 382 | 1.68 | 0.75 | 2.6E-12 | 2.04 | 1.03 | 6.4E-22 |
| Neurl2 | 115 | -1.48 | -0.57 | 5.3E-03 | -1.42 | -0.50 | 1.5E-02 |
| Neurl3 | 151 | 2.73 | 1.45 | 3.4E-12 | 2.74 | 1.46 | 3.5E-12 |
| Neurod1 | 2403 | -1.43 | -0.52 | 1.4E-03 | -1.58 | -0.66 | 3.5E-05 |
| Neurog1 | 475 | -2.00 | -1.00 | 4.6E-03 | -2.49 | -1.32 | 1.5E-04 |
| Nexmif | 3129 | -2.12 | -1.09 | 1.7E-03 | -1.76 | -0.81 | 2.3E-02 |
| Nexn | 473 | -2.54 | -1.34 | 3.2E-05 | -2.00 | -1.00 | 2.5E-03 |
| Nf1 | 8492 | -1.34 | -0.42 | 1.2E-04 | -1.27 | -0.35 | 1.9E-03 |
| Nfam1 | 283 | -1.52 | -0.60 | 3.3E-04 | -1.63 | -0.71 | 2.2E-05 |
| Nfatc1 | 689 | -1.65 | -0.72 | 4.9E-03 | -1.60 | -0.68 | 8.5E-03 |
| Nfatc2 | 552 | 2.04 | 1.03 | 5.8E-07 | 2.24 | 1.17 | 1.4E-08 |
| Nfatc2ip | 2440 | 2.11 | 1.07 | 1.8E-20 | 1.90 | 0.93 | 1.7E-15 |
| Nfatc4 | 3812 | -2.04 | -1.03 | 9.2E-09 | -2.10 | -1.07 | 2.2E-09 |
| Nfe2 | 103 | -2.15 | -1.10 | 1.3E-02 | -2.51 | -1.33 | 2.6E-03 |
| Nfe2l2 | 2059 | 1.43 | 0.52 | 1.3E-06 | 1.48 | 0.56 | 1.7E-07 |
| Nfe2l3 | 803 | 4.19 | 2.07 | 8.5E-61 | 3.97 | 1.99 | 1.5E-56 |
| Nfia | 6750 | -1.71 | -0.78 | 1.4E-03 | -1.83 | -0.88 | 3.0E-04 |
| Nfib | 18590 | -2.04 | -1.03 | 5.5E-04 | -2.04 | -1.03 | 5.8E-04 |
| Nfic | 2525 | -1.51 | -0.60 | 7.9E-04 | -1.57 | -0.65 | 2.6E-04 |
| Nfix | 5859 | -2.03 | -1.02 | 4.0E-03 | -2.11 | -1.08 | 2.3E-03 |
| Nfkb1 | 2019 | -1.96 | -0.97 | 8.9E-27 | -2.00 | -1.00 | 2.0E-28 |
| Nfkb2 | 1194 | 3.15 | 1.66 | 1.6E-16 | 3.19 | 1.68 | 7.0E-17 |
| Nfkbia | 750 | -1.87 | -0.90 | 1.8E-23 | -2.02 | -1.02 | 2.5E-29 |
| Nfkbib | 867 | 1.41 | 0.50 | 1.5E-08 | 1.35 | 0.43 | 9.9E-07 |
| Nfkbie | 688 | 3.90 | 1.96 | 2.0E-69 | 3.57 | 1.83 | 1.0E-60 |
| Nfkbil1 | 565 | -1.27 | -0.35 | 3.8E-03 | -1.36 | -0.44 | 1.8E-04 |
| Nfkbiz | 500 | 1.35 | 0.44 | 5.2E-03 | 1.41 | 0.50 | 1.4E-03 |
| Nfu1 | 1593 | -1.40 | -0.48 | 2.2E-14 | -1.34 | -0.42 | 3.0E-11 |
| Nfyb | 4464 | -1.51 | -0.59 | 1.9E-12 | -1.53 | -0.62 | 1.7E-13 |
| Nfyc | 3064 | -2.16 | -1.11 | 3.0E-64 | -2.19 | -1.13 | 2.0E-66 |
| Ngdn | 2572 | 1.30 | 0.38 | 1.4E-04 | 1.32 | 0.41 | 3.7E-05 |

| | | | | | | | |
|---|---|---|---|---|---|---|---|
| Ngef | 397 | 2.08 | 1.06 | 3.1E-08 | 1.95 | 0.96 | 5.9E-07 |
| Ngly1 | 1682 | -1.47 | -0.55 | 1.1E-10 | -1.42 | -0.51 | 3.7E-09 |
| Nhej1 | 194 | -2.15 | -1.10 | 1.1E-16 | -2.40 | -1.26 | 3.4E-21 |
| Nhlh1 | 2918 | -1.63 | -0.71 | 8.1E-06 | -1.62 | -0.70 | 1.1E-05 |
| Nhlh2 | 5362 | -2.89 | -1.53 | 1.7E-16 | -2.74 | -1.46 | 4.9E-15 |
| Nhlrc1 | 586 | 1.95 | 0.96 | 1.1E-15 | 1.87 | 0.91 | 4.7E-14 |
| Nhlrc2 | 3753 | -1.68 | -0.75 | 5.2E-19 | -1.62 | -0.70 | 1.4E-16 |
| Nhp2 | 3319 | 1.37 | 0.45 | 7.1E-04 | 1.26 | 0.34 | 1.4E-02 |
| Nhsl2 | 1894 | -1.38 | -0.46 | 5.0E-04 | -1.27 | -0.34 | 1.2E-02 |
| Niban1 | 371 | -2.01 | -1.01 | 2.0E-04 | -1.68 | -0.75 | 7.0E-03 |
| Nif3l1 | 3005 | -1.35 | -0.43 | 4.7E-09 | -1.36 | -0.44 | 1.4E-09 |
| Nim1k | 551 | 1.63 | 0.70 | 9.0E-08 | 1.49 | 0.58 | 1.3E-05 |
| Nin | 4425 | -1.32 | -0.40 | 1.3E-03 | -1.20 | -0.26 | 4.2E-02 |
| Ninj1 | 704 | 2.63 | 1.39 | 9.9E-19 | 2.80 | 1.48 | 5.7E-21 |
| Nipa2 | 4612 | 1.32 | 0.40 | 9.0E-08 | 1.36 | 0.44 | 3.8E-09 |
| Nipal1 | 136 | 2.53 | 1.34 | 1.4E-06 | 2.55 | 1.35 | 1.2E-06 |
| Nipal2 | 164 | -3.20 | -1.68 | 1.0E-10 | -3.71 | -1.89 | 3.6E-13 |
| Nipal3 | 2856 | 2.27 | 1.18 | 3.2E-24 | 2.31 | 1.21 | 3.0E-25 |
| Nipsnap2 | 3417 | -1.72 | -0.78 | 1.0E-28 | -1.81 | -0.86 | 3.8E-34 |
| Nisch | 40348 | -1.52 | -0.61 | 2.4E-22 | -1.51 | -0.59 | 4.5E-21 |
| Nit1 | 2863 | 1.25 | 0.32 | 2.9E-06 | 1.17 | 0.23 | 1.4E-03 |
| Nkain4 | 1161 | -1.36 | -0.45 | 1.6E-04 | -1.33 | -0.42 | 4.7E-04 |
| Nkapl | 138 | 1.29 | 0.37 | 1.6E-02 | 1.31 | 0.39 | 1.1E-02 |
| Nkd1 | 5147 | 1.71 | 0.77 | 1.6E-11 | 1.58 | 0.66 | 1.0E-08 |
| Nkiras1 | 1723 | 1.97 | 0.98 | 8.1E-32 | 2.02 | 1.01 | 3.6E-34 |
| Nkiras2 | 4808 | 1.27 | 0.34 | 1.8E-06 | 1.17 | 0.22 | 2.5E-03 |
| Nlk | 6570 | 1.45 | 0.53 | 4.7E-07 | 1.35 | 0.43 | 5.0E-05 |
| Nln | 4003 | 1.61 | 0.69 | 2.8E-21 | 1.71 | 0.77 | 1.5E-26 |
| Nlrx1 | 296 | -2.67 | -1.41 | 5.2E-24 | -2.61 | -1.39 | 3.9E-23 |
| Nmd3 | 3367 | 1.28 | 0.36 | 6.0E-06 | 1.25 | 0.33 | 3.9E-05 |
| Nme4 | 974 | -1.45 | -0.53 | 1.6E-03 | -1.43 | -0.52 | 2.4E-03 |
| Nme5 | 229 | 1.61 | 0.69 | 6.9E-06 | 1.66 | 0.73 | 1.8E-06 |
| Nme6 | 1056 | -1.43 | -0.52 | 5.7E-06 | -1.63 | -0.71 | 3.7E-10 |
| Nmi | 175 | -2.77 | -1.47 | 1.1E-12 | -2.45 | -1.29 | 4.8E-10 |
| Nmnat1 | 462 | -3.63 | -1.86 | 4.6E-55 | -4.02 | -2.01 | 3.3E-64 |
| Nmnat3 | 168 | 2.21 | 1.14 | 6.9E-15 | 2.00 | 1.00 | 1.8E-11 |
| Nmrk1 | 448 | 1.46 | 0.55 | 2.2E-03 | 1.67 | 0.74 | 3.0E-05 |
| Nmt2 | 10711 | 1.52 | 0.61 | 3.0E-17 | 1.57 | 0.65 | 1.5E-19 |
| Noa1 | 2066 | 1.80 | 0.84 | 7.9E-27 | 1.68 | 0.75 | 2.7E-21 |
| Noc4l | 2059 | 1.39 | 0.48 | 6.6E-05 | 1.30 | 0.38 | 1.8E-03 |
| Noct | 2475 | 1.53 | 0.61 | 4.3E-04 | 1.44 | 0.52 | 2.8E-03 |
| Nod1 | 332 | -1.51 | -0.59 | 2.2E-03 | -1.47 | -0.55 | 4.6E-03 |
| Nod2 | 134 | 6.06 | 2.60 | 2.1E-35 | 5.68 | 2.51 | 9.9E-33 |
| Nog | 443 | -1.55 | -0.63 | 2.8E-06 | -1.53 | -0.62 | 4.6E-06 |
| Nol7 | 4572 | -1.34 | -0.42 | 4.2E-04 | -1.43 | -0.52 | 1.4E-05 |
| Nolc1 | 10912 | 1.47 | 0.56 | 1.7E-08 | 1.35 | 0.44 | 1.5E-05 |
| Nomo1 | 6606 | 1.70 | 0.76 | 2.5E-12 | 1.57 | 0.65 | 3.4E-09 |

| | | | | | | | |
|---|---|---|---|---|---|---|---|
| Nono | 51016 | -1.20 | -0.26 | 1.0E-02 | -1.26 | -0.34 | 7.0E-04 |
| Nop10 | 3193 | 1.49 | 0.58 | 1.3E-06 | 1.38 | 0.46 | 1.5E-04 |
| Nop14 | 4335 | 1.33 | 0.41 | 2.0E-08 | 1.29 | 0.37 | 4.4E-07 |
| Nop16 | 2542 | 1.36 | 0.45 | 1.6E-06 | 1.29 | 0.37 | 8.8E-05 |
| Nop53 | 4778 | -1.53 | -0.62 | 3.6E-12 | -1.67 | -0.74 | 5.0E-17 |
| Nop9 | 3755 | 1.48 | 0.57 | 2.3E-09 | 1.32 | 0.40 | 2.8E-05 |
| Nopchap1 | 9108 | 2.12 | 1.08 | 2.7E-36 | 2.07 | 1.05 | 6.9E-34 |
| Nos3 | 594 | -1.97 | -0.98 | 8.6E-15 | -2.23 | -1.16 | 3.7E-20 |
| Nosip | 1793 | -1.60 | -0.68 | 2.9E-24 | -1.65 | -0.72 | 2.3E-27 |
| Notch3 | 7321 | -1.95 | -0.96 | 1.0E-08 | -2.07 | -1.05 | 3.8E-10 |
| Notch4 | 758 | -2.88 | -1.52 | 1.2E-31 | -3.18 | -1.67 | 1.1E-37 |
| Nova1 | 10201 | -1.90 | -0.92 | 1.3E-05 | -1.83 | -0.87 | 4.2E-05 |
| Nox4 | 332 | -2.82 | -1.50 | 2.6E-27 | -2.71 | -1.44 | 5.6E-25 |
| Noxo1 | 262 | -1.28 | -0.35 | 1.3E-03 | -1.27 | -0.35 | 1.7E-03 |
| Noxred1 | 289 | 1.80 | 0.85 | 4.9E-10 | 1.86 | 0.90 | 4.6E-11 |
| Npas1 | 195 | 3.74 | 1.90 | 1.7E-10 | 3.38 | 1.76 | 4.0E-09 |
| Npat | 5328 | 1.47 | 0.55 | 8.5E-07 | 1.59 | 0.67 | 1.7E-09 |
| Npc1 | 5127 | 1.74 | 0.80 | 2.8E-25 | 1.78 | 0.83 | 2.4E-27 |
| Npepl1 | 1632 | -1.80 | -0.85 | 6.6E-17 | -2.01 | -1.01 | 4.3E-23 |
| Nphp1 | 1991 | -1.32 | -0.40 | 1.8E-03 | -1.40 | -0.49 | 9.7E-05 |
| Nphp3 | 1933 | 1.89 | 0.92 | 1.2E-26 | 1.93 | 0.95 | 5.3E-28 |
| Nphp4 | 1152 | 1.44 | 0.52 | 2.4E-04 | 1.35 | 0.43 | 2.7E-03 |
| Npl | 736 | 2.03 | 1.02 | 2.8E-11 | 1.96 | 0.97 | 2.1E-10 |
| Npm2 | 167 | 1.56 | 0.64 | 7.0E-06 | 1.67 | 0.74 | 2.2E-07 |
| Npr2 | 2976 | -1.77 | -0.82 | 4.2E-12 | -1.67 | -0.74 | 5.3E-10 |
| Npr3 | 1342 | -2.27 | -1.18 | 2.7E-06 | -1.91 | -0.94 | 2.6E-04 |
| Nprl2 | 2232 | 1.98 | 0.98 | 1.2E-34 | 1.96 | 0.97 | 5.2E-34 |
| Nprl3 | 1675 | -1.37 | -0.45 | 3.3E-11 | -1.29 | -0.37 | 1.2E-07 |
| Nptn | 7273 | 1.40 | 0.48 | 3.3E-05 | 1.51 | 0.59 | 2.5E-07 |
| Nptx2 | 633 | 2.73 | 1.45 | 1.4E-30 | 2.31 | 1.21 | 1.4E-21 |
| Nqo1 | 194 | -1.64 | -0.72 | 3.9E-03 | -1.59 | -0.67 | 7.6E-03 |
| Nqo2 | 832 | -1.51 | -0.59 | 7.4E-05 | -1.49 | -0.58 | 1.2E-04 |
| Nr1d1 | 234 | 2.42 | 1.27 | 2.9E-08 | 2.09 | 1.06 | 4.4E-06 |
| Nr2c1 | 1652 | -1.35 | -0.44 | 1.7E-10 | -1.45 | -0.53 | 3.8E-15 |
| Nr2c2 | 11778 | -1.37 | -0.46 | 3.7E-04 | -1.35 | -0.43 | 8.9E-04 |
| Nr2c2ap | 1179 | 1.88 | 0.91 | 4.7E-25 | 1.77 | 0.82 | 1.2E-20 |
| Nr2f1 | 15518 | -1.63 | -0.70 | 9.4E-06 | -1.93 | -0.95 | 1.2E-09 |
| Nr2f6 | 2061 | 1.42 | 0.50 | 1.7E-04 | 1.34 | 0.42 | 2.1E-03 |
| Nr3c2 | 254 | 2.04 | 1.03 | 7.0E-04 | 2.10 | 1.07 | 4.3E-04 |
| Nr4a3 | 749 | 1.52 | 0.61 | 1.5E-02 | 1.66 | 0.74 | 3.0E-03 |
| Nr5a2 | 292 | -1.71 | -0.77 | 8.8E-04 | -1.47 | -0.56 | 2.0E-02 |
| Nr6a1 | 1821 | 3.70 | 1.89 | 6.5E-75 | 3.87 | 1.95 | 1.0E-79 |
| Nradd | 814 | -1.49 | -0.57 | 5.4E-07 | -1.51 | -0.59 | 2.1E-07 |
| Nrarp | 4113 | -1.29 | -0.37 | 4.1E-02 | -1.46 | -0.55 | 1.8E-03 |
| Nras | 15777 | -1.36 | -0.44 | 4.5E-09 | -1.42 | -0.51 | 1.2E-11 |
| Nrde2 | 1295 | -4.22 | -2.08 | 2.7E-140 | -4.02 | -2.01 | 5.3E-131 |
| Nrep | 61680 | -1.33 | -0.41 | 3.0E-04 | -1.28 | -0.36 | 2.1E-03 |

| | | | | | | | |
|---|---|---|---|---|---|---|---|
| Nrg2 | 164 | 1.68 | 0.75 | 2.0E-03 | 1.55 | 0.63 | 9.6E-03 |
| Nrgn | 312 | 1.79 | 0.84 | 1.4E-04 | 2.09 | 1.06 | 1.3E-06 |
| Nrip3 | 979 | 6.35 | 2.67 | 4.5E-53 | 7.23 | 2.85 | 3.0E-60 |
| Nrn1 | 2888 | -1.93 | -0.95 | 3.8E-05 | -1.94 | -0.96 | 3.0E-05 |
| Nrp1 | 3191 | -2.00 | -1.00 | 2.8E-06 | -1.69 | -0.76 | 5.2E-04 |
| Nrros | 320 | -1.59 | -0.67 | 4.6E-07 | -1.60 | -0.67 | 3.8E-07 |
| Nrsn1 | 1282 | 1.57 | 0.65 | 7.9E-03 | 1.53 | 0.61 | 1.3E-02 |
| Nrtn | 152 | 3.30 | 1.72 | 1.8E-14 | 3.79 | 1.92 | 2.4E-17 |
| Nsd1 | 20486 | -1.74 | -0.80 | 2.9E-11 | -1.58 | -0.66 | 4.4E-08 |
| Nsd2 | 33101 | -1.33 | -0.41 | 1.0E-04 | -1.28 | -0.35 | 9.4E-04 |
| Nsd3 | 13338 | 1.48 | 0.56 | 1.8E-14 | 1.54 | 0.62 | 1.8E-17 |
| Nsdhl | 3385 | 1.40 | 0.48 | 2.0E-04 | 1.27 | 0.35 | 8.8E-03 |
| Nsf | 7264 | 1.50 | 0.58 | 5.2E-09 | 1.48 | 0.57 | 1.1E-08 |
| Nsg1 | 13587 | -2.14 | -1.10 | 3.4E-07 | -2.10 | -1.07 | 6.6E-07 |
| Nsmaf | 2891 | -2.15 | -1.10 | 4.1E-38 | -2.08 | -1.05 | 1.1E-34 |
| Nsmce1 | 1642 | -1.68 | -0.74 | 1.9E-16 | -1.72 | -0.78 | 3.9E-18 |
| Nsmce3 | 1585 | 1.44 | 0.53 | 1.8E-18 | 1.36 | 0.44 | 2.3E-13 |
| Nsmf | 6084 | 1.32 | 0.40 | 1.3E-03 | 1.34 | 0.43 | 6.8E-04 |
| Nsun3 | 954 | -1.42 | -0.50 | 2.3E-07 | -1.46 | -0.54 | 1.8E-08 |
| Nsun4 | 1118 | -1.41 | -0.50 | 7.2E-08 | -1.49 | -0.58 | 3.1E-10 |
| Nsun6 | 1295 | 1.22 | 0.29 | 1.4E-02 | 1.34 | 0.42 | 3.5E-04 |
| Nt5c | 1466 | -1.55 | -0.64 | 8.8E-15 | -1.60 | -0.68 | 1.2E-16 |
| Nt5c3b | 3341 | -1.40 | -0.49 | 1.4E-16 | -1.42 | -0.51 | 3.8E-18 |
| Nt5dc2 | 12248 | 1.42 | 0.51 | 2.6E-05 | 1.28 | 0.36 | 3.9E-03 |
| Nt5dc3 | 3561 | 2.26 | 1.18 | 1.8E-06 | 2.36 | 1.24 | 5.0E-07 |
| Nt5m | 950 | -1.98 | -0.99 | 1.8E-25 | -2.05 | -1.04 | 4.6E-28 |
| Ntan1 | 3319 | -1.28 | -0.35 | 2.5E-06 | -1.28 | -0.35 | 2.3E-06 |
| Nthl1 | 501 | -1.30 | -0.38 | 5.3E-03 | -1.57 | -0.65 | 9.7E-07 |
| Ntmt1 | 2942 | 1.58 | 0.66 | 2.7E-14 | 1.51 | 0.60 | 1.1E-11 |
| Ntn3 | 1021 | -1.57 | -0.65 | 1.1E-02 | -1.65 | -0.72 | 4.6E-03 |
| Ntn4 | 1222 | -1.88 | -0.91 | 2.1E-07 | -2.09 | -1.06 | 1.1E-09 |
| Nubp1 | 1694 | -2.10 | -1.07 | 7.4E-27 | -2.41 | -1.27 | 3.3E-37 |
| Nubp2 | 2616 | 1.91 | 0.94 | 2.0E-20 | 1.72 | 0.79 | 1.1E-14 |
| Nucb1 | 6531 | 1.42 | 0.51 | 9.0E-06 | 1.34 | 0.43 | 2.2E-04 |
| Nucb2 | 1637 | 1.21 | 0.28 | 1.0E-03 | 1.28 | 0.36 | 1.8E-05 |
| Nucks1 | 17329 | -1.39 | -0.47 | 2.4E-03 | -1.66 | -0.73 | 1.4E-06 |
| Nudcd1 | 4209 | 1.44 | 0.52 | 2.1E-07 | 1.42 | 0.51 | 4.4E-07 |
| Nudt1 | 677 | -2.08 | -1.05 | 4.1E-16 | -2.32 | -1.21 | 6.8E-21 |
| Nudt11 | 3096 | 1.40 | 0.48 | 2.3E-04 | 1.40 | 0.49 | 2.1E-04 |
| Nudt12 | 918 | 1.52 | 0.61 | 1.0E-11 | 1.58 | 0.66 | 8.9E-14 |
| Nudt14 | 674 | -1.22 | -0.29 | 4.8E-03 | -1.35 | -0.43 | 1.2E-05 |
| Nudt15 | 319 | -5.35 | -2.42 | 7.3E-57 | -5.27 | -2.40 | 1.1E-55 |
| Nudt16 | 567 | -1.23 | -0.30 | 3.6E-03 | -1.34 | -0.42 | 2.8E-05 |
| Nudt16l1 | 2640 | -1.28 | -0.36 | 1.4E-07 | -1.31 | -0.39 | 7.4E-09 |
| Nudt18 | 815 | 2.34 | 1.23 | 8.7E-53 | 2.16 | 1.11 | 2.0E-43 |
| Nudt21 | 15711 | 1.27 | 0.34 | 1.9E-04 | 1.26 | 0.34 | 2.5E-04 |
| Nudt22 | 893 | 2.70 | 1.43 | 9.1E-33 | 2.52 | 1.34 | 2.1E-28 |

| | | | | | | | |
|---|---|---|---|---|---|---|---|
| Nudt3 | 6450 | -1.81 | -0.85 | 9.7E-08 | -1.84 | -0.88 | 3.7E-08 |
| Nudt7 | 428 | 1.62 | 0.70 | 9.0E-07 | 1.65 | 0.73 | 3.0E-07 |
| Nuf2 | 4458 | -1.48 | -0.57 | 3.8E-03 | -1.65 | -0.72 | 1.9E-04 |
| Numa1 | 17622 | -1.79 | -0.84 | 9.1E-25 | -1.73 | -0.79 | 6.3E-22 |
| Numb | 4118 | 1.78 | 0.83 | 1.7E-17 | 1.69 | 0.76 | 1.6E-14 |
| Nup205 | 10236 | -1.54 | -0.62 | 2.4E-08 | -1.43 | -0.52 | 4.6E-06 |
| Nup210 | 3981 | -1.32 | -0.41 | 1.9E-03 | -1.46 | -0.55 | 2.2E-05 |
| Nup210l | 725 | 2.71 | 1.44 | 2.5E-37 | 2.93 | 1.55 | 7.6E-43 |
| Nup37 | 1515 | -1.35 | -0.43 | 1.4E-02 | -1.41 | -0.50 | 4.1E-03 |
| Nup62cl | 337 | 4.31 | 2.11 | 3.0E-24 | 4.03 | 2.01 | 2.6E-22 |
| Nup93 | 7692 | 1.62 | 0.70 | 6.5E-09 | 1.50 | 0.59 | 1.3E-06 |
| Nupl2 | 803 | -1.48 | -0.57 | 1.8E-19 | -1.54 | -0.63 | 2.0E-23 |
| Nwd1 | 819 | 1.83 | 0.87 | 4.2E-03 | 2.17 | 1.12 | 1.8E-04 |
| Nwd2 | 615 | -2.83 | -1.50 | 2.4E-05 | -2.31 | -1.21 | 8.6E-04 |
| Nxf1 | 17752 | 1.27 | 0.35 | 4.8E-08 | 1.26 | 0.34 | 1.6E-07 |
| Nxn | 6394 | 1.35 | 0.43 | 1.9E-03 | 1.25 | 0.33 | 2.2E-02 |
| Nxpe3 | 1146 | 1.93 | 0.95 | 1.3E-47 | 1.84 | 0.88 | 6.4E-41 |
| Nxph1 | 911 | -2.60 | -1.38 | 8.0E-07 | -2.48 | -1.31 | 3.3E-06 |
| Nxph3 | 459 | 2.53 | 1.34 | 4.6E-11 | 2.29 | 1.20 | 5.4E-09 |
| Oasl2 | 117 | -3.25 | -1.70 | 4.1E-14 | -2.21 | -1.14 | 3.9E-07 |
| Oaz2 | 10228 | 1.33 | 0.41 | 1.9E-24 | 1.32 | 0.40 | 3.6E-24 |
| Obscn | 125 | -3.39 | -1.76 | 1.6E-02 | -2.90 | -1.53 | 4.1E-02 |
| Obsl1 | 3857 | -2.18 | -1.13 | 4.5E-09 | -2.29 | -1.20 | 4.2E-10 |
| Ocel1 | 242 | -1.85 | -0.89 | 6.2E-05 | -1.78 | -0.83 | 1.9E-04 |
| Ociad2 | 772 | -1.48 | -0.56 | 4.8E-08 | -1.47 | -0.55 | 7.8E-08 |
| Odad1 | 464 | 1.32 | 0.40 | 2.5E-03 | 1.35 | 0.43 | 1.2E-03 |
| Odad3 | 507 | 1.42 | 0.51 | 6.9E-05 | 1.35 | 0.43 | 9.3E-04 |
| Odc1 | 24222 | 1.49 | 0.58 | 5.0E-06 | 1.43 | 0.52 | 5.3E-05 |
| Odf2 | 13484 | 1.55 | 0.63 | 2.7E-14 | 1.51 | 0.60 | 5.8E-13 |
| Odf2l | 2956 | -1.91 | -0.93 | 1.2E-26 | -1.80 | -0.85 | 3.6E-22 |
| Odr4 | 2639 | -1.84 | -0.88 | 1.4E-44 | -1.76 | -0.82 | 2.1E-38 |
| Ofd1 | 2004 | -1.35 | -0.44 | 8.8E-09 | -1.34 | -0.42 | 2.4E-08 |
| Ogfod1 | 3673 | 2.50 | 1.32 | 6.9E-101 | 2.53 | 1.34 | 1.6E-103 |
| Ogfod2 | 1349 | 2.09 | 1.07 | 3.6E-27 | 1.95 | 0.97 | 1.8E-22 |
| Ogfod3 | 349 | -1.89 | -0.92 | 2.9E-28 | -2.04 | -1.03 | 4.6E-36 |
| Ogfr | 5977 | -1.19 | -0.25 | 8.5E-04 | -1.30 | -0.38 | 3.5E-07 |
| Ogfrl1 | 2898 | 1.74 | 0.80 | 3.5E-28 | 1.73 | 0.79 | 1.1E-27 |
| Ogn | 2623 | -3.11 | -1.64 | 1.4E-04 | -2.74 | -1.45 | 8.3E-04 |
| Ogt | 39637 | -1.51 | -0.60 | 1.5E-06 | -1.30 | -0.38 | 2.8E-03 |
| Ola1 | 7123 | 1.44 | 0.53 | 4.7E-10 | 1.38 | 0.46 | 7.2E-08 |
| Olfml2a | 1014 | -2.01 | -1.01 | 6.4E-04 | -2.06 | -1.04 | 4.0E-04 |
| Olfml2b | 548 | -1.51 | -0.59 | 5.4E-03 | -1.37 | -0.45 | 3.9E-02 |
| Olfml3 | 1215 | -2.55 | -1.35 | 1.4E-05 | -2.47 | -1.31 | 2.9E-05 |
| Olig1 | 341 | 2.11 | 1.08 | 7.4E-05 | 1.49 | 0.58 | 4.5E-02 |
| Oma1 | 501 | -1.32 | -0.40 | 2.4E-05 | -1.42 | -0.50 | 6.0E-08 |
| Onecut2 | 6783 | 1.40 | 0.48 | 1.2E-02 | 1.56 | 0.64 | 7.2E-04 |
| Ooep | 292 | -1.33 | -0.41 | 1.5E-03 | -1.33 | -0.42 | 1.4E-03 |

| | | | | | | | |
|---|---|---|---|---|---|---|---|
| Opa3 | 3112 | -1.80 | -0.84 | 1.1E-19 | -2.00 | -1.00 | 5.1E-27 |
| Opn3 | 438 | 2.62 | 1.39 | 1.3E-19 | 2.42 | 1.28 | 1.1E-16 |
| Oprk1 | 186 | 2.83 | 1.50 | 4.7E-19 | 2.84 | 1.51 | 4.2E-19 |
| Oprl1 | 3489 | -1.91 | -0.94 | 4.2E-05 | -1.76 | -0.82 | 4.1E-04 |
| Optc | 245 | -1.82 | -0.86 | 1.3E-08 | -1.55 | -0.63 | 4.2E-05 |
| Optn | 910 | 3.27 | 1.71 | 3.7E-58 | 3.23 | 1.69 | 5.0E-57 |
| Orai1 | 674 | 2.23 | 1.16 | 3.1E-34 | 2.18 | 1.12 | 2.8E-32 |
| Orai2 | 2262 | 1.43 | 0.52 | 2.0E-04 | 1.47 | 0.55 | 7.7E-05 |
| Orai3 | 822 | 1.72 | 0.79 | 5.6E-10 | 1.73 | 0.79 | 3.8E-10 |
| Orc1 | 857 | 1.62 | 0.69 | 4.8E-03 | 1.52 | 0.60 | 1.6E-02 |
| Orc3 | 5370 | -1.49 | -0.57 | 1.0E-08 | -1.42 | -0.51 | 3.8E-07 |
| Orc4 | 7212 | 1.35 | 0.43 | 5.6E-09 | 1.40 | 0.48 | 7.6E-11 |
| Ormdl2 | 1450 | 2.52 | 1.33 | 3.2E-20 | 2.47 | 1.30 | 2.2E-19 |
| Ormdl3 | 5815 | 1.81 | 0.86 | 1.7E-28 | 1.69 | 0.76 | 2.0E-22 |
| Os9 | 10969 | 1.57 | 0.65 | 2.1E-16 | 1.56 | 0.64 | 7.7E-16 |
| Osbpl10 | 323 | -2.02 | -1.01 | 7.4E-12 | -2.16 | -1.11 | 5.9E-14 |
| Osbpl11 | 3981 | 1.29 | 0.36 | 2.9E-09 | 1.25 | 0.32 | 1.6E-07 |
| Osbpl2 | 4577 | 1.60 | 0.68 | 1.5E-22 | 1.54 | 0.62 | 2.8E-19 |
| Osbpl3 | 2059 | 1.82 | 0.86 | 3.8E-32 | 1.76 | 0.81 | 1.1E-28 |
| Osbpl5 | 2180 | -1.44 | -0.52 | 4.4E-05 | -1.36 | -0.45 | 6.1E-04 |
| Osbpl7 | 1261 | 1.29 | 0.36 | 1.2E-04 | 1.23 | 0.30 | 2.0E-03 |
| Oscar | 117 | 2.73 | 1.45 | 1.8E-17 | 2.83 | 1.50 | 4.9E-18 |
| Oser1 | 1651 | 1.31 | 0.39 | 1.8E-05 | 1.21 | 0.27 | 3.0E-03 |
| Osr1 | 634 | -2.97 | -1.57 | 1.0E-06 | -2.98 | -1.57 | 1.1E-06 |
| Osr2 | 404 | -3.54 | -1.83 | 1.7E-02 | -2.96 | -1.57 | 4.4E-02 |
| Ostf1 | 1558 | 1.82 | 0.87 | 5.3E-12 | 1.85 | 0.89 | 1.3E-12 |
| Ostm1 | 4094 | 2.45 | 1.29 | 1.2E-52 | 2.36 | 1.24 | 3.0E-48 |
| Otor | 240 | -2.88 | -1.52 | 3.1E-04 | -3.15 | -1.65 | 8.8E-05 |
| Otp | 1318 | -2.64 | -1.40 | 2.1E-08 | -2.31 | -1.21 | 1.8E-06 |
| Otub1 | 6816 | -1.40 | -0.49 | 2.4E-13 | -1.43 | -0.52 | 7.4E-15 |
| Otub2 | 1001 | -1.38 | -0.46 | 9.9E-10 | -1.39 | -0.48 | 2.0E-10 |
| Otud1 | 309 | 1.79 | 0.84 | 4.9E-11 | 1.67 | 0.74 | 8.2E-09 |
| Otud3 | 956 | 1.51 | 0.59 | 9.3E-08 | 1.60 | 0.67 | 1.0E-09 |
| Otud7b | 5002 | 1.25 | 0.32 | 2.1E-04 | 1.22 | 0.28 | 1.2E-03 |
| Otulin | 1414 | 1.37 | 0.46 | 4.6E-06 | 1.26 | 0.33 | 1.4E-03 |
| Otulinl | 587 | 1.90 | 0.93 | 4.9E-11 | 1.81 | 0.86 | 1.4E-09 |
| Ovgp1 | 794 | 2.58 | 1.37 | 1.7E-27 | 2.72 | 1.45 | 1.4E-30 |
| Oxld1 | 146 | -1.43 | -0.52 | 1.7E-04 | -1.29 | -0.37 | 9.1E-03 |
| Oxsm | 612 | -1.24 | -0.31 | 3.2E-04 | -1.28 | -0.35 | 3.0E-05 |
| Oxsr1 | 9488 | 1.67 | 0.74 | 1.9E-44 | 1.57 | 0.65 | 9.2E-35 |
| P2rx4 | 252 | -2.71 | -1.44 | 4.0E-20 | -2.71 | -1.44 | 6.0E-20 |
| P2rx7 | 210 | -2.26 | -1.18 | 6.8E-06 | -1.67 | -0.74 | 6.3E-03 |
| P2ry1 | 533 | 1.31 | 0.39 | 3.2E-04 | 1.32 | 0.41 | 1.9E-04 |
| P2ry12 | 179 | -1.99 | -1.00 | 2.4E-09 | -2.31 | -1.21 | 5.2E-13 |
| P3h1 | 2999 | 1.41 | 0.49 | 1.1E-02 | 1.35 | 0.43 | 2.7E-02 |
| P3h2 | 397 | -2.34 | -1.22 | 3.5E-08 | -2.28 | -1.19 | 9.9E-08 |
| P3h3 | 3650 | -1.85 | -0.89 | 3.1E-11 | -1.90 | -0.93 | 3.2E-12 |

| | | | | | | | |
|---|---|---|---|---|---|---|---|
| P4ha3 | 971 | -2.60 | -1.38 | 3.9E-07 | -2.45 | -1.29 | 2.0E-06 |
| P4htm | 556 | -1.52 | -0.60 | 3.7E-05 | -1.59 | -0.67 | 4.3E-06 |
| Pa2g4 | 14613 | 1.35 | 0.43 | 2.2E-05 | 1.22 | 0.29 | 5.8E-03 |
| Pabpc1l | 274 | 4.48 | 2.16 | 2.0E-53 | 3.41 | 1.77 | 5.4E-37 |
| Pabpc4l | 818 | -1.50 | -0.59 | 4.0E-03 | -1.38 | -0.46 | 2.7E-02 |
| Pabpn1 | 4363 | -1.30 | -0.38 | 6.8E-03 | -1.41 | -0.50 | 2.7E-04 |
| Pacsin2 | 4329 | 1.44 | 0.53 | 6.0E-07 | 1.34 | 0.42 | 8.1E-05 |
| Paf1 | 7865 | 2.08 | 1.06 | 2.0E-50 | 1.93 | 0.95 | 1.8E-40 |
| Pafah1b1 | 29765 | -1.30 | -0.38 | 1.1E-04 | -1.25 | -0.32 | 1.5E-03 |
| Pafah2 | 1333 | 1.56 | 0.64 | 3.2E-13 | 1.49 | 0.57 | 8.3E-11 |
| Pag1 | 4748 | 1.91 | 0.93 | 6.8E-09 | 2.03 | 1.02 | 1.5E-10 |
| Paics | 23426 | 1.29 | 0.37 | 1.4E-02 | 1.27 | 0.34 | 2.3E-02 |
| Paip1 | 12566 | -1.36 | -0.45 | 2.2E-03 | -1.61 | -0.69 | 1.2E-06 |
| Paip2 | 14026 | -1.31 | -0.39 | 7.4E-09 | -1.29 | -0.36 | 5.8E-08 |
| Paip2b | 12791 | 1.56 | 0.64 | 4.0E-28 | 1.56 | 0.64 | 1.8E-28 |
| Pak1ip1 | 3967 | 1.36 | 0.45 | 3.0E-05 | 1.34 | 0.42 | 9.0E-05 |
| Pak4 | 1502 | -1.70 | -0.76 | 7.4E-07 | -1.95 | -0.96 | 3.1E-10 |
| Pak5 | 2953 | -1.54 | -0.62 | 6.4E-04 | -1.45 | -0.53 | 3.8E-03 |
| Pakap | 139 | -2.79 | -1.48 | 1.9E-03 | -2.35 | -1.23 | 1.1E-02 |
| Pald1 | 3174 | 1.53 | 0.62 | 3.3E-07 | 1.50 | 0.58 | 1.5E-06 |
| Palm3 | 720 | 1.99 | 0.99 | 7.7E-06 | 1.89 | 0.92 | 4.2E-05 |
| Palmd | 688 | -2.10 | -1.07 | 2.4E-07 | -1.87 | -0.90 | 1.5E-05 |
| Pals1 | 5339 | 1.45 | 0.53 | 1.6E-11 | 1.60 | 0.68 | 5.5E-18 |
| Pam | 5537 | -1.30 | -0.38 | 2.4E-06 | -1.31 | -0.39 | 8.8E-07 |
| Pan2 | 5265 | -1.91 | -0.94 | 2.1E-30 | -2.04 | -1.03 | 2.9E-36 |
| Pan3 | 5259 | -1.26 | -0.33 | 1.0E-03 | -1.32 | -0.40 | 7.0E-05 |
| Pank1 | 1648 | -1.41 | -0.50 | 6.8E-09 | -1.54 | -0.63 | 1.2E-13 |
| Pank4 | 3261 | -1.31 | -0.39 | 1.8E-14 | -1.26 | -0.34 | 6.3E-11 |
| Panx1 | 6792 | -1.42 | -0.51 | 4.6E-03 | -1.42 | -0.51 | 4.7E-03 |
| Panx2 | 332 | -1.84 | -0.88 | 1.0E-03 | -1.81 | -0.85 | 1.5E-03 |
| Paox | 1233 | 1.77 | 0.83 | 9.4E-24 | 1.68 | 0.75 | 1.6E-19 |
| Pappa | 1006 | -2.45 | -1.29 | 3.1E-13 | -2.50 | -1.32 | 9.2E-14 |
| Papss1 | 8672 | 1.50 | 0.58 | 8.4E-30 | 1.44 | 0.53 | 1.7E-24 |
| Papss2 | 732 | -5.25 | -2.39 | 1.2E-05 | -5.18 | -2.37 | 1.5E-05 |
| Paqr3 | 2037 | 2.03 | 1.02 | 5.0E-23 | 2.12 | 1.08 | 1.6E-25 |
| Paqr4 | 2309 | 1.91 | 0.94 | 2.9E-28 | 1.79 | 0.84 | 8.0E-23 |
| Paqr8 | 2154 | -1.37 | -0.46 | 6.3E-04 | -1.35 | -0.43 | 1.3E-03 |
| Pard3 | 5859 | -1.41 | -0.50 | 8.8E-16 | -1.46 | -0.54 | 1.2E-18 |
| Pard6b | 1257 | 1.96 | 0.97 | 3.1E-13 | 1.69 | 0.76 | 2.0E-08 |
| Pard6g | 4629 | 1.25 | 0.32 | 1.2E-04 | 1.16 | 0.21 | 1.4E-02 |
| Parl | 2551 | -1.21 | -0.28 | 7.2E-03 | -1.27 | -0.35 | 6.7E-04 |
| Parn | 2294 | -1.63 | -0.70 | 1.2E-24 | -1.75 | -0.81 | 2.7E-32 |
| Parp11 | 4400 | -1.84 | -0.88 | 5.8E-21 | -1.80 | -0.85 | 1.7E-19 |
| Parp12 | 362 | -2.81 | -1.49 | 5.0E-23 | -2.74 | -1.46 | 7.7E-22 |
| Parp14 | 138 | -2.69 | -1.43 | 5.3E-09 | -2.03 | -1.02 | 3.7E-05 |
| Parp2 | 2664 | -1.60 | -0.68 | 3.0E-13 | -1.66 | -0.73 | 3.9E-15 |
| Parp8 | 1728 | -2.15 | -1.10 | 7.7E-17 | -1.99 | -0.99 | 7.2E-14 |

| | | | | | | | |
|---|---|---|---|---|---|---|---|
| Parp9 | 269 | -2.70 | -1.43 | 3.1E-26 | -3.01 | -1.59 | 8.6E-32 |
| Parva | 8020 | -1.52 | -0.60 | 1.3E-03 | -1.59 | -0.67 | 3.4E-04 |
| Pate2 | 225 | 1.31 | 0.39 | 5.5E-03 | 1.34 | 0.42 | 2.3E-03 |
| Patj | 1326 | -2.29 | -1.20 | 1.3E-31 | -2.20 | -1.14 | 1.1E-28 |
| Patz1 | 5862 | -2.10 | -1.07 | 2.1E-22 | -2.31 | -1.21 | 2.9E-28 |
| Pawr | 1151 | 2.37 | 1.25 | 4.8E-16 | 2.11 | 1.08 | 2.8E-12 |
| Pax7 | 1240 | -1.74 | -0.80 | 3.4E-03 | -1.68 | -0.75 | 7.0E-03 |
| Paxx | 871 | -1.37 | -0.45 | 2.8E-03 | -1.47 | -0.55 | 2.3E-04 |
| Pbld2 | 122 | -1.30 | -0.38 | 1.8E-02 | -1.28 | -0.35 | 2.7E-02 |
| Pbrm1 | 15819 | -2.07 | -1.05 | 4.6E-26 | -1.93 | -0.95 | 1.9E-21 |
| Pbx2 | 7483 | -1.51 | -0.60 | 1.4E-05 | -1.78 | -0.83 | 7.6E-10 |
| Pbx4 | 1111 | 2.22 | 1.15 | 1.2E-13 | 2.05 | 1.03 | 3.1E-11 |
| Pbxip1 | 2932 | 1.44 | 0.52 | 8.7E-04 | 1.41 | 0.50 | 1.6E-03 |
| Pcbd1 | 282 | -1.63 | -0.70 | 5.6E-07 | -1.51 | -0.60 | 2.6E-05 |
| Pcbd2 | 201 | -1.27 | -0.34 | 5.0E-02 | -1.31 | -0.39 | 2.5E-02 |
| Pcbp1 | 13191 | -1.58 | -0.66 | 3.0E-09 | -1.78 | -0.83 | 4.8E-14 |
| Pcbp2 | 24042 | -1.18 | -0.24 | 3.6E-02 | -1.30 | -0.38 | 6.7E-04 |
| Pcbp3 | 3242 | 2.03 | 1.02 | 6.0E-21 | 1.96 | 0.97 | 4.3E-19 |
| Pcca | 2377 | 1.50 | 0.59 | 4.9E-07 | 1.51 | 0.59 | 4.8E-07 |
| Pccb | 2484 | -1.67 | -0.74 | 1.1E-47 | -1.81 | -0.86 | 4.0E-63 |
| Pcdh10 | 8761 | 1.47 | 0.55 | 4.4E-03 | 1.54 | 0.62 | 1.4E-03 |
| Pcdh17 | 5315 | 1.73 | 0.79 | 7.1E-05 | 1.84 | 0.88 | 9.4E-06 |
| Pcdh18 | 7448 | -1.70 | -0.76 | 1.9E-06 | -1.57 | -0.65 | 5.6E-05 |
| Pcdh20 | 515 | 2.08 | 1.06 | 8.0E-08 | 2.36 | 1.24 | 2.5E-10 |
| Pcdha11 | 113 | -2.30 | -1.20 | 2.5E-04 | -1.94 | -0.95 | 4.1E-03 |
| Pcdha3 | 113 | -2.00 | -1.00 | 1.2E-03 | -1.97 | -0.98 | 1.7E-03 |
| Pcdha7 | 112 | -1.90 | -0.93 | 1.3E-02 | -2.01 | -1.01 | 6.8E-03 |
| Pcdhb10 | 103 | -2.97 | -1.57 | 1.4E-13 | -3.10 | -1.63 | 1.5E-14 |
| Pcdhb11 | 128 | -2.48 | -1.31 | 1.1E-04 | -2.58 | -1.36 | 5.5E-05 |
| Pcdhb14 | 113 | -2.68 | -1.42 | 1.7E-07 | -2.56 | -1.35 | 6.9E-07 |
| Pcdhb16 | 221 | -3.44 | -1.78 | 1.5E-07 | -2.48 | -1.31 | 1.5E-04 |
| Pcdhb17 | 469 | -3.05 | -1.61 | 6.5E-11 | -2.53 | -1.34 | 6.8E-08 |
| Pcdhb18 | 214 | -2.69 | -1.43 | 5.7E-06 | -2.31 | -1.21 | 1.5E-04 |
| Pcdhb19 | 430 | -3.22 | -1.69 | 5.0E-07 | -2.44 | -1.29 | 1.6E-04 |
| Pcdhb20 | 386 | -2.84 | -1.51 | 1.3E-07 | -2.54 | -1.34 | 2.7E-06 |
| Pcdhb21 | 150 | -2.89 | -1.53 | 3.0E-05 | -2.23 | -1.16 | 2.0E-03 |
| Pcdhb22 | 483 | -2.04 | -1.03 | 3.7E-06 | -2.04 | -1.03 | 3.7E-06 |
| Pcdhb3 | 192 | -3.13 | -1.65 | 3.4E-17 | -2.79 | -1.48 | 3.1E-14 |
| Pcdhb4 | 178 | -2.75 | -1.46 | 2.3E-10 | -2.71 | -1.44 | 5.2E-10 |
| Pcdhb5 | 218 | -2.75 | -1.46 | 2.6E-10 | -2.61 | -1.38 | 2.3E-09 |
| Pcdhb6 | 116 | -2.96 | -1.57 | 2.4E-11 | -2.78 | -1.47 | 3.5E-10 |
| Pcdhb7 | 405 | -2.66 | -1.41 | 1.3E-09 | -2.23 | -1.15 | 9.5E-07 |
| Pcdhb8 | 178 | -3.02 | -1.59 | 1.1E-13 | -2.75 | -1.46 | 8.5E-12 |
| Pcdhb9 | 192 | -3.24 | -1.69 | 2.5E-10 | -2.84 | -1.51 | 2.0E-08 |
| Pcdhga1 | 217 | -1.80 | -0.85 | 2.7E-06 | -1.54 | -0.62 | 8.7E-04 |
| Pcdhga10 | 517 | -1.61 | -0.69 | 3.6E-11 | -1.59 | -0.67 | 1.6E-10 |
| Pcdhga11 | 947 | -2.05 | -1.04 | 6.7E-14 | -2.11 | -1.08 | 7.7E-15 |

| | | | | | | | |
|---|---|---|---|---|---|---|---|
| Pcdhga12 | 709 | -1.58 | -0.66 | 2.0E-03 | -1.62 | -0.70 | 1.0E-03 |
| Pcdhga2 | 580 | -1.38 | -0.47 | 5.3E-03 | -1.35 | -0.44 | 1.0E-02 |
| Pcdhga3 | 664 | -1.38 | -0.46 | 1.8E-03 | -1.53 | -0.61 | 2.8E-05 |
| Pcdhga4 | 302 | -1.70 | -0.77 | 4.2E-06 | -1.56 | -0.64 | 1.4E-04 |
| Pcdhga5 | 473 | -1.31 | -0.39 | 4.4E-04 | -1.37 | -0.45 | 4.0E-05 |
| Pcdhga6 | 405 | -1.86 | -0.90 | 1.2E-08 | -1.85 | -0.89 | 1.9E-08 |
| Pcdhga7 | 584 | -1.60 | -0.68 | 3.0E-08 | -1.73 | -0.79 | 8.2E-11 |
| Pcdhga8 | 335 | -1.46 | -0.54 | 9.1E-04 | -1.32 | -0.41 | 1.5E-02 |
| Pcdhga9 | 322 | -1.58 | -0.66 | 2.1E-06 | -1.75 | -0.80 | 5.3E-09 |
| Pcdhgb1 | 291 | -1.71 | -0.78 | 8.0E-05 | -1.67 | -0.74 | 2.0E-04 |
| Pcdhgb2 | 424 | -1.61 | -0.69 | 6.2E-08 | -1.61 | -0.68 | 7.8E-08 |
| Pcdhgb4 | 351 | -1.55 | -0.63 | 2.1E-07 | -1.52 | -0.60 | 9.3E-07 |
| Pcdhgb5 | 462 | -1.62 | -0.69 | 2.2E-07 | -1.71 | -0.78 | 5.9E-09 |
| Pcdhgb6 | 1079 | -1.61 | -0.68 | 4.9E-06 | -1.64 | -0.71 | 2.1E-06 |
| Pcdhgb7 | 570 | -1.73 | -0.79 | 1.7E-11 | -1.69 | -0.76 | 1.5E-10 |
| Pcdhgc3 | 6382 | 1.33 | 0.41 | 6.0E-03 | 1.30 | 0.38 | 1.1E-02 |
| Pced1a | 1376 | -2.20 | -1.14 | 1.6E-91 | -2.24 | -1.16 | 3.9E-96 |
| Pced1b | 191 | -1.63 | -0.70 | 2.7E-04 | -1.76 | -0.81 | 2.0E-05 |
| Pcgf1 | 786 | -1.35 | -0.43 | 7.1E-14 | -1.33 | -0.41 | 1.2E-12 |
| Pcgf2 | 5286 | -1.69 | -0.76 | 5.8E-07 | -1.81 | -0.86 | 1.5E-08 |
| Pcgf3 | 10060 | -1.24 | -0.31 | 3.4E-06 | -1.28 | -0.36 | 6.0E-08 |
| Pcgf5 | 1863 | 1.36 | 0.44 | 4.0E-03 | 1.44 | 0.53 | 5.1E-04 |
| Pcid2 | 2970 | -1.19 | -0.25 | 4.4E-03 | -1.28 | -0.36 | 2.9E-05 |
| Pcif1 | 4387 | -1.45 | -0.54 | 1.7E-07 | -1.58 | -0.66 | 9.3E-11 |
| Pck2 | 1166 | -1.50 | -0.58 | 3.2E-07 | -1.60 | -0.67 | 2.5E-09 |
| Pcnt | 3373 | -2.29 | -1.19 | 1.0E-23 | -2.22 | -1.15 | 4.7E-22 |
| Pcnx4 | 3878 | 2.25 | 1.17 | 2.6E-65 | 2.40 | 1.26 | 2.7E-75 |
| Pcolce2 | 140 | -4.38 | -2.13 | 1.8E-19 | -3.59 | -1.85 | 6.6E-15 |
| Pcp4l1 | 1133 | 2.40 | 1.27 | 3.6E-10 | 2.43 | 1.28 | 2.3E-10 |
| Pcsk1 | 305 | 1.47 | 0.56 | 1.3E-02 | 1.62 | 0.70 | 1.9E-03 |
| Pcsk2 | 2274 | 1.36 | 0.45 | 3.5E-02 | 1.40 | 0.49 | 2.2E-02 |
| Pcsk4 | 207 | 2.36 | 1.24 | 1.3E-10 | 2.21 | 1.14 | 3.7E-09 |
| Pcsk6 | 655 | -1.79 | -0.84 | 6.4E-07 | -1.61 | -0.69 | 5.4E-05 |
| Pctp | 515 | 1.88 | 0.91 | 5.1E-04 | 1.57 | 0.65 | 1.6E-02 |
| Pcx | 2714 | 1.60 | 0.68 | 6.8E-10 | 1.52 | 0.60 | 5.7E-08 |
| Pcyox1 | 10636 | 1.30 | 0.37 | 5.3E-11 | 1.23 | 0.30 | 1.4E-07 |
| Pcyt1a | 7273 | 1.30 | 0.38 | 9.1E-09 | 1.30 | 0.37 | 1.3E-08 |
| Pcyt1b | 3430 | -2.10 | -1.07 | 1.2E-16 | -2.03 | -1.02 | 2.8E-15 |
| Pcyt2 | 4545 | 1.54 | 0.62 | 6.2E-12 | 1.42 | 0.51 | 2.8E-08 |
| Pdcd10 | 3779 | 1.22 | 0.28 | 2.8E-03 | 1.33 | 0.41 | 9.8E-06 |
| Pdcl3 | 4311 | 1.30 | 0.37 | 4.0E-07 | 1.26 | 0.34 | 5.5E-06 |
| Pde10a | 2543 | -1.55 | -0.63 | 4.6E-03 | -1.52 | -0.61 | 6.8E-03 |
| Pde3a | 1410 | -1.64 | -0.71 | 2.0E-04 | -1.64 | -0.71 | 2.3E-04 |
| Pde4a | 1571 | 1.58 | 0.66 | 1.0E-03 | 1.45 | 0.53 | 9.6E-03 |
| Pde5a | 2634 | -1.31 | -0.39 | 2.3E-05 | -1.24 | -0.31 | 8.0E-04 |
| Pde6d | 2492 | 1.31 | 0.39 | 3.9E-08 | 1.23 | 0.30 | 3.9E-05 |
| Pde7b | 561 | -2.00 | -1.00 | 9.6E-07 | -1.99 | -1.00 | 1.1E-06 |

| | | | | | | | |
|---|---|---|---|---|---|---|---|
| Pde8b | 219 | -1.66 | -0.73 | 1.4E-03 | -1.72 | -0.78 | 5.8E-04 |
| Pde9a | 2649 | 1.65 | 0.73 | 1.3E-25 | 1.51 | 0.60 | 1.4E-17 |
| Pdgfb | 601 | -1.22 | -0.29 | 2.3E-02 | -1.41 | -0.50 | 5.6E-05 |
| Pdgfc | 1326 | -2.05 | -1.03 | 6.5E-06 | -1.92 | -0.94 | 5.0E-05 |
| Pdgfd | 180 | -1.69 | -0.75 | 3.4E-03 | -1.87 | -0.90 | 4.4E-04 |
| Pdgfra | 6469 | -2.49 | -1.31 | 1.5E-04 | -2.10 | -1.07 | 2.5E-03 |
| Pdha1 | 12367 | 1.32 | 0.40 | 9.8E-07 | 1.26 | 0.34 | 4.9E-05 |
| Pdhx | 3130 | 1.25 | 0.33 | 3.0E-08 | 1.30 | 0.38 | 1.4E-10 |
| Pdia5 | 1044 | 1.31 | 0.39 | 5.6E-03 | 1.35 | 0.43 | 2.6E-03 |
| Pdk1 | 4064 | 1.24 | 0.31 | 1.1E-02 | 1.33 | 0.41 | 6.5E-04 |
| Pdk2 | 1859 | 2.00 | 1.00 | 1.8E-19 | 2.02 | 1.02 | 3.9E-20 |
| Pdk4 | 205 | 2.25 | 1.17 | 8.3E-12 | 2.53 | 1.34 | 7.2E-15 |
| Pdlim4 | 1133 | 1.66 | 0.73 | 3.5E-05 | 1.70 | 0.77 | 1.5E-05 |
| Pdlim5 | 4085 | 2.20 | 1.14 | 1.0E-29 | 2.15 | 1.11 | 3.0E-28 |
| Pdlim7 | 3519 | 1.46 | 0.55 | 8.5E-07 | 1.34 | 0.42 | 1.9E-04 |
| Pdp1 | 4889 | 1.33 | 0.41 | 2.7E-04 | 1.38 | 0.47 | 3.1E-05 |
| Pdp2 | 2326 | 1.30 | 0.37 | 2.1E-07 | 1.30 | 0.37 | 2.4E-07 |
| Pdpk1 | 9306 | 1.37 | 0.45 | 1.0E-06 | 1.39 | 0.47 | 3.2E-07 |
| Pdrg1 | 4504 | 1.38 | 0.47 | 6.8E-20 | 1.40 | 0.48 | 2.9E-21 |
| Pds5b | 13938 | -1.37 | -0.45 | 2.7E-06 | -1.22 | -0.29 | 4.1E-03 |
| Pdss1 | 1545 | 1.23 | 0.30 | 4.4E-03 | 1.29 | 0.37 | 3.6E-04 |
| Pdss2 | 593 | -1.38 | -0.46 | 6.2E-09 | -1.44 | -0.53 | 2.4E-11 |
| Pdxk | 2900 | 2.25 | 1.17 | 9.8E-35 | 2.03 | 1.02 | 8.0E-27 |
| Pdzd2 | 1144 | -1.89 | -0.92 | 1.7E-06 | -1.79 | -0.84 | 1.6E-05 |
| Pdzd7 | 386 | 2.14 | 1.10 | 6.2E-06 | 1.92 | 0.94 | 1.2E-04 |
| Pdzd8 | 8176 | 1.44 | 0.53 | 5.7E-22 | 1.50 | 0.58 | 1.2E-26 |
| Pdzk1ip1 | 173 | -1.77 | -0.83 | 6.6E-03 | -1.66 | -0.73 | 1.8E-02 |
| Pdzph1 | 221 | -1.94 | -0.95 | 2.0E-03 | -2.24 | -1.16 | 1.4E-04 |
| Pea15a | 27187 | 1.25 | 0.32 | 1.5E-04 | 1.22 | 0.29 | 8.5E-04 |
| Peak1 | 2739 | 1.64 | 0.71 | 1.6E-04 | 2.02 | 1.01 | 3.8E-08 |
| Pear1 | 972 | -1.40 | -0.48 | 4.6E-02 | -1.52 | -0.61 | 1.1E-02 |
| Pecam1 | 2748 | -1.22 | -0.28 | 1.0E-02 | -1.26 | -0.33 | 2.4E-03 |
| Pecr | 490 | 1.35 | 0.43 | 2.6E-04 | 1.39 | 0.48 | 4.2E-05 |
| Pef1 | 1858 | -1.67 | -0.74 | 1.2E-18 | -1.75 | -0.81 | 4.0E-22 |
| Peg12 | 1943 | -1.69 | -0.76 | 1.1E-03 | -2.01 | -1.00 | 1.2E-05 |
| Peli2 | 2869 | 1.31 | 0.39 | 1.1E-03 | 1.28 | 0.36 | 3.3E-03 |
| Peli3 | 973 | 2.09 | 1.07 | 4.4E-14 | 2.01 | 1.01 | 1.3E-12 |
| Penk | 1392 | -2.51 | -1.33 | 5.1E-03 | -1.98 | -0.98 | 4.4E-02 |
| Pepd | 1988 | 1.60 | 0.68 | 2.8E-10 | 1.56 | 0.64 | 2.2E-09 |
| Per1 | 3226 | 1.95 | 0.97 | 1.1E-11 | 1.77 | 0.83 | 8.2E-09 |
| Per2 | 1092 | 1.71 | 0.77 | 1.0E-05 | 1.68 | 0.75 | 2.2E-05 |
| Per3 | 1173 | 1.85 | 0.89 | 8.7E-21 | 1.92 | 0.94 | 5.5E-23 |
| Perm1 | 293 | 2.66 | 1.41 | 5.1E-19 | 2.96 | 1.57 | 3.4E-23 |
| Perp | 438 | -3.34 | -1.74 | 3.0E-03 | -2.66 | -1.41 | 1.9E-02 |
| Pex1 | 2115 | -1.26 | -0.33 | 1.5E-05 | -1.18 | -0.24 | 1.8E-03 |
| Pex11a | 846 | 1.90 | 0.93 | 2.3E-33 | 1.80 | 0.85 | 3.0E-28 |
| Pex11g | 212 | -2.01 | -1.01 | 9.0E-22 | -2.07 | -1.05 | 4.2E-23 |

| | | | | | | | |
|---|---|---|---|---|---|---|---|
| Pex12 | 481 | -1.39 | -0.47 | 1.6E-09 | -1.47 | -0.56 | 1.0E-12 |
| Pex13 | 4933 | 1.62 | 0.70 | 6.8E-20 | 1.67 | 0.74 | 5.9E-22 |
| Pex16 | 961 | 1.31 | 0.39 | 6.5E-06 | 1.23 | 0.30 | 5.3E-04 |
| Pex26 | 2397 | 1.85 | 0.89 | 1.9E-19 | 1.75 | 0.80 | 3.7E-16 |
| Pex5 | 6802 | 1.58 | 0.66 | 2.9E-12 | 1.46 | 0.55 | 7.0E-09 |
| Pex6 | 3890 | 1.39 | 0.48 | 4.6E-05 | 1.35 | 0.43 | 2.8E-04 |
| Pex7 | 836 | -2.40 | -1.27 | 3.4E-59 | -2.40 | -1.26 | 1.2E-58 |
| Pfdn1 | 3377 | -1.57 | -0.65 | 9.0E-08 | -1.59 | -0.67 | 4.1E-08 |
| Pfdn6 | 5364 | 1.51 | 0.59 | 4.3E-04 | 1.32 | 0.40 | 1.9E-02 |
| Pfkfb1 | 239 | -1.57 | -0.66 | 4.1E-07 | -1.68 | -0.75 | 8.9E-09 |
| Pfkfb2 | 2491 | 1.36 | 0.44 | 1.3E-05 | 1.45 | 0.53 | 1.2E-07 |
| Pfkfb3 | 3112 | -2.18 | -1.12 | 8.3E-18 | -2.10 | -1.07 | 2.2E-16 |
| Pfkp | 14468 | 1.88 | 0.91 | 3.6E-03 | 2.55 | 1.35 | 9.2E-06 |
| Pfn1 | 10690 | -1.27 | -0.34 | 8.9E-03 | -1.41 | -0.49 | 1.1E-04 |
| Pfn4 | 210 | -3.99 | -2.00 | 6.6E-25 | -3.65 | -1.87 | 5.1E-22 |
| Pgam1 | 12146 | 1.35 | 0.43 | 7.1E-06 | 1.23 | 0.30 | 2.4E-03 |
| Pgap1 | 15824 | 1.57 | 0.65 | 1.9E-04 | 1.85 | 0.88 | 2.8E-07 |
| Pgap3 | 354 | -4.11 | -2.04 | 9.6E-76 | -4.51 | -2.17 | 2.2E-85 |
| Pgbd1 | 590 | -1.47 | -0.56 | 1.3E-05 | -1.64 | -0.71 | 1.8E-08 |
| Pgbd5 | 1690 | 2.38 | 1.25 | 4.9E-21 | 2.25 | 1.17 | 1.4E-18 |
| Pgd | 9982 | 1.36 | 0.44 | 7.4E-06 | 1.23 | 0.30 | 3.5E-03 |
| Pgf | 348 | -1.27 | -0.35 | 1.5E-02 | -1.23 | -0.30 | 3.8E-02 |
| Pgghg | 1129 | -1.67 | -0.74 | 1.9E-06 | -1.75 | -0.81 | 1.7E-07 |
| Pgk1 | 25746 | 1.32 | 0.40 | 5.4E-04 | 1.25 | 0.32 | 7.0E-03 |
| Pgm2 | 3130 | 1.84 | 0.88 | 9.7E-59 | 1.84 | 0.88 | 5.0E-59 |
| Pgm3 | 2790 | 1.56 | 0.64 | 9.2E-14 | 1.61 | 0.69 | 1.2E-15 |
| Pgp | 2958 | 1.94 | 0.96 | 9.9E-33 | 1.76 | 0.82 | 5.2E-24 |
| Pgpep1 | 5687 | 1.57 | 0.65 | 1.6E-08 | 1.51 | 0.60 | 2.3E-07 |
| Pgr | 169 | 1.60 | 0.68 | 8.3E-03 | 1.68 | 0.75 | 3.6E-03 |
| Pgrmc2 | 5271 | 1.68 | 0.75 | 1.2E-16 | 1.51 | 0.59 | 6.8E-11 |
| Phactr2 | 3670 | -1.60 | -0.68 | 6.7E-07 | -1.48 | -0.57 | 4.2E-05 |
| Phactr3 | 2006 | -1.60 | -0.68 | 8.6E-03 | -1.64 | -0.71 | 6.1E-03 |
| Phactr4 | 4597 | -1.30 | -0.38 | 1.1E-02 | -1.40 | -0.49 | 8.0E-04 |
| Phaf1 | 2851 | 1.57 | 0.65 | 2.8E-36 | 1.52 | 0.61 | 1.9E-31 |
| Phb | 6881 | 1.29 | 0.37 | 2.3E-05 | 1.27 | 0.34 | 1.1E-04 |
| Pheta1 | 3116 | 1.51 | 0.60 | 2.3E-10 | 1.39 | 0.47 | 6.6E-07 |
| Pheta2 | 202 | -3.57 | -1.83 | 3.3E-20 | -3.54 | -1.83 | 9.5E-20 |
| Phf11c | 445 | 2.71 | 1.44 | 6.2E-50 | 2.84 | 1.50 | 8.2E-54 |
| Phf13 | 3205 | 1.64 | 0.71 | 1.2E-16 | 1.54 | 0.62 | 5.5E-13 |
| Phf14 | 11218 | -1.74 | -0.80 | 2.7E-18 | -1.77 | -0.82 | 2.0E-19 |
| Phf19 | 335 | -1.54 | -0.62 | 6.7E-06 | -1.78 | -0.83 | 1.3E-09 |
| Phf2 | 6735 | -1.85 | -0.88 | 5.0E-08 | -2.00 | -1.00 | 5.6E-10 |
| Phf20 | 8339 | 1.24 | 0.31 | 1.2E-06 | 1.29 | 0.36 | 5.4E-09 |
| Phka2 | 2953 | 1.20 | 0.27 | 1.4E-02 | 1.26 | 0.33 | 2.0E-03 |
| Phkb | 3910 | -1.34 | -0.42 | 3.3E-05 | -1.21 | -0.28 | 7.5E-03 |
| Phkg2 | 3609 | 1.26 | 0.34 | 4.5E-08 | 1.19 | 0.25 | 4.8E-05 |
| Phldb2 | 6885 | -1.49 | -0.58 | 6.4E-03 | -1.41 | -0.49 | 2.3E-02 |

| | | | | | | | |
|---|---|---|---|---|---|---|---|
| Phldb3 | 132 | 4.83 | 2.27 | 8.7E-07 | 4.89 | 2.29 | 7.8E-07 |
| Phospho2 | 2340 | 1.27 | 0.34 | 7.0E-05 | 1.35 | 0.44 | 2.4E-07 |
| Phtf1 | 4560 | -1.55 | -0.64 | 3.4E-19 | -1.56 | -0.64 | 2.2E-19 |
| Phtf2 | 3888 | 1.49 | 0.57 | 4.5E-16 | 1.53 | 0.62 | 3.0E-18 |
| Phyh | 1347 | -1.43 | -0.51 | 8.6E-20 | -1.39 | -0.47 | 8.3E-17 |
| Phyhip | 193 | 1.86 | 0.90 | 3.8E-07 | 1.70 | 0.77 | 1.8E-05 |
| Pi16 | 115 | -1.91 | -0.93 | 4.3E-10 | -2.48 | -1.31 | 8.6E-19 |
| Pi4k2a | 5780 | 2.15 | 1.10 | 2.4E-66 | 2.08 | 1.06 | 4.4E-61 |
| Pi4k2b | 2638 | 2.20 | 1.14 | 8.8E-18 | 2.18 | 1.12 | 2.0E-17 |
| Pias3 | 2259 | -1.48 | -0.57 | 1.7E-10 | -1.56 | -0.64 | 3.8E-13 |
| Pick1 | 1257 | -1.32 | -0.40 | 2.5E-03 | -1.38 | -0.46 | 4.7E-04 |
| Pid1 | 3366 | -1.56 | -0.64 | 8.0E-05 | -1.36 | -0.44 | 8.1E-03 |
| Piezo2 | 1515 | -2.22 | -1.15 | 1.5E-03 | -1.99 | -0.99 | 7.2E-03 |
| Piga | 2529 | 1.66 | 0.74 | 3.0E-10 | 1.62 | 0.70 | 3.0E-09 |
| Pigb | 1251 | -1.38 | -0.46 | 1.0E-13 | -1.34 | -0.42 | 1.3E-11 |
| Pigc | 1658 | -1.70 | -0.77 | 7.5E-54 | -1.64 | -0.71 | 2.7E-46 |
| Pigf | 1493 | 2.44 | 1.29 | 3.4E-32 | 2.43 | 1.28 | 1.0E-31 |
| Pigh | 1044 | -1.39 | -0.47 | 5.5E-07 | -1.38 | -0.46 | 8.7E-07 |
| Pigk | 2989 | -1.40 | -0.49 | 9.3E-06 | -1.40 | -0.48 | 1.2E-05 |
| Pigm | 4879 | 1.59 | 0.67 | 6.9E-08 | 1.45 | 0.53 | 2.3E-05 |
| Pign | 4913 | 1.50 | 0.58 | 1.6E-07 | 1.71 | 0.77 | 1.6E-12 |
| Pigo | 2063 | -1.51 | -0.60 | 1.5E-13 | -1.65 | -0.72 | 2.7E-19 |
| Pigp | 2067 | -1.62 | -0.69 | 3.5E-20 | -1.60 | -0.68 | 1.9E-19 |
| Pigq | 4021 | -1.44 | -0.53 | 4.1E-23 | -1.52 | -0.61 | 1.0E-29 |
| Pigs | 4860 | 1.86 | 0.90 | 1.4E-24 | 1.75 | 0.81 | 4.3E-20 |
| Pigu | 2333 | -1.36 | -0.44 | 2.6E-12 | -1.38 | -0.47 | 1.0E-13 |
| Pigv | 1408 | 1.38 | 0.46 | 4.5E-07 | 1.37 | 0.46 | 6.2E-07 |
| Pigw | 537 | -2.08 | -1.05 | 1.4E-28 | -2.16 | -1.11 | 1.0E-31 |
| Pigyl | 880 | 1.31 | 0.39 | 2.5E-04 | 1.40 | 0.49 | 3.5E-06 |
| Pih1d1 | 2553 | -1.72 | -0.78 | 1.2E-32 | -1.82 | -0.86 | 3.8E-39 |
| Pih1d2 | 225 | -1.42 | -0.50 | 7.2E-04 | -1.37 | -0.46 | 2.6E-03 |
| Pik3c2a | 7213 | 1.25 | 0.32 | 3.7E-02 | 1.49 | 0.57 | 1.2E-04 |
| Pik3c2b | 4083 | -1.65 | -0.72 | 7.3E-06 | -1.81 | -0.86 | 9.1E-08 |
| Pik3cb | 2150 | 1.73 | 0.79 | 3.6E-22 | 1.77 | 0.82 | 1.2E-23 |
| Pik3cg | 194 | -2.20 | -1.14 | 4.0E-09 | -2.41 | -1.27 | 6.5E-11 |
| Pik3ip1 | 790 | -1.41 | -0.50 | 5.9E-06 | -1.39 | -0.48 | 1.6E-05 |
| Pik3r3 | 11699 | -1.40 | -0.49 | 7.0E-05 | -1.43 | -0.52 | 2.4E-05 |
| Pik3r4 | 3800 | 1.26 | 0.34 | 3.3E-05 | 1.24 | 0.31 | 1.7E-04 |
| Pik3r5 | 200 | 2.58 | 1.36 | 3.6E-06 | 2.72 | 1.44 | 1.0E-06 |
| Pik3r6 | 177 | 1.29 | 0.36 | 7.9E-04 | 1.24 | 0.32 | 3.7E-03 |
| Pikfyve | 9674 | 2.16 | 1.11 | 2.3E-18 | 2.44 | 1.29 | 3.7E-24 |
| Pim1 | 2603 | 1.54 | 0.62 | 2.4E-04 | 1.36 | 0.45 | 1.0E-02 |
| Pim2 | 1367 | 2.05 | 1.04 | 7.2E-24 | 1.79 | 0.84 | 6.6E-16 |
| Pin1 | 3848 | 1.63 | 0.71 | 4.5E-19 | 1.54 | 0.62 | 5.9E-15 |
| Pip4k2a | 2860 | 1.92 | 0.94 | 1.1E-39 | 1.94 | 0.96 | 5.0E-41 |
| Pip4k2b | 14253 | 1.51 | 0.59 | 3.0E-08 | 1.46 | 0.54 | 4.2E-07 |
| Pip4k2c | 2598 | 1.39 | 0.48 | 1.4E-09 | 1.39 | 0.48 | 1.4E-09 |

| | | | | | | | |
|---|---|---|---|---|---|---|---|
| Pip4p1 | 4135 | 1.73 | 0.79 | 2.8E-39 | 1.61 | 0.69 | 1.8E-30 |
| Pip4p2 | 4716 | -1.55 | -0.63 | 1.1E-12 | -1.64 | -0.71 | 6.1E-16 |
| Pip5k1b | 282 | 1.38 | 0.46 | 1.1E-03 | 1.23 | 0.29 | 4.6E-02 |
| Pip5kl1 | 244 | 2.37 | 1.24 | 3.4E-21 | 2.22 | 1.15 | 2.3E-18 |
| Pipox | 593 | -2.01 | -1.00 | 5.0E-08 | -2.35 | -1.23 | 1.4E-11 |
| Pithd1 | 1798 | 1.41 | 0.50 | 2.6E-12 | 1.22 | 0.29 | 6.4E-05 |
| Pitpna | 11361 | 1.47 | 0.56 | 5.8E-28 | 1.38 | 0.47 | 6.4E-20 |
| Pitpnb | 9524 | 1.32 | 0.40 | 2.6E-08 | 1.18 | 0.24 | 1.0E-03 |
| Pitpnc1 | 6213 | -1.35 | -0.43 | 1.2E-05 | -1.37 | -0.45 | 4.9E-06 |
| Pitpnm2 | 3374 | -1.80 | -0.85 | 2.7E-08 | -1.82 | -0.87 | 1.5E-08 |
| Pitpnm3 | 349 | -1.89 | -0.92 | 9.2E-03 | -1.69 | -0.75 | 3.6E-02 |
| Pitx1 | 157 | -3.67 | -1.88 | 5.1E-03 | -5.34 | -2.42 | 2.6E-04 |
| Pja2 | 22678 | 1.22 | 0.28 | 2.0E-02 | 1.35 | 0.43 | 2.8E-04 |
| Pkd2 | 5420 | -2.73 | -1.45 | 3.3E-43 | -2.56 | -1.36 | 5.7E-38 |
| Pknox2 | 3828 | 1.26 | 0.34 | 1.6E-06 | 1.13 | 0.18 | 1.7E-02 |
| Pkp1 | 127 | -6.56 | -2.71 | 6.8E-03 | -6.69 | -2.74 | 6.6E-03 |
| Pkp2 | 686 | 5.35 | 2.42 | 1.6E-21 | 4.56 | 2.19 | 9.0E-18 |
| Pkp4 | 8350 | 1.75 | 0.81 | 9.1E-41 | 1.72 | 0.78 | 2.5E-38 |
| Pla2g4c | 631 | 1.50 | 0.59 | 1.6E-05 | 1.54 | 0.62 | 6.0E-06 |
| Plag1 | 2036 | 1.54 | 0.62 | 6.8E-06 | 1.86 | 0.90 | 5.8E-11 |
| Plat | 4185 | 1.56 | 0.64 | 2.9E-12 | 1.47 | 0.56 | 1.7E-09 |
| Plbd2 | 2702 | 1.64 | 0.71 | 1.2E-11 | 1.55 | 0.64 | 1.7E-09 |
| Plcb3 | 4809 | 2.01 | 1.01 | 8.8E-11 | 2.01 | 1.01 | 9.5E-11 |
| Plcb4 | 2810 | -1.46 | -0.55 | 3.6E-05 | -1.34 | -0.43 | 1.6E-03 |
| Plcd1 | 1522 | 1.32 | 0.41 | 1.7E-02 | 1.30 | 0.38 | 2.9E-02 |
| Plcd3 | 496 | 3.47 | 1.79 | 2.2E-17 | 3.64 | 1.86 | 1.5E-18 |
| Plce1 | 2831 | -1.54 | -0.62 | 8.1E-09 | -1.31 | -0.39 | 4.3E-04 |
| Plcg2 | 339 | 1.95 | 0.97 | 1.2E-07 | 1.74 | 0.80 | 1.6E-05 |
| Plch1 | 4955 | 1.53 | 0.61 | 7.1E-05 | 1.63 | 0.71 | 4.5E-06 |
| Plcl2 | 2119 | -1.75 | -0.81 | 8.2E-13 | -1.80 | -0.85 | 6.2E-14 |
| Plcxd2 | 3734 | 1.95 | 0.96 | 1.3E-10 | 2.26 | 1.18 | 2.9E-15 |
| Pld3 | 4215 | 1.49 | 0.58 | 1.6E-08 | 1.50 | 0.59 | 8.4E-09 |
| Pld5 | 145 | -3.74 | -1.90 | 9.8E-11 | -3.32 | -1.73 | 4.9E-09 |
| Plec | 7010 | 1.45 | 0.53 | 1.3E-03 | 1.49 | 0.58 | 4.4E-04 |
| Plek | 727 | -1.74 | -0.80 | 4.3E-05 | -1.68 | -0.75 | 1.3E-04 |
| Plekha3 | 2122 | 1.34 | 0.42 | 2.6E-13 | 1.28 | 0.35 | 1.4E-09 |
| Plekha4 | 180 | -2.70 | -1.43 | 3.1E-04 | -2.49 | -1.32 | 1.0E-03 |
| Plekha5 | 5411 | -1.93 | -0.95 | 1.5E-33 | -1.95 | -0.96 | 1.5E-34 |
| Plekha6 | 8305 | -1.51 | -0.59 | 5.5E-04 | -1.57 | -0.65 | 1.5E-04 |
| Plekha7 | 3088 | 2.13 | 1.09 | 5.6E-16 | 1.97 | 0.98 | 3.6E-13 |
| Plekha8 | 5867 | 1.38 | 0.46 | 3.2E-12 | 1.37 | 0.45 | 1.2E-11 |
| Plekhb2 | 3773 | 1.69 | 0.75 | 1.9E-40 | 1.64 | 0.72 | 1.0E-36 |
| Plekhf1 | 330 | 4.68 | 2.23 | 4.1E-40 | 4.80 | 2.26 | 7.0E-41 |
| Plekhg1 | 2486 | -2.09 | -1.06 | 1.0E-17 | -2.12 | -1.08 | 2.3E-18 |
| Plekhg2 | 6442 | -1.26 | -0.34 | 1.8E-02 | -1.34 | -0.42 | 3.2E-03 |
| Plekhg4 | 1159 | 3.99 | 2.00 | 9.4E-32 | 4.23 | 2.08 | 2.5E-34 |
| Plekhh2 | 1655 | 1.52 | 0.61 | 8.4E-04 | 1.76 | 0.81 | 5.2E-06 |

| | | | | | | | |
|---|---|---|---|---|---|---|---|
| Plekhh3 | 1070 | 2.69 | 1.43 | 4.8E-35 | 2.49 | 1.32 | 6.5E-30 |
| Plekhm1 | 2965 | 1.68 | 0.75 | 4.1E-12 | 1.57 | 0.65 | 2.4E-09 |
| Plekhm2 | 2297 | -1.46 | -0.54 | 1.3E-06 | -1.57 | -0.65 | 5.4E-09 |
| Plekho1 | 3810 | -1.79 | -0.84 | 8.9E-15 | -1.92 | -0.94 | 3.0E-18 |
| Plekho2 | 1761 | 2.46 | 1.30 | 6.5E-17 | 2.38 | 1.25 | 9.3E-16 |
| Plin2 | 2216 | -1.98 | -0.99 | 2.2E-12 | -2.00 | -1.00 | 1.1E-12 |
| Plk1 | 3447 | -1.54 | -0.62 | 5.4E-03 | -1.83 | -0.87 | 7.0E-05 |
| Plk2 | 7132 | -1.47 | -0.56 | 5.9E-07 | -1.48 | -0.57 | 3.5E-07 |
| Plod3 | 3582 | -1.33 | -0.41 | 2.0E-05 | -1.42 | -0.51 | 9.5E-08 |
| Plp1 | 1332 | -1.65 | -0.72 | 4.4E-10 | -1.54 | -0.62 | 9.2E-08 |
| Plp2 | 1038 | -1.99 | -0.99 | 7.4E-13 | -2.04 | -1.03 | 1.0E-13 |
| Plpbp | 3893 | 1.27 | 0.34 | 1.2E-07 | 1.33 | 0.42 | 9.8E-11 |
| Plpp3 | 6037 | -1.85 | -0.89 | 3.7E-30 | -1.85 | -0.89 | 5.4E-30 |
| Plpp5 | 1196 | -2.43 | -1.28 | 4.3E-46 | -2.38 | -1.25 | 5.6E-44 |
| Plpp7 | 262 | -1.87 | -0.90 | 2.2E-07 | -1.73 | -0.79 | 7.2E-06 |
| Plppr2 | 4711 | 1.41 | 0.50 | 9.0E-03 | 1.39 | 0.47 | 1.5E-02 |
| Plppr3 | 14124 | -1.44 | -0.52 | 7.1E-06 | -1.48 | -0.57 | 9.5E-07 |
| Plrg1 | 5822 | 1.27 | 0.34 | 4.5E-04 | 1.24 | 0.31 | 1.4E-03 |
| Pls1 | 725 | 2.38 | 1.25 | 1.1E-12 | 2.43 | 1.28 | 3.7E-13 |
| Pls3 | 7182 | 1.43 | 0.52 | 8.4E-05 | 1.47 | 0.55 | 2.2E-05 |
| Plscr3 | 2888 | -1.25 | -0.32 | 5.6E-03 | -1.32 | -0.40 | 5.2E-04 |
| Plscr4 | 146 | -2.18 | -1.12 | 6.2E-07 | -2.14 | -1.10 | 1.4E-06 |
| Plxna3 | 18041 | 1.31 | 0.39 | 1.0E-02 | 1.39 | 0.47 | 1.9E-03 |
| Plxnb1 | 17385 | -1.79 | -0.84 | 1.2E-08 | -1.91 | -0.93 | 2.6E-10 |
| Pm20d2 | 108 | -1.68 | -0.75 | 1.5E-03 | -2.26 | -1.18 | 5.4E-07 |
| Pmaip1 | 652 | 3.61 | 1.85 | 2.5E-10 | 3.13 | 1.65 | 2.2E-08 |
| Pmf1 | 1767 | -1.54 | -0.63 | 2.9E-04 | -1.90 | -0.92 | 5.0E-08 |
| Pmm1 | 1980 | 1.83 | 0.87 | 1.9E-26 | 1.78 | 0.83 | 4.4E-24 |
| Pmm2 | 2334 | -2.06 | -1.04 | 1.8E-59 | -2.10 | -1.07 | 6.3E-62 |
| Pmp22 | 1376 | -2.34 | -1.23 | 9.3E-06 | -2.21 | -1.14 | 4.2E-05 |
| Pmpcb | 4600 | -1.25 | -0.32 | 2.1E-04 | -1.27 | -0.34 | 5.9E-05 |
| Pms2 | 2321 | 1.32 | 0.40 | 3.1E-07 | 1.30 | 0.38 | 1.6E-06 |
| Pmvk | 2886 | 1.34 | 0.42 | 1.1E-06 | 1.28 | 0.36 | 5.1E-05 |
| Pnisr | 26071 | -1.95 | -0.96 | 1.5E-24 | -1.81 | -0.86 | 1.0E-19 |
| Pnkd | 3936 | 1.58 | 0.66 | 7.6E-13 | 1.52 | 0.61 | 6.3E-11 |
| Pnkp | 1824 | 1.32 | 0.40 | 5.0E-07 | 1.21 | 0.28 | 8.1E-04 |
| Pnldc1 | 246 | -1.36 | -0.44 | 4.5E-04 | -1.33 | -0.41 | 1.2E-03 |
| pnma1 | 143 | -1.48 | -0.57 | 1.3E-05 | -1.66 | -0.73 | 1.5E-08 |
| Pnma2 | 3038 | 2.48 | 1.31 | 2.4E-23 | 2.61 | 1.39 | 5.1E-26 |
| Pnma3 | 667 | -1.39 | -0.47 | 1.5E-02 | -1.52 | -0.61 | 1.4E-03 |
| Pnn | 20926 | -1.23 | -0.30 | 1.2E-02 | -1.31 | -0.38 | 1.0E-03 |
| Pno1 | 2529 | 1.48 | 0.57 | 5.4E-05 | 1.37 | 0.45 | 1.6E-03 |
| Pnoc | 559 | -3.08 | -1.62 | 2.0E-09 | -3.04 | -1.60 | 3.1E-09 |
| Pnpla2 | 1152 | 2.44 | 1.29 | 4.1E-28 | 2.33 | 1.22 | 2.1E-25 |
| Pnpla3 | 284 | 3.75 | 1.91 | 3.1E-58 | 3.37 | 1.75 | 7.2E-50 |
| Pnpla6 | 3170 | -1.28 | -0.35 | 1.2E-04 | -1.27 | -0.34 | 1.9E-04 |
| Pnpla7 | 498 | -1.27 | -0.34 | 2.0E-03 | -1.28 | -0.35 | 1.4E-03 |

| | | | | | | | |
|---|---|---|---|---|---|---|---|
| Pnpo | 1163 | 1.61 | 0.68 | 9.1E-09 | 1.58 | 0.66 | 3.5E-08 |
| Pnrc1 | 3055 | -1.28 | -0.36 | 2.7E-06 | -1.34 | -0.42 | 1.6E-08 |
| Pnrc2 | 7960 | -1.28 | -0.36 | 1.7E-03 | -1.32 | -0.40 | 3.3E-04 |
| Poc1a | 907 | 1.83 | 0.87 | 5.8E-06 | 1.46 | 0.54 | 6.6E-03 |
| Poc1b | 3050 | 1.33 | 0.42 | 1.1E-04 | 1.20 | 0.26 | 2.0E-02 |
| Poc5 | 3976 | 1.37 | 0.45 | 9.8E-14 | 1.37 | 0.46 | 1.0E-13 |
| Podn | 147 | -2.59 | -1.37 | 1.8E-05 | -2.21 | -1.14 | 4.4E-04 |
| Podxl | 7193 | 1.35 | 0.44 | 2.0E-08 | 1.27 | 0.34 | 1.7E-05 |
| Pofut1 | 4371 | -1.52 | -0.61 | 7.4E-16 | -1.57 | -0.65 | 5.9E-18 |
| Pogk | 15161 | -1.31 | -0.39 | 3.1E-05 | -1.27 | -0.35 | 2.4E-04 |
| Poglut2 | 2495 | 1.30 | 0.38 | 6.6E-06 | 1.33 | 0.41 | 1.2E-06 |
| Poglut3 | 777 | -3.15 | -1.66 | 3.9E-14 | -3.10 | -1.63 | 1.1E-13 |
| Pogz | 8242 | -2.18 | -1.12 | 2.3E-29 | -2.25 | -1.17 | 8.9E-32 |
| Pola2 | 4393 | 1.67 | 0.74 | 7.1E-08 | 1.57 | 0.65 | 3.1E-06 |
| Poldip3 | 8934 | -1.23 | -0.30 | 2.0E-07 | -1.31 | -0.39 | 3.8E-12 |
| Pole3 | 6772 | 1.48 | 0.57 | 9.6E-07 | 1.37 | 0.46 | 1.1E-04 |
| Pole4 | 2215 | -1.59 | -0.67 | 5.1E-22 | -1.66 | -0.73 | 7.2E-26 |
| Polg | 7065 | 1.91 | 0.93 | 4.5E-43 | 1.84 | 0.88 | 3.0E-38 |
| Polg2 | 704 | 1.30 | 0.38 | 8.0E-07 | 1.28 | 0.35 | 6.3E-06 |
| Polk | 1613 | 1.80 | 0.85 | 5.9E-20 | 2.05 | 1.03 | 9.1E-29 |
| Poln | 160 | 2.73 | 1.45 | 5.5E-13 | 2.82 | 1.49 | 1.6E-13 |
| Polr1d | 3958 | 1.47 | 0.56 | 4.9E-11 | 1.40 | 0.49 | 1.2E-08 |
| Polr1e | 2297 | 2.27 | 1.18 | 1.3E-43 | 2.22 | 1.15 | 3.5E-41 |
| Polr1f | 3119 | 1.30 | 0.38 | 5.8E-04 | 1.29 | 0.36 | 1.2E-03 |
| Polr2a | 20372 | -1.35 | -0.43 | 4.3E-04 | -1.38 | -0.47 | 1.2E-04 |
| Polr2b | 16783 | 1.74 | 0.80 | 7.9E-13 | 1.78 | 0.84 | 5.1E-14 |
| Polr2e | 3853 | 1.54 | 0.62 | 2.6E-06 | 1.40 | 0.49 | 3.3E-04 |
| Polr2h | 1978 | 1.31 | 0.39 | 1.6E-05 | 1.33 | 0.42 | 3.0E-06 |
| Polr2i | 980 | -1.25 | -0.32 | 1.6E-02 | -1.32 | -0.41 | 1.7E-03 |
| Polr2j | 2771 | 1.32 | 0.40 | 1.5E-04 | 1.23 | 0.30 | 5.3E-03 |
| Polr2l | 2673 | 1.59 | 0.67 | 5.6E-07 | 1.50 | 0.58 | 1.8E-05 |
| Polr2m | 16513 | -1.51 | -0.59 | 3.0E-09 | -1.69 | -0.76 | 2.6E-14 |
| Polr3b | 2647 | -1.64 | -0.71 | 4.9E-30 | -1.60 | -0.68 | 1.7E-27 |
| Polr3d | 1895 | -1.33 | -0.41 | 2.7E-05 | -1.47 | -0.55 | 7.3E-09 |
| Polr3g | 1463 | 2.19 | 1.13 | 2.2E-07 | 2.33 | 1.22 | 2.1E-08 |
| Polr3gl | 2913 | 1.77 | 0.83 | 5.7E-12 | 1.64 | 0.72 | 2.8E-09 |
| Polr3h | 2099 | 1.72 | 0.78 | 3.5E-35 | 1.68 | 0.74 | 4.0E-32 |
| Pomgnt1 | 3228 | -1.63 | -0.70 | 1.2E-31 | -1.69 | -0.76 | 7.8E-37 |
| Pomgnt2 | 1045 | 1.55 | 0.63 | 3.9E-07 | 1.51 | 0.59 | 2.6E-06 |
| Pomk | 1708 | -1.41 | -0.50 | 1.1E-24 | -1.45 | -0.54 | 2.0E-28 |
| Pomt1 | 2351 | -1.30 | -0.37 | 2.8E-07 | -1.29 | -0.36 | 7.1E-07 |
| Pomt2 | 2019 | -1.52 | -0.61 | 3.3E-42 | -1.51 | -0.59 | 7.2E-40 |
| Pon2 | 1369 | -1.72 | -0.79 | 2.0E-18 | -1.90 | -0.92 | 6.9E-25 |
| Pop5 | 1111 | 1.76 | 0.82 | 4.0E-17 | 1.61 | 0.69 | 2.2E-12 |
| Pop7 | 676 | -2.16 | -1.11 | 9.9E-31 | -2.24 | -1.17 | 9.1E-34 |
| Por | 7940 | 1.90 | 0.92 | 7.1E-16 | 1.71 | 0.77 | 2.3E-11 |
| Postn | 6560 | -3.00 | -1.59 | 5.1E-03 | -2.43 | -1.28 | 2.7E-02 |

| | | | | | | | |
|---|---|---|---|---|---|---|---|
| Pou6f1 | 6486 | -1.58 | -0.66 | 2.1E-03 | -1.62 | -0.69 | 1.4E-03 |
| Pou6f2 | 614 | -1.65 | -0.72 | 4.4E-03 | -1.48 | -0.56 | 3.0E-02 |
| Ppa2 | 1614 | -1.37 | -0.45 | 1.6E-09 | -1.43 | -0.51 | 7.3E-12 |
| Ppard | 2049 | 2.04 | 1.03 | 1.1E-12 | 1.97 | 0.98 | 1.2E-11 |
| Ppargc1a | 1179 | 1.65 | 0.72 | 2.5E-05 | 1.87 | 0.91 | 7.6E-08 |
| Ppargc1b | 174 | 2.17 | 1.12 | 5.2E-06 | 2.56 | 1.36 | 3.3E-08 |
| Ppat | 4071 | 1.75 | 0.81 | 5.0E-09 | 1.64 | 0.72 | 2.3E-07 |
| Ppcdc | 802 | -1.80 | -0.85 | 1.6E-41 | -1.78 | -0.83 | 9.3E-40 |
| Ppdpf | 3917 | -1.37 | -0.45 | 5.9E-07 | -1.52 | -0.60 | 1.7E-11 |
| Ppfia3 | 3569 | 1.46 | 0.54 | 2.3E-03 | 1.34 | 0.42 | 2.1E-02 |
| Ppfibp1 | 5507 | 1.50 | 0.59 | 1.5E-04 | 1.64 | 0.71 | 3.8E-06 |
| Ppfibp2 | 512 | -1.85 | -0.89 | 2.4E-03 | -1.70 | -0.77 | 9.5E-03 |
| Pphln1 | 6579 | 1.33 | 0.42 | 7.9E-06 | 1.28 | 0.35 | 2.0E-04 |
| Ppia | 52024 | -1.22 | -0.28 | 1.6E-02 | -1.26 | -0.34 | 3.9E-03 |
| Ppif | 2242 | 1.39 | 0.48 | 2.2E-07 | 1.32 | 0.40 | 2.3E-05 |
| Ppih | 1751 | -1.30 | -0.38 | 1.3E-02 | -1.36 | -0.44 | 3.5E-03 |
| Ppil3 | 1497 | -1.67 | -0.74 | 9.9E-22 | -1.71 | -0.78 | 6.1E-24 |
| Ppil6 | 216 | 3.62 | 1.86 | 5.0E-31 | 4.25 | 2.09 | 5.1E-38 |
| Ppm1b | 10545 | 1.42 | 0.50 | 1.1E-11 | 1.32 | 0.40 | 9.3E-08 |
| Ppm1d | 2918 | 1.33 | 0.41 | 2.6E-08 | 1.23 | 0.30 | 7.7E-05 |
| Ppm1e | 10602 | 1.50 | 0.59 | 3.3E-03 | 1.58 | 0.66 | 8.8E-04 |
| Ppm1f | 5811 | 1.26 | 0.34 | 1.6E-07 | 1.24 | 0.31 | 1.7E-06 |
| Ppm1g | 15251 | 1.68 | 0.75 | 3.1E-13 | 1.49 | 0.57 | 3.7E-08 |
| Ppm1j | 282 | 7.24 | 2.86 | 1.4E-36 | 7.60 | 2.93 | 4.4E-38 |
| Ppm1k | 2258 | 2.52 | 1.33 | 9.3E-39 | 2.37 | 1.25 | 7.1E-34 |
| Ppme1 | 7854 | 1.40 | 0.48 | 4.5E-17 | 1.33 | 0.42 | 6.4E-13 |
| Ppox | 4532 | 1.40 | 0.48 | 7.0E-15 | 1.33 | 0.41 | 7.5E-11 |
| Ppp1cb | 23071 | -1.40 | -0.48 | 4.3E-10 | -1.40 | -0.49 | 2.9E-10 |
| Ppp1r12a | 8968 | -1.31 | -0.39 | 4.3E-11 | -1.28 | -0.36 | 3.0E-09 |
| Ppp1r12c | 6838 | 1.61 | 0.69 | 1.3E-18 | 1.53 | 0.61 | 7.4E-15 |
| Ppp1r13l | 1015 | 3.23 | 1.69 | 3.2E-23 | 3.11 | 1.64 | 9.1E-22 |
| Ppp1r14b | 7110 | -1.22 | -0.28 | 2.4E-02 | -1.36 | -0.45 | 2.3E-04 |
| Ppp1r15a | 1282 | 2.01 | 1.01 | 8.8E-23 | 2.09 | 1.06 | 5.4E-25 |
| Ppp1r16a | 3598 | 1.53 | 0.61 | 1.4E-12 | 1.41 | 0.50 | 1.2E-08 |
| Ppp1r16b | 1142 | 2.33 | 1.22 | 5.8E-12 | 2.21 | 1.14 | 1.3E-10 |
| Ppp1r1a | 1826 | 1.60 | 0.68 | 5.5E-14 | 1.61 | 0.69 | 2.4E-14 |
| Ppp1r1b | 299 | 3.35 | 1.74 | 1.3E-04 | 4.05 | 2.02 | 8.5E-06 |
| Ppp1r21 | 4118 | 1.26 | 0.33 | 3.6E-04 | 1.27 | 0.34 | 2.4E-04 |
| Ppp1r26 | 2297 | -1.33 | -0.41 | 1.9E-04 | -1.29 | -0.37 | 9.1E-04 |
| Ppp1r35 | 1343 | 1.72 | 0.78 | 2.9E-10 | 1.60 | 0.68 | 5.1E-08 |
| Ppp1r37 | 5590 | 1.35 | 0.43 | 4.0E-04 | 1.28 | 0.35 | 4.2E-03 |
| Ppp1r3d | 343 | 2.40 | 1.26 | 1.8E-07 | 2.77 | 1.47 | 1.1E-09 |
| Ppp1r9a | 8368 | -1.91 | -0.94 | 1.5E-09 | -1.65 | -0.72 | 3.9E-06 |
| Ppp2r1b | 7788 | 1.54 | 0.62 | 2.2E-11 | 1.44 | 0.53 | 2.0E-08 |
| Ppp2r3c | 3139 | -1.70 | -0.76 | 1.4E-14 | -1.61 | -0.69 | 3.3E-12 |
| Ppp2r5a | 3114 | 1.63 | 0.71 | 1.6E-22 | 1.59 | 0.67 | 4.4E-20 |
| Ppp2r5d | 10594 | 1.79 | 0.84 | 5.1E-25 | 1.68 | 0.75 | 2.5E-20 |

| | | | | | | | |
|---|---|---|---|---|---|---|---|
| Ppp3cb | 14492 | -1.25 | -0.32 | 1.0E-02 | -1.29 | -0.37 | 3.0E-03 |
| Ppp3cc | 516 | -1.27 | -0.34 | 6.9E-03 | -1.25 | -0.32 | 1.2E-02 |
| Ppp3r1 | 9679 | 1.35 | 0.43 | 3.6E-04 | 1.31 | 0.39 | 1.2E-03 |
| Ppp4r1 | 5377 | -1.79 | -0.84 | 5.1E-42 | -1.90 | -0.93 | 1.3E-50 |
| Ppp4r2 | 9308 | -1.40 | -0.48 | 3.3E-09 | -1.44 | -0.53 | 7.2E-11 |
| Ppp4r3a | 11173 | -1.34 | -0.42 | 1.9E-05 | -1.39 | -0.47 | 1.5E-06 |
| Ppp4r3b | 9651 | -1.28 | -0.35 | 1.0E-03 | -1.19 | -0.25 | 2.4E-02 |
| Ppp4r4 | 1187 | 2.34 | 1.23 | 1.0E-59 | 2.33 | 1.22 | 3.2E-58 |
| Pprc1 | 3855 | -1.56 | -0.64 | 4.9E-09 | -1.65 | -0.72 | 3.3E-11 |
| Ppt2 | 682 | 1.28 | 0.36 | 6.5E-03 | 1.37 | 0.46 | 4.5E-04 |
| Pptc7 | 2419 | 1.48 | 0.57 | 2.8E-15 | 1.39 | 0.48 | 4.0E-11 |
| Pradc1 | 708 | 1.82 | 0.87 | 5.0E-13 | 1.62 | 0.69 | 1.1E-08 |
| Prdm10 | 2700 | -3.68 | -1.88 | 9.2E-56 | -3.74 | -1.90 | 5.9E-57 |
| Prdm11 | 631 | -2.67 | -1.42 | 9.1E-11 | -2.36 | -1.24 | 1.9E-08 |
| Prdm12 | 651 | -1.97 | -0.98 | 1.7E-04 | -1.97 | -0.98 | 1.9E-04 |
| Prdm15 | 2046 | -2.70 | -1.43 | 1.6E-103 | -2.59 | -1.37 | 4.0E-95 |
| Prdm16 | 2577 | -2.00 | -1.00 | 2.8E-12 | -1.88 | -0.91 | 2.5E-10 |
| Prdm5 | 617 | -2.13 | -1.09 | 7.5E-11 | -2.15 | -1.10 | 4.6E-11 |
| Prdm6 | 324 | -2.15 | -1.10 | 2.1E-04 | -1.94 | -0.95 | 1.6E-03 |
| Prdm9 | 940 | 2.64 | 1.40 | 4.6E-31 | 2.47 | 1.31 | 4.5E-27 |
| Preb | 8107 | 1.29 | 0.36 | 5.0E-11 | 1.22 | 0.28 | 5.4E-07 |
| Prelp | 424 | -2.69 | -1.43 | 1.5E-07 | -2.39 | -1.26 | 4.8E-06 |
| Prepl | 9915 | 1.22 | 0.29 | 7.9E-03 | 1.31 | 0.39 | 3.3E-04 |
| Prex1 | 6987 | 1.55 | 0.63 | 5.5E-08 | 1.62 | 0.69 | 2.4E-09 |
| Prex2 | 1360 | -1.96 | -0.97 | 1.3E-06 | -1.61 | -0.69 | 8.7E-04 |
| Prim2 | 3251 | 1.60 | 0.68 | 3.4E-07 | 1.58 | 0.66 | 9.0E-07 |
| Prkab2 | 3272 | 1.68 | 0.75 | 4.6E-33 | 1.73 | 0.79 | 4.9E-36 |
| Prkaca | 10570 | 1.80 | 0.84 | 2.7E-22 | 1.62 | 0.69 | 2.1E-15 |
| Prkag1 | 3170 | -1.26 | -0.33 | 3.6E-12 | -1.35 | -0.43 | 2.8E-20 |
| Prkar2a | 12648 | 1.30 | 0.38 | 6.2E-08 | 1.27 | 0.34 | 8.6E-07 |
| Prkcd | 4272 | 2.40 | 1.26 | 3.3E-53 | 2.53 | 1.34 | 1.2E-59 |
| Prkci | 8204 | 1.84 | 0.88 | 1.0E-59 | 1.87 | 0.91 | 4.1E-63 |
| Prkcq | 221 | -2.05 | -1.04 | 2.4E-05 | -1.89 | -0.91 | 2.3E-04 |
| Prkd1 | 1487 | 1.33 | 0.41 | 9.2E-03 | 1.33 | 0.42 | 8.9E-03 |
| Prkd2 | 1619 | 1.69 | 0.76 | 3.1E-11 | 1.65 | 0.72 | 3.4E-10 |
| Prkg2 | 504 | -1.90 | -0.93 | 1.5E-10 | -2.32 | -1.21 | 2.9E-17 |
| Prkx | 4490 | 1.46 | 0.55 | 2.0E-06 | 1.44 | 0.52 | 7.1E-06 |
| Prmt1 | 19553 | -1.20 | -0.27 | 5.6E-03 | -1.31 | -0.38 | 4.2E-05 |
| Prmt3 | 1694 | -1.61 | -0.69 | 1.0E-20 | -1.69 | -0.76 | 8.8E-25 |
| Prmt5 | 9545 | 1.27 | 0.34 | 3.3E-05 | 1.18 | 0.24 | 4.3E-03 |
| Prmt8 | 1839 | -1.47 | -0.56 | 1.3E-03 | -1.41 | -0.50 | 4.8E-03 |
| Prmt9 | 2548 | 1.22 | 0.28 | 5.7E-04 | 1.29 | 0.37 | 6.1E-06 |
| Prnp | 5047 | 1.51 | 0.60 | 1.0E-09 | 1.52 | 0.60 | 8.9E-10 |
| Prob1 | 989 | 2.01 | 1.01 | 3.1E-15 | 1.98 | 0.99 | 1.2E-14 |
| Procr | 127 | -1.53 | -0.62 | 2.2E-03 | -1.53 | -0.61 | 2.4E-03 |
| Prokr1 | 903 | 1.71 | 0.77 | 1.2E-08 | 1.87 | 0.91 | 1.7E-11 |
| Prorp | 940 | -1.55 | -0.63 | 7.2E-13 | -1.72 | -0.78 | 4.1E-19 |

| | | | | | | | |
|---|---|---|---|---|---|---|---|
| Prorsd1 | 386 | -1.60 | -0.68 | 1.3E-05 | -1.78 | -0.83 | 7.6E-08 |
| Pros1 | 1393 | -1.42 | -0.51 | 5.8E-03 | -1.41 | -0.50 | 7.3E-03 |
| Proser2 | 244 | 1.84 | 0.88 | 9.4E-07 | 2.07 | 1.05 | 4.0E-09 |
| Proser3 | 1137 | -2.04 | -1.03 | 7.1E-13 | -2.24 | -1.16 | 5.1E-16 |
| Prpf3 | 4753 | 1.26 | 0.34 | 4.9E-05 | 1.19 | 0.25 | 3.5E-03 |
| Prpf6 | 9328 | 1.36 | 0.44 | 3.8E-11 | 1.33 | 0.41 | 1.2E-09 |
| Prps1 | 3960 | 1.34 | 0.42 | 3.5E-06 | 1.31 | 0.39 | 1.6E-05 |
| Prr15 | 502 | 2.11 | 1.08 | 4.1E-05 | 1.62 | 0.70 | 9.9E-03 |
| Prr16 | 220 | -2.05 | -1.03 | 3.8E-04 | -1.56 | -0.64 | 3.5E-02 |
| Prr18 | 491 | -3.50 | -1.81 | 3.6E-09 | -3.61 | -1.85 | 1.5E-09 |
| Prr36 | 5899 | -1.89 | -0.92 | 2.0E-05 | -2.20 | -1.14 | 9.2E-08 |
| Prr5 | 802 | 2.25 | 1.17 | 7.6E-13 | 2.07 | 1.05 | 1.3E-10 |
| Prr5l | 656 | -2.22 | -1.15 | 3.9E-11 | -2.48 | -1.31 | 4.1E-14 |
| Prr7 | 478 | -1.68 | -0.75 | 8.2E-08 | -1.90 | -0.92 | 1.9E-11 |
| Prrg1 | 882 | -1.38 | -0.46 | 1.8E-05 | -1.30 | -0.38 | 5.3E-04 |
| Prrg2 | 447 | -2.32 | -1.21 | 1.3E-35 | -2.30 | -1.20 | 6.0E-35 |
| Prrg3 | 2698 | -1.36 | -0.44 | 5.9E-06 | -1.31 | -0.39 | 7.7E-05 |
| Prrg4 | 194 | 2.51 | 1.33 | 1.3E-05 | 3.23 | 1.69 | 2.4E-08 |
| Prrt3 | 270 | 1.56 | 0.64 | 6.3E-04 | 1.44 | 0.53 | 5.3E-03 |
| Prrx1 | 7465 | -3.31 | -1.73 | 2.9E-07 | -3.15 | -1.65 | 9.6E-07 |
| Prss23 | 1182 | -1.43 | -0.52 | 2.6E-02 | -1.52 | -0.60 | 9.2E-03 |
| Prss36 | 317 | -1.77 | -0.82 | 6.5E-25 | -1.65 | -0.72 | 1.7E-19 |
| Prune1 | 3326 | -1.33 | -0.42 | 9.5E-06 | -1.34 | -0.42 | 5.9E-06 |
| Prx | 614 | -1.27 | -0.35 | 1.0E-02 | -1.33 | -0.41 | 2.3E-03 |
| Prxl2b | 931 | 1.77 | 0.82 | 1.5E-15 | 1.70 | 0.77 | 1.2E-13 |
| Prxl2c | 1128 | 1.56 | 0.64 | 2.0E-07 | 1.56 | 0.64 | 2.3E-07 |
| Psat1 | 17607 | 1.37 | 0.45 | 1.1E-05 | 1.33 | 0.41 | 5.6E-05 |
| Psen2 | 693 | 1.96 | 0.97 | 1.6E-27 | 1.90 | 0.93 | 4.1E-25 |
| Pskh1 | 1945 | -1.35 | -0.43 | 5.9E-03 | -1.43 | -0.52 | 9.3E-04 |
| Psma2 | 10930 | 1.26 | 0.33 | 1.4E-04 | 1.26 | 0.33 | 1.8E-04 |
| Psma5 | 8361 | 1.25 | 0.33 | 5.4E-04 | 1.20 | 0.27 | 5.6E-03 |
| Psmb10 | 401 | -1.57 | -0.65 | 6.3E-14 | -1.48 | -0.57 | 1.1E-10 |
| Psmd2 | 29334 | 1.27 | 0.35 | 6.7E-05 | 1.19 | 0.25 | 4.2E-03 |
| Psmd4 | 9072 | 1.27 | 0.34 | 8.8E-05 | 1.19 | 0.25 | 5.5E-03 |
| Psmd5 | 5393 | 1.29 | 0.37 | 2.5E-04 | 1.24 | 0.31 | 2.7E-03 |
| Psmd9 | 2434 | 1.38 | 0.47 | 1.2E-11 | 1.35 | 0.43 | 4.4E-10 |
| Psme4 | 24645 | 1.39 | 0.48 | 1.1E-06 | 1.42 | 0.50 | 2.9E-07 |
| Pspc1 | 5966 | -1.49 | -0.58 | 3.5E-08 | -1.56 | -0.64 | 8.0E-10 |
| Ptbp2 | 26867 | -1.53 | -0.62 | 5.5E-08 | -1.56 | -0.65 | 1.3E-08 |
| Ptbp3 | 8517 | 1.28 | 0.36 | 2.0E-02 | 1.44 | 0.53 | 4.8E-04 |
| Ptch1 | 7873 | -1.21 | -0.27 | 4.8E-04 | -1.26 | -0.34 | 1.1E-05 |
| Ptdss1 | 5032 | 1.32 | 0.40 | 6.1E-07 | 1.38 | 0.46 | 8.6E-09 |
| Pten | 12976 | 1.37 | 0.45 | 3.9E-06 | 1.37 | 0.45 | 3.9E-06 |
| Ptger1 | 207 | 3.05 | 1.61 | 1.8E-10 | 3.58 | 1.84 | 2.1E-13 |
| Ptger4 | 131 | 2.02 | 1.01 | 5.4E-03 | 2.75 | 1.46 | 4.6E-05 |
| Ptges | 739 | 2.61 | 1.39 | 5.4E-06 | 2.58 | 1.37 | 7.6E-06 |
| Ptges2 | 1580 | 1.71 | 0.77 | 8.9E-27 | 1.68 | 0.75 | 4.3E-25 |

| | | | | | | | |
|---|---|---|---|---|---|---|---|
| Ptgfr | 113 | -5.61 | -2.49 | 9.9E-17 | -6.27 | -2.65 | 3.9E-18 |
| Pth1r | 1324 | -1.60 | -0.67 | 5.9E-03 | -1.55 | -0.63 | 1.1E-02 |
| Ptk2b | 117 | 1.75 | 0.80 | 1.6E-03 | 1.68 | 0.75 | 3.7E-03 |
| Ptk7 | 9855 | 1.29 | 0.36 | 1.8E-02 | 1.27 | 0.35 | 2.5E-02 |
| Ptms | 25036 | -1.32 | -0.40 | 2.7E-02 | -1.44 | -0.52 | 3.6E-03 |
| Ptn | 20272 | -1.55 | -0.63 | 9.6E-05 | -1.46 | -0.54 | 9.5E-04 |
| Ptp4a3 | 1474 | -1.60 | -0.68 | 1.3E-12 | -1.62 | -0.69 | 5.0E-13 |
| Ptpa | 9367 | -1.29 | -0.36 | 3.5E-08 | -1.39 | -0.48 | 2.4E-13 |
| Ptpn1 | 7485 | 1.59 | 0.67 | 8.2E-10 | 1.45 | 0.53 | 1.3E-06 |
| Ptpn11 | 16665 | 1.31 | 0.39 | 9.8E-16 | 1.27 | 0.35 | 8.7E-13 |
| Ptpn12 | 6041 | -1.37 | -0.45 | 2.4E-10 | -1.41 | -0.50 | 2.4E-12 |
| Ptpn13 | 8616 | 2.15 | 1.11 | 1.0E-18 | 2.34 | 1.23 | 8.8E-23 |
| Ptpn2 | 2285 | -1.75 | -0.81 | 1.3E-31 | -1.78 | -0.83 | 3.6E-33 |
| Ptpn3 | 1467 | 2.06 | 1.05 | 4.5E-04 | 2.66 | 1.41 | 1.4E-06 |
| Ptpn4 | 9452 | 1.32 | 0.41 | 1.6E-02 | 1.60 | 0.68 | 3.4E-05 |
| Ptpn5 | 4140 | -1.59 | -0.67 | 1.2E-02 | -1.49 | -0.57 | 3.7E-02 |
| Ptpn6 | 230 | -1.42 | -0.50 | 3.0E-02 | -1.58 | -0.66 | 4.0E-03 |
| Ptpn9 | 4302 | -1.82 | -0.87 | 2.3E-30 | -1.88 | -0.91 | 3.8E-33 |
| Ptprb | 1124 | -1.77 | -0.82 | 2.0E-08 | -1.83 | -0.87 | 2.6E-09 |
| Ptprn | 2345 | 1.94 | 0.95 | 5.4E-07 | 1.85 | 0.89 | 3.4E-06 |
| Ptpro | 3982 | -1.70 | -0.77 | 1.0E-02 | -1.67 | -0.74 | 1.4E-02 |
| Ptpru | 5216 | 1.56 | 0.65 | 3.2E-06 | 1.49 | 0.58 | 3.5E-05 |
| Ptrh1 | 274 | 2.08 | 1.05 | 4.2E-16 | 1.97 | 0.98 | 6.8E-14 |
| Ptrh2 | 3006 | -1.27 | -0.34 | 4.6E-04 | -1.33 | -0.41 | 2.1E-05 |
| Ptrhd1 | 365 | -3.12 | -1.64 | 2.2E-44 | -3.60 | -1.85 | 2.7E-55 |
| Pts | 2565 | -1.37 | -0.46 | 3.0E-12 | -1.39 | -0.48 | 3.5E-13 |
| Pttg1 | 1147 | 1.89 | 0.92 | 2.0E-16 | 1.92 | 0.94 | 5.6E-17 |
| Pttg1ip | 4768 | 1.30 | 0.38 | 1.4E-04 | 1.28 | 0.35 | 3.9E-04 |
| Ptx3 | 1640 | -3.37 | -1.75 | 2.4E-20 | -3.61 | -1.85 | 1.3E-22 |
| Puf60 | 8826 | -1.63 | -0.70 | 1.2E-15 | -1.79 | -0.84 | 8.1E-22 |
| Pum1 | 8711 | -1.32 | -0.40 | 1.7E-03 | -1.43 | -0.52 | 4.2E-05 |
| Pum2 | 25350 | -1.43 | -0.52 | 4.5E-08 | -1.39 | -0.47 | 6.8E-07 |
| Pura | 2394 | 1.32 | 0.40 | 2.7E-05 | 1.35 | 0.43 | 5.1E-06 |
| Purg | 3082 | 1.28 | 0.36 | 3.3E-03 | 1.22 | 0.29 | 1.9E-02 |
| Pus10 | 3922 | 1.28 | 0.35 | 1.6E-03 | 1.33 | 0.42 | 1.8E-04 |
| Pus3 | 1379 | -2.12 | -1.08 | 2.1E-27 | -2.09 | -1.06 | 2.2E-26 |
| Pus7 | 3393 | 1.79 | 0.84 | 1.7E-45 | 1.76 | 0.82 | 3.4E-43 |
| Pusl1 | 1434 | 1.60 | 0.67 | 1.1E-23 | 1.61 | 0.69 | 1.0E-24 |
| Pvr | 426 | -2.25 | -1.17 | 3.1E-12 | -2.10 | -1.07 | 2.4E-10 |
| Pwwp2a | 3963 | -1.25 | -0.33 | 4.5E-04 | -1.36 | -0.45 | 9.1E-07 |
| Pwwp3b | 3399 | 2.53 | 1.34 | 1.8E-23 | 2.67 | 1.42 | 3.2E-26 |
| Pxdn | 15773 | -1.48 | -0.57 | 4.5E-05 | -1.38 | -0.46 | 1.1E-03 |
| Pxk | 3371 | 1.63 | 0.70 | 6.0E-11 | 1.73 | 0.79 | 1.1E-13 |
| Pxmp2 | 307 | 2.17 | 1.12 | 5.6E-26 | 2.39 | 1.25 | 7.7E-32 |
| Pxmp4 | 1007 | 1.50 | 0.58 | 1.7E-07 | 1.43 | 0.52 | 3.8E-06 |
| Pxn | 6562 | 2.24 | 1.16 | 1.4E-25 | 2.20 | 1.14 | 1.2E-24 |
| Pxylp1 | 3755 | 1.40 | 0.49 | 2.0E-10 | 1.42 | 0.50 | 5.2E-11 |

| | | | | | | | |
|---|---|---|---|---|---|---|---|
| Pycard | 123 | -1.78 | -0.83 | 2.5E-03 | -1.53 | -0.62 | 2.9E-02 |
| Pygb | 14633 | 1.74 | 0.80 | 1.1E-27 | 1.76 | 0.82 | 1.1E-28 |
| Pygo1 | 5395 | -2.65 | -1.40 | 8.7E-22 | -2.40 | -1.27 | 6.8E-18 |
| Pym1 | 1562 | -1.86 | -0.89 | 7.4E-17 | -1.95 | -0.96 | 1.7E-19 |
| Pyroxd1 | 1521 | -1.68 | -0.74 | 3.7E-25 | -1.55 | -0.63 | 1.6E-18 |
| Pyroxd2 | 193 | -1.33 | -0.42 | 1.5E-02 | -1.31 | -0.39 | 2.7E-02 |
| Qars | 9485 | 1.42 | 0.51 | 6.9E-06 | 1.34 | 0.43 | 1.9E-04 |
| Qdpr | 4362 | 1.55 | 0.64 | 4.2E-11 | 1.50 | 0.58 | 1.8E-09 |
| Qk | 28560 | -1.64 | -0.71 | 2.5E-12 | -1.70 | -0.77 | 3.4E-14 |
| Qpctl | 1773 | 1.64 | 0.71 | 1.6E-28 | 1.58 | 0.66 | 1.3E-24 |
| Qrich1 | 7228 | -1.22 | -0.29 | 1.2E-05 | -1.33 | -0.41 | 2.3E-10 |
| Qrsl1 | 916 | -2.30 | -1.20 | 8.3E-54 | -2.15 | -1.10 | 3.2E-45 |
| Qser1 | 10610 | -1.36 | -0.44 | 1.8E-05 | -1.33 | -0.41 | 8.5E-05 |
| Qsox1 | 3638 | 1.72 | 0.78 | 1.6E-19 | 1.74 | 0.80 | 1.8E-20 |
| Qsox2 | 2118 | 1.41 | 0.50 | 3.4E-09 | 1.33 | 0.41 | 1.7E-06 |
| R3hdm2 | 7116 | -1.65 | -0.72 | 3.2E-09 | -1.76 | -0.81 | 2.3E-11 |
| Rab11b | 7660 | -2.06 | -1.05 | 2.8E-39 | -2.14 | -1.10 | 2.2E-43 |
| Rab11fip1 | 1229 | 4.58 | 2.20 | 8.2E-26 | 4.36 | 2.12 | 3.4E-24 |
| Rab11fip2 | 3691 | -2.07 | -1.05 | 1.4E-21 | -2.06 | -1.04 | 4.2E-21 |
| Rab11fip5 | 3422 | 2.08 | 1.05 | 6.9E-20 | 2.09 | 1.06 | 3.7E-20 |
| Rab12 | 5690 | 1.35 | 0.43 | 6.6E-07 | 1.28 | 0.36 | 4.0E-05 |
| Rab15 | 1617 | -1.75 | -0.81 | 2.0E-06 | -1.69 | -0.76 | 9.5E-06 |
| Rab1b | 6869 | -1.52 | -0.61 | 7.0E-18 | -1.59 | -0.67 | 7.7E-22 |
| Rab20 | 106 | 6.47 | 2.69 | 1.2E-23 | 6.84 | 2.77 | 1.0E-24 |
| Rab21 | 5093 | 1.43 | 0.51 | 6.9E-07 | 1.31 | 0.39 | 2.1E-04 |
| Rab26 | 128 | 8.08 | 3.01 | 1.4E-67 | 5.99 | 2.58 | 4.1E-53 |
| Rab27b | 323 | -1.67 | -0.74 | 2.1E-03 | -1.58 | -0.66 | 7.3E-03 |
| Rab29 | 307 | -1.29 | -0.37 | 3.9E-03 | -1.33 | -0.41 | 1.3E-03 |
| Rab2b | 2907 | -1.24 | -0.31 | 5.1E-06 | -1.28 | -0.35 | 1.1E-07 |
| Rab32 | 377 | -1.39 | -0.48 | 2.0E-02 | -1.37 | -0.45 | 3.0E-02 |
| Rab33a | 707 | -1.67 | -0.74 | 2.4E-05 | -1.63 | -0.71 | 5.0E-05 |
| Rab34 | 1804 | -2.39 | -1.26 | 9.4E-51 | -2.48 | -1.31 | 5.4E-55 |
| Rab37 | 184 | 2.33 | 1.22 | 5.5E-09 | 2.47 | 1.30 | 5.1E-10 |
| Rab39 | 642 | 3.61 | 1.85 | 2.3E-66 | 3.29 | 1.72 | 9.2E-58 |
| Rab3d | 2350 | 2.68 | 1.42 | 2.9E-75 | 2.54 | 1.34 | 2.4E-67 |
| Rab3gap1 | 2905 | -1.47 | -0.56 | 4.9E-34 | -1.40 | -0.48 | 1.2E-25 |
| Rab3gap2 | 5920 | 1.24 | 0.31 | 1.5E-04 | 1.31 | 0.39 | 1.1E-06 |
| Rab3il1 | 2078 | 1.36 | 0.44 | 1.2E-02 | 1.32 | 0.41 | 2.2E-02 |
| Rab3ip | 3408 | 2.08 | 1.05 | 6.1E-46 | 1.96 | 0.97 | 6.2E-39 |
| Rab40b | 500 | 2.63 | 1.39 | 5.6E-15 | 2.94 | 1.56 | 2.4E-18 |
| Rab40c | 3116 | 1.63 | 0.70 | 2.4E-13 | 1.53 | 0.62 | 1.7E-10 |
| Rab4a | 1257 | 1.46 | 0.55 | 6.4E-16 | 1.38 | 0.47 | 8.2E-12 |
| Rab5if | 4432 | 1.64 | 0.71 | 2.8E-21 | 1.59 | 0.67 | 8.2E-19 |
| Rab6a | 18719 | 1.25 | 0.33 | 2.4E-03 | 1.20 | 0.26 | 1.8E-02 |
| Rab8a | 6226 | 1.33 | 0.41 | 6.5E-07 | 1.22 | 0.29 | 6.1E-04 |
| Rab8b | 7534 | 1.33 | 0.41 | 8.0E-11 | 1.29 | 0.37 | 1.0E-08 |
| Rab9 | 2389 | 1.27 | 0.35 | 2.2E-05 | 1.29 | 0.37 | 8.5E-06 |

| | | | | | | | |
|---|---|---|---|---|---|---|---|
| Rab9b | 861 | -1.48 | -0.57 | 1.2E-02 | -1.49 | -0.57 | 1.1E-02 |
| Rabac1 | 2232 | 1.30 | 0.38 | 9.0E-04 | 1.39 | 0.47 | 3.5E-05 |
| Rabep1 | 11994 | 1.35 | 0.43 | 3.8E-05 | 1.42 | 0.51 | 1.1E-06 |
| Rabep2 | 905 | -1.39 | -0.47 | 1.1E-07 | -1.50 | -0.58 | 4.3E-11 |
| Rabepk | 1136 | 1.66 | 0.73 | 9.7E-18 | 1.68 | 0.75 | 2.1E-18 |
| Rabgap1 | 11993 | 1.24 | 0.31 | 7.8E-06 | 1.27 | 0.35 | 6.7E-07 |
| Rabgef1 | 2176 | 1.32 | 0.40 | 2.1E-08 | 1.30 | 0.38 | 1.5E-07 |
| Rabggta | 1174 | -1.70 | -0.76 | 2.2E-19 | -1.78 | -0.83 | 1.3E-22 |
| Rabif | 2051 | -1.70 | -0.76 | 1.9E-35 | -1.72 | -0.78 | 3.7E-37 |
| Rabl2 | 2848 | 1.43 | 0.52 | 8.9E-13 | 1.33 | 0.41 | 2.8E-08 |
| Rabl3 | 1138 | -2.47 | -1.31 | 7.6E-19 | -2.78 | -1.47 | 1.3E-23 |
| Rabl6 | 6117 | -1.21 | -0.28 | 2.3E-07 | -1.27 | -0.35 | 6.1E-11 |
| Rad21 | 24544 | -1.30 | -0.38 | 9.2E-06 | -1.37 | -0.46 | 7.5E-08 |
| Rad23a | 2592 | -1.25 | -0.33 | 2.1E-03 | -1.33 | -0.42 | 6.8E-05 |
| Rad51c | 899 | -1.27 | -0.35 | 1.0E-02 | -1.43 | -0.51 | 8.5E-05 |
| Rad54l | 3943 | 1.44 | 0.53 | 1.0E-02 | 1.36 | 0.44 | 3.4E-02 |
| Rad9b | 923 | -1.31 | -0.39 | 1.0E-04 | -1.22 | -0.29 | 5.1E-03 |
| Radil | 654 | 1.45 | 0.53 | 2.3E-03 | 1.32 | 0.40 | 2.6E-02 |
| Raet1d | 244 | -1.51 | -0.59 | 2.3E-02 | -1.53 | -0.62 | 1.8E-02 |
| Raf1 | 11657 | -1.62 | -0.69 | 2.3E-24 | -1.60 | -0.68 | 1.8E-23 |
| Rai1 | 7722 | -1.57 | -0.65 | 2.5E-07 | -1.56 | -0.64 | 3.3E-07 |
| Rai14 | 9557 | -1.43 | -0.52 | 4.4E-04 | -1.35 | -0.43 | 4.2E-03 |
| Ralb | 3292 | 1.43 | 0.52 | 3.4E-08 | 1.49 | 0.57 | 1.0E-09 |
| Ralgapa2 | 4401 | 1.38 | 0.46 | 5.3E-04 | 1.45 | 0.53 | 5.3E-05 |
| Raly | 16070 | 1.36 | 0.45 | 9.1E-05 | 1.24 | 0.31 | 7.2E-03 |
| Ramp2 | 694 | -2.41 | -1.27 | 5.4E-48 | -2.40 | -1.26 | 2.5E-47 |
| Ranbp17 | 928 | -1.59 | -0.67 | 3.6E-07 | -1.48 | -0.57 | 1.9E-05 |
| Ranbp3 | 6406 | -1.16 | -0.21 | 1.3E-02 | -1.27 | -0.34 | 2.8E-05 |
| Rangap1 | 18532 | 1.47 | 0.56 | 2.1E-06 | 1.34 | 0.42 | 4.4E-04 |
| Rangrf | 981 | 1.83 | 0.87 | 5.0E-09 | 1.62 | 0.69 | 4.5E-06 |
| Rap2a | 5118 | 1.39 | 0.47 | 1.4E-09 | 1.32 | 0.40 | 5.1E-07 |
| Rap2c | 7466 | -1.25 | -0.33 | 5.2E-06 | -1.24 | -0.31 | 1.8E-05 |
| Rapgef1 | 10436 | -1.42 | -0.50 | 2.9E-06 | -1.50 | -0.59 | 4.5E-08 |
| Rapgef5 | 5201 | -1.58 | -0.66 | 1.3E-03 | -1.44 | -0.53 | 1.1E-02 |
| Rapgefl1 | 1691 | 2.22 | 1.15 | 2.2E-13 | 2.22 | 1.15 | 1.9E-13 |
| Rarb | 1030 | -2.09 | -1.07 | 2.6E-05 | -2.27 | -1.18 | 3.0E-06 |
| Rarg | 1854 | -2.08 | -1.05 | 4.1E-04 | -2.10 | -1.07 | 3.5E-04 |
| Rars | 8480 | 1.33 | 0.41 | 7.7E-04 | 1.32 | 0.40 | 1.1E-03 |
| Rasa2 | 2217 | -1.84 | -0.88 | 3.1E-20 | -1.72 | -0.78 | 3.9E-16 |
| Rasa3 | 4266 | 1.67 | 0.74 | 2.3E-36 | 1.65 | 0.72 | 1.1E-34 |
| Rasa4 | 1701 | 2.43 | 1.28 | 1.7E-23 | 2.39 | 1.26 | 1.5E-22 |
| Rasd2 | 225 | 6.17 | 2.62 | 6.5E-09 | 6.48 | 2.70 | 2.6E-09 |
| Rasgef1c | 131 | -1.99 | -0.99 | 9.5E-03 | -1.77 | -0.82 | 3.6E-02 |
| Rasgrf1 | 381 | 3.65 | 1.87 | 1.8E-07 | 4.38 | 2.13 | 2.2E-09 |
| Rasgrp3 | 853 | -1.68 | -0.75 | 7.6E-15 | -1.58 | -0.66 | 9.9E-12 |
| Rasip1 | 598 | -2.20 | -1.14 | 8.0E-19 | -2.21 | -1.14 | 6.2E-19 |
| Rasl10b | 3257 | 1.71 | 0.77 | 8.4E-06 | 1.66 | 0.73 | 2.5E-05 |

| | | | | | | | |
|---|---|---|---|---|---|---|---|
| Rasl11a | 248 | -1.31 | -0.39 | 2.9E-02 | -1.44 | -0.53 | 2.8E-03 |
| Rasl12 | 406 | -1.26 | -0.33 | 3.1E-04 | -1.15 | -0.20 | 3.5E-02 |
| Rassf2 | 7103 | 1.57 | 0.66 | 1.4E-13 | 1.44 | 0.52 | 4.6E-09 |
| Rassf3 | 1707 | 1.53 | 0.61 | 3.3E-03 | 1.59 | 0.67 | 1.3E-03 |
| Rassf4 | 6296 | 2.38 | 1.25 | 5.6E-14 | 2.84 | 1.51 | 7.1E-20 |
| Rassf5 | 168 | 1.54 | 0.62 | 2.2E-02 | 1.66 | 0.74 | 6.4E-03 |
| Rassf7 | 1195 | 2.10 | 1.07 | 7.9E-19 | 1.96 | 0.97 | 1.4E-15 |
| Rassf8 | 2738 | 1.47 | 0.55 | 4.1E-08 | 1.49 | 0.58 | 9.4E-09 |
| Rassf9 | 175 | -2.44 | -1.29 | 2.7E-09 | -2.25 | -1.17 | 7.5E-08 |
| Raver2 | 2460 | 1.54 | 0.62 | 4.5E-07 | 1.63 | 0.70 | 8.9E-09 |
| Rb1 | 8425 | 1.44 | 0.53 | 4.1E-06 | 1.56 | 0.64 | 1.6E-08 |
| Rbbp8 | 3246 | -1.31 | -0.39 | 1.1E-02 | -1.33 | -0.41 | 7.2E-03 |
| Rbbp9 | 3562 | -1.56 | -0.64 | 5.5E-20 | -1.59 | -0.67 | 1.1E-21 |
| Rbfa | 1318 | 1.87 | 0.90 | 2.2E-22 | 1.69 | 0.76 | 3.3E-16 |
| Rbfox2 | 29853 | -1.30 | -0.38 | 1.0E-02 | -1.27 | -0.35 | 1.9E-02 |
| Rbl1 | 4163 | -1.38 | -0.46 | 8.9E-04 | -1.42 | -0.51 | 2.6E-04 |
| Rbl2 | 3008 | 1.45 | 0.54 | 8.8E-16 | 1.47 | 0.56 | 8.1E-17 |
| Rbm10 | 7185 | -1.51 | -0.60 | 1.5E-12 | -1.65 | -0.73 | 4.9E-18 |
| Rbm11 | 391 | 4.61 | 2.20 | 3.3E-54 | 4.74 | 2.25 | 3.2E-56 |
| Rbm12 | 3524 | -1.73 | -0.79 | 7.4E-15 | -1.85 | -0.88 | 2.0E-18 |
| Rbm19 | 1975 | -1.70 | -0.76 | 5.1E-12 | -1.89 | -0.92 | 6.2E-17 |
| Rbm20 | 667 | -1.83 | -0.88 | 6.6E-09 | -1.82 | -0.86 | 1.1E-08 |
| Rbm22 | 4983 | -1.33 | -0.42 | 6.2E-05 | -1.47 | -0.56 | 5.1E-08 |
| Rbm25 | 24288 | -1.46 | -0.54 | 1.8E-08 | -1.46 | -0.54 | 1.9E-08 |
| Rbm28 | 10895 | 1.36 | 0.44 | 1.1E-20 | 1.39 | 0.47 | 3.3E-23 |
| Rbm3 | 18751 | -1.37 | -0.46 | 4.9E-04 | -1.49 | -0.57 | 1.1E-05 |
| Rbm33 | 11486 | -1.50 | -0.59 | 3.8E-08 | -1.41 | -0.50 | 3.5E-06 |
| Rbm34 | 3179 | 1.35 | 0.44 | 3.3E-07 | 1.27 | 0.35 | 6.5E-05 |
| Rbm38 | 2168 | 1.82 | 0.86 | 5.9E-06 | 1.59 | 0.67 | 6.0E-04 |
| Rbm39 | 32934 | -1.94 | -0.95 | 4.7E-46 | -1.83 | -0.87 | 1.8E-38 |
| Rbm43 | 384 | -1.49 | -0.57 | 5.0E-05 | -1.49 | -0.57 | 5.5E-05 |
| Rbm46 | 346 | 1.48 | 0.56 | 8.2E-03 | 1.69 | 0.76 | 3.0E-04 |
| Rbm47 | 199 | -2.66 | -1.41 | 2.1E-12 | -2.81 | -1.49 | 1.0E-13 |
| Rbm4b | 5862 | -2.10 | -1.07 | 2.0E-22 | -2.03 | -1.02 | 2.6E-20 |
| Rbm5 | 23084 | -1.69 | -0.75 | 7.4E-24 | -1.61 | -0.69 | 6.4E-20 |
| Rbms1 | 4901 | -1.84 | -0.88 | 1.6E-05 | -1.70 | -0.76 | 2.2E-04 |
| Rbms2 | 3710 | 1.40 | 0.49 | 1.4E-04 | 1.42 | 0.50 | 8.1E-05 |
| Rbmx | 23217 | -1.48 | -0.57 | 1.3E-10 | -1.51 | -0.60 | 1.3E-11 |
| Rbpj | 11332 | -1.25 | -0.32 | 4.8E-06 | -1.29 | -0.37 | 1.5E-07 |
| Rbpms2 | 573 | 1.66 | 0.73 | 1.7E-03 | 1.43 | 0.52 | 3.1E-02 |
| Rbsn | 4618 | -1.35 | -0.43 | 1.0E-22 | -1.33 | -0.42 | 8.4E-21 |
| Rbx1 | 7621 | -1.51 | -0.60 | 1.4E-15 | -1.55 | -0.63 | 3.6E-17 |
| Rcan1 | 3802 | 1.67 | 0.74 | 4.1E-27 | 1.59 | 0.67 | 4.6E-22 |
| Rcbtb2 | 5810 | -3.41 | -1.77 | 4.0E-51 | -3.46 | -1.79 | 2.5E-52 |
| Rcc1l | 1129 | -1.21 | -0.28 | 2.7E-02 | -1.34 | -0.42 | 6.3E-04 |
| Rcc2 | 17690 | -1.39 | -0.47 | 5.7E-03 | -1.57 | -0.65 | 1.0E-04 |
| Rchy1 | 3161 | -1.30 | -0.38 | 2.9E-12 | -1.34 | -0.42 | 6.3E-15 |

| | | | | | | | |
|---|---|---|---|---|---|---|---|
| Rcl1 | 1965 | 2.16 | 1.11 | 3.3E-21 | 2.11 | 1.07 | 6.3E-20 |
| Rcn2 | 10029 | -1.39 | -0.48 | 2.0E-07 | -1.48 | -0.57 | 6.5E-10 |
| Rcor3 | 680 | -1.75 | -0.81 | 1.2E-08 | -1.52 | -0.60 | 3.3E-05 |
| Rcsd1 | 870 | -1.67 | -0.74 | 1.0E-06 | -1.67 | -0.74 | 9.3E-07 |
| Rdh11 | 3707 | 1.44 | 0.52 | 1.3E-13 | 1.45 | 0.53 | 3.5E-14 |
| Rdh12 | 110 | 1.33 | 0.42 | 1.3E-02 | 1.76 | 0.81 | 3.8E-07 |
| Rdh13 | 2792 | 1.34 | 0.42 | 6.4E-14 | 1.28 | 0.36 | 3.4E-10 |
| Rdh14 | 769 | -1.89 | -0.92 | 2.4E-20 | -1.92 | -0.94 | 2.2E-21 |
| Rdm1 | 1015 | -1.26 | -0.33 | 5.5E-03 | -1.34 | -0.42 | 3.5E-04 |
| Recql | 4098 | 1.34 | 0.43 | 2.5E-08 | 1.30 | 0.38 | 6.9E-07 |
| Reep3 | 6903 | -1.26 | -0.33 | 5.0E-10 | -1.26 | -0.33 | 5.5E-10 |
| Reep4 | 1519 | 1.61 | 0.69 | 2.2E-05 | 1.46 | 0.55 | 9.5E-04 |
| Reep6 | 623 | 6.74 | 2.75 | 3.5E-44 | 6.43 | 2.68 | 6.0E-42 |
| Rel | 493 | 2.35 | 1.23 | 7.1E-19 | 2.56 | 1.36 | 2.1E-22 |
| Rela | 1932 | -1.48 | -0.57 | 8.0E-09 | -1.63 | -0.70 | 4.3E-13 |
| Relch | 6621 | 1.42 | 0.50 | 4.6E-06 | 1.55 | 0.63 | 5.1E-09 |
| Rell1 | 2397 | 2.36 | 1.24 | 7.2E-47 | 2.29 | 1.20 | 7.3E-44 |
| Rell2 | 1180 | 1.72 | 0.79 | 1.9E-03 | 1.74 | 0.80 | 1.5E-03 |
| Relt | 367 | 2.54 | 1.34 | 2.0E-46 | 2.58 | 1.37 | 2.8E-47 |
| Resf1 | 3872 | -1.48 | -0.57 | 2.0E-21 | -1.44 | -0.53 | 2.4E-18 |
| Retreg3 | 7186 | 1.31 | 0.39 | 8.9E-17 | 1.25 | 0.32 | 1.0E-11 |
| Retsat | 307 | -1.42 | -0.51 | 1.1E-03 | -1.40 | -0.48 | 2.0E-03 |
| Rex1bd | 1297 | -1.27 | -0.34 | 1.4E-04 | -1.36 | -0.45 | 4.2E-07 |
| Rfc5 | 2500 | -1.38 | -0.46 | 1.0E-03 | -1.56 | -0.64 | 3.0E-06 |
| Rffl | 1871 | 1.99 | 0.99 | 1.5E-17 | 1.91 | 0.94 | 1.1E-15 |
| Rfk | 4664 | 1.52 | 0.60 | 4.2E-12 | 1.42 | 0.51 | 6.4E-09 |
| Rft1 | 805 | -1.94 | -0.96 | 6.4E-31 | -1.97 | -0.98 | 3.6E-32 |
| Rftn2 | 1295 | -1.42 | -0.50 | 1.5E-06 | -1.33 | -0.41 | 1.1E-04 |
| Rfx1 | 2234 | -1.48 | -0.56 | 7.1E-06 | -1.53 | -0.61 | 8.7E-07 |
| Rfx2 | 1128 | 2.57 | 1.36 | 5.9E-26 | 2.32 | 1.21 | 9.6E-21 |
| Rfx3 | 5786 | -1.47 | -0.56 | 5.0E-08 | -1.45 | -0.53 | 2.2E-07 |
| Rfx4 | 2512 | -1.73 | -0.79 | 7.1E-04 | -1.78 | -0.83 | 3.4E-04 |
| Rfxap | 1885 | 1.46 | 0.54 | 2.2E-16 | 1.41 | 0.49 | 8.7E-14 |
| Rgcc | 529 | -1.42 | -0.50 | 3.9E-03 | -1.35 | -0.44 | 1.3E-02 |
| Rgl1 | 3228 | 1.52 | 0.60 | 6.8E-18 | 1.44 | 0.53 | 5.9E-14 |
| Rgl3 | 463 | 4.98 | 2.32 | 1.2E-41 | 4.13 | 2.05 | 6.5E-33 |
| Rgma | 6011 | -1.29 | -0.37 | 3.3E-02 | -1.36 | -0.45 | 8.9E-03 |
| Rgs10 | 538 | 1.84 | 0.88 | 1.0E-06 | 1.74 | 0.80 | 9.7E-06 |
| Rgs12 | 7398 | -1.51 | -0.60 | 2.5E-09 | -1.68 | -0.75 | 5.5E-14 |
| Rgs13 | 112 | -1.71 | -0.78 | 2.0E-02 | -2.13 | -1.09 | 7.2E-04 |
| Rgs17 | 1821 | 1.36 | 0.45 | 4.5E-02 | 1.55 | 0.63 | 3.9E-03 |
| Rgs19 | 1294 | -1.42 | -0.50 | 7.4E-12 | -1.51 | -0.59 | 4.1E-16 |
| Rgs3 | 2881 | -1.74 | -0.80 | 1.1E-19 | -1.75 | -0.80 | 4.5E-20 |
| Rgs9bp | 472 | 8.35 | 3.06 | 4.3E-75 | 6.94 | 2.80 | 4.6E-63 |
| Rhbdd3 | 1075 | -1.59 | -0.67 | 9.8E-14 | -1.67 | -0.74 | 2.0E-16 |
| Rhbdf2 | 535 | 3.88 | 1.96 | 2.4E-17 | 3.98 | 1.99 | 7.1E-18 |
| Rhbdl1 | 202 | 2.57 | 1.36 | 2.7E-11 | 2.56 | 1.36 | 4.0E-11 |

| | | | | | | | |
|---|---|---|---|---|---|---|---|
| Rhebl1 | 1255 | 1.58 | 0.66 | 2.5E-14 | 1.51 | 0.60 | 4.2E-12 |
| Rhobtb1 | 1395 | -3.17 | -1.66 | 3.8E-53 | -3.37 | -1.75 | 1.2E-58 |
| Rhobtb3 | 9159 | -1.40 | -0.49 | 2.7E-05 | -1.44 | -0.53 | 4.8E-06 |
| Rhof | 450 | 2.70 | 1.44 | 1.9E-30 | 2.77 | 1.47 | 1.1E-31 |
| Rhoj | 1113 | -2.65 | -1.41 | 9.4E-18 | -2.44 | -1.28 | 5.7E-15 |
| Rhot2 | 1656 | 1.44 | 0.53 | 6.0E-05 | 1.27 | 0.35 | 1.1E-02 |
| Rhou | 2734 | -1.48 | -0.56 | 1.1E-07 | -1.63 | -0.70 | 2.5E-11 |
| Rhpn1 | 1963 | 2.82 | 1.49 | 1.1E-36 | 2.53 | 1.34 | 9.8E-30 |
| Rhpn2 | 911 | 6.67 | 2.74 | 5.4E-70 | 7.27 | 2.86 | 1.1E-75 |
| Ric1 | 8113 | 1.48 | 0.56 | 4.6E-07 | 1.67 | 0.74 | 2.3E-11 |
| Riiad1 | 398 | 1.92 | 0.94 | 1.0E-11 | 1.94 | 0.95 | 8.2E-12 |
| Rilpl1 | 1682 | 2.15 | 1.10 | 5.9E-24 | 2.05 | 1.03 | 4.1E-21 |
| Rilpl2 | 1121 | 2.40 | 1.26 | 1.8E-34 | 2.27 | 1.18 | 2.8E-30 |
| Rimbp3 | 433 | 2.31 | 1.21 | 1.9E-13 | 2.35 | 1.23 | 7.1E-14 |
| Rims2 | 3497 | -1.38 | -0.47 | 1.1E-02 | -1.48 | -0.57 | 1.7E-03 |
| Rims3 | 4156 | 1.95 | 0.97 | 2.0E-08 | 1.98 | 0.98 | 1.2E-08 |
| Rims4 | 1715 | 1.77 | 0.83 | 1.1E-03 | 1.61 | 0.69 | 7.8E-03 |
| Rin1 | 390 | 1.88 | 0.91 | 4.5E-13 | 1.64 | 0.71 | 2.2E-08 |
| Rin2 | 2072 | -1.58 | -0.66 | 1.1E-09 | -1.61 | -0.69 | 2.6E-10 |
| Rin3 | 463 | -2.83 | -1.50 | 2.1E-13 | -3.14 | -1.65 | 5.9E-16 |
| Ring1 | 2355 | -1.90 | -0.93 | 2.7E-07 | -2.18 | -1.13 | 2.9E-10 |
| Riok2 | 4907 | 1.42 | 0.51 | 2.7E-08 | 1.41 | 0.50 | 6.6E-08 |
| Riox1 | 1052 | 1.28 | 0.36 | 1.9E-04 | 1.26 | 0.33 | 6.8E-04 |
| Ripk1 | 1087 | -2.06 | -1.04 | 5.7E-34 | -1.98 | -0.99 | 2.6E-30 |
| Ripk2 | 1145 | 2.81 | 1.49 | 2.9E-85 | 2.85 | 1.51 | 6.4E-87 |
| Ripk3 | 526 | 1.96 | 0.97 | 1.5E-02 | 1.91 | 0.93 | 2.0E-02 |
| Ripor1 | 3295 | 1.36 | 0.44 | 2.1E-03 | 1.29 | 0.36 | 1.3E-02 |
| Ripor3 | 171 | -3.13 | -1.65 | 1.0E-05 | -3.43 | -1.78 | 1.9E-06 |
| Rit1 | 2411 | 1.63 | 0.71 | 3.8E-30 | 1.60 | 0.68 | 4.9E-28 |
| Rita1 | 1249 | 2.58 | 1.37 | 9.1E-42 | 2.63 | 1.40 | 2.3E-43 |
| Rmc1 | 2865 | 2.07 | 1.05 | 2.3E-31 | 2.20 | 1.14 | 1.7E-36 |
| Rmdn1 | 682 | -1.34 | -0.42 | 3.3E-04 | -1.23 | -0.30 | 1.4E-02 |
| Rmdn2 | 218 | -6.36 | -2.67 | 3.3E-77 | -7.53 | -2.91 | 1.8E-87 |
| Rmdn3 | 2672 | 1.32 | 0.40 | 1.4E-11 | 1.27 | 0.35 | 4.4E-09 |
| Rmi1 | 1833 | -1.41 | -0.49 | 5.7E-15 | -1.48 | -0.57 | 1.4E-19 |
| Rmnd5a | 10131 | 1.53 | 0.61 | 3.3E-27 | 1.52 | 0.61 | 6.3E-27 |
| Rmnd5b | 2194 | 1.35 | 0.44 | 3.1E-10 | 1.32 | 0.40 | 7.3E-09 |
| Rnaseh2a | 2685 | 1.85 | 0.89 | 1.7E-11 | 1.67 | 0.74 | 2.4E-08 |
| Rnaseh2b | 1694 | -1.96 | -0.97 | 9.3E-12 | -2.03 | -1.02 | 6.7E-13 |
| Rnaseh2c | 2154 | 1.38 | 0.46 | 4.4E-03 | 1.28 | 0.35 | 3.6E-02 |
| Rnasel | 5478 | -2.82 | -1.49 | 3.1E-27 | -2.68 | -1.42 | 8.1E-25 |
| Rnd1 | 1048 | 3.11 | 1.64 | 8.5E-31 | 2.84 | 1.51 | 3.3E-26 |
| Rnd3 | 17191 | -1.47 | -0.56 | 1.3E-07 | -1.44 | -0.53 | 6.9E-07 |
| Rnf10 | 22283 | 1.25 | 0.33 | 3.4E-04 | 1.18 | 0.24 | 1.2E-02 |
| Rnf113a2 | 1062 | 1.33 | 0.41 | 7.3E-06 | 1.27 | 0.35 | 1.5E-04 |
| Rnf115 | 5353 | 1.41 | 0.50 | 2.7E-10 | 1.35 | 0.43 | 5.9E-08 |
| Rnf122 | 3323 | -1.22 | -0.29 | 5.8E-05 | -1.32 | -0.40 | 1.3E-08 |

| | | | | | | | |
|---|---|---|---|---|---|---|---|
| Rnf128 | 202 | 1.59 | 0.67 | 1.6E-02 | 1.64 | 0.72 | 1.0E-02 |
| Rnf13 | 3567 | 1.32 | 0.40 | 6.4E-06 | 1.40 | 0.49 | 4.1E-08 |
| Rnf135 | 168 | -1.99 | -0.99 | 4.9E-08 | -1.97 | -0.98 | 8.1E-08 |
| Rnf144a | 5958 | -1.52 | -0.61 | 2.1E-15 | -1.55 | -0.63 | 1.3E-16 |
| Rnf150 | 2619 | -1.92 | -0.94 | 5.1E-17 | -1.74 | -0.80 | 2.1E-12 |
| Rnf166 | 1825 | 1.38 | 0.46 | 6.3E-11 | 1.33 | 0.41 | 1.1E-08 |
| Rnf169 | 3609 | -1.24 | -0.31 | 1.5E-04 | -1.25 | -0.33 | 7.5E-05 |
| Rnf180 | 865 | -2.14 | -1.10 | 3.9E-12 | -1.97 | -0.98 | 8.3E-10 |
| Rnf182 | 2274 | 1.38 | 0.46 | 5.8E-03 | 1.36 | 0.45 | 8.1E-03 |
| Rnf185 | 4524 | 1.62 | 0.70 | 4.4E-25 | 1.57 | 0.65 | 8.8E-22 |
| Rnf187 | 17783 | 1.31 | 0.39 | 1.1E-06 | 1.25 | 0.32 | 1.0E-04 |
| Rnf19b | 2770 | 1.38 | 0.47 | 7.2E-05 | 1.32 | 0.40 | 7.2E-04 |
| Rnf207 | 148 | -1.31 | -0.39 | 2.2E-02 | -1.29 | -0.36 | 3.5E-02 |
| Rnf215 | 2115 | -1.50 | -0.58 | 8.3E-12 | -1.54 | -0.62 | 3.1E-13 |
| Rnf217 | 1654 | -1.53 | -0.61 | 2.5E-04 | -1.37 | -0.45 | 7.7E-03 |
| Rnf223 | 282 | 2.78 | 1.47 | 8.3E-22 | 2.61 | 1.39 | 1.8E-19 |
| Rnf225 | 192 | 1.74 | 0.80 | 3.5E-10 | 1.60 | 0.68 | 1.7E-07 |
| Rnf227 | 4542 | 1.44 | 0.53 | 1.4E-02 | 1.54 | 0.62 | 3.2E-03 |
| Rnf31 | 2259 | 1.26 | 0.33 | 1.1E-03 | 1.25 | 0.32 | 1.7E-03 |
| Rnf34 | 4614 | 1.49 | 0.57 | 1.2E-13 | 1.50 | 0.59 | 3.6E-14 |
| Rnft1 | 1790 | 1.25 | 0.32 | 3.5E-03 | 1.27 | 0.35 | 1.6E-03 |
| Rngtt | 5063 | 1.79 | 0.84 | 2.0E-29 | 1.87 | 0.90 | 5.4E-34 |
| Rnh1 | 2002 | -1.90 | -0.92 | 2.6E-12 | -1.98 | -0.98 | 7.1E-14 |
| Rnmt | 8419 | 1.40 | 0.49 | 7.8E-10 | 1.47 | 0.55 | 2.0E-12 |
| Rnpc3 | 2700 | -1.59 | -0.67 | 3.2E-09 | -1.37 | -0.45 | 8.4E-05 |
| Rnpep | 3605 | -1.32 | -0.40 | 4.7E-09 | -1.42 | -0.50 | 1.5E-13 |
| Rnpepl1 | 2631 | -1.26 | -0.34 | 3.0E-03 | -1.36 | -0.44 | 7.3E-05 |
| Ro60 | 20892 | 1.83 | 0.87 | 5.0E-11 | 2.10 | 1.07 | 2.7E-16 |
| Robo4 | 695 | -2.11 | -1.08 | 1.5E-17 | -2.21 | -1.15 | 1.1E-19 |
| Rogdi | 2534 | 1.34 | 0.43 | 1.7E-08 | 1.29 | 0.37 | 1.2E-06 |
| Rom1 | 105 | -1.71 | -0.78 | 2.4E-06 | -1.67 | -0.74 | 8.5E-06 |
| Ror2 | 1379 | -1.42 | -0.51 | 2.0E-02 | -1.60 | -0.68 | 1.5E-03 |
| Rorc | 321 | 2.31 | 1.21 | 2.2E-08 | 2.39 | 1.25 | 6.1E-09 |
| Rp2 | 3024 | 1.59 | 0.67 | 2.8E-12 | 1.56 | 0.64 | 1.5E-11 |
| Rpa1 | 12109 | 1.92 | 0.94 | 3.3E-17 | 1.78 | 0.83 | 1.4E-13 |
| Rpa2 | 4235 | 1.53 | 0.61 | 7.9E-05 | 1.34 | 0.42 | 8.6E-03 |
| Rpa3 | 2669 | 1.30 | 0.38 | 4.2E-03 | 1.26 | 0.33 | 1.5E-02 |
| Rpain | 1629 | 1.37 | 0.45 | 5.6E-09 | 1.33 | 0.41 | 1.7E-07 |
| Rpap1 | 2974 | -1.93 | -0.95 | 1.2E-16 | -2.07 | -1.05 | 2.7E-20 |
| Rpap2 | 1284 | -1.25 | -0.32 | 1.2E-04 | -1.26 | -0.33 | 8.5E-05 |
| Rpf1 | 2309 | -1.45 | -0.54 | 1.5E-07 | -1.50 | -0.59 | 6.7E-09 |
| Rpgrip1l | 3644 | 1.47 | 0.55 | 2.0E-05 | 1.68 | 0.75 | 4.8E-09 |
| Rpn1 | 12140 | 1.23 | 0.29 | 1.3E-03 | 1.27 | 0.34 | 1.8E-04 |
| Rpp21 | 529 | -1.64 | -0.71 | 8.3E-18 | -1.79 | -0.84 | 1.0E-23 |
| Rpp38 | 340 | -1.69 | -0.76 | 2.7E-10 | -1.70 | -0.77 | 1.9E-10 |
| Rpp40 | 867 | 2.05 | 1.03 | 7.0E-15 | 1.95 | 0.96 | 4.4E-13 |
| Rprd1a | 7927 | 1.50 | 0.58 | 1.9E-17 | 1.51 | 0.60 | 2.9E-18 |

| | | | | | | | |
|---|---|---|---|---|---|---|---|
| Rprd2 | 4254 | -2.07 | -1.05 | 4.0E-20 | -2.15 | -1.11 | 3.3E-22 |
| Rprm | 2788 | -1.47 | -0.56 | 7.8E-04 | -1.37 | -0.45 | 7.6E-03 |
| Rps19bp1 | 1322 | 1.25 | 0.32 | 4.3E-05 | 1.19 | 0.25 | 1.3E-03 |
| Rps6ka1 | 2855 | 2.84 | 1.51 | 4.1E-37 | 2.60 | 1.38 | 3.2E-31 |
| Rps6ka2 | 5307 | 1.80 | 0.85 | 8.1E-20 | 1.80 | 0.85 | 8.6E-20 |
| Rps6ka3 | 11427 | 1.21 | 0.27 | 3.3E-03 | 1.29 | 0.37 | 3.3E-05 |
| Rps6ka5 | 2355 | 1.53 | 0.61 | 4.5E-05 | 1.63 | 0.71 | 2.1E-06 |
| Rpusd3 | 444 | -1.21 | -0.28 | 3.8E-02 | -1.46 | -0.54 | 2.2E-05 |
| Rragb | 2829 | 1.56 | 0.64 | 9.6E-10 | 1.54 | 0.62 | 3.0E-09 |
| Rras | 533 | 1.63 | 0.71 | 3.5E-05 | 1.45 | 0.53 | 2.3E-03 |
| Rras2 | 3016 | 1.79 | 0.84 | 3.7E-27 | 1.76 | 0.82 | 5.1E-26 |
| Rrm2 | 12342 | -1.61 | -0.69 | 4.7E-03 | -1.96 | -0.97 | 4.6E-05 |
| Rrm2b | 1684 | 1.51 | 0.59 | 2.6E-25 | 1.51 | 0.59 | 1.9E-25 |
| Rrp1 | 15728 | 1.30 | 0.38 | 8.1E-08 | 1.21 | 0.27 | 1.4E-04 |
| Rrp12 | 3528 | 1.64 | 0.71 | 2.0E-12 | 1.54 | 0.63 | 7.9E-10 |
| Rrp9 | 2358 | 2.03 | 1.02 | 2.3E-16 | 1.79 | 0.84 | 1.9E-11 |
| Rsbn1 | 8220 | 1.21 | 0.28 | 8.6E-03 | 1.30 | 0.38 | 2.0E-04 |
| Rsf1 | 8543 | -1.42 | -0.50 | 1.3E-06 | -1.27 | -0.34 | 1.2E-03 |
| Rskr | 107 | 2.22 | 1.15 | 2.3E-17 | 2.39 | 1.26 | 3.6E-20 |
| Rsph1 | 173 | 1.43 | 0.52 | 3.7E-03 | 1.53 | 0.61 | 5.0E-04 |
| Rsph10b | 544 | 2.03 | 1.02 | 8.3E-08 | 2.64 | 1.40 | 9.2E-14 |
| Rsph3b | 721 | 1.26 | 0.33 | 4.2E-07 | 1.12 | 0.17 | 1.5E-02 |
| Rsph4a | 231 | 1.58 | 0.66 | 1.1E-02 | 1.80 | 0.85 | 1.0E-03 |
| Rsph9 | 558 | 1.40 | 0.49 | 1.0E-05 | 1.50 | 0.58 | 9.6E-08 |
| Rspo2 | 1294 | -1.65 | -0.72 | 3.4E-03 | -2.12 | -1.09 | 6.2E-06 |
| Rspo3 | 1820 | -1.64 | -0.71 | 1.5E-04 | -2.02 | -1.01 | 4.5E-08 |
| Rsrc2 | 12072 | -1.68 | -0.75 | 9.5E-24 | -1.62 | -0.70 | 8.0E-21 |
| Rtbdn | 164 | 1.76 | 0.81 | 1.6E-03 | 1.86 | 0.90 | 4.9E-04 |
| Rtca | 4003 | 1.31 | 0.39 | 4.8E-07 | 1.24 | 0.31 | 9.7E-05 |
| Rtel1 | 4172 | -1.34 | -0.43 | 2.1E-07 | -1.39 | -0.48 | 7.2E-09 |
| Rtf1 | 12843 | 1.35 | 0.44 | 8.2E-13 | 1.27 | 0.34 | 2.3E-08 |
| Rtf2 | 5214 | 1.28 | 0.36 | 1.0E-06 | 1.20 | 0.26 | 5.6E-04 |
| Rtkn | 1660 | -2.02 | -1.01 | 1.7E-08 | -2.06 | -1.04 | 5.7E-09 |
| Rtl3 | 706 | -5.16 | -2.37 | 8.6E-06 | -4.48 | -2.16 | 5.5E-05 |
| Rtl6 | 4845 | -1.49 | -0.57 | 1.3E-05 | -1.58 | -0.66 | 4.2E-07 |
| Rtn4 | 30259 | -1.65 | -0.73 | 1.7E-12 | -1.73 | -0.79 | 1.0E-14 |
| Rtn4rl2 | 469 | 2.04 | 1.03 | 4.0E-09 | 2.01 | 1.01 | 7.6E-09 |
| Rtraf | 11323 | 1.29 | 0.36 | 1.3E-04 | 1.30 | 0.37 | 7.9E-05 |
| Rubcn | 5545 | 1.54 | 0.62 | 2.6E-12 | 1.56 | 0.64 | 5.0E-13 |
| Rufy1 | 1883 | -1.35 | -0.43 | 2.1E-09 | -1.32 | -0.40 | 1.8E-08 |
| Rundc1 | 2598 | 1.54 | 0.62 | 2.8E-22 | 1.46 | 0.55 | 2.5E-17 |
| Runx1 | 381 | -3.19 | -1.67 | 1.6E-04 | -2.98 | -1.57 | 4.3E-04 |
| Runx1t1 | 7184 | -1.62 | -0.70 | 4.1E-03 | -1.61 | -0.69 | 4.8E-03 |
| Runx2 | 782 | -3.89 | -1.96 | 1.5E-05 | -3.99 | -2.00 | 1.1E-05 |
| Rusc2 | 5884 | -1.98 | -0.99 | 4.9E-05 | -2.11 | -1.08 | 8.5E-06 |
| Rwdd2a | 435 | 7.14 | 2.84 | 3.5E-147 | 6.82 | 2.77 | 5.2E-143 |
| Rwdd2b | 467 | -3.44 | -1.78 | 7.3E-118 | -3.69 | -1.89 | 2.2E-128 |

| | | | | | | | |
|---|---|---|---|---|---|---|---|
| Rwdd3 | 521 | -1.40 | -0.48 | 5.4E-03 | -1.33 | -0.41 | 2.0E-02 |
| Rwdd4a | 5137 | 1.92 | 0.94 | 5.5E-29 | 1.87 | 0.90 | 6.3E-27 |
| Rxfp3 | 166 | -3.05 | -1.61 | 1.0E-10 | -3.54 | -1.82 | 3.1E-13 |
| Rxrb | 4358 | -1.27 | -0.35 | 5.6E-06 | -1.34 | -0.42 | 3.0E-08 |
| Rxylt1 | 2032 | 1.49 | 0.57 | 1.7E-13 | 1.51 | 0.59 | 2.6E-14 |
| S100a10 | 2920 | -1.48 | -0.57 | 2.5E-02 | -1.49 | -0.57 | 2.4E-02 |
| S100a13 | 157 | -1.30 | -0.38 | 2.4E-02 | -1.27 | -0.34 | 4.5E-02 |
| S100a16 | 596 | -2.17 | -1.12 | 3.0E-18 | -2.26 | -1.18 | 4.2E-20 |
| S100a6 | 439 | -1.29 | -0.37 | 6.2E-03 | -1.27 | -0.34 | 1.3E-02 |
| S1pr1 | 1963 | -1.25 | -0.32 | 2.4E-06 | -1.38 | -0.46 | 4.0E-12 |
| Sacm1l | 7648 | 1.55 | 0.64 | 7.9E-19 | 1.63 | 0.71 | 6.1E-23 |
| Safb | 14014 | -1.45 | -0.53 | 1.4E-04 | -1.58 | -0.66 | 2.2E-06 |
| Sag | 181 | 4.19 | 2.07 | 6.6E-33 | 4.68 | 2.23 | 8.3E-37 |
| Sall1 | 4174 | -1.84 | -0.88 | 1.4E-04 | -1.84 | -0.88 | 1.5E-04 |
| Sall2 | 8927 | -1.72 | -0.78 | 1.6E-12 | -1.92 | -0.94 | 1.6E-17 |
| Sall3 | 1797 | -1.48 | -0.56 | 4.1E-03 | -1.77 | -0.82 | 1.8E-05 |
| Sall4 | 546 | 2.37 | 1.24 | 8.3E-04 | 1.83 | 0.87 | 2.4E-02 |
| Samd1 | 2620 | -1.40 | -0.49 | 5.6E-05 | -1.55 | -0.63 | 1.5E-07 |
| Samd10 | 2579 | -2.14 | -1.09 | 1.5E-10 | -2.11 | -1.07 | 3.6E-10 |
| Samd11 | 170 | -1.90 | -0.92 | 1.4E-04 | -2.03 | -1.02 | 2.3E-05 |
| Samd14 | 10591 | -1.54 | -0.63 | 9.7E-05 | -1.59 | -0.67 | 2.8E-05 |
| Samd3 | 400 | -2.38 | -1.25 | 2.3E-04 | -2.84 | -1.51 | 7.6E-06 |
| Samd4b | 6947 | 1.45 | 0.54 | 2.9E-06 | 1.35 | 0.43 | 2.5E-04 |
| Samd5 | 2239 | -1.88 | -0.91 | 1.7E-20 | -1.94 | -0.95 | 3.0E-22 |
| Samd8 | 6348 | 1.73 | 0.79 | 2.1E-14 | 1.70 | 0.77 | 1.4E-13 |
| Samd9l | 228 | -1.98 | -0.98 | 3.5E-05 | -1.94 | -0.96 | 5.7E-05 |
| Samhd1 | 5052 | 1.53 | 0.61 | 2.2E-09 | 1.48 | 0.56 | 5.2E-08 |
| Samm50 | 4748 | -1.45 | -0.54 | 4.5E-12 | -1.56 | -0.64 | 8.3E-17 |
| Sap130 | 6963 | 1.36 | 0.45 | 2.1E-13 | 1.30 | 0.38 | 4.4E-10 |
| Sap30l | 1292 | 1.46 | 0.55 | 3.0E-15 | 1.46 | 0.55 | 3.7E-15 |
| Saraf | 8581 | 1.68 | 0.75 | 2.4E-20 | 1.72 | 0.78 | 1.1E-21 |
| Sardh | 489 | -1.91 | -0.94 | 3.8E-05 | -1.93 | -0.95 | 3.0E-05 |
| Sars2 | 811 | -2.53 | -1.34 | 4.7E-23 | -2.64 | -1.40 | 4.4E-25 |
| Sat1 | 2244 | 1.28 | 0.35 | 7.5E-05 | 1.32 | 0.40 | 4.6E-06 |
| Sat2 | 685 | 1.40 | 0.49 | 1.2E-10 | 1.41 | 0.50 | 6.7E-11 |
| Saxo2 | 296 | 1.31 | 0.39 | 3.0E-02 | 1.43 | 0.51 | 3.7E-03 |
| Saysd1 | 366 | -1.58 | -0.66 | 2.3E-10 | -1.66 | -0.73 | 3.3E-12 |
| Sbf1 | 12325 | 1.57 | 0.65 | 4.1E-08 | 1.53 | 0.61 | 2.8E-07 |
| Sbk3 | 118 | 2.93 | 1.55 | 6.2E-06 | 3.13 | 1.65 | 1.7E-06 |
| Sbno2 | 1513 | -2.10 | -1.07 | 3.2E-24 | -2.26 | -1.18 | 5.3E-29 |
| Sc5d | 7412 | 1.66 | 0.73 | 7.1E-30 | 1.66 | 0.73 | 1.9E-29 |
| Scaf4 | 5878 | 1.30 | 0.38 | 4.1E-04 | 1.26 | 0.34 | 1.8E-03 |
| Scaf8 | 5748 | -1.42 | -0.50 | 3.3E-15 | -1.43 | -0.52 | 6.4E-16 |
| Scamp1 | 8509 | 1.28 | 0.36 | 7.8E-03 | 1.32 | 0.40 | 3.2E-03 |
| Scamp2 | 2628 | -1.36 | -0.44 | 3.0E-19 | -1.42 | -0.51 | 8.1E-25 |
| Scand1 | 1287 | -1.28 | -0.35 | 7.5E-03 | -1.37 | -0.46 | 4.2E-04 |
| Scaper | 3960 | -1.65 | -0.73 | 4.3E-05 | -1.49 | -0.58 | 1.4E-03 |

| | | | | | | | |
|---|---|---|---|---|---|---|---|
| Scara3 | 922 | -3.02 | -1.59 | 6.1E-15 | -3.17 | -1.67 | 3.5E-16 |
| Scara5 | 424 | -2.38 | -1.25 | 1.9E-03 | -2.80 | -1.49 | 1.9E-04 |
| Scarb1 | 3129 | 2.56 | 1.35 | 9.4E-38 | 2.28 | 1.19 | 1.7E-29 |
| Scarf1 | 312 | -1.34 | -0.42 | 6.7E-04 | -1.51 | -0.59 | 1.2E-06 |
| Scarf2 | 1480 | -2.40 | -1.27 | 2.7E-04 | -2.52 | -1.33 | 1.3E-04 |
| Scd1 | 3523 | 2.10 | 1.07 | 6.7E-06 | 1.71 | 0.78 | 1.4E-03 |
| Scfd1 | 2748 | -2.26 | -1.18 | 1.7E-33 | -2.12 | -1.08 | 1.7E-28 |
| Sclt1 | 3048 | 1.26 | 0.34 | 2.5E-05 | 1.38 | 0.47 | 3.0E-09 |
| Scly | 1245 | 2.26 | 1.18 | 2.9E-34 | 2.15 | 1.11 | 2.4E-30 |
| Scml2 | 716 | -1.94 | -0.95 | 1.6E-23 | -1.85 | -0.89 | 2.1E-20 |
| Scn2b | 833 | -1.36 | -0.45 | 1.1E-02 | -1.36 | -0.45 | 1.2E-02 |
| Scn3a | 8413 | -2.01 | -1.00 | 1.6E-05 | -1.68 | -0.74 | 1.8E-03 |
| Scn5a | 1085 | 2.19 | 1.13 | 4.6E-11 | 2.23 | 1.16 | 2.0E-11 |
| Scnm1 | 2425 | -1.35 | -0.44 | 1.7E-06 | -1.34 | -0.43 | 3.0E-06 |
| Scpep1 | 3033 | 1.39 | 0.47 | 3.5E-04 | 1.28 | 0.36 | 7.9E-03 |
| Scrn2 | 719 | -1.60 | -0.68 | 2.3E-05 | -1.71 | -0.78 | 1.1E-06 |
| Scrn3 | 647 | -1.99 | -0.99 | 9.8E-26 | -1.97 | -0.98 | 7.4E-25 |
| Scrt1 | 3996 | -2.00 | -1.00 | 7.3E-05 | -2.10 | -1.07 | 2.0E-05 |
| Scrt2 | 3779 | -2.79 | -1.48 | 1.5E-11 | -2.73 | -1.45 | 4.5E-11 |
| Scube1 | 4902 | 1.46 | 0.54 | 5.4E-05 | 1.50 | 0.59 | 1.4E-05 |
| Scube3 | 5734 | -1.48 | -0.57 | 1.9E-03 | -1.65 | -0.73 | 5.0E-05 |
| Scx | 108 | -4.53 | -2.18 | 5.3E-08 | -3.56 | -1.83 | 6.1E-06 |
| Scyl2 | 7981 | 1.35 | 0.43 | 1.4E-11 | 1.42 | 0.51 | 1.4E-15 |
| Sdc3 | 11979 | -1.49 | -0.57 | 3.1E-03 | -1.52 | -0.61 | 1.7E-03 |
| Sdcbp | 16088 | 1.53 | 0.62 | 2.3E-09 | 1.63 | 0.71 | 5.2E-12 |
| Sdccag8 | 1430 | 1.51 | 0.59 | 2.7E-29 | 1.62 | 0.70 | 1.0E-39 |
| Sdf2 | 2595 | -1.29 | -0.36 | 2.6E-06 | -1.29 | -0.37 | 1.8E-06 |
| Sdf2l1 | 1277 | 1.98 | 0.99 | 4.4E-11 | 1.79 | 0.84 | 2.5E-08 |
| Sdf4 | 6880 | -1.33 | -0.42 | 5.4E-17 | -1.40 | -0.49 | 1.1E-22 |
| Sdr39u1 | 1162 | -1.45 | -0.54 | 3.7E-04 | -1.52 | -0.60 | 5.5E-05 |
| Sdsl | 200 | -1.28 | -0.35 | 1.7E-02 | -1.38 | -0.46 | 1.4E-03 |
| Sec11c | 2003 | 1.91 | 0.94 | 2.0E-26 | 1.98 | 0.99 | 3.6E-29 |
| Sec14l1 | 12601 | 1.83 | 0.88 | 2.7E-20 | 1.74 | 0.80 | 3.4E-17 |
| Sec14l2 | 403 | 1.32 | 0.40 | 4.4E-04 | 1.28 | 0.35 | 2.4E-03 |
| Sec14l5 | 793 | 4.60 | 2.20 | 3.8E-27 | 5.41 | 2.44 | 8.2E-33 |
| Sec16a | 12641 | 1.29 | 0.37 | 8.7E-05 | 1.36 | 0.45 | 1.5E-06 |
| Sec22c | 4964 | 1.33 | 0.41 | 7.3E-04 | 1.22 | 0.29 | 2.1E-02 |
| Sec23a | 12547 | 1.24 | 0.32 | 3.1E-05 | 1.32 | 0.40 | 8.5E-08 |
| Sec23b | 6991 | 1.46 | 0.55 | 9.7E-08 | 1.42 | 0.50 | 1.4E-06 |
| Sec24a | 3464 | 1.29 | 0.37 | 1.1E-04 | 1.30 | 0.38 | 8.0E-05 |
| Sec62 | 9104 | -1.25 | -0.32 | 8.2E-08 | -1.23 | -0.29 | 1.3E-06 |
| Secisbp2 | 1609 | -2.31 | -1.21 | 4.5E-48 | -2.49 | -1.31 | 9.0E-57 |
| Seh1l | 5055 | -2.36 | -1.24 | 1.3E-47 | -2.38 | -1.25 | 2.6E-48 |
| Sel1l | 12129 | 1.28 | 0.35 | 7.3E-06 | 1.33 | 0.41 | 1.1E-07 |
| Sel1l3 | 1613 | -1.51 | -0.59 | 9.2E-04 | -1.46 | -0.55 | 2.4E-03 |
| Selenoi | 7031 | 1.21 | 0.27 | 1.6E-03 | 1.28 | 0.36 | 1.9E-05 |
| Selenon | 4954 | -1.90 | -0.93 | 5.5E-17 | -1.98 | -0.99 | 4.4E-19 |

| | | | | | | | |
|---|---|---|---|---|---|---|---|
| Sema3a | 2749 | -2.24 | -1.16 | 2.5E-07 | -1.77 | -0.82 | 3.5E-04 |
| Sema3b | 220 | -1.51 | -0.59 | 6.9E-03 | -1.53 | -0.61 | 5.5E-03 |
| Sema3c | 3350 | -1.87 | -0.90 | 5.6E-07 | -1.80 | -0.85 | 2.9E-06 |
| Sema3d | 891 | -2.64 | -1.40 | 2.0E-03 | -1.91 | -0.93 | 4.7E-02 |
| Sema3g | 601 | -1.64 | -0.71 | 2.1E-08 | -1.75 | -0.81 | 1.6E-10 |
| Sema4a | 1676 | -1.54 | -0.63 | 2.2E-03 | -1.61 | -0.68 | 7.7E-04 |
| Sema4c | 4933 | -1.84 | -0.88 | 7.6E-13 | -2.10 | -1.07 | 1.5E-18 |
| Sema4d | 1612 | 1.90 | 0.93 | 4.8E-10 | 1.86 | 0.89 | 2.1E-09 |
| Sema6a | 5687 | -1.29 | -0.37 | 1.6E-03 | -1.37 | -0.45 | 7.8E-05 |
| Sema6b | 2249 | 1.88 | 0.91 | 4.5E-06 | 1.85 | 0.89 | 8.3E-06 |
| Sema6d | 5702 | -1.66 | -0.73 | 6.7E-04 | -1.42 | -0.51 | 2.2E-02 |
| Senp2 | 8528 | 1.53 | 0.62 | 3.2E-24 | 1.48 | 0.57 | 1.3E-20 |
| Senp3 | 8408 | 1.29 | 0.36 | 1.2E-19 | 1.24 | 0.31 | 2.0E-14 |
| Senp6 | 10112 | -2.39 | -1.25 | 8.2E-61 | -2.35 | -1.23 | 1.4E-58 |
| Sephs1 | 9449 | -2.11 | -1.07 | 6.2E-31 | -2.22 | -1.15 | 1.7E-35 |
| Sephs2 | 3275 | 1.62 | 0.70 | 1.4E-10 | 1.41 | 0.50 | 6.9E-06 |
| Sepsecs | 2467 | 1.31 | 0.39 | 3.5E-07 | 1.37 | 0.45 | 4.7E-09 |
| Septin1 | 808 | -1.45 | -0.54 | 1.8E-03 | -1.80 | -0.85 | 4.1E-07 |
| Septin4 | 792 | 2.13 | 1.09 | 2.3E-16 | 2.27 | 1.18 | 4.4E-19 |
| Septin7 | 20284 | -1.31 | -0.39 | 1.6E-05 | -1.27 | -0.35 | 1.2E-04 |
| Septin9 | 12761 | -1.25 | -0.32 | 6.5E-03 | -1.39 | -0.47 | 3.4E-05 |
| Serac1 | 2485 | 1.67 | 0.74 | 5.5E-11 | 1.88 | 0.91 | 4.7E-16 |
| Sergef | 851 | 1.27 | 0.34 | 4.5E-03 | 1.27 | 0.34 | 5.3E-03 |
| Serinc2 | 2587 | -2.01 | -1.00 | 1.0E-31 | -2.10 | -1.07 | 7.5E-36 |
| Serinc4 | 116 | -4.90 | -2.29 | 1.0E-28 | -4.52 | -2.18 | 4.8E-26 |
| Serinc5 | 5207 | 1.90 | 0.92 | 2.9E-13 | 2.12 | 1.09 | 5.8E-18 |
| Serp1 | 12830 | 1.26 | 0.34 | 2.6E-03 | 1.23 | 0.30 | 8.4E-03 |
| Serpinb6a | 2101 | -1.30 | -0.38 | 2.4E-02 | -1.29 | -0.37 | 2.9E-02 |
| Serping1 | 1024 | -1.99 | -0.99 | 5.7E-12 | -1.97 | -0.98 | 1.1E-11 |
| Serpini1 | 7446 | 1.35 | 0.43 | 4.5E-02 | 1.59 | 0.67 | 1.3E-03 |
| Sertad2 | 4072 | 1.53 | 0.61 | 2.0E-13 | 1.50 | 0.58 | 3.5E-12 |
| Sertad3 | 847 | 2.13 | 1.09 | 4.1E-24 | 1.95 | 0.96 | 4.7E-19 |
| Sertad4 | 3346 | -1.91 | -0.94 | 3.2E-19 | -1.95 | -0.97 | 2.7E-20 |
| Sertm1 | 1096 | 2.09 | 1.06 | 3.0E-09 | 2.03 | 1.02 | 1.2E-08 |
| Sesn1 | 3981 | 1.42 | 0.51 | 3.3E-10 | 1.56 | 0.64 | 1.0E-15 |
| Sesn2 | 1972 | 4.48 | 2.16 | 3.7E-70 | 4.10 | 2.04 | 3.1E-62 |
| Sesn3 | 17785 | 1.60 | 0.67 | 4.1E-17 | 1.74 | 0.80 | 1.8E-23 |
| Sestd1 | 12808 | -1.41 | -0.50 | 6.4E-04 | -1.32 | -0.40 | 8.0E-03 |
| Set | 42386 | -1.41 | -0.49 | 4.4E-04 | -1.59 | -0.66 | 1.3E-06 |
| Setbp1 | 9254 | -3.28 | -1.71 | 1.7E-29 | -3.22 | -1.69 | 1.2E-28 |
| Setd1b | 4177 | -2.04 | -1.03 | 1.6E-07 | -2.28 | -1.19 | 1.1E-09 |
| Setd3 | 10303 | 1.43 | 0.52 | 7.9E-41 | 1.44 | 0.53 | 2.1E-42 |
| Setd4 | 686 | -1.85 | -0.88 | 1.3E-19 | -1.92 | -0.94 | 5.7E-22 |
| Setd7 | 4188 | -1.53 | -0.61 | 6.4E-14 | -1.52 | -0.61 | 8.9E-14 |
| Setdb2 | 1135 | 1.56 | 0.64 | 1.1E-07 | 1.63 | 0.71 | 5.2E-09 |
| Setmar | 331 | -1.98 | -0.99 | 8.2E-17 | -1.87 | -0.90 | 2.4E-14 |
| Sf3a1 | 10723 | -1.39 | -0.47 | 4.3E-04 | -1.56 | -0.65 | 1.1E-06 |

| | | | | | | | |
|---|---|---|---|---|---|---|---|
| Sf3b5 | 1896 | 1.48 | 0.56 | 5.4E-06 | 1.33 | 0.41 | 1.4E-03 |
| Sf3b6 | 6240 | 1.30 | 0.38 | 7.3E-05 | 1.34 | 0.42 | 1.2E-05 |
| Sfi1 | 1449 | -1.52 | -0.60 | 2.5E-14 | -1.55 | -0.64 | 9.6E-16 |
| Sfmbt2 | 633 | 1.78 | 0.83 | 1.1E-08 | 1.94 | 0.95 | 5.7E-11 |
| Sfpq | 47502 | -1.58 | -0.66 | 1.8E-16 | -1.58 | -0.66 | 4.7E-16 |
| Sfrp1 | 17728 | 1.45 | 0.54 | 1.1E-03 | 1.37 | 0.46 | 6.2E-03 |
| Sfrp2 | 4464 | -1.80 | -0.84 | 4.0E-03 | -1.75 | -0.81 | 6.1E-03 |
| Sfrp4 | 264 | 1.98 | 0.98 | 1.3E-20 | 1.84 | 0.88 | 2.2E-16 |
| Sft2d2 | 3066 | 1.57 | 0.65 | 2.1E-06 | 1.52 | 0.60 | 1.1E-05 |
| Sft2d3 | 601 | -1.35 | -0.43 | 9.2E-07 | -1.37 | -0.45 | 2.5E-07 |
| Sfxn2 | 3160 | 1.92 | 0.94 | 4.0E-29 | 1.78 | 0.83 | 3.2E-23 |
| Sgcb | 2549 | 1.26 | 0.33 | 5.9E-04 | 1.29 | 0.36 | 1.4E-04 |
| Sgce | 1951 | -1.87 | -0.90 | 4.6E-34 | -1.94 | -0.96 | 2.3E-38 |
| Sgf29 | 862 | -1.84 | -0.88 | 3.1E-24 | -2.00 | -1.00 | 8.2E-31 |
| Sgk3 | 621 | -1.99 | -0.99 | 8.3E-07 | -1.83 | -0.87 | 1.8E-05 |
| Sgo1 | 2634 | -1.48 | -0.56 | 1.5E-03 | -1.63 | -0.71 | 5.6E-05 |
| Sgpl1 | 11314 | 1.43 | 0.52 | 1.0E-14 | 1.41 | 0.50 | 8.3E-14 |
| Sgpp1 | 1571 | 1.51 | 0.60 | 6.5E-10 | 1.60 | 0.68 | 2.2E-12 |
| Sgpp2 | 329 | 2.17 | 1.12 | 5.5E-04 | 2.28 | 1.19 | 2.5E-04 |
| Sgsm1 | 2744 | 1.70 | 0.77 | 2.4E-10 | 1.60 | 0.68 | 2.9E-08 |
| Sgsm2 | 4474 | 1.40 | 0.49 | 2.7E-04 | 1.34 | 0.42 | 1.8E-03 |
| Sgsm3 | 2524 | 1.76 | 0.81 | 8.5E-29 | 1.69 | 0.75 | 8.3E-25 |
| Sgtb | 5003 | 2.59 | 1.37 | 4.7E-61 | 2.73 | 1.45 | 1.2E-67 |
| Sh2b2 | 1643 | 1.33 | 0.41 | 1.7E-03 | 1.23 | 0.30 | 2.5E-02 |
| Sh2b3 | 2074 | 2.26 | 1.18 | 1.9E-20 | 2.06 | 1.04 | 3.6E-16 |
| Sh2d4b | 1118 | 3.11 | 1.64 | 3.7E-16 | 3.31 | 1.73 | 7.1E-18 |
| Sh2d5 | 969 | 2.34 | 1.23 | 1.0E-12 | 2.58 | 1.37 | 1.7E-15 |
| Sh3bgr | 135 | 1.74 | 0.80 | 1.1E-02 | 1.97 | 0.98 | 1.9E-03 |
| Sh3bgrl2 | 2180 | 2.48 | 1.31 | 5.2E-53 | 2.71 | 1.44 | 3.1E-63 |
| Sh3bp2 | 1827 | -1.93 | -0.95 | 2.7E-10 | -2.20 | -1.14 | 2.9E-14 |
| Sh3bp4 | 3448 | 1.62 | 0.69 | 6.8E-16 | 1.60 | 0.67 | 5.5E-15 |
| Sh3bp5 | 4917 | -1.96 | -0.97 | 6.8E-09 | -2.03 | -1.02 | 1.1E-09 |
| Sh3d19 | 1865 | -1.45 | -0.53 | 1.5E-03 | -1.45 | -0.54 | 1.4E-03 |
| Sh3glb2 | 5383 | 1.26 | 0.34 | 4.5E-03 | 1.29 | 0.36 | 2.1E-03 |
| Sh3pxd2a | 9733 | -1.41 | -0.50 | 5.9E-04 | -1.31 | -0.39 | 8.6E-03 |
| Sh3pxd2b | 8231 | -1.73 | -0.79 | 2.1E-11 | -1.78 | -0.83 | 1.2E-12 |
| Sh3tc1 | 266 | -1.45 | -0.54 | 3.6E-04 | -1.40 | -0.48 | 1.6E-03 |
| Sh3tc2 | 162 | -3.02 | -1.60 | 4.6E-11 | -2.85 | -1.51 | 6.3E-10 |
| Sh3yl1 | 1332 | 1.48 | 0.56 | 8.1E-07 | 1.47 | 0.56 | 9.6E-07 |
| Shank2 | 2071 | -2.21 | -1.14 | 1.8E-05 | -2.32 | -1.22 | 4.8E-06 |
| Shb | 1757 | 1.71 | 0.77 | 1.6E-04 | 1.55 | 0.63 | 2.7E-03 |
| Shc2 | 2467 | -1.42 | -0.51 | 1.9E-05 | -1.41 | -0.50 | 3.2E-05 |
| Shc4 | 325 | -1.69 | -0.75 | 1.4E-05 | -1.50 | -0.58 | 1.0E-03 |
| Shd | 2941 | -1.46 | -0.55 | 2.6E-06 | -1.48 | -0.56 | 1.3E-06 |
| She | 492 | -1.98 | -0.99 | 5.0E-16 | -1.95 | -0.96 | 3.2E-15 |
| Shf | 2446 | 1.58 | 0.66 | 2.1E-09 | 1.69 | 0.75 | 5.4E-12 |
| Shfl | 184 | 1.90 | 0.93 | 1.8E-05 | 1.88 | 0.91 | 2.9E-05 |

| | | | | | | | |
|---|---|---|---|---|---|---|---|
| Shisa3 | 1066 | -2.55 | -1.35 | 1.7E-06 | -2.31 | -1.21 | 2.2E-05 |
| Shisa4 | 1825 | -1.22 | -0.28 | 1.0E-02 | -1.36 | -0.44 | 4.1E-05 |
| Shisa5 | 1429 | -1.39 | -0.47 | 4.7E-04 | -1.45 | -0.54 | 6.1E-05 |
| Shisa6 | 651 | -2.13 | -1.09 | 1.8E-04 | -1.85 | -0.89 | 2.7E-03 |
| Shisal1 | 2229 | -1.91 | -0.93 | 1.5E-05 | -1.94 | -0.96 | 9.5E-06 |
| Shkbp1 | 2965 | 1.27 | 0.35 | 2.7E-05 | 1.14 | 0.19 | 3.2E-02 |
| Shld1 | 402 | -2.93 | -1.55 | 1.7E-49 | -2.73 | -1.45 | 1.7E-43 |
| Shld2 | 510 | -1.72 | -0.78 | 2.4E-14 | -1.58 | -0.66 | 1.5E-10 |
| Shoc2 | 9766 | 1.27 | 0.35 | 7.7E-06 | 1.26 | 0.34 | 1.7E-05 |
| Shox2 | 3795 | -1.95 | -0.97 | 6.0E-05 | -2.26 | -1.17 | 8.6E-07 |
| Shpk | 275 | 2.54 | 1.34 | 7.5E-22 | 2.51 | 1.33 | 3.7E-21 |
| Shprh | 9281 | 1.24 | 0.31 | 1.6E-02 | 1.42 | 0.51 | 5.7E-05 |
| Shq1 | 1162 | 2.00 | 1.00 | 3.4E-36 | 1.92 | 0.94 | 4.7E-32 |
| Shroom1 | 169 | -1.89 | -0.92 | 9.7E-05 | -1.89 | -0.92 | 1.1E-04 |
| Shroom3 | 3100 | 1.77 | 0.82 | 3.0E-06 | 1.55 | 0.63 | 4.4E-04 |
| Shroom4 | 1948 | -1.34 | -0.42 | 3.7E-03 | -1.26 | -0.33 | 2.6E-02 |
| Siae | 1336 | 1.34 | 0.42 | 4.0E-05 | 1.49 | 0.58 | 1.2E-08 |
| Siah1a | 2537 | -1.65 | -0.72 | 2.6E-15 | -1.80 | -0.85 | 5.3E-21 |
| Siah1b | 894 | -1.32 | -0.40 | 3.5E-02 | -1.49 | -0.57 | 1.9E-03 |
| Siah2 | 779 | 1.79 | 0.84 | 7.6E-24 | 1.77 | 0.82 | 8.8E-23 |
| Sidt2 | 5621 | 1.30 | 0.37 | 2.4E-05 | 1.26 | 0.34 | 1.6E-04 |
| Sigmar1 | 2518 | 1.53 | 0.62 | 8.3E-08 | 1.33 | 0.41 | 4.7E-04 |
| Sik1 | 2688 | 1.25 | 0.32 | 1.0E-02 | 1.26 | 0.33 | 8.0E-03 |
| Sik2 | 2420 | 1.63 | 0.70 | 7.3E-09 | 1.76 | 0.82 | 1.2E-11 |
| Sil1 | 709 | -1.39 | -0.48 | 2.8E-16 | -1.45 | -0.54 | 3.0E-20 |
| Sim2 | 224 | -2.06 | -1.04 | 9.2E-03 | -2.27 | -1.18 | 2.9E-03 |
| Sin3a | 12950 | 1.69 | 0.76 | 3.1E-27 | 1.62 | 0.69 | 8.5E-23 |
| Sipa1l2 | 9580 | 1.61 | 0.69 | 4.0E-10 | 1.63 | 0.70 | 1.7E-10 |
| Sipa1l3 | 3720 | 1.80 | 0.85 | 1.7E-08 | 1.71 | 0.78 | 2.8E-07 |
| Sirpa | 2545 | -1.49 | -0.58 | 1.1E-05 | -1.54 | -0.62 | 2.4E-06 |
| Sirt1 | 3366 | -1.49 | -0.57 | 5.2E-15 | -1.48 | -0.56 | 1.1E-14 |
| Sirt2 | 5935 | 1.39 | 0.47 | 1.3E-18 | 1.39 | 0.48 | 8.1E-19 |
| Sirt5 | 690 | -1.98 | -0.99 | 1.3E-31 | -1.75 | -0.81 | 1.9E-21 |
| Sirt6 | 2279 | -1.54 | -0.62 | 3.6E-06 | -1.74 | -0.80 | 2.3E-09 |
| Sirt7 | 2821 | 1.58 | 0.66 | 7.8E-19 | 1.48 | 0.56 | 8.0E-14 |
| Siva1 | 2373 | 1.54 | 0.62 | 5.6E-05 | 1.45 | 0.53 | 6.3E-04 |
| Six5 | 1478 | -1.83 | -0.87 | 1.2E-09 | -1.96 | -0.97 | 9.4E-12 |
| Ska1 | 908 | -1.47 | -0.56 | 2.3E-03 | -1.62 | -0.69 | 1.2E-04 |
| Ska2 | 6665 | -1.50 | -0.58 | 4.6E-07 | -1.59 | -0.67 | 6.7E-09 |
| Skap1 | 123 | 1.53 | 0.62 | 7.1E-05 | 1.35 | 0.43 | 6.5E-03 |
| Skida1 | 4816 | -1.33 | -0.41 | 2.1E-03 | -1.45 | -0.54 | 3.4E-05 |
| Sla | 178 | -2.38 | -1.25 | 1.7E-03 | -2.74 | -1.46 | 2.3E-04 |
| Slbp | 7176 | 1.36 | 0.44 | 7.3E-04 | 1.26 | 0.33 | 1.4E-02 |
| Slc10a4 | 450 | -1.75 | -0.80 | 5.3E-05 | -1.66 | -0.73 | 2.9E-04 |
| Slc11a2 | 3989 | 1.66 | 0.73 | 2.9E-36 | 1.68 | 0.75 | 2.9E-38 |
| Slc12a7 | 5929 | 1.32 | 0.40 | 1.4E-03 | 1.31 | 0.38 | 2.2E-03 |
| Slc12a9 | 2684 | 1.88 | 0.91 | 3.7E-15 | 1.80 | 0.85 | 2.8E-13 |

| | | | | | | | |
|---|---|---|---|---|---|---|---|
| Slc13a5 | 110 | 2.16 | 1.11 | 1.6E-05 | 2.41 | 1.27 | 8.8E-07 |
| Slc14a2 | 114 | -1.82 | -0.86 | 7.2E-06 | -1.98 | -0.99 | 2.4E-07 |
| Slc15a4 | 1493 | 1.31 | 0.39 | 4.4E-07 | 1.31 | 0.39 | 3.5E-07 |
| Slc16a1 | 3029 | 1.53 | 0.62 | 5.4E-04 | 1.42 | 0.51 | 5.0E-03 |
| Slc16a12 | 204 | -2.76 | -1.46 | 6.9E-20 | -2.47 | -1.31 | 4.7E-16 |
| Slc16a14 | 680 | -2.34 | -1.23 | 5.2E-10 | -2.44 | -1.29 | 7.2E-11 |
| Slc16a3 | 1490 | 2.13 | 1.09 | 2.0E-06 | 1.98 | 0.98 | 2.1E-05 |
| Slc16a4 | 131 | -2.45 | -1.29 | 5.1E-05 | -2.02 | -1.01 | 1.9E-03 |
| Slc16a9 | 214 | -1.69 | -0.76 | 4.7E-03 | -2.02 | -1.02 | 1.2E-04 |
| Slc17a5 | 1374 | 1.28 | 0.35 | 4.6E-07 | 1.26 | 0.34 | 1.6E-06 |
| Slc17a6 | 5910 | -2.41 | -1.27 | 2.9E-09 | -2.42 | -1.28 | 2.4E-09 |
| Slc17a7 | 847 | 1.63 | 0.70 | 1.3E-02 | 1.63 | 0.70 | 1.4E-02 |
| Slc17a8 | 404 | -1.72 | -0.78 | 1.1E-03 | -1.43 | -0.52 | 3.8E-02 |
| Slc17a9 | 221 | 5.87 | 2.55 | 1.2E-33 | 6.74 | 2.75 | 2.1E-38 |
| Slc18b1 | 1524 | -1.48 | -0.57 | 1.6E-08 | -1.51 | -0.60 | 2.8E-09 |
| Slc19a2 | 1507 | 2.19 | 1.13 | 2.1E-40 | 2.18 | 1.12 | 7.3E-40 |
| Slc1a4 | 5038 | 1.38 | 0.46 | 1.1E-09 | 1.30 | 0.38 | 7.5E-07 |
| Slc1a5 | 3713 | 3.00 | 1.59 | 9.5E-23 | 2.70 | 1.43 | 1.1E-18 |
| Slc1a6 | 272 | 1.82 | 0.87 | 8.9E-07 | 1.71 | 0.77 | 1.4E-05 |
| Slc20a1 | 6973 | -1.82 | -0.86 | 2.4E-26 | -1.81 | -0.86 | 4.7E-26 |
| Slc20a2 | 1726 | -1.76 | -0.82 | 2.7E-20 | -1.86 | -0.90 | 3.2E-24 |
| Slc22a15 | 1103 | -2.14 | -1.10 | 3.7E-39 | -1.90 | -0.93 | 6.7E-28 |
| Slc22a21 | 450 | 2.16 | 1.11 | 2.7E-15 | 2.41 | 1.27 | 1.5E-19 |
| Slc22a5 | 1147 | 1.38 | 0.47 | 1.8E-09 | 1.41 | 0.49 | 2.8E-10 |
| Slc22a6 | 363 | -2.08 | -1.05 | 4.2E-03 | -2.10 | -1.07 | 3.8E-03 |
| Slc22a8 | 124 | -2.13 | -1.09 | 9.8E-03 | -2.17 | -1.12 | 8.2E-03 |
| Slc23a4 | 121 | 1.37 | 0.45 | 5.1E-03 | 1.60 | 0.68 | 1.9E-05 |
| Slc24a3 | 797 | -1.55 | -0.63 | 3.3E-02 | -1.63 | -0.70 | 1.7E-02 |
| Slc25a1 | 4306 | 1.73 | 0.79 | 6.5E-25 | 1.60 | 0.68 | 1.2E-18 |
| Slc25a10 | 2480 | 2.44 | 1.29 | 1.6E-30 | 2.21 | 1.14 | 2.8E-24 |
| Slc25a12 | 5108 | 1.38 | 0.47 | 2.3E-10 | 1.35 | 0.43 | 6.1E-09 |
| Slc25a14 | 1611 | 1.30 | 0.37 | 3.8E-05 | 1.32 | 0.40 | 8.4E-06 |
| Slc25a15 | 1643 | 1.65 | 0.73 | 7.3E-17 | 1.60 | 0.67 | 1.3E-14 |
| Slc25a16 | 1638 | 1.61 | 0.69 | 2.1E-19 | 1.46 | 0.55 | 9.3E-13 |
| Slc25a19 | 1683 | 1.46 | 0.55 | 8.6E-08 | 1.42 | 0.51 | 9.2E-07 |
| Slc25a20 | 2698 | 1.49 | 0.58 | 2.6E-21 | 1.47 | 0.56 | 5.5E-20 |
| Slc25a22 | 4225 | 1.38 | 0.47 | 5.5E-04 | 1.32 | 0.40 | 3.0E-03 |
| Slc25a23 | 9786 | 1.77 | 0.82 | 4.4E-16 | 1.69 | 0.76 | 9.4E-14 |
| Slc25a24 | 2430 | -2.04 | -1.03 | 1.5E-04 | -2.07 | -1.05 | 1.1E-04 |
| Slc25a25 | 3581 | 1.47 | 0.56 | 7.0E-08 | 1.42 | 0.50 | 1.2E-06 |
| Slc25a26 | 336 | -3.95 | -1.98 | 1.5E-63 | -4.30 | -2.10 | 3.8E-70 |
| Slc25a28 | 2490 | 1.58 | 0.66 | 4.9E-14 | 1.42 | 0.50 | 2.0E-08 |
| Slc25a29 | 727 | 2.28 | 1.19 | 1.4E-19 | 2.01 | 1.01 | 1.8E-14 |
| Slc25a30 | 2817 | 1.93 | 0.95 | 2.1E-46 | 1.91 | 0.93 | 9.3E-45 |
| Slc25a33 | 723 | 1.63 | 0.70 | 4.3E-27 | 1.73 | 0.79 | 5.4E-34 |
| Slc25a35 | 349 | 2.48 | 1.31 | 3.3E-12 | 2.72 | 1.44 | 1.6E-14 |
| Slc25a38 | 1131 | -1.32 | -0.41 | 2.9E-09 | -1.39 | -0.48 | 1.5E-12 |

| | | | | | | | |
|---|---|---|---|---|---|---|---|
| Slc25a39 | 4509 | 1.70 | 0.77 | 1.8E-14 | 1.57 | 0.65 | 9.8E-11 |
| Slc25a42 | 1225 | 2.10 | 1.07 | 9.8E-12 | 2.06 | 1.04 | 3.7E-11 |
| Slc25a44 | 3891 | 1.56 | 0.65 | 3.2E-27 | 1.51 | 0.60 | 2.9E-23 |
| Slc25a45 | 122 | -2.49 | -1.32 | 1.7E-10 | -3.14 | -1.65 | 1.0E-15 |
| Slc25a53 | 1330 | -1.35 | -0.43 | 1.4E-07 | -1.35 | -0.44 | 1.4E-07 |
| Slc26a10 | 171 | 1.95 | 0.96 | 1.7E-07 | 1.57 | 0.65 | 6.5E-04 |
| Slc26a11 | 802 | 1.81 | 0.86 | 2.3E-19 | 1.70 | 0.77 | 1.6E-15 |
| Slc26a2 | 2342 | 1.34 | 0.42 | 7.0E-03 | 1.52 | 0.61 | 6.4E-05 |
| Slc27a1 | 5126 | 1.47 | 0.56 | 2.5E-04 | 1.50 | 0.58 | 1.3E-04 |
| Slc27a2 | 336 | -1.71 | -0.78 | 1.4E-03 | -2.16 | -1.11 | 3.1E-06 |
| Slc27a3 | 1164 | -2.13 | -1.09 | 2.8E-17 | -2.06 | -1.04 | 6.6E-16 |
| Slc29a1 | 2220 | 1.90 | 0.93 | 7.2E-26 | 1.75 | 0.81 | 4.3E-20 |
| Slc29a3 | 1740 | -1.36 | -0.44 | 5.8E-12 | -1.33 | -0.41 | 2.7E-10 |
| Slc29a4 | 3649 | 1.55 | 0.64 | 1.5E-05 | 1.54 | 0.62 | 2.4E-05 |
| Slc2a10 | 356 | -1.90 | -0.93 | 3.7E-08 | -1.92 | -0.94 | 2.5E-08 |
| Slc2a12 | 396 | -1.57 | -0.65 | 6.9E-04 | -1.44 | -0.53 | 7.1E-03 |
| Slc2a13 | 1621 | 1.64 | 0.71 | 9.7E-07 | 1.87 | 0.90 | 3.3E-10 |
| Slc2a3 | 5543 | 2.39 | 1.26 | 1.6E-15 | 2.54 | 1.34 | 1.4E-17 |
| Slc2a4 | 1131 | 1.97 | 0.98 | 5.7E-13 | 1.87 | 0.91 | 3.0E-11 |
| Slc2a6 | 206 | 2.18 | 1.13 | 1.1E-15 | 2.27 | 1.18 | 4.4E-17 |
| Slc2a8 | 1230 | 2.53 | 1.34 | 2.1E-46 | 2.46 | 1.30 | 1.6E-43 |
| Slc30a5 | 4959 | 1.43 | 0.51 | 4.0E-14 | 1.38 | 0.46 | 1.3E-11 |
| Slc30a7 | 4694 | 1.29 | 0.37 | 3.6E-04 | 1.43 | 0.52 | 2.8E-07 |
| Slc30a9 | 10518 | 1.40 | 0.49 | 2.4E-20 | 1.45 | 0.54 | 1.8E-24 |
| Slc31a1 | 7592 | 1.84 | 0.88 | 1.1E-30 | 1.84 | 0.88 | 1.3E-30 |
| Slc31a2 | 837 | 1.75 | 0.81 | 6.9E-18 | 1.71 | 0.77 | 1.7E-16 |
| Slc33a1 | 2313 | 1.59 | 0.67 | 5.1E-21 | 1.58 | 0.66 | 1.3E-20 |
| Slc35a1 | 2184 | -1.86 | -0.90 | 1.7E-29 | -1.83 | -0.87 | 7.3E-28 |
| Slc35a2 | 1860 | 1.42 | 0.51 | 3.9E-11 | 1.34 | 0.42 | 7.6E-08 |
| Slc35a4 | 4201 | -1.33 | -0.41 | 6.4E-05 | -1.43 | -0.51 | 4.3E-07 |
| Slc35b2 | 3654 | 1.25 | 0.33 | 5.3E-05 | 1.21 | 0.27 | 8.1E-04 |
| Slc35c1 | 1025 | 1.73 | 0.79 | 1.8E-16 | 1.66 | 0.73 | 3.0E-14 |
| Slc35c2 | 2757 | 1.44 | 0.53 | 1.1E-10 | 1.30 | 0.38 | 6.3E-06 |
| Slc35d1 | 1412 | -1.22 | -0.28 | 3.8E-02 | -1.36 | -0.44 | 9.1E-04 |
| Slc35e1 | 5103 | 1.48 | 0.56 | 2.7E-16 | 1.42 | 0.50 | 3.2E-13 |
| Slc35e2 | 5105 | 1.40 | 0.49 | 4.7E-12 | 1.33 | 0.42 | 4.8E-09 |
| Slc35e3 | 4828 | 1.30 | 0.38 | 3.1E-08 | 1.26 | 0.33 | 1.8E-06 |
| Slc35e4 | 826 | 3.00 | 1.59 | 9.4E-58 | 2.93 | 1.55 | 2.5E-55 |
| Slc35f5 | 2232 | 1.28 | 0.36 | 7.1E-05 | 1.30 | 0.38 | 2.4E-05 |
| Slc35f6 | 1727 | 1.79 | 0.84 | 4.6E-26 | 1.80 | 0.85 | 1.5E-26 |
| Slc35g2 | 423 | 2.42 | 1.28 | 1.2E-26 | 2.46 | 1.30 | 1.4E-27 |
| Slc36a1 | 2256 | 1.31 | 0.39 | 4.0E-03 | 1.29 | 0.36 | 8.3E-03 |
| Slc37a2 | 374 | 1.57 | 0.65 | 9.7E-11 | 1.41 | 0.49 | 1.3E-06 |
| Slc37a4 | 1452 | 1.45 | 0.54 | 2.4E-12 | 1.43 | 0.51 | 3.0E-11 |
| Slc38a1 | 18710 | -1.71 | -0.78 | 3.2E-11 | -1.74 | -0.80 | 9.4E-12 |
| Slc38a10 | 11004 | 1.62 | 0.69 | 9.3E-17 | 1.52 | 0.60 | 6.0E-13 |
| Slc38a2 | 22564 | -1.52 | -0.61 | 1.1E-19 | -1.53 | -0.61 | 6.5E-20 |

| | | | | | | | |
|---|---|---|---|---|---|---|---|
| Slc38a3 | 1852 | 1.48 | 0.57 | 4.8E-08 | 1.48 | 0.57 | 4.4E-08 |
| Slc38a7 | 2589 | 1.49 | 0.57 | 1.4E-11 | 1.42 | 0.50 | 3.8E-09 |
| Slc38a9 | 3959 | 1.72 | 0.78 | 4.6E-31 | 1.81 | 0.86 | 4.1E-37 |
| Slc39a10 | 10904 | -1.53 | -0.61 | 3.0E-13 | -1.44 | -0.53 | 3.1E-10 |
| Slc39a14 | 2360 | 1.39 | 0.47 | 3.9E-10 | 1.47 | 0.55 | 1.4E-13 |
| Slc39a2 | 894 | 1.61 | 0.69 | 1.2E-04 | 1.55 | 0.63 | 4.7E-04 |
| Slc39a3 | 2642 | -1.55 | -0.63 | 9.3E-06 | -1.63 | -0.70 | 6.1E-07 |
| Slc39a8 | 1275 | -1.24 | -0.31 | 3.4E-02 | -1.30 | -0.38 | 1.1E-02 |
| Slc3a1 | 459 | 4.93 | 2.30 | 2.0E-30 | 4.72 | 2.24 | 8.7E-29 |
| Slc41a2 | 228 | -2.52 | -1.33 | 1.4E-07 | -2.33 | -1.22 | 1.6E-06 |
| Slc41a3 | 222 | 1.54 | 0.63 | 5.8E-05 | 1.58 | 0.66 | 2.2E-05 |
| Slc43a2 | 2408 | -1.85 | -0.89 | 5.4E-06 | -1.79 | -0.84 | 1.9E-05 |
| Slc44a2 | 7853 | 1.77 | 0.82 | 1.6E-35 | 1.78 | 0.83 | 1.6E-36 |
| Slc45a3 | 744 | 1.97 | 0.98 | 8.6E-06 | 1.61 | 0.68 | 2.4E-03 |
| Slc45a4 | 3192 | 1.96 | 0.97 | 6.4E-33 | 1.86 | 0.89 | 6.2E-28 |
| Slc46a1 | 366 | -1.35 | -0.43 | 8.7E-06 | -1.47 | -0.55 | 5.6E-09 |
| Slc46a3 | 384 | 3.68 | 1.88 | 1.6E-63 | 3.33 | 1.73 | 2.9E-54 |
| Slc4a11 | 268 | 3.08 | 1.62 | 1.9E-33 | 2.68 | 1.42 | 1.0E-25 |
| Slc4a2 | 4977 | 1.60 | 0.67 | 2.8E-11 | 1.60 | 0.68 | 1.8E-11 |
| Slc4a3 | 2939 | -2.67 | -1.42 | 1.6E-33 | -2.62 | -1.39 | 2.2E-32 |
| Slc4a4 | 967 | -1.33 | -0.41 | 1.5E-03 | -1.29 | -0.37 | 5.0E-03 |
| Slc4a7 | 9417 | -1.25 | -0.32 | 1.5E-02 | -1.21 | -0.27 | 4.3E-02 |
| Slc52a2 | 1559 | 1.50 | 0.59 | 3.9E-16 | 1.47 | 0.56 | 1.7E-14 |
| Slc5a5 | 115 | 2.91 | 1.54 | 5.1E-15 | 2.21 | 1.15 | 8.4E-09 |
| Slc5a7 | 304 | -2.47 | -1.30 | 1.4E-05 | -2.05 | -1.03 | 7.3E-04 |
| Slc66a1 | 1131 | 2.06 | 1.04 | 1.1E-34 | 2.08 | 1.06 | 1.8E-35 |
| Slc66a2 | 1359 | 1.67 | 0.74 | 1.5E-09 | 1.70 | 0.76 | 4.3E-10 |
| Slc6a20a | 146 | -2.70 | -1.43 | 2.9E-08 | -2.54 | -1.34 | 1.7E-07 |
| Slc6a6 | 8286 | -1.25 | -0.32 | 1.6E-03 | -1.36 | -0.44 | 1.0E-05 |
| Slc6a8 | 5359 | 1.72 | 0.78 | 5.6E-29 | 1.62 | 0.69 | 4.3E-23 |
| Slc7a1 | 5217 | 1.31 | 0.39 | 5.3E-03 | 1.25 | 0.32 | 2.6E-02 |
| Slc7a11 | 667 | -3.25 | -1.70 | 5.4E-15 | -3.16 | -1.66 | 2.5E-14 |
| Slc7a2 | 1667 | 1.39 | 0.48 | 6.6E-03 | 1.61 | 0.69 | 6.6E-05 |
| Slc7a3 | 316 | 2.75 | 1.46 | 5.8E-38 | 2.79 | 1.48 | 4.9E-39 |
| Slc7a5 | 5758 | 1.57 | 0.65 | 2.8E-06 | 1.36 | 0.44 | 2.2E-03 |
| Slc7a7 | 293 | -1.48 | -0.57 | 5.4E-05 | -1.62 | -0.70 | 4.5E-07 |
| Slc7a8 | 851 | 1.33 | 0.41 | 1.0E-02 | 1.36 | 0.44 | 6.0E-03 |
| Slc8a1 | 4042 | -2.30 | -1.20 | 2.5E-05 | -1.79 | -0.84 | 4.2E-03 |
| Slc8a2 | 3842 | 1.99 | 0.99 | 4.1E-15 | 2.04 | 1.03 | 4.2E-16 |
| Slc8b1 | 870 | 4.20 | 2.07 | 3.8E-30 | 3.89 | 1.96 | 4.1E-27 |
| Slc9a3r1 | 2902 | 3.04 | 1.60 | 2.4E-31 | 2.94 | 1.55 | 1.9E-29 |
| Slc9a5 | 3399 | 1.71 | 0.77 | 8.5E-15 | 1.62 | 0.69 | 4.4E-12 |
| Slc9a6 | 7768 | 1.31 | 0.39 | 7.2E-03 | 1.39 | 0.47 | 1.1E-03 |
| Slc9a8 | 1811 | -3.13 | -1.65 | 4.3E-99 | -3.26 | -1.70 | 1.3E-105 |
| Slco1c1 | 419 | -2.22 | -1.15 | 9.3E-04 | -2.01 | -1.01 | 4.3E-03 |
| Slco2a1 | 211 | -2.43 | -1.28 | 2.4E-07 | -2.42 | -1.28 | 3.1E-07 |
| Slco2b1 | 431 | -1.78 | -0.84 | 2.5E-09 | -1.85 | -0.89 | 2.4E-10 |

| | | | | | | | |
|---|---|---|---|---|---|---|---|
| Slco3a1 | 1369 | -1.74 | -0.80 | 5.1E-09 | -1.55 | -0.63 | 5.4E-06 |
| Slco4a1 | 394 | 3.18 | 1.67 | 3.0E-19 | 3.17 | 1.67 | 4.2E-19 |
| Slco5a1 | 5386 | -2.26 | -1.17 | 5.5E-15 | -2.03 | -1.02 | 1.3E-11 |
| Slfn9 | 1857 | -1.43 | -0.51 | 1.5E-04 | -1.33 | -0.41 | 3.3E-03 |
| Slirp | 2316 | -1.31 | -0.38 | 1.3E-02 | -1.39 | -0.47 | 2.0E-03 |
| Slit2 | 10361 | 1.31 | 0.39 | 2.6E-04 | 1.46 | 0.54 | 2.2E-07 |
| Slitrk1 | 1872 | -1.44 | -0.53 | 8.6E-03 | -1.46 | -0.55 | 6.4E-03 |
| Slitrk4 | 1131 | 1.62 | 0.70 | 2.8E-05 | 1.72 | 0.78 | 2.3E-06 |
| Slitrk5 | 2183 | -1.56 | -0.64 | 6.1E-09 | -1.59 | -0.67 | 1.4E-09 |
| Slk | 10088 | 1.48 | 0.56 | 3.7E-13 | 1.61 | 0.68 | 4.7E-19 |
| Slmap | 6971 | 1.39 | 0.47 | 4.8E-23 | 1.37 | 0.45 | 7.7E-21 |
| Sltm | 10413 | -1.46 | -0.54 | 1.3E-13 | -1.48 | -0.56 | 1.8E-14 |
| Slx1b | 1699 | 1.28 | 0.36 | 3.6E-03 | 1.26 | 0.34 | 7.2E-03 |
| Slx4ip | 2428 | -1.63 | -0.71 | 9.7E-23 | -1.60 | -0.68 | 9.2E-21 |
| Smad1 | 3746 | -1.35 | -0.43 | 2.5E-06 | -1.39 | -0.48 | 1.6E-07 |
| Smad3 | 5247 | -1.64 | -0.71 | 2.2E-08 | -1.73 | -0.79 | 5.3E-10 |
| Smad5 | 15919 | -1.45 | -0.53 | 3.7E-20 | -1.42 | -0.51 | 1.4E-18 |
| Smagp | 124 | 1.42 | 0.50 | 2.2E-02 | 1.46 | 0.55 | 1.3E-02 |
| Smap2 | 6084 | 1.47 | 0.56 | 7.9E-06 | 1.44 | 0.52 | 3.4E-05 |
| Smarca2 | 10932 | 1.25 | 0.33 | 3.1E-02 | 1.28 | 0.35 | 2.0E-02 |
| Smarca4 | 29774 | -1.59 | -0.67 | 5.2E-06 | -1.69 | -0.75 | 2.5E-07 |
| Smarcad1 | 10508 | -1.89 | -0.92 | 2.4E-24 | -1.87 | -0.90 | 1.0E-23 |
| Smarcal1 | 3318 | -1.56 | -0.64 | 2.4E-22 | -1.52 | -0.60 | 1.8E-19 |
| Smarcb1 | 7275 | -1.64 | -0.72 | 2.2E-12 | -1.76 | -0.82 | 7.7E-16 |
| Smarcc1 | 21646 | -1.39 | -0.48 | 1.0E-09 | -1.46 | -0.55 | 2.6E-12 |
| Smarcc2 | 27817 | -1.55 | -0.64 | 1.9E-08 | -1.63 | -0.70 | 4.3E-10 |
| Smarcd1 | 10186 | -1.80 | -0.85 | 7.4E-08 | -2.05 | -1.03 | 4.3E-11 |
| Smarce1 | 23766 | -1.53 | -0.62 | 2.2E-12 | -1.63 | -0.70 | 1.1E-15 |
| Smc1a | 18407 | -1.37 | -0.45 | 2.0E-06 | -1.37 | -0.46 | 1.7E-06 |
| Smc5 | 12744 | 1.29 | 0.37 | 8.8E-05 | 1.37 | 0.45 | 1.3E-06 |
| Smchd1 | 15940 | 1.36 | 0.45 | 3.4E-05 | 1.51 | 0.59 | 2.6E-08 |
| Smco4 | 361 | -2.06 | -1.04 | 3.9E-09 | -2.29 | -1.20 | 1.2E-11 |
| Smg1 | 23470 | 1.46 | 0.55 | 3.4E-03 | 1.74 | 0.80 | 1.4E-05 |
| Smg5 | 9291 | -1.31 | -0.39 | 1.1E-07 | -1.39 | -0.47 | 1.4E-10 |
| Smg9 | 1076 | -2.35 | -1.23 | 2.7E-29 | -2.42 | -1.28 | 2.7E-31 |
| Smim10l1 | 4598 | -3.08 | -1.62 | 3.7E-40 | -3.04 | -1.60 | 2.8E-39 |
| Smim10l2a | 482 | -1.45 | -0.54 | 3.7E-03 | -1.50 | -0.59 | 1.4E-03 |
| Smim12 | 956 | 1.57 | 0.65 | 7.3E-09 | 1.43 | 0.51 | 6.2E-06 |
| Smim15 | 4907 | 1.29 | 0.37 | 6.1E-04 | 1.32 | 0.40 | 1.8E-04 |
| Smim18 | 289 | -3.71 | -1.89 | 5.4E-11 | -3.28 | -1.72 | 3.1E-09 |
| Smim27 | 109 | -1.69 | -0.75 | 3.2E-03 | -1.92 | -0.94 | 2.2E-04 |
| Smim3 | 322 | -1.46 | -0.55 | 1.6E-02 | -1.51 | -0.60 | 9.0E-03 |
| Smim4 | 522 | 1.62 | 0.69 | 1.4E-05 | 1.65 | 0.73 | 5.9E-06 |
| Smim8 | 791 | -1.45 | -0.53 | 4.0E-06 | -1.45 | -0.54 | 3.5E-06 |
| Smo | 7268 | -1.62 | -0.69 | 1.6E-06 | -1.83 | -0.87 | 1.2E-09 |
| Smoc1 | 3139 | 1.30 | 0.38 | 2.4E-02 | 1.30 | 0.38 | 2.6E-02 |
| Smox | 2171 | -1.19 | -0.25 | 1.0E-02 | -1.26 | -0.33 | 5.5E-04 |

| | | | | | | | |
|---|---|---|---|---|---|---|---|
| Smpd2 | 1965 | 1.50 | 0.59 | 1.3E-11 | 1.36 | 0.45 | 4.0E-07 |
| Smtnl2 | 508 | 2.19 | 1.13 | 5.2E-04 | 2.28 | 1.19 | 2.7E-04 |
| Smug1 | 2182 | -1.54 | -0.62 | 3.9E-12 | -1.55 | -0.63 | 2.8E-12 |
| Smyd2 | 1864 | -1.24 | -0.31 | 3.7E-04 | -1.28 | -0.35 | 4.3E-05 |
| Smyd3 | 1305 | 1.60 | 0.68 | 1.7E-15 | 1.49 | 0.58 | 1.4E-11 |
| Smyd4 | 899 | -1.23 | -0.30 | 1.1E-04 | -1.27 | -0.34 | 1.3E-05 |
| Smyd5 | 2332 | -1.42 | -0.51 | 1.7E-20 | -1.55 | -0.63 | 9.9E-31 |
| Snap23 | 4894 | 1.41 | 0.49 | 1.3E-04 | 1.31 | 0.39 | 2.8E-03 |
| Snap25 | 6289 | 1.64 | 0.71 | 8.5E-05 | 1.69 | 0.75 | 3.2E-05 |
| Snap29 | 4892 | 1.41 | 0.49 | 2.8E-14 | 1.42 | 0.50 | 7.4E-15 |
| Snapc1 | 4179 | 1.57 | 0.65 | 6.7E-15 | 1.65 | 0.72 | 6.5E-18 |
| Snapc4 | 4129 | 1.42 | 0.51 | 5.2E-12 | 1.41 | 0.50 | 8.9E-12 |
| Snapc5 | 1234 | -1.45 | -0.53 | 7.4E-03 | -1.65 | -0.72 | 2.4E-04 |
| Snapin | 2158 | -1.35 | -0.43 | 4.0E-10 | -1.39 | -0.47 | 8.1E-12 |
| Sncaip | 2577 | -2.06 | -1.04 | 1.2E-16 | -1.95 | -0.96 | 2.6E-14 |
| Sncb | 516 | 2.09 | 1.07 | 5.2E-03 | 2.00 | 1.00 | 9.2E-03 |
| Snd1 | 10450 | 1.27 | 0.35 | 3.4E-04 | 1.22 | 0.29 | 3.4E-03 |
| Sned1 | 4427 | -1.51 | -0.59 | 1.2E-02 | -1.44 | -0.53 | 2.7E-02 |
| Snn | 12252 | -1.92 | -0.94 | 3.5E-30 | -1.98 | -0.99 | 3.5E-33 |
| Snrnp25 | 1146 | 1.43 | 0.52 | 5.9E-04 | 1.34 | 0.42 | 6.6E-03 |
| Snrnp35 | 926 | -2.21 | -1.14 | 1.9E-12 | -2.30 | -1.20 | 1.2E-13 |
| Snrpa1 | 3122 | 1.29 | 0.37 | 2.5E-03 | 1.23 | 0.30 | 1.6E-02 |
| Snta1 | 726 | 2.74 | 1.45 | 1.9E-47 | 2.38 | 1.25 | 2.7E-35 |
| Sntb1 | 280 | -2.57 | -1.36 | 4.1E-06 | -2.36 | -1.24 | 3.3E-05 |
| Sntb2 | 2564 | 1.26 | 0.33 | 5.9E-03 | 1.32 | 0.40 | 9.3E-04 |
| Snu13 | 7699 | 1.27 | 0.35 | 3.0E-04 | 1.25 | 0.33 | 6.9E-04 |
| Snupn | 1253 | 1.41 | 0.50 | 9.8E-19 | 1.39 | 0.48 | 3.6E-17 |
| Snw1 | 8694 | -1.51 | -0.60 | 6.4E-12 | -1.58 | -0.66 | 3.1E-14 |
| Snx11 | 4124 | 1.46 | 0.55 | 6.9E-11 | 1.40 | 0.49 | 7.9E-09 |
| Snx14 | 3275 | 1.73 | 0.79 | 5.1E-49 | 1.81 | 0.85 | 1.9E-56 |
| Snx15 | 982 | -1.68 | -0.75 | 5.4E-12 | -1.65 | -0.72 | 2.9E-11 |
| Snx18 | 3658 | 1.30 | 0.38 | 6.0E-04 | 1.32 | 0.40 | 3.3E-04 |
| Snx19 | 4390 | -1.29 | -0.37 | 3.7E-13 | -1.31 | -0.38 | 2.8E-14 |
| Snx21 | 779 | 1.67 | 0.74 | 9.9E-14 | 1.59 | 0.66 | 3.2E-11 |
| Snx22 | 461 | 3.08 | 1.62 | 2.9E-29 | 3.19 | 1.67 | 8.0E-31 |
| Snx24 | 897 | -1.51 | -0.60 | 1.9E-03 | -1.63 | -0.70 | 2.5E-04 |
| Snx25 | 2779 | 2.10 | 1.07 | 2.2E-78 | 2.18 | 1.12 | 5.9E-86 |
| Snx30 | 5831 | 1.98 | 0.99 | 4.0E-30 | 2.09 | 1.06 | 1.2E-34 |
| Snx32 | 2821 | 1.35 | 0.44 | 4.4E-03 | 1.40 | 0.49 | 1.3E-03 |
| Snx33 | 834 | -1.42 | -0.50 | 8.2E-05 | -1.58 | -0.66 | 1.8E-07 |
| Snx8 | 2475 | 1.76 | 0.82 | 4.1E-24 | 1.64 | 0.71 | 1.9E-18 |
| Snx9 | 1323 | -2.74 | -1.45 | 2.4E-17 | -2.99 | -1.58 | 2.7E-20 |
| Soat1 | 4864 | 2.13 | 1.09 | 6.7E-35 | 1.98 | 0.98 | 1.9E-28 |
| Socs7 | 10486 | 1.47 | 0.55 | 4.7E-05 | 1.46 | 0.55 | 5.9E-05 |
| Sod2 | 6897 | 2.26 | 1.18 | 2.9E-152 | 2.29 | 1.20 | 1.3E-156 |
| Soga3 | 5035 | -1.90 | -0.93 | 8.4E-07 | -1.83 | -0.87 | 3.9E-06 |
| Sorbs2 | 9516 | -1.96 | -0.97 | 1.8E-12 | -1.80 | -0.84 | 1.1E-09 |

| | | | | | | | |
|---|---|---|---|---|---|---|---|
| Sorbs3 | 1351 | -1.48 | -0.57 | 1.9E-03 | -1.61 | -0.69 | 1.3E-04 |
| Sorcs2 | 2917 | -1.39 | -0.47 | 5.6E-06 | -1.46 | -0.55 | 1.2E-07 |
| Sord | 2502 | 1.89 | 0.92 | 1.0E-26 | 1.74 | 0.80 | 1.2E-20 |
| Sort1 | 17489 | 1.62 | 0.70 | 5.5E-18 | 1.65 | 0.72 | 4.4E-19 |
| Sos1 | 9101 | 1.25 | 0.32 | 6.5E-03 | 1.29 | 0.37 | 2.0E-03 |
| Sostdc1 | 486 | -2.37 | -1.24 | 2.2E-03 | -1.90 | -0.93 | 2.6E-02 |
| Sowahc | 2621 | 1.66 | 0.73 | 2.3E-49 | 1.65 | 0.73 | 1.6E-48 |
| Sox12 | 10023 | -1.54 | -0.62 | 3.9E-09 | -1.62 | -0.69 | 4.8E-11 |
| Sox17 | 356 | -1.53 | -0.62 | 3.3E-05 | -1.75 | -0.80 | 4.1E-08 |
| Sox18 | 409 | -1.93 | -0.95 | 1.1E-05 | -2.13 | -1.09 | 4.2E-07 |
| Sox4 | 50721 | -1.28 | -0.36 | 3.2E-02 | -1.38 | -0.46 | 4.9E-03 |
| Sox6 | 5597 | -1.29 | -0.37 | 2.3E-02 | -1.32 | -0.40 | 1.4E-02 |
| Sox7 | 428 | -1.74 | -0.80 | 3.7E-12 | -1.89 | -0.92 | 1.7E-15 |
| Sox9 | 5279 | -1.58 | -0.66 | 2.7E-04 | -1.79 | -0.84 | 2.6E-06 |
| Sp1 | 8616 | -1.26 | -0.33 | 7.3E-07 | -1.20 | -0.27 | 7.9E-05 |
| Sp2 | 1254 | -1.49 | -0.58 | 2.2E-03 | -1.63 | -0.71 | 1.5E-04 |
| Sp3 | 9744 | -1.51 | -0.59 | 3.1E-11 | -1.49 | -0.58 | 7.0E-11 |
| Sp7 | 157 | -2.92 | -1.55 | 1.8E-03 | -3.25 | -1.70 | 5.8E-04 |
| Spaca6 | 5327 | -3.74 | -1.90 | 2.9E-63 | -3.67 | -1.87 | 2.5E-61 |
| Spag4 | 580 | 1.58 | 0.66 | 1.2E-07 | 1.44 | 0.52 | 3.7E-05 |
| Spag6l | 438 | 1.71 | 0.78 | 5.5E-04 | 2.01 | 1.01 | 5.6E-06 |
| Spag7 | 2715 | -2.17 | -1.12 | 1.1E-27 | -2.56 | -1.36 | 2.9E-40 |
| Sparc | 25277 | -1.37 | -0.46 | 2.6E-02 | -1.40 | -0.49 | 1.8E-02 |
| Sparcl1 | 6073 | -1.59 | -0.67 | 2.4E-04 | -1.48 | -0.56 | 2.5E-03 |
| Spata1 | 653 | 1.32 | 0.40 | 1.7E-03 | 1.52 | 0.60 | 1.5E-06 |
| Spata13 | 9483 | 1.51 | 0.59 | 3.8E-04 | 1.48 | 0.56 | 7.9E-04 |
| Spata2 | 2931 | 1.71 | 0.78 | 6.8E-12 | 1.56 | 0.64 | 2.3E-08 |
| Spata24 | 187 | 1.37 | 0.46 | 4.2E-03 | 1.46 | 0.55 | 5.9E-04 |
| Spata2l | 191 | 2.90 | 1.53 | 1.5E-19 | 2.82 | 1.50 | 1.0E-18 |
| Spata33 | 590 | 1.55 | 0.63 | 4.5E-10 | 1.55 | 0.63 | 6.3E-10 |
| Spata5 | 2791 | 1.56 | 0.64 | 1.0E-12 | 1.46 | 0.55 | 1.4E-09 |
| Spata6 | 2035 | 2.13 | 1.09 | 1.5E-13 | 2.11 | 1.08 | 3.0E-13 |
| Spata7 | 1542 | 1.63 | 0.71 | 1.8E-29 | 1.65 | 0.72 | 7.9E-31 |
| Spats2 | 6748 | 1.45 | 0.53 | 4.8E-13 | 1.33 | 0.41 | 3.6E-08 |
| Spats2l | 1991 | -3.12 | -1.64 | 2.4E-26 | -2.88 | -1.53 | 6.7E-23 |
| Spc24 | 2030 | -1.51 | -0.60 | 8.3E-03 | -1.74 | -0.80 | 2.9E-04 |
| Specc1 | 5673 | 1.55 | 0.63 | 1.6E-15 | 1.52 | 0.60 | 3.7E-14 |
| Specc1l | 8424 | -1.83 | -0.87 | 1.5E-40 | -1.72 | -0.78 | 2.4E-32 |
| Spef1 | 3543 | 1.42 | 0.51 | 2.5E-03 | 1.34 | 0.42 | 1.5E-02 |
| Speg | 2616 | 1.43 | 0.52 | 1.5E-03 | 1.37 | 0.45 | 6.1E-03 |
| Spem1 | 137 | 3.39 | 1.76 | 2.0E-10 | 3.30 | 1.72 | 5.4E-10 |
| Spg11 | 4046 | -1.60 | -0.67 | 7.9E-08 | -1.49 | -0.57 | 6.3E-06 |
| Spg20 | 4046 | -1.41 | -0.49 | 3.6E-04 | -1.38 | -0.47 | 7.9E-04 |
| Spg7 | 3473 | -1.55 | -0.63 | 1.3E-14 | -1.60 | -0.68 | 6.8E-17 |
| Spin1 | 15490 | -1.79 | -0.84 | 1.8E-17 | -1.89 | -0.92 | 1.0E-20 |
| Spink10 | 796 | 2.26 | 1.18 | 2.4E-15 | 2.65 | 1.40 | 2.8E-21 |
| Spire1 | 10690 | 1.51 | 0.60 | 1.9E-11 | 1.53 | 0.62 | 4.1E-12 |

| | | | | | | | |
|---|---|---|---|---|---|---|---|
| Spon2 | 113 | -2.15 | -1.11 | 3.1E-06 | -2.72 | -1.44 | 8.9E-10 |
| Spopl | 2489 | -1.38 | -0.47 | 2.3E-06 | -1.35 | -0.43 | 1.4E-05 |
| Spout1 | 3364 | 1.38 | 0.46 | 1.4E-04 | 1.26 | 0.33 | 7.4E-03 |
| Sppl2a | 5520 | -1.51 | -0.59 | 1.6E-16 | -1.44 | -0.53 | 2.3E-13 |
| Sppl2b | 3055 | 1.44 | 0.52 | 2.4E-08 | 1.35 | 0.43 | 6.5E-06 |
| Spr | 1109 | -1.31 | -0.39 | 4.1E-03 | -1.52 | -0.61 | 5.2E-06 |
| Spring1 | 4743 | 1.79 | 0.84 | 1.7E-36 | 1.85 | 0.88 | 1.4E-40 |
| Sprn | 295 | -3.01 | -1.59 | 2.4E-07 | -3.07 | -1.62 | 1.5E-07 |
| Sprtn | 3559 | 1.34 | 0.42 | 5.2E-09 | 1.24 | 0.31 | 2.8E-05 |
| Spry1 | 1203 | -1.37 | -0.45 | 2.8E-03 | -1.54 | -0.63 | 2.4E-05 |
| Spryd7 | 1753 | 1.42 | 0.51 | 4.6E-05 | 1.53 | 0.61 | 8.0E-07 |
| Spsb1 | 650 | 1.32 | 0.40 | 2.2E-02 | 1.41 | 0.50 | 3.7E-03 |
| Spsb2 | 367 | 2.55 | 1.35 | 1.4E-23 | 2.40 | 1.26 | 7.1E-21 |
| Spsb3 | 2270 | 1.35 | 0.43 | 2.3E-08 | 1.26 | 0.34 | 1.5E-05 |
| Spsb4 | 2053 | -1.50 | -0.59 | 4.0E-03 | -1.78 | -0.83 | 2.9E-05 |
| Sptbn2 | 13417 | -1.76 | -0.81 | 2.4E-04 | -1.71 | -0.77 | 5.3E-04 |
| Sptbn4 | 1642 | 1.84 | 0.88 | 1.6E-02 | 2.01 | 1.01 | 5.7E-03 |
| Sptlc2 | 8005 | 1.96 | 0.97 | 2.0E-25 | 2.11 | 1.08 | 4.7E-31 |
| Spty2d1 | 2481 | 1.37 | 0.45 | 5.4E-06 | 1.35 | 0.44 | 1.2E-05 |
| Spx | 311 | 1.66 | 0.73 | 2.6E-05 | 1.95 | 0.96 | 1.9E-08 |
| Sqle | 13030 | 1.50 | 0.59 | 1.2E-12 | 1.40 | 0.49 | 4.2E-09 |
| Sqstm1 | 10091 | 1.55 | 0.63 | 6.2E-16 | 1.58 | 0.66 | 4.2E-17 |
| Sra1 | 2348 | -1.23 | -0.30 | 1.8E-09 | -1.30 | -0.37 | 6.3E-14 |
| Src | 5937 | -2.49 | -1.32 | 4.8E-21 | -2.70 | -1.44 | 7.6E-25 |
| Srcap | 371 | -3.68 | -1.88 | 8.6E-20 | -3.26 | -1.71 | 1.9E-16 |
| Srd5a1 | 1783 | 3.14 | 1.65 | 1.4E-92 | 3.17 | 1.66 | 2.6E-93 |
| Srd5a3 | 489 | -2.21 | -1.14 | 4.8E-52 | -2.18 | -1.13 | 1.3E-50 |
| Srek1 | 9527 | -2.85 | -1.51 | 4.7E-51 | -2.87 | -1.52 | 6.0E-52 |
| Srf | 2203 | 1.31 | 0.39 | 1.6E-03 | 1.20 | 0.27 | 3.7E-02 |
| Srgap1 | 6536 | -1.90 | -0.92 | 2.5E-05 | -1.91 | -0.93 | 2.0E-05 |
| Srgap2 | 10639 | 1.42 | 0.51 | 3.8E-08 | 1.46 | 0.55 | 2.2E-09 |
| Srprb | 4548 | 1.27 | 0.35 | 4.0E-09 | 1.20 | 0.27 | 8.5E-06 |
| Srpx | 451 | -2.16 | -1.11 | 1.2E-04 | -2.01 | -1.01 | 5.7E-04 |
| Srr | 4555 | 1.42 | 0.50 | 3.0E-06 | 1.52 | 0.61 | 1.6E-08 |
| Srsf1 | 1566 | -1.36 | -0.44 | 5.6E-03 | -1.33 | -0.41 | 1.2E-02 |
| Srsf3 | 47495 | -1.30 | -0.37 | 5.4E-05 | -1.29 | -0.36 | 1.0E-04 |
| Srsf4 | 7466 | -1.34 | -0.42 | 9.7E-04 | -1.46 | -0.55 | 1.4E-05 |
| Srsf5 | 35498 | -1.28 | -0.36 | 7.4E-05 | -1.33 | -0.41 | 3.5E-06 |
| Srsf6 | 34547 | -2.03 | -1.02 | 8.1E-29 | -2.20 | -1.14 | 1.6E-35 |
| Srsf7 | 25856 | -1.48 | -0.57 | 1.2E-12 | -1.47 | -0.56 | 2.2E-12 |
| Srxn1 | 1307 | 1.87 | 0.90 | 4.2E-19 | 1.70 | 0.77 | 4.2E-14 |
| Ss18l2 | 1282 | 1.52 | 0.60 | 4.2E-11 | 1.56 | 0.64 | 1.4E-12 |
| Ssbp2 | 8311 | 1.50 | 0.59 | 1.5E-11 | 1.59 | 0.67 | 1.3E-14 |
| Ssbp4 | 2812 | 1.45 | 0.54 | 3.3E-08 | 1.41 | 0.50 | 3.3E-07 |
| Ssc5d | 536 | -1.37 | -0.46 | 3.2E-02 | -1.50 | -0.59 | 5.1E-03 |
| Ssh1 | 2434 | -1.39 | -0.47 | 5.5E-06 | -1.52 | -0.61 | 3.8E-09 |
| Ssh3 | 1943 | 1.75 | 0.81 | 7.8E-07 | 1.89 | 0.92 | 2.0E-08 |

| | | | | | | | |
|---|---|---|---|---|---|---|---|
| Sspn | 646 | -2.26 | -1.18 | 8.9E-08 | -2.36 | -1.24 | 1.8E-08 |
| Sspo | 103 | -1.75 | -0.81 | 4.7E-04 | -1.49 | -0.58 | 1.5E-02 |
| Ssr2 | 6877 | 1.29 | 0.36 | 2.9E-03 | 1.26 | 0.34 | 5.9E-03 |
| Ssrp1 | 22251 | -1.31 | -0.39 | 7.7E-05 | -1.43 | -0.52 | 8.9E-08 |
| Sst | 803 | -1.79 | -0.84 | 1.0E-02 | -1.86 | -0.90 | 6.2E-03 |
| Sstr2 | 819 | -1.84 | -0.88 | 3.9E-05 | -1.53 | -0.61 | 5.4E-03 |
| Ssx2ip | 4403 | 1.86 | 0.89 | 4.9E-18 | 1.70 | 0.77 | 1.4E-13 |
| St18 | 3717 | -1.99 | -0.99 | 7.2E-10 | -1.98 | -0.99 | 9.8E-10 |
| St3gal4 | 965 | -1.39 | -0.48 | 4.1E-04 | -1.54 | -0.62 | 3.2E-06 |
| St3gal5 | 2703 | 1.36 | 0.45 | 9.4E-09 | 1.34 | 0.42 | 5.6E-08 |
| St3gal6 | 437 | -1.56 | -0.64 | 2.8E-07 | -1.68 | -0.74 | 2.4E-09 |
| St6gal1 | 3911 | -1.95 | -0.97 | 5.1E-62 | -2.04 | -1.03 | 2.1E-70 |
| St6galnac4 | 1628 | 2.17 | 1.12 | 4.5E-26 | 2.22 | 1.15 | 1.9E-27 |
| St6galnac6 | 1758 | 2.32 | 1.21 | 7.9E-31 | 2.30 | 1.20 | 3.2E-30 |
| St7 | 2111 | -1.57 | -0.65 | 5.2E-08 | -1.50 | -0.58 | 1.5E-06 |
| St7l | 1691 | -1.40 | -0.49 | 8.8E-12 | -1.31 | -0.39 | 8.5E-08 |
| St8sia1 | 4148 | -1.62 | -0.70 | 3.3E-04 | -1.45 | -0.54 | 7.1E-03 |
| St8sia2 | 9960 | -1.84 | -0.88 | 8.8E-05 | -1.91 | -0.94 | 3.1E-05 |
| St8sia3 | 1765 | -4.17 | -2.06 | 4.5E-18 | -4.14 | -2.05 | 7.0E-18 |
| St8sia5 | 157 | 2.54 | 1.35 | 4.2E-06 | 2.48 | 1.31 | 7.5E-06 |
| Stab1 | 4245 | -1.34 | -0.42 | 3.2E-02 | -1.41 | -0.49 | 1.0E-02 |
| Stac | 129 | -1.90 | -0.92 | 2.4E-04 | -1.91 | -0.94 | 2.1E-04 |
| Stac2 | 1114 | 5.32 | 2.41 | 1.8E-34 | 5.17 | 2.37 | 2.3E-33 |
| Stac3 | 490 | -1.44 | -0.52 | 1.6E-02 | -1.39 | -0.47 | 3.2E-02 |
| Stag1 | 8175 | -2.42 | -1.28 | 1.3E-51 | -2.34 | -1.23 | 7.8E-48 |
| Stag2 | 13202 | -1.55 | -0.63 | 5.0E-08 | -1.39 | -0.47 | 6.7E-05 |
| Stag3 | 965 | 1.65 | 0.72 | 4.9E-06 | 1.63 | 0.71 | 7.6E-06 |
| Stam2 | 5072 | 1.24 | 0.31 | 6.5E-06 | 1.26 | 0.34 | 1.3E-06 |
| Stambpl1 | 339 | 1.51 | 0.60 | 6.9E-04 | 1.72 | 0.78 | 7.4E-06 |
| Star | 329 | 2.69 | 1.43 | 1.1E-25 | 2.70 | 1.43 | 7.9E-26 |
| Stard10 | 671 | 1.64 | 0.72 | 4.9E-06 | 1.72 | 0.78 | 5.9E-07 |
| Stard4 | 5817 | 1.52 | 0.60 | 1.6E-07 | 1.51 | 0.59 | 2.4E-07 |
| Stard5 | 1341 | 2.68 | 1.42 | 1.0E-35 | 2.66 | 1.41 | 5.0E-35 |
| Stat1 | 690 | 1.88 | 0.91 | 2.1E-23 | 1.90 | 0.93 | 2.6E-24 |
| Stat2 | 3380 | 1.76 | 0.81 | 3.0E-31 | 1.80 | 0.85 | 3.9E-34 |
| Stat3 | 5850 | 2.72 | 1.45 | 1.2E-44 | 2.69 | 1.43 | 1.7E-43 |
| Stat6 | 763 | -1.75 | -0.80 | 4.6E-05 | -1.80 | -0.85 | 1.5E-05 |
| Stau2 | 4911 | -1.37 | -0.45 | 6.4E-05 | -1.41 | -0.49 | 1.4E-05 |
| Stbd1 | 629 | 2.08 | 1.06 | 4.7E-10 | 2.13 | 1.09 | 1.5E-10 |
| Stc1 | 534 | 1.35 | 0.43 | 8.9E-03 | 1.35 | 0.44 | 9.0E-03 |
| Stc2 | 494 | 2.83 | 1.50 | 2.1E-23 | 2.55 | 1.35 | 4.8E-19 |
| Steap1 | 140 | 3.02 | 1.59 | 7.9E-18 | 2.69 | 1.43 | 1.3E-14 |
| Stil | 4088 | 1.35 | 0.43 | 1.6E-02 | 1.30 | 0.38 | 3.5E-02 |
| Stim1 | 2915 | 2.24 | 1.16 | 1.2E-35 | 2.26 | 1.18 | 1.5E-36 |
| Stim2 | 3047 | 1.41 | 0.50 | 3.3E-08 | 1.39 | 0.48 | 1.1E-07 |
| Stimate | 1907 | 1.49 | 0.58 | 8.0E-22 | 1.44 | 0.53 | 1.9E-18 |
| Sting1 | 128 | -2.33 | -1.22 | 2.8E-06 | -2.14 | -1.10 | 2.9E-05 |

| | | | | | | | |
|---|---|---|---|---|---|---|---|
| Stip1 | 21805 | 1.39 | 0.47 | 2.3E-06 | 1.25 | 0.32 | 1.6E-03 |
| Stk10 | 845 | 1.81 | 0.85 | 9.9E-14 | 1.77 | 0.82 | 9.0E-13 |
| Stk17b | 1393 | 1.95 | 0.97 | 1.1E-15 | 1.98 | 0.99 | 2.4E-16 |
| Stk25 | 13624 | 1.25 | 0.32 | 1.3E-17 | 1.22 | 0.28 | 6.3E-14 |
| Stk38 | 13352 | 1.29 | 0.37 | 5.0E-10 | 1.23 | 0.30 | 7.9E-07 |
| Stk38l | 2637 | 1.94 | 0.96 | 6.0E-32 | 2.01 | 1.00 | 6.1E-35 |
| Stk4 | 8426 | 1.54 | 0.62 | 2.6E-16 | 1.52 | 0.60 | 1.9E-15 |
| Stk40 | 5297 | 1.73 | 0.79 | 4.3E-15 | 1.65 | 0.73 | 6.4E-13 |
| Stn1 | 1701 | -1.60 | -0.67 | 1.7E-13 | -1.65 | -0.72 | 3.5E-15 |
| Stom | 1955 | 2.89 | 1.53 | 7.3E-28 | 2.64 | 1.40 | 1.8E-23 |
| Ston2 | 4113 | 1.62 | 0.69 | 5.3E-07 | 1.67 | 0.74 | 7.6E-08 |
| Stra6 | 924 | -1.96 | -0.97 | 1.0E-07 | -2.12 | -1.09 | 1.9E-09 |
| Strip2 | 678 | 4.36 | 2.12 | 6.3E-41 | 4.77 | 2.25 | 1.1E-45 |
| Strn3 | 7456 | -1.57 | -0.65 | 5.6E-16 | -1.50 | -0.59 | 3.7E-13 |
| Strn4 | 10352 | -1.37 | -0.45 | 9.0E-06 | -1.42 | -0.50 | 8.7E-07 |
| Stub1 | 4959 | 1.25 | 0.32 | 1.0E-04 | 1.16 | 0.21 | 1.2E-02 |
| Stum | 563 | 2.23 | 1.16 | 1.7E-04 | 2.57 | 1.36 | 8.9E-06 |
| Stx11 | 282 | 2.71 | 1.44 | 1.8E-09 | 2.70 | 1.43 | 2.1E-09 |
| Stx16 | 7226 | -1.32 | -0.40 | 2.0E-05 | -1.36 | -0.45 | 2.0E-06 |
| Stx18 | 1175 | -1.59 | -0.67 | 1.1E-09 | -1.68 | -0.75 | 9.8E-12 |
| Stx1a | 2912 | 1.38 | 0.46 | 5.2E-03 | 1.36 | 0.45 | 7.0E-03 |
| Stx2 | 3037 | 1.41 | 0.50 | 1.4E-15 | 1.44 | 0.52 | 8.4E-17 |
| Stx3 | 1545 | 1.40 | 0.48 | 7.5E-05 | 1.38 | 0.46 | 1.6E-04 |
| Stx6 | 7454 | 1.45 | 0.53 | 7.1E-24 | 1.45 | 0.54 | 4.6E-24 |
| Stx7 | 9746 | 1.38 | 0.47 | 1.1E-04 | 1.40 | 0.48 | 6.2E-05 |
| Stxbp4 | 3633 | -1.54 | -0.62 | 2.9E-16 | -1.49 | -0.57 | 5.0E-14 |
| Stxbp5 | 4749 | 1.46 | 0.54 | 3.0E-06 | 1.50 | 0.58 | 4.8E-07 |
| Stxbp5l | 612 | 2.27 | 1.18 | 6.0E-04 | 2.96 | 1.56 | 3.9E-06 |
| Stxbp6 | 1105 | -1.50 | -0.58 | 4.1E-08 | -1.63 | -0.70 | 2.7E-11 |
| Styx | 2624 | 1.33 | 0.42 | 3.1E-05 | 1.41 | 0.50 | 4.9E-07 |
| Sub1 | 17595 | 1.26 | 0.33 | 7.5E-06 | 1.29 | 0.37 | 4.6E-07 |
| Suclg1 | 5144 | 1.45 | 0.53 | 6.0E-14 | 1.43 | 0.51 | 5.3E-13 |
| Suclg2 | 2080 | -1.39 | -0.48 | 3.0E-03 | -1.50 | -0.59 | 2.4E-04 |
| Suds3 | 10665 | 1.60 | 0.68 | 5.3E-27 | 1.55 | 0.64 | 7.9E-24 |
| Sulf2 | 12815 | -1.49 | -0.57 | 7.7E-03 | -1.50 | -0.59 | 6.3E-03 |
| Sult5a1 | 138 | -2.24 | -1.16 | 2.0E-02 | -2.08 | -1.06 | 3.7E-02 |
| Sumf1 | 1092 | -1.34 | -0.43 | 9.9E-05 | -1.45 | -0.53 | 8.5E-07 |
| Sumf2 | 1500 | -1.24 | -0.31 | 2.9E-03 | -1.25 | -0.33 | 1.5E-03 |
| Sumo3 | 6505 | -1.39 | -0.48 | 4.8E-09 | -1.51 | -0.59 | 3.1E-13 |
| Sun1 | 7751 | 1.68 | 0.75 | 3.6E-33 | 1.70 | 0.77 | 1.0E-34 |
| Suox | 1368 | 2.18 | 1.13 | 2.5E-27 | 2.08 | 1.05 | 4.9E-24 |
| Supt3 | 750 | -1.50 | -0.59 | 1.3E-07 | -1.59 | -0.67 | 1.7E-09 |
| Supt5 | 16134 | 1.42 | 0.51 | 6.9E-18 | 1.33 | 0.41 | 4.1E-12 |
| Supt6 | 16116 | 1.29 | 0.37 | 4.9E-08 | 1.29 | 0.37 | 6.9E-08 |
| Supt7l | 1799 | -1.72 | -0.78 | 1.8E-25 | -1.84 | -0.88 | 5.0E-32 |
| Supv3l1 | 1953 | -1.23 | -0.30 | 2.4E-05 | -1.29 | -0.37 | 1.8E-07 |
| Susd1 | 208 | -1.50 | -0.59 | 2.9E-03 | -1.54 | -0.62 | 1.7E-03 |

| | | | | | | | |
|---|---|---|---|---|---|---|---|
| Susd2 | 309 | 1.35 | 0.43 | 1.9E-03 | 1.35 | 0.43 | 1.9E-03 |
| Susd5 | 292 | -9.88 | -3.30 | 4.4E-05 | -7.69 | -2.94 | 3.2E-04 |
| Susd6 | 3279 | 1.35 | 0.43 | 3.3E-13 | 1.26 | 0.34 | 2.1E-08 |
| Svip | 2706 | 1.92 | 0.94 | 1.6E-23 | 2.02 | 1.02 | 1.8E-27 |
| Swap70 | 2701 | 1.35 | 0.43 | 9.4E-07 | 1.32 | 0.40 | 5.4E-06 |
| Swsap1 | 912 | -1.67 | -0.74 | 4.3E-16 | -1.93 | -0.95 | 4.7E-26 |
| Swt1 | 2572 | 1.71 | 0.78 | 9.7E-17 | 1.81 | 0.86 | 3.6E-20 |
| Syap1 | 4114 | 1.57 | 0.65 | 3.4E-09 | 1.54 | 0.62 | 1.7E-08 |
| Sybu | 2165 | -1.37 | -0.45 | 4.5E-02 | -1.44 | -0.53 | 1.7E-02 |
| Syce2 | 1215 | 1.57 | 0.65 | 4.0E-07 | 1.46 | 0.55 | 2.5E-05 |
| Syde1 | 1619 | -1.47 | -0.55 | 7.3E-04 | -1.56 | -0.64 | 8.3E-05 |
| Syde2 | 887 | 1.65 | 0.72 | 4.0E-23 | 1.77 | 0.83 | 8.3E-30 |
| Syn2 | 1795 | 1.63 | 0.70 | 3.8E-02 | 1.78 | 0.83 | 1.3E-02 |
| Syncrip | 22048 | -1.48 | -0.57 | 7.5E-10 | -1.51 | -0.60 | 7.8E-11 |
| Syndig1 | 346 | -1.37 | -0.46 | 1.4E-04 | -1.38 | -0.46 | 1.4E-04 |
| Syne3 | 316 | 2.76 | 1.47 | 2.0E-05 | 3.53 | 1.82 | 9.7E-08 |
| Syne4 | 892 | 1.47 | 0.56 | 1.5E-03 | 1.62 | 0.70 | 6.0E-05 |
| Syngr1 | 2628 | 2.29 | 1.20 | 3.0E-06 | 2.30 | 1.20 | 2.8E-06 |
| Syngr2 | 829 | 2.30 | 1.20 | 5.9E-14 | 2.12 | 1.08 | 1.6E-11 |
| Synj1 | 9528 | 1.46 | 0.55 | 1.1E-03 | 1.54 | 0.62 | 1.8E-04 |
| Synj2 | 310 | -1.81 | -0.86 | 6.6E-03 | -1.74 | -0.80 | 1.2E-02 |
| Synj2bp | 3736 | 1.39 | 0.48 | 3.3E-10 | 1.44 | 0.52 | 4.5E-12 |
| Synpo | 2894 | -2.02 | -1.01 | 7.6E-03 | -1.91 | -0.93 | 1.5E-02 |
| Synpo2l | 136 | -18.20 | -4.19 | 1.3E-06 | -8.63 | -3.11 | 4.2E-04 |
| Synrg | 6993 | -1.33 | -0.41 | 7.1E-04 | -1.33 | -0.41 | 8.1E-04 |
| Syt11 | 19138 | -1.32 | -0.40 | 1.2E-04 | -1.40 | -0.48 | 2.6E-06 |
| Syt14 | 4701 | 1.35 | 0.44 | 2.7E-04 | 1.47 | 0.55 | 2.9E-06 |
| Syt5 | 1215 | -1.84 | -0.88 | 9.1E-05 | -1.96 | -0.97 | 1.5E-05 |
| T2 | 116 | 3.23 | 1.69 | 5.5E-15 | 3.92 | 1.97 | 1.5E-19 |
| Tab1 | 2078 | -2.65 | -1.41 | 1.6E-92 | -2.86 | -1.52 | 7.8E-107 |
| Tab2 | 9166 | -1.37 | -0.45 | 2.1E-25 | -1.40 | -0.48 | 2.6E-28 |
| Tacc2 | 7124 | -1.50 | -0.58 | 5.9E-06 | -1.52 | -0.60 | 2.5E-06 |
| Tada1 | 3144 | -1.55 | -0.63 | 3.8E-14 | -1.51 | -0.59 | 1.5E-12 |
| Tada3 | 2064 | -2.02 | -1.01 | 9.4E-14 | -2.28 | -1.19 | 1.3E-18 |
| Taf1 | 11862 | 1.26 | 0.33 | 3.9E-04 | 1.36 | 0.44 | 1.7E-06 |
| Taf10 | 4280 | 1.68 | 0.75 | 2.2E-16 | 1.55 | 0.64 | 5.0E-12 |
| Taf12 | 2366 | 1.44 | 0.53 | 4.4E-09 | 1.30 | 0.38 | 3.9E-05 |
| Taf13 | 5379 | 1.74 | 0.80 | 1.2E-11 | 1.73 | 0.79 | 1.7E-11 |
| Taf3 | 2425 | 1.48 | 0.56 | 1.1E-09 | 1.40 | 0.48 | 2.1E-07 |
| Taf4 | 3798 | 1.37 | 0.45 | 4.0E-04 | 1.28 | 0.35 | 7.5E-03 |
| Taf4b | 421 | 2.47 | 1.30 | 4.9E-10 | 2.51 | 1.33 | 2.3E-10 |
| Taf5 | 2229 | 1.25 | 0.32 | 4.0E-05 | 1.18 | 0.24 | 2.7E-03 |
| Taf5l | 1612 | -1.82 | -0.86 | 3.2E-24 | -1.98 | -0.99 | 1.4E-31 |
| Taf6 | 5567 | 1.28 | 0.35 | 8.3E-05 | 1.18 | 0.24 | 1.1E-02 |
| Taf6l | 1731 | 1.65 | 0.72 | 4.3E-15 | 1.54 | 0.62 | 1.9E-11 |
| Taf8 | 2115 | -1.20 | -0.26 | 4.5E-03 | -1.29 | -0.37 | 4.4E-05 |
| Taf9b | 529 | -4.59 | -2.20 | 6.7E-91 | -4.96 | -2.31 | 7.1E-100 |

| | | | | | | | |
|---|---|---|---|---|---|---|---|
| Tafa4 | 313 | 2.52 | 1.33 | 2.3E-08 | 2.73 | 1.45 | 9.6E-10 |
| Tagap1 | 2327 | -1.27 | -0.34 | 7.2E-08 | -1.33 | -0.41 | 6.4E-11 |
| Tagln2 | 2484 | 1.26 | 0.34 | 1.4E-02 | 1.28 | 0.36 | 9.7E-03 |
| Tagln3 | 7661 | -1.92 | -0.94 | 4.5E-11 | -1.87 | -0.90 | 3.3E-10 |
| Tal1 | 2195 | -2.22 | -1.15 | 4.6E-05 | -2.33 | -1.22 | 1.5E-05 |
| Tal2 | 3441 | -2.40 | -1.26 | 2.2E-03 | -2.80 | -1.48 | 2.9E-04 |
| Taldo1 | 8784 | 1.27 | 0.35 | 4.4E-04 | 1.24 | 0.31 | 2.2E-03 |
| Tamalin | 845 | 1.62 | 0.70 | 1.9E-05 | 1.46 | 0.55 | 9.3E-04 |
| Tamm41 | 1643 | 1.25 | 0.33 | 4.8E-04 | 1.20 | 0.26 | 6.6E-03 |
| Tanc1 | 3209 | 1.30 | 0.38 | 4.3E-04 | 1.33 | 0.41 | 1.3E-04 |
| Tango6 | 546 | -3.09 | -1.63 | 1.7E-42 | -2.99 | -1.58 | 6.9E-40 |
| Tank | 2874 | 1.70 | 0.76 | 1.2E-23 | 1.82 | 0.87 | 4.5E-30 |
| Taok1 | 12512 | -1.49 | -0.57 | 4.8E-06 | -1.36 | -0.44 | 4.8E-04 |
| Taok3 | 3732 | 1.95 | 0.96 | 1.8E-14 | 1.89 | 0.91 | 3.9E-13 |
| Tap1 | 311 | 3.23 | 1.69 | 3.3E-21 | 3.06 | 1.61 | 2.6E-19 |
| Tap2 | 168 | 4.79 | 2.26 | 1.1E-28 | 4.79 | 2.26 | 1.2E-28 |
| Tapbpl | 112 | -2.39 | -1.26 | 6.0E-09 | -2.30 | -1.20 | 3.8E-08 |
| Tarbp2 | 2167 | -2.29 | -1.19 | 3.3E-44 | -2.53 | -1.34 | 2.9E-55 |
| Tars | 7300 | 1.40 | 0.48 | 1.0E-04 | 1.37 | 0.45 | 2.7E-04 |
| Tars2 | 2372 | -1.99 | -0.99 | 1.2E-45 | -2.02 | -1.01 | 2.2E-47 |
| Tarsl2 | 1149 | -1.48 | -0.57 | 1.9E-20 | -1.40 | -0.49 | 1.9E-15 |
| Tas1r3 | 123 | 1.94 | 0.95 | 2.4E-06 | 1.82 | 0.86 | 2.6E-05 |
| Tasor | 6356 | -1.51 | -0.59 | 1.8E-07 | -1.48 | -0.57 | 7.3E-07 |
| Tasor2 | 13914 | 1.22 | 0.28 | 4.3E-02 | 1.40 | 0.49 | 3.0E-04 |
| Tatdn2 | 3628 | 1.65 | 0.72 | 9.1E-20 | 1.64 | 0.71 | 1.7E-19 |
| Tatdn3 | 667 | -2.25 | -1.17 | 1.5E-30 | -2.13 | -1.09 | 1.2E-26 |
| Tax1bp3 | 1992 | -1.37 | -0.45 | 6.4E-10 | -1.43 | -0.51 | 1.7E-12 |
| Tbc1d1 | 4443 | 1.49 | 0.58 | 9.6E-16 | 1.53 | 0.61 | 2.9E-17 |
| Tbc1d10a | 1112 | 1.30 | 0.38 | 2.9E-04 | 1.28 | 0.36 | 8.5E-04 |
| Tbc1d10b | 6167 | 1.44 | 0.52 | 5.1E-06 | 1.37 | 0.46 | 8.4E-05 |
| Tbc1d12 | 2436 | 1.39 | 0.47 | 3.0E-06 | 1.51 | 0.60 | 2.3E-09 |
| Tbc1d15 | 4447 | 1.21 | 0.27 | 9.3E-03 | 1.26 | 0.34 | 1.1E-03 |
| Tbc1d16 | 25429 | 1.40 | 0.49 | 5.3E-04 | 1.35 | 0.43 | 2.6E-03 |
| Tbc1d2 | 253 | 2.95 | 1.56 | 1.4E-21 | 3.24 | 1.70 | 4.9E-25 |
| Tbc1d20 | 4949 | 1.41 | 0.50 | 4.8E-17 | 1.31 | 0.39 | 8.0E-11 |
| Tbc1d22a | 981 | -1.43 | -0.52 | 4.4E-12 | -1.40 | -0.49 | 1.1E-10 |
| Tbc1d24 | 9564 | 2.00 | 1.00 | 5.3E-12 | 2.11 | 1.08 | 7.6E-14 |
| Tbc1d2b | 4400 | 1.71 | 0.77 | 4.0E-09 | 1.77 | 0.82 | 4.1E-10 |
| Tbc1d31 | 3810 | 2.35 | 1.23 | 1.3E-34 | 2.40 | 1.26 | 3.0E-36 |
| Tbc1d32 | 1172 | -1.53 | -0.61 | 1.1E-09 | -1.43 | -0.51 | 4.0E-07 |
| Tbc1d4 | 1461 | -1.37 | -0.46 | 7.0E-05 | -1.30 | -0.38 | 1.1E-03 |
| Tbc1d5 | 1509 | -1.60 | -0.68 | 6.8E-26 | -1.59 | -0.67 | 4.1E-25 |
| Tbc1d7 | 1095 | -2.06 | -1.04 | 2.0E-54 | -2.07 | -1.05 | 1.1E-55 |
| Tbc1d8 | 1328 | 1.74 | 0.80 | 4.4E-09 | 1.72 | 0.78 | 9.4E-09 |
| Tbc1d8b | 1293 | 1.22 | 0.28 | 1.0E-02 | 1.37 | 0.45 | 2.6E-05 |
| Tbc1d9 | 7374 | -1.42 | -0.51 | 2.0E-03 | -1.42 | -0.50 | 2.3E-03 |
| Tbc1d9b | 4080 | -1.79 | -0.84 | 9.7E-33 | -1.77 | -0.83 | 1.8E-31 |

| | | | | | | | |
|---|---|---|---|---|---|---|---|
| Tbcd | 6672 | 1.34 | 0.43 | 4.7E-11 | 1.33 | 0.41 | 2.4E-10 |
| Tbce | 4473 | -2.27 | -1.18 | 7.2E-41 | -2.16 | -1.11 | 4.1E-36 |
| Tbkbp1 | 2381 | -2.03 | -1.02 | 4.5E-10 | -1.96 | -0.97 | 3.1E-09 |
| Tbl1x | 11400 | -1.64 | -0.72 | 2.6E-32 | -1.68 | -0.75 | 4.3E-35 |
| Tbl1xr1 | 4323 | -1.42 | -0.51 | 2.5E-07 | -1.57 | -0.65 | 2.4E-11 |
| Tbl2 | 4923 | 2.12 | 1.08 | 8.0E-42 | 2.10 | 1.07 | 8.8E-41 |
| Tbl3 | 4350 | 1.42 | 0.51 | 1.1E-05 | 1.29 | 0.37 | 1.7E-03 |
| Tbp | 4856 | 1.26 | 0.34 | 1.6E-06 | 1.28 | 0.36 | 2.3E-07 |
| Tbr1 | 3672 | -2.68 | -1.42 | 2.6E-05 | -2.85 | -1.51 | 7.8E-06 |
| Tbx18 | 1274 | -1.50 | -0.58 | 6.7E-04 | -1.54 | -0.62 | 2.8E-04 |
| Tbx2 | 608 | -2.76 | -1.46 | 4.7E-03 | -2.92 | -1.55 | 2.8E-03 |
| Tbx22 | 155 | -4.84 | -2.27 | 3.4E-02 | -8.10 | -3.02 | 4.5E-03 |
| Tbx6 | 134 | 1.73 | 0.79 | 1.4E-04 | 1.92 | 0.94 | 4.9E-06 |
| Tcaf1 | 29441 | -1.33 | -0.42 | 2.9E-05 | -1.31 | -0.39 | 1.0E-04 |
| Tcaf2 | 1145 | 2.14 | 1.10 | 3.5E-09 | 2.23 | 1.16 | 4.5E-10 |
| Tcea2 | 1056 | -1.42 | -0.51 | 3.6E-06 | -1.47 | -0.56 | 3.6E-07 |
| Tceal1 | 996 | -1.82 | -0.86 | 6.2E-13 | -1.66 | -0.73 | 1.7E-09 |
| Tceal3 | 1740 | 1.34 | 0.42 | 4.1E-04 | 1.35 | 0.43 | 2.8E-04 |
| Tceal5 | 1149 | 2.01 | 1.00 | 2.3E-11 | 2.03 | 1.02 | 1.0E-11 |
| Tceal6 | 893 | 2.08 | 1.06 | 9.2E-08 | 2.22 | 1.15 | 5.8E-09 |
| Tceal8 | 4977 | -1.53 | -0.62 | 1.7E-11 | -1.45 | -0.54 | 5.4E-09 |
| Tceanc | 839 | -1.32 | -0.41 | 2.9E-04 | -1.36 | -0.44 | 7.6E-05 |
| Tcf15 | 112 | 3.03 | 1.60 | 8.3E-13 | 2.79 | 1.48 | 5.0E-11 |
| Tcf19 | 3238 | 2.44 | 1.29 | 1.0E-17 | 2.27 | 1.19 | 3.7E-15 |
| Tcf3 | 10239 | -1.46 | -0.55 | 1.1E-04 | -1.65 | -0.72 | 2.5E-07 |
| Tcf4 | 24203 | -1.55 | -0.63 | 9.7E-07 | -1.48 | -0.56 | 1.4E-05 |
| Tcf7 | 1392 | 2.97 | 1.57 | 1.5E-35 | 2.65 | 1.41 | 8.8E-29 |
| Tcf7l2 | 10030 | -1.73 | -0.79 | 1.7E-02 | -2.47 | -1.30 | 5.4E-05 |
| Tcfl5 | 227 | 4.40 | 2.14 | 1.4E-37 | 4.61 | 2.21 | 6.2E-39 |
| Tchh | 280 | 3.37 | 1.75 | 2.3E-32 | 3.13 | 1.65 | 1.3E-28 |
| Tchp | 2927 | 1.64 | 0.72 | 4.3E-09 | 1.50 | 0.59 | 1.9E-06 |
| Tcirg1 | 1639 | 1.51 | 0.60 | 2.6E-03 | 1.58 | 0.66 | 7.7E-04 |
| Tcn2 | 1780 | 1.29 | 0.37 | 1.0E-02 | 1.30 | 0.38 | 9.7E-03 |
| Tcp11 | 255 | 3.60 | 1.85 | 5.0E-49 | 3.67 | 1.88 | 4.7E-50 |
| Tcp11l2 | 2118 | 2.96 | 1.56 | 4.7E-46 | 3.11 | 1.64 | 2.2E-50 |
| Tcta | 1161 | -1.60 | -0.67 | 1.6E-17 | -1.61 | -0.68 | 7.3E-18 |
| Tcte2 | 326 | -1.56 | -0.64 | 2.0E-06 | -1.46 | -0.55 | 5.4E-05 |
| Tctn1 | 4106 | 1.47 | 0.55 | 1.8E-08 | 1.41 | 0.49 | 6.7E-07 |
| Tctn3 | 1997 | 1.94 | 0.96 | 6.3E-22 | 1.99 | 1.00 | 1.9E-23 |
| Tdg | 9125 | -1.52 | -0.61 | 6.6E-09 | -1.65 | -0.73 | 2.4E-12 |
| Tdrd3 | 3615 | -1.74 | -0.80 | 2.8E-39 | -1.77 | -0.83 | 8.2E-42 |
| Tdrd6 | 284 | 5.62 | 2.49 | 7.3E-22 | 7.97 | 3.00 | 1.8E-30 |
| Tdrd7 | 2046 | 1.92 | 0.94 | 1.4E-30 | 1.93 | 0.95 | 8.3E-31 |
| Tdrd9 | 110 | -1.41 | -0.50 | 9.9E-03 | -1.41 | -0.49 | 1.0E-02 |
| Tdrkh | 6712 | 1.38 | 0.47 | 3.2E-05 | 1.39 | 0.48 | 2.2E-05 |
| Tdrp | 666 | 2.40 | 1.26 | 1.5E-32 | 2.01 | 1.01 | 3.0E-21 |
| Tead2 | 10451 | -2.19 | -1.13 | 2.5E-12 | -2.47 | -1.30 | 5.5E-16 |

| | | | | | | | |
|---|---|---|---|---|---|---|---|
| Tead3 | 3179 | 1.74 | 0.80 | 8.7E-10 | 1.69 | 0.75 | 6.8E-09 |
| Tec | 603 | 1.95 | 0.96 | 1.6E-09 | 2.03 | 1.02 | 1.4E-10 |
| Tecpr1 | 6097 | 1.52 | 0.60 | 1.1E-05 | 1.48 | 0.56 | 4.3E-05 |
| Tecpr2 | 3397 | 1.26 | 0.33 | 1.6E-03 | 1.31 | 0.39 | 1.4E-04 |
| Tedc1 | 2765 | 1.89 | 0.92 | 5.2E-09 | 1.60 | 0.68 | 2.2E-05 |
| Tedc2 | 1770 | 1.42 | 0.51 | 1.5E-05 | 1.33 | 0.41 | 5.9E-04 |
| Tef | 4931 | 1.35 | 0.43 | 6.2E-10 | 1.36 | 0.44 | 4.1E-10 |
| Tefm | 624 | -1.55 | -0.63 | 8.6E-09 | -1.60 | -0.68 | 3.7E-10 |
| Tek | 1451 | -1.54 | -0.62 | 3.8E-13 | -1.58 | -0.66 | 1.3E-14 |
| Tekt1 | 346 | 1.31 | 0.39 | 6.8E-04 | 1.61 | 0.69 | 8.4E-10 |
| Tekt2 | 414 | -1.39 | -0.48 | 1.4E-03 | -1.37 | -0.45 | 2.8E-03 |
| Telo2 | 2941 | -1.23 | -0.30 | 6.1E-05 | -1.28 | -0.35 | 1.2E-06 |
| Ten1 | 841 | -1.97 | -0.98 | 3.5E-44 | -2.02 | -1.01 | 5.6E-47 |
| Tent4b | 7568 | 1.90 | 0.93 | 1.3E-35 | 1.91 | 0.94 | 2.4E-36 |
| Tent5b | 139 | 8.14 | 3.03 | 5.0E-23 | 6.42 | 2.68 | 2.0E-18 |
| Tep1 | 820 | -1.91 | -0.94 | 1.1E-11 | -1.78 | -0.83 | 2.5E-09 |
| Tepsin | 1722 | 1.43 | 0.52 | 6.7E-07 | 1.31 | 0.39 | 2.4E-04 |
| Terf1 | 1692 | -1.23 | -0.30 | 1.7E-02 | -1.26 | -0.33 | 7.5E-03 |
| Terf2ip | 4521 | 1.56 | 0.64 | 1.5E-18 | 1.49 | 0.57 | 5.1E-15 |
| Tert | 1252 | -1.80 | -0.85 | 8.9E-21 | -1.83 | -0.87 | 9.5E-22 |
| Tesc | 162 | 2.90 | 1.54 | 9.1E-05 | 2.75 | 1.46 | 2.2E-04 |
| Tesk2 | 871 | 2.79 | 1.48 | 4.2E-47 | 2.62 | 1.39 | 1.4E-41 |
| Tet2 | 6298 | -1.41 | -0.50 | 3.8E-05 | -1.30 | -0.38 | 2.0E-03 |
| Tet3 | 11236 | -1.39 | -0.48 | 1.7E-03 | -1.50 | -0.59 | 8.5E-05 |
| Tex10 | 5225 | 1.42 | 0.51 | 8.4E-07 | 1.46 | 0.55 | 7.7E-08 |
| Tex2 | 3136 | 1.83 | 0.87 | 8.5E-11 | 1.87 | 0.90 | 1.7E-11 |
| Tex261 | 4276 | 1.26 | 0.33 | 2.1E-06 | 1.22 | 0.29 | 4.0E-05 |
| Tex264 | 1405 | 1.32 | 0.40 | 2.0E-03 | 1.41 | 0.50 | 1.1E-04 |
| Tex30 | 1244 | 1.80 | 0.85 | 3.6E-08 | 1.80 | 0.85 | 3.7E-08 |
| Tex9 | 5404 | 1.50 | 0.58 | 5.3E-08 | 1.53 | 0.62 | 7.7E-09 |
| Tfap2a | 954 | -1.85 | -0.89 | 2.8E-03 | -1.71 | -0.77 | 1.1E-02 |
| Tfap2b | 7460 | -2.81 | -1.49 | 7.0E-09 | -2.51 | -1.33 | 2.8E-07 |
| Tfap2d | 599 | -3.40 | -1.76 | 1.9E-07 | -3.22 | -1.69 | 6.4E-07 |
| Tfap4 | 800 | -1.69 | -0.76 | 1.8E-05 | -1.93 | -0.95 | 5.7E-08 |
| Tfcp2l1 | 356 | 5.44 | 2.44 | 1.6E-33 | 6.36 | 2.67 | 4.3E-39 |
| Tfe3 | 3803 | 1.90 | 0.92 | 1.7E-30 | 1.74 | 0.80 | 3.4E-23 |
| Tfeb | 402 | 4.50 | 2.17 | 1.7E-31 | 4.30 | 2.11 | 1.3E-29 |
| Tfr2 | 490 | 3.16 | 1.66 | 3.4E-13 | 3.77 | 1.91 | 4.0E-17 |
| Tfrc | 13483 | 1.24 | 0.31 | 3.9E-02 | 1.30 | 0.38 | 1.1E-02 |
| Tgfb1i1 | 1065 | -2.93 | -1.55 | 8.5E-11 | -3.25 | -1.70 | 9.1E-13 |
| Tgfb2 | 2234 | -2.52 | -1.33 | 4.1E-13 | -2.34 | -1.23 | 2.9E-11 |
| Tgfb3 | 1115 | -2.03 | -1.02 | 1.7E-04 | -1.96 | -0.97 | 4.0E-04 |
| Tgfbr2 | 1914 | -2.46 | -1.30 | 1.1E-11 | -2.46 | -1.30 | 1.1E-11 |
| Tgfbr3l | 409 | -1.45 | -0.54 | 2.0E-04 | -1.66 | -0.73 | 2.7E-07 |
| Tgif2 | 4559 | -1.21 | -0.28 | 3.3E-02 | -1.33 | -0.41 | 1.5E-03 |
| Tgm2 | 1296 | -1.33 | -0.41 | 2.9E-02 | -1.37 | -0.45 | 1.5E-02 |
| Tgoln1 | 7088 | 1.73 | 0.79 | 3.0E-12 | 1.82 | 0.87 | 1.4E-14 |

| | | | | | | | |
|---|---|---|---|---|---|---|---|
| Th | 476 | -1.42 | -0.50 | 5.1E-03 | -1.52 | -0.61 | 6.6E-04 |
| Tha1 | 235 | 1.77 | 0.82 | 1.8E-06 | 1.45 | 0.53 | 2.6E-03 |
| Thap1 | 750 | -1.48 | -0.57 | 1.9E-06 | -1.54 | -0.62 | 1.4E-07 |
| Thap11 | 2637 | 1.40 | 0.49 | 7.1E-08 | 1.32 | 0.40 | 1.4E-05 |
| Thap2 | 3393 | -1.27 | -0.35 | 6.7E-06 | -1.23 | -0.30 | 1.1E-04 |
| Thap3 | 1000 | -1.25 | -0.33 | 1.8E-02 | -1.39 | -0.48 | 3.8E-04 |
| Thap4 | 1759 | 1.35 | 0.44 | 3.3E-12 | 1.28 | 0.35 | 3.2E-08 |
| Thap7 | 780 | -1.63 | -0.70 | 2.5E-14 | -1.66 | -0.73 | 2.5E-15 |
| Thbd | 1311 | -1.57 | -0.65 | 2.5E-02 | -1.79 | -0.84 | 3.4E-03 |
| Thbs1 | 7121 | -1.91 | -0.93 | 1.8E-02 | -1.90 | -0.93 | 1.9E-02 |
| Thbs3 | 4764 | -1.30 | -0.38 | 3.1E-05 | -1.34 | -0.42 | 2.5E-06 |
| Them6 | 238 | 2.75 | 1.46 | 6.2E-27 | 2.55 | 1.35 | 5.8E-23 |
| Thg1l | 1015 | -1.94 | -0.96 | 1.6E-12 | -2.16 | -1.11 | 2.3E-16 |
| Thnsl1 | 2243 | 1.45 | 0.54 | 2.0E-07 | 1.52 | 0.60 | 5.9E-09 |
| Thnsl2 | 150 | -1.33 | -0.41 | 2.2E-02 | -1.32 | -0.40 | 2.5E-02 |
| Thoc1 | 7063 | -1.34 | -0.42 | 1.4E-07 | -1.29 | -0.36 | 7.2E-06 |
| Thoc6 | 1517 | -1.43 | -0.52 | 1.9E-06 | -1.56 | -0.64 | 2.8E-09 |
| Thoc7 | 5327 | -1.49 | -0.58 | 2.5E-09 | -1.58 | -0.66 | 1.1E-11 |
| Thop1 | 4068 | 1.56 | 0.64 | 2.3E-10 | 1.46 | 0.55 | 8.6E-08 |
| Thpo | 102 | 1.68 | 0.75 | 9.4E-07 | 1.62 | 0.70 | 7.5E-06 |
| Thsd4 | 2133 | -1.67 | -0.74 | 1.0E-07 | -1.47 | -0.56 | 7.3E-05 |
| Thtpa | 821 | -1.29 | -0.37 | 1.1E-04 | -1.40 | -0.49 | 2.7E-07 |
| Thumpd2 | 586 | -1.34 | -0.42 | 3.8E-06 | -1.38 | -0.47 | 2.5E-07 |
| Thy1 | 176 | 2.63 | 1.39 | 3.3E-02 | 3.47 | 1.79 | 5.3E-03 |
| Tia1 | 23494 | -2.17 | -1.12 | 1.4E-17 | -1.92 | -0.94 | 9.8E-13 |
| Ticam1 | 491 | 1.80 | 0.85 | 8.4E-08 | 1.65 | 0.72 | 6.5E-06 |
| Ticrr | 3962 | 1.53 | 0.62 | 3.3E-04 | 1.45 | 0.54 | 2.1E-03 |
| Tifa | 328 | 1.41 | 0.50 | 3.0E-04 | 1.42 | 0.50 | 2.6E-04 |
| Tigar | 2483 | 1.91 | 0.93 | 3.7E-22 | 1.79 | 0.84 | 2.6E-18 |
| Tigd5 | 857 | 1.51 | 0.59 | 7.7E-08 | 1.44 | 0.53 | 2.3E-06 |
| Timeless | 3945 | -1.52 | -0.60 | 4.4E-05 | -1.67 | -0.74 | 4.6E-07 |
| Timm10 | 972 | 1.27 | 0.35 | 2.4E-04 | 1.20 | 0.26 | 7.9E-03 |
| Timm13 | 2145 | -1.34 | -0.43 | 1.1E-05 | -1.40 | -0.49 | 4.6E-07 |
| Timm21 | 2194 | -1.69 | -0.76 | 1.4E-14 | -1.71 | -0.78 | 2.4E-15 |
| Timm44 | 3363 | -1.28 | -0.35 | 2.7E-09 | -1.36 | -0.44 | 5.5E-14 |
| Timm8a1 | 1048 | 1.38 | 0.47 | 5.1E-05 | 1.37 | 0.45 | 8.4E-05 |
| Timm9 | 2038 | 1.28 | 0.36 | 2.9E-07 | 1.28 | 0.36 | 4.2E-07 |
| Timmdc1 | 2330 | 1.26 | 0.33 | 1.1E-04 | 1.17 | 0.22 | 1.3E-02 |
| Timp1 | 160 | -1.60 | -0.68 | 1.7E-02 | -2.05 | -1.03 | 1.9E-04 |
| Timp2 | 5730 | -1.69 | -0.76 | 1.6E-08 | -1.77 | -0.83 | 7.1E-10 |
| Tincr | 434 | 2.42 | 1.28 | 1.4E-11 | 1.99 | 0.99 | 1.9E-07 |
| Tinf2 | 3256 | 1.45 | 0.54 | 1.4E-04 | 1.31 | 0.39 | 6.9E-03 |
| Tiparp | 3539 | 1.73 | 0.79 | 5.6E-25 | 1.81 | 0.86 | 2.5E-29 |
| Tiprl | 5126 | 1.29 | 0.37 | 2.0E-13 | 1.26 | 0.33 | 4.0E-11 |
| Tjp2 | 4312 | 1.53 | 0.62 | 3.6E-06 | 1.34 | 0.43 | 1.8E-03 |
| Tk1 | 2067 | -2.32 | -1.21 | 4.3E-13 | -2.54 | -1.35 | 7.3E-16 |
| Tk2 | 2477 | 1.46 | 0.54 | 3.4E-16 | 1.39 | 0.48 | 1.1E-12 |

| | | | | | | | |
|---|---|---|---|---|---|---|---|
| Tkfc | 2031 | -1.33 | -0.41 | 3.4E-05 | -1.41 | -0.49 | 7.2E-07 |
| Tlcd1 | 692 | 2.04 | 1.03 | 1.3E-22 | 2.07 | 1.05 | 2.3E-23 |
| Tlcd2 | 162 | -1.96 | -0.97 | 7.5E-04 | -2.22 | -1.15 | 5.9E-05 |
| Tlcd3a | 1830 | 1.33 | 0.41 | 1.2E-12 | 1.28 | 0.36 | 9.1E-10 |
| Tlcd5 | 2588 | 1.45 | 0.53 | 7.9E-17 | 1.38 | 0.46 | 4.8E-13 |
| f | 6244 | -1.29 | -0.37 | 1.1E-09 | -1.32 | -0.40 | 1.5E-11 |
| Tlk1 | 7690 | -1.45 | -0.53 | 1.3E-08 | -1.48 | -0.56 | 1.9E-09 |
| Tlk2 | 6144 | -1.42 | -0.50 | 1.2E-14 | -1.44 | -0.52 | 8.3E-16 |
| Tll1 | 321 | -1.80 | -0.85 | 2.7E-05 | -1.80 | -0.85 | 3.1E-05 |
| Tlr12 | 106 | -1.66 | -0.73 | 2.5E-03 | -1.54 | -0.62 | 1.1E-02 |
| Tlr3 | 167 | -1.75 | -0.81 | 1.2E-10 | -1.95 | -0.96 | 2.6E-14 |
| Tm2d1 | 1284 | -1.57 | -0.65 | 3.0E-10 | -1.54 | -0.63 | 1.7E-09 |
| Tm2d2 | 2855 | 1.50 | 0.59 | 3.6E-16 | 1.53 | 0.61 | 1.2E-17 |
| Tm4sf1 | 458 | -2.37 | -1.25 | 3.0E-11 | -2.28 | -1.19 | 2.3E-10 |
| Tm6sf2 | 365 | 1.78 | 0.83 | 4.1E-05 | 1.79 | 0.84 | 3.5E-05 |
| Tm9sf4 | 4866 | -1.34 | -0.42 | 2.6E-21 | -1.36 | -0.44 | 1.1E-23 |
| Tmbim1 | 479 | -2.36 | -1.24 | 2.4E-09 | -2.46 | -1.30 | 3.7E-10 |
| Tmbim4 | 1587 | 1.43 | 0.51 | 1.5E-07 | 1.39 | 0.48 | 1.4E-06 |
| Tmbim6 | 15557 | 1.28 | 0.36 | 3.2E-05 | 1.21 | 0.27 | 2.2E-03 |
| Tmc4 | 1438 | 3.08 | 1.62 | 2.6E-25 | 3.14 | 1.65 | 3.9E-26 |
| Tmc6 | 570 | 1.65 | 0.73 | 4.0E-03 | 1.65 | 0.72 | 4.3E-03 |
| Tmcc3 | 1229 | 1.53 | 0.61 | 4.4E-06 | 1.57 | 0.65 | 8.9E-07 |
| Tmco6 | 1031 | -1.23 | -0.30 | 1.8E-03 | -1.36 | -0.44 | 2.7E-06 |
| Tmed1 | 924 | -3.04 | -1.60 | 6.1E-38 | -3.00 | -1.59 | 3.3E-37 |
| Tmed2 | 8416 | 1.31 | 0.39 | 2.5E-06 | 1.26 | 0.34 | 6.9E-05 |
| Tmed3 | 1087 | -1.92 | -0.94 | 1.8E-16 | -1.85 | -0.89 | 9.4E-15 |
| Tmed5 | 3902 | 1.68 | 0.75 | 4.3E-11 | 1.63 | 0.70 | 6.5E-10 |
| Tmed8 | 4148 | 1.63 | 0.70 | 1.3E-14 | 1.81 | 0.86 | 4.7E-21 |
| Tmeff1 | 16520 | -1.31 | -0.39 | 9.2E-04 | -1.26 | -0.34 | 4.1E-03 |
| Tmem104 | 883 | 1.53 | 0.61 | 5.7E-08 | 1.44 | 0.52 | 4.0E-06 |
| Tmem106a | 133 | -1.43 | -0.52 | 6.2E-04 | -1.36 | -0.44 | 3.8E-03 |
| Tmem107 | 724 | 1.60 | 0.68 | 5.5E-09 | 1.49 | 0.57 | 1.1E-06 |
| Tmem108 | 1225 | 1.53 | 0.62 | 7.9E-06 | 1.37 | 0.45 | 1.4E-03 |
| Tmem109 | 1668 | -1.37 | -0.45 | 7.2E-05 | -1.49 | -0.58 | 3.1E-07 |
| Tmem115 | 2107 | 1.60 | 0.68 | 2.3E-14 | 1.56 | 0.64 | 7.2E-13 |
| Tmem119 | 865 | -2.30 | -1.20 | 1.1E-02 | -2.38 | -1.25 | 8.4E-03 |
| Tmem120a | 2053 | 2.25 | 1.17 | 4.1E-65 | 2.23 | 1.16 | 1.4E-63 |
| Tmem120b | 817 | 2.39 | 1.26 | 5.2E-32 | 2.54 | 1.35 | 1.6E-36 |
| Tmem121 | 963 | 2.43 | 1.28 | 1.5E-30 | 2.21 | 1.15 | 1.4E-24 |
| Tmem123 | 7806 | 1.55 | 0.63 | 1.6E-03 | 1.48 | 0.56 | 5.3E-03 |
| Tmem126a | 1475 | -1.29 | -0.36 | 1.2E-04 | -1.35 | -0.43 | 4.3E-06 |
| Tmem128 | 2360 | -1.28 | -0.35 | 3.9E-05 | -1.28 | -0.36 | 2.6E-05 |
| Tmem129 | 1905 | 1.58 | 0.66 | 1.7E-11 | 1.55 | 0.63 | 1.3E-10 |
| Tmem131l | 11569 | -1.30 | -0.37 | 9.3E-04 | -1.28 | -0.36 | 1.6E-03 |
| Tmem132a | 13999 | -1.26 | -0.33 | 2.6E-02 | -1.36 | -0.44 | 2.2E-03 |
| Tmem132e | 827 | 2.19 | 1.13 | 3.6E-07 | 2.41 | 1.27 | 1.0E-08 |
| Tmem135 | 2395 | 1.42 | 0.51 | 2.1E-13 | 1.41 | 0.50 | 5.7E-13 |

| | | | | | | | |
|---|---|---|---|---|---|---|---|
| Tmem138 | 1260 | -1.24 | -0.31 | 1.6E-03 | -1.32 | -0.40 | 4.3E-05 |
| Tmem141 | 325 | -1.53 | -0.61 | 1.2E-07 | -1.38 | -0.47 | 7.5E-05 |
| Tmem143 | 493 | -1.52 | -0.60 | 9.8E-08 | -1.44 | -0.53 | 3.9E-06 |
| Tmem144 | 224 | -1.37 | -0.45 | 2.0E-02 | -1.47 | -0.56 | 3.7E-03 |
| Tmem14a | 285 | -3.62 | -1.86 | 5.0E-41 | -2.84 | -1.50 | 6.4E-28 |
| Tmem14c | 3505 | 1.25 | 0.32 | 2.4E-04 | 1.19 | 0.25 | 5.4E-03 |
| Tmem150c | 916 | -1.64 | -0.72 | 3.2E-03 | -1.60 | -0.68 | 5.4E-03 |
| Tmem151a | 211 | 2.01 | 1.01 | 6.8E-05 | 1.86 | 0.89 | 4.7E-04 |
| Tmem151b | 3587 | -1.45 | -0.54 | 2.4E-03 | -1.53 | -0.61 | 5.1E-04 |
| Tmem159 | 1138 | 2.65 | 1.41 | 1.3E-22 | 2.58 | 1.37 | 2.1E-21 |
| Tmem160 | 587 | 1.71 | 0.77 | 6.2E-14 | 1.60 | 0.68 | 4.7E-11 |
| Tmem161b | 2984 | 1.42 | 0.50 | 1.9E-07 | 1.38 | 0.47 | 1.4E-06 |
| Tmem163 | 1496 | -1.62 | -0.69 | 1.4E-05 | -1.68 | -0.75 | 2.4E-06 |
| Tmem168 | 2511 | -1.59 | -0.67 | 2.6E-13 | -1.57 | -0.65 | 9.5E-13 |
| Tmem17 | 291 | -3.46 | -1.79 | 2.5E-33 | -3.28 | -1.71 | 9.2E-31 |
| Tmem170 | 1331 | 1.58 | 0.66 | 1.1E-11 | 1.48 | 0.57 | 6.0E-09 |
| Tmem170b | 5482 | 1.28 | 0.35 | 2.5E-02 | 1.27 | 0.35 | 2.7E-02 |
| Tmem175 | 1180 | -1.70 | -0.77 | 2.3E-23 | -1.68 | -0.75 | 4.0E-22 |
| Tmem176b | 2642 | -1.42 | -0.51 | 2.4E-07 | -1.38 | -0.47 | 2.6E-06 |
| Tmem178 | 919 | -1.79 | -0.84 | 4.5E-04 | -1.77 | -0.82 | 6.2E-04 |
| Tmem181a | 3817 | 1.38 | 0.46 | 1.7E-19 | 1.41 | 0.50 | 1.3E-22 |
| Tmem183a | 10585 | 1.26 | 0.33 | 2.0E-08 | 1.25 | 0.32 | 5.1E-08 |
| Tmem184b | 4161 | 1.52 | 0.61 | 1.6E-07 | 1.48 | 0.56 | 1.3E-06 |
| Tmem184c | 5364 | 1.57 | 0.65 | 8.6E-38 | 1.64 | 0.71 | 8.7E-46 |
| Tmem185b | 922 | -1.86 | -0.89 | 1.9E-37 | -1.92 | -0.94 | 2.4E-41 |
| Tmem186 | 2246 | -1.44 | -0.52 | 1.1E-14 | -1.49 | -0.57 | 2.1E-17 |
| Tmem189 | 1696 | 2.27 | 1.18 | 8.8E-66 | 2.28 | 1.19 | 5.1E-66 |
| Tmem19 | 2803 | 1.46 | 0.55 | 2.7E-36 | 1.38 | 0.47 | 1.4E-26 |
| Tmem192 | 490 | 1.63 | 0.70 | 1.3E-10 | 1.68 | 0.75 | 8.2E-12 |
| Tmem198 | 1269 | 1.90 | 0.93 | 5.2E-09 | 1.81 | 0.85 | 8.3E-08 |
| Tmem200a | 1466 | -2.26 | -1.17 | 3.1E-18 | -2.42 | -1.28 | 2.3E-21 |
| Tmem209 | 3624 | -1.62 | -0.69 | 1.1E-14 | -1.61 | -0.69 | 2.0E-14 |
| Tmem218 | 1080 | 1.54 | 0.62 | 1.0E-07 | 1.56 | 0.64 | 3.2E-08 |
| Tmem223 | 2143 | -1.94 | -0.95 | 1.8E-19 | -1.86 | -0.89 | 3.7E-17 |
| Tmem229a | 328 | -1.93 | -0.95 | 2.0E-08 | -1.84 | -0.88 | 2.0E-07 |
| Tmem230 | 1607 | -1.70 | -0.77 | 5.8E-28 | -1.66 | -0.73 | 2.8E-25 |
| Tmem231 | 1605 | 1.41 | 0.50 | 3.6E-08 | 1.29 | 0.37 | 6.4E-05 |
| Tmem241 | 646 | 1.49 | 0.57 | 1.3E-07 | 1.54 | 0.63 | 7.3E-09 |
| Tmem242 | 3130 | -1.26 | -0.34 | 9.9E-04 | -1.31 | -0.39 | 1.3E-04 |
| Tmem243 | 590 | 2.02 | 1.02 | 2.3E-14 | 2.02 | 1.01 | 3.2E-14 |
| Tmem245 | 7677 | 1.27 | 0.34 | 1.9E-02 | 1.47 | 0.56 | 9.3E-05 |
| Tmem25 | 503 | 1.33 | 0.41 | 1.9E-03 | 1.39 | 0.48 | 3.2E-04 |
| Tmem250-ps | 1407 | -1.87 | -0.91 | 8.9E-25 | -2.05 | -1.04 | 4.2E-32 |
| Tmem256 | 2625 | 1.31 | 0.39 | 3.3E-03 | 1.28 | 0.35 | 8.1E-03 |
| Tmem259 | 5207 | 1.25 | 0.32 | 7.9E-04 | 1.19 | 0.25 | 1.0E-02 |
| Tmem263 | 5944 | 1.54 | 0.62 | 1.4E-15 | 1.50 | 0.58 | 8.1E-14 |
| Tmem266 | 344 | 1.92 | 0.94 | 1.2E-09 | 1.79 | 0.84 | 7.9E-08 |

| | | | | | | | |
|---|---|---|---|---|---|---|---|
| Tmem269 | 149 | 3.84 | 1.94 | 1.4E-24 | 4.45 | 2.15 | 2.6E-29 |
| Tmem28 | 1042 | 2.08 | 1.06 | 1.3E-12 | 1.85 | 0.89 | 3.5E-09 |
| Tmem29 | 1659 | -1.77 | -0.82 | 1.8E-11 | -1.57 | -0.65 | 1.3E-07 |
| Tmem30a | 13736 | 1.29 | 0.36 | 6.8E-05 | 1.35 | 0.43 | 1.6E-06 |
| Tmem38a | 1566 | 1.67 | 0.74 | 2.7E-10 | 1.69 | 0.75 | 1.6E-10 |
| Tmem38b | 1178 | 1.89 | 0.92 | 1.0E-19 | 1.73 | 0.79 | 6.8E-15 |
| Tmem39a | 2452 | -1.44 | -0.52 | 2.8E-08 | -1.39 | -0.47 | 7.4E-07 |
| Tmem39b | 1217 | -2.03 | -1.02 | 9.6E-28 | -2.26 | -1.18 | 2.6E-36 |
| Tmem41a | 1127 | 2.05 | 1.03 | 5.9E-68 | 2.06 | 1.04 | 2.0E-68 |
| Tmem41b | 3997 | 1.41 | 0.50 | 4.2E-08 | 1.47 | 0.56 | 6.7E-10 |
| Tmem42 | 124 | -3.49 | -1.80 | 1.5E-32 | -4.20 | -2.07 | 5.9E-41 |
| Tmem43 | 4788 | 1.26 | 0.33 | 1.8E-06 | 1.16 | 0.21 | 3.4E-03 |
| Tmem44 | 5837 | -2.26 | -1.18 | 4.5E-09 | -2.18 | -1.12 | 2.7E-08 |
| Tmem45a | 330 | -1.94 | -0.95 | 1.5E-03 | -1.73 | -0.79 | 9.8E-03 |
| Tmem50a | 4115 | 1.35 | 0.43 | 1.2E-17 | 1.31 | 0.38 | 2.7E-14 |
| Tmem50b | 4482 | 2.93 | 1.55 | 1.0E-167 | 2.98 | 1.58 | 3.3E-173 |
| Tmem60 | 1396 | 1.44 | 0.52 | 8.9E-09 | 1.51 | 0.59 | 7.3E-11 |
| Tmem62 | 474 | -2.40 | -1.26 | 3.9E-18 | -2.11 | -1.08 | 1.4E-13 |
| Tmem64 | 3264 | 1.33 | 0.41 | 2.4E-02 | 1.36 | 0.44 | 1.4E-02 |
| Tmem68 | 2272 | -1.88 | -0.91 | 1.9E-24 | -1.83 | -0.87 | 1.6E-22 |
| Tmem69 | 1429 | -1.30 | -0.38 | 9.8E-05 | -1.28 | -0.35 | 2.8E-04 |
| Tmem74 | 805 | 1.44 | 0.52 | 5.6E-03 | 1.64 | 0.71 | 1.2E-04 |
| Tmem74b | 367 | -2.35 | -1.23 | 2.8E-05 | -2.31 | -1.21 | 4.3E-05 |
| Tmem80 | 1603 | -2.48 | -1.31 | 2.3E-41 | -2.66 | -1.41 | 1.1E-47 |
| Tmem94 | 3930 | -1.81 | -0.85 | 2.5E-12 | -1.94 | -0.96 | 2.7E-15 |
| Tmem98 | 1432 | -2.20 | -1.14 | 7.8E-12 | -2.45 | -1.29 | 7.2E-15 |
| Tmod3 | 11387 | 1.69 | 0.76 | 3.3E-27 | 1.75 | 0.80 | 2.4E-30 |
| Tmpo | 33304 | -1.49 | -0.57 | 3.8E-07 | -1.56 | -0.64 | 1.4E-08 |
| Tmppe | 504 | 1.43 | 0.51 | 2.4E-04 | 1.42 | 0.51 | 3.1E-04 |
| Tmprss5 | 115 | -1.93 | -0.95 | 1.0E-11 | -1.85 | -0.89 | 2.2E-10 |
| Tmprss7 | 133 | 1.33 | 0.41 | 2.4E-02 | 1.48 | 0.57 | 1.4E-03 |
| Tmtc3 | 4476 | 1.24 | 0.31 | 6.0E-03 | 1.31 | 0.39 | 3.9E-04 |
| Tmub1 | 805 | 1.24 | 0.31 | 4.1E-04 | 1.25 | 0.33 | 2.6E-04 |
| Tnc | 5966 | -2.21 | -1.14 | 2.4E-03 | -2.33 | -1.22 | 1.2E-03 |
| Tnfaip3 | 379 | 1.47 | 0.56 | 2.1E-04 | 1.59 | 0.67 | 7.2E-06 |
| Tnfaip8l1 | 579 | 1.50 | 0.59 | 8.8E-09 | 1.72 | 0.78 | 1.4E-14 |
| Tnfrsf10b | 720 | 2.16 | 1.11 | 2.0E-07 | 2.24 | 1.16 | 4.9E-08 |
| Tnfrsf12a | 297 | 4.65 | 2.22 | 1.9E-26 | 5.05 | 2.34 | 7.9E-29 |
| Tnfrsf1a | 1675 | 1.39 | 0.48 | 4.8E-03 | 1.36 | 0.44 | 9.7E-03 |
| Tnfrsf22 | 910 | 1.93 | 0.95 | 1.0E-12 | 1.78 | 0.83 | 4.5E-10 |
| Tnfrsf23 | 150 | 2.41 | 1.27 | 9.3E-10 | 2.73 | 1.45 | 2.6E-12 |
| Tnfrsf25 | 334 | 1.85 | 0.89 | 1.3E-05 | 1.91 | 0.93 | 4.3E-06 |
| Tnfsf9 | 235 | 1.61 | 0.69 | 1.6E-06 | 1.65 | 0.72 | 6.0E-07 |
| Tnip1 | 891 | -1.47 | -0.56 | 9.3E-06 | -1.42 | -0.51 | 6.2E-05 |
| Tnip2 | 620 | 1.74 | 0.80 | 3.4E-17 | 1.65 | 0.72 | 3.4E-14 |
| Tnk2 | 7612 | 1.43 | 0.52 | 1.6E-04 | 1.32 | 0.41 | 3.8E-03 |
| Tnks1bp1 | 7812 | 1.39 | 0.47 | 1.9E-06 | 1.38 | 0.46 | 3.4E-06 |

| | | | | | | | |
|---|---|---|---|---|---|---|---|
| Tnnc1 | 242 | -7.62 | -2.93 | 6.2E-04 | -5.15 | -2.37 | 6.8E-03 |
| Tnni1 | 196 | -15.77 | -3.98 | 3.2E-04 | -8.98 | -3.17 | 5.0E-03 |
| Tnni3 | 499 | 5.18 | 2.37 | 7.1E-32 | 4.15 | 2.05 | 4.3E-24 |
| Tnnt1 | 415 | -4.83 | -2.27 | 5.6E-04 | -3.03 | -1.60 | 1.8E-02 |
| Tnnt2 | 197 | -15.34 | -3.94 | 2.4E-04 | -9.35 | -3.22 | 3.1E-03 |
| Tnnt3 | 136 | -12.07 | -3.59 | 3.7E-05 | -7.48 | -2.90 | 1.1E-03 |
| Tnrc18 | 15512 | -2.32 | -1.21 | 4.4E-15 | -2.46 | -1.30 | 4.6E-17 |
| Tnrc6a | 14170 | -1.40 | -0.49 | 9.5E-08 | -1.26 | -0.34 | 3.3E-04 |
| Tnrc6b | 8717 | -1.41 | -0.49 | 1.8E-06 | -1.32 | -0.40 | 1.3E-04 |
| Tns1 | 4010 | -1.62 | -0.70 | 7.5E-06 | -1.54 | -0.63 | 6.4E-05 |
| Tns2 | 1379 | -3.34 | -1.74 | 4.5E-19 | -3.35 | -1.74 | 3.7E-19 |
| Tns3 | 4656 | -2.53 | -1.34 | 1.0E-13 | -2.40 | -1.26 | 2.6E-12 |
| Tob2 | 2108 | 1.45 | 0.54 | 2.1E-04 | 1.33 | 0.41 | 6.1E-03 |
| Toe1 | 1228 | -1.74 | -0.80 | 2.8E-21 | -1.88 | -0.91 | 2.7E-27 |
| Togaram1 | 4769 | 1.31 | 0.39 | 5.1E-06 | 1.44 | 0.53 | 6.0E-10 |
| Tom1 | 557 | 1.48 | 0.57 | 4.3E-06 | 1.40 | 0.49 | 9.4E-05 |
| Tom1l2 | 7710 | 1.84 | 0.88 | 3.2E-15 | 1.95 | 0.97 | 5.5E-18 |
| Tomm20 | 10877 | 1.27 | 0.34 | 1.2E-05 | 1.26 | 0.33 | 2.3E-05 |
| Tomm34 | 2832 | 1.40 | 0.49 | 1.7E-11 | 1.36 | 0.44 | 1.3E-09 |
| Tomm40 | 3723 | 1.34 | 0.42 | 9.8E-05 | 1.25 | 0.32 | 4.0E-03 |
| Tomm40l | 2031 | -1.30 | -0.38 | 1.1E-10 | -1.32 | -0.40 | 6.7E-12 |
| Top3a | 2357 | 1.71 | 0.77 | 1.3E-20 | 1.66 | 0.73 | 1.2E-18 |
| Top3b | 3624 | -1.49 | -0.58 | 1.6E-43 | -1.49 | -0.58 | 2.3E-43 |
| Topors | 4472 | -1.36 | -0.45 | 7.6E-12 | -1.38 | -0.47 | 8.7E-13 |
| Tor1aip1 | 2580 | -2.27 | -1.18 | 2.9E-46 | -2.28 | -1.19 | 6.6E-47 |
| Tor2a | 1529 | -1.47 | -0.56 | 9.6E-15 | -1.47 | -0.56 | 9.5E-15 |
| Tor3a | 472 | 2.04 | 1.03 | 6.1E-18 | 2.07 | 1.05 | 1.2E-18 |
| Tox | 2097 | -2.62 | -1.39 | 2.5E-31 | -2.82 | -1.50 | 3.1E-36 |
| Tox2 | 2504 | 2.01 | 1.01 | 1.9E-10 | 1.60 | 0.68 | 2.5E-05 |
| Tox3 | 4651 | -1.50 | -0.59 | 9.0E-04 | -1.67 | -0.74 | 2.1E-05 |
| Tox4 | 9426 | 1.40 | 0.49 | 6.6E-24 | 1.37 | 0.46 | 5.0E-21 |
| Tpcn1 | 4301 | 1.64 | 0.71 | 3.0E-08 | 1.63 | 0.71 | 4.1E-08 |
| Tpgs1 | 646 | -2.39 | -1.26 | 1.8E-26 | -2.51 | -1.33 | 4.0E-29 |
| Tpgs2 | 5489 | 1.27 | 0.34 | 6.3E-06 | 1.31 | 0.38 | 4.0E-07 |
| Tpk1 | 309 | -2.19 | -1.13 | 1.9E-08 | -2.07 | -1.05 | 2.2E-07 |
| Tpm4 | 20925 | -1.48 | -0.57 | 5.3E-04 | -1.57 | -0.65 | 7.1E-05 |
| Tpmt | 2079 | 1.82 | 0.86 | 3.8E-15 | 1.80 | 0.85 | 1.3E-14 |
| Tpp2 | 10807 | 1.47 | 0.55 | 1.8E-09 | 1.52 | 0.61 | 3.6E-11 |
| Tppp | 1479 | 2.03 | 1.02 | 2.9E-18 | 2.19 | 1.13 | 3.2E-22 |
| Tpra1 | 1797 | 1.36 | 0.44 | 1.3E-07 | 1.38 | 0.46 | 3.1E-08 |
| Tprkb | 1978 | -1.24 | -0.31 | 1.5E-05 | -1.32 | -0.40 | 2.0E-08 |
| Tprn | 719 | 1.84 | 0.88 | 2.4E-15 | 1.72 | 0.79 | 2.1E-12 |
| Tpst1 | 1950 | -1.28 | -0.36 | 1.0E-07 | -1.32 | -0.40 | 1.9E-09 |
| Tpst2 | 1187 | 2.20 | 1.14 | 9.6E-27 | 2.00 | 1.00 | 6.5E-21 |
| Tra2b | 21745 | -1.19 | -0.25 | 1.5E-02 | -1.25 | -0.33 | 1.3E-03 |
| Trabd | 3320 | 1.41 | 0.49 | 9.7E-07 | 1.30 | 0.38 | 2.5E-04 |
| Trabd2b | 672 | -2.05 | -1.04 | 1.1E-05 | -2.03 | -1.02 | 1.6E-05 |

| | | | | | | | |
|---|---|---|---|---|---|---|---|
| Tradd | 444 | 2.26 | 1.17 | 2.6E-21 | 1.99 | 0.99 | 1.4E-15 |
| Traf3 | 7313 | 1.40 | 0.49 | 1.2E-07 | 1.45 | 0.53 | 7.0E-09 |
| Traf3ip1 | 2398 | 1.34 | 0.43 | 1.1E-04 | 1.19 | 0.24 | 3.4E-02 |
| Traf4 | 5983 | -1.48 | -0.56 | 2.9E-14 | -1.54 | -0.63 | 2.0E-17 |
| Traf6 | 4656 | 1.46 | 0.55 | 2.5E-07 | 1.53 | 0.61 | 6.7E-09 |
| Traf7 | 9194 | 1.42 | 0.51 | 3.8E-11 | 1.32 | 0.41 | 1.8E-07 |
| Trafd1 | 7127 | 1.28 | 0.36 | 1.1E-07 | 1.24 | 0.31 | 4.3E-06 |
| Trak1 | 8329 | 1.32 | 0.40 | 8.7E-03 | 1.25 | 0.32 | 4.3E-02 |
| Tram2 | 2411 | 1.75 | 0.80 | 2.1E-08 | 1.77 | 0.83 | 8.3E-09 |
| Trappc1 | 5418 | 1.37 | 0.45 | 1.5E-06 | 1.37 | 0.45 | 1.5E-06 |
| Trappc11 | 7708 | 1.50 | 0.59 | 1.7E-15 | 1.55 | 0.63 | 1.0E-17 |
| Trappc12 | 3137 | -1.27 | -0.35 | 3.9E-12 | -1.26 | -0.33 | 4.2E-11 |
| Trappc14 | 4197 | 1.32 | 0.40 | 5.2E-04 | 1.25 | 0.32 | 6.1E-03 |
| Trappc6b | 2765 | -1.19 | -0.24 | 1.6E-02 | -1.27 | -0.34 | 6.3E-04 |
| Triap1 | 1459 | 1.50 | 0.59 | 2.5E-08 | 1.28 | 0.35 | 1.1E-03 |
| Trib2 | 8489 | -1.73 | -0.79 | 3.2E-07 | -1.73 | -0.79 | 3.4E-07 |
| Tril | 2635 | -2.05 | -1.03 | 2.2E-04 | -2.28 | -1.19 | 1.7E-05 |
| Trim11 | 2350 | 1.42 | 0.50 | 2.2E-08 | 1.32 | 0.40 | 1.3E-05 |
| Trim12a | 257 | -1.79 | -0.84 | 1.3E-03 | -1.55 | -0.63 | 1.9E-02 |
| Trim16 | 549 | -1.63 | -0.70 | 2.8E-05 | -1.44 | -0.53 | 2.1E-03 |
| Trim17 | 137 | 1.91 | 0.93 | 3.0E-03 | 2.14 | 1.10 | 4.6E-04 |
| Trim21 | 244 | -1.57 | -0.65 | 3.2E-04 | -1.57 | -0.65 | 3.9E-04 |
| Trim24 | 7933 | -1.57 | -0.65 | 2.7E-15 | -1.60 | -0.68 | 1.1E-16 |
| Trim27 | 5638 | 1.32 | 0.40 | 2.1E-05 | 1.22 | 0.29 | 2.8E-03 |
| Trim3 | 4883 | 1.35 | 0.43 | 3.0E-05 | 1.36 | 0.44 | 2.2E-05 |
| Trim30a | 122 | -1.75 | -0.80 | 1.9E-04 | -1.56 | -0.64 | 3.3E-03 |
| Trim33 | 11213 | -1.29 | -0.37 | 6.9E-05 | -1.24 | -0.31 | 9.5E-04 |
| Trim34a | 103 | -2.69 | -1.43 | 8.2E-12 | -2.83 | -1.50 | 3.5E-13 |
| Trim37 | 17170 | 1.40 | 0.48 | 1.0E-09 | 1.38 | 0.46 | 5.4E-09 |
| Trim39 | 4419 | -1.84 | -0.88 | 1.1E-26 | -1.83 | -0.87 | 3.2E-26 |
| Trim41 | 4917 | -1.25 | -0.32 | 2.7E-03 | -1.35 | -0.43 | 3.2E-05 |
| Trim56 | 1628 | -1.49 | -0.58 | 7.3E-08 | -1.45 | -0.54 | 5.6E-07 |
| Trim59 | 8176 | -1.32 | -0.40 | 8.7E-03 | -1.44 | -0.52 | 4.2E-04 |
| Trim6 | 158 | 2.30 | 1.20 | 7.4E-04 | 1.74 | 0.80 | 3.0E-02 |
| Trim62 | 3421 | 2.11 | 1.08 | 3.4E-23 | 2.13 | 1.09 | 8.0E-24 |
| Trim65 | 221 | -2.00 | -1.00 | 4.2E-09 | -2.39 | -1.26 | 1.4E-13 |
| Trim68 | 784 | -1.43 | -0.52 | 8.8E-05 | -1.35 | -0.44 | 1.2E-03 |
| Trim71 | 313 | 2.10 | 1.07 | 2.4E-03 | 1.66 | 0.73 | 4.5E-02 |
| Trim8 | 3106 | -1.76 | -0.82 | 4.6E-08 | -2.04 | -1.03 | 3.6E-12 |
| Trim9 | 4582 | 1.38 | 0.46 | 2.5E-02 | 1.44 | 0.53 | 9.5E-03 |
| Triobp | 5590 | 1.48 | 0.56 | 6.3E-06 | 1.39 | 0.47 | 1.8E-04 |
| Trip10 | 2161 | 2.56 | 1.36 | 3.9E-39 | 2.36 | 1.24 | 8.8E-33 |
| Trip11 | 4453 | -1.40 | -0.48 | 5.4E-06 | -1.32 | -0.40 | 2.3E-04 |
| Trip13 | 1751 | -1.35 | -0.43 | 3.6E-02 | -1.53 | -0.62 | 2.4E-03 |
| Trmo | 251 | -2.34 | -1.22 | 9.0E-22 | -2.81 | -1.49 | 1.3E-31 |
| Trmt1 | 4627 | 1.42 | 0.51 | 1.1E-13 | 1.40 | 0.49 | 1.4E-12 |
| Trmt10a | 1019 | -1.95 | -0.96 | 6.1E-16 | -2.11 | -1.07 | 1.8E-19 |

| | | | | | | | |
|---|---|---|---|---|---|---|---|
| Trmt112 | 2004 | -1.29 | -0.37 | 9.3E-03 | -1.45 | -0.54 | 9.6E-05 |
| Trmt12 | 1120 | 1.39 | 0.47 | 2.0E-09 | 1.37 | 0.45 | 1.1E-08 |
| Trmt2b | 1075 | 1.28 | 0.35 | 2.3E-04 | 1.31 | 0.39 | 6.1E-05 |
| Trmt9b | 450 | 1.87 | 0.91 | 2.1E-22 | 1.85 | 0.89 | 2.5E-21 |
| Trmu | 1510 | -1.21 | -0.27 | 1.5E-04 | -1.28 | -0.36 | 5.1E-07 |
| Trnp1 | 298 | 1.45 | 0.53 | 3.8E-02 | 1.55 | 0.63 | 1.4E-02 |
| Trp53bp2 | 6157 | 1.32 | 0.40 | 2.4E-07 | 1.29 | 0.37 | 1.4E-06 |
| Trp53i13 | 1053 | 1.89 | 0.92 | 3.2E-11 | 1.74 | 0.80 | 1.3E-08 |
| Trp53inp1 | 9855 | 1.38 | 0.46 | 3.8E-06 | 1.54 | 0.62 | 3.0E-10 |
| Trp53inp2 | 13213 | -1.55 | -0.64 | 3.1E-09 | -1.54 | -0.62 | 7.7E-09 |
| Trp53rkb | 706 | 1.37 | 0.46 | 1.4E-09 | 1.29 | 0.37 | 1.2E-06 |
| Trp73 | 821 | -1.48 | -0.56 | 7.6E-03 | -1.67 | -0.74 | 3.5E-04 |
| Trpc1 | 1254 | 1.72 | 0.78 | 8.0E-23 | 1.79 | 0.84 | 3.1E-26 |
| Trpc3 | 522 | -1.35 | -0.43 | 2.0E-03 | -1.48 | -0.56 | 4.7E-05 |
| Trpm3 | 2849 | -2.05 | -1.04 | 9.6E-05 | -1.72 | -0.78 | 4.1E-03 |
| Trpm4 | 1776 | 2.04 | 1.03 | 1.2E-15 | 2.11 | 1.08 | 5.0E-17 |
| Trpm5 | 237 | -6.18 | -2.63 | 7.6E-09 | -5.06 | -2.34 | 3.2E-07 |
| Trps1 | 3348 | -1.81 | -0.86 | 1.8E-06 | -1.65 | -0.72 | 7.2E-05 |
| Trpv2 | 487 | -1.91 | -0.93 | 7.6E-10 | -2.03 | -1.02 | 1.5E-11 |
| Trrap | 17379 | -1.49 | -0.57 | 2.6E-06 | -1.36 | -0.45 | 3.2E-04 |
| Trub2 | 2831 | -1.39 | -0.47 | 2.8E-06 | -1.37 | -0.45 | 7.9E-06 |
| Tsc2 | 11894 | 1.25 | 0.32 | 9.6E-04 | 1.25 | 0.32 | 8.7E-04 |
| Tsc22d1 | 30581 | -1.45 | -0.54 | 9.7E-26 | -1.48 | -0.56 | 6.7E-28 |
| Tsc22d2 | 6615 | 1.55 | 0.63 | 1.6E-08 | 1.48 | 0.57 | 4.7E-07 |
| Tsc22d3 | 2483 | -1.51 | -0.60 | 3.0E-11 | -1.51 | -0.59 | 3.5E-11 |
| Tsen2 | 1122 | -1.51 | -0.60 | 1.9E-17 | -1.53 | -0.61 | 5.9E-18 |
| Tsen34 | 5496 | 1.37 | 0.46 | 4.3E-10 | 1.29 | 0.37 | 5.9E-07 |
| Tsen54 | 1528 | 1.78 | 0.83 | 1.5E-28 | 1.64 | 0.71 | 4.9E-21 |
| Tsfm | 1548 | 1.38 | 0.46 | 9.7E-05 | 1.20 | 0.27 | 3.2E-02 |
| Tsga10 | 2471 | -1.50 | -0.59 | 1.0E-09 | -1.38 | -0.47 | 1.4E-06 |
| Tshr | 132 | 2.25 | 1.17 | 2.0E-04 | 2.49 | 1.32 | 2.8E-05 |
| Tshz3 | 1699 | 1.28 | 0.35 | 4.1E-02 | 1.43 | 0.52 | 2.1E-03 |
| Tspan11 | 1093 | -2.22 | -1.15 | 4.0E-17 | -2.03 | -1.02 | 1.3E-13 |
| Tspan15 | 1298 | 2.50 | 1.32 | 1.7E-98 | 2.40 | 1.26 | 4.1E-90 |
| Tspan17 | 1100 | 2.74 | 1.46 | 6.1E-25 | 2.80 | 1.49 | 7.2E-26 |
| Tspan18 | 4519 | -1.45 | -0.54 | 9.2E-06 | -1.61 | -0.69 | 1.1E-08 |
| Tspan2 | 1531 | 1.68 | 0.75 | 6.1E-16 | 1.73 | 0.79 | 1.7E-17 |
| Tspan33 | 946 | 3.61 | 1.85 | 1.8E-36 | 2.97 | 1.57 | 1.5E-26 |
| Tspan4 | 996 | 1.40 | 0.49 | 1.0E-03 | 1.36 | 0.45 | 2.8E-03 |
| Tspyl1 | 10650 | 1.57 | 0.65 | 3.3E-22 | 1.53 | 0.61 | 1.2E-19 |
| Tspyl3 | 916 | -1.76 | -0.81 | 1.3E-05 | -1.94 | -0.96 | 2.2E-07 |
| Tspyl4 | 13508 | 1.37 | 0.45 | 2.1E-03 | 1.46 | 0.54 | 1.8E-04 |
| Tspyl5 | 1666 | 1.34 | 0.43 | 5.8E-03 | 1.31 | 0.39 | 1.3E-02 |
| Tssk4 | 155 | 1.28 | 0.35 | 1.1E-02 | 1.39 | 0.47 | 7.0E-04 |
| Tst | 418 | 1.41 | 0.49 | 1.3E-03 | 1.48 | 0.56 | 2.0E-04 |
| Ttbk1 | 6304 | 1.51 | 0.59 | 4.6E-03 | 1.37 | 0.45 | 3.6E-02 |
| Ttbk2 | 10775 | 1.63 | 0.70 | 1.2E-07 | 1.75 | 0.81 | 1.1E-09 |

| | | | | | | | |
|---|---|---|---|---|---|---|---|
| Ttc1 | 3512 | 1.31 | 0.39 | 3.6E-12 | 1.31 | 0.38 | 6.8E-12 |
| Ttc12 | 481 | 1.63 | 0.71 | 1.0E-06 | 1.63 | 0.71 | 1.1E-06 |
| Ttc16 | 131 | 1.98 | 0.99 | 3.1E-07 | 2.19 | 1.13 | 4.2E-09 |
| Ttc17 | 2560 | -1.32 | -0.40 | 9.2E-06 | -1.27 | -0.35 | 1.4E-04 |
| Ttc19 | 2326 | 1.28 | 0.36 | 7.6E-03 | 1.45 | 0.54 | 3.3E-05 |
| Ttc21b | 3707 | 1.89 | 0.92 | 1.1E-29 | 2.01 | 1.01 | 1.4E-35 |
| Ttc26 | 2246 | 1.49 | 0.57 | 9.4E-13 | 1.52 | 0.61 | 4.3E-14 |
| Ttc30a2 | 100 | 1.68 | 0.75 | 1.2E-06 | 2.10 | 1.07 | 1.4E-12 |
| Ttc32 | 289 | -3.76 | -1.91 | 1.4E-43 | -3.36 | -1.75 | 2.4E-36 |
| Ttc37 | 4684 | 1.43 | 0.51 | 6.4E-06 | 1.67 | 0.74 | 5.4E-11 |
| Ttc39b | 1769 | 1.41 | 0.49 | 1.5E-02 | 1.63 | 0.71 | 3.8E-04 |
| Ttc39c | 964 | -1.34 | -0.42 | 8.9E-06 | -1.41 | -0.49 | 1.4E-07 |
| Ttc41 | 330 | -2.20 | -1.14 | 3.6E-24 | -1.99 | -0.99 | 1.4E-18 |
| Ttc6 | 196 | -1.49 | -0.58 | 2.6E-03 | -1.40 | -0.48 | 1.3E-02 |
| Ttc7 | 821 | 2.85 | 1.51 | 1.6E-25 | 2.84 | 1.50 | 2.6E-25 |
| Ttc7b | 1617 | -1.60 | -0.67 | 6.4E-26 | -1.75 | -0.81 | 1.3E-36 |
| Ttf2 | 3923 | 1.40 | 0.49 | 9.7E-04 | 1.32 | 0.40 | 8.2E-03 |
| Tti1 | 2683 | -1.45 | -0.54 | 8.8E-11 | -1.48 | -0.56 | 8.0E-12 |
| Ttll11 | 336 | 1.56 | 0.64 | 3.4E-08 | 1.55 | 0.64 | 6.5E-08 |
| Ttll12 | 4809 | 1.40 | 0.49 | 2.1E-09 | 1.36 | 0.44 | 6.7E-08 |
| Ttll3 | 1166 | 1.44 | 0.53 | 5.3E-05 | 1.47 | 0.56 | 2.0E-05 |
| Ttll7 | 4198 | 1.68 | 0.75 | 8.9E-05 | 2.23 | 1.16 | 6.0E-10 |
| Ttn | 914 | -15.26 | -3.93 | 1.8E-03 | -8.61 | -3.11 | 1.6E-02 |
| Ttpa | 169 | -1.77 | -0.82 | 4.3E-03 | -1.76 | -0.81 | 5.2E-03 |
| Ttpal | 5297 | -1.28 | -0.36 | 3.0E-16 | -1.30 | -0.38 | 1.3E-17 |
| Ttyh1 | 10943 | -1.45 | -0.54 | 4.9E-04 | -1.49 | -0.58 | 1.6E-04 |
| Ttyh2 | 1994 | 1.66 | 0.73 | 6.6E-10 | 1.73 | 0.79 | 2.0E-11 |
| Tuba4a | 873 | 2.84 | 1.50 | 3.5E-16 | 3.02 | 1.59 | 5.6E-18 |
| Tubb3 | 58234 | -1.25 | -0.33 | 1.8E-02 | -1.27 | -0.34 | 1.4E-02 |
| Tubb5 | 143831 | -1.19 | -0.25 | 6.0E-04 | -1.26 | -0.34 | 3.9E-06 |
| Tubgcp5 | 3696 | -1.29 | -0.36 | 2.7E-04 | -1.21 | -0.27 | 8.0E-03 |
| Tubgcp6 | 7162 | 1.68 | 0.75 | 1.2E-16 | 1.72 | 0.78 | 5.1E-18 |
| Tufm | 6414 | 1.38 | 0.47 | 1.0E-06 | 1.27 | 0.34 | 4.9E-04 |
| Tuft1 | 785 | 4.25 | 2.09 | 8.2E-53 | 4.13 | 2.04 | 1.3E-50 |
| Tug1 | 14599 | -1.19 | -0.25 | 1.6E-04 | -1.26 | -0.33 | 2.8E-07 |
| Tulp3 | 3921 | 1.49 | 0.57 | 3.0E-08 | 1.43 | 0.51 | 8.7E-07 |
| Tusc1 | 250 | -2.59 | -1.37 | 6.2E-27 | -2.52 | -1.33 | 3.0E-25 |
| Tusc2 | 3022 | 1.63 | 0.70 | 9.8E-20 | 1.55 | 0.63 | 3.6E-16 |
| Tut4 | 12929 | -1.33 | -0.41 | 9.6E-07 | -1.25 | -0.32 | 1.3E-04 |
| Twf2 | 1910 | 1.56 | 0.64 | 1.4E-10 | 1.58 | 0.66 | 3.3E-11 |
| Twsg1 | 7741 | -1.59 | -0.67 | 6.2E-15 | -1.58 | -0.66 | 9.0E-15 |
| Txlnb | 482 | -1.69 | -0.75 | 3.1E-03 | -1.62 | -0.70 | 6.9E-03 |
| Txlng | 3438 | 1.74 | 0.80 | 1.2E-32 | 1.75 | 0.81 | 6.7E-33 |
| Txndc15 | 2764 | -1.52 | -0.61 | 2.3E-16 | -1.46 | -0.55 | 1.2E-13 |
| Txndc16 | 7016 | 2.19 | 1.13 | 6.2E-32 | 2.40 | 1.26 | 9.1E-40 |
| Txndc17 | 3932 | 1.34 | 0.43 | 9.8E-06 | 1.27 | 0.35 | 3.9E-04 |
| Txndc9 | 5445 | -1.59 | -0.67 | 9.6E-18 | -1.57 | -0.65 | 9.8E-17 |

| | | | | | | | |
|---|---|---|---|---|---|---|---|
| Txnl4a | 2661 | 1.53 | 0.62 | 5.5E-13 | 1.52 | 0.61 | 1.6E-12 |
| Txnl4b | 505 | -2.43 | -1.28 | 6.4E-47 | -2.48 | -1.31 | 6.9E-49 |
| Txnrd3 | 1181 | 1.47 | 0.55 | 3.3E-10 | 1.31 | 0.39 | 1.1E-05 |
| Tysnd1 | 1631 | 1.53 | 0.61 | 7.8E-10 | 1.39 | 0.48 | 2.0E-06 |
| Tyw1 | 2171 | 1.27 | 0.35 | 6.7E-07 | 1.22 | 0.29 | 6.2E-05 |
| Tyw3 | 881 | -1.80 | -0.85 | 4.8E-22 | -1.96 | -0.97 | 1.2E-28 |
| Tyw5 | 681 | -2.12 | -1.09 | 3.3E-26 | -2.11 | -1.08 | 9.4E-26 |
| U2af2 | 18489 | -1.37 | -0.46 | 8.6E-05 | -1.54 | -0.63 | 4.9E-08 |
| Uap1 | 3684 | 1.53 | 0.61 | 3.0E-21 | 1.57 | 0.65 | 1.3E-23 |
| Uap1l1 | 256 | 1.74 | 0.80 | 2.8E-04 | 1.61 | 0.69 | 1.9E-03 |
| Uba3 | 8246 | -1.44 | -0.52 | 5.7E-13 | -1.39 | -0.48 | 5.2E-11 |
| Uba6 | 7911 | 1.57 | 0.65 | 3.3E-07 | 1.73 | 0.79 | 4.1E-10 |
| Ubald2 | 2110 | -1.46 | -0.54 | 2.5E-06 | -1.49 | -0.58 | 5.6E-07 |
| Ubap1l | 237 | 1.70 | 0.77 | 8.2E-03 | 1.81 | 0.85 | 3.1E-03 |
| Ubc | 10057 | -1.43 | -0.51 | 4.2E-05 | -1.48 | -0.57 | 5.3E-06 |
| Ube2a | 2741 | 1.40 | 0.48 | 2.9E-17 | 1.37 | 0.46 | 1.6E-15 |
| Ube2b | 11470 | 1.23 | 0.29 | 4.3E-05 | 1.30 | 0.38 | 9.8E-08 |
| Ube2c | 4290 | -1.50 | -0.59 | 7.6E-03 | -1.83 | -0.87 | 5.3E-05 |
| Ube2cbp | 200 | -3.11 | -1.64 | 4.5E-42 | -3.55 | -1.83 | 6.6E-51 |
| Ube2e1 | 7281 | -1.22 | -0.29 | 4.0E-04 | -1.27 | -0.34 | 3.0E-05 |
| Ube2e3 | 10363 | -1.66 | -0.73 | 1.5E-17 | -1.67 | -0.74 | 8.3E-18 |
| Ube2f | 3212 | 1.50 | 0.58 | 5.2E-11 | 1.47 | 0.56 | 3.4E-10 |
| Ube2g2 | 3007 | 1.26 | 0.33 | 7.3E-06 | 1.14 | 0.19 | 1.5E-02 |
| Ube2h | 11325 | 1.37 | 0.46 | 7.8E-08 | 1.29 | 0.37 | 1.7E-05 |
| Ube2o | 8269 | 1.43 | 0.51 | 1.4E-05 | 1.38 | 0.46 | 1.1E-04 |
| Ube2q2 | 4081 | -1.35 | -0.43 | 4.0E-10 | -1.35 | -0.43 | 4.3E-10 |
| Ube2r2 | 10444 | -1.71 | -0.77 | 6.9E-12 | -1.90 | -0.92 | 1.7E-16 |
| Ube2t | 1563 | 1.73 | 0.79 | 2.6E-09 | 1.79 | 0.84 | 2.4E-10 |
| Ube4a | 11712 | 1.28 | 0.36 | 3.1E-04 | 1.36 | 0.44 | 6.1E-06 |
| Ubfd1 | 7495 | -1.31 | -0.39 | 4.1E-12 | -1.40 | -0.49 | 7.8E-18 |
| Ubiad1 | 958 | -1.55 | -0.64 | 9.6E-14 | -1.57 | -0.65 | 2.8E-14 |
| Ubl3 | 3853 | 1.39 | 0.47 | 1.1E-07 | 1.29 | 0.37 | 3.5E-05 |
| Ubl4a | 2918 | -1.34 | -0.42 | 7.1E-10 | -1.46 | -0.55 | 5.3E-16 |
| Ubr1 | 5748 | -1.47 | -0.56 | 1.3E-06 | -1.37 | -0.46 | 9.7E-05 |
| Ubr2 | 9577 | 1.35 | 0.43 | 6.4E-06 | 1.42 | 0.51 | 1.0E-07 |
| Ubr7 | 8587 | 1.71 | 0.77 | 5.4E-13 | 1.62 | 0.70 | 8.9E-11 |
| Ubtd1 | 757 | 2.09 | 1.07 | 3.7E-16 | 2.12 | 1.09 | 1.3E-16 |
| Ubtf | 9942 | -1.99 | -0.99 | 8.7E-27 | -2.23 | -1.16 | 8.1E-36 |
| Ubxn2a | 2682 | 1.25 | 0.32 | 2.4E-02 | 1.33 | 0.41 | 2.9E-03 |
| Ubxn2b | 5075 | 1.54 | 0.62 | 2.9E-12 | 1.62 | 0.70 | 4.4E-15 |
| Ubxn6 | 2946 | -1.51 | -0.60 | 1.2E-11 | -1.49 | -0.58 | 5.6E-11 |
| Ubxn8 | 490 | -7.75 | -2.95 | 8.8E-112 | -7.39 | -2.88 | 1.2E-106 |
| Uchl3 | 4070 | 1.67 | 0.74 | 4.3E-19 | 1.67 | 0.74 | 4.6E-19 |
| Uchl4 | 105 | 1.87 | 0.90 | 1.2E-09 | 2.01 | 1.00 | 1.5E-11 |
| Uchl5 | 3391 | 1.57 | 0.65 | 3.6E-08 | 1.46 | 0.55 | 4.5E-06 |
| Uck1 | 2727 | 1.26 | 0.33 | 7.3E-06 | 1.13 | 0.18 | 2.0E-02 |
| Uckl1 | 1542 | -1.48 | -0.57 | 9.1E-16 | -1.49 | -0.58 | 3.9E-16 |

| | | | | | | | |
|---|---|---|---|---|---|---|---|
| Ucp2 | 2830 | 1.51 | 0.59 | 4.2E-03 | 1.42 | 0.50 | 1.7E-02 |
| Ufc1 | 3001 | -1.94 | -0.96 | 5.4E-36 | -2.04 | -1.03 | 1.3E-41 |
| Ufsp1 | 264 | 1.62 | 0.70 | 2.3E-09 | 1.68 | 0.75 | 1.6E-10 |
| Ugp2 | 6285 | 1.28 | 0.36 | 1.7E-05 | 1.29 | 0.37 | 9.7E-06 |
| Ugt8a | 813 | 2.52 | 1.33 | 4.0E-35 | 2.56 | 1.36 | 2.3E-36 |
| Uhrf1bp1 | 5571 | 1.70 | 0.76 | 1.2E-21 | 1.77 | 0.82 | 6.7E-25 |
| Uhrf1bp1l | 5392 | -1.39 | -0.48 | 3.5E-08 | -1.33 | -0.41 | 2.7E-06 |
| Ulk2 | 9553 | -1.51 | -0.59 | 1.5E-11 | -1.47 | -0.55 | 2.5E-10 |
| Umad1 | 1184 | 1.46 | 0.55 | 2.0E-07 | 1.45 | 0.54 | 3.1E-07 |
| Unc119b | 6746 | 1.58 | 0.66 | 2.0E-09 | 1.48 | 0.57 | 2.9E-07 |
| Unc13a | 10790 | 2.41 | 1.27 | 2.4E-07 | 2.38 | 1.25 | 4.1E-07 |
| Unc45a | 4861 | 1.35 | 0.43 | 4.8E-07 | 1.28 | 0.36 | 3.2E-05 |
| Unc5a | 1158 | 1.98 | 0.98 | 4.3E-05 | 1.85 | 0.89 | 2.5E-04 |
| Unc5b | 3707 | 1.69 | 0.75 | 4.9E-08 | 1.41 | 0.50 | 4.4E-04 |
| Unc5c | 4631 | -2.20 | -1.14 | 1.5E-06 | -1.86 | -0.89 | 2.1E-04 |
| Unc79 | 2448 | -2.25 | -1.17 | 1.4E-06 | -2.15 | -1.10 | 5.9E-06 |
| Unk | 3649 | -1.40 | -0.48 | 8.9E-05 | -1.49 | -0.57 | 2.8E-06 |
| Upf2 | 7646 | 1.31 | 0.39 | 1.6E-13 | 1.39 | 0.47 | 5.9E-19 |
| Upf3a | 2232 | -3.26 | -1.71 | 8.0E-86 | -3.40 | -1.77 | 6.6E-92 |
| Upk1a | 133 | 2.95 | 1.56 | 3.4E-19 | 2.50 | 1.32 | 4.4E-14 |
| Uqcc1 | 2275 | -1.52 | -0.60 | 4.2E-23 | -1.57 | -0.65 | 4.6E-27 |
| Uqcc2 | 2157 | -1.35 | -0.43 | 5.0E-05 | -1.37 | -0.46 | 1.6E-05 |
| Uqcc3 | 785 | -1.36 | -0.44 | 8.8E-07 | -1.40 | -0.48 | 7.0E-08 |
| Uqcrb | 3751 | -1.34 | -0.43 | 7.5E-03 | -1.42 | -0.51 | 1.4E-03 |
| Uqcrh | 9072 | 1.28 | 0.36 | 8.7E-05 | 1.25 | 0.32 | 4.4E-04 |
| Urb1 | 3093 | 1.42 | 0.51 | 3.4E-06 | 1.57 | 0.65 | 2.1E-09 |
| Urb2 | 3004 | 1.44 | 0.53 | 1.9E-06 | 1.41 | 0.50 | 8.9E-06 |
| Uri1 | 5183 | 1.50 | 0.58 | 1.2E-15 | 1.39 | 0.47 | 1.6E-10 |
| Urod | 3339 | 1.71 | 0.77 | 1.1E-11 | 1.61 | 0.69 | 1.6E-09 |
| Use1 | 2661 | -1.34 | -0.42 | 9.7E-08 | -1.34 | -0.43 | 6.6E-08 |
| Usf1 | 5916 | -1.47 | -0.56 | 1.8E-05 | -1.62 | -0.70 | 5.7E-08 |
| Usf2 | 2029 | -1.74 | -0.80 | 3.3E-12 | -1.88 | -0.91 | 1.9E-15 |
| Ush1c | 115 | -1.37 | -0.46 | 1.6E-02 | -1.34 | -0.43 | 2.8E-02 |
| Ushbp1 | 371 | -3.20 | -1.68 | 8.6E-47 | -3.17 | -1.66 | 6.6E-46 |
| Usp11 | 13046 | 1.26 | 0.34 | 5.7E-07 | 1.22 | 0.29 | 2.0E-05 |
| Usp13 | 1190 | -1.49 | -0.58 | 2.3E-06 | -1.35 | -0.44 | 4.6E-04 |
| Usp14 | 10421 | 1.31 | 0.39 | 1.2E-07 | 1.33 | 0.41 | 2.0E-08 |
| Usp15 | 8350 | 1.26 | 0.33 | 6.5E-04 | 1.37 | 0.46 | 1.2E-06 |
| Usp2 | 685 | 3.08 | 1.62 | 1.5E-34 | 2.78 | 1.47 | 1.2E-28 |
| Usp20 | 5012 | 2.30 | 1.20 | 1.5E-40 | 2.23 | 1.16 | 1.3E-37 |
| Usp21 | 4076 | -1.53 | -0.62 | 4.4E-09 | -1.66 | -0.73 | 1.7E-12 |
| Usp22 | 27317 | -1.36 | -0.45 | 5.6E-09 | -1.40 | -0.49 | 1.4E-10 |
| Usp24 | 8094 | -1.71 | -0.78 | 5.3E-10 | -1.50 | -0.59 | 3.4E-06 |
| Usp25 | 5073 | 2.37 | 1.24 | 2.8E-63 | 2.35 | 1.23 | 5.5E-62 |
| Usp28 | 6957 | 2.12 | 1.09 | 5.1E-79 | 2.10 | 1.07 | 4.8E-77 |
| Usp3 | 8060 | -1.23 | -0.30 | 6.0E-04 | -1.25 | -0.32 | 3.4E-04 |
| Usp30 | 2549 | 1.25 | 0.33 | 4.6E-09 | 1.15 | 0.20 | 3.5E-04 |

| | | | | | | | |
|---|---|---|---|---|---|---|---|
| Usp31 | 5042 | 1.47 | 0.56 | 1.8E-11 | 1.50 | 0.59 | 1.4E-12 |
| Usp32 | 9161 | 1.40 | 0.49 | 6.1E-04 | 1.53 | 0.62 | 1.1E-05 |
| Usp33 | 11847 | 1.48 | 0.57 | 4.7E-12 | 1.59 | 0.67 | 4.6E-16 |
| Usp35 | 1336 | 1.64 | 0.71 | 6.1E-10 | 1.60 | 0.68 | 3.8E-09 |
| Usp37 | 6619 | 1.24 | 0.31 | 1.9E-04 | 1.27 | 0.34 | 2.9E-05 |
| Usp38 | 5289 | 1.70 | 0.76 | 1.1E-20 | 1.66 | 0.73 | 7.2E-19 |
| Usp40 | 3424 | -2.32 | -1.21 | 7.0E-40 | -2.22 | -1.15 | 9.6E-36 |
| Usp42 | 4880 | -1.28 | -0.36 | 3.5E-07 | -1.30 | -0.38 | 7.6E-08 |
| Usp44 | 751 | 4.63 | 2.21 | 5.9E-11 | 3.59 | 1.84 | 6.3E-08 |
| Usp45 | 4065 | 1.41 | 0.50 | 1.6E-14 | 1.43 | 0.52 | 1.5E-15 |
| Usp46 | 4586 | -1.27 | -0.34 | 5.8E-06 | -1.27 | -0.34 | 8.0E-06 |
| Usp53 | 1386 | 1.88 | 0.91 | 2.2E-17 | 2.11 | 1.08 | 8.6E-24 |
| Usp6nl | 3130 | -2.07 | -1.05 | 1.3E-51 | -2.11 | -1.08 | 1.1E-54 |
| Ust | 474 | -1.35 | -0.43 | 7.4E-03 | -1.37 | -0.45 | 5.0E-03 |
| Utp11 | 3414 | 1.38 | 0.46 | 2.5E-06 | 1.28 | 0.36 | 3.6E-04 |
| Utp18 | 4018 | 1.37 | 0.45 | 4.1E-08 | 1.36 | 0.45 | 6.1E-08 |
| Utp20 | 4019 | -2.06 | -1.05 | 9.2E-13 | -1.93 | -0.95 | 1.1E-10 |
| Utp4 | 4066 | 1.86 | 0.90 | 2.3E-20 | 1.73 | 0.79 | 3.7E-16 |
| Uty | 1375 | -1.36 | -0.44 | 5.2E-03 | -13.29 | -3.73 | 3.2E-05 |
| Uvrag | 2289 | 1.82 | 0.86 | 2.5E-64 | 1.79 | 0.84 | 5.4E-60 |
| Uxt | 528 | -1.44 | -0.52 | 2.2E-04 | -1.55 | -0.63 | 7.9E-06 |
| Vamp1 | 1657 | 2.02 | 1.02 | 4.1E-65 | 2.02 | 1.01 | 7.9E-64 |
| Vamp3 | 6118 | 1.30 | 0.38 | 3.5E-03 | 1.24 | 0.31 | 1.9E-02 |
| Vamp4 | 3673 | 1.31 | 0.39 | 2.6E-09 | 1.35 | 0.43 | 4.3E-11 |
| Vash2 | 4450 | -1.60 | -0.68 | 4.1E-16 | -1.68 | -0.75 | 3.5E-19 |
| Vasp | 4288 | 1.59 | 0.67 | 2.0E-11 | 1.42 | 0.51 | 7.1E-07 |
| Vat1 | 12488 | 1.45 | 0.53 | 1.6E-10 | 1.38 | 0.47 | 2.5E-08 |
| Vat1l | 5770 | 1.53 | 0.61 | 6.7E-06 | 1.50 | 0.58 | 1.8E-05 |
| Vav1 | 117 | -1.38 | -0.46 | 3.5E-02 | -1.62 | -0.69 | 1.2E-03 |
| Vav2 | 6464 | 1.31 | 0.39 | 5.9E-03 | 1.24 | 0.31 | 3.0E-02 |
| Vav3 | 2170 | -1.51 | -0.60 | 3.7E-06 | -1.57 | -0.65 | 4.5E-07 |
| Vcam1 | 4798 | -1.63 | -0.71 | 1.6E-10 | -1.44 | -0.52 | 3.8E-06 |
| Vcan | 34687 | -2.25 | -1.17 | 5.8E-07 | -1.76 | -0.82 | 6.3E-04 |
| Vcl | 8570 | 2.09 | 1.06 | 1.1E-06 | 2.17 | 1.12 | 2.8E-07 |
| Vcpip1 | 8251 | 1.31 | 0.39 | 5.6E-07 | 1.40 | 0.48 | 4.3E-10 |
| Vdac2 | 11384 | 1.29 | 0.37 | 5.2E-10 | 1.26 | 0.34 | 1.6E-08 |
| Vdr | 170 | 6.19 | 2.63 | 1.5E-10 | 6.71 | 2.75 | 2.3E-11 |
| Vegfc | 373 | -1.52 | -0.60 | 4.0E-06 | -1.73 | -0.79 | 1.1E-09 |
| Vegfd | 111 | -2.21 | -1.14 | 2.6E-04 | -1.91 | -0.93 | 3.5E-03 |
| Veph1 | 1252 | -1.73 | -0.79 | 5.9E-03 | -1.74 | -0.80 | 5.8E-03 |
| Vgf | 190 | 1.87 | 0.91 | 1.4E-02 | 1.83 | 0.88 | 1.9E-02 |
| Vgll3 | 1101 | -2.49 | -1.31 | 1.0E-02 | -2.14 | -1.10 | 3.7E-02 |
| Vipr1 | 150 | 2.57 | 1.36 | 7.5E-14 | 2.99 | 1.58 | 5.9E-18 |
| Vipr2 | 289 | -1.86 | -0.90 | 1.9E-07 | -2.05 | -1.04 | 1.9E-09 |
| Vkorc1l1 | 7992 | 1.53 | 0.61 | 1.1E-25 | 1.45 | 0.53 | 1.1E-19 |
| Vmac | 1606 | 2.18 | 1.13 | 3.7E-66 | 2.08 | 1.06 | 5.6E-58 |
| Vmn2r1 | 117 | 2.46 | 1.30 | 4.1E-11 | 2.12 | 1.08 | 6.0E-08 |

| | | | | | | | |
|---|---|---|---|---|---|---|---|
| Vps13a | 5354 | 1.76 | 0.81 | 7.0E-05 | 2.20 | 1.14 | 1.5E-08 |
| Vps26a | 6362 | 1.62 | 0.70 | 6.2E-16 | 1.58 | 0.66 | 1.8E-14 |
| Vps26b | 9586 | 1.26 | 0.33 | 3.4E-03 | 1.24 | 0.31 | 7.0E-03 |
| Vps26c | 3950 | 1.43 | 0.51 | 1.7E-08 | 1.44 | 0.53 | 6.1E-09 |
| Vps29 | 6144 | 1.38 | 0.47 | 9.3E-24 | 1.38 | 0.46 | 3.1E-23 |
| Vps33b | 2467 | 1.31 | 0.39 | 5.2E-08 | 1.22 | 0.29 | 9.5E-05 |
| Vps35 | 18421 | 1.29 | 0.37 | 3.1E-04 | 1.40 | 0.48 | 2.1E-06 |
| Vps35l | 4929 | -1.46 | -0.54 | 8.7E-19 | -1.47 | -0.55 | 2.0E-19 |
| Vps36 | 4977 | -1.35 | -0.43 | 2.0E-10 | -1.37 | -0.45 | 2.6E-11 |
| Vps37a | 6721 | 1.35 | 0.44 | 2.1E-05 | 1.49 | 0.58 | 1.3E-08 |
| Vps37c | 1601 | -1.27 | -0.34 | 8.9E-03 | -1.31 | -0.39 | 3.0E-03 |
| Vps37d | 1406 | -1.31 | -0.39 | 5.7E-03 | -1.36 | -0.44 | 1.6E-03 |
| Vps51 | 1271 | -3.16 | -1.66 | 6.0E-135 | -3.40 | -1.76 | 3.2E-151 |
| Vps52 | 3466 | -1.62 | -0.70 | 1.6E-20 | -1.70 | -0.76 | 1.9E-24 |
| Vps54 | 4716 | -1.45 | -0.53 | 6.4E-15 | -1.48 | -0.56 | 1.7E-16 |
| Vps72 | 5730 | -1.33 | -0.41 | 5.0E-05 | -1.50 | -0.59 | 5.3E-09 |
| Vps9d1 | 2534 | -1.71 | -0.77 | 4.3E-12 | -1.72 | -0.79 | 2.0E-12 |
| Vrk2 | 424 | -1.79 | -0.84 | 8.5E-05 | -2.08 | -1.06 | 5.4E-07 |
| Vsig10l | 394 | 1.53 | 0.62 | 2.3E-06 | 1.60 | 0.68 | 1.9E-07 |
| Vsig2 | 161 | -1.76 | -0.81 | 2.3E-04 | -1.71 | -0.78 | 5.1E-04 |
| Vstm4 | 264 | -2.96 | -1.56 | 4.6E-12 | -2.83 | -1.50 | 3.9E-11 |
| Vstm5 | 205 | -4.50 | -2.17 | 2.4E-12 | -4.02 | -2.01 | 1.1E-10 |
| Vti1a | 2058 | -1.33 | -0.42 | 5.7E-14 | -1.40 | -0.49 | 6.7E-19 |
| Vwa7 | 164 | 2.50 | 1.32 | 6.6E-09 | 2.60 | 1.38 | 1.4E-09 |
| Vwa8 | 1662 | -1.47 | -0.56 | 2.9E-07 | -1.44 | -0.52 | 1.8E-06 |
| Vwc2 | 211 | -1.72 | -0.78 | 3.9E-03 | -1.68 | -0.75 | 5.7E-03 |
| Vxn | 1156 | -4.08 | -2.03 | 1.5E-19 | -2.83 | -1.50 | 3.7E-11 |
| Wac | 11365 | -1.34 | -0.42 | 1.4E-10 | -1.38 | -0.47 | 6.3E-13 |
| Wars2 | 727 | -2.89 | -1.53 | 7.0E-44 | -2.81 | -1.49 | 2.8E-41 |
| Wasf2 | 4611 | -1.24 | -0.31 | 1.9E-02 | -1.28 | -0.35 | 7.1E-03 |
| Wasf3 | 6105 | 1.63 | 0.70 | 2.9E-13 | 1.60 | 0.68 | 1.6E-12 |
| Washc1 | 2899 | 1.26 | 0.33 | 2.3E-05 | 1.22 | 0.29 | 2.5E-04 |
| Washc3 | 649 | -2.80 | -1.49 | 4.5E-86 | -2.60 | -1.38 | 7.7E-75 |
| Wbp1l | 5892 | 1.39 | 0.47 | 4.1E-06 | 1.25 | 0.32 | 2.6E-03 |
| Wbp2 | 3393 | -2.33 | -1.22 | 2.3E-16 | -2.43 | -1.28 | 4.9E-18 |
| Wdcp | 2022 | 2.28 | 1.19 | 2.9E-55 | 2.28 | 1.19 | 3.6E-55 |
| Wdfy2 | 5610 | 1.96 | 0.97 | 6.9E-52 | 1.94 | 0.96 | 4.1E-50 |
| Wdfy3 | 12067 | -2.06 | -1.04 | 5.2E-11 | -1.89 | -0.92 | 8.4E-09 |
| Wdhd1 | 6475 | 1.28 | 0.35 | 8.2E-03 | 1.25 | 0.32 | 1.9E-02 |
| Wdr11 | 4595 | -1.81 | -0.86 | 1.8E-46 | -1.82 | -0.87 | 1.7E-47 |
| Wdr12 | 4641 | 1.83 | 0.87 | 5.5E-35 | 1.92 | 0.94 | 3.0E-40 |
| Wdr13 | 8043 | -1.78 | -0.83 | 1.3E-07 | -1.71 | -0.78 | 9.4E-07 |
| Wdr17 | 1605 | 2.56 | 1.36 | 5.7E-20 | 2.70 | 1.43 | 4.0E-22 |
| Wdr19 | 3611 | 1.74 | 0.80 | 7.4E-16 | 1.89 | 0.92 | 1.1E-20 |
| Wdr25 | 391 | -1.55 | -0.64 | 3.9E-08 | -1.48 | -0.57 | 1.2E-06 |
| Wdr35 | 4986 | 1.52 | 0.61 | 9.1E-16 | 1.59 | 0.67 | 5.0E-19 |
| Wdr37 | 9100 | 1.39 | 0.48 | 1.9E-13 | 1.40 | 0.49 | 8.7E-14 |

| | | | | | | | |
|---|---|---|---|---|---|---|---|
| Wdr4 | 3027 | -1.43 | -0.52 | 2.4E-11 | -1.47 | -0.55 | 9.8E-13 |
| Wdr41 | 1513 | -2.70 | -1.43 | 1.1E-116 | -2.73 | -1.45 | 2.1E-118 |
| Wdr44 | 1568 | 1.53 | 0.61 | 3.7E-06 | 1.62 | 0.70 | 1.0E-07 |
| Wdr46 | 2803 | 1.68 | 0.75 | 1.6E-08 | 1.49 | 0.57 | 2.3E-05 |
| Wdr47 | 10290 | 1.40 | 0.48 | 3.9E-04 | 1.49 | 0.58 | 1.8E-05 |
| Wdr5 | 6808 | 1.51 | 0.59 | 2.2E-11 | 1.34 | 0.42 | 3.1E-06 |
| Wdr53 | 599 | -1.23 | -0.30 | 7.1E-04 | -1.36 | -0.45 | 2.2E-07 |
| Wdr54 | 840 | 1.93 | 0.95 | 1.5E-30 | 1.83 | 0.87 | 6.1E-26 |
| Wdr55 | 1636 | 1.26 | 0.33 | 3.2E-04 | 1.15 | 0.20 | 4.0E-02 |
| Wdr5b | 920 | -1.34 | -0.42 | 3.0E-07 | -1.33 | -0.41 | 4.2E-07 |
| Wdr61 | 3886 | 1.27 | 0.34 | 3.8E-04 | 1.22 | 0.28 | 3.6E-03 |
| Wdr62 | 2137 | 2.25 | 1.17 | 1.2E-11 | 2.09 | 1.06 | 1.1E-09 |
| Wdr70 | 3264 | 1.29 | 0.37 | 7.6E-05 | 1.20 | 0.26 | 6.2E-03 |
| Wdr73 | 2484 | -1.25 | -0.32 | 8.8E-07 | -1.28 | -0.35 | 8.9E-08 |
| Wdr81 | 2430 | 1.35 | 0.44 | 9.5E-04 | 1.29 | 0.36 | 6.8E-03 |
| Wdr89 | 276 | -1.35 | -0.43 | 1.2E-03 | -1.54 | -0.62 | 2.7E-06 |
| Wdr90 | 4465 | 1.25 | 0.32 | 1.3E-02 | 1.21 | 0.28 | 3.6E-02 |
| Wdr91 | 1831 | 1.32 | 0.40 | 1.1E-06 | 1.30 | 0.37 | 7.5E-06 |
| Wdtc1 | 5520 | 1.51 | 0.59 | 8.7E-10 | 1.48 | 0.56 | 5.4E-09 |
| Wfikkn2 | 650 | -2.09 | -1.06 | 3.8E-08 | -2.16 | -1.11 | 9.2E-09 |
| Wfs1 | 1305 | 2.41 | 1.27 | 1.1E-34 | 2.24 | 1.17 | 2.2E-29 |
| Whamm | 896 | 1.53 | 0.61 | 3.7E-14 | 1.49 | 0.57 | 2.4E-12 |
| Whrn | 3196 | 2.42 | 1.27 | 4.7E-23 | 2.30 | 1.20 | 1.4E-20 |
| Wipf1 | 1705 | -1.80 | -0.84 | 2.2E-07 | -1.74 | -0.80 | 1.1E-06 |
| Wipi1 | 3596 | 3.00 | 1.58 | 1.7E-38 | 3.18 | 1.67 | 1.4E-42 |
| Wipi2 | 9643 | 1.70 | 0.76 | 4.7E-49 | 1.64 | 0.72 | 3.4E-43 |
| Wiz | 7689 | -1.39 | -0.48 | 5.8E-04 | -1.54 | -0.63 | 4.7E-06 |
| Wnt10b | 116 | -3.73 | -1.90 | 2.4E-03 | -3.29 | -1.72 | 6.8E-03 |
| Wnt16 | 127 | 2.71 | 1.44 | 1.2E-03 | 3.13 | 1.65 | 2.0E-04 |
| Wnt3a | 207 | -3.64 | -1.87 | 1.6E-12 | -4.01 | -2.01 | 2.7E-14 |
| Wnt5a | 3387 | -2.78 | -1.47 | 5.2E-10 | -2.58 | -1.36 | 1.0E-08 |
| Wnt6 | 289 | -2.44 | -1.29 | 1.2E-03 | -2.42 | -1.27 | 1.5E-03 |
| Wnt7b | 4194 | -1.68 | -0.75 | 1.4E-06 | -1.97 | -0.98 | 1.5E-10 |
| Wrap53 | 1991 | 1.84 | 0.88 | 9.0E-26 | 1.71 | 0.78 | 4.0E-20 |
| Wrap73 | 1836 | 1.40 | 0.49 | 1.7E-09 | 1.22 | 0.29 | 5.2E-04 |
| Wrnip1 | 3529 | 1.26 | 0.33 | 2.6E-08 | 1.24 | 0.31 | 1.2E-07 |
| Wsb1 | 31033 | -1.42 | -0.50 | 6.3E-08 | -1.35 | -0.44 | 2.8E-06 |
| Wsb2 | 6503 | 1.26 | 0.33 | 2.9E-03 | 1.25 | 0.33 | 3.6E-03 |
| Wscd2 | 862 | -1.38 | -0.47 | 2.0E-04 | -1.46 | -0.55 | 1.0E-05 |
| Wtap | 8523 | -1.26 | -0.34 | 2.5E-07 | -1.29 | -0.37 | 1.5E-08 |
| Wtip | 1084 | -1.27 | -0.35 | 4.2E-05 | -1.27 | -0.34 | 7.8E-05 |
| Wwc1 | 4810 | 1.26 | 0.34 | 4.4E-03 | 1.19 | 0.25 | 3.7E-02 |
| Wwc2 | 4495 | 1.48 | 0.56 | 1.0E-12 | 1.56 | 0.64 | 4.6E-16 |
| Wwox | 842 | 1.32 | 0.40 | 4.9E-07 | 1.23 | 0.30 | 1.8E-04 |
| Wwp2 | 6670 | -2.23 | -1.16 | 1.8E-10 | -2.10 | -1.07 | 4.5E-09 |
| Wwtr1 | 5460 | 1.26 | 0.34 | 8.3E-04 | 1.28 | 0.36 | 3.5E-04 |
| Xab2 | 4342 | 1.25 | 0.32 | 4.8E-04 | 1.14 | 0.19 | 4.6E-02 |

| | | | | | | | |
|---|---|---|---|---|---|---|---|
| Xiap | 8307 | -2.01 | -1.01 | 1.3E-27 | -1.94 | -0.95 | 9.2E-25 |
| Xkr5 | 243 | -3.20 | -1.68 | 1.5E-12 | -3.41 | -1.77 | 8.9E-14 |
| Xkr8 | 646 | 1.50 | 0.59 | 9.8E-11 | 1.48 | 0.56 | 5.6E-10 |
| Xndc1 | 858 | -2.03 | -1.02 | 1.1E-16 | -2.04 | -1.03 | 7.6E-17 |
| Xpc | 3233 | 1.40 | 0.49 | 2.2E-17 | 1.41 | 0.49 | 1.2E-17 |
| Xpnpep3 | 1957 | 1.35 | 0.43 | 3.4E-11 | 1.38 | 0.47 | 3.8E-13 |
| Xpo5 | 13073 | 1.26 | 0.34 | 7.1E-05 | 1.23 | 0.30 | 4.9E-04 |
| Xpr1 | 27946 | 1.49 | 0.58 | 2.8E-03 | 1.53 | 0.61 | 1.6E-03 |
| Xrcc3 | 1217 | 2.47 | 1.31 | 1.3E-12 | 2.08 | 1.06 | 1.1E-08 |
| Xrcc4 | 489 | -2.16 | -1.11 | 6.9E-21 | -2.06 | -1.05 | 1.2E-18 |
| Xrcc5 | 2588 | -1.57 | -0.65 | 4.3E-05 | -1.59 | -0.66 | 2.8E-05 |
| Xrcc6 | 3705 | 1.36 | 0.45 | 7.6E-09 | 1.32 | 0.40 | 2.1E-07 |
| Xylb | 865 | 1.56 | 0.64 | 1.1E-06 | 1.42 | 0.51 | 1.4E-04 |
| Xylt1 | 2099 | -1.27 | -0.35 | 3.2E-03 | -1.23 | -0.30 | 1.2E-02 |
| Yap1 | 9174 | -1.70 | -0.77 | 7.7E-08 | -1.87 | -0.91 | 1.9E-10 |
| Yars | 8078 | 1.34 | 0.42 | 1.8E-12 | 1.25 | 0.32 | 1.3E-07 |
| Yars2 | 1086 | -1.42 | -0.51 | 6.4E-09 | -1.46 | -0.54 | 4.8E-10 |
| Ybey | 491 | -1.40 | -0.48 | 8.1E-06 | -1.41 | -0.49 | 6.0E-06 |
| Ybx2 | 3335 | 3.88 | 1.96 | 6.2E-95 | 3.51 | 1.81 | 1.6E-81 |
| Ydjc | 543 | 1.59 | 0.67 | 1.5E-04 | 1.71 | 0.78 | 1.0E-05 |
| Yeats2 | 8743 | -1.43 | -0.51 | 3.0E-18 | -1.38 | -0.47 | 3.4E-15 |
| Yeats4 | 4533 | -1.86 | -0.89 | 8.7E-17 | -2.01 | -1.01 | 3.6E-21 |
| Yes1 | 4825 | 1.51 | 0.59 | 1.1E-17 | 1.57 | 0.65 | 5.0E-21 |
| Yif1a | 1279 | 1.27 | 0.35 | 4.9E-03 | 1.23 | 0.30 | 1.6E-02 |
| Yif1b | 1218 | 1.31 | 0.39 | 1.2E-03 | 1.22 | 0.29 | 1.9E-02 |
| Yipf2 | 599 | -2.35 | -1.23 | 2.0E-69 | -2.61 | -1.38 | 1.0E-84 |
| Yju2 | 895 | -1.78 | -0.83 | 4.2E-13 | -2.05 | -1.04 | 7.8E-20 |
| Ykt6 | 11959 | 1.26 | 0.33 | 1.3E-11 | 1.18 | 0.23 | 2.1E-06 |
| Ypel1 | 3603 | -2.07 | -1.05 | 1.8E-23 | -2.22 | -1.15 | 6.7E-28 |
| Ypel2 | 2360 | 1.79 | 0.84 | 2.5E-16 | 1.70 | 0.77 | 8.8E-14 |
| Ypel4 | 1375 | -1.46 | -0.55 | 2.8E-03 | -1.47 | -0.56 | 2.4E-03 |
| Yrdc | 1773 | 1.36 | 0.45 | 3.4E-06 | 1.25 | 0.32 | 1.0E-03 |
| Ythdf1 | 7205 | 1.29 | 0.36 | 1.3E-08 | 1.17 | 0.23 | 4.9E-04 |
| Ythdf2 | 7904 | -1.24 | -0.31 | 9.2E-05 | -1.30 | -0.38 | 1.2E-06 |
| Ywhae | 73412 | -1.26 | -0.33 | 1.0E-04 | -1.36 | -0.44 | 1.2E-07 |
| Ywhah | 13565 | -1.43 | -0.51 | 6.0E-11 | -1.48 | -0.56 | 6.1E-13 |
| Ywhaz | 43502 | -1.35 | -0.43 | 1.8E-10 | -1.44 | -0.52 | 9.7E-15 |
| Zadh2 | 2087 | -1.20 | -0.26 | 2.4E-02 | -1.26 | -0.33 | 4.1E-03 |
| Zbed3 | 1723 | -1.61 | -0.69 | 4.8E-09 | -1.79 | -0.84 | 7.4E-13 |
| Zbtb1 | 2234 | -1.99 | -0.99 | 8.5E-51 | -1.95 | -0.96 | 2.8E-47 |
| Zbtb11 | 3024 | 1.25 | 0.33 | 1.9E-05 | 1.21 | 0.28 | 3.5E-04 |
| Zbtb14 | 2587 | -1.41 | -0.49 | 4.6E-08 | -1.42 | -0.50 | 2.8E-08 |
| Zbtb16 | 881 | 1.86 | 0.89 | 2.2E-04 | 1.70 | 0.77 | 1.7E-03 |
| Zbtb17 | 2743 | -1.51 | -0.59 | 6.5E-11 | -1.67 | -0.74 | 2.3E-16 |
| Zbtb2 | 1344 | -1.87 | -0.91 | 8.7E-24 | -1.93 | -0.95 | 6.9E-26 |
| Zbtb20 | 6592 | -2.52 | -1.34 | 2.0E-14 | -2.13 | -1.09 | 5.5E-10 |
| Zbtb22 | 1772 | -1.42 | -0.51 | 9.5E-09 | -1.61 | -0.68 | 5.0E-15 |

| | | | | | | | |
|---|---|---|---|---|---|---|---|
| Zbtb3 | 300 | -1.92 | -0.94 | 2.2E-18 | -2.02 | -1.02 | 6.3E-21 |
| Zbtb34 | 3174 | 1.49 | 0.57 | 2.2E-08 | 1.39 | 0.47 | 5.2E-06 |
| Zbtb38 | 1457 | -2.16 | -1.11 | 1.2E-07 | -2.01 | -1.00 | 1.9E-06 |
| Zbtb4 | 1452 | -1.33 | -0.41 | 1.2E-02 | -1.28 | -0.36 | 3.1E-02 |
| Zbtb40 | 1435 | -2.92 | -1.55 | 5.0E-38 | -2.90 | -1.54 | 1.7E-37 |
| Zbtb41 | 8934 | 1.24 | 0.31 | 1.4E-02 | 1.30 | 0.38 | 2.3E-03 |
| Zbtb42 | 494 | 2.41 | 1.27 | 7.3E-19 | 2.05 | 1.03 | 6.8E-13 |
| Zbtb43 | 3435 | 1.28 | 0.36 | 4.0E-10 | 1.30 | 0.37 | 4.7E-11 |
| Zbtb45 | 1207 | -1.77 | -0.82 | 2.6E-09 | -2.04 | -1.03 | 4.1E-14 |
| Zbtb46 | 2860 | 2.14 | 1.10 | 1.4E-36 | 2.06 | 1.04 | 3.8E-33 |
| Zbtb48 | 1459 | 1.44 | 0.52 | 3.3E-06 | 1.30 | 0.38 | 9.0E-04 |
| Zbtb7b | 537 | 3.27 | 1.71 | 8.3E-13 | 3.09 | 1.63 | 1.1E-11 |
| Zbtb8a | 612 | -1.85 | -0.89 | 9.5E-22 | -1.83 | -0.88 | 3.9E-21 |
| Zbtb8b | 664 | 1.84 | 0.88 | 4.7E-05 | 2.13 | 1.09 | 3.7E-07 |
| Zbtb9 | 266 | 2.03 | 1.02 | 2.2E-13 | 1.74 | 0.80 | 9.6E-09 |
| Zc2hc1a | 7340 | 1.36 | 0.44 | 1.8E-04 | 1.46 | 0.54 | 2.6E-06 |
| Zc2hc1c | 202 | 2.08 | 1.06 | 2.2E-19 | 2.14 | 1.10 | 9.6E-21 |
| Zc3h13 | 11899 | 1.31 | 0.39 | 7.1E-07 | 1.33 | 0.41 | 1.5E-07 |
| Zc3h14 | 13241 | 1.26 | 0.33 | 8.9E-04 | 1.27 | 0.35 | 4.2E-04 |
| Zc3h4 | 6993 | -2.62 | -1.39 | 3.8E-24 | -2.79 | -1.48 | 3.1E-27 |
| Zc3h7a | 9457 | 1.21 | 0.27 | 1.6E-05 | 1.31 | 0.39 | 2.5E-10 |
| Zc3h7b | 14482 | -1.61 | -0.69 | 1.3E-07 | -1.74 | -0.80 | 6.6E-10 |
| Zc3hav1l | 4136 | -2.46 | -1.30 | 1.6E-65 | -2.57 | -1.36 | 2.9E-72 |
| Zc4h2 | 3787 | -1.58 | -0.66 | 3.2E-08 | -1.49 | -0.58 | 1.5E-06 |
| Zcchc10 | 635 | -1.73 | -0.79 | 1.1E-09 | -1.94 | -0.96 | 1.1E-13 |
| Zcchc12 | 7495 | 1.59 | 0.67 | 3.2E-05 | 1.57 | 0.65 | 5.4E-05 |
| Zcchc17 | 3503 | 1.44 | 0.52 | 9.5E-19 | 1.43 | 0.52 | 2.3E-18 |
| Zcchc18 | 5879 | -1.24 | -0.31 | 4.3E-02 | -1.28 | -0.36 | 1.8E-02 |
| Zcchc2 | 2689 | 2.15 | 1.10 | 7.1E-52 | 2.10 | 1.07 | 8.3E-49 |
| Zcchc4 | 1442 | -1.78 | -0.83 | 1.9E-32 | -1.78 | -0.83 | 1.8E-32 |
| Zcchc9 | 1816 | -1.54 | -0.63 | 5.8E-21 | -1.63 | -0.71 | 2.2E-26 |
| Zdbf2 | 13706 | 1.60 | 0.68 | 8.1E-06 | 1.55 | 0.64 | 3.2E-05 |
| Zdhhc12 | 777 | 2.01 | 1.01 | 5.3E-18 | 1.79 | 0.84 | 8.5E-13 |
| Zdhhc13 | 2682 | 1.37 | 0.46 | 1.4E-08 | 1.41 | 0.50 | 5.4E-10 |
| Zdhhc17 | 6534 | -1.62 | -0.69 | 6.5E-10 | -1.51 | -0.59 | 1.7E-07 |
| Zdhhc20 | 8506 | 1.29 | 0.37 | 4.4E-11 | 1.35 | 0.44 | 5.1E-15 |
| Zdhhc23 | 462 | 2.03 | 1.02 | 5.3E-06 | 2.05 | 1.03 | 4.3E-06 |
| Zdhhc24 | 1564 | 1.37 | 0.46 | 5.8E-10 | 1.31 | 0.39 | 2.0E-07 |
| Zdhhc3 | 6586 | -1.23 | -0.29 | 1.3E-09 | -1.28 | -0.36 | 6.3E-14 |
| Zdhhc4 | 1491 | -1.24 | -0.31 | 1.4E-07 | -1.29 | -0.37 | 2.6E-10 |
| Zeb2 | 8459 | -2.12 | -1.08 | 3.6E-24 | -1.99 | -0.99 | 1.9E-20 |
| Zfand4 | 1432 | 1.58 | 0.66 | 1.7E-07 | 1.49 | 0.57 | 5.8E-06 |
| Zfand5 | 13265 | -1.26 | -0.33 | 5.6E-05 | -1.32 | -0.40 | 1.5E-06 |
| Zfand6 | 11669 | 1.41 | 0.50 | 1.4E-15 | 1.45 | 0.54 | 4.4E-18 |
| Zfat | 427 | -7.05 | -2.82 | 1.4E-91 | -7.47 | -2.90 | 5.4E-97 |
| Zfc3h1 | 13100 | 1.57 | 0.65 | 3.4E-09 | 1.78 | 0.83 | 2.6E-14 |
| Zfhx3 | 8229 | -1.96 | -0.97 | 5.0E-07 | -2.12 | -1.08 | 1.8E-08 |

| | | | | | | | |
|---|---|---|---|---|---|---|---|
| Zfhx4 | 12364 | -3.17 | -1.67 | 8.0E-49 | -2.88 | -1.53 | 4.2E-41 |
| Zfp1 | 1240 | -1.54 | -0.63 | 2.7E-13 | -1.60 | -0.67 | 3.5E-15 |
| Zfp105 | 1673 | -1.52 | -0.61 | 5.6E-11 | -1.74 | -0.80 | 4.1E-18 |
| Zfp111 | 1581 | -1.63 | -0.71 | 3.6E-24 | -1.70 | -0.76 | 1.3E-27 |
| Zfp113 | 3507 | -1.47 | -0.56 | 1.7E-12 | -1.48 | -0.56 | 8.7E-13 |
| Zfp114 | 275 | -2.22 | -1.15 | 4.5E-11 | -1.74 | -0.80 | 6.5E-06 |
| Zfp119b | 356 | -1.39 | -0.48 | 2.9E-06 | -1.46 | -0.55 | 6.0E-08 |
| Zfp142 | 3811 | 1.49 | 0.57 | 1.3E-10 | 1.50 | 0.59 | 3.5E-11 |
| Zfp146 | 7965 | -1.58 | -0.66 | 1.2E-20 | -1.58 | -0.66 | 1.6E-20 |
| Zfp157 | 6202 | -1.42 | -0.50 | 1.6E-07 | -1.34 | -0.42 | 1.6E-05 |
| Zfp169 | 1064 | 1.39 | 0.48 | 4.3E-02 | 1.59 | 0.67 | 3.8E-03 |
| Zfp174 | 639 | 1.29 | 0.37 | 1.3E-02 | 1.38 | 0.47 | 1.6E-03 |
| Zfp180 | 3261 | -1.35 | -0.44 | 2.5E-11 | -1.42 | -0.51 | 7.7E-15 |
| Zfp182 | 1697 | -1.34 | -0.42 | 1.1E-04 | -1.22 | -0.29 | 1.1E-02 |
| Zfp189 | 1767 | 1.27 | 0.35 | 8.8E-06 | 1.35 | 0.43 | 2.1E-08 |
| Zfp2 | 2785 | -1.82 | -0.87 | 4.0E-12 | -1.70 | -0.77 | 1.0E-09 |
| Zfp217 | 3397 | 1.49 | 0.58 | 4.7E-07 | 1.40 | 0.49 | 2.2E-05 |
| Zfp219 | 6817 | -1.67 | -0.74 | 9.0E-07 | -1.93 | -0.95 | 1.9E-10 |
| Zfp229 | 1688 | -1.42 | -0.51 | 6.5E-11 | -1.38 | -0.46 | 4.1E-09 |
| Zfp236 | 3882 | -2.20 | -1.14 | 2.2E-26 | -2.06 | -1.05 | 1.8E-22 |
| Zfp24 | 7979 | -1.61 | -0.69 | 3.4E-15 | -1.57 | -0.65 | 7.0E-14 |
| Zfp248 | 1254 | -1.37 | -0.46 | 3.9E-11 | -1.35 | -0.43 | 4.0E-10 |
| Zfp251 | 5453 | 1.39 | 0.48 | 2.0E-25 | 1.34 | 0.42 | 3.6E-20 |
| Zfp26 | 5885 | -1.37 | -0.46 | 9.3E-07 | -1.27 | -0.34 | 3.8E-04 |
| Zfp260 | 8170 | -1.84 | -0.88 | 2.3E-32 | -1.78 | -0.83 | 1.5E-28 |
| Zfp267 | 3366 | 1.24 | 0.32 | 3.6E-03 | 1.35 | 0.44 | 3.8E-05 |
| Zfp273 | 716 | -1.74 | -0.80 | 9.4E-11 | -1.62 | -0.70 | 2.0E-08 |
| Zfp276 | 2602 | 1.25 | 0.33 | 2.1E-05 | 1.23 | 0.30 | 1.2E-04 |
| Zfp277 | 3025 | 1.93 | 0.95 | 1.8E-21 | 1.97 | 0.97 | 1.1E-22 |
| Zfp280c | 4253 | -1.45 | -0.54 | 2.4E-07 | -1.42 | -0.51 | 1.3E-06 |
| Zfp280d | 6356 | -1.86 | -0.89 | 1.0E-25 | -1.89 | -0.92 | 4.0E-27 |
| Zfp282 | 3474 | -1.38 | -0.46 | 1.4E-07 | -1.46 | -0.54 | 5.9E-10 |
| Zfp286 | 2098 | -1.37 | -0.45 | 5.1E-07 | -1.28 | -0.35 | 1.2E-04 |
| Zfp292 | 13697 | -1.64 | -0.71 | 1.3E-05 | -1.38 | -0.46 | 6.2E-03 |
| Zfp317 | 5125 | -1.25 | -0.33 | 1.9E-08 | -1.19 | -0.25 | 1.6E-05 |
| Zfp324 | 2240 | 1.26 | 0.34 | 3.0E-07 | 1.32 | 0.40 | 8.7E-10 |
| Zfp326 | 7224 | -1.36 | -0.44 | 1.2E-07 | -1.41 | -0.49 | 2.3E-09 |
| Zfp334 | 5954 | -1.29 | -0.36 | 5.0E-06 | -1.35 | -0.43 | 5.6E-08 |
| Zfp335 | 2346 | -1.52 | -0.60 | 1.9E-09 | -1.57 | -0.65 | 8.3E-11 |
| Zfp341 | 517 | -4.29 | -2.10 | 1.6E-34 | -4.49 | -2.17 | 1.1E-36 |
| Zfp346 | 3584 | 1.60 | 0.68 | 1.1E-40 | 1.57 | 0.65 | 2.7E-37 |
| Zfp358 | 2672 | -1.56 | -0.64 | 1.6E-17 | -1.64 | -0.71 | 2.2E-21 |
| Zfp362 | 4782 | -2.00 | -1.00 | 1.5E-10 | -2.34 | -1.23 | 2.6E-15 |
| Zfp366 | 402 | -1.69 | -0.75 | 2.8E-07 | -1.69 | -0.76 | 2.2E-07 |
| Zfp36l1 | 11177 | -1.39 | -0.47 | 3.7E-03 | -1.58 | -0.66 | 3.6E-05 |
| Zfp36l2 | 2849 | -1.35 | -0.43 | 4.2E-03 | -1.44 | -0.53 | 4.3E-04 |
| Zfp37 | 2948 | -1.43 | -0.52 | 3.3E-09 | -1.43 | -0.52 | 3.7E-09 |

| | | | | | | | |
|---|---|---|---|---|---|---|---|
| Zfp383 | 3042 | -1.69 | -0.75 | 5.5E-32 | -1.69 | -0.76 | 1.5E-32 |
| Zfp384 | 5270 | -1.44 | -0.53 | 1.0E-24 | -1.46 | -0.54 | 1.4E-25 |
| Zfp385a | 3630 | 1.81 | 0.86 | 1.6E-10 | 1.59 | 0.66 | 1.0E-06 |
| Zfp395 | 2558 | 1.37 | 0.45 | 3.3E-03 | 1.33 | 0.41 | 8.1E-03 |
| Zfp407 | 2262 | -2.08 | -1.06 | 6.4E-20 | -1.88 | -0.91 | 5.1E-15 |
| Zfp414 | 804 | -2.07 | -1.05 | 3.5E-21 | -2.37 | -1.25 | 2.4E-29 |
| Zfp422 | 9862 | -1.87 | -0.90 | 4.6E-16 | -2.06 | -1.04 | 4.5E-21 |
| Zfp423 | 3023 | -1.96 | -0.97 | 2.5E-12 | -2.17 | -1.12 | 6.2E-16 |
| Zfp428 | 1831 | -2.44 | -1.28 | 4.2E-15 | -2.55 | -1.35 | 1.5E-16 |
| Zfp433 | 212 | -2.11 | -1.08 | 1.7E-12 | -1.79 | -0.84 | 4.9E-08 |
| Zfp438 | 263 | -1.94 | -0.95 | 2.0E-19 | -2.06 | -1.04 | 1.2E-22 |
| Zfp444 | 3554 | 1.57 | 0.65 | 4.9E-11 | 1.47 | 0.56 | 2.2E-08 |
| Zfp445 | 23451 | -1.41 | -0.50 | 5.0E-07 | -1.29 | -0.36 | 3.6E-04 |
| Zfp446 | 1546 | -1.44 | -0.53 | 1.4E-09 | -1.41 | -0.50 | 1.2E-08 |
| Zfp451 | 8614 | -1.33 | -0.41 | 1.6E-04 | -1.36 | -0.44 | 5.6E-05 |
| Zfp455 | 173 | -2.22 | -1.15 | 2.1E-08 | -2.01 | -1.01 | 1.1E-06 |
| Zfp459 | 134 | 1.53 | 0.62 | 3.2E-05 | 1.45 | 0.53 | 3.8E-04 |
| Zfp462 | 16096 | -2.88 | -1.53 | 2.8E-27 | -3.04 | -1.60 | 5.8E-30 |
| Zfp467 | 1555 | -3.18 | -1.67 | 1.1E-51 | -3.37 | -1.75 | 1.1E-56 |
| Zfp469 | 523 | -2.88 | -1.52 | 6.2E-06 | -3.13 | -1.65 | 1.0E-06 |
| Zfp493 | 542 | 1.32 | 0.40 | 1.4E-07 | 1.38 | 0.47 | 8.8E-10 |
| Zfp507 | 4725 | -1.34 | -0.42 | 2.2E-10 | -1.26 | -0.34 | 4.8E-07 |
| Zfp51 | 792 | -1.32 | -0.40 | 4.1E-04 | -1.22 | -0.29 | 1.4E-02 |
| Zfp511 | 1167 | -1.30 | -0.38 | 1.3E-05 | -1.42 | -0.51 | 3.1E-09 |
| Zfp512 | 8309 | -1.21 | -0.28 | 5.9E-05 | -1.25 | -0.32 | 5.1E-06 |
| Zfp512b | 10518 | -1.38 | -0.47 | 4.9E-05 | -1.41 | -0.50 | 1.5E-05 |
| Zfp513 | 1553 | -1.52 | -0.60 | 1.6E-09 | -1.63 | -0.71 | 1.1E-12 |
| Zfp516 | 6909 | 1.81 | 0.86 | 5.7E-26 | 1.83 | 0.87 | 5.7E-27 |
| Zfp518a | 3169 | -1.80 | -0.85 | 4.4E-17 | -1.61 | -0.69 | 1.4E-11 |
| Zfp518b | 8224 | -1.57 | -0.66 | 1.4E-10 | -1.62 | -0.69 | 1.2E-11 |
| Zfp52 | 451 | 1.84 | 0.88 | 1.7E-14 | 1.89 | 0.92 | 2.1E-15 |
| Zfp521 | 5820 | -1.89 | -0.92 | 2.9E-72 | -1.92 | -0.94 | 1.2E-75 |
| Zfp523 | 3484 | -1.37 | -0.45 | 2.9E-05 | -1.42 | -0.51 | 2.7E-06 |
| Zfp53 | 877 | 1.26 | 0.33 | 5.6E-07 | 1.31 | 0.39 | 6.7E-09 |
| Zfp532 | 8221 | -2.91 | -1.54 | 5.1E-64 | -2.92 | -1.54 | 4.1E-64 |
| Zfp551 | 412 | -1.71 | -0.77 | 3.3E-03 | -1.48 | -0.57 | 3.5E-02 |
| Zfp558 | 2187 | -1.50 | -0.59 | 1.6E-03 | -1.49 | -0.58 | 2.0E-03 |
| Zfp560 | 1304 | -1.89 | -0.92 | 6.8E-13 | -1.86 | -0.89 | 2.9E-12 |
| Zfp563 | 2200 | -1.41 | -0.50 | 3.2E-04 | -1.35 | -0.44 | 1.9E-03 |
| Zfp566 | 616 | -1.22 | -0.28 | 1.4E-03 | -1.29 | -0.37 | 2.2E-05 |
| Zfp57 | 12157 | -2.06 | -1.04 | 3.4E-10 | -2.10 | -1.07 | 9.9E-11 |
| Zfp583 | 623 | -2.03 | -1.02 | 3.2E-38 | -2.06 | -1.05 | 9.0E-40 |
| Zfp592 | 3247 | -1.59 | -0.67 | 1.7E-12 | -1.64 | -0.71 | 8.3E-14 |
| Zfp593 | 936 | 1.82 | 0.86 | 1.0E-13 | 1.63 | 0.70 | 1.8E-09 |
| Zfp595 | 469 | -1.71 | -0.78 | 4.6E-11 | -1.44 | -0.53 | 1.3E-05 |
| Zfp60 | 11655 | -1.35 | -0.44 | 1.9E-05 | -1.26 | -0.33 | 1.5E-03 |
| Zfp605 | 1602 | -1.51 | -0.59 | 1.2E-07 | -1.43 | -0.52 | 4.4E-06 |

| | | | | | | | |
|---|---|---|---|---|---|---|---|
| Zfp608 | 6053 | -1.44 | -0.53 | 3.6E-08 | -1.46 | -0.54 | 1.4E-08 |
| Zfp61 | 1983 | -1.36 | -0.44 | 8.8E-11 | -1.42 | -0.51 | 1.3E-13 |
| Zfp612 | 3339 | 1.82 | 0.86 | 3.2E-49 | 1.88 | 0.91 | 1.2E-54 |
| Zfp618 | 6158 | -4.16 | -2.06 | 8.9E-83 | -4.04 | -2.01 | 3.7E-79 |
| Zfp619 | 823 | -2.46 | -1.30 | 4.2E-14 | -2.08 | -1.06 | 1.2E-09 |
| Zfp62 | 8113 | -1.41 | -0.49 | 1.7E-10 | -1.31 | -0.39 | 6.5E-07 |
| Zfp622 | 2705 | 1.28 | 0.36 | 6.6E-08 | 1.22 | 0.29 | 1.4E-05 |
| Zfp628 | 968 | -2.31 | -1.21 | 7.2E-12 | -2.59 | -1.37 | 6.2E-15 |
| Zfp637 | 1956 | -2.07 | -1.05 | 6.2E-64 | -2.13 | -1.09 | 2.0E-68 |
| Zfp638 | 16010 | -1.68 | -0.75 | 1.4E-10 | -1.43 | -0.52 | 1.3E-05 |
| Zfp639 | 3869 | 1.22 | 0.29 | 5.1E-06 | 1.25 | 0.32 | 2.9E-07 |
| Zfp641 | 621 | -1.59 | -0.67 | 1.6E-03 | -1.82 | -0.87 | 3.8E-05 |
| Zfp644 | 6934 | -1.76 | -0.81 | 6.7E-22 | -1.61 | -0.68 | 8.8E-16 |
| Zfp646 | 5666 | -1.35 | -0.44 | 1.9E-06 | -1.37 | -0.45 | 8.0E-07 |
| Zfp647 | 1177 | -1.39 | -0.48 | 1.1E-11 | -1.58 | -0.66 | 1.7E-21 |
| Zfp651 | 2369 | -1.33 | -0.41 | 2.1E-03 | -1.35 | -0.44 | 1.0E-03 |
| Zfp653 | 843 | -1.90 | -0.92 | 2.3E-14 | -2.13 | -1.09 | 1.1E-19 |
| Zfp654 | 3734 | 1.28 | 0.36 | 7.1E-05 | 1.33 | 0.41 | 4.8E-06 |
| Zfp661 | 1390 | 1.36 | 0.44 | 4.4E-16 | 1.33 | 0.41 | 4.7E-14 |
| Zfp667 | 2531 | 2.25 | 1.17 | 3.6E-56 | 2.26 | 1.18 | 1.2E-56 |
| Zfp668 | 2202 | -1.35 | -0.44 | 6.5E-04 | -1.53 | -0.61 | 1.1E-06 |
| Zfp68 | 6844 | -1.32 | -0.40 | 3.6E-06 | -1.23 | -0.30 | 6.1E-04 |
| Zfp687 | 5934 | 1.69 | 0.76 | 6.6E-16 | 1.57 | 0.65 | 6.4E-12 |
| Zfp69 | 585 | 1.62 | 0.70 | 7.2E-18 | 1.67 | 0.74 | 1.3E-19 |
| Zfp697 | 2635 | -1.19 | -0.26 | 2.0E-02 | -1.25 | -0.32 | 2.8E-03 |
| Zfp7 | 1302 | 1.27 | 0.35 | 2.6E-06 | 1.20 | 0.26 | 6.0E-04 |
| Zfp710 | 2445 | -1.74 | -0.80 | 5.0E-06 | -2.06 | -1.04 | 1.5E-09 |
| Zfp711 | 4206 | -1.40 | -0.48 | 3.4E-04 | -1.42 | -0.50 | 1.8E-04 |
| Zfp715 | 3122 | 1.27 | 0.35 | 4.4E-04 | 1.35 | 0.43 | 1.0E-05 |
| Zfp729b | 752 | -1.26 | -0.34 | 9.7E-04 | -1.20 | -0.27 | 1.0E-02 |
| Zfp74 | 1344 | -2.08 | -1.05 | 1.5E-28 | -2.20 | -1.14 | 5.9E-33 |
| Zfp746 | 2619 | -1.31 | -0.39 | 1.1E-06 | -1.33 | -0.42 | 2.6E-07 |
| Zfp747 | 677 | 1.48 | 0.57 | 5.7E-12 | 1.35 | 0.44 | 1.9E-07 |
| Zfp759 | 686 | -1.42 | -0.51 | 6.6E-04 | -1.28 | -0.35 | 2.2E-02 |
| Zfp764 | 465 | -1.27 | -0.35 | 3.7E-04 | -1.24 | -0.31 | 1.6E-03 |
| Zfp772 | 1294 | 2.03 | 1.02 | 7.7E-35 | 1.83 | 0.87 | 1.4E-25 |
| Zfp773 | 1023 | -2.08 | -1.06 | 3.5E-07 | -2.12 | -1.08 | 1.9E-07 |
| Zfp775 | 967 | 1.25 | 0.33 | 1.2E-03 | 1.23 | 0.30 | 3.5E-03 |
| Zfp777 | 1941 | -1.19 | -0.25 | 1.8E-03 | -1.31 | -0.39 | 1.0E-06 |
| Zfp78 | 400 | 1.98 | 0.98 | 6.7E-18 | 1.97 | 0.98 | 1.5E-17 |
| Zfp781 | 182 | -1.30 | -0.38 | 1.6E-02 | -1.25 | -0.33 | 4.4E-02 |
| Zfp784 | 254 | -3.19 | -1.67 | 3.9E-49 | -3.24 | -1.70 | 3.3E-50 |
| Zfp787 | 1026 | -1.54 | -0.63 | 1.9E-10 | -1.56 | -0.65 | 5.6E-11 |
| Zfp788 | 3856 | -1.42 | -0.51 | 9.6E-06 | -1.40 | -0.48 | 3.2E-05 |
| Zfp800 | 1722 | 1.28 | 0.36 | 8.9E-05 | 1.28 | 0.36 | 8.5E-05 |
| Zfp804a | 553 | -1.88 | -0.91 | 3.6E-03 | -1.68 | -0.75 | 2.0E-02 |
| Zfp808 | 588 | -1.39 | -0.48 | 5.6E-04 | -1.41 | -0.49 | 4.0E-04 |

| | | | | | | | |
|---|---|---|---|---|---|---|---|
| Zfp809 | 728 | -1.57 | -0.65 | 2.0E-13 | -1.51 | -0.59 | 4.0E-11 |
| Zfp810 | 2704 | -1.94 | -0.95 | 1.4E-35 | -1.95 | -0.96 | 4.9E-36 |
| Zfp820 | 1083 | 1.44 | 0.52 | 2.2E-04 | 1.39 | 0.48 | 7.8E-04 |
| Zfp821 | 5509 | -1.59 | -0.67 | 2.1E-09 | -1.67 | -0.74 | 3.3E-11 |
| Zfp825 | 894 | -1.43 | -0.51 | 1.1E-08 | -1.41 | -0.49 | 3.7E-08 |
| Zfp827 | 3982 | -2.60 | -1.38 | 2.6E-31 | -2.52 | -1.34 | 1.6E-29 |
| Zfp846 | 3942 | -1.68 | -0.75 | 1.5E-05 | -1.44 | -0.53 | 2.7E-03 |
| Zfp85 | 847 | 1.32 | 0.40 | 8.2E-05 | 1.39 | 0.47 | 2.4E-06 |
| Zfp850 | 2057 | 1.44 | 0.52 | 1.5E-07 | 1.60 | 0.68 | 4.3E-12 |
| Zfp866 | 5327 | -1.57 | -0.65 | 1.5E-13 | -1.49 | -0.57 | 8.0E-11 |
| Zfp867 | 2057 | 1.41 | 0.50 | 5.4E-09 | 1.48 | 0.56 | 2.7E-11 |
| Zfp868 | 2268 | -1.31 | -0.39 | 1.1E-06 | -1.36 | -0.44 | 3.7E-08 |
| Zfp869 | 3460 | -1.50 | -0.58 | 4.6E-24 | -1.46 | -0.54 | 2.9E-21 |
| Zfp870 | 1672 | 1.28 | 0.35 | 2.6E-06 | 1.28 | 0.36 | 2.0E-06 |
| Zfp873 | 927 | -2.50 | -1.32 | 8.5E-34 | -2.37 | -1.24 | 7.0E-30 |
| Zfp879 | 122 | -2.23 | -1.15 | 1.1E-11 | -1.87 | -0.90 | 1.9E-07 |
| Zfp882 | 1714 | -1.63 | -0.70 | 8.3E-08 | -1.35 | -0.43 | 1.5E-03 |
| Zfp9 | 2178 | -1.28 | -0.36 | 1.2E-03 | -1.22 | -0.29 | 9.5E-03 |
| Zfp90 | 3209 | -1.96 | -0.97 | 9.3E-26 | -1.96 | -0.97 | 4.9E-26 |
| Zfp91 | 10023 | -1.38 | -0.47 | 1.6E-07 | -1.40 | -0.49 | 4.5E-08 |
| Zfp934 | 795 | -1.37 | -0.45 | 2.7E-04 | -1.48 | -0.57 | 4.3E-06 |
| Zfp935 | 244 | -1.27 | -0.34 | 2.3E-03 | -1.27 | -0.34 | 2.6E-03 |
| Zfp94 | 741 | -1.41 | -0.49 | 7.1E-11 | -1.40 | -0.49 | 8.6E-11 |
| Zfp941 | 1127 | 1.33 | 0.41 | 3.2E-03 | 1.38 | 0.46 | 7.1E-04 |
| Zfp942 | 2807 | 1.35 | 0.43 | 5.9E-07 | 1.42 | 0.51 | 2.4E-09 |
| Zfp944 | 2350 | 1.73 | 0.79 | 9.7E-23 | 1.73 | 0.79 | 4.0E-23 |
| Zfp946 | 4810 | 1.25 | 0.32 | 6.0E-03 | 1.28 | 0.36 | 2.4E-03 |
| Zfp948 | 2365 | 1.57 | 0.65 | 2.8E-19 | 1.60 | 0.68 | 1.4E-20 |
| Zfp949 | 1805 | 1.25 | 0.33 | 2.2E-04 | 1.32 | 0.40 | 4.4E-06 |
| Zfp951 | 326 | -1.64 | -0.72 | 5.1E-05 | -1.62 | -0.70 | 9.0E-05 |
| Zfp954 | 1227 | 1.42 | 0.51 | 2.6E-07 | 1.47 | 0.56 | 1.1E-08 |
| Zfp955b | 4021 | 1.26 | 0.33 | 6.7E-04 | 1.38 | 0.46 | 1.3E-06 |
| Zfp958 | 1727 | 1.44 | 0.53 | 3.0E-12 | 1.53 | 0.61 | 3.9E-16 |
| Zfp959 | 799 | -1.44 | -0.53 | 9.5E-09 | -1.45 | -0.54 | 6.7E-09 |
| Zfp963 | 386 | -1.27 | -0.34 | 2.0E-02 | -1.50 | -0.59 | 3.9E-05 |
| Zfp964 | 711 | 1.41 | 0.50 | 5.2E-07 | 1.44 | 0.53 | 8.9E-08 |
| Zfp979 | 188 | -2.59 | -1.38 | 3.4E-12 | -2.57 | -1.36 | 4.8E-12 |
| Zfp994 | 951 | -1.27 | -0.35 | 1.3E-03 | -1.22 | -0.28 | 9.5E-03 |
| Zfpm1 | 1297 | 2.56 | 1.36 | 1.2E-21 | 2.62 | 1.39 | 1.1E-22 |
| Zfr2 | 2469 | 1.87 | 0.90 | 1.2E-08 | 1.93 | 0.95 | 1.7E-09 |
| Zfta | 17016 | -2.73 | -1.45 | 7.0E-24 | -2.85 | -1.51 | 6.4E-26 |
| Zfx | 4466 | -2.49 | -1.32 | 3.9E-50 | -2.46 | -1.30 | 7.7E-49 |
| Zfyve21 | 1323 | 1.51 | 0.60 | 4.2E-05 | 1.43 | 0.52 | 4.1E-04 |
| Zfyve26 | 2417 | -1.46 | -0.55 | 1.9E-09 | -1.33 | -0.41 | 1.0E-05 |
| Zfyve27 | 6713 | 1.72 | 0.78 | 2.5E-15 | 1.79 | 0.84 | 1.8E-17 |
| Zfyve28 | 820 | 4.08 | 2.03 | 1.5E-44 | 4.12 | 2.04 | 6.1E-45 |
| Zfyve9 | 3888 | -2.14 | -1.09 | 3.4E-34 | -1.96 | -0.97 | 3.5E-27 |

| | | | | | | | |
|---|---|---|---|---|---|---|---|
| Zhx2 | 1383 | -1.93 | -0.95 | 2.4E-11 | -1.92 | -0.94 | 3.8E-11 |
| Zhx3 | 3313 | 1.35 | 0.44 | 3.7E-06 | 1.41 | 0.49 | 1.8E-07 |
| Zic4 | 6358 | -1.29 | -0.36 | 1.0E-02 | -1.48 | -0.56 | 4.5E-05 |
| Zkscan1 | 8027 | -1.25 | -0.32 | 1.9E-05 | -1.23 | -0.30 | 1.1E-04 |
| Zkscan14 | 596 | 1.34 | 0.43 | 1.1E-04 | 1.32 | 0.40 | 3.3E-04 |
| Zkscan16 | 668 | 1.83 | 0.87 | 1.3E-03 | 2.57 | 1.36 | 2.7E-07 |
| Zkscan6 | 1229 | 1.27 | 0.35 | 5.2E-06 | 1.23 | 0.30 | 8.1E-05 |
| Zkscan8 | 5228 | -1.59 | -0.67 | 3.9E-06 | -1.33 | -0.41 | 5.8E-03 |
| Zmat3 | 4663 | 1.66 | 0.73 | 2.0E-06 | 1.43 | 0.52 | 1.0E-03 |
| Zmiz1 | 19376 | -1.45 | -0.53 | 1.1E-03 | -1.53 | -0.61 | 1.8E-04 |
| Zmym1 | 2904 | 1.34 | 0.42 | 3.7E-07 | 1.40 | 0.48 | 4.0E-09 |
| Zmym2 | 16595 | -1.41 | -0.49 | 2.1E-06 | -1.28 | -0.36 | 6.7E-04 |
| Zmym3 | 9494 | -1.27 | -0.35 | 4.2E-03 | -1.22 | -0.29 | 1.9E-02 |
| Zmym4 | 12662 | -1.53 | -0.61 | 7.8E-14 | -1.45 | -0.54 | 5.7E-11 |
| Zmynd10 | 281 | 1.93 | 0.95 | 2.1E-08 | 1.80 | 0.85 | 6.3E-07 |
| Zmynd11 | 13113 | -1.87 | -0.90 | 7.8E-23 | -1.83 | -0.87 | 1.8E-21 |
| Zmynd19 | 5752 | 1.54 | 0.63 | 1.5E-18 | 1.44 | 0.52 | 2.8E-13 |
| Zmynd8 | 7708 | -1.94 | -0.96 | 1.1E-19 | -1.96 | -0.97 | 3.1E-20 |
| Znhit1 | 1396 | -1.81 | -0.86 | 2.4E-44 | -1.90 | -0.93 | 6.2E-52 |
| Znhit3 | 1370 | 1.36 | 0.45 | 8.8E-08 | 1.31 | 0.39 | 5.3E-06 |
| Znrf2 | 4891 | 1.56 | 0.65 | 1.0E-03 | 1.64 | 0.72 | 2.5E-04 |
| Znrf3 | 1959 | 1.37 | 0.46 | 8.3E-07 | 1.24 | 0.32 | 9.1E-04 |
| Zpr1 | 3319 | 1.27 | 0.34 | 1.2E-06 | 1.20 | 0.27 | 1.9E-04 |
| Zranb1 | 8021 | 1.25 | 0.33 | 4.9E-04 | 1.25 | 0.32 | 7.3E-04 |
| Zrsr2 | 3328 | -1.37 | -0.46 | 5.2E-13 | -1.40 | -0.49 | 7.6E-15 |
| Zscan2 | 1220 | -1.35 | -0.43 | 1.2E-03 | -1.34 | -0.42 | 1.5E-03 |
| Zscan25 | 830 | -1.50 | -0.58 | 2.2E-11 | -1.66 | -0.73 | 4.5E-17 |
| Zswim3 | 917 | 1.67 | 0.74 | 3.9E-17 | 1.58 | 0.66 | 5.6E-14 |
| Zswim7 | 190 | -1.80 | -0.85 | 3.1E-11 | -1.88 | -0.91 | 1.1E-12 |
| Zswim8 | 18251 | 1.37 | 0.45 | 5.0E-05 | 1.33 | 0.41 | 2.4E-04 |
| Zswim9 | 2176 | -1.44 | -0.52 | 9.9E-18 | -1.44 | -0.52 | 1.0E-17 |
| Zw10 | 4754 | 1.31 | 0.39 | 1.1E-05 | 1.30 | 0.38 | 1.6E-05 |
| Zxdb | 744 | -1.25 | -0.32 | 1.8E-02 | -1.26 | -0.34 | 1.3E-02 |
| Zxdc | 2298 | 1.29 | 0.37 | 1.6E-05 | 1.32 | 0.40 | 2.8E-06 |
| Zyx | 3763 | -1.29 | -0.37 | 2.0E-02 | -1.43 | -0.52 | 7.8E-04 |
| Zzz3 | 7137 | -2.00 | -1.00 | 1.4E-65 | -1.97 | -0.98 | 1.5E-62 |

| Gene | Counts | FC | LFC | padj |
|---|---|---|---|---|
| 2310061I04Rik | 1372 | -1.21 | -0.27 | 2.32E-02 |
| 2810408A11Rik | 1262 | -1.30 | -0.38 | 8.66E-03 |
| 4932438A13Rik | 11118 | 1.78 | 0.83 | 3.05E-03 |
| Aar2 | 2823 | -1.11 | -0.15 | 2.27E-03 |
| Acaca | 13162 | 1.26 | 0.34 | 7.94E-03 |
| Acad11 | 989 | 1.30 | 0.38 | 1.47E-02 |
| Acss3 | 1028 | 1.68 | 0.75 | 1.27E-02 |
| Adamts7 | 3381 | -1.27 | -0.35 | 1.67E-03 |
| Adarb2 | 1599 | 1.53 | 0.61 | 1.40E-02 |
| Adcy5 | 3439 | 1.30 | 0.37 | 2.27E-02 |
| Adprh | 5714 | -1.18 | -0.24 | 1.15E-02 |
| Adra1a | 356 | 1.98 | 0.99 | 1.12E-03 |
| Alg12 | 845 | -1.17 | -0.23 | 1.24E-02 |
| Alms1 | 5356 | 1.67 | 0.74 | 1.45E-03 |
| Amer1 | 4653 | 1.26 | 0.34 | 1.33E-02 |
| Amot | 9654 | 1.39 | 0.48 | 6.99E-03 |
| Anapc13 | 2092 | -1.46 | -0.54 | 1.14E-02 |
| Ankle1 | 1377 | -1.36 | -0.44 | 1.29E-02 |
| Ankrd40 | 4320 | -1.25 | -0.32 | 1.00E-02 |
| Ap1s1 | 5705 | -1.13 | -0.17 | 2.28E-02 |
| Apcdd1 | 10238 | -1.23 | -0.30 | 1.59E-03 |
| Apod | 1656 | -1.33 | -0.41 | 2.16E-02 |
| Arhgap12 | 7583 | 1.19 | 0.25 | 2.13E-03 |
| Arl6ip4 | 2097 | -1.21 | -0.27 | 2.40E-02 |
| Arpc1a | 8165 | -1.07 | -0.10 | 1.14E-02 |
| Asphd1 | 339 | -1.56 | -0.64 | 2.18E-02 |
| Atm | 6498 | 1.60 | 0.68 | 2.25E-02 |
| Atp11a | 8506 | 1.22 | 0.29 | 5.06E-03 |
| Atp2b1 | 13439 | 1.24 | 0.31 | 1.94E-02 |
| Atp5g2 | 5429 | -1.26 | -0.33 | 1.14E-02 |
| Atp7a | 4050 | 1.55 | 0.64 | 1.10E-02 |
| Atr | 6071 | 1.39 | 0.47 | 9.66E-03 |
| Atrn | 8254 | 1.27 | 0.35 | 4.53E-03 |
| Atxn2l | 14834 | -1.36 | -0.45 | 2.30E-03 |
| Axin1 | 4403 | -1.22 | -0.29 | 2.88E-03 |
| B230307C23Rik | 547 | 1.43 | 0.51 | 1.11E-03 |
| Babam2 | 1176 | -1.16 | -0.22 | 5.48E-03 |
| Bace2 | 192 | 1.49 | 0.58 | 1.98E-02 |
| Baz1a | 3933 | -1.26 | -0.33 | 1.48E-02 |
| Bbs7 | 1938 | 1.13 | 0.18 | 8.69E-03 |
| Bcar3 | 561 | -1.29 | -0.37 | 1.38E-02 |
| Bfar | 3491 | -1.11 | -0.15 | 1.03E-02 |
| Bloc1s5 | 1633 | -1.10 | -0.14 | 2.31E-02 |
| Bod1l | 10264 | 1.43 | 0.52 | 1.78E-04 |
| Bola3 | 578 | -1.45 | -0.54 | 2.16E-04 |
| Bsg | 12872 | -1.15 | -0.20 | 1.13E-02 |

| | | | | |
|---|---|---|---|---|
| Bud13 | 1542 | -1.16 | -0.22 | 2.42E-02 |
| Cacfd1 | 7691 | -1.32 | -0.40 | 2.60E-03 |
| Cage1 | 218 | 1.32 | 0.40 | 1.34E-02 |
| Cap2 | 1094 | -1.22 | -0.29 | 1.72E-02 |
| Capn15 | 3377 | -1.35 | -0.43 | 1.22E-02 |
| Ccdc71 | 2625 | -1.15 | -0.20 | 7.23E-03 |
| Ccdc74a | 187 | -1.40 | -0.49 | 4.05E-05 |
| Ccna2 | 12858 | -1.42 | -0.51 | 8.24E-03 |
| Cdadc1 | 2510 | 1.17 | 0.23 | 7.44E-03 |
| Cdc37l1 | 9737 | 1.43 | 0.52 | 2.16E-03 |
| Cdc42se1 | 5069 | -1.14 | -0.19 | 2.37E-02 |
| Cdc7 | 5452 | 1.11 | 0.15 | 1.46E-02 |
| Cds1 | 507 | 1.54 | 0.63 | 1.18E-02 |
| Cdv3 | 19643 | -1.20 | -0.26 | 1.17E-02 |
| Cenpt | 3556 | -1.20 | -0.26 | 1.22E-03 |
| Cep290 | 3038 | 1.38 | 0.46 | 5.84E-03 |
| Cep89 | 3001 | -1.22 | -0.29 | 6.35E-03 |
| Cerkl | 102 | 1.87 | 0.91 | 1.04E-04 |
| Cers6 | 6953 | 1.33 | 0.41 | 1.77E-02 |
| Cfap61 | 143 | 1.42 | 0.51 | 1.50E-02 |
| Cfap91 | 109 | -1.46 | -0.55 | 1.34E-02 |
| Chd1 | 9109 | 1.23 | 0.30 | 1.85E-03 |
| Chek1 | 2827 | -1.43 | -0.52 | 5.06E-03 |
| Chid1 | 2893 | 1.20 | 0.26 | 1.26E-02 |
| Chml | 2840 | 1.24 | 0.31 | 1.48E-02 |
| Chpt1 | 2383 | 1.24 | 0.31 | 1.84E-02 |
| Chtf8 | 106 | -1.29 | -0.37 | 1.86E-02 |
| Ciao3 | 1278 | -1.15 | -0.21 | 4.13E-03 |
| Cited2 | 4639 | -1.28 | -0.35 | 6.19E-03 |
| Cldn11 | 1825 | -1.38 | -0.47 | 1.61E-02 |
| Cnot3 | 5875 | -1.22 | -0.29 | 9.80E-03 |
| Cnot9 | 6788 | -1.23 | -0.29 | 2.15E-03 |
| Coa8 | 1080 | -1.17 | -0.23 | 1.93E-03 |
| Coasy | 2229 | -1.13 | -0.18 | 1.80E-02 |
| Cog8 | 1756 | -1.15 | -0.21 | 1.22E-03 |
| Col22a1 | 1457 | -1.23 | -0.30 | 2.31E-02 |
| Cops7b | 3301 | -1.13 | -0.18 | 2.11E-02 |
| Cops8 | 5467 | -1.13 | -0.17 | 2.14E-02 |
| Cox18 | 615 | -1.13 | -0.17 | 8.45E-03 |
| Cpne8 | 442 | -1.31 | -0.39 | 2.17E-02 |
| Crim1 | 3274 | 1.21 | 0.28 | 1.92E-02 |
| Csf2rb | 112 | -2.31 | -1.21 | 1.02E-03 |
| Csgalnact2 | 1421 | 1.21 | 0.28 | 4.87E-03 |
| Csk | 5914 | -1.28 | -0.35 | 2.15E-03 |
| Cspp1 | 7665 | 1.18 | 0.23 | 1.80E-02 |
| Ctnna1 | 14712 | -1.22 | -0.29 | 2.33E-02 |
| Cuedc1 | 200 | -2.81 | -1.49 | 4.97E-03 |

| | | | | |
|---|---|---|---|---|
| Cuedc2 | 5327 | -1.14 | -0.19 | 1.54E-02 |
| Cux1 | 12750 | -1.27 | -0.35 | 2.56E-03 |
| Cxxc4 | 8281 | -1.32 | -0.40 | 1.76E-02 |
| Cyp2d22 | 450 | 1.36 | 0.45 | 4.99E-03 |
| Dazap2 | 8566 | -1.18 | -0.24 | 2.05E-02 |
| Dbnl | 6405 | -1.15 | -0.20 | 1.78E-02 |
| Dcakd | 8709 | -1.26 | -0.34 | 8.18E-03 |
| Dcp2 | 12662 | 1.27 | 0.35 | 6.86E-04 |
| Dctd | 1384 | -1.27 | -0.35 | 1.33E-02 |
| Ddx23 | 8925 | -1.24 | -0.31 | 8.20E-04 |
| Ddx56 | 2894 | -1.23 | -0.30 | 4.78E-04 |
| Dennd1a | 3407 | -1.17 | -0.23 | 1.35E-02 |
| Diablo | 756 | 1.27 | 0.34 | 1.10E-02 |
| Dlgap5 | 3503 | -1.44 | -0.52 | 6.63E-03 |
| Dll1 | 4795 | -1.45 | -0.54 | 1.69E-02 |
| Dlx1 | 5903 | -2.29 | -1.19 | 2.02E-02 |
| Dmap1 | 2240 | -1.14 | -0.19 | 1.73E-02 |
| Dmd | 2433 | 1.24 | 0.31 | 2.01E-02 |
| Dmxl1 | 7691 | 1.56 | 0.65 | 1.04E-03 |
| Dmxl2 | 9776 | 1.49 | 0.57 | 2.21E-02 |
| Dnajc30 | 931 | -1.17 | -0.22 | 5.24E-04 |
| Dnttip1 | 2157 | -1.16 | -0.21 | 1.56E-02 |
| Dpp9 | 4589 | -1.30 | -0.38 | 5.91E-03 |
| Dram2 | 2528 | 1.19 | 0.25 | 1.36E-03 |
| Drosha | 11946 | -1.18 | -0.24 | 1.22E-02 |
| Dtx2 | 1336 | -1.20 | -0.27 | 1.27E-02 |
| Dusp12 | 1723 | -1.11 | -0.15 | 9.41E-03 |
| Dzip3 | 13882 | 1.21 | 0.27 | 1.98E-02 |
| E130309D02Rik | 2040 | -1.32 | -0.40 | 1.37E-03 |
| Ece2 | 235 | -1.48 | -0.57 | 2.30E-02 |
| Ecm1 | 556 | -1.38 | -0.46 | 4.06E-03 |
| Eef2kmt | 453 | -1.31 | -0.39 | 1.42E-03 |
| Eif2b1 | 2850 | -1.22 | -0.28 | 8.64E-03 |
| Eif2b4 | 2400 | -1.14 | -0.19 | 1.24E-02 |
| Eif3d | 10087 | -1.15 | -0.21 | 1.99E-02 |
| Eif4e2 | 7107 | -1.13 | -0.18 | 1.76E-03 |
| Eif4e3 | 838 | -1.26 | -0.33 | 1.41E-03 |
| Eif4enif1 | 7426 | -1.11 | -0.14 | 2.44E-02 |
| Eif4h | 24292 | -1.21 | -0.28 | 7.54E-03 |
| Elmod2 | 5783 | 1.21 | 0.28 | 9.14E-04 |
| Emb | 1344 | -1.48 | -0.57 | 8.14E-03 |
| Enkur | 209 | 1.36 | 0.45 | 5.38E-03 |
| Enpp2 | 3719 | 1.58 | 0.66 | 2.98E-03 |
| Epm2aip1 | 16521 | 1.27 | 0.35 | 1.57E-02 |
| Ercc1 | 1776 | -1.19 | -0.25 | 9.33E-03 |
| Ess2 | 2744 | -1.18 | -0.24 | 1.20E-03 |
| Etv6 | 2568 | -1.25 | -0.32 | 1.05E-02 |

| | | | | |
|---|---|---|---|---|
| Exoc3l | 371 | -1.28 | -0.36 | 5.03E-03 |
| Exosc4 | 1273 | -1.17 | -0.23 | 3.88E-03 |
| Exph5 | 106 | 2.79 | 1.48 | 5.45E-04 |
| Fam120c | 2059 | 1.52 | 0.61 | 1.43E-03 |
| Fam126b | 2939 | 1.47 | 0.55 | 2.34E-04 |
| Fam3c | 2442 | -1.18 | -0.24 | 5.94E-03 |
| Fam53b | 2458 | -1.23 | -0.30 | 2.19E-02 |
| Fam71e1 | 593 | -1.32 | -0.40 | 2.67E-03 |
| Fbll1 | 556 | 1.56 | 0.65 | 7.59E-03 |
| Fbxo5 | 2579 | -1.44 | -0.53 | 4.22E-03 |
| Fbxw5 | 3978 | -1.15 | -0.20 | 7.54E-03 |
| Fer | 2828 | 1.32 | 0.40 | 3.87E-03 |
| Fezf1 | 950 | -1.68 | -0.75 | 2.34E-02 |
| Fign | 2202 | -1.48 | -0.57 | 2.00E-03 |
| Fkbp10 | 2887 | -1.56 | -0.64 | 2.47E-02 |
| Fkbp15 | 3728 | 1.18 | 0.24 | 4.95E-03 |
| Flnb | 14801 | 1.35 | 0.43 | 1.69E-02 |
| Flrt1 | 1229 | 1.42 | 0.50 | 1.83E-02 |
| Fnbp4 | 11381 | -1.10 | -0.14 | 2.05E-02 |
| Frem2 | 2619 | 1.67 | 0.74 | 2.48E-02 |
| Fut8 | 3711 | -1.10 | -0.14 | 1.45E-02 |
| Fzd1 | 5399 | -1.29 | -0.36 | 3.56E-03 |
| Fzd4 | 1743 | 1.51 | 0.59 | 1.72E-02 |
| Gar1 | 2245 | -1.49 | -0.58 | 9.28E-03 |
| Garre1 | 5078 | 1.15 | 0.20 | 7.95E-03 |
| Gatc | 2143 | -1.17 | -0.23 | 5.02E-03 |
| Gbx2 | 808 | -1.60 | -0.68 | 7.24E-03 |
| Gfpt1 | 11453 | 1.24 | 0.31 | 3.27E-03 |
| Gimap9 | 145 | -1.53 | -0.61 | 2.12E-02 |
| Gls | 15431 | 1.22 | 0.28 | 1.43E-02 |
| Gm14226 | 120 | -5.48 | -2.45 | 5.46E-03 |
| Gm14295 | 427 | 1.60 | 0.68 | 8.80E-05 |
| Gm14305 | 808 | 1.35 | 0.43 | 2.06E-03 |
| Gm14434 | 151 | -6.98 | -2.80 | 7.78E-03 |
| Gm29666 | 380 | 1.23 | 0.30 | 9.35E-03 |
| Gnl3 | 4724 | -1.20 | -0.27 | 1.59E-02 |
| Gpatch4 | 1842 | -1.19 | -0.25 | 2.00E-02 |
| Gps2 | 2670 | -1.30 | -0.37 | 1.33E-03 |
| Grap | 295 | -1.23 | -0.30 | 4.01E-03 |
| Gripap1 | 6532 | -1.16 | -0.22 | 1.43E-02 |
| Grwd1 | 1719 | -1.25 | -0.32 | 8.77E-03 |
| Gstm7 | 2431 | -1.25 | -0.32 | 2.08E-02 |
| Gtf2h4 | 1146 | -1.19 | -0.26 | 2.05E-02 |
| Helt | 223 | -1.97 | -0.97 | 1.15E-02 |
| Herc1 | 24022 | 1.49 | 0.57 | 8.91E-03 |
| Herc2 | 19070 | 1.44 | 0.53 | 9.32E-03 |
| Hes6 | 5117 | -1.32 | -0.40 | 5.58E-03 |

| | | | | |
|---|---|---|---|---|
| Hey1 | 2557 | -1.29 | -0.37 | 6.62E-03 |
| Hmgn1 | 28667 | -1.19 | -0.25 | 2.02E-02 |
| Hnrnpa1 | 138907 | -1.23 | -0.29 | 1.69E-02 |
| Hoxa5 | 237 | -60.93 | -5.93 | 1.77E-03 |
| Hoxb5 | 211 | <-100 | -10.05 | 1.41E-03 |
| Hoxb8 | 170 | -78.71 | -6.30 | 7.01E-03 |
| Hoxc4 | 154 | -35.79 | -5.16 | 1.80E-03 |
| Hsd11b2 | 605 | -1.75 | -0.81 | 2.29E-03 |
| Hsp90ab1 | 180217 | -1.17 | -0.23 | 2.10E-02 |
| Hspa12a | 6876 | 1.49 | 0.57 | 2.28E-03 |
| Hspa4l | 361 | 1.75 | 0.81 | 7.11E-03 |
| Htra3 | 383 | -1.55 | -0.63 | 1.84E-03 |
| Ier5l | 941 | -1.23 | -0.30 | 1.47E-02 |
| Igdcc3 | 2075 | -1.90 | -0.93 | 2.22E-03 |
| Ikzf2 | 1166 | 1.33 | 0.41 | 1.52E-03 |
| Irf2bp2 | 2311 | -1.34 | -0.43 | 9.80E-04 |
| Irs1 | 5806 | -1.21 | -0.28 | 7.51E-03 |
| Irx6 | 135 | -1.46 | -0.55 | 2.06E-02 |
| Jak2 | 6870 | 1.22 | 0.29 | 2.49E-02 |
| Josd2 | 1870 | -1.20 | -0.26 | 1.04E-02 |
| Kank2 | 2603 | -1.34 | -0.42 | 1.23E-02 |
| Kbtbd3 | 582 | 1.24 | 0.31 | 1.25E-02 |
| Kcnip3 | 820 | -1.34 | -0.42 | 2.37E-02 |
| Kcnn2 | 633 | 1.39 | 0.47 | 3.12E-03 |
| Kctd2 | 2701 | -1.16 | -0.21 | 5.90E-04 |
| Kdm5a | 11814 | 1.23 | 0.30 | 1.26E-02 |
| Kif2c | 4226 | -1.49 | -0.57 | 2.84E-03 |
| Kif3a | 10207 | 1.20 | 0.27 | 2.22E-02 |
| Klf2 | 278 | -1.41 | -0.50 | 1.61E-02 |
| Krit1 | 9029 | 1.20 | 0.26 | 2.17E-03 |
| Laptm5 | 867 | -1.28 | -0.36 | 1.76E-02 |
| Lca5l | 269 | 1.18 | 0.24 | 2.19E-02 |
| Lcp1 | 2352 | -1.20 | -0.26 | 1.35E-02 |
| Lemd2 | 3723 | -1.23 | -0.30 | 2.73E-03 |
| Lifr | 2809 | 1.39 | 0.48 | 2.47E-02 |
| Lipg | 768 | 1.32 | 0.40 | 6.58E-03 |
| Lmnb1 | 28472 | -1.47 | -0.56 | 2.34E-03 |
| Lrig1 | 4799 | 1.21 | 0.27 | 9.75E-03 |
| Lrig2 | 5794 | 1.19 | 0.25 | 1.15E-02 |
| Lrrc57 | 1466 | -1.12 | -0.17 | 1.04E-02 |
| Lrrc8a | 739 | -1.47 | -0.56 | 3.37E-04 |
| Lrrtm2 | 1399 | 1.57 | 0.65 | 2.46E-02 |
| Lsm2 | 2197 | -1.33 | -0.41 | 1.52E-02 |
| Lsm4 | 1198 | -1.48 | -0.56 | 1.78E-03 |
| Lsm7 | 2165 | -1.23 | -0.30 | 2.04E-02 |
| Ltn1 | 7740 | 1.22 | 0.29 | 1.06E-02 |
| Lurap1l | 363 | 1.32 | 0.40 | 2.33E-03 |

| | | | | |
|---|---|---|---|---|
| Maf1 | 5688 | -1.18 | -0.24 | 5.51E-03 |
| Man1a | 1331 | 1.55 | 0.64 | 4.33E-03 |
| Mcm7 | 14806 | -1.35 | -0.43 | 7.63E-03 |
| Meak7 | 1038 | -1.15 | -0.21 | 5.38E-03 |
| Med15 | 3114 | -1.20 | -0.26 | 2.55E-03 |
| Mief1 | 3469 | -1.14 | -0.19 | 5.34E-03 |
| Mmp15 | 3767 | -1.30 | -0.37 | 1.17E-02 |
| Mmp28 | 545 | -1.29 | -0.37 | 1.44E-02 |
| Mrgbp | 2766 | -1.14 | -0.19 | 1.02E-02 |
| Mrpl1 | 1564 | -1.15 | -0.20 | 5.64E-03 |
| Mrps14 | 2050 | -1.22 | -0.29 | 1.70E-02 |
| Mrps21 | 2087 | -1.28 | -0.36 | 1.85E-02 |
| Mrps27 | 2039 | -1.23 | -0.30 | 7.24E-03 |
| Mst1r | 351 | 1.38 | 0.46 | 1.96E-02 |
| Msto1 | 1681 | -1.17 | -0.22 | 1.57E-02 |
| Msx3 | 432 | 3.03 | 1.60 | 1.58E-03 |
| Mto1 | 1246 | -1.18 | -0.24 | 1.77E-03 |
| Muc1 | 686 | -1.41 | -0.49 | 4.24E-03 |
| Mybbp1a | 13803 | -1.21 | -0.27 | 1.65E-02 |
| Mycbp2 | 22103 | 1.50 | 0.58 | 1.28E-02 |
| Mycl | 7030 | -1.15 | -0.21 | 2.43E-02 |
| Mynn | 3581 | 1.15 | 0.20 | 5.64E-03 |
| Myo9a | 12240 | 1.34 | 0.42 | 2.22E-02 |
| Naa10 | 1102 | -1.28 | -0.36 | 3.16E-03 |
| Naa12 | 5122 | -1.13 | -0.17 | 8.69E-03 |
| Naa25 | 6315 | 1.13 | 0.17 | 1.15E-02 |
| Naa38 | 767 | -1.35 | -0.43 | 1.44E-03 |
| Nacc1 | 8224 | -1.29 | -0.37 | 3.23E-03 |
| Nacc2 | 1438 | -1.24 | -0.31 | 1.05E-02 |
| Napb | 4916 | 1.37 | 0.46 | 8.70E-03 |
| Naxe | 2417 | -1.12 | -0.16 | 7.47E-03 |
| Ndufa4l2 | 123 | -1.50 | -0.58 | 1.65E-03 |
| Ndufab1 | 3977 | -1.16 | -0.22 | 1.45E-02 |
| Ndufb2 | 1044 | -1.17 | -0.23 | 2.46E-02 |
| Necab2 | 1384 | -1.18 | -0.24 | 2.49E-02 |
| Nell2 | 9177 | -1.29 | -0.37 | 1.08E-02 |
| Neurog2 | 6066 | -1.66 | -0.73 | 1.33E-02 |
| Nfat5 | 15255 | 1.34 | 0.42 | 6.02E-03 |
| Nktr | 23629 | 1.37 | 0.45 | 4.76E-03 |
| Nkx2-1 | 2086 | -2.46 | -1.30 | 8.65E-03 |
| Nkx2-2 | 355 | -1.77 | -0.82 | 1.63E-02 |
| Nkx2-4 | 120 | -2.42 | -1.28 | 8.69E-05 |
| Nob1 | 2867 | -1.16 | -0.21 | 6.33E-03 |
| Noc3l | 3626 | 1.15 | 0.21 | 2.05E-02 |
| Nr2f2 | 13726 | -1.61 | -0.69 | 1.85E-04 |
| Nsun5 | 1428 | -1.18 | -0.24 | 2.01E-03 |
| Nudt5 | 3718 | -1.26 | -0.33 | 7.21E-03 |

| | | | | |
|---|---|---|---|---|
| Nup153 | 12606 | 1.16 | 0.21 | 2.33E-02 |
| Nynrin | 11471 | -1.25 | -0.32 | 1.70E-02 |
| Oard1 | 461 | -2.32 | -1.22 | 1.80E-02 |
| Oxnad1 | 1334 | -1.13 | -0.18 | 1.70E-02 |
| Pacc1 | 1723 | -1.13 | -0.18 | 8.63E-03 |
| Pacrg | 255 | 1.26 | 0.34 | 2.09E-02 |
| Padi2 | 216 | 2.64 | 1.40 | 6.41E-04 |
| Pafah1b3 | 6793 | -1.17 | -0.22 | 2.37E-02 |
| Pak1 | 11100 | -1.24 | -0.31 | 6.32E-03 |
| Pbk | 4745 | -1.41 | -0.50 | 1.13E-02 |
| Pcmtd1 | 6739 | 1.24 | 0.32 | 7.12E-03 |
| Pcmtd2 | 6862 | 1.19 | 0.25 | 1.11E-02 |
| Pcolce | 1711 | -2.09 | -1.06 | 2.17E-02 |
| Pcyox1l | 1502 | -1.28 | -0.36 | 8.46E-03 |
| Pdcd6 | 3322 | 1.21 | 0.27 | 6.72E-03 |
| Pdgfrl | 232 | -1.61 | -0.69 | 1.62E-02 |
| Pdlim3 | 637 | -1.54 | -0.62 | 2.04E-03 |
| Pelp1 | 4608 | -1.30 | -0.37 | 7.65E-03 |
| Pex14 | 1641 | -1.15 | -0.20 | 1.05E-02 |
| Pgls | 2632 | -1.17 | -0.23 | 4.10E-03 |
| Phc1 | 9754 | -1.17 | -0.23 | 7.76E-03 |
| Phc3 | 6851 | 1.39 | 0.48 | 1.33E-03 |
| Piezo1 | 2339 | -1.48 | -0.57 | 2.32E-02 |
| Pik3ca | 10244 | 1.17 | 0.23 | 1.55E-02 |
| Pir | 156 | -1.50 | -0.58 | 1.62E-02 |
| Plekhj1 | 2296 | -1.24 | -0.31 | 7.14E-03 |
| Poll | 949 | -1.27 | -0.34 | 1.08E-02 |
| Polr3f | 4653 | 1.14 | 0.19 | 2.04E-02 |
| Pomc | 169 | -2.03 | -1.02 | 8.22E-05 |
| Pou2f2 | 4785 | -1.47 | -0.56 | 2.00E-02 |
| Ppfia4 | 1710 | 1.59 | 0.67 | 2.58E-03 |
| Ppip5k1 | 2203 | 1.18 | 0.24 | 2.15E-02 |
| Ppp1r18 | 5100 | -1.23 | -0.30 | 7.38E-03 |
| Ppp1r3c | 679 | 1.55 | 0.63 | 2.07E-02 |
| Ppp1r8 | 5668 | -1.18 | -0.24 | 1.21E-02 |
| Ppp2r1a | 24678 | -1.17 | -0.23 | 2.20E-03 |
| Ppp4c | 6694 | -1.18 | -0.24 | 1.55E-02 |
| Prmt7 | 3820 | -1.17 | -0.23 | 7.25E-04 |
| Prr12 | 7437 | -1.48 | -0.57 | 4.95E-03 |
| Prtn3 | 272 | -1.25 | -0.32 | 1.00E-02 |
| Psenen | 1893 | -1.22 | -0.29 | 1.38E-02 |
| Psmb6 | 8970 | -1.22 | -0.29 | 2.64E-03 |
| Psmc5 | 8309 | -1.25 | -0.32 | 2.05E-03 |
| Psmg3 | 334 | -1.26 | -0.34 | 1.95E-02 |
| Psmg4 | 775 | -1.38 | -0.47 | 1.07E-03 |
| Psph | 1024 | -1.25 | -0.32 | 2.10E-03 |
| Ptar1 | 7591 | 1.23 | 0.30 | 5.80E-03 |

| | | | | |
|---|---|---|---|---|
| Ptov1 | 17233 | -1.22 | -0.29 | 1.54E-02 |
| Pygl | 778 | 1.37 | 0.46 | 9.97E-03 |
| Qpct | 375 | 1.28 | 0.36 | 2.31E-02 |
| Qrfprl | 164 | -1.63 | -0.71 | 2.49E-02 |
| Rab30 | 4492 | 1.28 | 0.36 | 7.00E-03 |
| Rad54b | 1711 | -1.30 | -0.37 | 9.44E-03 |
| Rapgef6 | 5276 | 1.32 | 0.40 | 1.08E-02 |
| Rbm12b2 | 2733 | 1.15 | 0.20 | 2.00E-02 |
| Rbm6 | 14500 | -1.12 | -0.16 | 9.93E-03 |
| Rc3h1 | 6268 | 1.13 | 0.18 | 1.60E-02 |
| Rce1 | 1327 | -1.15 | -0.21 | 1.54E-02 |
| Rcor2 | 9542 | -1.48 | -0.56 | 2.03E-03 |
| Rexo5 | 1339 | 1.26 | 0.34 | 1.16E-02 |
| Rfc3 | 3153 | -1.29 | -0.37 | 3.77E-03 |
| Rflnb | 2048 | 1.41 | 0.50 | 1.98E-02 |
| Rho | 101 | 1.40 | 0.49 | 1.24E-02 |
| Rhob | 6351 | -1.25 | -0.32 | 4.54E-03 |
| Ric8b | 4244 | 1.09 | 0.12 | 2.50E-02 |
| Riok3 | 10080 | 1.11 | 0.15 | 2.49E-02 |
| Rmi2 | 515 | -1.42 | -0.50 | 2.31E-02 |
| Rnf19a | 5179 | 1.15 | 0.20 | 1.50E-02 |
| Rnf220 | 6619 | -1.19 | -0.25 | 1.33E-02 |
| Rnf25 | 2639 | -1.12 | -0.17 | 1.77E-03 |
| Rnf32 | 651 | 1.31 | 0.39 | 8.25E-04 |
| Rnf44 | 9449 | -1.40 | -0.49 | 2.21E-03 |
| Rock1 | 8388 | 1.17 | 0.22 | 4.44E-03 |
| Rock2 | 10892 | 1.28 | 0.36 | 1.74E-04 |
| Rpl13a | 1404 | -1.30 | -0.38 | 2.33E-02 |
| Rpl7l1 | 7184 | -1.29 | -0.36 | 7.48E-04 |
| Rpusd4 | 998 | -1.23 | -0.30 | 1.57E-03 |
| Rrad | 117 | 1.68 | 0.75 | 1.29E-02 |
| Rrnad1 | 3110 | -1.16 | -0.21 | 4.72E-03 |
| Ruvbl1 | 4144 | -1.21 | -0.28 | 2.17E-02 |
| Sacs | 9029 | 1.44 | 0.53 | 2.30E-02 |
| Sash1 | 3853 | 1.18 | 0.24 | 1.98E-02 |
| Sec22a | 2051 | 1.17 | 0.22 | 2.32E-02 |
| Sec61a2 | 6520 | 1.15 | 0.20 | 2.03E-02 |
| Secisbp2l | 9900 | 1.17 | 0.23 | 7.13E-03 |
| Selenot | 10483 | -1.18 | -0.24 | 5.51E-03 |
| Sema4b | 1776 | -1.24 | -0.31 | 7.90E-03 |
| Sema5b | 5244 | -1.44 | -0.52 | 8.05E-03 |
| Setx | 15624 | 1.25 | 0.33 | 2.47E-02 |
| Sf3a2 | 3635 | -1.48 | -0.57 | 2.44E-03 |
| Sf3a3 | 7195 | -1.25 | -0.33 | 1.47E-02 |
| Sh3gl3 | 1221 | 1.33 | 0.41 | 1.08E-03 |
| Sim1 | 1200 | -1.41 | -0.50 | 2.22E-02 |
| Sinhcaf | 7080 | -1.33 | -0.41 | 9.77E-03 |

| | | | | |
|---|---|---|---|---|
| Six4 | 1023 | 1.43 | 0.51 | 2.45E-02 |
| Skil | 8411 | 1.16 | 0.21 | 1.69E-02 |
| Slain2 | 5334 | -1.15 | -0.20 | 8.86E-03 |
| Slc16a10 | 949 | 1.45 | 0.54 | 1.40E-02 |
| Slc16a13 | 975 | -1.23 | -0.30 | 6.52E-04 |
| Slc1a1 | 879 | 1.70 | 0.77 | 2.18E-02 |
| Slc25a36 | 10791 | 1.23 | 0.29 | 3.37E-03 |
| Slc25a40 | 1936 | 1.23 | 0.30 | 4.00E-03 |
| Slc35b1 | 4144 | -1.13 | -0.17 | 1.35E-02 |
| Slc3a2 | 4594 | -1.16 | -0.22 | 2.81E-03 |
| Slc4a8 | 5501 | 1.48 | 0.56 | 1.28E-02 |
| Smad7 | 788 | -1.25 | -0.32 | 1.20E-02 |
| Smarcd3 | 5057 | -1.31 | -0.39 | 1.51E-02 |
| Smdt1 | 3229 | -1.24 | -0.31 | 5.15E-03 |
| Smndc1 | 4434 | -1.24 | -0.31 | 7.76E-03 |
| Snrnp70 | 33965 | -1.29 | -0.37 | 3.86E-03 |
| Snrpd1 | 9050 | -1.27 | -0.35 | 1.11E-02 |
| Sox11 | 64358 | -1.53 | -0.62 | 4.59E-03 |
| Sox2 | 10110 | -1.50 | -0.58 | 2.36E-02 |
| Spdl1 | 1584 | -1.38 | -0.47 | 1.71E-02 |
| Spred2 | 2422 | -1.28 | -0.35 | 4.05E-03 |
| Sptan1 | 69080 | 1.42 | 0.51 | 2.96E-03 |
| Srp68 | 6240 | -1.20 | -0.27 | 8.62E-04 |
| Srpr | 5578 | -1.19 | -0.25 | 1.83E-02 |
| Srrt | 15369 | -1.33 | -0.41 | 1.42E-03 |
| Stard9 | 3691 | 1.63 | 0.70 | 5.50E-03 |
| Stk24 | 3981 | -1.14 | -0.19 | 4.89E-03 |
| Strip1 | 10609 | -1.12 | -0.16 | 2.03E-02 |
| Styk1 | 115 | -1.58 | -0.66 | 7.09E-04 |
| Supt20 | 9397 | 1.07 | 0.10 | 1.08E-02 |
| Surf6 | 2709 | -1.13 | -0.17 | 1.68E-02 |
| Suv39h1 | 5175 | -1.24 | -0.31 | 2.05E-03 |
| Szrd1 | 6855 | -1.27 | -0.34 | 7.77E-03 |
| Tac1 | 479 | 1.44 | 0.53 | 6.44E-03 |
| Tbc1d23 | 4906 | 1.19 | 0.25 | 6.88E-03 |
| Tbc1d25 | 920 | 1.14 | 0.20 | 2.03E-02 |
| Tbpl1 | 6051 | -1.20 | -0.26 | 1.98E-03 |
| Tent5a | 1570 | -1.27 | -0.35 | 2.20E-02 |
| Tes | 3225 | 1.42 | 0.51 | 2.48E-02 |
| Tesk1 | 3448 | -1.24 | -0.31 | 8.72E-03 |
| Tex15 | 2197 | 1.99 | 0.99 | 5.54E-03 |
| Tfg | 7142 | -1.12 | -0.16 | 2.28E-02 |
| Thbs2 | 2507 | -2.51 | -1.33 | 2.10E-02 |
| Them4 | 1096 | 1.44 | 0.53 | 7.58E-03 |
| Thoc2l | 6347 | 1.26 | 0.34 | 1.63E-03 |
| Tie1 | 2207 | -1.31 | -0.39 | 5.20E-03 |
| Tigd3 | 703 | -1.25 | -0.32 | 9.34E-03 |

| | | | | |
|---|---|---|---|---|
| Tlcd4 | 600 | 1.50 | 0.59 | 4.26E-04 |
| Tle3 | 8820 | -1.37 | -0.46 | 4.75E-03 |
| Tmc3 | 117 | 2.71 | 1.44 | 2.01E-03 |
| Tmeff2 | 1739 | 1.71 | 0.77 | 9.80E-03 |
| Tmem101 | 1474 | -1.23 | -0.30 | 2.57E-03 |
| Tmem11 | 1619 | -1.19 | -0.25 | 1.34E-02 |
| Tmem177 | 1273 | 1.18 | 0.24 | 8.42E-03 |
| Tmem219 | 718 | -1.20 | -0.27 | 4.95E-03 |
| Tmem234 | 5705 | -1.20 | -0.26 | 1.13E-03 |
| Tmem67 | 2300 | 1.15 | 0.21 | 7.20E-03 |
| Tom1l1 | 1576 | 1.26 | 0.33 | 1.48E-02 |
| Tra2a | 28189 | 1.13 | 0.17 | 2.23E-02 |
| Trim28 | 31719 | -1.27 | -0.35 | 2.90E-03 |
| Trim46 | 3338 | -1.42 | -0.51 | 2.36E-02 |
| Trmt61a | 927 | -1.33 | -0.41 | 1.31E-02 |
| Trp53i11 | 11166 | -1.29 | -0.37 | 2.31E-02 |
| Tstd2 | 4425 | -1.11 | -0.15 | 1.40E-02 |
| Ttc8 | 1988 | -1.14 | -0.19 | 2.19E-02 |
| Tuba1b | 31678 | -1.33 | -0.41 | 5.86E-03 |
| Txn2 | 4601 | -1.20 | -0.26 | 1.65E-03 |
| Tyrobp | 214 | -1.36 | -0.45 | 1.20E-02 |
| Ubash3b | 2278 | 1.33 | 0.41 | 1.51E-02 |
| Ubb | 56162 | -1.14 | -0.19 | 2.41E-03 |
| Ube2q1 | 8249 | -1.16 | -0.21 | 1.80E-03 |
| Ubqln2 | 14660 | -1.30 | -0.38 | 1.29E-02 |
| Ubxn10 | 241 | 1.41 | 0.50 | 2.75E-03 |
| Usp16 | 4941 | 1.14 | 0.19 | 1.89E-02 |
| Usp19 | 12383 | -1.17 | -0.23 | 1.41E-02 |
| Vax1 | 273 | -2.24 | -1.16 | 1.35E-02 |
| Vgll4 | 4141 | -1.14 | -0.19 | 2.14E-02 |
| Vps13c | 5813 | 1.66 | 0.73 | 8.25E-04 |
| Vps4b | 6196 | 1.17 | 0.23 | 1.24E-02 |
| Vsig10 | 507 | -1.48 | -0.57 | 8.74E-04 |
| Vwf | 2227 | -1.44 | -0.52 | 1.62E-02 |
| Washc2 | 8354 | 1.11 | 0.14 | 1.16E-02 |
| Wbp11 | 7481 | -1.28 | -0.36 | 1.04E-02 |
| Wdr24 | 1815 | -1.15 | -0.20 | 1.22E-02 |
| Wdr33 | 7608 | -1.10 | -0.14 | 3.51E-03 |
| Wdr6 | 28810 | -1.29 | -0.37 | 2.10E-03 |
| Wdr66 | 293 | 1.41 | 0.49 | 1.19E-02 |
| Wdr83 | 2149 | -1.21 | -0.27 | 3.56E-04 |
| Wfdc2 | 300 | -1.60 | -0.67 | 9.48E-03 |
| Wnk1 | 29420 | 1.18 | 0.24 | 9.23E-03 |
| Wnt3 | 639 | -1.37 | -0.45 | 5.34E-03 |
| Wnt8b | 1393 | -2.15 | -1.10 | 4.50E-03 |
| Xpnpep2 | 212 | -1.96 | -0.97 | 5.20E-03 |
| Yae1d1 | 4139 | 1.25 | 0.32 | 3.26E-03 |

| | | | | |
|---|---|---|---|---|
| Ythdc1 | 10887 | -1.17 | -0.22 | 1.81E-02 |
| Zbtb49 | 1302 | 1.22 | 0.29 | 6.58E-04 |
| Zcchc3 | 2879 | -1.41 | -0.50 | 2.00E-03 |
| Zdhhc2 | 3149 | -1.30 | -0.37 | 4.56E-03 |
| Zfp106 | 19891 | 1.20 | 0.26 | 1.97E-02 |
| Zfp263 | 3439 | -1.09 | -0.12 | 9.68E-03 |
| Zfp354b | 247 | 1.31 | 0.39 | 1.74E-02 |
| Zfp385b | 211 | -1.56 | -0.64 | 7.35E-03 |
| Zfp398 | 4267 | 1.29 | 0.37 | 6.26E-03 |
| Zfp442 | 518 | 1.52 | 0.61 | 1.67E-02 |
| Zfp458 | 642 | 1.32 | 0.40 | 1.93E-02 |
| Zfp553 | 4181 | -1.40 | -0.49 | 2.60E-03 |
| Zfp58 | 536 | 1.13 | 0.18 | 1.22E-02 |
| Zfp607b | 1626 | 1.28 | 0.36 | 6.65E-03 |
| Zfp677 | 477 | 1.25 | 0.32 | 1.59E-02 |
| Zfp712 | 879 | 1.27 | 0.35 | 1.64E-03 |
| Zfp758 | 1147 | 1.19 | 0.25 | 3.34E-03 |
| Zfp768 | 2361 | -1.19 | -0.26 | 2.31E-02 |
| Zfp780b | 2358 | 1.16 | 0.22 | 8.06E-03 |
| Zfp790 | 2580 | 1.20 | 0.26 | 4.51E-03 |
| Zfp93 | 770 | -1.18 | -0.24 | 5.56E-03 |
| Zfp940 | 1471 | 1.18 | 0.23 | 1.11E-02 |
| Zfp976 | 813 | 1.24 | 0.30 | 5.20E-03 |
| Zic1 | 19643 | -1.33 | -0.41 | 8.79E-04 |
| Zic2 | 6033 | -1.38 | -0.47 | 1.67E-03 |
| Zkscan17 | 2963 | -1.31 | -0.39 | 3.64E-03 |
| Zkscan3 | 11708 | 1.15 | 0.21 | 1.76E-03 |
| Zkscan7 | 2134 | 1.09 | 0.13 | 1.60E-02 |
| Zzef1 | 6975 | 1.47 | 0.56 | 1.70E-03 |

| Gene | Counts | FC | LFC | padj |
|---|---|---|---|---|
| 1700029I15Rik | 104 | 1.32 | 0.40 | 4.0E-03 |
| 2610318N02Rik | 2168 | 1.46 | 0.55 | 2.4E-02 |
| 4933430I17Rik | 225 | 1.38 | 0.46 | 4.2E-03 |
| A630089N07Rik | 910 | -1.38 | -0.47 | 4.7E-04 |
| Abca1 | 5507 | -1.42 | -0.50 | 1.3E-02 |
| Adamts16 | 233 | -2.05 | -1.04 | 6.3E-03 |
| Adamts17 | 919 | -1.79 | -0.84 | 1.8E-02 |
| Adamts9 | 1803 | -1.76 | -0.82 | 4.7E-03 |
| Adcy2 | 689 | -1.49 | -0.58 | 7.9E-03 |
| Afap1 | 7205 | -1.26 | -0.33 | 2.4E-02 |
| Agbl3 | 727 | -1.32 | -0.40 | 2.2E-03 |
| Aif1 | 113 | -1.49 | -0.57 | 2.7E-04 |
| Akt3 | 13712 | -1.30 | -0.38 | 5.6E-03 |
| Alad | 1636 | 1.26 | 0.34 | 3.9E-03 |
| Alg6 | 112 | -1.80 | -0.85 | 1.4E-02 |
| Alkbh2 | 322 | 1.37 | 0.45 | 1.5E-04 |
| Alkbh7 | 518 | 1.25 | 0.32 | 2.2E-03 |
| Alox12 | 123 | -1.71 | -0.78 | 2.0E-02 |
| Anxa11 | 287 | 1.79 | 0.84 | 8.5E-03 |
| Ap4e1 | 2597 | -1.32 | -0.40 | 8.6E-04 |
| Arhgap5 | 12703 | -1.27 | -0.34 | 1.5E-02 |
| Arhgef6 | 855 | -1.49 | -0.58 | 3.7E-04 |
| Atf7 | 3263 | -1.26 | -0.34 | 4.8E-03 |
| Atp2a1 | 197 | -6.61 | -2.73 | 3.5E-03 |
| Begain | 1390 | -1.41 | -0.49 | 2.1E-02 |
| Btrc | 4169 | -1.32 | -0.40 | 1.2E-03 |
| C1ql3 | 219 | -2.20 | -1.14 | 1.2E-03 |
| Cacnb4 | 2101 | -1.68 | -0.75 | 3.9E-03 |
| Capn11 | 223 | 4.18 | 2.06 | 2.4E-02 |
| Car11 | 1095 | -1.38 | -0.46 | 1.8E-02 |
| Casq1 | 141 | -1.82 | -0.86 | 2.8E-03 |
| Cby1 | 2843 | 1.25 | 0.33 | 1.9E-03 |
| Ccdc120 | 1927 | 1.27 | 0.34 | 1.3E-02 |
| Ccdc141 | 708 | -1.65 | -0.73 | 6.9E-03 |
| Ccdc86 | 1231 | 1.25 | 0.32 | 2.4E-02 |
| Ccnd3 | 6546 | 1.34 | 0.42 | 2.0E-02 |
| Cdc25c | 3169 | 1.44 | 0.53 | 7.3E-03 |
| Cdc34 | 3392 | 1.30 | 0.38 | 2.1E-02 |
| Cdc6 | 2717 | 1.52 | 0.60 | 3.7E-03 |
| Cdh12 | 230 | -1.72 | -0.78 | 6.1E-03 |
| Cdh15 | 136 | -4.45 | -2.15 | 6.9E-03 |
| Cdh6 | 3723 | -1.32 | -0.40 | 1.6E-04 |
| Cdh7 | 1176 | -1.35 | -0.44 | 8.4E-03 |
| Cenph | 1962 | 1.40 | 0.49 | 2.0E-02 |
| Cep131 | 4405 | 1.32 | 0.40 | 9.3E-03 |
| Cfap57 | 159 | -1.44 | -0.53 | 2.7E-03 |

| | | | | |
|---|---|---|---|---|
| Chaf1a | 7134 | 1.39 | 0.48 | 1.0E-02 |
| Chl1 | 10632 | -1.81 | -0.86 | 1.7E-02 |
| Chm | 3554 | -1.32 | -0.40 | 1.9E-04 |
| Clp1 | 1725 | 1.31 | 0.39 | 2.2E-02 |
| Cox16 | 107 | -1.35 | -0.43 | 2.5E-02 |
| Cox20 | 140 | -3.59 | -1.84 | 1.0E-02 |
| Cplane1 | 6998 | -1.35 | -0.43 | 3.0E-03 |
| Crybg3 | 1826 | -1.65 | -0.73 | 8.6E-04 |
| Cubn | 187 | -1.49 | -0.57 | 1.2E-02 |
| D130040H23Rik | 637 | -1.50 | -0.58 | 2.2E-03 |
| Daxx | 3662 | 1.33 | 0.41 | 5.3E-04 |
| Dcx | 58900 | -1.61 | -0.69 | 4.6E-03 |
| Ddit4 | 5097 | 1.36 | 0.45 | 1.3E-02 |
| Ddx27 | 4143 | 1.26 | 0.33 | 2.5E-03 |
| Des | 251 | -4.09 | -2.03 | 1.1E-02 |
| Dnaaf5 | 1424 | 1.27 | 0.35 | 2.9E-03 |
| Dnah7b | 877 | -1.61 | -0.68 | 7.5E-03 |
| Dnajc9 | 9295 | 1.35 | 0.43 | 7.1E-03 |
| Dohh | 255 | 1.26 | 0.33 | 1.7E-02 |
| Dpf1 | 5973 | 1.32 | 0.40 | 2.5E-02 |
| Drc3 | 255 | -1.45 | -0.54 | 1.1E-03 |
| Dusp4 | 7143 | -1.40 | -0.49 | 2.5E-02 |
| Dusp9 | 430 | 1.71 | 0.77 | 1.6E-02 |
| Dxo | 1656 | 1.27 | 0.34 | 2.5E-03 |
| E2f4 | 3798 | 1.25 | 0.32 | 1.6E-02 |
| Efna4 | 1245 | 1.27 | 0.34 | 2.0E-02 |
| Egf | 243 | -1.29 | -0.36 | 1.3E-02 |
| Eif2ak2 | 1456 | -1.33 | -0.41 | 2.3E-02 |
| Erfl | 229 | -1.76 | -0.81 | 7.5E-03 |
| Esrra | 1994 | 1.31 | 0.39 | 3.2E-03 |
| Evx1 | 102 | -3.25 | -1.70 | 1.3E-02 |
| Fam162a | 2543 | 1.26 | 0.33 | 6.7E-03 |
| Fam72a | 340 | 1.42 | 0.51 | 4.9E-03 |
| Fbl | 7803 | 1.38 | 0.46 | 5.4E-03 |
| Fermt3 | 511 | 1.29 | 0.37 | 1.5E-02 |
| Fgf17 | 317 | -1.64 | -0.71 | 1.2E-02 |
| Fhad1 | 201 | -1.66 | -0.73 | 1.3E-03 |
| Fmn2 | 4236 | 1.28 | 0.35 | 1.4E-02 |
| Fndc1 | 331 | -2.11 | -1.08 | 1.8E-02 |
| Fndc5 | 378 | -2.33 | -1.22 | 2.6E-03 |
| Fosl2 | 765 | -1.54 | -0.62 | 4.3E-03 |
| Frrs1l | 2272 | -1.50 | -0.58 | 4.7E-03 |
| Gjb2 | 1061 | -2.23 | -1.16 | 1.5E-02 |
| Gldc | 3791 | 1.64 | 0.71 | 1.6E-03 |
| Gltp | 1893 | 1.33 | 0.41 | 4.5E-03 |
| Gm20604 | 518 | 1.32 | 0.40 | 9.6E-03 |
| Gon7 | 834 | 1.26 | 0.34 | 1.5E-02 |

| | | | | |
|---|---|---|---|---|
| Gpr146 | 193 | 1.40 | 0.48 | 1.9E-03 |
| Hapln3 | 143 | 1.33 | 0.42 | 1.2E-02 |
| Hivep2 | 2898 | -1.46 | -0.55 | 1.3E-02 |
| Hmgn5 | 6156 | 1.39 | 0.47 | 1.2E-02 |
| Homer2 | 4016 | -1.33 | -0.41 | 1.0E-03 |
| Hrob | 900 | 1.46 | 0.54 | 1.8E-02 |
| Htr2c | 500 | -1.84 | -0.88 | 1.9E-02 |
| Ifih1 | 206 | -1.32 | -0.40 | 1.3E-02 |
| Ifitm3 | 232 | -1.42 | -0.51 | 1.3E-02 |
| Ift74 | 5073 | 1.25 | 0.32 | 2.3E-02 |
| Igsf9 | 10216 | 1.28 | 0.36 | 7.1E-03 |
| Il4ra | 563 | 1.38 | 0.47 | 1.2E-02 |
| Inpp4b | 601 | -2.02 | -1.01 | 4.6E-04 |
| Iqub | 149 | -1.68 | -0.75 | 6.6E-03 |
| Irx4 | 161 | -3.36 | -1.75 | 5.1E-04 |
| Itga4 | 2591 | -1.43 | -0.51 | 1.2E-02 |
| Itgb3bp | 466 | 1.75 | 0.81 | 2.4E-02 |
| Kcnd3 | 3361 | -1.66 | -0.74 | 1.0E-02 |
| Kcnh2 | 2306 | 1.28 | 0.36 | 7.6E-03 |
| Kcnh6 | 115 | -2.06 | -1.05 | 2.4E-02 |
| Kcnmb2 | 329 | -2.11 | -1.07 | 5.5E-04 |
| Kcnq2 | 10691 | -1.57 | -0.65 | 1.6E-02 |
| Kcnu1 | 111 | -1.88 | -0.91 | 3.6E-03 |
| Kctd17 | 5151 | 1.26 | 0.34 | 4.8E-04 |
| Kctd4 | 230 | -2.25 | -1.17 | 1.5E-02 |
| Klf12 | 4189 | -1.61 | -0.69 | 8.8E-03 |
| Klhl41 | 168 | -3.53 | -1.82 | 2.5E-02 |
| L3mbtl2 | 2835 | 1.35 | 0.43 | 4.7E-05 |
| Lbr | 8479 | 1.34 | 0.42 | 1.9E-02 |
| Lcorl | 6197 | -1.30 | -0.38 | 1.7E-02 |
| Lepr | 1001 | -1.62 | -0.70 | 2.5E-02 |
| Letm2 | 1108 | -1.25 | -0.32 | 7.4E-03 |
| Lgals4 | 152 | -1.54 | -0.62 | 5.3E-05 |
| Lpl | 1617 | -1.29 | -0.37 | 1.6E-02 |
| Lrfn5 | 1079 | -1.89 | -0.92 | 1.3E-02 |
| Lrrc29 | 145 | -1.30 | -0.38 | 1.1E-02 |
| Lrrc46 | 169 | 1.38 | 0.47 | 1.1E-03 |
| Lrrcc1 | 4760 | -1.32 | -0.40 | 1.0E-03 |
| Lrrn3 | 5607 | -1.38 | -0.46 | 1.6E-02 |
| Luzp2 | 1366 | -1.76 | -0.81 | 1.0E-02 |
| Mab21l2 | 3352 | -1.66 | -0.73 | 2.1E-02 |
| Macf1 | 35179 | -1.48 | -0.56 | 2.9E-03 |
| Mafk | 879 | 1.29 | 0.36 | 6.4E-03 |
| Map1b | 128461 | -1.38 | -0.46 | 1.6E-02 |
| Mbnl1 | 4042 | -1.34 | -0.42 | 5.6E-03 |
| Mbnl3 | 708 | -1.65 | -0.72 | 1.8E-02 |
| Med18 | 558 | 1.30 | 0.38 | 1.1E-02 |

| | | | | |
|---|---|---|---|---|
| Med25 | 9144 | 1.31 | 0.39 | 2.8E-03 |
| Med26 | 933 | 1.29 | 0.37 | 3.9E-03 |
| Mif4gd | 647 | 1.28 | 0.35 | 9.1E-03 |
| Mme | 3257 | -2.42 | -1.28 | 7.8E-03 |
| Mns1 | 3659 | 1.42 | 0.50 | 7.0E-03 |
| Mtcl1 | 7105 | 1.33 | 0.41 | 1.6E-02 |
| Mutyh | 826 | 1.30 | 0.38 | 2.1E-02 |
| Mybpc1 | 594 | -3.16 | -1.66 | 7.6E-03 |
| Myh11 | 137 | -1.80 | -0.85 | 1.0E-02 |
| Mylpf | 550 | -3.52 | -1.81 | 2.4E-02 |
| Myo18b | 256 | -1.66 | -0.73 | 1.6E-02 |
| Myo5a | 17678 | -1.36 | -0.44 | 7.3E-03 |
| Naaladl1 | 385 | 1.51 | 0.59 | 9.9E-03 |
| Nalcn | 1502 | -1.52 | -0.60 | 5.2E-03 |
| Nectin1 | 3445 | 1.50 | 0.59 | 3.8E-03 |
| Nes | 67412 | 1.42 | 0.51 | 2.0E-02 |
| Nipa1 | 1722 | -1.26 | -0.33 | 1.6E-02 |
| Nmb | 185 | 1.35 | 0.44 | 1.1E-02 |
| Npm3 | 2109 | 1.29 | 0.36 | 1.0E-02 |
| Nr4a2 | 2860 | 1.35 | 0.43 | 1.7E-02 |
| Nrk | 2102 | -2.87 | -1.52 | 4.9E-03 |
| Nuak2 | 3379 | 1.60 | 0.68 | 1.3E-03 |
| Nupr1l | 113 | 1.36 | 0.45 | 1.6E-04 |
| Nxt1 | 1570 | 1.38 | 0.47 | 5.6E-03 |
| Ophn1 | 3837 | -1.28 | -0.35 | 1.2E-02 |
| Pacsin3 | 1373 | 1.27 | 0.34 | 1.8E-02 |
| Pamr1 | 305 | -2.30 | -1.20 | 1.4E-02 |
| Pappa2 | 1655 | -1.51 | -0.59 | 1.5E-02 |
| Parp16 | 499 | 1.26 | 0.33 | 9.7E-03 |
| Pcdha12 | 158 | -1.73 | -0.79 | 3.8E-03 |
| Pcm1 | 18035 | -1.37 | -0.45 | 3.5E-04 |
| Pde1c | 1638 | -1.46 | -0.54 | 1.2E-02 |
| Pdzrn4 | 1059 | -1.53 | -0.62 | 4.2E-03 |
| Peg10 | 26394 | 1.29 | 0.37 | 1.5E-02 |
| Pelo | 345 | 1.40 | 0.48 | 1.3E-03 |
| Pgm5 | 1022 | -2.39 | -1.25 | 4.4E-03 |
| Phip | 21455 | -1.44 | -0.53 | 1.2E-03 |
| Phlda1 | 1204 | -1.43 | -0.52 | 5.3E-03 |
| Pik3r2 | 356 | 1.91 | 0.93 | 1.2E-02 |
| Plekhb1 | 218 | -1.68 | -0.75 | 7.7E-03 |
| Plpp1 | 809 | -1.25 | -0.33 | 1.6E-02 |
| Plpp2 | 470 | 1.38 | 0.47 | 3.8E-03 |
| Polr2f | 2309 | 1.28 | 0.35 | 1.9E-03 |
| Pou2f1 | 7046 | -1.31 | -0.39 | 3.9E-03 |
| Pou3f1 | 1697 | 1.35 | 0.44 | 1.8E-02 |
| Ppp1r9b | 13672 | 1.28 | 0.35 | 5.3E-03 |
| Prrt1 | 1648 | 1.51 | 0.60 | 2.3E-02 |

| | | | | |
|---|---|---|---|---|
| Psmd10 | 148 | 1.91 | 0.93 | 2.0E-02 |
| Ptch2 | 878 | 1.27 | 0.35 | 1.2E-02 |
| Ptchd1 | 234 | -1.83 | -0.87 | 1.6E-02 |
| Pycr2 | 2525 | 1.27 | 0.35 | 7.9E-04 |
| Raet1e | 233 | 1.64 | 0.71 | 1.7E-02 |
| Rap1gap2 | 1651 | -1.35 | -0.43 | 2.5E-02 |
| Rbm41 | 1700 | -1.29 | -0.36 | 4.3E-03 |
| Recql4 | 1338 | 1.39 | 0.48 | 2.0E-02 |
| Rgs16 | 2796 | 1.54 | 0.63 | 4.1E-04 |
| Ribc1 | 173 | 1.46 | 0.54 | 9.3E-03 |
| Rlbp1 | 741 | -1.99 | -0.99 | 1.8E-02 |
| Rnf181 | 217 | 2.32 | 1.21 | 7.0E-03 |
| Rnf208 | 988 | 1.44 | 0.52 | 9.9E-03 |
| Robo3 | 6624 | -1.73 | -0.79 | 5.8E-03 |
| Rorb | 880 | 1.72 | 0.78 | 3.9E-04 |
| Rusc1 | 2256 | 1.36 | 0.45 | 1.5E-02 |
| Sanbr | 9330 | -1.28 | -0.36 | 1.8E-02 |
| Scn9a | 2003 | -2.28 | -1.19 | 2.2E-02 |
| Sdhaf1 | 1026 | 1.36 | 0.44 | 1.5E-04 |
| Sema5a | 8354 | -1.30 | -0.38 | 1.9E-02 |
| Sgcd | 199 | -1.98 | -0.98 | 2.4E-02 |
| Sh3d21 | 165 | -1.68 | -0.75 | 2.2E-02 |
| Shmt2 | 5253 | 1.33 | 0.41 | 2.5E-03 |
| Sidt1 | 116 | -2.00 | -1.00 | 2.1E-02 |
| Sipa1 | 2494 | 1.29 | 0.37 | 2.1E-02 |
| Slc35a5 | 2093 | -1.25 | -0.32 | 7.7E-03 |
| Slc35d2 | 139 | -1.67 | -0.74 | 3.9E-03 |
| Slc35g1 | 693 | 1.38 | 0.47 | 1.9E-03 |
| Slc44a5 | 1430 | -1.51 | -0.59 | 1.0E-02 |
| Slitrk6 | 563 | -2.63 | -1.40 | 6.2E-04 |
| Smarca1 | 5682 | -1.30 | -0.38 | 9.4E-04 |
| Smpdl3a | 661 | -1.39 | -0.47 | 9.2E-04 |
| Snapc3 | 4489 | -1.32 | -0.40 | 5.7E-03 |
| Son | 59465 | -1.32 | -0.41 | 3.8E-03 |
| Sowaha | 140 | 1.63 | 0.70 | 2.5E-02 |
| Sp5 | 441 | 1.72 | 0.78 | 2.5E-05 |
| Spag9 | 35116 | -1.32 | -0.40 | 1.6E-03 |
| Spire2 | 1474 | 1.30 | 0.37 | 1.4E-02 |
| Srebf1 | 4265 | 1.38 | 0.47 | 5.8E-04 |
| Srsf9 | 4304 | 1.29 | 0.36 | 1.6E-02 |
| Stap2 | 174 | -1.44 | -0.53 | 9.9E-04 |
| Stmn2 | 30680 | -1.38 | -0.46 | 1.4E-02 |
| Sulf1 | 10404 | -1.55 | -0.63 | 8.6E-03 |
| Sumo2 | 11981 | -1.34 | -0.42 | 2.4E-02 |
| Sun2 | 6345 | 1.26 | 0.33 | 9.5E-03 |
| Syne1 | 2080 | -1.70 | -0.76 | 1.4E-02 |
| Tbc1d22b | 3126 | 1.26 | 0.33 | 2.9E-03 |

| | | | | |
|---|---|---|---|---|
| Tcp11l1 | 1548 | -1.36 | -0.45 | 1.1E-02 |
| Tecta | 130 | -1.99 | -1.00 | 5.9E-03 |
| Thbs4 | 431 | -2.11 | -1.08 | 6.5E-03 |
| Thrb | 169 | -1.57 | -0.65 | 8.4E-04 |
| Thsd7a | 5730 | -1.62 | -0.69 | 5.0E-03 |
| Tmem106b | 6052 | -1.25 | -0.32 | 6.4E-03 |
| Tmem200c | 306 | 1.34 | 0.42 | 2.8E-03 |
| Tmem201 | 5359 | 1.27 | 0.34 | 1.2E-02 |
| Tmem37 | 156 | 1.64 | 0.71 | 1.2E-02 |
| Tmem81 | 220 | 1.28 | 0.36 | 1.7E-03 |
| Tmem86b | 1293 | 1.31 | 0.39 | 8.0E-03 |
| Tmod4 | 105 | 1.26 | 0.34 | 2.4E-02 |
| Tnfrsf11b | 126 | -1.40 | -0.48 | 6.4E-03 |
| Tnik | 8481 | -1.46 | -0.55 | 1.3E-02 |
| Tnnc2 | 110 | -6.75 | -2.75 | 8.9E-03 |
| Traip | 1831 | 1.32 | 0.40 | 1.0E-02 |
| Trim12c | 629 | -1.34 | -0.43 | 8.2E-03 |
| Trio | 14535 | -1.30 | -0.38 | 1.8E-02 |
| Ttc3 | 64621 | -1.35 | -0.43 | 5.1E-03 |
| Tubg1 | 3274 | 1.31 | 0.39 | 2.8E-03 |
| Ubr5 | 28114 | -1.28 | -0.36 | 3.3E-03 |
| Upp1 | 107 | 1.46 | 0.55 | 1.1E-02 |
| Usp34 | 19191 | -1.41 | -0.50 | 6.4E-03 |
| Vit | 1153 | -1.27 | -0.35 | 4.7E-03 |
| Vldlr | 4509 | -1.35 | -0.43 | 1.6E-02 |
| Vwa5b1 | 186 | -1.71 | -0.77 | 2.3E-02 |
| Vwc2l | 244 | -2.04 | -1.03 | 1.2E-02 |
| Wars | 6340 | 1.25 | 0.32 | 3.4E-03 |
| Wnt4 | 470 | -1.70 | -0.76 | 2.3E-02 |
| Xlr3b | 932 | 2.04 | 1.03 | 4.8E-03 |
| Yjefn3 | 201 | 1.45 | 0.54 | 4.2E-03 |
| Zbtb37 | 5534 | -1.28 | -0.36 | 5.5E-03 |
| Zc3h11a | 1059 | -1.58 | -0.66 | 2.2E-03 |
| Zc3h3 | 1682 | 1.37 | 0.45 | 5.2E-03 |
| Zdhhc18 | 2443 | 1.26 | 0.33 | 2.3E-02 |
| Zfp109 | 410 | -1.31 | -0.39 | 1.2E-05 |
| Zfp119a | 583 | -1.53 | -0.61 | 8.2E-04 |
| Zfp280b | 2585 | 1.28 | 0.35 | 1.9E-03 |
| Zfp367 | 2698 | 1.26 | 0.34 | 2.5E-02 |
| Zfp456 | 134 | -1.40 | -0.49 | 8.1E-03 |
| Zfp598 | 5072 | 1.28 | 0.35 | 1.1E-03 |
| Zfp729a | 1813 | -1.26 | -0.33 | 1.7E-02 |
| Zfp738 | 3449 | -1.27 | -0.35 | 6.2E-03 |
| Zfp937 | 565 | -1.34 | -0.42 | 2.9E-05 |
| Zfp950 | 3319 | -1.41 | -0.49 | 7.2E-04 |
| Zpld1 | 155 | -2.90 | -1.53 | 7.2E-03 |
