## Supplemental Table S3 for "Rapid effects of valproic acid on the fetal brain transcriptome: Implications for brain development and autism"

| | | Male | | Female | |
|---|---|---|---|---|---|
| **Gene** | **Counts** | **FC** | **$p_{FDR}$** | **FC** | **$p_{FDR}$** |
| *Abat* | 8490 | 1.58 | 2.4E-08 | 1.68 | 1.7E-10 |
| *Abca7* | 4344 | 2.59 | 3.7E-24 | 2.58 | 7.2E-24 |
| *Abce1* | 14484 | -1.25 | 2.2E-03 | -1.20 | 1.3E-02 |
| *Abl2* | 6428 | -1.33 | 1.3E-03 | -1.25 | 1.4E-02 |
| *Ache* | 892 | 1.73 | 1.1E-03 | 1.82 | 3.1E-04 |
| *Actb* | 107135 | -1.36 | 2.4E-08 | -1.48 | 4.5E-13 |
| *Actn4* | 15736 | 1.43 | 3.2E-05 | 1.37 | 2.6E-04 |
| *Acy1* | 1770 | 1.70 | 2.3E-17 | 1.68 | 1.2E-16 |
| *Ada* | 284 | 1.30 | 1.9E-02 | 1.35 | 7.7E-03 |
| *Adcy3* | 1057 | -1.69 | 1.4E-08 | -1.81 | 1.1E-10 |
| *Adk* | 3332 | 1.30 | 1.4E-02 | 1.30 | 1.4E-02 |
| *Adnp* | 233 | -1.63 | 4.0E-07 | -1.58 | 3.0E-06 |
| *Adora2a* | 296 | -1.25 | 4.9E-02 | -1.36 | 4.9E-03 |
| *Ago3* | 5603 | 1.60 | 3.5E-07 | 1.90 | 1.9E-12 |
| *Agtr2* | 720 | -6.44 | 2.7E-04 | -4.45 | 4.1E-03 |
| *Ahi1* | 10423 | 1.57 | 1.1E-04 | 1.60 | 5.6E-05 |
| *Ahnak* | 9132 | -3.07 | 1.2E-07 | -2.64 | 5.6E-06 |
| *Akap9* | 18776 | -2.30 | 5.3E-09 | -1.85 | 2.4E-05 |
| *Aldoa* | 19582 | 1.32 | 1.1E-03 | 1.25 | 1.2E-02 |
| *Ankrd17* | 2867 | -1.72 | 7.9E-13 | -1.66 | 2.6E-11 |
| *Anxa1* | 480 | -4.00 | 4.4E-05 | -3.16 | 8.3E-04 |
| *Ap1s2* | 5451 | 1.35 | 1.5E-03 | 1.33 | 2.7E-03 |
| *Apba2* | 6951 | -1.57 | 2.4E-04 | -1.59 | 1.3E-04 |
| *Aph1a* | 4926 | 1.43 | 2.2E-08 | 1.31 | 2.6E-05 |
| *Arf3* | 12109 | 1.27 | 5.6E-06 | 1.20 | 9.6E-04 |
| *Arhgef2* | 9820 | -1.95 | 1.2E-15 | -2.08 | 9.6E-19 |
| *Arid1b* | 5659 | -2.13 | 1.2E-16 | -2.17 | 2.7E-17 |
| *Arid2* | 11217 | -1.62 | 2.8E-15 | -1.59 | 4.7E-14 |
| *Arntl2* | 511 | 1.71 | 3.1E-06 | 2.17 | 1.2E-11 |
| *Ash1l* | 9264 | -1.69 | 1.4E-05 | -1.40 | 6.9E-03 |
| *Asxl3* | 5208 | -1.75 | 1.5E-10 | -1.57 | 2.7E-07 |
| *Atp10a* | 473 | -1.89 | 3.3E-16 | -1.89 | 4.2E-16 |
| *Atp6v0a2* | 6136 | 1.42 | 2.4E-14 | 1.40 | 2.6E-13 |
| *Atrx* | 29927 | -1.80 | 2.5E-06 | -1.44 | 5.0E-03 |
| *Bace1* | 6271 | 1.83 | 4.3E-22 | 1.90 | 2.0E-24 |
| *Baiap2l1* | 386 | 3.78 | 2.5E-09 | 3.63 | 8.1E-09 |
| *Bcas1* | 279 | -1.78 | 1.5E-03 | -1.52 | 2.4E-02 |
| *Bcl11a* | 10078 | -1.59 | 6.6E-05 | -1.70 | 5.2E-06 |
| *Bcorl1* | 4918 | 1.89 | 1.9E-10 | 1.80 | 5.5E-09 |
| *Brinp2* | 2925 | 2.82 | 2.7E-12 | 2.61 | 1.2E-10 |
| *Brsk2* | 7140 | -1.74 | 2.6E-04 | -1.86 | 4.1E-05 |
| *Brwd3* | 6365 | 1.42 | 2.2E-03 | 1.66 | 4.9E-06 |
| *Btaf1* | 10163 | 1.41 | 1.9E-02 | 1.62 | 6.9E-04 |
| *Cacna1b* | 6214 | 1.34 | 4.7E-02 | 1.42 | 1.7E-02 |
| *Cacna1c* | 3446 | -1.47 | 3.1E-03 | -1.38 | 1.5E-02 |
| *Cacna2d1* | 9393 | -1.84 | 2.7E-08 | -1.59 | 3.0E-05 |
| *Cacnb2* | 498 | -2.08 | 9.5E-13 | -1.80 | 1.2E-08 |
| *Cadm1* | 12690 | 1.25 | 1.3E-02 | 1.28 | 5.3E-03 |
| *Camk2a* | 4142 | 1.94 | 1.6E-07 | 1.84 | 1.4E-06 |
| *Caprin1* | 3184 | 1.25 | 1.4E-03 | 1.31 | 1.0E-04 |
| *Cask* | 10174 | -1.30 | 4.7E-03 | -1.24 | 2.3E-02 |
| *Casz1* | 1306 | 2.43 | 2.5E-08 | 3.00 | 3.2E-12 |

| | | Male | | Female | |
|---|---|---|---|---|---|
| **Gene** | **Counts** | **FC** | **$p_{FDR}$** | **FC** | **$p_{FDR}$** |
| *Cc2d1a* | 2014 | -1.38 | 3.6E-03 | -1.44 | 1.1E-03 |
| *Ccdc91* | 1888 | -1.95 | 2.9E-49 | -1.82 | 4.0E-40 |
| *Ccng1* | 5463 | 1.49 | 2.9E-08 | 1.58 | 1.4E-10 |
| *Cd276* | 3140 | -2.13 | 5.0E-36 | -2.18 | 2.4E-38 |
| *Cd38* | 232 | -1.84 | 8.7E-05 | -1.90 | 3.5E-05 |
| *Cdc42bpb* | 14607 | 1.34 | 2.8E-04 | 1.31 | 9.7E-04 |
| *Cdh11* | 12820 | -2.40 | 2.1E-06 | -2.37 | 3.3E-06 |
| *Cdk19* | 6534 | 1.17 | 1.4E-02 | 1.25 | 3.0E-04 |
| *Cdk5rap2* | 8016 | -1.37 | 6.2E-04 | -1.27 | 1.2E-02 |
| *Cdkl5* | 3056 | 1.71 | 5.9E-08 | 1.95 | 9.5E-12 |
| *Cdon* | 13736 | -1.37 | 4.4E-04 | -1.20 | 4.7E-02 |
| *Celf6* | 1504 | 1.51 | 1.1E-03 | 1.44 | 4.8E-03 |
| *Cep135* | 2358 | -1.51 | 7.3E-09 | -1.38 | 9.7E-06 |
| *Cep41* | 1580 | -1.91 | 2.3E-44 | -1.94 | 1.8E-46 |
| *Chd2* | 12282 | 1.21 | 2.5E-02 | 1.35 | 3.5E-04 |
| *Chd3* | 39510 | -1.98 | 9.7E-21 | -2.00 | 3.8E-21 |
| *Chd7* | 13012 | -1.35 | 1.4E-02 | -1.33 | 2.0E-02 |
| *Chrna7* | 407.96 | 1.42 | 1.9E-04 | 1.28 | 9.1E-03 |
| *Clcn4* | 10609 | -1.57 | 1.3E-06 | -1.48 | 3.5E-05 |
| *Cln8* | 2506 | 1.77 | 2.3E-31 | 1.68 | 4.8E-26 |
| *Cmpk2* | 785 | 1.26 | 2.5E-02 | 1.26 | 2.4E-02 |
| *Cnr1* | 6596 | -2.09 | 6.8E-05 | -1.84 | 1.2E-03 |
| *Cntnap2* | 11429 | 1.42 | 7.8E-04 | 1.26 | 3.6E-02 |
| *Cpeb4* | 8073 | 1.60 | 9.9E-05 | 1.56 | 2.5E-04 |
| *Cpsf7* | 11294 | -1.50 | 5.6E-09 | -1.61 | 5.6E-12 |
| *Cpt2* | 1271 | 1.38 | 4.1E-04 | 1.29 | 5.1E-03 |
| *Crebbp* | 5496 | -1.93 | 1.7E-10 | -2.06 | 2.0E-12 |
| *Csde1* | 31901 | -1.98 | 6.1E-40 | -2.01 | 3.0E-41 |
| *Csnk1e* | 14294 | -1.44 | 1.2E-06 | -1.54 | 6.5E-09 |
| *Csnk2a1* | 16382 | -1.27 | 5.8E-08 | -1.26 | 8.2E-08 |
| *Csnk2b* | 5717 | 1.39 | 1.0E-11 | 1.32 | 1.4E-08 |
| *Ctcf* | 9148 | -2.34 | 1.8E-38 | -2.56 | 6.6E-47 |
| *Ctr9* | 13165 | 1.54 | 1.2E-27 | 1.54 | 1.4E-27 |
| *Cul7* | 10139 | -1.69 | 4.5E-14 | -1.80 | 1.2E-17 |
| *Cx3cr1* | 687 | -2.05 | 1.2E-17 | -2.28 | 1.3E-22 |
| *Cyfip1* | 6506 | -1.45 | 1.4E-06 | -1.45 | 1.6E-06 |
| *Ddc* | 823 | -1.30 | 1.8E-02 | -1.35 | 6.1E-03 |
| *Ddx3x* | 52188 | -1.58 | 5.5E-09 | -1.63 | 4.1E-10 |
| *Deaf1* | 3240 | -1.47 | 4.7E-07 | -1.57 | 2.5E-09 |
| *Depdc5* | 6414 | 1.60 | 1.5E-25 | 1.66 | 5.1E-29 |
| *Dhcr7* | 4286 | 1.56 | 6.0E-08 | 1.38 | 1.2E-04 |
| *Dhx30* | 9045 | -1.55 | 3.6E-20 | -1.65 | 7.1E-26 |
| *Dip2c* | 3203 | -2.00 | 2.1E-11 | -1.94 | 1.9E-10 |
| *Disc1* | 150 | -2.17 | 8.3E-13 | -1.71 | 1.1E-06 |
| *Dixdc1* | 6724 | 1.41 | 1.4E-06 | 1.50 | 1.0E-08 |
| *Dmpk* | 2879 | 1.32 | 4.3E-03 | 1.26 | 2.2E-02 |
| *Dnah10* | 231 | 2.35 | 4.2E-06 | 2.49 | 8.6E-07 |
| *Dnah17* | 157 | 5.09 | 4.6E-23 | 5.86 | 7.3E-27 |
| *Dnmt3a* | 15809 | -2.22 | 4.4E-38 | -2.11 | 3.4E-33 |
| *Dock1* | 4066 | -1.49 | 7.5E-04 | -1.38 | 7.5E-03 |
| *Dock8* | 232 | -2.26 | 2.3E-04 | -2.08 | 1.0E-03 |
| *Dolk* | 1049 | 1.29 | 1.9E-04 | 1.22 | 5.4E-03 |

### Table S3 (cont)

| | | Male | | Female | |
|---|---|---|---|---|---|
| Gene | Counts | FC | $p_{FDR}$ | FC | $p_{FDR}$ |
| *Dpp4* | 163 | -1.87 | 5.5E-03 | -1.69 | 2.3E-02 |
| *Dpysl2* | 11973 | -1.40 | 2.7E-02 | -1.70 | 2.8E-04 |
| *Dpysl3* | 81721 | -1.46 | 4.0E-03 | -1.42 | 8.2E-03 |
| *Drd2* | 191 | 1.98 | 7.8E-06 | 1.92 | 2.1E-05 |
| *Dusp15* | 477 | 2.05 | 1.4E-05 | 2.24 | 9.4E-07 |
| *Dyrk1a* | 7472 | -1.44 | 2.6E-08 | -1.54 | 2.1E-11 |
| *Ebf3* | 12814 | -1.68 | 2.6E-04 | -1.60 | 1.1E-03 |
| *Ecpas* | 9645 | -1.62 | 2.0E-10 | -1.50 | 1.6E-07 |
| *Efr3a* | 5605 | 1.74 | 1.0E-15 | 1.91 | 5.7E-21 |
| *Ehmt1* | 11001 | -1.32 | 3.2E-07 | -1.44 | 5.5E-12 |
| *Ep300* | 10587 | -1.39 | 1.2E-05 | -1.31 | 4.0E-04 |
| *Epc2* | 5022 | -3.53 | 7.0E-57 | -3.45 | 8.2E-55 |
| *Ephb2* | 25546 | 1.44 | 1.4E-03 | 1.29 | 2.8E-02 |
| *Erbin* | 14418 | -2.02 | 2.6E-26 | -1.88 | 3.6E-21 |
| *Erg* | 1097 | -2.37 | 4.0E-09 | -2.44 | 1.2E-09 |
| *Exoc3* | 8530 | -1.26 | 7.8E-06 | -1.23 | 4.7E-05 |
| *Exoc6* | 1031 | -1.67 | 5.4E-12 | -1.60 | 2.1E-10 |
| *Ext1* | 5543 | 1.28 | 3.9E-04 | 1.29 | 2.6E-04 |
| *Fabp5* | 12171 | -1.45 | 1.2E-07 | -1.46 | 7.7E-08 |
| *Fam98c* | 976 | 2.59 | 4.5E-43 | 2.47 | 5.2E-39 |
| *Fbxo11* | 13248 | -1.37 | 3.5E-04 | -1.41 | 1.1E-04 |
| *Fbxo33* | 1584 | 1.59 | 1.5E-17 | 1.51 | 6.6E-14 |
| *Fezf2* | 1417 | -1.73 | 5.8E-03 | -1.99 | 4.1E-04 |
| *Foxp2* | 6899 | -1.49 | 2.2E-08 | -1.58 | 1.6E-10 |
| *Frk* | 338 | -3.44 | 2.6E-06 | -3.42 | 3.2E-06 |
| *Fxn* | 615 | -1.96 | 1.2E-17 | -2.04 | 1.4E-19 |
| *Gabbr2* | 570 | -2.47 | 7.9E-03 | -2.25 | 1.9E-02 |
| *Gabra4* | 739 | 2.46 | 3.0E-08 | 2.63 | 2.6E-09 |
| *Galnt10* | 4324 | 1.96 | 2.5E-16 | 1.91 | 4.3E-15 |
| *Galnt14* | 565 | -2.75 | 7.3E-21 | -2.53 | 1.2E-17 |
| *Gas2* | 1719 | -1.55 | 2.7E-04 | -1.45 | 2.1E-03 |
| *Gbe1* | 851 | -1.62 | 4.7E-07 | -1.67 | 7.3E-08 |
| *Gnai1* | 4191 | 1.42 | 4.4E-03 | 1.43 | 3.8E-03 |
| *Gnb1l* | 483 | 1.57 | 2.1E-15 | 1.60 | 1.3E-16 |
| *Grb10* | 45862 | -1.51 | 5.4E-03 | -1.49 | 6.7E-03 |
| *Grid2ip* | 169 | 2.70 | 6.3E-24 | 3.40 | 1.4E-34 |
| *Grik4* | 520 | -1.44 | 1.2E-03 | -1.38 | 4.7E-03 |
| *Grik5* | 7313 | -1.31 | 2.2E-02 | -1.40 | 3.6E-03 |
| *Grin2a* | 270 | -2.28 | 1.5E-08 | -2.27 | 2.0E-08 |
| *Gtf2i* | 23958 | -2.36 | 1.2E-34 | -2.54 | 8.0E-41 |
| *Hcfc1* | 17725 | -2.30 | 2.7E-29 | -2.29 | 4.5E-29 |
| *Hdlbp* | 25696 | -1.29 | 9.1E-03 | -1.25 | 2.5E-02 |
| *Hecw2* | 1165 | -2.41 | 4.0E-22 | -2.08 | 1.7E-15 |
| *Hivep2** | 2898 | -1.46 | 1.3E-02 | -1.32 | 8.0E-02 |
| *Hnrnpk* | 69304 | -1.30 | 9.0E-05 | -1.36 | 3.4E-06 |
| *Hnrnpu* | 3018 | -1.59 | 3.3E-04 | -1.59 | 3.4E-04 |
| *Homer1* | 5300 | 1.61 | 9.6E-20 | 1.63 | 7.6E-21 |
| *Ica1* | 848 | -1.41 | 2.3E-04 | -1.30 | 5.0E-03 |
| *Igf1* | 2781 | -3.30 | 8.9E-04 | -3.18 | 1.3E-03 |
| *Ikzf1* | 271 | -1.58 | 1.5E-05 | -1.59 | 1.2E-05 |
| *Inpp1* | 1129 | -2.01 | 6.8E-32 | -2.01 | 8.4E-32 |
| *Ints6* | 3398 | -1.53 | 3.8E-20 | -1.47 | 2.7E-16 |
| *Iqgap3* | 4143 | 1.40 | 1.9E-03 | 1.39 | 2.5E-03 |
| *Iqsec2* | 1627 | 1.47 | 8.2E-05 | 1.40 | 7.0E-04 |
| *Irf2bpl* | 5211 | -1.30 | 2.7E-02 | -1.52 | 3.4E-04 |
| *Itpr1* | 4428 | 1.77 | 6.2E-07 | 2.01 | 6.7E-10 |
| *Kansl1* | 8932 | -1.43 | 1.0E-14 | -1.47 | 3.6E-17 |
| *Kansl1l* | 2089 | -1.80 | 1.7E-19 | -1.61 | 4.1E-13 |
| *Kat2b* | 1923 | -1.71 | 1.0E-09 | -1.73 | 4.2E-10 |
| *Kat6b* | 7883 | -2.68 | 1.9E-23 | -2.43 | 3.8E-19 |
| *Katnal1* | 2730 | -1.97 | 2.7E-24 | -2.04 | 9.5E-27 |
| *Kcnb1* | 1903 | 1.44 | 4.3E-03 | 1.49 | 1.6E-03 |
| *Kcnc3* | 5043 | 5.14 | 3.4E-42 | 4.74 | 3.4E-38 |
| *Kcnj10* | 1277 | 2.72 | 9.7E-28 | 2.66 | 1.8E-26 |
| *Kcnq2** | 10691 | -1.57 | 1.6E-02 | -1.42 | 6.4E-02 |
| *Kdm1b* | 5673 | 2.26 | 3.4E-130 | 2.28 | 4.1E-133 |
| *Kdm5c* | 12138 | -1.40 | 1.1E-14 | -1.53 | 3.7E-23 |
| *Kdm6b* | 8176 | -1.30 | 4.8E-02 | -1.51 | 1.4E-03 |
| *Kif13b* | 3411 | 1.29 | 4.0E-03 | 1.48 | 4.3E-06 |
| *Kif14* | 2071 | -1.46 | 4.8E-04 | -1.48 | 2.9E-04 |
| *Klf16* | 1425 | 2.00 | 1.5E-16 | 1.78 | 8.3E-12 |
| *Klf7* | 6968 | -1.26 | 2.5E-02 | -1.27 | 2.3E-02 |
| *Kmt2e* | 18204 | -1.33 | 3.4E-04 | -1.36 | 8.9E-05 |
| *Kmt5b* | 6543 | -6.45 | 2.2E-62 | -6.59 | 9.4E-64 |
| *Kptn* | 941 | -2.00 | 4.4E-45 | -2.16 | 2.4E-55 |
| *Lamb1* | 8515 | -1.45 | 4.2E-03 | -1.39 | 1.3E-02 |
| *Las1l* | 6968 | 1.40 | 4.1E-24 | 1.37 | 1.2E-21 |
| *Ldb1* | 15935 | -1.30 | 6.7E-05 | -1.40 | 3.0E-07 |
| *Ldlr* | 2768 | -1.36 | 3.2E-02 | -1.57 | 1.1E-03 |
| *Lin7b* | 298 | 6.34 | 4.5E-69 | 6.45 | 1.5E-69 |
| *Lnpk* | 5745 | 1.28 | 3.6E-03 | 1.46 | 3.3E-06 |
| *Lrfn2* | 233 | -1.63 | 3.9E-02 | -1.67 | 3.1E-02 |
| *Lrp2* | 2856 | 1.43 | 3.2E-02 | 1.64 | 2.8E-03 |
| *Lrrc1* | 2446 | 1.64 | 7.0E-12 | 1.59 | 9.9E-11 |
| *Lrrc4* | 888 | -3.27 | 1.1E-12 | -3.64 | 8.2E-15 |
| *Lztr1* | 7298 | -1.32 | 3.1E-15 | -1.33 | 6.8E-16 |
| *Lzts2* | 3586 | -1.20 | 3.2E-02 | -1.33 | 5.7E-04 |
| *Maoa* | 3840 | 1.36 | 9.8E-05 | 1.39 | 2.0E-05 |
| *Map1a* | 10875 | 1.62 | 2.0E-04 | 1.61 | 2.3E-04 |
| *Map4k1* | 311 | -1.27 | 6.1E-03 | -1.37 | 3.2E-04 |
| *Mapk8ip1* | 12031 | 1.45 | 1.9E-07 | 1.37 | 1.0E-05 |
| *Mark1* | 5757 | -1.39 | 3.0E-03 | -1.42 | 1.7E-03 |
| *Mast3* | 433 | 2.89 | 1.1E-05 | 2.18 | 1.5E-03 |
| *Mbd3* | 6414 | -2.01 | 1.9E-13 | -2.27 | 3.7E-18 |
| *Mbd4* | 2576 | 1.30 | 1.3E-05 | 1.33 | 1.8E-06 |
| *Mboat7* | 5265 | 1.93 | 3.0E-26 | 1.91 | 2.5E-25 |
| *Mecp2* | 11258 | -1.59 | 2.3E-05 | -1.54 | 7.6E-05 |
| *Megf10* | 3621 | -1.96 | 6.1E-22 | -1.80 | 9.4E-17 |
| *Meis2* | 15231 | -1.49 | 9.6E-04 | -1.39 | 7.0E-03 |
| *Mink1* | 6672 | 1.33 | 6.4E-04 | 1.25 | 9.3E-03 |
| *Mrtfb* | 3074 | -1.45 | 6.7E-10 | -1.39 | 5.4E-08 |
| *Mtf1* | 1803 | -1.16 | 2.5E-02 | -1.25 | 5.6E-04 |
| *Mtor* | 10534 | 1.48 | 7.4E-11 | 1.56 | 1.1E-13 |
| *Myh9* | 16822 | 1.30 | 3.5E-02 | 1.29 | 4.4E-02 |

### Table S3 (cont)

| | | Male | | Female | |
|---|---|---|---|---|---|
| **Gene** | **Counts** | **FC** | **$p_{FDR}$** | **FC** | **$p_{FDR}$** |
| *Mylk* | 1785 | -1.20 | 4.4E-02 | -1.25 | 1.5E-02 |
| *Myo1e* | 3165 | 3.35 | 1.4E-64 | 3.28 | 2.9E-62 |
| *Myo5a** | 17678 | -1.36 | 7.3E-03 | -1.22 | 9.8E-02 |
| *Myo9b* | 12403 | 1.43 | 3.5E-08 | 1.52 | 5.1E-11 |
| *Nacc1* | 8224 | -1.15 | 1.4E-01 | -1.29 | 3.2E-03 |
| *Nexmif* | 3129 | -2.12 | 1.7E-03 | -1.76 | 2.3E-02 |
| *Nf1* | 8492 | -1.34 | 1.2E-04 | -1.27 | 1.9E-03 |
| *Nfe2l3* | 803 | 4.19 | 8.5E-61 | 3.97 | 1.5E-56 |
| *Nfia* | 6750 | -1.71 | 1.4E-03 | -1.83 | 3.0E-04 |
| *Nfib* | 18590 | -2.04 | 5.5E-04 | -2.04 | 5.8E-04 |
| *Nfix* | 5859 | -2.03 | 4.0E-03 | -2.11 | 2.3E-03 |
| *Nipa2* | 4612 | 1.32 | 9.0E-08 | 1.36 | 3.8E-09 |
| *Nkx2-2†* | 355 | -1.14 | 6.4E-01 | -1.77 | 1.6E-02 |
| *Nr1d1* | 234 | 2.42 | 2.9E-08 | 2.09 | 4.4E-06 |
| *Nr2f1* | 15518 | -1.63 | 9.4E-06 | -1.93 | 1.2E-09 |
| *Nr3c2* | 254 | 2.04 | 7.0E-04 | 2.10 | 4.3E-04 |
| *Nr4a2** | 2860 | 1.35 | 1.7E-02 | 1.26 | 7.4E-02 |
| *Nsd1* | 20486 | -1.74 | 2.9E-11 | -1.58 | 4.4E-08 |
| *Nsd2* | 33101 | -1.33 | 1.0E-04 | -1.28 | 9.4E-04 |
| *Nsmce3* | 1585 | 1.44 | 1.8E-18 | 1.36 | 2.3E-13 |
| *Nxf1* | 17752 | 1.27 | 4.8E-08 | 1.26 | 1.6E-07 |
| *Nxph1* | 911 | -2.60 | 8.0E-07 | -2.48 | 3.3E-06 |
| *Ofd1* | 2004 | -1.35 | 8.8E-09 | -1.34 | 2.4E-08 |
| *Ophn1** | 3837 | -1.28 | 1.2E-02 | -1.18 | 1.1E-01 |
| *Patj* | 1326 | -2.29 | 1.3E-31 | -2.20 | 1.1E-28 |
| *Pcca* | 2377 | 1.50 | 4.9E-07 | 1.51 | 4.8E-07 |
| *Pccb* | 2484 | -1.67 | 1.1E-47 | -1.81 | 4.0E-63 |
| *Pcdh10* | 8761 | 1.47 | 4.4E-03 | 1.54 | 1.4E-03 |
| *Pcdha11* | 113 | -2.30 | 2.5E-04 | -1.94 | 4.1E-03 |
| *Pcdha3* | 113 | -2.00 | 1.2E-03 | -1.97 | 1.7E-03 |
| *Pcdha7* | 112 | -1.90 | 1.3E-02 | -2.01 | 6.8E-03 |
| *Pdk2* | 1859 | 2.00 | 1.8E-19 | 2.02 | 3.9E-20 |
| *Pdzd8* | 8176 | 1.44 | 5.7E-22 | 1.50 | 1.2E-26 |
| *Per1* | 3226 | 1.95 | 1.1E-11 | 1.77 | 8.2E-09 |
| *Per2* | 1092 | 1.71 | 1.0E-05 | 1.68 | 2.2E-05 |
| *Pex7* | 836 | -2.40 | 3.4E-59 | -2.40 | 1.2E-58 |
| *Phb* | 6881 | 1.29 | 2.3E-05 | 1.27 | 1.1E-04 |
| *Phf2* | 6735 | -1.85 | 5.0E-08 | -2.00 | 5.6E-10 |
| *Phf14* | 11218 | -1.74 | 2.7E-18 | -1.77 | 2.0E-19 |
| *Phip** | 21455 | -1.44 | 1.2E-03 | -1.22 | 9.0E-02 |
| *Pik3cg* | 194 | -2.20 | 4.0E-09 | -2.41 | 6.5E-11 |
| *Pitx1* | 157 | -3.67 | 5.1E-03 | -5.34 | 2.6E-04 |
| *Plxnb1* | 17385 | -1.79 | 1.2E-08 | -1.91 | 2.6E-10 |
| *Pnpla7* | 498 | -1.27 | 2.0E-03 | -1.28 | 1.4E-03 |
| *Pogz* | 8242 | -2.18 | 2.3E-29 | -2.25 | 8.9E-32 |
| *Polr2a* | 20372 | -1.35 | 4.3E-04 | -1.38 | 1.2E-04 |
| *Pola2* | 4393 | 1.67 | 7.1E-08 | 1.57 | 3.1E-06 |
| *Pomgnt1* | 3228 | -1.63 | 1.2E-31 | -1.69 | 7.8E-37 |
| *Pomt1* | 2351 | -1.30 | 2.8E-07 | -1.29 | 7.1E-07 |
| *Ppm1d* | 2918 | 1.33 | 2.6E-08 | 1.23 | 7.7E-05 |
| *Ppp1r1b* | 299 | 3.35 | 1.3E-04 | 4.05 | 8.5E-06 |
| *Ppp1r9b** | 13672 | 1.28 | 5.3E-03 | 1.17 | 9.4E-02 |

| | | Male | | Female | |
|---|---|---|---|---|---|
| **Gene** | **Counts** | **FC** | **$p_{FDR}$** | **FC** | **$p_{FDR}$** |
| *Ppp2r1b* | 7788 | 1.54 | 2.2E-11 | 1.44 | 2.0E-08 |
| *Ppp2r5d* | 10594 | 1.79 | 5.1E-25 | 1.68 | 2.5E-20 |
| *Prex1* | 6987 | 1.55 | 5.5E-08 | 1.62 | 2.4E-09 |
| *Prkd1* | 1487 | 1.33 | 9.2E-03 | 1.33 | 8.9E-03 |
| *Prr12** | 7437 | -1.31 | 6.4E-02 | -1.48 | 4.9E-03 |
| *Ptbp2* | 26867 | -1.53 | 5.5E-08 | -1.56 | 1.3E-08 |
| *Ptchd1** | 234 | -1.83 | 1.6E-02 | -1.65 | 5.1E-02 |
| *Ptdss1* | 5032 | 1.32 | 6.1E-07 | 1.38 | 8.6E-09 |
| *Ptpn4* | 9452 | 1.32 | 1.6E-02 | 1.60 | 3.4E-05 |
| *Prkd2* | 1619 | 1.69 | 3.1E-11 | 1.65 | 3.4E-10 |
| *Pten* | 12976 | 1.37 | 3.9E-06 | 1.37 | 3.9E-06 |
| *Ptk7* | 9855 | 1.29 | 1.8E-02 | 1.27 | 2.5E-02 |
| *Ptpn11* | 16665 | 1.31 | 9.8E-16 | 1.27 | 8.7E-13 |
| *Ptprb* | 1124 | -1.77 | 2.0E-08 | -1.83 | 2.6E-09 |
| *Pxdn* | 15773 | -1.48 | 4.5E-05 | -1.38 | 1.1E-03 |
| *Rab11fip5* | 3422 | 2.08 | 6.9E-20 | 2.09 | 3.7E-20 |
| *Rad21* | 24544 | -1.30 | 9.2E-06 | -1.37 | 7.5E-08 |
| *Rai1* | 7722 | -1.57 | 2.5E-07 | -1.56 | 3.3E-07 |
| *Ranbp17* | 928 | -1.59 | 3.6E-07 | -1.48 | 1.9E-05 |
| *Rassf5* | 168 | 1.54 | 2.2E-02 | 1.66 | 6.4E-03 |
| *Reep3* | 6903 | -1.26 | 5.0E-10 | -1.26 | 5.5E-10 |
| *Rfx3* | 5786 | -1.47 | 5.0E-08 | -1.45 | 2.2E-07 |
| *Rfx4* | 2512 | -1.73 | 7.1E-04 | -1.78 | 3.4E-04 |
| *Rims2* | 3497 | -1.38 | 1.1E-02 | -1.48 | 1.7E-03 |
| *Rims3* | 4156 | 1.95 | 2.0E-08 | 1.98 | 1.2E-08 |
| *Rnf135* | 168 | -1.99 | 4.9E-08 | -1.97 | 8.1E-08 |
| *Rorb** | 880 | 1.72 | 3.9E-04 | 1.32 | 8.8E-02 |
| *Rps6ka2* | 5307 | 1.80 | 8.1E-20 | 1.80 | 8.6E-20 |
| *Rps6ka3* | 11427 | 1.21 | 3.3E-03 | 1.29 | 3.3E-05 |
| *Runx1t1* | 7184 | -1.62 | 4.1E-03 | -1.61 | 4.8E-03 |
| *Samd11* | 170 | -1.90 | 1.4E-04 | -2.03 | 2.3E-05 |
| *Sbf1* | 12325 | 1.57 | 4.1E-08 | 1.53 | 2.8E-07 |
| *Scaf4* | 5878 | 1.30 | 4.1E-04 | 1.26 | 1.8E-03 |
| *Scn9a*†* | 2003 | -2.28 | 2.2E-02 | -1.29 | 5.3E-01 |
| *Sdc2* | 2896 | -1.53 | 2.6E-02 | -1.49 | 3.8E-02 |
| *Sema5a** | 8354 | -1.30 | 1.9E-02 | -1.16 | 2.1E-01 |
| *Set* | 42386 | -1.41 | 4.4E-04 | -1.59 | 1.3E-06 |
| *Setbp1* | 9254 | -3.28 | 1.7E-29 | -3.22 | 1.2E-28 |
| *Setd1b* | 4177 | -2.04 | 1.6E-07 | -2.28 | 1.1E-09 |
| *Setdb2* | 1135 | 1.56 | 1.1E-07 | 1.63 | 5.2E-09 |
| *Sgsm3* | 2524 | 1.76 | 8.5E-29 | 1.69 | 8.3E-25 |
| *Shank2* | 2071 | -2.21 | 1.8E-05 | -2.32 | 4.8E-06 |
| *Sik1* | 2688 | 1.25 | 1.0E-02 | 1.26 | 8.0E-03 |
| *Sin3a* | 12950 | 1.69 | 3.1E-27 | 1.62 | 8.5E-23 |
| *Slc22a15* | 1103 | -2.14 | 3.7E-39 | -1.90 | 6.7E-28 |
| *Slc25a12* | 5108 | 1.38 | 2.3E-10 | 1.35 | 6.1E-09 |
| *Slc25a39* | 4509 | 1.70 | 1.8E-14 | 1.57 | 9.8E-11 |
| *Slc29a4* | 3649 | 1.55 | 1.5E-05 | 1.54 | 2.4E-05 |
| *Slc38a10* | 11004 | 1.62 | 9.3E-17 | 1.52 | 6.0E-13 |
| *Slc6a8* | 5359 | 1.72 | 5.6E-29 | 1.62 | 4.3E-23 |
| *Slc7a3* | 316 | 2.75 | 5.8E-38 | 2.79 | 4.9E-39 |
| *Slc7a5* | 5758 | 1.57 | 2.8E-06 | 1.36 | 2.2E-03 |

### Table S3 (cont)

| Gene | Counts | Male FC | Male pFDR | Female FC | Female pFDR |
|---|---|---|---|---|---|
| *Slc7a7* | 293 | -1.48 | 5.4E-05 | -1.62 | 4.5E-07 |
| *Slc9a6* | 7768 | 1.31 | 7.2E-03 | 1.39 | 1.1E-03 |
| *Slitrk5* | 2183 | -1.56 | 6.1E-09 | -1.59 | 1.4E-09 |
| *Smarca2* | 10932 | 1.25 | 3.1E-02 | 1.28 | 2.0E-02 |
| *Smarca4* | 29774 | -1.59 | 5.2E-06 | -1.69 | 2.5E-07 |
| *Smarcc2* | 27817 | -1.55 | 1.9E-08 | -1.63 | 4.3E-10 |
| *Smc1a* | 18407 | -1.37 | 2.0E-06 | -1.37 | 1.7E-06 |
| *Snap25* | 6289 | 1.64 | 8.5E-05 | 1.69 | 3.2E-05 |
| *Snd1* | 10450 | 1.27 | 3.4E-04 | 1.22 | 3.4E-03 |
| *Snx14* | 3275 | 1.73 | 5.1E-49 | 1.81 | 1.9E-56 |
| *Son** | 59465 | -1.32 | 3.8E-03 | -1.14 | 2.2E-01 |
| *Sox6* | 5597 | -1.29 | 2.3E-02 | -1.32 | 1.4E-02 |
| *Sparcl1* | 6073 | -1.59 | 2.4E-04 | -1.48 | 2.5E-03 |
| *Srcap* | 371 | -3.68 | 8.6E-20 | -3.26 | 1.9E-16 |
| *Ssrp1* | 22251 | -1.31 | 7.7E-05 | -1.43 | 8.9E-08 |
| *St7* | 2111 | -1.57 | 5.2E-08 | -1.50 | 1.5E-06 |
| *St8sia2* | 9960 | -1.84 | 8.8E-05 | -1.91 | 3.1E-05 |
| *Stag1* | 8175 | -2.42 | 1.3E-51 | -2.34 | 7.8E-48 |
| *Stx1a* | 2912 | 1.38 | 5.2E-03 | 1.36 | 7.0E-03 |
| *Stxbp5* | 4749 | 1.46 | 3.0E-06 | 1.50 | 4.8E-07 |
| *Syap1* | 4114 | 1.57 | 3.4E-09 | 1.54 | 1.7E-08 |
| *Sybu* | 2165 | -1.37 | 4.5E-02 | -1.44 | 1.7E-02 |
| *Syn2* | 1795 | 1.63 | 3.8E-02 | 1.78 | 1.3E-02 |
| *Syncrip* | 22048 | -1.48 | 7.5E-10 | -1.51 | 7.8E-11 |
| *Synj1* | 9528 | 1.46 | 1.1E-03 | 1.54 | 1.8E-04 |
| *Taf1* | 11862 | 1.26 | 3.9E-04 | 1.36 | 1.7E-06 |
| *Taf4* | 3798 | 1.37 | 4.0E-04 | 1.28 | 7.5E-03 |
| *Taf6* | 5567 | 1.28 | 8.3E-05 | 1.18 | 1.1E-02 |
| *Taok1* | 12512 | -1.49 | 4.8E-06 | -1.36 | 4.8E-04 |
| *Tbc1d31* | 3810 | 2.35 | 1.3E-34 | 2.40 | 3.0E-36 |
| *Tbc1d5* | 1509 | -1.60 | 6.8E-26 | -1.59 | 4.1E-25 |
| *Tbl1x* | 11400 | -1.64 | 2.6E-32 | -1.68 | 4.3E-35 |
| *Tbl1xr1* | 4323 | -1.42 | 2.5E-07 | -1.57 | 2.4E-11 |
| *Tbr1* | 3672 | -2.68 | 2.6E-05 | -2.85 | 7.8E-06 |
| *Tbx22* | 155 | -4.84 | 3.4E-02 | -8.10 | 4.5E-03 |
| *Tceal1* | 996 | -1.82 | 6.2E-13 | -1.66 | 1.7E-09 |
| *Tcf4* | 24203 | -1.55 | 9.7E-07 | -1.48 | 1.4E-05 |
| *Tcf7l2* | 10030 | -1.73 | 1.7E-02 | -2.47 | 5.4E-05 |
| *Tet2* | 6298 | -1.41 | 3.8E-05 | -1.30 | 2.0E-03 |
| *Tet3* | 11236 | -1.39 | 1.7E-03 | -1.50 | 8.5E-05 |
| *Thbs1* | 7121 | -1.91 | 1.8E-02 | -1.90 | 1.9E-02 |
| *Tlk2* | 6144 | -1.42 | 1.2E-14 | -1.44 | 8.3E-16 |
| *Tm9sf4* | 4866 | -1.34 | 2.6E-21 | -1.36 | 1.1E-23 |
| *Tmem39b* | 1217 | -2.03 | 9.6E-28 | -2.26 | 2.6E-36 |
| *Tnrc6b* | 8717 | -1.41 | 1.8E-06 | -1.32 | 1.3E-04 |

| Gene | Counts | Male FC | Male pFDR | Female FC | Female pFDR |
|---|---|---|---|---|---|
| *Tns2* | 1379 | -3.34 | 4.5E-19 | -3.35 | 3.7E-19 |
| *Top3b* | 3624 | -1.49 | 1.6E-43 | -1.49 | 2.3E-43 |
| *Traf7* | 9194 | 1.42 | 3.8E-11 | 1.32 | 1.8E-07 |
| *Trappc6b* | 2765 | -1.19 | 1.6E-02 | -1.27 | 6.3E-04 |
| *Trim33* | 11213 | -1.29 | 6.9E-05 | -1.24 | 9.5E-04 |
| *Trio** | 14535 | -1.30 | 1.8E-02 | -1.19 | 1.4E-01 |
| *Trrap* | 17379 | -1.49 | 2.6E-06 | -1.36 | 3.2E-04 |
| *Tsc2* | 11894 | 1.25 | 9.6E-04 | 1.25 | 8.7E-04 |
| *Tshz3* | 1699 | 1.28 | 4.1E-02 | 1.43 | 2.1E-03 |
| *Tspan17* | 1100 | 2.74 | 6.1E-25 | 2.80 | 7.2E-26 |
| *Tspan4* | 996 | 1.40 | 1.0E-03 | 1.36 | 2.8E-03 |
| *Ttn†* | 914 | -15.26 | 1.8E-03 | -8.61 | 1.6E-02 |
| *Tubgcp5* | 3696 | -1.29 | 2.7E-04 | -1.21 | 8.0E-03 |
| *Ube2h* | 11325 | 1.37 | 7.8E-08 | 1.29 | 1.7E-05 |
| *Ubr1* | 5748 | -1.47 | 1.3E-06 | -1.37 | 9.7E-05 |
| *Ubr5** | 28114 | -1.28 | 3.3E-03 | -1.18 | 5.5E-02 |
| *Unc13a* | 10790 | 2.41 | 2.4E-07 | 2.38 | 4.1E-07 |
| *Unc79* | 2448 | -2.25 | 1.4E-06 | -2.15 | 5.9E-06 |
| *Upf2* | 7646 | 1.31 | 1.6E-13 | 1.39 | 5.9E-19 |
| *Usp15* | 8350 | 1.26 | 6.5E-04 | 1.37 | 1.2E-06 |
| *Usp2* | 685 | 3.08 | 1.5E-34 | 2.78 | 1.2E-28 |
| *Usp30* | 2549 | 1.25 | 4.6E-09 | 1.15 | 3.5E-04 |
| *Usp45* | 4065 | 1.41 | 1.6E-14 | 1.43 | 1.5E-15 |
| *Vdr* | 170 | 6.19 | 1.5E-10 | 6.71 | 2.3E-11 |
| *Vezf1** | 398 | -1.31 | 4.8E-02 | -1.11 | 4.8E-01 |
| *Vwa7* | 164 | 2.50 | 6.6E-09 | 2.60 | 1.4E-09 |
| *Wac* | 11365 | -1.34 | 1.4E-10 | -1.38 | 6.3E-13 |
| *Wdfy3* | 12067 | -2.06 | 5.2E-11 | -1.89 | 8.4E-09 |
| *Wnt1* | 77 | -2.22 | 1.4E-02 | -2.23 | 1.4E-02 |
| *Xpc* | 3233 | 1.40 | 2.2E-17 | 1.41 | 1.2E-17 |
| *Xrcc6* | 3705 | 1.36 | 7.6E-09 | 1.32 | 2.1E-07 |
| *Yeats2* | 8743 | -1.43 | 3.0E-18 | -1.38 | 3.4E-15 |
| *Ywhae* | 73412 | -1.26 | 1.0E-04 | -1.36 | 1.2E-07 |
| *Ywhaz* | 43502 | -1.35 | 1.8E-10 | -1.44 | 9.7E-15 |
| *Zbtb16* | 881 | 1.86 | 2.2E-04 | 1.70 | 1.7E-03 |
| *Zbtb20* | 6592 | -2.52 | 2.0E-14 | -2.13 | 5.5E-10 |
| *Zc3h4* | 6993 | -2.62 | 3.8E-24 | -2.79 | 3.1E-27 |
| *Zfyve26* | 2417 | -1.46 | 1.9E-09 | -1.33 | 1.0E-05 |
| *Zmiz1* | 19376 | -1.45 | 1.1E-03 | -1.53 | 1.8E-04 |
| *Zmym2* | 16595 | -1.41 | 2.1E-06 | -1.28 | 6.7E-04 |
| *Zmynd11* | 13113 | -1.87 | 7.8E-23 | -1.83 | 1.8E-21 |
| *Zmynd8* | 7708 | -1.94 | 1.1E-19 | -1.96 | 3.1E-20 |

*Significant effect of VPA in one sex only

**Supplemental Table S3**. 399 genes that have been reported to be linked to autism by GWAS (SFARI list) that are also significantly up- or downregulated by VPA (from Table S2) in the fetal mouse brain.
