## Supplemental Table S4 for "Rapid effects of valproic acid on the fetal brain transcriptome: Implications for brain development and autism"

| | | Male | | Female | |
|---|---|---|---|---|---|
| Gene | Counts | FC | $p_{FDR}$ | FC | $p_{FDR}$ |
| Adra2a | 747 | -1.66 | 2.3E-09 | -2.01 | 1.1E-16 |
| Aif1* | 113 | -1.49 | 2.7E-04 | -1.15 | 2.4E-01 |
| Ak2 | 7116 | 1.41 | 2.2E-04 | 1.27 | 1.2E-02 |
| Akap8 | 17422 | -1.26 | 3.2E-05 | -1.29 | 6.1E-06 |
| Akap8l | 8525 | 1.60 | 5.9E-12 | 1.61 | 2.6E-12 |
| Akt1 | 15951 | -1.40 | 5.5E-09 | -1.45 | 4.7E-11 |
| Akt2 | 4906 | -2.03 | 2.9E-50 | -2.08 | 5.6E-54 |
| Amotl2 | 6480 | -1.60 | 8.2E-11 | -1.72 | 4.1E-14 |
| Apbb1ip | 142 | -1.73 | 1.1E-04 | -1.57 | 1.7E-03 |
| Apc2 | 25582 | -1.41 | 3.9E-02 | -1.48 | 1.7E-02 |
| Arc | 170 | 1.64 | 1.7E-04 | 1.66 | 1.4E-04 |
| Arntl | 1203 | 2.17 | 8.8E-45 | 2.07 | 1.6E-39 |
| Asxl1 | 9564 | -1.39 | 7.4E-06 | -1.42 | 1.8E-06 |
| Bcl10 | 2338 | 1.68 | 2.6E-19 | 1.59 | 1.4E-15 |
| Bcl11b | 7112 | -1.58 | 1.7E-02 | -1.67 | 8.1E-03 |
| Bdnf | 423 | 2.57 | 9.7E-12 | 3.10 | 3.3E-16 |
| Bmp2 | 291 | -1.49 | 3.6E-02 | -1.68 | 6.3E-03 |
| Bola2 | 2477 | 1.59 | 2.6E-05 | 1.48 | 5.2E-04 |
| Brca1 | 5043 | 1.43 | 4.0E-03 | 1.48 | 1.7E-03 |
| Brd1 | 7073 | -1.39 | 1.9E-04 | -1.47 | 1.5E-05 |
| Brd2 | 19021 | -1.26 | 1.3E-04 | -1.35 | 6.2E-07 |
| Brwd1 | 13743 | -1.37 | 5.2E-06 | -1.29 | 2.3E-04 |
| C1qtnf1 | 444 | -1.45 | 2.7E-04 | -1.43 | 4.5E-04 |
| C2cd5 | 6100 | -1.54 | 1.9E-20 | -1.50 | 3.3E-18 |
| C9orf72 | 4763 | 1.74 | 6.8E-17 | 1.79 | 2.4E-18 |
| Cacnb4* | 2101 | -1.68 | 3.9E-03 | -1.37 | 9.8E-02 |
| Calm1 | 52234 | 1.44 | 1.0E-23 | 1.44 | 2.4E-24 |
| Camk1 | 1971 | -1.61 | 5.9E-24 | -1.60 | 5.7E-23 |
| Camkk1 | 2077 | 2.66 | 7.2E-41 | 2.53 | 6.9E-37 |
| Camkv | 1980 | 2.50 | 3.4E-28 | 2.30 | 1.1E-23 |
| Carm1 | 6783 | -1.40 | 1.9E-06 | -1.52 | 1.4E-09 |
| Casq2 | 132 | -14.04 | 5.9E-04 | -7.48 | 1.1E-02 |
| Cbln2 | 2008 | -1.82 | 1.5E-05 | -1.71 | 1.2E-04 |
| Cbs | 736 | 5.02 | 2.1E-38 | 5.08 | 7.7E-39 |
| Cd200 | 4220 | -2.05 | 1.3E-04 | -2.05 | 1.2E-04 |
| Cdh7* | 13012 | -1.35 | 8.4E-03 | -1.24 | 6.8E-02 |
| Cdh15* | 136 | -4.45 | 6.9E-03 | -2.61 | 9.7E-02 |
| Clic6 | 1509 | -1.80 | 3.4E-04 | -1.58 | 6.3E-03 |
| Col6a6 | 212 | -4.15 | 3.8E-04 | -4.13 | 4.2E-04 |
| Creb3 | 2520 | 1.42 | 4.5E-15 | 1.43 | 1.2E-15 |
| Cry1 | 3590 | 2.57 | 1.2E-98 | 2.41 | 5.4E-86 |
| Cry2 | 3881 | 2.01 | 2.8E-27 | 1.94 | 1.1E-24 |
| Csf1r | 1501 | -1.39 | 8.8E-04 | -1.46 | 1.2E-04 |
| Csgalnact1 | 179 | -4.40 | 2.5E-08 | -3.40 | 5.4E-06 |
| Cux1* | 12750 | -1.17 | 5.2E-02 | -1.27 | 2.6E-03 |
| Cxcl14 | 1869 | -3.84 | 2.6E-04 | -3.31 | 1.3E-03 |
| Dab2 | 6440 | -1.69 | 1.0E-02 | -1.63 | 1.8E-02 |
| Dcbld2 | 9176 | 1.74 | 2.6E-25 | 1.68 | 3.3E-22 |
| Dcps | 2121 | -1.27 | 3.8E-03 | -1.34 | 3.2E-04 |
| Dhfr | 4614 | 1.56 | 3.7E-04 | 1.38 | 1.1E-02 |
| Dlg5 | 6522 | -2.95 | 3.9E-61 | -2.96 | 1.3E-61 |
| Dlx1* | 5903 | -1.77 | 1.2E-01 | -2.29 | 2.0E-02 |
| Dnajc12 | 1458 | 4.40 | 5.0E-179 | 4.66 | 1.6E-192 |
| Dop1b | 4122 | 1.41 | 4.7E-03 | 1.47 | 1.5E-03 |
| Draxin | 7093 | -1.46 | 1.0E-02 | -1.48 | 8.0E-03 |
| Dyrk1b | 1249 | -2.21 | 3.1E-12 | -2.12 | 3.7E-11 |
| Ebf1 | 6893 | -1.54 | 6.9E-05 | -1.34 | 9.0E-03 |
| Efna3 | 1500 | 2.61 | 1.2E-19 | 2.86 | 3.1E-23 |
| Egfr | 1567 | -1.89 | 1.1E-02 | -1.69 | 3.9E-02 |
| Egr1 | 400 | 1.74 | 2.8E-04 | 1.71 | 4.8E-04 |
| Elavl4 | 13824 | -1.58 | 1.1E-05 | -1.45 | 4.3E-04 |
| Emp2 | 457 | -1.49 | 4.9E-02 | -1.62 | 1.6E-02 |
| Eomes | 3938 | -1.81 | 1.6E-02 | -1.86 | 1.1E-02 |
| Ets2 | 2942 | -1.45 | 8.6E-03 | -1.42 | 1.5E-02 |
| Evx1* | 102 | -3.25 | 1.3E-02 | -1.42 | 5.1E-01 |
| Farp1 | 6684 | -1.72 | 5.0E-11 | -1.72 | 7.1E-11 |
| Fbxl21 | 373 | 3.18 | 2.4E-56 | 3.23 | 2.4E-57 |
| Fbxl3 | 4323 | -1.77 | 5.5E-43 | -1.70 | 2.2E-37 |
| Fezf1* | 950 | -1.20 | 4.6E-01 | -1.68 | 2.3E-02 |
| Fgf2 | 295 | -2.94 | 3.9E-15 | -2.79 | 1.1E-13 |
| Fgf17* | 317 | -1.64 | 1.2E-02 | -1.45 | 6.8E-02 |
| Fgfr1op2 | 6435 | 1.26 | 5.9E-05 | 1.31 | 3.4E-06 |
| Flrt2 | 7693 | -1.73 | 2.6E-08 | -1.52 | 3.0E-05 |
| Fos | 108 | 1.92 | 1.0E-03 | 2.49 | 3.2E-06 |
| Foxa1 | 549 | -1.47 | 2.2E-02 | -1.75 | 5.7E-04 |
| Foxc1 | 1824 | -1.79 | 6.9E-04 | -2.13 | 7.9E-06 |
| Foxo1 | 2595 | 1.84 | 2.8E-13 | 1.80 | 1.8E-12 |
| Foxo3 | 4755 | 1.66 | 1.9E-11 | 1.70 | 2.6E-12 |
| Foxo4 | 3776 | 1.37 | 1.8E-05 | 1.36 | 3.1E-05 |
| Foxo6 | 990 | 1.56 | 1.2E-02 | 1.50 | 2.3E-02 |
| Fzd5 | 608 | 2.35 | 1.0E-10 | 1.80 | 1.3E-05 |
| Gabra1 | 591 | -1.75 | 2.6E-03 | -1.68 | 5.9E-03 |
| Gad1 | 3462 | -2.50 | 1.0E-04 | -2.54 | 7.7E-05 |
| Gad2 | 5532 | -3.18 | 2.9E-08 | -3.38 | 4.9E-09 |
| Gas7 | 1662 | -1.64 | 6.9E-03 | -1.67 | 4.8E-03 |
| Gbx1 | 311 | -2.29 | 3.5E-03 | -2.39 | 2.2E-03 |
| Gch1 | 460 | 5.58 | 7.6E-65 | 5.25 | 1.8E-60 |
| Gfpt2 | 1177 | 1.67 | 1.2E-07 | 1.78 | 2.5E-09 |
| Gls2 | 319 | 4.06 | 2.3E-62 | 4.84 | 4.0E-75 |
| Grb2 | 9573 | 1.44 | 5.9E-08 | 1.37 | 3.4E-06 |
| Grm2 | 321 | -1.39 | 2.8E-02 | -1.49 | 7.1E-03 |
| Grm3 | 394 | -1.84 | 2.7E-03 | -1.59 | 2.7E-02 |
| Hapln1 | 1964 | -4.31 | 1.6E-05 | -3.45 | 3.0E-04 |
| Hapln4 | 275 | 27.66 | 3.9E-126 | 23.71 | 2.8E-119 |
| Hes1 | 1561 | -1.52 | 3.5E-06 | -1.70 | 2.6E-09 |
| Hnrnpab | 47049 | -1.53 | 2.7E-05 | -1.75 | 1.5E-08 |
| Hnrnph1 | 93936 | -1.60 | 9.0E-14 | -1.61 | 5.2E-14 |
| Homer3 | 3874 | -1.27 | 1.0E-02 | -1.43 | 9.4E-05 |
| Hrh3 | 823 | 2.58 | 5.1E-15 | 2.34 | 2.6E-12 |
| Hsp1a | 235 | 6.27 | 1.4E-31 | 5.59 | 6.7E-28 |
| Hsp1b | 412 | 5.93 | 3.1E-35 | 5.20 | 1.9E-30 |
| Hsp2 | 996 | 6.34 | 1.0E-96 | 6.09 | 1.8E-92 |
| Icam5 | 104 | 16.47 | 6.4E-39 | 16.73 | 3.0E-37 |
| Insyn1 | 1174 | -1.42 | 1.6E-02 | -1.44 | 1.2E-02 |

### Table S4 (cont)

| | | Male | | Female | |
|---|---|---|---|---|---|
| Gene | Counts | FC | pFDR | FC | pFDR |
| *Itpka* | 175 | 17.28 | 1.1E-58 | 10.83 | 3.3E-44 |
| *JunB* | 206 | 2.19 | 5.8E-10 | 2.20 | 5.4E-10 |
| *Kbtbd2* | 4187 | -1.24 | 1.2E-05 | -1.31 | 2.0E-08 |
| *Kcnmb2** | 329 | -2.11 | 5.5E-04 | -1.45 | 1.1E-01 |
| *Kif17* | 387 | 4.17 | 2.5E-27 | 4.83 | 1.1E-32 |
| *Klf4* | 882 | -1.56 | 1.3E-02 | -1.59 | 9.9E-03 |
| *Kras* | 16185 | 1.38 | 2.1E-07 | 1.32 | 6.2E-06 |
| *Lamp5* | 587 | -6.05 | 9.0E-04 | -4.36 | 7.7E-03 |
| *Lhx8* | 610 | -2.58 | 7.4E-03 | -2.81 | 3.5E-03 |
| *Lmx1a* | 1011 | -1.57 | 1.5E-03 | -1.81 | 1.9E-05 |
| *Lrp6** | 10654 | -1.26 | 1.0E-04 | -1.14 | 3.4E-02 |
| *Lrrk1* | 709 | -2.99 | 2.8E-13 | -2.85 | 3.1E-12 |
| *Lrrk2* | 647 | -2.40 | 1.1E-08 | -2.20 | 2.9E-07 |
| *Maf* | 3212 | -1.63 | 3.8E-06 | -1.64 | 2.8E-06 |
| *Mafb* | 999 | -1.57 | 8.5E-05 | -1.71 | 2.2E-06 |
| *Map2* | 68999 | -1.59 | 2.1E-04 | -1.41 | 7.8E-03 |
| *Map2k1* | 4241 | 1.70 | 2.3E-18 | 1.67 | 3.6E-17 |
| *Mapk1* | 20963 | 1.33 | 3.2E-15 | 1.29 | 7.0E-13 |
| *Mapk14* | 7022 | 1.55 | 4.6E-40 | 1.52 | 9.9E-36 |
| *Marcks* | 64056 | -2.10 | 3.1E-25 | -2.26 | 2.7E-30 |
| *Mcam* | 4295 | 1.79 | 4.6E-17 | 1.65 | 5.1E-13 |
| *Mef2d* | 6108 | 1.54 | 1.3E-05 | 1.47 | 1.1E-04 |
| *Metrn* | 4092 | 2.39 | 5.0E-10 | 2.07 | 2.6E-07 |
| *Mllt11* | 16031 | -1.56 | 2.5E-03 | -1.49 | 8.0E-03 |
| *Ncoa3* | 5325 | 1.35 | 1.9E-06 | 1.38 | 3.7E-07 |
| *Ndst3* | 2340 | -1.64 | 9.5E-05 | -1.47 | 3.0E-03 |
| *Nell2** | 9177 | -1.22 | 5.5E-02 | -1.29 | 1.1E-02 |
| *Neurod1* | 2403 | -1.43 | 1.4E-03 | -1.58 | 3.5E-05 |
| *Neurog1* | 475 | -2.00 | 4.6E-03 | -2.49 | 1.5E-04 |
| *Neurog2** | 6066 | -1.37 | 1.4E-01 | -1.66 | 1.3E-02 |
| *Nfatc1* | 283 | -1.52 | 3.3E-04 | -1.63 | 2.2E-05 |
| *Nfatc2* | 689 | -1.65 | 4.9E-03 | -1.60 | 8.5E-03 |
| *Nfatc4* | 3812 | -2.04 | 9.2E-09 | -2.10 | 2.2E-09 |
| *Nhej1* | 194 | -2.15 | 1.1E-16 | -2.40 | 3.4E-21 |
| *Nhlh1* | 2918 | -1.63 | 8.1E-06 | -1.62 | 1.1E-05 |
| *Nhlh2* | 5362 | -2.89 | 1.7E-16 | -2.74 | 4.9E-15 |
| *Nkx2-1** | 2086 | -1.74 | 1.2E-01 | -2.46 | 8.6E-03 |
| *Nkx2-4*†* | 120 | -1.30 | 2.9E-01 | -2.42 | 8.7E-05 |
| *Nprl3* | 1675 | -1.37 | 3.3E-11 | -1.29 | 1.2E-07 |
| *Nras* | 15777 | -1.36 | 4.5E-09 | -1.42 | 1.2E-11 |
| *Ntn4* | 1222 | -1.88 | 2.1E-07 | -2.09 | 1.1E-09 |
| *Odc1* | 24222 | 1.49 | 5.0E-06 | 1.43 | 5.3E-05 |
| *Ogt* | 39637 | -1.51 | 1.5E-06 | -1.30 | 2.8E-03 |
| *Olig1** | 341 | 2.11 | 7.4E-05 | 1.49 | 4.5E-02 |
| *Otp* | 1318 | -2.64 | 2.1E-08 | -2.31 | 1.8E-06 |
| *P2rx7* | 210 | -2.26 | 6.8E-06 | -1.67 | 6.3E-03 |
| *Palm3* | 720 | 1.99 | 7.7E-06 | 1.89 | 4.2E-05 |
| *Pax7* | 1240 | -1.74 | 3.4E-03 | -1.68 | 7.0E-03 |
| *Pdk1* | 4064 | 1.24 | 1.1E-02 | 1.33 | 6.5E-04 |
| *Per3* | 1173 | 1.85 | 8.7E-21 | 1.92 | 5.5E-23 |
| *Pigh* | 1044 | -1.39 | 5.5E-07 | -1.38 | 8.7E-07 |
| *Pigp* | 2067 | -1.62 | 3.5E-20 | -1.60 | 1.9E-19 |

| | | Male | | Female | |
|---|---|---|---|---|---|
| Gene | Counts | FC | pFDR | FC | pFDR |
| *Pik3cb* | 2150 | 1.73 | 3.6E-22 | 1.77 | 1.2E-23 |
| *Pik3r4* | 3800 | 1.26 | 3.3E-05 | 1.24 | 1.7E-04 |
| *Plcg2* | 339 | 1.95 | 1.2E-07 | 1.74 | 1.6E-05 |
| *Plk1* | 3447 | -1.54 | 5.4E-03 | -1.83 | 7.0E-05 |
| *Pnisr* | 26071 | -1.95 | 1.5E-24 | -1.81 | 1.0E-19 |
| *Ppm1j* | 282 | 7.24 | 1.4E-36 | 7.60 | 4.4E-38 |
| *Ppp3cb* | 14492 | -1.25 | 1.0E-02 | -1.29 | 3.0E-03 |
| *Ppp3cc* | 516 | -1.27 | 6.9E-03 | -1.25 | 1.2E-02 |
| *Ptk2b* | 117 | 1.75 | 1.6E-03 | 1.68 | 3.7E-03 |
| *Ptpn1* | 7485 | 1.59 | 8.2E-10 | 1.45 | 1.3E-06 |
| *Ptpn5* | 4140 | -1.59 | 1.2E-02 | -1.49 | 3.7E-02 |
| *Ptpro* | 3982 | -1.70 | 1.0E-02 | -1.67 | 1.4E-02 |
| *Pttg1ip* | 4768 | 1.30 | 1.4E-04 | 1.28 | 3.9E-04 |
| *Pura* | 2394 | 1.32 | 2.7E-05 | 1.35 | 5.1E-06 |
| *Raf1* | 11657 | -1.62 | 2.3E-24 | -1.60 | 1.8E-23 |
| *Rarb* | 1030 | -2.09 | 2.6E-05 | -2.27 | 3.0E-06 |
| *Rcan1* | 3802 | 1.67 | 4.1E-27 | 1.59 | 4.6E-22 |
| *Rcan1* | 3802 | 1.67 | 4.1E-27 | 1.59 | 4.6E-22 |
| *Rmnd5b* | 2194 | 1.35 | 3.1E-10 | 1.32 | 7.3E-09 |
| *Rnd3* | 17191 | -1.47 | 1.3E-07 | -1.44 | 6.9E-07 |
| *Rnf19b* | 2770 | 1.38 | 7.2E-05 | 1.32 | 7.2E-04 |
| *Robo3** | 6624 | -1.73 | 5.8E-03 | -1.23 | 3.4E-01 |
| *Robo4* | 695 | -2.11 | 1.5E-17 | -2.21 | 1.1E-19 |
| *Rtn4rl1* | 469 | 2.04 | 4.0E-09 | 2.01 | 7.6E-09 |
| *Runx1* | 381 | -3.19 | 1.6E-04 | -2.98 | 4.3E-04 |
| *Runx2* | 782 | -3.89 | 1.5E-05 | -3.99 | 1.1E-05 |
| *Rxrb* | 4358 | -1.27 | 5.6E-06 | -1.34 | 3.0E-08 |
| *S1pr1* | 1963 | -1.25 | 2.4E-06 | -1.38 | 4.0E-12 |
| *Sema3a* | 2749 | -2.24 | 2.5E-07 | -1.77 | 3.5E-04 |
| *Setd4* | 686 | -1.85 | 1.3E-19 | -1.92 | 5.7E-22 |
| *Sh3bgr* | 135 | 1.74 | 1.1E-02 | 1.97 | 1.9E-03 |
| *Shc2* | 2467 | -1.42 | 1.9E-05 | -1.41 | 3.2E-05 |
| *Shc4* | 325 | -1.69 | 1.4E-05 | -1.50 | 1.0E-03 |
| *Shox2* | 3795 | -1.95 | 6.0E-05 | -2.26 | 8.6E-07 |
| *Sim2* | 224 | -2.06 | 9.2E-03 | -2.27 | 2.9E-03 |
| *Sirt1* | 3366 | -1.49 | 5.2E-15 | -1.48 | 1.1E-14 |
| *Slc17a6* | 5910 | -2.41 | 2.9E-09 | -2.42 | 2.4E-09 |
| *Slc17a7* | 847 | 1.63 | 1.3E-02 | 1.63 | 1.4E-02 |
| *Slc17a8* | 404 | -1.72 | 1.1E-03 | -1.43 | 3.8E-02 |
| *Slc17a9* | 221 | 5.87 | 1.2E-33 | 6.74 | 2.1E-38 |
| *Slc35e1* | 5103 | 1.48 | 2.7E-16 | 1.42 | 3.2E-13 |
| *Slc8a1* | 4042 | -2.30 | 2.5E-05 | -1.79 | 4.2E-03 |
| *Slc8a2* | 3842 | 1.99 | 4.1E-15 | 2.04 | 4.2E-16 |
| *Slc9a3r1* | 2902 | 3.04 | 2.4E-31 | 2.94 | 1.9E-29 |
| *Slit2* | 10361 | 1.31 | 2.6E-04 | 1.46 | 2.2E-07 |
| *Slitrk1* | 1872 | -1.44 | 8.6E-03 | -1.46 | 6.4E-03 |
| *Slitrk4* | 1131 | 1.62 | 2.8E-05 | 1.72 | 2.3E-06 |
| *Smarcb1* | 7275 | -1.64 | 2.2E-12 | -1.76 | 7.7E-16 |
| *Smyd2* | 1864 | -1.24 | 3.7E-04 | -1.28 | 4.3E-05 |
| *Sos1* | 9101 | 1.25 | 6.5E-03 | 1.29 | 2.0E-03 |
| *Sox2** | 10110 | -1.21 | 3.3E-01 | -1.50 | 2.4E-02 |
| *Sp1* | 8616 | -1.26 | 7.3E-07 | -1.20 | 7.9E-05 |

### Table S4 (cont)

| | | Male | | Female | |
|---|---|---|---|---|---|
| Gene | Counts | FC | $p_{FDR}$ | FC | $p_{FDR}$ |
| *Sp2* | 1254 | -1.49 | 2.2E-03 | -1.63 | 1.5E-04 |
| *Sp3* | 9744 | -1.51 | 3.1E-11 | -1.49 | 7.0E-11 |
| *Sparc* | 25277 | -1.37 | 2.6E-02 | -1.40 | 1.8E-02 |
| *Src* | 5937 | -2.49 | 4.8E-21 | -2.70 | 7.6E-25 |
| *Srebf1** | 4265 | 1.38 | 5.8E-04 | 1.21 | 5.7E-02 |
| *Sst* | 803 | -1.79 | 1.0E-02 | -1.86 | 6.2E-03 |
| *Stat3* | 5850 | 2.72 | 1.2E-44 | 2.69 | 1.7E-43 |
| *Stc1* | 534 | 1.35 | 8.9E-03 | 1.35 | 9.0E-03 |
| *Stc2* | 494 | 2.83 | 2.1E-23 | 2.55 | 4.8E-19 |
| *Stx18* | 1175 | -1.59 | 1.1E-09 | -1.68 | 9.8E-12 |
| *Sumo3* | 6505 | -1.39 | 4.8E-09 | -1.51 | 3.1E-13 |
| *Tacc2* | 7124 | -1.50 | 5.9E-06 | -1.52 | 2.5E-06 |
| *Tctn1* | 4106 | 1.47 | 1.8E-08 | 1.41 | 6.7E-07 |
| *Tctn3* | 1997 | 1.94 | 6.3E-22 | 1.99 | 1.9E-23 |
| *Tfap2a* | 954 | -1.85 | 2.8E-03 | -1.71 | 1.1E-02 |
| *Tgfb2* | 2234 | -2.52 | 4.1E-13 | -2.34 | 2.9E-11 |
| *Tgfb3* | 1115 | -2.03 | 1.7E-04 | -1.96 | 4.0E-04 |
| *Th* | 476 | -1.42 | 5.1E-03 | -1.52 | 6.6E-04 |
| *Tlr3* | 167 | -1.75 | 1.2E-10 | -1.95 | 2.6E-14 |
| *Tmem108* | 1225 | 1.53 | 7.9E-06 | 1.37 | 1.4E-03 |
| *Tmem163* | 1496 | -1.62 | 1.4E-05 | -1.68 | 2.4E-06 |
| *Top3a* | 2357 | 1.71 | 1.3E-20 | 1.66 | 1.2E-18 |
| *Trib2* | 8489 | -1.73 | 3.2E-07 | -1.73 | 3.4E-07 |
| *Trnp1* | 298 | 1.45 | 3.8E-02 | 1.55 | 1.4E-02 |
| *Trpm4* | 1776 | 2.04 | 1.2E-15 | 2.11 | 5.0E-17 |
| *Tubb3* | 58234 | -1.25 | 1.8E-02 | -1.27 | 1.4E-02 |
| *Tug1* | 14599 | -1.19 | 1.6E-04 | -1.26 | 2.8E-07 |
| *Twf2* | 1910 | 1.56 | 1.4E-10 | 1.58 | 3.3E-11 |
| *Unc5b* | 3707 | 1.69 | 4.9E-08 | 1.41 | 4.4E-04 |
| *Unc5c* | 4631 | -2.20 | 1.5E-06 | -1.86 | 2.1E-04 |
| *Usp34** | 19191 | -1.41 | 6.4E-03 | -1.17 | 2.7E-01 |
| *Vegfc* | 373 | -1.52 | 4.0E-06 | -1.73 | 1.1E-09 |
| *Vegfd* | 111 | -2.21 | 2.6E-04 | -1.91 | 3.5E-03 |
| *Wdr4* | 3027 | -1.43 | 2.4E-11 | -1.47 | 9.8E-13 |
| *Wdr41* | 1513 | -2.70 | 1.1E-116 | -2.73 | 2.1E-118 |
| *Wnt10b* | 116 | -3.73 | 2.4E-03 | -3.29 | 6.8E-03 |
| *Wnt16* | 127 | 2.71 | 1.2E-03 | 3.13 | 2.0E-04 |
| *Wnt3** | 639 | -1.19 | 1.3E-01 | -1.37 | 5.3E-03 |
| *Wnt3a* | 207 | -3.64 | 1.6E-12 | -4.01 | 2.7E-14 |
| *Wnt4** | 470 | -1.70 | 2.3E-02 | -1.39 | 1.8E-01 |
| *Wnt7b* | 4194 | -1.68 | 1.4E-06 | -1.97 | 1.5E-10 |
| *Wsb1* | 31033 | -1.42 | 6.3E-08 | -1.35 | 2.8E-06 |
| *Zfp804a* | 553 | -1.88 | 3.6E-03 | -1.68 | 2.0E-02 |
| *Zmym4* | 12662 | -1.53 | 7.8E-14 | -1.45 | 5.7E-11 |

* Significant effect of VPA in one sex only

**Supplemental Table S4**. 252 genes involved in brain development and neurodevelopmental disorders that are dysregulated in the fetal brain by VPA (from Table S2) that have ***not*** been reported to be associated with autism in GWAS.
